## Supplemental Tables, Figures,and Datasets for "Identification of novel tail-anchored membrane proteins integrated by the bacterial twin-arginine translocase"

#### SUPPLEMENTARY INFORMATION

| NAME | SEQUENCE 5´→3´ [a] | USE[b] |
|---|---|---|
| JES3 | ACAGAGGAGGATCCATGTCACTCAGTCGGCGTCAG | RL |
| JES4 | TTGCGCTCGAGCTCCGGTACCGGATTGACCAACAG | RL |
| C1F | TATAGAGCTCCTGACCGACCCGCGTA | RL |
| C1R | TATAGCATGCTTAACCCCAGGTAGCA | RL |
| C2F | TATAGAGCTCACCGCTGCTCCGGACA | RL |
| C2R | TATAGCATGCTTATTAAGCCTGACCACCACC | RL |
| C6F | TATAGAGCTCTCTACCCTGCCGCTGG | RL |
| C6R | TATAGCATGCTTAAGAAGCACGAGCA | RL |
| C8F | TATAGAGCTCGCTCTGGGTCCGGCTG | RL |
| C8R | TATAGCATGCTTACGGACGACGACGTT | RL |
| SP1F | TATAGGATCCATGCGTCACGCTCG | RL |
| SP1R | TATATCTAGAAGCAACAGCGTTGGTA | RL |
| SP2F | TATAGGATCCATGACCACCTCTTCTC | RL |
| SP2R | TATATCTAGATAGCAGCAGCCTGGTCC | RL |
| SP4F | TATAGGATCCATGGGTAACGCTGTTT | RL |
| SP4R | TATATCTAGAAGCCTGAGCCGGAGCAGAAC | RL |
| SP8F | TATAGGATCCATGTCTCTGCGTCGTC | RL |
| SP8R | TATATCTAGAAGCCAGAGCAGCCGGAGCAGCCA | RL |
| SP9F | TATAGGATCCATGTCTGGTATGGC | RL |
| SP9R | TATATCTAGAAGCAACTTCCGGAGCC | RL |
| ESN102 | TGTAAAACGACGGCCAGT | RL |
| ESN103 | CAGGAAACAGCTATGAC | RL |
| ESN237 | AGCTGGCACACCACCTTCCAC | RL |
| ESN434 | GTGGAAGGTGGTGTGCCAGCT | RL |
| ESN526 | CGTATGTTGTGTGGAATTGTGAGC | SEQ |
| ESN527 | GGGTAACGCCAGGGTTTTCCC | SEQ |
| ESN538 | GTGACCTGGATCGAGAACAACG | SEQ |
| ESN539 | CGGACAAGTGGAGCTGCTTG | SEQ |
| ESN540 | CAGTGAGTGGATCC*ATG*ACCTCTCGTCGTGGTACCTTC | RL |
| ESN541 | CCTAGTTCCTCTAGAAGCCAGAGCCGGCGGAGCCAG | RL |
| ESN542 | CAGTGAGTGGATCC*ATG*CCGGCGACCACCGGTGATC | RL |
| ESN543 | CCTAGTTCCTCTAGACGCCGCCGCGGTGGTCGGCG | RL |
| ESN544 | CAGTGAGTGGATCC*ATG*GGTATCGCTGCTTCTGGTCGTC | RL |
| ESN545 | CCTAGTTCCTCTAGAAGCCAGAGCACGAACACCCAG | RL |
| ESN546 | CAGTGAGTGGATCC*ATG*AGCCGTCGTCGTACCGCGCTG | RL |
| ESN547 | CCTAGTTCCTCTAGACGCGCTCGCGCTGGTCGCC | RL |
| ESN550 | CAGTGAGTGAGCTCGCTGGTACCCCGCTGGCTGTTCTG | RL |
| ESN551 | CCTAGTTCCGCATGC*TTA*CAGACCCAGGGTACGCTGAACC | RL |
| ESN552 | CAGTGAGTGAGCTCCAGACCGGTGGTGACCTGGCTGC | RL |
| ESN553 | CCTAGTTCCGCATGC*TTA*ACCGTTACGAGCAGCCATACGACGAC | RL |
| ESN554 | CAGTGAGTGAGCTCGGTAGCGACGCGCCGGTTGCG | RL |
| ESN555 | CCTAGTTCCGCATGC*TTA*CGCCTGCGCATCACGCGCACC | RL |
| ESN556 | CAGTGAGTGAGCTCGACGATGCGACCCCGGCGGCG | RL |
| ESN557 | CCTAGTTCCGCATGC*TTA*CGCACCCGCCGCACGACGAC | RL |
| ESN565 | CAGTGAGTGAGCTCGTTGCTTCTCTGCCGTGGCTG | RL |
| ESN566 | CCTAGTTCCGCATGC*TTA*CGGCTGAGCTTCGGTCAGC | RL |
| ESN584 | CAGTGAGTGAGCTCCCGGAGCTGTATCACGGTC | RL |
| ESN585 | CCTAGTTCCGCATGC*TTA*CTCATGATGATCCTCCTCGTC | RL |
| ESN591 | GGTCAATCCGGTACCG*TAA*GCATGCAAGCTTGGCGTAATCATGG | QC |
| ESN592 | C*TTA*CGGTACCGGATTGACCAACAGTTGCCCAATCGAGCCAC | QC |
| ESN669 | AACGCAGCTACTCTGTGGTATC | SEQ |
| ESN675 | CAGAGCAGCCAGACCCAGAGC | KLD |

| | | |
|---|---|---|
| ESN676 | CTGGGTGGT**GCC**TCTGACGCTC | KLD |

**Table S1. Oligonucleotides used in this study.**

a = restriction sites used for restriction-ligation cloning of SP/CT-encoding PCR products are underlined (please note that, unlike the gBlocks for SP/CT 1-9, those for SP/CT 10-16 were synthesised to include these sites already and the target sequences were amplified from conserved extensions for the re-usable oligos ESN102, ESN103, ESN237, and ESN434). The start codons of SP-coding sequences and the stop codons of CT-coding sequences are in italics. The C24A mutant codon of SP9 is in bold.

b = molecular application of each primer. RL: restriction-ligation cloning; QC: QuickChange whole-plasmid PCR; KLD: whole-plasmid PCR circularised with NEB's KLD kit; SEQ: sequencing (ESN526 and ESN527: annealing to opposite ends of the multiple cloning site of the pSU40 backbone; ESN538: internal *amiA* primer; ESN539 and ESN669: internal *sufI* primers).

| NAME | SEQUENCE |
|---|---|
| SP1 | ATGCGTCACGCTCGTCGTCGTGTTGTTCGTCGTGTTACCCGTCTGGCTGCTGTTGGTGGTCTGCTGCTGGGTGGTGCTATGGTTACCAACGCTGTTGCT |
| SP2 | ATGACCACCTCTTCTCCGTCTGCTCCGCGTCGTCGTATCCTGCGTTCTGCTACCGCTCTGGTTGGTGGTGTTGCTCTGGCTGTTGCTGTTCCGCTGGCTGCTTCTGCTCACGTTCGTGTTTCTCCGGACCAGGCTGCTGCT |
| SP3 | ATGACCTCTCGTCGTGGTACCTTCCTGGCTGCTCTGGTTACCGCTTCTCTGATCCCGCTGGCTCCGCCGGCTCTGGCT |
| SP4 | ATGGGTAACGCTGTTTCTGGTCGTCGTACCCTGCTGTCTGGTACCGCTGTTCTGGCTGCTGTTGCTCTGATCGCTCTGGGTTCTGCTCCGGCTCAGGCT |
| SP5 | ATGCCGGCGACCACCGGTGATCGTAGCCGTCGTCGTCCGCTGGCGTTTGCGGCGGCGGTTGCGACCGCGGCGGCGATTGGCGCGGCGAGCTTGGCGGCGGCGCCGACCACCGCGGCGGCG |
| SP6 | ATGGGTATCGCTGCTTCTGGTCGTCGTACCCTGCTGTCTGCTACCGCTGTTTCTGCTACCGCTGCTCTGATCGCTCTGGGTGCTGCTCCGGCTCAGGCTGACGCTATCAAACCGGACCTGGGTGTTCGTGCTCTGGCT |
| SP7 | ATGAGCCGTCGTCGTACCGCGCTGCTGACCGCGCCGGCTGCTGCGGCGGCGGCGGCGCTGACCCTGGTGGGCGCGGCGACCAGCGCGAGCGCG |
| SP8 | ATGTCTCTGCGTCGTCGTGCTGCTCTGCTGGTTACCGCTGCTTCTACCCTGACCGCTCTGGCTGCTCCGGCTGCTCTGGCT |
| SP9 | ATGTCTGGTATGGCTGGTCGTCGTTCTCTGGTTGCTGCTGCTCTGGGTCTGGCTGCTCTGCTGGGTGGTTGCTCTGACGCTCGTCCGCCGGCTCCGGAAGTTGCT |
| SP10 | ATGACGCCTAACCGCCGTCCTGTGGTACTTGCCGCAGCGGTAGCAACCGCTTTGGGGTTAGGGGCTACTGCTCTTACGTTACCGGCTGTGGCG |
| SP11 | ATGTTAACACGCCGCAACGCTTTAAGCGCGCTTGCCTTAGGCGTCGTCGGAGTGTCGATCGCTCCAACAGCCTTGGCG |
| SP12 | ATGAAGAACCTTAAGCGCGTACCTTTGGCAGGGCGTCGCACAGCTGTCGCCGCGGCCGCCGCAGCCACTGCGGTGGCATTAGCAGGCCCTGCTTACGCG |
| SP13 | ATGAACAGTACAACCTCGCGTCCAAGTGCACGTCGCCGCACTATTGCCGCTGCTGTAGCCGCATTCGCTGCTGGTCCAGCTCTTCTTACTGCACCCGCCGCACATGCG |
| SP14 | ATGAGTCCTATTGGTCGCCGCCGCTGGTTACACGTGTCTTGTGGGGGATCAACTATCTTAAACTCTAATACCTTTCGCCTTGCTGCCCTTGCTGTAGCTGCTGCTCCTGTAGCCCTTCTGGTGGCCGCACCTGCCCATGCG |
| SP15 | ATGCCTGCGCCGAAGAAAGCGCTTCGCCGTTCTTTGACTGTTACCTCGGCCGCGTTAACAGCTTTGGCATTGCTGTCTCCTGCAGCTCTTGCG |
| SP16 | ATGCGCGACTCCTCTGGCCTTTCGCGCCGTGCAGCCTTGGGACTTGCAGCGGCCTTGCCCTTAGCTGCAGCAACGGCAACGACAGCTGATGCG |
| CT1 | CTGACCGACCCGCGTAACGTTGGTCCGGGTCTGCTGGTTATCGCTGGTTCTCTGGTTGCTCTGGTTGCTACCCGTTGGATCCGTGCTGAACAGGACCGTAAAGCTTACCGTCAGCACTACTCTGCTACCTGGGGTTAA |

| | |
|---|---|
| CT2 | ACCGCTGCTCCGGACACCACCGTTACCACCGCTGCTTCTGACACCTCTGCTACCTCTTCTGCTGTTGCTGTTGGTCTGGGTGTTGGTGGTCTGGCTCTGGGTGCTGTTGCTCTGGTTGTTGCTGTTTTCGCTCTGACCCGTGTTCGTCGTGAAGGTGGTGGTCAGGCTTAA |
| CT3 | GCTGGTACCCCGCTGGCTGTTCTGCTGGGTCTGTTCGCTGCTATCGCTGTTGCTGTTGGTGCTATCAAACCGCTGCACTCTTTCCTGCTGCAGGTTCAGCGTACCCTGGGTCTGTAA |
| CT4 | CAGACCGGTGGTGACCTGGCTGCTACCGGTTCTGACTCTACCCTGCCGGTTGCTGGTGCTGCTGGTGCTGCTCTGCTGGCTGGTGGTGGTCTGTTCTACGCTATGCGTCGTCGTATGGCTGCTCGTAACGGTTAA |
| CT5 | GGTAGCGACGCGCCGGTTGCGGCGCTGGGCACCGCGGCGGCGCTGGCGGTGGCGGCGGGTGCGGGCGTGGTTTTCGCGGTTCGTCGTCGTCGTGGTGCGCGTGATGCGCAGGCGTAA |
| CT6 | TCTACCCTGCCGCTGGCTGGTGCTGCTGGTGCTGCTCTGCTGGCTGGTGGTTCTATCGTTTGGGCTGTTCGTCGTCGTTCTGCTGCTCGTGCTTCTTAA |
| CT7 | GACGATGCGACCCCGGCGGCGCGTGCGCAAGTGCGTGCGAGCGCGCAAGAGCGTAGCGAAGTTCTGGCGGCGACCGGTGCGCGTACCGGCGTGCTGCTGGCGGCGGGTGCGCTGGCGCTGGGTCTGGGTGCGGGTCTGGTTACCTGGCGTCGTCGTCGTGCGGCGGGTGCGTAA |
| CT8 | GCTCTGGGTCCGGCTGCTACCAAACGTGCTACCGGTGCTGAACTGCGTTCTGACGACAAAAAAGACGACGGTCTGTCTTCTTCTGCTACCACCTGGATCATCGTTGGCGTTGTATTCGTCGCTTCTGCTGGTTTCGGTCTGCTGCTGTCTGGTCGTAAACGTCGTCGTCCGTAA |
| CT9 | GTTGCTTCTCTGCCGTGGCTGCCGGTTGCTGCTATCGGTCTGCTGCTGGTTCGTGTTCTGTGGCGTCTGCGTCGTCGTCGTCGTGCTCGTATCGGTCGTCCGCTGACCGAAGCTCAGCCGTAA |
| CT10 | GGCGCACTTGCGTCTACTGGTGCCGAAGTCAACGCTGGGGCCCTGGCAGGGGCTGCCGCATTAGTGGTCGCCGCGGGCGCTGGCGTGGTGGTGGCGGCGTAA |
| CT10* | GGCGCACTTGCGTCTACTGGTGCCGAAGTCAACGCTGGGGCCCTGGCAGGGGCTGCCGCATTAGTGGTCGCCGCGGGCGCTGGCGTGGTGGTGGCGGCGCGTCGCCGTAATGCTGCCGGGGATGCCGCTAGCTAA |
| CT11 | GGCGCGGACGCACTGTCCTTATTAGGAATCGCAGGGGCTGCTTCGGTAGCAGGTGCCGGGTTTATGTTATCACGTCGCAAAAACGACTCGGAATAA |
| CT12 | GGTAGTAGTGCGACTACCGTTGCAATGGGTGCAGCAGCGACAGGACTTGTCTTAGCAGGAGCCGGTGCTCTGTATGCTGTCCGCCGTCGTGCGAATAGCTAA |
| CT13 | GGTAGCTCGTTGACTCCATACTTGGCTGGAGGATCGTTGGCGCTTTTAGCTGGGGGAGCCGGCGCACTTGCGCTTACGCGCCGTGCCCGTGCGGCTCGCGCTCGTGGCTAA |
| CT14 | GGTAGCAGCACTACGCCCTACATTGCGGGCGGTGCCGCTTTGTTGTTAGCTTTAGGGGCTGGTGCAATGGTAGTAGCCCGCCGCCGTGGAGCCCAGGATTAA |
| CT15 | GGCGCCGGCGTCGCTGCTGGCATCGCGGCTGGGGTTTTAGCATTGTTAGCAGGAGGAGCTTTTCTGATTAACCGTAAATGGCCGTTGCCAGATTTGGTCCGCCGTCGTACCCGCCCGTAA |

| | |
|---|---|
| CT16 | GGCGCACCGTGGGGTTTAGTTGGAGGTGCAGGTGCTGCCGTCGCACTGCTTGCAGGCGGAGCACTGGCCTTGCGCCGTCGT<br>CGTCGCGCAGCAGGTGAAGCAAGTGAACCAGTAGATGCTGTAGAGGAAGGTGCTGCTGAACCTAGCGAAGCTAAGGGTCGG<br>CGTCGTCGTCGTTAA |

**Table S2.** DNA sequences of codon-optimised signal peptide and C-tail coding sequences used in this study.

| Accession | Organism | Genomic module |
| --- | --- | --- |
| WP_013506165.1 | *Desulfurispirillum indicum* | FeS-CytB |
| WP_014856242.1 | *Melioribacter roseus* | FeS-quinol oxidase-CytC3 |
| WP_049676148.1 | *Desulfocarbo indianensis* | FeS-quinol oxidase- octahaem CytC |
| WP_028325305.1 | *Desulfatirhabdium butyrativorans* | FeS-quinol oxidase- octahaem CytC |
| WP_054030243.1 | *Desulfatitalea tepidiphila* | FeS-quinol oxidase- CytC3- octahaem CytC |
| WP_020589710.1 | *Desulfobacter curvatus* | FeS-CytB |
| WP_004071579.1 | *Desulfobacter postgatei* | FeS-CytB |
| WP_084070237.1 | *Desulfobacterium vacuolatum* | FeS-quinol oxidase- octahaem CytC |
| WP_014957271.1 | *Desulfobacula toluolica* | FeS-CytB |
| WP_012173729.1 | *Desulfococcus oleovorans* | FeS-CytB |
| WP_054692440.1 | *Desulfosarcina cetonica* | FeS-quinol oxidase-CytC3-CytC3 |
| WP_022663865.1 | *Desulfospira joergensenii* | FeS-CytB |
| WP_015723347.1 | *Desulfobulbus propionicus* | CytC3-FeS-quinol oxidase |
| WP_027370080.1 | *Desulfovermiculus halophilus* | FeS-quinol oxidase- CytC3- octahaem CytC |
| WP_071544310.1 | *Desulfovibrio dechloracetivorans* | FeS-CytB |
| WP_071546240.1 | *Desulfovibrio dechloracetivorans* | FeS-quinol oxidase- octahaem CytC |
| WP_062251202.1 | *Desulfovibrio fairfieldensis* | FeS-CytB |
| WP_066806246.1 | *Desulfovibrio indicus* | FeS-quinol oxidase-octahaem CytC |
| WP_008683453.1 | *Desulfovibrio sp. 3_1_syn3* | FeS-CytB |
| WP_009302998.1 | *Desulfovibrio sp. 6_1_46AFAA* | FeS-CytB |
| WP_014259604.1 | *Desulfovibrio africanus* | FeS-CytB |
| WP_027368072.1 | *Desulfovibrio africanus* | FeS-CytB |
| WP_029459533.1 | *Desulfovibrio alcoholivorans* | FeS-CytB |
| WP_020886939.1 | *Desulfovibrio alkalitolerans* | FeS-CytB |
| WP_014323720.1 | *Desulfovibrio desulfuricans* | FeS-quinol oxidase-octahaem CytC |

| | | |
|---|---|---|
| WP_022659754.1 | *Desulfovibrio desulfuricans* | FeS-CytB |
| WP_024823537.1 | *Desulfovibrio magneticus* | FeS-CytB |
| WP_015862494.1 | *Desulfovibrio magneticus* | FeS-CytB |
| WP_015416321.1 | *Desulfovibrio piezophilus* | FeS-CytB |
| WP_027985824.1 | *delta proteobacterium PSCGC 5451* | FeS-CytB |
| WP_003398576.1 | *Anoxybacillus flavithermus* | FeS-CytB |
| WP_008409599.1 | *Desulfotomaculum hydrothermale* | FeS-CytB |
| WP_012447020.1 | *Natranaerobius thermophilus* | FeS-CytB |
| WP_011418038.1 | *Syntrophus aciditrophicus* | FeS-CytB |
| YP_002248525.1 | *Thermodesulfovibrio yellowstonii* | FeS-CytB |
| WP_051534117.1** | *Desulfitibacter alkalitolerans* DSM 16504 | FeS-CytB |
| WP_028318203.1** | *Desulfobulbus elongatus* DSM 2908 | CytC3-FeS-quinol oxidase |
| WP_028580723.1** | *Desulfogranum japonicum* DSM 18378 | CytC3-FeS-quinol oxidase |
| WP_035247752.1** | *Desulfogranum mediterraneum* DSM 13871 | CytC3-FeS-quinol oxidase |
| WP_020676593.1** | *Geopsychrobacter electrodiphilus DSM 16401* | CytC3-FeS-quinol oxidase |
| WP_004088007.1** | *Helicobacter bilis* | FeS-CytB |
| WP_072908093.1** | *Malonomonas rubra DSM 5091* | CytC3-FeS-quinol oxidase |
| WP_053334878.1** | *Thermoanaerobaculum aquaticum* | CytC3-FeS-quinol oxidase |

Table S3. 4Fe-4S tail-anchored ferredoxin proteins, and partner proteins with which they are co-encoded at the same genomic locus. ** Indicates proteins that were identified in dataset S2.

| | |
|---|---|
| FAD-containing proteins | |
| WP_056452081.1 | FAD-dependent oxidoreductase *Brevundimonas* sp. Root1279 |
| WP_082731613.1 | FAD-dependent oxidoreductase, *Sphingomonas* |
| YP_001636932.1 | FAD-dependent pyridine nucleotide-disulfide oxidoreductase, *Chloroflexus aurantiacus* J-10-fl |
| WP_082690100.1 | FAD-binding monooxygenase, *Leptolyngbya* sp. NIES-2104 |
| WP_052700213.1** | FAD/NAD(P)-binding oxidoreductase, *Methylocucumis oryzae* |
| WP_020186848.1** | NAD(P)/FAD-dependent oxidoreductase, *Methylopila* sp. 73B |
| NAD-containing proteins | |
| WP_046090413.1 | NAD-dependent dehydratase, *Streptomyces antioxidans* |
| WP_030280942.1 | NAD-dependent dehydratase, *Streptomyces catenulae* |
| WP_052860708.1 | NAD-dependent dehydratase, *Streptomyces celluloflavus* |
| WP_062046107.1 | NAD-dependent dehydratase, *Streptomyces ciscaucasicus* |
| WP_078643469.1 | NAD-dependent dehydratase, *Streptomyces hygroscopicus* |
| WP_044570187.1 | NAD-dependent dehydratase, *Streptomyces iranensis* |
| WP_079429679.1 | NAD-dependent dehydratase, *Streptomyces katrae* |
| WP_067236815.1 | NAD-dependent dehydratase, *Streptomyces longwoodensis* |
| WP_055470700.1 | NAD-dependent dehydratase, *Streptomyces pathocidini* |
| WP_086886330.1 | NAD-dependent dehydratase, *Streptomyces rhizosphaericus* |
| WP_051779201.1 | MULTISPECIES: NAD-dependent dehydratase, *Streptomyces* |
| WP_065964212.1 | NAD-dependent dehydratase, *Streptomyces sparsogenes* |
| WP_053757466.1 | NAD-dependent dehydratase, *Streptomyces* sp. AS58 |
| WP_037799844.1 | MULTISPECIES: NAD-dependent dehydratase, *Streptomyces* |
| WP_069765881.1 | NAD-dependent dehydratase, *Streptomyces* sp. LUP47B |
| WP_073773307.1 | NAD-dependent dehydratase, *Streptomyces* sp. MJM1172 |
| WP_055547987.1 | NAD-dependent dehydratase, *Streptomyces* sp. NBRC 110028 |
| WP_031076788.1 | NAD-dependent dehydratase, *Streptomyces* sp. NRRL S-118 |
| WP_030711692.1 | NAD-dependent dehydratase, *Streptomyces* sp. NRRL S-237 |
| WP_037952333.1 | NAD-dependent dehydratase, *Streptomyces* sp. PRh5 |
| WP_057612235.1 | NAD-dependent dehydratase, *Streptomyces* sp. Root369 |
| WP_009715199.1 | MULTISPECIES: NAD-dependent dehydratase, *Streptomyces* |
| WP_069815624.1 | NAD-dependent dehydratase, *Streptomyces* sp. TP-A0874 |
| WP_079154649.1 | NAD-dependent dehydratase, *Streptomyces subrutilus* |
| WP_030878525.1 | NAD-dependent dehydratase, *Streptomyces varsoviensis* |
| WP_059145201.1 | NAD-dependent dehydratase, *Streptomyces violaceusniger* |
| WP_051763101.1 | NAD-dependent dehydratase, *Streptomyces virginiae* |

| | |
|---|---|
| WP_053167991.1 | NAD-dependent dehydratase, *Streptomyces virginiae* |
| WP_014176621.1 | NAD-dependent dehydratase, *Streptomyces bingchenggensis* |
| WP_030225883.1 | NAD-dependent dehydratase, *Streptomyces lavendulae* |
| WP_065964212.1 | NAD-dependent dehydratase, *Streptomyces sparsogenes* |
| WP_014054769.1 | NAD-dependent dehydratase, *Streptomyces violaceusniger* |
| WP_087353490.1 | NAD-dependent protein deacylase, *Olsenella* sp. An188 |
| Copper-containing proteins | |
| WP_026166134.1 | copper resistance protein, *Corynebacterium mastitidis* |
| WP_085235053.1 | copper resistance protein CopC, *Mycobacterium conspicuum* |
| WP_066814184.1 | copper resistance protein CopC, *Mycobacterium* sp. 1100029.7 |
| WP_077078633.1 | copper resistance protein CopC, *Mycobacterium* sp. AB215 |
| WP_005248910.1 | MULTISPECIES: copper resistance protein C, *Rhodococcus* |
| WP_039951718.1 | copper resistance protein CopC, *Rhodococcus imtechensis* |
| WP_041811727.1 | copper resistance protein CopC, *Rhodococcus jostii* |
| WP_012691057.1 | copper resistance protein C, *Rhodococcus opacus* |
| WP_005254749.1 | copper resistance protein, *Rhodococcus opacus* |
| WP_037228321.1 | copper resistance protein CopC, *Rhodococcus wratislaviensis* |
| WP_006540442.1 | copper resistance protein CopC, *Frankia* sp. EUN1f |
| WP_006594284.1 | copper resistance protein C, *Kineosphaera limosa* |
| WP_052129147.1, WP_053773647.1, WP_081235111.1 | copper resistance protein CopC, *Clavibacter michiganensis* |
| WP_048809232.1 | MULTISPECIES: Copper resistance protein C precursor, *Microbacterium* |
| WP_067311181.1 | copper resistance protein CopC, *Micromonospora rifamycinica* |
| WP_083706190.1** | copper resistance protein CopC, *Arthrobacter* sp. QXT-31 |
| WP_051866792.1** | copper resistance protein CopC, *Corynebacterium atypicum* |
| WP_018023019.1** | copper resistance protein CopC, *Corynebacterium doosanense* CAU |
| WP_083352005.1** | copper resistance protein CopC, *Leucobacter chromiiresistens* DSM 22788 |
| WP_058594659.1** | copper resistance protein CopC, *Leucobacter chromiiresistens* NS354 |
| WP_082334822.1** WP_076688294.1** | copper resistance protein CopC, *Microbacterium aurantiacum* |
| WP_058232366.1** | copper resistance protein CopC, *Microbacterium enclense* |
| WP_082784215.1** | copper resistance protein CopC, *Microbacterium laevaniformans* |
| WP_084344888.1** | copper resistance protein CopC, *Microbacterium resistens* NBRC 103078 |
| WP_029266231.1** | copper resistance protein CopC, *Microbacterium* sp. Cr-K32 |

| | |
|---|---|
| WP_082516518.1** | copper resistance protein CopC, *Microbacterium* sp. Leaf436 |
| WP_054678997.1** | copper resistance protein CopC, *Microbacterium* sp. No. 7 |
| WP_076674855.1** | copper resistance protein CopC, *Microbacterium* sp. RU1D |
| WP_016463500.1** | copper resistance protein CopC, *Microbacterium* sp. oral taxon 186 str. F0373 |
| WP_067359874.1** | copper resistance protein CopC. *Micromonospora rosaria* |
| WP_079705109.1** | copper resistance protein CopC, *Plantibacter cousiniae* |
| WP_079001510.1** | copper resistance protein CopC, *Plantibacter flavus* |
| WP_085511799.1** | copper resistance protein CopC, *Plantibacter flavus* DSM 14012 |
| WP_064294246.1** | copper resistance protein CopC, *Plantibacter* sp. H53 |
| WP_056779050.1** | copper resistance protein CopC, *Plantibacter* sp. Leaf171 |
| WP_056008599.1** | copper resistance protein CopC, *Plantibacter* sp. Leaf314 |
| WP_071259775.1** | copper resistance protein CopC, *Plantibacter* sp. MMLR14_011 |
| WP_086474591.1** | copper resistance protein CopC, *Plantibacter* sp. VKM Ac-1784 |
| WP_040269875.1** | copper resistance protein CopC, *Streptomonospora alba* |
| WP_051814695.1** | copper resistance protein CopC, *Streptomyces iakyrus* |

Table S4. Candidate tail-anchored Tat substrates that are predicted to bind FAD, NAD or copper cofactors. ** Indicates proteins that were identified in dataset S2.

.

| Accession | Organism | Further analysis |
|---|---|---|
| 'Tat pathway protein signal' annotation | | |
| WP_012680923.1 | *Acidobacterium capsulatum* | Has N-terminal pyridoxal phosphate-dependent aminotransferase (PRK09105) domain. The twin arginines form part of this domain and are therefore unlikely to form part of a signal sequence. |
| WP_050369614.1 | *Streptomyces acidiscabies* | alginate lyase family protein. The C-tail is conserved but the twin arginine motif is not conserved across close homologues |
| WP_067001745.1 | *Streptomyces cellostaticus* | alginate lyase family protein. The C-tail is conserved but the twin arginine motif is not conserved across close homologues |
| WP_046705711.1 | *Streptomyces europaeiscabiei* | alginate lyase family protein. The C-tail is conserved but the twin arginine motif is not conserved across close homologues |
| WP_086800221.1<br>WP_079024514.1<br>WP_086747795.1<br>WP_086754741.1 | *Streptomyces scabiei* | alginate lyase family protein. The C-tail is conserved but the twin arginine motif is not conserved across close homologues |
| WP_070699546.1 | *Atopobium* sp. HMSC064B08 | Protein with an N-terminal peptidoglycan hydrolase CwlO domain and a C-terminal NLPC_P60 domain. The NLPC_P60 domain overlaps with the hydrophobic stretch and it is therefore unlikely to be a C-tail. |
| 'Twin-arginine translocation annotation' | | |
| WP_088345824.1 | *Rhodomicrobium vannielii* | ABC transporter substrate-binding protein, FhuD family. The hydrophobic nature of the C-terminal region is not conserved across close homologues |
| WP_035708011.1 | *Haematobacter massiliensis* | Has a C-terminal lipid-binding SYLF domain that covers the hydrophobic 'tail' region and is therefore unlikely to be a TMH. |
| WP_008328674.1 | *Maritimibacter alkaliphilus* | Has a C-terminal lipid-binding SYLF domain that covers the hydrophobic 'tail' region and is therefore unlikely to be a TMH. |
| WP_072295392.1 | *Paracoccus* sp. SM22M-07 | Has a C-terminal lipid-binding SYLF domain that covers the hydrophobic 'tail' region and is therefore unlikely to be a TMH. |
| WP_068297290.1 | *Rhodobacter* sp. CCB-MM2 | Has a C-terminal lipid-binding SYLF domain that covers the hydrophobic 'tail' region and is therefore unlikely to be a TMH |
| WP_007290909.1<br>WP_007293159.1 | delta proteobacterium MLMS-1 | 4Fe-4S dicluster domain-containing protein. These protein records were suppressed because they are no longer annotated on any genome and Blast analysis fails to identify any close homologues. |
| WP_057494184.1<br>WP_074726554.1 | *Stenotrophomonas pavanii* | Gluconate 2-dehydrogenase subunit 3. The hydrophobic tail region is part of the gluconate_2-dh3 fold and unlikely to be a TMH. |
| WP_061479189.1 | *Stenotrophomonas* sp. DDT-1 | Gluconate 2-dehydrogenase subunit 3. The hydrophobic tail region is part of the gluconate_2-dh3 fold and unlikely to be a TMH. |
| WP_087921816.1 | *Stenotrophomonas* sp. WZN-1 | Gluconate 2-dehydrogenase subunit 3. The hydrophobic tail region is part of the gluconate_2-dh3 fold and unlikely to be a TMH. |

| | | |
|---|---|---|
| WP_049467359.1<br>WP_049404954.1<br>WP_046429410.1<br>WP_065196568.1<br>WP_065186014.1<br>WP_058980795.1<br>WP_072168650.1<br>WP_057496275.1<br>WP_065198061.1<br>WP_064237859.1<br>WP_049434537.1<br>WP_087802949.1<br>WP_088496559.1<br>WP_053462412.1<br>WP_053505877.1<br>WP_053442649.1<br>WP_054171204.1<br>WP_062606679.1<br>WP_049403241.1 | *Stenotrophomonas maltophilia* | Gluconate 2-dehydrogenase subunit 3. The hydrophobic tail region is part of the gluconate_2-dh3 fold and unlikely to be a TMH. |
| WP_035353858.1** | *Edaphobacter aggregans* DSM 19364 | No homologues in the database. |
| WP_083145723.1** | *Mycolicibacterium parafortuitum* | RR conserved. However Alphafold model suggests that the hydrophobic region is part of the protein fold. |
| WP_083090608.1** | *Rothia nasimurium* | Homologue of WP_083145723.1 |

Table S5. Protein accessions with Tat pathway protein signal or twin-arginine translocation pathway in their annotation. ** Indicates proteins that were identified in dataset S2.

.

| Accession | Organism |
| --- | --- |
| WP_043259400.1 | *Streptomyces hokutonensis* |
| WP_043435093.1 | *Streptomyces pluripotens* |
| WP_006380203.1 | *Streptomyces turgidiscabies* |
| WP_037801963.1 | *Streptomyces* sp. LaPpAH-108 |
| WP_053849945.1 | *Streptomyces* sp. NRRL B-24085 |
| WP_029393839.1 | *Streptomyces* sp. TOR3209 |
| WP_040902940.1 | *Streptomyces chartreusis* |
| WP_015655752.1 | *Streptomyces davawensis* |
| WP_039830567.1<br>WP_037666917.1 | *Streptomyces* |
| WP_014676716.1 | *Streptomyces hygroscopicus* |
| WP_026151595.1 | *Streptomyces prunicolor* |
| WP_007386666.1 | *Streptomyces sviceus* |
| WP_003994962.1<br>WP_037885301.1 | *Streptomyces viridochromogenes* |
| WP_043733548.1 | *Streptomyces zinciresistens* |
| WP_086565138.1[#] | *Streptomyces africanus* |
| WP_079047754.1[#] | *Streptomyces cyanoalbus* |
| WP_086855524.1[#] | *Streptomyces daghestanicus* |
| WP_075028213.1[#] | *Streptomyces mirabilis* |
| WP_070389912.1[#] | *Streptomyces pactum* |
| WP_079040672.1[#] | *Streptomyces prasinus* |
| WP_078878315.1[#] | *Streptomyces* |
| WP_073890878.1[#] | *Streptomyces* sp. CB01883 |
| WP_073934166.1[#] | *Streptomyces* sp. CB02400 |
| WP_073728708.1[#] | *Streptomyces* sp. CB02414 |
| WP_087804880.1[#] | *Streptomyces* sp. CS113 |
| WP_087788440.1[#] | *Streptomyces* sp. CS159 |
| WP_079308928.1[#] | *Streptomyces* sp. GKU 895 |
| WP_079130289.1[#] | *Streptomyces* sp. H-KF8 |
| WP_076082650.1[#] | *Streptomyces* sp. IMTB 2501 |
| WP_077797205.1[#] | *Streptomyces* sp. JHA26 |
| WP_076972742.1[#] | *Streptomyces* sp. M1013 |
| WP_081220707.1[#] | *Streptomyces* sp. M41(2017) |
| WP_071381920.1[#] | *Streptomyces* sp. MUSC 1 |
| WP_071371441.1[#] | *Streptomyces* sp. MUSC 14 |
| WP_086602166.1[#] | *Streptomyces swartbergensis* |
| WP_086698134.1[#] | *Streptomyces tricolor* |
| WP_086867759.1[#] | *Streptomyces viridochromogenes* |
| WP_081238835.1[#] | *Streptomyces viridosporus* |
| WP_016699903.1** | *Actinoalloteichus spitiensis* |
| WP_058266926.1** | *Arthrobacter enclensis* |
| WP_016325237.1** | *Streptomyces lividans* |

Table S6. Protein accessions for proteins of the S1 peptidase family. Accessions marked ♯ were annotated as serine protease in dataset S1 but were assigned as S1 peptidase based on current annotation; ** indicates proteins that were identified in dataset S2.

.

| | |
|---|---|
| WP_027241321.1 | *Pseudophaeobacter arcticus* |
| WP_049167940.1<br>WP_049150213.1 | *Corynebacterium propinquum* |
| WP_075811183.1 | *Corynebacterium* sp. CNJ-954 |
| WP_064834573.1 | *Corynebacterium* sp. EPI-003-04-2554_SCH2473622 |
| WP_006821438.1 | *Corynebacterium casei* |
| WP_038546560.1 | *Corynebacterium glyciniphilum* |
| WP_027017577.1 | *Corynebacterium pseudodiphtheriticum* |
| WP_043687850.1 | *Nocardia abscessus* |
| WP_039795807.1 | *Nocardia araoensis* |
| WP_014981882.1 | *Nocardia brasiliensis* |
| WP_036533687.1 | *Nocardia cyriacigeorgica* |
| WP_040865004.1 | *Nocardia exalbida* |
| WP_022597221.1<br>WP_005518562.1<br>WP_013416807.1<br>WP_064076622.1 | *Rhodococcus hoagii* |
| WP_013416807.1 | *Rhodococcus equi* |
| WP_069856650.1 | *Rhodococcus* |
| WP_051901016.1 | *Streptomyces aureus* |
| WP_010987315.1 | *Streptomyces avermitilis* |
| WP_059415102.1 | *Streptomyces azureus* |
| WP_086765287.1 | *Streptomyces bobili* |
| WP_066996028.1 | *Streptomyces cellostaticus* |
| WP_086838583.1 | *Streptomyces daghestanicus* |
| WP_030597466.1 | *Streptomyces fulvoviolaceus* |
| WP_071657364.1 | *Streptomyces gilvigriseus* |
| WP_067299969.1 | *Streptomyces griseochromogenes* |
| WP_078967395.1<br>WP_037656023.1<br>WP_085566213.1<br>WP_085568484.1 | *Streptomyces griseofuscus* |
| WP_037702946.1 | *Streptomyces lavenduligriseus* |
| WP_046417906.1 | *Streptomyces malaysiense* |
| WP_075026534.1 | *Streptomyces mirabilis* |
| WP_079172327.1<br>WP_031119869.1<br>WP_039657545.1<br>WP_051707667.1<br>WP_030994093.1 | *Streptomyces* |
| WP_086806635.1 | *Streptomyces murinus* |
| WP_052454075.1 | *Streptomyces nodosus* |
| WP_079064839.1 | *Streptomyces olivochromogenes* |
| WP_079101010.1 | *Streptomyces pathocidini* |
| WP_069779106.1<br>WP_079161659.1<br>WP_079161534.1 | *Streptomyces puniciscabiei* |
| WP_062702374.1 | *Streptomyces regalis* |
| WP_059249036.1<br>WP_059252215.1 | *Streptomyces reticuli* |
| WP_078629026.1 | *Streptomyces* sp. 142MFCol3.1 |
| WP_079662723.1 | *Streptomyces* sp. 3124.6 |

| | |
|---|---|
| WP_019981628.1 | *Streptomyces* sp. Amel2xE9 |
| WP_081546686.1 | *Streptomyces* sp. B9173 |
| WP_079192310.1 | *Streptomyces* sp. CB01883 |
| WP_043261201.1 | *Streptomyces* sp. e14 |
| WP_067039987.1 | *Streptomyces* sp. FXJ1.172 |
| WP_037801489.1 | *Streptomyces* sp. LaPpAH-108 |
| WP_081216934.1 | *Streptomyces* sp. M41(2017) |
| WP_053657310.1 | *Streptomyces* sp. MMG1121 |
| WP_079168661.1 | *Streptomyces* sp. MUSC 93 |
| WP_078890368.1 | *Streptomyces* sp. NBRC 110468 |
| WP_053707807.1 | *Streptomyces* sp. NRRL B-3648 |
| WP_079050130.1 | *Streptomyces* sp. NRRL F-5122 |
| WP_033281943.1 | *Streptomyces* sp. NRRL F-525 |
| WP_030345652.1 | *Streptomyces* sp. NRRL S-1022 |
| WP_037861009.1 | *Streptomyces* sp. NRRL S-340 |
| WP_030842128.1 | *Streptomyces* sp. NRRL S-475 |
| WP_030228728.1 | *Streptomyces* sp. NRRL WC-3626 |
| WP_057600964.1 | *Streptomyces* sp. Root1310 |
| WP_079041571.1 | *Streptomyces* sp. Root264 |
| WP_046261706.1 | *Streptomyces* sp. WM6386 |
| WP_086697472.1 | *Streptomyces tricolor* |
| WP_053196823.1 | *Streptomyces viridochromogenes* |
| WP_048587024.1 | *Streptomyces viridochromogenes* |
| WP_067119775.1 | *Streptomyces yokosukanensis* |
| WP_051754093.1 | *Streptomyces achromogenes* |
| WP_020273774.1 | *Streptomyces afghaniensis* |
| WP_054230871.1 | *Actinobacteria bacterium OK006* |
| WP_054220105.1 | *Actinobacteria bacterium OK074* |
| WP_054238031.1 | *Actinobacteria bacterium OV320* |
| WP_018121428.1** | *Corynebacterium propinquum* |
| WP_023019056.1** | *Corynebacterium* |

Table S7. Protein accessions for proteins annotated as D-alanyl-D-alanine carboxypeptidase. ** Indicates proteins that were identified in dataset S2.

| Accession | Organism | Comments/further analysis |
|---|---|---|
| WP_034515529.1 | *Actinomyces slackii* | Choice-of-anchor L domain-containing protein. Predicted to be a Sec signal peptide by SignalP 6.0 |
| WP_023019151.1 | *Corynebacterium* | LPXTG cell wall anchor domain-containing protein. Predicted to be a Sec signal peptide by SignalP 6.0. RR not conserved across closely related protein sequences |
| WP_016464758.1 | *Microbacterium* sp. oral taxon 186 | Tyrosine protein phosphatase. Misannotated previously |
| WP_051814589.1 | *Streptomyces iakyrus* | LPXTG cell wall anchor domain-containing protein. Potentially wrong start codon assigned in this accession. RR not conserved. |
| WP_046502582.1 | *Streptomyces* sp. NRRL B-24891 | LAETG motif-containing sortase-dependent surface protein. This particular accession predicted to be Tat dependent by SignalP 6.0, although the RR not conserved across closely related protein sequences |
| WP_010637519.1 | *Streptomyces* sp. S4 | HtaA domain-containing protein, partial. This particular accession predicted to be Tat dependent by SignalP 6.0. Lots of closely related/identical proteins by Blast. |
| WP_015611441.1 | *Streptomyces fulvissimus* | LPXTG cell wall anchor domain-containing protein. Strongly predicted Tat dependent by SignalP 6.0. RR motif not always conserved |
| WP_051840821.1 | *Streptomyces lavendulae* | Annotated as LPXTG in dataset S1 but current annotation indicates it is an LAETG motif-containing sortase-dependent surface protein. The RR motif not conserved across close homologues |
| WP_013151425.1 | *Nocardiopsis dassonvillei* | LPXTG cell wall anchor domain-containing protein. The RR motif is not conserved across homologous proteins |
| WP_016115629.1 | *Bacillus cereus* | M4 family metallopeptidase. The RR motif not conserved across close homologues |
| WP_016133109.1 | *Bacillus cereus* | M4 family metallopeptidase. Identical protein to WP_016115629.1 |
| WP_007790986.1 | *Peptostreptococcus stomatis* | Ig-like domain-containing protein. This bacterium lacks *tatA* or *tatC* genes so cannot be a Tat substrate |
| WP_067783620.1** | *Actinomyces vulturis* | LPXTG cell wall anchor domain-containing protein. Strongly predicted to be Tat dependent by SignalP 6.0. Very few related proteins, RR not conserved. |
| WP_034625822.1** | *Cellulomonas cellasea* | LPXTG cell wall anchor domain-containing protein. Weak prediction for Tat dependence by SignalP 6.0. Homologues generally also have paired Arg but also with weak prediction for Tat dependence |
| WP_087509670.1** | *Cellulomonas iranensis* | LPXTG cell wall anchor domain-containing protein. Strong prediction for Tat dependence. twin-arginine reasonably well conserved across homologous proteins |
| WP_070320500.1** | *Cellulomonas iranensis* NBRC 101100 | LPXTG cell wall anchor domain-containing protein. Very close homologue of WP_087509670.1 |
| WP_046529179.1** | *Cellulomonas* sp. FA1 | LPXTG cell wall anchor domain-containing protein. Very close homologue of WP_087509670.1 |
| WP_005209628.1** | *Clostridium celatum* DSM 1785 | LPXTG cell wall anchor domain-containing protein. Not recognised as a Tat signal peptide by SignalP 6.0. Very few homologues |
| WP_087014056.1** | *Leucobacter* sp. 7(1) | SpaH/EbpB family LPXTG-anchored major pilin. Close homologues predicted to be Sec dependent |
| WP_057873001.1** | *Loigolactobacillus rennini* DSM 20253 | LPXTG cell wall anchor domain-containing protein. This bacterium lacks *tatA* or *tatC* genes so cannot be a Tat substrate |
| WP_049350406.1** | *Rothia mucilaginosa* | LPXTG cell wall anchor domain-containing protein. The RR motif not conserved across close homologues |
| WP_037602190.1** | *Streptacidiphilus rugosus* AM-16 | LPXTG cell wall anchor domain-containing protein. Partial conservation of RR motif in homologous proteins |
| WP_037580257.1** | *Streptococcus equi* subsp. *ruminatorum* CECT 5772 | LPXTG cell wall anchor domain-containing protein. This bacterium lacks *tatA* or *tatC* genes so cannot be a Tat substrate |
| WP_060178439.1** | *Streptomyces* sp. IMTB 1903 | LPXTG cell wall anchor domain-containing protein. RR motif well conserved across homologous proteins |

| | | |
|---|---|---|
| WP_030231344.1** | *Streptomyces* sp. NRRL S-350 | LPXTG cell wall anchor domain-containing protein. RR motif not conserved across homologous proteins |
| WP_018509850.1** | *Streptomyces* sp. SID5594 | LPXTG cell wall anchor domain-containing protein. RR motif well conserved across homologous proteins |
| WP_046259605.1** | *Streptomyces* sp. WM6386 | LPXTG cell wall anchor domain-containing protein. RR motif well conserved across homologous proteins |
| WP_086778847.1 | *Streptomyces fimicarius* | Annotated as a peptidase in dataset 1 but current annotation is LAETG motif-containing sortase-dependent surface protein |
| WP_076967992.1 | *Streptomyces sp. IB2014 011-1* | Annotated as a peptidase in dataset 1 but current annotation is LAETG motif-containing sortase-dependent surface protein |
| WP_030568954.1 | *Streptomyces cyaneofuscatus* | Annotated as a peptidase in dataset 1 but current annotation is LAETG motif-containing sortase-dependent surface protein |
| WP_084749454.1 | *Streptomyces albolongus* | Annotated as a peptidase in dataset 1 but current annotation is LAETG motif-containing sortase-dependent surface protein |
| WP_032758621.1 | *Streptomyces alboviridis* | Annotated as a peptidase in dataset 1 but current annotation is LAETG motif-containing sortase-dependent surface protein |
| WP_031122887.1 | *Streptomyces sp. NRRL S-623* | Annotated as a peptidase in dataset 1 but current annotation is LAETG motif-containing sortase-dependent surface protein |
| WP_055558699.1 | *Streptomyces luridiscabiei* | Annotated as a peptidase in dataset 1 but current annotation is LAETG motif-containing sortase-dependent surface protein |
| WP_070013327.1** | *Streptomyces abyssalis* | SCO1860 family LAETG-anchored protein |
| WP_004943253.1** | *Streptomyces mobaraensis* NBRC 13819 | SCO1860 family LAETG-anchored protein |
| WP_070014799.1** | *Streptomyces nanshensis* | SCO1860 family LAETG-anchored protein |
| WP_064073155.1** | *Streptomyces noursei* | SCO1860 family LAETG-anchored protein |
| WP_055573436.1** | *Streptomyces prasinopilosus* CGMCC 4.3504 | SCO1860 family LAETG-anchored protein |
| WP_055694013.1** | *Streptomyces prasinopilosus* NRRL B-2711 | SCO1860 family LAETG-anchored protein |
| WP_055606274.1** | *Streptomyces prasinus* ATCC 13879 | SCO1860 family LAETG-anchored protein |
| WP_055688174.1** | *Streptomyces prasinus* NRRL B-2712 | SCO1860 family LAETG-anchored protein |
| WP_073752756.1** | *Streptomyces* sp. CB03234 | SCO1860 family LAETG-anchored protein |
| WP_065002969.1** | *Streptomyces* sp. H-KF8 | SCO1860 family LAETG-anchored protein |
| WP_030791129.1** | *Streptomyces* sp. NRRL S-920 | SCO1860 family LAETG-anchored protein |
| WP_078852820.1** | *Streptomyces* sp. NRRL S-1824 | SCO1860 family LAETG-anchored protein |
| WP_069464636.1** | *Actinacidiphila rubida* | LAETG motif-containing sortase-dependent surface protein. RR motif not conserved across homologous proteins |
| WP_030079189.1** | *Streptomyces baarnensis* | LAETG motif-containing sortase-dependent surface protein. RR motif not conserved across homologous proteins |
| WP_044384405.1** | *Streptomyces cyaneogriseus* subsp. *noncyanogenus* | LAETG motif-containing sortase-dependent surface protein. RR in wrong position relative to h-region |
| WP_029385521.1** | *Streptomyces leeuwenhoekii* C34 | LAETG motif-containing sortase-dependent surface protein. Almost identical to WP_044384405.1 |
| WP_048572334.1** | *Streptomyces leeuwenhoekii* C58 | LAETG motif-containing sortase-dependent surface protein. Almost identical to WP_044384405.1 |
| WP_059130410.1** | *Streptomyces* sp. NRRL F-5122 | LAETG motif-containing sortase-dependent surface protein. RR motif not conserved across homologous proteins |
| WP_053642605.1** | *Streptomyces* sp. NRRL F-6491 | LAETG motif-containing sortase-dependent surface protein. RR motif not conserved across homologous proteins |

Table S8. Protein accessions with LPXTG or LAETG in their annotation. ** indicates the proteins were identified in dataset S2.

**Fig S1.** A. Multiple sequence alignment of seven S1 peptidase sequences from *Streptomyces*. The red box indicates the likely signal peptide region (predicted using SignalP-6.0) and the yellow the C-tail region (identified from DeepTMHMM). B. AlphaFold structure prediction for Uniprot entry D9XEL1, an S1-peptidase from *Streptomyces viridochromogenes*, coloured according to confidence (from dark blue which is very high through to red which is very low).

**Fig S2.** Multiple sequence alignment of six streptomycete D-alanyl-D-alanine carboxypeptidases from dataset S1 alongside S. coelicolor DacC (CAB9066.1). The twin arginines of the signal peptide are indicated in red. The C-tail regions of each protein (identified from DeepTMHMM) are shown in yellow.

**Fig S3.** A. Multiple sequence alignment of seven LPXTG/LAETG-motif containing peptidase sequences from *Streptomyces*. The red box indicates the likely signal peptide region (predicted using SignalP-6.0) and the yellow the C-tail region (identified from DeepTMHMM). The asterisks indicate the position of the sortase recognition motif.

**Fig S4.** A. Multiple sequence alignment of SCO1860 family LAETG-anchored proteins identified in dataset S2. The twin arginines of the signal peptide are indicated in red. peptidase sequences from *Streptomyces*. The yellow box indicates the C-tail region (identified from DeepTMHMM). The asterisks indicate the position of the sortase recognition motif.

**Fig S5.** No detectable export of AmiA by signal peptides of candidates 3, 5, 7, 8, 9, 14 and 16. Plasmids encoding the indicated signal peptide (SP) - AmiA fusions were introduced into *E. coli* strains MC4100 Δ*amiA* Δ*amiC* (circles) and MC4100 Δ*amiA* Δ*amiC* Δ*tatABC* (triangles and labelled tat-) and grown for 20 h in LB in the presence of 0.5% SDS without shaking in a plate reader. Growth curves correspond to an averaged triplicate set ± one SD (standard deviation). For clarity, growth curves are displayed either in groups of 2 or individually alongside the positive ("SP_TorA") and negative ("Vector", i.e., pUniAmiA) controls in each plot.

```
WP_043259400.1   1 MRHARRRVVRRVTRLTAVGGLLLGGAMVTQAAMASDTPARDAATLSAPQTPADKGSALVAKLGSARTAGSWVDSGGRTVV
WP_006380203.1   1 MRHARRRIVRRATRLAAVGGVILGGLMVTQAVAS-EPPE-VNLPFSSAQAAGTTGSELVTELGTDRTAGSWIAADGKPVV
WP_043435093.1   1 MRHARRRVVRRLTRLAAVGGLLLGCTMVARAAVASEPPPASAR---SLSSAGNTGAAWVARLGTARTAGTWIGSDGRPVV
WP_016325237.1   1 MRHARRRIVRRVARLAAVGGLLLGGTMVTRAVAS-EPPDASAAPRTFAQTPSGAGGDLVSRLGTGRTAGHWVDSEGRPVV
WP_079047754.1   1 MRHARRRVVRRVTRLAAVGGLLLGATMVTRAVAS-EPPAPDPVPYAAAEEDTGPGSALVARLGASRTAGSWLGEDGRTVV
WP_086565138.1   1 MRHARRRVVRRVTRLAAVGGLLLGGAMVTNAVAS-ESPDPSIGVPYSARPATSMGTDLVSRLGTSRTAGSWVDSAGRPVV
WP_086855524.1   1 MRHARRRVVRRVTRLAAVGGLLLGGAMVTRAVAS-EPPDPTIGVPYSAQPATSMGTDLVSRLGTSRTAGSWVDSAGRPVV

WP_043259400.1  81 AVTDKATAAEVTKAGAEAKVVPYSMNQLKSAATTLSSAPKIAGTAWAMDYRNNKVLVEGDSSVSATSWAAMTKVAKSIGA
WP_006380203.1  79 AVTDAEAATEVEKAGARAKIVDHSMNDLKSATKTLRSAPRVAGTAWAVDYTKNEVVVQADSTVSAADWAKLSAVAKSIGG
WP_043435093.1  78 AVTDDAAARAVRRTGAKAKIVRNSMNDLTSATATLRAAPRVAGTAWAVDYRTNTVKVQADRTVSAANWSRISQVAKKIGG
WP_016325237.1  80 AVTDERAAEAVRRAGATAKRVRHSMSELRSAASTLRSAPRVAGTAWALDYRSNEVVVRGDRTVSGSDWSRLTAVAEGIGG
WP_079047754.1  80 AVTDEKAAAEVREAGADPKVVRHSMRDLKSATKTLGSAPRVPGTAWAMDYRSNEVVVRGDSTVSADDWSDLTRVAKGAGG
WP_086565138.1  80 AVTDEKAADEVERAGARAKMVRHSMSDLKSATATLRSAPRVPGTAWALDYKSNEVVVRGDSTVSASDWSRLTDVAEGIGG
WP_086855524.1  80 AVTDEKAADEVERAGAQAKMVRHSMSDLKSATATLRSAPRVPGTAWALDYKSNEVVVRGDRTVSASDWSRLTDVAEGIGG

WP_043259400.1 161 SVEMQRTQGKFTTRINGAQPILSTGGRCSAGYNVTNGQTDFILTAGHCGPVGSIWFADNQGDQQLGKTVSDNFPGSDFSL
WP_006380203.1 159 SVRMERSEGTFTTRINAAQPILSTGGRCSAGFNVTNGSADFILTAGHCGPNGSVWFADTGGSTEIGKTISSSFPINDYSL
WP_043435093.1 158 FVHMERTKGAFTTRLNGALPILSAGGRCSAGFNVTNGSTDFILTAGHCGPAGSTWFADNQGNQQLGRTVNSVFPGHDFSL
WP_016325237.1 160 FVRMERTEGAFTTRLNGAEPILSTAGRCSAGFNVTDGTSDFILTAGHCGPTGSVWFGDRPGDGQVGRTVAGSFPGDDFSL
WP_079047754.1 160 FVRMERTEGAFTTRINGAEPILSTAGRCSAGYNVTDGKREFILTAGHCGPDGSVWFADDGGRKKVGTTVGGSFPGDDYSL
WP_086565138.1 160 FVRMERTEGTFTTRVNGAQPILSTAGRCSAGFNVTDGTTDYILTAGHCGPDGSVWFADDRGRKQVGSSLKGSFPGDDYSL
WP_086855524.1 160 FVRMERTEGTFTTRINGAEPILSTAGRCSAGFNVTDGKTDFILTAGHCGPDGSVWFADDRGREQVGTSLKGSFPGDDYSL

WP_043259400.1 241 VQYASGSAGEGADVVAIGNGSGVTITGLADAAVGEKVFRSGSTSGLHDGEVTALDATVNYPEGTVTGLVQTTVCAEPGDS
WP_006380203.1 239 VQYNGGSAGDGADVVAIGNGRGVRITGAADPTVGQKVFRSGSTTGLRDGKVTGLNATVNYPEGTVSGLIETTVCAEAGDS
WP_043435093.1 238 VQYSSGQAGKGADVVAIGGGKGVRITGAADPAVGQRVFRSGSTSGLHDGQVTALNATVNYPEGTVSGLIETTVCAEPGDS
WP_016325237.1 240 VEYANGKAGDGADVVAVGDGKGVRITGAGEPAVGQRVFRSGSTSGLRDGRVTALDATVNYPEGTVTGLIDTDVCAEPGDS
WP_079047754.1 240 IEYDGGRAGEGAGVVSVGNGQGIRITGVGDPTVGQRVFRSGSTSGLRDGEVTALNATVNYPEGTVSGLIETNVCAEPGDS
WP_086565138.1 240 VKYDSGRAGEGADVVAVGGGKGVRITGVADAEVGQKIFRSGSTSGLREGEVTALNATVNYPEGTVTGLIETNVCAEPGDS
WP_086855524.1 240 VKYDSGRAGEGADVVAVGGGKGVRITGVADAEVGQKIFRSGSTSGLREGEVTALNATVNYPEGTVTGLIETNVCAEPGDS

WP_043259400.1 321 GGPLFSNGIALGVTSGGSGDCT-AGGTTFFQPLTKALAALNVKLLVAPKTA-----ASPAPSASAAGSAAAGTGSVTPGT
WP_006380203.1 319 GGPLFSEGIALGITSGGNGDCK-TGGTTFFEPVTEALTALNVRLIGQAGQ-----AAAPAPSG-AATQSAVAPGEAQPGS
WP_043435093.1 318 GGPLFSEGVALGVTSGGNGDCT-TGGTTFFQPVTKAMTALGVRLIVSAQNAGGTQSGAPSPGPSTPPQRAIAPSAASPGS
WP_016325237.1 320 GGPMFSEGVALGVTSGGSGDCA-KGGTTFFQPLPEAMASLGVRLIVPGREGAAAGTSAPAGQA-P---GTRAPGASGAGA
WP_079047754.1 320 GGPMFSEGIALGVTSGGSGDCRQAGGTTFFQPVTKALAQLRVDLLVAPKAGGGADTAAPAPSAST---GAATPDAASP--
WP_086565138.1 320 GGPMFADGIALGVTSGGSGDCT-AGGTTFFQPVTKALTELGVKLIVAKQAG-G-TGASPAPAASQ---GAASPGSSAP--
WP_086855524.1 320 GGPMFADGVALGLTSGGSGDCA-AGGTTFFQPVTKALTELGVKLIVAKQAG-GATGASPAPSASQ---GAASPGSSAP--

WP_043259400.1 395 SAPATNTTAGTTTLASRLTDPK VGPGLLVIAGSLIALVATRWIRAEQDRKAFQRHYSAMWG
WP_006380203.1 392 VAPVTGGSAG-EALLDKLADPK VGPGLLIAGGSLIALVATRWFRVEQDRRDYRRSYSATWS
WP_043435093.1 397 SAPVTGTAQG-GALLARLTDAR IGPGLLVIAGSLIALVATRYIRAEQERRAYQQHYSATWG
WP_016325237.1 395 PAPVAGV-DG-STLLARLTDRR IGPGLLVVAGSLVALVATRWIREEQDRKAYRRRYSATWG
WP_079047754.1 395 GSSAAGI-NA-STLLSRLADPR VGPGMLVIGGSLIALVATRWIRAEQDRKAYQRYYSATWS
WP_086565138.1 392 ---VTGE-DV-SGLLTRLTDPR VGPGLLVIAGSLVALVATRWIRAEQDRKAYRQHYSATWG
WP_086855524.1 393 ---VTGE-DV-SGLLTRLTDPR VGPGLLVIAGSLVALVATRWIRAEQDRKAYRQYYSATWG
```

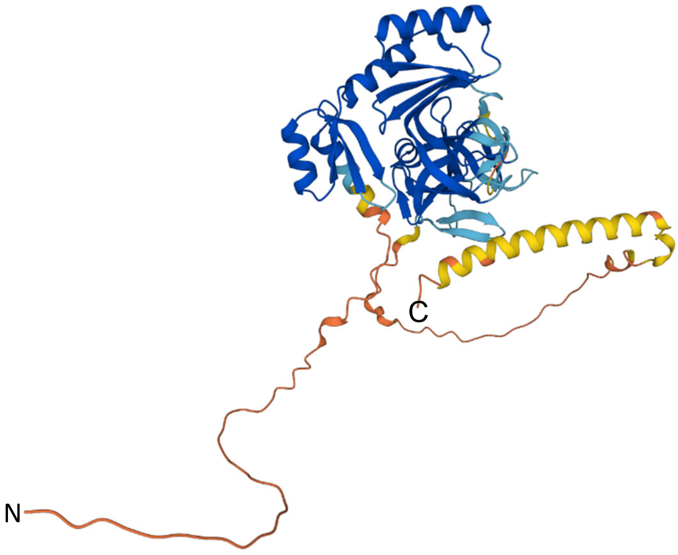


Fig S1

```
CAB89066.1       1 ----MPAP---KKALRRSLTVTSAALTALALLSPAALAAPSPSTSPSATPPASMSGVGGARLGKPGTQVDLAGGAPVLPK
WP_067119775.1   1 ----MPAP---KPTGRRSLLVTSATLLSLSATAPAALAAPPPGASPTATPPARMSTVGGERLGQPGTQVNLAPGVPVLPK
WP_086806635.1   1 ----MPAP---KNTARRSLLISAATLLSLAVTAPAALAAPAAGKAPAATPPAHMSTLGGARLGLPGTQVNLAPGVPVLPK
WP_053707807.1   1 ----MPAP---KTTGRRSLLVTSATLLSLTVTAPAALAAQAPSPSPTATPPAGMSTLGGARLGQPGTQVNLAPGVPVLPK
WP_078967395.1   1 MR-------ESTRLSRRAVLGLTAAALPTSAVLPLTAAA------P-ARAASPATAIGGERLARGGLQVDGAAAVPT--K
WP_051901016.1   1 MR-------DSSGLSRRAALGLAA-------ALPLAAAT------A-T-TADATSTIGGERLARAGIQVSGASGLPK--H
WP_079064839.1   1 MRDSSPVPPRAAQLSRRAALGLAA-------AVPLAAAS------P-A--SAATPVIGGERLARAGVQVGAAAGLPK--R

CAB89066.1      74 DLTARSWIVADAESGDVLAAHNAHWELAPASTLKMLFADTLLPKWPKTTKRKVQVSDLAGIGSGSSMVGIKEEETYTVHD
WP_067119775.1  74 DITARSWIVTDAESGDVLAAHNAHWQLPPASTLKMLFADTLLPRFPRTMEHKVVPSDLAGMGVGSSVVGIKEGRTYNVRD
WP_086806635.1  74 DLTARSWIVSDAQSGEVLASHNAHWALPPASTMKMLFADTLLPRFPRTTEHKVAAKDLAGMGEGSSVVGIKPGKTYSVHD
WP_053707807.1  74 ELTARSWIVSDAESGEVLAAHNAHWRLPPASTLKMLFADTVLPRFPKGTLHKVAPSDLKGMGAGSSVVGIKEGGTYSVHD
WP_078967395.1  65 LLTARAWLVADGTTGEVLASFNAHQRLAPASTLKMLFADTVLKKFDRTEKYTVKDSDISDVPAGSSLVGVKAGITYTVEQ
WP_051901016.1  57 -LTARSWLVADLTSGEVLASYNAHLRLPPASTLKMLFADAVLKKFERTETYKVVDADLADVPAGSSLVGVKPGITYTVHQ
WP_079064839.1  63 -LTARSWLVADHQSGEVLASYHAHTHLAPASTLKMLFADTVLKNFERTERHRVTDADLADIPAGSSMVGVKPGITYTVEQ

CAB89066.1     154 LWLGVFLRSGNDAVHVLSAMNDGVEKTVADMNAHAKELQAGDTTVVSPDGYDMPGQVSSAYDLTLFARSGLQKKDFREYA
WP_067119775.1 154 LWLGVFLRSGNDAVHVLSAMNDGVAQTVKDMQAHADELQARDTHVISPDGYDAPGQVSSAYDLSLIARSGLQKKDFRDYC
WP_086806635.1 154 LWLGVFLRSGNDAVHVLSAMNGGVAQTVQDMQHHADELQALDTHVVSPDGYDEPHQVSSAYDLTLVARSGLQKKDFRDYC
WP_053707807.1 154 LWLGVFLRSGNDAVHVLSAMNGGVAQTVQDMQDHAEELQALDTHVVSPDGYDAPEQVSSAYDLTLFARSGLQKRDFRDYC
WP_078967395.1 145 LWQGVFLRSGNDAVHVLAHMNGGIAKTVAEMQARAQDLQALDTHVVSPDGFDHPGQLSSAYDLTLFARHGLADPDFRGYC
WP_051901016.1 136 LWQGVFLRSGNDAVHVLSHMNGGLPATVAEMQALAKDLQALDTHVVSPDGFDHKGQLSSAYDLTLFARHGLADADFRGYC
WP_079064839.1 142 LWLGVFLRSGNDAVHVLSHMNGGLARTVAQMQAKAQDLQALDTHVVSPDGFDHKGQVSSAYDLTLFARHGLADADFRAYC

CAB89066.1     234 STVRAKFPGETKKDKKGKTSREPFEIQNTNRLLAGDADVEVYPGIAGVKNGYTTNAGNTFTGVAEQGGKVLLVTVMHPAE
WP_067119775.1 234 STVHAKFGD--------------AEIQNTNRLLSGDADVPVYQGIAGVKNGNTTNAGATFTGVAERNGRVLLVTAMNPEK
WP_086806635.1 234 STVSADFGG--------------QEIRNTNRLLSGDTDVPVYQGIAGVKNGNTTHAGATFTGVAQRGKQVLLVTVMNPQK
WP_053707807.1 234 STVSATFGS--------------TEIRNTNRLLSGDSDVPVYPGIAGVKNGNTTHAGATFTGVAERGGKVLLVTVMNPEK
WP_078967395.1 225 GTRIANFPAGGKK---------TFQIQNTDRLLTGAWGLKTYDGLIGVKNGYTSHAGNTFTGAATRDGRTLLVTVMHPAT
WP_051901016.1 216 STKTADFPAGGKK---------TFQIQNTDRLLTGAWGVKPYEGLIGVKNGYTSHAGNTFTGAATRDGRTLLVTVMHPKS
WP_079064839.1 222 HTKTANFPAGGKK---------TFQIQNTDRLLTGAWGVPTYDGLIGVKNGYTSNAGNTFTGAATRGGRTLLVTVMHPKS

CAB89066.1     314 DKHNEVYKETARLFDWGFRAAGKVRPVGELVRPRSATAASPSAEA-----------SGEAGGTGKGGAGTKPVAGAAAGD
WP_067119775.1 300 HDHNEVYKETAKLFDWGFRAAGKVNPVGELVPPKSALPSPG-------------ASASAGQSAGPHGSGSKPVADATAES
WP_086806635.1 300 QEHNEVYKETAKLFDWGFQAAGKVEPVGELVPPRGAAQTGTAPSGSPQASA--PAGRA-----------GSK--PVAAAAT
WP_053707807.1 300 HEHNEVYKETAKLLDWGFQAAGKVEPVGELVPPRSAATAQPDPSASAGRDAAQAAGRAAGRGSGTGGPGVKAAASATTPS
WP_078967395.1 296 -GSNAVYEQTAALLDWGFTHGRTARPVGALVKPLSEGGAAASPAPAGGQAPAGA------------------PAK-----
WP_051901016.1 287 -GGSGVYEETAQLLDWGFGKGAQAQPVGTLVGPLSEQGASKAPTSAAKAAAGAP------------------ASAGAAKS
WP_079064839.1 293 -GGSGVYEETAALLDWGFKEGAKARSVGTLVDPLSEGGASAAPTRKAARAAAGA------------------PAAAADGG

CAB89066.1     383 GAGGAGVAAGIAAGVLALLAGGAFLINRKWPLPDLVRRR-----TRP---------------------------------
WP_067119775.1 367 GSGGVGIAAAIAGGVLVLLAGGVFLINRRWPLPDLMRRR-----PRP---------------------------------
WP_086806635.1 366 GSRGIGTALAVAGGLLVLLAGGVFLVNRRWPLPDLMRRR-----PRA---------------------------------
WP_053707807.1 380 GTSGAGIALAVAGGLLALLGGGAFLVNRRWPLPDLMRRR-----ARP---------------------------------
WP_078967395.1 352 GPSAARLVEGAAGTAALLAAAGGFLLRRRRRAVEAREAPGDHRPPAPRAAEPDPQEPPAPT-DPSGGGRHRR--------
WP_051901016.1 348 DGAPWGLVGG-AGAAVALLAGGALALRRRRRAAGEASEPVDAV-------EEGAAEPSEAKGRRRRR-------------
WP_079064839.1 354 GSSSWGLVGG-AGGAVALLAGGAYAWRRRRRVAGDEAPDGRQE-------RHDGQD--RRDGRDRRDGRDRRDGQERQER

CAB89066.1         ---
WP_067119775.1     ---
WP_086806635.1     ---
WP_053707807.1     ---
WP_078967395.1     ---
WP_051901016.1     ---
WP_079064839.1 424 HQG
```

Fig S2

```
WP_086778847.1   1 MRITPSRRRTLRGSTAAVAATALVALG-AAPALADEPEPDLGVGTIAPIKGVQAGSAFSTPLTFLNKGTEEVPVSYVTYA
WP_076967992.1   1 MGIAASGRRTLLSATAVSATAALIALG-AAPAQADAIKPDLGVRALAPVTGIAAGSGFGLPVSLLNKGTEELPKVWVSYT
WP_030568954.1   1 MGIAASGRRTLLSATAVSATAALIALG-AAPAQADAIKPDLGVRALAPVTGIAAGSGFGLPVSLLNKGTEELPKVWVSYT
WP_084749454.1   1 MGIAVSGRRTPLSGGAVLASAALIALTATAPAQAAGTGPDLGVGALTPVTGIAPGGTFGLPVSVLNKGTEEVPKVYLSYS
WP_032758621.1   1 MGNAVSGRRTLLSGTAVLAAVALIALG-SAPAQADETEPDLGVRGLTPVTGIAPGSGFGLPVSLLNKGTEELPKAYVTYS
WP_031122887.1   1 MGNAVSGRRTLLSGTAVLAAVALIALG-SAPAQADETEPDLGVRGLTPVTGITPGSGFGLPVSLLNKGTEELPKAYVTYS
WP_055558699.1   1 MGNAVSGRRTLLSGTAVLAAVALIALG-SAPAQADETEPDLGVRGLTPVTGITPGSGFGLPVSLLNKGTEELPKAYVTYS

WP_086778847.1  80 VSPGLQADETYKNCSYYTIPSHDEIPDEQVAACKIDQPLKPGVVYGTEKPLSLKALDSALNDDLRVTIGVTEPDPDSGDT
WP_076967992.1  80 YSSGIAPAETYSNCVYYTLSPGGEWPEEHRADCTVDQPLKPGVLYGTEQPIALTARANALYDEVGLGLLAEEPVPHP-DI
WP_030568954.1  80 YSSGIAPAETYSNCVYYTLSPGGEWPKEHRADCTVDQPLKPGVLYGTEQPIALTARANALYDEVGLGLLAEEPVPHP-DI
WP_084749454.1  81 FSVGLETAETYSNCVYFRINSYDERPERNEALCTLDQPLKPGVLYGTEKPLGLKAGADALYDRAGYSVSADKPAPDD--G
WP_032758621.1  80 FSSGLATAETYSNCLYSTIGSYDEMPERHRAVCTVDQPLKPGVLYRTEKPIGLKAGANALYDTAGLGLWASPPGPDD--G
WP_031122887.1  80 FSSGLATAETYSNCLYSTIGSYDEMPERHRAVCTVDQPLKPGVLYGTEKPIGLKAGANALYDTAGLGLWASPPGPDD--G
WP_055558699.1  80 FSSGLATAETYSNCLYSTIGSYDEMPERHRAVCTVDQPLRPGVLYGTEKPIGLKAGANALYDTAGLGLWASPPGPDD--G

WP_086778847.1 160 GADEPVPGTGDPLTLVEKGPATDADRANHPESAASADVTAANTADFGLTGAELKGEPGEKVTATVKFTNAGPARVQGARD
WP_076967992.1 159 HHGEPEPGTGGPLKLVERGEATAADRAAHAEPGSRTEVTAVNTADYALTGAELQGKVGDKVTALVTFTNEGPARVYRDQG
WP_030568954.1 159 HHGEPEPGTGGPLKLVERGEATAADRAAHAEPGSRTEVTAVNTADYALTGAELQGKVGDKVTALVTFTNEGPARVYRDQG
WP_084749454.1 159 GESEHVPGTGAPLKLVERGEATEADRAAHPDSSVGADVTAVNTADFALTGAELKGKVGDKVTAQVKFTNKGPAWVMGERD
WP_032758621.1 158 GETDPVPGTGAPLTLVERGEATEADRAAHAESGSWTDVTAVNTADYALTGAELKGKVGDKVTAQVKFTNKGPAWVYRELG
WP_031122887.1 158 GETDPVPGTGAPLTLVERGEATEADRAAHAESGARTDVTAVNTADYALTGAELRGKVGDKVTAQVKFTNKGPAWVYRELG
WP_055558699.1 158 GETDPVPGTGAPLTLVERGEATEADRAAHAESGARTDVTAVNTADYALTGAELKGKVGDKVTAQVKFTNKGPAWVYRELG

WP_086778847.1 240 QSVATVDVHIPKGTSVVKAHGYCDAVTKTHYRCGTSQSWVDVNGGESYDFVLRIDKAVGRTTGEVSFGGPERPFDRNAAN
WP_076967992.1 239 TSAANVDVHIPAGTTVTKAHGFCRPVIQTHYRCGTSQSWVEAKGGETYSFVLRIDKAVGRTTGRMSFGGAARPFDKNAAN
WP_030568954.1 239 TSAANVDVHIPAGTTVTKAHGFCRPVIQTHYRCGTSQSWVEAKGGETYSFVLRIDKAVGRTTGRMSFGGAARPFDKNAAN
WP_084749454.1 239 TSVTTVDVRIPAGVTVTKANGFCTPVTKTHYTCGTSQAWVDPTDGATYPFVLRIDKAVGRTTGKVSFAGPARPFDKNAAN
WP_032758621.1 238 TGAASVDVRIPAGTTVTKANGYCSKVTKTHYRCGTSQSWVDEEGGETYPFVLRIDKAVGRTTGKVSFGGQSRPFDRNPAN
WP_031122887.1 238 TGAASVDVRIPAGTTVTKANGYCSKVTKTHYRCGTSQSWVDEEGGETYSFVLRIDKAVGRTTGKVSFGGQSRPFDRNPAN
WP_055558699.1 238 TGAASVDVRIPAGTTVTKANGYCSKVTKTHYRCGTSQSWVDEEGGETYSFVLRIDKAVGRTTGKVSFGGQSRPFDRNPAN

WP_086778847.1 320 DTAQIVIDTGKGSDTSGSSGSS--GS---------TG-GTGGSSSTGGSD-------TTGGTSGTGGSGSSGGSAGTGGT
WP_076967992.1 319 DTAEIVVDTPEPPTDGTESTGGTSSTG-G--SAGPTGSTDGGTGSTGGSTSGSSGSASSGSTGGSDTGTTGGSATSGSGS
WP_030568954.1 319 DTAEIVVDTPEPPTDGTESTGGTSSTG-G--SAGPTGSTDGGTGSTGGSTSGSSGSASSGSTGGSDTGTTGGSATSGSGS
WP_084749454.1 319 DTAEIVVETGEPATTGGTASTGGSGTSGGSASSGGSTGSDGGTGSTGGSTSGSSGSTATG------GTTGSDSTTSGSGS
WP_032758621.1 318 DTAEIVVETGKAGSTGGTTSGGS-----------ANGGSDGGTGSTGGSTSGSSGSTGGGSTN---TTGGSGSATSGGGS
WP_031122887.1 318 DTAEIVVETGKAGSTGGTTSGGS-----------ANGGSDGGTGSTGGSTSGSSGSTGGGSTD---TTGGSGSATSGGGS
WP_055558699.1 318 DTAEIVVETGKAGSTGGTTSGGS-----------ANGGSDGGTGSTGGSTSGSSGSTGGGSTD---TTGGSGSATSGGGS

WP_086778847.1 381 TTGGATPQTGGNLAATGSDSTAPLAGMAAAAVVAGGIVWAVRRRSAARGN
WP_076967992.1 396 TSGGTTPQTGGGLAATGSDSTLPLAGAAGAALLAGGSIVWAVRRRSAARAS
WP_030568954.1 396 TSGGTTPQTGGGLAATGSDSTLPLAGAAGAALLAGGSIVWAVRRRSAARAS
WP_084749454.1 393 TTGGTTPQTGGNLAATGSDSTLPVAGAAGAAVLAGAGLFYAMRRRTAARKG
WP_032758621.1 384 TTGGATPQTAGDLAATGSDSTLPVAGAAGAALLAGGGLFYAMRRRTAARNG
WP_031122887.1 384 TAGGATPQTGGDLAATGSDSTLPVAGAAGAALLAGGGLFYAMRRRMAARNG
WP_055558699.1 384 TAGGATPQTGGDLAATGSDSTLPVAGAAGAALLAGGGLFYAMRRRTAARNG
                               *****
```

Fig S3

```
WP_070013327.1   1 --------------------MITHV-----LSSRARRSV--LLSAATVMV-GGAMVSVP--AHAASADGHSDSTGNATAVA
WP_070014799.1   1 --------------------MITHV-----SGSRARRSA--LLGAAAVLA-GGTFLATP--AHAAPAHGHSGSKGKAGATA
WP_004943253.1   1 --------------------MSSTT-RSSFGMPVSRRAV---VPSAAALAAALTALSAS-PASAAPAT-GDHRTGAADAVV
WP_055573436.1   1 --------------------MNSTNSRV-----LARRRT--VVAAAAVLAAGSAALAGAGTAQATGE------PGSASAAV
WP_055694013.1   1 --------------------MNSTNSRV-----PARRRT--VVAAAAVLAAGSAALAGAGTAQATGE------PGSASAAV
WP_055606274.1   1 --------------------MNSTDSRA-----SARRRA--VVATATVLAAGPVALAGAGAAQATGE------HGSASAAV
WP_055688174.1   1 --------------------MNSTDSRA-----SARRRA--VVATVTVLAAGPVALAGAGAAQATGE------HGSASAAV
WP_065002969.1   1 --------------------MNSTDSRA-----SARRRA--VVATVTVLAAGPAALAGAGAAQATGE------HGSASAAV
WP_078852820.1   1 MSPIGRRRWLHVSCGGSTILNSNTFRLAALA----------------VAAAPVALLVAAPAHATGAT-TTTGGGKASAVV
WP_030791129.1   1 --------------------MNSFTFRMPACRRHRRR--LVVVATATALAAGPAALAGAGSARATGG--DHDRGGRADAVV
WP_064073155.1   1 --------------------MNSTT-----SRPSARRRTI--AAAVAAFAAGPALLTAPAA-HAT-G-----GEGRASAVV
WP_073752756.1   1 --------------------MNSNTVRLPARLPAARRRALSTAAAAVALAAGPALLAAPMA-HATGG-----GEGRASAVV

WP_070013327.1  52 ARADLQVSVAD-AAKVPVKASLNSVSAPEDAEETLLSAEVKGANKGQPLKLVQADVARSGAKADARRSTGHVKLAGLRAF
WP_070014799.1  52 ARADLQVSVAD-AAKVPVRASLNEVSAPRSADRTLLTAEVNGANKGRPLKLVQAEVARSAAKTGADRSSGHVKLAGLRAF
WP_004943253.1  56 LRAGLDVSLLDKTVRVPVNASLNDVHAPGNAARTALSVRVGGAEQGRPVELLRADAATARATADEGTATGRAELVRAKVH
WP_055573436.1  49 LRTGLDVALLDETVNVPLTVSLNEVHAPRSAHRTALTADLEGVDGGRPFSVLRAEVADARATVEGNRAEASTTLAEARIH
WP_055694013.1  49 LRTGLDVALLDGTVNVPLTVSLNEVHAPRSAHRTALTADLEGVDGGRPFSVLRAEVADARATVEGNRAEASTTLAEARIH
WP_055606274.1  49 LRTGLDVALLDRTVNVPLTVSLNEVHAPRSAHRTALTADLEGVDRGRPFPVLRAEAADARATVEGNRAEASTTLAKARIH
WP_055688174.1  49 LRTGLDVALLDRTVNVPLTVSLNEVHAPRSARRTALTADLEGVDRGRPFPVLRAEVADARATVEGNRAEASTTLAKARIH
WP_065002969.1  49 LRTGLDVALLDRTVNVPLTVSLNEVHAPRSARRTALTADLEGVDRGRPFPVLRAEVADARATVEGNRAEASTTLAKARIH
WP_078852820.1  64 LRTALDVSLLSKTVDVPLKTTLNEVHAPASAEQTALRVRLDGVNHGRPFSVLRADVATANATVDKHRAEGYSNLAKARVH
WP_030791129.1  58 LRTGLDVSLLNKTVNVPLKVSLNEVQAPASAEKTALTARLDGVDGGQPFSVLRADVATARATVKGGRAEGYSNIARAEMH
WP_064073155.1  48 LRTDLDVALLNKAVHVPLTATLNEVQAPASARRTALTVTLDGVEQGRPVSVLMADVATAQATADGRRGEGRTELARAKVH
WP_073752756.1  56 LRTDLDVSLLDKTVHVPLKATLNEVQAPASARKTALTVTLDGVEQGRPVSVLTADVATAKATADARRAEGHTNLAHATLH

WP_070013327.1 131 APGLLGKPLVSADLLTATATCKAGEKPMAKADL-ENVSVLGKPVTVNGVGSQQVKVPGVGTVDVALEQETTKAASGAATA
WP_070014799.1 131 APGLLGKPLVSADLLTATATCEAGAKPTAQADL-ENVSVLGKPVNVNAPGGQQIKLPGVGTVDVSLEQETTKTASGAATA
WP_004943253.1 136 VPGLPLLSLIEVRQVTSEAVCRAGQRPTATSNLLGPVTVLGKKITLSTGGTTDVDVPGVGRVRLDLSKTATTSHTGAATA
WP_055573436.1 129 LPGLPLLPLVELEAAGSRAVCETGRTPLASATLPAAVTVFGKRVGLTASGPTEVEVPGVGEVRLELSTTETTSRTAAATA
WP_055694013.1 129 LPGLPLLPLVELEAAGSRAVCETGRTPLASATLPAAVTVFGKRVGLTASGPTEVEVPGVGEVRLDLSTTETTSRTAAATA
WP_055606274.1 129 LPGLPLLSLVELEAVSSKAVCEMGRTPLASATLPAAVTVLGKRVRLTASGPTEVEVPGVGEVRLDLSKTETTSRTAAATA
WP_055688174.1 129 LPGLPLLSLVELEAVSSKAVCETGRTPLASATLPAAVTVLGKRVRLTASGPTEVEVPGVGEVRLDLSKTETTSRTAAATA
WP_065002969.1 129 LPGLPLLSLVELEAVSSKAVCETGRTPLASATLPAAVTVLGKRVRLTASGPTEVEVPGVGEVRLDLSKTETTSRTAAATA
WP_078852820.1 144 LPGLPLLSLIEVEKVTSKAVCEVGRRPVAESNVLGHVSVLGKRVTLTSGGTTRVDVPGVGEVRLDLSKTRTTSRMAAATA
WP_030791129.1 138 VPGLPLLSLIEVEKVTSKAVCEAGKRPVAEANLLGGVTVLGKKVTLTAGGTTEVKVPGVGEVTLALSKRRTTSRTAVAAA
WP_064073155.1 128 VPGLPLLSLIELEKVTARALCEAGGQPVAEANVQGRVTVLGKRVTLTAGGPTKVEVPGVGEVTLTLSDTRTTSRTAAATA
WP_073752756.1 136 VPGLPLLSLIEVEKVTSRAVCEAGEQPVAEANLLGRVTVLGKRVTLTAGGPTKVDVPGVGEVTLDLSTTRTTSRTAAATA

WP_070013327.1 210 LKLSYEVNPAKLNIVKAAGEIVLSEATCRTPAGGGEEGTGGTGGNEPGPQTGE---------------------------
WP_070014799.1 210 LKLSYEVNPAKLNIVKATGDIVLSEATCETPKGGSD-------GNEPGPQTGGGD-------------------------
WP_004943253.1 216 LELAVDVNPLKLGVAEVHGRVTLVRAGCETPAGAGKPAEQPPAPARQAPA----K-------------EATRTSGDGIKP
WP_055573436.1 209 LRLEVSVDPLELNVAGVDGTVTLAEATCEAPGGDAAPA---PGGQREPERREEPQEQGRERERADAVPPA--GSGTGVEP
WP_055694013.1 209 LRLEVSVDPLKLNVAEVDGTVTLAEATCEAPGGDAAPA---PGGQREPERREEPQEQGRERERADAVPPA--GSGTGVEP
WP_055606274.1 209 LRLEVSVDPLKLNVAEVNGTVTLAEATCEAPGGDAAPAPAAPGGQREPEPQEEGKGQKEEERADAAAPPAAGGTDTGVEP
WP_055688174.1 209 LRLEVSVDPLKLNVAEVNGTVTLAEATCEAPGGDAAPAPAAPGGQREPEPQEEGKGQKE-ERADAAAPPAASGTDTGVEP
WP_065002969.1 209 LRLEVSVDPLKLNVAEVNGTVTLAEATCEAPGGDAAPAPAAPGGQREPEPQEEGKGQKEEERADAAAPPATSGTDTGVEP
WP_078852820.1 224 LRLEVSVNPLKLNVADVKGEVTLAEATCETPKGPKPTKPAG--GSDGGTA------G-----------GDNGDGGTDVKP
WP_030791129.1 218 LELKVSVNPLKLNVAEVEGTVTLAGAGCEAPRAERERTGGG--TKPQG----------------------KPAKPV----
WP_064073155.1 208 LALTVSVNPLKANVADVTGTITLAEATCESPKAPAAAAAPA--APTAP----------------------AAAKPADVRP
WP_073752756.1 216 LALTVSVNPLKANVADVTGTVTLAEATCESPTAP-----------PKP----------------------AEARSAGVQP

WP_070013327.1 263 ---------DGGDGLAETGGSSATPFIAGAGALLVAGGGAMYLI[illegible]RRGARSTS--
WP_070014799.1 258 -GGDAGSNNGDGGGLAETGGDATTPIIAATGALLVAGGTTMYLV[illegible]RRAARNS---
WP_004943253.1 279 QTV------ADKRDLAETGGSSATPWIAGAAGLLVAGGGAVIA[illegible]RRRAAARGRG
WP_055573436.1 284 RGARAGRAA---PGLAETGGDPAAPYVAGGALALLAAGGAAVLL[illegible]RRGRRS----
WP_055694013.1 284 RGARAGRAA---PGLAETGGDPAAPYVAGGALALLAAGGAAVLL[illegible]RRGRRS----
WP_055606274.1 289 QGARAERAAPARTGLAETGGDSTTPYVAGGALALLATGGAAVLL[illegible]RRGRRT----
WP_055688174.1 288 QGARAERAAPARTGLAETGGDSTTPYVAGGALALLAAGGAAVLL[illegible]RRGRRT----
WP_065002969.1 289 QGTRAERAAPTRTGLAETGGDSTTPYVAGGALALLAAGGAAVLL[illegible]RRGRRT----
WP_078852820.1 285 QTGTGHAPAAVDTSLAETGGSSTTPYIAGGAALLLALGAGAMVV[illegible]RRRGAQD---
WP_030791129.1 270 --G----HKATEANLAETGGNSTTPYIAGGAVVLLIAGGGAVAM[illegible]RRSRSRG---
WP_064073155.1 264 QTA----GEPPEANLAETGGSSLTPYLAGGSLALLAGGAGALAL[illegible]RRARAARARG
WP_073752756.1 263 QSA----GEPQQANLAETGGSSLTPYLAGGSLALLTAGAATFAL[illegible]RRARTTRERG
                                  *****
```

Fig S4

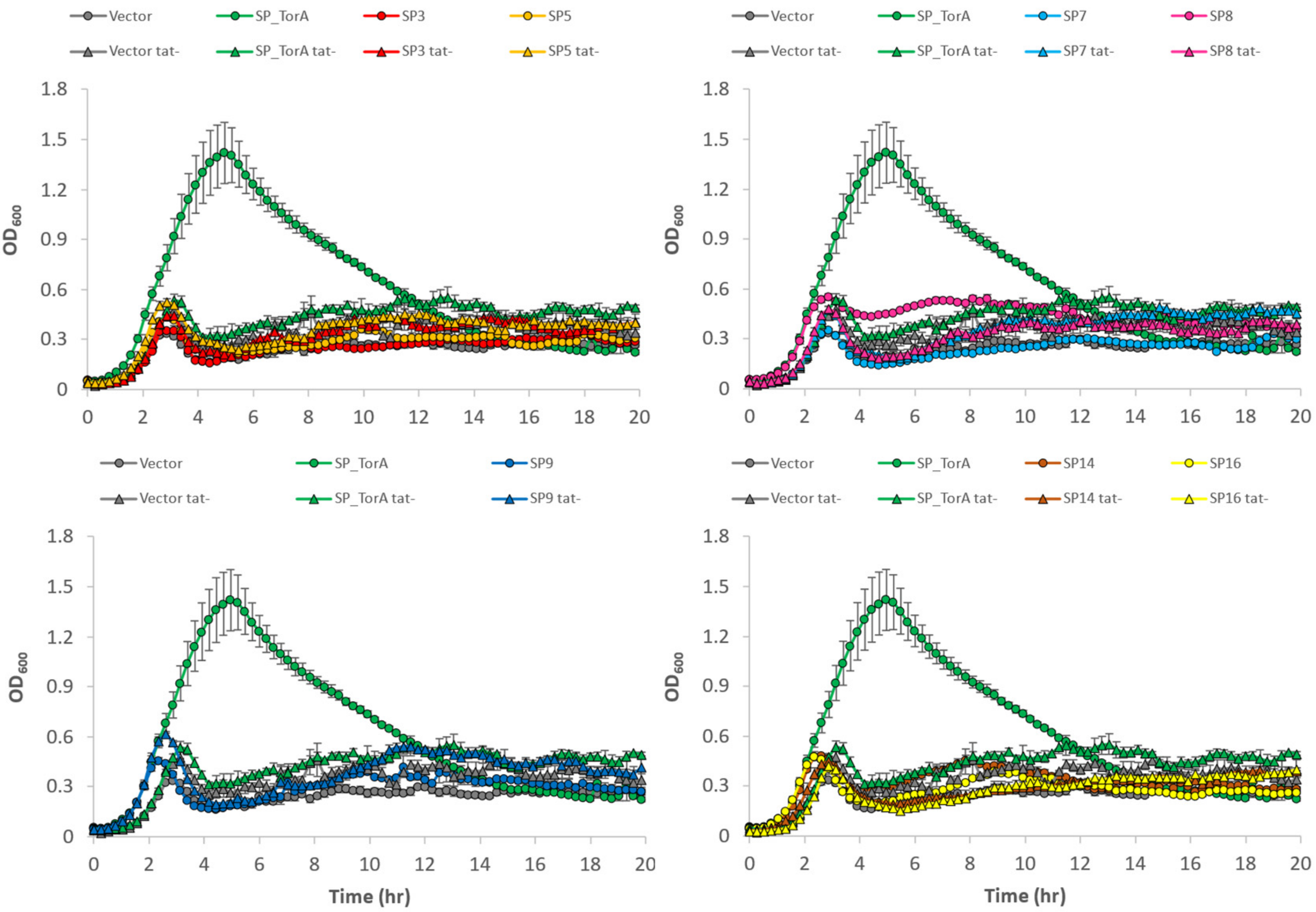
Vector
SP_TorA
SP3
SP5
Vector tat-
SP_TorA tat-
SP3 tat-
SP5 tat-
SP7
SP8
SP7 tat-
SP8 tat-
SP9
SP9 tat-
SP14
SP16
SP14 tat-
SP16 tat-
OD600
Time (hr)

Fig S5

**Dataset S1**

##### Column descriptions ###

# #

### 1. Serial:

### 2. Accession: Genome accession number.

### 3. Organism: Name of organism.

### 4. Lineage: Taxonomic classification of the organism.

### 5. ProteinId: Protein Identifier for this protein in the genome.

### 6. First60AminoAcids: First 60 amino acids of the protein.

### 7. Product: Annotated product for the protein.

### 8. TMHMMoutput: Output produced by TMHMM version 2.0. TMHMM produced

### multi-line output. In this column the lines in TMHMM output have been

### separated by the string "\n". Note that these are two characters not

### the single newline character programmers are used to.

Serial AccessionOrganism Lineage First60AminoAcids ProteinId Product TMHMMoutput

1 GCF_000688455.1_ASM68845v1 Acidobacterium ailaaui Acidobacteria; Acidobacteriia; Acidobacteriales; Acidobacteriaceae; Acidobacterium MSRRTFVSSATAGLAALGALSSAAEGHAQLVWTSKNWKLAEFETLLREPARIRQVYDVTQ WP_026442391.1 hypothetical protein [Acidobacterium ailaaui] Length: 233\nNumber of predicted TMHs: 1\nExp number of AAs in TMHs: 21.25002\nExp number, first 60 AAs: 1.35114\nTotal prob of N-in: 0.67991\ninside 1 201\nTMhelix 202 224\noutside 225 233

2 GCF_000022565.1_ASM2256v1 Acidobacterium capsulatum ATCC 51196 Acidobacteria; Acidobacteriia; Acidobacteriales; Acidobacteriaceae; Acidobacterium; Acidobacterium capsulatum MKSISRRSFVTTAAAGMAALGSLGPALPAAQGQAVEMASDWDISSFNQLAQSPARVKQLF WP_012680923.1 Tat pathway signal sequence domain-containing protein [Acidobacterium capsulatum] Length: 237\nNumber of predicted TMHs: 1\nExp number of AAs in TMHs: 31.62059\nExp number, first 60 AAs: 5.92535\nTotal prob of N-in: 0.86701\ninside 1 205\nTMhelix 206 228\noutside 229 237

3 GCF_000014005.1_ASM1400v1 Candidatus Koribacter versatilis Ellin345 Acidobacteria; Acidobacteriia; Acidobacteriales; Acidobacteriaceae; Candidatus Koribacter; Candidatus Koribacter versatilis MGEKALMSKKPTIEEHLKATGVTRRSFVQLCGMLMAAAPIGLSLTSKASAQEVAKVVGKA WP_011525036.1 hydrogenase 2 small subunit [Candidatus Koribacter versatilis] Length: 401\nNumber of predicted TMHs: 1\nExp number of AAs in TMHs: 19.93057\nExp number, first 60 AAs: 2.05251\nTotal prob of N-in: 0.15168\noutside 1 344\nTMhelix 345 367\ninside 368 401

4 GCF_000014005.1_ASM1400v1 Candidatus Koribacter versatilis Ellin345 Acidobacteria; Acidobacteriia; Acidobacteriales; Acidobacteriaceae; Candidatus Koribacter; Candidatus Koribacter versatilis MPSDDRLAASVDVTNAVRVSSRLDRLAIGWERRGAIVKAAVIGVIVSTTLAFVIPKQYES WP_011521694.1 LPS biosynthesis protein [Candidatus Koribacter versatilis] Length: 428\nNumber of predicted TMHs: 1\nExp number of AAs in TMHs: 44.11157\nExp number, first 60 AAs: 17.46214\nTotal prob of N-in: 0.61372\nPOSSIBLE N-term signal sequence\noutside 1 363\nTMhelix 364 383\ninside 384 428

5 GCF_000745965.1_ASM74596v1 Edaphobacter aggregans DSM 19364 Acidobacteria; Acidobacteriia; Acidobacteriales; Acidobacteriaceae; Edaphobacter; Edaphobacter aggregans MTSRRTFLTAGGAVGLVAALPSPIQATPSTELPKKSTSILFDTITTMSLNTLKELATPGP WP_035353858.1 hypothetical protein [Edaphobacter aggregans] Length: 348\nNumber of predicted TMHs: 1\nExp number of AAs in TMHs:

31.25686\nExp number, first 60 AAs: 7.24535\nTotal prob of N-in: 0.38342\noutside 1 324\nTMhelix 325 347\ninside 348 348

6 GCF_000745965.1_ASM74596v1 Edaphobacter aggregans DSM 19364 Acidobacteria; Acidobacteriia; Acidobacteriales; Acidobacteriaceae; Edaphobacter; Edaphobacter aggregans MLDFLKCLRMVVPSEPNTVRTHNNRSRRGWLTSFALVLLLSRVASAQSHTVTTLTATATG WP_035347944.1 hypothetical protein [Edaphobacter aggregans] Length: 1490\nNumber of predicted TMHs: 2\nExp number of AAs in TMHs: 41.43765\nExp number, first 60 AAs: 0.79926\nTotal prob of N-in: 0.04323\noutside 1 1398\nTMhelix 1399 1418\ninside 1419 1430\nTMhelix 1431 1453\noutside 1454 1490

7 GCF_000178955.2_ASM17895v2 Granulicella mallensis MP5ACTX8 Acidobacteria; Acidobacteriia; Acidobacteriales; Acidobacteriaceae; Granulicella; Granulicella mallensis MSSTDHISTPVATKPTANWVTNATFLWTRRRTLFRVGVVSLVLSGIIAFTIPKQYQSTAR WP_014264939.1 LPS biosynthesis protein [Granulicella mallensis] Length: 429\nNumber of predicted TMHs: 2\nExp number of AAs in TMHs: 46.7979399999999\nExp number, first 60 AAs: 19.74729\nTotal prob of N-in: 0.93126\nPOSSIBLE N-term signal sequence\ninside 1 32\nTMhelix 33 52\noutside 53 372\nTMhelix 373 395\ninside 396 429

8 GCF_000421065.1_ASM42106v1 Acidobacteriaceae bacterium TAA166 Acidobacteria; Acidobacteriia; Acidobacteriales; Acidobacteriaceae; unclassified Acidobacteriaceae MSGGFEQAHEHLEHTSHGHGSDPHNRRAAILIAIMAAALSITEFASKDAQTEYLTNHIAAWP_022843598.1 hypothetical protein [Acidobacteriaceae bacterium TAA166] Length: 190\nNumber of predicted TMHs: 2\nExp number of AAs in TMHs: 56.45437\nExp number, first 60 AAs: 14.1371\nTotal prob of N-in: 0.19605\nPOSSIBLE N-term signal sequence\ninside 1 140\nTMhelix 141 163\noutside 164 166\nTMhelix 167 189\ninside 190 190

9 GCF_001482755.1_ASM148275v1 Chloracidobacterium thermophilum Acidobacteria; Blastocatellia; Chloracidobacterium MTQSASRRKHIVVVGAGFGGVAFCQSFPEGLADITLVDRNNYHLFQPLLYQVATADLSPA WP_058867355.1 pyridine nucleotide-disulfide oxidoreductase [Chloracidobacterium thermophilum] Length: 429\nNumber of predicted TMHs: 1\nExp number of AAs in TMHs: 27.77766\nExp number, first 60 AAs: 5.33717\nTotal prob of N-in: 0.41544\noutside 1 376\nTMhelix 377 399\ninside 400 429

10 GCF_000226295.1_ASM22629v1 Chloracidobacterium thermophilum B Acidobacteria; Blastocatellia; Chloracidobacterium; Chloracidobacterium thermophilum MRGQGLRRRQFLRYATSGACGGVFGLLAAGLPNTGLAEAGVLPATDEGKLPKGFREPTLT WP_014099504.1 hypothetical protein [Chloracidobacterium thermophilum] Length: 194\nNumber of predicted TMHs: 2\nExp number of AAs in TMHs: 36.77337\nExp number, first 60 AAs: 18.25984\nTotal prob of N-in: 0.90896\nPOSSIBLE N-term signal sequence\ninside 1 19\nTMhelix 20 42\noutside 43 147\nTMhelix 148 170\ninside 171 194

11 GCF_000226295.1_ASM22629v1 Chloracidobacterium thermophilum B Acidobacteria; Blastocatellia; Chloracidobacterium; Chloracidobacterium thermophilum MTQSASRRKHIVVVGAGFGGVAFCQAFPEGLADITLVDRNNYHLFQPLLYQVATADLSPS WP_014099596.1 pyridine nucleotide-disulfide oxidoreductase [Chloracidobacterium thermophilum] Length: 429\nNumber of predicted TMHs: 1\nExp number of AAs in TMHs: 32.44365\nExp number, first 60 AAs: 10.56253\nTotal prob of N-in: 0.59020\nPOSSIBLE N-term signal sequence\noutside 1 376\nTMhelix 377 399\ninside 400 429

12 GCF_000820845.2_K22 Pyrinomonas methylaliphatogenes Acidobacteria; Blastocatellia; Pyrinomonas MSSPRESRATVATAHRAQNYNAMNSPMKREDPTARDSRRRFVRALHVWVGLITALPILAW WP_083437566.1 hypothetical protein [Pyrinomonas methylaliphatogenes] Length: 207\nNumber of predicted TMHs: 2\nExp number of AAs in TMHs: 44.77042\nExp number, first 60 AAs: 15.77322\nTotal prob of N-in: 0.98379\nPOSSIBLE N-term signal sequence\ninside 1 46\nTMhelix 47 69\noutside 70 165\nTMhelix 166 188\ninside 189 207

13 GCF_000820845.2_K22 Pyrinomonas methylaliphatogenes Acidobacteria; Blastocatellia; Pyrinomonas MSAQPIIWQTLAQDEGARAGILRTRRATIETPVFMPVGTLGSVKGVRFEALETELDARII WP_041977479.1 tRNA-guanine(34) transglycosylase [Pyrinomonas methylaliphatogenes] Length: 392\nNumber of predicted TMHs: 1\nExp number of AAs in TMHs: 17.7233\nExp number, first 60 AAs: 0.01064\nTotal prob of N-in: 0.09285\noutside 1 338\nTMhelix 339 361\ninside 362 392

14 GCF_000014905.1_ASM1490v1 Candidatus Solibacter usitatus Ellin6076 Acidobacteria; Solibacteres; Solibacterales; Solibacteraceae; Candidatus Solibacter; Candidatus Solibacter usitatus MIAGRRKLMTRAHFCLASFALFCGTAQASILLTANLTNAGENPPTHPTLVSGAPRPASFGWP_083783854.1 hypothetical protein [Candidatus Solibacter usitatus] Length: 216\nNumber of predicted TMHs: 2\nExp number of AAs in

TMHs: 74.32572\nExp number, first 60 AAs: 22.67883\nTotal prob of N-in: 0.98292\nPOSSIBLE N-term signal sequence\ninside 1 11\nTMhelix 12 34\noutside 35 187\nTMhelix 188 210\ninside 211 216

15 GCF_000687145.1_TheAqu1.0 Thermoanaerobaculum aquaticum Acidobacteria; unclassified Acidobacteria; Acidobacteria subdivision 23; Thermoanaerobaculum MNRRTFLTLAASAAAAAASRTAGAVTFQREEREEQEFVGVLMDTTRCIGCRQCEVACAEV WP_053334878.1 hypothetical protein [Thermoanaerobaculum aquaticum] Length: 284\nNumber of predicted TMHs: 1\nExp number of AAs in TMHs: 19.95718\nExp number, first 60 AAs: 0.46448\nTotal prob of N-in: 0.34451\noutside 1 255\nTMhelix 256 275\ninside 276 284

16 GCF_000171895.1_ASM17189v1 Hydrogenivirga sp. 128-5-R1-1 Aquificae; Aquificae; Aquificales; Aquificaceae; Hydrogenivirga MQTLPEVFKAKGYSRRDFLKFATLTTAILGLPPSMLPKVVHALETKPKPVVIWLEFQDCA WP_008287443.1 Ni/Fe hydrogenase [Hydrogenivirga sp. 128-5-R1-1] Length: 344\nNumber of predicted TMHs: 1\nExp number of AAs in TMHs: 16.61373\nExp number, first 60 AAs: 1.34194\nTotal prob of N-in: 0.08914\noutside 1 318\nTMhelix 319 336\ninside 337 344

17 GCF_000010785.1_ASM1078v1 Hydrogenobacter thermophilus TK-6 Aquificae; Aquificae; Aquificales; Aquificaceae; Hydrogenobacter; Hydrogenobacter thermophilus MESLYHIYRRRGYTRRDFLKSVLLLSAVLGFPPTSFGQAVKALELKPKPVVIWLEFQDCA WP_012962742.1 Ni/Fe hydrogenase [Hydrogenobacter thermophilus] Length: 345\nNumber of predicted TMHs: 2\nExp number of AAs in TMHs: 34.60353\nExp number, first 60 AAs: 16.4103\nTotal prob of N-in: 0.84750\nPOSSIBLE N-term signal sequence\ninside 1 20\nTMhelix 21 40\noutside 41 315\nTMhelix 316 338\ninside 339 345

18 GCF_000164905.1_ASM16490v1 Hydrogenobacter thermophilus TK-6 Aquificae; Aquificae; Aquificales; Aquificaceae; Hydrogenobacter; Hydrogenobacter thermophilus MESLYHIYRRRGYTRRDFLKSVLLLSAVLGFPPTSFGQAVKALELKPKPVVIWLEFQDCA WP_012962742.1 Ni/Fe hydrogenase [Hydrogenobacter thermophilus] Length: 345\nNumber of predicted TMHs: 2\nExp number of AAs in TMHs: 34.60353\nExp number, first 60 AAs: 16.4103\nTotal prob of N-in: 0.84750\nPOSSIBLE N-term signal sequence\ninside 1 20\nTMhelix 21 40\noutside 41 315\nTMhelix 316 338\ninside 339 345

19 GCF_000025605.1_ASM2560v1 Thermocrinis albus DSM 14484 Aquificae; Aquificae; Aquificales; Aquificaceae; Thermocrinis; Thermocrinis albus METVWDLFRRRGYSRREFLSFVSVTASLLALPANLLMSFGRALELREKPVVIWLEFQDCA WP_012992454.1 Ni/Fe hydrogenase [Thermocrinis albus] Length: 348\nNumber of predicted TMHs: 2\nExp number of AAs in TMHs: 39.3859099999999\nExp number, first 60 AAs: 20.03198\nTotal prob of N-in: 0.94334\nPOSSIBLE N-term signal sequence\ninside 1 20\nTMhelix 21 43\noutside 44 319\nTMhelix 320 339\ninside 340 348

20 GCF_000177635.2_ASM17763v2 Desulfurispirillum indicum S5 Chrysiogenetes; Chrysiogenetes; Chrysiogenales; Chrysiogenaceae; Desulfurispirillum; Desulfurispirillum indicum MQFSRRSFLAGMAAAATATAVPTKAKAARRYSGDISPDSYATLIDLTKCDGCPGKDMAAC WP_013506165.1 4Fe-4S ferredoxin iron-sulfur-binding domain-containing protein [Desulfurispirillum indicum] Length: 349\nNumber of predicted TMHs: 1\nExp number of AAs in TMHs: 21.28963\nExp number, first 60 AAs: 0.24015\nTotal prob of N-in: 0.08382\noutside 1 300\nTMhelix 301 323\ninside 324 349

21 GCF_000024845.1_ASM2484v1 Rhodothermus marinus DSM 4252 FCB group; Bacteroidetes/Chlorobi group; Bacteroidetes; Bacteroidetes Order II. Incertae sedis; Rhodothermaceae; Rhodothermus; Rhodothermus marinus MLPEQSLTTPLPATATAAPARRAAVLGVGAALPAHREPSTETERRLGLPPGWIARRTGIR WP_012842626.1 ketoacyl-ACP synthase III [Rhodothermus marinus] Length: 343\nNumber of predicted TMHs: 1\nExp number of AAs in TMHs: 17.12196\nExp number, first 60 AAs: 0.00231\nTotal prob of N-in: 0.01032\noutside 1 318\nTMhelix 319 341\ninside 342 343

22 GCF_000565305.1_ASM56530v1 Rhodothermus marinus SG0.5JP17-171 FCB group; Bacteroidetes/Chlorobi group; Bacteroidetes; Bacteroidetes Order II. Incertae sedis; Rhodothermaceae; Rhodothermus; Rhodothermus marinus MLPEQSLTTPLPATTTAAPARRAAVLGVGAALPAHREPSAETERRLGLPPGWIARRTGIR WP_014065834.1 ketoacyl-ACP synthase III [Rhodothermus marinus] Length: 343\nNumber of predicted TMHs: 1\nExp number of AAs in TMHs: 17.40927\nExp number, first 60 AAs: 0.00143\nTotal prob of N-in: 0.01106\noutside 1 318\nTMhelix 319 341\ninside 342 343

23 GCF_000224745.1_ASM22474v1 Rhodothermus marinus SG0.5JP17-172 FCB group; Bacteroidetes/Chlorobi group; Bacteroidetes; Bacteroidetes Order II. Incertae sedis; Rhodothermaceae; Rhodothermus; Rhodothermus marinus MLPEQSLTTPLPATTTAAPARRAAVLGVGAALPAHREPSAETERRLGLPPGWIARRTGIR WP_014065834.1 ketoacyl-ACP

synthase III [Rhodothermus marinus] Length: 343\nNumber of predicted TMHs: 1\nExp number of AAs in TMHs: 17.40927\nExp number, first 60 AAs: 0.00143\nTotal prob of N-in: 0.01106\noutside 1 318\nTMhelix 319 341\ninside 342 343

24 GCF_000013045.1_ASM1304v1 Salinibacter ruber DSM 13855 FCB group; Bacteroidetes/Chlorobi group; Bacteroidetes; Bacteroidetes Order II. Incertae sedis; Rhodothermaceae; Salinibacter; Salinibacter ruber MSNAFAGFLANRREALLQVLLIAGSLAALTYTANQFVLRADLTANNRYTLAGASHDIAQT YP_445373.1 hypothetical protein SRU_1249 [Salinibacter ruber DSM 13855] Length: 529\nNumber of predicted TMHs: 1\nExp number of AAs in TMHs: 29.6311599999999\nExp number, first 60 AAs: 8.43014\nTotal prob of N-in: 0.36953\noutside 1 489\nTMhelix 490 512\ninside 513 529

25 GCF_000090405.2_ASM9040v2 Salinibacter ruber M8 FCB group; Bacteroidetes/Chlorobi group; Bacteroidetes; Bacteroidetes Order II. Incertae sedis; Rhodothermaceae; Salinibacter; Salinibacter ruber MSNAFAGFLANRREALLQVLLIAGSLAALTYTANQFVLRADLTANNRYTLAGASHDIAQT WP_013061775.1 hypothetical protein [Salinibacter ruber] Length: 529\nNumber of predicted TMHs: 1\nExp number of AAs in TMHs: 29.62681\nExp number, first 60 AAs: 8.42821\nTotal prob of N-in: 0.36958\noutside 1 489\nTMhelix 490 512\ninside 513 529

26 GCF_001518995.2_ASM151899v2 Rhodothermaceae bacterium RA FCB group; Bacteroidetes/Chlorobi group; Bacteroidetes; Bacteroidetes Order II. Incertae sedis; Rhodothermaceae; unclassified Rhodothermaceae MDVQPTSSPPSGPIDERRARAEQTFWEMAAVLYRRRRFILGTTMAVAVLSVVLSLMMPVW WP_068125319.1 hypothetical protein [Rhodothermaceae bacterium RA] Length: 403\nNumber of predicted TMHs: 2\nExp number of AAs in TMHs: 46.07167\nExp number, first 60 AAs: 22.7586\nTotal prob of N-in: 0.97419\nPOSSIBLE N-term signal sequence\ninside 1 37\nTMhelix 38 60\noutside 61 360\nTMhelix 361 380\ninside 381 403

27 GCF_000613985.1_ASM61398v1 Prevotella nanceiensis DSM 19126 = JCM 15639 FCB group; Bacteroidetes/Chlorobi group; Bacteroidetes; Bacteroidia; Bacteroidales; Prevotellaceae; Prevotella; Prevotella nanceiensis MNEGELLYREMNEEDPWYMGDHDEEDEDNKDYSGEEEETPRRNAFVLFWNPTNSNYTIED WP_018362232.1 hypothetical protein [Prevotella nanceiensis] Length: 232\nNumber of predicted TMHs: 1\nExp number of AAs in TMHs: 19.19451\nExp number, first 60 AAs: 0.00035\nTotal prob of N-in: 0.91716\ninside 1 213\nTMhelix 214 231\noutside 232 232

28 GCF_000379965.1_ASM37996v1 Prevotella nanceiensis DSM 19126 = JCM 15639 FCB group; Bacteroidetes/Chlorobi group; Bacteroidetes; Bacteroidia; Bacteroidales; Prevotellaceae; Prevotella; Prevotella nanceiensis MNEGELLYREMNEEDPWYMGDHDEEDEDNKDYSGEEEETPRRNAFVLFWNPTNSNYTIED WP_018362232.1 hypothetical protein [Prevotella nanceiensis] Length: 232\nNumber of predicted TMHs: 1\nExp number of AAs in TMHs: 19.19451\nExp number, first 60 AAs: 0.00035\nTotal prob of N-in: 0.91716\ninside 1 213\nTMhelix 214 231\noutside 232 232

29 GCF_000312445.1_ASM31244v1 Phocaeicola abscessus CCUG 55929 FCB group; Bacteroidetes/Chlorobi group; Bacteroidetes; Bacteroidia; Bacteroidales; unclassified Bacteroidales; Phocaeicola; Phocaeicola abscessus MSQTHTPPIKETERQYEEQEIDLTEIVRKLWQRRKLIYKVCGGAALFGLLIAFSIPKEYT WP_019208887.1 hypothetical protein [Phocaeicola abscessus] Length: 378\nNumber of predicted TMHs: 2\nExp number of AAs in TMHs: 41.53124\nExp number, first 60 AAs: 19.40515\nTotal prob of N-in: 0.95571\nPOSSIBLE N-term signal sequence\ninside 1 35\nTMhelix 36 55\noutside 56 334\nTMhelix 335 354\ninside 355 378

30 GCF_002127285.1_ASM212728v1 Chitinophagaceae bacterium IBVUCB1 FCB group; Bacteroidetes/Chlorobi group; Bacteroidetes; Chitinophagia; Chitinophagales; Chitinophagaceae; unclassified Chitinophagaceae MAYHERRKNFIFSHYSISHPQHMSLQNQQANQTNNTNTDGLYIAIIRLLSSCIKKIKNSI WP_086099802.1 hypothetical protein [Chitinophagaceae bacterium IBVUCB1] Length: 316\nNumber of predicted TMHs: 2\nExp number of AAs in TMHs: 44.25117\nExp number, first 60 AAs: 1.60904\nTotal prob of N-in: 0.82368\ninside 1 59\nTMhelix 60 79\noutside 80 286\nTMhelix 287 309\ninside 310 316

31 GCF_001424405.1_Leaf189 Dyadobacter sp. Leaf189 FCB group; Bacteroidetes/Chlorobi group; Bacteroidetes; Cytophagia; Cytophagales; Cytophagaceae; Dyadobacter MRKTKRYKRRNLLTSPGTLTYVGPEIALKTTVRRIQYNEKMIKDEAVKSLKDCLPAGEGE WP_056283350.1 magnesium and cobalt transport protein CorA [Dyadobacter sp. Leaf189] Length: 358\nNumber of predicted TMHs: 2\nExp number of AAs in TMHs: 41.65788\nExp number, first 60 AAs: 0.0032\nTotal prob of N-in: 0.99366\ninside 1 300\nTMhelix 301 320\noutside 321 329\nTMhelix 330 352\ninside 353 358

32 GCF_000701505.1_ASM70150v1 Dyadobacter crusticola DSM 16708 FCB group; Bacteroidetes/Chlorobi group; Bacteroidetes; Cytophagia; Cytophagales; Cytophagaceae; Dyadobacter; Dyadobacter crusticola
MKKTKRYKRRNLLTSPGTLTYVGPDIALKTTVRRIQYNEKMIKDEPVKSLKDCLPAERGE WP_031525750.1 magnesium and cobalt transport protein CorA [Dyadobacter crusticola] Length: 358\nNumber of predicted TMHs: 2\nExp number of AAs in TMHs: 41.72572\nExp number, first 60 AAs: 0.00317\nTotal prob of N-in: 0.99317\ninside 1 300\nTMhelix 301 320\noutside 321 329\nTMhelix 330 352\ninside 353 358

33 GCF_002108455.1_ASM210845v1 Fibrella sp. ES10-3-2-2 FCB group; Bacteroidetes/Chlorobi group; Bacteroidetes; Cytophagia; Cytophagales; Cytophagaceae; Fibrella
MSTIQPANRVTDSDEDVIEIRLSDIVSFLKRSRRAMVLGAVVGGVLGALYAFSLPNQYTS WP_085411212.1 lipopolysaccharide biosynthesis protein [Fibrella sp. ES10-3-2-2] Length: 355\nNumber of predicted TMHs: 2\nExp number of AAs in TMHs: 46.96366\nExp number, first 60 AAs: 19.69321\nTotal prob of N-in: 0.89022\nPOSSIBLE N-term signal sequence\ninside 1 34\nTMhelix 35 54\noutside 55 330\nTMhelix 331 350\ninside 351 355

34 GCF_000383955.1_ASM38395v1 Rudanella lutea DSM 19387 FCB group; Bacteroidetes/Chlorobi group; Bacteroidetes; Cytophagia; Cytophagales; Cytophagaceae; Rudanella; Rudanella lutea
MVIIGPTCLTHMLETPDFELATSPLASRPVWATLWQARRRLLGSTLAGLLLGLILAFVLP WP_083940528.1 hypothetical protein [Rudanella lutea] Length: 360\nNumber of predicted TMHs: 2\nExp number of AAs in TMHs: 41.47953\nExp number, first 60 AAs: 18.9382\nTotal prob of N-in: 0.57810\nPOSSIBLE N-term signal sequence\ninside 1 40\nTMhelix 41 60\noutside 61 334\nTMhelix 335 357\ninside 358 360

35 GCF_001988955.1_ASM198895v1 Spirosoma montaniterrae FCB group; Bacteroidetes/Chlorobi group; Bacteroidetes; Cytophagia; Cytophagales; Cytophagaceae; Spirosoma
MSTTTLSQPTIQPDRDPDEIEIRVSDIIQFVKDSRRAMFLWGAALAIVGAIYAFTQPNEY WP_077133151.1 lipopolysaccharide biosynthesis protein [Spirosoma montaniterrae] Length: 360\nNumber of predicted TMHs: 2\nExp number of AAs in TMHs: 39.51387\nExp number, first 60 AAs: 18.69954\nTotal prob of N-in: 0.56777\nPOSSIBLE N-term signal sequence\ninside 1 36\nTMhelix 37 56\noutside 57 332\nTMhelix 333 352\ninside 353 360

36 GCF_001988955.1_ASM198895v1 Spirosoma montaniterrae FCB group; Bacteroidetes/Chlorobi group; Bacteroidetes; Cytophagia; Cytophagales; Cytophagaceae; Spirosoma
MNKSRRDVLKRMSLAALAGFTPLYGFSDATKKRIVTGEDMYRGIFFAEGAFAQLIPEIQS WP_077132096.1 hypothetical protein [Spirosoma montaniterrae] Length: 205\nNumber of predicted TMHs: 1\nExp number of AAs in TMHs: 23.06755\nExp number, first 60 AAs: 1.52134\nTotal prob of N-in: 0.89716\ninside 1 151\nTMhelix 152 174\noutside 175 205

37 GCF_000974425.1_ASM97442v1 Spirosoma radiotolerans FCB group; Bacteroidetes/Chlorobi group; Bacteroidetes; Cytophagia; Cytophagales; Cytophagaceae; Spirosoma
MKTSRRSALKQLSFALLAGFSPLAGFGSSEKNRLFTGEEMYRGIFFAEGQFARLIPEIQS WP_046574408.1 hypothetical protein [Spirosoma radiotolerans] Length: 204\nNumber of predicted TMHs: 1\nExp number of AAs in TMHs: 24.23603\nExp number, first 60 AAs: 2.46925\nTotal prob of N-in: 0.92828\ninside 1 150\nTMhelix 151 173\noutside 174 204

38 GCF_002067135.1_ASM206713v1 Spirosoma rigui FCB group; Bacteroidetes/Chlorobi group; Bacteroidetes; Cytophagia; Cytophagales; Cytophagaceae; Spirosoma
MSVTEGKSTEPKSHEIEIRLSDIVQFLKSSRRTVISWALGGLVVGGVYAAFQPNQFTSYV WP_080237806.1 lipopolysaccharide biosynthesis protein [Spirosoma rigui] Length: 362\nNumber of predicted TMHs: 2\nExp number of AAs in TMHs: 42.51158\nExp number, first 60 AAs: 17.87224\nTotal prob of N-in: 0.70684\nPOSSIBLE N-term signal sequence\ninside 1 33\nTMhelix 34 51\noutside 52 336\nTMhelix 337 359\ninside 360 362

39 GCF_000715495.1_ASM71549v1 Hymenobacter sp. IS2118 FCB group; Bacteroidetes/Chlorobi group; Bacteroidetes; Cytophagia; Cytophagales; Hymenobacteraceae; Hymenobacter
PAMPPPAPRRAAPAPRQLDAVPVAPPDTTRRATVATPLPGPMAAPAPVYAPTPAGQLPGT WP_035564895.1 hypothetical protein, partial [Hymenobacter sp. IS2118] Length: 291\nNumber of predicted TMHs: 2\nExp number of AAs in TMHs: 51.292\nExp number, first 60 AAs: 0.015\nTotal prob of N-in: 0.38387\ninside 1 227\nTMhelix 228 245\noutside 246 254\nTMhelix 255 272\ninside 273 291

40 GCF_900119185.1_IMG-taxon_2596583572_annotated_assembly Sinomicrobium oceani FCB group; Bacteroidetes/Chlorobi group; Bacteroidetes; Flavobacteriia; Flavobacteriales; Flavobacteriaceae; Sinomicrobium
MKKKTRPRKRRRKNLGSVPGTMTYTGSKTEQEFYIEVIDFSKEQCSRTVYNDVKDVFRYAWP_072315592.1 magnesium and cobalt transport protein CorA [Sinomicrobium oceani] Length: 355\nNumber of predicted TMHs: 2\nExp number of

AAs in TMHs: 41.83398\nExp number, first 60 AAs: 0\nTotal prob of N-in: 0.99727\ninside 1 293\nTMhelix 294 316\noutside 317 330\nTMhelix 331 349\ninside 350 355

41 GCF_000442105.1_BactbactF0290v01 Bacteroidetes bacterium oral taxon 272 str. F0290 FCB group; Bacteroidetes/Chlorobi group; Bacteroidetes; unclassified Bacteroidetes; unclassified Bacteroidetes (miscellaneous); Bacteroidetes bacterium oral taxon 272
MSQTHTPPIKETERQYEEQEIDLTEIVRKLWQRRKLIYKVCGGAALFGLLIAFSIPKEYT WP_019208887.1 hypothetical protein [Phocaeicola abscessus] Length: 378\nNumber of predicted TMHs: 2\nExp number of AAs in TMHs: 41.53124\nExp number, first 60 AAs: 19.40515\nTotal prob of N-in: 0.95571\nPOSSIBLE N-term signal sequence\ninside 1 35\nTMhelix 36 55\noutside 56 334\nTMhelix 335 354\ninside 355 378

42 GCF_000279145.1_ASM27914v1 Melioribacter roseus P3M-2 FCB group; Bacteroidetes/Chlorobi group; Ignavibacteriae; Ignavibacteria; Ignavibacteriales; Melioribacteraceae; Melioribacter; Melioribacter roseus
MSKQTRRNFIKTAAVAGSVIAGLPQKAKAAPKNILSDDRMGVLVDTTVCVGCRKCEWACK WP_014856242.1 4Fe-4S ferredoxin [Melioribacter roseus] Length: 300\nNumber of predicted TMHs: 1\nExp number of AAs in TMHs: 25.11422\nExp number, first 60 AAs: 0.13205\nTotal prob of N-in: 0.36501\noutside 1 266\nTMhelix 267 289\ninside 290 300

43 GCF_000695095.2_ASM69509v2 Gemmatimonas phototrophica FCB group; Gemmatimonadetes; Gemmatimonadetes; Gemmatimonadales; Gemmatimonadaceae; Gemmatimonas
MSHSSAPAHRQRRAAVPTIALAAWTIALPIMGCYSNVPVATGVTPTPGEGVVVLTTDGTS WP_026850174.1 hypothetical protein [Gemmatimonas phototrophica] Length: 162\nNumber of predicted TMHs: 2\nExp number of AAs in TMHs: 42.8737\nExp number, first 60 AAs: 20.43253\nTotal prob of N-in: 0.09416\nPOSSIBLE N-term signal sequence\noutside 1 14\nTMhelix 15 37\ninside 38 127\nTMhelix 128 150\noutside 151 162

44 GCF_000010305.1_ASM1030v1 Gemmatimonas aurantiaca T-27 FCB group; Gemmatimonadetes; Gemmatimonadetes; Gemmatimonadales; Gemmatimonadaceae; Gemmatimonas; Gemmatimonas aurantiaca
MRALWQSQSFRLPTRRRAWVALFGATAALGMAGCGGEESGAMTADLERDLQMAVNAQRPR WP_041265614.1 hypothetical protein [Gemmatimonas aurantiaca] Length: 241\nNumber of predicted TMHs: 1\nExp number of AAs in TMHs: 33.37214\nExp number, first 60 AAs: 11.07004\nTotal prob of N-in: 0.55783\nPOSSIBLE N-term signal sequence\noutside 1 209\nTMhelix 210 232\ninside 233 241

45 GCF_000196815.1_ASM19681v1 Nitrospira defluvii Nitrospirae; Nitrospira; Nitrospirales; Nitrospiraceae; Nitrospira MSRLLDTPQVVMTPSRGHGANLLSRDGMVPQDLLSVVSRRRGLVATVLMVSLLGAFAVSA WP_013247481.1 LPS biosynthesis protein [Nitrospira defluvii] Length: 408\nNumber of predicted TMHs: 2\nExp number of AAs in TMHs: 55.6903900000001\nExp number, first 60 AAs: 18.57896\nTotal prob of N-in: 0.62484\nPOSSIBLE N-term signal sequence\noutside 1 41\nTMhelix 42 64\ninside 65 373\nTMhelix 374 396\noutside 397 408

46 GCF_001273775.1_ASM127377v1 Nitrospira moscoviensis Nitrospirae; Nitrospira; Nitrospirales; Nitrospiraceae; Nitrospira MWGARRNALAVAVAAGVSLSSPVVLLATHEADHRFTVEGFVCGADGKGRANVDVLIKDTR WP_053378551.1 hypothetical protein [Nitrospira moscoviensis] Length: 174\nNumber of predicted TMHs: 2\nExp number of AAs in TMHs: 43.63376\nExp number, first 60 AAs: 21.41021\nTotal prob of N-in: 0.98134\nPOSSIBLE N-term signal sequence\ninside 1 6\nTMhelix 7 29\noutside 30 132\nTMhelix 133 155\ninside 156 174

47 GCF_900170025.1_NSND_Contig Nitrospira sp. ND1 Nitrospirae; Nitrospira; Nitrospirales; Nitrospiraceae; Nitrospira MSRVLDSPQVVMTPPRGHGANLTSRDGMGPQDLLSVVSRRRGLVATIIAASLLGAFAVSA WP_080878775.1 hypothetical protein [Nitrospira sp. ND1] Length: 408\nNumber of predicted TMHs: 2\nExp number of AAs in TMHs: 49.11103\nExp number, first 60 AAs: 18.41976\nTotal prob of N-in: 0.52726\nPOSSIBLE N-term signal sequence\ninside 1 41\nTMhelix 42 64\noutside 65 373\nTMhelix 374 396\ninside 397 408

48 GCF_001514535.1_ASM151453v1 Thermodesulfovibrio aggregans Nitrospirae; Nitrospira; Nitrospirales; Nitrospiraceae; Thermodesulfovibrio
MNISRRGFLALSGAVCGSLILGNEVEAREGKYATLIDLTKCDGCKDEPIPRCVKACRDYN WP_059176917.1 oxidoreductase [Thermodesulfovibrio aggregans]Length: 324\nNumber of predicted TMHs: 1\nExp number of AAs in TMHs: 18.68824\nExp number, first 60 AAs: 0.63701\nTotal prob of N-in: 0.21349\noutside 1 293\nTMhelix 294 313\ninside 314 324

49 GCF_000482825.1_ASM48282v1 Thermodesulfovibrio islandicus DSM 12570 Nitrospirae; Nitrospira; Nitrospirales; Nitrospiraceae; Thermodesulfovibrio; Thermodesulfovibrio islandicus
MEISRRGFLALSSAIGASLLLGNNAEAVEGKYATLIDLTKCDGCKDEPIPRCVKACREYN WP_012545259.1 MULTISPECIES:

oxidoreductase [Thermodesulfovibrio] Length: 325\nNumber of predicted TMHs: 1\nExp number of AAs in TMHs: 17.96531\nExp number, first 60 AAs: 0.6169\nTotal prob of N-in: 0.07244\noutside 1 293\nTMhelix 294 313\ninside 314 325

50 GCF_000020985.1_ASM2098v1 Thermodesulfovibrio yellowstonii DSM 11347 Nitrospirae; Nitrospira; Nitrospirales; Nitrospiraceae; Thermodesulfovibrio; Thermodesulfovibrio yellowstonii
MEISRRGFLALSSAIGASLLLGNNAEAVEGKYATLIDLTKCDGCKDEPIPRCVKACREYN YP_002248525.1 iron-sulfur protein [Thermodesulfovibrio yellowstonii DSM 11347] Length: 325\nNumber of predicted TMHs: 1\nExp number of AAs in TMHs: 17.96531\nExp number, first 60 AAs: 0.6169\nTotal prob of N-in: 0.07244\noutside 1 293\nTMhelix 294 313\ninside 314 325

51 GCF_001705755.1_ASM170575v1 Acidithiobacillus thiooxidans Proteobacteria; Acidithiobacillia; Acidithiobacillales; Acidithiobacillaceae; Acidithiobacillus
LATAGFATILLWMLRESHSRRALLGGVSALLVLSGVALAAGSVSFTHPVGHKLRVSLVQG WP_065966182.1 apolipoprotein N-acyltransferase, partial [Acidithiobacillus thiooxidans] Length: 334\nNumber of predicted TMHs: 2\nExp number of AAs in TMHs: 43.1001399999999\nExp number, first 60 AAs: 21.78244\nTotal prob of N-in: 0.90299\nPOSSIBLE N-term signal sequence\ninside 1 20\nTMhelix 21 43\noutside 44 305\nTMhelix 306 325\ninside 326 334

52 GCF_000813765.1_ASM81376v1 Brevundimonas nasdae Proteobacteria; Alphaproteobacteria; Caulobacterales; Caulobacteraceae; Brevundimonas
MSKIILSRRRLLVAAGALSLAGCQSVPPSEDTPTGSRTVGDYELDSGDKIRLSVFGEEEL WP_039245948.1 polysaccharide biosynthesis protein [Brevundimonas nasdae] Length: 191\nNumber of predicted TMHs: 1\nExp number of AAs in TMHs: 18.13226\nExp number, first 60 AAs: 0.2933\nTotal prob of N-in: 0.20568\noutside 1 127\nTMhelix 128 150\ninside 151 191

53 GCF_001425945.1_Root1279 Brevundimonas sp. Root1279 Proteobacteria; Alphaproteobacteria; Caulobacterales; Caulobacteraceae; Brevundimonas
MARRTVVIVGAGFGGLAAAKALKHANVDVVLIDRTNHHLFQPLLYQVATAALSPADIATA WP_056452081.1 FAD-dependent oxidoreductase [Brevundimonas sp. Root1279] Length: 414\nNumber of predicted TMHs: 1\nExp number of AAs in TMHs: 29.09579\nExp number, first 60 AAs: 2.7821\nTotal prob of N-in: 0.41979\noutside 1 360\nTMhelix 361 383\ninside 384 414

54 GCF_000421705.1_ASM42170v1 Brevundimonas naejangsanensis DSM 23858 Proteobacteria; Alphaproteobacteria; Caulobacterales; Caulobacteraceae; Brevundimonas; Brevundimonas naejangsanensis
MEKPRRRRRSPEEARREALASARILLLSGGPNAVTLAAVAAEIGVTHANLIHHFGSAAGL WP_029673426.1 TetR family transcriptional regulator [Brevundimonas naejangsanensis] Length: 202\nNumber of predicted TMHs: 2\nExp number of AAs in TMHs: 36.46862\nExp number, first 60 AAs: 17.65938\nTotal prob of N-in: 0.94839\nPOSSIBLE N-term signal sequence\ninside 1 23\nTMhelix 24 46\noutside 47 153\nTMhelix 154 176\ninside 177 202

55 GCF_000281955.1_Caulobacter.strAP07_v1.0 Caulobacter sp. AP07 Proteobacteria; Alphaproteobacteria; Caulobacterales; Caulobacteraceae; Caulobacter
MVILGRRDLYGSGKAPRSWSPSLCSRRRAAFGWLAFSFGGAVSRKILIINGHPDASAERL WP_081497709.1 hypothetical protein [Caulobacter sp. AP07] Length: 227\nNumber of predicted TMHs: 1\nExp number of AAs in TMHs: 26.61804\nExp number, first 60 AAs: 1.63673\nTotal prob of N-in: 0.41179\noutside 1 163\nTMhelix 164 186\ninside 187 227

56 GCF_001449105.1_ASM144910v1 Caulobacter vibrioides Proteobacteria; Alphaproteobacteria; Caulobacterales; Caulobacteraceae; Caulobacter
MSFLESIRTTTSRRPIITAVTGVAVLATAFAALYFFALRKPYGVLFTDLRTMDAATIVAD WP_058346279.1 flagellar M-ring protein FliF [Caulobacter vibrioides] Length: 410\nNumber of predicted TMHs: 2\nExp number of AAs in TMHs: 45.2303799999999\nExp number, first 60 AAs: 22.32074\nTotal prob of N-in: 0.89608\nPOSSIBLE N-term signal sequence\ninside 1 15\nTMhelix 16 38\noutside 39 353\nTMhelix 354 376\ninside 377 410

57 GCF_001463885.1_ASM146388v1 Aureimonas altamirensis Proteobacteria; Alphaproteobacteria; Rhizobiales; Aurantimonadaceae; Aureimonas
MTHRAPQAATPSAPVASRRRSAVVAACGALLTLGFVSLGVWQVERLYWKLDLIERVDARI WP_060606411.1
Surfeit locus 1 family protein [Aureimonas altamirensis] Length: 273\nNumber of predicted TMHs: 2\nExp number of AAs in TMHs: 50.95838\nExp number, first 60 AAs: 22.63591\nTotal prob of N-in: 0.99568\nPOSSIBLE N-term signal sequence\ninside 1 20\nTMhelix 21 43\noutside 44 222\nTMhelix 223 245\ninside 246 273

58 GCF_000800175.1_ASM80017v1 Aureimonas altamirensis Proteobacteria; Alphaproteobacteria; Rhizobiales; Aurantimonadaceae; Aureimonas MGHVPPQHQHDADGRGQQHAVTHRAPQAGPSSAPVASRRRSAVVAACGALLTLGFVSLGV WP_082016008.1 hypothetical protein [Aureimonas altamirensis] Length: 293\nNumber of predicted TMHs: 2\nExp number of AAs in TMHs: 52.18068\nExp number, first 60 AAs: 18.75251\nTotal prob of N-in: 0.84697\nPOSSIBLE N-term signal sequence\ninside 1 40\nTMhelix 41 63\noutside 64 242\nTMhelix 243 265\ninside 266 293

59 GCF_001423245.1_Leaf324 Aureimonas sp. Leaf324 Proteobacteria; Alphaproteobacteria; Rhizobiales; Aurantimonadaceae; Aureimonas MRARRSLLAAALVILLPAAALGQDSPAPLAETFEIGLSTETISVGTNFGGSRLVVFGALD WP_055888441.1 hypothetical protein [Aureimonas sp. Leaf324] Length: 265\nNumber of predicted TMHs: 1\nExp number of AAs in TMHs: 32.28833\nExp number, first 60 AAs: 9.98297\nTotal prob of N-in: 0.67364\noutside 1 239\nTMhelix 240 262\ninside 263 265

60 GCF_900141975.1_IMG-taxon_2585427611_annotated_assembly Aureimonas altamirensis DSM 21988 Proteobacteria; Alphaproteobacteria; Rhizobiales; Aurantimonadaceae; Aureimonas; Aureimonas altamirensis MTHRAPQAATPSAPVASRRRSAVVAACGALLTLGFVSLGVWQVERLYWKLDLIERVDARI WP_060606411.1 Surfeit locus 1 family protein [Aureimonas altamirensis] Length: 273\nNumber of predicted TMHs: 2\nExp number of AAs in TMHs: 50.95838\nExp number, first 60 AAs: 22.63591\nTotal prob of N-in: 0.99568\nPOSSIBLE N-term signal sequence\ninside 1 20\nTMhelix 21 43\noutside 44 222\nTMhelix 223 245\ninside 246 273

61 GCF_900176465.1_IMG-taxon_2718217637_annotated_assembly Fulvimarina manganoxydans Proteobacteria; Alphaproteobacteria; Rhizobiales; Aurantimonadaceae; Fulvimarina MQVNKAMESPLPQSATASPSSSRRSYIPFIAVMAGGVIDLLTKEWAQTVLQPYQPPIAIL WP_084410011.1 signal peptidase II [Fulvimarina manganoxydans] Length: 180\nNumber of predicted TMHs: 2\nExp number of AAs in TMHs: 53.13664\nExp number, first 60 AAs: 6.19815\nTotal prob of N-in: 0.76052\ninside 1 87\nTMhelix 88 105\noutside 106 147\nTMhelix 148 170\ninside 171 180

62 GCF_001006325.1_ASM100632v1 Afipia massiliensis Proteobacteria; Alphaproteobacteria; Rhizobiales; Bradyrhizobiaceae; Afipia MAISESDRHDAYLPRIQAPALDVDGIAAMLRRRAAFITAVTLACAGICLAWILLSAPKYV WP_046829270.1 succinoglycan biosynthesis transporter [Afipia massiliensis]Length: 467\nNumber of predicted TMHs: 2\nExp number of AAs in TMHs: 44.26059\nExp number, first 60 AAs: 21.82262\nTotal prob of N-in: 0.89152\nPOSSIBLE N-term signal sequence\ninside 1 33\nTMhelix 34 56\noutside 57 431\nTMhelix 432 454\ninside 455 467

63 GCF_000497635.1_OHSU_J-C4 Afipia sp. OHSU_I-C4 Proteobacteria; Alphaproteobacteria; Rhizobiales; Bradyrhizobiaceae; Afipia MAISENDRRDAYLSRIQVPALDVHGFATMLRRRAAFIGAVAAICAAISLAWILLSAPKYV WP_024573716.1 MULTISPECIES: succinoglycan biosynthesis transporter [Afipia] Length: 459\nNumber of predicted TMHs: 2\nExp number of AAs in TMHs: 56.0569000000001\nExp number, first 60 AAs: 21.67303\nTotal prob of N-in: 0.98414\nPOSSIBLE N-term signal sequence\ninside 1 33\nTMhelix 34 56\noutside 57 421\nTMhelix 422 444\ninside 445 459

64 GCF_000497655.1_OHSU_J-C6 Afipia sp. OHSU_I-C6 Proteobacteria; Alphaproteobacteria; Rhizobiales; Bradyrhizobiaceae; Afipia MAISENDRRDAYLSRIQVPALDVHGFATMLRRRAAFIGAVAAICAAISLAWILLSAPKYV WP_024573716.1 MULTISPECIES: succinoglycan biosynthesis transporter [Afipia] Length: 459\nNumber of predicted TMHs: 2\nExp number of AAs in TMHs: 56.0569000000001\nExp number, first 60 AAs: 21.67303\nTotal prob of N-in: 0.98414\nPOSSIBLE N-term signal sequence\ninside 1 33\nTMhelix 34 56\noutside 57 421\nTMhelix 422 444\ninside 445 459

65 GCF_000497575.1_OHSU_S-C1 Afipia sp. OHSU_II-C1 Proteobacteria; Alphaproteobacteria; Rhizobiales; Bradyrhizobiaceae; Afipia MAISENDRRDAYLSRIQVPALDVHGFATMLRRRAAFIGAVAAICAAISLAWILLSAPKYV WP_024573716.1 MULTISPECIES: succinoglycan biosynthesis transporter [Afipia] Length: 459\nNumber of predicted TMHs: 2\nExp number of AAs in TMHs: 56.0569000000001\nExp number, first 60 AAs: 21.67303\nTotal prob of N-in: 0.98414\nPOSSIBLE N-term signal sequence\ninside 1 33\nTMhelix 34 56\noutside 57 421\nTMhelix 422 444\ninside 445 459

66 GCF_000497595.1_OHSU_S-C2 Afipia sp. OHSU_II-C2 Proteobacteria; Alphaproteobacteria; Rhizobiales; Bradyrhizobiaceae; Afipia MAISENDRRDAYLSRIQVPALDVHGFATMLRRRAAFIGAVAAICAAISLAWILLSAPKYV WP_024573716.1 MULTISPECIES: succinoglycan biosynthesis transporter [Afipia] Length: 459\nNumber of predicted TMHs: 2\nExp number of AAs in TMHs: 56.0569000000001\nExp number, first 60 AAs: 21.67303\nTotal prob of N-in: 0.98414\nPOSSIBLE N-term signal sequence\ninside 1 33\nTMhelix 34 56\noutside 57 421\nTMhelix 422 444\ninside 445 459

67 GCF_000497555.1_OHSU_uncloned Afipia sp. OHSU_II-uncloned Proteobacteria; Alphaproteobacteria; Rhizobiales; Bradyrhizobiaceae; Afipia MAISENDRRDAYLSRIQVPALDVHGFATMLRRRAAFIGAVAAICAAISLAWILLSAPKYV WP_024573716.1 MULTISPECIES: succinoglycan biosynthesis transporter [Afipia] Length: 459\nNumber of predicted TMHs: 2\nExp number of AAs in TMHs: 56.0569000000001\nExp number, first 60 AAs: 21.67303\nTotal prob of N-in: 0.98414\nPOSSIBLE N-term signal sequence\ninside 1 33\nTMhelix 34 56\noutside 57 421\nTMhelix 422 444\ninside 445 459

68 GCF_000497615.1_OHSU_J_uncloned Afipia sp. OHSU_I-uncloned Proteobacteria; Alphaproteobacteria; Rhizobiales; Bradyrhizobiaceae; Afipia MAISENDRRDAYLSRIQVPALDVHGFATMLRRRAAFIGAVAAICAAISLAWILLSAPKYV WP_024573716.1 MULTISPECIES: succinoglycan biosynthesis transporter [Afipia] Length: 459\nNumber of predicted TMHs: 2\nExp number of AAs in TMHs: 56.0569000000001\nExp number, first 60 AAs: 21.67303\nTotal prob of N-in: 0.98414\nPOSSIBLE N-term signal sequence\ninside 1 33\nTMhelix 34 56\noutside 57 421\nTMhelix 422 444\ninside 445 459

69 GCF_000516555.1_P52-10 Afipia sp. P52-10 Proteobacteria; Alphaproteobacteria; Rhizobiales; Bradyrhizobiaceae; Afipia MSRRGALAIALAALLLAAGAGRAQAERLIASISNHRVTITPEYSGEELLLFGSIERDADT WP_034464293.1 membrane protein [Afipia sp. P52-10] Length: 261\nNumber of predicted TMHs: 1\nExp number of AAs in TMHs: 33.25907\nExp number, first 60 AAs: 10.6142\nTotal prob of N-in: 0.54172\nPOSSIBLE N-term signal sequence\ninside 1 235\nTMhelix 236 258\noutside 259 261

70 GCF_000308295.2_Ab_34632 Afipia birgiae 34632 Proteobacteria; Alphaproteobacteria; Rhizobiales; Bradyrhizobiaceae; Afipia; Afipia birgiae MAISEGDRRDAYLPRIQAPALDVDGIASMLRRRAAFIAAVTLACVGICLAWILLSAPRYI WP_019196192.1 MULTISPECIES: hypothetical protein [Bradyrhizobiaceae] Length: 467\nNumber of predicted TMHs: 2\nExp number of AAs in TMHs: 43.10182\nExp number, first 60 AAs: 21.87111\nTotal prob of N-in: 0.98975\nPOSSIBLE N-term signal sequence\ninside 1 33\nTMhelix 34 56\noutside 57 431\nTMhelix 432 454\ninside 455 467

71 GCF_001295925.1_ASM129592v1 Bosea sp. AAP35 Proteobacteria; Alphaproteobacteria; Rhizobiales; Bradyrhizobiaceae; Bosea MTHTRRSLIKGSLALAGAGLASLASPALAQQTYSESRSFSQNELVNSGHQFFGNVSRGLA WP_054144656.1 hypothetical protein [Bosea sp. AAP35] Length: 196\nNumber of predicted TMHs: 1\nExp number of AAs in TMHs: 22.59999\nExp number, first 60 AAs: 6.78389\nTotal prob of N-in: 0.84879\ninside 1 147\nTMhelix 148 170\noutside 171 196

72 GCF_001748145.1_ASM174814v1 Bosea sp. BIWAKO-01 Proteobacteria; Alphaproteobacteria; Rhizobiales; Bradyrhizobiaceae; Bosea MSHSSADATRRSMLLAAASGLAAAATPARAAPRDLKLTDLQKEADVACVYHCDFGDPQRF WP_069879344.1 hypothetical protein [Bosea sp. BIWAKO-01] Length: 174\nNumber of predicted TMHs: 1\nExp number of AAs in TMHs: 23.66863\nExp number, first 60 AAs: 4.76267\nTotal prob of N-in: 0.82108\ninside 1 147\nTMhelix 148 165\noutside 166 174

73 GCF_001426225.1_Leaf344 Bosea sp. Leaf344 Proteobacteria; Alphaproteobacteria; Rhizobiales; Bradyrhizobiaceae; Bosea MSAVETGERRRTLLWPTLLTIAGTIVLVMLGTWQLMRMSEKRAFIDRLAAQASGPPAAMP WP_056707620.1 hypothetical protein [Bosea sp. Leaf344] Length: 253\nNumber of predicted TMHs: 2\nExp number of AAs in TMHs: 44.2637699999999\nExp number, first 60 AAs: 22.75782\nTotal prob of N-in: 0.98369\nPOSSIBLE N-term signal sequence\ninside 1 12\nTMhelix 13 35\noutside 36 224\nTMhelix 225 247\ninside 248 253

74 GCF_001426225.1_Leaf344 Bosea sp. Leaf344 Proteobacteria; Alphaproteobacteria; Rhizobiales; Bradyrhizobiaceae; Bosea MIVTTRRTLIRGAAFALVALSQLPGVARAQSDGWFDIAGDDGKPMANTRLPVELTSEIEA WP_082540354.1 hypothetical protein [Bosea sp. Leaf344] Length: 234\nNumber of predicted TMHs: 1\nExp number of AAs in TMHs: 16.31398\nExp number, first 60 AAs: 1.11133\nTotal prob of N-in: 0.71837\ninside 1 188\nTMhelix 189 211\noutside 212 234

75 GCF_001429395.1_Root381 Bosea sp. Root381 Proteobacteria; Alphaproteobacteria; Rhizobiales; Bradyrhizobiaceae; Bosea MNGDAMAHMISAQTDRAVLEASDVRLGHLRARRRSFLKRYGWSWLVVALPTLLALVYYGL WP_082609718.1 hypothetical protein [Bosea sp. Root381] Length: 401\nNumber of predicted TMHs: 2\nExp number of AAs in TMHs: 44.99317\nExp number, first 60 AAs: 20.70059\nTotal prob of N-in: 0.83161\nPOSSIBLE N-term signal sequence\ninside 1 39\nTMhelix 40 62\noutside 63 367\nTMhelix 368 390\ninside 391 401

76 GCF_001429415.1_Root670 Bosea sp. Root670 Proteobacteria; Alphaproteobacteria; Rhizobiales; Bradyrhizobiaceae; Bosea MTQTRRNLIKAGLAMAGASVSGLPLGARAQNYQNQQSFSQNELVGSGHKFFGNVSRGLAL WP_082617556.1 hypothetical protein [Bosea sp. Root670] Length: 195\nNumber of predicted TMHs:

1\nExp number of AAs in TMHs: 18.76981\nExp number, first 60 AAs: 1.01737\nTotal prob of N-in: 0.85910\ninside 1 146\nTMhelix 147 169\noutside 170 195

77 GCF_000745475.1_ASM74547v1 Bosea sp. UNC402CLCol Proteobacteria; Alphaproteobacteria; Rhizobiales; Bradyrhizobiaceae; Bosea MIQTRRSLIKGGLAVAGTGLAGLPIPAIAQAQNYDNQRSFSQNELVGSGHKFFGNVSRGL WP_038359302.1 hypothetical protein [Bosea sp. UNC402CLCol] Length: 197\nNumber of predicted TMHs: 1\nExp number of AAs in TMHs: 33.42113\nExp number, first 60 AAs: 10.67496\nTotal prob of N-in: 0.95962\nPOSSIBLE N-term signal sequence\ninside 1 148\nTMhelix 149 171\noutside 172 197

78 GCF_900168195.1_IMG-taxon_2595698224_annotated_assembly Bosea thiooxidans Proteobacteria; Alphaproteobacteria; Rhizobiales; Bradyrhizobiaceae; Bosea MNQTRRSLIKGGLAVAGTGLTGLSVPALAQQQNYGDQRSFSQNELVSSGHKFFGNVSRGL WP_055729429.1 hypothetical protein [Bosea thiooxidans] Length: 197\nNumber of predicted TMHs: 1\nExp number of AAs in TMHs: 24.28801\nExp number, first 60 AAs: 8.40336\nTotal prob of N-in: 0.86901\ninside 1 148\nTMhelix 149 171\noutside 172 197

79 GCF_001420315.1_ASM142031v1 Bosea thiooxidans Proteobacteria; Alphaproteobacteria; Rhizobiales; Bradyrhizobiaceae; Bosea MNQTRRSLIKGGLAVAGTGLTGLSVPALAQQQNYGDQRSFSQNELVSSGHKFFGNVSRGL WP_055729429.1 hypothetical protein [Bosea thiooxidans] Length: 197\nNumber of predicted TMHs: 1\nExp number of AAs in TMHs: 24.28801\nExp number, first 60 AAs: 8.40336\nTotal prob of N-in: 0.86901\ninside 1 148\nTMhelix 149 171\noutside 172 197

80 GCF_001693385.1_ASM169338v1 Bradyrhizobium icense Proteobacteria; Alphaproteobacteria; Rhizobiales; Bradyrhizobiaceae; Bradyrhizobium MTGLSSRRPAVAGFAIFTLLMVAVFAGLGIWQLQRRVEKHALIAMLNERLAAAPEALPAQ WP_065726590.1 surfeit 1 protein [Bradyrhizobium icense] Length: 255\nNumber of predicted TMHs: 2\nExp number of AAs in TMHs: 43.57728\nExp number, first 60 AAs: 22.079\nTotal prob of N-in: 0.63731\nPOSSIBLE N-term signal sequence\ninside 1 8\nTMhelix 9 31\noutside 32 229\nTMhelix 230 249\ninside 250 255

81 GCF_001440395.1_ASM144039v1 Bradyrhizobium jicamae Proteobacteria; Alphaproteobacteria; Rhizobiales; Bradyrhizobiaceae; Bradyrhizobium MSALSSRRPAVTGFAIFTLLMVMAFSGLGIWQLQRRVEKHALIARLNERLAAVPEALPAA WP_057837174.1 surfeit 1 protein [Bradyrhizobium jicamae] Length: 255\nNumber of predicted TMHs: 2\nExp number of AAs in TMHs: 44.19113\nExp number, first 60 AAs: 21.71068\nTotal prob of N-in: 0.74698\nPOSSIBLE N-term signal sequence\ninside 1 8\nTMhelix 9 31\noutside 32 229\nTMhelix 230 249\ninside 250 255

82 GCF_001440475.1_ASM144047v1 Bradyrhizobium lablabi Proteobacteria; Alphaproteobacteria; Rhizobiales; Bradyrhizobiaceae; Bradyrhizobium MSALSSRRPAVAGFAIFTLLMVTAFSGLGIWQLQRRVEKHALIARLNERLAAAPEALPAA WP_057862396.1 surfeit 1 protein [Bradyrhizobium lablabi] Length: 255\nNumber of predicted TMHs: 2\nExp number of AAs in TMHs: 45.02028\nExp number, first 60 AAs: 22.00244\nTotal prob of N-in: 0.73028\nPOSSIBLE N-term signal sequence\ninside 1 8\nTMhelix 9 31\noutside 32 229\nTMhelix 230 249\ninside 250 255

83 GCF_900141755.1_IMG-taxon_2698536816_annotated_assembly Bradyrhizobium lablabi Proteobacteria; Alphaproteobacteria; Rhizobiales; Bradyrhizobiaceae; Bradyrhizobium MTGIEARRRNVAGFAIFTSVMLMVFIGLGVWQLQRRVEKHALIAALTERLAAAPAPLPTP WP_079541884.1 surfeit 1 protein [Bradyrhizobium lablabi] Length: 251\nNumber of predicted TMHs: 2\nExp number of AAs in TMHs: 42.3515099999999\nExp number, first 60 AAs: 21.77148\nTotal prob of N-in: 0.98599\nPOSSIBLE N-term signal sequence\ninside 1 11\nTMhelix 12 34\noutside 35 225\nTMhelix 226 245\ninside 246 251

84 GCF_001440035.1_ABySSBR3351v1 Bradyrhizobium manausense Proteobacteria; Alphaproteobacteria; Rhizobiales; Bradyrhizobiaceae; Bradyrhizobium MGAATETFYGVIRRQGITRRNFLKFCSLTAASFGLGPLAVSRIANALETRPRIPVIWMHG WP_057742222.1 uptake hydrogenase small subunit [Bradyrhizobium manausense] Length: 363\nNumber of predicted TMHs: 2\nExp number of AAs in TMHs: 37.57562\nExp number, first 60 AAs: 17.20676\nTotal prob of N-in: 0.79574\nPOSSIBLE N-term signal sequence\ninside 1 20\nTMhelix 21 43\noutside 44 326\nTMhelix 327 349\ninside 350 363

85 GCF_001693515.1_ASM169351v1 Bradyrhizobium paxllaeri Proteobacteria; Alphaproteobacteria; Rhizobiales; Bradyrhizobiaceae; Bradyrhizobium MSALSSRRPAVVGFATFTLLMVTAFSGLGIWQLQRRVEKHALIARLNERLAAAPEALPAA WP_065752960.1 surfeit 1 protein [Bradyrhizobium paxllaeri] Length: 255\nNumber of predicted TMHs: 2\nExp number of

AAs in TMHs: 45.14926\nExp number, first 60 AAs: 22.1285\nTotal prob of N-in: 0.57303\nPOSSIBLE N-term signal sequence\ninside 1 8\nTMhelix 9 31\noutside 32 229\nTMhelix 230 249\ninside 250 255

86 GCF_001556255.1_ASM155625v1 Bradyrhizobium sp. CCH10-C7 Proteobacteria; Alphaproteobacteria; Rhizobiales; Bradyrhizobiaceae; Bradyrhizobium MAISEGDRRDAYLPRIQAPALDVDGIASMLRRRAAFIAAVTLACVGICLAWILLSAPRYI WP_019196192.1 MULTISPECIES: hypothetical protein [Bradyrhizobiaceae] Length: 467\nNumber of predicted TMHs: 2\nExp number of AAs in TMHs: 43.10182\nExp number, first 60 AAs: 21.87111\nTotal prob of N-in: 0.98975\nPOSSIBLE N-term signal sequence\ninside 1 33\nTMhelix 34 56\noutside 57 431\nTMhelix 432 454\ninside 455 467

87 GCF_001556245.1_ASM155624v1 Bradyrhizobium sp. CCH4-A6 Proteobacteria; Alphaproteobacteria; Rhizobiales; Bradyrhizobiaceae; Bradyrhizobium MAISEGDRRDAYLPRIQAPALDVDGIASMLRRRAAFIAAVTLACVGICLAWILLSAPRYI WP_019196192.1 MULTISPECIES: hypothetical protein [Bradyrhizobiaceae] Length: 467\nNumber of predicted TMHs: 2\nExp number of AAs in TMHs: 43.10182\nExp number, first 60 AAs: 21.87111\nTotal prob of N-in: 0.98975\nPOSSIBLE N-term signal sequence\ninside 1 33\nTMhelix 34 56\noutside 57 431\nTMhelix 432 454\ninside 455 467

88 GCF_001556045.1_ASM155604v1 Bradyrhizobium sp. CCH5-F6 Proteobacteria; Alphaproteobacteria; Rhizobiales; Bradyrhizobiaceae; Bradyrhizobium MPQSRDRICDHNRLARRAFLVRLAALALLAVSVPVTPSLAQASGQVRVKIVKAGLLVGGG WP_061024826.1 hypothetical protein [Bradyrhizobium sp. CCH5-F6] Length: 159\nNumber of predicted TMHs: 2\nExp number of AAs in TMHs: 72.3524800000001\nExp number, first 60 AAs: 31.68164\nTotal prob of N-in: 0.99189\nPOSSIBLE N-term signal sequence\ninside 1 19\nTMhelix 20 42\noutside 43 105\nTMhelix 106 128\ninside 129 159

89 GCF_000472925.1_ASM47292v1 Bradyrhizobium sp. Ec3.3 Proteobacteria; Alphaproteobacteria; Rhizobiales; Bradyrhizobiaceae; Bradyrhizobium MNGTATRRRGVVGFGLFTLAMIAAFIALGVWQLQRRVAKHELIAALTERLAAPPVALPPP WP_027521434.1 hypothetical protein [Bradyrhizobium sp. Ec3.3] Length: 251\nNumber of predicted TMHs: 2\nExp number of AAs in TMHs: 51.18275\nExp number, first 60 AAs: 21.92627\nTotal prob of N-in: 0.59365\nPOSSIBLE N-term signal sequence\ninside 1 11\nTMhelix 12 34\noutside 35 225\nTMhelix 226 245\ninside 246 251

90 GCF_000938235.1_ASM93823v1 Bradyrhizobium sp. LTSPM299 Proteobacteria; Alphaproteobacteria; Rhizobiales; Bradyrhizobiaceae; Bradyrhizobium MRRRALLAIMGLLAATLAAAPADAERLIVSVSNHRVTVTPNYSGGELVLFGSVEKDASTP WP_044589585.1 membrane protein [Bradyrhizobium sp. LTSPM299] Length: 260\nNumber of predicted TMHs: 1\nExp number of AAs in TMHs: 25.76387\nExp number, first 60 AAs: 3.17836\nTotal prob of N-in: 0.76827\ninside 1 234\nTMhelix 235 257\noutside 258 260

91 GCF_900176205.1_BRAD285_PRJEB202261_v1 Bradyrhizobium sp. ORS 285 Proteobacteria; Alphaproteobacteria; Rhizobiales; Bradyrhizobiaceae; Bradyrhizobium MFDHHFLGGLAHDQIALLFMAAAAKAPRRSAVRMSTRVAGISAAILAIATIVMACYIAHM WP_006613759.1 DUF3592 domain-containing protein [Bradyrhizobium sp. ORS 285] Length: 193\nNumber of predicted TMHs: 2\nExp number of AAs in TMHs: 49.51449\nExp number, first 60 AAs: 29.05025\nTotal prob of N-in: 0.53299\nPOSSIBLE N-term signal sequence\ninside 1 37\nTMhelix 38 60\noutside 61 140\nTMhelix 141 160\ninside 161 193

92 GCF_000239755.1_ASM23975v2 Bradyrhizobium sp. ORS 285 Proteobacteria; Alphaproteobacteria; Rhizobiales; Bradyrhizobiaceae; Bradyrhizobium MFDHHFLGGLAHDQIALLFMAAAAKAPRRSAVRMSTRVAGISAAILAIATIVMACYIAHM WP_006613759.1 DUF3592 domain-containing protein [Bradyrhizobium sp. ORS 285] Length: 193\nNumber of predicted TMHs: 2\nExp number of AAs in TMHs: 49.51449\nExp number, first 60 AAs: 29.05025\nTotal prob of N-in: 0.53299\nPOSSIBLE N-term signal sequence\ninside 1 37\nTMhelix 38 60\noutside 61 140\nTMhelix 141 160\ninside 161 193

93 GCF_000472425.1_ASM47242v1 Bradyrhizobium sp. Tv2a-2 Proteobacteria; Alphaproteobacteria; Rhizobiales; Bradyrhizobiaceae; Bradyrhizobium MINPYRVFRRGEPARRRRHIGDGGSLAAAGLLIACLAAQPALAWDYWGGDAGGTRFSSIA WP_024518011.1 quinoprotein glucose dehydrogenase [Bradyrhizobium sp. Tv2a-2] Length: 691\nNumber of predicted TMHs: 2\nExp number of AAs in TMHs: 50.03864\nExp number, first 60 AAs: 18.76246\nTotal prob of N-in: 0.84264\nPOSSIBLE N-term signal sequence\ninside 1 24\nTMhelix 25 47\noutside 48 666\nTMhelix 667 686\ninside 687 691

94 GCF_000701345.1_ASM70134v1 Bradyrhizobium sp. URHD0069 Proteobacteria; Alphaproteobacteria; Rhizobiales; Bradyrhizobiaceae; Bradyrhizobium
MLTDMLPILWRRRRVIYAALVLTSGAAFLFYNTVGERYEAYTLMRVGQGIKDRSAGAGSG WP_051677293.1 hypothetical protein [Bradyrhizobium sp. URHD0069] Length: 485\nNumber of predicted TMHs: 2\nExp number of AAs in TMHs: 43.74465\nExp number, first 60 AAs: 21.53652\nTotal prob of N-in: 0.93900\nPOSSIBLE N-term signal sequence\ninside 1 12\nTMhelix 13 35\noutside 36 424\nTMhelix 425 447\ninside 448 485

95 GCF_001440385.1_ASM144038v1 Bradyrhizobium valentinum Proteobacteria; Alphaproteobacteria; Rhizobiales; Bradyrhizobiaceae; Bradyrhizobium
MTGLLSRRPAVTGFAIFTLVMVAVFASLGIWQMQRRVEKHALIARLNERLAAAPGALPQQ WP_057849324.1 surfeit 1 protein [Bradyrhizobium valentinum] Length: 255\nNumber of predicted TMHs: 2\nExp number of AAs in TMHs: 42.86168\nExp number, first 60 AAs: 21.66479\nTotal prob of N-in: 0.46332\nPOSSIBLE N-term signal sequence\noutside 1 9\nTMhelix 10 32\ninside 33 229\nTMhelix 230 249\noutside 250 255

96 GCF_001440405.1_ASM144040v1 Bradyrhizobium valentinum Proteobacteria; Alphaproteobacteria; Rhizobiales; Bradyrhizobiaceae; Bradyrhizobium
MTGLLSRRPAVTGFAIFTLVMVAVFASLGIWQMQRRVEKHALIARLNERLAAAPGALPQQ WP_057849324.1 surfeit 1 protein [Bradyrhizobium valentinum] Length: 255\nNumber of predicted TMHs: 2\nExp number of AAs in TMHs: 42.86168\nExp number, first 60 AAs: 21.66479\nTotal prob of N-in: 0.46332\nPOSSIBLE N-term signal sequence\noutside 1 9\nTMhelix 10 32\ninside 33 229\nTMhelix 230 249\noutside 250 255

97 GCF_001295845.1_ASM129584v1 Rhodopseudomonas sp. AAP120 Proteobacteria; Alphaproteobacteria; Rhizobiales; Bradyrhizobiaceae; Rhodopseudomonas
MTIVSRRAALALLASVAFIVPASAQQQPMPPRASAGPNTYGPDELTAAGHRFFGNVSRGL WP_054163953.1 hypothetical protein [Rhodopseudomonas sp. AAP120] Length: 197\nNumber of predicted TMHs: 1\nExp number of AAs in TMHs: 22.3687\nExp number, first 60 AAs: 3.25108\nTotal prob of N-in: 0.81426\ninside 1 148\nTMhelix 149 171\noutside 172 197

98 GCF_000013685.1_ASM1368v1 Rhodopseudomonas palustris BisB5 Proteobacteria; Alphaproteobacteria; Rhizobiales; Bradyrhizobiaceae; Rhodopseudomonas; Rhodopseudomonas palustris
MTIVSRRAALALFASAAFIVPAAAQQQQPLPPRAQASPSTYGPEELTNAGHRFFGNVSRG WP_011504069.1 hypothetical protein [Rhodopseudomonas palustris] Length: 198\nNumber of predicted TMHs: 1\nExp number of AAs in TMHs: 23.34464\nExp number, first 60 AAs: 2.98532\nTotal prob of N-in: 0.85034\ninside 1 149\nTMhelix 150 172\noutside 173 198

99 GCF_000526135.1_ASM52613v1 Hyphomicrobium sp. 802 Proteobacteria; Alphaproteobacteria; Rhizobiales; Hyphomicrobiaceae; Hyphomicrobium
MTFFGIEFDRRTALLAGALALTIAAMIGFHLVRNQRIADVLAIIDITGSMNTRDMGDPRG WP_013946138.1 MULTISPECIES: VWA domain-containing protein [Hyphomicrobium] Length: 341\nNumber of predicted TMHs: 2\nExp number of AAs in TMHs: 41.54814\nExp number, first 60 AAs: 20.29518\nTotal prob of N-in: 0.48706\nPOSSIBLE N-term signal sequence\ninside 1 12\nTMhelix 13 32\noutside 33 293\nTMhelix 294 316\ninside 317 341

100 GCF_000253295.1_ASM25329v1 Hyphomicrobium sp. MC1 Proteobacteria; Alphaproteobacteria; Rhizobiales; Hyphomicrobiaceae; Hyphomicrobium
MTFFGIEFDRRTALLAGALALTIAAMIGFHLVRNQRIADVLAIIDITGSMNTRDMGDPRG WP_013946138.1 MULTISPECIES: VWA domain-containing protein [Hyphomicrobium] Length: 341\nNumber of predicted TMHs: 2\nExp number of AAs in TMHs: 41.54814\nExp number, first 60 AAs: 20.29518\nTotal prob of N-in: 0.48706\nPOSSIBLE N-term signal sequence\ninside 1 12\nTMhelix 13 32\noutside 33 293\nTMhelix 294 316\ninside 317 341

101 GCF_002198715.1_ASM219871v1 Rhodomicrobium vannielii Proteobacteria; Alphaproteobacteria; Rhizobiales; Hyphomicrobiaceae; Rhodomicrobium
MRMTRRAALASLAACLAAPADSASGIRFTHAYGETVLPRPARRVVSLGYTTQDALLALGV WP_088345824.1 twin-arginine translocation pathway signal protein [Rhodomicrobium vannielii] Length: 328\nNumber of predicted TMHs: 1\nExp number of AAs in TMHs: 20.96153\nExp number, first 60 AAs: 0.43492\nTotal prob of N-in: 0.83865\ninside 1 276\nTMhelix 277 299\noutside 300 328

102 GCF_002198685.1_ASM219868v1 Rhodomicrobium vannielii Proteobacteria; Alphaproteobacteria; Rhizobiales; Hyphomicrobiaceae; Rhodomicrobium
MRMTRRAALASLAACLAAPADSASGIRFTHAYGETVLPRPARRVVSLGYTTQDALLALGV WP_088345824.1 twin-arginine translocation pathway signal protein [Rhodomicrobium vannielii] Length: 328\nNumber of predicted TMHs:

1\nExp number of AAs in TMHs: 20.96153\nExp number, first 60 AAs: 0.43492\nTotal prob of N-in: 0.83865\ninside 1 276\nTMhelix 277 299\noutside 300 328

103 GCF_000166055.1_ASM16605v1 Rhodomicrobium vannielii ATCC 17100 Proteobacteria; Alphaproteobacteria; Rhizobiales; Hyphomicrobiaceae; Rhodomicrobium; Rhodomicrobium vannielii MTLRSCTAEKARGEVSFMLHRRSVLSIGVALGAAPLLSSAVEAQSADRSRDVILAVADAK WP_041787646.1 hypothetical protein [Rhodomicrobium vannielii] Length: 272\nNumber of predicted TMHs: 1\nExp number of AAs in TMHs: 23.2314\nExp number, first 60 AAs: 3.21616\nTotal prob of N-in: 0.46150\noutside 1 233\nTMhelix 234 256\ninside 257 272

104 GCF_001043915.1_ASM104391v1 Methylobacterium aquaticum Proteobacteria; Alphaproteobacteria; Rhizobiales; Methylobacteriaceae; Methylobacterium MTVPDPAGRRAAIRDLVAPALASLVCLAILIGLGVWQLERKAWKEALIDRIVARSRIEPP WP_048462209.1 surfeit 1 [Methylobacterium aquaticum] Length: 266\nNumber of predicted TMHs: 2\nExp number of AAs in TMHs: 44.78158\nExp number, first 60 AAs: 21.53878\nTotal prob of N-in: 0.73186\nPOSSIBLE N-term signal sequence\ninside 1 15\nTMhelix 16 38\noutside 39 222\nTMhelix 223 245\ninside 246 266

105 GCF_001477085.1_ASM147708v1 Methylobacterium aquaticum Proteobacteria; Alphaproteobacteria; Rhizobiales; Methylobacteriaceae; Methylobacterium MTAPAPGAGAAERRAALRDLVAPALASLVCLAILIGLGVWQLERKAWKEALIDRIIARSR WP_058618654.1 surfeit 1 [Methylobacterium aquaticum] Length: 280\nNumber of predicted TMHs: 2\nExp number of AAs in TMHs: 44.68611\nExp number, first 60 AAs: 22.3317\nTotal prob of N-in: 0.51012\nPOSSIBLE N-term signal sequence\ninside 1 19\nTMhelix 20 42\noutside 43 231\nTMhelix 232 254\ninside 255 280

106 GCF_001476215.1_ASM147621v1 Methylobacterium aquaticum Proteobacteria; Alphaproteobacteria; Rhizobiales; Methylobacteriaceae; Methylobacterium MTAPAPGAGAAERRAALRDLVAPALASLVCLAILIGLGVWQLERKAWKEALIDRIIARSR WP_058618654.1 surfeit 1 [Methylobacterium aquaticum] Length: 280\nNumber of predicted TMHs: 2\nExp number of AAs in TMHs: 44.68611\nExp number, first 60 AAs: 22.3317\nTotal prob of N-in: 0.51012\nPOSSIBLE N-term signal sequence\ninside 1 19\nTMhelix 20 42\noutside 43 231\nTMhelix 232 254\ninside 255 280

107 GCF_001548015.1_ASM154801v1 Methylobacterium aquaticum Proteobacteria; Alphaproteobacteria; Rhizobiales; Methylobacteriaceae; Methylobacterium MTAPAPGPGAAERRAALRDLVAPALASLVCLGILVGLGVWQLERKAWKEALIDRIVARSR WP_060850032.1 surfeit 1 [Methylobacterium aquaticum] Length: 279\nNumber of predicted TMHs: 2\nExp number of AAs in TMHs: 44.51131\nExp number, first 60 AAs: 22.18531\nTotal prob of N-in: 0.31725\nPOSSIBLE N-term signal sequence\noutside 1 19\nTMhelix 20 42\ninside 43 231\nTMhelix 232 254\noutside 255 279

108 GCF_001548015.1_ASM154801v1 Methylobacterium aquaticum Proteobacteria; Alphaproteobacteria; Rhizobiales; Methylobacteriaceae; Methylobacterium MTDETTQEPYACGCAGAPLPRRSLLAAWLALLLAPADAFAAGEPGNAANASTVKAPNPKA WP_060849731.1 hypothetical protein [Methylobacterium aquaticum] Length: 215\nNumber of predicted TMHs: 1\nExp number of AAs in TMHs: 17.82078\nExp number, first 60 AAs: 1.0831\nTotal prob of N-in: 0.62622\ninside 1 185\nTMhelix 186 208\noutside 209 215

109 GCF_001548015.1_ASM154801v1 Methylobacterium aquaticum Proteobacteria; Alphaproteobacteria; Rhizobiales; Methylobacteriaceae; Methylobacterium MPQLGDYILRLVPLRSHLSVLVLARRPYLRADHFSGGGSVQGVGFLRVLRTLRLLRTYHL WP_082743019.1 Ion transporter [Methylobacterium aquaticum] Length: 197\nNumber of predicted TMHs: 2\nExp number of AAs in TMHs: 44.67191\nExp number, first 60 AAs: 0.99799\nTotal prob of N-in: 0.93043\ninside 1 77\nTMhelix 78 100\noutside 101 131\nTMhelix 132 154\ninside 155 197

110 GCF_001476615.1_ASM147661v1 Methylobacterium aquaticum Proteobacteria; Alphaproteobacteria; Rhizobiales; Methylobacteriaceae; Methylobacterium MTAPAPGAGAAERRAALRDLVAPALASLVCLAILIGLGVWQLERKAWKEALIDRIIARSR WP_058618654.1 surfeit 1 [Methylobacterium aquaticum] Length: 280\nNumber of predicted TMHs: 2\nExp number of AAs in TMHs: 44.68611\nExp number, first 60 AAs: 22.3317\nTotal prob of N-in: 0.51012\nPOSSIBLE N-term signal sequence\ninside 1 19\nTMhelix 20 42\noutside 43 231\nTMhelix 232 254\ninside 255 280

111 GCF_001043895.1_ASM104389v1 Methylobacterium platani Proteobacteria; Alphaproteobacteria; Rhizobiales; Methylobacteriaceae; Methylobacterium

MTAPAPGAGAAERRAALRDLVAPALASLVCLAILIGLGVWQLERKAWKEALIDRIVARSR WP_048428092.1 surfeit 1 [Methylobacterium platani] Length: 280\nNumber of predicted TMHs: 2\nExp number of AAs in TMHs: 45.33593\nExp number, first 60 AAs: 22.32788\nTotal prob of N-in: 0.49438\nPOSSIBLE N-term signal sequence\ninside 1 19\nTMhelix 20 42\noutside 43 231\nTMhelix 232 254\ninside 255 280

112 GCF_001043875.1_ASM104387v1 Methylobacterium platani Proteobacteria; Alphaproteobacteria; Rhizobiales; Methylobacteriaceae; Methylobacterium
MTAPAPGAGAAERRAALRDLVAPALASLVCLAILIGLGVWQLERKAWKEALIDRIVARSR WP_048428092.1 surfeit 1 [Methylobacterium platani] Length: 280\nNumber of predicted TMHs: 2\nExp number of AAs in TMHs: 45.33593\nExp number, first 60 AAs: 22.32788\nTotal prob of N-in: 0.49438\nPOSSIBLE N-term signal sequence\ninside 1 19\nTMhelix 20 42\noutside 43 231\nTMhelix 232 254\ninside 255 280

113 GCF_001653715.1_ASM165371v1 Methylobacterium platani Proteobacteria; Alphaproteobacteria; Rhizobiales; Methylobacteriaceae; Methylobacterium
MSAPLTASERRAAIRDLVAPALATLVCLAILVGLGVWQLERKAWKEALIDRIVARSRIDP WP_048431791.1 surfeit 1 [Methylobacterium platani] Length: 267\nNumber of predicted TMHs: 2\nExp number of AAs in TMHs: 44.80971\nExp number, first 60 AAs: 21.95878\nTotal prob of N-in: 0.62344\nPOSSIBLE N-term signal sequence\ninside 1 16\nTMhelix 17 39\noutside 40 223\nTMhelix 224 246\ninside 247 267

114 GCF_001653715.1_ASM165371v1 Methylobacterium platani Proteobacteria; Alphaproteobacteria; Rhizobiales; Methylobacteriaceae; Methylobacterium
MHTLSANAAGTERRGIFATLPAMLAWPAVAASDAAEAFERVTAPDPVFAAIETHRAEQVA WP_048435058.1 hypothetical protein [Methylobacterium platani] Length: 188\nNumber of predicted TMHs: 1\nExp number of AAs in TMHs: 22.85436\nExp number, first 60 AAs: 1.55214\nTotal prob of N-in: 0.29690\noutside 1 162\nTMhelix 163 185\ninside 186 188

115 GCF_001643095.1_ASM164309v1 Methylobacterium populi Proteobacteria; Alphaproteobacteria; Rhizobiales; Methylobacteriaceae; Methylobacterium
MPHDPIRPAATTRRAALLGLSAGLSSGLLGTVVPAAAYQDDGVRPDSFRASEVVENGHRF WP_012453627.1 hypothetical protein [Methylobacterium populi] Length: 205\nNumber of predicted TMHs: 2\nExp number of AAs in TMHs: 39.69176\nExp number, first 60 AAs: 19.987\nTotal prob of N-in: 0.59493\nPOSSIBLE N-term signal sequence\ninside 1 15\nTMhelix 16 38\noutside 39 156\nTMhelix 157 179\ninside 180 205

116 GCF_000372825.1_ASM37282v1 Methylobacterium sp. 77 Proteobacteria; Alphaproteobacteria; Rhizobiales; Methylobacteriaceae; Methylobacterium
MVRQVLNGGRRSVLAAFGLACLATIGGCADSRPRNPESGASESRTMARLNPVAQKTDAKL WP_081631254.1 hypothetical protein [Methylobacterium sp. 77] Length: 302\nNumber of predicted TMHs: 1\nExp number of AAs in TMHs: 28.49457\nExp number, first 60 AAs: 5.98221\nTotal prob of N-in: 0.32246\noutside 1 260\nTMhelix 261 283\ninside 284 302

117 GCF_001542815.1_ASM154281v1 Methylobacterium sp. AMS5 Proteobacteria; Alphaproteobacteria; Rhizobiales; Methylobacteriaceae; Methylobacterium
MPHDPIRTGGTTRRAALLGLSAGLSGGLLAGAAPALAYQDDGVRPDSFRAAEVVENGHRF WP_060769419.1 hypothetical protein [Methylobacterium sp. AMS5] Length: 205\nNumber of predicted TMHs: 2\nExp number of AAs in TMHs: 40.2427\nExp number, first 60 AAs: 21.06824\nTotal prob of N-in: 0.61328\nPOSSIBLE N-term signal sequence\ninside 1 15\nTMhelix 16 38\noutside 39 156\nTMhelix 157 179\ninside 180 205

118 GCF_001556475.1_ASM155647v1 Methylobacterium sp. CCH5-D2 Proteobacteria; Alphaproteobacteria; Rhizobiales; Methylobacteriaceae; Methylobacterium
MSARARRRILAAVAACGLLAACQEQGRQDEAATAAPQVRPAAKQASQDGGAPDRASRAKL WP_066924169.1 hypothetical protein [Methylobacterium sp. CCH5-D2] Length: 299\nNumber of predicted TMHs: 1\nExp number of AAs in TMHs: 23.28233\nExp number, first 60 AAs: 0.02425\nTotal prob of N-in: 0.02428\noutside 1 256\nTMhelix 257 279\ninside 280 299

119 GCF_001556475.1_ASM155647v1 Methylobacterium sp. CCH5-D2 Proteobacteria; Alphaproteobacteria; Rhizobiales; Methylobacteriaceae; Methylobacterium
MLGGTTRRPPLARQVVVVTGASSGIGLATARMAAARGARVVLAARSADALAEIRAGIEAA WP_066922328.1 short-chain dehydrogenase [Methylobacterium sp. CCH5-D2] Length: 341\nNumber of predicted TMHs: 1\nExp number of AAs in TMHs: 25.18644\nExp number, first 60 AAs: 3.05348\nTotal prob of N-in: 0.12265\noutside 1 311\nTMhelix 312 334\ninside 335 341

120 GCF_001423285.1_Leaf111 Methylobacterium sp. Leaf111 Proteobacteria; Alphaproteobacteria; Rhizobiales; Methylobacteriaceae; Methylobacterium
MASPRPGARRSIFALAALAVTGLILVAALAALGTWQVQRRIWKHDLIARVEARVHADPVA WP_056492945.1
Surfeit locus 1 family protein [Methylobacterium sp. Leaf111] Length: 251\nNumber of predicted TMHs: 2\nExp number of AAs in TMHs: 45.39455\nExp number, first 60 AAs: 22.72625\nTotal prob of N-in: 0.81272\nPOSSIBLE N-term signal sequence\ninside 1 12\nTMhelix 13 35\noutside 36 217\nTMhelix 218 240\ninside 241 251

121 GCF_001423325.1_Leaf113 Methylobacterium sp. Leaf113 Proteobacteria; Alphaproteobacteria; Rhizobiales; Methylobacteriaceae; Methylobacterium
MSRAPARRAAVALALLLAATGGARAVQPDEVLKDPVMEARARDISSGLRCLVCQNQSIDD WP_056178514.1
cytochrome c biogenesis protein [Methylobacterium sp. Leaf113] Length: 156\nNumber of predicted TMHs: 2\nExp number of AAs in TMHs: 36.80708\nExp number, first 60 AAs: 14.25074\nTotal prob of N-in: 0.80772\nPOSSIBLE N-term signal sequence\ninside 1 8\nTMhelix 9 26\noutside 27 103\nTMhelix 104 126\ninside 127 156

122 GCF_001422985.1_Leaf117 Methylobacterium sp. Leaf117 Proteobacteria; Alphaproteobacteria; Rhizobiales; Methylobacteriaceae; Methylobacterium
MSRAPARRAAVALALSLAATGGARAVQPDEVLKDPVMEARARDISSGLRCLVCQNQSIDD WP_056483469.1
cytochrome c biogenesis protein [Methylobacterium sp. Leaf117] Length: 156\nNumber of predicted TMHs: 1\nExp number of AAs in TMHs: 29.47166\nExp number, first 60 AAs: 6.96857\nTotal prob of N-in: 0.51782\noutside 1 102\nTMhelix 103 125\ninside 126 156

123 GCF_001422985.1_Leaf117 Methylobacterium sp. Leaf117 Proteobacteria; Alphaproteobacteria; Rhizobiales; Methylobacteriaceae; Methylobacterium
MSRASSTGRRQTVVVTGASAGVGRAIAHEFARHRWNVAVLARGEAGLAGTVRDIERAGGR WP_056487710.1
short-chain dehydrogenase [Methylobacterium sp. Leaf117] Length: 344\nNumber of predicted TMHs: 1\nExp number of AAs in TMHs: 31.10899\nExp number, first 60 AAs: 0.34135\nTotal prob of N-in: 0.47219\noutside 1 315\nTMhelix 316 335\ninside 336 344

124 GCF_001422935.1_Leaf119 Methylobacterium sp. Leaf119 Proteobacteria; Alphaproteobacteria; Rhizobiales; Methylobacteriaceae; Methylobacterium
MPHDPIRTGGTTRRAALLGLSAGLSGGLLAGAVPALAYQDDGVRPDSFRAAEVVENGHRF WP_003597644.1
MULTISPECIES: hypothetical protein [Methylobacterium] Length: 205\nNumber of predicted TMHs: 2\nExp number of AAs in TMHs: 42.38169\nExp number, first 60 AAs: 21.87362\nTotal prob of N-in: 0.54505\nPOSSIBLE N-term signal sequence\noutside 1 14\nTMhelix 15 37\ninside 38 156\nTMhelix 157 179\noutside 180 205

125 GCF_001423385.1_Leaf121 Methylobacterium sp. Leaf121 Proteobacteria; Alphaproteobacteria; Rhizobiales; Methylobacteriaceae; Methylobacterium
MPHDPIRTGGTTRRAALLGLSAGLSGGLLAGATPALAYQDDGVRPDSFRAAEVVENGHRF WP_015822348.1
MULTISPECIES: hypothetical protein [Methylobacterium] Length: 205\nNumber of predicted TMHs: 2\nExp number of AAs in TMHs: 39.86733\nExp number, first 60 AAs: 20.08937\nTotal prob of N-in: 0.62081\nPOSSIBLE N-term signal sequence\ninside 1 15\nTMhelix 16 38\noutside 39 156\nTMhelix 157 179\ninside 180 205

126 GCF_001423385.1_Leaf121 Methylobacterium sp. Leaf121 Proteobacteria; Alphaproteobacteria; Rhizobiales; Methylobacteriaceae; Methylobacterium
MTTYTATRGAIPLSSYDRPTASAGQSPSIAPAQLLAMLKRRRRLIALSMVLGTLGGVVVV WP_056499635.1
lipopolysaccharide biosynthesis protein [Methylobacterium sp. Leaf121] Length: 491\nNumber of predicted TMHs: 2\nExp number of AAs in TMHs: 41.40682\nExp number, first 60 AAs: 16.7865\nTotal prob of N-in: 0.97165\nPOSSIBLE N-term signal sequence\ninside 1 43\nTMhelix 44 61\noutside 62 448\nTMhelix 449 471\ninside 472 491

127 GCF_001423025.1_Leaf122 Methylobacterium sp. Leaf122 Proteobacteria; Alphaproteobacteria; Rhizobiales; Methylobacteriaceae; Methylobacterium
MTTYTATRGAIPLASYDRPAASAGQSPSIAPAQLLAMLRHRRRLIALSMILGTLGGVVVI WP_056115908.1
lipopolysaccharide biosynthesis protein [Methylobacterium sp. Leaf122] Length: 491\nNumber of predicted TMHs: 2\nExp number of AAs in TMHs: 41.43065\nExp number, first 60 AAs: 16.82503\nTotal prob of N-in: 0.96832\nPOSSIBLE N-term signal sequence\ninside 1 43\nTMhelix 44 61\noutside 62 448\nTMhelix 449 471\ninside 472 491

128 GCF_001423025.1_Leaf122 Methylobacterium sp. Leaf122 Proteobacteria; Alphaproteobacteria; Rhizobiales; Methylobacteriaceae; Methylobacterium
MPHDPIRTGGTTRRAALLGLSAGLSGGLLAGAVPALAYQDDGVRPDSFRAAEVVENGHRF WP_003597644.1

MULTISPECIES: hypothetical protein [Methylobacterium] Length: 205\nNumber of predicted TMHs: 2\nExp number of AAs in TMHs: 42.38169\nExp number, first 60 AAs: 21.87362\nTotal prob of N-in: 0.54505\nPOSSIBLE N-term signal sequence\noutside 1 14\nTMhelix 15 37\ninside 38 156\nTMhelix 157 179\noutside 180 205

129 GCF_001423405.1_Leaf123 Methylobacterium sp. Leaf123 Proteobacteria; Alphaproteobacteria; Rhizobiales; Methylobacteriaceae; Methylobacterium
MTTYTATRGAIPLSSYDRPAAATGHAPSITPTQLLAMLKRRRQLIVLSMILGTLGGVVVV WP_056199558.1
lipopolysaccharide biosynthesis protein [Methylobacterium sp. Leaf123] Length: 491\nNumber of predicted TMHs: 2\nExp number of AAs in TMHs: 41.4639\nExp number, first 60 AAs: 16.88528\nTotal prob of N-in: 0.96262\nPOSSIBLE N-term signal sequence\ninside 1 43\nTMhelix 44 61\noutside 62 448\nTMhelix 449 471\ninside 472 491

130 GCF_001423405.1_Leaf123 Methylobacterium sp. Leaf123 Proteobacteria; Alphaproteobacteria; Rhizobiales; Methylobacteriaceae; Methylobacterium
MPHDPIRTGGTTRRAALLGLSAGLSGGLLTGAVPALAYQDDGVRPDSFRASEVVENGHRF WP_056204758.1
hypothetical protein [Methylobacterium sp. Leaf123] Length: 205\nNumber of predicted TMHs: 2\nExp number of AAs in TMHs: 41.68973\nExp number, first 60 AAs: 21.37962\nTotal prob of N-in: 0.54286\nPOSSIBLE N-term signal sequence\noutside 1 14\nTMhelix 15 37\ninside 38 156\nTMhelix 157 179\noutside 180 205

131 GCF_001425465.1_Leaf456 Methylobacterium sp. Leaf456 Proteobacteria; Alphaproteobacteria; Rhizobiales; Methylobacteriaceae; Methylobacterium
MPHESGDEFRATRRAAVLGLAAGLIAAAVPAGAVDDMGRPESFRAAEVVENGHRFFGSVS WP_056240996.1
hypothetical protein [Methylobacterium sp. Leaf456] Length: 200\nNumber of predicted TMHs: 2\nExp number of AAs in TMHs: 38.30474\nExp number, first 60 AAs: 19.28273\nTotal prob of N-in: 0.28621\nPOSSIBLE N-term signal sequence\noutside 1 14\nTMhelix 15 34\ninside 35 151\nTMhelix 152 174\noutside 175 200

132 GCF_001424705.1_Leaf465 Methylobacterium sp. Leaf465 Proteobacteria; Alphaproteobacteria; Rhizobiales; Methylobacteriaceae; Methylobacterium
MASPRPGARRSIFALAALAVTGLILVAALAALGTWQVQRRTWKHDLIARVEARVHADPVA WP_056222799.1
MULTISPECIES: Surfeit locus 1 family protein [Methylobacterium] Length: 251\nNumber of predicted TMHs: 2\nExp number of AAs in TMHs: 45.4297699999999\nExp number, first 60 AAs: 22.71752\nTotal prob of N-in: 0.76260\nPOSSIBLE N-term signal sequence\ninside 1 12\nTMhelix 13 35\noutside 36 217\nTMhelix 218 240\ninside 241 251

133 GCF_001422795.1_Leaf88 Methylobacterium sp. Leaf88 Proteobacteria; Alphaproteobacteria; Rhizobiales; Methylobacteriaceae; Methylobacterium
MASPRPGARRSTFALAALAVTGLILVAALAALGTWQVQRRTWKHDLIARVEARVHADPIA WP_056274816.1
Surfeit locus 1 family protein [Methylobacterium sp. Leaf88] Length: 251\nNumber of predicted TMHs: 2\nExp number of AAs in TMHs: 45.41381\nExp number, first 60 AAs: 22.66975\nTotal prob of N-in: 0.73204\nPOSSIBLE N-term signal sequence\ninside 1 12\nTMhelix 13 35\noutside 36 217\nTMhelix 218 240\ninside 241 251

134 GCF_001422215.1_Leaf89 Methylobacterium sp. Leaf89 Proteobacteria; Alphaproteobacteria; Rhizobiales; Methylobacteriaceae; Methylobacterium
MASPRPGARRSTFALAALAVTGLILVAALAALGTWQVQRRTWKHDLIARVEARVHADPVA WP_056351180.1
Surfeit locus 1 family protein [Methylobacterium sp. Leaf89] Length: 251\nNumber of predicted TMHs: 2\nExp number of AAs in TMHs: 45.37744\nExp number, first 60 AAs: 22.67005\nTotal prob of N-in: 0.73319\nPOSSIBLE N-term signal sequence\ninside 1 12\nTMhelix 13 35\noutside 36 217\nTMhelix 218 240\ninside 241 251

135 GCF_001422265.1_Leaf90 Methylobacterium sp. Leaf90 Proteobacteria; Alphaproteobacteria; Rhizobiales; Methylobacteriaceae; Methylobacterium
MPHDPIRTGGTTRRAALLGLSAGLSGGLLAGATPALAYQDDGVRPDSFRAAEVVENGHRF WP_015822348.1
MULTISPECIES: hypothetical protein [Methylobacterium] Length: 205\nNumber of predicted TMHs: 2\nExp number of AAs in TMHs: 39.86733\nExp number, first 60 AAs: 20.08937\nTotal prob of N-in: 0.62081\nPOSSIBLE N-term signal sequence\ninside 1 15\nTMhelix 16 38\noutside 39 156\nTMhelix 157 179\ninside 180 205

136 GCF_001422305.1_Leaf92 Methylobacterium sp. Leaf92 Proteobacteria; Alphaproteobacteria; Rhizobiales; Methylobacteriaceae; Methylobacterium
MTTYTATRGAIPLASYDRPAASAGQSPSIAPAQLLAMLRHRRRLIALSMILGTLGGVVVI WP_056153970.1
lipopolysaccharide biosynthesis protein [Methylobacterium sp. Leaf92] Length: 491\nNumber of predicted TMHs: 2\nExp number of AAs in TMHs: 41.2493\nExp number, first 60 AAs: 16.76807\nTotal prob of N-in:

0.94177\nPOSSIBLE N-term signal sequence\ninside 1 43\nTMhelix 44 61\noutside 62 448\nTMhelix 449 471\ninside 472 491

137 GCF_001422305.1_Leaf92 Methylobacterium sp. Leaf92 Proteobacteria; Alphaproteobacteria; Rhizobiales; Methylobacteriaceae; Methylobacterium
MPHDPIRTGGTTRRAALLGLSAGLSGGLLAGATPALAYQDDGVRPDSFRAAEVVENGHRF WP_015822348.1
MULTISPECIES: hypothetical protein [Methylobacterium] Length: 205\nNumber of predicted TMHs: 2\nExp number of AAs in TMHs: 39.86733\nExp number, first 60 AAs: 20.08937\nTotal prob of N-in: 0.62081\nPOSSIBLE N-term signal sequence\ninside 1 15\nTMhelix 16 38\noutside 39 156\nTMhelix 157 179\ninside 180 205

138 GCF_001426045.1_Leaf94 Methylobacterium sp. Leaf94 Proteobacteria; Alphaproteobacteria; Rhizobiales; Methylobacteriaceae; Methylobacterium
MASPRPGARRSIFALAALAVTGLILVAALAALGTWQVQRRTWKHDLIARVEARVHADPVA WP_056222799.1
MULTISPECIES: Surfeit locus 1 family protein [Methylobacterium] Length: 251\nNumber of predicted TMHs: 2\nExp number of AAs in TMHs: 45.4297699999999\nExp number, first 60 AAs: 22.71752\nTotal prob of N-in: 0.76260\nPOSSIBLE N-term signal sequence\ninside 1 12\nTMhelix 13 35\noutside 36 217\nTMhelix 218 240\ninside 241 251

139 GCF_001422375.1_Leaf99 Methylobacterium sp. Leaf99 Proteobacteria; Alphaproteobacteria; Rhizobiales; Methylobacteriaceae; Methylobacterium
MSGMAGRRSLVAAALGLAALLGGCSDARPPAPEVASPLPSMQANRSAAAKLAFTHDLTLG WP_056088981.1
hypothetical protein [Methylobacterium sp. Leaf99] Length: 293\nNumber of predicted TMHs: 1\nExp number of AAs in TMHs: 22.64914\nExp number, first 60 AAs: 0.17927\nTotal prob of N-in: 0.01018\noutside 1 251\nTMhelix 252 274\ninside 275 293

140 GCF_000333655.1_Methylobacterium_sp._MB200_genome_sequence_1.0 Methylobacterium sp. MB200 Proteobacteria; Alphaproteobacteria; Rhizobiales; Methylobacteriaceae; Methylobacterium
MPHDPIRTATTRRAALLGLSAGLSGGLLASAVPALAYQDDGVRPDSFRASEVVENGHRFF WP_017485181.1
hypothetical protein [Methylobacterium sp. MB200] Length: 204\nNumber of predicted TMHs: 2\nExp number of AAs in TMHs: 41.47998\nExp number, first 60 AAs: 21.06658\nTotal prob of N-in: 0.63558\nPOSSIBLE N-term signal sequence\ninside 1 12\nTMhelix 13 35\noutside 36 155\nTMhelix 156 178\ninside 179 204

141 GCF_000333655.1_Methylobacterium_sp._MB200_genome_sequence_1.0 Methylobacterium sp. MB200 Proteobacteria; Alphaproteobacteria; Rhizobiales; Methylobacteriaceae; Methylobacterium
MNRGEAGRRGRCGSRTGSARRRLLGLLLGLLPALALLAGCSDNRRELRQGGERADAAAPA WP_017484839.1
DUF4349 domain-containing protein [Methylobacterium sp. MB200] Length: 311\nNumber of predicted TMHs: 2\nExp number of AAs in TMHs: 38.91081\nExp number, first 60 AAs: 17.47421\nTotal prob of N-in: 0.92393\nPOSSIBLE N-term signal sequence\ninside 1 20\nTMhelix 21 38\noutside 39 272\nTMhelix 273 295\ninside 296 311

142 GCF_000800935.1_ZNC0032.1 Methylobacterium sp. ZNC0032 Proteobacteria; Alphaproteobacteria; Rhizobiales; Methylobacteriaceae; Methylobacterium
MSPMTQTRRNLITGGLAVAGASLTGAPLSALAQNYQNQQSFSQNELVGSGHKFFGNVSRG WP_047576354.1
hypothetical protein [Methylobacterium sp. ZNC0032] Length: 198\nNumber of predicted TMHs: 1\nExp number of AAs in TMHs: 28.50211\nExp number, first 60 AAs: 10.5136\nTotal prob of N-in: 0.59334\nPOSSIBLE N-term signal sequence\ninside 1 149\nTMhelix 150 172\noutside 173 198

143 GCF_001043975.1_ASM104397v1 Methylobacterium variabile Proteobacteria; Alphaproteobacteria; Rhizobiales; Methylobacteriaceae; Methylobacterium
MTGGRRSALRDLLAPGLASLVCLAILIGLGTWQLERKAWKEALIDRIVARSRIEPPAPLP WP_048445522.1 surfeit 1 [Methylobacterium variabile] Length: 263\nNumber of predicted TMHs: 2\nExp number of AAs in TMHs: 43.82783\nExp number, first 60 AAs: 22.24961\nTotal prob of N-in: 0.96683\nPOSSIBLE N-term signal sequence\ninside 1 11\nTMhelix 12 34\noutside 35 218\nTMhelix 219 241\ninside 242 263

144 GCF_002117815.1_ASM211781v1 Methylobacterium zatmanii Proteobacteria; Alphaproteobacteria; Rhizobiales; Methylobacteriaceae; Methylobacterium
MPHDPIRTGGTTRRAALLGLSAGLSGGLLAGATPALAYQDDGVRPDSFRAAEVVENGHRF WP_015822348.1
MULTISPECIES: hypothetical protein [Methylobacterium] Length: 205\nNumber of predicted TMHs: 2\nExp number of AAs in TMHs: 39.86733\nExp number, first 60 AAs: 20.08937\nTotal prob of N-in: 0.62081\nPOSSIBLE N-term signal sequence\ninside 1 15\nTMhelix 16 38\noutside 39 156\nTMhelix 157 179\ninside 180 205

145 GCF_002117815.1_ASM211781v1 Methylobacterium zatmanii Proteobacteria; Alphaproteobacteria; Rhizobiales; Methylobacteriaceae; Methylobacterium MTTYTATRGAIPLSSYDRPTASAGQSPSIAPAQLLAMLKRRRRLIALSMVLGTLGGVVVA WP_085856269.1 lipopolysaccharide biosynthesis protein [Methylobacterium zatmanii] Length: 490\nNumber of predicted TMHs: 2\nExp number of AAs in TMHs: 39.50423\nExp number, first 60 AAs: 15.24783\nTotal prob of N-in: 0.89692\nPOSSIBLE N-term signal sequence\ninside 1 43\nTMhelix 44 61\noutside 62 447\nTMhelix 448 470\ninside 471 490

146 GCF_001854495.1_ASM185449v1 Methylobacterium extorquens Proteobacteria; Alphaproteobacteria; Rhizobiales; Methylobacteriaceae; Methylobacterium; Methylobacterium extorquens group MPHDPIRTRGTTRRAALLGLSAGLSGGLLAGAVPALAYQDDGVRPDSFRAAEVVENGHRF WP_082291284.1 hypothetical protein [Methylobacterium extorquens] Length: 205\nNumber of predicted TMHs: 2\nExp number of AAs in TMHs: 42.25638\nExp number, first 60 AAs: 21.93083\nTotal prob of N-in: 0.81986\nPOSSIBLE N-term signal sequence\ninside 1 15\nTMhelix 16 38\noutside 39 156\nTMhelix 157 179\ninside 180 205

147 GCF_001854495.1_ASM185449v1 Methylobacterium extorquens Proteobacteria; Alphaproteobacteria; Rhizobiales; Methylobacteriaceae; Methylobacterium; Methylobacterium extorquens group MTTYTATRGAIPLSSYDRPAASAGQSPSIAPAQLLAMLRHRRRLIALSMILGTLGGVVVV WP_082290253.1 lipopolysaccharide biosynthesis protein [Methylobacterium extorquens] Length: 491\nNumber of predicted TMHs: 2\nExp number of AAs in TMHs: 41.28213\nExp number, first 60 AAs: 16.68735\nTotal prob of N-in: 0.95649\nPOSSIBLE N-term signal sequence\ninside 1 43\nTMhelix 44 61\noutside 62 448\nTMhelix 449 471\ninside 472 491

148 GCF_001971665.1_ASM197166v1 Methylobacterium extorquens Proteobacteria; Alphaproteobacteria; Rhizobiales; Methylobacteriaceae; Methylobacterium; Methylobacterium extorquens group MPHDPIRTGGTTRRAALLGLSAGLSGGLLAGATPALAYQDDGVRPDSFRAAEVVENGHRF WP_015822348.1 MULTISPECIES: hypothetical protein [Methylobacterium] Length: 205\nNumber of predicted TMHs: 2\nExp number of AAs in TMHs: 39.86733\nExp number, first 60 AAs: 20.08937\nTotal prob of N-in: 0.62081\nPOSSIBLE N-term signal sequence\ninside 1 15\nTMhelix 16 38\noutside 39 156\nTMhelix 157 179\ninside 180 205

149 GCF_001971665.1_ASM197166v1 Methylobacterium extorquens Proteobacteria; Alphaproteobacteria; Rhizobiales; Methylobacteriaceae; Methylobacterium; Methylobacterium extorquens group MTTYTVTRGAIPLSSYDRPAASAGQSPSIAPAQLLAMLRHRRRLIALSMILGTLGGVVVI WP_076642097.1 lipopolysaccharide biosynthesis protein [Methylobacterium extorquens] Length: 491\nNumber of predicted TMHs: 2\nExp number of AAs in TMHs: 41.39425\nExp number, first 60 AAs: 16.79938\nTotal prob of N-in: 0.96133\nPOSSIBLE N-term signal sequence\ninside 1 43\nTMhelix 44 61\noutside 62 448\nTMhelix 449 471\ninside 472 491

150 GCF_000022685.1_ASM2268v1 Methylobacterium extorquens AM1 Proteobacteria; Alphaproteobacteria; Rhizobiales; Methylobacteriaceae; Methylobacterium; Methylobacterium extorquens group; Methylobacterium extorquens MPHDPIRTGGTTRRAALLGLSAGLSGGLLAGAVPALAYQDDGVRPDSFRAAEVVENGHRF WP_003597644.1 MULTISPECIES: hypothetical protein [Methylobacterium] Length: 205\nNumber of predicted TMHs: 2\nExp number of AAs in TMHs: 42.38169\nExp number, first 60 AAs: 21.87362\nTotal prob of N-in: 0.54505\nPOSSIBLE N-term signal sequence\noutside 1 14\nTMhelix 15 37\ninside 38 156\nTMhelix 157 179\noutside 180 205

151 GCF_000022685.1_ASM2268v1 Methylobacterium extorquens AM1 Proteobacteria; Alphaproteobacteria; Rhizobiales; Methylobacteriaceae; Methylobacterium; Methylobacterium extorquens group; Methylobacterium extorquens MTTYTATRGAIPLSSYDRPTASAGQSPSIAPAQLLAMLKRRRRLIALSMILGTLGGVVVV WP_003601398.1 LPS biosynthesis protein [Methylobacterium extorquens] Length: 491\nNumber of predicted TMHs: 2\nExp number of AAs in TMHs: 41.50849\nExp number, first 60 AAs: 16.88745\nTotal prob of N-in: 0.97611\nPOSSIBLE N-term signal sequence\ninside 1 43\nTMhelix 44 61\noutside 62 448\nTMhelix 449 471\ninside 472 491

152 GCF_000021845.1_ASM2184v1 Methylobacterium extorquens CM4 Proteobacteria; Alphaproteobacteria; Rhizobiales; Methylobacteriaceae; Methylobacterium; Methylobacterium extorquens group; Methylobacterium extorquens MPHDPIRTGGTTRRAALLGLSAGLSGGLLAGAVPALAYQDDGVRPDSFRAAEVVENGHRF WP_003597644.1 MULTISPECIES: hypothetical protein [Methylobacterium] Length: 205\nNumber of predicted TMHs: 2\nExp number of AAs in TMHs: 42.38169\nExp number, first 60 AAs: 21.87362\nTotal prob of N-in: 0.54505\nPOSSIBLE N-term signal sequence\noutside 1 14\nTMhelix 15 37\ninside 38 156\nTMhelix 157 179\noutside 180 205

153 GCF_000021845.1_ASM2184v1 Methylobacterium extorquens CM4 Proteobacteria; Alphaproteobacteria; Rhizobiales; Methylobacteriaceae; Methylobacterium; Methylobacterium extorquens group; Methylobacterium extorquens MTTYTATRGAIPLSSYDRSVASAGQSPSIAPAQLLAMLRHRRRLIALSMILGTLGGVLVV WP_015952381.1 LPS biosynthesis protein [Methylobacterium extorquens] Length: 491\nNumber of predicted TMHs: 2\nExp number of AAs in TMHs: 41.1931\nExp number, first 60 AAs: 16.70109\nTotal prob of N-in: 0.94931\nPOSSIBLE N-term signal sequence\ninside 1 43\nTMhelix 44 61\noutside 62 448\nTMhelix 449 471\ninside 472 491

154 GCF_000083545.1_ASM8354v1 Methylobacterium extorquens DM4 Proteobacteria; Alphaproteobacteria; Rhizobiales; Methylobacteriaceae; Methylobacterium; Methylobacterium extorquens group; Methylobacterium extorquens MPHDPIRTGGTTRRAALLGLSAGLSGGLLAGATPALAYQDDGVRPDSFRAAEVVENGHRF WP_015822348.1 MULTISPECIES: hypothetical protein [Methylobacterium] Length: 205\nNumber of predicted TMHs: 2\nExp number of AAs in TMHs: 39.86733\nExp number, first 60 AAs: 20.08937\nTotal prob of N-in: 0.62081\nPOSSIBLE N-term signal sequence\ninside 1 15\nTMhelix 16 38\noutside 39 156\nTMhelix 157 179\ninside 180 205

155 GCF_000243435.1_ASM24343v2 Methylobacterium extorquens DSM 13060 Proteobacteria; Alphaproteobacteria; Rhizobiales; Methylobacteriaceae; Methylobacterium; Methylobacterium extorquens group; Methylobacterium extorquens MPHDPIRTGGTTRRAALLGLSAGLSGGLLAGAVPALAYQDDGVRPDSFRAAEVVENGHRF WP_003597644.1 MULTISPECIES: hypothetical protein [Methylobacterium] Length: 205\nNumber of predicted TMHs: 2\nExp number of AAs in TMHs: 42.38169\nExp number, first 60 AAs: 21.87362\nTotal prob of N-in: 0.54505\nPOSSIBLE N-term signal sequence\noutside 1 14\nTMhelix 15 37\ninside 38 156\nTMhelix 157 179\noutside 180 205

156 GCF_000243435.1_ASM24343v2 Methylobacterium extorquens DSM 13060 Proteobacteria; Alphaproteobacteria; Rhizobiales; Methylobacteriaceae; Methylobacterium; Methylobacterium extorquens group; Methylobacterium extorquens MTTYTATRGAIPLSSYDRPTASAGQSPSIAPAQLLAMLKRRRRLIALSMILGTLGGVVVV WP_003601398.1 LPS biosynthesis protein [Methylobacterium extorquens] Length: 491\nNumber of predicted TMHs: 2\nExp number of AAs in TMHs: 41.50849\nExp number, first 60 AAs: 16.88745\nTotal prob of N-in: 0.97611\nPOSSIBLE N-term signal sequence\ninside 1 43\nTMhelix 44 61\noutside 62 448\nTMhelix 449 471\ninside 472 491

157 GCF_000018845.1_ASM1884v1 Methylobacterium extorquens PA1 Proteobacteria; Alphaproteobacteria; Rhizobiales; Methylobacteriaceae; Methylobacterium; Methylobacterium extorquens group; Methylobacterium extorquens MPHDPIRTGGTTRRAALLGLSAGLSGGLLAGAVPALAYQDDGVRPDSFRAAEVVENGHRF WP_003597644.1 MULTISPECIES: hypothetical protein [Methylobacterium] Length: 205\nNumber of predicted TMHs: 2\nExp number of AAs in TMHs: 42.38169\nExp number, first 60 AAs: 21.87362\nTotal prob of N-in: 0.54505\nPOSSIBLE N-term signal sequence\noutside 1 14\nTMhelix 15 37\ninside 38 156\nTMhelix 157 179\noutside 180 205

158 GCF_000364445.1_ASM36444v1 Methylobacterium mesophilicum SR1.6/6 Proteobacteria; Alphaproteobacteria; Rhizobiales; Methylobacteriaceae; Methylobacterium; Methylobacterium mesophilicum MAARARRTVIAAIALIASIPAHAVQPDEVLKDPVLEHRAREISAELRCLVCQNQSIDDSD WP_039893440.1 cytochrome c-type biogenesis protein CcmH [Methylobacterium mesophilicum] Length: 154\nNumber of predicted TMHs: 2\nExp number of AAs in TMHs: 37.14593\nExp number, first 60 AAs: 14.41781\nTotal prob of N-in: 0.85703\nPOSSIBLE N-term signal sequence\ninside 1 6\nTMhelix 7 24\noutside 25 105\nTMhelix 106 128\ninside 129 154

159 GCF_001043885.1_ASM104388v1 Methylobacterium platani JCM 14648 Proteobacteria; Alphaproteobacteria; Rhizobiales; Methylobacteriaceae; Methylobacterium; Methylobacterium platani MHTLSANAAGTERRGIFATLPAMLAWPAVAASDAAEAFERVTAPDPVFAAIETHRAEQVA WP_048435058.1 hypothetical protein [Methylobacterium platani] Length: 188\nNumber of predicted TMHs: 1\nExp number of AAs in TMHs: 22.85436\nExp number, first 60 AAs: 1.55214\nTotal prob of N-in: 0.29690\noutside 1 162\nTMhelix 163 185\ninside 186 188

160 GCF_001043885.1_ASM104388v1 Methylobacterium platani JCM 14648 Proteobacteria; Alphaproteobacteria; Rhizobiales; Methylobacteriaceae; Methylobacterium; Methylobacterium platani MSAPLTASERRAAIRDLVAPALATLVCLAILVGLGVWQLERKAWKEALIDRIVARSRIDP WP_048431791.1 surfeit 1 [Methylobacterium platani] Length: 267\nNumber of predicted TMHs: 2\nExp number of AAs in TMHs: 44.80971\nExp number, first 60 AAs: 21.95878\nTotal prob of N-in: 0.62344\nPOSSIBLE N-term signal sequence\ninside 1 16\nTMhelix 17 39\noutside 40 223\nTMhelix 224 246\ninside 247 267

161 GCF_000019945.1_ASM1994v1 Methylobacterium populi BJ001 Proteobacteria; Alphaproteobacteria; Rhizobiales; Methylobacteriaceae; Methylobacterium; Methylobacterium populi
MPHDPIRPAATTRRAALLGLSAGLSSGLLGTVVPAAAYQDDGVRPDSFRASEVVENGHRF WP_012453627.1
hypothetical protein [Methylobacterium populi] Length: 205\nNumber of predicted TMHs: 2\nExp number of AAs in TMHs: 39.69176\nExp number, first 60 AAs: 19.987\nTotal prob of N-in: 0.59493\nPOSSIBLE N-term signal sequence\ninside 1 15\nTMhelix 16 38\noutside 39 156\nTMhelix 157 179\ninside 180 205

162 GCF_002117405.1_ASM211740v1 Methylocystis bryophila Proteobacteria; Alphaproteobacteria; Rhizobiales; Methylocystaceae; Methylocystis
MVSFGQSAAGPRLRERGGGFRRRAGRRRLLALALLAGLVGPAFAVDPGEKLADPALESRA WP_085770606.1
cytochrome C biogenesis protein [Methylocystis bryophila] Length: 181\nNumber of predicted TMHs: 2\nExp number of AAs in TMHs: 38.6472199999999\nExp number, first 60 AAs: 17.72034\nTotal prob of N-in: 0.96926\nPOSSIBLE N-term signal sequence\ninside 1 27\nTMhelix 28 45\noutside 46 126\nTMhelix 127 149\ninside 150 181

163 GCF_000685825.1_ASM68582v1 Methylocystis sp. LW5 Proteobacteria; Alphaproteobacteria; Rhizobiales; Methylocystaceae; Methylocystis
MDTRSSRRNIFALSFLFALATAGSAFAHGGVKLEQDECVTTVGPVKVHFIGYQRKGEPEE WP_026598801.1 hypothetical protein [Methylocystis sp. LW5] Length: 216\nNumber of predicted TMHs: 2\nExp number of AAs in TMHs: 42.33461\nExp number, first 60 AAs: 20.44294\nTotal prob of N-in: 0.93701\nPOSSIBLE N-term signal sequence\ninside 1 8\nTMhelix 9 31\noutside 32 178\nTMhelix 179 201\ninside 202 216

164 GCF_000283235.1_MetPar_1.0 Methylocystis parvus OBBP Proteobacteria; Alphaproteobacteria; Rhizobiales; Methylocystaceae; Methylocystis; Methylocystis parvus
MRASIGFVCAFVAKKERRGNVFAKLLMAAAVSMVGFIAQAEAHGGVSLDQGQCIMKIGPD WP_016920776.1
hypothetical protein [Methylocystis parvus] Length: 206\nNumber of predicted TMHs: 2\nExp number of AAs in TMHs: 40.27208\nExp number, first 60 AAs: 20.0164\nTotal prob of N-in: 0.98868\nPOSSIBLE N-term signal sequence\ninside 1 20\nTMhelix 21 40\noutside 41 179\nTMhelix 180 198\ninside 199 206

165 GCF_000526315.1_ASM52631v1 Methylopila sp. 73B Proteobacteria; Alphaproteobacteria; Rhizobiales; Methylocystaceae; Methylopila
MTDTSLADTPKARRKVVVVGAGFAGLSVARGLAKADVDVTVLDRHNYHCFQPLLYQVATA WP_020186848.1
hypothetical protein [Methylopila sp. 73B] Length: 427\nNumber of predicted TMHs: 1\nExp number of AAs in TMHs: 25.6728\nExp number, first 60 AAs: 2.44931\nTotal prob of N-in: 0.15662\noutside 1 373\nTMhelix 374 396\ninside 397 427

166 GCF_001675455.1_ASM167545v1 Methylosinus sp. 3S-1 Proteobacteria; Alphaproteobacteria; Rhizobiales; Methylocystaceae; Methylosinus
MDTRSSRRRIFLLAFLSGIATAGYAFAHDGVKLEQDTCVLKVGPVKVHFIGYQNKGEPQE WP_003610832.1 MULTISPECIES: hypothetical protein [Methylosinus] Length: 219\nNumber of predicted TMHs: 2\nExp number of AAs in TMHs: 40.07186\nExp number, first 60 AAs: 18.44045\nTotal prob of N-in: 0.92262\nPOSSIBLE N-term signal sequence\ninside 1 8\nTMhelix 9 28\noutside 29 177\nTMhelix 178 200\ninside 201 219

167 GCF_000527115.1_ASM52711v1 Methylosinus sp. LW3 Proteobacteria; Alphaproteobacteria; Rhizobiales; Methylocystaceae; Methylosinus
MDTRSSRRNIFALSFLFALATAGSAFAHGGVKLEQDECVTTVGPVKVHFIGYQRKGEPEE WP_024880159.1 hypothetical protein [Methylosinus sp. LW3] Length: 216\nNumber of predicted TMHs: 2\nExp number of AAs in TMHs: 42.32664\nExp number, first 60 AAs: 20.42674\nTotal prob of N-in: 0.93534\nPOSSIBLE N-term signal sequence\ninside 1 8\nTMhelix 9 31\noutside 32 178\nTMhelix 179 201\ninside 202 216

168 GCF_001644125.1_ASM164412v1 Methylosinus sp. R-45379 Proteobacteria; Alphaproteobacteria; Rhizobiales; Methylocystaceae; Methylosinus
MDTRSSRRNIFALSFLFALATAGSAFAHGGVKLEQDECVTTVGPVKVHFIGYQRKGEPEE WP_064031969.1 hypothetical protein [Methylosinus sp. R-45379] Length: 216\nNumber of predicted TMHs: 2\nExp number of AAs in TMHs: 42.33486\nExp number, first 60 AAs: 20.44277\nTotal prob of N-in: 0.93700\nPOSSIBLE N-term signal sequence\ninside 1 8\nTMhelix 9 31\noutside 32 178\nTMhelix 179 201\ninside 202 216

169 GCF_000178815.2_ASM17881v2 Methylosinus trichosporium OB3b Proteobacteria; Alphaproteobacteria; Rhizobiales; Methylocystaceae; Methylosinus; Methylosinus trichosporium
MDTRSSRRRIFLLAFLSGIATAGYAFAHDGVKLEQDTCVLKVGPVKVHFIGYQNKGEPQE WP_003610832.1 MULTISPECIES: hypothetical protein [Methylosinus] Length: 219\nNumber of predicted TMHs: 2\nExp number of AAs in TMHs:

40.07186\nExp number, first 60 AAs: 18.44045\nTotal prob of N-in: 0.92262\nPOSSIBLE N-term signal sequence\ninside 1 8\nTMhelix 9 28\noutside 29 177\nTMhelix 178 200\ninside 201 219

170 GCF_000585625.1_Aqudef_v9 Aquamicrobium defluvii Proteobacteria; Alphaproteobacteria; Rhizobiales; Phyllobacteriaceae; Aquamicrobium MKRRAPLPVRAALLVAISAACLASPSEAHRRINPGEIQGIPMASLSHGQMAVIADYRSEI WP_035022923.1 hypothetical protein [Aquamicrobium defluvii] Length: 209\nNumber of predicted TMHs: 1\nExp number of AAs in TMHs: 23.55342\nExp number, first 60 AAs: 2.23898\nTotal prob of N-in: 0.37098\noutside 1 175\nTMhelix 176 198\ninside 199 209

171 GCF_000798645.1_ASM79864v1 Mesorhizobium sp. F7 Proteobacteria; Alphaproteobacteria; Rhizobiales; Phyllobacteriaceae; Mesorhizobium MASSSPASARRVGRRSPLATTALLVVGLLATGGAYALATTTATAETTAASAQLIDEGGKL WP_047566605.1 cystathionine beta-lyase [Mesorhizobium sp. F7] Length: 266\nNumber of predicted TMHs: 2\nExp number of AAs in TMHs: 42.99993\nExp number, first 60 AAs: 21.50354\nTotal prob of N-in: 0.81518\nPOSSIBLE N-term signal sequence\ninside 1 20\nTMhelix 21 43\noutside 44 243\nTMhelix 244 263\ninside 264 266

172 GCF_001014615.1_ASM101461v1 Mesorhizobium sp. LC103 Proteobacteria; Alphaproteobacteria; Rhizobiales; Phyllobacteriaceae; Mesorhizobium MDAGWAERFFSRRRVLVTGGSSGIGAAIAAGFAAAGAEVTATGVTEVEIVAAAGEERLSG WP_047147864.1 2-deoxy-D-gluconate 3-dehydrogenase [Mesorhizobium sp. LC103] Length: 248\nNumber of predicted TMHs: 2\nExp number of AAs in TMHs: 52.32931\nExp number, first 60 AAs: 20.19119\nTotal prob of N-in: 0.31131\nPOSSIBLE N-term signal sequence\noutside 1 14\nTMhelix 15 37\ninside 38 218\nTMhelix 219 241\noutside 242 248

173 GCF_000502695.1_Mloti_LSHC426A00 Mesorhizobium sp. LSHC426A00 Proteobacteria; Alphaproteobacteria; Rhizobiales; Phyllobacteriaceae; Mesorhizobium MSRSHPPRRPTLAAMQPTSFFCAKASWRSRWRWTWRLGRRGWAGWKARTGGEGKHGYGGT WP_084561162.1 hypothetical protein [Mesorhizobium sp. LSHC426A00] Length: 255\nNumber of predicted TMHs: 1\nExp number of AAs in TMHs: 26.82341\nExp number, first 60 AAs: 0.12519\nTotal prob of N-in: 0.77932\ninside 1 189\nTMhelix 190 212\noutside 213 255

174 GCF_000502835.1_Mloti_LSJC280B00 Mesorhizobium sp. LSJC280B00 Proteobacteria; Alphaproteobacteria; Rhizobiales; Phyllobacteriaceae; Mesorhizobium MSAGSGTPPISATASNNSAFERIANARRRALLQQHGPVAALSSAPVTPQPPEPVATVTLD WP_031211881.1 LPS biosynthesis protein [Mesorhizobium sp. LSJC280B00] Length: 516\nNumber of predicted TMHs: 2\nExp number of AAs in TMHs: 45.64555\nExp number, first 60 AAs: 0.0094\nTotal prob of N-in: 0.37715\noutside 1 78\nTMhelix 79 101\ninside 102 476\nTMhelix 477 499\noutside 500 516

175 GCF_001427385.1_Root157 Mesorhizobium sp. Root157 Proteobacteria; Alphaproteobacteria; Rhizobiales; Phyllobacteriaceae; Mesorhizobium MTSSSGSVGAPSRRRTLIAAVLGLAGIAVLLALGTWQVQRLHWKEAILQTIDQRMHAAPL WP_056104246.1 cytochrome oxidase [Mesorhizobium sp. Root157] Length: 253\nNumber of predicted TMHs: 2\nExp number of AAs in TMHs: 50.51325\nExp number, first 60 AAs: 21.3244\nTotal prob of N-in: 0.52390\nPOSSIBLE N-term signal sequence\ninside 1 16\nTMhelix 17 39\noutside 40 224\nTMhelix 225 244\ninside 245 253

176 GCF_001427285.1_Root552 Mesorhizobium sp. Root552 Proteobacteria; Alphaproteobacteria; Rhizobiales; Phyllobacteriaceae; Mesorhizobium MTSQPVSVGAPSRRRTIAALVLGLAGIAILLALGTWQVQRLHWKEAILQTIDQRMHAPAQ WP_056245182.1 cytochrome oxidase [Mesorhizobium sp. Root552] Length: 251\nNumber of predicted TMHs: 2\nExp number of AAs in TMHs: 42.60315\nExp number, first 60 AAs: 21.48432\nTotal prob of N-in: 0.92518\nPOSSIBLE N-term signal sequence\ninside 1 16\nTMhelix 17 39\noutside 40 224\nTMhelix 225 244\ninside 245 251

177 GCF_001686985.1_ASM168698v1 Mesorhizobium amorphae CCNWGS0123 Proteobacteria; Alphaproteobacteria; Rhizobiales; Phyllobacteriaceae; Mesorhizobium; Mesorhizobium amorphae MRKTETFYDVIRRQGITRRSFIKFCSLTAASLGFGPDAKTAMAEALETKERVPVIWMHGL WP_065518353.1 uptake hydrogenase small subunit [Mesorhizobium amorphae] Length: 360\nNumber of predicted TMHs: 1\nExp number of AAs in TMHs: 22.58413\nExp number, first 60 AAs: 3.30493\nTotal prob of N-in: 0.22543\noutside 1 325\nTMhelix 326 348\ninside 349 360

178 GCF_000350085.1_ASM35008v1 Mesorhizobium metallidurans STM 2683 Proteobacteria; Alphaproteobacteria; Rhizobiales; Phyllobacteriaceae; Mesorhizobium; Mesorhizobium metallidurans

MPAASGPSSPSAADSDISALERIASARRRAMAQQGAPATAHAPTGAPTTARVAAPVVPPH WP_008876244.1 polysaccharide biosynthesis, transporter [Mesorhizobium metallidurans] Length: 529\nNumber of predicted TMHs: 2\nExp number of AAs in TMHs: 45.81695\nExp number, first 60 AAs: 0.00222\nTotal prob of N-in: 0.04002\noutside 1 89\nTMhelix 90 112\ninside 113 488\nTMhelix 489 511\noutside 512 529

179 GCF_000733725.1_ASM73372v1 Nitratireductor basaltis Proteobacteria; Alphaproteobacteria; Rhizobiales; Phyllobacteriaceae; Nitratireductor
MRASRRANLILLSLALPALVVLLMLGTWQVKRLQWKETLVARIEERINAEPQPLERLDDL WP_036481955.1 hypothetical protein [Nitratireductor basaltis] Length: 244\nNumber of predicted TMHs: 2\nExp number of AAs in TMHs: 43.28783\nExp number, first 60 AAs: 22.01838\nTotal prob of N-in: 0.99521\nPOSSIBLE N-term signal sequence\ninside 1 6\nTMhelix 7 29\noutside 30 213\nTMhelix 214 236\ninside 237 244

180 GCF_000282595.1_Phyllobacterium.strYR531_v1.0 Phyllobacterium sp. YR531 Proteobacteria; Alphaproteobacteria; Rhizobiales; Phyllobacteriaceae; Phyllobacterium
MIDSSKAARRGTFPWITLILGTVVFLTLMALGTWQVERLQWKEQLLAEIEQRTHAAPASL WP_008128946.1 SURF1 family protein [Phyllobacterium sp. YR531] Length: 262\nNumber of predicted TMHs: 2\nExp number of AAs in TMHs: 41.83747\nExp number, first 60 AAs: 22.32501\nTotal prob of N-in: 0.92632\nPOSSIBLE N-term signal sequence\ninside 1 12\nTMhelix 13 35\noutside 36 218\nTMhelix 219 241\ninside 242 262

181 GCF_000745735.1_ASM74573v1 Agrobacterium rhizogenes Proteobacteria; Alphaproteobacteria; Rhizobiales; Rhizobiaceae; Rhizobium/Agrobacterium group; Agrobacterium
MRKIFKQNNPAKRRAVIFFALSAVSLGIGRVEAQTNELQALVEKGRGIATAADCMACHTV WP_051983093.1 cytochrome c [Agrobacterium rhizogenes] Length: 485\nNumber of predicted TMHs: 1\nExp number of AAs in TMHs: 26.9022\nExp number, first 60 AAs: 4.30508999999999\nTotal prob of N-in: 0.23156\noutside 1 452\nTMhelix 453 475\ninside 476 485

182 GCF_900013505.1_AHY_PRJEB12192_v1 Agrobacterium genomosp. 7 str. RV3 Proteobacteria; Alphaproteobacteria; Rhizobiales; Rhizobiaceae; Rhizobium/Agrobacterium group; Agrobacterium; Agrobacterium tumefaciens complex; Agrobacterium genomosp. 7
MSVVLLPSFLCLSQESSRRASARRRTPLSPRTWAGWIPVTSTGMRERVARLALLLTLVFA WP_080856138.1 cytochrome c-type biogenesis protein CcmH [Agrobacterium genomosp. 7] Length: 198\nNumber of predicted TMHs: 1\nExp number of AAs in TMHs: 36.00344\nExp number, first 60 AAs: 7.89985\nTotal prob of N-in: 0.69760\noutside 1 149\nTMhelix 150 172\ninside 173 198

183 GCF_001297245.1_ASM129724v1 Rhizobium acidisoli Proteobacteria; Alphaproteobacteria; Rhizobiales; Rhizobiaceae; Rhizobium/Agrobacterium group; Rhizobium
MSDNVYDVIVIGGGHAGSEAASAAARLGAKTALVTHRRDTIGVMSCNPAIGGLGKGHLVR WP_054185485.1 tRNA uridine-5-carboxymethylaminomethyl(34) synthesis enzyme MnmG [Rhizobium acidisoli] Length: 625\nNumber of predicted TMHs: 1\nExp number of AAs in TMHs: 19.92105\nExp number, first 60 AAs: 0.11745\nTotal prob of N-in: 0.07171\noutside 1 591\nTMhelix 592 614\ninside 615 625

184 GCF_900094555.1_IMG-taxon_2615840609_annotated_assembly Rhizobium hainanense Proteobacteria; Alphaproteobacteria; Rhizobiales; Rhizobiaceae; Rhizobium/Agrobacterium group; Rhizobium
MRRRRAVKRFLTTVTVTALSALLAVTSFDPAAAAPIQPLIQPAIQQTGDQPAGLTEVQYR WP_075855358.1 hypothetical protein [Rhizobium hainanense] Length: 163\nNumber of predicted TMHs: 1\nExp number of AAs in TMHs: 25.12958\nExp number, first 60 AAs: 8.44288\nTotal prob of N-in: 0.96530\ninside 1 104\nTMhelix 105 124\noutside 125 163

185 GCF_000799975.1_ASM79997v1 Rhizobium leguminosarum Proteobacteria; Alphaproteobacteria; Rhizobiales; Rhizobiaceae; Rhizobium/Agrobacterium group; Rhizobium
MRKIFKQNNPAKRRAVIFFALSAVSLGIGRVEAQTNELQALVEKGRGIATAADCMACHTV WP_052212217.1 cytochrome c [Rhizobium leguminosarum] Length: 485\nNumber of predicted TMHs: 1\nExp number of AAs in TMHs: 26.90345\nExp number, first 60 AAs: 4.30659999999999\nTotal prob of N-in: 0.23117\noutside 1 452\nTMhelix 453 475\ninside 476 485

186 GCF_001938945.1_ASM193894v1 Rhizobium rhizosphaerae Proteobacteria; Alphaproteobacteria; Rhizobiales; Rhizobiaceae; Rhizobium/Agrobacterium group; Rhizobium
MPPDARVRRRTVVTLLLAVLVTAGLLSLGTWQVKRLFWKQDLIARVEARVHAPPVPVPPA WP_075636314.1 Surfeit locus 1 family protein [Rhizobium rhizosphaerae] Length: 242\nNumber of predicted TMHs: 2\nExp number of AAs in TMHs: 42.89807\nExp number, first 60 AAs: 20.76893\nTotal prob of N-in: 0.99615\nPOSSIBLE N-term signal sequence\ninside 1 11\nTMhelix 12 31\noutside 32 214\nTMhelix 215 237\ninside 238 242

187 GCF_002078095.1_ASM207809v1 Rhizobium rhizosphaerae Proteobacteria; Alphaproteobacteria; Rhizobiales; Rhizobiaceae; Rhizobium/Agrobacterium group; Rhizobium
MPPDARVRRRTVVTLLLAVLVTAGLLSLGTWQVKRLFWKQDLIARVEARVHAPPVPVPPP WP_081176957.1
Surfeit locus 1 family protein [Rhizobium rhizosphaerae] Length: 242\nNumber of predicted TMHs: 2\nExp number of AAs in TMHs: 42.89807\nExp number, first 60 AAs: 20.76893\nTotal prob of N-in: 0.99615\nPOSSIBLE N-term signal sequence\ninside 1 11\nTMhelix 12 31\noutside 32 214\nTMhelix 215 237\ninside 238 242

188 GCF_000746265.1_ASM74626v1 Rhizobium sp. CF394 Proteobacteria; Alphaproteobacteria; Rhizobiales; Rhizobiaceae; Rhizobium/Agrobacterium group; Rhizobium
MTEFEISAPRRRRSVLTGIAVLVALAILLSLGTWQVERLHWKERLLSDIAERRATPAASL WP_037135714.1 cytochrome oxidase [Rhizobium sp. CF394] Length: 249\nNumber of predicted TMHs: 2\nExp number of AAs in TMHs: 42.66405\nExp number, first 60 AAs: 22.20222\nTotal prob of N-in: 0.95966\nPOSSIBLE N-term signal sequence\ninside 1 12\nTMhelix 13 35\noutside 36 222\nTMhelix 223 242\ninside 243 249

189 GCF_001854865.1_ASM185486v1 Rhizobium sp. LCM 4573 Proteobacteria; Alphaproteobacteria; Rhizobiales; Rhizobiaceae; Rhizobium/Agrobacterium group; Rhizobium
MLTRRTVLTVSLAAAAASTLGSRLSFAQQLTKEDVFFDKDAPVLGNPKGDVTIVEFFDYQ WP_071007428.1 disulfide bond formation protein DsbA [Rhizobium sp. LCM 4573] Length: 203\nNumber of predicted TMHs: 1\nExp number of AAs in TMHs: 21.14923\nExp number, first 60 AAs: 4.33496\nTotal prob of N-in: 0.71016\ninside 1 165\nTMhelix 166 188\noutside 189 203

190 GCF_001424245.1_Leaf155 Rhizobium sp. Leaf155 Proteobacteria; Alphaproteobacteria; Rhizobiales; Rhizobiaceae; Rhizobium/Agrobacterium group; Rhizobium
MDASVYNFLTAHRHFSATARRELVKLTFLPVLSYSAAISQRRKVSKGAIVKSRILDIFGR WP_082479029.1 hypothetical protein [Rhizobium sp. Leaf155] Length: 312\nNumber of predicted TMHs: 2\nExp number of AAs in TMHs: 46.39423\nExp number, first 60 AAs: 1.29682\nTotal prob of N-in: 0.11889\noutside 1 60\nTMhelix 61 83\ninside 84 275\nTMhelix 276 298\noutside 299 312

191 GCF_001424325.1_Leaf341 Rhizobium sp. Leaf341 Proteobacteria; Alphaproteobacteria; Rhizobiales; Rhizobiaceae; Rhizobium/Agrobacterium group; Rhizobium
MSPDPSRRPDTAAGSTRRAGTNDALAPPGAAVSRGRSRRGLVVFGVASAFVTVVLLALGI WP_082475616.1
Surfeit locus 1 family protein [Rhizobium sp. Leaf341] Length: 278\nNumber of predicted TMHs: 2\nExp number of AAs in TMHs: 44.59037\nExp number, first 60 AAs: 20.24173\nTotal prob of N-in: 0.93980\nPOSSIBLE N-term signal sequence\ninside 1 40\nTMhelix 41 63\noutside 64 242\nTMhelix 243 265\ninside 266 278

192 GCF_001424045.1_Leaf371 Rhizobium sp. Leaf371 Proteobacteria; Alphaproteobacteria; Rhizobiales; Rhizobiaceae; Rhizobium/Agrobacterium group; Rhizobium
MLETADPALTTPRRRVVIIGAGFGGLACAGALGDTDVDVIVIDRRNHNLFQPLLYQVATA WP_062595595.1
pyridine nucleotide-disulfide oxidoreductase [Rhizobium sp. Leaf371] Length: 434\nNumber of predicted TMHs: 1\nExp number of AAs in TMHs: 24.3759\nExp number, first 60 AAs: 4.54116\nTotal prob of N-in: 0.19458\noutside 1 378\nTMhelix 379 396\ninside 397 434

193 GCF_001425345.1_Leaf383 Rhizobium sp. Leaf383 Proteobacteria; Alphaproteobacteria; Rhizobiales; Rhizobiaceae; Rhizobium/Agrobacterium group; Rhizobium
MNVLEEADLARALPRRRVVIIGAGFGGLACAGALGHAEVDVVVIDRRNHNLFQPLLYQVA WP_062471281.1
MULTISPECIES: pyridine nucleotide-disulfide oxidoreductase [Rhizobium] Length: 436\nNumber of predicted TMHs: 1\nExp number of AAs in TMHs: 33.04413\nExp number, first 60 AAs: 13.41147\nTotal prob of N-in: 0.59501\nPOSSIBLE N-term signal sequence\noutside 1 380\nTMhelix 381 398\ninside 399 436

194 GCF_001425345.1_Leaf383 Rhizobium sp. Leaf383 Proteobacteria; Alphaproteobacteria; Rhizobiales; Rhizobiaceae; Rhizobium/Agrobacterium group; Rhizobium
MSPDPSRRPDTAAGSTRKTGTDDALAPPGAAVSRGRSRRGLVVFGVASTFVTVVLLALGI WP_082480194.1
MULTISPECIES: Surfeit locus 1 family protein [Rhizobium] Length: 278\nNumber of predicted TMHs: 2\nExp number of AAs in TMHs: 44.51364\nExp number, first 60 AAs: 20.24205\nTotal prob of N-in: 0.93594\nPOSSIBLE N-term signal sequence\ninside 1 40\nTMhelix 41 63\noutside 64 242\nTMhelix 243 265\ninside 266 278

195 GCF_001424065.1_Leaf384 Rhizobium sp. Leaf384 Proteobacteria; Alphaproteobacteria; Rhizobiales; Rhizobiaceae; Rhizobium/Agrobacterium group; Rhizobium
MSPDPSRRPDTAAGSTRKTGTDDALAPPGAAVSRGRSRRGLVVFGVASTFVTVVLLALGI WP_082480194.1
MULTISPECIES: Surfeit locus 1 family protein [Rhizobium] Length: 278\nNumber of predicted TMHs: 2\nExp

number of AAs in TMHs: 44.51364\nExp number, first 60 AAs: 20.24205\nTotal prob of N-in: 0.93594\nPOSSIBLE N-term signal sequence\ninside 1 40\nTMhelix 41 63\noutside 64 242\nTMhelix 243 265\ninside 266 278

196 GCF_001424065.1_Leaf384 Rhizobium sp. Leaf384 Proteobacteria; Alphaproteobacteria; Rhizobiales; Rhizobiaceae; Rhizobium/Agrobacterium group; Rhizobium MNVLEEADLARALPRRRVVIIGAGFGGLACAGALGHAEVDVVVIDRRNHNLFQPLLYQVA WP_062471281.1 MULTISPECIES: pyridine nucleotide-disulfide oxidoreductase [Rhizobium] Length: 436\nNumber of predicted TMHs: 1\nExp number of AAs in TMHs: 33.04413\nExp number, first 60 AAs: 13.41147\nTotal prob of N-in: 0.59501\nPOSSIBLE N-term signal sequence\noutside 1 380\nTMhelix 381 398\ninside 399 436

197 GCF_000247475.1_ASM24747v2 Rhizobium sp. PDO1-076 Proteobacteria; Alphaproteobacteria; Rhizobiales; Rhizobiaceae; Rhizobium/Agrobacterium group; Rhizobium MNTRTKNPVELDCATSHRRSRRPAIKAVGITTLAVTGLVAALTFGASAETPLAPRLPTTQ WP_007608427.1 hypothetical protein [Rhizobium sp. PDO1-076] Length: 157\nNumber of predicted TMHs: 2\nExp number of AAs in TMHs: 43.22282\nExp number, first 60 AAs: 21.88133\nTotal prob of N-in: 0.99842\nPOSSIBLE N-term signal sequence\ninside 1 23\nTMhelix 24 46\noutside 47 118\nTMhelix 119 141\ninside 142 157

198 GCF_000799775.1_ASM79977v1 Rhizobium sp. YR528 Proteobacteria; Alphaproteobacteria; Rhizobiales; Rhizobiaceae; Rhizobium/Agrobacterium group; Rhizobium MTGTMDGRRQRRPLRFKTLTNSPADQAPKRRSPIALTIFCGFLVVVLCAVFALGVWQVER WP_047504179.1 hypothetical protein [Rhizobium sp. YR528] Length: 268\nNumber of predicted TMHs: 2\nExp number of AAs in TMHs: 60.81005\nExp number, first 60 AAs: 22.83892\nTotal prob of N-in: 0.26784\nPOSSIBLE N-term signal sequence\ninside 1 33\nTMhelix 34 56\noutside 57 240\nTMhelix 241 260\ninside 261 268

199 GCF_000698845.1_ASM69884v1 Rhizobium etli bv. mimosae str. IE4771 Proteobacteria; Alphaproteobacteria; Rhizobiales; Rhizobiaceae; Rhizobium/Agrobacterium group; Rhizobium; Rhizobium etli; Rhizobium etli bv. mimosae MTGNVYDVIVIGGGHAGTEAASAAARLGAKTALVTHRRDTIGVMSCNPAIGGLGKGHLVR WP_038692108.1 tRNA uridine-5-carboxymethylaminomethyl(34) synthesis enzyme MnmG [Rhizobium etli] Length: 625\nNumber of predicted TMHs: 1\nExp number of AAs in TMHs: 17.90373\nExp number, first 60 AAs: 0.0964\nTotal prob of N-in: 0.01000\noutside 1 591\nTMhelix 592 614\ninside 615 625

200 GCF_000021345.1_ASM2134v1 Rhizobium leguminosarum bv. trifolii WSM2304 Proteobacteria; Alphaproteobacteria; Rhizobiales; Rhizobiaceae; Rhizobium/Agrobacterium group; Rhizobium; Rhizobium leguminosarum; Rhizobium leguminosarum bv. trifolii MTDKVYDVIVIGGGHAGSEAASAAARLGAKTALVTHRRDTIGVMSCNPAIGGLGKGHLVR WP_012559386.1 tRNA uridine-5-carboxymethylaminomethyl(34) synthesis enzyme MnmG [Rhizobium leguminosarum] Length: 625\nNumber of predicted TMHs: 1\nExp number of AAs in TMHs: 20.9423\nExp number, first 60 AAs: 0.12893\nTotal prob of N-in: 0.12246\noutside 1 591\nTMhelix 592 614\ninside 615 625

201 GCF_000271785.1_ASM27178v1 Rhizobium leguminosarum bv. trifolii WSM597 Proteobacteria; Alphaproteobacteria; Rhizobiales; Rhizobiaceae; Rhizobium/Agrobacterium group; Rhizobium; Rhizobium leguminosarum; Rhizobium leguminosarum bv. trifolii MSDNVYDVIVIGGGHAGSEAASAAARLGAKIALVTHRRDTIGVMSCNPAIGGLGKGHLVR WP_003589539.1 tRNA uridine-5-carboxymethylaminomethyl(34) synthesis enzyme MnmG [Rhizobium leguminosarum] Length: 625\nNumber of predicted TMHs: 1\nExp number of AAs in TMHs: 19.94344\nExp number, first 60 AAs: 0.12718\nTotal prob of N-in: 0.07295\noutside 1 591\nTMhelix 592 614\ninside 615 625

202 GCF_000518785.1_ASM51878v1 Rhizobium selenitireducens ATCC BAA-1503 Proteobacteria; Alphaproteobacteria; Rhizobiales; Rhizobiaceae; Rhizobium/Agrobacterium group; Rhizobium; Rhizobium selenitireducens MAETTRRGPVRLVFTVVAVAAAMAILLSLGTWQVKRLHWKEALIADIEVRRNAAPLDLAG WP_028736919.1 cytochrome oxidase [Rhizobium selenitireducens] Length: 244\nNumber of predicted TMHs: 2\nExp number of AAs in TMHs: 43.49493\nExp number, first 60 AAs: 22.53278\nTotal prob of N-in: 0.99034\nPOSSIBLE N-term signal sequence\ninside 1 11\nTMhelix 12 34\noutside 35 220\nTMhelix 221 240\ninside 241 244

203 GCF_000697965.2_ASM69796v2 Ensifer adhaerens Proteobacteria; Alphaproteobacteria; Rhizobiales; Rhizobiaceae; Sinorhizobium/Ensifer group; Ensifer MFTRRTVLTATVATAAAVLVGGRSSFAQQLTKEEVFFDKDAPVLGNPKGDVTIVEFFDYQ WP_034794695.1 MULTISPECIES: DSBA oxidoreductase [Ensifer] Length: 202\nNumber of predicted TMHs: 1\nExp number of AAs in TMHs: 22.55708\nExp number, first 60 AAs: 5.51195999999999\nTotal prob of N-in: 0.68987\ninside 1 165\nTMhelix 166 188\noutside 189 202

204 GCF_002007205.1_ASM200720v1 Ensifer adhaerens Proteobacteria; Alphaproteobacteria; Rhizobiales; Rhizobiaceae; Sinorhizobium/Ensifer group; Ensifer

MLTRRTVLTTTVATAAAVLVGGRPSFAQQLTKEEVFFDKDAPVLGNPKGDVTVVEFFDYQ WP_077966187.1 disulfide bond formation protein DsbA [Ensifer adhaerens] Length: 202\nNumber of predicted TMHs: 1\nExp number of AAs in TMHs: 19.67916\nExp number, first 60 AAs: 2.06494\nTotal prob of N-in: 0.76623\ninside 1 165\nTMhelix 166 188\noutside 189 202

205 GCF_002078505.1_ASM207850v1 Ensifer aridi Proteobacteria; Alphaproteobacteria; Rhizobiales; Rhizobiaceae; Sinorhizobium/Ensifer group; Ensifer MPSRSSPFTSAVTSGTIASLVTTAALALFAGLERRSIFQPTNATSHWLHGENAGHVRVADWP_081160173.1 hypothetical protein [Ensifer aridi] Length: 162\nNumber of predicted TMHs: 2\nExp number of AAs in TMHs: 54.38751\nExp number, first 60 AAs: 22.2793\nTotal prob of N-in: 0.03008\nPOSSIBLE N-term signal sequence\noutside 1 9\nTMhelix 10 32\ninside 33 132\nTMhelix 133 150\noutside 151 162

206 GCF_002093435.1_ASM209343v1 Ensifer aridi Proteobacteria; Alphaproteobacteria; Rhizobiales; Rhizobiaceae; Sinorhizobium/Ensifer group; Ensifer MPSRSSPFTSAVTSGTIASLVTTAALALFAGLERRSIFQPTNATSHWLHGENAGHVRVADWP_085032700.1 hypothetical protein [Ensifer aridi] Length: 162\nNumber of predicted TMHs: 2\nExp number of AAs in TMHs: 55.45865\nExp number, first 60 AAs: 22.30075\nTotal prob of N-in: 0.02631\nPOSSIBLE N-term signal sequence\noutside 1 9\nTMhelix 10 32\ninside 33 127\nTMhelix 128 150\noutside 151 162

207 GCF_001426365.1_Root1298 Ensifer sp. Root1298 Proteobacteria; Alphaproteobacteria; Rhizobiales; Rhizobiaceae; Sinorhizobium/Ensifer group; Ensifer MFTRRTVLTATVATAAAVLVGGRSSFAQQLTKEEVFFDKDAPVLGNPKGDVTIVEFFDYQ WP_060516608.1 MULTISPECIES: disulfide bond formation protein DsbA [Ensifer] Length: 202\nNumber of predicted TMHs: 1\nExp number of AAs in TMHs: 22.55637\nExp number, first 60 AAs: 5.51192999999999\nTotal prob of N-in: 0.68987\ninside 1 165\nTMhelix 166 188\noutside 189 202

208 GCF_001426465.1_Root1312 Ensifer sp. Root1312 Proteobacteria; Alphaproteobacteria; Rhizobiales; Rhizobiaceae; Sinorhizobium/Ensifer group; Ensifer MFTRRTVLTATVATAAAVLVGGRSSFAQQLTKEEVFFDKDAPVLGNPKGDVTIVEFFDYQ WP_060516608.1 MULTISPECIES: disulfide bond formation protein DsbA [Ensifer] Length: 202\nNumber of predicted TMHs: 1\nExp number of AAs in TMHs: 22.55637\nExp number, first 60 AAs: 5.51192999999999\nTotal prob of N-in: 0.68987\ninside 1 165\nTMhelix 166 188\noutside 189 202

209 GCF_001426275.1_Root423 Ensifer sp. Root423 Proteobacteria; Alphaproteobacteria; Rhizobiales; Rhizobiaceae; Sinorhizobium/Ensifer group; Ensifer MFTRRTVLTATVATAAAVLVGGRSSFAQQLTKEEVFFDKDAPVLGNPKGDVTIVEFFDYQ WP_034794695.1 MULTISPECIES: DSBA oxidoreductase [Ensifer] Length: 202\nNumber of predicted TMHs: 1\nExp number of AAs in TMHs: 22.55708\nExp number, first 60 AAs: 5.51195999999999\nTotal prob of N-in: 0.68987\ninside 1 165\nTMhelix 166 188\noutside 189 202

210 GCF_001427045.1_Root558 Ensifer sp. Root558 Proteobacteria; Alphaproteobacteria; Rhizobiales; Rhizobiaceae; Sinorhizobium/Ensifer group; Ensifer MFTRRTVLTATVATAAAVLVGGRSSFAQQLTKEEVFFDKDAPVLGNPKGDVTIVEFFDYQ WP_060530531.1 disulfide bond formation protein DsbA [Ensifer sp. Root558] Length: 202\nNumber of predicted TMHs: 1\nExp number of AAs in TMHs: 22.59136\nExp number, first 60 AAs: 5.29216\nTotal prob of N-in: 0.70210\ninside 1 165\nTMhelix 166 188\noutside 189 202

211 GCF_001429125.1_Root74 Ensifer sp. Root74 Proteobacteria; Alphaproteobacteria; Rhizobiales; Rhizobiaceae; Sinorhizobium/Ensifer group; Ensifer MFTRRTVLTATVATAAAVLVGGRSSFAQQLTKEEVFFDKDAPVLGNPKGDVTIVEFFDYQ WP_060516608.1 MULTISPECIES: disulfide bond formation protein DsbA [Ensifer] Length: 202\nNumber of predicted TMHs: 1\nExp number of AAs in TMHs: 22.55637\nExp number, first 60 AAs: 5.51192999999999\nTotal prob of N-in: 0.68987\ninside 1 165\nTMhelix 166 188\noutside 189 202

212 GCF_001429005.1_Root954 Ensifer sp. Root954 Proteobacteria; Alphaproteobacteria; Rhizobiales; Rhizobiaceae; Sinorhizobium/Ensifer group; Ensifer MFTRRTVLTATVATAAAVLVGGRSSFAQQLTKEEVFFDKDAPVLGNPKGDVTIVEFFDYQ WP_060516608.1 MULTISPECIES: disulfide bond formation protein DsbA [Ensifer] Length: 202\nNumber of predicted TMHs: 1\nExp number of AAs in TMHs: 22.55637\nExp number, first 60 AAs: 5.51192999999999\nTotal prob of N-in: 0.68987\ninside 1 165\nTMhelix 166 188\noutside 189 202

213 GCF_000799055.1_ZNC0028.1 Ensifer sp. ZNC0028 Proteobacteria; Alphaproteobacteria; Rhizobiales; Rhizobiaceae; Sinorhizobium/Ensifer group; Ensifer
MLTRRTVLTATVATAAAVLVGGRPSFAQQLTKEEVFFDKDAPVLGNPKGDVTIVEFFDYQ WP_043620617.1
DSBA oxidoreductase [Ensifer sp. ZNC0028] Length: 202\nNumber of predicted TMHs: 1\nExp number of AAs in TMHs: 20.5091\nExp number, first 60 AAs: 4.12810999999999\nTotal prob of N-in: 0.71812\ninside 1 165\nTMhelix 166 188\noutside 189 202

214 GCF_002002725.1_ASM200272v1 Sinorhizobium sp. A49 Proteobacteria; Alphaproteobacteria; Rhizobiales; Rhizobiaceae; Sinorhizobium/Ensifer group; Sinorhizobium
MLTRRTVLTTSFATAAAMLVGDRPSFAQQLTKEEVFFDKDAPVLGNPNGDVTVVEFFDYQ WP_077498189.1
disulfide bond formation protein DsbA [Sinorhizobium sp. A49] Length: 207\nNumber of predicted TMHs: 1\nExp number of AAs in TMHs: 17.56029\nExp number, first 60 AAs: 0.22325\nTotal prob of N-in: 0.74516\ninside 1 165\nTMhelix 166 188\noutside 189 207

215 GCF_001461765.1_ASM146176v1 Sinorhizobium sp. GW3 Proteobacteria; Alphaproteobacteria; Rhizobiales; Rhizobiaceae; Sinorhizobium/Ensifer group; Sinorhizobium
MFTRRTVLTTTVATAAAVLVGGRSSFAQQLTKEEVFFDKDAPVLGNPKGDVTIVEFFDYQ WP_060607982.1
disulfide bond formation protein DsbA [Sinorhizobium sp. GW3] Length: 202\nNumber of predicted TMHs: 1\nExp number of AAs in TMHs: 20.45155\nExp number, first 60 AAs: 2.85367\nTotal prob of N-in: 0.73769\ninside 1 165\nTMhelix 166 188\noutside 189 202

216 GCF_000261945.1_83666 Sinorhizobium fredii CCBAU 83666 Proteobacteria; Alphaproteobacteria; Rhizobiales; Rhizobiaceae; Sinorhizobium/Ensifer group; Sinorhizobium; Sinorhizobium fredii group; Sinorhizobium fredii
MRPRRAAVTLPNTWLPGADWADRYEVIVFTSRMTAAEAAWRALGSAPRWVRNLLSFRNRL WP_037435650.1
hypothetical protein [Sinorhizobium fredii] Length: 154\nNumber of predicted TMHs: 1\nExp number of AAs in TMHs: 22.50255\nExp number, first 60 AAs: 0.00201\nTotal prob of N-in: 0.97386\ninside 1 128\nTMhelix 129 151\noutside 152 154

217 GCF_000374525.1_ASM37452v1 Amorphus coralli DSM 19760 Proteobacteria; Alphaproteobacteria; Rhizobiales; Rhodobiaceae; Amorphus; Amorphus coralli
MEAARQQRGRTHRRSLIRGLVAFGLMLGCAVPAAAEKLVASLSAPRVEISSNFAGAEISVWP_018698701.1 hypothetical protein [Amorphus coralli] Length: 268\nNumber of predicted TMHs: 2\nExp number of AAs in TMHs: 42.16243\nExp number, first 60 AAs: 18.8632\nTotal prob of N-in: 0.96187\nPOSSIBLE N-term signal sequence\ninside 1 15\nTMhelix 16 35\noutside 36 242\nTMhelix 243 265\ninside 266 268

218 GCF_001927285.1_ASM192728v1 Rhizobiales bacterium MIMtkB18 Proteobacteria; Alphaproteobacteria; Rhizobiales; unclassified Rhizobiales; unclassified Rhizobiales (miscellaneous)
MKANSDDGRRPPRRSLIPLTIAALVALAVLLGLGTWQVERLGWKEDLIARVNARITAEPG WP_075217005.1
hypothetical protein [Rhizobiales bacterium MIMtkB18] Length: 256\nNumber of predicted TMHs: 2\nExp number of AAs in TMHs: 42.3177\nExp number, first 60 AAs: 21.97327\nTotal prob of N-in: 0.94450\nPOSSIBLE N-term signal sequence\ninside 1 15\nTMhelix 16 38\noutside 39 228\nTMhelix 229 248\ninside 249 256

219 GCF_001676615.1_ASM167661v1 Labrys sp. WJW Proteobacteria; Alphaproteobacteria; Rhizobiales; Xanthobacteraceae; Labrys MNDILPRRKALTMMAGMALSATGFVSPSLALAQSERPERISSNELVNSGHRFFGTVSRDL
WP_068286233.1 hypothetical protein [Labrys sp. WJW] Length: 197\nNumber of predicted TMHs: 2\nExp number of AAs in TMHs: 41.10332\nExp number, first 60 AAs: 18.72915\nTotal prob of N-in: 0.16518\nPOSSIBLE N-term signal sequence\noutside 1 9\nTMhelix 10 32\ninside 33 138\nTMhelix 139 161\noutside 162 197

220 GCF_000420265.1_ASM42026v1 Oceanicaulis alexandrii DSM 11625 Proteobacteria; Alphaproteobacteria; Rhodobacterales; Hyphomonadaceae; Oceanicaulis; Oceanicaulis alexandrii
MKPSLISTRSRLSVWARGRIPPVPIPTRRNVMARFRPYPVLTLLMLPALALLLWLGSWQLWP_084683472.1 hypothetical protein [Oceanicaulis alexandrii] Length: 267\nNumber of predicted TMHs: 2\nExp number of AAs in TMHs: 43.43189\nExp number, first 60 AAs: 21.31391\nTotal prob of N-in: 0.69591\nPOSSIBLE N-term signal sequence\ninside 1 37\nTMhelix 38 60\noutside 61 234\nTMhelix 235 257\ninside 258 267

221 GCF_001620265.1_ASM162026v1 Defluviimonas alba Proteobacteria; Alphaproteobacteria; Rhodobacterales; Rhodobacteraceae; Defluviimonas
MSQIETFYEVMRRQGITRRSFMKYCSLTASALGLGPAFVPKIAHAMETKPRTPVIWVNGL WP_066811897.1
uptake hydrogenase small subunit [Defluviimonas alba] Length: 361\nNumber of predicted TMHs: 2\nExp number of AAs in TMHs: 34.77662\nExp number, first 60 AAs: 17.01123\nTotal prob of N-in: 0.83565\nPOSSIBLE N-term signal sequence\ninside 1 19\nTMhelix 20 39\noutside 40 325\nTMhelix 326 348\ninside 349 361

222 GCF_000687435.1_SOAPdenovo_for_Version_1.05 Defluviimonas sp. 20V17 Proteobacteria; Alphaproteobacteria; Rhodobacterales; Rhodobacteraceae; Defluviimonas
MGAKMLIWVKQWHPRRSAVAFFMAAALSLGATAGAAAPTPSSRLSGLPETVGMGLRHLGD WP_035841397.1 PEP-CTERM domain protein [Defluviimonas sp. 20V17] Length: 267\nNumber of predicted TMHs: 2\nExp number of AAs in TMHs: 59.69228\nExp number, first 60 AAs: 19.79106\nTotal prob of N-in: 0.94460\nPOSSIBLE N-term signal sequence\ninside 1 16\nTMhelix 17 39\noutside 40 237\nTMhelix 238 260\ninside 261 267

223 GCF_002196955.1_ASM219695v1 Haematobacter massiliensis Proteobacteria; Alphaproteobacteria; Rhodobacterales; Rhodobacteraceae; Haematobacter
MVQISRRGFFIATGGAALGLTAACGNGVGTANSQRIDARVDATRDFLFSRYPGTVDLAAK WP_035708011.1 twin-arginine translocation pathway signal [Haematobacter massiliensis] Length: 187\nNumber of predicted TMHs: 2\nExp number of AAs in TMHs: 51.84043\nExp number, first 60 AAs: 17.00967\nTotal prob of N-in: 0.47391\nPOSSIBLE N-term signal sequence\ninside 1 6\nTMhelix 7 29\noutside 30 156\nTMhelix 157 179\ninside 180 187

224 GCF_000740795.1_ASM74079v1 Haematobacter massiliensis Proteobacteria; Alphaproteobacteria; Rhodobacterales; Rhodobacteraceae; Haematobacter
MVQISRRGFFIATGGAALGLTAACGNGVGTANSQRIDARVDATRDFLFSRYPGTVDLAAK WP_035708011.1 twin-arginine translocation pathway signal [Haematobacter massiliensis] Length: 187\nNumber of predicted TMHs: 2\nExp number of AAs in TMHs: 51.84043\nExp number, first 60 AAs: 17.00967\nTotal prob of N-in: 0.47391\nPOSSIBLE N-term signal sequence\ninside 1 6\nTMhelix 7 29\noutside 30 156\nTMhelix 157 179\ninside 180 187

225 GCF_002196885.1_ASM219688v1 Haematobacter massiliensis Proteobacteria; Alphaproteobacteria; Rhodobacterales; Rhodobacteraceae; Haematobacter
MVQISRRGFFIATGGAALGLTAACGNGVGTANSQRIDARVDATRDFLFSRYPGTVDLAAK WP_035708011.1 twin-arginine translocation pathway signal [Haematobacter massiliensis] Length: 187\nNumber of predicted TMHs: 2\nExp number of AAs in TMHs: 51.84043\nExp number, first 60 AAs: 17.00967\nTotal prob of N-in: 0.47391\nPOSSIBLE N-term signal sequence\ninside 1 6\nTMhelix 7 29\noutside 30 156\nTMhelix 157 179\ninside 180 187

226 GCF_000152805.1_ASM15280v1 Maritimibacter alkaliphilus HTCC2654 Proteobacteria; Alphaproteobacteria; Rhodobacterales; Rhodobacteraceae; Maritimibacter; Maritimibacter alkaliphilus
MKTYTRRGVLLAAGGAGASLALAGCDNGMGSNAAGTIDARVDSTLNYLFSTYPEAQDLRS WP_008328674.1 twin-arginine translocation pathway signal sequence domain protein [Maritimibacter alkaliphilus] Length: 188\nNumber of predicted TMHs: 1\nExp number of AAs in TMHs: 44.96086\nExp number, first 60 AAs: 6.71621\nTotal prob of N-in: 0.91543\ninside 1 155\nTMhelix 156 178\noutside 179 188

227 GCF_001879715.1_ASM187971v1 Nioella nitratireducens Proteobacteria; Alphaproteobacteria; Rhodobacterales; Rhodobacteraceae; Nioella
MRRRNFVLAGLAGVGVTAIGARGSRLLAQETQGIAPIAAFATDMAGGRVVVRAYPAEGRG WP_071675328.1 hypothetical protein [Nioella nitratireducens] Length: 221\nNumber of predicted TMHs: 1\nExp number of AAs in TMHs: 39.95733\nExp number, first 60 AAs: 17.3785\nTotal prob of N-in: 0.82685\nPOSSIBLE N-term signal sequence\noutside 1 151\nTMhelix 152 174\ninside 175 221

228 GCF_001639985.1_ASM163998v1 Paenirhodobacter sp. MME-103 Proteobacteria; Alphaproteobacteria; Rhodobacterales; Rhodobacteraceae; Paenirhodobacter
MIRRSQPLSERLEARGISRRSFLKFCTTTASLLALSPALAPRIAEALESPRRPSVIWLSF WP_068767481.1 Ni/Fe hydrogenase [Paenirhodobacter sp. MME-103] Length: 361\nNumber of predicted TMHs: 1\nExp number of AAs in TMHs: 43.08287\nExp number, first 60 AAs: 5.7731\nTotal prob of N-in: 0.55489\noutside 1 317\nTMhelix 318 340\ninside 341 361

229 GCF_001517385.1_JGI_assembly Pannonibacter indicus Proteobacteria; Alphaproteobacteria; Rhodobacterales; Rhodobacteraceae; Pannonibacter
MADSKASGTSSSEGRPISRRALMGFMLGSVASAALMGRSGAALAQGSPPGGSYNQQEILE WP_055456381.1 hypothetical protein [Pannonibacter indicus] Length: 210\nNumber of predicted TMHs: 1\nExp number of AAs in TMHs: 32.53059\nExp number, first 60 AAs: 10.97294\nTotal prob of N-in: 0.80366\nPOSSIBLE N-term signal sequence\ninside 1 161\nTMhelix 162 184\noutside 185 210

230 GCF_001418225.1_ASM141822v1 Pannonibacter indicus Proteobacteria; Alphaproteobacteria; Rhodobacterales; Rhodobacteraceae; Pannonibacter
MADSKASGTSSSEGRPISRRALMGFMLGSVASAALMGRSGAALAQGSPPGGSYNQQEILE WP_055456381.1 hypothetical protein [Pannonibacter indicus] Length: 210\nNumber of predicted TMHs: 1\nExp number of

AAs in TMHs: 32.53059\nExp number, first 60 AAs: 10.97294\nTotal prob of N-in: 0.80366\nPOSSIBLE N-term signal sequence\ninside 1 161\nTMhelix 162 184\noutside 185 210

231 GCF_001484065.1_ASM148406v1 Pannonibacter phragmitetus Proteobacteria; Alphaproteobacteria; Rhodobacterales; Rhodobacteraceae; Pannonibacter MADNKASGTSSSEGRPISRRALMGFMLGSVASAALMGRSGAALAQGSPPGGSYNQQEILD WP_050472495.1 hypothetical protein [Pannonibacter phragmitetus] Length: 210\nNumber of predicted TMHs: 1\nExp number of AAs in TMHs: 33.07507\nExp number, first 60 AAs: 11.17652\nTotal prob of N-in: 0.85418\nPOSSIBLE N-term signal sequence\ninside 1 161\nTMhelix 162 184\noutside 185 210

232 GCF_001484065.1_ASM148406v1 Pannonibacter phragmitetus Proteobacteria; Alphaproteobacteria; Rhodobacterales; Rhodobacteraceae; Pannonibacter MSQVETFYDVMRRQGITRRSFMKYCSLTAAALGLGPSFVPKIAHAMETKPRTPVIWIHGL WP_050474643.1 uptake hydrogenase small subunit [Pannonibacter phragmitetus] Length: 361\nNumber of predicted TMHs: 2\nExp number of AAs in TMHs: 32.48973\nExp number, first 60 AAs: 17.57341\nTotal prob of N-in: 0.86257\nPOSSIBLE N-term signal sequence\ninside 1 19\nTMhelix 20 39\noutside 40 325\nTMhelix 326 348\ninside 349 361

233 GCF_001188555.1_ASM118855v1 Pannonibacter phragmitetus Proteobacteria; Alphaproteobacteria; Rhodobacterales; Rhodobacteraceae; Pannonibacter MSQVETFYDVMRRQGITRRSFMKYCSLTAAALGLGPSFVPKIAHAMETKPRTPVIWIHGL WP_050474643.1 uptake hydrogenase small subunit [Pannonibacter phragmitetus] Length: 361\nNumber of predicted TMHs: 2\nExp number of AAs in TMHs: 32.48973\nExp number, first 60 AAs: 17.57341\nTotal prob of N-in: 0.86257\nPOSSIBLE N-term signal sequence\ninside 1 19\nTMhelix 20 39\noutside 40 325\nTMhelix 326 348\ninside 349 361

234 GCF_001188555.1_ASM118855v1 Pannonibacter phragmitetus Proteobacteria; Alphaproteobacteria; Rhodobacterales; Rhodobacteraceae; Pannonibacter MADNKASGTSSSEGRPISRRALMGFMLGSVASAALMGRSGAALAQGSPPGGSYNQQEILD WP_050472495.1 hypothetical protein [Pannonibacter phragmitetus] Length: 210\nNumber of predicted TMHs: 1\nExp number of AAs in TMHs: 33.07507\nExp number, first 60 AAs: 11.17652\nTotal prob of N-in: 0.85418\nPOSSIBLE N-term signal sequence\ninside 1 161\nTMhelix 162 184\noutside 185 210

235 GCF_000382365.1_ASM38236v1 Pannonibacter phragmitetus DSM 14782 Proteobacteria; Alphaproteobacteria; Rhodobacterales; Rhodobacteraceae; Pannonibacter; Pannonibacter phragmitetus MADNKASGTSSSESRPISRRALMGFMLGSVASAALMGRSGAAQAQGSPPGGSYNQQEILD WP_019965077.1 hypothetical protein [Pannonibacter phragmitetus] Length: 210\nNumber of predicted TMHs: 1\nExp number of AAs in TMHs: 28.89844\nExp number, first 60 AAs: 7.59725\nTotal prob of N-in: 0.78847\ninside 1 161\nTMhelix 162 184\noutside 185 210

236 GCF_900142875.1_IMG-taxon_2695420926_annotated_assembly Paracoccus solventivorans Proteobacteria; Alphaproteobacteria; Rhodobacterales; Rhodobacteraceae; Paracoccus MTINRRKMLGAGAAGLMLAGIGAVALRTRGTSPDAMTIDDVLHDPDNPVLGNPGGRLTIV WP_073063818.1 disulfide bond formation protein DsbA [Paracoccus solventivorans] Length: 210\nNumber of predicted TMHs: 2\nExp number of AAs in TMHs: 43.9033899999999\nExp number, first 60 AAs: 16.45804\nTotal prob of N-in: 0.90631\nPOSSIBLE N-term signal sequence\ninside 1 8\nTMhelix 9 26\noutside 27 171\nTMhelix 172 194\ninside 195 210

237 GCF_001887735.1_ASM188773v1 Paracoccus sp. SM22M-07 Proteobacteria; Alphaproteobacteria; Rhodobacterales; Rhodobacteraceae; Paracoccus MSISTKLSRRTLLASGGAALLAAGCTNAVGTNGPAQLDGRVDQTHQYLVSTYPASAPLLQ WP_072295392.1 twin-arginine translocation pathway signal [Paracoccus sp. SM22M-07] Length: 208\nNumber of predicted TMHs: 2\nExp number of AAs in TMHs: 62.65962\nExp number, first 60 AAs: 10.64541\nTotal prob of N-in: 0.95175\nPOSSIBLE N-term signal sequence\ninside 1 129\nTMhelix 130 152\noutside 153 155\nTMhelix 156 178\ninside 179 208

238 GCF_002087335.1_ASM208733v1 Pseudophaeobacter leonis Proteobacteria; Alphaproteobacteria; Rhodobacterales; Rhodobacteraceae; Pseudophaeobacter MGPIFSLADLFDMLRRRAAVILYVTVLGSILSIWVALNQQHVYPSVEVIQVTRPVIADDL WP_083097746.1 DUF874 domain-containing protein [Pseudophaeobacter leonis] Length: 426\nNumber of predicted TMHs: 2\nExp number of AAs in TMHs: 40.84844\nExp number, first 60 AAs: 19.04112\nTotal prob of N-in: 0.74484\nPOSSIBLE N-term signal sequence\ninside 1 19\nTMhelix 20 37\noutside 38 355\nTMhelix 356 378\ninside 379 426

239 GCF_000473205.1_ASM47320v1 Pseudophaeobacter arcticus DSM 23566 Proteobacteria; Alphaproteobacteria; Rhodobacterales; Rhodobacteraceae; Pseudophaeobacter; Pseudophaeobacter arcticus MFSRRSFLTAAAAALAAPAALANAPQTSLRPQARAAAGLGSGAGVTQASTTQASAATSLE WP_027241321.1 D-alanyl-D-alanine carboxypeptidase [Pseudophaeobacter arcticus] Length: 507\nNumber of predicted TMHs: 1\nExp number of AAs in TMHs: 22.50392\nExp number, first 60 AAs: 3.16219\nTotal prob of N-in: 0.16363\noutside 1 441\nTMhelix 442 464\ninside 465 507

240 GCF_001509585.1_ASM150958v1 Pseudoruegeria marinistellae Proteobacteria; Alphaproteobacteria; Rhodobacterales; Rhodobacteraceae; Pseudoruegeria MSQIETFYDVMRRQGITRRSFMKYCSLTAAALGLGPAYVPKIAHAMETKPRTPVIWVHGL WP_068114235.1 uptake hydrogenase small subunit [Pseudoruegeria marinistellae] Length: 365\nNumber of predicted TMHs: 2\nExp number of AAs in TMHs: 38.20853\nExp number, first 60 AAs: 18.53106\nTotal prob of N-in: 0.91667\nPOSSIBLE N-term signal sequence\ninside 1 19\nTMhelix 20 39\noutside 40 326\nTMhelix 327 349\ninside 350 365

241 GCF_900110875.1_IMG-taxon_2684622848_annotated_assembly Pseudovibrio axinellae Proteobacteria; Alphaproteobacteria; Rhodobacterales; Rhodobacteraceae; Pseudovibrio MAERRKLVWIIGASSGIGSSLSRLYARSGWRVVTSARSRQKLQDMAACFELMKVVPLDVT WP_068006864.1 short-chain dehydrogenase [Pseudovibrio axinellae] Length: 250\nNumber of predicted TMHs: 1\nExp number of AAs in TMHs: 29.1115\nExp number, first 60 AAs: 5.07134\nTotal prob of N-in: 0.53337\noutside 1 220\nTMhelix 221 243\ninside 244 250

242 GCF_001623255.1_ASM162325v1 Pseudovibrio axinellae Proteobacteria; Alphaproteobacteria; Rhodobacterales; Rhodobacteraceae; Pseudovibrio MAERRKLVWIIGASSGIGSSLSRLYARSGWRVVTSARSRQKLQDMAACFELMKVVPLDVT WP_068006864.1 short-chain dehydrogenase [Pseudovibrio axinellae] Length: 250\nNumber of predicted TMHs: 1\nExp number of AAs in TMHs: 29.1115\nExp number, first 60 AAs: 5.07134\nTotal prob of N-in: 0.53337\noutside 1 220\nTMhelix 221 243\ninside 244 250

243 GCF_001941685.1_ASM194168v1 Pseudovibrio sp. Ab134 Proteobacteria; Alphaproteobacteria; Rhodobacterales; Rhodobacteraceae; Pseudovibrio MAVIKVAILLMVQIMEEEVGDLTPARIMAVLMPLLAQTKTLARRRQILVQLTIAVQQVIS WP_083646242.1 hypothetical protein [Pseudovibrio sp. Ab134] Length: 160\nNumber of predicted TMHs: 2\nExp number of AAs in TMHs: 43.98225\nExp number, first 60 AAs: 10.96557\nTotal prob of N-in: 0.84335\nPOSSIBLE N-term signal sequence\ninside 1 46\nTMhelix 47 69\noutside 70 100\nTMhelix 101 123\ninside 124 160

244 GCF_001720585.1_ASM172058v1 Rhodobacter johrii Proteobacteria; Alphaproteobacteria; Rhodobacterales; Rhodobacteraceae; Rhodobacter MRMSRRPFLPALAALTLLTLAPAAAMAKPEACLTSLEGFPTALAYDTKDEEVRANRSIRE WP_069330809.1 hypothetical protein [Rhodobacter johrii] Length: 208\nNumber of predicted TMHs: 1\nExp number of AAs in TMHs: 38.35533\nExp number, first 60 AAs: 13.16276\nTotal prob of N-in: 0.79491\nPOSSIBLE N-term signal sequence\noutside 1 182\nTMhelix 183 205\ninside 206 208

245 GCF_000327525.1_Rhodobacter_sp._AKP1 Rhodobacter sp. AKP1 Proteobacteria; Alphaproteobacteria; Rhodobacterales; Rhodobacteraceae; Rhodobacter MRMSRRPFLPALAPLILLTLAPAAAMAKPEACLTSLEGFPTALAYDTGDEEVRANRSIRE WP_009561837.1 hypothetical protein [Rhodobacter sp. AKP1] Length: 208\nNumber of predicted TMHs: 1\nExp number of AAs in TMHs: 38.60733\nExp number, first 60 AAs: 12.54701\nTotal prob of N-in: 0.78611\nPOSSIBLE N-term signal sequence\noutside 1 182\nTMhelix 183 205\ninside 206 208

246 GCF_001691415.1_ASM169141v1 Rhodobacter sp. CCB-MM2 Proteobacteria; Alphaproteobacteria; Rhodobacterales; Rhodobacteraceae; Rhodobacter MTQDLRATRPVSPAVSPQISRRGLLVGGAATSLLAACGNPVGNFNAERLDARVDATRDYL WP_068297290.1 twin-arginine translocation pathway signal [Rhodobacter sp. CCB-MM2] Length: 200\nNumber of predicted TMHs: 2\nExp number of AAs in TMHs: 45.64345\nExp number, first 60 AAs: 4.14028\nTotal prob of N-in: 0.74650\ninside 1 66\nTMhelix 67 89\noutside 90 170\nTMhelix 171 193\ninside 194 200

247 GCF_001576595.1_ASM157659v1 Rhodobacter sphaeroides Proteobacteria; Alphaproteobacteria; Rhodobacterales; Rhodobacteraceae; Rhodobacter MRMSRRPFLPALAALALLTLAPAAAMAKPEACLTSLEGFPTALAYDTGDEEVRANRSIRE WP_017139944.1 hypothetical protein [Rhodobacter sphaeroides] Length: 208\nNumber of predicted TMHs: 2\nExp number of AAs in TMHs:

40.28641\nExp number, first 60 AAs: 15.17633\nTotal prob of N-in: 0.85847\nPOSSIBLE N-term signal sequence\ninside 1 6\nTMhelix 7 26\noutside 27 182\nTMhelix 183 205\ninside 206 208

248 GCF_001685625.1_ASM168562v1 Rhodobacter sphaeroides Proteobacteria; Alphaproteobacteria; Rhodobacterales; Rhodobacteraceae; Rhodobacter
MRMSRRPFLPALAALALLTLAPAAAMAKPEACLTSLEGFPTALAYDTGDEEVRANRSIRE WP_017139944.1 hypothetical protein [Rhodobacter sphaeroides] Length: 208\nNumber of predicted TMHs: 2\nExp number of AAs in TMHs: 40.28641\nExp number, first 60 AAs: 15.17633\nTotal prob of N-in: 0.85847\nPOSSIBLE N-term signal sequence\ninside 1 6\nTMhelix 7 26\noutside 27 182\nTMhelix 183 205\ninside 206 208

249 GCF_001856665.1_ASM185666v1 Rhodobacter sp. LPB0142 Proteobacteria; Alphaproteobacteria; Rhodobacterales; Rhodobacteraceae; Rhodobacter
MIRRSQPLSERLEARGISRRSFLKFCTTTASLLALSPALAPRIAEALESPRRPSVIWLSF WP_071165778.1 Ni/Fe hydrogenase [Rhodobacter sp. LPB0142] Length: 361\nNumber of predicted TMHs: 1\nExp number of AAs in TMHs: 47.0999\nExp number, first 60 AAs: 5.87293\nTotal prob of N-in: 0.57631\noutside 1 317\nTMhelix 318 340\ninside 341 361

250 GCF_000176015.1_ASM17601v1 Rhodobacter sp. SW2 Proteobacteria; Alphaproteobacteria; Rhodobacterales; Rhodobacteraceae; Rhodobacter
MSDRGTIGEAMQRQGISRRSLLKYTGYLASILALPPMASRAMADALASAGRQSVIWLSFQ WP_008030250.1 Ni/Fe hydrogenase [Rhodobacter sp. SW2] Length: 358\nNumber of predicted TMHs: 1\nExp number of AAs in TMHs: 39.72509\nExp number, first 60 AAs: 15.74169\nTotal prob of N-in: 0.80532\nPOSSIBLE N-term signal sequence\noutside 1 317\nTMhelix 318 340\ninside 341 358

251 GCF_000176015.1_ASM17601v1 Rhodobacter sp. SW2 Proteobacteria; Alphaproteobacteria; Rhodobacterales; Rhodobacteraceae; Rhodobacter
MSQIETFYDVMRRQGITRRSFMKYCSLTAAALGLGPAFVPKIAHAMETKPRTPVIWVHGL WP_008030829.1 hydrogenase (NiFe) small subunit HydA [Rhodobacter sp. SW2] Length: 361\nNumber of predicted TMHs: 2\nExp number of AAs in TMHs: 39.35316\nExp number, first 60 AAs: 19.29413\nTotal prob of N-in: 0.94633\nPOSSIBLE N-term signal sequence\ninside 1 19\nTMhelix 20 39\noutside 40 325\nTMhelix 326 348\ninside 349 361

252 GCF_000273405.1_Rhod_Spha_2_4_1_V1 Rhodobacter sphaeroides 2.4.1 Proteobacteria; Alphaproteobacteria; Rhodobacterales; Rhodobacteraceae; Rhodobacter; Rhodobacter sphaeroides
MRMSRRPFLPALAALALLTLAPAAAMAKPEACLTSLEGFPTALAYDTGDEEVRANRSIRE WP_017139944.1 hypothetical protein [Rhodobacter sphaeroides] Length: 208\nNumber of predicted TMHs: 2\nExp number of AAs in TMHs: 40.28641\nExp number, first 60 AAs: 15.17633\nTotal prob of N-in: 0.85847\nPOSSIBLE N-term signal sequence\ninside 1 6\nTMhelix 7 26\noutside 27 182\nTMhelix 183 205\ninside 206 208

253 GCF_000269625.1_PB_Rhod_Spha_2_4_1_V1 Rhodobacter sphaeroides 2.4.1 Proteobacteria; Alphaproteobacteria; Rhodobacterales; Rhodobacteraceae; Rhodobacter; Rhodobacter sphaeroides
MRMSRRPFLPALAALALLTLAPAAAMAKPEACLTSLEGFPTALAYDTGDEEVRANRSIRE WP_017139944.1 hypothetical protein [Rhodobacter sphaeroides] Length: 208\nNumber of predicted TMHs: 2\nExp number of AAs in TMHs: 40.28641\nExp number, first 60 AAs: 15.17633\nTotal prob of N-in: 0.85847\nPOSSIBLE N-term signal sequence\ninside 1 6\nTMhelix 7 26\noutside 27 182\nTMhelix 183 205\ninside 206 208

254 GCF_000012905.2_ASM1290v2 Rhodobacter sphaeroides 2.4.1 Proteobacteria; Alphaproteobacteria; Rhodobacterales; Rhodobacteraceae; Rhodobacter; Rhodobacter sphaeroides
MRMSRRPFLPALAALALLTLAPAAAMAKPEACLTSLEGFPTALAYDTGDEEVRANRSIRE YP_353758.2 hypothetical protein RSP_0681 [Rhodobacter sphaeroides 2.4.1] Length: 208\nNumber of predicted TMHs: 2\nExp number of AAs in TMHs: 40.28641\nExp number, first 60 AAs: 15.17633\nTotal prob of N-in: 0.85847\nPOSSIBLE N-term signal sequence\ninside 1 6\nTMhelix 7 26\noutside 27 182\nTMhelix 183 205\ninside 206 208

255 GCF_000016405.1_ASM1640v1 Rhodobacter sphaeroides ATCC 17025 Proteobacteria; Alphaproteobacteria; Rhodobacterales; Rhodobacteraceae; Rhodobacter; Rhodobacter sphaeroides
MPQIETFYDVMRRQGITRRSFMKYCSLTAAALGLGPSFVPKIAHAMETKPRTPVIWVHGL WP_011910314.1 hypothetical protein [Rhodobacter sphaeroides] Length: 368\nNumber of predicted TMHs: 2\nExp number of AAs in TMHs: 35.15752\nExp number, first 60 AAs: 16.24211\nTotal prob of N-in: 0.79807\nPOSSIBLE N-term signal sequence\ninside 1 19\nTMhelix 20 39\noutside 40 325\nTMhelix 326 348\ninside 349 368

256 GCF_000015985.1_ASM1598v1 Rhodobacter sphaeroides ATCC 17029 Proteobacteria; Alphaproteobacteria; Rhodobacterales; Rhodobacteraceae; Rhodobacter; Rhodobacter sphaeroides

MRMSRRPFLPALAALALLTLAPAAAMAKPEACLTSLEGFPTALAYDTGDEEVRANRSIRE WP_011841602.1 hypothetical protein [Rhodobacter sphaeroides] Length: 208\nNumber of predicted TMHs: 2\nExp number of AAs in TMHs: 40.28757\nExp number, first 60 AAs: 15.17641\nTotal prob of N-in: 0.85847\nPOSSIBLE N-term signal sequence\ninside 1 6\nTMhelix 7 26\noutside 27 182\nTMhelix 183 205\ninside 206 208

257 GCF_000021005.1_ASM2100v1 Rhodobacter sphaeroides KD131 Proteobacteria; Alphaproteobacteria; Rhodobacterales; Rhodobacteraceae; Rhodobacter; Rhodobacter sphaeroides
MRMSRRPFLLALAALALLTLAPAAAMAKPEACLTSLEGFPTALAYDTGDEEVRANRSIRE WP_015921152.1 hypothetical protein [Rhodobacter sphaeroides] Length: 208\nNumber of predicted TMHs: 2\nExp number of AAs in TMHs: 42.83982\nExp number, first 60 AAs: 19.29354\nTotal prob of N-in: 0.97437\nPOSSIBLE N-term signal sequence\ninside 1 6\nTMhelix 7 26\noutside 27 182\nTMhelix 183 205\ninside 206 208

258 GCF_000212605.1_ASM21260v1 Rhodobacter sphaeroides WS8N Proteobacteria; Alphaproteobacteria; Rhodobacterales; Rhodobacteraceae; Rhodobacter; Rhodobacter sphaeroides
MRMSRRPFLPALAALALLTLAPAAAMAKPEACLTSLEGFPTALAYDTGDEEVRANRSIRE WP_002720863.1 hypothetical protein [Rhodobacter sphaeroides] Length: 208\nNumber of predicted TMHs: 2\nExp number of AAs in TMHs: 39.70623\nExp number, first 60 AAs: 15.61495\nTotal prob of N-in: 0.85655\nPOSSIBLE N-term signal sequence\ninside 1 6\nTMhelix 7 26\noutside 27 182\nTMhelix 183 205\ninside 206 208

259 GCF_002079305.1_ASM207930v1 Rhodovulum sp. P5 Proteobacteria; Alphaproteobacteria; Rhodobacterales; Rhodobacteraceae; Rhodovulum
MSHHETIGEGLARRGISRRALLRYTAWLASLMALPPTASRVMAETLAGQPRQPVIWLSFQ WP_081531658.1 Ni/Fe hydrogenase [Rhodovulum sp. P5] Length: 362\nNumber of predicted TMHs: 1\nExp number of AAs in TMHs: 31.90729\nExp number, first 60 AAs: 11.57757\nTotal prob of N-in: 0.56537\nPOSSIBLE N-term signal sequence\noutside 1 319\nTMhelix 320 342\ninside 343 362

260 GCF_000293785.1_ASM29378v1 Rhodovulum sp. PH10 Proteobacteria; Alphaproteobacteria; Rhodobacterales; Rhodobacteraceae; Rhodovulum
MRSVEPAAAPRRRLLVPAVMSLVGVALLIGLGVWQLQRREWKHALIATLDERLSAAPVAL WP_008386842.1 SURF1 family protein [Rhodovulum sp. PH10] Length: 261\nNumber of predicted TMHs: 2\nExp number of AAs in TMHs: 58.6019200000001\nExp number, first 60 AAs: 21.58913\nTotal prob of N-in: 0.48061\nPOSSIBLE N-term signal sequence\ninside 1 12\nTMhelix 13 35\noutside 36 227\nTMhelix 228 250\ninside 251 261

261 GCF_001548075.1_ASM154807v1 Rhodovulum sulfidophilum Proteobacteria; Alphaproteobacteria; Rhodobacterales; Rhodobacteraceae; Rhodovulum
MPRRALIALLLALSLTPVLSGPLRAVEPNEVLADPAEEARAREISKEIRCLVCRNESIDD WP_042464940.1 cytochrome c biogenesis protein CcdA [Rhodovulum sulfidophilum] Length: 158\nNumber of predicted TMHs: 2\nExp number of AAs in TMHs: 37.50402\nExp number, first 60 AAs: 15.27651\nTotal prob of N-in: 0.78914\nPOSSIBLE N-term signal sequence\ninside 1 6\nTMhelix 7 26\noutside 27 107\nTMhelix 108 130\ninside 131 158

262 GCF_001941695.1_ASM194169v1 Rhodovulum sulfidophilum Proteobacteria; Alphaproteobacteria; Rhodobacterales; Rhodobacteraceae; Rhodovulum
MPRRALIALLLALSLTPVLSGPLRAVEPNEVLADPAEEARAREISKEIRCLVCRNESIDD WP_042464940.1 cytochrome c biogenesis protein CcdA [Rhodovulum sulfidophilum] Length: 158\nNumber of predicted TMHs: 2\nExp number of AAs in TMHs: 37.50402\nExp number, first 60 AAs: 15.27651\nTotal prob of N-in: 0.78914\nPOSSIBLE N-term signal sequence\ninside 1 6\nTMhelix 7 26\noutside 27 107\nTMhelix 108 130\ninside 131 158

263 GCF_001633145.1_ASM163314v1 Rhodovulum sulfidophilum Proteobacteria; Alphaproteobacteria; Rhodobacterales; Rhodobacteraceae; Rhodovulum
MPRRALIALLLALSLTPVLSGPLRAVEPNEVLADPAEEARAREISKEIRCLVCRNESIDD WP_042464940.1 cytochrome c biogenesis protein CcdA [Rhodovulum sulfidophilum] Length: 158\nNumber of predicted TMHs: 2\nExp number of AAs in TMHs: 37.50402\nExp number, first 60 AAs: 15.27651\nTotal prob of N-in: 0.78914\nPOSSIBLE N-term signal sequence\ninside 1 6\nTMhelix 7 26\noutside 27 107\nTMhelix 108 130\ninside 131 158

264 GCF_001941745.1_ASM194174v1 Rhodovulum sulfidophilum Proteobacteria; Alphaproteobacteria; Rhodobacterales; Rhodobacteraceae; Rhodovulum
MPRRALIALLLALSLTPVLSGPLRAVEPNEVLADPAEEARAREISKEIRCLVCRNESIDD WP_042464940.1 cytochrome c biogenesis protein CcdA [Rhodovulum sulfidophilum] Length: 158\nNumber of predicted TMHs: 2\nExp number of AAs in TMHs: 37.50402\nExp number, first 60 AAs: 15.27651\nTotal prob of N-in: 0.78914\nPOSSIBLE N-term signal sequence\ninside 1 6\nTMhelix 7 26\noutside 27 107\nTMhelix 108 130\ninside 131 158

265 GCF_000520135.2_ASM52013v2 Rhodovulum sulfidophilum DSM 1374 Proteobacteria; Alphaproteobacteria; Rhodobacterales; Rhodobacteraceae; Rhodovulum; Rhodovulum sulfidophilum
MPRRALIALLLALSLTPVLSGPLRAVEPNEVLADPAEEARAREISKEIRCLVCRNESIDD WP_042464940.1 cytochrome c biogenesis protein CcdA [Rhodovulum sulfidophilum] Length: 158\nNumber of predicted TMHs: 2\nExp number of AAs in TMHs: 37.50402\nExp number, first 60 AAs: 15.27651\nTotal prob of N-in: 0.78914\nPOSSIBLE N-term signal sequence\ninside 1 6\nTMhelix 7 26\noutside 27 107\nTMhelix 108 130\ninside 131 158

266 GCF_001633165.1_ASM163316v1 Rhodovulum sulfidophilum DSM 1374 Proteobacteria; Alphaproteobacteria; Rhodobacterales; Rhodobacteraceae; Rhodovulum; Rhodovulum sulfidophilum
MPRRALIALLLALSLTPVLSGPLRAVEPNEVLADPAEEARAREISKEIRCLVCRNESIDD WP_042464940.1 cytochrome c biogenesis protein CcdA [Rhodovulum sulfidophilum] Length: 158\nNumber of predicted TMHs: 2\nExp number of AAs in TMHs: 37.50402\nExp number, first 60 AAs: 15.27651\nTotal prob of N-in: 0.78914\nPOSSIBLE N-term signal sequence\ninside 1 6\nTMhelix 7 26\noutside 27 107\nTMhelix 108 130\ninside 131 158

267 GCF_900172355.1_R.antarcticus_CECT7023_Spades_Prokka Roseisalinus antarcticus Proteobacteria; Alphaproteobacteria; Rhodobacterales; Rhodobacteraceae; Roseisalinus
MDIPHTPSFRLDGRRALVTGASSGIGLGAAVALAEVGAHVVCAARGADRLNDAVAAIRAS WP_085880936.1 3-oxoacyl-ACP reductase [Roseisalinus antarcticus] Length: 253\nNumber of predicted TMHs: 2\nExp number of AAs in TMHs: 33.64248\nExp number, first 60 AAs: 14.27939\nTotal prob of N-in: 0.13913\nPOSSIBLE N-term signal sequence\noutside 1 19\nTMhelix 20 42\ninside 43 223\nTMhelix 224 246\noutside 247 253

268 GCF_000521865.1_SOAPdenovo_v1.05Roseivivax isoporae LMG 25204 Proteobacteria; Alphaproteobacteria; Rhodobacterales; Rhodobacteraceae; Roseivivax; Roseivivax isoporae
MSRRAAFRRAGAALVLTCALALPALAVQPDEMLGDPGLESRAREISEGLRCLVCQNESID WP_043765875.1 cytochrome c biogenesis protein CcdA [Roseivivax isoporae] Length: 159\nNumber of predicted TMHs: 2\nExp number of AAs in TMHs: 43.55586\nExp number, first 60 AAs: 21.18863\nTotal prob of N-in: 0.99154\nPOSSIBLE N-term signal sequence\ninside 1 4\nTMhelix 5 27\noutside 28 108\nTMhelix 109 131\ninside 132 159

269 GCF_001650975.1_ASM165097v1 Roseovarius indicusProteobacteria; Alphaproteobacteria; Rhodobacterales; Rhodobacteraceae; Roseovarius
MFGLLDEIIDRLADILELTDGRIEGVSRGLFGNGLETEVADRRSALATLGQAGALVSDVR WP_082917107.1 hypothetical protein [Roseovarius indicus] Length: 188\nNumber of predicted TMHs: 2\nExp number of AAs in TMHs: 44.53491\nExp number, first 60 AAs: 0.01893\nTotal prob of N-in: 0.88706\ninside 1 132\nTMhelix 133 155\noutside 156 159\nTMhelix 160 182\ninside 183 188

270 GCF_000152625.1_ASM15262v1 Roseovarius nubinhibens ISMProteobacteria; Alphaproteobacteria; Rhodobacterales; Rhodobacteraceae; Roseovarius; Roseovarius nubinhibens
MPHLTRRRLLGTLSATGLTTPALPLFAQSYDAPSCKAQLSHGDWLIKTQFVARYANDSFI WP_009814865.1 hypothetical protein [Roseovarius nubinhibens] Length: 384\nNumber of predicted TMHs: 1\nExp number of AAs in TMHs: 21.37458\nExp number, first 60 AAs: 0.21423\nTotal prob of N-in: 0.01574\noutside 1 342\nTMhelix 343 365\ninside 366 384

271 GCF_000714535.1_SOAPdenovo_v1.05Thioclava pacifica DSM 10166 Proteobacteria; Alphaproteobacteria; Rhodobacterales; Rhodobacteraceae; Thioclava; Thioclava pacifica
MPAIETFYDVMRRQGITRRSFMKYCSLTAAALGLGPAYVPKIAEAMETKPRTPVIWVHGL WP_084713737.1 uptake hydrogenase small subunit [Thioclava pacifica] Length: 358\nNumber of predicted TMHs: 2\nExp number of AAs in TMHs: 30.58511\nExp number, first 60 AAs: 16.57981\nTotal prob of N-in: 0.82125\nPOSSIBLE N-term signal sequence\ninside 1 19\nTMhelix 20 39\noutside 40 325\nTMhelix 326 348\ninside 349 358

272 GCF_002166865.1_ASM216686v1 Rhodobacteraceae bacterium WFHF2C18 Proteobacteria; Alphaproteobacteria; Rhodobacterales; Rhodobacteraceae; unclassified Rhodobacteraceae
MNTRHIPLIVNALGLTITRKKLHRIERMTVLIDWHRVRLSERERTRRDPVSGVAVPYFIA WP_087594503.1 hypothetical protein [Rhodobacteraceae bacterium WFHF2C18] Length: 161\nNumber of predicted TMHs: 1\nExp number of AAs in TMHs: 41.07853\nExp number, first 60 AAs: 5.13811\nTotal prob of N-in: 0.88374\ninside 1 119\nTMhelix 120 137\noutside 138 161

273 GCF_002153515.1_ASM215351v1 Acetobacter indonesiensis Proteobacteria; Alphaproteobacteria; Rhodospirillales; Acetobacteraceae; Acetobacter
MSNFDPKELKILSDILALVLEDQPGQSSSALEAIKNRARRDNVTGGALKNLFQAIAPNPS WP_086613831.1 hypothetical protein [Acetobacter indonesiensis] Length: 179\nNumber of predicted TMHs: 1\nExp number of AAs in TMHs:

20.06534\nExp number, first 60 AAs: 0.00234\nTotal prob of N-in: 0.68142\ninside 1 135\nTMhelix 136 158\noutside 159 179

274 GCF_002153545.1_ASM215354v1 Acetobacter indonesiensis Proteobacteria; Alphaproteobacteria; Rhodospirillales; Acetobacteraceae; Acetobacter MSNFDPKELKILSDILALVLEDQPGQSSSALEAIKNRARRDNVTGGALKNLFQAIAPNPS WP_048847868.1 hypothetical protein [Acetobacter indonesiensis] Length: 175\nNumber of predicted TMHs: 1\nExp number of AAs in TMHs: 21.18261\nExp number, first 60 AAs: 0.00185\nTotal prob of N-in: 0.75946\ninside 1 135\nTMhelix 136 158\noutside 159 175

275 GCF_002153485.1_ASM215348v1 Acetobacter sp. DmW_043 Proteobacteria; Alphaproteobacteria; Rhodospirillales; Acetobacteraceae; Acetobacter MPHLNDKQHLPACPTRRKILAAGIAGLAGTTLIKQHDALAANSENVSPPSVISTPARVWG WP_086554872.1 gluconolactonase [Acetobacter sp. DmW_043] Length: 409\nNumber of predicted TMHs: 1\nExp number of AAs in TMHs: 18.42401\nExp number, first 60 AAs: 0.05423\nTotal prob of N-in: 0.02970\noutside 1 340\nTMhelix 341 363\ninside 364 409

276 GCF_002153695.1_ASM215369v1 Acetobacter sp. DsW_059 Proteobacteria; Alphaproteobacteria; Rhodospirillales; Acetobacteraceae; Acetobacter MPHLNDKQHLPACTTRRKVLAAGIAGLAGTTLIKQHDALAANSENVSPPSVISTPARVWG WP_086635223.1 gluconolactonase [Acetobacter sp. DsW_059] Length: 409\nNumber of predicted TMHs: 1\nExp number of AAs in TMHs: 18.45668\nExp number, first 60 AAs: 0.08769\nTotal prob of N-in: 0.03157\noutside 1 340\nTMhelix 341 363\ninside 364 409

277 GCF_002153745.1_ASM215374v1 Acetobacter sp. DsW_063 Proteobacteria; Alphaproteobacteria; Rhodospirillales; Acetobacteraceae; Acetobacter MNAQTSPDDEARRALIRQKRMATGMLGGMAALTLAGYILPARGWLSESLWIDTLRAGGRA WP_086643025.1 DUF445 domain-containing protein [Acetobacter sp. DsW_063] Length: 430\nNumber of predicted TMHs: 2\nExp number of AAs in TMHs: 44.00197\nExp number, first 60 AAs: 21.30591\nTotal prob of N-in: 0.96985\nPOSSIBLE N-term signal sequence\ninside 1 20\nTMhelix 21 40\noutside 41 406\nTMhelix 407 429\ninside 430 430

278 GCF_000963945.1_ASM96394v1 Acetobacter indonesiensis 5H-1 Proteobacteria; Alphaproteobacteria; Rhodospirillales; Acetobacteraceae; Acetobacter; Acetobacter indonesiensis MSNFDPKELKILSDILALVLEDQPGQSSSALEAIKNRARRDNVTGGALKNLFQAIAPNPS WP_048847868.1 hypothetical protein [Acetobacter indonesiensis] Length: 175\nNumber of predicted TMHs: 1\nExp number of AAs in TMHs: 21.18261\nExp number, first 60 AAs: 0.00185\nTotal prob of N-in: 0.75946\ninside 1 135\nTMhelix 136 158\noutside 159 175

279 GCF_000429165.1_ASM42916v1 Acetobacter nitrogenifigens DSM 23921 = LMG 23498 Proteobacteria; Alphaproteobacteria; Rhodospirillales; Acetobacteraceae; Acetobacter; Acetobacter nitrogenifigens MNAQTSPDDDARRALIRQKRMATGMLGGMAVVTLAGYILPARGWLSESLWIDTLRAGGRA WP_026396542.1 DUF445 domain-containing protein [Acetobacter nitrogenifigens] Length: 430\nNumber of predicted TMHs: 2\nExp number of AAs in TMHs: 44.14883\nExp number, first 60 AAs: 21.38187\nTotal prob of N-in: 0.96766\nPOSSIBLE N-term signal sequence\ninside 1 20\nTMhelix 21 40\noutside 41 406\nTMhelix 407 429\ninside 430 430

280 GCF_900156265.1_IMG-taxon_2681812815_annotated_assembly Acidiphilium rubrum Proteobacteria; Alphaproteobacteria; Rhodospirillales; Acetobacteraceae; Acidiphilium MKRRSAIAASAAAFCAFPAASARAGSGSALDASVVRMKLQKGITPDVAVDCLKLRANLRN WP_029313089.1 MULTISPECIES: hypothetical protein [Acidiphilium] Length: 166\nNumber of predicted TMHs: 2\nExp number of AAs in TMHs: 29.18964\nExp number, first 60 AAs: 2.54444\nTotal prob of N-in: 0.80468\ninside 1 85\nTMhelix 86 108\noutside 109 122\nTMhelix 123 145\ninside 146 166

281 GCF_000701585.1_ASM70158v1 Acidiphilium angustum ATCC 35903 Proteobacteria; Alphaproteobacteria; Rhodospirillales; Acetobacteraceae; Acidiphilium; Acidiphilium angustum MKRRSAIAASAAAFCAFPAASARAGSGSALDASVVRMKLQKGITPDVAVDCLKLRANLRN WP_029313089.1 MULTISPECIES: hypothetical protein [Acidiphilium] Length: 166\nNumber of predicted TMHs: 2\nExp number of AAs in TMHs: 29.18964\nExp number, first 60 AAs: 2.54444\nTotal prob of N-in: 0.80468\ninside 1 85\nTMhelix 86 108\noutside 109 122\nTMhelix 123 145\ninside 146 166

282 GCF_000067045.1_ASM6704v1 Gluconacetobacter diazotrophicus PA1 5 Proteobacteria; Alphaproteobacteria; Rhodospirillales; Acetobacteraceae; Gluconacetobacter; Gluconacetobacter diazotrophicus MTRVTETRNDMERNGFDRLTERLARYLAGRSSRRGALARLGGWAASVPLFPLLPVWRGDA WP_012225357.1 methylamine dehydrogenase (amicyanin) light chain [Gluconacetobacter diazotrophicus] Length: 194\nNumber of predicted TMHs: 1\nExp number of AAs in TMHs: 16.56111\nExp number, first 60 AAs: 3.53842\nTotal prob of N-in: 0.83729\ninside 1 170\nTMhelix 171 193\noutside 194 194

283 GCF_000021325.1_ASM2132v1 Gluconacetobacter diazotrophicus PA1 5 Proteobacteria; Alphaproteobacteria; Rhodospirillales; Acetobacteraceae; Gluconacetobacter; Gluconacetobacter diazotrophicus MTRVTETRNDMERNGFDRLTERLARYLAGRSSRRGALARLGGWAASVPLFPLLPVWRGDA WP_012225357.1 methylamine dehydrogenase (amicyanin) light chain [Gluconacetobacter diazotrophicus] Length: 194\nNumber of predicted TMHs: 1\nExp number of AAs in TMHs: 16.56111\nExp number, first 60 AAs: 3.53842\nTotal prob of N-in: 0.83729\ninside 1 170\nTMhelix 171 193\noutside 194 194

284 GCF_001580715.1_ASM158071v1 Gluconobacter japonicus Proteobacteria; Alphaproteobacteria; Rhodospirillales; Acetobacteraceae; Gluconobacter MKPQSRVPTALCEVPRRRLLLGGLAFGSMSLLASHALTNTPKASDITDFLILSEFLTGRK WP_062015704.1 hypothetical protein [Gluconobacter japonicus] Length: 181\nNumber of predicted TMHs: 1\nExp number of AAs in TMHs: 36.68426\nExp number, first 60 AAs: 13.77237\nTotal prob of N-in: 0.88349\nPOSSIBLE N-term signal sequence\ninside 1 142\nTMhelix 143 165\noutside 166 181

285 GCF_001580785.1_ASM158078v1 Gluconobacter japonicus Proteobacteria; Alphaproteobacteria; Rhodospirillales; Acetobacteraceae; Gluconobacter MKPQSRVPTALCEVPRRRLLLGGLAFGSMSLLASHALTNTPKASDITDFLILSEFLTGRK WP_082785218.1 hypothetical protein [Gluconobacter japonicus] Length: 181\nNumber of predicted TMHs: 1\nExp number of AAs in TMHs: 37.84386\nExp number, first 60 AAs: 13.38674\nTotal prob of N-in: 0.88889\nPOSSIBLE N-term signal sequence\ninside 1 141\nTMhelix 142 164\noutside 165 181

286 GCF_001580675.1_ASM158067v1 Gluconobacter oxydans Proteobacteria; Alphaproteobacteria; Rhodospirillales; Acetobacteraceae; Gluconobacter MHQTMSWIRRRSVFLGVVVLPTVLAGGYYGLVASPQYQSEAEFVVRGQPSQSPGALAGLL WP_062494212.1 hypothetical protein [Gluconobacter oxydans] Length: 367\nNumber of predicted TMHs: 2\nExp number of AAs in TMHs: 44.18222\nExp number, first 60 AAs: 21.59432\nTotal prob of N-in: 0.99158\nPOSSIBLE N-term signal sequence\ninside 1 11\nTMhelix 12 34\noutside 35 336\nTMhelix 337 359\ninside 360 367

287 GCF_001580675.1_ASM158067v1 Gluconobacter oxydans Proteobacteria; Alphaproteobacteria; Rhodospirillales; Acetobacteraceae; Gluconobacter MPDAAEHGARMTGKRRQTILKMKTLIGVMLPALLATGVPARSATLHPLAPIAQAEMGNST WP_062495313.1 hypothetical protein [Gluconobacter oxydans] Length: 467\nNumber of predicted TMHs: 2\nExp number of AAs in TMHs: 56.91205\nExp number, first 60 AAs: 14.32906\nTotal prob of N-in: 0.70009\nPOSSIBLE N-term signal sequence\noutside 1 406\nTMhelix 407 426\ninside 427 437\nTMhelix 438 460\noutside 461 467

288 GCF_001580635.1_ASM158063v1 Gluconobacter oxydans Proteobacteria; Alphaproteobacteria; Rhodospirillales; Acetobacteraceae; Gluconobacter MPDAAEHGARMTGKRRQTILKMKTLIGVMLPALLVTGVPARSATLHPLAPIAQAEMGNSI WP_062446384.1 hypothetical protein [Gluconobacter oxydans] Length: 467\nNumber of predicted TMHs: 2\nExp number of AAs in TMHs: 59.1253\nExp number, first 60 AAs: 16.36295\nTotal prob of N-in: 0.79315\nPOSSIBLE N-term signal sequence\noutside 1 406\nTMhelix 407 426\ninside 427 437\nTMhelix 438 460\noutside 461 467

289 GCF_001580625.1_ASM158062v1 Gluconobacter oxydans Proteobacteria; Alphaproteobacteria; Rhodospirillales; Acetobacteraceae; Gluconobacter MPDAAEHGARMTGKRRQTILKMKTLIGVMLPALLATGVPARSATLHPLAPIAQAEMGTST WP_052327572.1 hypothetical protein [Gluconobacter oxydans] Length: 467\nNumber of predicted TMHs: 2\nExp number of AAs in TMHs: 57.1693\nExp number, first 60 AAs: 14.42903\nTotal prob of N-in: 0.70287\nPOSSIBLE N-term signal sequence\noutside 1 406\nTMhelix 407 426\ninside 427 437\nTMhelix 438 460\noutside 461 467

290 GCF_001580705.1_ASM158070v1 Gluconobacter oxydans Proteobacteria; Alphaproteobacteria; Rhodospirillales; Acetobacteraceae; Gluconobacter MLLLAKDKNSASSIGVTSGSRDMPDTAEHRARMTGKRRQTILKMKTLIGVMLPALLATGV WP_081105739.1 hypothetical protein [Gluconobacter oxydans] Length: 489\nNumber of predicted TMHs: 2\nExp number of

AAs in TMHs: 55.1244499999999\nExp number, first 60 AAs: 10.84295\nTotal prob of N-in: 0.59480\nPOSSIBLE N-term signal sequence\noutside 1 428\nTMhelix 429 448\ninside 449 459\nTMhelix 460 482\noutside 483 489

291 GCF_001581045.1_ASM158104v1 Gluconobacter oxydans Proteobacteria; Alphaproteobacteria; Rhodospirillales; Acetobacteraceae; Gluconobacter
MPDAAEHRTRVTGKRRQTILKMKTLIGVMLPALLATGVPARSATLHPLAPIAQAEMGNST WP_062455509.1
hypothetical protein [Gluconobacter oxydans] Length: 467\nNumber of predicted TMHs: 2\nExp number of AAs in TMHs: 59.05422\nExp number, first 60 AAs: 16.38618\nTotal prob of N-in: 0.81452\nPOSSIBLE N-term signal sequence\noutside 1 406\nTMhelix 407 426\ninside 427 437\nTMhelix 438 460\noutside 461 467

292 GCF_001580625.1_ASM158062v1 Gluconobacter oxydans Proteobacteria; Alphaproteobacteria; Rhodospirillales; Acetobacteraceae; Gluconobacter
MHQTMSWIRRRSVFLGVVVLPTVLAGGYYGLVASPQYQSEAEFVVRGQPSQSPGALAGLL WP_052327497.1
hypothetical protein [Gluconobacter oxydans] Length: 367\nNumber of predicted TMHs: 2\nExp number of AAs in TMHs: 44.18222\nExp number, first 60 AAs: 21.59432\nTotal prob of N-in: 0.99158\nPOSSIBLE N-term signal sequence\ninside 1 11\nTMhelix 12 34\noutside 35 336\nTMhelix 337 359\ninside 360 367

293 GCF_000583855.1_ASM58385v1 Gluconobacter oxydans DSM 3504 Proteobacteria; Alphaproteobacteria; Rhodospirillales; Acetobacteraceae; Gluconobacter; Gluconobacter oxydans
MHQTMSWIRRRSVFLGVVVLPTVLAGGYYGLVASPQYQSEAEFVVRGQPSQSPGALAGLL WP_052327497.1
hypothetical protein [Gluconobacter oxydans] Length: 367\nNumber of predicted TMHs: 2\nExp number of AAs in TMHs: 44.18222\nExp number, first 60 AAs: 21.59432\nTotal prob of N-in: 0.99158\nPOSSIBLE N-term signal sequence\ninside 1 11\nTMhelix 12 34\noutside 35 336\nTMhelix 337 359\ninside 360 367

294 GCF_000583855.1_ASM58385v1 Gluconobacter oxydans DSM 3504 Proteobacteria; Alphaproteobacteria; Rhodospirillales; Acetobacteraceae; Gluconobacter; Gluconobacter oxydans
MPDAAEHGARMTGKRRQTILKMKTLIGVMLPALLATGVPARSATLHPLAPIAQAEMGTST WP_052327572.1
hypothetical protein [Gluconobacter oxydans] Length: 467\nNumber of predicted TMHs: 2\nExp number of AAs in TMHs: 57.1693\nExp number, first 60 AAs: 14.42903\nTotal prob of N-in: 0.70287\nPOSSIBLE N-term signal sequence\noutside 1 406\nTMhelix 407 426\ninside 427 437\nTMhelix 438 460\noutside 461 467

295 GCF_002009295.1_ASM200929v1 Komagataeibacter nataicola Proteobacteria; Alphaproteobacteria; Rhodospirillales; Acetobacteraceae; Komagataeibacter
MNGIPQTDQDARRTLLRYKRSATGLLAAMAGLTVAGYALPQMGWLADRPWLEILRSGTKA WP_078524424.1
DUF445 domain-containing protein [Komagataeibacter nataicola] Length: 431\nNumber of predicted TMHs: 2\nExp number of AAs in TMHs: 51.46243\nExp number, first 60 AAs: 23.67005\nTotal prob of N-in: 0.98333\nPOSSIBLE N-term signal sequence\ninside 1 20\nTMhelix 21 43\noutside 44 407\nTMhelix 408 430\ninside 431 431

296 GCF_000817255.2_ASM81725v2 Komagataeibacter intermedius AF2 Proteobacteria; Alphaproteobacteria; Rhodospirillales; Acetobacteraceae; Komagataeibacter; Komagataeibacter intermedius
MNGIPQTDQDARRTLLRHKRSATGLLAVMAGLTVAGYALPQMGWLAERPWLEILRSGTKA WP_039734618.1
DUF445 domain-containing protein [Komagataeibacter intermedius] Length: 431\nNumber of predicted TMHs: 2\nExp number of AAs in TMHs: 49.32245\nExp number, first 60 AAs: 23.0803\nTotal prob of N-in: 0.98268\nPOSSIBLE N-term signal sequence\ninside 1 20\nTMhelix 21 43\noutside 44 407\nTMhelix 408 430\ninside 431 431

297 GCF_000964425.1_ASM96442v1 Komagataeibacter intermedius TF2 Proteobacteria; Alphaproteobacteria; Rhodospirillales; Acetobacteraceae; Komagataeibacter; Komagataeibacter intermedius
MNGIPQTDQDARRTLLRHKRSATGLLAVMAGLTVAGYALPQMGWLAERPWLEILRSGTKA WP_039734618.1
DUF445 domain-containing protein [Komagataeibacter intermedius] Length: 431\nNumber of predicted TMHs: 2\nExp number of AAs in TMHs: 49.32245\nExp number, first 60 AAs: 23.0803\nTotal prob of N-in: 0.98268\nPOSSIBLE N-term signal sequence\ninside 1 20\nTMhelix 21 43\noutside 44 407\nTMhelix 408 430\ninside 431 431

298 GCF_000550765.1_ASM55076v1 Komagataeibacter xylinus E25 Proteobacteria; Alphaproteobacteria; Rhodospirillales; Acetobacteraceae; Komagataeibacter; Komagataeibacter xylinus
MNAIPQTDQDARRTLLRYKRSATGLLAAMAGLTVAGYALPQMGWLAERPWLEILRSGTKA WP_025438929.1
DUF445 domain-containing protein [Komagataeibacter xylinus] Length: 431\nNumber of predicted TMHs: 2\nExp number of AAs in TMHs: 49.1127400000001\nExp number, first 60 AAs: 23.11667\nTotal prob of N-in: 0.98845\nPOSSIBLE N-term signal sequence\ninside 1 20\nTMhelix 21 43\noutside 44 407\nTMhelix 408 430\ninside 431 431

299 GCF_001571345.1_ASM157134v1 Komagataeibacter xylinus NBRC 15237 Proteobacteria; Alphaproteobacteria; Rhodospirillales; Acetobacteraceae; Komagataeibacter; Komagataeibacter xylinus
MHEPGHQQRVAQRRRIMLRRGLLLGGLLALLPLGSGARAQPHDYREPPPRHWSFQGPLGH WP_061272509.1
ubiquinol-cytochrome C reductase [Komagataeibacter xylinus] Length: 272\nNumber of predicted TMHs: 2\nExp number of AAs in TMHs: 38.13899\nExp number, first 60 AAs: 18.52286\nTotal prob of N-in: 0.96172\nPOSSIBLE N-term signal sequence\ninside 1 20\nTMhelix 21 40\noutside 41 244\nTMhelix 245 263\ninside 264 272

300 GCF_000711725.1_ASM71172v1 Roseomonas aerilata DSM 19363 Proteobacteria; Alphaproteobacteria; Rhodospirillales; Acetobacteraceae; Roseomonas; Roseomonas aerilata
MDLGMRGRRALVIGGSSGIGLATAGLLLKEGAEVTIAGRDSGRLDAARAQLQAETGTAPA WP_043830793.1
short-chain dehydrogenase [Roseomonas aerilata] Length: 260\nNumber of predicted TMHs: 1\nExp number of AAs in TMHs: 31.23198\nExp number, first 60 AAs: 6.26751\nTotal prob of N-in: 0.65014\ninside 1 229\nTMhelix 230 252\noutside 253 260

301 GCF_000963885.1_ASM96388v1 Tanticharoenia sakaeratensis NBRC 103193 Proteobacteria; Alphaproteobacteria; Rhodospirillales; Acetobacteraceae; Tanticharoenia; Tanticharoenia sakaeratensis
MVGVLAWVRRRATFLGVVVFPTLVAAGYYGLVASPQYMSEAEFVVRGQTSQTPGMLAGLL WP_053053819.1
capsule biosynthesis protein [Tanticharoenia sakaeratensis] Length: 367\nNumber of predicted TMHs: 2\nExp number of AAs in TMHs: 43.50007\nExp number, first 60 AAs: 21.66614\nTotal prob of N-in: 0.97596\nPOSSIBLE N-term signal sequence\ninside 1 11\nTMhelix 12 34\noutside 35 336\nTMhelix 337 359\ninside 360 367

302 GCF_002027385.1_ASM202738v1 Azospirillum brasilense Proteobacteria; Alphaproteobacteria; Rhodospirillales; Rhodospirillaceae; Azospirillum
MGDLDEYEGALAGLSRREFLAYCTGVAATLGLSPAFGVKIANAAVAAPRPTVIWLSGQAC WP_051140592.1
hydrogenase 2 small subunit [Azospirillum brasilense] Length: 375\nNumber of predicted TMHs: 1\nExp number of AAs in TMHs: 23.41947\nExp number, first 60 AAs: 3.5638\nTotal prob of N-in: 0.13436\noutside 1 335\nTMhelix 336 358\ninside 359 375

303 GCF_000632475.1_ASM63247v2 Azospirillum brasilense Proteobacteria; Alphaproteobacteria; Rhodospirillales; Rhodospirillaceae; Azospirillum
MGDLDEYEGALAGLSRREFLTYCTGVAATLGLSPAFGVKIANAAVAAPRPTVIWLSGQAC WP_051658414.1
hydrogenase 2 small subunit [Azospirillum brasilense] Length: 375\nNumber of predicted TMHs: 1\nExp number of AAs in TMHs: 21.83645\nExp number, first 60 AAs: 2.02467\nTotal prob of N-in: 0.07516\noutside 1 335\nTMhelix 336 358\ninside 359 375

304 GCF_002027445.1_ASM202744v1 Azospirillum brasilense Proteobacteria; Alphaproteobacteria; Rhodospirillales; Rhodospirillaceae; Azospirillum
MGDLDEYEGALAGLSRREFLAYCTGVAATLGLSPAFGVKIANAAVAAPRPTVIWLSGQAC WP_051140592.1
hydrogenase 2 small subunit [Azospirillum brasilense] Length: 375\nNumber of predicted TMHs: 1\nExp number of AAs in TMHs: 23.41947\nExp number, first 60 AAs: 3.5638\nTotal prob of N-in: 0.13436\noutside 1 335\nTMhelix 336 358\ninside 359 375

305 GCF_001315015.1_ASM131501v1 Azospirillum brasilense Proteobacteria; Alphaproteobacteria; Rhodospirillales; Rhodospirillaceae; Azospirillum
MGDLDEYEGALAGLSRREFLAYCTGVAATLGLSPAFGVKIANAAVAAPRPTVIWLSGQAC WP_051140592.1
hydrogenase 2 small subunit [Azospirillum brasilense] Length: 375\nNumber of predicted TMHs: 1\nExp number of AAs in TMHs: 23.41947\nExp number, first 60 AAs: 3.5638\nTotal prob of N-in: 0.13436\noutside 1 335\nTMhelix 336 358\ninside 359 375

306 GCF_000404045.1_ASM40404v1 Azospirillum brasilense FP2 Proteobacteria; Alphaproteobacteria; Rhodospirillales; Rhodospirillaceae; Azospirillum; Azospirillum brasilense
MGDLDEYEGALAGLSRREFLAYCTGVAATLGLSPAFGVKIANAAVAAPRPTVIWLSGQAC WP_051140592.1
hydrogenase 2 small subunit [Azospirillum brasilense] Length: 375\nNumber of predicted TMHs: 1\nExp number of AAs in TMHs: 23.41947\nExp number, first 60 AAs: 3.5638\nTotal prob of N-in: 0.13436\noutside 1 335\nTMhelix 336 358\ninside 359 375

307 GCF_000237365.1_ASM23736v1 Azospirillum brasilense Sp245 Proteobacteria; Alphaproteobacteria; Rhodospirillales; Rhodospirillaceae; Azospirillum; Azospirillum brasilense
MPDGHVDLDAEHIGDLDEYEGALAGLSRREFLTYCTGVAATLGLSPAFGVKIANAAVAAP WP_014198381.1
hydrogenase 2 small subunit [Azospirillum brasilense] Length: 387\nNumber of predicted TMHs: 1\nExp

number of AAs in TMHs: 21.91434\nExp number, first 60 AAs: 1.08647\nTotal prob of N-in: 0.02974\noutside 1 347\nTMhelix 348 370\ninside 371 387

308 GCF_000429625.1_ASM42962v1 Azospirillum halopraeferens DSM 3675 Proteobacteria; Alphaproteobacteria; Rhodospirillales; Rhodospirillaceae; Azospirillum; Azospirillum halopraeferens MRAPMIGGGMEGPMREAVRTPTYGGAGDRRRSIIALAGTLIAVFVVNSLIGLFAIRYAGD WP_029007384.1 hypothetical protein [Azospirillum halopraeferens] Length: 244\nNumber of predicted TMHs: 2\nExp number of AAs in TMHs: 44.9462\nExp number, first 60 AAs: 22.85903\nTotal prob of N-in: 0.16763\nPOSSIBLE N-term signal sequence\noutside 1 32\nTMhelix 33 55\ninside 56 218\nTMhelix 219 241\noutside 242 244

309 GCF_000968135.1_ASM96813v1 Magnetospira sp. QH-2 Proteobacteria; Alphaproteobacteria; Rhodospirillales; Rhodospirillaceae; Magnetospira MGNNKTLTEVLQSRGVSRRGFLKYCAGLASIMALPPAMAPRIAAALESQRRPSVIWLSFQ WP_046020110.1 Ni/Fe hydrogenase [Magnetospira sp. QH-2] Length: 363\nNumber of predicted TMHs: 1\nExp number of AAs in TMHs: 37.18445\nExp number, first 60 AAs: 16.30761\nTotal prob of N-in: 0.80423\nPOSSIBLE N-term signal sequence\noutside 1 318\nTMhelix 319 341\ninside 342 363

310 GCF_000968135.1_ASM96813v1 Magnetospira sp. QH-2 Proteobacteria; Alphaproteobacteria; Rhodospirillales; Rhodospirillaceae; Magnetospira MALLAPAFRHAERVNCSLSPRQTEPRGLAMPRHPDFHAPHAISHRRRSPLPGILLAMGTV WP_082085472.1 marine proteobacterial sortase target protein [Magnetospira sp. QH-2] Length: 761\nNumber of predicted TMHs: 2\nExp number of AAs in TMHs: 40.94963\nExp number, first 60 AAs: 10.00881\nTotal prob of N-in: 0.96543\nPOSSIBLE N-term signal sequence\ninside 1 49\nTMhelix 50 72\noutside 73 730\nTMhelix 731 753\ninside 754 761

311 GCF_000342045.1_MagSO-1_1.0 Magnetospirillum caucaseumProteobacteria; Alphaproteobacteria; Rhodospirillales; Rhodospirillaceae; Magnetospirillum MTITRRNFLRRAAAGGTAAVAACATLAPAGAEARENLKVPDNAVGLLFDGTLCIGCKACM WP_008620031.1 hydrogenase 2 protein HybA [Magnetospirillum caucaseum] Length: 339\nNumber of predicted TMHs: 1\nExp number of AAs in TMHs: 23.65835\nExp number, first 60 AAs: 4.39345\nTotal prob of N-in: 0.79845\ninside 1 295\nTMhelix 296 318\noutside 319 339

312 GCF_002105535.1_ASM210553v1 Magnetospirillum sp. ME-1 Proteobacteria; Alphaproteobacteria; Rhodospirillales; Rhodospirillaceae; Magnetospirillum MTITRRKFLGGTAGGAAAVAACSALAPSPAEARESHKVPENAVGLLFDGTLCIGCKACMS WP_085375359.1 hydrogenase 2 protein HybA [Magnetospirillum sp. ME-1] Length: 338\nNumber of predicted TMHs: 1\nExp number of AAs in TMHs: 18.84483\nExp number, first 60 AAs: 0.46107\nTotal prob of N-in: 0.70486\ninside 1 294\nTMhelix 295 317\noutside 318 338

313 GCF_001511835.1_XM-1 Magnetospirillum sp. XM-1 Proteobacteria; Alphaproteobacteria; Rhodospirillales; Rhodospirillaceae; Magnetospirillum MTITRRNFLRRAAAGGTAAAAACATLAPAGAEARESHKVPENAVGLLFDGTLCIGCKACM WP_068435929.1 hydrogenase 2 protein HybA [Magnetospirillum sp. XM-1] Length: 339\nNumber of predicted TMHs: 1\nExp number of AAs in TMHs: 19.90629\nExp number, first 60 AAs: 1.68441\nTotal prob of N-in: 0.78856\ninside 1 296\nTMhelix 297 316\noutside 317 339

314 GCF_000225995.1_ASM22599v1 Nitrospirillum amazonense Y2 Proteobacteria; Alphaproteobacteria; Rhodospirillales; Rhodospirillaceae; Nitrospirillum; Nitrospirillum amazonense MGRTRRSLVVTAVALLALAGCDNQESTGVPVPPPPPPVIAARAPVADKAGTEEIVVTGAQ WP_050898663.1 DUF4349 domain-containing protein [Nitrospirillum amazonense] Length: 290\nNumber of predicted TMHs: 1\nExp number of AAs in TMHs: 23.09537\nExp number, first 60 AAs: 0.09104\nTotal prob of N-in: 0.22995\noutside 1 257\nTMhelix 258 280\ninside 281 290

315 GCF_000429645.1_ASM42964v1 Niveispirillum irakense DSM 11586 Proteobacteria; Alphaproteobacteria; Rhodospirillales; Rhodospirillaceae; Niveispirillum; Niveispirillum irakense MKRFFCSLKRRSAIGLTTLALASLLSAPAFAVMPDEKLSDPVMEARARDISKELRCLVCQ WP_051329943.1 cytochrome c-type biogenesis protein CcmH [Niveispirillum irakense] Length: 177\nNumber of predicted TMHs: 2\nExp number of AAs in TMHs: 44.32701\nExp number, first 60 AAs: 21.63879\nTotal prob of N-in: 0.99854\nPOSSIBLE N-term signal sequence\ninside 1 11\nTMhelix 12 34\noutside 35 113\nTMhelix 114 136\ninside 137 177

316 GCF_001939945.1_ASM193994v1 Rhodocista sp. MIMtkB3 Proteobacteria; Alphaproteobacteria; Rhodospirillales; Rhodospirillaceae; Rhodocista
MTAPPPPVPPLSRRALLGAGAGLALAGLAPGGTWAEAGPRRGGRIRVAGIASSTADTLDP WP_075772809.1 ABC transporter substrate-binding protein [Rhodocista sp. MIMtkB3] Length: 518\nNumber of predicted TMHs: 2\nExp number of AAs in TMHs: 39.11268\nExp number, first 60 AAs: 5.25068\nTotal prob of N-in: 0.09040\noutside 1 469\nTMhelix 470 487\ninside 488 493\nTMhelix 494 516\noutside 517 518

317 GCF_000013085.1_ASM1308v1 Rhodospirillum rubrum ATCC 11170 Proteobacteria; Alphaproteobacteria; Rhodospirillales; Rhodospirillaceae; Rhodospirillum; Rhodospirillum rubrum
MGETETFYEVIRRQGISRRGFLKFCGVTAAGLGLGAGGAARIAQALETKPRVPVIWLHGLYP_426249.1 Ni-Fe hydrogenase, small subunit [Rhodospirillum rubrum ATCC 11170] Length: 361\nNumber of predicted TMHs: 2\nExp number of AAs in TMHs: 37.0095\nExp number, first 60 AAs: 21.46728\nTotal prob of N-in: 0.96564\nPOSSIBLE N-term signal sequence\ninside 1 20\nTMhelix 21 43\noutside 44 325\nTMhelix 326 348\ninside 349 361

318 GCF_000225955.1_ASM22595v1 Rhodospirillum rubrum F11 Proteobacteria; Alphaproteobacteria; Rhodospirillales; Rhodospirillaceae; Rhodospirillum; Rhodospirillum rubrum
MGETETFYEVIRRQGISRRGFLKFCGVTAAGLGLGAGGAARIAQALETKPRVPVIWLHGLWP_011388916.1 Ni-Fe hydrogenase small subunit [Rhodospirillum rubrum] Length: 361\nNumber of predicted TMHs: 2\nExp number of AAs in TMHs: 37.0095\nExp number, first 60 AAs: 21.46728\nTotal prob of N-in: 0.96564\nPOSSIBLE N-term signal sequence\ninside 1 20\nTMhelix 21 43\noutside 44 325\nTMhelix 326 348\ninside 349 361

319 GCF_000264455.2_ASM26445v2 Tistrella mobilis KA081020-065 Proteobacteria; Alphaproteobacteria; Rhodospirillales; Rhodospirillaceae; Tistrella; Tistrella mobilis
MAIHPSFTAPARARARRSHIRLLALGALALQLGAAPAMAQSQGPVDLKLWGQTPADAGGA WP_014744601.1 cellulose synthase subunit B [Tistrella mobilis] Length: 929\nNumber of predicted TMHs: 1\nExp number of AAs in TMHs: 38.50699\nExp number, first 60 AAs: 14.2731\nTotal prob of N-in: 0.70861\nPOSSIBLE N-term signal sequence\noutside 1 898\nTMhelix 899 921\ninside 922 929

320 GCF_900167455.1_IMG-taxon_2582581318_annotated_assembly Enhydrobacter aerosaccus Proteobacteria; Alphaproteobacteria; Rhodospirillales; unclassified Rhodospirillales; Enhydrobacter
MIDSIALARTCHSPTSCAICGGRRTFVGLVNCMLASLDRLVRSSYLNERVPFRVNVVARR WP_085937527.1 hypothetical protein [Enhydrobacter aerosaccus] Length: 418\nNumber of predicted TMHs: 2\nExp number of AAs in TMHs: 45.0428599999999\nExp number, first 60 AAs: 0.23521\nTotal prob of N-in: 0.92965\ninside 1 60\nTMhelix 61 83\noutside 84 388\nTMhelix 389 411\ninside 412 418

321 GCF_001634625.1_ASM163462v1 Croceicoccus sp. H4 Proteobacteria; Alphaproteobacteria; Sphingomonadales; Erythrobacteraceae; Croceicoccus
MGGRRKAIGGLIALALAAGMAAPALPARAEDASREREVAAIIAGSDRTAAPVARNVDKLS WP_066555293.1 hypothetical protein [Croceicoccus sp. H4] Length: 405\nNumber of predicted TMHs: 2\nExp number of AAs in TMHs: 38.32568\nExp number, first 60 AAs: 18.92806\nTotal prob of N-in: 0.93756\nPOSSIBLE N-term signal sequence\ninside 1 6\nTMhelix 7 26\noutside 27 365\nTMhelix 366 384\ninside 385 405

322 GCF_001077815.2_ASM107781v2 Erythrobacter atlanticus Proteobacteria; Alphaproteobacteria; Sphingomonadales; Erythrobacteraceae; Erythrobacter
MDFSQFWDAGGAAIVLGGTLLATVLASGRREMVAALGSIGQLGQKRFDYEKTRAEIAYDV WP_048884997.1 hypothetical protein [Erythrobacter atlanticus] Length: 217\nNumber of predicted TMHs: 2\nExp number of AAs in TMHs: 51.38337\nExp number, first 60 AAs: 20.92785\nTotal prob of N-in: 0.02361\nPOSSIBLE N-term signal sequence\noutside 1 4\nTMhelix 5 27\ninside 28 150\nTMhelix 151 173\noutside 174 217

323 GCF_001021555.1_ASM102155v1 Erythrobacter marinus Proteobacteria; Alphaproteobacteria; Sphingomonadales; Erythrobacteraceae; Erythrobacter
MQQQLFSLAFSRRAALRSGAMLGAGAAFSALPFGPAAFAQDLPAAHRAASRWPAVSRFIR WP_047093296.1 hypothetical protein [Erythrobacter marinus] Length: 449\nNumber of predicted TMHs: 1\nExp number of AAs in TMHs: 28.11231\nExp number, first 60 AAs: 6.65914\nTotal prob of N-in: 0.31362\noutside 1 387\nTMhelix 388 410\ninside 411 449

324 GCF_001013305.1_ASM101330v1 Erythrobacter marinus Proteobacteria; Alphaproteobacteria; Sphingomonadales; Erythrobacteraceae; Erythrobacter
MQQQLFSLAFSRRAALRSGAMLGAGAAFSALPFGPAAFAQDLPAAHRAASRWPAVSRFIR WP_047093296.1 hypothetical protein [Erythrobacter marinus] Length: 449\nNumber of predicted TMHs: 1\nExp number of

AAs in TMHs: 28.11231\nExp number, first 60 AAs: 6.65914\nTotal prob of N-in: 0.31362\noutside 1 387\nTMhelix 388 410\ninside 411 449

325 GCF_002198665.1_ASM219866v1 Novosphingobium sp. B 225 Proteobacteria; Alphaproteobacteria; Sphingomonadales; Sphingomonadaceae; Novosphingobium MAETRSLTAADGRRQMLLLGGGALLLALVITAVWYLFIHTPYVAAFTQLKTDDAALIVDE WP_088307329.1 flagellar M-ring protein FliF [Novosphingobium sp. B 225] Length: 420\nNumber of predicted TMHs: 2\nExp number of AAs in TMHs: 45.23758\nExp number, first 60 AAs: 22.56689\nTotal prob of N-in: 0.99395\nPOSSIBLE N-term signal sequence\ninside 1 15\nTMhelix 16 38\noutside 39 365\nTMhelix 366 388\ninside 389 420

326 GCF_000445125.1_ASM44512v1 Novosphingobium lindaniclasticum LE124 Proteobacteria; Alphaproteobacteria; Sphingomonadales; Sphingomonadaceae; Novosphingobium; Novosphingobium lindaniclasticum MPHTPPSRRTLIPAMVLALAAPAAIVVHAASAVADETASNDAEAEDGPAAGTLLLRGQGV WP_031342145.1 marine proteobacterial sortase target protein [Novosphingobium lindaniclasticum] Length: 722\nNumber of predicted TMHs: 1\nExp number of AAs in TMHs: 33.04069\nExp number, first 60 AAs: 10.14087\nTotal prob of N-in: 0.45739\nPOSSIBLE N-term signal sequence\noutside 1 694\nTMhelix 695 717\ninside 718 722

327 GCF_000375445.1_ASM37544v1 Novosphingobium nitrogenifigens DSM 19370 Proteobacteria; Alphaproteobacteria; Sphingomonadales; Sphingomonadaceae; Novosphingobium; Novosphingobium nitrogenifigens MNRIDQTHAFALRVMRETRMRRRAYLGTGLVLALLTAFPQPYVARARIVPQDASSIGLNS WP_008068685.1 capsule polysaccharide export protein [Novosphingobium nitrogenifigens] Length: 374\nNumber of predicted TMHs: 2\nExp number of AAs in TMHs: 46.11014\nExp number, first 60 AAs: 20.96586\nTotal prob of N-in: 0.98024\nPOSSIBLE N-term signal sequence\ninside 1 23\nTMhelix 24 43\noutside 44 337\nTMhelix 338 360\ninside 361 374

328 GCF_000375445.1_ASM37544v1 Novosphingobium nitrogenifigens DSM 19370 Proteobacteria; Alphaproteobacteria; Sphingomonadales; Sphingomonadaceae; Novosphingobium; Novosphingobium nitrogenifigens MSFVQIIRILLARRKLLGWTVLGCFLCGLAAIALIPTKYEATSRVLLDLVKPDPVTQQVL WP_008068854.1 chain length determinant protein [Novosphingobium nitrogenifigens] Length: 423\nNumber of predicted TMHs: 2\nExp number of AAs in TMHs: 42.572\nExp number, first 60 AAs: 20.24334\nTotal prob of N-in: 0.97938\nPOSSIBLE N-term signal sequence\ninside 1 15\nTMhelix 16 35\noutside 36 372\nTMhelix 373 395\ninside 396 423

329 GCF_000192575.1_NovNit_2.0 Novosphingobium nitrogenifigens DSM 19370 Proteobacteria; Alphaproteobacteria; Sphingomonadales; Sphingomonadaceae; Novosphingobium; Novosphingobium nitrogenifigens MSFVQIIRILLARRKLLGWTVLGCFLCGLAAIALIPTKYEATSRVLLDLVKPDPVTQQVL WP_008068854.1 chain length determinant protein [Novosphingobium nitrogenifigens] Length: 423\nNumber of predicted TMHs: 2\nExp number of AAs in TMHs: 42.572\nExp number, first 60 AAs: 20.24334\nTotal prob of N-in: 0.97938\nPOSSIBLE N-term signal sequence\ninside 1 15\nTMhelix 16 35\noutside 36 372\nTMhelix 373 395\ninside 396 423

330 GCF_000192575.1_NovNit_2.0 Novosphingobium nitrogenifigens DSM 19370 Proteobacteria; Alphaproteobacteria; Sphingomonadales; Sphingomonadaceae; Novosphingobium; Novosphingobium nitrogenifigens MNRIDQTHAFALRVMRETRMRRRAYLGTGLVLALLTAFPQPYVARARIVPQDASSIGLNS WP_008068685.1 capsule polysaccharide export protein [Novosphingobium nitrogenifigens] Length: 374\nNumber of predicted TMHs: 2\nExp number of AAs in TMHs: 46.11014\nExp number, first 60 AAs: 20.96586\nTotal prob of N-in: 0.98024\nPOSSIBLE N-term signal sequence\ninside 1 23\nTMhelix 24 43\noutside 44 337\nTMhelix 338 360\ninside 361 374

331 GCF_001598555.1_ASM159855v1 Novosphingobium rosa NBRC 15208 Proteobacteria; Alphaproteobacteria; Sphingomonadales; Sphingomonadaceae; Novosphingobium; Novosphingobium rosa MTEQYEANRRTILGAVAGVAGLAYAGPVLAQGTSAAEQTPRQAWATKDGAVRIGPNAAPV WP_068086363.1 hypothetical protein [Novosphingobium rosa] Length: 203\nNumber of predicted TMHs: 1\nExp number of AAs in TMHs: 47.79399\nExp number, first 60 AAs: 20.41172\nTotal prob of N-in: 0.92187\nPOSSIBLE N-term signal sequence\ninside 1 151\nTMhelix 152 174\noutside 175 203

332 GCF_000466945.1_ASM46694v1 Novosphingobium tardaugens NBRC 16725 Proteobacteria; Alphaproteobacteria; Sphingomonadales; Sphingomonadaceae; Novosphingobium; Novosphingobium tardaugens MTSARHPRSPRRRAVIAGVAILVAAGLVALGLWQLERREEKHALIAAVAERAYNPPVAAP WP_021688355.1 hypothetical protein [Novosphingobium tardaugens] Length: 248\nNumber of predicted TMHs: 2\nExp number of AAs in TMHs: 63.9590499999999\nExp number, first 60 AAs: 21.69389\nTotal prob of N-in: 0.66547\nPOSSIBLE N-term signal sequence\ninside 1 12\nTMhelix 13 35\noutside 36 213\nTMhelix 214 236\ninside 237 248

333 GCF_000722875.1_Schl_MUMv1 Sphingobium chlorophenolicum Proteobacteria; Alphaproteobacteria; Sphingomonadales; Sphingomonadaceae; Sphingobium
MRRRALLALTPALLLLCAADEPMLVPDVSQRDVEIQYSFTGADLLLFGAIVYPDGRRPKK WP_037451034.1 hypothetical protein [Sphingobium chlorophenolicum] Length: 253\nNumber of predicted TMHs: 1\nExp number of AAs in TMHs: 24.95438\nExp number, first 60 AAs: 2.36555\nTotal prob of N-in: 0.25984\noutside 1 226\nTMhelix 227 249\ninside 250 253

334 GCF_001005725.1_ASM100572v1 Sphingobium chungbukense Proteobacteria; Alphaproteobacteria; Sphingomonadales; Sphingomonadaceae; Sphingobium
MKRRALLALPPAMLLLTAADEPMLVPDVSQREVEIQYSFTGADLLLFGAIVYPDGRRPKT WP_046763191.1 hypothetical protein [Sphingobium chungbukense] Length: 253\nNumber of predicted TMHs: 1\nExp number of AAs in TMHs: 22.92438\nExp number, first 60 AAs: 0.39235\nTotal prob of N-in: 0.13143\noutside 1 226\nTMhelix 227 249\ninside 250 253

335 GCF_001580035.1_ASM158003v1 Sphingobium sp. 22B Proteobacteria; Alphaproteobacteria; Sphingomonadales; Sphingomonadaceae; Sphingobium
MRRRALLALPPALLLLTAADEPMLVPDVSQREVEIQYSFTGADLLLFGAIVYPDERRPSK WP_061939000.1 MULTISPECIES: hypothetical protein [Sphingomonadaceae] Length: 253\nNumber of predicted TMHs: 1\nExp number of AAs in TMHs: 22.68767\nExp number, first 60 AAs: 0.18927\nTotal prob of N-in: 0.10493\noutside 1 226\nTMhelix 227 249\ninside 250 253

336 GCF_001550165.1_ASM155016v1 Sphingobium sp. AM Proteobacteria; Alphaproteobacteria; Sphingomonadales; Sphingomonadaceae; Sphingobium
MRRRALLALPPALLLLTAADEPMLVPDVSQREVEIQYSFTGADLLLFGAIVYPDERRPSK WP_061939000.1 MULTISPECIES: hypothetical protein [Sphingomonadaceae] Length: 253\nNumber of predicted TMHs: 1\nExp number of AAs in TMHs: 22.68767\nExp number, first 60 AAs: 0.18927\nTotal prob of N-in: 0.10493\noutside 1 226\nTMhelix 227 249\ninside 250 253

337 GCF_000722555.1_ASM72255v1 Sphingobium sp. DC-2 Proteobacteria; Alphaproteobacteria; Sphingomonadales; Sphingomonadaceae; Sphingobium
MSKSKRARNRRGFLIAAAVALLACALLSLMARPPICTCGHIELWHGALDSGNSQHIADWY WP_030541714.1 hypothetical protein [Sphingobium sp. DC-2] Length: 190\nNumber of predicted TMHs: 2\nExp number of AAs in TMHs: 70.44107\nExp number, first 60 AAs: 24.7934\nTotal prob of N-in: 0.99978\nPOSSIBLE N-term signal sequence\ninside 1 11\nTMhelix 12 31\noutside 32 142\nTMhelix 143 165\ninside 166 190

338 GCF_000445085.1_ASM44508v1 Sphingobium sp. HDIP04 Proteobacteria; Alphaproteobacteria; Sphingomonadales; Sphingomonadaceae; Sphingobium
MRRRALLALPPALLLLTAADEPMLVPDVSQRDVEIQYSFTGADLLLFGAIVYPDGRRPSK WP_013039593.1 MULTISPECIES: hypothetical protein [Sphingobium] Length: 253\nNumber of predicted TMHs: 1\nExp number of AAs in TMHs: 22.8541\nExp number, first 60 AAs: 0.38394\nTotal prob of N-in: 0.14523\noutside 1 226\nTMhelix 227 249\ninside 250 253

339 GCF_001699185.1_ASM169918v1 Sphingobium sp. Ndbn-10 Proteobacteria; Alphaproteobacteria; Sphingomonadales; Sphingomonadaceae; Sphingobium
MRRRALLALPPALLLLTAADEPMLVPDVSQREVEIQYSFTGADLLLFGAIVYPDGRRPSK WP_025550228.1 MULTISPECIES: hypothetical protein [Sphingobium] Length: 253\nNumber of predicted TMHs: 1\nExp number of AAs in TMHs: 22.73123\nExp number, first 60 AAs: 0.2328\nTotal prob of N-in: 0.10612\noutside 1 226\nTMhelix 227 249\ninside 250 253

340 GCF_001699185.1_ASM169918v1 Sphingobium sp. Ndbn-10 Proteobacteria; Alphaproteobacteria; Sphingomonadales; Sphingomonadaceae; Sphingobium
MKSNRRPVIVVTGASAGVGRAVVRRFAQHGARIALIARGKDGLEATGREVEEAGGEALKL WP_065846115.1 short-chain dehydrogenase [Sphingobium sp. Ndbn-10] Length: 334\nNumber of predicted TMHs: 1\nExp number of AAs in TMHs: 20.82983\nExp number, first 60 AAs: 1.31122\nTotal prob of N-in: 0.20498\noutside 1 310\nTMhelix 311 330\ninside 331 334

341 GCF_000943805.1_ASM94380v1 Sphingobium sp. YBL2 Proteobacteria; Alphaproteobacteria; Sphingomonadales; Sphingomonadaceae; Sphingobium
MRRRALLALPPALLLLTAADEPMLVPDVSQREVEIQYSFTGADLLLFGAIVYPDGRRPSK WP_025550228.1 MULTISPECIES: hypothetical protein [Sphingobium] Length: 253\nNumber of predicted TMHs: 1\nExp number of AAs in TMHs:

22.73123\nExp number, first 60 AAs: 0.2328\nTotal prob of N-in: 0.10612\noutside 1 226\nTMhelix 227 249\ninside 250 253

342 GCF_900013425.1_ASM90001342v1 Sphingobium sp. Z007 Proteobacteria; Alphaproteobacteria; Sphingomonadales; Sphingomonadaceae; Sphingobium
MRRRRRTNRRGFIVAALIALAAGVILFLMHRPPICTCGYVELWHGALDNGNSQHIADWYS WP_088184295.1 hypothetical protein [Sphingobium sp. Z007] Length: 189\nNumber of predicted TMHs: 2\nExp number of AAs in TMHs: 65.1597800000001\nExp number, first 60 AAs: 23.81076\nTotal prob of N-in: 0.99984\nPOSSIBLE N-term signal sequence\ninside 1 11\nTMhelix 12 34\noutside 35 141\nTMhelix 142 164\ninside 165 189

343 GCF_900013445.1_Sphingobium_sp_Z007_rep_A_with_predator Sphingobium sp. Z007 Proteobacteria; Alphaproteobacteria; Sphingomonadales; Sphingomonadaceae; Sphingobium
MRRRRRTNRRGFIVAALIALAAGVILFLMHRPPICTCGYVELWHGALDNGNSQHIADWYS WP_088184295.1 hypothetical protein [Sphingobium sp. Z007] Length: 189\nNumber of predicted TMHs: 2\nExp number of AAs in TMHs: 65.1597800000001\nExp number, first 60 AAs: 23.81076\nTotal prob of N-in: 0.99984\nPOSSIBLE N-term signal sequence\ninside 1 11\nTMhelix 12 34\noutside 35 141\nTMhelix 142 164\ninside 165 189

344 GCF_900013455.1_Sphingobium_sp_Z007_rep_C_with_predator Sphingobium sp. Z007 Proteobacteria; Alphaproteobacteria; Sphingomonadales; Sphingomonadaceae; Sphingobium
MRRRRRTNRRGFIVAALIALAAGVILFLMHRPPICTCGYVELWHGALDNGNSQHIADWYS WP_088184295.1 hypothetical protein [Sphingobium sp. Z007] Length: 189\nNumber of predicted TMHs: 2\nExp number of AAs in TMHs: 65.1597800000001\nExp number, first 60 AAs: 23.81076\nTotal prob of N-in: 0.99984\nPOSSIBLE N-term signal sequence\ninside 1 11\nTMhelix 12 34\noutside 35 141\nTMhelix 142 164\ninside 165 189

345 GCF_900013435.1_Sphingobium_sp_Z007_rep_B_with_predator Sphingobium sp. Z007 Proteobacteria; Alphaproteobacteria; Sphingomonadales; Sphingomonadaceae; Sphingobium
MRRRRRTNRRGFIVAALIALAAGVILFLMHRPPICTCGYVELWHGALDNGNSQHIADWYS WP_088184295.1 hypothetical protein [Sphingobium sp. Z007] Length: 189\nNumber of predicted TMHs: 2\nExp number of AAs in TMHs: 65.1597800000001\nExp number, first 60 AAs: 23.81076\nTotal prob of N-in: 0.99984\nPOSSIBLE N-term signal sequence\ninside 1 11\nTMhelix 12 34\noutside 35 141\nTMhelix 142 164\ninside 165 189

346 GCF_900012545.1_Sphingobium_sp_Z007_rep_A_without_predator Sphingobium sp. Z007 Proteobacteria; Alphaproteobacteria; Sphingomonadales; Sphingomonadaceae; Sphingobium
MRRRRRTNRRGFIVAALIALAAGVILFLMHRPPICTCGYVELWHGALDNGNSQHIADWYS WP_088184295.1 hypothetical protein [Sphingobium sp. Z007] Length: 189\nNumber of predicted TMHs: 2\nExp number of AAs in TMHs: 65.1597800000001\nExp number, first 60 AAs: 23.81076\nTotal prob of N-in: 0.99984\nPOSSIBLE N-term signal sequence\ninside 1 11\nTMhelix 12 34\noutside 35 141\nTMhelix 142 164\ninside 165 189

347 GCF_900010525.1_Sphingobium_sp_Z007_rep_B_without_predator Sphingobium sp. Z007 Proteobacteria; Alphaproteobacteria; Sphingomonadales; Sphingomonadaceae; Sphingobium
MRRRRRTNRRGFIVAALIALAAGVILFLMHRPPICTCGYVELWHGALDNGNSQHIADWYS WP_088184295.1 hypothetical protein [Sphingobium sp. Z007] Length: 189\nNumber of predicted TMHs: 2\nExp number of AAs in TMHs: 65.1597800000001\nExp number, first 60 AAs: 23.81076\nTotal prob of N-in: 0.99984\nPOSSIBLE N-term signal sequence\ninside 1 11\nTMhelix 12 34\noutside 35 141\nTMhelix 142 164\ninside 165 189

348 GCF_000421925.1_ABySS Sphingobium chinhatense IP26 Proteobacteria; Alphaproteobacteria; Sphingomonadales; Sphingomonadaceae; Sphingobium; Sphingobium chinhatense
MRRRALLALPPALLLLTAADEPMLVPDVSQRDVEIQYSFTGADLLLFGAIVYPDGRRPSK WP_013039593.1 MULTISPECIES: hypothetical protein [Sphingobium] Length: 253\nNumber of predicted TMHs: 1\nExp number of AAs in TMHs: 22.8541\nExp number, first 60 AAs: 0.38394\nTotal prob of N-in: 0.14523\noutside 1 226\nTMhelix 227 249\ninside 250 253

349 GCF_000147835.2_ASM14783v3 Sphingobium chlorophenolicum L-1 Proteobacteria; Alphaproteobacteria; Sphingomonadales; Sphingomonadaceae; Sphingobium; Sphingobium chlorophenolicum
MRRRALLALTPALLLLCAADEPVLVPDVSQRDVEIQYSFTGADLLLFGAIVYPDGRRPKK WP_013846095.1 hypothetical protein [Sphingobium chlorophenolicum] Length: 253\nNumber of predicted TMHs: 1\nExp number of AAs in TMHs: 24.57785\nExp number, first 60 AAs: 1.9899\nTotal prob of N-in: 0.25166\noutside 1 226\nTMhelix 227 249\ninside 250 253

350 GCF_001598515.1_ASM159851v1 Sphingobium chlorophenolicum NBRC 16172 Proteobacteria; Alphaproteobacteria; Sphingomonadales; Sphingomonadaceae; Sphingobium; Sphingobium chlorophenolicum

MRRRALLALTPALLLLCAADEPMLVPDVSQRDVEIQYSFTGADLLLFGAIVYPDGRRPKK WP_037451034.1 hypothetical protein [Sphingobium chlorophenolicum] Length: 253\nNumber of predicted TMHs: 1\nExp number of AAs in TMHs: 24.95438\nExp number, first 60 AAs: 2.36555\nTotal prob of N-in: 0.25984\noutside 1 226\nTMhelix 227 249\ninside 250 253

351 GCF_001046645.1_S.czeLL01 Sphingobium czechense LL01 Proteobacteria; Alphaproteobacteria; Sphingomonadales; Sphingomonadaceae; Sphingobium; Sphingobium czechense MRRTQRINRRGLIAAALIALAACVILFLMRRPPICTCGYVELWHGALDSGNSQHIADWYS WP_066606327.1 hypothetical protein [Sphingobium czechense] Length: 189\nNumber of predicted TMHs: 2\nExp number of AAs in TMHs: 68.92185\nExp number, first 60 AAs: 23.93377\nTotal prob of N-in: 0.99897\nPOSSIBLE N-term signal sequence\ninside 1 11\nTMhelix 12 29\noutside 30 141\nTMhelix 142 164\ninside 165 189

352 GCF_000264945.2_ASM26494v2 Sphingobium indicum B90A Proteobacteria; Alphaproteobacteria; Sphingomonadales; Sphingomonadaceae; Sphingobium; Sphingobium indicum MRRRALLALPPALLLLTAADEPILVPDVSQRDVEIQYSFTGADLLLFGAIVYPDGRRPSK WP_007684696.1 MULTISPECIES: hypothetical protein [Sphingomonadaceae] Length: 253\nNumber of predicted TMHs: 1\nExp number of AAs in TMHs: 22.86686\nExp number, first 60 AAs: 0.3967\nTotal prob of N-in: 0.14573\noutside 1 226\nTMhelix 227 249\ninside 250 253

353 GCF_000091125.1_ASM9112v1 Sphingobium japonicum UT26S Proteobacteria; Alphaproteobacteria; Sphingomonadales; Sphingomonadaceae; Sphingobium; Sphingobium japonicum MRRRALLALPPALLLLTAADEPMLVPDVSQRDVEIQYSFTGADLLLFGAIVYPDGRRPSK WP_013039593.1 MULTISPECIES: hypothetical protein [Sphingobium] Length: 253\nNumber of predicted TMHs: 1\nExp number of AAs in TMHs: 22.8541\nExp number, first 60 AAs: 0.38394\nTotal prob of N-in: 0.14523\noutside 1 226\nTMhelix 227 249\ninside 250 253

354 GCF_000633575.2_de_novo_sequence_assembly Sphingobium lucknowense F2 Proteobacteria; Alphaproteobacteria; Sphingomonadales; Sphingomonadaceae; Sphingobium; Sphingobium lucknowense MRRRALLALPPALLLLTAADEPMLVPDVSQRDVEIQYSFTGADLLLFGAIVYPDGRRPSK WP_013039593.1 MULTISPECIES: hypothetical protein [Sphingobium] Length: 253\nNumber of predicted TMHs: 1\nExp number of AAs in TMHs: 22.8541\nExp number, first 60 AAs: 0.38394\nTotal prob of N-in: 0.14523\noutside 1 226\nTMhelix 227 249\ninside 250 253

355 GCF_000445065.1_ASM44506v1 Sphingobium quisquiliarum P25 Proteobacteria; Alphaproteobacteria; Sphingomonadales; Sphingomonadaceae; Sphingobium; Sphingobium quisquiliarum MSKSKRARNRRGFLIAAAVALLACALLFLMARPPICTCGHIELWHGALDSGNSQHIADWY WP_021237981.1 MULTISPECIES: membrane protein [Sphingomonadaceae] Length: 190\nNumber of predicted TMHs: 2\nExp number of AAs in TMHs: 70.36498\nExp number, first 60 AAs: 24.71952\nTotal prob of N-in: 0.99979\nPOSSIBLE N-term signal sequence\ninside 1 11\nTMhelix 12 31\noutside 32 142\nTMhelix 143 165\ninside 166 190

356 GCF_001651765.1_ASM165176v1 Sphingomonas paucimobilis Proteobacteria; Alphaproteobacteria; Sphingomonadales; Sphingomonadaceae; Sphingomonas MRRRALLALPPALLLLTAADEPMLVPDVSQREVEIQYSFTGADLLLFGAIVYPDERRPSK WP_061939000.1 MULTISPECIES: hypothetical protein [Sphingomonadaceae] Length: 253\nNumber of predicted TMHs: 1\nExp number of AAs in TMHs: 22.68767\nExp number, first 60 AAs: 0.18927\nTotal prob of N-in: 0.10493\noutside 1 226\nTMhelix 227 249\ninside 250 253

357 GCF_001029575.1_ASM102957v1 Sphingomonas paucimobilis Proteobacteria; Alphaproteobacteria; Sphingomonadales; Sphingomonadaceae; Sphingomonas MALDRRQMLQLGTLIAGGWAVSSVLKRTAPIGRDVANPEALAAIFDDRRSPASGPPTASL WP_047867254.1 MULTISPECIES: DSBA oxidoreductase [Sphingomonas] Length: 214\nNumber of predicted TMHs: 1\nExp number of AAs in TMHs: 21.55624\nExp number, first 60 AAs: 0.17312\nTotal prob of N-in: 0.48488\ninside 1 174\nTMhelix 175 197\noutside 198 214

358 GCF_000632085.1_Spau505_v1 Sphingomonas paucimobilis Proteobacteria; Alphaproteobacteria; Sphingomonadales; Sphingomonadaceae; Sphingomonas MSKSKRARNRRGFLIAAAVALLACALLFLMARPPICTCGHIELWHGALDSGNSQHIADWY WP_021237981.1 MULTISPECIES: membrane protein [Sphingomonadaceae] Length: 190\nNumber of predicted TMHs: 2\nExp number of AAs in TMHs: 70.36498\nExp number, first 60 AAs: 24.71952\nTotal prob of N-in: 0.99979\nPOSSIBLE N-term signal sequence\ninside 1 11\nTMhelix 12 31\noutside 32 142\nTMhelix 143 165\ninside 166 190

359 GCF_000783375.1_ASM78337v1 Sphingomonas sp. Ant20 Proteobacteria; Alphaproteobacteria; Sphingomonadales; Sphingomonadaceae; Sphingomonas
MPAPLTRRQIMQLGALVVGGWAASMVLRRTVPIGRDIGRSSALAKILDGKGFPMSGPADA WP_037527302.1
DSBA oxidoreductase [Sphingomonas sp. Ant20] Length: 214\nNumber of predicted TMHs: 2\nExp number of AAs in TMHs: 26.60212\nExp number, first 60 AAs: 12.09221\nTotal prob of N-in: 0.18174\nPOSSIBLE N-term signal sequence\noutside 1 9\nTMhelix 10 27\ninside 28 176\nTMhelix 177 199\noutside 200 214

360 GCF_000797515.1_ASM79751v1 Sphingomonas sp. Ant H11 Proteobacteria; Alphaproteobacteria; Sphingomonadales; Sphingomonadaceae; Sphingomonas
MRLGDASAPVLNQRKVIIDYVDANGEASRRTIVPFLIAGSFAGGRLDVLYLEARCLTANA WP_033966416.1 WYL domain-containing protein [Sphingomonas sp. Ant H11] Length: 153\nNumber of predicted TMHs: 1\nExp number of AAs in TMHs: 24.95493\nExp number, first 60 AAs: 4.12096\nTotal prob of N-in: 0.55719\ninside 1 118\nTMhelix 119 138\noutside 139 153

361 GCF_000576455.2_ASM57645v2 Sphingomonas sp. BHC-A Proteobacteria; Alphaproteobacteria; Sphingomonadales; Sphingomonadaceae; Sphingomonas
MRRRALLALPPALLLLTAADEPILVPDVSQRDVEIQYSFTGADLLLFGAIVYPDGRRPSK WP_007684696.1 MULTISPECIES: hypothetical protein [Sphingomonadaceae] Length: 253\nNumber of predicted TMHs: 1\nExp number of AAs in TMHs: 22.86686\nExp number, first 60 AAs: 0.3967\nTotal prob of N-in: 0.14573\noutside 1 226\nTMhelix 227 249\ninside 250 253

362 GCF_001557105.1_ASM155710v1 Sphingomonas sp. CCH13-B11 Proteobacteria; Alphaproteobacteria; Sphingomonadales; Sphingomonadaceae; Sphingomonas
MQRQVIEHTSVVPAHAGETVIADRDGPDRRPHIVIVGAGFGGLSLVRSLRRTDARITVID WP_066482177.1 MULTISPECIES: pyridine nucleotide-disulfide oxidoreductase [Sphingomonas] Length: 462\nNumber of predicted TMHs: 1\nExp number of AAs in TMHs: 21.24968\nExp number, first 60 AAs: 1.16228\nTotal prob of N-in: 0.07512\noutside 1 394\nTMhelix 395 417\ninside 418 462

363 GCF_001557005.1_ASM155700v1 Sphingomonas sp. CCH15-F11 Proteobacteria; Alphaproteobacteria; Sphingomonadales; Sphingomonadaceae; Sphingomonas
MQRQVIEHTSVAPAPAGETVIADRKGPDRRPHIVIVGAGFGGLSLVRSLRRTDARITVID WP_082731613.1 MULTISPECIES: FAD-dependent oxidoreductase [Sphingomonas] Length: 462\nNumber of predicted TMHs: 1\nExp number of AAs in TMHs: 22.18402\nExp number, first 60 AAs: 2.09564\nTotal prob of N-in: 0.12407\noutside 1 394\nTMhelix 395 417\ninside 418 462

364 GCF_001556095.1_ASM155609v1 Sphingomonas sp. CCH16-B10 Proteobacteria; Alphaproteobacteria; Sphingomonadales; Sphingomonadaceae; Sphingomonas
MQRQVIEHTSVVPAHAGETVIADRDGPDRRPHIVIVGAGFGGLSLVRSLRRTDARITVID WP_066482177.1 MULTISPECIES: pyridine nucleotide-disulfide oxidoreductase [Sphingomonas] Length: 462\nNumber of predicted TMHs: 1\nExp number of AAs in TMHs: 21.24968\nExp number, first 60 AAs: 1.16228\nTotal prob of N-in: 0.07512\noutside 1 394\nTMhelix 395 417\ninside 418 462

365 GCF_001557405.1_ASM155740v1 Sphingomonas sp. CCH18-H6 Proteobacteria; Alphaproteobacteria; Sphingomonadales; Sphingomonadaceae; Sphingomonas
MQRQVIEHTSVAPAPAGETVIADRKGPDRRPHIVIVGAGFGGLSLVRSLRRTDARITVID WP_082731613.1 MULTISPECIES: FAD-dependent oxidoreductase [Sphingomonas] Length: 462\nNumber of predicted TMHs: 1\nExp number of AAs in TMHs: 22.18402\nExp number, first 60 AAs: 2.09564\nTotal prob of N-in: 0.12407\noutside 1 394\nTMhelix 395 417\ninside 418 462

366 GCF_001556975.1_ASM155697v1 Sphingomonas sp. CCH19-C6 Proteobacteria; Alphaproteobacteria; Sphingomonadales; Sphingomonadaceae; Sphingomonas
MQRQVIEHTSVVPAHAGETVIADRDGPDRRPHIVIVGAGFGGLSLVRSLRRTDARITVID WP_066482177.1 MULTISPECIES: pyridine nucleotide-disulfide oxidoreductase [Sphingomonas] Length: 462\nNumber of predicted TMHs: 1\nExp number of AAs in TMHs: 21.24968\nExp number, first 60 AAs: 1.16228\nTotal prob of N-in: 0.07512\noutside 1 394\nTMhelix 395 417\ninside 418 462

367 GCF_001556565.1_ASM155656v1 Sphingomonas sp. CCH20-B6 Proteobacteria; Alphaproteobacteria; Sphingomonadales; Sphingomonadaceae; Sphingomonas
MALDRRQMLQLGTLIAGGWAVSSVLKRTAPIGRDVANPEALAAIFDDRRSPASGPPTASL WP_066760183.1
disulfide bond formation protein DsbA [Sphingomonas sp. CCH20-B6] Length: 214\nNumber of

predicted TMHs: 1\nExp number of AAs in TMHs: 21.55581\nExp number, first 60 AAs: 0.17312\nTotal prob of N-in: 0.48486\ninside 1 174\nTMhelix 175 197\noutside 198 214

368 GCF_001556565.1_ASM155656v1 Sphingomonas sp. CCH20-B6 Proteobacteria; Alphaproteobacteria; Sphingomonadales; Sphingomonadaceae; Sphingomonas
MQRQVIEHTSVVPAHAGETVIADRDGPDRRPHIVIVGAGFGGLSLVRSLRRTDARITVID WP_066482177.1 MULTISPECIES: pyridine nucleotide-disulfide oxidoreductase [Sphingomonas] Length: 462\nNumber of predicted TMHs: 1\nExp number of AAs in TMHs: 21.24968\nExp number, first 60 AAs: 1.16228\nTotal prob of N-in: 0.07512\noutside 1 394\nTMhelix 395 417\ninside 418 462

369 GCF_001555965.1_ASM155596v1 Sphingomonas sp. CCH21-G11 Proteobacteria; Alphaproteobacteria; Sphingomonadales; Sphingomonadaceae; Sphingomonas
MQRQVIEHTSVVPAHAGETVIADRDGPDRRPHIVIVGAGFGGLSLVRSLRRTDARITVID WP_066482177.1 MULTISPECIES: pyridine nucleotide-disulfide oxidoreductase [Sphingomonas] Length: 462\nNumber of predicted TMHs: 1\nExp number of AAs in TMHs: 21.24968\nExp number, first 60 AAs: 1.16228\nTotal prob of N-in: 0.07512\noutside 1 394\nTMhelix 395 417\ninside 418 462

370 GCF_001555785.1_ASM155578v1 Sphingomonas sp. CCH5-A5 Proteobacteria; Alphaproteobacteria; Sphingomonadales; Sphingomonadaceae; Sphingomonas
MQRQVIEHTSVVPAHAGETVIADRDGPDRRPHIVIVGAGFGGLSLVRSLRRTDARITVID WP_066482177.1 MULTISPECIES: pyridine nucleotide-disulfide oxidoreductase [Sphingomonas] Length: 462\nNumber of predicted TMHs: 1\nExp number of AAs in TMHs: 21.24968\nExp number, first 60 AAs: 1.16228\nTotal prob of N-in: 0.07512\noutside 1 394\nTMhelix 395 417\ninside 418 462

371 GCF_002127205.1_ASM212720v1 Sphingomonas sp. IBVSS1 Proteobacteria; Alphaproteobacteria; Sphingomonadales; Sphingomonadaceae; Sphingomonas
MKRRAAILLAASALAPAAAQTPATPRPNIVLIIADDWGFSDPGRYGSEIATPNIDALADT WP_086117650.1 hypothetical protein [Sphingomonas sp. IBVSS1] Length: 575\nNumber of predicted TMHs: 1\nExp number of AAs in TMHs: 20.92638\nExp number, first 60 AAs: 1.6731\nTotal prob of N-in: 0.08954\noutside 1 551\nTMhelix 552 569\ninside 570 575

372 GCF_001421415.1_Leaf208 Sphingomonas sp. Leaf208 Proteobacteria; Alphaproteobacteria; Sphingomonadales; Sphingomonadaceae; Sphingomonas
MLRMGMVAIWLLAPKAALWITQEARFRSRARRTFLFRVLAASNGQMALSPNYYSVLGVPS WP_082452852.1 hypothetical protein [Sphingomonas sp. Leaf208]Length: 223\nNumber of predicted TMHs: 1\nExp number of AAs in TMHs: 32.2934500000001\nExp number, first 60 AAs: 10.21782\nTotal prob of N-in: 0.38773\nPOSSIBLE N-term signal sequence\noutside 1 159\nTMhelix 160 182\ninside 183 223

373 GCF_001421785.1_Leaf33 Sphingomonas sp. Leaf33 Proteobacteria; Alphaproteobacteria; Sphingomonadales; Sphingomonadaceae; Sphingomonas
MTTRRPVLALVAALVTAGLIALGVWQLERRAWKLDLIARVDARIHAASAAFPWSGGAPQD WP_056427832.1 Surfeit locus 1 family protein [Sphingomonas sp. Leaf33] Length: 230\nNumber of predicted TMHs: 2\nExp number of AAs in TMHs: 43.90401\nExp number, first 60 AAs: 20.38312\nTotal prob of N-in: 0.95549\nPOSSIBLE N-term signal sequence\ninside 1 6\nTMhelix 7 26\noutside 27 202\nTMhelix 203 225\ninside 226 230

374 GCF_001421785.1_Leaf33 Sphingomonas sp. Leaf33 Proteobacteria; Alphaproteobacteria; Sphingomonadales; Sphingomonadaceae; Sphingomonas
MIALDSSRRRQWLLIGGVFAAVVALLAGAYYLFLSGGFTVLADHVRPGEAAAIVAELDKR WP_056427251.1 hypothetical protein [Sphingomonas sp. Leaf33] Length: 417\nNumber of predicted TMHs: 2\nExp number of AAs in TMHs: 42.11615\nExp number, first 60 AAs: 22.68705\nTotal prob of N-in: 0.99988\nPOSSIBLE N-term signal sequence\ninside 1 11\nTMhelix 12 34\noutside 35 360\nTMhelix 361 380\ninside 381 417

375 GCF_001426195.1_Leaf339 Sphingomonas sp. Leaf339 Proteobacteria; Alphaproteobacteria; Sphingomonadales; Sphingomonadaceae; Sphingomonas
MNRREMLVGGVSVGALTTADRALARAPATMAASGLPMPAPVRDLPKALPVADPGRTLLER WP_056530765.1 beta-galactosidase [Sphingomonas sp. Leaf339] Length: 963\nNumber of predicted TMHs: 1\nExp number of AAs in TMHs: 14.49514\nExp number, first 60 AAs: 0.4048\nTotal prob of N-in: 0.02113\noutside 1 939\nTMhelix 940 962\ninside 963 963

376 GCF_000293195.2_ASM29319v2 Sphingomonas sp. LH128 Proteobacteria; Alphaproteobacteria; Sphingomonadales; Sphingomonadaceae; Sphingomonas

MPDIDMIVRRLQRRSLVVSACILGLVLTACAVTGASLLQPMLLVALGLPIWDRIWWHRKR WP_008833422.1 hypothetical protein [Sphingomonas sp. LH128] Length: 182\nNumber of predicted TMHs: 2\nExp number of AAs in TMHs: 67.88088\nExp number, first 60 AAs: 27.68801\nTotal prob of N-in: 0.86576\nPOSSIBLE N-term signal sequence\ninside 1 15\nTMhelix 16 38\noutside 39 132\nTMhelix 133 155\ninside 156 182

377 GCF_001047295.1_ASM104729v1 Sphingomonas sp. MEA3-1 Proteobacteria; Alphaproteobacteria; Sphingomonadales; Sphingomonadaceae; Sphingomonas
MRRRALLALPPALLLLTAAEEPMLVPDVSQREVEIQYSFTGADLLLFGAIVYPDGRRPSK WP_048577909.1 transmembrane protein [Sphingomonas sp. MEA3-1] Length: 253\nNumber of predicted TMHs: 1\nExp number of AAs in TMHs: 22.73256\nExp number, first 60 AAs: 0.23413\nTotal prob of N-in: 0.10612\noutside 1 226\nTMhelix 227 249\ninside 250 253

378 GCF_000242835.1_ASM24283v1 Sphingomonas sp. PAMC 26617 Proteobacteria; Alphaproteobacteria; Sphingomonadales; Sphingomonadaceae; Sphingomonas
MSTPRRGFVRGAVTVVWLAVIAALIGLGTWQVQRRAWKLDLIARVDARVHAAPVAAPATA WP_037478759.1 SURF1 family protein [Sphingomonas sp. PAMC 26617] Length: 233\nNumber of predicted TMHs: 2\nExp number of AAs in TMHs: 43.91707\nExp number, first 60 AAs: 21.65393\nTotal prob of N-in: 0.97992\nPOSSIBLE N-term signal sequence\ninside 1 6\nTMhelix 7 29\noutside 30 199\nTMhelix 200 222\ninside 223 233

379 GCF_001428865.1_Root241 Sphingomonas sp. Root241 Proteobacteria; Alphaproteobacteria; Sphingomonadales; Sphingomonadaceae; Sphingomonas
MTSGARRNWFLAFALLAGAGFTALGVWQIERRAWKLDLIARVDARIHAPAAPLPARADWV WP_082596605.1 Surfeit locus 1 family protein [Sphingomonas sp. Root241] Length: 234\nNumber of predicted TMHs: 2\nExp number of AAs in TMHs: 44.04567\nExp number, first 60 AAs: 21.85345\nTotal prob of N-in: 0.97949\nPOSSIBLE N-term signal sequence\ninside 1 6\nTMhelix 7 29\noutside 30 205\nTMhelix 206 228\ninside 229 234

380 GCF_000153545.1_ASM15354v1 Sphingomonas sp. SKA58 Proteobacteria; Alphaproteobacteria; Sphingomonadales; Sphingomonadaceae; Sphingomonas
MTASHRRRNRRGFLLAALIAAMACGLLFLMDRPPICACGRVEVWHGALDSGNSQHIADWY WP_037541996.1 hypothetical protein [Sphingomonas sp. SKA58] Length: 190\nNumber of predicted TMHs: 2\nExp number of AAs in TMHs: 64.33612\nExp number, first 60 AAs: 21.65891\nTotal prob of N-in: 0.99951\nPOSSIBLE N-term signal sequence\ninside 1 12\nTMhelix 13 30\noutside 31 142\nTMhelix 143 165\ninside 166 190

381 GCF_001650735.1_ASM165073v1 Sphingomonas sp. TDK1 Proteobacteria; Alphaproteobacteria; Sphingomonadales; Sphingomonadaceae; Sphingomonas
MALDRRQMLQLGTLIAGGWAVSSVLKRTAPIGRDVANPEALAAIFDDRRSPASGPPTASL WP_035386738.1 MULTISPECIES: DSBA oxidoreductase [Bacteria] Length: 214\nNumber of predicted TMHs: 1\nExp number of AAs in TMHs: 21.55582\nExp number, first 60 AAs: 0.17312\nTotal prob of N-in: 0.48486\ninside 1 174\nTMhelix 175 197\noutside 198 214

382 GCF_001592345.1_ASM159234v1 Sphingomonas adhaesiva NBRC 15099 Proteobacteria; Alphaproteobacteria; Sphingomonadales; Sphingomonadaceae; Sphingomonas; Sphingomonas adhaesiva
MALDRRQMLQLGTLIAGGWAVSSVLKRTAPIGRDVANPEALAAIFDDRRSPASGPPTASL WP_047867254.1 MULTISPECIES: DSBA oxidoreductase [Sphingomonas] Length: 214\nNumber of predicted TMHs: 1\nExp number of AAs in TMHs: 21.55624\nExp number, first 60 AAs: 0.17312\nTotal prob of N-in: 0.48488\ninside 1 174\nTMhelix 175 197\noutside 198 214

383 GCF_000974765.1_ASM97476v1 Sphingomonas changbaiensis NBRC 104936 Proteobacteria; Alphaproteobacteria; Sphingomonadales; Sphingomonadaceae; Sphingomonas; Sphingomonas changbaiensis
MPGEPIVSGPGKPRRRTLMMLVLLAGTMLFIALGLWQVERRSEKLALIAAVQARVHAAPV WP_084689619.1 Surfeit locus 1 family protein [Sphingomonas changbaiensis] Length: 243\nNumber of predicted TMHs: 2\nExp number of AAs in TMHs: 38.57206\nExp number, first 60 AAs: 19.05317\nTotal prob of N-in: 0.95617\nPOSSIBLE N-term signal sequence\ninside 1 20\nTMhelix 21 38\noutside 39 217\nTMhelix 218 237\ninside 238 243

384 GCF_001591025.1_ASM159102v1 Sphingomonas soli NBRC 100801 Proteobacteria; Alphaproteobacteria; Sphingomonadales; Sphingomonadaceae; Sphingomonas; Sphingomonas soli
MKRRAILLLAPLLMGQSQPGQKPVLVPDVSQRDIEIAYSFTGAELLLFGAILYPGGRVPG WP_066799781.1 hypothetical protein [Sphingomonas soli] Length: 257\nNumber of predicted TMHs: 1\nExp number of AAs in TMHs: 24.11897\nExp number, first 60 AAs: 1.56887\nTotal prob of N-in: 0.40003\noutside 1 231\nTMhelix 232 254\ninside 255 257

385 GCF_000803645.1_ASM80364v1 Sphingopyxis fribergensis Proteobacteria; Alphaproteobacteria; Sphingomonadales; Sphingomonadaceae; Sphingopyxis
MTRTRRGAFAAAALIAAMGLAALGVWQVERRAWKHALVAAVDARIHAAPAAAPGPDAWGR WP_039574822.1
Surfeit locus 1 family protein [Sphingopyxis fribergensis] Length: 233\nNumber of predicted TMHs: 2\nExp number of AAs in TMHs: 46.75144\nExp number, first 60 AAs: 21.17189\nTotal prob of N-in: 0.98539\nPOSSIBLE N-term signal sequence\ninside 1 6\nTMhelix 7 26\noutside 27 206\nTMhelix 207 229\ninside 230 233

386 GCF_001956775.1_ASM195677v1 Sphingopyxis granuli Proteobacteria; Alphaproteobacteria; Sphingomonadales; Sphingomonadaceae; Sphingopyxis
MTATRRAVLAAAALLVTACLLALGVWQLERRVWKHALIAAVETRLHADPVAAPGPAAWPA WP_076076135.1
Surfeit locus 1 family protein [Sphingopyxis granuli] Length: 235\nNumber of predicted TMHs: 2\nExp number of AAs in TMHs: 45.53349\nExp number, first 60 AAs: 21.18833\nTotal prob of N-in: 0.95701\nPOSSIBLE N-term signal sequence\ninside 1 6\nTMhelix 7 26\noutside 27 206\nTMhelix 207 229\ninside 230 235

387 GCF_001559015.1_ASM155901v1 Sphingopyxis granuli Proteobacteria; Alphaproteobacteria; Sphingomonadales; Sphingomonadaceae; Sphingopyxis
MTATRRAVLAAAALLVTACLLALGVWQLERRVWKHALIAAVETRLQTDPVAAPGPAAWPA WP_067184963.1
Surfeit locus 1 family protein [Sphingopyxis granuli] Length: 235\nNumber of predicted TMHs: 2\nExp number of AAs in TMHs: 44.71161\nExp number, first 60 AAs: 21.1699\nTotal prob of N-in: 0.96885\nPOSSIBLE N-term signal sequence\ninside 1 6\nTMhelix 7 26\noutside 27 206\nTMhelix 207 229\ninside 230 235

388 GCF_001468495.1_ASM146849v1 Sphingopyxis sp. A083 Proteobacteria; Alphaproteobacteria; Sphingomonadales; Sphingomonadaceae; Sphingopyxis
MTRPRRAAFAAAALVAAALLAALGVWQLERREWKHALVAAVDARIHAAPVPAPPPAAWRG WP_058812601.1
Surfeit locus 1 family protein [Sphingopyxis sp. A083] Length: 241\nNumber of predicted TMHs: 2\nExp number of AAs in TMHs: 56.30971\nExp number, first 60 AAs: 21.00501\nTotal prob of N-in: 0.94403\nPOSSIBLE N-term signal sequence\ninside 1 6\nTMhelix 7 26\noutside 27 206\nTMhelix 207 229\ninside 230 241

389 GCF_001047015.1_ASM104701v1 Sphingopyxis sp. C-1 Proteobacteria; Alphaproteobacteria; Sphingomonadales; Sphingomonadaceae; Sphingopyxis
MPGRLGGSRRAAIIAIGALTALAPAQAQPVKPVPKHPNIVILLADDWGFSDVGSFGSEIA WP_083435958.1 arylsulfatase [Sphingopyxis sp. C-1] Length: 613\nNumber of predicted TMHs: 1\nExp number of AAs in TMHs: 20.98782\nExp number, first 60 AAs: 0.63877\nTotal prob of N-in: 0.03706\noutside 1 572\nTMhelix 573 592\ninside 593 613

390 GCF_001047015.1_ASM104701v1 Sphingopyxis sp. C-1 Proteobacteria; Alphaproteobacteria; Sphingomonadales; Sphingomonadaceae; Sphingopyxis
MKRTRRAAFAAAALIAAIGLAALGVWQLERRVWKHELVAAVEARIHAAPVAAPGPGAWDA WP_062179790.1
Surfeit locus 1 family protein [Sphingopyxis sp. C-1] Length: 233\nNumber of predicted TMHs: 2\nExp number of AAs in TMHs: 42.95983\nExp number, first 60 AAs: 20.94164\nTotal prob of N-in: 0.99246\nPOSSIBLE N-term signal sequence\ninside 1 6\nTMhelix 7 26\noutside 27 206\nTMhelix 207 229\ninside 230 233

391 GCF_001467395.1_ASM146739v1 Sphingopyxis sp. H005 Proteobacteria; Alphaproteobacteria; Sphingomonadales; Sphingomonadaceae; Sphingopyxis
MTRTRRAAFAAAALIAAVALAALGVWQLERRVWKHELVAAVEARIHAAPVAAPWPSAWDA WP_058536255.1
MULTISPECIES: Surfeit locus 1 family protein [Sphingopyxis] Length: 233\nNumber of predicted TMHs: 2\nExp number of AAs in TMHs: 48.28018\nExp number, first 60 AAs: 20.95142\nTotal prob of N-in: 0.99129\nPOSSIBLE N-term signal sequence\ninside 1 6\nTMhelix 7 26\noutside 27 206\nTMhelix 207 229\ninside 230 233

392 GCF_001468235.1_ASM146823v1 Sphingopyxis sp. H012 Proteobacteria; Alphaproteobacteria; Sphingomonadales; Sphingomonadaceae; Sphingopyxis
MTRTRRAAFAAAALIAAVALAALGVWQLERRVWKHELVAAVEARIHAAPVAAPWPSAWDA WP_058536255.1
MULTISPECIES: Surfeit locus 1 family protein [Sphingopyxis] Length: 233\nNumber of predicted TMHs: 2\nExp number of AAs in TMHs: 48.28018\nExp number, first 60 AAs: 20.95142\nTotal prob of N-in: 0.99129\nPOSSIBLE N-term signal sequence\ninside 1 6\nTMhelix 7 26\noutside 27 206\nTMhelix 207 229\ninside 230 233

393 GCF_001467405.1_ASM146740v1 Sphingopyxis sp. H038 Proteobacteria; Alphaproteobacteria; Sphingomonadales; Sphingomonadaceae; Sphingopyxis
MTRTRRAAFAAAALIAAVALAALGVWQLERRVWKHELVAAVEARIHAAPVAAPWPSAWDA WP_058536255.1
MULTISPECIES: Surfeit locus 1 family protein [Sphingopyxis] Length: 233\nNumber of predicted TMHs: 2\nExp number of AAs in TMHs: 48.28018\nExp number, first 60 AAs: 20.95142\nTotal prob of N-in: 0.99129\nPOSSIBLE N-term signal sequence\ninside 1 6\nTMhelix 7 26\noutside 27 206\nTMhelix 207 229\ninside 230 233

394 GCF_001468225.1_ASM146822v1 Sphingopyxis sp. H053 Proteobacteria; Alphaproteobacteria; Sphingomonadales; Sphingomonadaceae; Sphingopyxis
MTRTRRAAFAAAALIAAVALAALGVWQLERRVWKHELVAAVEARIHAAPVAAPWPSAWDA WP_058536255.1
MULTISPECIES: Surfeit locus 1 family protein [Sphingopyxis] Length: 233\nNumber of predicted TMHs: 2\nExp number of AAs in TMHs: 48.28018\nExp number, first 60 AAs: 20.95142\nTotal prob of N-in: 0.99129\nPOSSIBLE N-term signal sequence\ninside 1 6\nTMhelix 7 26\noutside 27 206\nTMhelix 207 229\ninside 230 233

395 GCF_001468315.1_ASM146831v1 Sphingopyxis sp. H057 Proteobacteria; Alphaproteobacteria; Sphingomonadales; Sphingomonadaceae; Sphingopyxis
MTRPRRAAFAAAALVAAACLAALGIWQVERRAWKHELVAAVDARIHAPPVAAPGPGRWPT WP_058454724.1
MULTISPECIES: Surfeit locus 1 family protein [Sphingopyxis] Length: 233\nNumber of predicted TMHs: 2\nExp number of AAs in TMHs: 42.81905\nExp number, first 60 AAs: 21.12953\nTotal prob of N-in: 0.99193\nPOSSIBLE N-term signal sequence\ninside 1 6\nTMhelix 7 26\noutside 27 206\nTMhelix 207 229\ninside 230 233

396 GCF_001468345.1_ASM146834v1 Sphingopyxis sp. H067 Proteobacteria; Alphaproteobacteria; Sphingomonadales; Sphingomonadaceae; Sphingopyxis
MTRPRRAAFAAAALVAAACLAALGIWQVERRAWKHELVAAVDARIHAPPVAAPGPGRWPT WP_058454724.1
MULTISPECIES: Surfeit locus 1 family protein [Sphingopyxis] Length: 233\nNumber of predicted TMHs: 2\nExp number of AAs in TMHs: 42.81905\nExp number, first 60 AAs: 21.12953\nTotal prob of N-in: 0.99193\nPOSSIBLE N-term signal sequence\ninside 1 6\nTMhelix 7 26\noutside 27 206\nTMhelix 207 229\ninside 230 233

397 GCF_001468365.1_ASM146836v1 Sphingopyxis sp. H071 Proteobacteria; Alphaproteobacteria; Sphingomonadales; Sphingomonadaceae; Sphingopyxis
MTRPRRAAFAAAALVAAACLAALGIWQVERRAWKHELVAAVDARIHAPPVAAPGPGRWPT WP_058454724.1
MULTISPECIES: Surfeit locus 1 family protein [Sphingopyxis] Length: 233\nNumber of predicted TMHs: 2\nExp number of AAs in TMHs: 42.81905\nExp number, first 60 AAs: 21.12953\nTotal prob of N-in: 0.99193\nPOSSIBLE N-term signal sequence\ninside 1 6\nTMhelix 7 26\noutside 27 206\nTMhelix 207 229\ninside 230 233

398 GCF_001468395.1_ASM146839v1 Sphingopyxis sp. H073 Proteobacteria; Alphaproteobacteria; Sphingomonadales; Sphingomonadaceae; Sphingopyxis
MTRPRRAAFAAAALVAAACLAALGIWQVERRAWKHELVAAVDARIHAPPVAAPGPGRWPT WP_058454724.1
MULTISPECIES: Surfeit locus 1 family protein [Sphingopyxis] Length: 233\nNumber of predicted TMHs: 2\nExp number of AAs in TMHs: 42.81905\nExp number, first 60 AAs: 21.12953\nTotal prob of N-in: 0.99193\nPOSSIBLE N-term signal sequence\ninside 1 6\nTMhelix 7 26\noutside 27 206\nTMhelix 207 229\ninside 230 233

399 GCF_001467445.1_ASM146744v1 Sphingopyxis sp. H077 Proteobacteria; Alphaproteobacteria; Sphingomonadales; Sphingomonadaceae; Sphingopyxis
MTRTRRAAFAAAALIAAVALAALGVWQLERRVWKHELVAAVEARIHAAPVAAPWPSAWDA WP_058536255.1
MULTISPECIES: Surfeit locus 1 family protein [Sphingopyxis] Length: 233\nNumber of predicted TMHs: 2\nExp number of AAs in TMHs: 48.28018\nExp number, first 60 AAs: 20.95142\nTotal prob of N-in: 0.99129\nPOSSIBLE N-term signal sequence\ninside 1 6\nTMhelix 7 26\noutside 27 206\nTMhelix 207 229\ninside 230 233

400 GCF_001467455.1_ASM146745v1 Sphingopyxis sp. H080 Proteobacteria; Alphaproteobacteria; Sphingomonadales; Sphingomonadaceae; Sphingopyxis
MTRTRRAAFAAAALIAAVALAALGVWQLERRVWKHELVAAVEARIHAAPVAAPWPSAWDA WP_058536255.1
MULTISPECIES: Surfeit locus 1 family protein [Sphingopyxis] Length: 233\nNumber of predicted TMHs: 2\nExp number of AAs in TMHs: 48.28018\nExp number, first 60 AAs: 20.95142\nTotal prob of N-in: 0.99129\nPOSSIBLE N-term signal sequence\ninside 1 6\nTMhelix 7 26\noutside 27 206\nTMhelix 207 229\ninside 230 233

401 GCF_001468385.1_ASM146838v1 Sphingopyxis sp. H081 Proteobacteria; Alphaproteobacteria; Sphingomonadales; Sphingomonadaceae; Sphingopyxis
MTRPRRAAFAAAALVAAACLAALGIWQVERRAWKHELVAAVDARIHAPPVAAPGPGRWPT WP_058454724.1
MULTISPECIES: Surfeit locus 1 family protein [Sphingopyxis] Length: 233\nNumber of predicted TMHs: 2\nExp number of AAs in TMHs: 42.81905\nExp number, first 60 AAs: 21.12953\nTotal prob of N-in: 0.99193\nPOSSIBLE N-term signal sequence\ninside 1 6\nTMhelix 7 26\noutside 27 206\nTMhelix 207 229\ninside 230 233

402 GCF_001467385.1_ASM146738v1 Sphingopyxis sp. H085 Proteobacteria; Alphaproteobacteria; Sphingomonadales; Sphingomonadaceae; Sphingopyxis
MTRTRRAAFAAAALIAAVALAALGVWQLERRVWKHELVAAVEARIHAAPVAAPWPSAWDA WP_058536255.1
MULTISPECIES: Surfeit locus 1 family protein [Sphingopyxis] Length: 233\nNumber of predicted TMHs:

2\nExp number of AAs in TMHs: 48.28018\nExp number, first 60 AAs: 20.95142\nTotal prob of N-in: 0.99129\nPOSSIBLE N-term signal sequence\ninside 1 6\nTMhelix 7 26\noutside 27 206\nTMhelix 207 229\ninside 230 233

403 GCF_001467475.1_ASM146747v1 Sphingopyxis sp. H093 Proteobacteria; Alphaproteobacteria; Sphingomonadales; Sphingomonadaceae; Sphingopyxis
MTRTRRAAFAAAALIAAVALAALGVWQLERRVWKHELVAAVEARIHAAPVAAPWPSAWDA WP_058536255.1
MULTISPECIES: Surfeit locus 1 family protein [Sphingopyxis] Length: 233\nNumber of predicted TMHs: 2\nExp number of AAs in TMHs: 48.28018\nExp number, first 60 AAs: 20.95142\nTotal prob of N-in: 0.99129\nPOSSIBLE N-term signal sequence\ninside 1 6\nTMhelix 7 26\noutside 27 206\nTMhelix 207 229\ninside 230 233

404 GCF_001468425.1_ASM146842v1 Sphingopyxis sp. H100 Proteobacteria; Alphaproteobacteria; Sphingomonadales; Sphingomonadaceae; Sphingopyxis
MTRPRRAAFAAAALVAAACLAALGIWQVERRAWKHELVAAVDARIHAPPVAAPGPGRWPT WP_058454724.1
MULTISPECIES: Surfeit locus 1 family protein [Sphingopyxis] Length: 233\nNumber of predicted TMHs: 2\nExp number of AAs in TMHs: 42.81905\nExp number, first 60 AAs: 21.12953\nTotal prob of N-in: 0.99193\nPOSSIBLE N-term signal sequence\ninside 1 6\nTMhelix 7 26\noutside 27 206\nTMhelix 207 229\ninside 230 233

405 GCF_001468475.1_ASM146847v1 Sphingopyxis sp. H107 Proteobacteria; Alphaproteobacteria; Sphingomonadales; Sphingomonadaceae; Sphingopyxis
MTRPRRAAFAAAALVAAACLAALGIWQVERRAWKHELVAAVDARIHAPPVAAPGPGRWPT WP_058454724.1
MULTISPECIES: Surfeit locus 1 family protein [Sphingopyxis] Length: 233\nNumber of predicted TMHs: 2\nExp number of AAs in TMHs: 42.81905\nExp number, first 60 AAs: 21.12953\nTotal prob of N-in: 0.99193\nPOSSIBLE N-term signal sequence\ninside 1 6\nTMhelix 7 26\noutside 27 206\nTMhelix 207 229\ninside 230 233

406 GCF_001468465.1_ASM146846v1 Sphingopyxis sp. H115 Proteobacteria; Alphaproteobacteria; Sphingomonadales; Sphingomonadaceae; Sphingopyxis
MTRTRRAAFAAAALIAALGLAALGVWQVERRAWKHELIAAVDARVHAAPAKAPRPDRWDA WP_058803118.1
Surfeit locus 1 family protein [Sphingopyxis sp. H115] Length: 233\nNumber of predicted TMHs: 2\nExp number of AAs in TMHs: 44.92482\nExp number, first 60 AAs: 21.26646\nTotal prob of N-in: 0.98729\nPOSSIBLE N-term signal sequence\ninside 1 6\nTMhelix 7 29\noutside 30 206\nTMhelix 207 229\ninside 230 233

407 GCF_001468465.1_ASM146846v1 Sphingopyxis sp. H115 Proteobacteria; Alphaproteobacteria; Sphingomonadales; Sphingomonadaceae; Sphingopyxis
MPGRLGGSRRAAIIATGAMIALLPVHARPAEPAPKQPNIVILLADDWGFSDVGSFGSEMA WP_058803291.1
arylsulfatase [Sphingopyxis sp. H115] Length: 613\nNumber of predicted TMHs: 1\nExp number of AAs in TMHs: 21.22031\nExp number, first 60 AAs: 0.845289999999999\nTotal prob of N-in: 0.04400\noutside 1 571\nTMhelix 572 594\ninside 595 613

408 GCF_000756385.1_ASM75638v1 Sphingopyxis sp. LC363 Proteobacteria; Alphaproteobacteria; Sphingomonadales; Sphingomonadaceae; Sphingopyxis
MTPARRAAFAAAALIAAIGLAALGAWQIERRAWKHELIAAVDARIHAPPVAAPGPDGWNA WP_037553521.1
Surfeit locus 1 family protein [Sphingopyxis sp. LC363] Length: 233\nNumber of predicted TMHs: 2\nExp number of AAs in TMHs: 43.59221\nExp number, first 60 AAs: 21.3735\nTotal prob of N-in: 0.92555\nPOSSIBLE N-term signal sequence\ninside 1 6\nTMhelix 7 29\noutside 30 206\nTMhelix 207 229\ninside 230 233

409 GCF_000756375.1_ASM75637v1 Sphingopyxis sp. LC81 Proteobacteria; Alphaproteobacteria; Sphingomonadales; Sphingomonadaceae; Sphingopyxis
MTRSRRAAFAAAALIAAIGLAALGVWQLERRVWKHELVAAVEARIHAAPVAAPGPGAWDA WP_037511434.1
Surfeit locus 1 family protein [Sphingopyxis sp. LC81] Length: 233\nNumber of predicted TMHs: 2\nExp number of AAs in TMHs: 58.48123\nExp number, first 60 AAs: 20.97118\nTotal prob of N-in: 0.97774\nPOSSIBLE N-term signal sequence\ninside 1 6\nTMhelix 7 26\noutside 27 206\nTMhelix 207 229\ninside 230 233

410 GCF_000371385.1_Sphingopyxis_sp._MC1 Sphingopyxis sp. MC1 Proteobacteria; Alphaproteobacteria; Sphingomonadales; Sphingomonadaceae; Sphingopyxis
MGDAPPQHQYDADAARHEPVAVTRPRRAAFAAAALVVAALLAALGVWQLERREWKHALVA WP_081611131.1
Surfeit locus 1 family protein [Sphingopyxis sp. MC1] Length: 262\nNumber of predicted TMHs: 2\nExp number of AAs in TMHs: 44.16228\nExp number, first 60 AAs: 20.35941\nTotal prob of N-in: 0.10632\nPOSSIBLE N-term signal sequence\noutside 1 27\nTMhelix 28 47\ninside 48 227\nTMhelix 228 250\noutside 251 262

411 GCF_001427105.1_Root154 Sphingopyxis sp. Root154 Proteobacteria; Alphaproteobacteria; Sphingomonadales; Sphingomonadaceae; Sphingopyxis

MTHPRRAAFAAAALIAAIGLAALGIWQLERRVWKHELVEAVEARIHAAPVAAPGPGAWDA WP_056369465.1 MULTISPECIES: Surfeit locus 1 family protein [Sphingopyxis] Length: 233\nNumber of predicted TMHs: 2\nExp number of AAs in TMHs: 50.95432\nExp number, first 60 AAs: 20.85505\nTotal prob of N-in: 0.97474\nPOSSIBLE N-term signal sequence\ninside 1 6\nTMhelix 7 26\noutside 27 206\nTMhelix 207 229\ninside 230 233

412 GCF_001429105.1_Root214 Sphingopyxis sp. Root214 Proteobacteria; Alphaproteobacteria; Sphingomonadales; Sphingomonadaceae; Sphingopyxis MTHPRRAAFAAAALIAAIGLAALGIWQLERRVWKHELVEAVEARIHAAPVAAPGPGAWDA WP_056369465.1 MULTISPECIES: Surfeit locus 1 family protein [Sphingopyxis] Length: 233\nNumber of predicted TMHs: 2\nExp number of AAs in TMHs: 50.95432\nExp number, first 60 AAs: 20.85505\nTotal prob of N-in: 0.97474\nPOSSIBLE N-term signal sequence\ninside 1 6\nTMhelix 7 26\noutside 27 206\nTMhelix 207 229\ninside 230 233

413 GCF_900177755.1_IMG-taxon_2708742469_annotated_assembly Sphingopyxis ummariensis Proteobacteria; Alphaproteobacteria; Sphingomonadales; Sphingomonadaceae; Sphingopyxis MTRPRRAAFAAAALVVAALLAALGVWQLERREWKHALVAAVDARIHAAPVPAPPPPAWRG WP_086457152.1 Surfeit locus 1 family protein [Sphingopyxis ummariensis] Length: 241\nNumber of predicted TMHs: 2\nExp number of AAs in TMHs: 56.27941\nExp number, first 60 AAs: 21.01528\nTotal prob of N-in: 0.94423\nPOSSIBLE N-term signal sequence\ninside 1 6\nTMhelix 7 26\noutside 27 206\nTMhelix 207 229\ninside 230 241

414 GCF_000013985.1_ASM1398v1 Sphingopyxis alaskensis RB2256 Proteobacteria; Alphaproteobacteria; Sphingomonadales; Sphingomonadaceae; Sphingopyxis; Sphingopyxis alaskensis MAAGWRTTRRGRTGASRRAAIIALACLLGSTSGLAQSRAAPPRQPNIVILLADDWGFSDV WP_011540613.1 arylsulfatase [Sphingopyxis alaskensis] Length: 609\nNumber of predicted TMHs: 1\nExp number of AAs in TMHs: 33.96543\nExp number, first 60 AAs: 15.00873\nTotal prob of N-in: 0.75540\nPOSSIBLE N-term signal sequence\noutside 1 584\nTMhelix 585 602\ninside 603 609

415 GCF_001591045.1_ASM159104v1 Sphingopyxis granuli NBRC 100800 Proteobacteria; Alphaproteobacteria; Sphingomonadales; Sphingomonadaceae; Sphingopyxis; Sphingopyxis granuli MTATRRAVLAAAALLVTACLLALGVWQLERRVWKHALIAAVGTRLHADPVAAPGPAAWPA WP_067107604.1 Surfeit locus 1 family protein [Sphingopyxis granuli] Length: 235\nNumber of predicted TMHs: 2\nExp number of AAs in TMHs: 42.85327\nExp number, first 60 AAs: 21.27767\nTotal prob of N-in: 0.95408\nPOSSIBLE N-term signal sequence\ninside 1 6\nTMhelix 7 26\noutside 27 206\nTMhelix 207 229\ninside 230 235

416 GCF_001598815.1_ASM159881v1 Sphingopyxis terrae NBRC 15098 Proteobacteria; Alphaproteobacteria; Sphingomonadales; Sphingomonadaceae; Sphingopyxis; Sphingopyxis terrae MTRPRRAAFAAAALVVAALLAALGVWQLERREWKHALVAAVDARIHAAPVAAPPPAAWRG WP_062770644.1 Surfeit locus 1 family protein [Sphingopyxis terrae] Length: 241\nNumber of predicted TMHs: 2\nExp number of AAs in TMHs: 55.70143\nExp number, first 60 AAs: 21.10054\nTotal prob of N-in: 0.94691\nPOSSIBLE N-term signal sequence\ninside 1 6\nTMhelix 7 26\noutside 27 206\nTMhelix 207 229\ninside 230 241

417 GCF_000024465.1_ASM2446v1 Candidatus Puniceispirillum marinum IMCC1322 Proteobacteria; Alphaproteobacteria; unclassified Alphaproteobacteria; SAR116 cluster; Candidatus Puniceispirillum; Candidatus Puniceispirillum marinum MATKQKPHNKKVIGHDSQLVLASASPRRRELLAQIGLSPAIMVADIDETPHKREKPQPYA WP_013046185.1 septum formation protein Maf [Candidatus Puniceispirillum marinum] Length: 214\nNumber of predicted TMHs: 1\nExp number of AAs in TMHs: 14.98364\nExp number, first 60 AAs: 0.00087\nTotal prob of N-in: 0.79472\ninside 1 182\nTMhelix 183 205\noutside 206 214

418 GCF_002209535.1_ASM220953v1 Achromobacter marplatensis Proteobacteria; Betaproteobacteria; Burkholderiales; Alcaligenaceae; Achromobacter MYNRRSIIRLTLLACAVGAGIAVACGPDFPAQLLDDRTGTLRSTPANSFSFEVTRLVAPA WP_088590309.1 hypothetical protein [Achromobacter marplatensis] Length: 836\nNumber of predicted TMHs: 2\nExp number of AAs in TMHs: 39.29622\nExp number, first 60 AAs: 18.38551\nTotal prob of N-in: 0.91193\nPOSSIBLE N-term signal sequence\ninside 1 6\nTMhelix 7 24\noutside 25 801\nTMhelix 802 821\ninside 822 836

419 GCF_001298915.1_11861_6_5 Achromobacter sp. 2789STDY5608611 Proteobacteria; Betaproteobacteria; Burkholderiales; Alcaligenaceae; Achromobacter MNPDILRAPRPAPRRAALAAGLRAACGGVALALAIAARATAGEAPAAAPDPALEAHVQAF WP_081322336.1 MULTISPECIES: hypothetical protein [Achromobacter] Length: 187\nNumber of predicted TMHs: 2\nExp number of AAs in TMHs: 40.06129\nExp number, first 60 AAs: 18.98302\nTotal prob of N-in: 0.86077\nPOSSIBLE N-term signal sequence\ninside 1 19\nTMhelix 20 42\noutside 43 123\nTMhelix 124 146\ninside 147 187

420 GCF_001298935.1_11861_6_23 Achromobacter sp. 2789STDY5608630 Proteobacteria; Betaproteobacteria; Burkholderiales; Alcaligenaceae; Achromobacter
MNPDILRAPRPAPRRAALAAGLRAACGGVALALAIAARATAGEAPAAAPDPALEAHVQAF WP_081322336.1
MULTISPECIES: hypothetical protein [Achromobacter] Length: 187\nNumber of predicted TMHs: 2\nExp number of AAs in TMHs: 40.06129\nExp number, first 60 AAs: 18.98302\nTotal prob of N-in: 0.86077\nPOSSIBLE N-term signal sequence\ninside 1 19\nTMhelix 20 42\noutside 43 123\nTMhelix 124 146\ninside 147 187

421 GCF_001299015.1_11861_6_24 Achromobacter sp. 2789STDY5608632 Proteobacteria; Betaproteobacteria; Burkholderiales; Alcaligenaceae; Achromobacter
MNPDILRAPRPAPRRAALAAGLRAACGGVALALAIAARATAGEAPAAAPDPALEAHVQAF WP_080982266.1
MULTISPECIES: hypothetical protein [Achromobacter] Length: 187\nNumber of predicted TMHs: 2\nExp number of AAs in TMHs: 40.06132\nExp number, first 60 AAs: 18.98302\nTotal prob of N-in: 0.86077\nPOSSIBLE N-term signal sequence\ninside 1 19\nTMhelix 20 42\noutside 43 123\nTMhelix 124 146\ninside 147 187

422 GCF_001299215.1_11861_6_72 Achromobacter sp. 2789STDY5663447 Proteobacteria; Betaproteobacteria; Burkholderiales; Alcaligenaceae; Achromobacter
MNPDILRAPRPAPRRAALAAGLRAACGGVALALAIAARATAGEAPAAAPDPALEAHVQAF WP_080982266.1
MULTISPECIES: hypothetical protein [Achromobacter] Length: 187\nNumber of predicted TMHs: 2\nExp number of AAs in TMHs: 40.06132\nExp number, first 60 AAs: 18.98302\nTotal prob of N-in: 0.86077\nPOSSIBLE N-term signal sequence\ninside 1 19\nTMhelix 20 42\noutside 43 123\nTMhelix 124 146\ninside 147 187

423 GCF_001299315.1_11861_6_73 Achromobacter sp. ATCC35328 Proteobacteria; Betaproteobacteria; Burkholderiales; Alcaligenaceae; Achromobacter
MNNEETFYQAMRRQGVTRRSFLKYCSLAATSLGLGAGMAPKIAWALENKPRIPVVWIHGL WP_001058323.1
MULTISPECIES: hydrogenase-1 small chain [Proteobacteria] Length: 372\nNumber of predicted TMHs: 2\nExp number of AAs in TMHs: 37.2608499999999\nExp number, first 60 AAs: 19.52923\nTotal prob of N-in: 0.94351\nPOSSIBLE N-term signal sequence\ninside 1 19\nTMhelix 20 39\noutside 40 325\nTMhelix 326 348\ninside 349 372

424 GCF_001299315.1_11861_6_73 Achromobacter sp. ATCC35328 Proteobacteria; Betaproteobacteria; Burkholderiales; Alcaligenaceae; Achromobacter
MTGDNTLIHSHGINRRDFMKLCAALAATMGLSSKAAAEMAESVTNPQRPPVIWIGAQECT WP_000145410.1
MULTISPECIES: hydrogenase-2 small chain [Proteobacteria] Length: 372\nNumber of predicted TMHs: 1\nExp number of AAs in TMHs: 23.72226\nExp number, first 60 AAs: 0.22038\nTotal prob of N-in: 0.12994\noutside 1 331\nTMhelix 332 354\ninside 355 372

425 GCF_002209495.1_ASM220949v1 Achromobacter sp. HZ28 Proteobacteria; Betaproteobacteria; Burkholderiales; Alcaligenaceae; Achromobacter
MLKNASPTTKPTMQKPVSPAPHRRRRPTLARLSLALLTSCSLVAQAAEVGAATAGTVPPA WP_088596650.1
cellulose synthase regulator BcsB [Achromobacter sp. HZ28] Length: 785\nNumber of predicted TMHs: 1\nExp number of AAs in TMHs: 24.8247\nExp number, first 60 AAs: 2.32423\nTotal prob of N-in: 0.10703\noutside 1 754\nTMhelix 755 777\ninside 778 785

426 GCF_002209485.1_ASM220948v1 Achromobacter sp. HZ34 Proteobacteria; Betaproteobacteria; Burkholderiales; Alcaligenaceae; Achromobacter
MLKNASPTTKPTMQKPVSPAPHRRRRPTLARLSLALLTSCSLVAQAAEVGAATAGTVPPA WP_088596650.1
cellulose synthase regulator BcsB [Achromobacter sp. HZ28] Length: 785\nNumber of predicted TMHs: 1\nExp number of AAs in TMHs: 24.8247\nExp number, first 60 AAs: 2.32423\nTotal prob of N-in: 0.10703\noutside 1 754\nTMhelix 755 777\ninside 778 785

427 GCF_001471535.1_ASM147153v1 Achromobacter xylosoxidans Proteobacteria; Betaproteobacteria; Burkholderiales; Alcaligenaceae; Achromobacter
MNPDILRAPRPAPRRAALAAGLRAACGGVALALAIAARATAGEAPAAAPDPALEAHVQAF WP_081322336.1
MULTISPECIES: hypothetical protein [Achromobacter] Length: 187\nNumber of predicted TMHs: 2\nExp number of AAs in TMHs: 40.06129\nExp number, first 60 AAs: 18.98302\nTotal prob of N-in: 0.86077\nPOSSIBLE N-term signal sequence\ninside 1 19\nTMhelix 20 42\noutside 43 123\nTMhelix 124 146\ninside 147 187

428 GCF_001457475.1_NCTC10807 Achromobacter xylosoxidans Proteobacteria; Betaproteobacteria; Burkholderiales; Alcaligenaceae; Achromobacter
MNPDILRAPRPAPRRAALAAGLRAACGGVALALAIAARATAGEAPAAAPDPALEARVQAF WP_081247826.1
hypothetical protein [Achromobacter xylosoxidans] Length: 187\nNumber of predicted TMHs: 2\nExp

number of AAs in TMHs: 40.0597\nExp number, first 60 AAs: 18.98153\nTotal prob of N-in: 0.86072\nPOSSIBLE N-term signal sequence\ninside 1 19\nTMhelix 20 42\noutside 43 123\nTMhelix 124 146\ninside 147 187

429 GCF_001057595.1_ASM105759v1 Achromobacter xylosoxidans Proteobacteria; Betaproteobacteria; Burkholderiales; Alcaligenaceae; Achromobacter MNPDILRAPRPAPRRAALAAGLRAACGGVALALAIAARATAGEAPAAAPDPALEAHVQAF WP_080982266.1 MULTISPECIES: hypothetical protein [Achromobacter] Length: 187\nNumber of predicted TMHs: 2\nExp number of AAs in TMHs: 40.06132\nExp number, first 60 AAs: 18.98302\nTotal prob of N-in: 0.86077\nPOSSIBLE N-term signal sequence\ninside 1 19\nTMhelix 20 42\noutside 43 123\nTMhelix 124 146\ninside 147 187

430 GCF_001038045.1_ASM103804v1 Achromobacter xylosoxidans Proteobacteria; Betaproteobacteria; Burkholderiales; Alcaligenaceae; Achromobacter MNPDILRAPRPTPRPAPRRAALAAGLRAACGGVALALAIAARATAGEAPAAAPDPALEAH WP_080969392.1 hypothetical protein [Achromobacter xylosoxidans] Length: 191\nNumber of predicted TMHs: 2\nExp number of AAs in TMHs: 39.60172\nExp number, first 60 AAs: 18.54883\nTotal prob of N-in: 0.85130\nPOSSIBLE N-term signal sequence\ninside 1 20\nTMhelix 21 43\noutside 44 127\nTMhelix 128 150\ninside 151 191

431 GCF_000286415.1_ASM28641v1 Achromobacter piechaudii HLE Proteobacteria; Betaproteobacteria; Burkholderiales; Alcaligenaceae; Achromobacter; Achromobacter piechaudii MYNRRSIIRLTLLACAVGAGIAVACGPDFPAQLLDDRAGTLRSTPANSFSFEVTRLVAPA WP_006226706.1 hypothetical protein [Achromobacter piechaudii] Length: 836\nNumber of predicted TMHs: 2\nExp number of AAs in TMHs: 40.2832799999999\nExp number, first 60 AAs: 18.43682\nTotal prob of N-in: 0.91306\nPOSSIBLE N-term signal sequence\ninside 1 6\nTMhelix 7 24\noutside 25 798\nTMhelix 799 821\ninside 822 836

432 GCF_001598595.1_ASM159859v1 Achromobacter xylosoxidans NBRC 15126 = ATCC 27061 Proteobacteria; Betaproteobacteria; Burkholderiales; Alcaligenaceae; Achromobacter; Achromobacter xylosoxidans MNPDILRAPRPAPRRAALAAGLRAACGGVALALAIAARATAGEAPAAAPDPALEARVQAF WP_081247826.1 hypothetical protein [Achromobacter xylosoxidans] Length: 187\nNumber of predicted TMHs: 2\nExp number of AAs in TMHs: 40.0597\nExp number, first 60 AAs: 18.98153\nTotal prob of N-in: 0.86072\nPOSSIBLE N-term signal sequence\ninside 1 19\nTMhelix 20 42\noutside 43 123\nTMhelix 124 146\ninside 147 187

433 GCF_001571085.1_ASM157108v1 Azohydromonas lata NBRC 102462 Proteobacteria; Betaproteobacteria; Burkholderiales; Alcaligenaceae; Azohydromonas; Azohydromonas lata MLSLRARRTVVALAAVLGTSLTARLGWWQLDRAAQKESLQQAVDAHAALPPLPASGLART WP_066337416.1 transmembrane cytochrome oxidase [Azohydromonas lata] Length: 246\nNumber of predicted TMHs: 1\nExp number of AAs in TMHs: 28.3475\nExp number, first 60 AAs: 7.27408\nTotal prob of N-in: 0.38659\noutside 1 216\nTMhelix 217 239\ninside 240 246

434 GCF_900078315.1_7642_7_30 Bordetella ansorpii Proteobacteria; Betaproteobacteria; Burkholderiales; Alcaligenaceae; Bordetella MSADTAHRRRAFLADTTALILFFTTTGALNERYIAGMTWDQVLHARLLGAALMVPVARPY WP_066408845.1 alanine transporter [Bordetella ansorpii] Length: 151\nNumber of predicted TMHs: 2\nExp number of AAs in TMHs: 73.64719\nExp number, first 60 AAs: 29.05964\nTotal prob of N-in: 0.93675\nPOSSIBLE N-term signal sequence\ninside 1 75\nTMhelix 76 98\noutside 99 107\nTMhelix 108 130\ninside 131 151

435 GCF_001078275.1_ASM107827v1 Bordetella hinzii Proteobacteria; Betaproteobacteria; Burkholderiales; Alcaligenaceae; Bordetella MKKNSLSTQTPQARVWPRRRALSRLSLVLLTSCSLLAQAAETPAPAPEPAPAPPPFKADQ WP_029577391.1 cellulose synthase BcsB subunit [Bordetella hinzii] Length: 770\nNumber of predicted TMHs: 1\nExp number of AAs in TMHs: 28.82266\nExp number, first 60 AAs: 6.4729\nTotal prob of N-in: 0.32498\noutside 1 739\nTMhelix 740 762\ninside 763 770

436 GCF_001078295.1_ASM107829v1 Bordetella hinzii Proteobacteria; Betaproteobacteria; Burkholderiales; Alcaligenaceae; Bordetella MKKNSLSTQTPQARVWPRRRALSRLSLVLLTSCSLLAQAAETPAPAPEPAPAPPPFKADQ WP_029577391.1 cellulose synthase BcsB subunit [Bordetella hinzii] Length: 770\nNumber of predicted TMHs: 1\nExp number of AAs in TMHs: 28.82266\nExp number, first 60 AAs: 6.4729\nTotal prob of N-in: 0.32498\noutside 1 739\nTMhelix 740 762\ninside 763 770

437 GCF_000657695.1_gbhv01 Bordetella hinzii 1277 Proteobacteria; Betaproteobacteria; Burkholderiales; Alcaligenaceae; Bordetella; Bordetella hinzii MKKNSLSTQTPQARVWPRRRALSRLSLVLLTSCSLLAQAAETPAPAPEPAPAPPPFKADQ WP_029577391.1 cellulose synthase BcsB subunit [Bordetella hinzii] Length: 770\nNumber of predicted TMHs: 1\nExp

number of AAs in TMHs: 28.82266\nExp number, first 60 AAs: 6.4729\nTotal prob of N-in: 0.32498\noutside 1 739\nTMhelix 740 762\ninside 763 770

438 GCF_000657775.1_gbh01v01 Bordetella hinzii 4161 Proteobacteria; Betaproteobacteria; Burkholderiales; Alcaligenaceae; Bordetella; Bordetella hinzii MKKNSLSTQTPQARVWPRRRALSRLSLVLLTSCSLLAQAAETPAPAPEPAPAPPPFKADQ WP_029577391.1 cellulose synthase BcsB subunit [Bordetella hinzii] Length: 770\nNumber of predicted TMHs: 1\nExp number of AAs in TMHs: 28.82266\nExp number, first 60 AAs: 6.4729\nTotal prob of N-in: 0.32498\noutside 1 739\nTMhelix 740 762\ninside 763 770

439 GCF_000657755.1_gbh02v01 Bordetella hinzii 5132 Proteobacteria; Betaproteobacteria; Burkholderiales; Alcaligenaceae; Bordetella; Bordetella hinzii MKKNSLSTQTPQARVWPRRRALSRLSLVLLTSCSLLAQAAETPAPAPEPAPAPPPFKADQ WP_029577391.1 cellulose synthase BcsB subunit [Bordetella hinzii] Length: 770\nNumber of predicted TMHs: 1\nExp number of AAs in TMHs: 28.82266\nExp number, first 60 AAs: 6.4729\nTotal prob of N-in: 0.32498\noutside 1 739\nTMhelix 740 762\ninside 763 770

440 GCF_000471685.1_Bhinzii1.0 Bordetella hinzii ATCC 51730 Proteobacteria; Betaproteobacteria; Burkholderiales; Alcaligenaceae; Bordetella; Bordetella hinzii MKKNSLSTQTPQARVWPRRRALSRLSLVLLTSCSLLAQAAETPAPAPEPAPAPPPFKADQ WP_029577391.1 cellulose synthase BcsB subunit [Bordetella hinzii] Length: 770\nNumber of predicted TMHs: 1\nExp number of AAs in TMHs: 28.82266\nExp number, first 60 AAs: 6.4729\nTotal prob of N-in: 0.32498\noutside 1 739\nTMhelix 740 762\ninside 763 770

441 GCF_000657815.1_gbh04v01 Bordetella hinzii CA90 BAL1384 Proteobacteria; Betaproteobacteria; Burkholderiales; Alcaligenaceae; Bordetella; Bordetella hinzii MKKNSLSTQTPQARVWPRRRALSRLSLVLLTSCSLLAQAAETPAPAPEPAPAPPPFKADQ WP_029577391.1 cellulose synthase BcsB subunit [Bordetella hinzii] Length: 770\nNumber of predicted TMHs: 1\nExp number of AAs in TMHs: 28.82266\nExp number, first 60 AAs: 6.4729\nTotal prob of N-in: 0.32498\noutside 1 739\nTMhelix 740 762\ninside 763 770

442 GCF_000657715.1_gbh06v01 Bordetella hinzii L60 Proteobacteria; Betaproteobacteria; Burkholderiales; Alcaligenaceae; Bordetella; Bordetella hinzii MKKNSLSTQTPQARVWPRRRALSRLSLVLLTSCSLLAQAAETPAPAPEPAPAPPPFKADQ WP_029577391.1 cellulose synthase BcsB subunit [Bordetella hinzii] Length: 770\nNumber of predicted TMHs: 1\nExp number of AAs in TMHs: 28.82266\nExp number, first 60 AAs: 6.4729\nTotal prob of N-in: 0.32498\noutside 1 739\nTMhelix 740 762\ninside 763 770

443 GCF_001548475.1_ASM154847v1 Bordetella hinzii LMG 13501 Proteobacteria; Betaproteobacteria; Burkholderiales; Alcaligenaceae; Bordetella; Bordetella hinzii MKKNSLSTQTPQARVWPRRRALSRLSLVLLTSCSLLAQAAETPAPAPEPAPAPPPFKADQ WP_029577391.1 cellulose synthase BcsB subunit [Bordetella hinzii] Length: 770\nNumber of predicted TMHs: 1\nExp number of AAs in TMHs: 28.82266\nExp number, first 60 AAs: 6.4729\nTotal prob of N-in: 0.32498\noutside 1 739\nTMhelix 740 762\ninside 763 770

444 GCF_000657735.1_gbh07v01 Bordetella hinzii OH87 BAL007II Proteobacteria; Betaproteobacteria; Burkholderiales; Alcaligenaceae; Bordetella; Bordetella hinzii MKKNSLSTQTPQARVWPRRRALSRLSLVLLTSCSLLAQAAETPAPAPEPAPAPPPFKADQ WP_029577391.1 cellulose synthase BcsB subunit [Bordetella hinzii] Length: 770\nNumber of predicted TMHs: 1\nExp number of AAs in TMHs: 28.82266\nExp number, first 60 AAs: 6.4729\nTotal prob of N-in: 0.32498\noutside 1 739\nTMhelix 740 762\ninside 763 770

445 GCF_000612685.1_scf7180000000004 Castellaniella defragrans 65Phen Proteobacteria; Betaproteobacteria; Burkholderiales; Alcaligenaceae; Castellaniella; Castellaniella defragrans MQNVPSEPGFLSLRQIHAMLAARRALILNVAGSLFFLTVMLTLLLPRTWTASSSLYINYR WP_043684137.1 hypothetical protein [Castellaniella defragrans] Length: 465\nNumber of predicted TMHs: 2\nExp number of AAs in TMHs: 43.80579\nExp number, first 60 AAs: 21.43023\nTotal prob of N-in: 0.95826\nPOSSIBLE N-term signal sequence\ninside 1 25\nTMhelix 26 45\noutside 46 410\nTMhelix 411 433\ninside 434 465

446 GCF_002105195.1_ASM210519v1 Derxia lacustris Proteobacteria; Betaproteobacteria; Burkholderiales; Alcaligenaceae; Derxia MSAYLPDVIGIDYDRSRRLTLREFLVIAFRDRRKILTGFFLTFTLAVSVSFVPKPSYTAD

WP_085318242.1 hypothetical protein [Derxia lacustris] Length: 495\nNumber of predicted TMHs: 1\nExp number of AAs in TMHs: 34.43941\nExp number, first 60 AAs: 12.13407\nTotal prob of N-in: 0.46833\nPOSSIBLE N-term signal sequence\noutside 1 441\nTMhelix 442 464\ninside 465 495

447 GCF_000482785.1_ASM48278v1 Derxia gummosa DSM 723 Proteobacteria; Betaproteobacteria; Burkholderiales; Alcaligenaceae; Derxia; Derxia gummosa MSAYLPDVISIDTDTRRQLTLREFLVIAFRDRRKILTGFFLTLTLGVSASFLPKPSYTAD WP_028311354.1 hypothetical protein [Derxia gummosa] Length: 494\nNumber of predicted TMHs: 1\nExp number of AAs in TMHs: 29.92567\nExp number, first 60 AAs: 7.77266\nTotal prob of N-in: 0.34377\noutside 1 445\nTMhelix 446 468\ninside 469 494

448 GCF_000482785.1_ASM48278v1 Derxia gummosa DSM 723 Proteobacteria; Betaproteobacteria; Burkholderiales; Alcaligenaceae; Derxia; Derxia gummosa MLTTNLPAVFRARRRTLFVTSALVALLCAVALAVLPRKYVASAEILYDVKASDPIVGAML WP_028311763.1 hypothetical protein [Derxia gummosa] Length: 457\nNumber of predicted TMHs: 2\nExp number of AAs in TMHs: 41.9446000000001\nExp number, first 60 AAs: 20.13666\nTotal prob of N-in: 0.95445\nPOSSIBLE N-term signal sequence\ninside 1 16\nTMhelix 17 36\noutside 37 395\nTMhelix 396 418\ninside 419 457

449 GCF_000482785.1_ASM48278v1 Derxia gummosa DSM 723 Proteobacteria; Betaproteobacteria; Burkholderiales; Alcaligenaceae; Derxia; Derxia gummosa MSDGPAPRRLLAIIRAPRRSGGLIRRGQYLNQQELAEISKMRRRAFLMTGMALGGSQLGG WP_028312690.1 hypothetical protein [Derxia gummosa]Length: 240\nNumber of predicted TMHs: 1\nExp number of AAs in TMHs: 30.45691\nExp number, first 60 AAs: 6.78327\nTotal prob of N-in: 0.14809\noutside 1 215\nTMhelix 216 238\ninside 239 240

450 GCF_000988095.1_ASM98809v1 Kerstersia gyiorum Proteobacteria; Betaproteobacteria; Burkholderiales; Alcaligenaceae; Kerstersia MSQASSPRKQLRALAEARRGIIVPGAFNALSARVIADLGFEAIYITGAGVTNMSFGLPDQ WP_068366698.1 carboxyvinyl-carboxyphosphonate phosphorylmutase [Kerstersia gyiorum] Length: 290\nNumber of predicted TMHs: 1\nExp number of AAs in TMHs: 23.44439\nExp number, first 60 AAs: 2.91724\nTotal prob of N-in: 0.40540\noutside 1 217\nTMhelix 218 240\ninside 241 290

451 GCF_001729305.1_ASM172930v1 Burkholderia sp. A2 Proteobacteria; Betaproteobacteria; Burkholderiales; Burkholderiaceae; Burkholderia MSLSITRRRLLAAGVAASSLALSGCFTPKLYKNDAYSEHVSAFMMTEDGKKLVVLGTRYH WP_069617352.1 hypothetical protein [Burkholderia sp. A2] Length: 212\nNumber of predicted TMHs: 1\nExp number of AAs in TMHs: 33.22626\nExp number, first 60 AAs: 1.40446\nTotal prob of N-in: 0.71273\ninside 1 186\nTMhelix 187 209\noutside 210 212

452 GCF_001719665.1_ASM171966v1 Burkholderia sp. CQ001 Proteobacteria; Betaproteobacteria; Burkholderiales; Burkholderiaceae; Burkholderia MNVSITRRRLLAAGVAASSLALSGCFTPKLYKDDGYSEQVSGFMMTEDGKKLVVLGTDYH WP_039366460.1 MULTISPECIES: hypothetical protein [Burkholderia] Length: 212\nNumber of predicted TMHs: 1\nExp number of AAs in TMHs: 25.9561\nExp number, first 60 AAs: 0.849399999999999\nTotal prob of N-in: 0.56566\noutside 1 180\nTMhelix 181 203\ninside 204 212

453 GCF_001719665.1_ASM171966v1 Burkholderia sp. CQ001 Proteobacteria; Betaproteobacteria; Burkholderiales; Burkholderiaceae; Burkholderia MKRRTFLALPAVAASGMLAGCSDANIGPITNRMLKDPEYRETLSAFLISADGKSLVVIGK WP_039354933.1 MULTISPECIES: 5-formyltetrahydrofolate cyclo-ligase [Burkholderia] Length: 212\nNumber of predicted TMHs: 1\nExp number of AAs in TMHs: 24.954\nExp number, first 60 AAs: 4.26972\nTotal prob of N-in: 0.39654\noutside 1 188\nTMhelix 189 211\ninside 212 212

454 GCF_001806955.1_ASM180695v1 Burkholderia sp. HMSC10F09 Proteobacteria; Betaproteobacteria; Burkholderiales; Burkholderiaceae; Burkholderia MSDDIDNGGGHGGFTRRDMMRVMAAGGMMAAGGGALLMPGQAAFAAPAPKRGGKIRVANE WP_059873413.1 MULTISPECIES: ABC transporter substrate-binding protein [Burkholderia] Length: 528\nNumber of predicted TMHs: 2\nExp number of AAs in TMHs: 36.36756\nExp number, first 60 AAs: 21.42525\nTotal prob of N-in: 0.89413\nPOSSIBLE N-term signal sequence\ninside 1 20\nTMhelix 21 43\noutside 44 503\nTMhelix 504 526\ninside 527 528

455 GCF_001043615.1_ASM104361v1 Burkholderia sp. LK4 Proteobacteria; Betaproteobacteria; Burkholderiales; Burkholderiaceae; Burkholderia MKRRTFLALPAVAASGMLAGCSNFGPVTKAWMSDSKYDETLSTFLITADGKQLVVLGKKY WP_027787377.1

MULTISPECIES: hypothetical protein [Burkholderia] Length: 210\nNumber of predicted TMHs: 1\nExp number of AAs in TMHs: 34.58187\nExp number, first 60 AAs: 12.86351\nTotal prob of N-in: 0.46668\nPOSSIBLE N-term signal sequence\noutside 1 186\nTMhelix 187 209\ninside 210 210

456 GCF_900119135.1_IMG-taxon_2599185355_annotated_assembly Burkholderia sp. NFACC33-1 Proteobacteria; Betaproteobacteria; Burkholderiales; Burkholderiaceae; Burkholderia MSISIARRRLLAAGMAASSLALSGCFTPKLYKDDAYSENVSGFMMTEDGKKLVVLGTRYH WP_072441791.1 MULTISPECIES: hypothetical protein [Burkholderia] Length: 212\nNumber of predicted TMHs: 1\nExp number of AAs in TMHs: 25.41302\nExp number, first 60 AAs: 0.48578\nTotal prob of N-in: 0.31164\noutside 1 180\nTMhelix 181 203\ninside 204 212

457 GCF_900119325.1_IMG-taxon_2599185240_annotated_assembly Burkholderia sp. NFPP32 Proteobacteria; Betaproteobacteria; Burkholderiales; Burkholderiaceae; Burkholderia MSISIARRRLLAAGMAASSLALSGCFTPKLYKDDAYSENVSGFMMTEDGKKLVVLGTRYH WP_072441791.1 MULTISPECIES: hypothetical protein [Burkholderia] Length: 212\nNumber of predicted TMHs: 1\nExp number of AAs in TMHs: 25.41302\nExp number, first 60 AAs: 0.48578\nTotal prob of N-in: 0.31164\noutside 1 180\nTMhelix 181 203\ninside 204 212

458 GCF_001577465.1_ASM157746v1 Burkholderia sp. PAMC 28687 Proteobacteria; Betaproteobacteria; Burkholderiales; Burkholderiaceae; Burkholderia MESGMSGFVMYFTDGSRRRQFLLAGALALLLCGVMAAAWFVLHPRYQVLFSDLRPQDASS WP_061998645.1 flagellar M-ring protein FliF [Burkholderia sp. PAMC 28687]Length: 508\nNumber of predicted TMHs: 2\nExp number of AAs in TMHs: 44.46051\nExp number, first 60 AAs: 22.6129\nTotal prob of N-in: 0.99770\nPOSSIBLE N-term signal sequence\ninside 1 20\nTMhelix 21 43\noutside 44 448\nTMhelix 449 471\ninside 472 508

459 GCF_001190015.1_ASM119001v1 Candidatus Burkholderia crenata Proteobacteria; Betaproteobacteria; Burkholderiales; Burkholderiaceae; Burkholderia MQRTAQGWLVLTQLTHHNAAAVGVSHGAPVRPASSLAAFHRLSDRRKLLIGTQAVLGTFA WP_082191787.1 hypothetical protein [Candidatus Burkholderia crenata] Length: 170\nNumber of predicted TMHs: 2\nExp number of AAs in TMHs: 45.84469\nExp number, first 60 AAs: 8.85908\nTotal prob of N-in: 0.89827\ninside 1 53\nTMhelix 54 76\noutside 77 122\nTMhelix 123 145\ninside 146 170

460 GCF_001527785.1_ASM152778v1 Burkholderia anthina Proteobacteria; Betaproteobacteria; Burkholderiales; Burkholderiaceae; Burkholderia; Burkholderia cepacia complex MKRRTFLALPAMAASGMLAGCADFGPVTKEWMKDSKYDETLSTFLITSDGKQLVVLGKRY WP_059585348.1 hypothetical protein [Burkholderia anthina] Length: 210\nNumber of predicted TMHs: 1\nExp number of AAs in TMHs: 32.06835\nExp number, first 60 AAs: 10.06597\nTotal prob of N-in: 0.54569\nPOSSIBLE N-term signal sequence\noutside 1 184\nTMhelix 185 207\ninside 208 210

461 GCF_001524825.1_ASM152482v1 Burkholderia anthina Proteobacteria; Betaproteobacteria; Burkholderiales; Burkholderiaceae; Burkholderia; Burkholderia cepacia complex MKRRTFLALPAMAASGMLAGCADFGPVTKEWMKDSKYDETLSTFLITSDGKQLVVLGKRY WP_059585348.1 hypothetical protein [Burkholderia anthina] Length: 210\nNumber of predicted TMHs: 1\nExp number of AAs in TMHs: 32.06835\nExp number, first 60 AAs: 10.06597\nTotal prob of N-in: 0.54569\nPOSSIBLE N-term signal sequence\noutside 1 184\nTMhelix 185 207\ninside 208 210

462 GCF_001530695.1_ASM153069v1 Burkholderia anthina Proteobacteria; Betaproteobacteria; Burkholderiales; Burkholderiaceae; Burkholderia; Burkholderia cepacia complex MKRRTFLALPAMAASGMLAGCADFGPVTKEWMKDSKYDETLSTFLITSDGKQLVVLGKRY WP_059585348.1 hypothetical protein [Burkholderia anthina] Length: 210\nNumber of predicted TMHs: 1\nExp number of AAs in TMHs: 32.06835\nExp number, first 60 AAs: 10.06597\nTotal prob of N-in: 0.54569\nPOSSIBLE N-term signal sequence\noutside 1 184\nTMhelix 185 207\ninside 208 210

463 GCF_001547525.1_ASM154752v1 Burkholderia anthina Proteobacteria; Betaproteobacteria; Burkholderiales; Burkholderiaceae; Burkholderia; Burkholderia cepacia complex MKRRTFLALPAMAASGMLAGCADFGPVTKEWMKDSKYDETLSTFLITSDGKQLVVLGKRY WP_059585348.1 hypothetical protein [Burkholderia anthina] Length: 210\nNumber of predicted TMHs: 1\nExp number of AAs in TMHs: 32.06835\nExp number, first 60 AAs: 10.06597\nTotal prob of N-in: 0.54569\nPOSSIBLE N-term signal sequence\noutside 1 184\nTMhelix 185 207\ninside 208 210

464	GCF_001524815.1_ASM152481v1	Burkholderia anthina	Proteobacteria; Betaproteobacteria; Burkholderiales; Burkholderiaceae; Burkholderia; Burkholderia cepacia complex
	MKRRTFLALPAMAASGMLAGCADFGPVTKEWMKDSKYDETLSTFLITSDGKQLVVLGKRY	WP_059585348.1	hypothetical protein [Burkholderia anthina]	Length: 210\nNumber of predicted TMHs: 1\nExp number of AAs in TMHs: 32.06835\nExp number, first 60 AAs: 10.06597\nTotal prob of N-in: 0.54569\nPOSSIBLE N-term signal sequence\noutside 1 184\nTMhelix 185 207\ninside 208 210

465	GCF_001527555.1_ASM152755v1	Burkholderia anthina	Proteobacteria; Betaproteobacteria; Burkholderiales; Burkholderiaceae; Burkholderia; Burkholderia cepacia complex
	MKRRTFLALPAMAASGMLAGCADFGPVTKEWMKDSKYDETLSTFLITSDGKQLVVLGKRY	WP_059585348.1	hypothetical protein [Burkholderia anthina]	Length: 210\nNumber of predicted TMHs: 1\nExp number of AAs in TMHs: 32.06835\nExp number, first 60 AAs: 10.06597\nTotal prob of N-in: 0.54569\nPOSSIBLE N-term signal sequence\noutside 1 184\nTMhelix 185 207\ninside 208 210

466	GCF_001991315.1_ASM199131v1	Burkholderia cenocepacia	Proteobacteria; Betaproteobacteria; Burkholderiales; Burkholderiaceae; Burkholderia; Burkholderia cepacia complex
	MSTSITRRRLLAAGVAASSLALSGCFTPKLYKNDAYSEHVSAFMMTEDGKKLVVLGTRYH	WP_077021267.1	hypothetical protein [Burkholderia cenocepacia]	Length: 212\nNumber of predicted TMHs: 1\nExp number of AAs in TMHs: 24.17376\nExp number, first 60 AAs: 0.92388\nTotal prob of N-in: 0.37951\noutside 1 180\nTMhelix 181 203\ninside 204 212

467	GCF_001991315.1_ASM199131v1	Burkholderia cenocepacia	Proteobacteria; Betaproteobacteria; Burkholderiales; Burkholderiaceae; Burkholderia; Burkholderia cepacia complex
	METLLTRIRASFADASPRRQLISVAVLVALVASLIAAGYFALRPRYQVLFRDLKPQDAAT	WP_077020516.1	flagellar M-ring protein FliF [Burkholderia cenocepacia]	Length: 491\nNumber of predicted TMHs: 2\nExp number of AAs in TMHs: 41.34236\nExp number, first 60 AAs: 22.1383\nTotal prob of N-in: 0.99080\nPOSSIBLE N-term signal sequence\ninside 1 20\nTMhelix 21 43\noutside 44 427\nTMhelix 428 450\ninside 451 491

468	GCF_001991325.1_ASM199132v1	Burkholderia cenocepacia	Proteobacteria; Betaproteobacteria; Burkholderiales; Burkholderiaceae; Burkholderia; Burkholderia cepacia complex
	MKRRTFLALPAVAASGMLAGCSSADLGPVTHRLMKDPEYRETLSAFLISADGKKLVVIGKWP_006483583.1	MULTISPECIES: hypothetical protein [Burkholderia cepacia complex]	Length: 212\nNumber of predicted TMHs: 1\nExp number of AAs in TMHs: 28.03134\nExp number, first 60 AAs: 6.30523\nTotal prob of N-in: 0.48947\noutside 1 187\nTMhelix 188 210\ninside 211 212

469	GCF_001991325.1_ASM199132v1	Burkholderia cenocepacia	Proteobacteria; Betaproteobacteria; Burkholderiales; Burkholderiaceae; Burkholderia; Burkholderia cepacia complex
	MSDDIDNGGKGGFTRRDMMRVMAASGMMAVGGGGLLTSAQSAFAAPAPKRGGKIRVANES	WP_006482802.1	MULTISPECIES: ABC transporter substrate-binding protein [Burkholderia cepacia complex]	Length: 527\nNumber of predicted TMHs: 2\nExp number of AAs in TMHs: 36.7303599999999\nExp number, first 60 AAs: 22.08937\nTotal prob of N-in: 0.97312\nPOSSIBLE N-term signal sequence\ninside 1 20\nTMhelix 21 43\noutside 44 502\nTMhelix 503 525\ninside 526 527

470	GCF_001991315.1_ASM199131v1	Burkholderia cenocepacia	Proteobacteria; Betaproteobacteria; Burkholderiales; Burkholderiaceae; Burkholderia; Burkholderia cepacia complex
	MKRRTFLALPAVAASGMLAGCSSADLGPVTHRLMKDPEYRETLSAFLISADGKKLVVIGKWP_006483583.1	MULTISPECIES: hypothetical protein [Burkholderia cepacia complex]	Length: 212\nNumber of predicted TMHs: 1\nExp number of AAs in TMHs: 28.03134\nExp number, first 60 AAs: 6.30523\nTotal prob of N-in: 0.48947\noutside 1 187\nTMhelix 188 210\ninside 211 212

471	GCF_001984375.1_ASM198437v1	Burkholderia cenocepacia	Proteobacteria; Betaproteobacteria; Burkholderiales; Burkholderiaceae; Burkholderia; Burkholderia cepacia complex
	MSDDIDNGGKGGFTRRDMMRVMAASGMMAVGGGGLLTSAQSAFAAPAPKRGGKIRVANES	WP_006482802.1	MULTISPECIES: ABC transporter substrate-binding protein [Burkholderia cepacia complex]	Length: 527\nNumber of predicted TMHs: 2\nExp number of AAs in TMHs: 36.7303599999999\nExp number, first 60 AAs: 22.08937\nTotal prob of N-in: 0.97312\nPOSSIBLE N-term signal sequence\ninside 1 20\nTMhelix 21 43\noutside 44 502\nTMhelix 503 525\ninside 526 527

472	GCF_001606135.1_ASM160613v1	Burkholderia cenocepacia	Proteobacteria; Betaproteobacteria; Burkholderiales; Burkholderiaceae; Burkholderia; Burkholderia cepacia complex
	MKRRTFLALPAVAASGMLAGCSSADLGPVTHRLMKDPEYRETLSAFLISADGKKLVVIGKWP_034201760.1	MULTISPECIES:

hypothetical protein [Burkholderia cepacia complex] Length: 212\nNumber of predicted TMHs: 1\nExp number of AAs in TMHs: 28.11606\nExp number, first 60 AAs: 6.33522\nTotal prob of N-in: 0.49088\noutside 1 187\nTMhelix 188 210\ninside 211 212

473 GCF_001984325.1_ASM198432v1 Burkholderia cenocepacia Proteobacteria; Betaproteobacteria; Burkholderiales; Burkholderiaceae; Burkholderia; Burkholderia cepacia complex MSTSITRRRLLAAGVAASSLALSGCFTPKLYKNDAYSEHVSAFMMTEDGKKLVVLGTRYH WP_077021267.1 hypothetical protein [Burkholderia cenocepacia] Length: 212\nNumber of predicted TMHs: 1\nExp number of AAs in TMHs: 24.17376\nExp number, first 60 AAs: 0.92388\nTotal prob of N-in: 0.37951\noutside 1 180\nTMhelix 181 203\ninside 204 212

474 GCF_001984325.1_ASM198432v1 Burkholderia cenocepacia Proteobacteria; Betaproteobacteria; Burkholderiales; Burkholderiaceae; Burkholderia; Burkholderia cepacia complex METLLTRIRASFADASPRRQLISVAVLVALVASLIAAGYFALRPRYQVLFRDLKPQDAAT WP_077020516.1 flagellar M-ring protein FliF [Burkholderia cenocepacia] Length: 491\nNumber of predicted TMHs: 2\nExp number of AAs in TMHs: 41.34236\nExp number, first 60 AAs: 22.1383\nTotal prob of N-in: 0.99080\nPOSSIBLE N-term signal sequence\ninside 1 20\nTMhelix 21 43\noutside 44 427\nTMhelix 428 450\ninside 451 491

475 GCF_001071255.1_ASM107125v1 Burkholderia cenocepacia Proteobacteria; Betaproteobacteria; Burkholderiales; Burkholderiaceae; Burkholderia; Burkholderia cepacia complex MKRRTFLALPAVAASGMLAGCSSADLGPVTHRLMKDPEYQETLSAFLISADGKKLVVIGK WP_048986526.1 hypothetical protein [Burkholderia cenocepacia] Length: 212\nNumber of predicted TMHs: 1\nExp number of AAs in TMHs: 28.09653\nExp number, first 60 AAs: 6.31597\nTotal prob of N-in: 0.49053\noutside 1 187\nTMhelix 188 210\ninside 211 212

476 GCF_001984325.1_ASM198432v1 Burkholderia cenocepacia Proteobacteria; Betaproteobacteria; Burkholderiales; Burkholderiaceae; Burkholderia; Burkholderia cepacia complex MKRRTFLALPAVAASGMLAGCSSADLGPVTHRLMKDPEYRETLSAFLISADGKKLVVIGKWP_006483583.1 MULTISPECIES: hypothetical protein [Burkholderia cepacia complex] Length: 212\nNumber of predicted TMHs: 1\nExp number of AAs in TMHs: 28.03134\nExp number, first 60 AAs: 6.30523\nTotal prob of N-in: 0.48947\noutside 1 187\nTMhelix 188 210\ninside 211 212

477 GCF_001541445.1_ASM154144v1 Burkholderia cenocepacia Proteobacteria; Betaproteobacteria; Burkholderiales; Burkholderiaceae; Burkholderia; Burkholderia cepacia complex MSLSITRRRLLAAGVAASSVALSGCFTPKLYKNDAYSERVSAFMMTEDGKKLVVLGTRYH WP_050012404.1 hypothetical protein [Burkholderia cenocepacia] Length: 212\nNumber of predicted TMHs: 1\nExp number of AAs in TMHs: 28.40203\nExp number, first 60 AAs: 1.09227\nTotal prob of N-in: 0.42249\noutside 1 180\nTMhelix 181 203\ninside 204 212

478 GCF_001072455.1_ASM107245v1 Burkholderia cenocepacia Proteobacteria; Betaproteobacteria; Burkholderiales; Burkholderiaceae; Burkholderia; Burkholderia cepacia complex MSDDIDNGGKGGFTRRDMMRVMAASGMMAVGGGGLLTSAQSAFAAPAPKRGGKIRVANES WP_006482802.1 MULTISPECIES: ABC transporter substrate-binding protein [Burkholderia cepacia complex] Length: 527\nNumber of predicted TMHs: 2\nExp number of AAs in TMHs: 36.7303599999999\nExp number, first 60 AAs: 22.08937\nTotal prob of N-in: 0.97312\nPOSSIBLE N-term signal sequence\ninside 1 20\nTMhelix 21 43\noutside 44 502\nTMhelix 503 525\ninside 526 527

479 GCF_000755725.1_ASM75572v1 Burkholderia cenocepacia Proteobacteria; Betaproteobacteria; Burkholderiales; Burkholderiaceae; Burkholderia; Burkholderia cepacia complex MTISTTRRRWLGAALAASSVMLSGCFTPKLYEEGGYHEDVSGFMVTEDGKKLVVLGARYH WP_040131189.1 hypothetical protein [Burkholderia cenocepacia] Length: 213\nNumber of predicted TMHs: 1\nExp number of AAs in TMHs: 22.93041\nExp number, first 60 AAs: 0.53063\nTotal prob of N-in: 0.73121\ninside 1 180\nTMhelix 181 203\noutside 204 213

480 GCF_001059305.1_ASM105930v1 Burkholderia cenocepacia Proteobacteria; Betaproteobacteria; Burkholderiales; Burkholderiaceae; Burkholderia; Burkholderia cepacia complex MKRRTFLALPAVAASGMLAGCSSADLGPVTHRLMKDPEYQETLSAFLISADGKKLVVIGK WP_048986526.1 hypothetical protein [Burkholderia cenocepacia] Length: 212\nNumber of predicted TMHs: 1\nExp number of AAs in TMHs: 28.09653\nExp number, first 60 AAs: 6.31597\nTotal prob of N-in: 0.49053\noutside 1 187\nTMhelix 188 210\ninside 211 212

481	GCF_001056435.1_ASM105643v1	Burkholderia cenocepacia	Proteobacteria; Betaproteobacteria; Burkholderiales; Burkholderiaceae; Burkholderia; Burkholderia cepacia complex
	MSDDIDNGGKGGFTRRDMMRVMAASGMMAVGGGGLLTSAQSAFAAPAPKRGGKIRVANES	WP_049106171.1	ABC transporter substrate-binding protein [Burkholderia cenocepacia]	Length: 527\nNumber of predicted TMHs:  2\nExp number of AAs in TMHs: 36.7303599999999\nExp number, first 60 AAs:  22.08937\nTotal prob of N-in:  0.97312\nPOSSIBLE N-term signal sequence\ninside   1   20\nTMhelix   21   43\noutside   44   502\nTMhelix   503   525\ninside   526   527

482	GCF_001056435.1_ASM105643v1	Burkholderia cenocepacia	Proteobacteria; Betaproteobacteria; Burkholderiales; Burkholderiaceae; Burkholderia; Burkholderia cepacia complex
	MKRRTFLALPAVAASGMLAGCSSADLGPVTHRLMKDPEYQETLSAFLISADGKKLVVIGK	WP_048986526.1	hypothetical protein [Burkholderia cenocepacia]	Length: 212\nNumber of predicted TMHs:  1\nExp number of AAs in TMHs: 28.09653\nExp number, first 60 AAs:  6.31597\nTotal prob of N-in:  0.49053\noutside   1   187\nTMhelix   188   210\ninside   211   212

483	GCF_001059305.1_ASM105930v1	Burkholderia cenocepacia	Proteobacteria; Betaproteobacteria; Burkholderiales; Burkholderiaceae; Burkholderia; Burkholderia cepacia complex
	MSDDIDNGGKGGFTRRDMMRVMAASGMMAVGGGGLLTSAQSAFAAPAPKRGGKIRVANES	WP_006482802.1	MULTISPECIES: ABC transporter substrate-binding protein [Burkholderia cepacia complex]	Length: 527\nNumber of predicted TMHs:  2\nExp number of AAs in TMHs: 36.7303599999999\nExp number, first 60 AAs:  22.08937\nTotal prob of N-in:  0.97312\nPOSSIBLE N-term signal sequence\ninside   1   20\nTMhelix   21   43\noutside   44   502\nTMhelix   503   525\ninside   526   527

484	GCF_001991415.1_ASM199141v1	Burkholderia cenocepacia	Proteobacteria; Betaproteobacteria; Burkholderiales; Burkholderiaceae; Burkholderia; Burkholderia cepacia complex
	MSTSITRRRLLAAGVAASSLALSGCFTPKLYKNDAYSEHVSAFMMTEDGKKLVVLGTRYH	WP_077021267.1	hypothetical protein [Burkholderia cenocepacia]	Length: 212\nNumber of predicted TMHs:  1\nExp number of AAs in TMHs: 24.17376\nExp number, first 60 AAs:  0.92388\nTotal prob of N-in:  0.37951\noutside   1   180\nTMhelix   181   203\ninside   204   212

485	GCF_002071955.1_ASM207195v1	Burkholderia cenocepacia	Proteobacteria; Betaproteobacteria; Burkholderiales; Burkholderiaceae; Burkholderia; Burkholderia cepacia complex
	MKRRTFLALPAVAASGMLAGCSSADLGPVTHRLMKDPEYRETLSAFLISADGKKLVVIGKWP_062884514.1	hypothetical protein [Burkholderia cenocepacia]	Length: 212\nNumber of predicted TMHs:  1\nExp number of AAs in TMHs: 28.03134\nExp number, first 60 AAs:  6.30523\nTotal prob of N-in:  0.48947\noutside   1   187\nTMhelix   188   210\ninside   211   212

486	GCF_002071955.1_ASM207195v1	Burkholderia cenocepacia	Proteobacteria; Betaproteobacteria; Burkholderiales; Burkholderiaceae; Burkholderia; Burkholderia cepacia complex
	METLLTRIRASFADASPRRQLISVAVLVALVASLIAAGYFALRPRYQVLFRDLKPQDAAT	WP_060214257.1	flagellar M-ring protein FliF [Burkholderia cenocepacia]	Length: 491\nNumber of predicted TMHs:  2\nExp number of AAs in TMHs: 41.34235\nExp number, first 60 AAs:  22.1383\nTotal prob of N-in:  0.99080\nPOSSIBLE N-term signal sequence\ninside   1   20\nTMhelix   21   43\noutside   44   427\nTMhelix   428   450\ninside   451   491

487	GCF_001994655.1_ASM199465v1	Burkholderia cenocepacia	Proteobacteria; Betaproteobacteria; Burkholderiales; Burkholderiaceae; Burkholderia; Burkholderia cepacia complex
	MSTSITRRRLLAAGVAASSLALSGCFTPKLYKNDAYSEHVSAFMMTEDGKKLVVLGTRYH	WP_077021267.1	hypothetical protein [Burkholderia cenocepacia]	Length: 212\nNumber of predicted TMHs:  1\nExp number of AAs in TMHs: 24.17376\nExp number, first 60 AAs:  0.92388\nTotal prob of N-in:  0.37951\noutside   1   180\nTMhelix   181   203\ninside   204   212

488	GCF_002071885.1_ASM207188v1	Burkholderia cenocepacia	Proteobacteria; Betaproteobacteria; Burkholderiales; Burkholderiaceae; Burkholderia; Burkholderia cepacia complex
	MSDDIDNGGKGGFTRRDMMRVMAASGMMAVGGGGLLTSAQSAFAAPAPKRGGKIRVANES	WP_006482802.1	MULTISPECIES: ABC transporter substrate-binding protein [Burkholderia cepacia complex]	Length: 527\nNumber of predicted TMHs:  2\nExp number of AAs in TMHs: 36.7303599999999\nExp number, first 60 AAs:  22.08937\nTotal prob of N-in:  0.97312\nPOSSIBLE N-term signal sequence\ninside   1   20\nTMhelix   21   43\noutside   44   502\nTMhelix   503   525\ninside   526   527

489	GCF_001994655.1_ASM199465v1	Burkholderia cenocepacia	Proteobacteria; Betaproteobacteria; Burkholderiales; Burkholderiaceae; Burkholderia; Burkholderia cepacia complex

METLLTRIRASFADASPRRQLISVAVLVALVASLIAAGYFALRPRYQVLFRDLKPQDAAT WP_077020516.1 flagellar M-ring protein FliF [Burkholderia cenocepacia] Length: 491\nNumber of predicted TMHs: 2\nExp number of AAs in TMHs: 41.34236\nExp number, first 60 AAs: 22.1383\nTotal prob of N-in: 0.99080\nPOSSIBLE N-term signal sequence\ninside 1 20\nTMhelix 21 43\noutside 44 427\nTMhelix 428 450\ninside 451 491

490 GCF_001994695.1_ASM199469v1 Burkholderia cenocepacia Proteobacteria; Betaproteobacteria; Burkholderiales; Burkholderiaceae; Burkholderia; Burkholderia cepacia complex
MSDDIDNGGKGGFTRRDMMRVMAASGMMAVGGGGLLTSAQSAFAAPAPKRGGKIRVANES WP_077187164.1 ABC transporter substrate-binding protein [Burkholderia cenocepacia] Length: 527\nNumber of predicted TMHs: 2\nExp number of AAs in TMHs: 36.7300399999999\nExp number, first 60 AAs: 22.0894\nTotal prob of N-in: 0.97312\nPOSSIBLE N-term signal sequence\ninside 1 20\nTMhelix 21 43\noutside 44 502\nTMhelix 503 525\ninside 526 527

491 GCF_001994695.1_ASM199469v1 Burkholderia cenocepacia Proteobacteria; Betaproteobacteria; Burkholderiales; Burkholderiaceae; Burkholderia; Burkholderia cepacia complex
MSTSITRRRLLAAGVAASSLALSGCFTPKLYKNDAYSEHVSAFMMTEDGKKLVVLGTRYH WP_077021267.1
hypothetical protein [Burkholderia cenocepacia] Length: 212\nNumber of predicted TMHs: 1\nExp number of AAs in TMHs: 24.17376\nExp number, first 60 AAs: 0.92388\nTotal prob of N-in: 0.37951\noutside 1 180\nTMhelix 181 203\ninside 204 212

492 GCF_001994695.1_ASM199469v1 Burkholderia cenocepacia Proteobacteria; Betaproteobacteria; Burkholderiales; Burkholderiaceae; Burkholderia; Burkholderia cepacia complex
METLLTRIRASFADASPRRQLISVAVLVALVASLIAAGYFALRPRYQVLFRDLKPQDAAT WP_077020516.1 flagellar M-ring protein FliF [Burkholderia cenocepacia] Length: 491\nNumber of predicted TMHs: 2\nExp number of AAs in TMHs: 41.34236\nExp number, first 60 AAs: 22.1383\nTotal prob of N-in: 0.99080\nPOSSIBLE N-term signal sequence\ninside 1 20\nTMhelix 21 43\noutside 44 427\nTMhelix 428 450\ninside 451 491

493 GCF_001994735.1_ASM199473v1 Burkholderia cenocepacia Proteobacteria; Betaproteobacteria; Burkholderiales; Burkholderiaceae; Burkholderia; Burkholderia cepacia complex
MKRRTFLALPAVAASGMLAGCSSADLGPVTHRLMKDPEYRETLSAFLISADGKKLVVIGKWP_006483583.1 MULTISPECIES: hypothetical protein [Burkholderia cepacia complex] Length: 212\nNumber of predicted TMHs: 1\nExp number of AAs in TMHs: 28.03134\nExp number, first 60 AAs: 6.30523\nTotal prob of N-in: 0.48947\noutside 1 187\nTMhelix 188 210\ninside 211 212

494 GCF_001994735.1_ASM199473v1 Burkholderia cenocepacia Proteobacteria; Betaproteobacteria; Burkholderiales; Burkholderiaceae; Burkholderia; Burkholderia cepacia complex
MSDDIDNGGKGGFTRRDMMRVMAASGMMAVGGGGLLTSAQSAFAAPAPKRGGKIRVANES WP_006482802.1
MULTISPECIES: ABC transporter substrate-binding protein [Burkholderia cepacia complex] Length: 527\nNumber of predicted TMHs: 2\nExp number of AAs in TMHs: 36.7303599999999\nExp number, first 60 AAs: 22.08937\nTotal prob of N-in: 0.97312\nPOSSIBLE N-term signal sequence\ninside 1 20\nTMhelix 21 43\noutside 44 502\nTMhelix 503 525\ninside 526 527

495 GCF_001994695.1_ASM199469v1 Burkholderia cenocepacia Proteobacteria; Betaproteobacteria; Burkholderiales; Burkholderiaceae; Burkholderia; Burkholderia cepacia complex
MKRRTFLALPAVAASGMLAGCSSADLGPVTHRLMKDPEYRETLSAFLISADGKKLVVIGKWP_006483583.1 MULTISPECIES: hypothetical protein [Burkholderia cepacia complex] Length: 212\nNumber of predicted TMHs: 1\nExp number of AAs in TMHs: 28.03134\nExp number, first 60 AAs: 6.30523\nTotal prob of N-in: 0.48947\noutside 1 187\nTMhelix 188 210\ninside 211 212

496 GCF_001994655.1_ASM199465v1 Burkholderia cenocepacia Proteobacteria; Betaproteobacteria; Burkholderiales; Burkholderiaceae; Burkholderia; Burkholderia cepacia complex
MKRRTFLALPAVAASGMLAGCSSADLGPVTHRLMKDPEYRETLSAFLISADGKKLVVIGKWP_006483583.1 MULTISPECIES: hypothetical protein [Burkholderia cepacia complex] Length: 212\nNumber of predicted TMHs: 1\nExp number of AAs in TMHs: 28.03134\nExp number, first 60 AAs: 6.30523\nTotal prob of N-in: 0.48947\noutside 1 187\nTMhelix 188 210\ninside 211 212

497 GCF_001994655.1_ASM199465v1 Burkholderia cenocepacia Proteobacteria; Betaproteobacteria; Burkholderiales; Burkholderiaceae; Burkholderia; Burkholderia cepacia complex
MSDDIDNGGKGGFTRRDMMRVMAASGMMAVGGGGLLTSAQSAFAAPAPKRGGKIRVANES WP_006482802.1
MULTISPECIES: ABC transporter substrate-binding protein [Burkholderia cepacia complex] Length: 527\nNumber of predicted TMHs: 2\nExp number of AAs in TMHs: 36.7303599999999\nExp number, first 60 AAs:

22.08937\nTotal prob of N-in: 0.97312\nPOSSIBLE N-term signal sequence\ninside 1 20\nTMhelix 21 43\noutside 44 502\nTMhelix 503 525\ninside 526 527

498 GCF_001994615.1_ASM199461v1 Burkholderia cenocepacia Proteobacteria; Betaproteobacteria; Burkholderiales; Burkholderiaceae; Burkholderia; Burkholderia cepacia complex MKRRTFLALPAVAASGMLAGCSSADLGPVTHRLMKDPEYRETLSAFLISADGKKLVVIGKWP_006483583.1 MULTISPECIES: hypothetical protein [Burkholderia cepacia complex] Length: 212\nNumber of predicted TMHs: 1\nExp number of AAs in TMHs: 28.03134\nExp number, first 60 AAs: 6.30523\nTotal prob of N-in: 0.48947\noutside 1 187\nTMhelix 188 210\ninside 211 212

499 GCF_001994615.1_ASM199461v1 Burkholderia cenocepacia Proteobacteria; Betaproteobacteria; Burkholderiales; Burkholderiaceae; Burkholderia; Burkholderia cepacia complex MSDDIDNGGKGGFTRRDMMRVMAASGMMAVGGGGLLTSAQSAFAAPAPKRGGKIRVANES WP_006482802.1 MULTISPECIES: ABC transporter substrate-binding protein [Burkholderia cepacia complex] Length: 527\nNumber of predicted TMHs: 2\nExp number of AAs in TMHs: 36.7303599999999\nExp number, first 60 AAs: 22.08937\nTotal prob of N-in: 0.97312\nPOSSIBLE N-term signal sequence\ninside 1 20\nTMhelix 21 43\noutside 44 502\nTMhelix 503 525\ninside 526 527

500 GCF_001994375.1_ASM199437v1 Burkholderia cenocepacia Proteobacteria; Betaproteobacteria; Burkholderiales; Burkholderiaceae; Burkholderia; Burkholderia cepacia complex MKRRTFLALPAVAASGMLAGCSSADLGPVTHRLMKDPEYRETLSAFLISADGKKLVVIGKWP_034201760.1 MULTISPECIES: hypothetical protein [Burkholderia cepacia complex] Length: 212\nNumber of predicted TMHs: 1\nExp number of AAs in TMHs: 28.11606\nExp number, first 60 AAs: 6.33522\nTotal prob of N-in: 0.49088\noutside 1 187\nTMhelix 188 210\ninside 211 212

501 GCF_001994335.1_ASM199433v1 Burkholderia cenocepacia Proteobacteria; Betaproteobacteria; Burkholderiales; Burkholderiaceae; Burkholderia; Burkholderia cepacia complex MKRRTFLALPAVAASGMLAGCSSADLGPVTHRLMKDPEYRETLSAFLISADGKKLVVIGKWP_034201760.1 MULTISPECIES: hypothetical protein [Burkholderia cepacia complex] Length: 212\nNumber of predicted TMHs: 1\nExp number of AAs in TMHs: 28.11606\nExp number, first 60 AAs: 6.33522\nTotal prob of N-in: 0.49088\noutside 1 187\nTMhelix 188 210\ninside 211 212

502 GCF_001994405.1_ASM199440v1 Burkholderia cenocepacia Proteobacteria; Betaproteobacteria; Burkholderiales; Burkholderiaceae; Burkholderia; Burkholderia cepacia complex MSDDIDNGGKGGFTRRDMMRVMAASGMMAVGGGGLLTSAQSAFAAPAPKRGGKIRVANES WP_006482802.1 MULTISPECIES: ABC transporter substrate-binding protein [Burkholderia cepacia complex] Length: 527\nNumber of predicted TMHs: 2\nExp number of AAs in TMHs: 36.7303599999999\nExp number, first 60 AAs: 22.08937\nTotal prob of N-in: 0.97312\nPOSSIBLE N-term signal sequence\ninside 1 20\nTMhelix 21 43\noutside 44 502\nTMhelix 503 525\ninside 526 527

503 GCF_001994375.1_ASM199437v1 Burkholderia cenocepacia Proteobacteria; Betaproteobacteria; Burkholderiales; Burkholderiaceae; Burkholderia; Burkholderia cepacia complex MSDDIDNGGKGGFTRRDMMRVMAASGMMAVGGGGLLTSAQSAFAAPAPKRGGKIRVANES WP_006482802.1 MULTISPECIES: ABC transporter substrate-binding protein [Burkholderia cepacia complex] Length: 527\nNumber of predicted TMHs: 2\nExp number of AAs in TMHs: 36.7303599999999\nExp number, first 60 AAs: 22.08937\nTotal prob of N-in: 0.97312\nPOSSIBLE N-term signal sequence\ninside 1 20\nTMhelix 21 43\noutside 44 502\nTMhelix 503 525\ninside 526 527

504 GCF_001994335.1_ASM199433v1 Burkholderia cenocepacia Proteobacteria; Betaproteobacteria; Burkholderiales; Burkholderiaceae; Burkholderia; Burkholderia cepacia complex MSDDIDNGGKGGFTRRDMMRVMAASGMMAVGGGGLLTSAQSAFAAPAPKRGGKIRVANES WP_006482802.1 MULTISPECIES: ABC transporter substrate-binding protein [Burkholderia cepacia complex] Length: 527\nNumber of predicted TMHs: 2\nExp number of AAs in TMHs: 36.7303599999999\nExp number, first 60 AAs: 22.08937\nTotal prob of N-in: 0.97312\nPOSSIBLE N-term signal sequence\ninside 1 20\nTMhelix 21 43\noutside 44 502\nTMhelix 503 525\ninside 526 527

505 GCF_001994285.1_ASM199428v1 Burkholderia cenocepacia Proteobacteria; Betaproteobacteria; Burkholderiales; Burkholderiaceae; Burkholderia; Burkholderia cepacia complex MSDDIDNGGKGGFTRRDMMRVMAASGMMAVGGGGLLTSAQSAFAAPAPKRGGKIRVANES WP_006482802.1 MULTISPECIES: ABC transporter substrate-binding protein [Burkholderia cepacia complex] Length: 527\nNumber of predicted TMHs: 2\nExp number of AAs in TMHs: 36.7303599999999\nExp number, first 60 AAs:

22.08937\nTotal prob of N-in: 0.97312\nPOSSIBLE N-term signal sequence\ninside 1 20\nTMhelix 21 43\noutside 44 502\nTMhelix 503 525\ninside 526 527

506 GCF_001994195.1_ASM199419v1 Burkholderia cenocepacia Proteobacteria; Betaproteobacteria; Burkholderiales; Burkholderiaceae; Burkholderia; Burkholderia cepacia complex
MKRRTFLALPAVAASGMLAGCSSADLGPVTHRLMKDPEYRETLSAFLISADGKKLVVIGKWP_034201760.1 MULTISPECIES: hypothetical protein [Burkholderia cepacia complex] Length: 212\nNumber of predicted TMHs: 1\nExp number of AAs in TMHs: 28.11606\nExp number, first 60 AAs: 6.33522\nTotal prob of N-in: 0.49088\noutside 1 187\nTMhelix 188 210\ninside 211 212

507 GCF_001993825.1_ASM199382v1 Burkholderia cenocepacia Proteobacteria; Betaproteobacteria; Burkholderiales; Burkholderiaceae; Burkholderia; Burkholderia cepacia complex
MKRRTFLALPAVAASGMLAGCSSADLGPVTHRLMKDPEYRETLSAFLISADGKKLVVIGKWP_034201760.1 MULTISPECIES: hypothetical protein [Burkholderia cepacia complex] Length: 212\nNumber of predicted TMHs: 1\nExp number of AAs in TMHs: 28.11606\nExp number, first 60 AAs: 6.33522\nTotal prob of N-in: 0.49088\noutside 1 187\nTMhelix 188 210\ninside 211 212

508 GCF_001993925.1_ASM199392v1 Burkholderia cenocepacia Proteobacteria; Betaproteobacteria; Burkholderiales; Burkholderiaceae; Burkholderia; Burkholderia cepacia complex
MSDDIDNGGKGGFTRRDMMRVMAASGMMAVGGGGLLTSAQSAFAAPAPKRGGKIRVANES WP_006482802.1
MULTISPECIES: ABC transporter substrate-binding protein [Burkholderia cepacia complex] Length: 527\nNumber of predicted TMHs: 2\nExp number of AAs in TMHs: 36.7303599999999\nExp number, first 60 AAs: 22.08937\nTotal prob of N-in: 0.97312\nPOSSIBLE N-term signal sequence\ninside 1 20\nTMhelix 21 43\noutside 44 502\nTMhelix 503 525\ninside 526 527

509 GCF_001993895.1_ASM199389v1 Burkholderia cenocepacia Proteobacteria; Betaproteobacteria; Burkholderiales; Burkholderiaceae; Burkholderia; Burkholderia cepacia complex
MSDDIDNGGKGGFTRRDMMRVMAASGMMAVGGGGLLTSAQSAFAAPAPKRGGKIRVANES WP_006482802.1
MULTISPECIES: ABC transporter substrate-binding protein [Burkholderia cepacia complex] Length: 527\nNumber of predicted TMHs: 2\nExp number of AAs in TMHs: 36.7303599999999\nExp number, first 60 AAs: 22.08937\nTotal prob of N-in: 0.97312\nPOSSIBLE N-term signal sequence\ninside 1 20\nTMhelix 21 43\noutside 44 502\nTMhelix 503 525\ninside 526 527

510 GCF_001993855.1_ASM199385v1 Burkholderia cenocepacia Proteobacteria; Betaproteobacteria; Burkholderiales; Burkholderiaceae; Burkholderia; Burkholderia cepacia complex
MSDDIDNGGKGGFTRRDMMRVMAASGMMAVGGGGLLTSAQSAFAAPAPKRGGKIRVANES WP_006482802.1
MULTISPECIES: ABC transporter substrate-binding protein [Burkholderia cepacia complex] Length: 527\nNumber of predicted TMHs: 2\nExp number of AAs in TMHs: 36.7303599999999\nExp number, first 60 AAs: 22.08937\nTotal prob of N-in: 0.97312\nPOSSIBLE N-term signal sequence\ninside 1 20\nTMhelix 21 43\noutside 44 502\nTMhelix 503 525\ninside 526 527

511 GCF_001993795.1_ASM199379v1 Burkholderia cenocepacia Proteobacteria; Betaproteobacteria; Burkholderiales; Burkholderiaceae; Burkholderia; Burkholderia cepacia complex
MKRRTFLALPAVAASGMLAGCSSADLGPVTHRLMKDPEYRETLSAFLISADGKKLVVIGKWP_006483583.1 MULTISPECIES: hypothetical protein [Burkholderia cepacia complex] Length: 212\nNumber of predicted TMHs: 1\nExp number of AAs in TMHs: 28.03134\nExp number, first 60 AAs: 6.30523\nTotal prob of N-in: 0.48947\noutside 1 187\nTMhelix 188 210\ninside 211 212

512 GCF_001993495.1_ASM199349v1 Burkholderia cenocepacia Proteobacteria; Betaproteobacteria; Burkholderiales; Burkholderiaceae; Burkholderia; Burkholderia cepacia complex
MSTSITRRRLLAAGVAASSLALSGCFTPKLYKNDAYSEHVSAFMMTEDGKKLVVLGTRYH WP_077021267.1
hypothetical protein [Burkholderia cenocepacia] Length: 212\nNumber of predicted TMHs: 1\nExp number of AAs in TMHs: 24.17376\nExp number, first 60 AAs: 0.92388\nTotal prob of N-in: 0.37951\noutside 1 180\nTMhelix 181 203\ninside 204 212

513 GCF_001993825.1_ASM199382v1 Burkholderia cenocepacia Proteobacteria; Betaproteobacteria; Burkholderiales; Burkholderiaceae; Burkholderia; Burkholderia cepacia complex
MSDDIDNGGKGGFTRRDMMRVMAASGMMAVGGGGLLTSAQSAFAAPAPKRGGKIRVANES WP_006482802.1
MULTISPECIES: ABC transporter substrate-binding protein [Burkholderia cepacia complex] Length: 527\nNumber of predicted TMHs: 2\nExp number of AAs in TMHs: 36.7303599999999\nExp number, first 60 AAs:

22.08937\nTotal prob of N-in: 0.97312\nPOSSIBLE N-term signal sequence\ninside 1 20\nTMhelix 21 43\noutside 44 502\nTMhelix 503 525\ninside 526 527

514 GCF_001993255.1_ASM199325v1 Burkholderia cenocepacia Proteobacteria; Betaproteobacteria; Burkholderiales; Burkholderiaceae; Burkholderia; Burkholderia cepacia complex MSTSITRRRLLAAGVAASSLALSGCFTPKLYKNDAYSEHVSAFMMTEDGKKLVVLGTRYH WP_077239688.1 hypothetical protein [Burkholderia cenocepacia] Length: 212\nNumber of predicted TMHs: 1\nExp number of AAs in TMHs: 24.13365\nExp number, first 60 AAs: 0.92254\nTotal prob of N-in: 0.37835\noutside 1 180\nTMhelix 181 203\ninside 204 212

515 GCF_001993255.1_ASM199325v1 Burkholderia cenocepacia Proteobacteria; Betaproteobacteria; Burkholderiales; Burkholderiaceae; Burkholderia; Burkholderia cepacia complex MSDDIDNGGKGGFTRRDMMRVMAASGMMAVGGGGLLTSAQSAFAAPAPKRGGKIRVANES WP_077187164.1 ABC transporter substrate-binding protein [Burkholderia cenocepacia] Length: 527\nNumber of predicted TMHs: 2\nExp number of AAs in TMHs: 36.7300399999999\nExp number, first 60 AAs: 22.0894\nTotal prob of N-in: 0.97312\nPOSSIBLE N-term signal sequence\ninside 1 20\nTMhelix 21 43\noutside 44 502\nTMhelix 503 525\ninside 526 527

516 GCF_001993335.1_ASM199333v1 Burkholderia cenocepacia Proteobacteria; Betaproteobacteria; Burkholderiales; Burkholderiaceae; Burkholderia; Burkholderia cepacia complex MSDDIDNGGKGGFTRRDMMRVMAASGMMAVGGGGLLTSAQSAFAAPAPKRGGKIRVANES WP_077187164.1 ABC transporter substrate-binding protein [Burkholderia cenocepacia] Length: 527\nNumber of predicted TMHs: 2\nExp number of AAs in TMHs: 36.7300399999999\nExp number, first 60 AAs: 22.0894\nTotal prob of N-in: 0.97312\nPOSSIBLE N-term signal sequence\ninside 1 20\nTMhelix 21 43\noutside 44 502\nTMhelix 503 525\ninside 526 527

517 GCF_001993335.1_ASM199333v1 Burkholderia cenocepacia Proteobacteria; Betaproteobacteria; Burkholderiales; Burkholderiaceae; Burkholderia; Burkholderia cepacia complex MSTSITRRRLLAAGVAASSLALSGCFTPKLYKNDAYSEHVSAFMMTEDGKKLVVLGTRYH WP_077021267.1 hypothetical protein [Burkholderia cenocepacia] Length: 212\nNumber of predicted TMHs: 1\nExp number of AAs in TMHs: 24.17376\nExp number, first 60 AAs: 0.92388\nTotal prob of N-in: 0.37951\noutside 1 180\nTMhelix 181 203\ninside 204 212

518 GCF_001993335.1_ASM199333v1 Burkholderia cenocepacia Proteobacteria; Betaproteobacteria; Burkholderiales; Burkholderiaceae; Burkholderia; Burkholderia cepacia complex METLLTRIRASFADASPRRQLISVAVLVALVASLIAAGYFALRPRYQVLFRDLKPQDAAT WP_077020516.1 flagellar M-ring protein FliF [Burkholderia cenocepacia] Length: 491\nNumber of predicted TMHs: 2\nExp number of AAs in TMHs: 41.34236\nExp number, first 60 AAs: 22.1383\nTotal prob of N-in: 0.99080\nPOSSIBLE N-term signal sequence\ninside 1 20\nTMhelix 21 43\noutside 44 427\nTMhelix 428 450\ninside 451 491

519 GCF_001993295.1_ASM199329v1 Burkholderia cenocepacia Proteobacteria; Betaproteobacteria; Burkholderiales; Burkholderiaceae; Burkholderia; Burkholderia cepacia complex MSDDIDNGGKGGFTRRDMMRVMAASGMMAVGGGGLLTSAQSAFAAPAPKRGGKIRVANES WP_077187164.1 ABC transporter substrate-binding protein [Burkholderia cenocepacia] Length: 527\nNumber of predicted TMHs: 2\nExp number of AAs in TMHs: 36.7300399999999\nExp number, first 60 AAs: 22.0894\nTotal prob of N-in: 0.97312\nPOSSIBLE N-term signal sequence\ninside 1 20\nTMhelix 21 43\noutside 44 502\nTMhelix 503 525\ninside 526 527

520 GCF_001993295.1_ASM199329v1 Burkholderia cenocepacia Proteobacteria; Betaproteobacteria; Burkholderiales; Burkholderiaceae; Burkholderia; Burkholderia cepacia complex MSTSITRRRLLAAGVAASSLALSGCFTPKLYKNDAYSEHVSAFMMTEDGKKLVVLGTRYH WP_077021267.1 hypothetical protein [Burkholderia cenocepacia] Length: 212\nNumber of predicted TMHs: 1\nExp number of AAs in TMHs: 24.17376\nExp number, first 60 AAs: 0.92388\nTotal prob of N-in: 0.37951\noutside 1 180\nTMhelix 181 203\ninside 204 212

521 GCF_001993435.1_ASM199343v1 Burkholderia cenocepacia Proteobacteria; Betaproteobacteria; Burkholderiales; Burkholderiaceae; Burkholderia; Burkholderia cepacia complex MKRRTFLALPAVAASGMLAGCSSADLGPVTHRLMKDPEYRETLSAFLISADGKKLVVIGKWP_006483583.1 MULTISPECIES: hypothetical protein [Burkholderia cepacia complex] Length: 212\nNumber of predicted TMHs: 1\nExp number of AAs in TMHs: 28.03134\nExp number, first 60 AAs: 6.30523\nTotal prob of N-in: 0.48947\noutside 1 187\nTMhelix 188 210\ninside 211 212

522 GCF_001993435.1_ASM199343v1 Burkholderia cenocepacia Proteobacteria; Betaproteobacteria; Burkholderiales; Burkholderiaceae; Burkholderia; Burkholderia cepacia complex

MSDDIDNGGKGGFTRRDMMRVMAASGMMAVGGGGLLTSAQSAFAAPAPKRGGKIRVANES WP_006482802.1 MULTISPECIES: ABC transporter substrate-binding protein [Burkholderia cepacia complex] Length: 527\nNumber of predicted TMHs: 2\nExp number of AAs in TMHs: 36.7303599999999\nExp number, first 60 AAs: 22.08937\nTotal prob of N-in: 0.97312\nPOSSIBLE N-term signal sequence\ninside 1 20\nTMhelix 21 43\noutside 44 502\nTMhelix 503 525\ninside 526 527

523 GCF_001993395.1_ASM199339v1 Burkholderia cenocepacia Proteobacteria; Betaproteobacteria; Burkholderiales; Burkholderiaceae; Burkholderia; Burkholderia cepacia complex MSDDIDNGGKGGFTRRDMMRVMAASGMMAVGGGGLLTSAQSAFAAPAPKRGGKIRVANES WP_006482802.1 MULTISPECIES: ABC transporter substrate-binding protein [Burkholderia cepacia complex] Length: 527\nNumber of predicted TMHs: 2\nExp number of AAs in TMHs: 36.7303599999999\nExp number, first 60 AAs: 22.08937\nTotal prob of N-in: 0.97312\nPOSSIBLE N-term signal sequence\ninside 1 20\nTMhelix 21 43\noutside 44 502\nTMhelix 503 525\ninside 526 527

524 GCF_001993335.1_ASM199333v1 Burkholderia cenocepacia Proteobacteria; Betaproteobacteria; Burkholderiales; Burkholderiaceae; Burkholderia; Burkholderia cepacia complex MKRRTFLALPAVAASGMLAGCSSADLGPVTHRLMKDPEYRETLSAFLISADGKKLVVIGKWP_006483583.1 MULTISPECIES: hypothetical protein [Burkholderia cepacia complex] Length: 212\nNumber of predicted TMHs: 1\nExp number of AAs in TMHs: 28.03134\nExp number, first 60 AAs: 6.30523\nTotal prob of N-in: 0.48947\noutside 1 187\nTMhelix 188 210\ninside 211 212

525 GCF_001992925.1_ASM199292v1 Burkholderia cenocepacia Proteobacteria; Betaproteobacteria; Burkholderiales; Burkholderiaceae; Burkholderia; Burkholderia cepacia complex MKRRTFLALPAVAASGMLAGCSSADLGPVTHRLMKAPEYRETLSAFLISADGKKLVVIGKWP_077234160.1 5-formyltetrahydrofolate cyclo-ligase [Burkholderia cenocepacia] Length: 212\nNumber of predicted TMHs: 1\nExp number of AAs in TMHs: 28.26337\nExp number, first 60 AAs: 6.55359\nTotal prob of N-in: 0.49573\noutside 1 187\nTMhelix 188 210\ninside 211 212

526 GCF_001992925.1_ASM199292v1 Burkholderia cenocepacia Proteobacteria; Betaproteobacteria; Burkholderiales; Burkholderiaceae; Burkholderia; Burkholderia cepacia complex MSDDIDNGGKGGFTRRDMMRVMAASGMMAVGGGGLLTSAQSAFAAPAPKRGGKIRVANES WP_077199690.1 ABC transporter substrate-binding protein [Burkholderia cenocepacia] Length: 527\nNumber of predicted TMHs: 2\nExp number of AAs in TMHs: 36.77039\nExp number, first 60 AAs: 22.08962\nTotal prob of N-in: 0.97295\nPOSSIBLE N-term signal sequence\ninside 1 20\nTMhelix 21 43\noutside 44 502\nTMhelix 503 525\ninside 526 527

527 GCF_001993295.1_ASM199329v1 Burkholderia cenocepacia Proteobacteria; Betaproteobacteria; Burkholderiales; Burkholderiaceae; Burkholderia; Burkholderia cepacia complex MKRRTFLALPAVAASGMLAGCSSADLGPVTHRLMKDPEYRETLSAFLISADGKKLVVIGKWP_006483583.1 MULTISPECIES: hypothetical protein [Burkholderia cepacia complex] Length: 212\nNumber of predicted TMHs: 1\nExp number of AAs in TMHs: 28.03134\nExp number, first 60 AAs: 6.30523\nTotal prob of N-in: 0.48947\noutside 1 187\nTMhelix 188 210\ninside 211 212

528 GCF_001993055.1_ASM199305v1 Burkholderia cenocepacia Proteobacteria; Betaproteobacteria; Burkholderiales; Burkholderiaceae; Burkholderia; Burkholderia cepacia complex MKRRTFLALPAVAASGMLAGCSSADLGPVTHRLMKDPEYRETLSAFLISADGKKLVVIGKWP_034201760.1 MULTISPECIES: hypothetical protein [Burkholderia cepacia complex] Length: 212\nNumber of predicted TMHs: 1\nExp number of AAs in TMHs: 28.11606\nExp number, first 60 AAs: 6.33522\nTotal prob of N-in: 0.49088\noutside 1 187\nTMhelix 188 210\ninside 211 212

529 GCF_001993075.1_ASM199307v1 Burkholderia cenocepacia Proteobacteria; Betaproteobacteria; Burkholderiales; Burkholderiaceae; Burkholderia; Burkholderia cepacia complex MSDDIDNGGKGGFTRRDMMRVMAASGMMAVGGGGLLTSAQSAFAAPAPKRGGKIRVANES WP_006482802.1 MULTISPECIES: ABC transporter substrate-binding protein [Burkholderia cepacia complex] Length: 527\nNumber of predicted TMHs: 2\nExp number of AAs in TMHs: 36.7303599999999\nExp number, first 60 AAs: 22.08937\nTotal prob of N-in: 0.97312\nPOSSIBLE N-term signal sequence\ninside 1 20\nTMhelix 21 43\noutside 44 502\nTMhelix 503 525\ninside 526 527

530 GCF_001993005.1_ASM199300v1 Burkholderia cenocepacia Proteobacteria; Betaproteobacteria; Burkholderiales; Burkholderiaceae; Burkholderia; Burkholderia cepacia complex MKRRTFLALPAVAASGMLAGCSSADLGPVTHRLMKDPEYRETLSAFLISADGKKLVVIGKWP_006483583.1 MULTISPECIES: hypothetical protein [Burkholderia cepacia complex] Length: 212\nNumber of predicted TMHs: 1\nExp number of

AAs in TMHs: 28.03134\nExp number, first 60 AAs: 6.30523\nTotal prob of N-in: 0.48947\noutside 1 187\nTMhelix 188 210\ninside 211 212

531 GCF_001992965.1_ASM199296v1 Burkholderia cenocepacia Proteobacteria; Betaproteobacteria; Burkholderiales; Burkholderiaceae; Burkholderia; Burkholderia cepacia complex MKRRTFLALPAVAASGMLAGCSSADLGPVTHRLMKDPEYRETLSAFLISADGKKLVVIGKWP_006483583.1 MULTISPECIES: hypothetical protein [Burkholderia cepacia complex] Length: 212\nNumber of predicted TMHs: 1\nExp number of AAs in TMHs: 28.03134\nExp number, first 60 AAs: 6.30523\nTotal prob of N-in: 0.48947\noutside 1 187\nTMhelix 188 210\ninside 211 212

532 GCF_001993005.1_ASM199300v1 Burkholderia cenocepacia Proteobacteria; Betaproteobacteria; Burkholderiales; Burkholderiaceae; Burkholderia; Burkholderia cepacia complex MSDDIDNGGKGGFTRRDMMRVMAASGMMAVGGGGLLTSAQSAFAAPAPKRGGKIRVANES WP_006482802.1 MULTISPECIES: ABC transporter substrate-binding protein [Burkholderia cepacia complex] Length: 527\nNumber of predicted TMHs: 2\nExp number of AAs in TMHs: 36.7303599999999\nExp number, first 60 AAs: 22.08937\nTotal prob of N-in: 0.97312\nPOSSIBLE N-term signal sequence\ninside 1 20\nTMhelix 21 43\noutside 44 502\nTMhelix 503 525\ninside 526 527

533 GCF_001992965.1_ASM199296v1 Burkholderia cenocepacia Proteobacteria; Betaproteobacteria; Burkholderiales; Burkholderiaceae; Burkholderia; Burkholderia cepacia complex MSDDIDNGGKGGFTRRDMMRVMAASGMMAVGGGGLLTSAQSAFAAPAPKRGGKIRVANES WP_006482802.1 MULTISPECIES: ABC transporter substrate-binding protein [Burkholderia cepacia complex] Length: 527\nNumber of predicted TMHs: 2\nExp number of AAs in TMHs: 36.7303599999999\nExp number, first 60 AAs: 22.08937\nTotal prob of N-in: 0.97312\nPOSSIBLE N-term signal sequence\ninside 1 20\nTMhelix 21 43\noutside 44 502\nTMhelix 503 525\ninside 526 527

534 GCF_001993055.1_ASM199305v1 Burkholderia cenocepacia Proteobacteria; Betaproteobacteria; Burkholderiales; Burkholderiaceae; Burkholderia; Burkholderia cepacia complex MSDDIDNGGKGGFTRRDMMRVMAASGMMAVGGGGLLTSAQSAFAAPAPKRGGKIRVANES WP_006482802.1 MULTISPECIES: ABC transporter substrate-binding protein [Burkholderia cepacia complex] Length: 527\nNumber of predicted TMHs: 2\nExp number of AAs in TMHs: 36.7303599999999\nExp number, first 60 AAs: 22.08937\nTotal prob of N-in: 0.97312\nPOSSIBLE N-term signal sequence\ninside 1 20\nTMhelix 21 43\noutside 44 502\nTMhelix 503 525\ninside 526 527

535 GCF_001992475.1_ASM199247v1 Burkholderia cenocepacia Proteobacteria; Betaproteobacteria; Burkholderiales; Burkholderiaceae; Burkholderia; Burkholderia cepacia complex MSTSITRRRLLAAGVAASSLALSGCFTPKLYKNDAYSEHVSAFMMTEDGKKLVVLGTRYH WP_077021267.1 hypothetical protein [Burkholderia cenocepacia] Length: 212\nNumber of predicted TMHs: 1\nExp number of AAs in TMHs: 24.17376\nExp number, first 60 AAs: 0.92388\nTotal prob of N-in: 0.37951\noutside 1 180\nTMhelix 181 203\ninside 204 212

536 GCF_001992195.1_ASM199219v1 Burkholderia cenocepacia Proteobacteria; Betaproteobacteria; Burkholderiales; Burkholderiaceae; Burkholderia; Burkholderia cepacia complex MSDDIDNGGKGGFTRRDMMRVMAASGMMAVGGGGLLTSAQSAFAAPAPKRGGKIRVANES WP_077187164.1 ABC transporter substrate-binding protein [Burkholderia cenocepacia] Length: 527\nNumber of predicted TMHs: 2\nExp number of AAs in TMHs: 36.7300399999999\nExp number, first 60 AAs: 22.0894\nTotal prob of N-in: 0.97312\nPOSSIBLE N-term signal sequence\ninside 1 20\nTMhelix 21 43\noutside 44 502\nTMhelix 503 525\ninside 526 527

537 GCF_001992195.1_ASM199219v1 Burkholderia cenocepacia Proteobacteria; Betaproteobacteria; Burkholderiales; Burkholderiaceae; Burkholderia; Burkholderia cepacia complex MSTSITRRRLLAAGVAASSLALSGCFTPKLYKNDAYSEHVSAFMMTEDGKKLVVLGTRYH WP_077021267.1 hypothetical protein [Burkholderia cenocepacia] Length: 212\nNumber of predicted TMHs: 1\nExp number of AAs in TMHs: 24.17376\nExp number, first 60 AAs: 0.92388\nTotal prob of N-in: 0.37951\noutside 1 180\nTMhelix 181 203\ninside 204 212

538 GCF_001992475.1_ASM199247v1 Burkholderia cenocepacia Proteobacteria; Betaproteobacteria; Burkholderiales; Burkholderiaceae; Burkholderia; Burkholderia cepacia complex MKRRTFLALPAVAASGMLAGCSSADLGPVTHRLMKDPEYRETLSAFLISADGKKLVVIGKWP_006483583.1 MULTISPECIES: hypothetical protein [Burkholderia cepacia complex] Length: 212\nNumber of predicted TMHs: 1\nExp number of AAs in TMHs: 28.03134\nExp number, first 60 AAs: 6.30523\nTotal prob of N-in: 0.48947\noutside 1 187\nTMhelix 188 210\ninside 211 212

539 GCF_001992475.1_ASM199247v1 Burkholderia cenocepacia Proteobacteria; Betaproteobacteria; Burkholderiales; Burkholderiaceae; Burkholderia; Burkholderia cepacia complex
MSDDIDNGGKGGFTRRDMMRVMAASGMMAVGGGGLLTSAQSAFAAPAPKRGGKIRVANES WP_006482802.1
MULTISPECIES: ABC transporter substrate-binding protein [Burkholderia cepacia complex] Length: 527\nNumber of predicted TMHs: 2\nExp number of AAs in TMHs: 36.7303599999999\nExp number, first 60 AAs: 22.08937\nTotal prob of N-in: 0.97312\nPOSSIBLE N-term signal sequence\ninside 1 20\nTMhelix 21 43\noutside 44 502\nTMhelix 503 525\ninside 526 527

540 GCF_001992195.1_ASM199219v1 Burkholderia cenocepacia Proteobacteria; Betaproteobacteria; Burkholderiales; Burkholderiaceae; Burkholderia; Burkholderia cepacia complex
METLLTRIRASFADASPRRQLISVAVLVALVASLIAAGYFALRPRYQVLFRDLKPQDAAT WP_077020516.1 flagellar M-ring protein FliF [Burkholderia cenocepacia] Length: 491\nNumber of predicted TMHs: 2\nExp number of AAs in TMHs: 41.34236\nExp number, first 60 AAs: 22.1383\nTotal prob of N-in: 0.99080\nPOSSIBLE N-term signal sequence\ninside 1 20\nTMhelix 21 43\noutside 44 427\nTMhelix 428 450\ninside 451 491

541 GCF_001992445.1_ASM199244v1 Burkholderia cenocepacia Proteobacteria; Betaproteobacteria; Burkholderiales; Burkholderiaceae; Burkholderia; Burkholderia cepacia complex
MSTSITRRRLLAAGVAASSLALSGCFTPKLYKNDAYSEHVSAFMMTEDGKKLVVLGTRYH WP_077021267.1
hypothetical protein [Burkholderia cenocepacia] Length: 212\nNumber of predicted TMHs: 1\nExp number of AAs in TMHs: 24.17376\nExp number, first 60 AAs: 0.92388\nTotal prob of N-in: 0.37951\noutside 1 180\nTMhelix 181 203\ninside 204 212

542 GCF_001992335.1_ASM199233v1 Burkholderia cenocepacia Proteobacteria; Betaproteobacteria; Burkholderiales; Burkholderiaceae; Burkholderia; Burkholderia cepacia complex
METLLTRIRASFADASPRRQLISVAVLVALVASLIAAGYFALRPRYQVLFRDLKPQDAAT WP_077020516.1 flagellar M-ring protein FliF [Burkholderia cenocepacia] Length: 491\nNumber of predicted TMHs: 2\nExp number of AAs in TMHs: 41.34236\nExp number, first 60 AAs: 22.1383\nTotal prob of N-in: 0.99080\nPOSSIBLE N-term signal sequence\ninside 1 20\nTMhelix 21 43\noutside 44 427\nTMhelix 428 450\ninside 451 491

543 GCF_001992375.1_ASM199237v1 Burkholderia cenocepacia Proteobacteria; Betaproteobacteria; Burkholderiales; Burkholderiaceae; Burkholderia; Burkholderia cepacia complex
MKRRTFLALPAVAASGMLAGCSSADLGPVTHRLMKDPEYRETLSAFLISADGKKLVVIGKWP_006483583.1 MULTISPECIES: hypothetical protein [Burkholderia cepacia complex] Length: 212\nNumber of predicted TMHs: 1\nExp number of AAs in TMHs: 28.03134\nExp number, first 60 AAs: 6.30523\nTotal prob of N-in: 0.48947\noutside 1 187\nTMhelix 188 210\ninside 211 212

544 GCF_001992335.1_ASM199233v1 Burkholderia cenocepacia Proteobacteria; Betaproteobacteria; Burkholderiales; Burkholderiaceae; Burkholderia; Burkholderia cepacia complex
MKRRTFLALPAVAASGMLAGCSSADLGPVTHRLMKDPEYRETLSAFLISADGKKLVVIGKWP_006483583.1 MULTISPECIES: hypothetical protein [Burkholderia cepacia complex] Length: 212\nNumber of predicted TMHs: 1\nExp number of AAs in TMHs: 28.03134\nExp number, first 60 AAs: 6.30523\nTotal prob of N-in: 0.48947\noutside 1 187\nTMhelix 188 210\ninside 211 212

545 GCF_001992335.1_ASM199233v1 Burkholderia cenocepacia Proteobacteria; Betaproteobacteria; Burkholderiales; Burkholderiaceae; Burkholderia; Burkholderia cepacia complex
MSDDIDNGGKGGFTRRDMMRVMAASGMMAVGGGGLLTSAQSAFAAPAPKRGGKIRVANES WP_006482802.1
MULTISPECIES: ABC transporter substrate-binding protein [Burkholderia cepacia complex] Length: 527\nNumber of predicted TMHs: 2\nExp number of AAs in TMHs: 36.7303599999999\nExp number, first 60 AAs: 22.08937\nTotal prob of N-in: 0.97312\nPOSSIBLE N-term signal sequence\ninside 1 20\nTMhelix 21 43\noutside 44 502\nTMhelix 503 525\ninside 526 527

546 GCF_001992285.1_ASM199228v1 Burkholderia cenocepacia Proteobacteria; Betaproteobacteria; Burkholderiales; Burkholderiaceae; Burkholderia; Burkholderia cepacia complex
MSDDIDNGGKGGFTRRDMMRVMAASGMMAVGGGGLLTSAQSAFAAPAPKRGGKIRVANES WP_006482802.1
MULTISPECIES: ABC transporter substrate-binding protein [Burkholderia cepacia complex] Length: 527\nNumber of predicted TMHs: 2\nExp number of AAs in TMHs: 36.7303599999999\nExp number, first 60 AAs: 22.08937\nTotal prob of N-in: 0.97312\nPOSSIBLE N-term signal sequence\ninside 1 20\nTMhelix 21 43\noutside 44 502\nTMhelix 503 525\ninside 526 527

547 GCF_001992225.1_ASM199222v1 Burkholderia cenocepacia Proteobacteria; Betaproteobacteria; Burkholderiales; Burkholderiaceae; Burkholderia; Burkholderia cepacia complex

MSDDIDNGGKGGFTRRDMMRVMAASGMMAVGGGGLLTSAQSAFAAPAPKRGGKIRVANES WP_006482802.1 MULTISPECIES: ABC transporter substrate-binding protein [Burkholderia cepacia complex] Length: 527\nNumber of predicted TMHs: 2\nExp number of AAs in TMHs: 36.7303599999999\nExp number, first 60 AAs: 22.08937\nTotal prob of N-in: 0.97312\nPOSSIBLE N-term signal sequence\ninside 1 20\nTMhelix 21 43\noutside 44 502\nTMhelix 503 525\ninside 526 527

548 GCF_001991835.1_ASM199183v1 Burkholderia cenocepacia Proteobacteria; Betaproteobacteria; Burkholderiales; Burkholderiaceae; Burkholderia; Burkholderia cepacia complex MSTSITRRRLLAAGVAASSLALSGCFTPKLYKNDAYSEHVSAFMMTEDGKKLVVLGTRYH WP_077021267.1 hypothetical protein [Burkholderia cenocepacia] Length: 212\nNumber of predicted TMHs: 1\nExp number of AAs in TMHs: 24.17376\nExp number, first 60 AAs: 0.92388\nTotal prob of N-in: 0.37951\noutside 1 180\nTMhelix 181 203\ninside 204 212

549 GCF_001991975.1_ASM199197v1 Burkholderia cenocepacia Proteobacteria; Betaproteobacteria; Burkholderiales; Burkholderiaceae; Burkholderia; Burkholderia cepacia complex MKRRTFLALPAVAASGMLAGCSSADLGPVTHRLMKDPEYRETLSAFLISADGKKLVVIGKWP_006483583.1 MULTISPECIES: hypothetical protein [Burkholderia cepacia complex] Length: 212\nNumber of predicted TMHs: 1\nExp number of AAs in TMHs: 28.03134\nExp number, first 60 AAs: 6.30523\nTotal prob of N-in: 0.48947\noutside 1 187\nTMhelix 188 210\ninside 211 212

550 GCF_001991695.1_ASM199169v1 Burkholderia cenocepacia Proteobacteria; Betaproteobacteria; Burkholderiales; Burkholderiaceae; Burkholderia; Burkholderia cepacia complex METLLTRIRASFADASPRRQLISVAVLVALVASLIAAGYFALRPRYQVLFRDLKPQDAAT WP_077020516.1 flagellar M-ring protein FliF [Burkholderia cenocepacia] Length: 491\nNumber of predicted TMHs: 2\nExp number of AAs in TMHs: 41.34236\nExp number, first 60 AAs: 22.1383\nTotal prob of N-in: 0.99080\nPOSSIBLE N-term signal sequence\ninside 1 20\nTMhelix 21 43\noutside 44 427\nTMhelix 428 450\ninside 451 491

551 GCF_001991665.1_ASM199166v1 Burkholderia cenocepacia Proteobacteria; Betaproteobacteria; Burkholderiales; Burkholderiaceae; Burkholderia; Burkholderia cepacia complex MSTSITRRRLLAAGVAASSLALSGCFTPKLYKNDAYSEHVSAFMMTEDGKKLVVLGTRYH WP_077021267.1 hypothetical protein [Burkholderia cenocepacia] Length: 212\nNumber of predicted TMHs: 1\nExp number of AAs in TMHs: 24.17376\nExp number, first 60 AAs: 0.92388\nTotal prob of N-in: 0.37951\noutside 1 180\nTMhelix 181 203\ninside 204 212

552 GCF_001991665.1_ASM199166v1 Burkholderia cenocepacia Proteobacteria; Betaproteobacteria; Burkholderiales; Burkholderiaceae; Burkholderia; Burkholderia cepacia complex METLLTRIRASFADASPRRQLISVAVLVALVASLIAAGYFALRPRYQVLFRDLKPQDAAT WP_077020516.1 flagellar M-ring protein FliF [Burkholderia cenocepacia] Length: 491\nNumber of predicted TMHs: 2\nExp number of AAs in TMHs: 41.34236\nExp number, first 60 AAs: 22.1383\nTotal prob of N-in: 0.99080\nPOSSIBLE N-term signal sequence\ninside 1 20\nTMhelix 21 43\noutside 44 427\nTMhelix 428 450\ninside 451 491

553 GCF_001991555.1_ASM199155v1 Burkholderia cenocepacia Proteobacteria; Betaproteobacteria; Burkholderiales; Burkholderiaceae; Burkholderia; Burkholderia cepacia complex MSTSITRRRLLAAGVAASSLALSGCFTPKLYKNDAYSEHVSAFMMTEDGKKLVVLGTRYH WP_077021267.1 hypothetical protein [Burkholderia cenocepacia] Length: 212\nNumber of predicted TMHs: 1\nExp number of AAs in TMHs: 24.17376\nExp number, first 60 AAs: 0.92388\nTotal prob of N-in: 0.37951\noutside 1 180\nTMhelix 181 203\ninside 204 212

554 GCF_001991625.1_ASM199162v1 Burkholderia cenocepacia Proteobacteria; Betaproteobacteria; Burkholderiales; Burkholderiaceae; Burkholderia; Burkholderia cepacia complex MSTSITRRRLLAAGVAASSLALSGCFTPKLYKNDAYSEHVSAFMMTEDGKKLVVLGTRYH WP_077021267.1 hypothetical protein [Burkholderia cenocepacia] Length: 212\nNumber of predicted TMHs: 1\nExp number of AAs in TMHs: 24.17376\nExp number, first 60 AAs: 0.92388\nTotal prob of N-in: 0.37951\noutside 1 180\nTMhelix 181 203\ninside 204 212

555 GCF_001991625.1_ASM199162v1 Burkholderia cenocepacia Proteobacteria; Betaproteobacteria; Burkholderiales; Burkholderiaceae; Burkholderia; Burkholderia cepacia complex METLLTRIRASFADASPRRQLISVAVLVALVASLIAAGYFALRPRYQVLFRDLKPQDAAT WP_077020516.1 flagellar M-ring protein FliF [Burkholderia cenocepacia] Length: 491\nNumber of predicted TMHs: 2\nExp number of AAs in TMHs: 41.34236\nExp number, first 60 AAs: 22.1383\nTotal prob of N-in: 0.99080\nPOSSIBLE N-term signal sequence\ninside 1 20\nTMhelix 21 43\noutside 44 427\nTMhelix 428 450\ninside 451 491

556 GCF_001991755.1_ASM199175v1 Burkholderia cenocepacia Proteobacteria; Betaproteobacteria; Burkholderiales; Burkholderiaceae; Burkholderia; Burkholderia cepacia complex
MKRRTFLALPAVAASGMLAGCSSADLGPVTHRLMKDPEYRETLSAFLISADGKKLVVIGKWP_006483583.1 MULTISPECIES: hypothetical protein [Burkholderia cepacia complex] Length: 212\nNumber of predicted TMHs: 1\nExp number of AAs in TMHs: 28.03134\nExp number, first 60 AAs: 6.30523\nTotal prob of N-in: 0.48947\noutside 1 187\nTMhelix 188 210\ninside 211 212

557 GCF_001991755.1_ASM199175v1 Burkholderia cenocepacia Proteobacteria; Betaproteobacteria; Burkholderiales; Burkholderiaceae; Burkholderia; Burkholderia cepacia complex
MSDDIDNGGKGGFTRRDMMRVMAASGMMAVGGGGLLTSAQSAFAAPAPKRGGKIRVANES WP_006482802.1
MULTISPECIES: ABC transporter substrate-binding protein [Burkholderia cepacia complex] Length: 527\nNumber of predicted TMHs: 2\nExp number of AAs in TMHs: 36.7303599999999\nExp number, first 60 AAs: 22.08937\nTotal prob of N-in: 0.97312\nPOSSIBLE N-term signal sequence\ninside 1 20\nTMhelix 21 43\noutside 44 502\nTMhelix 503 525\ninside 526 527

558 GCF_001991495.1_ASM199149v1 Burkholderia cenocepacia Proteobacteria; Betaproteobacteria; Burkholderiales; Burkholderiaceae; Burkholderia; Burkholderia cepacia complex
METLLTRIRASFADASPRRQLISVAVLVALVASLIAAGYFALRPRYQVLFRDLKPQDAAT WP_077020516.1 flagellar M-ring protein FliF [Burkholderia cenocepacia] Length: 491\nNumber of predicted TMHs: 2\nExp number of AAs in TMHs: 41.34236\nExp number, first 60 AAs: 22.1383\nTotal prob of N-in: 0.99080\nPOSSIBLE N-term signal sequence\ninside 1 20\nTMhelix 21 43\noutside 44 427\nTMhelix 428 450\ninside 451 491

559 GCF_001991735.1_ASM199173v1 Burkholderia cenocepacia Proteobacteria; Betaproteobacteria; Burkholderiales; Burkholderiaceae; Burkholderia; Burkholderia cepacia complex
MSTSITRRRLLAAGVAASSLALSGCFTPKLYKNDAYSEHVSAFMMTEDGKKLVVLGTRYH WP_077021267.1
hypothetical protein [Burkholderia cenocepacia] Length: 212\nNumber of predicted TMHs: 1\nExp number of AAs in TMHs: 24.17376\nExp number, first 60 AAs: 0.92388\nTotal prob of N-in: 0.37951\noutside 1 180\nTMhelix 181 203\ninside 204 212

560 GCF_001991735.1_ASM199173v1 Burkholderia cenocepacia Proteobacteria; Betaproteobacteria; Burkholderiales; Burkholderiaceae; Burkholderia; Burkholderia cepacia complex
METLLTRIRASFADASPRRQLISVAVLVALVASLIAAGYFALRPRYQVLFRDLKPQDAAT WP_077020516.1 flagellar M-ring protein FliF [Burkholderia cenocepacia] Length: 491\nNumber of predicted TMHs: 2\nExp number of AAs in TMHs: 41.34236\nExp number, first 60 AAs: 22.1383\nTotal prob of N-in: 0.99080\nPOSSIBLE N-term signal sequence\ninside 1 20\nTMhelix 21 43\noutside 44 427\nTMhelix 428 450\ninside 451 491

561 GCF_001991735.1_ASM199173v1 Burkholderia cenocepacia Proteobacteria; Betaproteobacteria; Burkholderiales; Burkholderiaceae; Burkholderia; Burkholderia cepacia complex
MKRRTFLALPAVAASGMLAGCSSADLGPVTHRLMKDPEYRETLSAFLISADGKKLVVIGKWP_006483583.1 MULTISPECIES: hypothetical protein [Burkholderia cepacia complex] Length: 212\nNumber of predicted TMHs: 1\nExp number of AAs in TMHs: 28.03134\nExp number, first 60 AAs: 6.30523\nTotal prob of N-in: 0.48947\noutside 1 187\nTMhelix 188 210\ninside 211 212

562 GCF_001991555.1_ASM199155v1 Burkholderia cenocepacia Proteobacteria; Betaproteobacteria; Burkholderiales; Burkholderiaceae; Burkholderia; Burkholderia cepacia complex
MSDDIDNGGKGGFTRRDMMRVMAASGMMAVGGGGLLTSAQSAFAAPAPKRGGKIRVANES WP_006482802.1
MULTISPECIES: ABC transporter substrate-binding protein [Burkholderia cepacia complex] Length: 527\nNumber of predicted TMHs: 2\nExp number of AAs in TMHs: 36.7303599999999\nExp number, first 60 AAs: 22.08937\nTotal prob of N-in: 0.97312\nPOSSIBLE N-term signal sequence\ninside 1 20\nTMhelix 21 43\noutside 44 502\nTMhelix 503 525\ninside 526 527

563 GCF_001991515.1_ASM199151v1 Burkholderia cenocepacia Proteobacteria; Betaproteobacteria; Burkholderiales; Burkholderiaceae; Burkholderia; Burkholderia cepacia complex
MKRRTFLALPAVAASGMLAGCSSADLGPVTHRLMKDPEYRETLSAFLISADGKKLVVIGKWP_006483583.1 MULTISPECIES: hypothetical protein [Burkholderia cepacia complex] Length: 212\nNumber of predicted TMHs: 1\nExp number of AAs in TMHs: 28.03134\nExp number, first 60 AAs: 6.30523\nTotal prob of N-in: 0.48947\noutside 1 187\nTMhelix 188 210\ninside 211 212

564 GCF_001991515.1_ASM199151v1 Burkholderia cenocepacia Proteobacteria; Betaproteobacteria; Burkholderiales; Burkholderiaceae; Burkholderia; Burkholderia cepacia complex
MSDDIDNGGKGGFTRRDMMRVMAASGMMAVGGGGLLTSAQSAFAAPAPKRGGKIRVANES WP_006482802.1

MULTISPECIES: ABC transporter substrate-binding protein [Burkholderia cepacia complex] Length: 527\nNumber of predicted TMHs: 2\nExp number of AAs in TMHs: 36.7303599999999\nExp number, first 60 AAs: 22.08937\nTotal prob of N-in: 0.97312\nPOSSIBLE N-term signal sequence\ninside 1 20\nTMhelix 21 43\noutside 44 502\nTMhelix 503 525\ninside 526 527

565 GCF_001991495.1_ASM199149v1 Burkholderia cenocepacia Proteobacteria; Betaproteobacteria; Burkholderiales; Burkholderiaceae; Burkholderia; Burkholderia cepacia complex MKRRTFLALPAVAASGMLAGCSSADLGPVTHRLMKDPEYRETLSAFLISADGKKLVVIGKWP_006483583.1 MULTISPECIES: hypothetical protein [Burkholderia cepacia complex] Length: 212\nNumber of predicted TMHs: 1\nExp number of AAs in TMHs: 28.03134\nExp number, first 60 AAs: 6.30523\nTotal prob of N-in: 0.48947\noutside 1 187\nTMhelix 188 210\ninside 211 212

566 GCF_001991495.1_ASM199149v1 Burkholderia cenocepacia Proteobacteria; Betaproteobacteria; Burkholderiales; Burkholderiaceae; Burkholderia; Burkholderia cepacia complex MSDDIDNGGKGGFTRRDMMRVMAASGMMAVGGGGLLTSAQSAFAAPAPKRGGKIRVANES WP_006482802.1 MULTISPECIES: ABC transporter substrate-binding protein [Burkholderia cepacia complex] Length: 527\nNumber of predicted TMHs: 2\nExp number of AAs in TMHs: 36.7303599999999\nExp number, first 60 AAs: 22.08937\nTotal prob of N-in: 0.97312\nPOSSIBLE N-term signal sequence\ninside 1 20\nTMhelix 21 43\noutside 44 502\nTMhelix 503 525\ninside 526 527

567 GCF_001991455.1_ASM199145v1 Burkholderia cenocepacia Proteobacteria; Betaproteobacteria; Burkholderiales; Burkholderiaceae; Burkholderia; Burkholderia cepacia complex MSDDIDNGGKGGFTRRDMMRVMAASGMMAVGGGGLLTSAQSAFAAPAPKRGGKIRVANES WP_006482802.1 MULTISPECIES: ABC transporter substrate-binding protein [Burkholderia cepacia complex] Length: 527\nNumber of predicted TMHs: 2\nExp number of AAs in TMHs: 36.7303599999999\nExp number, first 60 AAs: 22.08937\nTotal prob of N-in: 0.97312\nPOSSIBLE N-term signal sequence\ninside 1 20\nTMhelix 21 43\noutside 44 502\nTMhelix 503 525\ninside 526 527

568 GCF_001991415.1_ASM199141v1 Burkholderia cenocepacia Proteobacteria; Betaproteobacteria; Burkholderiales; Burkholderiaceae; Burkholderia; Burkholderia cepacia complex MKRRTFLALPAVAASGMLAGCSSADLGPVTHRLMKDPEYRETLSAFLISADGKKLVVIGKWP_006483583.1 MULTISPECIES: hypothetical protein [Burkholderia cepacia complex] Length: 212\nNumber of predicted TMHs: 1\nExp number of AAs in TMHs: 28.03134\nExp number, first 60 AAs: 6.30523\nTotal prob of N-in: 0.48947\noutside 1 187\nTMhelix 188 210\ninside 211 212

569 GCF_001991195.1_ASM199119v1 Burkholderia cenocepacia Proteobacteria; Betaproteobacteria; Burkholderiales; Burkholderiaceae; Burkholderia; Burkholderia cepacia complex MKRRTFLALPAVAASGMLAGCSSADLGPVTHRLMKDPEYRETLSAFLISADGKKLVVIGKWP_034201760.1 MULTISPECIES: hypothetical protein [Burkholderia cepacia complex] Length: 212\nNumber of predicted TMHs: 1\nExp number of AAs in TMHs: 28.11606\nExp number, first 60 AAs: 6.33522\nTotal prob of N-in: 0.49088\noutside 1 187\nTMhelix 188 210\ninside 211 212

570 GCF_001991325.1_ASM199132v1 Burkholderia cenocepacia Proteobacteria; Betaproteobacteria; Burkholderiales; Burkholderiaceae; Burkholderia; Burkholderia cepacia complex MSTSITRRRLLAAGVAASSLALSGCFTPKLYKNDAYSEHVSAFMMTEDGKKLVVLGTRYH WP_077021267.1 hypothetical protein [Burkholderia cenocepacia] Length: 212\nNumber of predicted TMHs: 1\nExp number of AAs in TMHs: 24.17376\nExp number, first 60 AAs: 0.92388\nTotal prob of N-in: 0.37951\noutside 1 180\nTMhelix 181 203\ninside 204 212

571 GCF_001991325.1_ASM199132v1 Burkholderia cenocepacia Proteobacteria; Betaproteobacteria; Burkholderiales; Burkholderiaceae; Burkholderia; Burkholderia cepacia complex METLLTRIRASFADASPRRQLISVAVLVALVASLIAAGYFALRPRYQVLFRDLKPQDAAT WP_077020516.1 flagellar M-ring protein FliF [Burkholderia cenocepacia] Length: 491\nNumber of predicted TMHs: 2\nExp number of AAs in TMHs: 41.34236\nExp number, first 60 AAs: 22.1383\nTotal prob of N-in: 0.99080\nPOSSIBLE N-term signal sequence\ninside 1 20\nTMhelix 21 43\noutside 44 427\nTMhelix 428 450\ninside 451 491

572 GCF_001993675.1_ASM199367v1 Burkholderia cenocepacia Proteobacteria; Betaproteobacteria; Burkholderiales; Burkholderiaceae; Burkholderia; Burkholderia cepacia complex MKRRTFLALPAVAASGMLAGCSSADLGPVTHRLMKDPEYRETLSAFLISADGKKLVVIGKWP_034201760.1 MULTISPECIES: hypothetical protein [Burkholderia cepacia complex] Length: 212\nNumber of predicted TMHs: 1\nExp number of

AAs in TMHs: 28.11606\nExp number, first 60 AAs: 6.33522\nTotal prob of N-in: 0.49088\noutside 1 187\nTMhelix 188 210\ninside 211 212

573 GCF_001993435.1_ASM199343v1 Burkholderia cenocepacia Proteobacteria; Betaproteobacteria; Burkholderiales; Burkholderiaceae; Burkholderia; Burkholderia cepacia complex METLLTRIRASFADASPRRQLISVAVLVALVASLIAAGYFALRPRYQVLFRDLKPQDAAT WP_077020516.1 flagellar M-ring protein FliF [Burkholderia cenocepacia] Length: 491\nNumber of predicted TMHs: 2\nExp number of AAs in TMHs: 41.34236\nExp number, first 60 AAs: 22.1383\nTotal prob of N-in: 0.99080\nPOSSIBLE N-term signal sequence\ninside 1 20\nTMhelix 21 43\noutside 44 427\nTMhelix 428 450\ninside 451 491

574 GCF_001993715.1_ASM199371v1 Burkholderia cenocepacia Proteobacteria; Betaproteobacteria; Burkholderiales; Burkholderiaceae; Burkholderia; Burkholderia cepacia complex MKRRTFLALPAVAASGMLAGCSSADLGPVTHRLMKDPEYRETLSAFLISADGKKLVVIGKWP_034201760.1 MULTISPECIES: hypothetical protein [Burkholderia cepacia complex] Length: 212\nNumber of predicted TMHs: 1\nExp number of AAs in TMHs: 28.11606\nExp number, first 60 AAs: 6.33522\nTotal prob of N-in: 0.49088\noutside 1 187\nTMhelix 188 210\ninside 211 212

575 GCF_001993715.1_ASM199371v1 Burkholderia cenocepacia Proteobacteria; Betaproteobacteria; Burkholderiales; Burkholderiaceae; Burkholderia; Burkholderia cepacia complex MSDDIDNGGKGGFTRRDMMRVMAASGMMAVGGGGLLTSAQSAFAAPAPKRGGKIRVANES WP_006482802.1 MULTISPECIES: ABC transporter substrate-binding protein [Burkholderia cepacia complex] Length: 527\nNumber of predicted TMHs: 2\nExp number of AAs in TMHs: 36.7303599999999\nExp number, first 60 AAs: 22.08937\nTotal prob of N-in: 0.97312\nPOSSIBLE N-term signal sequence\ninside 1 20\nTMhelix 21 43\noutside 44 502\nTMhelix 503 525\ninside 526 527

576 GCF_001993675.1_ASM199367v1 Burkholderia cenocepacia Proteobacteria; Betaproteobacteria; Burkholderiales; Burkholderiaceae; Burkholderia; Burkholderia cepacia complex MSDDIDNGGKGGFTRRDMMRVMAASGMMAVGGGGLLTSAQSAFAAPAPKRGGKIRVANES WP_006482802.1 MULTISPECIES: ABC transporter substrate-binding protein [Burkholderia cepacia complex] Length: 527\nNumber of predicted TMHs: 2\nExp number of AAs in TMHs: 36.7303599999999\nExp number, first 60 AAs: 22.08937\nTotal prob of N-in: 0.97312\nPOSSIBLE N-term signal sequence\ninside 1 20\nTMhelix 21 43\noutside 44 502\nTMhelix 503 525\ninside 526 527

577 GCF_001993535.1_ASM199353v1 Burkholderia cenocepacia Proteobacteria; Betaproteobacteria; Burkholderiales; Burkholderiaceae; Burkholderia; Burkholderia cepacia complex MSTSITRRRLLAAGVAASSLALSGCFTPKLYKNDAYSEHVSAFMMTEDGKKLVVLGTRYH WP_077021267.1 hypothetical protein [Burkholderia cenocepacia] Length: 212\nNumber of predicted TMHs: 1\nExp number of AAs in TMHs: 24.17376\nExp number, first 60 AAs: 0.92388\nTotal prob of N-in: 0.37951\noutside 1 180\nTMhelix 181 203\ninside 204 212

578 GCF_001993565.1_ASM199356v1 Burkholderia cenocepacia Proteobacteria; Betaproteobacteria; Burkholderiales; Burkholderiaceae; Burkholderia; Burkholderia cepacia complex MKRRTFLALPAVAASGMLAGCSSADLGPVTHRLMKDPEYRETLSAFLISADGKKLVVIGKWP_006483583.1 MULTISPECIES: hypothetical protein [Burkholderia cepacia complex] Length: 212\nNumber of predicted TMHs: 1\nExp number of AAs in TMHs: 28.03134\nExp number, first 60 AAs: 6.30523\nTotal prob of N-in: 0.48947\noutside 1 187\nTMhelix 188 210\ninside 211 212

579 GCF_001993535.1_ASM199353v1 Burkholderia cenocepacia Proteobacteria; Betaproteobacteria; Burkholderiales; Burkholderiaceae; Burkholderia; Burkholderia cepacia complex MKRRTFLALPAVAASGMLAGCSSADLGPVTHRLMKDPEYRETLSAFLISADGKKLVVIGKWP_006483583.1 MULTISPECIES: hypothetical protein [Burkholderia cepacia complex] Length: 212\nNumber of predicted TMHs: 1\nExp number of AAs in TMHs: 28.03134\nExp number, first 60 AAs: 6.30523\nTotal prob of N-in: 0.48947\noutside 1 187\nTMhelix 188 210\ninside 211 212

580 GCF_001993495.1_ASM199349v1 Burkholderia cenocepacia Proteobacteria; Betaproteobacteria; Burkholderiales; Burkholderiaceae; Burkholderia; Burkholderia cepacia complex MKRRTFLALPAVAASGMLAGCSSADLGPVTHRLMKDPEYRETLSAFLISADGKKLVVIGKWP_006483583.1 MULTISPECIES: hypothetical protein [Burkholderia cepacia complex] Length: 212\nNumber of predicted TMHs: 1\nExp number of AAs in TMHs: 28.03134\nExp number, first 60 AAs: 6.30523\nTotal prob of N-in: 0.48947\noutside 1 187\nTMhelix 188 210\ninside 211 212

581 GCF_001993535.1_ASM199353v1 Burkholderia cenocepacia Proteobacteria; Betaproteobacteria; Burkholderiales; Burkholderiaceae; Burkholderia; Burkholderia cepacia complex
MSDDIDNGGKGGFTRRDMMRVMAASGMMAVGGGGLLTSAQSAFAAPAPKRGGKIRVANES WP_006482802.1 MULTISPECIES: ABC transporter substrate-binding protein [Burkholderia cepacia complex] Length: 527\nNumber of predicted TMHs: 2\nExp number of AAs in TMHs: 36.7303599999999\nExp number, first 60 AAs: 22.08937\nTotal prob of N-in: 0.97312\nPOSSIBLE N-term signal sequence\ninside 1 20\nTMhelix 21 43\noutside 44 502\nTMhelix 503 525\ninside 526 527

582 GCF_001993475.1_ASM199347v1 Burkholderia cenocepacia Proteobacteria; Betaproteobacteria; Burkholderiales; Burkholderiaceae; Burkholderia; Burkholderia cepacia complex
MKRRTFLALPAVAASGMLAGCSSADLGPVTHRLMKDPEYRETLSAFLISADGKKLVVIGKWP_006483583.1 MULTISPECIES: hypothetical protein [Burkholderia cepacia complex] Length: 212\nNumber of predicted TMHs: 1\nExp number of AAs in TMHs: 28.03134\nExp number, first 60 AAs: 6.30523\nTotal prob of N-in: 0.48947\noutside 1 187\nTMhelix 188 210\ninside 211 212

583 GCF_001993495.1_ASM199349v1 Burkholderia cenocepacia Proteobacteria; Betaproteobacteria; Burkholderiales; Burkholderiaceae; Burkholderia; Burkholderia cepacia complex
MSDDIDNGGKGGFTRRDMMRVMAASGMMAVGGGGLLTSAQSAFAAPAPKRGGKIRVANES WP_006482802.1 MULTISPECIES: ABC transporter substrate-binding protein [Burkholderia cepacia complex] Length: 527\nNumber of predicted TMHs: 2\nExp number of AAs in TMHs: 36.7303599999999\nExp number, first 60 AAs: 22.08937\nTotal prob of N-in: 0.97312\nPOSSIBLE N-term signal sequence\ninside 1 20\nTMhelix 21 43\noutside 44 502\nTMhelix 503 525\ninside 526 527

584 GCF_001993475.1_ASM199347v1 Burkholderia cenocepacia Proteobacteria; Betaproteobacteria; Burkholderiales; Burkholderiaceae; Burkholderia; Burkholderia cepacia complex
MSDDIDNGGKGGFTRRDMMRVMAASGMMAVGGGGLLTSAQSAFAAPAPKRGGKIRVANES WP_006482802.1 MULTISPECIES: ABC transporter substrate-binding protein [Burkholderia cepacia complex] Length: 527\nNumber of predicted TMHs: 2\nExp number of AAs in TMHs: 36.7303599999999\nExp number, first 60 AAs: 22.08937\nTotal prob of N-in: 0.97312\nPOSSIBLE N-term signal sequence\ninside 1 20\nTMhelix 21 43\noutside 44 502\nTMhelix 503 525\ninside 526 527

585 GCF_001993075.1_ASM199307v1 Burkholderia cenocepacia Proteobacteria; Betaproteobacteria; Burkholderiales; Burkholderiaceae; Burkholderia; Burkholderia cepacia complex
MKRRTFLALPAVAASGMLAGCSSADLGPVTHRLMKDPEYRETLSAFLISADGKKLVVIGKWP_034201760.1 MULTISPECIES: hypothetical protein [Burkholderia cepacia complex] Length: 212\nNumber of predicted TMHs: 1\nExp number of AAs in TMHs: 28.11606\nExp number, first 60 AAs: 6.33522\nTotal prob of N-in: 0.49088\noutside 1 187\nTMhelix 188 210\ninside 211 212

586 GCF_001993185.1_ASM199318v1 Burkholderia cenocepacia Proteobacteria; Betaproteobacteria; Burkholderiales; Burkholderiaceae; Burkholderia; Burkholderia cepacia complex
MSDDIDNGGKGGFTRRDMMRVMAASGMMAVGGGGLLTSAQSAFAAPAPKRGGKIRVANES WP_077180776.1 ABC transporter substrate-binding protein [Burkholderia cenocepacia] Length: 527\nNumber of predicted TMHs: 2\nExp number of AAs in TMHs: 36.7312299999999\nExp number, first 60 AAs: 22.0894\nTotal prob of N-in: 0.97312\nPOSSIBLE N-term signal sequence\ninside 1 20\nTMhelix 21 43\noutside 44 502\nTMhelix 503 525\ninside 526 527

587 GCF_001993155.1_ASM199315v1 Burkholderia cenocepacia Proteobacteria; Betaproteobacteria; Burkholderiales; Burkholderiaceae; Burkholderia; Burkholderia cepacia complex
MSDDIDNGGKGGFTRRDMMRVMAASGMMAVGGGGLLTSAQSAFAAPAPKRGGKIRVANES WP_077180776.1 ABC transporter substrate-binding protein [Burkholderia cenocepacia] Length: 527\nNumber of predicted TMHs: 2\nExp number of AAs in TMHs: 36.7312299999999\nExp number, first 60 AAs: 22.0894\nTotal prob of N-in: 0.97312\nPOSSIBLE N-term signal sequence\ninside 1 20\nTMhelix 21 43\noutside 44 502\nTMhelix 503 525\ninside 526 527

588 GCF_001993185.1_ASM199318v1 Burkholderia cenocepacia Proteobacteria; Betaproteobacteria; Burkholderiales; Burkholderiaceae; Burkholderia; Burkholderia cepacia complex
MKRRTFLALPAVAASGMLAGCSSADLGPVTHRLMKDPEYRETLSAFLISADGKKLVVIGKWP_006483583.1 MULTISPECIES: hypothetical protein [Burkholderia cepacia complex] Length: 212\nNumber of predicted TMHs: 1\nExp number of AAs in TMHs: 28.03134\nExp number, first 60 AAs: 6.30523\nTotal prob of N-in: 0.48947\noutside 1 187\nTMhelix 188 210\ninside 211 212

589 GCF_001993155.1_ASM199315v1 Burkholderia cenocepacia Proteobacteria; Betaproteobacteria; Burkholderiales; Burkholderiaceae; Burkholderia; Burkholderia cepacia complex

MKRRTFLALPAVAASGMLAGCSSADLGPVTHRLMKDPEYRETLSAFLISADGKKLVVIGKWP_006483583.1 MULTISPECIES: hypothetical protein [Burkholderia cepacia complex] Length: 212\nNumber of predicted TMHs: 1\nExp number of AAs in TMHs: 28.03134\nExp number, first 60 AAs: 6.30523\nTotal prob of N-in: 0.48947\noutside 1 187\nTMhelix 188 210\ninside 211 212

590 GCF_001993115.1_ASM199311v1 Burkholderia cenocepacia Proteobacteria; Betaproteobacteria; Burkholderiales; Burkholderiaceae; Burkholderia; Burkholderia cepacia complex
MKRRTFLALPAVAASGMLAGCSSADLGPVTHRLMKDPEYRETLSAFLISADGKKLVVIGKWP_006483583.1 MULTISPECIES: hypothetical protein [Burkholderia cepacia complex] Length: 212\nNumber of predicted TMHs: 1\nExp number of AAs in TMHs: 28.03134\nExp number, first 60 AAs: 6.30523\nTotal prob of N-in: 0.48947\noutside 1 187\nTMhelix 188 210\ninside 211 212

591 GCF_001993115.1_ASM199311v1 Burkholderia cenocepacia Proteobacteria; Betaproteobacteria; Burkholderiales; Burkholderiaceae; Burkholderia; Burkholderia cepacia complex
MSDDIDNGGKGGFTRRDMMRVMAASGMMAVGGGGLLTSAQSAFAAPAPKRGGKIRVANES WP_006482802.1
MULTISPECIES: ABC transporter substrate-binding protein [Burkholderia cepacia complex] Length: 527\nNumber of predicted TMHs: 2\nExp number of AAs in TMHs: 36.7303599999999\nExp number, first 60 AAs: 22.08937\nTotal prob of N-in: 0.97312\nPOSSIBLE N-term signal sequence\ninside 1 20\nTMhelix 21 43\noutside 44 502\nTMhelix 503 525\ninside 526 527

592 GCF_001992825.1_ASM199282v1 Burkholderia cenocepacia Proteobacteria; Betaproteobacteria; Burkholderiales; Burkholderiaceae; Burkholderia; Burkholderia cepacia complex
MKRRTFLALPAVAASGMLAGCSSADLGPVTHRLMKDPEYRETLSAFLISADGKKLVVIGKWP_034201760.1 MULTISPECIES: hypothetical protein [Burkholderia cepacia complex] Length: 212\nNumber of predicted TMHs: 1\nExp number of AAs in TMHs: 28.11606\nExp number, first 60 AAs: 6.33522\nTotal prob of N-in: 0.49088\noutside 1 187\nTMhelix 188 210\ninside 211 212

593 GCF_001992895.1_ASM199289v1 Burkholderia cenocepacia Proteobacteria; Betaproteobacteria; Burkholderiales; Burkholderiaceae; Burkholderia; Burkholderia cepacia complex
MKRRTFLALPAVAASGMLAGCSSADLGPVTHRLMKDPEYRETLSAFLISADGKKLVVIGKWP_006483583.1 MULTISPECIES: hypothetical protein [Burkholderia cepacia complex] Length: 212\nNumber of predicted TMHs: 1\nExp number of AAs in TMHs: 28.03134\nExp number, first 60 AAs: 6.30523\nTotal prob of N-in: 0.48947\noutside 1 187\nTMhelix 188 210\ninside 211 212

594 GCF_001992875.1_ASM199287v1 Burkholderia cenocepacia Proteobacteria; Betaproteobacteria; Burkholderiales; Burkholderiaceae; Burkholderia; Burkholderia cepacia complex
MKRRTFLALPAVAASGMLAGCSSADLGPVTHRLMKDPEYRETLSAFLISADGKKLVVIGKWP_006483583.1 MULTISPECIES: hypothetical protein [Burkholderia cepacia complex] Length: 212\nNumber of predicted TMHs: 1\nExp number of AAs in TMHs: 28.03134\nExp number, first 60 AAs: 6.30523\nTotal prob of N-in: 0.48947\noutside 1 187\nTMhelix 188 210\ninside 211 212

595 GCF_001992785.1_ASM199278v1 Burkholderia cenocepacia Proteobacteria; Betaproteobacteria; Burkholderiales; Burkholderiaceae; Burkholderia; Burkholderia cepacia complex
MKRRTFLALPAVAASGMLAGCSSADLGPVTHRLMKDPEYRETLSAFLISADGKKLVVIGKWP_034201760.1 MULTISPECIES: hypothetical protein [Burkholderia cepacia complex] Length: 212\nNumber of predicted TMHs: 1\nExp number of AAs in TMHs: 28.11606\nExp number, first 60 AAs: 6.33522\nTotal prob of N-in: 0.49088\noutside 1 187\nTMhelix 188 210\ninside 211 212

596 GCF_001992875.1_ASM199287v1 Burkholderia cenocepacia Proteobacteria; Betaproteobacteria; Burkholderiales; Burkholderiaceae; Burkholderia; Burkholderia cepacia complex
MSDDIDNGGKGGFTRRDMMRVMAASGMMAVGGGGLLTSAQSAFAAPAPKRGGKIRVANES WP_006482802.1
MULTISPECIES: ABC transporter substrate-binding protein [Burkholderia cepacia complex] Length: 527\nNumber of predicted TMHs: 2\nExp number of AAs in TMHs: 36.7303599999999\nExp number, first 60 AAs: 22.08937\nTotal prob of N-in: 0.97312\nPOSSIBLE N-term signal sequence\ninside 1 20\nTMhelix 21 43\noutside 44 502\nTMhelix 503 525\ninside 526 527

597 GCF_001992825.1_ASM199282v1 Burkholderia cenocepacia Proteobacteria; Betaproteobacteria; Burkholderiales; Burkholderiaceae; Burkholderia; Burkholderia cepacia complex
MSDDIDNGGKGGFTRRDMMRVMAASGMMAVGGGGLLTSAQSAFAAPAPKRGGKIRVANES WP_006482802.1
MULTISPECIES: ABC transporter substrate-binding protein [Burkholderia cepacia complex] Length: 527\nNumber of predicted TMHs: 2\nExp number of AAs in TMHs: 36.7303599999999\nExp number, first 60 AAs:

22.08937\nTotal prob of N-in: 0.97312\nPOSSIBLE N-term signal sequence\ninside 1 20\nTMhelix 21 43\noutside 44 502\nTMhelix 503 525\ninside 526 527

598 GCF_001992785.1_ASM199278v1 Burkholderia cenocepacia Proteobacteria; Betaproteobacteria; Burkholderiales; Burkholderiaceae; Burkholderia; Burkholderia cepacia complex MSDDIDNGGKGGFTRRDMMRVMAASGMMAVGGGGLLTSAQSAFAAPAPKRGGKIRVANES WP_006482802.1 MULTISPECIES: ABC transporter substrate-binding protein [Burkholderia cepacia complex] Length: 527\nNumber of predicted TMHs: 2\nExp number of AAs in TMHs: 36.7303599999999\nExp number, first 60 AAs: 22.08937\nTotal prob of N-in: 0.97312\nPOSSIBLE N-term signal sequence\ninside 1 20\nTMhelix 21 43\noutside 44 502\nTMhelix 503 525\ninside 526 527

599 GCF_001992445.1_ASM199244v1 Burkholderia cenocepacia Proteobacteria; Betaproteobacteria; Burkholderiales; Burkholderiaceae; Burkholderia; Burkholderia cepacia complex MKRRTFLALPAVAASGMLAGCSSADLGPVTHRLMKDPEYRETLSAFLISADGKKLVVIGKWP_006483583.1 MULTISPECIES: hypothetical protein [Burkholderia cepacia complex] Length: 212\nNumber of predicted TMHs: 1\nExp number of AAs in TMHs: 28.03134\nExp number, first 60 AAs: 6.30523\nTotal prob of N-in: 0.48947\noutside 1 187\nTMhelix 188 210\ninside 211 212

600 GCF_001992445.1_ASM199244v1 Burkholderia cenocepacia Proteobacteria; Betaproteobacteria; Burkholderiales; Burkholderiaceae; Burkholderia; Burkholderia cepacia complex MSDDIDNGGKGGFTRRDMMRVMAASGMMAVGGGGLLTSAQSAFAAPAPKRGGKIRVANES WP_006482802.1 MULTISPECIES: ABC transporter substrate-binding protein [Burkholderia cepacia complex] Length: 527\nNumber of predicted TMHs: 2\nExp number of AAs in TMHs: 36.7303599999999\nExp number, first 60 AAs: 22.08937\nTotal prob of N-in: 0.97312\nPOSSIBLE N-term signal sequence\ninside 1 20\nTMhelix 21 43\noutside 44 502\nTMhelix 503 525\ninside 526 527

601 GCF_001992285.1_ASM199228v1 Burkholderia cenocepacia Proteobacteria; Betaproteobacteria; Burkholderiales; Burkholderiaceae; Burkholderia; Burkholderia cepacia complex MKRRTFLALPAVAASGMLAGCSSADLGPVTHRLMKDPEYRETLSAFLISADGKKLVVIGKWP_034201760.1 MULTISPECIES: hypothetical protein [Burkholderia cepacia complex] Length: 212\nNumber of predicted TMHs: 1\nExp number of AAs in TMHs: 28.11606\nExp number, first 60 AAs: 6.33522\nTotal prob of N-in: 0.49088\noutside 1 187\nTMhelix 188 210\ninside 211 212

602 GCF_001992095.1_ASM199209v1 Burkholderia cenocepacia Proteobacteria; Betaproteobacteria; Burkholderiales; Burkholderiaceae; Burkholderia; Burkholderia cepacia complex MSDDIDNGGKGGFTRRDMMRVMAASGMMAVGGGGLLTSAQSAFAAPAPKRGGKIRVANES WP_077180776.1 ABC transporter substrate-binding protein [Burkholderia cenocepacia] Length: 527\nNumber of predicted TMHs: 2\nExp number of AAs in TMHs: 36.7312299999999\nExp number, first 60 AAs: 22.0894\nTotal prob of N-in: 0.97312\nPOSSIBLE N-term signal sequence\ninside 1 20\nTMhelix 21 43\noutside 44 502\nTMhelix 503 525\ninside 526 527

603 GCF_001992395.1_ASM199239v1 Burkholderia cenocepacia Proteobacteria; Betaproteobacteria; Burkholderiales; Burkholderiaceae; Burkholderia; Burkholderia cepacia complex MSTSITRRRLLAAGVAASSLALSGCFTPKLYKNDAYSEHVSAFMMTEDGKKLVVLGTRYH WP_077021267.1 hypothetical protein [Burkholderia cenocepacia] Length: 212\nNumber of predicted TMHs: 1\nExp number of AAs in TMHs: 24.17376\nExp number, first 60 AAs: 0.92388\nTotal prob of N-in: 0.37951\noutside 1 180\nTMhelix 181 203\ninside 204 212

604 GCF_001992395.1_ASM199239v1 Burkholderia cenocepacia Proteobacteria; Betaproteobacteria; Burkholderiales; Burkholderiaceae; Burkholderia; Burkholderia cepacia complex METLLTRIRASFADASPRRQLISVAVLVALVASLIAAGYFALRPRYQVLFRDLKPQDAAT WP_077020516.1 flagellar M-ring protein FliF [Burkholderia cenocepacia] Length: 491\nNumber of predicted TMHs: 2\nExp number of AAs in TMHs: 41.34236\nExp number, first 60 AAs: 22.1383\nTotal prob of N-in: 0.99080\nPOSSIBLE N-term signal sequence\ninside 1 20\nTMhelix 21 43\noutside 44 427\nTMhelix 428 450\ninside 451 491

605 GCF_001992395.1_ASM199239v1 Burkholderia cenocepacia Proteobacteria; Betaproteobacteria; Burkholderiales; Burkholderiaceae; Burkholderia; Burkholderia cepacia complex MKRRTFLALPAVAASGMLAGCSSADLGPVTHRLMKDPEYRETLSAFLISADGKKLVVIGKWP_006483583.1 MULTISPECIES: hypothetical protein [Burkholderia cepacia complex] Length: 212\nNumber of predicted TMHs: 1\nExp number of AAs in TMHs: 28.03134\nExp number, first 60 AAs: 6.30523\nTotal prob of N-in: 0.48947\noutside 1 187\nTMhelix 188 210\ninside 211 212

606 GCF_001992195.1_ASM199219v1 Burkholderia cenocepacia Proteobacteria; Betaproteobacteria; Burkholderiales; Burkholderiaceae; Burkholderia; Burkholderia cepacia complex
MKRRTFLALPAVAASGMLAGCSSADLGPVTHRLMKDPEYRETLSAFLISADGKKLVVIGKWP_006483583.1 MULTISPECIES: hypothetical protein [Burkholderia cepacia complex] Length: 212\nNumber of predicted TMHs: 1\nExp number of AAs in TMHs: 28.03134\nExp number, first 60 AAs: 6.30523\nTotal prob of N-in: 0.48947\noutside 1 187\nTMhelix 188 210\ninside 211 212

607 GCF_001992115.1_ASM199211v1 Burkholderia cenocepacia Proteobacteria; Betaproteobacteria; Burkholderiales; Burkholderiaceae; Burkholderia; Burkholderia cepacia complex
MKRRTFLALPAVAASGMLAGCSSADLGPVTHRLMKDPEYRETLSAFLISADGKKLVVIGKWP_006483583.1 MULTISPECIES: hypothetical protein [Burkholderia cepacia complex] Length: 212\nNumber of predicted TMHs: 1\nExp number of AAs in TMHs: 28.03134\nExp number, first 60 AAs: 6.30523\nTotal prob of N-in: 0.48947\noutside 1 187\nTMhelix 188 210\ninside 211 212

608 GCF_001991835.1_ASM199183v1 Burkholderia cenocepacia Proteobacteria; Betaproteobacteria; Burkholderiales; Burkholderiaceae; Burkholderia; Burkholderia cepacia complex
METLLTRIRASFADASPRRQLISVAVLVALVASLIAAGYFALRPRYQVLFRDLKPQDAAT WP_077020516.1 flagellar M-ring protein FliF [Burkholderia cenocepacia] Length: 491\nNumber of predicted TMHs: 2\nExp number of AAs in TMHs: 41.34236\nExp number, first 60 AAs: 22.1383\nTotal prob of N-in: 0.99080\nPOSSIBLE N-term signal sequence\ninside 1 20\nTMhelix 21 43\noutside 44 427\nTMhelix 428 450\ninside 451 491

609 GCF_001992155.1_ASM199215v1 Burkholderia cenocepacia Proteobacteria; Betaproteobacteria; Burkholderiales; Burkholderiaceae; Burkholderia; Burkholderia cepacia complex
MSDDIDNGGKGGFTRRDMMRVMAASGMMAVGGGGLLTSAQSAFAAPAPKRGGKIRVANES WP_077187164.1 ABC transporter substrate-binding protein [Burkholderia cenocepacia] Length: 527\nNumber of predicted TMHs: 2\nExp number of AAs in TMHs: 36.7300399999999\nExp number, first 60 AAs: 22.0894\nTotal prob of N-in: 0.97312\nPOSSIBLE N-term signal sequence\ninside 1 20\nTMhelix 21 43\noutside 44 502\nTMhelix 503 525\ninside 526 527

610 GCF_001992155.1_ASM199215v1 Burkholderia cenocepacia Proteobacteria; Betaproteobacteria; Burkholderiales; Burkholderiaceae; Burkholderia; Burkholderia cepacia complex
MSTSITRRRLLAAGVAASSLALSGCFTPKLYKNDAYSEHVSAFMMTEDGKKLVVLGTRYH WP_077021267.1
hypothetical protein [Burkholderia cenocepacia] Length: 212\nNumber of predicted TMHs: 1\nExp number of AAs in TMHs: 24.17376\nExp number, first 60 AAs: 0.92388\nTotal prob of N-in: 0.37951\noutside 1 180\nTMhelix 181 203\ninside 204 212

611 GCF_001992155.1_ASM199215v1 Burkholderia cenocepacia Proteobacteria; Betaproteobacteria; Burkholderiales; Burkholderiaceae; Burkholderia; Burkholderia cepacia complex
METLLTRIRASFADASPRRQLISVAVLVALVASLIAAGYFALRPRYQVLFRDLKPQDAAT WP_077020516.1 flagellar M-ring protein FliF [Burkholderia cenocepacia] Length: 491\nNumber of predicted TMHs: 2\nExp number of AAs in TMHs: 41.34236\nExp number, first 60 AAs: 22.1383\nTotal prob of N-in: 0.99080\nPOSSIBLE N-term signal sequence\ninside 1 20\nTMhelix 21 43\noutside 44 427\nTMhelix 428 450\ninside 451 491

612 GCF_001991755.1_ASM199175v1 Burkholderia cenocepacia Proteobacteria; Betaproteobacteria; Burkholderiales; Burkholderiaceae; Burkholderia; Burkholderia cepacia complex
MSTSITRRRLLAAGVAASSLALSGCFTPKLYKNDAYSEHVSAFMMTEDGKKLVVLGTRYH WP_077021267.1
hypothetical protein [Burkholderia cenocepacia] Length: 212\nNumber of predicted TMHs: 1\nExp number of AAs in TMHs: 24.17376\nExp number, first 60 AAs: 0.92388\nTotal prob of N-in: 0.37951\noutside 1 180\nTMhelix 181 203\ninside 204 212

613 GCF_001991935.1_ASM199193v1 Burkholderia cenocepacia Proteobacteria; Betaproteobacteria; Burkholderiales; Burkholderiaceae; Burkholderia; Burkholderia cepacia complex
MSTSITRRRLLAAGVAASSLALSGCFTPKLYKNDAYSEHVSAFMMTEDGKKLVVLGTRYH WP_077021267.1
hypothetical protein [Burkholderia cenocepacia] Length: 212\nNumber of predicted TMHs: 1\nExp number of AAs in TMHs: 24.17376\nExp number, first 60 AAs: 0.92388\nTotal prob of N-in: 0.37951\noutside 1 180\nTMhelix 181 203\ninside 204 212

614 GCF_001991915.1_ASM199191v1 Burkholderia cenocepacia Proteobacteria; Betaproteobacteria; Burkholderiales; Burkholderiaceae; Burkholderia; Burkholderia cepacia complex
MSDDIDNGGKGGFTRRDMMRVMAASGMMAVGGGGLLTSAQSAFAAPAPKRGGKIRVANES WP_077180776.1 ABC transporter substrate-binding protein [Burkholderia cenocepacia] Length: 527\nNumber of predicted TMHs: 2\nExp

number of AAs in TMHs: 36.7312299999999\nExp number, first 60 AAs: 22.0894\nTotal prob of N-in: 0.97312\nPOSSIBLE N-term signal sequence\ninside 1 20\nTMhelix 21 43\noutside 44 502\nTMhelix 503 525\ninside 526 527

615 GCF_001991935.1_ASM199193v1 Burkholderia cenocepacia Proteobacteria; Betaproteobacteria; Burkholderiales; Burkholderiaceae; Burkholderia; Burkholderia cepacia complex METLLTRIRASFADASPRRQLISVAVLVALVASLIAAGYFALRPRYQVLFRDLKPQDAAT WP_077020516.1 flagellar M-ring protein FliF [Burkholderia cenocepacia] Length: 491\nNumber of predicted TMHs: 2\nExp number of AAs in TMHs: 41.34236\nExp number, first 60 AAs: 22.1383\nTotal prob of N-in: 0.99080\nPOSSIBLE N-term signal sequence\ninside 1 20\nTMhelix 21 43\noutside 44 427\nTMhelix 428 450\ninside 451 491

616 GCF_001991935.1_ASM199193v1 Burkholderia cenocepacia Proteobacteria; Betaproteobacteria; Burkholderiales; Burkholderiaceae; Burkholderia; Burkholderia cepacia complex MKRRTFLALPAVAASGMLAGCSSADLGPVTHRLMKDPEYRETLSAFLISADGKKLVVIGKWP_006483583.1 MULTISPECIES: hypothetical protein [Burkholderia cepacia complex] Length: 212\nNumber of predicted TMHs: 1\nExp number of AAs in TMHs: 28.03134\nExp number, first 60 AAs: 6.30523\nTotal prob of N-in: 0.48947\noutside 1 187\nTMhelix 188 210\ninside 211 212

617 GCF_001991915.1_ASM199191v1 Burkholderia cenocepacia Proteobacteria; Betaproteobacteria; Burkholderiales; Burkholderiaceae; Burkholderia; Burkholderia cepacia complex MKRRTFLALPAVAASGMLAGCSSADLGPVTHRLMKDPEYRETLSAFLISADGKKLVVIGKWP_006483583.1 MULTISPECIES: hypothetical protein [Burkholderia cepacia complex] Length: 212\nNumber of predicted TMHs: 1\nExp number of AAs in TMHs: 28.03134\nExp number, first 60 AAs: 6.30523\nTotal prob of N-in: 0.48947\noutside 1 187\nTMhelix 188 210\ninside 211 212

618 GCF_001991935.1_ASM199193v1 Burkholderia cenocepacia Proteobacteria; Betaproteobacteria; Burkholderiales; Burkholderiaceae; Burkholderia; Burkholderia cepacia complex MSDDIDNGGKGGFTRRDMMRVMAASGMMAVGGGGLLTSAQSAFAAPAPKRGGKIRVANES WP_006482802.1 MULTISPECIES: ABC transporter substrate-binding protein [Burkholderia cepacia complex] Length: 527\nNumber of predicted TMHs: 2\nExp number of AAs in TMHs: 36.7303599999999\nExp number, first 60 AAs: 22.08937\nTotal prob of N-in: 0.97312\nPOSSIBLE N-term signal sequence\ninside 1 20\nTMhelix 21 43\noutside 44 502\nTMhelix 503 525\ninside 526 527

619 GCF_001991515.1_ASM199151v1 Burkholderia cenocepacia Proteobacteria; Betaproteobacteria; Burkholderiales; Burkholderiaceae; Burkholderia; Burkholderia cepacia complex MSTSITRRRLLAAGVAASSLALSGCFTPKLYKNDAYSEHVSAFMMTEDGKKLVVLGTRYH WP_077021267.1 hypothetical protein [Burkholderia cenocepacia] Length: 212\nNumber of predicted TMHs: 1\nExp number of AAs in TMHs: 24.17376\nExp number, first 60 AAs: 0.92388\nTotal prob of N-in: 0.37951\noutside 1 180\nTMhelix 181 203\ninside 204 212

620 GCF_001991735.1_ASM199173v1 Burkholderia cenocepacia Proteobacteria; Betaproteobacteria; Burkholderiales; Burkholderiaceae; Burkholderia; Burkholderia cepacia complex MSDDIDNGGKGGFTRRDMMRVMAASGMMAVGGGGLLTSAQSAFAAPAPKRGGKIRVANES WP_006482802.1 MULTISPECIES: ABC transporter substrate-binding protein [Burkholderia cepacia complex] Length: 527\nNumber of predicted TMHs: 2\nExp number of AAs in TMHs: 36.7303599999999\nExp number, first 60 AAs: 22.08937\nTotal prob of N-in: 0.97312\nPOSSIBLE N-term signal sequence\ninside 1 20\nTMhelix 21 43\noutside 44 502\nTMhelix 503 525\ninside 526 527

621 GCF_001991415.1_ASM199141v1 Burkholderia cenocepacia Proteobacteria; Betaproteobacteria; Burkholderiales; Burkholderiaceae; Burkholderia; Burkholderia cepacia complex METLLTRIRASFADASPRRQLISVAVLVALVASLIAAGYFALRPRYQVLFRDLKPQDAAT WP_077020516.1 flagellar M-ring protein FliF [Burkholderia cenocepacia] Length: 491\nNumber of predicted TMHs: 2\nExp number of AAs in TMHs: 41.34236\nExp number, first 60 AAs: 22.1383\nTotal prob of N-in: 0.99080\nPOSSIBLE N-term signal sequence\ninside 1 20\nTMhelix 21 43\noutside 44 427\nTMhelix 428 450\ninside 451 491

622 GCF_001991695.1_ASM199169v1 Burkholderia cenocepacia Proteobacteria; Betaproteobacteria; Burkholderiales; Burkholderiaceae; Burkholderia; Burkholderia cepacia complex MKRRTFLALPAVAASGMLAGCSSADLGPVTHRLMKDPEYRETLSAFLISADGKKLVVIGKWP_006483583.1 MULTISPECIES: hypothetical protein [Burkholderia cepacia complex] Length: 212\nNumber of predicted TMHs: 1\nExp number of AAs in TMHs: 28.03134\nExp number, first 60 AAs: 6.30523\nTotal prob of N-in: 0.48947\noutside 1 187\nTMhelix 188 210\ninside 211 212

623 GCF_001991665.1_ASM199166v1 Burkholderia cenocepacia Proteobacteria; Betaproteobacteria; Burkholderiales; Burkholderiaceae; Burkholderia; Burkholderia cepacia complex
MKRRTFLALPAVAASGMLAGCSSADLGPVTHRLMKDPEYRETLSAFLISADGKKLVVIGKWP_006483583.1 MULTISPECIES: hypothetical protein [Burkholderia cepacia complex] Length: 212\nNumber of predicted TMHs: 1\nExp number of AAs in TMHs: 28.03134\nExp number, first 60 AAs: 6.30523\nTotal prob of N-in: 0.48947\noutside 1 187\nTMhelix 188 210\ninside 211 212

624 GCF_001991695.1_ASM199169v1 Burkholderia cenocepacia Proteobacteria; Betaproteobacteria; Burkholderiales; Burkholderiaceae; Burkholderia; Burkholderia cepacia complex
MSDDIDNGGKGGFTRRDMMRVMAASGMMAVGGGGLLTSAQSAFAAPAPKRGGKIRVANES WP_006482802.1 MULTISPECIES: ABC transporter substrate-binding protein [Burkholderia cepacia complex] Length: 527\nNumber of predicted TMHs: 2\nExp number of AAs in TMHs: 36.7303599999999\nExp number, first 60 AAs: 22.08937\nTotal prob of N-in: 0.97312\nPOSSIBLE N-term signal sequence\ninside 1 20\nTMhelix 21 43\noutside 44 502\nTMhelix 503 525\ninside 526 527

625 GCF_001991665.1_ASM199166v1 Burkholderia cenocepacia Proteobacteria; Betaproteobacteria; Burkholderiales; Burkholderiaceae; Burkholderia; Burkholderia cepacia complex
MSDDIDNGGKGGFTRRDMMRVMAASGMMAVGGGGLLTSAQSAFAAPAPKRGGKIRVANES WP_006482802.1 MULTISPECIES: ABC transporter substrate-binding protein [Burkholderia cepacia complex] Length: 527\nNumber of predicted TMHs: 2\nExp number of AAs in TMHs: 36.7303599999999\nExp number, first 60 AAs: 22.08937\nTotal prob of N-in: 0.97312\nPOSSIBLE N-term signal sequence\ninside 1 20\nTMhelix 21 43\noutside 44 502\nTMhelix 503 525\ninside 526 527

626 GCF_001991625.1_ASM199162v1 Burkholderia cenocepacia Proteobacteria; Betaproteobacteria; Burkholderiales; Burkholderiaceae; Burkholderia; Burkholderia cepacia complex
MKRRTFLALPAVAASGMLAGCSSADLGPVTHRLMKDPEYRETLSAFLISADGKKLVVIGKWP_006483583.1 MULTISPECIES: hypothetical protein [Burkholderia cepacia complex] Length: 212\nNumber of predicted TMHs: 1\nExp number of AAs in TMHs: 28.03134\nExp number, first 60 AAs: 6.30523\nTotal prob of N-in: 0.48947\noutside 1 187\nTMhelix 188 210\ninside 211 212

627 GCF_001991415.1_ASM199141v1 Burkholderia cenocepacia Proteobacteria; Betaproteobacteria; Burkholderiales; Burkholderiaceae; Burkholderia; Burkholderia cepacia complex
MSDDIDNGGKGGFTRRDMMRVMAASGMMAVGGGGLLTSAQSAFAAPAPKRGGKIRVANES WP_006482802.1 MULTISPECIES: ABC transporter substrate-binding protein [Burkholderia cepacia complex] Length: 527\nNumber of predicted TMHs: 2\nExp number of AAs in TMHs: 36.7303599999999\nExp number, first 60 AAs: 22.08937\nTotal prob of N-in: 0.97312\nPOSSIBLE N-term signal sequence\ninside 1 20\nTMhelix 21 43\noutside 44 502\nTMhelix 503 525\ninside 526 527

628 GCF_001991275.1_ASM199127v1 Burkholderia cenocepacia Proteobacteria; Betaproteobacteria; Burkholderiales; Burkholderiaceae; Burkholderia; Burkholderia cepacia complex
MKRRTFLALPAVAASGMLAGCSSADLGPVTHRLMKDPEYRETLSAFLISADGKKLVVIGKWP_034201760.1 MULTISPECIES: hypothetical protein [Burkholderia cepacia complex] Length: 212\nNumber of predicted TMHs: 1\nExp number of AAs in TMHs: 28.11606\nExp number, first 60 AAs: 6.33522\nTotal prob of N-in: 0.49088\noutside 1 187\nTMhelix 188 210\ninside 211 212

629 GCF_001991375.1_ASM199137v1 Burkholderia cenocepacia Proteobacteria; Betaproteobacteria; Burkholderiales; Burkholderiaceae; Burkholderia; Burkholderia cepacia complex
MSTSITRRRLLAAGVAASSLALSGCFTPKLYKNDAYSEHVSAFMMTEDGKKLVVLGTRYH WP_077021267.1 hypothetical protein [Burkholderia cenocepacia] Length: 212\nNumber of predicted TMHs: 1\nExp number of AAs in TMHs: 24.17376\nExp number, first 60 AAs: 0.92388\nTotal prob of N-in: 0.37951\noutside 1 180\nTMhelix 181 203\ninside 204 212

630 GCF_001991375.1_ASM199137v1 Burkholderia cenocepacia Proteobacteria; Betaproteobacteria; Burkholderiales; Burkholderiaceae; Burkholderia; Burkholderia cepacia complex
METLLTRIRASFADASPRRQLISVAVLVALVASLIAAGYFALRPRYQVLFRDLKPQDAAT WP_077020516.1 flagellar M-ring protein FliF [Burkholderia cenocepacia] Length: 491\nNumber of predicted TMHs: 2\nExp number of AAs in TMHs: 41.34236\nExp number, first 60 AAs: 22.1383\nTotal prob of N-in: 0.99080\nPOSSIBLE N-term signal sequence\ninside 1 20\nTMhelix 21 43\noutside 44 427\nTMhelix 428 450\ninside 451 491

631 GCF_001991375.1_ASM199137v1 Burkholderia cenocepacia Proteobacteria; Betaproteobacteria; Burkholderiales; Burkholderiaceae; Burkholderia; Burkholderia cepacia complex

MKRRTFLALPAVAASGMLAGCSSADLGPVTHRLMKDPEYRETLSAFLISADGKKLVVIGKWP_006483583.1 MULTISPECIES: hypothetical protein [Burkholderia cepacia complex] Length: 212\nNumber of predicted TMHs: 1\nExp number of AAs in TMHs: 28.03134\nExp number, first 60 AAs: 6.30523\nTotal prob of N-in: 0.48947\noutside 1 187\nTMhelix 188 210\ninside 211 212

632 GCF_001991375.1_ASM199137v1 Burkholderia cenocepacia Proteobacteria; Betaproteobacteria; Burkholderiales; Burkholderiaceae; Burkholderia; Burkholderia cepacia complex MSDDIDNGGKGGFTRRDMMRVMAASGMMAVGGGGLLTSAQSAFAAPAPKRGGKIRVANES WP_006482802.1 MULTISPECIES: ABC transporter substrate-binding protein [Burkholderia cepacia complex] Length: 527\nNumber of predicted TMHs: 2\nExp number of AAs in TMHs: 36.7303599999999\nExp number, first 60 AAs: 22.08937\nTotal prob of N-in: 0.97312\nPOSSIBLE N-term signal sequence\ninside 1 20\nTMhelix 21 43\noutside 44 502\nTMhelix 503 525\ninside 526 527

633 GCF_001991235.1_ASM199123v1 Burkholderia cenocepacia Proteobacteria; Betaproteobacteria; Burkholderiales; Burkholderiaceae; Burkholderia; Burkholderia cepacia complex MKRRTFLALPAVAASGMLAGCSSADLGPVTHRLMKDPEYRETLSAFLISADGKKLVVIGKWP_034201760.1 MULTISPECIES: hypothetical protein [Burkholderia cepacia complex] Length: 212\nNumber of predicted TMHs: 1\nExp number of AAs in TMHs: 28.11606\nExp number, first 60 AAs: 6.33522\nTotal prob of N-in: 0.49088\noutside 1 187\nTMhelix 188 210\ninside 211 212

634 GCF_001991195.1_ASM199119v1 Burkholderia cenocepacia Proteobacteria; Betaproteobacteria; Burkholderiales; Burkholderiaceae; Burkholderia; Burkholderia cepacia complex MSDDIDNGGKGGFTRRDMMRVMAASGMMAVGGGGLLTSAQSAFAAPAPKRGGKIRVANES WP_006482802.1 MULTISPECIES: ABC transporter substrate-binding protein [Burkholderia cepacia complex] Length: 527\nNumber of predicted TMHs: 2\nExp number of AAs in TMHs: 36.7303599999999\nExp number, first 60 AAs: 22.08937\nTotal prob of N-in: 0.97312\nPOSSIBLE N-term signal sequence\ninside 1 20\nTMhelix 21 43\noutside 44 502\nTMhelix 503 525\ninside 526 527

635 GCF_001720525.1_ASM172052v1 Burkholderia cenocepacia Proteobacteria; Betaproteobacteria; Burkholderiales; Burkholderiaceae; Burkholderia; Burkholderia cepacia complex MKRRTFLALPAVAASGMLAGCSSADLGPVTHRLMKDPEYRETLSAFLISADGKKLVVIGKWP_069352077.1 5-formyltetrahydrofolate cyclo-ligase [Burkholderia cenocepacia] Length: 212\nNumber of predicted TMHs: 1\nExp number of AAs in TMHs: 28.13384\nExp number, first 60 AAs: 6.35524\nTotal prob of N-in: 0.49086\noutside 1 187\nTMhelix 188 210\ninside 211 212

636 GCF_001991155.1_ASM199115v1 Burkholderia cenocepacia Proteobacteria; Betaproteobacteria; Burkholderiales; Burkholderiaceae; Burkholderia; Burkholderia cepacia complex MKRRTFLALPAVAASGMLAGCSSADLGPVTHRLMKDPEYRETLSAFLISADGKKLVVIGKWP_034201760.1 MULTISPECIES: hypothetical protein [Burkholderia cepacia complex] Length: 212\nNumber of predicted TMHs: 1\nExp number of AAs in TMHs: 28.11606\nExp number, first 60 AAs: 6.33522\nTotal prob of N-in: 0.49088\noutside 1 187\nTMhelix 188 210\ninside 211 212

637 GCF_001991135.1_ASM199113v1 Burkholderia cenocepacia Proteobacteria; Betaproteobacteria; Burkholderiales; Burkholderiaceae; Burkholderia; Burkholderia cepacia complex MKRRTFLALPAVAASGMLAGCSSADLGPVTHRLMKDPEYRETLSAFLISADGKKLVVIGKWP_034201760.1 MULTISPECIES: hypothetical protein [Burkholderia cepacia complex] Length: 212\nNumber of predicted TMHs: 1\nExp number of AAs in TMHs: 28.11606\nExp number, first 60 AAs: 6.33522\nTotal prob of N-in: 0.49088\noutside 1 187\nTMhelix 188 210\ninside 211 212

638 GCF_001984375.1_ASM198437v1 Burkholderia cenocepacia Proteobacteria; Betaproteobacteria; Burkholderiales; Burkholderiaceae; Burkholderia; Burkholderia cepacia complex MKRRTFLALPAVAASGMLAGCSSADLGPVTHRLMKDPEYRETLSAFLISADGKKLVVIGKWP_034201760.1 MULTISPECIES: hypothetical protein [Burkholderia cepacia complex] Length: 212\nNumber of predicted TMHs: 1\nExp number of AAs in TMHs: 28.11606\nExp number, first 60 AAs: 6.33522\nTotal prob of N-in: 0.49088\noutside 1 187\nTMhelix 188 210\ninside 211 212

639 GCF_001991155.1_ASM199115v1 Burkholderia cenocepacia Proteobacteria; Betaproteobacteria; Burkholderiales; Burkholderiaceae; Burkholderia; Burkholderia cepacia complex MSDDIDNGGKGGFTRRDMMRVMAASGMMAVGGGGLLTSAQSAFAAPAPKRGGKIRVANES WP_006482802.1 MULTISPECIES: ABC transporter substrate-binding protein [Burkholderia cepacia complex] Length: 527\nNumber of predicted TMHs: 2\nExp number of AAs in TMHs: 36.7303599999999\nExp number, first 60 AAs:

22.08937\nTotal prob of N-in: 0.97312\nPOSSIBLE N-term signal sequence\ninside 1 20\nTMhelix 21 43\noutside 44 502\nTMhelix 503 525\ninside 526 527

640 GCF_001991135.1_ASM199113v1 Burkholderia cenocepacia Proteobacteria; Betaproteobacteria; Burkholderiales; Burkholderiaceae; Burkholderia; Burkholderia cepacia complex MSDDIDNGGKGGFTRRDMMRVMAASGMMAVGGGGLLTSAQSAFAAPAPKRGGKIRVANES WP_006482802.1 MULTISPECIES: ABC transporter substrate-binding protein [Burkholderia cepacia complex] Length: 527\nNumber of predicted TMHs: 2\nExp number of AAs in TMHs: 36.7303599999999\nExp number, first 60 AAs: 22.08937\nTotal prob of N-in: 0.97312\nPOSSIBLE N-term signal sequence\ninside 1 20\nTMhelix 21 43\noutside 44 502\nTMhelix 503 525\ninside 526 527

641 GCF_001531965.1_ASM153196v1 Burkholderia cenocepacia Proteobacteria; Betaproteobacteria; Burkholderiales; Burkholderiaceae; Burkholderia; Burkholderia cepacia complex METLLTRIRASFADASPRRQLISVAVLVALVASLIAAGYFALRPRYQVLFRDLKPQDAAT WP_060214257.1 flagellar M-ring protein FliF [Burkholderia cenocepacia] Length: 491\nNumber of predicted TMHs: 2\nExp number of AAs in TMHs: 41.34235\nExp number, first 60 AAs: 22.1383\nTotal prob of N-in: 0.99080\nPOSSIBLE N-term signal sequence\ninside 1 20\nTMhelix 21 43\noutside 44 427\nTMhelix 428 450\ninside 451 491

642 GCF_001484665.1_ASM148466v1 Burkholderia cenocepacia Proteobacteria; Betaproteobacteria; Burkholderiales; Burkholderiaceae; Burkholderia; Burkholderia cepacia complex MKRRTFLALPAVAASGMLAGCSSADLGPVTHRLMKDPEYRETLSAFLISADGKKLVVIGKWP_034201760.1 MULTISPECIES: hypothetical protein [Burkholderia cepacia complex] Length: 212\nNumber of predicted TMHs: 1\nExp number of AAs in TMHs: 28.11606\nExp number, first 60 AAs: 6.33522\nTotal prob of N-in: 0.49088\noutside 1 187\nTMhelix 188 210\ninside 211 212

643 GCF_001531965.1_ASM153196v1 Burkholderia cenocepacia Proteobacteria; Betaproteobacteria; Burkholderiales; Burkholderiaceae; Burkholderia; Burkholderia cepacia complex MKRRTFLALPAVAASGMLAGCSSADLGPVTHRLMKDPEYRETLSAFLISADGKKLVVIGKWP_006483583.1 MULTISPECIES: hypothetical protein [Burkholderia cepacia complex] Length: 212\nNumber of predicted TMHs: 1\nExp number of AAs in TMHs: 28.03134\nExp number, first 60 AAs: 6.30523\nTotal prob of N-in: 0.48947\noutside 1 187\nTMhelix 188 210\ninside 211 212

644 GCF_001484665.1_ASM148466v1 Burkholderia cenocepacia Proteobacteria; Betaproteobacteria; Burkholderiales; Burkholderiaceae; Burkholderia; Burkholderia cepacia complex MSDDIDNGGKGGFTRRDMMRVMAASGMMAVGGGGLLTSAQSAFAAPAPKRGGKIRVANES WP_006482802.1 MULTISPECIES: ABC transporter substrate-binding protein [Burkholderia cepacia complex] Length: 527\nNumber of predicted TMHs: 2\nExp number of AAs in TMHs: 36.7303599999999\nExp number, first 60 AAs: 22.08937\nTotal prob of N-in: 0.97312\nPOSSIBLE N-term signal sequence\ninside 1 20\nTMhelix 21 43\noutside 44 502\nTMhelix 503 525\ninside 526 527

645 GCF_001531965.1_ASM153196v1 Burkholderia cenocepacia Proteobacteria; Betaproteobacteria; Burkholderiales; Burkholderiaceae; Burkholderia; Burkholderia cepacia complex MSDDIDNGGKGGFTRRDMMRVMAASGMMAVGGGGLLTSAQSAFAAPAPKRGGKIRVANES WP_006482802.1 MULTISPECIES: ABC transporter substrate-binding protein [Burkholderia cepacia complex] Length: 527\nNumber of predicted TMHs: 2\nExp number of AAs in TMHs: 36.7303599999999\nExp number, first 60 AAs: 22.08937\nTotal prob of N-in: 0.97312\nPOSSIBLE N-term signal sequence\ninside 1 20\nTMhelix 21 43\noutside 44 502\nTMhelix 503 525\ninside 526 527

646 GCF_001055325.1_ASM105532v1 Burkholderia cenocepacia Proteobacteria; Betaproteobacteria; Burkholderiales; Burkholderiaceae; Burkholderia; Burkholderia cepacia complex MKRRTFLALPAVAASGMLAGCSSADLGPVTHRLMKDPEYQETLSAFLISADGKKLVVIGK WP_048986526.1 hypothetical protein [Burkholderia cenocepacia] Length: 212\nNumber of predicted TMHs: 1\nExp number of AAs in TMHs: 28.09653\nExp number, first 60 AAs: 6.31597\nTotal prob of N-in: 0.49053\noutside 1 187\nTMhelix 188 210\ninside 211 212

647 GCF_001072455.1_ASM107245v1 Burkholderia cenocepacia Proteobacteria; Betaproteobacteria; Burkholderiales; Burkholderiaceae; Burkholderia; Burkholderia cepacia complex MKRRTFLALPAVAASGMLAGCSSADLGPVTHRLMKDPEYQETLSAFLISADGKKLVVIGK WP_048986526.1 hypothetical protein [Burkholderia cenocepacia] Length: 212\nNumber of predicted TMHs: 1\nExp number of AAs in TMHs: 28.09653\nExp number, first 60 AAs: 6.31597\nTotal prob of N-in: 0.49053\noutside 1 187\nTMhelix 188 210\ninside 211 212

648 GCF_001994835.1_ASM199483v1 Burkholderia cenocepacia Proteobacteria; Betaproteobacteria; Burkholderiales; Burkholderiaceae; Burkholderia; Burkholderia cepacia complex
MSTSITRRRLLAAGVAASSLALSGCFTPKLYKNDAYSEHVSAFMMTEDGKKLVVLGTRYH WP_077021267.1 hypothetical protein [Burkholderia cenocepacia] Length: 212\nNumber of predicted TMHs: 1\nExp number of AAs in TMHs: 24.17376\nExp number, first 60 AAs: 0.92388\nTotal prob of N-in: 0.37951\noutside 1 180\nTMhelix 181 203\ninside 204 212

649 GCF_002071955.1_ASM207195v1 Burkholderia cenocepacia Proteobacteria; Betaproteobacteria; Burkholderiales; Burkholderiaceae; Burkholderia; Burkholderia cepacia complex
MSDDIDNGGKGGFTRRDMMRVMAASGMMAVGGGGLLTSAQSAFAAPAPKRGGKIRVANES WP_006482802.1 MULTISPECIES: ABC transporter substrate-binding protein [Burkholderia cepacia complex] Length: 527\nNumber of predicted TMHs: 2\nExp number of AAs in TMHs: 36.7303599999999\nExp number, first 60 AAs: 22.08937\nTotal prob of N-in: 0.97312\nPOSSIBLE N-term signal sequence\ninside 1 20\nTMhelix 21 43\noutside 44 502\nTMhelix 503 525\ninside 526 527

650 GCF_002071935.1_ASM207193v1 Burkholderia cenocepacia Proteobacteria; Betaproteobacteria; Burkholderiales; Burkholderiaceae; Burkholderia; Burkholderia cepacia complex
MSDDIDNGGKGGFTRRDMMRVMAASGMMAVGGGGLLTSAQSAFAAPAPKRGGKIRVANES WP_006482802.1 MULTISPECIES: ABC transporter substrate-binding protein [Burkholderia cepacia complex] Length: 527\nNumber of predicted TMHs: 2\nExp number of AAs in TMHs: 36.7303599999999\nExp number, first 60 AAs: 22.08937\nTotal prob of N-in: 0.97312\nPOSSIBLE N-term signal sequence\ninside 1 20\nTMhelix 21 43\noutside 44 502\nTMhelix 503 525\ninside 526 527

651 GCF_001994835.1_ASM199483v1 Burkholderia cenocepacia Proteobacteria; Betaproteobacteria; Burkholderiales; Burkholderiaceae; Burkholderia; Burkholderia cepacia complex
METLLTRIRASFADASPRRQLISVAVLVALVASLIAAGYFALRPRYQVLFRDLKPQDAAT WP_077020516.1 flagellar M-ring protein FliF [Burkholderia cenocepacia] Length: 491\nNumber of predicted TMHs: 2\nExp number of AAs in TMHs: 41.34236\nExp number, first 60 AAs: 22.1383\nTotal prob of N-in: 0.99080\nPOSSIBLE N-term signal sequence\ninside 1 20\nTMhelix 21 43\noutside 44 427\nTMhelix 428 450\ninside 451 491

652 GCF_002071885.1_ASM207188v1 Burkholderia cenocepacia Proteobacteria; Betaproteobacteria; Burkholderiales; Burkholderiaceae; Burkholderia; Burkholderia cepacia complex
MKRRTFLALPAVAASGMLAGCSSADLGPVTHRLMKDPEYRETLSAFLISADGKKLVVIGKWP_062884514.1 hypothetical protein [Burkholderia cenocepacia] Length: 212\nNumber of predicted TMHs: 1\nExp number of AAs in TMHs: 28.03134\nExp number, first 60 AAs: 6.30523\nTotal prob of N-in: 0.48947\noutside 1 187\nTMhelix 188 210\ninside 211 212

653 GCF_002071885.1_ASM207188v1 Burkholderia cenocepacia Proteobacteria; Betaproteobacteria; Burkholderiales; Burkholderiaceae; Burkholderia; Burkholderia cepacia complex
METLLTRIRASFADASPRRQLISVAVLVALVASLIAAGYFALRPRYQVLFRDLKPQDAAT WP_060214257.1 flagellar M-ring protein FliF [Burkholderia cenocepacia] Length: 491\nNumber of predicted TMHs: 2\nExp number of AAs in TMHs: 41.34235\nExp number, first 60 AAs: 22.1383\nTotal prob of N-in: 0.99080\nPOSSIBLE N-term signal sequence\ninside 1 20\nTMhelix 21 43\noutside 44 427\nTMhelix 428 450\ninside 451 491

654 GCF_001994615.1_ASM199461v1 Burkholderia cenocepacia Proteobacteria; Betaproteobacteria; Burkholderiales; Burkholderiaceae; Burkholderia; Burkholderia cepacia complex
METLLTRIRASFADASPRRQLISVAVLVALVASLIAAGYFALRPRYQVLFRDLKPQDAAT WP_077020516.1 flagellar M-ring protein FliF [Burkholderia cenocepacia] Length: 491\nNumber of predicted TMHs: 2\nExp number of AAs in TMHs: 41.34236\nExp number, first 60 AAs: 22.1383\nTotal prob of N-in: 0.99080\nPOSSIBLE N-term signal sequence\ninside 1 20\nTMhelix 21 43\noutside 44 427\nTMhelix 428 450\ninside 451 491

655 GCF_001994775.1_ASM199477v1 Burkholderia cenocepacia Proteobacteria; Betaproteobacteria; Burkholderiales; Burkholderiaceae; Burkholderia; Burkholderia cepacia complex
MKRRTFLALPAVAASGMLAGCSSADLGPVTHRLMKDPEYRETLSAFLISADGKKLVVIGKWP_034201760.1 MULTISPECIES: hypothetical protein [Burkholderia cepacia complex] Length: 212\nNumber of predicted TMHs: 1\nExp number of AAs in TMHs: 28.11606\nExp number, first 60 AAs: 6.33522\nTotal prob of N-in: 0.49088\noutside 1 187\nTMhelix 188 210\ninside 211 212

656 GCF_001994875.1_ASM199487v1 Burkholderia cenocepacia Proteobacteria; Betaproteobacteria; Burkholderiales; Burkholderiaceae; Burkholderia; Burkholderia cepacia complex
MKRRTFLALPAVAASGMLAGCSSADLGPVTHRLMKDPEYRETLSAFLISADGKKLVVIGKWP_006483583.1 MULTISPECIES:

hypothetical protein [Burkholderia cepacia complex] Length: 212\nNumber of predicted TMHs: 1\nExp number of AAs in TMHs: 28.03134\nExp number, first 60 AAs: 6.30523\nTotal prob of N-in: 0.48947\noutside 1 187\nTMhelix 188 210\ninside 211 212

657 GCF_001994875.1_ASM199487v1 Burkholderia cenocepacia Proteobacteria; Betaproteobacteria; Burkholderiales; Burkholderiaceae; Burkholderia; Burkholderia cepacia complex MSDDIDNGGKGGFTRRDMMRVMAASGMMAVGGGGLLTSAQSAFAAPAPKRGGKIRVANES WP_006482802.1 MULTISPECIES: ABC transporter substrate-binding protein [Burkholderia cepacia complex] Length: 527\nNumber of predicted TMHs: 2\nExp number of AAs in TMHs: 36.7303599999999\nExp number, first 60 AAs: 22.08937\nTotal prob of N-in: 0.97312\nPOSSIBLE N-term signal sequence\ninside 1 20\nTMhelix 21 43\noutside 44 502\nTMhelix 503 525\ninside 526 527

658 GCF_001994835.1_ASM199483v1 Burkholderia cenocepacia Proteobacteria; Betaproteobacteria; Burkholderiales; Burkholderiaceae; Burkholderia; Burkholderia cepacia complex MKRRTFLALPAVAASGMLAGCSSADLGPVTHRLMKDPEYRETLSAFLISADGKKLVVIGKWP_006483583.1 MULTISPECIES: hypothetical protein [Burkholderia cepacia complex] Length: 212\nNumber of predicted TMHs: 1\nExp number of AAs in TMHs: 28.03134\nExp number, first 60 AAs: 6.30523\nTotal prob of N-in: 0.48947\noutside 1 187\nTMhelix 188 210\ninside 211 212

659 GCF_001994815.1_ASM199481v1 Burkholderia cenocepacia Proteobacteria; Betaproteobacteria; Burkholderiales; Burkholderiaceae; Burkholderia; Burkholderia cepacia complex MSDDIDNGGKGGFTRRDMMRVMAASGMMAVGGGGLLTSAQSAFAAPAPKRGGKIRVANES WP_006482802.1 MULTISPECIES: ABC transporter substrate-binding protein [Burkholderia cepacia complex] Length: 527\nNumber of predicted TMHs: 2\nExp number of AAs in TMHs: 36.7303599999999\nExp number, first 60 AAs: 22.08937\nTotal prob of N-in: 0.97312\nPOSSIBLE N-term signal sequence\ninside 1 20\nTMhelix 21 43\noutside 44 502\nTMhelix 503 525\ninside 526 527

660 GCF_001994775.1_ASM199477v1 Burkholderia cenocepacia Proteobacteria; Betaproteobacteria; Burkholderiales; Burkholderiaceae; Burkholderia; Burkholderia cepacia complex MSDDIDNGGKGGFTRRDMMRVMAASGMMAVGGGGLLTSAQSAFAAPAPKRGGKIRVANES WP_006482802.1 MULTISPECIES: ABC transporter substrate-binding protein [Burkholderia cepacia complex] Length: 527\nNumber of predicted TMHs: 2\nExp number of AAs in TMHs: 36.7303599999999\nExp number, first 60 AAs: 22.08937\nTotal prob of N-in: 0.97312\nPOSSIBLE N-term signal sequence\ninside 1 20\nTMhelix 21 43\noutside 44 502\nTMhelix 503 525\ninside 526 527

661 GCF_001994445.1_ASM199444v1 Burkholderia cenocepacia Proteobacteria; Betaproteobacteria; Burkholderiales; Burkholderiaceae; Burkholderia; Burkholderia cepacia complex MKRRTFLALPAVAASGMLAGCSSADLGPVTHRLMKDPEYRETLSAFLISADGKKLVVIGKWP_034201760.1 MULTISPECIES: hypothetical protein [Burkholderia cepacia complex] Length: 212\nNumber of predicted TMHs: 1\nExp number of AAs in TMHs: 28.11606\nExp number, first 60 AAs: 6.33522\nTotal prob of N-in: 0.49088\noutside 1 187\nTMhelix 188 210\ninside 211 212

662 GCF_001994505.1_ASM199450v1 Burkholderia cenocepacia Proteobacteria; Betaproteobacteria; Burkholderiales; Burkholderiaceae; Burkholderia; Burkholderia cepacia complex MSDDIDNGGKGGFTRRDMMRVMAASGMMAVGGGGLLTSAQSAFAAPAPKRGGKIRVANES WP_006482802.1 MULTISPECIES: ABC transporter substrate-binding protein [Burkholderia cepacia complex] Length: 527\nNumber of predicted TMHs: 2\nExp number of AAs in TMHs: 36.7303599999999\nExp number, first 60 AAs: 22.08937\nTotal prob of N-in: 0.97312\nPOSSIBLE N-term signal sequence\ninside 1 20\nTMhelix 21 43\noutside 44 502\nTMhelix 503 525\ninside 526 527

663 GCF_001994475.1_ASM199447v1 Burkholderia cenocepacia Proteobacteria; Betaproteobacteria; Burkholderiales; Burkholderiaceae; Burkholderia; Burkholderia cepacia complex MSDDIDNGGKGGFTRRDMMRVMAASGMMAVGGGGLLTSAQSAFAAPAPKRGGKIRVANES WP_006482802.1 MULTISPECIES: ABC transporter substrate-binding protein [Burkholderia cepacia complex] Length: 527\nNumber of predicted TMHs: 2\nExp number of AAs in TMHs: 36.7303599999999\nExp number, first 60 AAs: 22.08937\nTotal prob of N-in: 0.97312\nPOSSIBLE N-term signal sequence\ninside 1 20\nTMhelix 21 43\noutside 44 502\nTMhelix 503 525\ninside 526 527

664 GCF_001994265.1_ASM199426v1 Burkholderia cenocepacia Proteobacteria; Betaproteobacteria; Burkholderiales; Burkholderiaceae; Burkholderia; Burkholderia cepacia complex MKRRTFLALPAVAASGMLAGCSSADLGPVTHRLMKDPEYRETLSAFLISADGKKLVVIGKWP_034201760.1 MULTISPECIES:

hypothetical protein [Burkholderia cepacia complex]	Length: 212\nNumber of predicted TMHs: 1\nExp number of AAs in TMHs: 28.11606\nExp number, first 60 AAs: 6.33522\nTotal prob of N-in: 0.49088\noutside 1 187\nTMhelix 188 210\ninside 211 212

665	GCF_001994445.1_ASM199444v1	Burkholderia cenocepacia	Proteobacteria; Betaproteobacteria; Burkholderiales; Burkholderiaceae; Burkholderia; Burkholderia cepacia complex
MSDDIDNGGKGGFTRRDMMRVMAASGMMAVGGGGLLTSAQSAFAAPAPKRGGKIRVANES	WP_006482802.1
MULTISPECIES: ABC transporter substrate-binding protein [Burkholderia cepacia complex]	Length: 527\nNumber of predicted TMHs: 2\nExp number of AAs in TMHs: 36.7303599999999\nExp number, first 60 AAs: 22.08937\nTotal prob of N-in: 0.97312\nPOSSIBLE N-term signal sequence\ninside 1 20\nTMhelix 21 43\noutside 44 502\nTMhelix 503 525\ninside 526 527

666	GCF_001994215.1_ASM199421v1	Burkholderia cenocepacia	Proteobacteria; Betaproteobacteria; Burkholderiales; Burkholderiaceae; Burkholderia; Burkholderia cepacia complex
MKRRTFLALPAVAASGMLAGCSSADLGPVTHRLMKDPEYRETLSAFLISADGKKLVVIGKWP_034201760.1	MULTISPECIES: hypothetical protein [Burkholderia cepacia complex]	Length: 212\nNumber of predicted TMHs: 1\nExp number of AAs in TMHs: 28.11606\nExp number, first 60 AAs: 6.33522\nTotal prob of N-in: 0.49088\noutside 1 187\nTMhelix 188 210\ninside 211 212

667	GCF_001994405.1_ASM199440v1	Burkholderia cenocepacia	Proteobacteria; Betaproteobacteria; Burkholderiales; Burkholderiaceae; Burkholderia; Burkholderia cepacia complex
MKRRTFLALPAVAASGMLAGCSSADLGPVTHRLMKDPEYRETLSAFLISADGKKLVVIGKWP_034201760.1	MULTISPECIES: hypothetical protein [Burkholderia cepacia complex]	Length: 212\nNumber of predicted TMHs: 1\nExp number of AAs in TMHs: 28.11606\nExp number, first 60 AAs: 6.33522\nTotal prob of N-in: 0.49088\noutside 1 187\nTMhelix 188 210\ninside 211 212

668	GCF_001994025.1_ASM199402v1	Burkholderia cenocepacia	Proteobacteria; Betaproteobacteria; Burkholderiales; Burkholderiaceae; Burkholderia; Burkholderia cepacia complex
MKRRTFLALPAVAASGMLAGCSSADLGPVTHRLMKDPEYRETLSAFLISADGKKLVVIGKWP_006483583.1	MULTISPECIES: hypothetical protein [Burkholderia cepacia complex]	Length: 212\nNumber of predicted TMHs: 1\nExp number of AAs in TMHs: 28.03134\nExp number, first 60 AAs: 6.30523\nTotal prob of N-in: 0.48947\noutside 1 187\nTMhelix 188 210\ninside 211 212

669	GCF_001994035.1_ASM199403v1	Burkholderia cenocepacia	Proteobacteria; Betaproteobacteria; Burkholderiales; Burkholderiaceae; Burkholderia; Burkholderia cepacia complex
MSDDIDNGGKGGFTRRDMMRVMAASGMMAVGGGGLLTSAQSAFAAPAPKRGGKIRVANES	WP_006482802.1
MULTISPECIES: ABC transporter substrate-binding protein [Burkholderia cepacia complex]	Length: 527\nNumber of predicted TMHs: 2\nExp number of AAs in TMHs: 36.7303599999999\nExp number, first 60 AAs: 22.08937\nTotal prob of N-in: 0.97312\nPOSSIBLE N-term signal sequence\ninside 1 20\nTMhelix 21 43\noutside 44 502\nTMhelix 503 525\ninside 526 527

670	GCF_001994025.1_ASM199402v1	Burkholderia cenocepacia	Proteobacteria; Betaproteobacteria; Burkholderiales; Burkholderiaceae; Burkholderia; Burkholderia cepacia complex
MSDDIDNGGKGGFTRRDMMRVMAASGMMAVGGGGLLTSAQSAFAAPAPKRGGKIRVANES	WP_006482802.1
MULTISPECIES: ABC transporter substrate-binding protein [Burkholderia cepacia complex]	Length: 527\nNumber of predicted TMHs: 2\nExp number of AAs in TMHs: 36.7303599999999\nExp number, first 60 AAs: 22.08937\nTotal prob of N-in: 0.97312\nPOSSIBLE N-term signal sequence\ninside 1 20\nTMhelix 21 43\noutside 44 502\nTMhelix 503 525\ninside 526 527

671	GCF_001993975.1_ASM199397v1	Burkholderia cenocepacia	Proteobacteria; Betaproteobacteria; Burkholderiales; Burkholderiaceae; Burkholderia; Burkholderia cepacia complex
MKRRTFLALPAVAASGMLAGCSSADLGPVTHRLMKDPEYRETLSAFLISADGKKLVVIGKWP_006483583.1	MULTISPECIES: hypothetical protein [Burkholderia cepacia complex]	Length: 212\nNumber of predicted TMHs: 1\nExp number of AAs in TMHs: 28.03134\nExp number, first 60 AAs: 6.30523\nTotal prob of N-in: 0.48947\noutside 1 187\nTMhelix 188 210\ninside 211 212

672	GCF_001993925.1_ASM199392v1	Burkholderia cenocepacia	Proteobacteria; Betaproteobacteria; Burkholderiales; Burkholderiaceae; Burkholderia; Burkholderia cepacia complex
MKRRTFLALPAVAASGMLAGCSSADLGPVTHRLMKDPEYRETLSAFLISADGKKLVVIGKWP_006483583.1	MULTISPECIES: hypothetical protein [Burkholderia cepacia complex]	Length: 212\nNumber of predicted TMHs: 1\nExp number of

AAs in TMHs: 28.03134\nExp number, first 60 AAs: 6.30523\nTotal prob of N-in: 0.48947\noutside 1 187\nTMhelix 188 210\ninside 211 212

673 GCF_001993895.1_ASM199389v1 Burkholderia cenocepacia Proteobacteria; Betaproteobacteria; Burkholderiales; Burkholderiaceae; Burkholderia; Burkholderia cepacia complex MKRRTFLALPAVAASGMLAGCSSADLGPVTHRLMKDPEYRETLSAFLISADGKKLVVIGKWP_006483583.1 MULTISPECIES: hypothetical protein [Burkholderia cepacia complex] Length: 212\nNumber of predicted TMHs: 1\nExp number of AAs in TMHs: 28.03134\nExp number, first 60 AAs: 6.30523\nTotal prob of N-in: 0.48947\noutside 1 187\nTMhelix 188 210\ninside 211 212

674 GCF_001993975.1_ASM199397v1 Burkholderia cenocepacia Proteobacteria; Betaproteobacteria; Burkholderiales; Burkholderiaceae; Burkholderia; Burkholderia cepacia complex MSDDIDNGGKGGFTRRDMMRVMAASGMMAVGGGGLLTSAQSAFAAPAPKRGGKIRVANES WP_006482802.1 MULTISPECIES: ABC transporter substrate-binding protein [Burkholderia cepacia complex] Length: 527\nNumber of predicted TMHs: 2\nExp number of AAs in TMHs: 36.7303599999999\nExp number, first 60 AAs: 22.08937\nTotal prob of N-in: 0.97312\nPOSSIBLE N-term signal sequence\ninside 1 20\nTMhelix 21 43\noutside 44 502\nTMhelix 503 525\ninside 526 527

675 GCF_001993435.1_ASM199343v1 Burkholderia cenocepacia Proteobacteria; Betaproteobacteria; Burkholderiales; Burkholderiaceae; Burkholderia; Burkholderia cepacia complex MSTSITRRRLLAAGVAASSLALSGCFTPKLYKNDAYSEHVSAFMMTEDGKKLVVLGTRYH WP_077021267.1 hypothetical protein [Burkholderia cenocepacia] Length: 212\nNumber of predicted TMHs: 1\nExp number of AAs in TMHs: 24.17376\nExp number, first 60 AAs: 0.92388\nTotal prob of N-in: 0.37951\noutside 1 180\nTMhelix 181 203\ninside 204 212

676 GCF_001991355.1_ASM199135v1 Burkholderia cenocepacia Proteobacteria; Betaproteobacteria; Burkholderiales; Burkholderiaceae; Burkholderia; Burkholderia cepacia complex MSTSITRRRLLAAGVAASSLALSGCFTPKLYKNDAYSEHVSAFMMTEDGKKLVVLGTRYH WP_077021267.1 hypothetical protein [Burkholderia cenocepacia] Length: 212\nNumber of predicted TMHs: 1\nExp number of AAs in TMHs: 24.17376\nExp number, first 60 AAs: 0.92388\nTotal prob of N-in: 0.37951\noutside 1 180\nTMhelix 181 203\ninside 204 212

677 GCF_001992135.1_ASM199213v1 Burkholderia cenocepacia Proteobacteria; Betaproteobacteria; Burkholderiales; Burkholderiaceae; Burkholderia; Burkholderia cepacia complex METLLTRIRASFADASPRRQLISVAVLVALVASLIAAGYFALRPRYQVLFRDLKPQDAAT WP_077020516.1 flagellar M-ring protein FliF [Burkholderia cenocepacia] Length: 491\nNumber of predicted TMHs: 2\nExp number of AAs in TMHs: 41.34236\nExp number, first 60 AAs: 22.1383\nTotal prob of N-in: 0.99080\nPOSSIBLE N-term signal sequence\ninside 1 20\nTMhelix 21 43\noutside 44 427\nTMhelix 428 450\ninside 451 491

678 GCF_001992035.1_ASM199203v1 Burkholderia cenocepacia Proteobacteria; Betaproteobacteria; Burkholderiales; Burkholderiaceae; Burkholderia; Burkholderia cepacia complex METLLTRIRASFADASPRRQLISVAVLVALVASLIAAGYFALRPRYQVLFRDLKPQDAAT WP_077020516.1 flagellar M-ring protein FliF [Burkholderia cenocepacia] Length: 491\nNumber of predicted TMHs: 2\nExp number of AAs in TMHs: 41.34236\nExp number, first 60 AAs: 22.1383\nTotal prob of N-in: 0.99080\nPOSSIBLE N-term signal sequence\ninside 1 20\nTMhelix 21 43\noutside 44 427\nTMhelix 428 450\ninside 451 491

679 GCF_001992075.1_ASM199207v1 Burkholderia cenocepacia Proteobacteria; Betaproteobacteria; Burkholderiales; Burkholderiaceae; Burkholderia; Burkholderia cepacia complex MKRRTFLALPAVAASGMLAGCSSADLGPVTHRLMKDPEYRETLSAFLISADGKKLVVIGKWP_006483583.1 MULTISPECIES: hypothetical protein [Burkholderia cepacia complex] Length: 212\nNumber of predicted TMHs: 1\nExp number of AAs in TMHs: 28.03134\nExp number, first 60 AAs: 6.30523\nTotal prob of N-in: 0.48947\noutside 1 187\nTMhelix 188 210\ninside 211 212

680 GCF_001992135.1_ASM199213v1 Burkholderia cenocepacia Proteobacteria; Betaproteobacteria; Burkholderiales; Burkholderiaceae; Burkholderia; Burkholderia cepacia complex MKRRTFLALPAVAASGMLAGCSSADLGPVTHRLMKDPEYRETLSAFLISADGKKLVVIGKWP_006483583.1 MULTISPECIES: hypothetical protein [Burkholderia cepacia complex] Length: 212\nNumber of predicted TMHs: 1\nExp number of AAs in TMHs: 28.03134\nExp number, first 60 AAs: 6.30523\nTotal prob of N-in: 0.48947\noutside 1 187\nTMhelix 188 210\ninside 211 212

681 GCF_001992035.1_ASM199203v1 Burkholderia cenocepacia Proteobacteria; Betaproteobacteria; Burkholderiales; Burkholderiaceae; Burkholderia; Burkholderia cepacia complex
MKRRTFLALPAVAASGMLAGCSSADLGPVTHRLMKDPEYRETLSAFLISADGKKLVVIGKWP_006483583.1 MULTISPECIES: hypothetical protein [Burkholderia cepacia complex] Length: 212\nNumber of predicted TMHs: 1\nExp number of AAs in TMHs: 28.03134\nExp number, first 60 AAs: 6.30523\nTotal prob of N-in: 0.48947\noutside 1 187\nTMhelix 188 210\ninside 211 212

682 GCF_001992035.1_ASM199203v1 Burkholderia cenocepacia Proteobacteria; Betaproteobacteria; Burkholderiales; Burkholderiaceae; Burkholderia; Burkholderia cepacia complex
MSDDIDNGGKGGFTRRDMMRVMAASGMMAVGGGGLLTSAQSAFAAPAPKRGGKIRVANES WP_006482802.1 MULTISPECIES: ABC transporter substrate-binding protein [Burkholderia cepacia complex] Length: 527\nNumber of predicted TMHs: 2\nExp number of AAs in TMHs: 36.7303599999999\nExp number, first 60 AAs: 22.08937\nTotal prob of N-in: 0.97312\nPOSSIBLE N-term signal sequence\ninside 1 20\nTMhelix 21 43\noutside 44 502\nTMhelix 503 525\ninside 526 527

683 GCF_001991655.1_ASM199165v1 Burkholderia cenocepacia Proteobacteria; Betaproteobacteria; Burkholderiales; Burkholderiaceae; Burkholderia; Burkholderia cepacia complex
METLLTRIRASFADASPRRQLISVAVLVALVASLIAAGYFALRPRYQVLFRDLKPQDAAT WP_077020516.1 flagellar M-ring protein FliF [Burkholderia cenocepacia] Length: 491\nNumber of predicted TMHs: 2\nExp number of AAs in TMHs: 41.34236\nExp number, first 60 AAs: 22.1383\nTotal prob of N-in: 0.99080\nPOSSIBLE N-term signal sequence\ninside 1 20\nTMhelix 21 43\noutside 44 427\nTMhelix 428 450\ninside 451 491

684 GCF_001991615.1_ASM199161v1 Burkholderia cenocepacia Proteobacteria; Betaproteobacteria; Burkholderiales; Burkholderiaceae; Burkholderia; Burkholderia cepacia complex
MSTSITRRRLLAAGVAASSLALSGCFTPKLYKNDAYSEHVSAFMMTEDGKKLVVLGTRYH WP_077021267.1 hypothetical protein [Burkholderia cenocepacia] Length: 212\nNumber of predicted TMHs: 1\nExp number of AAs in TMHs: 24.17376\nExp number, first 60 AAs: 0.92388\nTotal prob of N-in: 0.37951\noutside 1 180\nTMhelix 181 203\ninside 204 212

685 GCF_001991615.1_ASM199161v1 Burkholderia cenocepacia Proteobacteria; Betaproteobacteria; Burkholderiales; Burkholderiaceae; Burkholderia; Burkholderia cepacia complex
METLLTRIRASFADASPRRQLISVAVLVALVASLIAAGYFALRPRYQVLFRDLKPQDAAT WP_077020516.1 flagellar M-ring protein FliF [Burkholderia cenocepacia] Length: 491\nNumber of predicted TMHs: 2\nExp number of AAs in TMHs: 41.34236\nExp number, first 60 AAs: 22.1383\nTotal prob of N-in: 0.99080\nPOSSIBLE N-term signal sequence\ninside 1 20\nTMhelix 21 43\noutside 44 427\nTMhelix 428 450\ninside 451 491

686 GCF_001992205.1_ASM199220v1 Burkholderia cenocepacia Proteobacteria; Betaproteobacteria; Burkholderiales; Burkholderiaceae; Burkholderia; Burkholderia cepacia complex
MKRRTFLALPAVAASGMLAGCSSADLGPVTHRLMKDPEYRETLSAFLISADGKKLVVIGKWP_006483583.1 MULTISPECIES: hypothetical protein [Burkholderia cepacia complex] Length: 212\nNumber of predicted TMHs: 1\nExp number of AAs in TMHs: 28.03134\nExp number, first 60 AAs: 6.30523\nTotal prob of N-in: 0.48947\noutside 1 187\nTMhelix 188 210\ninside 211 212

687 GCF_001992495.1_ASM199249v1 Burkholderia cenocepacia Proteobacteria; Betaproteobacteria; Burkholderiales; Burkholderiaceae; Burkholderia; Burkholderia cepacia complex
MKRRTFLALPAVAASGMLAGCSSADLGPVTHRLMKDPEYRETLSAFLISADGKKLVVIGKWP_006483583.1 MULTISPECIES: hypothetical protein [Burkholderia cepacia complex] Length: 212\nNumber of predicted TMHs: 1\nExp number of AAs in TMHs: 28.03134\nExp number, first 60 AAs: 6.30523\nTotal prob of N-in: 0.48947\noutside 1 187\nTMhelix 188 210\ninside 211 212

688 GCF_001991845.1_ASM199184v1 Burkholderia cenocepacia Proteobacteria; Betaproteobacteria; Burkholderiales; Burkholderiaceae; Burkholderia; Burkholderia cepacia complex
MSTSITRRRLLAAGVAASSLALSGCFTPKLYKNDAYSEHVSAFMMTEDGKKLVVLGTRYH WP_077021267.1 hypothetical protein [Burkholderia cenocepacia] Length: 212\nNumber of predicted TMHs: 1\nExp number of AAs in TMHs: 24.17376\nExp number, first 60 AAs: 0.92388\nTotal prob of N-in: 0.37951\noutside 1 180\nTMhelix 181 203\ninside 204 212

689 GCF_001991885.1_ASM199188v1 Burkholderia cenocepacia Proteobacteria; Betaproteobacteria; Burkholderiales; Burkholderiaceae; Burkholderia; Burkholderia cepacia complex
MSDDIDNGGKGGFTRRDMMRVMAASGMMAVGGGGLLTSAQSAFAAPAPKRGGKIRVANES WP_077180776.1 ABC transporter substrate-binding protein [Burkholderia cenocepacia] Length: 527\nNumber of predicted TMHs: 2\nExp

number of AAs in TMHs: 36.7312299999999\nExp number, first 60 AAs: 22.0894\nTotal prob of N-in: 0.97312\nPOSSIBLE N-term signal sequence\ninside 1 20\nTMhelix 21 43\noutside 44 502\nTMhelix 503 525\ninside 526 527

690 GCF_001991845.1_ASM199184v1 Burkholderia cenocepacia Proteobacteria; Betaproteobacteria; Burkholderiales; Burkholderiaceae; Burkholderia; Burkholderia cepacia complex METLLTRIRASFADASPRRQLISVAVLVALVASLIAAGYFALRPRYQVLFRDLKPQDAAT WP_077020516.1 flagellar M-ring protein FliF [Burkholderia cenocepacia] Length: 491\nNumber of predicted TMHs: 2\nExp number of AAs in TMHs: 41.34236\nExp number, first 60 AAs: 22.1383\nTotal prob of N-in: 0.99080\nPOSSIBLE N-term signal sequence\ninside 1 20\nTMhelix 21 43\noutside 44 427\nTMhelix 428 450\ninside 451 491

691 GCF_001991955.1_ASM199195v1 Burkholderia cenocepacia Proteobacteria; Betaproteobacteria; Burkholderiales; Burkholderiaceae; Burkholderia; Burkholderia cepacia complex MKRRTFLALPAVAASGMLAGCSSADLGPVTHRLMKDPEYRETLSAFLISADGKKLVVIGKWP_006483583.1 MULTISPECIES: hypothetical protein [Burkholderia cepacia complex] Length: 212\nNumber of predicted TMHs: 1\nExp number of AAs in TMHs: 28.03134\nExp number, first 60 AAs: 6.30523\nTotal prob of N-in: 0.48947\noutside 1 187\nTMhelix 188 210\ninside 211 212

692 GCF_001991995.1_ASM199199v1 Burkholderia cenocepacia Proteobacteria; Betaproteobacteria; Burkholderiales; Burkholderiaceae; Burkholderia; Burkholderia cepacia complex MKRRTFLALPAVAASGMLAGCSSADLGPVTHRLMKDPEYRETLSAFLISADGKKLVVIGKWP_006483583.1 MULTISPECIES: hypothetical protein [Burkholderia cepacia complex] Length: 212\nNumber of predicted TMHs: 1\nExp number of AAs in TMHs: 28.03134\nExp number, first 60 AAs: 6.30523\nTotal prob of N-in: 0.48947\noutside 1 187\nTMhelix 188 210\ninside 211 212

693 GCF_001991885.1_ASM199188v1 Burkholderia cenocepacia Proteobacteria; Betaproteobacteria; Burkholderiales; Burkholderiaceae; Burkholderia; Burkholderia cepacia complex MKRRTFLALPAVAASGMLAGCSSADLGPVTHRLMKDPEYRETLSAFLISADGKKLVVIGKWP_006483583.1 MULTISPECIES: hypothetical protein [Burkholderia cepacia complex] Length: 212\nNumber of predicted TMHs: 1\nExp number of AAs in TMHs: 28.03134\nExp number, first 60 AAs: 6.30523\nTotal prob of N-in: 0.48947\noutside 1 187\nTMhelix 188 210\ninside 211 212

694 GCF_001991845.1_ASM199184v1 Burkholderia cenocepacia Proteobacteria; Betaproteobacteria; Burkholderiales; Burkholderiaceae; Burkholderia; Burkholderia cepacia complex MSDDIDNGGKGGFTRRDMMRVMAASGMMAVGGGGLLTSAQSAFAAPAPKRGGKIRVANES WP_006482802.1 MULTISPECIES: ABC transporter substrate-binding protein [Burkholderia cepacia complex] Length: 527\nNumber of predicted TMHs: 2\nExp number of AAs in TMHs: 36.7303599999999\nExp number, first 60 AAs: 22.08937\nTotal prob of N-in: 0.97312\nPOSSIBLE N-term signal sequence\ninside 1 20\nTMhelix 21 43\noutside 44 502\nTMhelix 503 525\ninside 526 527

695 GCF_001991815.1_ASM199181v1 Burkholderia cenocepacia Proteobacteria; Betaproteobacteria; Burkholderiales; Burkholderiaceae; Burkholderia; Burkholderia cepacia complex MKRRTFLALPAVAASGMLAGCSSADLGPVTHRLMKDPEYRETLSAFLISADGKKLVVIGKWP_006483583.1 MULTISPECIES: hypothetical protein [Burkholderia cepacia complex] Length: 212\nNumber of predicted TMHs: 1\nExp number of AAs in TMHs: 28.03134\nExp number, first 60 AAs: 6.30523\nTotal prob of N-in: 0.48947\noutside 1 187\nTMhelix 188 210\ninside 211 212

696 GCF_001991385.1_ASM199138v1 Burkholderia cenocepacia Proteobacteria; Betaproteobacteria; Burkholderiales; Burkholderiaceae; Burkholderia; Burkholderia cepacia complex MSTSITRRRLLAAGVAASSLALSGCFTPKLYKNDAYSEHVSAFMMTEDGKKLVVLGTRYH WP_077021267.1 hypothetical protein [Burkholderia cenocepacia] Length: 212\nNumber of predicted TMHs: 1\nExp number of AAs in TMHs: 24.17376\nExp number, first 60 AAs: 0.92388\nTotal prob of N-in: 0.37951\noutside 1 180\nTMhelix 181 203\ninside 204 212

697 GCF_001991815.1_ASM199181v1 Burkholderia cenocepacia Proteobacteria; Betaproteobacteria; Burkholderiales; Burkholderiaceae; Burkholderia; Burkholderia cepacia complex MSDDIDNGGKGGFTRRDMMRVMAASGMMAVGGGGLLTSAQSAFAAPAPKRGGKIRVANES WP_006482802.1 MULTISPECIES: ABC transporter substrate-binding protein [Burkholderia cepacia complex] Length: 527\nNumber of predicted TMHs: 2\nExp number of AAs in TMHs: 36.7303599999999\nExp number, first 60 AAs: 22.08937\nTotal prob of N-in: 0.97312\nPOSSIBLE N-term signal sequence\ninside 1 20\nTMhelix 21 43\noutside 44 502\nTMhelix 503 525\ninside 526 527

698 GCF_001991795.1_ASM199179v1 Burkholderia cenocepacia Proteobacteria; Betaproteobacteria; Burkholderiales; Burkholderiaceae; Burkholderia; Burkholderia cepacia complex
MKRRTFLALPAVAASGMLAGCSSADLGPVTHRLMKDPEYRETLSAFLISADGKKLVVIGKWP_006483583.1 MULTISPECIES: hypothetical protein [Burkholderia cepacia complex] Length: 212\nNumber of predicted TMHs: 1\nExp number of AAs in TMHs: 28.03134\nExp number, first 60 AAs: 6.30523\nTotal prob of N-in: 0.48947\noutside 1 187\nTMhelix 188 210\ninside 211 212

699 GCF_001991795.1_ASM199179v1 Burkholderia cenocepacia Proteobacteria; Betaproteobacteria; Burkholderiales; Burkholderiaceae; Burkholderia; Burkholderia cepacia complex
MSDDIDNGGKGGFTRRDMMRVMAASGMMAVGGGGLLTSAQSAFAAPAPKRGGKIRVANES WP_006482802.1 MULTISPECIES: ABC transporter substrate-binding protein [Burkholderia cepacia complex] Length: 527\nNumber of predicted TMHs: 2\nExp number of AAs in TMHs: 36.7303599999999\nExp number, first 60 AAs: 22.08937\nTotal prob of N-in: 0.97312\nPOSSIBLE N-term signal sequence\ninside 1 20\nTMhelix 21 43\noutside 44 502\nTMhelix 503 525\ninside 526 527

700 GCF_001991385.1_ASM199138v1 Burkholderia cenocepacia Proteobacteria; Betaproteobacteria; Burkholderiales; Burkholderiaceae; Burkholderia; Burkholderia cepacia complex
METLLTRIRASFADASPRRQLISVAVLVALVASLIAAGYFALRPRYQVLFRDLKPQDAAT WP_077020516.1 flagellar M-ring protein FliF [Burkholderia cenocepacia] Length: 491\nNumber of predicted TMHs: 2\nExp number of AAs in TMHs: 41.34236\nExp number, first 60 AAs: 22.1383\nTotal prob of N-in: 0.99080\nPOSSIBLE N-term signal sequence\ninside 1 20\nTMhelix 21 43\noutside 44 427\nTMhelix 428 450\ninside 451 491

701 GCF_001991845.1_ASM199184v1 Burkholderia cenocepacia Proteobacteria; Betaproteobacteria; Burkholderiales; Burkholderiaceae; Burkholderia; Burkholderia cepacia complex
MKRRTFLALPAVAASGMLAGCSSADLGPVTHRLMKDPEYRETLSAFLISADGKKLVVIGKWP_006483583.1 MULTISPECIES: hypothetical protein [Burkholderia cepacia complex] Length: 212\nNumber of predicted TMHs: 1\nExp number of AAs in TMHs: 28.03134\nExp number, first 60 AAs: 6.30523\nTotal prob of N-in: 0.48947\noutside 1 187\nTMhelix 188 210\ninside 211 212

702 GCF_001991745.1_ASM199174v1 Burkholderia cenocepacia Proteobacteria; Betaproteobacteria; Burkholderiales; Burkholderiaceae; Burkholderia; Burkholderia cepacia complex
MSTSITRRRLLAAGVAASSLALSGCFTPKLYKNDAYSEHVSAFMMTEDGKKLVVLGTRYH WP_077021267.1 hypothetical protein [Burkholderia cenocepacia] Length: 212\nNumber of predicted TMHs: 1\nExp number of AAs in TMHs: 24.17376\nExp number, first 60 AAs: 0.92388\nTotal prob of N-in: 0.37951\noutside 1 180\nTMhelix 181 203\ninside 204 212

703 GCF_001991745.1_ASM199174v1 Burkholderia cenocepacia Proteobacteria; Betaproteobacteria; Burkholderiales; Burkholderiaceae; Burkholderia; Burkholderia cepacia complex
METLLTRIRASFADASPRRQLISVAVLVALVASLIAAGYFALRPRYQVLFRDLKPQDAAT WP_077020516.1 flagellar M-ring protein FliF [Burkholderia cenocepacia] Length: 491\nNumber of predicted TMHs: 2\nExp number of AAs in TMHs: 41.34236\nExp number, first 60 AAs: 22.1383\nTotal prob of N-in: 0.99080\nPOSSIBLE N-term signal sequence\ninside 1 20\nTMhelix 21 43\noutside 44 427\nTMhelix 428 450\ninside 451 491

704 GCF_001991745.1_ASM199174v1 Burkholderia cenocepacia Proteobacteria; Betaproteobacteria; Burkholderiales; Burkholderiaceae; Burkholderia; Burkholderia cepacia complex
MKRRTFLALPAVAASGMLAGCSSADLGPVTHRLMKDPEYRETLSAFLISADGKKLVVIGKWP_006483583.1 MULTISPECIES: hypothetical protein [Burkholderia cepacia complex] Length: 212\nNumber of predicted TMHs: 1\nExp number of AAs in TMHs: 28.03134\nExp number, first 60 AAs: 6.30523\nTotal prob of N-in: 0.48947\noutside 1 187\nTMhelix 188 210\ninside 211 212

705 GCF_001991715.1_ASM199171v1 Burkholderia cenocepacia Proteobacteria; Betaproteobacteria; Burkholderiales; Burkholderiaceae; Burkholderia; Burkholderia cepacia complex
MSTSITRRRLLAAGVAASSLALSGCFTPKLYKNDAYSEHVSAFMMTEDGKKLVVLGTRYH WP_077021267.1 hypothetical protein [Burkholderia cenocepacia] Length: 212\nNumber of predicted TMHs: 1\nExp number of AAs in TMHs: 24.17376\nExp number, first 60 AAs: 0.92388\nTotal prob of N-in: 0.37951\noutside 1 180\nTMhelix 181 203\ninside 204 212

706 GCF_001991715.1_ASM199171v1 Burkholderia cenocepacia Proteobacteria; Betaproteobacteria; Burkholderiales; Burkholderiaceae; Burkholderia; Burkholderia cepacia complex
METLLTRIRASFADASPRRQLISVAVLVALVASLIAAGYFALRPRYQVLFRDLKPQDAAT WP_077020516.1 flagellar M-ring protein FliF [Burkholderia cenocepacia] Length: 491\nNumber of predicted TMHs: 2\nExp number of AAs in

TMHs: 41.34236\nExp number, first 60 AAs:  22.1383\nTotal prob of N-in:   0.99080\nPOSSIBLE N-term signal sequence\ninside   1   20\nTMhelix   21   43\noutside   44   427\nTMhelix   428   450\ninside   451   491

707 GCF_001991715.1_ASM199171v1 Burkholderia cenocepacia Proteobacteria; Betaproteobacteria; Burkholderiales; Burkholderiaceae; Burkholderia; Burkholderia cepacia complex MKRRTFLALPAVAASGMLAGCSSADLGPVTHRLMKDPEYRETLSAFLISADGKKLVVIGKWP_006483583.1 MULTISPECIES: hypothetical protein [Burkholderia cepacia complex] Length: 212\nNumber of predicted TMHs:  1\nExp number of AAs in TMHs: 28.03134\nExp number, first 60 AAs:  6.30523\nTotal prob of N-in:   0.48947\noutside   1   187\nTMhelix   188   210\ninside   211   212

708 GCF_001991655.1_ASM199165v1 Burkholderia cenocepacia Proteobacteria; Betaproteobacteria; Burkholderiales; Burkholderiaceae; Burkholderia; Burkholderia cepacia complex MKRRTFLALPAVAASGMLAGCSSADLGPVTHRLMKDPEYRETLSAFLISADGKKLVVIGKWP_006483583.1 MULTISPECIES: hypothetical protein [Burkholderia cepacia complex] Length: 212\nNumber of predicted TMHs:  1\nExp number of AAs in TMHs: 28.03134\nExp number, first 60 AAs:  6.30523\nTotal prob of N-in:   0.48947\noutside   1   187\nTMhelix   188   210\ninside   211   212

709 GCF_001991745.1_ASM199174v1 Burkholderia cenocepacia Proteobacteria; Betaproteobacteria; Burkholderiales; Burkholderiaceae; Burkholderia; Burkholderia cepacia complex MSDDIDNGGKGGFTRRDMMRVMAASGMMAVGGGGLLTSAQSAFAAPAPKRGGKIRVANES WP_006482802.1 MULTISPECIES: ABC transporter substrate-binding protein [Burkholderia cepacia complex] Length: 527\nNumber of predicted TMHs:  2\nExp number of AAs in TMHs: 36.7303599999999\nExp number, first 60 AAs: 22.08937\nTotal prob of N-in:   0.97312\nPOSSIBLE N-term signal sequence\ninside   1   20\nTMhelix   21   43\noutside   44   502\nTMhelix   503   525\ninside   526   527

710 GCF_001991715.1_ASM199171v1 Burkholderia cenocepacia Proteobacteria; Betaproteobacteria; Burkholderiales; Burkholderiaceae; Burkholderia; Burkholderia cepacia complex MSDDIDNGGKGGFTRRDMMRVMAASGMMAVGGGGLLTSAQSAFAAPAPKRGGKIRVANES WP_006482802.1 MULTISPECIES: ABC transporter substrate-binding protein [Burkholderia cepacia complex] Length: 527\nNumber of predicted TMHs:  2\nExp number of AAs in TMHs: 36.7303599999999\nExp number, first 60 AAs: 22.08937\nTotal prob of N-in:   0.97312\nPOSSIBLE N-term signal sequence\ninside   1   20\nTMhelix   21   43\noutside   44   502\nTMhelix   503   525\ninside   526   527

711 GCF_001991655.1_ASM199165v1 Burkholderia cenocepacia Proteobacteria; Betaproteobacteria; Burkholderiales; Burkholderiaceae; Burkholderia; Burkholderia cepacia complex MSDDIDNGGKGGFTRRDMMRVMAASGMMAVGGGGLLTSAQSAFAAPAPKRGGKIRVANES WP_006482802.1 MULTISPECIES: ABC transporter substrate-binding protein [Burkholderia cepacia complex] Length: 527\nNumber of predicted TMHs:  2\nExp number of AAs in TMHs: 36.7303599999999\nExp number, first 60 AAs: 22.08937\nTotal prob of N-in:   0.97312\nPOSSIBLE N-term signal sequence\ninside   1   20\nTMhelix   21   43\noutside   44   502\nTMhelix   503   525\ninside   526   527

712 GCF_001991615.1_ASM199161v1 Burkholderia cenocepacia Proteobacteria; Betaproteobacteria; Burkholderiales; Burkholderiaceae; Burkholderia; Burkholderia cepacia complex MKRRTFLALPAVAASGMLAGCSSADLGPVTHRLMKDPEYRETLSAFLISADGKKLVVIGKWP_006483583.1 MULTISPECIES: hypothetical protein [Burkholderia cepacia complex] Length: 212\nNumber of predicted TMHs:  1\nExp number of AAs in TMHs: 28.03134\nExp number, first 60 AAs:  6.30523\nTotal prob of N-in:   0.48947\noutside   1   187\nTMhelix   188   210\ninside   211   212

713 GCF_001991615.1_ASM199161v1 Burkholderia cenocepacia Proteobacteria; Betaproteobacteria; Burkholderiales; Burkholderiaceae; Burkholderia; Burkholderia cepacia complex MSDDIDNGGKGGFTRRDMMRVMAASGMMAVGGGGLLTSAQSAFAAPAPKRGGKIRVANES WP_006482802.1 MULTISPECIES: ABC transporter substrate-binding protein [Burkholderia cepacia complex] Length: 527\nNumber of predicted TMHs:  2\nExp number of AAs in TMHs: 36.7303599999999\nExp number, first 60 AAs: 22.08937\nTotal prob of N-in:   0.97312\nPOSSIBLE N-term signal sequence\ninside   1   20\nTMhelix   21   43\noutside   44   502\nTMhelix   503   525\ninside   526   527

714 GCF_001991475.1_ASM199147v1 Burkholderia cenocepacia Proteobacteria; Betaproteobacteria; Burkholderiales; Burkholderiaceae; Burkholderia; Burkholderia cepacia complex MKRRTFLALPAVAASGMLAGCSSADLGPVTHRLMKDPEYRETLSAFLISADGKKLVVIGKWP_034201760.1 MULTISPECIES: hypothetical protein [Burkholderia cepacia complex] Length: 212\nNumber of predicted TMHs:  1\nExp number of

AAs in TMHs: 28.11606\nExp number, first 60 AAs: 6.33522\nTotal prob of N-in: 0.49088\noutside 1 187\nTMhelix 188 210\ninside 211 212

715 GCF_001991535.1_ASM199153v1 Burkholderia cenocepacia Proteobacteria; Betaproteobacteria; Burkholderiales; Burkholderiaceae; Burkholderia; Burkholderia cepacia complex MSTSITRRRLLAAGVAASSLALSGCFTPKLYKNDAYSEHVSAFMMTEDGKKLVVLGTRYH WP_077021267.1 hypothetical protein [Burkholderia cenocepacia] Length: 212\nNumber of predicted TMHs: 1\nExp number of AAs in TMHs: 24.17376\nExp number, first 60 AAs: 0.92388\nTotal prob of N-in: 0.37951\noutside 1 180\nTMhelix 181 203\ninside 204 212

716 GCF_001991535.1_ASM199153v1 Burkholderia cenocepacia Proteobacteria; Betaproteobacteria; Burkholderiales; Burkholderiaceae; Burkholderia; Burkholderia cepacia complex METLLTRIRASFADASPRRQLISVAVLVALVASLIAAGYFALRPRYQVLFRDLKPQDAAT WP_077020516.1 flagellar M-ring protein FliF [Burkholderia cenocepacia] Length: 491\nNumber of predicted TMHs: 2\nExp number of AAs in TMHs: 41.34236\nExp number, first 60 AAs: 22.1383\nTotal prob of N-in: 0.99080\nPOSSIBLE N-term signal sequence\ninside 1 20\nTMhelix 21 43\noutside 44 427\nTMhelix 428 450\ninside 451 491

717 GCF_001991535.1_ASM199153v1 Burkholderia cenocepacia Proteobacteria; Betaproteobacteria; Burkholderiales; Burkholderiaceae; Burkholderia; Burkholderia cepacia complex MKRRTFLALPAVAASGMLAGCSSADLGPVTHRLMKDPEYRETLSAFLISADGKKLVVIGKWP_006483583.1 MULTISPECIES: hypothetical protein [Burkholderia cepacia complex] Length: 212\nNumber of predicted TMHs: 1\nExp number of AAs in TMHs: 28.03134\nExp number, first 60 AAs: 6.30523\nTotal prob of N-in: 0.48947\noutside 1 187\nTMhelix 188 210\ninside 211 212

718 GCF_001991565.1_ASM199156v1 Burkholderia cenocepacia Proteobacteria; Betaproteobacteria; Burkholderiales; Burkholderiaceae; Burkholderia; Burkholderia cepacia complex MKRRTFLALPAVAASGMLAGCSSADLGPVTHRLMKDPEYRETLSAFLISADGKKLVVIGKWP_034201760.1 MULTISPECIES: hypothetical protein [Burkholderia cepacia complex] Length: 212\nNumber of predicted TMHs: 1\nExp number of AAs in TMHs: 28.11606\nExp number, first 60 AAs: 6.33522\nTotal prob of N-in: 0.49088\noutside 1 187\nTMhelix 188 210\ninside 211 212

719 GCF_001991565.1_ASM199156v1 Burkholderia cenocepacia Proteobacteria; Betaproteobacteria; Burkholderiales; Burkholderiaceae; Burkholderia; Burkholderia cepacia complex MSDDIDNGGKGGFTRRDMMRVMAASGMMAVGGGGLLTSAQSAFAAPAPKRGGKIRVANES WP_006482802.1 MULTISPECIES: ABC transporter substrate-binding protein [Burkholderia cepacia complex] Length: 527\nNumber of predicted TMHs: 2\nExp number of AAs in TMHs: 36.7303599999999\nExp number, first 60 AAs: 22.08937\nTotal prob of N-in: 0.97312\nPOSSIBLE N-term signal sequence\ninside 1 20\nTMhelix 21 43\noutside 44 502\nTMhelix 503 525\ninside 526 527

720 GCF_001991535.1_ASM199153v1 Burkholderia cenocepacia Proteobacteria; Betaproteobacteria; Burkholderiales; Burkholderiaceae; Burkholderia; Burkholderia cepacia complex MSDDIDNGGKGGFTRRDMMRVMAASGMMAVGGGGLLTSAQSAFAAPAPKRGGKIRVANES WP_006482802.1 MULTISPECIES: ABC transporter substrate-binding protein [Burkholderia cepacia complex] Length: 527\nNumber of predicted TMHs: 2\nExp number of AAs in TMHs: 36.7303599999999\nExp number, first 60 AAs: 22.08937\nTotal prob of N-in: 0.97312\nPOSSIBLE N-term signal sequence\ninside 1 20\nTMhelix 21 43\noutside 44 502\nTMhelix 503 525\ninside 526 527

721 GCF_001991435.1_ASM199143v1 Burkholderia cenocepacia Proteobacteria; Betaproteobacteria; Burkholderiales; Burkholderiaceae; Burkholderia; Burkholderia cepacia complex MKRRTFLALPAVAASGMLAGCSSADLGPVTHRLMKDPEYRETLSAFLISADGKKLVVIGKWP_034201760.1 MULTISPECIES: hypothetical protein [Burkholderia cepacia complex] Length: 212\nNumber of predicted TMHs: 1\nExp number of AAs in TMHs: 28.11606\nExp number, first 60 AAs: 6.33522\nTotal prob of N-in: 0.49088\noutside 1 187\nTMhelix 188 210\ninside 211 212

722 GCF_001991475.1_ASM199147v1 Burkholderia cenocepacia Proteobacteria; Betaproteobacteria; Burkholderiales; Burkholderiaceae; Burkholderia; Burkholderia cepacia complex MSDDIDNGGKGGFTRRDMMRVMAASGMMAVGGGGLLTSAQSAFAAPAPKRGGKIRVANES WP_006482802.1 MULTISPECIES: ABC transporter substrate-binding protein [Burkholderia cepacia complex] Length: 527\nNumber of predicted TMHs: 2\nExp number of AAs in TMHs: 36.7303599999999\nExp number, first 60 AAs: 22.08937\nTotal prob of N-in: 0.97312\nPOSSIBLE N-term signal sequence\ninside 1 20\nTMhelix 21 43\noutside 44 502\nTMhelix 503 525\ninside 526 527

723	GCF_001991385.1_ASM199138v1	Burkholderia cenocepacia	Proteobacteria; Betaproteobacteria; Burkholderiales; Burkholderiaceae; Burkholderia; Burkholderia cepacia complex
	MKRRTFLALPAVAASGMLAGCSSADLGPVTHRLMKDPEYRETLSAFLISADGKKLVVIGKWP_006483583.1	MULTISPECIES: hypothetical protein [Burkholderia cepacia complex]	Length: 212\nNumber of predicted TMHs: 1\nExp number of AAs in TMHs: 28.03134\nExp number, first 60 AAs: 6.30523\nTotal prob of N-in: 0.48947\noutside 1 187\nTMhelix 188 210\ninside 211 212

724	GCF_001991435.1_ASM199143v1	Burkholderia cenocepacia	Proteobacteria; Betaproteobacteria; Burkholderiales; Burkholderiaceae; Burkholderia; Burkholderia cepacia complex
	MSDDIDNGGKGGFTRRDMMRVMAASGMMAVGGGGLLTSAQSAFAAPAPKRGGKIRVANES	WP_006482802.1	MULTISPECIES: ABC transporter substrate-binding protein [Burkholderia cepacia complex]	Length: 527\nNumber of predicted TMHs: 2\nExp number of AAs in TMHs: 36.7303599999999\nExp number, first 60 AAs: 22.08937\nTotal prob of N-in: 0.97312\nPOSSIBLE N-term signal sequence\ninside 1 20\nTMhelix 21 43\noutside 44 502\nTMhelix 503 525\ninside 526 527

725	GCF_001991385.1_ASM199138v1	Burkholderia cenocepacia	Proteobacteria; Betaproteobacteria; Burkholderiales; Burkholderiaceae; Burkholderia; Burkholderia cepacia complex
	MSDDIDNGGKGGFTRRDMMRVMAASGMMAVGGGGLLTSAQSAFAAPAPKRGGKIRVANES	WP_006482802.1	MULTISPECIES: ABC transporter substrate-binding protein [Burkholderia cepacia complex]	Length: 527\nNumber of predicted TMHs: 2\nExp number of AAs in TMHs: 36.7303599999999\nExp number, first 60 AAs: 22.08937\nTotal prob of N-in: 0.97312\nPOSSIBLE N-term signal sequence\ninside 1 20\nTMhelix 21 43\noutside 44 502\nTMhelix 503 525\ninside 526 527

726	GCF_001991215.1_ASM199121v1	Burkholderia cenocepacia	Proteobacteria; Betaproteobacteria; Burkholderiales; Burkholderiaceae; Burkholderia; Burkholderia cepacia complex
	MKRRTFLALPAVAASGMLAGCSSADLGPVTHRLMKDPEYRETLSAFLISADGKKLVVIGKWP_034201760.1	MULTISPECIES: hypothetical protein [Burkholderia cepacia complex]	Length: 212\nNumber of predicted TMHs: 1\nExp number of AAs in TMHs: 28.11606\nExp number, first 60 AAs: 6.33522\nTotal prob of N-in: 0.49088\noutside 1 187\nTMhelix 188 210\ninside 211 212

727	GCF_001991355.1_ASM199135v1	Burkholderia cenocepacia	Proteobacteria; Betaproteobacteria; Burkholderiales; Burkholderiaceae; Burkholderia; Burkholderia cepacia complex
	METLLTRIRASFADASPRRQLISVAVLVALVASLIAAGYFALRPRYQVLFRDLKPQDAAT	WP_077020516.1	flagellar M-ring protein FliF [Burkholderia cenocepacia]	Length: 491\nNumber of predicted TMHs: 2\nExp number of AAs in TMHs: 41.34236\nExp number, first 60 AAs: 22.1383\nTotal prob of N-in: 0.99080\nPOSSIBLE N-term signal sequence\ninside 1 20\nTMhelix 21 43\noutside 44 427\nTMhelix 428 450\ninside 451 491

728	GCF_001991355.1_ASM199135v1	Burkholderia cenocepacia	Proteobacteria; Betaproteobacteria; Burkholderiales; Burkholderiaceae; Burkholderia; Burkholderia cepacia complex
	MKRRTFLALPAVAASGMLAGCSSADLGPVTHRLMKDPEYRETLSAFLISADGKKLVVIGKWP_006483583.1	MULTISPECIES: hypothetical protein [Burkholderia cepacia complex]	Length: 212\nNumber of predicted TMHs: 1\nExp number of AAs in TMHs: 28.03134\nExp number, first 60 AAs: 6.30523\nTotal prob of N-in: 0.48947\noutside 1 187\nTMhelix 188 210\ninside 211 212

729	GCF_001991355.1_ASM199135v1	Burkholderia cenocepacia	Proteobacteria; Betaproteobacteria; Burkholderiales; Burkholderiaceae; Burkholderia; Burkholderia cepacia complex
	MSDDIDNGGKGGFTRRDMMRVMAASGMMAVGGGGLLTSAQSAFAAPAPKRGGKIRVANES	WP_006482802.1	MULTISPECIES: ABC transporter substrate-binding protein [Burkholderia cepacia complex]	Length: 527\nNumber of predicted TMHs: 2\nExp number of AAs in TMHs: 36.7303599999999\nExp number, first 60 AAs: 22.08937\nTotal prob of N-in: 0.97312\nPOSSIBLE N-term signal sequence\ninside 1 20\nTMhelix 21 43\noutside 44 502\nTMhelix 503 525\ninside 526 527

730	GCF_001991175.1_ASM199117v1	Burkholderia cenocepacia	Proteobacteria; Betaproteobacteria; Burkholderiales; Burkholderiaceae; Burkholderia; Burkholderia cepacia complex
	MKRRTFLALPAVAASGMLAGCSSADLGPVTHRLMKDPEYRETLSAFLISADGKKLVVIGKWP_034201760.1	MULTISPECIES: hypothetical protein [Burkholderia cepacia complex]	Length: 212\nNumber of predicted TMHs: 1\nExp number of AAs in TMHs: 28.11606\nExp number, first 60 AAs: 6.33522\nTotal prob of N-in: 0.49088\noutside 1 187\nTMhelix 188 210\ninside 211 212

731	GCF_001991115.1_ASM199111v1	Burkholderia cenocepacia	Proteobacteria; Betaproteobacteria; Burkholderiales; Burkholderiaceae; Burkholderia; Burkholderia cepacia complex

MKRRTFLALPAVAASGMLAGCSSADLGPVTHRLMKDPEYRETLSAFLISADGKKLVVIGKWP_034201760.1 MULTISPECIES: hypothetical protein [Burkholderia cepacia complex] Length: 212\nNumber of predicted TMHs: 1\nExp number of AAs in TMHs: 28.11606\nExp number, first 60 AAs: 6.33522\nTotal prob of N-in: 0.49088\noutside 1 187\nTMhelix 188 210\ninside 211 212

732 GCF_001991255.1_ASM199125v1 Burkholderia cenocepacia Proteobacteria; Betaproteobacteria; Burkholderiales; Burkholderiaceae; Burkholderia; Burkholderia cepacia complex MSTSITRRRLLAAGVAASSLALSGCFTPKLYKNDAYSEHVSAFMMTEDGKKLVVLGTRYH WP_077021267.1 hypothetical protein [Burkholderia cenocepacia] Length: 212\nNumber of predicted TMHs: 1\nExp number of AAs in TMHs: 24.17376\nExp number, first 60 AAs: 0.92388\nTotal prob of N-in: 0.37951\noutside 1 180\nTMhelix 181 203\ninside 204 212

733 GCF_001991255.1_ASM199125v1 Burkholderia cenocepacia Proteobacteria; Betaproteobacteria; Burkholderiales; Burkholderiaceae; Burkholderia; Burkholderia cepacia complex METLLTRIRASFADASPRRQLISVAVLVALVASLIAAGYFALRPRYQVLFRDLKPQDAAT WP_077020516.1 flagellar M-ring protein FliF [Burkholderia cenocepacia] Length: 491\nNumber of predicted TMHs: 2\nExp number of AAs in TMHs: 41.34236\nExp number, first 60 AAs: 22.1383\nTotal prob of N-in: 0.99080\nPOSSIBLE N-term signal sequence\ninside 1 20\nTMhelix 21 43\noutside 44 427\nTMhelix 428 450\ninside 451 491

734 GCF_001991295.1_ASM199129v1 Burkholderia cenocepacia Proteobacteria; Betaproteobacteria; Burkholderiales; Burkholderiaceae; Burkholderia; Burkholderia cepacia complex MKRRTFLALPAVAASGMLAGCSSADLGPVTHRLMKDPEYRETLSAFLISADGKKLVVIGKWP_034201760.1 MULTISPECIES: hypothetical protein [Burkholderia cepacia complex] Length: 212\nNumber of predicted TMHs: 1\nExp number of AAs in TMHs: 28.11606\nExp number, first 60 AAs: 6.33522\nTotal prob of N-in: 0.49088\noutside 1 187\nTMhelix 188 210\ninside 211 212

735 GCF_001991295.1_ASM199129v1 Burkholderia cenocepacia Proteobacteria; Betaproteobacteria; Burkholderiales; Burkholderiaceae; Burkholderia; Burkholderia cepacia complex MSDDIDNGGKGGFTRRDMMRVMAASGMMAVGGGGLLTSAQSAFAAPAPKRGGKIRVANES WP_006482802.1 MULTISPECIES: ABC transporter substrate-binding protein [Burkholderia cepacia complex] Length: 527\nNumber of predicted TMHs: 2\nExp number of AAs in TMHs: 36.7303599999999\nExp number, first 60 AAs: 22.08937\nTotal prob of N-in: 0.97312\nPOSSIBLE N-term signal sequence\ninside 1 20\nTMhelix 21 43\noutside 44 502\nTMhelix 503 525\ninside 526 527

736 GCF_001991255.1_ASM199125v1 Burkholderia cenocepacia Proteobacteria; Betaproteobacteria; Burkholderiales; Burkholderiaceae; Burkholderia; Burkholderia cepacia complex MKRRTFLALPAVAASGMLAGCSSADLGPVTHRLMKDPEYRETLSAFLISADGKKLVVIGKWP_006483583.1 MULTISPECIES: hypothetical protein [Burkholderia cepacia complex] Length: 212\nNumber of predicted TMHs: 1\nExp number of AAs in TMHs: 28.03134\nExp number, first 60 AAs: 6.30523\nTotal prob of N-in: 0.48947\noutside 1 187\nTMhelix 188 210\ninside 211 212

737 GCF_001991255.1_ASM199125v1 Burkholderia cenocepacia Proteobacteria; Betaproteobacteria; Burkholderiales; Burkholderiaceae; Burkholderia; Burkholderia cepacia complex MSDDIDNGGKGGFTRRDMMRVMAASGMMAVGGGGLLTSAQSAFAAPAPKRGGKIRVANES WP_006482802.1 MULTISPECIES: ABC transporter substrate-binding protein [Burkholderia cepacia complex] Length: 527\nNumber of predicted TMHs: 2\nExp number of AAs in TMHs: 36.7303599999999\nExp number, first 60 AAs: 22.08937\nTotal prob of N-in: 0.97312\nPOSSIBLE N-term signal sequence\ninside 1 20\nTMhelix 21 43\noutside 44 502\nTMhelix 503 525\ninside 526 527

738 GCF_001984355.1_ASM198435v1 Burkholderia cenocepacia Proteobacteria; Betaproteobacteria; Burkholderiales; Burkholderiaceae; Burkholderia; Burkholderia cepacia complex MKRRTFLALPAVAASGMLAGCSSADLGPVTHRLMKDPEYRETLSAFLISADGKKLVVIGKWP_034201760.1 MULTISPECIES: hypothetical protein [Burkholderia cepacia complex] Length: 212\nNumber of predicted TMHs: 1\nExp number of AAs in TMHs: 28.11606\nExp number, first 60 AAs: 6.33522\nTotal prob of N-in: 0.49088\noutside 1 187\nTMhelix 188 210\ninside 211 212

739 GCF_001984315.1_ASM198431v1 Burkholderia cenocepacia Proteobacteria; Betaproteobacteria; Burkholderiales; Burkholderiaceae; Burkholderia; Burkholderia cepacia complex MKRRTFLALPAVAASGMLAGCSSADLGPVTHRLMKDPEYRETLSAFLISADGKKLVVIGKWP_034201760.1 MULTISPECIES: hypothetical protein [Burkholderia cepacia complex] Length: 212\nNumber of predicted TMHs: 1\nExp number of

AAs in TMHs: 28.11606\nExp number, first 60 AAs: 6.33522\nTotal prob of N-in: 0.49088\noutside 1 187\nTMhelix 188 210\ninside 211 212

740 GCF_001991215.1_ASM199121v1 Burkholderia cenocepacia Proteobacteria; Betaproteobacteria; Burkholderiales; Burkholderiaceae; Burkholderia; Burkholderia cepacia complex MSDDIDNGGKGGFTRRDMMRVMAASGMMAVGGGGLLTSAQSAFAAPAPKRGGKIRVANES WP_006482802.1 MULTISPECIES: ABC transporter substrate-binding protein [Burkholderia cepacia complex] Length: 527\nNumber of predicted TMHs: 2\nExp number of AAs in TMHs: 36.7303599999999\nExp number, first 60 AAs: 22.08937\nTotal prob of N-in: 0.97312\nPOSSIBLE N-term signal sequence\ninside 1 20\nTMhelix 21 43\noutside 44 502\nTMhelix 503 525\ninside 526 527

741 GCF_001991175.1_ASM199117v1 Burkholderia cenocepacia Proteobacteria; Betaproteobacteria; Burkholderiales; Burkholderiaceae; Burkholderia; Burkholderia cepacia complex MSDDIDNGGKGGFTRRDMMRVMAASGMMAVGGGGLLTSAQSAFAAPAPKRGGKIRVANES WP_006482802.1 MULTISPECIES: ABC transporter substrate-binding protein [Burkholderia cepacia complex] Length: 527\nNumber of predicted TMHs: 2\nExp number of AAs in TMHs: 36.7303599999999\nExp number, first 60 AAs: 22.08937\nTotal prob of N-in: 0.97312\nPOSSIBLE N-term signal sequence\ninside 1 20\nTMhelix 21 43\noutside 44 502\nTMhelix 503 525\ninside 526 527

742 GCF_001991115.1_ASM199111v1 Burkholderia cenocepacia Proteobacteria; Betaproteobacteria; Burkholderiales; Burkholderiaceae; Burkholderia; Burkholderia cepacia complex MSDDIDNGGKGGFTRRDMMRVMAASGMMAVGGGGLLTSAQSAFAAPAPKRGGKIRVANES WP_006482802.1 MULTISPECIES: ABC transporter substrate-binding protein [Burkholderia cepacia complex] Length: 527\nNumber of predicted TMHs: 2\nExp number of AAs in TMHs: 36.7303599999999\nExp number, first 60 AAs: 22.08937\nTotal prob of N-in: 0.97312\nPOSSIBLE N-term signal sequence\ninside 1 20\nTMhelix 21 43\noutside 44 502\nTMhelix 503 525\ninside 526 527

743 GCF_001984315.1_ASM198431v1 Burkholderia cenocepacia Proteobacteria; Betaproteobacteria; Burkholderiales; Burkholderiaceae; Burkholderia; Burkholderia cepacia complex MSDDIDNGGKGGFTRRDMMRVMAASGMMAVGGGGLLTSAQSAFAAPAPKRGGKIRVANES WP_006482802.1 MULTISPECIES: ABC transporter substrate-binding protein [Burkholderia cepacia complex] Length: 527\nNumber of predicted TMHs: 2\nExp number of AAs in TMHs: 36.7303599999999\nExp number, first 60 AAs: 22.08937\nTotal prob of N-in: 0.97312\nPOSSIBLE N-term signal sequence\ninside 1 20\nTMhelix 21 43\noutside 44 502\nTMhelix 503 525\ninside 526 527

744 GCF_001059335.1_ASM105933v1 Burkholderia cenocepacia Proteobacteria; Betaproteobacteria; Burkholderiales; Burkholderiaceae; Burkholderia; Burkholderia cepacia complex MKRRTFLALPAVAASGMLAGCSSADLGPVTHRLMKDPEYQETLSAFLISADGKKLVVIGK WP_048986526.1 hypothetical protein [Burkholderia cenocepacia] Length: 212\nNumber of predicted TMHs: 1\nExp number of AAs in TMHs: 28.09653\nExp number, first 60 AAs: 6.31597\nTotal prob of N-in: 0.49053\noutside 1 187\nTMhelix 188 210\ninside 211 212

745 GCF_001984285.1_ASM198428v1 Burkholderia cenocepacia Proteobacteria; Betaproteobacteria; Burkholderiales; Burkholderiaceae; Burkholderia; Burkholderia cepacia complex MKRRTFLALPAVAASGMLAGCSSADLGPVTHRLMKDPEYRETLSAFLISADGKKLVVIGKWP_034201760.1 MULTISPECIES: hypothetical protein [Burkholderia cepacia complex] Length: 212\nNumber of predicted TMHs: 1\nExp number of AAs in TMHs: 28.11606\nExp number, first 60 AAs: 6.33522\nTotal prob of N-in: 0.49088\noutside 1 187\nTMhelix 188 210\ninside 211 212

746 GCF_001984285.1_ASM198428v1 Burkholderia cenocepacia Proteobacteria; Betaproteobacteria; Burkholderiales; Burkholderiaceae; Burkholderia; Burkholderia cepacia complex MSDDIDNGGKGGFTRRDMMRVMAASGMMAVGGGGLLTSAQSAFAAPAPKRGGKIRVANES WP_006482802.1 MULTISPECIES: ABC transporter substrate-binding protein [Burkholderia cepacia complex] Length: 527\nNumber of predicted TMHs: 2\nExp number of AAs in TMHs: 36.7303599999999\nExp number, first 60 AAs: 22.08937\nTotal prob of N-in: 0.97312\nPOSSIBLE N-term signal sequence\ninside 1 20\nTMhelix 21 43\noutside 44 502\nTMhelix 503 525\ninside 526 527

747 GCF_001071265.1_ASM107126v1 Burkholderia cenocepacia Proteobacteria; Betaproteobacteria; Burkholderiales; Burkholderiaceae; Burkholderia; Burkholderia cepacia complex MKRRTFLALPAVAASGMLAGCSSADLGPVTHRLMKDPEYQETLSAFLISADGKKLVVIGK WP_048986526.1 hypothetical protein [Burkholderia cenocepacia] Length: 212\nNumber of predicted TMHs: 1\nExp number of

AAs in TMHs: 28.09653\nExp number, first 60 AAs: 6.31597\nTotal prob of N-in: 0.49053\noutside 1 187\nTMhelix 188 210\ninside 211 212

748 GCF_001057175.1_ASM105717v1 Burkholderia cenocepacia Proteobacteria; Betaproteobacteria; Burkholderiales; Burkholderiaceae; Burkholderia; Burkholderia cepacia complex MKRRTFLALPAVAASGMLAGCSSADLGPVTHRLMKDPEYQETLSAFLISADGKKLVVIGK WP_048986526.1 hypothetical protein [Burkholderia cenocepacia] Length: 212\nNumber of predicted TMHs: 1\nExp number of AAs in TMHs: 28.09653\nExp number, first 60 AAs: 6.31597\nTotal prob of N-in: 0.49053\noutside 1 187\nTMhelix 188 210\ninside 211 212

749 GCF_001718895.1_ASM171889v1 Burkholderia cenocepacia Proteobacteria; Betaproteobacteria; Burkholderiales; Burkholderiaceae; Burkholderia; Burkholderia cepacia complex MKRRTFLALPAVAASGMLAGCSSADLGPVTHRLMKDPEYRETLSAFLISADGKKLVVIGKWP_060268096.1 hypothetical protein [Burkholderia cenocepacia] Length: 212\nNumber of predicted TMHs: 1\nExp number of AAs in TMHs: 27.67564\nExp number, first 60 AAs: 6.28086\nTotal prob of N-in: 0.50388\noutside 1 188\nTMhelix 189 211\ninside 212 212

750 GCF_001984275.1_ASM198427v1 Burkholderia cenocepacia Proteobacteria; Betaproteobacteria; Burkholderiales; Burkholderiaceae; Burkholderia; Burkholderia cepacia complex MSTSITRRRLLAAGVAASSLALSGCFTPKLYKNDAYSEHVSAFMMTEDGKKLVVLGTRYH WP_077021267.1 hypothetical protein [Burkholderia cenocepacia] Length: 212\nNumber of predicted TMHs: 1\nExp number of AAs in TMHs: 24.17376\nExp number, first 60 AAs: 0.92388\nTotal prob of N-in: 0.37951\noutside 1 180\nTMhelix 181 203\ninside 204 212

751 GCF_001984275.1_ASM198427v1 Burkholderia cenocepacia Proteobacteria; Betaproteobacteria; Burkholderiales; Burkholderiaceae; Burkholderia; Burkholderia cepacia complex MKRRTFLALPAVAASGMLAGCSSADLGPVTHRLMKDPEYRETLSAFLISADGKKLVVIGKWP_006483583.1 MULTISPECIES: hypothetical protein [Burkholderia cepacia complex] Length: 212\nNumber of predicted TMHs: 1\nExp number of AAs in TMHs: 28.03134\nExp number, first 60 AAs: 6.30523\nTotal prob of N-in: 0.48947\noutside 1 187\nTMhelix 188 210\ninside 211 212

752 GCF_001718895.1_ASM171889v1 Burkholderia cenocepacia Proteobacteria; Betaproteobacteria; Burkholderiales; Burkholderiaceae; Burkholderia; Burkholderia cepacia complex MSDDIDNGGKGGFTRRDMMRVMAASGMMAVGGGGLLTSAQSAFAAPAPKRGGKIRVANES WP_006482802.1 MULTISPECIES: ABC transporter substrate-binding protein [Burkholderia cepacia complex] Length: 527\nNumber of predicted TMHs: 2\nExp number of AAs in TMHs: 36.7303599999999\nExp number, first 60 AAs: 22.08937\nTotal prob of N-in: 0.97312\nPOSSIBLE N-term signal sequence\ninside 1 20\nTMhelix 21 43\noutside 44 502\nTMhelix 503 525\ninside 526 527

753 GCF_001056655.1_ASM105665v1 Burkholderia cenocepacia Proteobacteria; Betaproteobacteria; Burkholderiales; Burkholderiaceae; Burkholderia; Burkholderia cepacia complex MKRRTFLALPAVAASGMLAGCSSADLGPVTHRLMKDPEYQETLSAFLISADGKKLVVIGK WP_048986526.1 hypothetical protein [Burkholderia cenocepacia] Length: 212\nNumber of predicted TMHs: 1\nExp number of AAs in TMHs: 28.09653\nExp number, first 60 AAs: 6.31597\nTotal prob of N-in: 0.49053\noutside 1 187\nTMhelix 188 210\ninside 211 212

754 GCF_001606115.1_ASM160611v1 Burkholderia cenocepacia Proteobacteria; Betaproteobacteria; Burkholderiales; Burkholderiaceae; Burkholderia; Burkholderia cepacia complex METLLTRIRASFADASPRRQLISVAVLVALVASLIAAGYFALRPRYQVLFRDLKPQDAAT WP_062886032.1 flagellar M-ring protein FliF [Burkholderia cenocepacia] Length: 491\nNumber of predicted TMHs: 2\nExp number of AAs in TMHs: 41.3427699999999\nExp number, first 60 AAs: 22.1383\nTotal prob of N-in: 0.99079\nPOSSIBLE N-term signal sequence\ninside 1 20\nTMhelix 21 43\noutside 44 427\nTMhelix 428 450\ninside 451 491

755 GCF_001533105.1_ASM153310v1 Burkholderia cenocepacia Proteobacteria; Betaproteobacteria; Burkholderiales; Burkholderiaceae; Burkholderia; Burkholderia cepacia complex MKRRTFLALPAVAASGMLAGCSSADLGPVTHRLMKDPEYRETLSAFLISADGKKLVVIGKWP_060268096.1 hypothetical protein [Burkholderia cenocepacia] Length: 212\nNumber of predicted TMHs: 1\nExp number of AAs in TMHs: 27.67564\nExp number, first 60 AAs: 6.28086\nTotal prob of N-in: 0.50388\noutside 1 188\nTMhelix 189 211\ninside 212 212

756 GCF_001606115.1_ASM160611v1 Burkholderia cenocepacia Proteobacteria; Betaproteobacteria; Burkholderiales; Burkholderiaceae; Burkholderia; Burkholderia cepacia complex
MKRRTFLALPAVAASGMLAGCSSADLGPVTHRLMKDPEYRETLSAFLISADGKKLVVIGKWP_062884514.1 hypothetical protein [Burkholderia cenocepacia] Length: 212\nNumber of predicted TMHs: 1\nExp number of AAs in TMHs: 28.03134\nExp number, first 60 AAs: 6.30523\nTotal prob of N-in: 0.48947\noutside 1 187\nTMhelix 188 210\ninside 211 212

757 GCF_001606115.1_ASM160611v1 Burkholderia cenocepacia Proteobacteria; Betaproteobacteria; Burkholderiales; Burkholderiaceae; Burkholderia; Burkholderia cepacia complex
MSDDIDNGGKGGFTRRDMMRVMAASGMMAVGGGGLLTSAQSAFAAPAPKRGGKIRVANES WP_006482802.1 MULTISPECIES: ABC transporter substrate-binding protein [Burkholderia cepacia complex] Length: 527\nNumber of predicted TMHs: 2\nExp number of AAs in TMHs: 36.7303599999999\nExp number, first 60 AAs: 22.08937\nTotal prob of N-in: 0.97312\nPOSSIBLE N-term signal sequence\ninside 1 20\nTMhelix 21 43\noutside 44 502\nTMhelix 503 525\ninside 526 527

758 GCF_001533105.1_ASM153310v1 Burkholderia cenocepacia Proteobacteria; Betaproteobacteria; Burkholderiales; Burkholderiaceae; Burkholderia; Burkholderia cepacia complex
MSDDIDNGGKGGFTRRDMMRVMAASGMMAVGGGGLLTSAQSAFAAPAPKRGGKIRVANES WP_006482802.1 MULTISPECIES: ABC transporter substrate-binding protein [Burkholderia cepacia complex] Length: 527\nNumber of predicted TMHs: 2\nExp number of AAs in TMHs: 36.7303599999999\nExp number, first 60 AAs: 22.08937\nTotal prob of N-in: 0.97312\nPOSSIBLE N-term signal sequence\ninside 1 20\nTMhelix 21 43\noutside 44 502\nTMhelix 503 525\ninside 526 527

759 GCF_001056365.1_ASM105636v1 Burkholderia cenocepacia Proteobacteria; Betaproteobacteria; Burkholderiales; Burkholderiaceae; Burkholderia; Burkholderia cepacia complex
MKRRTFLALPAVAASGMLAGCSSADLGPVTHRLMKDPEYQETLSAFLISADGKKLVVIGK WP_048986526.1 hypothetical protein [Burkholderia cenocepacia] Length: 212\nNumber of predicted TMHs: 1\nExp number of AAs in TMHs: 28.09653\nExp number, first 60 AAs: 6.31597\nTotal prob of N-in: 0.49053\noutside 1 187\nTMhelix 188 210\ninside 211 212

760 GCF_001524105.1_ASM152410v1 Burkholderia cenocepacia Proteobacteria; Betaproteobacteria; Burkholderiales; Burkholderiaceae; Burkholderia; Burkholderia cepacia complex
MSDDIDNGGKGGFTRRDMMRVMAASGMMAVGGGGLLTSAQSAFAAPAPKRGGKIRVANES WP_059723004.1 ABC transporter substrate-binding protein [Burkholderia cenocepacia] Length: 527\nNumber of predicted TMHs: 2\nExp number of AAs in TMHs: 36.4044299999999\nExp number, first 60 AAs: 22.08933\nTotal prob of N-in: 0.97308\nPOSSIBLE N-term signal sequence\ninside 1 20\nTMhelix 21 43\noutside 44 502\nTMhelix 503 525\ninside 526 527

761 GCF_001524105.1_ASM152410v1 Burkholderia cenocepacia Proteobacteria; Betaproteobacteria; Burkholderiales; Burkholderiaceae; Burkholderia; Burkholderia cepacia complex
MKRRTFLALPAVAASGMLAGCSSADLGPVTHRLMKDPEYRETLSAFLISADGKKLVVIGKWP_006483583.1 MULTISPECIES: hypothetical protein [Burkholderia cepacia complex] Length: 212\nNumber of predicted TMHs: 1\nExp number of AAs in TMHs: 28.03134\nExp number, first 60 AAs: 6.30523\nTotal prob of N-in: 0.48947\noutside 1 187\nTMhelix 188 210\ninside 211 212

762 GCF_001071265.1_ASM107126v1 Burkholderia cenocepacia Proteobacteria; Betaproteobacteria; Burkholderiales; Burkholderiaceae; Burkholderia; Burkholderia cepacia complex
MSDDIDNGGKGGFTRRDMMRVMAASGMMAVGGGGLLTSAQSAFAAPAPKRGGKIRVANES WP_006482802.1 MULTISPECIES: ABC transporter substrate-binding protein [Burkholderia cepacia complex] Length: 527\nNumber of predicted TMHs: 2\nExp number of AAs in TMHs: 36.7303599999999\nExp number, first 60 AAs: 22.08937\nTotal prob of N-in: 0.97312\nPOSSIBLE N-term signal sequence\ninside 1 20\nTMhelix 21 43\noutside 44 502\nTMhelix 503 525\ninside 526 527

763 GCF_001059335.1_ASM105933v1 Burkholderia cenocepacia Proteobacteria; Betaproteobacteria; Burkholderiales; Burkholderiaceae; Burkholderia; Burkholderia cepacia complex
MSDDIDNGGKGGFTRRDMMRVMAASGMMAVGGGGLLTSAQSAFAAPAPKRGGKIRVANES WP_006482802.1 MULTISPECIES: ABC transporter substrate-binding protein [Burkholderia cepacia complex] Length: 527\nNumber of predicted TMHs: 2\nExp number of AAs in TMHs: 36.7303599999999\nExp number, first 60 AAs: 22.08937\nTotal prob of N-in: 0.97312\nPOSSIBLE N-term signal sequence\ninside 1 20\nTMhelix 21 43\noutside 44 502\nTMhelix 503 525\ninside 526 527

764 GCF_001057175.1_ASM105717v1 Burkholderia cenocepacia Proteobacteria; Betaproteobacteria; Burkholderiales; Burkholderiaceae; Burkholderia; Burkholderia cepacia complex
MSDDIDNGGKGGFTRRDMMRVMAASGMMAVGGGGLLTSAQSAFAAPAPKRGGKIRVANES WP_006482802.1
MULTISPECIES: ABC transporter substrate-binding protein [Burkholderia cepacia complex] Length: 527\nNumber of predicted TMHs: 2\nExp number of AAs in TMHs: 36.7303599999999\nExp number, first 60 AAs: 22.08937\nTotal prob of N-in: 0.97312\nPOSSIBLE N-term signal sequence\ninside 1 20\nTMhelix 21 43\noutside 44 502\nTMhelix 503 525\ninside 526 527

765 GCF_001056655.1_ASM105665v1 Burkholderia cenocepacia Proteobacteria; Betaproteobacteria; Burkholderiales; Burkholderiaceae; Burkholderia; Burkholderia cepacia complex
MSDDIDNGGKGGFTRRDMMRVMAASGMMAVGGGGLLTSAQSAFAAPAPKRGGKIRVANES WP_006482802.1
MULTISPECIES: ABC transporter substrate-binding protein [Burkholderia cepacia complex] Length: 527\nNumber of predicted TMHs: 2\nExp number of AAs in TMHs: 36.7303599999999\nExp number, first 60 AAs: 22.08937\nTotal prob of N-in: 0.97312\nPOSSIBLE N-term signal sequence\ninside 1 20\nTMhelix 21 43\noutside 44 502\nTMhelix 503 525\ninside 526 527

766 GCF_001056365.1_ASM105636v1 Burkholderia cenocepacia Proteobacteria; Betaproteobacteria; Burkholderiales; Burkholderiaceae; Burkholderia; Burkholderia cepacia complex
MSDDIDNGGKGGFTRRDMMRVMAASGMMAVGGGGLLTSAQSAFAAPAPKRGGKIRVANES WP_006482802.1
MULTISPECIES: ABC transporter substrate-binding protein [Burkholderia cepacia complex] Length: 527\nNumber of predicted TMHs: 2\nExp number of AAs in TMHs: 36.7303599999999\nExp number, first 60 AAs: 22.08937\nTotal prob of N-in: 0.97312\nPOSSIBLE N-term signal sequence\ninside 1 20\nTMhelix 21 43\noutside 44 502\nTMhelix 503 525\ninside 526 527

767 GCF_000705535.1_BukCp6.0 Burkholderia cenocepacia Proteobacteria; Betaproteobacteria; Burkholderiales; Burkholderiaceae; Burkholderia; Burkholderia cepacia complex
MSTSITRRRLLAAAMAASSLALSGCFTPKLYKDDAYSEYVTSFMITQDGKKLVVLGTRYH WP_034189316.1 MULTISPECIES: hypothetical protein [Burkholderia cepacia complex] Length: 212\nNumber of predicted TMHs: 1\nExp number of AAs in TMHs: 25.28913\nExp number, first 60 AAs: 1.00711\nTotal prob of N-in: 0.41198\noutside 1 180\nTMhelix 181 203\ninside 204 212

768 GCF_000981305.1_ASM98130v1 Burkholderia cenocepacia Proteobacteria; Betaproteobacteria; Burkholderiales; Burkholderiaceae; Burkholderia; Burkholderia cepacia complex
MKRRTFLALPAVAASGMLAGCSSADLGPVTHRLMKDPEYRETLSAFLISADGKKLVVIGKWP_006483583.1 MULTISPECIES: hypothetical protein [Burkholderia cepacia complex] Length: 212\nNumber of predicted TMHs: 1\nExp number of AAs in TMHs: 28.03134\nExp number, first 60 AAs: 6.30523\nTotal prob of N-in: 0.48947\noutside 1 187\nTMhelix 188 210\ninside 211 212

769 GCF_000981305.1_ASM98130v1 Burkholderia cenocepacia Proteobacteria; Betaproteobacteria; Burkholderiales; Burkholderiaceae; Burkholderia; Burkholderia cepacia complex
MSDDIDNGGKGGFTRRDMMRVMAASGMMAVGGGGLLTSAQSAFAAPAPKRGGKIRVANES WP_006482802.1
MULTISPECIES: ABC transporter substrate-binding protein [Burkholderia cepacia complex] Length: 527\nNumber of predicted TMHs: 2\nExp number of AAs in TMHs: 36.7303599999999\nExp number, first 60 AAs: 22.08937\nTotal prob of N-in: 0.97312\nPOSSIBLE N-term signal sequence\ninside 1 20\nTMhelix 21 43\noutside 44 502\nTMhelix 503 525\ninside 526 527

770 GCF_000705535.1_BukCp6.0 Burkholderia cenocepacia Proteobacteria; Betaproteobacteria; Burkholderiales; Burkholderiaceae; Burkholderia; Burkholderia cepacia complex
MKRRTFLALPAVAAAGMLAGCTDVSIGPVTDRMMKDPEYRETLSAFLISADGKKLVVIGK WP_034186868.1
hypothetical protein [Burkholderia cenocepacia] Length: 212\nNumber of predicted TMHs: 2\nExp number of AAs in TMHs: 40.99654\nExp number, first 60 AAs: 18.36165\nTotal prob of N-in: 0.80910\nPOSSIBLE N-term signal sequence\ninside 1 4\nTMhelix 5 27\noutside 28 187\nTMhelix 188 210\ninside 211 212

771 GCF_002071945.1_ASM207194v1 Burkholderia cenocepacia Proteobacteria; Betaproteobacteria; Burkholderiales; Burkholderiaceae; Burkholderia; Burkholderia cepacia complex
MKRRTFLALPAVAASGMLAGCSSADLGPVTHRLMKDPEYRETLSAFLISADGKKLVVIGKWP_062884514.1 hypothetical protein [Burkholderia cenocepacia] Length: 212\nNumber of predicted TMHs: 1\nExp number of AAs in TMHs: 28.03134\nExp number, first 60 AAs: 6.30523\nTotal prob of N-in: 0.48947\noutside 1 187\nTMhelix 188 210\ninside 211 212

772 GCF_002071945.1_ASM207194v1 Burkholderia cenocepacia Proteobacteria; Betaproteobacteria; Burkholderiales; Burkholderiaceae; Burkholderia; Burkholderia cepacia complex
METLLTRIRASFADASPRRQLISVAVLVALVASLIAAGYFALRPRYQVLFRDLKPQDAAT WP_060214257.1 flagellar M-ring protein FliF [Burkholderia cenocepacia] Length: 491\nNumber of predicted TMHs: 2\nExp number of AAs in TMHs: 41.34235\nExp number, first 60 AAs: 22.1383\nTotal prob of N-in: 0.99080\nPOSSIBLE N-term signal sequence\ninside 1 20\nTMhelix 21 43\noutside 44 427\nTMhelix 428 450\ninside 451 491

773 GCF_002071905.1_ASM207190v1 Burkholderia cenocepacia Proteobacteria; Betaproteobacteria; Burkholderiales; Burkholderiaceae; Burkholderia; Burkholderia cepacia complex
MKRRTFLALPAVAASGMLAGCSSADLGPVTHRLMKDPEYRETLSAFLISADGKKLVVIGKWP_062884514.1 hypothetical protein [Burkholderia cenocepacia] Length: 212\nNumber of predicted TMHs: 1\nExp number of AAs in TMHs: 28.03134\nExp number, first 60 AAs: 6.30523\nTotal prob of N-in: 0.48947\noutside 1 187\nTMhelix 188 210\ninside 211 212

774 GCF_002071905.1_ASM207190v1 Burkholderia cenocepacia Proteobacteria; Betaproteobacteria; Burkholderiales; Burkholderiaceae; Burkholderia; Burkholderia cepacia complex
METLLTRIRASFADASPRRQLISVAVLVALVASLIAAGYFALRPRYQVLFRDLKPQDAAT WP_060214257.1 flagellar M-ring protein FliF [Burkholderia cenocepacia] Length: 491\nNumber of predicted TMHs: 2\nExp number of AAs in TMHs: 41.34235\nExp number, first 60 AAs: 22.1383\nTotal prob of N-in: 0.99080\nPOSSIBLE N-term signal sequence\ninside 1 20\nTMhelix 21 43\noutside 44 427\nTMhelix 428 450\ninside 451 491

775 GCF_002071875.1_ASM207187v1 Burkholderia cenocepacia Proteobacteria; Betaproteobacteria; Burkholderiales; Burkholderiaceae; Burkholderia; Burkholderia cepacia complex
MKRRTFLALPAVAASGMLAGCSSADLGPVTHRLMKDPEYRETLSAFLISADGKKLVVIGKWP_062884514.1 hypothetical protein [Burkholderia cenocepacia] Length: 212\nNumber of predicted TMHs: 1\nExp number of AAs in TMHs: 28.03134\nExp number, first 60 AAs: 6.30523\nTotal prob of N-in: 0.48947\noutside 1 187\nTMhelix 188 210\ninside 211 212

776 GCF_002071875.1_ASM207187v1 Burkholderia cenocepacia Proteobacteria; Betaproteobacteria; Burkholderiales; Burkholderiaceae; Burkholderia; Burkholderia cepacia complex
METLLTRIRASFADASPRRQLISVAVLVALVASLIAAGYFALRPRYQVLFRDLKPQDAAT WP_060214257.1 flagellar M-ring protein FliF [Burkholderia cenocepacia] Length: 491\nNumber of predicted TMHs: 2\nExp number of AAs in TMHs: 41.34235\nExp number, first 60 AAs: 22.1383\nTotal prob of N-in: 0.99080\nPOSSIBLE N-term signal sequence\ninside 1 20\nTMhelix 21 43\noutside 44 427\nTMhelix 428 450\ninside 451 491

777 GCF_001999885.1_ASM199988v1 Burkholderia cenocepacia Proteobacteria; Betaproteobacteria; Burkholderiales; Burkholderiaceae; Burkholderia; Burkholderia cepacia complex
MSTSITRRRLLAAGVAASSLALSGCFTPKLYKNDAYSEHVSAFMMTEDGKKLVVLGTRYH WP_077021267.1
hypothetical protein [Burkholderia cenocepacia] Length: 212\nNumber of predicted TMHs: 1\nExp number of AAs in TMHs: 24.17376\nExp number, first 60 AAs: 0.92388\nTotal prob of N-in: 0.37951\noutside 1 180\nTMhelix 181 203\ninside 204 212

778 GCF_002071995.1_ASM207199v1 Burkholderia cenocepacia Proteobacteria; Betaproteobacteria; Burkholderiales; Burkholderiaceae; Burkholderia; Burkholderia cepacia complex
MKRRTFLALPAVAASGMLAGCSSADLGPVTHRLMKDPEYRETLSAFLISADGKKLVVIGKWP_034201760.1 MULTISPECIES: hypothetical protein [Burkholderia cepacia complex] Length: 212\nNumber of predicted TMHs: 1\nExp number of AAs in TMHs: 28.11606\nExp number, first 60 AAs: 6.33522\nTotal prob of N-in: 0.49088\noutside 1 187\nTMhelix 188 210\ninside 211 212

779 GCF_002071995.1_ASM207199v1 Burkholderia cenocepacia Proteobacteria; Betaproteobacteria; Burkholderiales; Burkholderiaceae; Burkholderia; Burkholderia cepacia complex
MSDDIDNGGKGGFTRRDMMRVMAASGMMAVGGGGLLTSAQSAFAAPAPKRGGKIRVANES WP_006482802.1
MULTISPECIES: ABC transporter substrate-binding protein [Burkholderia cepacia complex] Length: 527\nNumber of predicted TMHs: 2\nExp number of AAs in TMHs: 36.7303599999999\nExp number, first 60 AAs: 22.08937\nTotal prob of N-in: 0.97312\nPOSSIBLE N-term signal sequence\ninside 1 20\nTMhelix 21 43\noutside 44 502\nTMhelix 503 525\ninside 526 527

780 GCF_002071945.1_ASM207194v1 Burkholderia cenocepacia Proteobacteria; Betaproteobacteria; Burkholderiales; Burkholderiaceae; Burkholderia; Burkholderia cepacia complex
MSDDIDNGGKGGFTRRDMMRVMAASGMMAVGGGGLLTSAQSAFAAPAPKRGGKIRVANES WP_006482802.1
MULTISPECIES: ABC transporter substrate-binding protein [Burkholderia cepacia complex] Length:

527\nNumber of predicted TMHs:  2\nExp number of AAs in TMHs: 36.7303599999999\nExp number, first 60 AAs:  22.08937\nTotal prob of N-in:   0.97312\nPOSSIBLE N-term signal sequence\ninside   1   20\nTMhelix   21   43\noutside   44   502\nTMhelix   503   525\ninside   526   527

781 GCF_002071905.1_ASM207190v1 Burkholderia cenocepacia Proteobacteria; Betaproteobacteria; Burkholderiales; Burkholderiaceae; Burkholderia; Burkholderia cepacia complex MSDDIDNGGKGGFTRRDMMRVMAASGMMAVGGGGLLTSAQSAFAAPAPKRGGKIRVANES WP_006482802.1 MULTISPECIES: ABC transporter substrate-binding protein [Burkholderia cepacia complex] Length: 527\nNumber of predicted TMHs:  2\nExp number of AAs in TMHs: 36.7303599999999\nExp number, first 60 AAs:  22.08937\nTotal prob of N-in:   0.97312\nPOSSIBLE N-term signal sequence\ninside   1   20\nTMhelix   21   43\noutside   44   502\nTMhelix   503   525\ninside   526   527

782 GCF_002071875.1_ASM207187v1 Burkholderia cenocepacia Proteobacteria; Betaproteobacteria; Burkholderiales; Burkholderiaceae; Burkholderia; Burkholderia cepacia complex MSDDIDNGGKGGFTRRDMMRVMAASGMMAVGGGGLLTSAQSAFAAPAPKRGGKIRVANES WP_006482802.1 MULTISPECIES: ABC transporter substrate-binding protein [Burkholderia cepacia complex] Length: 527\nNumber of predicted TMHs:  2\nExp number of AAs in TMHs: 36.7303599999999\nExp number, first 60 AAs:  22.08937\nTotal prob of N-in:   0.97312\nPOSSIBLE N-term signal sequence\ninside   1   20\nTMhelix   21   43\noutside   44   502\nTMhelix   503   525\ninside   526   527

783 GCF_001994665.1_ASM199466v1 Burkholderia cenocepacia Proteobacteria; Betaproteobacteria; Burkholderiales; Burkholderiaceae; Burkholderia; Burkholderia cepacia complex MSDDIDNGGKGGFTRRDMMRVMAASGMMAVGGGGLLTSAQSAFAAPAPKRGGKIRVANES WP_077180776.1 ABC transporter substrate-binding protein [Burkholderia cenocepacia] Length: 527\nNumber of predicted TMHs:  2\nExp number of AAs in TMHs: 36.7312299999999\nExp number, first 60 AAs:  22.0894\nTotal prob of N-in:   0.97312\nPOSSIBLE N-term signal sequence\ninside   1   20\nTMhelix   21   43\noutside   44   502\nTMhelix   503   525\ninside   526   527

784 GCF_001994895.1_ASM199489v1 Burkholderia cenocepacia Proteobacteria; Betaproteobacteria; Burkholderiales; Burkholderiaceae; Burkholderia; Burkholderia cepacia complex MKRRTFLALPAVAASGMLAGCSSADLGPVTHRLMKDPEYRETLSAFLISADGKKLVVIGKWP_034201760.1 MULTISPECIES: hypothetical protein [Burkholderia cepacia complex] Length: 212\nNumber of predicted TMHs:  1\nExp number of AAs in TMHs: 28.11606\nExp number, first 60 AAs:  6.33522\nTotal prob of N-in:   0.49088\noutside   1   187\nTMhelix   188   210\ninside   211   212

785 GCF_001994605.1_ASM199460v1 Burkholderia cenocepacia Proteobacteria; Betaproteobacteria; Burkholderiales; Burkholderiaceae; Burkholderia; Burkholderia cepacia complex MSTSITRRRLLAAGVAASSLALSGCFTPKLYKNDAYSEHVSAFMMTEDGKKLVVLGTRYH WP_077021267.1 hypothetical protein [Burkholderia cenocepacia] Length: 212\nNumber of predicted TMHs:  1\nExp number of AAs in TMHs: 24.17376\nExp number, first 60 AAs:  0.92388\nTotal prob of N-in:   0.37951\noutside   1   180\nTMhelix   181   203\ninside   204   212

786 GCF_001999885.1_ASM199988v1 Burkholderia cenocepacia Proteobacteria; Betaproteobacteria; Burkholderiales; Burkholderiaceae; Burkholderia; Burkholderia cepacia complex MKRRTFLALPAVAASGMLAGCSSADLGPVTHRLMKDPEYRETLSAFLISADGKKLVVIGKWP_006483583.1 MULTISPECIES: hypothetical protein [Burkholderia cepacia complex] Length: 212\nNumber of predicted TMHs:  1\nExp number of AAs in TMHs: 28.03134\nExp number, first 60 AAs:  6.30523\nTotal prob of N-in:   0.48947\noutside   1   187\nTMhelix   188   210\ninside   211   212

787 GCF_001999885.1_ASM199988v1 Burkholderia cenocepacia Proteobacteria; Betaproteobacteria; Burkholderiales; Burkholderiaceae; Burkholderia; Burkholderia cepacia complex MSDDIDNGGKGGFTRRDMMRVMAASGMMAVGGGGLLTSAQSAFAAPAPKRGGKIRVANES WP_006482802.1 MULTISPECIES: ABC transporter substrate-binding protein [Burkholderia cepacia complex] Length: 527\nNumber of predicted TMHs:  2\nExp number of AAs in TMHs: 36.7303599999999\nExp number, first 60 AAs:  22.08937\nTotal prob of N-in:   0.97312\nPOSSIBLE N-term signal sequence\ninside   1   20\nTMhelix   21   43\noutside   44   502\nTMhelix   503   525\ninside   526   527

788 GCF_001999805.1_ASM199980v1 Burkholderia cenocepacia Proteobacteria; Betaproteobacteria; Burkholderiales; Burkholderiaceae; Burkholderia; Burkholderia cepacia complex MSDDIDNGGKGGFTRRDMMRVMAASGMMAVGGGGLLTSAQSAFAAPAPKRGGKIRVANES WP_006482802.1 MULTISPECIES: ABC transporter substrate-binding protein [Burkholderia cepacia complex] Length: 527\nNumber of predicted TMHs:  2\nExp number of AAs in TMHs: 36.7303599999999\nExp number, first 60 AAs:

22.08937\nTotal prob of N-in: 0.97312\nPOSSIBLE N-term signal sequence\ninside 1 20\nTMhelix 21 43\noutside 44 502\nTMhelix 503 525\ninside 526 527

789 GCF_001994895.1_ASM199489v1 Burkholderia cenocepacia Proteobacteria; Betaproteobacteria; Burkholderiales; Burkholderiaceae; Burkholderia; Burkholderia cepacia complex MSDDIDNGGKGGFTRRDMMRVMAASGMMAVGGGGLLTSAQSAFAAPAPKRGGKIRVANES WP_006482802.1 MULTISPECIES: ABC transporter substrate-binding protein [Burkholderia cepacia complex] Length: 527\nNumber of predicted TMHs: 2\nExp number of AAs in TMHs: 36.7303599999999\nExp number, first 60 AAs: 22.08937\nTotal prob of N-in: 0.97312\nPOSSIBLE N-term signal sequence\ninside 1 20\nTMhelix 21 43\noutside 44 502\nTMhelix 503 525\ninside 526 527

790 GCF_001994605.1_ASM199460v1 Burkholderia cenocepacia Proteobacteria; Betaproteobacteria; Burkholderiales; Burkholderiaceae; Burkholderia; Burkholderia cepacia complex METLLTRIRASFADASPRRQLISVAVLVALVASLIAAGYFALRPRYQVLFRDLKPQDAAT WP_077020516.1 flagellar M-ring protein FliF [Burkholderia cenocepacia] Length: 491\nNumber of predicted TMHs: 2\nExp number of AAs in TMHs: 41.34236\nExp number, first 60 AAs: 22.1383\nTotal prob of N-in: 0.99080\nPOSSIBLE N-term signal sequence\ninside 1 20\nTMhelix 21 43\noutside 44 427\nTMhelix 428 450\ninside 451 491

791 GCF_001994715.1_ASM199471v1 Burkholderia cenocepacia Proteobacteria; Betaproteobacteria; Burkholderiales; Burkholderiaceae; Burkholderia; Burkholderia cepacia complex MSTSITRRRLLAAGVAASSLALSGCFTPKLYKNDAYSEHVSAFMMTEDGKKLVVLGTRYH WP_077021267.1 hypothetical protein [Burkholderia cenocepacia] Length: 212\nNumber of predicted TMHs: 1\nExp number of AAs in TMHs: 24.17376\nExp number, first 60 AAs: 0.92388\nTotal prob of N-in: 0.37951\noutside 1 180\nTMhelix 181 203\ninside 204 212

792 GCF_001994855.1_ASM199485v1 Burkholderia cenocepacia Proteobacteria; Betaproteobacteria; Burkholderiales; Burkholderiaceae; Burkholderia; Burkholderia cepacia complex MSTSITRRRLLAAGVAASSLALSGCFTPKLYKNDAYSEHVSAFMMTEDGKKLVVLGTRYH WP_077021267.1 hypothetical protein [Burkholderia cenocepacia] Length: 212\nNumber of predicted TMHs: 1\nExp number of AAs in TMHs: 24.17376\nExp number, first 60 AAs: 0.92388\nTotal prob of N-in: 0.37951\noutside 1 180\nTMhelix 181 203\ninside 204 212

793 GCF_001994855.1_ASM199485v1 Burkholderia cenocepacia Proteobacteria; Betaproteobacteria; Burkholderiales; Burkholderiaceae; Burkholderia; Burkholderia cepacia complex METLLTRIRASFADASPRRQLISVAVLVALVASLIAAGYFALRPRYQVLFRDLKPQDAAT WP_077020516.1 flagellar M-ring protein FliF [Burkholderia cenocepacia] Length: 491\nNumber of predicted TMHs: 2\nExp number of AAs in TMHs: 41.34236\nExp number, first 60 AAs: 22.1383\nTotal prob of N-in: 0.99080\nPOSSIBLE N-term signal sequence\ninside 1 20\nTMhelix 21 43\noutside 44 427\nTMhelix 428 450\ninside 451 491

794 GCF_001994795.1_ASM199479v1 Burkholderia cenocepacia Proteobacteria; Betaproteobacteria; Burkholderiales; Burkholderiaceae; Burkholderia; Burkholderia cepacia complex MKRRTFLALPAVAASGMLAGCSSADLGPVTHRLMKDPEYRETLSAFLISADGKKLVVIGKWP_034201760.1 MULTISPECIES: hypothetical protein [Burkholderia cepacia complex] Length: 212\nNumber of predicted TMHs: 1\nExp number of AAs in TMHs: 28.11606\nExp number, first 60 AAs: 6.33522\nTotal prob of N-in: 0.49088\noutside 1 187\nTMhelix 188 210\ninside 211 212

795 GCF_001994855.1_ASM199485v1 Burkholderia cenocepacia Proteobacteria; Betaproteobacteria; Burkholderiales; Burkholderiaceae; Burkholderia; Burkholderia cepacia complex MKRRTFLALPAVAASGMLAGCSSADLGPVTHRLMKDPEYRETLSAFLISADGKKLVVIGKWP_006483583.1 MULTISPECIES: hypothetical protein [Burkholderia cepacia complex] Length: 212\nNumber of predicted TMHs: 1\nExp number of AAs in TMHs: 28.03134\nExp number, first 60 AAs: 6.30523\nTotal prob of N-in: 0.48947\noutside 1 187\nTMhelix 188 210\ninside 211 212

796 GCF_001994855.1_ASM199485v1 Burkholderia cenocepacia Proteobacteria; Betaproteobacteria; Burkholderiales; Burkholderiaceae; Burkholderia; Burkholderia cepacia complex MSDDIDNGGKGGFTRRDMMRVMAASGMMAVGGGGLLTSAQSAFAAPAPKRGGKIRVANES WP_006482802.1 MULTISPECIES: ABC transporter substrate-binding protein [Burkholderia cepacia complex] Length: 527\nNumber of predicted TMHs: 2\nExp number of AAs in TMHs: 36.7303599999999\nExp number, first 60 AAs: 22.08937\nTotal prob of N-in: 0.97312\nPOSSIBLE N-term signal sequence\ninside 1 20\nTMhelix 21 43\noutside 44 502\nTMhelix 503 525\ninside 526 527

797 GCF_001994795.1_ASM199479v1 Burkholderia cenocepacia Proteobacteria; Betaproteobacteria; Burkholderiales; Burkholderiaceae; Burkholderia; Burkholderia cepacia complex
MSDDIDNGGKGGFTRRDMMRVMAASGMMAVGGGGLLTSAQSAFAAPAPKRGGKIRVANES WP_006482802.1
MULTISPECIES: ABC transporter substrate-binding protein [Burkholderia cepacia complex] Length: 527\nNumber of predicted TMHs: 2\nExp number of AAs in TMHs: 36.7303599999999\nExp number, first 60 AAs: 22.08937\nTotal prob of N-in: 0.97312\nPOSSIBLE N-term signal sequence\ninside 1 20\nTMhelix 21 43\noutside 44 502\nTMhelix 503 525\ninside 526 527

798 GCF_001994755.1_ASM199475v1 Burkholderia cenocepacia Proteobacteria; Betaproteobacteria; Burkholderiales; Burkholderiaceae; Burkholderia; Burkholderia cepacia complex
MKRRTFLALPAVAASGMLAGCSSADLGPVTHRLMKDPEYRETLSAFLISADGKKLVVIGKWP_006483583.1 MULTISPECIES: hypothetical protein [Burkholderia cepacia complex] Length: 212\nNumber of predicted TMHs: 1\nExp number of AAs in TMHs: 28.03134\nExp number, first 60 AAs: 6.30523\nTotal prob of N-in: 0.48947\noutside 1 187\nTMhelix 188 210\ninside 211 212

799 GCF_001994755.1_ASM199475v1 Burkholderia cenocepacia Proteobacteria; Betaproteobacteria; Burkholderiales; Burkholderiaceae; Burkholderia; Burkholderia cepacia complex
MSDDIDNGGKGGFTRRDMMRVMAASGMMAVGGGGLLTSAQSAFAAPAPKRGGKIRVANES WP_006482802.1
MULTISPECIES: ABC transporter substrate-binding protein [Burkholderia cepacia complex] Length: 527\nNumber of predicted TMHs: 2\nExp number of AAs in TMHs: 36.7303599999999\nExp number, first 60 AAs: 22.08937\nTotal prob of N-in: 0.97312\nPOSSIBLE N-term signal sequence\ninside 1 20\nTMhelix 21 43\noutside 44 502\nTMhelix 503 525\ninside 526 527

800 GCF_001994715.1_ASM199471v1 Burkholderia cenocepacia Proteobacteria; Betaproteobacteria; Burkholderiales; Burkholderiaceae; Burkholderia; Burkholderia cepacia complex
MKRRTFLALPAVAASGMLAGCSSADLGPVTHRLMKDPEYRETLSAFLISADGKKLVVIGKWP_006483583.1 MULTISPECIES: hypothetical protein [Burkholderia cepacia complex] Length: 212\nNumber of predicted TMHs: 1\nExp number of AAs in TMHs: 28.03134\nExp number, first 60 AAs: 6.30523\nTotal prob of N-in: 0.48947\noutside 1 187\nTMhelix 188 210\ninside 211 212

801 GCF_001994715.1_ASM199471v1 Burkholderia cenocepacia Proteobacteria; Betaproteobacteria; Burkholderiales; Burkholderiaceae; Burkholderia; Burkholderia cepacia complex
MSDDIDNGGKGGFTRRDMMRVMAASGMMAVGGGGLLTSAQSAFAAPAPKRGGKIRVANES WP_006482802.1
MULTISPECIES: ABC transporter substrate-binding protein [Burkholderia cepacia complex] Length: 527\nNumber of predicted TMHs: 2\nExp number of AAs in TMHs: 36.7303599999999\nExp number, first 60 AAs: 22.08937\nTotal prob of N-in: 0.97312\nPOSSIBLE N-term signal sequence\ninside 1 20\nTMhelix 21 43\noutside 44 502\nTMhelix 503 525\ninside 526 527

802 GCF_001994665.1_ASM199466v1 Burkholderia cenocepacia Proteobacteria; Betaproteobacteria; Burkholderiales; Burkholderiaceae; Burkholderia; Burkholderia cepacia complex
MKRRTFLALPAVAASGMLAGCSSADLGPVTHRLMKDPEYRETLSAFLISADGKKLVVIGKWP_006483583.1 MULTISPECIES: hypothetical protein [Burkholderia cepacia complex] Length: 212\nNumber of predicted TMHs: 1\nExp number of AAs in TMHs: 28.03134\nExp number, first 60 AAs: 6.30523\nTotal prob of N-in: 0.48947\noutside 1 187\nTMhelix 188 210\ninside 211 212

803 GCF_001994605.1_ASM199460v1 Burkholderia cenocepacia Proteobacteria; Betaproteobacteria; Burkholderiales; Burkholderiaceae; Burkholderia; Burkholderia cepacia complex
MKRRTFLALPAVAASGMLAGCSSADLGPVTHRLMKDPEYRETLSAFLISADGKKLVVIGKWP_006483583.1 MULTISPECIES: hypothetical protein [Burkholderia cepacia complex] Length: 212\nNumber of predicted TMHs: 1\nExp number of AAs in TMHs: 28.03134\nExp number, first 60 AAs: 6.30523\nTotal prob of N-in: 0.48947\noutside 1 187\nTMhelix 188 210\ninside 211 212

804 GCF_001994605.1_ASM199460v1 Burkholderia cenocepacia Proteobacteria; Betaproteobacteria; Burkholderiales; Burkholderiaceae; Burkholderia; Burkholderia cepacia complex
MSDDIDNGGKGGFTRRDMMRVMAASGMMAVGGGGLLTSAQSAFAAPAPKRGGKIRVANES WP_006482802.1
MULTISPECIES: ABC transporter substrate-binding protein [Burkholderia cepacia complex] Length: 527\nNumber of predicted TMHs: 2\nExp number of AAs in TMHs: 36.7303599999999\nExp number, first 60 AAs: 22.08937\nTotal prob of N-in: 0.97312\nPOSSIBLE N-term signal sequence\ninside 1 20\nTMhelix 21 43\noutside 44 502\nTMhelix 503 525\ninside 526 527

805 GCF_001999805.1_ASM199980v1 Burkholderia cenocepacia Proteobacteria; Betaproteobacteria; Burkholderiales; Burkholderiaceae; Burkholderia; Burkholderia cepacia complex
MKRRTFLALPAVAASGMLAGCSSADLGPVTHRLMKDPEYRETLSAFLISADGKKLVVIGKWP_034201760.1 MULTISPECIES: hypothetical protein [Burkholderia cepacia complex] Length: 212\nNumber of predicted TMHs: 1\nExp number of AAs in TMHs: 28.11606\nExp number, first 60 AAs: 6.33522\nTotal prob of N-in: 0.49088\noutside 1 187\nTMhelix 188 210\ninside 211 212

806 GCF_001994395.1_ASM199439v1 Burkholderia cenocepacia Proteobacteria; Betaproteobacteria; Burkholderiales; Burkholderiaceae; Burkholderia; Burkholderia cepacia complex
MKRRTFLALPAVAASGMLAGCSSADLGPVTHRLMKDPEYRETLSAFLISADGKKLVVIGKWP_034201760.1 MULTISPECIES: hypothetical protein [Burkholderia cepacia complex] Length: 212\nNumber of predicted TMHs: 1\nExp number of AAs in TMHs: 28.11606\nExp number, first 60 AAs: 6.33522\nTotal prob of N-in: 0.49088\noutside 1 187\nTMhelix 188 210\ninside 211 212

807 GCF_001994555.1_ASM199455v1 Burkholderia cenocepacia Proteobacteria; Betaproteobacteria; Burkholderiales; Burkholderiaceae; Burkholderia; Burkholderia cepacia complex
MKRRTFLALPAVAASGMLAGCSSADLGPVTHRLMKDPEYRETLSAFLISADGKKLVVIGKWP_034201760.1 MULTISPECIES: hypothetical protein [Burkholderia cepacia complex] Length: 212\nNumber of predicted TMHs: 1\nExp number of AAs in TMHs: 28.11606\nExp number, first 60 AAs: 6.33522\nTotal prob of N-in: 0.49088\noutside 1 187\nTMhelix 188 210\ninside 211 212

808 GCF_001994555.1_ASM199455v1 Burkholderia cenocepacia Proteobacteria; Betaproteobacteria; Burkholderiales; Burkholderiaceae; Burkholderia; Burkholderia cepacia complex
MSDDIDNGGKGGFTRRDMMRVMAASGMMAVGGGGLLTSAQSAFAAPAPKRGGKIRVANES WP_006482802.1 MULTISPECIES: ABC transporter substrate-binding protein [Burkholderia cepacia complex] Length: 527\nNumber of predicted TMHs: 2\nExp number of AAs in TMHs: 36.7303599999999\nExp number, first 60 AAs: 22.08937\nTotal prob of N-in: 0.97312\nPOSSIBLE N-term signal sequence\ninside 1 20\nTMhelix 21 43\noutside 44 502\nTMhelix 503 525\ninside 526 527

809 GCF_001994345.1_ASM199434v1 Burkholderia cenocepacia Proteobacteria; Betaproteobacteria; Burkholderiales; Burkholderiaceae; Burkholderia; Burkholderia cepacia complex
MKRRTFLALPAVAASGMLAGCSSADLGPVTHRLMKDPEYRETLSAFLISADGKKLVVIGKWP_034201760.1 MULTISPECIES: hypothetical protein [Burkholderia cepacia complex] Length: 212\nNumber of predicted TMHs: 1\nExp number of AAs in TMHs: 28.11606\nExp number, first 60 AAs: 6.33522\nTotal prob of N-in: 0.49088\noutside 1 187\nTMhelix 188 210\ninside 211 212

810 GCF_001994495.1_ASM199449v1 Burkholderia cenocepacia Proteobacteria; Betaproteobacteria; Burkholderiales; Burkholderiaceae; Burkholderia; Burkholderia cepacia complex
MKRRTFLALPAVAASGMLAGCSSADLGPVTHRLMKDPEYRETLSAFLISADGKKLVVIGKWP_034201760.1 MULTISPECIES: hypothetical protein [Burkholderia cepacia complex] Length: 212\nNumber of predicted TMHs: 1\nExp number of AAs in TMHs: 28.11606\nExp number, first 60 AAs: 6.33522\nTotal prob of N-in: 0.49088\noutside 1 187\nTMhelix 188 210\ninside 211 212

811 GCF_001994535.1_ASM199453v1 Burkholderia cenocepacia Proteobacteria; Betaproteobacteria; Burkholderiales; Burkholderiaceae; Burkholderia; Burkholderia cepacia complex
MKRRTFLALPAVAASGMLAGCSSADLGPVTHRLMKDPEYRETLSAFLISADGKKLVVIGKWP_034201760.1 MULTISPECIES: hypothetical protein [Burkholderia cepacia complex] Length: 212\nNumber of predicted TMHs: 1\nExp number of AAs in TMHs: 28.11606\nExp number, first 60 AAs: 6.33522\nTotal prob of N-in: 0.49088\noutside 1 187\nTMhelix 188 210\ninside 211 212

812 GCF_001994535.1_ASM199453v1 Burkholderia cenocepacia Proteobacteria; Betaproteobacteria; Burkholderiales; Burkholderiaceae; Burkholderia; Burkholderia cepacia complex
MSDDIDNGGKGGFTRRDMMRVMAASGMMAVGGGGLLTSAQSAFAAPAPKRGGKIRVANES WP_006482802.1 MULTISPECIES: ABC transporter substrate-binding protein [Burkholderia cepacia complex] Length: 527\nNumber of predicted TMHs: 2\nExp number of AAs in TMHs: 36.7303599999999\nExp number, first 60 AAs: 22.08937\nTotal prob of N-in: 0.97312\nPOSSIBLE N-term signal sequence\ninside 1 20\nTMhelix 21 43\noutside 44 502\nTMhelix 503 525\ninside 526 527

813 GCF_001994495.1_ASM199449v1 Burkholderia cenocepacia Proteobacteria; Betaproteobacteria; Burkholderiales; Burkholderiaceae; Burkholderia; Burkholderia cepacia complex
MSDDIDNGGKGGFTRRDMMRVMAASGMMAVGGGGLLTSAQSAFAAPAPKRGGKIRVANES WP_006482802.1

MULTISPECIES: ABC transporter substrate-binding protein [Burkholderia cepacia complex] Length: 527\nNumber of predicted TMHs: 2\nExp number of AAs in TMHs: 36.7303599999999\nExp number, first 60 AAs: 22.08937\nTotal prob of N-in: 0.97312\nPOSSIBLE N-term signal sequence\ninside 1 20\nTMhelix 21 43\noutside 44 502\nTMhelix 503 525\ninside 526 527

814 GCF_001994435.1_ASM199443v1 Burkholderia cenocepacia Proteobacteria; Betaproteobacteria; Burkholderiales; Burkholderiaceae; Burkholderia; Burkholderia cepacia complex MKRRTFLALPAVAASGMLAGCSSADLGPVTHRLMKDPEYRETLSAFLISADGKKLVVIGKWP_006483583.1 MULTISPECIES: hypothetical protein [Burkholderia cepacia complex] Length: 212\nNumber of predicted TMHs: 1\nExp number of AAs in TMHs: 28.03134\nExp number, first 60 AAs: 6.30523\nTotal prob of N-in: 0.48947\noutside 1 187\nTMhelix 188 210\ninside 211 212

815 GCF_001994435.1_ASM199443v1 Burkholderia cenocepacia Proteobacteria; Betaproteobacteria; Burkholderiales; Burkholderiaceae; Burkholderia; Burkholderia cepacia complex MSDDIDNGGKGGFTRRDMMRVMAASGMMAVGGGGLLTSAQSAFAAPAPKRGGKIRVANES WP_006482802.1 MULTISPECIES: ABC transporter substrate-binding protein [Burkholderia cepacia complex] Length: 527\nNumber of predicted TMHs: 2\nExp number of AAs in TMHs: 36.7303599999999\nExp number, first 60 AAs: 22.08937\nTotal prob of N-in: 0.97312\nPOSSIBLE N-term signal sequence\ninside 1 20\nTMhelix 21 43\noutside 44 502\nTMhelix 503 525\ninside 526 527

816 GCF_001994315.1_ASM199431v1 Burkholderia cenocepacia Proteobacteria; Betaproteobacteria; Burkholderiales; Burkholderiaceae; Burkholderia; Burkholderia cepacia complex MKRRTFLALPAVAASGMLAGCSSADLGPVTHRLMKDPEYRETLSAFLISADGKKLVVIGKWP_034201760.1 MULTISPECIES: hypothetical protein [Burkholderia cepacia complex] Length: 212\nNumber of predicted TMHs: 1\nExp number of AAs in TMHs: 28.11606\nExp number, first 60 AAs: 6.33522\nTotal prob of N-in: 0.49088\noutside 1 187\nTMhelix 188 210\ninside 211 212

817 GCF_001994255.1_ASM199425v1 Burkholderia cenocepacia Proteobacteria; Betaproteobacteria; Burkholderiales; Burkholderiaceae; Burkholderia; Burkholderia cepacia complex MKRRTFLALPAVAASGMLAGCSSADLGPVTHRLMKDPEYRETLSAFLISADGKKLVVIGKWP_034201760.1 MULTISPECIES: hypothetical protein [Burkholderia cepacia complex] Length: 212\nNumber of predicted TMHs: 1\nExp number of AAs in TMHs: 28.11606\nExp number, first 60 AAs: 6.33522\nTotal prob of N-in: 0.49088\noutside 1 187\nTMhelix 188 210\ninside 211 212

818 GCF_001994395.1_ASM199439v1 Burkholderia cenocepacia Proteobacteria; Betaproteobacteria; Burkholderiales; Burkholderiaceae; Burkholderia; Burkholderia cepacia complex MSDDIDNGGKGGFTRRDMMRVMAASGMMAVGGGGLLTSAQSAFAAPAPKRGGKIRVANES WP_006482802.1 MULTISPECIES: ABC transporter substrate-binding protein [Burkholderia cepacia complex] Length: 527\nNumber of predicted TMHs: 2\nExp number of AAs in TMHs: 36.7303599999999\nExp number, first 60 AAs: 22.08937\nTotal prob of N-in: 0.97312\nPOSSIBLE N-term signal sequence\ninside 1 20\nTMhelix 21 43\noutside 44 502\nTMhelix 503 525\ninside 526 527

819 GCF_001994345.1_ASM199434v1 Burkholderia cenocepacia Proteobacteria; Betaproteobacteria; Burkholderiales; Burkholderiaceae; Burkholderia; Burkholderia cepacia complex MSDDIDNGGKGGFTRRDMMRVMAASGMMAVGGGGLLTSAQSAFAAPAPKRGGKIRVANES WP_006482802.1 MULTISPECIES: ABC transporter substrate-binding protein [Burkholderia cepacia complex] Length: 527\nNumber of predicted TMHs: 2\nExp number of AAs in TMHs: 36.7303599999999\nExp number, first 60 AAs: 22.08937\nTotal prob of N-in: 0.97312\nPOSSIBLE N-term signal sequence\ninside 1 20\nTMhelix 21 43\noutside 44 502\nTMhelix 503 525\ninside 526 527

820 GCF_001994315.1_ASM199431v1 Burkholderia cenocepacia Proteobacteria; Betaproteobacteria; Burkholderiales; Burkholderiaceae; Burkholderia; Burkholderia cepacia complex MSDDIDNGGKGGFTRRDMMRVMAASGMMAVGGGGLLTSAQSAFAAPAPKRGGKIRVANES WP_006482802.1 MULTISPECIES: ABC transporter substrate-binding protein [Burkholderia cepacia complex] Length: 527\nNumber of predicted TMHs: 2\nExp number of AAs in TMHs: 36.7303599999999\nExp number, first 60 AAs: 22.08937\nTotal prob of N-in: 0.97312\nPOSSIBLE N-term signal sequence\ninside 1 20\nTMhelix 21 43\noutside 44 502\nTMhelix 503 525\ninside 526 527

821 GCF_001994255.1_ASM199425v1 Burkholderia cenocepacia Proteobacteria; Betaproteobacteria; Burkholderiales; Burkholderiaceae; Burkholderia; Burkholderia cepacia complex MSDDIDNGGKGGFTRRDMMRVMAASGMMAVGGGGLLTSAQSAFAAPAPKRGGKIRVANES WP_006482802.1

MULTISPECIES: ABC transporter substrate-binding protein [Burkholderia cepacia complex] Length: 527\nNumber of predicted TMHs: 2\nExp number of AAs in TMHs: 36.7303599999999\nExp number, first 60 AAs: 22.08937\nTotal prob of N-in: 0.97312\nPOSSIBLE N-term signal sequence\ninside 1 20\nTMhelix 21 43\noutside 44 502\nTMhelix 503 525\ninside 526 527

822 GCF_001994115.1_ASM199411v1 Burkholderia cenocepacia Proteobacteria; Betaproteobacteria; Burkholderiales; Burkholderiaceae; Burkholderia; Burkholderia cepacia complex
MKRRTFLALPAVAASGMLAGCSSADLGPVTHRLMKDPEYRETLSAFLISADGKKLVVIGKWP_034201760.1 MULTISPECIES: hypothetical protein [Burkholderia cepacia complex] Length: 212\nNumber of predicted TMHs: 1\nExp number of AAs in TMHs: 28.11606\nExp number, first 60 AAs: 6.33522\nTotal prob of N-in: 0.49088\noutside 1 187\nTMhelix 188 210\ninside 211 212

823 GCF_001994205.1_ASM199420v1 Burkholderia cenocepacia Proteobacteria; Betaproteobacteria; Burkholderiales; Burkholderiaceae; Burkholderia; Burkholderia cepacia complex
MKRRTFLALPAVAASGMLAGCSSADLGPVTHRLMKDPEYRETLSAFLISADGKKLVVIGKWP_034201760.1 MULTISPECIES: hypothetical protein [Burkholderia cepacia complex] Length: 212\nNumber of predicted TMHs: 1\nExp number of AAs in TMHs: 28.11606\nExp number, first 60 AAs: 6.33522\nTotal prob of N-in: 0.49088\noutside 1 187\nTMhelix 188 210\ninside 211 212

824 GCF_001994165.1_ASM199416v1 Burkholderia cenocepacia Proteobacteria; Betaproteobacteria; Burkholderiales; Burkholderiaceae; Burkholderia; Burkholderia cepacia complex
MKRRTFLALPAVAASGMLAGCSSADLGPVTHRLMKDPEYRETLSAFLISADGKKLVVIGKWP_034201760.1 MULTISPECIES: hypothetical protein [Burkholderia cepacia complex] Length: 212\nNumber of predicted TMHs: 1\nExp number of AAs in TMHs: 28.11606\nExp number, first 60 AAs: 6.33522\nTotal prob of N-in: 0.49088\noutside 1 187\nTMhelix 188 210\ninside 211 212

825 GCF_001994205.1_ASM199420v1 Burkholderia cenocepacia Proteobacteria; Betaproteobacteria; Burkholderiales; Burkholderiaceae; Burkholderia; Burkholderia cepacia complex
MSDDIDNGGKGGFTRRDMMRVMAASGMMAVGGGGLLTSAQSAFAAPAPKRGGKIRVANES WP_006482802.1
MULTISPECIES: ABC transporter substrate-binding protein [Burkholderia cepacia complex] Length: 527\nNumber of predicted TMHs: 2\nExp number of AAs in TMHs: 36.7303599999999\nExp number, first 60 AAs: 22.08937\nTotal prob of N-in: 0.97312\nPOSSIBLE N-term signal sequence\ninside 1 20\nTMhelix 21 43\noutside 44 502\nTMhelix 503 525\ninside 526 527

826 GCF_001994165.1_ASM199416v1 Burkholderia cenocepacia Proteobacteria; Betaproteobacteria; Burkholderiales; Burkholderiaceae; Burkholderia; Burkholderia cepacia complex
MSDDIDNGGKGGFTRRDMMRVMAASGMMAVGGGGLLTSAQSAFAAPAPKRGGKIRVANES WP_006482802.1
MULTISPECIES: ABC transporter substrate-binding protein [Burkholderia cepacia complex] Length: 527\nNumber of predicted TMHs: 2\nExp number of AAs in TMHs: 36.7303599999999\nExp number, first 60 AAs: 22.08937\nTotal prob of N-in: 0.97312\nPOSSIBLE N-term signal sequence\ninside 1 20\nTMhelix 21 43\noutside 44 502\nTMhelix 503 525\ninside 526 527

827 GCF_001994145.1_ASM199414v1 Burkholderia cenocepacia Proteobacteria; Betaproteobacteria; Burkholderiales; Burkholderiaceae; Burkholderia; Burkholderia cepacia complex
MKRRTFLALPAVAASGMLAGCSSADLGPVTHRLMKDPEYRETLSAFLISADGKKLVVIGKWP_034201760.1 MULTISPECIES: hypothetical protein [Burkholderia cepacia complex] Length: 212\nNumber of predicted TMHs: 1\nExp number of AAs in TMHs: 28.11606\nExp number, first 60 AAs: 6.33522\nTotal prob of N-in: 0.49088\noutside 1 187\nTMhelix 188 210\ninside 211 212

828 GCF_001994145.1_ASM199414v1 Burkholderia cenocepacia Proteobacteria; Betaproteobacteria; Burkholderiales; Burkholderiaceae; Burkholderia; Burkholderia cepacia complex
MSDDIDNGGKGGFTRRDMMRVMAASGMMAVGGGGLLTSAQSAFAAPAPKRGGKIRVANES WP_006482802.1
MULTISPECIES: ABC transporter substrate-binding protein [Burkholderia cepacia complex] Length: 527\nNumber of predicted TMHs: 2\nExp number of AAs in TMHs: 36.7303599999999\nExp number, first 60 AAs: 22.08937\nTotal prob of N-in: 0.97312\nPOSSIBLE N-term signal sequence\ninside 1 20\nTMhelix 21 43\noutside 44 502\nTMhelix 503 525\ninside 526 527

829 GCF_001994075.1_ASM199407v1 Burkholderia cenocepacia Proteobacteria; Betaproteobacteria; Burkholderiales; Burkholderiaceae; Burkholderia; Burkholderia cepacia complex
MSDDIDNGGKGGFTRRDMMRVMAASGMMAVGGGGLLTSAQSAFAAPAPKRGGKIRVANES WP_006482802.1
MULTISPECIES: ABC transporter substrate-binding protein [Burkholderia cepacia complex] Length:

527\nNumber of predicted TMHs:  2\nExp number of AAs in TMHs: 36.7303599999999\nExp number, first 60 AAs:  22.08937\nTotal prob of N-in:   0.97312\nPOSSIBLE N-term signal sequence\ninside   1   20\nTMhelix   21   43\noutside   44   502\nTMhelix   503   525\ninside   526   527

830 GCF_001993985.1_ASM199398v1 Burkholderia cenocepacia Proteobacteria; Betaproteobacteria; Burkholderiales; Burkholderiaceae; Burkholderia; Burkholderia cepacia complex MKRRTFLALPAVAASGMLAGCSSADLGPVTHRLMKDPEYRETLSAFLISADGKKLVVIGKWP_006483583.1 MULTISPECIES: hypothetical protein [Burkholderia cepacia complex] Length: 212\nNumber of predicted TMHs:  1\nExp number of AAs in TMHs: 28.03134\nExp number, first 60 AAs:  6.30523\nTotal prob of N-in:   0.48947\noutside   1   187\nTMhelix   188   210\ninside   211   212

831 GCF_001994115.1_ASM199411v1 Burkholderia cenocepacia Proteobacteria; Betaproteobacteria; Burkholderiales; Burkholderiaceae; Burkholderia; Burkholderia cepacia complex MSDDIDNGGKGGFTRRDMMRVMAASGMMAVGGGGLLTSAQSAFAAPAPKRGGKIRVANES WP_006482802.1 MULTISPECIES: ABC transporter substrate-binding protein [Burkholderia cepacia complex] Length: 527\nNumber of predicted TMHs:  2\nExp number of AAs in TMHs: 36.7303599999999\nExp number, first 60 AAs:  22.08937\nTotal prob of N-in:   0.97312\nPOSSIBLE N-term signal sequence\ninside   1   20\nTMhelix   21   43\noutside   44   502\nTMhelix   503   525\ninside   526   527

832 GCF_001993945.1_ASM199394v1 Burkholderia cenocepacia Proteobacteria; Betaproteobacteria; Burkholderiales; Burkholderiaceae; Burkholderia; Burkholderia cepacia complex MKRRTFLALPAVAASGMLAGCSSADLGPVTHRLMKDPEYRETLSAFLISADGKKLVVIGKWP_006483583.1 MULTISPECIES: hypothetical protein [Burkholderia cepacia complex] Length: 212\nNumber of predicted TMHs:  1\nExp number of AAs in TMHs: 28.03134\nExp number, first 60 AAs:  6.30523\nTotal prob of N-in:   0.48947\noutside   1   187\nTMhelix   188   210\ninside   211   212

833 GCF_001993985.1_ASM199398v1 Burkholderia cenocepacia Proteobacteria; Betaproteobacteria; Burkholderiales; Burkholderiaceae; Burkholderia; Burkholderia cepacia complex MSDDIDNGGKGGFTRRDMMRVMAASGMMAVGGGGLLTSAQSAFAAPAPKRGGKIRVANES WP_006482802.1 MULTISPECIES: ABC transporter substrate-binding protein [Burkholderia cepacia complex] Length: 527\nNumber of predicted TMHs:  2\nExp number of AAs in TMHs: 36.7303599999999\nExp number, first 60 AAs:  22.08937\nTotal prob of N-in:   0.97312\nPOSSIBLE N-term signal sequence\ninside   1   20\nTMhelix   21   43\noutside   44   502\nTMhelix   503   525\ninside   526   527

834 GCF_001993945.1_ASM199394v1 Burkholderia cenocepacia Proteobacteria; Betaproteobacteria; Burkholderiales; Burkholderiaceae; Burkholderia; Burkholderia cepacia complex MSDDIDNGGKGGFTRRDMMRVMAASGMMAVGGGGLLTSAQSAFAAPAPKRGGKIRVANES WP_006482802.1 MULTISPECIES: ABC transporter substrate-binding protein [Burkholderia cepacia complex] Length: 527\nNumber of predicted TMHs:  2\nExp number of AAs in TMHs: 36.7303599999999\nExp number, first 60 AAs:  22.08937\nTotal prob of N-in:   0.97312\nPOSSIBLE N-term signal sequence\ninside   1   20\nTMhelix   21   43\noutside   44   502\nTMhelix   503   525\ninside   526   527

835 GCF_001993905.1_ASM199390v1 Burkholderia cenocepacia Proteobacteria; Betaproteobacteria; Burkholderiales; Burkholderiaceae; Burkholderia; Burkholderia cepacia complex MKRRTFLALPAVAASGMLAGCSSADLGPVTHRLMKDPEYRETLSAFLISADGKKLVVIGKWP_006483583.1 MULTISPECIES: hypothetical protein [Burkholderia cepacia complex] Length: 212\nNumber of predicted TMHs:  1\nExp number of AAs in TMHs: 28.03134\nExp number, first 60 AAs:  6.30523\nTotal prob of N-in:   0.48947\noutside   1   187\nTMhelix   188   210\ninside   211   212

836 GCF_001993635.1_ASM199363v1 Burkholderia cenocepacia Proteobacteria; Betaproteobacteria; Burkholderiales; Burkholderiaceae; Burkholderia; Burkholderia cepacia complex MSDDIDNGGKGGFTRRDMMRVMAASGMMAVGGGGLLTSAQSAFAAPAPKRGGKIRVANES WP_077187164.1 ABC transporter substrate-binding protein [Burkholderia cenocepacia] Length: 527\nNumber of predicted TMHs:  2\nExp number of AAs in TMHs: 36.7300399999999\nExp number, first 60 AAs:  22.0894\nTotal prob of N-in:   0.97312\nPOSSIBLE N-term signal sequence\ninside   1   20\nTMhelix   21   43\noutside   44   502\nTMhelix   503   525\ninside   526   527

837 GCF_001993635.1_ASM199363v1 Burkholderia cenocepacia Proteobacteria; Betaproteobacteria; Burkholderiales; Burkholderiaceae; Burkholderia; Burkholderia cepacia complex MSTSITRRRLLAAGVAASSLALSGCFTPKLYKNDAYSEHVSAFMMTEDGKKLVVLGTRYH WP_077021267.1 hypothetical protein [Burkholderia cenocepacia] Length: 212\nNumber of predicted TMHs:  1\nExp number of

AAs in TMHs: 24.17376\nExp number, first 60 AAs: 0.92388\nTotal prob of N-in: 0.37951\noutside 1 180\nTMhelix 181 203\ninside 204 212

838 GCF_001993635.1_ASM199363v1 Burkholderia cenocepacia Proteobacteria; Betaproteobacteria; Burkholderiales; Burkholderiaceae; Burkholderia; Burkholderia cepacia complex METLLTRIRASFADASPRRQLISVAVLVALVASLIAAGYFALRPRYQVLFRDLKPQDAAT WP_077020516.1 flagellar M-ring protein FliF [Burkholderia cenocepacia] Length: 491\nNumber of predicted TMHs: 2\nExp number of AAs in TMHs: 41.34236\nExp number, first 60 AAs: 22.1383\nTotal prob of N-in: 0.99080\nPOSSIBLE N-term signal sequence\ninside 1 20\nTMhelix 21 43\noutside 44 427\nTMhelix 428 450\ninside 451 491

839 GCF_001993455.1_ASM199345v1 Burkholderia cenocepacia Proteobacteria; Betaproteobacteria; Burkholderiales; Burkholderiaceae; Burkholderia; Burkholderia cepacia complex MSTSITRRRLLAAGVAASSLALSGCFTPKLYKNDAYSEHVSAFMMTEDGKKLVVLGTRYH WP_077021267.1 hypothetical protein [Burkholderia cenocepacia] Length: 212\nNumber of predicted TMHs: 1\nExp number of AAs in TMHs: 24.17376\nExp number, first 60 AAs: 0.92388\nTotal prob of N-in: 0.37951\noutside 1 180\nTMhelix 181 203\ninside 204 212

840 GCF_001993905.1_ASM199390v1 Burkholderia cenocepacia Proteobacteria; Betaproteobacteria; Burkholderiales; Burkholderiaceae; Burkholderia; Burkholderia cepacia complex MSDDIDNGGKGGFTRRDMMRVMAASGMMAVGGGGLLTSAQSAFAAPAPKRGGKIRVANES WP_006482802.1 MULTISPECIES: ABC transporter substrate-binding protein [Burkholderia cepacia complex] Length: 527\nNumber of predicted TMHs: 2\nExp number of AAs in TMHs: 36.7303599999999\nExp number, first 60 AAs: 22.08937\nTotal prob of N-in: 0.97312\nPOSSIBLE N-term signal sequence\ninside 1 20\nTMhelix 21 43\noutside 44 502\nTMhelix 503 525\ninside 526 527

841 GCF_001993875.1_ASM199387v1 Burkholderia cenocepacia Proteobacteria; Betaproteobacteria; Burkholderiales; Burkholderiaceae; Burkholderia; Burkholderia cepacia complex MSDDIDNGGKGGFTRRDMMRVMAASGMMAVGGGGLLTSAQSAFAAPAPKRGGKIRVANES WP_006482802.1 MULTISPECIES: ABC transporter substrate-binding protein [Burkholderia cepacia complex] Length: 527\nNumber of predicted TMHs: 2\nExp number of AAs in TMHs: 36.7303599999999\nExp number, first 60 AAs: 22.08937\nTotal prob of N-in: 0.97312\nPOSSIBLE N-term signal sequence\ninside 1 20\nTMhelix 21 43\noutside 44 502\nTMhelix 503 525\ninside 526 527

842 GCF_001993805.1_ASM199380v1 Burkholderia cenocepacia Proteobacteria; Betaproteobacteria; Burkholderiales; Burkholderiaceae; Burkholderia; Burkholderia cepacia complex MSDDIDNGGKGGFTRRDMMRVMAASGMMAVGGGGLLTSAQSAFAAPAPKRGGKIRVANES WP_006482802.1 MULTISPECIES: ABC transporter substrate-binding protein [Burkholderia cepacia complex] Length: 527\nNumber of predicted TMHs: 2\nExp number of AAs in TMHs: 36.7303599999999\nExp number, first 60 AAs: 22.08937\nTotal prob of N-in: 0.97312\nPOSSIBLE N-term signal sequence\ninside 1 20\nTMhelix 21 43\noutside 44 502\nTMhelix 503 525\ninside 526 527

843 GCF_001993455.1_ASM199345v1 Burkholderia cenocepacia Proteobacteria; Betaproteobacteria; Burkholderiales; Burkholderiaceae; Burkholderia; Burkholderia cepacia complex METLLTRIRASFADASPRRQLISVAVLVALVASLIAAGYFALRPRYQVLFRDLKPQDAAT WP_077020516.1 flagellar M-ring protein FliF [Burkholderia cenocepacia] Length: 491\nNumber of predicted TMHs: 2\nExp number of AAs in TMHs: 41.34236\nExp number, first 60 AAs: 22.1383\nTotal prob of N-in: 0.99080\nPOSSIBLE N-term signal sequence\ninside 1 20\nTMhelix 21 43\noutside 44 427\nTMhelix 428 450\ninside 451 491

844 GCF_001993765.1_ASM199376v1 Burkholderia cenocepacia Proteobacteria; Betaproteobacteria; Burkholderiales; Burkholderiaceae; Burkholderia; Burkholderia cepacia complex MKRRTFLALPAVAASGMLAGCSSADLGPVTHRLMKDPEYRETLSAFLISADGKKLVVIGKWP_006483583.1 MULTISPECIES: hypothetical protein [Burkholderia cepacia complex] Length: 212\nNumber of predicted TMHs: 1\nExp number of AAs in TMHs: 28.03134\nExp number, first 60 AAs: 6.30523\nTotal prob of N-in: 0.48947\noutside 1 187\nTMhelix 188 210\ninside 211 212

845 GCF_001993685.1_ASM199368v1 Burkholderia cenocepacia Proteobacteria; Betaproteobacteria; Burkholderiales; Burkholderiaceae; Burkholderia; Burkholderia cepacia complex MKRRTFLALPAVAASGMLAGCSSADLGPVTHRLMKDPEYRETLSAFLISADGKKLVVIGKWP_034201760.1 MULTISPECIES: hypothetical protein [Burkholderia cepacia complex] Length: 212\nNumber of predicted TMHs: 1\nExp number of AAs in TMHs: 28.11606\nExp number, first 60 AAs: 6.33522\nTotal prob of N-in: 0.49088\noutside 1 187\nTMhelix 188 210\ninside 211 212

846 GCF_001993725.1_ASM199372v1 Burkholderia cenocepacia Proteobacteria; Betaproteobacteria; Burkholderiales; Burkholderiaceae; Burkholderia; Burkholderia cepacia complex MKRRTFLALPAVAASGMLAGCSSADLGPVTHRLMKDPEYRETLSAFLISADGKKLVVIGKWP_006483583.1 MULTISPECIES: hypothetical protein [Burkholderia cepacia complex] Length: 212\nNumber of predicted TMHs: 1\nExp number of AAs in TMHs: 28.03134\nExp number, first 60 AAs: 6.30523\nTotal prob of N-in: 0.48947\noutside 1 187\nTMhelix 188 210\ninside 211 212

847 GCF_001993765.1_ASM199376v1 Burkholderia cenocepacia Proteobacteria; Betaproteobacteria; Burkholderiales; Burkholderiaceae; Burkholderia; Burkholderia cepacia complex MSDDIDNGGKGGFTRRDMMRVMAASGMMAVGGGGLLTSAQSAFAAPAPKRGGKIRVANES WP_006482802.1 MULTISPECIES: ABC transporter substrate-binding protein [Burkholderia cepacia complex] Length: 527\nNumber of predicted TMHs: 2\nExp number of AAs in TMHs: 36.7303599999999\nExp number, first 60 AAs: 22.08937\nTotal prob of N-in: 0.97312\nPOSSIBLE N-term signal sequence\ninside 1 20\nTMhelix 21 43\noutside 44 502\nTMhelix 503 525\ninside 526 527

848 GCF_001993725.1_ASM199372v1 Burkholderia cenocepacia Proteobacteria; Betaproteobacteria; Burkholderiales; Burkholderiaceae; Burkholderia; Burkholderia cepacia complex MSDDIDNGGKGGFTRRDMMRVMAASGMMAVGGGGLLTSAQSAFAAPAPKRGGKIRVANES WP_006482802.1 MULTISPECIES: ABC transporter substrate-binding protein [Burkholderia cepacia complex] Length: 527\nNumber of predicted TMHs: 2\nExp number of AAs in TMHs: 36.7303599999999\nExp number, first 60 AAs: 22.08937\nTotal prob of N-in: 0.97312\nPOSSIBLE N-term signal sequence\ninside 1 20\nTMhelix 21 43\noutside 44 502\nTMhelix 503 525\ninside 526 527

849 GCF_001993685.1_ASM199368v1 Burkholderia cenocepacia Proteobacteria; Betaproteobacteria; Burkholderiales; Burkholderiaceae; Burkholderia; Burkholderia cepacia complex MSDDIDNGGKGGFTRRDMMRVMAASGMMAVGGGGLLTSAQSAFAAPAPKRGGKIRVANES WP_006482802.1 MULTISPECIES: ABC transporter substrate-binding protein [Burkholderia cepacia complex] Length: 527\nNumber of predicted TMHs: 2\nExp number of AAs in TMHs: 36.7303599999999\nExp number, first 60 AAs: 22.08937\nTotal prob of N-in: 0.97312\nPOSSIBLE N-term signal sequence\ninside 1 20\nTMhelix 21 43\noutside 44 502\nTMhelix 503 525\ninside 526 527

850 GCF_001993635.1_ASM199363v1 Burkholderia cenocepacia Proteobacteria; Betaproteobacteria; Burkholderiales; Burkholderiaceae; Burkholderia; Burkholderia cepacia complex MKRRTFLALPAVAASGMLAGCSSADLGPVTHRLMKDPEYRETLSAFLISADGKKLVVIGKWP_006483583.1 MULTISPECIES: hypothetical protein [Burkholderia cepacia complex] Length: 212\nNumber of predicted TMHs: 1\nExp number of AAs in TMHs: 28.03134\nExp number, first 60 AAs: 6.30523\nTotal prob of N-in: 0.48947\noutside 1 187\nTMhelix 188 210\ninside 211 212

851 GCF_001993415.1_ASM199341v1 Burkholderia cenocepacia Proteobacteria; Betaproteobacteria; Burkholderiales; Burkholderiaceae; Burkholderia; Burkholderia cepacia complex MKRRTFLALPAVAASGMLAGCSSADLGPVTHRLMKDPEYRETLSAFLISADGKKLVVIGKWP_034201760.1 MULTISPECIES: hypothetical protein [Burkholderia cepacia complex] Length: 212\nNumber of predicted TMHs: 1\nExp number of AAs in TMHs: 28.11606\nExp number, first 60 AAs: 6.33522\nTotal prob of N-in: 0.49088\noutside 1 187\nTMhelix 188 210\ninside 211 212

852 GCF_001993275.1_ASM199327v1 Burkholderia cenocepacia Proteobacteria; Betaproteobacteria; Burkholderiales; Burkholderiaceae; Burkholderia; Burkholderia cepacia complex MSDDIDNGGKGGFTRRDMMRVMAASGMMAVGGGGLLTSAQSAFAAPAPKRGGKIRVANES WP_077187164.1 ABC transporter substrate-binding protein [Burkholderia cenocepacia] Length: 527\nNumber of predicted TMHs: 2\nExp number of AAs in TMHs: 36.7300399999999\nExp number, first 60 AAs: 22.0894\nTotal prob of N-in: 0.97312\nPOSSIBLE N-term signal sequence\ninside 1 20\nTMhelix 21 43\noutside 44 502\nTMhelix 503 525\ninside 526 527

853 GCF_001993275.1_ASM199327v1 Burkholderia cenocepacia Proteobacteria; Betaproteobacteria; Burkholderiales; Burkholderiaceae; Burkholderia; Burkholderia cepacia complex MSTSITRRRLLAAGVAASSLALSGCFTPKLYKNDAYSEHVSAFMMTEDGKKLVVLGTRYH WP_077021267.1 hypothetical protein [Burkholderia cenocepacia] Length: 212\nNumber of predicted TMHs: 1\nExp number of AAs in TMHs: 24.17376\nExp number, first 60 AAs: 0.92388\nTotal prob of N-in: 0.37951\noutside 1 180\nTMhelix 181 203\ninside 204 212

854 GCF_001993275.1_ASM199327v1 Burkholderia cenocepacia Proteobacteria; Betaproteobacteria; Burkholderiales; Burkholderiaceae; Burkholderia; Burkholderia cepacia complex

METLLTRIRASFADASPRRQLISVAVLVALVASLIAAGYFALRPRYQVLFRDLKPQDAAT WP_077020516.1 flagellar M-ring protein FliF [Burkholderia cenocepacia] Length: 491\nNumber of predicted TMHs: 2\nExp number of AAs in TMHs: 41.34236\nExp number, first 60 AAs: 22.1383\nTotal prob of N-in: 0.99080\nPOSSIBLE N-term signal sequence\ninside 1 20\nTMhelix 21 43\noutside 44 427\nTMhelix 428 450\ninside 451 491

855 GCF_001994075.1_ASM199407v1 Burkholderia cenocepacia Proteobacteria; Betaproteobacteria; Burkholderiales; Burkholderiaceae; Burkholderia; Burkholderia cepacia complex
MKRRTFLALPAVAASGMLAGCSSADLGPVTHRLMKDPEYRETLSAFLISADGKKLVVIGKWP_006483583.1 MULTISPECIES: hypothetical protein [Burkholderia cepacia complex] Length: 212\nNumber of predicted TMHs: 1\nExp number of AAs in TMHs: 28.03134\nExp number, first 60 AAs: 6.30523\nTotal prob of N-in: 0.48947\noutside 1 187\nTMhelix 188 210\ninside 211 212

856 GCF_001993135.1_ASM199313v1 Burkholderia cenocepacia Proteobacteria; Betaproteobacteria; Burkholderiales; Burkholderiaceae; Burkholderia; Burkholderia cepacia complex
MSDDIDNGGKGGFTRRDMMRVMAASGMMAVGGGGLLTSAQSAFAAPAPKRGGKIRVANES WP_077180776.1 ABC transporter substrate-binding protein [Burkholderia cenocepacia] Length: 527\nNumber of predicted TMHs: 2\nExp number of AAs in TMHs: 36.7312299999999\nExp number, first 60 AAs: 22.0894\nTotal prob of N-in: 0.97312\nPOSSIBLE N-term signal sequence\ninside 1 20\nTMhelix 21 43\noutside 44 502\nTMhelix 503 525\ninside 526 527

857 GCF_001993375.1_ASM199337v1 Burkholderia cenocepacia Proteobacteria; Betaproteobacteria; Burkholderiales; Burkholderiaceae; Burkholderia; Burkholderia cepacia complex
MKRRTFLALPAVAASGMLAGCSSADLGPVTHRLMKDPEYRETLSAFLISADGKKLVVIGKWP_034201760.1 MULTISPECIES: hypothetical protein [Burkholderia cepacia complex] Length: 212\nNumber of predicted TMHs: 1\nExp number of AAs in TMHs: 28.11606\nExp number, first 60 AAs: 6.33522\nTotal prob of N-in: 0.49088\noutside 1 187\nTMhelix 188 210\ninside 211 212

858 GCF_001993215.1_ASM199321v1 Burkholderia cenocepacia Proteobacteria; Betaproteobacteria; Burkholderiales; Burkholderiaceae; Burkholderia; Burkholderia cepacia complex
MSDDIDNGGKGGFTRRDMMRVMAASGMMAVGGGGLLTSAQSAFAAPAPKRGGKIRVANES WP_077187164.1 ABC transporter substrate-binding protein [Burkholderia cenocepacia] Length: 527\nNumber of predicted TMHs: 2\nExp number of AAs in TMHs: 36.7300399999999\nExp number, first 60 AAs: 22.0894\nTotal prob of N-in: 0.97312\nPOSSIBLE N-term signal sequence\ninside 1 20\nTMhelix 21 43\noutside 44 502\nTMhelix 503 525\ninside 526 527

859 GCF_001993215.1_ASM199321v1 Burkholderia cenocepacia Proteobacteria; Betaproteobacteria; Burkholderiales; Burkholderiaceae; Burkholderia; Burkholderia cepacia complex
MSTSITRRRLLAAGVAASSLALSGCFTPKLYKNDAYSEHVSAFMMTEDGKKLVVLGTRYH WP_077021267.1
hypothetical protein [Burkholderia cenocepacia] Length: 212\nNumber of predicted TMHs: 1\nExp number of AAs in TMHs: 24.17376\nExp number, first 60 AAs: 0.92388\nTotal prob of N-in: 0.37951\noutside 1 180\nTMhelix 181 203\ninside 204 212

860 GCF_001993215.1_ASM199321v1 Burkholderia cenocepacia Proteobacteria; Betaproteobacteria; Burkholderiales; Burkholderiaceae; Burkholderia; Burkholderia cepacia complex
METLLTRIRASFADASPRRQLISVAVLVALVASLIAAGYFALRPRYQVLFRDLKPQDAAT WP_077020516.1 flagellar M-ring protein FliF [Burkholderia cenocepacia] Length: 491\nNumber of predicted TMHs: 2\nExp number of AAs in TMHs: 41.34236\nExp number, first 60 AAs: 22.1383\nTotal prob of N-in: 0.99080\nPOSSIBLE N-term signal sequence\ninside 1 20\nTMhelix 21 43\noutside 44 427\nTMhelix 428 450\ninside 451 491

861 GCF_001993175.1_ASM199317v1 Burkholderia cenocepacia Proteobacteria; Betaproteobacteria; Burkholderiales; Burkholderiaceae; Burkholderia; Burkholderia cepacia complex
MSDDIDNGGKGGFTRRDMMRVMAASGMMAVGGGGLLTSAQSAFAAPAPKRGGKIRVANES WP_077180776.1 ABC transporter substrate-binding protein [Burkholderia cenocepacia] Length: 527\nNumber of predicted TMHs: 2\nExp number of AAs in TMHs: 36.7312299999999\nExp number, first 60 AAs: 22.0894\nTotal prob of N-in: 0.97312\nPOSSIBLE N-term signal sequence\ninside 1 20\nTMhelix 21 43\noutside 44 502\nTMhelix 503 525\ninside 526 527

862 GCF_001993805.1_ASM199380v1 Burkholderia cenocepacia Proteobacteria; Betaproteobacteria; Burkholderiales; Burkholderiaceae; Burkholderia; Burkholderia cepacia complex
MKRRTFLALPAVAASGMLAGCSSADLGPVTHRLMKDPEYRETLSAFLISADGKKLVVIGKWP_006483583.1 MULTISPECIES: hypothetical protein [Burkholderia cepacia complex] Length: 212\nNumber of predicted TMHs: 1\nExp number of AAs in TMHs: 28.03134\nExp number, first 60 AAs: 6.30523\nTotal prob of N-in: 0.48947\noutside 1 187\nTMhelix 188 210\ninside 211 212

863 GCF_001993555.1_ASM199355v1 Burkholderia cenocepacia Proteobacteria; Betaproteobacteria; Burkholderiales; Burkholderiaceae; Burkholderia; Burkholderia cepacia complex MSTSITRRRLLAAGVAASSLALSGCFTPKLYKNDAYSEHVSAFMMTEDGKKLVVLGTRYH WP_077021267.1 hypothetical protein [Burkholderia cenocepacia] Length: 212\nNumber of predicted TMHs: 1\nExp number of AAs in TMHs: 24.17376\nExp number, first 60 AAs: 0.92388\nTotal prob of N-in: 0.37951\noutside 1 180\nTMhelix 181 203\ninside 204 212

864 GCF_001993555.1_ASM199355v1 Burkholderia cenocepacia Proteobacteria; Betaproteobacteria; Burkholderiales; Burkholderiaceae; Burkholderia; Burkholderia cepacia complex MKRRTFLALPAVAASGMLAGCSSADLGPVTHRLMKDPEYRETLSAFLISADGKKLVVIGKWP_006483583.1 MULTISPECIES: hypothetical protein [Burkholderia cepacia complex] Length: 212\nNumber of predicted TMHs: 1\nExp number of AAs in TMHs: 28.03134\nExp number, first 60 AAs: 6.30523\nTotal prob of N-in: 0.48947\noutside 1 187\nTMhelix 188 210\ninside 211 212

865 GCF_001993515.1_ASM199351v1 Burkholderia cenocepacia Proteobacteria; Betaproteobacteria; Burkholderiales; Burkholderiaceae; Burkholderia; Burkholderia cepacia complex MSTSITRRRLLAAGVAASSLALSGCFTPKLYKNDAYSEHVSAFMMTEDGKKLVVLGTRYH WP_077021267.1 hypothetical protein [Burkholderia cenocepacia] Length: 212\nNumber of predicted TMHs: 1\nExp number of AAs in TMHs: 24.17376\nExp number, first 60 AAs: 0.92388\nTotal prob of N-in: 0.37951\noutside 1 180\nTMhelix 181 203\ninside 204 212

866 GCF_001993515.1_ASM199351v1 Burkholderia cenocepacia Proteobacteria; Betaproteobacteria; Burkholderiales; Burkholderiaceae; Burkholderia; Burkholderia cepacia complex MKRRTFLALPAVAASGMLAGCSSADLGPVTHRLMKDPEYRETLSAFLISADGKKLVVIGKWP_006483583.1 MULTISPECIES: hypothetical protein [Burkholderia cepacia complex] Length: 212\nNumber of predicted TMHs: 1\nExp number of AAs in TMHs: 28.03134\nExp number, first 60 AAs: 6.30523\nTotal prob of N-in: 0.48947\noutside 1 187\nTMhelix 188 210\ninside 211 212

867 GCF_001993355.1_ASM199335v1 Burkholderia cenocepacia Proteobacteria; Betaproteobacteria; Burkholderiales; Burkholderiaceae; Burkholderia; Burkholderia cepacia complex MKRRTFLALPAVAASGMLAGCSSADLGPVTHRLMKDPEYRETLSAFLISADGKKLVVIGKWP_034201760.1 MULTISPECIES: hypothetical protein [Burkholderia cepacia complex] Length: 212\nNumber of predicted TMHs: 1\nExp number of AAs in TMHs: 28.11606\nExp number, first 60 AAs: 6.33522\nTotal prob of N-in: 0.49088\noutside 1 187\nTMhelix 188 210\ninside 211 212

868 GCF_001993555.1_ASM199355v1 Burkholderia cenocepacia Proteobacteria; Betaproteobacteria; Burkholderiales; Burkholderiaceae; Burkholderia; Burkholderia cepacia complex MSDDIDNGGKGGFTRRDMMRVMAASGMMAVGGGGLLTSAQSAFAAPAPKRGGKIRVANES WP_006482802.1 MULTISPECIES: ABC transporter substrate-binding protein [Burkholderia cepacia complex] Length: 527\nNumber of predicted TMHs: 2\nExp number of AAs in TMHs: 36.7303599999999\nExp number, first 60 AAs: 22.08937\nTotal prob of N-in: 0.97312\nPOSSIBLE N-term signal sequence\ninside 1 20\nTMhelix 21 43\noutside 44 502\nTMhelix 503 525\ninside 526 527

869 GCF_001993515.1_ASM199351v1 Burkholderia cenocepacia Proteobacteria; Betaproteobacteria; Burkholderiales; Burkholderiaceae; Burkholderia; Burkholderia cepacia complex MSDDIDNGGKGGFTRRDMMRVMAASGMMAVGGGGLLTSAQSAFAAPAPKRGGKIRVANES WP_006482802.1 MULTISPECIES: ABC transporter substrate-binding protein [Burkholderia cepacia complex] Length: 527\nNumber of predicted TMHs: 2\nExp number of AAs in TMHs: 36.7303599999999\nExp number, first 60 AAs: 22.08937\nTotal prob of N-in: 0.97312\nPOSSIBLE N-term signal sequence\ninside 1 20\nTMhelix 21 43\noutside 44 502\nTMhelix 503 525\ninside 526 527

870 GCF_001993875.1_ASM199387v1 Burkholderia cenocepacia Proteobacteria; Betaproteobacteria; Burkholderiales; Burkholderiaceae; Burkholderia; Burkholderia cepacia complex MKRRTFLALPAVAASGMLAGCSSADLGPVTHRLMKDPEYRETLSAFLISADGKKLVVIGKWP_006483583.1 MULTISPECIES: hypothetical protein [Burkholderia cepacia complex] Length: 212\nNumber of predicted TMHs: 1\nExp number of AAs in TMHs: 28.03134\nExp number, first 60 AAs: 6.30523\nTotal prob of N-in: 0.48947\noutside 1 187\nTMhelix 188 210\ninside 211 212

871 GCF_001993455.1_ASM199345v1 Burkholderia cenocepacia Proteobacteria; Betaproteobacteria; Burkholderiales; Burkholderiaceae; Burkholderia; Burkholderia cepacia complex MKRRTFLALPAVAASGMLAGCSSADLGPVTHRLMKDPEYRETLSAFLISADGKKLVVIGKWP_006483583.1 MULTISPECIES:

hypothetical protein [Burkholderia cepacia complex] Length: 212\nNumber of predicted TMHs: 1\nExp number of AAs in TMHs: 28.03134\nExp number, first 60 AAs: 6.30523\nTotal prob of N-in: 0.48947\noutside 1 187\nTMhelix 188 210\ninside 211 212

872 GCF_001993455.1_ASM199345v1 Burkholderia cenocepacia Proteobacteria; Betaproteobacteria; Burkholderiales; Burkholderiaceae; Burkholderia; Burkholderia cepacia complex MSDDIDNGGKGGFTRRDMMRVMAASGMMAVGGGGLLTSAQSAFAAPAPKRGGKIRVANES WP_006482802.1 MULTISPECIES: ABC transporter substrate-binding protein [Burkholderia cepacia complex] Length: 527\nNumber of predicted TMHs: 2\nExp number of AAs in TMHs: 36.7303599999999\nExp number, first 60 AAs: 22.08937\nTotal prob of N-in: 0.97312\nPOSSIBLE N-term signal sequence\ninside 1 20\nTMhelix 21 43\noutside 44 502\nTMhelix 503 525\ninside 526 527

873 GCF_001993415.1_ASM199341v1 Burkholderia cenocepacia Proteobacteria; Betaproteobacteria; Burkholderiales; Burkholderiaceae; Burkholderia; Burkholderia cepacia complex MSDDIDNGGKGGFTRRDMMRVMAASGMMAVGGGGLLTSAQSAFAAPAPKRGGKIRVANES WP_006482802.1 MULTISPECIES: ABC transporter substrate-binding protein [Burkholderia cepacia complex] Length: 527\nNumber of predicted TMHs: 2\nExp number of AAs in TMHs: 36.7303599999999\nExp number, first 60 AAs: 22.08937\nTotal prob of N-in: 0.97312\nPOSSIBLE N-term signal sequence\ninside 1 20\nTMhelix 21 43\noutside 44 502\nTMhelix 503 525\ninside 526 527

874 GCF_001993375.1_ASM199337v1 Burkholderia cenocepacia Proteobacteria; Betaproteobacteria; Burkholderiales; Burkholderiaceae; Burkholderia; Burkholderia cepacia complex MSDDIDNGGKGGFTRRDMMRVMAASGMMAVGGGGLLTSAQSAFAAPAPKRGGKIRVANES WP_006482802.1 MULTISPECIES: ABC transporter substrate-binding protein [Burkholderia cepacia complex] Length: 527\nNumber of predicted TMHs: 2\nExp number of AAs in TMHs: 36.7303599999999\nExp number, first 60 AAs: 22.08937\nTotal prob of N-in: 0.97312\nPOSSIBLE N-term signal sequence\ninside 1 20\nTMhelix 21 43\noutside 44 502\nTMhelix 503 525\ninside 526 527

875 GCF_001993315.1_ASM199331v1 Burkholderia cenocepacia Proteobacteria; Betaproteobacteria; Burkholderiales; Burkholderiaceae; Burkholderia; Burkholderia cepacia complex MKRRTFLALPAVAASGMLAGCSSADLGPVTHRLMKDPEYRETLSAFLISADGKKLVVIGKWP_006483583.1 MULTISPECIES: hypothetical protein [Burkholderia cepacia complex] Length: 212\nNumber of predicted TMHs: 1\nExp number of AAs in TMHs: 28.03134\nExp number, first 60 AAs: 6.30523\nTotal prob of N-in: 0.48947\noutside 1 187\nTMhelix 188 210\ninside 211 212

876 GCF_001993355.1_ASM199335v1 Burkholderia cenocepacia Proteobacteria; Betaproteobacteria; Burkholderiales; Burkholderiaceae; Burkholderia; Burkholderia cepacia complex MSDDIDNGGKGGFTRRDMMRVMAASGMMAVGGGGLLTSAQSAFAAPAPKRGGKIRVANES WP_006482802.1 MULTISPECIES: ABC transporter substrate-binding protein [Burkholderia cepacia complex] Length: 527\nNumber of predicted TMHs: 2\nExp number of AAs in TMHs: 36.7303599999999\nExp number, first 60 AAs: 22.08937\nTotal prob of N-in: 0.97312\nPOSSIBLE N-term signal sequence\ninside 1 20\nTMhelix 21 43\noutside 44 502\nTMhelix 503 525\ninside 526 527

877 GCF_001993315.1_ASM199331v1 Burkholderia cenocepacia Proteobacteria; Betaproteobacteria; Burkholderiales; Burkholderiaceae; Burkholderia; Burkholderia cepacia complex MSDDIDNGGKGGFTRRDMMRVMAASGMMAVGGGGLLTSAQSAFAAPAPKRGGKIRVANES WP_006482802.1 MULTISPECIES: ABC transporter substrate-binding protein [Burkholderia cepacia complex] Length: 527\nNumber of predicted TMHs: 2\nExp number of AAs in TMHs: 36.7303599999999\nExp number, first 60 AAs: 22.08937\nTotal prob of N-in: 0.97312\nPOSSIBLE N-term signal sequence\ninside 1 20\nTMhelix 21 43\noutside 44 502\nTMhelix 503 525\ninside 526 527

878 GCF_001993275.1_ASM199327v1 Burkholderia cenocepacia Proteobacteria; Betaproteobacteria; Burkholderiales; Burkholderiaceae; Burkholderia; Burkholderia cepacia complex MKRRTFLALPAVAASGMLAGCSSADLGPVTHRLMKDPEYRETLSAFLISADGKKLVVIGKWP_006483583.1 MULTISPECIES: hypothetical protein [Burkholderia cepacia complex] Length: 212\nNumber of predicted TMHs: 1\nExp number of AAs in TMHs: 28.03134\nExp number, first 60 AAs: 6.30523\nTotal prob of N-in: 0.48947\noutside 1 187\nTMhelix 188 210\ninside 211 212

879 GCF_001993215.1_ASM199321v1 Burkholderia cenocepacia Proteobacteria; Betaproteobacteria; Burkholderiales; Burkholderiaceae; Burkholderia; Burkholderia cepacia complex MKRRTFLALPAVAASGMLAGCSSADLGPVTHRLMKDPEYRETLSAFLISADGKKLVVIGKWP_006483583.1 MULTISPECIES:

hypothetical protein [Burkholderia cepacia complex] Length: 212\nNumber of predicted TMHs: 1\nExp number of AAs in TMHs: 28.03134\nExp number, first 60 AAs: 6.30523\nTotal prob of N-in: 0.48947\noutside 1 187\nTMhelix 188 210\ninside 211 212

880 GCF_001993175.1_ASM199317v1 Burkholderia cenocepacia Proteobacteria; Betaproteobacteria; Burkholderiales; Burkholderiaceae; Burkholderia; Burkholderia cepacia complex MKRRTFLALPAVAASGMLAGCSSADLGPVTHRLMKDPEYRETLSAFLISADGKKLVVIGKWP_006483583.1 MULTISPECIES: hypothetical protein [Burkholderia cepacia complex] Length: 212\nNumber of predicted TMHs: 1\nExp number of AAs in TMHs: 28.03134\nExp number, first 60 AAs: 6.30523\nTotal prob of N-in: 0.48947\noutside 1 187\nTMhelix 188 210\ninside 211 212

881 GCF_001993135.1_ASM199313v1 Burkholderia cenocepacia Proteobacteria; Betaproteobacteria; Burkholderiales; Burkholderiaceae; Burkholderia; Burkholderia cepacia complex MKRRTFLALPAVAASGMLAGCSSADLGPVTHRLMKDPEYRETLSAFLISADGKKLVVIGKWP_006483583.1 MULTISPECIES: hypothetical protein [Burkholderia cepacia complex] Length: 212\nNumber of predicted TMHs: 1\nExp number of AAs in TMHs: 28.03134\nExp number, first 60 AAs: 6.30523\nTotal prob of N-in: 0.48947\noutside 1 187\nTMhelix 188 210\ninside 211 212

882 GCF_001992855.1_ASM199285v1 Burkholderia cenocepacia Proteobacteria; Betaproteobacteria; Burkholderiales; Burkholderiaceae; Burkholderia; Burkholderia cepacia complex MKRRTFLALPAVAASGMLAGCSSADLGPVTHRLMKDPEYRETLSAFLISADGKKLVVIGKWP_034201760.1 MULTISPECIES: hypothetical protein [Burkholderia cepacia complex] Length: 212\nNumber of predicted TMHs: 1\nExp number of AAs in TMHs: 28.11606\nExp number, first 60 AAs: 6.33522\nTotal prob of N-in: 0.49088\noutside 1 187\nTMhelix 188 210\ninside 211 212

883 GCF_001993095.1_ASM199309v1 Burkholderia cenocepacia Proteobacteria; Betaproteobacteria; Burkholderiales; Burkholderiaceae; Burkholderia; Burkholderia cepacia complex MKRRTFLALPAVAASGMLAGCSSADLGPVTHRLMKDPEYRETLSAFLISADGKKLVVIGKWP_034201760.1 MULTISPECIES: hypothetical protein [Burkholderia cepacia complex] Length: 212\nNumber of predicted TMHs: 1\nExp number of AAs in TMHs: 28.11606\nExp number, first 60 AAs: 6.33522\nTotal prob of N-in: 0.49088\noutside 1 187\nTMhelix 188 210\ninside 211 212

884 GCF_001993095.1_ASM199309v1 Burkholderia cenocepacia Proteobacteria; Betaproteobacteria; Burkholderiales; Burkholderiaceae; Burkholderia; Burkholderia cepacia complex MSDDIDNGGKGGFTRRDMMRVMAASGMMAVGGGGLLTSAQSAFAAPAPKRGGKIRVANES WP_006482802.1 MULTISPECIES: ABC transporter substrate-binding protein [Burkholderia cepacia complex] Length: 527\nNumber of predicted TMHs: 2\nExp number of AAs in TMHs: 36.7303599999999\nExp number, first 60 AAs: 22.08937\nTotal prob of N-in: 0.97312\nPOSSIBLE N-term signal sequence\ninside 1 20\nTMhelix 21 43\noutside 44 502\nTMhelix 503 525\ninside 526 527

885 GCF_001992955.1_ASM199295v1 Burkholderia cenocepacia Proteobacteria; Betaproteobacteria; Burkholderiales; Burkholderiaceae; Burkholderia; Burkholderia cepacia complex MKRRTFLALPAVAASGMLAGCSSADLGPVTHRLMKDPEYRETLSAFLISADGKKLVVIGKWP_006483583.1 MULTISPECIES: hypothetical protein [Burkholderia cepacia complex] Length: 212\nNumber of predicted TMHs: 1\nExp number of AAs in TMHs: 28.03134\nExp number, first 60 AAs: 6.30523\nTotal prob of N-in: 0.48947\noutside 1 187\nTMhelix 188 210\ninside 211 212

886 GCF_001993035.1_ASM199303v1 Burkholderia cenocepacia Proteobacteria; Betaproteobacteria; Burkholderiales; Burkholderiaceae; Burkholderia; Burkholderia cepacia complex MKRRTFLALPAVAASGMLAGCSSADLGPVTHRLMKDPEYRETLSAFLISADGKKLVVIGKWP_006483583.1 MULTISPECIES: hypothetical protein [Burkholderia cepacia complex] Length: 212\nNumber of predicted TMHs: 1\nExp number of AAs in TMHs: 28.03134\nExp number, first 60 AAs: 6.30523\nTotal prob of N-in: 0.48947\noutside 1 187\nTMhelix 188 210\ninside 211 212

887 GCF_001993035.1_ASM199303v1 Burkholderia cenocepacia Proteobacteria; Betaproteobacteria; Burkholderiales; Burkholderiaceae; Burkholderia; Burkholderia cepacia complex MSDDIDNGGKGGFTRRDMMRVMAASGMMAVGGGGLLTSAQSAFAAPAPKRGGKIRVANES WP_006482802.1 MULTISPECIES: ABC transporter substrate-binding protein [Burkholderia cepacia complex] Length: 527\nNumber of predicted TMHs: 2\nExp number of AAs in TMHs: 36.7303599999999\nExp number, first 60 AAs: 22.08937\nTotal prob of N-in: 0.97312\nPOSSIBLE N-term signal sequence\ninside 1 20\nTMhelix 21 43\noutside 44 502\nTMhelix 503 525\ninside 526 527

888 GCF_001992915.1_ASM199291v1 Burkholderia cenocepacia Proteobacteria; Betaproteobacteria; Burkholderiales; Burkholderiaceae; Burkholderia; Burkholderia cepacia complex
MKRRTFLALPAVAASGMLAGCSSADLGPVTHRLMKDPEYRETLSAFLISADGKKLVVIGKWP_006483583.1 MULTISPECIES: hypothetical protein [Burkholderia cepacia complex] Length: 212\nNumber of predicted TMHs: 1\nExp number of AAs in TMHs: 28.03134\nExp number, first 60 AAs: 6.30523\nTotal prob of N-in: 0.48947\noutside 1 187\nTMhelix 188 210\ninside 211 212

889 GCF_001992995.1_ASM199299v1 Burkholderia cenocepacia Proteobacteria; Betaproteobacteria; Burkholderiales; Burkholderiaceae; Burkholderia; Burkholderia cepacia complex
MKRRTFLALPAVAASGMLAGCSSADLGPVTHRLMKDPEYRETLSAFLISADGKKLVVIGKWP_006483583.1 MULTISPECIES: hypothetical protein [Burkholderia cepacia complex] Length: 212\nNumber of predicted TMHs: 1\nExp number of AAs in TMHs: 28.03134\nExp number, first 60 AAs: 6.30523\nTotal prob of N-in: 0.48947\noutside 1 187\nTMhelix 188 210\ninside 211 212

890 GCF_001992995.1_ASM199299v1 Burkholderia cenocepacia Proteobacteria; Betaproteobacteria; Burkholderiales; Burkholderiaceae; Burkholderia; Burkholderia cepacia complex
MSDDIDNGGKGGFTRRDMMRVMAASGMMAVGGGGLLTSAQSAFAAPAPKRGGKIRVANES WP_006482802.1
MULTISPECIES: ABC transporter substrate-binding protein [Burkholderia cepacia complex] Length: 527\nNumber of predicted TMHs: 2\nExp number of AAs in TMHs: 36.7303599999999\nExp number, first 60 AAs: 22.08937\nTotal prob of N-in: 0.97312\nPOSSIBLE N-term signal sequence\ninside 1 20\nTMhelix 21 43\noutside 44 502\nTMhelix 503 525\ninside 526 527

891 GCF_001992955.1_ASM199295v1 Burkholderia cenocepacia Proteobacteria; Betaproteobacteria; Burkholderiales; Burkholderiaceae; Burkholderia; Burkholderia cepacia complex
MSDDIDNGGKGGFTRRDMMRVMAASGMMAVGGGGLLTSAQSAFAAPAPKRGGKIRVANES WP_006482802.1
MULTISPECIES: ABC transporter substrate-binding protein [Burkholderia cepacia complex] Length: 527\nNumber of predicted TMHs: 2\nExp number of AAs in TMHs: 36.7303599999999\nExp number, first 60 AAs: 22.08937\nTotal prob of N-in: 0.97312\nPOSSIBLE N-term signal sequence\ninside 1 20\nTMhelix 21 43\noutside 44 502\nTMhelix 503 525\ninside 526 527

892 GCF_001992815.1_ASM199281v1 Burkholderia cenocepacia Proteobacteria; Betaproteobacteria; Burkholderiales; Burkholderiaceae; Burkholderia; Burkholderia cepacia complex
MKRRTFLALPAVAASGMLAGCSSADLGPVTHRLMKDPEYRETLSAFLISADGKKLVVIGKWP_034201760.1 MULTISPECIES: hypothetical protein [Burkholderia cepacia complex] Length: 212\nNumber of predicted TMHs: 1\nExp number of AAs in TMHs: 28.11606\nExp number, first 60 AAs: 6.33522\nTotal prob of N-in: 0.49088\noutside 1 187\nTMhelix 188 210\ninside 211 212

893 GCF_001992915.1_ASM199291v1 Burkholderia cenocepacia Proteobacteria; Betaproteobacteria; Burkholderiales; Burkholderiaceae; Burkholderia; Burkholderia cepacia complex
MSDDIDNGGKGGFTRRDMMRVMAASGMMAVGGGGLLTSAQSAFAAPAPKRGGKIRVANES WP_006482802.1
MULTISPECIES: ABC transporter substrate-binding protein [Burkholderia cepacia complex] Length: 527\nNumber of predicted TMHs: 2\nExp number of AAs in TMHs: 36.7303599999999\nExp number, first 60 AAs: 22.08937\nTotal prob of N-in: 0.97312\nPOSSIBLE N-term signal sequence\ninside 1 20\nTMhelix 21 43\noutside 44 502\nTMhelix 503 525\ninside 526 527

894 GCF_001992855.1_ASM199285v1 Burkholderia cenocepacia Proteobacteria; Betaproteobacteria; Burkholderiales; Burkholderiaceae; Burkholderia; Burkholderia cepacia complex
MSDDIDNGGKGGFTRRDMMRVMAASGMMAVGGGGLLTSAQSAFAAPAPKRGGKIRVANES WP_006482802.1
MULTISPECIES: ABC transporter substrate-binding protein [Burkholderia cepacia complex] Length: 527\nNumber of predicted TMHs: 2\nExp number of AAs in TMHs: 36.7303599999999\nExp number, first 60 AAs: 22.08937\nTotal prob of N-in: 0.97312\nPOSSIBLE N-term signal sequence\ninside 1 20\nTMhelix 21 43\noutside 44 502\nTMhelix 503 525\ninside 526 527

895 GCF_001992385.1_ASM199238v1 Burkholderia cenocepacia Proteobacteria; Betaproteobacteria; Burkholderiales; Burkholderiaceae; Burkholderia; Burkholderia cepacia complex
MSTSITRRRLLAAGVAASSLALSGCFTPKLYKNDAYSEHVSAFMMTEDGKKLVVLGTRYH WP_077021267.1
hypothetical protein [Burkholderia cenocepacia] Length: 212\nNumber of predicted TMHs: 1\nExp number of AAs in TMHs: 24.17376\nExp number, first 60 AAs: 0.92388\nTotal prob of N-in: 0.37951\noutside 1 180\nTMhelix 181 203\ninside 204 212

896 GCF_001992815.1_ASM199281v1 Burkholderia cenocepacia Proteobacteria; Betaproteobacteria; Burkholderiales; Burkholderiaceae; Burkholderia; Burkholderia cepacia complex
MSDDIDNGGKGGFTRRDMMRVMAASGMMAVGGGGLLTSAQSAFAAPAPKRGGKIRVANES WP_006482802.1
MULTISPECIES: ABC transporter substrate-binding protein [Burkholderia cepacia complex] Length: 527\nNumber of predicted TMHs: 2\nExp number of AAs in TMHs: 36.7303599999999\nExp number, first 60 AAs: 22.08937\nTotal prob of N-in: 0.97312\nPOSSIBLE N-term signal sequence\ninside 1 20\nTMhelix 21 43\noutside 44 502\nTMhelix 503 525\ninside 526 527

897 GCF_001992385.1_ASM199238v1 Burkholderia cenocepacia Proteobacteria; Betaproteobacteria; Burkholderiales; Burkholderiaceae; Burkholderia; Burkholderia cepacia complex
METLLTRIRASFADASPRRQLISVAVLVALVASLIAAGYFALRPRYQVLFRDLKPQDAAT WP_077020516.1 flagellar M-ring protein FliF [Burkholderia cenocepacia] Length: 491\nNumber of predicted TMHs: 2\nExp number of AAs in TMHs: 41.34236\nExp number, first 60 AAs: 22.1383\nTotal prob of N-in: 0.99080\nPOSSIBLE N-term signal sequence\ninside 1 20\nTMhelix 21 43\noutside 44 427\nTMhelix 428 450\ninside 451 491

898 GCF_001992765.1_ASM199276v1 Burkholderia cenocepacia Proteobacteria; Betaproteobacteria; Burkholderiales; Burkholderiaceae; Burkholderia; Burkholderia cepacia complex
MKRRTFLALPAVAASGMLAGCSSADLGPVTHRLMKDPEYRETLSAFLISADGKKLVVIGKWP_034201760.1 MULTISPECIES: hypothetical protein [Burkholderia cepacia complex] Length: 212\nNumber of predicted TMHs: 1\nExp number of AAs in TMHs: 28.11606\nExp number, first 60 AAs: 6.33522\nTotal prob of N-in: 0.49088\noutside 1 187\nTMhelix 188 210\ninside 211 212

899 GCF_001992345.1_ASM199234v1 Burkholderia cenocepacia Proteobacteria; Betaproteobacteria; Burkholderiales; Burkholderiaceae; Burkholderia; Burkholderia cepacia complex
MSTSITRRRLLAAGVAASSLALSGCFTPKLYKNDAYSEHVSAFMMTEDGKKLVVLGTRYH WP_077021267.1
hypothetical protein [Burkholderia cenocepacia] Length: 212\nNumber of predicted TMHs: 1\nExp number of AAs in TMHs: 24.17376\nExp number, first 60 AAs: 0.92388\nTotal prob of N-in: 0.37951\noutside 1 180\nTMhelix 181 203\ninside 204 212

900 GCF_001992765.1_ASM199276v1 Burkholderia cenocepacia Proteobacteria; Betaproteobacteria; Burkholderiales; Burkholderiaceae; Burkholderia; Burkholderia cepacia complex
MSDDIDNGGKGGFTRRDMMRVMAASGMMAVGGGGLLTSAQSAFAAPAPKRGGKIRVANES WP_006482802.1
MULTISPECIES: ABC transporter substrate-binding protein [Burkholderia cepacia complex] Length: 527\nNumber of predicted TMHs: 2\nExp number of AAs in TMHs: 36.7303599999999\nExp number, first 60 AAs: 22.08937\nTotal prob of N-in: 0.97312\nPOSSIBLE N-term signal sequence\ninside 1 20\nTMhelix 21 43\noutside 44 502\nTMhelix 503 525\ninside 526 527

901 GCF_001992345.1_ASM199234v1 Burkholderia cenocepacia Proteobacteria; Betaproteobacteria; Burkholderiales; Burkholderiaceae; Burkholderia; Burkholderia cepacia complex
METLLTRIRASFADASPRRQLISVAVLVALVASLIAAGYFALRPRYQVLFRDLKPQDAAT WP_077020516.1 flagellar M-ring protein FliF [Burkholderia cenocepacia] Length: 491\nNumber of predicted TMHs: 2\nExp number of AAs in TMHs: 41.34236\nExp number, first 60 AAs: 22.1383\nTotal prob of N-in: 0.99080\nPOSSIBLE N-term signal sequence\ninside 1 20\nTMhelix 21 43\noutside 44 427\nTMhelix 428 450\ninside 451 491

902 GCF_001992205.1_ASM199220v1 Burkholderia cenocepacia Proteobacteria; Betaproteobacteria; Burkholderiales; Burkholderiaceae; Burkholderia; Burkholderia cepacia complex
MSDDIDNGGKGGFTRRDMMRVMAASGMMAVGGGGLLTSAQSAFAAPAPKRGGKIRVANES WP_077187164.1 ABC transporter substrate-binding protein [Burkholderia cenocepacia] Length: 527\nNumber of predicted TMHs: 2\nExp number of AAs in TMHs: 36.7300399999999\nExp number, first 60 AAs: 22.0894\nTotal prob of N-in: 0.97312\nPOSSIBLE N-term signal sequence\ninside 1 20\nTMhelix 21 43\noutside 44 502\nTMhelix 503 525\ninside 526 527

903 GCF_001992205.1_ASM199220v1 Burkholderia cenocepacia Proteobacteria; Betaproteobacteria; Burkholderiales; Burkholderiaceae; Burkholderia; Burkholderia cepacia complex
MSTSITRRRLLAAGVAASSLALSGCFTPKLYKNDAYSEHVSAFMMTEDGKKLVVLGTRYH WP_077021267.1
hypothetical protein [Burkholderia cenocepacia] Length: 212\nNumber of predicted TMHs: 1\nExp number of AAs in TMHs: 24.17376\nExp number, first 60 AAs: 0.92388\nTotal prob of N-in: 0.37951\noutside 1 180\nTMhelix 181 203\ninside 204 212

904 GCF_001992495.1_ASM199249v1 Burkholderia cenocepacia Proteobacteria; Betaproteobacteria; Burkholderiales; Burkholderiaceae; Burkholderia; Burkholderia cepacia complex
MSTSITRRRLLAAGVAASSLALSGCFTPKLYKNDAYSEHVSAFMMTEDGKKLVVLGTRYH WP_077021267.1

hypothetical protein [Burkholderia cenocepacia] Length: 212\nNumber of predicted TMHs: 1\nExp number of AAs in TMHs: 24.17376\nExp number, first 60 AAs: 0.92388\nTotal prob of N-in: 0.37951\noutside 1 180\nTMhelix 181 203\ninside 204 212

905 GCF_001992165.1_ASM199216v1 Burkholderia cenocepacia Proteobacteria; Betaproteobacteria; Burkholderiales; Burkholderiaceae; Burkholderia; Burkholderia cepacia complex MSDDIDNGGKGGFTRRDMMRVMAASGMMAVGGGGLLTSAQSAFAAPAPKRGGKIRVANES WP_077180776.1 ABC transporter substrate-binding protein [Burkholderia cenocepacia] Length: 527\nNumber of predicted TMHs: 2\nExp number of AAs in TMHs: 36.7312299999999\nExp number, first 60 AAs: 22.0894\nTotal prob of N-in: 0.97312\nPOSSIBLE N-term signal sequence\ninside 1 20\nTMhelix 21 43\noutside 44 502\nTMhelix 503 525\ninside 526 527

906 GCF_001992595.1_ASM199259v1 Burkholderia cenocepacia Proteobacteria; Betaproteobacteria; Burkholderiales; Burkholderiaceae; Burkholderia; Burkholderia cepacia complex MSTSITRRRLLAAGVAASSLALSGCFTPKLYKNDAYSEHVSAFMMTEDGKKLVVLGTRYH WP_077021267.1 hypothetical protein [Burkholderia cenocepacia] Length: 212\nNumber of predicted TMHs: 1\nExp number of AAs in TMHs: 24.17376\nExp number, first 60 AAs: 0.92388\nTotal prob of N-in: 0.37951\noutside 1 180\nTMhelix 181 203\ninside 204 212

907 GCF_001992565.1_ASM199256v1 Burkholderia cenocepacia Proteobacteria; Betaproteobacteria; Burkholderiales; Burkholderiaceae; Burkholderia; Burkholderia cepacia complex MKRRTFLALPAVAASGMLAGCSSADLGPVTHRLMKDPEYRETLSAFLISADGKKLVVIGKWP_034201760.1 MULTISPECIES: hypothetical protein [Burkholderia cepacia complex] Length: 212\nNumber of predicted TMHs: 1\nExp number of AAs in TMHs: 28.11606\nExp number, first 60 AAs: 6.33522\nTotal prob of N-in: 0.49088\noutside 1 187\nTMhelix 188 210\ninside 211 212

908 GCF_001992565.1_ASM199256v1 Burkholderia cenocepacia Proteobacteria; Betaproteobacteria; Burkholderiales; Burkholderiaceae; Burkholderia; Burkholderia cepacia complex MSDDIDNGGKGGFTRRDMMRVMAASGMMAVGGGGLLTSAQSAFAAPAPKRGGKIRVANES WP_006482802.1 MULTISPECIES: ABC transporter substrate-binding protein [Burkholderia cepacia complex] Length: 527\nNumber of predicted TMHs: 2\nExp number of AAs in TMHs: 36.7303599999999\nExp number, first 60 AAs: 22.08937\nTotal prob of N-in: 0.97312\nPOSSIBLE N-term signal sequence\ninside 1 20\nTMhelix 21 43\noutside 44 502\nTMhelix 503 525\ninside 526 527

909 GCF_001992595.1_ASM199259v1 Burkholderia cenocepacia Proteobacteria; Betaproteobacteria; Burkholderiales; Burkholderiaceae; Burkholderia; Burkholderia cepacia complex MSDDIDNGGKGGFTRRDMMRVMAASGMMAVGGGGLLTSAQSAFAAPAPKRGGKIRVANES WP_006482802.1 MULTISPECIES: ABC transporter substrate-binding protein [Burkholderia cepacia complex] Length: 527\nNumber of predicted TMHs: 2\nExp number of AAs in TMHs: 36.7303599999999\nExp number, first 60 AAs: 22.08937\nTotal prob of N-in: 0.97312\nPOSSIBLE N-term signal sequence\ninside 1 20\nTMhelix 21 43\noutside 44 502\nTMhelix 503 525\ninside 526 527

910 GCF_001992435.1_ASM199243v1 Burkholderia cenocepacia Proteobacteria; Betaproteobacteria; Burkholderiales; Burkholderiaceae; Burkholderia; Burkholderia cepacia complex MSTSITRRRLLAAGVAASSLALSGCFTPKLYKNDAYSEHVSAFMMTEDGKKLVVLGTRYH WP_077021267.1 hypothetical protein [Burkholderia cenocepacia] Length: 212\nNumber of predicted TMHs: 1\nExp number of AAs in TMHs: 24.17376\nExp number, first 60 AAs: 0.92388\nTotal prob of N-in: 0.37951\noutside 1 180\nTMhelix 181 203\ninside 204 212

911 GCF_001992135.1_ASM199213v1 Burkholderia cenocepacia Proteobacteria; Betaproteobacteria; Burkholderiales; Burkholderiaceae; Burkholderia; Burkholderia cepacia complex MSDDIDNGGKGGFTRRDMMRVMAASGMMAVGGGGLLTSAQSAFAAPAPKRGGKIRVANES WP_077187164.1 ABC transporter substrate-binding protein [Burkholderia cenocepacia] Length: 527\nNumber of predicted TMHs: 2\nExp number of AAs in TMHs: 36.7300399999999\nExp number, first 60 AAs: 22.0894\nTotal prob of N-in: 0.97312\nPOSSIBLE N-term signal sequence\ninside 1 20\nTMhelix 21 43\noutside 44 502\nTMhelix 503 525\ninside 526 527

912 GCF_001992135.1_ASM199213v1 Burkholderia cenocepacia Proteobacteria; Betaproteobacteria; Burkholderiales; Burkholderiaceae; Burkholderia; Burkholderia cepacia complex MSTSITRRRLLAAGVAASSLALSGCFTPKLYKNDAYSEHVSAFMMTEDGKKLVVLGTRYH WP_077021267.1 hypothetical protein [Burkholderia cenocepacia] Length: 212\nNumber of predicted TMHs: 1\nExp number of AAs in TMHs: 24.17376\nExp number, first 60 AAs: 0.92388\nTotal prob of N-in: 0.37951\noutside 1 180\nTMhelix 181 203\ninside 204 212

913 GCF_001992075.1_ASM199207v1 Burkholderia cenocepacia Proteobacteria; Betaproteobacteria; Burkholderiales; Burkholderiaceae; Burkholderia; Burkholderia cepacia complex
MSDDIDNGGKGGFTRRDMMRVMAASGMMAVGGGGLLTSAQSAFAAPAPKRGGKIRVANES WP_077180776.1 ABC transporter substrate-binding protein [Burkholderia cenocepacia] Length: 527\nNumber of predicted TMHs: 2\nExp number of AAs in TMHs: 36.7312299999999\nExp number, first 60 AAs: 22.0894\nTotal prob of N-in: 0.97312\nPOSSIBLE N-term signal sequence\ninside 1 20\nTMhelix 21 43\noutside 44 502\nTMhelix 503 525\ninside 526 527

914 GCF_001992535.1_ASM199253v1 Burkholderia cenocepacia Proteobacteria; Betaproteobacteria; Burkholderiales; Burkholderiaceae; Burkholderia; Burkholderia cepacia complex
MSTSITRRRLLAAGVAASSLALSGCFTPKLYKNDAYSEHVSAFMMTEDGKKLVVLGTRYH WP_077021267.1
hypothetical protein [Burkholderia cenocepacia] Length: 212\nNumber of predicted TMHs: 1\nExp number of AAs in TMHs: 24.17376\nExp number, first 60 AAs: 0.92388\nTotal prob of N-in: 0.37951\noutside 1 180\nTMhelix 181 203\ninside 204 212

915 GCF_001992535.1_ASM199253v1 Burkholderia cenocepacia Proteobacteria; Betaproteobacteria; Burkholderiales; Burkholderiaceae; Burkholderia; Burkholderia cepacia complex
MKRRTFLALPAVAASGMLAGCSSADLGPVTHRLMKDPEYRETLSAFLISADGKKLVVIGKWP_006483583.1 MULTISPECIES: hypothetical protein [Burkholderia cepacia complex] Length: 212\nNumber of predicted TMHs: 1\nExp number of AAs in TMHs: 28.03134\nExp number, first 60 AAs: 6.30523\nTotal prob of N-in: 0.48947\noutside 1 187\nTMhelix 188 210\ninside 211 212

916 GCF_001992535.1_ASM199253v1 Burkholderia cenocepacia Proteobacteria; Betaproteobacteria; Burkholderiales; Burkholderiaceae; Burkholderia; Burkholderia cepacia complex
MSDDIDNGGKGGFTRRDMMRVMAASGMMAVGGGGLLTSAQSAFAAPAPKRGGKIRVANES WP_006482802.1
MULTISPECIES: ABC transporter substrate-binding protein [Burkholderia cepacia complex] Length: 527\nNumber of predicted TMHs: 2\nExp number of AAs in TMHs: 36.7303599999999\nExp number, first 60 AAs: 22.08937\nTotal prob of N-in: 0.97312\nPOSSIBLE N-term signal sequence\ninside 1 20\nTMhelix 21 43\noutside 44 502\nTMhelix 503 525\ninside 526 527

917 GCF_001992385.1_ASM199238v1 Burkholderia cenocepacia Proteobacteria; Betaproteobacteria; Burkholderiales; Burkholderiaceae; Burkholderia; Burkholderia cepacia complex
MKRRTFLALPAVAASGMLAGCSSADLGPVTHRLMKDPEYRETLSAFLISADGKKLVVIGKWP_006483583.1 MULTISPECIES: hypothetical protein [Burkholderia cepacia complex] Length: 212\nNumber of predicted TMHs: 1\nExp number of AAs in TMHs: 28.03134\nExp number, first 60 AAs: 6.30523\nTotal prob of N-in: 0.48947\noutside 1 187\nTMhelix 188 210\ninside 211 212

918 GCF_001992435.1_ASM199243v1 Burkholderia cenocepacia Proteobacteria; Betaproteobacteria; Burkholderiales; Burkholderiaceae; Burkholderia; Burkholderia cepacia complex
MKRRTFLALPAVAASGMLAGCSSADLGPVTHRLMKDPEYRETLSAFLISADGKKLVVIGKWP_006483583.1 MULTISPECIES: hypothetical protein [Burkholderia cepacia complex] Length: 212\nNumber of predicted TMHs: 1\nExp number of AAs in TMHs: 28.03134\nExp number, first 60 AAs: 6.30523\nTotal prob of N-in: 0.48947\noutside 1 187\nTMhelix 188 210\ninside 211 212

919 GCF_001992495.1_ASM199249v1 Burkholderia cenocepacia Proteobacteria; Betaproteobacteria; Burkholderiales; Burkholderiaceae; Burkholderia; Burkholderia cepacia complex
MSDDIDNGGKGGFTRRDMMRVMAASGMMAVGGGGLLTSAQSAFAAPAPKRGGKIRVANES WP_006482802.1
MULTISPECIES: ABC transporter substrate-binding protein [Burkholderia cepacia complex] Length: 527\nNumber of predicted TMHs: 2\nExp number of AAs in TMHs: 36.7303599999999\nExp number, first 60 AAs: 22.08937\nTotal prob of N-in: 0.97312\nPOSSIBLE N-term signal sequence\ninside 1 20\nTMhelix 21 43\noutside 44 502\nTMhelix 503 525\ninside 526 527

920 GCF_001992035.1_ASM199203v1 Burkholderia cenocepacia Proteobacteria; Betaproteobacteria; Burkholderiales; Burkholderiaceae; Burkholderia; Burkholderia cepacia complex
MSTSITRRRLLAAGVAASSLALSGCFTPKLYKNDAYSEHVSAFMMTEDGKKLVVLGTRYH WP_077021267.1
hypothetical protein [Burkholderia cenocepacia] Length: 212\nNumber of predicted TMHs: 1\nExp number of AAs in TMHs: 24.17376\nExp number, first 60 AAs: 0.92388\nTotal prob of N-in: 0.37951\noutside 1 180\nTMhelix 181 203\ninside 204 212

921 GCF_001992385.1_ASM199238v1 Burkholderia cenocepacia Proteobacteria; Betaproteobacteria; Burkholderiales; Burkholderiaceae; Burkholderia; Burkholderia cepacia complex
MSDDIDNGGKGGFTRRDMMRVMAASGMMAVGGGGLLTSAQSAFAAPAPKRGGKIRVANES WP_006482802.1

MULTISPECIES: ABC transporter substrate-binding protein [Burkholderia cepacia complex] Length: 527\nNumber of predicted TMHs: 2\nExp number of AAs in TMHs: 36.7303599999999\nExp number, first 60 AAs: 22.08937\nTotal prob of N-in: 0.97312\nPOSSIBLE N-term signal sequence\ninside 1 20\nTMhelix 21 43\noutside 44 502\nTMhelix 503 525\ninside 526 527

922 GCF_001992345.1_ASM199234v1 Burkholderia cenocepacia Proteobacteria; Betaproteobacteria; Burkholderiales; Burkholderiaceae; Burkholderia; Burkholderia cepacia complex MKRRTFLALPAVAASGMLAGCSSADLGPVTHRLMKDPEYRETLSAFLISADGKKLVVIGKWP_006483583.1 MULTISPECIES: hypothetical protein [Burkholderia cepacia complex] Length: 212\nNumber of predicted TMHs: 1\nExp number of AAs in TMHs: 28.03134\nExp number, first 60 AAs: 6.30523\nTotal prob of N-in: 0.48947\noutside 1 187\nTMhelix 188 210\ninside 211 212

923 GCF_001992345.1_ASM199234v1 Burkholderia cenocepacia Proteobacteria; Betaproteobacteria; Burkholderiales; Burkholderiaceae; Burkholderia; Burkholderia cepacia complex MSDDIDNGGKGGFTRRDMMRVMAASGMMAVGGGGLLTSAQSAFAAPAPKRGGKIRVANES WP_006482802.1 MULTISPECIES: ABC transporter substrate-binding protein [Burkholderia cepacia complex] Length: 527\nNumber of predicted TMHs: 2\nExp number of AAs in TMHs: 36.7303599999999\nExp number, first 60 AAs: 22.08937\nTotal prob of N-in: 0.97312\nPOSSIBLE N-term signal sequence\ninside 1 20\nTMhelix 21 43\noutside 44 502\nTMhelix 503 525\ninside 526 527

924 GCF_001991955.1_ASM199195v1 Burkholderia cenocepacia Proteobacteria; Betaproteobacteria; Burkholderiales; Burkholderiaceae; Burkholderia; Burkholderia cepacia complex MSDDIDNGGKGGFTRRDMMRVMAASGMMAVGGGGLLTSAQSAFAAPAPKRGGKIRVANES WP_077180776.1 ABC transporter substrate-binding protein [Burkholderia cenocepacia] Length: 527\nNumber of predicted TMHs: 2\nExp number of AAs in TMHs: 36.7312299999999\nExp number, first 60 AAs: 22.0894\nTotal prob of N-in: 0.97312\nPOSSIBLE N-term signal sequence\ninside 1 20\nTMhelix 21 43\noutside 44 502\nTMhelix 503 525\ninside 526 527

925 GCF_001992315.1_ASM199231v1 Burkholderia cenocepacia Proteobacteria; Betaproteobacteria; Burkholderiales; Burkholderiaceae; Burkholderia; Burkholderia cepacia complex MSTSITRRRLLAAGVAASSLALSGCFTPKLYKNDAYSEHVSAFMMTEDGKKLVVLGTRYH WP_077021267.1 hypothetical protein [Burkholderia cenocepacia] Length: 212\nNumber of predicted TMHs: 1\nExp number of AAs in TMHs: 24.17376\nExp number, first 60 AAs: 0.92388\nTotal prob of N-in: 0.37951\noutside 1 180\nTMhelix 181 203\ninside 204 212

926 GCF_001992315.1_ASM199231v1 Burkholderia cenocepacia Proteobacteria; Betaproteobacteria; Burkholderiales; Burkholderiaceae; Burkholderia; Burkholderia cepacia complex METLLTRIRASFADASPRRQLISVAVLVALVASLIAAGYFALRPRYQVLFRDLKPQDAAT WP_077020516.1 flagellar M-ring protein FliF [Burkholderia cenocepacia] Length: 491\nNumber of predicted TMHs: 2\nExp number of AAs in TMHs: 41.34236\nExp number, first 60 AAs: 22.1383\nTotal prob of N-in: 0.99080\nPOSSIBLE N-term signal sequence\ninside 1 20\nTMhelix 21 43\noutside 44 427\nTMhelix 428 450\ninside 451 491

927 GCF_001992315.1_ASM199231v1 Burkholderia cenocepacia Proteobacteria; Betaproteobacteria; Burkholderiales; Burkholderiaceae; Burkholderia; Burkholderia cepacia complex MKRRTFLALPAVAASGMLAGCSSADLGPVTHRLMKDPEYRETLSAFLISADGKKLVVIGKWP_006483583.1 MULTISPECIES: hypothetical protein [Burkholderia cepacia complex] Length: 212\nNumber of predicted TMHs: 1\nExp number of AAs in TMHs: 28.03134\nExp number, first 60 AAs: 6.30523\nTotal prob of N-in: 0.48947\noutside 1 187\nTMhelix 188 210\ninside 211 212

928 GCF_001992315.1_ASM199231v1 Burkholderia cenocepacia Proteobacteria; Betaproteobacteria; Burkholderiales; Burkholderiaceae; Burkholderia; Burkholderia cepacia complex MSDDIDNGGKGGFTRRDMMRVMAASGMMAVGGGGLLTSAQSAFAAPAPKRGGKIRVANES WP_006482802.1 MULTISPECIES: ABC transporter substrate-binding protein [Burkholderia cepacia complex] Length: 527\nNumber of predicted TMHs: 2\nExp number of AAs in TMHs: 36.7303599999999\nExp number, first 60 AAs: 22.08937\nTotal prob of N-in: 0.97312\nPOSSIBLE N-term signal sequence\ninside 1 20\nTMhelix 21 43\noutside 44 502\nTMhelix 503 525\ninside 526 527

929 GCF_001991995.1_ASM199199v1 Burkholderia cenocepacia Proteobacteria; Betaproteobacteria; Burkholderiales; Burkholderiaceae; Burkholderia; Burkholderia cepacia complex MSDDIDNGGKGGFTRRDMMRVMAASGMMAVGGGGLLTSAQSAFAAPAPKRGGKIRVANES WP_077180776.1 ABC transporter substrate-binding protein [Burkholderia cenocepacia] Length: 527\nNumber of predicted TMHs: 2\nExp

number of AAs in TMHs: 36.7312299999999\nExp number, first 60 AAs: 22.0894\nTotal prob of N-in: 0.97312\nPOSSIBLE N-term signal sequence\ninside 1 20\nTMhelix 21 43\noutside 44 502\nTMhelix 503 525\ninside 526 527

930 GCF_001992205.1_ASM199220v1 Burkholderia cenocepacia Proteobacteria; Betaproteobacteria; Burkholderiales; Burkholderiaceae; Burkholderia; Burkholderia cepacia complex METLLTRIRASFADASPRRQLISVAVLVALVASLIAAGYFALRPRYQVLFRDLKPQDAAT WP_077020516.1 flagellar M-ring protein FliF [Burkholderia cenocepacia] Length: 491\nNumber of predicted TMHs: 2\nExp number of AAs in TMHs: 41.34236\nExp number, first 60 AAs: 22.1383\nTotal prob of N-in: 0.99080\nPOSSIBLE N-term signal sequence\ninside 1 20\nTMhelix 21 43\noutside 44 427\nTMhelix 428 450\ninside 451 491

931 GCF_001992255.1_ASM199225v1 Burkholderia cenocepacia Proteobacteria; Betaproteobacteria; Burkholderiales; Burkholderiaceae; Burkholderia; Burkholderia cepacia complex MKRRTFLALPAVAASGMLAGCSSADLGPVTHRLMKDPEYRETLSAFLISADGKKLVVIGKWP_034201760.1 MULTISPECIES: hypothetical protein [Burkholderia cepacia complex] Length: 212\nNumber of predicted TMHs: 1\nExp number of AAs in TMHs: 28.11606\nExp number, first 60 AAs: 6.33522\nTotal prob of N-in: 0.49088\noutside 1 187\nTMhelix 188 210\ninside 211 212

932 GCF_001991815.1_ASM199181v1 Burkholderia cenocepacia Proteobacteria; Betaproteobacteria; Burkholderiales; Burkholderiaceae; Burkholderia; Burkholderia cepacia complex MSTSITRRRLLAAGVAASSLALSGCFTPKLYKNDAYSEHVSAFMMTEDGKKLVVLGTRYH WP_077021267.1 hypothetical protein [Burkholderia cenocepacia] Length: 212\nNumber of predicted TMHs: 1\nExp number of AAs in TMHs: 24.17376\nExp number, first 60 AAs: 0.92388\nTotal prob of N-in: 0.37951\noutside 1 180\nTMhelix 181 203\ninside 204 212

933 GCF_001992165.1_ASM199216v1 Burkholderia cenocepacia Proteobacteria; Betaproteobacteria; Burkholderiales; Burkholderiaceae; Burkholderia; Burkholderia cepacia complex MKRRTFLALPAVAASGMLAGCSSADLGPVTHRLMKDPEYRETLSAFLISADGKKLVVIGKWP_006483583.1 MULTISPECIES: hypothetical protein [Burkholderia cepacia complex] Length: 212\nNumber of predicted TMHs: 1\nExp number of AAs in TMHs: 28.03134\nExp number, first 60 AAs: 6.30523\nTotal prob of N-in: 0.48947\noutside 1 187\nTMhelix 188 210\ninside 211 212

934 GCF_001992255.1_ASM199225v1 Burkholderia cenocepacia Proteobacteria; Betaproteobacteria; Burkholderiales; Burkholderiaceae; Burkholderia; Burkholderia cepacia complex MSDDIDNGGKGGFTRRDMMRVMAASGMMAVGGGGLLTSAQSAFAAPAPKRGGKIRVANES WP_006482802.1 MULTISPECIES: ABC transporter substrate-binding protein [Burkholderia cepacia complex] Length: 527\nNumber of predicted TMHs: 2\nExp number of AAs in TMHs: 36.7303599999999\nExp number, first 60 AAs: 22.08937\nTotal prob of N-in: 0.97312\nPOSSIBLE N-term signal sequence\ninside 1 20\nTMhelix 21 43\noutside 44 502\nTMhelix 503 525\ninside 526 527

935 GCF_001991815.1_ASM199181v1 Burkholderia cenocepacia Proteobacteria; Betaproteobacteria; Burkholderiales; Burkholderiaceae; Burkholderia; Burkholderia cepacia complex METLLTRIRASFADASPRRQLISVAVLVALVASLIAAGYFALRPRYQVLFRDLKPQDAAT WP_077020516.1 flagellar M-ring protein FliF [Burkholderia cenocepacia] Length: 491\nNumber of predicted TMHs: 2\nExp number of AAs in TMHs: 41.34236\nExp number, first 60 AAs: 22.1383\nTotal prob of N-in: 0.99080\nPOSSIBLE N-term signal sequence\ninside 1 20\nTMhelix 21 43\noutside 44 427\nTMhelix 428 450\ninside 451 491

936 GCF_001991795.1_ASM199179v1 Burkholderia cenocepacia Proteobacteria; Betaproteobacteria; Burkholderiales; Burkholderiaceae; Burkholderia; Burkholderia cepacia complex MSTSITRRRLLAAGVAASSLALSGCFTPKLYKNDAYSEHVSAFMMTEDGKKLVVLGTRYH WP_077021267.1 hypothetical protein [Burkholderia cenocepacia] Length: 212\nNumber of predicted TMHs: 1\nExp number of AAs in TMHs: 24.17376\nExp number, first 60 AAs: 0.92388\nTotal prob of N-in: 0.37951\noutside 1 180\nTMhelix 181 203\ninside 204 212

937 GCF_001991795.1_ASM199179v1 Burkholderia cenocepacia Proteobacteria; Betaproteobacteria; Burkholderiales; Burkholderiaceae; Burkholderia; Burkholderia cepacia complex METLLTRIRASFADASPRRQLISVAVLVALVASLIAAGYFALRPRYQVLFRDLKPQDAAT WP_077020516.1 flagellar M-ring protein FliF [Burkholderia cenocepacia] Length: 491\nNumber of predicted TMHs: 2\nExp number of AAs in TMHs: 41.34236\nExp number, first 60 AAs: 22.1383\nTotal prob of N-in: 0.99080\nPOSSIBLE N-term signal sequence\ninside 1 20\nTMhelix 21 43\noutside 44 427\nTMhelix 428 450\ninside 451 491

938 GCF_001993755.1_ASM199375v1 Burkholderia cenocepacia Proteobacteria; Betaproteobacteria; Burkholderiales; Burkholderiaceae; Burkholderia; Burkholderia cepacia complex
MKRRTFLALPAVAASGMLAGCSSADLGPVTHRLMKDPEYRETLSAFLISADGKKLVVIGKWP_006483583.1 MULTISPECIES: hypothetical protein [Burkholderia cepacia complex] Length: 212\nNumber of predicted TMHs: 1\nExp number of AAs in TMHs: 28.03134\nExp number, first 60 AAs: 6.30523\nTotal prob of N-in: 0.48947\noutside 1 187\nTMhelix 188 210\ninside 211 212

939 GCF_001992895.1_ASM199289v1 Burkholderia cenocepacia Proteobacteria; Betaproteobacteria; Burkholderiales; Burkholderiaceae; Burkholderia; Burkholderia cepacia complex
MSDDIDNGGKGGFTRRDMMRVMAASGMMAVGGGGLLTSAQSAFAAPAPKRGGKIRVANES WP_077199690.1 ABC transporter substrate-binding protein [Burkholderia cenocepacia] Length: 527\nNumber of predicted TMHs: 2\nExp number of AAs in TMHs: 36.77039\nExp number, first 60 AAs: 22.08962\nTotal prob of N-in: 0.97295\nPOSSIBLE N-term signal sequence\ninside 1 20\nTMhelix 21 43\noutside 44 502\nTMhelix 503 525\ninside 526 527

940 GCF_001993235.1_ASM199323v1 Burkholderia cenocepacia Proteobacteria; Betaproteobacteria; Burkholderiales; Burkholderiaceae; Burkholderia; Burkholderia cepacia complex
MSDDIDNGGKGGFTRRDMMRVMAASGMMAVGGGGLLTSAQSAFAAPAPKRGGKIRVANES WP_077187164.1 ABC transporter substrate-binding protein [Burkholderia cenocepacia] Length: 527\nNumber of predicted TMHs: 2\nExp number of AAs in TMHs: 36.7300399999999\nExp number, first 60 AAs: 22.0894\nTotal prob of N-in: 0.97312\nPOSSIBLE N-term signal sequence\ninside 1 20\nTMhelix 21 43\noutside 44 502\nTMhelix 503 525\ninside 526 527

941 GCF_001993235.1_ASM199323v1 Burkholderia cenocepacia Proteobacteria; Betaproteobacteria; Burkholderiales; Burkholderiaceae; Burkholderia; Burkholderia cepacia complex
MSTSITRRRLLAAGVAASSLALSGCFTPKLYKNDAYSEHVSAFMMTEDGKKLVVLGTRYH WP_077021267.1
hypothetical protein [Burkholderia cenocepacia] Length: 212\nNumber of predicted TMHs: 1\nExp number of AAs in TMHs: 24.17376\nExp number, first 60 AAs: 0.92388\nTotal prob of N-in: 0.37951\noutside 1 180\nTMhelix 181 203\ninside 204 212

942 GCF_001993255.1_ASM199325v1 Burkholderia cenocepacia Proteobacteria; Betaproteobacteria; Burkholderiales; Burkholderiaceae; Burkholderia; Burkholderia cepacia complex
METLLTRIRASFADASPRRQLISVAVLVALVASLIAAGYFALRPRYQVLFRDLKPQDAAT WP_077020516.1 flagellar M-ring protein FliF [Burkholderia cenocepacia] Length: 491\nNumber of predicted TMHs: 2\nExp number of AAs in TMHs: 41.34236\nExp number, first 60 AAs: 22.1383\nTotal prob of N-in: 0.99080\nPOSSIBLE N-term signal sequence\ninside 1 20\nTMhelix 21 43\noutside 44 427\nTMhelix 428 450\ninside 451 491

943 GCF_001720525.1_ASM172052v1 Burkholderia cenocepacia Proteobacteria; Betaproteobacteria; Burkholderiales; Burkholderiaceae; Burkholderia; Burkholderia cepacia complex
MSDDIDNGGKGGFTRRDMMRVMAASGMMAVGGGGLLTSAQSAFAAPAPKRGGKIRVANES WP_006482802.1
MULTISPECIES: ABC transporter substrate-binding protein [Burkholderia cepacia complex] Length: 527\nNumber of predicted TMHs: 2\nExp number of AAs in TMHs: 36.7303599999999\nExp number, first 60 AAs: 22.08937\nTotal prob of N-in: 0.97312\nPOSSIBLE N-term signal sequence\ninside 1 20\nTMhelix 21 43\noutside 44 502\nTMhelix 503 525\ninside 526 527

944 GCF_001606135.1_ASM160613v1 Burkholderia cenocepacia Proteobacteria; Betaproteobacteria; Burkholderiales; Burkholderiaceae; Burkholderia; Burkholderia cepacia complex
MSDDIDNGGKGGFTRRDMMRVMAASGMMAVGGGGLLTSAQSAFAAPAPKRGGKIRVANES WP_006482802.1
MULTISPECIES: ABC transporter substrate-binding protein [Burkholderia cepacia complex] Length: 527\nNumber of predicted TMHs: 2\nExp number of AAs in TMHs: 36.7303599999999\nExp number, first 60 AAs: 22.08937\nTotal prob of N-in: 0.97312\nPOSSIBLE N-term signal sequence\ninside 1 20\nTMhelix 21 43\noutside 44 502\nTMhelix 503 525\ninside 526 527

945 GCF_001056705.1_ASM105670v1 Burkholderia cenocepacia Proteobacteria; Betaproteobacteria; Burkholderiales; Burkholderiaceae; Burkholderia; Burkholderia cepacia complex
MKRRTFLALPAVAASGMLAGCSSADLGPVTHRLMKDPEYQETLSAFLISADGKKLVVIGK WP_048986526.1
hypothetical protein [Burkholderia cenocepacia] Length: 212\nNumber of predicted TMHs: 1\nExp number of AAs in TMHs: 28.09653\nExp number, first 60 AAs: 6.31597\nTotal prob of N-in: 0.49053\noutside 1 187\nTMhelix 188 210\ninside 211 212

946 GCF_001993235.1_ASM199323v1 Burkholderia cenocepacia Proteobacteria; Betaproteobacteria; Burkholderiales; Burkholderiaceae; Burkholderia; Burkholderia cepacia complex
METLLTRIRASFADASPRRQLISVAVLVALVASLIAAGYFALRPRYQVLFRDLKPQDAAT WP_077020516.1 flagellar M-

ring protein FliF [Burkholderia cenocepacia] Length: 491\nNumber of predicted TMHs: 2\nExp number of AAs in TMHs: 41.34236\nExp number, first 60 AAs: 22.1383\nTotal prob of N-in: 0.99080\nPOSSIBLE N-term signal sequence\ninside 1 20\nTMhelix 21 43\noutside 44 427\nTMhelix 428 450\ninside 451 491

947 GCF_001993255.1_ASM199325v1 Burkholderia cenocepacia Proteobacteria; Betaproteobacteria; Burkholderiales; Burkholderiaceae; Burkholderia; Burkholderia cepacia complex MKRRTFLALPAVAASGMLAGCSSADLGPVTHRLMKDPEYRETLSAFLISADGKKLVVIGKWP_006483583.1 MULTISPECIES: hypothetical protein [Burkholderia cepacia complex] Length: 212\nNumber of predicted TMHs: 1\nExp number of AAs in TMHs: 28.03134\nExp number, first 60 AAs: 6.30523\nTotal prob of N-in: 0.48947\noutside 1 187\nTMhelix 188 210\ninside 211 212

948 GCF_001056375.1_ASM105637v1 Burkholderia cenocepacia Proteobacteria; Betaproteobacteria; Burkholderiales; Burkholderiaceae; Burkholderia; Burkholderia cepacia complex MKRRTFLALPAVAASGMLAGCSSADLGPVTHRLMKDPEYQETLSAFLISADGKKLVVIGK WP_048986526.1 hypothetical protein [Burkholderia cenocepacia] Length: 212\nNumber of predicted TMHs: 1\nExp number of AAs in TMHs: 28.09653\nExp number, first 60 AAs: 6.31597\nTotal prob of N-in: 0.49053\noutside 1 187\nTMhelix 188 210\ninside 211 212

949 GCF_001993235.1_ASM199323v1 Burkholderia cenocepacia Proteobacteria; Betaproteobacteria; Burkholderiales; Burkholderiaceae; Burkholderia; Burkholderia cepacia complex MKRRTFLALPAVAASGMLAGCSSADLGPVTHRLMKDPEYRETLSAFLISADGKKLVVIGKWP_006483583.1 MULTISPECIES: hypothetical protein [Burkholderia cepacia complex] Length: 212\nNumber of predicted TMHs: 1\nExp number of AAs in TMHs: 28.03134\nExp number, first 60 AAs: 6.30523\nTotal prob of N-in: 0.48947\noutside 1 187\nTMhelix 188 210\ninside 211 212

950 GCF_001992555.1_ASM199255v1 Burkholderia cenocepacia Proteobacteria; Betaproteobacteria; Burkholderiales; Burkholderiaceae; Burkholderia; Burkholderia cepacia complex MSTSITRRRLLAAGVAASSLALSGCFTPKLYKNDAYSEHVSAFMMTEDGKKLVVLGTRYH WP_077021267.1 hypothetical protein [Burkholderia cenocepacia] Length: 212\nNumber of predicted TMHs: 1\nExp number of AAs in TMHs: 24.17376\nExp number, first 60 AAs: 0.92388\nTotal prob of N-in: 0.37951\noutside 1 180\nTMhelix 181 203\ninside 204 212

951 GCF_001992505.1_ASM199250v1 Burkholderia cenocepacia Proteobacteria; Betaproteobacteria; Burkholderiales; Burkholderiaceae; Burkholderia; Burkholderia cepacia complex MSTSITRRRLLAAGVAASSLALSGCFTPKLYKNDAYSEHVSAFMMTEDGKKLVVLGTRYH WP_077021267.1 hypothetical protein [Burkholderia cenocepacia] Length: 212\nNumber of predicted TMHs: 1\nExp number of AAs in TMHs: 24.17376\nExp number, first 60 AAs: 0.92388\nTotal prob of N-in: 0.37951\noutside 1 180\nTMhelix 181 203\ninside 204 212

952 GCF_001992555.1_ASM199255v1 Burkholderia cenocepacia Proteobacteria; Betaproteobacteria; Burkholderiales; Burkholderiaceae; Burkholderia; Burkholderia cepacia complex MKRRTFLALPAVAASGMLAGCSSADLGPVTHRLMKDPEYRETLSAFLISADGKKLVVIGKWP_006483583.1 MULTISPECIES: hypothetical protein [Burkholderia cepacia complex] Length: 212\nNumber of predicted TMHs: 1\nExp number of AAs in TMHs: 28.03134\nExp number, first 60 AAs: 6.30523\nTotal prob of N-in: 0.48947\noutside 1 187\nTMhelix 188 210\ninside 211 212

953 GCF_001992555.1_ASM199255v1 Burkholderia cenocepacia Proteobacteria; Betaproteobacteria; Burkholderiales; Burkholderiaceae; Burkholderia; Burkholderia cepacia complex MSDDIDNGGKGGFTRRDMMRVMAASGMMAVGGGGLLTSAQSAFAAPAPKRGGKIRVANES WP_006482802.1 MULTISPECIES: ABC transporter substrate-binding protein [Burkholderia cepacia complex] Length: 527\nNumber of predicted TMHs: 2\nExp number of AAs in TMHs: 36.7303599999999\nExp number, first 60 AAs: 22.08937\nTotal prob of N-in: 0.97312\nPOSSIBLE N-term signal sequence\ninside 1 20\nTMhelix 21 43\noutside 44 502\nTMhelix 503 525\ninside 526 527

954 GCF_001992505.1_ASM199250v1 Burkholderia cenocepacia Proteobacteria; Betaproteobacteria; Burkholderiales; Burkholderiaceae; Burkholderia; Burkholderia cepacia complex MKRRTFLALPAVAASGMLAGCSSADLGPVTHRLMKDPEYRETLSAFLISADGKKLVVIGKWP_006483583.1 MULTISPECIES: hypothetical protein [Burkholderia cepacia complex] Length: 212\nNumber of predicted TMHs: 1\nExp number of AAs in TMHs: 28.03134\nExp number, first 60 AAs: 6.30523\nTotal prob of N-in: 0.48947\noutside 1 187\nTMhelix 188 210\ninside 211 212

955 GCF_001992505.1_ASM199250v1 Burkholderia cenocepacia Proteobacteria; Betaproteobacteria; Burkholderiales; Burkholderiaceae; Burkholderia; Burkholderia cepacia complex
MSDDIDNGGKGGFTRRDMMRVMAASGMMAVGGGGLLTSAQSAFAAPAPKRGGKIRVANES WP_006482802.1 MULTISPECIES: ABC transporter substrate-binding protein [Burkholderia cepacia complex] Length: 527\nNumber of predicted TMHs: 2\nExp number of AAs in TMHs: 36.7303599999999\nExp number, first 60 AAs: 22.08937\nTotal prob of N-in: 0.97312\nPOSSIBLE N-term signal sequence\ninside 1 20\nTMhelix 21 43\noutside 44 502\nTMhelix 503 525\ninside 526 527

956 GCF_001992115.1_ASM199211v1 Burkholderia cenocepacia Proteobacteria; Betaproteobacteria; Burkholderiales; Burkholderiaceae; Burkholderia; Burkholderia cepacia complex
MSDDIDNGGKGGFTRRDMMRVMAASGMMAVGGGGLLTSAQSAFAAPAPKRGGKIRVANES WP_077180776.1 ABC transporter substrate-binding protein [Burkholderia cenocepacia] Length: 527\nNumber of predicted TMHs: 2\nExp number of AAs in TMHs: 36.7312299999999\nExp number, first 60 AAs: 22.0894\nTotal prob of N-in: 0.97312\nPOSSIBLE N-term signal sequence\ninside 1 20\nTMhelix 21 43\noutside 44 502\nTMhelix 503 525\ninside 526 527

957 GCF_000818575.1_BcenoS511.01 Burkholderia cenocepacia Proteobacteria; Betaproteobacteria; Burkholderiales; Burkholderiaceae; Burkholderia; Burkholderia cepacia complex
MSLSITRRRLLAAGVAASSVALSGCFTPKLYKNDAYSERVSAFMMTEDGKKLVVLGTRYH WP_050012404.1 hypothetical protein [Burkholderia cenocepacia] Length: 212\nNumber of predicted TMHs: 1\nExp number of AAs in TMHs: 28.40203\nExp number, first 60 AAs: 1.09227\nTotal prob of N-in: 0.42249\noutside 1 180\nTMhelix 181 203\ninside 204 212

958 GCF_001056705.1_ASM105670v1 Burkholderia cenocepacia Proteobacteria; Betaproteobacteria; Burkholderiales; Burkholderiaceae; Burkholderia; Burkholderia cepacia complex
MSDDIDNGGKGGFTRRDMMRVMAASGMMAVGGGGLLTSAQSAFAAPAPKRGGKIRVANES WP_006482802.1 MULTISPECIES: ABC transporter substrate-binding protein [Burkholderia cepacia complex] Length: 527\nNumber of predicted TMHs: 2\nExp number of AAs in TMHs: 36.7303599999999\nExp number, first 60 AAs: 22.08937\nTotal prob of N-in: 0.97312\nPOSSIBLE N-term signal sequence\ninside 1 20\nTMhelix 21 43\noutside 44 502\nTMhelix 503 525\ninside 526 527

959 GCF_000755725.1_ASM75572v1 Burkholderia cenocepacia Proteobacteria; Betaproteobacteria; Burkholderiales; Burkholderiaceae; Burkholderia; Burkholderia cepacia complex
MKRRTLLRASAIAVTAGLAGCTSMLFEDGRYDETVDRFLVSEDGKKFVVLGQQYHYIFDM WP_040127356.1 MULTISPECIES: hypothetical protein [Burkholderia cepacia complex] Length: 209\nNumber of predicted TMHs: 1\nExp number of AAs in TMHs: 28.96048\nExp number, first 60 AAs: 5.85843999999999\nTotal prob of N-in: 0.86015\ninside 1 182\nTMhelix 183 205\noutside 206 209

960 GCF_001994195.1_ASM199419v1 Burkholderia cenocepacia Proteobacteria; Betaproteobacteria; Burkholderiales; Burkholderiaceae; Burkholderia; Burkholderia cepacia complex
MSDDIDNGGKGGFTRRDMMRVMAASGMMAVGGGGLLTSAQSAFAAPAPKRGGKIRVANES WP_006482802.1 MULTISPECIES: ABC transporter substrate-binding protein [Burkholderia cepacia complex] Length: 527\nNumber of predicted TMHs: 2\nExp number of AAs in TMHs: 36.7303599999999\nExp number, first 60 AAs: 22.08937\nTotal prob of N-in: 0.97312\nPOSSIBLE N-term signal sequence\ninside 1 20\nTMhelix 21 43\noutside 44 502\nTMhelix 503 525\ninside 526 527

961 GCF_001992275.1_ASM199227v1 Burkholderia cenocepacia Proteobacteria; Betaproteobacteria; Burkholderiales; Burkholderiaceae; Burkholderia; Burkholderia cepacia complex
MKRRTFLALPAVAASGMLAGCSSADLGPVTHRLMKDPEYRETLSAFLISADGKKLVVIGKWP_034201760.1 MULTISPECIES: hypothetical protein [Burkholderia cepacia complex] Length: 212\nNumber of predicted TMHs: 1\nExp number of AAs in TMHs: 28.11606\nExp number, first 60 AAs: 6.33522\nTotal prob of N-in: 0.49088\noutside 1 187\nTMhelix 188 210\ninside 211 212

962 GCF_001056375.1_ASM105637v1 Burkholderia cenocepacia Proteobacteria; Betaproteobacteria; Burkholderiales; Burkholderiaceae; Burkholderia; Burkholderia cepacia complex
MSDDIDNGGKGGFTRRDMMRVMAASGMMAVGGGGLLTSAQSAFAAPAPKRGGKIRVANES WP_006482802.1 MULTISPECIES: ABC transporter substrate-binding protein [Burkholderia cepacia complex] Length: 527\nNumber of predicted TMHs: 2\nExp number of AAs in TMHs: 36.7303599999999\nExp number, first 60 AAs: 22.08937\nTotal prob of N-in: 0.97312\nPOSSIBLE N-term signal sequence\ninside 1 20\nTMhelix 21 43\noutside 44 502\nTMhelix 503 525\ninside 526 527

963 GCF_001055325.1_ASM105532v1 Burkholderia cenocepacia Proteobacteria; Betaproteobacteria; Burkholderiales; Burkholderiaceae; Burkholderia; Burkholderia cepacia complex
MSDDIDNGGKGGFTRRDMMRVMAASGMMAVGGGGLLTSAQSAFAAPAPKRGGKIRVANES WP_006482802.1
MULTISPECIES: ABC transporter substrate-binding protein [Burkholderia cepacia complex] Length: 527\nNumber of predicted TMHs: 2\nExp number of AAs in TMHs: 36.7303599999999\nExp number, first 60 AAs: 22.08937\nTotal prob of N-in: 0.97312\nPOSSIBLE N-term signal sequence\ninside 1 20\nTMhelix 21 43\noutside 44 502\nTMhelix 503 525\ninside 526 527

964 GCF_001994135.1_ASM199413v1 Burkholderia cenocepacia Proteobacteria; Betaproteobacteria; Burkholderiales; Burkholderiaceae; Burkholderia; Burkholderia cepacia complex
MKRRTFLALPAVAASGMLAGCSSADLGPVTHRLMKDPEYRETLSAFLISADGKKLVVIGKWP_034201760.1 MULTISPECIES: hypothetical protein [Burkholderia cepacia complex] Length: 212\nNumber of predicted TMHs: 1\nExp number of AAs in TMHs: 28.11606\nExp number, first 60 AAs: 6.33522\nTotal prob of N-in: 0.49088\noutside 1 187\nTMhelix 188 210\ninside 211 212

965 GCF_001994215.1_ASM199421v1 Burkholderia cenocepacia Proteobacteria; Betaproteobacteria; Burkholderiales; Burkholderiaceae; Burkholderia; Burkholderia cepacia complex
MSDDIDNGGKGGFTRRDMMRVMAASGMMAVGGGGLLTSAQSAFAAPAPKRGGKIRVANES WP_006482802.1
MULTISPECIES: ABC transporter substrate-binding protein [Burkholderia cepacia complex] Length: 527\nNumber of predicted TMHs: 2\nExp number of AAs in TMHs: 36.7303599999999\nExp number, first 60 AAs: 22.08937\nTotal prob of N-in: 0.97312\nPOSSIBLE N-term signal sequence\ninside 1 20\nTMhelix 21 43\noutside 44 502\nTMhelix 503 525\ninside 526 527

966 GCF_001994475.1_ASM199447v1 Burkholderia cenocepacia Proteobacteria; Betaproteobacteria; Burkholderiales; Burkholderiaceae; Burkholderia; Burkholderia cepacia complex
MKRRTFLALPAVAASGMLAGCSSADLGPVTHRLMKDPEYRETLSAFLISADGKKLVVIGKWP_034201760.1 MULTISPECIES: hypothetical protein [Burkholderia cepacia complex] Length: 212\nNumber of predicted TMHs: 1\nExp number of AAs in TMHs: 28.11606\nExp number, first 60 AAs: 6.33522\nTotal prob of N-in: 0.49088\noutside 1 187\nTMhelix 188 210\ninside 211 212

967 GCF_001994285.1_ASM199428v1 Burkholderia cenocepacia Proteobacteria; Betaproteobacteria; Burkholderiales; Burkholderiaceae; Burkholderia; Burkholderia cepacia complex
MKRRTFLALPAVAASGMLAGCSSADLGPVTHRLMKDPEYRETLSAFLISADGKKLVVIGKWP_034201760.1 MULTISPECIES: hypothetical protein [Burkholderia cepacia complex] Length: 212\nNumber of predicted TMHs: 1\nExp number of AAs in TMHs: 28.11606\nExp number, first 60 AAs: 6.33522\nTotal prob of N-in: 0.49088\noutside 1 187\nTMhelix 188 210\ninside 211 212

968 GCF_001992395.1_ASM199239v1 Burkholderia cenocepacia Proteobacteria; Betaproteobacteria; Burkholderiales; Burkholderiaceae; Burkholderia; Burkholderia cepacia complex
MSDDIDNGGKGGFTRRDMMRVMAASGMMAVGGGGLLTSAQSAFAAPAPKRGGKIRVANES WP_006482802.1
MULTISPECIES: ABC transporter substrate-binding protein [Burkholderia cepacia complex] Length: 527\nNumber of predicted TMHs: 2\nExp number of AAs in TMHs: 36.7303599999999\nExp number, first 60 AAs: 22.08937\nTotal prob of N-in: 0.97312\nPOSSIBLE N-term signal sequence\ninside 1 20\nTMhelix 21 43\noutside 44 502\nTMhelix 503 525\ninside 526 527

969 GCF_001994545.1_ASM199454v1 Burkholderia cenocepacia Proteobacteria; Betaproteobacteria; Burkholderiales; Burkholderiaceae; Burkholderia; Burkholderia cepacia complex
MSDDIDNGGKGGFTRRDMMRVMAASGMMAVGGGGLLTSAQSAFAAPAPKRGGKIRVANES WP_006482802.1
MULTISPECIES: ABC transporter substrate-binding protein [Burkholderia cepacia complex] Length: 527\nNumber of predicted TMHs: 2\nExp number of AAs in TMHs: 36.7303599999999\nExp number, first 60 AAs: 22.08937\nTotal prob of N-in: 0.97312\nPOSSIBLE N-term signal sequence\ninside 1 20\nTMhelix 21 43\noutside 44 502\nTMhelix 503 525\ninside 526 527

970 GCF_001992045.1_ASM199204v1 Burkholderia cenocepacia Proteobacteria; Betaproteobacteria; Burkholderiales; Burkholderiaceae; Burkholderia; Burkholderia cepacia complex
MSDDIDNGGKGGFTRRDMMRVMAASGMMAVGGGGLLTSAQSAFAAPAPKRGGKIRVANES WP_077180776.1 ABC transporter substrate-binding protein [Burkholderia cenocepacia] Length: 527\nNumber of predicted TMHs: 2\nExp number of AAs in TMHs: 36.7312299999999\nExp number, first 60 AAs: 22.0894\nTotal prob of N-in: 0.97312\nPOSSIBLE N-term signal sequence\ninside 1 20\nTMhelix 21 43\noutside 44 502\nTMhelix 503 525\ninside 526 527

971	GCF_001994595.1_ASM199459v1	Burkholderia cenocepacia	Proteobacteria; Betaproteobacteria; Burkholderiales; Burkholderiaceae; Burkholderia; Burkholderia cepacia complex
MSDDIDNGGKGGFTRRDMMRVMAASGMMAVGGGGLLTSAQSAFAAPAPKRGGKIRVANES	WP_006482802.1	MULTISPECIES: ABC transporter substrate-binding protein [Burkholderia cepacia complex]	Length: 527\nNumber of predicted TMHs: 2\nExp number of AAs in TMHs: 36.7303599999999\nExp number, first 60 AAs: 22.08937\nTotal prob of N-in: 0.97312\nPOSSIBLE N-term signal sequence\ninside 1 20\nTMhelix 21 43\noutside 44 502\nTMhelix 503 525\ninside 526 527

972	GCF_001992005.1_ASM199200v1	Burkholderia cenocepacia	Proteobacteria; Betaproteobacteria; Burkholderiales; Burkholderiaceae; Burkholderia; Burkholderia cepacia complex
MSDDIDNGGKGGFTRRDMMRVMAASGMMAVGGGGLLTSAQSAFAAPAPKRGGKIRVANES	WP_077180776.1	ABC transporter substrate-binding protein [Burkholderia cenocepacia]	Length: 527\nNumber of predicted TMHs: 2\nExp number of AAs in TMHs: 36.7312299999999\nExp number, first 60 AAs: 22.0894\nTotal prob of N-in: 0.97312\nPOSSIBLE N-term signal sequence\ninside 1 20\nTMhelix 21 43\noutside 44 502\nTMhelix 503 525\ninside 526 527

973	GCF_001992375.1_ASM199237v1	Burkholderia cenocepacia	Proteobacteria; Betaproteobacteria; Burkholderiales; Burkholderiaceae; Burkholderia; Burkholderia cepacia complex
MSTSITRRRLLAAGVAASSLALSGCFTPKLYKNDAYSEHVSAFMMTEDGKKLVVLGTRYH	WP_077021267.1	hypothetical protein [Burkholderia cenocepacia]	Length: 212\nNumber of predicted TMHs: 1\nExp number of AAs in TMHs: 24.17376\nExp number, first 60 AAs: 0.92388\nTotal prob of N-in: 0.37951\noutside 1 180\nTMhelix 181 203\ninside 204 212

974	GCF_001992335.1_ASM199233v1	Burkholderia cenocepacia	Proteobacteria; Betaproteobacteria; Burkholderiales; Burkholderiaceae; Burkholderia; Burkholderia cepacia complex
MSTSITRRRLLAAGVAASSLALSGCFTPKLYKNDAYSEHVSAFMMTEDGKKLVVLGTRYH	WP_077021267.1	hypothetical protein [Burkholderia cenocepacia]	Length: 212\nNumber of predicted TMHs: 1\nExp number of AAs in TMHs: 24.17376\nExp number, first 60 AAs: 0.92388\nTotal prob of N-in: 0.37951\noutside 1 180\nTMhelix 181 203\ninside 204 212

975	GCF_001993855.1_ASM199385v1	Burkholderia cenocepacia	Proteobacteria; Betaproteobacteria; Burkholderiales; Burkholderiaceae; Burkholderia; Burkholderia cepacia complex
MKRRTFLALPAVAASGMLAGCSSADLGPVTHRLMKDPEYRETLSAFLISADGKKLVVIGKWP_034201760.1	MULTISPECIES: hypothetical protein [Burkholderia cepacia complex]	Length: 212\nNumber of predicted TMHs: 1\nExp number of AAs in TMHs: 28.11606\nExp number, first 60 AAs: 6.33522\nTotal prob of N-in: 0.49088\noutside 1 187\nTMhelix 188 210\ninside 211 212

976	GCF_001994545.1_ASM199454v1	Burkholderia cenocepacia	Proteobacteria; Betaproteobacteria; Burkholderiales; Burkholderiaceae; Burkholderia; Burkholderia cepacia complex
MKRRTFLALPAVAASGMLAGCSSADLGPVTHRLMKDPEYRETLSAFLISADGKKLVVIGKWP_034201760.1	MULTISPECIES: hypothetical protein [Burkholderia cepacia complex]	Length: 212\nNumber of predicted TMHs: 1\nExp number of AAs in TMHs: 28.11606\nExp number, first 60 AAs: 6.33522\nTotal prob of N-in: 0.49088\noutside 1 187\nTMhelix 188 210\ninside 211 212

977	GCF_001994595.1_ASM199459v1	Burkholderia cenocepacia	Proteobacteria; Betaproteobacteria; Burkholderiales; Burkholderiaceae; Burkholderia; Burkholderia cepacia complex
MKRRTFLALPAVAASGMLAGCSSADLGPVTHRLMKDPEYRETLSAFLISADGKKLVVIGKWP_034201760.1	MULTISPECIES: hypothetical protein [Burkholderia cepacia complex]	Length: 212\nNumber of predicted TMHs: 1\nExp number of AAs in TMHs: 28.11606\nExp number, first 60 AAs: 6.33522\nTotal prob of N-in: 0.49088\noutside 1 187\nTMhelix 188 210\ninside 211 212

978	GCF_001992375.1_ASM199237v1	Burkholderia cenocepacia	Proteobacteria; Betaproteobacteria; Burkholderiales; Burkholderiaceae; Burkholderia; Burkholderia cepacia complex
METLLTRIRASFADASPRRQLISVAVLVALVASLIAAGYFALRPRYQVLFRDLKPQDAAT	WP_077020516.1	flagellar M-ring protein FliF [Burkholderia cenocepacia]	Length: 491\nNumber of predicted TMHs: 2\nExp number of AAs in TMHs: 41.34236\nExp number, first 60 AAs: 22.1383\nTotal prob of N-in: 0.99080\nPOSSIBLE N-term signal sequence\ninside 1 20\nTMhelix 21 43\noutside 44 427\nTMhelix 428 450\ninside 451 491

979	GCF_001992155.1_ASM199215v1	Burkholderia cenocepacia	Proteobacteria; Betaproteobacteria; Burkholderiales; Burkholderiaceae; Burkholderia; Burkholderia cepacia complex
MKRRTFLALPAVAASGMLAGCSSADLGPVTHRLMKDPEYRETLSAFLISADGKKLVVIGKWP_006483583.1	MULTISPECIES: hypothetical protein [Burkholderia cepacia complex]	Length: 212\nNumber of predicted TMHs: 1\nExp number of

AAs in TMHs: 28.03134\nExp number, first 60 AAs: 6.30523\nTotal prob of N-in: 0.48947\noutside 1 187\nTMhelix 188 210\ninside 211 212

980 GCF_001991755.1_ASM199175v1 Burkholderia cenocepacia Proteobacteria; Betaproteobacteria; Burkholderiales; Burkholderiaceae; Burkholderia; Burkholderia cepacia complex METLLTRIRASFADASPRRQLISVAVLVALVASLIAAGYFALRPRYQVLFRDLKPQDAAT WP_077020516.1 flagellar M-ring protein FliF [Burkholderia cenocepacia] Length: 491\nNumber of predicted TMHs: 2\nExp number of AAs in TMHs: 41.34236\nExp number, first 60 AAs: 22.1383\nTotal prob of N-in: 0.99080\nPOSSIBLE N-term signal sequence\ninside 1 20\nTMhelix 21 43\noutside 44 427\nTMhelix 428 450\ninside 451 491

981 GCF_001992225.1_ASM199222v1 Burkholderia cenocepacia Proteobacteria; Betaproteobacteria; Burkholderiales; Burkholderiaceae; Burkholderia; Burkholderia cepacia complex MKRRTFLALPAVAASGMLAGCSSADLGPVTHRLMKDPEYRETLSAFLISADGKKLVVIGKWP_034201760.1 MULTISPECIES: hypothetical protein [Burkholderia cepacia complex] Length: 212\nNumber of predicted TMHs: 1\nExp number of AAs in TMHs: 28.11606\nExp number, first 60 AAs: 6.33522\nTotal prob of N-in: 0.49088\noutside 1 187\nTMhelix 188 210\ninside 211 212

982 GCF_001991975.1_ASM199197v1 Burkholderia cenocepacia Proteobacteria; Betaproteobacteria; Burkholderiales; Burkholderiaceae; Burkholderia; Burkholderia cepacia complex MSDDIDNGGKGGFTRRDMMRVMAASGMMAVGGGGLLTSAQSAFAAPAPKRGGKIRVANES WP_077180776.1 ABC transporter substrate-binding protein [Burkholderia cenocepacia] Length: 527\nNumber of predicted TMHs: 2\nExp number of AAs in TMHs: 36.7312299999999\nExp number, first 60 AAs: 22.0894\nTotal prob of N-in: 0.97312\nPOSSIBLE N-term signal sequence\ninside 1 20\nTMhelix 21 43\noutside 44 502\nTMhelix 503 525\ninside 526 527

983 GCF_001992095.1_ASM199209v1 Burkholderia cenocepacia Proteobacteria; Betaproteobacteria; Burkholderiales; Burkholderiaceae; Burkholderia; Burkholderia cepacia complex MKRRTFLALPAVAASGMLAGCSSADLGPVTHRLMKDPEYRETLSAFLISADGKKLVVIGKWP_006483583.1 MULTISPECIES: hypothetical protein [Burkholderia cepacia complex] Length: 212\nNumber of predicted TMHs: 1\nExp number of AAs in TMHs: 28.03134\nExp number, first 60 AAs: 6.30523\nTotal prob of N-in: 0.48947\noutside 1 187\nTMhelix 188 210\ninside 211 212

984 GCF_002071935.1_ASM207193v1 Burkholderia cenocepacia Proteobacteria; Betaproteobacteria; Burkholderiales; Burkholderiaceae; Burkholderia; Burkholderia cepacia complex MKRRTFLALPAVAASGMLAGCSSADLGPVTHRLMKDPEYRETLSAFLISADGKKLVVIGKWP_062884514.1 hypothetical protein [Burkholderia cenocepacia] Length: 212\nNumber of predicted TMHs: 1\nExp number of AAs in TMHs: 28.03134\nExp number, first 60 AAs: 6.30523\nTotal prob of N-in: 0.48947\noutside 1 187\nTMhelix 188 210\ninside 211 212

985 GCF_002071935.1_ASM207193v1 Burkholderia cenocepacia Proteobacteria; Betaproteobacteria; Burkholderiales; Burkholderiaceae; Burkholderia; Burkholderia cepacia complex METLLTRIRASFADASPRRQLISVAVLVALVASLIAAGYFALRPRYQVLFRDLKPQDAAT WP_060214257.1 flagellar M-ring protein FliF [Burkholderia cenocepacia] Length: 491\nNumber of predicted TMHs: 2\nExp number of AAs in TMHs: 41.34235\nExp number, first 60 AAs: 22.1383\nTotal prob of N-in: 0.99080\nPOSSIBLE N-term signal sequence\ninside 1 20\nTMhelix 21 43\noutside 44 427\nTMhelix 428 450\ninside 451 491

986 GCF_002083015.1_ASM208301v1 Burkholderia cenocepacia Proteobacteria; Betaproteobacteria; Burkholderiales; Burkholderiaceae; Burkholderia; Burkholderia cepacia complex MKRRTFLALPAVAASGMLAGCSSADLGPVTHRLMKDPEYRETLSAFLISADGKKLVVIGKWP_034201760.1 MULTISPECIES: hypothetical protein [Burkholderia cepacia complex] Length: 212\nNumber of predicted TMHs: 1\nExp number of AAs in TMHs: 28.11606\nExp number, first 60 AAs: 6.33522\nTotal prob of N-in: 0.49088\noutside 1 187\nTMhelix 188 210\ninside 211 212

987 GCF_001994835.1_ASM199483v1 Burkholderia cenocepacia Proteobacteria; Betaproteobacteria; Burkholderiales; Burkholderiaceae; Burkholderia; Burkholderia cepacia complex MSDDIDNGGKGGFTRRDMMRVMAASGMMAVGGGGLLTSAQSAFAAPAPKRGGKIRVANES WP_077187164.1 ABC transporter substrate-binding protein [Burkholderia cenocepacia] Length: 527\nNumber of predicted TMHs: 2\nExp number of AAs in TMHs: 36.7300399999999\nExp number, first 60 AAs: 22.0894\nTotal prob of N-in: 0.97312\nPOSSIBLE N-term signal sequence\ninside 1 20\nTMhelix 21 43\noutside 44 502\nTMhelix 503 525\ninside 526 527

988 GCF_001994815.1_ASM199481v1 Burkholderia cenocepacia Proteobacteria; Betaproteobacteria; Burkholderiales; Burkholderiaceae; Burkholderia; Burkholderia cepacia complex

MKRRTFLALPAVAASGMLAGCSSADLGPVTHRLMKDPEYRETLSAFLISADGKKLVVIGKWP_034201760.1 MULTISPECIES: hypothetical protein [Burkholderia cepacia complex] Length: 212\nNumber of predicted TMHs: 1\nExp number of AAs in TMHs: 28.11606\nExp number, first 60 AAs: 6.33522\nTotal prob of N-in: 0.49088\noutside 1 187\nTMhelix 188 210\ninside 211 212

989 GCF_001994735.1_ASM199473v1 Burkholderia cenocepacia Proteobacteria; Betaproteobacteria; Burkholderiales; Burkholderiaceae; Burkholderia; Burkholderia cepacia complex
MSTSITRRRLLAAGVAASSLALSGCFTPKLYKNDAYSEHVSAFMMTEDGKKLVVLGTRYH WP_077021267.1
hypothetical protein [Burkholderia cenocepacia] Length: 212\nNumber of predicted TMHs: 1\nExp number of AAs in TMHs: 24.17376\nExp number, first 60 AAs: 0.92388\nTotal prob of N-in: 0.37951\noutside 1 180\nTMhelix 181 203\ninside 204 212

990 GCF_001999925.1_ASM199992v1 Burkholderia cenocepacia Proteobacteria; Betaproteobacteria; Burkholderiales; Burkholderiaceae; Burkholderia; Burkholderia cepacia complex
MKRRTFLALPAVAASGMLAGCSSADLGPVTHRLMKDPEYRETLSAFLISADGKKLVVIGKWP_034201760.1 MULTISPECIES: hypothetical protein [Burkholderia cepacia complex] Length: 212\nNumber of predicted TMHs: 1\nExp number of AAs in TMHs: 28.11606\nExp number, first 60 AAs: 6.33522\nTotal prob of N-in: 0.49088\noutside 1 187\nTMhelix 188 210\ninside 211 212

991 GCF_001994615.1_ASM199461v1 Burkholderia cenocepacia Proteobacteria; Betaproteobacteria; Burkholderiales; Burkholderiaceae; Burkholderia; Burkholderia cepacia complex
MSTSITRRRLLAAGVAASSLALSGCFTPKLYKNDAYSEHVSAFMMTEDGKKLVVLGTRYH WP_077021267.1
hypothetical protein [Burkholderia cenocepacia] Length: 212\nNumber of predicted TMHs: 1\nExp number of AAs in TMHs: 24.17376\nExp number, first 60 AAs: 0.92388\nTotal prob of N-in: 0.37951\noutside 1 180\nTMhelix 181 203\ninside 204 212

992 GCF_001994915.1_ASM199491v1 Burkholderia cenocepacia Proteobacteria; Betaproteobacteria; Burkholderiales; Burkholderiaceae; Burkholderia; Burkholderia cepacia complex
MKRRTFLALPAVAASGMLAGCSSADLGPVTHRLMKDPEYRETLSAFLISADGKKLVVIGKWP_006483583.1 MULTISPECIES: hypothetical protein [Burkholderia cepacia complex] Length: 212\nNumber of predicted TMHs: 1\nExp number of AAs in TMHs: 28.03134\nExp number, first 60 AAs: 6.30523\nTotal prob of N-in: 0.48947\noutside 1 187\nTMhelix 188 210\ninside 211 212

993 GCF_001999925.1_ASM199992v1 Burkholderia cenocepacia Proteobacteria; Betaproteobacteria; Burkholderiales; Burkholderiaceae; Burkholderia; Burkholderia cepacia complex
MSDDIDNGGKGGFTRRDMMRVMAASGMMAVGGGGLLTSAQSAFAAPAPKRGGKIRVANES WP_006482802.1
MULTISPECIES: ABC transporter substrate-binding protein [Burkholderia cepacia complex] Length: 527\nNumber of predicted TMHs: 2\nExp number of AAs in TMHs: 36.7303599999999\nExp number, first 60 AAs: 22.08937\nTotal prob of N-in: 0.97312\nPOSSIBLE N-term signal sequence\ninside 1 20\nTMhelix 21 43\noutside 44 502\nTMhelix 503 525\ninside 526 527

994 GCF_001994915.1_ASM199491v1 Burkholderia cenocepacia Proteobacteria; Betaproteobacteria; Burkholderiales; Burkholderiaceae; Burkholderia; Burkholderia cepacia complex
MSDDIDNGGKGGFTRRDMMRVMAASGMMAVGGGGLLTSAQSAFAAPAPKRGGKIRVANES WP_006482802.1
MULTISPECIES: ABC transporter substrate-binding protein [Burkholderia cepacia complex] Length: 527\nNumber of predicted TMHs: 2\nExp number of AAs in TMHs: 36.7303599999999\nExp number, first 60 AAs: 22.08937\nTotal prob of N-in: 0.97312\nPOSSIBLE N-term signal sequence\ninside 1 20\nTMhelix 21 43\noutside 44 502\nTMhelix 503 525\ninside 526 527

995 GCF_001994505.1_ASM199450v1 Burkholderia cenocepacia Proteobacteria; Betaproteobacteria; Burkholderiales; Burkholderiaceae; Burkholderia; Burkholderia cepacia complex
MKRRTFLALPAVAASGMLAGCSSADLGPVTHRLMKDPEYRETLSAFLISADGKKLVVIGKWP_034201760.1 MULTISPECIES: hypothetical protein [Burkholderia cepacia complex] Length: 212\nNumber of predicted TMHs: 1\nExp number of AAs in TMHs: 28.11606\nExp number, first 60 AAs: 6.33522\nTotal prob of N-in: 0.49088\noutside 1 187\nTMhelix 188 210\ninside 211 212

996 GCF_001991695.1_ASM199169v1 Burkholderia cenocepacia Proteobacteria; Betaproteobacteria; Burkholderiales; Burkholderiaceae; Burkholderia; Burkholderia cepacia complex
MSTSITRRRLLAAGVAASSLALSGCFTPKLYKNDAYSEHVSAFMMTEDGKKLVVLGTRYH WP_077021267.1
hypothetical protein [Burkholderia cenocepacia] Length: 212\nNumber of predicted TMHs: 1\nExp number of

AAs in TMHs: 24.17376\nExp number, first 60 AAs: 0.92388\nTotal prob of N-in: 0.37951\noutside 1 180\nTMhelix 181 203\ninside 204 212

997 GCF_001992045.1_ASM199204v1 Burkholderia cenocepacia Proteobacteria; Betaproteobacteria; Burkholderiales; Burkholderiaceae; Burkholderia; Burkholderia cepacia complex MKRRTFLALPAVAASGMLAGCSSADLGPVTHRLMKDPEYRETLSAFLISADGKKLVVIGKWP_006483583.1 MULTISPECIES: hypothetical protein [Burkholderia cepacia complex] Length: 212\nNumber of predicted TMHs: 1\nExp number of AAs in TMHs: 28.03134\nExp number, first 60 AAs: 6.30523\nTotal prob of N-in: 0.48947\noutside 1 187\nTMhelix 188 210\ninside 211 212

998 GCF_001992005.1_ASM199200v1 Burkholderia cenocepacia Proteobacteria; Betaproteobacteria; Burkholderiales; Burkholderiaceae; Burkholderia; Burkholderia cepacia complex MKRRTFLALPAVAASGMLAGCSSADLGPVTHRLMKDPEYRETLSAFLISADGKKLVVIGKWP_006483583.1 MULTISPECIES: hypothetical protein [Burkholderia cepacia complex] Length: 212\nNumber of predicted TMHs: 1\nExp number of AAs in TMHs: 28.03134\nExp number, first 60 AAs: 6.30523\nTotal prob of N-in: 0.48947\noutside 1 187\nTMhelix 188 210\ninside 211 212

999 GCF_001991875.1_ASM199187v1 Burkholderia cenocepacia Proteobacteria; Betaproteobacteria; Burkholderiales; Burkholderiaceae; Burkholderia; Burkholderia cepacia complex MSTSITRRRLLAAGVAASSLALSGCFTPKLYKNDAYSEHVSAFMMTEDGKKLVVLGTRYH WP_077021267.1 hypothetical protein [Burkholderia cenocepacia] Length: 212\nNumber of predicted TMHs: 1\nExp number of AAs in TMHs: 24.17376\nExp number, first 60 AAs: 0.92388\nTotal prob of N-in: 0.37951\noutside 1 180\nTMhelix 181 203\ninside 204 212

1000 GCF_001991875.1_ASM199187v1 Burkholderia cenocepacia Proteobacteria; Betaproteobacteria; Burkholderiales; Burkholderiaceae; Burkholderia; Burkholderia cepacia complex METLLTRIRASFADASPRRQLISVAVLVALVASLIAAGYFALRPRYQVLFRDLKPQDAAT WP_077020516.1 flagellar M-ring protein FliF [Burkholderia cenocepacia] Length: 491\nNumber of predicted TMHs: 2\nExp number of AAs in TMHs: 41.34236\nExp number, first 60 AAs: 22.1383\nTotal prob of N-in: 0.99080\nPOSSIBLE N-term signal sequence\ninside 1 20\nTMhelix 21 43\noutside 44 427\nTMhelix 428 450\ninside 451 491

1001 GCF_001991875.1_ASM199187v1 Burkholderia cenocepacia Proteobacteria; Betaproteobacteria; Burkholderiales; Burkholderiaceae; Burkholderia; Burkholderia cepacia complex MKRRTFLALPAVAASGMLAGCSSADLGPVTHRLMKDPEYRETLSAFLISADGKKLVVIGKWP_006483583.1 MULTISPECIES: hypothetical protein [Burkholderia cepacia complex] Length: 212\nNumber of predicted TMHs: 1\nExp number of AAs in TMHs: 28.03134\nExp number, first 60 AAs: 6.30523\nTotal prob of N-in: 0.48947\noutside 1 187\nTMhelix 188 210\ninside 211 212

1002 GCF_001991835.1_ASM199183v1 Burkholderia cenocepacia Proteobacteria; Betaproteobacteria; Burkholderiales; Burkholderiaceae; Burkholderia; Burkholderia cepacia complex MKRRTFLALPAVAASGMLAGCSSADLGPVTHRLMKDPEYRETLSAFLISADGKKLVVIGKWP_006483583.1 MULTISPECIES: hypothetical protein [Burkholderia cepacia complex] Length: 212\nNumber of predicted TMHs: 1\nExp number of AAs in TMHs: 28.03134\nExp number, first 60 AAs: 6.30523\nTotal prob of N-in: 0.48947\noutside 1 187\nTMhelix 188 210\ninside 211 212

1003 GCF_001991495.1_ASM199149v1 Burkholderia cenocepacia Proteobacteria; Betaproteobacteria; Burkholderiales; Burkholderiaceae; Burkholderia; Burkholderia cepacia complex MSTSITRRRLLAAGVAASSLALSGCFTPKLYKNDAYSEHVSAFMMTEDGKKLVVLGTRYH WP_077021267.1 hypothetical protein [Burkholderia cenocepacia] Length: 212\nNumber of predicted TMHs: 1\nExp number of AAs in TMHs: 24.17376\nExp number, first 60 AAs: 0.92388\nTotal prob of N-in: 0.37951\noutside 1 180\nTMhelix 181 203\ninside 204 212

1004 GCF_001991875.1_ASM199187v1 Burkholderia cenocepacia Proteobacteria; Betaproteobacteria; Burkholderiales; Burkholderiaceae; Burkholderia; Burkholderia cepacia complex MSDDIDNGGKGGFTRRDMMRVMAASGMMAVGGGGLLTSAQSAFAAPAPKRGGKIRVANES WP_006482802.1 MULTISPECIES: ABC transporter substrate-binding protein [Burkholderia cepacia complex] Length: 527\nNumber of predicted TMHs: 2\nExp number of AAs in TMHs: 36.7303599999999\nExp number, first 60 AAs: 22.08937\nTotal prob of N-in: 0.97312\nPOSSIBLE N-term signal sequence\ninside 1 20\nTMhelix 21 43\noutside 44 502\nTMhelix 503 525\ninside 526 527

1005 GCF_001991835.1_ASM199183v1 Burkholderia cenocepacia Proteobacteria; Betaproteobacteria; Burkholderiales; Burkholderiaceae; Burkholderia; Burkholderia cepacia complex
MSDDIDNGGKGGFTRRDMMRVMAASGMMAVGGGGLLTSAQSAFAAPAPKRGGKIRVANES WP_006482802.1
MULTISPECIES: ABC transporter substrate-binding protein [Burkholderia cepacia complex] Length: 527\nNumber of predicted TMHs: 2\nExp number of AAs in TMHs: 36.7303599999999\nExp number, first 60 AAs: 22.08937\nTotal prob of N-in: 0.97312\nPOSSIBLE N-term signal sequence\ninside 1 20\nTMhelix 21 43\noutside 44 502\nTMhelix 503 525\ninside 526 527

1006 GCF_001991625.1_ASM199162v1 Burkholderia cenocepacia Proteobacteria; Betaproteobacteria; Burkholderiales; Burkholderiaceae; Burkholderia; Burkholderia cepacia complex
MSDDIDNGGKGGFTRRDMMRVMAASGMMAVGGGGLLTSAQSAFAAPAPKRGGKIRVANES WP_006482802.1
MULTISPECIES: ABC transporter substrate-binding protein [Burkholderia cepacia complex] Length: 527\nNumber of predicted TMHs: 2\nExp number of AAs in TMHs: 36.7303599999999\nExp number, first 60 AAs: 22.08937\nTotal prob of N-in: 0.97312\nPOSSIBLE N-term signal sequence\ninside 1 20\nTMhelix 21 43\noutside 44 502\nTMhelix 503 525\ninside 526 527

1007 GCF_001991455.1_ASM199145v1 Burkholderia cenocepacia Proteobacteria; Betaproteobacteria; Burkholderiales; Burkholderiaceae; Burkholderia; Burkholderia cepacia complex
MKRRTFLALPAVAASGMLAGCSSADLGPVTHRLMKDPEYRETLSAFLISADGKKLVVIGKWP_034201760.1 MULTISPECIES: hypothetical protein [Burkholderia cepacia complex] Length: 212\nNumber of predicted TMHs: 1\nExp number of AAs in TMHs: 28.11606\nExp number, first 60 AAs: 6.33522\nTotal prob of N-in: 0.49088\noutside 1 187\nTMhelix 188 210\ninside 211 212

1008 GCF_001991555.1_ASM199155v1 Burkholderia cenocepacia Proteobacteria; Betaproteobacteria; Burkholderiales; Burkholderiaceae; Burkholderia; Burkholderia cepacia complex
METLLTRIRASFADASPRRQLISVAVLVALVASLIAAGYFALRPRYQVLFRDLKPQDAAT WP_077020516.1 flagellar M-ring protein FliF [Burkholderia cenocepacia] Length: 491\nNumber of predicted TMHs: 2\nExp number of AAs in TMHs: 41.34236\nExp number, first 60 AAs: 22.1383\nTotal prob of N-in: 0.99080\nPOSSIBLE N-term signal sequence\ninside 1 20\nTMhelix 21 43\noutside 44 427\nTMhelix 428 450\ninside 451 491

1009 GCF_001991515.1_ASM199151v1 Burkholderia cenocepacia Proteobacteria; Betaproteobacteria; Burkholderiales; Burkholderiaceae; Burkholderia; Burkholderia cepacia complex
METLLTRIRASFADASPRRQLISVAVLVALVASLIAAGYFALRPRYQVLFRDLKPQDAAT WP_077020516.1 flagellar M-ring protein FliF [Burkholderia cenocepacia] Length: 491\nNumber of predicted TMHs: 2\nExp number of AAs in TMHs: 41.34236\nExp number, first 60 AAs: 22.1383\nTotal prob of N-in: 0.99080\nPOSSIBLE N-term signal sequence\ninside 1 20\nTMhelix 21 43\noutside 44 427\nTMhelix 428 450\ninside 451 491

1010 GCF_001991595.1_ASM199159v1 Burkholderia cenocepacia Proteobacteria; Betaproteobacteria; Burkholderiales; Burkholderiaceae; Burkholderia; Burkholderia cepacia complex
MKRRTFLALPAVAASGMLAGCSSADLGPVTHRLMKDPEYRETLSAFLISADGKKLVVIGKWP_034201760.1 MULTISPECIES: hypothetical protein [Burkholderia cepacia complex] Length: 212\nNumber of predicted TMHs: 1\nExp number of AAs in TMHs: 28.11606\nExp number, first 60 AAs: 6.33522\nTotal prob of N-in: 0.49088\noutside 1 187\nTMhelix 188 210\ninside 211 212

1011 GCF_001991595.1_ASM199159v1 Burkholderia cenocepacia Proteobacteria; Betaproteobacteria; Burkholderiales; Burkholderiaceae; Burkholderia; Burkholderia cepacia complex
MSDDIDNGGKGGFTRRDMMRVMAASGMMAVGGGGLLTSAQSAFAAPAPKRGGKIRVANES WP_006482802.1
MULTISPECIES: ABC transporter substrate-binding protein [Burkholderia cepacia complex] Length: 527\nNumber of predicted TMHs: 2\nExp number of AAs in TMHs: 36.7303599999999\nExp number, first 60 AAs: 22.08937\nTotal prob of N-in: 0.97312\nPOSSIBLE N-term signal sequence\ninside 1 20\nTMhelix 21 43\noutside 44 502\nTMhelix 503 525\ninside 526 527

1012 GCF_001991555.1_ASM199155v1 Burkholderia cenocepacia Proteobacteria; Betaproteobacteria; Burkholderiales; Burkholderiaceae; Burkholderia; Burkholderia cepacia complex
MKRRTFLALPAVAASGMLAGCSSADLGPVTHRLMKDPEYRETLSAFLISADGKKLVVIGKWP_006483583.1 MULTISPECIES: hypothetical protein [Burkholderia cepacia complex] Length: 212\nNumber of predicted TMHs: 1\nExp number of AAs in TMHs: 28.03134\nExp number, first 60 AAs: 6.30523\nTotal prob of N-in: 0.48947\noutside 1 187\nTMhelix 188 210\ninside 211 212

1013 GCF_001720575.1_ASM172057v1 Burkholderia cenocepacia Proteobacteria; Betaproteobacteria; Burkholderiales; Burkholderiaceae; Burkholderia; Burkholderia cepacia complex

METLLTRIRASFADASPRRQLISVAVLVALVASLIAAGYFALRPRYQVLFRDLKPQDAAT WP_069352339.1 flagellar M-ring protein FliF [Burkholderia cenocepacia] Length: 491\nNumber of predicted TMHs: 2\nExp number of AAs in TMHs: 41.3425099999999\nExp number, first 60 AAs: 22.1383\nTotal prob of N-in: 0.99079\nPOSSIBLE N-term signal sequence\ninside 1 20\nTMhelix 21 43\noutside 44 427\nTMhelix 428 450\ninside 451 491

1014 GCF_001720575.1_ASM172057v1 Burkholderia cenocepacia Proteobacteria; Betaproteobacteria; Burkholderiales; Burkholderiaceae; Burkholderia; Burkholderia cepacia complex MKRRTFLALPAVAASGMLAGCSSADLGPVTHRLMKDPEYRETLSAFLISADGKKLVVIGKWP_069352077.1 5-formyltetrahydrofolate cyclo-ligase [Burkholderia cenocepacia] Length: 212\nNumber of predicted TMHs: 1\nExp number of AAs in TMHs: 28.13384\nExp number, first 60 AAs: 6.35524\nTotal prob of N-in: 0.49086\noutside 1 187\nTMhelix 188 210\ninside 211 212

1015 GCF_001991315.1_ASM199131v1 Burkholderia cenocepacia Proteobacteria; Betaproteobacteria; Burkholderiales; Burkholderiaceae; Burkholderia; Burkholderia cepacia complex MSDDIDNGGKGGFTRRDMMRVMAASGMMAVGGGGLLTSAQSAFAAPAPKRGGKIRVANES WP_006482802.1 MULTISPECIES: ABC transporter substrate-binding protein [Burkholderia cepacia complex] Length: 527\nNumber of predicted TMHs: 2\nExp number of AAs in TMHs: 36.7303599999999\nExp number, first 60 AAs: 22.08937\nTotal prob of N-in: 0.97312\nPOSSIBLE N-term signal sequence\ninside 1 20\nTMhelix 21 43\noutside 44 502\nTMhelix 503 525\ninside 526 527

1016 GCF_001991275.1_ASM199127v1 Burkholderia cenocepacia Proteobacteria; Betaproteobacteria; Burkholderiales; Burkholderiaceae; Burkholderia; Burkholderia cepacia complex MSDDIDNGGKGGFTRRDMMRVMAASGMMAVGGGGLLTSAQSAFAAPAPKRGGKIRVANES WP_006482802.1 MULTISPECIES: ABC transporter substrate-binding protein [Burkholderia cepacia complex] Length: 527\nNumber of predicted TMHs: 2\nExp number of AAs in TMHs: 36.7303599999999\nExp number, first 60 AAs: 22.08937\nTotal prob of N-in: 0.97312\nPOSSIBLE N-term signal sequence\ninside 1 20\nTMhelix 21 43\noutside 44 502\nTMhelix 503 525\ninside 526 527

1017 GCF_001991235.1_ASM199123v1 Burkholderia cenocepacia Proteobacteria; Betaproteobacteria; Burkholderiales; Burkholderiaceae; Burkholderia; Burkholderia cepacia complex MSDDIDNGGKGGFTRRDMMRVMAASGMMAVGGGGLLTSAQSAFAAPAPKRGGKIRVANES WP_006482802.1 MULTISPECIES: ABC transporter substrate-binding protein [Burkholderia cepacia complex] Length: 527\nNumber of predicted TMHs: 2\nExp number of AAs in TMHs: 36.7303599999999\nExp number, first 60 AAs: 22.08937\nTotal prob of N-in: 0.97312\nPOSSIBLE N-term signal sequence\ninside 1 20\nTMhelix 21 43\noutside 44 502\nTMhelix 503 525\ninside 526 527

1018 GCF_001994085.1_ASM199408v1 Burkholderia cenocepacia Proteobacteria; Betaproteobacteria; Burkholderiales; Burkholderiaceae; Burkholderia; Burkholderia cepacia complex MKRRTFLALPAVAASGMLAGCSSADLGPVTHRLMKDPEYRETLSAFLISADGKKLVVIGKWP_034201760.1 MULTISPECIES: hypothetical protein [Burkholderia cepacia complex] Length: 212\nNumber of predicted TMHs: 1\nExp number of AAs in TMHs: 28.11606\nExp number, first 60 AAs: 6.33522\nTotal prob of N-in: 0.49088\noutside 1 187\nTMhelix 188 210\ninside 211 212

1019 GCF_001994085.1_ASM199408v1 Burkholderia cenocepacia Proteobacteria; Betaproteobacteria; Burkholderiales; Burkholderiaceae; Burkholderia; Burkholderia cepacia complex MSDDIDNGGKGGFTRRDMMRVMAASGMMAVGGGGLLTSAQSAFAAPAPKRGGKIRVANES WP_006482802.1 MULTISPECIES: ABC transporter substrate-binding protein [Burkholderia cepacia complex] Length: 527\nNumber of predicted TMHs: 2\nExp number of AAs in TMHs: 36.7303599999999\nExp number, first 60 AAs: 22.08937\nTotal prob of N-in: 0.97312\nPOSSIBLE N-term signal sequence\ninside 1 20\nTMhelix 21 43\noutside 44 502\nTMhelix 503 525\ninside 526 527

1020 GCF_001720525.1_ASM172052v1 Burkholderia cenocepacia Proteobacteria; Betaproteobacteria; Burkholderiales; Burkholderiaceae; Burkholderia; Burkholderia cepacia complex METLLTRIRASFADASPRRQLISVAVLVALVASLIAAGYFALRPRYQVLFRDLKPQDAAT WP_069352339.1 flagellar M-ring protein FliF [Burkholderia cenocepacia] Length: 491\nNumber of predicted TMHs: 2\nExp number of AAs in TMHs: 41.3425099999999\nExp number, first 60 AAs: 22.1383\nTotal prob of N-in: 0.99079\nPOSSIBLE N-term signal sequence\ninside 1 20\nTMhelix 21 43\noutside 44 427\nTMhelix 428 450\ninside 451 491

1021 GCF_001984325.1_ASM198432v1 Burkholderia cenocepacia Proteobacteria; Betaproteobacteria; Burkholderiales; Burkholderiaceae; Burkholderia; Burkholderia cepacia complex MSDDIDNGGKGGFTRRDMMRVMAASGMMAVGGGGLLTSAQSAFAAPAPKRGGKIRVANES WP_006482802.1

MULTISPECIES: ABC transporter substrate-binding protein [Burkholderia cepacia complex] Length: 527\nNumber of predicted TMHs: 2\nExp number of AAs in TMHs: 36.7303599999999\nExp number, first 60 AAs: 22.08937\nTotal prob of N-in: 0.97312\nPOSSIBLE N-term signal sequence\ninside 1 20\nTMhelix 21 43\noutside 44 502\nTMhelix 503 525\ninside 526 527

1022 GCF_001720575.1_ASM172057v1 Burkholderia cenocepacia Proteobacteria; Betaproteobacteria; Burkholderiales; Burkholderiaceae; Burkholderia; Burkholderia cepacia complex
MSDDIDNGGKGGFTRRDMMRVMAASGMMAVGGGGLLTSAQSAFAAPAPKRGGKIRVANES WP_006482802.1 MULTISPECIES: ABC transporter substrate-binding protein [Burkholderia cepacia complex] Length: 527\nNumber of predicted TMHs: 2\nExp number of AAs in TMHs: 36.7303599999999\nExp number, first 60 AAs: 22.08937\nTotal prob of N-in: 0.97312\nPOSSIBLE N-term signal sequence\ninside 1 20\nTMhelix 21 43\noutside 44 502\nTMhelix 503 525\ninside 526 527

1023 GCF_001994035.1_ASM199403v1 Burkholderia cenocepacia Proteobacteria; Betaproteobacteria; Burkholderiales; Burkholderiaceae; Burkholderia; Burkholderia cepacia complex
MKRRTFLALPAVAASGMLAGCSSADLGPVTHRLMKDPEYRETLSAFLISADGKKLVVIGKWP_006483583.1 MULTISPECIES: hypothetical protein [Burkholderia cepacia complex] Length: 212\nNumber of predicted TMHs: 1\nExp number of AAs in TMHs: 28.03134\nExp number, first 60 AAs: 6.30523\nTotal prob of N-in: 0.48947\noutside 1 187\nTMhelix 188 210\ninside 211 212

1024 GCF_001993795.1_ASM199379v1 Burkholderia cenocepacia Proteobacteria; Betaproteobacteria; Burkholderiales; Burkholderiaceae; Burkholderia; Burkholderia cepacia complex
MSDDIDNGGKGGFTRRDMMRVMAASGMMAVGGGGLLTSAQSAFAAPAPKRGGKIRVANES WP_006482802.1 MULTISPECIES: ABC transporter substrate-binding protein [Burkholderia cepacia complex] Length: 527\nNumber of predicted TMHs: 2\nExp number of AAs in TMHs: 36.7303599999999\nExp number, first 60 AAs: 22.08937\nTotal prob of N-in: 0.97312\nPOSSIBLE N-term signal sequence\ninside 1 20\nTMhelix 21 43\noutside 44 502\nTMhelix 503 525\ninside 526 527

1025 GCF_001993755.1_ASM199375v1 Burkholderia cenocepacia Proteobacteria; Betaproteobacteria; Burkholderiales; Burkholderiaceae; Burkholderia; Burkholderia cepacia complex
MSDDIDNGGKGGFTRRDMMRVMAASGMMAVGGGGLLTSAQSAFAAPAPKRGGKIRVANES WP_006482802.1 MULTISPECIES: ABC transporter substrate-binding protein [Burkholderia cepacia complex] Length: 527\nNumber of predicted TMHs: 2\nExp number of AAs in TMHs: 36.7303599999999\nExp number, first 60 AAs: 22.08937\nTotal prob of N-in: 0.97312\nPOSSIBLE N-term signal sequence\ninside 1 20\nTMhelix 21 43\noutside 44 502\nTMhelix 503 525\ninside 526 527

1026 GCF_001993495.1_ASM199349v1 Burkholderia cenocepacia Proteobacteria; Betaproteobacteria; Burkholderiales; Burkholderiaceae; Burkholderia; Burkholderia cepacia complex
METLLTRIRASFADASPRRQLISVAVLVALVASLIAAGYFALRPRYQVLFRDLKPQDAAT WP_077020516.1 flagellar M-ring protein FliF [Burkholderia cenocepacia] Length: 491\nNumber of predicted TMHs: 2\nExp number of AAs in TMHs: 41.34236\nExp number, first 60 AAs: 22.1383\nTotal prob of N-in: 0.99080\nPOSSIBLE N-term signal sequence\ninside 1 20\nTMhelix 21 43\noutside 44 427\nTMhelix 428 450\ninside 451 491

1027 GCF_001993475.1_ASM199347v1 Burkholderia cenocepacia Proteobacteria; Betaproteobacteria; Burkholderiales; Burkholderiaceae; Burkholderia; Burkholderia cepacia complex
MSTSITRRRLLAAGVAASSLALSGCFTPKLYKNDAYSEHVSAFMMTEDGKKLVVLGTRYH WP_077021267.1 hypothetical protein [Burkholderia cenocepacia] Length: 212\nNumber of predicted TMHs: 1\nExp number of AAs in TMHs: 24.17376\nExp number, first 60 AAs: 0.92388\nTotal prob of N-in: 0.37951\noutside 1 180\nTMhelix 181 203\ninside 204 212

1028 GCF_001993475.1_ASM199347v1 Burkholderia cenocepacia Proteobacteria; Betaproteobacteria; Burkholderiales; Burkholderiaceae; Burkholderia; Burkholderia cepacia complex
METLLTRIRASFADASPRRQLISVAVLVALVASLIAAGYFALRPRYQVLFRDLKPQDAAT WP_077020516.1 flagellar M-ring protein FliF [Burkholderia cenocepacia] Length: 491\nNumber of predicted TMHs: 2\nExp number of AAs in TMHs: 41.34236\nExp number, first 60 AAs: 22.1383\nTotal prob of N-in: 0.99080\nPOSSIBLE N-term signal sequence\ninside 1 20\nTMhelix 21 43\noutside 44 427\nTMhelix 428 450\ninside 451 491

1029 GCF_001993295.1_ASM199329v1 Burkholderia cenocepacia Proteobacteria; Betaproteobacteria; Burkholderiales; Burkholderiaceae; Burkholderia; Burkholderia cepacia complex
METLLTRIRASFADASPRRQLISVAVLVALVASLIAAGYFALRPRYQVLFRDLKPQDAAT WP_077020516.1 flagellar M-ring protein FliF [Burkholderia cenocepacia] Length: 491\nNumber of predicted TMHs: 2\nExp number of AAs in

TMHs: 41.34236\nExp number, first 60 AAs: 22.1383\nTotal prob of N-in: 0.99080\nPOSSIBLE N-term signal sequence\ninside 1 20\nTMhelix 21 43\noutside 44 427\nTMhelix 428 450\ninside 451 491

1030 GCF_001993645.1_ASM199364v1 Burkholderia cenocepacia Proteobacteria; Betaproteobacteria; Burkholderiales; Burkholderiaceae; Burkholderia; Burkholderia cepacia complex MKRRTFLALPAVAASGMLAGCSSADLGPVTHRLMKDPEYRETLSAFLISADGKKLVVIGKWP_034201760.1 MULTISPECIES: hypothetical protein [Burkholderia cepacia complex] Length: 212\nNumber of predicted TMHs: 1\nExp number of AAs in TMHs: 28.11606\nExp number, first 60 AAs: 6.33522\nTotal prob of N-in: 0.49088\noutside 1 187\nTMhelix 188 210\ninside 211 212

1031 GCF_001993645.1_ASM199364v1 Burkholderia cenocepacia Proteobacteria; Betaproteobacteria; Burkholderiales; Burkholderiaceae; Burkholderia; Burkholderia cepacia complex MSDDIDNGGKGGFTRRDMMRVMAASGMMAVGGGGLLTSAQSAFAAPAPKRGGKIRVANES WP_006482802.1 MULTISPECIES: ABC transporter substrate-binding protein [Burkholderia cepacia complex] Length: 527\nNumber of predicted TMHs: 2\nExp number of AAs in TMHs: 36.7303599999999\nExp number, first 60 AAs: 22.08937\nTotal prob of N-in: 0.97312\nPOSSIBLE N-term signal sequence\ninside 1 20\nTMhelix 21 43\noutside 44 502\nTMhelix 503 525\ninside 526 527

1032 GCF_001993395.1_ASM199339v1 Burkholderia cenocepacia Proteobacteria; Betaproteobacteria; Burkholderiales; Burkholderiaceae; Burkholderia; Burkholderia cepacia complex MKRRTFLALPAVAASGMLAGCSSADLGPVTHRLMKDPEYRETLSAFLISADGKKLVVIGKWP_034201760.1 MULTISPECIES: hypothetical protein [Burkholderia cepacia complex] Length: 212\nNumber of predicted TMHs: 1\nExp number of AAs in TMHs: 28.11606\nExp number, first 60 AAs: 6.33522\nTotal prob of N-in: 0.49088\noutside 1 187\nTMhelix 188 210\ninside 211 212

1033 GCF_001073275.1_ASM107327v1 Burkholderia cenocepacia Proteobacteria; Betaproteobacteria; Burkholderiales; Burkholderiaceae; Burkholderia; Burkholderia cepacia complex MKRRTFLALPAVAASGMLAGCSSADLGPVTHRLMKDPEYQETLSAFLISADGKKLVVIGK WP_048986526.1 hypothetical protein [Burkholderia cenocepacia] Length: 212\nNumber of predicted TMHs: 1\nExp number of AAs in TMHs: 28.09653\nExp number, first 60 AAs: 6.31597\nTotal prob of N-in: 0.49053\noutside 1 187\nTMhelix 188 210\ninside 211 212

1034 GCF_001993565.1_ASM199356v1 Burkholderia cenocepacia Proteobacteria; Betaproteobacteria; Burkholderiales; Burkholderiaceae; Burkholderia; Burkholderia cepacia complex MSDDIDNGGKGGFTRRDMMRVMAASGMMAVGGGGLLTSAQSAFAAPAPKRGGKIRVANES WP_077187164.1 ABC transporter substrate-binding protein [Burkholderia cenocepacia] Length: 527\nNumber of predicted TMHs: 2\nExp number of AAs in TMHs: 36.7300399999999\nExp number, first 60 AAs: 22.0894\nTotal prob of N-in: 0.97312\nPOSSIBLE N-term signal sequence\ninside 1 20\nTMhelix 21 43\noutside 44 502\nTMhelix 503 525\ninside 526 527

1035 GCF_001994135.1_ASM199413v1 Burkholderia cenocepacia Proteobacteria; Betaproteobacteria; Burkholderiales; Burkholderiaceae; Burkholderia; Burkholderia cepacia complex MSDDIDNGGKGGFTRRDMMRVMAASGMMAVGGGGLLTSAQSAFAAPAPKRGGKIRVANES WP_006482802.1 MULTISPECIES: ABC transporter substrate-binding protein [Burkholderia cepacia complex] Length: 527\nNumber of predicted TMHs: 2\nExp number of AAs in TMHs: 36.7303599999999\nExp number, first 60 AAs: 22.08937\nTotal prob of N-in: 0.97312\nPOSSIBLE N-term signal sequence\ninside 1 20\nTMhelix 21 43\noutside 44 502\nTMhelix 503 525\ninside 526 527

1036 GCF_001993565.1_ASM199356v1 Burkholderia cenocepacia Proteobacteria; Betaproteobacteria; Burkholderiales; Burkholderiaceae; Burkholderia; Burkholderia cepacia complex MSTSITRRRLLAAGVAASSLALSGCFTPKLYKNDAYSEHVSAFMMTEDGKKLVVLGTRYH WP_077021267.1 hypothetical protein [Burkholderia cenocepacia] Length: 212\nNumber of predicted TMHs: 1\nExp number of AAs in TMHs: 24.17376\nExp number, first 60 AAs: 0.92388\nTotal prob of N-in: 0.37951\noutside 1 180\nTMhelix 181 203\ninside 204 212

1037 GCF_001993565.1_ASM199356v1 Burkholderia cenocepacia Proteobacteria; Betaproteobacteria; Burkholderiales; Burkholderiaceae; Burkholderia; Burkholderia cepacia complex METLLTRIRASFADASPRRQLISVAVLVALVASLIAAGYFALRPRYQVLFRDLKPQDAAT WP_077020516.1 flagellar M-ring protein FliF [Burkholderia cenocepacia] Length: 491\nNumber of predicted TMHs: 2\nExp number of AAs in TMHs: 41.34236\nExp number, first 60 AAs: 22.1383\nTotal prob of N-in: 0.99080\nPOSSIBLE N-term signal sequence\ninside 1 20\nTMhelix 21 43\noutside 44 427\nTMhelix 428 450\ninside 451 491

1038 GCF_001521915.1_ASM152191v1 Burkholderia cepacia Proteobacteria; Betaproteobacteria; Burkholderiales; Burkholderiaceae; Burkholderia; Burkholderia cepacia complex
MKRRALLTAGVIVVAAGLTGCTAMLFEDGRYEETVDRFLVSEDGKKFVVLGEKYHYIFDM WP_059496097.1
hypothetical protein [Burkholderia cepacia] Length: 208\nNumber of predicted TMHs: 2\nExp number of AAs in TMHs: 44.13865\nExp number, first 60 AAs: 21.11648\nTotal prob of N-in: 0.75846\nPOSSIBLE N-term signal sequence\ninside 1 6\nTMhelix 7 26\noutside 27 183\nTMhelix 184 206\ninside 207 208

1039 GCF_001522335.1_ASM152233v1 Burkholderia cepacia Proteobacteria; Betaproteobacteria; Burkholderiales; Burkholderiaceae; Burkholderia; Burkholderia cepacia complex
MKRRTFLALPAVAASGMLAGCSNFGPVTKAWMSDSKYDETLSTFLITADGKQLVVLGKKY WP_027787377.1
MULTISPECIES: hypothetical protein [Burkholderia] Length: 210\nNumber of predicted TMHs: 1\nExp number of AAs in TMHs: 34.58187\nExp number, first 60 AAs: 12.86351\nTotal prob of N-in: 0.46668\nPOSSIBLE N-term signal sequence\noutside 1 186\nTMhelix 187 209\ninside 210 210

1040 GCF_001522335.1_ASM152233v1 Burkholderia cepacia Proteobacteria; Betaproteobacteria; Burkholderiales; Burkholderiaceae; Burkholderia; Burkholderia cepacia complex
MKRRALLTAGVIVVAAGLTGCTAMLFEDGRYEETVDRFLVSEDGKKFVVLGEKYHYIFDM WP_059677781.1
hypothetical protein [Burkholderia cepacia] Length: 208\nNumber of predicted TMHs: 2\nExp number of AAs in TMHs: 44.88033\nExp number, first 60 AAs: 21.13087\nTotal prob of N-in: 0.76656\nPOSSIBLE N-term signal sequence\ninside 1 6\nTMhelix 7 26\noutside 27 183\nTMhelix 184 206\ninside 207 208

1041 GCF_000757365.1_BHX_1 Burkholderia cepacia Proteobacteria; Betaproteobacteria; Burkholderiales; Burkholderiaceae; Burkholderia; Burkholderia cepacia complex
MKRRTFLALPAVAASGMLAGCSSADLGPVTHRLMKDPEYRETLSAFLISADGKKLVVIGKWP_034201760.1 MULTISPECIES: hypothetical protein [Burkholderia cepacia complex] Length: 212\nNumber of predicted TMHs: 1\nExp number of AAs in TMHs: 28.11606\nExp number, first 60 AAs: 6.33522\nTotal prob of N-in: 0.49088\noutside 1 187\nTMhelix 188 210\ninside 211 212

1042 GCF_000755805.1_ASM75580v1 Burkholderia cepacia Proteobacteria; Betaproteobacteria; Burkholderiales; Burkholderiaceae; Burkholderia; Burkholderia cepacia complex
MKRRTFLALPAVAASGMLAGCSSADLGPVTHRLMKDPEYRETLSAFLISADGKKLVVIGKWP_034201760.1 MULTISPECIES: hypothetical protein [Burkholderia cepacia complex] Length: 212\nNumber of predicted TMHs: 1\nExp number of AAs in TMHs: 28.11606\nExp number, first 60 AAs: 6.33522\nTotal prob of N-in: 0.49088\noutside 1 187\nTMhelix 188 210\ninside 211 212

1043 GCF_001521975.1_ASM152197v1 Burkholderia cepacia Proteobacteria; Betaproteobacteria; Burkholderiales; Burkholderiaceae; Burkholderia; Burkholderia cepacia complex
MKRRTFLALPAVAASGMLAGCSNFGPVTKAWMSDSKYDETLSTFLITADGKQLVVLGKKY WP_059488505.1
hypothetical protein [Burkholderia cepacia] Length: 210\nNumber of predicted TMHs: 1\nExp number of AAs in TMHs: 34.5304\nExp number, first 60 AAs: 12.84424\nTotal prob of N-in: 0.46568\nPOSSIBLE N-term signal sequence\noutside 1 186\nTMhelix 187 209\ninside 210 210

1044 GCF_000724975.1_ASM72497v1 Burkholderia cepacia Proteobacteria; Betaproteobacteria; Burkholderiales; Burkholderiaceae; Burkholderia; Burkholderia cepacia complex
MSTSITRRRLLAAGVAASSLALSGCFTRALYENHDEPYVEHVSAFMITKDGKKLVVLGER WP_043180153.1 hypothetical protein [Burkholderia cepacia] Length: 213\nNumber of predicted TMHs: 1\nExp number of AAs in TMHs: 23.81691\nExp number, first 60 AAs: 1.71629\nTotal prob of N-in: 0.64617\noutside 1 182\nTMhelix 183 205\ninside 206 213

1045 GCF_001522785.1_ASM152278v1 Burkholderia cepacia Proteobacteria; Betaproteobacteria; Burkholderiales; Burkholderiaceae; Burkholderia; Burkholderia cepacia complex
MKRRTFLALPAVAASGMLAGCSNFGPVTKAWMSDSKYDETLSTFLITADGKQLVVLGKKY WP_059488505.1
hypothetical protein [Burkholderia cepacia] Length: 210\nNumber of predicted TMHs: 1\nExp number of AAs in TMHs: 34.5304\nExp number, first 60 AAs: 12.84424\nTotal prob of N-in: 0.46568\nPOSSIBLE N-term signal sequence\noutside 1 186\nTMhelix 187 209\ninside 210 210

1046 GCF_001522785.1_ASM152278v1 Burkholderia cepacia Proteobacteria; Betaproteobacteria; Burkholderiales; Burkholderiaceae; Burkholderia; Burkholderia cepacia complex
MKRRALLTAGVIVVAAGLTGCTAMLFEDGRYEETVDRFLVSEDGKKFVVLGEKYHYIFDM WP_059496097.1
hypothetical protein [Burkholderia cepacia] Length: 208\nNumber of predicted TMHs: 2\nExp number of

AAs in TMHs: 44.13865\nExp number, first 60 AAs: 21.11648\nTotal prob of N-in: 0.75846\nPOSSIBLE N-term signal sequence\ninside 1 6\nTMhelix 7 26\noutside 27 183\nTMhelix 184 206\ninside 207 208

1047 GCF_001522625.1_ASM152262v1 Burkholderia cepacia Proteobacteria; Betaproteobacteria; Burkholderiales; Burkholderiaceae; Burkholderia; Burkholderia cepacia complex MKTSITRRRLLAAAMAASTLALSGCFTPKLYENDAYSEYVTSFMITQDGKKLVVLGTNYH WP_059485162.1 hypothetical protein [Burkholderia cepacia] Length: 212\nNumber of predicted TMHs: 1\nExp number of AAs in TMHs: 24.29452\nExp number, first 60 AAs: 1.02778\nTotal prob of N-in: 0.64928\noutside 1 180\nTMhelix 181 203\ninside 204 212

1048 GCF_001522625.1_ASM152262v1 Burkholderia cepacia Proteobacteria; Betaproteobacteria; Burkholderiales; Burkholderiaceae; Burkholderia; Burkholderia cepacia complex MKRRTFLALPAVAASGMLAGCSNFGPVTKAWMSDSKYDETLSTFLITADGKQLVVLGKKY WP_059488505.1 hypothetical protein [Burkholderia cepacia] Length: 210\nNumber of predicted TMHs: 1\nExp number of AAs in TMHs: 34.5304\nExp number, first 60 AAs: 12.84424\nTotal prob of N-in: 0.46568\nPOSSIBLE N-term signal sequence\noutside 1 186\nTMhelix 187 209\ninside 210 210

1049 GCF_001525765.1_ASM152576v1 Burkholderia cepacia Proteobacteria; Betaproteobacteria; Burkholderiales; Burkholderiaceae; Burkholderia; Burkholderia cepacia complex MKTSITRRRLLAAAMAASTLALSGCFTPKLYENDAYSEYVTSFMITQDGKKLVVLGTNYH WP_059485162.1 hypothetical protein [Burkholderia cepacia] Length: 212\nNumber of predicted TMHs: 1\nExp number of AAs in TMHs: 24.29452\nExp number, first 60 AAs: 1.02778\nTotal prob of N-in: 0.64928\noutside 1 180\nTMhelix 181 203\ninside 204 212

1050 GCF_001525765.1_ASM152576v1 Burkholderia cepacia Proteobacteria; Betaproteobacteria; Burkholderiales; Burkholderiaceae; Burkholderia; Burkholderia cepacia complex MKRRTFLALPAVAASGMLAGCSNFGPVTKAWMSDSKYDETLSTFLITADGKQLVVLGKKY WP_059488505.1 hypothetical protein [Burkholderia cepacia] Length: 210\nNumber of predicted TMHs: 1\nExp number of AAs in TMHs: 34.5304\nExp number, first 60 AAs: 12.84424\nTotal prob of N-in: 0.46568\nPOSSIBLE N-term signal sequence\noutside 1 186\nTMhelix 187 209\ninside 210 210

1051 GCF_001524765.1_ASM152476v1 Burkholderia cepacia Proteobacteria; Betaproteobacteria; Burkholderiales; Burkholderiaceae; Burkholderia; Burkholderia cepacia complex MKRRALLTAGAIVVAAGLTGCTAMLFENGRYEETVDRFLVSEDGKKFVVLGEKYHYIFDM WP_059682727.1 hypothetical protein [Burkholderia cepacia] Length: 208\nNumber of predicted TMHs: 2\nExp number of AAs in TMHs: 44.24426\nExp number, first 60 AAs: 20.97809\nTotal prob of N-in: 0.67235\nPOSSIBLE N-term signal sequence\noutside 1 4\nTMhelix 5 27\ninside 28 184\nTMhelix 185 207\noutside 208 208

1052 GCF_001525765.1_ASM152576v1 Burkholderia cepacia Proteobacteria; Betaproteobacteria; Burkholderiales; Burkholderiaceae; Burkholderia; Burkholderia cepacia complex MKRRALLTAGVIVVAAGLTGCTAMLFEDGRYEETVDRFLVSEDGKKFVVLGEKYHYIFDM WP_059496097.1 hypothetical protein [Burkholderia cepacia] Length: 208\nNumber of predicted TMHs: 2\nExp number of AAs in TMHs: 44.13865\nExp number, first 60 AAs: 21.11648\nTotal prob of N-in: 0.75846\nPOSSIBLE N-term signal sequence\ninside 1 6\nTMhelix 7 26\noutside 27 183\nTMhelix 184 206\ninside 207 208

1053 GCF_001522925.1_ASM152292v1 Burkholderia cepacia Proteobacteria; Betaproteobacteria; Burkholderiales; Burkholderiaceae; Burkholderia; Burkholderia cepacia complex MKTSITRRRLLAAAMAASTLALSGCFTPKLYENDAYSEYVTSFMITQDGKKLVVLGTNYH WP_059485162.1 hypothetical protein [Burkholderia cepacia] Length: 212\nNumber of predicted TMHs: 1\nExp number of AAs in TMHs: 24.29452\nExp number, first 60 AAs: 1.02778\nTotal prob of N-in: 0.64928\noutside 1 180\nTMhelix 181 203\ninside 204 212

1054 GCF_001522925.1_ASM152292v1 Burkholderia cepacia Proteobacteria; Betaproteobacteria; Burkholderiales; Burkholderiaceae; Burkholderia; Burkholderia cepacia complex MKRRTFLALPAVAASGMLAGCSNFGPVTKAWMSDSKYDETLSTFLITADGKQLVVLGKKY WP_059488505.1 hypothetical protein [Burkholderia cepacia] Length: 210\nNumber of predicted TMHs: 1\nExp number of AAs in TMHs: 34.5304\nExp number, first 60 AAs: 12.84424\nTotal prob of N-in: 0.46568\nPOSSIBLE N-term signal sequence\noutside 1 186\nTMhelix 187 209\ninside 210 210

1055 GCF_001522925.1_ASM152292v1 Burkholderia cepacia Proteobacteria; Betaproteobacteria; Burkholderiales; Burkholderiaceae; Burkholderia; Burkholderia cepacia complex

MKRRALLTAGVIVVAAGLTGCTAMLFEDGRYEETVDRFLVSEDGKKFVVLGEKYHYIFDM WP_059496097.1 hypothetical protein [Burkholderia cepacia] Length: 208\nNumber of predicted TMHs: 2\nExp number of AAs in TMHs: 44.13865\nExp number, first 60 AAs: 21.11648\nTotal prob of N-in: 0.75846\nPOSSIBLE N-term signal sequence\ninside 1 6\nTMhelix 7 26\noutside 27 183\nTMhelix 184 206\ninside 207 208

1056 GCF_001522965.1_ASM152296v1 Burkholderia cepacia Proteobacteria; Betaproteobacteria; Burkholderiales; Burkholderiaceae; Burkholderia; Burkholderia cepacia complex MKTSITRRRLLAAAMAASTLALSGCFTPKLYENDAYSEYVTSFMITQDGKKLVVLGTNYH WP_059485162.1 hypothetical protein [Burkholderia cepacia] Length: 212\nNumber of predicted TMHs: 1\nExp number of AAs in TMHs: 24.29452\nExp number, first 60 AAs: 1.02778\nTotal prob of N-in: 0.64928\noutside 1 180\nTMhelix 181 203\ninside 204 212

1057 GCF_001525895.1_ASM152589v1 Burkholderia cepacia Proteobacteria; Betaproteobacteria; Burkholderiales; Burkholderiaceae; Burkholderia; Burkholderia cepacia complex MKTSITRRRLLAAAMAASTLALSGCFTPKLYENDAYSEYVTSFMITQDGKKLVVLGTNYH WP_059485162.1 hypothetical protein [Burkholderia cepacia] Length: 212\nNumber of predicted TMHs: 1\nExp number of AAs in TMHs: 24.29452\nExp number, first 60 AAs: 1.02778\nTotal prob of N-in: 0.64928\noutside 1 180\nTMhelix 181 203\ninside 204 212

1058 GCF_001525785.1_ASM152578v1 Burkholderia cepacia Proteobacteria; Betaproteobacteria; Burkholderiales; Burkholderiaceae; Burkholderia; Burkholderia cepacia complex MKRRTFLALPAVAASGMLAGCSNFGPVTKAWMSDSKYDETLSTFLITADGKQLVVLGKKY WP_059488505.1 hypothetical protein [Burkholderia cepacia] Length: 210\nNumber of predicted TMHs: 1\nExp number of AAs in TMHs: 34.5304\nExp number, first 60 AAs: 12.84424\nTotal prob of N-in: 0.46568\nPOSSIBLE N-term signal sequence\noutside 1 186\nTMhelix 187 209\ninside 210 210

1059 GCF_001529525.1_ASM152952v1 Burkholderia cepacia Proteobacteria; Betaproteobacteria; Burkholderiales; Burkholderiaceae; Burkholderia; Burkholderia cepacia complex MKRRTFLALPAVAASGMLAGCSNFGPVTKAWMSDSKYDETLSTFLITADGKQLVVLGKKY WP_059488505.1 hypothetical protein [Burkholderia cepacia] Length: 210\nNumber of predicted TMHs: 1\nExp number of AAs in TMHs: 34.5304\nExp number, first 60 AAs: 12.84424\nTotal prob of N-in: 0.46568\nPOSSIBLE N-term signal sequence\noutside 1 186\nTMhelix 187 209\ninside 210 210

1060 GCF_001530045.1_ASM153004v1 Burkholderia cepacia Proteobacteria; Betaproteobacteria; Burkholderiales; Burkholderiaceae; Burkholderia; Burkholderia cepacia complex MKRRTFLALPAVAASGMLAGCSNFGPVTKAWMSDSKYDETLSTFLITADGKQLVVLGKKY WP_059488505.1 hypothetical protein [Burkholderia cepacia] Length: 210\nNumber of predicted TMHs: 1\nExp number of AAs in TMHs: 34.5304\nExp number, first 60 AAs: 12.84424\nTotal prob of N-in: 0.46568\nPOSSIBLE N-term signal sequence\noutside 1 186\nTMhelix 187 209\ninside 210 210

1061 GCF_001525895.1_ASM152589v1 Burkholderia cepacia Proteobacteria; Betaproteobacteria; Burkholderiales; Burkholderiaceae; Burkholderia; Burkholderia cepacia complex MKRRTFLALPAVAASGMLAGCSNFGPVTKAWMSDSKYDETLSTFLITADGKQLVVLGKKY WP_059488505.1 hypothetical protein [Burkholderia cepacia] Length: 210\nNumber of predicted TMHs: 1\nExp number of AAs in TMHs: 34.5304\nExp number, first 60 AAs: 12.84424\nTotal prob of N-in: 0.46568\nPOSSIBLE N-term signal sequence\noutside 1 186\nTMhelix 187 209\ninside 210 210

1062 GCF_001525785.1_ASM152578v1 Burkholderia cepacia Proteobacteria; Betaproteobacteria; Burkholderiales; Burkholderiaceae; Burkholderia; Burkholderia cepacia complex MKRRALLTAGVIVVAAGLTGCTAMLFEDGRYEETVDRFLVSEDGKKFVVLGEKYHYIFDM WP_059496097.1 hypothetical protein [Burkholderia cepacia] Length: 208\nNumber of predicted TMHs: 2\nExp number of AAs in TMHs: 44.13865\nExp number, first 60 AAs: 21.11648\nTotal prob of N-in: 0.75846\nPOSSIBLE N-term signal sequence\ninside 1 6\nTMhelix 7 26\noutside 27 183\nTMhelix 184 206\ninside 207 208

1063 GCF_001530045.1_ASM153004v1 Burkholderia cepacia Proteobacteria; Betaproteobacteria; Burkholderiales; Burkholderiaceae; Burkholderia; Burkholderia cepacia complex MKTSITRRRLLAAAMAASTLALSGCFTPKLYENDAYSEYVTSFMITQDGKKLVVLGTNYH WP_060050342.1 hypothetical protein [Burkholderia cepacia] Length: 212\nNumber of predicted TMHs: 1\nExp number of AAs in TMHs: 24.23033\nExp number, first 60 AAs: 1.03058\nTotal prob of N-in: 0.64819\noutside 1 180\nTMhelix 181 203\ninside 204 212

1064 GCF_001522815.1_ASM152281v1 Burkholderia cepacia Proteobacteria; Betaproteobacteria; Burkholderiales; Burkholderiaceae; Burkholderia; Burkholderia cepacia complex
MKRRALLTAGVIVVAAGLTGCTAMLFEDGRYEETVDRFLVSEDGKKFVVLGEKYHYIFDM WP_059496097.1
hypothetical protein [Burkholderia cepacia] Length: 208\nNumber of predicted TMHs: 2\nExp number of AAs in TMHs: 44.13865\nExp number, first 60 AAs: 21.11648\nTotal prob of N-in: 0.75846\nPOSSIBLE N-term signal sequence\ninside 1 6\nTMhelix 7 26\noutside 27 183\nTMhelix 184 206\ninside 207 208

1065 GCF_001524125.1_ASM152412v1 Burkholderia cepacia Proteobacteria; Betaproteobacteria; Burkholderiales; Burkholderiaceae; Burkholderia; Burkholderia cepacia complex
MKRRSFLALPAVAASGMLAGCSNFGPVTKAWMSDSKYDETLSTFLITADGKQLVVLGKKY WP_059621060.1
hypothetical protein [Burkholderia cepacia] Length: 210\nNumber of predicted TMHs: 1\nExp number of AAs in TMHs: 34.37239\nExp number, first 60 AAs: 12.58285\nTotal prob of N-in: 0.49163\nPOSSIBLE N-term signal sequence\noutside 1 186\nTMhelix 187 209\ninside 210 210

1066 GCF_001528485.1_ASM152848v1 Burkholderia cepacia Proteobacteria; Betaproteobacteria; Burkholderiales; Burkholderiaceae; Burkholderia; Burkholderia cepacia complex
MKRRTFLALPAVAASGMLAGCSNFGPVTKAWMSDSKYDETLSTFLITADGKQLVVLGKKY WP_027787377.1
MULTISPECIES: hypothetical protein [Burkholderia] Length: 210\nNumber of predicted TMHs: 1\nExp number of AAs in TMHs: 34.58187\nExp number, first 60 AAs: 12.86351\nTotal prob of N-in: 0.46668\nPOSSIBLE N-term signal sequence\noutside 1 186\nTMhelix 187 209\ninside 210 210

1067 GCF_001526785.1_ASM152678v1 Burkholderia cepacia Proteobacteria; Betaproteobacteria; Burkholderiales; Burkholderiaceae; Burkholderia; Burkholderia cepacia complex
MKRRTFLALPAVAASGMLAGCSNFGPVTKAWMSDSKYDETLSTFLITADGKQLVVLGKKY WP_027787377.1
MULTISPECIES: hypothetical protein [Burkholderia] Length: 210\nNumber of predicted TMHs: 1\nExp number of AAs in TMHs: 34.58187\nExp number, first 60 AAs: 12.86351\nTotal prob of N-in: 0.46668\nPOSSIBLE N-term signal sequence\noutside 1 186\nTMhelix 187 209\ninside 210 210

1068 GCF_001528485.1_ASM152848v1 Burkholderia cepacia Proteobacteria; Betaproteobacteria; Burkholderiales; Burkholderiaceae; Burkholderia; Burkholderia cepacia complex
MKTSITRRRLLAAAMAASTLALSGCFTPKLYENDAYSEYVTSFMITQDGKKLVVLGTNYH WP_059485162.1 hypothetical protein [Burkholderia cepacia] Length: 212\nNumber of predicted TMHs: 1\nExp number of AAs in TMHs: 24.29452\nExp number, first 60 AAs: 1.02778\nTotal prob of N-in: 0.64928\noutside 1 180\nTMhelix 181 203\ninside 204 212

1069 GCF_001528485.1_ASM152848v1 Burkholderia cepacia Proteobacteria; Betaproteobacteria; Burkholderiales; Burkholderiaceae; Burkholderia; Burkholderia cepacia complex
MPDRKRRALLTAGVIVVAAGLTGCTAMLFEDGRYEETVDRFLVSEDGKKFVVLGEKYHYI WP_060056353.1
hypothetical protein [Burkholderia cepacia] Length: 211\nNumber of predicted TMHs: 2\nExp number of AAs in TMHs: 45.99861\nExp number, first 60 AAs: 21.9561\nTotal prob of N-in: 0.94962\nPOSSIBLE N-term signal sequence\ninside 1 6\nTMhelix 7 29\noutside 30 186\nTMhelix 187 209\ninside 210 211

1070 GCF_001526785.1_ASM152678v1 Burkholderia cepacia Proteobacteria; Betaproteobacteria; Burkholderiales; Burkholderiaceae; Burkholderia; Burkholderia cepacia complex
MKRRALLTAGVIVVAAGLTGCTAMLFEDGRYEETVDRFLVSEDGKKLVVLGEKYHYIFDM WP_059861788.1
hypothetical protein [Burkholderia cepacia] Length: 208\nNumber of predicted TMHs: 2\nExp number of AAs in TMHs: 44.86644\nExp number, first 60 AAs: 21.12438\nTotal prob of N-in: 0.76640\nPOSSIBLE N-term signal sequence\ninside 1 6\nTMhelix 7 26\noutside 27 183\nTMhelix 184 206\ninside 207 208

1071 GCF_001526445.1_ASM152644v1 Burkholderia cepacia Proteobacteria; Betaproteobacteria; Burkholderiales; Burkholderiaceae; Burkholderia; Burkholderia cepacia complex
MKRRTFLALPAVAASGMLAGCSNFGPVTKAWMSDSKYDETLSTFLITADGKQLVVLGKRY WP_059526018.1
hypothetical protein [Burkholderia cepacia] Length: 210\nNumber of predicted TMHs: 1\nExp number of AAs in TMHs: 34.61771\nExp number, first 60 AAs: 12.90764\nTotal prob of N-in: 0.46822\nPOSSIBLE N-term signal sequence\noutside 1 186\nTMhelix 187 209\ninside 210 210

1072 GCF_001528845.1_ASM152884v1 Burkholderia cepacia Proteobacteria; Betaproteobacteria; Burkholderiales; Burkholderiaceae; Burkholderia; Burkholderia cepacia complex
MKRRTFLALPAVAASGMLAGCSNFGPVTKAWMSDSKYDETLSTFLITADGKQLVVLGKKY WP_027787377.1
MULTISPECIES: hypothetical protein [Burkholderia] Length: 210\nNumber of predicted TMHs: 1\nExp

number of AAs in TMHs: 34.58187\nExp number, first 60 AAs: 12.86351\nTotal prob of N-in: 0.46668\nPOSSIBLE N-term signal sequence\noutside 1 186\nTMhelix 187 209\ninside 210 210

1073 GCF_001525945.1_ASM152594v1 Burkholderia cepacia Proteobacteria; Betaproteobacteria; Burkholderiales; Burkholderiaceae; Burkholderia; Burkholderia cepacia complex MKRRALLTAGVIVVAAGLTGCTAMLFEDGRYEETVDRFLVSEDGKKFVVLGEKYHYIFDM WP_059496097.1 hypothetical protein [Burkholderia cepacia] Length: 208\nNumber of predicted TMHs: 2\nExp number of AAs in TMHs: 44.13865\nExp number, first 60 AAs: 21.11648\nTotal prob of N-in: 0.75846\nPOSSIBLE N-term signal sequence\ninside 1 6\nTMhelix 7 26\noutside 27 183\nTMhelix 184 206\ninside 207 208

1074 GCF_000757365.1_BHX_1 Burkholderia cepacia Proteobacteria; Betaproteobacteria; Burkholderiales; Burkholderiaceae; Burkholderia; Burkholderia cepacia complex MSDDIDNGGKGGFTRRDMMRVMAASGMMAVGGGGLLTSAQSAFAAPAPKRGGKIRVANES WP_006482802.1 MULTISPECIES: ABC transporter substrate-binding protein [Burkholderia cepacia complex] Length: 527\nNumber of predicted TMHs: 2\nExp number of AAs in TMHs: 36.7303599999999\nExp number, first 60 AAs: 22.08937\nTotal prob of N-in: 0.97312\nPOSSIBLE N-term signal sequence\ninside 1 20\nTMhelix 21 43\noutside 44 502\nTMhelix 503 525\ninside 526 527

1075 GCF_001522785.1_ASM152278v1 Burkholderia cepacia Proteobacteria; Betaproteobacteria; Burkholderiales; Burkholderiaceae; Burkholderia; Burkholderia cepacia complex MKTSITRRRLLAAAMAASTLALSGCFTPKLYENDAYSEYVTSFMITQDGKKLVVLGTNYH WP_059485162.1 hypothetical protein [Burkholderia cepacia] Length: 212\nNumber of predicted TMHs: 1\nExp number of AAs in TMHs: 24.29452\nExp number, first 60 AAs: 1.02778\nTotal prob of N-in: 0.64928\noutside 1 180\nTMhelix 181 203\ninside 204 212

1076 GCF_000755805.1_ASM75580v1 Burkholderia cepacia Proteobacteria; Betaproteobacteria; Burkholderiales; Burkholderiaceae; Burkholderia; Burkholderia cepacia complex MSDDIDNGGKGGFTRRDMMRVMAASGMMAVGGGGLLTSAQSAFAAPAPKRGGKIRVANES WP_006482802.1 MULTISPECIES: ABC transporter substrate-binding protein [Burkholderia cepacia complex] Length: 527\nNumber of predicted TMHs: 2\nExp number of AAs in TMHs: 36.7303599999999\nExp number, first 60 AAs: 22.08937\nTotal prob of N-in: 0.97312\nPOSSIBLE N-term signal sequence\ninside 1 20\nTMhelix 21 43\noutside 44 502\nTMhelix 503 525\ninside 526 527

1077 GCF_001530055.1_ASM153005v1 Burkholderia cepacia Proteobacteria; Betaproteobacteria; Burkholderiales; Burkholderiaceae; Burkholderia; Burkholderia cepacia complex MKRRTFLALPAVAASGMLAGCSNFGPVTKAWMSDSKYDETLSTFLITADGKQLVVLGKKY WP_059488505.1 hypothetical protein [Burkholderia cepacia] Length: 210\nNumber of predicted TMHs: 1\nExp number of AAs in TMHs: 34.5304\nExp number, first 60 AAs: 12.84424\nTotal prob of N-in: 0.46568\nPOSSIBLE N-term signal sequence\noutside 1 186\nTMhelix 187 209\ninside 210 210

1078 GCF_001530105.1_ASM153010v1 Burkholderia cepacia Proteobacteria; Betaproteobacteria; Burkholderiales; Burkholderiaceae; Burkholderia; Burkholderia cepacia complex MKRRTFLALPAVAASGMLAGCSNFGPVTKAWMSDSKYDETLSTFLITADGKQLVVLGKKY WP_059488505.1 hypothetical protein [Burkholderia cepacia] Length: 210\nNumber of predicted TMHs: 1\nExp number of AAs in TMHs: 34.5304\nExp number, first 60 AAs: 12.84424\nTotal prob of N-in: 0.46568\nPOSSIBLE N-term signal sequence\noutside 1 186\nTMhelix 187 209\ninside 210 210

1079 GCF_000724975.1_ASM72497v1 Burkholderia cepacia Proteobacteria; Betaproteobacteria; Burkholderiales; Burkholderiaceae; Burkholderia; Burkholderia cepacia complex MKRRAALLGLSLLASVAAADGTPGAGRGAQRFSLDLDGAAAYYQLTVPQPVYAASRRDDL WP_043180671.1 membrane protein [Burkholderia cepacia] Length: 452\nNumber of predicted TMHs: 1\nExp number of AAs in TMHs: 24.81887\nExp number, first 60 AAs: 3.37091\nTotal prob of N-in: 0.33853\noutside 1 418\nTMhelix 419 441\ninside 442 452

1080 GCF_000724975.1_ASM72497v1 Burkholderia cepacia Proteobacteria; Betaproteobacteria; Burkholderiales; Burkholderiaceae; Burkholderia; Burkholderia cepacia complex MKRRTLLTAGAIIVAAGLTGCTAMLFEDGRYEETVDRFLVSEDGKKFVVLGKKYHYIFDM WP_043182701.1 hypothetical protein [Burkholderia cepacia] Length: 208\nNumber of predicted TMHs: 2\nExp number of AAs in TMHs: 43.33739\nExp number, first 60 AAs: 20.04151\nTotal prob of N-in: 0.74735\nPOSSIBLE N-term signal sequence\ninside 1 6\nTMhelix 7 26\noutside 27 183\nTMhelix 184 206\ninside 207 208

1081 GCF_001039005.1_ASM103900v1 Burkholderia cepacia Proteobacteria; Betaproteobacteria; Burkholderiales; Burkholderiaceae; Burkholderia; Burkholderia cepacia complex MKRRTFLALPAVAASGMLAGCSDFGPVTNRMMKDPEYRETLSAFLISADGKKLVVLGKQY WP_048243193.1 hypothetical protein [Burkholderia cepacia] Length: 210\nNumber of predicted TMHs: 1\nExp number of AAs in TMHs: 29.86055\nExp number, first 60 AAs: 8.74784000000001\nTotal prob of N-in: 0.51411\noutside 1 186\nTMhelix 187 209\ninside 210 210

1082 GCF_001530805.1_ASM153080v1 Burkholderia cepacia Proteobacteria; Betaproteobacteria; Burkholderiales; Burkholderiaceae; Burkholderia; Burkholderia cepacia complex MKRRTFLALPAVAASGMLAGCSNFGPVTKAWMSDSKYDETLSTFLITADGKQLVVLGKKY WP_059488505.1 hypothetical protein [Burkholderia cepacia] Length: 210\nNumber of predicted TMHs: 1\nExp number of AAs in TMHs: 34.5304\nExp number, first 60 AAs: 12.84424\nTotal prob of N-in: 0.46568\nPOSSIBLE N-term signal sequence\noutside 1 186\nTMhelix 187 209\ninside 210 210

1083 GCF_001530105.1_ASM153010v1 Burkholderia cepacia Proteobacteria; Betaproteobacteria; Burkholderiales; Burkholderiaceae; Burkholderia; Burkholderia cepacia complex MKTSITRRRLLAAAMAASTLALSGCFTPKLYENDAYSEYVTSFMITQDGKKLVVLGTNYH WP_060050342.1 hypothetical protein [Burkholderia cepacia] Length: 212\nNumber of predicted TMHs: 1\nExp number of AAs in TMHs: 24.23033\nExp number, first 60 AAs: 1.03058\nTotal prob of N-in: 0.64819\noutside 1 180\nTMhelix 181 203\ninside 204 212

1084 GCF_001530805.1_ASM153080v1 Burkholderia cepacia Proteobacteria; Betaproteobacteria; Burkholderiales; Burkholderiaceae; Burkholderia; Burkholderia cepacia complex MKTSITRRRLLAAAMAASTLALSGCFTPKLYENDAYSEYVTSFMITQDGKKLVVLGTNYH WP_060050342.1 hypothetical protein [Burkholderia cepacia] Length: 212\nNumber of predicted TMHs: 1\nExp number of AAs in TMHs: 24.23033\nExp number, first 60 AAs: 1.03058\nTotal prob of N-in: 0.64819\noutside 1 180\nTMhelix 181 203\ninside 204 212

1085 GCF_001530055.1_ASM153005v1 Burkholderia cepacia Proteobacteria; Betaproteobacteria; Burkholderiales; Burkholderiaceae; Burkholderia; Burkholderia cepacia complex MKTSITRRRLLAAAMAASTLALSGCFTPKLYENDAYSEYVTSFMITQDGKKLVVLGTNYH WP_060050342.1 hypothetical protein [Burkholderia cepacia] Length: 212\nNumber of predicted TMHs: 1\nExp number of AAs in TMHs: 24.23033\nExp number, first 60 AAs: 1.03058\nTotal prob of N-in: 0.64819\noutside 1 180\nTMhelix 181 203\ninside 204 212

1086 GCF_001531875.1_ASM153187v1 Burkholderia cepacia Proteobacteria; Betaproteobacteria; Burkholderiales; Burkholderiaceae; Burkholderia; Burkholderia cepacia complex MKRRTFLALPAVAASGMLAGCSNFGPVTKAWMSDSKYDETLSTFLITADGKQLVVLGKKY WP_027787377.1 MULTISPECIES: hypothetical protein [Burkholderia] Length: 210\nNumber of predicted TMHs: 1\nExp number of AAs in TMHs: 34.58187\nExp number, first 60 AAs: 12.86351\nTotal prob of N-in: 0.46668\nPOSSIBLE N-term signal sequence\noutside 1 186\nTMhelix 187 209\ninside 210 210

1087 GCF_001531505.1_ASM153150v1 Burkholderia cepacia Proteobacteria; Betaproteobacteria; Burkholderiales; Burkholderiaceae; Burkholderia; Burkholderia cepacia complex MKRRTFLALPAVAASGMLAGCSNFGPVTKAWMSDSKYDETLSTFLITADGKQLVVLGKKY WP_059488505.1 hypothetical protein [Burkholderia cepacia] Length: 210\nNumber of predicted TMHs: 1\nExp number of AAs in TMHs: 34.5304\nExp number, first 60 AAs: 12.84424\nTotal prob of N-in: 0.46568\nPOSSIBLE N-term signal sequence\noutside 1 186\nTMhelix 187 209\ninside 210 210

1088 GCF_001531915.1_ASM153191v1 Burkholderia cepacia Proteobacteria; Betaproteobacteria; Burkholderiales; Burkholderiaceae; Burkholderia; Burkholderia cepacia complex MKRRALLTAGAIVVAAGLTGCTAMLFEDGRYEETVDRFLVSEDGKKFVVLGEKYHYIFDM WP_059700133.1 hypothetical protein [Burkholderia cepacia] Length: 208\nNumber of predicted TMHs: 2\nExp number of AAs in TMHs: 43.66178\nExp number, first 60 AAs: 20.68411\nTotal prob of N-in: 0.74282\nPOSSIBLE N-term signal sequence\noutside 1 4\nTMhelix 5 27\ninside 28 184\nTMhelix 185 207\noutside 208 208

1089 GCF_001531915.1_ASM153191v1 Burkholderia cepacia Proteobacteria; Betaproteobacteria; Burkholderiales; Burkholderiaceae; Burkholderia; Burkholderia cepacia complex MKRRTFLALPAVAASGMLAGCSNFGPVTKAWMSDSKYDETLSTFLITADGKQLVVLGKKY WP_059700302.1 hypothetical protein [Burkholderia cepacia] Length: 210\nNumber of predicted TMHs: 1\nExp number of

AAs in TMHs: 34.58208\nExp number, first 60 AAs: 12.86353\nTotal prob of N-in: 0.46668\nPOSSIBLE N-term signal sequence\noutside 1 186\nTMhelix 187 209\ninside 210 210

1090 GCF_001533295.1_ASM153329v1 Burkholderia cepacia Proteobacteria; Betaproteobacteria; Burkholderiales; Burkholderiaceae; Burkholderia; Burkholderia cepacia complex MKRRTFLALPAVAASGMLAGCSNFGPVTKAWMSDSKYDETLSTFLITADGKQLVVLGKKY WP_027787377.1 MULTISPECIES: hypothetical protein [Burkholderia] Length: 210\nNumber of predicted TMHs: 1\nExp number of AAs in TMHs: 34.58187\nExp number, first 60 AAs: 12.86351\nTotal prob of N-in: 0.46668\nPOSSIBLE N-term signal sequence\noutside 1 186\nTMhelix 187 209\ninside 210 210

1091 GCF_001525925.1_ASM152592v1 Burkholderia cepacia Proteobacteria; Betaproteobacteria; Burkholderiales; Burkholderiaceae; Burkholderia; Burkholderia cepacia complex MKRRALLTAGVIVVAAGLTGCTAMLFEDGRYEETVDRFLVSEDGKKFVVLGEKYHYIFDM WP_059496097.1 hypothetical protein [Burkholderia cepacia] Length: 208\nNumber of predicted TMHs: 2\nExp number of AAs in TMHs: 44.13865\nExp number, first 60 AAs: 21.11648\nTotal prob of N-in: 0.75846\nPOSSIBLE N-term signal sequence\ninside 1 6\nTMhelix 7 26\noutside 27 183\nTMhelix 184 206\ninside 207 208

1092 GCF_000756845.1_BHW_1 Burkholderia cepacia Proteobacteria; Betaproteobacteria; Burkholderiales; Burkholderiaceae; Burkholderia; Burkholderia cepacia complex MKRRAALLGLSLLASFAAADGAPNAGRAMQRFALDLDGSAAYYQLTVPQPVYAASRRDDL WP_034204171.1 MULTISPECIES: membrane protein [Burkholderia cepacia complex] Length: 452\nNumber of predicted TMHs: 1\nExp number of AAs in TMHs: 26.99094\nExp number, first 60 AAs: 5.49432\nTotal prob of N-in: 0.35405\noutside 1 418\nTMhelix 419 441\ninside 442 452

1093 GCF_000877315.1_BIG Burkholderia cepacia Proteobacteria; Betaproteobacteria; Burkholderiales; Burkholderiaceae; Burkholderia; Burkholderia cepacia complex MSDDIDNGGKGGFTRRDMMRVMAASGMMAVGGGGLLTSAQSAFAAPAPKRGGKIRVANES WP_006482802.1 MULTISPECIES: ABC transporter substrate-binding protein [Burkholderia cepacia complex] Length: 527\nNumber of predicted TMHs: 2\nExp number of AAs in TMHs: 36.7303599999999\nExp number, first 60 AAs: 22.08937\nTotal prob of N-in: 0.97312\nPOSSIBLE N-term signal sequence\ninside 1 20\nTMhelix 21 43\noutside 44 502\nTMhelix 503 525\ninside 526 527

1094 GCF_001052915.1_ASM105291v1 Burkholderia cepacia Proteobacteria; Betaproteobacteria; Burkholderiales; Burkholderiaceae; Burkholderia; Burkholderia cepacia complex MKRRTFLALPAVAASGMLAGCSNFGPVTKAWMSDSKYDETLSTFLITADGKQLVVLGKKY WP_027787377.1 MULTISPECIES: hypothetical protein [Burkholderia] Length: 210\nNumber of predicted TMHs: 1\nExp number of AAs in TMHs: 34.58187\nExp number, first 60 AAs: 12.86351\nTotal prob of N-in: 0.46668\nPOSSIBLE N-term signal sequence\noutside 1 186\nTMhelix 187 209\ninside 210 210

1095 GCF_001052915.1_ASM105291v1 Burkholderia cepacia Proteobacteria; Betaproteobacteria; Burkholderiales; Burkholderiaceae; Burkholderia; Burkholderia cepacia complex MKRRALLTAGAIVAAGLTGCTAMLFEDGRYEETVDRFLVSEDGKKFVVLGEKYHYIFDMP WP_027787499.1 hypothetical protein [Burkholderia cepacia] Length: 207\nNumber of predicted TMHs: 1\nExp number of AAs in TMHs: 38.38341\nExp number, first 60 AAs: 15.45171\nTotal prob of N-in: 0.76513\nPOSSIBLE N-term signal sequence\ninside 1 183\nTMhelix 184 206\noutside 207 207

1096 GCF_000877315.1_BIG Burkholderia cepacia Proteobacteria; Betaproteobacteria; Burkholderiales; Burkholderiaceae; Burkholderia; Burkholderia cepacia complex MKRRTFLALPAVAASGMLAGCSSADLGPVTHRLMKDPEYRETLSAFLISADGKKLVVIGKWP_006483583.1 MULTISPECIES: hypothetical protein [Burkholderia cepacia complex] Length: 212\nNumber of predicted TMHs: 1\nExp number of AAs in TMHs: 28.03134\nExp number, first 60 AAs: 6.30523\nTotal prob of N-in: 0.48947\noutside 1 187\nTMhelix 188 210\ninside 211 212

1097 GCF_001522215.1_ASM152221v1 Burkholderia cepacia Proteobacteria; Betaproteobacteria; Burkholderiales; Burkholderiaceae; Burkholderia; Burkholderia cepacia complex MKRRTFLALPAVAASGMLAGCSNFGPVTKAWMSDSKYDETLSTFLITADGKQLVVLGKKY WP_027787377.1 MULTISPECIES: hypothetical protein [Burkholderia] Length: 210\nNumber of predicted TMHs: 1\nExp number of AAs in TMHs: 34.58187\nExp number, first 60 AAs: 12.86351\nTotal prob of N-in: 0.46668\nPOSSIBLE N-term signal sequence\noutside 1 186\nTMhelix 187 209\ninside 210 210

1098 GCF_001522215.1_ASM152221v1 Burkholderia cepacia Proteobacteria; Betaproteobacteria; Burkholderiales; Burkholderiaceae; Burkholderia; Burkholderia cepacia complex MKRRALLTAGVIVVAAGLTGCTAMLFEDGRYEETVDRFLVSEDGKKFVVLGEKYHYIFDM WP_059677781.1 hypothetical protein [Burkholderia cepacia] Length: 208\nNumber of predicted TMHs: 2\nExp number of AAs in TMHs: 44.88033\nExp number, first 60 AAs: 21.13087\nTotal prob of N-in: 0.76656\nPOSSIBLE N-term signal sequence\ninside 1 6\nTMhelix 7 26\noutside 27 183\nTMhelix 184 206\ninside 207 208

1099 GCF_001522305.1_ASM152230v1 Burkholderia cepacia Proteobacteria; Betaproteobacteria; Burkholderiales; Burkholderiaceae; Burkholderia; Burkholderia cepacia complex MKTSITRRRLLAAAMAASTLALSGCFTPKLYENDAYSEYVTSFMITQDGKKLVVLGTNYH WP_059485162.1 hypothetical protein [Burkholderia cepacia] Length: 212\nNumber of predicted TMHs: 1\nExp number of AAs in TMHs: 24.29452\nExp number, first 60 AAs: 1.02778\nTotal prob of N-in: 0.64928\noutside 1 180\nTMhelix 181 203\ninside 204 212

1100 GCF_001522305.1_ASM152230v1 Burkholderia cepacia Proteobacteria; Betaproteobacteria; Burkholderiales; Burkholderiaceae; Burkholderia; Burkholderia cepacia complex MKRRTFLALPAVAASGMLAGCSNFGPVTKAWMSDSKYDETLSTFLITADGKQLVVLGKKY WP_059488505.1 hypothetical protein [Burkholderia cepacia] Length: 210\nNumber of predicted TMHs: 1\nExp number of AAs in TMHs: 34.5304\nExp number, first 60 AAs: 12.84424\nTotal prob of N-in: 0.46568\nPOSSIBLE N-term signal sequence\noutside 1 186\nTMhelix 187 209\ninside 210 210

1101 GCF_001522525.1_ASM152252v1 Burkholderia cepacia Proteobacteria; Betaproteobacteria; Burkholderiales; Burkholderiaceae; Burkholderia; Burkholderia cepacia complex MKRRTFLALPAVAASGMLAGCSNFGPVTKAWMSDSKYDETLSTFLITADGKQLVVLGKKY WP_059488505.1 hypothetical protein [Burkholderia cepacia] Length: 210\nNumber of predicted TMHs: 1\nExp number of AAs in TMHs: 34.5304\nExp number, first 60 AAs: 12.84424\nTotal prob of N-in: 0.46568\nPOSSIBLE N-term signal sequence\noutside 1 186\nTMhelix 187 209\ninside 210 210

1102 GCF_001522695.1_ASM152269v1 Burkholderia cepacia Proteobacteria; Betaproteobacteria; Burkholderiales; Burkholderiaceae; Burkholderia; Burkholderia cepacia complex MKTSITRRRLLAAAMAASTLALSGCFTPKLYENDAYSEYVTSFMITQDGKKLVVLGTNYH WP_059485162.1 hypothetical protein [Burkholderia cepacia] Length: 212\nNumber of predicted TMHs: 1\nExp number of AAs in TMHs: 24.29452\nExp number, first 60 AAs: 1.02778\nTotal prob of N-in: 0.64928\noutside 1 180\nTMhelix 181 203\ninside 204 212

1103 GCF_001522695.1_ASM152269v1 Burkholderia cepacia Proteobacteria; Betaproteobacteria; Burkholderiales; Burkholderiaceae; Burkholderia; Burkholderia cepacia complex MKRRTFLALPAVAASGMLAGCSNFGPVTKAWMSDSKYDETLSTFLITADGKQLVVLGKKY WP_059488505.1 hypothetical protein [Burkholderia cepacia] Length: 210\nNumber of predicted TMHs: 1\nExp number of AAs in TMHs: 34.5304\nExp number, first 60 AAs: 12.84424\nTotal prob of N-in: 0.46568\nPOSSIBLE N-term signal sequence\noutside 1 186\nTMhelix 187 209\ninside 210 210

1104 GCF_001522695.1_ASM152269v1 Burkholderia cepacia Proteobacteria; Betaproteobacteria; Burkholderiales; Burkholderiaceae; Burkholderia; Burkholderia cepacia complex MKRRALLTAGVIVVAAGLTGCTAMLFEDGRYEETVDRFLVSEDGKKFVVLGEKYHYIFDM WP_059496097.1 hypothetical protein [Burkholderia cepacia] Length: 208\nNumber of predicted TMHs: 2\nExp number of AAs in TMHs: 44.13865\nExp number, first 60 AAs: 21.11648\nTotal prob of N-in: 0.75846\nPOSSIBLE N-term signal sequence\ninside 1 6\nTMhelix 7 26\noutside 27 183\nTMhelix 184 206\ninside 207 208

1105 GCF_000974835.2_ASM97483v2 Burkholderia cepacia Proteobacteria; Betaproteobacteria; Burkholderiales; Burkholderiaceae; Burkholderia; Burkholderia cepacia complex MSDDIDNGGGHGGFTRRDMMRVMAAGGLMAAGSGALLMPAQSAFAAPAPKRGGKIRVANE WP_035975758.1 MULTISPECIES: ABC transporter substrate-binding protein [Burkholderia cepacia complex] Length: 528\nNumber of predicted TMHs: 2\nExp number of AAs in TMHs: 35.70904\nExp number, first 60 AAs: 19.87611\nTotal prob of N-in: 0.85006\nPOSSIBLE N-term signal sequence\ninside 1 23\nTMhelix 24 46\noutside 47 503\nTMhelix 504 526\ninside 527 528

1106 GCF_001522525.1_ASM152252v1 Burkholderia cepacia Proteobacteria; Betaproteobacteria; Burkholderiales; Burkholderiaceae; Burkholderia; Burkholderia cepacia complex MKRRALLTAGVIVVAAGLTGCTAMLFEDGRYEETVDRFLVSEDGKKFVVLGEKYHYIFDM WP_059496097.1 hypothetical protein [Burkholderia cepacia] Length: 208\nNumber of predicted TMHs: 2\nExp number of

AAs in TMHs: 44.13865\nExp number, first 60 AAs: 21.11648\nTotal prob of N-in: 0.75846\nPOSSIBLE N-term signal sequence\ninside 1 6\nTMhelix 7 26\noutside 27 183\nTMhelix 184 206\ninside 207 208

1107 GCF_001522805.1_ASM152280v1 Burkholderia cepacia Proteobacteria; Betaproteobacteria; Burkholderiales; Burkholderiaceae; Burkholderia; Burkholderia cepacia complex MKTSITRRRLLAAAMAASTLALSGCFTPKLYENDAYSEYVTSFMITQDGKKLVVLGTNYH WP_059485162.1 hypothetical protein [Burkholderia cepacia] Length: 212\nNumber of predicted TMHs: 1\nExp number of AAs in TMHs: 24.29452\nExp number, first 60 AAs: 1.02778\nTotal prob of N-in: 0.64928\noutside 1 180\nTMhelix 181 203\ninside 204 212

1108 GCF_001522805.1_ASM152280v1 Burkholderia cepacia Proteobacteria; Betaproteobacteria; Burkholderiales; Burkholderiaceae; Burkholderia; Burkholderia cepacia complex MKRRTFLALPAVAASGMLAGCSNFGPVTKAWMSDSKYDETLSTFLITADGKQLVVLGKKY WP_059488505.1 hypothetical protein [Burkholderia cepacia] Length: 210\nNumber of predicted TMHs: 1\nExp number of AAs in TMHs: 34.5304\nExp number, first 60 AAs: 12.84424\nTotal prob of N-in: 0.46568\nPOSSIBLE N-term signal sequence\noutside 1 186\nTMhelix 187 209\ninside 210 210

1109 GCF_001522875.1_ASM152287v1 Burkholderia cepacia Proteobacteria; Betaproteobacteria; Burkholderiales; Burkholderiaceae; Burkholderia; Burkholderia cepacia complex MKTSITRRRLLAAAMAASTLALSGCFTPKLYENDAYSEYVTSFMITQDGKKLVVLGTNYH WP_059485162.1 hypothetical protein [Burkholderia cepacia] Length: 212\nNumber of predicted TMHs: 1\nExp number of AAs in TMHs: 24.29452\nExp number, first 60 AAs: 1.02778\nTotal prob of N-in: 0.64928\noutside 1 180\nTMhelix 181 203\ninside 204 212

1110 GCF_001522875.1_ASM152287v1 Burkholderia cepacia Proteobacteria; Betaproteobacteria; Burkholderiales; Burkholderiaceae; Burkholderia; Burkholderia cepacia complex MKRRTFLALPAVAASGMLAGCSNFGPVTKAWMSDSKYDETLSTFLITADGKQLVVLGKKY WP_059488505.1 hypothetical protein [Burkholderia cepacia] Length: 210\nNumber of predicted TMHs: 1\nExp number of AAs in TMHs: 34.5304\nExp number, first 60 AAs: 12.84424\nTotal prob of N-in: 0.46568\nPOSSIBLE N-term signal sequence\noutside 1 186\nTMhelix 187 209\ninside 210 210

1111 GCF_001522805.1_ASM152280v1 Burkholderia cepacia Proteobacteria; Betaproteobacteria; Burkholderiales; Burkholderiaceae; Burkholderia; Burkholderia cepacia complex MKRRALLTAGVIVVAAGLTGCTAMLFEDGRYEETVDRFLVSEDGKKFVVLGEKYHYIFDM WP_059496097.1 hypothetical protein [Burkholderia cepacia] Length: 208\nNumber of predicted TMHs: 2\nExp number of AAs in TMHs: 44.13865\nExp number, first 60 AAs: 21.11648\nTotal prob of N-in: 0.75846\nPOSSIBLE N-term signal sequence\ninside 1 6\nTMhelix 7 26\noutside 27 183\nTMhelix 184 206\ninside 207 208

1112 GCF_001522875.1_ASM152287v1 Burkholderia cepacia Proteobacteria; Betaproteobacteria; Burkholderiales; Burkholderiaceae; Burkholderia; Burkholderia cepacia complex MKRRALLTAGVIVVAAGLTGCTAMLFEDGRYEETVDRFLVSEDGKKFVVLGEKYHYIFDM WP_059496097.1 hypothetical protein [Burkholderia cepacia] Length: 208\nNumber of predicted TMHs: 2\nExp number of AAs in TMHs: 44.13865\nExp number, first 60 AAs: 21.11648\nTotal prob of N-in: 0.75846\nPOSSIBLE N-term signal sequence\ninside 1 6\nTMhelix 7 26\noutside 27 183\nTMhelix 184 206\ninside 207 208

1113 GCF_001522305.1_ASM152230v1 Burkholderia cepacia Proteobacteria; Betaproteobacteria; Burkholderiales; Burkholderiaceae; Burkholderia; Burkholderia cepacia complex MKRRALLTAGVIVVAAGLTGCTAMLFEDGRYEETVDRFLVSEDGKKFVVLGEKYHYIFDM WP_059496097.1 hypothetical protein [Burkholderia cepacia] Length: 208\nNumber of predicted TMHs: 2\nExp number of AAs in TMHs: 44.13865\nExp number, first 60 AAs: 21.11648\nTotal prob of N-in: 0.75846\nPOSSIBLE N-term signal sequence\ninside 1 6\nTMhelix 7 26\noutside 27 183\nTMhelix 184 206\ninside 207 208

1114 GCF_001522945.1_ASM152294v1 Burkholderia cepacia Proteobacteria; Betaproteobacteria; Burkholderiales; Burkholderiaceae; Burkholderia; Burkholderia cepacia complex MKTSITRRRLLAAAMAASTLALSGCFTPKLYENDAYSEYVTSFMITQDGKKLVVLGTNYH WP_059485162.1 hypothetical protein [Burkholderia cepacia] Length: 212\nNumber of predicted TMHs: 1\nExp number of AAs in TMHs: 24.29452\nExp number, first 60 AAs: 1.02778\nTotal prob of N-in: 0.64928\noutside 1 180\nTMhelix 181 203\ninside 204 212

1115 GCF_001522985.1_ASM152298v1 Burkholderia cepacia Proteobacteria; Betaproteobacteria; Burkholderiales; Burkholderiaceae; Burkholderia; Burkholderia cepacia complex

MKTSITRRRLLAAAMAASTLALSGCFTPKLYENDAYSEYVTSFMITQDGKKLVVLGTNYH WP_059485162.1 hypothetical protein [Burkholderia cepacia] Length: 212\nNumber of predicted TMHs: 1\nExp number of AAs in TMHs: 24.29452\nExp number, first 60 AAs: 1.02778\nTotal prob of N-in: 0.64928\noutside 1 180\nTMhelix 181 203\ninside 204 212

1116 GCF_001522945.1_ASM152294v1 Burkholderia cepacia Proteobacteria; Betaproteobacteria; Burkholderiales; Burkholderiaceae; Burkholderia; Burkholderia cepacia complex MKRRTFLALPAVAASGMLAGCSNFGPVTKAWMSDSKYDETLSTFLITADGKQLVVLGKKY WP_059488505.1 hypothetical protein [Burkholderia cepacia] Length: 210\nNumber of predicted TMHs: 1\nExp number of AAs in TMHs: 34.5304\nExp number, first 60 AAs: 12.84424\nTotal prob of N-in: 0.46568\nPOSSIBLE N-term signal sequence\noutside 1 186\nTMhelix 187 209\ninside 210 210

1117 GCF_001522945.1_ASM152294v1 Burkholderia cepacia Proteobacteria; Betaproteobacteria; Burkholderiales; Burkholderiaceae; Burkholderia; Burkholderia cepacia complex MKRRALLTAGVIVVAAGLTGCTAMLFEDGRYEETVDRFLVSEDGKKFVVLGEKYHYIFDM WP_059496097.1 hypothetical protein [Burkholderia cepacia] Length: 208\nNumber of predicted TMHs: 2\nExp number of AAs in TMHs: 44.13865\nExp number, first 60 AAs: 21.11648\nTotal prob of N-in: 0.75846\nPOSSIBLE N-term signal sequence\ninside 1 6\nTMhelix 7 26\noutside 27 183\nTMhelix 184 206\ninside 207 208

1118 GCF_001525775.1_ASM152577v1 Burkholderia cepacia Proteobacteria; Betaproteobacteria; Burkholderiales; Burkholderiaceae; Burkholderia; Burkholderia cepacia complex MKRRTFLALPAVAASGMLAGCSNFGPVTKAWMSDSKYDETLSTFLITADGKQLVVLGKKY WP_059488505.1 hypothetical protein [Burkholderia cepacia] Length: 210\nNumber of predicted TMHs: 1\nExp number of AAs in TMHs: 34.5304\nExp number, first 60 AAs: 12.84424\nTotal prob of N-in: 0.46568\nPOSSIBLE N-term signal sequence\noutside 1 186\nTMhelix 187 209\ninside 210 210

1119 GCF_001526755.1_ASM152675v1 Burkholderia cepacia Proteobacteria; Betaproteobacteria; Burkholderiales; Burkholderiaceae; Burkholderia; Burkholderia cepacia complex MKRRTFLALPAVAASGMLAGCSNFGPVTKAWMSDSKYDETLSTFLITADGKQLVVLGKKY WP_059488505.1 hypothetical protein [Burkholderia cepacia] Length: 210\nNumber of predicted TMHs: 1\nExp number of AAs in TMHs: 34.5304\nExp number, first 60 AAs: 12.84424\nTotal prob of N-in: 0.46568\nPOSSIBLE N-term signal sequence\noutside 1 186\nTMhelix 187 209\ninside 210 210

1120 GCF_001530665.1_ASM153066v1 Burkholderia cepacia Proteobacteria; Betaproteobacteria; Burkholderiales; Burkholderiaceae; Burkholderia; Burkholderia cepacia complex MKTSITRRRLLAAGMAASTLALSGCFTPKLYENDAYSEYVTSFMITQDGKKLVVLGTNYH WP_060086226.1 hypothetical protein [Burkholderia cepacia] Length: 212\nNumber of predicted TMHs: 1\nExp number of AAs in TMHs: 23.88551\nExp number, first 60 AAs: 0.71364\nTotal prob of N-in: 0.64707\noutside 1 180\nTMhelix 181 203\ninside 204 212

1121 GCF_001525775.1_ASM152577v1 Burkholderia cepacia Proteobacteria; Betaproteobacteria; Burkholderiales; Burkholderiaceae; Burkholderia; Burkholderia cepacia complex MKRRALLTAGVIVVAAGLTGCTAMLFEDGRYEETVDRFLVSEDGKKFVVLGEKYHYIFDM WP_059496097.1 hypothetical protein [Burkholderia cepacia] Length: 208\nNumber of predicted TMHs: 2\nExp number of AAs in TMHs: 44.13865\nExp number, first 60 AAs: 21.11648\nTotal prob of N-in: 0.75846\nPOSSIBLE N-term signal sequence\ninside 1 6\nTMhelix 7 26\noutside 27 183\nTMhelix 184 206\ninside 207 208

1122 GCF_001526755.1_ASM152675v1 Burkholderia cepacia Proteobacteria; Betaproteobacteria; Burkholderiales; Burkholderiaceae; Burkholderia; Burkholderia cepacia complex MKRRALLTAGVIVVAAGLTGCTAMLFEDGRYEETVDRFLVSEDGKKFVVLGEKYHYIFDM WP_059858336.1 hypothetical protein [Burkholderia cepacia] Length: 208\nNumber of predicted TMHs: 2\nExp number of AAs in TMHs: 44.87971\nExp number, first 60 AAs: 21.13087\nTotal prob of N-in: 0.76655\nPOSSIBLE N-term signal sequence\ninside 1 6\nTMhelix 7 26\noutside 27 183\nTMhelix 184 206\ninside 207 208

1123 GCF_001526255.1_ASM152625v1 Burkholderia cepacia Proteobacteria; Betaproteobacteria; Burkholderiales; Burkholderiaceae; Burkholderia; Burkholderia cepacia complex MKASITRRRLLAAGVAASSLALSGCFTPKLYKDDGYHEYVSGFMITQDGKKLVVLGTDYH WP_059525490.1 hypothetical protein [Burkholderia cepacia] Length: 212\nNumber of predicted TMHs: 1\nExp number of AAs in TMHs: 25.1207\nExp number, first 60 AAs: 0.80309\nTotal prob of N-in: 0.62748\noutside 1 180\nTMhelix 181 203\ninside 204 212

1124 GCF_001526255.1_ASM152625v1 Burkholderia cepacia Proteobacteria; Betaproteobacteria; Burkholderiales; Burkholderiaceae; Burkholderia; Burkholderia cepacia complex MKRRTFLALPAVAASGMLAGCSNFGPVTKAWMSDSKYDETLSTFLITADGKQLVVLGKRY WP_059526018.1 hypothetical protein [Burkholderia cepacia] Length: 210\nNumber of predicted TMHs: 1\nExp number of AAs in TMHs: 34.61771\nExp number, first 60 AAs: 12.90764\nTotal prob of N-in: 0.46822\nPOSSIBLE N-term signal sequence\noutside 1 186\nTMhelix 187 209\ninside 210 210

1125 GCF_001526905.1_ASM152690v1 Burkholderia cepacia Proteobacteria; Betaproteobacteria; Burkholderiales; Burkholderiaceae; Burkholderia; Burkholderia cepacia complex MKRRTFLALPAVAASGMLAGCSNFGPVTKAWMSDSKYDETLSTFLITADGKQLVVLGKKY WP_059488505.1 hypothetical protein [Burkholderia cepacia] Length: 210\nNumber of predicted TMHs: 1\nExp number of AAs in TMHs: 34.5304\nExp number, first 60 AAs: 12.84424\nTotal prob of N-in: 0.46568\nPOSSIBLE N-term signal sequence\noutside 1 186\nTMhelix 187 209\ninside 210 210

1126 GCF_001524665.1_ASM152466v1 Burkholderia cepacia Proteobacteria; Betaproteobacteria; Burkholderiales; Burkholderiaceae; Burkholderia; Burkholderia cepacia complex MKASITRRRLLAAGVAASSLALSGCFTPKLYKDDGYHEYVSGFMITQDGKKLVVLGTDYH WP_059525490.1 hypothetical protein [Burkholderia cepacia] Length: 212\nNumber of predicted TMHs: 1\nExp number of AAs in TMHs: 25.1207\nExp number, first 60 AAs: 0.80309\nTotal prob of N-in: 0.62748\noutside 1 180\nTMhelix 181 203\ninside 204 212

1127 GCF_001524665.1_ASM152466v1 Burkholderia cepacia Proteobacteria; Betaproteobacteria; Burkholderiales; Burkholderiaceae; Burkholderia; Burkholderia cepacia complex MKRRTFLALPAVAASGMLAGCSNFGPVTKAWMSDSKYDETLSTFLITADGKQLVVLGKRY WP_059526018.1 hypothetical protein [Burkholderia cepacia] Length: 210\nNumber of predicted TMHs: 1\nExp number of AAs in TMHs: 34.61771\nExp number, first 60 AAs: 12.90764\nTotal prob of N-in: 0.46822\nPOSSIBLE N-term signal sequence\noutside 1 186\nTMhelix 187 209\ninside 210 210

1128 GCF_001522985.1_ASM152298v1 Burkholderia cepacia Proteobacteria; Betaproteobacteria; Burkholderiales; Burkholderiaceae; Burkholderia; Burkholderia cepacia complex MKRRALLTAGVIVVAAGLTGCTAMLFEDGRYEETVDRFLVSEDGKKFVVLGEKYHYIFDM WP_059496097.1 hypothetical protein [Burkholderia cepacia] Length: 208\nNumber of predicted TMHs: 2\nExp number of AAs in TMHs: 44.13865\nExp number, first 60 AAs: 21.11648\nTotal prob of N-in: 0.75846\nPOSSIBLE N-term signal sequence\ninside 1 6\nTMhelix 7 26\noutside 27 183\nTMhelix 184 206\ninside 207 208

1129 GCF_001528675.1_ASM152867v1 Burkholderia cepacia Proteobacteria; Betaproteobacteria; Burkholderiales; Burkholderiaceae; Burkholderia; Burkholderia cepacia complex MKRRTFLALPAVAASGMLAGCSNFGPVTKAWMSDSKYDETLSTFLITADGKQLVVLGKKY WP_027787377.1 MULTISPECIES: hypothetical protein [Burkholderia] Length: 210\nNumber of predicted TMHs: 1\nExp number of AAs in TMHs: 34.58187\nExp number, first 60 AAs: 12.86351\nTotal prob of N-in: 0.46668\nPOSSIBLE N-term signal sequence\noutside 1 186\nTMhelix 187 209\ninside 210 210

1130 GCF_001528675.1_ASM152867v1 Burkholderia cepacia Proteobacteria; Betaproteobacteria; Burkholderiales; Burkholderiaceae; Burkholderia; Burkholderia cepacia complex MKRRALLTAGAIVVAAGLTGCTAMLFEDGRYEETVDRFLISEDGKKFVVLGEKYHYIFDM WP_060062973.1 hypothetical protein [Burkholderia cepacia] Length: 208\nNumber of predicted TMHs: 2\nExp number of AAs in TMHs: 44.02748\nExp number, first 60 AAs: 20.69721\nTotal prob of N-in: 0.74685\nPOSSIBLE N-term signal sequence\noutside 1 4\nTMhelix 5 27\ninside 28 184\nTMhelix 185 207\noutside 208 208

1131 GCF_001524835.1_ASM152483v1 Burkholderia cepacia Proteobacteria; Betaproteobacteria; Burkholderiales; Burkholderiaceae; Burkholderia; Burkholderia cepacia complex MKRRTFIALPAVAASGMLAGCSNFGPVTKAWMSDSKYDETLSTFLITADGKQLVVLGKKY WP_059653140.1 hypothetical protein [Burkholderia cepacia] Length: 210\nNumber of predicted TMHs: 1\nExp number of AAs in TMHs: 35.71481\nExp number, first 60 AAs: 14.07468\nTotal prob of N-in: 0.47915\nPOSSIBLE N-term signal sequence\noutside 1 186\nTMhelix 187 209\ninside 210 210

1132 GCF_001526345.1_ASM152634v1 Burkholderia cepacia Proteobacteria; Betaproteobacteria; Burkholderiales; Burkholderiaceae; Burkholderia; Burkholderia cepacia complex MKRRALLTAGVIVVAAGLTGCTAMLFEDGRYEETVDRFLVSEDGKKLVVLGEKYHYIFDM WP_059861788.1 hypothetical protein [Burkholderia cepacia] Length: 208\nNumber of predicted TMHs: 2\nExp number of

AAs in TMHs: 44.86644\nExp number, first 60 AAs:  21.12438\nTotal prob of N-in:   0.76640\nPOSSIBLE N-term signal sequence\ninside   1   6\nTMhelix   7   26\noutside   27   183\nTMhelix   184   206\ninside   207   208

1133 GCF_001530665.1_ASM153066v1 Burkholderia cepacia Proteobacteria; Betaproteobacteria; Burkholderiales; Burkholderiaceae; Burkholderia; Burkholderia cepacia complex
MKRRALLTAGAIVVAAGLTGCTAMLFEDGRYEETVDRFLVSEDGKKFVVLGEKYHYIFDM WP_059700133.1
hypothetical protein [Burkholderia cepacia] Length: 208\nNumber of predicted TMHs:  2\nExp number of AAs in TMHs: 43.66178\nExp number, first 60 AAs:  20.68411\nTotal prob of N-in:   0.74282\nPOSSIBLE N-term signal sequence\noutside   1   4\nTMhelix   5   27\ninside   28   184\nTMhelix   185   207\noutside   208   208

1134 GCF_001530665.1_ASM153066v1 Burkholderia cepacia Proteobacteria; Betaproteobacteria; Burkholderiales; Burkholderiaceae; Burkholderia; Burkholderia cepacia complex
MKRRTFLALPAVAASGMLAGCSNFGPVTKAWMSDSKYDETLSTFLITADGKQLVVLGKKY WP_027787377.1
MULTISPECIES: hypothetical protein [Burkholderia] Length: 210\nNumber of predicted TMHs:  1\nExp number of AAs in TMHs: 34.58187\nExp number, first 60 AAs:  12.86351\nTotal prob of N-in:   0.46668\nPOSSIBLE N-term signal sequence\noutside   1   186\nTMhelix   187   209\ninside   210   210

1135 GCF_001529685.1_ASM152968v1 Burkholderia cepacia Proteobacteria; Betaproteobacteria; Burkholderiales; Burkholderiaceae; Burkholderia; Burkholderia cepacia complex
MKRRTFLALPAVAASGMLAGCSNFGPVTKAWMSDSKYDETLSTFLITADGKQLVVLGKKY WP_059488505.1
hypothetical protein [Burkholderia cepacia] Length: 210\nNumber of predicted TMHs:  1\nExp number of AAs in TMHs: 34.5304\nExp number, first 60 AAs:  12.84424\nTotal prob of N-in:   0.46568\nPOSSIBLE N-term signal sequence\noutside   1   186\nTMhelix   187   209\ninside   210   210

1136 GCF_001529145.1_ASM152914v1 Burkholderia cepacia Proteobacteria; Betaproteobacteria; Burkholderiales; Burkholderiaceae; Burkholderia; Burkholderia cepacia complex
MKRRALLTAGAIVVAAGLTGCTAMLFEDGRYEETVDRFLVSEDGKKFVVLGEKYHYIFDM WP_059700133.1
hypothetical protein [Burkholderia cepacia] Length: 208\nNumber of predicted TMHs:  2\nExp number of AAs in TMHs: 43.66178\nExp number, first 60 AAs:  20.68411\nTotal prob of N-in:   0.74282\nPOSSIBLE N-term signal sequence\noutside   1   4\nTMhelix   5   27\ninside   28   184\nTMhelix   185   207\noutside   208   208

1137 GCF_001525965.1_ASM152596v1 Burkholderia cepacia Proteobacteria; Betaproteobacteria; Burkholderiales; Burkholderiaceae; Burkholderia; Burkholderia cepacia complex
MKRRALLTAGVIVVAAGLTGCTAMLFEDGRYEETVDRFLVSEDGKKFVVLGEKYHYIFDM WP_059496097.1
hypothetical protein [Burkholderia cepacia] Length: 208\nNumber of predicted TMHs:  2\nExp number of AAs in TMHs: 44.13865\nExp number, first 60 AAs:  21.11648\nTotal prob of N-in:   0.75846\nPOSSIBLE N-term signal sequence\ninside   1   6\nTMhelix   7   26\noutside   27   183\nTMhelix   184   206\ninside   207   208

1138 GCF_001532885.1_ASM153288v1 Burkholderia cepacia Proteobacteria; Betaproteobacteria; Burkholderiales; Burkholderiaceae; Burkholderia; Burkholderia cepacia complex
MKRRTFLALPAVAASGMLAGCSNFGPVTKAWMSDSKYDETLSTFLITADGKQLVVLGKKY WP_060343570.1
hypothetical protein [Burkholderia cepacia] Length: 210\nNumber of predicted TMHs:  1\nExp number of AAs in TMHs: 34.53037\nExp number, first 60 AAs:  12.84423\nTotal prob of N-in:   0.46568\nPOSSIBLE N-term signal sequence\noutside   1   186\nTMhelix   187   209\ninside   210   210

1139 GCF_001532885.1_ASM153288v1 Burkholderia cepacia Proteobacteria; Betaproteobacteria; Burkholderiales; Burkholderiaceae; Burkholderia; Burkholderia cepacia complex
MKRRALLTAGAIVVAAGLTGCTAMLFEDGRYEETVDRFLVSEDGKKFVVLGEKYHYIFDM WP_060343506.1
hypothetical protein [Burkholderia cepacia] Length: 208\nNumber of predicted TMHs:  2\nExp number of AAs in TMHs: 43.89503\nExp number, first 60 AAs:  20.69402\nTotal prob of N-in:   0.74543\nPOSSIBLE N-term signal sequence\noutside   1   4\nTMhelix   5   27\ninside   28   184\nTMhelix   185   207\noutside   208   208

1140 GCF_001532125.1_ASM153212v1 Burkholderia cepacia Proteobacteria; Betaproteobacteria; Burkholderiales; Burkholderiaceae; Burkholderia; Burkholderia cepacia complex
MKTSTTRRRLLAAAMAASTLALSGCFTPKLYENDAYSEYVTSFMITQDGKKLVVLGTNYH WP_060280409.1
hypothetical protein [Burkholderia cepacia] Length: 212\nNumber of predicted TMHs:  1\nExp number of AAs in TMHs: 24.45717\nExp number, first 60 AAs:  1.10603\nTotal prob of N-in:   0.72957\ninside   1   180\nTMhelix   181   203\noutside   204   212

1141 GCF_001533445.1_ASM153344v1 Burkholderia cepacia Proteobacteria; Betaproteobacteria; Burkholderiales; Burkholderiaceae; Burkholderia; Burkholderia cepacia complex

MPDRKRRALLTAGVIVVAAGLTGCTAMLFEDGRYEETVDRFLVSEDGKKFVVLGEKYHYI WP_060330709.1 hypothetical protein [Burkholderia cepacia] Length: 211\nNumber of predicted TMHs: 2\nExp number of AAs in TMHs: 45.99641\nExp number, first 60 AAs: 21.95605\nTotal prob of N-in: 0.94961\nPOSSIBLE N-term signal sequence\ninside 1 6\nTMhelix 7 29\noutside 30 186\nTMhelix 187 209\ninside 210 211

1142 GCF_002209085.1_ASM220908v1 Burkholderia cepacia Proteobacteria; Betaproteobacteria; Burkholderiales; Burkholderiaceae; Burkholderia; Burkholderia cepacia complex MKRRALLTAGAIVAAGLTGCTAMLFEDGRYEETVDRFLVSEDGKKFVVLGEKYHYIFDMP WP_027787499.1 hypothetical protein [Burkholderia cepacia] Length: 207\nNumber of predicted TMHs: 1\nExp number of AAs in TMHs: 38.38341\nExp number, first 60 AAs: 15.45171\nTotal prob of N-in: 0.76513\nPOSSIBLE N-term signal sequence\ninside 1 183\nTMhelix 184 206\noutside 207 207

1143 GCF_002209085.1_ASM220908v1 Burkholderia cepacia Proteobacteria; Betaproteobacteria; Burkholderiales; Burkholderiaceae; Burkholderia; Burkholderia cepacia complex MKRRTFLALPAVAASGMLAGCSNFGPVTKAWMSDSKYDETLSTFLITADGKQLVVLGKKY WP_027787377.1 MULTISPECIES: hypothetical protein [Burkholderia] Length: 210\nNumber of predicted TMHs: 1\nExp number of AAs in TMHs: 34.58187\nExp number, first 60 AAs: 12.86351\nTotal prob of N-in: 0.46668\nPOSSIBLE N-term signal sequence\noutside 1 186\nTMhelix 187 209\ninside 210 210

1144 GCF_001533445.1_ASM153344v1 Burkholderia cepacia Proteobacteria; Betaproteobacteria; Burkholderiales; Burkholderiaceae; Burkholderia; Burkholderia cepacia complex MKRRTFLALPAVAASGMLAGCSNFGPVTKAWMSDSKYDETLSTFLITADGKQLVVLGKKY WP_027787377.1 MULTISPECIES: hypothetical protein [Burkholderia] Length: 210\nNumber of predicted TMHs: 1\nExp number of AAs in TMHs: 34.58187\nExp number, first 60 AAs: 12.86351\nTotal prob of N-in: 0.46668\nPOSSIBLE N-term signal sequence\noutside 1 186\nTMhelix 187 209\ninside 210 210

1145 GCF_001536795.1_ASM153679v1 Burkholderia cepacia Proteobacteria; Betaproteobacteria; Burkholderiales; Burkholderiaceae; Burkholderia; Burkholderia cepacia complex MKRRALLTAGVIVVAAGLTGCTAMLFEDGRYEETVDRFLVSEDGKKLVVLGEKYHYIFDM WP_059861788.1 hypothetical protein [Burkholderia cepacia] Length: 208\nNumber of predicted TMHs: 2\nExp number of AAs in TMHs: 44.86644\nExp number, first 60 AAs: 21.12438\nTotal prob of N-in: 0.76640\nPOSSIBLE N-term signal sequence\ninside 1 6\nTMhelix 7 26\noutside 27 183\nTMhelix 184 206\ninside 207 208

1146 GCF_001536795.1_ASM153679v1 Burkholderia cepacia Proteobacteria; Betaproteobacteria; Burkholderiales; Burkholderiaceae; Burkholderia; Burkholderia cepacia complex MKRRTFLALPAVAASGMLAGCSNFGPVTKAWMSDSKYDETLSTFLITADGKQLVVLGKKY WP_027787377.1 MULTISPECIES: hypothetical protein [Burkholderia] Length: 210\nNumber of predicted TMHs: 1\nExp number of AAs in TMHs: 34.58187\nExp number, first 60 AAs: 12.86351\nTotal prob of N-in: 0.46668\nPOSSIBLE N-term signal sequence\noutside 1 186\nTMhelix 187 209\ninside 210 210

1147 GCF_001532555.1_ASM153255v1 Burkholderia cepacia Proteobacteria; Betaproteobacteria; Burkholderiales; Burkholderiaceae; Burkholderia; Burkholderia cepacia complex MKTSITRRRLLAAGMAASTLALSGCFTPKLYENDAYSEYVTSFMITQDGKKLVVLGTNYH WP_060086226.1 hypothetical protein [Burkholderia cepacia] Length: 212\nNumber of predicted TMHs: 1\nExp number of AAs in TMHs: 23.88551\nExp number, first 60 AAs: 0.71364\nTotal prob of N-in: 0.64707\noutside 1 180\nTMhelix 181 203\ninside 204 212

1148 GCF_001532555.1_ASM153255v1 Burkholderia cepacia Proteobacteria; Betaproteobacteria; Burkholderiales; Burkholderiaceae; Burkholderia; Burkholderia cepacia complex MKRRALLTAGAIVVAAGLTGCTAMLFEDGRYEETVDRFLVSEDGKKFVVLGEKYHYIFDM WP_059700133.1 hypothetical protein [Burkholderia cepacia] Length: 208\nNumber of predicted TMHs: 2\nExp number of AAs in TMHs: 43.66178\nExp number, first 60 AAs: 20.68411\nTotal prob of N-in: 0.74282\nPOSSIBLE N-term signal sequence\noutside 1 4\nTMhelix 5 27\ninside 28 184\nTMhelix 185 207\noutside 208 208

1149 GCF_001532955.1_ASM153295v1 Burkholderia cepacia Proteobacteria; Betaproteobacteria; Burkholderiales; Burkholderiaceae; Burkholderia; Burkholderia cepacia complex MKRRALLTAGAIVAAGLTGCTAMLFEDGRYEETVDRFLVSEDGKKFVVLGEKYHYIFDMP WP_027787499.1 hypothetical protein [Burkholderia cepacia] Length: 207\nNumber of predicted TMHs: 1\nExp number of AAs in TMHs: 38.38341\nExp number, first 60 AAs: 15.45171\nTotal prob of N-in: 0.76513\nPOSSIBLE N-term signal sequence\ninside 1 183\nTMhelix 184 206\noutside 207 207

1150 GCF_001532955.1_ASM153295v1 Burkholderia cepacia Proteobacteria; Betaproteobacteria; Burkholderiales; Burkholderiaceae; Burkholderia; Burkholderia cepacia complex
MKRRTFLALPAVAASGMLAGCSNFGPVTKAWMSDSKYDETLSTFLITADGKQLVVLGKKY WP_027787377.1
MULTISPECIES: hypothetical protein [Burkholderia] Length: 210\nNumber of predicted TMHs: 1\nExp number of AAs in TMHs: 34.58187\nExp number, first 60 AAs: 12.86351\nTotal prob of N-in: 0.46668\nPOSSIBLE N-term signal sequence\noutside 1 186\nTMhelix 187 209\ninside 210 210

1151 GCF_001532125.1_ASM153212v1 Burkholderia cepacia Proteobacteria; Betaproteobacteria; Burkholderiales; Burkholderiaceae; Burkholderia; Burkholderia cepacia complex
MKRRTFLALPAVAASGMLAGCSNFGPVTKAWMSDSKYDETLSTFLITADGKQLVVLGKKY WP_027787377.1
MULTISPECIES: hypothetical protein [Burkholderia] Length: 210\nNumber of predicted TMHs: 1\nExp number of AAs in TMHs: 34.58187\nExp number, first 60 AAs: 12.86351\nTotal prob of N-in: 0.46668\nPOSSIBLE N-term signal sequence\noutside 1 186\nTMhelix 187 209\ninside 210 210

1152 GCF_001532555.1_ASM153255v1 Burkholderia cepacia Proteobacteria; Betaproteobacteria; Burkholderiales; Burkholderiaceae; Burkholderia; Burkholderia cepacia complex
MKRRTFLALPAVAASGMLAGCSNFGPVTKAWMSDSKYDETLSTFLITADGKQLVVLGKKY WP_027787377.1
MULTISPECIES: hypothetical protein [Burkholderia] Length: 210\nNumber of predicted TMHs: 1\nExp number of AAs in TMHs: 34.58187\nExp number, first 60 AAs: 12.86351\nTotal prob of N-in: 0.46668\nPOSSIBLE N-term signal sequence\noutside 1 186\nTMhelix 187 209\ninside 210 210

1153 GCF_001529145.1_ASM152914v1 Burkholderia cepacia Proteobacteria; Betaproteobacteria; Burkholderiales; Burkholderiaceae; Burkholderia; Burkholderia cepacia complex
MKTSITRRRLLAAGMAASTLALSGCFTPKLYENDAYSEYVTSFMITQDGKKLVVLGTNYH WP_060086226.1 hypothetical protein [Burkholderia cepacia] Length: 212\nNumber of predicted TMHs: 1\nExp number of AAs in TMHs: 23.88551\nExp number, first 60 AAs: 0.71364\nTotal prob of N-in: 0.64707\noutside 1 180\nTMhelix 181 203\ninside 204 212

1154 GCF_001529145.1_ASM152914v1 Burkholderia cepacia Proteobacteria; Betaproteobacteria; Burkholderiales; Burkholderiaceae; Burkholderia; Burkholderia cepacia complex
MKRRTFLALPAVAASGMLAGCSNFGPVTKAWMSDSKYDETLSTFLITADGKQLVVLGKKY WP_060084896.1
hypothetical protein [Burkholderia cepacia] Length: 210\nNumber of predicted TMHs: 1\nExp number of AAs in TMHs: 34.63096\nExp number, first 60 AAs: 12.87279\nTotal prob of N-in: 0.46689\nPOSSIBLE N-term signal sequence\noutside 1 186\nTMhelix 187 209\ninside 210 210

1155 GCF_001530135.1_ASM153013v1 Burkholderia cepacia Proteobacteria; Betaproteobacteria; Burkholderiales; Burkholderiaceae; Burkholderia; Burkholderia cepacia complex
MKTSITRRRLLAAAMAASTLALSGCFTPKLYENDAYSEYVTSFMITQDGKKLVVLGTNYH WP_060050342.1 hypothetical protein [Burkholderia cepacia] Length: 212\nNumber of predicted TMHs: 1\nExp number of AAs in TMHs: 24.23033\nExp number, first 60 AAs: 1.03058\nTotal prob of N-in: 0.64819\noutside 1 180\nTMhelix 181 203\ninside 204 212

1156 GCF_001530135.1_ASM153013v1 Burkholderia cepacia Proteobacteria; Betaproteobacteria; Burkholderiales; Burkholderiaceae; Burkholderia; Burkholderia cepacia complex
MKRRTFLALPAVAASGMLAGCSNFGPVTKAWMSDSKYDETLSTFLITADGKQLVVLGKKY WP_059488505.1
hypothetical protein [Burkholderia cepacia] Length: 210\nNumber of predicted TMHs: 1\nExp number of AAs in TMHs: 34.5304\nExp number, first 60 AAs: 12.84424\nTotal prob of N-in: 0.46568\nPOSSIBLE N-term signal sequence\noutside 1 186\nTMhelix 187 209\ninside 210 210

1157 GCF_001531905.1_ASM153190v1 Burkholderia cepacia Proteobacteria; Betaproteobacteria; Burkholderiales; Burkholderiaceae; Burkholderia; Burkholderia cepacia complex
MKRRALLTAGVIVVAAGLTGCTAMLFEDGRYEETVDRFLVSEDGKKFVVLGEKYHYIFDM WP_060230611.1
hypothetical protein [Burkholderia cepacia] Length: 208\nNumber of predicted TMHs: 2\nExp number of AAs in TMHs: 44.88016\nExp number, first 60 AAs: 21.13087\nTotal prob of N-in: 0.76656\nPOSSIBLE N-term signal sequence\ninside 1 6\nTMhelix 7 26\noutside 27 183\nTMhelix 184 206\ninside 207 208

1158 GCF_001531905.1_ASM153190v1 Burkholderia cepacia Proteobacteria; Betaproteobacteria; Burkholderiales; Burkholderiaceae; Burkholderia; Burkholderia cepacia complex
MKRRTFLALPAVAASGMLAGCSNFGPVTKAWMSDSKYDETLSTFLITADGKQLVVLGKKY WP_027787377.1
MULTISPECIES: hypothetical protein [Burkholderia] Length: 210\nNumber of predicted TMHs: 1\nExp

number of AAs in TMHs: 34.58187\nExp number, first 60 AAs: 12.86351\nTotal prob of N-in: 0.46668\nPOSSIBLE N-term signal sequence\noutside 1 186\nTMhelix 187 209\ninside 210 210

1159 GCF_001531525.1_ASM153152v1 Burkholderia cepacia Proteobacteria; Betaproteobacteria; Burkholderiales; Burkholderiaceae; Burkholderia; Burkholderia cepacia complex MKRRTFLALPAVAASGMLAGCSNFGPVTKAWMSDSKYDETLSTFLITADGKQLVVLGKKY WP_027787377.1 MULTISPECIES: hypothetical protein [Burkholderia] Length: 210\nNumber of predicted TMHs: 1\nExp number of AAs in TMHs: 34.58187\nExp number, first 60 AAs: 12.86351\nTotal prob of N-in: 0.46668\nPOSSIBLE N-term signal sequence\noutside 1 186\nTMhelix 187 209\ninside 210 210

1160 GCF_001529685.1_ASM152968v1 Burkholderia cepacia Proteobacteria; Betaproteobacteria; Burkholderiales; Burkholderiaceae; Burkholderia; Burkholderia cepacia complex MKTSITRRRLLAAAMAASTLALSGCFTPKLYENDAYSEYVTSFMITQDGKKLVVLGTNYH WP_060050342.1 hypothetical protein [Burkholderia cepacia] Length: 212\nNumber of predicted TMHs: 1\nExp number of AAs in TMHs: 24.23033\nExp number, first 60 AAs: 1.03058\nTotal prob of N-in: 0.64819\noutside 1 180\nTMhelix 181 203\ninside 204 212

1161 GCF_001530085.1_ASM153008v1 Burkholderia cepacia Proteobacteria; Betaproteobacteria; Burkholderiales; Burkholderiaceae; Burkholderia; Burkholderia cepacia complex MKTSITRRRLLAAAMAASTLALSGCFTPKLYENDAYSEYVTSFMITQDGKKLVVLGTNYH WP_060050342.1 hypothetical protein [Burkholderia cepacia] Length: 212\nNumber of predicted TMHs: 1\nExp number of AAs in TMHs: 24.23033\nExp number, first 60 AAs: 1.03058\nTotal prob of N-in: 0.64819\noutside 1 180\nTMhelix 181 203\ninside 204 212

1162 GCF_001530085.1_ASM153008v1 Burkholderia cepacia Proteobacteria; Betaproteobacteria; Burkholderiales; Burkholderiaceae; Burkholderia; Burkholderia cepacia complex MKRRTFLALPAVAASGMLAGCSNFGPVTKAWMSDSKYDETLSTFLITADGKQLVVLGKKY WP_059488505.1 hypothetical protein [Burkholderia cepacia] Length: 210\nNumber of predicted TMHs: 1\nExp number of AAs in TMHs: 34.5304\nExp number, first 60 AAs: 12.84424\nTotal prob of N-in: 0.46568\nPOSSIBLE N-term signal sequence\noutside 1 186\nTMhelix 187 209\ninside 210 210

1163 GCF_001530905.1_ASM153090v1 Burkholderia cepacia Proteobacteria; Betaproteobacteria; Burkholderiales; Burkholderiaceae; Burkholderia; Burkholderia cepacia complex MKRRALLTAGVIVVAAGLTGRTAMLFEDGRYEETVDRFLVSEDGKKFVVLGEKYHYIFDM WP_060216934.1 hypothetical protein [Burkholderia cepacia] Length: 208\nNumber of predicted TMHs: 1\nExp number of AAs in TMHs: 36.926\nExp number, first 60 AAs: 13.53423\nTotal prob of N-in: 0.80331\nPOSSIBLE N-term signal sequence\ninside 1 184\nTMhelix 185 207\noutside 208 208

1164 GCF_001529825.1_ASM152982v1 Burkholderia cepacia Proteobacteria; Betaproteobacteria; Burkholderiales; Burkholderiaceae; Burkholderia; Burkholderia cepacia complex MKTSITRRRLLAAAMAASTLALSGCFTPKLYENDAYSEYVTSFMITQDGKKLVVLGTNYH WP_060050342.1 hypothetical protein [Burkholderia cepacia] Length: 212\nNumber of predicted TMHs: 1\nExp number of AAs in TMHs: 24.23033\nExp number, first 60 AAs: 1.03058\nTotal prob of N-in: 0.64819\noutside 1 180\nTMhelix 181 203\ninside 204 212

1165 GCF_001530905.1_ASM153090v1 Burkholderia cepacia Proteobacteria; Betaproteobacteria; Burkholderiales; Burkholderiaceae; Burkholderia; Burkholderia cepacia complex MKRRTFLALPAVAASGMLAGCSNFGPVTKAWMSDSKYDETLSTFLITADGKQLVVLGKKY WP_027787377.1 MULTISPECIES: hypothetical protein [Burkholderia] Length: 210\nNumber of predicted TMHs: 1\nExp number of AAs in TMHs: 34.58187\nExp number, first 60 AAs: 12.86351\nTotal prob of N-in: 0.46668\nPOSSIBLE N-term signal sequence\noutside 1 186\nTMhelix 187 209\ninside 210 210

1166 GCF_001529825.1_ASM152982v1 Burkholderia cepacia Proteobacteria; Betaproteobacteria; Burkholderiales; Burkholderiaceae; Burkholderia; Burkholderia cepacia complex MKRRTFLALPAVAASGMLAGCSNFGPVTKAWMSDSKYDETLSTFLITADGKQLVVLGKKY WP_059488505.1 hypothetical protein [Burkholderia cepacia] Length: 210\nNumber of predicted TMHs: 1\nExp number of AAs in TMHs: 34.5304\nExp number, first 60 AAs: 12.84424\nTotal prob of N-in: 0.46568\nPOSSIBLE N-term signal sequence\noutside 1 186\nTMhelix 187 209\ninside 210 210

1167 GCF_001530205.1_ASM153020v1 Burkholderia cepacia Proteobacteria; Betaproteobacteria; Burkholderiales; Burkholderiaceae; Burkholderia; Burkholderia cepacia complex

MKTSTTRRRLLAAAMAASTLALSGCFTPKLYENDAYSEYVTSFMITQDGKKLVVLGTNYH WP_060222271.1 hypothetical protein [Burkholderia cepacia] Length: 212\nNumber of predicted TMHs: 1\nExp number of AAs in TMHs: 24.38513\nExp number, first 60 AAs: 1.10944\nTotal prob of N-in: 0.72864\ninside 1 180\nTMhelix 181 203\noutside 204 212

1168 GCF_001522985.1_ASM152298v1 Burkholderia cepacia Proteobacteria; Betaproteobacteria; Burkholderiales; Burkholderiaceae; Burkholderia; Burkholderia cepacia complex MKRRTFLALPAVAASGMLAGCSNFGPVTKAWMSDSKYDETLSTFLITADGKQLVVLGKKY WP_059488505.1 hypothetical protein [Burkholderia cepacia] Length: 210\nNumber of predicted TMHs: 1\nExp number of AAs in TMHs: 34.5304\nExp number, first 60 AAs: 12.84424\nTotal prob of N-in: 0.46568\nPOSSIBLE N-term signal sequence\noutside 1 186\nTMhelix 187 209\ninside 210 210

1169 GCF_001530205.1_ASM153020v1 Burkholderia cepacia Proteobacteria; Betaproteobacteria; Burkholderiales; Burkholderiaceae; Burkholderia; Burkholderia cepacia complex MKRRTFLALPAVAASGMLAGCSNFGPVTKAWMSDSKYDETLSTFLITADGKQLVVLGKKY WP_060221786.1 hypothetical protein [Burkholderia cepacia] Length: 210\nNumber of predicted TMHs: 1\nExp number of AAs in TMHs: 34.5304\nExp number, first 60 AAs: 12.84424\nTotal prob of N-in: 0.46568\nPOSSIBLE N-term signal sequence\noutside 1 186\nTMhelix 187 209\ninside 210 210

1170 GCF_001530205.1_ASM153020v1 Burkholderia cepacia Proteobacteria; Betaproteobacteria; Burkholderiales; Burkholderiaceae; Burkholderia; Burkholderia cepacia complex MKRRALLTAGVIVVAAGLTGRTAMLFEDGRYEETVDRFLVSEDGKKFVVLGEKYHYIFDM WP_060216934.1 hypothetical protein [Burkholderia cepacia] Length: 208\nNumber of predicted TMHs: 1\nExp number of AAs in TMHs: 36.926\nExp number, first 60 AAs: 13.53423\nTotal prob of N-in: 0.80331\nPOSSIBLE N-term signal sequence\ninside 1 184\nTMhelix 185 207\noutside 208 208

1171 GCF_001530005.1_ASM153000v1 Burkholderia cepacia Proteobacteria; Betaproteobacteria; Burkholderiales; Burkholderiaceae; Burkholderia; Burkholderia cepacia complex MKTSITRRRLLAAAMAASTLALSGCFTPKLYENDAYSEYVTSFMITQDGKKLVVLGTNYH WP_060050342.1 hypothetical protein [Burkholderia cepacia] Length: 212\nNumber of predicted TMHs: 1\nExp number of AAs in TMHs: 24.23033\nExp number, first 60 AAs: 1.03058\nTotal prob of N-in: 0.64819\noutside 1 180\nTMhelix 181 203\ninside 204 212

1172 GCF_001526345.1_ASM152634v1 Burkholderia cepacia Proteobacteria; Betaproteobacteria; Burkholderiales; Burkholderiaceae; Burkholderia; Burkholderia cepacia complex MKRRTFLALPAVAASGMLAGCSNFGPVTKAWMSDSKYDETLSTFLITADGKQLVVLGKKY WP_027787377.1 MULTISPECIES: hypothetical protein [Burkholderia] Length: 210\nNumber of predicted TMHs: 1\nExp number of AAs in TMHs: 34.58187\nExp number, first 60 AAs: 12.86351\nTotal prob of N-in: 0.46668\nPOSSIBLE N-term signal sequence\noutside 1 186\nTMhelix 187 209\ninside 210 210

1173 GCF_001525925.1_ASM152592v1 Burkholderia cepacia Proteobacteria; Betaproteobacteria; Burkholderiales; Burkholderiaceae; Burkholderia; Burkholderia cepacia complex MKTSITRRRLLAAAMAASTLALSGCFTPKLYENDAYSEYVTSFMITQDGKKLVVLGTNYH WP_059485162.1 hypothetical protein [Burkholderia cepacia] Length: 212\nNumber of predicted TMHs: 1\nExp number of AAs in TMHs: 24.29452\nExp number, first 60 AAs: 1.02778\nTotal prob of N-in: 0.64928\noutside 1 180\nTMhelix 181 203\ninside 204 212

1174 GCF_001530005.1_ASM153000v1 Burkholderia cepacia Proteobacteria; Betaproteobacteria; Burkholderiales; Burkholderiaceae; Burkholderia; Burkholderia cepacia complex MKRRTFLALPAVAASGMLAGCSNFGPVTKAWMSDSKYDETLSTFLITADGKQLVVLGKKY WP_059488505.1 hypothetical protein [Burkholderia cepacia] Length: 210\nNumber of predicted TMHs: 1\nExp number of AAs in TMHs: 34.5304\nExp number, first 60 AAs: 12.84424\nTotal prob of N-in: 0.46568\nPOSSIBLE N-term signal sequence\noutside 1 186\nTMhelix 187 209\ninside 210 210

1175 GCF_001525965.1_ASM152596v1 Burkholderia cepacia Proteobacteria; Betaproteobacteria; Burkholderiales; Burkholderiaceae; Burkholderia; Burkholderia cepacia complex MKTSITRRRLLAAAMAASTLALSGCFTPKLYENDAYSEYVTSFMITQDGKKLVVLGTNYH WP_059485162.1 hypothetical protein [Burkholderia cepacia] Length: 212\nNumber of predicted TMHs: 1\nExp number of AAs in TMHs: 24.29452\nExp number, first 60 AAs: 1.02778\nTotal prob of N-in: 0.64928\noutside 1 180\nTMhelix 181 203\ninside 204 212

1176 GCF_001525925.1_ASM152592v1 Burkholderia cepacia Proteobacteria; Betaproteobacteria; Burkholderiales; Burkholderiaceae; Burkholderia; Burkholderia cepacia complex MKRRTFLALPAVAASGMLAGCSNFGPVTKAWMSDSKYDETLSTFLITADGKQLVVLGKKY WP_059488505.1 hypothetical protein [Burkholderia cepacia] Length: 210\nNumber of predicted TMHs: 1\nExp number of AAs in TMHs: 34.5304\nExp number, first 60 AAs: 12.84424\nTotal prob of N-in: 0.46568\nPOSSIBLE N-term signal sequence\noutside 1 186\nTMhelix 187 209\ninside 210 210

1177 GCF_001525965.1_ASM152596v1 Burkholderia cepacia Proteobacteria; Betaproteobacteria; Burkholderiales; Burkholderiaceae; Burkholderia; Burkholderia cepacia complex MKRRTFLALPAVAASGMLAGCSNFGPVTKAWMSDSKYDETLSTFLITADGKQLVVLGKKY WP_059488505.1 hypothetical protein [Burkholderia cepacia] Length: 210\nNumber of predicted TMHs: 1\nExp number of AAs in TMHs: 34.5304\nExp number, first 60 AAs: 12.84424\nTotal prob of N-in: 0.46568\nPOSSIBLE N-term signal sequence\noutside 1 186\nTMhelix 187 209\ninside 210 210

1178 GCF_001525775.1_ASM152577v1 Burkholderia cepacia Proteobacteria; Betaproteobacteria; Burkholderiales; Burkholderiaceae; Burkholderia; Burkholderia cepacia complex MKTSITRRRLLAAAMAASTLALSGCFTPKLYENDAYSEYVTSFMITQDGKKLVVLGTNYH WP_059485162.1 hypothetical protein [Burkholderia cepacia] Length: 212\nNumber of predicted TMHs: 1\nExp number of AAs in TMHs: 24.29452\nExp number, first 60 AAs: 1.02778\nTotal prob of N-in: 0.64928\noutside 1 180\nTMhelix 181 203\ninside 204 212

1179 GCF_001530185.1_ASM153018v1 Burkholderia cepacia Proteobacteria; Betaproteobacteria; Burkholderiales; Burkholderiaceae; Burkholderia; Burkholderia cepacia complex MKRRALLTAGAIVVAAGLTGCTAMLFEDGRYEETVDRFLVSEDGKKFVVLGEKYHYIFDM WP_059700133.1 hypothetical protein [Burkholderia cepacia] Length: 208\nNumber of predicted TMHs: 2\nExp number of AAs in TMHs: 43.66178\nExp number, first 60 AAs: 20.68411\nTotal prob of N-in: 0.74282\nPOSSIBLE N-term signal sequence\noutside 1 4\nTMhelix 5 27\ninside 28 184\nTMhelix 185 207\noutside 208 208

1180 GCF_001530185.1_ASM153018v1 Burkholderia cepacia Proteobacteria; Betaproteobacteria; Burkholderiales; Burkholderiaceae; Burkholderia; Burkholderia cepacia complex MKTSITRRRLLAAGMAASTLALSGCFTPKLYENDAYSEYVTSFMITQDGKKLVVLGTNYH WP_060086226.1 hypothetical protein [Burkholderia cepacia] Length: 212\nNumber of predicted TMHs: 1\nExp number of AAs in TMHs: 23.88551\nExp number, first 60 AAs: 0.71364\nTotal prob of N-in: 0.64707\noutside 1 180\nTMhelix 181 203\ninside 204 212

1181 GCF_001530415.1_ASM153041v1 Burkholderia cepacia Proteobacteria; Betaproteobacteria; Burkholderiales; Burkholderiaceae; Burkholderia; Burkholderia cepacia complex MKTSITRRRLLAAGMAASTLALSGCFTPKLYENDAYSEYVTSFMITQDGKKLVVLGTNYH WP_060086226.1 hypothetical protein [Burkholderia cepacia] Length: 212\nNumber of predicted TMHs: 1\nExp number of AAs in TMHs: 23.88551\nExp number, first 60 AAs: 0.71364\nTotal prob of N-in: 0.64707\noutside 1 180\nTMhelix 181 203\ninside 204 212

1182 GCF_001530185.1_ASM153018v1 Burkholderia cepacia Proteobacteria; Betaproteobacteria; Burkholderiales; Burkholderiaceae; Burkholderia; Burkholderia cepacia complex MKRRTFLALPVVAASGMLAGCSNFGPVTKAWMSDSKYDETLSTFLITADGKQLVVLGKKY WP_060172271.1 hypothetical protein [Burkholderia cepacia] Length: 210\nNumber of predicted TMHs: 1\nExp number of AAs in TMHs: 37.62368\nExp number, first 60 AAs: 15.87108\nTotal prob of N-in: 0.50888\nPOSSIBLE N-term signal sequence\noutside 1 186\nTMhelix 187 209\ninside 210 210

1183 GCF_001530415.1_ASM153041v1 Burkholderia cepacia Proteobacteria; Betaproteobacteria; Burkholderiales; Burkholderiaceae; Burkholderia; Burkholderia cepacia complex MKRRTFLALPVVAASGMLAGCSNFGPVTKAWMSDSKYDETLSTFLITADGKQLVVLGKKY WP_060172271.1 hypothetical protein [Burkholderia cepacia] Length: 210\nNumber of predicted TMHs: 1\nExp number of AAs in TMHs: 37.62368\nExp number, first 60 AAs: 15.87108\nTotal prob of N-in: 0.50888\nPOSSIBLE N-term signal sequence\noutside 1 186\nTMhelix 187 209\ninside 210 210

1184 GCF_001531875.1_ASM153187v1 Burkholderia cepacia Proteobacteria; Betaproteobacteria; Burkholderiales; Burkholderiaceae; Burkholderia; Burkholderia cepacia complex MKRRALLTAGAIVVAAGLTGCTAMLFENGRYEETVDRFLVSEDGKKFVVLGEKYHYIFDM WP_059682727.1 hypothetical protein [Burkholderia cepacia] Length: 208\nNumber of predicted TMHs: 2\nExp number of

AAs in TMHs: 44.24426\nExp number, first 60 AAs: 20.97809\nTotal prob of N-in: 0.67235\nPOSSIBLE N-term signal sequence\noutside 1 4\nTMhelix 5 27\ninside 28 184\nTMhelix 185 207\noutside 208 208

1185 GCF_001530785.1_ASM153078v1 Burkholderia cepacia Proteobacteria; Betaproteobacteria; Burkholderiales; Burkholderiaceae; Burkholderia; Burkholderia cepacia complex MKRRTFLALPAVAASGMLAGCSNFGPVTKAWMSDSKYDETLSTFLITADGKQLVVLGKKY WP_059488505.1 hypothetical protein [Burkholderia cepacia] Length: 210\nNumber of predicted TMHs: 1\nExp number of AAs in TMHs: 34.5304\nExp number, first 60 AAs: 12.84424\nTotal prob of N-in: 0.46568\nPOSSIBLE N-term signal sequence\noutside 1 186\nTMhelix 187 209\ninside 210 210

1186 GCF_001532525.1_ASM153252v1 Burkholderia cepacia Proteobacteria; Betaproteobacteria; Burkholderiales; Burkholderiaceae; Burkholderia; Burkholderia cepacia complex MKRRTFLALPAVAASGMLAGCSNFGPVTKAWMSDSKYDETLSTFLITADGKQLVVLGKKY WP_027787377.1 MULTISPECIES: hypothetical protein [Burkholderia] Length: 210\nNumber of predicted TMHs: 1\nExp number of AAs in TMHs: 34.58187\nExp number, first 60 AAs: 12.86351\nTotal prob of N-in: 0.46668\nPOSSIBLE N-term signal sequence\noutside 1 186\nTMhelix 187 209\ninside 210 210

1187 GCF_001532235.1_ASM153223v1 Burkholderia cepacia Proteobacteria; Betaproteobacteria; Burkholderiales; Burkholderiaceae; Burkholderia; Burkholderia cepacia complex MKRRTFLALPAVAASGMLAGCSNFGPVTKAWMSDSKYDETLSTFLITADGKQLVVLGKKY WP_027787377.1 MULTISPECIES: hypothetical protein [Burkholderia] Length: 210\nNumber of predicted TMHs: 1\nExp number of AAs in TMHs: 34.58187\nExp number, first 60 AAs: 12.86351\nTotal prob of N-in: 0.46668\nPOSSIBLE N-term signal sequence\noutside 1 186\nTMhelix 187 209\ninside 210 210

1188 GCF_001530785.1_ASM153078v1 Burkholderia cepacia Proteobacteria; Betaproteobacteria; Burkholderiales; Burkholderiaceae; Burkholderia; Burkholderia cepacia complex MKTSITRRRLLAAAMAASTLALSGCFTPKLYENDAYSEYVTSFMITQDGKKLVVLGTNYH WP_060050342.1 hypothetical protein [Burkholderia cepacia] Length: 212\nNumber of predicted TMHs: 1\nExp number of AAs in TMHs: 24.23033\nExp number, first 60 AAs: 1.03058\nTotal prob of N-in: 0.64819\noutside 1 180\nTMhelix 181 203\ninside 204 212

1189 GCF_001532525.1_ASM153252v1 Burkholderia cepacia Proteobacteria; Betaproteobacteria; Burkholderiales; Burkholderiaceae; Burkholderia; Burkholderia cepacia complex MKRRALLTAGAIVVAAGLTGCTAMLFEDGRYEETVDRFLVSEDGKKFVVLGEKYHYIFDM WP_059700133.1 hypothetical protein [Burkholderia cepacia] Length: 208\nNumber of predicted TMHs: 2\nExp number of AAs in TMHs: 43.66178\nExp number, first 60 AAs: 20.68411\nTotal prob of N-in: 0.74282\nPOSSIBLE N-term signal sequence\noutside 1 4\nTMhelix 5 27\ninside 28 184\nTMhelix 185 207\noutside 208 208

1190 GCF_001532705.1_ASM153270v1 Burkholderia cepacia Proteobacteria; Betaproteobacteria; Burkholderiales; Burkholderiaceae; Burkholderia; Burkholderia cepacia complex MKRRTFLALPAVAASGMLAGCSNFGPVTKAWMSDSKYDETLSTFLITADGKQLVVLGKKY WP_027787377.1 MULTISPECIES: hypothetical protein [Burkholderia] Length: 210\nNumber of predicted TMHs: 1\nExp number of AAs in TMHs: 34.58187\nExp number, first 60 AAs: 12.86351\nTotal prob of N-in: 0.46668\nPOSSIBLE N-term signal sequence\noutside 1 186\nTMhelix 187 209\ninside 210 210

1191 GCF_001532525.1_ASM153252v1 Burkholderia cepacia Proteobacteria; Betaproteobacteria; Burkholderiales; Burkholderiaceae; Burkholderia; Burkholderia cepacia complex MKTSITRRRLLAAGMAASTLALSGCFTPKLYENDAYSEYVTSFMITQDGKKLVVLGTNYH WP_060086226.1 hypothetical protein [Burkholderia cepacia] Length: 212\nNumber of predicted TMHs: 1\nExp number of AAs in TMHs: 23.88551\nExp number, first 60 AAs: 0.71364\nTotal prob of N-in: 0.64707\noutside 1 180\nTMhelix 181 203\ninside 204 212

1192 GCF_001718395.1_ASM171839v1 Burkholderia cepacia Proteobacteria; Betaproteobacteria; Burkholderiales; Burkholderiaceae; Burkholderia; Burkholderia cepacia complex MKRRTFLALPAVAASGMLAGCSNFGPVTKAWMSDSKYDETLSTFLITADGKQLVVLGKKY WP_027787377.1 MULTISPECIES: hypothetical protein [Burkholderia] Length: 210\nNumber of predicted TMHs: 1\nExp number of AAs in TMHs: 34.58187\nExp number, first 60 AAs: 12.86351\nTotal prob of N-in: 0.46668\nPOSSIBLE N-term signal sequence\noutside 1 186\nTMhelix 187 209\ninside 210 210

1193 GCF_001534085.1_ASM153408v1 Burkholderia cepacia Proteobacteria; Betaproteobacteria; Burkholderiales; Burkholderiaceae; Burkholderia; Burkholderia cepacia complex

MKRRTFLALPAVAASGMLAGCSNFGPVTKAWMSDSKYDETLSTFLITADGKQLVVLGKKY WP_027787377.1 MULTISPECIES: hypothetical protein [Burkholderia] Length: 210\nNumber of predicted TMHs: 1\nExp number of AAs in TMHs: 34.58187\nExp number, first 60 AAs: 12.86351\nTotal prob of N-in: 0.46668\nPOSSIBLE N-term signal sequence\noutside 1 186\nTMhelix 187 209\ninside 210 210

1194 GCF_001533295.1_ASM153329v1 Burkholderia cepacia Proteobacteria; Betaproteobacteria; Burkholderiales; Burkholderiaceae; Burkholderia; Burkholderia cepacia complex MKTSITRRRLLAAGMAASTLALSGCFTPKLYENDAYSEYVTSFMITQDGKKLVVLGTNYH WP_060086226.1 hypothetical protein [Burkholderia cepacia] Length: 212\nNumber of predicted TMHs: 1\nExp number of AAs in TMHs: 23.88551\nExp number, first 60 AAs: 0.71364\nTotal prob of N-in: 0.64707\noutside 1 180\nTMhelix 181 203\ninside 204 212

1195 GCF_001532235.1_ASM153223v1 Burkholderia cepacia Proteobacteria; Betaproteobacteria; Burkholderiales; Burkholderiaceae; Burkholderia; Burkholderia cepacia complex MKTSITRRRLLAAAMAASTLALSGCFTPKLYENDAYSEYVTSFMITQDGKKLVVLGTNYH WP_060308855.1 hypothetical protein [Burkholderia cepacia] Length: 212\nNumber of predicted TMHs: 1\nExp number of AAs in TMHs: 24.99192\nExp number, first 60 AAs: 0.96967\nTotal prob of N-in: 0.69859\noutside 1 180\nTMhelix 181 203\ninside 204 212

1196 GCF_001533295.1_ASM153329v1 Burkholderia cepacia Proteobacteria; Betaproteobacteria; Burkholderiales; Burkholderiaceae; Burkholderia; Burkholderia cepacia complex MKRRALLTAGAIVVAAGLTGCTAMLFEDGRYEETVDRFLVSEDGKKFVVLGEKYHYIFDM WP_060372671.1 hypothetical protein [Burkholderia cepacia] Length: 208\nNumber of predicted TMHs: 2\nExp number of AAs in TMHs: 43.68346\nExp number, first 60 AAs: 20.68525\nTotal prob of N-in: 0.74306\nPOSSIBLE N-term signal sequence\noutside 1 4\nTMhelix 5 27\ninside 28 184\nTMhelix 185 207\noutside 208 208

1197 GCF_001525945.1_ASM152594v1 Burkholderia cepacia Proteobacteria; Betaproteobacteria; Burkholderiales; Burkholderiaceae; Burkholderia; Burkholderia cepacia complex MKRRTFLALPAVAASGMLAGCSNFGPVTKAWMSDSKYDETLSTFLITADGKQLVVLGKKY WP_059488505.1 hypothetical protein [Burkholderia cepacia] Length: 210\nNumber of predicted TMHs: 1\nExp number of AAs in TMHs: 34.5304\nExp number, first 60 AAs: 12.84424\nTotal prob of N-in: 0.46568\nPOSSIBLE N-term signal sequence\noutside 1 186\nTMhelix 187 209\ninside 210 210

1198 GCF_001526205.1_ASM152620v1 Burkholderia cepacia Proteobacteria; Betaproteobacteria; Burkholderiales; Burkholderiaceae; Burkholderia; Burkholderia cepacia complex MKASITRRRLLAAGVAASSLALSGCFTPKLYKDDGYHEYVSGFMITQDGKKLVVLGTDYH WP_059525490.1 hypothetical protein [Burkholderia cepacia] Length: 212\nNumber of predicted TMHs: 1\nExp number of AAs in TMHs: 25.1207\nExp number, first 60 AAs: 0.80309\nTotal prob of N-in: 0.62748\noutside 1 180\nTMhelix 181 203\ninside 204 212

1199 GCF_001526205.1_ASM152620v1 Burkholderia cepacia Proteobacteria; Betaproteobacteria; Burkholderiales; Burkholderiaceae; Burkholderia; Burkholderia cepacia complex MKRRTFLALPAVAASGMLAGCSNFGPVTKAWMSDSKYDETLSTFLITADGKQLVVLGKRY WP_059526018.1 hypothetical protein [Burkholderia cepacia] Length: 210\nNumber of predicted TMHs: 1\nExp number of AAs in TMHs: 34.61771\nExp number, first 60 AAs: 12.90764\nTotal prob of N-in: 0.46822\nPOSSIBLE N-term signal sequence\noutside 1 186\nTMhelix 187 209\ninside 210 210

1200 GCF_001528845.1_ASM152884v1 Burkholderia cepacia Proteobacteria; Betaproteobacteria; Burkholderiales; Burkholderiaceae; Burkholderia; Burkholderia cepacia complex MKRRALLTAGAIVAAGLTGCTAMLFEDGRYEETVDRFLVSEDGKKFVVLGEKYHYIFDMP WP_027787499.1 hypothetical protein [Burkholderia cepacia] Length: 207\nNumber of predicted TMHs: 1\nExp number of AAs in TMHs: 38.38341\nExp number, first 60 AAs: 15.45171\nTotal prob of N-in: 0.76513\nPOSSIBLE N-term signal sequence\ninside 1 183\nTMhelix 184 206\noutside 207 207

1201 GCF_001528685.1_ASM152868v1 Burkholderia cepacia Proteobacteria; Betaproteobacteria; Burkholderiales; Burkholderiaceae; Burkholderia; Burkholderia cepacia complex MKRRALLTAGVIVVAAGLTGCTAMLFEDGRYEETVDRFLVSEDGKKFVVLGEKYHYIFDM WP_060069881.1 hypothetical protein [Burkholderia cepacia] Length: 208\nNumber of predicted TMHs: 2\nExp number of AAs in TMHs: 44.67702\nExp number, first 60 AAs: 21.12768\nTotal prob of N-in: 0.76365\nPOSSIBLE N-term signal sequence\ninside 1 6\nTMhelix 7 26\noutside 27 183\nTMhelix 184 206\ninside 207 208

1202 GCF_001526445.1_ASM152644v1 Burkholderia cepacia Proteobacteria; Betaproteobacteria; Burkholderiales; Burkholderiaceae; Burkholderia; Burkholderia cepacia complex
MKASITRRRLLAAGVAASSLALSGCFTPKLYKDDGYHEYVSGFMITQDGKKLIVLGTDYH WP_059904810.1 hypothetical protein [Burkholderia cepacia] Length: 212\nNumber of predicted TMHs: 1\nExp number of AAs in TMHs: 25.12914\nExp number, first 60 AAs: 0.80813\nTotal prob of N-in: 0.62758\noutside 1 180\nTMhelix 181 203\ninside 204 212

1203 GCF_001525895.1_ASM152589v1 Burkholderia cepacia Proteobacteria; Betaproteobacteria; Burkholderiales; Burkholderiaceae; Burkholderia; Burkholderia cepacia complex
MKRRALLTAGVIVVAAGLTGCTAMLFEDGRYEETVDRFLVSEDGKKFVVLGEKYHYIFDM WP_059496097.1 hypothetical protein [Burkholderia cepacia] Length: 208\nNumber of predicted TMHs: 2\nExp number of AAs in TMHs: 44.13865\nExp number, first 60 AAs: 21.11648\nTotal prob of N-in: 0.75846\nPOSSIBLE N-term signal sequence\ninside 1 6\nTMhelix 7 26\noutside 27 183\nTMhelix 184 206\ninside 207 208

1204 GCF_001529885.1_ASM152988v1 Burkholderia cepacia Proteobacteria; Betaproteobacteria; Burkholderiales; Burkholderiaceae; Burkholderia; Burkholderia cepacia complex
MKRRTFLALPAVAASGMLAGCSNFGPVTKAWMSDSKYDETLSTFLITADGKQLVVLGKKY WP_027787377.1 MULTISPECIES: hypothetical protein [Burkholderia] Length: 210\nNumber of predicted TMHs: 1\nExp number of AAs in TMHs: 34.58187\nExp number, first 60 AAs: 12.86351\nTotal prob of N-in: 0.46668\nPOSSIBLE N-term signal sequence\noutside 1 186\nTMhelix 187 209\ninside 210 210

1205 GCF_001529755.1_ASM152975v1 Burkholderia cepacia Proteobacteria; Betaproteobacteria; Burkholderiales; Burkholderiaceae; Burkholderia; Burkholderia cepacia complex
MKRRALLTAGVIVVAAGLTGCTAMLFEDGRYEETVDRFLVSEDGKKFVVLGEKYHYIFDM WP_060069881.1 hypothetical protein [Burkholderia cepacia] Length: 208\nNumber of predicted TMHs: 2\nExp number of AAs in TMHs: 44.67702\nExp number, first 60 AAs: 21.12768\nTotal prob of N-in: 0.76365\nPOSSIBLE N-term signal sequence\ninside 1 6\nTMhelix 7 26\noutside 27 183\nTMhelix 184 206\ninside 207 208

1206 GCF_001530415.1_ASM153041v1 Burkholderia cepacia Proteobacteria; Betaproteobacteria; Burkholderiales; Burkholderiaceae; Burkholderia; Burkholderia cepacia complex
MKRRALLTAGAIVVAAGLTGCTAMLFEDGRYEETVDRFLVSEDGKKFVVLGEKYHYIFDM WP_059700133.1 hypothetical protein [Burkholderia cepacia] Length: 208\nNumber of predicted TMHs: 2\nExp number of AAs in TMHs: 43.66178\nExp number, first 60 AAs: 20.68411\nTotal prob of N-in: 0.74282\nPOSSIBLE N-term signal sequence\noutside 1 4\nTMhelix 5 27\ninside 28 184\nTMhelix 185 207\noutside 208 208

1207 GCF_000724975.1_ASM72497v1 Burkholderia cepacia Proteobacteria; Betaproteobacteria; Burkholderiales; Burkholderiaceae; Burkholderia; Burkholderia cepacia complex
MKRRTFLALPAVAASGMLAGCADFGPATKAWMKDSEYDETLSTFLITADGKQLVVLGQKY WP_043182909.1 hypothetical protein [Burkholderia cepacia] Length: 210\nNumber of predicted TMHs: 1\nExp number of AAs in TMHs: 29.62266\nExp number, first 60 AAs: 7.72237\nTotal prob of N-in: 0.56326\noutside 1 184\nTMhelix 185 207\ninside 208 210

1208 GCF_001039005.1_ASM103900v1 Burkholderia cepacia Proteobacteria; Betaproteobacteria; Burkholderiales; Burkholderiaceae; Burkholderia; Burkholderia cepacia complex
MSTSITRRRLLAASVAASSLALSGCFTPKLYKDDAYSEYVSGFMISEDGKKLVVLGMRYH WP_048243592.1 hypothetical protein [Burkholderia cepacia] Length: 212\nNumber of predicted TMHs: 1\nExp number of AAs in TMHs: 24.37992\nExp number, first 60 AAs: 0.57929\nTotal prob of N-in: 0.56787\noutside 1 180\nTMhelix 181 203\ninside 204 212

1209 GCF_001053975.1_ASM105397v1 Burkholderia cepacia Proteobacteria; Betaproteobacteria; Burkholderiales; Burkholderiaceae; Burkholderia; Burkholderia cepacia complex
MKRRTFLALPAVAASGMLAGCSNFGPVTKAWMSDSKYDETLSTFLITADGKQLVVLGKKY WP_027787377.1 MULTISPECIES: hypothetical protein [Burkholderia] Length: 210\nNumber of predicted TMHs: 1\nExp number of AAs in TMHs: 34.58187\nExp number, first 60 AAs: 12.86351\nTotal prob of N-in: 0.46668\nPOSSIBLE N-term signal sequence\noutside 1 186\nTMhelix 187 209\ninside 210 210

1210 GCF_001053975.1_ASM105397v1 Burkholderia cepacia Proteobacteria; Betaproteobacteria; Burkholderiales; Burkholderiaceae; Burkholderia; Burkholderia cepacia complex
MKRRALLTAGAIVAAGLTGCTAMLFEDGRYEETVDRFLVSEDGKKFVVLGEKYHYIFDMP WP_027787499.1 hypothetical protein [Burkholderia cepacia] Length: 207\nNumber of predicted TMHs: 1\nExp number of

AAs in TMHs: 38.38341\nExp number, first 60 AAs: 15.45171\nTotal prob of N-in: 0.76513\nPOSSIBLE N-term signal sequence\ninside 1 183\nTMhelix 184 206\noutside 207 207

1211 GCF_001521915.1_ASM152191v1 Burkholderia cepacia Proteobacteria; Betaproteobacteria; Burkholderiales; Burkholderiaceae; Burkholderia; Burkholderia cepacia complex MKTSITRRRLLAAAMAASTLALSGCFTPKLYENDAYSEYVTSFMITQDGKKLVVLGTNYH WP_059485162.1 hypothetical protein [Burkholderia cepacia] Length: 212\nNumber of predicted TMHs: 1\nExp number of AAs in TMHs: 24.29452\nExp number, first 60 AAs: 1.02778\nTotal prob of N-in: 0.64928\noutside 1 180\nTMhelix 181 203\ninside 204 212

1212 GCF_001521915.1_ASM152191v1 Burkholderia cepacia Proteobacteria; Betaproteobacteria; Burkholderiales; Burkholderiaceae; Burkholderia; Burkholderia cepacia complex MKRRTFLALPAVAASGMLAGCSNFGPVTKAWMSDSKYDETLSTFLITADGKQLVVLGKKY WP_059488505.1 hypothetical protein [Burkholderia cepacia] Length: 210\nNumber of predicted TMHs: 1\nExp number of AAs in TMHs: 34.5304\nExp number, first 60 AAs: 12.84424\nTotal prob of N-in: 0.46568\nPOSSIBLE N-term signal sequence\noutside 1 186\nTMhelix 187 209\ninside 210 210

1213 GCF_001521975.1_ASM152197v1 Burkholderia cepacia Proteobacteria; Betaproteobacteria; Burkholderiales; Burkholderiaceae; Burkholderia; Burkholderia cepacia complex MKTSITRRRLLAAAMAASTLALSGCFTPKLYENDAYSEYVTSFMITQDGKKLVVLGTNYH WP_059485162.1 hypothetical protein [Burkholderia cepacia] Length: 212\nNumber of predicted TMHs: 1\nExp number of AAs in TMHs: 24.29452\nExp number, first 60 AAs: 1.02778\nTotal prob of N-in: 0.64928\noutside 1 180\nTMhelix 181 203\ninside 204 212

1214 GCF_001522295.1_ASM152229v1 Burkholderia cepacia Proteobacteria; Betaproteobacteria; Burkholderiales; Burkholderiaceae; Burkholderia; Burkholderia cepacia complex MKRRALLTAGVIVVAAGLTGCTAMLFEDGRYEETVDRFLVSEDGKKFVVLGEKYHYIFDM WP_059588168.1 hypothetical protein [Burkholderia cepacia] Length: 208\nNumber of predicted TMHs: 2\nExp number of AAs in TMHs: 44.91369\nExp number, first 60 AAs: 21.12905\nTotal prob of N-in: 0.76333\nPOSSIBLE N-term signal sequence\ninside 1 6\nTMhelix 7 26\noutside 27 183\nTMhelix 184 206\ninside 207 208

1215 GCF_001522295.1_ASM152229v1 Burkholderia cepacia Proteobacteria; Betaproteobacteria; Burkholderiales; Burkholderiaceae; Burkholderia; Burkholderia cepacia complex MKRRTFLALPAVAASGMLTGCSNFGPVTKAWMSDSKYDETLSTFLITADGKQLVVLGKKY WP_059588290.1 hypothetical protein [Burkholderia cepacia] Length: 210\nNumber of predicted TMHs: 1\nExp number of AAs in TMHs: 31.54854\nExp number, first 60 AAs: 9.85597\nTotal prob of N-in: 0.43345\noutside 1 186\nTMhelix 187 209\ninside 210 210

1216 GCF_001522575.1_ASM152257v1 Burkholderia cepacia Proteobacteria; Betaproteobacteria; Burkholderiales; Burkholderiaceae; Burkholderia; Burkholderia cepacia complex MKTSITRRRLLAAAMAASTLALSGCFTPKLYENDAYSEYVTSFMITQDGKKLVVLGTNYH WP_059485162.1 hypothetical protein [Burkholderia cepacia] Length: 212\nNumber of predicted TMHs: 1\nExp number of AAs in TMHs: 24.29452\nExp number, first 60 AAs: 1.02778\nTotal prob of N-in: 0.64928\noutside 1 180\nTMhelix 181 203\ninside 204 212

1217 GCF_001522575.1_ASM152257v1 Burkholderia cepacia Proteobacteria; Betaproteobacteria; Burkholderiales; Burkholderiaceae; Burkholderia; Burkholderia cepacia complex MKRRTFLALPAVAASGMLAGCSNFGPVTKAWMSDSKYDETLSTFLITADGKQLVVLGKKY WP_059488505.1 hypothetical protein [Burkholderia cepacia] Length: 210\nNumber of predicted TMHs: 1\nExp number of AAs in TMHs: 34.5304\nExp number, first 60 AAs: 12.84424\nTotal prob of N-in: 0.46568\nPOSSIBLE N-term signal sequence\noutside 1 186\nTMhelix 187 209\ninside 210 210

1218 GCF_001522815.1_ASM152281v1 Burkholderia cepacia Proteobacteria; Betaproteobacteria; Burkholderiales; Burkholderiaceae; Burkholderia; Burkholderia cepacia complex MKTSITRRRLLAAAMAASTLALSGCFTPKLYENDAYSEYVTSFMITQDGKKLVVLGTNYH WP_059485162.1 hypothetical protein [Burkholderia cepacia] Length: 212\nNumber of predicted TMHs: 1\nExp number of AAs in TMHs: 24.29452\nExp number, first 60 AAs: 1.02778\nTotal prob of N-in: 0.64928\noutside 1 180\nTMhelix 181 203\ninside 204 212

1219 GCF_001522815.1_ASM152281v1 Burkholderia cepacia Proteobacteria; Betaproteobacteria; Burkholderiales; Burkholderiaceae; Burkholderia; Burkholderia cepacia complex

MKRRTFLALPAVAASGMLAGCSNFGPVTKAWMSDSKYDETLSTFLITADGKQLVVLGKKY WP_059488505.1 hypothetical protein [Burkholderia cepacia] Length: 210\nNumber of predicted TMHs: 1\nExp number of AAs in TMHs: 34.5304\nExp number, first 60 AAs: 12.84424\nTotal prob of N-in: 0.46568\nPOSSIBLE N-term signal sequence\noutside 1 186\nTMhelix 187 209\ninside 210 210

1220 GCF_001522575.1_ASM152257v1 Burkholderia cepacia Proteobacteria; Betaproteobacteria; Burkholderiales; Burkholderiaceae; Burkholderia; Burkholderia cepacia complex MKRRALLTAGVIVVAAGLTGCTAMLFEDGRYEETVDRFLVSEDGKKFVVLGEKYHYIFDM WP_059496097.1 hypothetical protein [Burkholderia cepacia] Length: 208\nNumber of predicted TMHs: 2\nExp number of AAs in TMHs: 44.13865\nExp number, first 60 AAs: 21.11648\nTotal prob of N-in: 0.75846\nPOSSIBLE N-term signal sequence\ninside 1 6\nTMhelix 7 26\noutside 27 183\nTMhelix 184 206\ninside 207 208

1221 GCF_001523685.1_ASM152368v1 Burkholderia cepacia Proteobacteria; Betaproteobacteria; Burkholderiales; Burkholderiaceae; Burkholderia; Burkholderia cepacia complex MKRRTFLALPAVAASGMLAGCSNFGPVTKAWMSDSKYDETLSTFLITADGKQLVVLGKKY WP_027787377.1 MULTISPECIES: hypothetical protein [Burkholderia] Length: 210\nNumber of predicted TMHs: 1\nExp number of AAs in TMHs: 34.58187\nExp number, first 60 AAs: 12.86351\nTotal prob of N-in: 0.46668\nPOSSIBLE N-term signal sequence\noutside 1 186\nTMhelix 187 209\ninside 210 210

1222 GCF_001524765.1_ASM152476v1 Burkholderia cepacia Proteobacteria; Betaproteobacteria; Burkholderiales; Burkholderiaceae; Burkholderia; Burkholderia cepacia complex MKRRTFLALPAVAASGMLAGCSNFGPVTKAWMSDSKYDETLSTFLITADGKQLVVLGKKY WP_027787377.1 MULTISPECIES: hypothetical protein [Burkholderia] Length: 210\nNumber of predicted TMHs: 1\nExp number of AAs in TMHs: 34.58187\nExp number, first 60 AAs: 12.86351\nTotal prob of N-in: 0.46668\nPOSSIBLE N-term signal sequence\noutside 1 186\nTMhelix 187 209\ninside 210 210

1223 GCF_001522965.1_ASM152296v1 Burkholderia cepacia Proteobacteria; Betaproteobacteria; Burkholderiales; Burkholderiaceae; Burkholderia; Burkholderia cepacia complex MKRRTFLALPAVAASGMLAGCSNFGPVTKAWMSDSKYDETLSTFLITADGKQLVVLGKKY WP_059488505.1 hypothetical protein [Burkholderia cepacia] Length: 210\nNumber of predicted TMHs: 1\nExp number of AAs in TMHs: 34.5304\nExp number, first 60 AAs: 12.84424\nTotal prob of N-in: 0.46568\nPOSSIBLE N-term signal sequence\noutside 1 186\nTMhelix 187 209\ninside 210 210

1224 GCF_001522965.1_ASM152296v1 Burkholderia cepacia Proteobacteria; Betaproteobacteria; Burkholderiales; Burkholderiaceae; Burkholderia; Burkholderia cepacia complex MKRRALLTAGVIVVAAGLTGCTAMLFEDGRYEETVDRFLVSEDGKKFVVLGEKYHYIFDM WP_059496097.1 hypothetical protein [Burkholderia cepacia] Length: 208\nNumber of predicted TMHs: 2\nExp number of AAs in TMHs: 44.13865\nExp number, first 60 AAs: 21.11648\nTotal prob of N-in: 0.75846\nPOSSIBLE N-term signal sequence\ninside 1 6\nTMhelix 7 26\noutside 27 183\nTMhelix 184 206\ninside 207 208

1225 GCF_001523085.1_ASM152308v1 Burkholderia cepacia Proteobacteria; Betaproteobacteria; Burkholderiales; Burkholderiaceae; Burkholderia; Burkholderia cepacia complex MKRRTFLALPAVAASGMLAGCSNFGPVTKAWMSDSKYDETLSTFLITADGKQLVVLGKKY WP_059488505.1 hypothetical protein [Burkholderia cepacia] Length: 210\nNumber of predicted TMHs: 1\nExp number of AAs in TMHs: 34.5304\nExp number, first 60 AAs: 12.84424\nTotal prob of N-in: 0.46568\nPOSSIBLE N-term signal sequence\noutside 1 186\nTMhelix 187 209\ninside 210 210

1226 GCF_001523085.1_ASM152308v1 Burkholderia cepacia Proteobacteria; Betaproteobacteria; Burkholderiales; Burkholderiaceae; Burkholderia; Burkholderia cepacia complex MKRRALLTAGVIVVAAGLTGCTAMLFEDGRYEETVDRFLVSEDGKKFVVLGEKYHYIFDM WP_059496097.1 hypothetical protein [Burkholderia cepacia] Length: 208\nNumber of predicted TMHs: 2\nExp number of AAs in TMHs: 44.13865\nExp number, first 60 AAs: 21.11648\nTotal prob of N-in: 0.75846\nPOSSIBLE N-term signal sequence\ninside 1 6\nTMhelix 7 26\noutside 27 183\nTMhelix 184 206\ninside 207 208

1227 GCF_001522625.1_ASM152262v1 Burkholderia cepacia Proteobacteria; Betaproteobacteria; Burkholderiales; Burkholderiaceae; Burkholderia; Burkholderia cepacia complex MKRRALLTAGVIVVAAGLTGCTAMLFEDGRYEETVDRFLVSEDGKKFVVLGEKYHYIFDM WP_059496097.1 hypothetical protein [Burkholderia cepacia] Length: 208\nNumber of predicted TMHs: 2\nExp number of AAs in TMHs: 44.13865\nExp number, first 60 AAs: 21.11648\nTotal prob of N-in: 0.75846\nPOSSIBLE N-term signal sequence\ninside 1 6\nTMhelix 7 26\noutside 27 183\nTMhelix 184 206\ninside 207 208

1228 GCF_001525785.1_ASM152578v1 Burkholderia cepacia Proteobacteria; Betaproteobacteria; Burkholderiales; Burkholderiaceae; Burkholderia; Burkholderia cepacia complex MKTSITRRRLLAAAMAASTLALSGCFTPKLYENDAYSEYVTSFMITQDGKKLVVLGTNYH WP_059485162.1 hypothetical protein [Burkholderia cepacia] Length: 212\nNumber of predicted TMHs: 1\nExp number of AAs in TMHs: 24.29452\nExp number, first 60 AAs: 1.02778\nTotal prob of N-in: 0.64928\noutside 1 180\nTMhelix 181 203\ninside 204 212

1229 GCF_001525945.1_ASM152594v1 Burkholderia cepacia Proteobacteria; Betaproteobacteria; Burkholderiales; Burkholderiaceae; Burkholderia; Burkholderia cepacia complex MKTSITRRRLLAAAMAASTLALSGCFTPKLYENDAYSEYVTSFMITQDGKKLVVLGTNYH WP_059485162.1 hypothetical protein [Burkholderia cepacia] Length: 212\nNumber of predicted TMHs: 1\nExp number of AAs in TMHs: 24.29452\nExp number, first 60 AAs: 1.02778\nTotal prob of N-in: 0.64928\noutside 1 180\nTMhelix 181 203\ninside 204 212

1230 GCF_001720405.1_ASM172040v1 Burkholderia contaminans Proteobacteria; Betaproteobacteria; Burkholderiales; Burkholderiaceae; Burkholderia; Burkholderia cepacia complex MNVSITRRRLLAAGVAASSLALSGCFTPKLYKDDGYSEQVSGFMMTEDGKKLVVLGTDYH WP_039366460.1 MULTISPECIES: hypothetical protein [Burkholderia] Length: 212\nNumber of predicted TMHs: 1\nExp number of AAs in TMHs: 25.9561\nExp number, first 60 AAs: 0.849399999999999\nTotal prob of N-in: 0.56566\noutside 1 180\nTMhelix 181 203\ninside 204 212

1231 GCF_001865715.1_ASM186571v1 Burkholderia contaminans Proteobacteria; Betaproteobacteria; Burkholderiales; Burkholderiaceae; Burkholderia; Burkholderia cepacia complex MNVSITRRRLLAAGVAASSLALSGCFTPKLYKDDGYSEQVSGFMMTEDGKKLVVLGTDYH WP_071332150.1 hypothetical protein [Burkholderia contaminans] Length: 212\nNumber of predicted TMHs: 1\nExp number of AAs in TMHs: 25.96473\nExp number, first 60 AAs: 0.849319999999999\nTotal prob of N-in: 0.56583\noutside 1 180\nTMhelix 181 203\ninside 204 212

1232 GCF_001720405.1_ASM172040v1 Burkholderia contaminans Proteobacteria; Betaproteobacteria; Burkholderiales; Burkholderiaceae; Burkholderia; Burkholderia cepacia complex MKRRTFLALPAVAASGMLAGCSDANIGPITNRMLKDPEYRETLSAFLISADGKSLVVIGK WP_039354933.1 MULTISPECIES: 5-formyltetrahydrofolate cyclo-ligase [Burkholderia] Length: 212\nNumber of predicted TMHs: 1\nExp number of AAs in TMHs: 24.954\nExp number, first 60 AAs: 4.26972\nTotal prob of N-in: 0.39654\noutside 1 188\nTMhelix 189 211\ninside 212 212

1233 GCF_001029145.1_ASM102914v1 Burkholderia contaminans Proteobacteria; Betaproteobacteria; Burkholderiales; Burkholderiaceae; Burkholderia; Burkholderia cepacia complex MNVSITRRRLLAAGVAASSLALSGCFTPKLYKDDGYSEQVSGFMMTEDGKKLVVLGTDYH WP_046548430.1 hypothetical protein [Burkholderia contaminans] Length: 212\nNumber of predicted TMHs: 1\nExp number of AAs in TMHs: 25.30439\nExp number, first 60 AAs: 0.83557\nTotal prob of N-in: 0.55248\noutside 1 180\nTMhelix 181 203\ninside 204 212

1234 GCF_001758385.2_ASM175838v2 Burkholderia contaminans Proteobacteria; Betaproteobacteria; Burkholderiales; Burkholderiaceae; Burkholderia; Burkholderia cepacia complex MKRRTFLALPAVAASGMLAGCSDANIGPITNRMLKDPEYRETLSAFLISADGKSLVVIGK WP_039354933.1 MULTISPECIES: 5-formyltetrahydrofolate cyclo-ligase [Burkholderia] Length: 212\nNumber of predicted TMHs: 1\nExp number of AAs in TMHs: 24.954\nExp number, first 60 AAs: 4.26972\nTotal prob of N-in: 0.39654\noutside 1 188\nTMhelix 189 211\ninside 212 212

1235 GCF_001865715.1_ASM186571v1 Burkholderia contaminans Proteobacteria; Betaproteobacteria; Burkholderiales; Burkholderiaceae; Burkholderia; Burkholderia cepacia complex MKRRTLLTAGAVIVAAGLAGCTAMLFEDGHYEETVDRFLVSEDGKKFVVLGKKYHYIFDM WP_071334406.1 hypothetical protein [Burkholderia contaminans] Length: 208\nNumber of predicted TMHs: 2\nExp number of AAs in TMHs: 43.02465\nExp number, first 60 AAs: 19.94061\nTotal prob of N-in: 0.89156\nPOSSIBLE N-term signal sequence\ninside 1 6\nTMhelix 7 26\noutside 27 183\nTMhelix 184 206\ninside 207 208

1236 GCF_001865715.1_ASM186571v1 Burkholderia contaminans Proteobacteria; Betaproteobacteria; Burkholderiales; Burkholderiaceae; Burkholderia; Burkholderia cepacia complex MKRRTFIALPAVAASGMLAGCSDANIGPITNRMLKDPEYRETLSAFLISADGKSLVVIGK WP_071334492.1 5-formyltetrahydrofolate cyclo-ligase [Burkholderia contaminans] Length: 212\nNumber of predicted TMHs: 1\nExp

number of AAs in TMHs: 25.58861\nExp number, first 60 AAs: 4.80471\nTotal prob of N-in: 0.48605\noutside 1 188\nTMhelix 189 211\ninside 212 212

1237 GCF_001029145.1_ASM102914v1 Burkholderia contaminans Proteobacteria; Betaproteobacteria; Burkholderiales; Burkholderiaceae; Burkholderia; Burkholderia cepacia complex
MKRRTFLALPAVAASGMLAGCSNFGPVTKAWMKDSEYDETLSTFLITANGKQLVVLGKKY WP_047850866.1
hypothetical protein [Burkholderia contaminans] Length: 210\nNumber of predicted TMHs: 1\nExp number of AAs in TMHs: 33.66129\nExp number, first 60 AAs: 11.93749\nTotal prob of N-in: 0.44319\nPOSSIBLE N-term signal sequence\noutside 1 186\nTMhelix 187 209\ninside 210 210

1238 GCF_001029145.1_ASM102914v1 Burkholderia contaminans Proteobacteria; Betaproteobacteria; Burkholderiales; Burkholderiaceae; Burkholderia; Burkholderia cepacia complex
MKRRTLLTAGAVIVAAGLTGCTAMLFEDGQYEETVDRFLVSEDGKKFVVLGTKYHYILDM WP_047850963.1
hypothetical protein [Burkholderia contaminans] Length: 208\nNumber of predicted TMHs: 2\nExp number of AAs in TMHs: 42.39032\nExp number, first 60 AAs: 19.00032\nTotal prob of N-in: 0.85259\nPOSSIBLE N-term signal sequence\ninside 1 6\nTMhelix 7 26\noutside 27 183\nTMhelix 184 206\ninside 207 208

1239 GCF_001758385.2_ASM175838v2 Burkholderia contaminans Proteobacteria; Betaproteobacteria; Burkholderiales; Burkholderiaceae; Burkholderia; Burkholderia cepacia complex
MNVSITRRRLLAAGVAASSLALSGCFTPKLYKDDGYSEQVSGFMMTEDGKKLVVLGTDYH WP_039366460.1
MULTISPECIES: hypothetical protein [Burkholderia] Length: 212\nNumber of predicted TMHs: 1\nExp number of AAs in TMHs: 25.9561\nExp number, first 60 AAs: 0.849399999999999\nTotal prob of N-in: 0.56566\noutside 1 180\nTMhelix 181 203\ninside 204 212

1240 GCF_001523065.1_ASM152306v1 Burkholderia diffusa Proteobacteria; Betaproteobacteria; Burkholderiales; Burkholderiaceae; Burkholderia; Burkholderia cepacia complex
MSDDIDNGGKGGFTRRDMMRVMAASGMMAVGGGGLLTGARSAFAAPAPKRGGKIRVANES WP_059543332.1 ABC transporter substrate-binding protein [Burkholderia diffusa] Length: 527\nNumber of predicted TMHs: 2\nExp number of AAs in TMHs: 37.70292\nExp number, first 60 AAs: 21.84645\nTotal prob of N-in: 0.97213\nPOSSIBLE N-term signal sequence\ninside 1 20\nTMhelix 21 43\noutside 44 502\nTMhelix 503 525\ninside 526 527

1241 GCF_000012945.1_ASM1294v1 Burkholderia lata Proteobacteria; Betaproteobacteria; Burkholderiales; Burkholderiaceae; Burkholderia; Burkholderia cepacia complex
MSTSITRRRLLAAGMAASSLALSGCFTPALYENHDETYVEHVSAFMITKDGRKLVVLGER WP_011354338.1 hypothetical protein [Burkholderia lata] Length: 213\nNumber of predicted TMHs: 1\nExp number of AAs in TMHs: 24.80102\nExp number, first 60 AAs: 2.69778\nTotal prob of N-in: 0.60154\noutside 1 182\nTMhelix 183 205\ninside 206 213

1242 GCF_000012945.1_ASM1294v1 Burkholderia lata Proteobacteria; Betaproteobacteria; Burkholderiales; Burkholderiaceae; Burkholderia; Burkholderia cepacia complex
MKRRTFLALPAVAASGMLAGCSNFGPVTKAWMEDSEYDETLSTFLITADGKQLVVLGKKY WP_011352018.1
hypothetical protein [Burkholderia lata] Length: 210\nNumber of predicted TMHs: 1\nExp number of AAs in TMHs: 29.12587\nExp number, first 60 AAs: 7.38469\nTotal prob of N-in: 0.44274\noutside 1 186\nTMhelix 187 209\ninside 210 210

1243 GCF_000012945.1_ASM1294v1 Burkholderia lata Proteobacteria; Betaproteobacteria; Burkholderiales; Burkholderiaceae; Burkholderia; Burkholderia cepacia complex
MKRRTLLTAGAIIVAAGLTGCTAMLFEDSHYEETVDRFLVSEDGKKFVVLGKKYHYIFDM WP_011351870.1 hypothetical protein [Burkholderia lata] Length: 208\nNumber of predicted TMHs: 2\nExp number of AAs in TMHs: 43.47055\nExp number, first 60 AAs: 19.56622\nTotal prob of N-in: 0.86157\nPOSSIBLE N-term signal sequence\ninside 1 6\nTMhelix 7 26\noutside 27 183\nTMhelix 184 206\ninside 207 208

1244 GCF_001718575.1_ASM171857v1 Burkholderia lata Proteobacteria; Betaproteobacteria; Burkholderiales; Burkholderiaceae; Burkholderia; Burkholderia cepacia complex
MKRRTFLALPAVAASAMLAGCADADIGPITNRMMKDPEYRETLSAFLISADGKSLVVIGK WP_069247994.1 5-formyltetrahydrofolate cyclo-ligase [Burkholderia lata] Length: 212\nNumber of predicted TMHs: 1\nExp number of AAs in TMHs: 29.24129\nExp number, first 60 AAs: 8.20568\nTotal prob of N-in: 0.54502\noutside 1 188\nTMhelix 189 211\ninside 212 212

1245 GCF_001718575.1_ASM171857v1 Burkholderia lata Proteobacteria; Betaproteobacteria; Burkholderiales; Burkholderiaceae; Burkholderia; Burkholderia cepacia complex
MKRRTLLTTGAIVVAAGLAGCTAMLFEDGHYEETVDRFLVSEDGKQFVVLGKKYHYIFDM WP_069247875.1

hypothetical protein [Burkholderia lata] Length: 208\nNumber of predicted TMHs: 2\nExp number of AAs in TMHs: 40.93368\nExp number, first 60 AAs: 18.02997\nTotal prob of N-in: 0.84906\nPOSSIBLE N-term signal sequence\ninside 1 6\nTMhelix 7 26\noutside 27 183\nTMhelix 184 206\ninside 207 208

1246 GCF_001038835.1_ASM103883v1 Burkholderia lata Proteobacteria; Betaproteobacteria; Burkholderiales; Burkholderiaceae; Burkholderia; Burkholderia cepacia complex MKRRTFLALPAVAASGMLAGCSNFGPVTKAWMSDSKYDETLSTFLITADGKQLVVLGKKY WP_027787377.1 MULTISPECIES: hypothetical protein [Burkholderia] Length: 210\nNumber of predicted TMHs: 1\nExp number of AAs in TMHs: 34.58187\nExp number, first 60 AAs: 12.86351\nTotal prob of N-in: 0.46668\nPOSSIBLE N-term signal sequence\noutside 1 186\nTMhelix 187 209\ninside 210 210

1247 GCF_001038745.1_ASM103874v1 Burkholderia lata Proteobacteria; Betaproteobacteria; Burkholderiales; Burkholderiaceae; Burkholderia; Burkholderia cepacia complex MKRRTFLALPAVAASGMLAGCSNFGPVTKAWMSDSKYDETLSTFLITADGKQLVVLGKKY WP_027787377.1 MULTISPECIES: hypothetical protein [Burkholderia] Length: 210\nNumber of predicted TMHs: 1\nExp number of AAs in TMHs: 34.58187\nExp number, first 60 AAs: 12.86351\nTotal prob of N-in: 0.46668\nPOSSIBLE N-term signal sequence\noutside 1 186\nTMhelix 187 209\ninside 210 210

1248 GCF_001718555.1_ASM171855v1 Burkholderia metallica Proteobacteria; Betaproteobacteria; Burkholderiales; Burkholderiaceae; Burkholderia; Burkholderia cepacia complex MSLSITRRRLLAAGIAASSLALSGCFTPALYENRDETYIEHVSAFMITKDGKKLVVLGER WP_069260495.1 hypothetical protein [Burkholderia metallica] Length: 213\nNumber of predicted TMHs: 1\nExp number of AAs in TMHs: 26.81248\nExp number, first 60 AAs: 4.9294\nTotal prob of N-in: 0.48611\noutside 1 182\nTMhelix 183 205\ninside 206 213

1249 GCF_000756965.1_BHV_1 Burkholderia multivorans Proteobacteria; Betaproteobacteria; Burkholderiales; Burkholderiaceae; Burkholderia; Burkholderia cepacia complex MSDDIDNGGGHGGFTRRDMMRVMAAGGMMAAGGGALLMPGQAAFAAPAPKRGGKIRVANE WP_035956625.1 ABC transporter substrate-binding protein [Burkholderia multivorans] Length: 528\nNumber of predicted TMHs: 2\nExp number of AAs in TMHs: 36.36737\nExp number, first 60 AAs: 21.42524\nTotal prob of N-in: 0.89413\nPOSSIBLE N-term signal sequence\ninside 1 20\nTMhelix 21 43\noutside 44 503\nTMhelix 504 526\ninside 527 528

1250 GCF_001528045.1_ASM152804v1 Burkholderia multivorans Proteobacteria; Betaproteobacteria; Burkholderiales; Burkholderiaceae; Burkholderia; Burkholderia cepacia complex MSDDIDNGGGHGGFTRRDMMRVMAAGGMMAAGGGALLMPGQAAFAAPAPKRGGKIRVANE WP_035956625.1 ABC transporter substrate-binding protein [Burkholderia multivorans] Length: 528\nNumber of predicted TMHs: 2\nExp number of AAs in TMHs: 36.36737\nExp number, first 60 AAs: 21.42524\nTotal prob of N-in: 0.89413\nPOSSIBLE N-term signal sequence\ninside 1 20\nTMhelix 21 43\noutside 44 503\nTMhelix 504 526\ninside 527 528

1251 GCF_001302465.1_BMUNKI379.1 Burkholderia multivorans Proteobacteria; Betaproteobacteria; Burkholderiales; Burkholderiaceae; Burkholderia; Burkholderia cepacia complex MSDDIDNGGGHGGFTRRDMMRVMAAGGMMAAGGGALLMPGQAAFAAPAPKRGGKIRVANE WP_054315074.1 ABC transporter substrate-binding protein [Burkholderia multivorans] Length: 528\nNumber of predicted TMHs: 2\nExp number of AAs in TMHs: 36.36756\nExp number, first 60 AAs: 21.42525\nTotal prob of N-in: 0.89413\nPOSSIBLE N-term signal sequence\ninside 1 20\nTMhelix 21 43\noutside 44 503\nTMhelix 504 526\ninside 527 528

1252 GCF_001530485.1_ASM153048v1 Burkholderia multivorans Proteobacteria; Betaproteobacteria; Burkholderiales; Burkholderiaceae; Burkholderia; Burkholderia cepacia complex MSDDIDNGGGHGGFTRRDMMRVMAAGGMMAAGGGALLMPGQAAFAAPAPKRGGKIRVANE WP_060110928.1 ABC transporter substrate-binding protein [Burkholderia multivorans] Length: 528\nNumber of predicted TMHs: 2\nExp number of AAs in TMHs: 36.43565\nExp number, first 60 AAs: 21.42523\nTotal prob of N-in: 0.89414\nPOSSIBLE N-term signal sequence\ninside 1 20\nTMhelix 21 43\noutside 44 503\nTMhelix 504 526\ninside 527 528

1253 GCF_001532145.1_ASM153214v1 Burkholderia multivorans Proteobacteria; Betaproteobacteria; Burkholderiales; Burkholderiaceae; Burkholderia; Burkholderia cepacia complex MSDDIDNGGGHGGFTRRDMMRVMAAGGMMAAGGGALLMPGQAAFAAPAPKRGGKIRVANE WP_060145365.1 ABC transporter substrate-binding protein [Burkholderia multivorans] Length: 528\nNumber of predicted TMHs: 2\nExp number of AAs in TMHs: 36.36871\nExp number, first 60 AAs: 21.42528\nTotal prob of N-in: 0.89412\nPOSSIBLE N-term signal sequence\ninside 1 20\nTMhelix 21 43\noutside 44 503\nTMhelix 504 526\ninside 527 528

1254 GCF_001531955.1_ASM153195v1 Burkholderia multivorans Proteobacteria; Betaproteobacteria; Burkholderiales; Burkholderiaceae; Burkholderia; Burkholderia cepacia complex
MSDDIDNGGGHGGFTRRDMMRVMAAGGMMAAGGGALLMPGQAAFAAPAPKRGGKIRVANE WP_060181816.1 ABC transporter substrate-binding protein [Burkholderia multivorans] Length: 528\nNumber of predicted TMHs: 2\nExp number of AAs in TMHs: 36.36755\nExp number, first 60 AAs: 21.42524\nTotal prob of N-in: 0.89413\nPOSSIBLE N-term signal sequence\ninside 1 20\nTMhelix 21 43\noutside 44 503\nTMhelix 504 526\ninside 527 528

1255 GCF_001534105.1_ASM153410v1 Burkholderia multivorans Proteobacteria; Betaproteobacteria; Burkholderiales; Burkholderiaceae; Burkholderia; Burkholderia cepacia complex
MSDDIDNGGGHGGFTRRDMMRVMAAGGMMAAGGGALLMPGQAAFAAPAPKRGGKIRVANE WP_060181816.1 ABC transporter substrate-binding protein [Burkholderia multivorans] Length: 528\nNumber of predicted TMHs: 2\nExp number of AAs in TMHs: 36.36755\nExp number, first 60 AAs: 21.42524\nTotal prob of N-in: 0.89413\nPOSSIBLE N-term signal sequence\ninside 1 20\nTMhelix 21 43\noutside 44 503\nTMhelix 504 526\ninside 527 528

1256 GCF_001528425.1_ASM152842v1 Burkholderia multivorans Proteobacteria; Betaproteobacteria; Burkholderiales; Burkholderiaceae; Burkholderia; Burkholderia cepacia complex
MSDDIDNGGGHGGFTRRDMMRVMAAGGMMAAGGGALLMPGQAAFAAPAPKRGGKIRVANE WP_059873413.1
MULTISPECIES: ABC transporter substrate-binding protein [Burkholderia] Length: 528\nNumber of predicted TMHs: 2\nExp number of AAs in TMHs: 36.36756\nExp number, first 60 AAs: 21.42525\nTotal prob of N-in: 0.89413\nPOSSIBLE N-term signal sequence\ninside 1 20\nTMhelix 21 43\noutside 44 503\nTMhelix 504 526\ninside 527 528

1257 GCF_001526715.1_ASM152671v1 Burkholderia multivorans Proteobacteria; Betaproteobacteria; Burkholderiales; Burkholderiaceae; Burkholderia; Burkholderia cepacia complex
MSDDIDNGGGHGGFTRRDMMRVMAAGGMMAAGGGALLMPGQAAFAAPAPKRGGKIRVANE WP_059785739.1 ABC transporter substrate-binding protein [Burkholderia multivorans] Length: 528\nNumber of predicted TMHs: 2\nExp number of AAs in TMHs: 36.36735\nExp number, first 60 AAs: 21.42522\nTotal prob of N-in: 0.89415\nPOSSIBLE N-term signal sequence\ninside 1 20\nTMhelix 21 43\noutside 44 503\nTMhelix 504 526\ninside 527 528

1258 GCF_000807815.1_D2214.1 Burkholderia multivorans Proteobacteria; Betaproteobacteria; Burkholderiales; Burkholderiaceae; Burkholderia; Burkholderia cepacia complex
MSDDIDNGGGHGGFTRRDMMRVMAAGGMMAAGGGALLMPGQAAFAAPAPKRGGKIRVANE WP_039223870.1 ABC transporter substrate-binding protein [Burkholderia multivorans] Length: 528\nNumber of predicted TMHs: 2\nExp number of AAs in TMHs: 36.36755\nExp number, first 60 AAs: 21.42524\nTotal prob of N-in: 0.89413\nPOSSIBLE N-term signal sequence\ninside 1 20\nTMhelix 21 43\noutside 44 503\nTMhelix 504 526\ninside 527 528

1259 GCF_000807825.1_D2095.1 Burkholderia multivorans Proteobacteria; Betaproteobacteria; Burkholderiales; Burkholderiaceae; Burkholderia; Burkholderia cepacia complex
MSDDIDNGGGHGGFTRRDMMRVMAAGGMMAAGGGALLMPGQAAFAAPAPKRGGKIRVANE WP_039223870.1 ABC transporter substrate-binding protein [Burkholderia multivorans] Length: 528\nNumber of predicted TMHs: 2\nExp number of AAs in TMHs: 36.36755\nExp number, first 60 AAs: 21.42524\nTotal prob of N-in: 0.89413\nPOSSIBLE N-term signal sequence\ninside 1 20\nTMhelix 21 43\noutside 44 503\nTMhelix 504 526\ninside 527 528

1260 GCF_001058025.1_ASM105802v1 Burkholderia multivorans Proteobacteria; Betaproteobacteria; Burkholderiales; Burkholderiaceae; Burkholderia; Burkholderia cepacia complex
MNAPETIDPLHADLPREGVAMLHPAARPLTDAELAARRSRSRRATFIKWLRKVHGWVGLW WP_049018021.1
peptidase, partial [Burkholderia multivorans] Length: 216\nNumber of predicted TMHs: 2\nExp number of AAs in TMHs: 44.4077\nExp number, first 60 AAs: 6.75104\nTotal prob of N-in: 0.25127\noutside 1 52\nTMhelix 53 75\ninside 76 192\nTMhelix 193 215\noutside 216 216

1261 GCF_001058025.1_ASM105802v1 Burkholderia multivorans Proteobacteria; Betaproteobacteria; Burkholderiales; Burkholderiaceae; Burkholderia; Burkholderia cepacia complex
MSVIDSVEIAATPVPRTAPGARTGSARSRRGTFLQWLRKIHGWVGLWGAALGLLFGTTGF WP_049018213.1
peptidase, partial [Burkholderia multivorans] Length: 205\nNumber of predicted TMHs: 2\nExp number of AAs in TMHs: 44.74785\nExp number, first 60 AAs: 19.8795\nTotal prob of N-in: 0.14783\nPOSSIBLE N-term signal sequence\noutside 1 39\nTMhelix 40 62\ninside 63 181\nTMhelix 182 204\noutside 205 205

1262 GCF_001532085.1_ASM153208v1 Burkholderia pseudomultivorans Proteobacteria; Betaproteobacteria; Burkholderiales; Burkholderiaceae; Burkholderia; Burkholderia cepacia complex
MKRRTLLRASAIAVTAGLAGCTSMLFEDGRYDETVDRFLVSEDGKKFVVLGQQYHYIFDM WP_060251299.1
hypothetical protein [Burkholderia pseudomultivorans] Length: 209\nNumber of predicted TMHs: 1\nExp

number of AAs in TMHs: 28.98884\nExp number, first 60 AAs: 5.86933\nTotal prob of N-in: 0.86025\ninside 1 182\nTMhelix 183 205\noutside 206 209

1263 GCF_001532085.1_ASM153208v1 Burkholderia pseudomultivorans Proteobacteria; Betaproteobacteria; Burkholderiales; Burkholderiaceae; Burkholderia; Burkholderia cepacia complex
MTISTTRRRWLGAALAASSVMLSGCFTPKLYEEGGYHEDVSGFMVTEDGKKLVVLGARYH WP_060254451.1
hypothetical protein [Burkholderia pseudomultivorans] Length: 213\nNumber of predicted TMHs: 1\nExp number of AAs in TMHs: 22.92992\nExp number, first 60 AAs: 0.5306\nTotal prob of N-in: 0.73121\ninside 1 180\nTMhelix 181 203\noutside 204 213

1264 GCF_001524505.1_ASM152450v1 Burkholderia pseudomultivorans Proteobacteria; Betaproteobacteria; Burkholderiales; Burkholderiaceae; Burkholderia; Burkholderia cepacia complex
MTISTTRRRWLGAALAASSVMLSGCFTPKLYEEGGYHEDVSGFMVTEDGKKLVVLGARYH WP_059481399.1
hypothetical protein [Burkholderia pseudomultivorans] Length: 213\nNumber of predicted TMHs: 1\nExp number of AAs in TMHs: 22.92992\nExp number, first 60 AAs: 0.5306\nTotal prob of N-in: 0.73121\ninside 1 180\nTMhelix 181 203\noutside 204 213

1265 GCF_001532035.1_ASM153203v1 Burkholderia pseudomultivorans Proteobacteria; Betaproteobacteria; Burkholderiales; Burkholderiaceae; Burkholderia; Burkholderia cepacia complex
MRISTTRRRWLGAALAASSIMLSGCFTPKLYEEGGYHEDVSGFMVTEDGKKLVVLGARYH WP_060245633.1
hypothetical protein [Burkholderia pseudomultivorans] Length: 213\nNumber of predicted TMHs: 1\nExp number of AAs in TMHs: 23.17873\nExp number, first 60 AAs: 0.7519\nTotal prob of N-in: 0.78114\ninside 1 180\nTMhelix 181 203\noutside 204 213

1266 GCF_001523105.1_ASM152310v1 Burkholderia pseudomultivorans Proteobacteria; Betaproteobacteria; Burkholderiales; Burkholderiaceae; Burkholderia; Burkholderia cepacia complex
MTISTTRRRWLGAALAASSVMLSGCFTPKLYEEGGYHEDVSGFMVTEDGKKLVVLGARYH WP_059517016.1
hypothetical protein [Burkholderia pseudomultivorans] Length: 213\nNumber of predicted TMHs: 1\nExp number of AAs in TMHs: 23.39939\nExp number, first 60 AAs: 0.57121\nTotal prob of N-in: 0.69940\ninside 1 180\nTMhelix 181 203\noutside 204 213

1267 GCF_001523105.1_ASM152310v1 Burkholderia pseudomultivorans Proteobacteria; Betaproteobacteria; Burkholderiales; Burkholderiaceae; Burkholderia; Burkholderia cepacia complex
MKRRTLLRASAIAVTAGLAGCTSMLFEDGRYDETVDRFLVSEDGKKFVVLGQQYHYIFDM WP_059514641.1
hypothetical protein [Burkholderia pseudomultivorans] Length: 209\nNumber of predicted TMHs: 1\nExp number of AAs in TMHs: 28.98898\nExp number, first 60 AAs: 5.86933\nTotal prob of N-in: 0.86026\ninside 1 182\nTMhelix 183 205\noutside 206 209

1268 GCF_001524505.1_ASM152450v1 Burkholderia pseudomultivorans Proteobacteria; Betaproteobacteria; Burkholderiales; Burkholderiaceae; Burkholderia; Burkholderia cepacia complex
MKRRTLLRASAIAVTAGLAGCTSMLFEDGRYDETVDRFLVSEDGKKFVVLGRQYHYIFDM WP_059479327.1
hypothetical protein [Burkholderia pseudomultivorans] Length: 209\nNumber of predicted TMHs: 1\nExp number of AAs in TMHs: 28.95561\nExp number, first 60 AAs: 5.85426\nTotal prob of N-in: 0.86014\ninside 1 182\nTMhelix 183 205\noutside 206 209

1269 GCF_001718415.1_ASM171841v1 Burkholderia pseudomultivorans Proteobacteria; Betaproteobacteria; Burkholderiales; Burkholderiaceae; Burkholderia; Burkholderia cepacia complex
MTISTTRRRWLGAALAASSVMLSGCFTPKLYEEGGYHEDVSGFMVTEDGKKLVVLGARYH WP_069243740.1
hypothetical protein [Burkholderia pseudomultivorans] Length: 213\nNumber of predicted TMHs: 1\nExp number of AAs in TMHs: 22.92897\nExp number, first 60 AAs: 0.53068\nTotal prob of N-in: 0.73119\ninside 1 180\nTMhelix 181 203\noutside 204 213

1270 GCF_001718415.1_ASM171841v1 Burkholderia pseudomultivorans Proteobacteria; Betaproteobacteria; Burkholderiales; Burkholderiaceae; Burkholderia; Burkholderia cepacia complex
MKRRTLLRASAIAVTAGLAGCTSMLFEDGRYDETVDRFLVSEDGKKFVVLGQQYHYIFDM WP_059514641.1
hypothetical protein [Burkholderia pseudomultivorans] Length: 209\nNumber of predicted TMHs: 1\nExp number of AAs in TMHs: 28.98898\nExp number, first 60 AAs: 5.86933\nTotal prob of N-in: 0.86026\ninside 1 182\nTMhelix 183 205\noutside 206 209

1271 GCF_001532065.1_ASM153206v1 Burkholderia pseudomultivorans Proteobacteria; Betaproteobacteria; Burkholderiales; Burkholderiaceae; Burkholderia; Burkholderia cepacia complex

MKRRTLLRASAIAVTAGLAGCTSMLFEDGRYDETVDRFLVSEDGKKFVVLGRQYHYIFDM WP_059479327.1 hypothetical protein [Burkholderia pseudomultivorans] Length: 209\nNumber of predicted TMHs: 1\nExp number of AAs in TMHs: 28.95561\nExp number, first 60 AAs: 5.85426\nTotal prob of N-in: 0.86014\ninside 1 182\nTMhelix 183 205\noutside 206 209

1272 GCF_001533465.1_ASM153346v1 Burkholderia pseudomultivorans Proteobacteria; Betaproteobacteria; Burkholderiales; Burkholderiaceae; Burkholderia; Burkholderia cepacia complex MKRRTLLRASAIAVTAGLAGCTSMLFEDGRYDETVDRFLVSEDGKKFVVLGQQYHYIFDM WP_040127356.1 MULTISPECIES: hypothetical protein [Burkholderia cepacia complex] Length: 209\nNumber of predicted TMHs: 1\nExp number of AAs in TMHs: 28.96048\nExp number, first 60 AAs: 5.85843999999999\nTotal prob of N-in: 0.86015\ninside 1 182\nTMhelix 183 205\noutside 206 209

1273 GCF_001525825.1_ASM152582v1 Burkholderia pseudomultivorans Proteobacteria; Betaproteobacteria; Burkholderiales; Burkholderiaceae; Burkholderia; Burkholderia cepacia complex MTISTTRRRWLGAALAASSVMLSGCFTPKLYEEGGYHEDVSGFMVTEDGKKLVVLGARYH WP_059605889.1 hypothetical protein [Burkholderia pseudomultivorans] Length: 213\nNumber of predicted TMHs: 1\nExp number of AAs in TMHs: 22.93041\nExp number, first 60 AAs: 0.53063\nTotal prob of N-in: 0.73121\ninside 1 180\nTMhelix 181 203\noutside 204 213

1274 GCF_001523165.1_ASM152316v1 Burkholderia pseudomultivorans Proteobacteria; Betaproteobacteria; Burkholderiales; Burkholderiaceae; Burkholderia; Burkholderia cepacia complex MTISTTRRRWLGAALAASSVMLSGCFTPKLYEEGGYHEDVSGFMVTEDGKKLVVLGARYH WP_059517016.1 hypothetical protein [Burkholderia pseudomultivorans] Length: 213\nNumber of predicted TMHs: 1\nExp number of AAs in TMHs: 23.39939\nExp number, first 60 AAs: 0.57121\nTotal prob of N-in: 0.69940\ninside 1 180\nTMhelix 181 203\noutside 204 213

1275 GCF_001533465.1_ASM153346v1 Burkholderia pseudomultivorans Proteobacteria; Betaproteobacteria; Burkholderiales; Burkholderiaceae; Burkholderia; Burkholderia cepacia complex MTISTTRRRWLGAALAASSVMLSGCFTPKLYEEGGYHEDVSAFMVTEDGKKLVVLGARYH WP_060335132.1 hypothetical protein [Burkholderia pseudomultivorans] Length: 213\nNumber of predicted TMHs: 1\nExp number of AAs in TMHs: 22.92886\nExp number, first 60 AAs: 0.52906\nTotal prob of N-in: 0.73108\ninside 1 180\nTMhelix 181 203\noutside 204 213

1276 GCF_001532065.1_ASM153206v1 Burkholderia pseudomultivorans Proteobacteria; Betaproteobacteria; Burkholderiales; Burkholderiaceae; Burkholderia; Burkholderia cepacia complex MTISTTRRRWLGAALAASSVMLSGCFTPKLYEEGGYHEDVSGFMVTEDGKKLVVLGARYH WP_060299571.1 hypothetical protein [Burkholderia pseudomultivorans] Length: 213\nNumber of predicted TMHs: 1\nExp number of AAs in TMHs: 22.95776\nExp number, first 60 AAs: 0.474210000000001\nTotal prob of N-in: 0.76966\ninside 1 180\nTMhelix 181 203\noutside 204 213

1277 GCF_001525825.1_ASM152582v1 Burkholderia pseudomultivorans Proteobacteria; Betaproteobacteria; Burkholderiales; Burkholderiaceae; Burkholderia; Burkholderia cepacia complex MKRRTLLRASAIAVTAGLAGCTSMLFEDGRYDETVDRFLVSEDGKKFVVLGQQYHYIFDM WP_059514641.1 hypothetical protein [Burkholderia pseudomultivorans] Length: 209\nNumber of predicted TMHs: 1\nExp number of AAs in TMHs: 28.98898\nExp number, first 60 AAs: 5.86933\nTotal prob of N-in: 0.86026\ninside 1 182\nTMhelix 183 205\noutside 206 209

1278 GCF_001523165.1_ASM152316v1 Burkholderia pseudomultivorans Proteobacteria; Betaproteobacteria; Burkholderiales; Burkholderiaceae; Burkholderia; Burkholderia cepacia complex MKRRTLLRASAIAVTAGLAGCTSMLFEDGRYDETVDRFLVSEDGKKFVVLGQQYHYIFDM WP_059514641.1 hypothetical protein [Burkholderia pseudomultivorans] Length: 209\nNumber of predicted TMHs: 1\nExp number of AAs in TMHs: 28.98898\nExp number, first 60 AAs: 5.86933\nTotal prob of N-in: 0.86026\ninside 1 182\nTMhelix 183 205\noutside 206 209

1279 GCF_001028665.1_ASM102866v1 Burkholderia pyrrocinia Proteobacteria; Betaproteobacteria; Burkholderiales; Burkholderiaceae; Burkholderia; Burkholderia cepacia complex MSISITRRRLLAAGLAASSLALSGCFTPKLYKDDAYSEYVSGFMISEDGKKLVVLGMRYH WP_047902623.1 MULTISPECIES: hypothetical protein [Burkholderia cepacia complex] Length: 212\nNumber of predicted TMHs: 1\nExp number of AAs in TMHs: 23.99618\nExp number, first 60 AAs: 0.75736\nTotal prob of N-in: 0.41515\noutside 1 180\nTMhelix 181 203\ninside 204 212

1280 GCF_001028665.1_ASM102866v1 Burkholderia pyrrocinia Proteobacteria; Betaproteobacteria; Burkholderiales; Burkholderiaceae; Burkholderia; Burkholderia cepacia complex
MKRRTFLALPAVAASGMLAGCSDFGPVTNHMMKDPEYRETLTAFLISADGKKLVVLGKRY WP_047900707.1
hypothetical protein [Burkholderia pyrrocinia] Length: 216\nNumber of predicted TMHs: 1\nExp number of AAs in TMHs: 30.2816\nExp number, first 60 AAs: 8.09011000000001\nTotal prob of N-in: 0.44832\noutside 1 184\nTMhelix 185 207\ninside 208 216

1281 GCF_000743695.1_LYC2_genome1.0 Burkholderia pyrrocinia Proteobacteria; Betaproteobacteria; Burkholderiales; Burkholderiaceae; Burkholderia; Burkholderia cepacia complex
MSISITRRRLLAAGIAASSVTLSGCFTPKLYKDDAYSEEVSGFMITEDGKKLVVLGTRYH WP_034180619.1 hypothetical protein [Burkholderia pyrrocinia] Length: 212\nNumber of predicted TMHs: 1\nExp number of AAs in TMHs: 24.43089\nExp number, first 60 AAs: 0.44099\nTotal prob of N-in: 0.31905\noutside 1 180\nTMhelix 181 203\ninside 204 212

1282 GCF_001524085.1_ASM152408v1 Burkholderia seminalis Proteobacteria; Betaproteobacteria; Burkholderiales; Burkholderiaceae; Burkholderia; Burkholderia cepacia complex
MKRRTFLALPAVAAAGMLAGCTDVSIGPVTDRMMKDPEYRETLSAFLISADGKKLVVIGK WP_059557670.1
hypothetical protein [Burkholderia seminalis] Length: 212\nNumber of predicted TMHs: 2\nExp number of AAs in TMHs: 40.47155\nExp number, first 60 AAs: 18.26817\nTotal prob of N-in: 0.80532\nPOSSIBLE N-term signal sequence\ninside 1 4\nTMhelix 5 27\noutside 28 187\nTMhelix 188 210\ninside 211 212

1283 GCF_001718535.1_ASM171853v1 Burkholderia seminalis Proteobacteria; Betaproteobacteria; Burkholderiales; Burkholderiaceae; Burkholderia; Burkholderia cepacia complex
MKRRTFLALPAVAAAGMLAGCTDVSIGPVTDRMMKDPEYRETLSAFLISADGKKLVVIGK WP_059557670.1
hypothetical protein [Burkholderia seminalis] Length: 212\nNumber of predicted TMHs: 2\nExp number of AAs in TMHs: 40.47155\nExp number, first 60 AAs: 18.26817\nTotal prob of N-in: 0.80532\nPOSSIBLE N-term signal sequence\ninside 1 4\nTMhelix 5 27\noutside 28 187\nTMhelix 188 210\ninside 211 212

1284 GCF_001524085.1_ASM152408v1 Burkholderia seminalis Proteobacteria; Betaproteobacteria; Burkholderiales; Burkholderiaceae; Burkholderia; Burkholderia cepacia complex
MSTSITRRRLLAAAMAASSLALSGCFTPKLYKDDAYSEYVTSFMITQDGKKLVVLGTRYH WP_034189316.1 MULTISPECIES: hypothetical protein [Burkholderia cepacia complex] Length: 212\nNumber of predicted TMHs: 1\nExp number of AAs in TMHs: 25.28913\nExp number, first 60 AAs: 1.00711\nTotal prob of N-in: 0.41198\noutside 1 180\nTMhelix 181 203\ninside 204 212

1285 GCF_001718535.1_ASM171853v1 Burkholderia seminalis Proteobacteria; Betaproteobacteria; Burkholderiales; Burkholderiaceae; Burkholderia; Burkholderia cepacia complex
MSTSITRRRLLAAAMAASSLALSGCFTPKLYKDDAYSEYVTSFMITQDGKKLVVLGTRYH WP_034189316.1 MULTISPECIES: hypothetical protein [Burkholderia cepacia complex] Length: 212\nNumber of predicted TMHs: 1\nExp number of AAs in TMHs: 25.28913\nExp number, first 60 AAs: 1.00711\nTotal prob of N-in: 0.41198\noutside 1 180\nTMhelix 181 203\ninside 204 212

1286 GCF_001742345.1_ASM174234v1 Burkholderia seminalis Proteobacteria; Betaproteobacteria; Burkholderiales; Burkholderiaceae; Burkholderia; Burkholderia cepacia complex
MSTSITRRRLLAAAMAASSLALSGCFTPKLYKDDAYSEYVTSFMITQDGKKLVVLGTRYH WP_034189316.1 MULTISPECIES: hypothetical protein [Burkholderia cepacia complex] Length: 212\nNumber of predicted TMHs: 1\nExp number of AAs in TMHs: 25.28913\nExp number, first 60 AAs: 1.00711\nTotal prob of N-in: 0.41198\noutside 1 180\nTMhelix 181 203\ninside 204 212

1287 GCF_000466765.1_Burkholderia_vietnamiensis Burkholderia sp. AU4i Proteobacteria; Betaproteobacteria; Burkholderiales; Burkholderiaceae; Burkholderia; Burkholderia cepacia complex
MKRRTFLALPAVAASGMLVGCSNFGPVTKAWMSDSKYDETLSTFLITADGKQLVVLGKKY WP_021159606.1
hypothetical protein [Burkholderia sp. AU4i] Length: 210\nNumber of predicted TMHs: 1\nExp number of AAs in TMHs: 37.46363\nExp number, first 60 AAs: 15.76001\nTotal prob of N-in: 0.49698\nPOSSIBLE N-term signal sequence\noutside 1 186\nTMhelix 187 209\ninside 210 210

1288 GCF_002099195.1_ASM209919v1 Burkholderia sp. CAMPA 1040 Proteobacteria; Betaproteobacteria; Burkholderiales; Burkholderiaceae; Burkholderia; Burkholderia cepacia complex
MKRRTFLALPAVAASGMLAGCANFGPVTKAWMKDSKYDETLSTFLITADGKQLVVLGKKY WP_085039069.1 5-formyltetrahydrofolate cyclo-ligase [Burkholderia sp. CAMPA 1040] Length: 210\nNumber of predicted TMHs: 2\nExp

number of AAs in TMHs: 41.40762\nExp number, first 60 AAs: 19.71784\nTotal prob of N-in: 0.40351\nPOSSIBLE N-term signal sequence\noutside 1 4\nTMhelix 5 27\ninside 28 189\nTMhelix 190 209\noutside 210 210

1289 GCF_002099195.1_ASM209919v1 Burkholderia sp. CAMPA 1040 Proteobacteria; Betaproteobacteria; Burkholderiales; Burkholderiaceae; Burkholderia; Burkholderia cepacia complex MSTSITRRRLLAAGMAASSLALSGCFTPALYENHDETYVEHVSAFMITKDGKKLVVLGER WP_085038676.1 hypothetical protein [Burkholderia sp. CAMPA 1040]Length: 213\nNumber of predicted TMHs: 1\nExp number of AAs in TMHs: 24.80625\nExp number, first 60 AAs: 2.62355\nTotal prob of N-in: 0.56102\noutside 1 182\nTMhelix 183 205\ninside 206 213

1290 GCF_001524725.1_ASM152472v1 Burkholderia sp. MSMB1072 Proteobacteria; Betaproteobacteria; Burkholderiales; Burkholderiaceae; Burkholderia; Burkholderia cepacia complex MKRRAALLGLSLLASFAAADGAPNAGRAMQRFALDLDGSAAYYQLTVPQPVYAASRRDDL WP_059789976.1 hypothetical protein [Burkholderia sp. MSMB1072] Length: 451\nNumber of predicted TMHs: 1\nExp number of AAs in TMHs: 26.3808\nExp number, first 60 AAs: 5.47341\nTotal prob of N-in: 0.35715\noutside 1 417\nTMhelix 418 440\ninside 441 451

1291 GCF_001528895.1_ASM152889v1 Burkholderia sp. MSMB1078WGS Proteobacteria; Betaproteobacteria; Burkholderiales; Burkholderiaceae; Burkholderia; Burkholderia cepacia complex MKRRAALLGLSLLASFAAADGAPNAGRAMQRFALDLDGSAAYYQLTVPQPVYAASRRDDL WP_060030825.1 hypothetical protein [Burkholderia sp. MSMB1078WGS] Length: 451\nNumber of predicted TMHs: 1\nExp number of AAs in TMHs: 26.22521\nExp number, first 60 AAs: 5.00445\nTotal prob of N-in: 0.42314\noutside 1 417\nTMhelix 418 440\ninside 441 451

1292 GCF_001528895.1_ASM152889v1 Burkholderia sp. MSMB1078WGS Proteobacteria; Betaproteobacteria; Burkholderiales; Burkholderiaceae; Burkholderia; Burkholderia cepacia complex MESLLTRIRASFADASPRRQLISVAVLVALVASLITAGYFALRPRYQVLFRDLKPQDAAT WP_060036044.1 MULTISPECIES: flagellar M-ring protein FliF [Burkholderia cepacia complex] Length: 491\nNumber of predicted TMHs: 2\nExp number of AAs in TMHs: 42.71388\nExp number, first 60 AAs: 22.14192\nTotal prob of N-in: 0.99321\nPOSSIBLE N-term signal sequence\ninside 1 20\nTMhelix 21 43\noutside 44 428\nTMhelix 429 451\ninside 452 491

1293 GCF_001534535.1_ASM153453v1 Burkholderia sp. MSMB1459WGS Proteobacteria; Betaproteobacteria; Burkholderiales; Burkholderiaceae; Burkholderia; Burkholderia cepacia complex MESLLTRIRASFADASPRRQLISVAVLVALVASLITAGYFALRPRYQVLFRDLKPQDAAT WP_060036044.1 MULTISPECIES: flagellar M-ring protein FliF [Burkholderia cepacia complex] Length: 491\nNumber of predicted TMHs: 2\nExp number of AAs in TMHs: 42.71388\nExp number, first 60 AAs: 22.14192\nTotal prob of N-in: 0.99321\nPOSSIBLE N-term signal sequence\ninside 1 20\nTMhelix 21 43\noutside 44 428\nTMhelix 429 451\ninside 452 491

1294 GCF_001534535.1_ASM153453v1 Burkholderia sp. MSMB1459WGS Proteobacteria; Betaproteobacteria; Burkholderiales; Burkholderiaceae; Burkholderia; Burkholderia cepacia complex MKRRAALLGLSLLASFAAADGAPNAGRAMQRFALDLDGSAAYYQLTVPQPVYAASRRDDL WP_060353807.1 hypothetical protein [Burkholderia sp. MSMB1459WGS] Length: 451\nNumber of predicted TMHs: 1\nExp number of AAs in TMHs: 26.2388\nExp number, first 60 AAs: 5.00141000000001\nTotal prob of N-in: 0.42361\noutside 1 417\nTMhelix 418 440\ninside 441 451

1295 GCF_001526325.1_ASM152632v1 Burkholderia sp. MSMB1826 Proteobacteria; Betaproteobacteria; Burkholderiales; Burkholderiaceae; Burkholderia; Burkholderia cepacia complex MKRRAALLGLSLLASFAAADGAPNAGRAMQRFALDLDGSAAYYQLTVPQPVYAASRRDDL WP_059823442.1 hypothetical protein [Burkholderia sp. MSMB1826] Length: 452\nNumber of predicted TMHs: 1\nExp number of AAs in TMHs: 27.09136\nExp number, first 60 AAs: 5.48732\nTotal prob of N-in: 0.35503\noutside 1 418\nTMhelix 419 441\ninside 442 452

1296 GCF_001526365.1_ASM152636v1 Burkholderia sp. MSMB1835 Proteobacteria; Betaproteobacteria; Burkholderiales; Burkholderiaceae; Burkholderia; Burkholderia cepacia complex MKRRAALLGLSLLASFAAADGAPNAGRAMQRFALDLDGSAAYYQLTVPQPVYAASRRDDL WP_059835798.1 hypothetical protein [Burkholderia sp. MSMB1835] Length: 452\nNumber of predicted TMHs: 1\nExp number of AAs in TMHs: 25.65671\nExp number, first 60 AAs: 5.57342\nTotal prob of N-in: 0.34296\noutside 1 418\nTMhelix 419 441\ninside 442 452

1297 GCF_001532755.1_ASM153275v1 Burkholderia sp. MSMB2157WGS Proteobacteria; Betaproteobacteria; Burkholderiales; Burkholderiaceae; Burkholderia; Burkholderia cepacia complex

MKRRAALLGLSLLASFAAADGAPNAGRAMQRFALDLDGSAAYYQLTVPQPVYAASRRDDL WP_034204171.1 MULTISPECIES: membrane protein [Burkholderia cepacia complex] Length: 452\nNumber of predicted TMHs: 1\nExp number of AAs in TMHs: 26.99094\nExp number, first 60 AAs: 5.49432\nTotal prob of N-in: 0.35405\noutside 1 418\nTMhelix 419 441\ninside 442 452

1298 GCF_001522585.1_ASM152258v1 Burkholderia sp. NRF60-BP8 Proteobacteria; Betaproteobacteria; Burkholderiales; Burkholderiaceae; Burkholderia; Burkholderia cepacia complex MKRRAALLGLSLLASFAAADGAPNAGRAMQRFALDLDGSAAYYQLTVPQPVYAASRRDDL WP_059781113.1 hypothetical protein [Burkholderia sp. NRF60-BP8] Length: 452\nNumber of predicted TMHs: 1\nExp number of AAs in TMHs: 27.28651\nExp number, first 60 AAs: 5.48904000000001\nTotal prob of N-in: 0.35479\noutside 1 418\nTMhelix 419 441\ninside 442 452

1299 GCF_001685505.1_ASM168550v1 Burkholderia stabilis Proteobacteria; Betaproteobacteria; Burkholderiales; Burkholderiaceae; Burkholderia; Burkholderia cepacia complex MSISITRRRLLAAGLAASSLALSGCFTPKLYKDDAYSEYVSGFMISEDGKKLVVLGMRYH WP_047902623.1 MULTISPECIES: hypothetical protein [Burkholderia cepacia complex] Length: 212\nNumber of predicted TMHs: 1\nExp number of AAs in TMHs: 23.99618\nExp number, first 60 AAs: 0.75736\nTotal prob of N-in: 0.41515\noutside 1 180\nTMhelix 181 203\ninside 204 212

1300 GCF_001742165.1_ASM174216v1 Burkholderia stabilis Proteobacteria; Betaproteobacteria; Burkholderiales; Burkholderiaceae; Burkholderia; Burkholderia cepacia complex MKRRTLLTAGAIATATAAGLTGCTAMLFEDGHYKETVDRFLVSEDGKKFVVLGKKYHYIF WP_069745444.1 hypothetical protein [Burkholderia stabilis] Length: 210\nNumber of predicted TMHs: 2\nExp number of AAs in TMHs: 40.70876\nExp number, first 60 AAs: 17.46609\nTotal prob of N-in: 0.81154\nPOSSIBLE N-term signal sequence\ninside 1 6\nTMhelix 7 29\noutside 30 185\nTMhelix 186 208\ninside 209 210

1301 GCF_001742165.1_ASM174216v1 Burkholderia stabilis Proteobacteria; Betaproteobacteria; Burkholderiales; Burkholderiaceae; Burkholderia; Burkholderia cepacia complex MNVSITRRRLLAASLAATSVALSGCFTPKLYKDDAYSEKVSGFMITQDGKKLVVLGTRYH WP_069748857.1 hypothetical protein [Burkholderia stabilis] Length: 211\nNumber of predicted TMHs: 1\nExp number of AAs in TMHs: 25.03255\nExp number, first 60 AAs: 1.1725\nTotal prob of N-in: 0.52423\noutside 1 179\nTMhelix 180 202\ninside 203 211

1302 GCF_001530395.1_ASM153039v1 Burkholderia stagnalis Proteobacteria; Betaproteobacteria; Burkholderiales; Burkholderiaceae; Burkholderia; Burkholderia cepacia complex MTTTITRRRPLGAGALAGSLALSGCFTPNLFDDKQYTEGVSRFMITEDGRKLVVLGERYH WP_059884723.1 hypothetical protein [Burkholderia stagnalis] Length: 213\nNumber of predicted TMHs: 1\nExp number of AAs in TMHs: 25.00217\nExp number, first 60 AAs: 2.88427\nTotal prob of N-in: 0.69349\noutside 1 180\nTMhelix 181 203\ninside 204 213

1303 GCF_001529545.1_ASM152954v1 Burkholderia stagnalis Proteobacteria; Betaproteobacteria; Burkholderiales; Burkholderiaceae; Burkholderia; Burkholderia cepacia complex MTTTITRRRPLGAGALAGSLALSGCFTPNLFDDKQYTEGVSRFMITEDGRKLVVLGERYH WP_059884723.1 hypothetical protein [Burkholderia stagnalis] Length: 213\nNumber of predicted TMHs: 1\nExp number of AAs in TMHs: 25.00217\nExp number, first 60 AAs: 2.88427\nTotal prob of N-in: 0.69349\noutside 1 180\nTMhelix 181 203\ninside 204 213

1304 GCF_001528085.1_ASM152808v1 Burkholderia stagnalis Proteobacteria; Betaproteobacteria; Burkholderiales; Burkholderiaceae; Burkholderia; Burkholderia cepacia complex MTTTITRRRPLGAGALAGSLALSGCFTPNLFDDKQYTEGVSRFMITEDGRKLVVLGERYH WP_059884723.1 hypothetical protein [Burkholderia stagnalis] Length: 213\nNumber of predicted TMHs: 1\nExp number of AAs in TMHs: 25.00217\nExp number, first 60 AAs: 2.88427\nTotal prob of N-in: 0.69349\noutside 1 180\nTMhelix 181 203\ninside 204 213

1305 GCF_001526995.1_ASM152699v1 Burkholderia stagnalis Proteobacteria; Betaproteobacteria; Burkholderiales; Burkholderiaceae; Burkholderia; Burkholderia cepacia complex MNRLHPPISASGRRRTVRLAVLLASAVLVAGCQQELYGGLAERDCNEMMAALLQNGVDAQ WP_059891627.1 EscJ/YscJ/HrcJ family type III secretion inner membrane ring protein [Burkholderia stagnalis] Length: 282\nNumber of predicted TMHs: 1\nExp number of AAs in TMHs: 28.52094\nExp number, first 60 AAs: 5.59939\nTotal prob of N-in: 0.31120\noutside 1 223\nTMhelix 224 246\ninside 247 282

1306 GCF_001527645.1_ASM152764v1 Burkholderia stagnalis Proteobacteria; Betaproteobacteria; Burkholderiales; Burkholderiaceae; Burkholderia; Burkholderia cepacia complex
MTTTITRRRPLGAGALAGSLALSGCFTPKLFDDKQYTEGVSRFMITEDGRKLVVLGDRYH WP_059921586.1 hypothetical protein [Burkholderia stagnalis] Length: 213\nNumber of predicted TMHs: 1\nExp number of AAs in TMHs: 22.88495\nExp number, first 60 AAs: 0.72184\nTotal prob of N-in: 0.70181\noutside 1 180\nTMhelix 181 203\ninside 204 213

1307 GCF_001533075.1_ASM153307v1 Burkholderia stagnalis Proteobacteria; Betaproteobacteria; Burkholderiales; Burkholderiaceae; Burkholderia; Burkholderia cepacia complex
MNRLHPPISASGRRRTVRLAVLLASAVLVAGCQQELYGGLAERDCNEMMAALLQNGVDAQ WP_059891627.1 EscJ/YscJ/HrcJ family type III secretion inner membrane ring protein [Burkholderia stagnalis] Length: 282\nNumber of predicted TMHs: 1\nExp number of AAs in TMHs: 28.52094\nExp number, first 60 AAs: 5.59939\nTotal prob of N-in: 0.31120\noutside 1 223\nTMhelix 224 246\ninside 247 282

1308 GCF_001534285.1_ASM153428v1 Burkholderia stagnalis Proteobacteria; Betaproteobacteria; Burkholderiales; Burkholderiaceae; Burkholderia; Burkholderia cepacia complex
MTTTITRRRPLGAGALAGSLALSGCFTPNLFDDKQYTEGVSRFMITEDGRKLVVLGERYH WP_059884723.1 hypothetical protein [Burkholderia stagnalis] Length: 213\nNumber of predicted TMHs: 1\nExp number of AAs in TMHs: 25.00217\nExp number, first 60 AAs: 2.88427\nTotal prob of N-in: 0.69349\noutside 1 180\nTMhelix 181 203\ninside 204 213

1309 GCF_001533205.1_ASM153320v1 Burkholderia stagnalis Proteobacteria; Betaproteobacteria; Burkholderiales; Burkholderiaceae; Burkholderia; Burkholderia cepacia complex
MTTTITRRRPLGAGALAGSLALSGCFTPNLFDDKQYTEGVSRFMITEDGRKLVVLGERYH WP_059884723.1 hypothetical protein [Burkholderia stagnalis] Length: 213\nNumber of predicted TMHs: 1\nExp number of AAs in TMHs: 25.00217\nExp number, first 60 AAs: 2.88427\nTotal prob of N-in: 0.69349\noutside 1 180\nTMhelix 181 203\ninside 204 213

1310 GCF_001533815.1_ASM153381v1 Burkholderia stagnalis Proteobacteria; Betaproteobacteria; Burkholderiales; Burkholderiaceae; Burkholderia; Burkholderia cepacia complex
MTTTITRRRPLGAGALAGSLALSGCFTPKLFDDKQYTEGVSRFMITEDGRKLVVLGERYH WP_059801629.1 hypothetical protein [Burkholderia stagnalis] Length: 213\nNumber of predicted TMHs: 1\nExp number of AAs in TMHs: 22.88495\nExp number, first 60 AAs: 0.72184\nTotal prob of N-in: 0.70181\noutside 1 180\nTMhelix 181 203\ninside 204 213

1311 GCF_001533575.1_ASM153357v1 Burkholderia stagnalis Proteobacteria; Betaproteobacteria; Burkholderiales; Burkholderiaceae; Burkholderia; Burkholderia cepacia complex
MTTTITRRRPLGAGALGGSLALSGCFMPKLFDDKQYTEGVSRFMITEDGRKLVVLGERYH WP_060363596.1 hypothetical protein [Burkholderia stagnalis] Length: 213\nNumber of predicted TMHs: 1\nExp number of AAs in TMHs: 23.11898\nExp number, first 60 AAs: 1.21631\nTotal prob of N-in: 0.74366\noutside 1 180\nTMhelix 181 203\ninside 204 213

1312 GCF_001523185.1_ASM152318v1 Burkholderia stagnalis Proteobacteria; Betaproteobacteria; Burkholderiales; Burkholderiaceae; Burkholderia; Burkholderia cepacia complex
MTTTITRRRPLGAGALAGSLALSGCFTPKLFDDKQYTEGVSRFMITEDGRKLVVLGERYH WP_059801629.1 hypothetical protein [Burkholderia stagnalis] Length: 213\nNumber of predicted TMHs: 1\nExp number of AAs in TMHs: 22.88495\nExp number, first 60 AAs: 0.72184\nTotal prob of N-in: 0.70181\noutside 1 180\nTMhelix 181 203\ninside 204 213

1313 GCF_001533065.1_ASM153306v1 Burkholderia stagnalis Proteobacteria; Betaproteobacteria; Burkholderiales; Burkholderiaceae; Burkholderia; Burkholderia cepacia complex
MTTTITRRRPLGAGALAGSLALSGCFTPNLFDDKQYTEGVSRFMITEDGRKLVVLGERYH WP_059884723.1 hypothetical protein [Burkholderia stagnalis] Length: 213\nNumber of predicted TMHs: 1\nExp number of AAs in TMHs: 25.00217\nExp number, first 60 AAs: 2.88427\nTotal prob of N-in: 0.69349\noutside 1 180\nTMhelix 181 203\ninside 204 213

1314 GCF_001533115.1_ASM153311v1 Burkholderia stagnalis Proteobacteria; Betaproteobacteria; Burkholderiales; Burkholderiaceae; Burkholderia; Burkholderia cepacia complex
MTTTITRRRPLGAGALAGSLALSGCFTPNLFDDKQYTEGVSRFMITEDGRKLVVLGERYH WP_059884723.1 hypothetical protein [Burkholderia stagnalis] Length: 213\nNumber of predicted TMHs: 1\nExp number of AAs in TMHs:

25.00217\nExp number, first 60 AAs: 2.88427\nTotal prob of N-in: 0.69349\noutside 1 180\nTMhelix 181 203\ninside 204 213

1315 GCF_001533235.1_ASM153323v1 Burkholderia stagnalis Proteobacteria; Betaproteobacteria; Burkholderiales; Burkholderiaceae; Burkholderia; Burkholderia cepacia complex MTTTITRRRPLGAGALAGSLALSGCFTPNLFDDKQYTEGVSRFMITEDGRKLVVLGERYH WP_059884723.1 hypothetical protein [Burkholderia stagnalis] Length: 213\nNumber of predicted TMHs: 1\nExp number of AAs in TMHs: 25.00217\nExp number, first 60 AAs: 2.88427\nTotal prob of N-in: 0.69349\noutside 1 180\nTMhelix 181 203\ninside 204 213

1316 GCF_001533485.1_ASM153348v1 Burkholderia stagnalis Proteobacteria; Betaproteobacteria; Burkholderiales; Burkholderiaceae; Burkholderia; Burkholderia cepacia complex MTTTITRRRPLGAGALAGSLALSGCFTPNLFDDKQYTEGVSRFMITEDGRKLVVLGERYH WP_060291916.1 hypothetical protein [Burkholderia stagnalis] Length: 213\nNumber of predicted TMHs: 1\nExp number of AAs in TMHs: 24.94682\nExp number, first 60 AAs: 2.73248\nTotal prob of N-in: 0.73040\noutside 1 180\nTMhelix 181 203\ninside 204 213

1317 GCF_001526815.1_ASM152681v1 Burkholderia stagnalis Proteobacteria; Betaproteobacteria; Burkholderiales; Burkholderiaceae; Burkholderia; Burkholderia cepacia complex MNRLHPPISASGRRRTVRLAVLLASAVLVAGCQQELYGGLAERDCNEMMAALLQNGVDAQ WP_059891627.1 EscJ/YscJ/HrcJ family type III secretion inner membrane ring protein [Burkholderia stagnalis] Length: 282\nNumber of predicted TMHs: 1\nExp number of AAs in TMHs: 28.52094\nExp number, first 60 AAs: 5.59939\nTotal prob of N-in: 0.31120\noutside 1 223\nTMhelix 224 246\ninside 247 282

1318 GCF_001526825.1_ASM152682v1 Burkholderia stagnalis Proteobacteria; Betaproteobacteria; Burkholderiales; Burkholderiaceae; Burkholderia; Burkholderia cepacia complex MTTTITRRRPLGAGALAGSLALSGCFTPNLFDDKQYTEGVSRFMITEDGRKLVVLGERYH WP_059884723.1 hypothetical protein [Burkholderia stagnalis] Length: 213\nNumber of predicted TMHs: 1\nExp number of AAs in TMHs: 25.00217\nExp number, first 60 AAs: 2.88427\nTotal prob of N-in: 0.69349\noutside 1 180\nTMhelix 181 203\ninside 204 213

1319 GCF_001527665.1_ASM152766v1 Burkholderia stagnalis Proteobacteria; Betaproteobacteria; Burkholderiales; Burkholderiaceae; Burkholderia; Burkholderia cepacia complex MTTTITRRRPLGAGALAGSLALSGCFTPKLFDDKQYTEGVSRFMITEDGRKLVVLGERYH WP_059801629.1 hypothetical protein [Burkholderia stagnalis] Length: 213\nNumber of predicted TMHs: 1\nExp number of AAs in TMHs: 22.88495\nExp number, first 60 AAs: 0.72184\nTotal prob of N-in: 0.70181\noutside 1 180\nTMhelix 181 203\ninside 204 213

1320 GCF_001532995.1_ASM153299v1 Burkholderia stagnalis Proteobacteria; Betaproteobacteria; Burkholderiales; Burkholderiaceae; Burkholderia; Burkholderia cepacia complex MTTTITRRRPLGAGALAGSLALSGCFTPNLFDDKQYTEGVSRFMITEDGRKLVVLGERYH WP_060291916.1 hypothetical protein [Burkholderia stagnalis] Length: 213\nNumber of predicted TMHs: 1\nExp number of AAs in TMHs: 24.94682\nExp number, first 60 AAs: 2.73248\nTotal prob of N-in: 0.73040\noutside 1 180\nTMhelix 181 203\ninside 204 213

1321 GCF_001529625.1_ASM152962v1 Burkholderia stagnalis Proteobacteria; Betaproteobacteria; Burkholderiales; Burkholderiaceae; Burkholderia; Burkholderia cepacia complex MNRLHPPISASGRRRTVRLAVLLASAVLVAGCQQELYGGLAERDCNEMMAALLQNGVDAQ WP_059891627.1 EscJ/YscJ/HrcJ family type III secretion inner membrane ring protein [Burkholderia stagnalis] Length: 282\nNumber of predicted TMHs: 1\nExp number of AAs in TMHs: 28.52094\nExp number, first 60 AAs: 5.59939\nTotal prob of N-in: 0.31120\noutside 1 223\nTMhelix 224 246\ninside 247 282

1322 GCF_001529505.1_ASM152950v1 Burkholderia stagnalis Proteobacteria; Betaproteobacteria; Burkholderiales; Burkholderiaceae; Burkholderia; Burkholderia cepacia complex MTTTITRRRPLGAGALAGSLALSGCFTPNLFDDKQYTEGVSRFMITEDGRKLVVLGERYH WP_059884723.1 hypothetical protein [Burkholderia stagnalis] Length: 213\nNumber of predicted TMHs: 1\nExp number of AAs in TMHs: 25.00217\nExp number, first 60 AAs: 2.88427\nTotal prob of N-in: 0.69349\noutside 1 180\nTMhelix 181 203\ninside 204 213

1323 GCF_001530645.1_ASM153064v1 Burkholderia stagnalis Proteobacteria; Betaproteobacteria; Burkholderiales; Burkholderiaceae; Burkholderia; Burkholderia cepacia complex

MTTTITRRRPLGTGALAGSLALSGCFTPKLFDDKQYTEGVSRFMITEDGRKLVVLGERYH WP_060157732.1 hypothetical protein [Burkholderia stagnalis] Length: 213\nNumber of predicted TMHs: 1\nExp number of AAs in TMHs: 22.48409\nExp number, first 60 AAs: 0.32855\nTotal prob of N-in: 0.69770\noutside 1 180\nTMhelix 181 203\ninside 204 213

1324 GCF_001533495.1_ASM153349v1 Burkholderia stagnalis Proteobacteria; Betaproteobacteria; Burkholderiales; Burkholderiaceae; Burkholderia; Burkholderia cepacia complex MTTTITRRRPLGAGALAGSLALSGCFTPKLFDDKQYTEGVSRFMITEDGRKLVVLGERYH WP_059801629.1 hypothetical protein [Burkholderia stagnalis] Length: 213\nNumber of predicted TMHs: 1\nExp number of AAs in TMHs: 22.88495\nExp number, first 60 AAs: 0.72184\nTotal prob of N-in: 0.70181\noutside 1 180\nTMhelix 181 203\ninside 204 213

1325 GCF_001533705.1_ASM153370v1 Burkholderia stagnalis Proteobacteria; Betaproteobacteria; Burkholderiales; Burkholderiaceae; Burkholderia; Burkholderia cepacia complex MTTTITRRRPLGAGALAGSLALSGCFTPKLFDDKQYTEGVSRFMITEDGRKLVVLGERYH WP_059801629.1 hypothetical protein [Burkholderia stagnalis] Length: 213\nNumber of predicted TMHs: 1\nExp number of AAs in TMHs: 22.88495\nExp number, first 60 AAs: 0.72184\nTotal prob of N-in: 0.70181\noutside 1 180\nTMhelix 181 203\ninside 204 213

1326 GCF_001533845.1_ASM153384v1 Burkholderia stagnalis Proteobacteria; Betaproteobacteria; Burkholderiales; Burkholderiaceae; Burkholderia; Burkholderia cepacia complex MTTTITRRRPLGAGALAGSLALSGCFTPKLFDDKQYTEGVSRFMITEDGRKLVVLGERYH WP_059801629.1 hypothetical protein [Burkholderia stagnalis] Length: 213\nNumber of predicted TMHs: 1\nExp number of AAs in TMHs: 22.88495\nExp number, first 60 AAs: 0.72184\nTotal prob of N-in: 0.70181\noutside 1 180\nTMhelix 181 203\ninside 204 213

1327 GCF_001528995.1_ASM152899v1 Burkholderia territorii Proteobacteria; Betaproteobacteria; Burkholderiales; Burkholderiaceae; Burkholderia; Burkholderia cepacia complex MESLLTRIRASFADASPRRQLISVAVLVALVASLIAAGYFVLRPRYQVLFRDLKPQDAAT WP_060023992.1 flagellar M-ring protein FliF [Burkholderia territorii] Length: 491\nNumber of predicted TMHs: 2\nExp number of AAs in TMHs: 43.9898799999999\nExp number, first 60 AAs: 22.26454\nTotal prob of N-in: 0.99321\nPOSSIBLE N-term signal sequence\ninside 1 20\nTMhelix 21 43\noutside 44 433\nTMhelix 434 456\ninside 457 491

1328 GCF_001522365.1_ASM152236v1 Burkholderia territorii Proteobacteria; Betaproteobacteria; Burkholderiales; Burkholderiaceae; Burkholderia; Burkholderia cepacia complex MTPGRRSALASLGAMCVQGTWATYANHAFGPWIAGRAAVVQGICSFGMTYAVTRLIEWSV WP_059542378.1 hypothetical protein [Burkholderia territorii] Length: 158\nNumber of predicted TMHs: 2\nExp number of AAs in TMHs: 77.02686\nExp number, first 60 AAs: 35.59335\nTotal prob of N-in: 0.82864\nPOSSIBLE N-term signal sequence\ninside 1 71\nTMhelix 72 94\noutside 95 97\nTMhelix 98 120\ninside 121 158

1329 GCF_001521925.1_ASM152192v1 Burkholderia territorii Proteobacteria; Betaproteobacteria; Burkholderiales; Burkholderiaceae; Burkholderia; Burkholderia cepacia complex MTPGRRSALASLGAMCVQGTWATYANHAFGPWIAGRAAVVQGICSFGMTYAVTRLIEWSV WP_059542378.1 hypothetical protein [Burkholderia territorii] Length: 158\nNumber of predicted TMHs: 2\nExp number of AAs in TMHs: 77.02686\nExp number, first 60 AAs: 35.59335\nTotal prob of N-in: 0.82864\nPOSSIBLE N-term signal sequence\ninside 1 71\nTMhelix 72 94\noutside 95 97\nTMhelix 98 120\ninside 121 158

1330 GCF_001533385.1_ASM153338v1 Burkholderia territorii Proteobacteria; Betaproteobacteria; Burkholderiales; Burkholderiaceae; Burkholderia; Burkholderia cepacia complex MESLLTRIRASFTDASPRRQLISVAVLVALVASLIAAGYFVLRPRYQVLFRDLKPQDAAA WP_060361870.1 flagellar M-ring protein FliF [Burkholderia territorii] Length: 491\nNumber of predicted TMHs: 2\nExp number of AAs in TMHs: 43.99179\nExp number, first 60 AAs: 22.26719\nTotal prob of N-in: 0.99403\nPOSSIBLE N-term signal sequence\ninside 1 20\nTMhelix 21 43\noutside 44 433\nTMhelix 434 456\ninside 457 491

1331 GCF_001531615.1_ASM153161v1 Burkholderia ubonensis Proteobacteria; Betaproteobacteria; Burkholderiales; Burkholderiaceae; Burkholderia; Burkholderia cepacia complex MSDDIDKGGGKGGFTRRDMMRVMAASGMMAVGGGGLLMDAQSAFAAPAPKRGGKIRVANE WP_060065546.1 ABC transporter substrate-binding protein [Burkholderia ubonensis] Length: 528\nNumber of predicted TMHs: 1\nExp number of AAs in TMHs: 31.88338\nExp number, first 60 AAs: 16.09709\nTotal prob of N-in: 0.73476\nPOSSIBLE N-term signal sequence\noutside 1 503\nTMhelix 504 526\ninside 527 528

1332 GCF_001527105.1_ASM152710v1 Burkholderia ubonensis Proteobacteria; Betaproteobacteria; Burkholderiales; Burkholderiaceae; Burkholderia; Burkholderia cepacia complex
MSDDIDKGGGKGGFTRRDMMRVMAASGMMAVGGGGLLMDAQSAFAAPAPKRGGKIRVANE WP_059972434.1 ABC transporter substrate-binding protein [Burkholderia ubonensis] Length: 528\nNumber of predicted TMHs: 1\nExp number of AAs in TMHs: 32.0963\nExp number, first 60 AAs: 16.09802\nTotal prob of N-in: 0.73456\nPOSSIBLE N-term signal sequence\noutside 1 503\nTMhelix 504 526\ninside 527 528

1333 GCF_001527095.1_ASM152709v1 Burkholderia ubonensis Proteobacteria; Betaproteobacteria; Burkholderiales; Burkholderiaceae; Burkholderia; Burkholderia cepacia complex
MSDDIDKGGGKGGFTRRDMMRVMAASGMMAVGGGGLLMDAQSAFAAPAPKRGGKIRVANE WP_059991183.1 ABC transporter substrate-binding protein [Burkholderia ubonensis] Length: 528\nNumber of predicted TMHs: 1\nExp number of AAs in TMHs: 31.88336\nExp number, first 60 AAs: 16.09709\nTotal prob of N-in: 0.73476\nPOSSIBLE N-term signal sequence\noutside 1 503\nTMhelix 504 526\ninside 527 528

1334 GCF_001528985.1_ASM152898v1 Burkholderia ubonensis Proteobacteria; Betaproteobacteria; Burkholderiales; Burkholderiaceae; Burkholderia; Burkholderia cepacia complex
MSDDIDNGGGKGGFTRRDMMRVMAASGMMAVGGGGLLMDAQSAFAAPAPKRGGKIRVANE WP_059946573.1 ABC transporter substrate-binding protein [Burkholderia ubonensis] Length: 528\nNumber of predicted TMHs: 1\nExp number of AAs in TMHs: 31.30297\nExp number, first 60 AAs: 15.42728\nTotal prob of N-in: 0.69081\nPOSSIBLE N-term signal sequence\noutside 1 503\nTMhelix 504 526\ninside 527 528

1335 GCF_001529225.1_ASM152922v1 Burkholderia ubonensis Proteobacteria; Betaproteobacteria; Burkholderiales; Burkholderiaceae; Burkholderia; Burkholderia cepacia complex
MSDDIDNGGGGKGGFTRRDMMRVMAASGMMAVGGGGLLMDAQSAFAAPAPKRGGKIRVAN WP_060045123.1 ABC transporter substrate-binding protein [Burkholderia ubonensis] Length: 529\nNumber of predicted TMHs: 1\nExp number of AAs in TMHs: 30.50215\nExp number, first 60 AAs: 14.12787\nTotal prob of N-in: 0.59108\nPOSSIBLE N-term signal sequence\noutside 1 504\nTMhelix 505 527\ninside 528 529

1336 GCF_001529105.1_ASM152910v1 Burkholderia ubonensis Proteobacteria; Betaproteobacteria; Burkholderiales; Burkholderiaceae; Burkholderia; Burkholderia cepacia complex
MSDDIDKGGGKGGFTRRDMMRVMAASGMMAVGGGGLLMDAQSAFAAPAPKRGGKIRVANE WP_060065546.1 ABC transporter substrate-binding protein [Burkholderia ubonensis] Length: 528\nNumber of predicted TMHs: 1\nExp number of AAs in TMHs: 31.88338\nExp number, first 60 AAs: 16.09709\nTotal prob of N-in: 0.73476\nPOSSIBLE N-term signal sequence\noutside 1 503\nTMhelix 504 526\ninside 527 528

1337 GCF_001530825.1_ASM153082v1 Burkholderia ubonensis Proteobacteria; Betaproteobacteria; Burkholderiales; Burkholderiaceae; Burkholderia; Burkholderia cepacia complex
MSDDIDKGGGKGGFTRRDMMRVMAASGMMAVGGGGLLMDAQSAFAAPAPKRGGKIRVANE WP_059979451.1 ABC transporter substrate-binding protein [Burkholderia ubonensis] Length: 528\nNumber of predicted TMHs: 1\nExp number of AAs in TMHs: 32.09636\nExp number, first 60 AAs: 16.09804\nTotal prob of N-in: 0.73456\nPOSSIBLE N-term signal sequence\noutside 1 503\nTMhelix 504 526\ninside 527 528

1338 GCF_001882635.1_ASM188263v1 Burkholderia ubonensis Proteobacteria; Betaproteobacteria; Burkholderiales; Burkholderiaceae; Burkholderia; Burkholderia cepacia complex
MSDDIDNGGKGGFTRRDMMRVMAASGMMAVGGGGLLMDAQSAFAAPAPKRGGKIRVANES WP_059849896.1 ABC transporter substrate-binding protein [Burkholderia ubonensis] Length: 527\nNumber of predicted TMHs: 2\nExp number of AAs in TMHs: 36.08404\nExp number, first 60 AAs: 20.24786\nTotal prob of N-in: 0.91899\nPOSSIBLE N-term signal sequence\ninside 1 20\nTMhelix 21 43\noutside 44 502\nTMhelix 503 525\ninside 526 527

1339 GCF_001882505.1_ASM188250v1 Burkholderia ubonensis Proteobacteria; Betaproteobacteria; Burkholderiales; Burkholderiaceae; Burkholderia; Burkholderia cepacia complex
MSDDIDNGGKGGFTRRDMMRVMAASGMMAVGGGGLLMDAQSAFAAPAPKRGGKIRVANES WP_059707130.1 ABC transporter substrate-binding protein [Burkholderia ubonensis] Length: 527\nNumber of predicted TMHs: 2\nExp number of AAs in TMHs: 35.87636\nExp number, first 60 AAs: 20.2481\nTotal prob of N-in: 0.91905\nPOSSIBLE N-term signal sequence\ninside 1 20\nTMhelix 21 43\noutside 44 502\nTMhelix 503 525\ninside 526 527

1340 GCF_001546975.1_ASM154697v1 Burkholderia ubonensis Proteobacteria; Betaproteobacteria; Burkholderiales; Burkholderiaceae; Burkholderia; Burkholderia cepacia complex
MSDDIDKGGGKGGFTRRDMMRVMAASGMMAVGGGGLLMDAQSAFAAPAPKRGGKIRVANE WP_060065546.1 ABC transporter substrate-binding protein [Burkholderia ubonensis] Length: 528\nNumber of predicted TMHs: 1\nExp

number of AAs in TMHs: 31.88338\nExp number, first 60 AAs: 16.09709\nTotal prob of N-in: 0.73476\nPOSSIBLE N-term signal sequence\noutside 1 503\nTMhelix 504 526\ninside 527 528

1341 GCF_001534445.1_ASM153444v1 Burkholderia ubonensis Proteobacteria; Betaproteobacteria; Burkholderiales; Burkholderiaceae; Burkholderia; Burkholderia cepacia complex MSDDIDNGGKGGFTRRDMMRVMAASGMMAVGGGGLLMDAQSAFAAPAPKRGGKIRVANES WP_059849896.1 ABC transporter substrate-binding protein [Burkholderia ubonensis] Length: 527\nNumber of predicted TMHs: 2\nExp number of AAs in TMHs: 36.08404\nExp number, first 60 AAs: 20.24786\nTotal prob of N-in: 0.91899\nPOSSIBLE N-term signal sequence\ninside 1 20\nTMhelix 21 43\noutside 44 502\nTMhelix 503 525\ninside 526 527

1342 GCF_001533945.1_ASM153394v1 Burkholderia ubonensis Proteobacteria; Betaproteobacteria; Burkholderiales; Burkholderiaceae; Burkholderia; Burkholderia cepacia complex MSDDIDKGGGKGGFTRRDMMRVMAASGMMAVGGGGLLMDAQSAFAAPAPKRGGKIRVANE WP_060294615.1 ABC transporter substrate-binding protein [Burkholderia ubonensis] Length: 528\nNumber of predicted TMHs: 1\nExp number of AAs in TMHs: 32.09636\nExp number, first 60 AAs: 16.09801\nTotal prob of N-in: 0.73456\nPOSSIBLE N-term signal sequence\noutside 1 503\nTMhelix 504 526\ninside 527 528

1343 GCF_001531595.1_ASM153159v1 Burkholderia ubonensis Proteobacteria; Betaproteobacteria; Burkholderiales; Burkholderiaceae; Burkholderia; Burkholderia cepacia complex MSDDIDKGGGKGGFTRRDMMRVMAASGMMAVGGGGLLMDAQSAFAAPAPKRGGKIRVANE WP_060164996.1 ABC transporter substrate-binding protein [Burkholderia ubonensis] Length: 528\nNumber of predicted TMHs: 1\nExp number of AAs in TMHs: 32.0777799999999\nExp number, first 60 AAs: 16.09563\nTotal prob of N-in: 0.73469\nPOSSIBLE N-term signal sequence\noutside 1 503\nTMhelix 504 526\ninside 527 528

1344 GCF_001527885.1_ASM152788v1 Burkholderia ubonensis Proteobacteria; Betaproteobacteria; Burkholderiales; Burkholderiaceae; Burkholderia; Burkholderia cepacia complex MSDDIDNGGGKGGFTRRDMMRVMAASGMMAVGGGGLLMDAQSAFAAPAPKRGGKIRVANE WP_059945475.1 ABC transporter substrate-binding protein [Burkholderia ubonensis] Length: 528\nNumber of predicted TMHs: 1\nExp number of AAs in TMHs: 32.28161\nExp number, first 60 AAs: 15.42808\nTotal prob of N-in: 0.69069\nPOSSIBLE N-term signal sequence\noutside 1 503\nTMhelix 504 526\ninside 527 528

1345 GCF_001882825.1_ASM188282v1 Burkholderia ubonensis Proteobacteria; Betaproteobacteria; Burkholderiales; Burkholderiaceae; Burkholderia; Burkholderia cepacia complex MSDDIDKGGGKGGFTRRDMMRVMAASGMMAVGGGGLLMDAQSAFAAPAPKRGGKIRVANE WP_071766125.1 ABC transporter substrate-binding protein [Burkholderia ubonensis] Length: 528\nNumber of predicted TMHs: 1\nExp number of AAs in TMHs: 32.0963\nExp number, first 60 AAs: 16.09802\nTotal prob of N-in: 0.73456\nPOSSIBLE N-term signal sequence\noutside 1 503\nTMhelix 504 526\ninside 527 528

1346 GCF_001539165.1_ASM153916v1 Burkholderia ubonensis Proteobacteria; Betaproteobacteria; Burkholderiales; Burkholderiaceae; Burkholderia; Burkholderia cepacia complex MSDDIDKGGGKGGFTRRDMMRVMAASGMMAVGGGGLLMDAQSAFAAPAPKRGGKIRVANE WP_060449921.1 ABC transporter substrate-binding protein [Burkholderia ubonensis] Length: 528\nNumber of predicted TMHs: 1\nExp number of AAs in TMHs: 31.88352\nExp number, first 60 AAs: 16.09709\nTotal prob of N-in: 0.73476\nPOSSIBLE N-term signal sequence\noutside 1 503\nTMhelix 504 526\ninside 527 528

1347 GCF_001534425.1_ASM153442v1 Burkholderia ubonensis Proteobacteria; Betaproteobacteria; Burkholderiales; Burkholderiaceae; Burkholderia; Burkholderia cepacia complex MSDDIDNGGKGGFTRRDMMRVMAASGMMAVGGGGLLMDAQSAFAAPAPKRGGKIRVANES WP_060378261.1 ABC transporter substrate-binding protein [Burkholderia ubonensis] Length: 527\nNumber of predicted TMHs: 2\nExp number of AAs in TMHs: 35.87653\nExp number, first 60 AAs: 20.2481\nTotal prob of N-in: 0.91905\nPOSSIBLE N-term signal sequence\ninside 1 20\nTMhelix 21 43\noutside 44 502\nTMhelix 503 525\ninside 526 527

1348 GCF_001534195.1_ASM153419v1 Burkholderia ubonensis Proteobacteria; Betaproteobacteria; Burkholderiales; Burkholderiaceae; Burkholderia; Burkholderia cepacia complex MSDDIDKGGGKGGFTRRDMMRVMAASGMMAVGGGGLLMDAQSAFAAPAPKRGGKIRVANE WP_060203251.1 ABC transporter substrate-binding protein [Burkholderia ubonensis] Length: 528\nNumber of predicted TMHs: 1\nExp number of AAs in TMHs: 31.88349\nExp number, first 60 AAs: 16.09709\nTotal prob of N-in: 0.73476\nPOSSIBLE N-term signal sequence\noutside 1 503\nTMhelix 504 526\ninside 527 528

1349 GCF_001531825.1_ASM153182v1 Burkholderia ubonensis Proteobacteria; Betaproteobacteria; Burkholderiales; Burkholderiaceae; Burkholderia; Burkholderia cepacia complex

MSDDIDNGGKGGFTRRDMMRVMAASGMMAVGGGGLLMDAQSAFAAPAPKRGGKIRVANES WP_045565706.1 ABC transporter substrate-binding protein [Burkholderia ubonensis] Length: 527\nNumber of predicted TMHs: 2\nExp number of AAs in TMHs: 36.08388\nExp number, first 60 AAs: 20.24786\nTotal prob of N-in: 0.91899\nPOSSIBLE N-term signal sequence\ninside 1 20\nTMhelix 21 43\noutside 44 502\nTMhelix 503 525\ninside 526 527

1350 GCF_001531045.1_ASM153104v1 Burkholderia ubonensis Proteobacteria; Betaproteobacteria; Burkholderiales; Burkholderiaceae; Burkholderia; Burkholderia cepacia complex
MSDDIDNGGKGGFTRRDMMRVMAASGMMAVGGGGLLMDAQSAFAAPAPKRGGKIRVANES WP_059653009.1 ABC transporter substrate-binding protein [Burkholderia ubonensis] Length: 527\nNumber of predicted TMHs: 2\nExp number of AAs in TMHs: 36.08388\nExp number, first 60 AAs: 20.24786\nTotal prob of N-in: 0.91899\nPOSSIBLE N-term signal sequence\ninside 1 20\nTMhelix 21 43\noutside 44 502\nTMhelix 503 525\ninside 526 527

1351 GCF_001531515.1_ASM153151v1 Burkholderia ubonensis Proteobacteria; Betaproteobacteria; Burkholderiales; Burkholderiaceae; Burkholderia; Burkholderia cepacia complex
MSDDIDNGGKGGFTRRDMMRVMAASGMMAVGGGGLLMDAQSAFAAPAPKRGGKIRVANES WP_045565706.1 ABC transporter substrate-binding protein [Burkholderia ubonensis] Length: 527\nNumber of predicted TMHs: 2\nExp number of AAs in TMHs: 36.08388\nExp number, first 60 AAs: 20.24786\nTotal prob of N-in: 0.91899\nPOSSIBLE N-term signal sequence\ninside 1 20\nTMhelix 21 43\noutside 44 502\nTMhelix 503 525\ninside 526 527

1352 GCF_001529865.1_ASM152986v1 Burkholderia ubonensis Proteobacteria; Betaproteobacteria; Burkholderiales; Burkholderiaceae; Burkholderia; Burkholderia cepacia complex
MNAPETIDTPQAGTPRAGVTVLPHPAAARRLTGDELAARRQRSRRATFIKWLRKVHGWVG WP_060088081.1
peptidase, partial [Burkholderia ubonensis] Length: 231\nNumber of predicted TMHs: 2\nExp number of AAs in TMHs: 45.25994\nExp number, first 60 AAs: 5.46724\nTotal prob of N-in: 0.94374\ninside 1 54\nTMhelix 55 77\noutside 78 196\nTMhelix 197 219\ninside 220 231

1353 GCF_001525885.1_ASM152588v1 Burkholderia ubonensis Proteobacteria; Betaproteobacteria; Burkholderiales; Burkholderiaceae; Burkholderia; Burkholderia cepacia complex
MSDDIDKGGGKGGFTRRDMMRVMAASGMMAVGGGGLLMDAQSAFAAPAPKRGGKIRVANE WP_059661917.1 ABC transporter substrate-binding protein [Burkholderia ubonensis] Length: 528\nNumber of predicted TMHs: 1\nExp number of AAs in TMHs: 32.09621\nExp number, first 60 AAs: 16.09802\nTotal prob of N-in: 0.73456\nPOSSIBLE N-term signal sequence\noutside 1 503\nTMhelix 504 526\ninside 527 528

1354 GCF_001526645.1_ASM152664v1 Burkholderia ubonensis Proteobacteria; Betaproteobacteria; Burkholderiales; Burkholderiaceae; Burkholderia; Burkholderia cepacia complex
MSDDIDKGGGKGGFTRRDMMRVMAASGMMAVGGGGLLMDAQSAFAAPAPKRGGKIRVANE WP_059811412.1 ABC transporter substrate-binding protein [Burkholderia ubonensis] Length: 528\nNumber of predicted TMHs: 1\nExp number of AAs in TMHs: 31.88329\nExp number, first 60 AAs: 16.09709\nTotal prob of N-in: 0.73476\nPOSSIBLE N-term signal sequence\noutside 1 503\nTMhelix 504 526\ninside 527 528

1355 GCF_001526955.1_ASM152695v1 Burkholderia ubonensis Proteobacteria; Betaproteobacteria; Burkholderiales; Burkholderiaceae; Burkholderia; Burkholderia cepacia complex
MSDDIDNGGKGGFTRRDMMRVMAASGMMAVGGGGLLMDAQSAFAAPAPKRGGKIRVANES WP_059849896.1 ABC transporter substrate-binding protein [Burkholderia ubonensis] Length: 527\nNumber of predicted TMHs: 2\nExp number of AAs in TMHs: 36.08404\nExp number, first 60 AAs: 20.24786\nTotal prob of N-in: 0.91899\nPOSSIBLE N-term signal sequence\ninside 1 20\nTMhelix 21 43\noutside 44 502\nTMhelix 503 525\ninside 526 527

1356 GCF_001527065.1_ASM152706v1 Burkholderia ubonensis Proteobacteria; Betaproteobacteria; Burkholderiales; Burkholderiaceae; Burkholderia; Burkholderia cepacia complex
MSDDIDNGGGKGGFTRRDMMRVMAASGMMAVGGGGLLMDAQSAFAAPAPKRGGKIRVANE WP_059924245.1 ABC transporter substrate-binding protein [Burkholderia ubonensis] Length: 528\nNumber of predicted TMHs: 1\nExp number of AAs in TMHs: 32.28118\nExp number, first 60 AAs: 15.42803\nTotal prob of N-in: 0.69070\nPOSSIBLE N-term signal sequence\noutside 1 503\nTMhelix 504 526\ninside 527 528

1357 GCF_001527525.1_ASM152752v1 Burkholderia ubonensis Proteobacteria; Betaproteobacteria; Burkholderiales; Burkholderiaceae; Burkholderia; Burkholderia cepacia complex
MSDDIDNGGKGGFTRRDMMRVMAASGMMAVGGGGLLMDAQSAFAAPAPKRGGKIRVANES WP_060009876.1 ABC transporter substrate-binding protein [Burkholderia ubonensis] Length: 527\nNumber of predicted TMHs: 2\nExp number of AAs in TMHs: 36.08074\nExp number, first 60 AAs: 20.24775\nTotal prob of N-in: 0.91899\nPOSSIBLE N-term signal sequence\ninside 1 20\nTMhelix 21 43\noutside 44 502\nTMhelix 503 525\ninside 526 527

1358 GCF_001529215.1_ASM152921v1 Burkholderia ubonensis Proteobacteria; Betaproteobacteria; Burkholderiales; Burkholderiaceae; Burkholderia; Burkholderia cepacia complex
MSDDIDNGGGKGGFTRRDMMRVMAASGMMAVGGGGLLMDAQSAFAAPAPKRGGKIRVANE WP_060039427.1 ABC transporter substrate-binding protein [Burkholderia ubonensis] Length: 528\nNumber of predicted TMHs: 1\nExp number of AAs in TMHs: 31.51879\nExp number, first 60 AAs: 15.42904\nTotal prob of N-in: 0.69053\nPOSSIBLE N-term signal sequence\noutside 1 503\nTMhelix 504 526\ninside 527 528

1359 GCF_001527085.1_ASM152708v1 Burkholderia ubonensis Proteobacteria; Betaproteobacteria; Burkholderiales; Burkholderiaceae; Burkholderia; Burkholderia cepacia complex
MSDDIDNGGGKGGFTRRDMMRVMAASGMMAVGGGGLLMDAQSAFAAPAPKRGGKIRVANE WP_059878170.1 ABC transporter substrate-binding protein [Burkholderia ubonensis] Length: 528\nNumber of predicted TMHs: 1\nExp number of AAs in TMHs: 31.51969\nExp number, first 60 AAs: 15.4292\nTotal prob of N-in: 0.69052\nPOSSIBLE N-term signal sequence\noutside 1 503\nTMhelix 504 526\ninside 527 528

1360 GCF_001526865.1_ASM152686v1 Burkholderia ubonensis Proteobacteria; Betaproteobacteria; Burkholderiales; Burkholderiaceae; Burkholderia; Burkholderia cepacia complex
MSDDIDKGGGKGGFTRRDMMRVMAASGMMAVGGGGLLMDAQSAFAAPAPKRGGKIRVANE WP_059489522.1 ABC transporter substrate-binding protein [Burkholderia ubonensis] Length: 528\nNumber of predicted TMHs: 1\nExp number of AAs in TMHs: 31.88352\nExp number, first 60 AAs: 16.09709\nTotal prob of N-in: 0.73476\nPOSSIBLE N-term signal sequence\noutside 1 503\nTMhelix 504 526\ninside 527 528

1361 GCF_001525705.1_ASM152570v1 Burkholderia ubonensis Proteobacteria; Betaproteobacteria; Burkholderiales; Burkholderiaceae; Burkholderia; Burkholderia cepacia complex
MSDDIDNGGKGGFTRRDMMRVMAASGMMAVGGGGLLMDAQSAFAAPAPKRGGKIRVANES WP_059849896.1 ABC transporter substrate-binding protein [Burkholderia ubonensis] Length: 527\nNumber of predicted TMHs: 2\nExp number of AAs in TMHs: 36.08404\nExp number, first 60 AAs: 20.24786\nTotal prob of N-in: 0.91899\nPOSSIBLE N-term signal sequence\ninside 1 20\nTMhelix 21 43\noutside 44 502\nTMhelix 503 525\ninside 526 527

1362 GCF_001524365.1_ASM152436v1 Burkholderia ubonensis Proteobacteria; Betaproteobacteria; Burkholderiales; Burkholderiaceae; Burkholderia; Burkholderia cepacia complex
MSDDIDNGGGKGDFTRRDMMRVMAASGMMAVGGGGLLMDAQSAFAAPAPKRGGKIRVANE WP_059744049.1 ABC transporter substrate-binding protein [Burkholderia ubonensis] Length: 528\nNumber of predicted TMHs: 1\nExp number of AAs in TMHs: 31.70915\nExp number, first 60 AAs: 15.83511\nTotal prob of N-in: 0.71046\nPOSSIBLE N-term signal sequence\noutside 1 503\nTMhelix 504 526\ninside 527 528

1363 GCF_001523825.1_ASM152382v1 Burkholderia ubonensis Proteobacteria; Betaproteobacteria; Burkholderiales; Burkholderiaceae; Burkholderia; Burkholderia cepacia complex
MSDDIDNGGKGGFTRRDMMRVMAASGMMAVGGGGLLMDAQSAFAAPAPKRGGKIRVANES WP_059766173.1 ABC transporter substrate-binding protein [Burkholderia ubonensis] Length: 527\nNumber of predicted TMHs: 2\nExp number of AAs in TMHs: 35.87653\nExp number, first 60 AAs: 20.2481\nTotal prob of N-in: 0.91905\nPOSSIBLE N-term signal sequence\ninside 1 20\nTMhelix 21 43\noutside 44 502\nTMhelix 503 525\ninside 526 527

1364 GCF_001524405.1_ASM152440v1 Burkholderia ubonensis Proteobacteria; Betaproteobacteria; Burkholderiales; Burkholderiaceae; Burkholderia; Burkholderia cepacia complex
MSDDIDNGGKGGFTRRDMMRVMAASGMMAVGGGGLLMDAQSAFAAPAPKRGGKIRVANES WP_059653009.1 ABC transporter substrate-binding protein [Burkholderia ubonensis] Length: 527\nNumber of predicted TMHs: 2\nExp number of AAs in TMHs: 36.08388\nExp number, first 60 AAs: 20.24786\nTotal prob of N-in: 0.91899\nPOSSIBLE N-term signal sequence\ninside 1 20\nTMhelix 21 43\noutside 44 502\nTMhelix 503 525\ninside 526 527

1365 GCF_001523485.1_ASM152348v1 Burkholderia ubonensis Proteobacteria; Betaproteobacteria; Burkholderiales; Burkholderiaceae; Burkholderia; Burkholderia cepacia complex
MSDDIDKGGGKGGFTRRDMMRVMAASGMMAVGGGGLLMDAQSAFAAPAPKRGGKIRVANE WP_059489522.1 ABC transporter substrate-binding protein [Burkholderia ubonensis] Length: 528\nNumber of predicted TMHs: 1\nExp number of AAs in TMHs: 31.88352\nExp number, first 60 AAs: 16.09709\nTotal prob of N-in: 0.73476\nPOSSIBLE N-term signal sequence\noutside 1 503\nTMhelix 504 526\ninside 527 528

1366 GCF_001528245.1_ASM152824v1 Burkholderia ubonensis Proteobacteria; Betaproteobacteria; Burkholderiales; Burkholderiaceae; Burkholderia; Burkholderia cepacia complex
MSDDIDNGGKGGFTRRDMMRVMAASGMMAVGGGGLLMDAQSAFAAPAPKRGGKIRVANES WP_045565706.1 ABC transporter substrate-binding protein [Burkholderia ubonensis] Length: 527\nNumber of predicted TMHs: 2\nExp

number of AAs in TMHs: 36.08388\nExp number, first 60 AAs: 20.24786\nTotal prob of N-in: 0.91899\nPOSSIBLE N-term signal sequence\ninside 1 20\nTMhelix 21 43\noutside 44 502\nTMhelix 503 525\ninside 526 527

1367 GCF_001529655.1_ASM152965v1 Burkholderia ubonensis Proteobacteria; Betaproteobacteria; Burkholderiales; Burkholderiaceae; Burkholderia; Burkholderia cepacia complex MSDDIDNGGKGGFTRRDMMRVMAASGMMAVGGGGLLMDAQSAFAAPAPKRGGKIRVANES WP_060096822.1 ABC transporter substrate-binding protein [Burkholderia ubonensis] Length: 527\nNumber of predicted TMHs: 2\nExp number of AAs in TMHs: 35.5658399999999\nExp number, first 60 AAs: 20.24797\nTotal prob of N-in: 0.91902\nPOSSIBLE N-term signal sequence\ninside 1 20\nTMhelix 21 43\noutside 44 502\nTMhelix 503 525\ninside 526 527

1368 GCF_001530335.1_ASM153033v1 Burkholderia ubonensis Proteobacteria; Betaproteobacteria; Burkholderiales; Burkholderiaceae; Burkholderia; Burkholderia cepacia complex MSDDIDNGGGGKGGFTRRDMMRVMAASGMMAVGGGGLLMDAQSAFAAPAPKRGGKIRVAN WP_060090452.1 ABC transporter substrate-binding protein [Burkholderia ubonensis] Length: 529\nNumber of predicted TMHs: 1\nExp number of AAs in TMHs: 30.50228\nExp number, first 60 AAs: 14.12788\nTotal prob of N-in: 0.59108\nPOSSIBLE N-term signal sequence\noutside 1 504\nTMhelix 505 527\ninside 528 529

1369 GCF_001530375.1_ASM153037v1 Burkholderia ubonensis Proteobacteria; Betaproteobacteria; Burkholderiales; Burkholderiaceae; Burkholderia; Burkholderia cepacia complex MSDDIDNGGKGGFTRRDMMRVMAASGMMAVGGGLLMDAQSAFAAPAPKRGGKIRVANESS WP_060124274.1 ABC transporter substrate-binding protein [Burkholderia ubonensis] Length: 526\nNumber of predicted TMHs: 2\nExp number of AAs in TMHs: 37.63637\nExp number, first 60 AAs: 21.0439\nTotal prob of N-in: 0.93977\nPOSSIBLE N-term signal sequence\ninside 1 20\nTMhelix 21 43\noutside 44 501\nTMhelix 502 524\ninside 525 526

1370 GCF_001530545.1_ASM153054v1 Burkholderia ubonensis Proteobacteria; Betaproteobacteria; Burkholderiales; Burkholderiaceae; Burkholderia; Burkholderia cepacia complex MSDDIDKGGGKGGFTRRDMMRVMAASGMMAVGGGGLLMDAQSAFAAPAPKRGGKIRVANE WP_059979451.1 ABC transporter substrate-binding protein [Burkholderia ubonensis] Length: 528\nNumber of predicted TMHs: 1\nExp number of AAs in TMHs: 32.09636\nExp number, first 60 AAs: 16.09804\nTotal prob of N-in: 0.73456\nPOSSIBLE N-term signal sequence\noutside 1 503\nTMhelix 504 526\ninside 527 528

1371 GCF_001531605.1_ASM153160v1 Burkholderia ubonensis Proteobacteria; Betaproteobacteria; Burkholderiales; Burkholderiaceae; Burkholderia; Burkholderia cepacia complex MSDDIDKGGGKGGFTRRDMMRVMAASGMMAVGGGGLLMDAQSAFAAPAPKRGGKIRVANE WP_060203251.1 ABC transporter substrate-binding protein [Burkholderia ubonensis] Length: 528\nNumber of predicted TMHs: 1\nExp number of AAs in TMHs: 31.88349\nExp number, first 60 AAs: 16.09709\nTotal prob of N-in: 0.73476\nPOSSIBLE N-term signal sequence\noutside 1 503\nTMhelix 504 526\ninside 527 528

1372 GCF_001528145.1_ASM152814v1 Burkholderia ubonensis Proteobacteria; Betaproteobacteria; Burkholderiales; Burkholderiaceae; Burkholderia; Burkholderia cepacia complex MSDDIDKGGGKGGFTRRDMMRVMAASGMMAVGGGGLLMDAQSAFAAPAPKRGGKIRVANE WP_059979451.1 ABC transporter substrate-binding protein [Burkholderia ubonensis] Length: 528\nNumber of predicted TMHs: 1\nExp number of AAs in TMHs: 32.09636\nExp number, first 60 AAs: 16.09804\nTotal prob of N-in: 0.73456\nPOSSIBLE N-term signal sequence\noutside 1 503\nTMhelix 504 526\ninside 527 528

1373 GCF_001532805.1_ASM153280v1 Burkholderia ubonensis Proteobacteria; Betaproteobacteria; Burkholderiales; Burkholderiaceae; Burkholderia; Burkholderia cepacia complex MSDDIDKGGGKGGFTRRDMMRVMAASGMMAVGGGGLLMDAQSAFAAPAPKRGGKIRVANE WP_060338303.1 ABC transporter substrate-binding protein [Burkholderia ubonensis] Length: 528\nNumber of predicted TMHs: 1\nExp number of AAs in TMHs: 31.88348\nExp number, first 60 AAs: 16.09709\nTotal prob of N-in: 0.73476\nPOSSIBLE N-term signal sequence\noutside 1 503\nTMhelix 504 526\ninside 527 528

1374 GCF_001533405.1_ASM153340v1 Burkholderia ubonensis Proteobacteria; Betaproteobacteria; Burkholderiales; Burkholderiaceae; Burkholderia; Burkholderia cepacia complex MSDDIDKGGGKGGFTRRDMMRVMAASGMMAVGGGGLLMDAQSAFAAPAPKRGGKIRVANE WP_059489522.1 ABC transporter substrate-binding protein [Burkholderia ubonensis] Length: 528\nNumber of predicted TMHs: 1\nExp number of AAs in TMHs: 31.88352\nExp number, first 60 AAs: 16.09709\nTotal prob of N-in: 0.73476\nPOSSIBLE N-term signal sequence\noutside 1 503\nTMhelix 504 526\ninside 527 528

1375 GCF_001533535.1_ASM153353v1 Burkholderia ubonensis Proteobacteria; Betaproteobacteria; Burkholderiales; Burkholderiaceae; Burkholderia; Burkholderia cepacia complex
MSDDIDKGGGKGGFTRRDMMRVMAASGMMAVGGGGLLMDAQSAFAAPAPKRGGKIRVANE WP_060203251.1 ABC transporter substrate-binding protein [Burkholderia ubonensis] Length: 528\nNumber of predicted TMHs: 1\nExp number of AAs in TMHs: 31.88349\nExp number, first 60 AAs: 16.09709\nTotal prob of N-in: 0.73476\nPOSSIBLE N-term signal sequence\noutside 1 503\nTMhelix 504 526\ninside 527 528

1376 GCF_001533625.1_ASM153362v1 Burkholderia ubonensis Proteobacteria; Betaproteobacteria; Burkholderiales; Burkholderiaceae; Burkholderia; Burkholderia cepacia complex
MSDDIDKGGGKGGFTRRDMMRVMAASGMMAVGGGGLLMDAQSAFAAPAPKRGGKIRVANE WP_060203251.1 ABC transporter substrate-binding protein [Burkholderia ubonensis] Length: 528\nNumber of predicted TMHs: 1\nExp number of AAs in TMHs: 31.88349\nExp number, first 60 AAs: 16.09709\nTotal prob of N-in: 0.73476\nPOSSIBLE N-term signal sequence\noutside 1 503\nTMhelix 504 526\ninside 527 528

1377 GCF_001536805.1_ASM153680v1 Burkholderia ubonensis Proteobacteria; Betaproteobacteria; Burkholderiales; Burkholderiaceae; Burkholderia; Burkholderia cepacia complex
MSDDIDKGGGKGGFTRRDMMRVMAASGMMAVGGGGLLMDAQSAFAAPAPKRGGKIRVANE WP_060065546.1 ABC transporter substrate-binding protein [Burkholderia ubonensis] Length: 528\nNumber of predicted TMHs: 1\nExp number of AAs in TMHs: 31.88338\nExp number, first 60 AAs: 16.09709\nTotal prob of N-in: 0.73476\nPOSSIBLE N-term signal sequence\noutside 1 503\nTMhelix 504 526\ninside 527 528

1378 GCF_001882515.1_ASM188251v1 Burkholderia ubonensis Proteobacteria; Betaproteobacteria; Burkholderiales; Burkholderiaceae; Burkholderia; Burkholderia cepacia complex
MSDDIDKGGGKGGFTRRDMMRVMAASGMMAVGGGGLLMDAQSAFAAPAPKRGGKIRVANE WP_059972434.1 ABC transporter substrate-binding protein [Burkholderia ubonensis] Length: 528\nNumber of predicted TMHs: 1\nExp number of AAs in TMHs: 32.0963\nExp number, first 60 AAs: 16.09802\nTotal prob of N-in: 0.73456\nPOSSIBLE N-term signal sequence\noutside 1 503\nTMhelix 504 526\ninside 527 528

1379 GCF_001901485.1_ASM190148v1 Burkholderia ubonensis Proteobacteria; Betaproteobacteria; Burkholderiales; Burkholderiaceae; Burkholderia; Burkholderia cepacia complex
MSDDIDKGGGKGGFTRRDMMRVMAASGMMAVGGGGLLMDAQSAFAAPAPKRGGKIRVANE WP_060203251.1 ABC transporter substrate-binding protein [Burkholderia ubonensis] Length: 528\nNumber of predicted TMHs: 1\nExp number of AAs in TMHs: 31.88349\nExp number, first 60 AAs: 16.09709\nTotal prob of N-in: 0.73476\nPOSSIBLE N-term signal sequence\noutside 1 503\nTMhelix 504 526\ninside 527 528

1380 GCF_001882555.1_ASM188255v1 Burkholderia ubonensis Proteobacteria; Betaproteobacteria; Burkholderiales; Burkholderiaceae; Burkholderia; Burkholderia cepacia complex
MSDDIDNGGGGKGGFTRRDMMRVMAASGMMAVGGGGLLMDAQSAFAAPAPKRGGKIRVAN WP_071761684.1 ABC transporter substrate-binding protein [Burkholderia ubonensis] Length: 529\nNumber of predicted TMHs: 1\nExp number of AAs in TMHs: 30.27644\nExp number, first 60 AAs: 14.12306\nTotal prob of N-in: 0.59155\nPOSSIBLE N-term signal sequence\noutside 1 504\nTMhelix 505 527\ninside 528 529

1381 GCF_001882475.1_ASM188247v1 Burkholderia ubonensis Proteobacteria; Betaproteobacteria; Burkholderiales; Burkholderiaceae; Burkholderia; Burkholderia cepacia complex
MSDDIDKGGGKGGFTRRDMMRVMAASGMMAVGGGGLLMDAQSAFAAPAPKRGGKIRVANE WP_060203251.1 ABC transporter substrate-binding protein [Burkholderia ubonensis] Length: 528\nNumber of predicted TMHs: 1\nExp number of AAs in TMHs: 31.88349\nExp number, first 60 AAs: 16.09709\nTotal prob of N-in: 0.73476\nPOSSIBLE N-term signal sequence\noutside 1 503\nTMhelix 504 526\ninside 527 528

1382 GCF_001529155.1_ASM152915v1 Burkholderia ubonensis Proteobacteria; Betaproteobacteria; Burkholderiales; Burkholderiaceae; Burkholderia; Burkholderia cepacia complex
MSDDIDNGGGKGGFTRRDMMRVMAASGMMAVGGGGLLMDAQSAFAAPAPKRGGKIRVANE WP_060039427.1 ABC transporter substrate-binding protein [Burkholderia ubonensis] Length: 528\nNumber of predicted TMHs: 1\nExp number of AAs in TMHs: 31.51879\nExp number, first 60 AAs: 15.42904\nTotal prob of N-in: 0.69053\nPOSSIBLE N-term signal sequence\noutside 1 503\nTMhelix 504 526\ninside 527 528

1383 GCF_001529125.1_ASM152912v1 Burkholderia ubonensis Proteobacteria; Betaproteobacteria; Burkholderiales; Burkholderiaceae; Burkholderia; Burkholderia cepacia complex
MSDDIDNGGGKGGFTRRDMMRVMAASGMMAVGGGGLLMDAQSAFAAPAPKRGGKIRVANE WP_060054458.1 ABC transporter substrate-binding protein [Burkholderia ubonensis] Length: 528\nNumber of predicted TMHs: 1\nExp

number of AAs in TMHs: 32.28118\nExp number, first 60 AAs:  15.42803\nTotal prob of N-in:   0.69070\nPOSSIBLE N-term signal sequence\noutside   1   503\nTMhelix   504   526\ninside   527   528

1384 GCF_001528205.1_ASM152820v1 Burkholderia ubonensis Proteobacteria; Betaproteobacteria; Burkholderiales; Burkholderiaceae; Burkholderia; Burkholderia cepacia complex MSDDIDNGGKGGKGGFTRRDMMRVMAASGMMAVGGGGLLMDAQSAFAAPAPKRGGKIRVA WP_060194953.1 ABC transporter substrate-binding protein [Burkholderia ubonensis] Length: 530\nNumber of predicted TMHs:  1\nExp number of AAs in TMHs: 32.02507\nExp number, first 60 AAs:  16.0547\nTotal prob of N-in:   0.72830\nPOSSIBLE N-term signal sequence\noutside   1   505\nTMhelix   506   528\ninside   529   530

1385 GCF_001530165.1_ASM153016v1 Burkholderia ubonensis Proteobacteria; Betaproteobacteria; Burkholderiales; Burkholderiaceae; Burkholderia; Burkholderia cepacia complex MSDDIDNGGGGKGGFTRRDMMRVMAASGMMAVGGGGLLMDAQSAFAAPAPKRGGKIRVAN WP_060090452.1 ABC transporter substrate-binding protein [Burkholderia ubonensis] Length: 529\nNumber of predicted TMHs:  1\nExp number of AAs in TMHs: 30.50228\nExp number, first 60 AAs:  14.12788\nTotal prob of N-in:   0.59108\nPOSSIBLE N-term signal sequence\noutside   1   504\nTMhelix   505   527\ninside   528   529

1386 GCF_001882865.1_ASM188286v1 Burkholderia ubonensis Proteobacteria; Betaproteobacteria; Burkholderiales; Burkholderiaceae; Burkholderia; Burkholderia cepacia complex MSDDIDNGGKGGFTRRDMMRVMAASGMMAVGGGGLLMDAQSAFAAPAPKRGGKIRVANES WP_059707130.1 ABC transporter substrate-binding protein [Burkholderia ubonensis] Length: 527\nNumber of predicted TMHs:  2\nExp number of AAs in TMHs: 35.87636\nExp number, first 60 AAs:  20.2481\nTotal prob of N-in:   0.91905\nPOSSIBLE N-term signal sequence\ninside   1   20\nTMhelix   21   43\noutside   44   502\nTMhelix   503   525\ninside   526   527

1387 GCF_001523775.1_ASM152377v1 Burkholderia ubonensis Proteobacteria; Betaproteobacteria; Burkholderiales; Burkholderiaceae; Burkholderia; Burkholderia cepacia complex MSDDIDNGGGKGGFTRRDMMRVMAASGMMAVGGGGLLMDAQSAFAAPAPKRGGKIRVANE WP_059838574.1 ABC transporter substrate-binding protein [Burkholderia ubonensis] Length: 528\nNumber of predicted TMHs:  1\nExp number of AAs in TMHs: 31.32024\nExp number, first 60 AAs:  15.42717\nTotal prob of N-in:   0.69083\nPOSSIBLE N-term signal sequence\noutside   1   503\nTMhelix   504   526\ninside   527   528

1388 GCF_001533035.1_ASM153303v1 Burkholderia ubonensis Proteobacteria; Betaproteobacteria; Burkholderiales; Burkholderiaceae; Burkholderia; Burkholderia cepacia complex MSDDIDKGGGKGGFTRRDMMRVMAASGMMAVGGGGLLMDAQSAFAAPAPKRGGKIRVANE WP_060203251.1 ABC transporter substrate-binding protein [Burkholderia ubonensis] Length: 528\nNumber of predicted TMHs:  1\nExp number of AAs in TMHs: 31.88349\nExp number, first 60 AAs:  16.09709\nTotal prob of N-in:   0.73476\nPOSSIBLE N-term signal sequence\noutside   1   503\nTMhelix   504   526\ninside   527   528

1389 GCF_001533285.1_ASM153328v1 Burkholderia ubonensis Proteobacteria; Betaproteobacteria; Burkholderiales; Burkholderiaceae; Burkholderia; Burkholderia cepacia complex MSDDIDNGGGGKGGFTRRDMMRVMAASGMMAVGGGGLLMDAQSAFAAPAPKRGGKIRVAN WP_060284860.1 ABC transporter substrate-binding protein [Burkholderia ubonensis] Length: 529\nNumber of predicted TMHs:  1\nExp number of AAs in TMHs: 30.50216\nExp number, first 60 AAs:  14.12788\nTotal prob of N-in:   0.59108\nPOSSIBLE N-term signal sequence\noutside   1   504\nTMhelix   505   527\ninside   528   529

1390 GCF_001533565.1_ASM153356v1 Burkholderia ubonensis Proteobacteria; Betaproteobacteria; Burkholderiales; Burkholderiaceae; Burkholderia; Burkholderia cepacia complex MSDDIDKGGGKGGFTRRDMMRVMAASGMMAVGGGGLLMDAQSAFAAPAPKRGGKIRVANE WP_060203251.1 ABC transporter substrate-binding protein [Burkholderia ubonensis] Length: 528\nNumber of predicted TMHs:  1\nExp number of AAs in TMHs: 31.88349\nExp number, first 60 AAs:  16.09709\nTotal prob of N-in:   0.73476\nPOSSIBLE N-term signal sequence\noutside   1   503\nTMhelix   504   526\ninside   527   528

1391 GCF_001533655.1_ASM153365v1 Burkholderia ubonensis Proteobacteria; Betaproteobacteria; Burkholderiales; Burkholderiaceae; Burkholderia; Burkholderia cepacia complex MSDDIDKGGGKGGFTRRDMMRVMAASGMMAVGGGGLLMDAQSAFAAPAPKRGGKIRVANE WP_060203251.1 ABC transporter substrate-binding protein [Burkholderia ubonensis] Length: 528\nNumber of predicted TMHs:  1\nExp number of AAs in TMHs: 31.88349\nExp number, first 60 AAs:  16.09709\nTotal prob of N-in:   0.73476\nPOSSIBLE N-term signal sequence\noutside   1   503\nTMhelix   504   526\ninside   527   528

1392 GCF_001882425.1_ASM188242v1 Burkholderia ubonensis Proteobacteria; Betaproteobacteria; Burkholderiales; Burkholderiaceae; Burkholderia; Burkholderia cepacia complex

MSDDIDNGGGKGDFTRRDMMRVMAASGMMAVGGGGLLMDAQSAFAAPAPKRGGKIRVANE WP_059744049.1 ABC transporter substrate-binding protein [Burkholderia ubonensis] Length: 528\nNumber of predicted TMHs: 1\nExp number of AAs in TMHs: 31.70915\nExp number, first 60 AAs: 15.83511\nTotal prob of N-in: 0.71046\nPOSSIBLE N-term signal sequence\noutside 1 503\nTMhelix 504 526\ninside 527 528

1393 GCF_001882585.1_ASM188258v1 Burkholderia ubonensis Proteobacteria; Betaproteobacteria; Burkholderiales; Burkholderiaceae; Burkholderia; Burkholderia cepacia complex MSDDIDNGGGGKGGFTRRDMMRVMAASGMMAVGGGGLLMDAQSAFAAPAPKRGGKIRVAN WP_071761684.1 ABC transporter substrate-binding protein [Burkholderia ubonensis] Length: 529\nNumber of predicted TMHs: 1\nExp number of AAs in TMHs: 30.27644\nExp number, first 60 AAs: 14.12306\nTotal prob of N-in: 0.59155\nPOSSIBLE N-term signal sequence\noutside 1 504\nTMhelix 505 527\ninside 528 529

1394 GCF_001882645.1_ASM188264v1 Burkholderia ubonensis Proteobacteria; Betaproteobacteria; Burkholderiales; Burkholderiaceae; Burkholderia; Burkholderia cepacia complex MSDDIDKGGGKGGFTRRDMMRVMAASGMMAVGGGGLLMDAQSAFAAPAPKRGGKIRVANE WP_071853290.1 ABC transporter substrate-binding protein [Burkholderia ubonensis] Length: 528\nNumber of predicted TMHs: 1\nExp number of AAs in TMHs: 31.88352\nExp number, first 60 AAs: 16.09709\nTotal prob of N-in: 0.73476\nPOSSIBLE N-term signal sequence\noutside 1 503\nTMhelix 504 526\ninside 527 528

1395 GCF_001882785.1_ASM188278v1 Burkholderia ubonensis Proteobacteria; Betaproteobacteria; Burkholderiales; Burkholderiaceae; Burkholderia; Burkholderia cepacia complex MSDDIDNGGGKGGFTRRDMMRVMAASGMMAVGGGGLLMDAQSAFAAPAPKRGGKIRVANE WP_060039427.1 ABC transporter substrate-binding protein [Burkholderia ubonensis] Length: 528\nNumber of predicted TMHs: 1\nExp number of AAs in TMHs: 31.51879\nExp number, first 60 AAs: 15.42904\nTotal prob of N-in: 0.69053\nPOSSIBLE N-term signal sequence\noutside 1 503\nTMhelix 504 526\ninside 527 528

1396 GCF_001523425.1_ASM152342v1 Burkholderia ubonensis Proteobacteria; Betaproteobacteria; Burkholderiales; Burkholderiaceae; Burkholderia; Burkholderia cepacia complex MSDDIDNGGKGGFTRRDMMRVMAASGMMAVGGGGLLMDAQSAFAAPAPKRGGKIRVANES WP_059707130.1 ABC transporter substrate-binding protein [Burkholderia ubonensis] Length: 527\nNumber of predicted TMHs: 2\nExp number of AAs in TMHs: 35.87636\nExp number, first 60 AAs: 20.2481\nTotal prob of N-in: 0.91905\nPOSSIBLE N-term signal sequence\ninside 1 20\nTMhelix 21 43\noutside 44 502\nTMhelix 503 525\ninside 526 527

1397 GCF_001528875.1_ASM152887v1 Burkholderia ubonensis Proteobacteria; Betaproteobacteria; Burkholderiales; Burkholderiaceae; Burkholderia; Burkholderia cepacia complex MSDDIDNGGKGGFTRRDMMRVMAASGMMAVGGGGLLMDAQSAFAAPAPKRGGKIRVANES WP_059925480.1 ABC transporter substrate-binding protein [Burkholderia ubonensis] Length: 527\nNumber of predicted TMHs: 2\nExp number of AAs in TMHs: 36.08403\nExp number, first 60 AAs: 20.24786\nTotal prob of N-in: 0.91899\nPOSSIBLE N-term signal sequence\ninside 1 20\nTMhelix 21 43\noutside 44 502\nTMhelix 503 525\ninside 526 527

1398 GCF_001529705.1_ASM152970v1 Burkholderia ubonensis Proteobacteria; Betaproteobacteria; Burkholderiales; Burkholderiaceae; Burkholderia; Burkholderia cepacia complex MSDDIDNGGGGKGGFTRRDMMRVMAASGMMAVGGGGLLMDAQSAFAAPAPKRGGKIRVAN WP_060012706.1 ABC transporter substrate-binding protein [Burkholderia ubonensis] Length: 529\nNumber of predicted TMHs: 1\nExp number of AAs in TMHs: 30.27493\nExp number, first 60 AAs: 14.1228\nTotal prob of N-in: 0.59153\nPOSSIBLE N-term signal sequence\noutside 1 504\nTMhelix 505 527\ninside 528 529

1399 GCF_001529605.1_ASM152960v1 Burkholderia ubonensis Proteobacteria; Betaproteobacteria; Burkholderiales; Burkholderiaceae; Burkholderia; Burkholderia cepacia complex MSDDIDKGGGKGGFTRRDMMRVMAASGMMAVGGGGLLMDAQSAFAAPAPKRGGKIRVANE WP_060034152.1 ABC transporter substrate-binding protein [Burkholderia ubonensis] Length: 528\nNumber of predicted TMHs: 1\nExp number of AAs in TMHs: 32.86358\nExp number, first 60 AAs: 16.09747\nTotal prob of N-in: 0.73468\nPOSSIBLE N-term signal sequence\noutside 1 503\nTMhelix 504 526\ninside 527 528

1400 GCF_001530565.1_ASM153056v1 Burkholderia ubonensis Proteobacteria; Betaproteobacteria; Burkholderiales; Burkholderiaceae; Burkholderia; Burkholderia cepacia complex MSDDIDNGGKGGFTRRDMMRVMAASGMMAVGGGGLLMDAQSAFAAPAPKRGGKIRVANES WP_045565706.1 ABC transporter substrate-binding protein [Burkholderia ubonensis] Length: 527\nNumber of predicted TMHs: 2\nExp number of AAs in TMHs: 36.08388\nExp number, first 60 AAs: 20.24786\nTotal prob of N-in: 0.91899\nPOSSIBLE N-term signal sequence\ninside 1 20\nTMhelix 21 43\noutside 44 502\nTMhelix 503 525\ninside 526 527

1401 GCF_001532865.1_ASM153286v1 Burkholderia ubonensis Proteobacteria; Betaproteobacteria; Burkholderiales; Burkholderiaceae; Burkholderia; Burkholderia cepacia complex MSDDIDKGGGKGGFTRRDMMRVMAASGMMAVGGGGLLMDAQSAFAAPAPKRGGKIRVANE WP_059991183.1 ABC transporter substrate-binding protein [Burkholderia ubonensis] Length: 528\nNumber of predicted TMHs: 1\nExp number of AAs in TMHs: 31.88336\nExp number, first 60 AAs: 16.09709\nTotal prob of N-in: 0.73476\nPOSSIBLE N-term signal sequence\noutside 1 503\nTMhelix 504 526\ninside 527 528

1402 GCF_000959545.1_ASM95954v1 Burkholderia ambifaria AMMD Proteobacteria; Betaproteobacteria; Burkholderiales; Burkholderiaceae; Burkholderia; Burkholderia cepacia complex; Burkholderia ambifaria MRISITRRRLLAAGMAAASLPLSGCFTPKLYKESAYHEQVSAFMITEDGKKLVVLGQDYH WP_011659356.1 hypothetical protein [Burkholderia ambifaria] Length: 215\nNumber of predicted TMHs: 1\nExp number of AAs in TMHs: 24.18844\nExp number, first 60 AAs: 0.24064\nTotal prob of N-in: 0.48889\noutside 1 183\nTMhelix 184 206\ninside 207 215

1403 GCF_000203915.1_ASM20391v1 Burkholderia ambifaria AMMD Proteobacteria; Betaproteobacteria; Burkholderiales; Burkholderiaceae; Burkholderia; Burkholderia cepacia complex; Burkholderia ambifaria MRISITRRRLLAAGMAAASLPLSGCFTPKLYKESAYHEQVSAFMITEDGKKLVVLGQDYH WP_011659356.1 hypothetical protein [Burkholderia ambifaria] Length: 215\nNumber of predicted TMHs: 1\nExp number of AAs in TMHs: 24.18844\nExp number, first 60 AAs: 0.24064\nTotal prob of N-in: 0.48889\noutside 1 183\nTMhelix 184 206\ninside 207 215

1404 GCF_000181975.1_ASM18197v1 Burkholderia ambifaria IOP40-10 Proteobacteria; Betaproteobacteria; Burkholderiales; Burkholderiaceae; Burkholderia; Burkholderia cepacia complex; Burkholderia ambifaria MRISITRRRLLAAGMAAASLPLSGCFTPKLYKDSWYYERVSAFMITEDGKKLVVLGQDYH WP_006752991.1 hypothetical protein [Burkholderia ambifaria] Length: 215\nNumber of predicted TMHs: 1\nExp number of AAs in TMHs: 24.09895\nExp number, first 60 AAs: 0.45514\nTotal prob of N-in: 0.60274\noutside 1 183\nTMhelix 184 206\ninside 207 215

1405 GCF_000019925.1_ASM1992v1 Burkholderia ambifaria MC40-6 Proteobacteria; Betaproteobacteria; Burkholderiales; Burkholderiaceae; Burkholderia; Burkholderia cepacia complex; Burkholderia ambifaria MRISITRRRLLAAGMAAASLPLSGCFTPKLYKESAYHEQVSAFMITEDGKKLVVLGRDYH WP_012366622.1 hypothetical protein [Burkholderia ambifaria] Length: 215\nNumber of predicted TMHs: 1\nExp number of AAs in TMHs: 24.28164\nExp number, first 60 AAs: 0.23704\nTotal prob of N-in: 0.53897\noutside 1 183\nTMhelix 184 206\ninside 207 215

1406 GCF_000182015.1_ASM18201v1 Burkholderia ambifaria MEX-5 Proteobacteria; Betaproteobacteria; Burkholderiales; Burkholderiaceae; Burkholderia; Burkholderia cepacia complex; Burkholderia ambifaria MRISITRRRLLAAGMAAASLPLSGCFTPKLYKDSWYYEQVSAFMITEDGKKLVVLGQDYH WP_006761406.1 hypothetical protein [Burkholderia ambifaria] Length: 215\nNumber of predicted TMHs: 1\nExp number of AAs in TMHs: 23.5305\nExp number, first 60 AAs: 0.45014\nTotal prob of N-in: 0.51907\noutside 1 183\nTMhelix 184 206\ninside 207 215

1407 GCF_000333135.2_gbccbc7v2.0 Burkholderia cenocepacia BC7 Proteobacteria; Betaproteobacteria; Burkholderiales; Burkholderiaceae; Burkholderia; Burkholderia cepacia complex; Burkholderia cenocepacia MSDDIDNGGKGGFTRRDMMRVMAASGMMAVGGGGLLTSAQSAFAAPAPKRGGKIRVANES WP_006482802.1 MULTISPECIES: ABC transporter substrate-binding protein [Burkholderia cepacia complex] Length: 527\nNumber of predicted TMHs: 2\nExp number of AAs in TMHs: 36.7303599999999\nExp number, first 60 AAs: 22.08937\nTotal prob of N-in: 0.97312\nPOSSIBLE N-term signal sequence\ninside 1 20\nTMhelix 21 43\noutside 44 502\nTMhelix 503 525\ninside 526 527

1408 GCF_000333135.2_gbccbc7v2.0 Burkholderia cenocepacia BC7 Proteobacteria; Betaproteobacteria; Burkholderiales; Burkholderiaceae; Burkholderia; Burkholderia cepacia complex; Burkholderia cenocepacia MKRRTFLALPAVAASGMLAGCSSADLGPVTHRLMKDPEYRETLSAFLISADGKKLVVIGK WP_006483583.1 MULTISPECIES: hypothetical protein [Burkholderia cepacia complex] Length: 212\nNumber of predicted TMHs: 1\nExp number of AAs in TMHs: 28.03134\nExp number, first 60 AAs: 6.30523\nTotal prob of N-in: 0.48947\noutside 1 187\nTMhelix 188 210\ninside 211 212

1409 GCF_000236215.2_ASM23621v4 Burkholderia cenocepacia H111 Proteobacteria; Betaproteobacteria; Burkholderiales; Burkholderiaceae; Burkholderia; Burkholderia cepacia complex; Burkholderia cenocepacia MSDDIDNGGKGGFTRRDMMRVMAASGMMAVGGGGLLTSAQSAFAAPAPKRGGKIRVANES WP_006482802.1 MULTISPECIES: ABC transporter substrate-binding protein [Burkholderia cepacia complex]

Length: 527\nNumber of predicted TMHs: 2\nExp number of AAs in TMHs: 36.7303599999999\nExp number, first 60 AAs: 22.08937\nTotal prob of N-in: 0.97312\nPOSSIBLE N-term signal sequence\ninside 1 20\nTMhelix 21 43\noutside 44 502\nTMhelix 503 525\ninside 526 527

1410 GCF_000236215.2_ASM23621v4 Burkholderia cenocepacia H111 Proteobacteria; Betaproteobacteria; Burkholderiales; Burkholderiaceae; Burkholderia; Burkholderia cepacia complex; Burkholderia cenocepacia MKRRTFLALPAVAASGMLAGCSSADLGPVTHRLMKDPEYRETLSAFLISADGKKLVVIGKWP_006483583.1 MULTISPECIES: hypothetical protein [Burkholderia cepacia complex] Length: 212\nNumber of predicted TMHs: 1\nExp number of AAs in TMHs: 28.03134\nExp number, first 60 AAs: 6.30523\nTotal prob of N-in: 0.48947\noutside 1 187\nTMhelix 188 210\ninside 211 212

1411 GCF_000009485.1_ASM948v1 Burkholderia cenocepacia J2315 Proteobacteria; Betaproteobacteria; Burkholderiales; Burkholderiaceae; Burkholderia; Burkholderia cepacia complex; Burkholderia cenocepacia MKRRTFLALPAVAASGMLAGCSSADLGPVTHRLMKDPEYRETLSAFLISADGKKLVVIGKWP_006483583.1 MULTISPECIES: hypothetical protein [Burkholderia cepacia complex] Length: 212\nNumber of predicted TMHs: 1\nExp number of AAs in TMHs: 28.03134\nExp number, first 60 AAs: 6.30523\nTotal prob of N-in: 0.48947\noutside 1 187\nTMhelix 188 210\ninside 211 212

1412 GCF_000009485.1_ASM948v1 Burkholderia cenocepacia J2315 Proteobacteria; Betaproteobacteria; Burkholderiales; Burkholderiaceae; Burkholderia; Burkholderia cepacia complex; Burkholderia cenocepacia MSDDIDNGGKGGFTRRDMMRVMAASGMMAVGGGGLLTSAQSAFAAPAPKRGGKIRVANES WP_006482802.1 MULTISPECIES: ABC transporter substrate-binding protein [Burkholderia cepacia complex] Length: 527\nNumber of predicted TMHs: 2\nExp number of AAs in TMHs: 36.7303599999999\nExp number, first 60 AAs: 22.08937\nTotal prob of N-in: 0.97312\nPOSSIBLE N-term signal sequence\ninside 1 20\nTMhelix 21 43\noutside 44 502\nTMhelix 503 525\ninside 526 527

1413 GCF_000333155.2_gbcckv2.0Burkholderia cenocepacia K56-2Valvano Proteobacteria; Betaproteobacteria; Burkholderiales; Burkholderiaceae; Burkholderia; Burkholderia cepacia complex; Burkholderia cenocepacia MSDDIDNGGKGGFTRRDMMRVMAASGMMAVGGGGLLTSAQSAFAAPAPKRGGKIRVANES WP_006482802.1 MULTISPECIES: ABC transporter substrate-binding protein [Burkholderia cepacia complex] Length: 527\nNumber of predicted TMHs: 2\nExp number of AAs in TMHs: 36.7303599999999\nExp number, first 60 AAs: 22.08937\nTotal prob of N-in: 0.97312\nPOSSIBLE N-term signal sequence\ninside 1 20\nTMhelix 21 43\noutside 44 502\nTMhelix 503 525\ninside 526 527

1414 GCF_000333155.2_gbcckv2.0Burkholderia cenocepacia K56-2Valvano Proteobacteria; Betaproteobacteria; Burkholderiales; Burkholderiaceae; Burkholderia; Burkholderia cepacia complex; Burkholderia cenocepacia MKRRTFLALPAVAASGMLAGCSSADLGPVTHRLMKDPEYRETLSAFLISADGKKLVVIGKWP_006483583.1 MULTISPECIES: hypothetical protein [Burkholderia cepacia complex] Length: 212\nNumber of predicted TMHs: 1\nExp number of AAs in TMHs: 28.03134\nExp number, first 60 AAs: 6.30523\nTotal prob of N-in: 0.48947\noutside 1 187\nTMhelix 188 210\ninside 211 212

1415 GCF_001411495.1_ASM141149v1 Burkholderia cepacia ATCC 25416 Proteobacteria; Betaproteobacteria; Burkholderiales; Burkholderiaceae; Burkholderia; Burkholderia cepacia complex; Burkholderia cepacia MKRRALLTAGAIVAAGLTGCTAMLFEDGRYEETVDRFLVSEDGKKFVVLGEKYHYIFDMP WP_027787499.1 hypothetical protein [Burkholderia cepacia] Length: 207\nNumber of predicted TMHs: 1\nExp number of AAs in TMHs: 38.38341\nExp number, first 60 AAs: 15.45171\nTotal prob of N-in: 0.76513\nPOSSIBLE N-term signal sequence\ninside 1 183\nTMhelix 184 206\noutside 207 207

1416 GCF_000473485.1_ASM47348v1 Burkholderia cepacia ATCC 25416 Proteobacteria; Betaproteobacteria; Burkholderiales; Burkholderiaceae; Burkholderia; Burkholderia cepacia complex; Burkholderia cepacia MKRRTFLALPAVAASGMLAGCSNFGPVTKAWMSDSKYDETLSTFLITADGKQLVVLGKKY WP_027787377.1 MULTISPECIES: hypothetical protein [Burkholderia] Length: 210\nNumber of predicted TMHs: 1\nExp number of AAs in TMHs: 34.58187\nExp number, first 60 AAs: 12.86351\nTotal prob of N-in: 0.46668\nPOSSIBLE N-term signal sequence\noutside 1 186\nTMhelix 187 209\ninside 210 210

1417 GCF_001411495.1_ASM141149v1 Burkholderia cepacia ATCC 25416 Proteobacteria; Betaproteobacteria; Burkholderiales; Burkholderiaceae; Burkholderia; Burkholderia cepacia complex; Burkholderia cepacia MKRRTFLALPAVAASGMLAGCSNFGPVTKAWMSDSKYDETLSTFLITADGKQLVVLGKKY WP_027787377.1 MULTISPECIES: hypothetical protein [Burkholderia] Length: 210\nNumber of predicted TMHs: 1\nExp number of AAs in TMHs: 34.58187\nExp number, first 60 AAs: 12.86351\nTotal prob of N-in: 0.46668\nPOSSIBLE N-term signal sequence\noutside 1 186\nTMhelix 187 209\ninside 210 210

1418 GCF_000473485.1_ASM47348v1 Burkholderia cepacia ATCC 25416 Proteobacteria; Betaproteobacteria; Burkholderiales; Burkholderiaceae; Burkholderia; Burkholderia cepacia complex; Burkholderia cepacia MKRRALLTAGAIVAAGLTGCTAMLFEDGRYEETVDRFLVSEDGKKFVVLGEKYHYIFDMP WP_027787499.1 hypothetical protein [Burkholderia cepacia] Length: 207\nNumber of predicted TMHs: 1\nExp number of AAs in TMHs: 38.38341\nExp number, first 60 AAs: 15.45171\nTotal prob of N-in: 0.76513\nPOSSIBLE N-term signal sequence\ninside 1 183\nTMhelix 184 206\noutside 207 207

1419 GCF_000504665.1_ASM50466v1 Burkholderia cepacia Bu72 Proteobacteria; Betaproteobacteria; Burkholderiales; Burkholderiaceae; Burkholderia; Burkholderia cepacia complex; Burkholderia cepacia MNGSITRRRMLAACIAASSLTLSGCFTPALYENRDEAYIEHVSAFLITKDGKKLVVLGER WP_027781201.1 hypothetical protein [Burkholderia cepacia] Length: 213\nNumber of predicted TMHs: 1\nExp number of AAs in TMHs: 23.49227\nExp number, first 60 AAs: 1.25519\nTotal prob of N-in: 0.79117\ninside 1 180\nTMhelix 181 203\noutside 204 213

1420 GCF_000504665.1_ASM50466v1 Burkholderia cepacia Bu72 Proteobacteria; Betaproteobacteria; Burkholderiales; Burkholderiaceae; Burkholderia; Burkholderia cepacia complex; Burkholderia cepacia MKRRTFLALPAVAASGMLAGCADFGPVTKEWMKDSKYDETLSTFLITSDGKQLVVLGKRY WP_027784469.1 hypothetical protein [Burkholderia cepacia] Length: 210\nNumber of predicted TMHs: 1\nExp number of AAs in TMHs: 34.60809\nExp number, first 60 AAs: 11.71623\nTotal prob of N-in: 0.58249\nPOSSIBLE N-term signal sequence\noutside 1 185\nTMhelix 186 208\ninside 209 210

1421 GCF_000292915.1_ASM29291v1 Burkholderia cepacia GG4 Proteobacteria; Betaproteobacteria; Burkholderiales; Burkholderiaceae; Burkholderia; Burkholderia cepacia complex; Burkholderia cepacia MKRRTLLTAGAIVAAASLAGCTAMLFEDGRYEETVERFLVSEDGKKFVVLGKKYHYIFDM WP_041490491.1 hypothetical protein [Burkholderia cepacia] Length: 208\nNumber of predicted TMHs: 2\nExp number of AAs in TMHs: 40.40878\nExp number, first 60 AAs: 16.73081\nTotal prob of N-in: 0.74733\nPOSSIBLE N-term signal sequence\ninside 1 6\nTMhelix 7 26\noutside 27 183\nTMhelix 184 206\ninside 207 208

1422 GCF_000701165.1_ASM70116v2 Burkholderia cepacia JBK9 Proteobacteria; Betaproteobacteria; Burkholderiales; Burkholderiaceae; Burkholderia; Burkholderia cepacia complex; Burkholderia cepacia MSTSITRRRLLAAGMAASSLALSGCFTPALYENHDETYIEHVSAFLITKDGKKLVVLGER WP_059239508.1 hypothetical protein [Burkholderia cepacia] Length: 213\nNumber of predicted TMHs: 1\nExp number of AAs in TMHs: 24.70562\nExp number, first 60 AAs: 2.43635\nTotal prob of N-in: 0.66908\ninside 1 180\nTMhelix 181 203\noutside 204 213

1423 GCF_000701165.1_ASM70116v2 Burkholderia cepacia JBK9 Proteobacteria; Betaproteobacteria; Burkholderiales; Burkholderiaceae; Burkholderia; Burkholderia cepacia complex; Burkholderia cepacia MNPARRRFVVAGAIAVGALSVGACAPHMTKDLFSDHPYTEQIETFAMTGDRSKLVVIGQT WP_059236981.1 hypothetical protein [Burkholderia cepacia] Length: 213\nNumber of predicted TMHs: 2\nExp number of AAs in TMHs: 55.1943000000001\nExp number, first 60 AAs: 21.94434\nTotal prob of N-in: 0.98672\nPOSSIBLE N-term signal sequence\ninside 1 6\nTMhelix 7 29\noutside 30 186\nTMhelix 187 209\ninside 210 213

1424 GCF_000701165.1_ASM70116v2 Burkholderia cepacia JBK9 Proteobacteria; Betaproteobacteria; Burkholderiales; Burkholderiaceae; Burkholderia; Burkholderia cepacia complex; Burkholderia cepacia MKRRTFLALPAVAASGMLAACSDVNIGPITNRMMKDPEYRETLSAFLISADGKKLVVIGK WP_059234152.1 hypothetical protein [Burkholderia cepacia] Length: 212\nNumber of predicted TMHs: 1\nExp number of AAs in TMHs: 32.9565\nExp number, first 60 AAs: 10.92677\nTotal prob of N-in: 0.56235\nPOSSIBLE N-term signal sequence\noutside 1 187\nTMhelix 188 210\ninside 211 212

1425 GCF_000987055.1_ASM98705v1 Burkholderia contaminans FFH2055 Proteobacteria; Betaproteobacteria; Burkholderiales; Burkholderiaceae; Burkholderia; Burkholderia cepacia complex; Burkholderia contaminans MKRRTFLALPAVAASGMLAGCSNFGPVTKAWMKDSEYDETLSTFLITADGKQLVVLGKKY WP_046545747.1 hypothetical protein [Burkholderia contaminans] Length: 210\nNumber of predicted TMHs: 1\nExp number of AAs in TMHs: 33.22614\nExp number, first 60 AAs: 11.49922\nTotal prob of N-in: 0.43506\nPOSSIBLE N-term signal sequence\noutside 1 186\nTMhelix 187 209\ninside 210 210

1426 GCF_000987055.1_ASM98705v1 Burkholderia contaminans FFH2055 Proteobacteria; Betaproteobacteria; Burkholderiales; Burkholderiaceae; Burkholderia; Burkholderia cepacia complex; Burkholderia contaminans MKRRTLLTAGAVIVAAGLTGCTAMLFEDGQYEETVDRFLVSEDGKKFVVLGTKYHYILDM WP_046545623.1 hypothetical protein [Burkholderia contaminans] Length: 208\nNumber of predicted TMHs:

2\nExp number of AAs in TMHs: 42.44834\nExp number, first 60 AAs: 19.01288\nTotal prob of N-in: 0.85365\nPOSSIBLE N-term signal sequence\ninside 1 6\nTMhelix 7 26\noutside 27 183\nTMhelix 184 206\ninside 207 208

1427 GCF_000987055.1_ASM98705v1 Burkholderia contaminans FFH2055 Proteobacteria; Betaproteobacteria; Burkholderiales; Burkholderiaceae; Burkholderia; Burkholderia cepacia complex; Burkholderia contaminans MNVSITRRRLLAAGVAASSLALSGCFTPKLYKDDGYSEQVSGFMMTEDGKKLVVLGTDYH WP_046548430.1 hypothetical protein [Burkholderia contaminans] Length: 212\nNumber of predicted TMHs: 1\nExp number of AAs in TMHs: 25.30439\nExp number, first 60 AAs: 0.83557\nTotal prob of N-in: 0.55248\noutside 1 180\nTMhelix 181 203\ninside 204 212

1428 GCF_000987075.1_ASM98707v1 Burkholderia contaminans LMG 23361 Proteobacteria; Betaproteobacteria; Burkholderiales; Burkholderiaceae; Burkholderia; Burkholderia cepacia complex; Burkholderia contaminans MNVSITRRRLLAAGVAASSLALSGCFTPKLYKDDGYSEQVSGFMMTEDGKKLVVLGTDYH WP_039366460.1 MULTISPECIES: hypothetical protein [Burkholderia] Length: 212\nNumber of predicted TMHs: 1\nExp number of AAs in TMHs: 25.9561\nExp number, first 60 AAs: 0.849399999999999\nTotal prob of N-in: 0.56566\noutside 1 180\nTMhelix 181 203\ninside 204 212

1429 GCF_000987075.1_ASM98707v1 Burkholderia contaminans LMG 23361 Proteobacteria; Betaproteobacteria; Burkholderiales; Burkholderiaceae; Burkholderia; Burkholderia cepacia complex; Burkholderia contaminans MKRRTFLALPAVAASGMLAGCSDANIGPITNRMLKDPEYRETLSAFLISADGKSLVVIGK WP_039354933.1 MULTISPECIES: 5-formyltetrahydrofolate cyclo-ligase [Burkholderia] Length: 212\nNumber of predicted TMHs: 1\nExp number of AAs in TMHs: 24.954\nExp number, first 60 AAs: 4.26972\nTotal prob of N-in: 0.39654\noutside 1 188\nTMhelix 189 211\ninside 212 212

1430 GCF_000959505.1_ASM95950v1 Burkholderia dolosa AU0158 Proteobacteria; Betaproteobacteria; Burkholderiales; Burkholderiaceae; Burkholderia; Burkholderia cepacia complex; Burkholderia dolosa MSDDIDNGGGHGGFTRRDMMRVMAAGGLMAAGSGALLMPAQSAFAAPAPKRGGKIRVANE WP_035975758.1 MULTISPECIES: ABC transporter substrate-binding protein [Burkholderia cepacia complex] Length: 528\nNumber of predicted TMHs: 2\nExp number of AAs in TMHs: 35.70904\nExp number, first 60 AAs: 19.87611\nTotal prob of N-in: 0.85006\nPOSSIBLE N-term signal sequence\ninside 1 23\nTMhelix 24 46\noutside 47 503\nTMhelix 504 526\ninside 527 528

1431 GCF_000497165.1_Bdolosa1.0 Burkholderia dolosa PC543 Proteobacteria; Betaproteobacteria; Burkholderiales; Burkholderiaceae; Burkholderia; Burkholderia cepacia complex; Burkholderia dolosa MSDDIDNGGGHGGFTRRDMMRVMAAGGLMAAGSGALLMPAQSAFAAPAPKRGGKIRVANE WP_035975758.1 MULTISPECIES: ABC transporter substrate-binding protein [Burkholderia cepacia complex] Length: 528\nNumber of predicted TMHs: 2\nExp number of AAs in TMHs: 35.70904\nExp number, first 60 AAs: 19.87611\nTotal prob of N-in: 0.85006\nPOSSIBLE N-term signal sequence\ninside 1 23\nTMhelix 24 46\noutside 47 503\nTMhelix 504 526\ninside 527 528

1432 GCF_000286555.1_gbmulcf1v1.0 Burkholderia multivorans ATCC BAA-247 Proteobacteria; Betaproteobacteria; Burkholderiales; Burkholderiaceae; Burkholderia; Burkholderia cepacia complex; Burkholderia multivorans MNAPETIDPLHADLPREGVAVLHPAARPLTDAELAARRSRSRRATFIKWLRKVHGWVGLW WP_006410297.1 hypothetical protein, partial [Burkholderia multivorans] Length: 235\nNumber of predicted TMHs: 2\nExp number of AAs in TMHs: 45.19864\nExp number, first 60 AAs: 7.29721\nTotal prob of N-in: 0.78336\ninside 1 52\nTMhelix 53 75\noutside 76 194\nTMhelix 195 217\ninside 218 235

1433 GCF_000959245.1_ASM95924v1 Burkholderia ubonensis MSMB22 Proteobacteria; Betaproteobacteria; Burkholderiales; Burkholderiaceae; Burkholderia; Burkholderia cepacia complex; Burkholderia ubonensis MSDDIDNGGKGGFTRRDMMRVMAASGMMAVGGGGLLMDAQSAFAAPAPKRGGKIRVANES WP_045565706.1 ABC transporter substrate-binding protein [Burkholderia ubonensis] Length: 527\nNumber of predicted TMHs: 2\nExp number of AAs in TMHs: 36.08388\nExp number, first 60 AAs: 20.24786\nTotal prob of N-in: 0.91899\nPOSSIBLE N-term signal sequence\ninside 1 20\nTMhelix 21 43\noutside 44 502\nTMhelix 503 525\ninside 526 527

1434 GCF_000773415.1_BCW_38 Burkholderia pseudomallei MSHR7500 Proteobacteria; Betaproteobacteria; Burkholderiales; Burkholderiaceae; Burkholderia; pseudomallei group; Burkholderia pseudomallei MIMKPLRLPISAAGARRAALVACVALFAGCRQELYGGLAERDCNEMMAALLQNGVDAQKK WP_080305681.1 EscJ/YscJ/HrcJ family type III secretion inner membrane ring protein [Burkholderia pseudomallei] Length: 280\nNumber of predicted TMHs: 1\nExp number of AAs in TMHs: 22.93604\nExp number, first 60 AAs: 0.28279\nTotal prob of N-in: 0.16078\noutside 1 221\nTMhelix 222 244\ninside 245 280

1435 GCF_000648335.1_Burkholderia_pseudomallei_assembly Burkholderia pseudomallei PB08298010 Proteobacteria; Betaproteobacteria; Burkholderiales; Burkholderiaceae; Burkholderia; pseudomallei group; Burkholderia pseudomallei MIMKPLRLPISAAGARRAARRAALVACVALFAGCRQELYGGLAERDCNEMMAALLQNGVD WP_080291690.1 EscJ/YscJ/HrcJ family type III secretion inner membrane ring protein, partial [Burkholderia pseudomallei] Length: 270\nNumber of predicted TMHs: 1\nExp number of AAs in TMHs: 22.93164\nExp number, first 60 AAs: 0.25806\nTotal prob of N-in: 0.17089\noutside 1 225\nTMhelix 226 248\ninside 249 270

1436 GCF_000959345.1_ASM95934v1 Burkholderia pseudomallei PB08298010 Proteobacteria; Betaproteobacteria; Burkholderiales; Burkholderiaceae; Burkholderia; pseudomallei group; Burkholderia pseudomallei MIMKPLRLPISAAGARRAARRAALVACVALFAGCRQELYGGLAERDCNEMMAALLQNGVD WP_080341075.1 EscJ/YscJ/HrcJ family type III secretion inner membrane ring protein [Burkholderia pseudomallei] Length: 284\nNumber of predicted TMHs: 1\nExp number of AAs in TMHs: 22.91062\nExp number, first 60 AAs: 0.25946\nTotal prob of N-in: 0.15668\noutside 1 225\nTMhelix 226 248\ninside 249 284

1437 GCF_000698595.1_SOAPdenovo_v1.05Caballeronia glathei Proteobacteria; Betaproteobacteria; Burkholderiales; Burkholderiaceae; Caballeronia MSSRRRVVVTGMGIVSCLGNTLDAVADALCAARPGIVRIDAWRERGFVSQVAGVASVANE WP_035926162.1 3-oxoacyl-ACP synthase [Caballeronia glathei] Length: 409\nNumber of predicted TMHs: 1\nExp number of AAs in TMHs: 19.88178\nExp number, first 60 AAs: 3.03101\nTotal prob of N-in: 0.80732\ninside 1 383\nTMhelix 384 403\noutside 404 409

1438 GCF_001544475.1_ASM1544475v2 Caballeronia humi Proteobacteria; Betaproteobacteria; Burkholderiales; Burkholderiaceae; Caballeronia MPTRRPARRRYLQAPLHAASAVHVSTFSKLAPTLLTNPELARSAEAGAVPLTREQVLATA WP_087667151.1 hypothetical protein [Caballeronia humi] Length: 194\nNumber of predicted TMHs: 1\nExp number of AAs in TMHs: 34.77245\nExp number, first 60 AAs: 0.02637\nTotal prob of N-in: 0.66076\noutside 1 147\nTMhelix 148 170\ninside 171 194

1439 GCF_000876015.1_ASM87601v1 Cupriavidus basilensis Proteobacteria; Betaproteobacteria; Burkholderiales; Burkholderiaceae; Cupriavidus MHAPCTAPPGALSRRGFLKVGLGFSAALACAGLLPALSGCAPAPTNPAAGMRFLRDGDVA WP_059410969.1 hypothetical protein [Cupriavidus basilensis] Length: 193\nNumber of predicted TMHs: 2\nExp number of AAs in TMHs: 44.72246\nExp number, first 60 AAs: 21.82958\nTotal prob of N-in: 0.05259\nPOSSIBLE N-term signal sequence\noutside 1 19\nTMhelix 20 42\ninside 43 149\nTMhelix 150 172\noutside 173 193

1440 GCF_001658125.1_ASM165812v1 Cupriavidus gilardii Proteobacteria; Betaproteobacteria; Burkholderiales; Burkholderiaceae; Cupriavidus MTRIDSTVATPRRRGAMLTTAWLLAAMALPPAHSQPASPNSAVEGPSSNGGATPGVSAPA WP_064576041.1 cellulose synthase regulator BcsB [Cupriavidus gilardii] Length: 780\nNumber of predicted TMHs: 1\nExp number of AAs in TMHs: 23.98513\nExp number, first 60 AAs: 1.17831\nTotal prob of N-in: 0.06614\noutside 1 748\nTMhelix 749 771\ninside 772 780

1441 GCF_000709025.1_assembly_v1_of_C._metallidurans_NA1 Cupriavidus metallidurans Proteobacteria; Betaproteobacteria; Burkholderiales; Burkholderiaceae; Cupriavidus MGQKEEFGQAVNWSPVAKSIVRRISANRRPFLLTTALASLALFALMSYLLRHYQSEGFLR WP_011517311.1 hypothetical protein [Cupriavidus metallidurans] Length: 398\nNumber of predicted TMHs: 2\nExp number of AAs in TMHs: 43.70294\nExp number, first 60 AAs: 20.7357\nTotal prob of N-in: 0.97486\nPOSSIBLE N-term signal sequence\ninside 1 30\nTMhelix 31 50\noutside 51 356\nTMhelix 357 379\ninside 380 398

1442 GCF_001598055.1_ASM159805v1 Cupriavidus nantongensis Proteobacteria; Betaproteobacteria; Burkholderiales; Burkholderiaceae; Cupriavidus MRPNPHHRRAVADADADAEALHALTRRGFLKVGLGFSAALACTALVPSLAGCSSADPAPQ WP_062796007.1 hypothetical protein [Cupriavidus nantongensis] Length: 205\nNumber of predicted TMHs: 1\nExp number of AAs in TMHs: 28.44297\nExp number, first 60 AAs: 6.59348\nTotal prob of N-in: 0.87420\ninside 1 164\nTMhelix 165 187\noutside 188 205

1443 GCF_001853325.1_ASM185332v1 Cupriavidus necator Proteobacteria; Betaproteobacteria; Burkholderiales; Burkholderiaceae; Cupriavidus MAPNHPHRRADADAEALHALTRRGFLKVGLGFSAALACTALVPSLAGCSSAHPAPQAGMT WP_011616288.1 hypothetical protein [Cupriavidus necator] Length: 201\nNumber of predicted TMHs: 1\nExp number of AAs in TMHs: 27.71099\nExp number, first 60 AAs: 6.30589\nTotal prob of N-in: 0.73802\ninside 1 160\nTMhelix 161 183\noutside 184 201

1444 GCF_001481455.1_ASM148145v1 Cupriavidus necator Proteobacteria; Betaproteobacteria; Burkholderiales; Burkholderiaceae; Cupriavidus MAPNHPHRRADADAEALHALTRRGFLKVGLGFSAALACTALVPSLAGCSSAHPAPQAGMT WP_011616288.1 hypothetical protein [Cupriavidus necator] Length: 201\nNumber of predicted TMHs: 1\nExp number of AAs in TMHs: 27.71099\nExp number, first 60 AAs: 6.30589\nTotal prob of N-in: 0.73802\ninside 1 160\nTMhelix 161 183\noutside 184 201

1445 GCF_002011925.1_ASM201192v1 Cupriavidus necator Proteobacteria; Betaproteobacteria; Burkholderiales; Burkholderiaceae; Cupriavidus MPPNHPHRRADADADALHALTRRGFLKVGIGFSAALACTALVPSLAGCSSADPAPQAGMK WP_078198379.1 hypothetical protein [Cupriavidus necator] Length: 201\nNumber of predicted TMHs: 1\nExp number of AAs in TMHs: 30.48805\nExp number, first 60 AAs: 10.67233\nTotal prob of N-in: 0.51928\nPOSSIBLE N-term signal sequence\ninside 1 160\nTMhelix 161 183\noutside 184 201

1446 GCF_000426345.1_ASM42634v1 Cupriavidus sp. amp6 Proteobacteria; Betaproteobacteria; Burkholderiales; Burkholderiaceae; Cupriavidus MSTHSQGAPHRRADADAGALHALSRRGFLKVGLGFSAMLACTALVPSLSGCGVADAAPQA WP_029046437.1 hypothetical protein [Cupriavidus sp. amp6] Length: 204\nNumber of predicted TMHs: 2\nExp number of AAs in TMHs: 38.69186\nExp number, first 60 AAs: 17.92277\nTotal prob of N-in: 0.16749\nPOSSIBLE N-term signal sequence\noutside 1 30\nTMhelix 31 53\ninside 54 163\nTMhelix 164 186\noutside 187 204

1447 GCF_001652915.1_ASM165291v1 Cupriavidus sp. D384 Proteobacteria; Betaproteobacteria; Burkholderiales; Burkholderiaceae; Cupriavidus MCMTATSQPDDRSACRAGRMPRRRLFVRRLYACLQASFQAFQSASDVSAMPAACQRLLPY WP_082926441.1 hypothetical protein [Cupriavidus sp. D384] Length: 1379\nNumber of predicted TMHs: 2\nExp number of AAs in TMHs: 42.9201199999999\nExp number, first 60 AAs: 2.18437\nTotal prob of N-in: 0.96488\ninside 1 59\nTMhelix 60 82\noutside 83 1351\nTMhelix 1352 1374\ninside 1375 1379

1448 GCF_000812465.1_ASM81246v1 Cupriavidus sp. IDO Proteobacteria; Betaproteobacteria; Burkholderiales; Burkholderiaceae; Cupriavidus MRLNPRRSLLLCLALSGTPALGAGPPCGTPATPIAEIQGSTPTSPLAGRKVEIEAVVTAD WP_039006796.1 endonuclease [Cupriavidus sp. IDO] Length: 655\nNumber of predicted TMHs: 1\nExp number of AAs in TMHs: 17.65815\nExp number, first 60 AAs: 0.23232\nTotal prob of N-in: 0.01172\noutside 1 625\nTMhelix 626 648\ninside 649 655

1449 GCF_001854325.1_ASM185432v1 Cupriavidus sp. USMAA1020 Proteobacteria; Betaproteobacteria; Burkholderiales; Burkholderiaceae; Cupriavidus MVNSRPSDAGLLSVNANQLGAILFARRRVIVGVALATLLATALLLAVRPRLWTASSDVYV WP_084545496.1 hypothetical protein [Cupriavidus sp. USMAA1020] Length: 468\nNumber of predicted TMHs: 2\nExp number of AAs in TMHs: 49.45046\nExp number, first 60 AAs: 19.72683\nTotal prob of N-in: 0.56528\nPOSSIBLE N-term signal sequence\ninside 1 28\nTMhelix 29 47\noutside 48 415\nTMhelix 416 438\ninside 439 468

1450 GCF_001854305.1_ASM185430v1 Cupriavidus sp. USMAA2-4 Proteobacteria; Betaproteobacteria; Burkholderiales; Burkholderiaceae; Cupriavidus MPPSAGRYQRGCCVVNSRPSDAGLLSVNANQLGAILFARRRVIVGVALATLLATALLLAV WP_084084527.1 hypothetical protein [Cupriavidus sp. USMAA2-4]Length: 481\nNumber of predicted TMHs: 2\nExp number of AAs in TMHs: 44.83514\nExp number, first 60 AAs: 18.40858\nTotal prob of N-in: 0.77483\nPOSSIBLE N-term signal sequence\ninside 1 41\nTMhelix 42 60\noutside 61 428\nTMhelix 429 451\ninside 452 481

1451 GCF_001854285.1_ASM185428v1 Cupriavidus sp. USMAHM13 Proteobacteria; Betaproteobacteria; Burkholderiales; Burkholderiaceae; Cupriavidus MVNSRPSDAGLLSVNANQLGAILFARRRVIVGVALATLLATALLLAVRPRLWTASSDVYV WP_083383982.1 hypothetical protein [Cupriavidus sp. USMAHM13] Length: 468\nNumber of predicted TMHs: 2\nExp number of AAs in TMHs: 49.44813\nExp number, first 60 AAs: 19.72701\nTotal prob of N-in: 0.56513\nPOSSIBLE N-term signal sequence\ninside 1 28\nTMhelix 29 47\noutside 48 415\nTMhelix 416 438\ninside 439 468

1452 GCF_000395345.1_CupWS1.0 Cupriavidus sp. WS Proteobacteria; Betaproteobacteria; Burkholderiales; Burkholderiaceae; Cupriavidus MHAPCTAPPGALSRRGFLKVGLGFSAALACAGLLPALSGCAPAPTNPTAGMRFLRDGDVA WP_020206616.1 hypothetical protein [Cupriavidus sp. WS] Length: 193\nNumber of predicted TMHs: 2\nExp number of AAs in TMHs: 44.85854\nExp number, first 60 AAs: 21.95983\nTotal prob of N-in: 0.04444\nPOSSIBLE N-term signal sequence\noutside 1 19\nTMhelix 20 42\ninside 43 149\nTMhelix 150 172\noutside 173 193

1453 GCF_000243095.1_ASM24309v2 Cupriavidus basilensis OR16 Proteobacteria; Betaproteobacteria; Burkholderiales; Burkholderiaceae; Cupriavidus; Cupriavidus basilensis
MMESVVTEIFYRLSTAFFWPVALALLVLFVLSLADLGGLIVQSRRRAVAPRSDLPALANA WP_006158490.1 hypothetical protein [Cupriavidus basilensis] Length: 205\nNumber of predicted TMHs: 2\nExp number of AAs in TMHs: 45.6688600000001\nExp number, first 60 AAs: 22.60352\nTotal prob of N-in: 0.68463\nPOSSIBLE N-term signal sequence\ninside 1 8\nTMhelix 9 31\noutside 32 137\nTMhelix 138 160\ninside 161 205

1454 GCF_000243095.1_ASM24309v2 Cupriavidus basilensis OR16 Proteobacteria; Betaproteobacteria; Burkholderiales; Burkholderiaceae; Cupriavidus; Cupriavidus basilensis
MHRRSFLAAGTLATLALGGCATAALYDEIEKKEYGDYKEIISEILISQDGKSIVILGDKY WP_006159236.1 hypothetical protein [Cupriavidus basilensis] Length: 214\nNumber of predicted TMHs: 2\nExp number of AAs in TMHs: 39.92289\nExp number, first 60 AAs: 18.03034\nTotal prob of N-in: 0.82666\nPOSSIBLE N-term signal sequence\ninside 1 6\nTMhelix 7 26\noutside 27 189\nTMhelix 190 212\ninside 213 214

1455 GCF_001281465.1_ASM128146v1 Cupriavidus gilardii CR3 Proteobacteria; Betaproteobacteria; Burkholderiales; Burkholderiaceae; Cupriavidus; Cupriavidus gilardii
MTRIDSTVATPRRRGAMLTTAWLLAAMALPPAHSQPASPNSAVEGPSSQGSAASGVGAAP WP_053822122.1 cellulose synthase regulator BcsB [Cupriavidus gilardii] Length: 780\nNumber of predicted TMHs: 1\nExp number of AAs in TMHs: 23.90454\nExp number, first 60 AAs: 1.17857\nTotal prob of N-in: 0.06590\noutside 1 748\nTMhelix 749 771\ninside 772 780

1456 GCF_000196015.1_ASM19601v1 Cupriavidus metallidurans CH34 Proteobacteria; Betaproteobacteria; Burkholderiales; Burkholderiaceae; Cupriavidus; Cupriavidus metallidurans
MGQKEEFGQAVNWSPVAKSIVRRISANRRPFLLTTALASLALFALMSYLLRHYQSEGFLR WP_011517311.1 hypothetical protein [Cupriavidus metallidurans] Length: 398\nNumber of predicted TMHs: 2\nExp number of AAs in TMHs: 43.70294\nExp number, first 60 AAs: 20.7357\nTotal prob of N-in: 0.97486\nPOSSIBLE N-term signal sequence\ninside 1 30\nTMhelix 31 50\noutside 51 356\nTMhelix 357 379\ninside 380 398

1457 GCF_000744095.1_ASM74409v1 Cupriavidus necator A5-1 Proteobacteria; Betaproteobacteria; Burkholderiales; Burkholderiaceae; Cupriavidus; Cupriavidus necator
MPPDPSHAPSHAPHRSADAETLHALSRRGFLKVGFGFTAALACTALVPSLAGCSAGNPAP WP_042883642.1 hypothetical protein [Cupriavidus necator] Length: 206\nNumber of predicted TMHs: 2\nExp number of AAs in TMHs: 42.80237\nExp number, first 60 AAs: 21.2044\nTotal prob of N-in: 0.02563\nPOSSIBLE N-term signal sequence\noutside 1 32\nTMhelix 33 55\ninside 56 165\nTMhelix 166 188\noutside 189 206

1458 GCF_001598755.1_ASM159875v1 Cupriavidus necator NBRC 102504 Proteobacteria; Betaproteobacteria; Burkholderiales; Burkholderiaceae; Cupriavidus; Cupriavidus necator
MAPNHPHRRADADAEALHALTRRGFLKVGLGFSAALACTALVPSLAGCSSAHPAPQAGMT WP_011616288.1 hypothetical protein [Cupriavidus necator] Length: 201\nNumber of predicted TMHs: 1\nExp number of AAs in TMHs: 27.71099\nExp number, first 60 AAs: 6.30589\nTotal prob of N-in: 0.73802\ninside 1 160\nTMhelix 161 183\noutside 184 201

1459 GCF_000009285.1_ASM928v2 Ralstonia eutropha H16 Proteobacteria; Betaproteobacteria; Burkholderiales; Burkholderiaceae; Cupriavidus; Cupriavidus necator
MAPNHPHRRADADAEALHALTRRGFLKVGLGFSAALACTALVPSLAGCSSAHPAPQAGMT WP_011616288.1 hypothetical protein [Cupriavidus necator] Length: 201\nNumber of predicted TMHs: 1\nExp number of AAs in TMHs: 27.71099\nExp number, first 60 AAs: 6.30589\nTotal prob of N-in: 0.73802\ninside 1 160\nTMhelix 161 183\noutside 184 201

1460 GCF_001592245.1_ASM159224v1 Cupriavidus oxalaticus NBRC 13593 Proteobacteria; Betaproteobacteria; Burkholderiales; Burkholderiaceae; Cupriavidus; Cupriavidus oxalaticus
MTENVNLRRRALLQGGAGLALSSLVVHGAAQTWPARPIRLLTGYAPGGSSDITARAISAP WP_063239409.1 hypothetical protein [Cupriavidus oxalaticus] Length: 329\nNumber of predicted TMHs: 2\nExp number of AAs in TMHs: 41.5989700000001\nExp number, first 60 AAs: 2.44644\nTotal prob of N-in: 0.91682\ninside 1 187\nTMhelix 188 210\noutside 211 256\nTMhelix 257 279\ninside 280 329

1461 GCF_000203875.1_ASM20387v1 Ralstonia eutropha JMP134 Proteobacteria; Betaproteobacteria; Burkholderiales; Burkholderiaceae; Cupriavidus; Cupriavidus pinatubonensis
MNRKRPSAMPRSAYAQTPRRRLLSASLALMMLSAPTWPWHGVLAAPTSAATSTTAAAAPQ WP_011299652.1 cellulose synthase regulator BcsB [Cupriavidus pinatubonensis] Length: 781\nNumber of predicted TMHs:

1\nExp number of AAs in TMHs: 37.61556\nExp number, first 60 AAs: 14.91374\nTotal prob of N-in: 0.70287\nPOSSIBLE N-term signal sequence\noutside 1 746\nTMhelix 747 769\ninside 770 781

1462 GCF_000069785.1_ASM6978v1 Cupriavidus taiwanensis LMG 19424 Proteobacteria; Betaproteobacteria; Burkholderiales; Burkholderiaceae; Cupriavidus; Cupriavidus taiwanensis MRPNPHHRRAVAEADALHALTRRGFLKVGLGFSAALACTALVPSLAGCSSADPAPQAGMS WP_012354378.1 hypothetical protein [Cupriavidus taiwanensis] Length: 201\nNumber of predicted TMHs: 1\nExp number of AAs in TMHs: 24.31715\nExp number, first 60 AAs: 5.48385\nTotal prob of N-in: 0.85321\ninside 1 160\nTMhelix 161 183\noutside 184 201

1463 GCF_000472465.1_ASM47246v1 Cupriavidus taiwanensis STM 6018 Proteobacteria; Betaproteobacteria; Burkholderiales; Burkholderiaceae; Cupriavidus; Cupriavidus taiwanensis MRPNPHHRRAVADADALHALTRRGFLKVGLGFSAALACTALVPSLAGCSSADPAPQAGMS WP_025585582.1 hypothetical protein [Cupriavidus taiwanensis] Length: 201\nNumber of predicted TMHs: 1\nExp number of AAs in TMHs: 25.62847\nExp number, first 60 AAs: 6.11359\nTotal prob of N-in: 0.85716\ninside 1 160\nTMhelix 161 183\noutside 184 201

1464 GCF_000372525.1_ASM37252v1 Cupriavidus taiwanensis STM 6070 Proteobacteria; Betaproteobacteria; Burkholderiales; Burkholderiaceae; Cupriavidus; Cupriavidus taiwanensis MRPNPHHRRHAADADALHALTRRGFLKVGLGFSAALACSALVPSLAGCSFTDPAPQAGMS WP_018004969.1 hypothetical protein [Cupriavidus taiwanensis] Length: 201\nNumber of predicted TMHs: 1\nExp number of AAs in TMHs: 25.46165\nExp number, first 60 AAs: 8.10927\nTotal prob of N-in: 0.78778\ninside 1 157\nTMhelix 158 180\noutside 181 201

1465 GCF_001010785.1_ASM101078v1 Pandoraea apista Proteobacteria; Betaproteobacteria; Burkholderiales; Burkholderiaceae; Pandoraea MPALRRRSFMQARSHLIFGIGATWLMHRTGWAASVHAWPLTLVGALLPDIDHPKSMLGRR WP_082117624.1 hypothetical protein [Pandoraea apista] Length: 161\nNumber of predicted TMHs: 1\nExp number of AAs in TMHs: 63.3517\nExp number, first 60 AAs: 13.68286\nTotal prob of N-in: 0.84122\nPOSSIBLE N-term signal sequence\ninside 1 135\nTMhelix 136 158\noutside 159 161

1466 GCF_001027265.1_ASM102726v1 Pandoraea apista Proteobacteria; Betaproteobacteria; Burkholderiales; Burkholderiaceae; Pandoraea MPALRRRSFMQARSHLIFGIGATWLMHRTGWAASVHAWPLTLVGALLPDIDHPKSMLGRR WP_082117624.1 hypothetical protein [Pandoraea apista] Length: 161\nNumber of predicted TMHs: 1\nExp number of AAs in TMHs: 63.3517\nExp number, first 60 AAs: 13.68286\nTotal prob of N-in: 0.84122\nPOSSIBLE N-term signal sequence\ninside 1 135\nTMhelix 136 158\noutside 159 161

1467 GCF_001465595.2_ASM146559v2 Pandoraea apista Proteobacteria; Betaproteobacteria; Burkholderiales; Burkholderiaceae; Pandoraea MPALRRRSFMQARSHLIFGIGATWLMHRTGWAASVHAWPLTLVGALLPDIDHPKSMLGRR WP_082117624.1 hypothetical protein [Pandoraea apista] Length: 161\nNumber of predicted TMHs: 1\nExp number of AAs in TMHs: 63.3517\nExp number, first 60 AAs: 13.68286\nTotal prob of N-in: 0.84122\nPOSSIBLE N-term signal sequence\ninside 1 135\nTMhelix 136 158\noutside 159 161

1468 GCF_001049515.1_Pandoraea_assembly_1 Pandoraea apista Proteobacteria; Betaproteobacteria; Burkholderiales; Burkholderiaceae; Pandoraea MPALRRRSFMQARSHLIFGIGATWLMHRTGWAASVHAWPLTLVGALLPDIDHPKSMLGRR WP_082117624.1 hypothetical protein [Pandoraea apista] Length: 161\nNumber of predicted TMHs: 1\nExp number of AAs in TMHs: 63.3517\nExp number, first 60 AAs: 13.68286\nTotal prob of N-in: 0.84122\nPOSSIBLE N-term signal sequence\ninside 1 135\nTMhelix 136 158\noutside 159 161

1469 GCF_001699815.1_ASM169981v1 Ralstonia pickettii Proteobacteria; Betaproteobacteria; Burkholderiales; Burkholderiaceae; Ralstonia MKTQRRAFLHSVAAACLAAFCVSVQAEPDLLQPDEAFRVSARRVDARVVELEYIIAPGYH WP_009242072.1 MULTISPECIES: thiol:disulfide interchange protein [Burkholderiales] Length: 151\nNumber of predicted TMHs: 1\nExp number of AAs in TMHs: 19.90009\nExp number, first 60 AAs: 9.03290000000001\nTotal prob of N-in: 0.83681\ninside 1 115\nTMhelix 116 138\noutside 139 151

1470 GCF_001708525.1_ASM170852v1 Ralstonia solanacearum Proteobacteria; Betaproteobacteria; Burkholderiales; Burkholderiaceae; Ralstonia MKRRTLLTTGAFLACANLAGCATAKLFEDDKYQETVDRFLVSENGRKLVVLGKRYHYILD WP_023470008.1 hypothetical protein [Ralstonia solanacearum] Length: 209\nNumber of predicted TMHs: 1\nExp number of AAs in TMHs: 23.90156\nExp number, first 60 AAs: 2.19803\nTotal prob of N-in: 0.78605\ninside 1 172\nTMhelix 173 195\noutside 196 209

1471 GCF_000825845.1_Grenada91 Ralstonia solanacearum Proteobacteria; Betaproteobacteria; Burkholderiales; Burkholderiaceae; Ralstonia
MRSEVDTDTAQACRSRRTWLRGAGALAVGAGLGLPVAHAAASLEVGHPAPPLVLRTLDGH WP_013208365.1 alkyl hydroperoxide reductase [Ralstonia solanacearum] Length: 185\nNumber of predicted TMHs: 2\nExp number of AAs in TMHs: 36.74875\nExp number, first 60 AAs: 21.75705\nTotal prob of N-in: 0.98108\nPOSSIBLE N-term signal sequence\ninside 1 18\nTMhelix 19 41\noutside 42 122\nTMhelix 123 145\ninside 146 185

1472 GCF_001644855.1_ASM164485v1 Ralstonia solanacearum Proteobacteria; Betaproteobacteria; Burkholderiales; Burkholderiaceae; Ralstonia
MLDIALRNLIRNHRRSIVTAASIAIGSVAVTLFSAFAANVTNGLQTETVGRLGHLNILRK WP_064051878.1 hypothetical protein, partial [Ralstonia solanacearum] Length: 392\nNumber of predicted TMHs: 2\nExp number of AAs in TMHs: 40.42504\nExp number, first 60 AAs: 21.047\nTotal prob of N-in: 0.89564\nPOSSIBLE N-term signal sequence\ninside 1 15\nTMhelix 16 38\noutside 39 330\nTMhelix 331 353\ninside 354 392

1473 GCF_001373315.1_CFBP3858 Ralstonia solanacearum Proteobacteria; Betaproteobacteria; Burkholderiales; Burkholderiaceae; Ralstonia
MRSEVDTDTAQACRSRRTWLRGAGALAVGAGLGLPVAHAAASLEVGHPAPPLVLRTLDGH WP_013208365.1 alkyl hydroperoxide reductase [Ralstonia solanacearum] Length: 185\nNumber of predicted TMHs: 2\nExp number of AAs in TMHs: 36.74875\nExp number, first 60 AAs: 21.75705\nTotal prob of N-in: 0.98108\nPOSSIBLE N-term signal sequence\ninside 1 18\nTMhelix 19 41\noutside 42 122\nTMhelix 123 145\ninside 146 185

1474 GCF_001373275.1_IBSBF1900.fas.gz Ralstonia solanacearum Proteobacteria; Betaproteobacteria; Burkholderiales; Burkholderiaceae; Ralstonia
MRSEVDTDTAQACRSRRTWLRGAGALAVGAGLGLPVAHAAASLEVGHPAPPLVLRTLDGH WP_013208365.1 alkyl hydroperoxide reductase [Ralstonia solanacearum] Length: 185\nNumber of predicted TMHs: 2\nExp number of AAs in TMHs: 36.74875\nExp number, first 60 AAs: 21.75705\nTotal prob of N-in: 0.98108\nPOSSIBLE N-term signal sequence\ninside 1 18\nTMhelix 19 41\noutside 42 122\nTMhelix 123 145\ninside 146 185

1475 GCF_002162015.1_ASM216201v1 Ralstonia solanacearum Proteobacteria; Betaproteobacteria; Burkholderiales; Burkholderiaceae; Ralstonia
MPEIGNREDIIMKRRTLLTTGAFLASANLMGCTAMLFEDGRYKETVDRFLVSENGKKLVV WP_087451541.1 hypothetical protein [Ralstonia solanacearum] Length: 220\nNumber of predicted TMHs: 2\nExp number of AAs in TMHs: 41.17626\nExp number, first 60 AAs: 18.31127\nTotal prob of N-in: 0.32533\nPOSSIBLE N-term signal sequence\noutside 1 14\nTMhelix 15 37\ninside 38 193\nTMhelix 194 216\noutside 217 220

1476 GCF_000825785.1_B50 Ralstonia solanacearum Proteobacteria; Betaproteobacteria; Burkholderiales; Burkholderiaceae; Ralstonia MRSEVDTDTAQACRSRRTWLRGAGALAVGAGLGLPVAHAAASLEVGHPAPPLVLRTLDGH WP_080894115.1 alkyl hydroperoxide reductase, partial [Ralstonia solanacearum] Length: 179\nNumber of predicted TMHs: 2\nExp number of AAs in TMHs: 36.68618\nExp number, first 60 AAs: 21.75334\nTotal prob of N-in: 0.98113\nPOSSIBLE N-term signal sequence\ninside 1 18\nTMhelix 19 41\noutside 42 122\nTMhelix 123 145\ninside 146 179

1477 GCF_000227255.2_Ralstonia_sp_5_2_56FAA_V2 Ralstonia sp. 5_2_56FAA Proteobacteria; Betaproteobacteria; Burkholderiales; Burkholderiaceae; Ralstonia
MKTQRRAFLHSVAAACLAAFCVSVQAEPDLLQPDEAFRVSARRVDARVVELEYIIAPGYH WP_009242072.1 MULTISPECIES: thiol:disulfide interchange protein [Burkholderiales] Length: 151\nNumber of predicted TMHs: 1\nExp number of AAs in TMHs: 19.90009\nExp number, first 60 AAs: 9.03290000000001\nTotal prob of N-in: 0.83681\ninside 1 115\nTMhelix 116 138\noutside 139 151

1478 GCF_000165085.1_ASM16508v1 Ralstonia sp. 5_7_47FAA Proteobacteria; Betaproteobacteria; Burkholderiales; Burkholderiaceae; Ralstonia
MKTQRRAFLHSVAAACLAAFCVSVQAEPDLLQPDEAFRVSARRVDARVVELEYIIAPGYH WP_009242072.1 MULTISPECIES: thiol:disulfide interchange protein [Burkholderiales] Length: 151\nNumber of predicted TMHs: 1\nExp number of AAs in TMHs: 19.90009\nExp number, first 60 AAs: 9.03290000000001\nTotal prob of N-in: 0.83681\ninside 1 115\nTMhelix 116 138\noutside 139 151

1479 GCF_000389805.1_GA3-3_1.0 Ralstonia sp. GA3-3 Proteobacteria; Betaproteobacteria; Burkholderiales; Burkholderiaceae; Ralstonia MAPNHPHRRADADAEALHALTRRGFLKVGLGFSAALACTALVPSLAGCSSAHPAPQAGMT WP_010811364.1 hypothetical protein [Ralstonia sp. GA3-3] Length: 201\nNumber of predicted TMHs: 1\nExp number of AAs in TMHs: 27.71486\nExp number, first 60 AAs: 6.30745\nTotal prob of N-in: 0.73803\ninside 1 160\nTMhelix 161 183\noutside 184 201

1480 GCF_001887535.1_ASM188753v1 Ralstonia solanacearum FJAT-1458 Proteobacteria; Betaproteobacteria; Burkholderiales; Burkholderiaceae; Ralstonia; Ralstonia solanacearum
MKRRTLLTTGAFLACANLAGCATAKLFEDDKYQETVDRFLVSENGRKLVVLGKRYHYILD WP_023470008.1 hypothetical protein [Ralstonia solanacearum] Length: 209\nNumber of predicted TMHs: 1\nExp number of AAs in TMHs: 23.90156\nExp number, first 60 AAs: 2.19803\nTotal prob of N-in: 0.78605\ninside 1 172\nTMhelix 173 195\noutside 196 209

1481 GCF_000331895.1_FJAT-1458 Ralstonia solanacearum FJAT-1458 Proteobacteria; Betaproteobacteria; Burkholderiales; Burkholderiaceae; Ralstonia; Ralstonia solanacearum
MKRRTLLTTGAFLACANLAGCATAKLFEDDKYQETVDRFLVSENGRKLVVLGKRYHYILD WP_023470008.1 hypothetical protein [Ralstonia solanacearum] Length: 209\nNumber of predicted TMHs: 1\nExp number of AAs in TMHs: 23.90156\nExp number, first 60 AAs: 2.19803\nTotal prob of N-in: 0.78605\ninside 1 172\nTMhelix 173 195\noutside 196 209

1482 GCF_002155245.1_ASM215524v1 Ralstonia solanacearum FJAT-91 Proteobacteria; Betaproteobacteria; Burkholderiales; Burkholderiaceae; Ralstonia; Ralstonia solanacearum
MKRRTLLTTGAFLACANLAGCATAKLFEDDKYQETVDRFLVSENGRKLVVLGKRYHYILD WP_023470008.1 hypothetical protein [Ralstonia solanacearum] Length: 209\nNumber of predicted TMHs: 1\nExp number of AAs in TMHs: 23.90156\nExp number, first 60 AAs: 2.19803\nTotal prob of N-in: 0.78605\ninside 1 172\nTMhelix 173 195\noutside 196 209

1483 GCF_000331875.1_FJAT-91 Ralstonia solanacearum FJAT-91 Proteobacteria; Betaproteobacteria; Burkholderiales; Burkholderiaceae; Ralstonia; Ralstonia solanacearum
MKRRTLLTTGAFLACANLAGCATAKLFEDDKYQETVDRFLVSENGRKLVVLGKRYHYILD WP_023470008.1 hypothetical protein [Ralstonia solanacearum] Length: 209\nNumber of predicted TMHs: 1\nExp number of AAs in TMHs: 23.90156\nExp number, first 60 AAs: 2.19803\nTotal prob of N-in: 0.78605\ninside 1 172\nTMhelix 173 195\noutside 196 209

1484 GCF_000430925.2_RsSD54V2.0 Ralstonia solanacearum SD54Proteobacteria; Betaproteobacteria; Burkholderiales; Burkholderiaceae; Ralstonia; Ralstonia solanacearum
MKRRTLLTTGAFLACANLAGCATAKLFEDDKYQETVDRFLVSENGRKLVVLGKRYHYILD WP_023470008.1 hypothetical protein [Ralstonia solanacearum] Length: 209\nNumber of predicted TMHs: 1\nExp number of AAs in TMHs: 23.90156\nExp number, first 60 AAs: 2.19803\nTotal prob of N-in: 0.78605\ninside 1 172\nTMhelix 173 195\noutside 196 209

1485 GCF_000223115.1_ASM22311v2 Ralstonia solanacearum Y45 Proteobacteria; Betaproteobacteria; Burkholderiales; Burkholderiaceae; Ralstonia; Ralstonia solanacearum
MPEIGNREDIIMKRRTLLTTGAFLASANLMGCTAMLFEDGRYKETVDRFLVSENGKKLVV WP_019717437.1 hypothetical protein [Ralstonia solanacearum] Length: 220\nNumber of predicted TMHs: 2\nExp number of AAs in TMHs: 41.17583\nExp number, first 60 AAs: 18.31127\nTotal prob of N-in: 0.32532\nPOSSIBLE N-term signal sequence\noutside 1 14\nTMhelix 15 37\ninside 38 193\nTMhelix 194 216\noutside 217 220

1486 GCF_001428665.1_Root219 Acidovorax sp. Root219 Proteobacteria; Betaproteobacteria; Burkholderiales; Comamonadaceae; Acidovorax MGRTGGVQRMLPKPTPRPTTVSMPRRTALLAAAAATALAACGGGGSSSDEPPAEPVGQVI WP_082607805.1 hypothetical protein [Acidovorax sp. Root219] Length: 291\nNumber of predicted TMHs: 2\nExp number of AAs in TMHs: 43.68648\nExp number, first 60 AAs: 0.7908\nTotal prob of N-in: 0.96420\ninside 1 172\nTMhelix 173 195\noutside 196 229\nTMhelix 230 252\ninside 253 291

1487 GCF_000204645.1_ASM20464v1 Alicycliphilus denitrificans K601 Proteobacteria; Betaproteobacteria; Burkholderiales; Comamonadaceae; Alicycliphilus; Alicycliphilus denitrificans
MKTQRRAFLHSVAAACLAAFCVSVQAEPDLLQPDEAFRVSARRVDARVVELEYIIAPGYH WP_009242072.1 MULTISPECIES: thiol:disulfide interchange protein [Burkholderiales] Length: 151\nNumber of predicted TMHs: 1\nExp number of AAs in TMHs: 19.90009\nExp number, first 60 AAs: 9.03290000000001\nTotal prob of N-in: 0.83681\ninside 1 115\nTMhelix 116 138\noutside 139 151

1488 GCF_001044395.1_ASM104439v1 Caenimonas sp. SL110 Proteobacteria; Betaproteobacteria; Burkholderiales; Comamonadaceae; Caenimonas
MEESGFHVLLDGARVGPYDRRTIVGMRIKKTLTSGHVLVASNGAQLTVAELIGQPPAERF WP_048438794.1 hypothetical protein [Caenimonas sp. SL110] Length: 218\nNumber of predicted TMHs: 1\nExp number of AAs in TMHs: 32.24145\nExp number, first 60 AAs: 0.13224\nTotal prob of N-in: 0.47403\noutside 1 192\nTMhelix 193 215\ninside 216 218

1489 GCF_002025745.1_ASM202574v1 Hydrogenophaga sp. H7 Proteobacteria; Betaproteobacteria; Burkholderiales; Comamonadaceae; Hydrogenophaga
MTPRRALLWGAGALLCGLFLALGAWQVQRLAWKNDLVARVAARAHAEPAAAPGADTWPAL WP_079367090.1
Surfeit locus 1 family protein [Hydrogenophaga sp. H7] Length: 251\nNumber of predicted TMHs: 2\nExp number of AAs in TMHs: 44.1366\nExp number, first 60 AAs: 21.02305\nTotal prob of N-in: 0.88170\nPOSSIBLE N-term signal sequence\ninside 1 6\nTMhelix 7 26\noutside 27 212\nTMhelix 213 235\ninside 236 251

1490 GCF_000422885.1_ASM42288v1 Ottowia thiooxydans DSM 14619 Proteobacteria; Betaproteobacteria; Burkholderiales; Comamonadaceae; Ottowia; Ottowia thiooxydans
MNNASIQVSVVIPTHHRRDLLKRLLTSLQVQSLAPSQFEVLIVHNHTDDGTEIMAQQWCA WP_028604672.1
hypothetical protein [Ottowia thiooxydans] Length: 327\nNumber of predicted TMHs: 2\nExp number of AAs in TMHs: 41.31095\nExp number, first 60 AAs: 0.00163\nTotal prob of N-in: 0.20121\noutside 1 259\nTMhelix 260 282\ninside 283 293\nTMhelix 294 316\noutside 317 327

1491 GCF_001425705.1_Root1217 Pelomonas sp. Root1217 Proteobacteria; Betaproteobacteria; Burkholderiales; Comamonadaceae; Pelomonas MTTRRAVLSLSVAPLIGCGHPDWHREGYMESVNSLAITPAGDKLVVFGGTYHYVLNPPSA
WP_057298407.1 hypothetical protein [Pelomonas sp. Root1217] Length: 201\nNumber of predicted TMHs: 1\nExp number of AAs in TMHs: 21.22655\nExp number, first 60 AAs: 0.70739\nTotal prob of N-in: 0.33881\noutside 1 171\nTMhelix 172 194\ninside 195 201

1492 GCF_001425165.1_Root405 Pelomonas sp. Root405 Proteobacteria; Betaproteobacteria; Burkholderiales; Comamonadaceae; Pelomonas MLSRRGWLVLVAAVLSAALTARLGFWQLDRAAYKLELQSAIDAQAERPALGNAELDGAEL
WP_056263090.1 MULTISPECIES: transmembrane cytochrome oxidase [Pelomonas] Length: 235\nNumber of predicted TMHs: 2\nExp number of AAs in TMHs: 41.72521\nExp number, first 60 AAs: 19.81727\nTotal prob of N-in: 0.92566\nPOSSIBLE N-term signal sequence\ninside 1 6\nTMhelix 7 26\noutside 27 203\nTMhelix 204 226\ninside 227 235

1493 GCF_001427705.1_Root662 Pelomonas sp. Root662 Proteobacteria; Betaproteobacteria; Burkholderiales; Comamonadaceae; Pelomonas MLSRRGWLVLVAAVLSAALTARLGFWQLDRAAYKLELQSAIDAQAERPALGNAELDGAEL
WP_056263090.1 MULTISPECIES: transmembrane cytochrome oxidase [Pelomonas] Length: 235\nNumber of predicted TMHs: 2\nExp number of AAs in TMHs: 41.72521\nExp number, first 60 AAs: 19.81727\nTotal prob of N-in: 0.92566\nPOSSIBLE N-term signal sequence\ninside 1 6\nTMhelix 7 26\noutside 27 203\nTMhelix 204 226\ninside 227 235

1494 GCF_001580455.1_ASM158045v1 Ramlibacter tataouinensis Proteobacteria; Betaproteobacteria; Burkholderiales; Comamonadaceae; Ramlibacter
MPDSCTRRKALAMVAAGLAAPSAAADDAALSRELRVGAALLMRHTQTVPGVGDPPGWRLD WP_061501402.1
histidine phosphatase family protein [Ramlibacter tataouinensis] Length: 194\nNumber of predicted TMHs: 1\nExp number of AAs in TMHs: 17.64724\nExp number, first 60 AAs: 0.05642\nTotal prob of N-in: 0.82449\ninside 1 146\nTMhelix 147 169\noutside 170 194

1495 GCF_000215705.1_ASM21570v1 Ramlibacter tataouinensis TTB310 Proteobacteria; Betaproteobacteria; Burkholderiales; Comamonadaceae; Ramlibacter; Ramlibacter tataouinensis
MHPSVSASPSAETASRRQVVVITGASSGIGRAAALRFAQDGADLVLAARGTEALAYVAAE WP_013899373.1
short-chain dehydrogenase [Ramlibacter tataouinensis] Length: 334\nNumber of predicted TMHs: 1\nExp number of AAs in TMHs: 23.29243\nExp number, first 60 AAs: 0.10092\nTotal prob of N-in: 0.08318\noutside 1 305\nTMhelix 306 328\ninside 329 334

1496 GCF_000215705.1_ASM21570v1 Ramlibacter tataouinensis TTB310 Proteobacteria; Betaproteobacteria; Burkholderiales; Comamonadaceae; Ramlibacter; Ramlibacter tataouinensis
MSLTMTCAPSSARRRAMPSPMPPPAPVTMMDLFWTRMGVSWMLLEWMRVSATARSAQQQP WP_081466270.1
hypothetical protein [Ramlibacter tataouinensis] Length: 283\nNumber of predicted TMHs: 2\nExp number of AAs in TMHs: 44.64104\nExp number, first 60 AAs: 2.86822\nTotal prob of N-in: 0.52631\ninside 1 83\nTMhelix 84 106\noutside 107 216\nTMhelix 217 239\ninside 240 283

1497 GCF_002198735.1_ASM219873v1 Comamonadaceae bacterium A 288 Proteobacteria; Betaproteobacteria; Burkholderiales; Comamonadaceae; unclassified Comamonadaceae
MTEPLLLGDMLRERGVSRRAFLKYASYMSSLMALPPSAAAAMADGLAKARRQAVIWLSFQ WP_088282586.1
Ni/Fe hydrogenase [Comamonadaceae bacterium A 288] Length: 352\nNumber of predicted TMHs: 1\nExp number of AAs in TMHs: 21.74849\nExp number, first 60 AAs: 4.06492\nTotal prob of N-in: 0.17704\noutside 1 321\nTMhelix 322 341\ninside 342 352

1498 GCF_000745855.1_ASM74585v1 Xenophilus azovorans DSM 13620 Proteobacteria; Betaproteobacteria; Burkholderiales; Comamonadaceae; Xenophilus; Xenophilus azovorans MTVRAAPTGPAAASRGRRAALLAGVAVGIAGFVLLGNWQVQRLAWKRDLIARVDARIHAP WP_038217501.1 hypothetical protein [Xenophilus azovorans] Length: 254\nNumber of predicted TMHs: 2\nExp number of AAs in TMHs: 47.50944\nExp number, first 60 AAs: 20.58123\nTotal prob of N-in: 0.85674\nPOSSIBLE N-term signal sequence\ninside 1 20\nTMhelix 21 40\noutside 41 223\nTMhelix 224 246\ninside 247 254

1499 GCF_001189915.1_ASM118991v1 Herbaspirillum autotrophicum Proteobacteria; Betaproteobacteria; Burkholderiales; Oxalobacteraceae; Herbaspirillum MNFQTPESGLSSTQLAAMISARRGLIYKTLLATVALTVAVTLILPKTYTASSDVYLDYKG WP_050465065.1 hypothetical protein [Herbaspirillum autotrophicum] Length: 460\nNumber of predicted TMHs: 1\nExp number of AAs in TMHs: 35.09195\nExp number, first 60 AAs: 14.09006\nTotal prob of N-in: 0.66073\nPOSSIBLE N-term signal sequence\noutside 1 409\nTMhelix 410 432\ninside 433 460

1500 GCF_002025725.1_ASM202572v1 Herbaspirillum frisingense Proteobacteria; Betaproteobacteria; Burkholderiales; Oxalobacteraceae; Herbaspirillum MAPAQAASPAADALRKLADDTWIQRRETLSTLGFTTPLVLASNDSSREIYLPVPANVPLS WP_079217386.1 hypothetical protein [Herbaspirillum frisingense] Length: 724\nNumber of predicted TMHs: 1\nExp number of AAs in TMHs: 20.96493\nExp number, first 60 AAs: 0.00054\nTotal prob of N-in: 0.00061\noutside 1 681\nTMhelix 682 701\ninside 702 724

1501 GCF_001189965.1_ASM118996v1 Herbaspirillum rhizosphaeraeProteobacteria; Betaproteobacteria; Burkholderiales; Oxalobacteraceae; Herbaspirillum MNLQAPETGLSSKQLASMISARRSTIFKTLLATMAVTIVVTLLLPKTYTASSDVFLDYKG WP_050476765.1 hypothetical protein [Herbaspirillum rhizosphaerae] Length: 460\nNumber of predicted TMHs: 1\nExp number of AAs in TMHs: 34.71926\nExp number, first 60 AAs: 13.54724\nTotal prob of N-in: 0.63408\nPOSSIBLE N-term signal sequence\noutside 1 409\nTMhelix 410 432\ninside 433 460

1502 GCF_001189965.1_ASM118996v1 Herbaspirillum rhizosphaeraeProteobacteria; Betaproteobacteria; Burkholderiales; Oxalobacteraceae; Herbaspirillum MTHLPRRLSTHLSRRSAVIVLSALLYVAGAASVHAQEQSLEKPPAPPAAVAPTAPAAASA WP_083444747.1 cytochrome C [Herbaspirillum rhizosphaerae] Length: 508\nNumber of predicted TMHs: 1\nExp number of AAs in TMHs: 34.1564599999999\nExp number, first 60 AAs: 11.52651\nTotal prob of N-in: 0.54985\nPOSSIBLE N-term signal sequence\noutside 1 477\nTMhelix 478 500\ninside 501 508

1503 GCF_000282135.1_Herbaspirillum.strCF444_v1.0Herbaspirillum sp. CF444 Proteobacteria; Betaproteobacteria; Burkholderiales; Oxalobacteraceae; Herbaspirillum MNLQAPETGLSSKQLASMISARRSTIFKTLLATVAATIVVTLLLPKTYTASSDVYLDYKG WP_007882715.1 exopolysaccharide biosynthesis protein [Herbaspirillum sp. CF444] Length: 460\nNumber of predicted TMHs: 1\nExp number of AAs in TMHs: 31.68511\nExp number, first 60 AAs: 10.47441\nTotal prob of N-in: 0.49130\nPOSSIBLE N-term signal sequence\noutside 1 409\nTMhelix 410 432\ninside 433 460

1504 GCF_000577615.1_RV1423 Herbaspirillum sp. RV1423 Proteobacteria; Betaproteobacteria; Burkholderiales; Oxalobacteraceae; Herbaspirillum MNLQAPETGLSSRQLASMISARRGTIFKTLLATMAITIVVTLLLPKTYTASSDVFLDYKG WP_034291961.1 hypothetical protein [Herbaspirillum sp. RV1423] Length: 460\nNumber of predicted TMHs: 1\nExp number of AAs in TMHs: 37.67195\nExp number, first 60 AAs: 16.00063\nTotal prob of N-in: 0.73825\nPOSSIBLE N-term signal sequence\noutside 1 409\nTMhelix 410 432\ninside 433 460

1505 GCF_001267925.1_ASM126792v1 Herbaspirillum hiltneri N3 Proteobacteria; Betaproteobacteria; Burkholderiales; Oxalobacteraceae; Herbaspirillum; Herbaspirillum hiltneri MNLQAPETGLSSKQLASMISARRGTIFKTLMATMAVTIVVTLLLPKTYTASSDVFLDYKG WP_053198831.1 hypothetical protein [Herbaspirillum hiltneri] Length: 460\nNumber of predicted TMHs: 1\nExp number of AAs in TMHs: 33.64456\nExp number, first 60 AAs: 12.42944\nTotal prob of N-in: 0.57650\nPOSSIBLE N-term signal sequence\noutside 1 409\nTMhelix 410 432\ninside 433 460

1506 GCF_000256565.1_Asm2.0 Herbaspirillum lusitanum P6-12 Proteobacteria; Betaproteobacteria; Burkholderiales; Oxalobacteraceae; Herbaspirillum; Herbaspirillum lusitanum MNLQAPETGLSSKQLASMISARRSTIFKTLLATVAATIVVTLLLPKTYTASSDIYLDYKG WP_016834044.1 hypothetical protein [Herbaspirillum lusitanum] Length: 460\nNumber of predicted TMHs: 1\nExp number of AAs in TMHs: 32.92191\nExp number, first 60 AAs: 11.60922\nTotal prob of N-in: 0.54363\nPOSSIBLE N-term signal sequence\noutside 1 409\nTMhelix 410 432\ninside 433 460

1507 GCF_000300435.1_ASM30043v1 Herbaspirillum rubrisubalbicans Os34 Proteobacteria; Betaproteobacteria; Burkholderiales; Oxalobacteraceae; Herbaspirillum; Herbaspirillum rubrisubalbicans MQQDRRSVLKQLAAGSAALLTLPLQAAHGANPGNSLPATIRIGVAQLDRNWLLLVKSLSA WP_017453384.1 hypothetical protein [Herbaspirillum rubrisubalbicans] Length: 153\nNumber of predicted TMHs: 2\nExp number of AAs in TMHs: 47.52527\nExp number, first 60 AAs: 4.00532\nTotal prob of N-in: 0.90128\ninside 1 83\nTMhelix 84 106\noutside 107 115\nTMhelix 116 135\ninside 136 153

1508 GCF_000300415.1_ASM30041v1 Herbaspirillum rubrisubalbicans Os45 Proteobacteria; Betaproteobacteria; Burkholderiales; Oxalobacteraceae; Herbaspirillum; Herbaspirillum rubrisubalbicans MQQDRRSVLKQLAAGSAALLTLPLQAAHGANPGNSLPATIRIGVAQLDRNWLLLVKSLSA WP_017453384.1 hypothetical protein [Herbaspirillum rubrisubalbicans] Length: 153\nNumber of predicted TMHs: 2\nExp number of AAs in TMHs: 47.52527\nExp number, first 60 AAs: 4.00532\nTotal prob of N-in: 0.90128\ninside 1 83\nTMhelix 84 106\noutside 107 115\nTMhelix 116 135\ninside 136 153

1509 GCF_001445815.1_ASM144581v1 Janthinobacterium sp. Ant5-2-1 Proteobacteria; Betaproteobacteria; Burkholderiales; Oxalobacteraceae; Janthinobacterium MHNQNTQDNQDQQDDFEDDNATPFTDMLAIAWRSRRTIISVTIAAVVIGVGYTYAFPSYK WP_046686133.1 MULTISPECIES: LPS biosynthesis protein [Janthinobacterium] Length: 412\nNumber of predicted TMHs: 2\nExp number of AAs in TMHs: 43.08311\nExp number, first 60 AAs: 21.26952\nTotal prob of N-in: 0.99786\nPOSSIBLE N-term signal sequence\ninside 1 36\nTMhelix 37 59\noutside 60 380\nTMhelix 381 403\ninside 404 412

1510 GCF_000988085.1_ASM98808v1 Janthinobacterium sp. KBS0711 Proteobacteria; Betaproteobacteria; Burkholderiales; Oxalobacteraceae; Janthinobacterium MHNQNTQDNQDQQDDFEDDNATPFTDMLAIAWRSRRTIISVTIAAVVIGVGYTYAFPSYK WP_046686133.1 MULTISPECIES: LPS biosynthesis protein [Janthinobacterium] Length: 412\nNumber of predicted TMHs: 2\nExp number of AAs in TMHs: 43.08311\nExp number, first 60 AAs: 21.26952\nTotal prob of N-in: 0.99786\nPOSSIBLE N-term signal sequence\ninside 1 36\nTMhelix 37 59\noutside 60 380\nTMhelix 381 403\ninside 404 412

1511 GCF_001758645.1_ASM175864v1 Janthinobacterium sp. MP5059B Proteobacteria; Betaproteobacteria; Burkholderiales; Oxalobacteraceae; Janthinobacterium MHKEVKMIEGDEIDENILPVTDLLTTVWQGRRTLLAITLLAVAAGLAGRMLFSEYKSEGF WP_070281466.1 lipopolysaccharide biosynthesis protein [Janthinobacterium sp. MP5059B] Length: 407\nNumber of predicted TMHs: 2\nExp number of AAs in TMHs: 39.77553\nExp number, first 60 AAs: 18.01765\nTotal prob of N-in: 0.89939\nPOSSIBLE N-term signal sequence\ninside 1 33\nTMhelix 34 53\noutside 54 380\nTMhelix 381 400\ninside 401 407

1512 GCF_000745325.1_ASM74532v1 Janthinobacterium sp. RA13 Proteobacteria; Betaproteobacteria; Burkholderiales; Oxalobacteraceae; Janthinobacterium MHKEVKMIEGDEIDENILPVTDLLTTVWQGRRTLLAITLLAVAVGLAGRMLFSEYKSEGF WP_035821865.1 hypothetical protein [Janthinobacterium sp. RA13] Length: 407\nNumber of predicted TMHs: 2\nExp number of AAs in TMHs: 40.51585\nExp number, first 60 AAs: 18.75295\nTotal prob of N-in: 0.93526\nPOSSIBLE N-term signal sequence\ninside 1 33\nTMhelix 34 53\noutside 54 380\nTMhelix 381 400\ninside 401 407

1513 GCF_000723165.1_JAG1 Janthinobacterium agaricidamnosum NBRC 102515 = DSM 9628 Proteobacteria; Betaproteobacteria; Burkholderiales; Oxalobacteraceae; Janthinobacterium; Janthinobacterium agaricidamnosum MTSAFLQRRRLLQIASANCILLLFERPGFAQADTLDDASLAALISDFAKLRTLTYNEVRD WP_038490009.1 hypothetical protein [Janthinobacterium agaricidamnosum] Length: 192\nNumber of predicted TMHs: 1\nExp number of AAs in TMHs: 21.72251\nExp number, first 60 AAs: 0.04712\nTotal prob of N-in: 0.88989\ninside 1 140\nTMhelix 141 163\noutside 164 192

1514 GCF_000723165.1_JAG1 Janthinobacterium agaricidamnosum NBRC 102515 = DSM 9628 Proteobacteria; Betaproteobacteria; Burkholderiales; Oxalobacteraceae; Janthinobacterium; Janthinobacterium agaricidamnosum MSTSKTTDWTLIPGSKHEIEQVRERCRRMVRRRATMSAGVSAMPLPGIDVVSDVGLFTVL WP_038497525.1 hypothetical protein [Janthinobacterium agaricidamnosum] Length: 167\nNumber of predicted TMHs: 1\nExp number of AAs in TMHs: 38.45869\nExp number, first 60 AAs: 12.02463\nTotal prob of N-in: 0.86715\nPOSSIBLE N-term signal sequence\ninside 1 125\nTMhelix 126 148\noutside 149 167

1515 GCF_001571185.1_ASM157118v1 Janthinobacterium agaricidamnosum NBRC 102515 = DSM 9628 Proteobacteria; Betaproteobacteria; Burkholderiales; Oxalobacteraceae; Janthinobacterium; Janthinobacterium agaricidamnosum MTSAFLQRRRLLQIASANCILLLFERPGFAQADTLDDASLAALISDFAKLRTLTYNEVRD WP_038490009.1 hypothetical protein [Janthinobacterium agaricidamnosum] Length: 192\nNumber of predicted TMHs:

1\nExp number of AAs in TMHs: 21.72251\nExp number, first 60 AAs: 0.04712\nTotal prob of N-in: 0.88989\ninside 1 140\nTMhelix 141 163\noutside 164 192

1516 GCF_001571185.1_ASM157118v1 Janthinobacterium agaricidamnosum NBRC 102515 = DSM 9628 Proteobacteria; Betaproteobacteria; Burkholderiales; Oxalobacteraceae; Janthinobacterium; Janthinobacterium agaricidamnosum MSTSKTTDWTLIPGSKHEIEQVRERCRRMVRRRATMSAGVSAMPLPGIDVVSDVGLFTVL WP_038497525.1 hypothetical protein [Janthinobacterium agaricidamnosum] Length: 167\nNumber of predicted TMHs: 1\nExp number of AAs in TMHs: 38.45869\nExp number, first 60 AAs: 12.02463\nTotal prob of N-in: 0.86715\nPOSSIBLE N-term signal sequence\ninside 1 125\nTMhelix 126 148\noutside 149 167

1517 GCF_000759615.1_ASM75961v1 Massilia sp. JS1662 Proteobacteria; Betaproteobacteria; Burkholderiales; Oxalobacteraceae; Massilia MIAIVAVVIGLLAALAYAARRRWIDAVLALVAGVALAGTVGEFALPGEAGATLTVDAADP WP_036240134.1 hypothetical protein [Massilia sp. JS1662] Length: 504\nNumber of predicted TMHs: 2\nExp number of AAs in TMHs: 46.0662899999999\nExp number, first 60 AAs: 26.84887\nTotal prob of N-in: 0.48974\nPOSSIBLE N-term signal sequence\ninside 1 1\nTMhelix 2 19\noutside 20 481\nTMhelix 482 500\ninside 501 504

1518 GCF_001424165.1_Leaf139 Massilia sp. Leaf139 Proteobacteria; Betaproteobacteria; Burkholderiales; Oxalobacteraceae; Massilia MNLRQFLSLLRARRRLILSTLFATVLLALGWSLVQSKTYTATASVLLNYKGVDPLTGVTT WP_056339178.1 chain length determinant protein EpsF [Massilia sp. Leaf139] Length: 469\nNumber of predicted TMHs: 2\nExp number of AAs in TMHs: 40.03207\nExp number, first 60 AAs: 18.10426\nTotal prob of N-in: 0.77367\nPOSSIBLE N-term signal sequence\ninside 1 15\nTMhelix 16 35\noutside 36 397\nTMhelix 398 420\ninside 421 469

1519 GCF_001182045.1_Dakarella_massiliensis Dakarella massiliensis Proteobacteria; Betaproteobacteria; Burkholderiales; Sutterellaceae; Dakarella MAETQYEALRRQGVSRRSFLQFCSLTAASLGLGSAGAKDIATAMQTKPRVPVVWLHGLEC WP_049686423.1 hydrogenase [Dakarella massiliensis] Length: 367\nNumber of predicted TMHs: 1\nExp number of AAs in TMHs: 23.82984\nExp number, first 60 AAs: 2.2065\nTotal prob of N-in: 0.23084\noutside 1 327\nTMhelix 328 350\ninside 351 367

1520 GCF_900128485.1_PRJEB18048 Duodenibacillus massiliensis Proteobacteria; Betaproteobacteria; Burkholderiales; Sutterellaceae; Duodenibacillus MQNDDKKEVPMVTSVPEANRMPETVGSSAAGEPKLYKLKNWWRRRSVFEKEFCSFVALPT WP_084765547.1 hypothetical protein [Duodenibacillus massiliensis] Length: 407\nNumber of predicted TMHs: 2\nExp number of AAs in TMHs: 44.53419\nExp number, first 60 AAs: 8.38143\nTotal prob of N-in: 0.97169\ninside 1 51\nTMhelix 52 74\noutside 75 376\nTMhelix 377 399\ninside 400 407

1521 GCF_900128485.1_PRJEB18048 Duodenibacillus massiliensis Proteobacteria; Betaproteobacteria; Burkholderiales; Sutterellaceae; Duodenibacillus MMETQYEALRRQGISRRSFLQFCSLTAASLGLGSAGAKDIAEAMQTKPRTPVVWLHGLEC WP_022373966.1 hydrogenase-2 small chain [Duodenibacillus massiliensis] Length: 364\nNumber of predicted TMHs: 1\nExp number of AAs in TMHs: 20.35935\nExp number, first 60 AAs: 1.86988\nTotal prob of N-in: 0.17261\noutside 1 326\nTMhelix 327 349\ninside 350 364

1522 GCF_000205025.1_ASM20502v1 Parasutterella excrementihominis YIT 11859 Proteobacteria; Betaproteobacteria; Burkholderiales; Sutterellaceae; Parasutterella; Parasutterella excrementihominis MTETNYQLMRRQGVSRRSFLKFCSLTAASLGLGANGAADIAHAMETKPRTPVIWLHGLEC WP_008810871.1 nickel-iron hydrogenase small subunit [Parasutterella excrementihominis] Length: 366\nNumber of predicted TMHs: 1\nExp number of AAs in TMHs: 24.28046\nExp number, first 60 AAs: 6.6755\nTotal prob of N-in: 0.40992\noutside 1 328\nTMhelix 329 351\ninside 352 366

1523 GCF_000186505.1_Sutt_wads_3_1_45B_V1 Sutterella wadsworthensis 3_1_45B Proteobacteria; Betaproteobacteria; Burkholderiales; Sutterellaceae; Sutterella; Sutterella wadsworthensis MFETQYEALRRRGISRRSFLQFCSLTAASLGLGSAGAQEIAHAMETKPRTPVIWLHGLEC WP_005428750.1 ferredoxin hydrogenase [Sutterella wadsworthensis] Length: 364\nNumber of predicted TMHs: 1\nExp number of AAs in TMHs: 17.76306\nExp number, first 60 AAs: 4.02757\nTotal prob of N-in: 0.20351\noutside 1 327\nTMhelix 328 350\ninside 351 364

1524 GCF_000411515.1_Sutt_wads_HGA0223_V1 Sutterella wadsworthensis HGA0223 Proteobacteria; Betaproteobacteria; Burkholderiales; Sutterellaceae; Sutterella; Sutterella wadsworthensis

MFETQYEALRRRGISRRSFLQFCSLTAASLGLGSAGAQEIAHAMETKPRTPVIWLHGLEC WP_005428750.1 ferredoxin hydrogenase [Sutterella wadsworthensis] Length: 364\nNumber of predicted TMHs: 1\nExp number of AAs in TMHs: 17.76306\nExp number, first 60 AAs: 4.02757\nTotal prob of N-in: 0.20351\noutside 1 327\nTMhelix 328 350\ninside 351 364

1525 GCF_000799605.1_ASM79960v1 Methylibium sp. CF468 Proteobacteria; Betaproteobacteria; Burkholderiales; unclassified Burkholderiales; Burkholderiales Genera incertae sedis; Methylibium MRPRRAALTAAAALLFAGFAALGTWQVQRLSWKLDLIERVGQRVRAAPVEAPAGPVTAAA WP_082004703.1 Surfeit locus 1 family protein [Methylibium sp. CF468] Length: 245\nNumber of predicted TMHs: 2\nExp number of AAs in TMHs: 39.7317200000001\nExp number, first 60 AAs: 20.20864\nTotal prob of N-in: 0.87112\nPOSSIBLE N-term signal sequence\ninside 1 6\nTMhelix 7 26\noutside 27 195\nTMhelix 196 218\ninside 219 245

1526 GCF_000799605.1_ASM79960v1 Methylibium sp. CF468 Proteobacteria; Betaproteobacteria; Burkholderiales; unclassified Burkholderiales; Burkholderiales Genera incertae sedis; Methylibium MNPARRQLLTASSCLALAAPARALMAGTAPDSPDARIDANTPSSPWTSAVAVLTPSGIFS WP_047509482.1 hypothetical protein [Methylibium sp. CF468] Length: 322\nNumber of predicted TMHs: 1\nExp number of AAs in TMHs: 40.09819\nExp number, first 60 AAs: 8.64309\nTotal prob of N-in: 0.39403\noutside 1 290\nTMhelix 291 313\ninside 314 322

1527 GCF_000800095.1_ASM80009v1 Methylibium sp. YR605 Proteobacteria; Betaproteobacteria; Burkholderiales; unclassified Burkholderiales; Burkholderiales Genera incertae sedis; Methylibium MGDVPHEQQHDAHDARHASAAVRPRRAALTAAAALLFAGFAALGTWQVQRLSWKLDLIER WP_082011022.1 hypothetical protein [Methylibium sp. YR605] Length: 266\nNumber of predicted TMHs: 2\nExp number of AAs in TMHs: 40.7224599999999\nExp number, first 60 AAs: 20.19117\nTotal prob of N-in: 0.42212\nPOSSIBLE N-term signal sequence\noutside 1 26\nTMhelix 27 46\ninside 47 218\nTMhelix 219 241\noutside 242 266

1528 GCF_001653795.1_ASM165379v1 Mitsuaria sp. 7 Proteobacteria; Betaproteobacteria; Burkholderiales; unclassified Burkholderiales; Burkholderiales Genera incertae sedis; Mitsuaria MRIATRVYQAFAPRRSWITAPGLLAIALVASGSAATAATYSAESRSEARTLNLTAPVDAK WP_067273155.1 PEP-CTERM sorting domain-containing protein [Mitsuaria sp. 7] Length: 347\nNumber of predicted TMHs: 2\nExp number of AAs in TMHs: 35.2958\nExp number, first 60 AAs: 20.78531\nTotal prob of N-in: 0.93663\nPOSSIBLE N-term signal sequence\ninside 1 19\nTMhelix 20 42\noutside 43 317\nTMhelix 318 335\ninside 336 347

1529 GCF_001598255.1_ASM159825v1 Mitsuaria chitosanitabida NBRC 102408 Proteobacteria; Betaproteobacteria; Burkholderiales; unclassified Burkholderiales; Burkholderiales Genera incertae sedis; Mitsuaria; Mitsuaria chitosanitabida MAETTTLPAAPPRRGRRGVLLIAAVAGIAATVALGLWQLDRADQKKALQRAIDERALLSP WP_084452563.1 hypothetical protein [Mitsuaria chitosanitabida] Length: 285\nNumber of predicted TMHs: 2\nExp number of AAs in TMHs: 42.76596\nExp number, first 60 AAs: 21.24296\nTotal prob of N-in: 0.24468\nPOSSIBLE N-term signal sequence\noutside 1 14\nTMhelix 15 37\ninside 38 250\nTMhelix 251 273\noutside 274 285

1530 GCF_001477625.1_ASM147762v1 Paucibacter sp. KCTC 42545 Proteobacteria; Betaproteobacteria; Burkholderiales; unclassified Burkholderiales; Burkholderiales Genera incertae sedis; Paucibacter MTQLLLPRRQFLRLAGATPLALSGLATLGGCAESSSLKAPIKPYTETVKALLISRDGKHI WP_058720172.1 hypothetical protein [Paucibacter sp. KCTC 42545] Length: 220\nNumber of predicted TMHs: 1\nExp number of AAs in TMHs: 22.46103\nExp number, first 60 AAs: 2.27056\nTotal prob of N-in: 0.31512\noutside 1 194\nTMhelix 195 214\ninside 215 220

1531 GCF_001425785.1_Root1221 Rhizobacter sp. Root1221 Proteobacteria; Betaproteobacteria; Burkholderiales; unclassified Burkholderiales; Burkholderiales Genera incertae sedis; Rhizobacter MTDKNRNLVGQCVVLTGATSGIGRATAVQFARAGADLVLAARRETVLDEVAAQCQALGAQ WP_056669290.1 short-chain dehydrogenase [Rhizobacter sp. Root1221] Length: 328\nNumber of predicted TMHs: 1\nExp number of AAs in TMHs: 35.83491\nExp number, first 60 AAs: 1.63653\nTotal prob of N-in: 0.68874\noutside 1 302\nTMhelix 303 322\ninside 323 328

1532 GCF_001425825.1_Root1238 Rhizobacter sp. Root1238 Proteobacteria; Betaproteobacteria; Burkholderiales; unclassified Burkholderiales; Burkholderiales Genera incertae sedis; Rhizobacter MGDVPHEQQHDADDARHAGPAVRPRRAALIAAAALLFAGFAALGTWQVQRLSWKLDLIER WP_082541695.1 MULTISPECIES: hypothetical protein [Rhizobacter] Length: 270\nNumber of predicted TMHs: 2\nExp number of AAs in TMHs: 40.8798\nExp number, first 60 AAs: 20.22602\nTotal prob of N-in: 0.13947\nPOSSIBLE N-term signal sequence\noutside 1 26\nTMhelix 27 46\ninside 47 220\nTMhelix 221 243\noutside 244 270

1533 GCF_001428145.1_Root16D2 Rhizobacter sp. Root16D2 Proteobacteria; Betaproteobacteria; Burkholderiales; unclassified Burkholderiales; Burkholderiales Genera incertae sedis; Rhizobacter
MGDVPHEQQHDADDARHAGPAVRPRRAALIAAAALLFAGFAALGTWQVQRLSWKLDLIER WP_082541695.1
MULTISPECIES: hypothetical protein [Rhizobacter] Length: 270\nNumber of predicted TMHs: 2\nExp number of AAs in TMHs: 40.8798\nExp number, first 60 AAs: 20.22602\nTotal prob of N-in: 0.13947\nPOSSIBLE N-term signal sequence\noutside 1 26\nTMhelix 27 46\ninside 47 220\nTMhelix 221 243\noutside 244 270

1534 GCF_001424785.1_Root29 Rhizobacter sp. Root29 Proteobacteria; Betaproteobacteria; Burkholderiales; unclassified Burkholderiales; Burkholderiales Genera incertae sedis; Rhizobacter
MGDVPHEQQHDADDARHAGPAVRPRRAALIAAAALLFAGFAALGTWQVQRLSWKLDLIER WP_082541695.1
MULTISPECIES: hypothetical protein [Rhizobacter] Length: 270\nNumber of predicted TMHs: 2\nExp number of AAs in TMHs: 40.8798\nExp number, first 60 AAs: 20.22602\nTotal prob of N-in: 0.13947\nPOSSIBLE N-term signal sequence\noutside 1 26\nTMhelix 27 46\ninside 47 220\nTMhelix 221 243\noutside 244 270

1535 GCF_002205645.1_ASM220564v1 Roseateles aquatilis Proteobacteria; Betaproteobacteria; Burkholderiales; unclassified Burkholderiales; Burkholderiales Genera incertae sedis; Roseateles
MVEAGKTTASGRAATTGRRRGLLLVATVAGVAMTVSLALWQLSRADQKRALQRAIDERAL WP_088386249.1
transmembrane cytochrome oxidase [Roseateles aquatilis] Length: 288\nNumber of predicted TMHs: 2\nExp number of AAs in TMHs: 43.8763899999999\nExp number, first 60 AAs: 22.2053\nTotal prob of N-in: 0.99367\nPOSSIBLE N-term signal sequence\ninside 1 20\nTMhelix 21 43\noutside 44 253\nTMhelix 254 276\ninside 277 288

1536 GCF_001483865.1_ASM148386v1 Roseateles depolymerans Proteobacteria; Betaproteobacteria; Burkholderiales; unclassified Burkholderiales; Burkholderiales Genera incertae sedis; Roseateles
MNALLLIARGKRRGPFRMGWASALLVLALAGCNSSSELYAGLSERDANDITAALSDHGID WP_058936276.1
EscJ/YscJ/HrcJ family type III secretion inner membrane ring protein [Roseateles depolymerans] Length: 331\nNumber of predicted TMHs: 1\nExp number of AAs in TMHs: 26.72684\nExp number, first 60 AAs: 3.84743\nTotal prob of N-in: 0.16976\noutside 1 285\nTMhelix 286 308\ninside 309 331

1537 GCF_002205615.1_ASM220561v1 Roseateles terrae Proteobacteria; Betaproteobacteria; Burkholderiales; unclassified Burkholderiales; Burkholderiales Genera incertae sedis; Roseateles
MNALTPQRRSLLRGSRWLSTTALVLVLAGCNSSSELYGGLSERDANDITAALSDHGIQAS WP_088450548.1 EscJ/YscJ/HrcJ family type III secretion inner membrane ring protein [Roseateles terrae] Length: 338\nNumber of predicted TMHs: 1\nExp number of AAs in TMHs: 38.78414\nExp number, first 60 AAs: 0.45821\nTotal prob of N-in: 0.70766\noutside 1 279\nTMhelix 280 302\ninside 303 338

1538 GCF_000257145.1_ASM25714v1 Rubrivivax gelatinosus CBS Proteobacteria; Betaproteobacteria; Burkholderiales; unclassified Burkholderiales; Burkholderiales Genera incertae sedis; Rubrivivax; Rubrivivax gelatinosus
MTPLVGLAARSAWARRGTLVLVALSIALSTALLLTLERLREDVRASFSQAVSGTDLVVGA WP_043818688.1 ABC transporter permease, partial [Rubrivivax gelatinosus] Length: 338\nNumber of predicted TMHs: 2\nExp number of AAs in TMHs: 43.3180299999999\nExp number, first 60 AAs: 20.237\nTotal prob of N-in: 0.86789\nPOSSIBLE N-term signal sequence\ninside 1 11\nTMhelix 12 34\noutside 35 296\nTMhelix 297 319\ninside 320 338

1539 GCF_000284255.1_ASM28425v1 Rubrivivax gelatinosus IL144 Proteobacteria; Betaproteobacteria; Burkholderiales; unclassified Burkholderiales; Burkholderiales Genera incertae sedis; Rubrivivax; Rubrivivax gelatinosus
MNQDLRPEEHSSQAPGSDTETALTPADILAALRSRRKLLTLAPICTGIVVLGASFMIKPT WP_014430309.1 capsule polysaccharide transporter [Rubrivivax gelatinosus] Length: 377\nNumber of predicted TMHs: 2\nExp number of AAs in TMHs: 38.98661\nExp number, first 60 AAs: 19.54209\nTotal prob of N-in: 0.92166\nPOSSIBLE N-term signal sequence\ninside 1 37\nTMhelix 38 57\noutside 58 348\nTMhelix 349 366\ninside 367 377

1540 GCF_000144975.1_ASM14497v1 Burkholderiales bacterium 1_1_47 Proteobacteria; Betaproteobacteria; Burkholderiales; unclassified Burkholderiales; unclassified Burkholderiales (miscellaneous)
MTETNYQLMRRQGVSRRSFLKFCSLTAASLGLGANGAADIAHAMETKPRTPVIWLHGLEC WP_008810871.1
nickel-iron hydrogenase small subunit [Parasutterella excrementihominis] Length: 366\nNumber of predicted TMHs: 1\nExp number of AAs in TMHs: 24.28046\nExp number, first 60 AAs: 6.6755\nTotal prob of N-in: 0.40992\noutside 1 328\nTMhelix 329 351\ninside 352 366

1541 GCF_000244995.1_ASM24499v1 Burkholderiales bacterium JOSHI_001 Proteobacteria; Betaproteobacteria; Burkholderiales; unclassified Burkholderiales; unclassified Burkholderiales (miscellaneous)
MNPRGVAYTEFEADALRPCFSVTKPRREPMTMKRNLLASATVAVLALPMAAQAHLTFFQG WP_083839884.1
hypothetical protein [Burkholderiales bacterium JOSHI_001] Length: 225\nNumber of predicted TMHs:

2\nExp number of AAs in TMHs: 43.47704\nExp number, first 60 AAs: 21.65756\nTotal prob of N-in: 0.98449\nPOSSIBLE N-term signal sequence\ninside 1 35\nTMhelix 36 58\noutside 59 193\nTMhelix 194 216\ninside 217 225

1542 GCF_000244995.1_ASM24499v1 Burkholderiales bacterium JOSHI_001 Proteobacteria; Betaproteobacteria; Burkholderiales; unclassified Burkholderiales; unclassified Burkholderiales (miscellaneous) MSEPLLLGELLRQRGVSRRSLLKFAATMASALALPPHAAAAMATGLARARRQAVIWLSFQ WP_009551545.1 Ni/Fe hydrogenase [Burkholderiales bacterium JOSHI_001] Length: 368\nNumber of predicted TMHs: 1\nExp number of AAs in TMHs: 19.61402\nExp number, first 60 AAs: 4.66148\nTotal prob of N-in: 0.15508\noutside 1 321\nTMhelix 322 341\ninside 342 368

1543 GCF_001688905.2_ASM168890v2 Burkholderiales bacterium YL45 Proteobacteria; Betaproteobacteria; Burkholderiales; unclassified Burkholderiales; unclassified Burkholderiales (miscellaneous) MTETNYEMMRRQGVSRRSFLKFCSLTAASLGLGASGAADIAHAMETKPRTPVIWLHGLEC WP_066595802.1 hydrogenase [Burkholderiales bacterium YL45] Length: 366\nNumber of predicted TMHs: 1\nExp number of AAs in TMHs: 32.77943\nExp number, first 60 AAs: 11.9387\nTotal prob of N-in: 0.63508\nPOSSIBLE N-term signal sequence\noutside 1 328\nTMhelix 329 351\ninside 352 366

1544 GCF_000735045.1_Fv_myxofaciens_P3G_1.0 Ferrovum myxofaciens Proteobacteria; Betaproteobacteria; Ferrovales; Ferrovaceae; Ferrovum MKWFELTVGLRYTRAKRRNHFISFISLISMAGIALGVAALIVVLSVMNGFQKEMRERILG WP_035417212.1 lipoprotein-releasing system transmembrane subunit LolC, partial [Ferrovum myxofaciens] Length: 310\nNumber of predicted TMHs: 2\nExp number of AAs in TMHs: 47.37872\nExp number, first 60 AAs: 23.51011\nTotal prob of N-in: 0.95897\nPOSSIBLE N-term signal sequence\ninside 1 20\nTMhelix 21 43\noutside 44 272\nTMhelix 273 295\ninside 296 310

1545 GCF_001808185.1_ASM180818v1 Jeongeupia sp. USM3 Proteobacteria; Betaproteobacteria; Neisseriales; Chromobacteriaceae; Jeongeupia MARPSTSRMTSNNTMATSFMNAPAPVTLQPDVSTRRPALAGLAALVLLTLVLGAWQCRRG WP_083300840.1 hypothetical protein [Jeongeupia sp. USM3] Length: 242\nNumber of predicted TMHs: 2\nExp number of AAs in TMHs: 37.35233\nExp number, first 60 AAs: 17.7394\nTotal prob of N-in: 0.86211\nPOSSIBLE N-term signal sequence\ninside 1 37\nTMhelix 38 55\noutside 56 215\nTMhelix 216 235\ninside 236 242

1546 GCF_000226875.1_ASM22687v1 Neisseria shayeganii 871 Proteobacteria; Betaproteobacteria; Neisseriales; Neisseriaceae; Neisseria; Neisseria shayeganii MKLPFGWKLPKREEIFASRWAKPFAPLFDHPCFWTLNRRRAALSVAIGLFGGLMPGPTQM WP_009117763.1 hypothetical protein [Neisseria shayeganii] Length: 176\nNumber of predicted TMHs: 2\nExp number of AAs in TMHs: 63.75032\nExp number, first 60 AAs: 14.29957\nTotal prob of N-in: 0.21576\nPOSSIBLE N-term signal sequence\ninside 1 46\nTMhelix 47 69\noutside 70 138\nTMhelix 139 161\ninside 162 176

1547 GCF_000025705.1_ASM2570v1 Sideroxydans lithotrophicus ES-1 Proteobacteria; Betaproteobacteria; Nitrosomonadales; Gallionellaceae; Sideroxydans; Sideroxydans lithotrophicus MPGTVGEGLAQRGISRREFLQFCTSLAALLALPPSMASVMAEAISKARRQSVIWLSFQEC WP_041421154.1 Ni/Fe hydrogenase [Sideroxydans lithotrophicus] Length: 362\nNumber of predicted TMHs: 1\nExp number of AAs in TMHs: 27.51166\nExp number, first 60 AAs: 3.52622\nTotal prob of N-in: 0.12727\noutside 1 314\nTMhelix 315 337\ninside 338 362

1548 GCF_000971475.1_ASM97147v1 Sulfuricella sp. T08 Proteobacteria; Betaproteobacteria; Nitrosomonadales; Gallionellaceae; Sulfuricella MKRRDFLKGALAGGAMLASGSPAQARPNKTMPPEAVGLLYDSTLCIGCKACVAACKESNG WP_059418418.1 hydrogenase 2 protein HybA [Sulfuricella sp. T08] Length: 321\nNumber of predicted TMHs: 1\nExp number of AAs in TMHs: 19.23718\nExp number, first 60 AAs: 0.04698\nTotal prob of N-in: 0.38987\ninside 1 288\nTMhelix 289 308\noutside 309 321

1549 GCF_000297055.2_ASM29705v2 Sulfuricella denitrificans skB26 Proteobacteria; Betaproteobacteria; Nitrosomonadales; Gallionellaceae; Sulfuricella; Sulfuricella denitrificans MKRREFLKGALAGGALLATSSPAEARPNKTMPPEAVGLLYDSTLCIGCKACVAACKEAND WP_009206450.1 hydrogenase [Sulfuricella denitrificans] Length: 321\nNumber of predicted TMHs: 1\nExp number of AAs in TMHs: 18.68086\nExp number, first 60 AAs: 0.02311\nTotal prob of N-in: 0.37450\ninside 1 288\nTMhelix 289 308\noutside 309 321

1550 GCF_002162035.1_ASM216203v1 Sulfuriferula sp. AH1 Proteobacteria; Betaproteobacteria; Nitrosomonadales; Gallionellaceae; Sulfuriferula

MDEFDTLGAGLARSGISRRSFLKFCTAMASLMALPPSAAYAMADALKRAQRQSVIWLSFQ WP_087447741.1 Ni/Fe hydrogenase [Sulfuriferula sp. AH1] Length: 352\nNumber of predicted TMHs: 1\nExp number of AAs in TMHs: 37.66341\nExp number, first 60 AAs: 15.00688\nTotal prob of N-in: 0.56825\nPOSSIBLE N-term signal sequence\noutside 1 321\nTMhelix 322 341\ninside 342 352

1551 GCF_000175095.2_ASM17509v2 Nitrosomonas sp. AL212 Proteobacteria; Betaproteobacteria; Nitrosomonadales; Nitrosomonadaceae; Nitrosomonas MDFSRFFLIILARRTLILLTLVVTMSTTLLVSLLMSKSYKSTAVMVLTHKGADPVTGLIM WP_013647253.1 chain length determinant protein EpsF [Nitrosomonas sp. AL212] Length: 454\nNumber of predicted TMHs: 2\nExp number of AAs in TMHs: 42.8588399999999\nExp number, first 60 AAs: 20.83549\nTotal prob of N-in: 0.33194\nPOSSIBLE N-term signal sequence\noutside 1 14\nTMhelix 15 34\ninside 35 387\nTMhelix 388 410\noutside 411 454

1552 GCF_000376425.1_ASM37642v1 Thiobacillus denitrificans DSM 12475 Proteobacteria; Betaproteobacteria; Nitrosomonadales; Thiobacillaceae; Thiobacillus; Thiobacillus denitrificans MQGSLVDAFARQGVSRRSFLKYCATLASMMALPPVAGRAMAEAMAAARRPSVIWLPFQEC WP_018077114.1 Ni/Fe hydrogenase [Thiobacillus denitrificans] Length: 359\nNumber of predicted TMHs: 1\nExp number of AAs in TMHs: 26.48539\nExp number, first 60 AAs: 2.88431\nTotal prob of N-in: 0.15529\noutside 1 315\nTMhelix 316 338\ninside 339 359

1553 GCF_900156155.1_IMG-taxon_2681812808_annotated_assembly Azoarcus tolulyticus Proteobacteria; Betaproteobacteria; Rhodocyclales; Zoogloeaceae; Azoarcus MNTRRDFLKCALAGGAAVATAGAGPAQARGNAEISPDAVGLLFDSTLCIGCKACVAACKA WP_076602483.1 hydrogenase 2 protein HybA [Azoarcus tolulyticus] Length: 334\nNumber of predicted TMHs: 1\nExp number of AAs in TMHs: 19.77068\nExp number, first 60 AAs: 1.53305\nTotal prob of N-in: 0.53550\ninside 1 292\nTMhelix 293 312\noutside 313 334

1554 GCF_000373965.1_ASM37396v1 Uliginosibacterium gangwonense DSM 18521 Proteobacteria; Betaproteobacteria; Rhodocyclales; Zoogloeaceae; Uliginosibacterium; Uliginosibacterium gangwonense MITHYRIIFTGTHCGGRRRVMIRNQHLAQLALAVCLSGAALLALPAHAETGRVIVKFRSS WP_083917589.1 hypothetical protein [Uliginosibacterium gangwonense] Length: 631\nNumber of predicted TMHs: 2\nExp number of AAs in TMHs: 43.5748799999999\nExp number, first 60 AAs: 21.06316\nTotal prob of N-in: 0.95502\nPOSSIBLE N-term signal sequence\ninside 1 25\nTMhelix 26 48\noutside 49 606\nTMhelix 607 629\ninside 630 631

1555 GCF_000024165.1_ASM2416v1 Candidatus Accumulibacter phosphatis clade IIA str. UW-1 Proteobacteria; Betaproteobacteria; unclassified Betaproteobacteria; Candidatus Accumulibacter; Candidatus Accumulibacter phosphatis MANTSLLKLESDANIAVASKRLGMPRREFLQFCASVAASLGLPAGADAAVAEAVAAKKRP WP_015766771.1 hydrogenase 2 small subunit [Candidatus Accumulibacter phosphatis] Length: 395\nNumber of predicted TMHs: 1\nExp number of AAs in TMHs: 19.99184\nExp number, first 60 AAs: 0.58245\nTotal prob of N-in: 0.05311\noutside 1 353\nTMhelix 354 376\ninside 377 395

1556 GCF_001184205.1_ASM118420v1 Desulfocarbo indianensis Proteobacteria; delta/epsilon subdivisions; Deltaproteobacteria; Desulfarculales; Desulfarculaceae; Desulfocarbo MRFDRRKFLKLSTAAAAGTMLNPGPGQAKAAPPGPDSLGCLVDTTLCVGCRQCEAACNQR WP_049676148.1 4Fe-4S ferredoxin [Desulfocarbo indianensis] Length: 312\nNumber of predicted TMHs: 1\nExp number of AAs in TMHs: 22.35596\nExp number, first 60 AAs: 0.00208\nTotal prob of N-in: 0.51437\noutside 1 267\nTMhelix 268 290\ninside 291 312

1557 GCF_000429925.1_ASM42992v1 Desulfatirhabdium butyrativorans DSM 18734 Proteobacteria; delta/epsilon subdivisions; Deltaproteobacteria; Desulfobacterales; Desulfobacteraceae; Desulfatirhabdium; Desulfatirhabdium butyrativorans MKLDRRGFFKIVGIGASAAVAGAPKTSHAWQSKAPPDPYGCLVDLTRCVGCRKCELACNE WP_028325305.1 4Fe-4S ferredoxin [Desulfatirhabdium butyrativorans] Length: 299\nNumber of predicted TMHs: 1\nExp number of AAs in TMHs: 19.62587\nExp number, first 60 AAs: 0.56485\nTotal prob of N-in: 0.11569\noutside 1 267\nTMhelix 268 290\ninside 291 299

1558 GCF_001293685.1_ASM129368v1 Desulfatitalea tepidiphila Proteobacteria; delta/epsilon subdivisions; Deltaproteobacteria; Desulfobacterales; Desulfobacteraceae; Desulfatitalea MGISRRNFFKILSVAGAATATAQTPARAWQSQAPSEAYGCLVDLTRCIGCRKCEQACNEV WP_054030243.1 4Fe-4S ferredoxin [Desulfatitalea tepidiphila] Length: 306\nNumber of predicted TMHs: 1\nExp number of AAs in TMHs: 21.26268\nExp number, first 60 AAs: 0.05615\nTotal prob of N-in: 0.51312\ninside 1 268\nTMhelix 269 291\noutside 292 306

1559 GCF_000373985.1_ASM37398v1 Desulfobacter curvatus DSM 3379 Proteobacteria; delta/epsilon subdivisions; Deltaproteobacteria; Desulfobacterales; Desulfobacteraceae; Desulfobacter; Desulfobacter curvatus MTDGEKGGVSRRGFLKRVAAGGAAMLLPGTVSASRQGDELASLHDLSKCIGCGECVSACT WP_020589710.1 4Fe-4S ferredoxin [Desulfobacter curvatus] Length: 334\nNumber of predicted TMHs: 1\nExp number of AAs in TMHs: 22.61598\nExp number, first 60 AAs: 2.27967\nTotal prob of N-in: 0.19168\noutside 1 296\nTMhelix 297 319\ninside 320 334

1560 GCF_000373985.1_ASM37398v1 Desulfobacter curvatus DSM 3379 Proteobacteria; delta/epsilon subdivisions; Deltaproteobacteria; Desulfobacterales; Desulfobacteraceae; Desulfobacter; Desulfobacter curvatus MLRRMKYEQKGKHTMTADNHLNRRDFLISLGQGAAVCIGGFFLINHWPDCPPALSNPIIN WP_083927905.1 hypothetical protein [Desulfobacter curvatus] Length: 164\nNumber of predicted TMHs: 2\nExp number of AAs in TMHs: 38.20673\nExp number, first 60 AAs: 19.44357\nTotal prob of N-in: 0.98166\nPOSSIBLE N-term signal sequence\ninside 1 25\nTMhelix 26 48\noutside 49 133\nTMhelix 134 156\ninside 157 164

1561 GCF_000233695.2_ASM23369v3 Desulfobacter postgatei 2ac9 Proteobacteria; delta/epsilon subdivisions; Deltaproteobacteria; Desulfobacterales; Desulfobacteraceae; Desulfobacter; Desulfobacter postgatei MKDDKRVSVSRRGFLKGVAVGGASMLLPGTVRASRQGNALASLHDLSKCIGCGECVSACS WP_004071579.1 4Fe-4S ferredoxin [Desulfobacter postgatei]Length: 334\nNumber of predicted TMHs: 1\nExp number of AAs in TMHs: 30.07275\nExp number, first 60 AAs: 10.16048\nTotal prob of N-in: 0.73978\nPOSSIBLE N-term signal sequence\ninside 1 296\nTMhelix 297 319\noutside 320 334

1562 GCF_900176365.1_IMG-taxon_2681813565_annotated_assembly Desulfobacterium vacuolatum DSM 3385 Proteobacteria; delta/epsilon subdivisions; Deltaproteobacteria; Desulfobacterales; Desulfobacteraceae; Desulfobacterium; Desulfobacterium vacuolatum MNINRRGFFKVMGAAGATAMTPAAGARAWESAAPPDAFGCLVDLTRCIGCRKCEEACNRV WP_084070237.1 4Fe-4S ferredoxin [Desulfobacterium vacuolatum] Length: 307\nNumber of predicted TMHs: 1\nExp number of AAs in TMHs: 23.45689\nExp number, first 60 AAs: 3.74846\nTotal prob of N-in: 0.21526\noutside 1 264\nTMhelix 265 287\ninside 288 307

1563 GCF_000307105.1_ASM30710v1 Desulfobacula toluolica Tol2 Proteobacteria; delta/epsilon subdivisions; Deltaproteobacteria; Desulfobacterales; Desulfobacteraceae; Desulfobacula; Desulfobacula toluolica MTARKIRPKISRRAFLKGSITVAGSVAATASIARAARSLSGEKKEPLATLIDISKCIGCE WP_014957271.1 4Fe-4S ferredoxin [Desulfobacula toluolica] Length: 326\nNumber of predicted TMHs: 1\nExp number of AAs in TMHs: 24.63059\nExp number, first 60 AAs: 5.43439\nTotal prob of N-in: 0.69067\ninside 1 297\nTMhelix 298 320\noutside 321 326

1564 GCF_000018405.1_ASM1840v1 Desulfococcus oleovorans Hxd3 Proteobacteria; delta/epsilon subdivisions; Deltaproteobacteria; Desulfobacterales; Desulfobacteraceae; Desulfococcus; Desulfococcus oleovorans MPDTISRRNFLKAGLITAGAVAASGTGLASMAGAASGEPLCTLLDLSRCIGCEECVYACR WP_012173729.1 4Fe-4S ferredoxin [Desulfococcus oleovorans] Length: 328\nNumber of predicted TMHs: 2\nExp number of AAs in TMHs: 40.51029\nExp number, first 60 AAs: 18.1721\nTotal prob of N-in: 0.83298\nPOSSIBLE N-term signal sequence\ninside 1 12\nTMhelix 13 35\noutside 36 294\nTMhelix 295 317\ninside 318 328

1565 GCF_001311845.1_ASM131184v1 Desulfosarcina cetonica JCM 12296 Proteobacteria; delta/epsilon subdivisions; Deltaproteobacteria; Desulfobacterales; Desulfobacteraceae; Desulfosarcina; Desulfosarcina cetonica MGLNRRDFFKWLTVAGSAVAGSGRSAHAWQSRAPADPLGCLVDLTRCIGCRKCEEACNRV WP_054692440.1 4Fe-4S ferredoxin [Desulfosarcina cetonica] Length: 307\nNumber of predicted TMHs: 1\nExp number of AAs in TMHs: 20.30207\nExp number, first 60 AAs: 0.00654999999999999\nTotal prob of N-in: 0.51570\ninside 1 268\nTMhelix 269 291\noutside 292 307

1566 GCF_000420085.1_ASM42008v1 Desulfospira joergensenii DSM 10085 Proteobacteria; delta/epsilon subdivisions; Deltaproteobacteria; Desulfobacterales; Desulfobacteraceae; Desulfospira; Desulfospira joergensenii MTSEISRRAFLKSSLGAAGSVAASSGLARAAAAAETPEDPLATLIDVSKCIGCEECVHAC WP_022663865.1 4Fe-4S ferredoxin [Desulfospira joergensenii] Length: 319\nNumber of predicted TMHs: 1\nExp number of AAs in TMHs: 19.63546\nExp number, first 60 AAs: 0.13568\nTotal prob of N-in: 0.04593\noutside 1 288\nTMhelix 289 311\ninside 312 319

1567 GCF_000621145.1_ASM62114v1 Desulfobulbus elongatus DSM 2908 Proteobacteria; delta/epsilon subdivisions; Deltaproteobacteria; Desulfobacterales; Desulfobulbaceae; Desulfobulbus; Desulfobulbus elongatus MPTLIDRRRFLRGGLLASAAATVSLAPAQTEAASFEGYPNAMGVLVDLSRCVGCRSCEAA WP_028318203.1 hypothetical protein [Desulfobulbus elongatus] Length: 309\nNumber of predicted TMHs: 1\nExp number of

AAs in TMHs: 23.12625\nExp number, first 60 AAs: 0.706289999999999\nTotal prob of N-in: 0.09201\noutside 1 270\nTMhelix 271 293\ninside 294 309

1568 GCF_000429945.1_ASM42994v1 Desulfobulbus japonicus DSM 18378 Proteobacteria; delta/epsilon subdivisions; Deltaproteobacteria; Desulfobacterales; Desulfobulbaceae; Desulfobulbus; Desulfobulbus japonicus MKKAVSRRKFLKGGLAGSAALSLTTAKRSEAKSFEGYPDAMGVLVDLSRCVGCRSCEAAC WP_028580723.1 hypothetical protein [Desulfobulbus japonicus] Length: 308\nNumber of predicted TMHs: 1\nExp number of AAs in TMHs: 23.54981\nExp number, first 60 AAs: 0.01056\nTotal prob of N-in: 0.18363\noutside 1 269\nTMhelix 270 292\ninside 293 308

1569 GCF_000429945.1_ASM42994v1 Desulfobulbus japonicus DSM 18378 Proteobacteria; delta/epsilon subdivisions; Deltaproteobacteria; Desulfobacterales; Desulfobulbaceae; Desulfobulbus; Desulfobulbus japonicus MKRREFIKGAVGSLALASASGAGIANARTVQALPPGAVGILYDSTLCVGCNACMAACKKA WP_028579801.1 hypothetical protein [Desulfobulbus japonicus] Length: 329\nNumber of predicted TMHs: 1\nExp number of AAs in TMHs: 22.30953\nExp number, first 60 AAs: 1.57885\nTotal prob of N-in: 0.12392\noutside 1 285\nTMhelix 286 308\ninside 309 329

1570 GCF_000429965.1_ASM42996v1 Desulfobulbus mediterraneus DSM 13871 Proteobacteria; delta/epsilon subdivisions; Deltaproteobacteria; Desulfobacterales; Desulfobulbaceae; Desulfobulbus; Desulfobulbus mediterraneus MKIKKLLNRRTFLKGSLAGSAAAVSLSRSPEAKAATFEGYPDSMGVLVDLSRCVGCRSCE WP_035247752.1 hypothetical protein [Desulfobulbus mediterraneus] Length: 313\nNumber of predicted TMHs: 1\nExp number of AAs in TMHs: 22.4446\nExp number, first 60 AAs: 0.02691\nTotal prob of N-in: 0.18585\noutside 1 272\nTMhelix 273 295\ninside 296 313

1571 GCF_000186885.1_ASM18688v1 Desulfobulbus propionicus DSM 2032 Proteobacteria; delta/epsilon subdivisions; Deltaproteobacteria; Desulfobacterales; Desulfobulbaceae; Desulfobulbus; Desulfobulbus propionicus MKTLINRRRFLRGSLAASAAATVSLAKKTDAATFEGYPDAMGVLVDLSRCVGCRSCEAAC WP_015723347.1 4Fe-4S ferredoxin [Desulfobulbus propionicus] Length: 309\nNumber of predicted TMHs: 1\nExp number of AAs in TMHs: 22.78665\nExp number, first 60 AAs: 0.03002\nTotal prob of N-in: 0.10401\noutside 1 269\nTMhelix 270 292\ninside 293 309

1572 GCF_900130015.1_IMG-taxon_2585428080_annotated_assembly Desulfofustis glycolicus DSM 9705 Proteobacteria; delta/epsilon subdivisions; Deltaproteobacteria; Desulfobacterales; Desulfobulbaceae; Desulfofustis; Desulfofustis glycolicus MSRRNFLRTGLATFCATLIGTHAEKTAARDFEGYHQSKGVLVDLTRCIGCRSCEAACNKE WP_073377441.1 hypothetical protein [Desulfofustis glycolicus] Length: 312\nNumber of predicted TMHs: 1\nExp number of AAs in TMHs: 22.37439\nExp number, first 60 AAs: 0.19868\nTotal prob of N-in: 0.26481\noutside 1 266\nTMhelix 267 289\ninside 290 312

1573 GCF_000620765.1_ASM62076v1 Desulfovermiculus halophilus DSM 18834 Proteobacteria; delta/epsilon subdivisions; Deltaproteobacteria; Desulfovibrionales; Desulfohalobiaceae; Desulfovermiculus; Desulfovermiculus halophilus MDSTRRGFLKTLGAAGAATGATALTASNQARAWTSDAPDNPNACLVDLTRCIGCRLCEQA WP_027370080.1 4Fe-4S ferredoxin [Desulfovermiculus halophilus] Length: 311\nNumber of predicted TMHs: 1\nExp number of AAs in TMHs: 20.99481\nExp number, first 60 AAs: 0.32951\nTotal prob of N-in: 0.56379\ninside 1 273\nTMhelix 274 296\noutside 297 311

1574 GCF_000620765.1_ASM62076v1 Desulfovermiculus halophilus DSM 18834 Proteobacteria; delta/epsilon subdivisions; Deltaproteobacteria; Desulfovibrionales; Desulfohalobiaceae; Desulfovermiculus; Desulfovermiculus halophilus MNRRKFLILNGPNLGHLGIRQPDIYGQKTMDDLPRLIRATLGPERAAGIDWDMDQKNSEG WP_027369443.1 3-dehydroquinate dehydratase [Desulfovermiculus halophilus] Length: 153\nNumber of predicted TMHs: 1\nExp number of AAs in TMHs: 22.75091\nExp number, first 60 AAs: 0.00145\nTotal prob of N-in: 0.83905\ninside 1 121\nTMhelix 122 144\noutside 145 153

1575 GCF_000934745.1_ASM93474v1 Desulfonatronum thioautotrophicum Proteobacteria; delta/epsilon subdivisions; Deltaproteobacteria; Desulfovibrionales; Desulfonatronaceae; Desulfonatronum MEKVMGDSVTRQGRRAWFRLISALFGATVVAAALSFAATPHAQVGETAGVAKQVRFELNQ WP_084604463.1 hypothetical protein [Desulfonatronum thioautotrophicum] Length: 761\nNumber of predicted TMHs: 2\nExp number of AAs in TMHs: 44.37162\nExp number, first 60 AAs: 21.78135\nTotal prob of N-in: 0.99702\nPOSSIBLE N-term signal sequence\ninside 1 19\nTMhelix 20 42\noutside 43 720\nTMhelix 721 743\ninside 744 761

1576 GCF_001874525.1_ASM187452v1 Desulfovibrio dechloracetivorans Proteobacteria; delta/epsilon subdivisions; Deltaproteobacteria; Desulfovibrionales; Desulfovibrionaceae; Desulfovibrio

MSEKKYRAKAVSRRGFLKALGVGSAGALIPAAPTLAAQERAGAPSDGELATLLDLSKCIG WP_071544310.1 4Fe-4S ferredoxin [Desulfovibrio dechloracetivorans] Length: 354\nNumber of predicted TMHs: 1\nExp number of AAs in TMHs: 20.61627\nExp number, first 60 AAs: 2.40385\nTotal prob of N-in: 0.82202\ninside 1 319\nTMhelix 320 342\noutside 343 354

1577 GCF_001874525.1_ASM187452v1 Desulfovibrio dechloracetivorans Proteobacteria; delta/epsilon subdivisions; Deltaproteobacteria; Desulfovibrionales; Desulfovibrionaceae; Desulfovibrio MTITRRGFLGALGIAGAASTLPGNAQAWQSKAPPDPFGCLVDLTRCVGCRKCEQACNEVN WP_071546240.1 4Fe-4S ferredoxin [Desulfovibrio dechloracetivorans] Length: 302\nNumber of predicted TMHs: 1\nExp number of AAs in TMHs: 22.61861\nExp number, first 60 AAs: 2.36964\nTotal prob of N-in: 0.57353\ninside 1 267\nTMhelix 268 290\noutside 291 302

1578 GCF_001553605.1_ASM155360v1 Desulfovibrio fairfieldensis Proteobacteria; delta/epsilon subdivisions; Deltaproteobacteria; Desulfovibrionales; Desulfovibrionaceae; Desulfovibrio MGKNRFFQRRRFLQGAAGLLLGGTLPSALVPVAAKAGNLTGEPLTTLIDVDACDGCGACV WP_062251202.1 4Fe-4S ferredoxin [Desulfovibrio fairfieldensis] Length: 329\nNumber of predicted TMHs: 2\nExp number of AAs in TMHs: 39.90096\nExp number, first 60 AAs: 20.0665\nTotal prob of N-in: 0.89301\nPOSSIBLE N-term signal sequence\ninside 1 11\nTMhelix 12 34\noutside 35 300\nTMhelix 301 320\ninside 321 329

1579 GCF_001563225.1_ASM156322v1 Desulfovibrio indicus Proteobacteria; delta/epsilon subdivisions; Deltaproteobacteria; Desulfovibrionales; Desulfovibrionaceae; Desulfovibrio MPITRRSFLAALGIAGATAAAPKNAEAWQSKAPPDPYGCLVDLTRCVGCRKCEQACKEVN WP_066806246.1 4Fe-4S ferredoxin [Desulfovibrio indicus] Length: 302\nNumber of predicted TMHs: 1\nExp number of AAs in TMHs: 20.06391\nExp number, first 60 AAs: 0.07373\nTotal prob of N-in: 0.43639\ninside 1 267\nTMhelix 268 290\noutside 291 302

1580 GCF_000145315.2_Desulfovibrio_sp3_1_syn3_V2 Desulfovibrio sp. 3_1_syn3 Proteobacteria; delta/epsilon subdivisions; Deltaproteobacteria; Desulfovibrionales; Desulfovibrionaceae; Desulfovibrio MGKNRFFQRRRFLQGAAGLLLGGTLPSALAPVAAKAGNLTGEPLTTLIDVDACDGCGACV WP_008683453.1 4Fe-4S ferredoxin [Desulfovibrio sp. 3_1_syn3] Length: 329\nNumber of predicted TMHs: 2\nExp number of AAs in TMHs: 38.97466\nExp number, first 60 AAs: 19.13865\nTotal prob of N-in: 0.85513\nPOSSIBLE N-term signal sequence\ninside 1 11\nTMhelix 12 34\noutside 35 300\nTMhelix 301 320\ninside 321 329

1581 GCF_000224635.1_Desulfovibrio_sp_6_1_46AFAA_V1 Desulfovibrio sp. 6_1_46AFAA Proteobacteria; delta/epsilon subdivisions; Deltaproteobacteria; Desulfovibrionales; Desulfovibrionaceae; Desulfovibrio MGKNRFFQRRRFLQGAAGLLLGGTLPSALAPVAAKAGNLTGEPLTTLIDVDACDGCGACV WP_009302998.1 4Fe-4S ferredoxin [Desulfovibrio sp. 6_1_46AFAA] Length: 329\nNumber of predicted TMHs: 2\nExp number of AAs in TMHs: 38.97466\nExp number, first 60 AAs: 19.13865\nTotal prob of N-in: 0.85513\nPOSSIBLE N-term signal sequence\ninside 1 11\nTMhelix 12 34\noutside 35 300\nTMhelix 301 320\ninside 321 329

1582 GCF_000344315.1_ASM34431v1 Desulfovibrio africanus PCS Proteobacteria; delta/epsilon subdivisions; Deltaproteobacteria; Desulfovibrionales; Desulfovibrionaceae; Desulfovibrio; Desulfovibrio africanus MQTRQPLQLGSALTLEPESRPGSSLREMLAVLFRRRKAVLAFFLSVTFAAGLFTYFMPKT WP_005986650.1 hypothetical protein [Desulfovibrio africanus] Length: 537\nNumber of predicted TMHs: 2\nExp number of AAs in TMHs: 42.44994\nExp number, first 60 AAs: 19.8445\nTotal prob of N-in: 0.95717\nPOSSIBLE N-term signal sequence\ninside 1 37\nTMhelix 38 57\noutside 58 480\nTMhelix 481 503\ninside 504 537

1583 GCF_000344315.1_ASM34431v1 Desulfovibrio africanus PCS Proteobacteria; delta/epsilon subdivisions; Deltaproteobacteria; Desulfovibrionales; Desulfovibrionaceae; Desulfovibrio; Desulfovibrio africanus MQAPDNRSTVCPLRLPAPTDVSGDAALLRFLAVVRKRRRFLLAMPLAAGALTLLACLLII WP_005988823.1 hypothetical protein [Desulfovibrio africanus] Length: 326\nNumber of predicted TMHs: 2\nExp number of AAs in TMHs: 45.32532\nExp number, first 60 AAs: 19.67531\nTotal prob of N-in: 0.86752\nPOSSIBLE N-term signal sequence\ninside 1 39\nTMhelix 40 62\noutside 63 290\nTMhelix 291 313\ninside 314 326

1584 GCF_000212675.2_ASM21267v2 Desulfovibrio africanus str. Walvis Bay Proteobacteria; delta/epsilon subdivisions; Deltaproteobacteria; Desulfovibrionales; Desulfovibrionaceae; Desulfovibrio; Desulfovibrio africanus MNTKNKARFSRRAFLAAAAVGGAGLVLPGTARATAKGEPVATLLDLSKCIGCEACVLACQ WP_014259604.1 4Fe-4S ferredoxin [Desulfovibrio africanus] Length: 335\nNumber of predicted TMHs: 2\nExp number of AAs in TMHs: 42.01806\nExp number, first 60 AAs: 21.64995\nTotal prob of N-in: 0.99011\nPOSSIBLE N-term signal sequence\ninside 1 12\nTMhelix 13 35\noutside 36 302\nTMhelix 303 325\ninside 326 335

1585 GCF_000422545.1_ASM42254v1 Desulfovibrio africanus subsp. africanus DSM 2603 Proteobacteria; delta/epsilon subdivisions; Deltaproteobacteria; Desulfovibrionales; Desulfovibrionaceae; Desulfovibrio; Desulfovibrio africanus; Desulfovibrio africanus subsp. africanus MNTKNKARFSRRAFLAAAAVGGAGLVLPGTARAAAKGEPLATLLDLSKCIGCEACVLACQ WP_027368072.1 4Fe-4S ferredoxin [Desulfovibrio africanus] Length: 335\nNumber of predicted TMHs: 2\nExp number of AAs in TMHs: 42.81116\nExp number, first 60 AAs: 22.13873\nTotal prob of N-in: 0.99564\nPOSSIBLE N-term signal sequence\ninside 1 12\nTMhelix 13 35\noutside 36 302\nTMhelix 303 325\ninside 326 335

1586 GCF_000702665.1_ASM70266v1 Desulfovibrio alcoholivorans DSM 5433 Proteobacteria; delta/epsilon subdivisions; Deltaproteobacteria; Desulfovibrionales; Desulfovibrionaceae; Desulfovibrio; Desulfovibrio alcoholivorans MNRRAFLTALTTAAAAVAAPGTAKTVLARDAASAGEPLATLLDLSQCVGCGACVAACRDA WP_029459533.1 4Fe-4S ferredoxin [Desulfovibrio alcoholivorans] Length: 330\nNumber of predicted TMHs: 1\nExp number of AAs in TMHs: 18.78828\nExp number, first 60 AAs: 1.56724\nTotal prob of N-in: 0.08976\noutside 1 287\nTMhelix 288 310\ninside 311 330

1587 GCF_000422245.1_D_alkalitolerans_DSM16529_v1.0 Desulfovibrio alkalitolerans DSM 16529 Proteobacteria; delta/epsilon subdivisions; Deltaproteobacteria; Desulfovibrionales; Desulfovibrionaceae; Desulfovibrio; Desulfovibrio alkalitolerans MSKNPEETTPAKERGLSRRAFLGGLAAAGAACLAPGTAKAATASSAPRTIELATLIDLSR WP_020886939.1 4Fe-4S ferredoxin, iron-sulfur binding domain-containing protein [Desulfovibrio alkalitolerans] Length: 338\nNumber of predicted TMHs: 1\nExp number of AAs in TMHs: 37.09164\nExp number, first 60 AAs: 14.64247\nTotal prob of N-in: 0.70749\nPOSSIBLE N-term signal sequence\noutside 1 303\nTMhelix 304 326\ninside 327 338

1588 GCF_000189295.2_ASM18929v2 Desulfovibrio desulfuricans ND132 Proteobacteria; delta/epsilon subdivisions; Deltaproteobacteria; Desulfovibrionales; Desulfovibrionaceae; Desulfovibrio; Desulfovibrio desulfuricans MTISRRGFLGALGVAGAVSAVPGNAQAWESKAPPDPFGCLVDLSRCVGCRKCEQACKEVN WP_014323720.1 4Fe-4S ferredoxin [Desulfovibrio desulfuricans] Length: 302\nNumber of predicted TMHs: 1\nExp number of AAs in TMHs: 21.57854\nExp number, first 60 AAs: 2.08137\nTotal prob of N-in: 0.39694\noutside 1 263\nTMhelix 264 286\ninside 287 302

1589 GCF_000420465.1_ASM42046v1 Desulfovibrio desulfuricans subsp. desulfuricans DSM 642 Proteobacteria; delta/epsilon subdivisions; Deltaproteobacteria; Desulfovibrionales; Desulfovibrionaceae; Desulfovibrio; Desulfovibrio desulfuricans; Desulfovibrio desulfuricans subsp. desulfuricans MGNKLFLQRRSVIKAMAGTAGLSLAGATPSVASSLQNWVCGDDRATIIDVDLCNGCGACV WP_022659754.1 4Fe-4S ferredoxin [Desulfovibrio desulfuricans] Length: 329\nNumber of predicted TMHs: 1\nExp number of AAs in TMHs: 20.16323\nExp number, first 60 AAs: 0.28157\nTotal prob of N-in: 0.03318\noutside 1 297\nTMhelix 298 320\ninside 321 329

1590 GCF_000429285.1_ASM42928v1 Desulfovibrio gigas DSM 1382 = ATCC 19364 Proteobacteria; delta/epsilon subdivisions; Deltaproteobacteria; Desulfovibrionales; Desulfovibrionaceae; Desulfovibrio; Desulfovibrio gigas MPFPSSPVRVLARACARVFAREGGTRRRVLAGIGLCLAAALLLVLLASAWIPRAARPDAP WP_081696777.1 hypothetical protein [Desulfovibrio gigas] Length: 381\nNumber of predicted TMHs: 2\nExp number of AAs in TMHs: 46.5952499999999\nExp number, first 60 AAs: 22.69976\nTotal prob of N-in: 0.68669\nPOSSIBLE N-term signal sequence\ninside 1 28\nTMhelix 29 51\noutside 52 349\nTMhelix 350 372\ninside 373 381

1591 GCF_000468495.1_ASM46849v1 Desulfovibrio gigas DSM 1382 = ATCC 19364 Proteobacteria; delta/epsilon subdivisions; Deltaproteobacteria; Desulfovibrionales; Desulfovibrionaceae; Desulfovibrio; Desulfovibrio gigas MPFPSSPVRVLARACARVFAREGGTRRRVLAGIGLCLAAALLLVLLASAWIPRAARPDAP WP_081696777.1 hypothetical protein [Desulfovibrio gigas] Length: 381\nNumber of predicted TMHs: 2\nExp number of AAs in TMHs: 46.5952499999999\nExp number, first 60 AAs: 22.69976\nTotal prob of N-in: 0.68669\nPOSSIBLE N-term signal sequence\ninside 1 28\nTMhelix 29 51\noutside 52 349\nTMhelix 350 372\ninside 373 381

1592 GCF_000526455.1_ASM52645v1 Desulfovibrio cf. magneticus IFRC170 Proteobacteria; delta/epsilon subdivisions; Deltaproteobacteria; Desulfovibrionales; Desulfovibrionaceae; Desulfovibrio; Desulfovibrio magneticus MNRRAFLKTLGTGGAVLAAPTAATTLLARDASAASGEPLATLYDLSKCIGCGACVAACRD WP_024823537.1 4Fe-4S ferredoxin [Desulfovibrio magneticus] Length: 325\nNumber of predicted TMHs: 1\nExp number of AAs in TMHs: 18.09303\nExp number, first 60 AAs: 1.96934\nTotal prob of N-in: 0.12404\noutside 1 290\nTMhelix 291 310\ninside 311 325

1593 GCF_000010665.1_ASM1066v1 Desulfovibrio magneticus RS-1 Proteobacteria; delta/epsilon subdivisions; Deltaproteobacteria; Desulfovibrionales; Desulfovibrionaceae; Desulfovibrio; Desulfovibrio magneticus

MNRRAFLKSLGTAVAAAPVAAPLLARDAAAASGEQLATLFDLSKCVGCGACVAGCREAHA WP_015862494.1 4Fe-4S ferredoxin [Desulfovibrio magneticus] Length: 321\nNumber of predicted TMHs: 1\nExp number of AAs in TMHs: 19.24336\nExp number, first 60 AAs: 0.668399999999999\nTotal prob of N-in: 0.06738\noutside 1 286\nTMhelix 287 309\ninside 310 321

1594 GCF_000341895.1_ASM34189v1 Desulfovibrio piezophilus C1TLV30 Proteobacteria; delta/epsilon subdivisions; Deltaproteobacteria; Desulfovibrionales; Desulfovibrionaceae; Desulfovibrio; Desulfovibrio piezophilus MNKKRQSSVGVSRRGFLKAFGMGSAAALLPGNSLARERVSTASEGELATLLDLSKCIGCG WP_015416321.1 4Fe-4S ferredoxin iron-sulfur-binding domain protein [Desulfovibrio piezophilus] Length: 344\nNumber of predicted TMHs: 1\nExp number of AAs in TMHs: 18.00614\nExp number, first 60 AAs: 0.53065\nTotal prob of N-in: 0.21440\noutside 1 303\nTMhelix 304 326\ninside 327 344

1595 GCF_001611275.1_ASM161127v1 Desulfuromonas sp. DDH964 Proteobacteria; delta/epsilon subdivisions; Deltaproteobacteria; Desulfuromonadales; Desulfuromonadaceae; Desulfuromonas MDLSRRKFLKLGAVAGGSAVCNLARPAAAREMPDIDPDWYGMLNDSTRCIGCKACMVACK WP_066728980.1 hypothetical protein [Desulfuromonas sp. DDH964] Length: 309\nNumber of predicted TMHs: 1\nExp number of AAs in TMHs: 19.86472\nExp number, first 60 AAs: 0.06591\nTotal prob of N-in: 0.04176\noutside 1 274\nTMhelix 275 292\ninside 293 309

1596 GCF_900142125.1_IMG-taxon_2585428155_annotated_assembly Malonomonas rubra DSM 5091 Proteobacteria; delta/epsilon subdivisions; Deltaproteobacteria; Desulfuromonadales; Desulfuromonadaceae; Malonomonas; Malonomonas rubra MSKVSRRKFLAGSLAGGAAVVAAPAKKALAAGNFSGFPDGMGVLVDLTRCVGCRSCEAAC WP_072908093.1 hypothetical protein [Malonomonas rubra] Length: 310\nNumber of predicted TMHs: 1\nExp number of AAs in TMHs: 21.72637\nExp number, first 60 AAs: 1.11846\nTotal prob of N-in: 0.50594\ninside 1 271\nTMhelix 272 294\noutside 295 310

1597 GCF_000711225.1_ASM71122v1 Pelobacter seleniigenes DSM 18267 Proteobacteria; delta/epsilon subdivisions; Deltaproteobacteria; Desulfuromonadales; Desulfuromonadaceae; Pelobacter; Pelobacter seleniigenes MSKITRRKFLAGSVAGSAAVMMTPAKKALAAGSFEGFPDGMGVLVDLTRCIGCRTCEAAC WP_029914989.1 hypothetical protein [Pelobacter seleniigenes] Length: 313\nNumber of predicted TMHs: 1\nExp number of AAs in TMHs: 23.64494\nExp number, first 60 AAs: 0.35887\nTotal prob of N-in: 0.65351\ninside 1 269\nTMhelix 270 292\noutside 293 313

1598 GCF_000711225.1_ASM71122v1 Pelobacter seleniigenes DSM 18267 Proteobacteria; delta/epsilon subdivisions; Deltaproteobacteria; Desulfuromonadales; Desulfuromonadaceae; Pelobacter; Pelobacter seleniigenes MDISRRKFFKLTAVGGAAVCSAATPAAAQRQPVKADPDWYGMLNDSTRCIGCKACQVACK WP_029915213.1 hypothetical protein [Pelobacter seleniigenes] Length: 301\nNumber of predicted TMHs: 1\nExp number of AAs in TMHs: 21.67169\nExp number, first 60 AAs: 0.38848\nTotal prob of N-in: 0.04056\noutside 1 264\nTMhelix 265 287\ninside 288 301

1599 GCF_900103065.1_IMG-taxon_2599185148_annotated_assembly Geoalkalibacter ferrihydriticus Proteobacteria; delta/epsilon subdivisions; Deltaproteobacteria; Desulfuromonadales; Geobacteraceae; Geoalkalibacter MPISRRKFFKVAAATGAGACAALTAPAQASAAKLPDDPGYGMLNDSTRCIGCKACQAACK WP_040101249.1 hypothetical protein [Geoalkalibacter ferrihydriticus] Length: 302\nNumber of predicted TMHs: 1\nExp number of AAs in TMHs: 18.65243\nExp number, first 60 AAs: 0.52844\nTotal prob of N-in: 0.14948\noutside 1 266\nTMhelix 267 289\ninside 290 302

1600 GCF_000827125.1_ASM82712v1 Geoalkalibacter subterraneusProteobacteria; delta/epsilon subdivisions; Deltaproteobacteria; Desulfuromonadales; Geobacteraceae; Geoalkalibacter MKIDRRKFLKLSAAAGGTAACSMASPAAAKGPKAPDPNWYGMLNDSTRCIGCKACQVACK WP_040201369.1 hypothetical protein [Geoalkalibacter subterraneus] Length: 309\nNumber of predicted TMHs: 1\nExp number of AAs in TMHs: 19.93943\nExp number, first 60 AAs: 0.20977\nTotal prob of N-in: 0.11749\noutside 1 274\nTMhelix 275 292\ninside 293 309

1601 GCF_000820505.1_ASM82050v1 Geoalkalibacter ferrihydriticus DSM 17813 Proteobacteria; delta/epsilon subdivisions; Deltaproteobacteria; Desulfuromonadales; Geobacteraceae; Geoalkalibacter; Geoalkalibacter ferrihydriticus MPISRRKFFKVAAATGAGACAALTAPAQASAAKLPDDPGYGMLNDSTRCIGCKACQAACK WP_040101249.1 hypothetical protein [Geoalkalibacter ferrihydriticus] Length: 302\nNumber of predicted TMHs: 1\nExp number of AAs in TMHs: 18.65243\nExp number, first 60 AAs: 0.52844\nTotal prob of N-in: 0.14948\noutside 1 266\nTMhelix 267 289\ninside 290 302

1602 GCF_000023645.1_ASM2364v1 Geobacter sp. M21 Proteobacteria; delta/epsilon subdivisions; Deltaproteobacteria; Desulfuromonadales; Geobacteraceae; Geobacter MRFHDPQMLFLLALLVPLFVWIRRGESRRPALPLSAAFAGNPLPDTLRSRLARLLPYLRL WP_015837714.1 hypothetical protein [Geobacter sp. M21] Length: 331\nNumber of predicted TMHs: 1\nExp number of AAs in TMHs: 34.34913\nExp number, first 60 AAs: 10.21019\nTotal prob of N-in: 0.16991\nPOSSIBLE N-term signal sequence\noutside 1 307\nTMhelix 308 327\ninside 328 331

1603 GCF_900167465.1_IMG-taxon_2585428117_annotated_assembly Geobacter thiogenes Proteobacteria; delta/epsilon subdivisions; Deltaproteobacteria; Desulfuromonadales; Geobacteraceae; Geobacter MERRDFLKGCLACGVTAATVSVASGKAMAAGSFEGYPDAMGVLVDLTRCVGCRSCEAACN WP_078789847.1 hypothetical protein [Geobacter thiogenes] Length: 301\nNumber of predicted TMHs: 1\nExp number of AAs in TMHs: 19.40941\nExp number, first 60 AAs: 1.20956\nTotal prob of N-in: 0.45790\ninside 1 262\nTMhelix 263 285\noutside 286 301

1604 GCF_000020385.1_ASM2038v1 Geobacter lovleyi SZ Proteobacteria; delta/epsilon subdivisions; Deltaproteobacteria; Desulfuromonadales; Geobacteraceae; Geobacter; Geobacter lovleyi MERRDFLKGCLACGVTAATVSVASGKAMAAGSFEGYPDAMGVLVDLTRCVGCRSCEAACN WP_041243266.1 hypothetical protein [Geobacter lovleyi] Length: 301\nNumber of predicted TMHs: 1\nExp number of AAs in TMHs: 19.41177\nExp number, first 60 AAs: 1.2088\nTotal prob of N-in: 0.45830\ninside 1 262\nTMhelix 263 285\noutside 286 301

1605 GCF_000384395.1_ASM38439v1 Geopsychrobacter electrodiphilus DSM 16401 Proteobacteria; delta/epsilon subdivisions; Deltaproteobacteria; Desulfuromonadales; Geobacteraceae; Geopsychrobacter; Geopsychrobacter electrodiphilus MSKVSRRRFLAGGLASGAAVLLAPAKKVLAAGSFSGYPDGMGVLVDLTRCLGCRSCEAAC WP_020676593.1 hypothetical protein [Geopsychrobacter electrodiphilus] Length: 312\nNumber of predicted TMHs: 1\nExp number of AAs in TMHs: 23.6398\nExp number, first 60 AAs: 2.28212\nTotal prob of N-in: 0.49539\noutside 1 270\nTMhelix 271 293\ninside 294 312

1606 GCF_000017505.1_ASM1750v1 Anaeromyxobacter sp. Fw109-5 Proteobacteria; delta/epsilon subdivisions; Deltaproteobacteria; Myxococcales; Cystobacterineae; Anaeromyxobacteraceae; Anaeromyxobacter MSPRRALLAAALSTAPLFAAAADVWVAGATEKIRPDAQPRQTTEARIAAARNEFEAFQVV WP_011985708.1 hypothetical protein [Anaeromyxobacter sp. Fw109-5] Length: 604\nNumber of predicted TMHs: 2\nExp number of AAs in TMHs: 36.87372\nExp number, first 60 AAs: 15.66667\nTotal prob of N-in: 0.69895\nPOSSIBLE N-term signal sequence\ninside 1 6\nTMhelix 7 29\noutside 30 575\nTMhelix 576 598\ninside 599 604

1607 GCF_000017505.1_ASM1750v1 Anaeromyxobacter sp. Fw109-5 Proteobacteria; delta/epsilon subdivisions; Deltaproteobacteria; Myxococcales; Cystobacterineae; Anaeromyxobacteraceae; Anaeromyxobacter MPNVSRRTVLGTLGGAAALAAAPAPAEARAPKPPRDDDLGLLFDSTRCVGCRACTVRCRE WP_012098326.1 hydrogenase 2 protein HybA [Anaeromyxobacter sp. Fw109-5] Length: 293\nNumber of predicted TMHs: 2\nExp number of AAs in TMHs: 40.52893\nExp number, first 60 AAs: 0.21823\nTotal prob of N-in: 0.91132\ninside 1 223\nTMhelix 224 246\noutside 247 260\nTMhelix 261 280\ninside 281 293

1608 GCF_000017505.1_ASM1750v1 Anaeromyxobacter sp. Fw109-5 Proteobacteria; delta/epsilon subdivisions; Deltaproteobacteria; Myxococcales; Cystobacterineae; Anaeromyxobacteraceae; Anaeromyxobacter MNDMRSEVLGLGTAKGVSRRDFIKVCTTAAAAVGLPAWAGEKMAENVAKGRKPSVIWLHF WP_041448759.1 hydrogenase 2 small subunit [Anaeromyxobacter sp. Fw109-5] Length: 376\nNumber of predicted TMHs: 1\nExp number of AAs in TMHs: 18.83664\nExp number, first 60 AAs: 0.66492\nTotal prob of N-in: 0.07262\noutside 1 339\nTMhelix 340 362\ninside 363 376

1609 GCF_000020805.1_ASM2080v1 Anaeromyxobacter sp. K Proteobacteria; delta/epsilon subdivisions; Deltaproteobacteria; Myxococcales; Cystobacterineae; Anaeromyxobacteraceae; Anaeromyxobacter MKTRRRAFTAAASALALSLLPLAASAADAWVASATEKIRPDAKARPQTEAHLSAARNEFA WP_012525440.1 hypothetical protein [Anaeromyxobacter sp. K] Length: 609\nNumber of predicted TMHs: 1\nExp number of AAs in TMHs: 39.47811\nExp number, first 60 AAs: 17.42459\nTotal prob of N-in: 0.79771\nPOSSIBLE N-term signal sequence\noutside 1 570\nTMhelix 571 593\ninside 594 609

1610 GCF_000964525.1_ASM96452v1 Anaeromyxobacter sp. PSR-1 Proteobacteria; delta/epsilon subdivisions; Deltaproteobacteria; Myxococcales; Cystobacterineae; Anaeromyxobacteraceae; Anaeromyxobacter MKTRRRAFTAAAVALALSLLPLAASAAEAWVASATEKIRPDAKARPQTEAHLAAARNEFA WP_059439524.1 hypothetical protein [Anaeromyxobacter sp. PSR-1] Length: 607\nNumber of predicted TMHs: 2\nExp

number of AAs in TMHs: 40.08595\nExp number, first 60 AAs: 20.842\nTotal prob of N-in: 0.95021\nPOSSIBLE N-term signal sequence\ninside 1 6\nTMhelix 7 29\noutside 30 568\nTMhelix 569 591\ninside 592 607

1611 GCF_000022145.1_ASM2214v1 Anaeromyxobacter dehalogenans 2CP-1 Proteobacteria; delta/epsilon subdivisions; Deltaproteobacteria; Myxococcales; Cystobacterineae; Anaeromyxobacteraceae; Anaeromyxobacter; Anaeromyxobacter dehalogenans MKTRRRAFTAAASALALSLLPLTASAADAWVASATEKIRPDAKARPQTEAHLSAARNEFA WP_012632768.1 hypothetical protein [Anaeromyxobacter dehalogenans] Length: 609\nNumber of predicted TMHs: 1\nExp number of AAs in TMHs: 37.26036\nExp number, first 60 AAs: 15.23966\nTotal prob of N-in: 0.69799\nPOSSIBLE N-term signal sequence\noutside 1 570\nTMhelix 571 593\ninside 594 609

1612 GCF_000255295.1_ASM25529v1 Corallococcus coralloides DSM 2259 Proteobacteria; delta/epsilon subdivisions; Deltaproteobacteria; Myxococcales; Cystobacterineae; Myxococcaceae; Corallococcus; Corallococcus coralloides MTRRTPVFAAPLLALLFLTGCSIELQHELTEADANEIYVLLSKNGINAKKEKAEGGNEVR WP_014398427.1 type III secretion system protein [Corallococcus coralloides] Length: 272\nNumber of predicted TMHs: 1\nExp number of AAs in TMHs: 23.34225\nExp number, first 60 AAs: 1.30527\nTotal prob of N-in: 0.90075\ninside 1 225\nTMhelix 226 248\noutside 249 272

1613 GCF_900106535.1_IMG-taxon_2693429903_annotated_assembly Myxococcus xanthus Proteobacteria; delta/epsilon subdivisions; Deltaproteobacteria; Myxococcales; Cystobacterineae; Myxococcaceae; Myxococcus MARPVEPAPLEGPVRFGPYTLVRRIGAGGMGEVFLAREEAPRRACVVKKVLPQLMASPQF WP_020478435.1 serine/threonine protein kinase [Myxococcus xanthus] Length: 663\nNumber of predicted TMHs: 1\nExp number of AAs in TMHs: 18.35093\nExp number, first 60 AAs: 0.00078\nTotal prob of N-in: 0.00103\noutside 1 641\nTMhelix 642 662\ninside 663 663

1614 GCF_000219105.1_ASM21910v1 Myxococcus fulvus HW-1 Proteobacteria; delta/epsilon subdivisions; Deltaproteobacteria; Myxococcales; Cystobacterineae; Myxococcaceae; Myxococcus; Myxococcus fulvus MARPVEPAPLEGPVRFGPYTLARRIGAGGMGEVFLAREESPRRACVVKKVLPQLMASPQF WP_082207061.1 serine/threonine protein kinase [Myxococcus fulvus] Length: 684\nNumber of predicted TMHs: 1\nExp number of AAs in TMHs: 20.53272\nExp number, first 60 AAs: 0.00058\nTotal prob of N-in: 0.00295\noutside 1 660\nTMhelix 661 683\ninside 684 684

1615 GCF_000331735.1_ASM33173v1 Myxococcus stipitatus DSM 14675 Proteobacteria; delta/epsilon subdivisions; Deltaproteobacteria; Myxococcales; Cystobacterineae; Myxococcaceae; Myxococcus; Myxococcus stipitatus MARPVEPEPLAGPVRFGPYTLVRRIGAGGMGEVFLAREESPRRACVVKKVLPQLMQSPQF WP_015346040.1 serine/threonine protein kinase [Myxococcus stipitatus] Length: 598\nNumber of predicted TMHs: 1\nExp number of AAs in TMHs: 22.75862\nExp number, first 60 AAs: 0.0009\nTotal prob of N-in: 0.00057\noutside 1 574\nTMhelix 575 597\ninside 598 598

1616 GCF_000012685.1_ASM1268v1 Myxococcus xanthus DK 1622 Proteobacteria; delta/epsilon subdivisions; Deltaproteobacteria; Myxococcales; Cystobacterineae; Myxococcaceae; Myxococcus; Myxococcus xanthus MARPVEPAPLEGPVRFGPYTLVRRIGAGGMGEVFLAREEAPRRACVVKKVLPQLMASPQF WP_020478435.1 serine/threonine protein kinase [Myxococcus xanthus] Length: 663\nNumber of predicted TMHs: 1\nExp number of AAs in TMHs: 18.35093\nExp number, first 60 AAs: 0.00078\nTotal prob of N-in: 0.00103\noutside 1 641\nTMhelix 642 662\ninside 663 663

1617 GCF_000278585.1_ASM27858v2 Myxococcus xanthus DZ2 Proteobacteria; delta/epsilon subdivisions; Deltaproteobacteria; Myxococcales; Cystobacterineae; Myxococcaceae; Myxococcus; Myxococcus xanthus MARPVEPAPLEGPVRFGPYTLVRRIGAGGMGEVFLAREEAPRRACVVKKVLPQLMASPQF WP_020478435.1 serine/threonine protein kinase [Myxococcus xanthus] Length: 663\nNumber of predicted TMHs: 1\nExp number of AAs in TMHs: 18.35093\nExp number, first 60 AAs: 0.00078\nTotal prob of N-in: 0.00103\noutside 1 641\nTMhelix 642 662\ninside 663 663

1618 GCF_000340515.1_Myxococcus_xanthus_DZF1 Myxococcus xanthus DZF1 Proteobacteria; delta/epsilon subdivisions; Deltaproteobacteria; Myxococcales; Cystobacterineae; Myxococcaceae; Myxococcus; Myxococcus xanthus MARPVEPAPLEGPVRFGPYTLVRRIGAGGMGEVFLAREEAPRRACVVKKVLPQLMASPQF WP_020478435.1 serine/threonine protein kinase [Myxococcus xanthus] Length: 663\nNumber of predicted TMHs: 1\nExp number of AAs in TMHs: 18.35093\nExp number, first 60 AAs: 0.00078\nTotal prob of N-in: 0.00103\noutside 1 641\nTMhelix 642 662\ninside 663 663

1619 GCF_001189295.1_ASM118929v1 Chondromyces crocatus Proteobacteria; delta/epsilon subdivisions; Deltaproteobacteria; Myxococcales; Sorangiineae; Polyangiaceae; Chondromyces
MTEPTHPMTEPTPCRSERARESTRSRRGPFIAAALFTGLCLATVPAAAQPAERSATTRDP WP_050435163.1 hypothetical protein [Chondromyces crocatus] Length: 360\nNumber of predicted TMHs: 2\nExp number of AAs in TMHs: 61.70389\nExp number, first 60 AAs: 15.91983\nTotal prob of N-in: 0.39737\nPOSSIBLE N-term signal sequence\ninside 1 253\nTMhelix 254 276\noutside 277 309\nTMhelix 310 332\ninside 333 360

1620 GCF_001189295.1_ASM118929v1 Chondromyces crocatus Proteobacteria; delta/epsilon subdivisions; Deltaproteobacteria; Myxococcales; Sorangiineae; Polyangiaceae; Chondromyces
MRRNLALRFWRVHIASSSPLLHVVSTRRSPLGSRGALALALGLGVWLAPSGAEADVLPPL WP_050434613.1 hypothetical protein [Chondromyces crocatus] Length: 202\nNumber of predicted TMHs: 1\nExp number of AAs in TMHs: 25.60587\nExp number, first 60 AAs: 5.74653\nTotal prob of N-in: 0.35415\noutside 1 163\nTMhelix 164 186\ninside 187 202

1621 GCF_000601485.1_Chondromyces_apiculatus_DSM_436_genome_assembly Chondromyces apiculatus DSM 436 Proteobacteria; delta/epsilon subdivisions; Deltaproteobacteria; Myxococcales; Sorangiineae; Polyangiaceae; Chondromyces; Chondromyces apiculatus
MRRRAAVLAVLSLALLSLSSAHAQQPPPPPAAAAQPPAGAAQPSVDAAQPPPGELRVRAS WP_081864772.1 hypothetical protein [Chondromyces apiculatus] Length: 232\nNumber of predicted TMHs: 2\nExp number of AAs in TMHs: 50.35529\nExp number, first 60 AAs: 4.98433\nTotal prob of N-in: 0.84893\ninside 1 161\nTMhelix 162 184\noutside 185 193\nTMhelix 194 216\ninside 217 232

1622 GCF_000601485.1_Chondromyces_apiculatus_DSM_436_genome_assembly Chondromyces apiculatus DSM 436 Proteobacteria; delta/epsilon subdivisions; Deltaproteobacteria; Myxococcales; Sorangiineae; Polyangiaceae; Chondromyces; Chondromyces apiculatus
MSLPRRRPTIVAILVLSLATASTSGAMAQQPAVRAPSPADKALAEALFLDGRKRMDAGEI WP_052374220.1 hypothetical protein [Chondromyces apiculatus] Length: 405\nNumber of predicted TMHs: 2\nExp number of AAs in TMHs: 48.53656\nExp number, first 60 AAs: 3.02623\nTotal prob of N-in: 0.77669\ninside 1 294\nTMhelix 295 317\noutside 318 350\nTMhelix 351 373\ninside 374 405

1623 GCF_000601485.1_Chondromyces_apiculatus_DSM_436_genome_assembly Chondromyces apiculatus DSM 436 Proteobacteria; delta/epsilon subdivisions; Deltaproteobacteria; Myxococcales; Sorangiineae; Polyangiaceae; Chondromyces; Chondromyces apiculatus
MNTRRRLFQAATLLLAGVIGFFVMLRSPEPAHGQGIMARWGTVGIETEEERQLFWSLLCM WP_044245686.1 cytochrome c biogenesis protein [Chondromyces apiculatus] Length: 184\nNumber of predicted TMHs: 2\nExp number of AAs in TMHs: 40.43933\nExp number, first 60 AAs: 19.30881\nTotal prob of N-in: 0.99853\nPOSSIBLE N-term signal sequence\ninside 1 6\nTMhelix 7 25\noutside 26 115\nTMhelix 116 138\ninside 139 184

1624 GCF_000418325.1_ASM41832v1 Sorangium cellulosum So0157-2 Proteobacteria; delta/epsilon subdivisions; Deltaproteobacteria; Myxococcales; Sorangiineae; Polyangiaceae; Sorangium; Sorangium cellulosum
MMQAHFRCARPRAAAARRRVVAWALGLAAAAPAPSVAAEPPAPPAAAAPSAVAEPPAPSV WP_049949682.1 hypothetical protein [Sorangium cellulosum] Length: 263\nNumber of predicted TMHs: 2\nExp number of AAs in TMHs: 54.61175\nExp number, first 60 AAs: 0.77856\nTotal prob of N-in: 0.99840\ninside 1 161\nTMhelix 162 184\noutside 185 198\nTMhelix 199 221\ninside 222 263

1625 GCF_000067165.1_ASM6716v1 Sorangium cellulosum So ce56 Proteobacteria; delta/epsilon subdivisions; Deltaproteobacteria; Myxococcales; Sorangiineae; Polyangiaceae; Sorangium; Sorangium cellulosum
MLCHRRLSSPCPASGCAAAVSRAPLAGAGRRRAVALLSALAALASLAPAAGAAEPTREER WP_012237712.1 hypothetical protein [Sorangium cellulosum] Length: 386\nNumber of predicted TMHs: 2\nExp number of AAs in TMHs: 48.66073\nExp number, first 60 AAs: 3.22438\nTotal prob of N-in: 0.27017\noutside 1 265\nTMhelix 266 288\ninside 289 324\nTMhelix 325 347\noutside 348 386

1626 GCF_000737325.1_ASM73732v2 Sandaracinus amylolyticus Proteobacteria; delta/epsilon subdivisions; Deltaproteobacteria; Myxococcales; Sorangiineae; Sandaracinaceae; Sandaracinus
MPGSGARRSVLAAVVIACAAALPARASAADAIVLSIGGDAPDATAREARDAVIAALSGDG WP_053234517.1 hypothetical protein [Sandaracinus amylolyticus] Length: 298\nNumber of predicted TMHs: 2\nExp number of AAs in TMHs: 51.3128\nExp number, first 60 AAs: 5.98483\nTotal prob of N-in: 0.31983\noutside 1 206\nTMhelix 207 229\ninside 230 249\nTMhelix 250 272\noutside 273 298

1627 GCF_000737325.1_ASM73732v2 Sandaracinus amylolyticus Proteobacteria; delta/epsilon subdivisions; Deltaproteobacteria; Myxococcales; Sorangiineae; Sandaracinaceae; Sandaracinus

MAVADMERRGFLTTVALGATGIGTGAVWLPGCGGAPTRPTDLATAHAETLAERLDRGVAS WP_053231613.1 hypothetical protein [Sandaracinus amylolyticus] Length: 330\nNumber of predicted TMHs: 2\nExp number of AAs in TMHs: 60.03699\nExp number, first 60 AAs: 14.85398\nTotal prob of N-in: 0.45020\nPOSSIBLE N-term signal sequence\noutside 1 275\nTMhelix 276 298\ninside 299 304\nTMhelix 305 327\noutside 328 330

1628 GCF_000737325.1_ASM73732v2 Sandaracinus amylolyticus Proteobacteria; delta/epsilon subdivisions; Deltaproteobacteria; Myxococcales; Sorangiineae; Sandaracinaceae; Sandaracinus MDRRRRGTLLASALIAAACALAPTTRASAQPPQGGGGGSESMSQATTTEGDMNDERARGA WP_053230789.1 hypothetical protein [Sandaracinus amylolyticus] Length: 292\nNumber of predicted TMHs: 2\nExp number of AAs in TMHs: 46.887\nExp number, first 60 AAs: 3.68663\nTotal prob of N-in: 0.92486\ninside 1 183\nTMhelix 184 206\noutside 207 239\nTMhelix 240 262\ninside 263 292

1629 GCF_000737325.1_ASM73732v2 Sandaracinus amylolyticus Proteobacteria; delta/epsilon subdivisions; Deltaproteobacteria; Myxococcales; Sorangiineae; Sandaracinaceae; Sandaracinus MEVEMDTRRLGTRRDPVALTLLRVVAGVTMAAHGWQKVEGFDGWRDTVASMGVPAADVLA WP_053235340.1 hypothetical protein [Sandaracinus amylolyticus] Length: 162\nNumber of predicted TMHs: 2\nExp number of AAs in TMHs: 75.15936\nExp number, first 60 AAs: 20.56567\nTotal prob of N-in: 0.93302\nPOSSIBLE N-term signal sequence\ninside 1 71\nTMhelix 72 94\noutside 95 108\nTMhelix 109 131\ninside 132 162

1630 GCF_000266945.1_ASM26694v1 Desulfomonile tiedjei DSM 6799 Proteobacteria; delta/epsilon subdivisions; Deltaproteobacteria; Syntrophobacterales; Syntrophaceae; Desulfomonile; Desulfomonile tiedjei MNADRRNFLKMLAGTGMALGFGLEGNALADVASLRGYPDQFGVLVDTTVCIGCRRCEWAC WP_014813017.1 Fe-S-cluster-containing hydrogenase subunit [Desulfomonile tiedjei] Length: 312\nNumber of predicted TMHs: 1\nExp number of AAs in TMHs: 30.4455\nExp number, first 60 AAs: 3.05155\nTotal prob of N-in: 0.47444\noutside 1 267\nTMhelix 268 290\ninside 291 312

1631 GCF_000013405.1_ASM1340v1 Syntrophus aciditrophicus SB Proteobacteria; delta/epsilon subdivisions; Deltaproteobacteria; Syntrophobacterales; Syntrophaceae; Syntrophus; Syntrophus aciditrophicus MEAEKISKISRRSLLKASTGALAIPGLAFPGTVQADVKEQLATLIDLSRCNGCVDLAKPS WP_011418038.1 (Fe-S)-binding protein [Syntrophus aciditrophicus] Length: 336\nNumber of predicted TMHs: 1\nExp number of AAs in TMHs: 25.14155\nExp number, first 60 AAs: 5.6359\nTotal prob of N-in: 0.28614\noutside 1 301\nTMhelix 302 321\ninside 322 336

1632 GCF_900176285.1_IMG-taxon_2585428172_annotated_assembly Desulfacinum hydrothermale DSM 13146 Proteobacteria; delta/epsilon subdivisions; Deltaproteobacteria; Syntrophobacterales; Syntrophobacteraceae; Desulfacinum; Desulfacinum hydrothermale MVMPKQKKARRSLVYGSNALVSTLVFLAILVFVALIAQRHPLRVDLTESGEFSLSEQSRK WP_084057466.1 hypothetical protein [Desulfacinum hydrothermale] Length: 488\nNumber of predicted TMHs: 2\nExp number of AAs in TMHs: 44.80881\nExp number, first 60 AAs: 22.9426\nTotal prob of N-in: 0.99977\nPOSSIBLE N-term signal sequence\ninside 1 12\nTMhelix 13 35\noutside 36 459\nTMhelix 460 482\ninside 483 488

1633 GCF_900129305.1_IMG-taxon_2582580750_annotated_assembly Desulfacinum infernum DSM 9756 Proteobacteria; delta/epsilon subdivisions; Deltaproteobacteria; Syntrophobacterales; Syntrophobacteraceae; Desulfacinum; Desulfacinum infernum MVMPKQKRARRSLVYGSNTFVSILVFLAILVFVALIAERHPLRVDLTDSGEFSLSEQSRK WP_073038219.1 hypothetical protein [Desulfacinum infernum] Length: 494\nNumber of predicted TMHs: 2\nExp number of AAs in TMHs: 45.04904\nExp number, first 60 AAs: 22.77221\nTotal prob of N-in: 0.99993\nPOSSIBLE N-term signal sequence\ninside 1 12\nTMhelix 13 35\noutside 36 466\nTMhelix 467 489\ninside 490 494

1634 GCF_000168275.1_ASM16827v1 delta proteobacterium MLMS-1 Proteobacteria; delta/epsilon subdivisions; Deltaproteobacteria; unclassified Deltaproteobacteria; unclassified Deltaproteobacteria (miscellaneous) MDENKLNNQGRRDFLRKLAILPVAGVCLQAGEAEASRREYVKLIDTTKCIGCHRCTSACDWP_007290909.1 twin-arginine translocation pathway signal protein [delta proteobacterium MLMS-1] Length: 291\nNumber of predicted TMHs: 1\nExp number of AAs in TMHs: 32.1649\nExp number, first 60 AAs: 0.04976\nTotal prob of N-in: 0.83970\ninside 1 254\nTMhelix 255 277\noutside 278 291

1635 GCF_000168275.1_ASM16827v1 delta proteobacterium MLMS-1 Proteobacteria; delta/epsilon subdivisions; Deltaproteobacteria; unclassified Deltaproteobacteria; unclassified Deltaproteobacteria (miscellaneous) MDENKLNNQGRRDFLRKLAILPVAGVCLQAGEAEASRREYVKLIDTTKCIGCHRCTSACDWP_007293159.1 twin-arginine translocation pathway signal protein [delta proteobacterium MLMS-1] Length: 291\nNumber of predicted TMHs: 1\nExp number of AAs in TMHs: 32.1649\nExp number, first 60 AAs: 0.04976\nTotal prob of N-in: 0.83970\ninside 1 254\nTMhelix 255 277\noutside 278 291

1636 GCF_000483025.1_ASM48302v1 delta proteobacterium PSCGC 5451 Proteobacteria; delta/epsilon subdivisions; Deltaproteobacteria; unclassified Deltaproteobacteria; unclassified Deltaproteobacteria (miscellaneous) MAQQISRRSFLKGGMAAAAGVAASSLPGTEALAGPDESKQLATLIDISKCVGCEACVDAC WP_027985824.1 4Fe-4S ferredoxin [delta proteobacterium PSCGC 5451] Length: 319\nNumber of predicted TMHs: 1\nExp number of AAs in TMHs: 21.95468\nExp number, first 60 AAs: 0.23888\nTotal prob of N-in: 0.46524\noutside 1 294\nTMhelix 295 317\ninside 318 319

1637 GCF_001762225.1_BCW_6447 Campylobacter coli Proteobacteria; delta/epsilon subdivisions; Epsilonproteobacteria; Campylobacterales; Campylobacteraceae; Campylobacter MIDYNQIESRLSALEKLPSLNPENSIPKALERSGFTRRDFMKWAGAMTAFLALPASFTPMWP_070241983.1 Ni/Fe hydrogenase [Campylobacter coli] Length: 379\nNumber of predicted TMHs: 1\nExp number of AAs in TMHs: 30.2914\nExp number, first 60 AAs: 13.0589\nTotal prob of N-in: 0.71245\nPOSSIBLE N-term signal sequence\noutside 1 352\nTMhelix 353 372\ninside 373 379

1638 GCF_001762245.1_BCW_6448 Campylobacter coli Proteobacteria; delta/epsilon subdivisions; Epsilonproteobacteria; Campylobacterales; Campylobacteraceae; Campylobacter MIDYNQIESRLSALEKLPSLNPENSIPKALERSGFTRRDFMKWAGAMTAFLALPASFTPMWP_070241983.1 Ni/Fe hydrogenase [Campylobacter coli] Length: 379\nNumber of predicted TMHs: 1\nExp number of AAs in TMHs: 30.2914\nExp number, first 60 AAs: 13.0589\nTotal prob of N-in: 0.71245\nPOSSIBLE N-term signal sequence\noutside 1 352\nTMhelix 353 372\ninside 373 379

1639 GCF_001763995.1_ASM176399v1 Campylobacter coli Proteobacteria; delta/epsilon subdivisions; Epsilonproteobacteria; Campylobacterales; Campylobacteraceae; Campylobacter MIDYNQIESRLSALEKLPSLNPENSIPKALERSGFTRRDFMKWAGAMTAFLALPASFTPMWP_070241983.1 Ni/Fe hydrogenase [Campylobacter coli] Length: 379\nNumber of predicted TMHs: 1\nExp number of AAs in TMHs: 30.2914\nExp number, first 60 AAs: 13.0589\nTotal prob of N-in: 0.71245\nPOSSIBLE N-term signal sequence\noutside 1 352\nTMhelix 353 372\ninside 373 379

1640 GCF_001764045.1_ASM176404v1 Campylobacter coli Proteobacteria; delta/epsilon subdivisions; Epsilonproteobacteria; Campylobacterales; Campylobacteraceae; Campylobacter MIDYNQIESRLSALEKLPSLNPENSIPKALERSGFTRRDFMKWAGAMTAFLALPASFTPMWP_070241983.1 Ni/Fe hydrogenase [Campylobacter coli] Length: 379\nNumber of predicted TMHs: 1\nExp number of AAs in TMHs: 30.2914\nExp number, first 60 AAs: 13.0589\nTotal prob of N-in: 0.71245\nPOSSIBLE N-term signal sequence\noutside 1 352\nTMhelix 353 372\ninside 373 379

1641 GCF_002165825.1_ASM216582v1 Campylobacter concisus Proteobacteria; delta/epsilon subdivisions; Epsilonproteobacteria; Campylobacterales; Campylobacteraceae; Campylobacter MNNDLRQKINRRLSELSALPKMKNDTSIAQLLKDKGFTRRDFMKWAGAMTAFMALPSAMT WP_087581625.1 Ni/Fe hydrogenase [Campylobacter concisus] Length: 381\nNumber of predicted TMHs: 2\nExp number of AAs in TMHs: 37.50523\nExp number, first 60 AAs: 14.40614\nTotal prob of N-in: 0.87349\nPOSSIBLE N-term signal sequence\ninside 1 41\nTMhelix 42 64\noutside 65 354\nTMhelix 355 377\ninside 378 381

1642 GCF_002165785.1_ASM216578v1 Campylobacter concisus Proteobacteria; delta/epsilon subdivisions; Epsilonproteobacteria; Campylobacterales; Campylobacteraceae; Campylobacter MNNDLRQKIDRRLSELSALPKMKSDSSIAQLLKEKGFTRRDFMKWAGAMTAFMALPSAMT WP_087584544.1 Ni/Fe hydrogenase [Campylobacter concisus] Length: 381\nNumber of predicted TMHs: 2\nExp number of AAs in TMHs: 34.1642\nExp number, first 60 AAs: 12.83657\nTotal prob of N-in: 0.82515\nPOSSIBLE N-term signal sequence\ninside 1 41\nTMhelix 42 64\noutside 65 354\nTMhelix 355 377\ninside 378 381

1643 GCF_002165875.1_ASM216587v1 Campylobacter concisus Proteobacteria; delta/epsilon subdivisions; Epsilonproteobacteria; Campylobacterales; Campylobacteraceae; Campylobacter MNNDLRQKIDRRLSELSALPKMKSDSSIAQLLKEKGFTRRDFMKWAGAMTAFMALPSAMT WP_084041138.1 Ni/Fe hydrogenase [Campylobacter concisus] Length: 381\nNumber of predicted TMHs: 2\nExp number of AAs in TMHs: 34.18696\nExp number, first 60 AAs: 12.8353\nTotal prob of N-in: 0.82522\nPOSSIBLE N-term signal sequence\ninside 1 41\nTMhelix 42 64\noutside 65 354\nTMhelix 355 377\ninside 378 381

1644 GCF_001891085.1_ASM189108v1 Campylobacter concisus Proteobacteria; delta/epsilon subdivisions; Epsilonproteobacteria; Campylobacterales; Campylobacteraceae; Campylobacter MNNDLRQKIDRRLSELSALPKMKSDSSIAQLLKDKGFTRRDFMKWAGAMTAFMALPSAMT WP_072594279.1 Ni/Fe hydrogenase [Campylobacter concisus] Length: 381\nNumber of predicted TMHs: 2\nExp number of

AAs in TMHs: 34.71024\nExp number, first 60 AAs:  13.32009\nTotal prob of N-in:   0.83892\nPOSSIBLE N-term signal sequence\ninside   1   41\nTMhelix   42   64\noutside   65   354\nTMhelix   355   377\ninside   378   381

1645 GCF_002092855.1_ASM209285v1 Campylobacter concisus Proteobacteria; delta/epsilon subdivisions; Epsilonproteobacteria; Campylobacterales; Campylobacteraceae; Campylobacter MNNDLRQKIDRRLSELSALPKMKSDSSIAQLLKDKGFTRRDFMKWAGVMTAFMALPSAMT WP_084107888.1 Ni/Fe hydrogenase [Campylobacter concisus] Length: 381\nNumber of predicted TMHs:  2\nExp number of AAs in TMHs: 36.36419\nExp number, first 60 AAs:  14.86867\nTotal prob of N-in:   0.90101\nPOSSIBLE N-term signal sequence\ninside   1   41\nTMhelix   42   64\noutside   65   354\nTMhelix   355   377\ninside   378   381

1646 GCF_002165775.1_ASM216577v1 Campylobacter concisus Proteobacteria; delta/epsilon subdivisions; Epsilonproteobacteria; Campylobacterales; Campylobacteraceae; Campylobacter MNNDLRQKINRRLSELSALPKMKNDTSIAQLLKDKGFTRRDFMKWAGAMTAFMALPSAMT WP_087580127.1 Ni/Fe hydrogenase [Campylobacter concisus] Length: 381\nNumber of predicted TMHs:  2\nExp number of AAs in TMHs: 39.06839\nExp number, first 60 AAs:  13.87807\nTotal prob of N-in:   0.87899\nPOSSIBLE N-term signal sequence\ninside   1   41\nTMhelix   42   64\noutside   65   354\nTMhelix   355   377\ninside   378   381

1647 GCF_002165815.1_ASM216581v1 Campylobacter concisus Proteobacteria; delta/epsilon subdivisions; Epsilonproteobacteria; Campylobacterales; Campylobacteraceae; Campylobacter MNNDLRQKIDRRLSELSALPKMKNDTSIAQLLKDKGFTRRDFMKWAGAMTAFMALPSAMT WP_087583277.1 Ni/Fe hydrogenase [Campylobacter concisus] Length: 381\nNumber of predicted TMHs:  2\nExp number of AAs in TMHs: 36.98104\nExp number, first 60 AAs:  14.00642\nTotal prob of N-in:   0.84629\nPOSSIBLE N-term signal sequence\ninside   1   41\nTMhelix   42   64\noutside   65   354\nTMhelix   355   377\ninside   378   381

1648 GCF_002165895.1_ASM216589v1 Campylobacter concisus Proteobacteria; delta/epsilon subdivisions; Epsilonproteobacteria; Campylobacterales; Campylobacteraceae; Campylobacter MNNDLRQKIDRRLSELSALPKMKSDSSIAQLLKEKGFTRRDFMKWAGAMTAFMALPSAMT WP_087585940.1 Ni/Fe hydrogenase [Campylobacter concisus] Length: 381\nNumber of predicted TMHs:  2\nExp number of AAs in TMHs: 38.31154\nExp number, first 60 AAs:  12.62016\nTotal prob of N-in:   0.83976\nPOSSIBLE N-term signal sequence\ninside   1   41\nTMhelix   42   64\noutside   65   354\nTMhelix   355   377\ninside   378   381

1649 GCF_002092845.1_ASM209284v1 Campylobacter concisus Proteobacteria; delta/epsilon subdivisions; Epsilonproteobacteria; Campylobacterales; Campylobacteraceae; Campylobacter MNNDLRQKIDRRLSELSALPKMKSDSSIAQLLKEKGFTRRDFMKWAGAMTAFMALPSAMT WP_084041138.1 Ni/Fe hydrogenase [Campylobacter concisus] Length: 381\nNumber of predicted TMHs:  2\nExp number of AAs in TMHs: 34.18696\nExp number, first 60 AAs:  12.8353\nTotal prob of N-in:   0.82522\nPOSSIBLE N-term signal sequence\ninside   1   41\nTMhelix   42   64\noutside   65   354\nTMhelix   355   377\ninside   378   381

1650 GCF_001298465.1_ASM129846v1 Campylobacter concisus Proteobacteria; delta/epsilon subdivisions; Epsilonproteobacteria; Campylobacterales; Campylobacteraceae; Campylobacter MNNDLRQKIDRRLSELSALPKMKSDSSIAQLLKDKGFTRRDFMKWAGAMTAFMALPSAMT WP_021091342.1 hypothetical protein [Campylobacter concisus] Length: 381\nNumber of predicted TMHs:  2\nExp number of AAs in TMHs: 34.64315\nExp number, first 60 AAs:  13.34172\nTotal prob of N-in:   0.83884\nPOSSIBLE N-term signal sequence\ninside   1   41\nTMhelix   42   64\noutside   65   354\nTMhelix   355   377\ninside   378   381

1651 GCF_002114545.1_ASM211454v1 Campylobacter concisus Proteobacteria; delta/epsilon subdivisions; Epsilonproteobacteria; Campylobacterales; Campylobacteraceae; Campylobacter MNNDLRQKIDRRLSELSALPKMKSDSSIAQLLKEKGFTRRDFMKWAGAMTAFMALPSAMT WP_084041138.1 Ni/Fe hydrogenase [Campylobacter concisus] Length: 381\nNumber of predicted TMHs:  2\nExp number of AAs in TMHs: 34.18696\nExp number, first 60 AAs:  12.8353\nTotal prob of N-in:   0.82522\nPOSSIBLE N-term signal sequence\ninside   1   41\nTMhelix   42   64\noutside   65   354\nTMhelix   355   377\ninside   378   381

1652 GCF_002165855.1_ASM216585v1 Campylobacter concisus Proteobacteria; delta/epsilon subdivisions; Epsilonproteobacteria; Campylobacterales; Campylobacteraceae; Campylobacter MNNDLRQKIDRRLSELSALPKMKSDSSIAQLLKEKGFTRRDFMKWAGAMTAFMALPSAMT WP_084041138.1 Ni/Fe hydrogenase [Campylobacter concisus] Length: 381\nNumber of predicted TMHs:  2\nExp number of AAs in TMHs: 34.18696\nExp number, first 60 AAs:  12.8353\nTotal prob of N-in:   0.82522\nPOSSIBLE N-term signal sequence\ninside   1   41\nTMhelix   42   64\noutside   65   354\nTMhelix   355   377\ninside   378   381

1653 GCF_900116585.1_IMG-taxon_2687453784_annotated_assembly Campylobacter hyointestinalis Proteobacteria; delta/epsilon subdivisions; Epsilonproteobacteria; Campylobacterales; Campylobacteraceae;

Campylobacter MDRRSFFKFSAVAASSSLFATSNLEDKKIMSIIDLDLCDGCKNEPIPLCVKACRDKNEPN WP_064019789.1 oxidoreductase [Campylobacter hyointestinalis] Length: 310\nNumber of predicted TMHs: 1\nExp number of AAs in TMHs: 22.01747\nExp number, first 60 AAs: 0.07136\nTotal prob of N-in: 0.02459\noutside 1 281\nTMhelix 282 304\ninside 305 310

1654 GCF_001643995.1_ASM164399v1 Campylobacter iguaniorum Proteobacteria; delta/epsilon subdivisions; Epsilonproteobacteria; Campylobacterales; Campylobacteraceae; Campylobacter MVDHKLLNTIQDRLDALSKMPKIKENSSIMAVLKENGFTRRDFMKWAGMMTAAMALPASF WP_038454336.1 Ni/Fe hydrogenase [Campylobacter iguaniorum] Length: 383\nNumber of predicted TMHs: 1\nExp number of AAs in TMHs: 25.44254\nExp number, first 60 AAs: 3.95027\nTotal prob of N-in: 0.29028\noutside 1 356\nTMhelix 357 379\ninside 380 383

1655 GCF_000736415.1_ASM73641v1 Campylobacter iguaniorum Proteobacteria; delta/epsilon subdivisions; Epsilonproteobacteria; Campylobacterales; Campylobacteraceae; Campylobacter MVDHKLLNTIQDRLDALSKMPKIKENSSIMAVLKENGFTRRDFMKWAGMMTAAMALPASF WP_038454336.1 Ni/Fe hydrogenase [Campylobacter iguaniorum] Length: 383\nNumber of predicted TMHs: 1\nExp number of AAs in TMHs: 25.44254\nExp number, first 60 AAs: 3.95027\nTotal prob of N-in: 0.29028\noutside 1 356\nTMhelix 357 379\ninside 380 383

1656 GCF_001483985.1_ASM148398v1 Campylobacter iguaniorum Proteobacteria; delta/epsilon subdivisions; Epsilonproteobacteria; Campylobacterales; Campylobacteraceae; Campylobacter MVDHKLLNTIQDRLDALSKMPKIKENSSIMAVLKENGFTRRDFMKWAGMMTAAMALPASF WP_038454336.1 Ni/Fe hydrogenase [Campylobacter iguaniorum] Length: 383\nNumber of predicted TMHs: 1\nExp number of AAs in TMHs: 25.44254\nExp number, first 60 AAs: 3.95027\nTotal prob of N-in: 0.29028\noutside 1 356\nTMhelix 357 379\ninside 380 383

1657 GCF_001570705.1_ASM157070v1 Campylobacter jejuni Proteobacteria; delta/epsilon subdivisions; Epsilonproteobacteria; Campylobacterales; Campylobacteraceae; Campylobacter MIDYHQIESRLSALEKLPSLKDNDSIAKALEKSGFSRRDFMKWAGAMTAFLALPASFTPM WP_061210015.1 Ni/Fe hydrogenase [Campylobacter jejuni] Length: 379\nNumber of predicted TMHs: 1\nExp number of AAs in TMHs: 28.4713\nExp number, first 60 AAs: 9.84365\nTotal prob of N-in: 0.56300\noutside 1 352\nTMhelix 353 375\ninside 376 379

1658 GCF_000705255.1_Campylobacter_9_barcode Campylobacter mucosalis Proteobacteria; delta/epsilon subdivisions; Epsilonproteobacteria; Campylobacterales; Campylobacteraceae; Campylobacter MNQDVLIKINERLNWLSSLKSIDKGESITKALKNSGFSRRDFMKWAGAMTAFMALPASFT WP_034966984.1 Ni/Fe hydrogenase [Campylobacter mucosalis] Length: 382\nNumber of predicted TMHs: 1\nExp number of AAs in TMHs: 26.68071\nExp number, first 60 AAs: 7.81656\nTotal prob of N-in: 0.49285\noutside 1 354\nTMhelix 355 374\ninside 375 382

1659 GCF_000238755.1_Campylobacter_sp_10_1_50_V1 Campylobacter sp. 10_1_50 Proteobacteria; delta/epsilon subdivisions; Epsilonproteobacteria; Campylobacterales; Campylobacteraceae; Campylobacter MNNDLRQKINRRLSELSALPKMKNDTSIAQLLKDKGFTRRDFMKWAGAMTAFMALPSAMT WP_009293888.1 quinone-reactive Ni/Fe-hydrogenase small chain [Campylobacter sp. 10_1_50] Length: 381\nNumber of predicted TMHs: 2\nExp number of AAs in TMHs: 36.7937899999999\nExp number, first 60 AAs: 14.27418\nTotal prob of N-in: 0.87396\nPOSSIBLE N-term signal sequence\ninside 1 41\nTMhelix 42 64\noutside 65 354\nTMhelix 355 377\ninside 378 381

1660 GCF_000287855.1_CspFOBRC14v1.0 Campylobacter sp. FOBRC14 Proteobacteria; delta/epsilon subdivisions; Epsilonproteobacteria; Campylobacterales; Campylobacteraceae; Campylobacter MVHNIHEKANRRLNELSLLPKIKENGTIAKILEQNGFTRRDFMKWAGAMTAFMALPASMT WP_009651157.1 MULTISPECIES: quinone-reactive Ni/Fe-hydrogenase small chain [Campylobacter] Length: 381\nNumber of predicted TMHs: 1\nExp number of AAs in TMHs: 33.75286\nExp number, first 60 AAs: 9.47284\nTotal prob of N-in: 0.64572\noutside 1 354\nTMhelix 355 377\ninside 378 381

1661 GCF_000762855.1_ASM76285v1 Campylobacter sp. MIT 97-5078 Proteobacteria; delta/epsilon subdivisions; Epsilonproteobacteria; Campylobacterales; Campylobacteraceae; Campylobacter MTFKELEARLKELESLPSMKGNSSIPKALEEAGFSRRDFMKWAGAMTAFLALPASFAPMV WP_034905116.1 Ni/Fe hydrogenase [Campylobacter sp. MIT 97-5078] Length: 379\nNumber of predicted TMHs: 1\nExp number of AAs in TMHs: 23.36419\nExp number, first 60 AAs: 7.3363\nTotal prob of N-in: 0.27574\noutside 1 351\nTMhelix 352 371\ninside 372 379

1662 GCF_002139915.1_ASM213991v1 Campylobacter sp. NCTC 13003 Proteobacteria; delta/epsilon subdivisions; Epsilonproteobacteria; Campylobacterales; Campylobacteraceae; Campylobacter
MVDNHNITTIQNRLSELAKLPKMKANSSIMDVLNEHGFTRRDFMKWAGAMTALMALPASF WP_086302671.1
Ni/Fe hydrogenase [Campylobacter sp. NCTC 13003] Length: 383\nNumber of predicted TMHs: 2\nExp number of AAs in TMHs: 34.4757299999999\nExp number, first 60 AAs: 10.91641\nTotal prob of N-in: 0.76446\nPOSSIBLE N-term signal sequence\ninside 1 45\nTMhelix 46 68\noutside 69 356\nTMhelix 357 376\ninside 377 383

1663 GCF_002139465.1_ASM213946v1 Campylobacter sp. P0021 Proteobacteria; delta/epsilon subdivisions; Epsilonproteobacteria; Campylobacterales; Campylobacteraceae; Campylobacter
MVDNHNITTIQNRLSELAKLPKMKANSSIMDVLNEHGFTRRDFMKWAGAMTALMALPASF WP_086229493.1
Ni/Fe hydrogenase [Campylobacter sp. P0088] Length: 383\nNumber of predicted TMHs: 2\nExp number of AAs in TMHs: 34.4668299999999\nExp number, first 60 AAs: 10.91725\nTotal prob of N-in: 0.76446\nPOSSIBLE N-term signal sequence\ninside 1 45\nTMhelix 46 68\noutside 69 356\nTMhelix 357 376\ninside 377 383

1664 GCF_002139455.1_ASM213945v1 Campylobacter sp. P0023 Proteobacteria; delta/epsilon subdivisions; Epsilonproteobacteria; Campylobacterales; Campylobacteraceae; Campylobacter
MVDNHNITTIQNRLSELAKLPKMKANSSIMDVLNEHGFTRRDFMKWAGAMTALMALPASF WP_086229493.1
Ni/Fe hydrogenase [Campylobacter sp. P0088] Length: 383\nNumber of predicted TMHs: 2\nExp number of AAs in TMHs: 34.4668299999999\nExp number, first 60 AAs: 10.91725\nTotal prob of N-in: 0.76446\nPOSSIBLE N-term signal sequence\ninside 1 45\nTMhelix 46 68\noutside 69 356\nTMhelix 357 376\ninside 377 383

1665 GCF_002139775.1_ASM213977v1 Campylobacter sp. P0024 Proteobacteria; delta/epsilon subdivisions; Epsilonproteobacteria; Campylobacterales; Campylobacteraceae; Campylobacter
MVDNHNVTTIQNRLSELAKLPKMKANSSIMDVLNEHGFTRRDFMKWAGAMTALMALPASF WP_086293345.1
Ni/Fe hydrogenase [Campylobacter sp. P0024] Length: 383\nNumber of predicted TMHs: 2\nExp number of AAs in TMHs: 35.8555599999999\nExp number, first 60 AAs: 11.0779\nTotal prob of N-in: 0.76233\nPOSSIBLE N-term signal sequence\ninside 1 45\nTMhelix 46 68\noutside 69 356\nTMhelix 357 379\ninside 380 383

1666 GCF_002139075.1_ASM213907v1 Campylobacter sp. P0078 Proteobacteria; delta/epsilon subdivisions; Epsilonproteobacteria; Campylobacterales; Campylobacteraceae; Campylobacter
MVDNHNVTTIQNRLSELAKLPKMKSNSSIMEVLNEHGFTRRDFMKWAGAMTALMALPASF WP_086237247.1
Ni/Fe hydrogenase [Campylobacter sp. P0078] Length: 383\nNumber of predicted TMHs: 2\nExp number of AAs in TMHs: 34.40876\nExp number, first 60 AAs: 11.13559\nTotal prob of N-in: 0.76324\nPOSSIBLE N-term signal sequence\ninside 1 45\nTMhelix 46 68\noutside 69 356\nTMhelix 357 376\ninside 377 383

1667 GCF_002139495.1_ASM213949v1 Campylobacter sp. P0085 Proteobacteria; delta/epsilon subdivisions; Epsilonproteobacteria; Campylobacterales; Campylobacteraceae; Campylobacter
MVDNHNITTIQNRLSELAKLPKMKANSSIMDVLNEHGFTRRDFMKWAGAMTALMALPASF WP_086253088.1
Ni/Fe hydrogenase [Campylobacter sp. P091] Length: 383\nNumber of predicted TMHs: 2\nExp number of AAs in TMHs: 34.46748\nExp number, first 60 AAs: 10.91753\nTotal prob of N-in: 0.76445\nPOSSIBLE N-term signal sequence\ninside 1 45\nTMhelix 46 68\noutside 69 356\nTMhelix 357 376\ninside 377 383

1668 GCF_002139505.1_ASM213950v1 Campylobacter sp. P0087 Proteobacteria; delta/epsilon subdivisions; Epsilonproteobacteria; Campylobacterales; Campylobacteraceae; Campylobacter
MVDNHNITTIQNRLSELAKLPKMKANSSIMDVLNEHGFTRRDFMKWAGAMTALMALPASF WP_086282549.1
Ni/Fe hydrogenase [Campylobacter sp. P0087] Length: 383\nNumber of predicted TMHs: 2\nExp number of AAs in TMHs: 35.12412\nExp number, first 60 AAs: 10.8322\nTotal prob of N-in: 0.76531\nPOSSIBLE N-term signal sequence\ninside 1 45\nTMhelix 46 68\noutside 69 356\nTMhelix 357 376\ninside 377 383

1669 GCF_002138695.1_ASM213869v1 Campylobacter sp. P0088 Proteobacteria; delta/epsilon subdivisions; Epsilonproteobacteria; Campylobacterales; Campylobacteraceae; Campylobacter
MVDNHNITTIQNRLSELAKLPKMKANSSIMDVLNEHGFTRRDFMKWAGAMTALMALPASF WP_086229493.1
Ni/Fe hydrogenase [Campylobacter sp. P0088] Length: 383\nNumber of predicted TMHs: 2\nExp number of AAs in TMHs: 34.4668299999999\nExp number, first 60 AAs: 10.91725\nTotal prob of N-in: 0.76446\nPOSSIBLE N-term signal sequence\ninside 1 45\nTMhelix 46 68\noutside 69 356\nTMhelix 357 376\ninside 377 383

1670 GCF_002139535.1_ASM213953v1 Campylobacter sp. P0094 Proteobacteria; delta/epsilon subdivisions; Epsilonproteobacteria; Campylobacterales; Campylobacteraceae; Campylobacter
MVDNHNVTTIQNRLSELAKLPKMKANSSIMDVLNEHGFTRRDFMKWAGAMTALMALPASF WP_086277851.1
Ni/Fe hydrogenase [Campylobacter sp. P0094] Length: 383\nNumber of predicted TMHs: 1\nExp number of

AAs in TMHs: 32.5626\nExp number, first 60 AAs: 9.27044\nTotal prob of N-in: 0.68975\noutside 1 356\nTMhelix 357 376\ninside 377 383

1671 GCF_002138705.1_ASM213870v1 Campylobacter sp. P0098 Proteobacteria; delta/epsilon subdivisions; Epsilonproteobacteria; Campylobacterales; Campylobacteraceae; Campylobacter
MVDNHNITTIQNRLSELAKLPKMKANSSIMDVLNEHGFTRRDFMKWAGAMTALMALPASF WP_086224116.1
Ni/Fe hydrogenase [Campylobacter sp. P0098] Length: 383\nNumber of predicted TMHs: 2\nExp number of AAs in TMHs: 34.4668199999999\nExp number, first 60 AAs: 10.91724\nTotal prob of N-in: 0.76446\nPOSSIBLE N-term signal sequence\ninside 1 45\nTMhelix 46 68\noutside 69 356\nTMhelix 357 376\ninside 377 383

1672 GCF_002139545.1_ASM213954v1 Campylobacter sp. P0103 Proteobacteria; delta/epsilon subdivisions; Epsilonproteobacteria; Campylobacterales; Campylobacteraceae; Campylobacter
MVDNHNITTIQNRLSELAKLPKMKANSSIMDVLNEHGFTRRDFMKWAGAMTALMALPASF WP_086224116.1
Ni/Fe hydrogenase [Campylobacter sp. P0098] Length: 383\nNumber of predicted TMHs: 2\nExp number of AAs in TMHs: 34.4668199999999\nExp number, first 60 AAs: 10.91724\nTotal prob of N-in: 0.76446\nPOSSIBLE N-term signal sequence\ninside 1 45\nTMhelix 46 68\noutside 69 356\nTMhelix 357 376\ninside 377 383

1673 GCF_002139575.1_ASM213957v1 Campylobacter sp. P0106 Proteobacteria; delta/epsilon subdivisions; Epsilonproteobacteria; Campylobacterales; Campylobacteraceae; Campylobacter
MVDNHNITTIQNRLSELAKLPKMKANSSIMDVLNEHGFTRRDFMKWAGAMTALMALPASF WP_086227799.1
Ni/Fe hydrogenase [Campylobacter sp. P0109] Length: 383\nNumber of predicted TMHs: 2\nExp number of AAs in TMHs: 34.46784\nExp number, first 60 AAs: 10.9175\nTotal prob of N-in: 0.76445\nPOSSIBLE N-term signal sequence\ninside 1 45\nTMhelix 46 68\noutside 69 356\nTMhelix 357 376\ninside 377 383

1674 GCF_002139585.1_ASM213958v1 Campylobacter sp. P0107 Proteobacteria; delta/epsilon subdivisions; Epsilonproteobacteria; Campylobacterales; Campylobacteraceae; Campylobacter
MVDNHNITTIQNRLSELAKLPKMKANSSIMDVLNEHGFTRRDFMKWAGAMTALMALPASF WP_086229493.1
Ni/Fe hydrogenase [Campylobacter sp. P0088] Length: 383\nNumber of predicted TMHs: 2\nExp number of AAs in TMHs: 34.4668299999999\nExp number, first 60 AAs: 10.91725\nTotal prob of N-in: 0.76446\nPOSSIBLE N-term signal sequence\ninside 1 45\nTMhelix 46 68\noutside 69 356\nTMhelix 357 376\ninside 377 383

1675 GCF_002138715.1_ASM213871v1 Campylobacter sp. P0108 Proteobacteria; delta/epsilon subdivisions; Epsilonproteobacteria; Campylobacterales; Campylobacteraceae; Campylobacter
MVDNRNLSTIQNRLAELAKLPKMKANSSIMDVLNEHGFTRRDFMKWAGAMTALMALPASF WP_086226486.1
Ni/Fe hydrogenase [Campylobacter sp. P0108] Length: 383\nNumber of predicted TMHs: 1\nExp number of AAs in TMHs: 33.02325\nExp number, first 60 AAs: 10.03035\nTotal prob of N-in: 0.70017\nPOSSIBLE N-term signal sequence\noutside 1 356\nTMhelix 357 376\ninside 377 383

1676 GCF_002138755.1_ASM213875v1 Campylobacter sp. P0109 Proteobacteria; delta/epsilon subdivisions; Epsilonproteobacteria; Campylobacterales; Campylobacteraceae; Campylobacter
MVDNHNITTIQNRLSELAKLPKMKANSSIMDVLNEHGFTRRDFMKWAGAMTALMALPASF WP_086227799.1
Ni/Fe hydrogenase [Campylobacter sp. P0109] Length: 383\nNumber of predicted TMHs: 2\nExp number of AAs in TMHs: 34.46784\nExp number, first 60 AAs: 10.9175\nTotal prob of N-in: 0.76445\nPOSSIBLE N-term signal sequence\ninside 1 45\nTMhelix 46 68\noutside 69 356\nTMhelix 357 376\ninside 377 383

1677 GCF_002139615.1_ASM213961v1 Campylobacter sp. P0111 Proteobacteria; delta/epsilon subdivisions; Epsilonproteobacteria; Campylobacterales; Campylobacteraceae; Campylobacter
MVDNHNITTIQNRLSELAKLPKMKANSSIMDVLNEHGFTRRDFMKWAGAMTALMALPASF WP_086229493.1
Ni/Fe hydrogenase [Campylobacter sp. P0088] Length: 383\nNumber of predicted TMHs: 2\nExp number of AAs in TMHs: 34.4668299999999\nExp number, first 60 AAs: 10.91725\nTotal prob of N-in: 0.76446\nPOSSIBLE N-term signal sequence\ninside 1 45\nTMhelix 46 68\noutside 69 356\nTMhelix 357 376\ninside 377 383

1678 GCF_002139105.1_ASM213910v1 Campylobacter sp. P0124 Proteobacteria; delta/epsilon subdivisions; Epsilonproteobacteria; Campylobacterales; Campylobacteraceae; Campylobacter
MVDNHNVTTIQNRLSELAKLPKMKSNSSIMEVLNEHGFTRRDFMKWAGAMTALMALPASF WP_086237247.1
Ni/Fe hydrogenase [Campylobacter sp. P0078] Length: 383\nNumber of predicted TMHs: 2\nExp number of AAs in TMHs: 34.40876\nExp number, first 60 AAs: 11.13559\nTotal prob of N-in: 0.76324\nPOSSIBLE N-term signal sequence\ninside 1 45\nTMhelix 46 68\noutside 69 356\nTMhelix 357 376\ninside 377 383

1679 GCF_002139625.1_ASM213962v1 Campylobacter sp. P0132 Proteobacteria; delta/epsilon subdivisions; Epsilonproteobacteria; Campylobacterales; Campylobacteraceae; Campylobacter

MVDNHNITTIQNRLSELAKLPKMKANSSIMDVLNEHGFTRRDFMKWAGAMTALMALPASF WP_086234315.1 Ni/Fe hydrogenase [Campylobacter sp. P0140] Length: 383\nNumber of predicted TMHs: 2\nExp number of AAs in TMHs: 34.46783\nExp number, first 60 AAs: 10.9175\nTotal prob of N-in: 0.76445\nPOSSIBLE N-term signal sequence\ninside 1 45\nTMhelix 46 68\noutside 69 356\nTMhelix 357 376\ninside 377 383

1680 GCF_002139665.1_ASM213966v1 Campylobacter sp. P0134 Proteobacteria; delta/epsilon subdivisions; Epsilonproteobacteria; Campylobacterales; Campylobacteraceae; Campylobacter MVDNHNITTIQNRLSELAKLPKMKANSSIMDVLNEHGFTRRDFMKWAGAMTALMALPASF WP_086253088.1 Ni/Fe hydrogenase [Campylobacter sp. P091] Length: 383\nNumber of predicted TMHs: 2\nExp number of AAs in TMHs: 34.46748\nExp number, first 60 AAs: 10.91753\nTotal prob of N-in: 0.76445\nPOSSIBLE N-term signal sequence\ninside 1 45\nTMhelix 46 68\noutside 69 356\nTMhelix 357 376\ninside 377 383

1681 GCF_002138775.1_ASM213877v1 Campylobacter sp. P0135 Proteobacteria; delta/epsilon subdivisions; Epsilonproteobacteria; Campylobacterales; Campylobacteraceae; Campylobacter MVDNHNITTIQNRLSELAKLPKMKANSSIMDVLNEHGFTRRDFMKWAGAMTALMALPASF WP_086229493.1 Ni/Fe hydrogenase [Campylobacter sp. P0088] Length: 383\nNumber of predicted TMHs: 2\nExp number of AAs in TMHs: 34.4668299999999\nExp number, first 60 AAs: 10.91725\nTotal prob of N-in: 0.76446\nPOSSIBLE N-term signal sequence\ninside 1 45\nTMhelix 46 68\noutside 69 356\nTMhelix 357 376\ninside 377 383

1682 GCF_002138785.1_ASM213878v1 Campylobacter sp. P0136 Proteobacteria; delta/epsilon subdivisions; Epsilonproteobacteria; Campylobacterales; Campylobacteraceae; Campylobacter MVDNHNITTIQNRLSELAKLPKMKANSSIMDVLNEHGFTRRDFMKWAGAMTALMALPASF WP_086224116.1 Ni/Fe hydrogenase [Campylobacter sp. P0098] Length: 383\nNumber of predicted TMHs: 2\nExp number of AAs in TMHs: 34.4668199999999\nExp number, first 60 AAs: 10.91724\nTotal prob of N-in: 0.76446\nPOSSIBLE N-term signal sequence\ninside 1 45\nTMhelix 46 68\noutside 69 356\nTMhelix 357 376\ninside 377 383

1683 GCF_002138795.1_ASM213879v1 Campylobacter sp. P0138 Proteobacteria; delta/epsilon subdivisions; Epsilonproteobacteria; Campylobacterales; Campylobacteraceae; Campylobacter MVDNHNITTIQNRLSELAKLPKMKANSSIMDVLNEHGFTRRDFMKWAGAMTALMALPASF WP_086224116.1 Ni/Fe hydrogenase [Campylobacter sp. P0098] Length: 383\nNumber of predicted TMHs: 2\nExp number of AAs in TMHs: 34.4668199999999\nExp number, first 60 AAs: 10.91724\nTotal prob of N-in: 0.76446\nPOSSIBLE N-term signal sequence\ninside 1 45\nTMhelix 46 68\noutside 69 356\nTMhelix 357 376\ninside 377 383

1684 GCF_002139655.1_ASM213965v1 Campylobacter sp. P0139 Proteobacteria; delta/epsilon subdivisions; Epsilonproteobacteria; Campylobacterales; Campylobacteraceae; Campylobacter MVDNHNITTIQNRLSELAKLPKMKANSSIMDVLNEHGFTRRDFMKWAGAMTALMALPASF WP_086224116.1 Ni/Fe hydrogenase [Campylobacter sp. P0098] Length: 383\nNumber of predicted TMHs: 2\nExp number of AAs in TMHs: 34.4668199999999\nExp number, first 60 AAs: 10.91724\nTotal prob of N-in: 0.76446\nPOSSIBLE N-term signal sequence\ninside 1 45\nTMhelix 46 68\noutside 69 356\nTMhelix 357 376\ninside 377 383

1685 GCF_002138835.1_ASM213883v1 Campylobacter sp. P0140 Proteobacteria; delta/epsilon subdivisions; Epsilonproteobacteria; Campylobacterales; Campylobacteraceae; Campylobacter MVDNHNITTIQNRLSELAKLPKMKANSSIMDVLNEHGFTRRDFMKWAGAMTALMALPASF WP_086234315.1 Ni/Fe hydrogenase [Campylobacter sp. P0140] Length: 383\nNumber of predicted TMHs: 2\nExp number of AAs in TMHs: 34.46783\nExp number, first 60 AAs: 10.9175\nTotal prob of N-in: 0.76445\nPOSSIBLE N-term signal sequence\ninside 1 45\nTMhelix 46 68\noutside 69 356\nTMhelix 357 376\ninside 377 383

1686 GCF_002139695.1_ASM213969v1 Campylobacter sp. P0187 Proteobacteria; delta/epsilon subdivisions; Epsilonproteobacteria; Campylobacterales; Campylobacteraceae; Campylobacter MVDNHNITTIQNRLSELAKLPKMKANSSIMDVLNEHGFTRRDFMKWAGAMTALMALPASF WP_086224116.1 Ni/Fe hydrogenase [Campylobacter sp. P0098] Length: 383\nNumber of predicted TMHs: 2\nExp number of AAs in TMHs: 34.4668199999999\nExp number, first 60 AAs: 10.91724\nTotal prob of N-in: 0.76446\nPOSSIBLE N-term signal sequence\ninside 1 45\nTMhelix 46 68\noutside 69 356\nTMhelix 357 376\ninside 377 383

1687 GCF_002138895.1_ASM213889v1 Campylobacter sp. P0209 Proteobacteria; delta/epsilon subdivisions; Epsilonproteobacteria; Campylobacterales; Campylobacteraceae; Campylobacter MVDNHNITTIQNRLSELAKLPKMKANSSIMDVLNEHGFTRRDFMKWAGAMTALMALPASF WP_086224116.1 Ni/Fe hydrogenase [Campylobacter sp. P0098] Length: 383\nNumber of predicted TMHs: 2\nExp number of AAs in TMHs: 34.4668199999999\nExp number, first 60 AAs: 10.91724\nTotal prob of N-in: 0.76446\nPOSSIBLE N-term signal sequence\ninside 1 45\nTMhelix 46 68\noutside 69 356\nTMhelix 357 376\ninside 377 383

1688 GCF_002138855.1_ASM213885v1 Campylobacter sp. P0222 Proteobacteria; delta/epsilon subdivisions; Epsilonproteobacteria; Campylobacterales; Campylobacteraceae; Campylobacter
MVDNHNITTIQNRLSELAKLPKMKANSSIMDVLNEHGFTRRDFMKWAGAMTALMALPASF WP_086240242.1
Ni/Fe hydrogenase [Campylobacter sp. P0222] Length: 383\nNumber of predicted TMHs: 2\nExp number of AAs in TMHs: 34.70972\nExp number, first 60 AAs: 10.91593\nTotal prob of N-in: 0.76526\nPOSSIBLE N-term signal sequence\ninside 1 45\nTMhelix 46 68\noutside 69 356\nTMhelix 357 376\ninside 377 383

1689 GCF_002138865.1_ASM213886v1 Campylobacter sp. P0227 Proteobacteria; delta/epsilon subdivisions; Epsilonproteobacteria; Campylobacterales; Campylobacteraceae; Campylobacter
MVDNHNITTIQNRLSELAKLPKMKANSSIMDVLNEHGFTRRDFMKWAGAMTALMALPASF WP_086243030.1
Ni/Fe hydrogenase [Campylobacter sp. P0227] Length: 383\nNumber of predicted TMHs: 2\nExp number of AAs in TMHs: 34.4757499999999\nExp number, first 60 AAs: 10.91641\nTotal prob of N-in: 0.76446\nPOSSIBLE N-term signal sequence\ninside 1 45\nTMhelix 46 68\noutside 69 356\nTMhelix 357 376\ninside 377 383

1690 GCF_002138915.1_ASM213891v1 Campylobacter sp. P031 Proteobacteria; delta/epsilon subdivisions; Epsilonproteobacteria; Campylobacterales; Campylobacteraceae; Campylobacter
MVDNHNITTIQNRLSELAKLPKMKANSSIMDVLNEHGFTRRDFMKWAGAMTALMALPASF WP_086224116.1
Ni/Fe hydrogenase [Campylobacter sp. P0098] Length: 383\nNumber of predicted TMHs: 2\nExp number of AAs in TMHs: 34.4668199999999\nExp number, first 60 AAs: 10.91724\nTotal prob of N-in: 0.76446\nPOSSIBLE N-term signal sequence\ninside 1 45\nTMhelix 46 68\noutside 69 356\nTMhelix 357 376\ninside 377 383

1691 GCF_002139705.1_ASM213970v1 Campylobacter sp. P090 Proteobacteria; delta/epsilon subdivisions; Epsilonproteobacteria; Campylobacterales; Campylobacteraceae; Campylobacter
MVDNHNITTIQNRLSELAKLPKMKANSSIMDVLNEHGFTRRDFMKWAGAMTALMALPASF WP_086224116.1
Ni/Fe hydrogenase [Campylobacter sp. P0098] Length: 383\nNumber of predicted TMHs: 2\nExp number of AAs in TMHs: 34.4668199999999\nExp number, first 60 AAs: 10.91724\nTotal prob of N-in: 0.76446\nPOSSIBLE N-term signal sequence\ninside 1 45\nTMhelix 46 68\noutside 69 356\nTMhelix 357 376\ninside 377 383

1692 GCF_002138935.1_ASM213893v1 Campylobacter sp. P091 Proteobacteria; delta/epsilon subdivisions; Epsilonproteobacteria; Campylobacterales; Campylobacteraceae; Campylobacter
MVDNHNITTIQNRLSELAKLPKMKANSSIMDVLNEHGFTRRDFMKWAGAMTALMALPASF WP_086253088.1
Ni/Fe hydrogenase [Campylobacter sp. P091] Length: 383\nNumber of predicted TMHs: 2\nExp number of AAs in TMHs: 34.46748\nExp number, first 60 AAs: 10.91753\nTotal prob of N-in: 0.76445\nPOSSIBLE N-term signal sequence\ninside 1 45\nTMhelix 46 68\noutside 69 356\nTMhelix 357 376\ninside 377 383

1693 GCF_002138945.1_ASM213894v1 Campylobacter sp. P093 Proteobacteria; delta/epsilon subdivisions; Epsilonproteobacteria; Campylobacterales; Campylobacteraceae; Campylobacter
MVDNHNITTIQNRLSELAKLPKMKANSSIMDVLNEHGFTRRDFMKWAGAMTALMALPASF WP_086224116.1
Ni/Fe hydrogenase [Campylobacter sp. P0098] Length: 383\nNumber of predicted TMHs: 2\nExp number of AAs in TMHs: 34.4668199999999\nExp number, first 60 AAs: 10.91724\nTotal prob of N-in: 0.76446\nPOSSIBLE N-term signal sequence\ninside 1 45\nTMhelix 46 68\noutside 69 356\nTMhelix 357 376\ninside 377 383

1694 GCF_002138975.1_ASM213897v1 Campylobacter sp. P146 Proteobacteria; delta/epsilon subdivisions; Epsilonproteobacteria; Campylobacterales; Campylobacteraceae; Campylobacter
MVDNHNITTIQNRLSELAKLPKMKANSSIMDVLNEHGFTRRDFMKWAGAMTALMALPASF WP_086224116.1
Ni/Fe hydrogenase [Campylobacter sp. P0098] Length: 383\nNumber of predicted TMHs: 2\nExp number of AAs in TMHs: 34.4668199999999\nExp number, first 60 AAs: 10.91724\nTotal prob of N-in: 0.76446\nPOSSIBLE N-term signal sequence\ninside 1 45\nTMhelix 46 68\noutside 69 356\nTMhelix 357 376\ninside 377 383

1695 GCF_002138995.1_ASM213899v1 Campylobacter sp. P148 Proteobacteria; delta/epsilon subdivisions; Epsilonproteobacteria; Campylobacterales; Campylobacteraceae; Campylobacter
MVDNHNITTIQNRLSELAKLPKMKANSSIMDVLNEHGFTRRDFMKWAGAMTALMALPASF WP_086224116.1
Ni/Fe hydrogenase [Campylobacter sp. P0098] Length: 383\nNumber of predicted TMHs: 2\nExp number of AAs in TMHs: 34.4668199999999\nExp number, first 60 AAs: 10.91724\nTotal prob of N-in: 0.76446\nPOSSIBLE N-term signal sequence\ninside 1 45\nTMhelix 46 68\noutside 69 356\nTMhelix 357 376\ninside 377 383

1696 GCF_002139745.1_ASM213974v1 Campylobacter sp. P155 Proteobacteria; delta/epsilon subdivisions; Epsilonproteobacteria; Campylobacterales; Campylobacteraceae; Campylobacter
MVDNHNITTIQNRLSELAKLPKMKANSSIMDVLNEHGFTRRDFMKWAGAMTALMALPASF WP_086288076.1
Ni/Fe hydrogenase [Campylobacter sp. P155] Length: 383\nNumber of predicted TMHs: 2\nExp number of

AAs in TMHs: 33.99546\nExp number, first 60 AAs: 10.95993\nTotal prob of N-in: 0.76344\nPOSSIBLE N-term signal sequence\ninside 1 45\nTMhelix 46 68\noutside 69 356\nTMhelix 357 376\ninside 377 383

1697 GCF_002139015.1_ASM213901v1 Campylobacter sp. P157 Proteobacteria; delta/epsilon subdivisions; Epsilonproteobacteria; Campylobacterales; Campylobacteraceae; Campylobacter MVDNHNITTIQNRLSELAKLPKMKANSSIMDVLNEHGFTRRDFMKWAGAMTALMALPASF WP_086235800.1 Ni/Fe hydrogenase [Campylobacter sp. P160] Length: 383\nNumber of predicted TMHs: 2\nExp number of AAs in TMHs: 35.1235\nExp number, first 60 AAs: 10.83195\nTotal prob of N-in: 0.76531\nPOSSIBLE N-term signal sequence\ninside 1 45\nTMhelix 46 68\noutside 69 356\nTMhelix 357 376\ninside 377 383

1698 GCF_002139025.1_ASM213902v1 Campylobacter sp. P159 Proteobacteria; delta/epsilon subdivisions; Epsilonproteobacteria; Campylobacterales; Campylobacteraceae; Campylobacter MVDNHNITTIQNRLSELAKLPKMKANSSIMDVLNEHGFTRRDFMKWAGAMTALMALPASF WP_086229493.1 Ni/Fe hydrogenase [Campylobacter sp. P0088] Length: 383\nNumber of predicted TMHs: 2\nExp number of AAs in TMHs: 34.4668299999999\nExp number, first 60 AAs: 10.91725\nTotal prob of N-in: 0.76446\nPOSSIBLE N-term signal sequence\ninside 1 45\nTMhelix 46 68\noutside 69 356\nTMhelix 357 376\ninside 377 383

1699 GCF_002139055.1_ASM213905v1 Campylobacter sp. P160 Proteobacteria; delta/epsilon subdivisions; Epsilonproteobacteria; Campylobacterales; Campylobacteraceae; Campylobacter MVDNHNITTIQNRLSELAKLPKMKANSSIMDVLNEHGFTRRDFMKWAGAMTALMALPASF WP_086235800.1 Ni/Fe hydrogenase [Campylobacter sp. P160] Length: 383\nNumber of predicted TMHs: 2\nExp number of AAs in TMHs: 35.1235\nExp number, first 60 AAs: 10.83195\nTotal prob of N-in: 0.76531\nPOSSIBLE N-term signal sequence\ninside 1 45\nTMhelix 46 68\noutside 69 356\nTMhelix 357 376\ninside 377 383

1700 GCF_002139735.1_ASM213973v1 Campylobacter sp. P162 Proteobacteria; delta/epsilon subdivisions; Epsilonproteobacteria; Campylobacterales; Campylobacteraceae; Campylobacter MVDNHNITTIQNRLSELAKLPKMKANSSIMDVLNEHGFTRRDFMKWAGAMTALMALPASF WP_086234315.1 Ni/Fe hydrogenase [Campylobacter sp. P0140] Length: 383\nNumber of predicted TMHs: 2\nExp number of AAs in TMHs: 34.46783\nExp number, first 60 AAs: 10.9175\nTotal prob of N-in: 0.76445\nPOSSIBLE N-term signal sequence\ninside 1 45\nTMhelix 46 68\noutside 69 356\nTMhelix 357 376\ninside 377 383

1701 GCF_002139875.1_ASM213987v1 Campylobacter sp. RM12175 Proteobacteria; delta/epsilon subdivisions; Epsilonproteobacteria; Campylobacterales; Campylobacteraceae; Campylobacter MVDNRNLSTIQNRLAELAKLPKMKENSSIMEVLSEHGFTRRDFMKWAGAMTALMALPASF WP_086247137.1 Ni/Fe hydrogenase [Campylobacter sp. S0112] Length: 383\nNumber of predicted TMHs: 1\nExp number of AAs in TMHs: 32.32817\nExp number, first 60 AAs: 9.79613\nTotal prob of N-in: 0.68853\noutside 1 356\nTMhelix 357 376\ninside 377 383

1702 GCF_002139855.1_ASM213985v1 Campylobacter sp. RM6137 Proteobacteria; delta/epsilon subdivisions; Epsilonproteobacteria; Campylobacterales; Campylobacteraceae; Campylobacter MVDNHNVTTIQNRLSELAKLPKMKSNSSIMEVLNEHGFTRRDFMKWAGAMTALMALPASF WP_086297856.1 Ni/Fe hydrogenase [Campylobacter sp. RM6137] Length: 383\nNumber of predicted TMHs: 2\nExp number of AAs in TMHs: 34.40536\nExp number, first 60 AAs: 11.13575\nTotal prob of N-in: 0.76324\nPOSSIBLE N-term signal sequence\ninside 1 45\nTMhelix 46 68\noutside 69 356\nTMhelix 357 376\ninside 377 383

1703 GCF_002139095.1_ASM213909v1 Campylobacter sp. RM8835 Proteobacteria; delta/epsilon subdivisions; Epsilonproteobacteria; Campylobacterales; Campylobacteraceae; Campylobacter MVDNRNLSTIQNRLAELAKLPKMKENSSIMEVLSEHGFTRRDFMKWAGAMTALMALPASF WP_086247137.1 Ni/Fe hydrogenase [Campylobacter sp. S0112] Length: 383\nNumber of predicted TMHs: 1\nExp number of AAs in TMHs: 32.32817\nExp number, first 60 AAs: 9.79613\nTotal prob of N-in: 0.68853\noutside 1 356\nTMhelix 357 376\ninside 377 383

1704 GCF_002139895.1_ASM213989v1 Campylobacter sp. RM8964 Proteobacteria; delta/epsilon subdivisions; Epsilonproteobacteria; Campylobacterales; Campylobacteraceae; Campylobacter MVDNRNLSTIQNRLAELAKLPKMKENSSIMEVLSEHGFTRRDFMKWAGAMTALMALPASF WP_086333853.1 Ni/Fe hydrogenase [Campylobacter sp. RM8964] Length: 383\nNumber of predicted TMHs: 2\nExp number of AAs in TMHs: 34.60878\nExp number, first 60 AAs: 11.22558\nTotal prob of N-in: 0.75469\nPOSSIBLE N-term signal sequence\ninside 1 45\nTMhelix 46 68\noutside 69 356\nTMhelix 357 376\ninside 377 383

1705 GCF_002139135.1_ASM213913v1 Campylobacter sp. RM8965 Proteobacteria; delta/epsilon subdivisions; Epsilonproteobacteria; Campylobacterales; Campylobacteraceae; Campylobacter

MVDNRNLSTIQNRLAELAKLPKMKENSSIMEVLSEHGFTRRDFMKWAGAMTALMALPASF WP_086247137.1 Ni/Fe hydrogenase [Campylobacter sp. S0112] Length: 383\nNumber of predicted TMHs: 1\nExp number of AAs in TMHs: 32.32817\nExp number, first 60 AAs: 9.79613\nTotal prob of N-in: 0.68853\noutside 1 356\nTMhelix 357 376\ninside 377 383

1706 GCF_002139155.1_ASM213915v1 Campylobacter sp. RM8966 Proteobacteria; delta/epsilon subdivisions; Epsilonproteobacteria; Campylobacterales; Campylobacteraceae; Campylobacter MVDNRNLSTIQNRLAELAKLPKMKENSSIMEVLSEHGFTRRDFMKWAGAMTALMALPASF WP_086247137.1 Ni/Fe hydrogenase [Campylobacter sp. S0112] Length: 383\nNumber of predicted TMHs: 1\nExp number of AAs in TMHs: 32.32817\nExp number, first 60 AAs: 9.79613\nTotal prob of N-in: 0.68853\noutside 1 356\nTMhelix 357 376\ninside 377 383

1707 GCF_002139795.1_ASM213979v1 Campylobacter sp. RM8970 Proteobacteria; delta/epsilon subdivisions; Epsilonproteobacteria; Campylobacterales; Campylobacteraceae; Campylobacter MVDNRNLSTIQNRLAELAKLPKMKENSSIMEVLSEHGFTRRDFMKWAGAMTALMALPASF WP_086247137.1 Ni/Fe hydrogenase [Campylobacter sp. S0112] Length: 383\nNumber of predicted TMHs: 1\nExp number of AAs in TMHs: 32.32817\nExp number, first 60 AAs: 9.79613\nTotal prob of N-in: 0.68853\noutside 1 356\nTMhelix 357 376\ninside 377 383

1708 GCF_002139825.1_ASM213982v1 Campylobacter sp. RM9261 Proteobacteria; delta/epsilon subdivisions; Epsilonproteobacteria; Campylobacterales; Campylobacteraceae; Campylobacter MVDNRNLSTIQNRLAELAKLPKMKENSSIMEVLSEHGFTRRDFMKWAGAMTALMALPASF WP_086247137.1 Ni/Fe hydrogenase [Campylobacter sp. S0112] Length: 383\nNumber of predicted TMHs: 1\nExp number of AAs in TMHs: 32.32817\nExp number, first 60 AAs: 9.79613\nTotal prob of N-in: 0.68853\noutside 1 356\nTMhelix 357 376\ninside 377 383

1709 GCF_002139175.1_ASM213917v1 Campylobacter sp. RM9262 Proteobacteria; delta/epsilon subdivisions; Epsilonproteobacteria; Campylobacterales; Campylobacteraceae; Campylobacter MVDNRNLSTIQNRLAELAKLPKMKENSSIMEVLSEHGFTRRDFMKWAGAMTALMALPASF WP_086247137.1 Ni/Fe hydrogenase [Campylobacter sp. S0112] Length: 383\nNumber of predicted TMHs: 1\nExp number of AAs in TMHs: 32.32817\nExp number, first 60 AAs: 9.79613\nTotal prob of N-in: 0.68853\noutside 1 356\nTMhelix 357 376\ninside 377 383

1710 GCF_002139815.1_ASM213981v1 Campylobacter sp. RM9263 Proteobacteria; delta/epsilon subdivisions; Epsilonproteobacteria; Campylobacterales; Campylobacteraceae; Campylobacter MVDNRNLSTIQNRLAELAKLPKMKENSSIMEVLSEHGFTRRDFMKWAGAMTALMALPASF WP_086247137.1 Ni/Fe hydrogenase [Campylobacter sp. S0112] Length: 383\nNumber of predicted TMHs: 1\nExp number of AAs in TMHs: 32.32817\nExp number, first 60 AAs: 9.79613\nTotal prob of N-in: 0.68853\noutside 1 356\nTMhelix 357 376\ninside 377 383

1711 GCF_002139185.1_ASM213918v1 Campylobacter sp. S0112 Proteobacteria; delta/epsilon subdivisions; Epsilonproteobacteria; Campylobacterales; Campylobacteraceae; Campylobacter MVDNRNLSTIQNRLAELAKLPKMKENSSIMEVLSEHGFTRRDFMKWAGAMTALMALPASF WP_086247137.1 Ni/Fe hydrogenase [Campylobacter sp. S0112] Length: 383\nNumber of predicted TMHs: 1\nExp number of AAs in TMHs: 32.32817\nExp number, first 60 AAs: 9.79613\nTotal prob of N-in: 0.68853\noutside 1 356\nTMhelix 357 376\ninside 377 383

1712 GCF_000017725.2_ASM1772v2 Campylobacter concisus 13826 Proteobacteria; delta/epsilon subdivisions; Epsilonproteobacteria; Campylobacterales; Campylobacteraceae; Campylobacter; Campylobacter concisus MNNDLRQKINRRLSELSALSKMKNDTSIAQLLKDKGFTRRDFMKWAGAMTAFMALPSAMT WP_048809818.1 Ni/Fe hydrogenase [Campylobacter concisus] Length: 381\nNumber of predicted TMHs: 2\nExp number of AAs in TMHs: 37.4411399999999\nExp number, first 60 AAs: 14.24401\nTotal prob of N-in: 0.87262\nPOSSIBLE N-term signal sequence\ninside 1 41\nTMhelix 42 64\noutside 65 354\nTMhelix 355 377\ninside 378 381

1713 GCF_000466705.1_ATCC51561 Campylobacter concisus ATCC 51561 Proteobacteria; delta/epsilon subdivisions; Epsilonproteobacteria; Campylobacterales; Campylobacteraceae; Campylobacter; Campylobacter concisus MNNDLRQKINRRLSELSALPKMKNDTSIAQLLKDKGFTRRDFMKWAGAMTAFMALPSAMT WP_021085837.1 Quinone-reactive Ni [Campylobacter concisus] Length: 381\nNumber of predicted TMHs: 2\nExp number of AAs in TMHs: 37.4411399999999\nExp number, first 60 AAs: 14.24402\nTotal prob of N-in: 0.87262\nPOSSIBLE N-term signal sequence\ninside 1 41\nTMhelix 42 64\noutside 65 354\nTMhelix 355 377\ninside 378 381

1714 GCF_000466745.1_ATCC51562 Campylobacter concisus ATCC 51562 Proteobacteria; delta/epsilon subdivisions; Epsilonproteobacteria; Campylobacterales; Campylobacteraceae; Campylobacter; Campylobacter concisus MNNDLRQKIDRRLSELSALPKMKSDSSIAQLLKDKGFTRRDFMKWAGAMTAFMALPSAMT WP_021091342.1 hypothetical protein [Campylobacter concisus] Length: 381\nNumber of predicted TMHs: 2\nExp number of AAs in TMHs: 34.64315\nExp number, first 60 AAs: 13.34172\nTotal prob of N-in: 0.83884\nPOSSIBLE N-term signal sequence\ninside 1 41\nTMhelix 42 64\noutside 65 354\nTMhelix 355 377\ninside 378 381

1715 GCF_000466665.1_UNSW1 Campylobacter concisus UNSW1 Proteobacteria; delta/epsilon subdivisions; Epsilonproteobacteria; Campylobacterales; Campylobacteraceae; Campylobacter; Campylobacter concisus MNNDLRQKINRRLSELSALPKMKNDTSIAQLLKDKGFTRRDFMKWAGAMTAFMALPSAMT WP_021085837.1 Quinone-reactive Ni [Campylobacter concisus] Length: 381\nNumber of predicted TMHs: 2\nExp number of AAs in TMHs: 37.4411399999999\nExp number, first 60 AAs: 14.24402\nTotal prob of N-in: 0.87262\nPOSSIBLE N-term signal sequence\ninside 1 41\nTMhelix 42 64\noutside 65 354\nTMhelix 355 377\ninside 378 381

1716 GCF_000466725.1_UNSW2 Campylobacter concisus UNSW2 Proteobacteria; delta/epsilon subdivisions; Epsilonproteobacteria; Campylobacterales; Campylobacteraceae; Campylobacter; Campylobacter concisus MNNDLRQKINRRLSELSALPKMKNDTSIAQLLKDKGFTRRDFMKWAGAMTAFMALPSAMT WP_002940950.1 quinone-reactive Ni/Fe-hydrogenase small chain [Campylobacter concisus] Length: 381\nNumber of predicted TMHs: 2\nExp number of AAs in TMHs: 38.5527599999999\nExp number, first 60 AAs: 14.21245\nTotal prob of N-in: 0.87296\nPOSSIBLE N-term signal sequence\ninside 1 41\nTMhelix 42 64\noutside 65 354\nTMhelix 355 377\ninside 378 381

1717 GCF_000466645.1_UNSW3 Campylobacter concisus UNSW3 Proteobacteria; delta/epsilon subdivisions; Epsilonproteobacteria; Campylobacterales; Campylobacteraceae; Campylobacter; Campylobacter concisus MNNDLRQKINRRLSELSALPKMKNDTSIAQLLKDKGFTRRDFMKWAGAMTAFMALPSAMT WP_002940950.1 quinone-reactive Ni/Fe-hydrogenase small chain [Campylobacter concisus] Length: 381\nNumber of predicted TMHs: 2\nExp number of AAs in TMHs: 38.5527599999999\nExp number, first 60 AAs: 14.21245\nTotal prob of N-in: 0.87296\nPOSSIBLE N-term signal sequence\ninside 1 41\nTMhelix 42 64\noutside 65 354\nTMhelix 355 377\ninside 378 381

1718 GCF_000259315.1_ASM25931v1 Campylobacter concisus UNSWCD Proteobacteria; delta/epsilon subdivisions; Epsilonproteobacteria; Campylobacterales; Campylobacteraceae; Campylobacter; Campylobacter concisus MNNDLRQKINRRLSELSALPKMKNDTSIAQLLKDKGFTRRDFMKWAGAMTAFMALPSAMT WP_002940950.1 quinone-reactive Ni/Fe-hydrogenase small chain [Campylobacter concisus] Length: 381\nNumber of predicted TMHs: 2\nExp number of AAs in TMHs: 38.5527599999999\nExp number, first 60 AAs: 14.21245\nTotal prob of N-in: 0.87296\nPOSSIBLE N-term signal sequence\ninside 1 41\nTMhelix 42 64\noutside 65 354\nTMhelix 355 377\ninside 378 381

1719 GCF_000466685.1_UNSWCS Campylobacter concisus UNSWCS Proteobacteria; delta/epsilon subdivisions; Epsilonproteobacteria; Campylobacterales; Campylobacteraceae; Campylobacter; Campylobacter concisus MNNDLRQKINRRLSELSALPKMKNDTSIAQLLKDKGFTRRDFMKWAGAMTAFMALPSAMT WP_002940950.1 quinone-reactive Ni/Fe-hydrogenase small chain [Campylobacter concisus] Length: 381\nNumber of predicted TMHs: 2\nExp number of AAs in TMHs: 38.5527599999999\nExp number, first 60 AAs: 14.21245\nTotal prob of N-in: 0.87296\nPOSSIBLE N-term signal sequence\ninside 1 41\nTMhelix 42 64\noutside 65 354\nTMhelix 355 377\ninside 378 381

1720 GCF_000017465.2_ASM1746v2 Campylobacter curvus 525.92 Proteobacteria; delta/epsilon subdivisions; Epsilonproteobacteria; Campylobacterales; Campylobacteraceae; Campylobacter; Campylobacter curvus MVHNIHEKANRRLNELSLLPKIKENGTIAKILEQNGFTRRDFMKWAGAMTAFMALPASMT WP_009651157.1 MULTISPECIES: quinone-reactive Ni/Fe-hydrogenase small chain [Campylobacter] Length: 381\nNumber of predicted TMHs: 1\nExp number of AAs in TMHs: 33.75286\nExp number, first 60 AAs: 9.47284\nTotal prob of N-in: 0.64572\noutside 1 354\nTMhelix 355 377\ninside 378 381

1721 GCF_000376325.1_ASM37632v1 Campylobacter curvus DSM 6644 Proteobacteria; delta/epsilon subdivisions; Epsilonproteobacteria; Campylobacterales; Campylobacteraceae; Campylobacter; Campylobacter curvus MVQNIHEKANRRLNKLSLLPKIKENGTIAKILEQNGFTRRDFMKWAGAMTAFMALPASMT WP_018136330.1 quinone-reactive Ni/Fe-hydrogenase small chain [Campylobacter curvus] Length: 381\nNumber of predicted TMHs: 1\nExp number of AAs in TMHs: 34.17619\nExp number, first 60 AAs: 9.78484\nTotal prob of N-in: 0.69388\noutside 1 354\nTMhelix 355 377\ninside 378 381

1722 GCF_001487085.1_17150_1_75 Campylobacter fetus subsp. fetus Proteobacteria; delta/epsilon subdivisions; Epsilonproteobacteria; Campylobacterales; Campylobacteraceae; Campylobacter; Campylobacter fetus MVEHKLLNTIQDRLDALSKMPKVKKNSSIMAVLKENGFTRRDFMKWAGAMTAMMALPASF WP_059425527.1 Ni/Fe hydrogenase [Campylobacter fetus] Length: 383\nNumber of predicted TMHs: 1\nExp number of AAs in TMHs: 21.33594\nExp number, first 60 AAs: 2.35497\nTotal prob of N-in: 0.19219\noutside 1 356\nTMhelix 357 376\ninside 377 383

1723 GCF_001486865.1_17059_2_3 Campylobacter fetus subsp. fetus Proteobacteria; delta/epsilon subdivisions; Epsilonproteobacteria; Campylobacterales; Campylobacteraceae; Campylobacter; Campylobacter fetus MDRRSFFKFSAVAASSPLFATSNLEEKKIMSIIDLDLCDGCKNELMPLCVKACRDKNEPN WP_059426839.1 oxidoreductase [Campylobacter fetus] Length: 312\nNumber of predicted TMHs: 1\nExp number of AAs in TMHs: 22.31823\nExp number, first 60 AAs: 0.04111\nTotal prob of N-in: 0.01396\noutside 1 281\nTMhelix 282 304\ninside 305 312

1724 GCF_001486865.1_17059_2_3 Campylobacter fetus subsp. fetus Proteobacteria; delta/epsilon subdivisions; Epsilonproteobacteria; Campylobacterales; Campylobacteraceae; Campylobacter; Campylobacter fetus MVEHKLLNTIQDRLDALSKMPKVKKNSSIMAVLKENGFTRRDFMKWAGAMTAMMALPASF WP_059426924.1 Ni/Fe hydrogenase [Campylobacter fetus] Length: 383\nNumber of predicted TMHs: 1\nExp number of AAs in TMHs: 21.55052\nExp number, first 60 AAs: 2.3641\nTotal prob of N-in: 0.19410\noutside 1 356\nTMhelix 357 376\ninside 377 383

1725 GCF_001487005.1_17059_2_1 Campylobacter fetus subsp. fetus Proteobacteria; delta/epsilon subdivisions; Epsilonproteobacteria; Campylobacterales; Campylobacteraceae; Campylobacter; Campylobacter fetus MVEHKLLNTIQDRLDALSKMPKVKKNSSIMAVLKENGFTRRDFMKWAGAMTAMMALPASF WP_059426924.1 Ni/Fe hydrogenase [Campylobacter fetus] Length: 383\nNumber of predicted TMHs: 1\nExp number of AAs in TMHs: 21.55052\nExp number, first 60 AAs: 2.3641\nTotal prob of N-in: 0.19410\noutside 1 356\nTMhelix 357 376\ninside 377 383

1726 GCF_001486845.1_17150_1_81 Campylobacter fetus subsp. fetus Proteobacteria; delta/epsilon subdivisions; Epsilonproteobacteria; Campylobacterales; Campylobacteraceae; Campylobacter; Campylobacter fetus MVEHKLLNTIQDRLDALSKMPKVKKNSSIMAVLKENGFTRRDFMKWAGAMTAMMALPASF WP_059425527.1 Ni/Fe hydrogenase [Campylobacter fetus] Length: 383\nNumber of predicted TMHs: 1\nExp number of AAs in TMHs: 21.33594\nExp number, first 60 AAs: 2.35497\nTotal prob of N-in: 0.19219\noutside 1 356\nTMhelix 357 376\ninside 377 383

1727 GCF_001487005.1_17059_2_1 Campylobacter fetus subsp. fetus Proteobacteria; delta/epsilon subdivisions; Epsilonproteobacteria; Campylobacterales; Campylobacteraceae; Campylobacter; Campylobacter fetus MDRRSFFKFSAVAASSPLFATSNLEEKKIMSIIDLDLCDGCKNELMPLCVKACRDKNEPN WP_059426839.1 oxidoreductase [Campylobacter fetus] Length: 312\nNumber of predicted TMHs: 1\nExp number of AAs in TMHs: 22.31823\nExp number, first 60 AAs: 0.04111\nTotal prob of N-in: 0.01396\noutside 1 281\nTMhelix 282 304\ninside 305 312

1728 GCF_001486945.1_17150_1_76 Campylobacter fetus subsp. fetus Proteobacteria; delta/epsilon subdivisions; Epsilonproteobacteria; Campylobacterales; Campylobacteraceae; Campylobacter; Campylobacter fetus MVEHKLLNTIQDRLDALSKMPKVKKNSSIMAVLKENGFTRRDFMKWAGAMTAMMALPASF WP_059426924.1 Ni/Fe hydrogenase [Campylobacter fetus] Length: 383\nNumber of predicted TMHs: 1\nExp number of AAs in TMHs: 21.55052\nExp number, first 60 AAs: 2.3641\nTotal prob of N-in: 0.19410\noutside 1 356\nTMhelix 357 376\ninside 377 383

1729 GCF_001486945.1_17150_1_76 Campylobacter fetus subsp. fetus Proteobacteria; delta/epsilon subdivisions; Epsilonproteobacteria; Campylobacterales; Campylobacteraceae; Campylobacter; Campylobacter fetus MDRRSFFKFSAVAASSPLFATSNLEEKKIMSIIDLDLCDGCKNELMPLCVKACRDKNEPN WP_059426839.1 oxidoreductase [Campylobacter fetus] Length: 312\nNumber of predicted TMHs: 1\nExp number of AAs in TMHs: 22.31823\nExp number, first 60 AAs: 0.04111\nTotal prob of N-in: 0.01396\noutside 1 281\nTMhelix 282 304\ninside 305 312

1730 GCF_001643975.1_ASM164397v1 Campylobacter hyointestinalis subsp. lawsonii Proteobacteria; delta/epsilon subdivisions; Epsilonproteobacteria; Campylobacterales; Campylobacteraceae; Campylobacter; Campylobacter hyointestinalis MDRRSFFKFSAVAASSPLFATSNLEDKKIMSIIDLDLCDGCKNEPIPLCVKACRDKNEPN WP_063998339.1 oxidoreductase [Campylobacter hyointestinalis] Length: 310\nNumber of predicted TMHs:

1\nExp number of AAs in TMHs: 20.40362\nExp number, first 60 AAs: 0.04353\nTotal prob of N-in: 0.06262\noutside 1 281\nTMhelix 282 304\ninside 305 310

1731 GCF_001643975.1_ASM164397v1 Campylobacter hyointestinalis subsp. lawsonii Proteobacteria; delta/epsilon subdivisions; Epsilonproteobacteria; Campylobacterales; Campylobacteraceae; Campylobacter; Campylobacter hyointestinalis MVEHKLLNTIQDRLDALSKMPKFKKNSSIMSVLNEHGFTRRDFMKWAGAMTALMALPASF WP_063998141.1 Ni/Fe hydrogenase [Campylobacter hyointestinalis] Length: 383\nNumber of predicted TMHs: 1\nExp number of AAs in TMHs: 28.54262\nExp number, first 60 AAs: 7.17732\nTotal prob of N-in: 0.47958\noutside 1 356\nTMhelix 357 376\ninside 377 383

1732 GCF_001643955.1_ASM164395v1 Campylobacter hyointestinalis subsp. hyointestinalis LMG 9260 Proteobacteria; delta/epsilon subdivisions; Epsilonproteobacteria; Campylobacterales; Campylobacteraceae; Campylobacter; Campylobacter hyointestinalis; Campylobacter hyointestinalis subsp. hyointestinalis MDRRSFFKFSAVAASSSLFATSNLEDKKIMSIIDLDLCDGCKNEPIPLCVKACRDKNEPN WP_064019789.1 oxidoreductase [Campylobacter hyointestinalis] Length: 310\nNumber of predicted TMHs: 1\nExp number of AAs in TMHs: 22.01747\nExp number, first 60 AAs: 0.07136\nTotal prob of N-in: 0.02459\noutside 1 281\nTMhelix 282 304\ninside 305 310

1733 GCF_000344295.2_velvet Campylobacter showae CC57C Proteobacteria; delta/epsilon subdivisions; Epsilonproteobacteria; Campylobacterales; Campylobacteraceae; Campylobacter; Campylobacter showae MRDDVLVRINERLNVLAALPTLKNGSIGEALKKSGFSRRDFMKWAGAMTAFMALPASMTP WP_002952686.1 thymidylate kinase [Campylobacter showae] Length: 382\nNumber of predicted TMHs: 1\nExp number of AAs in TMHs: 28.73337\nExp number, first 60 AAs: 5.94567\nTotal prob of N-in: 0.42528\noutside 1 353\nTMhelix 354 376\ninside 377 382

1734 GCF_000313615.1_Velvet_assembly_of_Campylobacter_showae_CSUNSWCD Campylobacter showae CSUNSWCD Proteobacteria; delta/epsilon subdivisions; Epsilonproteobacteria; Campylobacterales; Campylobacteraceae; Campylobacter; Campylobacter showae MRDDVLVRINERLNVLAALPTLKNGSIGEALKKSGFSRRDFMKWAGAMTAFMALPASMTP WP_002952686.1 thymidylate kinase [Campylobacter showae] Length: 382\nNumber of predicted TMHs: 1\nExp number of AAs in TMHs: 28.73337\nExp number, first 60 AAs: 5.94567\nTotal prob of N-in: 0.42528\noutside 1 353\nTMhelix 354 376\ninside 377 382

1735 GCF_000175655.1_ASM17565v1 Campylobacter showae RM3277 Proteobacteria; delta/epsilon subdivisions; Epsilonproteobacteria; Campylobacterales; Campylobacteraceae; Campylobacter; Campylobacter showae MKDDVLVRINERLNVLATLPTFKNGSIGEALKKSGFSRRDFMKWAGAMTAFMALPASMAP WP_002947953.1 quinone-reactive Ni/Fe-hydrogenase small chain [Campylobacter showae] Length: 382\nNumber of predicted TMHs: 1\nExp number of AAs in TMHs: 31.29117\nExp number, first 60 AAs: 8.12759\nTotal prob of N-in: 0.50112\noutside 1 353\nTMhelix 354 376\ninside 377 382

1736 GCF_000788295.1_ASM78829v1 Campylobacter sputorum biovar sputorum Proteobacteria; delta/epsilon subdivisions; Epsilonproteobacteria; Campylobacterales; Campylobacteraceae; Campylobacter; Campylobacter sputorum MKQDNLAAKISARLNEVQNMPKMHNGSIMQKLQERGFTRRDFMKWAGAMTAMLSLPTSFI WP_033915906.1 Ni/Fe hydrogenase [Campylobacter sputorum] Length: 379\nNumber of predicted TMHs: 1\nExp number of AAs in TMHs: 24.01502\nExp number, first 60 AAs: 2.99413\nTotal prob of N-in: 0.39495\noutside 1 355\nTMhelix 356 375\ninside 376 379

1737 GCF_000765745.1_ASM76574v1 Helicobacter apodemus Proteobacteria; delta/epsilon subdivisions; Epsilonproteobacteria; Campylobacterales; Helicobacteraceae; Helicobacter MQREQVLLEKAQKRLEEIHKIPALKKGNSITKALRRNGISRRDFMKWAGAMTAMLSLPAS WP_034553322.1 Ni/Fe hydrogenase [Helicobacter apodemus] Length: 383\nNumber of predicted TMHs: 1\nExp number of AAs in TMHs: 24.9605\nExp number, first 60 AAs: 3.65683\nTotal prob of N-in: 0.25338\noutside 1 355\nTMhelix 356 378\ninside 379 383

1738 GCF_000765845.1_ASM76584v1 Helicobacter bilis Proteobacteria; delta/epsilon subdivisions; Epsilonproteobacteria; Campylobacterales; Helicobacteraceae; Helicobacter MQRRNFLKSLALTPVLVAGANAEPSDSTKEGKEETRSLIDSSEFSKTKGKASIINLDKCD WP_004088007.1 hypothetical protein [Helicobacter bilis] Length: 345\nNumber of predicted TMHs: 1\nExp number of AAs in TMHs: 18.27315\nExp number, first 60 AAs: 0.00939\nTotal prob of N-in: 0.10106\noutside 1 313\nTMhelix 314 336\ninside 337 345

1739 GCF_000765785.1_ASM76578v1 Helicobacter bilis Proteobacteria; delta/epsilon subdivisions; Epsilonproteobacteria; Campylobacterales; Helicobacteraceae; Helicobacter
MQRRNFLKSLALTPVLVAGANAEPSDSTKEGKEETRSLIDSSEFSKTKGKASIINLDKCD WP_004088007.1 hypothetical protein [Helicobacter bilis] Length: 345\nNumber of predicted TMHs: 1\nExp number of AAs in TMHs: 18.27315\nExp number, first 60 AAs: 0.00939\nTotal prob of N-in: 0.10106\noutside 1 313\nTMhelix 314 336\ninside 337 345

1740 GCF_001999985.1_ASM199998v1 Helicobacter bilis Proteobacteria; delta/epsilon subdivisions; Epsilonproteobacteria; Campylobacterales; Helicobacteraceae; Helicobacter
MQRRNFLKSLALTPVLVAGANAESSDSTKDGKKEKRSLIDSSEFSKTKGKASIINLDKCD WP_077387918.1 oxidoreductase [Helicobacter bilis] Length: 345\nNumber of predicted TMHs: 1\nExp number of AAs in TMHs: 18.27618\nExp number, first 60 AAs: 0.01242\nTotal prob of N-in: 0.10119\noutside 1 313\nTMhelix 314 336\ninside 337 345

1741 GCF_001602095.1_ASM160209v1 Helicobacter himalayensis Proteobacteria; delta/epsilon subdivisions; Epsilonproteobacteria; Campylobacterales; Helicobacteraceae; Helicobacter
MSDYTQEKLLERINKHLDELEKLPKIRENTSIMQTLQEHGFSRRDFMKWAGGMAAMLSLP WP_066387589.1 Ni/Fe hydrogenase [Helicobacter himalayensis] Length: 393\nNumber of predicted TMHs: 1\nExp number of AAs in TMHs: 32.91046\nExp number, first 60 AAs: 8.69508\nTotal prob of N-in: 0.49399\noutside 1 359\nTMhelix 360 382\ninside 383 393

1742 GCF_000763135.1_ASM76313v1 Helicobacter jaachi Proteobacteria; delta/epsilon subdivisions; Epsilonproteobacteria; Campylobacterales; Helicobacteraceae; Helicobacter
MNIEGAHEELFRKATQRLDELEKLPAHNANKNLFEALESKGFSRRDFMKWSGMMTTALAL WP_081946299.1 Ni/Fe hydrogenase [Helicobacter jaachi] Length: 386\nNumber of predicted TMHs: 1\nExp number of AAs in TMHs: 21.3299\nExp number, first 60 AAs: 1.12954\nTotal prob of N-in: 0.08877\noutside 1 357\nTMhelix 358 380\ninside 381 386

1743 GCF_000765825.1_ASM76582v1 Helicobacter magdeburgensis Proteobacteria; delta/epsilon subdivisions; Epsilonproteobacteria; Campylobacterales; Helicobacteraceae; Helicobacter
MQRRNFLKSLALTPVLVAGANAEPSDSTKEGKEETRSLIDSSEFSKTKGKASIINLDKCD WP_034587156.1 oxidoreductase [Helicobacter magdeburgensis] Length: 345\nNumber of predicted TMHs: 1\nExp number of AAs in TMHs: 18.27358\nExp number, first 60 AAs: 0.00939\nTotal prob of N-in: 0.10108\noutside 1 313\nTMhelix 314 336\ninside 337 345

1744 GCF_001698785.1_ASM169878v1 Helicobacter pullorum Proteobacteria; delta/epsilon subdivisions; Epsilonproteobacteria; Campylobacterales; Helicobacteraceae; Helicobacter
MEREQVLLQKAQERLKEINKFPALKKGNSIKKMLKENGISRRDFMKWAGAMTAMLSLPAS WP_005020544.1 quinone-reactive Ni/Fe-hydrogenase small chain [Helicobacter pullorum] Length: 385\nNumber of predicted TMHs: 1\nExp number of AAs in TMHs: 21.91577\nExp number, first 60 AAs: 0.64271\nTotal prob of N-in: 0.09800\noutside 1 355\nTMhelix 356 378\ninside 379 385

1745 GCF_001698675.1_ASM169867v1 Helicobacter pullorum Proteobacteria; delta/epsilon subdivisions; Epsilonproteobacteria; Campylobacterales; Helicobacteraceae; Helicobacter
MEREQVLLQKAQERLKEINKFPALKKGNSIKKMLKENGISRRDFMKWAGAMTAMLSLPAS WP_065827860.1 Ni/Fe hydrogenase [Helicobacter pullorum] Length: 385\nNumber of predicted TMHs: 1\nExp number of AAs in TMHs: 21.38773\nExp number, first 60 AAs: 0.64291\nTotal prob of N-in: 0.08711\noutside 1 355\nTMhelix 356 378\ninside 379 385

1746 GCF_001698755.1_ASM169875v1 Helicobacter pullorum Proteobacteria; delta/epsilon subdivisions; Epsilonproteobacteria; Campylobacterales; Helicobacteraceae; Helicobacter
MEREQVLLQKAQERLKEINKFPALKKGNSIKKMLKENGISRRDFMKWAGAMTAMLSLPAS WP_005020544.1 quinone-reactive Ni/Fe-hydrogenase small chain [Helicobacter pullorum] Length: 385\nNumber of predicted TMHs: 1\nExp number of AAs in TMHs: 21.91577\nExp number, first 60 AAs: 0.64271\nTotal prob of N-in: 0.09800\noutside 1 355\nTMhelix 356 378\ninside 379 385

1747 GCF_001698595.1_ASM169859v1 Helicobacter pullorum Proteobacteria; delta/epsilon subdivisions; Epsilonproteobacteria; Campylobacterales; Helicobacteraceae; Helicobacter
MEREQVLLQKAQERLKEINKFPALKKGNSIKKMLKENGISRRDFMKWAGAMTAMLSLPAS WP_065827860.1 Ni/Fe hydrogenase [Helicobacter pullorum] Length: 385\nNumber of predicted TMHs: 1\nExp number of AAs in TMHs: 21.38773\nExp number, first 60 AAs: 0.64291\nTotal prob of N-in: 0.08711\noutside 1 355\nTMhelix 356 378\ninside 379 385

1748 GCF_001298045.1_ASM129804v1 Helicobacter pullorum Proteobacteria; delta/epsilon subdivisions; Epsilonproteobacteria; Campylobacterales; Helicobacteraceae; Helicobacter
MEREQVLLQKAQERLKEINKFPALKKGNSIKKMLKENGISRRDFMKWAGAMTAMLSLPAS WP_005020544.1
quinone-reactive Ni/Fe-hydrogenase small chain [Helicobacter pullorum] Length: 385\nNumber of predicted TMHs: 1\nExp number of AAs in TMHs: 21.91577\nExp number, first 60 AAs: 0.64271\nTotal prob of N-in: 0.09800\noutside 1 355\nTMhelix 356 378\ninside 379 385

1749 GCF_001298065.1_ASM129806v1 Helicobacter pullorum Proteobacteria; delta/epsilon subdivisions; Epsilonproteobacteria; Campylobacterales; Helicobacteraceae; Helicobacter
MEREQVLLQKAQERLKEINKFPALKKGNSIKKMLKENGISRRDFMKWAGAMTAMLSLPAS WP_054194698.1
Ni/Fe hydrogenase [Helicobacter pullorum] Length: 385\nNumber of predicted TMHs: 1\nExp number of AAs in TMHs: 21.42489\nExp number, first 60 AAs: 0.64344\nTotal prob of N-in: 0.08825\noutside 1 355\nTMhelix 356 378\ninside 379 385

1750 GCF_001298055.1_ASM129805v1 Helicobacter pullorum Proteobacteria; delta/epsilon subdivisions; Epsilonproteobacteria; Campylobacterales; Helicobacteraceae; Helicobacter
MEREQVLLQKAQERLKEINKFPALKKGNSIKKMLKENGISRRDFMKWAGAMTAMLSLPAS WP_054194698.1
Ni/Fe hydrogenase [Helicobacter pullorum] Length: 385\nNumber of predicted TMHs: 1\nExp number of AAs in TMHs: 21.42489\nExp number, first 60 AAs: 0.64344\nTotal prob of N-in: 0.08825\noutside 1 355\nTMhelix 356 378\ninside 379 385

1751 GCF_001698605.1_ASM169860v1 Helicobacter pullorum Proteobacteria; delta/epsilon subdivisions; Epsilonproteobacteria; Campylobacterales; Helicobacteraceae; Helicobacter
MEREQVLLQKAQERLKEINKFPALKKGNSIKKMLKENGISRRDFMKWAGAMTAMLSLPAS WP_005020544.1
quinone-reactive Ni/Fe-hydrogenase small chain [Helicobacter pullorum] Length: 385\nNumber of predicted TMHs: 1\nExp number of AAs in TMHs: 21.91577\nExp number, first 60 AAs: 0.64271\nTotal prob of N-in: 0.09800\noutside 1 355\nTMhelix 356 378\ninside 379 385

1752 GCF_001698625.1_ASM169862v1 Helicobacter pullorum Proteobacteria; delta/epsilon subdivisions; Epsilonproteobacteria; Campylobacterales; Helicobacteraceae; Helicobacter
MEREQVLLQKAQERLKEINKFPALKKGNSIKKMLKENGISRRDFMKWAGAMTAMLSLPAS WP_060660962.1
Ni/Fe hydrogenase [Helicobacter pullorum] Length: 385\nNumber of predicted TMHs: 1\nExp number of AAs in TMHs: 22.13084\nExp number, first 60 AAs: 0.64285\nTotal prob of N-in: 0.09674\noutside 1 355\nTMhelix 356 378\ninside 379 385

1753 GCF_001698705.1_ASM169870v1 Helicobacter pullorum Proteobacteria; delta/epsilon subdivisions; Epsilonproteobacteria; Campylobacterales; Helicobacteraceae; Helicobacter
MEREQVLLQKAQERLKEINKFPALKKGNSIKKMLKENGISRRDFMKWAGAMTAMLSLPAS WP_060660962.1
Ni/Fe hydrogenase [Helicobacter pullorum] Length: 385\nNumber of predicted TMHs: 1\nExp number of AAs in TMHs: 22.13084\nExp number, first 60 AAs: 0.64285\nTotal prob of N-in: 0.09674\noutside 1 355\nTMhelix 356 378\ninside 379 385

1754 GCF_001698685.1_ASM169868v1 Helicobacter pullorum Proteobacteria; delta/epsilon subdivisions; Epsilonproteobacteria; Campylobacterales; Helicobacteraceae; Helicobacter
MEREQVLLQKAQERLKEINKFPALKKGNSIKKMLKENGISRRDFMKWAGAMTAMLSLPAS WP_065827860.1
Ni/Fe hydrogenase [Helicobacter pullorum] Length: 385\nNumber of predicted TMHs: 1\nExp number of AAs in TMHs: 21.38773\nExp number, first 60 AAs: 0.64291\nTotal prob of N-in: 0.08711\noutside 1 355\nTMhelix 356 378\ninside 379 385

1755 GCF_001698765.1_ASM169876v1 Helicobacter pullorum Proteobacteria; delta/epsilon subdivisions; Epsilonproteobacteria; Campylobacterales; Helicobacteraceae; Helicobacter
MEREQVLLQKAQERLKEINKFPALKKGNSIKKMLKENGISRRDFMKWAGAMTAMLSLPAS WP_054194698.1
Ni/Fe hydrogenase [Helicobacter pullorum] Length: 385\nNumber of predicted TMHs: 1\nExp number of AAs in TMHs: 21.42489\nExp number, first 60 AAs: 0.64344\nTotal prob of N-in: 0.08825\noutside 1 355\nTMhelix 356 378\ninside 379 385

1756 GCF_001299515.1_ASM129951v1 Helicobacter pullorum Proteobacteria; delta/epsilon subdivisions; Epsilonproteobacteria; Campylobacterales; Helicobacteraceae; Helicobacter
MEREQVLLQKAQERLKEINKFPALKKGNSIKKMLKENGISRRDFMKWAGAMTAMLSLPAS WP_060660962.1
Ni/Fe hydrogenase [Helicobacter pullorum] Length: 385\nNumber of predicted TMHs: 1\nExp number of

AAs in TMHs: 22.13084\nExp number, first 60 AAs: 0.64285\nTotal prob of N-in: 0.09674\noutside 1 355\nTMhelix 356 378\ninside 379 385

1757 GCF_001698695.1_ASM169869v1 Helicobacter pullorum Proteobacteria; delta/epsilon subdivisions; Epsilonproteobacteria; Campylobacterales; Helicobacteraceae; Helicobacter MEREQVLLQKAQERLKEINKFPALKKGNSIKKMLKENGISRRDFMKWAGAMTAMLSLPAS WP_005020544.1 quinone-reactive Ni/Fe-hydrogenase small chain [Helicobacter pullorum] Length: 385\nNumber of predicted TMHs: 1\nExp number of AAs in TMHs: 21.91577\nExp number, first 60 AAs: 0.64271\nTotal prob of N-in: 0.09800\noutside 1 355\nTMhelix 356 378\ninside 379 385

1758 GCF_001698615.1_ASM169861v1 Helicobacter pullorum Proteobacteria; delta/epsilon subdivisions; Epsilonproteobacteria; Campylobacterales; Helicobacteraceae; Helicobacter MEREQVLLQKAQERLKEINKFPALKKGNSIKKMLKENGISRRDFMKWAGAMTAMLSLPAS WP_065827860.1 Ni/Fe hydrogenase [Helicobacter pullorum] Length: 385\nNumber of predicted TMHs: 1\nExp number of AAs in TMHs: 21.38773\nExp number, first 60 AAs: 0.64291\nTotal prob of N-in: 0.08711\noutside 1 355\nTMhelix 356 378\ninside 379 385

1759 GCF_000762845.1_ASM76284v1 Helicobacter sp. MIT 01-6451 Proteobacteria; delta/epsilon subdivisions; Epsilonproteobacteria; Campylobacterales; Helicobacteraceae; Helicobacter MKIQDAHQELFQKASKRLERLSKVSAHNADKNLLQELKSKGFSRRDFIKWSGMMTAALAL WP_034363568.1 Ni/Fe hydrogenase [Helicobacter sp. MIT 01-6451] Length: 386\nNumber of predicted TMHs: 1\nExp number of AAs in TMHs: 27.93636\nExp number, first 60 AAs: 4.8463\nTotal prob of N-in: 0.45799\noutside 1 357\nTMhelix 358 380\ninside 381 386

1760 GCF_000762925.1_ASM76292v1 Helicobacter sp. MIT 03-1614 Proteobacteria; delta/epsilon subdivisions; Epsilonproteobacteria; Campylobacterales; Helicobacteraceae; Helicobacter MKTQDAYQELFKKASQRLDELSKLPAHNAEKDLLQELESKGFSRRDFMKWSGMMTAALAL WP_034367361.1 Ni/Fe hydrogenase [Helicobacter sp. MIT 03-1614] Length: 386\nNumber of predicted TMHs: 1\nExp number of AAs in TMHs: 26.27319\nExp number, first 60 AAs: 3.44018\nTotal prob of N-in: 0.40058\noutside 1 357\nTMhelix 358 380\ninside 381 386

1761 GCF_000765685.1_ASM76568v1 Helicobacter sp. MIT 03-1616 Proteobacteria; delta/epsilon subdivisions; Epsilonproteobacteria; Campylobacterales; Helicobacteraceae; Helicobacter MKTQDAHKELFGKASQRLEELSKLPAHNAERDLLTELESKGFSRRDFIKWSGMMTAALAL WP_034326141.1 MULTISPECIES: Ni/Fe hydrogenase [Helicobacter] Length: 386\nNumber of predicted TMHs: 1\nExp number of AAs in TMHs: 27.15789\nExp number, first 60 AAs: 3.71904\nTotal prob of N-in: 0.40327\noutside 1 357\nTMhelix 358 380\ninside 381 386

1762 GCF_000765665.1_ASM76566v1 Helicobacter sp. MIT 05-5293 Proteobacteria; delta/epsilon subdivisions; Epsilonproteobacteria; Campylobacterales; Helicobacteraceae; Helicobacter MQRRNFLKSLALTPVLVASANAESTDSTKKDTRSLIDSSKFSKTKGKASIIDLDKCDGCV WP_034369927.1 oxidoreductase [Helicobacter sp. MIT 05-5293] Length: 341\nNumber of predicted TMHs: 1\nExp number of AAs in TMHs: 18.34608\nExp number, first 60 AAs: 0.00738\nTotal prob of N-in: 0.11879\noutside 1 310\nTMhelix 311 333\ninside 334 341

1763 GCF_000765695.1_ASM76569v1 Helicobacter sp. MIT 11-5569 Proteobacteria; delta/epsilon subdivisions; Epsilonproteobacteria; Campylobacterales; Helicobacteraceae; Helicobacter MERENNLLQQVESRLAEIAKFPALKKGNSIKKMLKEKGISRRDFMKWAGAMTAMLSLPAS WP_034286826.1 Ni/Fe hydrogenase [Helicobacter sp. MIT 11-5569] Length: 383\nNumber of predicted TMHs: 1\nExp number of AAs in TMHs: 21.82971\nExp number, first 60 AAs: 1.96249\nTotal prob of N-in: 0.15566\noutside 1 355\nTMhelix 356 378\ninside 379 383

1764 GCF_000765695.1_ASM76569v1 Helicobacter sp. MIT 11-5569 Proteobacteria; delta/epsilon subdivisions; Epsilonproteobacteria; Campylobacterales; Helicobacteraceae; Helicobacter MNRRNFFKIIGIAGVGALSANAKEIPQNTQDSIKSPYKTASIIDVGLCDGCKDLGTPKCV WP_052083416.1 oxidoreductase [Helicobacter sp. MIT 11-5569] Length: 344\nNumber of predicted TMHs: 1\nExp number of AAs in TMHs: 19.21064\nExp number, first 60 AAs: 0.0782\nTotal prob of N-in: 0.11344\noutside 1 311\nTMhelix 312 334\ninside 335 344

1765 GCF_000765905.1_ASM76590v1 Helicobacter trogontum Proteobacteria; delta/epsilon subdivisions; Epsilonproteobacteria; Campylobacterales; Helicobacteraceae; Helicobacter

MQRRNFLKSLALTPMLVAGANAESSDSTKDGKKEKRSLIDSSEFSKTKGKASIINLDKCD WP_034344427.1 oxidoreductase [Helicobacter trogontum] Length: 348\nNumber of predicted TMHs: 1\nExp number of AAs in TMHs: 18.94377\nExp number, first 60 AAs: 0.01029\nTotal prob of N-in: 0.05606\noutside 1 313\nTMhelix 314 336\ninside 337 348

1766 GCF_001460635.1_htyphlonius_mit97-6810 Helicobacter typhlonius Proteobacteria; delta/epsilon subdivisions; Epsilonproteobacteria; Campylobacterales; Helicobacteraceae; Helicobacter MKTQDAHKELFGKASQRLEELSKLPAHNAERDLLTELESKGFSRRDFIKWSGMMTAALAL WP_034326141.1 MULTISPECIES: Ni/Fe hydrogenase [Helicobacter] Length: 386\nNumber of predicted TMHs: 1\nExp number of AAs in TMHs: 27.15789\nExp number, first 60 AAs: 3.71904\nTotal prob of N-in: 0.40327\noutside 1 357\nTMhelix 358 380\ninside 381 386

1767 GCF_000765765.1_ASM76576v1 Helicobacter typhlonius Proteobacteria; delta/epsilon subdivisions; Epsilonproteobacteria; Campylobacterales; Helicobacteraceae; Helicobacter MKTQDAHKELFGKASQRLEELSKLPAHNAERDLLTELESKGFSRRDFIKWSGMMTAALAL WP_034326141.1 MULTISPECIES: Ni/Fe hydrogenase [Helicobacter] Length: 386\nNumber of predicted TMHs: 1\nExp number of AAs in TMHs: 27.15789\nExp number, first 60 AAs: 3.71904\nTotal prob of N-in: 0.40327\noutside 1 357\nTMhelix 358 380\ninside 381 386

1768 GCF_000158435.2_Heli_bilis_ATCC43879_V2 Helicobacter bilis ATCC 43879 Proteobacteria; delta/epsilon subdivisions; Epsilonproteobacteria; Campylobacterales; Helicobacteraceae; Helicobacter; Helicobacter bilis MQRRNFLKSLALTPVLVAGANAESSDSTKEGKEKTRSLIDSSEFSKTKGKASIINLDKCD WP_005219174.1 oxidoreductase [Helicobacter bilis] Length: 345\nNumber of predicted TMHs: 1\nExp number of AAs in TMHs: 18.27428\nExp number, first 60 AAs: 0.01056\nTotal prob of N-in: 0.10111\noutside 1 313\nTMhelix 314 336\ninside 337 345

1769 GCF_000686565.1_ASM68656v1 Helicobacter bilis ATCC 51630 Proteobacteria; delta/epsilon subdivisions; Epsilonproteobacteria; Campylobacterales; Helicobacteraceae; Helicobacter; Helicobacter bilis MQRRNFLKSLALTPVLVAGANAEPSDSTKEGKEETRSLIDSSEFSKTKGKASIINLDKCD WP_004088007.1 hypothetical protein [Helicobacter bilis] Length: 345\nNumber of predicted TMHs: 1\nExp number of AAs in TMHs: 18.27315\nExp number, first 60 AAs: 0.00939\nTotal prob of N-in: 0.10106\noutside 1 313\nTMhelix 314 336\ninside 337 345

1770 GCF_000364285.1_Heli_bili_WiWa_V1 Helicobacter bilis WiWa Proteobacteria; delta/epsilon subdivisions; Epsilonproteobacteria; Campylobacterales; Helicobacteraceae; Helicobacter; Helicobacter bilis MQRRNFLKSLALTPVLVAGANAEPSDSTKEGKEETRSLIDSSEFSKTKGKASIINLDKCD WP_004088007.1 hypothetical protein [Helicobacter bilis] Length: 345\nNumber of predicted TMHs: 1\nExp number of AAs in TMHs: 18.27315\nExp number, first 60 AAs: 0.00939\nTotal prob of N-in: 0.10106\noutside 1 313\nTMhelix 314 336\ninside 337 345

1771 GCF_000155455.1_ASM15545v1 Helicobacter canadensis MIT 98-5491 Proteobacteria; delta/epsilon subdivisions; Epsilonproteobacteria; Campylobacterales; Helicobacteraceae; Helicobacter; Helicobacter canadensis MEREELLLQKAQKRLKEISKFPALKEGNSIKKMLQEKGVSRRDFLKWAGAMTAMLSLPAS WP_006655924.1 quinone-reactive Ni/Fe-hydrogenase small chain [Helicobacter canadensis] Length: 383\nNumber of predicted TMHs: 1\nExp number of AAs in TMHs: 21.90653\nExp number, first 60 AAs: 0.59315\nTotal prob of N-in: 0.07627\noutside 1 356\nTMhelix 357 376\ninside 377 383

1772 GCF_000162575.1_ASM16257v1 Helicobacter canadensis MIT 98-5491 Proteobacteria; delta/epsilon subdivisions; Epsilonproteobacteria; Campylobacterales; Helicobacteraceae; Helicobacter; Helicobacter canadensis MEREELLLQKAQKRLKEISKFPALKEGNSIKKMLQEKGVSRRDFLKWAGAMTAMLSLPAS WP_006655924.1 quinone-reactive Ni/Fe-hydrogenase small chain [Helicobacter canadensis] Length: 383\nNumber of predicted TMHs: 1\nExp number of AAs in TMHs: 21.90653\nExp number, first 60 AAs: 0.59315\nTotal prob of N-in: 0.07627\noutside 1 356\nTMhelix 357 376\ninside 377 383

1773 GCF_000507865.1_Heli_cani_NCTC_12740_V1 Helicobacter canis NCTC 12740 Proteobacteria; delta/epsilon subdivisions; Epsilonproteobacteria; Campylobacterales; Helicobacteraceae; Helicobacter; Helicobacter canis MAEVSARLDAIEKLPAHNGDKELIAELESKGISRRDFMKWAGAMTAALSLPASFAPLTAR WP_034578294.1 Ni/Fe hydrogenase [Helicobacter canis] Length: 376\nNumber of predicted TMHs: 1\nExp number of AAs in TMHs: 22.2099\nExp number, first 60 AAs: 2.28235\nTotal prob of N-in: 0.15417\noutside 1 350\nTMhelix 351 373\ninside 374 376

1774 GCF_000509365.1_ASM50936v1 Helicobacter fennelliae MRY12-0050 Proteobacteria; delta/epsilon subdivisions; Epsilonproteobacteria; Campylobacterales; Helicobacteraceae; Helicobacter; Helicobacter fennelliae

MEDKHAKHLFSMVSKRLKELEKMPIHNEHKNLMDELHKNGLTRRDFMKWAGAMTAALSLP WP_023949824.1 quinone-reactive Ni/Fe-hydrogenase small chain precursor [Helicobacter fennelliae] Length: 384\nNumber of predicted TMHs: 1\nExp number of AAs in TMHs: 28.48957\nExp number, first 60 AAs: 4.52893\nTotal prob of N-in: 0.23216\noutside 1 358\nTMhelix 359 381\ninside 382 384

1775 GCF_000007905.1_ASM790v1 Helicobacter hepaticus ATCC 51449 Proteobacteria; delta/epsilon subdivisions; Epsilonproteobacteria; Campylobacterales; Helicobacteraceae; Helicobacter; Helicobacter hepaticus MEETNQNNQTNTSRRNFLKFLGVASVSAVSVNAAQSTDSKQPQYKMASIIDLGLCDGCRD WP_011116318.1 oxidoreductase [Helicobacter hepaticus] Length: 345\nNumber of predicted TMHs: 1\nExp number of AAs in TMHs: 23.35456\nExp number, first 60 AAs: 3.01427\nTotal prob of N-in: 0.55498\ninside 1 313\nTMhelix 314 336\noutside 337 345

1776 GCF_000007905.1_ASM790v1 Helicobacter hepaticus ATCC 51449 Proteobacteria; delta/epsilon subdivisions; Epsilonproteobacteria; Campylobacterales; Helicobacteraceae; Helicobacter; Helicobacter hepaticus MKTQDAYQELFKKASQRLDELSKLPAHNAEKDLLQELESKGFSRRDFMKWSGMMTAALAL WP_011114899.1 quinone-reactive Ni/Fe-hydrogenase small chain [Helicobacter hepaticus] Length: 386\nNumber of predicted TMHs: 1\nExp number of AAs in TMHs: 26.27309\nExp number, first 60 AAs: 3.44018\nTotal prob of N-in: 0.40058\noutside 1 357\nTMhelix 358 380\ninside 381 386

1777 GCF_000091985.1_ASM9198v1 Helicobacter mustelae 12198 Proteobacteria; delta/epsilon subdivisions; Epsilonproteobacteria; Campylobacterales; Helicobacteraceae; Helicobacter; Helicobacter mustelae MKNEEHLFKKIAKRLDFLAELPACKEDKNLDAELSKKGIDRRDFMKWAGAMTATLALPAA WP_013023320.1 quinone-reactive Ni/Fe-hydrogenase small chain [Helicobacter mustelae] Length: 381\nNumber of predicted TMHs: 1\nExp number of AAs in TMHs: 42.9144\nExp number, first 60 AAs: 8.23453\nTotal prob of N-in: 0.60057\noutside 1 355\nTMhelix 356 375\ninside 376 381

1778 GCF_000155495.1_ASM15549v1 Helicobacter pullorum MIT 98-5489 Proteobacteria; delta/epsilon subdivisions; Epsilonproteobacteria; Campylobacterales; Helicobacteraceae; Helicobacter; Helicobacter pullorum MEREQVLLQKAQERLKEINKFPALKKGNSIKKMLKENGISRRDFMKWAGAMTAMLSLPAS WP_005020544.1 quinone-reactive Ni/Fe-hydrogenase small chain [Helicobacter pullorum] Length: 385\nNumber of predicted TMHs: 1\nExp number of AAs in TMHs: 21.91577\nExp number, first 60 AAs: 0.64271\nTotal prob of N-in: 0.09800\noutside 1 355\nTMhelix 356 378\ninside 379 385

1779 GCF_000687535.1_ASM68753v1 Helicobacter rodentium ATCC 700285 Proteobacteria; delta/epsilon subdivisions; Epsilonproteobacteria; Campylobacterales; Helicobacteraceae; Helicobacter; Helicobacter rodentium MEKEQTLLRQAEERLAKLAKFPALKKGDSIKKMLKEKGISRRDFMKWAGAMTAMLSLPAS WP_026944261.1 Ni/Fe hydrogenase [Helicobacter rodentium] Length: 383\nNumber of predicted TMHs: 1\nExp number of AAs in TMHs: 21.98335\nExp number, first 60 AAs: 1.41287\nTotal prob of N-in: 0.13250\noutside 1 355\nTMhelix 356 378\ninside 379 383

1780 GCF_000687535.1_ASM68753v1 Helicobacter rodentium ATCC 700285 Proteobacteria; delta/epsilon subdivisions; Epsilonproteobacteria; Campylobacterales; Helicobacteraceae; Helicobacter; Helicobacter rodentium MKETNRNNQANSSRRRFLKFLGVASVSAVSVNATQPTDSKVSSLAHNPQYKMASIIDLGL WP_026943105.1 oxidoreductase [Helicobacter rodentium] Length: 351\nNumber of predicted TMHs: 1\nExp number of AAs in TMHs: 19.85703\nExp number, first 60 AAs: 2.41082\nTotal prob of N-in: 0.40173\noutside 1 317\nTMhelix 318 340\ninside 341 351

1781 GCF_000158455.1_ASM15845v1 Helicobacter winghamensis ATCC BAA-430 Proteobacteria; delta/epsilon subdivisions; Epsilonproteobacteria; Campylobacterales; Helicobacteraceae; Helicobacter; Helicobacter winghamensis MEKDNLLLKQAEERLAKLAKFPALKKGNSIKKMLKEKGVSRRDFMKWAGAMTAMLSLPAS WP_083781382.1 Ni/Fe hydrogenase [Helicobacter winghamensis] Length: 383\nNumber of predicted TMHs: 1\nExp number of AAs in TMHs: 21.95985\nExp number, first 60 AAs: 1.02856\nTotal prob of N-in: 0.07495\noutside 1 355\nTMhelix 356 378\ninside 379 383

1782 GCF_000196135.1_ASM19613v1 Wolinella succinogenes DSM 1740 Proteobacteria; delta/epsilon subdivisions; Epsilonproteobacteria; Campylobacterales; Helicobacteraceae; Wolinella; Wolinella succinogenes MERTQELFKKASQRLQELSSLPAIREDKDIPQMLEEKGIERRDFMKWAGAMTAMLSLPAT WP_011139498.1 quinone-reactive Ni/Fe-hydrogenase small chain [Wolinella succinogenes] Length: 386\nNumber of predicted TMHs: 1\nExp number of AAs in TMHs: 23.00028\nExp number, first 60 AAs: 1.96035\nTotal prob of N-in: 0.30254\noutside 1 357\nTMhelix 358 380\ninside 381 386

1783 GCF_000196135.1_ASM19613v1 Wolinella succinogenes DSM 1740 Proteobacteria; delta/epsilon subdivisions; Epsilonproteobacteria; Campylobacterales; Helicobacteraceae; Wolinella; Wolinella succinogenes MENLSRRRFFKKVALASTLSLAPLSVQAKEEKTNQKRIASIIDLTLCDGCEGEAMPLCVK WP_011138510.1 oxidoreductase [Wolinella succinogenes] Length: 329\nNumber of predicted TMHs: 1\nExp number of AAs in TMHs: 19.85761\nExp number, first 60 AAs: 0.02132\nTotal prob of N-in: 0.03168\noutside 1 298\nTMhelix 299 321\ninside 322 329

1784 GCF_001705075.1_ASM170507v1 Acidiferrobacter thiooxydans Proteobacteria; Gammaproteobacteria; Acidiferrobacterales; Acidiferrobacteraceae; Acidiferrobacter MGGVLEKRGFTRRAFLQLCATVSATLALSPADAAEMARRLQYAPRPTVIYLSYQECTGCLWP_065970735.1 Ni/Fe hydrogenase [Acidiferrobacter thiooxydans] Length: 358\nNumber of predicted TMHs: 1\nExp number of AAs in TMHs: 37.32241\nExp number, first 60 AAs: 1.07037\nTotal prob of N-in: 0.16268\noutside 1 316\nTMhelix 317 336\ninside 337 358

1785 GCF_000820025.1_PRJEB7034 Aeromonas popoffii Proteobacteria; Gammaproteobacteria; Aeromonadales; Aeromonadaceae; Aeromonas MVEDNVLSAHGINRRDFMKLCAGMAATLGLSQHAVAKMATALTGPERPPVIWIGAQECTG WP_042034313.1 hydrogenase 2 small subunit [Aeromonas popoffii] Length: 377\nNumber of predicted TMHs: 1\nExp number of AAs in TMHs: 21.27685\nExp number, first 60 AAs: 0.8388\nTotal prob of N-in: 0.04675\noutside 1 334\nTMhelix 335 357\ninside 358 377

1786 GCF_900156095.1_IMG-taxon_2708742413_annotated_assembly Aeromonas sp. RU39B Proteobacteria; Gammaproteobacteria; Aeromonadales; Aeromonadaceae; Aeromonas MHINDSFLTRLGVSRRDFMKFCAGLAATAGLPVSAAEKVASALSAPERPPVIWIGAQECT WP_076577314.1 hydrogenase 2 small subunit [Aeromonas sp. RU39B] Length: 381\nNumber of predicted TMHs: 1\nExp number of AAs in TMHs: 18.51034\nExp number, first 60 AAs: 0.43854\nTotal prob of N-in: 0.03369\noutside 1 333\nTMhelix 334 356\ninside 357 381

1787 GCF_000315195.1_ASM31519v1 Aeromonas dhakensis AAK1 Proteobacteria; Gammaproteobacteria; Aeromonadales; Aeromonadaceae; Aeromonas; Aeromonas dhakensis MVEDTFLSAHGINRRDFMKLCAGMAATLGLSQHAVAKMATALTSPERPPVIWIGAQECTG WP_010633706.1 MULTISPECIES: hydrogenase 2 small subunit [Aeromonas] Length: 377\nNumber of predicted TMHs: 1\nExp number of AAs in TMHs: 18.75985\nExp number, first 60 AAs: 0.45554\nTotal prob of N-in: 0.02788\noutside 1 334\nTMhelix 335 357\ninside 358 377

1788 GCF_000612075.2_ASM61207v2 Aeromonas hydrophila YL17 Proteobacteria; Gammaproteobacteria; Aeromonadales; Aeromonadaceae; Aeromonas; Aeromonas hydrophila MVEDTFLSAHGINRRDFMKLCAGMAATLGLSQHAVAKMATALTSPERPPVIWIGAQECTG WP_010633706.1 MULTISPECIES: hydrogenase 2 small subunit [Aeromonas] Length: 377\nNumber of predicted TMHs: 1\nExp number of AAs in TMHs: 18.75985\nExp number, first 60 AAs: 0.45554\nTotal prob of N-in: 0.02788\noutside 1 334\nTMhelix 335 357\ninside 358 377

1789 GCF_001043175.1_ASM104317v1 Marinobacter psychrophilus Proteobacteria; Gammaproteobacteria; Alteromonadales; Alteromonadaceae; Marinobacter MEERRNQPINTYQPDYSDEISLVDLAATFIRRRRVFYAVFLLCTLGGLAYALLTPEKFDY WP_048386345.1 lipopolysaccharide biosynthesis protein [Marinobacter psychrophilus] Length: 260\nNumber of predicted TMHs: 2\nExp number of AAs in TMHs: 41.52921\nExp number, first 60 AAs: 19.40121\nTotal prob of N-in: 0.93493\nPOSSIBLE N-term signal sequence\ninside 1 34\nTMhelix 35 54\noutside 55 223\nTMhelix 224 246\ninside 247 260

1790 GCF_000283275.1_ASM28327v1 Marinobacter sp. BSs20148 Proteobacteria; Gammaproteobacteria; Alteromonadales; Alteromonadaceae; Marinobacter MEERRNQPNNTYQPEYSDEISLVDLAATFIRRRRVFYAVFLLCTLGGLAYALLTPEKFDY WP_014871804.1 hypothetical protein [Marinobacter sp. BSs20148] Length: 261\nNumber of predicted TMHs: 2\nExp number of AAs in TMHs: 41.37642\nExp number, first 60 AAs: 19.41605\nTotal prob of N-in: 0.95579\nPOSSIBLE N-term signal sequence\ninside 1 34\nTMhelix 35 54\noutside 55 224\nTMhelix 225 247\ninside 248 261

1791 GCF_000686085.1_ASM68608v1 Marinobacter sp. HL-58 Proteobacteria; Gammaproteobacteria; Alteromonadales; Alteromonadaceae; Marinobacter MDSTPERRQTYDDEISLVDLATTFIRRRRVFYVVFLATTLAGLAYALLLVPEKYEYVSLY WP_027832639.1 LPS biosynthesis protein [Marinobacter sp. HL-58] Length: 255\nNumber of predicted TMHs: 2\nExp number of AAs in

TMHs: 43.13888\nExp number, first 60 AAs: 21.04712\nTotal prob of N-in: 0.93218\nPOSSIBLE N-term signal sequence\ninside 1 29\nTMhelix 30 52\noutside 53 221\nTMhelix 222 244\ninside 245 255

1792 GCF_002104475.1_ASM210447v1 Colwellia mytili Proteobacteria; Gammaproteobacteria; Alteromonadales; Colwelliaceae; Colwellia
MRISMTSFICSKSSIKLSRRQIINNVHCAMTLFLVTFISSFTLSAAELNMLALVSDRSAA WP_085298870.1 cobalamin biosynthesis protein CobN [Colwellia mytili] Length: 1541\nNumber of predicted TMHs: 2\nExp number of AAs in TMHs: 41.55097\nExp number, first 60 AAs: 22.11339\nTotal prob of N-in: 0.98396\nPOSSIBLE N-term signal sequence\ninside 1 20\nTMhelix 21 43\noutside 44 1511\nTMhelix 1512 1530\ninside 1531 1541

1793 GCF_900129905.1_IMG-taxon_2588254304_annotated_assembly Ferrimonas marina Proteobacteria; Gammaproteobacteria; Alteromonadales; Ferrimonadaceae; Ferrimonas
MNKYDALLAQCRQRLDSLAEQPPLKQASLRDSMEAQGISRRDFMKWSATTATVLALPMQF WP_067656581.1 hydrogenase [Ferrimonas marina] Length: 381\nNumber of predicted TMHs: 1\nExp number of AAs in TMHs: 20.90269\nExp number, first 60 AAs: 0.57046\nTotal prob of N-in: 0.09810\noutside 1 345\nTMhelix 346 368\ninside 369 381

1794 GCF_000148645.1_ASM14864v1 Ferrimonas balearica DSM 9799 Proteobacteria; Gammaproteobacteria; Alteromonadales; Ferrimonadaceae; Ferrimonas; Ferrimonas balearica
MSDYQALMARCEARLEKLAAQAPVKTDSLQTRLDAAGISRRDFMKWSAVTTTALALPMQF WP_013346648.1 quinone-reactive Ni/Fe-hydrogenase small chain [Ferrimonas balearica] Length: 381\nNumber of predicted TMHs: 1\nExp number of AAs in TMHs: 22.2201\nExp number, first 60 AAs: 0.42977\nTotal prob of N-in: 0.16070\noutside 1 349\nTMhelix 350 372\ninside 373 381

1795 GCF_000422645.1_ASM42264v1 Ferrimonas futtsuensis DSM 18154 Proteobacteria; Gammaproteobacteria; Alteromonadales; Ferrimonadaceae; Ferrimonas; Ferrimonas futtsuensis
MGQYDALFSRGKKRLSELKKINAVKSTPLMQRLESQGISRREFMTWSAQVTAMLSLPLSF WP_028110914.1 hydrogenase [Ferrimonas futtsuensis] Length: 379\nNumber of predicted TMHs: 1\nExp number of AAs in TMHs: 20.82494\nExp number, first 60 AAs: 0.08447\nTotal prob of N-in: 0.07645\noutside 1 349\nTMhelix 350 372\ninside 373 379

1796 GCF_000425405.1_ASM42540v1 Ferrimonas kyonanensis DSM 18153 Proteobacteria; Gammaproteobacteria; Alteromonadales; Ferrimonadaceae; Ferrimonas; Ferrimonas kyonanensis
MGQYDALFSRGKKRLTELKNIDAVKSTPLMKRLESQGISRREFMTWSAQVTAMLSLPLSF WP_028112169.1 hydrogenase [Ferrimonas kyonanensis] Length: 379\nNumber of predicted TMHs: 1\nExp number of AAs in TMHs: 21.56189\nExp number, first 60 AAs: 0.09644\nTotal prob of N-in: 0.06057\noutside 1 349\nTMhelix 350 372\ninside 373 379

1797 GCF_001571125.1_ASM157112v1 Ferrimonas marina NBRC 102583 Proteobacteria; Gammaproteobacteria; Alteromonadales; Ferrimonadaceae; Ferrimonas; Ferrimonas marina
MNKYDALLAQCRQRLDSLAEQPPLKQASLRDSMEAQGISRRDFMKWSATTATVLALPMQF WP_067656581.1 hydrogenase [Ferrimonas marina] Length: 381\nNumber of predicted TMHs: 1\nExp number of AAs in TMHs: 20.90269\nExp number, first 60 AAs: 0.57046\nTotal prob of N-in: 0.09810\noutside 1 345\nTMhelix 346 368\ninside 369 381

1798 GCF_000422665.1_ASM42266v1 Ferrimonas senticii DSM 18821 Proteobacteria; Gammaproteobacteria; Alteromonadales; Ferrimonadaceae; Ferrimonas; Ferrimonas senticii
MANYDALLVTARTRIAELKAQAPQKQKSLDTRMEEGGVSRRDFMKWSATTAAALGLPLQF WP_028115082.1 hydrogenase [Ferrimonas senticii] Length: 381\nNumber of predicted TMHs: 1\nExp number of AAs in TMHs: 19.46287\nExp number, first 60 AAs: 0.10673\nTotal prob of N-in: 0.07602\noutside 1 350\nTMhelix 351 373\ninside 374 381

1799 GCF_000422665.1_ASM42266v1 Ferrimonas senticii DSM 18821 Proteobacteria; Gammaproteobacteria; Alteromonadales; Ferrimonadaceae; Ferrimonas; Ferrimonas senticii
MSLSPRRRRILLARLLQIAFAGALVALVVLVWQQQRLQVQGLLHQQGQRSVDQLLAQASV WP_028116374.1 hypothetical protein [Ferrimonas senticii] Length: 224\nNumber of predicted TMHs: 2\nExp number of AAs in TMHs: 42.88506\nExp number, first 60 AAs: 22.30309\nTotal prob of N-in: 0.59658\nPOSSIBLE N-term signal sequence\noutside 1 9\nTMhelix 10 32\ninside 33 165\nTMhelix 166 185\noutside 186 224

1800 GCF_000947195.1_Shewanella_algae_MARS_14 Shewanella algae Proteobacteria; Gammaproteobacteria; Alteromonadales; Shewanellaceae; Shewanella

MDTHAALYEQGRQRLEKLRQLPARQDKSLQERMLDHGLTRRDFMKWSASVTALLALPLPF WP_044734537.1 hydrogenase [Shewanella algae] Length: 378\nNumber of predicted TMHs: 1\nExp number of AAs in TMHs: 23.63941\nExp number, first 60 AAs: 1.5826\nTotal prob of N-in: 0.19886\noutside 1 349\nTMhelix 350 372\ninside 373 378

1801 GCF_001858195.1_ASM185819v1 Shewanella algae Proteobacteria; Gammaproteobacteria; Alteromonadales; Shewanellaceae; Shewanella
MDTHAALYEQGRQRLEKLRQLPARQDKSLQERMLDHGLTRRDFMKWSASVTAMLALPLPF WP_062793451.1 hydrogenase [Shewanella algae] Length: 378\nNumber of predicted TMHs: 1\nExp number of AAs in TMHs: 23.10927\nExp number, first 60 AAs: 1.24403\nTotal prob of N-in: 0.18447\noutside 1 349\nTMhelix 350 372\ninside 373 378

1802 GCF_001870495.1_ASM187049v1 Shewanella algae Proteobacteria; Gammaproteobacteria; Alteromonadales; Shewanellaceae; Shewanella
MDTHAALYEQGRQRLEKLRQLPARQDKSLQERMLELGLTRRDFMKWSASVTAMMALPLPF WP_039035909.1 MULTISPECIES: hydrogenase [Shewanella] Length: 378\nNumber of predicted TMHs: 1\nExp number of AAs in TMHs: 23.71306\nExp number, first 60 AAs: 0.88418\nTotal prob of N-in: 0.18963\noutside 1 349\nTMhelix 350 372\ninside 373 378

1803 GCF_001620325.1_ASM162032v1 Shewanella baltica Proteobacteria; Gammaproteobacteria; Alteromonadales; Shewanellaceae; Shewanella
MDTHAALYEQGKARLDALRQFAPRQQQSLIEKMQQHGISRRDFMKWSAMVTSMLALPLPF WP_006081384.1 MULTISPECIES: quinone-reactive Ni/Fe-hydrogenase small chain [Shewanella] Length: 378\nNumber of predicted TMHs: 1\nExp number of AAs in TMHs: 24.66445\nExp number, first 60 AAs: 1.48416\nTotal prob of N-in: 0.25198\noutside 1 349\nTMhelix 350 372\ninside 373 378

1804 GCF_001735525.1_ASM173552v1 Shewanella colwelliana Proteobacteria; Gammaproteobacteria; Alteromonadales; Shewanellaceae; Shewanella
MDTHAALYEQGKARLEMLRSLEPRHAHSLQTKMEEHGVTRRDFMKWSAAVTAMLALPLPF WP_069672046.1 hydrogenase [Shewanella colwelliana] Length: 378\nNumber of predicted TMHs: 1\nExp number of AAs in TMHs: 26.60579\nExp number, first 60 AAs: 5.48181\nTotal prob of N-in: 0.40422\noutside 1 349\nTMhelix 350 372\ninside 373 378

1805 GCF_000753795.1_ASM75379v1 Shewanella mangrovi Proteobacteria; Gammaproteobacteria; Alteromonadales; Shewanellaceae; Shewanella
METHAALFEQGRKRLEALRQMPPRQQESLKDKLLEHGISRRDFMKWSASVTALLALPLPF WP_037439697.1 hydrogenase [Shewanella mangrovi] Length: 378\nNumber of predicted TMHs: 1\nExp number of AAs in TMHs: 21.9702\nExp number, first 60 AAs: 0.42258\nTotal prob of N-in: 0.06421\noutside 1 349\nTMhelix 350 372\ninside 373 378

1806 GCF_000753795.1_ASM75379v1 Shewanella mangrovi Proteobacteria; Gammaproteobacteria; Alteromonadales; Shewanellaceae; Shewanella
MDTHSALFEQGRKRLQALRQLPPRQETSLQERMLKHGITRRDFMKWSASVTALLALPLPF WP_037439703.1 hydrogenase [Shewanella mangrovi] Length: 378\nNumber of predicted TMHs: 1\nExp number of AAs in TMHs: 21.98065\nExp number, first 60 AAs: 1.20849\nTotal prob of N-in: 0.11804\noutside 1 349\nTMhelix 350 372\ninside 373 378

1807 GCF_900156405.1_IMG-taxon_2681812898_annotated_assembly Shewanella morhuae Proteobacteria; Gammaproteobacteria; Alteromonadales; Shewanellaceae; Shewanella
MDTHAALYEQGKARLDALRQFAPRQQQSLIEKMQQHGISRRDFMKWSAMVTGMLALPLPF WP_076500176.1 hydrogenase [Shewanella morhuae] Length: 378\nNumber of predicted TMHs: 1\nExp number of AAs in TMHs: 28.55763\nExp number, first 60 AAs: 4.4504\nTotal prob of N-in: 0.34088\noutside 1 349\nTMhelix 350 372\ninside 373 378

1808 GCF_000518445.1_ASM51844v1 Shewanella sp. 38A_GOM-205m Proteobacteria; Gammaproteobacteria; Alteromonadales; Shewanellaceae; Shewanella
MDTHAALYEQGRQRLEKLRQLPARQDKSLQERMLDHGLTRRDFMKWSASVTAMLALPLPF WP_028779962.1 hydrogenase [Shewanella sp. 38A_GOM-205m] Length: 378\nNumber of predicted TMHs: 1\nExp number of AAs in TMHs: 24.85972\nExp number, first 60 AAs: 1.31631\nTotal prob of N-in: 0.21290\noutside 1 349\nTMhelix 350 372\ninside 373 378

1809 GCF_000832025.1_ASM83202v1 Shewanella sp. cp20 Proteobacteria; Gammaproteobacteria; Alteromonadales; Shewanellaceae; Shewanella
MDTHAALYQQGQARLEELRRLAPRHAQTLQDKMEEHGITRRDFMKWSAAVTAMLALPLPF WP_041511451.1 hydrogenase [Shewanella sp. cp20] Length: 378\nNumber of predicted TMHs: 1\nExp number of AAs in TMHs: 29.33845\nExp number, first 60 AAs: 7.34673\nTotal prob of N-in: 0.54191\noutside 1 349\nTMhelix 350 372\ninside 373 378

1810 GCF_000773485.1_ASM77348v1 Shewanella sp. ECSMB14101 Proteobacteria; Gammaproteobacteria; Alteromonadales; Shewanellaceae; Shewanella
MDTHAALYQQGQARLEELRRLAPRHAHTLQDKMEEHGISRRDFMKWSAAVTAMLALPLPF WP_033538297.1 hydrogenase [Shewanella sp. ECSMB14101] Length: 378\nNumber of predicted TMHs: 1\nExp number of AAs in TMHs: 28.88504\nExp number, first 60 AAs: 6.08921\nTotal prob of N-in: 0.47582\noutside 1 349\nTMhelix 350 372\ninside 373 378

1811 GCF_000773485.1_ASM77348v1 Shewanella sp. ECSMB14101 Proteobacteria; Gammaproteobacteria; Alteromonadales; Shewanellaceae; Shewanella
MKLSRRSLLLSALTAAIACSTHADDTKPLQTLIPGQEKLKLERADKQRNDGMYFVRFKES WP_033540164.1 peptidase S8 [Shewanella sp. ECSMB14101] Length: 1192\nNumber of predicted TMHs: 1\nExp number of AAs in TMHs: 15.81229\nExp number, first 60 AAs: 0.13151\nTotal prob of N-in: 0.00725\noutside 1 1166\nTMhelix 1167 1186\ninside 1187 1192

1812 GCF_000813075.1_ASM81307v1 Shewanella sp. ECSMB14102 Proteobacteria; Gammaproteobacteria; Alteromonadales; Shewanellaceae; Shewanella
MDTHAALYEQGRQRLEKLRQLPARQDKSLQERMLELGLTRRDFMKWSASVTAMMALPLPF WP_039035909.1 MULTISPECIES: hydrogenase [Shewanella] Length: 378\nNumber of predicted TMHs: 1\nExp number of AAs in TMHs: 23.71306\nExp number, first 60 AAs: 0.88418\nTotal prob of N-in: 0.18963\noutside 1 349\nTMhelix 350 372\ninside 373 378

1813 GCF_001887095.2_ASM188709v2 Shewanella sp. SACH Proteobacteria; Gammaproteobacteria; Alteromonadales; Shewanellaceae; Shewanella
MDTHAALYEQGKARLDALRQFAPRQQQSLIEKMQQHGISRRDFMKWSAMVTSMLALPLPF WP_006081384.1 MULTISPECIES: quinone-reactive Ni/Fe-hydrogenase small chain [Shewanella] Length: 378\nNumber of predicted TMHs: 1\nExp number of AAs in TMHs: 24.66445\nExp number, first 60 AAs: 1.48416\nTotal prob of N-in: 0.25198\noutside 1 349\nTMhelix 350 372\ninside 373 378

1814 GCF_000015185.1_ASM1518v1 Shewanella sp. W3-18-1 Proteobacteria; Gammaproteobacteria; Alteromonadales; Shewanellaceae; Shewanella
MDTHAALYEQGKARLEALRQLAPRQQQSLVEKMQQHGISRRDFMKWSAMVTGMLALPLPF WP_011789250.1 MULTISPECIES: quinone-reactive Ni/Fe-hydrogenase small chain [Shewanella] Length: 378\nNumber of predicted TMHs: 1\nExp number of AAs in TMHs: 28.13644\nExp number, first 60 AAs: 2.74758\nTotal prob of N-in: 0.30256\noutside 1 349\nTMhelix 350 372\ninside 373 378

1815 GCF_001598875.1_ASM159887v1 Shewanella algae JCM 21037 = NBRC 103173 Proteobacteria; Gammaproteobacteria; Alteromonadales; Shewanellaceae; Shewanella; Shewanella algae
MDTHAALYEQGRQRLEKLRQLPARQDKSLQERMLDHGLTRRDFMKWSASVTAMLALPLPF WP_062793451.1 hydrogenase [Shewanella algae] Length: 378\nNumber of predicted TMHs: 1\nExp number of AAs in TMHs: 23.10927\nExp number, first 60 AAs: 1.24403\nTotal prob of N-in: 0.18447\noutside 1 349\nTMhelix 350 372\ninside 373 378

1816 GCF_000015245.1_ASM1524v1 Shewanella amazonensis SB2B Proteobacteria; Gammaproteobacteria; Alteromonadales; Shewanellaceae; Shewanella; Shewanella amazonensis
MDTHAALYEQGKQRLETLRSLPRRHEETIFDKMAQHGITRRDFMKWSASVTAMLALPLPF WP_011759793.1 quinone-reactive Ni/Fe-hydrogenase small chain [Shewanella amazonensis] Length: 378\nNumber of predicted TMHs: 1\nExp number of AAs in TMHs: 27.38419\nExp number, first 60 AAs: 4.25081\nTotal prob of N-in: 0.32575\noutside 1 349\nTMhelix 350 372\ninside 373 378

1817 GCF_000147735.2_ASM14773v3 Shewanella baltica BA175 Proteobacteria; Gammaproteobacteria; Alteromonadales; Shewanellaceae; Shewanella; Shewanella baltica
MDTHAALYEQGKARLDALRQFAPRQQQSLIEKMQQHGISRRDFMKWSAMVTSMLALPLPF WP_006081384.1 MULTISPECIES: quinone-reactive Ni/Fe-hydrogenase small chain [Shewanella] Length: 378\nNumber of

predicted TMHs: 1\nExp number of AAs in TMHs: 24.66445\nExp number, first 60 AAs: 1.48416\nTotal prob of N-in: 0.25198\noutside 1 349\nTMhelix 350 372\ninside 373 378

1818 GCF_000215895.1_ASM21589v1 Shewanella baltica OS117 Proteobacteria; Gammaproteobacteria; Alteromonadales; Shewanellaceae; Shewanella; Shewanella baltica
MDTHAALYEQGKARLDALRQFAPRQQQSLIEKMQQHGISRRDFMKWSAMVTSMLALPLPF WP_011846647.1 quinone-reactive Ni/Fe-hydrogenase small chain [Shewanella baltica] Length: 378\nNumber of predicted TMHs: 1\nExp number of AAs in TMHs: 23.92704\nExp number, first 60 AAs: 1.47675\nTotal prob of N-in: 0.24104\noutside 1 349\nTMhelix 350 372\ninside 373 378

1819 GCF_000015845.1_ASM1584v1 Shewanella baltica OS155 Proteobacteria; Gammaproteobacteria; Alteromonadales; Shewanellaceae; Shewanella; Shewanella baltica
MDTHAALYEQGKARLDALRQFAPRQQQSLIEKMQQHGISRRDFMKWSAMVTSMLALPLPF WP_011846647.1 quinone-reactive Ni/Fe-hydrogenase small chain [Shewanella baltica] Length: 378\nNumber of predicted TMHs: 1\nExp number of AAs in TMHs: 23.92704\nExp number, first 60 AAs: 1.47675\nTotal prob of N-in: 0.24104\noutside 1 349\nTMhelix 350 372\ninside 373 378

1820 GCF_000179535.2_ASM17953v2 Shewanella baltica OS183 Proteobacteria; Gammaproteobacteria; Alteromonadales; Shewanellaceae; Shewanella; Shewanella baltica
MDTHAALYEQGKARLDALRQFAPRQQQSLIEKMQQHGISRRDFMKWSAMVTSMLALPLPF WP_006081384.1 MULTISPECIES: quinone-reactive Ni/Fe-hydrogenase small chain [Shewanella] Length: 378\nNumber of predicted TMHs: 1\nExp number of AAs in TMHs: 24.66445\nExp number, first 60 AAs: 1.48416\nTotal prob of N-in: 0.25198\noutside 1 349\nTMhelix 350 372\ninside 373 378

1821 GCF_000017325.1_ASM1732v1 Shewanella baltica OS185 Proteobacteria; Gammaproteobacteria; Alteromonadales; Shewanellaceae; Shewanella; Shewanella baltica
MDTHAALYEQGKARLDALRQFAPRQQQSLIEKMQQHGISRRDFMKWSAMVTSMLALPLPF WP_006081384.1 MULTISPECIES: quinone-reactive Ni/Fe-hydrogenase small chain [Shewanella] Length: 378\nNumber of predicted TMHs: 1\nExp number of AAs in TMHs: 24.66445\nExp number, first 60 AAs: 1.48416\nTotal prob of N-in: 0.25198\noutside 1 349\nTMhelix 350 372\ninside 373 378

1822 GCF_000018765.1_ASM1876v1 Shewanella baltica OS195 Proteobacteria; Gammaproteobacteria; Alteromonadales; Shewanellaceae; Shewanella; Shewanella baltica
MDTHAALYEQGKARLDALRQFAPRQQQSLIEKMQQHGISRRDFMKWSAMVTSMLALPLPF WP_006081384.1 MULTISPECIES: quinone-reactive Ni/Fe-hydrogenase small chain [Shewanella] Length: 378\nNumber of predicted TMHs: 1\nExp number of AAs in TMHs: 24.66445\nExp number, first 60 AAs: 1.48416\nTotal prob of N-in: 0.25198\noutside 1 349\nTMhelix 350 372\ninside 373 378

1823 GCF_000021665.1_ASM2166v1 Shewanella baltica OS223 Proteobacteria; Gammaproteobacteria; Alteromonadales; Shewanellaceae; Shewanella; Shewanella baltica
MDTHAALYEQGKARLDALRQFAPRQQQSLIEKMQQHGISRRDFMKWSAMVTSMLALPLPF WP_006081384.1 MULTISPECIES: quinone-reactive Ni/Fe-hydrogenase small chain [Shewanella] Length: 378\nNumber of predicted TMHs: 1\nExp number of AAs in TMHs: 24.66445\nExp number, first 60 AAs: 1.48416\nTotal prob of N-in: 0.25198\noutside 1 349\nTMhelix 350 372\ninside 373 378

1824 GCF_000231345.1_ASM23134v2 Shewanella baltica OS625 Proteobacteria; Gammaproteobacteria; Alteromonadales; Shewanellaceae; Shewanella; Shewanella baltica
MDTHAALYEQGKARLDALRQFAPRQQQSLIEKMQQHGISRRDFMKWSAMVTSMLALPLPF WP_006081384.1 MULTISPECIES: quinone-reactive Ni/Fe-hydrogenase small chain [Shewanella] Length: 378\nNumber of predicted TMHs: 1\nExp number of AAs in TMHs: 24.66445\nExp number, first 60 AAs: 1.48416\nTotal prob of N-in: 0.25198\noutside 1 349\nTMhelix 350 372\ninside 373 378

1825 GCF_000178875.2_ASM17887v2 Shewanella baltica OS678 Proteobacteria; Gammaproteobacteria; Alteromonadales; Shewanellaceae; Shewanella; Shewanella baltica
MDTHAALYEQGKARLDALRQFAPRQQQSLIEKMQQHGISRRDFMKWSAMVTSMLALPLPF WP_006081384.1 MULTISPECIES: quinone-reactive Ni/Fe-hydrogenase small chain [Shewanella] Length: 378\nNumber of predicted TMHs: 1\nExp number of AAs in TMHs: 24.66445\nExp number, first 60 AAs: 1.48416\nTotal prob of N-in: 0.25198\noutside 1 349\nTMhelix 350 372\ninside 373 378

1826 GCF_000518705.1_ASM51870v1 Shewanella colwelliana ATCC 39565 Proteobacteria; Gammaproteobacteria; Alteromonadales; Shewanellaceae; Shewanella; Shewanella colwelliana

MDTHAALYEQGKARLEMLRSLEPRHAHSLQTKMEEHGVTRRDFMKWSAAVTAMLALPLPF WP_028762125.1 hydrogenase [Shewanella colwelliana] Length: 378\nNumber of predicted TMHs: 1\nExp number of AAs in TMHs: 26.76957\nExp number, first 60 AAs: 5.68247\nTotal prob of N-in: 0.41249\noutside 1 349\nTMhelix 350 372\ninside 373 378

1827 GCF_000016065.1_ASM1606v1 Shewanella loihica PV-4 Proteobacteria; Gammaproteobacteria; Alteromonadales; Shewanellaceae; Shewanella; Shewanella loihica
MDTHAALYQQGQARLEELRRLAPRHAQTLQDKMEEHGITRRDFMKWSAAVTAMLALPLPF WP_011865563.1 quinone-reactive Ni/Fe-hydrogenase small chain [Shewanella loihica] Length: 378\nNumber of predicted TMHs: 1\nExp number of AAs in TMHs: 29.52983\nExp number, first 60 AAs: 7.3078\nTotal prob of N-in: 0.54348\noutside 1 349\nTMhelix 350 372\ninside 373 378

1828 GCF_000614975.1_ASM61497v1 Shewanella marina JCM 15074 Proteobacteria; Gammaproteobacteria; Alteromonadales; Shewanellaceae; Shewanella; Shewanella marina
MDTHAALYEQGLARIASLRKLAPRKSKTLQQVMEDHGITRRDFMKWSASISAMLALPLPF WP_025820305.1 hydrogenase [Shewanella marina] Length: 379\nNumber of predicted TMHs: 1\nExp number of AAs in TMHs: 35.4395\nExp number, first 60 AAs: 2.7214\nTotal prob of N-in: 0.41530\noutside 1 350\nTMhelix 351 373\ninside 374 379

1829 GCF_000169215.2_ASM16921v2 Shewanella putrefaciens 200 Proteobacteria; Gammaproteobacteria; Alteromonadales; Shewanellaceae; Shewanella; Shewanella putrefaciens
MDTHAALYEQGKARLEALRQLAPRQQQSLVEKMQQHGISRRDFMKWSAMVTGMLALPLPF WP_011789250.1 MULTISPECIES: quinone-reactive Ni/Fe-hydrogenase small chain [Shewanella] Length: 378\nNumber of predicted TMHs: 1\nExp number of AAs in TMHs: 28.13644\nExp number, first 60 AAs: 2.74758\nTotal prob of N-in: 0.30256\noutside 1 349\nTMhelix 350 372\ninside 373 378

1830 GCF_000016585.1_ASM1658v1 Shewanella putrefaciens CN-32 Proteobacteria; Gammaproteobacteria; Alteromonadales; Shewanellaceae; Shewanella; Shewanella putrefaciens
MDTHAALYEQGKARLEALRQLAPRQQQSLVEKMQQHGISRRDFMKWSAMVTGMLALPLPF WP_011789250.1 MULTISPECIES: quinone-reactive Ni/Fe-hydrogenase small chain [Shewanella] Length: 378\nNumber of predicted TMHs: 1\nExp number of AAs in TMHs: 28.13644\nExp number, first 60 AAs: 2.74758\nTotal prob of N-in: 0.30256\noutside 1 349\nTMhelix 350 372\ninside 373 378

1831 GCF_000519065.1_ASM51906v1 Shewanella putrefaciens HRCR-6 Proteobacteria; Gammaproteobacteria; Alteromonadales; Shewanellaceae; Shewanella; Shewanella putrefaciens
MDTHAALYEQGKARLDALRQFAPRQQQSLIEKMQQHGISRRDFMKWSAMVTSMLALPLPF WP_028760176.1 hydrogenase [Shewanella putrefaciens] Length: 378\nNumber of predicted TMHs: 1\nExp number of AAs in TMHs: 24.6609\nExp number, first 60 AAs: 1.4845\nTotal prob of N-in: 0.25189\noutside 1 349\nTMhelix 350 372\ninside 373 378

1832 GCF_001591325.1_ASM159132v1 Shewanella putrefaciens JCM 20190 = NBRC 3908 Proteobacteria; Gammaproteobacteria; Alteromonadales; Shewanellaceae; Shewanella; Shewanella putrefaciens
MDTHAALYEQGKARLEALRQLAPRQQQSLVEKMQQHGISRRDFMKWSAMVTGMLALPLPF WP_011789250.1 MULTISPECIES: quinone-reactive Ni/Fe-hydrogenase small chain [Shewanella] Length: 378\nNumber of predicted TMHs: 1\nExp number of AAs in TMHs: 28.13644\nExp number, first 60 AAs: 2.74758\nTotal prob of N-in: 0.30256\noutside 1 349\nTMhelix 350 372\ninside 373 378

1833 GCF_000518805.1_ASM51880v1 Shewanella waksmanii ATCC BAA-643 Proteobacteria; Gammaproteobacteria; Alteromonadales; Shewanellaceae; Shewanella; Shewanella waksmanii
MDTHAALYQQGQARLAKLRSLKPRHVDSLKDKMKQHGISRRDFMKWSAAVTSMLALPLPF WP_028774220.1 hydrogenase [Shewanella waksmanii] Length: 378\nNumber of predicted TMHs: 1\nExp number of AAs in TMHs: 26.61896\nExp number, first 60 AAs: 4.71386\nTotal prob of N-in: 0.39232\noutside 1 349\nTMhelix 350 372\ninside 373 378

1834 GCF_001187555.1_ASM118755v1 Gilvimarinus polysaccharolyticus Proteobacteria; Gammaproteobacteria; Cellvibrionales; Cellvibrionaceae; Gilvimarinus
MSYKLWRKAEKIARRAILSMLTITATVVWAEPDSGRLVLLADDDSVAGTTTEALHLGTHI WP_084005438.1 marine proteobacterial sortase target protein [Gilvimarinus polysaccharolyticus] Length: 702\nNumber of predicted TMHs: 2\nExp number of AAs in TMHs: 34.66936\nExp number, first 60 AAs: 13.92826\nTotal prob of N-in: 0.74333\nPOSSIBLE N-term signal sequence\ninside 1 11\nTMhelix 12 30\noutside 31 669\nTMhelix 670 692\ninside 693 702

1835 GCF_000025485.1_ASM2548v1 Allochromatium vinosum DSM 180 Proteobacteria; Gammaproteobacteria; Chromatiales; Chromatiaceae; Allochromatium; Allochromatium vinosum MRKRSDKTLGESLRDHGLSRRGFLKLCAATASLMALPPSMAPAIAAALEQARRPSVIWLS WP_012971233.1 Ni/Fe hydrogenase [Allochromatium vinosum] Length: 367\nNumber of predicted TMHs: 2\nExp number of AAs in TMHs: 50.63256\nExp number, first 60 AAs: 5.37019\nTotal prob of N-in: 0.76887\ninside 1 190\nTMhelix 191 213\noutside 214 320\nTMhelix 321 343\ninside 344 367

1836 GCF_000379525.1_ASM37952v1 Lamprocystis purpurea DSM 4197 Proteobacteria; Gammaproteobacteria; Chromatiales; Chromatiaceae; Lamprocystis; Lamprocystis purpurea MTNRRELLGAAALIPLALGLRSAMAAEGKPAHEKAGHEAPKQADFLFVQNAQGITYADGR WP_026199800.1 hypothetical protein [Lamprocystis purpurea] Length: 178\nNumber of predicted TMHs: 1\nExp number of AAs in TMHs: 31.67168\nExp number, first 60 AAs: 10.46293\nTotal prob of N-in: 0.59780\nPOSSIBLE N-term signal sequence\noutside 1 148\nTMhelix 149 171\ninside 172 178

1837 GCF_000379525.1_ASM37952v1 Lamprocystis purpurea DSM 4197 Proteobacteria; Gammaproteobacteria; Chromatiales; Chromatiaceae; Lamprocystis; Lamprocystis purpurea MNRSAASRREPALGPYLSDQLDVYGISRRRFIEFALGITATLGLPLGMSGQVLAALEDTK WP_020503333.1 hydrogenase 2 small subunit [Lamprocystis purpurea] Length: 402\nNumber of predicted TMHs: 2\nExp number of AAs in TMHs: 40.47169\nExp number, first 60 AAs: 19.29949\nTotal prob of N-in: 0.87744\nPOSSIBLE N-term signal sequence\ninside 1 33\nTMhelix 34 56\noutside 57 354\nTMhelix 355 377\ninside 378 402

1838 GCF_000223985.1_ASM22398v1 Thiocapsa marina 5811 Proteobacteria; Gammaproteobacteria; Chromatiales; Chromatiaceae; Thiocapsa; Thiocapsa marina MPSRRGTLRLAATIALSMLGSPALAHKLQVFAFAEGPRIGGSAYFAGGGAASGARIEVLDWP_007194980.1 hypothetical protein [Thiocapsa marina] Length: 210\nNumber of predicted TMHs: 1\nExp number of AAs in TMHs: 21.49261\nExp number, first 60 AAs: 2.60279\nTotal prob of N-in: 0.29101\noutside 1 183\nTMhelix 184 201\ninside 202 210

1839 GCF_000227745.2_ASM22774v3 Thiocystis violascens DSM 198 Proteobacteria; Gammaproteobacteria; Chromatiales; Chromatiaceae; Thiocystis; Thiocystis violascens MKDVDVFSNETAIGPYLADQLDIRGISRRGFLKFTLGITAAMGLPFGMSGQVLAALEQSQ WP_014780159.1 hydrogenase 2 small subunit [Thiocystis violascens] Length: 404\nNumber of predicted TMHs: 1\nExp number of AAs in TMHs: 34.61427\nExp number, first 60 AAs: 13.31243\nTotal prob of N-in: 0.58974\nPOSSIBLE N-term signal sequence\noutside 1 354\nTMhelix 355 377\ninside 378 404

1840 GCF_000227745.2_ASM22774v3 Thiocystis violascens DSM 198 Proteobacteria; Gammaproteobacteria; Chromatiales; Chromatiaceae; Thiocystis; Thiocystis violascens MPVRKTPDKTLGESLRDHGLSRRGFLKFCAATASMMALPPSMVPVIAAALEAARRPSVIW WP_014779051.1 Ni/Fe hydrogenase [Thiocystis violascens] Length: 369\nNumber of predicted TMHs: 1\nExp number of AAs in TMHs: 42.0651\nExp number, first 60 AAs: 8.68836\nTotal prob of N-in: 0.60655\noutside 1 322\nTMhelix 323 345\ninside 346 369

1841 GCF_000227745.2_ASM22774v3 Thiocystis violascens DSM 198 Proteobacteria; Gammaproteobacteria; Chromatiales; Chromatiaceae; Thiocystis; Thiocystis violascens MAKETTLGDLLRSQGVSRRAFLQFCTTTATMMALPAAVIPSMARALEQARRPSVIWLSFQ WP_014777803.1 Ni/Fe hydrogenase [Thiocystis violascens] Length: 354\nNumber of predicted TMHs: 2\nExp number of AAs in TMHs: 68.95645\nExp number, first 60 AAs: 20.67345\nTotal prob of N-in: 0.85382\nPOSSIBLE N-term signal sequence\ninside 1 20\nTMhelix 21 43\noutside 44 320\nTMhelix 321 338\ninside 339 354

1842 GCF_000327045.1_ASM32704v1 Thioflavicoccus mobilis 8321 Proteobacteria; Gammaproteobacteria; Chromatiales; Chromatiaceae; Thioflavicoccus; Thioflavicoccus mobilis MAATKTLGEMLRERGVSRRGFLKFCAATASAMALPPSMAPAIAAALERARRPSVVWLSFQ WP_015282143.1 hydrogenase (NiFe) small subunit HydA [Thioflavicoccus mobilis] Length: 365\nNumber of predicted TMHs: 1\nExp number of AAs in TMHs: 39.98671\nExp number, first 60 AAs: 15.94885\nTotal prob of N-in: 0.73112\nPOSSIBLE N-term signal sequence\noutside 1 318\nTMhelix 319 341\ninside 342 365

1843 GCF_000224065.1_ASM22406v1 Thiorhodococcus drewsii AZ1 Proteobacteria; Gammaproteobacteria; Chromatiales; Chromatiaceae; Thiorhodococcus; Thiorhodococcus drewsii MTTGETLGEFLGRQGVSRRAFLKFCSAMASMMALPAASVSALADALEQARRPSVIWLSFQ WP_007039030.1 Ni/Fe hydrogenase [Thiorhodococcus drewsii] Length: 354\nNumber of predicted TMHs: 1\nExp number of AAs in TMHs: 65.46378\nExp number, first 60 AAs: 17.16945\nTotal prob of N-in: 0.73522\nPOSSIBLE N-term signal sequence\noutside 1 320\nTMhelix 321 339\ninside 340 354

1844 GCF_000228725.2_ASM22872v3 Thiorhodovibrio sp. 970 Proteobacteria; Gammaproteobacteria; Chromatiales; Chromatiaceae; Thiorhodovibrio
MADRPTPNLQKTLDSPGGHHSMTLGETLERNGISRRGFLKFCVTTASLMALPPTAATAIA WP_009147065.1
Ni/Fe hydrogenase [Thiorhodovibrio sp. 970] Length: 382\nNumber of predicted TMHs: 1\nExp number of AAs in TMHs: 48.61162\nExp number, first 60 AAs: 5.43158\nTotal prob of N-in: 0.34753\noutside 1 336\nTMhelix 337 359\ninside 360 382

1845 GCF_001753245.1_ASM175324v1 Acidihalobacter prosperus Proteobacteria; Gammaproteobacteria; Chromatiales; Ectothiorhodospiraceae; Acidihalobacter
MAIETFYDVMRRQGISRRSFLKFCSLTATSLGLSPAFVPKIAWAMENKPRTPVLWMHGLE WP_070079895.1
uptake hydrogenase small subunit [Acidihalobacter prosperus] Length: 368\nNumber of predicted TMHs: 1\nExp number of AAs in TMHs: 24.60033\nExp number, first 60 AAs: 10.32866\nTotal prob of N-in: 0.50738\nPOSSIBLE N-term signal sequence\noutside 1 325\nTMhelix 326 348\ninside 349 368

1846 GCF_001753165.1_ASM175316v1 Acidihalobacter prosperus Proteobacteria; Gammaproteobacteria; Chromatiales; Ectothiorhodospiraceae; Acidihalobacter
METFYDVMRRQGISRRSFLKFCSLTATTLGLSPAFVPKIAWAMENTPRTPVLWMHGLECT WP_083251005.1
uptake hydrogenase small subunit [Acidihalobacter prosperus] Length: 366\nNumber of predicted TMHs: 1\nExp number of AAs in TMHs: 32.82965\nExp number, first 60 AAs: 12.17786\nTotal prob of N-in: 0.54951\nPOSSIBLE N-term signal sequence\noutside 1 322\nTMhelix 323 345\ninside 346 366

1847 GCF_001753165.1_ASM175316v1 Acidihalobacter prosperus Proteobacteria; Gammaproteobacteria; Chromatiales; Ectothiorhodospiraceae; Acidihalobacter
MDADDKNTIGDYLRSQGISRRAFLKLCAAMASALALPPGAALAMAETLPQKRRPSVIYMS WP_070072154.1
Ni/Fe hydrogenase [Acidihalobacter prosperus] Length: 355\nNumber of predicted TMHs: 2\nExp number of AAs in TMHs: 52.7135\nExp number, first 60 AAs: 19.94885\nTotal prob of N-in: 0.89653\nPOSSIBLE N-term signal sequence\ninside 1 20\nTMhelix 21 43\noutside 44 321\nTMhelix 322 344\ninside 345 355

1848 GCF_000321415.2_ASM32141v2 Thioalkalivibrio nitratireducens DSM 14787 Proteobacteria; Gammaproteobacteria; Chromatiales; Ectothiorhodospiraceae; Thioalkalivibrio; Thioalkalivibrio nitratireducens
MKEHEMTLGEELRRQGITRRSFLKFCAAVASSMAIPASMVPAMAASLAQARRQSVIWMPY WP_083499426.1
Ni/Fe hydrogenase [Thioalkalivibrio nitratireducens] Length: 353\nNumber of predicted TMHs: 2\nExp number of AAs in TMHs: 38.26707\nExp number, first 60 AAs: 20.66437\nTotal prob of N-in: 0.91607\nPOSSIBLE N-term signal sequence\ninside 1 20\nTMhelix 21 43\noutside 44 317\nTMhelix 318 340\ninside 341 353

1849 GCF_000321415.2_ASM32141v2 Thioalkalivibrio nitratireducens DSM 14787 Proteobacteria; Gammaproteobacteria; Chromatiales; Ectothiorhodospiraceae; Thioalkalivibrio; Thioalkalivibrio nitratireducens
MLRRRSALLAVSLLFVGVMLLSLYVHGTGVVHNDPERNIWIPERLTVPLQLQVAFNEEEI WP_015258557.1 hypothetical protein [Thioalkalivibrio nitratireducens] Length: 465\nNumber of predicted TMHs: 2\nExp number of AAs in TMHs: 44.78544\nExp number, first 60 AAs: 21.8918\nTotal prob of N-in: 0.99479\nPOSSIBLE N-term signal sequence\ninside 1 6\nTMhelix 7 29\noutside 30 437\nTMhelix 438 460\ninside 461 465

1850 GCF_000227685.2_ASM22768v3 Thioalkalivibrio paradoxus ARh 1 Proteobacteria; Gammaproteobacteria; Chromatiales; Ectothiorhodospiraceae; Thioalkalivibrio; Thioalkalivibrio paradoxus
MLRRRSALLAVSVLFVAVLLVSLVTHGTGVIQNDPARNIWVPGQLTLPLQLQVAFNDDEI WP_006747624.1
cytochrome c-552/DMSO reductase-like, heme-binding domain protein [Thioalkalivibrio paradoxus]
Length: 465\nNumber of predicted TMHs: 2\nExp number of AAs in TMHs: 45.17944\nExp number, first 60 AAs: 22.30918\nTotal prob of N-in: 0.99684\nPOSSIBLE N-term signal sequence\ninside 1 6\nTMhelix 7 29\noutside 30 437\nTMhelix 438 460\ninside 461 465

1851 GCF_000021985.1_ASM2198v1 Thioalkalivibrio sulfidiphilus HL-EbGr7 Proteobacteria; Gammaproteobacteria; Chromatiales; Ectothiorhodospiraceae; Thioalkalivibrio; Thioalkalivibrio sulfidiphilus
MKGSQSDEAEQTRGGSRREAAYYPDDEISLIDLWRTLVRRRSIIGFTVVGSLVLGIMFLI WP_041441629.1 hypothetical protein [Thioalkalivibrio sulfidiphilus] Length: 408\nNumber of predicted TMHs: 2\nExp number of AAs in TMHs: 41.81775\nExp number, first 60 AAs: 18.51989\nTotal prob of N-in: 0.99056\nPOSSIBLE N-term signal sequence\ninside 1 41\nTMhelix 42 61\noutside 62 377\nTMhelix 378 400\ninside 401 408

1852 GCF_000227725.1_ASM22772v1 Thiorhodospira sibirica ATCC 700588 Proteobacteria; Gammaproteobacteria; Chromatiales; Ectothiorhodospiraceae; Thiorhodospira; Thiorhodospira sibirica
MKKADTLGDELRRQGISRRTFLKFCAAMATSMALPASMAPVMADALAATRRQSVIWLSFQ WP_006786091.1
Ni/Fe hydrogenase [Thiorhodospira sibirica] Length: 352\nNumber of predicted TMHs: 2\nExp number of

AAs in TMHs: 47.08912\nExp number, first 60 AAs: 21.90288\nTotal prob of N-in: 0.97456\nPOSSIBLE N-term signal sequence\ninside 1 20\nTMhelix 21 43\noutside 44 317\nTMhelix 318 340\ninside 341 352

1853 GCF_000427805.1_ASM42780v1 Budvicia aquatica DSM 5075 = ATCC 35567 Proteobacteria; Gammaproteobacteria; Enterobacterales; Budviciaceae; Budvicia; Budvicia aquatica
MIEDNPLLPSHGINRRDFMKLCTALAATMGLSANAAAEIAESVSNPQRPPVIWIGAQECT WP_029095537.1
hydrogenase 2 small subunit [Budvicia aquatica] Length: 372\nNumber of predicted TMHs: 1\nExp number of AAs in TMHs: 22.03326\nExp number, first 60 AAs: 0.19218\nTotal prob of N-in: 0.02508\noutside 1 330\nTMhelix 331 353\ninside 354 372

1854 GCF_000248015.1_ASM24801v2 Atlantibacter hermannii NBRC 105704 Proteobacteria; Gammaproteobacteria; Enterobacterales; Enterobacteriaceae; Atlantibacter; Atlantibacter hermannii
MSEGQNAFYHNGISRRDFMTLCAALSATMGLTGNASAEMVEALASPSRPPVVWIGAQECT WP_002435487.1
hydrogenase 2 small subunit [Atlantibacter hermannii] Length: 376\nNumber of predicted TMHs: 1\nExp number of AAs in TMHs: 21.97162\nExp number, first 60 AAs: 0.6658\nTotal prob of N-in: 0.11559\noutside 1 330\nTMhelix 331 353\ninside 354 376

1855 GCF_001654925.1_Bbr51605_DRAFTv1 Buttiauxella brennerae ATCC 51605 Proteobacteria; Gammaproteobacteria; Enterobacterales; Enterobacteriaceae; Buttiauxella; Buttiauxella brennerae
MFKRRGLLTAVTVAISLSLFAGPALANPGNGNSGAHGNSGNGNSGNKGNSSAHSNGNDKS WP_064558476.1
hypothetical protein [Buttiauxella brennerae] Length: 165\nNumber of predicted TMHs: 2\nExp number of AAs in TMHs: 40.23663\nExp number, first 60 AAs: 19.81096\nTotal prob of N-in: 0.97252\nPOSSIBLE N-term signal sequence\ninside 1 6\nTMhelix 7 26\noutside 27 140\nTMhelix 141 163\ninside 164 165

1856 GCF_000936345.1_3e8A_assembly Citrobacter amalonaticus Proteobacteria; Gammaproteobacteria; Enterobacterales; Enterobacteriaceae; Citrobacter
MTGDNTLINSHGVNRRDFMKLCAALAATMGLSSKAAAEMAESVSRPQRPPVVWIGAQECT WP_042998143.1
MULTISPECIES: hydrogenase 2 small subunit [Gammaproteobacteria] Length: 372\nNumber of predicted TMHs: 1\nExp number of AAs in TMHs: 27.41754\nExp number, first 60 AAs: 0.28297\nTotal prob of N-in: 0.25533\noutside 1 331\nTMhelix 332 354\ninside 355 372

1857 GCF_001559075.1_ASM155907v1 Citrobacter amalonaticus Proteobacteria; Gammaproteobacteria; Enterobacterales; Enterobacteriaceae; Citrobacter
MTGDNTLINSHGVNRRDFMKLCAALAATMGLSSKAAAEMAESVSRPQRPPVVWIGAQECT WP_042998143.1
MULTISPECIES: hydrogenase 2 small subunit [Gammaproteobacteria] Length: 372\nNumber of predicted TMHs: 1\nExp number of AAs in TMHs: 27.41754\nExp number, first 60 AAs: 0.28297\nTotal prob of N-in: 0.25533\noutside 1 331\nTMhelix 332 354\ninside 355 372

1858 GCF_001373155.1_3e8B_assembly Citrobacter amalonaticus Proteobacteria; Gammaproteobacteria; Enterobacterales; Enterobacteriaceae; Citrobacter
MTGDNTLINSHGVNRRDFMKLCAALAATMGLSSKAAAEMAESVSRPQRPPVVWIGAQECT WP_042998143.1
MULTISPECIES: hydrogenase 2 small subunit [Gammaproteobacteria] Length: 372\nNumber of predicted TMHs: 1\nExp number of AAs in TMHs: 27.41754\nExp number, first 60 AAs: 0.28297\nTotal prob of N-in: 0.25533\noutside 1 331\nTMhelix 332 354\ninside 355 372

1859 GCF_001276105.1_ASM127610v1 Citrobacter amalonaticus Proteobacteria; Gammaproteobacteria; Enterobacterales; Enterobacteriaceae; Citrobacter
MTGDNTLINSHGVNRRDFMKLCAALAATMGLSSKAAAEMAESVSRPQRPPVVWIGAQECT WP_042998143.1
MULTISPECIES: hydrogenase 2 small subunit [Gammaproteobacteria] Length: 372\nNumber of predicted TMHs: 1\nExp number of AAs in TMHs: 27.41754\nExp number, first 60 AAs: 0.28297\nTotal prob of N-in: 0.25533\noutside 1 331\nTMhelix 332 354\ninside 355 372

1860 GCF_000972645.1_ASM97264v1 Citrobacter amalonaticus Proteobacteria; Gammaproteobacteria; Enterobacterales; Enterobacteriaceae; Citrobacter
MTGDNSLINSNGINRRDFMKLCAALAATMGLSSKAAAEMAESVSRPQRPPVIWIGAQECT WP_016154493.1
MULTISPECIES: hydrogenase-2 small chain [Enterobacteriaceae] Length: 372\nNumber of predicted TMHs: 1\nExp number of AAs in TMHs: 26.90274\nExp number, first 60 AAs: 0.32271\nTotal prob of N-in: 0.22239\noutside 1 331\nTMhelix 332 354\ninside 355 372

1861 GCF_001471655.1_ASM147165v1 Citrobacter amalonaticus Proteobacteria; Gammaproteobacteria; Enterobacterales; Enterobacteriaceae; Citrobacter

MTGDNTLINSHGVNRRDFMKLCAALAATMGLSSKAAAEMAESVSRPQRPPVVWIGAQECT WP_042998143.1 MULTISPECIES: hydrogenase 2 small subunit [Gammaproteobacteria] Length: 372\nNumber of predicted TMHs: 1\nExp number of AAs in TMHs: 27.41754\nExp number, first 60 AAs: 0.28297\nTotal prob of N-in: 0.25533\noutside 1 331\nTMhelix 332 354\ninside 355 372

1862 GCF_001276125.1_ASM127612v1 Citrobacter amalonaticus Proteobacteria; Gammaproteobacteria; Enterobacterales; Enterobacteriaceae; Citrobacter MTGDNTLINSHGVNRRDFMKLCAALAATMGLSSKAAAEMAESVSRPQRPPVVWIGAQECT WP_042998143.1 MULTISPECIES: hydrogenase 2 small subunit [Gammaproteobacteria] Length: 372\nNumber of predicted TMHs: 1\nExp number of AAs in TMHs: 27.41754\nExp number, first 60 AAs: 0.28297\nTotal prob of N-in: 0.25533\noutside 1 331\nTMhelix 332 354\ninside 355 372

1863 GCF_002151695.1_ASM215169v1 Citrobacter amalonaticus Proteobacteria; Gammaproteobacteria; Enterobacterales; Enterobacteriaceae; Citrobacter MTGDNTLINSHGVNRRDFMKLCAALAATMGLSSKAAAEMAESVSRPQRPPVVWIGAQECT WP_042998143.1 MULTISPECIES: hydrogenase 2 small subunit [Gammaproteobacteria] Length: 372\nNumber of predicted TMHs: 1\nExp number of AAs in TMHs: 27.41754\nExp number, first 60 AAs: 0.28297\nTotal prob of N-in: 0.25533\noutside 1 331\nTMhelix 332 354\ninside 355 372

1864 GCF_001558935.1_ASM155893v1 Citrobacter amalonaticus Proteobacteria; Gammaproteobacteria; Enterobacterales; Enterobacteriaceae; Citrobacter MTGDNTLINSHGVNRRDFMKLCAALAATMGLSSKAAAEMAESVSRPQRPPVVWIGAQECT WP_042998143.1 MULTISPECIES: hydrogenase 2 small subunit [Gammaproteobacteria] Length: 372\nNumber of predicted TMHs: 1\nExp number of AAs in TMHs: 27.41754\nExp number, first 60 AAs: 0.28297\nTotal prob of N-in: 0.25533\noutside 1 331\nTMhelix 332 354\ninside 355 372

1865 GCF_000731055.1_ASM73105v1 Citrobacter amalonaticus Proteobacteria; Gammaproteobacteria; Enterobacterales; Enterobacteriaceae; Citrobacter MTGDNTLINSHGVNRRDFMKLCAALAATMGLSSKAAAEMAESVSRPQRPPVVWIGAQECT WP_042998143.1 MULTISPECIES: hydrogenase 2 small subunit [Gammaproteobacteria] Length: 372\nNumber of predicted TMHs: 1\nExp number of AAs in TMHs: 27.41754\nExp number, first 60 AAs: 0.28297\nTotal prob of N-in: 0.25533\noutside 1 331\nTMhelix 332 354\ninside 355 372

1866 GCF_900079995.1_ASM90007999v1 Citrobacter europaeus Proteobacteria; Gammaproteobacteria; Enterobacterales; Enterobacteriaceae; Citrobacter MTGDNSLINSNGINRRDFMKLCAALAATMGLSSKAAAEMAESVSRPQRPPVIWIGAQECT WP_043017615.1 MULTISPECIES: hydrogenase 2 small subunit [Citrobacter freundii complex] Length: 372\nNumber of predicted TMHs: 1\nExp number of AAs in TMHs: 27.21963\nExp number, first 60 AAs: 0.31687\nTotal prob of N-in: 0.24323\noutside 1 331\nTMhelix 332 354\ninside 355 372

1867 GCF_900080005.1_ASM90008000v1 Citrobacter europaeus Proteobacteria; Gammaproteobacteria; Enterobacterales; Enterobacteriaceae; Citrobacter MTGDNSLINSNGINRRDFMKLCAALAATMGLSSKAAAEMAESVSRPQRPPVIWIGAQECT WP_043017615.1 MULTISPECIES: hydrogenase 2 small subunit [Citrobacter freundii complex] Length: 372\nNumber of predicted TMHs: 1\nExp number of AAs in TMHs: 27.21963\nExp number, first 60 AAs: 0.31687\nTotal prob of N-in: 0.24323\noutside 1 331\nTMhelix 332 354\ninside 355 372

1868 GCF_001546305.1_ASM154630v1 Citrobacter koseri Proteobacteria; Gammaproteobacteria; Enterobacterales; Enterobacteriaceae; Citrobacter MTGDNTLITSHGVNRRDFMKLCAALAATMGLSSKAAAEMAESVSRPQRPPVVWIGAQECT WP_012135130.1 MULTISPECIES: hydrogenase 2 small subunit [Enterobacteriaceae] Length: 372\nNumber of predicted TMHs: 1\nExp number of AAs in TMHs: 24.24593\nExp number, first 60 AAs: 0.31574\nTotal prob of N-in: 0.12382\noutside 1 331\nTMhelix 332 354\ninside 355 372

1869 GCF_001858105.1_ASM185810v1 Citrobacter koseri Proteobacteria; Gammaproteobacteria; Enterobacterales; Enterobacteriaceae; Citrobacter MTGDNTLITSHGVNRRDFMKLCAALAATMGLSSKAAAEMAESVSRPQRPPVVWIGAQECT WP_012135130.1 MULTISPECIES: hydrogenase 2 small subunit [Enterobacteriaceae] Length: 372\nNumber of predicted TMHs: 1\nExp number of AAs in TMHs: 24.24593\nExp number, first 60 AAs: 0.31574\nTotal prob of N-in: 0.12382\noutside 1 331\nTMhelix 332 354\ninside 355 372

1870 GCF_001546325.1_ASM154632v1 Citrobacter koseri Proteobacteria; Gammaproteobacteria; Enterobacterales; Enterobacteriaceae; Citrobacter
MTGDNTLITSHGVNRRDFMKLCAALAATMGLSSKAAAEMAESVSRPQRPPVVWIGAQECT WP_012135130.1
MULTISPECIES: hydrogenase 2 small subunit [Enterobacteriaceae] Length: 372\nNumber of predicted TMHs: 1\nExp number of AAs in TMHs: 24.24593\nExp number, first 60 AAs: 0.31574\nTotal prob of N-in: 0.12382\noutside 1 331\nTMhelix 332 354\ninside 355 372

1871 GCF_001552875.1_ASM155287v1 Citrobacter koseri Proteobacteria; Gammaproteobacteria; Enterobacterales; Enterobacteriaceae; Citrobacter
MTGDNTLITSHGVNRRDFMKLCAALAATMGLSSKAAAEMAESVSRPQRPPVVWIGAQECT WP_012135130.1
MULTISPECIES: hydrogenase 2 small subunit [Enterobacteriaceae] Length: 372\nNumber of predicted TMHs: 1\nExp number of AAs in TMHs: 24.24593\nExp number, first 60 AAs: 0.31574\nTotal prob of N-in: 0.12382\noutside 1 331\nTMhelix 332 354\ninside 355 372

1872 GCF_001057775.1_ASM105777v1 Citrobacter koseri Proteobacteria; Gammaproteobacteria; Enterobacterales; Enterobacteriaceae; Citrobacter
MTGDNTLITSHGVNRRDFMKLCAALAATMGLSSKAAAEMAESVSRPQRPPVVWIGAQECT WP_047461202.1
hydrogenase 2 small subunit [Citrobacter koseri] Length: 372\nNumber of predicted TMHs: 1\nExp number of AAs in TMHs: 23.67459\nExp number, first 60 AAs: 0.31664\nTotal prob of N-in: 0.12036\noutside 1 331\nTMhelix 332 354\ninside 355 372

1873 GCF_000783445.1_ASM78344v1 Citrobacter koseri Proteobacteria; Gammaproteobacteria; Enterobacterales; Enterobacteriaceae; Citrobacter
MTGDNTLITSHGVNRRDFMKLCAALAATMGLSSKAAAEMAESVSRPQRPPVVWIGAQECT WP_047461202.1
hydrogenase 2 small subunit [Citrobacter koseri] Length: 372\nNumber of predicted TMHs: 1\nExp number of AAs in TMHs: 23.67459\nExp number, first 60 AAs: 0.31664\nTotal prob of N-in: 0.12036\noutside 1 331\nTMhelix 332 354\ninside 355 372

1874 GCF_000826205.1_Citrobacter_pasteurii_55-13 Citrobacter pasteurii Proteobacteria; Gammaproteobacteria; Enterobacterales; Enterobacteriaceae; Citrobacter
MTGDNSLINSNGINRRDFMKLCAALAATMGLSSKAAAEMAESVSRPQRPPVIWIGAQECT WP_040233985.1
hydrogenase 2 small subunit [Citrobacter pasteurii] Length: 372\nNumber of predicted TMHs: 1\nExp number of AAs in TMHs: 27.21943\nExp number, first 60 AAs: 0.31687\nTotal prob of N-in: 0.24323\noutside 1 331\nTMhelix 332 354\ninside 355 372

1875 GCF_000835925.1_ASM83592v1 Citrobacter rodentium Proteobacteria; Gammaproteobacteria; Enterobacterales; Enterobacteriaceae; Citrobacter
MTGDNTLINSHGVNRRDFMKLCAALAATMGLSSKAAAEMAESVSRPQRPPVVWIGAQECT WP_012907591.1
hydrogenase 2 small subunit [Citrobacter rodentium] Length: 372\nNumber of predicted TMHs: 1\nExp number of AAs in TMHs: 24.50861\nExp number, first 60 AAs: 0.31328\nTotal prob of N-in: 0.14317\noutside 1 331\nTMhelix 332 354\ninside 355 372

1876 GCF_000158355.2_Citrobacter_sp_30_2_V2 Citrobacter sp. 30_2 Proteobacteria; Gammaproteobacteria; Enterobacterales; Enterobacteriaceae; Citrobacter
MTGDNSLINSSGINRRDFMKLCAALAATMGLSSKAAAEMAESVSRPQRPPVIWIGAQECT WP_003024486.1
MULTISPECIES: hydrogenase 2 small subunit [Enterobacteriaceae] Length: 372\nNumber of predicted TMHs: 1\nExp number of AAs in TMHs: 27.14621\nExp number, first 60 AAs: 0.22683\nTotal prob of N-in: 0.23982\noutside 1 331\nTMhelix 332 354\ninside 355 372

1877 GCF_001463265.1_ASM146326v1 Citrobacter sp. 50677481 Proteobacteria; Gammaproteobacteria; Enterobacterales; Enterobacteriaceae; Citrobacter
MTGDNTLINSHGVNRRDFMKLCAALAATMGLSSKAAAEMAESVSRPQRPPVIWIGAQECT WP_042289187.1
MULTISPECIES: hydrogenase 2 small subunit [Citrobacter] Length: 372\nNumber of predicted TMHs: 1\nExp number of AAs in TMHs: 24.58538\nExp number, first 60 AAs: 0.33179\nTotal prob of N-in: 0.14844\noutside 1 331\nTMhelix 332 354\ninside 355 372

1878 GCF_000277565.1_ASM27756v1 Citrobacter sp. A1 Proteobacteria; Gammaproteobacteria; Enterobacterales; Enterobacteriaceae; Citrobacter
MTGDNSLINSSGINRRDFMKLCAALAATMGLSSKAAAEMAESVSRPQRPPVIWIGAQECT WP_003024486.1
MULTISPECIES: hydrogenase 2 small subunit [Enterobacteriaceae] Length: 372\nNumber of predicted TMHs:

1\nExp number of AAs in TMHs: 27.14621\nExp number, first 60 AAs: 0.22683\nTotal prob of N-in: 0.23982\noutside 1 331\nTMhelix 332 354\ninside 355 372

1879 GCF_001721255.1_ASM172125v1 Citrobacter sp. CFSAN044567 Proteobacteria; Gammaproteobacteria; Enterobacterales; Enterobacteriaceae; Citrobacter
MTGDNSLINSSGINRRDFMKLCAALAATMGLSSKAAAEMAESVSRPQRPPVIWIGAQECT WP_003024486.1
MULTISPECIES: hydrogenase 2 small subunit [Enterobacteriaceae] Length: 372\nNumber of predicted TMHs: 1\nExp number of AAs in TMHs: 27.14621\nExp number, first 60 AAs: 0.22683\nTotal prob of N-in: 0.23982\noutside 1 331\nTMhelix 332 354\ninside 355 372

1880 GCF_001297795.1_ASM129779v1 Citrobacter sp. CtB7.12 Proteobacteria; Gammaproteobacteria; Enterobacterales; Enterobacteriaceae; Citrobacter
MTGDNTLINSHGVNRRDFMKLCAALAATMGLSSKAAAEMAESVSRPQRPPVVWIGAQECT WP_054177123.1
hydrogenase 2 small subunit [Citrobacter sp. CtB7.12] Length: 372\nNumber of predicted TMHs: 1\nExp number of AAs in TMHs: 27.41881\nExp number, first 60 AAs: 0.28297\nTotal prob of N-in: 0.25535\noutside 1 331\nTMhelix 332 354\ninside 355 372

1881 GCF_001559235.1_ASM155923v1 Citrobacter sp. FDAARGOS_156 Proteobacteria; Gammaproteobacteria; Enterobacterales; Enterobacteriaceae; Citrobacter
MTGDNSLINSNGINRRDFMKLCAALAATMGLSSKAAAEMAESVSRPQRPPVIWIGAQECT WP_016154493.1
MULTISPECIES: hydrogenase-2 small chain [Enterobacteriaceae] Length: 372\nNumber of predicted TMHs: 1\nExp number of AAs in TMHs: 26.90274\nExp number, first 60 AAs: 0.32271\nTotal prob of N-in: 0.22239\noutside 1 331\nTMhelix 332 354\ninside 355 372

1882 GCF_000809165.1_ASM80916v1 Citrobacter sp. JT3 Proteobacteria; Gammaproteobacteria; Enterobacterales; Enterobacteriaceae; Citrobacter
MTGDNTLINSHGVNRRDFMKLCAALAATMGLSSKAAAEMAESVSRPQRPPVVWIGAQECT WP_042998143.1
MULTISPECIES: hydrogenase 2 small subunit [Gammaproteobacteria] Length: 372\nNumber of predicted TMHs: 1\nExp number of AAs in TMHs: 27.41754\nExp number, first 60 AAs: 0.28297\nTotal prob of N-in: 0.25533\noutside 1 331\nTMhelix 332 354\ninside 355 372

1883 GCF_000398845.1_Esch_coli_KTE151_V1 Citrobacter sp. KTE151 Proteobacteria; Gammaproteobacteria; Enterobacterales; Enterobacteriaceae; Citrobacter
MTGDNSLINSNGINRRDFMKLCAALAATMGLSSKAAAEMAESVSRPQRPPVIWIGAQECT WP_016154493.1
MULTISPECIES: hydrogenase-2 small chain [Enterobacteriaceae] Length: 372\nNumber of predicted TMHs: 1\nExp number of AAs in TMHs: 26.90274\nExp number, first 60 AAs: 0.32271\nTotal prob of N-in: 0.22239\noutside 1 331\nTMhelix 332 354\ninside 355 372

1884 GCF_000398825.1_Esch_coli_KTE30_V1 Citrobacter sp. KTE30 Proteobacteria; Gammaproteobacteria; Enterobacterales; Enterobacteriaceae; Citrobacter
MTGDNSLINSSGINRRDFMKLCAALAATMGLSSKAAAEMAESVSRPQRPPVIWIGAQECT WP_003024486.1
MULTISPECIES: hydrogenase 2 small subunit [Enterobacteriaceae] Length: 372\nNumber of predicted TMHs: 1\nExp number of AAs in TMHs: 27.14621\nExp number, first 60 AAs: 0.22683\nTotal prob of N-in: 0.23982\noutside 1 331\nTMhelix 332 354\ninside 355 372

1885 GCF_000398865.1_Esch_coli_KTE32_V1 Citrobacter sp. KTE32 Proteobacteria; Gammaproteobacteria; Enterobacterales; Enterobacteriaceae; Citrobacter
MTGDNSLINSNGINRRDFMKLCAALAATMGLSSKAAAEMAESVSRPQRPPVIWIGAQECT WP_016154493.1
MULTISPECIES: hydrogenase-2 small chain [Enterobacteriaceae] Length: 372\nNumber of predicted TMHs: 1\nExp number of AAs in TMHs: 26.90274\nExp number, first 60 AAs: 0.32271\nTotal prob of N-in: 0.22239\noutside 1 331\nTMhelix 332 354\ninside 355 372

1886 GCF_000313895.1_L17dn Citrobacter sp. L17 Proteobacteria; Gammaproteobacteria; Enterobacterales; Enterobacteriaceae; Citrobacter MTGDNSLINSSGINRRDFMKLCAALAATMGLSSKAAAEMAESVSRPQRPPVIWIGAQECT
WP_003024486.1 MULTISPECIES: hydrogenase 2 small subunit [Enterobacteriaceae] Length: 372\nNumber of predicted TMHs: 1\nExp number of AAs in TMHs: 27.14621\nExp number, first 60 AAs: 0.22683\nTotal prob of N-in: 0.23982\noutside 1 331\nTMhelix 332 354\ninside 355 372

1887 GCF_000692135.1_Citr_sp_MGH_55_V1 Citrobacter sp. MGH 55 Proteobacteria; Gammaproteobacteria; Enterobacterales; Enterobacteriaceae; Citrobacter
MTGDNTLINSHGVNRRDFMKLCAALAATMGLSSKAAAEMAESVSRPQRPPVVWIGAQECT WP_042998143.1

MULTISPECIES: hydrogenase 2 small subunit [Gammaproteobacteria] Length: 372\nNumber of predicted TMHs: 1\nExp number of AAs in TMHs: 27.41754\nExp number, first 60 AAs: 0.28297\nTotal prob of N-in: 0.25533\noutside 1 331\nTMhelix 332 354\ninside 355 372

1888 GCF_000739675.1_ASM73967v1 Citrobacter sp. S-77Proteobacteria; Gammaproteobacteria; Enterobacterales; Enterobacteriaceae; Citrobacter
MTGDNSLINSNGINRRDFMKLCAALAATMGLSSKAAAEMAESVSRPQRPPVIWIGAQECT WP_045447522.1
MULTISPECIES: hydrogenase 2 small subunit [Citrobacter] Length: 372\nNumber of predicted TMHs: 1\nExp number of AAs in TMHs: 26.06135\nExp number, first 60 AAs: 0.32735\nTotal prob of N-in: 0.20887\noutside 1 331\nTMhelix 332 354\ninside 355 372

1889 GCF_000981805.1_ASM98180v1 Citrobacter amalonaticus Y19Proteobacteria; Gammaproteobacteria; Enterobacterales; Enterobacteriaceae; Citrobacter; Citrobacter amalonaticus
MTGDNTLINSHGVNRRDFMKLCAALAATMGLSSKAAAEMAESVSRPQRPPVVWIGAQECT WP_046497002.1
hydrogenase 2 small subunit [Citrobacter amalonaticus] Length: 372\nNumber of predicted TMHs: 1\nExp number of AAs in TMHs: 27.64581\nExp number, first 60 AAs: 0.27351\nTotal prob of N-in: 0.28342\noutside 1 331\nTMhelix 332 354\ninside 355 372

1890 GCF_000764735.1_ASM76473v1 Citrobacter farmeri GTC 1319Proteobacteria; Gammaproteobacteria; Enterobacterales; Enterobacteriaceae; Citrobacter; Citrobacter farmeri
MTGDNTLINSHGVNRRDFMKLCAALAATMGLSSKAAAEMAESVSRPQRPPVVWIGAQECT WP_042324189.1
hydrogenase 2 small subunit [Citrobacter farmeri] Length: 372\nNumber of predicted TMHs: 1\nExp number of AAs in TMHs: 27.41806\nExp number, first 60 AAs: 0.28299\nTotal prob of N-in: 0.25533\noutside 1 331\nTMhelix 332 354\ninside 355 372

1891 GCF_002075345.1_ASM207534v1 Citrobacter braakii Proteobacteria; Gammaproteobacteria; Enterobacterales; Enterobacteriaceae; Citrobacter; Citrobacter freundii complex
MTGDNSLINSNGINRRDFMKLCAALAATMGLSSKAAAEMAESVSRPQRPPVIWIGAQECT WP_016154493.1
MULTISPECIES: hydrogenase-2 small chain [Enterobacteriaceae] Length: 372\nNumber of predicted TMHs: 1\nExp number of AAs in TMHs: 26.90274\nExp number, first 60 AAs: 0.32271\nTotal prob of N-in: 0.22239\noutside 1 331\nTMhelix 332 354\ninside 355 372

1892 GCF_001062165.1_ASM106216v1 Citrobacter braakii Proteobacteria; Gammaproteobacteria; Enterobacterales; Enterobacteriaceae; Citrobacter; Citrobacter freundii complex
MTGDNSLINSNGINRRDFMKLCAALAATMGLSSKAAAEMAESVSRPQRPPVIWIGAQECT WP_016154493.1
MULTISPECIES: hydrogenase-2 small chain [Enterobacteriaceae] Length: 372\nNumber of predicted TMHs: 1\nExp number of AAs in TMHs: 26.90274\nExp number, first 60 AAs: 0.32271\nTotal prob of N-in: 0.22239\noutside 1 331\nTMhelix 332 354\ninside 355 372

1893 GCF_000786275.1_ASM78627v1 Citrobacter braakii Proteobacteria; Gammaproteobacteria; Enterobacterales; Enterobacteriaceae; Citrobacter; Citrobacter freundii complex
MTGDNSLINSNGINRRDFMKLCAALAATMGLSSKAAAEMAESVSRPQRPPVIWIGAQECT WP_016154493.1
MULTISPECIES: hydrogenase-2 small chain [Enterobacteriaceae] Length: 372\nNumber of predicted TMHs: 1\nExp number of AAs in TMHs: 26.90274\nExp number, first 60 AAs: 0.32271\nTotal prob of N-in: 0.22239\noutside 1 331\nTMhelix 332 354\ninside 355 372

1894 GCF_001067775.1_ASM106777v1 Citrobacter braakii Proteobacteria; Gammaproteobacteria; Enterobacterales; Enterobacteriaceae; Citrobacter; Citrobacter freundii complex
MTGDNSLINSNGINRRDFMKLCAALAATMGLSSKAAAEMAESVSRPQRPPVIWIGAQECT WP_016154493.1
MULTISPECIES: hydrogenase-2 small chain [Enterobacteriaceae] Length: 372\nNumber of predicted TMHs: 1\nExp number of AAs in TMHs: 26.90274\nExp number, first 60 AAs: 0.32271\nTotal prob of N-in: 0.22239\noutside 1 331\nTMhelix 332 354\ninside 355 372

1895 GCF_000786265.1_ASM78626v1 Citrobacter braakii Proteobacteria; Gammaproteobacteria; Enterobacterales; Enterobacteriaceae; Citrobacter; Citrobacter freundii complex
MTGDNSLINSNGINRRDFMKLCAALAATMGLSSKAAAEMAESVSRPQRPPVIWIGAQECT WP_016154493.1
MULTISPECIES: hydrogenase-2 small chain [Enterobacteriaceae] Length: 372\nNumber of predicted TMHs: 1\nExp number of AAs in TMHs: 26.90274\nExp number, first 60 AAs: 0.32271\nTotal prob of N-in: 0.22239\noutside 1 331\nTMhelix 332 354\ninside 355 372

1896 GCF_002073755.1_ASM207375v1 Citrobacter braakii Proteobacteria; Gammaproteobacteria; Enterobacterales; Enterobacteriaceae; Citrobacter; Citrobacter freundii complex
MTGDNSLINSNGINRRDFMKLCAALAATMGLSSKAAAEMAESVSRPQRPPVIWIGAQECT WP_043017615.1
MULTISPECIES: hydrogenase 2 small subunit [Citrobacter freundii complex] Length: 372\nNumber of predicted TMHs: 1\nExp number of AAs in TMHs: 27.21963\nExp number, first 60 AAs: 0.31687\nTotal prob of N-in: 0.24323\noutside 1 331\nTMhelix 332 354\ninside 355 372

1897 GCF_001059745.1_ASM105974v1 Citrobacter braakii Proteobacteria; Gammaproteobacteria; Enterobacterales; Enterobacteriaceae; Citrobacter; Citrobacter freundii complex
MTGDNSLINSNGINRRDFMKLCAALAATMGLSSKAAAEMAESVSRPQRPPVIWIGAQECT WP_016154493.1
MULTISPECIES: hydrogenase-2 small chain [Enterobacteriaceae] Length: 372\nNumber of predicted TMHs: 1\nExp number of AAs in TMHs: 26.90274\nExp number, first 60 AAs: 0.32271\nTotal prob of N-in: 0.22239\noutside 1 331\nTMhelix 332 354\ninside 355 372

1898 GCF_001065805.1_ASM106580v1 Citrobacter braakii Proteobacteria; Gammaproteobacteria; Enterobacterales; Enterobacteriaceae; Citrobacter; Citrobacter freundii complex
MTGDNSLINSNGINRRDFMKLCAALAATMGLSSKAAAEMAESVSRPQRPPVIWIGAQECT WP_049268745.1
hydrogenase 2 small subunit [Citrobacter braakii] Length: 372\nNumber of predicted TMHs: 1\nExp number of AAs in TMHs: 26.94462\nExp number, first 60 AAs: 0.32389\nTotal prob of N-in: 0.22173\noutside 1 332\nTMhelix 333 355\ninside 356 372

1899 GCF_002208845.1_ASM220884v1 Citrobacter braakii Proteobacteria; Gammaproteobacteria; Enterobacterales; Enterobacteriaceae; Citrobacter; Citrobacter freundii complex
MTGDNSLINSNGINRRDFMKLCAALAATMGLSSKAAAEMAESVSRPQRPPVIWIGAQECT WP_016154493.1
MULTISPECIES: hydrogenase-2 small chain [Enterobacteriaceae] Length: 372\nNumber of predicted TMHs: 1\nExp number of AAs in TMHs: 26.90274\nExp number, first 60 AAs: 0.32271\nTotal prob of N-in: 0.22239\noutside 1 331\nTMhelix 332 354\ninside 355 372

1900 GCF_001317155.2_ASM131715v2 Citrobacter freundiiProteobacteria; Gammaproteobacteria; Enterobacterales; Enterobacteriaceae; Citrobacter; Citrobacter freundii complex
MTGDNSLINSSGINRRDFMKLCAALAATMGLSSKAAAEMAESVSRPQRPPVIWIGAQECT WP_057068275.1
hydrogenase 2 small subunit [Citrobacter freundii] Length: 372\nNumber of predicted TMHs: 1\nExp number of AAs in TMHs: 27.15001\nExp number, first 60 AAs: 0.22682\nTotal prob of N-in: 0.23980\noutside 1 331\nTMhelix 332 354\ninside 355 372

1901 GCF_001413155.1_ASM141315v1 Citrobacter freundiiProteobacteria; Gammaproteobacteria; Enterobacterales; Enterobacteriaceae; Citrobacter; Citrobacter freundii complex
MTGDNSLINSSGINRRDFMKLCAALAATMGLSSKAAAEMAESVSRPQRPPVIWIGAQECT WP_003024486.1
MULTISPECIES: hydrogenase 2 small subunit [Enterobacteriaceae] Length: 372\nNumber of predicted TMHs: 1\nExp number of AAs in TMHs: 27.14621\nExp number, first 60 AAs: 0.22683\nTotal prob of N-in: 0.23982\noutside 1 331\nTMhelix 332 354\ninside 355 372

1902 GCF_001413115.1_ASM141311v1 Citrobacter freundiiProteobacteria; Gammaproteobacteria; Enterobacterales; Enterobacteriaceae; Citrobacter; Citrobacter freundii complex
MTGDNSLINSSGINRRDFMKLCAALAATMGLSSKAAAEMAESVSRPQRPPVIWIGAQECT WP_003024486.1
MULTISPECIES: hydrogenase 2 small subunit [Enterobacteriaceae] Length: 372\nNumber of predicted TMHs: 1\nExp number of AAs in TMHs: 27.14621\nExp number, first 60 AAs: 0.22683\nTotal prob of N-in: 0.23982\noutside 1 331\nTMhelix 332 354\ninside 355 372

1903 GCF_001413295.1_ASM141329v1 Citrobacter freundiiProteobacteria; Gammaproteobacteria; Enterobacterales; Enterobacteriaceae; Citrobacter; Citrobacter freundii complex
MTGDNSLINSSGINRRDFMKLCAALAATMGLSSKAAAEMAESVSRPQRPPVIWIGAQECT WP_003024486.1
MULTISPECIES: hydrogenase 2 small subunit [Enterobacteriaceae] Length: 372\nNumber of predicted TMHs: 1\nExp number of AAs in TMHs: 27.14621\nExp number, first 60 AAs: 0.22683\nTotal prob of N-in: 0.23982\noutside 1 331\nTMhelix 332 354\ninside 355 372

1904 GCF_001880865.1_ASM188086v1 Citrobacter freundiiProteobacteria; Gammaproteobacteria; Enterobacterales; Enterobacteriaceae; Citrobacter; Citrobacter freundii complex
MTGDNSLINSSGINRRDFMKLCAALAATMGLSSKAAAEMAESVSRPQRPPVIWIGAQECT WP_003024486.1
MULTISPECIES: hydrogenase 2 small subunit [Enterobacteriaceae] Length: 372\nNumber of predicted TMHs:

1\nExp number of AAs in TMHs: 27.14621\nExp number, first 60 AAs: 0.22683\nTotal prob of N-in: 0.23982\noutside 1 331\nTMhelix 332 354\ninside 355 372

1905 GCF_001413075.1_ASM141307v1 Citrobacter freundiiProteobacteria; Gammaproteobacteria; Enterobacterales; Enterobacteriaceae; Citrobacter; Citrobacter freundii complex
MTGDNSLINSSGINRRDFMKLCAALAATMGLSSKAAAEMAESVSRPQRPPVIWIGAQECT WP_003024486.1
MULTISPECIES: hydrogenase 2 small subunit [Enterobacteriaceae] Length: 372\nNumber of predicted TMHs: 1\nExp number of AAs in TMHs: 27.14621\nExp number, first 60 AAs: 0.22683\nTotal prob of N-in: 0.23982\noutside 1 331\nTMhelix 332 354\ninside 355 372

1906 GCF_001922445.1_ASM192244v1 Citrobacter freundiiProteobacteria; Gammaproteobacteria; Enterobacterales; Enterobacteriaceae; Citrobacter; Citrobacter freundii complex
MTGDNSLINSNGINRRDFMKLCAALAATMGLSSKAAAEMAESVSRPQRPPVIWIGAQECT WP_043017615.1
MULTISPECIES: hydrogenase 2 small subunit [Citrobacter freundii complex] Length: 372\nNumber of predicted TMHs: 1\nExp number of AAs in TMHs: 27.21963\nExp number, first 60 AAs: 0.31687\nTotal prob of N-in: 0.24323\noutside 1 331\nTMhelix 332 354\ninside 355 372

1907 GCF_002189125.1_ASM218912v1 Citrobacter freundiiProteobacteria; Gammaproteobacteria; Enterobacterales; Enterobacteriaceae; Citrobacter; Citrobacter freundii complex
MTGDNSLINSSGINRRDFMKLCAALAATMGLSSKAAAEMAESVSRPQRPPVIWIGAQECT WP_003024486.1
MULTISPECIES: hydrogenase 2 small subunit [Enterobacteriaceae] Length: 372\nNumber of predicted TMHs: 1\nExp number of AAs in TMHs: 27.14621\nExp number, first 60 AAs: 0.22683\nTotal prob of N-in: 0.23982\noutside 1 331\nTMhelix 332 354\ninside 355 372

1908 GCF_002151785.1_ASM215178v1 Citrobacter freundiiProteobacteria; Gammaproteobacteria; Enterobacterales; Enterobacteriaceae; Citrobacter; Citrobacter freundii complex
MTGDNSLINSSGINRRDFMKLCAALAATMGLSSKAAAEMAESVSRPQRPPVIWIGAQECT WP_003024486.1
MULTISPECIES: hydrogenase 2 small subunit [Enterobacteriaceae] Length: 372\nNumber of predicted TMHs: 1\nExp number of AAs in TMHs: 27.14621\nExp number, first 60 AAs: 0.22683\nTotal prob of N-in: 0.23982\noutside 1 331\nTMhelix 332 354\ninside 355 372

1909 GCF_002151755.1_ASM215175v1 Citrobacter freundiiProteobacteria; Gammaproteobacteria; Enterobacterales; Enterobacteriaceae; Citrobacter; Citrobacter freundii complex
MTGDNSLINSSGINRRDFMKLCAALAATMGLSSKAAAEMAESVSRPQRPPVIWIGAQECT WP_003024486.1
MULTISPECIES: hydrogenase 2 small subunit [Enterobacteriaceae] Length: 372\nNumber of predicted TMHs: 1\nExp number of AAs in TMHs: 27.14621\nExp number, first 60 AAs: 0.22683\nTotal prob of N-in: 0.23982\noutside 1 331\nTMhelix 332 354\ninside 355 372

1910 GCF_001880955.1_ASM188095v1 Citrobacter freundiiProteobacteria; Gammaproteobacteria; Enterobacterales; Enterobacteriaceae; Citrobacter; Citrobacter freundii complex
MTGDNSLINSSGINRRDFMKLCAALAATMGLSSKAAAEMAESVSRPQRPPVIWIGAQECT WP_003024486.1
MULTISPECIES: hydrogenase 2 small subunit [Enterobacteriaceae] Length: 372\nNumber of predicted TMHs: 1\nExp number of AAs in TMHs: 27.14621\nExp number, first 60 AAs: 0.22683\nTotal prob of N-in: 0.23982\noutside 1 331\nTMhelix 332 354\ninside 355 372

1911 GCF_001880915.1_ASM188091v1 Citrobacter freundiiProteobacteria; Gammaproteobacteria; Enterobacterales; Enterobacteriaceae; Citrobacter; Citrobacter freundii complex
MTGDNSLINSSGINRRDFMKLCAALAATMGLSSKAAAEMAESVSRPQRPPVIWIGAQECT WP_003024486.1
MULTISPECIES: hydrogenase 2 small subunit [Enterobacteriaceae] Length: 372\nNumber of predicted TMHs: 1\nExp number of AAs in TMHs: 27.14621\nExp number, first 60 AAs: 0.22683\nTotal prob of N-in: 0.23982\noutside 1 331\nTMhelix 332 354\ninside 355 372

1912 GCF_001867255.1_ASM186725v1 Citrobacter freundiiProteobacteria; Gammaproteobacteria; Enterobacterales; Enterobacteriaceae; Citrobacter; Citrobacter freundii complex
MRQNSTGEETRVMNTNNEETFYQAMRRKGVSRRSFLKYCSLAATSLGLGAGMTPRIAWAL WP_071444621.1
hydrogenase [Citrobacter freundii] Length: 387\nNumber of predicted TMHs: 1\nExp number of AAs in TMHs: 32.7962199999999\nExp number, first 60 AAs: 16.18145\nTotal prob of N-in: 0.76200\nPOSSIBLE N-term signal sequence\noutside 1 339\nTMhelix 340 362\ninside 363 387

1913 GCF_001718055.1_ASM171805v1 Citrobacter freundiiProteobacteria; Gammaproteobacteria; Enterobacterales; Enterobacteriaceae; Citrobacter; Citrobacter freundii complex

MTGDNSLINSSGINRRDFMKLCAALAATMGLSSKAAAEMAESVSRPQRPPVIWIGAQECT WP_003024486.1 MULTISPECIES: hydrogenase 2 small subunit [Enterobacteriaceae] Length: 372\nNumber of predicted TMHs: 1\nExp number of AAs in TMHs: 27.14621\nExp number, first 60 AAs: 0.22683\nTotal prob of N-in: 0.23982\noutside 1 331\nTMhelix 332 354\ninside 355 372

1914 GCF_001867255.1_ASM186725v1 Citrobacter freundiiProteobacteria; Gammaproteobacteria; Enterobacterales; Enterobacteriaceae; Citrobacter; Citrobacter freundii complex MTGDNSLINSSGINRRDFMKLCAALAATMGLSSKAAAEMAESVSRPQRPPVIWIGAQECT WP_003024486.1 MULTISPECIES: hydrogenase 2 small subunit [Enterobacteriaceae] Length: 372\nNumber of predicted TMHs: 1\nExp number of AAs in TMHs: 27.14621\nExp number, first 60 AAs: 0.22683\nTotal prob of N-in: 0.23982\noutside 1 331\nTMhelix 332 354\ninside 355 372

1915 GCF_001880795.1_ASM188079v1 Citrobacter freundiiProteobacteria; Gammaproteobacteria; Enterobacterales; Enterobacteriaceae; Citrobacter; Citrobacter freundii complex MTGDNSLINSSGINRRDFMKLCAALAATMGLSSKAAAEMAESVSRPQRPPVIWIGAQECT WP_003024486.1 MULTISPECIES: hydrogenase 2 small subunit [Enterobacteriaceae] Length: 372\nNumber of predicted TMHs: 1\nExp number of AAs in TMHs: 27.14621\nExp number, first 60 AAs: 0.22683\nTotal prob of N-in: 0.23982\noutside 1 331\nTMhelix 332 354\ninside 355 372

1916 GCF_001689745.1_ASM168974v1 Citrobacter freundiiProteobacteria; Gammaproteobacteria; Enterobacterales; Enterobacteriaceae; Citrobacter; Citrobacter freundii complex MTGDNSLINSNGINRRDFMKLCAALAATMGLSSKAAAEMAESVSRPQRPPVIWIGAQECT WP_016154493.1 MULTISPECIES: hydrogenase-2 small chain [Enterobacteriaceae] Length: 372\nNumber of predicted TMHs: 1\nExp number of AAs in TMHs: 26.90274\nExp number, first 60 AAs: 0.32271\nTotal prob of N-in: 0.22239\noutside 1 331\nTMhelix 332 354\ninside 355 372

1917 GCF_001718055.1_ASM171805v1 Citrobacter freundiiProteobacteria; Gammaproteobacteria; Enterobacterales; Enterobacteriaceae; Citrobacter; Citrobacter freundii complex MKGVLEKCTVEDFNYISATLDSYVSFTNDKRRKTLLSAYQNNPALHAELISLIDTQIKYF WP_001515206.1 MULTISPECIES: hypothetical protein [Enterobacteriaceae] Length: 249\nNumber of predicted TMHs: 2\nExp number of AAs in TMHs: 39.68696\nExp number, first 60 AAs: 0.01961\nTotal prob of N-in: 0.09053\noutside 1 157\nTMhelix 158 180\ninside 181 200\nTMhelix 201 220\noutside 221 249

1918 GCF_001482575.1_ASM148257v1 Citrobacter freundiiProteobacteria; Gammaproteobacteria; Enterobacterales; Enterobacteriaceae; Citrobacter; Citrobacter freundii complex MTGDNSLINSSGINRRDFMKLCAALAATMGLSSKAAAEMAESVSRPQRPPVIWIGAQECT WP_003024486.1 MULTISPECIES: hydrogenase 2 small subunit [Enterobacteriaceae] Length: 372\nNumber of predicted TMHs: 1\nExp number of AAs in TMHs: 27.14621\nExp number, first 60 AAs: 0.22683\nTotal prob of N-in: 0.23982\noutside 1 331\nTMhelix 332 354\ninside 355 372

1919 GCF_001880835.1_ASM188083v1 Citrobacter freundiiProteobacteria; Gammaproteobacteria; Enterobacterales; Enterobacteriaceae; Citrobacter; Citrobacter freundii complex MTGDNSLINSSGINRRDFMKLCAALAATMGLSSKAAAEMAESVSRPQRPPVIWIGAQECT WP_003024486.1 MULTISPECIES: hydrogenase 2 small subunit [Enterobacteriaceae] Length: 372\nNumber of predicted TMHs: 1\nExp number of AAs in TMHs: 27.14621\nExp number, first 60 AAs: 0.22683\nTotal prob of N-in: 0.23982\noutside 1 331\nTMhelix 332 354\ninside 355 372

1920 GCF_001413335.1_ASM141333v1 Citrobacter freundiiProteobacteria; Gammaproteobacteria; Enterobacterales; Enterobacteriaceae; Citrobacter; Citrobacter freundii complex MTGDNSLINSSGINRRDFMKLCAALAATMGLSSKAAAEMAESVSRPQRPPVIWIGAQECT WP_003024486.1 MULTISPECIES: hydrogenase 2 small subunit [Enterobacteriaceae] Length: 372\nNumber of predicted TMHs: 1\nExp number of AAs in TMHs: 27.14621\nExp number, first 60 AAs: 0.22683\nTotal prob of N-in: 0.23982\noutside 1 331\nTMhelix 332 354\ninside 355 372

1921 GCF_001413255.1_ASM141325v1 Citrobacter freundiiProteobacteria; Gammaproteobacteria; Enterobacterales; Enterobacteriaceae; Citrobacter; Citrobacter freundii complex MTGDNSLINSSGINRRDFMKLCAALAATMGLSSKAAAEMAESVSRPQRPPVIWIGAQECT WP_003024486.1 MULTISPECIES: hydrogenase 2 small subunit [Enterobacteriaceae] Length: 372\nNumber of predicted TMHs: 1\nExp number of AAs in TMHs: 27.14621\nExp number, first 60 AAs: 0.22683\nTotal prob of N-in: 0.23982\noutside 1 331\nTMhelix 332 354\ninside 355 372

1922 GCF_001413195.1_ASM141319v1 Citrobacter freundiiProteobacteria; Gammaproteobacteria; Enterobacterales; Enterobacteriaceae; Citrobacter; Citrobacter freundii complex
MTGDNSLINSSGINRRDFMKLCAALAATMGLSSKAAAEMAESVSRPQRPPVIWIGAQECT WP_003024486.1
MULTISPECIES: hydrogenase 2 small subunit [Enterobacteriaceae] Length: 372\nNumber of predicted TMHs: 1\nExp number of AAs in TMHs: 27.14621\nExp number, first 60 AAs: 0.22683\nTotal prob of N-in: 0.23982\noutside 1 331\nTMhelix 332 354\ninside 355 372

1923 GCF_001413235.1_ASM141323v1 Citrobacter freundiiProteobacteria; Gammaproteobacteria; Enterobacterales; Enterobacteriaceae; Citrobacter; Citrobacter freundii complex
MTGDNSLINSSGINRRDFMKLCAALAATMGLSSKAAAEMAESVSRPQRPPVIWIGAQECT WP_003024486.1
MULTISPECIES: hydrogenase 2 small subunit [Enterobacteriaceae] Length: 372\nNumber of predicted TMHs: 1\nExp number of AAs in TMHs: 27.14621\nExp number, first 60 AAs: 0.22683\nTotal prob of N-in: 0.23982\noutside 1 331\nTMhelix 332 354\ninside 355 372

1924 GCF_001412715.1_AssemblyName Citrobacter freundiiProteobacteria; Gammaproteobacteria; Enterobacterales; Enterobacteriaceae; Citrobacter; Citrobacter freundii complex
MTGDNSLINSSGINRRDFMKLCAALAATMGLSSKAAAEMAESVSRPQRPPVIWIGAQECT WP_003024486.1
MULTISPECIES: hydrogenase 2 small subunit [Enterobacteriaceae] Length: 372\nNumber of predicted TMHs: 1\nExp number of AAs in TMHs: 27.14621\nExp number, first 60 AAs: 0.22683\nTotal prob of N-in: 0.23982\noutside 1 331\nTMhelix 332 354\ninside 355 372

1925 GCF_001412735.1_ASM141273v1 Citrobacter freundiiProteobacteria; Gammaproteobacteria; Enterobacterales; Enterobacteriaceae; Citrobacter; Citrobacter freundii complex
MTGDNSLINSSGINRRDFMKLCAALAATMGLSSKAAAEMAESVSRPQRPPVIWIGAQECT WP_003024486.1
MULTISPECIES: hydrogenase 2 small subunit [Enterobacteriaceae] Length: 372\nNumber of predicted TMHs: 1\nExp number of AAs in TMHs: 27.14621\nExp number, first 60 AAs: 0.22683\nTotal prob of N-in: 0.23982\noutside 1 331\nTMhelix 332 354\ninside 355 372

1926 GCF_000937505.2_ASM93750v2 Citrobacter freundiiProteobacteria; Gammaproteobacteria; Enterobacterales; Enterobacteriaceae; Citrobacter; Citrobacter freundii complex
MTGDNSLINSSGINRRDFMKLCAALAATMGLSSKAAAEMAESVSRPQRPPVIWIGAQECT WP_044713741.1
hydrogenase 2 small subunit [Citrobacter freundii] Length: 372\nNumber of predicted TMHs: 1\nExp number of AAs in TMHs: 27.14708\nExp number, first 60 AAs: 0.22683\nTotal prob of N-in: 0.23983\noutside 1 331\nTMhelix 332 354\ninside 355 372

1927 GCF_001412795.1_ASM141279v1 Citrobacter freundiiProteobacteria; Gammaproteobacteria; Enterobacterales; Enterobacteriaceae; Citrobacter; Citrobacter freundii complex
MTGDNSLINSSGINRRDFMKLCAALAATMGLSSKAAAEMAESVSRPQRPPVIWIGAQECT WP_003024486.1
MULTISPECIES: hydrogenase 2 small subunit [Enterobacteriaceae] Length: 372\nNumber of predicted TMHs: 1\nExp number of AAs in TMHs: 27.14621\nExp number, first 60 AAs: 0.22683\nTotal prob of N-in: 0.23982\noutside 1 331\nTMhelix 332 354\ninside 355 372

1928 GCF_001412815.1_ASM141281v1 Citrobacter freundiiProteobacteria; Gammaproteobacteria; Enterobacterales; Enterobacteriaceae; Citrobacter; Citrobacter freundii complex
MTGDNSLINSSGINRRDFMKLCAALAATMGLSSKAAAEMAESVSRPQRPPVIWIGAQECT WP_003024486.1
MULTISPECIES: hydrogenase 2 small subunit [Enterobacteriaceae] Length: 372\nNumber of predicted TMHs: 1\nExp number of AAs in TMHs: 27.14621\nExp number, first 60 AAs: 0.22683\nTotal prob of N-in: 0.23982\noutside 1 331\nTMhelix 332 354\ninside 355 372

1929 GCF_001412855.1_ASM141285v1 Citrobacter freundiiProteobacteria; Gammaproteobacteria; Enterobacterales; Enterobacteriaceae; Citrobacter; Citrobacter freundii complex
MTGDNSLINSSGINRRDFMKLCAALAATMGLSSKAAAEMAESVSRPQRPPVIWIGAQECT WP_003024486.1
MULTISPECIES: hydrogenase 2 small subunit [Enterobacteriaceae] Length: 372\nNumber of predicted TMHs: 1\nExp number of AAs in TMHs: 27.14621\nExp number, first 60 AAs: 0.22683\nTotal prob of N-in: 0.23982\noutside 1 331\nTMhelix 332 354\ninside 355 372

1930 GCF_000783755.1_ASM78375v1 Citrobacter freundiiProteobacteria; Gammaproteobacteria; Enterobacterales; Enterobacteriaceae; Citrobacter; Citrobacter freundii complex
MTGDNSLINSSGINRRDFMKLCAALAATMGLSSKAAAEMAESVSRPQRPPVIWIGAQECT WP_003024486.1
MULTISPECIES: hydrogenase 2 small subunit [Enterobacteriaceae] Length: 372\nNumber of predicted TMHs:

1\nExp number of AAs in TMHs: 27.14621\nExp number, first 60 AAs: 0.22683\nTotal prob of N-in: 0.23982\noutside 1 331\nTMhelix 332 354\ninside 355 372

1931 GCF_000937455.2_ASM93745v2 Citrobacter freundiiProteobacteria; Gammaproteobacteria; Enterobacterales; Enterobacteriaceae; Citrobacter; Citrobacter freundii complex
MTGDNSLINSSGINRRDFMKLCAALAATMGLSSKAAAEMAESVSRPQRPPVIWIGAQECT WP_003024486.1
MULTISPECIES: hydrogenase 2 small subunit [Enterobacteriaceae] Length: 372\nNumber of predicted TMHs: 1\nExp number of AAs in TMHs: 27.14621\nExp number, first 60 AAs: 0.22683\nTotal prob of N-in: 0.23982\noutside 1 331\nTMhelix 332 354\ninside 355 372

1932 GCF_001055295.1_ASM105529v1 Citrobacter freundiiProteobacteria; Gammaproteobacteria; Enterobacterales; Enterobacteriaceae; Citrobacter; Citrobacter freundii complex
MTGDNSLINSSGINRRDFMKLCAALAATMGLSSKAAAEMAESVSRPQRPPVIWIGAQECT WP_003024486.1
MULTISPECIES: hydrogenase 2 small subunit [Enterobacteriaceae] Length: 372\nNumber of predicted TMHs: 1\nExp number of AAs in TMHs: 27.14621\nExp number, first 60 AAs: 0.22683\nTotal prob of N-in: 0.23982\noutside 1 331\nTMhelix 332 354\ninside 355 372

1933 GCF_001022155.1_ASM102215v1 Citrobacter freundiiProteobacteria; Gammaproteobacteria; Enterobacterales; Enterobacteriaceae; Citrobacter; Citrobacter freundii complex
MTGDNSLINSSGINRRDFMKLCAALAATMGLSSKAAAEMAESVSRPQRPPVIWIGAQECT WP_003024486.1
MULTISPECIES: hydrogenase 2 small subunit [Enterobacteriaceae] Length: 372\nNumber of predicted TMHs: 1\nExp number of AAs in TMHs: 27.14621\nExp number, first 60 AAs: 0.22683\nTotal prob of N-in: 0.23982\noutside 1 331\nTMhelix 332 354\ninside 355 372

1934 GCF_001057215.1_ASM105721v1 Citrobacter freundiiProteobacteria; Gammaproteobacteria; Enterobacterales; Enterobacteriaceae; Citrobacter; Citrobacter freundii complex
MTGDNSLINSSGINRRDFMKLCAALAATMGLSSKAAAEMAESVSRPQRPPVIWIGAQECT WP_003024486.1
MULTISPECIES: hydrogenase 2 small subunit [Enterobacteriaceae] Length: 372\nNumber of predicted TMHs: 1\nExp number of AAs in TMHs: 27.14621\nExp number, first 60 AAs: 0.22683\nTotal prob of N-in: 0.23982\noutside 1 331\nTMhelix 332 354\ninside 355 372

1935 GCF_001058675.1_ASM105867v1 Citrobacter freundiiProteobacteria; Gammaproteobacteria; Enterobacterales; Enterobacteriaceae; Citrobacter; Citrobacter freundii complex
MTGDNSLINSSGINRRDFMKLCAALAATMGLSSKAAAEMAESVSRPQRPPVIWIGAQECT WP_003024486.1
MULTISPECIES: hydrogenase 2 small subunit [Enterobacteriaceae] Length: 372\nNumber of predicted TMHs: 1\nExp number of AAs in TMHs: 27.14621\nExp number, first 60 AAs: 0.22683\nTotal prob of N-in: 0.23982\noutside 1 331\nTMhelix 332 354\ninside 355 372

1936 GCF_001059255.1_ASM105925v1 Citrobacter freundiiProteobacteria; Gammaproteobacteria; Enterobacterales; Enterobacteriaceae; Citrobacter; Citrobacter freundii complex
MTGDNSLINSSGINRRDFMKLCAALAATMGLSSKAAAEMAESVSRPQRPPVIWIGAQECT WP_003024486.1
MULTISPECIES: hydrogenase 2 small subunit [Enterobacteriaceae] Length: 372\nNumber of predicted TMHs: 1\nExp number of AAs in TMHs: 27.14621\nExp number, first 60 AAs: 0.22683\nTotal prob of N-in: 0.23982\noutside 1 331\nTMhelix 332 354\ninside 355 372

1937 GCF_001281005.1_ASM128100v1 Citrobacter freundiiProteobacteria; Gammaproteobacteria; Enterobacterales; Enterobacteriaceae; Citrobacter; Citrobacter freundii complex
MTGDNSLINSSGINRRDFMKLCAALAATMGLSSKAAAEMAESVSRPQRPPVIWIGAQECT WP_003024486.1
MULTISPECIES: hydrogenase 2 small subunit [Enterobacteriaceae] Length: 372\nNumber of predicted TMHs: 1\nExp number of AAs in TMHs: 27.14621\nExp number, first 60 AAs: 0.22683\nTotal prob of N-in: 0.23982\noutside 1 331\nTMhelix 332 354\ninside 355 372

1938 GCF_001316675.1_ASM131667v1 Citrobacter freundiiProteobacteria; Gammaproteobacteria; Enterobacterales; Enterobacteriaceae; Citrobacter; Citrobacter freundii complex
MTGDNSLINSSGINRRDFMKLCAALAATMGLSSKAAAEMAESVSRPQRPPVIWIGAQECT WP_003024486.1
MULTISPECIES: hydrogenase 2 small subunit [Enterobacteriaceae] Length: 372\nNumber of predicted TMHs: 1\nExp number of AAs in TMHs: 27.14621\nExp number, first 60 AAs: 0.22683\nTotal prob of N-in: 0.23982\noutside 1 331\nTMhelix 332 354\ninside 355 372

1939 GCF_001413045.1_ASM141304v1 Citrobacter freundiiProteobacteria; Gammaproteobacteria; Enterobacterales; Enterobacteriaceae; Citrobacter; Citrobacter freundii complex

MTGDNSLINSSGINRRDFMKLCAALAATMGLSSKAAAEMAESVSRPQRPPVIWIGAQECT WP_003024486.1 MULTISPECIES: hydrogenase 2 small subunit [Enterobacteriaceae] Length: 372\nNumber of predicted TMHs: 1\nExp number of AAs in TMHs: 27.14621\nExp number, first 60 AAs: 0.22683\nTotal prob of N-in: 0.23982\noutside 1 331\nTMhelix 332 354\ninside 355 372

1940 GCF_001413095.1_ASM141309v1 Citrobacter freundiiProteobacteria; Gammaproteobacteria; Enterobacterales; Enterobacteriaceae; Citrobacter; Citrobacter freundii complex MTGDNSLINSSGINRRDFMKLCAALAATMGLSSKAAAEMAESVSRPQRPPVIWIGAQECT WP_003024486.1 MULTISPECIES: hydrogenase 2 small subunit [Enterobacteriaceae] Length: 372\nNumber of predicted TMHs: 1\nExp number of AAs in TMHs: 27.14621\nExp number, first 60 AAs: 0.22683\nTotal prob of N-in: 0.23982\noutside 1 331\nTMhelix 332 354\ninside 355 372

1941 GCF_001413125.1_ASM141312v1 Citrobacter freundiiProteobacteria; Gammaproteobacteria; Enterobacterales; Enterobacteriaceae; Citrobacter; Citrobacter freundii complex MTGDNSLINSSGINRRDFMKLCAALAATMGLSSKAAAEMAESVSRPQRPPVIWIGAQECT WP_003024486.1 MULTISPECIES: hydrogenase 2 small subunit [Enterobacteriaceae] Length: 372\nNumber of predicted TMHs: 1\nExp number of AAs in TMHs: 27.14621\nExp number, first 60 AAs: 0.22683\nTotal prob of N-in: 0.23982\noutside 1 331\nTMhelix 332 354\ninside 355 372

1942 GCF_001413165.1_ASM141316v1 Citrobacter freundiiProteobacteria; Gammaproteobacteria; Enterobacterales; Enterobacteriaceae; Citrobacter; Citrobacter freundii complex MTGDNSLINSSGINRRDFMKLCAALAATMGLSSKAAAEMAESVSRPQRPPVIWIGAQECT WP_003024486.1 MULTISPECIES: hydrogenase 2 small subunit [Enterobacteriaceae] Length: 372\nNumber of predicted TMHs: 1\nExp number of AAs in TMHs: 27.14621\nExp number, first 60 AAs: 0.22683\nTotal prob of N-in: 0.23982\noutside 1 331\nTMhelix 332 354\ninside 355 372

1943 GCF_001413215.1_ASM141321v1 Citrobacter freundiiProteobacteria; Gammaproteobacteria; Enterobacterales; Enterobacteriaceae; Citrobacter; Citrobacter freundii complex MTGDNSLINSSGINRRDFMKLCAALAATMGLSSKAAAEMAESVSRPQRPPVIWIGAQECT WP_003024486.1 MULTISPECIES: hydrogenase 2 small subunit [Enterobacteriaceae] Length: 372\nNumber of predicted TMHs: 1\nExp number of AAs in TMHs: 27.14621\nExp number, first 60 AAs: 0.22683\nTotal prob of N-in: 0.23982\noutside 1 331\nTMhelix 332 354\ninside 355 372

1944 GCF_001413275.1_ASM141327v1 Citrobacter freundiiProteobacteria; Gammaproteobacteria; Enterobacterales; Enterobacteriaceae; Citrobacter; Citrobacter freundii complex MTGDNSLINSSGINRRDFMKLCAALAATMGLSSKAAAEMAESVSRPQRPPVIWIGAQECT WP_003024486.1 MULTISPECIES: hydrogenase 2 small subunit [Enterobacteriaceae] Length: 372\nNumber of predicted TMHs: 1\nExp number of AAs in TMHs: 27.14621\nExp number, first 60 AAs: 0.22683\nTotal prob of N-in: 0.23982\noutside 1 331\nTMhelix 332 354\ninside 355 372

1945 GCF_001413315.1_ASM141331v1 Citrobacter freundiiProteobacteria; Gammaproteobacteria; Enterobacterales; Enterobacteriaceae; Citrobacter; Citrobacter freundii complex MTGDNSLINSSGINRRDFMKLCAALAATMGLSSKAAAEMAESVSRPQRPPVIWIGAQECT WP_003024486.1 MULTISPECIES: hydrogenase 2 small subunit [Enterobacteriaceae] Length: 372\nNumber of predicted TMHs: 1\nExp number of AAs in TMHs: 27.14621\nExp number, first 60 AAs: 0.22683\nTotal prob of N-in: 0.23982\noutside 1 331\nTMhelix 332 354\ninside 355 372

1946 GCF_001482545.1_ASM148254v1 Citrobacter freundiiProteobacteria; Gammaproteobacteria; Enterobacterales; Enterobacteriaceae; Citrobacter; Citrobacter freundii complex MTGDNSLINSSGINRRDFMKLCAALAATMGLSSKAAAEMAESVSRPQRPPVIWIGAQECT WP_003024486.1 MULTISPECIES: hydrogenase 2 small subunit [Enterobacteriaceae] Length: 372\nNumber of predicted TMHs: 1\nExp number of AAs in TMHs: 27.14621\nExp number, first 60 AAs: 0.22683\nTotal prob of N-in: 0.23982\noutside 1 331\nTMhelix 332 354\ninside 355 372

1947 GCF_001546285.1_ASM154628v1 Citrobacter freundiiProteobacteria; Gammaproteobacteria; Enterobacterales; Enterobacteriaceae; Citrobacter; Citrobacter freundii complex MTGDNSLINSSGINRRDFMKLCAALAATMGLSSKAAAEMAESVSRPQRPPVIWIGAQECT WP_003024486.1 MULTISPECIES: hydrogenase 2 small subunit [Enterobacteriaceae] Length: 372\nNumber of predicted TMHs: 1\nExp number of AAs in TMHs: 27.14621\nExp number, first 60 AAs: 0.22683\nTotal prob of N-in: 0.23982\noutside 1 331\nTMhelix 332 354\ninside 355 372

1948 GCF_001880775.1_ASM188077v1 Citrobacter freundiiProteobacteria; Gammaproteobacteria; Enterobacterales; Enterobacteriaceae; Citrobacter; Citrobacter freundii complex
MTGDNSLINSSGINRRDFMKLCAALAATMGLSSKAAAEMAESVSRPQRPPVIWIGAQECT WP_003024486.1
MULTISPECIES: hydrogenase 2 small subunit [Enterobacteriaceae] Length: 372\nNumber of predicted TMHs: 1\nExp number of AAs in TMHs: 27.14621\nExp number, first 60 AAs: 0.22683\nTotal prob of N-in: 0.23982\noutside 1 331\nTMhelix 332 354\ninside 355 372

1949 GCF_001702455.1_ASM170245v1 Citrobacter freundiiProteobacteria; Gammaproteobacteria; Enterobacterales; Enterobacteriaceae; Citrobacter; Citrobacter freundii complex
MTGDNSLINSSGINRRDFMKLCAALAATMGLSSKAAAEMAESVSRPQRPPVIWIGAQECT WP_003024486.1
MULTISPECIES: hydrogenase 2 small subunit [Enterobacteriaceae] Length: 372\nNumber of predicted TMHs: 1\nExp number of AAs in TMHs: 27.14621\nExp number, first 60 AAs: 0.22683\nTotal prob of N-in: 0.23982\noutside 1 331\nTMhelix 332 354\ninside 355 372

1950 GCF_001880825.1_ASM188082v1 Citrobacter freundiiProteobacteria; Gammaproteobacteria; Enterobacterales; Enterobacteriaceae; Citrobacter; Citrobacter freundii complex
MTGDNSLINSSGINRRDFMKLCAALAATMGLSSKAAAEMAESVSRPQRPPVIWIGAQECT WP_003024486.1
MULTISPECIES: hydrogenase 2 small subunit [Enterobacteriaceae] Length: 372\nNumber of predicted TMHs: 1\nExp number of AAs in TMHs: 27.14621\nExp number, first 60 AAs: 0.22683\nTotal prob of N-in: 0.23982\noutside 1 331\nTMhelix 332 354\ninside 355 372

1951 GCF_001880845.1_ASM188084v1 Citrobacter freundiiProteobacteria; Gammaproteobacteria; Enterobacterales; Enterobacteriaceae; Citrobacter; Citrobacter freundii complex
MTGDNSLINSSGINRRDFMKLCAALAATMGLSSKAAAEMAESVSRPQRPPVIWIGAQECT WP_003024486.1
MULTISPECIES: hydrogenase 2 small subunit [Enterobacteriaceae] Length: 372\nNumber of predicted TMHs: 1\nExp number of AAs in TMHs: 27.14621\nExp number, first 60 AAs: 0.22683\nTotal prob of N-in: 0.23982\noutside 1 331\nTMhelix 332 354\ninside 355 372

1952 GCF_001880905.1_ASM188090v1 Citrobacter freundiiProteobacteria; Gammaproteobacteria; Enterobacterales; Enterobacteriaceae; Citrobacter; Citrobacter freundii complex
MTGDNSLINSSGINRRDFMKLCAALAATMGLSSKAAAEMAESVSRPQRPPVIWIGAQECT WP_003024486.1
MULTISPECIES: hydrogenase 2 small subunit [Enterobacteriaceae] Length: 372\nNumber of predicted TMHs: 1\nExp number of AAs in TMHs: 27.14621\nExp number, first 60 AAs: 0.22683\nTotal prob of N-in: 0.23982\noutside 1 331\nTMhelix 332 354\ninside 355 372

1953 GCF_002151735.1_ASM215173v1 Citrobacter freundiiProteobacteria; Gammaproteobacteria; Enterobacterales; Enterobacteriaceae; Citrobacter; Citrobacter freundii complex
MTGDNSLINSSGINRRDFMKLCAALAATMGLSSKAAAEMAESVSRPQRPPVIWIGAQECT WP_003024486.1
MULTISPECIES: hydrogenase 2 small subunit [Enterobacteriaceae] Length: 372\nNumber of predicted TMHs: 1\nExp number of AAs in TMHs: 27.14621\nExp number, first 60 AAs: 0.22683\nTotal prob of N-in: 0.23982\noutside 1 331\nTMhelix 332 354\ninside 355 372

1954 GCF_001880945.1_ASM188094v1 Citrobacter freundiiProteobacteria; Gammaproteobacteria; Enterobacterales; Enterobacteriaceae; Citrobacter; Citrobacter freundii complex
MTGDNSLINSSGINRRDFMKLCAALAATMGLSSKAAAEMAESVSRPQRPPVIWIGAQECT WP_003024486.1
MULTISPECIES: hydrogenase 2 small subunit [Enterobacteriaceae] Length: 372\nNumber of predicted TMHs: 1\nExp number of AAs in TMHs: 27.14621\nExp number, first 60 AAs: 0.22683\nTotal prob of N-in: 0.23982\noutside 1 331\nTMhelix 332 354\ninside 355 372

1955 GCF_002151815.1_ASM215181v1 Citrobacter freundiiProteobacteria; Gammaproteobacteria; Enterobacterales; Enterobacteriaceae; Citrobacter; Citrobacter freundii complex
MTGDNSLINSNGINRRDFMKLCAALAATMGLSSKAAAEMAESVSRPQRPPVIWIGAQECT WP_016154493.1
MULTISPECIES: hydrogenase-2 small chain [Enterobacteriaceae] Length: 372\nNumber of predicted TMHs: 1\nExp number of AAs in TMHs: 26.90274\nExp number, first 60 AAs: 0.32271\nTotal prob of N-in: 0.22239\noutside 1 331\nTMhelix 332 354\ninside 355 372

1956 GCF_000714305.1_ASM71430v1 Citrobacter freundiiProteobacteria; Gammaproteobacteria; Enterobacterales; Enterobacteriaceae; Citrobacter; Citrobacter freundii complex
MTGDNSLINSSGINRRDFMKLCAALAATMGLSSKAAAEMAESVSRPQRPPVIWIGAQECT WP_003024486.1
MULTISPECIES: hydrogenase 2 small subunit [Enterobacteriaceae] Length: 372\nNumber of predicted TMHs:

1\nExp number of AAs in TMHs: 27.14621\nExp number, first 60 AAs: 0.22683\nTotal prob of N-in: 0.23982\noutside 1 331\nTMhelix 332 354\ninside 355 372

1957 GCF_000982845.1_ASM98284v1 Citrobacter freundiiProteobacteria; Gammaproteobacteria; Enterobacterales; Enterobacteriaceae; Citrobacter; Citrobacter freundii complex
MTGDNSLINSSGINRRDFMKLCAALAATMGLSSKAAAEMAESVSRPQRPPVIWIGAQECT WP_003024486.1
MULTISPECIES: hydrogenase 2 small subunit [Enterobacteriaceae] Length: 372\nNumber of predicted TMHs: 1\nExp number of AAs in TMHs: 27.14621\nExp number, first 60 AAs: 0.22683\nTotal prob of N-in: 0.23982\noutside 1 331\nTMhelix 332 354\ninside 355 372

1958 GCF_000783995.1_ASM78399v1 Citrobacter freundiiProteobacteria; Gammaproteobacteria; Enterobacterales; Enterobacteriaceae; Citrobacter; Citrobacter freundii complex
MTGDNSLINSNGINRRDFMKLCAALAATMGLSSKAAAEMAESVSRPQRPPVIWIGAQECT WP_043017615.1
MULTISPECIES: hydrogenase 2 small subunit [Citrobacter freundii complex] Length: 372\nNumber of predicted TMHs: 1\nExp number of AAs in TMHs: 27.21963\nExp number, first 60 AAs: 0.31687\nTotal prob of N-in: 0.24323\noutside 1 331\nTMhelix 332 354\ninside 355 372

1959 GCF_001022275.1_ASM102227v1 Citrobacter freundiiProteobacteria; Gammaproteobacteria; Enterobacterales; Enterobacteriaceae; Citrobacter; Citrobacter freundii complex
MTGDNSLINSSGINRRDFMKLCAALAATMGLSSKAAAEMAESVSRPQRPPVIWIGAQECT WP_003024486.1
MULTISPECIES: hydrogenase 2 small subunit [Enterobacteriaceae] Length: 372\nNumber of predicted TMHs: 1\nExp number of AAs in TMHs: 27.14621\nExp number, first 60 AAs: 0.22683\nTotal prob of N-in: 0.23982\noutside 1 331\nTMhelix 332 354\ninside 355 372

1960 GCF_001055315.1_ASM105531v1 Citrobacter freundiiProteobacteria; Gammaproteobacteria; Enterobacterales; Enterobacteriaceae; Citrobacter; Citrobacter freundii complex
MTGDNSLINSSGINRRDFMKLCAALAATMGLSSKAAAEMAESVSRPQRPPVIWIGAQECT WP_003024486.1
MULTISPECIES: hydrogenase 2 small subunit [Enterobacteriaceae] Length: 372\nNumber of predicted TMHs: 1\nExp number of AAs in TMHs: 27.14621\nExp number, first 60 AAs: 0.22683\nTotal prob of N-in: 0.23982\noutside 1 331\nTMhelix 332 354\ninside 355 372

1961 GCF_001058505.1_ASM105850v1 Citrobacter freundiiProteobacteria; Gammaproteobacteria; Enterobacterales; Enterobacteriaceae; Citrobacter; Citrobacter freundii complex
MTGDNSLINSSGINRRDFMKLCAALAATMGLSSKAAAEMAESVSRPQRPPVIWIGAQECT WP_003024486.1
MULTISPECIES: hydrogenase 2 small subunit [Enterobacteriaceae] Length: 372\nNumber of predicted TMHs: 1\nExp number of AAs in TMHs: 27.14621\nExp number, first 60 AAs: 0.22683\nTotal prob of N-in: 0.23982\noutside 1 331\nTMhelix 332 354\ninside 355 372

1962 GCF_001306025.1_ASM130602v1 Citrobacter freundiiProteobacteria; Gammaproteobacteria; Enterobacterales; Enterobacteriaceae; Citrobacter; Citrobacter freundii complex
MTGDNSLINSSGINRRDFMKLCAALAATMGLSSKAAAEMAESVSRPQRPPVIWIGAQECT WP_003024486.1
MULTISPECIES: hydrogenase 2 small subunit [Enterobacteriaceae] Length: 372\nNumber of predicted TMHs: 1\nExp number of AAs in TMHs: 27.14621\nExp number, first 60 AAs: 0.22683\nTotal prob of N-in: 0.23982\noutside 1 331\nTMhelix 332 354\ninside 355 372

1963 GCF_001273815.1_ASM127381v1 Citrobacter freundiiProteobacteria; Gammaproteobacteria; Enterobacterales; Enterobacteriaceae; Citrobacter; Citrobacter freundii complex
MTGDNSLINSNGINRRDFMKLCAALAATMGLSSKAAAEMAESVSRPQRPPVIWIGAQECT WP_016154493.1
MULTISPECIES: hydrogenase-2 small chain [Enterobacteriaceae] Length: 372\nNumber of predicted TMHs: 1\nExp number of AAs in TMHs: 26.90274\nExp number, first 60 AAs: 0.32271\nTotal prob of N-in: 0.22239\noutside 1 331\nTMhelix 332 354\ninside 355 372

1964 GCF_001412725.1_ASM141272v1 Citrobacter freundiiProteobacteria; Gammaproteobacteria; Enterobacterales; Enterobacteriaceae; Citrobacter; Citrobacter freundii complex
MTGDNSLINSSGINRRDFMKLCAALAATMGLSSKAAAEMAESVSRPQRPPVIWIGAQECT WP_003024486.1
MULTISPECIES: hydrogenase 2 small subunit [Enterobacteriaceae] Length: 372\nNumber of predicted TMHs: 1\nExp number of AAs in TMHs: 27.14621\nExp number, first 60 AAs: 0.22683\nTotal prob of N-in: 0.23982\noutside 1 331\nTMhelix 332 354\ninside 355 372

1965 GCF_001411885.1_ASM141188v1 Citrobacter freundiiProteobacteria; Gammaproteobacteria; Enterobacterales; Enterobacteriaceae; Citrobacter; Citrobacter freundii complex

MTGDNSLINSSGINRRDFMKLCAALAATMGLSSKAAAEMAESVSRPQRPPVIWIGAQECT WP_003024486.1 MULTISPECIES: hydrogenase 2 small subunit [Enterobacteriaceae] Length: 372\nNumber of predicted TMHs: 1\nExp number of AAs in TMHs: 27.14621\nExp number, first 60 AAs: 0.22683\nTotal prob of N-in: 0.23982\noutside 1 331\nTMhelix 332 354\ninside 355 372

1966 GCF_001412755.1_ASM141275v1 Citrobacter freundiiProteobacteria; Gammaproteobacteria; Enterobacterales; Enterobacteriaceae; Citrobacter; Citrobacter freundii complex MTGDNSLINSSGINRRDFMKLCAALAATMGLSSKAAAEMAESVSRPQRPPVIWIGAQECT WP_003024486.1 MULTISPECIES: hydrogenase 2 small subunit [Enterobacteriaceae] Length: 372\nNumber of predicted TMHs: 1\nExp number of AAs in TMHs: 27.14621\nExp number, first 60 AAs: 0.22683\nTotal prob of N-in: 0.23982\noutside 1 331\nTMhelix 332 354\ninside 355 372

1967 GCF_001412825.1_ASM141282v1 Citrobacter freundiiProteobacteria; Gammaproteobacteria; Enterobacterales; Enterobacteriaceae; Citrobacter; Citrobacter freundii complex MTGDNSLINSSGINRRDFMKLCAALAATMGLSSKAAAEMAESVSRPQRPPVIWIGAQECT WP_003024486.1 MULTISPECIES: hydrogenase 2 small subunit [Enterobacteriaceae] Length: 372\nNumber of predicted TMHs: 1\nExp number of AAs in TMHs: 27.14621\nExp number, first 60 AAs: 0.22683\nTotal prob of N-in: 0.23982\noutside 1 331\nTMhelix 332 354\ninside 355 372

1968 GCF_001412875.1_ASM141287v1 Citrobacter freundiiProteobacteria; Gammaproteobacteria; Enterobacterales; Enterobacteriaceae; Citrobacter; Citrobacter freundii complex MTGDNSLINSSGINRRDFMKLCAALAATMGLSSKAAAEMAESVSRPQRPPVIWIGAQECT WP_003024486.1 MULTISPECIES: hydrogenase 2 small subunit [Enterobacteriaceae] Length: 372\nNumber of predicted TMHs: 1\nExp number of AAs in TMHs: 27.14621\nExp number, first 60 AAs: 0.22683\nTotal prob of N-in: 0.23982\noutside 1 331\nTMhelix 332 354\ninside 355 372

1969 GCF_001413035.1_ASM141303v1 Citrobacter freundiiProteobacteria; Gammaproteobacteria; Enterobacterales; Enterobacteriaceae; Citrobacter; Citrobacter freundii complex MTGDNSLINSSGINRRDFMKLCAALAATMGLSSKAAAEMAESVSRPQRPPVIWIGAQECT WP_003024486.1 MULTISPECIES: hydrogenase 2 small subunit [Enterobacteriaceae] Length: 372\nNumber of predicted TMHs: 1\nExp number of AAs in TMHs: 27.14621\nExp number, first 60 AAs: 0.22683\nTotal prob of N-in: 0.23982\noutside 1 331\nTMhelix 332 354\ninside 355 372

1970 GCF_001317135.2_48_S23v.02 Citrobacter freundiiProteobacteria; Gammaproteobacteria; Enterobacterales; Enterobacteriaceae; Citrobacter; Citrobacter freundii complex MTGDNSLINSSGINRRDFMKLCAALAATMGLSSKAAAEMAESVSRPQRPPVIWIGAQECT WP_057068275.1 hydrogenase 2 small subunit [Citrobacter freundii] Length: 372\nNumber of predicted TMHs: 1\nExp number of AAs in TMHs: 27.15001\nExp number, first 60 AAs: 0.22682\nTotal prob of N-in: 0.23980\noutside 1 331\nTMhelix 332 354\ninside 355 372

1971 GCF_002151775.1_ASM215177v1 Citrobacter freundiiProteobacteria; Gammaproteobacteria; Enterobacterales; Enterobacteriaceae; Citrobacter; Citrobacter freundii complex MTGDNSLINSSGINRRDFMKLCAALAATMGLSSKAAAEMAESVSRPQRPPVIWIGAQECT WP_003024486.1 MULTISPECIES: hydrogenase 2 small subunit [Enterobacteriaceae] Length: 372\nNumber of predicted TMHs: 1\nExp number of AAs in TMHs: 27.14621\nExp number, first 60 AAs: 0.22683\nTotal prob of N-in: 0.23982\noutside 1 331\nTMhelix 332 354\ninside 355 372

1972 GCF_002192675.1_ASM219267v1 Citrobacter freundiiProteobacteria; Gammaproteobacteria; Enterobacterales; Enterobacteriaceae; Citrobacter; Citrobacter freundii complex MTGDNSLINSSGINRRDFMKLCAALAATMGLSSKAAAEMAESVSRPQRPPVIWIGAQECT WP_003024486.1 MULTISPECIES: hydrogenase 2 small subunit [Enterobacteriaceae] Length: 372\nNumber of predicted TMHs: 1\nExp number of AAs in TMHs: 27.14621\nExp number, first 60 AAs: 0.22683\nTotal prob of N-in: 0.23982\noutside 1 331\nTMhelix 332 354\ninside 355 372

1973 GCF_002042945.1_ASM204294v1 Citrobacter sp. A316 Proteobacteria; Gammaproteobacteria; Enterobacterales; Enterobacteriaceae; Citrobacter; Citrobacter freundii complex MRQNSTGEETRVMNTNNEETFYQAMRRKGVSRRSFLKYCSLAATSLGLGAGMTPRIAWAL WP_079939359.1 [Ni/Fe] hydrogenase small subunit [Citrobacter sp. A316] Length: 387\nNumber of predicted TMHs: 1\nExp number of AAs in TMHs: 33.86716\nExp number, first 60 AAs: 16.1538\nTotal prob of N-in: 0.76283\nPOSSIBLE N-term signal sequence\noutside 1 339\nTMhelix 340 362\ninside 363 387

1974 GCF_002042945.1_ASM204294v1 Citrobacter sp. A316 Proteobacteria; Gammaproteobacteria; Enterobacterales; Enterobacteriaceae; Citrobacter; Citrobacter freundii complex
MTGDNSLINSSGINRRDFMKLCAALAATMGLSSKAAAEMAESVSRPQRPPVIWIGAQECT WP_003024486.1
MULTISPECIES: hydrogenase 2 small subunit [Enterobacteriaceae] Length: 372\nNumber of predicted TMHs: 1\nExp number of AAs in TMHs: 27.14621\nExp number, first 60 AAs: 0.22683\nTotal prob of N-in: 0.23982\noutside 1 331\nTMhelix 332 354\ninside 355 372

1975 GCF_002042885.1_ASM204288v1 Citrobacter sp. A60 Proteobacteria; Gammaproteobacteria; Enterobacterales; Enterobacteriaceae; Citrobacter; Citrobacter freundii complex
MTGDNSLINSSGINRRDFMKLCAALAATMGLSSKAAAEMAESVSRPQRPPVIWIGAQECT WP_003024486.1
MULTISPECIES: hydrogenase 2 small subunit [Enterobacteriaceae] Length: 372\nNumber of predicted TMHs: 1\nExp number of AAs in TMHs: 27.14621\nExp number, first 60 AAs: 0.22683\nTotal prob of N-in: 0.23982\noutside 1 331\nTMhelix 332 354\ninside 355 372

1976 GCF_001586835.1_ASM158683v1 Citrobacter sp. AATXQ Proteobacteria; Gammaproteobacteria; Enterobacterales; Enterobacteriaceae; Citrobacter; Citrobacter freundii complex
MTGDNSLINSSGINRRDFMKLCAALAATMGLSSKAAAEMAESVSRPQRPPVIWIGAQECT WP_003024486.1
MULTISPECIES: hydrogenase 2 small subunit [Enterobacteriaceae] Length: 372\nNumber of predicted TMHs: 1\nExp number of AAs in TMHs: 27.14621\nExp number, first 60 AAs: 0.22683\nTotal prob of N-in: 0.23982\noutside 1 331\nTMhelix 332 354\ninside 355 372

1977 GCF_001586845.1_ASM158684v1 Citrobacter sp. AATXR Proteobacteria; Gammaproteobacteria; Enterobacterales; Enterobacteriaceae; Citrobacter; Citrobacter freundii complex
MTGDNSLINSSGINRRDFMKLCAALAATMGLSSKAAAEMAESVSRPQRPPVIWIGAQECT WP_003024486.1
MULTISPECIES: hydrogenase 2 small subunit [Enterobacteriaceae] Length: 372\nNumber of predicted TMHs: 1\nExp number of AAs in TMHs: 27.14621\nExp number, first 60 AAs: 0.22683\nTotal prob of N-in: 0.23982\noutside 1 331\nTMhelix 332 354\ninside 355 372

1978 GCF_001037575.1_Citr_freu_complex_BIDMC107_V1 Citrobacter sp. BIDMC107 Proteobacteria; Gammaproteobacteria; Enterobacterales; Enterobacteriaceae; Citrobacter; Citrobacter freundii complex
MTGDNSLINSSGINRRDFMKLCAALAATMGLSSKAAAEMAESVSRPQRPPVIWIGAQECT WP_003024486.1
MULTISPECIES: hydrogenase 2 small subunit [Enterobacteriaceae] Length: 372\nNumber of predicted TMHs: 1\nExp number of AAs in TMHs: 27.14621\nExp number, first 60 AAs: 0.22683\nTotal prob of N-in: 0.23982\noutside 1 331\nTMhelix 332 354\ninside 355 372

1979 GCF_001037585.1_Citr_freu_complex_BIDMC108_V1 Citrobacter sp. BIDMC108 Proteobacteria; Gammaproteobacteria; Enterobacterales; Enterobacteriaceae; Citrobacter; Citrobacter freundii complex
MTGDNSLINSSGINRRDFMKLCAALAATMGLSSKAAAEMAESVSRPQRPPVIWIGAQECT WP_003024486.1
MULTISPECIES: hydrogenase 2 small subunit [Enterobacteriaceae] Length: 372\nNumber of predicted TMHs: 1\nExp number of AAs in TMHs: 27.14621\nExp number, first 60 AAs: 0.22683\nTotal prob of N-in: 0.23982\noutside 1 331\nTMhelix 332 354\ninside 355 372

1980 GCF_001034465.1_Citr_freu_complex_MGH100_V1 Citrobacter sp. MGH100 Proteobacteria; Gammaproteobacteria; Enterobacterales; Enterobacteriaceae; Citrobacter; Citrobacter freundii complex
MTGDNSLINSNGINRRDFMKLCAALAATMGLSSKAAAEMAESVSRPQRPPVIWIGAQECT WP_016154493.1
MULTISPECIES: hydrogenase-2 small chain [Enterobacteriaceae] Length: 372\nNumber of predicted TMHs: 1\nExp number of AAs in TMHs: 26.90274\nExp number, first 60 AAs: 0.32271\nTotal prob of N-in: 0.22239\noutside 1 331\nTMhelix 332 354\ninside 355 372

1981 GCF_001034475.1_Citr_freu_complex_MGH103_V1 Citrobacter sp. MGH103 Proteobacteria; Gammaproteobacteria; Enterobacterales; Enterobacteriaceae; Citrobacter; Citrobacter freundii complex
MTGDNSLINSSGINRRDFMKLCAALAATMGLSSKAAAEMAESVSRPQRPPVIWIGAQECT WP_003024486.1
MULTISPECIES: hydrogenase 2 small subunit [Enterobacteriaceae] Length: 372\nNumber of predicted TMHs: 1\nExp number of AAs in TMHs: 27.14621\nExp number, first 60 AAs: 0.22683\nTotal prob of N-in: 0.23982\noutside 1 331\nTMhelix 332 354\ninside 355 372

1982 GCF_001034485.1_Citr_freu_complex_MGH104_V1 Citrobacter sp. MGH104 Proteobacteria; Gammaproteobacteria; Enterobacterales; Enterobacteriaceae; Citrobacter; Citrobacter freundii complex
MTGDNSLINSSGINRRDFMKLCAALAATMGLSSKAAAEMAESVSRPQRPPVIWIGAQECT WP_003024486.1
MULTISPECIES: hydrogenase 2 small subunit [Enterobacteriaceae] Length: 372\nNumber of predicted TMHs:

1\nExp number of AAs in TMHs: 27.14621\nExp number, first 60 AAs: 0.22683\nTotal prob of N-in: 0.23982\noutside 1 331\nTMhelix 332 354\ninside 355 372

1983 GCF_001037485.1_Citr_freu_complex_MGH105_V1 Citrobacter sp. MGH105 Proteobacteria; Gammaproteobacteria; Enterobacterales; Enterobacteriaceae; Citrobacter; Citrobacter freundii complex MTGDNSLINSSGINRRDFMKLCAALAATMGLSSKAAAEMAESVSRPQRPPVIWIGAQECT WP_003024486.1 MULTISPECIES: hydrogenase 2 small subunit [Enterobacteriaceae] Length: 372\nNumber of predicted TMHs: 1\nExp number of AAs in TMHs: 27.14621\nExp number, first 60 AAs: 0.22683\nTotal prob of N-in: 0.23982\noutside 1 331\nTMhelix 332 354\ninside 355 372

1984 GCF_001037495.1_Citr_freu_complex_MGH106_V1 Citrobacter sp. MGH106 Proteobacteria; Gammaproteobacteria; Enterobacterales; Enterobacteriaceae; Citrobacter; Citrobacter freundii complex MTGDNSLINSNGINRRDFMKLCAALAATMGLSSKAAAEMAESVSRPQRPPVIWIGAQECT WP_045447522.1 MULTISPECIES: hydrogenase 2 small subunit [Citrobacter] Length: 372\nNumber of predicted TMHs: 1\nExp number of AAs in TMHs: 26.06135\nExp number, first 60 AAs: 0.32735\nTotal prob of N-in: 0.20887\noutside 1 331\nTMhelix 332 354\ninside 355 372

1985 GCF_001037505.1_Citr_freu_complex_MGH109_V1 Citrobacter sp. MGH109 Proteobacteria; Gammaproteobacteria; Enterobacterales; Enterobacteriaceae; Citrobacter; Citrobacter freundii complex MTGDNSLINSSGINRRDFMKLCAALAATMGLSSKAAAEMAESVSRPQRPPVIWIGAQECT WP_003024486.1 MULTISPECIES: hydrogenase 2 small subunit [Enterobacteriaceae] Length: 372\nNumber of predicted TMHs: 1\nExp number of AAs in TMHs: 27.14621\nExp number, first 60 AAs: 0.22683\nTotal prob of N-in: 0.23982\noutside 1 331\nTMhelix 332 354\ninside 355 372

1986 GCF_001037515.1_Citr_freu_complex_MGH110_V1 Citrobacter sp. MGH110 Proteobacteria; Gammaproteobacteria; Enterobacterales; Enterobacteriaceae; Citrobacter; Citrobacter freundii complex MTGDNSLINSSGINRRDFMKLCAALAATMGLSSKAAAEMAESVSRPQRPPVIWIGAQECT WP_003024486.1 MULTISPECIES: hydrogenase 2 small subunit [Enterobacteriaceae] Length: 372\nNumber of predicted TMHs: 1\nExp number of AAs in TMHs: 27.14621\nExp number, first 60 AAs: 0.22683\nTotal prob of N-in: 0.23982\noutside 1 331\nTMhelix 332 354\ninside 355 372

1987 GCF_001034445.1_Citr_freu_complex_MGH99_V1 Citrobacter sp. MGH99 Proteobacteria; Gammaproteobacteria; Enterobacterales; Enterobacteriaceae; Citrobacter; Citrobacter freundii complex MTGDNSLINSSGINRRDFMKLCAALAATMGLSSKAAAEMAESVSRPQRPPVIWIGAQECT WP_003024486.1 MULTISPECIES: hydrogenase 2 small subunit [Enterobacteriaceae] Length: 372\nNumber of predicted TMHs: 1\nExp number of AAs in TMHs: 27.14621\nExp number, first 60 AAs: 0.22683\nTotal prob of N-in: 0.23982\noutside 1 331\nTMhelix 332 354\ninside 355 372

1988 GCF_002099305.1_ASM209930v1 Citrobacter werkmanii Proteobacteria; Gammaproteobacteria; Enterobacterales; Enterobacteriaceae; Citrobacter; Citrobacter freundii complex MTGDNSLINSSGINRRDFMKLCAALAATMGLSSKAAAEMAESVSRPQRPPVIWIGAQECT WP_003024486.1 MULTISPECIES: hydrogenase 2 small subunit [Enterobacteriaceae] Length: 372\nNumber of predicted TMHs: 1\nExp number of AAs in TMHs: 27.14621\nExp number, first 60 AAs: 0.22683\nTotal prob of N-in: 0.23982\noutside 1 331\nTMhelix 332 354\ninside 355 372

1989 GCF_002185305.1_ASM218530v1 Citrobacter werkmanii Proteobacteria; Gammaproteobacteria; Enterobacterales; Enterobacteriaceae; Citrobacter; Citrobacter freundii complex MTGDNSLINSSGINRRDFMKLCAALAATMGLSSKAAAEMAESVSRPQRPPVIWIGAQECT WP_003024486.1 MULTISPECIES: hydrogenase 2 small subunit [Enterobacteriaceae] Length: 372\nNumber of predicted TMHs: 1\nExp number of AAs in TMHs: 27.14621\nExp number, first 60 AAs: 0.22683\nTotal prob of N-in: 0.23982\noutside 1 331\nTMhelix 332 354\ninside 355 372

1990 GCF_002025225.1_ASM202522v1 Citrobacter werkmanii Proteobacteria; Gammaproteobacteria; Enterobacterales; Enterobacteriaceae; Citrobacter; Citrobacter freundii complex MTGDNSLINSSGINRRDFMKLCAALAATMGLSSKAAAEMAESVSRPQRPPVIWIGAQECT WP_003024486.1 MULTISPECIES: hydrogenase 2 small subunit [Enterobacteriaceae] Length: 372\nNumber of predicted TMHs: 1\nExp number of AAs in TMHs: 27.14621\nExp number, first 60 AAs: 0.22683\nTotal prob of N-in: 0.23982\noutside 1 331\nTMhelix 332 354\ninside 355 372

1991 GCF_002114305.1_ASM211430v1 Citrobacter werkmanii Proteobacteria; Gammaproteobacteria; Enterobacterales; Enterobacteriaceae; Citrobacter; Citrobacter freundii complex

MTGDNSLINSSGINRRDFMKLCAALAATMGLSSKAAAEMAESVSRPQRPPVIWIGAQECT WP_003024486.1 MULTISPECIES: hydrogenase 2 small subunit [Enterobacteriaceae] Length: 372\nNumber of predicted TMHs: 1\nExp number of AAs in TMHs: 27.14621\nExp number, first 60 AAs: 0.22683\nTotal prob of N-in: 0.23982\noutside 1 331\nTMhelix 332 354\ninside 355 372

1992 GCF_000238735.1_Citr_freu_4_7_47CFAA_V1 Citrobacter freundii 4_7_47CFAA Proteobacteria; Gammaproteobacteria; Enterobacterales; Enterobacteriaceae; Citrobacter; Citrobacter freundii complex; Citrobacter freundii MTGDNSLINSSGINRRDFMKLCAALAATMGLSSKAAAEMAESVSRPQRPPVIWIGAQECT WP_003024486.1 MULTISPECIES: hydrogenase 2 small subunit [Enterobacteriaceae] Length: 372\nNumber of predicted TMHs: 1\nExp number of AAs in TMHs: 27.14621\nExp number, first 60 AAs: 0.22683\nTotal prob of N-in: 0.23982\noutside 1 331\nTMhelix 332 354\ninside 355 372

1993 GCF_000312465.1_ASM31246v1 Citrobacter freundii ATCC 8090 = MTCC 1658 Proteobacteria; Gammaproteobacteria; Enterobacterales; Enterobacteriaceae; Citrobacter; Citrobacter freundii complex; Citrobacter freundii MTGDNSLINSSGINRRDFMKLCAALAATMGLSSKAAAEMAESVSRPQRPPVIWIGAQECT WP_003024486.1 MULTISPECIES: hydrogenase 2 small subunit [Enterobacteriaceae] Length: 372\nNumber of predicted TMHs: 1\nExp number of AAs in TMHs: 27.14621\nExp number, first 60 AAs: 0.22683\nTotal prob of N-in: 0.23982\noutside 1 331\nTMhelix 332 354\ninside 355 372

1994 GCF_000734905.1_ASM73490v1 Citrobacter freundii ATCC 8090 = MTCC 1658 Proteobacteria; Gammaproteobacteria; Enterobacterales; Enterobacteriaceae; Citrobacter; Citrobacter freundii complex; Citrobacter freundii MTGDNSLINSSGINRRDFMKLCAALAATMGLSSKAAAEMAESVSRPQRPPVIWIGAQECT WP_003024486.1 MULTISPECIES: hydrogenase 2 small subunit [Enterobacteriaceae] Length: 372\nNumber of predicted TMHs: 1\nExp number of AAs in TMHs: 27.14621\nExp number, first 60 AAs: 0.22683\nTotal prob of N-in: 0.23982\noutside 1 331\nTMhelix 332 354\ninside 355 372

1995 GCF_000648515.1_ASM64851v1 Citrobacter freundii CFNIH1 Proteobacteria; Gammaproteobacteria; Enterobacterales; Enterobacteriaceae; Citrobacter; Citrobacter freundii complex; Citrobacter freundii MTGDNSLINSSGINRRDFMKLCAALAATMGLSSKAAAEMAESVSRPQRPPVIWIGAQECT WP_003024486.1 MULTISPECIES: hydrogenase 2 small subunit [Enterobacteriaceae] Length: 372\nNumber of predicted TMHs: 1\nExp number of AAs in TMHs: 27.14621\nExp number, first 60 AAs: 0.22683\nTotal prob of N-in: 0.23982\noutside 1 331\nTMhelix 332 354\ninside 355 372

1996 GCF_000342325.1_Whole_genome_assembly Citrobacter freundii GTC 09479 Proteobacteria; Gammaproteobacteria; Enterobacterales; Enterobacteriaceae; Citrobacter; Citrobacter freundii complex; Citrobacter freundii MTGDNSLINSSGINRRDFMKLCAALAATMGLSSKAAAEMAESVSRPQRPPVIWIGAQECT WP_003024486.1 MULTISPECIES: hydrogenase 2 small subunit [Enterobacteriaceae] Length: 372\nNumber of predicted TMHs: 1\nExp number of AAs in TMHs: 27.14621\nExp number, first 60 AAs: 0.22683\nTotal prob of N-in: 0.23982\noutside 1 331\nTMhelix 332 354\ninside 355 372

1997 GCF_000388155.1_Whole_genome_assembly Citrobacter freundii GTC 09629 Proteobacteria; Gammaproteobacteria; Enterobacterales; Enterobacteriaceae; Citrobacter; Citrobacter freundii complex; Citrobacter freundii MTGDNSLINSSGINRRDFMKLCAALAATMGLSSKAAAEMAESVSRPQRPPVIWIGAQECT WP_003024486.1 MULTISPECIES: hydrogenase 2 small subunit [Enterobacteriaceae] Length: 372\nNumber of predicted TMHs: 1\nExp number of AAs in TMHs: 27.14621\nExp number, first 60 AAs: 0.22683\nTotal prob of N-in: 0.23982\noutside 1 331\nTMhelix 332 354\ninside 355 372

1998 GCF_000692115.1_Citr_freu_MGH_56_V1 Citrobacter freundii MGH 56 Proteobacteria; Gammaproteobacteria; Enterobacterales; Enterobacteriaceae; Citrobacter; Citrobacter freundii complex; Citrobacter freundii MTGDNSLINSNGINRRDFMKLCAALAATMGLSSKAAAEMAESVSRPQRPPVIWIGAQECT WP_043017615.1 MULTISPECIES: hydrogenase 2 small subunit [Citrobacter freundii complex] Length: 372\nNumber of predicted TMHs: 1\nExp number of AAs in TMHs: 27.21963\nExp number, first 60 AAs: 0.31687\nTotal prob of N-in: 0.24323\noutside 1 331\nTMhelix 332 354\ninside 355 372

1999 GCF_000759735.1_ASM75973v1 Citrobacter freundii NBRC 12681 Proteobacteria; Gammaproteobacteria; Enterobacterales; Enterobacteriaceae; Citrobacter; Citrobacter freundii complex; Citrobacter freundii MTGDNSLINSSGINRRDFMKLCAALAATMGLSSKAAAEMAESVSRPQRPPVIWIGAQECT WP_003024486.1 MULTISPECIES: hydrogenase 2 small subunit [Enterobacteriaceae] Length: 372\nNumber of predicted TMHs: 1\nExp number of AAs in TMHs: 27.14621\nExp number, first 60 AAs: 0.22683\nTotal prob of N-in: 0.23982\noutside 1 331\nTMhelix 332 354\ninside 355 372

2000 GCF_000582615.1_Citrobacter_assembly Citrobacter freundii RLS1 Proteobacteria; Gammaproteobacteria; Enterobacterales; Enterobacteriaceae; Citrobacter; Citrobacter freundii complex; Citrobacter freundii MTGDNSLINSSGINRRDFMKLCAALAATMGLSSKAAAEMAESVSRPQRPPVIWIGAQECT WP_003024486.1 MULTISPECIES: hydrogenase 2 small subunit [Enterobacteriaceae] Length: 372\nNumber of predicted TMHs: 1\nExp number of AAs in TMHs: 27.14621\nExp number, first 60 AAs: 0.22683\nTotal prob of N-in: 0.23982\noutside 1 331\nTMhelix 332 354\ninside 355 372

2001 GCF_000208765.1_ASM20876v1 Citrobacter freundii str. ballerup 7851/39 Proteobacteria; Gammaproteobacteria; Enterobacterales; Enterobacteriaceae; Citrobacter; Citrobacter freundii complex; Citrobacter freundii MTGDNSLINSNGINRRDFMKLCAALAATMGLSSKAAAEMAESVSRPQRPPVIWIGAQECT WP_016154493.1 MULTISPECIES: hydrogenase-2 small chain [Enterobacteriaceae] Length: 372\nNumber of predicted TMHs: 1\nExp number of AAs in TMHs: 26.90274\nExp number, first 60 AAs: 0.32271\nTotal prob of N-in: 0.22239\noutside 1 331\nTMhelix 332 354\ninside 355 372

2002 GCF_000521965.1_Citr_freu_UCI_31_V2 Citrobacter freundii UCI 31 Proteobacteria; Gammaproteobacteria; Enterobacterales; Enterobacteriaceae; Citrobacter; Citrobacter freundii complex; Citrobacter freundii MTGDNSLINSSGINRRDFMKLCAALAATMGLSSKAAAEMAESVSRPQRPPVIWIGAQECT WP_003024486.1 MULTISPECIES: hydrogenase 2 small subunit [Enterobacteriaceae] Length: 372\nNumber of predicted TMHs: 1\nExp number of AAs in TMHs: 27.14621\nExp number, first 60 AAs: 0.22683\nTotal prob of N-in: 0.23982\noutside 1 331\nTMhelix 332 354\ninside 355 372

2003 GCF_000521945.1_Citr_freu_UCI_32_V1 Citrobacter freundii UCI 32 Proteobacteria; Gammaproteobacteria; Enterobacterales; Enterobacteriaceae; Citrobacter; Citrobacter freundii complex; Citrobacter freundii MTGDNSLINSSGINRRDFMKLCAALAATMGLSSKAAAEMAESVSRPQRPPVIWIGAQECT WP_003024486.1 MULTISPECIES: hydrogenase 2 small subunit [Enterobacteriaceae] Length: 372\nNumber of predicted TMHs: 1\nExp number of AAs in TMHs: 27.14621\nExp number, first 60 AAs: 0.22683\nTotal prob of N-in: 0.23982\noutside 1 331\nTMhelix 332 354\ninside 355 372

2004 GCF_000759835.1_ASM75983v1 Citrobacter sedlakii NBRC 105722 Proteobacteria; Gammaproteobacteria; Enterobacterales; Enterobacteriaceae; Citrobacter; Citrobacter freundii complex; Citrobacter sedlakii MTGDNTLINSHGVNRRDFMKLCAALAATMGLSSKAAAEMAESVSRPQRPPVIWIGAQECT WP_042289187.1 MULTISPECIES: hydrogenase 2 small subunit [Citrobacter] Length: 372\nNumber of predicted TMHs: 1\nExp number of AAs in TMHs: 24.58538\nExp number, first 60 AAs: 0.33179\nTotal prob of N-in: 0.14844\noutside 1 331\nTMhelix 332 354\ninside 355 372

2005 GCF_000759755.1_ASM75975v1 Citrobacter werkmanii NBRC 105721 Proteobacteria; Gammaproteobacteria; Enterobacterales; Enterobacteriaceae; Citrobacter; Citrobacter freundii complex; Citrobacter werkmanii MTGDNSLINSSGINRRDFMKLCAALAATMGLSSKAAAEMAESVSRPQRPPVIWIGAQECT WP_003024486.1 MULTISPECIES: hydrogenase 2 small subunit [Enterobacteriaceae] Length: 372\nNumber of predicted TMHs: 1\nExp number of AAs in TMHs: 27.14621\nExp number, first 60 AAs: 0.22683\nTotal prob of N-in: 0.23982\noutside 1 331\nTMhelix 332 354\ninside 355 372

2006 GCF_000155975.1_ASM15597v1 Citrobacter youngae ATCC 29220 Proteobacteria; Gammaproteobacteria; Enterobacterales; Enterobacteriaceae; Citrobacter; Citrobacter freundii complex; Citrobacter youngae MTGDNSLINSSGINRRDFMKLCAALAATMGLSSKAAAEMAESVSRPQRPPVIWIGAQECT WP_003024486.1 MULTISPECIES: hydrogenase 2 small subunit [Enterobacteriaceae] Length: 372\nNumber of predicted TMHs: 1\nExp number of AAs in TMHs: 27.14621\nExp number, first 60 AAs: 0.22683\nTotal prob of N-in: 0.23982\noutside 1 331\nTMhelix 332 354\ninside 355 372

2007 GCF_000018045.1_ASM1804v1 Citrobacter koseri ATCC BAA-895 Proteobacteria; Gammaproteobacteria; Enterobacterales; Enterobacteriaceae; Citrobacter; Citrobacter koseri MTGDNTLITSHGVNRRDFMKLCAALAATMGLSSKAAAEMAESVSRPQRPPVVWIGAQECT WP_012135130.1 MULTISPECIES: hydrogenase 2 small subunit [Enterobacteriaceae] Length: 372\nNumber of predicted TMHs: 1\nExp number of AAs in TMHs: 24.24593\nExp number, first 60 AAs: 0.31574\nTotal prob of N-in: 0.12382\noutside 1 331\nTMhelix 332 354\ninside 355 372

2008 GCF_000027085.1_ASM2708v1 Citrobacter rodentium ICC168 Proteobacteria; Gammaproteobacteria; Enterobacterales; Enterobacteriaceae; Citrobacter; Citrobacter rodentium MTGDNTLINSHGVNRRDFMKLCAALAATMGLSSKAAAEMAESVSRPQRPPVVWIGAQECT WP_012907591.1 hydrogenase 2 small subunit [Citrobacter rodentium] Length: 372\nNumber of predicted TMHs: 1\nExp

number of AAs in TMHs: 24.50861\nExp number, first 60 AAs: 0.31328\nTotal prob of N-in: 0.14317\noutside 1 331\nTMhelix 332 354\ninside 355 372

2009 GCF_000759815.1_ASM75981v1 Citrobacter rodentium NBRC 105723 Proteobacteria; Gammaproteobacteria; Enterobacterales; Enterobacteriaceae; Citrobacter; Citrobacter rodentium MTGDNTLINSHGVNRRDFMKLCAALAATMGLSSKAAAEMAESVSRPQRPPVVWIGAQECT WP_012907591.1 hydrogenase 2 small subunit [Citrobacter rodentium] Length: 372\nNumber of predicted TMHs: 1\nExp number of AAs in TMHs: 24.50861\nExp number, first 60 AAs: 0.31328\nTotal prob of N-in: 0.14317\noutside 1 331\nTMhelix 332 354\ninside 355 372

2010 GCF_000814905.1_ASM81490v1 Enterobacter sp. Bisph1 Proteobacteria; Gammaproteobacteria; Enterobacterales; Enterobacteriaceae; Enterobacter MAGENQHFAPQGVSRRDFMKLCAALAATMGLSSRAAAEMAEAVSHPQRPPVVWIGAQECT WP_039056976.1 hydrogenase 2 small subunit [Enterobacter sp. Bisph1] Length: 372\nNumber of predicted TMHs: 1\nExp number of AAs in TMHs: 22.06781\nExp number, first 60 AAs: 0.10311\nTotal prob of N-in: 0.01669\noutside 1 330\nTMhelix 331 353\ninside 354 372

2011 GCF_000814915.1_widad Enterobacter sp. Bisph2 Proteobacteria; Gammaproteobacteria; Enterobacterales; Enterobacteriaceae; Enterobacter MADEIPQLTSFTSCGINRRDFMKLCTALAATMGLSSTAAAKMAQAVTQSRRPPVVWIGAQ WP_052249210.1 hydrogenase 2 small subunit [Enterobacter sp. Bisph2] Length: 375\nNumber of predicted TMHs: 1\nExp number of AAs in TMHs: 24.64967\nExp number, first 60 AAs: 1.94828\nTotal prob of N-in: 0.15052\noutside 1 333\nTMhelix 334 356\ninside 357 375

2012 GCF_001582075.1_ASM158207v1 Enterobacter sp. FY-07 Proteobacteria; Gammaproteobacteria; Enterobacterales; Enterobacteriaceae; Enterobacter MAGENQHFVSQGVSRRDFMKLCAALAATMGLSGKAAAEMAEAVSHPQRPPVIWIGAQECT WP_061493156.1 hydrogenase 2 small subunit [Enterobacter sp. FY-07] Length: 372\nNumber of predicted TMHs: 1\nExp number of AAs in TMHs: 23.58979\nExp number, first 60 AAs: 0.32199\nTotal prob of N-in: 0.09013\noutside 1 330\nTMhelix 331 353\ninside 354 372

2013 GCF_001022685.1_ASM102268v1 Enterobacter sp. GN02600 Proteobacteria; Gammaproteobacteria; Enterobacterales; Enterobacteriaceae; Enterobacter MTGDNSLINSNGINRRDFMKLCAALAATMGLSSKAAAEMAESVSRPQRPPVIWIGAQECT WP_016154493.1 MULTISPECIES: hydrogenase-2 small chain [Enterobacteriaceae] Length: 372\nNumber of predicted TMHs: 1\nExp number of AAs in TMHs: 26.90274\nExp number, first 60 AAs: 0.32271\nTotal prob of N-in: 0.22239\noutside 1 331\nTMhelix 332 354\ninside 355 372

2014 GCF_900119515.1_IMG-taxon_2600255257_annotated_assembly Enterobacter sp. NFIX03 Proteobacteria; Gammaproteobacteria; Enterobacterales; Enterobacteriaceae; Enterobacter MAGDNLHLASQGVSRRDFMKLCAALAATMGLSGKAAAEMAEAISHPQRPPVVWIGAQECT WP_007370143.1 MULTISPECIES: hydrogenase 2 small subunit [Enterobacteriaceae] Length: 372\nNumber of predicted TMHs: 1\nExp number of AAs in TMHs: 24.18752\nExp number, first 60 AAs: 0.2398\nTotal prob of N-in: 0.11792\noutside 1 332\nTMhelix 333 355\ninside 356 372

2015 GCF_900168315.1_IMG-taxon_2609459761_annotated_assembly Enterobacter sp. NFR05 Proteobacteria; Gammaproteobacteria; Enterobacterales; Enterobacteriaceae; Enterobacter MADEIPQLTSFISCGINRRDFMKLCTALAATMGLSSTAAAKMAQAVTQSKRPPVVWIGAQ WP_079497107.1 hydrogenase 2 small subunit [Enterobacter sp. NFR05] Length: 375\nNumber of predicted TMHs: 1\nExp number of AAs in TMHs: 24.29869\nExp number, first 60 AAs: 1.59149\nTotal prob of N-in: 0.13544\noutside 1 333\nTMhelix 334 356\ninside 357 375

2016 GCF_000410515.1_ASM41051v1 Enterobacter sp. R4-368 Proteobacteria; Gammaproteobacteria; Enterobacterales; Enterobacteriaceae; Enterobacter MAGENQQFASHGVSRRDFMKLCAALAATMGLSGKAAAEMAEAVSHPQRPPVIWIGAQECT WP_020455253.1 hydrogenase 2 small subunit [Enterobacter sp. R4-368] Length: 372\nNumber of predicted TMHs: 1\nExp number of AAs in TMHs: 25.64425\nExp number, first 60 AAs: 0.31979\nTotal prob of N-in: 0.22011\noutside 1 330\nTMhelix 331 353\ninside 354 372

2017 GCF_001888805.1_ASM188880v1 Enterobacter sp. Sa187 Proteobacteria; Gammaproteobacteria; Enterobacterales; Enterobacteriaceae; Enterobacter

MVTKTSAIPPGITRRDFMTLCAALAATMGLSSQAAARMAGALSQPQRLPVIWIGAQECTG WP_072569773.1 hydrogenase 2 small subunit [Enterobacter sp. Sa187] Length: 371\nNumber of predicted TMHs: 1\nExp number of AAs in TMHs: 28.79508\nExp number, first 60 AAs: 4.70252\nTotal prob of N-in: 0.36502\noutside 1 329\nTMhelix 330 352\ninside 353 371

2018 GCF_001052135.1_ASM105213v1 Enterobacter asburiae Proteobacteria; Gammaproteobacteria; Enterobacterales; Enterobacteriaceae; Enterobacter; Enterobacter cloacae complex MKGVLEKCTVEDFNYISATLDSYVSFTNDKRRKTLLSTYQNNPALHAELISLIDTQIKYF WP_048979315.1 hypothetical protein [Enterobacter asburiae] Length: 249\nNumber of predicted TMHs: 2\nExp number of AAs in TMHs: 39.67787\nExp number, first 60 AAs: 0.01154\nTotal prob of N-in: 0.09019\noutside 1 157\nTMhelix 158 180\ninside 181 200\nTMhelix 201 220\noutside 221 249

2019 GCF_001052355.1_ASM105235v1 Enterobacter asburiae Proteobacteria; Gammaproteobacteria; Enterobacterales; Enterobacteriaceae; Enterobacter; Enterobacter cloacae complex MKGVLEKCTVEDFNYISATLDSYVSFTNDKRRKTLLSTYQNNPALHAELISLIDTQIKYF WP_048979315.1 hypothetical protein [Enterobacter asburiae] Length: 249\nNumber of predicted TMHs: 2\nExp number of AAs in TMHs: 39.67787\nExp number, first 60 AAs: 0.01154\nTotal prob of N-in: 0.09019\noutside 1 157\nTMhelix 158 180\ninside 181 200\nTMhelix 201 220\noutside 221 249

2020 GCF_900077125.1_12045_4_44 Enterobacter cloacae Proteobacteria; Gammaproteobacteria; Enterobacterales; Enterobacteriaceae; Enterobacter; Enterobacter cloacae complex MTGDNSLINSSGINRRDFMKLCAALAATMGLSSKAAAEMAESVSRPQRPPVIWIGAQECT WP_003024486.1 MULTISPECIES: hydrogenase 2 small subunit [Enterobacteriaceae] Length: 372\nNumber of predicted TMHs: 1\nExp number of AAs in TMHs: 27.14621\nExp number, first 60 AAs: 0.22683\nTotal prob of N-in: 0.23982\noutside 1 331\nTMhelix 332 354\ninside 355 372

2021 GCF_001514725.1_ASM151472v1 Escherichia albertii Proteobacteria; Gammaproteobacteria; Enterobacterales; Enterobacteriaceae; Escherichia MNNEETFYQSMRRQGVTRRSFLKYCSLAATSLGLGAGMAPKIAWALENKPRIPVVWIHGL WP_002461900.1 MULTISPECIES: hydrogenase-1 small chain [Escherichia] Length: 372\nNumber of predicted TMHs: 2\nExp number of AAs in TMHs: 37.1074899999999\nExp number, first 60 AAs: 19.37835\nTotal prob of N-in: 0.93614\nPOSSIBLE N-term signal sequence\ninside 1 19\nTMhelix 20 39\noutside 40 325\nTMhelix 326 348\ninside 349 372

2022 GCF_001514825.1_ASM151482v1 Escherichia albertii Proteobacteria; Gammaproteobacteria; Enterobacterales; Enterobacteriaceae; Escherichia MNNEETFYQSMRRQGVTRRSFLKYCSLAATSLGLGAGMAPKIAWALENKPRIPVVWIHGL WP_002461900.1 MULTISPECIES: hydrogenase-1 small chain [Escherichia] Length: 372\nNumber of predicted TMHs: 2\nExp number of AAs in TMHs: 37.1074899999999\nExp number, first 60 AAs: 19.37835\nTotal prob of N-in: 0.93614\nPOSSIBLE N-term signal sequence\ninside 1 19\nTMhelix 20 39\noutside 40 325\nTMhelix 326 348\ninside 349 372

2023 GCF_001514725.1_ASM151472v1 Escherichia albertii Proteobacteria; Gammaproteobacteria; Enterobacterales; Enterobacteriaceae; Escherichia MTGDNTLIHSHGINRRDFMKLCAALAATMGLSSKAAAEMAESVTNPQRPPVIWIGAQECT WP_000145410.1 MULTISPECIES: hydrogenase-2 small chain [Proteobacteria] Length: 372\nNumber of predicted TMHs: 1\nExp number of AAs in TMHs: 23.72226\nExp number, first 60 AAs: 0.22038\nTotal prob of N-in: 0.12994\noutside 1 331\nTMhelix 332 354\ninside 355 372

2024 GCF_001514575.1_ASM151457v1 Escherichia albertii Proteobacteria; Gammaproteobacteria; Enterobacterales; Enterobacteriaceae; Escherichia MNNEETFYQSMRRQGVTRRSFLKYCSLAATSLGLGAGMAPKIAWALENKPRIPVVWIHGL WP_002461900.1 MULTISPECIES: hydrogenase-1 small chain [Escherichia] Length: 372\nNumber of predicted TMHs: 2\nExp number of AAs in TMHs: 37.1074899999999\nExp number, first 60 AAs: 19.37835\nTotal prob of N-in: 0.93614\nPOSSIBLE N-term signal sequence\ninside 1 19\nTMhelix 20 39\noutside 40 325\nTMhelix 326 348\ninside 349 372

2025 GCF_001515025.1_ASM151502v1 Escherichia albertii Proteobacteria; Gammaproteobacteria; Enterobacterales; Enterobacteriaceae; Escherichia MTGDNTLIHSHGINRRDFMKLCAALAATMGLSSKAAAEMAESVTNPQRPPVIWIGAQECT WP_000145410.1 MULTISPECIES: hydrogenase-2 small chain [Proteobacteria] Length: 372\nNumber of predicted TMHs:

1\nExp number of AAs in TMHs: 23.72226\nExp number, first 60 AAs: 0.22038\nTotal prob of N-in: 0.12994\noutside 1 331\nTMhelix 332 354\ninside 355 372

2026 GCF_001514965.1_ASM151496v1 Escherichia albertii Proteobacteria; Gammaproteobacteria; Enterobacterales; Enterobacteriaceae; Escherichia
MNNEETFYQSMRRQGVTRRSFLKYCSLAATSLGLGAGMAPKIAWALENKPRIPVVWIHGL WP_044712053.1
MULTISPECIES: hydrogenase-1 small chain [Escherichia] Length: 372\nNumber of predicted TMHs: 2\nExp number of AAs in TMHs: 36.9963799999999\nExp number, first 60 AAs: 19.37682\nTotal prob of N-in: 0.93606\nPOSSIBLE N-term signal sequence\ninside 1 19\nTMhelix 20 39\noutside 40 325\nTMhelix 326 348\ninside 349 372

2027 GCF_001515065.1_ASM151506v1 Escherichia albertii Proteobacteria; Gammaproteobacteria; Enterobacterales; Enterobacteriaceae; Escherichia
MTGDNTLIHSHGINRRDFMKLCAALAATMGLSSKAAAEMAESVTNPQRPPVIWIGAQECT WP_000145410.1
MULTISPECIES: hydrogenase-2 small chain [Proteobacteria] Length: 372\nNumber of predicted TMHs: 1\nExp number of AAs in TMHs: 23.72226\nExp number, first 60 AAs: 0.22038\nTotal prob of N-in: 0.12994\noutside 1 331\nTMhelix 332 354\ninside 355 372

2028 GCF_001514645.1_ASM151464v1 Escherichia albertii Proteobacteria; Gammaproteobacteria; Enterobacterales; Enterobacteriaceae; Escherichia
MNNEETFYQSMRRQGVTRRSFLKYCSLAATSLGLGAGMAPKIAWALENKPRIPVVWIHGL WP_002461900.1
MULTISPECIES: hydrogenase-1 small chain [Escherichia] Length: 372\nNumber of predicted TMHs: 2\nExp number of AAs in TMHs: 37.1074899999999\nExp number, first 60 AAs: 19.37835\nTotal prob of N-in: 0.93614\nPOSSIBLE N-term signal sequence\ninside 1 19\nTMhelix 20 39\noutside 40 325\nTMhelix 326 348\ninside 349 372

2029 GCF_001514965.1_ASM151496v1 Escherichia albertii Proteobacteria; Gammaproteobacteria; Enterobacterales; Enterobacteriaceae; Escherichia
MTGDNTLIHSHGINRRDFMKLCAALAATMGLSSKAAAEMAESVTNPQRPPVIWIGAQECT WP_000145410.1
MULTISPECIES: hydrogenase-2 small chain [Proteobacteria] Length: 372\nNumber of predicted TMHs: 1\nExp number of AAs in TMHs: 23.72226\nExp number, first 60 AAs: 0.22038\nTotal prob of N-in: 0.12994\noutside 1 331\nTMhelix 332 354\ninside 355 372

2030 GCF_001514625.1_ASM151462v1 Escherichia albertii Proteobacteria; Gammaproteobacteria; Enterobacterales; Enterobacteriaceae; Escherichia
MNNEETFYQSMRRQGVTRRSFLKYCSLAATSLGLGAGMAPKIAWALENKPRIPVVWIHGL WP_002461900.1
MULTISPECIES: hydrogenase-1 small chain [Escherichia] Length: 372\nNumber of predicted TMHs: 2\nExp number of AAs in TMHs: 37.1074899999999\nExp number, first 60 AAs: 19.37835\nTotal prob of N-in: 0.93614\nPOSSIBLE N-term signal sequence\ninside 1 19\nTMhelix 20 39\noutside 40 325\nTMhelix 326 348\ninside 349 372

2031 GCF_001514685.1_ASM151468v1 Escherichia albertii Proteobacteria; Gammaproteobacteria; Enterobacterales; Enterobacteriaceae; Escherichia
MNNEETFYQSMRRQGVTRRSFLKYCSLAATSLGLGAGMAPKIAWALENKPRIPVVWIHGL WP_002461900.1
MULTISPECIES: hydrogenase-1 small chain [Escherichia] Length: 372\nNumber of predicted TMHs: 2\nExp number of AAs in TMHs: 37.1074899999999\nExp number, first 60 AAs: 19.37835\nTotal prob of N-in: 0.93614\nPOSSIBLE N-term signal sequence\ninside 1 19\nTMhelix 20 39\noutside 40 325\nTMhelix 326 348\ninside 349 372

2032 GCF_001514555.1_ASM151455v1 Escherichia albertii Proteobacteria; Gammaproteobacteria; Enterobacterales; Enterobacteriaceae; Escherichia
MTGDNTLIHSHGINRRDFMKLCAALAATMGLSSKAAAEMAESVTNPQRPPVIWIGAQECT WP_000145410.1
MULTISPECIES: hydrogenase-2 small chain [Proteobacteria] Length: 372\nNumber of predicted TMHs: 1\nExp number of AAs in TMHs: 23.72226\nExp number, first 60 AAs: 0.22038\nTotal prob of N-in: 0.12994\noutside 1 331\nTMhelix 332 354\ninside 355 372

2033 GCF_001514825.1_ASM151482v1 Escherichia albertii Proteobacteria; Gammaproteobacteria; Enterobacterales; Enterobacteriaceae; Escherichia
MTGDNTLIHSHGINRRDFMKLCAALAATMGLSSKAAAEMAESVTNPQRPPVIWIGAQECT WP_000145410.1
MULTISPECIES: hydrogenase-2 small chain [Proteobacteria] Length: 372\nNumber of predicted TMHs:

1\nExp number of AAs in TMHs: 23.72226\nExp number, first 60 AAs: 0.22038\nTotal prob of N-in: 0.12994\noutside 1 331\nTMhelix 332 354\ninside 355 372

2034 GCF_001514785.1_ASM151478v1 Escherichia albertii Proteobacteria; Gammaproteobacteria; Enterobacterales; Enterobacteriaceae; Escherichia MTGDNTLIHSHGINRRDFMKLCAALAATMGLSSKAAAEMAESVTNPQRPPVIWIGAQECT WP_025238349.1 hydrogenase 2 small subunit [Escherichia albertii] Length: 372\nNumber of predicted TMHs: 1\nExp number of AAs in TMHs: 23.72042\nExp number, first 60 AAs: 0.22025\nTotal prob of N-in: 0.12992\noutside 1 331\nTMhelix 332 354\ninside 355 372

2035 GCF_001514885.1_ASM151488v1 Escherichia albertii Proteobacteria; Gammaproteobacteria; Enterobacterales; Enterobacteriaceae; Escherichia MTGDNTLIHSHGINRRDFMKLCAALAATMGLSSKAAAEMAESVTNPQRPPVIWIGAQECT WP_000145410.1 MULTISPECIES: hydrogenase-2 small chain [Proteobacteria] Length: 372\nNumber of predicted TMHs: 1\nExp number of AAs in TMHs: 23.72226\nExp number, first 60 AAs: 0.22038\nTotal prob of N-in: 0.12994\noutside 1 331\nTMhelix 332 354\ninside 355 372

2036 GCF_001514785.1_ASM151478v1 Escherichia albertii Proteobacteria; Gammaproteobacteria; Enterobacterales; Enterobacteriaceae; Escherichia MNNEETFYQSMRRQGVTRRSFLKYCSLAATSLGLGAGMAPKIAWALENKPRIPVVWIHGL WP_059219811.1 hydrogenase [Escherichia albertii] Length: 372\nNumber of predicted TMHs: 2\nExp number of AAs in TMHs: 37.1074899999999\nExp number, first 60 AAs: 19.37835\nTotal prob of N-in: 0.93614\nPOSSIBLE N-term signal sequence\ninside 1 19\nTMhelix 20 39\noutside 40 325\nTMhelix 326 348\ninside 349 372

2037 GCF_001514745.1_ASM151474v1 Escherichia albertii Proteobacteria; Gammaproteobacteria; Enterobacterales; Enterobacteriaceae; Escherichia MNNEETFYQSMRRQGVTRRSFLKYCSLAATSLGLGAGMAPKIAWALENKPRIPVVWIHGL WP_059219811.1 hydrogenase [Escherichia albertii] Length: 372\nNumber of predicted TMHs: 2\nExp number of AAs in TMHs: 37.1074899999999\nExp number, first 60 AAs: 19.37835\nTotal prob of N-in: 0.93614\nPOSSIBLE N-term signal sequence\ninside 1 19\nTMhelix 20 39\noutside 40 325\nTMhelix 326 348\ninside 349 372

2038 GCF_001514745.1_ASM151474v1 Escherichia albertii Proteobacteria; Gammaproteobacteria; Enterobacterales; Enterobacteriaceae; Escherichia MTGDNTLIHSHGINRRDFMKLCAALAATMGLSSKAAAEMAESVTNPQRPPVIWIGAQECT WP_025238349.1 hydrogenase 2 small subunit [Escherichia albertii] Length: 372\nNumber of predicted TMHs: 1\nExp number of AAs in TMHs: 23.72042\nExp number, first 60 AAs: 0.22025\nTotal prob of N-in: 0.12992\noutside 1 331\nTMhelix 332 354\ninside 355 372

2039 GCF_001514575.1_ASM151457v1 Escherichia albertii Proteobacteria; Gammaproteobacteria; Enterobacterales; Enterobacteriaceae; Escherichia MTGDNTLIHSHGINRRDFMKLCAALAATMGLSSKAAAEMAESVTNPQRPPVIWIGAQECT WP_000145410.1 MULTISPECIES: hydrogenase-2 small chain [Proteobacteria] Length: 372\nNumber of predicted TMHs: 1\nExp number of AAs in TMHs: 23.72226\nExp number, first 60 AAs: 0.22038\nTotal prob of N-in: 0.12994\noutside 1 331\nTMhelix 332 354\ninside 355 372

2040 GCF_001514645.1_ASM151464v1 Escherichia albertii Proteobacteria; Gammaproteobacteria; Enterobacterales; Enterobacteriaceae; Escherichia MTGDNTLIHSHGINRRDFMKLCAALAATMGLSSKAAAEMAESVTNPQRPPVIWIGAQECT WP_000145410.1 MULTISPECIES: hydrogenase-2 small chain [Proteobacteria] Length: 372\nNumber of predicted TMHs: 1\nExp number of AAs in TMHs: 23.72226\nExp number, first 60 AAs: 0.22038\nTotal prob of N-in: 0.12994\noutside 1 331\nTMhelix 332 354\ninside 355 372

2041 GCF_001514685.1_ASM151468v1 Escherichia albertii Proteobacteria; Gammaproteobacteria; Enterobacterales; Enterobacteriaceae; Escherichia MTGDNTLIHSHGINRRDFMKLCAALAATMGLSSKAAAEMAESVTNPQRPPVIWIGAQECT WP_000145410.1 MULTISPECIES: hydrogenase-2 small chain [Proteobacteria] Length: 372\nNumber of predicted TMHs: 1\nExp number of AAs in TMHs: 23.72226\nExp number, first 60 AAs: 0.22038\nTotal prob of N-in: 0.12994\noutside 1 331\nTMhelix 332 354\ninside 355 372

2042 GCF_001514555.1_ASM151455v1 Escherichia albertii Proteobacteria; Gammaproteobacteria; Enterobacterales; Enterobacteriaceae; Escherichia

MNNEETFYQSMRRQGVTRRSFLKYCSLAATSLGLGAGMAPKIAWALENKPRIPVVWIHGL WP_059219811.1 hydrogenase [Escherichia albertii] Length: 372\nNumber of predicted TMHs: 2\nExp number of AAs in TMHs: 37.1074899999999\nExp number, first 60 AAs: 19.37835\nTotal prob of N-in: 0.93614\nPOSSIBLE N-term signal sequence\ninside 1 19\nTMhelix 20 39\noutside 40 325\nTMhelix 326 348\ninside 349 372

2043 GCF_001514845.1_ASM151484v1 Escherichia albertii Proteobacteria; Gammaproteobacteria; Enterobacterales; Enterobacteriaceae; Escherichia MTGDNTLIHSHGINRRDFMKLCAALAATMGLSNKAAAEMAESVTNPQRPPVIWIGAQECT WP_059268075.1 hydrogenase 2 small subunit [Escherichia albertii] Length: 372\nNumber of predicted TMHs: 1\nExp number of AAs in TMHs: 23.66196\nExp number, first 60 AAs: 0.09463\nTotal prob of N-in: 0.12684\noutside 1 331\nTMhelix 332 354\ninside 355 372

2044 GCF_001515025.1_ASM151502v1 Escherichia albertii Proteobacteria; Gammaproteobacteria; Enterobacterales; Enterobacteriaceae; Escherichia MNNEETFYQSMRRQGVTRRSFLKYCSLAATSLGLGAGMAPKIAWALENKPRIPVVWIHGL WP_044712053.1 MULTISPECIES: hydrogenase-1 small chain [Escherichia] Length: 372\nNumber of predicted TMHs: 2\nExp number of AAs in TMHs: 36.9963799999999\nExp number, first 60 AAs: 19.37682\nTotal prob of N-in: 0.93606\nPOSSIBLE N-term signal sequence\ninside 1 19\nTMhelix 20 39\noutside 40 325\nTMhelix 326 348\ninside 349 372

2045 GCF_001514885.1_ASM151488v1 Escherichia albertii Proteobacteria; Gammaproteobacteria; Enterobacterales; Enterobacteriaceae; Escherichia MNNEETFYQSMRRQGVTRRSFLKYCSLAATSLGLGAGMAPKIAWALENKPRIPVVWIHGL WP_002461900.1 MULTISPECIES: hydrogenase-1 small chain [Escherichia] Length: 372\nNumber of predicted TMHs: 2\nExp number of AAs in TMHs: 37.1074899999999\nExp number, first 60 AAs: 19.37835\nTotal prob of N-in: 0.93614\nPOSSIBLE N-term signal sequence\ninside 1 19\nTMhelix 20 39\noutside 40 325\nTMhelix 326 348\ninside 349 372

2046 GCF_001515005.1_ASM151500v1 Escherichia albertii Proteobacteria; Gammaproteobacteria; Enterobacterales; Enterobacteriaceae; Escherichia MNNEETFYQSMRRQGVTRRSFLKYCSLAATSLGLGAGMAPKIAWALENKPRIPVVWIHGL WP_059219811.1 hydrogenase [Escherichia albertii] Length: 372\nNumber of predicted TMHs: 2\nExp number of AAs in TMHs: 37.1074899999999\nExp number, first 60 AAs: 19.37835\nTotal prob of N-in: 0.93614\nPOSSIBLE N-term signal sequence\ninside 1 19\nTMhelix 20 39\noutside 40 325\nTMhelix 326 348\ninside 349 372

2047 GCF_001515065.1_ASM151506v1 Escherichia albertii Proteobacteria; Gammaproteobacteria; Enterobacterales; Enterobacteriaceae; Escherichia MNNEETFYQSMRRQGVTRRSFLKYCSLAATSLGLGAGMAPKIAWALENKPRIPVVWIHGL WP_002461900.1 MULTISPECIES: hydrogenase-1 small chain [Escherichia] Length: 372\nNumber of predicted TMHs: 2\nExp number of AAs in TMHs: 37.1074899999999\nExp number, first 60 AAs: 19.37835\nTotal prob of N-in: 0.93614\nPOSSIBLE N-term signal sequence\ninside 1 19\nTMhelix 20 39\noutside 40 325\nTMhelix 326 348\ninside 349 372

2048 GCF_001515005.1_ASM151500v1 Escherichia albertii Proteobacteria; Gammaproteobacteria; Enterobacterales; Enterobacteriaceae; Escherichia MTGDNTLIHSHGINRRDFMKLCAALAATMGLSSKAAAEMAESVTNPQRPPVIWIGAQECT WP_059258712.1 hydrogenase 2 small subunit [Escherichia albertii] Length: 372\nNumber of predicted TMHs: 1\nExp number of AAs in TMHs: 21.2336\nExp number, first 60 AAs: 0.22126\nTotal prob of N-in: 0.11592\noutside 1 332\nTMhelix 333 355\ninside 356 372

2049 GCF_001514625.1_ASM151462v1 Escherichia albertii Proteobacteria; Gammaproteobacteria; Enterobacterales; Enterobacteriaceae; Escherichia MTGDNTLIHSHGINRRDFMKLCAALAATMGLSSKAAAEMAESVTNPQRPPVIWIGAQECT WP_000145410.1 MULTISPECIES: hydrogenase-2 small chain [Proteobacteria] Length: 372\nNumber of predicted TMHs: 1\nExp number of AAs in TMHs: 23.72226\nExp number, first 60 AAs: 0.22038\nTotal prob of N-in: 0.12994\noutside 1 331\nTMhelix 332 354\ninside 355 372

2050 GCF_001514945.1_ASM151494v1 Escherichia albertii Proteobacteria; Gammaproteobacteria; Enterobacterales; Enterobacteriaceae; Escherichia MNNEETFYQSMRRQGVTRRSFLKYCSLAATSLGLGAGMAPKIAWALENKPRIPVVWIHGL WP_044712053.1 MULTISPECIES: hydrogenase-1 small chain [Escherichia] Length: 372\nNumber of predicted TMHs: 2\nExp

number of AAs in TMHs: 36.9963799999999\nExp number, first 60 AAs: 19.37682\nTotal prob of N-in: 0.93606\nPOSSIBLE N-term signal sequence\ninside 1 19\nTMhelix 20 39\noutside 40 325\nTMhelix 326 348\ninside 349 372

2051 GCF_001514865.1_ASM151486v1 Escherichia albertii Proteobacteria; Gammaproteobacteria; Enterobacterales; Enterobacteriaceae; Escherichia
MNNEETFYQSMRRQGVTRRSFLKYCSLAATSLGLGAGMAPKIAWALENKPRIPVVWIHGL WP_059272123.1
hydrogenase [Escherichia albertii] Length: 372\nNumber of predicted TMHs: 2\nExp number of AAs in TMHs: 38.51918\nExp number, first 60 AAs: 19.38068\nTotal prob of N-in: 0.93612\nPOSSIBLE N-term signal sequence\ninside 1 19\nTMhelix 20 39\noutside 40 325\nTMhelix 326 348\ninside 349 372

2052 GCF_001514665.1_ASM151466v1 Escherichia albertii Proteobacteria; Gammaproteobacteria; Enterobacterales; Enterobacteriaceae; Escherichia
MNNEETFYQSMRRQGVTRRSFLKYCSLAATSLGLGAGMAPKIAWALENKPRIPVVWIHGL WP_044712053.1
MULTISPECIES: hydrogenase-1 small chain [Escherichia] Length: 372\nNumber of predicted TMHs: 2\nExp number of AAs in TMHs: 36.9963799999999\nExp number, first 60 AAs: 19.37682\nTotal prob of N-in: 0.93606\nPOSSIBLE N-term signal sequence\ninside 1 19\nTMhelix 20 39\noutside 40 325\nTMhelix 326 348\ninside 349 372

2053 GCF_001514765.1_ASM151476v1 Escherichia albertii Proteobacteria; Gammaproteobacteria; Enterobacterales; Enterobacteriaceae; Escherichia
MTGDNTLIHSHGINRRDFMKLCAALAATMGLSSKAAAEMAESVTNPQRPPVIWIGAQECT WP_025238349.1
hydrogenase 2 small subunit [Escherichia albertii] Length: 372\nNumber of predicted TMHs: 1\nExp number of AAs in TMHs: 23.72042\nExp number, first 60 AAs: 0.22025\nTotal prob of N-in: 0.12992\noutside 1 331\nTMhelix 332 354\ninside 355 372

2054 GCF_001514665.1_ASM151466v1 Escherichia albertii Proteobacteria; Gammaproteobacteria; Enterobacterales; Enterobacteriaceae; Escherichia
MTGDNTLIHSHGINRRDFMKLCAALAATMGLSSKAAAEMAESVTNPQRPPVIWIGAQECT WP_000145410.1
MULTISPECIES: hydrogenase-2 small chain [Proteobacteria] Length: 372\nNumber of predicted TMHs: 1\nExp number of AAs in TMHs: 23.72226\nExp number, first 60 AAs: 0.22038\nTotal prob of N-in: 0.12994\noutside 1 331\nTMhelix 332 354\ninside 355 372

2055 GCF_001514705.1_ASM151470v1 Escherichia albertii Proteobacteria; Gammaproteobacteria; Enterobacterales; Enterobacteriaceae; Escherichia
MNNEETFYQSMRRQGVTRRSFLKYCSLAATSLGLGAGMAPKIAWALENKPRIPVVWIHGL WP_002461900.1
MULTISPECIES: hydrogenase-1 small chain [Escherichia] Length: 372\nNumber of predicted TMHs: 2\nExp number of AAs in TMHs: 37.1074899999999\nExp number, first 60 AAs: 19.37835\nTotal prob of N-in: 0.93614\nPOSSIBLE N-term signal sequence\ninside 1 19\nTMhelix 20 39\noutside 40 325\nTMhelix 326 348\ninside 349 372

2056 GCF_001515045.1_ASM151504v1 Escherichia albertii Proteobacteria; Gammaproteobacteria; Enterobacterales; Enterobacteriaceae; Escherichia
MNNEETFYQSMRRQGVTRRSFLKYCSLAATSLGLGAGMAPKIAWALENKPRIPVVWIHGL WP_059219811.1
hydrogenase [Escherichia albertii] Length: 372\nNumber of predicted TMHs: 2\nExp number of AAs in TMHs: 37.1074899999999\nExp number, first 60 AAs: 19.37835\nTotal prob of N-in: 0.93614\nPOSSIBLE N-term signal sequence\ninside 1 19\nTMhelix 20 39\noutside 40 325\nTMhelix 326 348\ninside 349 372

2057 GCF_001514595.1_ASM151459v1 Escherichia albertii Proteobacteria; Gammaproteobacteria; Enterobacterales; Enterobacteriaceae; Escherichia
MNNEETFYQSMRRQGVTRRSFLKYCSLAATSLGLGAGMAPKIAWALENKPRIPVVWIHGL WP_010318334.1
hydrogenase-1 small chain [Escherichia albertii] Length: 372\nNumber of predicted TMHs: 2\nExp number of AAs in TMHs: 36.9963899999999\nExp number, first 60 AAs: 19.37682\nTotal prob of N-in: 0.93606\nPOSSIBLE N-term signal sequence\ninside 1 19\nTMhelix 20 39\noutside 40 325\nTMhelix 326 348\ninside 349 372

2058 GCF_001514705.1_ASM151470v1 Escherichia albertii Proteobacteria; Gammaproteobacteria; Enterobacterales; Enterobacteriaceae; Escherichia
MTGDNTLIHSHGINRRDFMKLCAALAATMGLSSKAAAEMAESVTNPQRPPVIWIGAQECT WP_000145410.1
MULTISPECIES: hydrogenase-2 small chain [Proteobacteria] Length: 372\nNumber of predicted TMHs: 1\nExp number of AAs in TMHs: 23.72226\nExp number, first 60 AAs: 0.22038\nTotal prob of N-in: 0.12994\noutside 1 331\nTMhelix 332 354\ninside 355 372

2059 GCF_001514595.1_ASM151459v1 Escherichia albertii Proteobacteria; Gammaproteobacteria; Enterobacterales; Enterobacteriaceae; Escherichia
MTGDNTLIHSHGINRRDFMKLCAALAATMGLSSKAAAEMAESVTNPQRPPVIWIGAQECT WP_000145410.1
MULTISPECIES: hydrogenase-2 small chain [Proteobacteria] Length: 372\nNumber of predicted TMHs: 1\nExp number of AAs in TMHs: 23.72226\nExp number, first 60 AAs: 0.22038\nTotal prob of N-in: 0.12994\noutside 1 331\nTMhelix 332 354\ninside 355 372

2060 GCF_001515045.1_ASM151504v1 Escherichia albertii Proteobacteria; Gammaproteobacteria; Enterobacterales; Enterobacteriaceae; Escherichia
MTGDNTLIHSHGINRRDFMKLCAALAATMGLSSKAAAEMAESVTNPQRPPVIWIGAQECT WP_025238349.1
hydrogenase 2 small subunit [Escherichia albertii] Length: 372\nNumber of predicted TMHs: 1\nExp number of AAs in TMHs: 23.72042\nExp number, first 60 AAs: 0.22025\nTotal prob of N-in: 0.12992\noutside 1 331\nTMhelix 332 354\ninside 355 372

2061 GCF_001514985.1_ASM151498v1 Escherichia albertii Proteobacteria; Gammaproteobacteria; Enterobacterales; Enterobacteriaceae; Escherichia
MNNEETFYQSMRRQGVTRRSFLKYCSLAATSLGLGAGMAPKIAWALENKPRIPVVWIHGL WP_002461900.1
MULTISPECIES: hydrogenase-1 small chain [Escherichia] Length: 372\nNumber of predicted TMHs: 2\nExp number of AAs in TMHs: 37.1074899999999\nExp number, first 60 AAs: 19.37835\nTotal prob of N-in: 0.93614\nPOSSIBLE N-term signal sequence\ninside 1 19\nTMhelix 20 39\noutside 40 325\nTMhelix 326 348\ninside 349 372

2062 GCF_001514905.1_ASM151490v1 Escherichia albertii Proteobacteria; Gammaproteobacteria; Enterobacterales; Enterobacteriaceae; Escherichia
MNNEETFYQSMRRQGVTRRSFLKYCSLAATSLGLGAGMAPKIAWALENKPRIPVVWIHGL WP_059227719.1
hydrogenase [Escherichia albertii] Length: 372\nNumber of predicted TMHs: 2\nExp number of AAs in TMHs: 38.99311\nExp number, first 60 AAs: 19.37771\nTotal prob of N-in: 0.93622\nPOSSIBLE N-term signal sequence\ninside 1 19\nTMhelix 20 39\noutside 40 325\nTMhelix 326 348\ninside 349 372

2063 GCF_001549955.1_ASM154995v1 Escherichia albertii Proteobacteria; Gammaproteobacteria; Enterobacterales; Enterobacteriaceae; Escherichia
MTGDNTLIHSHGINRRDFMKLCAALAATMGLSSKAAAEMAESVTNPQRPPVIWIGAQECT WP_000145410.1
MULTISPECIES: hydrogenase-2 small chain [Proteobacteria] Length: 372\nNumber of predicted TMHs: 1\nExp number of AAs in TMHs: 23.72226\nExp number, first 60 AAs: 0.22038\nTotal prob of N-in: 0.12994\noutside 1 331\nTMhelix 332 354\ninside 355 372

2064 GCF_001549955.1_ASM154995v1 Escherichia albertii Proteobacteria; Gammaproteobacteria; Enterobacterales; Enterobacteriaceae; Escherichia
MNNEETFYQSMRRQGVTRRSFLKYCSLAATSLGLGAGMAPKIAWALENKPRIPVVWIHGL WP_002461900.1
MULTISPECIES: hydrogenase-1 small chain [Escherichia] Length: 372\nNumber of predicted TMHs: 2\nExp number of AAs in TMHs: 37.1074899999999\nExp number, first 60 AAs: 19.37835\nTotal prob of N-in: 0.93614\nPOSSIBLE N-term signal sequence\ninside 1 19\nTMhelix 20 39\noutside 40 325\nTMhelix 326 348\ninside 349 372

2065 GCF_001514765.1_ASM151476v1 Escherichia albertii Proteobacteria; Gammaproteobacteria; Enterobacterales; Enterobacteriaceae; Escherichia
MNNEETFYQSMRRQGVTRRSFLKYCSLAATSLGLGAGMAPKIAWALENKPRIPVVWIHGL WP_059219811.1
hydrogenase [Escherichia albertii] Length: 372\nNumber of predicted TMHs: 2\nExp number of AAs in TMHs: 37.1074899999999\nExp number, first 60 AAs: 19.37835\nTotal prob of N-in: 0.93614\nPOSSIBLE N-term signal sequence\ninside 1 19\nTMhelix 20 39\noutside 40 325\nTMhelix 326 348\ninside 349 372

2066 GCF_001514865.1_ASM151486v1 Escherichia albertii Proteobacteria; Gammaproteobacteria; Enterobacterales; Enterobacteriaceae; Escherichia
MTGDNTLIHSHGINRRDFMKLCAALAATMGLSSKAAAEMAESVTNPQRPPVIWIGAQECT WP_000145410.1
MULTISPECIES: hydrogenase-2 small chain [Proteobacteria] Length: 372\nNumber of predicted TMHs: 1\nExp number of AAs in TMHs: 23.72226\nExp number, first 60 AAs: 0.22038\nTotal prob of N-in: 0.12994\noutside 1 331\nTMhelix 332 354\ninside 355 372

2067 GCF_001514805.1_ASM151480v1 Escherichia albertii Proteobacteria; Gammaproteobacteria; Enterobacterales; Enterobacteriaceae; Escherichia
MTGDNTLIHSHGINRRDFMKLCAALAATMGLSSKAAAEMAESVTNPQRPPVIWIGAQECT WP_000145410.1

MULTISPECIES: hydrogenase-2 small chain [Proteobacteria] Length: 372\nNumber of predicted TMHs: 1\nExp number of AAs in TMHs: 23.72226\nExp number, first 60 AAs: 0.22038\nTotal prob of N-in: 0.12994\noutside 1 331\nTMhelix 332 354\ninside 355 372

2068 GCF_001514925.1_ASM151492v1 Escherichia albertii Proteobacteria; Gammaproteobacteria; Enterobacterales; Enterobacteriaceae; Escherichia MNNEETFYQSMRRQGVTRRSFLKYCSLAATSLGLGAGMAPKIAWALENKPRIPVVWIHGL WP_002461900.1 MULTISPECIES: hydrogenase-1 small chain [Escherichia] Length: 372\nNumber of predicted TMHs: 2\nExp number of AAs in TMHs: 37.1074899999999\nExp number, first 60 AAs: 19.37835\nTotal prob of N-in: 0.93614\nPOSSIBLE N-term signal sequence\ninside 1 19\nTMhelix 20 39\noutside 40 325\nTMhelix 326 348\ninside 349 372

2069 GCF_001514905.1_ASM151490v1 Escherichia albertii Proteobacteria; Gammaproteobacteria; Enterobacterales; Enterobacteriaceae; Escherichia MTGDNTLIHSHGINRRDFMKLCAALAATMGLSSKAAAEMAESVTNPQRPPVIWIGAQECT WP_000145410.1 MULTISPECIES: hydrogenase-2 small chain [Proteobacteria] Length: 372\nNumber of predicted TMHs: 1\nExp number of AAs in TMHs: 23.72226\nExp number, first 60 AAs: 0.22038\nTotal prob of N-in: 0.12994\noutside 1 331\nTMhelix 332 354\ninside 355 372

2070 GCF_001514925.1_ASM151492v1 Escherichia albertii Proteobacteria; Gammaproteobacteria; Enterobacterales; Enterobacteriaceae; Escherichia MTGDNTLIHSHGINRRDFMKLCAALAATMGLSSKAAAEMAESVTNPQRPPVIWIGAQECT WP_000145410.1 MULTISPECIES: hydrogenase-2 small chain [Proteobacteria] Length: 372\nNumber of predicted TMHs: 1\nExp number of AAs in TMHs: 23.72226\nExp number, first 60 AAs: 0.22038\nTotal prob of N-in: 0.12994\noutside 1 331\nTMhelix 332 354\ninside 355 372

2071 GCF_001514945.1_ASM151494v1 Escherichia albertii Proteobacteria; Gammaproteobacteria; Enterobacterales; Enterobacteriaceae; Escherichia MTGDNTLIHSHGINRRDFMKLCAALAATMGLSSKAAAEMAESVTNPQRPPVIWIGAQECT WP_000145410.1 MULTISPECIES: hydrogenase-2 small chain [Proteobacteria] Length: 372\nNumber of predicted TMHs: 1\nExp number of AAs in TMHs: 23.72226\nExp number, first 60 AAs: 0.22038\nTotal prob of N-in: 0.12994\noutside 1 331\nTMhelix 332 354\ninside 355 372

2072 GCF_001514805.1_ASM151480v1 Escherichia albertii Proteobacteria; Gammaproteobacteria; Enterobacterales; Enterobacteriaceae; Escherichia MNNEETFYQSMRRQGVTRRSFLKYCSLAATSLGLGAGMAPKIAWALENKPRIPVVWIHGL WP_044712053.1 MULTISPECIES: hydrogenase-1 small chain [Escherichia] Length: 372\nNumber of predicted TMHs: 2\nExp number of AAs in TMHs: 36.9963799999999\nExp number, first 60 AAs: 19.37682\nTotal prob of N-in: 0.93606\nPOSSIBLE N-term signal sequence\ninside 1 19\nTMhelix 20 39\noutside 40 325\nTMhelix 326 348\ninside 349 372

2073 GCF_001514985.1_ASM151498v1 Escherichia albertii Proteobacteria; Gammaproteobacteria; Enterobacterales; Enterobacteriaceae; Escherichia MTGDNTLIHSHGINRRDFMKLCAALAATMGLSSKAAAEMAESVTNPQRPPVIWIGAQECT WP_000145410.1 MULTISPECIES: hydrogenase-2 small chain [Proteobacteria] Length: 372\nNumber of predicted TMHs: 1\nExp number of AAs in TMHs: 23.72226\nExp number, first 60 AAs: 0.22038\nTotal prob of N-in: 0.12994\noutside 1 331\nTMhelix 332 354\ninside 355 372

2074 GCF_001286085.1_7790_1_78 Escherichia coli Proteobacteria; Gammaproteobacteria; Enterobacterales; Enterobacteriaceae; Escherichia MTGDNTLIHSHGINRRDFMKLCAALAATMGLSSKAAAEMAESVTNPQRPPVIWIGAQECT WP_000145410.1 MULTISPECIES: hydrogenase-2 small chain [Proteobacteria] Length: 372\nNumber of predicted TMHs: 1\nExp number of AAs in TMHs: 23.72226\nExp number, first 60 AAs: 0.22038\nTotal prob of N-in: 0.12994\noutside 1 331\nTMhelix 332 354\ninside 355 372

2075 GCF_000599805.2_a5_miseq_linux_20140113 Escherichia coli Proteobacteria; Gammaproteobacteria; Enterobacterales; Enterobacteriaceae; Escherichia MTGDNTLIHSHGINRRDFMKLCAALAATMGLSSKAAAEMAESVTNPQRPPVIWIGAQECT WP_000145410.1 MULTISPECIES: hydrogenase-2 small chain [Proteobacteria] Length: 372\nNumber of predicted TMHs: 1\nExp number of AAs in TMHs: 23.72226\nExp number, first 60 AAs: 0.22038\nTotal prob of N-in: 0.12994\noutside 1 331\nTMhelix 332 354\ninside 355 372

2076 GCF_000599705.1_ASM59970v1 Escherichia coli Proteobacteria; Gammaproteobacteria; Enterobacterales; Enterobacteriaceae; Escherichia MNNEETFYQAMRRQGVTRRSFLKYCSLAATSLGLGAGMAPKIAWALENKPRIPVVWIHGL WP_001058323.1 MULTISPECIES: hydrogenase-1 small chain [Proteobacteria] Length: 372\nNumber of predicted TMHs: 2\nExp number of AAs in TMHs: 37.2608499999999\nExp number, first 60 AAs: 19.52923\nTotal prob of N-in: 0.94351\nPOSSIBLE N-term signal sequence\ninside 1 19\nTMhelix 20 39\noutside 40 325\nTMhelix 326 348\ninside 349 372

2077 GCF_000599705.1_ASM59970v1 Escherichia coli Proteobacteria; Gammaproteobacteria; Enterobacterales; Enterobacteriaceae; Escherichia MTGDNTLIHSHGINRRDFMKLCAALAATMGLSSKAAAEMAESVTNPQRPPVIWIGAQECT WP_000145410.1 MULTISPECIES: hydrogenase-2 small chain [Proteobacteria] Length: 372\nNumber of predicted TMHs: 1\nExp number of AAs in TMHs: 23.72226\nExp number, first 60 AAs: 0.22038\nTotal prob of N-in: 0.12994\noutside 1 331\nTMhelix 332 354\ninside 355 372

2078 GCF_000599645.1_ASM59964v1 Escherichia coli Proteobacteria; Gammaproteobacteria; Enterobacterales; Enterobacteriaceae; Escherichia MNNEETFYQAMRRQGVTRRSFLKYCSLAATSLGLGAGMAPKIAWALENKPRIPVVWIHGL WP_001058323.1 MULTISPECIES: hydrogenase-1 small chain [Proteobacteria] Length: 372\nNumber of predicted TMHs: 2\nExp number of AAs in TMHs: 37.2608499999999\nExp number, first 60 AAs: 19.52923\nTotal prob of N-in: 0.94351\nPOSSIBLE N-term signal sequence\ninside 1 19\nTMhelix 20 39\noutside 40 325\nTMhelix 326 348\ninside 349 372

2079 GCF_000599645.1_ASM59964v1 Escherichia coli Proteobacteria; Gammaproteobacteria; Enterobacterales; Enterobacteriaceae; Escherichia MTGDNTLIHSHGINRRDFMKLCAALAATMGLSSKAAAEMAESVTNPQRPPVIWIGAQECT WP_000145410.1 MULTISPECIES: hydrogenase-2 small chain [Proteobacteria] Length: 372\nNumber of predicted TMHs: 1\nExp number of AAs in TMHs: 23.72226\nExp number, first 60 AAs: 0.22038\nTotal prob of N-in: 0.12994\noutside 1 331\nTMhelix 332 354\ninside 355 372

2080 GCF_001284925.1_7790_1_88 Escherichia coli Proteobacteria; Gammaproteobacteria; Enterobacterales; Enterobacteriaceae; Escherichia MTGDNTLIHSHGINRRDFMKLCAALAATMGLSSKAAAEMAESVTNPQRPPVIWIGAQECT WP_000145410.1 MULTISPECIES: hydrogenase-2 small chain [Proteobacteria] Length: 372\nNumber of predicted TMHs: 1\nExp number of AAs in TMHs: 23.72226\nExp number, first 60 AAs: 0.22038\nTotal prob of N-in: 0.12994\noutside 1 331\nTMhelix 332 354\ninside 355 372

2081 GCF_001285005.1_8205_3_22 Escherichia coli Proteobacteria; Gammaproteobacteria; Enterobacterales; Enterobacteriaceae; Escherichia MTGDNTLIHSHGINRRDFMKLCAALAATMGLSSKAAAEMAESVTNPQRPPVIWIGAQECT WP_000145410.1 MULTISPECIES: hydrogenase-2 small chain [Proteobacteria] Length: 372\nNumber of predicted TMHs: 1\nExp number of AAs in TMHs: 23.72226\nExp number, first 60 AAs: 0.22038\nTotal prob of N-in: 0.12994\noutside 1 331\nTMhelix 332 354\ninside 355 372

2082 GCF_000937275.1_FHI12 Escherichia coli Proteobacteria; Gammaproteobacteria; Enterobacterales; Enterobacteriaceae; Escherichia MNNEETFYQAMRRQGVTRRSFLKYCSLAATSLGLGAGMAPKIAWALENKPRIPVVWIHGL WP_001058323.1 MULTISPECIES: hydrogenase-1 small chain [Proteobacteria] Length: 372\nNumber of predicted TMHs: 2\nExp number of AAs in TMHs: 37.2608499999999\nExp number, first 60 AAs: 19.52923\nTotal prob of N-in: 0.94351\nPOSSIBLE N-term signal sequence\ninside 1 19\nTMhelix 20 39\noutside 40 325\nTMhelix 326 348\ninside 349 372

2083 GCF_000937275.1_FHI12 Escherichia coli Proteobacteria; Gammaproteobacteria; Enterobacterales; Enterobacteriaceae; Escherichia MTGDNTLIHSHGINRRDFMKLCAALAATMGLSSKAAAEMAESVTNPQRPPVIWIGAQECT WP_000145410.1 MULTISPECIES: hydrogenase-2 small chain [Proteobacteria] Length: 372\nNumber of predicted TMHs: 1\nExp number of AAs in TMHs: 23.72226\nExp number, first 60 AAs: 0.22038\nTotal prob of N-in: 0.12994\noutside 1 331\nTMhelix 332 354\ninside 355 372

2084 GCF_000936455.1_46A_S2_assembly Escherichia coli Proteobacteria; Gammaproteobacteria; Enterobacterales; Enterobacteriaceae; Escherichia MNNEETFYQAMRRQGVTRRSFLKYCSLAATSLGLGAGMAPKIAWALENKPRIPVVWIHGL WP_001058323.1 MULTISPECIES: hydrogenase-1 small chain [Proteobacteria] Length: 372\nNumber of predicted TMHs:

2\nExp number of AAs in TMHs: 37.2608499999999\nExp number, first 60 AAs: 19.52923\nTotal prob of N-in: 0.94351\nPOSSIBLE N-term signal sequence\ninside 1 19\nTMhelix 20 39\noutside 40 325\nTMhelix 326 348\ninside 349 372

2085 GCF_000936455.1_46A_S2_assembly Escherichia coli Proteobacteria; Gammaproteobacteria; Enterobacterales; Enterobacteriaceae; Escherichia MTGDNTLIHSHGINRRDFMKLCAALAATMGLSSKAAAEMAESVTNPQRPPVIWIGAQECT WP_000145410.1 MULTISPECIES: hydrogenase-2 small chain [Proteobacteria] Length: 372\nNumber of predicted TMHs: 1\nExp number of AAs in TMHs: 23.72226\nExp number, first 60 AAs: 0.22038\nTotal prob of N-in: 0.12994\noutside 1 331\nTMhelix 332 354\ninside 355 372

2086 GCF_000937715.2_FHI65 Escherichia coli Proteobacteria; Gammaproteobacteria; Enterobacterales; Enterobacteriaceae; Escherichia MTGDNTLIHSHGINRRDFMKLCAALAATMGLSSKAAAEMAESVTNPQRPPVIWIGAQECT WP_000145410.1 MULTISPECIES: hydrogenase-2 small chain [Proteobacteria] Length: 372\nNumber of predicted TMHs: 1\nExp number of AAs in TMHs: 23.72226\nExp number, first 60 AAs: 0.22038\nTotal prob of N-in: 0.12994\noutside 1 331\nTMhelix 332 354\ninside 355 372

2087 GCF_000935505.1_ASM93550v1 Escherichia coli Proteobacteria; Gammaproteobacteria; Enterobacterales; Enterobacteriaceae; Escherichia MNNEETFYQAMRRQGVTRRSFLKYCSLAATSLGLGAGMAPKIAWALENKPRIPVVWIHGL WP_001058323.1 MULTISPECIES: hydrogenase-1 small chain [Proteobacteria] Length: 372\nNumber of predicted TMHs: 2\nExp number of AAs in TMHs: 37.2608499999999\nExp number, first 60 AAs: 19.52923\nTotal prob of N-in: 0.94351\nPOSSIBLE N-term signal sequence\ninside 1 19\nTMhelix 20 39\noutside 40 325\nTMhelix 326 348\ninside 349 372

2088 GCF_000798315.1_Escherichia_coli_CVM_N33633PS_v._1.0 Escherichia coli Proteobacteria; Gammaproteobacteria; Enterobacterales; Enterobacteriaceae; Escherichia MTGDNTLIHSHGINRRDFMKLCAALAATMGLSSKAAAEMAESVTNPQRPPVIWIGAQECT WP_000145410.1 MULTISPECIES: hydrogenase-2 small chain [Proteobacteria] Length: 372\nNumber of predicted TMHs: 1\nExp number of AAs in TMHs: 23.72226\nExp number, first 60 AAs: 0.22038\nTotal prob of N-in: 0.12994\noutside 1 331\nTMhelix 332 354\ninside 355 372

2089 GCF_000935505.1_ASM93550v1 Escherichia coli Proteobacteria; Gammaproteobacteria; Enterobacterales; Enterobacteriaceae; Escherichia MTGDNTLIHSHGINRRDFMKLCAALAATMGLSSKAAAEMAESVTNPQRPPVIWIGAQECT WP_000145410.1 MULTISPECIES: hydrogenase-2 small chain [Proteobacteria] Length: 372\nNumber of predicted TMHs: 1\nExp number of AAs in TMHs: 23.72226\nExp number, first 60 AAs: 0.22038\nTotal prob of N-in: 0.12994\noutside 1 331\nTMhelix 332 354\ninside 355 372

2090 GCF_000798315.1_Escherichia_coli_CVM_N33633PS_v._1.0 Escherichia coli Proteobacteria; Gammaproteobacteria; Enterobacterales; Enterobacteriaceae; Escherichia MNNEETFYQAMRRQGVTRRSFLKYCSLAATSLGLGAGMAPKIAWALENKPRIPVVWIHGL WP_001295940.1 MULTISPECIES: hydrogenase [Enterobacteriaceae] Length: 372\nNumber of predicted TMHs: 2\nExp number of AAs in TMHs: 37.2608899999999\nExp number, first 60 AAs: 19.52923\nTotal prob of N-in: 0.94351\nPOSSIBLE N-term signal sequence\ninside 1 19\nTMhelix 20 39\noutside 40 325\nTMhelix 326 348\ninside 349 372

2091 GCF_000938995.1_FHI25 Escherichia coli Proteobacteria; Gammaproteobacteria; Enterobacterales; Enterobacteriaceae; Escherichia MTGDNTLIHSHGINRRDFMKLCAALAATMGLSSKAAAEMAESVTNPQRPPVIWIGAQECT WP_000145410.1 MULTISPECIES: hydrogenase-2 small chain [Proteobacteria] Length: 372\nNumber of predicted TMHs: 1\nExp number of AAs in TMHs: 23.72226\nExp number, first 60 AAs: 0.22038\nTotal prob of N-in: 0.12994\noutside 1 331\nTMhelix 332 354\ninside 355 372

2092 GCF_000938995.1_FHI25 Escherichia coli Proteobacteria; Gammaproteobacteria; Enterobacterales; Enterobacteriaceae; Escherichia MNNEETFYQAMRRQGVTRRSFLKYCSLAATSLGLGAGMAPKIAWALENKPRIPVVWIHGL WP_001058323.1 MULTISPECIES: hydrogenase-1 small chain [Proteobacteria] Length: 372\nNumber of predicted TMHs: 2\nExp number of AAs in TMHs: 37.2608499999999\nExp number, first 60 AAs: 19.52923\nTotal prob of N-in: 0.94351\nPOSSIBLE N-term signal sequence\ninside 1 19\nTMhelix 20 39\noutside 40 325\nTMhelix 326 348\ninside 349 372

2093 GCF_000798375.1_Escherichia_coli_CVM_N38826PS_v._1.0 Escherichia coli Proteobacteria; Gammaproteobacteria; Enterobacterales; Enterobacteriaceae; Escherichia
MTGDNTLIHSHGINRRDFMKLCAALAATMGLSSKAAAEMAESVTNPQRPPVIWIGAQECT WP_000145410.1
MULTISPECIES: hydrogenase-2 small chain [Proteobacteria] Length: 372\nNumber of predicted TMHs: 1\nExp number of AAs in TMHs: 23.72226\nExp number, first 60 AAs: 0.22038\nTotal prob of N-in: 0.12994\noutside 1 331\nTMhelix 332 354\ninside 355 372

2094 GCF_000798375.1_Escherichia_coli_CVM_N38826PS_v._1.0 Escherichia coli Proteobacteria; Gammaproteobacteria; Enterobacterales; Enterobacteriaceae; Escherichia
MNNEETFYQAMRRQGVTRRSFLKYCSLAATSLGLGAGMAPKIAWALENKPRIPVVWIHGL WP_001058323.1
MULTISPECIES: hydrogenase-1 small chain [Proteobacteria] Length: 372\nNumber of predicted TMHs: 2\nExp number of AAs in TMHs: 37.2608499999999\nExp number, first 60 AAs: 19.52923\nTotal prob of N-in: 0.94351\nPOSSIBLE N-term signal sequence\ninside 1 19\nTMhelix 20 39\noutside 40 325\nTMhelix 326 348\ninside 349 372

2095 GCF_000935375.1_ASM93537v1 Escherichia coli Proteobacteria; Gammaproteobacteria; Enterobacterales; Enterobacteriaceae; Escherichia
MTGDNTLIHSHGINRRDFMKLCAALAATMGLSSKAAAEMAESVTNPQRPPVIWIGAQECT WP_000145410.1
MULTISPECIES: hydrogenase-2 small chain [Proteobacteria] Length: 372\nNumber of predicted TMHs: 1\nExp number of AAs in TMHs: 23.72226\nExp number, first 60 AAs: 0.22038\nTotal prob of N-in: 0.12994\noutside 1 331\nTMhelix 332 354\ninside 355 372

2096 GCF_000935075.1_ASM93507v1 Escherichia coli Proteobacteria; Gammaproteobacteria; Enterobacterales; Enterobacteriaceae; Escherichia
MNNEETFYQAMRRQGVTRRSFLKYCSLAATSLGLGAGMAPKIAWALENKPRIPVVWIHGL WP_001301613.1
hydrogenase [Escherichia coli] Length: 372\nNumber of predicted TMHs: 2\nExp number of AAs in TMHs: 37.46598\nExp number, first 60 AAs: 19.53139\nTotal prob of N-in: 0.94364\nPOSSIBLE N-term signal sequence\ninside 1 19\nTMhelix 20 39\noutside 40 325\nTMhelix 326 348\ninside 349 372

2097 GCF_000935075.1_ASM93507v1 Escherichia coli Proteobacteria; Gammaproteobacteria; Enterobacterales; Enterobacteriaceae; Escherichia
MTGDNTLIHSHGINRRDFMKLCAALAATMGLSSKAAAEMAESVTNPQRPPVIWIGAQECT WP_000145410.1
MULTISPECIES: hydrogenase-2 small chain [Proteobacteria] Length: 372\nNumber of predicted TMHs: 1\nExp number of AAs in TMHs: 23.72226\nExp number, first 60 AAs: 0.22038\nTotal prob of N-in: 0.12994\noutside 1 331\nTMhelix 332 354\ninside 355 372

2098 GCF_000939255.1_FHI72 Escherichia coli Proteobacteria; Gammaproteobacteria; Enterobacterales; Enterobacteriaceae; Escherichia MTGDNTLIHSHGINRRDFMKLCAALAATMGLSSKAAAEMVESVTNPQRPPVIWIGAQECT
WP_000145423.1 hydrogenase 2 small subunit [Escherichia coli] Length: 372\nNumber of predicted TMHs: 1\nExp number of AAs in TMHs: 23.78201\nExp number, first 60 AAs: 0.28355\nTotal prob of N-in: 0.13257\noutside 1 331\nTMhelix 332 354\ninside 355 372

2099 GCF_000932215.1_ASM93221v1 Escherichia coli Proteobacteria; Gammaproteobacteria; Enterobacterales; Enterobacteriaceae; Escherichia
MTGDNTLIHSHGINRRDFMKLCAALAATMGLSSKAAAEMAESVTNPQRPPVIWIGAQECT WP_044374594.1
hydrogenase 2 small subunit [Escherichia coli] Length: 372\nNumber of predicted TMHs: 1\nExp number of AAs in TMHs: 27.15394\nExp number, first 60 AAs: 0.2094\nTotal prob of N-in: 0.25674\noutside 1 331\nTMhelix 332 354\ninside 355 372

2100 GCF_000798435.1_Escherichia_coli_CVM_N33719PS_v._1.0 Escherichia coli Proteobacteria; Gammaproteobacteria; Enterobacterales; Enterobacteriaceae; Escherichia
MTGDNTLIHSHGINRRDFMKLCAALAATMGLSSKAAAEMAESVTNPQRPPVIWIGAQECT WP_000145410.1
MULTISPECIES: hydrogenase-2 small chain [Proteobacteria] Length: 372\nNumber of predicted TMHs: 1\nExp number of AAs in TMHs: 23.72226\nExp number, first 60 AAs: 0.22038\nTotal prob of N-in: 0.12994\noutside 1 331\nTMhelix 332 354\ninside 355 372

2101 GCF_000798435.1_Escherichia_coli_CVM_N33719PS_v._1.0 Escherichia coli Proteobacteria; Gammaproteobacteria; Enterobacterales; Enterobacteriaceae; Escherichia
MNNEETFYQAMRRQGVTRRSFLKYCSLAATSLGLGAGMAPKIAWALENKPRIPVVWIHGL WP_001058323.1
MULTISPECIES: hydrogenase-1 small chain [Proteobacteria] Length: 372\nNumber of predicted TMHs: 2\nExp number of AAs in TMHs: 37.2608499999999\nExp number, first 60 AAs: 19.52923\nTotal prob of N-in:

0.94351\nPOSSIBLE N-term signal sequence\ninside 1 19\nTMhelix 20 39\noutside 40 325\nTMhelix 326 348\ninside 349 372

2102 GCF_001285105.1_7748_7_38 Escherichia coli Proteobacteria; Gammaproteobacteria; Enterobacterales; Enterobacteriaceae; Escherichia MTGDNTLIHSHGINRRDFMKLCAALAATMGLSSKAAAEMAESVTNPQRPPVIWIGAQECT WP_000145410.1 MULTISPECIES: hydrogenase-2 small chain [Proteobacteria] Length: 372\nNumber of predicted TMHs: 1\nExp number of AAs in TMHs: 23.72226\nExp number, first 60 AAs: 0.22038\nTotal prob of N-in: 0.12994\noutside 1 331\nTMhelix 332 354\ninside 355 372

2103 GCF_001285105.1_7748_7_38 Escherichia coli Proteobacteria; Gammaproteobacteria; Enterobacterales; Enterobacteriaceae; Escherichia MNNEETFYQAMRRQGVTRRSFLKYCSLAATSLGLGAGMAPKIAWALENKPRIPVVWIHGL WP_001295940.1 MULTISPECIES: hydrogenase [Enterobacteriaceae] Length: 372\nNumber of predicted TMHs: 2\nExp number of AAs in TMHs: 37.2608899999999\nExp number, first 60 AAs: 19.52923\nTotal prob of N-in: 0.94351\nPOSSIBLE N-term signal sequence\ninside 1 19\nTMhelix 20 39\noutside 40 325\nTMhelix 326 348\ninside 349 372

2104 GCF_000833145.1_ASM83314v1 Escherichia coli Proteobacteria; Gammaproteobacteria; Enterobacterales; Enterobacteriaceae; Escherichia MNNEETFYQAMRRQGVTRRSFLKYCSLAATSLGLGAGMAPKIAWALENKPRIPVVWIHGL WP_001058323.1 MULTISPECIES: hydrogenase-1 small chain [Proteobacteria] Length: 372\nNumber of predicted TMHs: 2\nExp number of AAs in TMHs: 37.2608499999999\nExp number, first 60 AAs: 19.52923\nTotal prob of N-in: 0.94351\nPOSSIBLE N-term signal sequence\ninside 1 19\nTMhelix 20 39\noutside 40 325\nTMhelix 326 348\ninside 349 372

2105 GCF_000940555.1_Ec46A_L2_MIRA_assembly Escherichia coli Proteobacteria; Gammaproteobacteria; Enterobacterales; Enterobacteriaceae; Escherichia MNNEETFYQAMRRQGVTRRSFLKYCSLAATSLGLGAGMAPKIAWALENKPRIPVVWIHGL WP_001058323.1 MULTISPECIES: hydrogenase-1 small chain [Proteobacteria] Length: 372\nNumber of predicted TMHs: 2\nExp number of AAs in TMHs: 37.2608499999999\nExp number, first 60 AAs: 19.52923\nTotal prob of N-in: 0.94351\nPOSSIBLE N-term signal sequence\ninside 1 19\nTMhelix 20 39\noutside 40 325\nTMhelix 326 348\ninside 349 372

2106 GCF_000941195.1_57A_A8_assembly Escherichia coli Proteobacteria; Gammaproteobacteria; Enterobacterales; Enterobacteriaceae; Escherichia MNNEETFYQAMRRQGVTRRSFLKYCSLAATSLGLGAGMAPKIAWALENKPRIPVVWIHGL WP_001058323.1 MULTISPECIES: hydrogenase-1 small chain [Proteobacteria] Length: 372\nNumber of predicted TMHs: 2\nExp number of AAs in TMHs: 37.2608499999999\nExp number, first 60 AAs: 19.52923\nTotal prob of N-in: 0.94351\nPOSSIBLE N-term signal sequence\ninside 1 19\nTMhelix 20 39\noutside 40 325\nTMhelix 326 348\ninside 349 372

2107 GCF_001297985.1_ASM129798v1 Escherichia coli Proteobacteria; Gammaproteobacteria; Enterobacterales; Enterobacteriaceae; Escherichia MNNEETFYQAMRRQGVTRRSFLKYCSLAATSLGLGAGMAPKIAWALENKPRIPVVWIHGL WP_054191816.1 hydrogenase [Escherichia coli] Length: 372\nNumber of predicted TMHs: 2\nExp number of AAs in TMHs: 37.74755\nExp number, first 60 AAs: 19.52988\nTotal prob of N-in: 0.94380\nPOSSIBLE N-term signal sequence\ninside 1 19\nTMhelix 20 39\noutside 40 325\nTMhelix 326 348\ninside 349 372

2108 GCF_001297965.1_ASM129796v1 Escherichia coli Proteobacteria; Gammaproteobacteria; Enterobacterales; Enterobacteriaceae; Escherichia MTGDNTLIHSHGINRRDFMKLCAALAATMGLSSKAAAEMAESVTNPQRPPVIWIGAQECT WP_000145410.1 MULTISPECIES: hydrogenase-2 small chain [Proteobacteria] Length: 372\nNumber of predicted TMHs: 1\nExp number of AAs in TMHs: 23.72226\nExp number, first 60 AAs: 0.22038\nTotal prob of N-in: 0.12994\noutside 1 331\nTMhelix 332 354\ninside 355 372

2109 GCF_000941575.1_FHI92 Escherichia coli Proteobacteria; Gammaproteobacteria; Enterobacterales; Enterobacteriaceae; Escherichia MTGDNTLIHSHGINRRDFMKLCAALAATMGLSSKAAAEMAESVTNPQRPPVIWIGAQECT WP_000145410.1 MULTISPECIES: hydrogenase-2 small chain [Proteobacteria] Length: 372\nNumber of predicted TMHs: 1\nExp number of AAs in TMHs: 23.72226\nExp number, first 60 AAs: 0.22038\nTotal prob of N-in: 0.12994\noutside 1 331\nTMhelix 332 354\ninside 355 372

2110 GCF_001286705.1_8205_8_76 Escherichia coli Proteobacteria; Gammaproteobacteria; Enterobacterales; Enterobacteriaceae; Escherichia MNNEETFYQAMRRQGVTRRSFLKYCSLAATSLGLGAGMAPKIAWALENKPRIPVVWIHGL WP_001058323.1 MULTISPECIES: hydrogenase-1 small chain [Proteobacteria] Length: 372\nNumber of predicted TMHs: 2\nExp number of AAs in TMHs: 37.2608499999999\nExp number, first 60 AAs: 19.52923\nTotal prob of N-in: 0.94351\nPOSSIBLE N-term signal sequence\ninside 1 19\nTMhelix 20 39\noutside 40 325\nTMhelix 326 348\ninside 349 372

2111 GCF_001297965.1_ASM129796v1 Escherichia coli Proteobacteria; Gammaproteobacteria; Enterobacterales; Enterobacteriaceae; Escherichia MNNEETFYQAMRRQGVTRRSFLKYCSLAATSLGLGAGMAPKIAWALENKPRIPVVWIHGL WP_001301613.1 hydrogenase [Escherichia coli] Length: 372\nNumber of predicted TMHs: 2\nExp number of AAs in TMHs: 37.46598\nExp number, first 60 AAs: 19.53139\nTotal prob of N-in: 0.94364\nPOSSIBLE N-term signal sequence\ninside 1 19\nTMhelix 20 39\noutside 40 325\nTMhelix 326 348\ninside 349 372

2112 GCF_000941575.1_FHI92 Escherichia coli Proteobacteria; Gammaproteobacteria; Enterobacterales; Enterobacteriaceae; Escherichia MNNEETFYQAMRRQGVTRRSFLKYCSLAATSLGLGAGMAPKIAWALENKPRIPVVWIHGL WP_001717971.1 hydrogenase-1 small chain [Escherichia coli] Length: 372\nNumber of predicted TMHs: 2\nExp number of AAs in TMHs: 38.00708\nExp number, first 60 AAs: 19.5286\nTotal prob of N-in: 0.94354\nPOSSIBLE N-term signal sequence\ninside 1 19\nTMhelix 20 39\noutside 40 325\nTMhelix 326 348\ninside 349 372

2113 GCF_000944015.1_FHI71 Escherichia coli Proteobacteria; Gammaproteobacteria; Enterobacterales; Enterobacteriaceae; Escherichia MTGDNTLIHSHGINRRDFMKLCAALAATMGLSSKAAAEMAESVTNPQRPPVIWIGAQECT WP_000145410.1 MULTISPECIES: hydrogenase-2 small chain [Proteobacteria] Length: 372\nNumber of predicted TMHs: 1\nExp number of AAs in TMHs: 23.72226\nExp number, first 60 AAs: 0.22038\nTotal prob of N-in: 0.12994\noutside 1 331\nTMhelix 332 354\ninside 355 372

2114 GCF_000944015.1_FHI71 Escherichia coli Proteobacteria; Gammaproteobacteria; Enterobacterales; Enterobacteriaceae; Escherichia MNNEETFYQAMRRQGVTRRSFLKYCSLAATSLGLGAGMAPKIAWALENKPRIPVVWIHGL WP_001058323.1 MULTISPECIES: hydrogenase-1 small chain [Proteobacteria] Length: 372\nNumber of predicted TMHs: 2\nExp number of AAs in TMHs: 37.2608499999999\nExp number, first 60 AAs: 19.52923\nTotal prob of N-in: 0.94351\nPOSSIBLE N-term signal sequence\ninside 1 19\nTMhelix 20 39\noutside 40 325\nTMhelix 326 348\ninside 349 372

2115 GCF_001286565.1_8205_3_43 Escherichia coli Proteobacteria; Gammaproteobacteria; Enterobacterales; Enterobacteriaceae; Escherichia MNNEETFYQAMRRQGVTRRSFLKYCSLAATSLGLGAGMAPKIAWALENKPRIPVVWIHGL WP_001058323.1 MULTISPECIES: hydrogenase-1 small chain [Proteobacteria] Length: 372\nNumber of predicted TMHs: 2\nExp number of AAs in TMHs: 37.2608499999999\nExp number, first 60 AAs: 19.52923\nTotal prob of N-in: 0.94351\nPOSSIBLE N-term signal sequence\ninside 1 19\nTMhelix 20 39\noutside 40 325\nTMhelix 326 348\ninside 349 372

2116 GCF_001286565.1_8205_3_43 Escherichia coli Proteobacteria; Gammaproteobacteria; Enterobacterales; Enterobacteriaceae; Escherichia MTGDNTLIHSHGINRRDFMKLCAALAATMGLSSKAAAEMAESVTNPQRPPVIWIGAQECT WP_000145410.1 MULTISPECIES: hydrogenase-2 small chain [Proteobacteria] Length: 372\nNumber of predicted TMHs: 1\nExp number of AAs in TMHs: 23.72226\nExp number, first 60 AAs: 0.22038\nTotal prob of N-in: 0.12994\noutside 1 331\nTMhelix 332 354\ninside 355 372

2117 GCF_001286385.1_7748_7_24 Escherichia coli Proteobacteria; Gammaproteobacteria; Enterobacterales; Enterobacteriaceae; Escherichia MNNEETFYQAMRRQGVTRRSFLKYCSLAATSLGLGAGMAPKIAWALENKPRIPVVWIHGL WP_001058323.1 MULTISPECIES: hydrogenase-1 small chain [Proteobacteria] Length: 372\nNumber of predicted TMHs: 2\nExp number of AAs in TMHs: 37.2608499999999\nExp number, first 60 AAs: 19.52923\nTotal prob of N-in: 0.94351\nPOSSIBLE N-term signal sequence\ninside 1 19\nTMhelix 20 39\noutside 40 325\nTMhelix 326 348\ninside 349 372

2118 GCF_001286385.1_7748_7_24 Escherichia coli Proteobacteria; Gammaproteobacteria; Enterobacterales; Enterobacteriaceae; Escherichia MTGDNTLIHSHGINRRDFMKLCAALAATMGLSSKAAAEMAESVTNPQRPPVIWIGAQECT WP_000145410.1 MULTISPECIES: hydrogenase-2 small chain [Proteobacteria] Length: 372\nNumber of predicted TMHs:

1\nExp number of AAs in TMHs: 23.72226\nExp number, first 60 AAs: 0.22038\nTotal prob of N-in: 0.12994\noutside 1 331\nTMhelix 332 354\ninside 355 372

2119 GCF_001286325.1_8205_3_58 Escherichia coli Proteobacteria; Gammaproteobacteria; Enterobacterales; Enterobacteriaceae; Escherichia MTGDNTLIHSHGINRRDFMKLCAALAATMGLSSKAAAEMAESVTNPQRPPVIWIGAQECT WP_000145410.1 MULTISPECIES: hydrogenase-2 small chain [Proteobacteria] Length: 372\nNumber of predicted TMHs: 1\nExp number of AAs in TMHs: 23.72226\nExp number, first 60 AAs: 0.22038\nTotal prob of N-in: 0.12994\noutside 1 331\nTMhelix 332 354\ninside 355 372

2120 GCF_001191045.1_CFSAN026843_01-NGE.clc Escherichia coli Proteobacteria; Gammaproteobacteria; Enterobacterales; Enterobacteriaceae; Escherichia MNNEETFYQAMRRQGVTRRSFLKYCSLAATSLGLGAGMAPKIAWALENKPRIPVVWIHGL WP_001058323.1 MULTISPECIES: hydrogenase-1 small chain [Proteobacteria] Length: 372\nNumber of predicted TMHs: 2\nExp number of AAs in TMHs: 37.2608499999999\nExp number, first 60 AAs: 19.52923\nTotal prob of N-in: 0.94351\nPOSSIBLE N-term signal sequence\ninside 1 19\nTMhelix 20 39\noutside 40 325\nTMhelix 326 348\ninside 349 372

2121 GCF_000948005.1_ASM94800v1 Escherichia coli Proteobacteria; Gammaproteobacteria; Enterobacterales; Enterobacteriaceae; Escherichia MTGDNTLIHSHGINRRDFMKLCAALAATMGLSSKAAAEMAESVTNPQRPPVIWIGAQECT WP_000145410.1 MULTISPECIES: hydrogenase-2 small chain [Proteobacteria] Length: 372\nNumber of predicted TMHs: 1\nExp number of AAs in TMHs: 23.72226\nExp number, first 60 AAs: 0.22038\nTotal prob of N-in: 0.12994\noutside 1 331\nTMhelix 332 354\ninside 355 372

2122 GCF_000819375.1_Escherichia_coli_CVM_N33429PS_v1.0 Escherichia coli Proteobacteria; Gammaproteobacteria; Enterobacterales; Enterobacteriaceae; Escherichia MNNEETFYQAMRRQGVTRRSFLKYCSLAATSLGLGAGMAPKIAWALENKPRIPVVWIHGL WP_001058323.1 MULTISPECIES: hydrogenase-1 small chain [Proteobacteria] Length: 372\nNumber of predicted TMHs: 2\nExp number of AAs in TMHs: 37.2608499999999\nExp number, first 60 AAs: 19.52923\nTotal prob of N-in: 0.94351\nPOSSIBLE N-term signal sequence\ninside 1 19\nTMhelix 20 39\noutside 40 325\nTMhelix 326 348\ninside 349 372

2123 GCF_000819375.1_Escherichia_coli_CVM_N33429PS_v1.0 Escherichia coli Proteobacteria; Gammaproteobacteria; Enterobacterales; Enterobacteriaceae; Escherichia MTGDNTLIHSHGINRRDFMKLCAALAATMGLSSKAAAEMAESVTNPQRPPVIWIGAQECT WP_000145410.1 MULTISPECIES: hydrogenase-2 small chain [Proteobacteria] Length: 372\nNumber of predicted TMHs: 1\nExp number of AAs in TMHs: 23.72226\nExp number, first 60 AAs: 0.22038\nTotal prob of N-in: 0.12994\noutside 1 331\nTMhelix 332 354\ninside 355 372

2124 GCF_001286205.1_7748_7_48 Escherichia coli Proteobacteria; Gammaproteobacteria; Enterobacterales; Enterobacteriaceae; Escherichia MNNEETFYQAMRRQGVTRRSFLKYCSLAATSLGLGAGMAPKIAWALENKPRIPVVWIHGL WP_001058323.1 MULTISPECIES: hydrogenase-1 small chain [Proteobacteria] Length: 372\nNumber of predicted TMHs: 2\nExp number of AAs in TMHs: 37.2608499999999\nExp number, first 60 AAs: 19.52923\nTotal prob of N-in: 0.94351\nPOSSIBLE N-term signal sequence\ninside 1 19\nTMhelix 20 39\noutside 40 325\nTMhelix 326 348\ninside 349 372

2125 GCF_001286205.1_7748_7_48 Escherichia coli Proteobacteria; Gammaproteobacteria; Enterobacterales; Enterobacteriaceae; Escherichia MTGDNTLIHSHGINRRDFMKLCAALAATMGLSSKAAAEMAESVTNPQRPPVIWIGAQECT WP_000145410.1 MULTISPECIES: hydrogenase-2 small chain [Proteobacteria] Length: 372\nNumber of predicted TMHs: 1\nExp number of AAs in TMHs: 23.72226\nExp number, first 60 AAs: 0.22038\nTotal prob of N-in: 0.12994\noutside 1 331\nTMhelix 332 354\ninside 355 372

2126 GCF_001285145.1_7748_7_7Escherichia coli Proteobacteria; Gammaproteobacteria; Enterobacterales; Enterobacteriaceae; Escherichia MTGDNTLIHSHGINRRDFMKLCAALAATMGLSSKAAAEMAESVTNPQRPPVIWIGAQECT WP_000145410.1 MULTISPECIES: hydrogenase-2 small chain [Proteobacteria] Length: 372\nNumber of predicted TMHs: 1\nExp number of AAs in TMHs: 23.72226\nExp number, first 60 AAs: 0.22038\nTotal prob of N-in: 0.12994\noutside 1 331\nTMhelix 332 354\ninside 355 372

2127 GCF_001285145.1_7748_7_7Escherichia coli Proteobacteria; Gammaproteobacteria; Enterobacterales; Enterobacteriaceae; Escherichia MNNEETFYQAMRRQGVTRRSFLKYCSLAATSLGLGAGMAPKIAWALENKPRIPVVWIHGL WP_001295940.1 MULTISPECIES: hydrogenase [Enterobacteriaceae] Length: 372\nNumber of predicted TMHs: 2\nExp number of AAs in TMHs: 37.2608899999999\nExp number, first 60 AAs: 19.52923\nTotal prob of N-in: 0.94351\nPOSSIBLE N-term signal sequence\ninside 1 19\nTMhelix 20 39\noutside 40 325\nTMhelix 326 348\ninside 349 372

2128 GCF_000948005.1_ASM94800v1 Escherichia coli Proteobacteria; Gammaproteobacteria; Enterobacterales; Enterobacteriaceae; Escherichia MNNEETFYQAMRRQGVTRRSFLKYCSLAATSLGLGAGMAPKIAWALENKPRIPVVWIHGL WP_001058323.1 MULTISPECIES: hydrogenase-1 small chain [Proteobacteria] Length: 372\nNumber of predicted TMHs: 2\nExp number of AAs in TMHs: 37.2608499999999\nExp number, first 60 AAs: 19.52923\nTotal prob of N-in: 0.94351\nPOSSIBLE N-term signal sequence\ninside 1 19\nTMhelix 20 39\noutside 40 325\nTMhelix 326 348\ninside 349 372

2129 GCF_001286145.1_7553_7_62 Escherichia coli Proteobacteria; Gammaproteobacteria; Enterobacterales; Enterobacteriaceae; Escherichia MNNEETFYQAMRRQGVTRRSFLKYCSLAATSLGLGAGMAPKIAWALENKPRIPVVWIHGL WP_001058323.1 MULTISPECIES: hydrogenase-1 small chain [Proteobacteria] Length: 372\nNumber of predicted TMHs: 2\nExp number of AAs in TMHs: 37.2608499999999\nExp number, first 60 AAs: 19.52923\nTotal prob of N-in: 0.94351\nPOSSIBLE N-term signal sequence\ninside 1 19\nTMhelix 20 39\noutside 40 325\nTMhelix 326 348\ninside 349 372

2130 GCF_001285165.1_8205_8_82 Escherichia coli Proteobacteria; Gammaproteobacteria; Enterobacterales; Enterobacteriaceae; Escherichia MNNEETFYQAMRRQGVTRRSFLKYCSLAATSLGLGAGMAPKIAWALENKPRIPVVWIHGL WP_001058323.1 MULTISPECIES: hydrogenase-1 small chain [Proteobacteria] Length: 372\nNumber of predicted TMHs: 2\nExp number of AAs in TMHs: 37.2608499999999\nExp number, first 60 AAs: 19.52923\nTotal prob of N-in: 0.94351\nPOSSIBLE N-term signal sequence\ninside 1 19\nTMhelix 20 39\noutside 40 325\nTMhelix 326 348\ninside 349 372

2131 GCF_001286145.1_7553_7_62 Escherichia coli Proteobacteria; Gammaproteobacteria; Enterobacterales; Enterobacteriaceae; Escherichia MTGDNTLIHSHGINRRDFMKLCAALAATMGLSSKAAAEMAESVTNPQRPPVIWIGAQECT WP_000145410.1 MULTISPECIES: hydrogenase-2 small chain [Proteobacteria] Length: 372\nNumber of predicted TMHs: 1\nExp number of AAs in TMHs: 23.72226\nExp number, first 60 AAs: 0.22038\nTotal prob of N-in: 0.12994\noutside 1 331\nTMhelix 332 354\ninside 355 372

2132 GCF_000781375.1_ASM78137v1 Escherichia coli Proteobacteria; Gammaproteobacteria; Enterobacterales; Enterobacteriaceae; Escherichia MNNEETFYQAMRRQGVTRRSFLKYCSLAATSLGLGAGMAPKIAWALENKPRIPVVWIHGL WP_001058323.1 MULTISPECIES: hydrogenase-1 small chain [Proteobacteria] Length: 372\nNumber of predicted TMHs: 2\nExp number of AAs in TMHs: 37.2608499999999\nExp number, first 60 AAs: 19.52923\nTotal prob of N-in: 0.94351\nPOSSIBLE N-term signal sequence\ninside 1 19\nTMhelix 20 39\noutside 40 325\nTMhelix 326 348\ninside 349 372

2133 GCF_000781375.1_ASM78137v1 Escherichia coli Proteobacteria; Gammaproteobacteria; Enterobacterales; Enterobacteriaceae; Escherichia MTGDNTLIHSHGINRRDFMKLCAALAATMGLSSKAAAEMAESVTNPQRPPVIWIGAQECT WP_000145410.1 MULTISPECIES: hydrogenase-2 small chain [Proteobacteria] Length: 372\nNumber of predicted TMHs: 1\nExp number of AAs in TMHs: 23.72226\nExp number, first 60 AAs: 0.22038\nTotal prob of N-in: 0.12994\noutside 1 331\nTMhelix 332 354\ninside 355 372

2134 GCF_000780875.1_ASM78087v1 Escherichia coli Proteobacteria; Gammaproteobacteria; Enterobacterales; Enterobacteriaceae; Escherichia MTGDNTLIHSHGINRRDFMKLCAALAATMGLSSKAAAEMAESVTNPQRPPVIWIGAQECT WP_000145410.1 MULTISPECIES: hydrogenase-2 small chain [Proteobacteria] Length: 372\nNumber of predicted TMHs: 1\nExp number of AAs in TMHs: 23.72226\nExp number, first 60 AAs: 0.22038\nTotal prob of N-in: 0.12994\noutside 1 331\nTMhelix 332 354\ninside 355 372

2135 GCF_000780845.1_ASM78084v1 Escherichia coli Proteobacteria; Gammaproteobacteria; Enterobacterales; Enterobacteriaceae; Escherichia
MTGDNTLIHSHGINRRDFMKLCAALAATMGLSSKAAAEMAESVTNPQRPPVIWIGAQECT WP_000145410.1
MULTISPECIES: hydrogenase-2 small chain [Proteobacteria] Length: 372\nNumber of predicted TMHs: 1\nExp number of AAs in TMHs: 23.72226\nExp number, first 60 AAs: 0.22038\nTotal prob of N-in: 0.12994\noutside 1 331\nTMhelix 332 354\ninside 355 372

2136 GCF_000780455.1_ASM78045v1 Escherichia coli Proteobacteria; Gammaproteobacteria; Enterobacterales; Enterobacteriaceae; Escherichia
MNNEETFYQAMRRQGVTRRSFLKYCSLAATSLGLGAGMAPKIAWALENKPRIPVVWIHGL WP_001295940.1
MULTISPECIES: hydrogenase [Enterobacteriaceae] Length: 372\nNumber of predicted TMHs: 2\nExp number of AAs in TMHs: 37.2608899999999\nExp number, first 60 AAs: 19.52923\nTotal prob of N-in: 0.94351\nPOSSIBLE N-term signal sequence\ninside 1 19\nTMhelix 20 39\noutside 40 325\nTMhelix 326 348\ninside 349 372

2137 GCF_000780635.1_ASM78063v1 Escherichia coli Proteobacteria; Gammaproteobacteria; Enterobacterales; Enterobacteriaceae; Escherichia
MNNEETFYQAMRRQGVTRRSFLKYCSLAATSLGLGAGMAPKIAWALENKPRIPVVWIHGL WP_001295940.1
MULTISPECIES: hydrogenase [Enterobacteriaceae] Length: 372\nNumber of predicted TMHs: 2\nExp number of AAs in TMHs: 37.2608899999999\nExp number, first 60 AAs: 19.52923\nTotal prob of N-in: 0.94351\nPOSSIBLE N-term signal sequence\ninside 1 19\nTMhelix 20 39\noutside 40 325\nTMhelix 326 348\ninside 349 372

2138 GCF_000780835.1_ASM78083v1 Escherichia coli Proteobacteria; Gammaproteobacteria; Enterobacterales; Enterobacteriaceae; Escherichia
MNNEETFYQAMRRQGVTRRSFLKYCSLAATSLGLGAGMAPKIAWALENKPRIPVVWIHGL WP_001058323.1
MULTISPECIES: hydrogenase-1 small chain [Proteobacteria] Length: 372\nNumber of predicted TMHs: 2\nExp number of AAs in TMHs: 37.2608499999999\nExp number, first 60 AAs: 19.52923\nTotal prob of N-in: 0.94351\nPOSSIBLE N-term signal sequence\ninside 1 19\nTMhelix 20 39\noutside 40 325\nTMhelix 326 348\ninside 349 372

2139 GCF_001285285.1_8205_3_42 Escherichia coli Proteobacteria; Gammaproteobacteria; Enterobacterales; Enterobacteriaceae; Escherichia
MNNEETFYQAMRRQGVTRRSFLKYCSLAATSLGLGAGMAPKIAWALENKPRIPVVWIHGL WP_001058323.1
MULTISPECIES: hydrogenase-1 small chain [Proteobacteria] Length: 372\nNumber of predicted TMHs: 2\nExp number of AAs in TMHs: 37.2608499999999\nExp number, first 60 AAs: 19.52923\nTotal prob of N-in: 0.94351\nPOSSIBLE N-term signal sequence\ninside 1 19\nTMhelix 20 39\noutside 40 325\nTMhelix 326 348\ninside 349 372

2140 GCF_000780835.1_ASM78083v1 Escherichia coli Proteobacteria; Gammaproteobacteria; Enterobacterales; Enterobacteriaceae; Escherichia
MTGDNTLIHSHGINRRDFMKLCAALAATMGLSSKAAAEMAESVTNPQRPPVIWIGAQECT WP_000145410.1
MULTISPECIES: hydrogenase-2 small chain [Proteobacteria] Length: 372\nNumber of predicted TMHs: 1\nExp number of AAs in TMHs: 23.72226\nExp number, first 60 AAs: 0.22038\nTotal prob of N-in: 0.12994\noutside 1 331\nTMhelix 332 354\ninside 355 372

2141 GCF_000780675.1_ASM78067v1 Escherichia coli Proteobacteria; Gammaproteobacteria; Enterobacterales; Enterobacteriaceae; Escherichia
MNNEETFYQAMRRQGVTRRSFLKYCSLAATSLGLGAGMAPKIAWALENKPRIPVVWIHGL WP_001058323.1
MULTISPECIES: hydrogenase-1 small chain [Proteobacteria] Length: 372\nNumber of predicted TMHs: 2\nExp number of AAs in TMHs: 37.2608499999999\nExp number, first 60 AAs: 19.52923\nTotal prob of N-in: 0.94351\nPOSSIBLE N-term signal sequence\ninside 1 19\nTMhelix 20 39\noutside 40 325\nTMhelix 326 348\ninside 349 372

2142 GCF_000780315.1_ASM78031v1 Escherichia coli Proteobacteria; Gammaproteobacteria; Enterobacterales; Enterobacteriaceae; Escherichia
MNNEETFYQAMRRQGVTRRSFLKYCSLAATSLGLGAGMAPKIAWALENKPRIPVVWIHGL WP_001058323.1
MULTISPECIES: hydrogenase-1 small chain [Proteobacteria] Length: 372\nNumber of predicted TMHs: 2\nExp number of AAs in TMHs: 37.2608499999999\nExp number, first 60 AAs: 19.52923\nTotal prob of N-in: 0.94351\nPOSSIBLE N-term signal sequence\ninside 1 19\nTMhelix 20 39\noutside 40 325\nTMhelix 326 348\ninside 349 372

2143 GCF_000780415.1_ASM78041v1 Escherichia coli Proteobacteria; Gammaproteobacteria; Enterobacterales; Enterobacteriaceae; Escherichia
MTGDNTLIHSHGINRRDFMKLCAALAATMGLSSKAAAEMVESVTNPQRPPVIWIGAQECT WP_000145423.1
hydrogenase 2 small subunit [Escherichia coli] Length: 372\nNumber of predicted TMHs: 1\nExp number of AAs in TMHs: 23.78201\nExp number, first 60 AAs: 0.28355\nTotal prob of N-in: 0.13257\noutside 1 331\nTMhelix 332 354\ninside 355 372

2144 GCF_000780355.1_ASM78035v1 Escherichia coli Proteobacteria; Gammaproteobacteria; Enterobacterales; Enterobacteriaceae; Escherichia
MTGDNTLIHSHGINRRDFMKLCAALAATMGLSSKAAAEMAESVTNPQRPPVIWIGAQECT WP_000145410.1
MULTISPECIES: hydrogenase-2 small chain [Proteobacteria] Length: 372\nNumber of predicted TMHs: 1\nExp number of AAs in TMHs: 23.72226\nExp number, first 60 AAs: 0.22038\nTotal prob of N-in: 0.12994\noutside 1 331\nTMhelix 332 354\ninside 355 372

2145 GCF_000780255.1_ASM78025v1 Escherichia coli Proteobacteria; Gammaproteobacteria; Enterobacterales; Enterobacteriaceae; Escherichia
MTGDNTLIHSHGINRRDFMKLCAALAATMGLSSKAAAEMVESVTNPQRPPVIWIGAQECT WP_000145423.1
hydrogenase 2 small subunit [Escherichia coli] Length: 372\nNumber of predicted TMHs: 1\nExp number of AAs in TMHs: 23.78201\nExp number, first 60 AAs: 0.28355\nTotal prob of N-in: 0.13257\noutside 1 331\nTMhelix 332 354\ninside 355 372

2146 GCF_000780315.1_ASM78031v1 Escherichia coli Proteobacteria; Gammaproteobacteria; Enterobacterales; Enterobacteriaceae; Escherichia
MTGDNTLIHSHGINRRDFMKLCAALAATMGLSSKAAAEMAESVTNPQRPPVIWIGAQECT WP_000145410.1
MULTISPECIES: hydrogenase-2 small chain [Proteobacteria] Length: 372\nNumber of predicted TMHs: 1\nExp number of AAs in TMHs: 23.72226\nExp number, first 60 AAs: 0.22038\nTotal prob of N-in: 0.12994\noutside 1 331\nTMhelix 332 354\ninside 355 372

2147 GCF_000780215.1_ASM78021v1 Escherichia coli Proteobacteria; Gammaproteobacteria; Enterobacterales; Enterobacteriaceae; Escherichia
MTGDNTLIHSHGINRRDFMKLCAALAATMGLSSKAAAEMAESVTNPQRPPVIWIGAQECT WP_000145410.1
MULTISPECIES: hydrogenase-2 small chain [Proteobacteria] Length: 372\nNumber of predicted TMHs: 1\nExp number of AAs in TMHs: 23.72226\nExp number, first 60 AAs: 0.22038\nTotal prob of N-in: 0.12994\noutside 1 331\nTMhelix 332 354\ninside 355 372

2148 GCF_000779775.1_ASM77977v1 Escherichia coli Proteobacteria; Gammaproteobacteria; Enterobacterales; Enterobacteriaceae; Escherichia
MNNEETFYQAMRRQGVTRRSFLKYCSLAATSLGLGAGMAPKIAWALENKPRIPVVWIHGL WP_001295940.1
MULTISPECIES: hydrogenase [Enterobacteriaceae] Length: 372\nNumber of predicted TMHs: 2\nExp number of AAs in TMHs: 37.2608899999999\nExp number, first 60 AAs: 19.52923\nTotal prob of N-in: 0.94351\nPOSSIBLE N-term signal sequence\ninside 1 19\nTMhelix 20 39\noutside 40 325\nTMhelix 326 348\ninside 349 372

2149 GCF_000779995.1_ASM77999v1 Escherichia coli Proteobacteria; Gammaproteobacteria; Enterobacterales; Enterobacteriaceae; Escherichia
MNNEETFYQAMRRQGVTRRSFLKYCSLAATSLGLGAGMAPKIAWALENKPRIPVVWIHGL WP_001058323.1
MULTISPECIES: hydrogenase-1 small chain [Proteobacteria] Length: 372\nNumber of predicted TMHs: 2\nExp number of AAs in TMHs: 37.2608499999999\nExp number, first 60 AAs: 19.52923\nTotal prob of N-in: 0.94351\nPOSSIBLE N-term signal sequence\ninside 1 19\nTMhelix 20 39\noutside 40 325\nTMhelix 326 348\ninside 349 372

2150 GCF_000779995.1_ASM77999v1 Escherichia coli Proteobacteria; Gammaproteobacteria; Enterobacterales; Enterobacteriaceae; Escherichia
MTGDNTLIHSHGINRRDFMKLCAALAATMGLSSKAAAEMAESVTNPQRPPVIWIGAQECT WP_000145410.1
MULTISPECIES: hydrogenase-2 small chain [Proteobacteria] Length: 372\nNumber of predicted TMHs: 1\nExp number of AAs in TMHs: 23.72226\nExp number, first 60 AAs: 0.22038\nTotal prob of N-in: 0.12994\noutside 1 331\nTMhelix 332 354\ninside 355 372

2151 GCF_000779935.1_ASM77993v1 Escherichia coli Proteobacteria; Gammaproteobacteria; Enterobacterales; Enterobacteriaceae; Escherichia
MTGDNTLIHSHGINRRDFMKLCAALAATMGLSSKAAAEMVESVTNPQRPPVIWIGAQECT WP_000145423.1

hydrogenase 2 small subunit [Escherichia coli] Length: 372\nNumber of predicted TMHs: 1\nExp number of AAs in TMHs: 23.78201\nExp number, first 60 AAs: 0.28355\nTotal prob of N-in: 0.13257\noutside 1 331\nTMhelix 332 354\ninside 355 372

2152 GCF_000779615.1_ASM77961v1 Escherichia coli Proteobacteria; Gammaproteobacteria; Enterobacterales; Enterobacteriaceae; Escherichia
MNNEETFYQAMRRQGVTRRSFLKYCSLAATSLGLGAGMAPKIAWALENKPRIPVVWIHGL WP_001058323.1
MULTISPECIES: hydrogenase-1 small chain [Proteobacteria] Length: 372\nNumber of predicted TMHs: 2\nExp number of AAs in TMHs: 37.2608499999999\nExp number, first 60 AAs: 19.52923\nTotal prob of N-in: 0.94351\nPOSSIBLE N-term signal sequence\ninside 1 19\nTMhelix 20 39\noutside 40 325\nTMhelix 326 348\ninside 349 372

2153 GCF_000779835.1_ASM77983v1 Escherichia coli Proteobacteria; Gammaproteobacteria; Enterobacterales; Enterobacteriaceae; Escherichia
MTGDNTLIHSHGINRRDFMKLCAALAATMGLSSKAAAEMAESVTNPQRPPVIWIGAQECT WP_000145418.1
hydrogenase 2 small subunit [Escherichia coli] Length: 372\nNumber of predicted TMHs: 1\nExp number of AAs in TMHs: 23.72313\nExp number, first 60 AAs: 0.22034\nTotal prob of N-in: 0.12997\noutside 1 331\nTMhelix 332 354\ninside 355 372

2154 GCF_000779565.1_ASM77956v1 Escherichia coli Proteobacteria; Gammaproteobacteria; Enterobacterales; Enterobacteriaceae; Escherichia
MNNEETFYQAMRRQGVTRRSFLKYCSLAATSLGLGAGMAPKIAWALENKPRIPVVWIHGL WP_001295940.1
MULTISPECIES: hydrogenase [Enterobacteriaceae] Length: 372\nNumber of predicted TMHs: 2\nExp number of AAs in TMHs: 37.2608899999999\nExp number, first 60 AAs: 19.52923\nTotal prob of N-in: 0.94351\nPOSSIBLE N-term signal sequence\ninside 1 19\nTMhelix 20 39\noutside 40 325\nTMhelix 326 348\ninside 349 372

2155 GCF_000779775.1_ASM77977v1 Escherichia coli Proteobacteria; Gammaproteobacteria; Enterobacterales; Enterobacteriaceae; Escherichia
MTGDNTLIHSHGINRRDFMKLCAALAATMGLSSKAAAEMVESVTNPQRPPVIWIGAQECT WP_000145423.1
hydrogenase 2 small subunit [Escherichia coli] Length: 372\nNumber of predicted TMHs: 1\nExp number of AAs in TMHs: 23.78201\nExp number, first 60 AAs: 0.28355\nTotal prob of N-in: 0.13257\noutside 1 331\nTMhelix 332 354\ninside 355 372

2156 GCF_000779365.1_ASM77936v1 Escherichia coli Proteobacteria; Gammaproteobacteria; Enterobacterales; Enterobacteriaceae; Escherichia
MNNEETFYQAMRRQGVTRRSFLKYCSLAATSLGLGAGMAPKIAWALENKPRIPVVWIHGL WP_001058323.1
MULTISPECIES: hydrogenase-1 small chain [Proteobacteria] Length: 372\nNumber of predicted TMHs: 2\nExp number of AAs in TMHs: 37.2608499999999\nExp number, first 60 AAs: 19.52923\nTotal prob of N-in: 0.94351\nPOSSIBLE N-term signal sequence\ninside 1 19\nTMhelix 20 39\noutside 40 325\nTMhelix 326 348\ninside 349 372

2157 GCF_000779255.1_ASM77925v1 Escherichia coli Proteobacteria; Gammaproteobacteria; Enterobacterales; Enterobacteriaceae; Escherichia
MTGDNTLIHSHGINRRDFMKLCAALAATMGLSSKAAAEMAESVTNPQRPPVIWIGAQECT WP_000145410.1
MULTISPECIES: hydrogenase-2 small chain [Proteobacteria] Length: 372\nNumber of predicted TMHs: 1\nExp number of AAs in TMHs: 23.72226\nExp number, first 60 AAs: 0.22038\nTotal prob of N-in: 0.12994\noutside 1 331\nTMhelix 332 354\ninside 355 372

2158 GCF_000779365.1_ASM77936v1 Escherichia coli Proteobacteria; Gammaproteobacteria; Enterobacterales; Enterobacteriaceae; Escherichia
MTGDNTLIHSHGINRRDFMKLCAALAATMGLSSKAAAEMAESVTNPQRPPVIWIGAQECT WP_000145410.1
MULTISPECIES: hydrogenase-2 small chain [Proteobacteria] Length: 372\nNumber of predicted TMHs: 1\nExp number of AAs in TMHs: 23.72226\nExp number, first 60 AAs: 0.22038\nTotal prob of N-in: 0.12994\noutside 1 331\nTMhelix 332 354\ninside 355 372

2159 GCF_000779175.1_ASM77917v1 Escherichia coli Proteobacteria; Gammaproteobacteria; Enterobacterales; Enterobacteriaceae; Escherichia
MTGDNTLIHSHGINRRDFMKLCAALAATMGLSSKAAAEMAESVTNPQRPPVIWIGAQECT WP_000145410.1
MULTISPECIES: hydrogenase-2 small chain [Proteobacteria] Length: 372\nNumber of predicted TMHs:

1\nExp number of AAs in TMHs: 23.72226\nExp number, first 60 AAs: 0.22038\nTotal prob of N-in: 0.12994\noutside 1 331\nTMhelix 332 354\ninside 355 372

2160 GCF_000779295.1_ASM77929v1 Escherichia coli Proteobacteria; Gammaproteobacteria; Enterobacterales; Enterobacteriaceae; Escherichia
MTGDNTLIHSHGINRRDFMKLCAALAATMGLSSKAAAEMAESVTNPQRPPVIWIGAQECT WP_000145410.1
MULTISPECIES: hydrogenase-2 small chain [Proteobacteria] Length: 372\nNumber of predicted TMHs: 1\nExp number of AAs in TMHs: 23.72226\nExp number, first 60 AAs: 0.22038\nTotal prob of N-in: 0.12994\noutside 1 331\nTMhelix 332 354\ninside 355 372

2161 GCF_000779075.1_ASM77907v1 Escherichia coli Proteobacteria; Gammaproteobacteria; Enterobacterales; Enterobacteriaceae; Escherichia
MTGDNTLIHSHGINRRDFMKLCAALAATMGLSSKAAAEMAESVTNPQRPPVIWIGAQECT WP_000145410.1
MULTISPECIES: hydrogenase-2 small chain [Proteobacteria] Length: 372\nNumber of predicted TMHs: 1\nExp number of AAs in TMHs: 23.72226\nExp number, first 60 AAs: 0.22038\nTotal prob of N-in: 0.12994\noutside 1 331\nTMhelix 332 354\ninside 355 372

2162 GCF_000779095.1_ASM77909v1 Escherichia coli Proteobacteria; Gammaproteobacteria; Enterobacterales; Enterobacteriaceae; Escherichia
MTGDNTLIHSHGINRRDFMKLCAALAATMGLSSKAAAEMAESVTNPQRPPVIWIGAQECT WP_000145410.1
MULTISPECIES: hydrogenase-2 small chain [Proteobacteria] Length: 372\nNumber of predicted TMHs: 1\nExp number of AAs in TMHs: 23.72226\nExp number, first 60 AAs: 0.22038\nTotal prob of N-in: 0.12994\noutside 1 331\nTMhelix 332 354\ninside 355 372

2163 GCF_000778175.1_ASM77817v1 Escherichia coli Proteobacteria; Gammaproteobacteria; Enterobacterales; Enterobacteriaceae; Escherichia
MNNEETFYQAMRRQGVTRRSFLKYCSLAATSLGLGAGMAPKIAWALENKPRIPVVWIHGL WP_001295940.1
MULTISPECIES: hydrogenase [Enterobacteriaceae] Length: 372\nNumber of predicted TMHs: 2\nExp number of AAs in TMHs: 37.2608899999999\nExp number, first 60 AAs: 19.52923\nTotal prob of N-in: 0.94351\nPOSSIBLE N-term signal sequence\ninside 1 19\nTMhelix 20 39\noutside 40 325\nTMhelix 326 348\ninside 349 372

2164 GCF_000778765.1_ASM77876v1 Escherichia coli Proteobacteria; Gammaproteobacteria; Enterobacterales; Enterobacteriaceae; Escherichia
MNNEETFYQAMRRQGVTRRSFLKYCSLAATSLGLGAGMAPKIAWALENKPRIPVVWIHGL WP_001058323.1
MULTISPECIES: hydrogenase-1 small chain [Proteobacteria] Length: 372\nNumber of predicted TMHs: 2\nExp number of AAs in TMHs: 37.2608499999999\nExp number, first 60 AAs: 19.52923\nTotal prob of N-in: 0.94351\nPOSSIBLE N-term signal sequence\ninside 1 19\nTMhelix 20 39\noutside 40 325\nTMhelix 326 348\ninside 349 372

2165 GCF_000778765.1_ASM77876v1 Escherichia coli Proteobacteria; Gammaproteobacteria; Enterobacterales; Enterobacteriaceae; Escherichia
MTGDNTLIHSHGINRRDFMKLCAALAATMGLSSKAAAEMAESVTNPQRPPVIWIGAQECT WP_000145410.1
MULTISPECIES: hydrogenase-2 small chain [Proteobacteria] Length: 372\nNumber of predicted TMHs: 1\nExp number of AAs in TMHs: 23.72226\nExp number, first 60 AAs: 0.22038\nTotal prob of N-in: 0.12994\noutside 1 331\nTMhelix 332 354\ninside 355 372

2166 GCF_000778415.1_ASM77841v1 Escherichia coli Proteobacteria; Gammaproteobacteria; Enterobacterales; Enterobacteriaceae; Escherichia
MNNEETFYQAMRRQGVTRRSFLKYCSLAATSLGLGAGMAPKIAWALENKPRIPVVWIHGL WP_001058323.1
MULTISPECIES: hydrogenase-1 small chain [Proteobacteria] Length: 372\nNumber of predicted TMHs: 2\nExp number of AAs in TMHs: 37.2608499999999\nExp number, first 60 AAs: 19.52923\nTotal prob of N-in: 0.94351\nPOSSIBLE N-term signal sequence\ninside 1 19\nTMhelix 20 39\noutside 40 325\nTMhelix 326 348\ninside 349 372

2167 GCF_000778705.1_ASM77870v1 Escherichia coli Proteobacteria; Gammaproteobacteria; Enterobacterales; Enterobacteriaceae; Escherichia
MNNEETFYQAMRRQGVTRRSFLKYCSLAATSLGLGAGMAPKIAWALENKPRIPVVWIHGL WP_001295940.1
MULTISPECIES: hydrogenase [Enterobacteriaceae] Length: 372\nNumber of predicted TMHs: 2\nExp number of AAs in TMHs: 37.2608899999999\nExp number, first 60 AAs: 19.52923\nTotal prob of N-in:

0.94351\nPOSSIBLE N-term signal sequence\ninside 1 19\nTMhelix 20 39\noutside 40 325\nTMhelix 326 348\ninside 349 372

2168 GCF_000778705.1_ASM77870v1 Escherichia coli Proteobacteria; Gammaproteobacteria; Enterobacterales; Enterobacteriaceae; Escherichia MTGDNTLIHSHGINRRDFMKLCAALAATMGLSSKAAAEMVESVTNPQRPPVIWIGAQECT WP_000145423.1 hydrogenase 2 small subunit [Escherichia coli] Length: 372\nNumber of predicted TMHs: 1\nExp number of AAs in TMHs: 23.78201\nExp number, first 60 AAs: 0.28355\nTotal prob of N-in: 0.13257\noutside 1 331\nTMhelix 332 354\ninside 355 372

2169 GCF_000778315.1_ASM77831v1 Escherichia coli Proteobacteria; Gammaproteobacteria; Enterobacterales; Enterobacteriaceae; Escherichia MNNEETFYQAMRRQGVTRRSFLKYCSLAATSLGLGAGMAPKIAWALENKPRIPVVWIHGL WP_001058323.1 MULTISPECIES: hydrogenase-1 small chain [Proteobacteria] Length: 372\nNumber of predicted TMHs: 2\nExp number of AAs in TMHs: 37.2608499999999\nExp number, first 60 AAs: 19.52923\nTotal prob of N-in: 0.94351\nPOSSIBLE N-term signal sequence\ninside 1 19\nTMhelix 20 39\noutside 40 325\nTMhelix 326 348\ninside 349 372

2170 GCF_000778035.1_ASM77803v1 Escherichia coli Proteobacteria; Gammaproteobacteria; Enterobacterales; Enterobacteriaceae; Escherichia MTGDNTLIHSHGINRRDFMKLCAALAATMGLSSKAAAEMAESVTNPQRPPVIWIGAQECT WP_000145410.1 MULTISPECIES: hydrogenase-2 small chain [Proteobacteria] Length: 372\nNumber of predicted TMHs: 1\nExp number of AAs in TMHs: 23.72226\nExp number, first 60 AAs: 0.22038\nTotal prob of N-in: 0.12994\noutside 1 331\nTMhelix 332 354\ninside 355 372

2171 GCF_000777975.1_ASM77797v1 Escherichia coli Proteobacteria; Gammaproteobacteria; Enterobacterales; Enterobacteriaceae; Escherichia MNNEETFYQAMRRQGVTRRSFLKYCSLAATSLGLGAGMAPKIAWALENKPRIPVVWIHGL WP_001538532.1 hydrogenase-1 small chain [Escherichia coli] Length: 372\nNumber of predicted TMHs: 2\nExp number of AAs in TMHs: 37.27041\nExp number, first 60 AAs: 19.52927\nTotal prob of N-in: 0.94352\nPOSSIBLE N-term signal sequence\ninside 1 19\nTMhelix 20 39\noutside 40 325\nTMhelix 326 348\ninside 349 372

2172 GCF_000778015.1_ASM77801v1 Escherichia coli Proteobacteria; Gammaproteobacteria; Enterobacterales; Enterobacteriaceae; Escherichia MNNEETFYQAMRRQGVTRRSFLKYCSLAATSLGLGAGMAPKIAWALENKPRIPVVWIHGL WP_001538532.1 hydrogenase-1 small chain [Escherichia coli] Length: 372\nNumber of predicted TMHs: 2\nExp number of AAs in TMHs: 37.27041\nExp number, first 60 AAs: 19.52927\nTotal prob of N-in: 0.94352\nPOSSIBLE N-term signal sequence\ninside 1 19\nTMhelix 20 39\noutside 40 325\nTMhelix 326 348\ninside 349 372

2173 GCF_000778015.1_ASM77801v1 Escherichia coli Proteobacteria; Gammaproteobacteria; Enterobacterales; Enterobacteriaceae; Escherichia MTGDNTLIHSHGINRRDFMKLCAALAATMGLSSKAAAEMVESVTNPQRPPVIWIGAQECT WP_000145423.1 hydrogenase 2 small subunit [Escherichia coli] Length: 372\nNumber of predicted TMHs: 1\nExp number of AAs in TMHs: 23.78201\nExp number, first 60 AAs: 0.28355\nTotal prob of N-in: 0.13257\noutside 1 331\nTMhelix 332 354\ninside 355 372

2174 GCF_000777455.1_ASM77745v1 Escherichia coli Proteobacteria; Gammaproteobacteria; Enterobacterales; Enterobacteriaceae; Escherichia MNNEETFYQAMRRQGVTRRSFLKYCSLAATSLGLGAGMAPKIAWALENKPRIPVVWIHGL WP_001058323.1 MULTISPECIES: hydrogenase-1 small chain [Proteobacteria] Length: 372\nNumber of predicted TMHs: 2\nExp number of AAs in TMHs: 37.2608499999999\nExp number, first 60 AAs: 19.52923\nTotal prob of N-in: 0.94351\nPOSSIBLE N-term signal sequence\ninside 1 19\nTMhelix 20 39\noutside 40 325\nTMhelix 326 348\ninside 349 372

2175 GCF_000777845.1_ASM77784v1 Escherichia coli Proteobacteria; Gammaproteobacteria; Enterobacterales; Enterobacteriaceae; Escherichia MNNEETFYQAMRRQGVTRRSFLKYCSLAATSLGLGAGMAPKIAWALENKPRIPVVWIHGL WP_001058323.1 MULTISPECIES: hydrogenase-1 small chain [Proteobacteria] Length: 372\nNumber of predicted TMHs: 2\nExp number of AAs in TMHs: 37.2608499999999\nExp number, first 60 AAs: 19.52923\nTotal prob of N-in: 0.94351\nPOSSIBLE N-term signal sequence\ninside 1 19\nTMhelix 20 39\noutside 40 325\nTMhelix 326 348\ninside 349 372

2176 GCF_000777815.1_ASM77781v1 Escherichia coli Proteobacteria; Gammaproteobacteria; Enterobacterales; Enterobacteriaceae; Escherichia
MNNEETFYQAMRRQGVTRRSFLKYCSLAATSLGLGAGMAPKIAWALENKPRIPVVWIHGL WP_001058323.1
MULTISPECIES: hydrogenase-1 small chain [Proteobacteria] Length: 372\nNumber of predicted TMHs: 2\nExp number of AAs in TMHs: 37.2608499999999\nExp number, first 60 AAs: 19.52923\nTotal prob of N-in: 0.94351\nPOSSIBLE N-term signal sequence\ninside 1 19\nTMhelix 20 39\noutside 40 325\nTMhelix 326 348\ninside 349 372

2177 GCF_000777215.1_ASM77721v1 Escherichia coli Proteobacteria; Gammaproteobacteria; Enterobacterales; Enterobacteriaceae; Escherichia
MTGDNTLIHSHGINRRDFMKLCAALAATMGLSSKAAAEMAESVTNPQRPPVIWIGAQECT WP_000145410.1
MULTISPECIES: hydrogenase-2 small chain [Proteobacteria] Length: 372\nNumber of predicted TMHs: 1\nExp number of AAs in TMHs: 23.72226\nExp number, first 60 AAs: 0.22038\nTotal prob of N-in: 0.12994\noutside 1 331\nTMhelix 332 354\ninside 355 372

2178 GCF_000777105.1_ASM77710v1 Escherichia coli Proteobacteria; Gammaproteobacteria; Enterobacterales; Enterobacteriaceae; Escherichia
MTGDNTLIHSHGINRRDFMKLCAALAATMGLSSKAAAEMVESVTNPQRPPVIWIGAQECT WP_000145423.1
hydrogenase 2 small subunit [Escherichia coli] Length: 372\nNumber of predicted TMHs: 1\nExp number of AAs in TMHs: 23.78201\nExp number, first 60 AAs: 0.28355\nTotal prob of N-in: 0.13257\noutside 1 331\nTMhelix 332 354\ninside 355 372

2179 GCF_000776995.1_ASM77699v1 Escherichia coli Proteobacteria; Gammaproteobacteria; Enterobacterales; Enterobacteriaceae; Escherichia
MTGDNTLIHSHGINRRDFMKLCAALAATMGLSSKAAAEMVESVTNPQRPPVIWIGAQECT WP_000145423.1
hydrogenase 2 small subunit [Escherichia coli] Length: 372\nNumber of predicted TMHs: 1\nExp number of AAs in TMHs: 23.78201\nExp number, first 60 AAs: 0.28355\nTotal prob of N-in: 0.13257\noutside 1 331\nTMhelix 332 354\ninside 355 372

2180 GCF_001285405.1_7790_1_74 Escherichia coli Proteobacteria; Gammaproteobacteria; Enterobacterales; Enterobacteriaceae; Escherichia
MNNEETFYQAMRRQGVTRRSFLKYCSLAATSLGLGAGMAPKIAWALENKPRIPVVWIHGL WP_001295940.1
MULTISPECIES: hydrogenase [Enterobacteriaceae] Length: 372\nNumber of predicted TMHs: 2\nExp number of AAs in TMHs: 37.2608899999999\nExp number, first 60 AAs: 19.52923\nTotal prob of N-in: 0.94351\nPOSSIBLE N-term signal sequence\ninside 1 19\nTMhelix 20 39\noutside 40 325\nTMhelix 326 348\ninside 349 372

2181 GCF_000776925.1_ASM77692v1 Escherichia coli Proteobacteria; Gammaproteobacteria; Enterobacterales; Enterobacteriaceae; Escherichia
MTGDNTLIHSHGINRRDFMKLCAALAATMGLSSKAAAEMAESVTNPQRPPVIWIGAQECT WP_000145410.1
MULTISPECIES: hydrogenase-2 small chain [Proteobacteria] Length: 372\nNumber of predicted TMHs: 1\nExp number of AAs in TMHs: 23.72226\nExp number, first 60 AAs: 0.22038\nTotal prob of N-in: 0.12994\noutside 1 331\nTMhelix 332 354\ninside 355 372

2182 GCF_000776965.1_ASM77696v1 Escherichia coli Proteobacteria; Gammaproteobacteria; Enterobacterales; Enterobacteriaceae; Escherichia
MTGDNTLIHSHGINRRDFMKLCAALAATMGLSSKAAAEMAESVTNPQRPPVIWIGAQECT WP_000145410.1
MULTISPECIES: hydrogenase-2 small chain [Proteobacteria] Length: 372\nNumber of predicted TMHs: 1\nExp number of AAs in TMHs: 23.72226\nExp number, first 60 AAs: 0.22038\nTotal prob of N-in: 0.12994\noutside 1 331\nTMhelix 332 354\ninside 355 372

2183 GCF_000777065.1_ASM77706v1 Escherichia coli Proteobacteria; Gammaproteobacteria; Enterobacterales; Enterobacteriaceae; Escherichia
MTGDNTLIHSHGINRRDFMKLCAALAATMGLSSKAAAEMVESVTNPQRPPVIWIGAQECT WP_000145423.1
hydrogenase 2 small subunit [Escherichia coli] Length: 372\nNumber of predicted TMHs: 1\nExp number of AAs in TMHs: 23.78201\nExp number, first 60 AAs: 0.28355\nTotal prob of N-in: 0.13257\noutside 1 331\nTMhelix 332 354\ninside 355 372

2184 GCF_000776835.1_ASM77683v1 Escherichia coli Proteobacteria; Gammaproteobacteria; Enterobacterales; Enterobacteriaceae; Escherichia
MTGDNTLIHSHGINRRDFMKLCAALAATMGLSSKAAAEMVESVTNPQRPPVIWIGAQECT WP_000145423.1

hydrogenase 2 small subunit [Escherichia coli] Length: 372\nNumber of predicted TMHs: 1\nExp number of AAs in TMHs: 23.78201\nExp number, first 60 AAs: 0.28355\nTotal prob of N-in: 0.13257\noutside 1 331\nTMhelix 332 354\ninside 355 372

2185 GCF_000776795.1_ASM77679v1 Escherichia coli Proteobacteria; Gammaproteobacteria; Enterobacterales; Enterobacteriaceae; Escherichia MTGDNTLIHSHGINRRDFMKLCAALAATMGLSSKAAAEMAESVTNPQRPPVIWIGAQECT WP_000145410.1 MULTISPECIES: hydrogenase-2 small chain [Proteobacteria] Length: 372\nNumber of predicted TMHs: 1\nExp number of AAs in TMHs: 23.72226\nExp number, first 60 AAs: 0.22038\nTotal prob of N-in: 0.12994\noutside 1 331\nTMhelix 332 354\ninside 355 372

2186 GCF_000776355.1_ASM77635v1 Escherichia coli Proteobacteria; Gammaproteobacteria; Enterobacterales; Enterobacteriaceae; Escherichia MNNEETFYQAMRRQGVTRRSFLKYCSLAATSLGLGAGMAPKIAWALENKPRIPVVWIHGL WP_001058323.1 MULTISPECIES: hydrogenase-1 small chain [Proteobacteria] Length: 372\nNumber of predicted TMHs: 2\nExp number of AAs in TMHs: 37.2608499999999\nExp number, first 60 AAs: 19.52923\nTotal prob of N-in: 0.94351\nPOSSIBLE N-term signal sequence\ninside 1 19\nTMhelix 20 39\noutside 40 325\nTMhelix 326 348\ninside 349 372

2187 GCF_000776285.1_ASM77628v1 Escherichia coli Proteobacteria; Gammaproteobacteria; Enterobacterales; Enterobacteriaceae; Escherichia MNNEETFYQAMRRQGVTRRSFLKYCSLAATSLGLGAGMAPKIAWALENKPRIPVVWIHGL WP_001058323.1 MULTISPECIES: hydrogenase-1 small chain [Proteobacteria] Length: 372\nNumber of predicted TMHs: 2\nExp number of AAs in TMHs: 37.2608499999999\nExp number, first 60 AAs: 19.52923\nTotal prob of N-in: 0.94351\nPOSSIBLE N-term signal sequence\ninside 1 19\nTMhelix 20 39\noutside 40 325\nTMhelix 326 348\ninside 349 372

2188 GCF_000776595.1_ASM77659v1 Escherichia coli Proteobacteria; Gammaproteobacteria; Enterobacterales; Enterobacteriaceae; Escherichia MNNEETFYQAMRRQGVTRRSFLKYCSLAATSLGLGAGMAPKIAWALENKPRIPVVWIHGL WP_001295940.1 MULTISPECIES: hydrogenase [Enterobacteriaceae] Length: 372\nNumber of predicted TMHs: 2\nExp number of AAs in TMHs: 37.2608899999999\nExp number, first 60 AAs: 19.52923\nTotal prob of N-in: 0.94351\nPOSSIBLE N-term signal sequence\ninside 1 19\nTMhelix 20 39\noutside 40 325\nTMhelix 326 348\ninside 349 372

2189 GCF_000776255.1_ASM77625v1 Escherichia coli Proteobacteria; Gammaproteobacteria; Enterobacterales; Enterobacteriaceae; Escherichia MTGDNTLIHSHGINRRDFMKLCAALATTMGLSSKAAAEMAESVTNPQRPPVIWIGAQECT WP_023149629.1 MULTISPECIES: hydrogenase-2 small chain [Enterobacteriaceae] Length: 372\nNumber of predicted TMHs: 1\nExp number of AAs in TMHs: 23.63603\nExp number, first 60 AAs: 0.12971\nTotal prob of N-in: 0.12608\noutside 1 331\nTMhelix 332 354\ninside 355 372

2190 GCF_000776635.1_ASM77663v1 Escherichia coli Proteobacteria; Gammaproteobacteria; Enterobacterales; Enterobacteriaceae; Escherichia MNNEETFYQAMRRQGVTRRSFLKYCSLAATSLGLGAGMAPKIAWALENKPRIPVVWIHGL WP_001058323.1 MULTISPECIES: hydrogenase-1 small chain [Proteobacteria] Length: 372\nNumber of predicted TMHs: 2\nExp number of AAs in TMHs: 37.2608499999999\nExp number, first 60 AAs: 19.52923\nTotal prob of N-in: 0.94351\nPOSSIBLE N-term signal sequence\ninside 1 19\nTMhelix 20 39\noutside 40 325\nTMhelix 326 348\ninside 349 372

2191 GCF_001285505.1_7790_1_81 Escherichia coli Proteobacteria; Gammaproteobacteria; Enterobacterales; Enterobacteriaceae; Escherichia MTGDNTLIHSHGINRRDFMKLCAALAATMGLSSKAAAEMAESVTNPQRPPVIWIGAQECT WP_000145410.1 MULTISPECIES: hydrogenase-2 small chain [Proteobacteria] Length: 372\nNumber of predicted TMHs: 1\nExp number of AAs in TMHs: 23.72226\nExp number, first 60 AAs: 0.22038\nTotal prob of N-in: 0.12994\noutside 1 331\nTMhelix 332 354\ninside 355 372

2192 GCF_000776255.1_ASM77625v1 Escherichia coli Proteobacteria; Gammaproteobacteria; Enterobacterales; Enterobacteriaceae; Escherichia MNNEETFYQAMRRQGVTRRSFLKYCSLAATSLGLGAGMAPKIAWALENKPRIPVVWIHGL WP_001058323.1 MULTISPECIES: hydrogenase-1 small chain [Proteobacteria] Length: 372\nNumber of predicted TMHs:

2\nExp number of AAs in TMHs: 37.2608499999999\nExp number, first 60 AAs: 19.52923\nTotal prob of N-in: 0.94351\nPOSSIBLE N-term signal sequence\ninside 1 19\nTMhelix 20 39\noutside 40 325\nTMhelix 326 348\ninside 349 372

2193 GCF_000776635.1_ASM77663v1 Escherichia coli Proteobacteria; Gammaproteobacteria; Enterobacterales; Enterobacteriaceae; Escherichia MTGDNTLIHSHGINRRDFMKLCAALAATMGLSSKAAAEMAESVTNPQRPPVIWIGAQECT WP_000145410.1 MULTISPECIES: hydrogenase-2 small chain [Proteobacteria] Length: 372\nNumber of predicted TMHs: 1\nExp number of AAs in TMHs: 23.72226\nExp number, first 60 AAs: 0.22038\nTotal prob of N-in: 0.12994\noutside 1 331\nTMhelix 332 354\ninside 355 372

2194 GCF_000770035.1_ASM77003v1 Escherichia coli Proteobacteria; Gammaproteobacteria; Enterobacterales; Enterobacteriaceae; Escherichia MNNEETFYQAMRRQGVTRRSFLKYCSLAATSLGLGAGMAPKIAWALENKPRIPVVWIHGL WP_001058323.1 MULTISPECIES: hydrogenase-1 small chain [Proteobacteria] Length: 372\nNumber of predicted TMHs: 2\nExp number of AAs in TMHs: 37.2608499999999\nExp number, first 60 AAs: 19.52923\nTotal prob of N-in: 0.94351\nPOSSIBLE N-term signal sequence\ninside 1 19\nTMhelix 20 39\noutside 40 325\nTMhelix 326 348\ninside 349 372

2195 GCF_000768125.1_ASM76812v1 Escherichia coli Proteobacteria; Gammaproteobacteria; Enterobacterales; Enterobacteriaceae; Escherichia MNNEETFYQAMRRQGVTRRSFLKYCSLAATSLGLGAGMAPKIAWALENKPRIPVVWIHGL WP_001058323.1 MULTISPECIES: hydrogenase-1 small chain [Proteobacteria] Length: 372\nNumber of predicted TMHs: 2\nExp number of AAs in TMHs: 37.2608499999999\nExp number, first 60 AAs: 19.52923\nTotal prob of N-in: 0.94351\nPOSSIBLE N-term signal sequence\ninside 1 19\nTMhelix 20 39\noutside 40 325\nTMhelix 326 348\ninside 349 372

2196 GCF_000768485.1_ASM76848v1 Escherichia coli Proteobacteria; Gammaproteobacteria; Enterobacterales; Enterobacteriaceae; Escherichia MNNEETFYQAMRRQGVTRRSFLKYCSLAATSLGLGAGMAPKIAWALENKPRIPVVWIHGL WP_001058323.1 MULTISPECIES: hydrogenase-1 small chain [Proteobacteria] Length: 372\nNumber of predicted TMHs: 2\nExp number of AAs in TMHs: 37.2608499999999\nExp number, first 60 AAs: 19.52923\nTotal prob of N-in: 0.94351\nPOSSIBLE N-term signal sequence\ninside 1 19\nTMhelix 20 39\noutside 40 325\nTMhelix 326 348\ninside 349 372

2197 GCF_000773575.1_ASM77357v1 Escherichia coli Proteobacteria; Gammaproteobacteria; Enterobacterales; Enterobacteriaceae; Escherichia MNNEETFYQAMRRQGVTRRSFLKYCSLAATSLGLGAGMAPKIAWALENKPRIPVVWIHGL WP_001058323.1 MULTISPECIES: hydrogenase-1 small chain [Proteobacteria] Length: 372\nNumber of predicted TMHs: 2\nExp number of AAs in TMHs: 37.2608499999999\nExp number, first 60 AAs: 19.52923\nTotal prob of N-in: 0.94351\nPOSSIBLE N-term signal sequence\ninside 1 19\nTMhelix 20 39\noutside 40 325\nTMhelix 326 348\ninside 349 372

2198 GCF_000776045.1_ASM77604v1 Escherichia coli Proteobacteria; Gammaproteobacteria; Enterobacterales; Enterobacteriaceae; Escherichia MNNEETFYQAMRRQGVTRRSFLKYCSLAATSLGLGAGMAPKIAWALENKPRIPVVWIHGL WP_001058323.1 MULTISPECIES: hydrogenase-1 small chain [Proteobacteria] Length: 372\nNumber of predicted TMHs: 2\nExp number of AAs in TMHs: 37.2608499999999\nExp number, first 60 AAs: 19.52923\nTotal prob of N-in: 0.94351\nPOSSIBLE N-term signal sequence\ninside 1 19\nTMhelix 20 39\noutside 40 325\nTMhelix 326 348\ninside 349 372

2199 GCF_000754845.1_ASM75484v1 Escherichia coli Proteobacteria; Gammaproteobacteria; Enterobacterales; Enterobacteriaceae; Escherichia MTGDNTLIHSHGINRRDFMKLCAALAATMGLSSKAAAEMAESVTNPQRPPVIWIGAQECT WP_000145410.1 MULTISPECIES: hydrogenase-2 small chain [Proteobacteria] Length: 372\nNumber of predicted TMHs: 1\nExp number of AAs in TMHs: 23.72226\nExp number, first 60 AAs: 0.22038\nTotal prob of N-in: 0.12994\noutside 1 331\nTMhelix 332 354\ninside 355 372

2200 GCF_000753135.1_FHI64 Escherichia coli Proteobacteria; Gammaproteobacteria; Enterobacterales; Enterobacteriaceae; Escherichia MNNEETFYQAMRRQGVTRRSFLKYCSLAATSLGLGAGMAPKIAWALENKPRIPVVWIHGL WP_001058323.1 MULTISPECIES: hydrogenase-1 small chain [Proteobacteria] Length: 372\nNumber of

predicted TMHs: 2\nExp number of AAs in TMHs: 37.2608499999999\nExp number, first 60 AAs: 19.52923\nTotal prob of N-in: 0.94351\nPOSSIBLE N-term signal sequence\ninside 1 19\nTMhelix 20 39\noutside 40 325\nTMhelix 326 348\ninside 349 372

2201 GCF_000752755.1_FHI22 Escherichia coli Proteobacteria; Gammaproteobacteria; Enterobacterales; Enterobacteriaceae; Escherichia MNNEETFYQAMRRQGVTRRSFLKYCSLAATSLGLGAGMAPKIAWALENKPRIPVVWIHGL WP_001058323.1 MULTISPECIES: hydrogenase-1 small chain [Proteobacteria] Length: 372\nNumber of predicted TMHs: 2\nExp number of AAs in TMHs: 37.2608499999999\nExp number, first 60 AAs: 19.52923\nTotal prob of N-in: 0.94351\nPOSSIBLE N-term signal sequence\ninside 1 19\nTMhelix 20 39\noutside 40 325\nTMhelix 326 348\ninside 349 372

2202 GCF_000752815.1_FHI37 Escherichia coli Proteobacteria; Gammaproteobacteria; Enterobacterales; Enterobacteriaceae; Escherichia MNNEETFYQAMRRQGVTRRSFLKYCSLAATSLGLGAGMAPKIAWALENKPRIPVVWIHGL WP_001058323.1 MULTISPECIES: hydrogenase-1 small chain [Proteobacteria] Length: 372\nNumber of predicted TMHs: 2\nExp number of AAs in TMHs: 37.2608499999999\nExp number, first 60 AAs: 19.52923\nTotal prob of N-in: 0.94351\nPOSSIBLE N-term signal sequence\ninside 1 19\nTMhelix 20 39\noutside 40 325\nTMhelix 326 348\ninside 349 372

2203 GCF_000753275.1_FHI95 Escherichia coli Proteobacteria; Gammaproteobacteria; Enterobacterales; Enterobacteriaceae; Escherichia MTGDNTLIHSHGINRRDFMKLCAALAATMGLSSKAAAEMAESVTNPQRPPVIWIGAQECT WP_000145410.1 MULTISPECIES: hydrogenase-2 small chain [Proteobacteria] Length: 372\nNumber of predicted TMHs: 1\nExp number of AAs in TMHs: 23.72226\nExp number, first 60 AAs: 0.22038\nTotal prob of N-in: 0.12994\noutside 1 331\nTMhelix 332 354\ninside 355 372

2204 GCF_000753415.1_FHI93 Escherichia coli Proteobacteria; Gammaproteobacteria; Enterobacterales; Enterobacteriaceae; Escherichia MTGDNTLIHSHGINRRDFMKLCAALAATMGLSSKAAAEMVESVTNPQRPPVIWIGAQECT WP_000145423.1 hydrogenase 2 small subunit [Escherichia coli] Length: 372\nNumber of predicted TMHs: 1\nExp number of AAs in TMHs: 23.78201\nExp number, first 60 AAs: 0.28355\nTotal prob of N-in: 0.13257\noutside 1 331\nTMhelix 332 354\ninside 355 372

2205 GCF_000752895.1_FHI101 Escherichia coli Proteobacteria; Gammaproteobacteria; Enterobacterales; Enterobacteriaceae; Escherichia MNNEETFYQAMRRQGVTRRSFLKYCSLAATSLGLGAGMAPKIAWALENKPRIPVVWIHGL WP_001058323.1 MULTISPECIES: hydrogenase-1 small chain [Proteobacteria] Length: 372\nNumber of predicted TMHs: 2\nExp number of AAs in TMHs: 37.2608499999999\nExp number, first 60 AAs: 19.52923\nTotal prob of N-in: 0.94351\nPOSSIBLE N-term signal sequence\ninside 1 19\nTMhelix 20 39\noutside 40 325\nTMhelix 326 348\ninside 349 372

2206 GCF_000749575.1_ASM74957v1 Escherichia coli Proteobacteria; Gammaproteobacteria; Enterobacterales; Enterobacteriaceae; Escherichia MNNEETFYQAMRRQGVTRRSFLKYCSLAATSLGLGAGMAPKIAWALENKPRIPVVWIHGL WP_001295940.1 MULTISPECIES: hydrogenase [Enterobacteriaceae] Length: 372\nNumber of predicted TMHs: 2\nExp number of AAs in TMHs: 37.2608899999999\nExp number, first 60 AAs: 19.52923\nTotal prob of N-in: 0.94351\nPOSSIBLE N-term signal sequence\ninside 1 19\nTMhelix 20 39\noutside 40 325\nTMhelix 326 348\ninside 349 372

2207 GCF_000749545.1_ASM74954v1 Escherichia coli Proteobacteria; Gammaproteobacteria; Enterobacterales; Enterobacteriaceae; Escherichia MNNEETFYQAMRRQGVTRRSFLKYCSLAATSLGLGAGMAPKIAWALENKPRIPVVWIHGL WP_001295940.1 MULTISPECIES: hydrogenase [Enterobacteriaceae] Length: 372\nNumber of predicted TMHs: 2\nExp number of AAs in TMHs: 37.2608899999999\nExp number, first 60 AAs: 19.52923\nTotal prob of N-in: 0.94351\nPOSSIBLE N-term signal sequence\ninside 1 19\nTMhelix 20 39\noutside 40 325\nTMhelix 326 348\ninside 349 372

2208 GCF_000752295.1_FHI78 Escherichia coli Proteobacteria; Gammaproteobacteria; Enterobacterales; Enterobacteriaceae; Escherichia MNNEETFYQAMRRQGVTRRSFLKYCSLAATSLGLGAGMAPKIAWALENKPRIPVVWIHGL WP_001058323.1 MULTISPECIES: hydrogenase-1 small chain [Proteobacteria] Length: 372\nNumber of predicted TMHs: 2\nExp number of AAs in TMHs: 37.2608499999999\nExp number, first 60 AAs: 19.52923\nTotal prob of N-in: 0.94351\nPOSSIBLE N-term signal sequence\ninside 1 19\nTMhelix 20 39\noutside 40 325\nTMhelix 326 348\ninside 349 372

2209 GCF_000752575.1_FHI81 Escherichia coli Proteobacteria; Gammaproteobacteria; Enterobacterales; Enterobacteriaceae; Escherichia MNNEETFYQAMRRQGVTRRSFLKYCSLAATSLGLGAGMAPKIAWALENKPRIPVVWIHGL WP_001058323.1 MULTISPECIES: hydrogenase-1 small chain [Proteobacteria] Length: 372\nNumber of predicted TMHs: 2\nExp number of AAs in TMHs: 37.2608499999999\nExp number, first 60 AAs: 19.52923\nTotal prob of N-in: 0.94351\nPOSSIBLE N-term signal sequence\ninside 1 19\nTMhelix 20 39\noutside 40 325\nTMhelix 326 348\ninside 349 372

2210 GCF_000752575.1_FHI81 Escherichia coli Proteobacteria; Gammaproteobacteria; Enterobacterales; Enterobacteriaceae; Escherichia MTGDNTLIHSHGINRRDFMKLCAALAATMGLSSKAAAEMAESVTNPQRPPVIWIGAQECT WP_000145410.1 MULTISPECIES: hydrogenase-2 small chain [Proteobacteria] Length: 372\nNumber of predicted TMHs: 1\nExp number of AAs in TMHs: 23.72226\nExp number, first 60 AAs: 0.22038\nTotal prob of N-in: 0.12994\noutside 1 331\nTMhelix 332 354\ninside 355 372

2211 GCF_000752115.1_FHI46 Escherichia coli Proteobacteria; Gammaproteobacteria; Enterobacterales; Enterobacteriaceae; Escherichia MNNEETFYQAMRRQGVTRRSFLKYCSLAATSLGLGAGMAPKIAWALENKPRIPVVWIHGL WP_001058323.1 MULTISPECIES: hydrogenase-1 small chain [Proteobacteria] Length: 372\nNumber of predicted TMHs: 2\nExp number of AAs in TMHs: 37.2608499999999\nExp number, first 60 AAs: 19.52923\nTotal prob of N-in: 0.94351\nPOSSIBLE N-term signal sequence\ninside 1 19\nTMhelix 20 39\noutside 40 325\nTMhelix 326 348\ninside 349 372

2212 GCF_000752195.1_FHI3 Escherichia coli Proteobacteria; Gammaproteobacteria; Enterobacterales; Enterobacteriaceae; Escherichia MNNEETFYQAMRRQGVTRRSFLKYCSLAATSLGLGAGMAPKIAWALENKPRIPVVWIHGL WP_001058323.1 MULTISPECIES: hydrogenase-1 small chain [Proteobacteria] Length: 372\nNumber of predicted TMHs: 2\nExp number of AAs in TMHs: 37.2608499999999\nExp number, first 60 AAs: 19.52923\nTotal prob of N-in: 0.94351\nPOSSIBLE N-term signal sequence\ninside 1 19\nTMhelix 20 39\noutside 40 325\nTMhelix 326 348\ninside 349 372

2213 GCF_001191135.1_CFSAN026793_01-NGE.clc Escherichia coli Proteobacteria; Gammaproteobacteria; Enterobacterales; Enterobacteriaceae; Escherichia MTGDNTLIHSHGINRRDFMKLCAALAATMGLSSKAAAEMAESVTNPQRPPVIWIGAQECT WP_000145410.1 MULTISPECIES: hydrogenase-2 small chain [Proteobacteria] Length: 372\nNumber of predicted TMHs: 1\nExp number of AAs in TMHs: 23.72226\nExp number, first 60 AAs: 0.22038\nTotal prob of N-in: 0.12994\noutside 1 331\nTMhelix 332 354\ninside 355 372

2214 GCF_001191135.1_CFSAN026793_01-NGE.clc Escherichia coli Proteobacteria; Gammaproteobacteria; Enterobacterales; Enterobacteriaceae; Escherichia MNNEETFYQAMRRQGVTRRSFLKYCSLAATSLGLGAGMAPKIAWALENKPRIPVVWIHGL WP_001058323.1 MULTISPECIES: hydrogenase-1 small chain [Proteobacteria] Length: 372\nNumber of predicted TMHs: 2\nExp number of AAs in TMHs: 37.2608499999999\nExp number, first 60 AAs: 19.52923\nTotal prob of N-in: 0.94351\nPOSSIBLE N-term signal sequence\ninside 1 19\nTMhelix 20 39\noutside 40 325\nTMhelix 326 348\ninside 349 372

2215 GCF_001285565.1_8205_3_74 Escherichia coli Proteobacteria; Gammaproteobacteria; Enterobacterales; Enterobacteriaceae; Escherichia MNNEETFYQAMRRQGVTRRSFLKYCSLAATSLGLGAGMAPKIAWALENKPRIPVVWIHGL WP_001058323.1 MULTISPECIES: hydrogenase-1 small chain [Proteobacteria] Length: 372\nNumber of predicted TMHs: 2\nExp number of AAs in TMHs: 37.2608499999999\nExp number, first 60 AAs: 19.52923\nTotal prob of N-in: 0.94351\nPOSSIBLE N-term signal sequence\ninside 1 19\nTMhelix 20 39\noutside 40 325\nTMhelix 326 348\ninside 349 372

2216 GCF_000714995.1_JA69_HGAP2 Escherichia coli Proteobacteria; Gammaproteobacteria; Enterobacterales; Enterobacteriaceae; Escherichia MNNEETFYQAMRRQGVTRRSFLKYCSLAATSLGLGAGMAPKIAWALENKPRIPVVWIHGL WP_001058323.1 MULTISPECIES: hydrogenase-1 small chain [Proteobacteria] Length: 372\nNumber of predicted TMHs: 2\nExp number of AAs in TMHs: 37.2608499999999\nExp number, first 60 AAs: 19.52923\nTotal prob of N-in: 0.94351\nPOSSIBLE N-term signal sequence\ninside 1 19\nTMhelix 20 39\noutside 40 325\nTMhelix 326 348\ninside 349 372

2217 GCF_001057995.1_ASM105799v1 Escherichia coli Proteobacteria; Gammaproteobacteria; Enterobacterales; Enterobacteriaceae; Escherichia MNNEETFYQAMRRQGVTRRSFLKYCSLAATSLGLGAGMAPKIAWALENKPRIPVVWIHGL WP_001058323.1

MULTISPECIES: hydrogenase-1 small chain [Proteobacteria] Length: 372\nNumber of predicted TMHs: 2\nExp number of AAs in TMHs: 37.2608499999999\nExp number, first 60 AAs: 19.52923\nTotal prob of N-in: 0.94351\nPOSSIBLE N-term signal sequence\ninside 1 19\nTMhelix 20 39\noutside 40 325\nTMhelix 326 348\ninside 349 372

2218 GCF_001057995.1_ASM105799v1 Escherichia coli Proteobacteria; Gammaproteobacteria; Enterobacterales; Enterobacteriaceae; Escherichia MTGDNTLIHSHGINRRDFMKLCAALAATMGLSSKAAAEMAESVTNPQRPPVIWIGAQECT WP_000145410.1 MULTISPECIES: hydrogenase-2 small chain [Proteobacteria] Length: 372\nNumber of predicted TMHs: 1\nExp number of AAs in TMHs: 23.72226\nExp number, first 60 AAs: 0.22038\nTotal prob of N-in: 0.12994\noutside 1 331\nTMhelix 332 354\ninside 355 372

2219 GCF_001057125.1_ASM105712v1 Escherichia coli Proteobacteria; Gammaproteobacteria; Enterobacterales; Enterobacteriaceae; Escherichia MNNEETFYQAMRRQGVTRRSFLKYCSLAATSLGLGAGMAPKIAWALENKPRIPVVWIHGL WP_001295940.1 MULTISPECIES: hydrogenase [Enterobacteriaceae] Length: 372\nNumber of predicted TMHs: 2\nExp number of AAs in TMHs: 37.2608899999999\nExp number, first 60 AAs: 19.52923\nTotal prob of N-in: 0.94351\nPOSSIBLE N-term signal sequence\ninside 1 19\nTMhelix 20 39\noutside 40 325\nTMhelix 326 348\ninside 349 372

2220 GCF_001057385.1_ASM105738v1 Escherichia coli Proteobacteria; Gammaproteobacteria; Enterobacterales; Enterobacteriaceae; Escherichia MTGDNTLIHSHGINRRDFMKLCAALAATMGLSSKAAAEMAESVTNPQRPPVIWIGAQECT WP_000145410.1 MULTISPECIES: hydrogenase-2 small chain [Proteobacteria] Length: 372\nNumber of predicted TMHs: 1\nExp number of AAs in TMHs: 23.72226\nExp number, first 60 AAs: 0.22038\nTotal prob of N-in: 0.12994\noutside 1 331\nTMhelix 332 354\ninside 355 372

2221 GCF_001057125.1_ASM105712v1 Escherichia coli Proteobacteria; Gammaproteobacteria; Enterobacterales; Enterobacteriaceae; Escherichia MTGDNTLIHSHGINRRDFMKLCAALAATMGLSSKAAAEMVESVTNPQRPPVIWIGAQECT WP_000145423.1 hydrogenase 2 small subunit [Escherichia coli] Length: 372\nNumber of predicted TMHs: 1\nExp number of AAs in TMHs: 23.78201\nExp number, first 60 AAs: 0.28355\nTotal prob of N-in: 0.13257\noutside 1 331\nTMhelix 332 354\ninside 355 372

2222 GCF_001056965.1_ASM105696v1 Escherichia coli Proteobacteria; Gammaproteobacteria; Enterobacterales; Enterobacteriaceae; Escherichia MNNEETFYQAMRRQGVTRRSFLKYCSLAATSLGLGAGMAPKIAWALENKPRIPVVWIHGL WP_001058323.1 MULTISPECIES: hydrogenase-1 small chain [Proteobacteria] Length: 372\nNumber of predicted TMHs: 2\nExp number of AAs in TMHs: 37.2608499999999\nExp number, first 60 AAs: 19.52923\nTotal prob of N-in: 0.94351\nPOSSIBLE N-term signal sequence\ninside 1 19\nTMhelix 20 39\noutside 40 325\nTMhelix 326 348\ninside 349 372

2223 GCF_001056965.1_ASM105696v1 Escherichia coli Proteobacteria; Gammaproteobacteria; Enterobacterales; Enterobacteriaceae; Escherichia MTGDNTLIHSHGINRRDFMKLCAALAATMGLSSKAAAEMAESVTNPQRPPVIWIGAQECT WP_000145410.1 MULTISPECIES: hydrogenase-2 small chain [Proteobacteria] Length: 372\nNumber of predicted TMHs: 1\nExp number of AAs in TMHs: 23.72226\nExp number, first 60 AAs: 0.22038\nTotal prob of N-in: 0.12994\noutside 1 331\nTMhelix 332 354\ninside 355 372

2224 GCF_001285645.1_8205_3_23 Escherichia coli Proteobacteria; Gammaproteobacteria; Enterobacterales; Enterobacteriaceae; Escherichia MTGDNTLIHSHGINRRDFMKLCAALAATMGLSSKAAAEMAESVTNPQRPPVIWIGAQECT WP_000145410.1 MULTISPECIES: hydrogenase-2 small chain [Proteobacteria] Length: 372\nNumber of predicted TMHs: 1\nExp number of AAs in TMHs: 23.72226\nExp number, first 60 AAs: 0.22038\nTotal prob of N-in: 0.12994\noutside 1 331\nTMhelix 332 354\ninside 355 372

2225 GCF_001191205.1_CFSAN026794_01-NGE.clc Escherichia coli Proteobacteria; Gammaproteobacteria; Enterobacterales; Enterobacteriaceae; Escherichia MTGDNTLIHSHGINRRDFMKLCAALAATMGLSSKAAAEMAESVTNPQRPPVIWIGAQECT WP_000145410.1 MULTISPECIES: hydrogenase-2 small chain [Proteobacteria] Length: 372\nNumber of predicted TMHs:

1\nExp number of AAs in TMHs: 23.72226\nExp number, first 60 AAs: 0.22038\nTotal prob of N-in: 0.12994\noutside 1 331\nTMhelix 332 354\ninside 355 372

2226 GCF_001056875.1_ASM105687v1 Escherichia coli Proteobacteria; Gammaproteobacteria; Enterobacterales; Enterobacteriaceae; Escherichia
MNNEETFYQAMRRQGVTRRSFLKYCSLAATSLGLGAGMAPKIAWALENKPRIPVVWIHGL WP_001058323.1
MULTISPECIES: hydrogenase-1 small chain [Proteobacteria] Length: 372\nNumber of predicted TMHs: 2\nExp number of AAs in TMHs: 37.2608499999999\nExp number, first 60 AAs: 19.52923\nTotal prob of N-in: 0.94351\nPOSSIBLE N-term signal sequence\ninside 1 19\nTMhelix 20 39\noutside 40 325\nTMhelix 326 348\ninside 349 372

2227 GCF_001285745.1_7748_7_35 Escherichia coli Proteobacteria; Gammaproteobacteria; Enterobacterales; Enterobacteriaceae; Escherichia
MTGDNTLIHSHGINRRDFMKLCAALAATMGLSSKAAAEMAESVTNPQRPPVIWIGAQECT WP_000145410.1
MULTISPECIES: hydrogenase-2 small chain [Proteobacteria] Length: 372\nNumber of predicted TMHs: 1\nExp number of AAs in TMHs: 23.72226\nExp number, first 60 AAs: 0.22038\nTotal prob of N-in: 0.12994\noutside 1 331\nTMhelix 332 354\ninside 355 372

2228 GCF_001191205.1_CFSAN026794_01-NGE.clc Escherichia coli Proteobacteria; Gammaproteobacteria; Enterobacterales; Enterobacteriaceae; Escherichia
MNNEETFYQAMRRQGVTRRSFLKYCSLAATSLGLGAGMAPKIAWALENKPRIPVVWIHGL WP_001058323.1
MULTISPECIES: hydrogenase-1 small chain [Proteobacteria] Length: 372\nNumber of predicted TMHs: 2\nExp number of AAs in TMHs: 37.2608499999999\nExp number, first 60 AAs: 19.52923\nTotal prob of N-in: 0.94351\nPOSSIBLE N-term signal sequence\ninside 1 19\nTMhelix 20 39\noutside 40 325\nTMhelix 326 348\ninside 349 372

2229 GCF_000798535.1_Escherichia_coli_CVM_N36837PS_v._1.0 Escherichia coli Proteobacteria; Gammaproteobacteria; Enterobacterales; Enterobacteriaceae; Escherichia
MTGDNTLIHSHGINRRDFMKLCAALAATMGLSSKAAAEMAESVTNPQRPPVIWIGAQECT WP_000145410.1
MULTISPECIES: hydrogenase-2 small chain [Proteobacteria] Length: 372\nNumber of predicted TMHs: 1\nExp number of AAs in TMHs: 23.72226\nExp number, first 60 AAs: 0.22038\nTotal prob of N-in: 0.12994\noutside 1 331\nTMhelix 332 354\ninside 355 372

2230 GCF_000798535.1_Escherichia_coli_CVM_N36837PS_v._1.0 Escherichia coli Proteobacteria; Gammaproteobacteria; Enterobacterales; Enterobacteriaceae; Escherichia
MNNEETFYQAMRRQGVTRRSFLKYCSLAATSLGLGAGMAPKIAWALENKPRIPVVWIHGL WP_001058323.1
MULTISPECIES: hydrogenase-1 small chain [Proteobacteria] Length: 372\nNumber of predicted TMHs: 2\nExp number of AAs in TMHs: 37.2608499999999\nExp number, first 60 AAs: 19.52923\nTotal prob of N-in: 0.94351\nPOSSIBLE N-term signal sequence\ninside 1 19\nTMhelix 20 39\noutside 40 325\nTMhelix 326 348\ninside 349 372

2231 GCF_001268525.1_1.EC2988.1 Escherichia coli Proteobacteria; Gammaproteobacteria; Enterobacterales; Enterobacteriaceae; Escherichia
MNNEETFYQAMRRQGVTRRSFLKYCSLAATSLGLGAGMAPKIAWALENKPRIPVVWIHGL WP_001058323.1
MULTISPECIES: hydrogenase-1 small chain [Proteobacteria] Length: 372\nNumber of predicted TMHs: 2\nExp number of AAs in TMHs: 37.2608499999999\nExp number, first 60 AAs: 19.52923\nTotal prob of N-in: 0.94351\nPOSSIBLE N-term signal sequence\ninside 1 19\nTMhelix 20 39\noutside 40 325\nTMhelix 326 348\ninside 349 372

2232 GCF_000647815.1_ASM64781v3 Escherichia coli Proteobacteria; Gammaproteobacteria; Enterobacterales; Enterobacteriaceae; Escherichia
MNNEETFYQAMRRQGVTRRSFLKYCSLAATSLGLGAGMAPKIAWALENKPRIPVVWIHGL WP_032161483.1
hydrogenase [Escherichia coli] Length: 372\nNumber of predicted TMHs: 2\nExp number of AAs in TMHs: 38.07604\nExp number, first 60 AAs: 19.52977\nTotal prob of N-in: 0.94353\nPOSSIBLE N-term signal sequence\ninside 1 19\nTMhelix 20 39\noutside 40 325\nTMhelix 326 348\ninside 349 372

2233 GCF_000969495.1_ASM96949v1 Escherichia coli Proteobacteria; Gammaproteobacteria; Enterobacterales; Enterobacteriaceae; Escherichia
MTGDNTLIHSHGINRRDFMKLCAALAATMGLSSKAAAEMAESVTNPQRPPVIWIGAQECT WP_000145410.1
MULTISPECIES: hydrogenase-2 small chain [Proteobacteria] Length: 372\nNumber of predicted TMHs:

1\nExp number of AAs in TMHs: 23.72226\nExp number, first 60 AAs: 0.22038\nTotal prob of N-in: 0.12994\noutside 1 331\nTMhelix 332 354\ninside 355 372

2234 GCF_000965705.1_ASM96570v1 Escherichia coli Proteobacteria; Gammaproteobacteria; Enterobacterales; Enterobacteriaceae; Escherichia MNNEETFYQAMRRQGVTRRSFLKYCSLAATSLGLGAGMAPKIAWALENKPRIPVVWIHGL WP_001058323.1 MULTISPECIES: hydrogenase-1 small chain [Proteobacteria] Length: 372\nNumber of predicted TMHs: 2\nExp number of AAs in TMHs: 37.2608499999999\nExp number, first 60 AAs: 19.52923\nTotal prob of N-in: 0.94351\nPOSSIBLE N-term signal sequence\ninside 1 19\nTMhelix 20 39\noutside 40 325\nTMhelix 326 348\ninside 349 372

2235 GCF_000965705.1_ASM96570v1 Escherichia coli Proteobacteria; Gammaproteobacteria; Enterobacterales; Enterobacteriaceae; Escherichia MTGDNTLIHSHGINRRDFMKLCAALAATMGLSSKAAAEMAESVTNPQRPPVIWIGAQECT WP_000145410.1 MULTISPECIES: hydrogenase-2 small chain [Proteobacteria] Length: 372\nNumber of predicted TMHs: 1\nExp number of AAs in TMHs: 23.72226\nExp number, first 60 AAs: 0.22038\nTotal prob of N-in: 0.12994\noutside 1 331\nTMhelix 332 354\ninside 355 372

2236 GCF_000965635.1_ASM96563v1 Escherichia coli Proteobacteria; Gammaproteobacteria; Enterobacterales; Enterobacteriaceae; Escherichia MNNEETFYQAMRRQGVTRRSFLKYCSLAATSLGLGAGMAPKIAWALENKPRIPVVWIHGL WP_044723380.1 hydrogenase [Escherichia coli] Length: 372\nNumber of predicted TMHs: 2\nExp number of AAs in TMHs: 37.2612299999999\nExp number, first 60 AAs: 19.52923\nTotal prob of N-in: 0.94351\nPOSSIBLE N-term signal sequence\ninside 1 19\nTMhelix 20 39\noutside 40 325\nTMhelix 326 348\ninside 349 372

2237 GCF_000965635.1_ASM96563v1 Escherichia coli Proteobacteria; Gammaproteobacteria; Enterobacterales; Enterobacteriaceae; Escherichia MTGDNTLIHSHGINRRDFMKLCAALAATMGLSSKAAAEMAESVTNPQRPPVIWIGAQECT WP_000145410.1 MULTISPECIES: hydrogenase-2 small chain [Proteobacteria] Length: 372\nNumber of predicted TMHs: 1\nExp number of AAs in TMHs: 23.72226\nExp number, first 60 AAs: 0.22038\nTotal prob of N-in: 0.12994\noutside 1 331\nTMhelix 332 354\ninside 355 372

2238 GCF_001076455.1_ASM107645v1 Escherichia coli Proteobacteria; Gammaproteobacteria; Enterobacterales; Enterobacteriaceae; Escherichia MTGDNTLIHSHGINRRDFMKLCAALAATMGLSSKAAAEMAESVTNPQRPPVIWIGAQECT WP_000145410.1 MULTISPECIES: hydrogenase-2 small chain [Proteobacteria] Length: 372\nNumber of predicted TMHs: 1\nExp number of AAs in TMHs: 23.72226\nExp number, first 60 AAs: 0.22038\nTotal prob of N-in: 0.12994\noutside 1 331\nTMhelix 332 354\ninside 355 372

2239 GCF_001076455.1_ASM107645v1 Escherichia coli Proteobacteria; Gammaproteobacteria; Enterobacterales; Enterobacteriaceae; Escherichia MNNEETFYQAMRRQGVTRRSFLKYCSLAATSLGLGAGMAPKIAWALENKPRIPVVWIHGL WP_001058323.1 MULTISPECIES: hydrogenase-1 small chain [Proteobacteria] Length: 372\nNumber of predicted TMHs: 2\nExp number of AAs in TMHs: 37.2608499999999\nExp number, first 60 AAs: 19.52923\nTotal prob of N-in: 0.94351\nPOSSIBLE N-term signal sequence\ninside 1 19\nTMhelix 20 39\noutside 40 325\nTMhelix 326 348\ninside 349 372

2240 GCF_001268425.1_1.EC2993.1 Escherichia coli Proteobacteria; Gammaproteobacteria; Enterobacterales; Enterobacteriaceae; Escherichia MTGDNTLIHSHGINRRDFMKLCAALAATMGLSSKAAAEMAESVTNPQRPPVIWIGAQECT WP_000145410.1 MULTISPECIES: hydrogenase-2 small chain [Proteobacteria] Length: 372\nNumber of predicted TMHs: 1\nExp number of AAs in TMHs: 23.72226\nExp number, first 60 AAs: 0.22038\nTotal prob of N-in: 0.12994\noutside 1 331\nTMhelix 332 354\ninside 355 372

2241 GCF_000690945.1_S7438 Escherichia coli Proteobacteria; Gammaproteobacteria; Enterobacterales; Enterobacteriaceae; Escherichia MTGDNTLIHSHGINRRDFMKLCAALAATMGLSSKAAAEMAESVTNPQRPPVIWIGAQECT WP_000145410.1 MULTISPECIES: hydrogenase-2 small chain [Proteobacteria] Length: 372\nNumber of predicted TMHs: 1\nExp number of AAs in TMHs: 23.72226\nExp number, first 60 AAs: 0.22038\nTotal prob of N-in: 0.12994\noutside 1 331\nTMhelix 332 354\ninside 355 372

2242 GCF_000690945.1_S7438 Escherichia coli Proteobacteria; Gammaproteobacteria; Enterobacterales; Enterobacteriaceae; Escherichia MNNEETFYQAMRRQGVTRRSFLKYCSLAATSLGLGAGMAPKIAWALENKPRIPVVWIHGL WP_001058323.1 MULTISPECIES: hydrogenase-1 small chain [Proteobacteria] Length: 372\nNumber of predicted TMHs: 2\nExp number of AAs in TMHs: 37.2608499999999\nExp number, first 60 AAs: 19.52923\nTotal prob of N-in: 0.94351\nPOSSIBLE N-term signal sequence\ninside 1 19\nTMhelix 20 39\noutside 40 325\nTMhelix 326 348\ninside 349 372

2243 GCF_001285845.1_7790_1_64 Escherichia coli Proteobacteria; Gammaproteobacteria; Enterobacterales; Enterobacteriaceae; Escherichia MTGDNTLIHSHGINRRDFMKLCAALAATMGLSSKAAAEMAESVTNPQRPPVIWIGAQECT WP_000145410.1 MULTISPECIES: hydrogenase-2 small chain [Proteobacteria] Length: 372\nNumber of predicted TMHs: 1\nExp number of AAs in TMHs: 23.72226\nExp number, first 60 AAs: 0.22038\nTotal prob of N-in: 0.12994\noutside 1 331\nTMhelix 332 354\ninside 355 372

2244 GCF_001285845.1_7790_1_64 Escherichia coli Proteobacteria; Gammaproteobacteria; Enterobacterales; Enterobacteriaceae; Escherichia MNNEETFYQAMRRQGVTRRSFLKYCSLAATSLGLGAGMAPKIAWALENKPRIPVVWIHGL WP_001295940.1 MULTISPECIES: hydrogenase [Enterobacteriaceae] Length: 372\nNumber of predicted TMHs: 2\nExp number of AAs in TMHs: 37.2608899999999\nExp number, first 60 AAs: 19.52923\nTotal prob of N-in: 0.94351\nPOSSIBLE N-term signal sequence\ninside 1 19\nTMhelix 20 39\noutside 40 325\nTMhelix 326 348\ninside 349 372

2245 GCF_000951895.1_FHI30 Escherichia coli Proteobacteria; Gammaproteobacteria; Enterobacterales; Enterobacteriaceae; Escherichia MNNEETFYQAMRRQGVTRRSFLKYCSLAATSLGLGAGMAPKIAWALENKPRIPVVWIHGL WP_001058323.1 MULTISPECIES: hydrogenase-1 small chain [Proteobacteria] Length: 372\nNumber of predicted TMHs: 2\nExp number of AAs in TMHs: 37.2608499999999\nExp number, first 60 AAs: 19.52923\nTotal prob of N-in: 0.94351\nPOSSIBLE N-term signal sequence\ninside 1 19\nTMhelix 20 39\noutside 40 325\nTMhelix 326 348\ninside 349 372

2246 GCF_000691045.1_S7380 Escherichia coli Proteobacteria; Gammaproteobacteria; Enterobacterales; Enterobacteriaceae; Escherichia MTGDNTLIHSHGINRRDFMKLCAALAATMGLSSKAAAEMAESVTNPQRPPVIWIGAQECT WP_000145410.1 MULTISPECIES: hydrogenase-2 small chain [Proteobacteria] Length: 372\nNumber of predicted TMHs: 1\nExp number of AAs in TMHs: 23.72226\nExp number, first 60 AAs: 0.22038\nTotal prob of N-in: 0.12994\noutside 1 331\nTMhelix 332 354\ninside 355 372

2247 GCF_001183645.1_ASM118364v1 Escherichia coli Proteobacteria; Gammaproteobacteria; Enterobacterales; Enterobacteriaceae; Escherichia MTGDNTLIHSHGINRRDFMKLCAALAATMGLSSKAAAEMAESVTNPQRPPVIWIGAQECT WP_000145410.1 MULTISPECIES: hydrogenase-2 small chain [Proteobacteria] Length: 372\nNumber of predicted TMHs: 1\nExp number of AAs in TMHs: 23.72226\nExp number, first 60 AAs: 0.22038\nTotal prob of N-in: 0.12994\noutside 1 331\nTMhelix 332 354\ninside 355 372

2248 GCF_001183645.1_ASM118364v1 Escherichia coli Proteobacteria; Gammaproteobacteria; Enterobacterales; Enterobacteriaceae; Escherichia MNNEETFYQAMRRQGVTRRSFLKYCSLAATSLGLGAGMAPKIAWALENKPRIPVVWIHGL WP_001058323.1 MULTISPECIES: hydrogenase-1 small chain [Proteobacteria] Length: 372\nNumber of predicted TMHs: 2\nExp number of AAs in TMHs: 37.2608499999999\nExp number, first 60 AAs: 19.52923\nTotal prob of N-in: 0.94351\nPOSSIBLE N-term signal sequence\ninside 1 19\nTMhelix 20 39\noutside 40 325\nTMhelix 326 348\ninside 349 372

2249 GCF_000696545.1_v1 Escherichia coli Proteobacteria; Gammaproteobacteria; Enterobacterales; Enterobacteriaceae; Escherichia MNNEETFYQAMRRQGVTRRSFLKYCSLAATSLGLGAGMAPKIAWALENKPRIPVVWIHGL WP_001058323.1 MULTISPECIES: hydrogenase-1 small chain [Proteobacteria] Length: 372\nNumber of predicted TMHs: 2\nExp number of AAs in TMHs: 37.2608499999999\nExp number, first 60 AAs: 19.52923\nTotal prob of N-in: 0.94351\nPOSSIBLE N-term signal sequence\ninside 1 19\nTMhelix 20 39\noutside 40 325\nTMhelix 326 348\ninside 349 372

2250 GCF_000948535.1_ASM94853v1 Escherichia coli Proteobacteria; Gammaproteobacteria; Enterobacterales; Enterobacteriaceae; Escherichia MTGDNTLIHSHGINRRDFMKLCAALAATMGLSSKAAAEMAESVTNPQRPPVIWIGAQECT WP_000145410.1 MULTISPECIES: hydrogenase-2 small chain [Proteobacteria] Length: 372\nNumber of predicted TMHs:

1\nExp number of AAs in TMHs: 23.72226\nExp number, first 60 AAs: 0.22038\nTotal prob of N-in: 0.12994\noutside 1 331\nTMhelix 332 354\ninside 355 372

2251 GCF_000800675.1_ASM80067v1 Escherichia coli Proteobacteria; Gammaproteobacteria; Enterobacterales; Enterobacteriaceae; Escherichia
MTGDNTLIHSHGINRRDFMKLCAALAATMGLSSKAAAEMAESVTNPQRPPVIWIGAQECT WP_000145410.1
MULTISPECIES: hydrogenase-2 small chain [Proteobacteria] Length: 372\nNumber of predicted TMHs: 1\nExp number of AAs in TMHs: 23.72226\nExp number, first 60 AAs: 0.22038\nTotal prob of N-in: 0.12994\noutside 1 331\nTMhelix 332 354\ninside 355 372

2252 GCF_001285885.1_7790_1_94 Escherichia coli Proteobacteria; Gammaproteobacteria; Enterobacterales; Enterobacteriaceae; Escherichia
MTGDNTLIHSHGINRRDFMKLCAALAATMGLSSKAAAEMAESVTNPQRPPVIWIGAQECT WP_000145410.1
MULTISPECIES: hydrogenase-2 small chain [Proteobacteria] Length: 372\nNumber of predicted TMHs: 1\nExp number of AAs in TMHs: 23.72226\nExp number, first 60 AAs: 0.22038\nTotal prob of N-in: 0.12994\noutside 1 331\nTMhelix 332 354\ninside 355 372

2253 GCF_001285885.1_7790_1_94 Escherichia coli Proteobacteria; Gammaproteobacteria; Enterobacterales; Enterobacteriaceae; Escherichia
MNNEETFYQAMRRQGVTRRSFLKYCSLAATSLGLGAGMAPKIAWALENKPRIPVVWIHGL WP_001058323.1
MULTISPECIES: hydrogenase-1 small chain [Proteobacteria] Length: 372\nNumber of predicted TMHs: 2\nExp number of AAs in TMHs: 37.2608499999999\nExp number, first 60 AAs: 19.52923\nTotal prob of N-in: 0.94351\nPOSSIBLE N-term signal sequence\ninside 1 19\nTMhelix 20 39\noutside 40 325\nTMhelix 326 348\ninside 349 372

2254 GCF_001285965.1_8205_3_10 Escherichia coli Proteobacteria; Gammaproteobacteria; Enterobacterales; Enterobacteriaceae; Escherichia
MTGDNTLIHSHGINRRDFMKLCAALAATMGLSSKAAAEMAESVTNPQRPPVIWIGAQECT WP_000145410.1
MULTISPECIES: hydrogenase-2 small chain [Proteobacteria] Length: 372\nNumber of predicted TMHs: 1\nExp number of AAs in TMHs: 23.72226\nExp number, first 60 AAs: 0.22038\nTotal prob of N-in: 0.12994\noutside 1 331\nTMhelix 332 354\ninside 355 372

2255 GCF_001285965.1_8205_3_10 Escherichia coli Proteobacteria; Gammaproteobacteria; Enterobacterales; Enterobacteriaceae; Escherichia
MNNEETFYQAMRRQGVTRRSFLKYCSLAATSLGLGAGMAPKIAWALENKPRIPVVWIHGL WP_001058323.1
MULTISPECIES: hydrogenase-1 small chain [Proteobacteria] Length: 372\nNumber of predicted TMHs: 2\nExp number of AAs in TMHs: 37.2608499999999\nExp number, first 60 AAs: 19.52923\nTotal prob of N-in: 0.94351\nPOSSIBLE N-term signal sequence\ninside 1 19\nTMhelix 20 39\noutside 40 325\nTMhelix 326 348\ninside 349 372

2256 GCF_000948675.1_ASM94867v1 Escherichia coli Proteobacteria; Gammaproteobacteria; Enterobacterales; Enterobacteriaceae; Escherichia
MNNEETFYQAMRRQGVTRRSFLKYCSLAATSLGLGAGMAPKIAWALENKPRIPVVWIHGL WP_001058323.1
MULTISPECIES: hydrogenase-1 small chain [Proteobacteria] Length: 372\nNumber of predicted TMHs: 2\nExp number of AAs in TMHs: 37.2608499999999\nExp number, first 60 AAs: 19.52923\nTotal prob of N-in: 0.94351\nPOSSIBLE N-term signal sequence\ninside 1 19\nTMhelix 20 39\noutside 40 325\nTMhelix 326 348\ninside 349 372

2257 GCF_000948895.1_ASM94889v1 Escherichia coli Proteobacteria; Gammaproteobacteria; Enterobacterales; Enterobacteriaceae; Escherichia
MNNEETFYQAMRRQGVTRRSFLKYCSLAATSLGLGAGMAPKIAWALENKPRIPVVWIHGL WP_001301613.1
hydrogenase [Escherichia coli] Length: 372\nNumber of predicted TMHs: 2\nExp number of AAs in TMHs: 37.46598\nExp number, first 60 AAs: 19.53139\nTotal prob of N-in: 0.94364\nPOSSIBLE N-term signal sequence\ninside 1 19\nTMhelix 20 39\noutside 40 325\nTMhelix 326 348\ninside 349 372

2258 GCF_000948895.1_ASM94889v1 Escherichia coli Proteobacteria; Gammaproteobacteria; Enterobacterales; Enterobacteriaceae; Escherichia
MTGDNTLIHSHGINRRDFMKLCAALAATMGLSSKAAAEMAESVTNPQRPPVIWIGAQECT WP_000145410.1
MULTISPECIES: hydrogenase-2 small chain [Proteobacteria] Length: 372\nNumber of predicted TMHs: 1\nExp number of AAs in TMHs: 23.72226\nExp number, first 60 AAs: 0.22038\nTotal prob of N-in: 0.12994\noutside 1 331\nTMhelix 332 354\ninside 355 372

2259 GCF_000948765.1_ASM94876v1 Escherichia coli Proteobacteria; Gammaproteobacteria; Enterobacterales; Enterobacteriaceae; Escherichia
MNNEETFYQAMRRQGVTRRSFLKYCSLAATSLGLGAGMAPKIAWALENKPRIPVVWIHGL WP_001301613.1
hydrogenase [Escherichia coli] Length: 372\nNumber of predicted TMHs: 2\nExp number of AAs in TMHs: 37.46598\nExp number, first 60 AAs: 19.53139\nTotal prob of N-in: 0.94364\nPOSSIBLE N-term signal sequence\ninside 1 19\nTMhelix 20 39\noutside 40 325\nTMhelix 326 348\ninside 349 372

2260 GCF_000948875.1_ASM94887v1 Escherichia coli Proteobacteria; Gammaproteobacteria; Enterobacterales; Enterobacteriaceae; Escherichia
MTGDNTLIHSHGINRRDFMKLCAALAATMGLSSKAAAEMAESVTNPQRPPVIWIGAQECT WP_000145410.1
MULTISPECIES: hydrogenase-2 small chain [Proteobacteria] Length: 372\nNumber of predicted TMHs: 1\nExp number of AAs in TMHs: 23.72226\nExp number, first 60 AAs: 0.22038\nTotal prob of N-in: 0.12994\noutside 1 331\nTMhelix 332 354\ninside 355 372

2261 GCF_001284805.1_7748_7_14 Escherichia coli Proteobacteria; Gammaproteobacteria; Enterobacterales; Enterobacteriaceae; Escherichia
MTGDNTLIHSHGINRRDFMKLCAALAATMGLSSKAAAEMAESVTNPQRPPVIWIGAQECT WP_000145410.1
MULTISPECIES: hydrogenase-2 small chain [Proteobacteria] Length: 372\nNumber of predicted TMHs: 1\nExp number of AAs in TMHs: 23.72226\nExp number, first 60 AAs: 0.22038\nTotal prob of N-in: 0.12994\noutside 1 331\nTMhelix 332 354\ninside 355 372

2262 GCF_000778955.1_ASM77895v1 Escherichia coli Proteobacteria; Gammaproteobacteria; Enterobacterales; Enterobacteriaceae; Escherichia
MNNEETFYQAMRRQGVTRRSFLKYCSLAATSLGLGAGMAPKIAWALENKPRIPVVWIHGL WP_001058323.1
MULTISPECIES: hydrogenase-1 small chain [Proteobacteria] Length: 372\nNumber of predicted TMHs: 2\nExp number of AAs in TMHs: 37.2608499999999\nExp number, first 60 AAs: 19.52923\nTotal prob of N-in: 0.94351\nPOSSIBLE N-term signal sequence\ninside 1 19\nTMhelix 20 39\noutside 40 325\nTMhelix 326 348\ninside 349 372

2263 GCF_000779125.1_ASM77912v1 Escherichia coli Proteobacteria; Gammaproteobacteria; Enterobacterales; Enterobacteriaceae; Escherichia
MTGDNTLIHSHGINRRDFMKLCAALAATMGLSSKAAAEMAESVTNPQRPPVIWIGAQECT WP_000145410.1
MULTISPECIES: hydrogenase-2 small chain [Proteobacteria] Length: 372\nNumber of predicted TMHs: 1\nExp number of AAs in TMHs: 23.72226\nExp number, first 60 AAs: 0.22038\nTotal prob of N-in: 0.12994\noutside 1 331\nTMhelix 332 354\ninside 355 372

2264 GCF_000779005.1_ASM77900v1 Escherichia coli Proteobacteria; Gammaproteobacteria; Enterobacterales; Enterobacteriaceae; Escherichia
MNNEETFYQAMRRQGVTRRSFLKYCSLAATSLGLGAGMAPKIAWALENKPRIPVVWIHGL WP_001058323.1
MULTISPECIES: hydrogenase-1 small chain [Proteobacteria] Length: 372\nNumber of predicted TMHs: 2\nExp number of AAs in TMHs: 37.2608499999999\nExp number, first 60 AAs: 19.52923\nTotal prob of N-in: 0.94351\nPOSSIBLE N-term signal sequence\ninside 1 19\nTMhelix 20 39\noutside 40 325\nTMhelix 326 348\ninside 349 372

2265 GCF_000779215.1_ASM77921v1 Escherichia coli Proteobacteria; Gammaproteobacteria; Enterobacterales; Enterobacteriaceae; Escherichia
MTGDNTLIHSHGINRRDFMKLCAALAATMGLSSKAAAEMAESVTNPQRPPVIWIGAQECT WP_000145418.1
hydrogenase 2 small subunit [Escherichia coli] Length: 372\nNumber of predicted TMHs: 1\nExp number of AAs in TMHs: 23.72313\nExp number, first 60 AAs: 0.22034\nTotal prob of N-in: 0.12997\noutside 1 331\nTMhelix 332 354\ninside 355 372

2266 GCF_000779155.1_ASM77915v1 Escherichia coli Proteobacteria; Gammaproteobacteria; Enterobacterales; Enterobacteriaceae; Escherichia
MTGDNTLIHSHGINRRDFMKLCAALAATMGLSSKAAAEMAESVTNPQRPPVIWIGAQECT WP_000145410.1
MULTISPECIES: hydrogenase-2 small chain [Proteobacteria] Length: 372\nNumber of predicted TMHs: 1\nExp number of AAs in TMHs: 23.72226\nExp number, first 60 AAs: 0.22038\nTotal prob of N-in: 0.12994\noutside 1 331\nTMhelix 332 354\ninside 355 372

2267 GCF_000779655.1_ASM77965v1 Escherichia coli Proteobacteria; Gammaproteobacteria; Enterobacterales; Enterobacteriaceae; Escherichia
MTGDNTLIHSHGINRRDFMKLCAALAATMGLSSKAAAEMAESVTNPQRPPVIWIGAQECT WP_000145410.1

MULTISPECIES: hydrogenase-2 small chain [Proteobacteria] Length: 372\nNumber of predicted TMHs: 1\nExp number of AAs in TMHs: 23.72226\nExp number, first 60 AAs: 0.22038\nTotal prob of N-in: 0.12994\noutside 1 331\nTMhelix 332 354\ninside 355 372

2268 GCF_000779655.1_ASM77965v1 Escherichia coli Proteobacteria; Gammaproteobacteria; Enterobacterales; Enterobacteriaceae; Escherichia
MNNEETFYQAMRRQGVTRRSFLKYCSLAATSLGLGAGMAPKIAWALENKPRIPVVWIHGL WP_001295940.1
MULTISPECIES: hydrogenase [Enterobacteriaceae] Length: 372\nNumber of predicted TMHs: 2\nExp number of AAs in TMHs: 37.2608899999999\nExp number, first 60 AAs: 19.52923\nTotal prob of N-in: 0.94351\nPOSSIBLE N-term signal sequence\ninside 1 19\nTMhelix 20 39\noutside 40 325\nTMhelix 326 348\ninside 349 372

2269 GCF_000779545.1_ASM77954v1 Escherichia coli Proteobacteria; Gammaproteobacteria; Enterobacterales; Enterobacteriaceae; Escherichia
MNNEETFYQAMRRQGVTRRSFLKYCSLAATSLGLGAGMAPKIAWALENKPRIPVVWIHGL WP_001058323.1
MULTISPECIES: hydrogenase-1 small chain [Proteobacteria] Length: 372\nNumber of predicted TMHs: 2\nExp number of AAs in TMHs: 37.2608499999999\nExp number, first 60 AAs: 19.52923\nTotal prob of N-in: 0.94351\nPOSSIBLE N-term signal sequence\ninside 1 19\nTMhelix 20 39\noutside 40 325\nTMhelix 326 348\ninside 349 372

2270 GCF_000779735.1_ASM77973v1 Escherichia coli Proteobacteria; Gammaproteobacteria; Enterobacterales; Enterobacteriaceae; Escherichia
MTGDNTLIHSHGINRRDFMKLCAALAATMGLSSKAAAEMAESVTNPQRPPVIWIGAQECT WP_000145410.1
MULTISPECIES: hydrogenase-2 small chain [Proteobacteria] Length: 372\nNumber of predicted TMHs: 1\nExp number of AAs in TMHs: 23.72226\nExp number, first 60 AAs: 0.22038\nTotal prob of N-in: 0.12994\noutside 1 331\nTMhelix 332 354\ninside 355 372

2271 GCF_000779695.1_ASM77969v1 Escherichia coli Proteobacteria; Gammaproteobacteria; Enterobacterales; Enterobacteriaceae; Escherichia
MTGDNTLIHSHGINRRDFMKLCAALAATMGLSSKAAAEMAESVTNPQRPPVIWIGAQECT WP_000145410.1
MULTISPECIES: hydrogenase-2 small chain [Proteobacteria] Length: 372\nNumber of predicted TMHs: 1\nExp number of AAs in TMHs: 23.72226\nExp number, first 60 AAs: 0.22038\nTotal prob of N-in: 0.12994\noutside 1 331\nTMhelix 332 354\ninside 355 372

2272 GCF_000779585.1_ASM77958v1 Escherichia coli Proteobacteria; Gammaproteobacteria; Enterobacterales; Enterobacteriaceae; Escherichia
MNNEETFYQAMRRQGVTRRSFLKYCSLAATSLGLGAGMAPKIAWALENKPRIPVVWIHGL WP_001058323.1
MULTISPECIES: hydrogenase-1 small chain [Proteobacteria] Length: 372\nNumber of predicted TMHs: 2\nExp number of AAs in TMHs: 37.2608499999999\nExp number, first 60 AAs: 19.52923\nTotal prob of N-in: 0.94351\nPOSSIBLE N-term signal sequence\ninside 1 19\nTMhelix 20 39\noutside 40 325\nTMhelix 326 348\ninside 349 372

2273 GCF_000779745.1_ASM77974v1 Escherichia coli Proteobacteria; Gammaproteobacteria; Enterobacterales; Enterobacteriaceae; Escherichia
MTGDNTLIHSHGINRRDFMKLCAALAATMGLSSKAAAEMVESVTNPQRPPVIWIGAQECT WP_000145423.1
hydrogenase 2 small subunit [Escherichia coli] Length: 372\nNumber of predicted TMHs: 1\nExp number of AAs in TMHs: 23.78201\nExp number, first 60 AAs: 0.28355\nTotal prob of N-in: 0.13257\noutside 1 331\nTMhelix 332 354\ninside 355 372

2274 GCF_000780375.1_ASM78037v1 Escherichia coli Proteobacteria; Gammaproteobacteria; Enterobacterales; Enterobacteriaceae; Escherichia
MNNEETFYQAMRRQGVTRRSFLKYCSLAATSLGLGAGMAPKIAWALENKPRIPVVWIHGL WP_001295940.1
MULTISPECIES: hydrogenase [Enterobacteriaceae] Length: 372\nNumber of predicted TMHs: 2\nExp number of AAs in TMHs: 37.2608899999999\nExp number, first 60 AAs: 19.52923\nTotal prob of N-in: 0.94351\nPOSSIBLE N-term signal sequence\ninside 1 19\nTMhelix 20 39\noutside 40 325\nTMhelix 326 348\ninside 349 372

2275 GCF_000780395.1_ASM78039v1 Escherichia coli Proteobacteria; Gammaproteobacteria; Enterobacterales; Enterobacteriaceae; Escherichia
MNNEETFYQAMRRQGVTRRSFLKYCSLAATSLGLGAGMAPKIAWALENKPRIPVVWIHGL WP_001295940.1
MULTISPECIES: hydrogenase [Enterobacteriaceae] Length: 372\nNumber of predicted TMHs: 2\nExp

number of AAs in TMHs: 37.2608899999999\nExp number, first 60 AAs: 19.52923\nTotal prob of N-in: 0.94351\nPOSSIBLE N-term signal sequence\ninside 1 19\nTMhelix 20 39\noutside 40 325\nTMhelix 326 348\ninside 349 372

2276 GCF_000780235.1_ASM78023v1 Escherichia coli Proteobacteria; Gammaproteobacteria; Enterobacterales; Enterobacteriaceae; Escherichia
MNNEETFYQAMRRQGVTRRSFLKYCSLAATSLGLGAGMAPKIAWALENKPRIPVVWIHGL WP_001058323.1
MULTISPECIES: hydrogenase-1 small chain [Proteobacteria] Length: 372\nNumber of predicted TMHs: 2\nExp number of AAs in TMHs: 37.2608499999999\nExp number, first 60 AAs: 19.52923\nTotal prob of N-in: 0.94351\nPOSSIBLE N-term signal sequence\ninside 1 19\nTMhelix 20 39\noutside 40 325\nTMhelix 326 348\ninside 349 372

2277 GCF_000780175.1_ASM78017v1 Escherichia coli Proteobacteria; Gammaproteobacteria; Enterobacterales; Enterobacteriaceae; Escherichia
MNNEETFYQAMRRQGVTRRSFLKYCSLAATSLGLGAGMAPKIAWALENKPRIPVVWIHGL WP_001295940.1
MULTISPECIES: hydrogenase [Enterobacteriaceae] Length: 372\nNumber of predicted TMHs: 2\nExp number of AAs in TMHs: 37.2608899999999\nExp number, first 60 AAs: 19.52923\nTotal prob of N-in: 0.94351\nPOSSIBLE N-term signal sequence\ninside 1 19\nTMhelix 20 39\noutside 40 325\nTMhelix 326 348\ninside 349 372

2278 GCF_000780435.1_ASM78043v1 Escherichia coli Proteobacteria; Gammaproteobacteria; Enterobacterales; Enterobacteriaceae; Escherichia
MTGDNTLIHSHGINRRDFMKLCAALAATMGLSSKAAAEMVESVTNPQRPPVIWIGAQECT WP_000145423.1
hydrogenase 2 small subunit [Escherichia coli] Length: 372\nNumber of predicted TMHs: 1\nExp number of AAs in TMHs: 23.78201\nExp number, first 60 AAs: 0.28355\nTotal prob of N-in: 0.13257\noutside 1 331\nTMhelix 332 354\ninside 355 372

2279 GCF_000780435.1_ASM78043v1 Escherichia coli Proteobacteria; Gammaproteobacteria; Enterobacterales; Enterobacteriaceae; Escherichia
MNNEETFYQAMRRQGVTRRSFLKYCSLAATSLGLGAGMAPKIAWALENKPRIPVVWIHGL WP_001295940.1
MULTISPECIES: hydrogenase [Enterobacteriaceae] Length: 372\nNumber of predicted TMHs: 2\nExp number of AAs in TMHs: 37.2608899999999\nExp number, first 60 AAs: 19.52923\nTotal prob of N-in: 0.94351\nPOSSIBLE N-term signal sequence\ninside 1 19\nTMhelix 20 39\noutside 40 325\nTMhelix 326 348\ninside 349 372

2280 GCF_000780295.1_ASM78029v1 Escherichia coli Proteobacteria; Gammaproteobacteria; Enterobacterales; Enterobacteriaceae; Escherichia
MNNEETFYQAMRRQGVTRRSFLKYCSLAATSLGLGAGMAPKIAWALENKPRIPVVWIHGL WP_001058323.1
MULTISPECIES: hydrogenase-1 small chain [Proteobacteria] Length: 372\nNumber of predicted TMHs: 2\nExp number of AAs in TMHs: 37.2608499999999\nExp number, first 60 AAs: 19.52923\nTotal prob of N-in: 0.94351\nPOSSIBLE N-term signal sequence\ninside 1 19\nTMhelix 20 39\noutside 40 325\nTMhelix 326 348\ninside 349 372

2281 GCF_000780615.1_ASM78061v1 Escherichia coli Proteobacteria; Gammaproteobacteria; Enterobacterales; Enterobacteriaceae; Escherichia
MNNEETFYQAMRRQGVTRRSFLKYCSLAATSLGLGAGMAPKIAWALENKPRIPVVWIHGL WP_001295940.1
MULTISPECIES: hydrogenase [Enterobacteriaceae] Length: 372\nNumber of predicted TMHs: 2\nExp number of AAs in TMHs: 37.2608899999999\nExp number, first 60 AAs: 19.52923\nTotal prob of N-in: 0.94351\nPOSSIBLE N-term signal sequence\ninside 1 19\nTMhelix 20 39\noutside 40 325\nTMhelix 326 348\ninside 349 372

2282 GCF_000780915.1_ASM78091v1 Escherichia coli Proteobacteria; Gammaproteobacteria; Enterobacterales; Enterobacteriaceae; Escherichia
MTGDNTLIHSHGINRRDFMKLCAALAATMGLSSKAAAEMAESVTNPQRPPVIWIGAQECT WP_000145418.1
hydrogenase 2 small subunit [Escherichia coli] Length: 372\nNumber of predicted TMHs: 1\nExp number of AAs in TMHs: 23.72313\nExp number, first 60 AAs: 0.22034\nTotal prob of N-in: 0.12997\noutside 1 331\nTMhelix 332 354\ninside 355 372

2283 GCF_000780895.1_ASM78089v1 Escherichia coli Proteobacteria; Gammaproteobacteria; Enterobacterales; Enterobacteriaceae; Escherichia
MTGDNTLIHSHGINRRDFMKLCAALAATMGLSSKAAAEMAESVTNPQRPPVIWIGAQECT WP_000145410.1

MULTISPECIES: hydrogenase-2 small chain [Proteobacteria] Length: 372\nNumber of predicted TMHs: 1\nExp number of AAs in TMHs: 23.72226\nExp number, first 60 AAs: 0.22038\nTotal prob of N-in: 0.12994\noutside 1 331\nTMhelix 332 354\ninside 355 372

2284 GCF_000780965.1_ASM78096v1 Escherichia coli Proteobacteria; Gammaproteobacteria; Enterobacterales; Enterobacteriaceae; Escherichia
MTGDNTLIHSHGINRRDFMKLCAALAATMGLSSKAAAEMAESVTNPQRPPVIWIGAQECT WP_000145410.1
MULTISPECIES: hydrogenase-2 small chain [Proteobacteria] Length: 372\nNumber of predicted TMHs: 1\nExp number of AAs in TMHs: 23.72226\nExp number, first 60 AAs: 0.22038\nTotal prob of N-in: 0.12994\noutside 1 331\nTMhelix 332 354\ninside 355 372

2285 GCF_000781005.1_ASM78100v1 Escherichia coli Proteobacteria; Gammaproteobacteria; Enterobacterales; Enterobacteriaceae; Escherichia
MTGDNTLIHSHGINRRDFMKLCAALAATMGLSSKAAAEMVESVTNPQRPPVIWIGAQECT WP_000145423.1
hydrogenase 2 small subunit [Escherichia coli] Length: 372\nNumber of predicted TMHs: 1\nExp number of AAs in TMHs: 23.78201\nExp number, first 60 AAs: 0.28355\nTotal prob of N-in: 0.13257\noutside 1 331\nTMhelix 332 354\ninside 355 372

2286 GCF_000781035.1_ASM78103v1 Escherichia coli Proteobacteria; Gammaproteobacteria; Enterobacterales; Enterobacteriaceae; Escherichia
MTGDNTLIHSHGINRRDFMKLCAALAATMGLSSKAAAEMAESVTNPQRPPVIWIGAQECT WP_000145410.1
MULTISPECIES: hydrogenase-2 small chain [Proteobacteria] Length: 372\nNumber of predicted TMHs: 1\nExp number of AAs in TMHs: 23.72226\nExp number, first 60 AAs: 0.22038\nTotal prob of N-in: 0.12994\noutside 1 331\nTMhelix 332 354\ninside 355 372

2287 GCF_000780915.1_ASM78091v1 Escherichia coli Proteobacteria; Gammaproteobacteria; Enterobacterales; Enterobacteriaceae; Escherichia
MNNEETFYQAMRRQGVTRRSFLKYCSLAATSLGLGAGMAPKIAWALENKPRIPVVWIHGL WP_001058323.1
MULTISPECIES: hydrogenase-1 small chain [Proteobacteria] Length: 372\nNumber of predicted TMHs: 2\nExp number of AAs in TMHs: 37.2608499999999\nExp number, first 60 AAs: 19.52923\nTotal prob of N-in: 0.94351\nPOSSIBLE N-term signal sequence\ninside 1 19\nTMhelix 20 39\noutside 40 325\nTMhelix 326 348\ninside 349 372

2288 GCF_000781575.1_ASM78157v1 Escherichia coli Proteobacteria; Gammaproteobacteria; Enterobacterales; Enterobacteriaceae; Escherichia
MTGDNTLIHSHGINRRDFMKLCAALAATMGLSSKAAAEMAESVTNPQRPPVIWIGAQECT WP_000145410.1
MULTISPECIES: hydrogenase-2 small chain [Proteobacteria] Length: 372\nNumber of predicted TMHs: 1\nExp number of AAs in TMHs: 23.72226\nExp number, first 60 AAs: 0.22038\nTotal prob of N-in: 0.12994\noutside 1 331\nTMhelix 332 354\ninside 355 372

2289 GCF_000781335.1_ASM78133v1 Escherichia coli Proteobacteria; Gammaproteobacteria; Enterobacterales; Enterobacteriaceae; Escherichia
MNNEETFYQAMRRQGVTRRSFLKYCSLAATSLGLGAGMAPKIAWALENKPRIPVVWIHGL WP_001295940.1
MULTISPECIES: hydrogenase [Enterobacteriaceae] Length: 372\nNumber of predicted TMHs: 2\nExp number of AAs in TMHs: 37.2608899999999\nExp number, first 60 AAs: 19.52923\nTotal prob of N-in: 0.94351\nPOSSIBLE N-term signal sequence\ninside 1 19\nTMhelix 20 39\noutside 40 325\nTMhelix 326 348\ninside 349 372

2290 GCF_000781605.1_ASM78160v1 Escherichia coli Proteobacteria; Gammaproteobacteria; Enterobacterales; Enterobacteriaceae; Escherichia
MTGDNTLIHSHGINRRDFMKLCAALAATMGLSSKAAAEMAESVTNPQRPPVIWIGAQECT WP_000145410.1
MULTISPECIES: hydrogenase-2 small chain [Proteobacteria] Length: 372\nNumber of predicted TMHs: 1\nExp number of AAs in TMHs: 23.72226\nExp number, first 60 AAs: 0.22038\nTotal prob of N-in: 0.12994\noutside 1 331\nTMhelix 332 354\ninside 355 372

2291 GCF_000781655.1_ASM78165v1 Escherichia coli Proteobacteria; Gammaproteobacteria; Enterobacterales; Enterobacteriaceae; Escherichia
MTGDNTLIHSHGINRRDFMKLCAALAATMGLSSKAAAEMAESVTNPQRPPVIWIGAQECT WP_000145410.1
MULTISPECIES: hydrogenase-2 small chain [Proteobacteria] Length: 372\nNumber of predicted TMHs: 1\nExp number of AAs in TMHs: 23.72226\nExp number, first 60 AAs: 0.22038\nTotal prob of N-in: 0.12994\noutside 1 331\nTMhelix 332 354\ninside 355 372

2292 GCF_000781675.1_ASM78167v1 Escherichia coli Proteobacteria; Gammaproteobacteria; Enterobacterales; Enterobacteriaceae; Escherichia
MTGDNTLIHSHGINRRDFMKLCAALAATMGLSSKAAAEMAESVTNPQRPPVIWIGAQECT WP_000145410.1
MULTISPECIES: hydrogenase-2 small chain [Proteobacteria] Length: 372\nNumber of predicted TMHs: 1\nExp number of AAs in TMHs: 23.72226\nExp number, first 60 AAs: 0.22038\nTotal prob of N-in: 0.12994\noutside 1 331\nTMhelix 332 354\ninside 355 372

2293 GCF_000781755.1_ASM78175v1 Escherichia coli Proteobacteria; Gammaproteobacteria; Enterobacterales; Enterobacteriaceae; Escherichia
MTGDNTLIHSHGINRRDFMKLCAALAATMGLSSKAAAEMAESVTNPQRPPVIWIGAQECT WP_000145410.1
MULTISPECIES: hydrogenase-2 small chain [Proteobacteria] Length: 372\nNumber of predicted TMHs: 1\nExp number of AAs in TMHs: 23.72226\nExp number, first 60 AAs: 0.22038\nTotal prob of N-in: 0.12994\noutside 1 331\nTMhelix 332 354\ninside 355 372

2294 GCF_000781805.1_ASM78180v1 Escherichia coli Proteobacteria; Gammaproteobacteria; Enterobacterales; Enterobacteriaceae; Escherichia
MTGDNTLIHSHGINRRDFMKLCAALAATMGLSSKAAAEMAESVTNPQRPPVIWIGAQECT WP_000145410.1
MULTISPECIES: hydrogenase-2 small chain [Proteobacteria] Length: 372\nNumber of predicted TMHs: 1\nExp number of AAs in TMHs: 23.72226\nExp number, first 60 AAs: 0.22038\nTotal prob of N-in: 0.12994\noutside 1 331\nTMhelix 332 354\ninside 355 372

2295 GCF_000781935.1_ASM78193v1 Escherichia coli Proteobacteria; Gammaproteobacteria; Enterobacterales; Enterobacteriaceae; Escherichia
MTGDNTLIHSHGINRRDFMKLCAALAATMGLSSKAAAEMAESVTNPQRPPVIWIGAQECT WP_000145410.1
MULTISPECIES: hydrogenase-2 small chain [Proteobacteria] Length: 372\nNumber of predicted TMHs: 1\nExp number of AAs in TMHs: 23.72226\nExp number, first 60 AAs: 0.22038\nTotal prob of N-in: 0.12994\noutside 1 331\nTMhelix 332 354\ninside 355 372

2296 GCF_000782135.1_ASM78213v1 Escherichia coli Proteobacteria; Gammaproteobacteria; Enterobacterales; Enterobacteriaceae; Escherichia
MTGDNTLIHSHGINRRDFMKLCAALAATMGLSSKAAAEMVESVTNPQRPPVIWIGAQECT WP_000145423.1
hydrogenase 2 small subunit [Escherichia coli] Length: 372\nNumber of predicted TMHs: 1\nExp number of AAs in TMHs: 23.78201\nExp number, first 60 AAs: 0.28355\nTotal prob of N-in: 0.13257\noutside 1 331\nTMhelix 332 354\ninside 355 372

2297 GCF_000782175.1_ASM78217v1 Escherichia coli Proteobacteria; Gammaproteobacteria; Enterobacterales; Enterobacteriaceae; Escherichia
MTGDNTLIHSHGINRRDFMKLCAALAATMGLSSKAAAEMVESVTNPQRPPVIWIGAQECT WP_000145423.1
hydrogenase 2 small subunit [Escherichia coli] Length: 372\nNumber of predicted TMHs: 1\nExp number of AAs in TMHs: 23.78201\nExp number, first 60 AAs: 0.28355\nTotal prob of N-in: 0.13257\noutside 1 331\nTMhelix 332 354\ninside 355 372

2298 GCF_000782195.1_ASM78219v1 Escherichia coli Proteobacteria; Gammaproteobacteria; Enterobacterales; Enterobacteriaceae; Escherichia
MTGDNTLIHSHGINRRDFMKLCAALAATMGLSSKAAAEMVESVTNPQRPPVIWIGAQECT WP_000145423.1
hydrogenase 2 small subunit [Escherichia coli] Length: 372\nNumber of predicted TMHs: 1\nExp number of AAs in TMHs: 23.78201\nExp number, first 60 AAs: 0.28355\nTotal prob of N-in: 0.13257\noutside 1 331\nTMhelix 332 354\ninside 355 372

2299 GCF_000782095.1_ASM78209v1 Escherichia coli Proteobacteria; Gammaproteobacteria; Enterobacterales; Enterobacteriaceae; Escherichia
MNNEETFYQAMRRQGVTRRSFLKYCSLAATSLGLGAGMAPKIAWALENKPRIPVVWIHGL WP_001058323.1
MULTISPECIES: hydrogenase-1 small chain [Proteobacteria] Length: 372\nNumber of predicted TMHs: 2\nExp number of AAs in TMHs: 37.2608499999999\nExp number, first 60 AAs: 19.52923\nTotal prob of N-in: 0.94351\nPOSSIBLE N-term signal sequence\ninside 1 19\nTMhelix 20 39\noutside 40 325\nTMhelix 326 348\ninside 349 372

2300 GCF_000782235.1_ASM78223v1 Escherichia coli Proteobacteria; Gammaproteobacteria; Enterobacterales; Enterobacteriaceae; Escherichia
MTGDNTLIHSHGINRRDFMKLCAALAATMGLSSKAAAEMAESVTNPQRPPVIWIGAQECT WP_000145410.1
MULTISPECIES: hydrogenase-2 small chain [Proteobacteria] Length: 372\nNumber of predicted TMHs:

1\nExp number of AAs in TMHs: 23.72226\nExp number, first 60 AAs: 0.22038\nTotal prob of N-in: 0.12994\noutside 1 331\nTMhelix 332 354\ninside 355 372

2301 GCF_000782335.1_ASM78233v1 Escherichia coli Proteobacteria; Gammaproteobacteria; Enterobacterales; Enterobacteriaceae; Escherichia
MTGDNTLIHSHGINRRDFMKLCAALAATMGLSSKAAAEMAESVTNPQRPPVIWIGAQECT WP_000145410.1
MULTISPECIES: hydrogenase-2 small chain [Proteobacteria] Length: 372\nNumber of predicted TMHs: 1\nExp number of AAs in TMHs: 23.72226\nExp number, first 60 AAs: 0.22038\nTotal prob of N-in: 0.12994\noutside 1 331\nTMhelix 332 354\ninside 355 372

2302 GCF_000782295.1_ASM78229v1 Escherichia coli Proteobacteria; Gammaproteobacteria; Enterobacterales; Enterobacteriaceae; Escherichia
MTGDNTLIHSHGINRRDFMKLCAALAATMGLSSKAAAEMVESVTNPQRPPVIWIGAQECT WP_000145423.1
hydrogenase 2 small subunit [Escherichia coli] Length: 372\nNumber of predicted TMHs: 1\nExp number of AAs in TMHs: 23.78201\nExp number, first 60 AAs: 0.28355\nTotal prob of N-in: 0.13257\noutside 1 331\nTMhelix 332 354\ninside 355 372

2303 GCF_000782335.1_ASM78233v1 Escherichia coli Proteobacteria; Gammaproteobacteria; Enterobacterales; Enterobacteriaceae; Escherichia
MNNEETFYQAMRRQGVTRRSFLKYCSLAATSLGLGAGMAPKIAWALENKPRIPVVWIHGL WP_001295940.1
MULTISPECIES: hydrogenase [Enterobacteriaceae] Length: 372\nNumber of predicted TMHs: 2\nExp number of AAs in TMHs: 37.2608899999999\nExp number, first 60 AAs: 19.52923\nTotal prob of N-in: 0.94351\nPOSSIBLE N-term signal sequence\ninside 1 19\nTMhelix 20 39\noutside 40 325\nTMhelix 326 348\ninside 349 372

2304 GCF_000782355.1_ASM78235v1 Escherichia coli Proteobacteria; Gammaproteobacteria; Enterobacterales; Enterobacteriaceae; Escherichia
MNNEETFYQAMRRQGVTRRSFLKYCSLAATSLGLGAGMAPKIAWALENKPRIPVVWIHGL WP_001295940.1
MULTISPECIES: hydrogenase [Enterobacteriaceae] Length: 372\nNumber of predicted TMHs: 2\nExp number of AAs in TMHs: 37.2608899999999\nExp number, first 60 AAs: 19.52923\nTotal prob of N-in: 0.94351\nPOSSIBLE N-term signal sequence\ninside 1 19\nTMhelix 20 39\noutside 40 325\nTMhelix 326 348\ninside 349 372

2305 GCF_000782635.1_ASM78263v1 Escherichia coli Proteobacteria; Gammaproteobacteria; Enterobacterales; Enterobacteriaceae; Escherichia
MTGDNTLIHSHGINRRDFMKLCAALAATMGLSSKAAAEMVESVTNPQRPPVIWIGAQECT WP_000145423.1
hydrogenase 2 small subunit [Escherichia coli] Length: 372\nNumber of predicted TMHs: 1\nExp number of AAs in TMHs: 23.78201\nExp number, first 60 AAs: 0.28355\nTotal prob of N-in: 0.13257\noutside 1 331\nTMhelix 332 354\ninside 355 372

2306 GCF_000782615.1_ASM78261v1 Escherichia coli Proteobacteria; Gammaproteobacteria; Enterobacterales; Enterobacteriaceae; Escherichia
MTGDNTLIHSHGINRRDFMKLCAALAATMGLSSKAAAEMVESVTNPQRPPVIWIGAQECT WP_000145423.1
hydrogenase 2 small subunit [Escherichia coli] Length: 372\nNumber of predicted TMHs: 1\nExp number of AAs in TMHs: 23.78201\nExp number, first 60 AAs: 0.28355\nTotal prob of N-in: 0.13257\noutside 1 331\nTMhelix 332 354\ninside 355 372

2307 GCF_000782675.1_ASM78267v1 Escherichia coli Proteobacteria; Gammaproteobacteria; Enterobacterales; Enterobacteriaceae; Escherichia
MTGDNTLIHSHGINRRDFMKLCAALAATMGLSSKAAAEMVESVTNPQRPPVIWIGAQECT WP_000145423.1
hydrogenase 2 small subunit [Escherichia coli] Length: 372\nNumber of predicted TMHs: 1\nExp number of AAs in TMHs: 23.78201\nExp number, first 60 AAs: 0.28355\nTotal prob of N-in: 0.13257\noutside 1 331\nTMhelix 332 354\ninside 355 372

2308 GCF_000782735.1_ASM78273v1 Escherichia coli Proteobacteria; Gammaproteobacteria; Enterobacterales; Enterobacteriaceae; Escherichia
MTGDNTLIHSHGINRRDFMKLCAALAATMGLSSKAAAEMAESVTNPQRPPVIWIGAQECT WP_000145410.1
MULTISPECIES: hydrogenase-2 small chain [Proteobacteria] Length: 372\nNumber of predicted TMHs: 1\nExp number of AAs in TMHs: 23.72226\nExp number, first 60 AAs: 0.22038\nTotal prob of N-in: 0.12994\noutside 1 331\nTMhelix 332 354\ninside 355 372

2309 GCF_000782715.1_ASM78271v1 Escherichia coli Proteobacteria; Gammaproteobacteria; Enterobacterales; Enterobacteriaceae; Escherichia
MTGDNTLIHSHGINRRDFMKLCAALAATMGLSSKAAAEMVESVTNPQRPPVIWIGAQECT WP_000145423.1
hydrogenase 2 small subunit [Escherichia coli] Length: 372\nNumber of predicted TMHs: 1\nExp number of AAs in TMHs: 23.78201\nExp number, first 60 AAs: 0.28355\nTotal prob of N-in: 0.13257\noutside 1 331\nTMhelix 332 354\ninside 355 372

2310 GCF_000797655.1_Escherichia_coli_CVM_N36401PS_v._1.0 Escherichia coli Proteobacteria; Gammaproteobacteria; Enterobacterales; Enterobacteriaceae; Escherichia
MTGDNTLIHSHGINRRDFMKLCAALAATMGLSSKAAAEMAESVTNPQRPPVIWIGAQECT WP_000145410.1
MULTISPECIES: hydrogenase-2 small chain [Proteobacteria] Length: 372\nNumber of predicted TMHs: 1\nExp number of AAs in TMHs: 23.72226\nExp number, first 60 AAs: 0.22038\nTotal prob of N-in: 0.12994\noutside 1 331\nTMhelix 332 354\ninside 355 372

2311 GCF_000797715.1_Escherichia_coli_CVM_N38795PS_v._1.0 Escherichia coli Proteobacteria; Gammaproteobacteria; Enterobacterales; Enterobacteriaceae; Escherichia
MTGDNTLIHSHGINRRDFMKLCAALAATMGLSSKAAAEMAESVTNPQRPPVIWIGAQECT WP_000145410.1
MULTISPECIES: hydrogenase-2 small chain [Proteobacteria] Length: 372\nNumber of predicted TMHs: 1\nExp number of AAs in TMHs: 23.72226\nExp number, first 60 AAs: 0.22038\nTotal prob of N-in: 0.12994\noutside 1 331\nTMhelix 332 354\ninside 355 372

2312 GCF_000797675.1_Escherichia_coli_CVM_N38663PS_v._1.0 Escherichia coli Proteobacteria; Gammaproteobacteria; Enterobacterales; Enterobacteriaceae; Escherichia
MTGDNTLIHSHGINRRDFMKLCAALAATMGLSSKAAAEMAESVTNPQRPPVIWIGAQECT WP_000145410.1
MULTISPECIES: hydrogenase-2 small chain [Proteobacteria] Length: 372\nNumber of predicted TMHs: 1\nExp number of AAs in TMHs: 23.72226\nExp number, first 60 AAs: 0.22038\nTotal prob of N-in: 0.12994\noutside 1 331\nTMhelix 332 354\ninside 355 372

2313 GCF_000797735.1_Escherichia_coli_CVM_N33552PS_v._1.0 Escherichia coli Proteobacteria; Gammaproteobacteria; Enterobacterales; Enterobacteriaceae; Escherichia
MTGDNTLIHSHGINRRDFMKLCAALAATMGLSSKAAAEMAESVTNPQRPPVIWIGAQECT WP_000145410.1
MULTISPECIES: hydrogenase-2 small chain [Proteobacteria] Length: 372\nNumber of predicted TMHs: 1\nExp number of AAs in TMHs: 23.72226\nExp number, first 60 AAs: 0.22038\nTotal prob of N-in: 0.12994\noutside 1 331\nTMhelix 332 354\ninside 355 372

2314 GCF_000797735.1_Escherichia_coli_CVM_N33552PS_v._1.0 Escherichia coli Proteobacteria; Gammaproteobacteria; Enterobacterales; Enterobacteriaceae; Escherichia
MNNEETFYQAMRRQGVTRRSFLKYCSLAATSLGLGAGMAPKIAWALENKPRIPVVWIHGL WP_001058323.1
MULTISPECIES: hydrogenase-1 small chain [Proteobacteria] Length: 372\nNumber of predicted TMHs: 2\nExp number of AAs in TMHs: 37.2608499999999\nExp number, first 60 AAs: 19.52923\nTotal prob of N-in: 0.94351\nPOSSIBLE N-term signal sequence\ninside 1 19\nTMhelix 20 39\noutside 40 325\nTMhelix 326 348\ninside 349 372

2315 GCF_000797715.1_Escherichia_coli_CVM_N38795PS_v._1.0 Escherichia coli Proteobacteria; Gammaproteobacteria; Enterobacterales; Enterobacteriaceae; Escherichia
MNNEETFYQAMRRQGVTRRSFLKYCSLAATSLGLGAGMAPKIAWALENKPRIPVVWIHGL WP_001058323.1
MULTISPECIES: hydrogenase-1 small chain [Proteobacteria] Length: 372\nNumber of predicted TMHs: 2\nExp number of AAs in TMHs: 37.2608499999999\nExp number, first 60 AAs: 19.52923\nTotal prob of N-in: 0.94351\nPOSSIBLE N-term signal sequence\ninside 1 19\nTMhelix 20 39\noutside 40 325\nTMhelix 326 348\ninside 349 372

2316 GCF_000798035.1_Escherichia_coli_CVM_N34352PS_v._1.0 Escherichia coli Proteobacteria; Gammaproteobacteria; Enterobacterales; Enterobacteriaceae; Escherichia
MTGDNTLIHSHGINRRDFMKLCAALAATMGLSSKAAAEMAESVTNPQRPPVIWIGAQECT WP_000145410.1
MULTISPECIES: hydrogenase-2 small chain [Proteobacteria] Length: 372\nNumber of predicted TMHs: 1\nExp number of AAs in TMHs: 23.72226\nExp number, first 60 AAs: 0.22038\nTotal prob of N-in: 0.12994\noutside 1 331\nTMhelix 332 354\ninside 355 372

2317 GCF_000798035.1_Escherichia_coli_CVM_N34352PS_v._1.0 Escherichia coli Proteobacteria; Gammaproteobacteria; Enterobacterales; Enterobacteriaceae; Escherichia
MNNEETFYQAMRRQGVTRRSFLKYCSLAATSLGLGAGMAPKIAWALENKPRIPVVWIHGL WP_001058323.1

MULTISPECIES: hydrogenase-1 small chain [Proteobacteria] Length: 372\nNumber of predicted TMHs: 2\nExp number of AAs in TMHs: 37.2608499999999\nExp number, first 60 AAs: 19.52923\nTotal prob of N-in: 0.94351\nPOSSIBLE N-term signal sequence\ninside 1 19\nTMhelix 20 39\noutside 40 325\nTMhelix 326 348\ninside 349 372

2318 GCF_000797955.1_Escherichia_coli_CVM_N36400PS_v._1.0 Escherichia coli Proteobacteria; Gammaproteobacteria; Enterobacterales; Enterobacteriaceae; Escherichia
MNNEETFYQAMRRQGVTRRSFLKYCSLAATSLGLGAGMAPKIAWALENKPRIPVVWIHGL WP_001058323.1
MULTISPECIES: hydrogenase-1 small chain [Proteobacteria] Length: 372\nNumber of predicted TMHs: 2\nExp number of AAs in TMHs: 37.2608499999999\nExp number, first 60 AAs: 19.52923\nTotal prob of N-in: 0.94351\nPOSSIBLE N-term signal sequence\ninside 1 19\nTMhelix 20 39\noutside 40 325\nTMhelix 326 348\ninside 349 372

2319 GCF_000797855.1_Escherichia_coli_CVM_N33806PS_v._1.0 Escherichia coli Proteobacteria; Gammaproteobacteria; Enterobacterales; Enterobacteriaceae; Escherichia
MNNEETFYQAMRRQGVTRRSFLKYCSLAATSLGLGAGMAPKIAWALENKPRIPVVWIHGL WP_001295940.1
MULTISPECIES: hydrogenase [Enterobacteriaceae] Length: 372\nNumber of predicted TMHs: 2\nExp number of AAs in TMHs: 37.2608899999999\nExp number, first 60 AAs: 19.52923\nTotal prob of N-in: 0.94351\nPOSSIBLE N-term signal sequence\ninside 1 19\nTMhelix 20 39\noutside 40 325\nTMhelix 326 348\ninside 349 372

2320 GCF_000798015.1_Escherichia_coli_CVM_N34351PS_v._1.0 Escherichia coli Proteobacteria; Gammaproteobacteria; Enterobacterales; Enterobacteriaceae; Escherichia
MTGDNTLIHSHGINRRDFMKLCAALAATMGLSSKAAAEMAESVTNPQRPPVIWIGAQECT WP_000145410.1
MULTISPECIES: hydrogenase-2 small chain [Proteobacteria] Length: 372\nNumber of predicted TMHs: 1\nExp number of AAs in TMHs: 23.72226\nExp number, first 60 AAs: 0.22038\nTotal prob of N-in: 0.12994\noutside 1 331\nTMhelix 332 354\ninside 355 372

2321 GCF_000798115.1_Escherichia_coli_CVM_N36393PS_v._1.0 Escherichia coli Proteobacteria; Gammaproteobacteria; Enterobacterales; Enterobacteriaceae; Escherichia
MTGDNTLIHSHGINRRDFMKLCAALAATMGLSSKAAAEMAESVTNPQRPPVIWIGAQECT WP_000145410.1
MULTISPECIES: hydrogenase-2 small chain [Proteobacteria] Length: 372\nNumber of predicted TMHs: 1\nExp number of AAs in TMHs: 23.72226\nExp number, first 60 AAs: 0.22038\nTotal prob of N-in: 0.12994\noutside 1 331\nTMhelix 332 354\ninside 355 372

2322 GCF_000798015.1_Escherichia_coli_CVM_N34351PS_v._1.0 Escherichia coli Proteobacteria; Gammaproteobacteria; Enterobacterales; Enterobacteriaceae; Escherichia
MNNEETFYQAMRRQGVTRRSFLKYCSLAATSLGLGAGMAPKIAWALENKPRIPVVWIHGL WP_001058323.1
MULTISPECIES: hydrogenase-1 small chain [Proteobacteria] Length: 372\nNumber of predicted TMHs: 2\nExp number of AAs in TMHs: 37.2608499999999\nExp number, first 60 AAs: 19.52923\nTotal prob of N-in: 0.94351\nPOSSIBLE N-term signal sequence\ninside 1 19\nTMhelix 20 39\noutside 40 325\nTMhelix 326 348\ninside 349 372

2323 GCF_000798235.1_Escherichia_coli_CVM_N36834PS_v._1.0 Escherichia coli Proteobacteria; Gammaproteobacteria; Enterobacterales; Enterobacteriaceae; Escherichia
MNNEETFYQAMRRQGVTRRSFLKYCSLAATSLGLGAGMAPKIAWALENKPRIPVVWIHGL WP_001295940.1
MULTISPECIES: hydrogenase [Enterobacteriaceae] Length: 372\nNumber of predicted TMHs: 2\nExp number of AAs in TMHs: 37.2608899999999\nExp number, first 60 AAs: 19.52923\nTotal prob of N-in: 0.94351\nPOSSIBLE N-term signal sequence\ninside 1 19\nTMhelix 20 39\noutside 40 325\nTMhelix 326 348\ninside 349 372

2324 GCF_000798415.1_Escherichia_coli_CVM_N34557PS_v._1.0 Escherichia coli Proteobacteria; Gammaproteobacteria; Enterobacterales; Enterobacteriaceae; Escherichia
MTGDNTLIHSHGINRRDFMKLCAALAATMGLSSKAAAEMAESVTNPQRPPVIWIGAQECT WP_000145410.1
MULTISPECIES: hydrogenase-2 small chain [Proteobacteria] Length: 372\nNumber of predicted TMHs: 1\nExp number of AAs in TMHs: 23.72226\nExp number, first 60 AAs: 0.22038\nTotal prob of N-in: 0.12994\noutside 1 331\nTMhelix 332 354\ninside 355 372

2325 GCF_000798455.1_Escherichia_coli_CVM_N36158PS_v._1.0 Escherichia coli Proteobacteria; Gammaproteobacteria; Enterobacterales; Enterobacteriaceae; Escherichia
MTGDNTLIHSHGINRRDFMKLCAALAATMGLSSKAAAEMAESVTNPQRPPVIWIGAQECT WP_000145410.1

MULTISPECIES: hydrogenase-2 small chain [Proteobacteria] Length: 372\nNumber of predicted TMHs: 1\nExp number of AAs in TMHs: 23.72226\nExp number, first 60 AAs: 0.22038\nTotal prob of N-in: 0.12994\noutside 1 331\nTMhelix 332 354\ninside 355 372

2326 GCF_000798495.1_Escherichia_coli_CVM_N36918PS_v._1.0 Escherichia coli Proteobacteria; Gammaproteobacteria; Enterobacterales; Enterobacteriaceae; Escherichia MTGDNTLIHSHGINRRDFMKLCAALAATMGLSSKAAAEMAESVTNPQRPPVIWIGAQECT WP_000145410.1 MULTISPECIES: hydrogenase-2 small chain [Proteobacteria] Length: 372\nNumber of predicted TMHs: 1\nExp number of AAs in TMHs: 23.72226\nExp number, first 60 AAs: 0.22038\nTotal prob of N-in: 0.12994\noutside 1 331\nTMhelix 332 354\ninside 355 372

2327 GCF_000798515.1_Escherichia_coli_CVM_N33561PS_v._1.0 Escherichia coli Proteobacteria; Gammaproteobacteria; Enterobacterales; Enterobacteriaceae; Escherichia MTGDNTLIHSHGINRRDFMKLCAALAATMGLSSKAAAEMAESVTNPQRPPVIWIGAQECT WP_000145410.1 MULTISPECIES: hydrogenase-2 small chain [Proteobacteria] Length: 372\nNumber of predicted TMHs: 1\nExp number of AAs in TMHs: 23.72226\nExp number, first 60 AAs: 0.22038\nTotal prob of N-in: 0.12994\noutside 1 331\nTMhelix 332 354\ninside 355 372

2328 GCF_000798575.1_Escherichia_coli_CVM_N36404PS_v._1.0 Escherichia coli Proteobacteria; Gammaproteobacteria; Enterobacterales; Enterobacteriaceae; Escherichia MTGDNTLIHSHGINRRDFMKLCAALAATMGLSSKAAAEMAESVTNPQRPPVIWIGAQECT WP_000145410.1 MULTISPECIES: hydrogenase-2 small chain [Proteobacteria] Length: 372\nNumber of predicted TMHs: 1\nExp number of AAs in TMHs: 23.72226\nExp number, first 60 AAs: 0.22038\nTotal prob of N-in: 0.12994\noutside 1 331\nTMhelix 332 354\ninside 355 372

2329 GCF_000798415.1_Escherichia_coli_CVM_N34557PS_v._1.0 Escherichia coli Proteobacteria; Gammaproteobacteria; Enterobacterales; Enterobacteriaceae; Escherichia MNNEETFYQAMRRQGVTRRSFLKYCSLAATSLGLGAGMAPKIAWALENKPRIPVVWIHGL WP_001058323.1 MULTISPECIES: hydrogenase-1 small chain [Proteobacteria] Length: 372\nNumber of predicted TMHs: 2\nExp number of AAs in TMHs: 37.2608499999999\nExp number, first 60 AAs: 19.52923\nTotal prob of N-in: 0.94351\nPOSSIBLE N-term signal sequence\ninside 1 19\nTMhelix 20 39\noutside 40 325\nTMhelix 326 348\ninside 349 372

2330 GCF_000805835.1_ASM80583v1 Escherichia coli Proteobacteria; Gammaproteobacteria; Enterobacterales; Enterobacteriaceae; Escherichia MTGDNTLIHSHGINRRDFMKLCAALAATMGLSSKAAAEMAESVTNPQRPPVIWIGAQECT WP_038339805.1 hydrogenase 2 small subunit [Escherichia coli] Length: 372\nNumber of predicted TMHs: 1\nExp number of AAs in TMHs: 23.86829\nExp number, first 60 AAs: 0.22037\nTotal prob of N-in: 0.12535\noutside 1 332\nTMhelix 333 355\ninside 356 372

2331 GCF_000807565.1_ASM80756v1 Escherichia coli Proteobacteria; Gammaproteobacteria; Enterobacterales; Enterobacteriaceae; Escherichia MTGDNTLIHSHGINRRDFMKLCAALAATMGLSSKAAAEMAESVTNPQRPPVIWIGAQECT WP_000145410.1 MULTISPECIES: hydrogenase-2 small chain [Proteobacteria] Length: 372\nNumber of predicted TMHs: 1\nExp number of AAs in TMHs: 23.72226\nExp number, first 60 AAs: 0.22038\nTotal prob of N-in: 0.12994\noutside 1 331\nTMhelix 332 354\ninside 355 372

2332 GCF_000807565.1_ASM80756v1 Escherichia coli Proteobacteria; Gammaproteobacteria; Enterobacterales; Enterobacteriaceae; Escherichia MNNEETFYQAMRRQGVTRRSFLKYCSLAATSLGLGAGMAPKIAWALENKPRIPVVWIHGL WP_001058323.1 MULTISPECIES: hydrogenase-1 small chain [Proteobacteria] Length: 372\nNumber of predicted TMHs: 2\nExp number of AAs in TMHs: 37.2608499999999\nExp number, first 60 AAs: 19.52923\nTotal prob of N-in: 0.94351\nPOSSIBLE N-term signal sequence\ninside 1 19\nTMhelix 20 39\noutside 40 325\nTMhelix 326 348\ninside 349 372

2333 GCF_000804325.1_ASM80432v1 Escherichia coli Proteobacteria; Gammaproteobacteria; Enterobacterales; Enterobacteriaceae; Escherichia MNNEETFYQAMRRQGVTRRSFLKYCSLAATSLGLGAGMAPKIAWALENKPRIPVVWIHGL WP_032161483.1 hydrogenase [Escherichia coli] Length: 372\nNumber of predicted TMHs: 2\nExp number of AAs in TMHs: 38.07604\nExp number, first 60 AAs: 19.52977\nTotal prob of N-in: 0.94353\nPOSSIBLE N-term signal sequence\ninside 1 19\nTMhelix 20 39\noutside 40 325\nTMhelix 326 348\ninside 349 372

2334 GCF_000807635.1_ASM80763v1 Escherichia coli Proteobacteria; Gammaproteobacteria; Enterobacterales; Enterobacteriaceae; Escherichia
MTGDNTLIHSHGINRRDFMKLCAALAATMGLSSKAAAEMAESVTNPQRPPVIWIGAQECT WP_000145410.1
MULTISPECIES: hydrogenase-2 small chain [Proteobacteria] Length: 372\nNumber of predicted TMHs: 1\nExp number of AAs in TMHs: 23.72226\nExp number, first 60 AAs: 0.22038\nTotal prob of N-in: 0.12994\noutside 1 331\nTMhelix 332 354\ninside 355 372

2335 GCF_000807635.1_ASM80763v1 Escherichia coli Proteobacteria; Gammaproteobacteria; Enterobacterales; Enterobacteriaceae; Escherichia
MNNEETFYQAMRRQGVTRRSFLKYCSLAATSLGLGAGMAPKIAWALENKPRIPVVWIHGL WP_001058323.1
MULTISPECIES: hydrogenase-1 small chain [Proteobacteria] Length: 372\nNumber of predicted TMHs: 2\nExp number of AAs in TMHs: 37.2608499999999\nExp number, first 60 AAs: 19.52923\nTotal prob of N-in: 0.94351\nPOSSIBLE N-term signal sequence\ninside 1 19\nTMhelix 20 39\noutside 40 325\nTMhelix 326 348\ninside 349 372

2336 GCF_000819105.1_Escherichia_coli_CVM_N33720PS_v1.0 Escherichia coli Proteobacteria; Gammaproteobacteria; Enterobacterales; Enterobacteriaceae; Escherichia
MTGDNTLIHSHGINRRDFMKLCAALAATMGLSSKAAAEMAESVTNPQRPPVIWIGAQECT WP_000145410.1
MULTISPECIES: hydrogenase-2 small chain [Proteobacteria] Length: 372\nNumber of predicted TMHs: 1\nExp number of AAs in TMHs: 23.72226\nExp number, first 60 AAs: 0.22038\nTotal prob of N-in: 0.12994\noutside 1 331\nTMhelix 332 354\ninside 355 372

2337 GCF_000819005.1_Escherichia_coli_CVM_N33825PS_v1.0 Escherichia coli Proteobacteria; Gammaproteobacteria; Enterobacterales; Enterobacteriaceae; Escherichia
MNNEETFYQAMRRQGVTRRSFLKYCSLAATSLGLGAGMAPKIAWALENKPRIPVVWIHGL WP_001058323.1
MULTISPECIES: hydrogenase-1 small chain [Proteobacteria] Length: 372\nNumber of predicted TMHs: 2\nExp number of AAs in TMHs: 37.2608499999999\nExp number, first 60 AAs: 19.52923\nTotal prob of N-in: 0.94351\nPOSSIBLE N-term signal sequence\ninside 1 19\nTMhelix 20 39\noutside 40 325\nTMhelix 326 348\ninside 349 372

2338 GCF_000819105.1_Escherichia_coli_CVM_N33720PS_v1.0 Escherichia coli Proteobacteria; Gammaproteobacteria; Enterobacterales; Enterobacteriaceae; Escherichia
MNNEETFYQAMRRQGVTRRSFLKYCSLAATSLGLGAGMAPKIAWALENKPRIPVVWIHGL WP_001058323.1
MULTISPECIES: hydrogenase-1 small chain [Proteobacteria] Length: 372\nNumber of predicted TMHs: 2\nExp number of AAs in TMHs: 37.2608499999999\nExp number, first 60 AAs: 19.52923\nTotal prob of N-in: 0.94351\nPOSSIBLE N-term signal sequence\ninside 1 19\nTMhelix 20 39\noutside 40 325\nTMhelix 326 348\ninside 349 372

2339 GCF_000819125.1_Escherichia_coli_CVM_N34564PS_v1.0 Escherichia coli Proteobacteria; Gammaproteobacteria; Enterobacterales; Enterobacteriaceae; Escherichia
MNNEETFYQAMRRQGVTRRSFLKYCSLAATSLGLGAGMAPKIAWALENKPRIPVVWIHGL WP_001058323.1
MULTISPECIES: hydrogenase-1 small chain [Proteobacteria] Length: 372\nNumber of predicted TMHs: 2\nExp number of AAs in TMHs: 37.2608499999999\nExp number, first 60 AAs: 19.52923\nTotal prob of N-in: 0.94351\nPOSSIBLE N-term signal sequence\ninside 1 19\nTMhelix 20 39\noutside 40 325\nTMhelix 326 348\ninside 349 372

2340 GCF_000819225.1_Escherichia_coli_CVM_N36601PS_v1.0 Escherichia coli Proteobacteria; Gammaproteobacteria; Enterobacterales; Enterobacteriaceae; Escherichia
MTGDNTLIHSHGINRRDFMKLCAALAATMGLSSKAAAEMAESVTNPQRPPVIWIGAQECT WP_000145410.1
MULTISPECIES: hydrogenase-2 small chain [Proteobacteria] Length: 372\nNumber of predicted TMHs: 1\nExp number of AAs in TMHs: 23.72226\nExp number, first 60 AAs: 0.22038\nTotal prob of N-in: 0.12994\noutside 1 331\nTMhelix 332 354\ninside 355 372

2341 GCF_000819185.1_Escherichia_coli_CVM_N36113PS_v1.0 Escherichia coli Proteobacteria; Gammaproteobacteria; Enterobacterales; Enterobacteriaceae; Escherichia
MTGDNTLIHSHGINRRDFMKLCAALAATMGLSSKAAAEMAESVTNPQRPPVIWIGAQECT WP_000145410.1
MULTISPECIES: hydrogenase-2 small chain [Proteobacteria] Length: 372\nNumber of predicted TMHs: 1\nExp number of AAs in TMHs: 23.72226\nExp number, first 60 AAs: 0.22038\nTotal prob of N-in: 0.12994\noutside 1 331\nTMhelix 332 354\ninside 355 372

2342 GCF_000819325.1_Escherichia_coli_CVM_N38381PS_v1.0 Escherichia coli Proteobacteria; Gammaproteobacteria; Enterobacterales; Enterobacteriaceae; Escherichia
MNNEETFYQAMRRQGVTRRSFLKYCSLAATSLGLGAGMAPKIAWALENKPRIPVVWIHGL WP_001058323.1
MULTISPECIES: hydrogenase-1 small chain [Proteobacteria] Length: 372\nNumber of predicted TMHs: 2\nExp number of AAs in TMHs: 37.2608499999999\nExp number, first 60 AAs: 19.52923\nTotal prob of N-in: 0.94351\nPOSSIBLE N-term signal sequence\ninside 1 19\nTMhelix 20 39\noutside 40 325\nTMhelix 326 348\ninside 349 372

2343 GCF_000819645.1_ASM81964v1 Escherichia coli Proteobacteria; Gammaproteobacteria; Enterobacterales; Enterobacteriaceae; Escherichia
MTGDNTLIHSHGINRRDFMKLCAALAATMGLSSKAAAEMAESVTNPQRPPVIWIGAQECT WP_000145410.1
MULTISPECIES: hydrogenase-2 small chain [Proteobacteria] Length: 372\nNumber of predicted TMHs: 1\nExp number of AAs in TMHs: 23.72226\nExp number, first 60 AAs: 0.22038\nTotal prob of N-in: 0.12994\noutside 1 331\nTMhelix 332 354\ninside 355 372

2344 GCF_000826805.1_ASM82680v1 Escherichia coli Proteobacteria; Gammaproteobacteria; Enterobacterales; Enterobacteriaceae; Escherichia
MTGDNTLIHSHGINRRDFMKLCAALAATMGLSSKAAAEMAESVTNPQRPPVIWIGAQECT WP_000145410.1
MULTISPECIES: hydrogenase-2 small chain [Proteobacteria] Length: 372\nNumber of predicted TMHs: 1\nExp number of AAs in TMHs: 23.72226\nExp number, first 60 AAs: 0.22038\nTotal prob of N-in: 0.12994\noutside 1 331\nTMhelix 332 354\ninside 355 372

2345 GCF_000833635.2_ASM83363v2 Escherichia coli Proteobacteria; Gammaproteobacteria; Enterobacterales; Enterobacteriaceae; Escherichia
MTGDNTLIHSHGINRRDFMKLCAALAATMGLSSKAAAEMAESVTNPQRPPVIWIGAQECT WP_000145410.1
MULTISPECIES: hydrogenase-2 small chain [Proteobacteria] Length: 372\nNumber of predicted TMHs: 1\nExp number of AAs in TMHs: 23.72226\nExp number, first 60 AAs: 0.22038\nTotal prob of N-in: 0.12994\noutside 1 331\nTMhelix 332 354\ninside 355 372

2346 GCF_000819645.1_ASM81964v1 Escherichia coli Proteobacteria; Gammaproteobacteria; Enterobacterales; Enterobacteriaceae; Escherichia
MNNEETFYQAMRRQGVTRRSFLKYCSLAATSLGLGAGMAPKIAWALENKPRIPVVWIHGL WP_001058320.1
hydrogenase [Escherichia coli] Length: 372\nNumber of predicted TMHs: 2\nExp number of AAs in TMHs: 37.26082\nExp number, first 60 AAs: 19.52923\nTotal prob of N-in: 0.94351\nPOSSIBLE N-term signal sequence\ninside 1 19\nTMhelix 20 39\noutside 40 325\nTMhelix 326 348\ninside 349 372

2347 GCF_000936105.1_Ec57A_A8_MIRA_assembly Escherichia coli Proteobacteria; Gammaproteobacteria; Enterobacterales; Enterobacteriaceae; Escherichia
MNNEETFYQAMRRQGVTRRSFLKYCSLAATSLGLGAGMAPKIAWALENKPRIPVVWIHGL WP_001058323.1
MULTISPECIES: hydrogenase-1 small chain [Proteobacteria] Length: 372\nNumber of predicted TMHs: 2\nExp number of AAs in TMHs: 37.2608499999999\nExp number, first 60 AAs: 19.52923\nTotal prob of N-in: 0.94351\nPOSSIBLE N-term signal sequence\ninside 1 19\nTMhelix 20 39\noutside 40 325\nTMhelix 326 348\ninside 349 372

2348 GCF_000936245.1_FHI7 Escherichia coli Proteobacteria; Gammaproteobacteria; Enterobacterales; Enterobacteriaceae; Escherichia MTGDNTLIHSHGINRRDFMKLCAALAATMGLSSKAAAEMAESVTNPQRPPVIWIGAQECT
WP_000145410.1 MULTISPECIES: hydrogenase-2 small chain [Proteobacteria] Length: 372\nNumber of predicted TMHs: 1\nExp number of AAs in TMHs: 23.72226\nExp number, first 60 AAs: 0.22038\nTotal prob of N-in: 0.12994\noutside 1 331\nTMhelix 332 354\ninside 355 372

2349 GCF_000936495.1_FHI102 Escherichia coli Proteobacteria; Gammaproteobacteria; Enterobacterales; Enterobacteriaceae; Escherichia MTGDNTLIHSHGINRRDFMKLCAALAATMGLSSKAAAEMAESVTNPQRPPVIWIGAQECT
WP_000145410.1 MULTISPECIES: hydrogenase-2 small chain [Proteobacteria] Length: 372\nNumber of predicted TMHs: 1\nExp number of AAs in TMHs: 23.72226\nExp number, first 60 AAs: 0.22038\nTotal prob of N-in: 0.12994\noutside 1 331\nTMhelix 332 354\ninside 355 372

2350 GCF_000935515.1_ASM93551v1 Escherichia coli Proteobacteria; Gammaproteobacteria; Enterobacterales; Enterobacteriaceae; Escherichia
MNNEETFYQAMRRQGVTRRSFLKYCSLAATSLGLGAGMAPKIAWALENKPRIPVVWIHGL WP_001058323.1
MULTISPECIES: hydrogenase-1 small chain [Proteobacteria] Length: 372\nNumber of predicted TMHs: 2\nExp number of AAs in TMHs: 37.2608499999999\nExp number, first 60 AAs: 19.52923\nTotal prob of N-in:

0.94351\nPOSSIBLE N-term signal sequence\ninside 1 19\nTMhelix 20 39\noutside 40 325\nTMhelix 326 348\ninside 349 372

2351 GCF_000935555.1_ASM93555v1 Escherichia coli Proteobacteria; Gammaproteobacteria; Enterobacterales; Enterobacteriaceae; Escherichia MNNEETFYQAMRRQGVTRRSFLKYCSLAATSLGLGAGMAPKIAWALENKPRIPVVWIHGL WP_001058323.1 MULTISPECIES: hydrogenase-1 small chain [Proteobacteria] Length: 372\nNumber of predicted TMHs: 2\nExp number of AAs in TMHs: 37.2608499999999\nExp number, first 60 AAs: 19.52923\nTotal prob of N-in: 0.94351\nPOSSIBLE N-term signal sequence\ninside 1 19\nTMhelix 20 39\noutside 40 325\nTMhelix 326 348\ninside 349 372

2352 GCF_000935905.1_FHI21 Escherichia coli Proteobacteria; Gammaproteobacteria; Enterobacterales; Enterobacteriaceae; Escherichia MNNEETFYQAMRRQGVTRRSFLKYCSLAATSLGLGAGMAPKIAWALENKPRIPVVWIHGL WP_001058323.1 MULTISPECIES: hydrogenase-1 small chain [Proteobacteria] Length: 372\nNumber of predicted TMHs: 2\nExp number of AAs in TMHs: 37.2608499999999\nExp number, first 60 AAs: 19.52923\nTotal prob of N-in: 0.94351\nPOSSIBLE N-term signal sequence\ninside 1 19\nTMhelix 20 39\noutside 40 325\nTMhelix 326 348\ninside 349 372

2353 GCF_000939335.1_Ec46A_S2_MIRA_assembly Escherichia coli Proteobacteria; Gammaproteobacteria; Enterobacterales; Enterobacteriaceae; Escherichia MTGDNTLIHSHGINRRDFMKLCAALAATMGLSSKAAAEMAESVTNPQRPPVIWIGAQECT WP_000145410.1 MULTISPECIES: hydrogenase-2 small chain [Proteobacteria] Length: 372\nNumber of predicted TMHs: 1\nExp number of AAs in TMHs: 23.72226\nExp number, first 60 AAs: 0.22038\nTotal prob of N-in: 0.12994\noutside 1 331\nTMhelix 332 354\ninside 355 372

2354 GCF_000939755.1_FHI43 Escherichia coli Proteobacteria; Gammaproteobacteria; Enterobacterales; Enterobacteriaceae; Escherichia MTGDNTLIHSHGINRRDFMKLCAALAATMGLSSKAAAEMAESVTNPQRPPVIWIGAQECT WP_000145410.1 MULTISPECIES: hydrogenase-2 small chain [Proteobacteria] Length: 372\nNumber of predicted TMHs: 1\nExp number of AAs in TMHs: 23.72226\nExp number, first 60 AAs: 0.22038\nTotal prob of N-in: 0.12994\noutside 1 331\nTMhelix 332 354\ninside 355 372

2355 GCF_000939955.1_FHI79 Escherichia coli Proteobacteria; Gammaproteobacteria; Enterobacterales; Enterobacteriaceae; Escherichia MTGDNTLIHSHGINRRDFMKLCAALAATMGLSSKAAAEMAESVTNPQRPPVIWIGAQECT WP_000145410.1 MULTISPECIES: hydrogenase-2 small chain [Proteobacteria] Length: 372\nNumber of predicted TMHs: 1\nExp number of AAs in TMHs: 23.72226\nExp number, first 60 AAs: 0.22038\nTotal prob of N-in: 0.12994\noutside 1 331\nTMhelix 332 354\ninside 355 372

2356 GCF_000939335.1_Ec46A_S2_MIRA_assembly Escherichia coli Proteobacteria; Gammaproteobacteria; Enterobacterales; Enterobacteriaceae; Escherichia MNNEETFYQAMRRQGVTRRSFLKYCSLAATSLGLGAGMAPKIAWALENKPRIPVVWIHGL WP_001058323.1 MULTISPECIES: hydrogenase-1 small chain [Proteobacteria] Length: 372\nNumber of predicted TMHs: 2\nExp number of AAs in TMHs: 37.2608499999999\nExp number, first 60 AAs: 19.52923\nTotal prob of N-in: 0.94351\nPOSSIBLE N-term signal sequence\ninside 1 19\nTMhelix 20 39\noutside 40 325\nTMhelix 326 348\ninside 349 372

2357 GCF_000937415.1_FHI98 Escherichia coli Proteobacteria; Gammaproteobacteria; Enterobacterales; Enterobacteriaceae; Escherichia MNNEETFYQAMRRQGVTRRSFLKYCSLAATSLGLGAGMAPKIAWALENKPRIPVVWIHGL WP_001295940.1 MULTISPECIES: hydrogenase [Enterobacteriaceae] Length: 372\nNumber of predicted TMHs: 2\nExp number of AAs in TMHs: 37.2608899999999\nExp number, first 60 AAs: 19.52923\nTotal prob of N-in: 0.94351\nPOSSIBLE N-term signal sequence\ninside 1 19\nTMhelix 20 39\noutside 40 325\nTMhelix 326 348\ninside 349 372

2358 GCF_000939035.1_Ec26A_L2_MIRA_assembly Escherichia coli Proteobacteria; Gammaproteobacteria; Enterobacterales; Enterobacteriaceae; Escherichia MNNEETFYQAMRRQGVTRRSFLKYCSLAATSLGLGAGMAPKIAWALENKPRIPVVWIHGL WP_001058320.1 hydrogenase [Escherichia coli] Length: 372\nNumber of predicted TMHs: 2\nExp number of AAs in TMHs: 37.26082\nExp number, first 60 AAs: 19.52923\nTotal prob of N-in: 0.94351\nPOSSIBLE N-term signal sequence\ninside 1 19\nTMhelix 20 39\noutside 40 325\nTMhelix 326 348\ninside 349 372

2359 GCF_000946275.1_FHI97 Escherichia coli Proteobacteria; Gammaproteobacteria; Enterobacterales; Enterobacteriaceae; Escherichia MNNEETFYQAMRRQGVTRRSFLKYCSLAATSLGLGAGMAPKIAWALENKPRIPVVWIHGL

WP_024246676.1 hydrogenase-1 small chain [Escherichia coli] Length: 372\nNumber of predicted TMHs: 2\nExp number of AAs in TMHs: 37.8214899999999\nExp number, first 60 AAs: 19.52326\nTotal prob of N-in: 0.94395\nPOSSIBLE N-term signal sequence\ninside 1 19\nTMhelix 20 39\noutside 40 325\nTMhelix 326 348\ninside 349 372

2360 GCF_000941895.1_FHI6 Escherichia coli Proteobacteria; Gammaproteobacteria; Enterobacterales; Enterobacteriaceae; Escherichia MNNEETFYQAMRRQGVTRRSFLKYCSLAATSLGLGAGMAPKIAWALENKPRIPVVWIHGL WP_001058323.1 MULTISPECIES: hydrogenase-1 small chain [Proteobacteria] Length: 372\nNumber of predicted TMHs: 2\nExp number of AAs in TMHs: 37.2608499999999\nExp number, first 60 AAs: 19.52923\nTotal prob of N-in: 0.94351\nPOSSIBLE N-term signal sequence\ninside 1 19\nTMhelix 20 39\noutside 40 325\nTMhelix 326 348\ninside 349 372

2361 GCF_000946475.1_FHI34 Escherichia coli Proteobacteria; Gammaproteobacteria; Enterobacterales; Enterobacteriaceae; Escherichia MTGDNTLIHSHGINRRDFMKLCAALAATMGLSSKAAAEMAESVTNPQRPPVIWIGAQECT WP_000145410.1 MULTISPECIES: hydrogenase-2 small chain [Proteobacteria] Length: 372\nNumber of predicted TMHs: 1\nExp number of AAs in TMHs: 23.72226\nExp number, first 60 AAs: 0.22038\nTotal prob of N-in: 0.12994\noutside 1 331\nTMhelix 332 354\ninside 355 372

2362 GCF_000946475.1_FHI34 Escherichia coli Proteobacteria; Gammaproteobacteria; Enterobacterales; Enterobacteriaceae; Escherichia MNNEETFYQAMRRQGVTRRSFLKYCSLAATSLGLGAGMAPKIAWALENKPRIPVVWIHGL WP_001058323.1 MULTISPECIES: hydrogenase-1 small chain [Proteobacteria] Length: 372\nNumber of predicted TMHs: 2\nExp number of AAs in TMHs: 37.2608499999999\nExp number, first 60 AAs: 19.52923\nTotal prob of N-in: 0.94351\nPOSSIBLE N-term signal sequence\ninside 1 19\nTMhelix 20 39\noutside 40 325\nTMhelix 326 348\ninside 349 372

2363 GCF_000943355.2_FHI42 Escherichia coli Proteobacteria; Gammaproteobacteria; Enterobacterales; Enterobacteriaceae; Escherichia MNNEETFYQAMRRQGVTRRSFLKYCSLAATSLGLGAGMAPKIAWALENKPRIPVVWIHGL WP_001058323.1 MULTISPECIES: hydrogenase-1 small chain [Proteobacteria] Length: 372\nNumber of predicted TMHs: 2\nExp number of AAs in TMHs: 37.2608499999999\nExp number, first 60 AAs: 19.52923\nTotal prob of N-in: 0.94351\nPOSSIBLE N-term signal sequence\ninside 1 19\nTMhelix 20 39\noutside 40 325\nTMhelix 326 348\ninside 349 372

2364 GCF_000947375.1_4e8A_assembly Escherichia coli Proteobacteria; Gammaproteobacteria; Enterobacterales; Enterobacteriaceae; Escherichia MNNEETFYQAMRRQGVTRRSFLKYCSLAATSLGLGAGMAPKIAWALENKPRIPVVWIHGL WP_001058323.1 MULTISPECIES: hydrogenase-1 small chain [Proteobacteria] Length: 372\nNumber of predicted TMHs: 2\nExp number of AAs in TMHs: 37.2608499999999\nExp number, first 60 AAs: 19.52923\nTotal prob of N-in: 0.94351\nPOSSIBLE N-term signal sequence\ninside 1 19\nTMhelix 20 39\noutside 40 325\nTMhelix 326 348\ninside 349 372

2365 GCF_000941995.1_FHI59 Escherichia coli Proteobacteria; Gammaproteobacteria; Enterobacterales; Enterobacteriaceae; Escherichia MNNEETFYQAMRRQGVTRRSFLKYCSLAATSLGLGAGMAPKIAWALENKPRIPVVWIHGL WP_001295940.1 MULTISPECIES: hydrogenase [Enterobacteriaceae] Length: 372\nNumber of predicted TMHs: 2\nExp number of AAs in TMHs: 37.2608899999999\nExp number, first 60 AAs: 19.52923\nTotal prob of N-in: 0.94351\nPOSSIBLE N-term signal sequence\ninside 1 19\nTMhelix 20 39\noutside 40 325\nTMhelix 326 348\ninside 349 372

2366 GCF_000948745.1_ASM94874v1 Escherichia coli Proteobacteria; Gammaproteobacteria; Enterobacterales; Enterobacteriaceae; Escherichia MNNEETFYQAMRRQGVTRRSFLKYCSLAATSLGLGAGMAPKIAWALENKPRIPVVWIHGL WP_001058323.1 MULTISPECIES: hydrogenase-1 small chain [Proteobacteria] Length: 372\nNumber of predicted TMHs: 2\nExp number of AAs in TMHs: 37.2608499999999\nExp number, first 60 AAs: 19.52923\nTotal prob of N-in: 0.94351\nPOSSIBLE N-term signal sequence\ninside 1 19\nTMhelix 20 39\noutside 40 325\nTMhelix 326 348\ninside 349 372

2367 GCF_000948435.1_ASM94843v1 Escherichia coli Proteobacteria; Gammaproteobacteria; Enterobacterales; Enterobacteriaceae; Escherichia MNNEETFYQAMRRQGVTRRSFLKYCSLAATSLGLGAGMAPKIAWALENKPRIPVVWIHGL WP_001295940.1 MULTISPECIES: hydrogenase [Enterobacteriaceae] Length: 372\nNumber of predicted TMHs: 2\nExp number of AAs in TMHs: 37.2608899999999\nExp number, first 60 AAs: 19.52923\nTotal prob of N-in:

0.94351\nPOSSIBLE N-term signal sequence\ninside 1 19\nTMhelix 20 39\noutside 40 325\nTMhelix 326 348\ninside 349 372

2368 GCF_000948595.1_ASM94859v1 Escherichia coli Proteobacteria; Gammaproteobacteria; Enterobacterales; Enterobacteriaceae; Escherichia MNNEETFYQAMRRQGVTRRSFLKYCSLAATSLGLGAGMAPKIAWALENKPRIPVVWIHGL WP_001058323.1 MULTISPECIES: hydrogenase-1 small chain [Proteobacteria] Length: 372\nNumber of predicted TMHs: 2\nExp number of AAs in TMHs: 37.2608499999999\nExp number, first 60 AAs: 19.52923\nTotal prob of N-in: 0.94351\nPOSSIBLE N-term signal sequence\ninside 1 19\nTMhelix 20 39\noutside 40 325\nTMhelix 326 348\ninside 349 372

2369 GCF_000948785.1_ASM94878v1 Escherichia coli Proteobacteria; Gammaproteobacteria; Enterobacterales; Enterobacteriaceae; Escherichia MTGDNTLIHSHGINRRDFMKLCAALAATMGLSSKAAAEMAESVTNPQRPPVIWIGAQECT WP_000145410.1 MULTISPECIES: hydrogenase-2 small chain [Proteobacteria] Length: 372\nNumber of predicted TMHs: 1\nExp number of AAs in TMHs: 23.72226\nExp number, first 60 AAs: 0.22038\nTotal prob of N-in: 0.12994\noutside 1 331\nTMhelix 332 354\ninside 355 372

2370 GCF_000948785.1_ASM94878v1 Escherichia coli Proteobacteria; Gammaproteobacteria; Enterobacterales; Enterobacteriaceae; Escherichia MNNEETFYQAMRRQGVTRRSFLKYCSLAATSLGLGAGMAPKIAWALENKPRIPVVWIHGL WP_001058323.1 MULTISPECIES: hydrogenase-1 small chain [Proteobacteria] Length: 372\nNumber of predicted TMHs: 2\nExp number of AAs in TMHs: 37.2608499999999\nExp number, first 60 AAs: 19.52923\nTotal prob of N-in: 0.94351\nPOSSIBLE N-term signal sequence\ninside 1 19\nTMhelix 20 39\noutside 40 325\nTMhelix 326 348\ninside 349 372

2371 GCF_000948555.1_ASM94855v1 Escherichia coli Proteobacteria; Gammaproteobacteria; Enterobacterales; Enterobacteriaceae; Escherichia MNNEETFYQAMRRQGVTRRSFLKYCSLAATSLGLGAGIAPKIAWALENKPRIPVVWIHGL WP_023982790.1 hydrogenase [Escherichia coli] Length: 372\nNumber of predicted TMHs: 2\nExp number of AAs in TMHs: 37.8310099999999\nExp number, first 60 AAs: 20.15722\nTotal prob of N-in: 0.95896\nPOSSIBLE N-term signal sequence\ninside 1 20\nTMhelix 21 43\noutside 44 325\nTMhelix 326 348\ninside 349 372

2372 GCF_000948835.1_ASM94883v1 Escherichia coli Proteobacteria; Gammaproteobacteria; Enterobacterales; Enterobacteriaceae; Escherichia MTGDNTLIHSHGINRRDFMKLCAALAATMGLSSKAAAEMVESVTNPQRPPVIWIGAQECT WP_000145423.1 hydrogenase 2 small subunit [Escherichia coli] Length: 372\nNumber of predicted TMHs: 1\nExp number of AAs in TMHs: 23.78201\nExp number, first 60 AAs: 0.28355\nTotal prob of N-in: 0.13257\noutside 1 331\nTMhelix 332 354\ninside 355 372

2373 GCF_000951745.1_ASM95174v1 Escherichia coli Proteobacteria; Gammaproteobacteria; Enterobacterales; Enterobacteriaceae; Escherichia MNNEETFYQAMRRQGVTRRSFLKYCSLAATSLGLGAGMAPKIAWALENKPRIPVVWIHGL WP_001058323.1 MULTISPECIES: hydrogenase-1 small chain [Proteobacteria] Length: 372\nNumber of predicted TMHs: 2\nExp number of AAs in TMHs: 37.2608499999999\nExp number, first 60 AAs: 19.52923\nTotal prob of N-in: 0.94351\nPOSSIBLE N-term signal sequence\ninside 1 19\nTMhelix 20 39\noutside 40 325\nTMhelix 326 348\ninside 349 372

2374 GCF_000951855.1_FHI14 Escherichia coli Proteobacteria; Gammaproteobacteria; Enterobacterales; Enterobacteriaceae; Escherichia MNNEETFYQAMRRQGVTRRSFLKYCSLAATSLGLGAGMAPKIAWALENKPRIPVVWIHGL WP_001058323.1 MULTISPECIES: hydrogenase-1 small chain [Proteobacteria] Length: 372\nNumber of predicted TMHs: 2\nExp number of AAs in TMHs: 37.2608499999999\nExp number, first 60 AAs: 19.52923\nTotal prob of N-in: 0.94351\nPOSSIBLE N-term signal sequence\ninside 1 19\nTMhelix 20 39\noutside 40 325\nTMhelix 326 348\ninside 349 372

2375 GCF_000948905.1_ASM94890v1 Escherichia coli Proteobacteria; Gammaproteobacteria; Enterobacterales; Enterobacteriaceae; Escherichia MNNEETFYQAMRRQGVTRRSFLKYCSLAATSLGLGAGMAPKIAWALENKPRIPVVWIHGL WP_001058323.1 MULTISPECIES: hydrogenase-1 small chain [Proteobacteria] Length: 372\nNumber of predicted TMHs: 2\nExp number of AAs in TMHs: 37.2608499999999\nExp number, first 60 AAs: 19.52923\nTotal prob of N-in:

0.94351\nPOSSIBLE N-term signal sequence\ninside 1 19\nTMhelix 20 39\noutside 40 325\nTMhelix 326 348\ninside 349 372

2376 GCF_000952305.1_ASM95230v1 Escherichia coli Proteobacteria; Gammaproteobacteria; Enterobacterales; Enterobacteriaceae; Escherichia MTGDNTLIHSHGINRRDFMKLCAALAATMGLSSKAAAEMVESVTNPQRPPVIWIGAQECT WP_000145423.1 hydrogenase 2 small subunit [Escherichia coli] Length: 372\nNumber of predicted TMHs: 1\nExp number of AAs in TMHs: 23.78201\nExp number, first 60 AAs: 0.28355\nTotal prob of N-in: 0.13257\noutside 1 331\nTMhelix 332 354\ninside 355 372

2377 GCF_000951955.1_FHI100 Escherichia coli Proteobacteria; Gammaproteobacteria; Enterobacterales; Enterobacteriaceae; Escherichia MTGDNTLIHSHGINRRDFMKLCAALAATMGLSSKAAAEMAESVTNPQRPPVIWIGAQECT WP_000145410.1 MULTISPECIES: hydrogenase-2 small chain [Proteobacteria] Length: 372\nNumber of predicted TMHs: 1\nExp number of AAs in TMHs: 23.72226\nExp number, first 60 AAs: 0.22038\nTotal prob of N-in: 0.12994\noutside 1 331\nTMhelix 332 354\ninside 355 372

2378 GCF_000952385.1_ASM95238v1 Escherichia coli Proteobacteria; Gammaproteobacteria; Enterobacterales; Enterobacteriaceae; Escherichia MTGDNTLIHSHGINRRDFMKLCAALAATMGLSSKAAAEMVESVTNPQRPPVIWIGAQECT WP_000145423.1 hydrogenase 2 small subunit [Escherichia coli] Length: 372\nNumber of predicted TMHs: 1\nExp number of AAs in TMHs: 23.78201\nExp number, first 60 AAs: 0.28355\nTotal prob of N-in: 0.13257\noutside 1 331\nTMhelix 332 354\ninside 355 372

2379 GCF_000951955.1_FHI100 Escherichia coli Proteobacteria; Gammaproteobacteria; Enterobacterales; Enterobacteriaceae; Escherichia MNNEETFYQAMRRQGVTRRSFLKYCSLAATSLGLGAGMAPKIAWALENKPRIPVVWIHGL WP_001058323.1 MULTISPECIES: hydrogenase-1 small chain [Proteobacteria] Length: 372\nNumber of predicted TMHs: 2\nExp number of AAs in TMHs: 37.2608499999999\nExp number, first 60 AAs: 19.52923\nTotal prob of N-in: 0.94351\nPOSSIBLE N-term signal sequence\ninside 1 19\nTMhelix 20 39\noutside 40 325\nTMhelix 326 348\ninside 349 372

2380 GCF_000954045.1_ASM95404v1 Escherichia coli Proteobacteria; Gammaproteobacteria; Enterobacterales; Enterobacteriaceae; Escherichia MTGDNTLIHSHGINRRDFMKLCAALAATMGLSSKAAAEMAESVTNPQRPPVIWIGAQECT WP_000145410.1 MULTISPECIES: hydrogenase-2 small chain [Proteobacteria] Length: 372\nNumber of predicted TMHs: 1\nExp number of AAs in TMHs: 23.72226\nExp number, first 60 AAs: 0.22038\nTotal prob of N-in: 0.12994\noutside 1 331\nTMhelix 332 354\ninside 355 372

2381 GCF_000965555.1_ASM96555v1 Escherichia coli Proteobacteria; Gammaproteobacteria; Enterobacterales; Enterobacteriaceae; Escherichia MTGDNTLIHSHGINRRDFMKLCAALAATMGLSSKAAAEMAESVTNPQRPPVIWIGAQECT WP_000145410.1 MULTISPECIES: hydrogenase-2 small chain [Proteobacteria] Length: 372\nNumber of predicted TMHs: 1\nExp number of AAs in TMHs: 23.72226\nExp number, first 60 AAs: 0.22038\nTotal prob of N-in: 0.12994\noutside 1 331\nTMhelix 332 354\ninside 355 372

2382 GCF_000965575.1_ASM96557v1 Escherichia coli Proteobacteria; Gammaproteobacteria; Enterobacterales; Enterobacteriaceae; Escherichia MTGDNTLIHSHGINRRDFMKLCAALAATMGLSSKAAAEMAESVTNPQRPPVIWIGAQECT WP_000145410.1 MULTISPECIES: hydrogenase-2 small chain [Proteobacteria] Length: 372\nNumber of predicted TMHs: 1\nExp number of AAs in TMHs: 23.72226\nExp number, first 60 AAs: 0.22038\nTotal prob of N-in: 0.12994\noutside 1 331\nTMhelix 332 354\ninside 355 372

2383 GCF_000965665.1_ASM96566v1 Escherichia coli Proteobacteria; Gammaproteobacteria; Enterobacterales; Enterobacteriaceae; Escherichia MTGDNTLIHSHGINRRDFMKLCAALAATMGLSSKAAAEMAESVTNPQRPPVIWIGAQECT WP_000145410.1 MULTISPECIES: hydrogenase-2 small chain [Proteobacteria] Length: 372\nNumber of predicted TMHs: 1\nExp number of AAs in TMHs: 23.72226\nExp number, first 60 AAs: 0.22038\nTotal prob of N-in: 0.12994\noutside 1 331\nTMhelix 332 354\ninside 355 372

2384 GCF_000952385.1_ASM95238v1 Escherichia coli Proteobacteria; Gammaproteobacteria; Enterobacterales; Enterobacteriaceae; Escherichia MNNEETFYQAMRRQGVTRRSFLKYCSLAATSLGLGAGMAPKIAWALENKPRIPVVWIHGL WP_001295940.1

MULTISPECIES: hydrogenase [Enterobacteriaceae] Length: 372\nNumber of predicted TMHs: 2\nExp number of AAs in TMHs: 37.2608899999999\nExp number, first 60 AAs: 19.52923\nTotal prob of N-in: 0.94351\nPOSSIBLE N-term signal sequence\ninside 1 19\nTMhelix 20 39\noutside 40 325\nTMhelix 326 348\ninside 349 372

2385 GCF_000965625.1_ASM96562v1 Escherichia coli Proteobacteria; Gammaproteobacteria; Enterobacterales; Enterobacteriaceae; Escherichia
MTGDNTLIHSHGINRRDFMKLCAALAATMGLSSKAAAEMAESVTNPQRPPVIWIGAQECT WP_000145410.1
MULTISPECIES: hydrogenase-2 small chain [Proteobacteria] Length: 372\nNumber of predicted TMHs: 1\nExp number of AAs in TMHs: 23.72226\nExp number, first 60 AAs: 0.22038\nTotal prob of N-in: 0.12994\noutside 1 331\nTMhelix 332 354\ninside 355 372

2386 GCF_000954045.1_ASM95404v1 Escherichia coli Proteobacteria; Gammaproteobacteria; Enterobacterales; Enterobacteriaceae; Escherichia
MNNEETFYQAMRRQGVTRRSFLKYCSLAATSLGLGAGMAPKIAWALENKPRIPVVWIHGL WP_001058323.1
MULTISPECIES: hydrogenase-1 small chain [Proteobacteria] Length: 372\nNumber of predicted TMHs: 2\nExp number of AAs in TMHs: 37.2608499999999\nExp number, first 60 AAs: 19.52923\nTotal prob of N-in: 0.94351\nPOSSIBLE N-term signal sequence\ninside 1 19\nTMhelix 20 39\noutside 40 325\nTMhelix 326 348\ninside 349 372

2387 GCF_000965665.1_ASM96566v1 Escherichia coli Proteobacteria; Gammaproteobacteria; Enterobacterales; Enterobacteriaceae; Escherichia
MNNEETFYQAMRRQGVTRRSFLKYCSLAATSLGLGAGMAPKIAWALENKPRIPVVWIHGL WP_001058323.1
MULTISPECIES: hydrogenase-1 small chain [Proteobacteria] Length: 372\nNumber of predicted TMHs: 2\nExp number of AAs in TMHs: 37.2608499999999\nExp number, first 60 AAs: 19.52923\nTotal prob of N-in: 0.94351\nPOSSIBLE N-term signal sequence\ninside 1 19\nTMhelix 20 39\noutside 40 325\nTMhelix 326 348\ninside 349 372

2388 GCF_000971615.1_ASM97161v1 Escherichia coli Proteobacteria; Gammaproteobacteria; Enterobacterales; Enterobacteriaceae; Escherichia
MTGDNTLIHSHGINRRDFMKLCAALAATMGLSSKAAAEMAESVTNPQRPPVIWIGAQECT WP_000145410.1
MULTISPECIES: hydrogenase-2 small chain [Proteobacteria] Length: 372\nNumber of predicted TMHs: 1\nExp number of AAs in TMHs: 23.72226\nExp number, first 60 AAs: 0.22038\nTotal prob of N-in: 0.12994\noutside 1 331\nTMhelix 332 354\ninside 355 372

2389 GCF_000969545.1_ASM96954v1 Escherichia coli Proteobacteria; Gammaproteobacteria; Enterobacterales; Enterobacteriaceae; Escherichia
MNNEETFYQAMRRQGVTRRSFLKYCSLAATSLGLGAGMAPKIAWALENKPRIPVVWIHGL WP_001058323.1
MULTISPECIES: hydrogenase-1 small chain [Proteobacteria] Length: 372\nNumber of predicted TMHs: 2\nExp number of AAs in TMHs: 37.2608499999999\nExp number, first 60 AAs: 19.52923\nTotal prob of N-in: 0.94351\nPOSSIBLE N-term signal sequence\ninside 1 19\nTMhelix 20 39\noutside 40 325\nTMhelix 326 348\ninside 349 372

2390 GCF_000971615.1_ASM97161v1 Escherichia coli Proteobacteria; Gammaproteobacteria; Enterobacterales; Enterobacteriaceae; Escherichia
MNNEETFYQAMRRQGVTRRSFLKYCSLAATSLGLGAGMAPKIAWALENKPRIPVVWIHGL WP_001058323.1
MULTISPECIES: hydrogenase-1 small chain [Proteobacteria] Length: 372\nNumber of predicted TMHs: 2\nExp number of AAs in TMHs: 37.2608499999999\nExp number, first 60 AAs: 19.52923\nTotal prob of N-in: 0.94351\nPOSSIBLE N-term signal sequence\ninside 1 19\nTMhelix 20 39\noutside 40 325\nTMhelix 326 348\ninside 349 372

2391 GCF_000969545.1_ASM96954v1 Escherichia coli Proteobacteria; Gammaproteobacteria; Enterobacterales; Enterobacteriaceae; Escherichia
MTGDNTLIHSHGINRRDFMKLCAALAATMGLSSKAAAEMAESVTNPQRPPVIWIGAQECT WP_046125440.1
hydrogenase 2 small subunit, partial [Escherichia coli] Length: 356\nNumber of predicted TMHs: 1\nExp number of AAs in TMHs: 23.4608\nExp number, first 60 AAs: 0.22207\nTotal prob of N-in: 0.19726\noutside 1 332\nTMhelix 333 355\ninside 356 356

2392 GCF_000965575.1_ASM96557v1 Escherichia coli Proteobacteria; Gammaproteobacteria; Enterobacterales; Enterobacteriaceae; Escherichia
MNNEETFYQAMRRQGVTRRSFLKYCSLAATSLGLGAGMAPKIAWALENKPRIPVVWIHGL WP_024246676.1

hydrogenase-1 small chain [Escherichia coli] Length: 372\nNumber of predicted TMHs: 2\nExp number of AAs in TMHs: 37.8214899999999\nExp number, first 60 AAs: 19.52326\nTotal prob of N-in: 0.94395\nPOSSIBLE N-term signal sequence\ninside 1 19\nTMhelix 20 39\noutside 40 325\nTMhelix 326 348\ninside 349 372

2393 GCF_000965625.1_ASM96562v1 Escherichia coli Proteobacteria; Gammaproteobacteria; Enterobacterales; Enterobacteriaceae; Escherichia MNNEETFYQAMRRQGVTRRSFLKYCSLAATSLGLGAGMAPKIAWALENKPRIPVVWIHGL WP_044723380.1 hydrogenase [Escherichia coli] Length: 372\nNumber of predicted TMHs: 2\nExp number of AAs in TMHs: 37.2612299999999\nExp number, first 60 AAs: 19.52923\nTotal prob of N-in: 0.94351\nPOSSIBLE N-term signal sequence\ninside 1 19\nTMhelix 20 39\noutside 40 325\nTMhelix 326 348\ninside 349 372

2394 GCF_000987525.1_ASM98752v1 Escherichia coli Proteobacteria; Gammaproteobacteria; Enterobacterales; Enterobacteriaceae; Escherichia MNNEETFYQAMRRQGVTRRSFLKYCSLAATSLGLGAGMAPKIAWALENKPRIPVVWIHGL WP_001058320.1 hydrogenase [Escherichia coli] Length: 372\nNumber of predicted TMHs: 2\nExp number of AAs in TMHs: 37.26082\nExp number, first 60 AAs: 19.52923\nTotal prob of N-in: 0.94351\nPOSSIBLE N-term signal sequence\ninside 1 19\nTMhelix 20 39\noutside 40 325\nTMhelix 326 348\ninside 349 372

2395 GCF_000981065.1_ASM98106v1 Escherichia coli Proteobacteria; Gammaproteobacteria; Enterobacterales; Enterobacteriaceae; Escherichia MNNEETFYQAMRRQGVTRRSFLKYCSLAATSLGLGAGMAPKIAWALENKPRIPVVWIHGL WP_001058323.1 MULTISPECIES: hydrogenase-1 small chain [Proteobacteria] Length: 372\nNumber of predicted TMHs: 2\nExp number of AAs in TMHs: 37.2608499999999\nExp number, first 60 AAs: 19.52923\nTotal prob of N-in: 0.94351\nPOSSIBLE N-term signal sequence\ninside 1 19\nTMhelix 20 39\noutside 40 325\nTMhelix 326 348\ninside 349 372

2396 GCF_000987535.1_ASM98753v1 Escherichia coli Proteobacteria; Gammaproteobacteria; Enterobacterales; Enterobacteriaceae; Escherichia MTGDNTLIHSHGINRRDFMKLCAALAATMGLSSKAAAEMAESVTNPQRPPVIWIGAQECT WP_000145410.1 MULTISPECIES: hydrogenase-2 small chain [Proteobacteria] Length: 372\nNumber of predicted TMHs: 1\nExp number of AAs in TMHs: 23.72226\nExp number, first 60 AAs: 0.22038\nTotal prob of N-in: 0.12994\noutside 1 331\nTMhelix 332 354\ninside 355 372

2397 GCF_000987535.1_ASM98753v1 Escherichia coli Proteobacteria; Gammaproteobacteria; Enterobacterales; Enterobacteriaceae; Escherichia MNNEETFYQAMRRQGVTRRSFLKYCSLAATSLGLGAGMAPKIAWALENKPRIPVVWIHGL WP_001058320.1 hydrogenase [Escherichia coli] Length: 372\nNumber of predicted TMHs: 2\nExp number of AAs in TMHs: 37.26082\nExp number, first 60 AAs: 19.52923\nTotal prob of N-in: 0.94351\nPOSSIBLE N-term signal sequence\ninside 1 19\nTMhelix 20 39\noutside 40 325\nTMhelix 326 348\ninside 349 372

2398 GCF_001005605.1_ASM100560v1 Escherichia coli Proteobacteria; Gammaproteobacteria; Enterobacterales; Enterobacteriaceae; Escherichia MNNEETFYQAMRRQGVTRRSFLKYCSLAATSLGLGAGMAPKIAWALENKPRIPVVWIHGL WP_001058323.1 MULTISPECIES: hydrogenase-1 small chain [Proteobacteria] Length: 372\nNumber of predicted TMHs: 2\nExp number of AAs in TMHs: 37.2608499999999\nExp number, first 60 AAs: 19.52923\nTotal prob of N-in: 0.94351\nPOSSIBLE N-term signal sequence\ninside 1 19\nTMhelix 20 39\noutside 40 325\nTMhelix 326 348\ninside 349 372

2399 GCF_001012105.1_CFSAN026778_02.0Escherichia coli Proteobacteria; Gammaproteobacteria; Enterobacterales; Enterobacteriaceae; Escherichia MTGDNTLIHSHGINRRDFMKLCAALAATMGLSSKAAAEMAESVTNPQRPPVIWIGAQECT WP_000145410.1 MULTISPECIES: hydrogenase-2 small chain [Proteobacteria] Length: 372\nNumber of predicted TMHs: 1\nExp number of AAs in TMHs: 23.72226\nExp number, first 60 AAs: 0.22038\nTotal prob of N-in: 0.12994\noutside 1 331\nTMhelix 332 354\ninside 355 372

2400 GCF_001012085.1_CFSAN026834_01.0Escherichia coli Proteobacteria; Gammaproteobacteria; Enterobacterales; Enterobacteriaceae; Escherichia MTGDNTLIHSHGINRRDFMKLCAALAATMGLSSKAAAEMAESVTNPQRPPVIWIGAQECT WP_000145410.1 MULTISPECIES: hydrogenase-2 small chain [Proteobacteria] Length: 372\nNumber of predicted TMHs: 1\nExp number of AAs in TMHs: 23.72226\nExp number, first 60 AAs: 0.22038\nTotal prob of N-in: 0.12994\noutside 1 331\nTMhelix 332 354\ninside 355 372

2401 GCF_001012155.1_CFSAN026779_02.0Escherichia coli Proteobacteria; Gammaproteobacteria; Enterobacterales; Enterobacteriaceae; Escherichia
MTGDNTLIHSHGINRRDFMKLCAALAATMGLSSKAAAEMAESVTNPQRPPVIWIGAQECT WP_000145410.1
MULTISPECIES: hydrogenase-2 small chain [Proteobacteria] Length: 372\nNumber of predicted TMHs: 1\nExp number of AAs in TMHs: 23.72226\nExp number, first 60 AAs: 0.22038\nTotal prob of N-in: 0.12994\noutside 1 331\nTMhelix 332 354\ninside 355 372

2402 GCF_001012085.1_CFSAN026834_01.0Escherichia coli Proteobacteria; Gammaproteobacteria; Enterobacterales; Enterobacteriaceae; Escherichia
MNNEETFYQAMRRQGVTRRSFLKYCSLAATSLGLGAGMAPKIAWALENKPRIPVVWIHGL WP_001058323.1
MULTISPECIES: hydrogenase-1 small chain [Proteobacteria] Length: 372\nNumber of predicted TMHs: 2\nExp number of AAs in TMHs: 37.2608499999999\nExp number, first 60 AAs: 19.52923\nTotal prob of N-in: 0.94351\nPOSSIBLE N-term signal sequence\ninside 1 19\nTMhelix 20 39\noutside 40 325\nTMhelix 326 348\ninside 349 372

2403 GCF_001012105.1_CFSAN026778_02.0Escherichia coli Proteobacteria; Gammaproteobacteria; Enterobacterales; Enterobacteriaceae; Escherichia
MNNEETFYQAMRRQGVTRRSFLKYCSLAATSLGLGAGMAPKIAWALENKPRIPVVWIHGL WP_001058323.1
MULTISPECIES: hydrogenase-1 small chain [Proteobacteria] Length: 372\nNumber of predicted TMHs: 2\nExp number of AAs in TMHs: 37.2608499999999\nExp number, first 60 AAs: 19.52923\nTotal prob of N-in: 0.94351\nPOSSIBLE N-term signal sequence\ninside 1 19\nTMhelix 20 39\noutside 40 325\nTMhelix 326 348\ninside 349 372

2404 GCF_001012155.1_CFSAN026779_02.0Escherichia coli Proteobacteria; Gammaproteobacteria; Enterobacterales; Enterobacteriaceae; Escherichia
MNNEETFYQAMRRQGVTRRSFLKYCSLAATSLGLGAGMAPKIAWALENKPRIPVVWIHGL WP_001058323.1
MULTISPECIES: hydrogenase-1 small chain [Proteobacteria] Length: 372\nNumber of predicted TMHs: 2\nExp number of AAs in TMHs: 37.2608499999999\nExp number, first 60 AAs: 19.52923\nTotal prob of N-in: 0.94351\nPOSSIBLE N-term signal sequence\ninside 1 19\nTMhelix 20 39\noutside 40 325\nTMhelix 326 348\ninside 349 372

2405 GCF_001012275.1_CFSAN026791_01.0Escherichia coli Proteobacteria; Gammaproteobacteria; Enterobacterales; Enterobacteriaceae; Escherichia
MTGDNTLIHSHGINRRDFMKLCAALAATMGLSSKAAAEMAESVTNPQRPPVIWIGAQECT WP_000145410.1
MULTISPECIES: hydrogenase-2 small chain [Proteobacteria] Length: 372\nNumber of predicted TMHs: 1\nExp number of AAs in TMHs: 23.72226\nExp number, first 60 AAs: 0.22038\nTotal prob of N-in: 0.12994\noutside 1 331\nTMhelix 332 354\ninside 355 372

2406 GCF_001012195.1_CFSAN026788_02.0Escherichia coli Proteobacteria; Gammaproteobacteria; Enterobacterales; Enterobacteriaceae; Escherichia
MNNEETFYQAMRRQGVTRRSFLKYCSLAATSLGLGAGMAPKIAWALENKPRIPVVWIHGL WP_001058323.1
MULTISPECIES: hydrogenase-1 small chain [Proteobacteria] Length: 372\nNumber of predicted TMHs: 2\nExp number of AAs in TMHs: 37.2608499999999\nExp number, first 60 AAs: 19.52923\nTotal prob of N-in: 0.94351\nPOSSIBLE N-term signal sequence\ninside 1 19\nTMhelix 20 39\noutside 40 325\nTMhelix 326 348\ninside 349 372

2407 GCF_001012395.1_CFSAN026813_01.0Escherichia coli Proteobacteria; Gammaproteobacteria; Enterobacterales; Enterobacteriaceae; Escherichia
MTGDNTLIHSHGINRRDFMKLCAALAATMGLSSKAAAEMAESVTNPQRPPVIWIGAQECT WP_000145410.1
MULTISPECIES: hydrogenase-2 small chain [Proteobacteria] Length: 372\nNumber of predicted TMHs: 1\nExp number of AAs in TMHs: 23.72226\nExp number, first 60 AAs: 0.22038\nTotal prob of N-in: 0.12994\noutside 1 331\nTMhelix 332 354\ninside 355 372

2408 GCF_001012175.1_CFSAN026787_02.0Escherichia coli Proteobacteria; Gammaproteobacteria; Enterobacterales; Enterobacteriaceae; Escherichia
MNNEETFYQAMRRQGVTRRSFLKYCSLAATSLGLGAGMAPKIAWALENKPRIPVVWIHGL WP_001295940.1
MULTISPECIES: hydrogenase [Enterobacteriaceae] Length: 372\nNumber of predicted TMHs: 2\nExp number of AAs in TMHs: 37.2608899999999\nExp number, first 60 AAs: 19.52923\nTotal prob of N-in: 0.94351\nPOSSIBLE N-term signal sequence\ninside 1 19\nTMhelix 20 39\noutside 40 325\nTMhelix 326 348\ninside 349 372

2409 GCF_001012315.1_CFSAN026801_01.0 Escherichia coli Proteobacteria; Gammaproteobacteria; Enterobacterales; Enterobacteriaceae; Escherichia
MTGDNTLIHSHGINRRDFMKLCAALAATMGLSSKAAAEMAESVTNPQRPPVIWIGAQECT WP_000145410.1
MULTISPECIES: hydrogenase-2 small chain [Proteobacteria] Length: 372\nNumber of predicted TMHs: 1\nExp number of AAs in TMHs: 23.72226\nExp number, first 60 AAs: 0.22038\nTotal prob of N-in: 0.12994\noutside 1 331\nTMhelix 332 354\ninside 355 372

2410 GCF_001012445.1_CFSAN026826_02.0 Escherichia coli Proteobacteria; Gammaproteobacteria; Enterobacterales; Enterobacteriaceae; Escherichia
MTGDNTLIHSHGINRRDFMKLCAALAATMGLSSKAAAEMAESVTNPQRPPVIWIGAQECT WP_000145410.1
MULTISPECIES: hydrogenase-2 small chain [Proteobacteria] Length: 372\nNumber of predicted TMHs: 1\nExp number of AAs in TMHs: 23.72226\nExp number, first 60 AAs: 0.22038\nTotal prob of N-in: 0.12994\noutside 1 331\nTMhelix 332 354\ninside 355 372

2411 GCF_001012405.1_CFSAN026815_01.0 Escherichia coli Proteobacteria; Gammaproteobacteria; Enterobacterales; Enterobacteriaceae; Escherichia
MTGDNTLIHSHGINRRDFMKLCAALAATMGLSSKAAAEMAESVTNPQRPPVIWIGAQECT WP_000145410.1
MULTISPECIES: hydrogenase-2 small chain [Proteobacteria] Length: 372\nNumber of predicted TMHs: 1\nExp number of AAs in TMHs: 23.72226\nExp number, first 60 AAs: 0.22038\nTotal prob of N-in: 0.12994\noutside 1 331\nTMhelix 332 354\ninside 355 372

2412 GCF_001012505.1_CFSAN026836_01.0 Escherichia coli Proteobacteria; Gammaproteobacteria; Enterobacterales; Enterobacteriaceae; Escherichia
MTGDNTLIHSHGINRRDFMKLCAALAATMGLSSKAAAEMAESVTNPQRPPVIWIGAQECT WP_000145410.1
MULTISPECIES: hydrogenase-2 small chain [Proteobacteria] Length: 372\nNumber of predicted TMHs: 1\nExp number of AAs in TMHs: 23.72226\nExp number, first 60 AAs: 0.22038\nTotal prob of N-in: 0.12994\noutside 1 331\nTMhelix 332 354\ninside 355 372

2413 GCF_001012495.1_CFSAN026835_01.0 Escherichia coli Proteobacteria; Gammaproteobacteria; Enterobacterales; Enterobacteriaceae; Escherichia
MNNEETFYQAMRRQGVTRRSFLKYCSLAATSLGLGAGMAPKIAWALENKPRIPVVWIHGL WP_001058323.1
MULTISPECIES: hydrogenase-1 small chain [Proteobacteria] Length: 372\nNumber of predicted TMHs: 2\nExp number of AAs in TMHs: 37.2608499999999\nExp number, first 60 AAs: 19.52923\nTotal prob of N-in: 0.94351\nPOSSIBLE N-term signal sequence\ninside 1 19\nTMhelix 20 39\noutside 40 325\nTMhelix 326 348\ninside 349 372

2414 GCF_001012505.1_CFSAN026836_01.0 Escherichia coli Proteobacteria; Gammaproteobacteria; Enterobacterales; Enterobacteriaceae; Escherichia
MNNEETFYQAMRRQGVTRRSFLKYCSLAATSLGLGAGMAPKIAWALENKPRIPVVWIHGL WP_001058323.1
MULTISPECIES: hydrogenase-1 small chain [Proteobacteria] Length: 372\nNumber of predicted TMHs: 2\nExp number of AAs in TMHs: 37.2608499999999\nExp number, first 60 AAs: 19.52923\nTotal prob of N-in: 0.94351\nPOSSIBLE N-term signal sequence\ninside 1 19\nTMhelix 20 39\noutside 40 325\nTMhelix 326 348\ninside 349 372

2415 GCF_001012345.1_CFSAN026809_01.0 Escherichia coli Proteobacteria; Gammaproteobacteria; Enterobacterales; Enterobacteriaceae; Escherichia
MNNEETFYQAMRRQGVTRRSFLKYCSLAATSLGLGAGMAPKIAWALENKPRIPVVWIHGL WP_001295940.1
MULTISPECIES: hydrogenase [Enterobacteriaceae] Length: 372\nNumber of predicted TMHs: 2\nExp number of AAs in TMHs: 37.2608899999999\nExp number, first 60 AAs: 19.52923\nTotal prob of N-in: 0.94351\nPOSSIBLE N-term signal sequence\ninside 1 19\nTMhelix 20 39\noutside 40 325\nTMhelix 326 348\ninside 349 372

2416 GCF_001012545.1_CFSAN026844_01.0 Escherichia coli Proteobacteria; Gammaproteobacteria; Enterobacterales; Enterobacteriaceae; Escherichia
MTGDNTLIHSHGINRRDFMKLCAALAATMGLSSKAAAEMAESVTNPQRPPVIWIGAQECT WP_000145410.1
MULTISPECIES: hydrogenase-2 small chain [Proteobacteria] Length: 372\nNumber of predicted TMHs: 1\nExp number of AAs in TMHs: 23.72226\nExp number, first 60 AAs: 0.22038\nTotal prob of N-in: 0.12994\noutside 1 331\nTMhelix 332 354\ninside 355 372

2417 GCF_001012545.1_CFSAN026844_01.0 Escherichia coli Proteobacteria; Gammaproteobacteria; Enterobacterales; Enterobacteriaceae; Escherichia

MNNEETFYQAMRRQGVTRRSFLKYCSLAATSLGLGAGMAPKIAWALENKPRIPVVWIHGL WP_001058323.1 MULTISPECIES: hydrogenase-1 small chain [Proteobacteria] Length: 372\nNumber of predicted TMHs: 2\nExp number of AAs in TMHs: 37.2608499999999\nExp number, first 60 AAs: 19.52923\nTotal prob of N-in: 0.94351\nPOSSIBLE N-term signal sequence\ninside 1 19\nTMhelix 20 39\noutside 40 325\nTMhelix 326 348\ninside 349 372

2418 GCF_001030325.1_Esch_coli_BWH61_V1 Escherichia coli Proteobacteria; Gammaproteobacteria; Enterobacterales; Enterobacteriaceae; Escherichia MNNEETFYQAMRRQGVTRRSFLKYCSLAATSLGLGAGMAPKIAWALENKPRIPVVWIHGL WP_001295940.1 MULTISPECIES: hydrogenase [Enterobacteriaceae] Length: 372\nNumber of predicted TMHs: 2\nExp number of AAs in TMHs: 37.2608899999999\nExp number, first 60 AAs: 19.52923\nTotal prob of N-in: 0.94351\nPOSSIBLE N-term signal sequence\ninside 1 19\nTMhelix 20 39\noutside 40 325\nTMhelix 326 348\ninside 349 372

2419 GCF_001030345.1_Esch_coli_MGH107_V1 Escherichia coli Proteobacteria; Gammaproteobacteria; Enterobacterales; Enterobacteriaceae; Escherichia MTGDNTLIHSHGINRRDFMKLCAALAATMGLSSKAAAEMVESVTNPQRPPVIWIGAQECT WP_000145423.1 hydrogenase 2 small subunit [Escherichia coli] Length: 372\nNumber of predicted TMHs: 1\nExp number of AAs in TMHs: 23.78201\nExp number, first 60 AAs: 0.28355\nTotal prob of N-in: 0.13257\noutside 1 331\nTMhelix 332 354\ninside 355 372

2420 GCF_001030365.1_Esch_coli_BWH55_V1 Escherichia coli Proteobacteria; Gammaproteobacteria; Enterobacterales; Enterobacteriaceae; Escherichia MTGDNTLIHSHGINRRDFMKLCAALAATMGLSSKAAAEMVESVTNPQRPPVIWIGAQECT WP_000145423.1 hydrogenase 2 small subunit [Escherichia coli] Length: 372\nNumber of predicted TMHs: 1\nExp number of AAs in TMHs: 23.78201\nExp number, first 60 AAs: 0.28355\nTotal prob of N-in: 0.13257\noutside 1 331\nTMhelix 332 354\ninside 355 372

2421 GCF_001030345.1_Esch_coli_MGH107_V1 Escherichia coli Proteobacteria; Gammaproteobacteria; Enterobacterales; Enterobacteriaceae; Escherichia MNNEETFYQAMRRQGVTRRSFLKYCSLAATSLGLGAGMAPKIAWALENKPRIPVVWIHGL WP_001295940.1 MULTISPECIES: hydrogenase [Enterobacteriaceae] Length: 372\nNumber of predicted TMHs: 2\nExp number of AAs in TMHs: 37.2608899999999\nExp number, first 60 AAs: 19.52923\nTotal prob of N-in: 0.94351\nPOSSIBLE N-term signal sequence\ninside 1 19\nTMhelix 20 39\noutside 40 325\nTMhelix 326 348\ninside 349 372

2422 GCF_001030365.1_Esch_coli_BWH55_V1 Escherichia coli Proteobacteria; Gammaproteobacteria; Enterobacterales; Enterobacteriaceae; Escherichia MNNEETFYQAMRRQGVTRRSFLKYCSLAATSLGLGAGMAPKIAWALENKPRIPVVWIHGL WP_001295940.1 MULTISPECIES: hydrogenase [Enterobacteriaceae] Length: 372\nNumber of predicted TMHs: 2\nExp number of AAs in TMHs: 37.2608899999999\nExp number, first 60 AAs: 19.52923\nTotal prob of N-in: 0.94351\nPOSSIBLE N-term signal sequence\ninside 1 19\nTMhelix 20 39\noutside 40 325\nTMhelix 326 348\ninside 349 372

2423 GCF_001030545.1_Esch_coli_BIDMC103_V1 Escherichia coli Proteobacteria; Gammaproteobacteria; Enterobacterales; Enterobacteriaceae; Escherichia MTGDNTLIHSHGINRRDFMKLCAALAATMGLSSKAAAEMAESVTNPQRPPVIWIGAQECT WP_000145410.1 MULTISPECIES: hydrogenase-2 small chain [Proteobacteria] Length: 372\nNumber of predicted TMHs: 1\nExp number of AAs in TMHs: 23.72226\nExp number, first 60 AAs: 0.22038\nTotal prob of N-in: 0.12994\noutside 1 331\nTMhelix 332 354\ninside 355 372

2424 GCF_001030585.1_Esch_coli_BIDMC105_V1 Escherichia coli Proteobacteria; Gammaproteobacteria; Enterobacterales; Enterobacteriaceae; Escherichia MTGDNTLIHSHGINRRDFMKLCAALAATMGLSSKAAAEMVESVTNPQRPPVIWIGAQECT WP_000145423.1 hydrogenase 2 small subunit [Escherichia coli] Length: 372\nNumber of predicted TMHs: 1\nExp number of AAs in TMHs: 23.78201\nExp number, first 60 AAs: 0.28355\nTotal prob of N-in: 0.13257\noutside 1 331\nTMhelix 332 354\ninside 355 372

2425 GCF_001030595.1_Esch_coli_BIDMC106_V1 Escherichia coli Proteobacteria; Gammaproteobacteria; Enterobacterales; Enterobacteriaceae; Escherichia MTGDNTLIHSHGINRRDFMKLCAALAATMGLSSKAAAEMAESVTNPQRPPVIWIGAQECT WP_000145410.1

MULTISPECIES: hydrogenase-2 small chain [Proteobacteria] Length: 372\nNumber of predicted TMHs: 1\nExp number of AAs in TMHs: 23.72226\nExp number, first 60 AAs: 0.22038\nTotal prob of N-in: 0.12994\noutside 1 331\nTMhelix 332 354\ninside 355 372

2426 GCF_001054175.1_ASM105417v1 Escherichia coli Proteobacteria; Gammaproteobacteria; Enterobacterales; Enterobacteriaceae; Escherichia
MTGDNTLIHSHGINRRDFMKLCAALAATMGLSSKAAAEMAESVTNPQRPPVIWIGAQECT WP_000145410.1
MULTISPECIES: hydrogenase-2 small chain [Proteobacteria] Length: 372\nNumber of predicted TMHs: 1\nExp number of AAs in TMHs: 23.72226\nExp number, first 60 AAs: 0.22038\nTotal prob of N-in: 0.12994\noutside 1 331\nTMhelix 332 354\ninside 355 372

2427 GCF_001012615.1_CFSAN026816_01.0 Escherichia coli Proteobacteria; Gammaproteobacteria; Enterobacterales; Enterobacteriaceae; Escherichia
MNNEETFYQAMRRQGVTRRSFLKYCSLAATSLGLGAGMAPKIAWALENKPRIPVVWIHGL WP_001058320.1
hydrogenase [Escherichia coli] Length: 372\nNumber of predicted TMHs: 2\nExp number of AAs in TMHs: 37.26082\nExp number, first 60 AAs: 19.52923\nTotal prob of N-in: 0.94351\nPOSSIBLE N-term signal sequence\ninside 1 19\nTMhelix 20 39\noutside 40 325\nTMhelix 326 348\ninside 349 372

2428 GCF_001030635.1_Esch_coli_BIDMC112_V1 Escherichia coli Proteobacteria; Gammaproteobacteria; Enterobacterales; Enterobacteriaceae; Escherichia
MTGDNTLIHSHGINRRDFMKLCAALAATMGLSSKAAAEMAESVTNPQRPPVIWIGAQECT WP_000145410.1
MULTISPECIES: hydrogenase-2 small chain [Proteobacteria] Length: 372\nNumber of predicted TMHs: 1\nExp number of AAs in TMHs: 23.72226\nExp number, first 60 AAs: 0.22038\nTotal prob of N-in: 0.12994\noutside 1 331\nTMhelix 332 354\ninside 355 372

2429 GCF_001030635.1_Esch_coli_BIDMC112_V1 Escherichia coli Proteobacteria; Gammaproteobacteria; Enterobacterales; Enterobacteriaceae; Escherichia
MNNEETFYQAMRRQGVTRRSFLKYCSLAATSLGLGAGMAPKIAWALENKPRIPVVWIHGL WP_021557704.1
hydrogenase-1 small chain [Escherichia coli] Length: 372\nNumber of predicted TMHs: 2\nExp number of AAs in TMHs: 37.2609699999999\nExp number, first 60 AAs: 19.52923\nTotal prob of N-in: 0.94351\nPOSSIBLE N-term signal sequence\ninside 1 19\nTMhelix 20 39\noutside 40 325\nTMhelix 326 348\ninside 349 372

2430 GCF_001030435.1_Esch_coli_MGH122_V1 Escherichia coli Proteobacteria; Gammaproteobacteria; Enterobacterales; Enterobacteriaceae; Escherichia
MNNEETFYQAMRRQGVTRRSFLKYCSLAATSLGLGAGMAPKIAWALENKPRIPVVWIHGL WP_001058323.1
MULTISPECIES: hydrogenase-1 small chain [Proteobacteria] Length: 372\nNumber of predicted TMHs: 2\nExp number of AAs in TMHs: 37.2608499999999\nExp number, first 60 AAs: 19.52923\nTotal prob of N-in: 0.94351\nPOSSIBLE N-term signal sequence\ninside 1 19\nTMhelix 20 39\noutside 40 325\nTMhelix 326 348\ninside 349 372

2431 GCF_001030595.1_Esch_coli_BIDMC106_V1 Escherichia coli Proteobacteria; Gammaproteobacteria; Enterobacterales; Enterobacteriaceae; Escherichia
MNNEETFYQAMRRQGVTRRSFLKYCSLAATSLGLGAGMAPKIAWALENKPRIPVVWIHGL WP_001058323.1
MULTISPECIES: hydrogenase-1 small chain [Proteobacteria] Length: 372\nNumber of predicted TMHs: 2\nExp number of AAs in TMHs: 37.2608499999999\nExp number, first 60 AAs: 19.52923\nTotal prob of N-in: 0.94351\nPOSSIBLE N-term signal sequence\ninside 1 19\nTMhelix 20 39\noutside 40 325\nTMhelix 326 348\ninside 349 372

2432 GCF_001030485.1_Esch_coli_BIDMC98_V1 Escherichia coli Proteobacteria; Gammaproteobacteria; Enterobacterales; Enterobacteriaceae; Escherichia
MNNEETFYQAMRRQGVTRRSFLKYCSLAATSLGLGAGMAPKIAWALENKPRIPVVWIHGL WP_001058323.1
MULTISPECIES: hydrogenase-1 small chain [Proteobacteria] Length: 372\nNumber of predicted TMHs: 2\nExp number of AAs in TMHs: 37.2608499999999\nExp number, first 60 AAs: 19.52923\nTotal prob of N-in: 0.94351\nPOSSIBLE N-term signal sequence\ninside 1 19\nTMhelix 20 39\noutside 40 325\nTMhelix 326 348\ninside 349 372

2433 GCF_001030485.1_Esch_coli_BIDMC98_V1 Escherichia coli Proteobacteria; Gammaproteobacteria; Enterobacterales; Enterobacteriaceae; Escherichia
MTGDNTLIHSHGINRRDFMKLCAALAATMGLSSKAAAEMAESVTNPQRPPVIWIGAQECT WP_032298447.1
hydrogenase 2 small subunit [Escherichia coli] Length: 372\nNumber of predicted TMHs: 1\nExp number of

AAs in TMHs: 23.69807\nExp number, first 60 AAs: 0.221\nTotal prob of N-in: 0.12853\noutside 1 331\nTMhelix 332 354\ninside 355 372

2434 GCF_001030505.1_Esch_coli_BIDMC101_V1 Escherichia coli Proteobacteria; Gammaproteobacteria; Enterobacterales; Enterobacteriaceae; Escherichia
MNNEETFYQAMRRQGVTRRSFLKYCSLAATSLGLGAGMAPKIAWALENKPRIPVVWIHGL WP_001058323.1
MULTISPECIES: hydrogenase-1 small chain [Proteobacteria] Length: 372\nNumber of predicted TMHs: 2\nExp number of AAs in TMHs: 37.2608499999999\nExp number, first 60 AAs: 19.52923\nTotal prob of N-in: 0.94351\nPOSSIBLE N-term signal sequence\ninside 1 19\nTMhelix 20 39\noutside 40 325\nTMhelix 326 348\ninside 349 372

2435 GCF_001030675.1_Esch_coli_BIDMC113_V1 Escherichia coli Proteobacteria; Gammaproteobacteria; Enterobacterales; Enterobacteriaceae; Escherichia
MTGDNTLIHSHGINRRDFMKLCAALAATMGLSSKAAAEMVESVTNPQRPPVIWIGAQECT WP_000145423.1
hydrogenase 2 small subunit [Escherichia coli] Length: 372\nNumber of predicted TMHs: 1\nExp number of AAs in TMHs: 23.78201\nExp number, first 60 AAs: 0.28355\nTotal prob of N-in: 0.13257\noutside 1 331\nTMhelix 332 354\ninside 355 372

2436 GCF_001030675.1_Esch_coli_BIDMC113_V1 Escherichia coli Proteobacteria; Gammaproteobacteria; Enterobacterales; Enterobacteriaceae; Escherichia
MNNEETFYQAMRRQGVTRRSFLKYCSLAATSLGLGAGMAPKIAWALENKPRIPVVWIHGL WP_001295940.1
MULTISPECIES: hydrogenase [Enterobacteriaceae] Length: 372\nNumber of predicted TMHs: 2\nExp number of AAs in TMHs: 37.2608899999999\nExp number, first 60 AAs: 19.52923\nTotal prob of N-in: 0.94351\nPOSSIBLE N-term signal sequence\ninside 1 19\nTMhelix 20 39\noutside 40 325\nTMhelix 326 348\ninside 349 372

2437 GCF_001039125.1_ASM103912v1 Escherichia coli Proteobacteria; Gammaproteobacteria; Enterobacterales; Enterobacteriaceae; Escherichia
MNNEETFYQAMRRQGVTRRSFLKYCSLAATSLGLGAGMAPKIAWALENKPRIPVVWIHGL WP_001058323.1
MULTISPECIES: hydrogenase-1 small chain [Proteobacteria] Length: 372\nNumber of predicted TMHs: 2\nExp number of AAs in TMHs: 37.2608499999999\nExp number, first 60 AAs: 19.52923\nTotal prob of N-in: 0.94351\nPOSSIBLE N-term signal sequence\ninside 1 19\nTMhelix 20 39\noutside 40 325\nTMhelix 326 348\ninside 349 372

2438 GCF_001039155.1_ASM103915v1 Escherichia coli Proteobacteria; Gammaproteobacteria; Enterobacterales; Enterobacteriaceae; Escherichia
MTGDNTLIHSHGINRRDFMKLCAALAATMGLSSKAAAEMAESVTNPQRPPVIWIGAQECT WP_000145410.1
MULTISPECIES: hydrogenase-2 small chain [Proteobacteria] Length: 372\nNumber of predicted TMHs: 1\nExp number of AAs in TMHs: 23.72226\nExp number, first 60 AAs: 0.22038\nTotal prob of N-in: 0.12994\noutside 1 331\nTMhelix 332 354\ninside 355 372

2439 GCF_001039215.1_ASM103921v1 Escherichia coli Proteobacteria; Gammaproteobacteria; Enterobacterales; Enterobacteriaceae; Escherichia
MTGDNTLIHSHGINRRDFMKLCAALAATMGLSSKAAAEMAESVTNPQRPPVIWIGAQECT WP_000145410.1
MULTISPECIES: hydrogenase-2 small chain [Proteobacteria] Length: 372\nNumber of predicted TMHs: 1\nExp number of AAs in TMHs: 23.72226\nExp number, first 60 AAs: 0.22038\nTotal prob of N-in: 0.12994\noutside 1 331\nTMhelix 332 354\ninside 355 372

2440 GCF_001039415.1_ASM103941v1 Escherichia coli Proteobacteria; Gammaproteobacteria; Enterobacterales; Enterobacteriaceae; Escherichia
MTGDNTLIHSHGINRRDFMKLCAALAATMGLSSKAAAEMAESVTNPQRPPVIWIGAQECT WP_000145410.1
MULTISPECIES: hydrogenase-2 small chain [Proteobacteria] Length: 372\nNumber of predicted TMHs: 1\nExp number of AAs in TMHs: 23.72226\nExp number, first 60 AAs: 0.22038\nTotal prob of N-in: 0.12994\noutside 1 331\nTMhelix 332 354\ninside 355 372

2441 GCF_001043215.1_ASM104321v1 Escherichia coli Proteobacteria; Gammaproteobacteria; Enterobacterales; Enterobacteriaceae; Escherichia
MTGDNTLIHSHGINRRDFMKLCAALAATMGLSSKAAAEMAESVTNPQRPPVIWIGAQECT WP_000145410.1
MULTISPECIES: hydrogenase-2 small chain [Proteobacteria] Length: 372\nNumber of predicted TMHs: 1\nExp number of AAs in TMHs: 23.72226\nExp number, first 60 AAs: 0.22038\nTotal prob of N-in: 0.12994\noutside 1 331\nTMhelix 332 354\ninside 355 372

2442 GCF_001052125.1_ASM105212v1 Escherichia coli Proteobacteria; Gammaproteobacteria; Enterobacterales; Enterobacteriaceae; Escherichia
MTGDNTLIHSHGINRRDFMKLCAALAATMGLSSKAAAEMAESVTNPQRPPVIWIGAQECT WP_000145410.1
MULTISPECIES: hydrogenase-2 small chain [Proteobacteria] Length: 372\nNumber of predicted TMHs: 1\nExp number of AAs in TMHs: 23.72226\nExp number, first 60 AAs: 0.22038\nTotal prob of N-in: 0.12994\noutside 1 331\nTMhelix 332 354\ninside 355 372

2443 GCF_001039215.1_ASM103921v1 Escherichia coli Proteobacteria; Gammaproteobacteria; Enterobacterales; Enterobacteriaceae; Escherichia
MNNEETFYQAMRRQGVTRRSFLKYCSLAATSLGLGAGMAPKIAWALENKPRIPVVWIHGL WP_001058323.1
MULTISPECIES: hydrogenase-1 small chain [Proteobacteria] Length: 372\nNumber of predicted TMHs: 2\nExp number of AAs in TMHs: 37.2608499999999\nExp number, first 60 AAs: 19.52923\nTotal prob of N-in: 0.94351\nPOSSIBLE N-term signal sequence\ninside 1 19\nTMhelix 20 39\noutside 40 325\nTMhelix 326 348\ninside 349 372

2444 GCF_001039415.1_ASM103941v1 Escherichia coli Proteobacteria; Gammaproteobacteria; Enterobacterales; Enterobacteriaceae; Escherichia
MNNEETFYQAMRRQGVTRRSFLKYCSLAATSLGLGAGMAPKIAWALENKPRIPVVWIHGL WP_001058323.1
MULTISPECIES: hydrogenase-1 small chain [Proteobacteria] Length: 372\nNumber of predicted TMHs: 2\nExp number of AAs in TMHs: 37.2608499999999\nExp number, first 60 AAs: 19.52923\nTotal prob of N-in: 0.94351\nPOSSIBLE N-term signal sequence\ninside 1 19\nTMhelix 20 39\noutside 40 325\nTMhelix 326 348\ninside 349 372

2445 GCF_001043215.1_ASM104321v1 Escherichia coli Proteobacteria; Gammaproteobacteria; Enterobacterales; Enterobacteriaceae; Escherichia
MNNEETFYQAMRRQGVTRRSFLKYCSLAATSLGLGAGMAPKIAWALENKPRIPVVWIHGL WP_001058323.1
MULTISPECIES: hydrogenase-1 small chain [Proteobacteria] Length: 372\nNumber of predicted TMHs: 2\nExp number of AAs in TMHs: 37.2608499999999\nExp number, first 60 AAs: 19.52923\nTotal prob of N-in: 0.94351\nPOSSIBLE N-term signal sequence\ninside 1 19\nTMhelix 20 39\noutside 40 325\nTMhelix 326 348\ninside 349 372

2446 GCF_001053245.1_ASM105324v1 Escherichia coli Proteobacteria; Gammaproteobacteria; Enterobacterales; Enterobacteriaceae; Escherichia
MTGDNTLIHSHGINRRDFMKLCAALAATMGLSSKAAAEMAESVTNPQRPPVIWIGAQECT WP_000145410.1
MULTISPECIES: hydrogenase-2 small chain [Proteobacteria] Length: 372\nNumber of predicted TMHs: 1\nExp number of AAs in TMHs: 23.72226\nExp number, first 60 AAs: 0.22038\nTotal prob of N-in: 0.12994\noutside 1 331\nTMhelix 332 354\ninside 355 372

2447 GCF_001053065.1_ASM105306v1 Escherichia coli Proteobacteria; Gammaproteobacteria; Enterobacterales; Enterobacteriaceae; Escherichia
MTGDNTLIHSHGINRRDFMKLCAALAATMGLSSKAAAEMAESVTNPQRPPVIWIGAQECT WP_000145410.1
MULTISPECIES: hydrogenase-2 small chain [Proteobacteria] Length: 372\nNumber of predicted TMHs: 1\nExp number of AAs in TMHs: 23.72226\nExp number, first 60 AAs: 0.22038\nTotal prob of N-in: 0.12994\noutside 1 331\nTMhelix 332 354\ninside 355 372

2448 GCF_001053305.1_ASM105330v1 Escherichia coli Proteobacteria; Gammaproteobacteria; Enterobacterales; Enterobacteriaceae; Escherichia
MTGDNTLIHSHGINRRDFMKLCAALAATMGLSSKAAAEMVESVTNPQRPPVIWIGAQECT WP_000145423.1
hydrogenase 2 small subunit [Escherichia coli] Length: 372\nNumber of predicted TMHs: 1\nExp number of AAs in TMHs: 23.78201\nExp number, first 60 AAs: 0.28355\nTotal prob of N-in: 0.13257\noutside 1 331\nTMhelix 332 354\ninside 355 372

2449 GCF_001054145.1_ASM105414v1 Escherichia coli Proteobacteria; Gammaproteobacteria; Enterobacterales; Enterobacteriaceae; Escherichia
MTGDNTLIHSHGINRRDFMKLCAALAATMGLSSKAAAEMAESVTNPQRPPVIWIGAQECT WP_000145410.1
MULTISPECIES: hydrogenase-2 small chain [Proteobacteria] Length: 372\nNumber of predicted TMHs: 1\nExp number of AAs in TMHs: 23.72226\nExp number, first 60 AAs: 0.22038\nTotal prob of N-in: 0.12994\noutside 1 331\nTMhelix 332 354\ninside 355 372

2450 GCF_001054815.1_ASM105481v1 Escherichia coli Proteobacteria; Gammaproteobacteria; Enterobacterales; Enterobacteriaceae; Escherichia

MNNEETFYQAMRRQGVTRRSFLKYCSLAATSLGLGAGMAPKIAWALENKPRIPVVWIHGL WP_001058323.1 MULTISPECIES: hydrogenase-1 small chain [Proteobacteria] Length: 372\nNumber of predicted TMHs: 2\nExp number of AAs in TMHs: 37.2608499999999\nExp number, first 60 AAs: 19.52923\nTotal prob of N-in: 0.94351\nPOSSIBLE N-term signal sequence\ninside 1 19\nTMhelix 20 39\noutside 40 325\nTMhelix 326 348\ninside 349 372

2451 GCF_001054845.1_ASM105484v1 Escherichia coli Proteobacteria; Gammaproteobacteria; Enterobacterales; Enterobacteriaceae; Escherichia MNNEETFYQAMRRQGVTRRSFLKYCSLAATSLGLGAGMAPKIAWALENKPRIPVVWIHGL WP_001058323.1 MULTISPECIES: hydrogenase-1 small chain [Proteobacteria] Length: 372\nNumber of predicted TMHs: 2\nExp number of AAs in TMHs: 37.2608499999999\nExp number, first 60 AAs: 19.52923\nTotal prob of N-in: 0.94351\nPOSSIBLE N-term signal sequence\ninside 1 19\nTMhelix 20 39\noutside 40 325\nTMhelix 326 348\ninside 349 372

2452 GCF_001053305.1_ASM105330v1 Escherichia coli Proteobacteria; Gammaproteobacteria; Enterobacterales; Enterobacteriaceae; Escherichia MNNEETFYQAMRRQGVTRRSFLKYCSLAATSLGLGAGMAPKIAWALENKPRIPVVWIHGL WP_001295940.1 MULTISPECIES: hydrogenase [Enterobacteriaceae] Length: 372\nNumber of predicted TMHs: 2\nExp number of AAs in TMHs: 37.2608899999999\nExp number, first 60 AAs: 19.52923\nTotal prob of N-in: 0.94351\nPOSSIBLE N-term signal sequence\ninside 1 19\nTMhelix 20 39\noutside 40 325\nTMhelix 326 348\ninside 349 372

2453 GCF_001054145.1_ASM105414v1 Escherichia coli Proteobacteria; Gammaproteobacteria; Enterobacterales; Enterobacteriaceae; Escherichia MNNEETFYQAMRRQGVTRRSFLKYCSLAATSLGLGAGMAPKIAWALENKPRIPVVWIHGL WP_001058323.1 MULTISPECIES: hydrogenase-1 small chain [Proteobacteria] Length: 372\nNumber of predicted TMHs: 2\nExp number of AAs in TMHs: 37.2608499999999\nExp number, first 60 AAs: 19.52923\nTotal prob of N-in: 0.94351\nPOSSIBLE N-term signal sequence\ninside 1 19\nTMhelix 20 39\noutside 40 325\nTMhelix 326 348\ninside 349 372

2454 GCF_001056195.1_ASM105619v1 Escherichia coli Proteobacteria; Gammaproteobacteria; Enterobacterales; Enterobacteriaceae; Escherichia MTGDNTLIHSHGINRRDFMKLCAALAATMGLSSKAAAEMAESVTNPQRPPVIWIGAQECT WP_000145410.1 MULTISPECIES: hydrogenase-2 small chain [Proteobacteria] Length: 372\nNumber of predicted TMHs: 1\nExp number of AAs in TMHs: 23.72226\nExp number, first 60 AAs: 0.22038\nTotal prob of N-in: 0.12994\noutside 1 331\nTMhelix 332 354\ninside 355 372

2455 GCF_001056195.1_ASM105619v1 Escherichia coli Proteobacteria; Gammaproteobacteria; Enterobacterales; Enterobacteriaceae; Escherichia MNNEETFYQAMRRQGVTRRSFLKYCSLAATSLGLGAGMAPKIAWALENKPRIPVVWIHGL WP_001058323.1 MULTISPECIES: hydrogenase-1 small chain [Proteobacteria] Length: 372\nNumber of predicted TMHs: 2\nExp number of AAs in TMHs: 37.2608499999999\nExp number, first 60 AAs: 19.52923\nTotal prob of N-in: 0.94351\nPOSSIBLE N-term signal sequence\ninside 1 19\nTMhelix 20 39\noutside 40 325\nTMhelix 326 348\ninside 349 372

2456 GCF_001054615.1_ASM105461v1 Escherichia coli Proteobacteria; Gammaproteobacteria; Enterobacterales; Enterobacteriaceae; Escherichia MNNEETFYQAMRRQGVTRRSFLKYCSLAATSLGLGAGMAPKIAWALENKPRIPVVWIHGL WP_001295940.1 MULTISPECIES: hydrogenase [Enterobacteriaceae] Length: 372\nNumber of predicted TMHs: 2\nExp number of AAs in TMHs: 37.2608899999999\nExp number, first 60 AAs: 19.52923\nTotal prob of N-in: 0.94351\nPOSSIBLE N-term signal sequence\ninside 1 19\nTMhelix 20 39\noutside 40 325\nTMhelix 326 348\ninside 349 372

2457 GCF_001055685.1_ASM105568v1 Escherichia coli Proteobacteria; Gammaproteobacteria; Enterobacterales; Enterobacteriaceae; Escherichia MNNEETFYQAMRRQGVTRRSFLKYCSLAATSLGLGAGMAPKIAWALENKPRIPVVWIHGL WP_001295940.1 MULTISPECIES: hydrogenase [Enterobacteriaceae] Length: 372\nNumber of predicted TMHs: 2\nExp number of AAs in TMHs: 37.2608899999999\nExp number, first 60 AAs: 19.52923\nTotal prob of N-in: 0.94351\nPOSSIBLE N-term signal sequence\ninside 1 19\nTMhelix 20 39\noutside 40 325\nTMhelix 326 348\ninside 349 372

2458 GCF_001056025.1_ASM105602v1 Escherichia coli Proteobacteria; Gammaproteobacteria; Enterobacterales; Enterobacteriaceae; Escherichia
MNNEETFYQAMRRQGVTRRSFLKYCSLAATSLGLGAGMAPKIAWALENKPRIPVVWIHGL WP_001295940.1
MULTISPECIES: hydrogenase [Enterobacteriaceae] Length: 372\nNumber of predicted TMHs: 2\nExp number of AAs in TMHs: 37.2608899999999\nExp number, first 60 AAs: 19.52923\nTotal prob of N-in: 0.94351\nPOSSIBLE N-term signal sequence\ninside 1 19\nTMhelix 20 39\noutside 40 325\nTMhelix 326 348\ninside 349 372

2459 GCF_001056975.1_ASM105697v1 Escherichia coli Proteobacteria; Gammaproteobacteria; Enterobacterales; Enterobacteriaceae; Escherichia
MTGDNTLIHSHGINRRDFMKLCAALAATMGLSSKAAAEMAESVTNPQRPPVIWIGAQECT WP_000145410.1
MULTISPECIES: hydrogenase-2 small chain [Proteobacteria] Length: 372\nNumber of predicted TMHs: 1\nExp number of AAs in TMHs: 23.72226\nExp number, first 60 AAs: 0.22038\nTotal prob of N-in: 0.12994\noutside 1 331\nTMhelix 332 354\ninside 355 372

2460 GCF_001056975.1_ASM105697v1 Escherichia coli Proteobacteria; Gammaproteobacteria; Enterobacterales; Enterobacteriaceae; Escherichia
MNNEETFYQAMRRQGVTRRSFLKYCSLAATSLGLGAGMAPKIAWALENKPRIPVVWIHGL WP_001058323.1
MULTISPECIES: hydrogenase-1 small chain [Proteobacteria] Length: 372\nNumber of predicted TMHs: 2\nExp number of AAs in TMHs: 37.2608499999999\nExp number, first 60 AAs: 19.52923\nTotal prob of N-in: 0.94351\nPOSSIBLE N-term signal sequence\ninside 1 19\nTMhelix 20 39\noutside 40 325\nTMhelix 326 348\ninside 349 372

2461 GCF_001056845.1_ASM105684v1 Escherichia coli Proteobacteria; Gammaproteobacteria; Enterobacterales; Enterobacteriaceae; Escherichia
MNNEETFYQAMRRQGVTRRSFLKYCSLAATSLGLGAGMAPKIAWALENKPRIPVVWIHGL WP_001058323.1
MULTISPECIES: hydrogenase-1 small chain [Proteobacteria] Length: 372\nNumber of predicted TMHs: 2\nExp number of AAs in TMHs: 37.2608499999999\nExp number, first 60 AAs: 19.52923\nTotal prob of N-in: 0.94351\nPOSSIBLE N-term signal sequence\ninside 1 19\nTMhelix 20 39\noutside 40 325\nTMhelix 326 348\ninside 349 372

2462 GCF_001056815.1_ASM105681v1 Escherichia coli Proteobacteria; Gammaproteobacteria; Enterobacterales; Enterobacteriaceae; Escherichia
MNNEETFYQAMRRQGVTRRSFLKYCSLAATSLGLGAGMAPKIAWALENKPRIPVVWIHGL WP_001058323.1
MULTISPECIES: hydrogenase-1 small chain [Proteobacteria] Length: 372\nNumber of predicted TMHs: 2\nExp number of AAs in TMHs: 37.2608499999999\nExp number, first 60 AAs: 19.52923\nTotal prob of N-in: 0.94351\nPOSSIBLE N-term signal sequence\ninside 1 19\nTMhelix 20 39\noutside 40 325\nTMhelix 326 348\ninside 349 372

2463 GCF_001057065.1_ASM105706v1 Escherichia coli Proteobacteria; Gammaproteobacteria; Enterobacterales; Enterobacteriaceae; Escherichia
MNNEETFYQAMRRQGVTRRSFLKYCSLAATSLGLGAGMAPKIAWALENKPRIPVVWIHGL WP_001058323.1
MULTISPECIES: hydrogenase-1 small chain [Proteobacteria] Length: 372\nNumber of predicted TMHs: 2\nExp number of AAs in TMHs: 37.2608499999999\nExp number, first 60 AAs: 19.52923\nTotal prob of N-in: 0.94351\nPOSSIBLE N-term signal sequence\ninside 1 19\nTMhelix 20 39\noutside 40 325\nTMhelix 326 348\ninside 349 372

2464 GCF_001058275.1_ASM105827v1 Escherichia coli Proteobacteria; Gammaproteobacteria; Enterobacterales; Enterobacteriaceae; Escherichia
MTGDNTLIHSHGINRRDFMKLCAALAATMGLSSKAAAEMVESVTNPQRPPVIWIGAQECT WP_000145423.1
hydrogenase 2 small subunit [Escherichia coli] Length: 372\nNumber of predicted TMHs: 1\nExp number of AAs in TMHs: 23.78201\nExp number, first 60 AAs: 0.28355\nTotal prob of N-in: 0.13257\noutside 1 331\nTMhelix 332 354\ninside 355 372

2465 GCF_001058275.1_ASM105827v1 Escherichia coli Proteobacteria; Gammaproteobacteria; Enterobacterales; Enterobacteriaceae; Escherichia
MNNEETFYQAMRRQGVTRRSFLKYCSLAATSLGLGAGMAPKIAWALENKPRIPVVWIHGL WP_001295940.1
MULTISPECIES: hydrogenase [Enterobacteriaceae] Length: 372\nNumber of predicted TMHs: 2\nExp number of AAs in TMHs: 37.2608899999999\nExp number, first 60 AAs: 19.52923\nTotal prob of N-in: 0.94351\nPOSSIBLE N-term signal sequence\ninside 1 19\nTMhelix 20 39\noutside 40 325\nTMhelix 326 348\ninside 349 372

2466 GCF_001057225.1_ASM105722v1 Escherichia coli Proteobacteria; Gammaproteobacteria; Enterobacterales; Enterobacteriaceae; Escherichia
MNNEETFYQAMRRQGVTRRSFLKYCSLAATSLGLGAGMAPKIAWALENKPRIPVVWIHGL WP_001538532.1
hydrogenase-1 small chain [Escherichia coli] Length: 372\nNumber of predicted TMHs: 2\nExp number of AAs in TMHs: 37.27041\nExp number, first 60 AAs: 19.52927\nTotal prob of N-in: 0.94352\nPOSSIBLE N-term signal sequence\ninside 1 19\nTMhelix 20 39\noutside 40 325\nTMhelix 326 348\ninside 349 372

2467 GCF_001057975.1_ASM105797v1 Escherichia coli Proteobacteria; Gammaproteobacteria; Enterobacterales; Enterobacteriaceae; Escherichia
MNNEETFYQAMRRQGVTRRSFLKYCSLAATSLGLGAGMAPKIAWALENKPRIPVVWIHGL WP_001058323.1
MULTISPECIES: hydrogenase-1 small chain [Proteobacteria] Length: 372\nNumber of predicted TMHs: 2\nExp number of AAs in TMHs: 37.2608499999999\nExp number, first 60 AAs: 19.52923\nTotal prob of N-in: 0.94351\nPOSSIBLE N-term signal sequence\ninside 1 19\nTMhelix 20 39\noutside 40 325\nTMhelix 326 348\ninside 349 372

2468 GCF_001058795.1_ASM105879v1 Escherichia coli Proteobacteria; Gammaproteobacteria; Enterobacterales; Enterobacteriaceae; Escherichia
MTGDNTLIHSHGINRRDFMKLCAALAATMGLSSKAAAEMAESVTNPQRPPVIWIGAQECT WP_000145410.1
MULTISPECIES: hydrogenase-2 small chain [Proteobacteria] Length: 372\nNumber of predicted TMHs: 1\nExp number of AAs in TMHs: 23.72226\nExp number, first 60 AAs: 0.22038\nTotal prob of N-in: 0.12994\noutside 1 331\nTMhelix 332 354\ninside 355 372

2469 GCF_001058375.1_ASM105837v1 Escherichia coli Proteobacteria; Gammaproteobacteria; Enterobacterales; Enterobacteriaceae; Escherichia
MTGDNTLIHSHGINRRDFMKLCAALAATMGLSSKAAAEMVESVTNPQRPPVIWIGAQECT WP_000145423.1
hydrogenase 2 small subunit [Escherichia coli] Length: 372\nNumber of predicted TMHs: 1\nExp number of AAs in TMHs: 23.78201\nExp number, first 60 AAs: 0.28355\nTotal prob of N-in: 0.13257\noutside 1 331\nTMhelix 332 354\ninside 355 372

2470 GCF_001058935.1_ASM105893v1 Escherichia coli Proteobacteria; Gammaproteobacteria; Enterobacterales; Enterobacteriaceae; Escherichia
MTGDNTLIHSHGINRRDFMKLCAALAATMGLSSKAAAEMAESVTNPQRPPVIWIGAQECT WP_000145410.1
MULTISPECIES: hydrogenase-2 small chain [Proteobacteria] Length: 372\nNumber of predicted TMHs: 1\nExp number of AAs in TMHs: 23.72226\nExp number, first 60 AAs: 0.22038\nTotal prob of N-in: 0.12994\noutside 1 331\nTMhelix 332 354\ninside 355 372

2471 GCF_001058935.1_ASM105893v1 Escherichia coli Proteobacteria; Gammaproteobacteria; Enterobacterales; Enterobacteriaceae; Escherichia
MNNEETFYQAMRRQGVTRRSFLKYCSLAATSLGLGAGMAPKIAWALENKPRIPVVWIHGL WP_001058323.1
MULTISPECIES: hydrogenase-1 small chain [Proteobacteria] Length: 372\nNumber of predicted TMHs: 2\nExp number of AAs in TMHs: 37.2608499999999\nExp number, first 60 AAs: 19.52923\nTotal prob of N-in: 0.94351\nPOSSIBLE N-term signal sequence\ninside 1 19\nTMhelix 20 39\noutside 40 325\nTMhelix 326 348\ninside 349 372

2472 GCF_001058255.1_ASM105825v1 Escherichia coli Proteobacteria; Gammaproteobacteria; Enterobacterales; Enterobacteriaceae; Escherichia
MNNEETFYQAMRRQGVTRRSFLKYCSLAATSLGLGAGMAPKIAWALENKPRIPVVWIHGL WP_001295940.1
MULTISPECIES: hydrogenase [Enterobacteriaceae] Length: 372\nNumber of predicted TMHs: 2\nExp number of AAs in TMHs: 37.2608899999999\nExp number, first 60 AAs: 19.52923\nTotal prob of N-in: 0.94351\nPOSSIBLE N-term signal sequence\ninside 1 19\nTMhelix 20 39\noutside 40 325\nTMhelix 326 348\ninside 349 372

2473 GCF_001059015.1_ASM105901v1 Escherichia coli Proteobacteria; Gammaproteobacteria; Enterobacterales; Enterobacteriaceae; Escherichia
MTGDNTLIHSHGINRRDFMKLCAALAATMGLSSKAAAEMAESVTNPQRPPVIWIGAQECT WP_000145410.1
MULTISPECIES: hydrogenase-2 small chain [Proteobacteria] Length: 372\nNumber of predicted TMHs: 1\nExp number of AAs in TMHs: 23.72226\nExp number, first 60 AAs: 0.22038\nTotal prob of N-in: 0.12994\noutside 1 331\nTMhelix 332 354\ninside 355 372

2474 GCF_001059155.1_ASM105915v1 Escherichia coli Proteobacteria; Gammaproteobacteria; Enterobacterales; Enterobacteriaceae; Escherichia

MTGDNTLIHSHGINRRDFMKLCAALAATMGLSSKAAAEMAESVTNPQRPPVIWIGAQECT WP_000145410.1 MULTISPECIES: hydrogenase-2 small chain [Proteobacteria] Length: 372\nNumber of predicted TMHs: 1\nExp number of AAs in TMHs: 23.72226\nExp number, first 60 AAs: 0.22038\nTotal prob of N-in: 0.12994\noutside 1 331\nTMhelix 332 354\ninside 355 372

2475 GCF_001059095.1_ASM105909v1 Escherichia coli Proteobacteria; Gammaproteobacteria; Enterobacterales; Enterobacteriaceae; Escherichia MTGDNTLIHSHGINRRDFMKLCAALAATMGLSSKAAAEMVESVTNPQRPPVIWIGAQECT WP_000145423.1 hydrogenase 2 small subunit [Escherichia coli] Length: 372\nNumber of predicted TMHs: 1\nExp number of AAs in TMHs: 23.78201\nExp number, first 60 AAs: 0.28355\nTotal prob of N-in: 0.13257\noutside 1 331\nTMhelix 332 354\ninside 355 372

2476 GCF_001059035.1_ASM105903v1 Escherichia coli Proteobacteria; Gammaproteobacteria; Enterobacterales; Enterobacteriaceae; Escherichia MTGDNTLIHSHGINRRDFMKLCAALAATMGLSSKAAAEMAESVTNPQRPPVIWIGAQECT WP_000145410.1 MULTISPECIES: hydrogenase-2 small chain [Proteobacteria] Length: 372\nNumber of predicted TMHs: 1\nExp number of AAs in TMHs: 23.72226\nExp number, first 60 AAs: 0.22038\nTotal prob of N-in: 0.12994\noutside 1 331\nTMhelix 332 354\ninside 355 372

2477 GCF_001059735.1_ASM105973v1 Escherichia coli Proteobacteria; Gammaproteobacteria; Enterobacterales; Enterobacteriaceae; Escherichia MNNEETFYQAMRRQGVTRRSFLKYCSLAATSLGLGAGMAPKIAWALENKPRIPVVWIHGL WP_001295940.1 MULTISPECIES: hydrogenase [Enterobacteriaceae] Length: 372\nNumber of predicted TMHs: 2\nExp number of AAs in TMHs: 37.2608899999999\nExp number, first 60 AAs: 19.52923\nTotal prob of N-in: 0.94351\nPOSSIBLE N-term signal sequence\ninside 1 19\nTMhelix 20 39\noutside 40 325\nTMhelix 326 348\ninside 349 372

2478 GCF_001059035.1_ASM105903v1 Escherichia coli Proteobacteria; Gammaproteobacteria; Enterobacterales; Enterobacteriaceae; Escherichia MNNEETFYQAMRRQGVTRRSFLKYCSLAATSLGLGAGMAPKIAWALENKPRIPVVWIHGL WP_001058323.1 MULTISPECIES: hydrogenase-1 small chain [Proteobacteria] Length: 372\nNumber of predicted TMHs: 2\nExp number of AAs in TMHs: 37.2608499999999\nExp number, first 60 AAs: 19.52923\nTotal prob of N-in: 0.94351\nPOSSIBLE N-term signal sequence\ninside 1 19\nTMhelix 20 39\noutside 40 325\nTMhelix 326 348\ninside 349 372

2479 GCF_001059155.1_ASM105915v1 Escherichia coli Proteobacteria; Gammaproteobacteria; Enterobacterales; Enterobacteriaceae; Escherichia MNNEETFYQAMRRQGVTRRSFLKYCSLAATSLGLGAGMAPKIAWALENKPRIPVVWIHGL WP_001058323.1 MULTISPECIES: hydrogenase-1 small chain [Proteobacteria] Length: 372\nNumber of predicted TMHs: 2\nExp number of AAs in TMHs: 37.2608499999999\nExp number, first 60 AAs: 19.52923\nTotal prob of N-in: 0.94351\nPOSSIBLE N-term signal sequence\ninside 1 19\nTMhelix 20 39\noutside 40 325\nTMhelix 326 348\ninside 349 372

2480 GCF_001059575.1_ASM105957v1 Escherichia coli Proteobacteria; Gammaproteobacteria; Enterobacterales; Enterobacteriaceae; Escherichia MNNEETFYQAMRRQGVTRRSFLKYCSLAATSLGLGAGMAPKIAWALENKPRIPVVWIHGL WP_001058323.1 MULTISPECIES: hydrogenase-1 small chain [Proteobacteria] Length: 372\nNumber of predicted TMHs: 2\nExp number of AAs in TMHs: 37.2608499999999\nExp number, first 60 AAs: 19.52923\nTotal prob of N-in: 0.94351\nPOSSIBLE N-term signal sequence\ninside 1 19\nTMhelix 20 39\noutside 40 325\nTMhelix 326 348\ninside 349 372

2481 GCF_001059455.1_ASM105945v1 Escherichia coli Proteobacteria; Gammaproteobacteria; Enterobacterales; Enterobacteriaceae; Escherichia MNNEETFYQAMRRQGVTRRSFLKYCSLAATSLGLGAGMAPKIAWALENKPRIPVVWIHGL WP_001058323.1 MULTISPECIES: hydrogenase-1 small chain [Proteobacteria] Length: 372\nNumber of predicted TMHs: 2\nExp number of AAs in TMHs: 37.2608499999999\nExp number, first 60 AAs: 19.52923\nTotal prob of N-in: 0.94351\nPOSSIBLE N-term signal sequence\ninside 1 19\nTMhelix 20 39\noutside 40 325\nTMhelix 326 348\ninside 349 372

2482 GCF_001062415.1_ASM106241v1 Escherichia coli Proteobacteria; Gammaproteobacteria; Enterobacterales; Enterobacteriaceae; Escherichia

MTGDNTLIHSHGINRRDFMKLCAALAATMGLSSKAAAEMVESVTNPQRPPVIWIGAQECT WP_000145423.1 hydrogenase 2 small subunit [Escherichia coli] Length: 372\nNumber of predicted TMHs: 1\nExp number of AAs in TMHs: 23.78201\nExp number, first 60 AAs: 0.28355\nTotal prob of N-in: 0.13257\noutside 1 331\nTMhelix 332 354\ninside 355 372

2483 GCF_001063685.1_ASM106368v1 Escherichia coli Proteobacteria; Gammaproteobacteria; Enterobacterales; Enterobacteriaceae; Escherichia MTGDNTLIHSHGINRRDFMKLCAALAATMGLSSKAAAEMVESVTNPQRPPVIWIGAQECT WP_000145423.1 hydrogenase 2 small subunit [Escherichia coli] Length: 372\nNumber of predicted TMHs: 1\nExp number of AAs in TMHs: 23.78201\nExp number, first 60 AAs: 0.28355\nTotal prob of N-in: 0.13257\noutside 1 331\nTMhelix 332 354\ninside 355 372

2484 GCF_001077865.1_ASM107786v1 Escherichia coli Proteobacteria; Gammaproteobacteria; Enterobacterales; Enterobacteriaceae; Escherichia MTGDNTLIHSHGINRRDFMKLCAALAATMGLSSKAAAEMAESVTNPQRPPVIWIGAQECT WP_000145410.1 MULTISPECIES: hydrogenase-2 small chain [Proteobacteria] Length: 372\nNumber of predicted TMHs: 1\nExp number of AAs in TMHs: 23.72226\nExp number, first 60 AAs: 0.22038\nTotal prob of N-in: 0.12994\noutside 1 331\nTMhelix 332 354\ninside 355 372

2485 GCF_001077935.1_ASM107793v1 Escherichia coli Proteobacteria; Gammaproteobacteria; Enterobacterales; Enterobacteriaceae; Escherichia MTGDNTLIHSHGINRRDFMKLCAALAATMGLSSKAAAEMAESVTNPQRPPVIWIGAQECT WP_000145410.1 MULTISPECIES: hydrogenase-2 small chain [Proteobacteria] Length: 372\nNumber of predicted TMHs: 1\nExp number of AAs in TMHs: 23.72226\nExp number, first 60 AAs: 0.22038\nTotal prob of N-in: 0.12994\noutside 1 331\nTMhelix 332 354\ninside 355 372

2486 GCF_001077945.1_ASM107794v1 Escherichia coli Proteobacteria; Gammaproteobacteria; Enterobacterales; Enterobacteriaceae; Escherichia MTGDNTLIHSHGINRRDFMKLCAALAATMGLSSKAAAEMAESVTNPQRPPVIWIGAQECT WP_000145410.1 MULTISPECIES: hydrogenase-2 small chain [Proteobacteria] Length: 372\nNumber of predicted TMHs: 1\nExp number of AAs in TMHs: 23.72226\nExp number, first 60 AAs: 0.22038\nTotal prob of N-in: 0.12994\noutside 1 331\nTMhelix 332 354\ninside 355 372

2487 GCF_001078015.1_ASM107801v1 Escherichia coli Proteobacteria; Gammaproteobacteria; Enterobacterales; Enterobacteriaceae; Escherichia MTGDNTLIHSHGINRRDFMKLCAALAATMGLSSKAAAEMAESVTNPQRPPVIWIGAQECT WP_000145410.1 MULTISPECIES: hydrogenase-2 small chain [Proteobacteria] Length: 372\nNumber of predicted TMHs: 1\nExp number of AAs in TMHs: 23.72226\nExp number, first 60 AAs: 0.22038\nTotal prob of N-in: 0.12994\noutside 1 331\nTMhelix 332 354\ninside 355 372

2488 GCF_001065165.1_ASM106516v1 Escherichia coli Proteobacteria; Gammaproteobacteria; Enterobacterales; Enterobacteriaceae; Escherichia MNNEETFYQAMRRQGVTRRSFLKYCSLAATSLGLGAGMAPKIAWALENKPRIPVVWIHGL WP_001058323.1 MULTISPECIES: hydrogenase-1 small chain [Proteobacteria] Length: 372\nNumber of predicted TMHs: 2\nExp number of AAs in TMHs: 37.2608499999999\nExp number, first 60 AAs: 19.52923\nTotal prob of N-in: 0.94351\nPOSSIBLE N-term signal sequence\ninside 1 19\nTMhelix 20 39\noutside 40 325\nTMhelix 326 348\ninside 349 372

2489 GCF_001182815.1_ASM118281v1 Escherichia coli Proteobacteria; Gammaproteobacteria; Enterobacterales; Enterobacteriaceae; Escherichia MTGDNTLIHSHGINRRDFMKLCAALAATMGLSSKAAAEMAESVTNPQRPPVIWIGAQECT WP_000145410.1 MULTISPECIES: hydrogenase-2 small chain [Proteobacteria] Length: 372\nNumber of predicted TMHs: 1\nExp number of AAs in TMHs: 23.72226\nExp number, first 60 AAs: 0.22038\nTotal prob of N-in: 0.12994\noutside 1 331\nTMhelix 332 354\ninside 355 372

2490 GCF_001183665.1_ASM118366v1 Escherichia coli Proteobacteria; Gammaproteobacteria; Enterobacterales; Enterobacteriaceae; Escherichia MTGDNTLIHSHGINRRDFMKLCAALAATMGLSSKAAAEMAESVTNPQRPPVIWIGAQECT WP_000145410.1 MULTISPECIES: hydrogenase-2 small chain [Proteobacteria] Length: 372\nNumber of predicted TMHs: 1\nExp number of AAs in TMHs: 23.72226\nExp number, first 60 AAs: 0.22038\nTotal prob of N-in: 0.12994\noutside 1 331\nTMhelix 332 354\ninside 355 372

2491 GCF_001183665.1_ASM118366v1 Escherichia coli Proteobacteria; Gammaproteobacteria; Enterobacterales; Enterobacteriaceae; Escherichia
MNNEETFYQAMRRQGVTRRSFLKYCSLAATSLGLGAGMAPKIAWALENKPRIPVVWIHGL WP_001058323.1
MULTISPECIES: hydrogenase-1 small chain [Proteobacteria] Length: 372\nNumber of predicted TMHs: 2\nExp number of AAs in TMHs: 37.2608499999999\nExp number, first 60 AAs: 19.52923\nTotal prob of N-in: 0.94351\nPOSSIBLE N-term signal sequence\ninside 1 19\nTMhelix 20 39\noutside 40 325\nTMhelix 326 348\ninside 349 372

2492 GCF_001077945.1_ASM107794v1 Escherichia coli Proteobacteria; Gammaproteobacteria; Enterobacterales; Enterobacteriaceae; Escherichia
MNNEETFYQAMRRQGVTRRSFLKYCSLAATSLGLGAGMAPKIAWALENKPRIPVVWIHGL WP_001058323.1
MULTISPECIES: hydrogenase-1 small chain [Proteobacteria] Length: 372\nNumber of predicted TMHs: 2\nExp number of AAs in TMHs: 37.2608499999999\nExp number, first 60 AAs: 19.52923\nTotal prob of N-in: 0.94351\nPOSSIBLE N-term signal sequence\ninside 1 19\nTMhelix 20 39\noutside 40 325\nTMhelix 326 348\ninside 349 372

2493 GCF_001078015.1_ASM107801v1 Escherichia coli Proteobacteria; Gammaproteobacteria; Enterobacterales; Enterobacteriaceae; Escherichia
MNNEETFYQAMRRQGVTRRSFLKYCSLAATSLGLGAGMAPKIAWALENKPRIPVVWIHGL WP_001058323.1
MULTISPECIES: hydrogenase-1 small chain [Proteobacteria] Length: 372\nNumber of predicted TMHs: 2\nExp number of AAs in TMHs: 37.2608499999999\nExp number, first 60 AAs: 19.52923\nTotal prob of N-in: 0.94351\nPOSSIBLE N-term signal sequence\ninside 1 19\nTMhelix 20 39\noutside 40 325\nTMhelix 326 348\ninside 349 372

2494 GCF_001191025.1_CFSAN026792_01-NGE.clc Escherichia coli Proteobacteria; Gammaproteobacteria; Enterobacterales; Enterobacteriaceae; Escherichia
MTGDNTLIHSHGINRRDFMKLCAALAATMGLSSKAAAEMAESVTNPQRPPVIWIGAQECT WP_000145410.1
MULTISPECIES: hydrogenase-2 small chain [Proteobacteria] Length: 372\nNumber of predicted TMHs: 1\nExp number of AAs in TMHs: 23.72226\nExp number, first 60 AAs: 0.22038\nTotal prob of N-in: 0.12994\noutside 1 331\nTMhelix 332 354\ninside 355 372

2495 GCF_001191035.1_CFSAN026821_01-NGE.clc Escherichia coli Proteobacteria; Gammaproteobacteria; Enterobacterales; Enterobacteriaceae; Escherichia
MTGDNTLIHSHGINRRDFMKLCAALAATMGLSSKAAAEMAESVTNPQRPPVIWIGAQECT WP_000145410.1
MULTISPECIES: hydrogenase-2 small chain [Proteobacteria] Length: 372\nNumber of predicted TMHs: 1\nExp number of AAs in TMHs: 23.72226\nExp number, first 60 AAs: 0.22038\nTotal prob of N-in: 0.12994\noutside 1 331\nTMhelix 332 354\ninside 355 372

2496 GCF_001191275.1_CFSAN026786_02-NGE.clc Escherichia coli Proteobacteria; Gammaproteobacteria; Enterobacterales; Enterobacteriaceae; Escherichia
MTGDNTLIHSHGINRRDFMKLCAALAATMGLSSKAAAEMAESVTNPQRPPVIWIGAQECT WP_000145410.1
MULTISPECIES: hydrogenase-2 small chain [Proteobacteria] Length: 372\nNumber of predicted TMHs: 1\nExp number of AAs in TMHs: 23.72226\nExp number, first 60 AAs: 0.22038\nTotal prob of N-in: 0.12994\noutside 1 331\nTMhelix 332 354\ninside 355 372

2497 GCF_001191295.1_CFSAN026833_01-NGE.clc Escherichia coli Proteobacteria; Gammaproteobacteria; Enterobacterales; Enterobacteriaceae; Escherichia
MTGDNTLIHSHGINRRDFMKLCAALAATMGLSSKAAAEMAESVTNPQRPPVIWIGAQECT WP_000145410.1
MULTISPECIES: hydrogenase-2 small chain [Proteobacteria] Length: 372\nNumber of predicted TMHs: 1\nExp number of AAs in TMHs: 23.72226\nExp number, first 60 AAs: 0.22038\nTotal prob of N-in: 0.12994\noutside 1 331\nTMhelix 332 354\ninside 355 372

2498 GCF_001191195.1_CFSAN026846_01-NGE.clc Escherichia coli Proteobacteria; Gammaproteobacteria; Enterobacterales; Enterobacteriaceae; Escherichia
MNNEETFYQAMRRQGVTRRSFLKYCSLAATSLGLGAGMAPKIAWALENKPRIPVVWIHGL WP_001058323.1
MULTISPECIES: hydrogenase-1 small chain [Proteobacteria] Length: 372\nNumber of predicted TMHs: 2\nExp number of AAs in TMHs: 37.2608499999999\nExp number, first 60 AAs: 19.52923\nTotal prob of N-in: 0.94351\nPOSSIBLE N-term signal sequence\ninside 1 19\nTMhelix 20 39\noutside 40 325\nTMhelix 326 348\ninside 349 372

2499 GCF_001191215.1_CFSAN026817_01-NGE.clc Escherichia coli Proteobacteria; Gammaproteobacteria; Enterobacterales; Enterobacteriaceae; Escherichia
MNNEETFYQAMRRQGVTRRSFLKYCSLAATSLGLGAGMAPKIAWALENKPRIPVVWIHGL WP_001058323.1
MULTISPECIES: hydrogenase-1 small chain [Proteobacteria] Length: 372\nNumber of predicted TMHs: 2\nExp number of AAs in TMHs: 37.2608499999999\nExp number, first 60 AAs: 19.52923\nTotal prob of N-in: 0.94351\nPOSSIBLE N-term signal sequence\ninside 1 19\nTMhelix 20 39\noutside 40 325\nTMhelix 326 348\ninside 349 372

2500 GCF_001191115.1_CFSAN026783_02-NGE.clc Escherichia coli Proteobacteria; Gammaproteobacteria; Enterobacterales; Enterobacteriaceae; Escherichia
MNNEETFYQAMRRQGVTRRSFLKYCSLAATSLGLGAGMAPKIAWALENKPRIPVVWIHGL WP_001058323.1
MULTISPECIES: hydrogenase-1 small chain [Proteobacteria] Length: 372\nNumber of predicted TMHs: 2\nExp number of AAs in TMHs: 37.2608499999999\nExp number, first 60 AAs: 19.52923\nTotal prob of N-in: 0.94351\nPOSSIBLE N-term signal sequence\ninside 1 19\nTMhelix 20 39\noutside 40 325\nTMhelix 326 348\ninside 349 372

2501 GCF_001191125.1_CFSAN026804_01-NGE.clc Escherichia coli Proteobacteria; Gammaproteobacteria; Enterobacterales; Enterobacteriaceae; Escherichia
MNNEETFYQAMRRQGVTRRSFLKYCSLAATSLGLGAGMAPKIAWALENKPRIPVVWIHGL WP_001058323.1
MULTISPECIES: hydrogenase-1 small chain [Proteobacteria] Length: 372\nNumber of predicted TMHs: 2\nExp number of AAs in TMHs: 37.2608499999999\nExp number, first 60 AAs: 19.52923\nTotal prob of N-in: 0.94351\nPOSSIBLE N-term signal sequence\ninside 1 19\nTMhelix 20 39\noutside 40 325\nTMhelix 326 348\ninside 349 372

2502 GCF_001191185.1_CFSAN026847_01-NGE.clc Escherichia coli Proteobacteria; Gammaproteobacteria; Enterobacterales; Enterobacteriaceae; Escherichia
MNNEETFYQAMRRQGVTRRSFLKYCSLAATSLGLGAGMAPKIAWALENKPRIPVVWIHGL WP_001058323.1
MULTISPECIES: hydrogenase-1 small chain [Proteobacteria] Length: 372\nNumber of predicted TMHs: 2\nExp number of AAs in TMHs: 37.2608499999999\nExp number, first 60 AAs: 19.52923\nTotal prob of N-in: 0.94351\nPOSSIBLE N-term signal sequence\ninside 1 19\nTMhelix 20 39\noutside 40 325\nTMhelix 326 348\ninside 349 372

2503 GCF_001191295.1_CFSAN026833_01-NGE.clc Escherichia coli Proteobacteria; Gammaproteobacteria; Enterobacterales; Enterobacteriaceae; Escherichia
MNNEETFYQAMRRQGVTRRSFLKYCSLAATSLGLGAGMAPKIAWALENKPRIPVVWIHGL WP_001058323.1
MULTISPECIES: hydrogenase-1 small chain [Proteobacteria] Length: 372\nNumber of predicted TMHs: 2\nExp number of AAs in TMHs: 37.2608499999999\nExp number, first 60 AAs: 19.52923\nTotal prob of N-in: 0.94351\nPOSSIBLE N-term signal sequence\ninside 1 19\nTMhelix 20 39\noutside 40 325\nTMhelix 326 348\ninside 349 372

2504 GCF_001191395.1_CFSAN026824_01-NGE.clc Escherichia coli Proteobacteria; Gammaproteobacteria; Enterobacterales; Enterobacteriaceae; Escherichia
MTGDNTLIHSHGINRRDFMKLCAALAATMGLSSKAAAEMAESVTNPQRPPVIWIGAQECT WP_000145410.1
MULTISPECIES: hydrogenase-2 small chain [Proteobacteria] Length: 372\nNumber of predicted TMHs: 1\nExp number of AAs in TMHs: 23.72226\nExp number, first 60 AAs: 0.22038\nTotal prob of N-in: 0.12994\noutside 1 331\nTMhelix 332 354\ninside 355 372

2505 GCF_001191435.1_CFSAN026820_01-NGE.clc Escherichia coli Proteobacteria; Gammaproteobacteria; Enterobacterales; Enterobacteriaceae; Escherichia
MTGDNTLIHSHGINRRDFMKLCAALAATMGLSSKAAAEMAESVTNPQRPPVIWIGAQECT WP_000145410.1
MULTISPECIES: hydrogenase-2 small chain [Proteobacteria] Length: 372\nNumber of predicted TMHs: 1\nExp number of AAs in TMHs: 23.72226\nExp number, first 60 AAs: 0.22038\nTotal prob of N-in: 0.12994\noutside 1 331\nTMhelix 332 354\ninside 355 372

2506 GCF_001191455.1_CFSAN026831_01-NGE.clc Escherichia coli Proteobacteria; Gammaproteobacteria; Enterobacterales; Enterobacteriaceae; Escherichia
MTGDNTLIHSHGINRRDFMKLCAALAATMGLSSKAAAEMAESVTNPQRPPVIWIGAQECT WP_000145410.1
MULTISPECIES: hydrogenase-2 small chain [Proteobacteria] Length: 372\nNumber of predicted TMHs: 1\nExp number of AAs in TMHs: 23.72226\nExp number, first 60 AAs: 0.22038\nTotal prob of N-in: 0.12994\noutside 1 331\nTMhelix 332 354\ninside 355 372

2507 GCF_001245225.1_E._coli_A192PP Escherichia coli Proteobacteria; Gammaproteobacteria; Enterobacterales; Enterobacteriaceae; Escherichia
MTGDNTLIHSHGINRRDFMKLCAALAATMGLSSKAAAEMAESVTNPQRPPVIWIGAQECT WP_000145410.1
MULTISPECIES: hydrogenase-2 small chain [Proteobacteria] Length: 372\nNumber of predicted TMHs: 1\nExp number of AAs in TMHs: 23.72226\nExp number, first 60 AAs: 0.22038\nTotal prob of N-in: 0.12994\noutside 1 331\nTMhelix 332 354\ninside 355 372

2508 GCF_001191505.1_CFSAN026800_01-NGE.clc Escherichia coli Proteobacteria; Gammaproteobacteria; Enterobacterales; Enterobacteriaceae; Escherichia
MTGDNTLIHSHGINRRDFMKLCAALAATMGLSSKAAAEMAESVTNPQRPPVIWIGAQECT WP_000145410.1
MULTISPECIES: hydrogenase-2 small chain [Proteobacteria] Length: 372\nNumber of predicted TMHs: 1\nExp number of AAs in TMHs: 23.72226\nExp number, first 60 AAs: 0.22038\nTotal prob of N-in: 0.12994\noutside 1 331\nTMhelix 332 354\ninside 355 372

2509 GCF_001262855.1_ASM126285v1 Escherichia coli Proteobacteria; Gammaproteobacteria; Enterobacterales; Enterobacteriaceae; Escherichia
MNNEETFYQAMRRQGVTRRSFLKYCSLAATSLGLGAGMAPKIAWALENKPRIPVVWIHGL WP_001058323.1
MULTISPECIES: hydrogenase-1 small chain [Proteobacteria] Length: 372\nNumber of predicted TMHs: 2\nExp number of AAs in TMHs: 37.2608499999999\nExp number, first 60 AAs: 19.52923\nTotal prob of N-in: 0.94351\nPOSSIBLE N-term signal sequence\ninside 1 19\nTMhelix 20 39\noutside 40 325\nTMhelix 326 348\ninside 349 372

2510 GCF_001191435.1_CFSAN026820_01-NGE.clc Escherichia coli Proteobacteria; Gammaproteobacteria; Enterobacterales; Enterobacteriaceae; Escherichia
MNNEETFYQAMRRQGVTRRSFLKYCSLAATSLGLGAGMAPKIAWALENKPRIPVVWIHGL WP_050946405.1
hydrogenase [Escherichia coli] Length: 372\nNumber of predicted TMHs: 2\nExp number of AAs in TMHs: 37.2615099999999\nExp number, first 60 AAs: 19.52921\nTotal prob of N-in: 0.94351\nPOSSIBLE N-term signal sequence\ninside 1 19\nTMhelix 20 39\noutside 40 325\nTMhelix 326 348\ninside 349 372

2511 GCF_001262865.1_6842-1 Escherichia coli Proteobacteria; Gammaproteobacteria; Enterobacterales; Enterobacteriaceae; Escherichia MTGDNTLIHSHGINRRDFMKLCAALAATMGLSSKAAAEMAESVTNPQRPPVIWIGAQECT
WP_000145410.1 MULTISPECIES: hydrogenase-2 small chain [Proteobacteria] Length: 372\nNumber of predicted TMHs: 1\nExp number of AAs in TMHs: 23.72226\nExp number, first 60 AAs: 0.22038\nTotal prob of N-in: 0.12994\noutside 1 331\nTMhelix 332 354\ninside 355 372

2512 GCF_001262905.1_5710-2 Escherichia coli Proteobacteria; Gammaproteobacteria; Enterobacterales; Enterobacteriaceae; Escherichia MTGDNTLIHSHGINRRDFMKLCAALAATMGLSSKAAAEMAESVTNPQRPPVIWIGAQECT
WP_000145410.1 MULTISPECIES: hydrogenase-2 small chain [Proteobacteria] Length: 372\nNumber of predicted TMHs: 1\nExp number of AAs in TMHs: 23.72226\nExp number, first 60 AAs: 0.22038\nTotal prob of N-in: 0.12994\noutside 1 331\nTMhelix 332 354\ninside 355 372

2513 GCF_001262865.1_6842-1 Escherichia coli Proteobacteria; Gammaproteobacteria; Enterobacterales; Enterobacteriaceae; Escherichia MNNEETFYQAMRRQGVTRRSFLKYCSLAATSLGLGAGMAPKIAWALENKPRIPVVWIHGL
WP_001058323.1 MULTISPECIES: hydrogenase-1 small chain [Proteobacteria] Length: 372\nNumber of predicted TMHs: 2\nExp number of AAs in TMHs: 37.2608499999999\nExp number, first 60 AAs: 19.52923\nTotal prob of N-in: 0.94351\nPOSSIBLE N-term signal sequence\ninside 1 19\nTMhelix 20 39\noutside 40 325\nTMhelix 326 348\ninside 349 372

2514 GCF_001262905.1_5710-2 Escherichia coli Proteobacteria; Gammaproteobacteria; Enterobacterales; Enterobacteriaceae; Escherichia MNNEETFYQAMRRQGVTRRSFLKYCSLAATSLGLGAGMAPKIAWALENKPRIPVVWIHGL
WP_001058323.1 MULTISPECIES: hydrogenase-1 small chain [Proteobacteria] Length: 372\nNumber of predicted TMHs: 2\nExp number of AAs in TMHs: 37.2608499999999\nExp number, first 60 AAs: 19.52923\nTotal prob of N-in: 0.94351\nPOSSIBLE N-term signal sequence\ninside 1 19\nTMhelix 20 39\noutside 40 325\nTMhelix 326 348\ninside 349 372

2515 GCF_001265165.1_ASM126516v1 Escherichia coli Proteobacteria; Gammaproteobacteria; Enterobacterales; Enterobacteriaceae; Escherichia
MTGDNTLIHSHGINRRDFMKLCAALAATMGLSSKAAAEMAESVTNPQRPPVIWIGAQECT WP_000145410.1
MULTISPECIES: hydrogenase-2 small chain [Proteobacteria] Length: 372\nNumber of predicted TMHs: 1\nExp number of AAs in TMHs: 23.72226\nExp number, first 60 AAs: 0.22038\nTotal prob of N-in: 0.12994\noutside 1 331\nTMhelix 332 354\ninside 355 372

2516 GCF_001265155.1_100414 Escherichia coli Proteobacteria; Gammaproteobacteria; Enterobacterales; Enterobacteriaceae; Escherichia MTGDNTLIHSHGINRRDFMKLCAALAATMGLSSKAAAEMAESVTNPQRPPVIWIGAQECT WP_000145410.1 MULTISPECIES: hydrogenase-2 small chain [Proteobacteria] Length: 372\nNumber of predicted TMHs: 1\nExp number of AAs in TMHs: 23.72226\nExp number, first 60 AAs: 0.22038\nTotal prob of N-in: 0.12994\noutside 1 331\nTMhelix 332 354\ninside 355 372

2517 GCF_001265155.1_100414 Escherichia coli Proteobacteria; Gammaproteobacteria; Enterobacterales; Enterobacteriaceae; Escherichia MNNEETFYQAMRRQGVTRRSFLKYCSLAATSLGLGAGMAPKIAWALENKPRIPVVWIHGL WP_001058323.1 MULTISPECIES: hydrogenase-1 small chain [Proteobacteria] Length: 372\nNumber of predicted TMHs: 2\nExp number of AAs in TMHs: 37.2608499999999\nExp number, first 60 AAs: 19.52923\nTotal prob of N-in: 0.94351\nPOSSIBLE N-term signal sequence\ninside 1 19\nTMhelix 20 39\noutside 40 325\nTMhelix 326 348\ninside 349 372

2518 GCF_001263065.1_16118-2 Escherichia coli Proteobacteria; Gammaproteobacteria; Enterobacterales; Enterobacteriaceae; Escherichia MNNEETFYQAMRRQGVTRRSFLKYCSLAATSLGLGAGMAPKIAWALENKPRIPVVWIHGL WP_001058323.1 MULTISPECIES: hydrogenase-1 small chain [Proteobacteria] Length: 372\nNumber of predicted TMHs: 2\nExp number of AAs in TMHs: 37.2608499999999\nExp number, first 60 AAs: 19.52923\nTotal prob of N-in: 0.94351\nPOSSIBLE N-term signal sequence\ninside 1 19\nTMhelix 20 39\noutside 40 325\nTMhelix 326 348\ninside 349 372

2519 GCF_001265165.1_ASM126516v1 Escherichia coli Proteobacteria; Gammaproteobacteria; Enterobacterales; Enterobacteriaceae; Escherichia MNNEETFYQAMRRQGVTRRSFLKYCSLAATSLGLGAGMAPKIAWALENKPRIPVVWIHGL WP_001058323.1 MULTISPECIES: hydrogenase-1 small chain [Proteobacteria] Length: 372\nNumber of predicted TMHs: 2\nExp number of AAs in TMHs: 37.2608499999999\nExp number, first 60 AAs: 19.52923\nTotal prob of N-in: 0.94351\nPOSSIBLE N-term signal sequence\ninside 1 19\nTMhelix 20 39\noutside 40 325\nTMhelix 326 348\ninside 349 372

2520 GCF_001262965.1_1223-4 Escherichia coli Proteobacteria; Gammaproteobacteria; Enterobacterales; Enterobacteriaceae; Escherichia MNNEETFYQAMRRQGVTRRSFLKYCSLAATSLGLGAGMAPKIAWALENKPRIPVVWIHGL WP_001058323.1 MULTISPECIES: hydrogenase-1 small chain [Proteobacteria] Length: 372\nNumber of predicted TMHs: 2\nExp number of AAs in TMHs: 37.2608499999999\nExp number, first 60 AAs: 19.52923\nTotal prob of N-in: 0.94351\nPOSSIBLE N-term signal sequence\ninside 1 19\nTMhelix 20 39\noutside 40 325\nTMhelix 326 348\ninside 349 372

2521 GCF_001263735.1_ASM126373v1 Escherichia coli Proteobacteria; Gammaproteobacteria; Enterobacterales; Enterobacteriaceae; Escherichia MNNEETFYQAMRRQGVTRRSFLKYCSLAATSLGLGAGMAPKIAWALENKPRIPVVWIHGL WP_001058323.1 MULTISPECIES: hydrogenase-1 small chain [Proteobacteria] Length: 372\nNumber of predicted TMHs: 2\nExp number of AAs in TMHs: 37.2608499999999\nExp number, first 60 AAs: 19.52923\nTotal prob of N-in: 0.94351\nPOSSIBLE N-term signal sequence\ninside 1 19\nTMhelix 20 39\noutside 40 325\nTMhelix 326 348\ninside 349 372

2522 GCF_001264195.1_ASM126419v1 Escherichia coli Proteobacteria; Gammaproteobacteria; Enterobacterales; Enterobacteriaceae; Escherichia MNNEETFYQAMRRQGVTRRSFLKYCSLAATSLGLGAGMAPKIAWALENKPRIPVVWIHGL WP_001353311.1 MULTISPECIES: hydrogenase [Escherichia] Length: 372\nNumber of predicted TMHs: 2\nExp number of AAs in TMHs: 37.2638699999999\nExp number, first 60 AAs: 19.52924\nTotal prob of N-in: 0.94351\nPOSSIBLE N-term signal sequence\ninside 1 19\nTMhelix 20 39\noutside 40 325\nTMhelix 326 348\ninside 349 372

2523 GCF_001265315.1_300262 Escherichia coli Proteobacteria; Gammaproteobacteria; Enterobacterales; Enterobacteriaceae; Escherichia MTGDNTLIHSHGINRRDFMKLCAALAATMGLSSKAAAEMAESVTNPQRPPVIWIGAQECT WP_000145410.1 MULTISPECIES: hydrogenase-2 small chain [Proteobacteria] Length: 372\nNumber of predicted TMHs: 1\nExp number of AAs in TMHs: 23.72226\nExp number, first 60 AAs: 0.22038\nTotal prob of N-in: 0.12994\noutside 1 331\nTMhelix 332 354\ninside 355 372

2524 GCF_001265315.1_300262 Escherichia coli Proteobacteria; Gammaproteobacteria; Enterobacterales; Enterobacteriaceae; Escherichia MNNEETFYQAMRRQGVTRRSFLKYCSLAATSLGLGAGMAPKIAWALENKPRIPVVWIHGL WP_001058323.1 MULTISPECIES: hydrogenase-1 small chain [Proteobacteria] Length: 372\nNumber of predicted TMHs: 2\nExp number of AAs in TMHs: 37.2608499999999\nExp number, first 60 AAs: 19.52923\nTotal prob of

N-in: 0.94351\nPOSSIBLE N-term signal sequence\ninside 1 19\nTMhelix 20 39\noutside 40 325\nTMhelix 326 348\ninside 349 372

2525 GCF_001265395.1_302275 Escherichia coli Proteobacteria; Gammaproteobacteria; Enterobacterales; Enterobacteriaceae; Escherichia MTGDNTLIHSHGINRRDFMKLCAALAATMGLSSKAAAEMAESVTNPQRPPVIWIGAQECT WP_000145410.1 MULTISPECIES: hydrogenase-2 small chain [Proteobacteria] Length: 372\nNumber of predicted TMHs: 1\nExp number of AAs in TMHs: 23.72226\nExp number, first 60 AAs: 0.22038\nTotal prob of N-in: 0.12994\noutside 1 331\nTMhelix 332 354\ninside 355 372

2526 GCF_001265435.1_400791 Escherichia coli Proteobacteria; Gammaproteobacteria; Enterobacterales; Enterobacteriaceae; Escherichia MTGDNTLIHSHGINRRDFMKLCAALAATMGLSSKAAAEMAESVTNPQRPPVIWIGAQECT WP_000145410.1 MULTISPECIES: hydrogenase-2 small chain [Proteobacteria] Length: 372\nNumber of predicted TMHs: 1\nExp number of AAs in TMHs: 23.72226\nExp number, first 60 AAs: 0.22038\nTotal prob of N-in: 0.12994\noutside 1 331\nTMhelix 332 354\ninside 355 372

2527 GCF_001265455.1_302662 Escherichia coli Proteobacteria; Gammaproteobacteria; Enterobacterales; Enterobacteriaceae; Escherichia MTGDNTLIHSHGINRRDFMKLCAALAATMGLSSKAAAEMAESVTNPQRPPVIWIGAQECT WP_000145410.1 MULTISPECIES: hydrogenase-2 small chain [Proteobacteria] Length: 372\nNumber of predicted TMHs: 1\nExp number of AAs in TMHs: 23.72226\nExp number, first 60 AAs: 0.22038\nTotal prob of N-in: 0.12994\noutside 1 331\nTMhelix 332 354\ninside 355 372

2528 GCF_001265435.1_400791 Escherichia coli Proteobacteria; Gammaproteobacteria; Enterobacterales; Enterobacteriaceae; Escherichia MNNEETFYQAMRRQGVTRRSFLKYCSLAATSLGLGAGMAPKIAWALENKPRIPVVWIHGL WP_001058323.1 MULTISPECIES: hydrogenase-1 small chain [Proteobacteria] Length: 372\nNumber of predicted TMHs: 2\nExp number of AAs in TMHs: 37.2608499999999\nExp number, first 60 AAs: 19.52923\nTotal prob of N-in: 0.94351\nPOSSIBLE N-term signal sequence\ninside 1 19\nTMhelix 20 39\noutside 40 325\nTMhelix 326 348\ninside 349 372

2529 GCF_001265675.1_102598 Escherichia coli Proteobacteria; Gammaproteobacteria; Enterobacterales; Enterobacteriaceae; Escherichia MTGDNTLIHSHGINRRDFMKLCAALAATMGLSSKAAAEMAESVTNPQRPPVIWIGAQECT WP_000145410.1 MULTISPECIES: hydrogenase-2 small chain [Proteobacteria] Length: 372\nNumber of predicted TMHs: 1\nExp number of AAs in TMHs: 23.72226\nExp number, first 60 AAs: 0.22038\nTotal prob of N-in: 0.12994\noutside 1 331\nTMhelix 332 354\ninside 355 372

2530 GCF_001265675.1_102598 Escherichia coli Proteobacteria; Gammaproteobacteria; Enterobacterales; Enterobacteriaceae; Escherichia MNNEETFYQAMRRQGVTRRSFLKYCSLAATSLGLGAGMAPKIAWALENKPRIPVVWIHGL WP_001058323.1 MULTISPECIES: hydrogenase-1 small chain [Proteobacteria] Length: 372\nNumber of predicted TMHs: 2\nExp number of AAs in TMHs: 37.2608499999999\nExp number, first 60 AAs: 19.52923\nTotal prob of N-in: 0.94351\nPOSSIBLE N-term signal sequence\ninside 1 19\nTMhelix 20 39\noutside 40 325\nTMhelix 326 348\ninside 349 372

2531 GCF_001265455.1_302662 Escherichia coli Proteobacteria; Gammaproteobacteria; Enterobacterales; Enterobacteriaceae; Escherichia MNNEETFYQAMRRQGVTRRSFLKYCSLAATSLGLGAGMAPKIAWALENKPRIPVVWIHGL WP_001058323.1 MULTISPECIES: hydrogenase-1 small chain [Proteobacteria] Length: 372\nNumber of predicted TMHs: 2\nExp number of AAs in TMHs: 37.2608499999999\nExp number, first 60 AAs: 19.52923\nTotal prob of N-in: 0.94351\nPOSSIBLE N-term signal sequence\ninside 1 19\nTMhelix 20 39\noutside 40 325\nTMhelix 326 348\ninside 349 372

2532 GCF_001265765.1_300214 Escherichia coli Proteobacteria; Gammaproteobacteria; Enterobacterales; Enterobacteriaceae; Escherichia MTGDNTLIHSHGINRRDFMKLCAALAATMGLSSKAAAEMAESVTNPQRPPVIWIGAQECT WP_000145410.1 MULTISPECIES: hydrogenase-2 small chain [Proteobacteria] Length: 372\nNumber of predicted TMHs: 1\nExp number of AAs in TMHs: 23.72226\nExp number, first 60 AAs: 0.22038\nTotal prob of N-in: 0.12994\noutside 1 331\nTMhelix 332 354\ninside 355 372

2533 GCF_001265685.1_102929 Escherichia coli Proteobacteria; Gammaproteobacteria; Enterobacterales; Enterobacteriaceae; Escherichia MTGDNTLIHSHGINRRDFMKLCAALAATMGLSSKAAAEMAESVTNPQRPPVIWIGAQECT WP_000145410.1 MULTISPECIES: hydrogenase-2 small chain [Proteobacteria] Length: 372\nNumber of predicted TMHs: 1\nExp number of AAs in TMHs: 23.72226\nExp number, first 60 AAs: 0.22038\nTotal prob of N-in: 0.12994\noutside 1 331\nTMhelix 332 354\ninside 355 372

2534 GCF_001265735.1_200146 Escherichia coli Proteobacteria; Gammaproteobacteria; Enterobacterales; Enterobacteriaceae; Escherichia MTGDNTLIHSHGINRRDFMKLCAALAATMGLSSKAAAEMAESVTNPQRPPVIWIGAQECT

WP_000145410.1 MULTISPECIES: hydrogenase-2 small chain [Proteobacteria] Length: 372\nNumber of predicted TMHs: 1\nExp number of AAs in TMHs: 23.72226\nExp number, first 60 AAs: 0.22038\nTotal prob of N-in: 0.12994\noutside 1 331\nTMhelix 332 354\ninside 355 372

2535 GCF_001265835.1_302687 Escherichia coli Proteobacteria; Gammaproteobacteria; Enterobacterales; Enterobacteriaceae; Escherichia MNNEETFYQAMRRQGVTRRSFLKYCSLAATSLGLGAGMAPKIAWALENKPRIPVVWIHGL WP_001058323.1 MULTISPECIES: hydrogenase-1 small chain [Proteobacteria] Length: 372\nNumber of predicted TMHs: 2\nExp number of AAs in TMHs: 37.2608499999999\nExp number, first 60 AAs: 19.52923\nTotal prob of N-in: 0.94351\nPOSSIBLE N-term signal sequence\ninside 1 19\nTMhelix 20 39\noutside 40 325\nTMhelix 326 348\ninside 349 372

2536 GCF_001265875.1_303145 Escherichia coli Proteobacteria; Gammaproteobacteria; Enterobacterales; Enterobacteriaceae; Escherichia MNNEETFYQAMRRQGVTRRSFLKYCSLAATSLGLGAGMAPKIAWALENKPRIPVVWIHGL WP_001295940.1 MULTISPECIES: hydrogenase [Enterobacteriaceae] Length: 372\nNumber of predicted TMHs: 2\nExp number of AAs in TMHs: 37.2608899999999\nExp number, first 60 AAs: 19.52923\nTotal prob of N-in: 0.94351\nPOSSIBLE N-term signal sequence\ninside 1 19\nTMhelix 20 39\noutside 40 325\nTMhelix 326 348\ninside 349 372

2537 GCF_001265685.1_102929 Escherichia coli Proteobacteria; Gammaproteobacteria; Enterobacterales; Enterobacteriaceae; Escherichia MNNEETFYQAMRRQGVTRRSFLKYCSLAATSLGLGAGMAPKIAWALENKPRIPVVWIHGL WP_001058323.1 MULTISPECIES: hydrogenase-1 small chain [Proteobacteria] Length: 372\nNumber of predicted TMHs: 2\nExp number of AAs in TMHs: 37.2608499999999\nExp number, first 60 AAs: 19.52923\nTotal prob of N-in: 0.94351\nPOSSIBLE N-term signal sequence\ninside 1 19\nTMhelix 20 39\noutside 40 325\nTMhelix 326 348\ninside 349 372

2538 GCF_001265915.1_401210 Escherichia coli Proteobacteria; Gammaproteobacteria; Enterobacterales; Enterobacteriaceae; Escherichia MTGDNTLIHSHGINRRDFMKLCAALAATMGLSSKAAAEMAESVTNPQRPPVIWIGAQECT WP_000145410.1 MULTISPECIES: hydrogenase-2 small chain [Proteobacteria] Length: 372\nNumber of predicted TMHs: 1\nExp number of AAs in TMHs: 23.72226\nExp number, first 60 AAs: 0.22038\nTotal prob of N-in: 0.12994\noutside 1 331\nTMhelix 332 354\ninside 355 372

2539 GCF_001265925.1_401588 Escherichia coli Proteobacteria; Gammaproteobacteria; Enterobacterales; Enterobacteriaceae; Escherichia MTGDNTLIHSHGINRRDFMKLCAALAATMGLSSKAAAEMAESVTNPQRPPVIWIGAQECT WP_000145410.1 MULTISPECIES: hydrogenase-2 small chain [Proteobacteria] Length: 372\nNumber of predicted TMHs: 1\nExp number of AAs in TMHs: 23.72226\nExp number, first 60 AAs: 0.22038\nTotal prob of N-in: 0.12994\noutside 1 331\nTMhelix 332 354\ninside 355 372

2540 GCF_001265975.1_402310 Escherichia coli Proteobacteria; Gammaproteobacteria; Enterobacterales; Enterobacteriaceae; Escherichia MTGDNTLIHSHGINRRDFMKLCAALAATMGLSSKAAAEMAESVTNPQRPPVIWIGAQECT WP_000145410.1 MULTISPECIES: hydrogenase-2 small chain [Proteobacteria] Length: 372\nNumber of predicted TMHs: 1\nExp number of AAs in TMHs: 23.72226\nExp number, first 60 AAs: 0.22038\nTotal prob of N-in: 0.12994\noutside 1 331\nTMhelix 332 354\ninside 355 372

2541 GCF_001265925.1_401588 Escherichia coli Proteobacteria; Gammaproteobacteria; Enterobacterales; Enterobacteriaceae; Escherichia MNNEETFYQAMRRQGVTRRSFLKYCSLAATSLGLGAGMAPKIAWALENKPRIPVVWIHGL WP_001058323.1 MULTISPECIES: hydrogenase-1 small chain [Proteobacteria] Length: 372\nNumber of predicted TMHs: 2\nExp number of AAs in TMHs: 37.2608499999999\nExp number, first 60 AAs: 19.52923\nTotal prob of N-in: 0.94351\nPOSSIBLE N-term signal sequence\ninside 1 19\nTMhelix 20 39\noutside 40 325\nTMhelix 326 348\ninside 349 372

2542 GCF_001266075.1_100175 Escherichia coli Proteobacteria; Gammaproteobacteria; Enterobacterales; Enterobacteriaceae; Escherichia MTGDNTLIHSHGINRRDFMKLCAALAATMGLSSKAAAEMAESVTNPQRPPVIWIGAQECT WP_000145410.1 MULTISPECIES: hydrogenase-2 small chain [Proteobacteria] Length: 372\nNumber of predicted TMHs: 1\nExp number of AAs in TMHs: 23.72226\nExp number, first 60 AAs: 0.22038\nTotal prob of N-in: 0.12994\noutside 1 331\nTMhelix 332 354\ninside 355 372

2543 GCF_001266175.1_200077 Escherichia coli Proteobacteria; Gammaproteobacteria; Enterobacterales; Enterobacteriaceae; Escherichia MTGDNTLIHSHGINRRDFMKLCAALAATMGLSSKAAAEMAESVTNPQRPPVIWIGAQECT WP_000145410.1 MULTISPECIES: hydrogenase-2 small chain [Proteobacteria] Length: 372\nNumber of predicted TMHs: 1\nExp number of AAs in TMHs: 23.72226\nExp number, first 60 AAs: 0.22038\nTotal prob of N-in: 0.12994\noutside 1 331\nTMhelix 332 354\ninside 355 372

2544 GCF_001266115.1_102535 Escherichia coli Proteobacteria; Gammaproteobacteria; Enterobacterales; Enterobacteriaceae; Escherichia MNNEETFYQAMRRQGVTRRSFLKYCSLAATSLGLGAGMAPKIAWALENKPRIPVVWIHGL WP_001058323.1 MULTISPECIES: hydrogenase-1 small chain [Proteobacteria] Length: 372\nNumber of predicted TMHs: 2\nExp number of AAs in TMHs: 37.2608499999999\nExp number, first 60 AAs: 19.52923\nTotal prob of N-in: 0.94351\nPOSSIBLE N-term signal sequence\ninside 1 19\nTMhelix 20 39\noutside 40 325\nTMhelix 326 348\ninside 349 372

2545 GCF_001266175.1_200077 Escherichia coli Proteobacteria; Gammaproteobacteria; Enterobacterales; Enterobacteriaceae; Escherichia MNNEETFYQAMRRQGVTRRSFLKYCSLAATSLGLGAGMAPKIAWALENKPRIPVVWIHGL WP_052905910.1 hydrogenase [Escherichia coli] Length: 372\nNumber of predicted TMHs: 2\nExp number of AAs in TMHs: 37.2614\nExp number, first 60 AAs: 19.52921\nTotal prob of N-in: 0.94351\nPOSSIBLE N-term signal sequence\ninside 1 19\nTMhelix 20 39\noutside 40 325\nTMhelix 326 348\ninside 349 372

2546 GCF_001266075.1_100175 Escherichia coli Proteobacteria; Gammaproteobacteria; Enterobacterales; Enterobacteriaceae; Escherichia MNNEETFYQAMRRQGVTRRSFLKYCSLAATSLGLGAGMAPKIAWALENKPRIPVVWIHGL WP_001295940.1 MULTISPECIES: hydrogenase [Enterobacteriaceae] Length: 372\nNumber of predicted TMHs: 2\nExp number of AAs in TMHs: 37.2608899999999\nExp number, first 60 AAs: 19.52923\nTotal prob of N-in: 0.94351\nPOSSIBLE N-term signal sequence\ninside 1 19\nTMhelix 20 39\noutside 40 325\nTMhelix 326 348\ninside 349 372

2547 GCF_001266015.1_100100 Escherichia coli Proteobacteria; Gammaproteobacteria; Enterobacterales; Enterobacteriaceae; Escherichia MNNEETFYQAMRRQGVTRRSFLKYCSLAATSLGLGAGMAPKIAWALENKPRIPVVWIHGL WP_001058323.1 MULTISPECIES: hydrogenase-1 small chain [Proteobacteria] Length: 372\nNumber of predicted TMHs: 2\nExp number of AAs in TMHs: 37.2608499999999\nExp number, first 60 AAs: 19.52923\nTotal prob of N-in: 0.94351\nPOSSIBLE N-term signal sequence\ninside 1 19\nTMhelix 20 39\noutside 40 325\nTMhelix 326 348\ninside 349 372

2548 GCF_001266215.1_300847 Escherichia coli Proteobacteria; Gammaproteobacteria; Enterobacterales; Enterobacteriaceae; Escherichia MTGDNTLIHSHGINRRDFMKLCAALAATMGLSSKAAAEMAESVTNPQRPPVIWIGAQECT WP_000145410.1 MULTISPECIES: hydrogenase-2 small chain [Proteobacteria] Length: 372\nNumber of predicted TMHs: 1\nExp number of AAs in TMHs: 23.72226\nExp number, first 60 AAs: 0.22038\nTotal prob of N-in: 0.12994\noutside 1 331\nTMhelix 332 354\ninside 355 372

2549 GCF_001266355.1_401195 Escherichia coli Proteobacteria; Gammaproteobacteria; Enterobacterales; Enterobacteriaceae; Escherichia MTGDNTLIHSHGINRRDFMKLCAALAATMGLSSKAAAEMAESVTNPQRPPVIWIGAQECT WP_000145408.1 hydrogenase 2 small subunit [Escherichia coli] Length: 372\nNumber of predicted TMHs: 1\nExp number of AAs in TMHs: 23.72255\nExp number, first 60 AAs: 0.22038\nTotal prob of N-in: 0.12995\noutside 1 331\nTMhelix 332 354\ninside 355 372

2550 GCF_001266445.1_700283 Escherichia coli Proteobacteria; Gammaproteobacteria; Enterobacterales; Enterobacteriaceae; Escherichia MTGDNTLIHSHGINRRDFMKLCAALAATMGLSSKAAAEMVESVTNPQRPPVIWIGAQECT WP_000145423.1 hydrogenase 2 small subunit [Escherichia coli] Length: 372\nNumber of predicted TMHs: 1\nExp number of AAs in TMHs: 23.78201\nExp number, first 60 AAs: 0.28355\nTotal prob of N-in: 0.13257\noutside 1 331\nTMhelix 332 354\ninside 355 372

2551 GCF_001266375.1_401675 Escherichia coli Proteobacteria; Gammaproteobacteria; Enterobacterales; Enterobacteriaceae; Escherichia MNNEETFYQAMRRQGVTRRSFLKYCSLAATSLGLGAGMAPKIAWALENKPRIPVVWIHGL WP_001058323.1 MULTISPECIES: hydrogenase-1 small chain [Proteobacteria] Length: 372\nNumber of predicted TMHs: 2\nExp number of AAs in TMHs: 37.2608499999999\nExp number, first 60 AAs: 19.52923\nTotal prob of N-in: 0.94351\nPOSSIBLE N-term signal sequence\ninside 1 19\nTMhelix 20 39\noutside 40 325\nTMhelix 326 348\ninside 349 372

2552 GCF_001266445.1_700283 Escherichia coli Proteobacteria; Gammaproteobacteria; Enterobacterales; Enterobacteriaceae; Escherichia MNNEETFYQAMRRQGVTRRSFLKYCSLAATSLGLGAGMAPKIAWALENKPRIPVVWIHGL WP_001058323.1 MULTISPECIES: hydrogenase-1 small chain [Proteobacteria] Length: 372\nNumber of predicted TMHs: 2\nExp number of AAs in TMHs: 37.2608499999999\nExp number, first 60 AAs: 19.52923\nTotal prob of N-in: 0.94351\nPOSSIBLE N-term signal sequence\ninside 1 19\nTMhelix 20 39\noutside 40 325\nTMhelix 326 348\ninside 349 372

2553 GCF_001266355.1_401195 Escherichia coli Proteobacteria; Gammaproteobacteria; Enterobacterales; Enterobacteriaceae; Escherichia MNNEETFYQAMRRQGVTRRSFLKYCSLAATSLGLGAGMAPKIAWALENKPRIPVVWIHGL

WP_001058323.1 MULTISPECIES: hydrogenase-1 small chain [Proteobacteria] Length: 372\nNumber of predicted TMHs: 2\nExp number of AAs in TMHs: 37.2608499999999\nExp number, first 60 AAs: 19.52923\nTotal prob of N-in: 0.94351\nPOSSIBLE N-term signal sequence\ninside 1 19\nTMhelix 20 39\noutside 40 325\nTMhelix 326 348\ninside 349 372

2554 GCF_001266275.1_303139 Escherichia coli Proteobacteria; Gammaproteobacteria; Enterobacterales; Enterobacteriaceae; Escherichia MNNEETFYQAMRRQGVTRRSFLKYCSLAATSLGLGAGMAPKIAWALENKPRIPVVWIHGL WP_001058323.1 MULTISPECIES: hydrogenase-1 small chain [Proteobacteria] Length: 372\nNumber of predicted TMHs: 2\nExp number of AAs in TMHs: 37.2608499999999\nExp number, first 60 AAs: 19.52923\nTotal prob of N-in: 0.94351\nPOSSIBLE N-term signal sequence\ninside 1 19\nTMhelix 20 39\noutside 40 325\nTMhelix 326 348\ninside 349 372

2555 GCF_001267335.1_403116 Escherichia coli Proteobacteria; Gammaproteobacteria; Enterobacterales; Enterobacteriaceae; Escherichia MTGDNTLIHSHGINRRDFMKLCAALAATMGLSSKAAAEMAESVTNPQRPPVIWIGAQECT WP_000145408.1 hydrogenase 2 small subunit [Escherichia coli] Length: 372\nNumber of predicted TMHs: 1\nExp number of AAs in TMHs: 23.72255\nExp number, first 60 AAs: 0.22038\nTotal prob of N-in: 0.12995\noutside 1 331\nTMhelix 332 354\ninside 355 372

2556 GCF_001268345.1_YE1 Escherichia coli Proteobacteria; Gammaproteobacteria; Enterobacterales; Enterobacteriaceae; Escherichia MTGDNTLIHSHGINRRDFMKLCAALAATMGLSSKAAAEMAESVTNPQRPPVIWIGAQECT WP_000145410.1 MULTISPECIES: hydrogenase-2 small chain [Proteobacteria] Length: 372\nNumber of predicted TMHs: 1\nExp number of AAs in TMHs: 23.72226\nExp number, first 60 AAs: 0.22038\nTotal prob of N-in: 0.12994\noutside 1 331\nTMhelix 332 354\ninside 355 372

2557 GCF_001268685.1_YE19 Escherichia coli Proteobacteria; Gammaproteobacteria; Enterobacterales; Enterobacteriaceae; Escherichia MTGDNTLIHSHGINRRDFMKLCAALAATMGLSSKAAAEMAESVTNPQRPPVIWIGAQECT WP_053263529.1 hydrogenase 2 small subunit [Escherichia coli] Length: 372\nNumber of predicted TMHs: 1\nExp number of AAs in TMHs: 23.79108\nExp number, first 60 AAs: 0.22018\nTotal prob of N-in: 0.12753\noutside 1 331\nTMhelix 332 354\ninside 355 372

2558 GCF_001267355.1_401091 Escherichia coli Proteobacteria; Gammaproteobacteria; Enterobacterales; Enterobacteriaceae; Escherichia MNNEETFYQAMRRQGVTRRSFLKYCSLAATSLGLGAGMAPKIAWALENKPRIPVVWIHGL WP_001058323.1 MULTISPECIES: hydrogenase-1 small chain [Proteobacteria] Length: 372\nNumber of predicted TMHs: 2\nExp number of AAs in TMHs: 37.2608499999999\nExp number, first 60 AAs: 19.52923\nTotal prob of N-in: 0.94351\nPOSSIBLE N-term signal sequence\ninside 1 19\nTMhelix 20 39\noutside 40 325\nTMhelix 326 348\ninside 349 372

2559 GCF_001268225.1_1.EC2990.1 Escherichia coli Proteobacteria; Gammaproteobacteria; Enterobacterales; Enterobacteriaceae; Escherichia MNNEETFYQAMRRQGVTRRSFLKYCSLAATSLGLGAGMAPKIAWALENKPRIPVVWIHGL WP_001058323.1 MULTISPECIES: hydrogenase-1 small chain [Proteobacteria] Length: 372\nNumber of predicted TMHs: 2\nExp number of AAs in TMHs: 37.2608499999999\nExp number, first 60 AAs: 19.52923\nTotal prob of N-in: 0.94351\nPOSSIBLE N-term signal sequence\ninside 1 19\nTMhelix 20 39\noutside 40 325\nTMhelix 326 348\ninside 349 372

2560 GCF_001268205.1_1.EC2733.1 Escherichia coli Proteobacteria; Gammaproteobacteria; Enterobacterales; Enterobacteriaceae; Escherichia MNNEETFYQAMRRQGVTRRSFLKYCSLAATSLGLGAGMAPKIAWALENKPRIPVVWIHGL WP_001058323.1 MULTISPECIES: hydrogenase-1 small chain [Proteobacteria] Length: 372\nNumber of predicted TMHs: 2\nExp number of AAs in TMHs: 37.2608499999999\nExp number, first 60 AAs: 19.52923\nTotal prob of N-in: 0.94351\nPOSSIBLE N-term signal sequence\ninside 1 19\nTMhelix 20 39\noutside 40 325\nTMhelix 326 348\ninside 349 372

2561 GCF_001268305.1_YE15 Escherichia coli Proteobacteria; Gammaproteobacteria; Enterobacterales; Enterobacteriaceae; Escherichia MNNEETFYQAMRRQGVTRRSFLKYCSLAATSLGLGAGMAPKIAWALENKPRIPVVWIHGL WP_001058323.1 MULTISPECIES: hydrogenase-1 small chain [Proteobacteria] Length: 372\nNumber of predicted TMHs: 2\nExp number of AAs in TMHs: 37.2608499999999\nExp number, first 60 AAs: 19.52923\nTotal prob of N-in: 0.94351\nPOSSIBLE N-term signal sequence\ninside 1 19\nTMhelix 20 39\noutside 40 325\nTMhelix 326 348\ninside 349 372

2562 GCF_001268365.1_YE30 Escherichia coli Proteobacteria; Gammaproteobacteria; Enterobacterales; Enterobacteriaceae; Escherichia MTGDNTLIHSHGINRRDFMKLCAALAATMGLSSKAAAEMAESVTNPQRPPVIWIGAQECT WP_000145410.1 MULTISPECIES: hydrogenase-2 small chain [Proteobacteria] Length: 372\nNumber of predicted TMHs: 1\nExp number of AAs in TMHs: 23.72226\nExp number, first 60 AAs: 0.22038\nTotal prob of N-in: 0.12994\noutside 1 331\nTMhelix 332 354\ninside 355 372

2563 GCF_001268485.1_YE24 Escherichia coli Proteobacteria; Gammaproteobacteria; Enterobacterales; Enterobacteriaceae; Escherichia MNNEETFYQAMRRQGVTRRSFLKYCSLAATSLGLGAGMAPKIAWALENKPRIPVVWIHGL WP_001058323.1 MULTISPECIES: hydrogenase-1 small chain [Proteobacteria] Length: 372\nNumber of predicted TMHs: 2\nExp number of AAs in TMHs: 37.2608499999999\nExp number, first 60 AAs: 19.52923\nTotal prob of N-in: 0.94351\nPOSSIBLE N-term signal sequence\ninside 1 19\nTMhelix 20 39\noutside 40 325\nTMhelix 326 348\ninside 349 372

2564 GCF_001268265.1_1.EC2996.1 Escherichia coli Proteobacteria; Gammaproteobacteria; Enterobacterales; Enterobacteriaceae; Escherichia MNNEETFYQAMRRQGVTRRSFLKYCSLAATSLGLGAGMAPKIAWALENKPRIPVVWIHGL WP_053264900.1 hydrogenase [Escherichia coli] Length: 372\nNumber of predicted TMHs: 2\nExp number of AAs in TMHs: 37.24714\nExp number, first 60 AAs: 19.52921\nTotal prob of N-in: 0.94351\nPOSSIBLE N-term signal sequence\ninside 1 19\nTMhelix 20 39\noutside 40 325\nTMhelix 326 348\ninside 349 372

2565 GCF_001268545.1_YE11 Escherichia coli Proteobacteria; Gammaproteobacteria; Enterobacterales; Enterobacteriaceae; Escherichia MTGDNTLIHSHGINRRDFMKLCAALAATMGLSSKAAAEMAESVTNPQRPPVIWIGAQECT WP_000145410.1 MULTISPECIES: hydrogenase-2 small chain [Proteobacteria] Length: 372\nNumber of predicted TMHs: 1\nExp number of AAs in TMHs: 23.72226\nExp number, first 60 AAs: 0.22038\nTotal prob of N-in: 0.12994\noutside 1 331\nTMhelix 332 354\ninside 355 372

2566 GCF_001268505.1_YE16 Escherichia coli Proteobacteria; Gammaproteobacteria; Enterobacterales; Enterobacteriaceae; Escherichia MNNEETFYQAMRRQGVTRRSFLKYCSLAATSLGLGAGMAPKIAWALENKPRIPVVWIHGL WP_001058323.1 MULTISPECIES: hydrogenase-1 small chain [Proteobacteria] Length: 372\nNumber of predicted TMHs: 2\nExp number of AAs in TMHs: 37.2608499999999\nExp number, first 60 AAs: 19.52923\nTotal prob of N-in: 0.94351\nPOSSIBLE N-term signal sequence\ninside 1 19\nTMhelix 20 39\noutside 40 325\nTMhelix 326 348\ninside 349 372

2567 GCF_001268585.1_YE14 Escherichia coli Proteobacteria; Gammaproteobacteria; Enterobacterales; Enterobacteriaceae; Escherichia MTGDNTLIHSHGINRRDFMKLCAALAATMGLSSKAAAEMAESVTNPQRPPVIWIGAQECT WP_000145410.1 MULTISPECIES: hydrogenase-2 small chain [Proteobacteria] Length: 372\nNumber of predicted TMHs: 1\nExp number of AAs in TMHs: 23.72226\nExp number, first 60 AAs: 0.22038\nTotal prob of N-in: 0.12994\noutside 1 331\nTMhelix 332 354\ninside 355 372

2568 GCF_001268605.1_YE29 Escherichia coli Proteobacteria; Gammaproteobacteria; Enterobacterales; Enterobacteriaceae; Escherichia MTGDNTLIHSHGINRRDFMKLCAALAATMGLSSKAAAEMAESVTNPQRPPVIWIGAQECT WP_000145410.1 MULTISPECIES: hydrogenase-2 small chain [Proteobacteria] Length: 372\nNumber of predicted TMHs: 1\nExp number of AAs in TMHs: 23.72226\nExp number, first 60 AAs: 0.22038\nTotal prob of N-in: 0.12994\noutside 1 331\nTMhelix 332 354\ninside 355 372

2569 GCF_001268645.1_YE2 Escherichia coli Proteobacteria; Gammaproteobacteria; Enterobacterales; Enterobacteriaceae; Escherichia MTGDNTLIHSHGINRRDFMKLCAALAATMGLSSKAAAEMAESVTNPQRPPVIWIGAQECT WP_000145410.1 MULTISPECIES: hydrogenase-2 small chain [Proteobacteria] Length: 372\nNumber of predicted TMHs: 1\nExp number of AAs in TMHs: 23.72226\nExp number, first 60 AAs: 0.22038\nTotal prob of N-in: 0.12994\noutside 1 331\nTMhelix 332 354\ninside 355 372

2570 GCF_001268745.1_YE9 Escherichia coli Proteobacteria; Gammaproteobacteria; Enterobacterales; Enterobacteriaceae; Escherichia MTGDNTLIHSHGINRRDFMKLCAALAATMGLSSKAAAEMAESVTNPQRPPVIWIGAQECT WP_000145410.1 MULTISPECIES: hydrogenase-2 small chain [Proteobacteria] Length: 372\nNumber of predicted TMHs: 1\nExp number of AAs in TMHs: 23.72226\nExp number, first 60 AAs: 0.22038\nTotal prob of N-in: 0.12994\noutside 1 331\nTMhelix 332 354\ninside 355 372

2571 GCF_001268825.1_YE8 Escherichia coli Proteobacteria; Gammaproteobacteria; Enterobacterales; Enterobacteriaceae; Escherichia MTGDNTLIHSHGINRRDFMKLCAALAATMGLSSKAAAEMAESVTNPQRPPVIWIGAQECT WP_000145410.1 MULTISPECIES: hydrogenase-2 small chain [Proteobacteria] Length: 372\nNumber of

predicted TMHs: 1\nExp number of AAs in TMHs: 23.72226\nExp number, first 60 AAs: 0.22038\nTotal prob of N-in: 0.12994\noutside 1 331\nTMhelix 332 354\ninside 355 372

2572 GCF_001268785.1_YE25 Escherichia coli Proteobacteria; Gammaproteobacteria; Enterobacterales; Enterobacteriaceae; Escherichia MTGDNTLIHSHGINRRDFMKLCAALAATMGLSSKAAAEMAESVTNPQRPPVIWIGAQECT WP_000145410.1 MULTISPECIES: hydrogenase-2 small chain [Proteobacteria] Length: 372\nNumber of predicted TMHs: 1\nExp number of AAs in TMHs: 23.72226\nExp number, first 60 AAs: 0.22038\nTotal prob of N-in: 0.12994\noutside 1 331\nTMhelix 332 354\ninside 355 372

2573 GCF_001268745.1_YE9 Escherichia coli Proteobacteria; Gammaproteobacteria; Enterobacterales; Enterobacteriaceae; Escherichia MNNEETFYQAMRRQGVTRRSFLKYCSLAATSLGLGAGMAPKIAWALENKPRIPVVWIHGL WP_001058323.1 MULTISPECIES: hydrogenase-1 small chain [Proteobacteria] Length: 372\nNumber of predicted TMHs: 2\nExp number of AAs in TMHs: 37.2608499999999\nExp number, first 60 AAs: 19.52923\nTotal prob of N-in: 0.94351\nPOSSIBLE N-term signal sequence\ninside 1 19\nTMhelix 20 39\noutside 40 325\nTMhelix 326 348\ninside 349 372

2574 GCF_001268825.1_YE8 Escherichia coli Proteobacteria; Gammaproteobacteria; Enterobacterales; Enterobacteriaceae; Escherichia MNNEETFYQAMRRQGVTRRSFLKYCSLAATSLGLGAGMAPKIAWALENKPRIPVVWIHGL WP_001058323.1 MULTISPECIES: hydrogenase-1 small chain [Proteobacteria] Length: 372\nNumber of predicted TMHs: 2\nExp number of AAs in TMHs: 37.2608499999999\nExp number, first 60 AAs: 19.52923\nTotal prob of N-in: 0.94351\nPOSSIBLE N-term signal sequence\ninside 1 19\nTMhelix 20 39\noutside 40 325\nTMhelix 326 348\ninside 349 372

2575 GCF_001268725.1_YE7 Escherichia coli Proteobacteria; Gammaproteobacteria; Enterobacterales; Enterobacteriaceae; Escherichia MNNEETFYQAMRRQGVTRRSFLKYCSLAATSLGLGAGMAPKIAWALENKPRIPVVWIHGL WP_001058323.1 MULTISPECIES: hydrogenase-1 small chain [Proteobacteria] Length: 372\nNumber of predicted TMHs: 2\nExp number of AAs in TMHs: 37.2608499999999\nExp number, first 60 AAs: 19.52923\nTotal prob of N-in: 0.94351\nPOSSIBLE N-term signal sequence\ninside 1 19\nTMhelix 20 39\noutside 40 325\nTMhelix 326 348\ninside 349 372

2576 GCF_001268865.1_1.EC2739.1 Escherichia coli Proteobacteria; Gammaproteobacteria; Enterobacterales; Enterobacteriaceae; Escherichia MTGDNTLIHSHGINRRDFMKLCAALAATMGLSSKAAAEMAESVTNPQRPPVIWIGAQECT WP_000145410.1 MULTISPECIES: hydrogenase-2 small chain [Proteobacteria] Length: 372\nNumber of predicted TMHs: 1\nExp number of AAs in TMHs: 23.72226\nExp number, first 60 AAs: 0.22038\nTotal prob of N-in: 0.12994\noutside 1 331\nTMhelix 332 354\ninside 355 372

2577 GCF_001269085.1_1.EC2991.1 Escherichia coli Proteobacteria; Gammaproteobacteria; Enterobacterales; Enterobacteriaceae; Escherichia MTGDNTLIHSHGINRRDFMKLCAALAATMGLSSKAAAEMAESVTNPQRPPVIWIGAQECT WP_000145410.1 MULTISPECIES: hydrogenase-2 small chain [Proteobacteria] Length: 372\nNumber of predicted TMHs: 1\nExp number of AAs in TMHs: 23.72226\nExp number, first 60 AAs: 0.22038\nTotal prob of N-in: 0.12994\noutside 1 331\nTMhelix 332 354\ninside 355 372

2578 GCF_001269085.1_1.EC2991.1 Escherichia coli Proteobacteria; Gammaproteobacteria; Enterobacterales; Enterobacteriaceae; Escherichia MNNEETFYQAMRRQGVTRRSFLKYCSLAATSLGLGAGMAPKIAWALENKPRIPVVWIHGL WP_001058323.1 MULTISPECIES: hydrogenase-1 small chain [Proteobacteria] Length: 372\nNumber of predicted TMHs: 2\nExp number of AAs in TMHs: 37.2608499999999\nExp number, first 60 AAs: 19.52923\nTotal prob of N-in: 0.94351\nPOSSIBLE N-term signal sequence\ninside 1 19\nTMhelix 20 39\noutside 40 325\nTMhelix 326 348\ninside 349 372

2579 GCF_001269065.1_1.EC2729.1 Escherichia coli Proteobacteria; Gammaproteobacteria; Enterobacterales; Enterobacteriaceae; Escherichia MNNEETFYQAMRRQGVTRRSFLKYCSLAATSLGLGAGMAPKIAWALENKPRIPVVWIHGL WP_001058323.1 MULTISPECIES: hydrogenase-1 small chain [Proteobacteria] Length: 372\nNumber of predicted TMHs: 2\nExp number of AAs in TMHs: 37.2608499999999\nExp number, first 60 AAs: 19.52923\nTotal prob of N-in: 0.94351\nPOSSIBLE N-term signal sequence\ninside 1 19\nTMhelix 20 39\noutside 40 325\nTMhelix 326 348\ninside 349 372

2580 GCF_001268905.1_YE10 Escherichia coli Proteobacteria; Gammaproteobacteria; Enterobacterales; Enterobacteriaceae; Escherichia MNNEETFYQAMRRQGVTRRSFLKYCSLAATSLGLGAGMAPKIAWALENKPRIPVVWIHGL WP_001058323.1 MULTISPECIES: hydrogenase-1 small chain [Proteobacteria] Length: 372\nNumber of predicted TMHs: 2\nExp number of AAs in TMHs: 37.2608499999999\nExp number, first 60 AAs: 19.52923\nTotal prob of N-in: 0.94351\nPOSSIBLE N-term signal sequence\ninside 1 19\nTMhelix 20 39\noutside 40 325\nTMhelix 326 348\ninside 349 372

2581 GCF_001269165.1_YE27 Escherichia coli Proteobacteria; Gammaproteobacteria; Enterobacterales; Enterobacteriaceae; Escherichia MTGDNTLIHSHGINRRDFMKLCAALAATMGLSSKAAAEMAESVTNPQRPPVIWIGAQECT WP_000145410.1 MULTISPECIES: hydrogenase-2 small chain [Proteobacteria] Length: 372\nNumber of predicted TMHs: 1\nExp number of AAs in TMHs: 23.72226\nExp number, first 60 AAs: 0.22038\nTotal prob of N-in: 0.12994\noutside 1 331\nTMhelix 332 354\ninside 355 372

2582 GCF_001269165.1_YE27 Escherichia coli Proteobacteria; Gammaproteobacteria; Enterobacterales; Enterobacteriaceae; Escherichia MNNEETFYQAMRRQGVTRRSFLKYCSLAATSLGLGAGMAPKIAWALENKPRIPVVWIHGL WP_001058323.1 MULTISPECIES: hydrogenase-1 small chain [Proteobacteria] Length: 372\nNumber of predicted TMHs: 2\nExp number of AAs in TMHs: 37.2608499999999\nExp number, first 60 AAs: 19.52923\nTotal prob of N-in: 0.94351\nPOSSIBLE N-term signal sequence\ninside 1 19\nTMhelix 20 39\noutside 40 325\nTMhelix 326 348\ninside 349 372

2583 GCF_001268985.1_1.ECB31.1 Escherichia coli Proteobacteria; Gammaproteobacteria; Enterobacterales; Enterobacteriaceae; Escherichia MNNEETFYQAMRRQGVTRRSFLKYCSLAATSLGLGAGMAPKIAWALENKPRIPVVWIHGL WP_001058323.1 MULTISPECIES: hydrogenase-1 small chain [Proteobacteria] Length: 372\nNumber of predicted TMHs: 2\nExp number of AAs in TMHs: 37.2608499999999\nExp number, first 60 AAs: 19.52923\nTotal prob of N-in: 0.94351\nPOSSIBLE N-term signal sequence\ninside 1 19\nTMhelix 20 39\noutside 40 325\nTMhelix 326 348\ninside 349 372

2584 GCF_001269305.1_1.ECAF83.1 Escherichia coli Proteobacteria; Gammaproteobacteria; Enterobacterales; Enterobacteriaceae; Escherichia MTGDNTLIHSHGINRRDFMKLCAALAATMGLSSKAAAEMAESVTNPQRPPVIWIGAQECT WP_000145410.1 MULTISPECIES: hydrogenase-2 small chain [Proteobacteria] Length: 372\nNumber of predicted TMHs: 1\nExp number of AAs in TMHs: 23.72226\nExp number, first 60 AAs: 0.22038\nTotal prob of N-in: 0.12994\noutside 1 331\nTMhelix 332 354\ninside 355 372

2585 GCF_001276265.1_ASM127626v1 Escherichia coli Proteobacteria; Gammaproteobacteria; Enterobacterales; Enterobacteriaceae; Escherichia MTGDNTLIHSHGINRRDFMKLCAALAATMGLSSKAAAEMAESVTNPQRPPVIWIGAQECT WP_000145410.1 MULTISPECIES: hydrogenase-2 small chain [Proteobacteria] Length: 372\nNumber of predicted TMHs: 1\nExp number of AAs in TMHs: 23.72226\nExp number, first 60 AAs: 0.22038\nTotal prob of N-in: 0.12994\noutside 1 331\nTMhelix 332 354\ninside 355 372

2586 GCF_001277395.1_7790_1_92 Escherichia coli Proteobacteria; Gammaproteobacteria; Enterobacterales; Enterobacteriaceae; Escherichia MTGDNTLIHSHGINRRDFMKLCAALAATMGLSSKAAAEMAESVTNPQRPPVIWIGAQECT WP_000145410.1 MULTISPECIES: hydrogenase-2 small chain [Proteobacteria] Length: 372\nNumber of predicted TMHs: 1\nExp number of AAs in TMHs: 23.72226\nExp number, first 60 AAs: 0.22038\nTotal prob of N-in: 0.12994\noutside 1 331\nTMhelix 332 354\ninside 355 372

2587 GCF_001277415.1_7748_7_42 Escherichia coli Proteobacteria; Gammaproteobacteria; Enterobacterales; Enterobacteriaceae; Escherichia MTGDNTLIHSHGINRRDFMKLCAALAATMGLSSKAAAEMAESVTNPQRPPVIWIGAQECT WP_000145410.1 MULTISPECIES: hydrogenase-2 small chain [Proteobacteria] Length: 372\nNumber of predicted TMHs: 1\nExp number of AAs in TMHs: 23.72226\nExp number, first 60 AAs: 0.22038\nTotal prob of N-in: 0.12994\noutside 1 331\nTMhelix 332 354\ninside 355 372

2588 GCF_001277415.1_7748_7_42 Escherichia coli Proteobacteria; Gammaproteobacteria; Enterobacterales; Enterobacteriaceae; Escherichia MNNEETFYQAMRRQGVTRRSFLKYCSLAATSLGLGAGMAPKIAWALENKPRIPVVWIHGL WP_001058323.1 MULTISPECIES: hydrogenase-1 small chain [Proteobacteria] Length: 372\nNumber of predicted TMHs: 2\nExp number of AAs in TMHs: 37.2608499999999\nExp number, first 60 AAs: 19.52923\nTotal prob of N-in:

0.94351\nPOSSIBLE N-term signal sequence\ninside 1 19\nTMhelix 20 39\noutside 40 325\nTMhelix 326 348\ninside 349 372

2589 GCF_001269125.1_YE26 Escherichia coli Proteobacteria; Gammaproteobacteria; Enterobacterales; Enterobacteriaceae; Escherichia MTGDNTLIHSHGINRRDFMKLCAALAATMGLSSKAAAEMAESVTNPQRPPVIWIGAQECT WP_032298447.1 hydrogenase 2 small subunit [Escherichia coli] Length: 372\nNumber of predicted TMHs: 1\nExp number of AAs in TMHs: 23.69807\nExp number, first 60 AAs: 0.221\nTotal prob of N-in: 0.12853\noutside 1 331\nTMhelix 332 354\ninside 355 372

2590 GCF_001269265.1_1.EC2723.1 Escherichia coli Proteobacteria; Gammaproteobacteria; Enterobacterales; Enterobacteriaceae; Escherichia MNNEETFYQAMRRQGVTRRSFLKYCSLAATSLGLGAGMAPKIAWALENKPRIPVVWIHGL WP_001058323.1 MULTISPECIES: hydrogenase-1 small chain [Proteobacteria] Length: 372\nNumber of predicted TMHs: 2\nExp number of AAs in TMHs: 37.2608499999999\nExp number, first 60 AAs: 19.52923\nTotal prob of N-in: 0.94351\nPOSSIBLE N-term signal sequence\ninside 1 19\nTMhelix 20 39\noutside 40 325\nTMhelix 326 348\ninside 349 372

2591 GCF_001277555.1_7748_7_39 Escherichia coli Proteobacteria; Gammaproteobacteria; Enterobacterales; Enterobacteriaceae; Escherichia MTGDNTLIHSHGINRRDFMKLCAALAATMGLSSKAAAEMAESVTNPQRPPVIWIGAQECT WP_000145410.1 MULTISPECIES: hydrogenase-2 small chain [Proteobacteria] Length: 372\nNumber of predicted TMHs: 1\nExp number of AAs in TMHs: 23.72226\nExp number, first 60 AAs: 0.22038\nTotal prob of N-in: 0.12994\noutside 1 331\nTMhelix 332 354\ninside 355 372

2592 GCF_001277555.1_7748_7_39 Escherichia coli Proteobacteria; Gammaproteobacteria; Enterobacterales; Enterobacteriaceae; Escherichia MNNEETFYQAMRRQGVTRRSFLKYCSLAATSLGLGAGMAPKIAWALENKPRIPVVWIHGL WP_001058323.1 MULTISPECIES: hydrogenase-1 small chain [Proteobacteria] Length: 372\nNumber of predicted TMHs: 2\nExp number of AAs in TMHs: 37.2608499999999\nExp number, first 60 AAs: 19.52923\nTotal prob of N-in: 0.94351\nPOSSIBLE N-term signal sequence\ninside 1 19\nTMhelix 20 39\noutside 40 325\nTMhelix 326 348\ninside 349 372

2593 GCF_001277515.1_7790_1_59 Escherichia coli Proteobacteria; Gammaproteobacteria; Enterobacterales; Enterobacteriaceae; Escherichia MNNEETFYQAMRRQGVTRRSFLKYCSLAATSLGLGAGMAPKIAWALENKPRIPVVWIHGL WP_001295940.1 MULTISPECIES: hydrogenase [Enterobacteriaceae] Length: 372\nNumber of predicted TMHs: 2\nExp number of AAs in TMHs: 37.2608899999999\nExp number, first 60 AAs: 19.52923\nTotal prob of N-in: 0.94351\nPOSSIBLE N-term signal sequence\ninside 1 19\nTMhelix 20 39\noutside 40 325\nTMhelix 326 348\ninside 349 372

2594 GCF_001269305.1_1.ECAF83.1 Escherichia coli Proteobacteria; Gammaproteobacteria; Enterobacterales; Enterobacteriaceae; Escherichia MNNEETFYQAMRRQGVTRRSFLKYCSLAATSLGLGAGMAPKIAWALENKPRIPVVWIHGL WP_001498899.1 hydrogenase-1 small chain [Escherichia coli] Length: 372\nNumber of predicted TMHs: 2\nExp number of AAs in TMHs: 39.34459\nExp number, first 60 AAs: 19.53328\nTotal prob of N-in: 0.94350\nPOSSIBLE N-term signal sequence\ninside 1 19\nTMhelix 20 39\noutside 40 325\nTMhelix 326 348\ninside 349 372

2595 GCF_001277595.1_7748_7_19 Escherichia coli Proteobacteria; Gammaproteobacteria; Enterobacterales; Enterobacteriaceae; Escherichia MTGDNTLIHSHGINRRDFMKLCAALAATMGLSSKAAAEMAESVTNPQRPPVIWIGAQECT WP_000145410.1 MULTISPECIES: hydrogenase-2 small chain [Proteobacteria] Length: 372\nNumber of predicted TMHs: 1\nExp number of AAs in TMHs: 23.72226\nExp number, first 60 AAs: 0.22038\nTotal prob of N-in: 0.12994\noutside 1 331\nTMhelix 332 354\ninside 355 372

2596 GCF_001277615.1_7790_1_70 Escherichia coli Proteobacteria; Gammaproteobacteria; Enterobacterales; Enterobacteriaceae; Escherichia MTGDNTLIHSHGINRRDFMKLCAALAATMGLSSKAAAEMAESVTNPQRPPVIWIGAQECT WP_000145410.1 MULTISPECIES: hydrogenase-2 small chain [Proteobacteria] Length: 372\nNumber of predicted TMHs: 1\nExp number of AAs in TMHs: 23.72226\nExp number, first 60 AAs: 0.22038\nTotal prob of N-in: 0.12994\noutside 1 331\nTMhelix 332 354\ninside 355 372

2597 GCF_001277595.1_7748_7_19 Escherichia coli Proteobacteria; Gammaproteobacteria; Enterobacterales; Enterobacteriaceae; Escherichia
MNNEETFYQAMRRQGVTRRSFLKYCSLAATSLGLGAGMAPKIAWALENKPRIPVVWIHGL WP_001058323.1
MULTISPECIES: hydrogenase-1 small chain [Proteobacteria] Length: 372\nNumber of predicted TMHs: 2\nExp number of AAs in TMHs: 37.2608499999999\nExp number, first 60 AAs: 19.52923\nTotal prob of N-in: 0.94351\nPOSSIBLE N-term signal sequence\ninside 1 19\nTMhelix 20 39\noutside 40 325\nTMhelix 326 348\ninside 349 372

2598 GCF_001277695.1_7748_7_33 Escherichia coli Proteobacteria; Gammaproteobacteria; Enterobacterales; Enterobacteriaceae; Escherichia
MNNEETFYQAMRRQGVTRRSFLKYCSLAATSLGLGAGMAPKIAWALENKPRIPVVWIHGL WP_001058323.1
MULTISPECIES: hydrogenase-1 small chain [Proteobacteria] Length: 372\nNumber of predicted TMHs: 2\nExp number of AAs in TMHs: 37.2608499999999\nExp number, first 60 AAs: 19.52923\nTotal prob of N-in: 0.94351\nPOSSIBLE N-term signal sequence\ninside 1 19\nTMhelix 20 39\noutside 40 325\nTMhelix 326 348\ninside 349 372

2599 GCF_001277655.1_7553_7_71 Escherichia coli Proteobacteria; Gammaproteobacteria; Enterobacterales; Enterobacteriaceae; Escherichia
MNNEETFYQAMRRQGVTRRSFLKYCSLAATSLGLGAGMAPKIAWALENKPRIPVVWIHGL WP_001058323.1
MULTISPECIES: hydrogenase-1 small chain [Proteobacteria] Length: 372\nNumber of predicted TMHs: 2\nExp number of AAs in TMHs: 37.2608499999999\nExp number, first 60 AAs: 19.52923\nTotal prob of N-in: 0.94351\nPOSSIBLE N-term signal sequence\ninside 1 19\nTMhelix 20 39\noutside 40 325\nTMhelix 326 348\ninside 349 372

2600 GCF_001277755.1_7790_1_90 Escherichia coli Proteobacteria; Gammaproteobacteria; Enterobacterales; Enterobacteriaceae; Escherichia
MTGDNTLIHSHGINRRDFMKLCAALAATMGLSSKAAAEMAESVTNPQRPPVIWIGAQECT WP_000145410.1
MULTISPECIES: hydrogenase-2 small chain [Proteobacteria] Length: 372\nNumber of predicted TMHs: 1\nExp number of AAs in TMHs: 23.72226\nExp number, first 60 AAs: 0.22038\nTotal prob of N-in: 0.12994\noutside 1 331\nTMhelix 332 354\ninside 355 372

2601 GCF_001280325.1_ASM128032v1 Escherichia coli Proteobacteria; Gammaproteobacteria; Enterobacterales; Enterobacteriaceae; Escherichia
MTGDNTLIHSHGINRRDFMKLCAALAATMGLSSKAAAEMAESVTNPQRPPVIWIGAQECT WP_000145410.1
MULTISPECIES: hydrogenase-2 small chain [Proteobacteria] Length: 372\nNumber of predicted TMHs: 1\nExp number of AAs in TMHs: 23.72226\nExp number, first 60 AAs: 0.22038\nTotal prob of N-in: 0.12994\noutside 1 331\nTMhelix 332 354\ninside 355 372

2602 GCF_001280385.1_ASM128038v1 Escherichia coli Proteobacteria; Gammaproteobacteria; Enterobacterales; Enterobacteriaceae; Escherichia
MTGDNTLIHSHGINRRDFMKLCAALAATMGLSSKAAAEMAESVTNPQRPPVIWIGAQECT WP_000145410.1
MULTISPECIES: hydrogenase-2 small chain [Proteobacteria] Length: 372\nNumber of predicted TMHs: 1\nExp number of AAs in TMHs: 23.72226\nExp number, first 60 AAs: 0.22038\nTotal prob of N-in: 0.12994\noutside 1 331\nTMhelix 332 354\ninside 355 372

2603 GCF_001281685.1_ASM128168v1 Escherichia coli Proteobacteria; Gammaproteobacteria; Enterobacterales; Enterobacteriaceae; Escherichia
MTGDNTLIHSHGINRRDFMKLCAALAATMGLSSKAAAEMAESVTNPQRPPVIWIGAQECT WP_000145410.1
MULTISPECIES: hydrogenase-2 small chain [Proteobacteria] Length: 372\nNumber of predicted TMHs: 1\nExp number of AAs in TMHs: 23.72226\nExp number, first 60 AAs: 0.22038\nTotal prob of N-in: 0.12994\noutside 1 331\nTMhelix 332 354\ninside 355 372

2604 GCF_001277755.1_7790_1_90 Escherichia coli Proteobacteria; Gammaproteobacteria; Enterobacterales; Enterobacteriaceae; Escherichia
MNNEETFYQAMRRQGVTRRSFLKYCSLAATSLGLGAGMAPKIAWALENKPRIPVVWIHGL WP_001058323.1
MULTISPECIES: hydrogenase-1 small chain [Proteobacteria] Length: 372\nNumber of predicted TMHs: 2\nExp number of AAs in TMHs: 37.2608499999999\nExp number, first 60 AAs: 19.52923\nTotal prob of N-in: 0.94351\nPOSSIBLE N-term signal sequence\ninside 1 19\nTMhelix 20 39\noutside 40 325\nTMhelix 326 348\ninside 349 372

2605 GCF_001281755.1_ASM128175v1 Escherichia coli Proteobacteria; Gammaproteobacteria; Enterobacterales; Enterobacteriaceae; Escherichia
MTGDNTLIHSHGINRRDFMKLCAALAATMGLSSKAAAEMAESVTNPQRPPVIWIGAQECT WP_000145410.1
MULTISPECIES: hydrogenase-2 small chain [Proteobacteria] Length: 372\nNumber of predicted TMHs: 1\nExp number of AAs in TMHs: 23.72226\nExp number, first 60 AAs: 0.22038\nTotal prob of N-in: 0.12994\noutside 1 331\nTMhelix 332 354\ninside 355 372

2606 GCF_001281775.1_ASM128177v1 Escherichia coli Proteobacteria; Gammaproteobacteria; Enterobacterales; Enterobacteriaceae; Escherichia
MTGDNTLIHSHGINRRDFMKLCAALAATMGLSSKAAAEMAESVTNPQRPPVIWIGAQECT WP_000145410.1
MULTISPECIES: hydrogenase-2 small chain [Proteobacteria] Length: 372\nNumber of predicted TMHs: 1\nExp number of AAs in TMHs: 23.72226\nExp number, first 60 AAs: 0.22038\nTotal prob of N-in: 0.12994\noutside 1 331\nTMhelix 332 354\ninside 355 372

2607 GCF_001281815.1_ASM128181v1 Escherichia coli Proteobacteria; Gammaproteobacteria; Enterobacterales; Enterobacteriaceae; Escherichia
MTGDNTLIHSHGINRRDFMKLCAALAATMGLSSKAAAEMAESVTNPQRPPVIWIGAQECT WP_000145410.1
MULTISPECIES: hydrogenase-2 small chain [Proteobacteria] Length: 372\nNumber of predicted TMHs: 1\nExp number of AAs in TMHs: 23.72226\nExp number, first 60 AAs: 0.22038\nTotal prob of N-in: 0.12994\noutside 1 331\nTMhelix 332 354\ninside 355 372

2608 GCF_001281855.1_ASM128185v1 Escherichia coli Proteobacteria; Gammaproteobacteria; Enterobacterales; Enterobacteriaceae; Escherichia
MTGDNTLIHSHGINRRDFMKLCAALAATMGLSSKAAAEMAESVTNPQRPPVIWIGAQECT WP_000145410.1
MULTISPECIES: hydrogenase-2 small chain [Proteobacteria] Length: 372\nNumber of predicted TMHs: 1\nExp number of AAs in TMHs: 23.72226\nExp number, first 60 AAs: 0.22038\nTotal prob of N-in: 0.12994\noutside 1 331\nTMhelix 332 354\ninside 355 372

2609 GCF_001281885.1_ASM128188v1 Escherichia coli Proteobacteria; Gammaproteobacteria; Enterobacterales; Enterobacteriaceae; Escherichia
MTGDNTLIHSHGINRRDFMKLCAALAATMGLSSKAAAEMAESVTNPQRPPVIWIGAQECT WP_000145410.1
MULTISPECIES: hydrogenase-2 small chain [Proteobacteria] Length: 372\nNumber of predicted TMHs: 1\nExp number of AAs in TMHs: 23.72226\nExp number, first 60 AAs: 0.22038\nTotal prob of N-in: 0.12994\noutside 1 331\nTMhelix 332 354\ninside 355 372

2610 GCF_001281915.1_ASM128191v1 Escherichia coli Proteobacteria; Gammaproteobacteria; Enterobacterales; Enterobacteriaceae; Escherichia
MTGDNTLIHSHGINRRDFMKLCAALAATMGLSSKAAAEMAESVTNPQRPPVIWIGAQECT WP_000145410.1
MULTISPECIES: hydrogenase-2 small chain [Proteobacteria] Length: 372\nNumber of predicted TMHs: 1\nExp number of AAs in TMHs: 23.72226\nExp number, first 60 AAs: 0.22038\nTotal prob of N-in: 0.12994\noutside 1 331\nTMhelix 332 354\ninside 355 372

2611 GCF_001281965.1_ASM128196v1 Escherichia coli Proteobacteria; Gammaproteobacteria; Enterobacterales; Enterobacteriaceae; Escherichia
MTGDNTLIHSHGINRRDFMKLCAALAATMGLSSKAAAEMAESVTNPQRPPVIWIGAQECT WP_000145410.1
MULTISPECIES: hydrogenase-2 small chain [Proteobacteria] Length: 372\nNumber of predicted TMHs: 1\nExp number of AAs in TMHs: 23.72226\nExp number, first 60 AAs: 0.22038\nTotal prob of N-in: 0.12994\noutside 1 331\nTMhelix 332 354\ninside 355 372

2612 GCF_001282065.1_ASM128206v1 Escherichia coli Proteobacteria; Gammaproteobacteria; Enterobacterales; Enterobacteriaceae; Escherichia
MTGDNTLIHSHGINRRDFMKLCAALAATMGLSSKAAAEMAESVTNPQRPPVIWIGAQECT WP_000145410.1
MULTISPECIES: hydrogenase-2 small chain [Proteobacteria] Length: 372\nNumber of predicted TMHs: 1\nExp number of AAs in TMHs: 23.72226\nExp number, first 60 AAs: 0.22038\nTotal prob of N-in: 0.12994\noutside 1 331\nTMhelix 332 354\ninside 355 372

2613 GCF_001282155.1_ASM128215v1 Escherichia coli Proteobacteria; Gammaproteobacteria; Enterobacterales; Enterobacteriaceae; Escherichia
MTGDNTLIHSHGINRRDFMKLCAALAATMGLSSKAAAEMAESVTNPQRPPVIWIGAQECT WP_000145410.1
MULTISPECIES: hydrogenase-2 small chain [Proteobacteria] Length: 372\nNumber of predicted TMHs:

1\nExp number of AAs in TMHs: 23.72226\nExp number, first 60 AAs: 0.22038\nTotal prob of N-in: 0.12994\noutside 1 331\nTMhelix 332 354\ninside 355 372

2614 GCF_001282155.1_ASM128215v1 Escherichia coli Proteobacteria; Gammaproteobacteria; Enterobacterales; Enterobacteriaceae; Escherichia MNNEETFYQAMRRQGVTRRSFLKYCSLAATSLGLGAGMAPKIAWALENKPRIPVVWIHGL WP_001058320.1 hydrogenase [Escherichia coli] Length: 372\nNumber of predicted TMHs: 2\nExp number of AAs in TMHs: 37.26082\nExp number, first 60 AAs: 19.52923\nTotal prob of N-in: 0.94351\nPOSSIBLE N-term signal sequence\ninside 1 19\nTMhelix 20 39\noutside 40 325\nTMhelix 326 348\ninside 349 372

2615 GCF_001282025.1_ASM128202v1 Escherichia coli Proteobacteria; Gammaproteobacteria; Enterobacterales; Enterobacteriaceae; Escherichia MNNEETFYQAMRRQGVTRRSFLKYCSLAATSLGLGAGMAPKIAWALENKPRIPVVWIHGL WP_001301613.1 hydrogenase [Escherichia coli] Length: 372\nNumber of predicted TMHs: 2\nExp number of AAs in TMHs: 37.46598\nExp number, first 60 AAs: 19.53139\nTotal prob of N-in: 0.94364\nPOSSIBLE N-term signal sequence\ninside 1 19\nTMhelix 20 39\noutside 40 325\nTMhelix 326 348\ninside 349 372

2616 GCF_001282065.1_ASM128206v1 Escherichia coli Proteobacteria; Gammaproteobacteria; Enterobacterales; Enterobacteriaceae; Escherichia MNNEETFYQAMRRQGVTRRSFLKYCSLAATSLGLGAGMAPKIAWALENKPRIPVVWIHGL WP_001301613.1 hydrogenase [Escherichia coli] Length: 372\nNumber of predicted TMHs: 2\nExp number of AAs in TMHs: 37.46598\nExp number, first 60 AAs: 19.53139\nTotal prob of N-in: 0.94364\nPOSSIBLE N-term signal sequence\ninside 1 19\nTMhelix 20 39\noutside 40 325\nTMhelix 326 348\ninside 349 372

2617 GCF_001281815.1_ASM128181v1 Escherichia coli Proteobacteria; Gammaproteobacteria; Enterobacterales; Enterobacteriaceae; Escherichia MNNEETFYQAMRRQGVTRRSFLKYCSLAATSLGLGAGMAPKIAWALENKPRIPVVWIHGL WP_001301613.1 hydrogenase [Escherichia coli] Length: 372\nNumber of predicted TMHs: 2\nExp number of AAs in TMHs: 37.46598\nExp number, first 60 AAs: 19.53139\nTotal prob of N-in: 0.94364\nPOSSIBLE N-term signal sequence\ninside 1 19\nTMhelix 20 39\noutside 40 325\nTMhelix 326 348\ninside 349 372

2618 GCF_001281855.1_ASM128185v1 Escherichia coli Proteobacteria; Gammaproteobacteria; Enterobacterales; Enterobacteriaceae; Escherichia MNNEETFYQAMRRQGVTRRSFLKYCSLAATSLGLGAGMAPKIAWALENKPRIPVVWIHGL WP_001301613.1 hydrogenase [Escherichia coli] Length: 372\nNumber of predicted TMHs: 2\nExp number of AAs in TMHs: 37.46598\nExp number, first 60 AAs: 19.53139\nTotal prob of N-in: 0.94364\nPOSSIBLE N-term signal sequence\ninside 1 19\nTMhelix 20 39\noutside 40 325\nTMhelix 326 348\ninside 349 372

2619 GCF_001282275.1_ASM128227v1 Escherichia coli Proteobacteria; Gammaproteobacteria; Enterobacterales; Enterobacteriaceae; Escherichia MTGDNTLIHSHGINRRDFMKLCAALAATMGLSSKAAAEMAESVTNPQRPPVIWIGAQECT WP_000145410.1 MULTISPECIES: hydrogenase-2 small chain [Proteobacteria] Length: 372\nNumber of predicted TMHs: 1\nExp number of AAs in TMHs: 23.72226\nExp number, first 60 AAs: 0.22038\nTotal prob of N-in: 0.12994\noutside 1 331\nTMhelix 332 354\ninside 355 372

2620 GCF_001282335.1_ASM128233v1 Escherichia coli Proteobacteria; Gammaproteobacteria; Enterobacterales; Enterobacteriaceae; Escherichia MTGDNTLIHSHGINRRDFMKLCAALAATMGLSSKAAAEMAESVTNPQRPPVIWIGAQECT WP_000145410.1 MULTISPECIES: hydrogenase-2 small chain [Proteobacteria] Length: 372\nNumber of predicted TMHs: 1\nExp number of AAs in TMHs: 23.72226\nExp number, first 60 AAs: 0.22038\nTotal prob of N-in: 0.12994\noutside 1 331\nTMhelix 332 354\ninside 355 372

2621 GCF_001282375.1_ASM128237v1 Escherichia coli Proteobacteria; Gammaproteobacteria; Enterobacterales; Enterobacteriaceae; Escherichia MTGDNTLIHSHGINRRDFMKLCAALAATMGLSSKAAAEMAESVTNPQRPPVIWIGAQECT WP_000145410.1 MULTISPECIES: hydrogenase-2 small chain [Proteobacteria] Length: 372\nNumber of predicted TMHs: 1\nExp number of AAs in TMHs: 23.72226\nExp number, first 60 AAs: 0.22038\nTotal prob of N-in: 0.12994\noutside 1 331\nTMhelix 332 354\ninside 355 372

2622 GCF_001283145.1_7748_7_13 Escherichia coli Proteobacteria; Gammaproteobacteria; Enterobacterales; Enterobacteriaceae; Escherichia

MTGDNTLIHSHGINRRDFMKLCAALAATMGLSSKAAAEMAESVTNPQRPPVIWIGAQECT WP_000145410.1 MULTISPECIES: hydrogenase-2 small chain [Proteobacteria] Length: 372\nNumber of predicted TMHs: 1\nExp number of AAs in TMHs: 23.72226\nExp number, first 60 AAs: 0.22038\nTotal prob of N-in: 0.12994\noutside 1 331\nTMhelix 332 354\ninside 355 372

2623 GCF_001283185.1_8205_8_77 Escherichia coli Proteobacteria; Gammaproteobacteria; Enterobacterales; Enterobacteriaceae; Escherichia MTGDNTLIHSHGINRRDFMKLCAALAATMGLSSKAAAEMAESVTNPQRPPVIWIGAQECT WP_000145410.1 MULTISPECIES: hydrogenase-2 small chain [Proteobacteria] Length: 372\nNumber of predicted TMHs: 1\nExp number of AAs in TMHs: 23.72226\nExp number, first 60 AAs: 0.22038\nTotal prob of N-in: 0.12994\noutside 1 331\nTMhelix 332 354\ninside 355 372

2624 GCF_001283285.1_8205_3_51 Escherichia coli Proteobacteria; Gammaproteobacteria; Enterobacterales; Enterobacteriaceae; Escherichia MTGDNTLIHSHGINRRDFMKLCAALAATMGLSSKAAAEMAESVTNPQRPPVIWIGAQECT WP_000145410.1 MULTISPECIES: hydrogenase-2 small chain [Proteobacteria] Length: 372\nNumber of predicted TMHs: 1\nExp number of AAs in TMHs: 23.72226\nExp number, first 60 AAs: 0.22038\nTotal prob of N-in: 0.12994\noutside 1 331\nTMhelix 332 354\ninside 355 372

2625 GCF_001283765.1_7790_1_80 Escherichia coli Proteobacteria; Gammaproteobacteria; Enterobacterales; Enterobacteriaceae; Escherichia MTGDNTLIHSHGINRRDFMKLCAALAATMGLSSKAAAEMAESVTNPQRPPVIWIGAQECT WP_000145410.1 MULTISPECIES: hydrogenase-2 small chain [Proteobacteria] Length: 372\nNumber of predicted TMHs: 1\nExp number of AAs in TMHs: 23.72226\nExp number, first 60 AAs: 0.22038\nTotal prob of N-in: 0.12994\noutside 1 331\nTMhelix 332 354\ninside 355 372

2626 GCF_001282275.1_ASM128227v1 Escherichia coli Proteobacteria; Gammaproteobacteria; Enterobacterales; Enterobacteriaceae; Escherichia MNNEETFYQAMRRQGVTRRSFLKYCSLAATSLGLGAGMAPKIAWALENKPRIPVVWIHGL WP_001058320.1 hydrogenase [Escherichia coli] Length: 372\nNumber of predicted TMHs: 2\nExp number of AAs in TMHs: 37.26082\nExp number, first 60 AAs: 19.52923\nTotal prob of N-in: 0.94351\nPOSSIBLE N-term signal sequence\ninside 1 19\nTMhelix 20 39\noutside 40 325\nTMhelix 326 348\ninside 349 372

2627 GCF_001283305.1_8205_3_48 Escherichia coli Proteobacteria; Gammaproteobacteria; Enterobacterales; Enterobacteriaceae; Escherichia MNNEETFYQAMRRQGVTRRSFLKYCSLAATSLGLGAGMAPKIAWALENKPRIPVVWIHGL WP_001058323.1 MULTISPECIES: hydrogenase-1 small chain [Proteobacteria] Length: 372\nNumber of predicted TMHs: 2\nExp number of AAs in TMHs: 37.2608499999999\nExp number, first 60 AAs: 19.52923\nTotal prob of N-in: 0.94351\nPOSSIBLE N-term signal sequence\ninside 1 19\nTMhelix 20 39\noutside 40 325\nTMhelix 326 348\ninside 349 372

2628 GCF_001283185.1_8205_8_77 Escherichia coli Proteobacteria; Gammaproteobacteria; Enterobacterales; Enterobacteriaceae; Escherichia MNNEETFYQAMRRQGVTRRSFLKYCSLAATSLGLGAGMAPKIAWALENKPRIPVVWIHGL WP_001058323.1 MULTISPECIES: hydrogenase-1 small chain [Proteobacteria] Length: 372\nNumber of predicted TMHs: 2\nExp number of AAs in TMHs: 37.2608499999999\nExp number, first 60 AAs: 19.52923\nTotal prob of N-in: 0.94351\nPOSSIBLE N-term signal sequence\ninside 1 19\nTMhelix 20 39\noutside 40 325\nTMhelix 326 348\ninside 349 372

2629 GCF_001283285.1_8205_3_51 Escherichia coli Proteobacteria; Gammaproteobacteria; Enterobacterales; Enterobacteriaceae; Escherichia MNNEETFYQAMRRQGVTRRSFLKYCSLAATSLGLGAGMAPKIAWALENKPRIPVVWIHGL WP_001058323.1 MULTISPECIES: hydrogenase-1 small chain [Proteobacteria] Length: 372\nNumber of predicted TMHs: 2\nExp number of AAs in TMHs: 37.2608499999999\nExp number, first 60 AAs: 19.52923\nTotal prob of N-in: 0.94351\nPOSSIBLE N-term signal sequence\ninside 1 19\nTMhelix 20 39\noutside 40 325\nTMhelix 326 348\ninside 349 372

2630 GCF_001283505.1_8205_3_24 Escherichia coli Proteobacteria; Gammaproteobacteria; Enterobacterales; Enterobacteriaceae; Escherichia MTGDNTLIHSHGINRRDFMKLCAALAATMGLSSKAAAEMAESVTNPQRPPVIWIGAQECT WP_000145410.1 MULTISPECIES: hydrogenase-2 small chain [Proteobacteria] Length: 372\nNumber of predicted TMHs:

1\nExp number of AAs in TMHs: 23.72226\nExp number, first 60 AAs: 0.22038\nTotal prob of N-in: 0.12994\noutside 1 331\nTMhelix 332 354\ninside 355 372

2631 GCF_001283505.1_8205_3_24 Escherichia coli Proteobacteria; Gammaproteobacteria; Enterobacterales; Enterobacteriaceae; Escherichia
MNNEETFYQAMRRQGVTRRSFLKYCSLAATSLGLGAGMAPKIAWALENKPRIPVVWIHGL WP_001058323.1
MULTISPECIES: hydrogenase-1 small chain [Proteobacteria] Length: 372\nNumber of predicted TMHs: 2\nExp number of AAs in TMHs: 37.2608499999999\nExp number, first 60 AAs: 19.52923\nTotal prob of N-in: 0.94351\nPOSSIBLE N-term signal sequence\ninside 1 19\nTMhelix 20 39\noutside 40 325\nTMhelix 326 348\ninside 349 372

2632 GCF_001283145.1_7748_7_13 Escherichia coli Proteobacteria; Gammaproteobacteria; Enterobacterales; Enterobacteriaceae; Escherichia
MNNEETFYQAMRRQGVTRRSFLKYCSLAATSLGLGAGMAPKIAWALENKPRIPVVWIHGL WP_001295940.1
MULTISPECIES: hydrogenase [Enterobacteriaceae] Length: 372\nNumber of predicted TMHs: 2\nExp number of AAs in TMHs: 37.2608899999999\nExp number, first 60 AAs: 19.52923\nTotal prob of N-in: 0.94351\nPOSSIBLE N-term signal sequence\ninside 1 19\nTMhelix 20 39\noutside 40 325\nTMhelix 326 348\ninside 349 372

2633 GCF_001283545.1_7790_1_82 Escherichia coli Proteobacteria; Gammaproteobacteria; Enterobacterales; Enterobacteriaceae; Escherichia
MTGDNTLIHSHGINRRDFMKLCAALAATMGLSSKAAAEMAESVTNPQRPPVIWIGAQECT WP_000145410.1
MULTISPECIES: hydrogenase-2 small chain [Proteobacteria] Length: 372\nNumber of predicted TMHs: 1\nExp number of AAs in TMHs: 23.72226\nExp number, first 60 AAs: 0.22038\nTotal prob of N-in: 0.12994\noutside 1 331\nTMhelix 332 354\ninside 355 372

2634 GCF_001283365.1_7790_1_96 Escherichia coli Proteobacteria; Gammaproteobacteria; Enterobacterales; Enterobacteriaceae; Escherichia
MTGDNTLIHSHGINRRDFMKLCAALAATMGLSSKAAAEMAESVTNPQRPPVIWIGAQECT WP_053918372.1
hydrogenase 2 small subunit [Escherichia coli] Length: 372\nNumber of predicted TMHs: 1\nExp number of AAs in TMHs: 23.73548\nExp number, first 60 AAs: 0.22027\nTotal prob of N-in: 0.13021\noutside 1 331\nTMhelix 332 354\ninside 355 372

2635 GCF_001283665.1_7790_1_75 Escherichia coli Proteobacteria; Gammaproteobacteria; Enterobacterales; Enterobacteriaceae; Escherichia
MNNEETFYQAMRRQGVTRRSFLKYCSLAATSLGLGAGMAPKIAWALENKPRIPVVWIHGL WP_001058323.1
MULTISPECIES: hydrogenase-1 small chain [Proteobacteria] Length: 372\nNumber of predicted TMHs: 2\nExp number of AAs in TMHs: 37.2608499999999\nExp number, first 60 AAs: 19.52923\nTotal prob of N-in: 0.94351\nPOSSIBLE N-term signal sequence\ninside 1 19\nTMhelix 20 39\noutside 40 325\nTMhelix 326 348\ninside 349 372

2636 GCF_001283825.1_7748_7_25 Escherichia coli Proteobacteria; Gammaproteobacteria; Enterobacterales; Enterobacteriaceae; Escherichia
MTGDNTLIHSHGINRRDFMKLCAALAATMGLSSKAAAEMAESVTNPQRPPVIWIGAQECT WP_000145410.1
MULTISPECIES: hydrogenase-2 small chain [Proteobacteria] Length: 372\nNumber of predicted TMHs: 1\nExp number of AAs in TMHs: 23.72226\nExp number, first 60 AAs: 0.22038\nTotal prob of N-in: 0.12994\noutside 1 331\nTMhelix 332 354\ninside 355 372

2637 GCF_001283805.1_7748_7_40 Escherichia coli Proteobacteria; Gammaproteobacteria; Enterobacterales; Enterobacteriaceae; Escherichia
MTGDNTLIHSHGINRRDFMKLCAALAATMGLSSKAAAEMAESVTNPQRPPVIWIGAQECT WP_000145410.1
MULTISPECIES: hydrogenase-2 small chain [Proteobacteria] Length: 372\nNumber of predicted TMHs: 1\nExp number of AAs in TMHs: 23.72226\nExp number, first 60 AAs: 0.22038\nTotal prob of N-in: 0.12994\noutside 1 331\nTMhelix 332 354\ninside 355 372

2638 GCF_001283865.1_8205_3_32 Escherichia coli Proteobacteria; Gammaproteobacteria; Enterobacterales; Enterobacteriaceae; Escherichia
MTGDNTLIHSHGINRRDFMKLCAALAATMGLSSKAAAEMAESVTNPQRPPVIWIGAQECT WP_000145410.1
MULTISPECIES: hydrogenase-2 small chain [Proteobacteria] Length: 372\nNumber of predicted TMHs: 1\nExp number of AAs in TMHs: 23.72226\nExp number, first 60 AAs: 0.22038\nTotal prob of N-in: 0.12994\noutside 1 331\nTMhelix 332 354\ninside 355 372

2639 GCF_001283825.1_7748_7_25 Escherichia coli Proteobacteria; Gammaproteobacteria; Enterobacterales; Enterobacteriaceae; Escherichia
MNNEETFYQAMRRQGVTRRSFLKYCSLAATSLGLGAGMAPKIAWALENKPRIPVVWIHGL WP_001058323.1
MULTISPECIES: hydrogenase-1 small chain [Proteobacteria] Length: 372\nNumber of predicted TMHs: 2\nExp number of AAs in TMHs: 37.2608499999999\nExp number, first 60 AAs: 19.52923\nTotal prob of N-in: 0.94351\nPOSSIBLE N-term signal sequence\ninside 1 19\nTMhelix 20 39\noutside 40 325\nTMhelix 326 348\ninside 349 372

2640 GCF_001283805.1_7748_7_40 Escherichia coli Proteobacteria; Gammaproteobacteria; Enterobacterales; Enterobacteriaceae; Escherichia
MNNEETFYQAMRRQGVTRRSFLKYCSLAATSLGLGAGMAPKIAWALENKPRIPVVWIHGL WP_001295940.1
MULTISPECIES: hydrogenase [Enterobacteriaceae] Length: 372\nNumber of predicted TMHs: 2\nExp number of AAs in TMHs: 37.2608899999999\nExp number, first 60 AAs: 19.52923\nTotal prob of N-in: 0.94351\nPOSSIBLE N-term signal sequence\ninside 1 19\nTMhelix 20 39\noutside 40 325\nTMhelix 326 348\ninside 349 372

2641 GCF_001283925.1_7748_7_31 Escherichia coli Proteobacteria; Gammaproteobacteria; Enterobacterales; Enterobacteriaceae; Escherichia
MNNEETFYQAMRRQGVTRRSFLKYCSLAATSLGLGAGMAPKIAWALENKPRIPVVWIHGL WP_001058323.1
MULTISPECIES: hydrogenase-1 small chain [Proteobacteria] Length: 372\nNumber of predicted TMHs: 2\nExp number of AAs in TMHs: 37.2608499999999\nExp number, first 60 AAs: 19.52923\nTotal prob of N-in: 0.94351\nPOSSIBLE N-term signal sequence\ninside 1 19\nTMhelix 20 39\noutside 40 325\nTMhelix 326 348\ninside 349 372

2642 GCF_001283945.1_7790_1_87 Escherichia coli Proteobacteria; Gammaproteobacteria; Enterobacterales; Enterobacteriaceae; Escherichia
MNNEETFYQAMRRQGVTRRSFLKYCSLAATSLGLGAGMAPKIAWALENKPRIPVVWIHGL WP_001058323.1
MULTISPECIES: hydrogenase-1 small chain [Proteobacteria] Length: 372\nNumber of predicted TMHs: 2\nExp number of AAs in TMHs: 37.2608499999999\nExp number, first 60 AAs: 19.52923\nTotal prob of N-in: 0.94351\nPOSSIBLE N-term signal sequence\ninside 1 19\nTMhelix 20 39\noutside 40 325\nTMhelix 326 348\ninside 349 372

2643 GCF_001283985.1_7553_7_67 Escherichia coli Proteobacteria; Gammaproteobacteria; Enterobacterales; Enterobacteriaceae; Escherichia
MNNEETFYQAMRRQGVTRRSFLKYCSLAATSLGLGAGMAPKIAWALENKPRIPVVWIHGL WP_001058323.1
MULTISPECIES: hydrogenase-1 small chain [Proteobacteria] Length: 372\nNumber of predicted TMHs: 2\nExp number of AAs in TMHs: 37.2608499999999\nExp number, first 60 AAs: 19.52923\nTotal prob of N-in: 0.94351\nPOSSIBLE N-term signal sequence\ninside 1 19\nTMhelix 20 39\noutside 40 325\nTMhelix 326 348\ninside 349 372

2644 GCF_001283705.1_7748_7_16 Escherichia coli Proteobacteria; Gammaproteobacteria; Enterobacterales; Enterobacteriaceae; Escherichia
MNNEETFYQAMRRQGVTRRSFLKYCSLAATSLGLGAGMAPKIAWALENKPRIPVVWIHGL WP_052935581.1
hydrogenase [Escherichia coli] Length: 372\nNumber of predicted TMHs: 2\nExp number of AAs in TMHs: 37.2610899999999\nExp number, first 60 AAs: 19.52923\nTotal prob of N-in: 0.94351\nPOSSIBLE N-term signal sequence\ninside 1 19\nTMhelix 20 39\noutside 40 325\nTMhelix 326 348\ninside 349 372

2645 GCF_001284025.1_7790_1_52 Escherichia coli Proteobacteria; Gammaproteobacteria; Enterobacterales; Enterobacteriaceae; Escherichia
MTGDNTLIHSHGINRRDFMKLCAALAATMGLSSKAAAEMAESVTNPQRPPVIWIGAQECT WP_000145410.1
MULTISPECIES: hydrogenase-2 small chain [Proteobacteria] Length: 372\nNumber of predicted TMHs: 1\nExp number of AAs in TMHs: 23.72226\nExp number, first 60 AAs: 0.22038\nTotal prob of N-in: 0.12994\noutside 1 331\nTMhelix 332 354\ninside 355 372

2646 GCF_001284045.1_8205_3_14 Escherichia coli Proteobacteria; Gammaproteobacteria; Enterobacterales; Enterobacteriaceae; Escherichia
MTGDNTLIHSHGINRRDFMKLCAALAATMGLSSKAAAEMAESVTNPQRPPVIWIGAQECT WP_000145410.1
MULTISPECIES: hydrogenase-2 small chain [Proteobacteria] Length: 372\nNumber of predicted TMHs: 1\nExp number of AAs in TMHs: 23.72226\nExp number, first 60 AAs: 0.22038\nTotal prob of N-in: 0.12994\noutside 1 331\nTMhelix 332 354\ninside 355 372

2647 GCF_001284085.1_7553_7_70 Escherichia coli Proteobacteria; Gammaproteobacteria; Enterobacterales; Enterobacteriaceae; Escherichia
MTGDNTLIHSHGINRRDFMKLCAALAATMGLSSKAAAEMAESVTNPQRPPVIWIGAQECT WP_000145410.1
MULTISPECIES: hydrogenase-2 small chain [Proteobacteria] Length: 372\nNumber of predicted TMHs: 1\nExp number of AAs in TMHs: 23.72226\nExp number, first 60 AAs: 0.22038\nTotal prob of N-in: 0.12994\noutside 1 331\nTMhelix 332 354\ninside 355 372

2648 GCF_001284205.1_7748_7_12 Escherichia coli Proteobacteria; Gammaproteobacteria; Enterobacterales; Enterobacteriaceae; Escherichia
MNNEETFYQAMRRQGVTRRSFLKYCSLAATSLGLGAGMAPKIAWALENKPRIPVVWIHGL WP_001295940.1
MULTISPECIES: hydrogenase [Enterobacteriaceae] Length: 372\nNumber of predicted TMHs: 2\nExp number of AAs in TMHs: 37.2608899999999\nExp number, first 60 AAs: 19.52923\nTotal prob of N-in: 0.94351\nPOSSIBLE N-term signal sequence\ninside 1 19\nTMhelix 20 39\noutside 40 325\nTMhelix 326 348\ninside 349 372

2649 GCF_001284265.1_7790_1_57 Escherichia coli Proteobacteria; Gammaproteobacteria; Enterobacterales; Enterobacteriaceae; Escherichia
MNNEETFYQAMRRQGVTRRSFLKYCSLAATSLGLGAGMAPKIAWALENKPRIPVVWIHGL WP_044723380.1
hydrogenase [Escherichia coli] Length: 372\nNumber of predicted TMHs: 2\nExp number of AAs in TMHs: 37.2612299999999\nExp number, first 60 AAs: 19.52923\nTotal prob of N-in: 0.94351\nPOSSIBLE N-term signal sequence\ninside 1 19\nTMhelix 20 39\noutside 40 325\nTMhelix 326 348\ninside 349 372

2650 GCF_001284085.1_7553_7_70 Escherichia coli Proteobacteria; Gammaproteobacteria; Enterobacterales; Enterobacteriaceae; Escherichia
MNNEETFYQAMRRQGVTRRSFLKYCSLAATSLGLGAGMAPKIAWALENKPRIPVVWIHGL WP_001058323.1
MULTISPECIES: hydrogenase-1 small chain [Proteobacteria] Length: 372\nNumber of predicted TMHs: 2\nExp number of AAs in TMHs: 37.2608499999999\nExp number, first 60 AAs: 19.52923\nTotal prob of N-in: 0.94351\nPOSSIBLE N-term signal sequence\ninside 1 19\nTMhelix 20 39\noutside 40 325\nTMhelix 326 348\ninside 349 372

2651 GCF_001284245.1_7790_1_93 Escherichia coli Proteobacteria; Gammaproteobacteria; Enterobacterales; Enterobacteriaceae; Escherichia
MTGDNTLIHSHGINRRDFMKLCAALAATMGLSSKAAAEMAESVTNPQRPPVIWIGAQECT WP_000145410.1
MULTISPECIES: hydrogenase-2 small chain [Proteobacteria] Length: 372\nNumber of predicted TMHs: 1\nExp number of AAs in TMHs: 23.72226\nExp number, first 60 AAs: 0.22038\nTotal prob of N-in: 0.12994\noutside 1 331\nTMhelix 332 354\ninside 355 372

2652 GCF_001284325.1_8205_8_80 Escherichia coli Proteobacteria; Gammaproteobacteria; Enterobacterales; Enterobacteriaceae; Escherichia
MTGDNTLIHSHGINRRDFMKLCAALAATMGLSSKAAAEMAESVTNPQRPPVIWIGAQECT WP_000145410.1
MULTISPECIES: hydrogenase-2 small chain [Proteobacteria] Length: 372\nNumber of predicted TMHs: 1\nExp number of AAs in TMHs: 23.72226\nExp number, first 60 AAs: 0.22038\nTotal prob of N-in: 0.12994\noutside 1 331\nTMhelix 332 354\ninside 355 372

2653 GCF_001284245.1_7790_1_93 Escherichia coli Proteobacteria; Gammaproteobacteria; Enterobacterales; Enterobacteriaceae; Escherichia
MNNEETFYQAMRRQGVTRRSFLKYCSLAATSLGLGAGMAPKIAWALENKPRIPVVWIHGL WP_001058323.1
MULTISPECIES: hydrogenase-1 small chain [Proteobacteria] Length: 372\nNumber of predicted TMHs: 2\nExp number of AAs in TMHs: 37.2608499999999\nExp number, first 60 AAs: 19.52923\nTotal prob of N-in: 0.94351\nPOSSIBLE N-term signal sequence\ninside 1 19\nTMhelix 20 39\noutside 40 325\nTMhelix 326 348\ninside 349 372

2654 GCF_001284365.1_8205_8_72 Escherichia coli Proteobacteria; Gammaproteobacteria; Enterobacterales; Enterobacteriaceae; Escherichia
MNNEETFYQAMRRQGVTRRSFLKYCSLAATSLGLGAGMAPKIAWALENKPRIPVVWIHGL WP_001058323.1
MULTISPECIES: hydrogenase-1 small chain [Proteobacteria] Length: 372\nNumber of predicted TMHs: 2\nExp number of AAs in TMHs: 37.2608499999999\nExp number, first 60 AAs: 19.52923\nTotal prob of N-in: 0.94351\nPOSSIBLE N-term signal sequence\ninside 1 19\nTMhelix 20 39\noutside 40 325\nTMhelix 326 348\ninside 349 372

2655	GCF_001284525.1_7790_1_72	Escherichia coli	Proteobacteria; Gammaproteobacteria; Enterobacterales; Enterobacteriaceae; Escherichia	MTGDNTLIHSHGINRRDFMKLCAALAATMGLSSKAAAEMAESVTNPQRPPVIWIGAQECT	WP_000145410.1	MULTISPECIES: hydrogenase-2 small chain [Proteobacteria]	Length: 372\nNumber of predicted TMHs: 1\nExp number of AAs in TMHs: 23.72226\nExp number, first 60 AAs: 0.22038\nTotal prob of N-in: 0.12994\noutside 1 331\nTMhelix 332 354\ninside 355 372

2656	GCF_001284585.1_8205_3_2Escherichia coli	Proteobacteria; Gammaproteobacteria; Enterobacterales; Enterobacteriaceae; Escherichia	MTGDNTLIHSHGINRRDFMKLCAALAATMGLSSKAAAEMAESVTNPQRPPVIWIGAQECT	WP_000145410.1	MULTISPECIES: hydrogenase-2 small chain [Proteobacteria]	Length: 372\nNumber of predicted TMHs: 1\nExp number of AAs in TMHs: 23.72226\nExp number, first 60 AAs: 0.22038\nTotal prob of N-in: 0.12994\noutside 1 331\nTMhelix 332 354\ninside 355 372

2657	GCF_001284565.1_7790_1_86	Escherichia coli	Proteobacteria; Gammaproteobacteria; Enterobacterales; Enterobacteriaceae; Escherichia	MTGDNTLIHSHGINRRDFMKLCAALAATMGLSSKAAAEMAESVTNPQRPPVIWIGAQECT	WP_000145410.1	MULTISPECIES: hydrogenase-2 small chain [Proteobacteria]	Length: 372\nNumber of predicted TMHs: 1\nExp number of AAs in TMHs: 23.72226\nExp number, first 60 AAs: 0.22038\nTotal prob of N-in: 0.12994\noutside 1 331\nTMhelix 332 354\ninside 355 372

2658	GCF_001284585.1_8205_3_2Escherichia coli	Proteobacteria; Gammaproteobacteria; Enterobacterales; Enterobacteriaceae; Escherichia	MNNEETFYQAMRRQGVTRRSFLKYCSLAATSLGLGAGMAPKIAWALENKPRIPVVWIHGL	WP_001058323.1	MULTISPECIES: hydrogenase-1 small chain [Proteobacteria]	Length: 372\nNumber of predicted TMHs: 2\nExp number of AAs in TMHs: 37.2608499999999\nExp number, first 60 AAs: 19.52923\nTotal prob of N-in: 0.94351\nPOSSIBLE N-term signal sequence\ninside 1 19\nTMhelix 20 39\noutside 40 325\nTMhelix 326 348\ninside 349 372

2659	GCF_001284565.1_7790_1_86	Escherichia coli	Proteobacteria; Gammaproteobacteria; Enterobacterales; Enterobacteriaceae; Escherichia	MNNEETFYQAMRRQGVTRRSFLKYCSLAATSLGLGAGMAPKIAWALENKPRIPVVWIHGL	WP_001058323.1	MULTISPECIES: hydrogenase-1 small chain [Proteobacteria]	Length: 372\nNumber of predicted TMHs: 2\nExp number of AAs in TMHs: 37.2608499999999\nExp number, first 60 AAs: 19.52923\nTotal prob of N-in: 0.94351\nPOSSIBLE N-term signal sequence\ninside 1 19\nTMhelix 20 39\noutside 40 325\nTMhelix 326 348\ninside 349 372

2660	GCF_001284765.1_7748_7_37	Escherichia coli	Proteobacteria; Gammaproteobacteria; Enterobacterales; Enterobacteriaceae; Escherichia	MTGDNTLIHSHGINRRDFMKLCAALAATMGLSSKAAAEMAESVTNPQRPPVIWIGAQECT	WP_000145410.1	MULTISPECIES: hydrogenase-2 small chain [Proteobacteria]	Length: 372\nNumber of predicted TMHs: 1\nExp number of AAs in TMHs: 23.72226\nExp number, first 60 AAs: 0.22038\nTotal prob of N-in: 0.12994\noutside 1 331\nTMhelix 332 354\ninside 355 372

2661	GCF_001284745.1_7790_1_91	Escherichia coli	Proteobacteria; Gammaproteobacteria; Enterobacterales; Enterobacteriaceae; Escherichia	MNNEETFYQAMRRQGVTRRSFLKYCSLAATSLGLGAGMAPKIAWALENKPRIPVVWIHGL	WP_001058323.1	MULTISPECIES: hydrogenase-1 small chain [Proteobacteria]	Length: 372\nNumber of predicted TMHs: 2\nExp number of AAs in TMHs: 37.2608499999999\nExp number, first 60 AAs: 19.52923\nTotal prob of N-in: 0.94351\nPOSSIBLE N-term signal sequence\ninside 1 19\nTMhelix 20 39\noutside 40 325\nTMhelix 326 348\ninside 349 372

2662	GCF_001284765.1_7748_7_37	Escherichia coli	Proteobacteria; Gammaproteobacteria; Enterobacterales; Enterobacteriaceae; Escherichia	MNNEETFYQAMRRQGVTRRSFLKYCSLAATSLGLGAGMAPKIAWALENKPRIPVVWIHGL	WP_001058323.1	MULTISPECIES: hydrogenase-1 small chain [Proteobacteria]	Length: 372\nNumber of predicted TMHs: 2\nExp number of AAs in TMHs: 37.2608499999999\nExp number, first 60 AAs: 19.52923\nTotal prob of N-in: 0.94351\nPOSSIBLE N-term signal sequence\ninside 1 19\nTMhelix 20 39\noutside 40 325\nTMhelix 326 348\ninside 349 372

2663	GCF_001284705.1_8205_3_17	Escherichia coli	Proteobacteria; Gammaproteobacteria; Enterobacterales; Enterobacteriaceae; Escherichia	MNNEETFYQAMRRQGVTRRSFLKYCSLAATSLGLGAGMAPKIAWALENKPRIPVVWIHGL	WP_053882050.1

hydrogenase [Escherichia coli] Length: 372\nNumber of predicted TMHs: 2\nExp number of AAs in TMHs: 37.25814\nExp number, first 60 AAs: 19.52926\nTotal prob of N-in: 0.94351\nPOSSIBLE N-term signal sequence\ninside 1 19\nTMhelix 20 39\noutside 40 325\nTMhelix 326 348\ninside 349 372

2664 GCF_001284525.1_7790_1_72 Escherichia coli Proteobacteria; Gammaproteobacteria; Enterobacterales; Enterobacteriaceae; Escherichia MNNEETFYQAMRRQGVTRRSFLKYCSLAATSLGLGAGMAPKIAWALENKPRIPVVWIHGL WP_001058323.1 MULTISPECIES: hydrogenase-1 small chain [Proteobacteria] Length: 372\nNumber of predicted TMHs: 2\nExp number of AAs in TMHs: 37.2608499999999\nExp number, first 60 AAs: 19.52923\nTotal prob of N-in: 0.94351\nPOSSIBLE N-term signal sequence\ninside 1 19\nTMhelix 20 39\noutside 40 325\nTMhelix 326 348\ninside 349 372

2665 GCF_001284485.1_8205_3_8Escherichia coli Proteobacteria; Gammaproteobacteria; Enterobacterales; Enterobacteriaceae; Escherichia MNNEETFYQAMRRQGVTRRSFLKYCSLAATSLGLGAGMAPKIAWALENKPRIPVVWIHGL WP_024246676.1 hydrogenase-1 small chain [Escherichia coli] Length: 372\nNumber of predicted TMHs: 2\nExp number of AAs in TMHs: 37.8214899999999\nExp number, first 60 AAs: 19.52326\nTotal prob of N-in: 0.94395\nPOSSIBLE N-term signal sequence\ninside 1 19\nTMhelix 20 39\noutside 40 325\nTMhelix 326 348\ninside 349 372

2666 GCF_001284985.1_8205_3_20 Escherichia coli Proteobacteria; Gammaproteobacteria; Enterobacterales; Enterobacteriaceae; Escherichia MTGDNTLIHSHGINRRDFMKLCAALAATMGLSSKAAAEMAESVTNPQRPPVIWIGAQECT WP_000145410.1 MULTISPECIES: hydrogenase-2 small chain [Proteobacteria] Length: 372\nNumber of predicted TMHs: 1\nExp number of AAs in TMHs: 23.72226\nExp number, first 60 AAs: 0.22038\nTotal prob of N-in: 0.12994\noutside 1 331\nTMhelix 332 354\ninside 355 372

2667 GCF_001284985.1_8205_3_20 Escherichia coli Proteobacteria; Gammaproteobacteria; Enterobacterales; Enterobacteriaceae; Escherichia MNNEETFYQAMRRQGVTRRSFLKYCSLAATSLGLGAGMAPKIAWALENKPRIPVVWIHGL WP_001058323.1 MULTISPECIES: hydrogenase-1 small chain [Proteobacteria] Length: 372\nNumber of predicted TMHs: 2\nExp number of AAs in TMHs: 37.2608499999999\nExp number, first 60 AAs: 19.52923\nTotal prob of N-in: 0.94351\nPOSSIBLE N-term signal sequence\ninside 1 19\nTMhelix 20 39\noutside 40 325\nTMhelix 326 348\ninside 349 372

2668 GCF_001284865.1_8205_3_72 Escherichia coli Proteobacteria; Gammaproteobacteria; Enterobacterales; Enterobacteriaceae; Escherichia MNNEETFYQAMRRQGVTRRSFLKYCSLAATSLGLGAGMAPKIAWALENKPRIPVVWIHGL WP_001058323.1 MULTISPECIES: hydrogenase-1 small chain [Proteobacteria] Length: 372\nNumber of predicted TMHs: 2\nExp number of AAs in TMHs: 37.2608499999999\nExp number, first 60 AAs: 19.52923\nTotal prob of N-in: 0.94351\nPOSSIBLE N-term signal sequence\ninside 1 19\nTMhelix 20 39\noutside 40 325\nTMhelix 326 348\ninside 349 372

2669 GCF_001284905.1_8205_3_31 Escherichia coli Proteobacteria; Gammaproteobacteria; Enterobacterales; Enterobacteriaceae; Escherichia MNNEETFYQAMRRQGVTRRSFLKYCSLAATSLGLGAGMAPKIAWALENKPRIPVVWIHGL WP_001058323.1 MULTISPECIES: hydrogenase-1 small chain [Proteobacteria] Length: 372\nNumber of predicted TMHs: 2\nExp number of AAs in TMHs: 37.2608499999999\nExp number, first 60 AAs: 19.52923\nTotal prob of N-in: 0.94351\nPOSSIBLE N-term signal sequence\ninside 1 19\nTMhelix 20 39\noutside 40 325\nTMhelix 326 348\ninside 349 372

2670 GCF_001285025.1_7748_7_46 Escherichia coli Proteobacteria; Gammaproteobacteria; Enterobacterales; Enterobacteriaceae; Escherichia MTGDNTLIHSHGINRRDFMKLCAALAATMGLSSKAAAEMAESVTNPQRPPVIWIGAQECT WP_000145410.1 MULTISPECIES: hydrogenase-2 small chain [Proteobacteria] Length: 372\nNumber of predicted TMHs: 1\nExp number of AAs in TMHs: 23.72226\nExp number, first 60 AAs: 0.22038\nTotal prob of N-in: 0.12994\noutside 1 331\nTMhelix 332 354\ninside 355 372

2671 GCF_001285085.1_8205_3_12 Escherichia coli Proteobacteria; Gammaproteobacteria; Enterobacterales; Enterobacteriaceae; Escherichia MTGDNTLIHSHGINRRDFMKLCAALAATMGLSSKAAAEMAESVTNPQRPPVIWIGAQECT WP_000145410.1 MULTISPECIES: hydrogenase-2 small chain [Proteobacteria] Length: 372\nNumber of predicted TMHs:

1\nExp number of AAs in TMHs: 23.72226\nExp number, first 60 AAs: 0.22038\nTotal prob of N-in: 0.12994\noutside 1 331\nTMhelix 332 354\ninside 355 372

2672 GCF_001285065.1_8205_3_27 Escherichia coli Proteobacteria; Gammaproteobacteria; Enterobacterales; Enterobacteriaceae; Escherichia MTGDNTLIHSHGINRRDFMKLCAALAATMGLSSKAAAEMAESVTNPQRPPVIWIGAQECT WP_000145410.1 MULTISPECIES: hydrogenase-2 small chain [Proteobacteria] Length: 372\nNumber of predicted TMHs: 1\nExp number of AAs in TMHs: 23.72226\nExp number, first 60 AAs: 0.22038\nTotal prob of N-in: 0.12994\noutside 1 331\nTMhelix 332 354\ninside 355 372

2673 GCF_001285265.1_7790_1_54 Escherichia coli Proteobacteria; Gammaproteobacteria; Enterobacterales; Enterobacteriaceae; Escherichia MTGDNTLIHSHGINRRDFMKLCAALAATMGLSSKAAAEMAESVTNPQRPPVIWIGAQECT WP_000145410.1 MULTISPECIES: hydrogenase-2 small chain [Proteobacteria] Length: 372\nNumber of predicted TMHs: 1\nExp number of AAs in TMHs: 23.72226\nExp number, first 60 AAs: 0.22038\nTotal prob of N-in: 0.12994\noutside 1 331\nTMhelix 332 354\ninside 355 372

2674 GCF_001285205.1_8205_3_78 Escherichia coli Proteobacteria; Gammaproteobacteria; Enterobacterales; Enterobacteriaceae; Escherichia MNNEETFYQAMRRQGVTRRSFLKYCSLAATSLGLGAGMAPKIAWALENKPRIPVVWIHGL WP_001058323.1 MULTISPECIES: hydrogenase-1 small chain [Proteobacteria] Length: 372\nNumber of predicted TMHs: 2\nExp number of AAs in TMHs: 37.2608499999999\nExp number, first 60 AAs: 19.52923\nTotal prob of N-in: 0.94351\nPOSSIBLE N-term signal sequence\ninside 1 19\nTMhelix 20 39\noutside 40 325\nTMhelix 326 348\ninside 349 372

2675 GCF_001285245.1_8205_3_18 Escherichia coli Proteobacteria; Gammaproteobacteria; Enterobacterales; Enterobacteriaceae; Escherichia MNNEETFYQAMRRQGVTRRSFLKYCSLAATSLGLGAGMAPKIAWALENKPRIPVVWIHGL WP_001058323.1 MULTISPECIES: hydrogenase-1 small chain [Proteobacteria] Length: 372\nNumber of predicted TMHs: 2\nExp number of AAs in TMHs: 37.2608499999999\nExp number, first 60 AAs: 19.52923\nTotal prob of N-in: 0.94351\nPOSSIBLE N-term signal sequence\ninside 1 19\nTMhelix 20 39\noutside 40 325\nTMhelix 326 348\ninside 349 372

2676 GCF_001285305.1_8205_3_50 Escherichia coli Proteobacteria; Gammaproteobacteria; Enterobacterales; Enterobacteriaceae; Escherichia MTGDNTLIHSHGINRRDFMKLCAALAATMGLSSKAAAEMAESVTNPQRPPVIWIGAQECT WP_000145410.1 MULTISPECIES: hydrogenase-2 small chain [Proteobacteria] Length: 372\nNumber of predicted TMHs: 1\nExp number of AAs in TMHs: 23.72226\nExp number, first 60 AAs: 0.22038\nTotal prob of N-in: 0.12994\noutside 1 331\nTMhelix 332 354\ninside 355 372

2677 GCF_001285125.1_7553_7_72 Escherichia coli Proteobacteria; Gammaproteobacteria; Enterobacterales; Enterobacteriaceae; Escherichia MNNEETFYQAMRRQGVTRRSFLKYCSLAATSLGLGAGMAPKIAWALENKPRIPVVWIHGL WP_001295940.1 MULTISPECIES: hydrogenase [Enterobacteriaceae] Length: 372\nNumber of predicted TMHs: 2\nExp number of AAs in TMHs: 37.2608899999999\nExp number, first 60 AAs: 19.52923\nTotal prob of N-in: 0.94351\nPOSSIBLE N-term signal sequence\ninside 1 19\nTMhelix 20 39\noutside 40 325\nTMhelix 326 348\ninside 349 372

2678 GCF_001285345.1_7748_7_8Escherichia coli Proteobacteria; Gammaproteobacteria; Enterobacterales; Enterobacteriaceae; Escherichia MTGDNTLIHSHGINRRDFMKLCAALAATMGLSSKAAAEMAESVTNPQRPPVIWIGAQECT WP_000145410.1 MULTISPECIES: hydrogenase-2 small chain [Proteobacteria] Length: 372\nNumber of predicted TMHs: 1\nExp number of AAs in TMHs: 23.72226\nExp number, first 60 AAs: 0.22038\nTotal prob of N-in: 0.12994\noutside 1 331\nTMhelix 332 354\ninside 355 372

2679 GCF_001285605.1_8205_8_74 Escherichia coli Proteobacteria; Gammaproteobacteria; Enterobacterales; Enterobacteriaceae; Escherichia MTGDNTLIHSHGINRRDFMKLCAALAATMGLSSKAAAEMAESVTNPQRPPVIWIGAQECT WP_000145410.1 MULTISPECIES: hydrogenase-2 small chain [Proteobacteria] Length: 372\nNumber of predicted TMHs: 1\nExp number of AAs in TMHs: 23.72226\nExp number, first 60 AAs: 0.22038\nTotal prob of N-in: 0.12994\noutside 1 331\nTMhelix 332 354\ninside 355 372

2680	GCF_001285265.1_7790_1_54	Escherichia coli	Proteobacteria; Gammaproteobacteria; Enterobacterales; Enterobacteriaceae; Escherichia	MNNEETFYQAMRRQGVTRRSFLKYCSLAATSLGLGAGMAPKIAWALENKPRIPVVWIHGL	WP_001058323.1	MULTISPECIES: hydrogenase-1 small chain [Proteobacteria]	Length: 372\nNumber of predicted TMHs: 2\nExp number of AAs in TMHs: 37.2608499999999\nExp number, first 60 AAs: 19.52923\nTotal prob of N-in: 0.94351\nPOSSIBLE N-term signal sequence\ninside 1 19\nTMhelix 20 39\noutside 40 325\nTMhelix 326 348\ninside 349 372

2681	GCF_001285365.1_8205_3_7Escherichia coli	Proteobacteria; Gammaproteobacteria; Enterobacterales; Enterobacteriaceae; Escherichia	MTGDNTLIHSHGINRRDFMKLCAALAATMGLSSKAAAEMAESVTNPQRPPVIWIGAQECT	WP_000145410.1	MULTISPECIES: hydrogenase-2 small chain [Proteobacteria]	Length: 372\nNumber of predicted TMHs: 1\nExp number of AAs in TMHs: 23.72226\nExp number, first 60 AAs: 0.22038\nTotal prob of N-in: 0.12994\noutside 1 331\nTMhelix 332 354\ninside 355 372

2682	GCF_001285425.1_7790_1_85	Escherichia coli	Proteobacteria; Gammaproteobacteria; Enterobacterales; Enterobacteriaceae; Escherichia	MTGDNTLIHSHGINRRDFMKLCAALAATMGLSSKAAAEMAESVTNPQRPPVIWIGAQECT	WP_000145410.1	MULTISPECIES: hydrogenase-2 small chain [Proteobacteria]	Length: 372\nNumber of predicted TMHs: 1\nExp number of AAs in TMHs: 23.72226\nExp number, first 60 AAs: 0.22038\nTotal prob of N-in: 0.12994\noutside 1 331\nTMhelix 332 354\ninside 355 372

2683	GCF_001285465.1_7790_1_63	Escherichia coli	Proteobacteria; Gammaproteobacteria; Enterobacterales; Enterobacteriaceae; Escherichia	MTGDNTLIHSHGINRRDFMKLCAALAATMGLSSKAAAEMAESVTNPQRPPVIWIGAQECT	WP_000145410.1	MULTISPECIES: hydrogenase-2 small chain [Proteobacteria]	Length: 372\nNumber of predicted TMHs: 1\nExp number of AAs in TMHs: 23.72226\nExp number, first 60 AAs: 0.22038\nTotal prob of N-in: 0.12994\noutside 1 331\nTMhelix 332 354\ninside 355 372

2684	GCF_001285345.1_7748_7_8Escherichia coli	Proteobacteria; Gammaproteobacteria; Enterobacterales; Enterobacteriaceae; Escherichia	MNNEETFYQAMRRQGVTRRSFLKYCSLAATSLGLGAGMAPKIAWALENKPRIPVVWIHGL	WP_001058323.1	MULTISPECIES: hydrogenase-1 small chain [Proteobacteria]	Length: 372\nNumber of predicted TMHs: 2\nExp number of AAs in TMHs: 37.2608499999999\nExp number, first 60 AAs: 19.52923\nTotal prob of N-in: 0.94351\nPOSSIBLE N-term signal sequence\ninside 1 19\nTMhelix 20 39\noutside 40 325\nTMhelix 326 348\ninside 349 372

2685	GCF_001285485.1_8205_8_84	Escherichia coli	Proteobacteria; Gammaproteobacteria; Enterobacterales; Enterobacteriaceae; Escherichia	MTGDNTLIHSHGINRRDFMKLCAALAATMGLSSKAAAEMAESVTNPQRPPVIWIGAQECT	WP_000145410.1	MULTISPECIES: hydrogenase-2 small chain [Proteobacteria]	Length: 372\nNumber of predicted TMHs: 1\nExp number of AAs in TMHs: 23.72226\nExp number, first 60 AAs: 0.22038\nTotal prob of N-in: 0.12994\noutside 1 331\nTMhelix 332 354\ninside 355 372

2686	GCF_001285525.1_7748_7_4Escherichia coli	Proteobacteria; Gammaproteobacteria; Enterobacterales; Enterobacteriaceae; Escherichia	MTGDNTLIHSHGINRRDFMKLCAALAATMGLSSKAAAEMAESVTNPQRPPVIWIGAQECT	WP_000145410.1	MULTISPECIES: hydrogenase-2 small chain [Proteobacteria]	Length: 372\nNumber of predicted TMHs: 1\nExp number of AAs in TMHs: 23.72226\nExp number, first 60 AAs: 0.22038\nTotal prob of N-in: 0.12994\noutside 1 331\nTMhelix 332 354\ninside 355 372

2687	GCF_001285665.1_7748_7_21	Escherichia coli	Proteobacteria; Gammaproteobacteria; Enterobacterales; Enterobacteriaceae; Escherichia	MTGDNTLIHSHGINRRDFMKLCAALAATMGLSSKAAAEMAESVTNPQRPPVIWIGAQECT	WP_000145410.1	MULTISPECIES: hydrogenase-2 small chain [Proteobacteria]	Length: 372\nNumber of predicted TMHs: 1\nExp number of AAs in TMHs: 23.72226\nExp number, first 60 AAs: 0.22038\nTotal prob of N-in: 0.12994\noutside 1 331\nTMhelix 332 354\ninside 355 372

2688	GCF_001285665.1_7748_7_21	Escherichia coli	Proteobacteria; Gammaproteobacteria; Enterobacterales; Enterobacteriaceae; Escherichia	MNNEETFYQAMRRQGVTRRSFLKYCSLAATSLGLGAGMAPKIAWALENKPRIPVVWIHGL	WP_001058320.1	hydrogenase [Escherichia coli]	Length: 372\nNumber of predicted TMHs: 2\nExp number of AAs in TMHs: 37.26082\nExp number, first 60 AAs: 19.52923\nTotal prob of N-in: 0.94351\nPOSSIBLE N-term signal sequence\ninside 1 19\nTMhelix 20 39\noutside 40 325\nTMhelix 326 348\ninside 349 372

2689 GCF_001285705.1_8205_3_56 Escherichia coli Proteobacteria; Gammaproteobacteria; Enterobacterales; Enterobacteriaceae; Escherichia
MNNEETFYQAMRRQGVTRRSFLKYCSLAATSLGLGAGMAPKIAWALENKPRIPVVWIHGL WP_001058323.1
MULTISPECIES: hydrogenase-1 small chain [Proteobacteria] Length: 372\nNumber of predicted TMHs: 2\nExp number of AAs in TMHs: 37.2608499999999\nExp number, first 60 AAs: 19.52923\nTotal prob of N-in: 0.94351\nPOSSIBLE N-term signal sequence\ninside 1 19\nTMhelix 20 39\noutside 40 325\nTMhelix 326 348\ninside 349 372

2690 GCF_001285485.1_8205_8_84 Escherichia coli Proteobacteria; Gammaproteobacteria; Enterobacterales; Enterobacteriaceae; Escherichia
MNNEETFYQAMRRQGVTRRSFLKYCSLAATSLGLGAGMAPKIAWALENKPRIPVVWIHGL WP_001058323.1
MULTISPECIES: hydrogenase-1 small chain [Proteobacteria] Length: 372\nNumber of predicted TMHs: 2\nExp number of AAs in TMHs: 37.2608499999999\nExp number, first 60 AAs: 19.52923\nTotal prob of N-in: 0.94351\nPOSSIBLE N-term signal sequence\ninside 1 19\nTMhelix 20 39\noutside 40 325\nTMhelix 326 348\ninside 349 372

2691 GCF_001285725.1_7790_1_71 Escherichia coli Proteobacteria; Gammaproteobacteria; Enterobacterales; Enterobacteriaceae; Escherichia
MTGDNTLIHSHGINRRDFMKLCAALAATMGLSSKAAAEMAESVTNPQRPPVIWIGAQECT WP_000145410.1
MULTISPECIES: hydrogenase-2 small chain [Proteobacteria] Length: 372\nNumber of predicted TMHs: 1\nExp number of AAs in TMHs: 23.72226\nExp number, first 60 AAs: 0.22038\nTotal prob of N-in: 0.12994\noutside 1 331\nTMhelix 332 354\ninside 355 372

2692 GCF_001285725.1_7790_1_71 Escherichia coli Proteobacteria; Gammaproteobacteria; Enterobacterales; Enterobacteriaceae; Escherichia
MNNEETFYQAMRRQGVTRRSFLKYCSLAATSLGLGAGMAPKIAWALENKPRIPVVWIHGL WP_001359719.1
hydrogenase [Escherichia coli] Length: 372\nNumber of predicted TMHs: 2\nExp number of AAs in TMHs: 37.2176\nExp number, first 60 AAs: 19.53448\nTotal prob of N-in: 0.94360\nPOSSIBLE N-term signal sequence\ninside 1 19\nTMhelix 20 39\noutside 40 325\nTMhelix 326 348\ninside 349 372

2693 GCF_001285625.1_7790_1_84 Escherichia coli Proteobacteria; Gammaproteobacteria; Enterobacterales; Enterobacteriaceae; Escherichia
MNNEETFYQAMRRQGVTRRSFLKYCSLAATSLGLGAGMAPKIAWALENKPRIPVVWIHGL WP_001058323.1
MULTISPECIES: hydrogenase-1 small chain [Proteobacteria] Length: 372\nNumber of predicted TMHs: 2\nExp number of AAs in TMHs: 37.2608499999999\nExp number, first 60 AAs: 19.52923\nTotal prob of N-in: 0.94351\nPOSSIBLE N-term signal sequence\ninside 1 19\nTMhelix 20 39\noutside 40 325\nTMhelix 326 348\ninside 349 372

2694 GCF_001285905.1_8205_3_79 Escherichia coli Proteobacteria; Gammaproteobacteria; Enterobacterales; Enterobacteriaceae; Escherichia
MTGDNTLIHSHGINRRDFMKLCAALAATMGLSSKAAAEMAESVTNPQRPPVIWIGAQECT WP_000145410.1
MULTISPECIES: hydrogenase-2 small chain [Proteobacteria] Length: 372\nNumber of predicted TMHs: 1\nExp number of AAs in TMHs: 23.72226\nExp number, first 60 AAs: 0.22038\nTotal prob of N-in: 0.12994\noutside 1 331\nTMhelix 332 354\ninside 355 372

2695 GCF_001285945.1_8205_3_9Escherichia coli Proteobacteria; Gammaproteobacteria; Enterobacterales; Enterobacteriaceae; Escherichia MTGDNTLIHSHGINRRDFMKLCAALAATMGLSSKAAAEMAESVTNPQRPPVIWIGAQECT
WP_000145410.1 MULTISPECIES: hydrogenase-2 small chain [Proteobacteria] Length: 372\nNumber of predicted TMHs: 1\nExp number of AAs in TMHs: 23.72226\nExp number, first 60 AAs: 0.22038\nTotal prob of N-in: 0.12994\noutside 1 331\nTMhelix 332 354\ninside 355 372

2696 GCF_001285825.1_7748_7_20 Escherichia coli Proteobacteria; Gammaproteobacteria; Enterobacterales; Enterobacteriaceae; Escherichia
MNNEETFYQAMRRQGVTRRSFLKYCSLAATSLGLGAGMAPKIAWALENKPRIPVVWIHGL WP_001295940.1
MULTISPECIES: hydrogenase [Enterobacteriaceae] Length: 372\nNumber of predicted TMHs: 2\nExp number of AAs in TMHs: 37.2608899999999\nExp number, first 60 AAs: 19.52923\nTotal prob of N-in: 0.94351\nPOSSIBLE N-term signal sequence\ninside 1 19\nTMhelix 20 39\noutside 40 325\nTMhelix 326 348\ninside 349 372

2697 GCF_001285985.1_7748_7_5Escherichia coli Proteobacteria; Gammaproteobacteria; Enterobacterales; Enterobacteriaceae; Escherichia MTGDNTLIHSHGINRRDFMKLCAALAATMGLSSKAAAEMAESVTNPQRPPVIWIGAQECT

WP_000145410.1 MULTISPECIES: hydrogenase-2 small chain [Proteobacteria] Length: 372\nNumber of predicted TMHs: 1\nExp number of AAs in TMHs: 23.72226\nExp number, first 60 AAs: 0.22038\nTotal prob of N-in: 0.12994\noutside 1 331\nTMhelix 332 354\ninside 355 372

2698 GCF_001285865.1_7553_7_64 Escherichia coli Proteobacteria; Gammaproteobacteria; Enterobacterales; Enterobacteriaceae; Escherichia MNNEETFYQAMRRQGVTRRSFLKYCSLAATSLGLGAGMAPKIAWALENKPRIPVVWIHGL WP_001058323.1 MULTISPECIES: hydrogenase-1 small chain [Proteobacteria] Length: 372\nNumber of predicted TMHs: 2\nExp number of AAs in TMHs: 37.2608499999999\nExp number, first 60 AAs: 19.52923\nTotal prob of N-in: 0.94351\nPOSSIBLE N-term signal sequence\ninside 1 19\nTMhelix 20 39\noutside 40 325\nTMhelix 326 348\ninside 349 372

2699 GCF_001286045.1_7790_1_68 Escherichia coli Proteobacteria; Gammaproteobacteria; Enterobacterales; Enterobacteriaceae; Escherichia MTGDNTLIHSHGINRRDFMKLCAALAATMGLSSKAAAEMAESVTNPQRPPVIWIGAQECT WP_000145410.1 MULTISPECIES: hydrogenase-2 small chain [Proteobacteria] Length: 372\nNumber of predicted TMHs: 1\nExp number of AAs in TMHs: 23.72226\nExp number, first 60 AAs: 0.22038\nTotal prob of N-in: 0.12994\noutside 1 331\nTMhelix 332 354\ninside 355 372

2700 GCF_001286305.1_8205_3_81 Escherichia coli Proteobacteria; Gammaproteobacteria; Enterobacterales; Enterobacteriaceae; Escherichia MTGDNTLIHSHGINRRDFMKLCAALAATMGLSSKAAAEMAESVTNPQRPPVIWIGAQECT WP_000145410.1 MULTISPECIES: hydrogenase-2 small chain [Proteobacteria] Length: 372\nNumber of predicted TMHs: 1\nExp number of AAs in TMHs: 23.72226\nExp number, first 60 AAs: 0.22038\nTotal prob of N-in: 0.12994\noutside 1 331\nTMhelix 332 354\ninside 355 372

2701 GCF_001286185.1_7790_1_77 Escherichia coli Proteobacteria; Gammaproteobacteria; Enterobacterales; Enterobacteriaceae; Escherichia MNNEETFYQAMRRQGVTRRSFLKYCSLAATSLGLGAGMAPKIAWALENKPRIPVVWIHGL WP_001058323.1 MULTISPECIES: hydrogenase-1 small chain [Proteobacteria] Length: 372\nNumber of predicted TMHs: 2\nExp number of AAs in TMHs: 37.2608499999999\nExp number, first 60 AAs: 19.52923\nTotal prob of N-in: 0.94351\nPOSSIBLE N-term signal sequence\ninside 1 19\nTMhelix 20 39\noutside 40 325\nTMhelix 326 348\ninside 349 372

2702 GCF_001285905.1_8205_3_79 Escherichia coli Proteobacteria; Gammaproteobacteria; Enterobacterales; Enterobacteriaceae; Escherichia MNNEETFYQAMRRQGVTRRSFLKYCSLAATSLGLGAGMAPKIAWALENKPRIPVVWIHGL WP_053893390.1 hydrogenase [Escherichia coli] Length: 372\nNumber of predicted TMHs: 2\nExp number of AAs in TMHs: 37.0700899999999\nExp number, first 60 AAs: 19.52405\nTotal prob of N-in: 0.94332\nPOSSIBLE N-term signal sequence\ninside 1 19\nTMhelix 20 39\noutside 40 325\nTMhelix 326 348\ninside 349 372

2703 GCF_001286225.1_7748_7_29 Escherichia coli Proteobacteria; Gammaproteobacteria; Enterobacterales; Enterobacteriaceae; Escherichia MTGDNTLIHSHGINRRDFMKLCAALAATMGLSSKAAAEMAESVTNPQRPPVIWIGAQECT WP_000145410.1 MULTISPECIES: hydrogenase-2 small chain [Proteobacteria] Length: 372\nNumber of predicted TMHs: 1\nExp number of AAs in TMHs: 23.72226\nExp number, first 60 AAs: 0.22038\nTotal prob of N-in: 0.12994\noutside 1 331\nTMhelix 332 354\ninside 355 372

2704 GCF_001286285.1_7790_1_69 Escherichia coli Proteobacteria; Gammaproteobacteria; Enterobacterales; Enterobacteriaceae; Escherichia MTGDNTLIHSHGINRRDFMKLCAALAATMGLSSKAAAEMAESVTNPQRPPVIWIGAQECT WP_000145410.1 MULTISPECIES: hydrogenase-2 small chain [Proteobacteria] Length: 372\nNumber of predicted TMHs: 1\nExp number of AAs in TMHs: 23.72226\nExp number, first 60 AAs: 0.22038\nTotal prob of N-in: 0.12994\noutside 1 331\nTMhelix 332 354\ninside 355 372

2705 GCF_001286225.1_7748_7_29 Escherichia coli Proteobacteria; Gammaproteobacteria; Enterobacterales; Enterobacteriaceae; Escherichia MNNEETFYQAMRRQGVTRRSFLKYCSLAATSLGLGAGMAPKIAWALENKPRIPVVWIHGL WP_001058323.1 MULTISPECIES: hydrogenase-1 small chain [Proteobacteria] Length: 372\nNumber of predicted TMHs: 2\nExp number of AAs in TMHs: 37.2608499999999\nExp number, first 60 AAs: 19.52923\nTotal prob of N-in:

0.94351\nPOSSIBLE N-term signal sequence\ninside 1 19\nTMhelix 20 39\noutside 40 325\nTMhelix 326 348\ninside 349 372

2706 GCF_001286285.1_7790_1_69 Escherichia coli Proteobacteria; Gammaproteobacteria; Enterobacterales; Enterobacteriaceae; Escherichia MNNEETFYQAMRRQGVTRRSFLKYCSLAATSLGLGAGMAPKIAWALENKPRIPVVWIHGL WP_001058323.1 MULTISPECIES: hydrogenase-1 small chain [Proteobacteria] Length: 372\nNumber of predicted TMHs: 2\nExp number of AAs in TMHs: 37.2608499999999\nExp number, first 60 AAs: 19.52923\nTotal prob of N-in: 0.94351\nPOSSIBLE N-term signal sequence\ninside 1 19\nTMhelix 20 39\noutside 40 325\nTMhelix 326 348\ninside 349 372

2707 GCF_001285985.1_7748_7_5Escherichia coli Proteobacteria; Gammaproteobacteria; Enterobacterales; Enterobacteriaceae; Escherichia MNNEETFYQAMRRQGVTRRSFLKYCSLAATSLGLGAGMAPKIAWALENKPRIPVVWIHGL WP_001295940.1 MULTISPECIES: hydrogenase [Enterobacteriaceae] Length: 372\nNumber of predicted TMHs: 2\nExp number of AAs in TMHs: 37.2608899999999\nExp number, first 60 AAs: 19.52923\nTotal prob of N-in: 0.94351\nPOSSIBLE N-term signal sequence\ninside 1 19\nTMhelix 20 39\noutside 40 325\nTMhelix 326 348\ninside 349 372

2708 GCF_001286425.1_7553_7_68 Escherichia coli Proteobacteria; Gammaproteobacteria; Enterobacterales; Enterobacteriaceae; Escherichia MTGDNTLIHSHGINRRDFMKLCAALAATMGLSSKAAAEMAESVTNPQRPPVIWIGAQECT WP_000145410.1 MULTISPECIES: hydrogenase-2 small chain [Proteobacteria] Length: 372\nNumber of predicted TMHs: 1\nExp number of AAs in TMHs: 23.72226\nExp number, first 60 AAs: 0.22038\nTotal prob of N-in: 0.12994\noutside 1 331\nTMhelix 332 354\ninside 355 372

2709 GCF_001286465.1_7748_7_3Escherichia coli Proteobacteria; Gammaproteobacteria; Enterobacterales; Enterobacteriaceae; Escherichia MNNEETFYQAMRRQGVTRRSFLKYCSLAATSLGLGAGMAPKIAWALENKPRIPVVWIHGL WP_001058323.1 MULTISPECIES: hydrogenase-1 small chain [Proteobacteria] Length: 372\nNumber of predicted TMHs: 2\nExp number of AAs in TMHs: 37.2608499999999\nExp number, first 60 AAs: 19.52923\nTotal prob of N-in: 0.94351\nPOSSIBLE N-term signal sequence\ninside 1 19\nTMhelix 20 39\noutside 40 325\nTMhelix 326 348\ninside 349 372

2710 GCF_001286545.1_8205_3_80 Escherichia coli Proteobacteria; Gammaproteobacteria; Enterobacterales; Enterobacteriaceae; Escherichia MTGDNTLIHSHGINRRDFMKLCAALAATMGLSSKAAAEMAESVTNPQRPPVIWIGAQECT WP_000145410.1 MULTISPECIES: hydrogenase-2 small chain [Proteobacteria] Length: 372\nNumber of predicted TMHs: 1\nExp number of AAs in TMHs: 23.72226\nExp number, first 60 AAs: 0.22038\nTotal prob of N-in: 0.12994\noutside 1 331\nTMhelix 332 354\ninside 355 372

2711 GCF_001286585.1_8205_3_68 Escherichia coli Proteobacteria; Gammaproteobacteria; Enterobacterales; Enterobacteriaceae; Escherichia MTGDNTLIHSHGINRRDFMKLCAALAATMGLSSKAAAEMAESVTNPQRPPVIWIGAQECT WP_000145410.1 MULTISPECIES: hydrogenase-2 small chain [Proteobacteria] Length: 372\nNumber of predicted TMHs: 1\nExp number of AAs in TMHs: 23.72226\nExp number, first 60 AAs: 0.22038\nTotal prob of N-in: 0.12994\noutside 1 331\nTMhelix 332 354\ninside 355 372

2712 GCF_001286425.1_7553_7_68 Escherichia coli Proteobacteria; Gammaproteobacteria; Enterobacterales; Enterobacteriaceae; Escherichia MNNEETFYQAMRRQGVTRRSFLKYCSLAATSLGLGAGMAPKIAWALENKPRIPVVWIHGL WP_001058323.1 MULTISPECIES: hydrogenase-1 small chain [Proteobacteria] Length: 372\nNumber of predicted TMHs: 2\nExp number of AAs in TMHs: 37.2608499999999\nExp number, first 60 AAs: 19.52923\nTotal prob of N-in: 0.94351\nPOSSIBLE N-term signal sequence\ninside 1 19\nTMhelix 20 39\noutside 40 325\nTMhelix 326 348\ninside 349 372

2713 GCF_001286625.1_8205_3_29 Escherichia coli Proteobacteria; Gammaproteobacteria; Enterobacterales; Enterobacteriaceae; Escherichia MTGDNTLIHSHGINRRDFMKLCAALAATMGLSSKAAAEMAESVTNPQRPPVIWIGAQECT WP_000145410.1 MULTISPECIES: hydrogenase-2 small chain [Proteobacteria] Length: 372\nNumber of predicted TMHs: 1\nExp number of AAs in TMHs: 23.72226\nExp number, first 60 AAs: 0.22038\nTotal prob of N-in: 0.12994\noutside 1 331\nTMhelix 332 354\ninside 355 372

2714 GCF_001286545.1_8205_3_80 Escherichia coli Proteobacteria; Gammaproteobacteria; Enterobacterales; Enterobacteriaceae; Escherichia
MNNEETFYQAMRRQGVTRRSFLKYCSLAATSLGLGAGMAPKIAWALENKPRIPVVWIHGL WP_001058323.1
MULTISPECIES: hydrogenase-1 small chain [Proteobacteria] Length: 372\nNumber of predicted TMHs: 2\nExp number of AAs in TMHs: 37.2608499999999\nExp number, first 60 AAs: 19.52923\nTotal prob of N-in: 0.94351\nPOSSIBLE N-term signal sequence\ninside 1 19\nTMhelix 20 39\noutside 40 325\nTMhelix 326 348\ninside 349 372

2715 GCF_001286585.1_8205_3_68 Escherichia coli Proteobacteria; Gammaproteobacteria; Enterobacterales; Enterobacteriaceae; Escherichia
MNNEETFYQAMRRQGVTRRSFLKYCSLAATSLGLGAGMAPKIAWALENKPRIPVVWIHGL WP_001295940.1
MULTISPECIES: hydrogenase [Enterobacteriaceae] Length: 372\nNumber of predicted TMHs: 2\nExp number of AAs in TMHs: 37.2608899999999\nExp number, first 60 AAs: 19.52923\nTotal prob of N-in: 0.94351\nPOSSIBLE N-term signal sequence\ninside 1 19\nTMhelix 20 39\noutside 40 325\nTMhelix 326 348\ninside 349 372

2716 GCF_001297725.1_Ecoli_assembly01 Escherichia coli Proteobacteria; Gammaproteobacteria; Enterobacterales; Enterobacteriaceae; Escherichia
MTGDNTLIHSHGINRRDFMKLCAALAATMGLSSKAAAEMAESVTNPQRPPVIWIGAQECT WP_000145410.1
MULTISPECIES: hydrogenase-2 small chain [Proteobacteria] Length: 372\nNumber of predicted TMHs: 1\nExp number of AAs in TMHs: 23.72226\nExp number, first 60 AAs: 0.22038\nTotal prob of N-in: 0.12994\noutside 1 331\nTMhelix 332 354\ninside 355 372

2717 GCF_001297975.1_ASM129797v1 Escherichia coli Proteobacteria; Gammaproteobacteria; Enterobacterales; Enterobacteriaceae; Escherichia
MTGDNTLIHSHGINRRDFMKLCAALAATMGLSSKAAAEMAESVTNPQRPPVIWIGAQECT WP_000145410.1
MULTISPECIES: hydrogenase-2 small chain [Proteobacteria] Length: 372\nNumber of predicted TMHs: 1\nExp number of AAs in TMHs: 23.72226\nExp number, first 60 AAs: 0.22038\nTotal prob of N-in: 0.12994\noutside 1 331\nTMhelix 332 354\ninside 355 372

2718 GCF_001297725.1_Ecoli_assembly01 Escherichia coli Proteobacteria; Gammaproteobacteria; Enterobacterales; Enterobacteriaceae; Escherichia
MNNEETFYQAMRRQGVTRRSFLKYCSLAATSLGLGAGMAPKIAWALENKPRIPVVWIHGL WP_001058323.1
MULTISPECIES: hydrogenase-1 small chain [Proteobacteria] Length: 372\nNumber of predicted TMHs: 2\nExp number of AAs in TMHs: 37.2608499999999\nExp number, first 60 AAs: 19.52923\nTotal prob of N-in: 0.94351\nPOSSIBLE N-term signal sequence\ninside 1 19\nTMhelix 20 39\noutside 40 325\nTMhelix 326 348\ninside 349 372

2719 GCF_001297975.1_ASM129797v1 Escherichia coli Proteobacteria; Gammaproteobacteria; Enterobacterales; Enterobacteriaceae; Escherichia
MNNEETFYQAMRRQGVTRRSFLKYCSLAATSLGLGAGMAPKIAWALENKPRIPVVWIHGL WP_001301613.1
hydrogenase [Escherichia coli] Length: 372\nNumber of predicted TMHs: 2\nExp number of AAs in TMHs: 37.46598\nExp number, first 60 AAs: 19.53139\nTotal prob of N-in: 0.94364\nPOSSIBLE N-term signal sequence\ninside 1 19\nTMhelix 20 39\noutside 40 325\nTMhelix 326 348\ninside 349 372

2720 GCF_001286625.1_8205_3_29 Escherichia coli Proteobacteria; Gammaproteobacteria; Enterobacterales; Enterobacteriaceae; Escherichia
MNNEETFYQAMRRQGVTRRSFLKYCSLAATSLGLGAGMAPKIAWALENKPRIPVVWIHGL WP_001359719.1
hydrogenase [Escherichia coli] Length: 372\nNumber of predicted TMHs: 2\nExp number of AAs in TMHs: 37.2176\nExp number, first 60 AAs: 19.53448\nTotal prob of N-in: 0.94360\nPOSSIBLE N-term signal sequence\ninside 1 19\nTMhelix 20 39\noutside 40 325\nTMhelix 326 348\ninside 349 372

2721 GCF_001308165.1_ASM130816v1 Escherichia coli Proteobacteria; Gammaproteobacteria; Enterobacterales; Enterobacteriaceae; Escherichia
MTGDNTLIHSHGINRRDFMKLCAALAATMGLSSKAAAEMAESVTNPQRPPVIWIGAQECT WP_000145410.1
MULTISPECIES: hydrogenase-2 small chain [Proteobacteria] Length: 372\nNumber of predicted TMHs: 1\nExp number of AAs in TMHs: 23.72226\nExp number, first 60 AAs: 0.22038\nTotal prob of N-in: 0.12994\noutside 1 331\nTMhelix 332 354\ninside 355 372

2722 GCF_001309485.1_ASM130948v1 Escherichia coli Proteobacteria; Gammaproteobacteria; Enterobacterales; Enterobacteriaceae; Escherichia

MTGDNTLIHSHGINRRDFMKLCAALAATMGLSSKAAAEMAESVTNPQRPPVIWIGAQECT WP_000145410.1 MULTISPECIES: hydrogenase-2 small chain [Proteobacteria] Length: 372\nNumber of predicted TMHs: 1\nExp number of AAs in TMHs: 23.72226\nExp number, first 60 AAs: 0.22038\nTotal prob of N-in: 0.12994\noutside 1 331\nTMhelix 332 354\ninside 355 372

2723 GCF_001309535.1_ASM130953v1 Escherichia coli Proteobacteria; Gammaproteobacteria; Enterobacterales; Enterobacteriaceae; Escherichia MTGDNTLIHSHGINRRDFMKLCAALAATMGLSSKAAAEMAESVTNPQRPPVIWIGAQECT WP_000145410.1 MULTISPECIES: hydrogenase-2 small chain [Proteobacteria] Length: 372\nNumber of predicted TMHs: 1\nExp number of AAs in TMHs: 23.72226\nExp number, first 60 AAs: 0.22038\nTotal prob of N-in: 0.12994\noutside 1 331\nTMhelix 332 354\ninside 355 372

2724 GCF_001309565.1_ASM130956v1 Escherichia coli Proteobacteria; Gammaproteobacteria; Enterobacterales; Enterobacteriaceae; Escherichia MTGDNTLIHSHGINRRDFMKLCAALAATMGLSSKAAAEMAESVTNPQRPPVIWIGAQECT WP_000145410.1 MULTISPECIES: hydrogenase-2 small chain [Proteobacteria] Length: 372\nNumber of predicted TMHs: 1\nExp number of AAs in TMHs: 23.72226\nExp number, first 60 AAs: 0.22038\nTotal prob of N-in: 0.12994\noutside 1 331\nTMhelix 332 354\ninside 355 372

2725 GCF_001309535.1_ASM130953v1 Escherichia coli Proteobacteria; Gammaproteobacteria; Enterobacterales; Enterobacteriaceae; Escherichia MNNEETFYQAMRRQGVTRRSFLKYCSLAATSLGLGAGMAPKIAWALENKPRIPVVWIHGL WP_001058323.1 MULTISPECIES: hydrogenase-1 small chain [Proteobacteria] Length: 372\nNumber of predicted TMHs: 2\nExp number of AAs in TMHs: 37.2608499999999\nExp number, first 60 AAs: 19.52923\nTotal prob of N-in: 0.94351\nPOSSIBLE N-term signal sequence\ninside 1 19\nTMhelix 20 39\noutside 40 325\nTMhelix 326 348\ninside 349 372

2726 GCF_001309485.1_ASM130948v1 Escherichia coli Proteobacteria; Gammaproteobacteria; Enterobacterales; Enterobacteriaceae; Escherichia MNNEETFYQAMRRQGVTRRSFLKYCSLAATSLGLGAGMAPKIAWALENKPRIPVVWIHGL WP_001058323.1 MULTISPECIES: hydrogenase-1 small chain [Proteobacteria] Length: 372\nNumber of predicted TMHs: 2\nExp number of AAs in TMHs: 37.2608499999999\nExp number, first 60 AAs: 19.52923\nTotal prob of N-in: 0.94351\nPOSSIBLE N-term signal sequence\ninside 1 19\nTMhelix 20 39\noutside 40 325\nTMhelix 326 348\ninside 349 372

2727 GCF_001297995.1_ASM129799v1 Escherichia coli Proteobacteria; Gammaproteobacteria; Enterobacterales; Enterobacteriaceae; Escherichia MNNEETFYQAMRRQGVTRRSFLKYCSLAATSLGLGAGMAPKIAWALENKPRIPVVWIHGL WP_001301613.1 hydrogenase [Escherichia coli] Length: 372\nNumber of predicted TMHs: 2\nExp number of AAs in TMHs: 37.46598\nExp number, first 60 AAs: 19.53139\nTotal prob of N-in: 0.94364\nPOSSIBLE N-term signal sequence\ninside 1 19\nTMhelix 20 39\noutside 40 325\nTMhelix 326 348\ninside 349 372

2728 GCF_001306655.1_JA16 Escherichia coli Proteobacteria; Gammaproteobacteria; Enterobacterales; Enterobacteriaceae; Escherichia MNNEETFYQAMRRQGVTRRSFLKYCSLAATSLGLGAGMAPKIAWALENKPRIPVVWIHGL WP_001295940.1 MULTISPECIES: hydrogenase [Enterobacteriaceae] Length: 372\nNumber of predicted TMHs: 2\nExp number of AAs in TMHs: 37.2608899999999\nExp number, first 60 AAs: 19.52923\nTotal prob of N-in: 0.94351\nPOSSIBLE N-term signal sequence\ninside 1 19\nTMhelix 20 39\noutside 40 325\nTMhelix 326 348\ninside 349 372

2729 GCF_001309685.1_ASM130968v1 Escherichia coli Proteobacteria; Gammaproteobacteria; Enterobacterales; Enterobacteriaceae; Escherichia MTGDNTLIHSHGINRRDFMKLCAALAATMGLSSKAAAEMAESVTNPQRPPVIWIGAQECT WP_000145410.1 MULTISPECIES: hydrogenase-2 small chain [Proteobacteria] Length: 372\nNumber of predicted TMHs: 1\nExp number of AAs in TMHs: 23.72226\nExp number, first 60 AAs: 0.22038\nTotal prob of N-in: 0.12994\noutside 1 331\nTMhelix 332 354\ninside 355 372

2730 GCF_001309635.1_ASM130963v1 Escherichia coli Proteobacteria; Gammaproteobacteria; Enterobacterales; Enterobacteriaceae; Escherichia MTGDNTLIHSHGINRRDFMKLCAALAATMGLSSKAAAEMAESVTNPQRPPVIWIGAQECT WP_000145410.1 MULTISPECIES: hydrogenase-2 small chain [Proteobacteria] Length: 372\nNumber of predicted TMHs:

1\nExp number of AAs in TMHs: 23.72226\nExp number, first 60 AAs: 0.22038\nTotal prob of N-in: 0.12994\noutside 1 331\nTMhelix 332 354\ninside 355 372

2731 GCF_001309775.1_ASM130977v1 Escherichia coli Proteobacteria; Gammaproteobacteria; Enterobacterales; Enterobacteriaceae; Escherichia
MTGDNTLIHSHGINRRDFMKLCAALAATMGLSSKAAAEMAESVTNPQRPPVIWIGAQECT WP_000145410.1
MULTISPECIES: hydrogenase-2 small chain [Proteobacteria] Length: 372\nNumber of predicted TMHs: 1\nExp number of AAs in TMHs: 23.72226\nExp number, first 60 AAs: 0.22038\nTotal prob of N-in: 0.12994\noutside 1 331\nTMhelix 332 354\ninside 355 372

2732 GCF_001309675.1_ASM130967v1 Escherichia coli Proteobacteria; Gammaproteobacteria; Enterobacterales; Enterobacteriaceae; Escherichia
MTGDNTLIHSHGINRRDFMKLCAALAATMGLSSKAAAEMAESVTNPQRPPVIWIGAQECT WP_000145410.1
MULTISPECIES: hydrogenase-2 small chain [Proteobacteria] Length: 372\nNumber of predicted TMHs: 1\nExp number of AAs in TMHs: 23.72226\nExp number, first 60 AAs: 0.22038\nTotal prob of N-in: 0.12994\noutside 1 331\nTMhelix 332 354\ninside 355 372

2733 GCF_001309595.1_ASM130959v1 Escherichia coli Proteobacteria; Gammaproteobacteria; Enterobacterales; Enterobacteriaceae; Escherichia
MTGDNTLIHSHGINRRDFMKLCAALAATMGLSNKAAAEMAESVTNPQRPPVIWIGAQECT WP_054632347.1
hydrogenase 2 small subunit [Escherichia coli] Length: 372\nNumber of predicted TMHs: 1\nExp number of AAs in TMHs: 23.60505\nExp number, first 60 AAs: 0.09567\nTotal prob of N-in: 0.12455\noutside 1 331\nTMhelix 332 354\ninside 355 372

2734 GCF_001309885.1_ASM130988v1 Escherichia coli Proteobacteria; Gammaproteobacteria; Enterobacterales; Enterobacteriaceae; Escherichia
MNNEETFYQAMRRQGVTRRSFLKYCSLAATSLGLGAGMAPKIAWALENKPRIPVVWIHGL WP_001301613.1
hydrogenase [Escherichia coli] Length: 372\nNumber of predicted TMHs: 2\nExp number of AAs in TMHs: 37.46598\nExp number, first 60 AAs: 19.53139\nTotal prob of N-in: 0.94364\nPOSSIBLE N-term signal sequence\ninside 1 19\nTMhelix 20 39\noutside 40 325\nTMhelix 326 348\ninside 349 372

2735 GCF_001309685.1_ASM130968v1 Escherichia coli Proteobacteria; Gammaproteobacteria; Enterobacterales; Enterobacteriaceae; Escherichia
MNNEETFYQAMRRQGVTRRSFLKYCSLAATSLGLGAGMAPKIAWALENKPRIPVVWIHGL WP_001058323.1
MULTISPECIES: hydrogenase-1 small chain [Proteobacteria] Length: 372\nNumber of predicted TMHs: 2\nExp number of AAs in TMHs: 37.2608499999999\nExp number, first 60 AAs: 19.52923\nTotal prob of N-in: 0.94351\nPOSSIBLE N-term signal sequence\ninside 1 19\nTMhelix 20 39\noutside 40 325\nTMhelix 326 348\ninside 349 372

2736 GCF_001309735.1_ASM130973v1 Escherichia coli Proteobacteria; Gammaproteobacteria; Enterobacterales; Enterobacteriaceae; Escherichia
MNNEETFYQAMRRQGVTRRSFLKYCSLAATSLGLGAGMAPKIAWALENKPRIPVVWIHGL WP_001058323.1
MULTISPECIES: hydrogenase-1 small chain [Proteobacteria] Length: 372\nNumber of predicted TMHs: 2\nExp number of AAs in TMHs: 37.2608499999999\nExp number, first 60 AAs: 19.52923\nTotal prob of N-in: 0.94351\nPOSSIBLE N-term signal sequence\ninside 1 19\nTMhelix 20 39\noutside 40 325\nTMhelix 326 348\ninside 349 372

2737 GCF_001309675.1_ASM130967v1 Escherichia coli Proteobacteria; Gammaproteobacteria; Enterobacterales; Enterobacteriaceae; Escherichia
MNNEETFYQAMRRQGVTRRSFLKYCSLAATSLGLGAGMAPKIAWALENKPRIPVVWIHGL WP_001058323.1
MULTISPECIES: hydrogenase-1 small chain [Proteobacteria] Length: 372\nNumber of predicted TMHs: 2\nExp number of AAs in TMHs: 37.2608499999999\nExp number, first 60 AAs: 19.52923\nTotal prob of N-in: 0.94351\nPOSSIBLE N-term signal sequence\ninside 1 19\nTMhelix 20 39\noutside 40 325\nTMhelix 326 348\ninside 349 372

2738 GCF_001309775.1_ASM130977v1 Escherichia coli Proteobacteria; Gammaproteobacteria; Enterobacterales; Enterobacteriaceae; Escherichia
MNNEETFYQAMRRQGVTRRSFLKYCSLAATSLGLGAGMAPKIAWALENKPRIPVVWIHGL WP_001058323.1
MULTISPECIES: hydrogenase-1 small chain [Proteobacteria] Length: 372\nNumber of predicted TMHs: 2\nExp number of AAs in TMHs: 37.2608499999999\nExp number, first 60 AAs: 19.52923\nTotal prob of N-in:

0.94351\nPOSSIBLE N-term signal sequence\ninside 1 19\nTMhelix 20 39\noutside 40 325\nTMhelix 326 348\ninside 349 372

2739 GCF_001402805.1_ASM140280v1 Escherichia coli Proteobacteria; Gammaproteobacteria; Enterobacterales; Enterobacteriaceae; Escherichia MTGDNTLIHSHGINRRDFMKLCAALAATMGLSSKAAAEMAESVTNPQRPPVIWIGAQECT WP_000145410.1 MULTISPECIES: hydrogenase-2 small chain [Proteobacteria] Length: 372\nNumber of predicted TMHs: 1\nExp number of AAs in TMHs: 23.72226\nExp number, first 60 AAs: 0.22038\nTotal prob of N-in: 0.12994\noutside 1 331\nTMhelix 332 354\ninside 355 372

2740 GCF_001412975.1_ASM141297v1 Escherichia coli Proteobacteria; Gammaproteobacteria; Enterobacterales; Enterobacteriaceae; Escherichia MTGDNTLIHSHGINRRDFMKLCAALAATMGLSSKAAAEMAESVTNPQRPPVIWIGAQECT WP_000145410.1 MULTISPECIES: hydrogenase-2 small chain [Proteobacteria] Length: 372\nNumber of predicted TMHs: 1\nExp number of AAs in TMHs: 23.72226\nExp number, first 60 AAs: 0.22038\nTotal prob of N-in: 0.12994\noutside 1 331\nTMhelix 332 354\ninside 355 372

2741 GCF_001413005.1_AssemblyName Escherichia coli Proteobacteria; Gammaproteobacteria; Enterobacterales; Enterobacteriaceae; Escherichia MTGDNTLIHSHGINRRDFMKLCAALAATMGLSSKAAAEMAESVTNPQRPPVIWIGAQECT WP_000145410.1 MULTISPECIES: hydrogenase-2 small chain [Proteobacteria] Length: 372\nNumber of predicted TMHs: 1\nExp number of AAs in TMHs: 23.72226\nExp number, first 60 AAs: 0.22038\nTotal prob of N-in: 0.12994\noutside 1 331\nTMhelix 332 354\ninside 355 372

2742 GCF_001413355.1_ASM141335v1 Escherichia coli Proteobacteria; Gammaproteobacteria; Enterobacterales; Enterobacteriaceae; Escherichia MTGDNTLIHSHGINRRDFMKLCAALAATMGLSSKAAAEMAESVTNPQRPPVIWIGAQECT WP_000145410.1 MULTISPECIES: hydrogenase-2 small chain [Proteobacteria] Length: 372\nNumber of predicted TMHs: 1\nExp number of AAs in TMHs: 23.72226\nExp number, first 60 AAs: 0.22038\nTotal prob of N-in: 0.12994\noutside 1 331\nTMhelix 332 354\ninside 355 372

2743 GCF_001413395.1_ASM141339v1 Escherichia coli Proteobacteria; Gammaproteobacteria; Enterobacterales; Enterobacteriaceae; Escherichia MTGDNTLIHSHGINRRDFMKLCAALAATMGLSSKAAAEMAESVTNPQRPPVIWIGAQECT WP_000145410.1 MULTISPECIES: hydrogenase-2 small chain [Proteobacteria] Length: 372\nNumber of predicted TMHs: 1\nExp number of AAs in TMHs: 23.72226\nExp number, first 60 AAs: 0.22038\nTotal prob of N-in: 0.12994\noutside 1 331\nTMhelix 332 354\ninside 355 372

2744 GCF_001413395.1_ASM141339v1 Escherichia coli Proteobacteria; Gammaproteobacteria; Enterobacterales; Enterobacteriaceae; Escherichia MNNEETFYQAMRRQGVTRRSFLKYCSLAATSLGLGAGMAPKIAWALENKPRIPVVWIHGL WP_001058323.1 MULTISPECIES: hydrogenase-1 small chain [Proteobacteria] Length: 372\nNumber of predicted TMHs: 2\nExp number of AAs in TMHs: 37.2608499999999\nExp number, first 60 AAs: 19.52923\nTotal prob of N-in: 0.94351\nPOSSIBLE N-term signal sequence\ninside 1 19\nTMhelix 20 39\noutside 40 325\nTMhelix 326 348\ninside 349 372

2745 GCF_001412895.1_ASM141289v1 Escherichia coli Proteobacteria; Gammaproteobacteria; Enterobacterales; Enterobacteriaceae; Escherichia MNNEETFYQAMRRQGVTRRSFLKYCSLAATSLGLGAGMAPKIAWALENKPRIPVVWIHGL WP_001058323.1 MULTISPECIES: hydrogenase-1 small chain [Proteobacteria] Length: 372\nNumber of predicted TMHs: 2\nExp number of AAs in TMHs: 37.2608499999999\nExp number, first 60 AAs: 19.52923\nTotal prob of N-in: 0.94351\nPOSSIBLE N-term signal sequence\ninside 1 19\nTMhelix 20 39\noutside 40 325\nTMhelix 326 348\ninside 349 372

2746 GCF_001413485.1_ASM141348v1 Escherichia coli Proteobacteria; Gammaproteobacteria; Enterobacterales; Enterobacteriaceae; Escherichia MNNEETFYQAMRRQGVTRRSFLKYCSLAATSLGLGAGMAPKIAWALENKPRIPVVWIHGL WP_001058323.1 MULTISPECIES: hydrogenase-1 small chain [Proteobacteria] Length: 372\nNumber of predicted TMHs: 2\nExp number of AAs in TMHs: 37.2608499999999\nExp number, first 60 AAs: 19.52923\nTotal prob of N-in: 0.94351\nPOSSIBLE N-term signal sequence\ninside 1 19\nTMhelix 20 39\noutside 40 325\nTMhelix 326 348\ninside 349 372

2747 GCF_001413005.1_AssemblyName Escherichia coli Proteobacteria; Gammaproteobacteria; Enterobacterales; Enterobacteriaceae; Escherichia
MNNEETFYQAMRRQGVTRRSFLKYCSLAATSLGLGAGMAPKIAWALENKPRIPVVWIHGL WP_001058323.1
MULTISPECIES: hydrogenase-1 small chain [Proteobacteria] Length: 372\nNumber of predicted TMHs: 2\nExp number of AAs in TMHs: 37.2608499999999\nExp number, first 60 AAs: 19.52923\nTotal prob of N-in: 0.94351\nPOSSIBLE N-term signal sequence\ninside 1 19\nTMhelix 20 39\noutside 40 325\nTMhelix 326 348\ninside 349 372

2748 GCF_001413355.1_ASM141335v1 Escherichia coli Proteobacteria; Gammaproteobacteria; Enterobacterales; Enterobacteriaceae; Escherichia
MNNEETFYQAMRRQGVTRRSFLKYCSLAATSLGLGAGMAPKIAWALENKPRIPVVWIHGL WP_001058323.1
MULTISPECIES: hydrogenase-1 small chain [Proteobacteria] Length: 372\nNumber of predicted TMHs: 2\nExp number of AAs in TMHs: 37.2608499999999\nExp number, first 60 AAs: 19.52923\nTotal prob of N-in: 0.94351\nPOSSIBLE N-term signal sequence\ninside 1 19\nTMhelix 20 39\noutside 40 325\nTMhelix 326 348\ninside 349 372

2749 GCF_001413535.1_ASM141353v1 Escherichia coli Proteobacteria; Gammaproteobacteria; Enterobacterales; Enterobacteriaceae; Escherichia
MTGDNTLIHSHGINRRDFMKLCAALAATMGLSSKAAAEMAESVTNPQRPPVIWIGAQECT WP_000145410.1
MULTISPECIES: hydrogenase-2 small chain [Proteobacteria] Length: 372\nNumber of predicted TMHs: 1\nExp number of AAs in TMHs: 23.72226\nExp number, first 60 AAs: 0.22038\nTotal prob of N-in: 0.12994\noutside 1 331\nTMhelix 332 354\ninside 355 372

2750 GCF_001413535.1_ASM141353v1 Escherichia coli Proteobacteria; Gammaproteobacteria; Enterobacterales; Enterobacteriaceae; Escherichia
MNNEETFYQAMRRQGVTRRSFLKYCSLAATSLGLGAGMAPKIAWALENKPRIPVVWIHGL WP_001058323.1
MULTISPECIES: hydrogenase-1 small chain [Proteobacteria] Length: 372\nNumber of predicted TMHs: 2\nExp number of AAs in TMHs: 37.2608499999999\nExp number, first 60 AAs: 19.52923\nTotal prob of N-in: 0.94351\nPOSSIBLE N-term signal sequence\ninside 1 19\nTMhelix 20 39\noutside 40 325\nTMhelix 326 348\ninside 349 372

2751 GCF_001402805.1_ASM140280v1 Escherichia coli Proteobacteria; Gammaproteobacteria; Enterobacterales; Enterobacteriaceae; Escherichia
MNNEETFYQAMRRQGVTRRSFLKYCSLAATSLGLGAGMAPKIAWALENKPRIPVVWIHGL WP_001295940.1
MULTISPECIES: hydrogenase [Enterobacteriaceae] Length: 372\nNumber of predicted TMHs: 2\nExp number of AAs in TMHs: 37.2608899999999\nExp number, first 60 AAs: 19.52923\nTotal prob of N-in: 0.94351\nPOSSIBLE N-term signal sequence\ninside 1 19\nTMhelix 20 39\noutside 40 325\nTMhelix 326 348\ninside 349 372

2752 GCF_001309995.1_ASM130999v1 Escherichia coli Proteobacteria; Gammaproteobacteria; Enterobacterales; Enterobacteriaceae; Escherichia
MNNEETFYQAMRRQGVTRRSFLKYCSLAATSLGLGAGMAPKIAWALENKPRIPVVWIHGL WP_001301613.1
hydrogenase [Escherichia coli] Length: 372\nNumber of predicted TMHs: 2\nExp number of AAs in TMHs: 37.46598\nExp number, first 60 AAs: 19.53139\nTotal prob of N-in: 0.94364\nPOSSIBLE N-term signal sequence\ninside 1 19\nTMhelix 20 39\noutside 40 325\nTMhelix 326 348\ninside 349 372

2753 GCF_001413745.1_ASM141374v1 Escherichia coli Proteobacteria; Gammaproteobacteria; Enterobacterales; Enterobacteriaceae; Escherichia
MTGDNTLIHSHGINRRDFMKLCAALAATMGLSSKAAAEMAESVTNPQRPPVIWIGAQECT WP_000145410.1
MULTISPECIES: hydrogenase-2 small chain [Proteobacteria] Length: 372\nNumber of predicted TMHs: 1\nExp number of AAs in TMHs: 23.72226\nExp number, first 60 AAs: 0.22038\nTotal prob of N-in: 0.12994\noutside 1 331\nTMhelix 332 354\ninside 355 372

2754 GCF_001413805.1_ASM141380v1 Escherichia coli Proteobacteria; Gammaproteobacteria; Enterobacterales; Enterobacteriaceae; Escherichia
MTGDNTLIHSHGINRRDFMKLCAALAATMGLSSKAAAEMAESVTNPQRPPVIWIGAQECT WP_000145410.1
MULTISPECIES: hydrogenase-2 small chain [Proteobacteria] Length: 372\nNumber of predicted TMHs: 1\nExp number of AAs in TMHs: 23.72226\nExp number, first 60 AAs: 0.22038\nTotal prob of N-in: 0.12994\noutside 1 331\nTMhelix 332 354\ninside 355 372

2755 GCF_001413735.1_ASM141373v1 Escherichia coli Proteobacteria; Gammaproteobacteria; Enterobacterales; Enterobacteriaceae; Escherichia
MNNEETFYQAMRRQGVTRRSFLKYCSLAATSLGLGAGMAPKIAWALENKPRIPVVWIHGL WP_001058323.1
MULTISPECIES: hydrogenase-1 small chain [Proteobacteria] Length: 372\nNumber of predicted TMHs: 2\nExp number of AAs in TMHs: 37.2608499999999\nExp number, first 60 AAs: 19.52923\nTotal prob of N-in: 0.94351\nPOSSIBLE N-term signal sequence\ninside 1 19\nTMhelix 20 39\noutside 40 325\nTMhelix 326 348\ninside 349 372

2756 GCF_001413745.1_ASM141374v1 Escherichia coli Proteobacteria; Gammaproteobacteria; Enterobacterales; Enterobacteriaceae; Escherichia
MNNEETFYQAMRRQGVTRRSFLKYCSLAATSLGLGAGMAPKIAWALENKPRIPVVWIHGL WP_001058323.1
MULTISPECIES: hydrogenase-1 small chain [Proteobacteria] Length: 372\nNumber of predicted TMHs: 2\nExp number of AAs in TMHs: 37.2608499999999\nExp number, first 60 AAs: 19.52923\nTotal prob of N-in: 0.94351\nPOSSIBLE N-term signal sequence\ninside 1 19\nTMhelix 20 39\noutside 40 325\nTMhelix 326 348\ninside 349 372

2757 GCF_001413805.1_ASM141380v1 Escherichia coli Proteobacteria; Gammaproteobacteria; Enterobacterales; Enterobacteriaceae; Escherichia
MNNEETFYQAMRRQGVTRRSFLKYCSLAATSLGLGAGMAPKIAWALENKPRIPVVWIHGL WP_001058323.1
MULTISPECIES: hydrogenase-1 small chain [Proteobacteria] Length: 372\nNumber of predicted TMHs: 2\nExp number of AAs in TMHs: 37.2608499999999\nExp number, first 60 AAs: 19.52923\nTotal prob of N-in: 0.94351\nPOSSIBLE N-term signal sequence\ninside 1 19\nTMhelix 20 39\noutside 40 325\nTMhelix 326 348\ninside 349 372

2758 GCF_001413635.1_ASM141363v1 Escherichia coli Proteobacteria; Gammaproteobacteria; Enterobacterales; Enterobacteriaceae; Escherichia
MNNEETFYQAMRRQGVTRRSFLKYCSLAATSLGLGAGMAPKIAWALENKPRIPVVWIHGL WP_001058323.1
MULTISPECIES: hydrogenase-1 small chain [Proteobacteria] Length: 372\nNumber of predicted TMHs: 2\nExp number of AAs in TMHs: 37.2608499999999\nExp number, first 60 AAs: 19.52923\nTotal prob of N-in: 0.94351\nPOSSIBLE N-term signal sequence\ninside 1 19\nTMhelix 20 39\noutside 40 325\nTMhelix 326 348\ninside 349 372

2759 GCF_001413595.1_ASM141359v1 Escherichia coli Proteobacteria; Gammaproteobacteria; Enterobacterales; Enterobacteriaceae; Escherichia
MNNEETFYQAMRRQGVTRRSFLKYCSLAATSLGLGAGMAPKIAWALENKPRIPVVWIHGL WP_001058323.1
MULTISPECIES: hydrogenase-1 small chain [Proteobacteria] Length: 372\nNumber of predicted TMHs: 2\nExp number of AAs in TMHs: 37.2608499999999\nExp number, first 60 AAs: 19.52923\nTotal prob of N-in: 0.94351\nPOSSIBLE N-term signal sequence\ninside 1 19\nTMhelix 20 39\noutside 40 325\nTMhelix 326 348\ninside 349 372

2760 GCF_001413875.1_ASM141387v1 Escherichia coli Proteobacteria; Gammaproteobacteria; Enterobacterales; Enterobacteriaceae; Escherichia
MNNEETFYQAMRRQGVTRRSFLKYCSLAATSLGLGAGMAPKIAWALENKPRIPVVWIHGL WP_001058323.1
MULTISPECIES: hydrogenase-1 small chain [Proteobacteria] Length: 372\nNumber of predicted TMHs: 2\nExp number of AAs in TMHs: 37.2608499999999\nExp number, first 60 AAs: 19.52923\nTotal prob of N-in: 0.94351\nPOSSIBLE N-term signal sequence\ninside 1 19\nTMhelix 20 39\noutside 40 325\nTMhelix 326 348\ninside 349 372

2761 GCF_001413905.1_ASM141390v1 Escherichia coli Proteobacteria; Gammaproteobacteria; Enterobacterales; Enterobacteriaceae; Escherichia
MNNEETFYQAMRRQGVTRRSFLKYCSLAATSLGLGAGMAPKIAWALENKPRIPVVWIHGL WP_001058323.1
MULTISPECIES: hydrogenase-1 small chain [Proteobacteria] Length: 372\nNumber of predicted TMHs: 2\nExp number of AAs in TMHs: 37.2608499999999\nExp number, first 60 AAs: 19.52923\nTotal prob of N-in: 0.94351\nPOSSIBLE N-term signal sequence\ninside 1 19\nTMhelix 20 39\noutside 40 325\nTMhelix 326 348\ninside 349 372

2762 GCF_001419785.1_ASM141978v1 Escherichia coli Proteobacteria; Gammaproteobacteria; Enterobacterales; Enterobacteriaceae; Escherichia
MNNEETFYQAMRRQGVTRRSFLKYCSLAATSLGLGAGMAPKIAWALENKPRIPVVWIHGL WP_001058323.1
MULTISPECIES: hydrogenase-1 small chain [Proteobacteria] Length: 372\nNumber of predicted TMHs: 2\nExp number of AAs in TMHs: 37.2608499999999\nExp number, first 60 AAs: 19.52923\nTotal prob of N-in:

0.94351\nPOSSIBLE N-term signal sequence\ninside 1 19\nTMhelix 20 39\noutside 40 325\nTMhelix 326 348\ninside 349 372

2763 GCF_001419825.1_ASM141982v1 Escherichia coli Proteobacteria; Gammaproteobacteria; Enterobacterales; Enterobacteriaceae; Escherichia MTGDNTLIHSHGINRRDFMKLCAALAATMGLSSKAAAEMAESVTNPQRPPVIWIGAQECT WP_000145410.1 MULTISPECIES: hydrogenase-2 small chain [Proteobacteria] Length: 372\nNumber of predicted TMHs: 1\nExp number of AAs in TMHs: 23.72226\nExp number, first 60 AAs: 0.22038\nTotal prob of N-in: 0.12994\noutside 1 331\nTMhelix 332 354\ninside 355 372

2764 GCF_001419895.1_ASM141989v1 Escherichia coli Proteobacteria; Gammaproteobacteria; Enterobacterales; Enterobacteriaceae; Escherichia MTGDNTLIHSHGINRRDFMKLCAALAATMGLSSKAAAEMAESVTNPQRPPVIWIGAQECT WP_000145410.1 MULTISPECIES: hydrogenase-2 small chain [Proteobacteria] Length: 372\nNumber of predicted TMHs: 1\nExp number of AAs in TMHs: 23.72226\nExp number, first 60 AAs: 0.22038\nTotal prob of N-in: 0.12994\noutside 1 331\nTMhelix 332 354\ninside 355 372

2765 GCF_001419865.1_ASM141986v1 Escherichia coli Proteobacteria; Gammaproteobacteria; Enterobacterales; Enterobacteriaceae; Escherichia MTGDNTLIHSHGINRRDFMKLCAALAATMGLSSKAAAEMAESVTNPQRPPVIWIGAQECT WP_000145410.1 MULTISPECIES: hydrogenase-2 small chain [Proteobacteria] Length: 372\nNumber of predicted TMHs: 1\nExp number of AAs in TMHs: 23.72226\nExp number, first 60 AAs: 0.22038\nTotal prob of N-in: 0.12994\noutside 1 331\nTMhelix 332 354\ninside 355 372

2766 GCF_001419925.1_ASM141992v1 Escherichia coli Proteobacteria; Gammaproteobacteria; Enterobacterales; Enterobacteriaceae; Escherichia MTGDNTLIHSHGINRRDFMKLCAALAATMGLSSKAAAEMAESVTNPQRPPVIWIGAQECT WP_000145410.1 MULTISPECIES: hydrogenase-2 small chain [Proteobacteria] Length: 372\nNumber of predicted TMHs: 1\nExp number of AAs in TMHs: 23.72226\nExp number, first 60 AAs: 0.22038\nTotal prob of N-in: 0.12994\noutside 1 331\nTMhelix 332 354\ninside 355 372

2767 GCF_001419825.1_ASM141982v1 Escherichia coli Proteobacteria; Gammaproteobacteria; Enterobacterales; Enterobacteriaceae; Escherichia MNNEETFYQAMRRQGVTRRSFLKYCSLAATSLGLGAGMAPKIAWALENKPRIPVVWIHGL WP_001058323.1 MULTISPECIES: hydrogenase-1 small chain [Proteobacteria] Length: 372\nNumber of predicted TMHs: 2\nExp number of AAs in TMHs: 37.2608499999999\nExp number, first 60 AAs: 19.52923\nTotal prob of N-in: 0.94351\nPOSSIBLE N-term signal sequence\ninside 1 19\nTMhelix 20 39\noutside 40 325\nTMhelix 326 348\ninside 349 372

2768 GCF_001421045.1_ASM142104v1 Escherichia coli Proteobacteria; Gammaproteobacteria; Enterobacterales; Enterobacteriaceae; Escherichia MTGDNTLIHSHGINRRDFMKLCAALAATMGLSSKAAAEMAESVTNPQRPPVIWIGAQECT WP_000145410.1 MULTISPECIES: hydrogenase-2 small chain [Proteobacteria] Length: 372\nNumber of predicted TMHs: 1\nExp number of AAs in TMHs: 23.72226\nExp number, first 60 AAs: 0.22038\nTotal prob of N-in: 0.12994\noutside 1 331\nTMhelix 332 354\ninside 355 372

2769 GCF_001421045.1_ASM142104v1 Escherichia coli Proteobacteria; Gammaproteobacteria; Enterobacterales; Enterobacteriaceae; Escherichia MNNEETFYQAMRRQGVTRRSFLKYCSLAATSLGLGAGMAPKIAWALENKPRIPVVWIHGL WP_001058320.1 hydrogenase [Escherichia coli] Length: 372\nNumber of predicted TMHs: 2\nExp number of AAs in TMHs: 37.26082\nExp number, first 60 AAs: 19.52923\nTotal prob of N-in: 0.94351\nPOSSIBLE N-term signal sequence\ninside 1 19\nTMhelix 20 39\noutside 40 325\nTMhelix 326 348\ninside 349 372

2770 GCF_001419925.1_ASM141992v1 Escherichia coli Proteobacteria; Gammaproteobacteria; Enterobacterales; Enterobacteriaceae; Escherichia MNNEETFYQAMRRQGVTRRSFLKYCSLAATSLGLGAGMAPKIAWALENKPRIPVVWIHGL WP_001058323.1 MULTISPECIES: hydrogenase-1 small chain [Proteobacteria] Length: 372\nNumber of predicted TMHs: 2\nExp number of AAs in TMHs: 37.2608499999999\nExp number, first 60 AAs: 19.52923\nTotal prob of N-in: 0.94351\nPOSSIBLE N-term signal sequence\ninside 1 19\nTMhelix 20 39\noutside 40 325\nTMhelix 326 348\ninside 349 372

2771 GCF_001420045.1_ASM142004v1 Escherichia coli Proteobacteria; Gammaproteobacteria; Enterobacterales; Enterobacteriaceae; Escherichia
MNNEETFYQAMRRQGVTRRSFLKYCSLAATSLGLGAGMAPKIAWALENKPRIPVVWIHGL WP_001058323.1
MULTISPECIES: hydrogenase-1 small chain [Proteobacteria] Length: 372\nNumber of predicted TMHs: 2\nExp number of AAs in TMHs: 37.2608499999999\nExp number, first 60 AAs: 19.52923\nTotal prob of N-in: 0.94351\nPOSSIBLE N-term signal sequence\ninside 1 19\nTMhelix 20 39\noutside 40 325\nTMhelix 326 348\ninside 349 372

2772 GCF_001440525.1_ASM144052v1 Escherichia coli Proteobacteria; Gammaproteobacteria; Enterobacterales; Enterobacteriaceae; Escherichia
MTGDNTLIHSHGINRRDFMKLCAALAATMGLSSKAAAEMAESVTNPQRPPVIWIGAQECT WP_000145410.1
MULTISPECIES: hydrogenase-2 small chain [Proteobacteria] Length: 372\nNumber of predicted TMHs: 1\nExp number of AAs in TMHs: 23.72226\nExp number, first 60 AAs: 0.22038\nTotal prob of N-in: 0.12994\noutside 1 331\nTMhelix 332 354\ninside 355 372

2773 GCF_001440645.1_ASM144064v1 Escherichia coli Proteobacteria; Gammaproteobacteria; Enterobacterales; Enterobacteriaceae; Escherichia
MTGDNTLIHSHGINRRDFMKLCAALAATMGLSSKAAAEMAESVTNPQRPPVIWIGAQECT WP_000145410.1
MULTISPECIES: hydrogenase-2 small chain [Proteobacteria] Length: 372\nNumber of predicted TMHs: 1\nExp number of AAs in TMHs: 23.72226\nExp number, first 60 AAs: 0.22038\nTotal prob of N-in: 0.12994\noutside 1 331\nTMhelix 332 354\ninside 355 372

2774 GCF_001440615.1_ASM144061v1 Escherichia coli Proteobacteria; Gammaproteobacteria; Enterobacterales; Enterobacteriaceae; Escherichia
MTGDNTLIHSHGINRRDFMKLCAALAATMGLSSKAAAEMAESVTNPQRPPVIWIGAQECT WP_000145410.1
MULTISPECIES: hydrogenase-2 small chain [Proteobacteria] Length: 372\nNumber of predicted TMHs: 1\nExp number of AAs in TMHs: 23.72226\nExp number, first 60 AAs: 0.22038\nTotal prob of N-in: 0.12994\noutside 1 331\nTMhelix 332 354\ninside 355 372

2775 GCF_001440665.1_ASM144066v1 Escherichia coli Proteobacteria; Gammaproteobacteria; Enterobacterales; Enterobacteriaceae; Escherichia
MTGDNTLIHSHGINRRDFMKLCAALAATMGLSSKAAAEMAESVTNPQRPPVIWIGAQECT WP_000145410.1
MULTISPECIES: hydrogenase-2 small chain [Proteobacteria] Length: 372\nNumber of predicted TMHs: 1\nExp number of AAs in TMHs: 23.72226\nExp number, first 60 AAs: 0.22038\nTotal prob of N-in: 0.12994\noutside 1 331\nTMhelix 332 354\ninside 355 372

2776 GCF_001440735.1_ASM144073v1 Escherichia coli Proteobacteria; Gammaproteobacteria; Enterobacterales; Enterobacteriaceae; Escherichia
MTGDNTLIHSHGINRRDFMKLCAALAATMGLSSKAAAEMAESVTNPQRPPVIWIGAQECT WP_000145410.1
MULTISPECIES: hydrogenase-2 small chain [Proteobacteria] Length: 372\nNumber of predicted TMHs: 1\nExp number of AAs in TMHs: 23.72226\nExp number, first 60 AAs: 0.22038\nTotal prob of N-in: 0.12994\noutside 1 331\nTMhelix 332 354\ninside 355 372

2777 GCF_001440725.1_ASM144072v1 Escherichia coli Proteobacteria; Gammaproteobacteria; Enterobacterales; Enterobacteriaceae; Escherichia
MTGDNTLIHSHGINRRDFMKLCAALAATMGLSSKAAAEMAESVTNPQRPPVIWIGAQECT WP_000145410.1
MULTISPECIES: hydrogenase-2 small chain [Proteobacteria] Length: 372\nNumber of predicted TMHs: 1\nExp number of AAs in TMHs: 23.72226\nExp number, first 60 AAs: 0.22038\nTotal prob of N-in: 0.12994\noutside 1 331\nTMhelix 332 354\ninside 355 372

2778 GCF_001440775.1_ASM144077v1 Escherichia coli Proteobacteria; Gammaproteobacteria; Enterobacterales; Enterobacteriaceae; Escherichia
MTGDNTLIHSHGINRRDFMKLCAALAATMGLSSKAAAEMAESVTNPQRPPVIWIGAQECT WP_000145410.1
MULTISPECIES: hydrogenase-2 small chain [Proteobacteria] Length: 372\nNumber of predicted TMHs: 1\nExp number of AAs in TMHs: 23.72226\nExp number, first 60 AAs: 0.22038\nTotal prob of N-in: 0.12994\noutside 1 331\nTMhelix 332 354\ninside 355 372

2779 GCF_001440725.1_ASM144072v1 Escherichia coli Proteobacteria; Gammaproteobacteria; Enterobacterales; Enterobacteriaceae; Escherichia
MNNEETFYQAMRRQGVTRRSFLKSCSLAATSLGLGAGMAPKIAWALENKPRIPVVWIHGL WP_001058318.1
hydrogenase [Escherichia coli] Length: 372\nNumber of predicted TMHs: 1\nExp number of AAs in

TMHs: 33.59637\nExp number, first 60 AAs:  15.81976\nTotal prob of N-in:   0.77394\nPOSSIBLE N-term signal sequence\noutside   1   325\nTMhelix   326   348\ninside   349   372

2780 GCF_001440775.1_ASM144077v1 Escherichia coli Proteobacteria; Gammaproteobacteria; Enterobacterales; Enterobacteriaceae; Escherichia MNNEETFYQAMRRQGVTRRSFLKSCSLAATSLGLGAGMAPKIAWALENKPRIPVVWIHGL WP_001058318.1 hydrogenase [Escherichia coli] Length: 372\nNumber of predicted TMHs:  1\nExp number of AAs in TMHs: 33.59637\nExp number, first 60 AAs:  15.81976\nTotal prob of N-in:   0.77394\nPOSSIBLE N-term signal sequence\noutside   1   325\nTMhelix   326   348\ninside   349   372

2781 GCF_001440575.1_ASM144057v1 Escherichia coli Proteobacteria; Gammaproteobacteria; Enterobacterales; Enterobacteriaceae; Escherichia MNNEETFYQAMRRQGVTRRSFLKSCSLAATSLGLGAGMAPKIAWALENKPRIPVVWIHGL WP_001058318.1 hydrogenase [Escherichia coli] Length: 372\nNumber of predicted TMHs:  1\nExp number of AAs in TMHs: 33.59637\nExp number, first 60 AAs:  15.81976\nTotal prob of N-in:   0.77394\nPOSSIBLE N-term signal sequence\noutside   1   325\nTMhelix   326   348\ninside   349   372

2782 GCF_001440525.1_ASM144052v1 Escherichia coli Proteobacteria; Gammaproteobacteria; Enterobacterales; Enterobacteriaceae; Escherichia MNNEETFYQAMRRQGVTRRSFLKYCSLAATSLGLGAGMAPKIAWALENKPRIPVVWIHGL WP_001058323.1 MULTISPECIES: hydrogenase-1 small chain [Proteobacteria] Length: 372\nNumber of predicted TMHs: 2\nExp number of AAs in TMHs: 37.2608499999999\nExp number, first 60 AAs:  19.52923\nTotal prob of N-in: 0.94351\nPOSSIBLE N-term signal sequence\ninside   1   19\nTMhelix   20   39\noutside   40   325\nTMhelix   326 348\ninside   349   372

2783 GCF_001445675.1_ASM144567v1 Escherichia coli Proteobacteria; Gammaproteobacteria; Enterobacterales; Enterobacteriaceae; Escherichia MTGDNTLIHSHGINRRDFMKLCAALAATMGLSSKAAAEMAESVTNPQRPPVIWIGAQECT WP_000145410.1 MULTISPECIES: hydrogenase-2 small chain [Proteobacteria] Length: 372\nNumber of predicted TMHs: 1\nExp number of AAs in TMHs: 23.72226\nExp number, first 60 AAs:  0.22038\nTotal prob of N-in:   0.12994\noutside   1 331\nTMhelix   332   354\ninside   355   372

2784 GCF_001446455.1_ASM144645v1 Escherichia coli Proteobacteria; Gammaproteobacteria; Enterobacterales; Enterobacteriaceae; Escherichia MTGDNTLIHSHGINRRDFMKLCAALAATMGLSSKAAAEMAESVTNPQRPPVIWIGAQECT WP_000145410.1 MULTISPECIES: hydrogenase-2 small chain [Proteobacteria] Length: 372\nNumber of predicted TMHs: 1\nExp number of AAs in TMHs: 23.72226\nExp number, first 60 AAs:  0.22038\nTotal prob of N-in:   0.12994\noutside   1 331\nTMhelix   332   354\ninside   355   372

2785 GCF_001455035.2_EC22593v3 Escherichia coli Proteobacteria; Gammaproteobacteria; Enterobacterales; Enterobacteriaceae; Escherichia MTGDNTLIHSHGINRRDFMKLCAALAATMGLSSKAAAEMAESVTNPQRPPVIWIGAQECT WP_000145410.1 MULTISPECIES: hydrogenase-2 small chain [Proteobacteria] Length: 372\nNumber of predicted TMHs: 1\nExp number of AAs in TMHs: 23.72226\nExp number, first 60 AAs:  0.22038\nTotal prob of N-in:   0.12994\noutside   1 331\nTMhelix   332   354\ninside   355   372

2786 GCF_001448025.1_ASM144802v1 Escherichia coli Proteobacteria; Gammaproteobacteria; Enterobacterales; Enterobacteriaceae; Escherichia MTGDNTLIHSHGINRRDFMKLCAALAATMGLSSKAAAEMAESVTNPQRPPVIWIGAQECT WP_000145410.1 MULTISPECIES: hydrogenase-2 small chain [Proteobacteria] Length: 372\nNumber of predicted TMHs: 1\nExp number of AAs in TMHs: 23.72226\nExp number, first 60 AAs:  0.22038\nTotal prob of N-in:   0.12994\noutside   1 331\nTMhelix   332   354\ninside   355   372

2787 GCF_001462975.1_ASM146297v1 Escherichia coli Proteobacteria; Gammaproteobacteria; Enterobacterales; Enterobacteriaceae; Escherichia MTGDNTLIHSHGINRRDFMKLCAALAATMGLSSKAAAEMAESVTNPQRPPVIWIGAQECT WP_000145410.1 MULTISPECIES: hydrogenase-2 small chain [Proteobacteria] Length: 372\nNumber of predicted TMHs: 1\nExp number of AAs in TMHs: 23.72226\nExp number, first 60 AAs:  0.22038\nTotal prob of N-in:   0.12994\noutside   1 331\nTMhelix   332   354\ninside   355   372

2788 GCF_001462715.1_ASM146271v1 Escherichia coli Proteobacteria; Gammaproteobacteria; Enterobacterales; Enterobacteriaceae; Escherichia
MTGDNTLIHSHGINRRDFMKLCAALAATMGLSSKAAAEMAESVTNPQRPPVIWIGAQECT WP_000145410.1
MULTISPECIES: hydrogenase-2 small chain [Proteobacteria] Length: 372\nNumber of predicted TMHs: 1\nExp number of AAs in TMHs: 23.72226\nExp number, first 60 AAs: 0.22038\nTotal prob of N-in: 0.12994\noutside 1 331\nTMhelix 332 354\ninside 355 372

2789 GCF_001445675.1_ASM144567v1 Escherichia coli Proteobacteria; Gammaproteobacteria; Enterobacterales; Enterobacteriaceae; Escherichia
MNNEETFYQAMRRQGVTRRSFLKSCSLAATSLGLGAGMAPKIAWALENKPRIPVVWIHGL WP_001058318.1
hydrogenase [Escherichia coli] Length: 372\nNumber of predicted TMHs: 1\nExp number of AAs in TMHs: 33.59637\nExp number, first 60 AAs: 15.81976\nTotal prob of N-in: 0.77394\nPOSSIBLE N-term signal sequence\noutside 1 325\nTMhelix 326 348\ninside 349 372

2790 GCF_001455035.2_EC22593v3 Escherichia coli Proteobacteria; Gammaproteobacteria; Enterobacterales; Enterobacteriaceae; Escherichia
MNNEETFYQAMRRQGVTRRSFLKYCSLAATSLGLGAGMAPKIAWALENKPRIPVVWIHGL WP_001058323.1
MULTISPECIES: hydrogenase-1 small chain [Proteobacteria] Length: 372\nNumber of predicted TMHs: 2\nExp number of AAs in TMHs: 37.2608499999999\nExp number, first 60 AAs: 19.52923\nTotal prob of N-in: 0.94351\nPOSSIBLE N-term signal sequence\ninside 1 19\nTMhelix 20 39\noutside 40 325\nTMhelix 326 348\ninside 349 372

2791 GCF_001463345.1_ASM146334v1 Escherichia coli Proteobacteria; Gammaproteobacteria; Enterobacterales; Enterobacteriaceae; Escherichia
MTGDNTLIHSHGINRRDFMKLCAALAATMGLSSKAAAEMAESVTNPQRPPVIWIGAQECT WP_000145410.1
MULTISPECIES: hydrogenase-2 small chain [Proteobacteria] Length: 372\nNumber of predicted TMHs: 1\nExp number of AAs in TMHs: 23.72226\nExp number, first 60 AAs: 0.22038\nTotal prob of N-in: 0.12994\noutside 1 331\nTMhelix 332 354\ninside 355 372

2792 GCF_001463345.1_ASM146334v1 Escherichia coli Proteobacteria; Gammaproteobacteria; Enterobacterales; Enterobacteriaceae; Escherichia
MNNEETFYQAMRRQGVTRRSFLKYCSLAATSLGLGAGMAPKIAWALENKPRIPVVWIHGL WP_001058323.1
MULTISPECIES: hydrogenase-1 small chain [Proteobacteria] Length: 372\nNumber of predicted TMHs: 2\nExp number of AAs in TMHs: 37.2608499999999\nExp number, first 60 AAs: 19.52923\nTotal prob of N-in: 0.94351\nPOSSIBLE N-term signal sequence\ninside 1 19\nTMhelix 20 39\noutside 40 325\nTMhelix 326 348\ninside 349 372

2793 GCF_001462715.1_ASM146271v1 Escherichia coli Proteobacteria; Gammaproteobacteria; Enterobacterales; Enterobacteriaceae; Escherichia
MNNEETFYQAMRRQGVTRRSFLKYCSLAATSLGLGAGMAPKIAWALENKPRIPVVWIHGL WP_001058320.1
hydrogenase [Escherichia coli] Length: 372\nNumber of predicted TMHs: 2\nExp number of AAs in TMHs: 37.26082\nExp number, first 60 AAs: 19.52923\nTotal prob of N-in: 0.94351\nPOSSIBLE N-term signal sequence\ninside 1 19\nTMhelix 20 39\noutside 40 325\nTMhelix 326 348\ninside 349 372

2794 GCF_001463385.1_ASM146338v1 Escherichia coli Proteobacteria; Gammaproteobacteria; Enterobacterales; Enterobacteriaceae; Escherichia
MTGDNTLIHSHGINRRDFMKLCAALAATMGLSSKAAAEMAESVTNPQRPPVIWIGAQECT WP_000145410.1
MULTISPECIES: hydrogenase-2 small chain [Proteobacteria] Length: 372\nNumber of predicted TMHs: 1\nExp number of AAs in TMHs: 23.72226\nExp number, first 60 AAs: 0.22038\nTotal prob of N-in: 0.12994\noutside 1 331\nTMhelix 332 354\ninside 355 372

2795 GCF_001463385.1_ASM146338v1 Escherichia coli Proteobacteria; Gammaproteobacteria; Enterobacterales; Enterobacteriaceae; Escherichia
MNNEETFYQAMRRQGVTRRSFLKYCSLAATSLGLGAGMAPKIAWALENKPRIPVVWIHGL WP_001058320.1
hydrogenase [Escherichia coli] Length: 372\nNumber of predicted TMHs: 2\nExp number of AAs in TMHs: 37.26082\nExp number, first 60 AAs: 19.52923\nTotal prob of N-in: 0.94351\nPOSSIBLE N-term signal sequence\ninside 1 19\nTMhelix 20 39\noutside 40 325\nTMhelix 326 348\ninside 349 372

2796 GCF_001446455.1_ASM144645v1 Escherichia coli Proteobacteria; Gammaproteobacteria; Enterobacterales; Enterobacteriaceae; Escherichia
MNNEETFYQAMRRQGVTRRSFLKYCSLAATSLGLGAGMAPKIAWALENKPRIPVVWIHGL WP_058055433.1

hydrogenase [Escherichia coli] Length: 372\nNumber of predicted TMHs: 2\nExp number of AAs in TMHs: 36.72011\nExp number, first 60 AAs: 19.53447\nTotal prob of N-in: 0.94308\nPOSSIBLE N-term signal sequence\ninside 1 19\nTMhelix 20 39\noutside 40 325\nTMhelix 326 348\ninside 349 372

2797 GCF_001484295.1_ASM148429v1 Escherichia coli Proteobacteria; Gammaproteobacteria; Enterobacterales; Enterobacteriaceae; Escherichia
MTGDNTLIHSHGINRRDFMKLCAALAATMGLSSKAAAEMAESVTNPQRPPVIWIGAQECT WP_000145410.1
MULTISPECIES: hydrogenase-2 small chain [Proteobacteria] Length: 372\nNumber of predicted TMHs: 1\nExp number of AAs in TMHs: 23.72226\nExp number, first 60 AAs: 0.22038\nTotal prob of N-in: 0.12994\noutside 1 331\nTMhelix 332 354\ninside 355 372

2798 GCF_001484255.1_ASM148425v1 Escherichia coli Proteobacteria; Gammaproteobacteria; Enterobacterales; Enterobacteriaceae; Escherichia
MTGDNTLIHSHGINRRDFMKLCAALAATMGLSSKAAAEMAESVTNPQRPPVIWIGAQECT WP_000145410.1
MULTISPECIES: hydrogenase-2 small chain [Proteobacteria] Length: 372\nNumber of predicted TMHs: 1\nExp number of AAs in TMHs: 23.72226\nExp number, first 60 AAs: 0.22038\nTotal prob of N-in: 0.12994\noutside 1 331\nTMhelix 332 354\ninside 355 372

2799 GCF_001484305.1_ASM148430v1 Escherichia coli Proteobacteria; Gammaproteobacteria; Enterobacterales; Enterobacteriaceae; Escherichia
MTGDNTLIHSHGINRRDFMKLCAALAATMGLSSKAAAEMAESVTNPQRPPVIWIGAQECT WP_000145410.1
MULTISPECIES: hydrogenase-2 small chain [Proteobacteria] Length: 372\nNumber of predicted TMHs: 1\nExp number of AAs in TMHs: 23.72226\nExp number, first 60 AAs: 0.22038\nTotal prob of N-in: 0.12994\noutside 1 331\nTMhelix 332 354\ninside 355 372

2800 GCF_001484295.1_ASM148429v1 Escherichia coli Proteobacteria; Gammaproteobacteria; Enterobacterales; Enterobacteriaceae; Escherichia
MNNEETFYQAMRRQGVTRRSFLKYCSLAATSLGLGAGMAPKIAWALENKPRIPVVWIHGL WP_001058323.1
MULTISPECIES: hydrogenase-1 small chain [Proteobacteria] Length: 372\nNumber of predicted TMHs: 2\nExp number of AAs in TMHs: 37.2608499999999\nExp number, first 60 AAs: 19.52923\nTotal prob of N-in: 0.94351\nPOSSIBLE N-term signal sequence\ninside 1 19\nTMhelix 20 39\noutside 40 325\nTMhelix 326 348\ninside 349 372

2801 GCF_001484255.1_ASM148425v1 Escherichia coli Proteobacteria; Gammaproteobacteria; Enterobacterales; Enterobacteriaceae; Escherichia
MNNEETFYQAMRRQGVTRRSFLKYCSLAATSLGLGAGMAPKIAWALENKPRIPVVWIHGL WP_001058320.1
hydrogenase [Escherichia coli] Length: 372\nNumber of predicted TMHs: 2\nExp number of AAs in TMHs: 37.26082\nExp number, first 60 AAs: 19.52923\nTotal prob of N-in: 0.94351\nPOSSIBLE N-term signal sequence\ninside 1 19\nTMhelix 20 39\noutside 40 325\nTMhelix 326 348\ninside 349 372

2802 GCF_001484305.1_ASM148430v1 Escherichia coli Proteobacteria; Gammaproteobacteria; Enterobacterales; Enterobacteriaceae; Escherichia
MNNEETFYQAMRRQGVTRRSFLKYCSLAATSLGLGAGMAPKIAWALENKPRIPVVWIHGL WP_001058323.1
MULTISPECIES: hydrogenase-1 small chain [Proteobacteria] Length: 372\nNumber of predicted TMHs: 2\nExp number of AAs in TMHs: 37.2608499999999\nExp number, first 60 AAs: 19.52923\nTotal prob of N-in: 0.94351\nPOSSIBLE N-term signal sequence\ninside 1 19\nTMhelix 20 39\noutside 40 325\nTMhelix 326 348\ninside 349 372

2803 GCF_001463455.1_ASM146345v1 Escherichia coli Proteobacteria; Gammaproteobacteria; Enterobacterales; Enterobacteriaceae; Escherichia
MNNEETFYQAMRRQGVTRRSFLKYCSLAATSLGLGAGMAPKIAWALENKPRIPVVWIHGL WP_001058323.1
MULTISPECIES: hydrogenase-1 small chain [Proteobacteria] Length: 372\nNumber of predicted TMHs: 2\nExp number of AAs in TMHs: 37.2608499999999\nExp number, first 60 AAs: 19.52923\nTotal prob of N-in: 0.94351\nPOSSIBLE N-term signal sequence\ninside 1 19\nTMhelix 20 39\noutside 40 325\nTMhelix 326 348\ninside 349 372

2804 GCF_001484405.1_ASM148440v1 Escherichia coli Proteobacteria; Gammaproteobacteria; Enterobacterales; Enterobacteriaceae; Escherichia
MNNEETFYQAMRRQGVTRRSFLKYCSLAATSLGLGAGMAPKIAWALENKPRIPVVWIHGL WP_001058323.1
MULTISPECIES: hydrogenase-1 small chain [Proteobacteria] Length: 372\nNumber of predicted TMHs: 2\nExp number of AAs in TMHs: 37.2608499999999\nExp number, first 60 AAs: 19.52923\nTotal prob of N-in:

0.94351\nPOSSIBLE N-term signal sequence\ninside 1 19\nTMhelix 20 39\noutside 40 325\nTMhelix 326 348\ninside 349 372

2805 GCF_001484395.1_ASM148439v1 Escherichia coli Proteobacteria; Gammaproteobacteria; Enterobacterales; Enterobacteriaceae; Escherichia MNNEETFYQAMRRQGVTRRSFLKYCSLAATSLGLGAGMAPKIAWALENKPRIPVVWIHGL WP_001058323.1 MULTISPECIES: hydrogenase-1 small chain [Proteobacteria] Length: 372\nNumber of predicted TMHs: 2\nExp number of AAs in TMHs: 37.2608499999999\nExp number, first 60 AAs: 19.52923\nTotal prob of N-in: 0.94351\nPOSSIBLE N-term signal sequence\ninside 1 19\nTMhelix 20 39\noutside 40 325\nTMhelix 326 348\ninside 349 372

2806 GCF_001469815.1_ASM146981v1 Escherichia coli Proteobacteria; Gammaproteobacteria; Enterobacterales; Enterobacteriaceae; Escherichia MNNEETFYQAMRRQGVTRRSFLKYCSLAATSLGLGAGMAPKIAWALENKPRIPVVWIHGL WP_001295940.1 MULTISPECIES: hydrogenase [Enterobacteriaceae] Length: 372\nNumber of predicted TMHs: 2\nExp number of AAs in TMHs: 37.2608899999999\nExp number, first 60 AAs: 19.52923\nTotal prob of N-in: 0.94351\nPOSSIBLE N-term signal sequence\ninside 1 19\nTMhelix 20 39\noutside 40 325\nTMhelix 326 348\ninside 349 372

2807 GCF_001484455.1_ASM148445v1 Escherichia coli Proteobacteria; Gammaproteobacteria; Enterobacterales; Enterobacteriaceae; Escherichia MTGDNTLIHSHGINRRDFMKLCAALAATMGLSSKAAAEMAESVTNPQRPPVIWIGAQECT WP_000145410.1 MULTISPECIES: hydrogenase-2 small chain [Proteobacteria] Length: 372\nNumber of predicted TMHs: 1\nExp number of AAs in TMHs: 23.72226\nExp number, first 60 AAs: 0.22038\nTotal prob of N-in: 0.12994\noutside 1 331\nTMhelix 332 354\ninside 355 372

2808 GCF_001484485.1_ASM148448v1 Escherichia coli Proteobacteria; Gammaproteobacteria; Enterobacterales; Enterobacteriaceae; Escherichia MTGDNTLIHSHGINRRDFMKLCAALAATMGLSSKAAAEMAESVTNPQRPPVIWIGAQECT WP_000145410.1 MULTISPECIES: hydrogenase-2 small chain [Proteobacteria] Length: 372\nNumber of predicted TMHs: 1\nExp number of AAs in TMHs: 23.72226\nExp number, first 60 AAs: 0.22038\nTotal prob of N-in: 0.12994\noutside 1 331\nTMhelix 332 354\ninside 355 372

2809 GCF_001509655.1_ASM150965v1 Escherichia coli Proteobacteria; Gammaproteobacteria; Enterobacterales; Enterobacteriaceae; Escherichia MTGDNTLIHSHGINRRDFMKLCAALAATMGLSSKAAAEMAESVTNPQRPPVIWIGAQECT WP_000145410.1 MULTISPECIES: hydrogenase-2 small chain [Proteobacteria] Length: 372\nNumber of predicted TMHs: 1\nExp number of AAs in TMHs: 23.72226\nExp number, first 60 AAs: 0.22038\nTotal prob of N-in: 0.12994\noutside 1 331\nTMhelix 332 354\ninside 355 372

2810 GCF_001509715.1_ASM150971v1 Escherichia coli Proteobacteria; Gammaproteobacteria; Enterobacterales; Enterobacteriaceae; Escherichia MTGDNTLIHSHGINRRDFMKLCAALAATMGLSSKAAAEMAESVTNPQRPPVIWIGAQECT WP_000145410.1 MULTISPECIES: hydrogenase-2 small chain [Proteobacteria] Length: 372\nNumber of predicted TMHs: 1\nExp number of AAs in TMHs: 23.72226\nExp number, first 60 AAs: 0.22038\nTotal prob of N-in: 0.12994\noutside 1 331\nTMhelix 332 354\ninside 355 372

2811 GCF_001515725.1_ASM151572v1 Escherichia coli Proteobacteria; Gammaproteobacteria; Enterobacterales; Enterobacteriaceae; Escherichia MTGDNTLIHSHGINRRDFMKLCAALAATMGLSSKAAAEMAESVTNPQRPPVIWIGAQECT WP_000145410.1 MULTISPECIES: hydrogenase-2 small chain [Proteobacteria] Length: 372\nNumber of predicted TMHs: 1\nExp number of AAs in TMHs: 23.72226\nExp number, first 60 AAs: 0.22038\nTotal prob of N-in: 0.12994\noutside 1 331\nTMhelix 332 354\ninside 355 372

2812 GCF_001515725.1_ASM151572v1 Escherichia coli Proteobacteria; Gammaproteobacteria; Enterobacterales; Enterobacteriaceae; Escherichia MNNEETFYQAMRRQGVTRRSFLKYCSLAATSLGLGAGMAPKIAWALENKPRIPVVWIHGL WP_001058320.1 hydrogenase [Escherichia coli] Length: 372\nNumber of predicted TMHs: 2\nExp number of AAs in TMHs: 37.26082\nExp number, first 60 AAs: 19.52923\nTotal prob of N-in: 0.94351\nPOSSIBLE N-term signal sequence\ninside 1 19\nTMhelix 20 39\noutside 40 325\nTMhelix 326 348\ninside 349 372

2813 GCF_001513635.1_ASM151363v1 Escherichia coli Proteobacteria; Gammaproteobacteria; Enterobacterales; Enterobacteriaceae; Escherichia
MNNEETFYQAMRRQGVTRRSFLKYCSLAATSLGLGAGMAPKIAWALENKPRIPVVWIHGL WP_001295940.1
MULTISPECIES: hydrogenase [Enterobacteriaceae] Length: 372\nNumber of predicted TMHs: 2\nExp number of AAs in TMHs: 37.2608899999999\nExp number, first 60 AAs: 19.52923\nTotal prob of N-in: 0.94351\nPOSSIBLE N-term signal sequence\ninside 1 19\nTMhelix 20 39\noutside 40 325\nTMhelix 326 348\ninside 349 372

2814 GCF_001517685.1_NG-7574_Bell_ERRESI Escherichia coli Proteobacteria; Gammaproteobacteria; Enterobacterales; Enterobacteriaceae; Escherichia
MNNEETFYQAMRRQGVTRRSFLKYCSLAATSLGLGAGMAPKIAWALENKPRIPVVWIHGL WP_001295940.1
MULTISPECIES: hydrogenase [Enterobacteriaceae] Length: 372\nNumber of predicted TMHs: 2\nExp number of AAs in TMHs: 37.2608899999999\nExp number, first 60 AAs: 19.52923\nTotal prob of N-in: 0.94351\nPOSSIBLE N-term signal sequence\ninside 1 19\nTMhelix 20 39\noutside 40 325\nTMhelix 326 348\ninside 349 372

2815 GCF_001509655.1_ASM150965v1 Escherichia coli Proteobacteria; Gammaproteobacteria; Enterobacterales; Enterobacteriaceae; Escherichia
MNNEETFYQAMRRQGVTRRSFLKYCSLAATSLGLGAGMAPKIAWALENKPRIPVVWIHGL WP_001058323.1
MULTISPECIES: hydrogenase-1 small chain [Proteobacteria] Length: 372\nNumber of predicted TMHs: 2\nExp number of AAs in TMHs: 37.2608499999999\nExp number, first 60 AAs: 19.52923\nTotal prob of N-in: 0.94351\nPOSSIBLE N-term signal sequence\ninside 1 19\nTMhelix 20 39\noutside 40 325\nTMhelix 326 348\ninside 349 372

2816 GCF_001509715.1_ASM150971v1 Escherichia coli Proteobacteria; Gammaproteobacteria; Enterobacterales; Enterobacteriaceae; Escherichia
MNNEETFYQAMRRQGVTRRSFLKYCSLAATSLGLGAGMAPKIAWALENKPRIPVVWIHGL WP_001058323.1
MULTISPECIES: hydrogenase-1 small chain [Proteobacteria] Length: 372\nNumber of predicted TMHs: 2\nExp number of AAs in TMHs: 37.2608499999999\nExp number, first 60 AAs: 19.52923\nTotal prob of N-in: 0.94351\nPOSSIBLE N-term signal sequence\ninside 1 19\nTMhelix 20 39\noutside 40 325\nTMhelix 326 348\ninside 349 372

2817 GCF_001509675.1_ASM150967v1 Escherichia coli Proteobacteria; Gammaproteobacteria; Enterobacterales; Enterobacteriaceae; Escherichia
MNNEETFYQAMRRQGVTRRSFLKYCSLAATSLGLGAGMAPKIAWALENKPRIPVVWIHGL WP_001058323.1
MULTISPECIES: hydrogenase-1 small chain [Proteobacteria] Length: 372\nNumber of predicted TMHs: 2\nExp number of AAs in TMHs: 37.2608499999999\nExp number, first 60 AAs: 19.52923\nTotal prob of N-in: 0.94351\nPOSSIBLE N-term signal sequence\ninside 1 19\nTMhelix 20 39\noutside 40 325\nTMhelix 326 348\ninside 349 372

2818 GCF_001519315.1_ASM151931v1 Escherichia coli Proteobacteria; Gammaproteobacteria; Enterobacterales; Enterobacteriaceae; Escherichia
MTGDNTLIHSHGINRRDFMKLCAALAATMGLSSKAAAEMAESVTNPQRPPVIWIGAQECT WP_000145410.1
MULTISPECIES: hydrogenase-2 small chain [Proteobacteria] Length: 372\nNumber of predicted TMHs: 1\nExp number of AAs in TMHs: 23.72226\nExp number, first 60 AAs: 0.22038\nTotal prob of N-in: 0.12994\noutside 1 331\nTMhelix 332 354\ninside 355 372

2819 GCF_001519285.1_ASM151928v1 Escherichia coli Proteobacteria; Gammaproteobacteria; Enterobacterales; Enterobacteriaceae; Escherichia
MTGDNTLIHSHGINRRDFMKLCAALAATMGLSSKAAAEMAESVTNPQRPPVIWIGAQECT WP_000145410.1
MULTISPECIES: hydrogenase-2 small chain [Proteobacteria] Length: 372\nNumber of predicted TMHs: 1\nExp number of AAs in TMHs: 23.72226\nExp number, first 60 AAs: 0.22038\nTotal prob of N-in: 0.12994\noutside 1 331\nTMhelix 332 354\ninside 355 372

2820 GCF_001519355.1_ASM151935v1 Escherichia coli Proteobacteria; Gammaproteobacteria; Enterobacterales; Enterobacteriaceae; Escherichia
MTGDNTLIHSHGINRRDFMKLCAALAATMGLSSKAAAEMAESVTNPQRPPVIWIGAQECT WP_000145410.1
MULTISPECIES: hydrogenase-2 small chain [Proteobacteria] Length: 372\nNumber of predicted TMHs: 1\nExp number of AAs in TMHs: 23.72226\nExp number, first 60 AAs: 0.22038\nTotal prob of N-in: 0.12994\noutside 1 331\nTMhelix 332 354\ninside 355 372

2821 GCF_001519315.1_ASM151931v1 Escherichia coli Proteobacteria; Gammaproteobacteria; Enterobacterales; Enterobacteriaceae; Escherichia
MNNEETFYQAMRRQGVTRRSFLKYCSLAATSLGLGAGMAPKIAWALENKPRIPVVWIHGL WP_001058323.1
MULTISPECIES: hydrogenase-1 small chain [Proteobacteria] Length: 372\nNumber of predicted TMHs: 2\nExp number of AAs in TMHs: 37.2608499999999\nExp number, first 60 AAs: 19.52923\nTotal prob of N-in: 0.94351\nPOSSIBLE N-term signal sequence\ninside 1 19\nTMhelix 20 39\noutside 40 325\nTMhelix 326 348\ninside 349 372

2822 GCF_001519355.1_ASM151935v1 Escherichia coli Proteobacteria; Gammaproteobacteria; Enterobacterales; Enterobacteriaceae; Escherichia
MNNEETFYQAMRRQGVTRRSFLKYCSLAATSLGLGAGMAPKIAWALENKPRIPVVWIHGL WP_001058323.1
MULTISPECIES: hydrogenase-1 small chain [Proteobacteria] Length: 372\nNumber of predicted TMHs: 2\nExp number of AAs in TMHs: 37.2608499999999\nExp number, first 60 AAs: 19.52923\nTotal prob of N-in: 0.94351\nPOSSIBLE N-term signal sequence\ninside 1 19\nTMhelix 20 39\noutside 40 325\nTMhelix 326 348\ninside 349 372

2823 GCF_001519245.1_ASM151924v1 Escherichia coli Proteobacteria; Gammaproteobacteria; Enterobacterales; Enterobacteriaceae; Escherichia
MNNEETFYQAMRRQGVTRRSFLKYCSLAATSLGLGAGMAPKIAWALENKPRIPVVWIHGL WP_001058323.1
MULTISPECIES: hydrogenase-1 small chain [Proteobacteria] Length: 372\nNumber of predicted TMHs: 2\nExp number of AAs in TMHs: 37.2608499999999\nExp number, first 60 AAs: 19.52923\nTotal prob of N-in: 0.94351\nPOSSIBLE N-term signal sequence\ninside 1 19\nTMhelix 20 39\noutside 40 325\nTMhelix 326 348\ninside 349 372

2824 GCF_001519195.1_ASM151919v1 Escherichia coli Proteobacteria; Gammaproteobacteria; Enterobacterales; Enterobacteriaceae; Escherichia
MNNEETFYQAMRRQGVTRRSFLKYCSLAATSLGLGAGMAPKIAWALENKPRIPVVWIHGL WP_001058323.1
MULTISPECIES: hydrogenase-1 small chain [Proteobacteria] Length: 372\nNumber of predicted TMHs: 2\nExp number of AAs in TMHs: 37.2608499999999\nExp number, first 60 AAs: 19.52923\nTotal prob of N-in: 0.94351\nPOSSIBLE N-term signal sequence\ninside 1 19\nTMhelix 20 39\noutside 40 325\nTMhelix 326 348\ninside 349 372

2825 GCF_001519515.1_ASM151951v1 Escherichia coli Proteobacteria; Gammaproteobacteria; Enterobacterales; Enterobacteriaceae; Escherichia
MTGDNTLIHSHGINRRDFMKLCAALAATMGLSSKAAAEMAESVTNPQRPPVIWIGAQECT WP_000145410.1
MULTISPECIES: hydrogenase-2 small chain [Proteobacteria] Length: 372\nNumber of predicted TMHs: 1\nExp number of AAs in TMHs: 23.72226\nExp number, first 60 AAs: 0.22038\nTotal prob of N-in: 0.12994\noutside 1 331\nTMhelix 332 354\ninside 355 372

2826 GCF_001519555.1_ASM151955v1 Escherichia coli Proteobacteria; Gammaproteobacteria; Enterobacterales; Enterobacteriaceae; Escherichia
MTGDNTLIHSHGINRRDFMKLCAALAATMGLSSKAAAEMAESVTNPQRPPVIWIGAQECT WP_000145410.1
MULTISPECIES: hydrogenase-2 small chain [Proteobacteria] Length: 372\nNumber of predicted TMHs: 1\nExp number of AAs in TMHs: 23.72226\nExp number, first 60 AAs: 0.22038\nTotal prob of N-in: 0.12994\noutside 1 331\nTMhelix 332 354\ninside 355 372

2827 GCF_001519515.1_ASM151951v1 Escherichia coli Proteobacteria; Gammaproteobacteria; Enterobacterales; Enterobacteriaceae; Escherichia
MNNEETFYQAMRRQGVTRRSFLKYCSLAATSLGLGAGMAPKIAWALENKPRIPVVWIHGL WP_001058323.1
MULTISPECIES: hydrogenase-1 small chain [Proteobacteria] Length: 372\nNumber of predicted TMHs: 2\nExp number of AAs in TMHs: 37.2608499999999\nExp number, first 60 AAs: 19.52923\nTotal prob of N-in: 0.94351\nPOSSIBLE N-term signal sequence\ninside 1 19\nTMhelix 20 39\noutside 40 325\nTMhelix 326 348\ninside 349 372

2828 GCF_001519445.1_ASM151944v1 Escherichia coli Proteobacteria; Gammaproteobacteria; Enterobacterales; Enterobacteriaceae; Escherichia
MNNEETFYQAMRRQGVTRRSFLKYCSLAATSLGLGAGMAPKIAWALENKPRIPVVWIHGL WP_001058323.1
MULTISPECIES: hydrogenase-1 small chain [Proteobacteria] Length: 372\nNumber of predicted TMHs: 2\nExp number of AAs in TMHs: 37.2608499999999\nExp number, first 60 AAs: 19.52923\nTotal prob of N-in: 0.94351\nPOSSIBLE N-term signal sequence\ninside 1 19\nTMhelix 20 39\noutside 40 325\nTMhelix 326 348\ninside 349 372

2829 GCF_001519485.1_ASM151948v1 Escherichia coli Proteobacteria; Gammaproteobacteria; Enterobacterales; Enterobacteriaceae; Escherichia
MNNEETFYQAMRRQGVTRRSFLKYCSLAATSLGLGAGMAPKIAWALENKPRIPVVWIHGL WP_001058323.1
MULTISPECIES: hydrogenase-1 small chain [Proteobacteria] Length: 372\nNumber of predicted TMHs: 2\nExp number of AAs in TMHs: 37.2608499999999\nExp number, first 60 AAs: 19.52923\nTotal prob of N-in: 0.94351\nPOSSIBLE N-term signal sequence\ninside 1 19\nTMhelix 20 39\noutside 40 325\nTMhelix 326 348\ninside 349 372

2830 GCF_001519595.1_ASM151959v1 Escherichia coli Proteobacteria; Gammaproteobacteria; Enterobacterales; Enterobacteriaceae; Escherichia
MTGDNTLIHSHGINRRDFMKLCAALAATMGLSSKAAAEMAESVTNPQRPPVIWIGAQECT WP_000145410.1
MULTISPECIES: hydrogenase-2 small chain [Proteobacteria] Length: 372\nNumber of predicted TMHs: 1\nExp number of AAs in TMHs: 23.72226\nExp number, first 60 AAs: 0.22038\nTotal prob of N-in: 0.12994\noutside 1 331\nTMhelix 332 354\ninside 355 372

2831 GCF_001519645.1_ASM151964v1 Escherichia coli Proteobacteria; Gammaproteobacteria; Enterobacterales; Enterobacteriaceae; Escherichia
MNNEETFYQAMRRQGVTRRSFLKYCSLAATSLGLGAGMAPKIAWALENKPRIPVVWIHGL WP_001058323.1
MULTISPECIES: hydrogenase-1 small chain [Proteobacteria] Length: 372\nNumber of predicted TMHs: 2\nExp number of AAs in TMHs: 37.2608499999999\nExp number, first 60 AAs: 19.52923\nTotal prob of N-in: 0.94351\nPOSSIBLE N-term signal sequence\ninside 1 19\nTMhelix 20 39\noutside 40 325\nTMhelix 326 348\ninside 349 372

2832 GCF_001519855.1_ASM151985v1 Escherichia coli Proteobacteria; Gammaproteobacteria; Enterobacterales; Enterobacteriaceae; Escherichia
MTGDNTLIHSHGINRRDFMKLCAALAATMGLSSKAAAEMAESVTNPQRPPVIWIGAQECT WP_000145418.1
hydrogenase 2 small subunit [Escherichia coli] Length: 372\nNumber of predicted TMHs: 1\nExp number of AAs in TMHs: 23.72313\nExp number, first 60 AAs: 0.22034\nTotal prob of N-in: 0.12997\noutside 1 331\nTMhelix 332 354\ninside 355 372

2833 GCF_001519795.1_ASM151979v1 Escherichia coli Proteobacteria; Gammaproteobacteria; Enterobacterales; Enterobacteriaceae; Escherichia
MTGDNTLIHSHGINRRDFMKLCAALAATMGLSSKAAAEMAESVTNPQRPPVIWIGAQECT WP_000145410.1
MULTISPECIES: hydrogenase-2 small chain [Proteobacteria] Length: 372\nNumber of predicted TMHs: 1\nExp number of AAs in TMHs: 23.72226\nExp number, first 60 AAs: 0.22038\nTotal prob of N-in: 0.12994\noutside 1 331\nTMhelix 332 354\ninside 355 372

2834 GCF_001519835.1_ASM151983v1 Escherichia coli Proteobacteria; Gammaproteobacteria; Enterobacterales; Enterobacteriaceae; Escherichia
MTGDNTLIHSHGINRRDFMKLCAALAATMGLSSKAAAEMAESVTNPQRPPVIWIGAQECT WP_000145410.1
MULTISPECIES: hydrogenase-2 small chain [Proteobacteria] Length: 372\nNumber of predicted TMHs: 1\nExp number of AAs in TMHs: 23.72226\nExp number, first 60 AAs: 0.22038\nTotal prob of N-in: 0.12994\noutside 1 331\nTMhelix 332 354\ninside 355 372

2835 GCF_001519835.1_ASM151983v1 Escherichia coli Proteobacteria; Gammaproteobacteria; Enterobacterales; Enterobacteriaceae; Escherichia
MNNEETFYQAMRRQGVTRRSFLKYCSLAATSLGLGAGMAPKIAWALENKPRIPVVWIHGL WP_001058323.1
MULTISPECIES: hydrogenase-1 small chain [Proteobacteria] Length: 372\nNumber of predicted TMHs: 2\nExp number of AAs in TMHs: 37.2608499999999\nExp number, first 60 AAs: 19.52923\nTotal prob of N-in: 0.94351\nPOSSIBLE N-term signal sequence\ninside 1 19\nTMhelix 20 39\noutside 40 325\nTMhelix 326 348\ninside 349 372

2836 GCF_001519795.1_ASM151979v1 Escherichia coli Proteobacteria; Gammaproteobacteria; Enterobacterales; Enterobacteriaceae; Escherichia
MNNEETFYQAMRRQGVTRRSFLKYCSLAATSLGLGAGMAPKIAWALENKPRIPVVWIHGL WP_001058323.1
MULTISPECIES: hydrogenase-1 small chain [Proteobacteria] Length: 372\nNumber of predicted TMHs: 2\nExp number of AAs in TMHs: 37.2608499999999\nExp number, first 60 AAs: 19.52923\nTotal prob of N-in: 0.94351\nPOSSIBLE N-term signal sequence\ninside 1 19\nTMhelix 20 39\noutside 40 325\nTMhelix 326 348\ninside 349 372

2837 GCF_001519855.1_ASM151985v1 Escherichia coli Proteobacteria; Gammaproteobacteria; Enterobacterales; Enterobacteriaceae; Escherichia MNNEETFYQAMRRQGVTRRSFLKYCSLAATSLGLGAGMAPKIAWALENKPRIPVVWIHGL WP_021520630.1 hydrogenase-1 small chain [Escherichia coli] Length: 372\nNumber of predicted TMHs: 2\nExp number of AAs in TMHs: 37.8197599999999\nExp number, first 60 AAs: 19.53295\nTotal prob of N-in: 0.94396\nPOSSIBLE N-term signal sequence\ninside 1 19\nTMhelix 20 39\noutside 40 325\nTMhelix 326 348\ninside 349 372

2838 GCF_001519945.1_ASM151994v1 Escherichia coli Proteobacteria; Gammaproteobacteria; Enterobacterales; Enterobacteriaceae; Escherichia MNNEETFYQAMRRQGVTRRSFLKYCSLAATSLGLGAGMAPKIAWALENKPRIPVVWIHGL WP_001058323.1 MULTISPECIES: hydrogenase-1 small chain [Proteobacteria] Length: 372\nNumber of predicted TMHs: 2\nExp number of AAs in TMHs: 37.2608499999999\nExp number, first 60 AAs: 19.52923\nTotal prob of N-in: 0.94351\nPOSSIBLE N-term signal sequence\ninside 1 19\nTMhelix 20 39\noutside 40 325\nTMhelix 326 348\ninside 349 372

2839 GCF_001520095.1_ASM152009v1 Escherichia coli Proteobacteria; Gammaproteobacteria; Enterobacterales; Enterobacteriaceae; Escherichia MTGDNTLIHSHGINRRDFMKLCAALAATMGLSSKAAAEMVESVTNPQRPPVIWIGAQECT WP_000145423.1 hydrogenase 2 small subunit [Escherichia coli] Length: 372\nNumber of predicted TMHs: 1\nExp number of AAs in TMHs: 23.78201\nExp number, first 60 AAs: 0.28355\nTotal prob of N-in: 0.13257\noutside 1 331\nTMhelix 332 354\ninside 355 372

2840 GCF_001520135.1_ASM152013v1 Escherichia coli Proteobacteria; Gammaproteobacteria; Enterobacterales; Enterobacteriaceae; Escherichia MTGDNTLIHSHGINRRDFMKLCAALAATMGLSSKAAAEMAESVTNPQRPPVIWIGAQECT WP_000145410.1 MULTISPECIES: hydrogenase-2 small chain [Proteobacteria] Length: 372\nNumber of predicted TMHs: 1\nExp number of AAs in TMHs: 23.72226\nExp number, first 60 AAs: 0.22038\nTotal prob of N-in: 0.12994\noutside 1 331\nTMhelix 332 354\ninside 355 372

2841 GCF_001520095.1_ASM152009v1 Escherichia coli Proteobacteria; Gammaproteobacteria; Enterobacterales; Enterobacteriaceae; Escherichia MNNEETFYQAMRRQGVTRRSFLKYCSLAATSLGLGAGMAPKIAWALENKPRIPVVWIHGL WP_001295940.1 MULTISPECIES: hydrogenase [Enterobacteriaceae] Length: 372\nNumber of predicted TMHs: 2\nExp number of AAs in TMHs: 37.2608899999999\nExp number, first 60 AAs: 19.52923\nTotal prob of N-in: 0.94351\nPOSSIBLE N-term signal sequence\ninside 1 19\nTMhelix 20 39\noutside 40 325\nTMhelix 326 348\ninside 349 372

2842 GCF_001520135.1_ASM152013v1 Escherichia coli Proteobacteria; Gammaproteobacteria; Enterobacterales; Enterobacteriaceae; Escherichia MNNEETFYQAMRRQGVTRRSFLKYCSLAATSLGLGAGMAPKIAWALENKPRIPVVWIHGL WP_001295940.1 MULTISPECIES: hydrogenase [Enterobacteriaceae] Length: 372\nNumber of predicted TMHs: 2\nExp number of AAs in TMHs: 37.2608899999999\nExp number, first 60 AAs: 19.52923\nTotal prob of N-in: 0.94351\nPOSSIBLE N-term signal sequence\ninside 1 19\nTMhelix 20 39\noutside 40 325\nTMhelix 326 348\ninside 349 372

2843 GCF_001519935.1_ASM151993v1 Escherichia coli Proteobacteria; Gammaproteobacteria; Enterobacterales; Enterobacteriaceae; Escherichia MNNEETFYQAMRRQGVTRRSFLKYCSLAATSLGLGAGMAPKIAWALENKPRIPVVWIHGL WP_001295940.1 MULTISPECIES: hydrogenase [Enterobacteriaceae] Length: 372\nNumber of predicted TMHs: 2\nExp number of AAs in TMHs: 37.2608899999999\nExp number, first 60 AAs: 19.52923\nTotal prob of N-in: 0.94351\nPOSSIBLE N-term signal sequence\ninside 1 19\nTMhelix 20 39\noutside 40 325\nTMhelix 326 348\ninside 349 372

2844 GCF_001520055.1_ASM152005v1 Escherichia coli Proteobacteria; Gammaproteobacteria; Enterobacterales; Enterobacteriaceae; Escherichia MNNEETFYQAMRRQGVTRRSFLKYCSLAATSLGLGAGMAPKIAWALENKPRIPVVWIHGL WP_001058323.1 MULTISPECIES: hydrogenase-1 small chain [Proteobacteria] Length: 372\nNumber of predicted TMHs: 2\nExp number of AAs in TMHs: 37.2608499999999\nExp number, first 60 AAs: 19.52923\nTotal prob of N-in: 0.94351\nPOSSIBLE N-term signal sequence\ninside 1 19\nTMhelix 20 39\noutside 40 325\nTMhelix 326 348\ninside 349 372

2845 GCF_001520295.1_ASM152029v1 Escherichia coli Proteobacteria; Gammaproteobacteria; Enterobacterales; Enterobacteriaceae; Escherichia
MTGDNTLIHSHGINRRDFMKLCAALAATMGLSSKAAAEMVESVTNPQRPPVIWIGAQECT WP_000145423.1
hydrogenase 2 small subunit [Escherichia coli] Length: 372\nNumber of predicted TMHs: 1\nExp number of AAs in TMHs: 23.78201\nExp number, first 60 AAs: 0.28355\nTotal prob of N-in: 0.13257\noutside 1 331\nTMhelix 332 354\ninside 355 372

2846 GCF_001520295.1_ASM152029v1 Escherichia coli Proteobacteria; Gammaproteobacteria; Enterobacterales; Enterobacteriaceae; Escherichia
MNNEETFYQAMRRQGVTRRSFLKYCSLAATSLGLGAGMAPKIAWALENKPRIPVVWIHGL WP_001295940.1
MULTISPECIES: hydrogenase [Enterobacteriaceae] Length: 372\nNumber of predicted TMHs: 2\nExp number of AAs in TMHs: 37.2608899999999\nExp number, first 60 AAs: 19.52923\nTotal prob of N-in: 0.94351\nPOSSIBLE N-term signal sequence\ninside 1 19\nTMhelix 20 39\noutside 40 325\nTMhelix 326 348\ninside 349 372

2847 GCF_001520035.1_ASM152003v1 Escherichia coli Proteobacteria; Gammaproteobacteria; Enterobacterales; Enterobacteriaceae; Escherichia
MNNEETFYQAMRRQGVTRRSFLKYCSLAATSLGLGAGMAPKIAWALENKPRIPVVWIHGL WP_001295940.1
MULTISPECIES: hydrogenase [Enterobacteriaceae] Length: 372\nNumber of predicted TMHs: 2\nExp number of AAs in TMHs: 37.2608899999999\nExp number, first 60 AAs: 19.52923\nTotal prob of N-in: 0.94351\nPOSSIBLE N-term signal sequence\ninside 1 19\nTMhelix 20 39\noutside 40 325\nTMhelix 326 348\ninside 349 372

2848 GCF_001519995.1_ASM151999v1 Escherichia coli Proteobacteria; Gammaproteobacteria; Enterobacterales; Enterobacteriaceae; Escherichia
MNNEETFYQAMRRQGVTRRSFLKYCSLAATSLGLGAGMAPKIAWALENKPRIPVVWIHGL WP_001295940.1
MULTISPECIES: hydrogenase [Enterobacteriaceae] Length: 372\nNumber of predicted TMHs: 2\nExp number of AAs in TMHs: 37.2608899999999\nExp number, first 60 AAs: 19.52923\nTotal prob of N-in: 0.94351\nPOSSIBLE N-term signal sequence\ninside 1 19\nTMhelix 20 39\noutside 40 325\nTMhelix 326 348\ninside 349 372

2849 GCF_001520335.1_ASM152033v1 Escherichia coli Proteobacteria; Gammaproteobacteria; Enterobacterales; Enterobacteriaceae; Escherichia
MNNEETFYQAMRRQGVTRRSFLKYCSLAATSLGLGAGMAPKIAWALENKPRIPVVWIHGL WP_001058323.1
MULTISPECIES: hydrogenase-1 small chain [Proteobacteria] Length: 372\nNumber of predicted TMHs: 2\nExp number of AAs in TMHs: 37.2608499999999\nExp number, first 60 AAs: 19.52923\nTotal prob of N-in: 0.94351\nPOSSIBLE N-term signal sequence\ninside 1 19\nTMhelix 20 39\noutside 40 325\nTMhelix 326 348\ninside 349 372

2850 GCF_001520255.1_ASM152025v1 Escherichia coli Proteobacteria; Gammaproteobacteria; Enterobacterales; Enterobacteriaceae; Escherichia
MNNEETFYQAMRRQGVTRRSFLKYCSLAATSLGLGAGMAPKIAWALENKPRIPVVWIHGL WP_001058323.1
MULTISPECIES: hydrogenase-1 small chain [Proteobacteria] Length: 372\nNumber of predicted TMHs: 2\nExp number of AAs in TMHs: 37.2608499999999\nExp number, first 60 AAs: 19.52923\nTotal prob of N-in: 0.94351\nPOSSIBLE N-term signal sequence\ninside 1 19\nTMhelix 20 39\noutside 40 325\nTMhelix 326 348\ninside 349 372

2851 GCF_001520495.1_ASM152049v1 Escherichia coli Proteobacteria; Gammaproteobacteria; Enterobacterales; Enterobacteriaceae; Escherichia
MNNEETFYQAMRRQGVTRRSFLKYCSLAATSLGLGAGMAPKIAWALENKPRIPVVWIHGL WP_001058323.1
MULTISPECIES: hydrogenase-1 small chain [Proteobacteria] Length: 372\nNumber of predicted TMHs: 2\nExp number of AAs in TMHs: 37.2608499999999\nExp number, first 60 AAs: 19.52923\nTotal prob of N-in: 0.94351\nPOSSIBLE N-term signal sequence\ninside 1 19\nTMhelix 20 39\noutside 40 325\nTMhelix 326 348\ninside 349 372

2852 GCF_001520455.1_ASM152045v1 Escherichia coli Proteobacteria; Gammaproteobacteria; Enterobacterales; Enterobacteriaceae; Escherichia
MNNEETFYQAMRRQGVTRRSFLKYCSLAATSLGLGAGMAPKIAWALENKPRIPVVWIHGL WP_001295940.1
MULTISPECIES: hydrogenase [Enterobacteriaceae] Length: 372\nNumber of predicted TMHs: 2\nExp number of AAs in TMHs: 37.2608899999999\nExp number, first 60 AAs: 19.52923\nTotal prob of N-in:

0.94351\nPOSSIBLE N-term signal sequence\ninside 1 19\nTMhelix 20 39\noutside 40 325\nTMhelix 326 348\ninside 349 372

2853 GCF_001520355.1_ASM152035v1 Escherichia coli Proteobacteria; Gammaproteobacteria; Enterobacterales; Enterobacteriaceae; Escherichia
MNNEETFYQAMRRQGVTRRSFLKYCSLAATSLGLGAGMAPKIAWALENKPRIPVVWIHGL WP_001295940.1
MULTISPECIES: hydrogenase [Enterobacteriaceae] Length: 372\nNumber of predicted TMHs: 2\nExp number of AAs in TMHs: 37.2608899999999\nExp number, first 60 AAs: 19.52923\nTotal prob of N-in: 0.94351\nPOSSIBLE N-term signal sequence\ninside 1 19\nTMhelix 20 39\noutside 40 325\nTMhelix 326 348\ninside 349 372

2854 GCF_001520655.1_ASM152065v1 Escherichia coli Proteobacteria; Gammaproteobacteria; Enterobacterales; Enterobacteriaceae; Escherichia
MTGDNTLIHSHGINRRDFMKLCAALAATMGLSSKAAAEMVESVTNPQRPPVIWIGAQECT WP_000145423.1
hydrogenase 2 small subunit [Escherichia coli] Length: 372\nNumber of predicted TMHs: 1\nExp number of AAs in TMHs: 23.78201\nExp number, first 60 AAs: 0.28355\nTotal prob of N-in: 0.13257\noutside 1 331\nTMhelix 332 354\ninside 355 372

2855 GCF_001520655.1_ASM152065v1 Escherichia coli Proteobacteria; Gammaproteobacteria; Enterobacterales; Enterobacteriaceae; Escherichia
MNNEETFYQAMRRQGVTRRSFLKYCSLAATSLGLGAGMAPKIAWALENKPRIPVVWIHGL WP_001295940.1
MULTISPECIES: hydrogenase [Enterobacteriaceae] Length: 372\nNumber of predicted TMHs: 2\nExp number of AAs in TMHs: 37.2608899999999\nExp number, first 60 AAs: 19.52923\nTotal prob of N-in: 0.94351\nPOSSIBLE N-term signal sequence\ninside 1 19\nTMhelix 20 39\noutside 40 325\nTMhelix 326 348\ninside 349 372

2856 GCF_001520395.1_ASM152039v1 Escherichia coli Proteobacteria; Gammaproteobacteria; Enterobacterales; Enterobacteriaceae; Escherichia
MNNEETFYQAMRRQGVTRRSFLKYCSLAATSLGLGAGMAPKIAWALENKPRIPVVWIHGL WP_001295940.1
MULTISPECIES: hydrogenase [Enterobacteriaceae] Length: 372\nNumber of predicted TMHs: 2\nExp number of AAs in TMHs: 37.2608899999999\nExp number, first 60 AAs: 19.52923\nTotal prob of N-in: 0.94351\nPOSSIBLE N-term signal sequence\ninside 1 19\nTMhelix 20 39\noutside 40 325\nTMhelix 326 348\ninside 349 372

2857 GCF_001520695.1_ASM152069v1 Escherichia coli Proteobacteria; Gammaproteobacteria; Enterobacterales; Enterobacteriaceae; Escherichia
MTGDNTLIHSHGINRRDFMKLCAALAATMGLSSKAAAEMAESVTNPQRPPVIWIGAQECT WP_000145410.1
MULTISPECIES: hydrogenase-2 small chain [Proteobacteria] Length: 372\nNumber of predicted TMHs: 1\nExp number of AAs in TMHs: 23.72226\nExp number, first 60 AAs: 0.22038\nTotal prob of N-in: 0.12994\noutside 1 331\nTMhelix 332 354\ninside 355 372

2858 GCF_001520845.1_ASM152084v1 Escherichia coli Proteobacteria; Gammaproteobacteria; Enterobacterales; Enterobacteriaceae; Escherichia
MTGDNTLIHSHGINRRDFMKLCAALAATMGLSSKAAAEMAESVTNPQRPPVIWIGAQECT WP_000145410.1
MULTISPECIES: hydrogenase-2 small chain [Proteobacteria] Length: 372\nNumber of predicted TMHs: 1\nExp number of AAs in TMHs: 23.72226\nExp number, first 60 AAs: 0.22038\nTotal prob of N-in: 0.12994\noutside 1 331\nTMhelix 332 354\ninside 355 372

2859 GCF_001520835.1_ASM152083v1 Escherichia coli Proteobacteria; Gammaproteobacteria; Enterobacterales; Enterobacteriaceae; Escherichia
MTGDNTLIHSHGINRRDFMKLCAALAATMGLSSKAAAEMAESVTNPQRPPVIWIGAQECT WP_000145410.1
MULTISPECIES: hydrogenase-2 small chain [Proteobacteria] Length: 372\nNumber of predicted TMHs: 1\nExp number of AAs in TMHs: 23.72226\nExp number, first 60 AAs: 0.22038\nTotal prob of N-in: 0.12994\noutside 1 331\nTMhelix 332 354\ninside 355 372

2860 GCF_001520845.1_ASM152084v1 Escherichia coli Proteobacteria; Gammaproteobacteria; Enterobacterales; Enterobacteriaceae; Escherichia
MNNEETFYQAMRRQGVTRRSFLKYCSLAATSLGLGAGMAPKIAWALENKPRIPVVWIHGL WP_001058323.1
MULTISPECIES: hydrogenase-1 small chain [Proteobacteria] Length: 372\nNumber of predicted TMHs: 2\nExp number of AAs in TMHs: 37.2608499999999\nExp number, first 60 AAs: 19.52923\nTotal prob of N-in:

0.94351\nPOSSIBLE N-term signal sequence\ninside 1 19\nTMhelix 20 39\noutside 40 325\nTMhelix 326 348\ninside 349 372

2861 GCF_001520835.1_ASM152083v1 Escherichia coli Proteobacteria; Gammaproteobacteria; Enterobacterales; Enterobacteriaceae; Escherichia
MNNEETFYQAMRRQGVTRRSFLKYCSLAATSLGLGAGMAPKIAWALENKPRIPVVWIHGL WP_001058323.1
MULTISPECIES: hydrogenase-1 small chain [Proteobacteria] Length: 372\nNumber of predicted TMHs: 2\nExp number of AAs in TMHs: 37.2608499999999\nExp number, first 60 AAs: 19.52923\nTotal prob of N-in: 0.94351\nPOSSIBLE N-term signal sequence\ninside 1 19\nTMhelix 20 39\noutside 40 325\nTMhelix 326 348\ninside 349 372

2862 GCF_001520595.1_ASM152059v1 Escherichia coli Proteobacteria; Gammaproteobacteria; Enterobacterales; Enterobacteriaceae; Escherichia
MNNEETFYQAMRRQGVTRRSFLKYCSLAATSLGLGAGMAPKIAWALENKPRIPVVWIHGL WP_001295940.1
MULTISPECIES: hydrogenase [Enterobacteriaceae] Length: 372\nNumber of predicted TMHs: 2\nExp number of AAs in TMHs: 37.2608899999999\nExp number, first 60 AAs: 19.52923\nTotal prob of N-in: 0.94351\nPOSSIBLE N-term signal sequence\ninside 1 19\nTMhelix 20 39\noutside 40 325\nTMhelix 326 348\ninside 349 372

2863 GCF_001520895.1_ASM152089v1 Escherichia coli Proteobacteria; Gammaproteobacteria; Enterobacterales; Enterobacteriaceae; Escherichia
MTGDNTLIHSHGINRRDFMKLCAALAATMGLSSKAAAEMAESVTNPQRPPVIWIGAQECT WP_000145410.1
MULTISPECIES: hydrogenase-2 small chain [Proteobacteria] Length: 372\nNumber of predicted TMHs: 1\nExp number of AAs in TMHs: 23.72226\nExp number, first 60 AAs: 0.22038\nTotal prob of N-in: 0.12994\noutside 1 331\nTMhelix 332 354\ninside 355 372

2864 GCF_001520935.1_ASM152093v1 Escherichia coli Proteobacteria; Gammaproteobacteria; Enterobacterales; Enterobacteriaceae; Escherichia
MTGDNTLIHSHGINRRDFMKLCAALAATMGLSSKAAAEMAESVTNPQRPPVIWIGAQECT WP_000145410.1
MULTISPECIES: hydrogenase-2 small chain [Proteobacteria] Length: 372\nNumber of predicted TMHs: 1\nExp number of AAs in TMHs: 23.72226\nExp number, first 60 AAs: 0.22038\nTotal prob of N-in: 0.12994\noutside 1 331\nTMhelix 332 354\ninside 355 372

2865 GCF_001521055.1_ASM152105v1 Escherichia coli Proteobacteria; Gammaproteobacteria; Enterobacterales; Enterobacteriaceae; Escherichia
MTGDNTLIHSHGINRRDFMKLCAALAATMGLSSKAAAEMVESVTNPQRPPVIWIGAQECT WP_000145423.1
hydrogenase 2 small subunit [Escherichia coli] Length: 372\nNumber of predicted TMHs: 1\nExp number of AAs in TMHs: 23.78201\nExp number, first 60 AAs: 0.28355\nTotal prob of N-in: 0.13257\noutside 1 331\nTMhelix 332 354\ninside 355 372

2866 GCF_001521055.1_ASM152105v1 Escherichia coli Proteobacteria; Gammaproteobacteria; Enterobacterales; Enterobacteriaceae; Escherichia
MNNEETFYQAMRRQGVTRRSFLKYCSLAATSLGLGAGMAPKIAWALENKPRIPVVWIHGL WP_001295940.1
MULTISPECIES: hydrogenase [Enterobacteriaceae] Length: 372\nNumber of predicted TMHs: 2\nExp number of AAs in TMHs: 37.2608899999999\nExp number, first 60 AAs: 19.52923\nTotal prob of N-in: 0.94351\nPOSSIBLE N-term signal sequence\ninside 1 19\nTMhelix 20 39\noutside 40 325\nTMhelix 326 348\ninside 349 372

2867 GCF_001520795.1_ASM152079v1 Escherichia coli Proteobacteria; Gammaproteobacteria; Enterobacterales; Enterobacteriaceae; Escherichia
MNNEETFYQAMRRQGVTRRSFLKYCSLAATSLGLGAGMAPKIAWALENKPRIPVVWIHGL WP_001295940.1
MULTISPECIES: hydrogenase [Enterobacteriaceae] Length: 372\nNumber of predicted TMHs: 2\nExp number of AAs in TMHs: 37.2608899999999\nExp number, first 60 AAs: 19.52923\nTotal prob of N-in: 0.94351\nPOSSIBLE N-term signal sequence\ninside 1 19\nTMhelix 20 39\noutside 40 325\nTMhelix 326 348\ninside 349 372

2868 GCF_001521095.1_ASM152109v1 Escherichia coli Proteobacteria; Gammaproteobacteria; Enterobacterales; Enterobacteriaceae; Escherichia
MTGDNTLIHSHGINRRDFMKLCAALAATMGLSSKAAAEMVESVTNPQRPPVIWIGAQECT WP_000145423.1
hydrogenase 2 small subunit [Escherichia coli] Length: 372\nNumber of predicted TMHs: 1\nExp number of

AAs in TMHs: 23.78201\nExp number, first 60 AAs:  0.28355\nTotal prob of N-in:   0.13257\noutside   1   331\nTMhelix   332   354\ninside   355   372

2869 GCF_001521095.1_ASM152109v1 Escherichia coli Proteobacteria; Gammaproteobacteria; Enterobacterales; Enterobacteriaceae; Escherichia
MNNEETFYQAMRRQGVTRRSFLKYCSLAATSLGLGAGMAPKIAWALENKPRIPVVWIHGL WP_001295940.1
MULTISPECIES: hydrogenase [Enterobacteriaceae] Length: 372\nNumber of predicted TMHs:  2\nExp number of AAs in TMHs: 37.2608899999999\nExp number, first 60 AAs:  19.52923\nTotal prob of N-in:   0.94351\nPOSSIBLE N-term signal sequence\ninside   1   19\nTMhelix   20   39\noutside   40   325\nTMhelix   326   348\ninside   349   372

2870 GCF_001520955.1_ASM152095v1 Escherichia coli Proteobacteria; Gammaproteobacteria; Enterobacterales; Enterobacteriaceae; Escherichia
MNNEETFYQAMRRQGVTRRSFLKYCSLAATSLGLGAGMAPKIAWALENKPRIPVVWIHGL WP_001058323.1
MULTISPECIES: hydrogenase-1 small chain [Proteobacteria] Length: 372\nNumber of predicted TMHs:  2\nExp number of AAs in TMHs: 37.2608499999999\nExp number, first 60 AAs:  19.52923\nTotal prob of N-in:   0.94351\nPOSSIBLE N-term signal sequence\ninside   1   19\nTMhelix   20   39\noutside   40   325\nTMhelix   326   348\ninside   349   372

2871 GCF_001521275.1_ASM152127v1 Escherichia coli Proteobacteria; Gammaproteobacteria; Enterobacterales; Enterobacteriaceae; Escherichia
MNNEETFYQAMRRQGVTRRSFLKYCSLAATSLGLGAGMAPKIAWALENKPRIPVVWIHGL WP_001058323.1
MULTISPECIES: hydrogenase-1 small chain [Proteobacteria] Length: 372\nNumber of predicted TMHs:  2\nExp number of AAs in TMHs: 37.2608499999999\nExp number, first 60 AAs:  19.52923\nTotal prob of N-in:   0.94351\nPOSSIBLE N-term signal sequence\ninside   1   19\nTMhelix   20   39\noutside   40   325\nTMhelix   326   348\ninside   349   372

2872 GCF_001521195.1_ASM152119v1 Escherichia coli Proteobacteria; Gammaproteobacteria; Enterobacterales; Enterobacteriaceae; Escherichia
MNNEETFYQAMRRQGVTRRSFLKYCSLAATSLGLGAGMAPKIAWALENKPRIPVVWIHGL WP_021520630.1
hydrogenase-1 small chain [Escherichia coli] Length: 372\nNumber of predicted TMHs:  2\nExp number of AAs in TMHs: 37.8197599999999\nExp number, first 60 AAs:  19.53295\nTotal prob of N-in:   0.94396\nPOSSIBLE N-term signal sequence\ninside   1   19\nTMhelix   20   39\noutside   40   325\nTMhelix   326   348\ninside   349   372

2873 GCF_001521025.1_ASM152102v1 Escherichia coli Proteobacteria; Gammaproteobacteria; Enterobacterales; Enterobacteriaceae; Escherichia
MNNEETFYQAMRRQGVTRRSFLKYCSLAATSLGLGAGMAPKIAWALENKPRIPVVWIHGL WP_001295940.1
MULTISPECIES: hydrogenase [Enterobacteriaceae] Length: 372\nNumber of predicted TMHs:  2\nExp number of AAs in TMHs: 37.2608899999999\nExp number, first 60 AAs:  19.52923\nTotal prob of N-in:   0.94351\nPOSSIBLE N-term signal sequence\ninside   1   19\nTMhelix   20   39\noutside   40   325\nTMhelix   326   348\ninside   349   372

2874 GCF_001521285.1_ASM152128v1 Escherichia coli Proteobacteria; Gammaproteobacteria; Enterobacterales; Enterobacteriaceae; Escherichia
MTGDNTLIHSHGINRRDFMKLCAALAATMGLSSKAAAEMAESVTNPQRPPVIWIGAQECT WP_000145410.1
MULTISPECIES: hydrogenase-2 small chain [Proteobacteria] Length: 372\nNumber of predicted TMHs:  1\nExp number of AAs in TMHs: 23.72226\nExp number, first 60 AAs:  0.22038\nTotal prob of N-in:   0.12994\noutside   1   331\nTMhelix   332   354\ninside   355   372

2875 GCF_001521335.1_ASM152133v1 Escherichia coli Proteobacteria; Gammaproteobacteria; Enterobacterales; Enterobacteriaceae; Escherichia
MTGDNTLIHSHGINRRDFMKLCAALAATMGLSSKAAAEMVESVTNPQRPPVIWIGAQECT WP_000145423.1
hydrogenase 2 small subunit [Escherichia coli] Length: 372\nNumber of predicted TMHs:  1\nExp number of AAs in TMHs: 23.78201\nExp number, first 60 AAs:  0.28355\nTotal prob of N-in:   0.13257\noutside   1   331\nTMhelix   332   354\ninside   355   372

2876 GCF_001521375.1_ASM152137v1 Escherichia coli Proteobacteria; Gammaproteobacteria; Enterobacterales; Enterobacteriaceae; Escherichia
MTGDNTLIHSHGINRRDFMKLCAALAATMGLSSKAAAEMAESVTNPQRPPVIWIGAQECT WP_000145410.1
MULTISPECIES: hydrogenase-2 small chain [Proteobacteria] Length: 372\nNumber of predicted TMHs:

1\nExp number of AAs in TMHs: 23.72226\nExp number, first 60 AAs: 0.22038\nTotal prob of N-in: 0.12994\noutside 1 331\nTMhelix 332 354\ninside 355 372

2877 GCF_001521435.1_ASM152143v1 Escherichia coli Proteobacteria; Gammaproteobacteria; Enterobacterales; Enterobacteriaceae; Escherichia
MTGDNTLIHSHGINRRDFMKLCAALAATMGLSSKAAAEMAESVTNPQRPPVIWIGAQECT WP_000145410.1
MULTISPECIES: hydrogenase-2 small chain [Proteobacteria] Length: 372\nNumber of predicted TMHs: 1\nExp number of AAs in TMHs: 23.72226\nExp number, first 60 AAs: 0.22038\nTotal prob of N-in: 0.12994\noutside 1 331\nTMhelix 332 354\ninside 355 372

2878 GCF_001521575.1_ASM152157v1 Escherichia coli Proteobacteria; Gammaproteobacteria; Enterobacterales; Enterobacteriaceae; Escherichia
MTGDNTLIHSHGINRRDFMKLCAALAATMGLSSKAAAEMAESVTNPQRPPVIWIGAQECT WP_000145410.1
MULTISPECIES: hydrogenase-2 small chain [Proteobacteria] Length: 372\nNumber of predicted TMHs: 1\nExp number of AAs in TMHs: 23.72226\nExp number, first 60 AAs: 0.22038\nTotal prob of N-in: 0.12994\noutside 1 331\nTMhelix 332 354\ninside 355 372

2879 GCF_001521495.1_ASM152149v1 Escherichia coli Proteobacteria; Gammaproteobacteria; Enterobacterales; Enterobacteriaceae; Escherichia
MNNEETFYQAMRRQGVTRRSFLKYCSLAATSLGLGAGMAPKIAWALENKPRIPVVWIHGL WP_001295940.1
MULTISPECIES: hydrogenase [Enterobacteriaceae] Length: 372\nNumber of predicted TMHs: 2\nExp number of AAs in TMHs: 37.2608899999999\nExp number, first 60 AAs: 19.52923\nTotal prob of N-in: 0.94351\nPOSSIBLE N-term signal sequence\ninside 1 19\nTMhelix 20 39\noutside 40 325\nTMhelix 326 348\ninside 349 372

2880 GCF_001521475.1_ASM152147v1 Escherichia coli Proteobacteria; Gammaproteobacteria; Enterobacterales; Enterobacteriaceae; Escherichia
MTGDNTLIHSHGINRRDFMKLCAALAATMGLSSKAAAEMAESVTNPQRPPVIWIGAQECT WP_059321935.1
hydrogenase 2 small subunit [Escherichia coli] Length: 372\nNumber of predicted TMHs: 1\nExp number of AAs in TMHs: 23.8325\nExp number, first 60 AAs: 0.21938\nTotal prob of N-in: 0.13453\noutside 1 331\nTMhelix 332 354\ninside 355 372

2881 GCF_001521335.1_ASM152133v1 Escherichia coli Proteobacteria; Gammaproteobacteria; Enterobacterales; Enterobacteriaceae; Escherichia
MNNEETFYQAMRRQGVTRRSFLKYCSLAATSLGLGAGMAPKIAWALENKPRIPVVWIHGL WP_001295940.1
MULTISPECIES: hydrogenase [Enterobacteriaceae] Length: 372\nNumber of predicted TMHs: 2\nExp number of AAs in TMHs: 37.2608899999999\nExp number, first 60 AAs: 19.52923\nTotal prob of N-in: 0.94351\nPOSSIBLE N-term signal sequence\ninside 1 19\nTMhelix 20 39\noutside 40 325\nTMhelix 326 348\ninside 349 372

2882 GCF_001521375.1_ASM152137v1 Escherichia coli Proteobacteria; Gammaproteobacteria; Enterobacterales; Enterobacteriaceae; Escherichia
MNNEETFYQAMRRQGVTRRSFLKYCSLAATSLGLGAGMAPKIAWALENKPRIPVVWIHGL WP_001295940.1
MULTISPECIES: hydrogenase [Enterobacteriaceae] Length: 372\nNumber of predicted TMHs: 2\nExp number of AAs in TMHs: 37.2608899999999\nExp number, first 60 AAs: 19.52923\nTotal prob of N-in: 0.94351\nPOSSIBLE N-term signal sequence\ninside 1 19\nTMhelix 20 39\noutside 40 325\nTMhelix 326 348\ninside 349 372

2883 GCF_001521595.1_ASM152159v1 Escherichia coli Proteobacteria; Gammaproteobacteria; Enterobacterales; Enterobacteriaceae; Escherichia
MTGDNTLIHSHGINRRDFMKLCAALAATMGLSSKAAAEMAESVTNPQRPPVIWIGAQECT WP_000145410.1
MULTISPECIES: hydrogenase-2 small chain [Proteobacteria] Length: 372\nNumber of predicted TMHs: 1\nExp number of AAs in TMHs: 23.72226\nExp number, first 60 AAs: 0.22038\nTotal prob of N-in: 0.12994\noutside 1 331\nTMhelix 332 354\ninside 355 372

2884 GCF_001521635.1_ASM152163v1 Escherichia coli Proteobacteria; Gammaproteobacteria; Enterobacterales; Enterobacteriaceae; Escherichia
MTGDNTLIHSHGINRRDFMKLCAALAATMGLSSKAAAEMAESVTNPQRPPVIWIGAQECT WP_000145410.1
MULTISPECIES: hydrogenase-2 small chain [Proteobacteria] Length: 372\nNumber of predicted TMHs: 1\nExp number of AAs in TMHs: 23.72226\nExp number, first 60 AAs: 0.22038\nTotal prob of N-in: 0.12994\noutside 1 331\nTMhelix 332 354\ninside 355 372

2885 GCF_001521695.1_ASM152169v1 Escherichia coli Proteobacteria; Gammaproteobacteria; Enterobacterales; Enterobacteriaceae; Escherichia
MNNEETFYQAMRRQGVTRRSFLKYCSLAATSLGLGAGMAPKIAWALENKPRIPVVWIHGL WP_001058323.1
MULTISPECIES: hydrogenase-1 small chain [Proteobacteria] Length: 372\nNumber of predicted TMHs: 2\nExp number of AAs in TMHs: 37.2608499999999\nExp number, first 60 AAs: 19.52923\nTotal prob of N-in: 0.94351\nPOSSIBLE N-term signal sequence\ninside 1 19\nTMhelix 20 39\noutside 40 325\nTMhelix 326 348\ninside 349 372

2886 GCF_001542545.1_ASM154254v1 Escherichia coli Proteobacteria; Gammaproteobacteria; Enterobacterales; Enterobacteriaceae; Escherichia
MNNEETFYQAMRRQGVTRRSFLKYCSLAATSLGLGAGMAPKIAWALENKPRIPVVWIHGL WP_001058323.1
MULTISPECIES: hydrogenase-1 small chain [Proteobacteria] Length: 372\nNumber of predicted TMHs: 2\nExp number of AAs in TMHs: 37.2608499999999\nExp number, first 60 AAs: 19.52923\nTotal prob of N-in: 0.94351\nPOSSIBLE N-term signal sequence\ninside 1 19\nTMhelix 20 39\noutside 40 325\nTMhelix 326 348\ninside 349 372

2887 GCF_001521535.1_ASM152153v1 Escherichia coli Proteobacteria; Gammaproteobacteria; Enterobacterales; Enterobacteriaceae; Escherichia
MNNEETFYQAMRRQGVTRRSFLKYCSLAATSLGLGAGMAPKIAWALENKPRIPVVWIHGL WP_001295940.1
MULTISPECIES: hydrogenase [Enterobacteriaceae] Length: 372\nNumber of predicted TMHs: 2\nExp number of AAs in TMHs: 37.2608899999999\nExp number, first 60 AAs: 19.52923\nTotal prob of N-in: 0.94351\nPOSSIBLE N-term signal sequence\ninside 1 19\nTMhelix 20 39\noutside 40 325\nTMhelix 326 348\ninside 349 372

2888 GCF_001550005.1_ASM155000v1 Escherichia coli Proteobacteria; Gammaproteobacteria; Enterobacterales; Enterobacteriaceae; Escherichia
MTGDNTLIHSHGINRRDFMKLCAALAATMGLSSKAAAEMAESVTNPQRPPVIWIGAQECT WP_000145410.1
MULTISPECIES: hydrogenase-2 small chain [Proteobacteria] Length: 372\nNumber of predicted TMHs: 1\nExp number of AAs in TMHs: 23.72226\nExp number, first 60 AAs: 0.22038\nTotal prob of N-in: 0.12994\noutside 1 331\nTMhelix 332 354\ninside 355 372

2889 GCF_001554295.1_ASM155429v1 Escherichia coli Proteobacteria; Gammaproteobacteria; Enterobacterales; Enterobacteriaceae; Escherichia
MTGDNTLIHSHGINRRDFMKLCAALAATMGLSSKAAAEMAESVTNPQRPPVIWIGAQECT WP_000145410.1
MULTISPECIES: hydrogenase-2 small chain [Proteobacteria] Length: 372\nNumber of predicted TMHs: 1\nExp number of AAs in TMHs: 23.72226\nExp number, first 60 AAs: 0.22038\nTotal prob of N-in: 0.12994\noutside 1 331\nTMhelix 332 354\ninside 355 372

2890 GCF_001555495.1_ASM155549v1 Escherichia coli Proteobacteria; Gammaproteobacteria; Enterobacterales; Enterobacteriaceae; Escherichia
MTGDNTLIHSHGINRRDFMKLCAALAATMGLSSKAAAEMVESVTNPQRPPVIWIGAQECT WP_000145423.1
hydrogenase 2 small subunit [Escherichia coli] Length: 372\nNumber of predicted TMHs: 1\nExp number of AAs in TMHs: 23.78201\nExp number, first 60 AAs: 0.28355\nTotal prob of N-in: 0.13257\noutside 1 331\nTMhelix 332 354\ninside 355 372

2891 GCF_001561205.1_ASM156120v1 Escherichia coli Proteobacteria; Gammaproteobacteria; Enterobacterales; Enterobacteriaceae; Escherichia
MTGDNTLIHSHGINRRDFMKLCAALAATMGLSSKAAAEMVESVTNPQRPPVIWIGAQECT WP_000145423.1
hydrogenase 2 small subunit [Escherichia coli] Length: 372\nNumber of predicted TMHs: 1\nExp number of AAs in TMHs: 23.78201\nExp number, first 60 AAs: 0.28355\nTotal prob of N-in: 0.13257\noutside 1 331\nTMhelix 332 354\ninside 355 372

2892 GCF_001561015.1_ASM156101v1 Escherichia coli Proteobacteria; Gammaproteobacteria; Enterobacterales; Enterobacteriaceae; Escherichia
MTGDNTLIHSHGINRRDFMKLCAALAATMGLSSKAAAEMVESVTNPQRPPVIWIGAQECT WP_000145423.1
hydrogenase 2 small subunit [Escherichia coli] Length: 372\nNumber of predicted TMHs: 1\nExp number of AAs in TMHs: 23.78201\nExp number, first 60 AAs: 0.28355\nTotal prob of N-in: 0.13257\noutside 1 331\nTMhelix 332 354\ninside 355 372

2893 GCF_001561045.1_ASM156104v1 Escherichia coli Proteobacteria; Gammaproteobacteria; Enterobacterales; Enterobacteriaceae; Escherichia

MTGDNTLIHSHGINRRDFMKLCAALAATMGLSSKAAAEMAESVTNPQRPPVIWIGAQECT WP_000145410.1 MULTISPECIES: hydrogenase-2 small chain [Proteobacteria] Length: 372\nNumber of predicted TMHs: 1\nExp number of AAs in TMHs: 23.72226\nExp number, first 60 AAs: 0.22038\nTotal prob of N-in: 0.12994\noutside 1 331\nTMhelix 332 354\ninside 355 372

2894 GCF_001561105.1_ASM156110v1 Escherichia coli Proteobacteria; Gammaproteobacteria; Enterobacterales; Enterobacteriaceae; Escherichia MTGDNTLIHSHGINRRDFMKLCAALAATMGLSSKAAAEMVESVTNPQRPPVIWIGAQECT WP_000145423.1 hydrogenase 2 small subunit [Escherichia coli] Length: 372\nNumber of predicted TMHs: 1\nExp number of AAs in TMHs: 23.78201\nExp number, first 60 AAs: 0.28355\nTotal prob of N-in: 0.13257\noutside 1 331\nTMhelix 332 354\ninside 355 372

2895 GCF_001561135.1_ASM156113v1 Escherichia coli Proteobacteria; Gammaproteobacteria; Enterobacterales; Enterobacteriaceae; Escherichia MTGDNTLIHSHGINRRDFMKLCAALAATMGLSSKAAAEMAESVTNPQRPPVIWIGAQECT WP_000145410.1 MULTISPECIES: hydrogenase-2 small chain [Proteobacteria] Length: 372\nNumber of predicted TMHs: 1\nExp number of AAs in TMHs: 23.72226\nExp number, first 60 AAs: 0.22038\nTotal prob of N-in: 0.12994\noutside 1 331\nTMhelix 332 354\ninside 355 372

2896 GCF_001561195.1_ASM156119v1 Escherichia coli Proteobacteria; Gammaproteobacteria; Enterobacterales; Enterobacteriaceae; Escherichia MTGDNTLIHSHGINRRDFMKLCAALAATMGLSSKAAAEMAESVTNPQRPPVIWIGAQECT WP_000145410.1 MULTISPECIES: hydrogenase-2 small chain [Proteobacteria] Length: 372\nNumber of predicted TMHs: 1\nExp number of AAs in TMHs: 23.72226\nExp number, first 60 AAs: 0.22038\nTotal prob of N-in: 0.12994\noutside 1 331\nTMhelix 332 354\ninside 355 372

2897 GCF_001561295.1_ASM156129v1 Escherichia coli Proteobacteria; Gammaproteobacteria; Enterobacterales; Enterobacteriaceae; Escherichia MTGDNTLIHSHGINRRDFMKLCAALAATMGLSSKAAAEMAESVTNPQRPPVIWIGAQECT WP_000145410.1 MULTISPECIES: hydrogenase-2 small chain [Proteobacteria] Length: 372\nNumber of predicted TMHs: 1\nExp number of AAs in TMHs: 23.72226\nExp number, first 60 AAs: 0.22038\nTotal prob of N-in: 0.12994\noutside 1 331\nTMhelix 332 354\ninside 355 372

2898 GCF_001561295.1_ASM156129v1 Escherichia coli Proteobacteria; Gammaproteobacteria; Enterobacterales; Enterobacteriaceae; Escherichia MNNEETFYQAMRRQGVTRRSFLKYCSLAATSLGLGAGMAPKIAWALENKPRIPVVWIHGL WP_001058323.1 MULTISPECIES: hydrogenase-1 small chain [Proteobacteria] Length: 372\nNumber of predicted TMHs: 2\nExp number of AAs in TMHs: 37.2608499999999\nExp number, first 60 AAs: 19.52923\nTotal prob of N-in: 0.94351\nPOSSIBLE N-term signal sequence\ninside 1 19\nTMhelix 20 39\noutside 40 325\nTMhelix 326 348\ninside 349 372

2899 GCF_001561045.1_ASM156104v1 Escherichia coli Proteobacteria; Gammaproteobacteria; Enterobacterales; Enterobacteriaceae; Escherichia MNNEETFYQAMRRQGVTRRSFLKYCSLAATSLGLGAGMAPKIAWALENKPRIPVVWIHGL WP_001295940.1 MULTISPECIES: hydrogenase [Enterobacteriaceae] Length: 372\nNumber of predicted TMHs: 2\nExp number of AAs in TMHs: 37.2608899999999\nExp number, first 60 AAs: 19.52923\nTotal prob of N-in: 0.94351\nPOSSIBLE N-term signal sequence\ninside 1 19\nTMhelix 20 39\noutside 40 325\nTMhelix 326 348\ninside 349 372

2900 GCF_001561105.1_ASM156110v1 Escherichia coli Proteobacteria; Gammaproteobacteria; Enterobacterales; Enterobacteriaceae; Escherichia MNNEETFYQAMRRQGVTRRSFLKYCSLAATSLGLGAGMAPKIAWALENKPRIPVVWIHGL WP_001295940.1 MULTISPECIES: hydrogenase [Enterobacteriaceae] Length: 372\nNumber of predicted TMHs: 2\nExp number of AAs in TMHs: 37.2608899999999\nExp number, first 60 AAs: 19.52923\nTotal prob of N-in: 0.94351\nPOSSIBLE N-term signal sequence\ninside 1 19\nTMhelix 20 39\noutside 40 325\nTMhelix 326 348\ninside 349 372

2901 GCF_001561135.1_ASM156113v1 Escherichia coli Proteobacteria; Gammaproteobacteria; Enterobacterales; Enterobacteriaceae; Escherichia MNNEETFYQAMRRQGVTRRSFLKYCSLAATSLGLGAGMAPKIAWALENKPRIPVVWIHGL WP_001295940.1 MULTISPECIES: hydrogenase [Enterobacteriaceae] Length: 372\nNumber of predicted TMHs: 2\nExp

number of AAs in TMHs: 37.2608899999999\nExp number, first 60 AAs:  19.52923\nTotal prob of N-in:  0.94351\nPOSSIBLE N-term signal sequence\ninside   1   19\nTMhelix   20   39\noutside   40   325\nTMhelix   326   348\ninside   349   372

2902 GCF_001561355.1_ASM156135v1 Escherichia coli Proteobacteria; Gammaproteobacteria; Enterobacterales; Enterobacteriaceae; Escherichia
MTGDNTLIHSHGINRRDFMKLCAALAATMGLSSKAAAEMAESVTNPQRPPVIWIGAQECT WP_000145410.1
MULTISPECIES: hydrogenase-2 small chain [Proteobacteria] Length: 372\nNumber of predicted TMHs:  1\nExp number of AAs in TMHs: 23.72226\nExp number, first 60 AAs:  0.22038\nTotal prob of N-in:   0.12994\noutside   1   331\nTMhelix   332   354\ninside   355   372

2903 GCF_001561375.1_ASM156137v1 Escherichia coli Proteobacteria; Gammaproteobacteria; Enterobacterales; Enterobacteriaceae; Escherichia
MTGDNTLIHSHGINRRDFMKLCAALAATMGLSSKAAAEMAESVTNPQRPPVIWIGAQECT WP_000145410.1
MULTISPECIES: hydrogenase-2 small chain [Proteobacteria] Length: 372\nNumber of predicted TMHs:  1\nExp number of AAs in TMHs: 23.72226\nExp number, first 60 AAs:  0.22038\nTotal prob of N-in:   0.12994\noutside   1   331\nTMhelix   332   354\ninside   355   372

2904 GCF_001561445.1_ASM156144v1 Escherichia coli Proteobacteria; Gammaproteobacteria; Enterobacterales; Enterobacteriaceae; Escherichia
MNNEETFYQAMRRQGVTRRSFLKYCSLAATSLGLGAGMAPKIAWALENKPRIPVVWIHGL WP_001295940.1
MULTISPECIES: hydrogenase [Enterobacteriaceae] Length: 372\nNumber of predicted TMHs:  2\nExp number of AAs in TMHs: 37.2608899999999\nExp number, first 60 AAs:  19.52923\nTotal prob of N-in:  0.94351\nPOSSIBLE N-term signal sequence\ninside   1   19\nTMhelix   20   39\noutside   40   325\nTMhelix   326   348\ninside   349   372

2905 GCF_001561495.1_ASM156149v1 Escherichia coli Proteobacteria; Gammaproteobacteria; Enterobacterales; Enterobacteriaceae; Escherichia
MNNEETFYQAMRRQGVTRRSFLKYCSLAATSLGLGAGMAPKIAWALENKPRIPVVWIHGL WP_001295940.1
MULTISPECIES: hydrogenase [Enterobacteriaceae] Length: 372\nNumber of predicted TMHs:  2\nExp number of AAs in TMHs: 37.2608899999999\nExp number, first 60 AAs:  19.52923\nTotal prob of N-in:  0.94351\nPOSSIBLE N-term signal sequence\ninside   1   19\nTMhelix   20   39\noutside   40   325\nTMhelix   326   348\ninside   349   372

2906 GCF_001561285.1_ASM156128v1 Escherichia coli Proteobacteria; Gammaproteobacteria; Enterobacterales; Enterobacteriaceae; Escherichia
MNNEETFYQAMRRQGVTRRSFLKYCSLAATSLGLGAGMAPKIAWALENKPRIPVVWIHGL WP_001295940.1
MULTISPECIES: hydrogenase [Enterobacteriaceae] Length: 372\nNumber of predicted TMHs:  2\nExp number of AAs in TMHs: 37.2608899999999\nExp number, first 60 AAs:  19.52923\nTotal prob of N-in:  0.94351\nPOSSIBLE N-term signal sequence\ninside   1   19\nTMhelix   20   39\noutside   40   325\nTMhelix   326   348\ninside   349   372

2907 GCF_001561515.1_ASM156151v1 Escherichia coli Proteobacteria; Gammaproteobacteria; Enterobacterales; Enterobacteriaceae; Escherichia
MTGDNTLIHSHGINRRDFMKLCAALAATMGLSSKAAAEMVESVTNPQRPPVIWIGAQECT WP_000145423.1
hydrogenase 2 small subunit [Escherichia coli] Length: 372\nNumber of predicted TMHs:  1\nExp number of AAs in TMHs: 23.78201\nExp number, first 60 AAs:  0.28355\nTotal prob of N-in:   0.13257\noutside   1   331\nTMhelix   332   354\ninside   355   372

2908 GCF_001561515.1_ASM156151v1 Escherichia coli Proteobacteria; Gammaproteobacteria; Enterobacterales; Enterobacteriaceae; Escherichia
MNNEETFYQAMRRQGVTRRSFLKYCSLAATSLGLGAGMAPKIAWALENKPRIPVVWIHGL WP_001295940.1
MULTISPECIES: hydrogenase [Enterobacteriaceae] Length: 372\nNumber of predicted TMHs:  2\nExp number of AAs in TMHs: 37.2608899999999\nExp number, first 60 AAs:  19.52923\nTotal prob of N-in:  0.94351\nPOSSIBLE N-term signal sequence\ninside   1   19\nTMhelix   20   39\noutside   40   325\nTMhelix   326   348\ninside   349   372

2909 GCF_001561255.1_ASM156125v1 Escherichia coli Proteobacteria; Gammaproteobacteria; Enterobacterales; Enterobacteriaceae; Escherichia
MNNEETFYQAMRRQGVTRRSFLKYCSLAATSLGLGAGMAPKIAWALENKPRIPVVWIHGL WP_001295940.1
MULTISPECIES: hydrogenase [Enterobacteriaceae] Length: 372\nNumber of predicted TMHs:  2\nExp

number of AAs in TMHs: 37.2608899999999\nExp number, first 60 AAs: 19.52923\nTotal prob of N-in: 0.94351\nPOSSIBLE N-term signal sequence\ninside 1 19\nTMhelix 20 39\noutside 40 325\nTMhelix 326 348\ninside 349 372

2910 GCF_001561555.1_ASM156155v1 Escherichia coli Proteobacteria; Gammaproteobacteria; Enterobacterales; Enterobacteriaceae; Escherichia
MTGDNTLIHSHGINRRDFMKLCAALAATMGLSSKAAAEMAESVTNPQRPPVIWIGAQECT WP_000145410.1
MULTISPECIES: hydrogenase-2 small chain [Proteobacteria] Length: 372\nNumber of predicted TMHs: 1\nExp number of AAs in TMHs: 23.72226\nExp number, first 60 AAs: 0.22038\nTotal prob of N-in: 0.12994\noutside 1 331\nTMhelix 332 354\ninside 355 372

2911 GCF_001561755.1_ASM156175v1 Escherichia coli Proteobacteria; Gammaproteobacteria; Enterobacterales; Enterobacteriaceae; Escherichia
MNNEETFYQAMRRQGVTRRSFLKYCSLAATSLGLGAGMAPKIAWALENKPRIPVVWIHGL WP_001295940.1
MULTISPECIES: hydrogenase [Enterobacteriaceae] Length: 372\nNumber of predicted TMHs: 2\nExp number of AAs in TMHs: 37.2608899999999\nExp number, first 60 AAs: 19.52923\nTotal prob of N-in: 0.94351\nPOSSIBLE N-term signal sequence\ninside 1 19\nTMhelix 20 39\noutside 40 325\nTMhelix 326 348\ninside 349 372

2912 GCF_001561555.1_ASM156155v1 Escherichia coli Proteobacteria; Gammaproteobacteria; Enterobacterales; Enterobacteriaceae; Escherichia
MNNEETFYQAMRRQGVTRRSFLKYCSLAATSLGLGAGMAPKIAWALENKPRIPVVWIHGL WP_001295940.1
MULTISPECIES: hydrogenase [Enterobacteriaceae] Length: 372\nNumber of predicted TMHs: 2\nExp number of AAs in TMHs: 37.2608899999999\nExp number, first 60 AAs: 19.52923\nTotal prob of N-in: 0.94351\nPOSSIBLE N-term signal sequence\ninside 1 19\nTMhelix 20 39\noutside 40 325\nTMhelix 326 348\ninside 349 372

2913 GCF_001561785.1_ASM156178v1 Escherichia coli Proteobacteria; Gammaproteobacteria; Enterobacterales; Enterobacteriaceae; Escherichia
MTGDNTLIHSHGINRRDFMKLCAALAATMGLSSKAAAEMVESVTNPQRPPVIWIGAQECT WP_000145423.1
hydrogenase 2 small subunit [Escherichia coli] Length: 372\nNumber of predicted TMHs: 1\nExp number of AAs in TMHs: 23.78201\nExp number, first 60 AAs: 0.28355\nTotal prob of N-in: 0.13257\noutside 1 331\nTMhelix 332 354\ninside 355 372

2914 GCF_001561835.1_ASM156183v1 Escherichia coli Proteobacteria; Gammaproteobacteria; Enterobacterales; Enterobacteriaceae; Escherichia
MTGDNTLIHSHGINRRDFMKLCAALAATMGLSSKAAAEMAESVTNPQRPPVIWIGAQECT WP_000145410.1
MULTISPECIES: hydrogenase-2 small chain [Proteobacteria] Length: 372\nNumber of predicted TMHs: 1\nExp number of AAs in TMHs: 23.72226\nExp number, first 60 AAs: 0.22038\nTotal prob of N-in: 0.12994\noutside 1 331\nTMhelix 332 354\ninside 355 372

2915 GCF_001561785.1_ASM156178v1 Escherichia coli Proteobacteria; Gammaproteobacteria; Enterobacterales; Enterobacteriaceae; Escherichia
MNNEETFYQAMRRQGVTRRSFLKYCSLAATSLGLGAGMAPKIAWALENKPRIPVVWIHGL WP_001295940.1
MULTISPECIES: hydrogenase [Enterobacteriaceae] Length: 372\nNumber of predicted TMHs: 2\nExp number of AAs in TMHs: 37.2608899999999\nExp number, first 60 AAs: 19.52923\nTotal prob of N-in: 0.94351\nPOSSIBLE N-term signal sequence\ninside 1 19\nTMhelix 20 39\noutside 40 325\nTMhelix 326 348\ninside 349 372

2916 GCF_001562635.1_ASM156263v1 Escherichia coli Proteobacteria; Gammaproteobacteria; Enterobacterales; Enterobacteriaceae; Escherichia
MTGDNTLIHSHGINRRDFMKLCAALAATMGLSSKAAAEMAESVTNPQRPPVIWIGAQECT WP_000145410.1
MULTISPECIES: hydrogenase-2 small chain [Proteobacteria] Length: 372\nNumber of predicted TMHs: 1\nExp number of AAs in TMHs: 23.72226\nExp number, first 60 AAs: 0.22038\nTotal prob of N-in: 0.12994\noutside 1 331\nTMhelix 332 354\ninside 355 372

2917 GCF_001562645.1_ASM156264v1 Escherichia coli Proteobacteria; Gammaproteobacteria; Enterobacterales; Enterobacteriaceae; Escherichia
MTGDNTLIHSHGINRRDFMKLCAALAATMGLSSKAAAEMAESVTNPQRPPVIWIGAQECT WP_000145410.1
MULTISPECIES: hydrogenase-2 small chain [Proteobacteria] Length: 372\nNumber of predicted TMHs:

1\nExp number of AAs in TMHs: 23.72226\nExp number, first 60 AAs: 0.22038\nTotal prob of N-in: 0.12994\noutside 1 331\nTMhelix 332 354\ninside 355 372

2918 GCF_001562695.1_ASM156269v1 Escherichia coli Proteobacteria; Gammaproteobacteria; Enterobacterales; Enterobacteriaceae; Escherichia
MTGDNTLIHSHGINRRDFMKLCAALAATMGLSSKAAAEMAESVTNPQRPPVIWIGAQECT WP_000145410.1
MULTISPECIES: hydrogenase-2 small chain [Proteobacteria] Length: 372\nNumber of predicted TMHs: 1\nExp number of AAs in TMHs: 23.72226\nExp number, first 60 AAs: 0.22038\nTotal prob of N-in: 0.12994\noutside 1 331\nTMhelix 332 354\ninside 355 372

2919 GCF_001562335.1_ASM156233v1 Escherichia coli Proteobacteria; Gammaproteobacteria; Enterobacterales; Enterobacteriaceae; Escherichia
MNNEETFYQAMRRQGVTRRSFLKYCSLAATSLGLGAGMAPKIAWALENKPRIPVVWIHGL WP_001058323.1
MULTISPECIES: hydrogenase-1 small chain [Proteobacteria] Length: 372\nNumber of predicted TMHs: 2\nExp number of AAs in TMHs: 37.2608499999999\nExp number, first 60 AAs: 19.52923\nTotal prob of N-in: 0.94351\nPOSSIBLE N-term signal sequence\ninside 1 19\nTMhelix 20 39\noutside 40 325\nTMhelix 326 348\ninside 349 372

2920 GCF_001562715.1_ASM156271v1 Escherichia coli Proteobacteria; Gammaproteobacteria; Enterobacterales; Enterobacteriaceae; Escherichia
MTGDNTLIHSHGINRRDFMKLCAALAATMGLSSKAAAEMAESVTNPQRPPVIWIGAQECT WP_000145410.1
MULTISPECIES: hydrogenase-2 small chain [Proteobacteria] Length: 372\nNumber of predicted TMHs: 1\nExp number of AAs in TMHs: 23.72226\nExp number, first 60 AAs: 0.22038\nTotal prob of N-in: 0.12994\noutside 1 331\nTMhelix 332 354\ninside 355 372

2921 GCF_001562375.1_ASM156237v1 Escherichia coli Proteobacteria; Gammaproteobacteria; Enterobacterales; Enterobacteriaceae; Escherichia
MNNEETFYQAMRRQGVTRRSFLKYCSLAATSLGLGAGMAPKIAWALENKPRIPVVWIHGL WP_001058323.1
MULTISPECIES: hydrogenase-1 small chain [Proteobacteria] Length: 372\nNumber of predicted TMHs: 2\nExp number of AAs in TMHs: 37.2608499999999\nExp number, first 60 AAs: 19.52923\nTotal prob of N-in: 0.94351\nPOSSIBLE N-term signal sequence\ninside 1 19\nTMhelix 20 39\noutside 40 325\nTMhelix 326 348\ninside 349 372

2922 GCF_001562855.1_ASM156285v1 Escherichia coli Proteobacteria; Gammaproteobacteria; Enterobacterales; Enterobacteriaceae; Escherichia
MTGDNTLIHSHGINRRDFMKLCAALAATMGLSSKAAAEMAESVTNPQRPPVIWIGAQECT WP_000145410.1
MULTISPECIES: hydrogenase-2 small chain [Proteobacteria] Length: 372\nNumber of predicted TMHs: 1\nExp number of AAs in TMHs: 23.72226\nExp number, first 60 AAs: 0.22038\nTotal prob of N-in: 0.12994\noutside 1 331\nTMhelix 332 354\ninside 355 372

2923 GCF_001562635.1_ASM156263v1 Escherichia coli Proteobacteria; Gammaproteobacteria; Enterobacterales; Enterobacteriaceae; Escherichia
MNNEETFYQAMRRQGVTRRSFLKYCSLAATSLGLGAGMAPKIAWALENKPRIPVVWIHGL WP_001058323.1
MULTISPECIES: hydrogenase-1 small chain [Proteobacteria] Length: 372\nNumber of predicted TMHs: 2\nExp number of AAs in TMHs: 37.2608499999999\nExp number, first 60 AAs: 19.52923\nTotal prob of N-in: 0.94351\nPOSSIBLE N-term signal sequence\ninside 1 19\nTMhelix 20 39\noutside 40 325\nTMhelix 326 348\ninside 349 372

2924 GCF_001562755.1_ASM156275v1 Escherichia coli Proteobacteria; Gammaproteobacteria; Enterobacterales; Enterobacteriaceae; Escherichia
MTGDNTLIHSHGINRRDFMKLCAALAATMGLSSKAAAEMAESVTNPQRPPVIWIGAQECT WP_000145410.1
MULTISPECIES: hydrogenase-2 small chain [Proteobacteria] Length: 372\nNumber of predicted TMHs: 1\nExp number of AAs in TMHs: 23.72226\nExp number, first 60 AAs: 0.22038\nTotal prob of N-in: 0.12994\noutside 1 331\nTMhelix 332 354\ninside 355 372

2925 GCF_001562755.1_ASM156275v1 Escherichia coli Proteobacteria; Gammaproteobacteria; Enterobacterales; Enterobacteriaceae; Escherichia
MNNEETFYQAMRRQGVTRRSFLKYCSLAATSLGLGAGMAPKIAWALENKPRIPVVWIHGL WP_001058323.1
MULTISPECIES: hydrogenase-1 small chain [Proteobacteria] Length: 372\nNumber of predicted TMHs: 2\nExp number of AAs in TMHs: 37.2608499999999\nExp number, first 60 AAs: 19.52923\nTotal prob of N-in:

0.94351\nPOSSIBLE N-term signal sequence\ninside 1 19\nTMhelix 20 39\noutside 40 325\nTMhelix 326 348\ninside 349 372

2926 GCF_001562645.1_ASM156264v1 Escherichia coli Proteobacteria; Gammaproteobacteria; Enterobacterales; Enterobacteriaceae; Escherichia
MNNEETFYQAMRRQGVTRRSFLKYCSLAATSLGLGAGMAPKIAWALENKPRIPVVWIHGL WP_001058323.1
MULTISPECIES: hydrogenase-1 small chain [Proteobacteria] Length: 372\nNumber of predicted TMHs: 2\nExp number of AAs in TMHs: 37.2608499999999\nExp number, first 60 AAs: 19.52923\nTotal prob of N-in: 0.94351\nPOSSIBLE N-term signal sequence\ninside 1 19\nTMhelix 20 39\noutside 40 325\nTMhelix 326 348\ninside 349 372

2927 GCF_001562815.1_ASM156281v1 Escherichia coli Proteobacteria; Gammaproteobacteria; Enterobacterales; Enterobacteriaceae; Escherichia
MTGDNTLIHSHGINRRDFMKLCAALAATMGLSSKAAAEMAESVTNPQRPPVIWIGAQECT WP_000145410.1
MULTISPECIES: hydrogenase-2 small chain [Proteobacteria] Length: 372\nNumber of predicted TMHs: 1\nExp number of AAs in TMHs: 23.72226\nExp number, first 60 AAs: 0.22038\nTotal prob of N-in: 0.12994\noutside 1 331\nTMhelix 332 354\ninside 355 372

2928 GCF_001562815.1_ASM156281v1 Escherichia coli Proteobacteria; Gammaproteobacteria; Enterobacterales; Enterobacteriaceae; Escherichia
MNNEETFYQAMRRQGVTRRSFLKYCSLAATSLGLGAGMAPKIAWALENKPRIPVVWIHGL WP_001058323.1
MULTISPECIES: hydrogenase-1 small chain [Proteobacteria] Length: 372\nNumber of predicted TMHs: 2\nExp number of AAs in TMHs: 37.2608499999999\nExp number, first 60 AAs: 19.52923\nTotal prob of N-in: 0.94351\nPOSSIBLE N-term signal sequence\ninside 1 19\nTMhelix 20 39\noutside 40 325\nTMhelix 326 348\ninside 349 372

2929 GCF_001562695.1_ASM156269v1 Escherichia coli Proteobacteria; Gammaproteobacteria; Enterobacterales; Enterobacteriaceae; Escherichia
MNNEETFYQAMRRQGVTRRSFLKYCSLAATSLGLGAGMAPKIAWALENKPRIPVVWIHGL WP_001058323.1
MULTISPECIES: hydrogenase-1 small chain [Proteobacteria] Length: 372\nNumber of predicted TMHs: 2\nExp number of AAs in TMHs: 37.2608499999999\nExp number, first 60 AAs: 19.52923\nTotal prob of N-in: 0.94351\nPOSSIBLE N-term signal sequence\ninside 1 19\nTMhelix 20 39\noutside 40 325\nTMhelix 326 348\ninside 349 372

2930 GCF_001563405.1_ASM156340v1 Escherichia coli Proteobacteria; Gammaproteobacteria; Enterobacterales; Enterobacteriaceae; Escherichia
MNNEETFYQAMRRQGVTRRSFLKYCSLAATSLGLGAGMAPKIAWALENKPRIPVVWIHGL WP_001058323.1
MULTISPECIES: hydrogenase-1 small chain [Proteobacteria] Length: 372\nNumber of predicted TMHs: 2\nExp number of AAs in TMHs: 37.2608499999999\nExp number, first 60 AAs: 19.52923\nTotal prob of N-in: 0.94351\nPOSSIBLE N-term signal sequence\ninside 1 19\nTMhelix 20 39\noutside 40 325\nTMhelix 326 348\ninside 349 372

2931 GCF_001566635.1_ASM156663v1 Escherichia coli Proteobacteria; Gammaproteobacteria; Enterobacterales; Enterobacteriaceae; Escherichia
MTGDNTLIHSHGINRRDFMKLCAALAATMGLSSKAAAEMAESVTNPQRPPVIWIGAQECT WP_000145410.1
MULTISPECIES: hydrogenase-2 small chain [Proteobacteria] Length: 372\nNumber of predicted TMHs: 1\nExp number of AAs in TMHs: 23.72226\nExp number, first 60 AAs: 0.22038\nTotal prob of N-in: 0.12994\noutside 1 331\nTMhelix 332 354\ninside 355 372

2932 GCF_001566635.1_ASM156663v1 Escherichia coli Proteobacteria; Gammaproteobacteria; Enterobacterales; Enterobacteriaceae; Escherichia
MNNEETFYQAMRRQGVTRRSFLKYCSLAATSLGLGAGMAPKIAWALENKPRIPVVWIHGL WP_001295940.1
MULTISPECIES: hydrogenase [Enterobacteriaceae] Length: 372\nNumber of predicted TMHs: 2\nExp number of AAs in TMHs: 37.2608899999999\nExp number, first 60 AAs: 19.52923\nTotal prob of N-in: 0.94351\nPOSSIBLE N-term signal sequence\ninside 1 19\nTMhelix 20 39\noutside 40 325\nTMhelix 326 348\ninside 349 372

2933 GCF_001563715.1_ASM156371v1 Escherichia coli Proteobacteria; Gammaproteobacteria; Enterobacterales; Enterobacteriaceae; Escherichia
MNNEETFYQAMRRQGVTRRSFLKYCSLAATSLGLGAGMAPKIAWALENKPRIPVVWIHGL WP_001058323.1
MULTISPECIES: hydrogenase-1 small chain [Proteobacteria] Length: 372\nNumber of predicted TMHs:

2\nExp number of AAs in TMHs: 37.2608499999999\nExp number, first 60 AAs: 19.52923\nTotal prob of N-in: 0.94351\nPOSSIBLE N-term signal sequence\ninside 1 19\nTMhelix 20 39\noutside 40 325\nTMhelix 326 348\ninside 349 372

2934 GCF_001571485.1_ASM157148v1 Escherichia coli Proteobacteria; Gammaproteobacteria; Enterobacterales; Enterobacteriaceae; Escherichia MTGDNTLIHSHGINRRDFMKLCAALAATMGLSSKAAAEMAESVTNPQRPPVIWIGAQECT WP_000145410.1 MULTISPECIES: hydrogenase-2 small chain [Proteobacteria] Length: 372\nNumber of predicted TMHs: 1\nExp number of AAs in TMHs: 23.72226\nExp number, first 60 AAs: 0.22038\nTotal prob of N-in: 0.12994\noutside 1 331\nTMhelix 332 354\ninside 355 372

2935 GCF_001571495.1_ASM157149v1 Escherichia coli Proteobacteria; Gammaproteobacteria; Enterobacterales; Enterobacteriaceae; Escherichia MTGDNTLIHSHGINRRDFMKLCAALAATMGLSSKAAAEMAESVTNPQRPPVIWIGAQECT WP_000145410.1 MULTISPECIES: hydrogenase-2 small chain [Proteobacteria] Length: 372\nNumber of predicted TMHs: 1\nExp number of AAs in TMHs: 23.72226\nExp number, first 60 AAs: 0.22038\nTotal prob of N-in: 0.12994\noutside 1 331\nTMhelix 332 354\ninside 355 372

2936 GCF_001571565.1_ASM157156v1 Escherichia coli Proteobacteria; Gammaproteobacteria; Enterobacterales; Enterobacteriaceae; Escherichia MTGDNTLIHSHGINRRDFMKLCAALAATMGLSSKAAAEMAESVTNPQRPPVIWIGAQECT WP_000145410.1 MULTISPECIES: hydrogenase-2 small chain [Proteobacteria] Length: 372\nNumber of predicted TMHs: 1\nExp number of AAs in TMHs: 23.72226\nExp number, first 60 AAs: 0.22038\nTotal prob of N-in: 0.12994\noutside 1 331\nTMhelix 332 354\ninside 355 372

2937 GCF_001571675.1_ASM157167v1 Escherichia coli Proteobacteria; Gammaproteobacteria; Enterobacterales; Enterobacteriaceae; Escherichia MTGDNTLIHSHGINRRDFMKLCAALAATMGLSSKAAAEMAESVTNPQRPPVIWIGAQECT WP_000145410.1 MULTISPECIES: hydrogenase-2 small chain [Proteobacteria] Length: 372\nNumber of predicted TMHs: 1\nExp number of AAs in TMHs: 23.72226\nExp number, first 60 AAs: 0.22038\nTotal prob of N-in: 0.12994\noutside 1 331\nTMhelix 332 354\ninside 355 372

2938 GCF_001571725.1_ASM157172v1 Escherichia coli Proteobacteria; Gammaproteobacteria; Enterobacterales; Enterobacteriaceae; Escherichia MNNEETFYQAMRRQGVTRRSFLKYCSLAATSLGLGAGMAPKIAWALENKPRIPVVWIHGL WP_001058323.1 MULTISPECIES: hydrogenase-1 small chain [Proteobacteria] Length: 372\nNumber of predicted TMHs: 2\nExp number of AAs in TMHs: 37.2608499999999\nExp number, first 60 AAs: 19.52923\nTotal prob of N-in: 0.94351\nPOSSIBLE N-term signal sequence\ninside 1 19\nTMhelix 20 39\noutside 40 325\nTMhelix 326 348\ninside 349 372

2939 GCF_001571675.1_ASM157167v1 Escherichia coli Proteobacteria; Gammaproteobacteria; Enterobacterales; Enterobacteriaceae; Escherichia MNNEETFYQAMRRQGVTRRSFLKYCSLAATSLGLGAGMAPKIAWALENKPRIPVVWIHGL WP_001058323.1 MULTISPECIES: hydrogenase-1 small chain [Proteobacteria] Length: 372\nNumber of predicted TMHs: 2\nExp number of AAs in TMHs: 37.2608499999999\nExp number, first 60 AAs: 19.52923\nTotal prob of N-in: 0.94351\nPOSSIBLE N-term signal sequence\ninside 1 19\nTMhelix 20 39\noutside 40 325\nTMhelix 326 348\ninside 349 372

2940 GCF_001571485.1_ASM157148v1 Escherichia coli Proteobacteria; Gammaproteobacteria; Enterobacterales; Enterobacteriaceae; Escherichia MNNEETFYQAMRRQGVTRRSFLKYCSLAATSLGLGAGMAPKIAWALENKPRIPVVWIHGL WP_001058323.1 MULTISPECIES: hydrogenase-1 small chain [Proteobacteria] Length: 372\nNumber of predicted TMHs: 2\nExp number of AAs in TMHs: 37.2608499999999\nExp number, first 60 AAs: 19.52923\nTotal prob of N-in: 0.94351\nPOSSIBLE N-term signal sequence\ninside 1 19\nTMhelix 20 39\noutside 40 325\nTMhelix 326 348\ninside 349 372

2941 GCF_001571805.1_ASM157180v1 Escherichia coli Proteobacteria; Gammaproteobacteria; Enterobacterales; Enterobacteriaceae; Escherichia MTGDNTLIHSHGINRRDFMKLCAALAATMGLSSKAAAEMAESVTNPQRPPVIWIGAQECT WP_000145410.1 MULTISPECIES: hydrogenase-2 small chain [Proteobacteria] Length: 372\nNumber of predicted TMHs:

1\nExp number of AAs in TMHs: 23.72226\nExp number, first 60 AAs: 0.22038\nTotal prob of N-in: 0.12994\noutside 1 331\nTMhelix 332 354\ninside 355 372

2942 GCF_001571805.1_ASM157180v1 Escherichia coli Proteobacteria; Gammaproteobacteria; Enterobacterales; Enterobacteriaceae; Escherichia
MNNEETFYQAMRRQGVTRRSFLKYCSLAATSLGLGAGMAPKIAWALENKPRIPVVWIHGL WP_001058323.1
MULTISPECIES: hydrogenase-1 small chain [Proteobacteria] Length: 372\nNumber of predicted TMHs: 2\nExp number of AAs in TMHs: 37.2608499999999\nExp number, first 60 AAs: 19.52923\nTotal prob of N-in: 0.94351\nPOSSIBLE N-term signal sequence\ninside 1 19\nTMhelix 20 39\noutside 40 325\nTMhelix 326 348\ninside 349 372

2943 GCF_001571925.1_ASM157192v1 Escherichia coli Proteobacteria; Gammaproteobacteria; Enterobacterales; Enterobacteriaceae; Escherichia
MNNEETFYQAMRRQGVTRRSFLKYCSLAATSLGLGAGMAPKIAWALENKPRIPVVWIHGL WP_001058323.1
MULTISPECIES: hydrogenase-1 small chain [Proteobacteria] Length: 372\nNumber of predicted TMHs: 2\nExp number of AAs in TMHs: 37.2608499999999\nExp number, first 60 AAs: 19.52923\nTotal prob of N-in: 0.94351\nPOSSIBLE N-term signal sequence\ninside 1 19\nTMhelix 20 39\noutside 40 325\nTMhelix 326 348\ninside 349 372

2944 GCF_001571965.1_ASM157196v1 Escherichia coli Proteobacteria; Gammaproteobacteria; Enterobacterales; Enterobacteriaceae; Escherichia
MNNEETFYQAMRRQGVTRRSFLKYCSLAATSLGLGAGMAPKIAWALENKPRIPVVWIHGL WP_001058323.1
MULTISPECIES: hydrogenase-1 small chain [Proteobacteria] Length: 372\nNumber of predicted TMHs: 2\nExp number of AAs in TMHs: 37.2608499999999\nExp number, first 60 AAs: 19.52923\nTotal prob of N-in: 0.94351\nPOSSIBLE N-term signal sequence\ninside 1 19\nTMhelix 20 39\noutside 40 325\nTMhelix 326 348\ninside 349 372

2945 GCF_001572245.1_ASM157224v1 Escherichia coli Proteobacteria; Gammaproteobacteria; Enterobacterales; Enterobacteriaceae; Escherichia
MTGDNTLIHSHGINRRDFMKLCAALAATMGLSSKAAAEMAESVTNPQRPPVIWIGAQECT WP_000145410.1
MULTISPECIES: hydrogenase-2 small chain [Proteobacteria] Length: 372\nNumber of predicted TMHs: 1\nExp number of AAs in TMHs: 23.72226\nExp number, first 60 AAs: 0.22038\nTotal prob of N-in: 0.12994\noutside 1 331\nTMhelix 332 354\ninside 355 372

2946 GCF_001572215.1_ASM157221v1 Escherichia coli Proteobacteria; Gammaproteobacteria; Enterobacterales; Enterobacteriaceae; Escherichia
MTGDNTLIHSHGINRRDFMKLCAALAATMGLSSKAAAEMAESVTNPQRPPVIWIGAQECT WP_000145410.1
MULTISPECIES: hydrogenase-2 small chain [Proteobacteria] Length: 372\nNumber of predicted TMHs: 1\nExp number of AAs in TMHs: 23.72226\nExp number, first 60 AAs: 0.22038\nTotal prob of N-in: 0.12994\noutside 1 331\nTMhelix 332 354\ninside 355 372

2947 GCF_001572285.1_ASM157228v1 Escherichia coli Proteobacteria; Gammaproteobacteria; Enterobacterales; Enterobacteriaceae; Escherichia
MTGDNTLIHSHGINRRDFMKLCAALAATMGLSSKAAAEMAESVTNPQRPPVIWIGAQECT WP_000145410.1
MULTISPECIES: hydrogenase-2 small chain [Proteobacteria] Length: 372\nNumber of predicted TMHs: 1\nExp number of AAs in TMHs: 23.72226\nExp number, first 60 AAs: 0.22038\nTotal prob of N-in: 0.12994\noutside 1 331\nTMhelix 332 354\ninside 355 372

2948 GCF_001572245.1_ASM157224v1 Escherichia coli Proteobacteria; Gammaproteobacteria; Enterobacterales; Enterobacteriaceae; Escherichia
MNNEETFYQAMRRQGVTRRSFLKYCSLAATSLGLGAGMAPKIAWALENKPRIPVVWIHGL WP_001058323.1
MULTISPECIES: hydrogenase-1 small chain [Proteobacteria] Length: 372\nNumber of predicted TMHs: 2\nExp number of AAs in TMHs: 37.2608499999999\nExp number, first 60 AAs: 19.52923\nTotal prob of N-in: 0.94351\nPOSSIBLE N-term signal sequence\ninside 1 19\nTMhelix 20 39\noutside 40 325\nTMhelix 326 348\ninside 349 372

2949 GCF_001572215.1_ASM157221v1 Escherichia coli Proteobacteria; Gammaproteobacteria; Enterobacterales; Enterobacteriaceae; Escherichia
MNNEETFYQAMRRQGVTRRSFLKYCSLAATSLGLGAGMAPKIAWALENKPRIPVVWIHGL WP_001058323.1
MULTISPECIES: hydrogenase-1 small chain [Proteobacteria] Length: 372\nNumber of predicted TMHs: 2\nExp number of AAs in TMHs: 37.2608499999999\nExp number, first 60 AAs: 19.52923\nTotal prob of N-in:

0.94351\nPOSSIBLE N-term signal sequence\ninside 1 19\nTMhelix 20 39\noutside 40 325\nTMhelix 326 348\ninside 349 372

2950 GCF_001572285.1_ASM157228v1 Escherichia coli Proteobacteria; Gammaproteobacteria; Enterobacterales; Enterobacteriaceae; Escherichia MNNEETFYQAMRRQGVTRRSFLKYCSLAATSLGLGAGMAPKIAWALENKPRIPVVWIHGL WP_001058323.1 MULTISPECIES: hydrogenase-1 small chain [Proteobacteria] Length: 372\nNumber of predicted TMHs: 2\nExp number of AAs in TMHs: 37.2608499999999\nExp number, first 60 AAs: 19.52923\nTotal prob of N-in: 0.94351\nPOSSIBLE N-term signal sequence\ninside 1 19\nTMhelix 20 39\noutside 40 325\nTMhelix 326 348\ninside 349 372

2951 GCF_001571995.1_ASM157199v1 Escherichia coli Proteobacteria; Gammaproteobacteria; Enterobacterales; Enterobacteriaceae; Escherichia MNNEETFYQAMRRQGVTRRSFLKYCSLAATSLGLGAGMAPKIAWALENKPRIPVVWIHGL WP_001058323.1 MULTISPECIES: hydrogenase-1 small chain [Proteobacteria] Length: 372\nNumber of predicted TMHs: 2\nExp number of AAs in TMHs: 37.2608499999999\nExp number, first 60 AAs: 19.52923\nTotal prob of N-in: 0.94351\nPOSSIBLE N-term signal sequence\ninside 1 19\nTMhelix 20 39\noutside 40 325\nTMhelix 326 348\ninside 349 372

2952 GCF_001572325.1_ASM157232v1 Escherichia coli Proteobacteria; Gammaproteobacteria; Enterobacterales; Enterobacteriaceae; Escherichia MTGDNTLIHSHGINRRDFMKLCAALAATMGLSSKAAAEMAESVTNPQRPPVIWIGAQECT WP_000145410.1 MULTISPECIES: hydrogenase-2 small chain [Proteobacteria] Length: 372\nNumber of predicted TMHs: 1\nExp number of AAs in TMHs: 23.72226\nExp number, first 60 AAs: 0.22038\nTotal prob of N-in: 0.12994\noutside 1 331\nTMhelix 332 354\ninside 355 372

2953 GCF_001572325.1_ASM157232v1 Escherichia coli Proteobacteria; Gammaproteobacteria; Enterobacterales; Enterobacteriaceae; Escherichia MNNEETFYQAMRRQGVTRRSFLKYCSLAATSLGLGAGMAPKIAWALENKPRIPVVWIHGL WP_001058323.1 MULTISPECIES: hydrogenase-1 small chain [Proteobacteria] Length: 372\nNumber of predicted TMHs: 2\nExp number of AAs in TMHs: 37.2608499999999\nExp number, first 60 AAs: 19.52923\nTotal prob of N-in: 0.94351\nPOSSIBLE N-term signal sequence\ninside 1 19\nTMhelix 20 39\noutside 40 325\nTMhelix 326 348\ninside 349 372

2954 GCF_001572045.1_ASM157204v1 Escherichia coli Proteobacteria; Gammaproteobacteria; Enterobacterales; Enterobacteriaceae; Escherichia MNNEETFYQAMRRQGVTRRSFLKYCSLAATSLGLGAGMAPKIAWALENKPRIPVVWIHGL WP_001058323.1 MULTISPECIES: hydrogenase-1 small chain [Proteobacteria] Length: 372\nNumber of predicted TMHs: 2\nExp number of AAs in TMHs: 37.2608499999999\nExp number, first 60 AAs: 19.52923\nTotal prob of N-in: 0.94351\nPOSSIBLE N-term signal sequence\ninside 1 19\nTMhelix 20 39\noutside 40 325\nTMhelix 326 348\ninside 349 372

2955 GCF_001572345.1_ASM157234v1 Escherichia coli Proteobacteria; Gammaproteobacteria; Enterobacterales; Enterobacteriaceae; Escherichia MTGDNTLIHSHGINRRDFMKLCAALAATMGLSSKAAAEMAESVTNPQRPPVIWIGAQECT WP_000145410.1 MULTISPECIES: hydrogenase-2 small chain [Proteobacteria] Length: 372\nNumber of predicted TMHs: 1\nExp number of AAs in TMHs: 23.72226\nExp number, first 60 AAs: 0.22038\nTotal prob of N-in: 0.12994\noutside 1 331\nTMhelix 332 354\ninside 355 372

2956 GCF_001572345.1_ASM157234v1 Escherichia coli Proteobacteria; Gammaproteobacteria; Enterobacterales; Enterobacteriaceae; Escherichia MNNEETFYQAMRRQGVTRRSFLKYCSLAATSLGLGAGMAPKIAWALENKPRIPVVWIHGL WP_001058323.1 MULTISPECIES: hydrogenase-1 small chain [Proteobacteria] Length: 372\nNumber of predicted TMHs: 2\nExp number of AAs in TMHs: 37.2608499999999\nExp number, first 60 AAs: 19.52923\nTotal prob of N-in: 0.94351\nPOSSIBLE N-term signal sequence\ninside 1 19\nTMhelix 20 39\noutside 40 325\nTMhelix 326 348\ninside 349 372

2957 GCF_001572525.1_ASM157252v1 Escherichia coli Proteobacteria; Gammaproteobacteria; Enterobacterales; Enterobacteriaceae; Escherichia MTGDNTLIHSHGINRRDFMKLCAALAATMGLSSKAAAEMAESVTNPQRPPVIWIGAQECT WP_000145410.1 MULTISPECIES: hydrogenase-2 small chain [Proteobacteria] Length: 372\nNumber of predicted TMHs:

1\nExp number of AAs in TMHs: 23.72226\nExp number, first 60 AAs: 0.22038\nTotal prob of N-in: 0.12994\noutside 1 331\nTMhelix 332 354\ninside 355 372

2958 GCF_001572425.1_ASM157242v1 Escherichia coli Proteobacteria; Gammaproteobacteria; Enterobacterales; Enterobacteriaceae; Escherichia
MNNEETFYQAMRRQGVTRRSFLKYCSLAATSLGLGAGMAPKIAWALENKPRIPVVWIHGL WP_001058323.1
MULTISPECIES: hydrogenase-1 small chain [Proteobacteria] Length: 372\nNumber of predicted TMHs: 2\nExp number of AAs in TMHs: 37.2608499999999\nExp number, first 60 AAs: 19.52923\nTotal prob of N-in: 0.94351\nPOSSIBLE N-term signal sequence\ninside 1 19\nTMhelix 20 39\noutside 40 325\nTMhelix 326 348\ninside 349 372

2959 GCF_001572585.1_ASM157258v1 Escherichia coli Proteobacteria; Gammaproteobacteria; Enterobacterales; Enterobacteriaceae; Escherichia
MTGDNTLIHSHGINRRDFMKLCAALAATMGLSSKAAAEMAESVTNPQRPPVIWIGAQECT WP_000145410.1
MULTISPECIES: hydrogenase-2 small chain [Proteobacteria] Length: 372\nNumber of predicted TMHs: 1\nExp number of AAs in TMHs: 23.72226\nExp number, first 60 AAs: 0.22038\nTotal prob of N-in: 0.12994\noutside 1 331\nTMhelix 332 354\ninside 355 372

2960 GCF_001572635.1_ASM157263v1 Escherichia coli Proteobacteria; Gammaproteobacteria; Enterobacterales; Enterobacteriaceae; Escherichia
MTGDNTLIHSHGINRRDFMKLCAALAATMGLSSKAAAEMAESVTNPQRPPVIWIGAQECT WP_000145410.1
MULTISPECIES: hydrogenase-2 small chain [Proteobacteria] Length: 372\nNumber of predicted TMHs: 1\nExp number of AAs in TMHs: 23.72226\nExp number, first 60 AAs: 0.22038\nTotal prob of N-in: 0.12994\noutside 1 331\nTMhelix 332 354\ninside 355 372

2961 GCF_001572585.1_ASM157258v1 Escherichia coli Proteobacteria; Gammaproteobacteria; Enterobacterales; Enterobacteriaceae; Escherichia
MNNEETFYQAMRRQGVTRRSFLKYCSLAATSLGLGAGMAPKIAWALENKPRIPVVWIHGL WP_001058323.1
MULTISPECIES: hydrogenase-1 small chain [Proteobacteria] Length: 372\nNumber of predicted TMHs: 2\nExp number of AAs in TMHs: 37.2608499999999\nExp number, first 60 AAs: 19.52923\nTotal prob of N-in: 0.94351\nPOSSIBLE N-term signal sequence\ninside 1 19\nTMhelix 20 39\noutside 40 325\nTMhelix 326 348\ninside 349 372

2962 GCF_001575635.1_ASM157563v1 Escherichia coli Proteobacteria; Gammaproteobacteria; Enterobacterales; Enterobacteriaceae; Escherichia
MTGDNTLIHSHGINRRDFMKLCAALAATMGLSSKAAAEMAESVTNPQRPPVIWIGAQECT WP_000145410.1
MULTISPECIES: hydrogenase-2 small chain [Proteobacteria] Length: 372\nNumber of predicted TMHs: 1\nExp number of AAs in TMHs: 23.72226\nExp number, first 60 AAs: 0.22038\nTotal prob of N-in: 0.12994\noutside 1 331\nTMhelix 332 354\ninside 355 372

2963 GCF_001572485.1_ASM157248v1 Escherichia coli Proteobacteria; Gammaproteobacteria; Enterobacterales; Enterobacteriaceae; Escherichia
MNNEETFYQAMRRQGVTRRSFLKYCSLAATSLGLGAGMAPKIAWALENKPRIPVVWIHGL WP_001058323.1
MULTISPECIES: hydrogenase-1 small chain [Proteobacteria] Length: 372\nNumber of predicted TMHs: 2\nExp number of AAs in TMHs: 37.2608499999999\nExp number, first 60 AAs: 19.52923\nTotal prob of N-in: 0.94351\nPOSSIBLE N-term signal sequence\ninside 1 19\nTMhelix 20 39\noutside 40 325\nTMhelix 326 348\ninside 349 372

2964 GCF_001572525.1_ASM157252v1 Escherichia coli Proteobacteria; Gammaproteobacteria; Enterobacterales; Enterobacteriaceae; Escherichia
MNNEETFYQAMRRQGVTRRSFLKYCSLAATSLGLGAGMAPKIAWALENKPRIPVVWIHGL WP_001058323.1
MULTISPECIES: hydrogenase-1 small chain [Proteobacteria] Length: 372\nNumber of predicted TMHs: 2\nExp number of AAs in TMHs: 37.2608499999999\nExp number, first 60 AAs: 19.52923\nTotal prob of N-in: 0.94351\nPOSSIBLE N-term signal sequence\ninside 1 19\nTMhelix 20 39\noutside 40 325\nTMhelix 326 348\ninside 349 372

2965 GCF_001572535.1_ASM157253v1 Escherichia coli Proteobacteria; Gammaproteobacteria; Enterobacterales; Enterobacteriaceae; Escherichia
MNNEETFYQAMRRQGVTRRSFLKYCSLAATSLGLGAGMAPKIAWALENKPRIPVVWIHGL WP_001058323.1
MULTISPECIES: hydrogenase-1 small chain [Proteobacteria] Length: 372\nNumber of predicted TMHs: 2\nExp number of AAs in TMHs: 37.2608499999999\nExp number, first 60 AAs: 19.52923\nTotal prob of N-in:

0.94351\nPOSSIBLE N-term signal sequence\ninside 1 19\nTMhelix 20 39\noutside 40 325\nTMhelix 326 348\ninside 349 372

2966 GCF_001575375.1_ASM157537v1 Escherichia coli Proteobacteria; Gammaproteobacteria; Enterobacterales; Enterobacteriaceae; Escherichia MTGDNTLIHSHGINRRDFMKLCAALAATMGLSSKAAAEMAESVTNPQRPPVIWIGAQECT WP_000145410.1 MULTISPECIES: hydrogenase-2 small chain [Proteobacteria] Length: 372\nNumber of predicted TMHs: 1\nExp number of AAs in TMHs: 23.72226\nExp number, first 60 AAs: 0.22038\nTotal prob of N-in: 0.12994\noutside 1 331\nTMhelix 332 354\ninside 355 372

2967 GCF_001575455.1_ASM157545v1 Escherichia coli Proteobacteria; Gammaproteobacteria; Enterobacterales; Enterobacteriaceae; Escherichia MTGDNTLIHSHGINRRDFMKLCAALAATMGLSSKAAAEMAESVTNPQRPPVIWIGAQECT WP_000145410.1 MULTISPECIES: hydrogenase-2 small chain [Proteobacteria] Length: 372\nNumber of predicted TMHs: 1\nExp number of AAs in TMHs: 23.72226\nExp number, first 60 AAs: 0.22038\nTotal prob of N-in: 0.12994\noutside 1 331\nTMhelix 332 354\ninside 355 372

2968 GCF_001575465.1_ASM157546v1 Escherichia coli Proteobacteria; Gammaproteobacteria; Enterobacterales; Enterobacteriaceae; Escherichia MTGDNTLIHSHGINRRDFMKLCAALAATMGLSSKAAAEMAESVTNPQRPPVIWIGAQECT WP_000145410.1 MULTISPECIES: hydrogenase-2 small chain [Proteobacteria] Length: 372\nNumber of predicted TMHs: 1\nExp number of AAs in TMHs: 23.72226\nExp number, first 60 AAs: 0.22038\nTotal prob of N-in: 0.12994\noutside 1 331\nTMhelix 332 354\ninside 355 372

2969 GCF_001575515.1_ASM157551v1 Escherichia coli Proteobacteria; Gammaproteobacteria; Enterobacterales; Enterobacteriaceae; Escherichia MTGDNTLIHSHGINRRDFMKLCAALAATMGLSSKAAAEMAESVTNPQRPPVIWIGAQECT WP_000145410.1 MULTISPECIES: hydrogenase-2 small chain [Proteobacteria] Length: 372\nNumber of predicted TMHs: 1\nExp number of AAs in TMHs: 23.72226\nExp number, first 60 AAs: 0.22038\nTotal prob of N-in: 0.12994\noutside 1 331\nTMhelix 332 354\ninside 355 372

2970 GCF_001575615.1_ASM157561v1 Escherichia coli Proteobacteria; Gammaproteobacteria; Enterobacterales; Enterobacteriaceae; Escherichia MNNEETFYQAMRRQGVTRRSFLKYCSLAATSLGLGAGMAPKIAWALENKPRIPVVWIHGL WP_001058323.1 MULTISPECIES: hydrogenase-1 small chain [Proteobacteria] Length: 372\nNumber of predicted TMHs: 2\nExp number of AAs in TMHs: 37.2608499999999\nExp number, first 60 AAs: 19.52923\nTotal prob of N-in: 0.94351\nPOSSIBLE N-term signal sequence\ninside 1 19\nTMhelix 20 39\noutside 40 325\nTMhelix 326 348\ninside 349 372

2971 GCF_001575515.1_ASM157551v1 Escherichia coli Proteobacteria; Gammaproteobacteria; Enterobacterales; Enterobacteriaceae; Escherichia MNNEETFYQAMRRQGVTRRSFLKYCSLAATSLGLGAGMAPKIAWALENKPRIPVVWIHGL WP_001058323.1 MULTISPECIES: hydrogenase-1 small chain [Proteobacteria] Length: 372\nNumber of predicted TMHs: 2\nExp number of AAs in TMHs: 37.2608499999999\nExp number, first 60 AAs: 19.52923\nTotal prob of N-in: 0.94351\nPOSSIBLE N-term signal sequence\ninside 1 19\nTMhelix 20 39\noutside 40 325\nTMhelix 326 348\ninside 349 372

2972 GCF_001575705.1_ASM157570v1 Escherichia coli Proteobacteria; Gammaproteobacteria; Enterobacterales; Enterobacteriaceae; Escherichia MTGDNTLIHSHGINRRDFMKLCAALAATMGLSSKAAAEMAESVTNPQRPPVIWIGAQECT WP_000145410.1 MULTISPECIES: hydrogenase-2 small chain [Proteobacteria] Length: 372\nNumber of predicted TMHs: 1\nExp number of AAs in TMHs: 23.72226\nExp number, first 60 AAs: 0.22038\nTotal prob of N-in: 0.12994\noutside 1 331\nTMhelix 332 354\ninside 355 372

2973 GCF_001575465.1_ASM157546v1 Escherichia coli Proteobacteria; Gammaproteobacteria; Enterobacterales; Enterobacteriaceae; Escherichia MNNEETFYQAMRRQGVTRRSFLKYCSLAATSLGLGAGMAPKIAWALENKPRIPVVWIHGL WP_001058323.1 MULTISPECIES: hydrogenase-1 small chain [Proteobacteria] Length: 372\nNumber of predicted TMHs: 2\nExp number of AAs in TMHs: 37.2608499999999\nExp number, first 60 AAs: 19.52923\nTotal prob of N-in: 0.94351\nPOSSIBLE N-term signal sequence\ninside 1 19\nTMhelix 20 39\noutside 40 325\nTMhelix 326 348\ninside 349 372

2974 GCF_001575755.1_ASM157575v1 Escherichia coli Proteobacteria; Gammaproteobacteria; Enterobacterales; Enterobacteriaceae; Escherichia
MTGDNTLIHSHGINRRDFMKLCAALAATMGLSSKAAAEMAESVTNPQRPPVIWIGAQECT WP_000145410.1
MULTISPECIES: hydrogenase-2 small chain [Proteobacteria] Length: 372\nNumber of predicted TMHs: 1\nExp number of AAs in TMHs: 23.72226\nExp number, first 60 AAs: 0.22038\nTotal prob of N-in: 0.12994\noutside 1 331\nTMhelix 332 354\ninside 355 372

2975 GCF_001575395.1_ASM157539v1 Escherichia coli Proteobacteria; Gammaproteobacteria; Enterobacterales; Enterobacteriaceae; Escherichia
MTGDNTLIHSHGINRRDFMKLCAALAATMGLSSKAAAEMAESVTNPQRPPVIWIGAQECT WP_061359937.1
hydrogenase 2 small subunit [Escherichia coli] Length: 372\nNumber of predicted TMHs: 1\nExp number of AAs in TMHs: 24.349\nExp number, first 60 AAs: 0.21646\nTotal prob of N-in: 0.15472\noutside 1 331\nTMhelix 332 354\ninside 355 372

2976 GCF_001575545.1_ASM157554v1 Escherichia coli Proteobacteria; Gammaproteobacteria; Enterobacterales; Enterobacteriaceae; Escherichia
MNNEETFYQAMRRQGVTRRSFLKYCSLAATSLGLGAGMAPKIAWALENKPRIPVVWIHGL WP_001058323.1
MULTISPECIES: hydrogenase-1 small chain [Proteobacteria] Length: 372\nNumber of predicted TMHs: 2\nExp number of AAs in TMHs: 37.2608499999999\nExp number, first 60 AAs: 19.52923\nTotal prob of N-in: 0.94351\nPOSSIBLE N-term signal sequence\ninside 1 19\nTMhelix 20 39\noutside 40 325\nTMhelix 326 348\ninside 349 372

2977 GCF_001575855.1_ASM157585v1 Escherichia coli Proteobacteria; Gammaproteobacteria; Enterobacterales; Enterobacteriaceae; Escherichia
MNNEETFYQAMRRQGVTRRSFLKYCSLAATSLGLGAGMAPKIAWALENKPRIPVVWIHGL WP_001058323.1
MULTISPECIES: hydrogenase-1 small chain [Proteobacteria] Length: 372\nNumber of predicted TMHs: 2\nExp number of AAs in TMHs: 37.2608499999999\nExp number, first 60 AAs: 19.52923\nTotal prob of N-in: 0.94351\nPOSSIBLE N-term signal sequence\ninside 1 19\nTMhelix 20 39\noutside 40 325\nTMhelix 326 348\ninside 349 372

2978 GCF_001575795.1_ASM157579v1 Escherichia coli Proteobacteria; Gammaproteobacteria; Enterobacterales; Enterobacteriaceae; Escherichia
MNNEETFYQAMRRQGVTRRSFLKYCSLAATSLGLGAGMAPKIAWALENKPRIPVVWIHGL WP_058055433.1
hydrogenase [Escherichia coli] Length: 372\nNumber of predicted TMHs: 2\nExp number of AAs in TMHs: 36.72011\nExp number, first 60 AAs: 19.53447\nTotal prob of N-in: 0.94308\nPOSSIBLE N-term signal sequence\ninside 1 19\nTMhelix 20 39\noutside 40 325\nTMhelix 326 348\ninside 349 372

2979 GCF_001575775.1_ASM157577v1 Escherichia coli Proteobacteria; Gammaproteobacteria; Enterobacterales; Enterobacteriaceae; Escherichia
MNNEETFYQAMRRQGVTRRSFLKYCSLAATSLGLGAGMAPKIAWALENKPRIPVVWIHGL WP_001058323.1
MULTISPECIES: hydrogenase-1 small chain [Proteobacteria] Length: 372\nNumber of predicted TMHs: 2\nExp number of AAs in TMHs: 37.2608499999999\nExp number, first 60 AAs: 19.52923\nTotal prob of N-in: 0.94351\nPOSSIBLE N-term signal sequence\ninside 1 19\nTMhelix 20 39\noutside 40 325\nTMhelix 326 348\ninside 349 372

2980 GCF_001575915.1_ASM157591v1 Escherichia coli Proteobacteria; Gammaproteobacteria; Enterobacterales; Enterobacteriaceae; Escherichia
MTGDNTLIHSHGINRRDFMKLCAALAATMGLSSKAAAEMAESVTNPQRPPVIWIGAQECT WP_000145410.1
MULTISPECIES: hydrogenase-2 small chain [Proteobacteria] Length: 372\nNumber of predicted TMHs: 1\nExp number of AAs in TMHs: 23.72226\nExp number, first 60 AAs: 0.22038\nTotal prob of N-in: 0.12994\noutside 1 331\nTMhelix 332 354\ninside 355 372

2981 GCF_001575675.1_ASM157567v1 Escherichia coli Proteobacteria; Gammaproteobacteria; Enterobacterales; Enterobacteriaceae; Escherichia
MNNEETFYQAMRRQGVTRRSFLKYCSLAATSLGLGAGMAPKIAWALENKPRIPVVWIHGL WP_061355971.1
hydrogenase [Escherichia coli] Length: 372\nNumber of predicted TMHs: 2\nExp number of AAs in TMHs: 39.12167\nExp number, first 60 AAs: 19.52599\nTotal prob of N-in: 0.94362\nPOSSIBLE N-term signal sequence\ninside 1 19\nTMhelix 20 39\noutside 40 325\nTMhelix 326 348\ninside 349 372

2982 GCF_001575865.1_ASM157586v1 Escherichia coli Proteobacteria; Gammaproteobacteria; Enterobacterales; Enterobacteriaceae; Escherichia

MTGDNTLIHSHGINRRDFMKLCAALAATMGLSSKAAAEMAESVTNPQRPPVIWIGAQECT WP_000145410.1 MULTISPECIES: hydrogenase-2 small chain [Proteobacteria] Length: 372\nNumber of predicted TMHs: 1\nExp number of AAs in TMHs: 23.72226\nExp number, first 60 AAs: 0.22038\nTotal prob of N-in: 0.12994\noutside 1 331\nTMhelix 332 354\ninside 355 372

2983 GCF_001575965.1_ASM157596v1 Escherichia coli Proteobacteria; Gammaproteobacteria; Enterobacterales; Enterobacteriaceae; Escherichia MTGDNTLIHSHGINRRDFMKLCAALAATMGLSSKAAAEMAESVTNPQRPPVIWIGAQECT WP_000145410.1 MULTISPECIES: hydrogenase-2 small chain [Proteobacteria] Length: 372\nNumber of predicted TMHs: 1\nExp number of AAs in TMHs: 23.72226\nExp number, first 60 AAs: 0.22038\nTotal prob of N-in: 0.12994\noutside 1 331\nTMhelix 332 354\ninside 355 372

2984 GCF_001576105.1_ASM157610v1 Escherichia coli Proteobacteria; Gammaproteobacteria; Enterobacterales; Enterobacteriaceae; Escherichia MNNEETFYQAMRRQGVTRRSFLKYCSLAATSLGLGAGMAPKIAWALENKPRIPVVWIHGL WP_001058323.1 MULTISPECIES: hydrogenase-1 small chain [Proteobacteria] Length: 372\nNumber of predicted TMHs: 2\nExp number of AAs in TMHs: 37.2608499999999\nExp number, first 60 AAs: 19.52923\nTotal prob of N-in: 0.94351\nPOSSIBLE N-term signal sequence\ninside 1 19\nTMhelix 20 39\noutside 40 325\nTMhelix 326 348\ninside 349 372

2985 GCF_001575865.1_ASM157586v1 Escherichia coli Proteobacteria; Gammaproteobacteria; Enterobacterales; Enterobacteriaceae; Escherichia MNNEETFYQAMRRQGVTRRSFLKYCSLAATSLGLGAGMAPKIAWALENKPRIPVVWIHGL WP_001058323.1 MULTISPECIES: hydrogenase-1 small chain [Proteobacteria] Length: 372\nNumber of predicted TMHs: 2\nExp number of AAs in TMHs: 37.2608499999999\nExp number, first 60 AAs: 19.52923\nTotal prob of N-in: 0.94351\nPOSSIBLE N-term signal sequence\ninside 1 19\nTMhelix 20 39\noutside 40 325\nTMhelix 326 348\ninside 349 372

2986 GCF_001576075.1_ASM157607v1 Escherichia coli Proteobacteria; Gammaproteobacteria; Enterobacterales; Enterobacteriaceae; Escherichia MTGDNTLIHSHGINRRDFMKLCAALAATMGLSSKAAAEMAESVTNPQRPPVIWIGAQECT WP_000145410.1 MULTISPECIES: hydrogenase-2 small chain [Proteobacteria] Length: 372\nNumber of predicted TMHs: 1\nExp number of AAs in TMHs: 23.72226\nExp number, first 60 AAs: 0.22038\nTotal prob of N-in: 0.12994\noutside 1 331\nTMhelix 332 354\ninside 355 372

2987 GCF_001576075.1_ASM157607v1 Escherichia coli Proteobacteria; Gammaproteobacteria; Enterobacterales; Enterobacteriaceae; Escherichia MNNEETFYQAMRRQGVTRRSFLKYCSLAATSLGLGAGMAPKIAWALENKPRIPVVWIHGL WP_001058323.1 MULTISPECIES: hydrogenase-1 small chain [Proteobacteria] Length: 372\nNumber of predicted TMHs: 2\nExp number of AAs in TMHs: 37.2608499999999\nExp number, first 60 AAs: 19.52923\nTotal prob of N-in: 0.94351\nPOSSIBLE N-term signal sequence\ninside 1 19\nTMhelix 20 39\noutside 40 325\nTMhelix 326 348\ninside 349 372

2988 GCF_001575965.1_ASM157596v1 Escherichia coli Proteobacteria; Gammaproteobacteria; Enterobacterales; Enterobacteriaceae; Escherichia MNNEETFYQAMRRQGVTRRSFLKYCSLAATSLGLGAGMAPKIAWALENKPRIPVVWIHGL WP_001058323.1 MULTISPECIES: hydrogenase-1 small chain [Proteobacteria] Length: 372\nNumber of predicted TMHs: 2\nExp number of AAs in TMHs: 37.2608499999999\nExp number, first 60 AAs: 19.52923\nTotal prob of N-in: 0.94351\nPOSSIBLE N-term signal sequence\ninside 1 19\nTMhelix 20 39\noutside 40 325\nTMhelix 326 348\ninside 349 372

2989 GCF_001575945.1_ASM157594v1 Escherichia coli Proteobacteria; Gammaproteobacteria; Enterobacterales; Enterobacteriaceae; Escherichia MNNEETFYQAMRRQGVTRRSFLKYCSLAATSLGLGAGMAPKIAWALENKPRIPVVWIHGL WP_001058323.1 MULTISPECIES: hydrogenase-1 small chain [Proteobacteria] Length: 372\nNumber of predicted TMHs: 2\nExp number of AAs in TMHs: 37.2608499999999\nExp number, first 60 AAs: 19.52923\nTotal prob of N-in: 0.94351\nPOSSIBLE N-term signal sequence\ninside 1 19\nTMhelix 20 39\noutside 40 325\nTMhelix 326 348\ninside 349 372

2990 GCF_001576035.1_ASM157603v1 Escherichia coli Proteobacteria; Gammaproteobacteria; Enterobacterales; Enterobacteriaceae; Escherichia

MNNEETFYQAMRRQGVTRRSFLKYCSLAATSLGLGAGMAPKIAWALENKPRIPVVWIHGL WP_001058323.1 MULTISPECIES: hydrogenase-1 small chain [Proteobacteria] Length: 372\nNumber of predicted TMHs: 2\nExp number of AAs in TMHs: 37.2608499999999\nExp number, first 60 AAs: 19.52923\nTotal prob of N-in: 0.94351\nPOSSIBLE N-term signal sequence\ninside 1 19\nTMhelix 20 39\noutside 40 325\nTMhelix 326 348\ninside 349 372

2991 GCF_001576495.1_ASM157649v1 Escherichia coli Proteobacteria; Gammaproteobacteria; Enterobacterales; Enterobacteriaceae; Escherichia MTGDNTLIHSHGINRRDFMKLCAALAATMGLSSKAAAEMAESVTNPQRPPVIWIGAQECT WP_000145410.1 MULTISPECIES: hydrogenase-2 small chain [Proteobacteria] Length: 372\nNumber of predicted TMHs: 1\nExp number of AAs in TMHs: 23.72226\nExp number, first 60 AAs: 0.22038\nTotal prob of N-in: 0.12994\noutside 1 331\nTMhelix 332 354\ninside 355 372

2992 GCF_001576545.1_ASM157654v1 Escherichia coli Proteobacteria; Gammaproteobacteria; Enterobacterales; Enterobacteriaceae; Escherichia MTGDNTLIHSHGINRRDFMKLCAALAATMGLSSKAAAEMAESVTNPQRPPVIWIGAQECT WP_000145410.1 MULTISPECIES: hydrogenase-2 small chain [Proteobacteria] Length: 372\nNumber of predicted TMHs: 1\nExp number of AAs in TMHs: 23.72226\nExp number, first 60 AAs: 0.22038\nTotal prob of N-in: 0.12994\noutside 1 331\nTMhelix 332 354\ninside 355 372

2993 GCF_001576385.1_ASM157638v1 Escherichia coli Proteobacteria; Gammaproteobacteria; Enterobacterales; Enterobacteriaceae; Escherichia MNNEETFYQAMRRQGVTRRSFLKYCSLAATSLGLGAGMAPKIAWALENKPRIPVVWIHGL WP_001058323.1 MULTISPECIES: hydrogenase-1 small chain [Proteobacteria] Length: 372\nNumber of predicted TMHs: 2\nExp number of AAs in TMHs: 37.2608499999999\nExp number, first 60 AAs: 19.52923\nTotal prob of N-in: 0.94351\nPOSSIBLE N-term signal sequence\ninside 1 19\nTMhelix 20 39\noutside 40 325\nTMhelix 326 348\ninside 349 372

2994 GCF_001576465.1_ASM157646v1 Escherichia coli Proteobacteria; Gammaproteobacteria; Enterobacterales; Enterobacteriaceae; Escherichia MNNEETFYQAMRRQGVTRRSFLKYCSLAATSLGLGAGMAPKIAWALENKPRIPVVWIHGL WP_001058323.1 MULTISPECIES: hydrogenase-1 small chain [Proteobacteria] Length: 372\nNumber of predicted TMHs: 2\nExp number of AAs in TMHs: 37.2608499999999\nExp number, first 60 AAs: 19.52923\nTotal prob of N-in: 0.94351\nPOSSIBLE N-term signal sequence\ninside 1 19\nTMhelix 20 39\noutside 40 325\nTMhelix 326 348\ninside 349 372

2995 GCF_001576545.1_ASM157654v1 Escherichia coli Proteobacteria; Gammaproteobacteria; Enterobacterales; Enterobacteriaceae; Escherichia MNNEETFYQAMRRQGVTRRSFLKYCSLAATSLGLGAGMAPKIAWALENKPRIPVVWIHGL WP_001058323.1 MULTISPECIES: hydrogenase-1 small chain [Proteobacteria] Length: 372\nNumber of predicted TMHs: 2\nExp number of AAs in TMHs: 37.2608499999999\nExp number, first 60 AAs: 19.52923\nTotal prob of N-in: 0.94351\nPOSSIBLE N-term signal sequence\ninside 1 19\nTMhelix 20 39\noutside 40 325\nTMhelix 326 348\ninside 349 372

2996 GCF_001576375.1_ASM157637v1 Escherichia coli Proteobacteria; Gammaproteobacteria; Enterobacterales; Enterobacteriaceae; Escherichia MNNEETFYQAMRRQGVTRRSFLKYCSLAATSLGLGAGMAPKIAWALENKPRIPVVWIHGL WP_001058323.1 MULTISPECIES: hydrogenase-1 small chain [Proteobacteria] Length: 372\nNumber of predicted TMHs: 2\nExp number of AAs in TMHs: 37.2608499999999\nExp number, first 60 AAs: 19.52923\nTotal prob of N-in: 0.94351\nPOSSIBLE N-term signal sequence\ninside 1 19\nTMhelix 20 39\noutside 40 325\nTMhelix 326 348\ninside 349 372

2997 GCF_001582895.1_ASM158289v1 Escherichia coli Proteobacteria; Gammaproteobacteria; Enterobacterales; Enterobacteriaceae; Escherichia MTGDNTLIHSHGINRRDFMKLCAALAATMGLSSKAAAEMAESVTNPQRPPVIWIGAQECT WP_000145410.1 MULTISPECIES: hydrogenase-2 small chain [Proteobacteria] Length: 372\nNumber of predicted TMHs: 1\nExp number of AAs in TMHs: 23.72226\nExp number, first 60 AAs: 0.22038\nTotal prob of N-in: 0.12994\noutside 1 331\nTMhelix 332 354\ninside 355 372

2998 GCF_001582905.1_ASM158290v1 Escherichia coli Proteobacteria; Gammaproteobacteria; Enterobacterales; Enterobacteriaceae; Escherichia

MTGDNTLIHSHGINRRDFMKLCAALAATMGLSSKAAAEMAESVTNPQRPPVIWIGAQECT WP_000145410.1 MULTISPECIES: hydrogenase-2 small chain [Proteobacteria] Length: 372\nNumber of predicted TMHs: 1\nExp number of AAs in TMHs: 23.72226\nExp number, first 60 AAs: 0.22038\nTotal prob of N-in: 0.12994\noutside 1 331\nTMhelix 332 354\ninside 355 372

2999 GCF_001582895.1_ASM158289v1 Escherichia coli Proteobacteria; Gammaproteobacteria; Enterobacterales; Enterobacteriaceae; Escherichia MNNEETFYQAMRRQGVTRRSFLKYCSLAATSLGLGAGMAPKIAWALENKPRIPVVWIHGL WP_001058323.1 MULTISPECIES: hydrogenase-1 small chain [Proteobacteria] Length: 372\nNumber of predicted TMHs: 2\nExp number of AAs in TMHs: 37.2608499999999\nExp number, first 60 AAs: 19.52923\nTotal prob of N-in: 0.94351\nPOSSIBLE N-term signal sequence\ninside 1 19\nTMhelix 20 39\noutside 40 325\nTMhelix 326 348\ninside 349 372

3000 GCF_001582905.1_ASM158290v1 Escherichia coli Proteobacteria; Gammaproteobacteria; Enterobacterales; Enterobacteriaceae; Escherichia MNNEETFYQAMRRQGVTRRSFLKYCSLAATSLGLGAGMAPKIAWALENKPRIPVVWIHGL WP_001058323.1 MULTISPECIES: hydrogenase-1 small chain [Proteobacteria] Length: 372\nNumber of predicted TMHs: 2\nExp number of AAs in TMHs: 37.2608499999999\nExp number, first 60 AAs: 19.52923\nTotal prob of N-in: 0.94351\nPOSSIBLE N-term signal sequence\ninside 1 19\nTMhelix 20 39\noutside 40 325\nTMhelix 326 348\ninside 349 372

3001 GCF_001576235.1_ASM157623v1 Escherichia coli Proteobacteria; Gammaproteobacteria; Enterobacterales; Enterobacteriaceae; Escherichia MTGDNTLIHSHGINRRDFMKLCAALAATMGLSSKAAAEMAESVTNPQRPPVIWIGAQECT WP_061357712.1 hydrogenase 2 small subunit [Escherichia coli] Length: 372\nNumber of predicted TMHs: 1\nExp number of AAs in TMHs: 23.72413\nExp number, first 60 AAs: 0.22038\nTotal prob of N-in: 0.12999\noutside 1 331\nTMhelix 332 354\ninside 355 372

3002 GCF_001576315.1_ASM157631v1 Escherichia coli Proteobacteria; Gammaproteobacteria; Enterobacterales; Enterobacteriaceae; Escherichia MNNEETFYQAMRRQGVTRRSFLKYCSLAATSLGLGAGMAPKIAWALENKPRIPVVWIHGL WP_001058323.1 MULTISPECIES: hydrogenase-1 small chain [Proteobacteria] Length: 372\nNumber of predicted TMHs: 2\nExp number of AAs in TMHs: 37.2608499999999\nExp number, first 60 AAs: 19.52923\nTotal prob of N-in: 0.94351\nPOSSIBLE N-term signal sequence\ninside 1 19\nTMhelix 20 39\noutside 40 325\nTMhelix 326 348\ninside 349 372

3003 GCF_001576435.1_ASM157643v1 Escherichia coli Proteobacteria; Gammaproteobacteria; Enterobacterales; Enterobacteriaceae; Escherichia MNNEETFYQAMRRQGVTRRSFLKYCSLAATSLGLGAGMAPKIAWALENKPRIPVVWIHGL WP_001058323.1 MULTISPECIES: hydrogenase-1 small chain [Proteobacteria] Length: 372\nNumber of predicted TMHs: 2\nExp number of AAs in TMHs: 37.2608499999999\nExp number, first 60 AAs: 19.52923\nTotal prob of N-in: 0.94351\nPOSSIBLE N-term signal sequence\ninside 1 19\nTMhelix 20 39\noutside 40 325\nTMhelix 326 348\ninside 349 372

3004 GCF_001605825.2_ASM160582v2 Escherichia coli Proteobacteria; Gammaproteobacteria; Enterobacterales; Enterobacteriaceae; Escherichia MTGDNTLIHSHGINRRDFMKLCAALAATMGLSSKAAAEMVESVTNPQRPPVIWIGAQECT WP_000145423.1 hydrogenase 2 small subunit [Escherichia coli] Length: 372\nNumber of predicted TMHs: 1\nExp number of AAs in TMHs: 23.78201\nExp number, first 60 AAs: 0.28355\nTotal prob of N-in: 0.13257\noutside 1 331\nTMhelix 332 354\ninside 355 372

3005 GCF_001606365.1_ASM160636v1 Escherichia coli Proteobacteria; Gammaproteobacteria; Enterobacterales; Enterobacteriaceae; Escherichia MTGDNTLIHSHGINRRDFMKLCAALAATMGLSSKAAAEMAESVTNPQRPPVIWIGAQECT WP_000145410.1 MULTISPECIES: hydrogenase-2 small chain [Proteobacteria] Length: 372\nNumber of predicted TMHs: 1\nExp number of AAs in TMHs: 23.72226\nExp number, first 60 AAs: 0.22038\nTotal prob of N-in: 0.12994\noutside 1 331\nTMhelix 332 354\ninside 355 372

3006 GCF_001606375.1_ASM160637v1 Escherichia coli Proteobacteria; Gammaproteobacteria; Enterobacterales; Enterobacteriaceae; Escherichia MTGDNTLIHSHGINRRDFMKLCAALAATMGLSSKAAAEMAESVTNPQRPPVIWIGAQECT WP_000145410.1

MULTISPECIES: hydrogenase-2 small chain [Proteobacteria] Length: 372\nNumber of predicted TMHs: 1\nExp number of AAs in TMHs: 23.72226\nExp number, first 60 AAs: 0.22038\nTotal prob of N-in: 0.12994\noutside 1 331\nTMhelix 332 354\ninside 355 372

3007 GCF_001606435.1_ASM160643v1 Escherichia coli Proteobacteria; Gammaproteobacteria; Enterobacterales; Enterobacteriaceae; Escherichia
MTGDNTLIHSHGINRRDFMKLCAALAATMGLSSKAAAEMAESVTNPQRPPVIWIGAQECT WP_000145410.1
MULTISPECIES: hydrogenase-2 small chain [Proteobacteria] Length: 372\nNumber of predicted TMHs: 1\nExp number of AAs in TMHs: 23.72226\nExp number, first 60 AAs: 0.22038\nTotal prob of N-in: 0.12994\noutside 1 331\nTMhelix 332 354\ninside 355 372

3008 GCF_001606465.1_ASM160646v1 Escherichia coli Proteobacteria; Gammaproteobacteria; Enterobacterales; Enterobacteriaceae; Escherichia
MTGDNTLIHSHGINRRDFMKLCAALAATMGLSSKAAAEMAESVTNPQRPPVIWIGAQECT WP_000145410.1
MULTISPECIES: hydrogenase-2 small chain [Proteobacteria] Length: 372\nNumber of predicted TMHs: 1\nExp number of AAs in TMHs: 23.72226\nExp number, first 60 AAs: 0.22038\nTotal prob of N-in: 0.12994\noutside 1 331\nTMhelix 332 354\ninside 355 372

3009 GCF_001606475.1_ASM160647v1 Escherichia coli Proteobacteria; Gammaproteobacteria; Enterobacterales; Enterobacteriaceae; Escherichia
MNNEETFYQAMRRQGVTRRSFLKYCSLAATSLGLGAGMAPKIAWALENKPRIPVVWIHGL WP_001058323.1
MULTISPECIES: hydrogenase-1 small chain [Proteobacteria] Length: 372\nNumber of predicted TMHs: 2\nExp number of AAs in TMHs: 37.2608499999999\nExp number, first 60 AAs: 19.52923\nTotal prob of N-in: 0.94351\nPOSSIBLE N-term signal sequence\ninside 1 19\nTMhelix 20 39\noutside 40 325\nTMhelix 326 348\ninside 349 372

3010 GCF_001606525.1_ASM160652v1 Escherichia coli Proteobacteria; Gammaproteobacteria; Enterobacterales; Enterobacteriaceae; Escherichia
MTGDNTLIHSHGINRRDFMKLCAALAATMGLSSKAAAEMAESVTNPQRPPVIWIGAQECT WP_028131934.1
hydrogenase-2 small chain [Escherichia coli] Length: 372\nNumber of predicted TMHs: 1\nExp number of AAs in TMHs: 24.01565\nExp number, first 60 AAs: 0.22022\nTotal prob of N-in: 0.13096\noutside 1 331\nTMhelix 332 354\ninside 355 372

3011 GCF_001606595.1_ASM160659v1 Escherichia coli Proteobacteria; Gammaproteobacteria; Enterobacterales; Enterobacteriaceae; Escherichia
MNNEETFYQAMRRQGVTRRSFLKSCSLAATSLGLGAGMAPKIAWALENKPRIPVVWIHGL WP_001058318.1
hydrogenase [Escherichia coli] Length: 372\nNumber of predicted TMHs: 1\nExp number of AAs in TMHs: 33.59637\nExp number, first 60 AAs: 15.81976\nTotal prob of N-in: 0.77394\nPOSSIBLE N-term signal sequence\noutside 1 325\nTMhelix 326 348\ninside 349 372

3012 GCF_001606365.1_ASM160636v1 Escherichia coli Proteobacteria; Gammaproteobacteria; Enterobacterales; Enterobacteriaceae; Escherichia
MNNEETFYQAMRRQGVTRRSFLKYCSLAATSLGLGAGMAPKIAWALENKPRIPVVWIHGL WP_001058323.1
MULTISPECIES: hydrogenase-1 small chain [Proteobacteria] Length: 372\nNumber of predicted TMHs: 2\nExp number of AAs in TMHs: 37.2608499999999\nExp number, first 60 AAs: 19.52923\nTotal prob of N-in: 0.94351\nPOSSIBLE N-term signal sequence\ninside 1 19\nTMhelix 20 39\noutside 40 325\nTMhelix 326 348\ninside 349 372

3013 GCF_001606465.1_ASM160646v1 Escherichia coli Proteobacteria; Gammaproteobacteria; Enterobacterales; Enterobacteriaceae; Escherichia
MNNEETFYQAMRRQGVTRRSFLKYCSLAATSLGLGAGMAPKIAWALENKPRIPVVWIHGL WP_001058323.1
MULTISPECIES: hydrogenase-1 small chain [Proteobacteria] Length: 372\nNumber of predicted TMHs: 2\nExp number of AAs in TMHs: 37.2608499999999\nExp number, first 60 AAs: 19.52923\nTotal prob of N-in: 0.94351\nPOSSIBLE N-term signal sequence\ninside 1 19\nTMhelix 20 39\noutside 40 325\nTMhelix 326 348\ninside 349 372

3014 GCF_001605825.2_ASM160582v2 Escherichia coli Proteobacteria; Gammaproteobacteria; Enterobacterales; Enterobacteriaceae; Escherichia
MNNEETFYQAMRRQGVTRRSFLKYCSLAATSLGLGAGMAPKIAWALENKPRIPVVWIHGL WP_001295940.1
MULTISPECIES: hydrogenase [Enterobacteriaceae] Length: 372\nNumber of predicted TMHs: 2\nExp number of AAs in TMHs: 37.2608899999999\nExp number, first 60 AAs: 19.52923\nTotal prob of N-in:

0.94351\nPOSSIBLE N-term signal sequence\ninside 1 19\nTMhelix 20 39\noutside 40 325\nTMhelix 326 348\ninside 349 372

3015 GCF_001606625.1_ASM160662v1 Escherichia coli Proteobacteria; Gammaproteobacteria; Enterobacterales; Enterobacteriaceae; Escherichia MTGDNTLIHSHGINRRDFMKLCAALAATMGLSSKAAAEMAESVTNPQRPPVIWIGAQECT WP_000145410.1 MULTISPECIES: hydrogenase-2 small chain [Proteobacteria] Length: 372\nNumber of predicted TMHs: 1\nExp number of AAs in TMHs: 23.72226\nExp number, first 60 AAs: 0.22038\nTotal prob of N-in: 0.12994\noutside 1 331\nTMhelix 332 354\ninside 355 372

3016 GCF_001606675.1_ASM160667v1 Escherichia coli Proteobacteria; Gammaproteobacteria; Enterobacterales; Enterobacteriaceae; Escherichia MTGDNTLIHSHGINRRDFMKLCAALAATMGLSSKAAAEMAESVTNPQRPPVIWIGAQECT WP_000145410.1 MULTISPECIES: hydrogenase-2 small chain [Proteobacteria] Length: 372\nNumber of predicted TMHs: 1\nExp number of AAs in TMHs: 23.72226\nExp number, first 60 AAs: 0.22038\nTotal prob of N-in: 0.12994\noutside 1 331\nTMhelix 332 354\ninside 355 372

3017 GCF_001606985.1_ASM160698v1 Escherichia coli Proteobacteria; Gammaproteobacteria; Enterobacterales; Enterobacteriaceae; Escherichia MTGDNTLIHSHGINRRDFMKLCAALAATMGLSSKAAAEMAESVTNPQRPPVIWIGAQECT WP_000145410.1 MULTISPECIES: hydrogenase-2 small chain [Proteobacteria] Length: 372\nNumber of predicted TMHs: 1\nExp number of AAs in TMHs: 23.72226\nExp number, first 60 AAs: 0.22038\nTotal prob of N-in: 0.12994\noutside 1 331\nTMhelix 332 354\ninside 355 372

3018 GCF_001606975.1_ASM160697v1 Escherichia coli Proteobacteria; Gammaproteobacteria; Enterobacterales; Enterobacteriaceae; Escherichia MNNEETFYQAMRRQGVTRRSFLKYCSLAATSLGLGAGMAPKIAWALENKPRIPVVWIHGL WP_001058323.1 MULTISPECIES: hydrogenase-1 small chain [Proteobacteria] Length: 372\nNumber of predicted TMHs: 2\nExp number of AAs in TMHs: 37.2608499999999\nExp number, first 60 AAs: 19.52923\nTotal prob of N-in: 0.94351\nPOSSIBLE N-term signal sequence\ninside 1 19\nTMhelix 20 39\noutside 40 325\nTMhelix 326 348\ninside 349 372

3019 GCF_001606985.1_ASM160698v1 Escherichia coli Proteobacteria; Gammaproteobacteria; Enterobacterales; Enterobacteriaceae; Escherichia MNNEETFYQAMRRQGVTRRSFLKYCSLAATSLGLGAGMAPKIAWALENKPRIPVVWIHGL WP_001058323.1 MULTISPECIES: hydrogenase-1 small chain [Proteobacteria] Length: 372\nNumber of predicted TMHs: 2\nExp number of AAs in TMHs: 37.2608499999999\nExp number, first 60 AAs: 19.52923\nTotal prob of N-in: 0.94351\nPOSSIBLE N-term signal sequence\ninside 1 19\nTMhelix 20 39\noutside 40 325\nTMhelix 326 348\ninside 349 372

3020 GCF_001606785.1_ASM160678v1 Escherichia coli Proteobacteria; Gammaproteobacteria; Enterobacterales; Enterobacteriaceae; Escherichia MNNEETFYQAMRRQGVTRRSFLKYCSLAATSLGLGAGMAPKIAWALENKPRIPVVWIHGL WP_001058323.1 MULTISPECIES: hydrogenase-1 small chain [Proteobacteria] Length: 372\nNumber of predicted TMHs: 2\nExp number of AAs in TMHs: 37.2608499999999\nExp number, first 60 AAs: 19.52923\nTotal prob of N-in: 0.94351\nPOSSIBLE N-term signal sequence\ninside 1 19\nTMhelix 20 39\noutside 40 325\nTMhelix 326 348\ninside 349 372

3021 GCF_001606865.1_ASM160686v1 Escherichia coli Proteobacteria; Gammaproteobacteria; Enterobacterales; Enterobacteriaceae; Escherichia MNNEETFYQAMRRQGVTRRSFLKYCSLAATSLGLGAGMAPKIAWALENKPRIPVVWIHGL WP_001058323.1 MULTISPECIES: hydrogenase-1 small chain [Proteobacteria] Length: 372\nNumber of predicted TMHs: 2\nExp number of AAs in TMHs: 37.2608499999999\nExp number, first 60 AAs: 19.52923\nTotal prob of N-in: 0.94351\nPOSSIBLE N-term signal sequence\ninside 1 19\nTMhelix 20 39\noutside 40 325\nTMhelix 326 348\ninside 349 372

3022 GCF_001606895.1_ASM160689v1 Escherichia coli Proteobacteria; Gammaproteobacteria; Enterobacterales; Enterobacteriaceae; Escherichia MNNEETFYQAMRRQGVTRRSFLKYCSLAATSLGLGAGMAPKIAWALENKPRIPVVWIHGL WP_001058323.1 MULTISPECIES: hydrogenase-1 small chain [Proteobacteria] Length: 372\nNumber of predicted TMHs: 2\nExp number of AAs in TMHs: 37.2608499999999\nExp number, first 60 AAs: 19.52923\nTotal prob of N-in:

0.94351\nPOSSIBLE N-term signal sequence\ninside 1 19\nTMhelix 20 39\noutside 40 325\nTMhelix 326 348\ninside 349 372

3023 GCF_001606695.1_ASM160669v1 Escherichia coli Proteobacteria; Gammaproteobacteria; Enterobacterales; Enterobacteriaceae; Escherichia MNNEETFYQAMRRQGVTRRSFLKYCSLAATSLGLGAGMAPKIAWALENKPRIPVVWIHGL WP_001295940.1 MULTISPECIES: hydrogenase [Enterobacteriaceae] Length: 372\nNumber of predicted TMHs: 2\nExp number of AAs in TMHs: 37.2608899999999\nExp number, first 60 AAs: 19.52923\nTotal prob of N-in: 0.94351\nPOSSIBLE N-term signal sequence\ninside 1 19\nTMhelix 20 39\noutside 40 325\nTMhelix 326 348\ninside 349 372

3024 GCF_001607185.1_ASM160718v1 Escherichia coli Proteobacteria; Gammaproteobacteria; Enterobacterales; Enterobacteriaceae; Escherichia MTGDNTLIHSHGINRRDFMKLCAALAATMGLSSKAAAEMAESVTNPQRPPVIWIGAQECT WP_000145410.1 MULTISPECIES: hydrogenase-2 small chain [Proteobacteria] Length: 372\nNumber of predicted TMHs: 1\nExp number of AAs in TMHs: 23.72226\nExp number, first 60 AAs: 0.22038\nTotal prob of N-in: 0.12994\noutside 1 331\nTMhelix 332 354\ninside 355 372

3025 GCF_001607225.1_ASM160722v1 Escherichia coli Proteobacteria; Gammaproteobacteria; Enterobacterales; Enterobacteriaceae; Escherichia MTGDNTLIHSHGINRRDFMKLCAALAATMGLSSKAAAEMVESVTNPQRPPVIWIGAQECT WP_000145423.1 hydrogenase 2 small subunit [Escherichia coli] Length: 372\nNumber of predicted TMHs: 1\nExp number of AAs in TMHs: 23.78201\nExp number, first 60 AAs: 0.28355\nTotal prob of N-in: 0.13257\noutside 1 331\nTMhelix 332 354\ninside 355 372

3026 GCF_001607175.1_ASM160717v1 Escherichia coli Proteobacteria; Gammaproteobacteria; Enterobacterales; Enterobacteriaceae; Escherichia MTGDNTLIHSHGINRRDFMKLCAALAATMGLSSKAAAEMAESVTNPQRPPVIWIGAQECT WP_000145410.1 MULTISPECIES: hydrogenase-2 small chain [Proteobacteria] Length: 372\nNumber of predicted TMHs: 1\nExp number of AAs in TMHs: 23.72226\nExp number, first 60 AAs: 0.22038\nTotal prob of N-in: 0.12994\noutside 1 331\nTMhelix 332 354\ninside 355 372

3027 GCF_001607765.1_ASM160776v1 Escherichia coli Proteobacteria; Gammaproteobacteria; Enterobacterales; Enterobacteriaceae; Escherichia MTGDNTLIHSHGINRRDFMKLCAALAATMGLSSKAAAEMAESVTNPQRPPVIWIGAQECT WP_000145410.1 MULTISPECIES: hydrogenase-2 small chain [Proteobacteria] Length: 372\nNumber of predicted TMHs: 1\nExp number of AAs in TMHs: 23.72226\nExp number, first 60 AAs: 0.22038\nTotal prob of N-in: 0.12994\noutside 1 331\nTMhelix 332 354\ninside 355 372

3028 GCF_001607075.1_ASM160707v1 Escherichia coli Proteobacteria; Gammaproteobacteria; Enterobacterales; Enterobacteriaceae; Escherichia MNNEETFYQAMRRQGVTRRSFLKYCSLAATSLGLGAGMAPKIAWALENKPRIPVVWIHGL WP_001058323.1 MULTISPECIES: hydrogenase-1 small chain [Proteobacteria] Length: 372\nNumber of predicted TMHs: 2\nExp number of AAs in TMHs: 37.2608499999999\nExp number, first 60 AAs: 19.52923\nTotal prob of N-in: 0.94351\nPOSSIBLE N-term signal sequence\ninside 1 19\nTMhelix 20 39\noutside 40 325\nTMhelix 326 348\ninside 349 372

3029 GCF_001607285.1_ASM160728v1 Escherichia coli Proteobacteria; Gammaproteobacteria; Enterobacterales; Enterobacteriaceae; Escherichia MTGDNTLIHSHGINRRDFMKLCAALAATMGLSSKAAAEMAESVTNPQRPPVIWIGAQECT WP_000145410.1 MULTISPECIES: hydrogenase-2 small chain [Proteobacteria] Length: 372\nNumber of predicted TMHs: 1\nExp number of AAs in TMHs: 23.72226\nExp number, first 60 AAs: 0.22038\nTotal prob of N-in: 0.12994\noutside 1 331\nTMhelix 332 354\ninside 355 372

3030 GCF_001607265.1_ASM160726v1 Escherichia coli Proteobacteria; Gammaproteobacteria; Enterobacterales; Enterobacteriaceae; Escherichia MTGDNTLIHSHGINRRDFMKLCAALAATMGLSSKAAAEMAESVTNPQRPPVIWIGAQECT WP_000145410.1 MULTISPECIES: hydrogenase-2 small chain [Proteobacteria] Length: 372\nNumber of predicted TMHs: 1\nExp number of AAs in TMHs: 23.72226\nExp number, first 60 AAs: 0.22038\nTotal prob of N-in: 0.12994\noutside 1 331\nTMhelix 332 354\ninside 355 372

3031 GCF_001607125.1_ASM160712v1 Escherichia coli Proteobacteria; Gammaproteobacteria; Enterobacterales; Enterobacteriaceae; Escherichia
MNNEETFYQAMRRQGVTRRSFLKYCSLAATSLGLGAGMAPKIAWALENKPRIPVVWIHGL WP_001058323.1
MULTISPECIES: hydrogenase-1 small chain [Proteobacteria] Length: 372\nNumber of predicted TMHs: 2\nExp number of AAs in TMHs: 37.2608499999999\nExp number, first 60 AAs: 19.52923\nTotal prob of N-in: 0.94351\nPOSSIBLE N-term signal sequence\ninside 1 19\nTMhelix 20 39\noutside 40 325\nTMhelix 326 348\ninside 349 372

3032 GCF_001607185.1_ASM160718v1 Escherichia coli Proteobacteria; Gammaproteobacteria; Enterobacterales; Enterobacteriaceae; Escherichia
MNNEETFYQAMRRQGVTRRSFLKYCSLAATSLGLGAGMAPKIAWALENKPRIPVVWIHGL WP_001058323.1
MULTISPECIES: hydrogenase-1 small chain [Proteobacteria] Length: 372\nNumber of predicted TMHs: 2\nExp number of AAs in TMHs: 37.2608499999999\nExp number, first 60 AAs: 19.52923\nTotal prob of N-in: 0.94351\nPOSSIBLE N-term signal sequence\ninside 1 19\nTMhelix 20 39\noutside 40 325\nTMhelix 326 348\ninside 349 372

3033 GCF_001607385.1_ASM160738v1 Escherichia coli Proteobacteria; Gammaproteobacteria; Enterobacterales; Enterobacteriaceae; Escherichia
MTGDNTLIHSHGINRRDFMKLCAALAATMGLSSKAAAEMAESVTNPQRPPVIWIGAQECT WP_000145410.1
MULTISPECIES: hydrogenase-2 small chain [Proteobacteria] Length: 372\nNumber of predicted TMHs: 1\nExp number of AAs in TMHs: 23.72226\nExp number, first 60 AAs: 0.22038\nTotal prob of N-in: 0.12994\noutside 1 331\nTMhelix 332 354\ninside 355 372

3034 GCF_001607435.1_ASM160743v1 Escherichia coli Proteobacteria; Gammaproteobacteria; Enterobacterales; Enterobacteriaceae; Escherichia
MTGDNTLIHSHGINRRDFMKLCAALAATMGLSSKAAAEMAESVTNPQRPPVIWIGAQECT WP_000145410.1
MULTISPECIES: hydrogenase-2 small chain [Proteobacteria] Length: 372\nNumber of predicted TMHs: 1\nExp number of AAs in TMHs: 23.72226\nExp number, first 60 AAs: 0.22038\nTotal prob of N-in: 0.12994\noutside 1 331\nTMhelix 332 354\ninside 355 372

3035 GCF_001607385.1_ASM160738v1 Escherichia coli Proteobacteria; Gammaproteobacteria; Enterobacterales; Enterobacteriaceae; Escherichia
MNNEETFYQAMRRQGVTRRSFLKYCSLAATSLGLGAGMAPKIAWALENKPRIPVVWIHGL WP_001058323.1
MULTISPECIES: hydrogenase-1 small chain [Proteobacteria] Length: 372\nNumber of predicted TMHs: 2\nExp number of AAs in TMHs: 37.2608499999999\nExp number, first 60 AAs: 19.52923\nTotal prob of N-in: 0.94351\nPOSSIBLE N-term signal sequence\ninside 1 19\nTMhelix 20 39\noutside 40 325\nTMhelix 326 348\ninside 349 372

3036 GCF_001607435.1_ASM160743v1 Escherichia coli Proteobacteria; Gammaproteobacteria; Enterobacterales; Enterobacteriaceae; Escherichia
MNNEETFYQAMRRQGVTRRSFLKYCSLAATSLGLGAGMAPKIAWALENKPRIPVVWIHGL WP_001058323.1
MULTISPECIES: hydrogenase-1 small chain [Proteobacteria] Length: 372\nNumber of predicted TMHs: 2\nExp number of AAs in TMHs: 37.2608499999999\nExp number, first 60 AAs: 19.52923\nTotal prob of N-in: 0.94351\nPOSSIBLE N-term signal sequence\ninside 1 19\nTMhelix 20 39\noutside 40 325\nTMhelix 326 348\ninside 349 372

3037 GCF_001607175.1_ASM160717v1 Escherichia coli Proteobacteria; Gammaproteobacteria; Enterobacterales; Enterobacteriaceae; Escherichia
MNNEETFYQAMRRQGVTRRSFLKYCSLAATSLGLGAGMAPKIAWALENKPRIPVVWIHGL WP_001295940.1
MULTISPECIES: hydrogenase [Enterobacteriaceae] Length: 372\nNumber of predicted TMHs: 2\nExp number of AAs in TMHs: 37.2608899999999\nExp number, first 60 AAs: 19.52923\nTotal prob of N-in: 0.94351\nPOSSIBLE N-term signal sequence\ninside 1 19\nTMhelix 20 39\noutside 40 325\nTMhelix 326 348\ninside 349 372

3038 GCF_001607495.1_ASM160749v1 Escherichia coli Proteobacteria; Gammaproteobacteria; Enterobacterales; Enterobacteriaceae; Escherichia
MNNEETFYQAMRRQGVTRRSFLKYCSLAATSLGLGAGMAPKIAWALENKPRIPVVWIHGL WP_001058323.1
MULTISPECIES: hydrogenase-1 small chain [Proteobacteria] Length: 372\nNumber of predicted TMHs: 2\nExp number of AAs in TMHs: 37.2608499999999\nExp number, first 60 AAs: 19.52923\nTotal prob of N-in: 0.94351\nPOSSIBLE N-term signal sequence\ninside 1 19\nTMhelix 20 39\noutside 40 325\nTMhelix 326 348\ninside 349 372

3039 GCF_001607675.1_ASM160767v1 Escherichia coli Proteobacteria; Gammaproteobacteria; Enterobacterales; Enterobacteriaceae; Escherichia
MTGDNTLIHSHGINRRDFMKLCAALAATMGLSSKAAAEMAESVTNPQRPPVIWIGAQECT WP_000145410.1
MULTISPECIES: hydrogenase-2 small chain [Proteobacteria] Length: 372\nNumber of predicted TMHs: 1\nExp number of AAs in TMHs: 23.72226\nExp number, first 60 AAs: 0.22038\nTotal prob of N-in: 0.12994\noutside 1 331\nTMhelix 332 354\ninside 355 372

3040 GCF_001607735.1_ASM160773v1 Escherichia coli Proteobacteria; Gammaproteobacteria; Enterobacterales; Enterobacteriaceae; Escherichia
MTGDNTLIHSHGINRRDFMKLCAALAATMGLSSKAAAEMAESVTNPQRPPVIWIGAQECT WP_000145410.1
MULTISPECIES: hydrogenase-2 small chain [Proteobacteria] Length: 372\nNumber of predicted TMHs: 1\nExp number of AAs in TMHs: 23.72226\nExp number, first 60 AAs: 0.22038\nTotal prob of N-in: 0.12994\noutside 1 331\nTMhelix 332 354\ninside 355 372

3041 GCF_001607795.1_ASM160779v1 Escherichia coli Proteobacteria; Gammaproteobacteria; Enterobacterales; Enterobacteriaceae; Escherichia
MTGDNTLIHSHGINRRDFMKLCAALAATMGLSSKAAAEMAESVTNPQRPPVIWIGAQECT WP_000145410.1
MULTISPECIES: hydrogenase-2 small chain [Proteobacteria] Length: 372\nNumber of predicted TMHs: 1\nExp number of AAs in TMHs: 23.72226\nExp number, first 60 AAs: 0.22038\nTotal prob of N-in: 0.12994\noutside 1 331\nTMhelix 332 354\ninside 355 372

3042 GCF_001607735.1_ASM160773v1 Escherichia coli Proteobacteria; Gammaproteobacteria; Enterobacterales; Enterobacteriaceae; Escherichia
MNNEETFYQAMRRQGVTRRSFLKYCSLAATSLGLGAGMAPKIAWALENKPRIPVVWIHGL WP_001301613.1
hydrogenase [Escherichia coli] Length: 372\nNumber of predicted TMHs: 2\nExp number of AAs in TMHs: 37.46598\nExp number, first 60 AAs: 19.53139\nTotal prob of N-in: 0.94364\nPOSSIBLE N-term signal sequence\ninside 1 19\nTMhelix 20 39\noutside 40 325\nTMhelix 326 348\ninside 349 372

3043 GCF_001607675.1_ASM160767v1 Escherichia coli Proteobacteria; Gammaproteobacteria; Enterobacterales; Enterobacteriaceae; Escherichia
MNNEETFYQAMRRQGVTRRSFLKYCSLAATSLGLGAGMAPKIAWALENKPRIPVVWIHGL WP_001295940.1
MULTISPECIES: hydrogenase [Enterobacteriaceae] Length: 372\nNumber of predicted TMHs: 2\nExp number of AAs in TMHs: 37.2608899999999\nExp number, first 60 AAs: 19.52923\nTotal prob of N-in: 0.94351\nPOSSIBLE N-term signal sequence\ninside 1 19\nTMhelix 20 39\noutside 40 325\nTMhelix 326 348\ninside 349 372

3044 GCF_001607795.1_ASM160779v1 Escherichia coli Proteobacteria; Gammaproteobacteria; Enterobacterales; Enterobacteriaceae; Escherichia
MNNEETFYQAMRRQGVTRRSFLKYCSLAATSLGLGAGMAPKIAWALENKPRIPVVWIHGL WP_062876341.1
hydrogenase [Escherichia coli] Length: 372\nNumber of predicted TMHs: 2\nExp number of AAs in TMHs: 37.43626\nExp number, first 60 AAs: 19.52875\nTotal prob of N-in: 0.94361\nPOSSIBLE N-term signal sequence\ninside 1 19\nTMhelix 20 39\noutside 40 325\nTMhelix 326 348\ninside 349 372

3045 GCF_001607835.1_ASM160783v1 Escherichia coli Proteobacteria; Gammaproteobacteria; Enterobacterales; Enterobacteriaceae; Escherichia
MNNEETFYQAMRRQGVTRRSFLKYCSLAATSLGLGAGMAPKIAWALENKPRIPVVWIHGL WP_001058323.1
MULTISPECIES: hydrogenase-1 small chain [Proteobacteria] Length: 372\nNumber of predicted TMHs: 2\nExp number of AAs in TMHs: 37.2608499999999\nExp number, first 60 AAs: 19.52923\nTotal prob of N-in: 0.94351\nPOSSIBLE N-term signal sequence\ninside 1 19\nTMhelix 20 39\noutside 40 325\nTMhelix 326 348\ninside 349 372

3046 GCF_001607875.1_ASM160787v1 Escherichia coli Proteobacteria; Gammaproteobacteria; Enterobacterales; Enterobacteriaceae; Escherichia
MTGDNTLIHSHGINRRDFMKLCAALAATMGLSSKAAAEMAESVTNPQRPPVIWIGAQECT WP_000145410.1
MULTISPECIES: hydrogenase-2 small chain [Proteobacteria] Length: 372\nNumber of predicted TMHs: 1\nExp number of AAs in TMHs: 23.72226\nExp number, first 60 AAs: 0.22038\nTotal prob of N-in: 0.12994\noutside 1 331\nTMhelix 332 354\ninside 355 372

3047 GCF_001607925.1_ASM160792v1 Escherichia coli Proteobacteria; Gammaproteobacteria; Enterobacterales; Enterobacteriaceae; Escherichia
MTGDNTLIHSHGINRRDFMKLCAALAATMGLSSKAAAEMAESVTNPQRPPVIWIGAQECT WP_000145410.1

MULTISPECIES: hydrogenase-2 small chain [Proteobacteria] Length: 372\nNumber of predicted TMHs: 1\nExp number of AAs in TMHs: 23.72226\nExp number, first 60 AAs: 0.22038\nTotal prob of N-in: 0.12994\noutside 1 331\nTMhelix 332 354\ninside 355 372

3048 GCF_001607895.1_ASM160789v1 Escherichia coli Proteobacteria; Gammaproteobacteria; Enterobacterales; Enterobacteriaceae; Escherichia MTGDNTLIHSHGINRRDFMKLCAALAATMGLSSKAAAEMAESVTNPQRPPVIWIGAQECT WP_000145410.1 MULTISPECIES: hydrogenase-2 small chain [Proteobacteria] Length: 372\nNumber of predicted TMHs: 1\nExp number of AAs in TMHs: 23.72226\nExp number, first 60 AAs: 0.22038\nTotal prob of N-in: 0.12994\noutside 1 331\nTMhelix 332 354\ninside 355 372

3049 GCF_001607975.1_ASM160797v1 Escherichia coli Proteobacteria; Gammaproteobacteria; Enterobacterales; Enterobacteriaceae; Escherichia MTGDNTLIHSHGINRRDFMKLCAALAATMGLSSKAAAEMAESVTNPQRPPVIWIGAQECT WP_000145410.1 MULTISPECIES: hydrogenase-2 small chain [Proteobacteria] Length: 372\nNumber of predicted TMHs: 1\nExp number of AAs in TMHs: 23.72226\nExp number, first 60 AAs: 0.22038\nTotal prob of N-in: 0.12994\noutside 1 331\nTMhelix 332 354\ninside 355 372

3050 GCF_001608015.1_ASM160801v1 Escherichia coli Proteobacteria; Gammaproteobacteria; Enterobacterales; Enterobacteriaceae; Escherichia MTGDNTLIHSHGINRRDFMKLCAALAATMGLSSKAAAEMAESVTNPQRPPVIWIGAQECT WP_000145410.1 MULTISPECIES: hydrogenase-2 small chain [Proteobacteria] Length: 372\nNumber of predicted TMHs: 1\nExp number of AAs in TMHs: 23.72226\nExp number, first 60 AAs: 0.22038\nTotal prob of N-in: 0.12994\noutside 1 331\nTMhelix 332 354\ninside 355 372

3051 GCF_001607995.1_ASM160799v1 Escherichia coli Proteobacteria; Gammaproteobacteria; Enterobacterales; Enterobacteriaceae; Escherichia MTGDNTLIHSHGINRRDFMKLCAALAATMGLSSKAAAEMAESVTNPQRPPVIWIGAQECT WP_000145410.1 MULTISPECIES: hydrogenase-2 small chain [Proteobacteria] Length: 372\nNumber of predicted TMHs: 1\nExp number of AAs in TMHs: 23.72226\nExp number, first 60 AAs: 0.22038\nTotal prob of N-in: 0.12994\noutside 1 331\nTMhelix 332 354\ninside 355 372

3052 GCF_001608125.1_ASM160812v1 Escherichia coli Proteobacteria; Gammaproteobacteria; Enterobacterales; Enterobacteriaceae; Escherichia MTGDNTLIHSHGINRRDFMKLCAALAATMGLSSKAAAEMAESVTNPQRPPVIWIGAQECT WP_000145410.1 MULTISPECIES: hydrogenase-2 small chain [Proteobacteria] Length: 372\nNumber of predicted TMHs: 1\nExp number of AAs in TMHs: 23.72226\nExp number, first 60 AAs: 0.22038\nTotal prob of N-in: 0.12994\noutside 1 331\nTMhelix 332 354\ninside 355 372

3053 GCF_001608125.1_ASM160812v1 Escherichia coli Proteobacteria; Gammaproteobacteria; Enterobacterales; Enterobacteriaceae; Escherichia MNNEETFYQAMRRQGVTRRSFLKYCSLAATSLGLGAGMAPKIAWALENKPRIPVVWIHGL WP_001058323.1 MULTISPECIES: hydrogenase-1 small chain [Proteobacteria] Length: 372\nNumber of predicted TMHs: 2\nExp number of AAs in TMHs: 37.2608499999999\nExp number, first 60 AAs: 19.52923\nTotal prob of N-in: 0.94351\nPOSSIBLE N-term signal sequence\ninside 1 19\nTMhelix 20 39\noutside 40 325\nTMhelix 326 348\ninside 349 372

3054 GCF_001607895.1_ASM160789v1 Escherichia coli Proteobacteria; Gammaproteobacteria; Enterobacterales; Enterobacteriaceae; Escherichia MNNEETFYQAMRRQGVTRRSFLKYCSLAATSLGLGAGMAPKIAWALENKPRIPVVWIHGL WP_001058323.1 MULTISPECIES: hydrogenase-1 small chain [Proteobacteria] Length: 372\nNumber of predicted TMHs: 2\nExp number of AAs in TMHs: 37.2608499999999\nExp number, first 60 AAs: 19.52923\nTotal prob of N-in: 0.94351\nPOSSIBLE N-term signal sequence\ninside 1 19\nTMhelix 20 39\noutside 40 325\nTMhelix 326 348\ninside 349 372

3055 GCF_001608185.1_ASM160818v1 Escherichia coli Proteobacteria; Gammaproteobacteria; Enterobacterales; Enterobacteriaceae; Escherichia MTGDNTLIHSHGINRRDFMKLCAALAATMGLSSKAAAEMAESVTNPQRPPVIWIGAQECT WP_000145410.1 MULTISPECIES: hydrogenase-2 small chain [Proteobacteria] Length: 372\nNumber of predicted TMHs: 1\nExp number of AAs in TMHs: 23.72226\nExp number, first 60 AAs: 0.22038\nTotal prob of N-in: 0.12994\noutside 1 331\nTMhelix 332 354\ninside 355 372

3056 GCF_001608185.1_ASM160818v1 Escherichia coli Proteobacteria; Gammaproteobacteria; Enterobacterales; Enterobacteriaceae; Escherichia
MNNEETFYQAMRRQGVTRRSFLKYCSLAATSLGLGAGMAPKIAWALENKPRIPVVWIHGL WP_001058323.1
MULTISPECIES: hydrogenase-1 small chain [Proteobacteria] Length: 372\nNumber of predicted TMHs: 2\nExp number of AAs in TMHs: 37.2608499999999\nExp number, first 60 AAs: 19.52923\nTotal prob of N-in: 0.94351\nPOSSIBLE N-term signal sequence\ninside 1 19\nTMhelix 20 39\noutside 40 325\nTMhelix 326 348\ninside 349 372

3057 GCF_001607975.1_ASM160797v1 Escherichia coli Proteobacteria; Gammaproteobacteria; Enterobacterales; Enterobacteriaceae; Escherichia
MNNEETFYQAMRRQGVTRRSFLKYCSLAATSLGLGAGMAPKIAWALENKPRIPVVWIHGL WP_001058323.1
MULTISPECIES: hydrogenase-1 small chain [Proteobacteria] Length: 372\nNumber of predicted TMHs: 2\nExp number of AAs in TMHs: 37.2608499999999\nExp number, first 60 AAs: 19.52923\nTotal prob of N-in: 0.94351\nPOSSIBLE N-term signal sequence\ninside 1 19\nTMhelix 20 39\noutside 40 325\nTMhelix 326 348\ninside 349 372

3058 GCF_001608235.1_ASM160823v1 Escherichia coli Proteobacteria; Gammaproteobacteria; Enterobacterales; Enterobacteriaceae; Escherichia
MTGDNTLIHSHGINRRDFMKLCAALAATMGLSSKAAAEMVESVTNPQRPPVIWIGAQECT WP_000145423.1
hydrogenase 2 small subunit [Escherichia coli] Length: 372\nNumber of predicted TMHs: 1\nExp number of AAs in TMHs: 23.78201\nExp number, first 60 AAs: 0.28355\nTotal prob of N-in: 0.13257\noutside 1 331\nTMhelix 332 354\ninside 355 372

3059 GCF_001608075.1_ASM160807v1 Escherichia coli Proteobacteria; Gammaproteobacteria; Enterobacterales; Enterobacteriaceae; Escherichia
MNNEETFYQAMRRQGVTRRSFLKYCSLAATSLGLGAGMAPKIAWALENKPRIPVVWIHGL WP_001058323.1
MULTISPECIES: hydrogenase-1 small chain [Proteobacteria] Length: 372\nNumber of predicted TMHs: 2\nExp number of AAs in TMHs: 37.2608499999999\nExp number, first 60 AAs: 19.52923\nTotal prob of N-in: 0.94351\nPOSSIBLE N-term signal sequence\ninside 1 19\nTMhelix 20 39\noutside 40 325\nTMhelix 326 348\ninside 349 372

3060 GCF_001608235.1_ASM160823v1 Escherichia coli Proteobacteria; Gammaproteobacteria; Enterobacterales; Enterobacteriaceae; Escherichia
MNNEETFYQAMRRQGVTRRSFLKYCSLAATSLGLGAGMAPKIAWALENKPRIPVVWIHGL WP_001295940.1
MULTISPECIES: hydrogenase [Enterobacteriaceae] Length: 372\nNumber of predicted TMHs: 2\nExp number of AAs in TMHs: 37.2608899999999\nExp number, first 60 AAs: 19.52923\nTotal prob of N-in: 0.94351\nPOSSIBLE N-term signal sequence\ninside 1 19\nTMhelix 20 39\noutside 40 325\nTMhelix 326 348\ninside 349 372

3061 GCF_001608265.1_ASM160826v1 Escherichia coli Proteobacteria; Gammaproteobacteria; Enterobacterales; Enterobacteriaceae; Escherichia
MTGDNTLIHSHGINRRDFMKLCAALAATMGLSSKAAAEMAESVTNPQRPPVIWIGAQECT WP_000145410.1
MULTISPECIES: hydrogenase-2 small chain [Proteobacteria] Length: 372\nNumber of predicted TMHs: 1\nExp number of AAs in TMHs: 23.72226\nExp number, first 60 AAs: 0.22038\nTotal prob of N-in: 0.12994\noutside 1 331\nTMhelix 332 354\ninside 355 372

3062 GCF_001608285.1_ASM160828v1 Escherichia coli Proteobacteria; Gammaproteobacteria; Enterobacterales; Enterobacteriaceae; Escherichia
MTGDNTLIHSHGINRRDFMKLCAALAATMGLSSKAAAEMAESVTNPQRPPVIWIGAQECT WP_000145410.1
MULTISPECIES: hydrogenase-2 small chain [Proteobacteria] Length: 372\nNumber of predicted TMHs: 1\nExp number of AAs in TMHs: 23.72226\nExp number, first 60 AAs: 0.22038\nTotal prob of N-in: 0.12994\noutside 1 331\nTMhelix 332 354\ninside 355 372

3063 GCF_001608075.1_ASM160807v1 Escherichia coli Proteobacteria; Gammaproteobacteria; Enterobacterales; Enterobacteriaceae; Escherichia
MHSHGINRRDFMKLCAALAATMGLSSKAAAEMAESVTNPQRPPVIWIGAQECTGCTESLL WP_064769412.1
hydrogenase 2 small subunit [Escherichia coli] Length: 365\nNumber of predicted TMHs: 1\nExp number of AAs in TMHs: 26.84905\nExp number, first 60 AAs: 0.03931\nTotal prob of N-in: 0.32128\noutside 1 324\nTMhelix 325 347\ninside 348 365

3064 GCF_001609215.1_ASM160921v1 Escherichia coli Proteobacteria; Gammaproteobacteria; Enterobacterales; Enterobacteriaceae; Escherichia
MTGDNTLIHSHGINRRDFMKLCAALAATMGLSSKAAAEMAESVTNPQRPPVIWIGAQECT WP_000145410.1
MULTISPECIES: hydrogenase-2 small chain [Proteobacteria] Length: 372\nNumber of predicted TMHs: 1\nExp number of AAs in TMHs: 23.72226\nExp number, first 60 AAs: 0.22038\nTotal prob of N-in: 0.12994\noutside 1 331\nTMhelix 332 354\ninside 355 372

3065 GCF_001609265.1_ASM160926v1 Escherichia coli Proteobacteria; Gammaproteobacteria; Enterobacterales; Enterobacteriaceae; Escherichia
MNNEETFYQAMRRQGVTRRSFLKYCSLAATSLGLGAGMAPKIAWALENKPRIPVVWIHGL WP_001058323.1
MULTISPECIES: hydrogenase-1 small chain [Proteobacteria] Length: 372\nNumber of predicted TMHs: 2\nExp number of AAs in TMHs: 37.2608499999999\nExp number, first 60 AAs: 19.52923\nTotal prob of N-in: 0.94351\nPOSSIBLE N-term signal sequence\ninside 1 19\nTMhelix 20 39\noutside 40 325\nTMhelix 326 348\ninside 349 372

3066 GCF_001609315.1_ASM160931v1 Escherichia coli Proteobacteria; Gammaproteobacteria; Enterobacterales; Enterobacteriaceae; Escherichia
MNNEETFYQAMRRQGVTRRSFLKYCSLAATSLGLGAGMAPKIAWALENKPRIPVVWIHGL WP_001058323.1
MULTISPECIES: hydrogenase-1 small chain [Proteobacteria] Length: 372\nNumber of predicted TMHs: 2\nExp number of AAs in TMHs: 37.2608499999999\nExp number, first 60 AAs: 19.52923\nTotal prob of N-in: 0.94351\nPOSSIBLE N-term signal sequence\ninside 1 19\nTMhelix 20 39\noutside 40 325\nTMhelix 326 348\ninside 349 372

3067 GCF_001609735.1_ASM160973v1 Escherichia coli Proteobacteria; Gammaproteobacteria; Enterobacterales; Enterobacteriaceae; Escherichia
MNNEETFYQAMRRQGVTRRSFLKYCSLAATSLGLGAGMAPKIAWALENKPRIPVVWIHGL WP_001058323.1
MULTISPECIES: hydrogenase-1 small chain [Proteobacteria] Length: 372\nNumber of predicted TMHs: 2\nExp number of AAs in TMHs: 37.2608499999999\nExp number, first 60 AAs: 19.52923\nTotal prob of N-in: 0.94351\nPOSSIBLE N-term signal sequence\ninside 1 19\nTMhelix 20 39\noutside 40 325\nTMhelix 326 348\ninside 349 372

3068 GCF_001608285.1_ASM160828v1 Escherichia coli Proteobacteria; Gammaproteobacteria; Enterobacterales; Enterobacteriaceae; Escherichia
MNNEETFYQAMRRQGVTRRSFLKYCSLAATSLGLGAGMAPKIAWALENKPRIPVVWIHGL WP_001058323.1
MULTISPECIES: hydrogenase-1 small chain [Proteobacteria] Length: 372\nNumber of predicted TMHs: 2\nExp number of AAs in TMHs: 37.2608499999999\nExp number, first 60 AAs: 19.52923\nTotal prob of N-in: 0.94351\nPOSSIBLE N-term signal sequence\ninside 1 19\nTMhelix 20 39\noutside 40 325\nTMhelix 326 348\ninside 349 372

3069 GCF_001609745.1_ASM160974v1 Escherichia coli Proteobacteria; Gammaproteobacteria; Enterobacterales; Enterobacteriaceae; Escherichia
MTGDNTLIHSHGINRRDFMKLCAALAATMGLSSKAAAEMAESVTNPQRPPVIWIGAQECT WP_000145410.1
MULTISPECIES: hydrogenase-2 small chain [Proteobacteria] Length: 372\nNumber of predicted TMHs: 1\nExp number of AAs in TMHs: 23.72226\nExp number, first 60 AAs: 0.22038\nTotal prob of N-in: 0.12994\noutside 1 331\nTMhelix 332 354\ninside 355 372

3070 GCF_001609795.1_ASM160979v1 Escherichia coli Proteobacteria; Gammaproteobacteria; Enterobacterales; Enterobacteriaceae; Escherichia
MTGDNTLIHSHGINRRDFMKLCAALAATMGLSSKAAAEMAESVTNPQRPPVIWIGAQECT WP_000145410.1
MULTISPECIES: hydrogenase-2 small chain [Proteobacteria] Length: 372\nNumber of predicted TMHs: 1\nExp number of AAs in TMHs: 23.72226\nExp number, first 60 AAs: 0.22038\nTotal prob of N-in: 0.12994\noutside 1 331\nTMhelix 332 354\ninside 355 372

3071 GCF_001609835.1_ASM160983v1 Escherichia coli Proteobacteria; Gammaproteobacteria; Enterobacterales; Enterobacteriaceae; Escherichia
MTGDNTLIHSHGINRRDFMKLCAALAATMGLSSKAAAEMAESVTNPQRPPVIWIGAQECT WP_000145410.1
MULTISPECIES: hydrogenase-2 small chain [Proteobacteria] Length: 372\nNumber of predicted TMHs: 1\nExp number of AAs in TMHs: 23.72226\nExp number, first 60 AAs: 0.22038\nTotal prob of N-in: 0.12994\noutside 1 331\nTMhelix 332 354\ninside 355 372

3072 GCF_001612475.1_ASM161247v1 Escherichia coli Proteobacteria; Gammaproteobacteria; Enterobacterales; Enterobacteriaceae; Escherichia
MNNEETFYQAMRRQGVTRRSFLKYCSLAATSLGLGAGMAPKIAWALENKPRIPVVWIHGL WP_001301613.1
hydrogenase [Escherichia coli] Length: 372\nNumber of predicted TMHs: 2\nExp number of AAs in TMHs: 37.46598\nExp number, first 60 AAs: 19.53139\nTotal prob of N-in: 0.94364\nPOSSIBLE N-term signal sequence\ninside 1 19\nTMhelix 20 39\noutside 40 325\nTMhelix 326 348\ninside 349 372

3073 GCF_001614455.1_ASM161445v1 Escherichia coli Proteobacteria; Gammaproteobacteria; Enterobacterales; Enterobacteriaceae; Escherichia
MTGDNTLIHSHGINRRDFMKLCAALAATMGLSSKAAAEMAESVTNPQRPPVIWIGAQECT WP_000145410.1
MULTISPECIES: hydrogenase-2 small chain [Proteobacteria] Length: 372\nNumber of predicted TMHs: 1\nExp number of AAs in TMHs: 23.72226\nExp number, first 60 AAs: 0.22038\nTotal prob of N-in: 0.12994\noutside 1 331\nTMhelix 332 354\ninside 355 372

3074 GCF_001612475.1_ASM161247v1 Escherichia coli Proteobacteria; Gammaproteobacteria; Enterobacterales; Enterobacteriaceae; Escherichia
MTGDNTLIHSHGINRRDFMKLCAALAATMGLSSKAAAEMAESVTNPQRPPVIWIGAQECT WP_038339805.1
hydrogenase 2 small subunit [Escherichia coli] Length: 372\nNumber of predicted TMHs: 1\nExp number of AAs in TMHs: 23.86829\nExp number, first 60 AAs: 0.22037\nTotal prob of N-in: 0.12535\noutside 1 332\nTMhelix 333 355\ninside 356 372

3075 GCF_001614395.1_ASM161439v1 Escherichia coli Proteobacteria; Gammaproteobacteria; Enterobacterales; Enterobacteriaceae; Escherichia
MNNEETFYQAMRRQGVTRRSFLKYCSLAATSLGLGAGMAPKIAWALENKPRIPVVWIHGL WP_001058323.1
MULTISPECIES: hydrogenase-1 small chain [Proteobacteria] Length: 372\nNumber of predicted TMHs: 2\nExp number of AAs in TMHs: 37.2608499999999\nExp number, first 60 AAs: 19.52923\nTotal prob of N-in: 0.94351\nPOSSIBLE N-term signal sequence\ninside 1 19\nTMhelix 20 39\noutside 40 325\nTMhelix 326 348\ninside 349 372

3076 GCF_001614455.1_ASM161445v1 Escherichia coli Proteobacteria; Gammaproteobacteria; Enterobacterales; Enterobacteriaceae; Escherichia
MNNEETFYQAMRRQGVTRRSFLKYCSLAATSLGLGAGMAPKIAWALENKPRIPVVWIHGL WP_001058323.1
MULTISPECIES: hydrogenase-1 small chain [Proteobacteria] Length: 372\nNumber of predicted TMHs: 2\nExp number of AAs in TMHs: 37.2608499999999\nExp number, first 60 AAs: 19.52923\nTotal prob of N-in: 0.94351\nPOSSIBLE N-term signal sequence\ninside 1 19\nTMhelix 20 39\noutside 40 325\nTMhelix 326 348\ninside 349 372

3077 GCF_001614325.1_ASM161432v1 Escherichia coli Proteobacteria; Gammaproteobacteria; Enterobacterales; Enterobacteriaceae; Escherichia
MNNEETFYQAMRRQGVTRRSFLKYCSLAATSLGLGAGMAPKIAWALENKPRIPVVWIHGL WP_062876341.1
hydrogenase [Escherichia coli] Length: 372\nNumber of predicted TMHs: 2\nExp number of AAs in TMHs: 37.43626\nExp number, first 60 AAs: 19.52875\nTotal prob of N-in: 0.94361\nPOSSIBLE N-term signal sequence\ninside 1 19\nTMhelix 20 39\noutside 40 325\nTMhelix 326 348\ninside 349 372

3078 GCF_001614475.1_ASM161447v1 Escherichia coli Proteobacteria; Gammaproteobacteria; Enterobacterales; Enterobacteriaceae; Escherichia
MTGDNTLIHSHGINRRDFMKLCAALAATMGLSSKAAAEMAESVTNPQRPPVIWIGAQECT WP_000145410.1
MULTISPECIES: hydrogenase-2 small chain [Proteobacteria] Length: 372\nNumber of predicted TMHs: 1\nExp number of AAs in TMHs: 23.72226\nExp number, first 60 AAs: 0.22038\nTotal prob of N-in: 0.12994\noutside 1 331\nTMhelix 332 354\ninside 355 372

3079 GCF_001614565.1_ASM161456v1 Escherichia coli Proteobacteria; Gammaproteobacteria; Enterobacterales; Enterobacteriaceae; Escherichia
MNNEETFYQAMRRQGVTRRSFLKYCSLAATSLGLGAGMAPKIAWALENKPRIPVVWIHGL WP_001058323.1
MULTISPECIES: hydrogenase-1 small chain [Proteobacteria] Length: 372\nNumber of predicted TMHs: 2\nExp number of AAs in TMHs: 37.2608499999999\nExp number, first 60 AAs: 19.52923\nTotal prob of N-in: 0.94351\nPOSSIBLE N-term signal sequence\ninside 1 19\nTMhelix 20 39\noutside 40 325\nTMhelix 326 348\ninside 349 372

3080 GCF_001614665.1_ASM161466v1 Escherichia coli Proteobacteria; Gammaproteobacteria; Enterobacterales; Enterobacteriaceae; Escherichia

MNNEETFYQAMRRQGVTRRSFLKYCSLAATSLGLGAGMAPKIAWALENKPRIPVVWIHGL WP_001058323.1 MULTISPECIES: hydrogenase-1 small chain [Proteobacteria] Length: 372\nNumber of predicted TMHs: 2\nExp number of AAs in TMHs: 37.2608499999999\nExp number, first 60 AAs: 19.52923\nTotal prob of N-in: 0.94351\nPOSSIBLE N-term signal sequence\ninside 1 19\nTMhelix 20 39\noutside 40 325\nTMhelix 326 348\ninside 349 372

3081 GCF_001614475.1_ASM161447v1 Escherichia coli Proteobacteria; Gammaproteobacteria; Enterobacterales; Enterobacteriaceae; Escherichia MNNEETFYQAMRRQGVTRRSFLKYCSLAATSLGLGAGMAPKIAWALENKPRIPVVWIHGL WP_001058323.1 MULTISPECIES: hydrogenase-1 small chain [Proteobacteria] Length: 372\nNumber of predicted TMHs: 2\nExp number of AAs in TMHs: 37.2608499999999\nExp number, first 60 AAs: 19.52923\nTotal prob of N-in: 0.94351\nPOSSIBLE N-term signal sequence\ninside 1 19\nTMhelix 20 39\noutside 40 325\nTMhelix 326 348\ninside 349 372

3082 GCF_001614495.1_ASM161449v1 Escherichia coli Proteobacteria; Gammaproteobacteria; Enterobacterales; Enterobacteriaceae; Escherichia MNNEETFYQAMRRQGVTRRSFLKYCSLAATSLGLGAGMAPKIAWALENKPRIPVVWIHGL WP_001058323.1 MULTISPECIES: hydrogenase-1 small chain [Proteobacteria] Length: 372\nNumber of predicted TMHs: 2\nExp number of AAs in TMHs: 37.2608499999999\nExp number, first 60 AAs: 19.52923\nTotal prob of N-in: 0.94351\nPOSSIBLE N-term signal sequence\ninside 1 19\nTMhelix 20 39\noutside 40 325\nTMhelix 326 348\ninside 349 372

3083 GCF_001614755.1_ASM161475v1 Escherichia coli Proteobacteria; Gammaproteobacteria; Enterobacterales; Enterobacteriaceae; Escherichia MTGDNTLIHSHGINRRDFMKLCAALAATMGLSSKAAAEMAESVTNPQRPPVIWIGAQECT WP_000145410.1 MULTISPECIES: hydrogenase-2 small chain [Proteobacteria] Length: 372\nNumber of predicted TMHs: 1\nExp number of AAs in TMHs: 23.72226\nExp number, first 60 AAs: 0.22038\nTotal prob of N-in: 0.12994\noutside 1 331\nTMhelix 332 354\ninside 355 372

3084 GCF_001614755.1_ASM161475v1 Escherichia coli Proteobacteria; Gammaproteobacteria; Enterobacterales; Enterobacteriaceae; Escherichia MNNEETFYQAMRRQGVTRRSFLKYCSLAATSLGLGAGMAPKIAWALENKPRIPVVWIHGL WP_001058323.1 MULTISPECIES: hydrogenase-1 small chain [Proteobacteria] Length: 372\nNumber of predicted TMHs: 2\nExp number of AAs in TMHs: 37.2608499999999\nExp number, first 60 AAs: 19.52923\nTotal prob of N-in: 0.94351\nPOSSIBLE N-term signal sequence\ninside 1 19\nTMhelix 20 39\noutside 40 325\nTMhelix 326 348\ninside 349 372

3085 GCF_001614615.1_ASM161461v1 Escherichia coli Proteobacteria; Gammaproteobacteria; Enterobacterales; Enterobacteriaceae; Escherichia MNNEETFYQAMRRQGVTRRSFLKYCSLAATSLGLGAGMAPKIAWALENKPRIPVVWIHGL WP_001058323.1 MULTISPECIES: hydrogenase-1 small chain [Proteobacteria] Length: 372\nNumber of predicted TMHs: 2\nExp number of AAs in TMHs: 37.2608499999999\nExp number, first 60 AAs: 19.52923\nTotal prob of N-in: 0.94351\nPOSSIBLE N-term signal sequence\ninside 1 19\nTMhelix 20 39\noutside 40 325\nTMhelix 326 348\ninside 349 372

3086 GCF_001614715.1_ASM161471v1 Escherichia coli Proteobacteria; Gammaproteobacteria; Enterobacterales; Enterobacteriaceae; Escherichia MTGDNTLIHSHGINRRDFMKLCAALAATMGLSSKAAAEMAESVTNPQRPPVIWIGAQECT WP_029487472.1 hydrogenase 2 small subunit [Escherichia coli] Length: 372\nNumber of predicted TMHs: 1\nExp number of AAs in TMHs: 22.56987\nExp number, first 60 AAs: 0.23233\nTotal prob of N-in: 0.08221\noutside 1 331\nTMhelix 332 354\ninside 355 372

3087 GCF_001614975.1_ASM161497v1 Escherichia coli Proteobacteria; Gammaproteobacteria; Enterobacterales; Enterobacteriaceae; Escherichia MTGDNTLIHSHGINRRDFMKLCAALAATMGLSSKAAAEMAESVTNPQRPPVIWIGAQECT WP_000145410.1 MULTISPECIES: hydrogenase-2 small chain [Proteobacteria] Length: 372\nNumber of predicted TMHs: 1\nExp number of AAs in TMHs: 23.72226\nExp number, first 60 AAs: 0.22038\nTotal prob of N-in: 0.12994\noutside 1 331\nTMhelix 332 354\ninside 355 372

3088 GCF_001614905.1_ASM161490v1 Escherichia coli Proteobacteria; Gammaproteobacteria; Enterobacterales; Enterobacteriaceae; Escherichia

MTGDNTLIHSHGINRRDFMKLCAALAATMGLSSKAAAEMAESVTNPQRPPVIWIGAQECT WP_000145410.1 MULTISPECIES: hydrogenase-2 small chain [Proteobacteria] Length: 372\nNumber of predicted TMHs: 1\nExp number of AAs in TMHs: 23.72226\nExp number, first 60 AAs: 0.22038\nTotal prob of N-in: 0.12994\noutside 1 331\nTMhelix 332 354\ninside 355 372

3089 GCF_001614905.1_ASM161490v1 Escherichia coli Proteobacteria; Gammaproteobacteria; Enterobacterales; Enterobacteriaceae; Escherichia MNNEETFYQAMRRQGVTRRSFLKYCSLAATSLGLGAGMAPKIAWALENKPRIPVVWIHGL WP_001058323.1 MULTISPECIES: hydrogenase-1 small chain [Proteobacteria] Length: 372\nNumber of predicted TMHs: 2\nExp number of AAs in TMHs: 37.2608499999999\nExp number, first 60 AAs: 19.52923\nTotal prob of N-in: 0.94351\nPOSSIBLE N-term signal sequence\ninside 1 19\nTMhelix 20 39\noutside 40 325\nTMhelix 326 348\ninside 349 372

3090 GCF_001614975.1_ASM161497v1 Escherichia coli Proteobacteria; Gammaproteobacteria; Enterobacterales; Enterobacteriaceae; Escherichia MNNEETFYQAMRRQGVTRRSFLKYCSLAATSLGLGAGMAPKIAWALENKPRIPVVWIHGL WP_001295940.1 MULTISPECIES: hydrogenase [Enterobacteriaceae] Length: 372\nNumber of predicted TMHs: 2\nExp number of AAs in TMHs: 37.2608899999999\nExp number, first 60 AAs: 19.52923\nTotal prob of N-in: 0.94351\nPOSSIBLE N-term signal sequence\ninside 1 19\nTMhelix 20 39\noutside 40 325\nTMhelix 326 348\ninside 349 372

3091 GCF_001614775.1_ASM161477v1 Escherichia coli Proteobacteria; Gammaproteobacteria; Enterobacterales; Enterobacteriaceae; Escherichia MNNEETFYQAMRRQGVTRRSFLKYCSLAATSLGLGAGMAPKIAWALENKPRIPVVWIHGL WP_001058323.1 MULTISPECIES: hydrogenase-1 small chain [Proteobacteria] Length: 372\nNumber of predicted TMHs: 2\nExp number of AAs in TMHs: 37.2608499999999\nExp number, first 60 AAs: 19.52923\nTotal prob of N-in: 0.94351\nPOSSIBLE N-term signal sequence\ninside 1 19\nTMhelix 20 39\noutside 40 325\nTMhelix 326 348\ninside 349 372

3092 GCF_001615065.1_ASM161506v1 Escherichia coli Proteobacteria; Gammaproteobacteria; Enterobacterales; Enterobacteriaceae; Escherichia MTGDNTLIHSHGINRRDFMKLCAALAATMGLSSKAAAEMAESVTNPQRPPVIWIGAQECT WP_000145410.1 MULTISPECIES: hydrogenase-2 small chain [Proteobacteria] Length: 372\nNumber of predicted TMHs: 1\nExp number of AAs in TMHs: 23.72226\nExp number, first 60 AAs: 0.22038\nTotal prob of N-in: 0.12994\noutside 1 331\nTMhelix 332 354\ninside 355 372

3093 GCF_001615125.1_ASM161512v1 Escherichia coli Proteobacteria; Gammaproteobacteria; Enterobacterales; Enterobacteriaceae; Escherichia MNNEETFYQAMRRQGVTRRSFLKYCSLAATSLGLGAGMAPKIAWALENKPRIPVVWIHGL WP_001058320.1 hydrogenase [Escherichia coli] Length: 372\nNumber of predicted TMHs: 2\nExp number of AAs in TMHs: 37.26082\nExp number, first 60 AAs: 19.52923\nTotal prob of N-in: 0.94351\nPOSSIBLE N-term signal sequence\ninside 1 19\nTMhelix 20 39\noutside 40 325\nTMhelix 326 348\ninside 349 372

3094 GCF_001614805.1_ASM161480v1 Escherichia coli Proteobacteria; Gammaproteobacteria; Enterobacterales; Enterobacteriaceae; Escherichia MNNEETFYQAMRRQGVTRRSFLKYCSLAATSLGLGAGMAPKIAWALENKPRIPVVWIHGL WP_063501981.1 hydrogenase [Escherichia coli] Length: 372\nNumber of predicted TMHs: 2\nExp number of AAs in TMHs: 37.06322\nExp number, first 60 AAs: 19.52932\nTotal prob of N-in: 0.94340\nPOSSIBLE N-term signal sequence\ninside 1 19\nTMhelix 20 39\noutside 40 325\nTMhelix 326 348\ninside 349 372

3095 GCF_001615195.1_ASM161519v1 Escherichia coli Proteobacteria; Gammaproteobacteria; Enterobacterales; Enterobacteriaceae; Escherichia MTGDNTLIHSHGINRRDFMKLCAALAATMGLSSKAAAEMAESVTNPQRPPVIWIGAQECT WP_000145410.1 MULTISPECIES: hydrogenase-2 small chain [Proteobacteria] Length: 372\nNumber of predicted TMHs: 1\nExp number of AAs in TMHs: 23.72226\nExp number, first 60 AAs: 0.22038\nTotal prob of N-in: 0.12994\noutside 1 331\nTMhelix 332 354\ninside 355 372

3096 GCF_001615175.1_ASM161517v1 Escherichia coli Proteobacteria; Gammaproteobacteria; Enterobacterales; Enterobacteriaceae; Escherichia MNNEETFYQAMRRQGVTRRSFLKYCSLAATSLGLGAGMAPKIAWALENKPRIPVVWIHGL WP_001058323.1 MULTISPECIES: hydrogenase-1 small chain [Proteobacteria] Length: 372\nNumber of predicted TMHs:

2\nExp number of AAs in TMHs: 37.2608499999999\nExp number, first 60 AAs: 19.52923\nTotal prob of N-in: 0.94351\nPOSSIBLE N-term signal sequence\ninside 1 19\nTMhelix 20 39\noutside 40 325\nTMhelix 326 348\ninside 349 372

3097 GCF_001615275.1_ASM161527v1 Escherichia coli Proteobacteria; Gammaproteobacteria; Enterobacterales; Enterobacteriaceae; Escherichia MTGDNTLIHSHGINRRDFMKLCAALAATMGLSSKAAAEMAESVTNPQRPPVIWIGAQECT WP_000145410.1 MULTISPECIES: hydrogenase-2 small chain [Proteobacteria] Length: 372\nNumber of predicted TMHs: 1\nExp number of AAs in TMHs: 23.72226\nExp number, first 60 AAs: 0.22038\nTotal prob of N-in: 0.12994\noutside 1 331\nTMhelix 332 354\ninside 355 372

3098 GCF_001615295.1_ASM161529v1 Escherichia coli Proteobacteria; Gammaproteobacteria; Enterobacterales; Enterobacteriaceae; Escherichia MTGDNTLIHSHGINRRDFMKLCAALAATMGLSSKAAAEMAESVTNPQRPPVIWIGAQECT WP_000145410.1 MULTISPECIES: hydrogenase-2 small chain [Proteobacteria] Length: 372\nNumber of predicted TMHs: 1\nExp number of AAs in TMHs: 23.72226\nExp number, first 60 AAs: 0.22038\nTotal prob of N-in: 0.12994\noutside 1 331\nTMhelix 332 354\ninside 355 372

3099 GCF_001615225.1_ASM161522v1 Escherichia coli Proteobacteria; Gammaproteobacteria; Enterobacterales; Enterobacteriaceae; Escherichia MTGDNTLIHSHGINRRDFMKLCAALAATMGLSSKAAAEMAESVTNPQRPPVIWIGAQECT WP_000145410.1 MULTISPECIES: hydrogenase-2 small chain [Proteobacteria] Length: 372\nNumber of predicted TMHs: 1\nExp number of AAs in TMHs: 23.72226\nExp number, first 60 AAs: 0.22038\nTotal prob of N-in: 0.12994\noutside 1 331\nTMhelix 332 354\ninside 355 372

3100 GCF_001615325.1_ASM161532v1 Escherichia coli Proteobacteria; Gammaproteobacteria; Enterobacterales; Enterobacteriaceae; Escherichia MNNEETFYQAMRRQGVTRRSFLKYCSLAATSLGLGAGMAPKIAWALENKPRIPVVWIHGL WP_001058323.1 MULTISPECIES: hydrogenase-1 small chain [Proteobacteria] Length: 372\nNumber of predicted TMHs: 2\nExp number of AAs in TMHs: 37.2608499999999\nExp number, first 60 AAs: 19.52923\nTotal prob of N-in: 0.94351\nPOSSIBLE N-term signal sequence\ninside 1 19\nTMhelix 20 39\noutside 40 325\nTMhelix 326 348\ninside 349 372

3101 GCF_001615175.1_ASM161517v1 Escherichia coli Proteobacteria; Gammaproteobacteria; Enterobacterales; Enterobacteriaceae; Escherichia MTGDNTLIHSHGINRRDFMKLCAALAATMGLSSKAAAEMAESVTNPQRPPVIWIGAQECT WP_063088068.1 hydrogenase 2 small subunit [Escherichia coli] Length: 372\nNumber of predicted TMHs: 1\nExp number of AAs in TMHs: 23.73081\nExp number, first 60 AAs: 0.22055\nTotal prob of N-in: 0.13010\noutside 1 331\nTMhelix 332 354\ninside 355 372

3102 GCF_001615485.1_ASM161548v1 Escherichia coli Proteobacteria; Gammaproteobacteria; Enterobacterales; Enterobacteriaceae; Escherichia MTGDNTLIHSHGINRRDFMKLCAALAATMGLSSKAAAEMAESVTNPQRPPVIWIGAQECT WP_000145410.1 MULTISPECIES: hydrogenase-2 small chain [Proteobacteria] Length: 372\nNumber of predicted TMHs: 1\nExp number of AAs in TMHs: 23.72226\nExp number, first 60 AAs: 0.22038\nTotal prob of N-in: 0.12994\noutside 1 331\nTMhelix 332 354\ninside 355 372

3103 GCF_001615485.1_ASM161548v1 Escherichia coli Proteobacteria; Gammaproteobacteria; Enterobacterales; Enterobacteriaceae; Escherichia MNNEETFYQAMRRQGVTRRSFLKYCSLAATSLGLGAGMAPKIAWALENKPRIPVVWIHGL WP_001058323.1 MULTISPECIES: hydrogenase-1 small chain [Proteobacteria] Length: 372\nNumber of predicted TMHs: 2\nExp number of AAs in TMHs: 37.2608499999999\nExp number, first 60 AAs: 19.52923\nTotal prob of N-in: 0.94351\nPOSSIBLE N-term signal sequence\ninside 1 19\nTMhelix 20 39\noutside 40 325\nTMhelix 326 348\ninside 349 372

3104 GCF_001615225.1_ASM161522v1 Escherichia coli Proteobacteria; Gammaproteobacteria; Enterobacterales; Enterobacteriaceae; Escherichia MNNEETFYQAMRRQGVTRRSFLKYCSLAATSLGLGAGMAPKIAWALENKPRIPVVWIHGL WP_001058323.1 MULTISPECIES: hydrogenase-1 small chain [Proteobacteria] Length: 372\nNumber of predicted TMHs: 2\nExp number of AAs in TMHs: 37.2608499999999\nExp number, first 60 AAs: 19.52923\nTotal prob of N-in:

0.94351\nPOSSIBLE N-term signal sequence\ninside 1 19\nTMhelix 20 39\noutside 40 325\nTMhelix 326 348\ninside 349 372

3105 GCF_001615195.1_ASM161519v1 Escherichia coli Proteobacteria; Gammaproteobacteria; Enterobacterales; Enterobacteriaceae; Escherichia MNNEETFYQAMRRQGVTRRSFLKYCSLAATSLGLGAGMAPKIAWALENKPRIPVVWIHGL WP_001295940.1 MULTISPECIES: hydrogenase [Enterobacteriaceae] Length: 372\nNumber of predicted TMHs: 2\nExp number of AAs in TMHs: 37.2608899999999\nExp number, first 60 AAs: 19.52923\nTotal prob of N-in: 0.94351\nPOSSIBLE N-term signal sequence\ninside 1 19\nTMhelix 20 39\noutside 40 325\nTMhelix 326 348\ninside 349 372

3106 GCF_001615695.1_ASM161569v1 Escherichia coli Proteobacteria; Gammaproteobacteria; Enterobacterales; Enterobacteriaceae; Escherichia MNNEETFYQAMRRQGVTRRSFLKYCSLAATSLGLGAGMAPKIAWALENKPRIPVVWIHGL WP_001058323.1 MULTISPECIES: hydrogenase-1 small chain [Proteobacteria] Length: 372\nNumber of predicted TMHs: 2\nExp number of AAs in TMHs: 37.2608499999999\nExp number, first 60 AAs: 19.52923\nTotal prob of N-in: 0.94351\nPOSSIBLE N-term signal sequence\ninside 1 19\nTMhelix 20 39\noutside 40 325\nTMhelix 326 348\ninside 349 372

3107 GCF_001615565.1_ASM161556v1 Escherichia coli Proteobacteria; Gammaproteobacteria; Enterobacterales; Enterobacteriaceae; Escherichia MTGDNTLIHSHGINRRDFMKLCAALAATMGLSSKAAAEMAESVTNPQRPPVIWIGAQECT WP_000145410.1 MULTISPECIES: hydrogenase-2 small chain [Proteobacteria] Length: 372\nNumber of predicted TMHs: 1\nExp number of AAs in TMHs: 23.72226\nExp number, first 60 AAs: 0.22038\nTotal prob of N-in: 0.12994\noutside 1 331\nTMhelix 332 354\ninside 355 372

3108 GCF_001615975.1_ASM161597v1 Escherichia coli Proteobacteria; Gammaproteobacteria; Enterobacterales; Enterobacteriaceae; Escherichia MTGDNTLIHSHGINRRDFMKLCAALAATMGLSSKAAAEMAESVTNPQRPPVIWIGAQECT WP_000145410.1 MULTISPECIES: hydrogenase-2 small chain [Proteobacteria] Length: 372\nNumber of predicted TMHs: 1\nExp number of AAs in TMHs: 23.72226\nExp number, first 60 AAs: 0.22038\nTotal prob of N-in: 0.12994\noutside 1 331\nTMhelix 332 354\ninside 355 372

3109 GCF_001615925.1_ASM161592v1 Escherichia coli Proteobacteria; Gammaproteobacteria; Enterobacterales; Enterobacteriaceae; Escherichia MTGDNTLIHSHGINRRDFMKLCAALAATMGLSSKAAAEMAESVTNPQRPPVIWIGAQECT WP_000145410.1 MULTISPECIES: hydrogenase-2 small chain [Proteobacteria] Length: 372\nNumber of predicted TMHs: 1\nExp number of AAs in TMHs: 23.72226\nExp number, first 60 AAs: 0.22038\nTotal prob of N-in: 0.12994\noutside 1 331\nTMhelix 332 354\ninside 355 372

3110 GCF_001616035.1_ASM161603v1 Escherichia coli Proteobacteria; Gammaproteobacteria; Enterobacterales; Enterobacteriaceae; Escherichia MTGDNTLIHSHGINRRDFMKLCAALAATMGLSSKAAAEMAESVTNPQRPPVIWIGAQECT WP_000145410.1 MULTISPECIES: hydrogenase-2 small chain [Proteobacteria] Length: 372\nNumber of predicted TMHs: 1\nExp number of AAs in TMHs: 23.72226\nExp number, first 60 AAs: 0.22038\nTotal prob of N-in: 0.12994\noutside 1 331\nTMhelix 332 354\ninside 355 372

3111 GCF_001615985.1_ASM161598v1 Escherichia coli Proteobacteria; Gammaproteobacteria; Enterobacterales; Enterobacteriaceae; Escherichia MTGDNTLIHSHGINRRDFMKLCAALAATMGLSSKAAAEMAESVTNPQRPPVIWIGAQECT WP_000145410.1 MULTISPECIES: hydrogenase-2 small chain [Proteobacteria] Length: 372\nNumber of predicted TMHs: 1\nExp number of AAs in TMHs: 23.72226\nExp number, first 60 AAs: 0.22038\nTotal prob of N-in: 0.12994\noutside 1 331\nTMhelix 332 354\ninside 355 372

3112 GCF_001616055.1_ASM161605v1 Escherichia coli Proteobacteria; Gammaproteobacteria; Enterobacterales; Enterobacteriaceae; Escherichia MTGDNTLIHSHGINRRDFMKLCAALAATMGLSSKAAAEMAESVTNPQRPPVIWIGAQECT WP_000145410.1 MULTISPECIES: hydrogenase-2 small chain [Proteobacteria] Length: 372\nNumber of predicted TMHs: 1\nExp number of AAs in TMHs: 23.72226\nExp number, first 60 AAs: 0.22038\nTotal prob of N-in: 0.12994\noutside 1 331\nTMhelix 332 354\ninside 355 372

3113 GCF_001616095.1_ASM161609v1 Escherichia coli Proteobacteria; Gammaproteobacteria; Enterobacterales; Enterobacteriaceae; Escherichia
MTGDNTLIHSHGINRRDFMKLCAALAATMGLSSKAAAEMAESVTNPQRPPVIWIGAQECT WP_000145410.1
MULTISPECIES: hydrogenase-2 small chain [Proteobacteria] Length: 372\nNumber of predicted TMHs: 1\nExp number of AAs in TMHs: 23.72226\nExp number, first 60 AAs: 0.22038\nTotal prob of N-in: 0.12994\noutside 1 331\nTMhelix 332 354\ninside 355 372

3114 GCF_001616095.1_ASM161609v1 Escherichia coli Proteobacteria; Gammaproteobacteria; Enterobacterales; Enterobacteriaceae; Escherichia
MNNEETFYQAMRRQGVTRRSFLKYCSLAATSLGLGAGMAPKIAWALENKPRIPVVWIHGL WP_001058323.1
MULTISPECIES: hydrogenase-1 small chain [Proteobacteria] Length: 372\nNumber of predicted TMHs: 2\nExp number of AAs in TMHs: 37.2608499999999\nExp number, first 60 AAs: 19.52923\nTotal prob of N-in: 0.94351\nPOSSIBLE N-term signal sequence\ninside 1 19\nTMhelix 20 39\noutside 40 325\nTMhelix 326 348\ninside 349 372

3115 GCF_001616285.1_ASM161628v1 Escherichia coli Proteobacteria; Gammaproteobacteria; Enterobacterales; Enterobacteriaceae; Escherichia
MTGDNTLIHSHGINRRDFMKLCAALAATMGLSSKAAAEMAESVTNPQRPPVIWIGAQECT WP_000145410.1
MULTISPECIES: hydrogenase-2 small chain [Proteobacteria] Length: 372\nNumber of predicted TMHs: 1\nExp number of AAs in TMHs: 23.72226\nExp number, first 60 AAs: 0.22038\nTotal prob of N-in: 0.12994\noutside 1 331\nTMhelix 332 354\ninside 355 372

3116 GCF_001616335.1_ASM161633v1 Escherichia coli Proteobacteria; Gammaproteobacteria; Enterobacterales; Enterobacteriaceae; Escherichia
MTGDNTLIHSHGINRRDFMKLCAALAATMGLSSKAAAEMAESVTNPQRPPVIWIGAQECT WP_000145410.1
MULTISPECIES: hydrogenase-2 small chain [Proteobacteria] Length: 372\nNumber of predicted TMHs: 1\nExp number of AAs in TMHs: 23.72226\nExp number, first 60 AAs: 0.22038\nTotal prob of N-in: 0.12994\noutside 1 331\nTMhelix 332 354\ninside 355 372

3117 GCF_001616375.1_ASM161637v1 Escherichia coli Proteobacteria; Gammaproteobacteria; Enterobacterales; Enterobacteriaceae; Escherichia
MTGDNTLIHSHGINRRDFMKLCAALAATMGLSSKAAAEMAESVTNPQRPPVIWIGAQECT WP_000145410.1
MULTISPECIES: hydrogenase-2 small chain [Proteobacteria] Length: 372\nNumber of predicted TMHs: 1\nExp number of AAs in TMHs: 23.72226\nExp number, first 60 AAs: 0.22038\nTotal prob of N-in: 0.12994\noutside 1 331\nTMhelix 332 354\ninside 355 372

3118 GCF_001616285.1_ASM161628v1 Escherichia coli Proteobacteria; Gammaproteobacteria; Enterobacterales; Enterobacteriaceae; Escherichia
MNNEETFYQAMRRQGVTRRSFLKYCSLAATSLGLGAGMAPKIAWALENKPRIPVVWIHGL WP_001058323.1
MULTISPECIES: hydrogenase-1 small chain [Proteobacteria] Length: 372\nNumber of predicted TMHs: 2\nExp number of AAs in TMHs: 37.2608499999999\nExp number, first 60 AAs: 19.52923\nTotal prob of N-in: 0.94351\nPOSSIBLE N-term signal sequence\ninside 1 19\nTMhelix 20 39\noutside 40 325\nTMhelix 326 348\ninside 349 372

3119 GCF_001616385.1_ASM161638v1 Escherichia coli Proteobacteria; Gammaproteobacteria; Enterobacterales; Enterobacteriaceae; Escherichia
MTGDNTLIHSHGINRRDFMKLCAALAATMGLSSKAAAEMAESVTNPQRPPVIWIGAQECT WP_000145410.1
MULTISPECIES: hydrogenase-2 small chain [Proteobacteria] Length: 372\nNumber of predicted TMHs: 1\nExp number of AAs in TMHs: 23.72226\nExp number, first 60 AAs: 0.22038\nTotal prob of N-in: 0.12994\noutside 1 331\nTMhelix 332 354\ninside 355 372

3120 GCF_001616335.1_ASM161633v1 Escherichia coli Proteobacteria; Gammaproteobacteria; Enterobacterales; Enterobacteriaceae; Escherichia
MNNEETFYQAMRRQGVTRRSFLKYCSLAATSLGLGAGMAPKIAWALENKPRIPVVWIHGL WP_001058323.1
MULTISPECIES: hydrogenase-1 small chain [Proteobacteria] Length: 372\nNumber of predicted TMHs: 2\nExp number of AAs in TMHs: 37.2608499999999\nExp number, first 60 AAs: 19.52923\nTotal prob of N-in: 0.94351\nPOSSIBLE N-term signal sequence\ninside 1 19\nTMhelix 20 39\noutside 40 325\nTMhelix 326 348\ninside 349 372

3121 GCF_001616635.1_ASM161663v1 Escherichia coli Proteobacteria; Gammaproteobacteria; Enterobacterales; Enterobacteriaceae; Escherichia

MTGDNTLIHSHGINRRDFMKLCAALAATMGLSSKAAAEMAESVTNPQRPPVIWIGAQECT WP_000145410.1 MULTISPECIES: hydrogenase-2 small chain [Proteobacteria] Length: 372\nNumber of predicted TMHs: 1\nExp number of AAs in TMHs: 23.72226\nExp number, first 60 AAs: 0.22038\nTotal prob of N-in: 0.12994\noutside 1 331\nTMhelix 332 354\ninside 355 372

3122 GCF_001616635.1_ASM161663v1 Escherichia coli Proteobacteria; Gammaproteobacteria; Enterobacterales; Enterobacteriaceae; Escherichia MNNEETFYQAMRRQGVTRRSFLKYCSLAATSLGLGAGMAPKIAWALENKPRIPVVWIHGL WP_001058323.1 MULTISPECIES: hydrogenase-1 small chain [Proteobacteria] Length: 372\nNumber of predicted TMHs: 2\nExp number of AAs in TMHs: 37.2608499999999\nExp number, first 60 AAs: 19.52923\nTotal prob of N-in: 0.94351\nPOSSIBLE N-term signal sequence\ninside 1 19\nTMhelix 20 39\noutside 40 325\nTMhelix 326 348\ninside 349 372

3123 GCF_001616375.1_ASM161637v1 Escherichia coli Proteobacteria; Gammaproteobacteria; Enterobacterales; Enterobacteriaceae; Escherichia MNNEETFYQAMRRQGVTRRSFLKYCSLAATSLGLGAGMAPKIAWALENKPRIPVVWIHGL WP_001058323.1 MULTISPECIES: hydrogenase-1 small chain [Proteobacteria] Length: 372\nNumber of predicted TMHs: 2\nExp number of AAs in TMHs: 37.2608499999999\nExp number, first 60 AAs: 19.52923\nTotal prob of N-in: 0.94351\nPOSSIBLE N-term signal sequence\ninside 1 19\nTMhelix 20 39\noutside 40 325\nTMhelix 326 348\ninside 349 372

3124 GCF_001616385.1_ASM161638v1 Escherichia coli Proteobacteria; Gammaproteobacteria; Enterobacterales; Enterobacteriaceae; Escherichia MNNEETFYQAMRRQGVTRRSFLKYCSLAATSLGLGAGMAPKIAWALENKPRIPVVWIHGL WP_001058320.1 hydrogenase [Escherichia coli] Length: 372\nNumber of predicted TMHs: 2\nExp number of AAs in TMHs: 37.26082\nExp number, first 60 AAs: 19.52923\nTotal prob of N-in: 0.94351\nPOSSIBLE N-term signal sequence\ninside 1 19\nTMhelix 20 39\noutside 40 325\nTMhelix 326 348\ninside 349 372

3125 GCF_001616495.1_ASM161649v1 Escherichia coli Proteobacteria; Gammaproteobacteria; Enterobacterales; Enterobacteriaceae; Escherichia MNNEETFYQAMRRQGVTRRSFLKYCSLAATSLGLGAGMAPKIAWALENKPRIPVVWIHGL WP_001058323.1 MULTISPECIES: hydrogenase-1 small chain [Proteobacteria] Length: 372\nNumber of predicted TMHs: 2\nExp number of AAs in TMHs: 37.2608499999999\nExp number, first 60 AAs: 19.52923\nTotal prob of N-in: 0.94351\nPOSSIBLE N-term signal sequence\ninside 1 19\nTMhelix 20 39\noutside 40 325\nTMhelix 326 348\ninside 349 372

3126 GCF_001616575.1_ASM161657v1 Escherichia coli Proteobacteria; Gammaproteobacteria; Enterobacterales; Enterobacteriaceae; Escherichia MNNEETFYQAMRRQGVTRRSFLKYCSLAATSLGLGAGMAPKIAWALENKPRIPVVWIHGL WP_001058323.1 MULTISPECIES: hydrogenase-1 small chain [Proteobacteria] Length: 372\nNumber of predicted TMHs: 2\nExp number of AAs in TMHs: 37.2608499999999\nExp number, first 60 AAs: 19.52923\nTotal prob of N-in: 0.94351\nPOSSIBLE N-term signal sequence\ninside 1 19\nTMhelix 20 39\noutside 40 325\nTMhelix 326 348\ninside 349 372

3127 GCF_001616895.1_ASM161689v1 Escherichia coli Proteobacteria; Gammaproteobacteria; Enterobacterales; Enterobacteriaceae; Escherichia MNNEETFYQAMRRQGVTRRSFLKYCSLAATSLGLGAGMAPKIAWALENKPRIPVVWIHGL WP_001058323.1 MULTISPECIES: hydrogenase-1 small chain [Proteobacteria] Length: 372\nNumber of predicted TMHs: 2\nExp number of AAs in TMHs: 37.2608499999999\nExp number, first 60 AAs: 19.52923\nTotal prob of N-in: 0.94351\nPOSSIBLE N-term signal sequence\ninside 1 19\nTMhelix 20 39\noutside 40 325\nTMhelix 326 348\ninside 349 372

3128 GCF_001616665.1_ASM161666v1 Escherichia coli Proteobacteria; Gammaproteobacteria; Enterobacterales; Enterobacteriaceae; Escherichia MNNEETFYQAMRRQGVTRRSFLKYCSLAATSLGLGAGMAPKIAWALENKPRIPVVWIHGL WP_063106937.1 hydrogenase [Escherichia coli] Length: 372\nNumber of predicted TMHs: 2\nExp number of AAs in TMHs: 36.54338\nExp number, first 60 AAs: 19.53679\nTotal prob of N-in: 0.94294\nPOSSIBLE N-term signal sequence\ninside 1 19\nTMhelix 20 39\noutside 40 325\nTMhelix 326 348\ninside 349 372

3129 GCF_001616775.1_ASM161677v1 Escherichia coli Proteobacteria; Gammaproteobacteria; Enterobacterales; Enterobacteriaceae; Escherichia

MNNEETFYQAMRRQGVTRRSFLKYCSLAATSLGLGAGMAPKIAWALENKPRIPVVWIHGL WP_024246676.1 hydrogenase-1 small chain [Escherichia coli] Length: 372\nNumber of predicted TMHs: 2\nExp number of AAs in TMHs: 37.8214899999999\nExp number, first 60 AAs: 19.52326\nTotal prob of N-in: 0.94395\nPOSSIBLE N-term signal sequence\ninside 1 19\nTMhelix 20 39\noutside 40 325\nTMhelix 326 348\ninside 349 372

3130 GCF_001617135.1_ASM161713v1 Escherichia coli Proteobacteria; Gammaproteobacteria; Enterobacterales; Enterobacteriaceae; Escherichia MTGDNTLIHSHGINRRDFMKLCAALAATMGLSSKAAAEMAESVTNPQRPPVIWIGAQECT WP_000145410.1 MULTISPECIES: hydrogenase-2 small chain [Proteobacteria] Length: 372\nNumber of predicted TMHs: 1\nExp number of AAs in TMHs: 23.72226\nExp number, first 60 AAs: 0.22038\nTotal prob of N-in: 0.12994\noutside 1 331\nTMhelix 332 354\ninside 355 372

3131 GCF_001617115.1_ASM161711v1 Escherichia coli Proteobacteria; Gammaproteobacteria; Enterobacterales; Enterobacteriaceae; Escherichia MTGDNTLIHSHGINRRDFMKLCAALAATMGLSSKAAAEMAESVTNPQRPPVIWIGAQECT WP_000145410.1 MULTISPECIES: hydrogenase-2 small chain [Proteobacteria] Length: 372\nNumber of predicted TMHs: 1\nExp number of AAs in TMHs: 23.72226\nExp number, first 60 AAs: 0.22038\nTotal prob of N-in: 0.12994\noutside 1 331\nTMhelix 332 354\ninside 355 372

3132 GCF_001617175.1_ASM161717v1 Escherichia coli Proteobacteria; Gammaproteobacteria; Enterobacterales; Enterobacteriaceae; Escherichia MTGDNTLIHSHGINRRDFMKLCAALAATMGLSSKAAAEMAESVTNPQRPPVIWIGAQECT WP_000145410.1 MULTISPECIES: hydrogenase-2 small chain [Proteobacteria] Length: 372\nNumber of predicted TMHs: 1\nExp number of AAs in TMHs: 23.72226\nExp number, first 60 AAs: 0.22038\nTotal prob of N-in: 0.12994\noutside 1 331\nTMhelix 332 354\ninside 355 372

3133 GCF_001617175.1_ASM161717v1 Escherichia coli Proteobacteria; Gammaproteobacteria; Enterobacterales; Enterobacteriaceae; Escherichia MNNEETFYQAMRRQGVTRRSFLKYCSLAATSLGLGAGMAPKIAWALENKPRIPVVWIHGL WP_001058323.1 MULTISPECIES: hydrogenase-1 small chain [Proteobacteria] Length: 372\nNumber of predicted TMHs: 2\nExp number of AAs in TMHs: 37.2608499999999\nExp number, first 60 AAs: 19.52923\nTotal prob of N-in: 0.94351\nPOSSIBLE N-term signal sequence\ninside 1 19\nTMhelix 20 39\noutside 40 325\nTMhelix 326 348\ninside 349 372

3134 GCF_001617265.1_ASM161726v1 Escherichia coli Proteobacteria; Gammaproteobacteria; Enterobacterales; Enterobacteriaceae; Escherichia MNNEETFYQAMRRQGVTRRSFLKYCSLAATSLGLGAGMAPKIAWALENKPRIPVVWIHGL WP_001058323.1 MULTISPECIES: hydrogenase-1 small chain [Proteobacteria] Length: 372\nNumber of predicted TMHs: 2\nExp number of AAs in TMHs: 37.2608499999999\nExp number, first 60 AAs: 19.52923\nTotal prob of N-in: 0.94351\nPOSSIBLE N-term signal sequence\ninside 1 19\nTMhelix 20 39\noutside 40 325\nTMhelix 326 348\ninside 349 372

3135 GCF_001619145.1_ASM161914v1 Escherichia coli Proteobacteria; Gammaproteobacteria; Enterobacterales; Enterobacteriaceae; Escherichia MTGDNTLIHSHGINRRDFMKLCAALAATMGLSSKAAAEMAESVTNPQRPPVIWIGAQECT WP_000145410.1 MULTISPECIES: hydrogenase-2 small chain [Proteobacteria] Length: 372\nNumber of predicted TMHs: 1\nExp number of AAs in TMHs: 23.72226\nExp number, first 60 AAs: 0.22038\nTotal prob of N-in: 0.12994\noutside 1 331\nTMhelix 332 354\ninside 355 372

3136 GCF_001619145.1_ASM161914v1 Escherichia coli Proteobacteria; Gammaproteobacteria; Enterobacterales; Enterobacteriaceae; Escherichia MNNEETFYQAMRRQGVTRRSFLKYCSLAATSLGLGAGMAPKIAWALENKPRIPVVWIHGL WP_001301613.1 hydrogenase [Escherichia coli] Length: 372\nNumber of predicted TMHs: 2\nExp number of AAs in TMHs: 37.46598\nExp number, first 60 AAs: 19.53139\nTotal prob of N-in: 0.94364\nPOSSIBLE N-term signal sequence\ninside 1 19\nTMhelix 20 39\noutside 40 325\nTMhelix 326 348\ninside 349 372

3137 GCF_001617215.1_ASM161721v1 Escherichia coli Proteobacteria; Gammaproteobacteria; Enterobacterales; Enterobacteriaceae; Escherichia MNNEETFYQAMRRQGVTRRSFLKYCSLAATSLGLGAGMAPKIAWALENKPRIPVVWIHGL WP_024246676.1 hydrogenase-1 small chain [Escherichia coli] Length: 372\nNumber of predicted TMHs: 2\nExp number of

AAs in TMHs: 37.8214899999999\nExp number, first 60 AAs: 19.52326\nTotal prob of N-in: 0.94395\nPOSSIBLE N-term signal sequence\ninside 1 19\nTMhelix 20 39\noutside 40 325\nTMhelix 326 348\ninside 349 372

3138 GCF_001619195.1_ASM161919v1 Escherichia coli Proteobacteria; Gammaproteobacteria; Enterobacterales; Enterobacteriaceae; Escherichia
MTGDNTLIHSHGINRRDFMKLCAALAATMGLSSKAAAEMAESVTNPQRPPVIWIGAQECT WP_000145410.1
MULTISPECIES: hydrogenase-2 small chain [Proteobacteria] Length: 372\nNumber of predicted TMHs: 1\nExp number of AAs in TMHs: 23.72226\nExp number, first 60 AAs: 0.22038\nTotal prob of N-in: 0.12994\noutside 1 331\nTMhelix 332 354\ninside 355 372

3139 GCF_001619095.1_ASM161909v1 Escherichia coli Proteobacteria; Gammaproteobacteria; Enterobacterales; Enterobacteriaceae; Escherichia
MNNEETFYQAMRRQGVTRRSFLKYCSLAATSLGLGAGMAPKIAWALENKPRIPVVWIHGL WP_001301613.1
hydrogenase [Escherichia coli] Length: 372\nNumber of predicted TMHs: 2\nExp number of AAs in TMHs: 37.46598\nExp number, first 60 AAs: 19.53139\nTotal prob of N-in: 0.94364\nPOSSIBLE N-term signal sequence\ninside 1 19\nTMhelix 20 39\noutside 40 325\nTMhelix 326 348\ninside 349 372

3140 GCF_001621035.1_ASM162103v1 Escherichia coli Proteobacteria; Gammaproteobacteria; Enterobacterales; Enterobacteriaceae; Escherichia
MNNEETFYQAMRRQGVTRRSFLKYCSLAATSLGLGAGMAPKIAWALENKPRIPVVWIHGL WP_001295940.1
MULTISPECIES: hydrogenase [Enterobacteriaceae] Length: 372\nNumber of predicted TMHs: 2\nExp number of AAs in TMHs: 37.2608899999999\nExp number, first 60 AAs: 19.52923\nTotal prob of N-in: 0.94351\nPOSSIBLE N-term signal sequence\ninside 1 19\nTMhelix 20 39\noutside 40 325\nTMhelix 326 348\ninside 349 372

3141 GCF_001621145.1_ASM162114v1 Escherichia coli Proteobacteria; Gammaproteobacteria; Enterobacterales; Enterobacteriaceae; Escherichia
MNNEETFYQAMRRQGVTRRSFLKYCSLAATSLGLGAGMAPKIAWALENKPRIPVVWIHGL WP_001058323.1
MULTISPECIES: hydrogenase-1 small chain [Proteobacteria] Length: 372\nNumber of predicted TMHs: 2\nExp number of AAs in TMHs: 37.2608499999999\nExp number, first 60 AAs: 19.52923\nTotal prob of N-in: 0.94351\nPOSSIBLE N-term signal sequence\ninside 1 19\nTMhelix 20 39\noutside 40 325\nTMhelix 326 348\ninside 349 372

3142 GCF_001621145.1_ASM162114v1 Escherichia coli Proteobacteria; Gammaproteobacteria; Enterobacterales; Enterobacteriaceae; Escherichia
MTGDNTLIHSHGINRRDFMKLCAALAATMGLSSKAAAEMAESVTNPQRPPVIWIGAQECT WP_059321935.1
hydrogenase 2 small subunit [Escherichia coli] Length: 372\nNumber of predicted TMHs: 1\nExp number of AAs in TMHs: 23.8325\nExp number, first 60 AAs: 0.21938\nTotal prob of N-in: 0.13453\noutside 1 331\nTMhelix 332 354\ninside 355 372

3143 GCF_001620985.1_ASM162098v1 Escherichia coli Proteobacteria; Gammaproteobacteria; Enterobacterales; Enterobacteriaceae; Escherichia
MNNEETFYQAMRRQGVTRRSFLKYCSLAATSLGLGAGMAPKIAWALENKPRIPVVWIHGL WP_001058323.1
MULTISPECIES: hydrogenase-1 small chain [Proteobacteria] Length: 372\nNumber of predicted TMHs: 2\nExp number of AAs in TMHs: 37.2608499999999\nExp number, first 60 AAs: 19.52923\nTotal prob of N-in: 0.94351\nPOSSIBLE N-term signal sequence\ninside 1 19\nTMhelix 20 39\noutside 40 325\nTMhelix 326 348\ninside 349 372

3144 GCF_001619195.1_ASM161919v1 Escherichia coli Proteobacteria; Gammaproteobacteria; Enterobacterales; Enterobacteriaceae; Escherichia
MNNEETFYQAMRRQGVTRRSFLKYCSLAATSLGLGAGMAPKIAWALENKPRIPVVWIHGL WP_001301613.1
hydrogenase [Escherichia coli] Length: 372\nNumber of predicted TMHs: 2\nExp number of AAs in TMHs: 37.46598\nExp number, first 60 AAs: 19.53139\nTotal prob of N-in: 0.94364\nPOSSIBLE N-term signal sequence\ninside 1 19\nTMhelix 20 39\noutside 40 325\nTMhelix 326 348\ninside 349 372

3145 GCF_001619205.1_ASM161920v1 Escherichia coli Proteobacteria; Gammaproteobacteria; Enterobacterales; Enterobacteriaceae; Escherichia
MNNEETFYQAMRRQGVTRRSFLKYCSLAATSLGLGAGMAPKIAWALENKPRIPVVWIHGL WP_001301613.1
hydrogenase [Escherichia coli] Length: 372\nNumber of predicted TMHs: 2\nExp number of AAs in TMHs: 37.46598\nExp number, first 60 AAs: 19.53139\nTotal prob of N-in: 0.94364\nPOSSIBLE N-term signal sequence\ninside 1 19\nTMhelix 20 39\noutside 40 325\nTMhelix 326 348\ninside 349 372

3146 GCF_001621305.1_ASM162130v1 Escherichia coli Proteobacteria; Gammaproteobacteria; Enterobacterales; Enterobacteriaceae; Escherichia
MNNEETFYQAMRRQGVTRRSFLKYCSLAATSLGLGAGMAPKIAWALENKPRIPVVWIHGL WP_001295940.1
MULTISPECIES: hydrogenase [Enterobacteriaceae] Length: 372\nNumber of predicted TMHs: 2\nExp number of AAs in TMHs: 37.2608899999999\nExp number, first 60 AAs: 19.52923\nTotal prob of N-in: 0.94351\nPOSSIBLE N-term signal sequence\ninside 1 19\nTMhelix 20 39\noutside 40 325\nTMhelix 326 348\ninside 349 372

3147 GCF_001621575.1_ASM162157v1 Escherichia coli Proteobacteria; Gammaproteobacteria; Enterobacterales; Enterobacteriaceae; Escherichia
MTGDNTLIHSHGINRRDFMKLCAALAATMGLSSKAAAEMVESVTNPQRPPVIWIGAQECT WP_000145423.1
hydrogenase 2 small subunit [Escherichia coli] Length: 372\nNumber of predicted TMHs: 1\nExp number of AAs in TMHs: 23.78201\nExp number, first 60 AAs: 0.28355\nTotal prob of N-in: 0.13257\noutside 1 331\nTMhelix 332 354\ninside 355 372

3148 GCF_001621585.1_ASM162158v1 Escherichia coli Proteobacteria; Gammaproteobacteria; Enterobacterales; Enterobacteriaceae; Escherichia
MTGDNTLIHSHGINRRDFMKLCAALAATMGLSSKAAAEMAESVTNPQRPPVIWIGAQECT WP_000145410.1
MULTISPECIES: hydrogenase-2 small chain [Proteobacteria] Length: 372\nNumber of predicted TMHs: 1\nExp number of AAs in TMHs: 23.72226\nExp number, first 60 AAs: 0.22038\nTotal prob of N-in: 0.12994\noutside 1 331\nTMhelix 332 354\ninside 355 372

3149 GCF_001621675.1_ASM162167v1 Escherichia coli Proteobacteria; Gammaproteobacteria; Enterobacterales; Enterobacteriaceae; Escherichia
MTGDNTLIHSHGINRRDFMKLCAALAATMGLSSKAAAEMAESVTNPQRPPVIWIGAQECT WP_000145410.1
MULTISPECIES: hydrogenase-2 small chain [Proteobacteria] Length: 372\nNumber of predicted TMHs: 1\nExp number of AAs in TMHs: 23.72226\nExp number, first 60 AAs: 0.22038\nTotal prob of N-in: 0.12994\noutside 1 331\nTMhelix 332 354\ninside 355 372

3150 GCF_001621645.1_ASM162164v1 Escherichia coli Proteobacteria; Gammaproteobacteria; Enterobacterales; Enterobacteriaceae; Escherichia
MTGDNTLIHSHGINRRDFMKLCAALAATMGLSSKAAAEMAESVTNPQRPPVIWIGAQECT WP_000145410.1
MULTISPECIES: hydrogenase-2 small chain [Proteobacteria] Length: 372\nNumber of predicted TMHs: 1\nExp number of AAs in TMHs: 23.72226\nExp number, first 60 AAs: 0.22038\nTotal prob of N-in: 0.12994\noutside 1 331\nTMhelix 332 354\ninside 355 372

3151 GCF_001621705.1_ASM162170v1 Escherichia coli Proteobacteria; Gammaproteobacteria; Enterobacterales; Enterobacteriaceae; Escherichia
MTGDNTLIHSHGINRRDFMKLCAALAATMGLSSKAAAEMVESVTNPQRPPVIWIGAQECT WP_000145423.1
hydrogenase 2 small subunit [Escherichia coli] Length: 372\nNumber of predicted TMHs: 1\nExp number of AAs in TMHs: 23.78201\nExp number, first 60 AAs: 0.28355\nTotal prob of N-in: 0.13257\noutside 1 331\nTMhelix 332 354\ninside 355 372

3152 GCF_001621745.1_ASM162174v1 Escherichia coli Proteobacteria; Gammaproteobacteria; Enterobacterales; Enterobacteriaceae; Escherichia
MTGDNTLIHSHGINRRDFMKLCAALAATMGLSSKAAAEMAESVTNPQRPPVIWIGAQECT WP_000145410.1
MULTISPECIES: hydrogenase-2 small chain [Proteobacteria] Length: 372\nNumber of predicted TMHs: 1\nExp number of AAs in TMHs: 23.72226\nExp number, first 60 AAs: 0.22038\nTotal prob of N-in: 0.12994\noutside 1 331\nTMhelix 332 354\ninside 355 372

3153 GCF_001621805.1_ASM162180v1 Escherichia coli Proteobacteria; Gammaproteobacteria; Enterobacterales; Enterobacteriaceae; Escherichia
MTGDNTLIHSHGINRRDFMKLCAALAATMGLSSKAAAEMVESVTNPQRPPVIWIGAQECT WP_000145423.1
hydrogenase 2 small subunit [Escherichia coli] Length: 372\nNumber of predicted TMHs: 1\nExp number of AAs in TMHs: 23.78201\nExp number, first 60 AAs: 0.28355\nTotal prob of N-in: 0.13257\noutside 1 331\nTMhelix 332 354\ninside 355 372

3154 GCF_001621805.1_ASM162180v1 Escherichia coli Proteobacteria; Gammaproteobacteria; Enterobacterales; Enterobacteriaceae; Escherichia
MNNEETFYQAMRRQGVTRRSFLKYCSLAATSLGLGAGMAPKIAWALENKPRIPVVWIHGL WP_001295940.1
MULTISPECIES: hydrogenase [Enterobacteriaceae] Length: 372\nNumber of predicted TMHs: 2\nExp

number of AAs in TMHs: 37.2608899999999\nExp number, first 60 AAs: 19.52923\nTotal prob of N-in: 0.94351\nPOSSIBLE N-term signal sequence\ninside 1 19\nTMhelix 20 39\noutside 40 325\nTMhelix 326 348\ninside 349 372

3155 GCF_001622025.1_ASM162202v1 Escherichia coli Proteobacteria; Gammaproteobacteria; Enterobacterales; Enterobacteriaceae; Escherichia
MTGDNTLIHSHGINRRDFMKLCAALAATMGLSSKAAAEMAESVTNPQRPPVIWIGAQECT WP_000145410.1
MULTISPECIES: hydrogenase-2 small chain [Proteobacteria] Length: 372\nNumber of predicted TMHs: 1\nExp number of AAs in TMHs: 23.72226\nExp number, first 60 AAs: 0.22038\nTotal prob of N-in: 0.12994\noutside 1 331\nTMhelix 332 354\ninside 355 372

3156 GCF_001622025.1_ASM162202v1 Escherichia coli Proteobacteria; Gammaproteobacteria; Enterobacterales; Enterobacteriaceae; Escherichia
MNNEETFYQAMRRQGVTRRSFLKYCSLAATSLGLGAGMAPKIAWALENKPRIPVVWIHGL WP_001058323.1
MULTISPECIES: hydrogenase-1 small chain [Proteobacteria] Length: 372\nNumber of predicted TMHs: 2\nExp number of AAs in TMHs: 37.2608499999999\nExp number, first 60 AAs: 19.52923\nTotal prob of N-in: 0.94351\nPOSSIBLE N-term signal sequence\ninside 1 19\nTMhelix 20 39\noutside 40 325\nTMhelix 326 348\ninside 349 372

3157 GCF_001621835.1_ASM162183v1 Escherichia coli Proteobacteria; Gammaproteobacteria; Enterobacterales; Enterobacteriaceae; Escherichia
MNNEETFYQAMRRQGVTRRSFLKYCSLAATSLGLGAGMAPKIAWALENKPRIPVVWIHGL WP_001295940.1
MULTISPECIES: hydrogenase [Enterobacteriaceae] Length: 372\nNumber of predicted TMHs: 2\nExp number of AAs in TMHs: 37.2608899999999\nExp number, first 60 AAs: 19.52923\nTotal prob of N-in: 0.94351\nPOSSIBLE N-term signal sequence\ninside 1 19\nTMhelix 20 39\noutside 40 325\nTMhelix 326 348\ninside 349 372

3158 GCF_001622055.1_ASM162205v1 Escherichia coli Proteobacteria; Gammaproteobacteria; Enterobacterales; Enterobacteriaceae; Escherichia
MTGDNTLIHSHGINRRDFMKLCAALAATMGLSSKAAAEMAESVTNPQRPPVIWIGAQECT WP_000145410.1
MULTISPECIES: hydrogenase-2 small chain [Proteobacteria] Length: 372\nNumber of predicted TMHs: 1\nExp number of AAs in TMHs: 23.72226\nExp number, first 60 AAs: 0.22038\nTotal prob of N-in: 0.12994\noutside 1 331\nTMhelix 332 354\ninside 355 372

3159 GCF_001622075.1_ASM162207v1 Escherichia coli Proteobacteria; Gammaproteobacteria; Enterobacterales; Enterobacteriaceae; Escherichia
MTGDNTLIHSHGINRRDFMKLCAALAATMGLSSKAAAEMAESVTNPQRPPVIWIGAQECT WP_000145410.1
MULTISPECIES: hydrogenase-2 small chain [Proteobacteria] Length: 372\nNumber of predicted TMHs: 1\nExp number of AAs in TMHs: 23.72226\nExp number, first 60 AAs: 0.22038\nTotal prob of N-in: 0.12994\noutside 1 331\nTMhelix 332 354\ninside 355 372

3160 GCF_001622115.1_ASM162211v1 Escherichia coli Proteobacteria; Gammaproteobacteria; Enterobacterales; Enterobacteriaceae; Escherichia
MTGDNTLIHSHGINRRDFMKLCAALAATMGLSSKAAAEMVESVTNPQRPPVIWIGAQECT WP_000145423.1
hydrogenase 2 small subunit [Escherichia coli] Length: 372\nNumber of predicted TMHs: 1\nExp number of AAs in TMHs: 23.78201\nExp number, first 60 AAs: 0.28355\nTotal prob of N-in: 0.13257\noutside 1 331\nTMhelix 332 354\ninside 355 372

3161 GCF_001637845.1_ASM163784v1 Escherichia coli Proteobacteria; Gammaproteobacteria; Enterobacterales; Enterobacteriaceae; Escherichia
MNNEETFYQAMRRQGVTRRSFLKYCSLAATSLGLGAGMAPKIAWALENKPRIPVVWIHGL WP_001295940.1
MULTISPECIES: hydrogenase [Enterobacteriaceae] Length: 372\nNumber of predicted TMHs: 2\nExp number of AAs in TMHs: 37.2608899999999\nExp number, first 60 AAs: 19.52923\nTotal prob of N-in: 0.94351\nPOSSIBLE N-term signal sequence\ninside 1 19\nTMhelix 20 39\noutside 40 325\nTMhelix 326 348\ninside 349 372

3162 GCF_001637635.1_ASM163763v1 Escherichia coli Proteobacteria; Gammaproteobacteria; Enterobacterales; Enterobacteriaceae; Escherichia
MNNEETFYQAMRRQGVTRRSFLKYCSLAATSLGLGAGMAPKIAWALENKPRIPVVWIHGL WP_001058323.1
MULTISPECIES: hydrogenase-1 small chain [Proteobacteria] Length: 372\nNumber of predicted TMHs: 2\nExp number of AAs in TMHs: 37.2608499999999\nExp number, first 60 AAs: 19.52923\nTotal prob of N-in:

0.94351\nPOSSIBLE N-term signal sequence\ninside 1 19\nTMhelix 20 39\noutside 40 325\nTMhelix 326 348\ninside 349 372

3163 GCF_001637715.1_ASM163771v1 Escherichia coli Proteobacteria; Gammaproteobacteria; Enterobacterales; Enterobacteriaceae; Escherichia
MNNEETFYQAMRRQGVTRRSFLKYCSLAATSLGLGAGMAPKIAWALENKPRIPVVWIHGL WP_001058323.1
MULTISPECIES: hydrogenase-1 small chain [Proteobacteria] Length: 372\nNumber of predicted TMHs: 2\nExp number of AAs in TMHs: 37.2608499999999\nExp number, first 60 AAs: 19.52923\nTotal prob of N-in: 0.94351\nPOSSIBLE N-term signal sequence\ninside 1 19\nTMhelix 20 39\noutside 40 325\nTMhelix 326 348\ninside 349 372

3164 GCF_001637735.1_ASM163773v1 Escherichia coli Proteobacteria; Gammaproteobacteria; Enterobacterales; Enterobacteriaceae; Escherichia
MNNEETFYQAMRRQGVTRRSFLKYCSLAATSLGLGAGMAPKIAWALENKPRIPVVWIHGL WP_001058323.1
MULTISPECIES: hydrogenase-1 small chain [Proteobacteria] Length: 372\nNumber of predicted TMHs: 2\nExp number of AAs in TMHs: 37.2608499999999\nExp number, first 60 AAs: 19.52923\nTotal prob of N-in: 0.94351\nPOSSIBLE N-term signal sequence\ninside 1 19\nTMhelix 20 39\noutside 40 325\nTMhelix 326 348\ninside 349 372

3165 GCF_001637625.1_ASM163762v1 Escherichia coli Proteobacteria; Gammaproteobacteria; Enterobacterales; Enterobacteriaceae; Escherichia
MNNEETFYQAMRRQGVTRRSFLKYCSLAATSLGLGAGMAPKIAWALENKPRIPVVWIHGL WP_001295940.1
MULTISPECIES: hydrogenase [Enterobacteriaceae] Length: 372\nNumber of predicted TMHs: 2\nExp number of AAs in TMHs: 37.2608899999999\nExp number, first 60 AAs: 19.52923\nTotal prob of N-in: 0.94351\nPOSSIBLE N-term signal sequence\ninside 1 19\nTMhelix 20 39\noutside 40 325\nTMhelix 326 348\ninside 349 372

3166 GCF_001637885.1_ASM163788v1 Escherichia coli Proteobacteria; Gammaproteobacteria; Enterobacterales; Enterobacteriaceae; Escherichia
MTGDNTLIHSHGINRRDFMKLCAALAATMGLSSKAAAEMVESVTNPQRPPVIWIGAQECT WP_000145423.1
hydrogenase 2 small subunit [Escherichia coli] Length: 372\nNumber of predicted TMHs: 1\nExp number of AAs in TMHs: 23.78201\nExp number, first 60 AAs: 0.28355\nTotal prob of N-in: 0.13257\noutside 1 331\nTMhelix 332 354\ninside 355 372

3167 GCF_001637885.1_ASM163788v1 Escherichia coli Proteobacteria; Gammaproteobacteria; Enterobacterales; Enterobacteriaceae; Escherichia
MNNEETFYQAMRRQGVTRRSFLKYCSLAATSLGLGAGMAPKIAWALENKPRIPVVWIHGL WP_001295940.1
MULTISPECIES: hydrogenase [Enterobacteriaceae] Length: 372\nNumber of predicted TMHs: 2\nExp number of AAs in TMHs: 37.2608899999999\nExp number, first 60 AAs: 19.52923\nTotal prob of N-in: 0.94351\nPOSSIBLE N-term signal sequence\ninside 1 19\nTMhelix 20 39\noutside 40 325\nTMhelix 326 348\ninside 349 372

3168 GCF_001637945.1_ASM163794v1 Escherichia coli Proteobacteria; Gammaproteobacteria; Enterobacterales; Enterobacteriaceae; Escherichia
MNNEETFYQAMRRQGVTRRSFLKYCSLAATSLGLGAGMAPKIAWALENKPRIPVVWIHGL WP_001295940.1
MULTISPECIES: hydrogenase [Enterobacteriaceae] Length: 372\nNumber of predicted TMHs: 2\nExp number of AAs in TMHs: 37.2608899999999\nExp number, first 60 AAs: 19.52923\nTotal prob of N-in: 0.94351\nPOSSIBLE N-term signal sequence\ninside 1 19\nTMhelix 20 39\noutside 40 325\nTMhelix 326 348\ninside 349 372

3169 GCF_001637985.1_ASM163798v1 Escherichia coli Proteobacteria; Gammaproteobacteria; Enterobacterales; Enterobacteriaceae; Escherichia
MNNEETFYQAMRRQGVTRRSFLKYCSLAATSLGLGAGMAPKIAWALENKPRIPVVWIHGL WP_001295940.1
MULTISPECIES: hydrogenase [Enterobacteriaceae] Length: 372\nNumber of predicted TMHs: 2\nExp number of AAs in TMHs: 37.2608899999999\nExp number, first 60 AAs: 19.52923\nTotal prob of N-in: 0.94351\nPOSSIBLE N-term signal sequence\ninside 1 19\nTMhelix 20 39\noutside 40 325\nTMhelix 326 348\ninside 349 372

3170 GCF_001637905.1_ASM163790v1 Escherichia coli Proteobacteria; Gammaproteobacteria; Enterobacterales; Enterobacteriaceae; Escherichia
MNNEETFYQAMRRQGVTRRSFLKYCSLAATSLGLGAGMAPKIAWALENKPRIPVVWIHGL WP_001295940.1

MULTISPECIES: hydrogenase [Enterobacteriaceae] Length: 372\nNumber of predicted TMHs: 2\nExp number of AAs in TMHs: 37.2608899999999\nExp number, first 60 AAs: 19.52923\nTotal prob of N-in: 0.94351\nPOSSIBLE N-term signal sequence\ninside 1 19\nTMhelix 20 39\noutside 40 325\nTMhelix 326 348\ninside 349 372

3171 GCF_001638025.1_ASM163802v1 Escherichia coli Proteobacteria; Gammaproteobacteria; Enterobacterales; Enterobacteriaceae; Escherichia
MNNEETFYQAMRRQGVTRRSFLKYCSLAATSLGLGAGMAPKIAWALENKPRIPVVWIHGL WP_001295940.1
MULTISPECIES: hydrogenase [Enterobacteriaceae] Length: 372\nNumber of predicted TMHs: 2\nExp number of AAs in TMHs: 37.2608899999999\nExp number, first 60 AAs: 19.52923\nTotal prob of N-in: 0.94351\nPOSSIBLE N-term signal sequence\ninside 1 19\nTMhelix 20 39\noutside 40 325\nTMhelix 326 348\ninside 349 372

3172 GCF_001638045.1_ASM163804v1 Escherichia coli Proteobacteria; Gammaproteobacteria; Enterobacterales; Enterobacteriaceae; Escherichia
MNNEETFYQAMRRQGVTRRSFLKYCSLAATSLGLGAGMAPKIAWALENKPRIPVVWIHGL WP_001295940.1
MULTISPECIES: hydrogenase [Enterobacteriaceae] Length: 372\nNumber of predicted TMHs: 2\nExp number of AAs in TMHs: 37.2608899999999\nExp number, first 60 AAs: 19.52923\nTotal prob of N-in: 0.94351\nPOSSIBLE N-term signal sequence\ninside 1 19\nTMhelix 20 39\noutside 40 325\nTMhelix 326 348\ninside 349 372

3173 GCF_001641955.1_ASM164195v1 Escherichia coli Proteobacteria; Gammaproteobacteria; Enterobacterales; Enterobacteriaceae; Escherichia
MNNEETFYQAMRRQGVTRRSFLKYCSLAATSLGLGAGMAPKIAWALENKPRIPVVWIHGL WP_001058323.1
MULTISPECIES: hydrogenase-1 small chain [Proteobacteria] Length: 372\nNumber of predicted TMHs: 2\nExp number of AAs in TMHs: 37.2608499999999\nExp number, first 60 AAs: 19.52923\nTotal prob of N-in: 0.94351\nPOSSIBLE N-term signal sequence\ninside 1 19\nTMhelix 20 39\noutside 40 325\nTMhelix 326 348\ninside 349 372

3174 GCF_001647325.1_ASM164732v1 Escherichia coli Proteobacteria; Gammaproteobacteria; Enterobacterales; Enterobacteriaceae; Escherichia
MTGDNTLIHSHGINRRDFMKLCAALAATMGLSSKAAAEMVESVTNPQRPPVIWIGAQECT WP_000145423.1
hydrogenase 2 small subunit [Escherichia coli] Length: 372\nNumber of predicted TMHs: 1\nExp number of AAs in TMHs: 23.78201\nExp number, first 60 AAs: 0.28355\nTotal prob of N-in: 0.13257\noutside 1 331\nTMhelix 332 354\ninside 355 372

3175 GCF_001647345.1_ASM164734v1 Escherichia coli Proteobacteria; Gammaproteobacteria; Enterobacterales; Enterobacteriaceae; Escherichia
MTGDNTLIHSHGINRRDFMKLCAALAATMGLSSKAAAEMVESVTNPQRPPVIWIGAQECT WP_000145423.1
hydrogenase 2 small subunit [Escherichia coli] Length: 372\nNumber of predicted TMHs: 1\nExp number of AAs in TMHs: 23.78201\nExp number, first 60 AAs: 0.28355\nTotal prob of N-in: 0.13257\noutside 1 331\nTMhelix 332 354\ninside 355 372

3176 GCF_001647325.1_ASM164732v1 Escherichia coli Proteobacteria; Gammaproteobacteria; Enterobacterales; Enterobacteriaceae; Escherichia
MNNEETFYQAMRRQGVTRRSFLKYCSLAATSLGLGAGMAPKIAWALENKPRIPVVWIHGL WP_001295940.1
MULTISPECIES: hydrogenase [Enterobacteriaceae] Length: 372\nNumber of predicted TMHs: 2\nExp number of AAs in TMHs: 37.2608899999999\nExp number, first 60 AAs: 19.52923\nTotal prob of N-in: 0.94351\nPOSSIBLE N-term signal sequence\ninside 1 19\nTMhelix 20 39\noutside 40 325\nTMhelix 326 348\ninside 349 372

3177 GCF_001647345.1_ASM164734v1 Escherichia coli Proteobacteria; Gammaproteobacteria; Enterobacterales; Enterobacteriaceae; Escherichia
MNNEETFYQAMRRQGVTRRSFLKYCSLAATSLGLGAGMAPKIAWALENKPRIPVVWIHGL WP_001295940.1
MULTISPECIES: hydrogenase [Enterobacteriaceae] Length: 372\nNumber of predicted TMHs: 2\nExp number of AAs in TMHs: 37.2608899999999\nExp number, first 60 AAs: 19.52923\nTotal prob of N-in: 0.94351\nPOSSIBLE N-term signal sequence\ninside 1 19\nTMhelix 20 39\noutside 40 325\nTMhelix 326 348\ninside 349 372

3178 GCF_001644175.1_ASM164417v1 Escherichia coli Proteobacteria; Gammaproteobacteria; Enterobacterales; Enterobacteriaceae; Escherichia

MNNEETFYQAMRRQGVTRRSFLKYCSLAATSLGLGAGMAPKIAWALENKPRIPVVWIHGL WP_001295940.1 MULTISPECIES: hydrogenase [Enterobacteriaceae] Length: 372\nNumber of predicted TMHs: 2\nExp number of AAs in TMHs: 37.2608899999999\nExp number, first 60 AAs: 19.52923\nTotal prob of N-in: 0.94351\nPOSSIBLE N-term signal sequence\ninside 1 19\nTMhelix 20 39\noutside 40 325\nTMhelix 326 348\ninside 349 372

3179 GCF_001647395.1_ASM164739v1 Escherichia coli Proteobacteria; Gammaproteobacteria; Enterobacterales; Enterobacteriaceae; Escherichia MTGDNTLIHSHGINRRDFMKLCAALAATMGLSSKAAAEMVESVTNPQRPPVIWIGAQECT WP_000145423.1 hydrogenase 2 small subunit [Escherichia coli] Length: 372\nNumber of predicted TMHs: 1\nExp number of AAs in TMHs: 23.78201\nExp number, first 60 AAs: 0.28355\nTotal prob of N-in: 0.13257\noutside 1 331\nTMhelix 332 354\ninside 355 372

3180 GCF_001650565.1_ASM165056v1 Escherichia coli Proteobacteria; Gammaproteobacteria; Enterobacterales; Enterobacteriaceae; Escherichia MNNEETFYQAMRRQGVTRRSFLKYCSLAATSLGLGAGMAPKIAWALENKPRIPVVWIHGL WP_001058323.1 MULTISPECIES: hydrogenase-1 small chain [Proteobacteria] Length: 372\nNumber of predicted TMHs: 2\nExp number of AAs in TMHs: 37.2608499999999\nExp number, first 60 AAs: 19.52923\nTotal prob of N-in: 0.94351\nPOSSIBLE N-term signal sequence\ninside 1 19\nTMhelix 20 39\noutside 40 325\nTMhelix 326 348\ninside 349 372

3181 GCF_001647395.1_ASM164739v1 Escherichia coli Proteobacteria; Gammaproteobacteria; Enterobacterales; Enterobacteriaceae; Escherichia MNNEETFYQAMRRQGVTRRSFLKYCSLAATSLGLGAGMAPKIAWALENKPRIPVVWIHGL WP_001295940.1 MULTISPECIES: hydrogenase [Enterobacteriaceae] Length: 372\nNumber of predicted TMHs: 2\nExp number of AAs in TMHs: 37.2608899999999\nExp number, first 60 AAs: 19.52923\nTotal prob of N-in: 0.94351\nPOSSIBLE N-term signal sequence\ninside 1 19\nTMhelix 20 39\noutside 40 325\nTMhelix 326 348\ninside 349 372

3182 GCF_001651695.1_ASM165169v1 Escherichia coli Proteobacteria; Gammaproteobacteria; Enterobacterales; Enterobacteriaceae; Escherichia MTGDNTLIHSHGINRRDFMKLCAALAATMGLSSKAAAEMAESVTNPQRPPVIWIGAQECT WP_000145410.1 MULTISPECIES: hydrogenase-2 small chain [Proteobacteria] Length: 372\nNumber of predicted TMHs: 1\nExp number of AAs in TMHs: 23.72226\nExp number, first 60 AAs: 0.22038\nTotal prob of N-in: 0.12994\noutside 1 331\nTMhelix 332 354\ninside 355 372

3183 GCF_001651695.1_ASM165169v1 Escherichia coli Proteobacteria; Gammaproteobacteria; Enterobacterales; Enterobacteriaceae; Escherichia MNNEETFYQAMRRQGVTRRSFLKYCSLAATSLGLGAGMAPKIAWALENKPRIPVVWIHGL WP_001058323.1 MULTISPECIES: hydrogenase-1 small chain [Proteobacteria] Length: 372\nNumber of predicted TMHs: 2\nExp number of AAs in TMHs: 37.2608499999999\nExp number, first 60 AAs: 19.52923\nTotal prob of N-in: 0.94351\nPOSSIBLE N-term signal sequence\ninside 1 19\nTMhelix 20 39\noutside 40 325\nTMhelix 326 348\ninside 349 372

3184 GCF_001648715.1_ASM164871v1 Escherichia coli Proteobacteria; Gammaproteobacteria; Enterobacterales; Enterobacteriaceae; Escherichia MNNEETFYQAMRRQGVTRRSFLKYCSLAATSLGLGAGMAPKIAWALENKPRIPVVWIHGL WP_001295940.1 MULTISPECIES: hydrogenase [Enterobacteriaceae] Length: 372\nNumber of predicted TMHs: 2\nExp number of AAs in TMHs: 37.2608899999999\nExp number, first 60 AAs: 19.52923\nTotal prob of N-in: 0.94351\nPOSSIBLE N-term signal sequence\ninside 1 19\nTMhelix 20 39\noutside 40 325\nTMhelix 326 348\ninside 349 372

3185 GCF_001652805.1_ASM165280v1 Escherichia coli Proteobacteria; Gammaproteobacteria; Enterobacterales; Enterobacteriaceae; Escherichia MNNEETFYQAMRRQGVTRRSFLKYCSLAATSLGLGAGMAPKIAWALENKPRIPVVWIHGL WP_001538532.1 hydrogenase-1 small chain [Escherichia coli] Length: 372\nNumber of predicted TMHs: 2\nExp number of AAs in TMHs: 37.27041\nExp number, first 60 AAs: 19.52927\nTotal prob of N-in: 0.94352\nPOSSIBLE N-term signal sequence\ninside 1 19\nTMhelix 20 39\noutside 40 325\nTMhelix 326 348\ninside 349 372

3186 GCF_001657975.1_ASM165797v1 Escherichia coli Proteobacteria; Gammaproteobacteria; Enterobacterales; Enterobacteriaceae; Escherichia

MNNEETFYQAMRRQGVTRRSFLKYCSLAATSLGLGAGMAPKIAWALENKPRIPVVWIHGL WP_001058323.1 MULTISPECIES: hydrogenase-1 small chain [Proteobacteria] Length: 372\nNumber of predicted TMHs: 2\nExp number of AAs in TMHs: 37.2608499999999\nExp number, first 60 AAs: 19.52923\nTotal prob of N-in: 0.94351\nPOSSIBLE N-term signal sequence\ninside 1 19\nTMhelix 20 39\noutside 40 325\nTMhelix 326 348\ninside 349 372

3187 GCF_001653635.1_ASM165363v1 Escherichia coli Proteobacteria; Gammaproteobacteria; Enterobacterales; Enterobacteriaceae; Escherichia MNNEETFYQAMRRQGVTRRSFLKYCSLAATSLGLGAGMAPKIAWALENKPRIPVVWIHGL WP_001058323.1 MULTISPECIES: hydrogenase-1 small chain [Proteobacteria] Length: 372\nNumber of predicted TMHs: 2\nExp number of AAs in TMHs: 37.2608499999999\nExp number, first 60 AAs: 19.52923\nTotal prob of N-in: 0.94351\nPOSSIBLE N-term signal sequence\ninside 1 19\nTMhelix 20 39\noutside 40 325\nTMhelix 326 348\ninside 349 372

3188 GCF_001660195.1_ASM166019v1 Escherichia coli Proteobacteria; Gammaproteobacteria; Enterobacterales; Enterobacteriaceae; Escherichia MTGDNTLIHSHGINRRDFMKLCAALAATMGLSSKAAAEMAESVTNPQRPPVIWIGAQECT WP_000145410.1 MULTISPECIES: hydrogenase-2 small chain [Proteobacteria] Length: 372\nNumber of predicted TMHs: 1\nExp number of AAs in TMHs: 23.72226\nExp number, first 60 AAs: 0.22038\nTotal prob of N-in: 0.12994\noutside 1 331\nTMhelix 332 354\ninside 355 372

3189 GCF_001660195.1_ASM166019v1 Escherichia coli Proteobacteria; Gammaproteobacteria; Enterobacterales; Enterobacteriaceae; Escherichia MNNEETFYQAMRRQGVTRRSFLKYCSLAATSLGLGAGMAPKIAWALENKPRIPVVWIHGL WP_001295940.1 MULTISPECIES: hydrogenase [Enterobacteriaceae] Length: 372\nNumber of predicted TMHs: 2\nExp number of AAs in TMHs: 37.2608899999999\nExp number, first 60 AAs: 19.52923\nTotal prob of N-in: 0.94351\nPOSSIBLE N-term signal sequence\ninside 1 19\nTMhelix 20 39\noutside 40 325\nTMhelix 326 348\ninside 349 372

3190 GCF_001653555.1_ASM165355v1 Escherichia coli Proteobacteria; Gammaproteobacteria; Enterobacterales; Enterobacteriaceae; Escherichia MNNEETFYQAMRRQGVTRRSFLKYCSLAATSLGLGAGMAPKIAWALENKPRIPVVWIHGL WP_001295940.1 MULTISPECIES: hydrogenase [Enterobacteriaceae] Length: 372\nNumber of predicted TMHs: 2\nExp number of AAs in TMHs: 37.2608899999999\nExp number, first 60 AAs: 19.52923\nTotal prob of N-in: 0.94351\nPOSSIBLE N-term signal sequence\ninside 1 19\nTMhelix 20 39\noutside 40 325\nTMhelix 326 348\ninside 349 372

3191 GCF_001660545.2_ASM166054v3 Escherichia coli Proteobacteria; Gammaproteobacteria; Enterobacterales; Enterobacteriaceae; Escherichia MNNEETFYQAMRRQGVTRRSFLKYCSLAATSLGLGAGMAPKIAWALENKPRIPVVWIHGL WP_001295940.1 MULTISPECIES: hydrogenase [Enterobacteriaceae] Length: 372\nNumber of predicted TMHs: 2\nExp number of AAs in TMHs: 37.2608899999999\nExp number, first 60 AAs: 19.52923\nTotal prob of N-in: 0.94351\nPOSSIBLE N-term signal sequence\ninside 1 19\nTMhelix 20 39\noutside 40 325\nTMhelix 326 348\ninside 349 372

3192 GCF_001663075.1_ASM166307v1 Escherichia coli Proteobacteria; Gammaproteobacteria; Enterobacterales; Enterobacteriaceae; Escherichia MNNEETFYQAMRRQGVTRRSFLKYCSLAATSLGLGAGMAPKIAWALENKPRIPVVWIHGL WP_001058323.1 MULTISPECIES: hydrogenase-1 small chain [Proteobacteria] Length: 372\nNumber of predicted TMHs: 2\nExp number of AAs in TMHs: 37.2608499999999\nExp number, first 60 AAs: 19.52923\nTotal prob of N-in: 0.94351\nPOSSIBLE N-term signal sequence\ninside 1 19\nTMhelix 20 39\noutside 40 325\nTMhelix 326 348\ninside 349 372

3193 GCF_001660345.1_ASM166034v1 Escherichia coli Proteobacteria; Gammaproteobacteria; Enterobacterales; Enterobacteriaceae; Escherichia MNNEETFYQAMRRQGVTRRSFLKYCSLAATSLGLGAGMAPKIAWALENKPRIPVVWIHGL WP_001295940.1 MULTISPECIES: hydrogenase [Enterobacteriaceae] Length: 372\nNumber of predicted TMHs: 2\nExp number of AAs in TMHs: 37.2608899999999\nExp number, first 60 AAs: 19.52923\nTotal prob of N-in: 0.94351\nPOSSIBLE N-term signal sequence\ninside 1 19\nTMhelix 20 39\noutside 40 325\nTMhelix 326 348\ninside 349 372

3194 GCF_001675145.1_ASM167514v1 Escherichia coli Proteobacteria; Gammaproteobacteria; Enterobacterales; Enterobacteriaceae; Escherichia
MNNEETFYQAMRRQGVTRRSFLKYCSLAATSLGLGAGMAPKIAWALENKPRIPVVWIHGL WP_001058323.1
MULTISPECIES: hydrogenase-1 small chain [Proteobacteria] Length: 372\nNumber of predicted TMHs: 2\nExp number of AAs in TMHs: 37.2608499999999\nExp number, first 60 AAs: 19.52923\nTotal prob of N-in: 0.94351\nPOSSIBLE N-term signal sequence\ninside 1 19\nTMhelix 20 39\noutside 40 325\nTMhelix 326 348\ninside 349 372

3195 GCF_001677515.1_ASM167751v1 Escherichia coli Proteobacteria; Gammaproteobacteria; Enterobacterales; Enterobacteriaceae; Escherichia
MTGDNTLIHSHGINRRDFMKLCAALAATMGLSSKAAAEMAESVTNPQRPPVIWIGAQECT WP_000145410.1
MULTISPECIES: hydrogenase-2 small chain [Proteobacteria] Length: 372\nNumber of predicted TMHs: 1\nExp number of AAs in TMHs: 23.72226\nExp number, first 60 AAs: 0.22038\nTotal prob of N-in: 0.12994\noutside 1 331\nTMhelix 332 354\ninside 355 372

3196 GCF_001677535.2_ASM167753v2 Escherichia coli Proteobacteria; Gammaproteobacteria; Enterobacterales; Enterobacteriaceae; Escherichia
MTGDNTLIHSHGINRRDFMKLCAALAATMGLSSKAAAEMAESVTNPQRPPVIWIGAQECT WP_000145410.1
MULTISPECIES: hydrogenase-2 small chain [Proteobacteria] Length: 372\nNumber of predicted TMHs: 1\nExp number of AAs in TMHs: 23.72226\nExp number, first 60 AAs: 0.22038\nTotal prob of N-in: 0.12994\noutside 1 331\nTMhelix 332 354\ninside 355 372

3197 GCF_001677555.2_ASM167755v2 Escherichia coli Proteobacteria; Gammaproteobacteria; Enterobacterales; Enterobacteriaceae; Escherichia
MTGDNTLIHSHGINRRDFMKLCAALAATMGLSSKAAAEMAESVTNPQRPPVIWIGAQECT WP_000145410.1
MULTISPECIES: hydrogenase-2 small chain [Proteobacteria] Length: 372\nNumber of predicted TMHs: 1\nExp number of AAs in TMHs: 23.72226\nExp number, first 60 AAs: 0.22038\nTotal prob of N-in: 0.12994\noutside 1 331\nTMhelix 332 354\ninside 355 372

3198 GCF_001677615.2_ASM167761v2 Escherichia coli Proteobacteria; Gammaproteobacteria; Enterobacterales; Enterobacteriaceae; Escherichia
MNNEETFYQAMRRQGVTRRSFLKYCSLAATSLGLGAGMAPKIAWALENKPRIPVVWIHGL WP_001301613.1
hydrogenase [Escherichia coli] Length: 372\nNumber of predicted TMHs: 2\nExp number of AAs in TMHs: 37.46598\nExp number, first 60 AAs: 19.53139\nTotal prob of N-in: 0.94364\nPOSSIBLE N-term signal sequence\ninside 1 19\nTMhelix 20 39\noutside 40 325\nTMhelix 326 348\ninside 349 372

3199 GCF_001677645.2_ASM167764v2 Escherichia coli Proteobacteria; Gammaproteobacteria; Enterobacterales; Enterobacteriaceae; Escherichia
MNNEETFYQAMRRQGVTRRSFLKYCSLAATSLGLGAGMAPKIAWALENKPRIPVVWIHGL WP_001301613.1
hydrogenase [Escherichia coli] Length: 372\nNumber of predicted TMHs: 2\nExp number of AAs in TMHs: 37.46598\nExp number, first 60 AAs: 19.53139\nTotal prob of N-in: 0.94364\nPOSSIBLE N-term signal sequence\ninside 1 19\nTMhelix 20 39\noutside 40 325\nTMhelix 326 348\ninside 349 372

3200 GCF_001677775.2_ASM167777v2 Escherichia coli Proteobacteria; Gammaproteobacteria; Enterobacterales; Enterobacteriaceae; Escherichia
MNNEETFYQAMRRQGVTRRSFLKYCSLAATSLGLGAGMAPKIAWALENKPRIPVVWIHGL WP_001301613.1
hydrogenase [Escherichia coli] Length: 372\nNumber of predicted TMHs: 2\nExp number of AAs in TMHs: 37.46598\nExp number, first 60 AAs: 19.53139\nTotal prob of N-in: 0.94364\nPOSSIBLE N-term signal sequence\ninside 1 19\nTMhelix 20 39\noutside 40 325\nTMhelix 326 348\ninside 349 372

3201 GCF_001677705.2_ASM167770v2 Escherichia coli Proteobacteria; Gammaproteobacteria; Enterobacterales; Enterobacteriaceae; Escherichia
MNNEETFYQAMRRQGVTRRSFLKYCSLAATSLGLGAGMAPKIAWALENKPRIPVVWIHGL WP_001301613.1
hydrogenase [Escherichia coli] Length: 372\nNumber of predicted TMHs: 2\nExp number of AAs in TMHs: 37.46598\nExp number, first 60 AAs: 19.53139\nTotal prob of N-in: 0.94364\nPOSSIBLE N-term signal sequence\ninside 1 19\nTMhelix 20 39\noutside 40 325\nTMhelix 326 348\ninside 349 372

3202 GCF_001678925.1_ASM167892v1 Escherichia coli Proteobacteria; Gammaproteobacteria; Enterobacterales; Enterobacteriaceae; Escherichia
MTGDNTLIHSHGINRRDFMKLCAALAATMGLSSKAAAEMAESVTNPQRPPVIWIGAQECT WP_000145410.1
MULTISPECIES: hydrogenase-2 small chain [Proteobacteria] Length: 372\nNumber of predicted TMHs:

1\nExp number of AAs in TMHs: 23.72226\nExp number, first 60 AAs: 0.22038\nTotal prob of N-in: 0.12994\noutside 1 331\nTMhelix 332 354\ninside 355 372

3203 GCF_001678925.1_ASM167892v1 Escherichia coli Proteobacteria; Gammaproteobacteria; Enterobacterales; Enterobacteriaceae; Escherichia
MNNEETFYQAMRRQGVTRRSFLKYCSLAATSLGLGAGMAPKIAWALENKPRIPVVWIHGL WP_001058323.1
MULTISPECIES: hydrogenase-1 small chain [Proteobacteria] Length: 372\nNumber of predicted TMHs: 2\nExp number of AAs in TMHs: 37.2608499999999\nExp number, first 60 AAs: 19.52923\nTotal prob of N-in: 0.94351\nPOSSIBLE N-term signal sequence\ninside 1 19\nTMhelix 20 39\noutside 40 325\nTMhelix 326 348\ninside 349 372

3204 GCF_001677795.2_ASM167779v2 Escherichia coli Proteobacteria; Gammaproteobacteria; Enterobacterales; Enterobacteriaceae; Escherichia
MNNEETFYQAMRRQGVTRRSFLKYCSLAATSLGLGAGMAPKIAWALENKPRIPVVWIHGL WP_001301613.1
hydrogenase [Escherichia coli] Length: 372\nNumber of predicted TMHs: 2\nExp number of AAs in TMHs: 37.46598\nExp number, first 60 AAs: 19.53139\nTotal prob of N-in: 0.94364\nPOSSIBLE N-term signal sequence\ninside 1 19\nTMhelix 20 39\noutside 40 325\nTMhelix 326 348\ninside 349 372

3205 GCF_001692795.1_ASM169279v1 Escherichia coli Proteobacteria; Gammaproteobacteria; Enterobacterales; Enterobacteriaceae; Escherichia
MNNEETFYQAMRRQGVTRRSFLKYCSLAATSLGLGAGMAPKIAWALENKPRIPVVWIHGL WP_001058323.1
MULTISPECIES: hydrogenase-1 small chain [Proteobacteria] Length: 372\nNumber of predicted TMHs: 2\nExp number of AAs in TMHs: 37.2608499999999\nExp number, first 60 AAs: 19.52923\nTotal prob of N-in: 0.94351\nPOSSIBLE N-term signal sequence\ninside 1 19\nTMhelix 20 39\noutside 40 325\nTMhelix 326 348\ninside 349 372

3206 GCF_001692775.1_ASM169277v1 Escherichia coli Proteobacteria; Gammaproteobacteria; Enterobacterales; Enterobacteriaceae; Escherichia
MNNEETFYQAMRRQGVTRRSFLKYCSLAATSLGLGAGMAPKIAWALENKPRIPVVWIHGL WP_001058323.1
MULTISPECIES: hydrogenase-1 small chain [Proteobacteria] Length: 372\nNumber of predicted TMHs: 2\nExp number of AAs in TMHs: 37.2608499999999\nExp number, first 60 AAs: 19.52923\nTotal prob of N-in: 0.94351\nPOSSIBLE N-term signal sequence\ninside 1 19\nTMhelix 20 39\noutside 40 325\nTMhelix 326 348\ninside 349 372

3207 GCF_001700035.1_ASM170003v1 Escherichia coli Proteobacteria; Gammaproteobacteria; Enterobacterales; Enterobacteriaceae; Escherichia
MNNEETFYQAMRRQGVTRRSFLKYCSLAATSLGLGAGMAPKIAWALENKPRIPVVWIHGL WP_001058323.1
MULTISPECIES: hydrogenase-1 small chain [Proteobacteria] Length: 372\nNumber of predicted TMHs: 2\nExp number of AAs in TMHs: 37.2608499999999\nExp number, first 60 AAs: 19.52923\nTotal prob of N-in: 0.94351\nPOSSIBLE N-term signal sequence\ninside 1 19\nTMhelix 20 39\noutside 40 325\nTMhelix 326 348\ninside 349 372

3208 GCF_001709145.1_ASM170914v1 Escherichia coli Proteobacteria; Gammaproteobacteria; Enterobacterales; Enterobacteriaceae; Escherichia
MTGDNTLIHSHGINRRDFMKLCAALAATMGLSSKAAAEMAESVTNPQRPPVIWIGAQECT WP_000145410.1
MULTISPECIES: hydrogenase-2 small chain [Proteobacteria] Length: 372\nNumber of predicted TMHs: 1\nExp number of AAs in TMHs: 23.72226\nExp number, first 60 AAs: 0.22038\nTotal prob of N-in: 0.12994\noutside 1 331\nTMhelix 332 354\ninside 355 372

3209 GCF_001709145.1_ASM170914v1 Escherichia coli Proteobacteria; Gammaproteobacteria; Enterobacterales; Enterobacteriaceae; Escherichia
MNNEETFYQAMRRQGVTRRSFLKYCSLAATSLGLGAGMAPKIAWALENKPRIPVVWIHGL WP_001058320.1
hydrogenase [Escherichia coli] Length: 372\nNumber of predicted TMHs: 2\nExp number of AAs in TMHs: 37.26082\nExp number, first 60 AAs: 19.52923\nTotal prob of N-in: 0.94351\nPOSSIBLE N-term signal sequence\ninside 1 19\nTMhelix 20 39\noutside 40 325\nTMhelix 326 348\ninside 349 372

3210 GCF_001704035.1_ASM170403v1 Escherichia coli Proteobacteria; Gammaproteobacteria; Enterobacterales; Enterobacteriaceae; Escherichia
MNNEETFYQAMRRQGVTRRSFLKYCSLAATSLGLGAGMAPKIAWALENKPRIPVVWIHGL WP_001058323.1
MULTISPECIES: hydrogenase-1 small chain [Proteobacteria] Length: 372\nNumber of predicted TMHs: 2\nExp number of AAs in TMHs: 37.2608499999999\nExp number, first 60 AAs: 19.52923\nTotal prob of N-in:

0.94351\nPOSSIBLE N-term signal sequence\ninside 1 19\nTMhelix 20 39\noutside 40 325\nTMhelix 326 348\ninside 349 372

3211 GCF_001700105.1_ASM170010v1 Escherichia coli Proteobacteria; Gammaproteobacteria; Enterobacterales; Enterobacteriaceae; Escherichia MNNEETFYQAMRRQGVTRRSFLKYCSLAATSLGLGAGMAPKIAWALENKPRIPVVWIHGL WP_001058323.1 MULTISPECIES: hydrogenase-1 small chain [Proteobacteria] Length: 372\nNumber of predicted TMHs: 2\nExp number of AAs in TMHs: 37.2608499999999\nExp number, first 60 AAs: 19.52923\nTotal prob of N-in: 0.94351\nPOSSIBLE N-term signal sequence\ninside 1 19\nTMhelix 20 39\noutside 40 325\nTMhelix 326 348\ninside 349 372

3212 GCF_001713545.1_ASM171354v1 Escherichia coli Proteobacteria; Gammaproteobacteria; Enterobacterales; Enterobacteriaceae; Escherichia MTGDNTLIHSHGINRRDFMKLCAALAATMGLSSKAAAEMAESVTNPQRPPVIWIGAQECT WP_000145410.1 MULTISPECIES: hydrogenase-2 small chain [Proteobacteria] Length: 372\nNumber of predicted TMHs: 1\nExp number of AAs in TMHs: 23.72226\nExp number, first 60 AAs: 0.22038\nTotal prob of N-in: 0.12994\noutside 1 331\nTMhelix 332 354\ninside 355 372

3213 GCF_001713575.1_ASM171357v1 Escherichia coli Proteobacteria; Gammaproteobacteria; Enterobacterales; Enterobacteriaceae; Escherichia MTGDNTLIHSHGINRRDFMKLCAALAATMGLSSKAAAEMAESVTNPQRPPVIWIGAQECT WP_000145410.1 MULTISPECIES: hydrogenase-2 small chain [Proteobacteria] Length: 372\nNumber of predicted TMHs: 1\nExp number of AAs in TMHs: 23.72226\nExp number, first 60 AAs: 0.22038\nTotal prob of N-in: 0.12994\noutside 1 331\nTMhelix 332 354\ninside 355 372

3214 GCF_001742555.1_ASM174255v1 Escherichia coli Proteobacteria; Gammaproteobacteria; Enterobacterales; Enterobacteriaceae; Escherichia MNNEETFYQAMRRQGVTRRSFLKYCSLAATSLGLGAGMAPKIAWALENKPRIPVVWIHGL WP_001058323.1 MULTISPECIES: hydrogenase-1 small chain [Proteobacteria] Length: 372\nNumber of predicted TMHs: 2\nExp number of AAs in TMHs: 37.2608499999999\nExp number, first 60 AAs: 19.52923\nTotal prob of N-in: 0.94351\nPOSSIBLE N-term signal sequence\ninside 1 19\nTMhelix 20 39\noutside 40 325\nTMhelix 326 348\ninside 349 372

3215 GCF_001742625.1_ASM174262v1 Escherichia coli Proteobacteria; Gammaproteobacteria; Enterobacterales; Enterobacteriaceae; Escherichia MNNEETFYQAMRRQGVTRRSFLKYCSLAATSLGLGAGMAPKIAWALENKPRIPVVWIHGL WP_001058323.1 MULTISPECIES: hydrogenase-1 small chain [Proteobacteria] Length: 372\nNumber of predicted TMHs: 2\nExp number of AAs in TMHs: 37.2608499999999\nExp number, first 60 AAs: 19.52923\nTotal prob of N-in: 0.94351\nPOSSIBLE N-term signal sequence\ninside 1 19\nTMhelix 20 39\noutside 40 325\nTMhelix 326 348\ninside 349 372

3216 GCF_001713585.1_ASM171358v1 Escherichia coli Proteobacteria; Gammaproteobacteria; Enterobacterales; Enterobacteriaceae; Escherichia MNNEETFYQAMRRQGVTRRSFLKYCSLAATSLGLGAGMAPKIAWALENKPRIPVVWIHGL WP_069067259.1 hydrogenase [Escherichia coli] Length: 372\nNumber of predicted TMHs: 2\nExp number of AAs in TMHs: 37.25908\nExp number, first 60 AAs: 19.52926\nTotal prob of N-in: 0.94351\nPOSSIBLE N-term signal sequence\ninside 1 19\nTMhelix 20 39\noutside 40 325\nTMhelix 326 348\ninside 349 372

3217 GCF_001728785.1_ASM172878v1 Escherichia coli Proteobacteria; Gammaproteobacteria; Enterobacterales; Enterobacteriaceae; Escherichia MNNEETFYQAMRRQGVTRRSFLKYCSLAATSLGLGAGMAPKIAWALENKPRIPVVWIHGL WP_001301613.1 hydrogenase [Escherichia coli] Length: 372\nNumber of predicted TMHs: 2\nExp number of AAs in TMHs: 37.46598\nExp number, first 60 AAs: 19.53139\nTotal prob of N-in: 0.94364\nPOSSIBLE N-term signal sequence\ninside 1 19\nTMhelix 20 39\noutside 40 325\nTMhelix 326 348\ninside 349 372

3218 GCF_001730745.1_ASM173074v1 Escherichia coli Proteobacteria; Gammaproteobacteria; Enterobacterales; Enterobacteriaceae; Escherichia MNNEETFYQAMRRQGVTRRSFLKYCSLAATSLGLGAGMAPKIAWALENKPRIPVVWIHGL WP_001301613.1 hydrogenase [Escherichia coli] Length: 372\nNumber of predicted TMHs: 2\nExp number of AAs in TMHs: 37.46598\nExp number, first 60 AAs: 19.53139\nTotal prob of N-in: 0.94364\nPOSSIBLE N-term signal sequence\ninside 1 19\nTMhelix 20 39\noutside 40 325\nTMhelix 326 348\ninside 349 372

3219 GCF_001742705.1_ASM174270v1 Escherichia coli Proteobacteria; Gammaproteobacteria; Enterobacterales; Enterobacteriaceae; Escherichia
MNNEETFYQAMRRQGVTRRSFLKYCSLAATSLGLGAGMAPKIAWALENKPRIPVVWIHGL WP_001058323.1
MULTISPECIES: hydrogenase-1 small chain [Proteobacteria] Length: 372\nNumber of predicted TMHs: 2\nExp number of AAs in TMHs: 37.2608499999999\nExp number, first 60 AAs: 19.52923\nTotal prob of N-in: 0.94351\nPOSSIBLE N-term signal sequence\ninside 1 19\nTMhelix 20 39\noutside 40 325\nTMhelix 326 348\ninside 349 372

3220 GCF_001742725.1_ASM174272v1 Escherichia coli Proteobacteria; Gammaproteobacteria; Enterobacterales; Enterobacteriaceae; Escherichia
MTGDNTLIHSHGINRRDFMKLCAALAATMGLSSKAAAEMAESVTNPQRPPVIWIGAQECT WP_000145410.1
MULTISPECIES: hydrogenase-2 small chain [Proteobacteria] Length: 372\nNumber of predicted TMHs: 1\nExp number of AAs in TMHs: 23.72226\nExp number, first 60 AAs: 0.22038\nTotal prob of N-in: 0.12994\noutside 1 331\nTMhelix 332 354\ninside 355 372

3221 GCF_001748825.1_ASM174882v1 Escherichia coli Proteobacteria; Gammaproteobacteria; Enterobacterales; Enterobacteriaceae; Escherichia
MTGDNTLIHSHGINRRDFMKLCAALAATMGLSSKAAAEMAESVTNPQRPPVIWIGAQECT WP_000145410.1
MULTISPECIES: hydrogenase-2 small chain [Proteobacteria] Length: 372\nNumber of predicted TMHs: 1\nExp number of AAs in TMHs: 23.72226\nExp number, first 60 AAs: 0.22038\nTotal prob of N-in: 0.12994\noutside 1 331\nTMhelix 332 354\ninside 355 372

3222 GCF_001748715.1_ASM174871v1 Escherichia coli Proteobacteria; Gammaproteobacteria; Enterobacterales; Enterobacteriaceae; Escherichia
MNNEETFYQAMRRQGVTRRSFLKYCSLAATSLGLGAGMAPKIAWALENKPRIPVVWIHGL WP_001058323.1
MULTISPECIES: hydrogenase-1 small chain [Proteobacteria] Length: 372\nNumber of predicted TMHs: 2\nExp number of AAs in TMHs: 37.2608499999999\nExp number, first 60 AAs: 19.52923\nTotal prob of N-in: 0.94351\nPOSSIBLE N-term signal sequence\ninside 1 19\nTMhelix 20 39\noutside 40 325\nTMhelix 326 348\ninside 349 372

3223 GCF_001748805.1_ASM174880v1 Escherichia coli Proteobacteria; Gammaproteobacteria; Enterobacterales; Enterobacteriaceae; Escherichia
MNNEETFYQAMRRQGVTRRSFLKYCSLAATSLGLGAGMAPKIAWALENKPRIPVVWIHGL WP_001058323.1
MULTISPECIES: hydrogenase-1 small chain [Proteobacteria] Length: 372\nNumber of predicted TMHs: 2\nExp number of AAs in TMHs: 37.2608499999999\nExp number, first 60 AAs: 19.52923\nTotal prob of N-in: 0.94351\nPOSSIBLE N-term signal sequence\ninside 1 19\nTMhelix 20 39\noutside 40 325\nTMhelix 326 348\ninside 349 372

3224 GCF_001748825.1_ASM174882v1 Escherichia coli Proteobacteria; Gammaproteobacteria; Enterobacterales; Enterobacteriaceae; Escherichia
MNNEETFYQAMRRQGVTRRSFLKYCSLAATSLGLGAGMAPKIAWALENKPRIPVVWIHGL WP_001058323.1
MULTISPECIES: hydrogenase-1 small chain [Proteobacteria] Length: 372\nNumber of predicted TMHs: 2\nExp number of AAs in TMHs: 37.2608499999999\nExp number, first 60 AAs: 19.52923\nTotal prob of N-in: 0.94351\nPOSSIBLE N-term signal sequence\ninside 1 19\nTMhelix 20 39\noutside 40 325\nTMhelix 326 348\ninside 349 372

3225 GCF_001748715.1_ASM174871v1 Escherichia coli Proteobacteria; Gammaproteobacteria; Enterobacterales; Enterobacteriaceae; Escherichia
MTGDNTLIHSHGINRRDFMKLCAALAATMGLSSKAAAEMAESVTNPQRPPVIWIGAQECT WP_069914839.1
hydrogenase 2 small subunit [Escherichia coli] Length: 372\nNumber of predicted TMHs: 1\nExp number of AAs in TMHs: 24.17006\nExp number, first 60 AAs: 0.22021\nTotal prob of N-in: 0.14360\noutside 1 331\nTMhelix 332 354\ninside 355 372

3226 GCF_001748525.1_ASM174852v1 Escherichia coli Proteobacteria; Gammaproteobacteria; Enterobacterales; Enterobacteriaceae; Escherichia
MNNEETFYQAMRRQGVTRRSFLKYCSLAATSLGLGAGMAPKIAWALENKPRIPVVWIHGL WP_001295940.1
MULTISPECIES: hydrogenase [Enterobacteriaceae] Length: 372\nNumber of predicted TMHs: 2\nExp number of AAs in TMHs: 37.2608899999999\nExp number, first 60 AAs: 19.52923\nTotal prob of N-in: 0.94351\nPOSSIBLE N-term signal sequence\ninside 1 19\nTMhelix 20 39\noutside 40 325\nTMhelix 326 348\ninside 349 372

3227 GCF_001748885.1_ASM174888v1 Escherichia coli Proteobacteria; Gammaproteobacteria; Enterobacterales; Enterobacteriaceae; Escherichia
MTGDNTLIHSHGINRRDFMKLCAALAATMGLSSKAAAEMAESVTNPQRPPVIWIGAQECT WP_000145410.1
MULTISPECIES: hydrogenase-2 small chain [Proteobacteria] Length: 372\nNumber of predicted TMHs: 1\nExp number of AAs in TMHs: 23.72226\nExp number, first 60 AAs: 0.22038\nTotal prob of N-in: 0.12994\noutside 1 331\nTMhelix 332 354\ninside 355 372

3228 GCF_001748975.1_ASM174897v1 Escherichia coli Proteobacteria; Gammaproteobacteria; Enterobacterales; Enterobacteriaceae; Escherichia
MNNEETFYQAMRRQGVTRRSFLKYCSLAATSLGLGAGMAPKIAWALENKPRIPVVWIHGL WP_001058323.1
MULTISPECIES: hydrogenase-1 small chain [Proteobacteria] Length: 372\nNumber of predicted TMHs: 2\nExp number of AAs in TMHs: 37.2608499999999\nExp number, first 60 AAs: 19.52923\nTotal prob of N-in: 0.94351\nPOSSIBLE N-term signal sequence\ninside 1 19\nTMhelix 20 39\noutside 40 325\nTMhelix 326 348\ninside 349 372

3229 GCF_001749225.1_ASM174922v1 Escherichia coli Proteobacteria; Gammaproteobacteria; Enterobacterales; Enterobacteriaceae; Escherichia
MTGDNTLIHSHGINRRDFMKLCAALAATMGLSSKAAAEMAESVTNPQRPPVIWIGAQECT WP_000145410.1
MULTISPECIES: hydrogenase-2 small chain [Proteobacteria] Length: 372\nNumber of predicted TMHs: 1\nExp number of AAs in TMHs: 23.72226\nExp number, first 60 AAs: 0.22038\nTotal prob of N-in: 0.12994\noutside 1 331\nTMhelix 332 354\ninside 355 372

3230 GCF_001749225.1_ASM174922v1 Escherichia coli Proteobacteria; Gammaproteobacteria; Enterobacterales; Enterobacteriaceae; Escherichia
MNNEETFYQAMRRQGVTRRSFLKYCSLAATSLGLGAGMAPKIAWALENKPRIPVVWIHGL WP_001058323.1
MULTISPECIES: hydrogenase-1 small chain [Proteobacteria] Length: 372\nNumber of predicted TMHs: 2\nExp number of AAs in TMHs: 37.2608499999999\nExp number, first 60 AAs: 19.52923\nTotal prob of N-in: 0.94351\nPOSSIBLE N-term signal sequence\ninside 1 19\nTMhelix 20 39\noutside 40 325\nTMhelix 326 348\ninside 349 372

3231 GCF_001748915.1_ASM174891v1 Escherichia coli Proteobacteria; Gammaproteobacteria; Enterobacterales; Enterobacteriaceae; Escherichia
MNNEETFYQAMRRQGVTRRSFLKYCSLAATSLGLGAGMAPKIAWALENKPRIPVVWIHGL WP_001058323.1
MULTISPECIES: hydrogenase-1 small chain [Proteobacteria] Length: 372\nNumber of predicted TMHs: 2\nExp number of AAs in TMHs: 37.2608499999999\nExp number, first 60 AAs: 19.52923\nTotal prob of N-in: 0.94351\nPOSSIBLE N-term signal sequence\ninside 1 19\nTMhelix 20 39\noutside 40 325\nTMhelix 326 348\ninside 349 372

3232 GCF_001749215.1_ASM174921v1 Escherichia coli Proteobacteria; Gammaproteobacteria; Enterobacterales; Enterobacteriaceae; Escherichia
MTGDNTLIHSHGINRRDFMKLCAALAATMGLSSKAAAEMAESVTNPQRPPVIWIGAQECT WP_000145410.1
MULTISPECIES: hydrogenase-2 small chain [Proteobacteria] Length: 372\nNumber of predicted TMHs: 1\nExp number of AAs in TMHs: 23.72226\nExp number, first 60 AAs: 0.22038\nTotal prob of N-in: 0.12994\noutside 1 331\nTMhelix 332 354\ninside 355 372

3233 GCF_001749215.1_ASM174921v1 Escherichia coli Proteobacteria; Gammaproteobacteria; Enterobacterales; Enterobacteriaceae; Escherichia
MNNEETFYQAMRRQGVTRRSFLKYCSLAATSLGLGAGMAPKIAWALENKPRIPVVWIHGL WP_001058323.1
MULTISPECIES: hydrogenase-1 small chain [Proteobacteria] Length: 372\nNumber of predicted TMHs: 2\nExp number of AAs in TMHs: 37.2608499999999\nExp number, first 60 AAs: 19.52923\nTotal prob of N-in: 0.94351\nPOSSIBLE N-term signal sequence\ninside 1 19\nTMhelix 20 39\noutside 40 325\nTMhelix 326 348\ninside 349 372

3234 GCF_001749125.1_ASM174912v1 Escherichia coli Proteobacteria; Gammaproteobacteria; Enterobacterales; Enterobacteriaceae; Escherichia
MNNEETFYQAMRRQGVTRRSFLKYCSLAATSLGLGAGMAPKIAWALENKPRIPVVWIHGL WP_001058323.1
MULTISPECIES: hydrogenase-1 small chain [Proteobacteria] Length: 372\nNumber of predicted TMHs: 2\nExp number of AAs in TMHs: 37.2608499999999\nExp number, first 60 AAs: 19.52923\nTotal prob of N-in: 0.94351\nPOSSIBLE N-term signal sequence\ninside 1 19\nTMhelix 20 39\noutside 40 325\nTMhelix 326 348\ninside 349 372

3235 GCF_001749375.1_ASM174937v1 Escherichia coli Proteobacteria; Gammaproteobacteria; Enterobacterales; Enterobacteriaceae; Escherichia
MNNEETFYQAMRRQGVTRRSFLKYCSLAATSLGLGAGMAPKIAWALENKPRIPVVWIHGL WP_001058323.1
MULTISPECIES: hydrogenase-1 small chain [Proteobacteria] Length: 372\nNumber of predicted TMHs: 2\nExp number of AAs in TMHs: 37.2608499999999\nExp number, first 60 AAs: 19.52923\nTotal prob of N-in: 0.94351\nPOSSIBLE N-term signal sequence\ninside 1 19\nTMhelix 20 39\noutside 40 325\nTMhelix 326 348\ninside 349 372

3236 GCF_001749365.1_ASM174936v1 Escherichia coli Proteobacteria; Gammaproteobacteria; Enterobacterales; Enterobacteriaceae; Escherichia
MTGDNTLIHSHGINRRDFMKLCAALAATMGLSSKAAAEMAESVTNPQRPPVIWIGAQECT WP_028131934.1
hydrogenase-2 small chain [Escherichia coli] Length: 372\nNumber of predicted TMHs: 1\nExp number of AAs in TMHs: 24.01565\nExp number, first 60 AAs: 0.22022\nTotal prob of N-in: 0.13096\noutside 1 331\nTMhelix 332 354\ninside 355 372

3237 GCF_001749365.1_ASM174936v1 Escherichia coli Proteobacteria; Gammaproteobacteria; Enterobacterales; Enterobacteriaceae; Escherichia
MNNEETFYQAMRRQGVTRRSFLKYCSLAATSLGLGAGMAPKIAWALENKPRIPVVWIHGL WP_059321607.1
hydrogenase [Escherichia coli] Length: 372\nNumber of predicted TMHs: 2\nExp number of AAs in TMHs: 37.2608299999999\nExp number, first 60 AAs: 19.52923\nTotal prob of N-in: 0.94351\nPOSSIBLE N-term signal sequence\ninside 1 19\nTMhelix 20 39\noutside 40 325\nTMhelix 326 348\ninside 349 372

3238 GCF_001749415.1_ASM174941v1 Escherichia coli Proteobacteria; Gammaproteobacteria; Enterobacterales; Enterobacteriaceae; Escherichia
MNNEETFYQAMRRQGVTRRSFLKYCSLAATSLGLGAGMAPKIAWALENKPRIPVVWIHGL WP_001058323.1
MULTISPECIES: hydrogenase-1 small chain [Proteobacteria] Length: 372\nNumber of predicted TMHs: 2\nExp number of AAs in TMHs: 37.2608499999999\nExp number, first 60 AAs: 19.52923\nTotal prob of N-in: 0.94351\nPOSSIBLE N-term signal sequence\ninside 1 19\nTMhelix 20 39\noutside 40 325\nTMhelix 326 348\ninside 349 372

3239 GCF_001749465.1_ASM174946v1 Escherichia coli Proteobacteria; Gammaproteobacteria; Enterobacterales; Enterobacteriaceae; Escherichia
MNNEETFYQAMRRQGVTRRSFLKYCSLAATSLGLGAGMAPKIAWALENKPRIPVVWIHGL WP_001058323.1
MULTISPECIES: hydrogenase-1 small chain [Proteobacteria] Length: 372\nNumber of predicted TMHs: 2\nExp number of AAs in TMHs: 37.2608499999999\nExp number, first 60 AAs: 19.52923\nTotal prob of N-in: 0.94351\nPOSSIBLE N-term signal sequence\ninside 1 19\nTMhelix 20 39\noutside 40 325\nTMhelix 326 348\ninside 349 372

3240 GCF_001749585.1_ASM174958v1 Escherichia coli Proteobacteria; Gammaproteobacteria; Enterobacterales; Enterobacteriaceae; Escherichia
MTGDNTLIHSHGINRRDFMKLCAALAATMGLSSKAAAEMAESVTNPQRPPVIWIGAQECT WP_000145410.1
MULTISPECIES: hydrogenase-2 small chain [Proteobacteria] Length: 372\nNumber of predicted TMHs: 1\nExp number of AAs in TMHs: 23.72226\nExp number, first 60 AAs: 0.22038\nTotal prob of N-in: 0.12994\noutside 1 331\nTMhelix 332 354\ninside 355 372

3241 GCF_001749465.1_ASM174946v1 Escherichia coli Proteobacteria; Gammaproteobacteria; Enterobacterales; Enterobacteriaceae; Escherichia
MTGDNTLIHSHGINRRDFMKLCAALAATMGLSSKAAAEMAESVTNPQRPPVIWIGAQECT WP_069916757.1
hydrogenase 2 small subunit [Escherichia coli] Length: 372\nNumber of predicted TMHs: 1\nExp number of AAs in TMHs: 22.4747\nExp number, first 60 AAs: 0.22626\nTotal prob of N-in: 0.08094\noutside 1 331\nTMhelix 332 354\ninside 355 372

3242 GCF_001816285.1_ASM181628v1 Escherichia coli Proteobacteria; Gammaproteobacteria; Enterobacterales; Enterobacteriaceae; Escherichia
MTGDNTLIHSHGINRRDFMKLCAALAATMGLSSKAAAEMAESVTNPQRPPVIWIGAQECT WP_000145410.1
MULTISPECIES: hydrogenase-2 small chain [Proteobacteria] Length: 372\nNumber of predicted TMHs: 1\nExp number of AAs in TMHs: 23.72226\nExp number, first 60 AAs: 0.22038\nTotal prob of N-in: 0.12994\noutside 1 331\nTMhelix 332 354\ninside 355 372

3243 GCF_001854565.1_ASM185456v1 Escherichia coli Proteobacteria; Gammaproteobacteria; Enterobacterales; Enterobacteriaceae; Escherichia

MTGDNTLIHSHGINRRDFMKLCAALAATMGLSSKAAAEMAESVTNPQRPPVIWIGAQECT WP_000145410.1 MULTISPECIES: hydrogenase-2 small chain [Proteobacteria] Length: 372\nNumber of predicted TMHs: 1\nExp number of AAs in TMHs: 23.72226\nExp number, first 60 AAs: 0.22038\nTotal prob of N-in: 0.12994\noutside 1 331\nTMhelix 332 354\ninside 355 372

3244 GCF_001816295.1_ASM181629v1 Escherichia coli Proteobacteria; Gammaproteobacteria; Enterobacterales; Enterobacteriaceae; Escherichia MTGDNTLIHSHGINRRDFMKLCAALAATMGLSSKAAAEMAESVTNPQRPPVIWIGAQECT WP_000145410.1 MULTISPECIES: hydrogenase-2 small chain [Proteobacteria] Length: 372\nNumber of predicted TMHs: 1\nExp number of AAs in TMHs: 23.72226\nExp number, first 60 AAs: 0.22038\nTotal prob of N-in: 0.12994\noutside 1 331\nTMhelix 332 354\ninside 355 372

3245 GCF_001816285.1_ASM181628v1 Escherichia coli Proteobacteria; Gammaproteobacteria; Enterobacterales; Enterobacteriaceae; Escherichia MNNEETFYQAMRRQGVTRRSFLKYCSLAATSLGLGAGMAPKIAWALENKPRIPVVWIHGL WP_001058323.1 MULTISPECIES: hydrogenase-1 small chain [Proteobacteria] Length: 372\nNumber of predicted TMHs: 2\nExp number of AAs in TMHs: 37.2608499999999\nExp number, first 60 AAs: 19.52923\nTotal prob of N-in: 0.94351\nPOSSIBLE N-term signal sequence\ninside 1 19\nTMhelix 20 39\noutside 40 325\nTMhelix 326 348\ninside 349 372

3246 GCF_001816295.1_ASM181629v1 Escherichia coli Proteobacteria; Gammaproteobacteria; Enterobacterales; Enterobacteriaceae; Escherichia MNNEETFYQAMRRQGVTRRSFLKYCSLAATSLGLGAGMAPKIAWALENKPRIPVVWIHGL WP_001058323.1 MULTISPECIES: hydrogenase-1 small chain [Proteobacteria] Length: 372\nNumber of predicted TMHs: 2\nExp number of AAs in TMHs: 37.2608499999999\nExp number, first 60 AAs: 19.52923\nTotal prob of N-in: 0.94351\nPOSSIBLE N-term signal sequence\ninside 1 19\nTMhelix 20 39\noutside 40 325\nTMhelix 326 348\ninside 349 372

3247 GCF_001750845.1_ASM175084v1 Escherichia coli Proteobacteria; Gammaproteobacteria; Enterobacterales; Enterobacteriaceae; Escherichia MNNEETFYQAMRRQGVTRRSFLKYCSLAATSLGLGAGMAPKIAWALENKPRIPVVWIHGL WP_001058323.1 MULTISPECIES: hydrogenase-1 small chain [Proteobacteria] Length: 372\nNumber of predicted TMHs: 2\nExp number of AAs in TMHs: 37.2608499999999\nExp number, first 60 AAs: 19.52923\nTotal prob of N-in: 0.94351\nPOSSIBLE N-term signal sequence\ninside 1 19\nTMhelix 20 39\noutside 40 325\nTMhelix 326 348\ninside 349 372

3248 GCF_001865985.1_ASM186598v1 Escherichia coli Proteobacteria; Gammaproteobacteria; Enterobacterales; Enterobacteriaceae; Escherichia MTGDNTLIHSHGINRRDFMKLCAALAATMGLSSKAAAEMAESVTNPQRPPVIWIGAQECT WP_000145410.1 MULTISPECIES: hydrogenase-2 small chain [Proteobacteria] Length: 372\nNumber of predicted TMHs: 1\nExp number of AAs in TMHs: 23.72226\nExp number, first 60 AAs: 0.22038\nTotal prob of N-in: 0.12994\noutside 1 331\nTMhelix 332 354\ninside 355 372

3249 GCF_001874785.1_ASM187478v1 Escherichia coli Proteobacteria; Gammaproteobacteria; Enterobacterales; Enterobacteriaceae; Escherichia MTGDNTLIHSHGINRRDFMKLCAALAATMGLSSKAAAEMVESVTNPQRPPVIWIGAQECT WP_000145423.1 hydrogenase 2 small subunit [Escherichia coli] Length: 372\nNumber of predicted TMHs: 1\nExp number of AAs in TMHs: 23.78201\nExp number, first 60 AAs: 0.28355\nTotal prob of N-in: 0.13257\noutside 1 331\nTMhelix 332 354\ninside 355 372

3250 GCF_001865895.1_ASM186589v1 Escherichia coli Proteobacteria; Gammaproteobacteria; Enterobacterales; Enterobacteriaceae; Escherichia MNNEETFYQAMRRQGVTRRSFLKYCSLAATSLGLGAGMAPKIAWALENKPRIPVVWIHGL WP_001058323.1 MULTISPECIES: hydrogenase-1 small chain [Proteobacteria] Length: 372\nNumber of predicted TMHs: 2\nExp number of AAs in TMHs: 37.2608499999999\nExp number, first 60 AAs: 19.52923\nTotal prob of N-in: 0.94351\nPOSSIBLE N-term signal sequence\ninside 1 19\nTMhelix 20 39\noutside 40 325\nTMhelix 326 348\ninside 349 372

3251 GCF_001877815.1_ASM187781v1 Escherichia coli Proteobacteria; Gammaproteobacteria; Enterobacterales; Enterobacteriaceae; Escherichia MTGDNTLIHSHGINRRDFMKLCAALAATMGLSSKAAAEMAESVTNPQRPPVIWIGAQECT WP_000145410.1

MULTISPECIES: hydrogenase-2 small chain [Proteobacteria] Length: 372\nNumber of predicted TMHs: 1\nExp number of AAs in TMHs: 23.72226\nExp number, first 60 AAs: 0.22038\nTotal prob of N-in: 0.12994\noutside 1 331\nTMhelix 332 354\ninside 355 372

3252 GCF_001881055.1_ASM188105v1 Escherichia coli Proteobacteria; Gammaproteobacteria; Enterobacterales; Enterobacteriaceae; Escherichia MTGDNTLIHSHGINRRDFMKLCAALAATMGLSSKAAAEMAESVTNPQRPPVIWIGAQECT WP_000145410.1 MULTISPECIES: hydrogenase-2 small chain [Proteobacteria] Length: 372\nNumber of predicted TMHs: 1\nExp number of AAs in TMHs: 23.72226\nExp number, first 60 AAs: 0.22038\nTotal prob of N-in: 0.12994\noutside 1 331\nTMhelix 332 354\ninside 355 372

3253 GCF_001881045.1_ASM188104v1 Escherichia coli Proteobacteria; Gammaproteobacteria; Enterobacterales; Enterobacteriaceae; Escherichia MTGDNTLIHSHGINRRDFMKLCAALAATMGLSSKAAAEMAESVTNPQRPPVIWIGAQECT WP_000145410.1 MULTISPECIES: hydrogenase-2 small chain [Proteobacteria] Length: 372\nNumber of predicted TMHs: 1\nExp number of AAs in TMHs: 23.72226\nExp number, first 60 AAs: 0.22038\nTotal prob of N-in: 0.12994\noutside 1 331\nTMhelix 332 354\ninside 355 372

3254 GCF_001874845.1_ASM187484v1 Escherichia coli Proteobacteria; Gammaproteobacteria; Enterobacterales; Enterobacteriaceae; Escherichia MTGDNTLIHSHGINRRDFMKLCAALAATMGLSSKAAAEMAESVTNPQRPPVIWIGAQECT WP_000145410.1 MULTISPECIES: hydrogenase-2 small chain [Proteobacteria] Length: 372\nNumber of predicted TMHs: 1\nExp number of AAs in TMHs: 23.72226\nExp number, first 60 AAs: 0.22038\nTotal prob of N-in: 0.12994\noutside 1 331\nTMhelix 332 354\ninside 355 372

3255 GCF_001874845.1_ASM187484v1 Escherichia coli Proteobacteria; Gammaproteobacteria; Enterobacterales; Enterobacteriaceae; Escherichia MNNEETFYQAMRRQGVTRRSFLKYCSLAATSLGLGAGMAPKIAWALENKPRIPVVWIHGL WP_001295940.1 MULTISPECIES: hydrogenase [Enterobacteriaceae] Length: 372\nNumber of predicted TMHs: 2\nExp number of AAs in TMHs: 37.2608899999999\nExp number, first 60 AAs: 19.52923\nTotal prob of N-in: 0.94351\nPOSSIBLE N-term signal sequence\ninside 1 19\nTMhelix 20 39\noutside 40 325\nTMhelix 326 348\ninside 349 372

3256 GCF_001881465.1_ASM188146v1 Escherichia coli Proteobacteria; Gammaproteobacteria; Enterobacterales; Enterobacteriaceae; Escherichia MTGDNTLIHSHGINRRDFMKLCAALAATMGLSSKAAAEMAESVTNPQRPPVIWIGAQECT WP_000145410.1 MULTISPECIES: hydrogenase-2 small chain [Proteobacteria] Length: 372\nNumber of predicted TMHs: 1\nExp number of AAs in TMHs: 23.72226\nExp number, first 60 AAs: 0.22038\nTotal prob of N-in: 0.12994\noutside 1 331\nTMhelix 332 354\ninside 355 372

3257 GCF_001881515.1_ASM188151v1 Escherichia coli Proteobacteria; Gammaproteobacteria; Enterobacterales; Enterobacteriaceae; Escherichia MTGDNTLIHSHGINRRDFMKLCAALAATMGLSSKAAAEMAESVTNPQRPPVIWIGAQECT WP_000145410.1 MULTISPECIES: hydrogenase-2 small chain [Proteobacteria] Length: 372\nNumber of predicted TMHs: 1\nExp number of AAs in TMHs: 23.72226\nExp number, first 60 AAs: 0.22038\nTotal prob of N-in: 0.12994\noutside 1 331\nTMhelix 332 354\ninside 355 372

3258 GCF_001881665.1_ASM188166v1 Escherichia coli Proteobacteria; Gammaproteobacteria; Enterobacterales; Enterobacteriaceae; Escherichia MTGDNTLIHSHGINRRDFMKLCAALAATMGLSSKAAAEMAESVTNPQRPPVIWIGAQECT WP_000145410.1 MULTISPECIES: hydrogenase-2 small chain [Proteobacteria] Length: 372\nNumber of predicted TMHs: 1\nExp number of AAs in TMHs: 23.72226\nExp number, first 60 AAs: 0.22038\nTotal prob of N-in: 0.12994\noutside 1 331\nTMhelix 332 354\ninside 355 372

3259 GCF_001881515.1_ASM188151v1 Escherichia coli Proteobacteria; Gammaproteobacteria; Enterobacterales; Enterobacteriaceae; Escherichia MNNEETFYQAMRRQGVTRRSFLKYCSLAATSLGLGAGMAPKIAWALENKPRIPVVWIHGL WP_001058323.1 MULTISPECIES: hydrogenase-1 small chain [Proteobacteria] Length: 372\nNumber of predicted TMHs: 2\nExp number of AAs in TMHs: 37.2608499999999\nExp number, first 60 AAs: 19.52923\nTotal prob of N-in: 0.94351\nPOSSIBLE N-term signal sequence\ninside 1 19\nTMhelix 20 39\noutside 40 325\nTMhelix 326 348\ninside 349 372

3260 GCF_001881665.1_ASM188166v1 Escherichia coli Proteobacteria; Gammaproteobacteria; Enterobacterales; Enterobacteriaceae; Escherichia
MNNEETFYQAMRRQGVTRRSFLKYCSLAATSLGLGAGMAPKIAWALENKPRIPVVWIHGL WP_001058323.1
MULTISPECIES: hydrogenase-1 small chain [Proteobacteria] Length: 372\nNumber of predicted TMHs: 2\nExp number of AAs in TMHs: 37.2608499999999\nExp number, first 60 AAs: 19.52923\nTotal prob of N-in: 0.94351\nPOSSIBLE N-term signal sequence\ninside 1 19\nTMhelix 20 39\noutside 40 325\nTMhelix 326 348\ninside 349 372

3261 GCF_001884865.1_ASM188486v1 Escherichia coli Proteobacteria; Gammaproteobacteria; Enterobacterales; Enterobacteriaceae; Escherichia
MNNEETFYQAMRRQGVTRRSFLKYCSLAATSLGLGAGMAPKIAWALENKPRIPVVWIHGL WP_001301613.1
hydrogenase [Escherichia coli] Length: 372\nNumber of predicted TMHs: 2\nExp number of AAs in TMHs: 37.46598\nExp number, first 60 AAs: 19.53139\nTotal prob of N-in: 0.94364\nPOSSIBLE N-term signal sequence\ninside 1 19\nTMhelix 20 39\noutside 40 325\nTMhelix 326 348\ninside 349 372

3262 GCF_001884955.1_ASM188495v1 Escherichia coli Proteobacteria; Gammaproteobacteria; Enterobacterales; Enterobacteriaceae; Escherichia
MTGDNTLIHSHGINRRDFMKLCAALAATMGLSSKAAAEMAESVTNPQRPPVIWIGAQECS WP_061069243.1
hydrogenase 2 small subunit [Escherichia coli] Length: 372\nNumber of predicted TMHs: 1\nExp number of AAs in TMHs: 23.72219\nExp number, first 60 AAs: 0.22034\nTotal prob of N-in: 0.12994\noutside 1 331\nTMhelix 332 354\ninside 355 372

3263 GCF_001890285.1_ASM189028v1 Escherichia coli Proteobacteria; Gammaproteobacteria; Enterobacterales; Enterobacteriaceae; Escherichia
MTGDNTLIHSHGINRRDFMKLCAALAATMGLSSKAAAEMAESVTNPQRPPVIWIGAQECT WP_000145410.1
MULTISPECIES: hydrogenase-2 small chain [Proteobacteria] Length: 372\nNumber of predicted TMHs: 1\nExp number of AAs in TMHs: 23.72226\nExp number, first 60 AAs: 0.22038\nTotal prob of N-in: 0.12994\noutside 1 331\nTMhelix 332 354\ninside 355 372

3264 GCF_001884965.1_ASM188496v1 Escherichia coli Proteobacteria; Gammaproteobacteria; Enterobacterales; Enterobacteriaceae; Escherichia
MTGDNTLIHSHGINRRDFMKLCAALAATMGLSSKAAAEMAESVTNPQRPPVIWIGAQECS WP_061069243.1
hydrogenase 2 small subunit [Escherichia coli] Length: 372\nNumber of predicted TMHs: 1\nExp number of AAs in TMHs: 23.72219\nExp number, first 60 AAs: 0.22034\nTotal prob of N-in: 0.12994\noutside 1 331\nTMhelix 332 354\ninside 355 372

3265 GCF_001885025.1_ASM188502v1 Escherichia coli Proteobacteria; Gammaproteobacteria; Enterobacterales; Enterobacteriaceae; Escherichia
MTGDNTLIHSHGINRRDFMKLCAALAATMGLSSKAAAEMAESVTNPQRPPVIWIGAQECS WP_061069243.1
hydrogenase 2 small subunit [Escherichia coli] Length: 372\nNumber of predicted TMHs: 1\nExp number of AAs in TMHs: 23.72219\nExp number, first 60 AAs: 0.22034\nTotal prob of N-in: 0.12994\noutside 1 331\nTMhelix 332 354\ninside 355 372

3266 GCF_001886535.1_ASM188653v1 Escherichia coli Proteobacteria; Gammaproteobacteria; Enterobacterales; Enterobacteriaceae; Escherichia
MTGDNTLIHSHGINRRDFMKLCAALATTMGLSSKAAAEMAESVTNPQRPPVIWIGAQECT WP_023149629.1
MULTISPECIES: hydrogenase-2 small chain [Enterobacteriaceae] Length: 372\nNumber of predicted TMHs: 1\nExp number of AAs in TMHs: 23.63603\nExp number, first 60 AAs: 0.12971\nTotal prob of N-in: 0.12608\noutside 1 331\nTMhelix 332 354\ninside 355 372

3267 GCF_001886555.1_ASM188655v1 Escherichia coli Proteobacteria; Gammaproteobacteria; Enterobacterales; Enterobacteriaceae; Escherichia
MTGDNTLIHSHGINRRDFMKLCAALATTMGLSSKAAAEMAESVTNPQRPPVIWIGAQECT WP_023149629.1
MULTISPECIES: hydrogenase-2 small chain [Enterobacteriaceae] Length: 372\nNumber of predicted TMHs: 1\nExp number of AAs in TMHs: 23.63603\nExp number, first 60 AAs: 0.12971\nTotal prob of N-in: 0.12608\noutside 1 331\nTMhelix 332 354\ninside 355 372

3268 GCF_001890305.1_ASM189030v1 Escherichia coli Proteobacteria; Gammaproteobacteria; Enterobacterales; Enterobacteriaceae; Escherichia
MTGDNTLIHSHGINRRDFMKLCAALAATMGLSSKAAAEMAESVTNPQRPPVIWIGAQECT WP_000145410.1
MULTISPECIES: hydrogenase-2 small chain [Proteobacteria] Length: 372\nNumber of predicted TMHs:

1\nExp number of AAs in TMHs: 23.72226\nExp number, first 60 AAs: 0.22038\nTotal prob of N-in: 0.12994\noutside 1 331\nTMhelix 332 354\ninside 355 372

3269 GCF_001890285.1_ASM189028v1 Escherichia coli Proteobacteria; Gammaproteobacteria; Enterobacterales; Enterobacteriaceae; Escherichia MNNEETFYQAMRRQGVTRRSFLKYCSLAATSLGLGAGMAPKIAWALENKPRIPVVWIHGL WP_001301613.1 hydrogenase [Escherichia coli] Length: 372\nNumber of predicted TMHs: 2\nExp number of AAs in TMHs: 37.46598\nExp number, first 60 AAs: 19.53139\nTotal prob of N-in: 0.94364\nPOSSIBLE N-term signal sequence\ninside 1 19\nTMhelix 20 39\noutside 40 325\nTMhelix 326 348\ninside 349 372

3270 GCF_001891445.1_ASM189144v1 Escherichia coli Proteobacteria; Gammaproteobacteria; Enterobacterales; Enterobacteriaceae; Escherichia MTGDNTLIHSHGINRRDFMKLCAALAATMGLSSKAAAEMAESVTNPQRPPVIWIGAQECT WP_000145410.1 MULTISPECIES: hydrogenase-2 small chain [Proteobacteria] Length: 372\nNumber of predicted TMHs: 1\nExp number of AAs in TMHs: 23.72226\nExp number, first 60 AAs: 0.22038\nTotal prob of N-in: 0.12994\noutside 1 331\nTMhelix 332 354\ninside 355 372

3271 GCF_001891475.1_ASM189147v1 Escherichia coli Proteobacteria; Gammaproteobacteria; Enterobacterales; Enterobacteriaceae; Escherichia MTGDNTLIHSHGINRRDFMKLCAALAATMGLSSKAAAEMAESVTNPQRPPVIWIGAQECT WP_000145410.1 MULTISPECIES: hydrogenase-2 small chain [Proteobacteria] Length: 372\nNumber of predicted TMHs: 1\nExp number of AAs in TMHs: 23.72226\nExp number, first 60 AAs: 0.22038\nTotal prob of N-in: 0.12994\noutside 1 331\nTMhelix 332 354\ninside 355 372

3272 GCF_001891475.1_ASM189147v1 Escherichia coli Proteobacteria; Gammaproteobacteria; Enterobacterales; Enterobacteriaceae; Escherichia MNNEETFYQAMRRQGVTRRSFLKYCSLAATSLGLGAGMAPKIAWALENKPRIPVVWIHGL WP_001058323.1 MULTISPECIES: hydrogenase-1 small chain [Proteobacteria] Length: 372\nNumber of predicted TMHs: 2\nExp number of AAs in TMHs: 37.2608499999999\nExp number, first 60 AAs: 19.52923\nTotal prob of N-in: 0.94351\nPOSSIBLE N-term signal sequence\ninside 1 19\nTMhelix 20 39\noutside 40 325\nTMhelix 326 348\ninside 349 372

3273 GCF_001891445.1_ASM189144v1 Escherichia coli Proteobacteria; Gammaproteobacteria; Enterobacterales; Enterobacteriaceae; Escherichia MNNEETFYQAMRRQGVTRRSFLKYCSLAATSLGLGAGMAPKIAWALENKPRIPVVWIHGL WP_001295940.1 MULTISPECIES: hydrogenase [Enterobacteriaceae] Length: 372\nNumber of predicted TMHs: 2\nExp number of AAs in TMHs: 37.2608899999999\nExp number, first 60 AAs: 19.52923\nTotal prob of N-in: 0.94351\nPOSSIBLE N-term signal sequence\ninside 1 19\nTMhelix 20 39\noutside 40 325\nTMhelix 326 348\ninside 349 372

3274 GCF_001891225.1_ASM189122v1 Escherichia coli Proteobacteria; Gammaproteobacteria; Enterobacterales; Enterobacteriaceae; Escherichia MNNEETFYQAMRRQGVTRRSFLKYCSLAATSLGLGAGMAPKIAWALENKPRIPVVWIHGL WP_001295940.1 MULTISPECIES: hydrogenase [Enterobacteriaceae] Length: 372\nNumber of predicted TMHs: 2\nExp number of AAs in TMHs: 37.2608899999999\nExp number, first 60 AAs: 19.52923\nTotal prob of N-in: 0.94351\nPOSSIBLE N-term signal sequence\ninside 1 19\nTMhelix 20 39\noutside 40 325\nTMhelix 326 348\ninside 349 372

3275 GCF_001890345.1_ASM189034v1 Escherichia coli Proteobacteria; Gammaproteobacteria; Enterobacterales; Enterobacteriaceae; Escherichia MNNEETFYQAMRRQGVTRRSFLKYCSLAATSLGLGAGMAPKIAWALENKPRIPVVWIHGL WP_001301613.1 hydrogenase [Escherichia coli] Length: 372\nNumber of predicted TMHs: 2\nExp number of AAs in TMHs: 37.46598\nExp number, first 60 AAs: 19.53139\nTotal prob of N-in: 0.94364\nPOSSIBLE N-term signal sequence\ninside 1 19\nTMhelix 20 39\noutside 40 325\nTMhelix 326 348\ninside 349 372

3276 GCF_001891835.1_ASM189183v1 Escherichia coli Proteobacteria; Gammaproteobacteria; Enterobacterales; Enterobacteriaceae; Escherichia MNNEETFYQAMRRQGVTRRSFLKYCSLAATSLGLGAGMAPKIAWALENKPRIPVVWIHGL WP_001058323.1 MULTISPECIES: hydrogenase-1 small chain [Proteobacteria] Length: 372\nNumber of predicted TMHs: 2\nExp number of AAs in TMHs: 37.2608499999999\nExp number, first 60 AAs: 19.52923\nTotal prob of N-in:

0.94351\nPOSSIBLE N-term signal sequence\ninside 1 19\nTMhelix 20 39\noutside 40 325\nTMhelix 326 348\ninside 349 372

3277 GCF_001891695.1_ASM189169v1 Escherichia coli Proteobacteria; Gammaproteobacteria; Enterobacterales; Enterobacteriaceae; Escherichia
MNNEETFYQAMRRQGVTRRSFLKYCSLAATSLGLGAGMAPKIAWALENKPRIPVVWIHGL WP_001058323.1
MULTISPECIES: hydrogenase-1 small chain [Proteobacteria] Length: 372\nNumber of predicted TMHs: 2\nExp number of AAs in TMHs: 37.2608499999999\nExp number, first 60 AAs: 19.52923\nTotal prob of N-in: 0.94351\nPOSSIBLE N-term signal sequence\ninside 1 19\nTMhelix 20 39\noutside 40 325\nTMhelix 326 348\ninside 349 372

3278 GCF_001891545.1_ASM189154v1 Escherichia coli Proteobacteria; Gammaproteobacteria; Enterobacterales; Enterobacteriaceae; Escherichia
MNNEETFYQAMRRQGVTRRSFLKYCSLAATSLGLGAGMAPKIAWALENKPRIPVVWIHGL WP_001295940.1
MULTISPECIES: hydrogenase [Enterobacteriaceae] Length: 372\nNumber of predicted TMHs: 2\nExp number of AAs in TMHs: 37.2608899999999\nExp number, first 60 AAs: 19.52923\nTotal prob of N-in: 0.94351\nPOSSIBLE N-term signal sequence\ninside 1 19\nTMhelix 20 39\noutside 40 325\nTMhelix 326 348\ninside 349 372

3279 GCF_001891765.1_ASM189176v1 Escherichia coli Proteobacteria; Gammaproteobacteria; Enterobacterales; Enterobacteriaceae; Escherichia
MNNEETFYQAMRRQGVTRRSFLKYCSLAATSLGLGAGMAPKIAWALENKPRIPVVWIHGL WP_001058323.1
MULTISPECIES: hydrogenase-1 small chain [Proteobacteria] Length: 372\nNumber of predicted TMHs: 2\nExp number of AAs in TMHs: 37.2608499999999\nExp number, first 60 AAs: 19.52923\nTotal prob of N-in: 0.94351\nPOSSIBLE N-term signal sequence\ninside 1 19\nTMhelix 20 39\noutside 40 325\nTMhelix 326 348\ninside 349 372

3280 GCF_001891885.1_ASM189188v1 Escherichia coli Proteobacteria; Gammaproteobacteria; Enterobacterales; Enterobacteriaceae; Escherichia
MTGDNTLIHSHGINRRDFMKLCAALAATMGLSSKAAAEMAESVTNPQRPPVIWIGAQECT WP_000145410.1
MULTISPECIES: hydrogenase-2 small chain [Proteobacteria] Length: 372\nNumber of predicted TMHs: 1\nExp number of AAs in TMHs: 23.72226\nExp number, first 60 AAs: 0.22038\nTotal prob of N-in: 0.12994\noutside 1 331\nTMhelix 332 354\ninside 355 372

3281 GCF_001891865.1_ASM189186v1 Escherichia coli Proteobacteria; Gammaproteobacteria; Enterobacterales; Enterobacteriaceae; Escherichia
MTGDNTLIHSHGINRRDFMKLCAALAATMGLSSKAAAEMAESVTNPQRPPVIWIGAQECT WP_000145410.1
MULTISPECIES: hydrogenase-2 small chain [Proteobacteria] Length: 372\nNumber of predicted TMHs: 1\nExp number of AAs in TMHs: 23.72226\nExp number, first 60 AAs: 0.22038\nTotal prob of N-in: 0.12994\noutside 1 331\nTMhelix 332 354\ninside 355 372

3282 GCF_001891865.1_ASM189186v1 Escherichia coli Proteobacteria; Gammaproteobacteria; Enterobacterales; Enterobacteriaceae; Escherichia
MNNEETFYQAMRRQGVTRRSFLKYCSLAATSLGLGAGMAPKIAWALENKPRIPVVWIHGL WP_001058323.1
MULTISPECIES: hydrogenase-1 small chain [Proteobacteria] Length: 372\nNumber of predicted TMHs: 2\nExp number of AAs in TMHs: 37.2608499999999\nExp number, first 60 AAs: 19.52923\nTotal prob of N-in: 0.94351\nPOSSIBLE N-term signal sequence\ninside 1 19\nTMhelix 20 39\noutside 40 325\nTMhelix 326 348\ninside 349 372

3283 GCF_001891995.1_ASM189199v1 Escherichia coli Proteobacteria; Gammaproteobacteria; Enterobacterales; Enterobacteriaceae; Escherichia
MNNEETFYQAMRRQGVTRRSFLKYCSLAATSLGLGAGMAPKIAWALENKPRIPVVWIHGL WP_001295940.1
MULTISPECIES: hydrogenase [Enterobacteriaceae] Length: 372\nNumber of predicted TMHs: 2\nExp number of AAs in TMHs: 37.2608899999999\nExp number, first 60 AAs: 19.52923\nTotal prob of N-in: 0.94351\nPOSSIBLE N-term signal sequence\ninside 1 19\nTMhelix 20 39\noutside 40 325\nTMhelix 326 348\ninside 349 372

3284 GCF_001892135.1_ASM189213v1 Escherichia coli Proteobacteria; Gammaproteobacteria; Enterobacterales; Enterobacteriaceae; Escherichia
MNNEETFYQAMRRQGVTRRSFLKYCSLAATSLGLGAGMAPKIAWALENKPRIPVVWIHGL WP_001295940.1
MULTISPECIES: hydrogenase [Enterobacteriaceae] Length: 372\nNumber of predicted TMHs: 2\nExp

number of AAs in TMHs: 37.2608899999999\nExp number, first 60 AAs: 19.52923\nTotal prob of N-in: 0.94351\nPOSSIBLE N-term signal sequence\ninside 1 19\nTMhelix 20 39\noutside 40 325\nTMhelix 326 348\ninside 349 372

3285 GCF_001892105.1_ASM189210v1 Escherichia coli Proteobacteria; Gammaproteobacteria; Enterobacterales; Enterobacteriaceae; Escherichia
MNNEETFYQAMRRQGVTRRSFLKYCSLAATSLGLGAGMAPKIAWALENKPRIPVVWIHGL WP_001295940.1
MULTISPECIES: hydrogenase [Enterobacteriaceae] Length: 372\nNumber of predicted TMHs: 2\nExp number of AAs in TMHs: 37.2608899999999\nExp number, first 60 AAs: 19.52923\nTotal prob of N-in: 0.94351\nPOSSIBLE N-term signal sequence\ninside 1 19\nTMhelix 20 39\noutside 40 325\nTMhelix 326 348\ninside 349 372

3286 GCF_001892065.1_ASM189206v1 Escherichia coli Proteobacteria; Gammaproteobacteria; Enterobacterales; Enterobacteriaceae; Escherichia
MNNEETFYQAMRRQGVTRRSFLKYCSLAATSLGLGAGMAPKIAWALENKPRIPVVWIHGL WP_001295940.1
MULTISPECIES: hydrogenase [Enterobacteriaceae] Length: 372\nNumber of predicted TMHs: 2\nExp number of AAs in TMHs: 37.2608899999999\nExp number, first 60 AAs: 19.52923\nTotal prob of N-in: 0.94351\nPOSSIBLE N-term signal sequence\ninside 1 19\nTMhelix 20 39\noutside 40 325\nTMhelix 326 348\ninside 349 372

3287 GCF_001892185.1_ASM189218v1 Escherichia coli Proteobacteria; Gammaproteobacteria; Enterobacterales; Enterobacteriaceae; Escherichia
MNNEETFYQAMRRQGVTRRSFLKYCSLAATSLGLGAGMAPKIAWALENKPRIPVVWIHGL WP_001295940.1
MULTISPECIES: hydrogenase [Enterobacteriaceae] Length: 372\nNumber of predicted TMHs: 2\nExp number of AAs in TMHs: 37.2608899999999\nExp number, first 60 AAs: 19.52923\nTotal prob of N-in: 0.94351\nPOSSIBLE N-term signal sequence\ninside 1 19\nTMhelix 20 39\noutside 40 325\nTMhelix 326 348\ninside 349 372

3288 GCF_001892355.1_ASM189235v1 Escherichia coli Proteobacteria; Gammaproteobacteria; Enterobacterales; Enterobacteriaceae; Escherichia
MTGDNTLIHSHGINRRDFMKLCAALAATMGLSSKAAAEMAESVTNPQRPPVIWIGAQECT WP_000145410.1
MULTISPECIES: hydrogenase-2 small chain [Proteobacteria] Length: 372\nNumber of predicted TMHs: 1\nExp number of AAs in TMHs: 23.72226\nExp number, first 60 AAs: 0.22038\nTotal prob of N-in: 0.12994\noutside 1 331\nTMhelix 332 354\ninside 355 372

3289 GCF_001892335.1_ASM189233v1 Escherichia coli Proteobacteria; Gammaproteobacteria; Enterobacterales; Enterobacteriaceae; Escherichia
MNNEETFYQAMRRQGVTRRSFLKYCSLAATSLGLGAGMAPKIAWALENKPRIPVVWIHGL WP_001058323.1
MULTISPECIES: hydrogenase-1 small chain [Proteobacteria] Length: 372\nNumber of predicted TMHs: 2\nExp number of AAs in TMHs: 37.2608499999999\nExp number, first 60 AAs: 19.52923\nTotal prob of N-in: 0.94351\nPOSSIBLE N-term signal sequence\ninside 1 19\nTMhelix 20 39\noutside 40 325\nTMhelix 326 348\ninside 349 372

3290 GCF_001892645.1_ASM189264v1 Escherichia coli Proteobacteria; Gammaproteobacteria; Enterobacterales; Enterobacteriaceae; Escherichia
MNNEETFYQAMRRQGVTRRSFLKYCSLAATSLGLGAGMAPKIAWALENKPRIPVVWIHGL WP_001295940.1
MULTISPECIES: hydrogenase [Enterobacteriaceae] Length: 372\nNumber of predicted TMHs: 2\nExp number of AAs in TMHs: 37.2608899999999\nExp number, first 60 AAs: 19.52923\nTotal prob of N-in: 0.94351\nPOSSIBLE N-term signal sequence\ninside 1 19\nTMhelix 20 39\noutside 40 325\nTMhelix 326 348\ninside 349 372

3291 GCF_001892675.1_ASM189267v1 Escherichia coli Proteobacteria; Gammaproteobacteria; Enterobacterales; Enterobacteriaceae; Escherichia
MNNEETFYQAMRRQGVTRRSFLKYCSLAATSLGLGAGMAPKIAWALENKPRIPVVWIHGL WP_001295940.1
MULTISPECIES: hydrogenase [Enterobacteriaceae] Length: 372\nNumber of predicted TMHs: 2\nExp number of AAs in TMHs: 37.2608899999999\nExp number, first 60 AAs: 19.52923\nTotal prob of N-in: 0.94351\nPOSSIBLE N-term signal sequence\ninside 1 19\nTMhelix 20 39\noutside 40 325\nTMhelix 326 348\ninside 349 372

3292 GCF_001892745.1_ASM189274v1 Escherichia coli Proteobacteria; Gammaproteobacteria; Enterobacterales; Enterobacteriaceae; Escherichia

MNNEETFYQAMRRQGVTRRSFLKYCSLAATSLGLGAGMAPKIAWALENKPRIPVVWIHGL WP_001058323.1 MULTISPECIES: hydrogenase-1 small chain [Proteobacteria] Length: 372\nNumber of predicted TMHs: 2\nExp number of AAs in TMHs: 37.2608499999999\nExp number, first 60 AAs: 19.52923\nTotal prob of N-in: 0.94351\nPOSSIBLE N-term signal sequence\ninside 1 19\nTMhelix 20 39\noutside 40 325\nTMhelix 326 348\ninside 349 372

3293 GCF_001892765.1_ASM189276v1 Escherichia coli Proteobacteria; Gammaproteobacteria; Enterobacterales; Enterobacteriaceae; Escherichia MTGDNTLIHSHGINRRDFMKLCAALAATMGLSSKAAAEMAESVTNPQRPPVIWIGAQECT WP_000145410.1 MULTISPECIES: hydrogenase-2 small chain [Proteobacteria] Length: 372\nNumber of predicted TMHs: 1\nExp number of AAs in TMHs: 23.72226\nExp number, first 60 AAs: 0.22038\nTotal prob of N-in: 0.12994\noutside 1 331\nTMhelix 332 354\ninside 355 372

3294 GCF_001892825.1_ASM189282v1 Escherichia coli Proteobacteria; Gammaproteobacteria; Enterobacterales; Enterobacteriaceae; Escherichia MTGDNTLIHSHGINRRDFMKLCAALAATMGLSSKAAAEMAESVTNPQRPPVIWIGAQECT WP_000145410.1 MULTISPECIES: hydrogenase-2 small chain [Proteobacteria] Length: 372\nNumber of predicted TMHs: 1\nExp number of AAs in TMHs: 23.72226\nExp number, first 60 AAs: 0.22038\nTotal prob of N-in: 0.12994\noutside 1 331\nTMhelix 332 354\ninside 355 372

3295 GCF_001892865.1_ASM189286v1 Escherichia coli Proteobacteria; Gammaproteobacteria; Enterobacterales; Enterobacteriaceae; Escherichia MTGDNTLIHSHGINRRDFMKLCAALAATMGLSSKAAAEMAESVTNPQRPPVIWIGAQECT WP_000145410.1 MULTISPECIES: hydrogenase-2 small chain [Proteobacteria] Length: 372\nNumber of predicted TMHs: 1\nExp number of AAs in TMHs: 23.72226\nExp number, first 60 AAs: 0.22038\nTotal prob of N-in: 0.12994\noutside 1 331\nTMhelix 332 354\ninside 355 372

3296 GCF_001892835.1_ASM189283v1 Escherichia coli Proteobacteria; Gammaproteobacteria; Enterobacterales; Enterobacteriaceae; Escherichia MTGDNTLIHSHGINRRDFMKLCAALAATMGLSSKAAAEMAESVTNPQRPPVIWIGAQECT WP_000145410.1 MULTISPECIES: hydrogenase-2 small chain [Proteobacteria] Length: 372\nNumber of predicted TMHs: 1\nExp number of AAs in TMHs: 23.72226\nExp number, first 60 AAs: 0.22038\nTotal prob of N-in: 0.12994\noutside 1 331\nTMhelix 332 354\ninside 355 372

3297 GCF_001892865.1_ASM189286v1 Escherichia coli Proteobacteria; Gammaproteobacteria; Enterobacterales; Enterobacteriaceae; Escherichia MNNEETFYQAMRRQGVTRRSFLKYCSLAATSLGLGAGMAPKIAWALENKPRIPVVWIHGL WP_001058323.1 MULTISPECIES: hydrogenase-1 small chain [Proteobacteria] Length: 372\nNumber of predicted TMHs: 2\nExp number of AAs in TMHs: 37.2608499999999\nExp number, first 60 AAs: 19.52923\nTotal prob of N-in: 0.94351\nPOSSIBLE N-term signal sequence\ninside 1 19\nTMhelix 20 39\noutside 40 325\nTMhelix 326 348\ninside 349 372

3298 GCF_001892935.1_ASM189293v1 Escherichia coli Proteobacteria; Gammaproteobacteria; Enterobacterales; Enterobacteriaceae; Escherichia MNNEETFYQAMRRQGVTRRSFLKYCSLAATSLGLGAGMAPKIAWALENKPRIPVVWIHGL WP_001058323.1 MULTISPECIES: hydrogenase-1 small chain [Proteobacteria] Length: 372\nNumber of predicted TMHs: 2\nExp number of AAs in TMHs: 37.2608499999999\nExp number, first 60 AAs: 19.52923\nTotal prob of N-in: 0.94351\nPOSSIBLE N-term signal sequence\ninside 1 19\nTMhelix 20 39\noutside 40 325\nTMhelix 326 348\ninside 349 372

3299 GCF_001893155.1_ASM189315v1 Escherichia coli Proteobacteria; Gammaproteobacteria; Enterobacterales; Enterobacteriaceae; Escherichia MTGDNTLIHSHGINRRDFMKLCAALAATMGLSSKAAAEMAESVTNPQRPPVIWIGAQECT WP_000145410.1 MULTISPECIES: hydrogenase-2 small chain [Proteobacteria] Length: 372\nNumber of predicted TMHs: 1\nExp number of AAs in TMHs: 23.72226\nExp number, first 60 AAs: 0.22038\nTotal prob of N-in: 0.12994\noutside 1 331\nTMhelix 332 354\ninside 355 372

3300 GCF_001893155.1_ASM189315v1 Escherichia coli Proteobacteria; Gammaproteobacteria; Enterobacterales; Enterobacteriaceae; Escherichia MNNEETFYQAMRRQGVTRRSFLKYCSLAATSLGLGAGMAPKIAWALENKPRIPVVWIHGL WP_001295940.1 MULTISPECIES: hydrogenase [Enterobacteriaceae] Length: 372\nNumber of predicted TMHs: 2\nExp

number of AAs in TMHs: 37.2608899999999\nExp number, first 60 AAs: 19.52923\nTotal prob of N-in: 0.94351\nPOSSIBLE N-term signal sequence\ninside 1 19\nTMhelix 20 39\noutside 40 325\nTMhelix 326 348\ninside 349 372

3301 GCF_001892835.1_ASM189283v1 Escherichia coli Proteobacteria; Gammaproteobacteria; Enterobacterales; Enterobacteriaceae; Escherichia MNNEETFYQAMRRQGVTRRSFLKYCSLAATSLGLGAGMAPKIAWALENKPRIPVVWIHGL WP_001295940.1 MULTISPECIES: hydrogenase [Enterobacteriaceae] Length: 372\nNumber of predicted TMHs: 2\nExp number of AAs in TMHs: 37.2608899999999\nExp number, first 60 AAs: 19.52923\nTotal prob of N-in: 0.94351\nPOSSIBLE N-term signal sequence\ninside 1 19\nTMhelix 20 39\noutside 40 325\nTMhelix 326 348\ninside 349 372

3302 GCF_001893215.1_ASM189321v1 Escherichia coli Proteobacteria; Gammaproteobacteria; Enterobacterales; Enterobacteriaceae; Escherichia MTGDNTLIHSHGINRRDFMKLCAALAATMGLSSKAAAEMAESVTNPQRPPVIWIGAQECT WP_000145410.1 MULTISPECIES: hydrogenase-2 small chain [Proteobacteria] Length: 372\nNumber of predicted TMHs: 1\nExp number of AAs in TMHs: 23.72226\nExp number, first 60 AAs: 0.22038\nTotal prob of N-in: 0.12994\noutside 1 331\nTMhelix 332 354\ninside 355 372

3303 GCF_001893225.1_ASM189322v1 Escherichia coli Proteobacteria; Gammaproteobacteria; Enterobacterales; Enterobacteriaceae; Escherichia MTGDNTLIHSHGINRRDFMKLCAALAATMGLSSKAAAEMAESVTNPQRPPVIWIGAQECT WP_000145410.1 MULTISPECIES: hydrogenase-2 small chain [Proteobacteria] Length: 372\nNumber of predicted TMHs: 1\nExp number of AAs in TMHs: 23.72226\nExp number, first 60 AAs: 0.22038\nTotal prob of N-in: 0.12994\noutside 1 331\nTMhelix 332 354\ninside 355 372

3304 GCF_001893525.1_ASM189352v1 Escherichia coli Proteobacteria; Gammaproteobacteria; Enterobacterales; Enterobacteriaceae; Escherichia MTGDNTLIHSHGINRRDFMKLCAALAATMGLSSKAAAEMAESVTNPQRPPVIWIGAQECT WP_000145410.1 MULTISPECIES: hydrogenase-2 small chain [Proteobacteria] Length: 372\nNumber of predicted TMHs: 1\nExp number of AAs in TMHs: 23.72226\nExp number, first 60 AAs: 0.22038\nTotal prob of N-in: 0.12994\noutside 1 331\nTMhelix 332 354\ninside 355 372

3305 GCF_001893545.1_ASM189354v1 Escherichia coli Proteobacteria; Gammaproteobacteria; Enterobacterales; Enterobacteriaceae; Escherichia MNNEETFYQAMRRQGVTRRSFLKYCSLAATSLGLGAGMAPKIAWALENKPRIPVVWIHGL WP_001058323.1 MULTISPECIES: hydrogenase-1 small chain [Proteobacteria] Length: 372\nNumber of predicted TMHs: 2\nExp number of AAs in TMHs: 37.2608499999999\nExp number, first 60 AAs: 19.52923\nTotal prob of N-in: 0.94351\nPOSSIBLE N-term signal sequence\ninside 1 19\nTMhelix 20 39\noutside 40 325\nTMhelix 326 348\ninside 349 372

3306 GCF_001893525.1_ASM189352v1 Escherichia coli Proteobacteria; Gammaproteobacteria; Enterobacterales; Enterobacteriaceae; Escherichia MNNEETFYQAMRRQGVTRRSFLKYCSLAATSLGLGAGMAPKIAWALENKPRIPVVWIHGL WP_001058323.1 MULTISPECIES: hydrogenase-1 small chain [Proteobacteria] Length: 372\nNumber of predicted TMHs: 2\nExp number of AAs in TMHs: 37.2608499999999\nExp number, first 60 AAs: 19.52923\nTotal prob of N-in: 0.94351\nPOSSIBLE N-term signal sequence\ninside 1 19\nTMhelix 20 39\noutside 40 325\nTMhelix 326 348\ninside 349 372

3307 GCF_001893225.1_ASM189322v1 Escherichia coli Proteobacteria; Gammaproteobacteria; Enterobacterales; Enterobacteriaceae; Escherichia MNNEETFYQAMRRQGVTRRSFLKYCSLAATSLGLGAGMAPKIAWALENKPRIPVVWIHGL WP_001058323.1 MULTISPECIES: hydrogenase-1 small chain [Proteobacteria] Length: 372\nNumber of predicted TMHs: 2\nExp number of AAs in TMHs: 37.2608499999999\nExp number, first 60 AAs: 19.52923\nTotal prob of N-in: 0.94351\nPOSSIBLE N-term signal sequence\ninside 1 19\nTMhelix 20 39\noutside 40 325\nTMhelix 326 348\ninside 349 372

3308 GCF_001893315.1_ASM189331v1 Escherichia coli Proteobacteria; Gammaproteobacteria; Enterobacterales; Enterobacteriaceae; Escherichia MNNEETFYQAMRRQGVTRRSFLKYCSLAATSLGLGAGMAPKIAWALENKSRIPVVWIHGL WP_049595080.1 hydrogenase [Escherichia coli] Length: 372\nNumber of predicted TMHs: 2\nExp number of AAs in

TMHs: 37.1898399999999\nExp number, first 60 AAs:  19.44607\nTotal prob of N-in:   0.94159\nPOSSIBLE N-term signal sequence\ninside   1   19\nTMhelix   20   39\noutside   40   325\nTMhelix   326   348\ninside   349   372

3309 GCF_001893555.1_ASM189355v1 Escherichia coli Proteobacteria; Gammaproteobacteria; Enterobacterales; Enterobacteriaceae; Escherichia MTGDNTLIHSHGINRRDFMKLCAALAATMGLSSKAAAEMAESVTNPQRPPVIWIGAQECT WP_000145410.1 MULTISPECIES: hydrogenase-2 small chain [Proteobacteria] Length: 372\nNumber of predicted TMHs: 1\nExp number of AAs in TMHs: 23.72226\nExp number, first 60 AAs:  0.22038\nTotal prob of N-in:   0.12994\noutside   1   331\nTMhelix   332   354\ninside   355   372

3310 GCF_001893615.1_ASM189361v1 Escherichia coli Proteobacteria; Gammaproteobacteria; Enterobacterales; Enterobacteriaceae; Escherichia MTGDNTLIHSHGINRRDFMKLCAALAATMGLSSKAAAEMAESVTNPQRPPVIWIGAQECT WP_000145410.1 MULTISPECIES: hydrogenase-2 small chain [Proteobacteria] Length: 372\nNumber of predicted TMHs: 1\nExp number of AAs in TMHs: 23.72226\nExp number, first 60 AAs:  0.22038\nTotal prob of N-in:   0.12994\noutside   1   331\nTMhelix   332   354\ninside   355   372

3311 GCF_001893845.1_ASM189384v1 Escherichia coli Proteobacteria; Gammaproteobacteria; Enterobacterales; Enterobacteriaceae; Escherichia MNNEETFYQAMRRQGVTRRSFLKYCSLAATSLGLGAGMAPKIAWALENKPRIPVVWIHGL WP_001058323.1 MULTISPECIES: hydrogenase-1 small chain [Proteobacteria] Length: 372\nNumber of predicted TMHs: 2\nExp number of AAs in TMHs: 37.2608499999999\nExp number, first 60 AAs:  19.52923\nTotal prob of N-in:   0.94351\nPOSSIBLE N-term signal sequence\ninside   1   19\nTMhelix   20   39\noutside   40   325\nTMhelix   326   348\ninside   349   372

3312 GCF_001893935.1_ASM189393v1 Escherichia coli Proteobacteria; Gammaproteobacteria; Enterobacterales; Enterobacteriaceae; Escherichia MNNEETFYQAMRRQGVTRRSFLKYCSLAATSLGLGAGMAPKIAWALENKPRIPVVWIHGL WP_001058323.1 MULTISPECIES: hydrogenase-1 small chain [Proteobacteria] Length: 372\nNumber of predicted TMHs: 2\nExp number of AAs in TMHs: 37.2608499999999\nExp number, first 60 AAs:  19.52923\nTotal prob of N-in:   0.94351\nPOSSIBLE N-term signal sequence\ninside   1   19\nTMhelix   20   39\noutside   40   325\nTMhelix   326   348\ninside   349   372

3313 GCF_001893685.1_ASM189368v1 Escherichia coli Proteobacteria; Gammaproteobacteria; Enterobacterales; Enterobacteriaceae; Escherichia MNNEETFYQAMRRQGVTRRSFLKYCSLAATSLGLGAGMAPKIAWALENKPRIPVVWIHGL WP_001058323.1 MULTISPECIES: hydrogenase-1 small chain [Proteobacteria] Length: 372\nNumber of predicted TMHs: 2\nExp number of AAs in TMHs: 37.2608499999999\nExp number, first 60 AAs:  19.52923\nTotal prob of N-in:   0.94351\nPOSSIBLE N-term signal sequence\ninside   1   19\nTMhelix   20   39\noutside   40   325\nTMhelix   326   348\ninside   349   372

3314 GCF_001893705.1_ASM189370v1 Escherichia coli Proteobacteria; Gammaproteobacteria; Enterobacterales; Enterobacteriaceae; Escherichia MNNEETFYQAMRRQGVTRRSFLKYCSLAATSLGLGAGMAPKIAWALENKPRIPVVWIHGL WP_001058323.1 MULTISPECIES: hydrogenase-1 small chain [Proteobacteria] Length: 372\nNumber of predicted TMHs: 2\nExp number of AAs in TMHs: 37.2608499999999\nExp number, first 60 AAs:  19.52923\nTotal prob of N-in:   0.94351\nPOSSIBLE N-term signal sequence\ninside   1   19\nTMhelix   20   39\noutside   40   325\nTMhelix   326   348\ninside   349   372

3315 GCF_001893915.1_ASM189391v1 Escherichia coli Proteobacteria; Gammaproteobacteria; Enterobacterales; Enterobacteriaceae; Escherichia MTGDNTLIHSHGINRRDFMKLCAALAATMGLSSKAAAEMAESVTNPQRPPVIWIGAQECT WP_000145410.1 MULTISPECIES: hydrogenase-2 small chain [Proteobacteria] Length: 372\nNumber of predicted TMHs: 1\nExp number of AAs in TMHs: 23.72226\nExp number, first 60 AAs:  0.22038\nTotal prob of N-in:   0.12994\noutside   1   331\nTMhelix   332   354\ninside   355   372

3316 GCF_001893985.1_ASM189398v1 Escherichia coli Proteobacteria; Gammaproteobacteria; Enterobacterales; Enterobacteriaceae; Escherichia MTGDNTLIHSHGINRRDFMKLCAALAATMGLSSKAAAEMAESVTNPQRPPVIWIGAQECT WP_000145410.1 MULTISPECIES: hydrogenase-2 small chain [Proteobacteria] Length: 372\nNumber of predicted TMHs:

1\nExp number of AAs in TMHs: 23.72226\nExp number, first 60 AAs: 0.22038\nTotal prob of N-in: 0.12994\noutside 1 331\nTMhelix 332 354\ninside 355 372

3317 GCF_001893985.1_ASM189398v1 Escherichia coli Proteobacteria; Gammaproteobacteria; Enterobacterales; Enterobacteriaceae; Escherichia
MNNEETFYQAMRRQGVTRRSFLKYCSLAATSLGLGAGMAPKIAWALENKPRIPVVWIHGL WP_001058323.1
MULTISPECIES: hydrogenase-1 small chain [Proteobacteria] Length: 372\nNumber of predicted TMHs: 2\nExp number of AAs in TMHs: 37.2608499999999\nExp number, first 60 AAs: 19.52923\nTotal prob of N-in: 0.94351\nPOSSIBLE N-term signal sequence\ninside 1 19\nTMhelix 20 39\noutside 40 325\nTMhelix 326 348\ninside 349 372

3318 GCF_001894075.1_ASM189407v1 Escherichia coli Proteobacteria; Gammaproteobacteria; Enterobacterales; Enterobacteriaceae; Escherichia
MNNEETFYQAMRRQGVTRRSFLKYCSLAATSLGLGAGMAPKIAWALENKPRIPVVWIHGL WP_001295940.1
MULTISPECIES: hydrogenase [Enterobacteriaceae] Length: 372\nNumber of predicted TMHs: 2\nExp number of AAs in TMHs: 37.2608899999999\nExp number, first 60 AAs: 19.52923\nTotal prob of N-in: 0.94351\nPOSSIBLE N-term signal sequence\ninside 1 19\nTMhelix 20 39\noutside 40 325\nTMhelix 326 348\ninside 349 372

3319 GCF_001894315.1_ASM189431v1 Escherichia coli Proteobacteria; Gammaproteobacteria; Enterobacterales; Enterobacteriaceae; Escherichia
MTGDNTLIHSHGINRRDFMKLCAALAATMGLSSKAAAEMAESVTNPQRPPVIWIGAQECT WP_000145410.1
MULTISPECIES: hydrogenase-2 small chain [Proteobacteria] Length: 372\nNumber of predicted TMHs: 1\nExp number of AAs in TMHs: 23.72226\nExp number, first 60 AAs: 0.22038\nTotal prob of N-in: 0.12994\noutside 1 331\nTMhelix 332 354\ninside 355 372

3320 GCF_001894325.1_ASM189432v1 Escherichia coli Proteobacteria; Gammaproteobacteria; Enterobacterales; Enterobacteriaceae; Escherichia
MTGDNTLIHSHGINRRDFMKLCAALAATMGLSSKAAAEMAESVTNPQRPPVIWIGAQECT WP_000145410.1
MULTISPECIES: hydrogenase-2 small chain [Proteobacteria] Length: 372\nNumber of predicted TMHs: 1\nExp number of AAs in TMHs: 23.72226\nExp number, first 60 AAs: 0.22038\nTotal prob of N-in: 0.12994\noutside 1 331\nTMhelix 332 354\ninside 355 372

3321 GCF_001894385.1_ASM189438v1 Escherichia coli Proteobacteria; Gammaproteobacteria; Enterobacterales; Enterobacteriaceae; Escherichia
MTGDNTLIHSHGINRRDFMKLCAALAATMGLSSKAAAEMAESVTNPQRPPVIWIGAQECT WP_000145410.1
MULTISPECIES: hydrogenase-2 small chain [Proteobacteria] Length: 372\nNumber of predicted TMHs: 1\nExp number of AAs in TMHs: 23.72226\nExp number, first 60 AAs: 0.22038\nTotal prob of N-in: 0.12994\noutside 1 331\nTMhelix 332 354\ninside 355 372

3322 GCF_001894325.1_ASM189432v1 Escherichia coli Proteobacteria; Gammaproteobacteria; Enterobacterales; Enterobacteriaceae; Escherichia
MNNEETFYQAMRRQGVTRRSFLKYCSLAATSLGLGAGMAPKIAWALENKPRIPVVWIHGL WP_001058323.1
MULTISPECIES: hydrogenase-1 small chain [Proteobacteria] Length: 372\nNumber of predicted TMHs: 2\nExp number of AAs in TMHs: 37.2608499999999\nExp number, first 60 AAs: 19.52923\nTotal prob of N-in: 0.94351\nPOSSIBLE N-term signal sequence\ninside 1 19\nTMhelix 20 39\noutside 40 325\nTMhelix 326 348\ninside 349 372

3323 GCF_001894315.1_ASM189431v1 Escherichia coli Proteobacteria; Gammaproteobacteria; Enterobacterales; Enterobacteriaceae; Escherichia
MNNEETFYQAMRRQGVTRRSFLKYCSLAATSLGLGAGMAPKIAWALENKPRIPVVWIHGL WP_001058323.1
MULTISPECIES: hydrogenase-1 small chain [Proteobacteria] Length: 372\nNumber of predicted TMHs: 2\nExp number of AAs in TMHs: 37.2608499999999\nExp number, first 60 AAs: 19.52923\nTotal prob of N-in: 0.94351\nPOSSIBLE N-term signal sequence\ninside 1 19\nTMhelix 20 39\noutside 40 325\nTMhelix 326 348\ninside 349 372

3324 GCF_001894385.1_ASM189438v1 Escherichia coli Proteobacteria; Gammaproteobacteria; Enterobacterales; Enterobacteriaceae; Escherichia
MNNEETFYQAMRRQGVTRRSFLKYCSLAATSLGLGAGMAPKIAWALENKPRIPVVWIHGL WP_001058323.1
MULTISPECIES: hydrogenase-1 small chain [Proteobacteria] Length: 372\nNumber of predicted TMHs: 2\nExp number of AAs in TMHs: 37.2608499999999\nExp number, first 60 AAs: 19.52923\nTotal prob of N-in:

0.94351\nPOSSIBLE N-term signal sequence\ninside 1 19\nTMhelix 20 39\noutside 40 325\nTMhelix 326 348\ninside 349 372

3325 GCF_001900335.1_ASM190033v1 Escherichia coli Proteobacteria; Gammaproteobacteria; Enterobacterales; Enterobacteriaceae; Escherichia MTGDNTLIHSHGINRRDFMKLCAALAATMGLSSKAAAEMAESVTNPQRPPVIWIGAQECT WP_000145410.1 MULTISPECIES: hydrogenase-2 small chain [Proteobacteria] Length: 372\nNumber of predicted TMHs: 1\nExp number of AAs in TMHs: 23.72226\nExp number, first 60 AAs: 0.22038\nTotal prob of N-in: 0.12994\noutside 1 331\nTMhelix 332 354\ninside 355 372

3326 GCF_001900355.1_ASM190035v1 Escherichia coli Proteobacteria; Gammaproteobacteria; Enterobacterales; Enterobacteriaceae; Escherichia MTGDNTLIHSHGINRRDFMKLCAALAATMGLSSKAAAEMAESVTNPQRPPVIWIGAQECT WP_000145410.1 MULTISPECIES: hydrogenase-2 small chain [Proteobacteria] Length: 372\nNumber of predicted TMHs: 1\nExp number of AAs in TMHs: 23.72226\nExp number, first 60 AAs: 0.22038\nTotal prob of N-in: 0.12994\noutside 1 331\nTMhelix 332 354\ninside 355 372

3327 GCF_001900295.1_ASM190029v1 Escherichia coli Proteobacteria; Gammaproteobacteria; Enterobacterales; Enterobacteriaceae; Escherichia MNNEETFYQAMRRQGVTRRSFLKYCSLAATSLGLGAGMAPKIAWALENKPRIPVVWIHGL WP_001058323.1 MULTISPECIES: hydrogenase-1 small chain [Proteobacteria] Length: 372\nNumber of predicted TMHs: 2\nExp number of AAs in TMHs: 37.2608499999999\nExp number, first 60 AAs: 19.52923\nTotal prob of N-in: 0.94351\nPOSSIBLE N-term signal sequence\ninside 1 19\nTMhelix 20 39\noutside 40 325\nTMhelix 326 348\ninside 349 372

3328 GCF_001900355.1_ASM190035v1 Escherichia coli Proteobacteria; Gammaproteobacteria; Enterobacterales; Enterobacteriaceae; Escherichia MNNEETFYQAMRRQGVTRRSFLKYCSLAATSLGLGAGMAPKIAWALENKPRIPVVWIHGL WP_001058323.1 MULTISPECIES: hydrogenase-1 small chain [Proteobacteria] Length: 372\nNumber of predicted TMHs: 2\nExp number of AAs in TMHs: 37.2608499999999\nExp number, first 60 AAs: 19.52923\nTotal prob of N-in: 0.94351\nPOSSIBLE N-term signal sequence\ninside 1 19\nTMhelix 20 39\noutside 40 325\nTMhelix 326 348\ninside 349 372

3329 GCF_001894505.1_ASM189450v1 Escherichia coli Proteobacteria; Gammaproteobacteria; Enterobacterales; Enterobacteriaceae; Escherichia MNNEETFYQAMRRQGVTRRSFLKYCSLAATSLGLGAGMAPKIAWALENKPRIPVVWIHGL WP_001058323.1 MULTISPECIES: hydrogenase-1 small chain [Proteobacteria] Length: 372\nNumber of predicted TMHs: 2\nExp number of AAs in TMHs: 37.2608499999999\nExp number, first 60 AAs: 19.52923\nTotal prob of N-in: 0.94351\nPOSSIBLE N-term signal sequence\ninside 1 19\nTMhelix 20 39\noutside 40 325\nTMhelix 326 348\ninside 349 372

3330 GCF_001900395.1_ASM190039v1 Escherichia coli Proteobacteria; Gammaproteobacteria; Enterobacterales; Enterobacteriaceae; Escherichia MTGDNTLIHSHGINRRDFMKLCAALAATMGLSSKAAAEMAESVTNPQRPPVIWIGAQECT WP_000145410.1 MULTISPECIES: hydrogenase-2 small chain [Proteobacteria] Length: 372\nNumber of predicted TMHs: 1\nExp number of AAs in TMHs: 23.72226\nExp number, first 60 AAs: 0.22038\nTotal prob of N-in: 0.12994\noutside 1 331\nTMhelix 332 354\ninside 355 372

3331 GCF_001900575.1_ASM190057v1 Escherichia coli Proteobacteria; Gammaproteobacteria; Enterobacterales; Enterobacteriaceae; Escherichia MNNEETFYQAMRRQGVTRRSFLKYCSLAATSLGLGAGMAPKIAWALENKPRIPVVWIHGL WP_001058323.1 MULTISPECIES: hydrogenase-1 small chain [Proteobacteria] Length: 372\nNumber of predicted TMHs: 2\nExp number of AAs in TMHs: 37.2608499999999\nExp number, first 60 AAs: 19.52923\nTotal prob of N-in: 0.94351\nPOSSIBLE N-term signal sequence\ninside 1 19\nTMhelix 20 39\noutside 40 325\nTMhelix 326 348\ninside 349 372

3332 GCF_001900435.1_ASM190043v1 Escherichia coli Proteobacteria; Gammaproteobacteria; Enterobacterales; Enterobacteriaceae; Escherichia MNNEETFYQAMRRQGVTRRSFLKYCSLAATSLGLGAGMAPKIAWALENKPRIPVVWIHGL WP_001058323.1 MULTISPECIES: hydrogenase-1 small chain [Proteobacteria] Length: 372\nNumber of predicted TMHs: 2\nExp number of AAs in TMHs: 37.2608499999999\nExp number, first 60 AAs: 19.52923\nTotal prob of N-in:

0.94351\nPOSSIBLE N-term signal sequence\ninside 1 19\nTMhelix 20 39\noutside 40 325\nTMhelix 326 348\ninside 349 372

3333 GCF_001900715.1_ASM190071v1 Escherichia coli Proteobacteria; Gammaproteobacteria; Enterobacterales; Enterobacteriaceae; Escherichia MTGDNTLIHSHGINRRDFMKLCAALAATMGLSSKAAAEMAESVTNPQRPPVIWIGAQECT WP_000145410.1 MULTISPECIES: hydrogenase-2 small chain [Proteobacteria] Length: 372\nNumber of predicted TMHs: 1\nExp number of AAs in TMHs: 23.72226\nExp number, first 60 AAs: 0.22038\nTotal prob of N-in: 0.12994\noutside 1 331\nTMhelix 332 354\ninside 355 372

3334 GCF_001900735.1_ASM190073v1 Escherichia coli Proteobacteria; Gammaproteobacteria; Enterobacterales; Enterobacteriaceae; Escherichia MTGDNTLIHSHGINRRDFMKLCAALAATMGLSSKAAAEMAESVTNPQRPPVIWIGAQECT WP_000145410.1 MULTISPECIES: hydrogenase-2 small chain [Proteobacteria] Length: 372\nNumber of predicted TMHs: 1\nExp number of AAs in TMHs: 23.72226\nExp number, first 60 AAs: 0.22038\nTotal prob of N-in: 0.12994\noutside 1 331\nTMhelix 332 354\ninside 355 372

3335 GCF_001900715.1_ASM190071v1 Escherichia coli Proteobacteria; Gammaproteobacteria; Enterobacterales; Enterobacteriaceae; Escherichia MNNEETFYQAMRRQGVTRRSFLKYCSLAATSLGLGAGMAPKIAWALENKPRIPVVWIHGL WP_001058323.1 MULTISPECIES: hydrogenase-1 small chain [Proteobacteria] Length: 372\nNumber of predicted TMHs: 2\nExp number of AAs in TMHs: 37.2608499999999\nExp number, first 60 AAs: 19.52923\nTotal prob of N-in: 0.94351\nPOSSIBLE N-term signal sequence\ninside 1 19\nTMhelix 20 39\noutside 40 325\nTMhelix 326 348\ninside 349 372

3336 GCF_001900735.1_ASM190073v1 Escherichia coli Proteobacteria; Gammaproteobacteria; Enterobacterales; Enterobacteriaceae; Escherichia MNNEETFYQAMRRQGVTRRSFLKYCSLAATSLGLGAGMAPKIAWALENKPRIPVVWIHGL WP_001058323.1 MULTISPECIES: hydrogenase-1 small chain [Proteobacteria] Length: 372\nNumber of predicted TMHs: 2\nExp number of AAs in TMHs: 37.2608499999999\nExp number, first 60 AAs: 19.52923\nTotal prob of N-in: 0.94351\nPOSSIBLE N-term signal sequence\ninside 1 19\nTMhelix 20 39\noutside 40 325\nTMhelix 326 348\ninside 349 372

3337 GCF_001900535.1_ASM190053v1 Escherichia coli Proteobacteria; Gammaproteobacteria; Enterobacterales; Enterobacteriaceae; Escherichia MNNEETFYQAMRRQGVTRRSFLKYCSLAATSLGLGAGMAPKIAWALENKPRIPVVWIHGL WP_001058323.1 MULTISPECIES: hydrogenase-1 small chain [Proteobacteria] Length: 372\nNumber of predicted TMHs: 2\nExp number of AAs in TMHs: 37.2608499999999\nExp number, first 60 AAs: 19.52923\nTotal prob of N-in: 0.94351\nPOSSIBLE N-term signal sequence\ninside 1 19\nTMhelix 20 39\noutside 40 325\nTMhelix 326 348\ninside 349 372

3338 GCF_001901025.1_ASM190102v1 Escherichia coli Proteobacteria; Gammaproteobacteria; Enterobacterales; Enterobacteriaceae; Escherichia MNNEETFYQAMRRQGVTRRSFLKYCSLAATSLGLGAGMAPKIAWALENKPRIPVVWIHGL WP_053264900.1 hydrogenase [Escherichia coli] Length: 372\nNumber of predicted TMHs: 2\nExp number of AAs in TMHs: 37.24714\nExp number, first 60 AAs: 19.52921\nTotal prob of N-in: 0.94351\nPOSSIBLE N-term signal sequence\ninside 1 19\nTMhelix 20 39\noutside 40 325\nTMhelix 326 348\ninside 349 372

3339 GCF_001900795.1_ASM190079v1 Escherichia coli Proteobacteria; Gammaproteobacteria; Enterobacterales; Enterobacteriaceae; Escherichia MNNEETFYQAMRRQGVTRRSFLKYCSLAATSLGLGAGMAPKIAWALENKPRIPVVWIHGL WP_073511893.1 hydrogenase [Escherichia coli] Length: 372\nNumber of predicted TMHs: 2\nExp number of AAs in TMHs: 37.2607899999999\nExp number, first 60 AAs: 19.52923\nTotal prob of N-in: 0.94351\nPOSSIBLE N-term signal sequence\ninside 1 19\nTMhelix 20 39\noutside 40 325\nTMhelix 326 348\ninside 349 372

3340 GCF_001901125.1_ASM190112v1 Escherichia coli Proteobacteria; Gammaproteobacteria; Enterobacterales; Enterobacteriaceae; Escherichia MTGDNTLIHSHGINRRDFMKLCAALAATMGLSSKAAAEMAESVTNPQRPPVIWIGAQECT WP_000145410.1 MULTISPECIES: hydrogenase-2 small chain [Proteobacteria] Length: 372\nNumber of predicted TMHs: 1\nExp number of AAs in TMHs: 23.72226\nExp number, first 60 AAs: 0.22038\nTotal prob of N-in: 0.12994\noutside 1 331\nTMhelix 332 354\ninside 355 372

3341 GCF_001901085.1_ASM190108v1 Escherichia coli Proteobacteria; Gammaproteobacteria; Enterobacterales; Enterobacteriaceae; Escherichia
MTGDNTLIHSHGINRRDFMKLCAALAATMGLSSKAAAEMAESVTNPQRPPVIWIGAQECT WP_000145410.1
MULTISPECIES: hydrogenase-2 small chain [Proteobacteria] Length: 372\nNumber of predicted TMHs: 1\nExp number of AAs in TMHs: 23.72226\nExp number, first 60 AAs: 0.22038\nTotal prob of N-in: 0.12994\noutside 1 331\nTMhelix 332 354\ninside 355 372

3342 GCF_001901165.1_ASM190116v1 Escherichia coli Proteobacteria; Gammaproteobacteria; Enterobacterales; Enterobacteriaceae; Escherichia
MTGDNTLIHSHGINRRDFMKLCAALAATMGLSSKAAAEMAESVTNPQRPPVIWIGAQECT WP_000145410.1
MULTISPECIES: hydrogenase-2 small chain [Proteobacteria] Length: 372\nNumber of predicted TMHs: 1\nExp number of AAs in TMHs: 23.72226\nExp number, first 60 AAs: 0.22038\nTotal prob of N-in: 0.12994\noutside 1 331\nTMhelix 332 354\ninside 355 372

3343 GCF_001901085.1_ASM190108v1 Escherichia coli Proteobacteria; Gammaproteobacteria; Enterobacterales; Enterobacteriaceae; Escherichia
MNNEETFYQAMRRQGVTRRSFLKYCSLAATSLGLGAGMAPKIAWALENKPRIPVVWIHGL WP_001058323.1
MULTISPECIES: hydrogenase-1 small chain [Proteobacteria] Length: 372\nNumber of predicted TMHs: 2\nExp number of AAs in TMHs: 37.2608499999999\nExp number, first 60 AAs: 19.52923\nTotal prob of N-in: 0.94351\nPOSSIBLE N-term signal sequence\ninside 1 19\nTMhelix 20 39\noutside 40 325\nTMhelix 326 348\ninside 349 372

3344 GCF_001901125.1_ASM190112v1 Escherichia coli Proteobacteria; Gammaproteobacteria; Enterobacterales; Enterobacteriaceae; Escherichia
MNNEETFYQAMRRQGVTRRSFLKYCSLAATSLGLGAGMAPKIAWALENKPRIPVVWIHGL WP_001058323.1
MULTISPECIES: hydrogenase-1 small chain [Proteobacteria] Length: 372\nNumber of predicted TMHs: 2\nExp number of AAs in TMHs: 37.2608499999999\nExp number, first 60 AAs: 19.52923\nTotal prob of N-in: 0.94351\nPOSSIBLE N-term signal sequence\ninside 1 19\nTMhelix 20 39\noutside 40 325\nTMhelix 326 348\ninside 349 372

3345 GCF_001902735.1_ASM190273v1 Escherichia coli Proteobacteria; Gammaproteobacteria; Enterobacterales; Enterobacteriaceae; Escherichia
MTGDNTLIHSHGINRRDFMKLCAALAATMGLSSKAAAEMAESVTNPQRPPVIWIGAQECT WP_000145410.1
MULTISPECIES: hydrogenase-2 small chain [Proteobacteria] Length: 372\nNumber of predicted TMHs: 1\nExp number of AAs in TMHs: 23.72226\nExp number, first 60 AAs: 0.22038\nTotal prob of N-in: 0.12994\noutside 1 331\nTMhelix 332 354\ninside 355 372

3346 GCF_001902675.1_ASM190267v1 Escherichia coli Proteobacteria; Gammaproteobacteria; Enterobacterales; Enterobacteriaceae; Escherichia
MTGDNTLIHSHGINRRDFMKLCAALAATMGLSSKAAAEMAESVTNPQRPPVIWIGAQECT WP_000145410.1
MULTISPECIES: hydrogenase-2 small chain [Proteobacteria] Length: 372\nNumber of predicted TMHs: 1\nExp number of AAs in TMHs: 23.72226\nExp number, first 60 AAs: 0.22038\nTotal prob of N-in: 0.12994\noutside 1 331\nTMhelix 332 354\ninside 355 372

3347 GCF_001907325.1_ASM190732v1 Escherichia coli Proteobacteria; Gammaproteobacteria; Enterobacterales; Enterobacteriaceae; Escherichia
MTGDNTLIHSHGINRRDFMKLCAALAATMGLSSKAAAEMAESVTNPQRPPVIWIGAQECT WP_000145410.1
MULTISPECIES: hydrogenase-2 small chain [Proteobacteria] Length: 372\nNumber of predicted TMHs: 1\nExp number of AAs in TMHs: 23.72226\nExp number, first 60 AAs: 0.22038\nTotal prob of N-in: 0.12994\noutside 1 331\nTMhelix 332 354\ninside 355 372

3348 GCF_001907335.1_ASM190733v1 Escherichia coli Proteobacteria; Gammaproteobacteria; Enterobacterales; Enterobacteriaceae; Escherichia
MTGDNTLIHSHGINRRDFMKLCAALAATMGLSSKAAAEMAESVTNPQRPPVIWIGAQECT WP_000145410.1
MULTISPECIES: hydrogenase-2 small chain [Proteobacteria] Length: 372\nNumber of predicted TMHs: 1\nExp number of AAs in TMHs: 23.72226\nExp number, first 60 AAs: 0.22038\nTotal prob of N-in: 0.12994\noutside 1 331\nTMhelix 332 354\ninside 355 372

3349 GCF_001907435.1_ASM190743v1 Escherichia coli Proteobacteria; Gammaproteobacteria; Enterobacterales; Enterobacteriaceae; Escherichia
MTGDNTLIHSHGINRRDFMKLCAALAATMGLSSKAAAEMAESVTNPQRPPVIWIGAQECT WP_000145410.1

MULTISPECIES: hydrogenase-2 small chain [Proteobacteria] Length: 372\nNumber of predicted TMHs: 1\nExp number of AAs in TMHs: 23.72226\nExp number, first 60 AAs: 0.22038\nTotal prob of N-in: 0.12994\noutside 1 331\nTMhelix 332 354\ninside 355 372

3350 GCF_001901465.1_ASM190146v1 Escherichia coli Proteobacteria; Gammaproteobacteria; Enterobacterales; Enterobacteriaceae; Escherichia
MNNEETFYQAMRRQGVTRRSFLKYCSLAATSLGLGAGMAPKIAWALENKPRIPVVWIHGL WP_001058323.1
MULTISPECIES: hydrogenase-1 small chain [Proteobacteria] Length: 372\nNumber of predicted TMHs: 2\nExp number of AAs in TMHs: 37.2608499999999\nExp number, first 60 AAs: 19.52923\nTotal prob of N-in: 0.94351\nPOSSIBLE N-term signal sequence\ninside 1 19\nTMhelix 20 39\noutside 40 325\nTMhelix 326 348\ninside 349 372

3351 GCF_001907385.1_ASM190738v1 Escherichia coli Proteobacteria; Gammaproteobacteria; Enterobacterales; Enterobacteriaceae; Escherichia
MNNEETFYQAMRRQGVTRRSFLKYCSLAATSLGLGAGMAPKIAWALENKPRIPVVWIHGL WP_001058323.1
MULTISPECIES: hydrogenase-1 small chain [Proteobacteria] Length: 372\nNumber of predicted TMHs: 2\nExp number of AAs in TMHs: 37.2608499999999\nExp number, first 60 AAs: 19.52923\nTotal prob of N-in: 0.94351\nPOSSIBLE N-term signal sequence\ninside 1 19\nTMhelix 20 39\noutside 40 325\nTMhelix 326 348\ninside 349 372

3352 GCF_001902655.1_ASM190265v1 Escherichia coli Proteobacteria; Gammaproteobacteria; Enterobacterales; Enterobacteriaceae; Escherichia
MNNEETFYQAMRRQGVTRRSFLKYCSLAATSLGLGAGMAPKIAWALENKPRIPVVWIHGL WP_001563066.1
hydrogenase-1 small chain [Escherichia coli] Length: 372\nNumber of predicted TMHs: 2\nExp number of AAs in TMHs: 37.39588\nExp number, first 60 AAs: 19.52908\nTotal prob of N-in: 0.94358\nPOSSIBLE N-term signal sequence\ninside 1 19\nTMhelix 20 39\noutside 40 325\nTMhelix 326 348\ninside 349 372

3353 GCF_001901425.1_ASM190142v1 Escherichia coli Proteobacteria; Gammaproteobacteria; Enterobacterales; Enterobacteriaceae; Escherichia
MNNEETFYQAMRRQGVTRRSFLKYCSLAATSLGLGAGMAPKIAWALENKPRIPVVWIHGL WP_001295940.1
MULTISPECIES: hydrogenase [Enterobacteriaceae] Length: 372\nNumber of predicted TMHs: 2\nExp number of AAs in TMHs: 37.2608899999999\nExp number, first 60 AAs: 19.52923\nTotal prob of N-in: 0.94351\nPOSSIBLE N-term signal sequence\ninside 1 19\nTMhelix 20 39\noutside 40 325\nTMhelix 326 348\ninside 349 372

3354 GCF_001910475.1_ASM191047v1 Escherichia coli Proteobacteria; Gammaproteobacteria; Enterobacterales; Enterobacteriaceae; Escherichia
MTGDNTLIHSHGINRRDFMKLCAALAATMGLSSKAAAEMAESVTNPQRPPVIWIGAQECT WP_000145410.1
MULTISPECIES: hydrogenase-2 small chain [Proteobacteria] Length: 372\nNumber of predicted TMHs: 1\nExp number of AAs in TMHs: 23.72226\nExp number, first 60 AAs: 0.22038\nTotal prob of N-in: 0.12994\noutside 1 331\nTMhelix 332 354\ninside 355 372

3355 GCF_001907935.1_ASM190793v1 Escherichia coli Proteobacteria; Gammaproteobacteria; Enterobacterales; Enterobacteriaceae; Escherichia
MTGDNTLIHSHGINRRDFMKLCAALAATMGLSSKAAAEMAESVTNPQRPPVIWIGAQECT WP_000145410.1
MULTISPECIES: hydrogenase-2 small chain [Proteobacteria] Length: 372\nNumber of predicted TMHs: 1\nExp number of AAs in TMHs: 23.72226\nExp number, first 60 AAs: 0.22038\nTotal prob of N-in: 0.12994\noutside 1 331\nTMhelix 332 354\ninside 355 372

3356 GCF_001907545.1_ASM190754v1 Escherichia coli Proteobacteria; Gammaproteobacteria; Enterobacterales; Enterobacteriaceae; Escherichia
MNNEETFYQAMRRQGVTRRSFLKYCSLAATSLGLGAGMAPKIAWALENKPRIPVVWIHGL WP_001058323.1
MULTISPECIES: hydrogenase-1 small chain [Proteobacteria] Length: 372\nNumber of predicted TMHs: 2\nExp number of AAs in TMHs: 37.2608499999999\nExp number, first 60 AAs: 19.52923\nTotal prob of N-in: 0.94351\nPOSSIBLE N-term signal sequence\ninside 1 19\nTMhelix 20 39\noutside 40 325\nTMhelix 326 348\ninside 349 372

3357 GCF_001910845.1_ASM191084v1 Escherichia coli Proteobacteria; Gammaproteobacteria; Enterobacterales; Enterobacteriaceae; Escherichia
MTGDNTLIHSHGINRRDFMKLCAALAATMGLSSKAAAEMAESVTNPQRPPVIWIGAQECT WP_000145410.1
MULTISPECIES: hydrogenase-2 small chain [Proteobacteria] Length: 372\nNumber of predicted TMHs:

1\nExp number of AAs in TMHs: 23.72226\nExp number, first 60 AAs: 0.22038\nTotal prob of N-in: 0.12994\noutside 1 331\nTMhelix 332 354\ninside 355 372

3358 GCF_001910825.1_ASM191082v1 Escherichia coli Proteobacteria; Gammaproteobacteria; Enterobacterales; Enterobacteriaceae; Escherichia
MTGDNTLIHSHGINRRDFMKLCAALAATMGLSSKAAAEMAESVTNPQRPPVIWIGAQECT WP_000145410.1
MULTISPECIES: hydrogenase-2 small chain [Proteobacteria] Length: 372\nNumber of predicted TMHs: 1\nExp number of AAs in TMHs: 23.72226\nExp number, first 60 AAs: 0.22038\nTotal prob of N-in: 0.12994\noutside 1 331\nTMhelix 332 354\ninside 355 372

3359 GCF_001911245.1_ASM191124v1 Escherichia coli Proteobacteria; Gammaproteobacteria; Enterobacterales; Enterobacteriaceae; Escherichia
MNNEETFYQAMRRQGVTRRSFLKYCSLAATSLGLGAGMAPKIAWALENKPRIPVVWIHGL WP_001058323.1
MULTISPECIES: hydrogenase-1 small chain [Proteobacteria] Length: 372\nNumber of predicted TMHs: 2\nExp number of AAs in TMHs: 37.2608499999999\nExp number, first 60 AAs: 19.52923\nTotal prob of N-in: 0.94351\nPOSSIBLE N-term signal sequence\ninside 1 19\nTMhelix 20 39\noutside 40 325\nTMhelix 326 348\ninside 349 372

3360 GCF_001911305.1_ASM191130v1 Escherichia coli Proteobacteria; Gammaproteobacteria; Enterobacterales; Enterobacteriaceae; Escherichia
MNNEETFYQAMRRQGVTRRSFLKYCSLAATSLGLGAGMAPKIAWALENKPRIPVVWIHGL WP_001058320.1
hydrogenase [Escherichia coli] Length: 372\nNumber of predicted TMHs: 2\nExp number of AAs in TMHs: 37.26082\nExp number, first 60 AAs: 19.52923\nTotal prob of N-in: 0.94351\nPOSSIBLE N-term signal sequence\ninside 1 19\nTMhelix 20 39\noutside 40 325\nTMhelix 326 348\ninside 349 372

3361 GCF_001911095.1_ASM191109v1 Escherichia coli Proteobacteria; Gammaproteobacteria; Enterobacterales; Enterobacteriaceae; Escherichia
MNNEETFYQAMRRQGVTRRSFLKYCSLAATSLGLGAGMAPKIAWALENKPRIPVVWIHGL WP_001058323.1
MULTISPECIES: hydrogenase-1 small chain [Proteobacteria] Length: 372\nNumber of predicted TMHs: 2\nExp number of AAs in TMHs: 37.2608499999999\nExp number, first 60 AAs: 19.52923\nTotal prob of N-in: 0.94351\nPOSSIBLE N-term signal sequence\ninside 1 19\nTMhelix 20 39\noutside 40 325\nTMhelix 326 348\ninside 349 372

3362 GCF_001911405.1_ASM191140v1 Escherichia coli Proteobacteria; Gammaproteobacteria; Enterobacterales; Enterobacteriaceae; Escherichia
MTGDNTLIHSHGINRRDFMKLCAALAATMGLSSKAAAEMAESVTNPQRPPVIWIGAQECT WP_000145410.1
MULTISPECIES: hydrogenase-2 small chain [Proteobacteria] Length: 372\nNumber of predicted TMHs: 1\nExp number of AAs in TMHs: 23.72226\nExp number, first 60 AAs: 0.22038\nTotal prob of N-in: 0.12994\noutside 1 331\nTMhelix 332 354\ninside 355 372

3363 GCF_001911405.1_ASM191140v1 Escherichia coli Proteobacteria; Gammaproteobacteria; Enterobacterales; Enterobacteriaceae; Escherichia
MNNEETFYQAMRRQGVTRRSFLKYCSLAATSLGLGAGMAPKIAWALENKPRIPVVWIHGL WP_001353311.1
MULTISPECIES: hydrogenase [Escherichia] Length: 372\nNumber of predicted TMHs: 2\nExp number of AAs in TMHs: 37.2638699999999\nExp number, first 60 AAs: 19.52924\nTotal prob of N-in: 0.94351\nPOSSIBLE N-term signal sequence\ninside 1 19\nTMhelix 20 39\noutside 40 325\nTMhelix 326 348\ninside 349 372

3364 GCF_001911225.1_ASM191122v1 Escherichia coli Proteobacteria; Gammaproteobacteria; Enterobacterales; Enterobacteriaceae; Escherichia
MNNEETFYQAMRRQGVTRRSFLKYCSLAATSLGLGAGMAPKIAWALENKPRIPVVWIHGL WP_001058323.1
MULTISPECIES: hydrogenase-1 small chain [Proteobacteria] Length: 372\nNumber of predicted TMHs: 2\nExp number of AAs in TMHs: 37.2608499999999\nExp number, first 60 AAs: 19.52923\nTotal prob of N-in: 0.94351\nPOSSIBLE N-term signal sequence\ninside 1 19\nTMhelix 20 39\noutside 40 325\nTMhelix 326 348\ninside 349 372

3365 GCF_001911175.1_ASM191117v1 Escherichia coli Proteobacteria; Gammaproteobacteria; Enterobacterales; Enterobacteriaceae; Escherichia
MNNEETFYQAMRRQGVTRRSFLKYCSLAATSLGLGAGMAPKIAWALENKPRIPVVWIHGL WP_001058320.1
hydrogenase [Escherichia coli] Length: 372\nNumber of predicted TMHs: 2\nExp number of AAs in TMHs: 37.26082\nExp number, first 60 AAs: 19.52923\nTotal prob of N-in: 0.94351\nPOSSIBLE N-term signal sequence\ninside 1 19\nTMhelix 20 39\noutside 40 325\nTMhelix 326 348\ninside 349 372

3366 GCF_001911805.1_ASM191180v1 Escherichia coli Proteobacteria; Gammaproteobacteria; Enterobacterales; Enterobacteriaceae; Escherichia MTGDNTLIHSHGINRRDFMKLCAALAATMGLSSKAAAEMAESVTNPQRPPVIWIGAQECT WP_000145410.1 MULTISPECIES: hydrogenase-2 small chain [Proteobacteria] Length: 372\nNumber of predicted TMHs: 1\nExp number of AAs in TMHs: 23.72226\nExp number, first 60 AAs: 0.22038\nTotal prob of N-in: 0.12994\noutside 1 331\nTMhelix 332 354\ninside 355 372

3367 GCF_001911665.1_ASM191166v1 Escherichia coli Proteobacteria; Gammaproteobacteria; Enterobacterales; Enterobacteriaceae; Escherichia MNNEETFYQAMRRQGVTRRSFLKYCSLAATSLGLGAGMAPKIAWALENKPRIPVVWIHGL WP_001058323.1 MULTISPECIES: hydrogenase-1 small chain [Proteobacteria] Length: 372\nNumber of predicted TMHs: 2\nExp number of AAs in TMHs: 37.2608499999999\nExp number, first 60 AAs: 19.52923\nTotal prob of N-in: 0.94351\nPOSSIBLE N-term signal sequence\ninside 1 19\nTMhelix 20 39\noutside 40 325\nTMhelix 326 348\ninside 349 372

3368 GCF_001911755.1_ASM191175v1 Escherichia coli Proteobacteria; Gammaproteobacteria; Enterobacterales; Enterobacteriaceae; Escherichia MNNEETFYQAMRRQGVTRRSFLKYCSLAATSLGLGAGMAPKIAWALENKPRIPVVWIHGL WP_001058323.1 MULTISPECIES: hydrogenase-1 small chain [Proteobacteria] Length: 372\nNumber of predicted TMHs: 2\nExp number of AAs in TMHs: 37.2608499999999\nExp number, first 60 AAs: 19.52923\nTotal prob of N-in: 0.94351\nPOSSIBLE N-term signal sequence\ninside 1 19\nTMhelix 20 39\noutside 40 325\nTMhelix 326 348\ninside 349 372

3369 GCF_001911495.1_ASM191149v1 Escherichia coli Proteobacteria; Gammaproteobacteria; Enterobacterales; Enterobacteriaceae; Escherichia MNNEETFYQAMRRQGVTRRSFLKYCSLAATSLGLGAGMAPKIAWALENKPRIPVVWIHGL WP_001058323.1 MULTISPECIES: hydrogenase-1 small chain [Proteobacteria] Length: 372\nNumber of predicted TMHs: 2\nExp number of AAs in TMHs: 37.2608499999999\nExp number, first 60 AAs: 19.52923\nTotal prob of N-in: 0.94351\nPOSSIBLE N-term signal sequence\ninside 1 19\nTMhelix 20 39\noutside 40 325\nTMhelix 326 348\ninside 349 372

3370 GCF_002156845.1_ASM215684v1 Escherichia coli Proteobacteria; Gammaproteobacteria; Enterobacterales; Enterobacteriaceae; Escherichia MTGDNTLIHSHGINRRDFMKLCAALAATMGLSSKAAAEMAESVTNPQRPPVIWIGAQECT WP_000145410.1 MULTISPECIES: hydrogenase-2 small chain [Proteobacteria] Length: 372\nNumber of predicted TMHs: 1\nExp number of AAs in TMHs: 23.72226\nExp number, first 60 AAs: 0.22038\nTotal prob of N-in: 0.12994\noutside 1 331\nTMhelix 332 354\ninside 355 372

3371 GCF_002157245.1_ASM215724v1 Escherichia coli Proteobacteria; Gammaproteobacteria; Enterobacterales; Enterobacteriaceae; Escherichia MTGDNTLIHSHGINRRDFMKLCAALAATMGLSSKAAAEMVESVTNPQRPPVIWIGAQECT WP_000145423.1 hydrogenase 2 small subunit [Escherichia coli] Length: 372\nNumber of predicted TMHs: 1\nExp number of AAs in TMHs: 23.78201\nExp number, first 60 AAs: 0.28355\nTotal prob of N-in: 0.13257\noutside 1 331\nTMhelix 332 354\ninside 355 372

3372 GCF_002155835.1_ASM215583v1 Escherichia coli Proteobacteria; Gammaproteobacteria; Enterobacterales; Enterobacteriaceae; Escherichia MNNEETFYQAMRRQGVTRRSFLKYCSLAATSLGLGAGMAPKIAWALENKPRIPVVWIHGL WP_001058323.1 MULTISPECIES: hydrogenase-1 small chain [Proteobacteria] Length: 372\nNumber of predicted TMHs: 2\nExp number of AAs in TMHs: 37.2608499999999\nExp number, first 60 AAs: 19.52923\nTotal prob of N-in: 0.94351\nPOSSIBLE N-term signal sequence\ninside 1 19\nTMhelix 20 39\noutside 40 325\nTMhelix 326 348\ninside 349 372

3373 GCF_002156845.1_ASM215684v1 Escherichia coli Proteobacteria; Gammaproteobacteria; Enterobacterales; Enterobacteriaceae; Escherichia MNNEETFYQAMRRQGVTRRSFLKYCSLAATSLGLGAGMAPKIAWALENKPRIPVVWIHGL WP_001058323.1 MULTISPECIES: hydrogenase-1 small chain [Proteobacteria] Length: 372\nNumber of predicted TMHs: 2\nExp number of AAs in TMHs: 37.2608499999999\nExp number, first 60 AAs: 19.52923\nTotal prob of N-in: 0.94351\nPOSSIBLE N-term signal sequence\ninside 1 19\nTMhelix 20 39\noutside 40 325\nTMhelix 326 348\ninside 349 372

3374 GCF_002157245.1_ASM215724v1 Escherichia coli Proteobacteria; Gammaproteobacteria; Enterobacterales; Enterobacteriaceae; Escherichia
MNNEETFYQAMRRQGVTRRSFLKYCSLAATSLGLGAGMAPKIAWALENKPRIPVVWIHGL WP_001295940.1
MULTISPECIES: hydrogenase [Enterobacteriaceae] Length: 372\nNumber of predicted TMHs: 2\nExp number of AAs in TMHs: 37.2608899999999\nExp number, first 60 AAs: 19.52923\nTotal prob of N-in: 0.94351\nPOSSIBLE N-term signal sequence\ninside 1 19\nTMhelix 20 39\noutside 40 325\nTMhelix 326 348\ninside 349 372

3375 GCF_002155845.1_ASM215584v1 Escherichia coli Proteobacteria; Gammaproteobacteria; Enterobacterales; Enterobacteriaceae; Escherichia
MNNEETFYQAMRRQGVTRRSFLKYCSLAATSLGLGAGMAPKIAWALENKPRIPVVWIHGL WP_001058323.1
MULTISPECIES: hydrogenase-1 small chain [Proteobacteria] Length: 372\nNumber of predicted TMHs: 2\nExp number of AAs in TMHs: 37.2608499999999\nExp number, first 60 AAs: 19.52923\nTotal prob of N-in: 0.94351\nPOSSIBLE N-term signal sequence\ninside 1 19\nTMhelix 20 39\noutside 40 325\nTMhelix 326 348\ninside 349 372

3376 GCF_002160185.1_ASM216018v1 Escherichia coli Proteobacteria; Gammaproteobacteria; Enterobacterales; Enterobacteriaceae; Escherichia
MTGDNTLIHSHGINRRDFMKLCAALAATMGLSSKAAAEMAESVTNPQRPPVIWIGAQECT WP_000145410.1
MULTISPECIES: hydrogenase-2 small chain [Proteobacteria] Length: 372\nNumber of predicted TMHs: 1\nExp number of AAs in TMHs: 23.72226\nExp number, first 60 AAs: 0.22038\nTotal prob of N-in: 0.12994\noutside 1 331\nTMhelix 332 354\ninside 355 372

3377 GCF_002164095.1_ASM216409v1 Escherichia coli Proteobacteria; Gammaproteobacteria; Enterobacterales; Enterobacteriaceae; Escherichia
MTGDNTLIHSHGINRRDFMKLCAALAATMGLSSKAAAEMAESVTNPQRPPVIWIGAQECT WP_000145410.1
MULTISPECIES: hydrogenase-2 small chain [Proteobacteria] Length: 372\nNumber of predicted TMHs: 1\nExp number of AAs in TMHs: 23.72226\nExp number, first 60 AAs: 0.22038\nTotal prob of N-in: 0.12994\noutside 1 331\nTMhelix 332 354\ninside 355 372

3378 GCF_002164185.1_ASM216418v1 Escherichia coli Proteobacteria; Gammaproteobacteria; Enterobacterales; Enterobacteriaceae; Escherichia
MTGDNTLIHSHGINRRDFMKLCAALAATMGLSSKAAAEMAESVTNPQRPPVIWIGAQECT WP_000145410.1
MULTISPECIES: hydrogenase-2 small chain [Proteobacteria] Length: 372\nNumber of predicted TMHs: 1\nExp number of AAs in TMHs: 23.72226\nExp number, first 60 AAs: 0.22038\nTotal prob of N-in: 0.12994\noutside 1 331\nTMhelix 332 354\ninside 355 372

3379 GCF_002164105.1_ASM216410v1 Escherichia coli Proteobacteria; Gammaproteobacteria; Enterobacterales; Enterobacteriaceae; Escherichia
MTGDNTLIHSHGINRRDFMKLCAALAATMGLSSKAAAEMAESVTNPQRPPVIWIGAQECT WP_000145410.1
MULTISPECIES: hydrogenase-2 small chain [Proteobacteria] Length: 372\nNumber of predicted TMHs: 1\nExp number of AAs in TMHs: 23.72226\nExp number, first 60 AAs: 0.22038\nTotal prob of N-in: 0.12994\noutside 1 331\nTMhelix 332 354\ninside 355 372

3380 GCF_002164125.1_ASM216412v1 Escherichia coli Proteobacteria; Gammaproteobacteria; Enterobacterales; Enterobacteriaceae; Escherichia
MTGDNTLIHSHGINRRDFMKLCAALAATMGLSSKAAAEMAESVTNPQRPPVIWIGAQECT WP_000145410.1
MULTISPECIES: hydrogenase-2 small chain [Proteobacteria] Length: 372\nNumber of predicted TMHs: 1\nExp number of AAs in TMHs: 23.72226\nExp number, first 60 AAs: 0.22038\nTotal prob of N-in: 0.12994\noutside 1 331\nTMhelix 332 354\ninside 355 372

3381 GCF_002164195.1_ASM216419v1 Escherichia coli Proteobacteria; Gammaproteobacteria; Enterobacterales; Enterobacteriaceae; Escherichia
MTGDNTLIHSHGINRRDFMKLCAALAATMGLSSKAAAEMAESVTNPQRPPVIWIGAQECT WP_000145410.1
MULTISPECIES: hydrogenase-2 small chain [Proteobacteria] Length: 372\nNumber of predicted TMHs: 1\nExp number of AAs in TMHs: 23.72226\nExp number, first 60 AAs: 0.22038\nTotal prob of N-in: 0.12994\noutside 1 331\nTMhelix 332 354\ninside 355 372

3382 GCF_002164095.1_ASM216409v1 Escherichia coli Proteobacteria; Gammaproteobacteria; Enterobacterales; Enterobacteriaceae; Escherichia
MNNEETFYQAMRRQGVTRRSFLKYCSLAATSLGLGAGMAPKIAWALENKPRIPVVWIHGL WP_001301613.1

hydrogenase [Escherichia coli] Length: 372\nNumber of predicted TMHs: 2\nExp number of AAs in TMHs: 37.46598\nExp number, first 60 AAs: 19.53139\nTotal prob of N-in: 0.94364\nPOSSIBLE N-term signal sequence\ninside 1 19\nTMhelix 20 39\noutside 40 325\nTMhelix 326 348\ninside 349 372

3383 GCF_002164125.1_ASM216412v1 Escherichia coli Proteobacteria; Gammaproteobacteria; Enterobacterales; Enterobacteriaceae; Escherichia MNNEETFYQAMRRQGVTRRSFLKYCSLAATSLGLGAGMAPKIAWALENKPRIPVVWIHGL WP_001301613.1 hydrogenase [Escherichia coli] Length: 372\nNumber of predicted TMHs: 2\nExp number of AAs in TMHs: 37.46598\nExp number, first 60 AAs: 19.53139\nTotal prob of N-in: 0.94364\nPOSSIBLE N-term signal sequence\ninside 1 19\nTMhelix 20 39\noutside 40 325\nTMhelix 326 348\ninside 349 372

3384 GCF_002164415.1_ASM216441v1 Escherichia coli Proteobacteria; Gammaproteobacteria; Enterobacterales; Enterobacteriaceae; Escherichia MNNEETFYQAMRRQGVTRRSFLKYCSLAATSLGLGAGMAPKIAWALENKPRIPVVWIHGL WP_001301613.1 hydrogenase [Escherichia coli] Length: 372\nNumber of predicted TMHs: 2\nExp number of AAs in TMHs: 37.46598\nExp number, first 60 AAs: 19.53139\nTotal prob of N-in: 0.94364\nPOSSIBLE N-term signal sequence\ninside 1 19\nTMhelix 20 39\noutside 40 325\nTMhelix 326 348\ninside 349 372

3385 GCF_002164195.1_ASM216419v1 Escherichia coli Proteobacteria; Gammaproteobacteria; Enterobacterales; Enterobacteriaceae; Escherichia MNNEETFYQAMRRQGVTRRSFLKYCSLAATSLGLGAGMAPKIAWALENKPRIPVVWIHGL WP_001301613.1 hydrogenase [Escherichia coli] Length: 372\nNumber of predicted TMHs: 2\nExp number of AAs in TMHs: 37.46598\nExp number, first 60 AAs: 19.53139\nTotal prob of N-in: 0.94364\nPOSSIBLE N-term signal sequence\ninside 1 19\nTMhelix 20 39\noutside 40 325\nTMhelix 326 348\ninside 349 372

3386 GCF_002164375.1_ASM216437v1 Escherichia coli Proteobacteria; Gammaproteobacteria; Enterobacterales; Enterobacteriaceae; Escherichia MTGDNTLIHSHGINRRDFMKLCAALAATMGLSSKAAAEMAESVTNPQRPPVIWIGAQECT WP_000145410.1 MULTISPECIES: hydrogenase-2 small chain [Proteobacteria] Length: 372\nNumber of predicted TMHs: 1\nExp number of AAs in TMHs: 23.72226\nExp number, first 60 AAs: 0.22038\nTotal prob of N-in: 0.12994\noutside 1 331\nTMhelix 332 354\ninside 355 372

3387 GCF_002164455.1_ASM216445v1 Escherichia coli Proteobacteria; Gammaproteobacteria; Enterobacterales; Enterobacteriaceae; Escherichia MTGDNTLIHSHGINRRDFMKLCAALAATMGLSSKAAAEMAESVTNPQRPPVIWIGAQECT WP_000145410.1 MULTISPECIES: hydrogenase-2 small chain [Proteobacteria] Length: 372\nNumber of predicted TMHs: 1\nExp number of AAs in TMHs: 23.72226\nExp number, first 60 AAs: 0.22038\nTotal prob of N-in: 0.12994\noutside 1 331\nTMhelix 332 354\ninside 355 372

3388 GCF_002164635.1_ASM216463v1 Escherichia coli Proteobacteria; Gammaproteobacteria; Enterobacterales; Enterobacteriaceae; Escherichia MTGDNTLIHSHGINRRDFMKLCAALAATMGLSSKAAAEMAESVTNPQRPPVIWIGAQECT WP_000145410.1 MULTISPECIES: hydrogenase-2 small chain [Proteobacteria] Length: 372\nNumber of predicted TMHs: 1\nExp number of AAs in TMHs: 23.72226\nExp number, first 60 AAs: 0.22038\nTotal prob of N-in: 0.12994\noutside 1 331\nTMhelix 332 354\ninside 355 372

3389 GCF_002164515.1_ASM216451v1 Escherichia coli Proteobacteria; Gammaproteobacteria; Enterobacterales; Enterobacteriaceae; Escherichia MTGDNTLIHSHGINRRDFMKLCAALAATMGLSSKAAAEMAESVTNPQRPPVIWIGAQECT WP_000145410.1 MULTISPECIES: hydrogenase-2 small chain [Proteobacteria] Length: 372\nNumber of predicted TMHs: 1\nExp number of AAs in TMHs: 23.72226\nExp number, first 60 AAs: 0.22038\nTotal prob of N-in: 0.12994\noutside 1 331\nTMhelix 332 354\ninside 355 372

3390 GCF_002164545.1_ASM216454v1 Escherichia coli Proteobacteria; Gammaproteobacteria; Enterobacterales; Enterobacteriaceae; Escherichia MTGDNTLIHSHGINRRDFMKLCAALAATMGLSSKAAAEMAESVTNPQRPPVIWIGAQECT WP_000145410.1 MULTISPECIES: hydrogenase-2 small chain [Proteobacteria] Length: 372\nNumber of predicted TMHs: 1\nExp number of AAs in TMHs: 23.72226\nExp number, first 60 AAs: 0.22038\nTotal prob of N-in: 0.12994\noutside 1 331\nTMhelix 332 354\ninside 355 372

3391 GCF_002164515.1_ASM216451v1 Escherichia coli Proteobacteria; Gammaproteobacteria; Enterobacterales; Enterobacteriaceae; Escherichia
MNNEETFYQAMRRQGVTRRSFLKYCSLAATSLGLGAGMAPKIAWALENKPRIPVVWIHGL WP_001301613.1
hydrogenase [Escherichia coli] Length: 372\nNumber of predicted TMHs: 2\nExp number of AAs in TMHs: 37.46598\nExp number, first 60 AAs: 19.53139\nTotal prob of N-in: 0.94364\nPOSSIBLE N-term signal sequence\ninside 1 19\nTMhelix 20 39\noutside 40 325\nTMhelix 326 348\ninside 349 372

3392 GCF_002164545.1_ASM216454v1 Escherichia coli Proteobacteria; Gammaproteobacteria; Enterobacterales; Enterobacteriaceae; Escherichia
MNNEETFYQAMRRQGVTRRSFLKYCSLAATSLGLGAGMAPKIAWALENKPRIPVVWIHGL WP_001301613.1
hydrogenase [Escherichia coli] Length: 372\nNumber of predicted TMHs: 2\nExp number of AAs in TMHs: 37.46598\nExp number, first 60 AAs: 19.53139\nTotal prob of N-in: 0.94364\nPOSSIBLE N-term signal sequence\ninside 1 19\nTMhelix 20 39\noutside 40 325\nTMhelix 326 348\ninside 349 372

3393 GCF_002165005.1_ASM216500v1 Escherichia coli Proteobacteria; Gammaproteobacteria; Enterobacterales; Enterobacteriaceae; Escherichia
MTGDNTLIHSHGINRRDFMKLCAALAATMGLSSKAAAEMAESVTNPQRPPVIWIGAQECT WP_000145410.1
MULTISPECIES: hydrogenase-2 small chain [Proteobacteria] Length: 372\nNumber of predicted TMHs: 1\nExp number of AAs in TMHs: 23.72226\nExp number, first 60 AAs: 0.22038\nTotal prob of N-in: 0.12994\noutside 1 331\nTMhelix 332 354\ninside 355 372

3394 GCF_002165005.1_ASM216500v1 Escherichia coli Proteobacteria; Gammaproteobacteria; Enterobacterales; Enterobacteriaceae; Escherichia
MNNEETFYQAMRRQGVTRRSFLKYCSLAATSLGLGAGMAPKIAWALENKPRIPVVWIHGL WP_001295940.1
MULTISPECIES: hydrogenase [Enterobacteriaceae] Length: 372\nNumber of predicted TMHs: 2\nExp number of AAs in TMHs: 37.2608899999999\nExp number, first 60 AAs: 19.52923\nTotal prob of N-in: 0.94351\nPOSSIBLE N-term signal sequence\ninside 1 19\nTMhelix 20 39\noutside 40 325\nTMhelix 326 348\ninside 349 372

3395 GCF_002164535.1_ASM216453v1 Escherichia coli Proteobacteria; Gammaproteobacteria; Enterobacterales; Enterobacteriaceae; Escherichia
MNNEETFYQAMRRQGVTRRSFLKYCSLAATSLGLGAGMAPKIAWALENKPRIPVVWIHGL WP_001301613.1
hydrogenase [Escherichia coli] Length: 372\nNumber of predicted TMHs: 2\nExp number of AAs in TMHs: 37.46598\nExp number, first 60 AAs: 19.53139\nTotal prob of N-in: 0.94364\nPOSSIBLE N-term signal sequence\ninside 1 19\nTMhelix 20 39\noutside 40 325\nTMhelix 326 348\ninside 349 372

3396 GCF_002165095.1_ASM216509v1 Escherichia coli Proteobacteria; Gammaproteobacteria; Enterobacterales; Enterobacteriaceae; Escherichia
MTGDNTLIHSHGINRRDFMKLCAALAATMGLSSKAAAEMAESVTNPQRPPVIWIGAQECT WP_000145410.1
MULTISPECIES: hydrogenase-2 small chain [Proteobacteria] Length: 372\nNumber of predicted TMHs: 1\nExp number of AAs in TMHs: 23.72226\nExp number, first 60 AAs: 0.22038\nTotal prob of N-in: 0.12994\noutside 1 331\nTMhelix 332 354\ninside 355 372

3397 GCF_002165915.1_ASM216591v1 Escherichia coli Proteobacteria; Gammaproteobacteria; Enterobacterales; Enterobacteriaceae; Escherichia
MTGDNTLIHSHGINRRDFMKLCAALAATMGLSSKAAAEMAESVTNPQRPPVIWIGAQECT WP_000145410.1
MULTISPECIES: hydrogenase-2 small chain [Proteobacteria] Length: 372\nNumber of predicted TMHs: 1\nExp number of AAs in TMHs: 23.72226\nExp number, first 60 AAs: 0.22038\nTotal prob of N-in: 0.12994\noutside 1 331\nTMhelix 332 354\ninside 355 372

3398 GCF_002166115.1_ASM216611v1 Escherichia coli Proteobacteria; Gammaproteobacteria; Enterobacterales; Enterobacteriaceae; Escherichia
MNNEETFYQAMRRQGVTRRSFLKYCSLAATSLGLGAGMAPKIAWALENKPRIPVVWIHGL WP_001058323.1
MULTISPECIES: hydrogenase-1 small chain [Proteobacteria] Length: 372\nNumber of predicted TMHs: 2\nExp number of AAs in TMHs: 37.2608499999999\nExp number, first 60 AAs: 19.52923\nTotal prob of N-in: 0.94351\nPOSSIBLE N-term signal sequence\ninside 1 19\nTMhelix 20 39\noutside 40 325\nTMhelix 326 348\ninside 349 372

3399 GCF_002166075.1_ASM216607v1 Escherichia coli Proteobacteria; Gammaproteobacteria; Enterobacterales; Enterobacteriaceae; Escherichia
MNNEETFYQAMRRQGVTRRSFLKYCSLAATSLGLGAGMAPKIAWALENKPRIPVVWIHGL WP_001058323.1

MULTISPECIES: hydrogenase-1 small chain [Proteobacteria] Length: 372\nNumber of predicted TMHs: 2\nExp number of AAs in TMHs: 37.2608499999999\nExp number, first 60 AAs: 19.52923\nTotal prob of N-in: 0.94351\nPOSSIBLE N-term signal sequence\ninside 1 19\nTMhelix 20 39\noutside 40 325\nTMhelix 326 348\ninside 349 372

3400 GCF_002166135.1_ASM216613v1 Escherichia coli Proteobacteria; Gammaproteobacteria; Enterobacterales; Enterobacteriaceae; Escherichia
MTGDNTLIHSHGINRRDFMKLCAALAATMGLSSKAAAEMAESVTNPQRPPVIWIGAQECT WP_000145410.1
MULTISPECIES: hydrogenase-2 small chain [Proteobacteria] Length: 372\nNumber of predicted TMHs: 1\nExp number of AAs in TMHs: 23.72226\nExp number, first 60 AAs: 0.22038\nTotal prob of N-in: 0.12994\noutside 1 331\nTMhelix 332 354\ninside 355 372

3401 GCF_002166175.1_ASM216617v1 Escherichia coli Proteobacteria; Gammaproteobacteria; Enterobacterales; Enterobacteriaceae; Escherichia
MTGDNTLIHSHGINRRDFMKLCAALAATMGLSSKAAAEMAESVTNPQRPPVIWIGAQECT WP_000145410.1
MULTISPECIES: hydrogenase-2 small chain [Proteobacteria] Length: 372\nNumber of predicted TMHs: 1\nExp number of AAs in TMHs: 23.72226\nExp number, first 60 AAs: 0.22038\nTotal prob of N-in: 0.12994\noutside 1 331\nTMhelix 332 354\ninside 355 372

3402 GCF_002166195.1_ASM216619v1 Escherichia coli Proteobacteria; Gammaproteobacteria; Enterobacterales; Enterobacteriaceae; Escherichia
MTGDNTLIHSHGINRRDFMKLCAALAATMGLSSKAAAEMAESVTNPQRPPVIWIGAQECT WP_000145410.1
MULTISPECIES: hydrogenase-2 small chain [Proteobacteria] Length: 372\nNumber of predicted TMHs: 1\nExp number of AAs in TMHs: 23.72226\nExp number, first 60 AAs: 0.22038\nTotal prob of N-in: 0.12994\noutside 1 331\nTMhelix 332 354\ninside 355 372

3403 GCF_002166285.1_ASM216628v1 Escherichia coli Proteobacteria; Gammaproteobacteria; Enterobacterales; Enterobacteriaceae; Escherichia
MTGDNTLIHSHGINRRDFMKLCAALAATMGLSSKAAAEMVESVTNPQRPPVIWIGAQECT WP_000145423.1
hydrogenase 2 small subunit [Escherichia coli] Length: 372\nNumber of predicted TMHs: 1\nExp number of AAs in TMHs: 23.78201\nExp number, first 60 AAs: 0.28355\nTotal prob of N-in: 0.13257\noutside 1 331\nTMhelix 332 354\ninside 355 372

3404 GCF_002166285.1_ASM216628v1 Escherichia coli Proteobacteria; Gammaproteobacteria; Enterobacterales; Enterobacteriaceae; Escherichia
MNNEETFYQAMRRQGVTRRSFLKYCSLAATSLGLGAGMAPKIAWALENKPRIPVVWIHGL WP_001058323.1
MULTISPECIES: hydrogenase-1 small chain [Proteobacteria] Length: 372\nNumber of predicted TMHs: 2\nExp number of AAs in TMHs: 37.2608499999999\nExp number, first 60 AAs: 19.52923\nTotal prob of N-in: 0.94351\nPOSSIBLE N-term signal sequence\ninside 1 19\nTMhelix 20 39\noutside 40 325\nTMhelix 326 348\ninside 349 372

3405 GCF_002166525.1_ASM216652v1 Escherichia coli Proteobacteria; Gammaproteobacteria; Enterobacterales; Enterobacteriaceae; Escherichia
MTGDNTLIHSHGINRRDFMKLCAALAATMGLSSKAAAEMVESVTNPQRPPVIWIGAQECT WP_000145423.1
hydrogenase 2 small subunit [Escherichia coli] Length: 372\nNumber of predicted TMHs: 1\nExp number of AAs in TMHs: 23.78201\nExp number, first 60 AAs: 0.28355\nTotal prob of N-in: 0.13257\noutside 1 331\nTMhelix 332 354\ninside 355 372

3406 GCF_002166565.1_ASM216656v1 Escherichia coli Proteobacteria; Gammaproteobacteria; Enterobacterales; Enterobacteriaceae; Escherichia
MTGDNTLIHSHGINRRDFMKLCAALAATMGLSSKAAAEMAESVTNPQRPPVIWIGAQECT WP_000145410.1
MULTISPECIES: hydrogenase-2 small chain [Proteobacteria] Length: 372\nNumber of predicted TMHs: 1\nExp number of AAs in TMHs: 23.72226\nExp number, first 60 AAs: 0.22038\nTotal prob of N-in: 0.12994\noutside 1 331\nTMhelix 332 354\ninside 355 372

3407 GCF_002166525.1_ASM216652v1 Escherichia coli Proteobacteria; Gammaproteobacteria; Enterobacterales; Enterobacteriaceae; Escherichia
MNNEETFYQAMRRQGVTRRSFLKYCSLAATSLGLGAGMAPKIAWALENKPRIPVVWIHGL WP_001058323.1
MULTISPECIES: hydrogenase-1 small chain [Proteobacteria] Length: 372\nNumber of predicted TMHs: 2\nExp number of AAs in TMHs: 37.2608499999999\nExp number, first 60 AAs: 19.52923\nTotal prob of N-in:

0.94351\nPOSSIBLE N-term signal sequence\ninside 1 19\nTMhelix 20 39\noutside 40 325\nTMhelix 326 348\ninside 349 372

3408 GCF_002166595.1_ASM216659v1 Escherichia coli Proteobacteria; Gammaproteobacteria; Enterobacterales; Enterobacteriaceae; Escherichia MTGDNTLIHSHGINRRDFMKLCAALAATMGLSSKAAAEMAESVTNPQRPPVIWIGAQECT WP_000145410.1 MULTISPECIES: hydrogenase-2 small chain [Proteobacteria] Length: 372\nNumber of predicted TMHs: 1\nExp number of AAs in TMHs: 23.72226\nExp number, first 60 AAs: 0.22038\nTotal prob of N-in: 0.12994\noutside 1 331\nTMhelix 332 354\ninside 355 372

3409 GCF_002166625.1_ASM216662v1 Escherichia coli Proteobacteria; Gammaproteobacteria; Enterobacterales; Enterobacteriaceae; Escherichia MTGDNTLIHSHGINRRDFMKLCAALAATMGLSSKAAAEMAESVTNPQRPPVIWIGAQECT WP_000145410.1 MULTISPECIES: hydrogenase-2 small chain [Proteobacteria] Length: 372\nNumber of predicted TMHs: 1\nExp number of AAs in TMHs: 23.72226\nExp number, first 60 AAs: 0.22038\nTotal prob of N-in: 0.12994\noutside 1 331\nTMhelix 332 354\ninside 355 372

3410 GCF_002166675.1_ASM216667v1 Escherichia coli Proteobacteria; Gammaproteobacteria; Enterobacterales; Enterobacteriaceae; Escherichia MTGDNTLIHSHGINRRDFMKLCAALAATMGLSSKAAAEMAESVTNPQRPPVIWIGAQECT WP_000145410.1 MULTISPECIES: hydrogenase-2 small chain [Proteobacteria] Length: 372\nNumber of predicted TMHs: 1\nExp number of AAs in TMHs: 23.72226\nExp number, first 60 AAs: 0.22038\nTotal prob of N-in: 0.12994\noutside 1 331\nTMhelix 332 354\ninside 355 372

3411 GCF_002173145.1_ASM217314v1 Escherichia coli Proteobacteria; Gammaproteobacteria; Enterobacterales; Enterobacteriaceae; Escherichia MNNEETFYQAMRRQGVTRRSFLKYCSLAATSLGLGAGMAPKIAWALENKPRIPVVWIHGL WP_001058323.1 MULTISPECIES: hydrogenase-1 small chain [Proteobacteria] Length: 372\nNumber of predicted TMHs: 2\nExp number of AAs in TMHs: 37.2608499999999\nExp number, first 60 AAs: 19.52923\nTotal prob of N-in: 0.94351\nPOSSIBLE N-term signal sequence\ninside 1 19\nTMhelix 20 39\noutside 40 325\nTMhelix 326 348\ninside 349 372

3412 GCF_002173215.1_ASM217321v1 Escherichia coli Proteobacteria; Gammaproteobacteria; Enterobacterales; Enterobacteriaceae; Escherichia MNNEETFYQAMRRQGVTRRSFLKYCSLAATSLGLGAGMAPKIAWALENKPRIPVVWIHGL WP_001058323.1 MULTISPECIES: hydrogenase-1 small chain [Proteobacteria] Length: 372\nNumber of predicted TMHs: 2\nExp number of AAs in TMHs: 37.2608499999999\nExp number, first 60 AAs: 19.52923\nTotal prob of N-in: 0.94351\nPOSSIBLE N-term signal sequence\ninside 1 19\nTMhelix 20 39\noutside 40 325\nTMhelix 326 348\ninside 349 372

3413 GCF_002173275.1_ASM217327v1 Escherichia coli Proteobacteria; Gammaproteobacteria; Enterobacterales; Enterobacteriaceae; Escherichia MTGDNTLIHSHGINRRDFMKLCAALAATMGLSSKAAAEMAESVTNPQRPPVIWIGAQECT WP_000145410.1 MULTISPECIES: hydrogenase-2 small chain [Proteobacteria] Length: 372\nNumber of predicted TMHs: 1\nExp number of AAs in TMHs: 23.72226\nExp number, first 60 AAs: 0.22038\nTotal prob of N-in: 0.12994\noutside 1 331\nTMhelix 332 354\ninside 355 372

3414 GCF_002173275.1_ASM217327v1 Escherichia coli Proteobacteria; Gammaproteobacteria; Enterobacterales; Enterobacteriaceae; Escherichia MNNEETFYQAMRRQGVTRRSFLKYCSLAATSLGLGAGMAPKIAWALENKPRIPVVWIHGL WP_001058323.1 MULTISPECIES: hydrogenase-1 small chain [Proteobacteria] Length: 372\nNumber of predicted TMHs: 2\nExp number of AAs in TMHs: 37.2608499999999\nExp number, first 60 AAs: 19.52923\nTotal prob of N-in: 0.94351\nPOSSIBLE N-term signal sequence\ninside 1 19\nTMhelix 20 39\noutside 40 325\nTMhelix 326 348\ninside 349 372

3415 GCF_002173255.1_ASM217325v1 Escherichia coli Proteobacteria; Gammaproteobacteria; Enterobacterales; Enterobacteriaceae; Escherichia MNNEETFYQAMRRQGVTRRSFLKYCSLAATSLGLGAGMAPKIAWALENKPRIPVVWIHGL WP_001058323.1 MULTISPECIES: hydrogenase-1 small chain [Proteobacteria] Length: 372\nNumber of predicted TMHs: 2\nExp number of AAs in TMHs: 37.2608499999999\nExp number, first 60 AAs: 19.52923\nTotal prob of N-in:

0.94351\nPOSSIBLE N-term signal sequence\ninside 1 19\nTMhelix 20 39\noutside 40 325\nTMhelix 326 348\ninside 349 372

3416 GCF_002173335.1_ASM217333v1 Escherichia coli Proteobacteria; Gammaproteobacteria; Enterobacterales; Enterobacteriaceae; Escherichia MTGDNTLIHSHGINRRDFMKLCAALAATMGLSSKAAAEMAESVTNPQRPPVIWIGAQECT WP_000145410.1 MULTISPECIES: hydrogenase-2 small chain [Proteobacteria] Length: 372\nNumber of predicted TMHs: 1\nExp number of AAs in TMHs: 23.72226\nExp number, first 60 AAs: 0.22038\nTotal prob of N-in: 0.12994\noutside 1 331\nTMhelix 332 354\ninside 355 372

3417 GCF_002173345.1_ASM217334v1 Escherichia coli Proteobacteria; Gammaproteobacteria; Enterobacterales; Enterobacteriaceae; Escherichia MTGDNTLIHSHGINRRDFMKLCAALAATMGLSSKAAAEMAESVTNPQRPPVIWIGAQECT WP_000145410.1 MULTISPECIES: hydrogenase-2 small chain [Proteobacteria] Length: 372\nNumber of predicted TMHs: 1\nExp number of AAs in TMHs: 23.72226\nExp number, first 60 AAs: 0.22038\nTotal prob of N-in: 0.12994\noutside 1 331\nTMhelix 332 354\ninside 355 372

3418 GCF_002175555.1_ASM217555v1 Escherichia coli Proteobacteria; Gammaproteobacteria; Enterobacterales; Enterobacteriaceae; Escherichia MTGDNTLIHSHGINRRDFMKLCAALAATMGLSSKAAAEMAESVTNPQRPPVIWIGAQECT WP_000145410.1 MULTISPECIES: hydrogenase-2 small chain [Proteobacteria] Length: 372\nNumber of predicted TMHs: 1\nExp number of AAs in TMHs: 23.72226\nExp number, first 60 AAs: 0.22038\nTotal prob of N-in: 0.12994\noutside 1 331\nTMhelix 332 354\ninside 355 372

3419 GCF_002175615.1_ASM217561v1 Escherichia coli Proteobacteria; Gammaproteobacteria; Enterobacterales; Enterobacteriaceae; Escherichia MTGDNTLIHSHGINRRDFMKLCAALAATMGLSSKAAAEMAESVTNPQRPPVIWIGAQECT WP_000145410.1 MULTISPECIES: hydrogenase-2 small chain [Proteobacteria] Length: 372\nNumber of predicted TMHs: 1\nExp number of AAs in TMHs: 23.72226\nExp number, first 60 AAs: 0.22038\nTotal prob of N-in: 0.12994\noutside 1 331\nTMhelix 332 354\ninside 355 372

3420 GCF_002175405.1_ASM217540v1 Escherichia coli Proteobacteria; Gammaproteobacteria; Enterobacterales; Enterobacteriaceae; Escherichia MNNEETFYQAMRRQGVTRRSFLKYCSLAATSLGLGAGMAPKIAWALENKPRIPVVWIHGL WP_001301613.1 hydrogenase [Escherichia coli] Length: 372\nNumber of predicted TMHs: 2\nExp number of AAs in TMHs: 37.46598\nExp number, first 60 AAs: 19.53139\nTotal prob of N-in: 0.94364\nPOSSIBLE N-term signal sequence\ninside 1 19\nTMhelix 20 39\noutside 40 325\nTMhelix 326 348\ninside 349 372

3421 GCF_002175395.1_ASM217539v1 Escherichia coli Proteobacteria; Gammaproteobacteria; Enterobacterales; Enterobacteriaceae; Escherichia MNNEETFYQAMRRQGVTRRSFLKYCSLAATSLGLGAGMAPKIAWALENKPRIPVVWIHGL WP_001301613.1 hydrogenase [Escherichia coli] Length: 372\nNumber of predicted TMHs: 2\nExp number of AAs in TMHs: 37.46598\nExp number, first 60 AAs: 19.53139\nTotal prob of N-in: 0.94364\nPOSSIBLE N-term signal sequence\ninside 1 19\nTMhelix 20 39\noutside 40 325\nTMhelix 326 348\ninside 349 372

3422 GCF_002175625.1_ASM217562v1 Escherichia coli Proteobacteria; Gammaproteobacteria; Enterobacterales; Enterobacteriaceae; Escherichia MTGDNTLIHSHGINRRDFMKLCAALAATMGLSSKAAAEMAESVTNPQRPPVIWIGAQECT WP_000145410.1 MULTISPECIES: hydrogenase-2 small chain [Proteobacteria] Length: 372\nNumber of predicted TMHs: 1\nExp number of AAs in TMHs: 23.72226\nExp number, first 60 AAs: 0.22038\nTotal prob of N-in: 0.12994\noutside 1 331\nTMhelix 332 354\ninside 355 372

3423 GCF_002175655.1_ASM217565v1 Escherichia coli Proteobacteria; Gammaproteobacteria; Enterobacterales; Enterobacteriaceae; Escherichia MTGDNTLIHSHGINRRDFMKLCAALAATMGLSSKAAAEMAESVTNPQRPPVIWIGAQECT WP_000145410.1 MULTISPECIES: hydrogenase-2 small chain [Proteobacteria] Length: 372\nNumber of predicted TMHs: 1\nExp number of AAs in TMHs: 23.72226\nExp number, first 60 AAs: 0.22038\nTotal prob of N-in: 0.12994\noutside 1 331\nTMhelix 332 354\ninside 355 372

3424 GCF_002175655.1_ASM217565v1 Escherichia coli Proteobacteria; Gammaproteobacteria; Enterobacterales; Enterobacteriaceae; Escherichia

MNNEETFYQAMRRQGVTRRSFLKYCSLAATSLGLGAGMAPKIAWALENKPRIPVVWIHGL WP_001301613.1 hydrogenase [Escherichia coli] Length: 372\nNumber of predicted TMHs: 2\nExp number of AAs in TMHs: 37.46598\nExp number, first 60 AAs: 19.53139\nTotal prob of N-in: 0.94364\nPOSSIBLE N-term signal sequence\ninside 1 19\nTMhelix 20 39\noutside 40 325\nTMhelix 326 348\ninside 349 372

3425 GCF_002175885.1_ASM217588v1 Escherichia coli Proteobacteria; Gammaproteobacteria; Enterobacterales; Enterobacteriaceae; Escherichia MTGDNTLIHSHGINRRDFMKLCAALAATMGLSSKAAAEMAESVTNPQRPPVIWIGAQECT WP_000145410.1 MULTISPECIES: hydrogenase-2 small chain [Proteobacteria] Length: 372\nNumber of predicted TMHs: 1\nExp number of AAs in TMHs: 23.72226\nExp number, first 60 AAs: 0.22038\nTotal prob of N-in: 0.12994\noutside 1 331\nTMhelix 332 354\ninside 355 372

3426 GCF_002175935.1_ASM217593v1 Escherichia coli Proteobacteria; Gammaproteobacteria; Enterobacterales; Enterobacteriaceae; Escherichia MTGDNTLIHSHGINRRDFMKLCAALAATMGLSSKAAAEMAESVTNPQRPPVIWIGAQECT WP_000145410.1 MULTISPECIES: hydrogenase-2 small chain [Proteobacteria] Length: 372\nNumber of predicted TMHs: 1\nExp number of AAs in TMHs: 23.72226\nExp number, first 60 AAs: 0.22038\nTotal prob of N-in: 0.12994\noutside 1 331\nTMhelix 332 354\ninside 355 372

3427 GCF_002175805.1_ASM217580v1 Escherichia coli Proteobacteria; Gammaproteobacteria; Enterobacterales; Enterobacteriaceae; Escherichia MNNEETFYQAMRRQGVTRRSFLKYCSLAATSLGLGAGMAPKIAWALENKPRIPVVWIHGL WP_001301613.1 hydrogenase [Escherichia coli] Length: 372\nNumber of predicted TMHs: 2\nExp number of AAs in TMHs: 37.46598\nExp number, first 60 AAs: 19.53139\nTotal prob of N-in: 0.94364\nPOSSIBLE N-term signal sequence\ninside 1 19\nTMhelix 20 39\noutside 40 325\nTMhelix 326 348\ninside 349 372

3428 GCF_002175615.1_ASM217561v1 Escherichia coli Proteobacteria; Gammaproteobacteria; Enterobacterales; Enterobacteriaceae; Escherichia MNNEETFYQAMRRQGVTRRSFLKYCSLAATSLGLGAGMAPKIAWALENKPRIPVVWIHGL WP_001301613.1 hydrogenase [Escherichia coli] Length: 372\nNumber of predicted TMHs: 2\nExp number of AAs in TMHs: 37.46598\nExp number, first 60 AAs: 19.53139\nTotal prob of N-in: 0.94364\nPOSSIBLE N-term signal sequence\ninside 1 19\nTMhelix 20 39\noutside 40 325\nTMhelix 326 348\ninside 349 372

3429 GCF_002175955.1_ASM217595v1 Escherichia coli Proteobacteria; Gammaproteobacteria; Enterobacterales; Enterobacteriaceae; Escherichia MTGDNTLIHSHGINRRDFMKLCAALAATMGLSSKAAAEMAESVTNPQRPPVIWIGAQECT WP_000145410.1 MULTISPECIES: hydrogenase-2 small chain [Proteobacteria] Length: 372\nNumber of predicted TMHs: 1\nExp number of AAs in TMHs: 23.72226\nExp number, first 60 AAs: 0.22038\nTotal prob of N-in: 0.12994\noutside 1 331\nTMhelix 332 354\ninside 355 372

3430 GCF_002176015.1_ASM217601v1 Escherichia coli Proteobacteria; Gammaproteobacteria; Enterobacterales; Enterobacteriaceae; Escherichia MTGDNTLIHSHGINRRDFMKLCAALAATMGLSSKAAAEMAESVTNPQRPPVIWIGAQECT WP_000145410.1 MULTISPECIES: hydrogenase-2 small chain [Proteobacteria] Length: 372\nNumber of predicted TMHs: 1\nExp number of AAs in TMHs: 23.72226\nExp number, first 60 AAs: 0.22038\nTotal prob of N-in: 0.12994\noutside 1 331\nTMhelix 332 354\ninside 355 372

3431 GCF_002176025.1_ASM217602v1 Escherichia coli Proteobacteria; Gammaproteobacteria; Enterobacterales; Enterobacteriaceae; Escherichia MNNEETFYQAMRRQGVTRRSFLKYCSLAATSLGLGAGMAPKIAWALENKPRIPVVWIHGL WP_001301613.1 hydrogenase [Escherichia coli] Length: 372\nNumber of predicted TMHs: 2\nExp number of AAs in TMHs: 37.46598\nExp number, first 60 AAs: 19.53139\nTotal prob of N-in: 0.94364\nPOSSIBLE N-term signal sequence\ninside 1 19\nTMhelix 20 39\noutside 40 325\nTMhelix 326 348\ninside 349 372

3432 GCF_002176045.1_ASM217604v1 Escherichia coli Proteobacteria; Gammaproteobacteria; Enterobacterales; Enterobacteriaceae; Escherichia MNNEETFYQAMRRQGVTRRSFLKYCSLAATSLGLGAGMAPKIAWALENKPRIPVVWIHGL WP_001301613.1 hydrogenase [Escherichia coli] Length: 372\nNumber of predicted TMHs: 2\nExp number of AAs in TMHs: 37.46598\nExp number, first 60 AAs: 19.53139\nTotal prob of N-in: 0.94364\nPOSSIBLE N-term signal sequence\ninside 1 19\nTMhelix 20 39\noutside 40 325\nTMhelix 326 348\ninside 349 372

3433 GCF_002176355.1_ASM217635v1 Escherichia coli Proteobacteria; Gammaproteobacteria; Enterobacterales; Enterobacteriaceae; Escherichia
MTGDNTLIHSHGINRRDFMKLCAALAATMGLSSKAAAEMAESVTNPQRPPVIWIGAQECT WP_000145410.1
MULTISPECIES: hydrogenase-2 small chain [Proteobacteria] Length: 372\nNumber of predicted TMHs: 1\nExp number of AAs in TMHs: 23.72226\nExp number, first 60 AAs: 0.22038\nTotal prob of N-in: 0.12994\noutside 1 331\nTMhelix 332 354\ninside 355 372

3434 GCF_002176115.1_ASM217611v1 Escherichia coli Proteobacteria; Gammaproteobacteria; Enterobacterales; Enterobacteriaceae; Escherichia
MNNEETFYQAMRRQGVTRRSFLKYCSLAATSLGLGAGMAPKIAWALENKPRIPVVWIHGL WP_001301613.1
hydrogenase [Escherichia coli] Length: 372\nNumber of predicted TMHs: 2\nExp number of AAs in TMHs: 37.46598\nExp number, first 60 AAs: 19.53139\nTotal prob of N-in: 0.94364\nPOSSIBLE N-term signal sequence\ninside 1 19\nTMhelix 20 39\noutside 40 325\nTMhelix 326 348\ninside 349 372

3435 GCF_002176415.1_ASM217641v1 Escherichia coli Proteobacteria; Gammaproteobacteria; Enterobacterales; Enterobacteriaceae; Escherichia
MTGDNTLIHSHGINRRDFMKLCAALAATMGLSSKAAAEMAESVTNPQRPPVIWIGAQECT WP_000145410.1
MULTISPECIES: hydrogenase-2 small chain [Proteobacteria] Length: 372\nNumber of predicted TMHs: 1\nExp number of AAs in TMHs: 23.72226\nExp number, first 60 AAs: 0.22038\nTotal prob of N-in: 0.12994\noutside 1 331\nTMhelix 332 354\ninside 355 372

3436 GCF_002176425.1_ASM217642v1 Escherichia coli Proteobacteria; Gammaproteobacteria; Enterobacterales; Enterobacteriaceae; Escherichia
MTGDNTLIHSHGINRRDFMKLCAALAATMGLSSKAAAEMAESVTNPQRPPVIWIGAQECT WP_000145410.1
MULTISPECIES: hydrogenase-2 small chain [Proteobacteria] Length: 372\nNumber of predicted TMHs: 1\nExp number of AAs in TMHs: 23.72226\nExp number, first 60 AAs: 0.22038\nTotal prob of N-in: 0.12994\noutside 1 331\nTMhelix 332 354\ninside 355 372

3437 GCF_002180815.1_ASM218081v1 Escherichia coli Proteobacteria; Gammaproteobacteria; Enterobacterales; Enterobacteriaceae; Escherichia
MTGDNTLIHSHGINRRDFMKLCAALAATMGLSSKAAAEMVESVTNPQRPPVIWIGAQECT WP_000145423.1
hydrogenase 2 small subunit [Escherichia coli] Length: 372\nNumber of predicted TMHs: 1\nExp number of AAs in TMHs: 23.78201\nExp number, first 60 AAs: 0.28355\nTotal prob of N-in: 0.13257\noutside 1 331\nTMhelix 332 354\ninside 355 372

3438 GCF_002179755.1_ASM217975v1 Escherichia coli Proteobacteria; Gammaproteobacteria; Enterobacterales; Enterobacteriaceae; Escherichia
MTGDNTLIHSHGINRRDFMKLCAALAATMGLSSKAAAEMAESVTNPQRPPVIWIGAQECT WP_000145410.1
MULTISPECIES: hydrogenase-2 small chain [Proteobacteria] Length: 372\nNumber of predicted TMHs: 1\nExp number of AAs in TMHs: 23.72226\nExp number, first 60 AAs: 0.22038\nTotal prob of N-in: 0.12994\noutside 1 331\nTMhelix 332 354\ninside 355 372

3439 GCF_002181625.1_ASM218162v1 Escherichia coli Proteobacteria; Gammaproteobacteria; Enterobacterales; Enterobacteriaceae; Escherichia
MTGDNTLIHSHGINRRDFMKLCAALAATMGLSSKAAAEMVESVTNPQRPPVIWIGAQECT WP_000145423.1
hydrogenase 2 small subunit [Escherichia coli] Length: 372\nNumber of predicted TMHs: 1\nExp number of AAs in TMHs: 23.78201\nExp number, first 60 AAs: 0.28355\nTotal prob of N-in: 0.13257\noutside 1 331\nTMhelix 332 354\ninside 355 372

3440 GCF_002191055.1_ASM219105v1 Escherichia coli Proteobacteria; Gammaproteobacteria; Enterobacterales; Enterobacteriaceae; Escherichia
MTGDNTLIHSHGINRRDFMKLCAALAATMGLSSKAAAEMAESVTNPQRPPVIWIGAQECT WP_000145410.1
MULTISPECIES: hydrogenase-2 small chain [Proteobacteria] Length: 372\nNumber of predicted TMHs: 1\nExp number of AAs in TMHs: 23.72226\nExp number, first 60 AAs: 0.22038\nTotal prob of N-in: 0.12994\noutside 1 331\nTMhelix 332 354\ninside 355 372

3441 GCF_002176425.1_ASM217642v1 Escherichia coli Proteobacteria; Gammaproteobacteria; Enterobacterales; Enterobacteriaceae; Escherichia
MNNEETFYQAMRRQGVTRRSFLKYCSLAATSLGLGAGMAPKIAWALENKPRIPVVWIHGL WP_001301613.1
hydrogenase [Escherichia coli] Length: 372\nNumber of predicted TMHs: 2\nExp number of AAs in

TMHs: 37.46598\nExp number, first 60 AAs: 19.53139\nTotal prob of N-in: 0.94364\nPOSSIBLE N-term signal sequence\ninside 1 19\nTMhelix 20 39\noutside 40 325\nTMhelix 326 348\ninside 349 372

3442 GCF_002176415.1_ASM217641v1 Escherichia coli Proteobacteria; Gammaproteobacteria; Enterobacterales; Enterobacteriaceae; Escherichia
MNNEETFYQAMRRQGVTRRSFLKYCSLAATSLGLGAGMAPKIAWALENKPRIPVVWIHGL WP_001301613.1
hydrogenase [Escherichia coli] Length: 372\nNumber of predicted TMHs: 2\nExp number of AAs in TMHs: 37.46598\nExp number, first 60 AAs: 19.53139\nTotal prob of N-in: 0.94364\nPOSSIBLE N-term signal sequence\ninside 1 19\nTMhelix 20 39\noutside 40 325\nTMhelix 326 348\ninside 349 372

3443 GCF_002176475.1_ASM217647v1 Escherichia coli Proteobacteria; Gammaproteobacteria; Enterobacterales; Enterobacteriaceae; Escherichia
MNNEETFYQAMRRQGVTRRSFLKYCSLAATSLGLGAGMAPKIAWALENKPRIPVVWIHGL WP_001301613.1
hydrogenase [Escherichia coli] Length: 372\nNumber of predicted TMHs: 2\nExp number of AAs in TMHs: 37.46598\nExp number, first 60 AAs: 19.53139\nTotal prob of N-in: 0.94364\nPOSSIBLE N-term signal sequence\ninside 1 19\nTMhelix 20 39\noutside 40 325\nTMhelix 326 348\ninside 349 372

3444 GCF_002189985.1_ASM218998v1 Escherichia coli Proteobacteria; Gammaproteobacteria; Enterobacterales; Enterobacteriaceae; Escherichia
MTGDNTLIHSHGINRRDFMKLCAALAATMGLSSKAAAEMAESVTNPQRPPVIWIGAQECT WP_000145410.1
MULTISPECIES: hydrogenase-2 small chain [Proteobacteria] Length: 372\nNumber of predicted TMHs: 1\nExp number of AAs in TMHs: 23.72226\nExp number, first 60 AAs: 0.22038\nTotal prob of N-in: 0.12994\noutside 1 331\nTMhelix 332 354\ninside 355 372

3445 GCF_002189935.1_ASM218993v1 Escherichia coli Proteobacteria; Gammaproteobacteria; Enterobacterales; Enterobacteriaceae; Escherichia
MTGDNTLIHSHGINRRDFMKLCAALAATMGLSSKAAAEMAESVTNPQRPPVIWIGAQECT WP_000145410.1
MULTISPECIES: hydrogenase-2 small chain [Proteobacteria] Length: 372\nNumber of predicted TMHs: 1\nExp number of AAs in TMHs: 23.72226\nExp number, first 60 AAs: 0.22038\nTotal prob of N-in: 0.12994\noutside 1 331\nTMhelix 332 354\ninside 355 372

3446 GCF_002190025.1_ASM219002v1 Escherichia coli Proteobacteria; Gammaproteobacteria; Enterobacterales; Enterobacteriaceae; Escherichia
MTGDNTLIHSHGINRRDFMKLCAALAATMGLSSKAAAEMAESVTNPQRPPVIWIGAQECT WP_000145410.1
MULTISPECIES: hydrogenase-2 small chain [Proteobacteria] Length: 372\nNumber of predicted TMHs: 1\nExp number of AAs in TMHs: 23.72226\nExp number, first 60 AAs: 0.22038\nTotal prob of N-in: 0.12994\noutside 1 331\nTMhelix 332 354\ninside 355 372

3447 GCF_002190035.1_ASM219003v1 Escherichia coli Proteobacteria; Gammaproteobacteria; Enterobacterales; Enterobacteriaceae; Escherichia
MTGDNTLIHSHGINRRDFMKLCAALAATMGLSSKAAAEMAESVTNPQRPPVIWIGAQECT WP_000145410.1
MULTISPECIES: hydrogenase-2 small chain [Proteobacteria] Length: 372\nNumber of predicted TMHs: 1\nExp number of AAs in TMHs: 23.72226\nExp number, first 60 AAs: 0.22038\nTotal prob of N-in: 0.12994\noutside 1 331\nTMhelix 332 354\ninside 355 372

3448 GCF_002190085.1_ASM219008v1 Escherichia coli Proteobacteria; Gammaproteobacteria; Enterobacterales; Enterobacteriaceae; Escherichia
MTGDNTLIHSHGINRRDFMKLCAALAATMGLSSKAAAEMAESVTNPQRPPVIWIGAQECT WP_000145410.1
MULTISPECIES: hydrogenase-2 small chain [Proteobacteria] Length: 372\nNumber of predicted TMHs: 1\nExp number of AAs in TMHs: 23.72226\nExp number, first 60 AAs: 0.22038\nTotal prob of N-in: 0.12994\noutside 1 331\nTMhelix 332 354\ninside 355 372

3449 GCF_002189935.1_ASM218993v1 Escherichia coli Proteobacteria; Gammaproteobacteria; Enterobacterales; Enterobacteriaceae; Escherichia
MNNEETFYQAMRRQGVTRRSFLKYCSLAATSLGLGAGMAPKIAWALENKPRIPVVWIHGL WP_001058323.1
MULTISPECIES: hydrogenase-1 small chain [Proteobacteria] Length: 372\nNumber of predicted TMHs: 2\nExp number of AAs in TMHs: 37.2608499999999\nExp number, first 60 AAs: 19.52923\nTotal prob of N-in: 0.94351\nPOSSIBLE N-term signal sequence\ninside 1 19\nTMhelix 20 39\noutside 40 325\nTMhelix 326 348\ninside 349 372

3450 GCF_002190025.1_ASM219002v1 Escherichia coli Proteobacteria; Gammaproteobacteria; Enterobacterales; Enterobacteriaceae; Escherichia
MNNEETFYQAMRRQGVTRRSFLKYCSLAATSLGLGAGMAPKIAWALENKPRIPVVWIHGL WP_001058323.1
MULTISPECIES: hydrogenase-1 small chain [Proteobacteria] Length: 372\nNumber of predicted TMHs: 2\nExp number of AAs in TMHs: 37.2608499999999\nExp number, first 60 AAs: 19.52923\nTotal prob of N-in: 0.94351\nPOSSIBLE N-term signal sequence\ninside 1 19\nTMhelix 20 39\noutside 40 325\nTMhelix 326 348\ninside 349 372

3451 GCF_002190205.1_ASM219020v1 Escherichia coli Proteobacteria; Gammaproteobacteria; Enterobacterales; Enterobacteriaceae; Escherichia
MNNEETFYQAMRRQGVTRRSFLKYCSLAATSLGLGAGMAPKIAWALENKPRIPVVWIHGL WP_001058323.1
MULTISPECIES: hydrogenase-1 small chain [Proteobacteria] Length: 372\nNumber of predicted TMHs: 2\nExp number of AAs in TMHs: 37.2608499999999\nExp number, first 60 AAs: 19.52923\nTotal prob of N-in: 0.94351\nPOSSIBLE N-term signal sequence\ninside 1 19\nTMhelix 20 39\noutside 40 325\nTMhelix 326 348\ninside 349 372

3452 GCF_002190295.1_ASM219029v1 Escherichia coli Proteobacteria; Gammaproteobacteria; Enterobacterales; Enterobacteriaceae; Escherichia
MTGDNTLIHSHGINRRDFMKLCAALAATMGLSSKAAAEMAESVTNPQRPPVIWIGAQECT WP_000145410.1
MULTISPECIES: hydrogenase-2 small chain [Proteobacteria] Length: 372\nNumber of predicted TMHs: 1\nExp number of AAs in TMHs: 23.72226\nExp number, first 60 AAs: 0.22038\nTotal prob of N-in: 0.12994\noutside 1 331\nTMhelix 332 354\ninside 355 372

3453 GCF_002190355.1_ASM219035v1 Escherichia coli Proteobacteria; Gammaproteobacteria; Enterobacterales; Enterobacteriaceae; Escherichia
MTGDNTLIHSHGINRRDFMKLCAALAATMGLSSKAAAEMAESVTNPQRPPVIWIGAQECT WP_000145410.1
MULTISPECIES: hydrogenase-2 small chain [Proteobacteria] Length: 372\nNumber of predicted TMHs: 1\nExp number of AAs in TMHs: 23.72226\nExp number, first 60 AAs: 0.22038\nTotal prob of N-in: 0.12994\noutside 1 331\nTMhelix 332 354\ninside 355 372

3454 GCF_002190365.1_ASM219036v1 Escherichia coli Proteobacteria; Gammaproteobacteria; Enterobacterales; Enterobacteriaceae; Escherichia
MTGDNTLIHSHGINRRDFMKLCAALAATMGLSSKAAAEMAESVTNPQRPPVIWIGAQECT WP_000145410.1
MULTISPECIES: hydrogenase-2 small chain [Proteobacteria] Length: 372\nNumber of predicted TMHs: 1\nExp number of AAs in TMHs: 23.72226\nExp number, first 60 AAs: 0.22038\nTotal prob of N-in: 0.12994\noutside 1 331\nTMhelix 332 354\ninside 355 372

3455 GCF_002190395.1_ASM219039v1 Escherichia coli Proteobacteria; Gammaproteobacteria; Enterobacterales; Enterobacteriaceae; Escherichia
MTGDNTLIHSHGINRRDFMKLCAALAATMGLSSKAAAEMAESVTNPQRPPVIWIGAQECT WP_000145410.1
MULTISPECIES: hydrogenase-2 small chain [Proteobacteria] Length: 372\nNumber of predicted TMHs: 1\nExp number of AAs in TMHs: 23.72226\nExp number, first 60 AAs: 0.22038\nTotal prob of N-in: 0.12994\noutside 1 331\nTMhelix 332 354\ninside 355 372

3456 GCF_002190455.1_ASM219045v1 Escherichia coli Proteobacteria; Gammaproteobacteria; Enterobacterales; Enterobacteriaceae; Escherichia
MTGDNTLIHSHGINRRDFMKLCAALAATMGLSSKAAAEMAESVTNPQRPPVIWIGAQECT WP_000145410.1
MULTISPECIES: hydrogenase-2 small chain [Proteobacteria] Length: 372\nNumber of predicted TMHs: 1\nExp number of AAs in TMHs: 23.72226\nExp number, first 60 AAs: 0.22038\nTotal prob of N-in: 0.12994\noutside 1 331\nTMhelix 332 354\ninside 355 372

3457 GCF_002190465.1_ASM219046v1 Escherichia coli Proteobacteria; Gammaproteobacteria; Enterobacterales; Enterobacteriaceae; Escherichia
MTGDNTLIHSHGINRRDFMKLCAALAATMGLSSKAAAEMAESVTNPQRPPVIWIGAQECT WP_000145410.1
MULTISPECIES: hydrogenase-2 small chain [Proteobacteria] Length: 372\nNumber of predicted TMHs: 1\nExp number of AAs in TMHs: 23.72226\nExp number, first 60 AAs: 0.22038\nTotal prob of N-in: 0.12994\noutside 1 331\nTMhelix 332 354\ninside 355 372

3458 GCF_002191075.1_ASM219107v1 Escherichia coli Proteobacteria; Gammaproteobacteria; Enterobacterales; Enterobacteriaceae; Escherichia
MTGDNTLIHSHGINRRDFMKLCAALAATMGLSSKAAAEMAESVTNPQRPPVIWIGAQECT WP_000145410.1

MULTISPECIES: hydrogenase-2 small chain [Proteobacteria] Length: 372\nNumber of predicted TMHs: 1\nExp number of AAs in TMHs: 23.72226\nExp number, first 60 AAs: 0.22038\nTotal prob of N-in: 0.12994\noutside 1 331\nTMhelix 332 354\ninside 355 372

3459 GCF_002190825.1_ASM219082v1 Escherichia coli Proteobacteria; Gammaproteobacteria; Enterobacterales; Enterobacteriaceae; Escherichia
MTGDNTLIHSHGINRRDFMKLCAALAATMGLSSKAAAEMAESVTNPQRPPVIWIGAQECT WP_000145410.1
MULTISPECIES: hydrogenase-2 small chain [Proteobacteria] Length: 372\nNumber of predicted TMHs: 1\nExp number of AAs in TMHs: 23.72226\nExp number, first 60 AAs: 0.22038\nTotal prob of N-in: 0.12994\noutside 1 331\nTMhelix 332 354\ninside 355 372

3460 GCF_002190825.1_ASM219082v1 Escherichia coli Proteobacteria; Gammaproteobacteria; Enterobacterales; Enterobacteriaceae; Escherichia
MNNEETFYQAMRRQGVTRRSFLKYCSLAATSLGLGAGMAPKIAWALENKPRIPVVWIHGL WP_001058323.1
MULTISPECIES: hydrogenase-1 small chain [Proteobacteria] Length: 372\nNumber of predicted TMHs: 2\nExp number of AAs in TMHs: 37.2608499999999\nExp number, first 60 AAs: 19.52923\nTotal prob of N-in: 0.94351\nPOSSIBLE N-term signal sequence\ninside 1 19\nTMhelix 20 39\noutside 40 325\nTMhelix 326 348\ninside 349 372

3461 GCF_002190785.1_ASM219078v1 Escherichia coli Proteobacteria; Gammaproteobacteria; Enterobacterales; Enterobacteriaceae; Escherichia
MNNEETFYQAMRRQGVTRRSFLKYCSLAATSLGLGAGMAPKIAWALENKPRIPVVWIHGL WP_001058323.1
MULTISPECIES: hydrogenase-1 small chain [Proteobacteria] Length: 372\nNumber of predicted TMHs: 2\nExp number of AAs in TMHs: 37.2608499999999\nExp number, first 60 AAs: 19.52923\nTotal prob of N-in: 0.94351\nPOSSIBLE N-term signal sequence\ninside 1 19\nTMhelix 20 39\noutside 40 325\nTMhelix 326 348\ninside 349 372

3462 GCF_002190865.1_ASM219086v1 Escherichia coli Proteobacteria; Gammaproteobacteria; Enterobacterales; Enterobacteriaceae; Escherichia
MNNEETFYQAMRRQGVTRRSFLKYCSLAATSLGLGAGMAPKIAWALENKPRIPVVWIHGL WP_001295940.1
MULTISPECIES: hydrogenase [Enterobacteriaceae] Length: 372\nNumber of predicted TMHs: 2\nExp number of AAs in TMHs: 37.2608899999999\nExp number, first 60 AAs: 19.52923\nTotal prob of N-in: 0.94351\nPOSSIBLE N-term signal sequence\ninside 1 19\nTMhelix 20 39\noutside 40 325\nTMhelix 326 348\ninside 349 372

3463 GCF_002190715.1_ASM219071v1 Escherichia coli Proteobacteria; Gammaproteobacteria; Enterobacterales; Enterobacteriaceae; Escherichia
MNNEETFYQAMRRQGVTRRSFLKYCSLAATSLGLGAGMAPKIAWALENKPRIPVVWIHGL WP_001058323.1
MULTISPECIES: hydrogenase-1 small chain [Proteobacteria] Length: 372\nNumber of predicted TMHs: 2\nExp number of AAs in TMHs: 37.2608499999999\nExp number, first 60 AAs: 19.52923\nTotal prob of N-in: 0.94351\nPOSSIBLE N-term signal sequence\ninside 1 19\nTMhelix 20 39\noutside 40 325\nTMhelix 326 348\ninside 349 372

3464 GCF_002190885.1_ASM219088v1 Escherichia coli Proteobacteria; Gammaproteobacteria; Enterobacterales; Enterobacteriaceae; Escherichia
MTGDNTLIHSHGINRRDFMKLCAALAATMGLSSKAAAEMAESVTNPQRPPVIWIGAQECT WP_000145410.1
MULTISPECIES: hydrogenase-2 small chain [Proteobacteria] Length: 372\nNumber of predicted TMHs: 1\nExp number of AAs in TMHs: 23.72226\nExp number, first 60 AAs: 0.22038\nTotal prob of N-in: 0.12994\noutside 1 331\nTMhelix 332 354\ninside 355 372

3465 GCF_002190945.1_ASM219094v1 Escherichia coli Proteobacteria; Gammaproteobacteria; Enterobacterales; Enterobacteriaceae; Escherichia
MTGDNTLIHSHGINRRDFMKLCAALAATMGLSSKAAAEMAESVTNPQRPPVIWIGAQECT WP_000145410.1
MULTISPECIES: hydrogenase-2 small chain [Proteobacteria] Length: 372\nNumber of predicted TMHs: 1\nExp number of AAs in TMHs: 23.72226\nExp number, first 60 AAs: 0.22038\nTotal prob of N-in: 0.12994\noutside 1 331\nTMhelix 332 354\ninside 355 372

3466 GCF_002193295.1_ASM219329v1 Escherichia coli Proteobacteria; Gammaproteobacteria; Enterobacterales; Enterobacteriaceae; Escherichia
MTGDNTLIHSHGINRRDFMKLCAALAATMGLSSKAAAEMAESVTNPQRPPVIWIGAQECT WP_000145410.1
MULTISPECIES: hydrogenase-2 small chain [Proteobacteria] Length: 372\nNumber of predicted TMHs:

1\nExp number of AAs in TMHs: 23.72226\nExp number, first 60 AAs: 0.22038\nTotal prob of N-in: 0.12994\noutside 1 331\nTMhelix 332 354\ninside 355 372

3467 GCF_002193295.1_ASM219329v1 Escherichia coli Proteobacteria; Gammaproteobacteria; Enterobacterales; Enterobacteriaceae; Escherichia
MNNEETFYQAMRRQGVTRRSFLKYCSLAATSLGLGAGMAPKIAWALENKPRIPVVWIHGL WP_001058323.1
MULTISPECIES: hydrogenase-1 small chain [Proteobacteria] Length: 372\nNumber of predicted TMHs: 2\nExp number of AAs in TMHs: 37.2608499999999\nExp number, first 60 AAs: 19.52923\nTotal prob of N-in: 0.94351\nPOSSIBLE N-term signal sequence\ninside 1 19\nTMhelix 20 39\noutside 40 325\nTMhelix 326 348\ninside 349 372

3468 GCF_002191055.1_ASM219105v1 Escherichia coli Proteobacteria; Gammaproteobacteria; Enterobacterales; Enterobacteriaceae; Escherichia
MNNEETFYQAMRRQGVTRRSFLKYCSLAATSLGLGAGMAPKIAWALENKPRIPVVWIHGL WP_001295940.1
MULTISPECIES: hydrogenase [Enterobacteriaceae] Length: 372\nNumber of predicted TMHs: 2\nExp number of AAs in TMHs: 37.2608899999999\nExp number, first 60 AAs: 19.52923\nTotal prob of N-in: 0.94351\nPOSSIBLE N-term signal sequence\ninside 1 19\nTMhelix 20 39\noutside 40 325\nTMhelix 326 348\ninside 349 372

3469 GCF_002193355.1_ASM219335v1 Escherichia coli Proteobacteria; Gammaproteobacteria; Enterobacterales; Enterobacteriaceae; Escherichia
MTGDNTLIHSHGINRRDFMKLCAALAATMGLSSKAAAEMAESVTNPQRPPVIWIGAQECT WP_000145410.1
MULTISPECIES: hydrogenase-2 small chain [Proteobacteria] Length: 372\nNumber of predicted TMHs: 1\nExp number of AAs in TMHs: 23.72226\nExp number, first 60 AAs: 0.22038\nTotal prob of N-in: 0.12994\noutside 1 331\nTMhelix 332 354\ninside 355 372

3470 GCF_002193355.1_ASM219335v1 Escherichia coli Proteobacteria; Gammaproteobacteria; Enterobacterales; Enterobacteriaceae; Escherichia
MNNEETFYQAMRRQGVTRRSFLKYCSLAATSLGLGAGMAPKIAWALENKPRIPVVWIHGL WP_001058323.1
MULTISPECIES: hydrogenase-1 small chain [Proteobacteria] Length: 372\nNumber of predicted TMHs: 2\nExp number of AAs in TMHs: 37.2608499999999\nExp number, first 60 AAs: 19.52923\nTotal prob of N-in: 0.94351\nPOSSIBLE N-term signal sequence\ninside 1 19\nTMhelix 20 39\noutside 40 325\nTMhelix 326 348\ninside 349 372

3471 GCF_002193245.1_ASM219324v1 Escherichia coli Proteobacteria; Gammaproteobacteria; Enterobacterales; Enterobacteriaceae; Escherichia
MNNEETFYQAMRRQGVTRRSFLKYCSLAATSLGLGAGMAPKIAWALENKPRIPVVWIHGL WP_001058323.1
MULTISPECIES: hydrogenase-1 small chain [Proteobacteria] Length: 372\nNumber of predicted TMHs: 2\nExp number of AAs in TMHs: 37.2608499999999\nExp number, first 60 AAs: 19.52923\nTotal prob of N-in: 0.94351\nPOSSIBLE N-term signal sequence\ninside 1 19\nTMhelix 20 39\noutside 40 325\nTMhelix 326 348\ninside 349 372

3472 GCF_002192275.1_ASM219227v1 Escherichia coli Proteobacteria; Gammaproteobacteria; Enterobacterales; Enterobacteriaceae; Escherichia
MTGDNTLIHSHGINRRDFMKLCAALATTMGLSSKAAAEMAESVTNPQRPPVIWIGAQECT WP_023149629.1
MULTISPECIES: hydrogenase-2 small chain [Enterobacteriaceae] Length: 372\nNumber of predicted TMHs: 1\nExp number of AAs in TMHs: 23.63603\nExp number, first 60 AAs: 0.12971\nTotal prob of N-in: 0.12608\noutside 1 331\nTMhelix 332 354\ninside 355 372

3473 GCF_002194625.1_ASM219462v1 Escherichia coli Proteobacteria; Gammaproteobacteria; Enterobacterales; Enterobacteriaceae; Escherichia
MTGDNTLIHSHGINRRDFMKLCAALAATMGLSSKAAAEMAESVTNPQRPPVIWIGAQECT WP_000145410.1
MULTISPECIES: hydrogenase-2 small chain [Proteobacteria] Length: 372\nNumber of predicted TMHs: 1\nExp number of AAs in TMHs: 23.72226\nExp number, first 60 AAs: 0.22038\nTotal prob of N-in: 0.12994\noutside 1 331\nTMhelix 332 354\ninside 355 372

3474 GCF_002194605.1_ASM219460v1 Escherichia coli Proteobacteria; Gammaproteobacteria; Enterobacterales; Enterobacteriaceae; Escherichia
MTGDNTLIHSHGINRRDFMKLCAALAATMGLSSKAAAEMAESVTNPQRPPVIWIGAQECT WP_000145410.1
MULTISPECIES: hydrogenase-2 small chain [Proteobacteria] Length: 372\nNumber of predicted TMHs:

1\nExp number of AAs in TMHs: 23.72226\nExp number, first 60 AAs: 0.22038\nTotal prob of N-in: 0.12994\noutside 1 331\nTMhelix 332 354\ninside 355 372

3475 GCF_002194675.1_ASM219467v1 Escherichia coli Proteobacteria; Gammaproteobacteria; Enterobacterales; Enterobacteriaceae; Escherichia
MTGDNTLIHSHGINRRDFMKLCAALAATMGLSSKAAAEMAESVTNPQRPPVIWIGAQECT WP_000145410.1
MULTISPECIES: hydrogenase-2 small chain [Proteobacteria] Length: 372\nNumber of predicted TMHs: 1\nExp number of AAs in TMHs: 23.72226\nExp number, first 60 AAs: 0.22038\nTotal prob of N-in: 0.12994\noutside 1 331\nTMhelix 332 354\ninside 355 372

3476 GCF_002194705.1_ASM219470v1 Escherichia coli Proteobacteria; Gammaproteobacteria; Enterobacterales; Enterobacteriaceae; Escherichia
MTGDNTLIHSHGINRRDFMKLCAALAATMGLSSKAAAEMAESVTNPQRPPVIWIGAQECT WP_000145410.1
MULTISPECIES: hydrogenase-2 small chain [Proteobacteria] Length: 372\nNumber of predicted TMHs: 1\nExp number of AAs in TMHs: 23.72226\nExp number, first 60 AAs: 0.22038\nTotal prob of N-in: 0.12994\noutside 1 331\nTMhelix 332 354\ninside 355 372

3477 GCF_002194715.1_ASM219471v1 Escherichia coli Proteobacteria; Gammaproteobacteria; Enterobacterales; Enterobacteriaceae; Escherichia
MTGDNTLIHSHGINRRDFMKLCAALAATMGLSSKAAAEMAESVTNPQRPPVIWIGAQECT WP_000145410.1
MULTISPECIES: hydrogenase-2 small chain [Proteobacteria] Length: 372\nNumber of predicted TMHs: 1\nExp number of AAs in TMHs: 23.72226\nExp number, first 60 AAs: 0.22038\nTotal prob of N-in: 0.12994\noutside 1 331\nTMhelix 332 354\ninside 355 372

3478 GCF_002194765.1_ASM219476v1 Escherichia coli Proteobacteria; Gammaproteobacteria; Enterobacterales; Enterobacteriaceae; Escherichia
MTGDNTLIHSHGINRRDFMKLCAALAATMGLSSKAAAEMAESVTNPQRPPVIWIGAQECT WP_000145410.1
MULTISPECIES: hydrogenase-2 small chain [Proteobacteria] Length: 372\nNumber of predicted TMHs: 1\nExp number of AAs in TMHs: 23.72226\nExp number, first 60 AAs: 0.22038\nTotal prob of N-in: 0.12994\noutside 1 331\nTMhelix 332 354\ninside 355 372

3479 GCF_002194775.1_ASM219477v1 Escherichia coli Proteobacteria; Gammaproteobacteria; Enterobacterales; Enterobacteriaceae; Escherichia
MTGDNTLIHSHGINRRDFMKLCAALAATMGLSSKAAAEMAESVTNPQRPPVIWIGAQECT WP_000145410.1
MULTISPECIES: hydrogenase-2 small chain [Proteobacteria] Length: 372\nNumber of predicted TMHs: 1\nExp number of AAs in TMHs: 23.72226\nExp number, first 60 AAs: 0.22038\nTotal prob of N-in: 0.12994\noutside 1 331\nTMhelix 332 354\ninside 355 372

3480 GCF_002194835.1_ASM219483v1 Escherichia coli Proteobacteria; Gammaproteobacteria; Enterobacterales; Enterobacteriaceae; Escherichia
MTGDNTLIHSHGINRRDFMKLCAALAATMGLSSKAAAEMAESVTNPQRPPVIWIGAQECT WP_000145410.1
MULTISPECIES: hydrogenase-2 small chain [Proteobacteria] Length: 372\nNumber of predicted TMHs: 1\nExp number of AAs in TMHs: 23.72226\nExp number, first 60 AAs: 0.22038\nTotal prob of N-in: 0.12994\noutside 1 331\nTMhelix 332 354\ninside 355 372

3481 GCF_002195185.1_ASM219518v1 Escherichia coli Proteobacteria; Gammaproteobacteria; Enterobacterales; Enterobacteriaceae; Escherichia
MTGDNTLIHSHGINRRDFMKLCAALAATMGLSSKAAAEMAESVTNPQRPPVIWIGAQECT WP_000145410.1
MULTISPECIES: hydrogenase-2 small chain [Proteobacteria] Length: 372\nNumber of predicted TMHs: 1\nExp number of AAs in TMHs: 23.72226\nExp number, first 60 AAs: 0.22038\nTotal prob of N-in: 0.12994\noutside 1 331\nTMhelix 332 354\ninside 355 372

3482 GCF_002195235.1_ASM219523v1 Escherichia coli Proteobacteria; Gammaproteobacteria; Enterobacterales; Enterobacteriaceae; Escherichia
MTGDNTLIHSHGINRRDFMKLCAALAATMGLSSKAAAEMAESVTNPQRPPVIWIGAQECT WP_000145410.1
MULTISPECIES: hydrogenase-2 small chain [Proteobacteria] Length: 372\nNumber of predicted TMHs: 1\nExp number of AAs in TMHs: 23.72226\nExp number, first 60 AAs: 0.22038\nTotal prob of N-in: 0.12994\noutside 1 331\nTMhelix 332 354\ninside 355 372

3483 GCF_002195235.1_ASM219523v1 Escherichia coli Proteobacteria; Gammaproteobacteria; Enterobacterales; Enterobacteriaceae; Escherichia

MNNEETFYQAMRRQGVTRRSFLKYCSLAATSLGLGAGMAPKIAWALENKPRIPVVWIHGL WP_001301613.1 hydrogenase [Escherichia coli] Length: 372\nNumber of predicted TMHs: 2\nExp number of AAs in TMHs: 37.46598\nExp number, first 60 AAs: 19.53139\nTotal prob of N-in: 0.94364\nPOSSIBLE N-term signal sequence\ninside 1 19\nTMhelix 20 39\noutside 40 325\nTMhelix 326 348\ninside 349 372

3484 GCF_002194935.1_ASM219493v1 Escherichia coli Proteobacteria; Gammaproteobacteria; Enterobacterales; Enterobacteriaceae; Escherichia MNNEETFYQAMRRQGVTRRSFLKYCSLAATSLGLGAGMAPKIAWALENKPRIPVVWIHGL WP_001301613.1 hydrogenase [Escherichia coli] Length: 372\nNumber of predicted TMHs: 2\nExp number of AAs in TMHs: 37.46598\nExp number, first 60 AAs: 19.53139\nTotal prob of N-in: 0.94364\nPOSSIBLE N-term signal sequence\ninside 1 19\nTMhelix 20 39\noutside 40 325\nTMhelix 326 348\ninside 349 372

3485 GCF_002195245.1_ASM219524v1 Escherichia coli Proteobacteria; Gammaproteobacteria; Enterobacterales; Enterobacteriaceae; Escherichia MTGDNTLIHSHGINRRDFMKLCAALAATMGLSSKAAAEMAESVTNPQRPPVIWIGAQECT WP_000145410.1 MULTISPECIES: hydrogenase-2 small chain [Proteobacteria] Length: 372\nNumber of predicted TMHs: 1\nExp number of AAs in TMHs: 23.72226\nExp number, first 60 AAs: 0.22038\nTotal prob of N-in: 0.12994\noutside 1 331\nTMhelix 332 354\ninside 355 372

3486 GCF_002195265.1_ASM219526v1 Escherichia coli Proteobacteria; Gammaproteobacteria; Enterobacterales; Enterobacteriaceae; Escherichia MTGDNTLIHSHGINRRDFMKLCAALAATMGLSSKAAAEMAESVTNPQRPPVIWIGAQECT WP_000145410.1 MULTISPECIES: hydrogenase-2 small chain [Proteobacteria] Length: 372\nNumber of predicted TMHs: 1\nExp number of AAs in TMHs: 23.72226\nExp number, first 60 AAs: 0.22038\nTotal prob of N-in: 0.12994\noutside 1 331\nTMhelix 332 354\ninside 355 372

3487 GCF_002195335.1_ASM219533v1 Escherichia coli Proteobacteria; Gammaproteobacteria; Enterobacterales; Enterobacteriaceae; Escherichia MTGDNTLIHSHGINRRDFMKLCAALAATMGLSSKAAAEMAESVTNPQRPPVIWIGAQECT WP_000145410.1 MULTISPECIES: hydrogenase-2 small chain [Proteobacteria] Length: 372\nNumber of predicted TMHs: 1\nExp number of AAs in TMHs: 23.72226\nExp number, first 60 AAs: 0.22038\nTotal prob of N-in: 0.12994\noutside 1 331\nTMhelix 332 354\ninside 355 372

3488 GCF_002195615.1_ASM219561v1 Escherichia coli Proteobacteria; Gammaproteobacteria; Enterobacterales; Enterobacteriaceae; Escherichia MNNEETFYQAMRRQGVTRRSFLKYCSLAATSLGLGAGMAPKIAWALENKPRIPVVWIHGL WP_001301613.1 hydrogenase [Escherichia coli] Length: 372\nNumber of predicted TMHs: 2\nExp number of AAs in TMHs: 37.46598\nExp number, first 60 AAs: 19.53139\nTotal prob of N-in: 0.94364\nPOSSIBLE N-term signal sequence\ninside 1 19\nTMhelix 20 39\noutside 40 325\nTMhelix 326 348\ninside 349 372

3489 GCF_002195675.1_ASM219567v1 Escherichia coli Proteobacteria; Gammaproteobacteria; Enterobacterales; Enterobacteriaceae; Escherichia MNNEETFYQAMRRQGVTRRSFLKYCSLAATSLGLGAGMAPKIAWALENKPRIPVVWIHGL WP_001301613.1 hydrogenase [Escherichia coli] Length: 372\nNumber of predicted TMHs: 2\nExp number of AAs in TMHs: 37.46598\nExp number, first 60 AAs: 19.53139\nTotal prob of N-in: 0.94364\nPOSSIBLE N-term signal sequence\ninside 1 19\nTMhelix 20 39\noutside 40 325\nTMhelix 326 348\ninside 349 372

3490 GCF_002195705.1_ASM219570v1 Escherichia coli Proteobacteria; Gammaproteobacteria; Enterobacterales; Enterobacteriaceae; Escherichia MNNEETFYQAMRRQGVTRRSFLKYCSLAATSLGLGAGMAPKIAWALENKPRIPVVWIHGL WP_001301613.1 hydrogenase [Escherichia coli] Length: 372\nNumber of predicted TMHs: 2\nExp number of AAs in TMHs: 37.46598\nExp number, first 60 AAs: 19.53139\nTotal prob of N-in: 0.94364\nPOSSIBLE N-term signal sequence\ninside 1 19\nTMhelix 20 39\noutside 40 325\nTMhelix 326 348\ninside 349 372

3491 GCF_002195405.1_ASM219540v1 Escherichia coli Proteobacteria; Gammaproteobacteria; Enterobacterales; Enterobacteriaceae; Escherichia MNNEETFYQAMRRQGVTRRSFLKYCSLAATSLGLGAGMAPKIAWALENKPRIPVVWIHGL WP_001301613.1 hydrogenase [Escherichia coli] Length: 372\nNumber of predicted TMHs: 2\nExp number of AAs in TMHs: 37.46598\nExp number, first 60 AAs: 19.53139\nTotal prob of N-in: 0.94364\nPOSSIBLE N-term signal sequence\ninside 1 19\nTMhelix 20 39\noutside 40 325\nTMhelix 326 348\ninside 349 372

3492 GCF_002195725.1_ASM219572v1 Escherichia coli Proteobacteria; Gammaproteobacteria; Enterobacterales; Enterobacteriaceae; Escherichia
MTGDNTLIHSHGINRRDFMKLCAALAATMGLSSKAAAEMAESVTNPQRPPVIWIGAQECT WP_000145410.1
MULTISPECIES: hydrogenase-2 small chain [Proteobacteria] Length: 372\nNumber of predicted TMHs: 1\nExp number of AAs in TMHs: 23.72226\nExp number, first 60 AAs: 0.22038\nTotal prob of N-in: 0.12994\noutside 1 331\nTMhelix 332 354\ninside 355 372

3493 GCF_002195775.1_ASM219577v1 Escherichia coli Proteobacteria; Gammaproteobacteria; Enterobacterales; Enterobacteriaceae; Escherichia
MTGDNTLIHSHGINRRDFMKLCAALAATMGLSSKAAAEMAESVTNPQRPPVIWIGAQECT WP_000145410.1
MULTISPECIES: hydrogenase-2 small chain [Proteobacteria] Length: 372\nNumber of predicted TMHs: 1\nExp number of AAs in TMHs: 23.72226\nExp number, first 60 AAs: 0.22038\nTotal prob of N-in: 0.12994\noutside 1 331\nTMhelix 332 354\ninside 355 372

3494 GCF_002195725.1_ASM219572v1 Escherichia coli Proteobacteria; Gammaproteobacteria; Enterobacterales; Enterobacteriaceae; Escherichia
MNNEETFYQAMRRQGVTRRSFLKYCSLAATSLGLGAGMAPKIAWALENKPRIPVVWIHGL WP_001301613.1
hydrogenase [Escherichia coli] Length: 372\nNumber of predicted TMHs: 2\nExp number of AAs in TMHs: 37.46598\nExp number, first 60 AAs: 19.53139\nTotal prob of N-in: 0.94364\nPOSSIBLE N-term signal sequence\ninside 1 19\nTMhelix 20 39\noutside 40 325\nTMhelix 326 348\ninside 349 372

3495 GCF_002196635.1_ASM219663v1 Escherichia coli Proteobacteria; Gammaproteobacteria; Enterobacterales; Enterobacteriaceae; Escherichia
MNNEETFYQAMRRQGVTRRSFLKYCSLAATSLGLGAGMAPKIAWALENKPRIPVVWIHGL WP_001058323.1
MULTISPECIES: hydrogenase-1 small chain [Proteobacteria] Length: 372\nNumber of predicted TMHs: 2\nExp number of AAs in TMHs: 37.2608499999999\nExp number, first 60 AAs: 19.52923\nTotal prob of N-in: 0.94351\nPOSSIBLE N-term signal sequence\ninside 1 19\nTMhelix 20 39\noutside 40 325\nTMhelix 326 348\ninside 349 372

3496 GCF_002196495.1_ASM219649v1 Escherichia coli Proteobacteria; Gammaproteobacteria; Enterobacterales; Enterobacteriaceae; Escherichia
MNNEETFYQAMRRQGVTRRSFLKYCSLAATSLGLGAGMAPKIAWALENKPRIPVVWIHGL WP_001058323.1
MULTISPECIES: hydrogenase-1 small chain [Proteobacteria] Length: 372\nNumber of predicted TMHs: 2\nExp number of AAs in TMHs: 37.2608499999999\nExp number, first 60 AAs: 19.52923\nTotal prob of N-in: 0.94351\nPOSSIBLE N-term signal sequence\ninside 1 19\nTMhelix 20 39\noutside 40 325\nTMhelix 326 348\ninside 349 372

3497 GCF_002196735.1_ASM219673v1 Escherichia coli Proteobacteria; Gammaproteobacteria; Enterobacterales; Enterobacteriaceae; Escherichia
MTGDNTLIHSHGINRRDFMKLCAALAATMGLSSKAAAEMAESVTNPQRPPVIWIGAQECT WP_000145410.1
MULTISPECIES: hydrogenase-2 small chain [Proteobacteria] Length: 372\nNumber of predicted TMHs: 1\nExp number of AAs in TMHs: 23.72226\nExp number, first 60 AAs: 0.22038\nTotal prob of N-in: 0.12994\noutside 1 331\nTMhelix 332 354\ninside 355 372

3498 GCF_002197975.1_ASM219797v1 Escherichia coli Proteobacteria; Gammaproteobacteria; Enterobacterales; Enterobacteriaceae; Escherichia
MTGDNTLIHSHGINRRDFMKLCAALAATMGLSSKAAAEMAESVTNPQRPPVIWIGAQECT WP_000145410.1
MULTISPECIES: hydrogenase-2 small chain [Proteobacteria] Length: 372\nNumber of predicted TMHs: 1\nExp number of AAs in TMHs: 23.72226\nExp number, first 60 AAs: 0.22038\nTotal prob of N-in: 0.12994\noutside 1 331\nTMhelix 332 354\ninside 355 372

3499 GCF_002196735.1_ASM219673v1 Escherichia coli Proteobacteria; Gammaproteobacteria; Enterobacterales; Enterobacteriaceae; Escherichia
MNNEETFYQAMRRQGVTRRSFLKYCSLAATSLGLGAGMAPKIAWALENKPRIPVVWIHGL WP_001058323.1
MULTISPECIES: hydrogenase-1 small chain [Proteobacteria] Length: 372\nNumber of predicted TMHs: 2\nExp number of AAs in TMHs: 37.2608499999999\nExp number, first 60 AAs: 19.52923\nTotal prob of N-in: 0.94351\nPOSSIBLE N-term signal sequence\ninside 1 19\nTMhelix 20 39\noutside 40 325\nTMhelix 326 348\ninside 349 372

3500 GCF_002197975.1_ASM219797v1 Escherichia coli Proteobacteria; Gammaproteobacteria; Enterobacterales; Enterobacteriaceae; Escherichia

MNNEETFYQAMRRQGVTRRSFLKYCSLAATSLGLGAGMAPKIAWALENKPRIPVVWIHGL WP_001058323.1 MULTISPECIES: hydrogenase-1 small chain [Proteobacteria] Length: 372\nNumber of predicted TMHs: 2\nExp number of AAs in TMHs: 37.2608499999999\nExp number, first 60 AAs: 19.52923\nTotal prob of N-in: 0.94351\nPOSSIBLE N-term signal sequence\ninside 1 19\nTMhelix 20 39\noutside 40 325\nTMhelix 326 348\ninside 349 372

3501 GCF_900007725.1_M885_contigs Escherichia coli Proteobacteria; Gammaproteobacteria; Enterobacterales; Enterobacteriaceae; Escherichia MTGDNTLIHSHGINRRDFMKLCAALAATMGLSSKAAAEMAESVTNPQRPPVIWIGAQECT WP_000145410.1 MULTISPECIES: hydrogenase-2 small chain [Proteobacteria] Length: 372\nNumber of predicted TMHs: 1\nExp number of AAs in TMHs: 23.72226\nExp number, first 60 AAs: 0.22038\nTotal prob of N-in: 0.12994\noutside 1 331\nTMhelix 332 354\ninside 355 372

3502 GCF_900007315.1_VAP_LM33 Escherichia coli Proteobacteria; Gammaproteobacteria; Enterobacterales; Enterobacteriaceae; Escherichia MTGDNTLIHSHGINRRDFMKLCAALAATMGLSSKAAAEMAESVTNPQRPPVIWIGAQECT WP_000145410.1 MULTISPECIES: hydrogenase-2 small chain [Proteobacteria] Length: 372\nNumber of predicted TMHs: 1\nExp number of AAs in TMHs: 23.72226\nExp number, first 60 AAs: 0.22038\nTotal prob of N-in: 0.12994\noutside 1 331\nTMhelix 332 354\ninside 355 372

3503 GCF_900007735.1_M900_contigs Escherichia coli Proteobacteria; Gammaproteobacteria; Enterobacterales; Enterobacteriaceae; Escherichia MTGDNTLIHSHGINRRDFMKLCAALAATMGLSSKAAAEMAESVTNPQRPPVIWIGAQECT WP_000145410.1 MULTISPECIES: hydrogenase-2 small chain [Proteobacteria] Length: 372\nNumber of predicted TMHs: 1\nExp number of AAs in TMHs: 23.72226\nExp number, first 60 AAs: 0.22038\nTotal prob of N-in: 0.12994\noutside 1 331\nTMhelix 332 354\ninside 355 372

3504 GCF_900007725.1_M885_contigs Escherichia coli Proteobacteria; Gammaproteobacteria; Enterobacterales; Enterobacteriaceae; Escherichia MNNEETFYQAMRRQGVTRRSFLKYCSLAATSLGLGAGMAPKIAWALENKPRIPVVWIHGL WP_001058323.1 MULTISPECIES: hydrogenase-1 small chain [Proteobacteria] Length: 372\nNumber of predicted TMHs: 2\nExp number of AAs in TMHs: 37.2608499999999\nExp number, first 60 AAs: 19.52923\nTotal prob of N-in: 0.94351\nPOSSIBLE N-term signal sequence\ninside 1 19\nTMhelix 20 39\noutside 40 325\nTMhelix 326 348\ninside 349 372

3505 GCF_900007735.1_M900_contigs Escherichia coli Proteobacteria; Gammaproteobacteria; Enterobacterales; Enterobacteriaceae; Escherichia MNNEETFYQAMRRQGVTRRSFLKYCSLAATSLGLGAGMAPKIAWALENKPRIPVVWIHGL WP_001058323.1 MULTISPECIES: hydrogenase-1 small chain [Proteobacteria] Length: 372\nNumber of predicted TMHs: 2\nExp number of AAs in TMHs: 37.2608499999999\nExp number, first 60 AAs: 19.52923\nTotal prob of N-in: 0.94351\nPOSSIBLE N-term signal sequence\ninside 1 19\nTMhelix 20 39\noutside 40 325\nTMhelix 326 348\ninside 349 372

3506 GCF_900007315.1_VAP_LM33 Escherichia coli Proteobacteria; Gammaproteobacteria; Enterobacterales; Enterobacteriaceae; Escherichia MNNEETFYQAMRRQGVTRRSFLKYCSLAATSLGLGAGMAPKIAWALENKPRIPVVWIHGL WP_001295940.1 MULTISPECIES: hydrogenase [Enterobacteriaceae] Length: 372\nNumber of predicted TMHs: 2\nExp number of AAs in TMHs: 37.2608899999999\nExp number, first 60 AAs: 19.52923\nTotal prob of N-in: 0.94351\nPOSSIBLE N-term signal sequence\ninside 1 19\nTMhelix 20 39\noutside 40 325\nTMhelix 326 348\ninside 349 372

3507 GCF_002207985.1_ASM220798v1 Escherichia coli Proteobacteria; Gammaproteobacteria; Enterobacterales; Enterobacteriaceae; Escherichia MNNEETFYQAMRRQGVTRRSFLKYCSLAATSLGLGAGMAPKIAWALENKPRIPVVWIHGL WP_001058323.1 MULTISPECIES: hydrogenase-1 small chain [Proteobacteria] Length: 372\nNumber of predicted TMHs: 2\nExp number of AAs in TMHs: 37.2608499999999\nExp number, first 60 AAs: 19.52923\nTotal prob of N-in: 0.94351\nPOSSIBLE N-term signal sequence\ninside 1 19\nTMhelix 20 39\noutside 40 325\nTMhelix 326 348\ninside 349 372

3508 GCF_900015735.1_ED178_contigs Escherichia coli Proteobacteria; Gammaproteobacteria; Enterobacterales; Enterobacteriaceae; Escherichia

MTGDNTLIHSHGINRRDFMKLCAALAATMGLSSKAAAEMAESVTNPQRPPVIWIGAQECT WP_000145410.1 MULTISPECIES: hydrogenase-2 small chain [Proteobacteria] Length: 372\nNumber of predicted TMHs: 1\nExp number of AAs in TMHs: 23.72226\nExp number, first 60 AAs: 0.22038\nTotal prob of N-in: 0.12994\noutside 1 331\nTMhelix 332 354\ninside 355 372

3509 GCF_900015735.1_ED178_contigs Escherichia coli Proteobacteria; Gammaproteobacteria; Enterobacterales; Enterobacteriaceae; Escherichia MNNEETFYQAMRRQGVTRRSFLKYCSLAATSLGLGAGMAPKIAWALENKPRIPVVWIHGL WP_001058323.1 MULTISPECIES: hydrogenase-1 small chain [Proteobacteria] Length: 372\nNumber of predicted TMHs: 2\nExp number of AAs in TMHs: 37.2608499999999\nExp number, first 60 AAs: 19.52923\nTotal prob of N-in: 0.94351\nPOSSIBLE N-term signal sequence\ninside 1 19\nTMhelix 20 39\noutside 40 325\nTMhelix 326 348\ninside 349 372

3510 GCF_900015715.1_ED449_contigs Escherichia coli Proteobacteria; Gammaproteobacteria; Enterobacterales; Enterobacteriaceae; Escherichia MNNEETFYQAMRRQGVTRRSFLKYCSLAATSLGLGAGMAPKIAWALENKPRIPVVWIHGL WP_001301613.1 hydrogenase [Escherichia coli] Length: 372\nNumber of predicted TMHs: 2\nExp number of AAs in TMHs: 37.46598\nExp number, first 60 AAs: 19.53139\nTotal prob of N-in: 0.94364\nPOSSIBLE N-term signal sequence\ninside 1 19\nTMhelix 20 39\noutside 40 325\nTMhelix 326 348\ninside 349 372

3511 GCF_900014325.1_ED657_contigs Escherichia coli Proteobacteria; Gammaproteobacteria; Enterobacterales; Enterobacteriaceae; Escherichia MNNEETFYQAMRRQGVTRRSFLKYCSLAATSLGLGAGMAPKIAWALENKPRIPVVWIHGL WP_001058323.1 MULTISPECIES: hydrogenase-1 small chain [Proteobacteria] Length: 372\nNumber of predicted TMHs: 2\nExp number of AAs in TMHs: 37.2608499999999\nExp number, first 60 AAs: 19.52923\nTotal prob of N-in: 0.94351\nPOSSIBLE N-term signal sequence\ninside 1 19\nTMhelix 20 39\noutside 40 325\nTMhelix 326 348\ninside 349 372

3512 GCF_900015765.1_ED414_contigs Escherichia coli Proteobacteria; Gammaproteobacteria; Enterobacterales; Enterobacteriaceae; Escherichia MTGDNTLIHSHGINRRDFMKLCAALAATMGLSSKAAAEMAESVTNPQRPPVIWIGAQECT WP_000145410.1 MULTISPECIES: hydrogenase-2 small chain [Proteobacteria] Length: 372\nNumber of predicted TMHs: 1\nExp number of AAs in TMHs: 23.72226\nExp number, first 60 AAs: 0.22038\nTotal prob of N-in: 0.12994\noutside 1 331\nTMhelix 332 354\ninside 355 372

3513 GCF_900015785.1_ED073_contigs Escherichia coli Proteobacteria; Gammaproteobacteria; Enterobacterales; Enterobacteriaceae; Escherichia MTGDNTLIHSHGINRRDFMKLCAALAATMGLSSKAAAEMAESVTNPQRPPVIWIGAQECT WP_000145410.1 MULTISPECIES: hydrogenase-2 small chain [Proteobacteria] Length: 372\nNumber of predicted TMHs: 1\nExp number of AAs in TMHs: 23.72226\nExp number, first 60 AAs: 0.22038\nTotal prob of N-in: 0.12994\noutside 1 331\nTMhelix 332 354\ninside 355 372

3514 GCF_900015805.1_ED287_contigs Escherichia coli Proteobacteria; Gammaproteobacteria; Enterobacterales; Enterobacteriaceae; Escherichia MTGDNTLIHSHGINRRDFMKLCAALAATMGLSSKAAAEMAESVTNPQRPPVIWIGAQECT WP_000145410.1 MULTISPECIES: hydrogenase-2 small chain [Proteobacteria] Length: 372\nNumber of predicted TMHs: 1\nExp number of AAs in TMHs: 23.72226\nExp number, first 60 AAs: 0.22038\nTotal prob of N-in: 0.12994\noutside 1 331\nTMhelix 332 354\ninside 355 372

3515 GCF_900015815.1_ED603_contigs Escherichia coli Proteobacteria; Gammaproteobacteria; Enterobacterales; Enterobacteriaceae; Escherichia MTGDNTLIHSHGINRRDFMKLCAALAATMGLSSKAAAEMAESVTNPQRPPVIWIGAQECT WP_000145410.1 MULTISPECIES: hydrogenase-2 small chain [Proteobacteria] Length: 372\nNumber of predicted TMHs: 1\nExp number of AAs in TMHs: 23.72226\nExp number, first 60 AAs: 0.22038\nTotal prob of N-in: 0.12994\noutside 1 331\nTMhelix 332 354\ninside 355 372

3516 GCF_900015855.1_ED180_contigs Escherichia coli Proteobacteria; Gammaproteobacteria; Enterobacterales; Enterobacteriaceae; Escherichia MTGDNTLIHSHGINRRDFMKLCAALAATMGLSSKAAAEMAESVTNPQRPPVIWIGAQECT WP_000145410.1 MULTISPECIES: hydrogenase-2 small chain [Proteobacteria] Length: 372\nNumber of predicted TMHs:

1\nExp number of AAs in TMHs: 23.72226\nExp number, first 60 AAs: 0.22038\nTotal prob of N-in: 0.12994\noutside 1 331\nTMhelix 332 354\ninside 355 372

3517 GCF_900015865.1_ED411_contigs Escherichia coli Proteobacteria; Gammaproteobacteria; Enterobacterales; Enterobacteriaceae; Escherichia
MTGDNTLIHSHGINRRDFMKLCAALAATMGLSSKAAAEMAESVTNPQRPPVIWIGAQECT WP_000145410.1
MULTISPECIES: hydrogenase-2 small chain [Proteobacteria] Length: 372\nNumber of predicted TMHs: 1\nExp number of AAs in TMHs: 23.72226\nExp number, first 60 AAs: 0.22038\nTotal prob of N-in: 0.12994\noutside 1 331\nTMhelix 332 354\ninside 355 372

3518 GCF_900015885.1_ED729_contigs Escherichia coli Proteobacteria; Gammaproteobacteria; Enterobacterales; Enterobacteriaceae; Escherichia
MTGDNTLIHSHGINRRDFMKLCAALAATMGLSSKAAAEMAESVTNPQRPPVIWIGAQECT WP_000145410.1
MULTISPECIES: hydrogenase-2 small chain [Proteobacteria] Length: 372\nNumber of predicted TMHs: 1\nExp number of AAs in TMHs: 23.72226\nExp number, first 60 AAs: 0.22038\nTotal prob of N-in: 0.12994\noutside 1 331\nTMhelix 332 354\ninside 355 372

3519 GCF_900015905.1_ED679_contigs Escherichia coli Proteobacteria; Gammaproteobacteria; Enterobacterales; Enterobacteriaceae; Escherichia
MTGDNTLIHSHGINRRDFMKLCAALAATMGLSSKAAAEMAESVTNPQRPPVIWIGAQECT WP_000145410.1
MULTISPECIES: hydrogenase-2 small chain [Proteobacteria] Length: 372\nNumber of predicted TMHs: 1\nExp number of AAs in TMHs: 23.72226\nExp number, first 60 AAs: 0.22038\nTotal prob of N-in: 0.12994\noutside 1 331\nTMhelix 332 354\ninside 355 372

3520 GCF_900015985.1_M856_contigs Escherichia coli Proteobacteria; Gammaproteobacteria; Enterobacterales; Enterobacteriaceae; Escherichia
MTGDNTLIHSHGINRRDFMKLCAALAATMGLSSKAAAEMVESVTNPQRPPVIWIGAQECT WP_000145423.1
hydrogenase 2 small subunit [Escherichia coli] Length: 372\nNumber of predicted TMHs: 1\nExp number of AAs in TMHs: 23.78201\nExp number, first 60 AAs: 0.28355\nTotal prob of N-in: 0.13257\noutside 1 331\nTMhelix 332 354\ninside 355 372

3521 GCF_900015945.1_ED636_contigs Escherichia coli Proteobacteria; Gammaproteobacteria; Enterobacterales; Enterobacteriaceae; Escherichia
MTGDNTLIHSHGINRRDFMKLCAALAATMGLSSKAAAEMAESVTNPQRPPVIWIGAQECT WP_000145410.1
MULTISPECIES: hydrogenase-2 small chain [Proteobacteria] Length: 372\nNumber of predicted TMHs: 1\nExp number of AAs in TMHs: 23.72226\nExp number, first 60 AAs: 0.22038\nTotal prob of N-in: 0.12994\noutside 1 331\nTMhelix 332 354\ninside 355 372

3522 GCF_900016055.1_BCW5717_contigs Escherichia coli Proteobacteria; Gammaproteobacteria; Enterobacterales; Enterobacteriaceae; Escherichia
MNNEETFYQAMRRQGVTRRSFLKYCSLAATSLGLGAGMAPKIAWALENKPRIPVVWIHGL WP_001295940.1
MULTISPECIES: hydrogenase [Enterobacteriaceae] Length: 372\nNumber of predicted TMHs: 2\nExp number of AAs in TMHs: 37.2608899999999\nExp number, first 60 AAs: 19.52923\nTotal prob of N-in: 0.94351\nPOSSIBLE N-term signal sequence\ninside 1 19\nTMhelix 20 39\noutside 40 325\nTMhelix 326 348\ninside 349 372

3523 GCF_900015945.1_ED636_contigs Escherichia coli Proteobacteria; Gammaproteobacteria; Enterobacterales; Enterobacteriaceae; Escherichia
MNNEETFYQAMRRQGVTRRSFLKYCSLAATSLGLGAGMAPKIAWALENKPRIPVVWIHGL WP_001301613.1
hydrogenase [Escherichia coli] Length: 372\nNumber of predicted TMHs: 2\nExp number of AAs in TMHs: 37.46598\nExp number, first 60 AAs: 19.53139\nTotal prob of N-in: 0.94364\nPOSSIBLE N-term signal sequence\ninside 1 19\nTMhelix 20 39\noutside 40 325\nTMhelix 326 348\ninside 349 372

3524 GCF_900016085.1_ED366_contigs Escherichia coli Proteobacteria; Gammaproteobacteria; Enterobacterales; Enterobacteriaceae; Escherichia
MTGDNTLIHSHGINRRDFMKLCAALAATMGLSSKAAAEMAESVTNPQRPPVIWIGAQECT WP_000145410.1
MULTISPECIES: hydrogenase-2 small chain [Proteobacteria] Length: 372\nNumber of predicted TMHs: 1\nExp number of AAs in TMHs: 23.72226\nExp number, first 60 AAs: 0.22038\nTotal prob of N-in: 0.12994\noutside 1 331\nTMhelix 332 354\ninside 355 372

3525 GCF_900016115.1_ED430_contigs Escherichia coli Proteobacteria; Gammaproteobacteria; Enterobacterales; Enterobacteriaceae; Escherichia MTGDNTLIHSHGINRRDFMKLCAALAATMGLSSKAAAEMAESVTNPQRPPVIWIGAQECT WP_000145410.1 MULTISPECIES: hydrogenase-2 small chain [Proteobacteria] Length: 372\nNumber of predicted TMHs: 1\nExp number of AAs in TMHs: 23.72226\nExp number, first 60 AAs: 0.22038\nTotal prob of N-in: 0.12994\noutside 1 331\nTMhelix 332 354\ninside 355 372

3526 GCF_900042775.1_F1B2 Escherichia coli Proteobacteria; Gammaproteobacteria; Enterobacterales; Enterobacteriaceae; Escherichia MTGDNTLIHSHGINRRDFMKLCAALAATMGLSSKAAAEMAESVTNPQRPPVIWIGAQECT WP_000145410.1 MULTISPECIES: hydrogenase-2 small chain [Proteobacteria] Length: 372\nNumber of predicted TMHs: 1\nExp number of AAs in TMHs: 23.72226\nExp number, first 60 AAs: 0.22038\nTotal prob of N-in: 0.12994\noutside 1 331\nTMhelix 332 354\ninside 355 372

3527 GCF_900039535.1_F13P5 Escherichia coli Proteobacteria; Gammaproteobacteria; Enterobacterales; Enterobacteriaceae; Escherichia MTGDNTLIHSHGINRRDFMKLCAALAATMGLSSKAAAEMAESVTNPQRPPVIWIGAQECT WP_000145410.1 MULTISPECIES: hydrogenase-2 small chain [Proteobacteria] Length: 372\nNumber of predicted TMHs: 1\nExp number of AAs in TMHs: 23.72226\nExp number, first 60 AAs: 0.22038\nTotal prob of N-in: 0.12994\noutside 1 331\nTMhelix 332 354\ninside 355 372

3528 GCF_900016135.1_ED444_contigs Escherichia coli Proteobacteria; Gammaproteobacteria; Enterobacterales; Enterobacteriaceae; Escherichia MTGDNTLIHSHGINRRDFMKLCAALAATMGLSSKAAAEMAESVTNPQRPPVIWIGAQECT WP_000145410.1 MULTISPECIES: hydrogenase-2 small chain [Proteobacteria] Length: 372\nNumber of predicted TMHs: 1\nExp number of AAs in TMHs: 23.72226\nExp number, first 60 AAs: 0.22038\nTotal prob of N-in: 0.12994\noutside 1 331\nTMhelix 332 354\ninside 355 372

3529 GCF_900089795.1_CQ09 Escherichia coli Proteobacteria; Gammaproteobacteria; Enterobacterales; Enterobacteriaceae; Escherichia MTGDNTLIHSHGINRRDFMKLCAALAATMGLSSKAAAEMAESVTNPQRPPVIWIGAQECT WP_000145410.1 MULTISPECIES: hydrogenase-2 small chain [Proteobacteria] Length: 372\nNumber of predicted TMHs: 1\nExp number of AAs in TMHs: 23.72226\nExp number, first 60 AAs: 0.22038\nTotal prob of N-in: 0.12994\noutside 1 331\nTMhelix 332 354\ninside 355 372

3530 GCF_900093825.1_scaffolds_20160015 Escherichia coli Proteobacteria; Gammaproteobacteria; Enterobacterales; Enterobacteriaceae; Escherichia MTGDNTLIHSHGINRRDFMKLCAALAATMGLSSKAAAEMAESVTNPQRPPVIWIGAQECT WP_000145410.1 MULTISPECIES: hydrogenase-2 small chain [Proteobacteria] Length: 372\nNumber of predicted TMHs: 1\nExp number of AAs in TMHs: 23.72226\nExp number, first 60 AAs: 0.22038\nTotal prob of N-in: 0.12994\noutside 1 331\nTMhelix 332 354\ninside 355 372

3531 GCF_900093835.1_scaffolds_20151360 Escherichia coli Proteobacteria; Gammaproteobacteria; Enterobacterales; Enterobacteriaceae; Escherichia MTGDNTLIHSHGINRRDFMKLCAALAATMGLSSKAAAEMAESVTNPQRPPVIWIGAQECT WP_000145410.1 MULTISPECIES: hydrogenase-2 small chain [Proteobacteria] Length: 372\nNumber of predicted TMHs: 1\nExp number of AAs in TMHs: 23.72226\nExp number, first 60 AAs: 0.22038\nTotal prob of N-in: 0.12994\noutside 1 331\nTMhelix 332 354\ninside 355 372

3532 GCF_900093875.1_scaffolds_C333 Escherichia coli Proteobacteria; Gammaproteobacteria; Enterobacterales; Enterobacteriaceae; Escherichia MTGDNTLIHSHGINRRDFMKLCAALAATMGLSSKAAAEMAESVTNPQRPPVIWIGAQECT WP_000145410.1 MULTISPECIES: hydrogenase-2 small chain [Proteobacteria] Length: 372\nNumber of predicted TMHs: 1\nExp number of AAs in TMHs: 23.72226\nExp number, first 60 AAs: 0.22038\nTotal prob of N-in: 0.12994\noutside 1 331\nTMhelix 332 354\ninside 355 372

3533 GCF_900093855.1_scaffolds_20160007 Escherichia coli Proteobacteria; Gammaproteobacteria; Enterobacterales; Enterobacteriaceae; Escherichia MTGDNTLIHSHGINRRDFMKLCAALAATMGLSSKAAAEMAESVTNPQRPPVIWIGAQECT WP_000145410.1 MULTISPECIES: hydrogenase-2 small chain [Proteobacteria] Length: 372\nNumber of predicted TMHs: 1\nExp number of AAs in TMHs: 23.72226\nExp number, first 60 AAs: 0.22038\nTotal prob of N-in: 0.12994\noutside 1 331\nTMhelix 332 354\ninside 355 372

3534 GCF_900088965.1_ASM90008896v1 Escherichia coli Proteobacteria; Gammaproteobacteria; Enterobacterales; Enterobacteriaceae; Escherichia
MNNEETFYQAMRRQGVTRRSFLKYCSLAATSLGLGAGMAPKIAWALENKPRIPVVWIHGL WP_001058323.1
MULTISPECIES: hydrogenase-1 small chain [Proteobacteria] Length: 372\nNumber of predicted TMHs: 2\nExp number of AAs in TMHs: 37.2608499999999\nExp number, first 60 AAs: 19.52923\nTotal prob of N-in: 0.94351\nPOSSIBLE N-term signal sequence\ninside 1 19\nTMhelix 20 39\noutside 40 325\nTMhelix 326 348\ninside 349 372

3535 GCF_900096815.1_Ecoli_AG100_Sample2_M9_Assembly Escherichia coli Proteobacteria; Gammaproteobacteria; Enterobacterales; Enterobacteriaceae; Escherichia
MTGDNTLIHSHGINRRDFMKLCAALAATMGLSSKAAAEMAESVTNPQRPPVIWIGAQECT WP_000145410.1
MULTISPECIES: hydrogenase-2 small chain [Proteobacteria] Length: 372\nNumber of predicted TMHs: 1\nExp number of AAs in TMHs: 23.72226\nExp number, first 60 AAs: 0.22038\nTotal prob of N-in: 0.12994\noutside 1 331\nTMhelix 332 354\ninside 355 372

3536 GCF_900093835.1_scaffolds_20151360 Escherichia coli Proteobacteria; Gammaproteobacteria; Enterobacterales; Enterobacteriaceae; Escherichia
MNNEETFYQAMRRQGVTRRSFLKYCSLAATSLGLGAGMAPKIAWALENKPRIPVVWIHGL WP_001295940.1
MULTISPECIES: hydrogenase [Enterobacteriaceae] Length: 372\nNumber of predicted TMHs: 2\nExp number of AAs in TMHs: 37.2608899999999\nExp number, first 60 AAs: 19.52923\nTotal prob of N-in: 0.94351\nPOSSIBLE N-term signal sequence\ninside 1 19\nTMhelix 20 39\noutside 40 325\nTMhelix 326 348\ninside 349 372

3537 GCF_900093985.1_scaffolds_20160017 Escherichia coli Proteobacteria; Gammaproteobacteria; Enterobacterales; Enterobacteriaceae; Escherichia
MTGDNTLIHSHGINRRDFMKLCAALAATMGLSSKAAAEMAESVTNPQRPPVIWIGAQECT WP_000145410.1
MULTISPECIES: hydrogenase-2 small chain [Proteobacteria] Length: 372\nNumber of predicted TMHs: 1\nExp number of AAs in TMHs: 23.72226\nExp number, first 60 AAs: 0.22038\nTotal prob of N-in: 0.12994\noutside 1 331\nTMhelix 332 354\ninside 355 372

3538 GCF_900094005.1_scaffolds_20160006 Escherichia coli Proteobacteria; Gammaproteobacteria; Enterobacterales; Enterobacteriaceae; Escherichia
MTGDNTLIHSHGINRRDFMKLCAALAATMGLSSKAAAEMAESVTNPQRPPVIWIGAQECT WP_000145410.1
MULTISPECIES: hydrogenase-2 small chain [Proteobacteria] Length: 372\nNumber of predicted TMHs: 1\nExp number of AAs in TMHs: 23.72226\nExp number, first 60 AAs: 0.22038\nTotal prob of N-in: 0.12994\noutside 1 331\nTMhelix 332 354\ninside 355 372

3539 GCF_900094015.1_scaffolds_20151012 Escherichia coli Proteobacteria; Gammaproteobacteria; Enterobacterales; Enterobacteriaceae; Escherichia
MTGDNTLIHSHGINRRDFMKLCAALAATMGLSSKAAAEMAESVTNPQRPPVIWIGAQECT WP_000145410.1
MULTISPECIES: hydrogenase-2 small chain [Proteobacteria] Length: 372\nNumber of predicted TMHs: 1\nExp number of AAs in TMHs: 23.72226\nExp number, first 60 AAs: 0.22038\nTotal prob of N-in: 0.12994\noutside 1 331\nTMhelix 332 354\ninside 355 372

3540 GCF_900093985.1_scaffolds_20160017 Escherichia coli Proteobacteria; Gammaproteobacteria; Enterobacterales; Enterobacteriaceae; Escherichia
MNNEETFYQAMRRQGVTRRSFLKYCSLAATSLGLGAGMAPKIAWALENKPRIPVVWIHGL WP_001058323.1
MULTISPECIES: hydrogenase-1 small chain [Proteobacteria] Length: 372\nNumber of predicted TMHs: 2\nExp number of AAs in TMHs: 37.2608499999999\nExp number, first 60 AAs: 19.52923\nTotal prob of N-in: 0.94351\nPOSSIBLE N-term signal sequence\ninside 1 19\nTMhelix 20 39\noutside 40 325\nTMhelix 326 348\ninside 349 372

3541 GCF_900096785.1_Ecoli_AG100_Sample2_Doxycycline_Assembly Escherichia coli Proteobacteria; Gammaproteobacteria; Enterobacterales; Enterobacteriaceae; Escherichia
MTGDNTLIHSHGINRRDFMKLCAALAATMGLSSKAAAEMAESVTNPQRPPVIWIGAQECT WP_000145410.1
MULTISPECIES: hydrogenase-2 small chain [Proteobacteria] Length: 372\nNumber of predicted TMHs: 1\nExp number of AAs in TMHs: 23.72226\nExp number, first 60 AAs: 0.22038\nTotal prob of N-in: 0.12994\noutside 1 331\nTMhelix 332 354\ninside 355 372

3542 GCF_900096805.1_Ecoli_AG100_Sample1_Wildtype_Assembly Escherichia coli Proteobacteria; Gammaproteobacteria; Enterobacterales; Enterobacteriaceae; Escherichia

MTGDNTLIHSHGINRRDFMKLCAALAATMGLSSKAAAEMAESVTNPQRPPVIWIGAQECT WP_000145410.1 MULTISPECIES: hydrogenase-2 small chain [Proteobacteria] Length: 372\nNumber of predicted TMHs: 1\nExp number of AAs in TMHs: 23.72226\nExp number, first 60 AAs: 0.22038\nTotal prob of N-in: 0.12994\noutside 1 331\nTMhelix 332 354\ninside 355 372

3543 GCF_900128765.1_ASM90012876v1 Escherichia coli Proteobacteria; Gammaproteobacteria; Enterobacterales; Enterobacteriaceae; Escherichia MNNEETFYQAMRRQGVTRRSFLKYCSLAATSLGLGAGMAPKIAWALENKPRIPVVWIHGL WP_001058323.1 MULTISPECIES: hydrogenase-1 small chain [Proteobacteria] Length: 372\nNumber of predicted TMHs: 2\nExp number of AAs in TMHs: 37.2608499999999\nExp number, first 60 AAs: 19.52923\nTotal prob of N-in: 0.94351\nPOSSIBLE N-term signal sequence\ninside 1 19\nTMhelix 20 39\noutside 40 325\nTMhelix 326 348\ninside 349 372

3544 GCF_900166685.1_Isolate_2 Escherichia coli Proteobacteria; Gammaproteobacteria; Enterobacterales; Enterobacteriaceae; Escherichia MNNEETFYQAMRRQGVTRRSFLKYCSLAATSLGLGAGMAPKIAWALENKPRIPVVWIHGL WP_001295940.1 MULTISPECIES: hydrogenase [Enterobacteriaceae] Length: 372\nNumber of predicted TMHs: 2\nExp number of AAs in TMHs: 37.2608899999999\nExp number, first 60 AAs: 19.52923\nTotal prob of N-in: 0.94351\nPOSSIBLE N-term signal sequence\ninside 1 19\nTMhelix 20 39\noutside 40 325\nTMhelix 326 348\ninside 349 372

3545 GCF_900096805.1_Ecoli_AG100_Sample1_Wildtype_Assembly Escherichia coli Proteobacteria; Gammaproteobacteria; Enterobacterales; Enterobacteriaceae; Escherichia MNNEETFYQAMRRQGVTRRSFLKYCSLAATSLGLGAGMAPKIAWALENKPRIPVVWIHGL WP_001058323.1 MULTISPECIES: hydrogenase-1 small chain [Proteobacteria] Length: 372\nNumber of predicted TMHs: 2\nExp number of AAs in TMHs: 37.2608499999999\nExp number, first 60 AAs: 19.52923\nTotal prob of N-in: 0.94351\nPOSSIBLE N-term signal sequence\ninside 1 19\nTMhelix 20 39\noutside 40 325\nTMhelix 326 348\ninside 349 372

3546 GCF_900166695.1_Isolate_3 Escherichia coli Proteobacteria; Gammaproteobacteria; Enterobacterales; Enterobacteriaceae; Escherichia MTGDNTLIHSHGINRRDFMKLCAALAATMGLSSKAAAEMVESVTNPQRPPVIWIGAQECT WP_000145423.1 hydrogenase 2 small subunit [Escherichia coli] Length: 372\nNumber of predicted TMHs: 1\nExp number of AAs in TMHs: 23.78201\nExp number, first 60 AAs: 0.28355\nTotal prob of N-in: 0.13257\noutside 1 331\nTMhelix 332 354\ninside 355 372

3547 GCF_900166715.1_Isolate_16 Escherichia coli Proteobacteria; Gammaproteobacteria; Enterobacterales; Enterobacteriaceae; Escherichia MTGDNTLIHSHGINRRDFMKLCAALAATMGLSSKAAAEMVESVTNPQRPPVIWIGAQECT WP_000145423.1 hydrogenase 2 small subunit [Escherichia coli] Length: 372\nNumber of predicted TMHs: 1\nExp number of AAs in TMHs: 23.78201\nExp number, first 60 AAs: 0.28355\nTotal prob of N-in: 0.13257\noutside 1 331\nTMhelix 332 354\ninside 355 372

3548 GCF_900166745.1_Isolate_14 Escherichia coli Proteobacteria; Gammaproteobacteria; Enterobacterales; Enterobacteriaceae; Escherichia MTGDNTLIHSHGINRRDFMKLCAALAATMGLSSKAAAEMVESVTNPQRPPVIWIGAQECT WP_000145423.1 hydrogenase 2 small subunit [Escherichia coli] Length: 372\nNumber of predicted TMHs: 1\nExp number of AAs in TMHs: 23.78201\nExp number, first 60 AAs: 0.28355\nTotal prob of N-in: 0.13257\noutside 1 331\nTMhelix 332 354\ninside 355 372

3549 GCF_900166755.1_Isolate_17 Escherichia coli Proteobacteria; Gammaproteobacteria; Enterobacterales; Enterobacteriaceae; Escherichia MTGDNTLIHSHGINRRDFMKLCAALAATMGLSSKAAAEMAESVTNPQRPPVIWIGAQECT WP_000145410.1 MULTISPECIES: hydrogenase-2 small chain [Proteobacteria] Length: 372\nNumber of predicted TMHs: 1\nExp number of AAs in TMHs: 23.72226\nExp number, first 60 AAs: 0.22038\nTotal prob of N-in: 0.12994\noutside 1 331\nTMhelix 332 354\ninside 355 372

3550 GCF_900166745.1_Isolate_14 Escherichia coli Proteobacteria; Gammaproteobacteria; Enterobacterales; Enterobacteriaceae; Escherichia MNNEETFYQAMRRQGVTRRSFLKYCSLAATSLGLGAGMAPKIAWALENKPRIPVVWIHGL WP_001295940.1 MULTISPECIES: hydrogenase [Enterobacteriaceae] Length: 372\nNumber of predicted TMHs: 2\nExp number of AAs in TMHs: 37.2608899999999\nExp number, first 60 AAs: 19.52923\nTotal prob of N-in:

0.94351\nPOSSIBLE N-term signal sequence\ninside 1 19\nTMhelix 20 39\noutside 40 325\nTMhelix 326 348\ninside 349 372

3551 GCF_900166785.1_Isolate_4 Escherichia coli Proteobacteria; Gammaproteobacteria; Enterobacterales; Enterobacteriaceae; Escherichia MNNEETFYQAMRRQGVTRRSFLKYCSLAATSLGLGAGMAPKIAWALENKPRIPVVWIHGL WP_001295940.1 MULTISPECIES: hydrogenase [Enterobacteriaceae] Length: 372\nNumber of predicted TMHs: 2\nExp number of AAs in TMHs: 37.2608899999999\nExp number, first 60 AAs: 19.52923\nTotal prob of N-in: 0.94351\nPOSSIBLE N-term signal sequence\ninside 1 19\nTMhelix 20 39\noutside 40 325\nTMhelix 326 348\ninside 349 372

3552 GCF_900166805.1_Isolate_7 Escherichia coli Proteobacteria; Gammaproteobacteria; Enterobacterales; Enterobacteriaceae; Escherichia MNNEETFYQAMRRQGVTRRSFLKYCSLAATSLGLGAGMAPKIAWALENKPRIPVVWIHGL WP_001295940.1 MULTISPECIES: hydrogenase [Enterobacteriaceae] Length: 372\nNumber of predicted TMHs: 2\nExp number of AAs in TMHs: 37.2608899999999\nExp number, first 60 AAs: 19.52923\nTotal prob of N-in: 0.94351\nPOSSIBLE N-term signal sequence\ninside 1 19\nTMhelix 20 39\noutside 40 325\nTMhelix 326 348\ninside 349 372

3553 GCF_000599725.2_a5_miseq_linux_20140113 Escherichia coli Proteobacteria; Gammaproteobacteria; Enterobacterales; Enterobacteriaceae; Escherichia MTGDNTLIHSHGINRRDFMKLCAALAATMGLSSKAAAEMAESVTNPQRPPVIWIGAQECT WP_000145410.1 MULTISPECIES: hydrogenase-2 small chain [Proteobacteria] Length: 372\nNumber of predicted TMHs: 1\nExp number of AAs in TMHs: 23.72226\nExp number, first 60 AAs: 0.22038\nTotal prob of N-in: 0.12994\noutside 1 331\nTMhelix 332 354\ninside 355 372

3554 GCF_000599785.2_a5_miseq_linux_20140113 Escherichia coli Proteobacteria; Gammaproteobacteria; Enterobacterales; Enterobacteriaceae; Escherichia MTGDNTLIHSHGINRRDFMKLCAALAATMGLSSKAAAEMVESVTNPQRPPVIWIGAQECT WP_000145423.1 hydrogenase 2 small subunit [Escherichia coli] Length: 372\nNumber of predicted TMHs: 1\nExp number of AAs in TMHs: 23.78201\nExp number, first 60 AAs: 0.28355\nTotal prob of N-in: 0.13257\noutside 1 331\nTMhelix 332 354\ninside 355 372

3555 GCF_000599825.1_a5_miseq_linux_20140113 Escherichia coli Proteobacteria; Gammaproteobacteria; Enterobacterales; Enterobacteriaceae; Escherichia MTGDNTLIHSHGINRRDFMKLCAALAATMGLSSKAAAEMAESVTNPQRPPVIWIGAQECT WP_000145410.1 MULTISPECIES: hydrogenase-2 small chain [Proteobacteria] Length: 372\nNumber of predicted TMHs: 1\nExp number of AAs in TMHs: 23.72226\nExp number, first 60 AAs: 0.22038\nTotal prob of N-in: 0.12994\noutside 1 331\nTMhelix 332 354\ninside 355 372

3556 GCF_000599725.2_a5_miseq_linux_20140113 Escherichia coli Proteobacteria; Gammaproteobacteria; Enterobacterales; Enterobacteriaceae; Escherichia MNNEETFYQAMRRQGVTRRSFLKYCSLAATSLGLGAGMAPKIAWALENKPRIPVVWIHGL WP_001058320.1 hydrogenase [Escherichia coli] Length: 372\nNumber of predicted TMHs: 2\nExp number of AAs in TMHs: 37.26082\nExp number, first 60 AAs: 19.52923\nTotal prob of N-in: 0.94351\nPOSSIBLE N-term signal sequence\ninside 1 19\nTMhelix 20 39\noutside 40 325\nTMhelix 326 348\ninside 349 372

3557 GCF_000599765.2_a5_miseq_linux_20140113 Escherichia coli Proteobacteria; Gammaproteobacteria; Enterobacterales; Enterobacteriaceae; Escherichia MNNEETFYQAMRRQGVTRRSFLKYCSLAATSLGLGAGMAPKIAWALENKPRIPVVWIHGL WP_001058323.1 MULTISPECIES: hydrogenase-1 small chain [Proteobacteria] Length: 372\nNumber of predicted TMHs: 2\nExp number of AAs in TMHs: 37.2608499999999\nExp number, first 60 AAs: 19.52923\nTotal prob of N-in: 0.94351\nPOSSIBLE N-term signal sequence\ninside 1 19\nTMhelix 20 39\noutside 40 325\nTMhelix 326 348\ninside 349 372

3558 GCF_000736735.1_ASM73673v1 Escherichia coli Proteobacteria; Gammaproteobacteria; Enterobacterales; Enterobacteriaceae; Escherichia MTGDNTLIHSHGINRRDFMKLCAALAATMGLSSKAAAEMAESVTNPQRPPVIWIGAQECT WP_028131934.1 hydrogenase-2 small chain [Escherichia coli] Length: 372\nNumber of predicted TMHs: 1\nExp number of AAs in TMHs: 24.01565\nExp number, first 60 AAs: 0.22022\nTotal prob of N-in: 0.13096\noutside 1 331\nTMhelix 332 354\ninside 355 372

3559 GCF_000690965.1_S6400 Escherichia coli Proteobacteria; Gammaproteobacteria; Enterobacterales; Enterobacteriaceae; Escherichia MTGDNTLIHSHGINRRDFMKLCAALAATMGLSSKAAAEMAESVTNPQRPPVIWIGAQECT WP_000145410.1 MULTISPECIES: hydrogenase-2 small chain [Proteobacteria] Length: 372\nNumber of predicted TMHs: 1\nExp number of AAs in TMHs: 23.72226\nExp number, first 60 AAs: 0.22038\nTotal prob of N-in: 0.12994\noutside 1 331\nTMhelix 332 354\ninside 355 372

3560 GCF_000690965.1_S6400 Escherichia coli Proteobacteria; Gammaproteobacteria; Enterobacterales; Enterobacteriaceae; Escherichia MNNEETFYQAMRRQGVTRRSFLKYCSLAATSLGLGAGMAPKIAWALENKPRIPVVWIHGL WP_001058323.1 MULTISPECIES: hydrogenase-1 small chain [Proteobacteria] Length: 372\nNumber of predicted TMHs: 2\nExp number of AAs in TMHs: 37.2608499999999\nExp number, first 60 AAs: 19.52923\nTotal prob of N-in: 0.94351\nPOSSIBLE N-term signal sequence\ninside 1 19\nTMhelix 20 39\noutside 40 325\nTMhelix 326 348\ninside 349 372

3561 GCF_000708165.1_E_coli_EC2.contigs Escherichia coli Proteobacteria; Gammaproteobacteria; Enterobacterales; Enterobacteriaceae; Escherichia MTGDNTLIHSHGINRRDFMKLCAALAATMGLSSKAAAEMAESVTNPQRPPVIWIGAQECT WP_000145410.1 MULTISPECIES: hydrogenase-2 small chain [Proteobacteria] Length: 372\nNumber of predicted TMHs: 1\nExp number of AAs in TMHs: 23.72226\nExp number, first 60 AAs: 0.22038\nTotal prob of N-in: 0.12994\noutside 1 331\nTMhelix 332 354\ninside 355 372

3562 GCF_000714975.1_JA65_HGAP2 Escherichia coli Proteobacteria; Gammaproteobacteria; Enterobacterales; Enterobacteriaceae; Escherichia MTGDNTLIHSHGINRRDFMKLCAALAATMGLSSKAAAEMAESVTNPQRPPVIWIGAQECT WP_000145410.1 MULTISPECIES: hydrogenase-2 small chain [Proteobacteria] Length: 372\nNumber of predicted TMHs: 1\nExp number of AAs in TMHs: 23.72226\nExp number, first 60 AAs: 0.22038\nTotal prob of N-in: 0.12994\noutside 1 331\nTMhelix 332 354\ninside 355 372

3563 GCF_000714975.1_JA65_HGAP2 Escherichia coli Proteobacteria; Gammaproteobacteria; Enterobacterales; Enterobacteriaceae; Escherichia MNNEETFYQAMRRQGVTRRSFLKYCSLAATSLGLGAGMAPKIAWALENKPRIPVVWIHGL WP_001058323.1 MULTISPECIES: hydrogenase-1 small chain [Proteobacteria] Length: 372\nNumber of predicted TMHs: 2\nExp number of AAs in TMHs: 37.2608499999999\nExp number, first 60 AAs: 19.52923\nTotal prob of N-in: 0.94351\nPOSSIBLE N-term signal sequence\ninside 1 19\nTMhelix 20 39\noutside 40 325\nTMhelix 326 348\ninside 349 372

3564 GCF_000714895.1_JA03_HGAP2 Escherichia coli Proteobacteria; Gammaproteobacteria; Enterobacterales; Enterobacteriaceae; Escherichia MTGDNTLIHSHGINRRDFMKLCAALAATMGLSSKAAAEMAESVTNPQRPPVIWIGAQECT WP_000145410.1 MULTISPECIES: hydrogenase-2 small chain [Proteobacteria] Length: 372\nNumber of predicted TMHs: 1\nExp number of AAs in TMHs: 23.72226\nExp number, first 60 AAs: 0.22038\nTotal prob of N-in: 0.12994\noutside 1 331\nTMhelix 332 354\ninside 355 372

3565 GCF_000714895.1_JA03_HGAP2 Escherichia coli Proteobacteria; Gammaproteobacteria; Enterobacterales; Enterobacteriaceae; Escherichia MNNEETFYQAMRRQGVTRRSFLKYCSLAATSLGLGAGMAPKIAWALENKPRIPVVWIHGL WP_001058320.1 hydrogenase [Escherichia coli] Length: 372\nNumber of predicted TMHs: 2\nExp number of AAs in TMHs: 37.26082\nExp number, first 60 AAs: 19.52923\nTotal prob of N-in: 0.94351\nPOSSIBLE N-term signal sequence\ninside 1 19\nTMhelix 20 39\noutside 40 325\nTMhelix 326 348\ninside 349 372

3566 GCF_000752175.1_FHI41 Escherichia coli Proteobacteria; Gammaproteobacteria; Enterobacterales; Enterobacteriaceae; Escherichia MTGDNTLIHSHGINRRDFMKLCAALAATMGLSSKAAAEMAESVTNPQRPPVIWIGAQECT WP_000145410.1 MULTISPECIES: hydrogenase-2 small chain [Proteobacteria] Length: 372\nNumber of predicted TMHs: 1\nExp number of AAs in TMHs: 23.72226\nExp number, first 60 AAs: 0.22038\nTotal prob of N-in: 0.12994\noutside 1 331\nTMhelix 332 354\ninside 355 372

3567 GCF_000749565.1_ASM74956v1 Escherichia coli Proteobacteria; Gammaproteobacteria; Enterobacterales; Enterobacteriaceae; Escherichia MNNEETFYQAMRRQGVTRRSFLKYCSLAATSLGLGAGMAPKIAWALENKPRIPVVWIHGL WP_001295940.1 MULTISPECIES: hydrogenase [Enterobacteriaceae] Length: 372\nNumber of predicted TMHs: 2\nExp number of AAs in TMHs: 37.2608899999999\nExp number, first 60 AAs: 19.52923\nTotal prob of N-in:

0.94351\nPOSSIBLE N-term signal sequence\ninside 1 19\nTMhelix 20 39\noutside 40 325\nTMhelix 326 348\ninside 349 372

3568 GCF_000752235.1_FHI69 Escherichia coli Proteobacteria; Gammaproteobacteria; Enterobacterales; Enterobacteriaceae; Escherichia MTGDNTLIHSHGINRRDFMKLCAALAATMGLSSKAAAEMAESVTNPQRPPVIWIGAQECT WP_000145410.1 MULTISPECIES: hydrogenase-2 small chain [Proteobacteria] Length: 372\nNumber of predicted TMHs: 1\nExp number of AAs in TMHs: 23.72226\nExp number, first 60 AAs: 0.22038\nTotal prob of N-in: 0.12994\noutside 1 331\nTMhelix 332 354\ninside 355 372

3569 GCF_000752255.1_FHI60 Escherichia coli Proteobacteria; Gammaproteobacteria; Enterobacterales; Enterobacteriaceae; Escherichia MTGDNTLIHSHGINRRDFMKLCAALAATMGLSSKAAAEMAESVTNPQRPPVIWIGAQECT WP_000145410.1 MULTISPECIES: hydrogenase-2 small chain [Proteobacteria] Length: 372\nNumber of predicted TMHs: 1\nExp number of AAs in TMHs: 23.72226\nExp number, first 60 AAs: 0.22038\nTotal prob of N-in: 0.12994\noutside 1 331\nTMhelix 332 354\ninside 355 372

3570 GCF_000752175.1_FHI41 Escherichia coli Proteobacteria; Gammaproteobacteria; Enterobacterales; Enterobacteriaceae; Escherichia MNNEETFYQAMRRQGVTRRSFLKYCSLAATSLGLGAGMAPKIAWALENKPRIPVVWIHGL WP_001058323.1 MULTISPECIES: hydrogenase-1 small chain [Proteobacteria] Length: 372\nNumber of predicted TMHs: 2\nExp number of AAs in TMHs: 37.2608499999999\nExp number, first 60 AAs: 19.52923\nTotal prob of N-in: 0.94351\nPOSSIBLE N-term signal sequence\ninside 1 19\nTMhelix 20 39\noutside 40 325\nTMhelix 326 348\ninside 349 372

3571 GCF_000752235.1_FHI69 Escherichia coli Proteobacteria; Gammaproteobacteria; Enterobacterales; Enterobacteriaceae; Escherichia MNNEETFYQAMRRQGVTRRSFLKYCSLAATSLGLGAGMAPKIAWALENKPRIPVVWIHGL WP_001058323.1 MULTISPECIES: hydrogenase-1 small chain [Proteobacteria] Length: 372\nNumber of predicted TMHs: 2\nExp number of AAs in TMHs: 37.2608499999999\nExp number, first 60 AAs: 19.52923\nTotal prob of N-in: 0.94351\nPOSSIBLE N-term signal sequence\ninside 1 19\nTMhelix 20 39\noutside 40 325\nTMhelix 326 348\ninside 349 372

3572 GCF_000752655.1_FHI47 Escherichia coli Proteobacteria; Gammaproteobacteria; Enterobacterales; Enterobacteriaceae; Escherichia MNNEETFYQAMRRQGVTRRSFLKYCSLAATSLGLGAGMAPKIAWALENKPRIPVVWIHGL WP_001058323.1 MULTISPECIES: hydrogenase-1 small chain [Proteobacteria] Length: 372\nNumber of predicted TMHs: 2\nExp number of AAs in TMHs: 37.2608499999999\nExp number, first 60 AAs: 19.52923\nTotal prob of N-in: 0.94351\nPOSSIBLE N-term signal sequence\ninside 1 19\nTMhelix 20 39\noutside 40 325\nTMhelix 326 348\ninside 349 372

3573 GCF_000752875.1_FHI39 Escherichia coli Proteobacteria; Gammaproteobacteria; Enterobacterales; Enterobacteriaceae; Escherichia MTGDNTLIHSHGINRRDFMKLCAALAATMGLSSKAAAEMAESVTNPQRPPVIWIGAQECT WP_000145410.1 MULTISPECIES: hydrogenase-2 small chain [Proteobacteria] Length: 372\nNumber of predicted TMHs: 1\nExp number of AAs in TMHs: 23.72226\nExp number, first 60 AAs: 0.22038\nTotal prob of N-in: 0.12994\noutside 1 331\nTMhelix 332 354\ninside 355 372

3574 GCF_000752915.1_FHI77 Escherichia coli Proteobacteria; Gammaproteobacteria; Enterobacterales; Enterobacteriaceae; Escherichia MTGDNTLIHSHGINRRDFMKLCAALAATMGLSSKAAAEMAESVTNPQRPPVIWIGAQECT WP_000145410.1 MULTISPECIES: hydrogenase-2 small chain [Proteobacteria] Length: 372\nNumber of predicted TMHs: 1\nExp number of AAs in TMHs: 23.72226\nExp number, first 60 AAs: 0.22038\nTotal prob of N-in: 0.12994\noutside 1 331\nTMhelix 332 354\ninside 355 372

3575 GCF_000752955.1_FHI91 Escherichia coli Proteobacteria; Gammaproteobacteria; Enterobacterales; Enterobacteriaceae; Escherichia MTGDNTLIHSHGINRRDFMKLCAALAATMGLSSKAAAEMAESVTNPQRPPVIWIGAQECT WP_000145410.1 MULTISPECIES: hydrogenase-2 small chain [Proteobacteria] Length: 372\nNumber of predicted TMHs: 1\nExp number of AAs in TMHs: 23.72226\nExp number, first 60 AAs: 0.22038\nTotal prob of N-in: 0.12994\noutside 1 331\nTMhelix 332 354\ninside 355 372

3576 GCF_000753075.1_FHI73 Escherichia coli Proteobacteria; Gammaproteobacteria; Enterobacterales; Enterobacteriaceae; Escherichia MTGDNTLIHSHGINRRDFMKLCAALAATMGLSSKAAAEMAESVTNPQRPPVIWIGAQECT WP_000145410.1 MULTISPECIES: hydrogenase-2 small chain [Proteobacteria] Length: 372\nNumber of predicted TMHs: 1\nExp number of AAs in TMHs: 23.72226\nExp number, first 60 AAs: 0.22038\nTotal prob of N-in: 0.12994\noutside 1 331\nTMhelix 332 354\ninside 355 372

3577 GCF_000752975.1_StOlav172 Escherichia coli Proteobacteria; Gammaproteobacteria; Enterobacterales; Enterobacteriaceae; Escherichia

MTGDNTLIHSHGINRRDFMKLCAALAATMGLSSKAAAEMAESVTNPQRPPVIWIGAQECT WP_000145410.1 MULTISPECIES: hydrogenase-2 small chain [Proteobacteria] Length: 372\nNumber of predicted TMHs: 1\nExp number of AAs in TMHs: 23.72226\nExp number, first 60 AAs: 0.22038\nTotal prob of N-in: 0.12994\noutside 1 331\nTMhelix 332 354\ninside 355 372

3578 GCF_000753315.1_FHI35 Escherichia coli Proteobacteria; Gammaproteobacteria; Enterobacterales; Enterobacteriaceae; Escherichia MNNEETFYQAMRRQGVTRRSFLKYCSLAATSLGLGAGMAPKIAWALENKPRIPVVWIHGL WP_001058323.1 MULTISPECIES: hydrogenase-1 small chain [Proteobacteria] Length: 372\nNumber of predicted TMHs: 2\nExp number of AAs in TMHs: 37.2608499999999\nExp number, first 60 AAs: 19.52923\nTotal prob of N-in: 0.94351\nPOSSIBLE N-term signal sequence\ninside 1 19\nTMhelix 20 39\noutside 40 325\nTMhelix 326 348\ninside 349 372

3579 GCF_000768405.1_ASM76840v1 Escherichia coli Proteobacteria; Gammaproteobacteria; Enterobacterales; Enterobacteriaceae; Escherichia MTGDNTLIHSHGINRRDFMKLCAALAATMGLSSKAAAEMAESVTNPQRPPVIWIGAQECT WP_000145410.1 MULTISPECIES: hydrogenase-2 small chain [Proteobacteria] Length: 372\nNumber of predicted TMHs: 1\nExp number of AAs in TMHs: 23.72226\nExp number, first 60 AAs: 0.22038\nTotal prob of N-in: 0.12994\noutside 1 331\nTMhelix 332 354\ninside 355 372

3580 GCF_000768405.1_ASM76840v1 Escherichia coli Proteobacteria; Gammaproteobacteria; Enterobacterales; Enterobacteriaceae; Escherichia MNNEETFYQAMRRQGVTRRSFLKYCSLAATSLGLGAGMAPKIAWALENKPRIPVVWIHGL WP_001058323.1 MULTISPECIES: hydrogenase-1 small chain [Proteobacteria] Length: 372\nNumber of predicted TMHs: 2\nExp number of AAs in TMHs: 37.2608499999999\nExp number, first 60 AAs: 19.52923\nTotal prob of N-in: 0.94351\nPOSSIBLE N-term signal sequence\ninside 1 19\nTMhelix 20 39\noutside 40 325\nTMhelix 326 348\ninside 349 372

3581 GCF_000753515.1_FHI13 Escherichia coli Proteobacteria; Gammaproteobacteria; Enterobacterales; Enterobacteriaceae; Escherichia MNNEETFYQAMRRQGVTRRSFLKYCSLAATSLGLGAGMAPKIAWALENKPRIPVVWIHGL WP_001058323.1 MULTISPECIES: hydrogenase-1 small chain [Proteobacteria] Length: 372\nNumber of predicted TMHs: 2\nExp number of AAs in TMHs: 37.2608499999999\nExp number, first 60 AAs: 19.52923\nTotal prob of N-in: 0.94351\nPOSSIBLE N-term signal sequence\ninside 1 19\nTMhelix 20 39\noutside 40 325\nTMhelix 326 348\ninside 349 372

3582 GCF_000754855.1_ASM75485v1 Escherichia coli Proteobacteria; Gammaproteobacteria; Enterobacterales; Enterobacteriaceae; Escherichia MNNEETFYQAMRRQGVTRRSFLKYCSLAATSLGLGAGMAPKIAWALENKPRIPVVWIHGL WP_001058323.1 MULTISPECIES: hydrogenase-1 small chain [Proteobacteria] Length: 372\nNumber of predicted TMHs: 2\nExp number of AAs in TMHs: 37.2608499999999\nExp number, first 60 AAs: 19.52923\nTotal prob of N-in: 0.94351\nPOSSIBLE N-term signal sequence\ninside 1 19\nTMhelix 20 39\noutside 40 325\nTMhelix 326 348\ninside 349 372

3583 GCF_000755445.1_ASM75544v1 Escherichia coli Proteobacteria; Gammaproteobacteria; Enterobacterales; Enterobacteriaceae; Escherichia MNNEETFYQAMRRQGVTRRSFLKYCSLAATSLGLGAGMAPKIAWALENKPRIPVVWIHGL WP_001058323.1 MULTISPECIES: hydrogenase-1 small chain [Proteobacteria] Length: 372\nNumber of predicted TMHs: 2\nExp number of AAs in TMHs: 37.2608499999999\nExp number, first 60 AAs: 19.52923\nTotal prob of N-in: 0.94351\nPOSSIBLE N-term signal sequence\ninside 1 19\nTMhelix 20 39\noutside 40 325\nTMhelix 326 348\ninside 349 372

3584 GCF_000776135.1_ASM77613v1 Escherichia coli Proteobacteria; Gammaproteobacteria; Enterobacterales; Enterobacteriaceae; Escherichia MNNEETFYQAMRRQGVTRRSFLKYCSLAATSLGLGAGMAPKIAWALENKPRIPVVWIHGL WP_001058323.1 MULTISPECIES: hydrogenase-1 small chain [Proteobacteria] Length: 372\nNumber of predicted TMHs: 2\nExp number of AAs in TMHs: 37.2608499999999\nExp number, first 60 AAs: 19.52923\nTotal prob of N-in: 0.94351\nPOSSIBLE N-term signal sequence\ninside 1 19\nTMhelix 20 39\noutside 40 325\nTMhelix 326 348\ninside 349 372

3585 GCF_000773435.1_ASM77343v1 Escherichia coli Proteobacteria; Gammaproteobacteria; Enterobacterales; Enterobacteriaceae; Escherichia MNNEETFYQAMRRQGVTRRSFLKYCSLAATSLGLGAGMAPKIAWALENKPRIPVVWIHGL WP_047603468.1

hydrogenase [Escherichia coli] Length: 372\nNumber of predicted TMHs: 2\nExp number of AAs in TMHs: 39.28018\nExp number, first 60 AAs: 19.5238\nTotal prob of N-in: 0.94367\nPOSSIBLE N-term signal sequence\ninside 1 19\nTMhelix 20 39\noutside 40 326\nTMhelix 327 349\ninside 350 372

3586 GCF_000776235.1_ASM77623v1 Escherichia coli Proteobacteria; Gammaproteobacteria; Enterobacterales; Enterobacteriaceae; Escherichia MTGDNTLIHSHGINRRDFMKLCAALAATMGLSSKAAAEMAESVTNPQRPPVIWIGAQECT WP_000145410.1 MULTISPECIES: hydrogenase-2 small chain [Proteobacteria] Length: 372\nNumber of predicted TMHs: 1\nExp number of AAs in TMHs: 23.72226\nExp number, first 60 AAs: 0.22038\nTotal prob of N-in: 0.12994\noutside 1 331\nTMhelix 332 354\ninside 355 372

3587 GCF_000776315.1_ASM77631v1 Escherichia coli Proteobacteria; Gammaproteobacteria; Enterobacterales; Enterobacteriaceae; Escherichia MTGDNTLIHSHGINRRDFMKLCAALAATMGLSSKAAAEMAESVTNPQRPPVIWIGAQECT WP_000145410.1 MULTISPECIES: hydrogenase-2 small chain [Proteobacteria] Length: 372\nNumber of predicted TMHs: 1\nExp number of AAs in TMHs: 23.72226\nExp number, first 60 AAs: 0.22038\nTotal prob of N-in: 0.12994\noutside 1 331\nTMhelix 332 354\ninside 355 372

3588 GCF_000776215.1_ASM77621v1 Escherichia coli Proteobacteria; Gammaproteobacteria; Enterobacterales; Enterobacteriaceae; Escherichia MNNEETFYQAMRRQGVTRRSFLKYCSLAATSLGLGAGMAPKIAWALENKPRIPVVWIHGL WP_001058323.1 MULTISPECIES: hydrogenase-1 small chain [Proteobacteria] Length: 372\nNumber of predicted TMHs: 2\nExp number of AAs in TMHs: 37.2608499999999\nExp number, first 60 AAs: 19.52923\nTotal prob of N-in: 0.94351\nPOSSIBLE N-term signal sequence\ninside 1 19\nTMhelix 20 39\noutside 40 325\nTMhelix 326 348\ninside 349 372

3589 GCF_000776375.1_ASM77637v1 Escherichia coli Proteobacteria; Gammaproteobacteria; Enterobacterales; Enterobacteriaceae; Escherichia MTGDNTLIHSHGINRRDFMKLCAALAATMGLSSKAAAEMAESVTNPQRPPVIWIGAQECT WP_000145410.1 MULTISPECIES: hydrogenase-2 small chain [Proteobacteria] Length: 372\nNumber of predicted TMHs: 1\nExp number of AAs in TMHs: 23.72226\nExp number, first 60 AAs: 0.22038\nTotal prob of N-in: 0.12994\noutside 1 331\nTMhelix 332 354\ninside 355 372

3590 GCF_000776335.1_ASM77633v1 Escherichia coli Proteobacteria; Gammaproteobacteria; Enterobacterales; Enterobacteriaceae; Escherichia MTGDNTLIHSHGINRRDFMKLCAALAATMGLSSKAAAEMAESVTNPQRPPVIWIGAQECT WP_000145410.1 MULTISPECIES: hydrogenase-2 small chain [Proteobacteria] Length: 372\nNumber of predicted TMHs: 1\nExp number of AAs in TMHs: 23.72226\nExp number, first 60 AAs: 0.22038\nTotal prob of N-in: 0.12994\noutside 1 331\nTMhelix 332 354\ninside 355 372

3591 GCF_000776715.1_ASM77671v1 Escherichia coli Proteobacteria; Gammaproteobacteria; Enterobacterales; Enterobacteriaceae; Escherichia MNNEETFYQAMRRQGVTRRSFLKYCSLAATSLGLGAGMAPKIAWALENKPRIPVVWIHGL WP_001058323.1 MULTISPECIES: hydrogenase-1 small chain [Proteobacteria] Length: 372\nNumber of predicted TMHs: 2\nExp number of AAs in TMHs: 37.2608499999999\nExp number, first 60 AAs: 19.52923\nTotal prob of N-in: 0.94351\nPOSSIBLE N-term signal sequence\ninside 1 19\nTMhelix 20 39\noutside 40 325\nTMhelix 326 348\ninside 349 372

3592 GCF_000776775.1_ASM77677v1 Escherichia coli Proteobacteria; Gammaproteobacteria; Enterobacterales; Enterobacteriaceae; Escherichia MNNEETFYQAMRRQGVTRRSFLKYCSLAATSLGLGAGMAPKIAWALENKPRIPVVWIHGL WP_001058323.1 MULTISPECIES: hydrogenase-1 small chain [Proteobacteria] Length: 372\nNumber of predicted TMHs: 2\nExp number of AAs in TMHs: 37.2608499999999\nExp number, first 60 AAs: 19.52923\nTotal prob of N-in: 0.94351\nPOSSIBLE N-term signal sequence\ninside 1 19\nTMhelix 20 39\noutside 40 325\nTMhelix 326 348\ninside 349 372

3593 GCF_000776935.1_ASM77693v1 Escherichia coli Proteobacteria; Gammaproteobacteria; Enterobacterales; Enterobacteriaceae; Escherichia MNNEETFYQAMRRQGVTRRSFLKYCSLAATSLGLGAGMAPKIAWALENKPRIPVVWIHGL WP_001058323.1 MULTISPECIES: hydrogenase-1 small chain [Proteobacteria] Length: 372\nNumber of predicted TMHs: 2\nExp number of AAs in TMHs: 37.2608499999999\nExp number, first 60 AAs: 19.52923\nTotal prob of N-in:

0.94351\nPOSSIBLE N-term signal sequence\ninside 1 19\nTMhelix 20 39\noutside 40 325\nTMhelix 326 348\ninside 349 372

3594 GCF_000776815.1_ASM77681v1 Escherichia coli Proteobacteria; Gammaproteobacteria; Enterobacterales; Enterobacteriaceae; Escherichia
MNNEETFYQAMRRQGVTRRSFLKYCSLAATSLGLGAGMAPKIAWALENKPRIPVVWIHGL WP_001058323.1
MULTISPECIES: hydrogenase-1 small chain [Proteobacteria] Length: 372\nNumber of predicted TMHs: 2\nExp number of AAs in TMHs: 37.2608499999999\nExp number, first 60 AAs: 19.52923\nTotal prob of N-in: 0.94351\nPOSSIBLE N-term signal sequence\ninside 1 19\nTMhelix 20 39\noutside 40 325\nTMhelix 326 348\ninside 349 372

3595 GCF_000777025.1_ASM77702v1 Escherichia coli Proteobacteria; Gammaproteobacteria; Enterobacterales; Enterobacteriaceae; Escherichia
MTGDNTLIHSHGINRRDFMKLCAALAATMGLSSKAAAEMAESVTNPQRPPVIWIGAQECT WP_000145410.1
MULTISPECIES: hydrogenase-2 small chain [Proteobacteria] Length: 372\nNumber of predicted TMHs: 1\nExp number of AAs in TMHs: 23.72226\nExp number, first 60 AAs: 0.22038\nTotal prob of N-in: 0.12994\noutside 1 331\nTMhelix 332 354\ninside 355 372

3596 GCF_000776855.1_ASM77685v1 Escherichia coli Proteobacteria; Gammaproteobacteria; Enterobacterales; Enterobacteriaceae; Escherichia
MNNEETFYQAMRRQGVTRRSFLKYCSLAATSLGLGAGMAPKIAWALENKPRIPVVWIHGL WP_001058323.1
MULTISPECIES: hydrogenase-1 small chain [Proteobacteria] Length: 372\nNumber of predicted TMHs: 2\nExp number of AAs in TMHs: 37.2608499999999\nExp number, first 60 AAs: 19.52923\nTotal prob of N-in: 0.94351\nPOSSIBLE N-term signal sequence\ninside 1 19\nTMhelix 20 39\noutside 40 325\nTMhelix 326 348\ninside 349 372

3597 GCF_000776915.1_ASM77691v1 Escherichia coli Proteobacteria; Gammaproteobacteria; Enterobacterales; Enterobacteriaceae; Escherichia
MNNEETFYQAMRRQGVTRRSFLKYCSLAATSLGLGAGMAPKIAWALENKPRIPVVWIHGL WP_001295940.1
MULTISPECIES: hydrogenase [Enterobacteriaceae] Length: 372\nNumber of predicted TMHs: 2\nExp number of AAs in TMHs: 37.2608899999999\nExp number, first 60 AAs: 19.52923\nTotal prob of N-in: 0.94351\nPOSSIBLE N-term signal sequence\ninside 1 19\nTMhelix 20 39\noutside 40 325\nTMhelix 326 348\ninside 349 372

3598 GCF_000777495.1_ASM77749v1 Escherichia coli Proteobacteria; Gammaproteobacteria; Enterobacterales; Enterobacteriaceae; Escherichia
MTGDNTLIHSHGINRRDFMKLCAALAATMGLSSKAAAEMAESVTNPQRPPVIWIGAQECT WP_000145410.1
MULTISPECIES: hydrogenase-2 small chain [Proteobacteria] Length: 372\nNumber of predicted TMHs: 1\nExp number of AAs in TMHs: 23.72226\nExp number, first 60 AAs: 0.22038\nTotal prob of N-in: 0.12994\noutside 1 331\nTMhelix 332 354\ninside 355 372

3599 GCF_000777585.1_ASM77758v1 Escherichia coli Proteobacteria; Gammaproteobacteria; Enterobacterales; Enterobacteriaceae; Escherichia
MTGDNTLIHSHGINRRDFMKLCAALAATMGLSSKAAAEMVESVTNPQRPPVIWIGAQECT WP_000145423.1
hydrogenase 2 small subunit [Escherichia coli] Length: 372\nNumber of predicted TMHs: 1\nExp number of AAs in TMHs: 23.78201\nExp number, first 60 AAs: 0.28355\nTotal prob of N-in: 0.13257\noutside 1 331\nTMhelix 332 354\ninside 355 372

3600 GCF_000777655.1_ASM77765v1 Escherichia coli Proteobacteria; Gammaproteobacteria; Enterobacterales; Enterobacteriaceae; Escherichia
MTGDNTLIHSHGINRRDFMKLCAALAATMGLSSKAAAEMAESVTNPQRPPVIWIGAQECT WP_000145410.1
MULTISPECIES: hydrogenase-2 small chain [Proteobacteria] Length: 372\nNumber of predicted TMHs: 1\nExp number of AAs in TMHs: 23.72226\nExp number, first 60 AAs: 0.22038\nTotal prob of N-in: 0.12994\noutside 1 331\nTMhelix 332 354\ninside 355 372

3601 GCF_000777605.1_ASM77760v1 Escherichia coli Proteobacteria; Gammaproteobacteria; Enterobacterales; Enterobacteriaceae; Escherichia
MTGDNTLIHSHGINRRDFMKLCAALAATMGLSSKAAAEMAESVTNPQRPPVIWIGAQECT WP_000145410.1
MULTISPECIES: hydrogenase-2 small chain [Proteobacteria] Length: 372\nNumber of predicted TMHs: 1\nExp number of AAs in TMHs: 23.72226\nExp number, first 60 AAs: 0.22038\nTotal prob of N-in: 0.12994\noutside 1 331\nTMhelix 332 354\ninside 355 372

3602 GCF_000777375.1_ASM77737v1 Escherichia coli Proteobacteria; Gammaproteobacteria; Enterobacterales; Enterobacteriaceae; Escherichia
MNNEETFYQAMRRQGVTRRSFLKYCSLAATSLGLGAGMAPKIAWALENKPRIPVVWIHGL WP_001295940.1
MULTISPECIES: hydrogenase [Enterobacteriaceae] Length: 372\nNumber of predicted TMHs: 2\nExp number of AAs in TMHs: 37.2608899999999\nExp number, first 60 AAs: 19.52923\nTotal prob of N-in: 0.94351\nPOSSIBLE N-term signal sequence\ninside 1 19\nTMhelix 20 39\noutside 40 325\nTMhelix 326 348\ninside 349 372

3603 GCF_000777585.1_ASM77758v1 Escherichia coli Proteobacteria; Gammaproteobacteria; Enterobacterales; Enterobacteriaceae; Escherichia
MNNEETFYQAMRRQGVTRRSFLKYCSLAATSLGLGAGMAPKIAWALENKPRIPVVWIHGL WP_001058323.1
MULTISPECIES: hydrogenase-1 small chain [Proteobacteria] Length: 372\nNumber of predicted TMHs: 2\nExp number of AAs in TMHs: 37.2608499999999\nExp number, first 60 AAs: 19.52923\nTotal prob of N-in: 0.94351\nPOSSIBLE N-term signal sequence\ninside 1 19\nTMhelix 20 39\noutside 40 325\nTMhelix 326 348\ninside 349 372

3604 GCF_000777945.1_ASM77794v1 Escherichia coli Proteobacteria; Gammaproteobacteria; Enterobacterales; Enterobacteriaceae; Escherichia
MTGDNTLIHSHGINRRDFMKLCAALAATMGLSSKAAAEMVESVTNPQRPPVIWIGAQECT WP_000145423.1
hydrogenase 2 small subunit [Escherichia coli] Length: 372\nNumber of predicted TMHs: 1\nExp number of AAs in TMHs: 23.78201\nExp number, first 60 AAs: 0.28355\nTotal prob of N-in: 0.13257\noutside 1 331\nTMhelix 332 354\ninside 355 372

3605 GCF_000778235.1_ASM77823v1 Escherichia coli Proteobacteria; Gammaproteobacteria; Enterobacterales; Enterobacteriaceae; Escherichia
MNNEETFYQAMRRQGVTRRSFLKYCSLAATSLGLGAGMAPKIAWALENKPRIPVVWIHGL WP_001058323.1
MULTISPECIES: hydrogenase-1 small chain [Proteobacteria] Length: 372\nNumber of predicted TMHs: 2\nExp number of AAs in TMHs: 37.2608499999999\nExp number, first 60 AAs: 19.52923\nTotal prob of N-in: 0.94351\nPOSSIBLE N-term signal sequence\ninside 1 19\nTMhelix 20 39\noutside 40 325\nTMhelix 326 348\ninside 349 372

3606 GCF_000778215.1_ASM77821v1 Escherichia coli Proteobacteria; Gammaproteobacteria; Enterobacterales; Enterobacteriaceae; Escherichia
MNNEETFYQAMRRQGVTRRSFLKYCSLAATSLGLGAGMAPKIAWALENKPRIPVVWIHGL WP_001295940.1
MULTISPECIES: hydrogenase [Enterobacteriaceae] Length: 372\nNumber of predicted TMHs: 2\nExp number of AAs in TMHs: 37.2608899999999\nExp number, first 60 AAs: 19.52923\nTotal prob of N-in: 0.94351\nPOSSIBLE N-term signal sequence\ninside 1 19\nTMhelix 20 39\noutside 40 325\nTMhelix 326 348\ninside 349 372

3607 GCF_000777945.1_ASM77794v1 Escherichia coli Proteobacteria; Gammaproteobacteria; Enterobacterales; Enterobacteriaceae; Escherichia
MNNEETFYQAMRRQGVTRRSFLKYCSLAATSLGLGAGMAPKIAWALENKPRIPVVWIHGL WP_001295940.1
MULTISPECIES: hydrogenase [Enterobacteriaceae] Length: 372\nNumber of predicted TMHs: 2\nExp number of AAs in TMHs: 37.2608899999999\nExp number, first 60 AAs: 19.52923\nTotal prob of N-in: 0.94351\nPOSSIBLE N-term signal sequence\ninside 1 19\nTMhelix 20 39\noutside 40 325\nTMhelix 326 348\ninside 349 372

3608 GCF_000778285.1_ASM77828v1 Escherichia coli Proteobacteria; Gammaproteobacteria; Enterobacterales; Enterobacteriaceae; Escherichia
MTGDNTLIHSHGINRRDFMKLCAALAATMGLSSKAAAEMAESVTNPQRPPVIWIGAQECT WP_000145410.1
MULTISPECIES: hydrogenase-2 small chain [Proteobacteria] Length: 372\nNumber of predicted TMHs: 1\nExp number of AAs in TMHs: 23.72226\nExp number, first 60 AAs: 0.22038\nTotal prob of N-in: 0.12994\noutside 1 331\nTMhelix 332 354\ninside 355 372

3609 GCF_000778285.1_ASM77828v1 Escherichia coli Proteobacteria; Gammaproteobacteria; Enterobacterales; Enterobacteriaceae; Escherichia
MNNEETFYQAMRRQGVTRRSFLKYCSLAATSLGLGAGMAPKIAWALENKPRIPVVWIHGL WP_001058323.1
MULTISPECIES: hydrogenase-1 small chain [Proteobacteria] Length: 372\nNumber of predicted TMHs: 2\nExp number of AAs in TMHs: 37.2608499999999\nExp number, first 60 AAs: 19.52923\nTotal prob of N-in: 0.94351\nPOSSIBLE N-term signal sequence\ninside 1 19\nTMhelix 20 39\noutside 40 325\nTMhelix 326 348\ninside 349 372

3610 GCF_000778075.1_ASM77807v1 Escherichia coli Proteobacteria; Gammaproteobacteria; Enterobacterales; Enterobacteriaceae; Escherichia
MNNEETFYQAMRRQGVTRRSFLKYCSLAATSLGLGAGMAPKIAWALENKPRIPVVWIHGL WP_001058323.1
MULTISPECIES: hydrogenase-1 small chain [Proteobacteria] Length: 372\nNumber of predicted TMHs: 2\nExp number of AAs in TMHs: 37.2608499999999\nExp number, first 60 AAs: 19.52923\nTotal prob of N-in: 0.94351\nPOSSIBLE N-term signal sequence\ninside 1 19\nTMhelix 20 39\noutside 40 325\nTMhelix 326 348\ninside 349 372

3611 GCF_000778095.1_ASM77809v1 Escherichia coli Proteobacteria; Gammaproteobacteria; Enterobacterales; Enterobacteriaceae; Escherichia
MNNEETFYQAMRRQGVTRRSFLKYCSLAATSLGLGAGMAPKIAWALENKPRIPVVWIHGL WP_001058323.1
MULTISPECIES: hydrogenase-1 small chain [Proteobacteria] Length: 372\nNumber of predicted TMHs: 2\nExp number of AAs in TMHs: 37.2608499999999\nExp number, first 60 AAs: 19.52923\nTotal prob of N-in: 0.94351\nPOSSIBLE N-term signal sequence\ninside 1 19\nTMhelix 20 39\noutside 40 325\nTMhelix 326 348\ninside 349 372

3612 GCF_000779005.1_ASM77900v1 Escherichia coli Proteobacteria; Gammaproteobacteria; Enterobacterales; Enterobacteriaceae; Escherichia
MTGDNTLIHSHGINRRDFMKLCAALAATMGLSSKAAAEMAESVTNPQRPPVIWIGAQECT WP_000145410.1
MULTISPECIES: hydrogenase-2 small chain [Proteobacteria] Length: 372\nNumber of predicted TMHs: 1\nExp number of AAs in TMHs: 23.72226\nExp number, first 60 AAs: 0.22038\nTotal prob of N-in: 0.12994\noutside 1 331\nTMhelix 332 354\ninside 355 372

3613 GCF_000779025.1_ASM77902v1 Escherichia coli Proteobacteria; Gammaproteobacteria; Enterobacterales; Enterobacteriaceae; Escherichia
MTGDNTLIHSHGINRRDFMKLCAALAATMGLSSKAAAEMAESVTNPQRPPVIWIGAQECT WP_000145410.1
MULTISPECIES: hydrogenase-2 small chain [Proteobacteria] Length: 372\nNumber of predicted TMHs: 1\nExp number of AAs in TMHs: 23.72226\nExp number, first 60 AAs: 0.22038\nTotal prob of N-in: 0.12994\noutside 1 331\nTMhelix 332 354\ninside 355 372

3614 GCF_001284505.1_7748_7_26 Escherichia coli Proteobacteria; Gammaproteobacteria; Enterobacterales; Enterobacteriaceae; Escherichia
MTGDNTLIHSHGINRRDFMKLCAALAATMGLSSKAAAEMAESVTNPQRPPVIWIGAQECT WP_024228528.1
hydrogenase 2 small subunit [Escherichia coli] Length: 372\nNumber of predicted TMHs: 1\nExp number of AAs in TMHs: 23.61909\nExp number, first 60 AAs: 0.22137\nTotal prob of N-in: 0.12572\noutside 1 331\nTMhelix 332 354\ninside 355 372

3615 GCF_001284105.1_8205_3_63 Escherichia coli Proteobacteria; Gammaproteobacteria; Enterobacterales; Enterobacteriaceae; Escherichia
MNNEETFYQAMRRQGVTRRSFLKYCSLAATSLGLGAGMAPKIAWALENKPRIPVVWIHGL WP_001058323.1
MULTISPECIES: hydrogenase-1 small chain [Proteobacteria] Length: 372\nNumber of predicted TMHs: 2\nExp number of AAs in TMHs: 37.2608499999999\nExp number, first 60 AAs: 19.52923\nTotal prob of N-in: 0.94351\nPOSSIBLE N-term signal sequence\ninside 1 19\nTMhelix 20 39\noutside 40 325\nTMhelix 326 348\ninside 349 372

3616 GCF_000798055.1_Escherichia_coli_CVM_N33922PS_v._1.0 Escherichia coli Proteobacteria; Gammaproteobacteria; Enterobacterales; Enterobacteriaceae; Escherichia
MTGDNTLIHSHGINRRDFMKLCAALAATMGLSSKAAAEMAESVTNPQRPPVIWIGAQECT WP_000145410.1
MULTISPECIES: hydrogenase-2 small chain [Proteobacteria] Length: 372\nNumber of predicted TMHs: 1\nExp number of AAs in TMHs: 23.72226\nExp number, first 60 AAs: 0.22038\nTotal prob of N-in: 0.12994\noutside 1 331\nTMhelix 332 354\ninside 355 372

3617 GCF_001276585.2_ASM127658v2 Escherichia coli Proteobacteria; Gammaproteobacteria; Enterobacterales; Enterobacteriaceae; Escherichia
MNNEETFYQAMRRQGVTRRSFLKYCSLAATSLGLGAGMAPKIAWALENKPRIPVVWIHGL WP_001058323.1
MULTISPECIES: hydrogenase-1 small chain [Proteobacteria] Length: 372\nNumber of predicted TMHs: 2\nExp number of AAs in TMHs: 37.2608499999999\nExp number, first 60 AAs: 19.52923\nTotal prob of N-in: 0.94351\nPOSSIBLE N-term signal sequence\ninside 1 19\nTMhelix 20 39\noutside 40 325\nTMhelix 326 348\ninside 349 372

3618 GCF_000798095.1_Escherichia_coli_CVM_N36312PS_v._1.0 Escherichia coli Proteobacteria; Gammaproteobacteria; Enterobacterales; Enterobacteriaceae; Escherichia
MTGDNTLIHSHGINRRDFMKLCAALAATMGLSSKAAAEMAESVTNPQRPPVIWIGAQECT WP_000145410.1
MULTISPECIES: hydrogenase-2 small chain [Proteobacteria] Length: 372\nNumber of predicted TMHs: 1\nExp number of AAs in TMHs: 23.72226\nExp number, first 60 AAs: 0.22038\nTotal prob of N-in: 0.12994\noutside 1 331\nTMhelix 332 354\ninside 355 372

3619 GCF_001284345.1_8205_3_61 Escherichia coli Proteobacteria; Gammaproteobacteria; Enterobacterales; Enterobacteriaceae; Escherichia
MTGDNTLIHSHGINRRDFMKLCAALAATMGLSSKAAAEMAESVTNPQRPPVIWIGAQECT WP_000145410.1
MULTISPECIES: hydrogenase-2 small chain [Proteobacteria] Length: 372\nNumber of predicted TMHs: 1\nExp number of AAs in TMHs: 23.72226\nExp number, first 60 AAs: 0.22038\nTotal prob of N-in: 0.12994\noutside 1 331\nTMhelix 332 354\ninside 355 372

3620 GCF_001284445.1_7790_1_76 Escherichia coli Proteobacteria; Gammaproteobacteria; Enterobacterales; Enterobacteriaceae; Escherichia
MTGDNTLIHSHGINRRDFMKLCAALAATMGLSSKAAAEMAESVTNPQRPPVIWIGAQECT WP_000145410.1
MULTISPECIES: hydrogenase-2 small chain [Proteobacteria] Length: 372\nNumber of predicted TMHs: 1\nExp number of AAs in TMHs: 23.72226\nExp number, first 60 AAs: 0.22038\nTotal prob of N-in: 0.12994\noutside 1 331\nTMhelix 332 354\ninside 355 372

3621 GCF_000797835.1_Escherichia_coli_CVM_N33804PS_v._1.0 Escherichia coli Proteobacteria; Gammaproteobacteria; Enterobacterales; Enterobacteriaceae; Escherichia
MTGDNTLIHSHGINRRDFMKLCAALAATMGLSSKAAAEMAESVTNPQRPPVIWIGAQECT WP_000145410.1
MULTISPECIES: hydrogenase-2 small chain [Proteobacteria] Length: 372\nNumber of predicted TMHs: 1\nExp number of AAs in TMHs: 23.72226\nExp number, first 60 AAs: 0.22038\nTotal prob of N-in: 0.12994\noutside 1 331\nTMhelix 332 354\ninside 355 372

3622 GCF_001284005.1_7748_7_41 Escherichia coli Proteobacteria; Gammaproteobacteria; Enterobacterales; Enterobacteriaceae; Escherichia
MNNEETFYQAMRRQGVTRRSFLKYCSLAATSLGLGAGMAPKIAWALENKPRIPVVWIHGL WP_001058323.1
MULTISPECIES: hydrogenase-1 small chain [Proteobacteria] Length: 372\nNumber of predicted TMHs: 2\nExp number of AAs in TMHs: 37.2608499999999\nExp number, first 60 AAs: 19.52923\nTotal prob of N-in: 0.94351\nPOSSIBLE N-term signal sequence\ninside 1 19\nTMhelix 20 39\noutside 40 325\nTMhelix 326 348\ninside 349 372

3623 GCF_001269285.1_1.EC2992.1 Escherichia coli Proteobacteria; Gammaproteobacteria; Enterobacterales; Enterobacteriaceae; Escherichia
MNNEETFYQAMRRQGVTRRSFLKYCSLAATSLGLGAGMAPKIAWALENKPRIPVVWIHGL WP_001058323.1
MULTISPECIES: hydrogenase-1 small chain [Proteobacteria] Length: 372\nNumber of predicted TMHs: 2\nExp number of AAs in TMHs: 37.2608499999999\nExp number, first 60 AAs: 19.52923\nTotal prob of N-in: 0.94351\nPOSSIBLE N-term signal sequence\ninside 1 19\nTMhelix 20 39\noutside 40 325\nTMhelix 326 348\ninside 349 372

3624 GCF_001284185.1_8205_3_69 Escherichia coli Proteobacteria; Gammaproteobacteria; Enterobacterales; Enterobacteriaceae; Escherichia
MNNEETFYQAMRRQGVTRRSFLKYCSLAATSLGLGAGMAPKIAWALENKPRIPVVWIHGL WP_001058323.1
MULTISPECIES: hydrogenase-1 small chain [Proteobacteria] Length: 372\nNumber of predicted TMHs: 2\nExp number of AAs in TMHs: 37.2608499999999\nExp number, first 60 AAs: 19.52923\nTotal prob of N-in: 0.94351\nPOSSIBLE N-term signal sequence\ninside 1 19\nTMhelix 20 39\noutside 40 325\nTMhelix 326 348\ninside 349 372

3625 GCF_001269285.1_1.EC2992.1 Escherichia coli Proteobacteria; Gammaproteobacteria; Enterobacterales; Enterobacteriaceae; Escherichia
MTGDNTLIHSHGINRRDFMKLCAALAATMGLSSKAAAEMVESVTNPQRPPVIWIGAQECT WP_000145423.1
hydrogenase 2 small subunit [Escherichia coli] Length: 372\nNumber of predicted TMHs: 1\nExp number of AAs in TMHs: 23.78201\nExp number, first 60 AAs: 0.28355\nTotal prob of N-in: 0.13257\noutside 1 331\nTMhelix 332 354\ninside 355 372

3626 GCF_001277435.1_7748_7_2Escherichia coli Proteobacteria; Gammaproteobacteria; Enterobacterales; Enterobacteriaceae; Escherichia MTGDNTLIHSHGINRRDFMKLCAALAATMGLSSKAAAEMAESVTNPQRPPVIWIGAQECT

WP_000145410.1 MULTISPECIES: hydrogenase-2 small chain [Proteobacteria] Length: 372\nNumber of predicted TMHs: 1\nExp number of AAs in TMHs: 23.72226\nExp number, first 60 AAs: 0.22038\nTotal prob of N-in: 0.12994\noutside 1 331\nTMhelix 332 354\ninside 355 372

3627 GCF_001269185.1_YE13 Escherichia coli Proteobacteria; Gammaproteobacteria; Enterobacterales; Enterobacteriaceae; Escherichia MTGDNTLIHSHGINRRDFMKLCAALAATMGLSSKAAAEMAESVTNPQRPPVIWIGAQECT WP_000145410.1 MULTISPECIES: hydrogenase-2 small chain [Proteobacteria] Length: 372\nNumber of predicted TMHs: 1\nExp number of AAs in TMHs: 23.72226\nExp number, first 60 AAs: 0.22038\nTotal prob of N-in: 0.12994\noutside 1 331\nTMhelix 332 354\ninside 355 372

3628 GCF_001284225.1_8205_8_70 Escherichia coli Proteobacteria; Gammaproteobacteria; Enterobacterales; Enterobacteriaceae; Escherichia MNNEETFYQAMRRQGVTRRSFLKYCSLAATSLGLGAGMAPKIAWALENKPRIPVVWIHGL WP_001058320.1 hydrogenase [Escherichia coli] Length: 372\nNumber of predicted TMHs: 2\nExp number of AAs in TMHs: 37.26082\nExp number, first 60 AAs: 19.52923\nTotal prob of N-in: 0.94351\nPOSSIBLE N-term signal sequence\ninside 1 19\nTMhelix 20 39\noutside 40 325\nTMhelix 326 348\ninside 349 372

3629 GCF_001269325.1_YE6 Escherichia coli Proteobacteria; Gammaproteobacteria; Enterobacterales; Enterobacteriaceae; Escherichia MTGDNTLIHSHGINRRDFMKLCAALAATMGLSSKAAAEMAESVTNPQRPPVIWIGAQECT WP_000145410.1 MULTISPECIES: hydrogenase-2 small chain [Proteobacteria] Length: 372\nNumber of predicted TMHs: 1\nExp number of AAs in TMHs: 23.72226\nExp number, first 60 AAs: 0.22038\nTotal prob of N-in: 0.12994\noutside 1 331\nTMhelix 332 354\ninside 355 372

3630 GCF_001284665.1_7748_7_34 Escherichia coli Proteobacteria; Gammaproteobacteria; Enterobacterales; Enterobacteriaceae; Escherichia MNNEETFYQAMRRQGVTRRSFLKYCSLAATSLGLGAGMAPKIAWALENKPRIPVVWIHGL WP_001058323.1 MULTISPECIES: hydrogenase-1 small chain [Proteobacteria] Length: 372\nNumber of predicted TMHs: 2\nExp number of AAs in TMHs: 37.2608499999999\nExp number, first 60 AAs: 19.52923\nTotal prob of N-in: 0.94351\nPOSSIBLE N-term signal sequence\ninside 1 19\nTMhelix 20 39\noutside 40 325\nTMhelix 326 348\ninside 349 372

3631 GCF_001277675.1_7748_7_1Escherichia coli Proteobacteria; Gammaproteobacteria; Enterobacterales; Enterobacteriaceae; Escherichia MTGDNTLIHSHGINRRDFMKLCAALAATMGLSSKAAAEMAESVTNPQRPPVIWIGAQECT WP_000145410.1 MULTISPECIES: hydrogenase-2 small chain [Proteobacteria] Length: 372\nNumber of predicted TMHs: 1\nExp number of AAs in TMHs: 23.72226\nExp number, first 60 AAs: 0.22038\nTotal prob of N-in: 0.12994\noutside 1 331\nTMhelix 332 354\ninside 355 372

3632 GCF_001284825.1_7790_1_50 Escherichia coli Proteobacteria; Gammaproteobacteria; Enterobacterales; Enterobacteriaceae; Escherichia MNNEETFYQAMRRQGITRRSFLKYCSLAATSLGLGAGMAPKIAWALENKPRIPVVWIHGL WP_053897536.1 hydrogenase [Escherichia coli] Length: 372\nNumber of predicted TMHs: 2\nExp number of AAs in TMHs: 37.4535599999999\nExp number, first 60 AAs: 19.77343\nTotal prob of N-in: 0.95460\nPOSSIBLE N-term signal sequence\ninside 1 19\nTMhelix 20 39\noutside 40 325\nTMhelix 326 348\ninside 349 372

3633 GCF_001284605.1_7748_7_27 Escherichia coli Proteobacteria; Gammaproteobacteria; Enterobacterales; Enterobacteriaceae; Escherichia MTGDNTLIHSHGINRRDFMKLCAALAATMGLSSKAAAEMAESVTNPQRPPVIWIGAQECT WP_053881429.1 hydrogenase 2 small subunit [Escherichia coli] Length: 372\nNumber of predicted TMHs: 1\nExp number of AAs in TMHs: 21.67161\nExp number, first 60 AAs: 0.23069\nTotal prob of N-in: 0.04884\noutside 1 331\nTMhelix 332 354\ninside 355 372

3634 GCF_001284545.1_7790_1_53 Escherichia coli Proteobacteria; Gammaproteobacteria; Enterobacterales; Enterobacteriaceae; Escherichia MTGDNTLIHSHGINRRDFMKLCAALAATMGLSSKAAAEMAESVTNPQRPPVIWIGAQECT WP_000145410.1 MULTISPECIES: hydrogenase-2 small chain [Proteobacteria] Length: 372\nNumber of predicted TMHs: 1\nExp number of AAs in TMHs: 23.72226\nExp number, first 60 AAs: 0.22038\nTotal prob of N-in: 0.12994\noutside 1 331\nTMhelix 332 354\ninside 355 372

3635 GCF_000797895.1_Escherichia_coli_CVM_N38834PS_v._1.0 Escherichia coli Proteobacteria; Gammaproteobacteria; Enterobacterales; Enterobacteriaceae; Escherichia MTGDNTLIHSHGINRRDFMKLCAALAATMGLSSKAAAEMAESVTNPQRPPVIWIGAQECT WP_000145410.1

MULTISPECIES: hydrogenase-2 small chain [Proteobacteria] Length: 372\nNumber of predicted TMHs: 1\nExp number of AAs in TMHs: 23.72226\nExp number, first 60 AAs: 0.22038\nTotal prob of N-in: 0.12994\noutside 1 331\nTMhelix 332 354\ninside 355 372

3636 GCF_000797935.1_Escherichia_coli_CVM_N36396PS_v._1.0 Escherichia coli Proteobacteria; Gammaproteobacteria; Enterobacterales; Enterobacteriaceae; Escherichia MNNEETFYQAMRRQGVTRRSFLKYCSLAATSLGLGAGMAPKIAWALENKPRIPVVWIHGL WP_001058323.1 MULTISPECIES: hydrogenase-1 small chain [Proteobacteria] Length: 372\nNumber of predicted TMHs: 2\nExp number of AAs in TMHs: 37.2608499999999\nExp number, first 60 AAs: 19.52923\nTotal prob of N-in: 0.94351\nPOSSIBLE N-term signal sequence\ninside 1 19\nTMhelix 20 39\noutside 40 325\nTMhelix 326 348\ninside 349 372

3637 GCF_001277735.1_7748_7_15 Escherichia coli Proteobacteria; Gammaproteobacteria; Enterobacterales; Enterobacteriaceae; Escherichia MTGDNTLIHSHGINRRDFMKLCAALAATMGLSSKAAAEMAESVTNPQRPPVIWIGAQECT WP_000145410.1 MULTISPECIES: hydrogenase-2 small chain [Proteobacteria] Length: 372\nNumber of predicted TMHs: 1\nExp number of AAs in TMHs: 23.72226\nExp number, first 60 AAs: 0.22038\nTotal prob of N-in: 0.12994\noutside 1 331\nTMhelix 332 354\ninside 355 372

3638 GCF_001284885.1_8205_3_19 Escherichia coli Proteobacteria; Gammaproteobacteria; Enterobacterales; Enterobacteriaceae; Escherichia MTGDNTLIHSHGINRRDFMKLCAALAATMGLSSKAAAEMAESVTNPQRPPVIWIGAQECT WP_000145410.1 MULTISPECIES: hydrogenase-2 small chain [Proteobacteria] Length: 372\nNumber of predicted TMHs: 1\nExp number of AAs in TMHs: 23.72226\nExp number, first 60 AAs: 0.22038\nTotal prob of N-in: 0.12994\noutside 1 331\nTMhelix 332 354\ninside 355 372

3639 GCF_001284225.1_8205_8_70 Escherichia coli Proteobacteria; Gammaproteobacteria; Enterobacterales; Enterobacteriaceae; Escherichia MTGDNTLIHSHGINRRDFMKLCAALAATMGLSSKAAAEMAESVTNPQRPPVIWIGAQECT WP_000145410.1 MULTISPECIES: hydrogenase-2 small chain [Proteobacteria] Length: 372\nNumber of predicted TMHs: 1\nExp number of AAs in TMHs: 23.72226\nExp number, first 60 AAs: 0.22038\nTotal prob of N-in: 0.12994\noutside 1 331\nTMhelix 332 354\ninside 355 372

3640 GCF_001283785.1_8205_3_70 Escherichia coli Proteobacteria; Gammaproteobacteria; Enterobacterales; Enterobacteriaceae; Escherichia MNNEETFYQAMRRQGVTRRSFLKYCSLAATSLGLGAGMAPKIAWALENKPRIPVVWIHGL WP_001058323.1 MULTISPECIES: hydrogenase-1 small chain [Proteobacteria] Length: 372\nNumber of predicted TMHs: 2\nExp number of AAs in TMHs: 37.2608499999999\nExp number, first 60 AAs: 19.52923\nTotal prob of N-in: 0.94351\nPOSSIBLE N-term signal sequence\ninside 1 19\nTMhelix 20 39\noutside 40 325\nTMhelix 326 348\ninside 349 372

3641 GCF_001277775.1_7790_1_60 Escherichia coli Proteobacteria; Gammaproteobacteria; Enterobacterales; Enterobacteriaceae; Escherichia MTGDNTLIHSHGINRRDFMKLCAALAATMGLSSKAAAEMVESVTNPQRPPVIWIGAQECT WP_000145423.1 hydrogenase 2 small subunit [Escherichia coli] Length: 372\nNumber of predicted TMHs: 1\nExp number of AAs in TMHs: 23.78201\nExp number, first 60 AAs: 0.28355\nTotal prob of N-in: 0.13257\noutside 1 331\nTMhelix 332 354\ninside 355 372

3642 GCF_001277675.1_7748_7_1Escherichia coli Proteobacteria; Gammaproteobacteria; Enterobacterales; Enterobacteriaceae; Escherichia MNNEETFYQAMRRQGVTRRSFLKYCSLAATSLGLGAGMAPKIAWALENKPRIPVVWIHGL WP_001058323.1 MULTISPECIES: hydrogenase-1 small chain [Proteobacteria] Length: 372\nNumber of predicted TMHs: 2\nExp number of AAs in TMHs: 37.2608499999999\nExp number, first 60 AAs: 19.52923\nTotal prob of N-in: 0.94351\nPOSSIBLE N-term signal sequence\ninside 1 19\nTMhelix 20 39\noutside 40 325\nTMhelix 326 348\ninside 349 372

3643 GCF_001277635.1_7790_1_49 Escherichia coli Proteobacteria; Gammaproteobacteria; Enterobacterales; Enterobacteriaceae; Escherichia MTGDNTLIHSHGINRRDFMKLCAALAATMGLSSKAAAEMAESVTNPQRPPVIWIGAQECT WP_000145410.1 MULTISPECIES: hydrogenase-2 small chain [Proteobacteria] Length: 372\nNumber of predicted TMHs: 1\nExp number of AAs in TMHs: 23.72226\nExp number, first 60 AAs: 0.22038\nTotal prob of N-in: 0.12994\noutside 1 331\nTMhelix 332 354\ninside 355 372

3644 GCF_000599745.2_a5_miseq_linux_20140113 Escherichia coli Proteobacteria; Gammaproteobacteria; Enterobacterales; Enterobacteriaceae; Escherichia
MTGDNTLIHSHGINRRDFMKLCAALAATMGLSSKAAAEMAESVTNPQRPPVIWIGAQECT WP_000145410.1
MULTISPECIES: hydrogenase-2 small chain [Proteobacteria] Length: 372\nNumber of predicted TMHs: 1\nExp number of AAs in TMHs: 23.72226\nExp number, first 60 AAs: 0.22038\nTotal prob of N-in: 0.12994\noutside 1 331\nTMhelix 332 354\ninside 355 372

3645 GCF_000798475.1_Escherichia_coli_CVM_N36204PS_v._1.0 Escherichia coli Proteobacteria; Gammaproteobacteria; Enterobacterales; Enterobacteriaceae; Escherichia
MTGDNTLIHSHGINRRDFMKLCAALAATMGLSSKAAAEMAESVTNPQRPPVIWIGAQECT WP_000145410.1
MULTISPECIES: hydrogenase-2 small chain [Proteobacteria] Length: 372\nNumber of predicted TMHs: 1\nExp number of AAs in TMHs: 23.72226\nExp number, first 60 AAs: 0.22038\nTotal prob of N-in: 0.12994\noutside 1 331\nTMhelix 332 354\ninside 355 372

3646 GCF_000798475.1_Escherichia_coli_CVM_N36204PS_v._1.0 Escherichia coli Proteobacteria; Gammaproteobacteria; Enterobacterales; Enterobacteriaceae; Escherichia
MNNEETFYQAMRRQGVTRRSFLKYCSLAATSLGLGAGMAPKIAWALENKPRIPVVWIHGL WP_001058323.1
MULTISPECIES: hydrogenase-1 small chain [Proteobacteria] Length: 372\nNumber of predicted TMHs: 2\nExp number of AAs in TMHs: 37.2608499999999\nExp number, first 60 AAs: 19.52923\nTotal prob of N-in: 0.94351\nPOSSIBLE N-term signal sequence\ninside 1 19\nTMhelix 20 39\noutside 40 325\nTMhelix 326 348\ninside 349 372

3647 GCF_000696545.1_v1 Escherichia coli Proteobacteria; Gammaproteobacteria; Enterobacterales; Enterobacteriaceae; Escherichia MTGDNTLIHSHGINRRDFMKLCAALAATMGLSSKAAAEMAESVTNPQRPPVIWIGAQECT
WP_000145410.1 MULTISPECIES: hydrogenase-2 small chain [Proteobacteria] Length: 372\nNumber of predicted TMHs: 1\nExp number of AAs in TMHs: 23.72226\nExp number, first 60 AAs: 0.22038\nTotal prob of N-in: 0.12994\noutside 1 331\nTMhelix 332 354\ninside 355 372

3648 GCF_000948675.1_ASM94867v1 Escherichia coli Proteobacteria; Gammaproteobacteria; Enterobacterales; Enterobacteriaceae; Escherichia
MTGDNTLIHSHGINRRDFMKLCAALAATMGLSSKAAAEMAESVTNPQRPPVIWIGAQECT WP_000145410.1
MULTISPECIES: hydrogenase-2 small chain [Proteobacteria] Length: 372\nNumber of predicted TMHs: 1\nExp number of AAs in TMHs: 23.72226\nExp number, first 60 AAs: 0.22038\nTotal prob of N-in: 0.12994\noutside 1 331\nTMhelix 332 354\ninside 355 372

3649 GCF_000948615.1_ASM94861v1 Escherichia coli Proteobacteria; Gammaproteobacteria; Enterobacterales; Enterobacteriaceae; Escherichia
MTGDNTLIHSHGINRRDFMKLCAALAATMGLSSKAAAEMAESVTNPQRPPVIWIGAQECT WP_000145410.1
MULTISPECIES: hydrogenase-2 small chain [Proteobacteria] Length: 372\nNumber of predicted TMHs: 1\nExp number of AAs in TMHs: 23.72226\nExp number, first 60 AAs: 0.22038\nTotal prob of N-in: 0.12994\noutside 1 331\nTMhelix 332 354\ninside 355 372

3650 GCF_000948565.1_ASM94856v1 Escherichia coli Proteobacteria; Gammaproteobacteria; Enterobacterales; Enterobacteriaceae; Escherichia
MNNEETFYQAMRRQGVTRRSFLKYCSLAATSLGLGAGMAPKIAWALENKPRIPVVWIHGL WP_001058323.1
MULTISPECIES: hydrogenase-1 small chain [Proteobacteria] Length: 372\nNumber of predicted TMHs: 2\nExp number of AAs in TMHs: 37.2608499999999\nExp number, first 60 AAs: 19.52923\nTotal prob of N-in: 0.94351\nPOSSIBLE N-term signal sequence\ninside 1 19\nTMhelix 20 39\noutside 40 325\nTMhelix 326 348\ninside 349 372

3651 GCF_000948565.1_ASM94856v1 Escherichia coli Proteobacteria; Gammaproteobacteria; Enterobacterales; Enterobacteriaceae; Escherichia
MTGDNTLIHSHGINRRDFMKLCAALAATMGLSSKAAAEMAESVTNPQRPPVIWIGAQECT WP_000145410.1
MULTISPECIES: hydrogenase-2 small chain [Proteobacteria] Length: 372\nNumber of predicted TMHs: 1\nExp number of AAs in TMHs: 23.72226\nExp number, first 60 AAs: 0.22038\nTotal prob of N-in: 0.12994\noutside 1 331\nTMhelix 332 354\ninside 355 372

3652 GCF_000952225.1_ASM95222v1 Escherichia coli Proteobacteria; Gammaproteobacteria; Enterobacterales; Enterobacteriaceae; Escherichia
MTGDNTLIHSHGINRRDFMKLCAALAATMGLSSKAAAEMVESVTNPQRPPVIWIGAQECT WP_000145423.1
hydrogenase 2 small subunit [Escherichia coli] Length: 372\nNumber of predicted TMHs: 1\nExp number of

AAs in TMHs: 23.78201\nExp number, first 60 AAs: 0.28355\nTotal prob of N-in: 0.13257\noutside 1 331\nTMhelix 332 354\ninside 355 372

3653 GCF_000952225.1_ASM95222v1 Escherichia coli Proteobacteria; Gammaproteobacteria; Enterobacterales; Enterobacteriaceae; Escherichia MNNEETFYQAMRRQGVTRRSFLKYCSLAATSLGLGAGMAPKIAWALENKPRIPVVWIHGL WP_001295940.1 MULTISPECIES: hydrogenase [Enterobacteriaceae] Length: 372\nNumber of predicted TMHs: 2\nExp number of AAs in TMHs: 37.2608899999999\nExp number, first 60 AAs: 19.52923\nTotal prob of N-in: 0.94351\nPOSSIBLE N-term signal sequence\ninside 1 19\nTMhelix 20 39\noutside 40 325\nTMhelix 326 348\ninside 349 372

3654 GCF_000691105.1_S6685 Escherichia coli Proteobacteria; Gammaproteobacteria; Enterobacterales; Enterobacteriaceae; Escherichia MNNEETFYQAMRRQGVTRRSFLKYCSLAATSLGLGAGMAPKIAWALENKPRIPVVWIHGL WP_001295940.1 MULTISPECIES: hydrogenase [Enterobacteriaceae] Length: 372\nNumber of predicted TMHs: 2\nExp number of AAs in TMHs: 37.2608899999999\nExp number, first 60 AAs: 19.52923\nTotal prob of N-in: 0.94351\nPOSSIBLE N-term signal sequence\ninside 1 19\nTMhelix 20 39\noutside 40 325\nTMhelix 326 348\ninside 349 372

3655 GCF_000953765.1_ASM95376v1 Escherichia coli Proteobacteria; Gammaproteobacteria; Enterobacterales; Enterobacteriaceae; Escherichia MTGDNTLIHSHGINRRDFMKLCAALAATMGLSSKAAAEMAESVTNPQRPPVIWIGAQECT WP_000145410.1 MULTISPECIES: hydrogenase-2 small chain [Proteobacteria] Length: 372\nNumber of predicted TMHs: 1\nExp number of AAs in TMHs: 23.72226\nExp number, first 60 AAs: 0.22038\nTotal prob of N-in: 0.12994\noutside 1 331\nTMhelix 332 354\ninside 355 372

3656 GCF_000965545.1_ASM96554v1 Escherichia coli Proteobacteria; Gammaproteobacteria; Enterobacterales; Enterobacteriaceae; Escherichia MNNEETFYQAMRRQGVTRRSFLKYCSLAATSLGLGAGMAPKIAWALENKPRIPVVWIHGL WP_024246676.1 hydrogenase-1 small chain [Escherichia coli] Length: 372\nNumber of predicted TMHs: 2\nExp number of AAs in TMHs: 37.8214899999999\nExp number, first 60 AAs: 19.52326\nTotal prob of N-in: 0.94395\nPOSSIBLE N-term signal sequence\ninside 1 19\nTMhelix 20 39\noutside 40 325\nTMhelix 326 348\ninside 349 372

3657 GCF_001285185.1_7748_7_6Escherichia coli Proteobacteria; Gammaproteobacteria; Enterobacterales; Enterobacteriaceae; Escherichia MTGDNTLIHSHGINRRDFMKLCAALAATMGLSSKAAAEMAESVTNPQRPPVIWIGAQECT WP_000145410.1 MULTISPECIES: hydrogenase-2 small chain [Proteobacteria] Length: 372\nNumber of predicted TMHs: 1\nExp number of AAs in TMHs: 23.72226\nExp number, first 60 AAs: 0.22038\nTotal prob of N-in: 0.12994\noutside 1 331\nTMhelix 332 354\ninside 355 372

3658 GCF_000948515.1_ASM94851v1 Escherichia coli Proteobacteria; Gammaproteobacteria; Enterobacterales; Enterobacteriaceae; Escherichia MTGDNTLIHSHGINRRDFMKLCAALAATMGLSSKAAAEMAESVTNPQRPPVIWIGAQECT WP_000145410.1 MULTISPECIES: hydrogenase-2 small chain [Proteobacteria] Length: 372\nNumber of predicted TMHs: 1\nExp number of AAs in TMHs: 23.72226\nExp number, first 60 AAs: 0.22038\nTotal prob of N-in: 0.12994\noutside 1 331\nTMhelix 332 354\ninside 355 372

3659 GCF_001285225.1_7748_7_11 Escherichia coli Proteobacteria; Gammaproteobacteria; Enterobacterales; Enterobacteriaceae; Escherichia MTGDNTLIHSHGINRRDFMKLCAALAATMGLSSKAAAEMAESVTNPQRPPVIWIGAQECT WP_000145410.1 MULTISPECIES: hydrogenase-2 small chain [Proteobacteria] Length: 372\nNumber of predicted TMHs: 1\nExp number of AAs in TMHs: 23.72226\nExp number, first 60 AAs: 0.22038\nTotal prob of N-in: 0.12994\noutside 1 331\nTMhelix 332 354\ninside 355 372

3660 GCF_001285225.1_7748_7_11 Escherichia coli Proteobacteria; Gammaproteobacteria; Enterobacterales; Enterobacteriaceae; Escherichia MNNEETFYQAMRRQGVTRRSFLKYCSLAATSLGLGAGMAPKIAWALENKPRIPVVWIHGL WP_001295940.1 MULTISPECIES: hydrogenase [Enterobacteriaceae] Length: 372\nNumber of predicted TMHs: 2\nExp number of AAs in TMHs: 37.2608899999999\nExp number, first 60 AAs: 19.52923\nTotal prob of N-in: 0.94351\nPOSSIBLE N-term signal sequence\ninside 1 19\nTMhelix 20 39\noutside 40 325\nTMhelix 326 348\ninside 349 372

3661 GCF_000948475.1_ASM94847v1 Escherichia coli Proteobacteria; Gammaproteobacteria; Enterobacterales; Enterobacteriaceae; Escherichia
MNNEETFYQAMRRQGVTRRSFLKYCSLAATSLGLGAGMAPKIAWALENKPRIPVVWIHGL WP_001295940.1
MULTISPECIES: hydrogenase [Enterobacteriaceae] Length: 372\nNumber of predicted TMHs: 2\nExp number of AAs in TMHs: 37.2608899999999\nExp number, first 60 AAs: 19.52923\nTotal prob of N-in: 0.94351\nPOSSIBLE N-term signal sequence\ninside 1 19\nTMhelix 20 39\noutside 40 325\nTMhelix 326 348\ninside 349 372

3662 GCF_000948475.1_ASM94847v1 Escherichia coli Proteobacteria; Gammaproteobacteria; Enterobacterales; Enterobacteriaceae; Escherichia
MTGDNTLIHSHGINRRDFMKLCAALAATMGLSSKAAAEMAESVTNPQRPPVIWIGAQECT WP_000145410.1
MULTISPECIES: hydrogenase-2 small chain [Proteobacteria] Length: 372\nNumber of predicted TMHs: 1\nExp number of AAs in TMHs: 23.72226\nExp number, first 60 AAs: 0.22038\nTotal prob of N-in: 0.12994\noutside 1 331\nTMhelix 332 354\ninside 355 372

3663 GCF_000965565.1_ASM96556v1 Escherichia coli Proteobacteria; Gammaproteobacteria; Enterobacterales; Enterobacteriaceae; Escherichia
MTGDNTLIHSHGINRRDFMKLCAALAATMGLSSKAAAEMAESVTNPQRPPVIWIGAQECT WP_000145410.1
MULTISPECIES: hydrogenase-2 small chain [Proteobacteria] Length: 372\nNumber of predicted TMHs: 1\nExp number of AAs in TMHs: 23.72226\nExp number, first 60 AAs: 0.22038\nTotal prob of N-in: 0.12994\noutside 1 331\nTMhelix 332 354\ninside 355 372

3664 GCF_001285285.1_8205_3_42 Escherichia coli Proteobacteria; Gammaproteobacteria; Enterobacterales; Enterobacteriaceae; Escherichia
MTGDNTLIHSHGINRRDFMKLCAALAATMGLSSKAAAEMAESVTNPQRPPVIWIGAQECT WP_000145410.1
MULTISPECIES: hydrogenase-2 small chain [Proteobacteria] Length: 372\nNumber of predicted TMHs: 1\nExp number of AAs in TMHs: 23.72226\nExp number, first 60 AAs: 0.22038\nTotal prob of N-in: 0.12994\noutside 1 331\nTMhelix 332 354\ninside 355 372

3665 GCF_000965565.1_ASM96556v1 Escherichia coli Proteobacteria; Gammaproteobacteria; Enterobacterales; Enterobacteriaceae; Escherichia
MNNEETFYQAMRRQGVTRRSFLKYCSLAATSLGLGAGMAPKIAWALENKPRIPVVWIHGL WP_001058323.1
MULTISPECIES: hydrogenase-1 small chain [Proteobacteria] Length: 372\nNumber of predicted TMHs: 2\nExp number of AAs in TMHs: 37.2608499999999\nExp number, first 60 AAs: 19.52923\nTotal prob of N-in: 0.94351\nPOSSIBLE N-term signal sequence\ninside 1 19\nTMhelix 20 39\noutside 40 325\nTMhelix 326 348\ninside 349 372

3666 GCF_000967155.2_HUSEC2011CHR1 Escherichia coli Proteobacteria; Gammaproteobacteria; Enterobacterales; Enterobacteriaceae; Escherichia
MTGDNTLIHSHGINRRDFMKLCAALAATMGLSSKAAAEMAESVTNPQRPPVIWIGAQECT WP_000145410.1
MULTISPECIES: hydrogenase-2 small chain [Proteobacteria] Length: 372\nNumber of predicted TMHs: 1\nExp number of AAs in TMHs: 23.72226\nExp number, first 60 AAs: 0.22038\nTotal prob of N-in: 0.12994\noutside 1 331\nTMhelix 332 354\ninside 355 372

3667 GCF_000967155.2_HUSEC2011CHR1 Escherichia coli Proteobacteria; Gammaproteobacteria; Enterobacterales; Enterobacteriaceae; Escherichia
MNNEETFYQAMRRQGVTRRSFLKYCSLAATSLGLGAGMAPKIAWALENKPRIPVVWIHGL WP_001058323.1
MULTISPECIES: hydrogenase-1 small chain [Proteobacteria] Length: 372\nNumber of predicted TMHs: 2\nExp number of AAs in TMHs: 37.2608499999999\nExp number, first 60 AAs: 19.52923\nTotal prob of N-in: 0.94351\nPOSSIBLE N-term signal sequence\ninside 1 19\nTMhelix 20 39\noutside 40 325\nTMhelix 326 348\ninside 349 372

3668 GCF_001268525.1_1.EC2988.1 Escherichia coli Proteobacteria; Gammaproteobacteria; Enterobacterales; Enterobacteriaceae; Escherichia
MTGDNTLIHSHGINRRDFMKLCAALAATMGLSSKAAAEMAESVTNPQRPPVIWIGAQECT WP_000145410.1
MULTISPECIES: hydrogenase-2 small chain [Proteobacteria] Length: 372\nNumber of predicted TMHs: 1\nExp number of AAs in TMHs: 23.72226\nExp number, first 60 AAs: 0.22038\nTotal prob of N-in: 0.12994\noutside 1 331\nTMhelix 332 354\ninside 355 372

3669 GCF_001268565.1_1.EC2923.1 Escherichia coli Proteobacteria; Gammaproteobacteria; Enterobacterales; Enterobacteriaceae; Escherichia

MNNEETFYQAMRRQGVTRRSFLKYCSLAATSLGLGAGMAPKIAWALENKPRIPVVWIHGL WP_001058323.1 MULTISPECIES: hydrogenase-1 small chain [Proteobacteria] Length: 372\nNumber of predicted TMHs: 2\nExp number of AAs in TMHs: 37.2608499999999\nExp number, first 60 AAs: 19.52923\nTotal prob of N-in: 0.94351\nPOSSIBLE N-term signal sequence\ninside 1 19\nTMhelix 20 39\noutside 40 325\nTMhelix 326 348\ninside 349 372

3670 GCF_001285325.1_7748_7_23 Escherichia coli Proteobacteria; Gammaproteobacteria; Enterobacterales; Enterobacteriaceae; Escherichia MNNEETFYQAMRRQGVTRRSFLKYCSLAATSLGLGAGMAPKIAWALENKPRIPVVWIHGL WP_001295940.1 MULTISPECIES: hydrogenase [Enterobacteriaceae] Length: 372\nNumber of predicted TMHs: 2\nExp number of AAs in TMHs: 37.2608899999999\nExp number, first 60 AAs: 19.52923\nTotal prob of N-in: 0.94351\nPOSSIBLE N-term signal sequence\ninside 1 19\nTMhelix 20 39\noutside 40 325\nTMhelix 326 348\ninside 349 372

3671 GCF_000708145.1_E_coli_EC1.contigs Escherichia coli Proteobacteria; Gammaproteobacteria; Enterobacterales; Enterobacteriaceae; Escherichia MTGDNTLIHSHGINRRDFMKLCAALAATMGLSSKAAAEMAESVTNPQRPPVIWIGAQECT WP_000145410.1 MULTISPECIES: hydrogenase-2 small chain [Proteobacteria] Length: 372\nNumber of predicted TMHs: 1\nExp number of AAs in TMHs: 23.72226\nExp number, first 60 AAs: 0.22038\nTotal prob of N-in: 0.12994\noutside 1 331\nTMhelix 332 354\ninside 355 372

3672 GCF_000708145.1_E_coli_EC1.contigs Escherichia coli Proteobacteria; Gammaproteobacteria; Enterobacterales; Enterobacteriaceae; Escherichia MNNEETFYQAMRRQGVTRRSFLKYCSLAATSLGLGAGMAPKIAWALENKPRIPVVWIHGL WP_001058323.1 MULTISPECIES: hydrogenase-1 small chain [Proteobacteria] Length: 372\nNumber of predicted TMHs: 2\nExp number of AAs in TMHs: 37.2608499999999\nExp number, first 60 AAs: 19.52923\nTotal prob of N-in: 0.94351\nPOSSIBLE N-term signal sequence\ninside 1 19\nTMhelix 20 39\noutside 40 325\nTMhelix 326 348\ninside 349 372

3673 GCF_001056665.1_ASM105666v1 Escherichia coli Proteobacteria; Gammaproteobacteria; Enterobacterales; Enterobacteriaceae; Escherichia MTGDNTLIHSHGINRRDFMKLCAALAATMGLSSKAAAEMAESVTNPQRPPVIWIGAQECT WP_000145410.1 MULTISPECIES: hydrogenase-2 small chain [Proteobacteria] Length: 372\nNumber of predicted TMHs: 1\nExp number of AAs in TMHs: 23.72226\nExp number, first 60 AAs: 0.22038\nTotal prob of N-in: 0.12994\noutside 1 331\nTMhelix 332 354\ninside 355 372

3674 GCF_001056875.1_ASM105687v1 Escherichia coli Proteobacteria; Gammaproteobacteria; Enterobacterales; Enterobacteriaceae; Escherichia MTGDNTLIHSHGINRRDFMKLCAALAATMGLSSKAAAEMAESVTNPQRPPVIWIGAQECT WP_000145410.1 MULTISPECIES: hydrogenase-2 small chain [Proteobacteria] Length: 372\nNumber of predicted TMHs: 1\nExp number of AAs in TMHs: 23.72226\nExp number, first 60 AAs: 0.22038\nTotal prob of N-in: 0.12994\noutside 1 331\nTMhelix 332 354\ninside 355 372

3675 GCF_001285445.1_8205_3_65 Escherichia coli Proteobacteria; Gammaproteobacteria; Enterobacterales; Enterobacteriaceae; Escherichia MNNEETFYQAMRRQGVTRRSFLKYCSLAATSLGLGAGMAPKIAWALENKPRIPVVWIHGL WP_001058320.1 hydrogenase [Escherichia coli] Length: 372\nNumber of predicted TMHs: 2\nExp number of AAs in TMHs: 37.26082\nExp number, first 60 AAs: 19.52923\nTotal prob of N-in: 0.94351\nPOSSIBLE N-term signal sequence\ninside 1 19\nTMhelix 20 39\noutside 40 325\nTMhelix 326 348\ninside 349 372

3676 GCF_001057385.1_ASM105738v1 Escherichia coli Proteobacteria; Gammaproteobacteria; Enterobacterales; Enterobacteriaceae; Escherichia MNNEETFYQAMRRQGVTRRSFLKYCSLAATSLGLGAGMAPKIAWALENKPRIPVVWIHGL WP_001058323.1 MULTISPECIES: hydrogenase-1 small chain [Proteobacteria] Length: 372\nNumber of predicted TMHs: 2\nExp number of AAs in TMHs: 37.2608499999999\nExp number, first 60 AAs: 19.52923\nTotal prob of N-in: 0.94351\nPOSSIBLE N-term signal sequence\ninside 1 19\nTMhelix 20 39\noutside 40 325\nTMhelix 326 348\ninside 349 372

3677 GCF_001285505.1_7790_1_81 Escherichia coli Proteobacteria; Gammaproteobacteria; Enterobacterales; Enterobacteriaceae; Escherichia MNNEETFYQAMRRQGVTRRSFLKYCSLAATSLGLGAGMAPKIAWALENKPRIPVVWIHGL WP_001295940.1

MULTISPECIES: hydrogenase [Enterobacteriaceae] Length: 372\nNumber of predicted TMHs: 2\nExp number of AAs in TMHs: 37.2608899999999\nExp number, first 60 AAs: 19.52923\nTotal prob of N-in: 0.94351\nPOSSIBLE N-term signal sequence\ninside 1 19\nTMhelix 20 39\noutside 40 325\nTMhelix 326 348\ninside 349 372

3678 GCF_001057895.1_ASM105789v1 Escherichia coli Proteobacteria; Gammaproteobacteria; Enterobacterales; Enterobacteriaceae; Escherichia
MTGDNTLIHSHGINRRDFMKLCAALAATMGLSSKAAAEMVESVTNPQRPPVIWIGAQECT WP_000145423.1
hydrogenase 2 small subunit [Escherichia coli] Length: 372\nNumber of predicted TMHs: 1\nExp number of AAs in TMHs: 23.78201\nExp number, first 60 AAs: 0.28355\nTotal prob of N-in: 0.13257\noutside 1 331\nTMhelix 332 354\ninside 355 372

3679 GCF_001057895.1_ASM105789v1 Escherichia coli Proteobacteria; Gammaproteobacteria; Enterobacterales; Enterobacteriaceae; Escherichia
MNNEETFYQAMRRQGVTRRSFLKYCSLAATSLGLGAGMAPKIAWALENKPRIPVVWIHGL WP_001295940.1
MULTISPECIES: hydrogenase [Enterobacteriaceae] Length: 372\nNumber of predicted TMHs: 2\nExp number of AAs in TMHs: 37.2608899999999\nExp number, first 60 AAs: 19.52923\nTotal prob of N-in: 0.94351\nPOSSIBLE N-term signal sequence\ninside 1 19\nTMhelix 20 39\noutside 40 325\nTMhelix 326 348\ninside 349 372

3680 GCF_001058175.1_ASM105817v1 Escherichia coli Proteobacteria; Gammaproteobacteria; Enterobacterales; Enterobacteriaceae; Escherichia
MNNEETFYQAMRRQGVTRRSFLKYCSLAATSLGLGAGMAPKIAWALENKPRIPVVWIHGL WP_001295940.1
MULTISPECIES: hydrogenase [Enterobacteriaceae] Length: 372\nNumber of predicted TMHs: 2\nExp number of AAs in TMHs: 37.2608899999999\nExp number, first 60 AAs: 19.52923\nTotal prob of N-in: 0.94351\nPOSSIBLE N-term signal sequence\ninside 1 19\nTMhelix 20 39\noutside 40 325\nTMhelix 326 348\ninside 349 372

3681 GCF_001191105.1_CFSAN026819_01-NGE.clc Escherichia coli Proteobacteria; Gammaproteobacteria; Enterobacterales; Enterobacteriaceae; Escherichia
MNNEETFYQAMRRQGVTRRSFLKYCSLAATSLGLGAGMAPKIAWALENKPRIPVVWIHGL WP_001295940.1
MULTISPECIES: hydrogenase [Enterobacteriaceae] Length: 372\nNumber of predicted TMHs: 2\nExp number of AAs in TMHs: 37.2608899999999\nExp number, first 60 AAs: 19.52923\nTotal prob of N-in: 0.94351\nPOSSIBLE N-term signal sequence\ninside 1 19\nTMhelix 20 39\noutside 40 325\nTMhelix 326 348\ninside 349 372

3682 GCF_001191105.1_CFSAN026819_01-NGE.clc Escherichia coli Proteobacteria; Gammaproteobacteria; Enterobacterales; Enterobacteriaceae; Escherichia
MTGDNTLIHSHGINRRDFMKLCAALAATMGLSSKAAAEMAESVTNPQRPPVIWIGAQECT WP_000145410.1
MULTISPECIES: hydrogenase-2 small chain [Proteobacteria] Length: 372\nNumber of predicted TMHs: 1\nExp number of AAs in TMHs: 23.72226\nExp number, first 60 AAs: 0.22038\nTotal prob of N-in: 0.12994\noutside 1 331\nTMhelix 332 354\ninside 355 372

3683 GCF_001285565.1_8205_3_74 Escherichia coli Proteobacteria; Gammaproteobacteria; Enterobacterales; Enterobacteriaceae; Escherichia
MTGDNTLIHSHGINRRDFMKLCAALAATMGLSSKAAAEMAESVTNPQRPPVIWIGAQECT WP_000145410.1
MULTISPECIES: hydrogenase-2 small chain [Proteobacteria] Length: 372\nNumber of predicted TMHs: 1\nExp number of AAs in TMHs: 23.72226\nExp number, first 60 AAs: 0.22038\nTotal prob of N-in: 0.12994\noutside 1 331\nTMhelix 332 354\ninside 355 372

3684 GCF_000714915.1_UCD_JA17_pb Escherichia coli Proteobacteria; Gammaproteobacteria; Enterobacterales; Enterobacteriaceae; Escherichia
MTGDNTLIHSHGINRRDFMKLCAALAATMGLSSKAAAEMAESVTNPQRPPVIWIGAQECT WP_000145410.1
MULTISPECIES: hydrogenase-2 small chain [Proteobacteria] Length: 372\nNumber of predicted TMHs: 1\nExp number of AAs in TMHs: 23.72226\nExp number, first 60 AAs: 0.22038\nTotal prob of N-in: 0.12994\noutside 1 331\nTMhelix 332 354\ninside 355 372

3685 GCF_000714995.1_JA69_HGAP2 Escherichia coli Proteobacteria; Gammaproteobacteria; Enterobacterales; Enterobacteriaceae; Escherichia
MTGDNTLIHSHGINRRDFMKLCAALAATMGLSSKAAAEMAESVTNPQRPPVIWIGAQECT WP_000145410.1
MULTISPECIES: hydrogenase-2 small chain [Proteobacteria] Length: 372\nNumber of predicted TMHs:

1\nExp number of AAs in TMHs: 23.72226\nExp number, first 60 AAs: 0.22038\nTotal prob of N-in: 0.12994\noutside 1 331\nTMhelix 332 354\ninside 355 372

3686 GCF_000752295.1_FHI78 Escherichia coli Proteobacteria; Gammaproteobacteria; Enterobacterales; Enterobacteriaceae; Escherichia MTGDNTLIHSHGINRRDFMKLCAALAATMGLSSKAAAEMAESVTNPQRPPVIWIGAQECT WP_000145410.1 MULTISPECIES: hydrogenase-2 small chain [Proteobacteria] Length: 372\nNumber of predicted TMHs: 1\nExp number of AAs in TMHs: 23.72226\nExp number, first 60 AAs: 0.22038\nTotal prob of N-in: 0.12994\noutside 1 331\nTMhelix 332 354\ninside 355 372

3687 GCF_000947985.1_ASM94798v1 Escherichia coli Proteobacteria; Gammaproteobacteria; Enterobacterales; Enterobacteriaceae; Escherichia MNNEETFYQAMRRQGVTRRSFLKYCSLAATSLGLGAGMAPKIAWALENKPRIPVVWIHGL WP_001058323.1 MULTISPECIES: hydrogenase-1 small chain [Proteobacteria] Length: 372\nNumber of predicted TMHs: 2\nExp number of AAs in TMHs: 37.2608499999999\nExp number, first 60 AAs: 19.52923\nTotal prob of N-in: 0.94351\nPOSSIBLE N-term signal sequence\ninside 1 19\nTMhelix 20 39\noutside 40 325\nTMhelix 326 348\ninside 349 372

3688 GCF_000752355.1_FHI50 Escherichia coli Proteobacteria; Gammaproteobacteria; Enterobacterales; Enterobacteriaceae; Escherichia MTGDNTLIHSHGINRRDFMKLCAALAATMGLSSKAAAEMAESVTNPQRPPVIWIGAQECT WP_000145410.1 MULTISPECIES: hydrogenase-2 small chain [Proteobacteria] Length: 372\nNumber of predicted TMHs: 1\nExp number of AAs in TMHs: 23.72226\nExp number, first 60 AAs: 0.22038\nTotal prob of N-in: 0.12994\noutside 1 331\nTMhelix 332 354\ninside 355 372

3689 GCF_000947935.1_ASM94793v1 Escherichia coli Proteobacteria; Gammaproteobacteria; Enterobacterales; Enterobacteriaceae; Escherichia MNNEETFYQAMRRQGVTRRSFLKYCSLAATSLGLGAGMAPKIAWALENKPRIPVVWIHGL WP_001058323.1 MULTISPECIES: hydrogenase-1 small chain [Proteobacteria] Length: 372\nNumber of predicted TMHs: 2\nExp number of AAs in TMHs: 37.2608499999999\nExp number, first 60 AAs: 19.52923\nTotal prob of N-in: 0.94351\nPOSSIBLE N-term signal sequence\ninside 1 19\nTMhelix 20 39\noutside 40 325\nTMhelix 326 348\ninside 349 372

3690 GCF_000752455.1_FHI31 Escherichia coli Proteobacteria; Gammaproteobacteria; Enterobacterales; Enterobacteriaceae; Escherichia MTGDNTLIHSHGINRRDFMKLCAALAATMGLSSKAAAEMAESVTNPQRPPVIWIGAQECT WP_000145410.1 MULTISPECIES: hydrogenase-2 small chain [Proteobacteria] Length: 372\nNumber of predicted TMHs: 1\nExp number of AAs in TMHs: 23.72226\nExp number, first 60 AAs: 0.22038\nTotal prob of N-in: 0.12994\noutside 1 331\nTMhelix 332 354\ninside 355 372

3691 GCF_000947935.1_ASM94793v1 Escherichia coli Proteobacteria; Gammaproteobacteria; Enterobacterales; Enterobacteriaceae; Escherichia MTGDNTLIHSHGINRRDFMKLCAALAATMGLSSKAAAEMAESVTNPQRPPVIWIGAQECT WP_000145410.1 MULTISPECIES: hydrogenase-2 small chain [Proteobacteria] Length: 372\nNumber of predicted TMHs: 1\nExp number of AAs in TMHs: 23.72226\nExp number, first 60 AAs: 0.22038\nTotal prob of N-in: 0.12994\noutside 1 331\nTMhelix 332 354\ninside 355 372

3692 GCF_000752515.1_FHI61 Escherichia coli Proteobacteria; Gammaproteobacteria; Enterobacterales; Enterobacteriaceae; Escherichia MTGDNTLIHSHGINRRDFMKLCAALAATMGLSSKAAAEMAESVTNPQRPPVIWIGAQECT WP_000145410.1 MULTISPECIES: hydrogenase-2 small chain [Proteobacteria] Length: 372\nNumber of predicted TMHs: 1\nExp number of AAs in TMHs: 23.72226\nExp number, first 60 AAs: 0.22038\nTotal prob of N-in: 0.12994\noutside 1 331\nTMhelix 332 354\ninside 355 372

3693 GCF_000752355.1_FHI50 Escherichia coli Proteobacteria; Gammaproteobacteria; Enterobacterales; Enterobacteriaceae; Escherichia MNNEETFYQAMRRQGVTRRSFLKYCSLAATSLGLGAGMAPKIAWALENKPRIPVVWIHGL WP_001058323.1 MULTISPECIES: hydrogenase-1 small chain [Proteobacteria] Length: 372\nNumber of predicted TMHs: 2\nExp number of AAs in TMHs: 37.2608499999999\nExp number, first 60 AAs: 19.52923\nTotal prob of N-in: 0.94351\nPOSSIBLE N-term signal sequence\ninside 1 19\nTMhelix 20 39\noutside 40 325\nTMhelix 326 348\ninside 349 372

3694 GCF_000752455.1_FHI31 Escherichia coli Proteobacteria; Gammaproteobacteria; Enterobacterales; Enterobacteriaceae; Escherichia MNNEETFYQAMRRQGVTRRSFLKYCSLAATSLGLGAGMAPKIAWALENKPRIPVVWIHGL WP_001058323.1 MULTISPECIES: hydrogenase-1 small chain [Proteobacteria] Length: 372\nNumber of predicted TMHs: 2\nExp number of AAs in TMHs: 37.2608499999999\nExp number, first 60 AAs: 19.52923\nTotal prob of

N-in: 0.94351\nPOSSIBLE N-term signal sequence\ninside 1 19\nTMhelix 20 39\noutside 40 325\nTMhelix 326 348\ninside 349 372

3695 GCF_000752515.1_FHI61 Escherichia coli Proteobacteria; Gammaproteobacteria; Enterobacterales; Enterobacteriaceae; Escherichia MNNEETFYQAMRRQGVTRRSFLKYCSLAATSLGLGAGMAPKIAWALENKPRIPVVWIHGL WP_001058323.1 MULTISPECIES: hydrogenase-1 small chain [Proteobacteria] Length: 372\nNumber of predicted TMHs: 2\nExp number of AAs in TMHs: 37.2608499999999\nExp number, first 60 AAs: 19.52923\nTotal prob of N-in: 0.94351\nPOSSIBLE N-term signal sequence\ninside 1 19\nTMhelix 20 39\noutside 40 325\nTMhelix 326 348\ninside 349 372

3696 GCF_000946855.1_2d11B_assembly Escherichia coli Proteobacteria; Gammaproteobacteria; Enterobacterales; Enterobacteriaceae; Escherichia MNNEETFYQAMRRQGVTRRSFLKYCSLAATSLGLGAGMAPKIAWALENKPRIPVVWIHGL WP_032280669.1 hydrogenase [Escherichia coli] Length: 372\nNumber of predicted TMHs: 2\nExp number of AAs in TMHs: 37.2609299999999\nExp number, first 60 AAs: 19.52923\nTotal prob of N-in: 0.94351\nPOSSIBLE N-term signal sequence\ninside 1 19\nTMhelix 20 39\noutside 40 325\nTMhelix 326 348\ninside 349 372

3697 GCF_000752155.1_FHI52 Escherichia coli Proteobacteria; Gammaproteobacteria; Enterobacterales; Enterobacteriaceae; Escherichia MNNEETFYQAMRRQGVTRRSFLKYCSLAATSLGLGAGMAPKIAWALENKPRIPVVWIHGL WP_001058323.1 MULTISPECIES: hydrogenase-1 small chain [Proteobacteria] Length: 372\nNumber of predicted TMHs: 2\nExp number of AAs in TMHs: 37.2608499999999\nExp number, first 60 AAs: 19.52923\nTotal prob of N-in: 0.94351\nPOSSIBLE N-term signal sequence\ninside 1 19\nTMhelix 20 39\noutside 40 325\nTMhelix 326 348\ninside 349 372

3698 GCF_000753035.1_FHI38 Escherichia coli Proteobacteria; Gammaproteobacteria; Enterobacterales; Enterobacteriaceae; Escherichia MTGDNTLIHSHGINRRDFMKLCAALAATMGLSSKAAAEMAESVTNPQRPPVIWIGAQECT WP_000145410.1 MULTISPECIES: hydrogenase-2 small chain [Proteobacteria] Length: 372\nNumber of predicted TMHs: 1\nExp number of AAs in TMHs: 23.72226\nExp number, first 60 AAs: 0.22038\nTotal prob of N-in: 0.12994\noutside 1 331\nTMhelix 332 354\ninside 355 372

3699 GCF_000946855.1_2d11B_assembly Escherichia coli Proteobacteria; Gammaproteobacteria; Enterobacterales; Enterobacteriaceae; Escherichia MTGDNTLIHSHGINRRDFMKLCAALAATMGLSSKAAAEMAESVTNPQRPPVIWIGAQECT WP_000145410.1 MULTISPECIES: hydrogenase-2 small chain [Proteobacteria] Length: 372\nNumber of predicted TMHs: 1\nExp number of AAs in TMHs: 23.72226\nExp number, first 60 AAs: 0.22038\nTotal prob of N-in: 0.12994\noutside 1 331\nTMhelix 332 354\ninside 355 372

3700 GCF_000752935.1_FHI49 Escherichia coli Proteobacteria; Gammaproteobacteria; Enterobacterales; Enterobacteriaceae; Escherichia MTGDNTLIHSHGINRRDFMKLCAALAATMGLSSKAAAEMAESVTNPQRPPVIWIGAQECT WP_000145410.1 MULTISPECIES: hydrogenase-2 small chain [Proteobacteria] Length: 372\nNumber of predicted TMHs: 1\nExp number of AAs in TMHs: 23.72226\nExp number, first 60 AAs: 0.22038\nTotal prob of N-in: 0.12994\noutside 1 331\nTMhelix 332 354\ninside 355 372

3701 GCF_000753135.1_FHI64 Escherichia coli Proteobacteria; Gammaproteobacteria; Enterobacterales; Enterobacteriaceae; Escherichia MTGDNTLIHSHGINRRDFMKLCAALAATMGLSSKAAAEMAESVTNPQRPPVIWIGAQECT WP_000145410.1 MULTISPECIES: hydrogenase-2 small chain [Proteobacteria] Length: 372\nNumber of predicted TMHs: 1\nExp number of AAs in TMHs: 23.72226\nExp number, first 60 AAs: 0.22038\nTotal prob of N-in: 0.12994\noutside 1 331\nTMhelix 332 354\ninside 355 372

3702 GCF_000753115.1_FHI45 Escherichia coli Proteobacteria; Gammaproteobacteria; Enterobacterales; Enterobacteriaceae; Escherichia MTGDNTLIHSHGINRRDFMKLCAALAATMGLSSKAAAEMAESVTNPQRPPVIWIGAQECT WP_000145410.1 MULTISPECIES: hydrogenase-2 small chain [Proteobacteria] Length: 372\nNumber of predicted TMHs: 1\nExp number of AAs in TMHs: 23.72226\nExp number, first 60 AAs: 0.22038\nTotal prob of N-in: 0.12994\noutside 1 331\nTMhelix 332 354\ninside 355 372

3703 GCF_000753195.1_FHI17 Escherichia coli Proteobacteria; Gammaproteobacteria; Enterobacterales; Enterobacteriaceae; Escherichia MTGDNTLIHSHGINRRDFMKLCAALAATMGLSSKAAAEMVESVTNPQRPPVIWIGAQECT WP_000145423.1 hydrogenase 2 small subunit [Escherichia coli] Length: 372\nNumber of predicted TMHs: 1\nExp number of AAs in TMHs: 23.78201\nExp number, first 60 AAs: 0.28355\nTotal prob of N-in: 0.13257\noutside 1 331\nTMhelix 332 354\ninside 355 372

3704 GCF_000753035.1_FHI38 Escherichia coli Proteobacteria; Gammaproteobacteria; Enterobacterales; Enterobacteriaceae; Escherichia MNNEETFYQAMRRQGVTRRSFLKYCSLAATSLGLGAGMAPKIAWALENKPRIPVVWIHGL WP_001058323.1 MULTISPECIES: hydrogenase-1 small chain [Proteobacteria] Length: 372\nNumber of predicted TMHs: 2\nExp number of AAs in TMHs: 37.2608499999999\nExp number, first 60 AAs: 19.52923\nTotal prob of N-in: 0.94351\nPOSSIBLE N-term signal sequence\ninside 1 19\nTMhelix 20 39\noutside 40 325\nTMhelix 326 348\ninside 349 372

3705 GCF_000770255.1_ASM77025v1 Escherichia coli Proteobacteria; Gammaproteobacteria; Enterobacterales; Enterobacteriaceae; Escherichia MTGDNTLIHSHGINRRDFMKLCAALAATMGLSSKAAAEMAESVTNPQRPPVIWIGAQECT WP_000145410.1 MULTISPECIES: hydrogenase-2 small chain [Proteobacteria] Length: 372\nNumber of predicted TMHs: 1\nExp number of AAs in TMHs: 23.72226\nExp number, first 60 AAs: 0.22038\nTotal prob of N-in: 0.12994\noutside 1 331\nTMhelix 332 354\ninside 355 372

3706 GCF_000768485.1_ASM76848v1 Escherichia coli Proteobacteria; Gammaproteobacteria; Enterobacterales; Enterobacteriaceae; Escherichia MTGDNTLIHSHGINRRDFMKLCAALAATMGLSSKAAAEMAESVTNPQRPPVIWIGAQECT WP_000145410.1 MULTISPECIES: hydrogenase-2 small chain [Proteobacteria] Length: 372\nNumber of predicted TMHs: 1\nExp number of AAs in TMHs: 23.72226\nExp number, first 60 AAs: 0.22038\nTotal prob of N-in: 0.12994\noutside 1 331\nTMhelix 332 354\ninside 355 372

3707 GCF_000945955.1_FHI85 Escherichia coli Proteobacteria; Gammaproteobacteria; Enterobacterales; Enterobacteriaceae; Escherichia MTGDNTLIHSHGINRRDFMKLCAALAATMGLSSKAAAEMAESVTNPQRPPVIWIGAQECT WP_000145410.1 MULTISPECIES: hydrogenase-2 small chain [Proteobacteria] Length: 372\nNumber of predicted TMHs: 1\nExp number of AAs in TMHs: 23.72226\nExp number, first 60 AAs: 0.22038\nTotal prob of N-in: 0.12994\noutside 1 331\nTMhelix 332 354\ninside 355 372

3708 GCF_000770035.1_ASM77003v1 Escherichia coli Proteobacteria; Gammaproteobacteria; Enterobacterales; Enterobacteriaceae; Escherichia MTGDNTLIHSHGINRRDFMKLCAALAATMGLSSKAAAEMAESVTNPQRPPVIWIGAQECT WP_000145410.1 MULTISPECIES: hydrogenase-2 small chain [Proteobacteria] Length: 372\nNumber of predicted TMHs: 1\nExp number of AAs in TMHs: 23.72226\nExp number, first 60 AAs: 0.22038\nTotal prob of N-in: 0.12994\noutside 1 331\nTMhelix 332 354\ninside 355 372

3709 GCF_000770285.1_ASM77028v1 Escherichia coli Proteobacteria; Gammaproteobacteria; Enterobacterales; Enterobacteriaceae; Escherichia MTGDNTLIHSHGINRRDFMKLCAALAATMGLSSKAAAEMVESVTNPQRPPVIWIGAQECT WP_000145423.1 hydrogenase 2 small subunit [Escherichia coli] Length: 372\nNumber of predicted TMHs: 1\nExp number of AAs in TMHs: 23.78201\nExp number, first 60 AAs: 0.28355\nTotal prob of N-in: 0.13257\noutside 1 331\nTMhelix 332 354\ninside 355 372

3710 GCF_000773535.1_ASM77353v1 Escherichia coli Proteobacteria; Gammaproteobacteria; Enterobacterales; Enterobacteriaceae; Escherichia MTGDNTLIHSHGINRRDFMKLCAALAATMGLSSKAAAEMAESVTNPQRPPVIWIGAQECT WP_000145410.1 MULTISPECIES: hydrogenase-2 small chain [Proteobacteria] Length: 372\nNumber of predicted TMHs: 1\nExp number of AAs in TMHs: 23.72226\nExp number, first 60 AAs: 0.22038\nTotal prob of N-in: 0.12994\noutside 1 331\nTMhelix 332 354\ninside 355 372

3711 GCF_000776045.1_ASM77604v1 Escherichia coli Proteobacteria; Gammaproteobacteria; Enterobacterales; Enterobacteriaceae; Escherichia MTGDNTLIHSHGINRRDFMKLCAALAATMGLSSKAAAEMAESVTNPQRPPVIWIGAQECT WP_000145410.1 MULTISPECIES: hydrogenase-2 small chain [Proteobacteria] Length: 372\nNumber of predicted TMHs: 1\nExp number of AAs in TMHs: 23.72226\nExp number, first 60 AAs: 0.22038\nTotal prob of N-in: 0.12994\noutside 1 331\nTMhelix 332 354\ninside 355 372

3712 GCF_000773575.1_ASM77357v1 Escherichia coli Proteobacteria; Gammaproteobacteria; Enterobacterales; Enterobacteriaceae; Escherichia MTGDNTLIHSHGINRRDFMKLCAALAATMGLSSKAAAEMAESVTNPQRPPVIWIGAQECT WP_000145410.1 MULTISPECIES: hydrogenase-2 small chain [Proteobacteria] Length: 372\nNumber of predicted TMHs: 1\nExp number of AAs in TMHs: 23.72226\nExp number, first 60 AAs: 0.22038\nTotal prob of N-in: 0.12994\noutside 1 331\nTMhelix 332 354\ninside 355 372

3713 GCF_000773535.1_ASM77353v1 Escherichia coli Proteobacteria; Gammaproteobacteria; Enterobacterales; Enterobacteriaceae; Escherichia MNNEETFYQAMRRQGVTRRSFLKYCSLAATSLGLGAGMAPKIAWALENKPRIPVVWIHGL WP_001058323.1 MULTISPECIES: hydrogenase-1 small chain [Proteobacteria] Length: 372\nNumber of predicted TMHs: 2\nExp number of AAs in TMHs: 37.2608499999999\nExp number, first 60 AAs: 19.52923\nTotal prob of N-in: 0.94351\nPOSSIBLE N-term signal sequence\ninside 1 19\nTMhelix 20 39\noutside 40 325\nTMhelix 326 348\ninside 349 372

3714 GCF_000776155.1_ASM77615v1 Escherichia coli Proteobacteria; Gammaproteobacteria; Enterobacterales; Enterobacteriaceae; Escherichia MTGDNTLIHSHGINRRDFMKLCAALAATMGLSSKAAAEMVESVTNPQRPPVIWIGAQECT WP_000145423.1 hydrogenase 2 small subunit [Escherichia coli] Length: 372\nNumber of predicted TMHs: 1\nExp number of AAs in TMHs: 23.78201\nExp number, first 60 AAs: 0.28355\nTotal prob of N-in: 0.13257\noutside 1 331\nTMhelix 332 354\ninside 355 372

3715 GCF_000776195.1_ASM77619v1 Escherichia coli Proteobacteria; Gammaproteobacteria; Enterobacterales; Enterobacteriaceae; Escherichia MTGDNTLIHSHGINRRDFMKLCAALAATMGLSSKAAAEMAESVTNPQRPPVIWIGAQECT WP_000145410.1 MULTISPECIES: hydrogenase-2 small chain [Proteobacteria] Length: 372\nNumber of predicted TMHs: 1\nExp number of AAs in TMHs: 23.72226\nExp number, first 60 AAs: 0.22038\nTotal prob of N-in: 0.12994\noutside 1 331\nTMhelix 332 354\ninside 355 372

3716 GCF_000776285.1_ASM77628v1 Escherichia coli Proteobacteria; Gammaproteobacteria; Enterobacterales; Enterobacteriaceae; Escherichia MTGDNTLIHSHGINRRDFMKLCAALAATMGLSSKAAAEMAESVTNPQRPPVIWIGAQECT WP_000145410.1 MULTISPECIES: hydrogenase-2 small chain [Proteobacteria] Length: 372\nNumber of predicted TMHs: 1\nExp number of AAs in TMHs: 23.72226\nExp number, first 60 AAs: 0.22038\nTotal prob of N-in: 0.12994\noutside 1 331\nTMhelix 332 354\ninside 355 372

3717 GCF_000776355.1_ASM77635v1 Escherichia coli Proteobacteria; Gammaproteobacteria; Enterobacterales; Enterobacteriaceae; Escherichia MTGDNTLIHSHGINRRDFMKLCAALAATMGLSSKAAAEMAESVTNPQRPPVIWIGAQECT WP_000145410.1 MULTISPECIES: hydrogenase-2 small chain [Proteobacteria] Length: 372\nNumber of predicted TMHs: 1\nExp number of AAs in TMHs: 23.72226\nExp number, first 60 AAs: 0.22038\nTotal prob of N-in: 0.12994\noutside 1 331\nTMhelix 332 354\ninside 355 372

3718 GCF_000776395.1_ASM77639v1 Escherichia coli Proteobacteria; Gammaproteobacteria; Enterobacterales; Enterobacteriaceae; Escherichia MTGDNTLIHSHGINRRDFMKLCAALAATMGLSSKAAAEMVESVTNPQRPPVIWIGAQECT WP_000145423.1 hydrogenase 2 small subunit [Escherichia coli] Length: 372\nNumber of predicted TMHs: 1\nExp number of AAs in TMHs: 23.78201\nExp number, first 60 AAs: 0.28355\nTotal prob of N-in: 0.13257\noutside 1 331\nTMhelix 332 354\ninside 355 372

3719 GCF_000776505.1_ASM77650v1 Escherichia coli Proteobacteria; Gammaproteobacteria; Enterobacterales; Enterobacteriaceae; Escherichia MTGDNTLIHSHGINRRDFMKLCAALAATMGLSSKAAAEMAESVTNPQRPPVIWIGAQECT WP_000145410.1 MULTISPECIES: hydrogenase-2 small chain [Proteobacteria] Length: 372\nNumber of predicted TMHs: 1\nExp number of AAs in TMHs: 23.72226\nExp number, first 60 AAs: 0.22038\nTotal prob of N-in: 0.12994\noutside 1 331\nTMhelix 332 354\ninside 355 372

3720 GCF_000776455.1_ASM77645v1 Escherichia coli Proteobacteria; Gammaproteobacteria; Enterobacterales; Enterobacteriaceae; Escherichia MTGDNTLIHSHGINRRDFMKLCAALAATMGLSSKAAAEMAESVTNPQRPPVIWIGAQECT WP_000145418.1 hydrogenase 2 small subunit [Escherichia coli] Length: 372\nNumber of predicted TMHs: 1\nExp number of AAs in TMHs: 23.72313\nExp number, first 60 AAs: 0.22034\nTotal prob of N-in: 0.12997\noutside 1 331\nTMhelix 332 354\ninside 355 372

3721 GCF_000942915.1_FHI29 Escherichia coli Proteobacteria; Gammaproteobacteria; Enterobacterales; Enterobacteriaceae; Escherichia MNNEETFYQAMRRQGVTRRSFLKYCSLAATSLGLGAGMAPKIAWALENKPRIPVVWIHGL WP_001058323.1 MULTISPECIES: hydrogenase-1 small chain [Proteobacteria] Length: 372\nNumber of predicted TMHs: 2\nExp number of AAs in TMHs: 37.2608499999999\nExp number, first 60 AAs: 19.52923\nTotal prob of

N-in: 0.94351\nPOSSIBLE N-term signal sequence\ninside 1 19\nTMhelix 20 39\noutside 40 325\nTMhelix 326 348\ninside 349 372

3722 GCF_000942915.1_FHI29 Escherichia coli Proteobacteria; Gammaproteobacteria; Enterobacterales; Enterobacteriaceae; Escherichia MTGDNTLIHSHGINRRDFMKLCAALAATMGLSSKAAAEMAESVTNPQRPPVIWIGAQECT WP_000145410.1 MULTISPECIES: hydrogenase-2 small chain [Proteobacteria] Length: 372\nNumber of predicted TMHs: 1\nExp number of AAs in TMHs: 23.72226\nExp number, first 60 AAs: 0.22038\nTotal prob of N-in: 0.12994\noutside 1 331\nTMhelix 332 354\ninside 355 372

3723 GCF_000776595.1_ASM77659v1 Escherichia coli Proteobacteria; Gammaproteobacteria; Enterobacterales; Enterobacteriaceae; Escherichia MTGDNTLIHSHGINRRDFMKLCAALAATMGLSSKAAAEMVESVTNPQRPPVIWIGAQECT WP_000145423.1 hydrogenase 2 small subunit [Escherichia coli] Length: 372\nNumber of predicted TMHs: 1\nExp number of AAs in TMHs: 23.78201\nExp number, first 60 AAs: 0.28355\nTotal prob of N-in: 0.13257\noutside 1 331\nTMhelix 332 354\ninside 355 372

3724 GCF_000777165.1_ASM77716v1 Escherichia coli Proteobacteria; Gammaproteobacteria; Enterobacterales; Enterobacteriaceae; Escherichia MTGDNTLIHSHGINRRDFMKLCAALAATMGLSSKAAAEMAESVTNPQRPPVIWIGAQECT WP_000145410.1 MULTISPECIES: hydrogenase-2 small chain [Proteobacteria] Length: 372\nNumber of predicted TMHs: 1\nExp number of AAs in TMHs: 23.72226\nExp number, first 60 AAs: 0.22038\nTotal prob of N-in: 0.12994\noutside 1 331\nTMhelix 332 354\ninside 355 372

3725 GCF_000777165.1_ASM77716v1 Escherichia coli Proteobacteria; Gammaproteobacteria; Enterobacterales; Enterobacteriaceae; Escherichia MNNEETFYQAMRRQGVTRRSFLKYCSLAATSLGLGAGMAPKIAWALENKPRIPVVWIHGL WP_001058323.1 MULTISPECIES: hydrogenase-1 small chain [Proteobacteria] Length: 372\nNumber of predicted TMHs: 2\nExp number of AAs in TMHs: 37.2608499999999\nExp number, first 60 AAs: 19.52923\nTotal prob of N-in: 0.94351\nPOSSIBLE N-term signal sequence\ninside 1 19\nTMhelix 20 39\noutside 40 325\nTMhelix 326 348\ninside 349 372

3726 GCF_000777215.1_ASM77721v1 Escherichia coli Proteobacteria; Gammaproteobacteria; Enterobacterales; Enterobacteriaceae; Escherichia MNNEETFYQAMRRQGVTRRSFLKYCSLAATSLGLGAGMAPKIAWALENKPRIPVVWIHGL WP_001058323.1 MULTISPECIES: hydrogenase-1 small chain [Proteobacteria] Length: 372\nNumber of predicted TMHs: 2\nExp number of AAs in TMHs: 37.2608499999999\nExp number, first 60 AAs: 19.52923\nTotal prob of N-in: 0.94351\nPOSSIBLE N-term signal sequence\ninside 1 19\nTMhelix 20 39\noutside 40 325\nTMhelix 326 348\ninside 349 372

3727 GCF_000776795.1_ASM77679v1 Escherichia coli Proteobacteria; Gammaproteobacteria; Enterobacterales; Enterobacteriaceae; Escherichia MNNEETFYQAMRRQGVTRRSFLKYCSLAATSLGLGAGMAPKIAWALENKPRIPVVWIHGL WP_001058323.1 MULTISPECIES: hydrogenase-1 small chain [Proteobacteria] Length: 372\nNumber of predicted TMHs: 2\nExp number of AAs in TMHs: 37.2608499999999\nExp number, first 60 AAs: 19.52923\nTotal prob of N-in: 0.94351\nPOSSIBLE N-term signal sequence\ninside 1 19\nTMhelix 20 39\noutside 40 325\nTMhelix 326 348\ninside 349 372

3728 GCF_000777065.1_ASM77706v1 Escherichia coli Proteobacteria; Gammaproteobacteria; Enterobacterales; Enterobacteriaceae; Escherichia MNNEETFYQAMRRQGVTRRSFLKYCSLAATSLGLGAGMAPKIAWALENKPRIPVVWIHGL WP_001058323.1 MULTISPECIES: hydrogenase-1 small chain [Proteobacteria] Length: 372\nNumber of predicted TMHs: 2\nExp number of AAs in TMHs: 37.2608499999999\nExp number, first 60 AAs: 19.52923\nTotal prob of N-in: 0.94351\nPOSSIBLE N-term signal sequence\ninside 1 19\nTMhelix 20 39\noutside 40 325\nTMhelix 326 348\ninside 349 372

3729 GCF_000777285.1_ASM77728v1 Escherichia coli Proteobacteria; Gammaproteobacteria; Enterobacterales; Enterobacteriaceae; Escherichia MTGDNTLIHSHGINRRDFMKLCAALAATMGLSSKAAAEMAESVTNPQRPPVIWIGAQECT WP_000145410.1 MULTISPECIES: hydrogenase-2 small chain [Proteobacteria] Length: 372\nNumber of predicted TMHs: 1\nExp number of AAs in TMHs: 23.72226\nExp number, first 60 AAs: 0.22038\nTotal prob of N-in: 0.12994\noutside 1 331\nTMhelix 332 354\ninside 355 372

3730 GCF_000777285.1_ASM77728v1 Escherichia coli Proteobacteria; Gammaproteobacteria; Enterobacterales; Enterobacteriaceae; Escherichia
MNNEETFYQAMRRQGVTRRSFLKYCSLAATSLGLGAGMAPKIAWALENKPRIPVVWIHGL WP_001058323.1
MULTISPECIES: hydrogenase-1 small chain [Proteobacteria] Length: 372\nNumber of predicted TMHs: 2\nExp number of AAs in TMHs: 37.2608499999999\nExp number, first 60 AAs: 19.52923\nTotal prob of N-in: 0.94351\nPOSSIBLE N-term signal sequence\ninside 1 19\nTMhelix 20 39\noutside 40 325\nTMhelix 326 348\ninside 349 372

3731 GCF_000777455.1_ASM77745v1 Escherichia coli Proteobacteria; Gammaproteobacteria; Enterobacterales; Enterobacteriaceae; Escherichia
MTGDNTLIHSHGINRRDFMKLCAALAATMGLSSKAAAEMAESVTNPQRPPVIWIGAQECT WP_000145410.1
MULTISPECIES: hydrogenase-2 small chain [Proteobacteria] Length: 372\nNumber of predicted TMHs: 1\nExp number of AAs in TMHs: 23.72226\nExp number, first 60 AAs: 0.22038\nTotal prob of N-in: 0.12994\noutside 1 331\nTMhelix 332 354\ninside 355 372

3732 GCF_000777555.1_ASM77755v1 Escherichia coli Proteobacteria; Gammaproteobacteria; Enterobacterales; Enterobacteriaceae; Escherichia
MTGDNTLIHSHGINRRDFMKLCAALAATMGLSSKAAAEMVESVTNPQRPPVIWIGAQECT WP_000145423.1
hydrogenase 2 small subunit [Escherichia coli] Length: 372\nNumber of predicted TMHs: 1\nExp number of AAs in TMHs: 23.78201\nExp number, first 60 AAs: 0.28355\nTotal prob of N-in: 0.13257\noutside 1 331\nTMhelix 332 354\ninside 355 372

3733 GCF_000777515.1_ASM77751v1 Escherichia coli Proteobacteria; Gammaproteobacteria; Enterobacterales; Enterobacteriaceae; Escherichia
MTGDNTLIHSHGINRRDFMKLCAALAATMGLSSKAAAEMAESVTNPQRPPVIWIGAQECT WP_000145410.1
MULTISPECIES: hydrogenase-2 small chain [Proteobacteria] Length: 372\nNumber of predicted TMHs: 1\nExp number of AAs in TMHs: 23.72226\nExp number, first 60 AAs: 0.22038\nTotal prob of N-in: 0.12994\noutside 1 331\nTMhelix 332 354\ninside 355 372

3734 GCF_000777975.1_ASM77797v1 Escherichia coli Proteobacteria; Gammaproteobacteria; Enterobacterales; Enterobacteriaceae; Escherichia
MTGDNTLIHSHGINRRDFMKLCAALAATMGLSSKAAAEMVESVTNPQRPPVIWIGAQECT WP_000145423.1
hydrogenase 2 small subunit [Escherichia coli] Length: 372\nNumber of predicted TMHs: 1\nExp number of AAs in TMHs: 23.78201\nExp number, first 60 AAs: 0.28355\nTotal prob of N-in: 0.13257\noutside 1 331\nTMhelix 332 354\ninside 355 372

3735 GCF_000777625.1_ASM77762v1 Escherichia coli Proteobacteria; Gammaproteobacteria; Enterobacterales; Enterobacteriaceae; Escherichia
MTGDNTLIHSHGINRRDFMKLCAALAATMGLSSKAAAEMAESVTNPQRPPVIWIGAQECT WP_000145410.1
MULTISPECIES: hydrogenase-2 small chain [Proteobacteria] Length: 372\nNumber of predicted TMHs: 1\nExp number of AAs in TMHs: 23.72226\nExp number, first 60 AAs: 0.22038\nTotal prob of N-in: 0.12994\noutside 1 331\nTMhelix 332 354\ninside 355 372

3736 GCF_000777675.1_ASM77767v1 Escherichia coli Proteobacteria; Gammaproteobacteria; Enterobacterales; Enterobacteriaceae; Escherichia
MTGDNTLIHSHGINRRDFMKLCAALAATMGLSSKAAAEMAESVTNPQRPPVIWIGAQECT WP_000145410.1
MULTISPECIES: hydrogenase-2 small chain [Proteobacteria] Length: 372\nNumber of predicted TMHs: 1\nExp number of AAs in TMHs: 23.72226\nExp number, first 60 AAs: 0.22038\nTotal prob of N-in: 0.12994\noutside 1 331\nTMhelix 332 354\ninside 355 372

3737 GCF_000777845.1_ASM77784v1 Escherichia coli Proteobacteria; Gammaproteobacteria; Enterobacterales; Enterobacteriaceae; Escherichia
MTGDNTLIHSHGINRRDFMKLCAALAATMGLSSKAAAEMAESVTNPQRPPVIWIGAQECT WP_000145410.1
MULTISPECIES: hydrogenase-2 small chain [Proteobacteria] Length: 372\nNumber of predicted TMHs: 1\nExp number of AAs in TMHs: 23.72226\nExp number, first 60 AAs: 0.22038\nTotal prob of N-in: 0.12994\noutside 1 331\nTMhelix 332 354\ninside 355 372

3738 GCF_000777815.1_ASM77781v1 Escherichia coli Proteobacteria; Gammaproteobacteria; Enterobacterales; Enterobacteriaceae; Escherichia
MTGDNTLIHSHGINRRDFMKLCAALAATMGLSSKAAAEMAESVTNPQRPPVIWIGAQECT WP_000145410.1
MULTISPECIES: hydrogenase-2 small chain [Proteobacteria] Length: 372\nNumber of predicted TMHs:

1\nExp number of AAs in TMHs: 23.72226\nExp number, first 60 AAs: 0.22038\nTotal prob of N-in: 0.12994\noutside 1 331\nTMhelix 332 354\ninside 355 372

3739 GCF_000778275.1_ASM77827v1 Escherichia coli Proteobacteria; Gammaproteobacteria; Enterobacterales; Enterobacteriaceae; Escherichia
MTGDNTLIHSHGINRRDFMKLCAALAATMGLSSKAAAEMAESVTNPQRPPVIWIGAQECT WP_000145418.1
hydrogenase 2 small subunit [Escherichia coli] Length: 372\nNumber of predicted TMHs: 1\nExp number of AAs in TMHs: 23.72313\nExp number, first 60 AAs: 0.22034\nTotal prob of N-in: 0.12997\noutside 1 331\nTMhelix 332 354\ninside 355 372

3740 GCF_000778175.1_ASM77817v1 Escherichia coli Proteobacteria; Gammaproteobacteria; Enterobacterales; Enterobacteriaceae; Escherichia
MTGDNTLIHSHGINRRDFMKLCAALAATMGLSSKAAAEMVESVTNPQRPPVIWIGAQECT WP_000145423.1
hydrogenase 2 small subunit [Escherichia coli] Length: 372\nNumber of predicted TMHs: 1\nExp number of AAs in TMHs: 23.78201\nExp number, first 60 AAs: 0.28355\nTotal prob of N-in: 0.13257\noutside 1 331\nTMhelix 332 354\ninside 355 372

3741 GCF_000778315.1_ASM77831v1 Escherichia coli Proteobacteria; Gammaproteobacteria; Enterobacterales; Enterobacteriaceae; Escherichia
MTGDNTLIHSHGINRRDFMKLCAALAATMGLSSKAAAEMAESVTNPQRPPVIWIGAQECT WP_000145410.1
MULTISPECIES: hydrogenase-2 small chain [Proteobacteria] Length: 372\nNumber of predicted TMHs: 1\nExp number of AAs in TMHs: 23.72226\nExp number, first 60 AAs: 0.22038\nTotal prob of N-in: 0.12994\noutside 1 331\nTMhelix 332 354\ninside 355 372

3742 GCF_000941195.1_57A_A8_assembly Escherichia coli Proteobacteria; Gammaproteobacteria; Enterobacterales; Enterobacteriaceae; Escherichia
MTGDNTLIHSHGINRRDFMKLCAALAATMGLSSKAAAEMVESVTNPQRPPVIWIGAQECT WP_000145423.1
hydrogenase 2 small subunit [Escherichia coli] Length: 372\nNumber of predicted TMHs: 1\nExp number of AAs in TMHs: 23.78201\nExp number, first 60 AAs: 0.28355\nTotal prob of N-in: 0.13257\noutside 1 331\nTMhelix 332 354\ninside 355 372

3743 GCF_000778375.1_ASM77837v1 Escherichia coli Proteobacteria; Gammaproteobacteria; Enterobacterales; Enterobacteriaceae; Escherichia
MTGDNTLIHSHGINRRDFMKLCAALAATMGLSSKAAAEMAESVTNPQRPPVIWIGAQECT WP_000145410.1
MULTISPECIES: hydrogenase-2 small chain [Proteobacteria] Length: 372\nNumber of predicted TMHs: 1\nExp number of AAs in TMHs: 23.72226\nExp number, first 60 AAs: 0.22038\nTotal prob of N-in: 0.12994\noutside 1 331\nTMhelix 332 354\ninside 355 372

3744 GCF_000778415.1_ASM77841v1 Escherichia coli Proteobacteria; Gammaproteobacteria; Enterobacterales; Enterobacteriaceae; Escherichia
MTGDNTLIHSHGINRRDFMKLCAALAATMGLSSKAAAEMAESVTNPQRPPVIWIGAQECT WP_000145410.1
MULTISPECIES: hydrogenase-2 small chain [Proteobacteria] Length: 372\nNumber of predicted TMHs: 1\nExp number of AAs in TMHs: 23.72226\nExp number, first 60 AAs: 0.22038\nTotal prob of N-in: 0.12994\noutside 1 331\nTMhelix 332 354\ninside 355 372

3745 GCF_000778475.1_ASM77847v1 Escherichia coli Proteobacteria; Gammaproteobacteria; Enterobacterales; Enterobacteriaceae; Escherichia
MTGDNTLIHSHGINRRDFMKLCAALAATMGLSSKAAAEMVESVTNPQRPPVIWIGAQECT WP_000145423.1
hydrogenase 2 small subunit [Escherichia coli] Length: 372\nNumber of predicted TMHs: 1\nExp number of AAs in TMHs: 23.78201\nExp number, first 60 AAs: 0.28355\nTotal prob of N-in: 0.13257\noutside 1 331\nTMhelix 332 354\ninside 355 372

3746 GCF_000940555.1_Ec46A_L2_MIRA_assembly Escherichia coli Proteobacteria; Gammaproteobacteria; Enterobacterales; Enterobacteriaceae; Escherichia
MTGDNTLIHSHGINRRDFMKLCAALAATMGLSSKAAAEMAESVTNPQRPPVIWIGAQECT WP_000145410.1
MULTISPECIES: hydrogenase-2 small chain [Proteobacteria] Length: 372\nNumber of predicted TMHs: 1\nExp number of AAs in TMHs: 23.72226\nExp number, first 60 AAs: 0.22038\nTotal prob of N-in: 0.12994\noutside 1 331\nTMhelix 332 354\ninside 355 372

3747 GCF_000778545.1_ASM77854v1 Escherichia coli Proteobacteria; Gammaproteobacteria; Enterobacterales; Enterobacteriaceae; Escherichia

MTGDNTLIHSHGINRRDFMKLCAALAATMGLSSKAAAEMAESVTNPQRPPVIWIGAQECT WP_000145410.1 MULTISPECIES: hydrogenase-2 small chain [Proteobacteria] Length: 372\nNumber of predicted TMHs: 1\nExp number of AAs in TMHs: 23.72226\nExp number, first 60 AAs: 0.22038\nTotal prob of N-in: 0.12994\noutside 1 331\nTMhelix 332 354\ninside 355 372

3748 GCF_000778545.1_ASM77854v1 Escherichia coli Proteobacteria; Gammaproteobacteria; Enterobacterales; Enterobacteriaceae; Escherichia MNNEETFYQAMRRQGVTRRSFLKYCSLAATSLGLGAGMAPKIAWALENKPRIPVVWIHGL WP_001058323.1 MULTISPECIES: hydrogenase-1 small chain [Proteobacteria] Length: 372\nNumber of predicted TMHs: 2\nExp number of AAs in TMHs: 37.2608499999999\nExp number, first 60 AAs: 19.52923\nTotal prob of N-in: 0.94351\nPOSSIBLE N-term signal sequence\ninside 1 19\nTMhelix 20 39\noutside 40 325\nTMhelix 326 348\ninside 349 372

3749 GCF_000779455.1_ASM77945v1 Escherichia coli Proteobacteria; Gammaproteobacteria; Enterobacterales; Enterobacteriaceae; Escherichia MNNEETFYQAMRRQGVTRRSFLKYCSLAATSLGLGAGMAPKIAWALENKPRIPVVWIHGL WP_001058323.1 MULTISPECIES: hydrogenase-1 small chain [Proteobacteria] Length: 372\nNumber of predicted TMHs: 2\nExp number of AAs in TMHs: 37.2608499999999\nExp number, first 60 AAs: 19.52923\nTotal prob of N-in: 0.94351\nPOSSIBLE N-term signal sequence\ninside 1 19\nTMhelix 20 39\noutside 40 325\nTMhelix 326 348\ninside 349 372

3750 GCF_000779175.1_ASM77917v1 Escherichia coli Proteobacteria; Gammaproteobacteria; Enterobacterales; Enterobacteriaceae; Escherichia MNNEETFYQAMRRQGVTRRSFLKYCSLAATSLGLGAGMAPKIAWALENKPRIPVVWIHGL WP_001058323.1 MULTISPECIES: hydrogenase-1 small chain [Proteobacteria] Length: 372\nNumber of predicted TMHs: 2\nExp number of AAs in TMHs: 37.2608499999999\nExp number, first 60 AAs: 19.52923\nTotal prob of N-in: 0.94351\nPOSSIBLE N-term signal sequence\ninside 1 19\nTMhelix 20 39\noutside 40 325\nTMhelix 326 348\ninside 349 372

3751 GCF_000779495.1_ASM77949v1 Escherichia coli Proteobacteria; Gammaproteobacteria; Enterobacterales; Enterobacteriaceae; Escherichia MTGDNTLIHSHGINRRDFMKLCAALAATMGLSSKAAAEMAESVTNPQRPPVIWIGAQECT WP_000145410.1 MULTISPECIES: hydrogenase-2 small chain [Proteobacteria] Length: 372\nNumber of predicted TMHs: 1\nExp number of AAs in TMHs: 23.72226\nExp number, first 60 AAs: 0.22038\nTotal prob of N-in: 0.12994\noutside 1 331\nTMhelix 332 354\ninside 355 372

3752 GCF_000779075.1_ASM77907v1 Escherichia coli Proteobacteria; Gammaproteobacteria; Enterobacterales; Enterobacteriaceae; Escherichia MNNEETFYQAMRRQGVTRRSFLKYCSLAATSLGLGAGMAPKIAWALENKPRIPVVWIHGL WP_042004834.1 hydrogenase [Escherichia coli] Length: 372\nNumber of predicted TMHs: 2\nExp number of AAs in TMHs: 37.26875\nExp number, first 60 AAs: 19.52896\nTotal prob of N-in: 0.94352\nPOSSIBLE N-term signal sequence\ninside 1 19\nTMhelix 20 39\noutside 40 325\nTMhelix 326 348\ninside 349 372

3753 GCF_000779615.1_ASM77961v1 Escherichia coli Proteobacteria; Gammaproteobacteria; Enterobacterales; Enterobacteriaceae; Escherichia MTGDNTLIHSHGINRRDFMKLCAALAATMGLSSKAAAEMAESVTNPQRPPVIWIGAQECT WP_000145410.1 MULTISPECIES: hydrogenase-2 small chain [Proteobacteria] Length: 372\nNumber of predicted TMHs: 1\nExp number of AAs in TMHs: 23.72226\nExp number, first 60 AAs: 0.22038\nTotal prob of N-in: 0.12994\noutside 1 331\nTMhelix 332 354\ninside 355 372

3754 GCF_000779675.1_ASM77967v1 Escherichia coli Proteobacteria; Gammaproteobacteria; Enterobacterales; Enterobacteriaceae; Escherichia MTGDNTLIHSHGINRRDFMKLCAALAATMGLSSKAAAEMAESVTNPQRPPVIWIGAQECT WP_000145410.1 MULTISPECIES: hydrogenase-2 small chain [Proteobacteria] Length: 372\nNumber of predicted TMHs: 1\nExp number of AAs in TMHs: 23.72226\nExp number, first 60 AAs: 0.22038\nTotal prob of N-in: 0.12994\noutside 1 331\nTMhelix 332 354\ninside 355 372

3755 GCF_000779715.1_ASM77971v1 Escherichia coli Proteobacteria; Gammaproteobacteria; Enterobacterales; Enterobacteriaceae; Escherichia MTGDNTLIHSHGINRRDFMKLCAALAATMGLSSKAAAEMAESVTNPQRPPVIWIGAQECT WP_000145418.1 hydrogenase 2 small subunit [Escherichia coli] Length: 372\nNumber of predicted TMHs: 1\nExp number of

AAs in TMHs: 23.72313\nExp number, first 60 AAs: 0.22034\nTotal prob of N-in: 0.12997\noutside 1 331\nTMhelix 332 354\ninside 355 372

3756 GCF_000780015.1_ASM78001v1 Escherichia coli Proteobacteria; Gammaproteobacteria; Enterobacterales; Enterobacteriaceae; Escherichia MNNEETFYQAMRRQGVTRRSFLKYCSLAATSLGLGAGMAPKIAWALENKPRIPVVWIHGL WP_001295940.1 MULTISPECIES: hydrogenase [Enterobacteriaceae] Length: 372\nNumber of predicted TMHs: 2\nExp number of AAs in TMHs: 37.2608899999999\nExp number, first 60 AAs: 19.52923\nTotal prob of N-in: 0.94351\nPOSSIBLE N-term signal sequence\ninside 1 19\nTMhelix 20 39\noutside 40 325\nTMhelix 326 348\ninside 349 372

3757 GCF_000779835.1_ASM77983v1 Escherichia coli Proteobacteria; Gammaproteobacteria; Enterobacterales; Enterobacteriaceae; Escherichia MNNEETFYQAMRRQGVTRRSFLKYCSLAATSLGLGAGMAPKIAWALENKPRIPVVWIHGL WP_001058323.1 MULTISPECIES: hydrogenase-1 small chain [Proteobacteria] Length: 372\nNumber of predicted TMHs: 2\nExp number of AAs in TMHs: 37.2608499999999\nExp number, first 60 AAs: 19.52923\nTotal prob of N-in: 0.94351\nPOSSIBLE N-term signal sequence\ninside 1 19\nTMhelix 20 39\noutside 40 325\nTMhelix 326 348\ninside 349 372

3758 GCF_000780115.1_ASM78011v1 Escherichia coli Proteobacteria; Gammaproteobacteria; Enterobacterales; Enterobacteriaceae; Escherichia MTGDNTLIHSHGINRRDFMKLCAALAATMGLSSKAAAEMAESVTNPQRPPVIWIGAQECT WP_000145410.1 MULTISPECIES: hydrogenase-2 small chain [Proteobacteria] Length: 372\nNumber of predicted TMHs: 1\nExp number of AAs in TMHs: 23.72226\nExp number, first 60 AAs: 0.22038\nTotal prob of N-in: 0.12994\noutside 1 331\nTMhelix 332 354\ninside 355 372

3759 GCF_000940255.1_FHI15 Escherichia coli Proteobacteria; Gammaproteobacteria; Enterobacterales; Enterobacteriaceae; Escherichia MNNEETFYQAMRRQGVTRRSFLKYCSLAATSLGLGAGMAPKIAWALENKPRIPVVWIHGL WP_001058323.1 MULTISPECIES: hydrogenase-1 small chain [Proteobacteria] Length: 372\nNumber of predicted TMHs: 2\nExp number of AAs in TMHs: 37.2608499999999\nExp number, first 60 AAs: 19.52923\nTotal prob of N-in: 0.94351\nPOSSIBLE N-term signal sequence\ninside 1 19\nTMhelix 20 39\noutside 40 325\nTMhelix 326 348\ninside 349 372

3760 GCF_000940255.1_FHI15 Escherichia coli Proteobacteria; Gammaproteobacteria; Enterobacterales; Enterobacteriaceae; Escherichia MTGDNTLIHSHGINRRDFMKLCAALAATMGLSSKAAAEMAESVTNPQRPPVIWIGAQECT WP_000145410.1 MULTISPECIES: hydrogenase-2 small chain [Proteobacteria] Length: 372\nNumber of predicted TMHs: 1\nExp number of AAs in TMHs: 23.72226\nExp number, first 60 AAs: 0.22038\nTotal prob of N-in: 0.12994\noutside 1 331\nTMhelix 332 354\ninside 355 372

3761 GCF_000940035.2_FHI89 Escherichia coli Proteobacteria; Gammaproteobacteria; Enterobacterales; Enterobacteriaceae; Escherichia MNNEETFYQAMRRQGVTRRSFLKYCSLAATSLGLGAGMAPKIAWALENKPRIPVVWIHGL WP_001058323.1 MULTISPECIES: hydrogenase-1 small chain [Proteobacteria] Length: 372\nNumber of predicted TMHs: 2\nExp number of AAs in TMHs: 37.2608499999999\nExp number, first 60 AAs: 19.52923\nTotal prob of N-in: 0.94351\nPOSSIBLE N-term signal sequence\ninside 1 19\nTMhelix 20 39\noutside 40 325\nTMhelix 326 348\ninside 349 372

3762 GCF_000780155.1_ASM78015v1 Escherichia coli Proteobacteria; Gammaproteobacteria; Enterobacterales; Enterobacteriaceae; Escherichia MTGDNTLIHSHGINRRDFMKLCAALAATMGLSSKAAAEMAESVTNPQRPPVIWIGAQECT WP_000145410.1 MULTISPECIES: hydrogenase-2 small chain [Proteobacteria] Length: 372\nNumber of predicted TMHs: 1\nExp number of AAs in TMHs: 23.72226\nExp number, first 60 AAs: 0.22038\nTotal prob of N-in: 0.12994\noutside 1 331\nTMhelix 332 354\ninside 355 372

3763 GCF_000780115.1_ASM78011v1 Escherichia coli Proteobacteria; Gammaproteobacteria; Enterobacterales; Enterobacteriaceae; Escherichia MNNEETFYQAMRRQGVTRRSFLKYCSLAATSLGLGAGMAPKIAWALENKPRIPVVWIHGL WP_001058323.1 MULTISPECIES: hydrogenase-1 small chain [Proteobacteria] Length: 372\nNumber of predicted TMHs: 2\nExp number of AAs in TMHs: 37.2608499999999\nExp number, first 60 AAs: 19.52923\nTotal prob of N-in: 0.94351\nPOSSIBLE N-term signal sequence\ninside 1 19\nTMhelix 20 39\noutside 40 325\nTMhelix 326 348\ninside 349 372

3764 GCF_000780155.1_ASM78015v1 Escherichia coli Proteobacteria; Gammaproteobacteria; Enterobacterales; Enterobacteriaceae; Escherichia
MNNEETFYQAMRRQGVTRRSFLKYCSLAATSLGLGAGMAPKIAWALENKPRIPVVWIHGL WP_001058323.1
MULTISPECIES: hydrogenase-1 small chain [Proteobacteria] Length: 372\nNumber of predicted TMHs: 2\nExp number of AAs in TMHs: 37.2608499999999\nExp number, first 60 AAs: 19.52923\nTotal prob of N-in: 0.94351\nPOSSIBLE N-term signal sequence\ninside 1 19\nTMhelix 20 39\noutside 40 325\nTMhelix 326 348\ninside 349 372

3765 GCF_000779715.1_ASM77971v1 Escherichia coli Proteobacteria; Gammaproteobacteria; Enterobacterales; Enterobacteriaceae; Escherichia
MNNEETFYQAMRRQGVTRRSFLKYCSLAATSLGLGAGMAPKIAWALENKPRIPVVWIHGL WP_021520630.1
hydrogenase-1 small chain [Escherichia coli] Length: 372\nNumber of predicted TMHs: 2\nExp number of AAs in TMHs: 37.8197599999999\nExp number, first 60 AAs: 19.53295\nTotal prob of N-in: 0.94396\nPOSSIBLE N-term signal sequence\ninside 1 19\nTMhelix 20 39\noutside 40 325\nTMhelix 326 348\ninside 349 372

3766 GCF_000780415.1_ASM78041v1 Escherichia coli Proteobacteria; Gammaproteobacteria; Enterobacterales; Enterobacteriaceae; Escherichia
MNNEETFYQAMRRQGVTRRSFLKYCSLAATSLGLGAGMAPKIAWALENKPRIPVVWIHGL WP_001058323.1
MULTISPECIES: hydrogenase-1 small chain [Proteobacteria] Length: 372\nNumber of predicted TMHs: 2\nExp number of AAs in TMHs: 37.2608499999999\nExp number, first 60 AAs: 19.52923\nTotal prob of N-in: 0.94351\nPOSSIBLE N-term signal sequence\ninside 1 19\nTMhelix 20 39\noutside 40 325\nTMhelix 326 348\ninside 349 372

3767 GCF_000780355.1_ASM78035v1 Escherichia coli Proteobacteria; Gammaproteobacteria; Enterobacterales; Enterobacteriaceae; Escherichia
MNNEETFYQAMRRQGVTRRSFLKYCSLAATSLGLGAGMAPKIAWALENKPRIPVVWIHGL WP_001058323.1
MULTISPECIES: hydrogenase-1 small chain [Proteobacteria] Length: 372\nNumber of predicted TMHs: 2\nExp number of AAs in TMHs: 37.2608499999999\nExp number, first 60 AAs: 19.52923\nTotal prob of N-in: 0.94351\nPOSSIBLE N-term signal sequence\ninside 1 19\nTMhelix 20 39\noutside 40 325\nTMhelix 326 348\ninside 349 372

3768 GCF_000780735.1_ASM78073v1 Escherichia coli Proteobacteria; Gammaproteobacteria; Enterobacterales; Enterobacteriaceae; Escherichia
MTGDNTLIHSHGINRRDFMKLCAALAATMGLSSKAAAEMAESVTNPQRPPVIWIGAQECT WP_000145410.1
MULTISPECIES: hydrogenase-2 small chain [Proteobacteria] Length: 372\nNumber of predicted TMHs: 1\nExp number of AAs in TMHs: 23.72226\nExp number, first 60 AAs: 0.22038\nTotal prob of N-in: 0.12994\noutside 1 331\nTMhelix 332 354\ninside 355 372

3769 GCF_000780775.1_ASM78077v1 Escherichia coli Proteobacteria; Gammaproteobacteria; Enterobacterales; Enterobacteriaceae; Escherichia
MTGDNTLIHSHGINRRDFMKLCAALAATMGLSSKAAAEMAESVTNPQRPPVIWIGAQECT WP_000145410.1
MULTISPECIES: hydrogenase-2 small chain [Proteobacteria] Length: 372\nNumber of predicted TMHs: 1\nExp number of AAs in TMHs: 23.72226\nExp number, first 60 AAs: 0.22038\nTotal prob of N-in: 0.12994\noutside 1 331\nTMhelix 332 354\ninside 355 372

3770 GCF_000780735.1_ASM78073v1 Escherichia coli Proteobacteria; Gammaproteobacteria; Enterobacterales; Enterobacteriaceae; Escherichia
MNNEETFYQAMRRQGVTRRSFLKYCSLAATSLGLGAGMAPKIAWALENKPRIPVVWIHGL WP_001058323.1
MULTISPECIES: hydrogenase-1 small chain [Proteobacteria] Length: 372\nNumber of predicted TMHs: 2\nExp number of AAs in TMHs: 37.2608499999999\nExp number, first 60 AAs: 19.52923\nTotal prob of N-in: 0.94351\nPOSSIBLE N-term signal sequence\ninside 1 19\nTMhelix 20 39\noutside 40 325\nTMhelix 326 348\ninside 349 372

3771 GCF_000780775.1_ASM78077v1 Escherichia coli Proteobacteria; Gammaproteobacteria; Enterobacterales; Enterobacteriaceae; Escherichia
MNNEETFYQAMRRQGVTRRSFLKYCSLAATSLGLGAGMAPKIAWALENKPRIPVVWIHGL WP_001058323.1
MULTISPECIES: hydrogenase-1 small chain [Proteobacteria] Length: 372\nNumber of predicted TMHs: 2\nExp number of AAs in TMHs: 37.2608499999999\nExp number, first 60 AAs: 19.52923\nTotal prob of N-in: 0.94351\nPOSSIBLE N-term signal sequence\ninside 1 19\nTMhelix 20 39\noutside 40 325\nTMhelix 326 348\ninside 349 372

3772 GCF_000780255.1_ASM78025v1 Escherichia coli Proteobacteria; Gammaproteobacteria; Enterobacterales; Enterobacteriaceae; Escherichia MNNEETFYQAMRRQGVTRRSFLKYCSLAATSLGLGAGMAPKIAWALENKPRIPVVWIHGL WP_001295940.1 MULTISPECIES: hydrogenase [Enterobacteriaceae] Length: 372\nNumber of predicted TMHs: 2\nExp number of AAs in TMHs: 37.2608899999999\nExp number, first 60 AAs: 19.52923\nTotal prob of N-in: 0.94351\nPOSSIBLE N-term signal sequence\ninside 1 19\nTMhelix 20 39\noutside 40 325\nTMhelix 326 348\ninside 349 372

3773 GCF_000939255.1_FHI72 Escherichia coli Proteobacteria; Gammaproteobacteria; Enterobacterales; Enterobacteriaceae; Escherichia MNNEETFYQAMRRQGVTRRSFLKYCSLAATSLGLGAGMAPKIAWALENKPRIPVVWIHGL WP_001058323.1 MULTISPECIES: hydrogenase-1 small chain [Proteobacteria] Length: 372\nNumber of predicted TMHs: 2\nExp number of AAs in TMHs: 37.2608499999999\nExp number, first 60 AAs: 19.52923\nTotal prob of N-in: 0.94351\nPOSSIBLE N-term signal sequence\ninside 1 19\nTMhelix 20 39\noutside 40 325\nTMhelix 326 348\ninside 349 372

3774 GCF_001077955.1_ASM107795v1 Escherichia coli Proteobacteria; Gammaproteobacteria; Enterobacterales; Enterobacteriaceae; Escherichia MTGDNTLIHSHGINRRDFMKLCAALAATMGLSSKAAAEMAESVTNPQRPPVIWIGAQECT WP_000145410.1 MULTISPECIES: hydrogenase-2 small chain [Proteobacteria] Length: 372\nNumber of predicted TMHs: 1\nExp number of AAs in TMHs: 23.72226\nExp number, first 60 AAs: 0.22038\nTotal prob of N-in: 0.12994\noutside 1 331\nTMhelix 332 354\ninside 355 372

3775 GCF_000781175.1_ASM78117v1 Escherichia coli Proteobacteria; Gammaproteobacteria; Enterobacterales; Enterobacteriaceae; Escherichia MNNEETFYQAMRRQGVTRRSFLKYCSLAATSLGLGAGMAPKIAWALENKPRIPVVWIHGL WP_001058323.1 MULTISPECIES: hydrogenase-1 small chain [Proteobacteria] Length: 372\nNumber of predicted TMHs: 2\nExp number of AAs in TMHs: 37.2608499999999\nExp number, first 60 AAs: 19.52923\nTotal prob of N-in: 0.94351\nPOSSIBLE N-term signal sequence\ninside 1 19\nTMhelix 20 39\noutside 40 325\nTMhelix 326 348\ninside 349 372

3776 GCF_000780995.1_ASM78099v1 Escherichia coli Proteobacteria; Gammaproteobacteria; Enterobacterales; Enterobacteriaceae; Escherichia MNNEETFYQAMRRQGVTRRSFLKYCSLAATSLGLGAGMAPKIAWALENKPRIPVVWIHGL WP_001058323.1 MULTISPECIES: hydrogenase-1 small chain [Proteobacteria] Length: 372\nNumber of predicted TMHs: 2\nExp number of AAs in TMHs: 37.2608499999999\nExp number, first 60 AAs: 19.52923\nTotal prob of N-in: 0.94351\nPOSSIBLE N-term signal sequence\ninside 1 19\nTMhelix 20 39\noutside 40 325\nTMhelix 326 348\ninside 349 372

3777 GCF_000781275.1_ASM78127v1 Escherichia coli Proteobacteria; Gammaproteobacteria; Enterobacterales; Enterobacteriaceae; Escherichia MTGDNTLIHSHGINRRDFMKLCAALAATMGLSSKAAAEMAESVTNPQRPPVIWIGAQECT WP_000145410.1 MULTISPECIES: hydrogenase-2 small chain [Proteobacteria] Length: 372\nNumber of predicted TMHs: 1\nExp number of AAs in TMHs: 23.72226\nExp number, first 60 AAs: 0.22038\nTotal prob of N-in: 0.12994\noutside 1 331\nTMhelix 332 354\ninside 355 372

3778 GCF_000781315.1_ASM78131v1 Escherichia coli Proteobacteria; Gammaproteobacteria; Enterobacterales; Enterobacteriaceae; Escherichia MTGDNTLIHSHGINRRDFMKLCAALAATMGLSSKAAAEMAESVTNPQRPPVIWIGAQECT WP_000145410.1 MULTISPECIES: hydrogenase-2 small chain [Proteobacteria] Length: 372\nNumber of predicted TMHs: 1\nExp number of AAs in TMHs: 23.72226\nExp number, first 60 AAs: 0.22038\nTotal prob of N-in: 0.12994\noutside 1 331\nTMhelix 332 354\ninside 355 372

3779 GCF_000938175.1_Ec46A_L1_MIRA_assembly Escherichia coli Proteobacteria; Gammaproteobacteria; Enterobacterales; Enterobacteriaceae; Escherichia MNNEETFYQAMRRQGVTRRSFLKYCSLAATSLGLGAGMAPKIAWALENKPRIPVVWIHGL WP_001058323.1 MULTISPECIES: hydrogenase-1 small chain [Proteobacteria] Length: 372\nNumber of predicted TMHs: 2\nExp number of AAs in TMHs: 37.2608499999999\nExp number, first 60 AAs: 19.52923\nTotal prob of N-in: 0.94351\nPOSSIBLE N-term signal sequence\ninside 1 19\nTMhelix 20 39\noutside 40 325\nTMhelix 326 348\ninside 349 372

3780 GCF_000938175.1_Ec46A_L1_MIRA_assembly Escherichia coli Proteobacteria; Gammaproteobacteria; Enterobacterales; Enterobacteriaceae; Escherichia
MTGDNTLIHSHGINRRDFMKLCAALAATMGLSSKAAAEMAESVTNPQRPPVIWIGAQECT WP_000145410.1
MULTISPECIES: hydrogenase-2 small chain [Proteobacteria] Length: 372\nNumber of predicted TMHs: 1\nExp number of AAs in TMHs: 23.72226\nExp number, first 60 AAs: 0.22038\nTotal prob of N-in: 0.12994\noutside 1 331\nTMhelix 332 354\ninside 355 372

3781 GCF_000781275.1_ASM78127v1 Escherichia coli Proteobacteria; Gammaproteobacteria; Enterobacterales; Enterobacteriaceae; Escherichia
MNNEETFYQAMRRQGVTRRSFLKYCSLAATSLGLGAGMAPKIAWALENKPRIPVVWIHGL WP_001058323.1
MULTISPECIES: hydrogenase-1 small chain [Proteobacteria] Length: 372\nNumber of predicted TMHs: 2\nExp number of AAs in TMHs: 37.2608499999999\nExp number, first 60 AAs: 19.52923\nTotal prob of N-in: 0.94351\nPOSSIBLE N-term signal sequence\ninside 1 19\nTMhelix 20 39\noutside 40 325\nTMhelix 326 348\ninside 349 372

3782 GCF_000781315.1_ASM78131v1 Escherichia coli Proteobacteria; Gammaproteobacteria; Enterobacterales; Enterobacteriaceae; Escherichia
MNNEETFYQAMRRQGVTRRSFLKYCSLAATSLGLGAGMAPKIAWALENKPRIPVVWIHGL WP_001295940.1
MULTISPECIES: hydrogenase [Enterobacteriaceae] Length: 372\nNumber of predicted TMHs: 2\nExp number of AAs in TMHs: 37.2608899999999\nExp number, first 60 AAs: 19.52923\nTotal prob of N-in: 0.94351\nPOSSIBLE N-term signal sequence\ninside 1 19\nTMhelix 20 39\noutside 40 325\nTMhelix 326 348\ninside 349 372

3783 GCF_000781045.1_ASM78104v1 Escherichia coli Proteobacteria; Gammaproteobacteria; Enterobacterales; Enterobacteriaceae; Escherichia
MNNEETFYQAMRRQGVTRRSFLKYCSLAATSLGLGAGMAPKIAWALENKPRIPVVWIHGL WP_001058323.1
MULTISPECIES: hydrogenase-1 small chain [Proteobacteria] Length: 372\nNumber of predicted TMHs: 2\nExp number of AAs in TMHs: 37.2608499999999\nExp number, first 60 AAs: 19.52923\nTotal prob of N-in: 0.94351\nPOSSIBLE N-term signal sequence\ninside 1 19\nTMhelix 20 39\noutside 40 325\nTMhelix 326 348\ninside 349 372

3784 GCF_000599745.2_a5_miseq_linux_20140113 Escherichia coli Proteobacteria; Gammaproteobacteria; Enterobacterales; Enterobacteriaceae; Escherichia
MNNEETFYQAMRRQGVTRRSFLKYCSLAATSLGLGAGMAPKIAWALENKPRIPVVWIHGL WP_001058323.1
MULTISPECIES: hydrogenase-1 small chain [Proteobacteria] Length: 372\nNumber of predicted TMHs: 2\nExp number of AAs in TMHs: 37.2608499999999\nExp number, first 60 AAs: 19.52923\nTotal prob of N-in: 0.94351\nPOSSIBLE N-term signal sequence\ninside 1 19\nTMhelix 20 39\noutside 40 325\nTMhelix 326 348\ninside 349 372

3785 GCF_000599805.2_a5_miseq_linux_20140113 Escherichia coli Proteobacteria; Gammaproteobacteria; Enterobacterales; Enterobacteriaceae; Escherichia
MNNEETFYQAMRRQGVTRRSFLKYCSLAATSLGLGAGMAPKIAWALENKPRIPVVWIHGL WP_001058323.1
MULTISPECIES: hydrogenase-1 small chain [Proteobacteria] Length: 372\nNumber of predicted TMHs: 2\nExp number of AAs in TMHs: 37.2608499999999\nExp number, first 60 AAs: 19.52923\nTotal prob of N-in: 0.94351\nPOSSIBLE N-term signal sequence\ninside 1 19\nTMhelix 20 39\noutside 40 325\nTMhelix 326 348\ninside 349 372

3786 GCF_001286005.1_7790_1_62 Escherichia coli Proteobacteria; Gammaproteobacteria; Enterobacterales; Enterobacteriaceae; Escherichia
MTGDNTLIHSHGINRRDFMKLCAALAATMGLSSKAAAEMAESVTNPQRPPVIWIGAQECT WP_000145410.1
MULTISPECIES: hydrogenase-2 small chain [Proteobacteria] Length: 372\nNumber of predicted TMHs: 1\nExp number of AAs in TMHs: 23.72226\nExp number, first 60 AAs: 0.22038\nTotal prob of N-in: 0.12994\noutside 1 331\nTMhelix 332 354\ninside 355 372

3787 GCF_001286005.1_7790_1_62 Escherichia coli Proteobacteria; Gammaproteobacteria; Enterobacterales; Enterobacteriaceae; Escherichia
MNNEETFYQAMRRQGVTRRSFLKYCSLAATSLGLGAGMAPKIAWALENKPRIPVVWIHGL WP_001058323.1
MULTISPECIES: hydrogenase-1 small chain [Proteobacteria] Length: 372\nNumber of predicted TMHs: 2\nExp number of AAs in TMHs: 37.2608499999999\nExp number, first 60 AAs: 19.52923\nTotal prob of N-in: 0.94351\nPOSSIBLE N-term signal sequence\ninside 1 19\nTMhelix 20 39\noutside 40 325\nTMhelix 326 348\ninside 349 372

3788 GCF_000937095.2_FHI87 Escherichia coli Proteobacteria; Gammaproteobacteria; Enterobacterales; Enterobacteriaceae; Escherichia MNNEETFYQAMRRQGVTRRSFLKYCSLAATSLGLGAGMAPKIAWALENKPRIPVVWIHGL WP_001058323.1 MULTISPECIES: hydrogenase-1 small chain [Proteobacteria] Length: 372\nNumber of predicted TMHs: 2\nExp number of AAs in TMHs: 37.2608499999999\nExp number, first 60 AAs: 19.52923\nTotal prob of N-in: 0.94351\nPOSSIBLE N-term signal sequence\ninside 1 19\nTMhelix 20 39\noutside 40 325\nTMhelix 326 348\ninside 349 372

3789 GCF_000936125.1_FHI75 Escherichia coli Proteobacteria; Gammaproteobacteria; Enterobacterales; Enterobacteriaceae; Escherichia MNNEETFYQAMRRQGVTRRSFLKYCSLAATSLGLGAGMAPKIAWALENKPRIPVVWIHGL WP_001295940.1 MULTISPECIES: hydrogenase [Enterobacteriaceae] Length: 372\nNumber of predicted TMHs: 2\nExp number of AAs in TMHs: 37.2608899999999\nExp number, first 60 AAs: 19.52923\nTotal prob of N-in: 0.94351\nPOSSIBLE N-term signal sequence\ninside 1 19\nTMhelix 20 39\noutside 40 325\nTMhelix 326 348\ninside 349 372

3790 GCF_000936125.1_FHI75 Escherichia coli Proteobacteria; Gammaproteobacteria; Enterobacterales; Enterobacteriaceae; Escherichia MTGDNTLIHSHGINRRDFMKLCAALAATMGLSSKAAAEMAESVTNPQRPPVIWIGAQECT WP_000145410.1 MULTISPECIES: hydrogenase-2 small chain [Proteobacteria] Length: 372\nNumber of predicted TMHs: 1\nExp number of AAs in TMHs: 23.72226\nExp number, first 60 AAs: 0.22038\nTotal prob of N-in: 0.12994\noutside 1 331\nTMhelix 332 354\ninside 355 372

3791 GCF_001286085.1_7790_1_78 Escherichia coli Proteobacteria; Gammaproteobacteria; Enterobacterales; Enterobacteriaceae; Escherichia MNNEETFYQSMRRQGVTRRSFLKYCSLAATSLGLGAGMAPKIAWALENKPRIPVVWIHGL WP_002461900.1 MULTISPECIES: hydrogenase-1 small chain [Escherichia] Length: 372\nNumber of predicted TMHs: 2\nExp number of AAs in TMHs: 37.1074899999999\nExp number, first 60 AAs: 19.37835\nTotal prob of N-in: 0.93614\nPOSSIBLE N-term signal sequence\ninside 1 19\nTMhelix 20 39\noutside 40 325\nTMhelix 326 348\ninside 349 372

3792 GCF_000935525.1_ASM93552v1 Escherichia coli Proteobacteria; Gammaproteobacteria; Enterobacterales; Enterobacteriaceae; Escherichia MNNEETFYQAMRRQGVTRRSFLKYCSLAATSLGLGAGMAPKIAWALENKPRIPVVWIHGL WP_001058323.1 MULTISPECIES: hydrogenase-1 small chain [Proteobacteria] Length: 372\nNumber of predicted TMHs: 2\nExp number of AAs in TMHs: 37.2608499999999\nExp number, first 60 AAs: 19.52923\nTotal prob of N-in: 0.94351\nPOSSIBLE N-term signal sequence\ninside 1 19\nTMhelix 20 39\noutside 40 325\nTMhelix 326 348\ninside 349 372

3793 GCF_000935525.1_ASM93552v1 Escherichia coli Proteobacteria; Gammaproteobacteria; Enterobacterales; Enterobacteriaceae; Escherichia MTGDNTLIHSHGINRRDFMKLCAALAATMGLSSKAAAEMAESVTNPQRPPVIWIGAQECT WP_000145410.1 MULTISPECIES: hydrogenase-2 small chain [Proteobacteria] Length: 372\nNumber of predicted TMHs: 1\nExp number of AAs in TMHs: 23.72226\nExp number, first 60 AAs: 0.22038\nTotal prob of N-in: 0.12994\noutside 1 331\nTMhelix 332 354\ninside 355 372

3794 GCF_000935435.1_ASM93543v1 Escherichia coli Proteobacteria; Gammaproteobacteria; Enterobacterales; Enterobacteriaceae; Escherichia MTGDNTLIHSHGINRRDFMKLCAALAATMGLSSKAAAEMAESVTNPQRPPVIWIGAQECT WP_000145410.1 MULTISPECIES: hydrogenase-2 small chain [Proteobacteria] Length: 372\nNumber of predicted TMHs: 1\nExp number of AAs in TMHs: 23.72226\nExp number, first 60 AAs: 0.22038\nTotal prob of N-in: 0.12994\noutside 1 331\nTMhelix 332 354\ninside 355 372

3795 GCF_001306585.1_ASM130658v1 Escherichia coli Proteobacteria; Gammaproteobacteria; Enterobacterales; Enterobacteriaceae; Escherichia MTGDNTLIHSHGINRRDFMKLCAALAATMGLSSKAAAEMVESVTNPQRPPVIWIGAQECT WP_000145423.1 hydrogenase 2 small subunit [Escherichia coli] Length: 372\nNumber of predicted TMHs: 1\nExp number of AAs in TMHs: 23.78201\nExp number, first 60 AAs: 0.28355\nTotal prob of N-in: 0.13257\noutside 1 331\nTMhelix 332 354\ninside 355 372

3796 GCF_001286665.1_7553_7_65 Escherichia coli Proteobacteria; Gammaproteobacteria; Enterobacterales; Enterobacteriaceae; Escherichia MNNEETFYQAMRRQGVTRRSFLKYCSLAATSLGLGAGMAPKIAWALENKPRIPVVWIHGL WP_001058323.1 MULTISPECIES: hydrogenase-1 small chain [Proteobacteria] Length: 372\nNumber of predicted TMHs:

2\nExp number of AAs in TMHs: 37.2608499999999\nExp number, first 60 AAs: 19.52923\nTotal prob of N-in: 0.94351\nPOSSIBLE N-term signal sequence\ninside 1 19\nTMhelix 20 39\noutside 40 325\nTMhelix 326 348\ninside 349 372

3797 GCF_001286665.1_7553_7_65 Escherichia coli Proteobacteria; Gammaproteobacteria; Enterobacterales; Enterobacteriaceae; Escherichia MTGDNTLIHSHGINRRDFMKLCAALAATMGLSSKAAAEMAESVTNPQRPPVIWIGAQECT WP_000145406.1 hydrogenase-2 small chain [Escherichia coli] Length: 372\nNumber of predicted TMHs: 1\nExp number of AAs in TMHs: 23.72251\nExp number, first 60 AAs: 0.22027\nTotal prob of N-in: 0.13006\noutside 1 331\nTMhelix 332 354\ninside 355 372

3798 GCF_001286605.1_8205_3_1Escherichia coli Proteobacteria; Gammaproteobacteria; Enterobacterales; Enterobacteriaceae; Escherichia MNNEETFYQAMRRQGVTRRSFLKYCSLAATSLGLGAGMAPKIAWALENKPRIPVVWIHGL WP_001058323.1 MULTISPECIES: hydrogenase-1 small chain [Proteobacteria] Length: 372\nNumber of predicted TMHs: 2\nExp number of AAs in TMHs: 37.2608499999999\nExp number, first 60 AAs: 19.52923\nTotal prob of N-in: 0.94351\nPOSSIBLE N-term signal sequence\ninside 1 19\nTMhelix 20 39\noutside 40 325\nTMhelix 326 348\ninside 349 372

3799 GCF_001286605.1_8205_3_1Escherichia coli Proteobacteria; Gammaproteobacteria; Enterobacterales; Enterobacteriaceae; Escherichia MTGDNTLIHSHGINRRDFMKLCAALAATMGLSSKAAAEMAESVTNPQRPPVIWIGAQECT WP_000145410.1 MULTISPECIES: hydrogenase-2 small chain [Proteobacteria] Length: 372\nNumber of predicted TMHs: 1\nExp number of AAs in TMHs: 23.72226\nExp number, first 60 AAs: 0.22038\nTotal prob of N-in: 0.12994\noutside 1 331\nTMhelix 332 354\ninside 355 372

3800 GCF_001286485.1_8205_3_40 Escherichia coli Proteobacteria; Gammaproteobacteria; Enterobacterales; Enterobacteriaceae; Escherichia MNNEETFYQAMRRQGVTRRSFLKYCSLAATSLGLGAGMAPKIAWALENKPRIPVVWIHGL WP_001058323.1 MULTISPECIES: hydrogenase-1 small chain [Proteobacteria] Length: 372\nNumber of predicted TMHs: 2\nExp number of AAs in TMHs: 37.2608499999999\nExp number, first 60 AAs: 19.52923\nTotal prob of N-in: 0.94351\nPOSSIBLE N-term signal sequence\ninside 1 19\nTMhelix 20 39\noutside 40 325\nTMhelix 326 348\ninside 349 372

3801 GCF_001286485.1_8205_3_40 Escherichia coli Proteobacteria; Gammaproteobacteria; Enterobacterales; Enterobacteriaceae; Escherichia MTGDNTLIHSHGINRRDFMKLCAALAATMGLSSKAAAEMAESVTNPQRPPVIWIGAQECT WP_000145410.1 MULTISPECIES: hydrogenase-2 small chain [Proteobacteria] Length: 372\nNumber of predicted TMHs: 1\nExp number of AAs in TMHs: 23.72226\nExp number, first 60 AAs: 0.22038\nTotal prob of N-in: 0.12994\noutside 1 331\nTMhelix 332 354\ninside 355 372

3802 GCF_001286445.1_8205_3_60 Escherichia coli Proteobacteria; Gammaproteobacteria; Enterobacterales; Enterobacteriaceae; Escherichia MNNEETFYQAMRRQGVTRRSFLKYCSLAATSLGLGAGMAPKIAWALENKPRIPVVWIHGL WP_001058323.1 MULTISPECIES: hydrogenase-1 small chain [Proteobacteria] Length: 372\nNumber of predicted TMHs: 2\nExp number of AAs in TMHs: 37.2608499999999\nExp number, first 60 AAs: 19.52923\nTotal prob of N-in: 0.94351\nPOSSIBLE N-term signal sequence\ninside 1 19\nTMhelix 20 39\noutside 40 325\nTMhelix 326 348\ninside 349 372

3803 GCF_001286445.1_8205_3_60 Escherichia coli Proteobacteria; Gammaproteobacteria; Enterobacterales; Enterobacteriaceae; Escherichia MTGDNTLIHSHGINRRDFMKLCAALAATMGLSSKAAAEMAESVTNPQRPPVIWIGAQECT WP_000145410.1 MULTISPECIES: hydrogenase-2 small chain [Proteobacteria] Length: 372\nNumber of predicted TMHs: 1\nExp number of AAs in TMHs: 23.72226\nExp number, first 60 AAs: 0.22038\nTotal prob of N-in: 0.12994\noutside 1 331\nTMhelix 332 354\ninside 355 372

3804 GCF_001187545.1_ASM118754v1 Escherichia coli Proteobacteria; Gammaproteobacteria; Enterobacterales; Enterobacteriaceae; Escherichia MNNEETFYQAMRRQGVTRRSFLKYCSLAATSLGLGAGMAPKIAWALENKPRIPVVWIHGL WP_001058323.1 MULTISPECIES: hydrogenase-1 small chain [Proteobacteria] Length: 372\nNumber of predicted TMHs: 2\nExp number of AAs in TMHs: 37.2608499999999\nExp number, first 60 AAs: 19.52923\nTotal prob of N-in: 0.94351\nPOSSIBLE N-term signal sequence\ninside 1 19\nTMhelix 20 39\noutside 40 325\nTMhelix 326 348\ninside 349 372

3805 GCF_001286245.1_8205_8_71 Escherichia coli Proteobacteria; Gammaproteobacteria; Enterobacterales; Enterobacteriaceae; Escherichia
MNNEETFYQAMRRQGVTRRSFLKYCSLAATSLGLGAGMAPKIAWALENKPRIPVVWIHGL WP_001058323.1
MULTISPECIES: hydrogenase-1 small chain [Proteobacteria] Length: 372\nNumber of predicted TMHs: 2\nExp number of AAs in TMHs: 37.2608499999999\nExp number, first 60 AAs: 19.52923\nTotal prob of N-in: 0.94351\nPOSSIBLE N-term signal sequence\ninside 1 19\nTMhelix 20 39\noutside 40 325\nTMhelix 326 348\ninside 349 372

3806 GCF_001183685.1_ASM118368v1 Escherichia coli Proteobacteria; Gammaproteobacteria; Enterobacterales; Enterobacteriaceae; Escherichia
MTGDNTLIHSHGINRRDFMKLCAALAATMGLSSKAAAEMAESVTNPQRPPVIWIGAQECT WP_000145410.1
MULTISPECIES: hydrogenase-2 small chain [Proteobacteria] Length: 372\nNumber of predicted TMHs: 1\nExp number of AAs in TMHs: 23.72226\nExp number, first 60 AAs: 0.22038\nTotal prob of N-in: 0.12994\noutside 1 331\nTMhelix 332 354\ninside 355 372

3807 GCF_001891365.1_ASM189136v1 Escherichia coli Proteobacteria; Gammaproteobacteria; Enterobacterales; Enterobacteriaceae; Escherichia
MNNEETFYQAMRRQGVTRRSFLKYCSLAATSLGLGAGMAPKIAWALENKPRIPVVWIHGL WP_001295940.1
MULTISPECIES: hydrogenase [Enterobacteriaceae] Length: 372\nNumber of predicted TMHs: 2\nExp number of AAs in TMHs: 37.2608899999999\nExp number, first 60 AAs: 19.52923\nTotal prob of N-in: 0.94351\nPOSSIBLE N-term signal sequence\ninside 1 19\nTMhelix 20 39\noutside 40 325\nTMhelix 326 348\ninside 349 372

3808 GCF_001891295.1_ASM189129v1 Escherichia coli Proteobacteria; Gammaproteobacteria; Enterobacterales; Enterobacteriaceae; Escherichia
MNNEETFYQAMRRQGVTRRSFLKYCSLAATSLGLGAGMAPKIAWALENKPRIPVVWIHGL WP_001295940.1
MULTISPECIES: hydrogenase [Enterobacteriaceae] Length: 372\nNumber of predicted TMHs: 2\nExp number of AAs in TMHs: 37.2608899999999\nExp number, first 60 AAs: 19.52923\nTotal prob of N-in: 0.94351\nPOSSIBLE N-term signal sequence\ninside 1 19\nTMhelix 20 39\noutside 40 325\nTMhelix 326 348\ninside 349 372

3809 GCF_001891615.1_ASM189161v1 Escherichia coli Proteobacteria; Gammaproteobacteria; Enterobacterales; Enterobacteriaceae; Escherichia
MTGDNTLIHSHGINRRDFMKLCAALAATMGLSSKAAAEMAESVTNPQRPPVIWIGAQECT WP_000145410.1
MULTISPECIES: hydrogenase-2 small chain [Proteobacteria] Length: 372\nNumber of predicted TMHs: 1\nExp number of AAs in TMHs: 23.72226\nExp number, first 60 AAs: 0.22038\nTotal prob of N-in: 0.12994\noutside 1 331\nTMhelix 332 354\ninside 355 372

3810 GCF_001891685.1_ASM189168v1 Escherichia coli Proteobacteria; Gammaproteobacteria; Enterobacterales; Enterobacteriaceae; Escherichia
MTGDNTLIHSHGINRRDFMKLCAALAATMGLSSKAAAEMAESVTNPQRPPVIWIGAQECT WP_000145410.1
MULTISPECIES: hydrogenase-2 small chain [Proteobacteria] Length: 372\nNumber of predicted TMHs: 1\nExp number of AAs in TMHs: 23.72226\nExp number, first 60 AAs: 0.22038\nTotal prob of N-in: 0.12994\noutside 1 331\nTMhelix 332 354\ninside 355 372

3811 GCF_001891555.1_ASM189155v1 Escherichia coli Proteobacteria; Gammaproteobacteria; Enterobacterales; Enterobacteriaceae; Escherichia
MNNEETFYQAMRRQGVTRRSFLKYCSLAATSLGLGAGMAPKIAWALENKPRIPVVWIHGL WP_001295940.1
MULTISPECIES: hydrogenase [Enterobacteriaceae] Length: 372\nNumber of predicted TMHs: 2\nExp number of AAs in TMHs: 37.2608899999999\nExp number, first 60 AAs: 19.52923\nTotal prob of N-in: 0.94351\nPOSSIBLE N-term signal sequence\ninside 1 19\nTMhelix 20 39\noutside 40 325\nTMhelix 326 348\ninside 349 372

3812 GCF_001891785.1_ASM189178v1 Escherichia coli Proteobacteria; Gammaproteobacteria; Enterobacterales; Enterobacteriaceae; Escherichia
MNNEETFYQAMRRQGVTRRSFLKYCSLAATSLGLGAGMAPKIAWALENKPRIPVVWIHGL WP_001058323.1
MULTISPECIES: hydrogenase-1 small chain [Proteobacteria] Length: 372\nNumber of predicted TMHs: 2\nExp number of AAs in TMHs: 37.2608499999999\nExp number, first 60 AAs: 19.52923\nTotal prob of N-in: 0.94351\nPOSSIBLE N-term signal sequence\ninside 1 19\nTMhelix 20 39\noutside 40 325\nTMhelix 326 348\ninside 349 372

3813 GCF_001891925.1_ASM189192v1 Escherichia coli Proteobacteria; Gammaproteobacteria; Enterobacterales; Enterobacteriaceae; Escherichia
MTGDNTLIHSHGINRRDFMKLCAALAATMGLSSKAAAEMAESVTNPQRPPVIWIGAQECT WP_000145410.1
MULTISPECIES: hydrogenase-2 small chain [Proteobacteria] Length: 372\nNumber of predicted TMHs: 1\nExp number of AAs in TMHs: 23.72226\nExp number, first 60 AAs: 0.22038\nTotal prob of N-in: 0.12994\noutside 1 331\nTMhelix 332 354\ninside 355 372

3814 GCF_001891875.1_ASM189187v1 Escherichia coli Proteobacteria; Gammaproteobacteria; Enterobacterales; Enterobacteriaceae; Escherichia
MTGDNTLIHSHGINRRDFMKLCAALAATMGLSSKAAAEMAESVTNPQRPPVIWIGAQECT WP_000145410.1
MULTISPECIES: hydrogenase-2 small chain [Proteobacteria] Length: 372\nNumber of predicted TMHs: 1\nExp number of AAs in TMHs: 23.72226\nExp number, first 60 AAs: 0.22038\nTotal prob of N-in: 0.12994\noutside 1 331\nTMhelix 332 354\ninside 355 372

3815 GCF_001891975.1_ASM189197v1 Escherichia coli Proteobacteria; Gammaproteobacteria; Enterobacterales; Enterobacteriaceae; Escherichia
MTGDNTLIHSHGINRRDFMKLCAALAATMGLSSKAAAEMAESVTNPQRPPVIWIGAQECT WP_000145410.1
MULTISPECIES: hydrogenase-2 small chain [Proteobacteria] Length: 372\nNumber of predicted TMHs: 1\nExp number of AAs in TMHs: 23.72226\nExp number, first 60 AAs: 0.22038\nTotal prob of N-in: 0.12994\noutside 1 331\nTMhelix 332 354\ninside 355 372

3816 GCF_001891925.1_ASM189192v1 Escherichia coli Proteobacteria; Gammaproteobacteria; Enterobacterales; Enterobacteriaceae; Escherichia
MNNEETFYQAMRRQGVTRRSFLKYCSLAATSLGLGAGMAPKIAWALENKPRIPVVWIHGL WP_001058323.1
MULTISPECIES: hydrogenase-1 small chain [Proteobacteria] Length: 372\nNumber of predicted TMHs: 2\nExp number of AAs in TMHs: 37.2608499999999\nExp number, first 60 AAs: 19.52923\nTotal prob of N-in: 0.94351\nPOSSIBLE N-term signal sequence\ninside 1 19\nTMhelix 20 39\noutside 40 325\nTMhelix 326 348\ninside 349 372

3817 GCF_001891875.1_ASM189187v1 Escherichia coli Proteobacteria; Gammaproteobacteria; Enterobacterales; Enterobacteriaceae; Escherichia
MNNEETFYQAMRRQGVTRRSFLKYCSLAATSLGLGAGMAPKIAWALENKPRIPVVWIHGL WP_001058323.1
MULTISPECIES: hydrogenase-1 small chain [Proteobacteria] Length: 372\nNumber of predicted TMHs: 2\nExp number of AAs in TMHs: 37.2608499999999\nExp number, first 60 AAs: 19.52923\nTotal prob of N-in: 0.94351\nPOSSIBLE N-term signal sequence\ninside 1 19\nTMhelix 20 39\noutside 40 325\nTMhelix 326 348\ninside 349 372

3818 GCF_001892055.1_ASM189205v1 Escherichia coli Proteobacteria; Gammaproteobacteria; Enterobacterales; Enterobacteriaceae; Escherichia
MNNEETFYQAMRRQGVTRRSFLKYCSLAATSLGLGAGMAPKIAWALENKPRIPVVWIHGL WP_001058323.1
MULTISPECIES: hydrogenase-1 small chain [Proteobacteria] Length: 372\nNumber of predicted TMHs: 2\nExp number of AAs in TMHs: 37.2608499999999\nExp number, first 60 AAs: 19.52923\nTotal prob of N-in: 0.94351\nPOSSIBLE N-term signal sequence\ninside 1 19\nTMhelix 20 39\noutside 40 325\nTMhelix 326 348\ninside 349 372

3819 GCF_001892255.1_ASM189225v1 Escherichia coli Proteobacteria; Gammaproteobacteria; Enterobacterales; Enterobacteriaceae; Escherichia
MTGDNTLIHSHGINRRDFMKLCAALAATMGLSSKAAAEMAESVTNPQRPPVIWIGAQECT WP_000145410.1
MULTISPECIES: hydrogenase-2 small chain [Proteobacteria] Length: 372\nNumber of predicted TMHs: 1\nExp number of AAs in TMHs: 23.72226\nExp number, first 60 AAs: 0.22038\nTotal prob of N-in: 0.12994\noutside 1 331\nTMhelix 332 354\ninside 355 372

3820 GCF_001892295.1_ASM189229v1 Escherichia coli Proteobacteria; Gammaproteobacteria; Enterobacterales; Enterobacteriaceae; Escherichia
MTGDNTLIHSHGINRRDFMKLCAALAATMGLSSKAAAEMAESVTNPQRPPVIWIGAQECT WP_000145410.1
MULTISPECIES: hydrogenase-2 small chain [Proteobacteria] Length: 372\nNumber of predicted TMHs: 1\nExp number of AAs in TMHs: 23.72226\nExp number, first 60 AAs: 0.22038\nTotal prob of N-in: 0.12994\noutside 1 331\nTMhelix 332 354\ninside 355 372

3821 GCF_001892295.1_ASM189229v1 Escherichia coli Proteobacteria; Gammaproteobacteria; Enterobacterales; Enterobacteriaceae; Escherichia

MNNEETFYQAMRRQGVTRRSFLKYCSLAATSLGLGAGMAPKIAWALENKPRIPVVWIHGL WP_001058323.1 MULTISPECIES: hydrogenase-1 small chain [Proteobacteria] Length: 372\nNumber of predicted TMHs: 2\nExp number of AAs in TMHs: 37.2608499999999\nExp number, first 60 AAs: 19.52923\nTotal prob of N-in: 0.94351\nPOSSIBLE N-term signal sequence\ninside 1 19\nTMhelix 20 39\noutside 40 325\nTMhelix 326 348\ninside 349 372

3822 GCF_001892255.1_ASM189225v1 Escherichia coli Proteobacteria; Gammaproteobacteria; Enterobacterales; Enterobacteriaceae; Escherichia MNNEETFYQAMRRQGVTRRSFLKYCSLAATSLGLGAGMAPKIAWALENKPRIPVVWIHGL WP_001295940.1 MULTISPECIES: hydrogenase [Enterobacteriaceae] Length: 372\nNumber of predicted TMHs: 2\nExp number of AAs in TMHs: 37.2608899999999\nExp number, first 60 AAs: 19.52923\nTotal prob of N-in: 0.94351\nPOSSIBLE N-term signal sequence\ninside 1 19\nTMhelix 20 39\noutside 40 325\nTMhelix 326 348\ninside 349 372

3823 GCF_001892025.1_ASM189202v1 Escherichia coli Proteobacteria; Gammaproteobacteria; Enterobacterales; Enterobacteriaceae; Escherichia MNNEETFYQAMRRQGVTRRSFLKYCSLAATSLGLGAGMAPKIAWALENKPRIPVVWIHGL WP_001295940.1 MULTISPECIES: hydrogenase [Enterobacteriaceae] Length: 372\nNumber of predicted TMHs: 2\nExp number of AAs in TMHs: 37.2608899999999\nExp number, first 60 AAs: 19.52923\nTotal prob of N-in: 0.94351\nPOSSIBLE N-term signal sequence\ninside 1 19\nTMhelix 20 39\noutside 40 325\nTMhelix 326 348\ninside 349 372

3824 GCF_001892125.1_ASM189212v1 Escherichia coli Proteobacteria; Gammaproteobacteria; Enterobacterales; Enterobacteriaceae; Escherichia MNNEETFYQAMRRQGVTRRSFLKYCSLAATSLGLGAGMAPKIAWALENKPRIPVVWIHGL WP_001295940.1 MULTISPECIES: hydrogenase [Enterobacteriaceae] Length: 372\nNumber of predicted TMHs: 2\nExp number of AAs in TMHs: 37.2608899999999\nExp number, first 60 AAs: 19.52923\nTotal prob of N-in: 0.94351\nPOSSIBLE N-term signal sequence\ninside 1 19\nTMhelix 20 39\noutside 40 325\nTMhelix 326 348\ninside 349 372

3825 GCF_001892445.1_ASM189244v1 Escherichia coli Proteobacteria; Gammaproteobacteria; Enterobacterales; Enterobacteriaceae; Escherichia MTGDNTLIHSHGINRRDFMKLCAALAATMGLSSKAAAEMAESVTNPQRPPVIWIGAQECT WP_000145410.1 MULTISPECIES: hydrogenase-2 small chain [Proteobacteria] Length: 372\nNumber of predicted TMHs: 1\nExp number of AAs in TMHs: 23.72226\nExp number, first 60 AAs: 0.22038\nTotal prob of N-in: 0.12994\noutside 1 331\nTMhelix 332 354\ninside 355 372

3826 GCF_001892565.1_ASM189256v1 Escherichia coli Proteobacteria; Gammaproteobacteria; Enterobacterales; Enterobacteriaceae; Escherichia MTGDNTLIHSHGINRRDFMKLCAALAATMGLSSKAAAEMAESVTNPQRPPVIWIGAQECT WP_000145410.1 MULTISPECIES: hydrogenase-2 small chain [Proteobacteria] Length: 372\nNumber of predicted TMHs: 1\nExp number of AAs in TMHs: 23.72226\nExp number, first 60 AAs: 0.22038\nTotal prob of N-in: 0.12994\noutside 1 331\nTMhelix 332 354\ninside 355 372

3827 GCF_001892605.1_ASM189260v1 Escherichia coli Proteobacteria; Gammaproteobacteria; Enterobacterales; Enterobacteriaceae; Escherichia MTGDNTLIHSHGINRRDFMKLCAALAATMGLSSKAAAEMAESVTNPQRPPVIWIGAQECT WP_000145410.1 MULTISPECIES: hydrogenase-2 small chain [Proteobacteria] Length: 372\nNumber of predicted TMHs: 1\nExp number of AAs in TMHs: 23.72226\nExp number, first 60 AAs: 0.22038\nTotal prob of N-in: 0.12994\noutside 1 331\nTMhelix 332 354\ninside 355 372

3828 GCF_001892665.1_ASM189266v1 Escherichia coli Proteobacteria; Gammaproteobacteria; Enterobacterales; Enterobacteriaceae; Escherichia MTGDNTLIHSHGINRRDFMKLCAALAATMGLSSKAAAEMAESVTNPQRPPVIWIGAQECT WP_000145410.1 MULTISPECIES: hydrogenase-2 small chain [Proteobacteria] Length: 372\nNumber of predicted TMHs: 1\nExp number of AAs in TMHs: 23.72226\nExp number, first 60 AAs: 0.22038\nTotal prob of N-in: 0.12994\noutside 1 331\nTMhelix 332 354\ninside 355 372

3829 GCF_001892665.1_ASM189266v1 Escherichia coli Proteobacteria; Gammaproteobacteria; Enterobacterales; Enterobacteriaceae; Escherichia MNNEETFYQAMRRQGVTRRSFLKYCSLAATSLGLGAGMAPKIAWALENKPRIPVVWIHGL WP_001058323.1

MULTISPECIES: hydrogenase-1 small chain [Proteobacteria] Length: 372\nNumber of predicted TMHs: 2\nExp number of AAs in TMHs: 37.2608499999999\nExp number, first 60 AAs: 19.52923\nTotal prob of N-in: 0.94351\nPOSSIBLE N-term signal sequence\ninside 1 19\nTMhelix 20 39\noutside 40 325\nTMhelix 326 348\ninside 349 372

3830 GCF_001892325.1_ASM189232v1 Escherichia coli Proteobacteria; Gammaproteobacteria; Enterobacterales; Enterobacteriaceae; Escherichia
MTGDNTLIHSHGINRRDFMKLCAALATTMGLSSKAAAEMAESVTNPQRPPVIWIGAQECT WP_023149629.1
MULTISPECIES: hydrogenase-2 small chain [Enterobacteriaceae] Length: 372\nNumber of predicted TMHs: 1\nExp number of AAs in TMHs: 23.63603\nExp number, first 60 AAs: 0.12971\nTotal prob of N-in: 0.12608\noutside 1 331\nTMhelix 332 354\ninside 355 372

3831 GCF_001892725.1_ASM189272v1 Escherichia coli Proteobacteria; Gammaproteobacteria; Enterobacterales; Enterobacteriaceae; Escherichia
MTGDNTLIHSHGINRRDFMKLCAALAATMGLSSKAAAEMAESVTNPQRPPVIWIGAQECT WP_000145410.1
MULTISPECIES: hydrogenase-2 small chain [Proteobacteria] Length: 372\nNumber of predicted TMHs: 1\nExp number of AAs in TMHs: 23.72226\nExp number, first 60 AAs: 0.22038\nTotal prob of N-in: 0.12994\noutside 1 331\nTMhelix 332 354\ninside 355 372

3832 GCF_001892755.1_ASM189275v1 Escherichia coli Proteobacteria; Gammaproteobacteria; Enterobacterales; Enterobacteriaceae; Escherichia
MTGDNTLIHSHGINRRDFMKLCAALAATMGLSSKAAAEMAESVTNPQRPPVIWIGAQECT WP_000145410.1
MULTISPECIES: hydrogenase-2 small chain [Proteobacteria] Length: 372\nNumber of predicted TMHs: 1\nExp number of AAs in TMHs: 23.72226\nExp number, first 60 AAs: 0.22038\nTotal prob of N-in: 0.12994\noutside 1 331\nTMhelix 332 354\ninside 355 372

3833 GCF_001892905.1_ASM189290v1 Escherichia coli Proteobacteria; Gammaproteobacteria; Enterobacterales; Enterobacteriaceae; Escherichia
MTGDNTLIHSHGINRRDFMKLCAALAATMGLSSKAAAEMAESVTNPQRPPVIWIGAQECT WP_000145410.1
MULTISPECIES: hydrogenase-2 small chain [Proteobacteria] Length: 372\nNumber of predicted TMHs: 1\nExp number of AAs in TMHs: 23.72226\nExp number, first 60 AAs: 0.22038\nTotal prob of N-in: 0.12994\noutside 1 331\nTMhelix 332 354\ninside 355 372

3834 GCF_001892845.1_ASM189284v1 Escherichia coli Proteobacteria; Gammaproteobacteria; Enterobacterales; Enterobacteriaceae; Escherichia
MTGDNTLIHSHGINRRDFMKLCAALAATMGLSSKAAAEMAESVTNPQRPPVIWIGAQECT WP_000145410.1
MULTISPECIES: hydrogenase-2 small chain [Proteobacteria] Length: 372\nNumber of predicted TMHs: 1\nExp number of AAs in TMHs: 23.72226\nExp number, first 60 AAs: 0.22038\nTotal prob of N-in: 0.12994\noutside 1 331\nTMhelix 332 354\ninside 355 372

3835 GCF_001892795.1_ASM189279v1 Escherichia coli Proteobacteria; Gammaproteobacteria; Enterobacterales; Enterobacteriaceae; Escherichia
MNNEETFYQAMRRQGVTRRSFLKYCSLAATSLGLGAGMAPKIAWALENKPRIPVVWIHGL WP_001058323.1
MULTISPECIES: hydrogenase-1 small chain [Proteobacteria] Length: 372\nNumber of predicted TMHs: 2\nExp number of AAs in TMHs: 37.2608499999999\nExp number, first 60 AAs: 19.52923\nTotal prob of N-in: 0.94351\nPOSSIBLE N-term signal sequence\ninside 1 19\nTMhelix 20 39\noutside 40 325\nTMhelix 326 348\ninside 349 372

3836 GCF_001892905.1_ASM189290v1 Escherichia coli Proteobacteria; Gammaproteobacteria; Enterobacterales; Enterobacteriaceae; Escherichia
MNNEETFYQAMRRQGVTRRSFLKYCSLAATSLGLGAGMAPKIAWALENKPRIPVVWIHGL WP_001058323.1
MULTISPECIES: hydrogenase-1 small chain [Proteobacteria] Length: 372\nNumber of predicted TMHs: 2\nExp number of AAs in TMHs: 37.2608499999999\nExp number, first 60 AAs: 19.52923\nTotal prob of N-in: 0.94351\nPOSSIBLE N-term signal sequence\ninside 1 19\nTMhelix 20 39\noutside 40 325\nTMhelix 326 348\ninside 349 372

3837 GCF_001892845.1_ASM189284v1 Escherichia coli Proteobacteria; Gammaproteobacteria; Enterobacterales; Enterobacteriaceae; Escherichia
MNNEETFYQAMRRQGVTRRSFLKYCSLAATSLGLGAGMAPKIAWALENKPRIPVVWIHGL WP_001295940.1
MULTISPECIES: hydrogenase [Enterobacteriaceae] Length: 372\nNumber of predicted TMHs: 2\nExp number of AAs in TMHs: 37.2608899999999\nExp number, first 60 AAs: 19.52923\nTotal prob of N-in:

0.94351\nPOSSIBLE N-term signal sequence\ninside 1 19\nTMhelix 20 39\noutside 40 325\nTMhelix 326 348\ninside 349 372

3838 GCF_001892605.1_ASM189260v1 Escherichia coli Proteobacteria; Gammaproteobacteria; Enterobacterales; Enterobacteriaceae; Escherichia MNNEETFYQAMRRQGVTRRSFLKYCSLAATSLGLGAGMAPKIAWALENKPRIPVVWIHGL WP_001295940.1 MULTISPECIES: hydrogenase [Enterobacteriaceae] Length: 372\nNumber of predicted TMHs: 2\nExp number of AAs in TMHs: 37.2608899999999\nExp number, first 60 AAs: 19.52923\nTotal prob of N-in: 0.94351\nPOSSIBLE N-term signal sequence\ninside 1 19\nTMhelix 20 39\noutside 40 325\nTMhelix 326 348\ninside 349 372

3839 GCF_001893005.1_ASM189300v1 Escherichia coli Proteobacteria; Gammaproteobacteria; Enterobacterales; Enterobacteriaceae; Escherichia MTGDNTLIHSHGINRRDFMKLCAALAATMGLSSKAAAEMAESVTNPQRPPVIWIGAQECT WP_000145410.1 MULTISPECIES: hydrogenase-2 small chain [Proteobacteria] Length: 372\nNumber of predicted TMHs: 1\nExp number of AAs in TMHs: 23.72226\nExp number, first 60 AAs: 0.22038\nTotal prob of N-in: 0.12994\noutside 1 331\nTMhelix 332 354\ninside 355 372

3840 GCF_001893205.1_ASM189320v1 Escherichia coli Proteobacteria; Gammaproteobacteria; Enterobacterales; Enterobacteriaceae; Escherichia MNNEETFYQAMRRQGVTRRSFLKYCSLAATSLGLGAGMAPKIAWALENKPRIPVVWIHGL WP_001058323.1 MULTISPECIES: hydrogenase-1 small chain [Proteobacteria] Length: 372\nNumber of predicted TMHs: 2\nExp number of AAs in TMHs: 37.2608499999999\nExp number, first 60 AAs: 19.52923\nTotal prob of N-in: 0.94351\nPOSSIBLE N-term signal sequence\ninside 1 19\nTMhelix 20 39\noutside 40 325\nTMhelix 326 348\ninside 349 372

3841 GCF_001893245.1_ASM189324v1 Escherichia coli Proteobacteria; Gammaproteobacteria; Enterobacterales; Enterobacteriaceae; Escherichia MNNEETFYQAMRRQGVTRRSFLKYCSLAATSLGLGAGMAPKIAWALENKPRIPVVWIHGL WP_063501981.1 hydrogenase [Escherichia coli] Length: 372\nNumber of predicted TMHs: 2\nExp number of AAs in TMHs: 37.06322\nExp number, first 60 AAs: 19.52932\nTotal prob of N-in: 0.94340\nPOSSIBLE N-term signal sequence\ninside 1 19\nTMhelix 20 39\noutside 40 325\nTMhelix 326 348\ninside 349 372

3842 GCF_001893005.1_ASM189300v1 Escherichia coli Proteobacteria; Gammaproteobacteria; Enterobacterales; Enterobacteriaceae; Escherichia MNNEETFYQAMRRQGVTRRSFLKYCSLAATSLGLGAGMAPKIAWALENKPRIPVVWIHGL WP_001295940.1 MULTISPECIES: hydrogenase [Enterobacteriaceae] Length: 372\nNumber of predicted TMHs: 2\nExp number of AAs in TMHs: 37.2608899999999\nExp number, first 60 AAs: 19.52923\nTotal prob of N-in: 0.94351\nPOSSIBLE N-term signal sequence\ninside 1 19\nTMhelix 20 39\noutside 40 325\nTMhelix 326 348\ninside 349 372

3843 GCF_001893305.1_ASM189330v1 Escherichia coli Proteobacteria; Gammaproteobacteria; Enterobacterales; Enterobacteriaceae; Escherichia MTGDNTLIHSHGINRRDFMKLCAALAATMGLSSKAAAEMAESVTNPQRPPVIWIGAQECT WP_000145410.1 MULTISPECIES: hydrogenase-2 small chain [Proteobacteria] Length: 372\nNumber of predicted TMHs: 1\nExp number of AAs in TMHs: 23.72226\nExp number, first 60 AAs: 0.22038\nTotal prob of N-in: 0.12994\noutside 1 331\nTMhelix 332 354\ninside 355 372

3844 GCF_001893305.1_ASM189330v1 Escherichia coli Proteobacteria; Gammaproteobacteria; Enterobacterales; Enterobacteriaceae; Escherichia MNNEETFYQAMRRQGVTRRSFLKYCSLAATSLGLGAGMAPKIAWALENKPRIPVVWIHGL WP_001058323.1 MULTISPECIES: hydrogenase-1 small chain [Proteobacteria] Length: 372\nNumber of predicted TMHs: 2\nExp number of AAs in TMHs: 37.2608499999999\nExp number, first 60 AAs: 19.52923\nTotal prob of N-in: 0.94351\nPOSSIBLE N-term signal sequence\ninside 1 19\nTMhelix 20 39\noutside 40 325\nTMhelix 326 348\ninside 349 372

3845 GCF_001893025.1_ASM189302v1 Escherichia coli Proteobacteria; Gammaproteobacteria; Enterobacterales; Enterobacteriaceae; Escherichia MNNEETFYQAMRRQGVTRRSFLKYCSLAATSLGLGAGMAPKIAWALENKPRIPVVWIHGL WP_001058323.1 MULTISPECIES: hydrogenase-1 small chain [Proteobacteria] Length: 372\nNumber of predicted TMHs: 2\nExp number of AAs in TMHs: 37.2608499999999\nExp number, first 60 AAs: 19.52923\nTotal prob of N-in:

0.94351\nPOSSIBLE N-term signal sequence\ninside 1 19\nTMhelix 20 39\noutside 40 325\nTMhelix 326 348\ninside 349 372

3846 GCF_001893345.1_ASM189334v1 Escherichia coli Proteobacteria; Gammaproteobacteria; Enterobacterales; Enterobacteriaceae; Escherichia MTGDNTLIHSHGINRRDFMKLCAALAATMGLSSKAAAEMAESVTNPQRPPVIWIGAQECT WP_000145410.1 MULTISPECIES: hydrogenase-2 small chain [Proteobacteria] Length: 372\nNumber of predicted TMHs: 1\nExp number of AAs in TMHs: 23.72226\nExp number, first 60 AAs: 0.22038\nTotal prob of N-in: 0.12994\noutside 1 331\nTMhelix 332 354\ninside 355 372

3847 GCF_001893445.1_ASM189344v1 Escherichia coli Proteobacteria; Gammaproteobacteria; Enterobacterales; Enterobacteriaceae; Escherichia MNNEETFYQAMRRQGVTRRSFLKYCSLAATSLGLGAGMAPKIAWALENKPRIPVVWIHGL WP_001058323.1 MULTISPECIES: hydrogenase-1 small chain [Proteobacteria] Length: 372\nNumber of predicted TMHs: 2\nExp number of AAs in TMHs: 37.2608499999999\nExp number, first 60 AAs: 19.52923\nTotal prob of N-in: 0.94351\nPOSSIBLE N-term signal sequence\ninside 1 19\nTMhelix 20 39\noutside 40 325\nTMhelix 326 348\ninside 349 372

3848 GCF_001893165.1_ASM189316v1 Escherichia coli Proteobacteria; Gammaproteobacteria; Enterobacterales; Enterobacteriaceae; Escherichia MNNEETFYQAMRRQGVTRRSFLKYCSLAATSLGLGAGMAPKIAWALENKPRIPVVWIHGL WP_001295940.1 MULTISPECIES: hydrogenase [Enterobacteriaceae] Length: 372\nNumber of predicted TMHs: 2\nExp number of AAs in TMHs: 37.2608899999999\nExp number, first 60 AAs: 19.52923\nTotal prob of N-in: 0.94351\nPOSSIBLE N-term signal sequence\ninside 1 19\nTMhelix 20 39\noutside 40 325\nTMhelix 326 348\ninside 349 372

3849 GCF_001893625.1_ASM189362v1 Escherichia coli Proteobacteria; Gammaproteobacteria; Enterobacterales; Enterobacteriaceae; Escherichia MTGDNTLIHSHGINRRDFMKLCAALAATMGLSSKAAAEMAESVTNPQRPPVIWIGAQECT WP_000145410.1 MULTISPECIES: hydrogenase-2 small chain [Proteobacteria] Length: 372\nNumber of predicted TMHs: 1\nExp number of AAs in TMHs: 23.72226\nExp number, first 60 AAs: 0.22038\nTotal prob of N-in: 0.12994\noutside 1 331\nTMhelix 332 354\ninside 355 372

3850 GCF_001893605.1_ASM189360v1 Escherichia coli Proteobacteria; Gammaproteobacteria; Enterobacterales; Enterobacteriaceae; Escherichia MTGDNTLIHSHGINRRDFMKLCAALAATMGLSSKAAAEMAESVTNPQRPPVIWIGAQECT WP_000145410.1 MULTISPECIES: hydrogenase-2 small chain [Proteobacteria] Length: 372\nNumber of predicted TMHs: 1\nExp number of AAs in TMHs: 23.72226\nExp number, first 60 AAs: 0.22038\nTotal prob of N-in: 0.12994\noutside 1 331\nTMhelix 332 354\ninside 355 372

3851 GCF_001893695.1_ASM189369v1 Escherichia coli Proteobacteria; Gammaproteobacteria; Enterobacterales; Enterobacteriaceae; Escherichia MTGDNTLIHSHGINRRDFMKLCAALAATMGLSSKAAAEMAESVTNPQRPPVIWIGAQECT WP_000145410.1 MULTISPECIES: hydrogenase-2 small chain [Proteobacteria] Length: 372\nNumber of predicted TMHs: 1\nExp number of AAs in TMHs: 23.72226\nExp number, first 60 AAs: 0.22038\nTotal prob of N-in: 0.12994\noutside 1 331\nTMhelix 332 354\ninside 355 372

3852 GCF_001893715.1_ASM189371v1 Escherichia coli Proteobacteria; Gammaproteobacteria; Enterobacterales; Enterobacteriaceae; Escherichia MTGDNTLIHSHGINRRDFMKLCAALAATMGLSSKAAAEMAESVTNPQRPPVIWIGAQECT WP_000145410.1 MULTISPECIES: hydrogenase-2 small chain [Proteobacteria] Length: 372\nNumber of predicted TMHs: 1\nExp number of AAs in TMHs: 23.72226\nExp number, first 60 AAs: 0.22038\nTotal prob of N-in: 0.12994\noutside 1 331\nTMhelix 332 354\ninside 355 372

3853 GCF_001893625.1_ASM189362v1 Escherichia coli Proteobacteria; Gammaproteobacteria; Enterobacterales; Enterobacteriaceae; Escherichia MNNEETFYQAMRRQGVTRRSFLKYCSLAATSLGLGAGMAPKIAWALENKPRIPVVWIHGL WP_001058323.1 MULTISPECIES: hydrogenase-1 small chain [Proteobacteria] Length: 372\nNumber of predicted TMHs: 2\nExp number of AAs in TMHs: 37.2608499999999\nExp number, first 60 AAs: 19.52923\nTotal prob of N-in: 0.94351\nPOSSIBLE N-term signal sequence\ninside 1 19\nTMhelix 20 39\noutside 40 325\nTMhelix 326 348\ninside 349 372

3854 GCF_001893955.1_ASM189395v1 Escherichia coli Proteobacteria; Gammaproteobacteria; Enterobacterales; Enterobacteriaceae; Escherichia
MNNEETFYQAMRRQGVTRRSFLKYCSLAATSLGLGAGMAPKIAWALENKPRIPVVWIHGL WP_001058323.1
MULTISPECIES: hydrogenase-1 small chain [Proteobacteria] Length: 372\nNumber of predicted TMHs: 2\nExp number of AAs in TMHs: 37.2608499999999\nExp number, first 60 AAs: 19.52923\nTotal prob of N-in: 0.94351\nPOSSIBLE N-term signal sequence\ninside 1 19\nTMhelix 20 39\noutside 40 325\nTMhelix 326 348\ninside 349 372

3855 GCF_001893715.1_ASM189371v1 Escherichia coli Proteobacteria; Gammaproteobacteria; Enterobacterales; Enterobacteriaceae; Escherichia
MNNEETFYQAMRRQGVTRRSFLKYCSLAATSLGLGAGMAPKIAWALENKPRIPVVWIHGL WP_001058323.1
MULTISPECIES: hydrogenase-1 small chain [Proteobacteria] Length: 372\nNumber of predicted TMHs: 2\nExp number of AAs in TMHs: 37.2608499999999\nExp number, first 60 AAs: 19.52923\nTotal prob of N-in: 0.94351\nPOSSIBLE N-term signal sequence\ninside 1 19\nTMhelix 20 39\noutside 40 325\nTMhelix 326 348\ninside 349 372

3856 GCF_001893775.1_ASM189377v1 Escherichia coli Proteobacteria; Gammaproteobacteria; Enterobacterales; Enterobacteriaceae; Escherichia
MNNEETFYQAMRRQGVTRRSFLKYCSLAATSLGLGAGMAPKIAWALENKPRIPVVWIHGL WP_001058323.1
MULTISPECIES: hydrogenase-1 small chain [Proteobacteria] Length: 372\nNumber of predicted TMHs: 2\nExp number of AAs in TMHs: 37.2608499999999\nExp number, first 60 AAs: 19.52923\nTotal prob of N-in: 0.94351\nPOSSIBLE N-term signal sequence\ninside 1 19\nTMhelix 20 39\noutside 40 325\nTMhelix 326 348\ninside 349 372

3857 GCF_001893695.1_ASM189369v1 Escherichia coli Proteobacteria; Gammaproteobacteria; Enterobacterales; Enterobacteriaceae; Escherichia
MNNEETFYQAMRRQGVTRRSFLKYCSLAATSLGLGAGMAPKIAWALENKPRIPVVWIHGL WP_001295940.1
MULTISPECIES: hydrogenase [Enterobacteriaceae] Length: 372\nNumber of predicted TMHs: 2\nExp number of AAs in TMHs: 37.2608899999999\nExp number, first 60 AAs: 19.52923\nTotal prob of N-in: 0.94351\nPOSSIBLE N-term signal sequence\ninside 1 19\nTMhelix 20 39\noutside 40 325\nTMhelix 326 348\ninside 349 372

3858 GCF_001893995.1_ASM189399v1 Escherichia coli Proteobacteria; Gammaproteobacteria; Enterobacterales; Enterobacteriaceae; Escherichia
MTGDNTLIHSHGINRRDFMKLCAALAATMGLSSKAAAEMAESVTNPQRPPVIWIGAQECT WP_000145410.1
MULTISPECIES: hydrogenase-2 small chain [Proteobacteria] Length: 372\nNumber of predicted TMHs: 1\nExp number of AAs in TMHs: 23.72226\nExp number, first 60 AAs: 0.22038\nTotal prob of N-in: 0.12994\noutside 1 331\nTMhelix 332 354\ninside 355 372

3859 GCF_001894035.1_ASM189403v1 Escherichia coli Proteobacteria; Gammaproteobacteria; Enterobacterales; Enterobacteriaceae; Escherichia
MTGDNTLIHSHGINRRDFMKLCAALAATMGLSSKAAAEMAESVTNPQRPPVIWIGAQECT WP_000145410.1
MULTISPECIES: hydrogenase-2 small chain [Proteobacteria] Length: 372\nNumber of predicted TMHs: 1\nExp number of AAs in TMHs: 23.72226\nExp number, first 60 AAs: 0.22038\nTotal prob of N-in: 0.12994\noutside 1 331\nTMhelix 332 354\ninside 355 372

3860 GCF_001894085.1_ASM189408v1 Escherichia coli Proteobacteria; Gammaproteobacteria; Enterobacterales; Enterobacteriaceae; Escherichia
MTGDNTLIHSHGINRRDFMKLCAALAATMGLSSKAAAEMAESVTNPQRPPVIWIGAQECT WP_000145410.1
MULTISPECIES: hydrogenase-2 small chain [Proteobacteria] Length: 372\nNumber of predicted TMHs: 1\nExp number of AAs in TMHs: 23.72226\nExp number, first 60 AAs: 0.22038\nTotal prob of N-in: 0.12994\noutside 1 331\nTMhelix 332 354\ninside 355 372

3861 GCF_001894035.1_ASM189403v1 Escherichia coli Proteobacteria; Gammaproteobacteria; Enterobacterales; Enterobacteriaceae; Escherichia
MNNEETFYQAMRRQGVTRRSFLKYCSLAATSLGLGAGMAPKIAWALENKPRIPVVWIHGL WP_001058323.1
MULTISPECIES: hydrogenase-1 small chain [Proteobacteria] Length: 372\nNumber of predicted TMHs: 2\nExp number of AAs in TMHs: 37.2608499999999\nExp number, first 60 AAs: 19.52923\nTotal prob of N-in: 0.94351\nPOSSIBLE N-term signal sequence\ninside 1 19\nTMhelix 20 39\noutside 40 325\nTMhelix 326 348\ninside 349 372

3862 GCF_001894085.1_ASM189408v1 Escherichia coli Proteobacteria; Gammaproteobacteria; Enterobacterales; Enterobacteriaceae; Escherichia
MNNEETFYQAMRRQGVTRRSFLKYCSLAATSLGLGAGMAPKIAWALENKPRIPVVWIHGL WP_001058323.1
MULTISPECIES: hydrogenase-1 small chain [Proteobacteria] Length: 372\nNumber of predicted TMHs: 2\nExp number of AAs in TMHs: 37.2608499999999\nExp number, first 60 AAs: 19.52923\nTotal prob of N-in: 0.94351\nPOSSIBLE N-term signal sequence\ninside 1 19\nTMhelix 20 39\noutside 40 325\nTMhelix 326 348\ninside 349 372

3863 GCF_001894065.1_ASM189406v1 Escherichia coli Proteobacteria; Gammaproteobacteria; Enterobacterales; Enterobacteriaceae; Escherichia
MNNEETFYQAMRRQGVTRRSFLKYCSLAATSLGLGAGMAPKIAWALENKPRIPVVWIHGL WP_001058323.1
MULTISPECIES: hydrogenase-1 small chain [Proteobacteria] Length: 372\nNumber of predicted TMHs: 2\nExp number of AAs in TMHs: 37.2608499999999\nExp number, first 60 AAs: 19.52923\nTotal prob of N-in: 0.94351\nPOSSIBLE N-term signal sequence\ninside 1 19\nTMhelix 20 39\noutside 40 325\nTMhelix 326 348\ninside 349 372

3864 GCF_001894305.1_ASM189430v1 Escherichia coli Proteobacteria; Gammaproteobacteria; Enterobacterales; Enterobacteriaceae; Escherichia
MTGDNTLIHSHGINRRDFMKLCAALAATMGLSSKAAAEMAESVTNPQRPPVIWIGAQECT WP_000145410.1
MULTISPECIES: hydrogenase-2 small chain [Proteobacteria] Length: 372\nNumber of predicted TMHs: 1\nExp number of AAs in TMHs: 23.72226\nExp number, first 60 AAs: 0.22038\nTotal prob of N-in: 0.12994\noutside 1 331\nTMhelix 332 354\ninside 355 372

3865 GCF_001894335.1_ASM189433v1 Escherichia coli Proteobacteria; Gammaproteobacteria; Enterobacterales; Enterobacteriaceae; Escherichia
MTGDNTLIHSHGINRRDFMKLCAALAATMGLSSKAAAEMAESVTNPQRPPVIWIGAQECT WP_000145410.1
MULTISPECIES: hydrogenase-2 small chain [Proteobacteria] Length: 372\nNumber of predicted TMHs: 1\nExp number of AAs in TMHs: 23.72226\nExp number, first 60 AAs: 0.22038\nTotal prob of N-in: 0.12994\noutside 1 331\nTMhelix 332 354\ninside 355 372

3866 GCF_001894395.1_ASM189439v1 Escherichia coli Proteobacteria; Gammaproteobacteria; Enterobacterales; Enterobacteriaceae; Escherichia
MTGDNTLIHSHGINRRDFMKLCAALAATMGLSSKAAAEMAESVTNPQRPPVIWIGAQECT WP_000145410.1
MULTISPECIES: hydrogenase-2 small chain [Proteobacteria] Length: 372\nNumber of predicted TMHs: 1\nExp number of AAs in TMHs: 23.72226\nExp number, first 60 AAs: 0.22038\nTotal prob of N-in: 0.12994\noutside 1 331\nTMhelix 332 354\ninside 355 372

3867 GCF_001894425.1_ASM189442v1 Escherichia coli Proteobacteria; Gammaproteobacteria; Enterobacterales; Enterobacteriaceae; Escherichia
MTGDNTLIHSHGINRRDFMKLCAALAATMGLSSKAAAEMAESVTNPQRPPVIWIGAQECT WP_000145410.1
MULTISPECIES: hydrogenase-2 small chain [Proteobacteria] Length: 372\nNumber of predicted TMHs: 1\nExp number of AAs in TMHs: 23.72226\nExp number, first 60 AAs: 0.22038\nTotal prob of N-in: 0.12994\noutside 1 331\nTMhelix 332 354\ninside 355 372

3868 GCF_001894425.1_ASM189442v1 Escherichia coli Proteobacteria; Gammaproteobacteria; Enterobacterales; Enterobacteriaceae; Escherichia
MNNEETFYQAMRRQGVTRRSFLKYCSLAATSLGLGAGMAPKIAWALENKPRIPVVWIHGL WP_001058323.1
MULTISPECIES: hydrogenase-1 small chain [Proteobacteria] Length: 372\nNumber of predicted TMHs: 2\nExp number of AAs in TMHs: 37.2608499999999\nExp number, first 60 AAs: 19.52923\nTotal prob of N-in: 0.94351\nPOSSIBLE N-term signal sequence\ninside 1 19\nTMhelix 20 39\noutside 40 325\nTMhelix 326 348\ninside 349 372

3869 GCF_001894585.1_ASM189458v1 Escherichia coli Proteobacteria; Gammaproteobacteria; Enterobacterales; Enterobacteriaceae; Escherichia
MTGDNTLIHSHGINRRDFMKLCAALAATMGLSSKAAAEMAESVTNPQRPPVIWIGAQECT WP_000145410.1
MULTISPECIES: hydrogenase-2 small chain [Proteobacteria] Length: 372\nNumber of predicted TMHs: 1\nExp number of AAs in TMHs: 23.72226\nExp number, first 60 AAs: 0.22038\nTotal prob of N-in: 0.12994\noutside 1 331\nTMhelix 332 354\ninside 355 372

3870 GCF_001894545.1_ASM189454v1 Escherichia coli Proteobacteria; Gammaproteobacteria; Enterobacterales; Enterobacteriaceae; Escherichia

MTGDNTLIHSHGINRRDFMKLCAALAATMGLSSKAAAEMAESVTNPQRPPVIWIGAQECT WP_000145410.1 MULTISPECIES: hydrogenase-2 small chain [Proteobacteria] Length: 372\nNumber of predicted TMHs: 1\nExp number of AAs in TMHs: 23.72226\nExp number, first 60 AAs: 0.22038\nTotal prob of N-in: 0.12994\noutside 1 331\nTMhelix 332 354\ninside 355 372

3871 GCF_001894635.1_ASM189463v1 Escherichia coli Proteobacteria; Gammaproteobacteria; Enterobacterales; Enterobacteriaceae; Escherichia MTGDNTLIHSHGINRRDFMKLCAALAATMGLSSKAAAEMAESVTNPQRPPVIWIGAQECT WP_000145410.1 MULTISPECIES: hydrogenase-2 small chain [Proteobacteria] Length: 372\nNumber of predicted TMHs: 1\nExp number of AAs in TMHs: 23.72226\nExp number, first 60 AAs: 0.22038\nTotal prob of N-in: 0.12994\noutside 1 331\nTMhelix 332 354\ninside 355 372

3872 GCF_001894705.1_ASM189470v1 Escherichia coli Proteobacteria; Gammaproteobacteria; Enterobacterales; Enterobacteriaceae; Escherichia MTGDNTLIHSHGINRRDFMKLCAALAATMGLSSKAAAEMAESVTNPQRPPVIWIGAQECT WP_000145410.1 MULTISPECIES: hydrogenase-2 small chain [Proteobacteria] Length: 372\nNumber of predicted TMHs: 1\nExp number of AAs in TMHs: 23.72226\nExp number, first 60 AAs: 0.22038\nTotal prob of N-in: 0.12994\noutside 1 331\nTMhelix 332 354\ninside 355 372

3873 GCF_001894545.1_ASM189454v1 Escherichia coli Proteobacteria; Gammaproteobacteria; Enterobacterales; Enterobacteriaceae; Escherichia MNNEETFYQAMRRQGVTRRSFLKYCSLAATSLGLGAGMAPKIAWALENKPRIPVVWIHGL WP_001058323.1 MULTISPECIES: hydrogenase-1 small chain [Proteobacteria] Length: 372\nNumber of predicted TMHs: 2\nExp number of AAs in TMHs: 37.2608499999999\nExp number, first 60 AAs: 19.52923\nTotal prob of N-in: 0.94351\nPOSSIBLE N-term signal sequence\ninside 1 19\nTMhelix 20 39\noutside 40 325\nTMhelix 326 348\ninside 349 372

3874 GCF_001899625.1_ASM189962v1 Escherichia coli Proteobacteria; Gammaproteobacteria; Enterobacterales; Enterobacteriaceae; Escherichia MTGDNTLIHSHGINRRDFMKLCAALAATMGLSSKAAAEMAESVTNPQRPPVIWIGAQECT WP_000145410.1 MULTISPECIES: hydrogenase-2 small chain [Proteobacteria] Length: 372\nNumber of predicted TMHs: 1\nExp number of AAs in TMHs: 23.72226\nExp number, first 60 AAs: 0.22038\nTotal prob of N-in: 0.12994\noutside 1 331\nTMhelix 332 354\ninside 355 372

3875 GCF_001894585.1_ASM189458v1 Escherichia coli Proteobacteria; Gammaproteobacteria; Enterobacterales; Enterobacteriaceae; Escherichia MNNEETFYQAMRRQGVTRRSFLKYCSLAATSLGLGAGMAPKIAWALENKPRIPVVWIHGL WP_001295940.1 MULTISPECIES: hydrogenase [Enterobacteriaceae] Length: 372\nNumber of predicted TMHs: 2\nExp number of AAs in TMHs: 37.2608899999999\nExp number, first 60 AAs: 19.52923\nTotal prob of N-in: 0.94351\nPOSSIBLE N-term signal sequence\ninside 1 19\nTMhelix 20 39\noutside 40 325\nTMhelix 326 348\ninside 349 372

3876 GCF_001900495.1_ASM190049v1 Escherichia coli Proteobacteria; Gammaproteobacteria; Enterobacterales; Enterobacteriaceae; Escherichia MTGDNTLIHSHGINRRDFMKLCAALAATMGLSSKAAAEMAESVTNPQRPPVIWIGAQECT WP_000145410.1 MULTISPECIES: hydrogenase-2 small chain [Proteobacteria] Length: 372\nNumber of predicted TMHs: 1\nExp number of AAs in TMHs: 23.72226\nExp number, first 60 AAs: 0.22038\nTotal prob of N-in: 0.12994\noutside 1 331\nTMhelix 332 354\ninside 355 372

3877 GCF_001900495.1_ASM190049v1 Escherichia coli Proteobacteria; Gammaproteobacteria; Enterobacterales; Enterobacteriaceae; Escherichia MNNEETFYQAMRRQGVTRRSFLKYCSLAATSLGLGAGMAPKIAWALENKPRIPVVWIHGL WP_001058323.1 MULTISPECIES: hydrogenase-1 small chain [Proteobacteria] Length: 372\nNumber of predicted TMHs: 2\nExp number of AAs in TMHs: 37.2608499999999\nExp number, first 60 AAs: 19.52923\nTotal prob of N-in: 0.94351\nPOSSIBLE N-term signal sequence\ninside 1 19\nTMhelix 20 39\noutside 40 325\nTMhelix 326 348\ninside 349 372

3878 GCF_001894705.1_ASM189470v1 Escherichia coli Proteobacteria; Gammaproteobacteria; Enterobacterales; Enterobacteriaceae; Escherichia MNNEETFYQAMRRQGVTRRSFLKYCSLAATSLGLGAGMAPKIAWALENKPRIPVVWIHGL WP_001058323.1 MULTISPECIES: hydrogenase-1 small chain [Proteobacteria] Length: 372\nNumber of predicted TMHs:

2\nExp number of AAs in TMHs: 37.2608499999999\nExp number, first 60 AAs: 19.52923\nTotal prob of N-in: 0.94351\nPOSSIBLE N-term signal sequence\ninside 1 19\nTMhelix 20 39\noutside 40 325\nTMhelix 326 348\ninside 349 372

3879 GCF_001900555.1_ASM190055v1 Escherichia coli Proteobacteria; Gammaproteobacteria; Enterobacterales; Enterobacteriaceae; Escherichia
MTGDNTLIHSHGINRRDFMKLCAALAATMGLSSKAAAEMAESVTNPQRPPVIWIGAQECT WP_000145410.1
MULTISPECIES: hydrogenase-2 small chain [Proteobacteria] Length: 372\nNumber of predicted TMHs: 1\nExp number of AAs in TMHs: 23.72226\nExp number, first 60 AAs: 0.22038\nTotal prob of N-in: 0.12994\noutside 1 331\nTMhelix 332 354\ninside 355 372

3880 GCF_001900555.1_ASM190055v1 Escherichia coli Proteobacteria; Gammaproteobacteria; Enterobacterales; Enterobacteriaceae; Escherichia
MNNEETFYQAMRRQGVTRRSFLKYCSLAATSLGLGAGMAPKIAWALENKPRIPVVWIHGL WP_001058320.1
hydrogenase [Escherichia coli] Length: 372\nNumber of predicted TMHs: 2\nExp number of AAs in TMHs: 37.26082\nExp number, first 60 AAs: 19.52923\nTotal prob of N-in: 0.94351\nPOSSIBLE N-term signal sequence\ninside 1 19\nTMhelix 20 39\noutside 40 325\nTMhelix 326 348\ninside 349 372

3881 GCF_001900315.1_ASM190031v1 Escherichia coli Proteobacteria; Gammaproteobacteria; Enterobacterales; Enterobacteriaceae; Escherichia
MNNEETFYQAMRRQGVTRRSFLKYCSLAATSLGLGAGMAPKIAWALENKPRIPVVWIHGL WP_001058323.1
MULTISPECIES: hydrogenase-1 small chain [Proteobacteria] Length: 372\nNumber of predicted TMHs: 2\nExp number of AAs in TMHs: 37.2608499999999\nExp number, first 60 AAs: 19.52923\nTotal prob of N-in: 0.94351\nPOSSIBLE N-term signal sequence\ninside 1 19\nTMhelix 20 39\noutside 40 325\nTMhelix 326 348\ninside 349 372

3882 GCF_001900655.1_ASM190065v1 Escherichia coli Proteobacteria; Gammaproteobacteria; Enterobacterales; Enterobacteriaceae; Escherichia
MNNEETFYQAMRRQGVTRRSFLKYCSLAATSLGLGAGMAPKIAWALENKPRIPVVWIHGL WP_001058323.1
MULTISPECIES: hydrogenase-1 small chain [Proteobacteria] Length: 372\nNumber of predicted TMHs: 2\nExp number of AAs in TMHs: 37.2608499999999\nExp number, first 60 AAs: 19.52923\nTotal prob of N-in: 0.94351\nPOSSIBLE N-term signal sequence\ninside 1 19\nTMhelix 20 39\noutside 40 325\nTMhelix 326 348\ninside 349 372

3883 GCF_001900905.1_ASM190090v1 Escherichia coli Proteobacteria; Gammaproteobacteria; Enterobacterales; Enterobacteriaceae; Escherichia
MTGDNTLIHSHGINRRDFMKLCAALAATMGLSSKAAAEMAESVTNPQRPPVIWIGAQECT WP_000145410.1
MULTISPECIES: hydrogenase-2 small chain [Proteobacteria] Length: 372\nNumber of predicted TMHs: 1\nExp number of AAs in TMHs: 23.72226\nExp number, first 60 AAs: 0.22038\nTotal prob of N-in: 0.12994\noutside 1 331\nTMhelix 332 354\ninside 355 372

3884 GCF_001900905.1_ASM190090v1 Escherichia coli Proteobacteria; Gammaproteobacteria; Enterobacterales; Enterobacteriaceae; Escherichia
MNNEETFYQAMRRQGVTRRSFLKYCSLAATSLGLGAGMAPKIAWALENKPRIPVVWIHGL WP_001058323.1
MULTISPECIES: hydrogenase-1 small chain [Proteobacteria] Length: 372\nNumber of predicted TMHs: 2\nExp number of AAs in TMHs: 37.2608499999999\nExp number, first 60 AAs: 19.52923\nTotal prob of N-in: 0.94351\nPOSSIBLE N-term signal sequence\ninside 1 19\nTMhelix 20 39\noutside 40 325\nTMhelix 326 348\ninside 349 372

3885 GCF_001900775.1_ASM190077v1 Escherichia coli Proteobacteria; Gammaproteobacteria; Enterobacterales; Enterobacteriaceae; Escherichia
MNNEETFYQAMRRQGVTRRSFLKYCSLAATSLGLGAGMAPKIAWALENKPRIPVVWIHGL WP_001058323.1
MULTISPECIES: hydrogenase-1 small chain [Proteobacteria] Length: 372\nNumber of predicted TMHs: 2\nExp number of AAs in TMHs: 37.2608499999999\nExp number, first 60 AAs: 19.52923\nTotal prob of N-in: 0.94351\nPOSSIBLE N-term signal sequence\ninside 1 19\nTMhelix 20 39\noutside 40 325\nTMhelix 326 348\ninside 349 372

3886 GCF_001900945.1_ASM190094v1 Escherichia coli Proteobacteria; Gammaproteobacteria; Enterobacterales; Enterobacteriaceae; Escherichia
MTGDNTLIHSHGINRRDFMKLCAALAATMGLSSKAAAEMAESVTNPQRPPVIWIGAQECT WP_000145410.1
MULTISPECIES: hydrogenase-2 small chain [Proteobacteria] Length: 372\nNumber of predicted TMHs:

1\nExp number of AAs in TMHs: 23.72226\nExp number, first 60 AAs: 0.22038\nTotal prob of N-in: 0.12994\noutside 1 331\nTMhelix 332 354\ninside 355 372

3887 GCF_001900945.1_ASM190094v1 Escherichia coli Proteobacteria; Gammaproteobacteria; Enterobacterales; Enterobacteriaceae; Escherichia MNNEETFYQAMRRQGVTRRSFLKYCSLAATSLGLGAGMAPKIAWALENKPRIPVVWIHGL WP_001295940.1 MULTISPECIES: hydrogenase [Enterobacteriaceae] Length: 372\nNumber of predicted TMHs: 2\nExp number of AAs in TMHs: 37.2608899999999\nExp number, first 60 AAs: 19.52923\nTotal prob of N-in: 0.94351\nPOSSIBLE N-term signal sequence\ninside 1 19\nTMhelix 20 39\noutside 40 325\nTMhelix 326 348\ninside 349 372

3888 GCF_001900695.1_ASM190069v1 Escherichia coli Proteobacteria; Gammaproteobacteria; Enterobacterales; Enterobacteriaceae; Escherichia MNNEETFYQAMRRQGVTRRSFLKYCSLAATSLGLGAGMAPKIAWALENKPRIPVVWIHGL WP_001058323.1 MULTISPECIES: hydrogenase-1 small chain [Proteobacteria] Length: 372\nNumber of predicted TMHs: 2\nExp number of AAs in TMHs: 37.2608499999999\nExp number, first 60 AAs: 19.52923\nTotal prob of N-in: 0.94351\nPOSSIBLE N-term signal sequence\ninside 1 19\nTMhelix 20 39\noutside 40 325\nTMhelix 326 348\ninside 349 372

3889 GCF_001901105.1_ASM190110v1 Escherichia coli Proteobacteria; Gammaproteobacteria; Enterobacterales; Enterobacteriaceae; Escherichia MNNEETFYQAMRRQGVTRRSFLKYCSLAATSLGLGAGMAPKIAWALENKPRIPVVWIHGL WP_001058323.1 MULTISPECIES: hydrogenase-1 small chain [Proteobacteria] Length: 372\nNumber of predicted TMHs: 2\nExp number of AAs in TMHs: 37.2608499999999\nExp number, first 60 AAs: 19.52923\nTotal prob of N-in: 0.94351\nPOSSIBLE N-term signal sequence\ninside 1 19\nTMhelix 20 39\noutside 40 325\nTMhelix 326 348\ninside 349 372

3890 GCF_001901365.1_ASM190136v1 Escherichia coli Proteobacteria; Gammaproteobacteria; Enterobacterales; Enterobacteriaceae; Escherichia MTGDNTLIHSHGINRRDFMKLCAALAATMGLSSKAAAEMAESVTNPQRPPVIWIGAQECT WP_000145410.1 MULTISPECIES: hydrogenase-2 small chain [Proteobacteria] Length: 372\nNumber of predicted TMHs: 1\nExp number of AAs in TMHs: 23.72226\nExp number, first 60 AAs: 0.22038\nTotal prob of N-in: 0.12994\noutside 1 331\nTMhelix 332 354\ninside 355 372

3891 GCF_001901365.1_ASM190136v1 Escherichia coli Proteobacteria; Gammaproteobacteria; Enterobacterales; Enterobacteriaceae; Escherichia MNNEETFYQAMRRQGVTRRSFLKYCSLAATSLGLGAGMAPKIAWALENKPRIPVVWIHGL WP_001058323.1 MULTISPECIES: hydrogenase-1 small chain [Proteobacteria] Length: 372\nNumber of predicted TMHs: 2\nExp number of AAs in TMHs: 37.2608499999999\nExp number, first 60 AAs: 19.52923\nTotal prob of N-in: 0.94351\nPOSSIBLE N-term signal sequence\ninside 1 19\nTMhelix 20 39\noutside 40 325\nTMhelix 326 348\ninside 349 372

3892 GCF_001901215.1_ASM190121v1 Escherichia coli Proteobacteria; Gammaproteobacteria; Enterobacterales; Enterobacteriaceae; Escherichia MNNEETFYQAMRRQGVTRRSFLKYCSLAATSLGLGAGMAPKIAWALENKPRIPVVWIHGL WP_001058323.1 MULTISPECIES: hydrogenase-1 small chain [Proteobacteria] Length: 372\nNumber of predicted TMHs: 2\nExp number of AAs in TMHs: 37.2608499999999\nExp number, first 60 AAs: 19.52923\nTotal prob of N-in: 0.94351\nPOSSIBLE N-term signal sequence\ninside 1 19\nTMhelix 20 39\noutside 40 325\nTMhelix 326 348\ninside 349 372

3893 GCF_001901405.1_ASM190140v1 Escherichia coli Proteobacteria; Gammaproteobacteria; Enterobacterales; Enterobacteriaceae; Escherichia MTGDNTLIHSHGINRRDFMKLCAALAATMGLSSKAAAEMAESVTNPQRPPVIWIGAQECT WP_000145410.1 MULTISPECIES: hydrogenase-2 small chain [Proteobacteria] Length: 372\nNumber of predicted TMHs: 1\nExp number of AAs in TMHs: 23.72226\nExp number, first 60 AAs: 0.22038\nTotal prob of N-in: 0.12994\noutside 1 331\nTMhelix 332 354\ninside 355 372

3894 GCF_001901445.1_ASM190144v1 Escherichia coli Proteobacteria; Gammaproteobacteria; Enterobacterales; Enterobacteriaceae; Escherichia MTGDNTLIHSHGINRRDFMKLCAALAATMGLSSKAAAEMAESVTNPQRPPVIWIGAQECT WP_000145410.1 MULTISPECIES: hydrogenase-2 small chain [Proteobacteria] Length: 372\nNumber of predicted TMHs:

1\nExp number of AAs in TMHs: 23.72226\nExp number, first 60 AAs: 0.22038\nTotal prob of N-in: 0.12994\noutside 1 331\nTMhelix 332 354\ninside 355 372

3895 GCF_001901545.1_ASM190154v1 Escherichia coli Proteobacteria; Gammaproteobacteria; Enterobacterales; Enterobacteriaceae; Escherichia
MTGDNTLIHSHGINRRDFMKLCAALAATMGLSSKAAAEMAESVTNPQRPPVIWIGAQECT WP_000145410.1
MULTISPECIES: hydrogenase-2 small chain [Proteobacteria] Length: 372\nNumber of predicted TMHs: 1\nExp number of AAs in TMHs: 23.72226\nExp number, first 60 AAs: 0.22038\nTotal prob of N-in: 0.12994\noutside 1 331\nTMhelix 332 354\ninside 355 372

3896 GCF_001907395.1_ASM190739v1 Escherichia coli Proteobacteria; Gammaproteobacteria; Enterobacterales; Enterobacteriaceae; Escherichia
MNNEETFYQAMRRQGVTRRSFLKYCSLAATSLGLGAGMAPKIAWALENKPRIPVVWIHGL WP_001058323.1
MULTISPECIES: hydrogenase-1 small chain [Proteobacteria] Length: 372\nNumber of predicted TMHs: 2\nExp number of AAs in TMHs: 37.2608499999999\nExp number, first 60 AAs: 19.52923\nTotal prob of N-in: 0.94351\nPOSSIBLE N-term signal sequence\ninside 1 19\nTMhelix 20 39\noutside 40 325\nTMhelix 326 348\ninside 349 372

3897 GCF_001907505.1_ASM190750v1 Escherichia coli Proteobacteria; Gammaproteobacteria; Enterobacterales; Enterobacteriaceae; Escherichia
MTGDNTLIHSHGINRRDFMKLCAALAATMGLSSKAAAEMAESVTNPQRPPVIWIGAQECT WP_000145410.1
MULTISPECIES: hydrogenase-2 small chain [Proteobacteria] Length: 372\nNumber of predicted TMHs: 1\nExp number of AAs in TMHs: 23.72226\nExp number, first 60 AAs: 0.22038\nTotal prob of N-in: 0.12994\noutside 1 331\nTMhelix 332 354\ninside 355 372

3898 GCF_001902685.1_ASM190268v1 Escherichia coli Proteobacteria; Gammaproteobacteria; Enterobacterales; Enterobacteriaceae; Escherichia
MNNEETFYQAMRRQGVTRRSFLKYCSLAATSLGLGAGMAPKIAWALENKPRIPVVWIHGL WP_001058323.1
MULTISPECIES: hydrogenase-1 small chain [Proteobacteria] Length: 372\nNumber of predicted TMHs: 2\nExp number of AAs in TMHs: 37.2608499999999\nExp number, first 60 AAs: 19.52923\nTotal prob of N-in: 0.94351\nPOSSIBLE N-term signal sequence\ninside 1 19\nTMhelix 20 39\noutside 40 325\nTMhelix 326 348\ninside 349 372

3899 GCF_001907375.1_ASM190737v1 Escherichia coli Proteobacteria; Gammaproteobacteria; Enterobacterales; Enterobacteriaceae; Escherichia
MNNEETFYQAMRRQGVTRRSFLKYCSLAATSLGLGAGMAPKIAWALENKPRIPVVWIHGL WP_001058323.1
MULTISPECIES: hydrogenase-1 small chain [Proteobacteria] Length: 372\nNumber of predicted TMHs: 2\nExp number of AAs in TMHs: 37.2608499999999\nExp number, first 60 AAs: 19.52923\nTotal prob of N-in: 0.94351\nPOSSIBLE N-term signal sequence\ninside 1 19\nTMhelix 20 39\noutside 40 325\nTMhelix 326 348\ninside 349 372

3900 GCF_001907315.1_ASM190731v1 Escherichia coli Proteobacteria; Gammaproteobacteria; Enterobacterales; Enterobacteriaceae; Escherichia
MNNEETFYQAMRRQGVTRRSFLKYCSLAATSLGLGAGMAPKIAWALENKPRIPVVWIHGL WP_001058323.1
MULTISPECIES: hydrogenase-1 small chain [Proteobacteria] Length: 372\nNumber of predicted TMHs: 2\nExp number of AAs in TMHs: 37.2608499999999\nExp number, first 60 AAs: 19.52923\nTotal prob of N-in: 0.94351\nPOSSIBLE N-term signal sequence\ninside 1 19\nTMhelix 20 39\noutside 40 325\nTMhelix 326 348\ninside 349 372

3901 GCF_001907465.1_ASM190746v1 Escherichia coli Proteobacteria; Gammaproteobacteria; Enterobacterales; Enterobacteriaceae; Escherichia
MNNEETFYQAMRRQGVTRRSFLKYCSLAATSLGLGAGMAPKIAWALENKPRIPVVWIHGL WP_001058323.1
MULTISPECIES: hydrogenase-1 small chain [Proteobacteria] Length: 372\nNumber of predicted TMHs: 2\nExp number of AAs in TMHs: 37.2608499999999\nExp number, first 60 AAs: 19.52923\nTotal prob of N-in: 0.94351\nPOSSIBLE N-term signal sequence\ninside 1 19\nTMhelix 20 39\noutside 40 325\nTMhelix 326 348\ninside 349 372

3902 GCF_001907565.1_ASM190756v1 Escherichia coli Proteobacteria; Gammaproteobacteria; Enterobacterales; Enterobacteriaceae; Escherichia
MNNEETFYQAMRRQGVTRRSFLKYCSLAATSLGLGAGMAPKIAWALENKPRIPVVWIHGL WP_001058323.1
MULTISPECIES: hydrogenase-1 small chain [Proteobacteria] Length: 372\nNumber of predicted TMHs:

2\nExp number of AAs in TMHs: 37.2608499999999\nExp number, first 60 AAs: 19.52923\nTotal prob of N-in: 0.94351\nPOSSIBLE N-term signal sequence\ninside 1 19\nTMhelix 20 39\noutside 40 325\nTMhelix 326 348\ninside 349 372

3903 GCF_001910925.1_ASM191092v1 Escherichia coli Proteobacteria; Gammaproteobacteria; Enterobacterales; Enterobacteriaceae; Escherichia
MTGDNTLIHSHGINRRDFMKLCAALAATMGLSSKAAAEMAESVTNPQRPPVIWIGAQECT WP_000145410.1
MULTISPECIES: hydrogenase-2 small chain [Proteobacteria] Length: 372\nNumber of predicted TMHs: 1\nExp number of AAs in TMHs: 23.72226\nExp number, first 60 AAs: 0.22038\nTotal prob of N-in: 0.12994\noutside 1 331\nTMhelix 332 354\ninside 355 372

3904 GCF_001910835.1_ASM191083v1 Escherichia coli Proteobacteria; Gammaproteobacteria; Enterobacterales; Enterobacteriaceae; Escherichia
MNNEETFYQAMRRQGVTRRSFLKYCSLAATSLGLGAGMAPKIAWALENKPRIPVVWIHGL WP_001058323.1
MULTISPECIES: hydrogenase-1 small chain [Proteobacteria] Length: 372\nNumber of predicted TMHs: 2\nExp number of AAs in TMHs: 37.2608499999999\nExp number, first 60 AAs: 19.52923\nTotal prob of N-in: 0.94351\nPOSSIBLE N-term signal sequence\ninside 1 19\nTMhelix 20 39\noutside 40 325\nTMhelix 326 348\ninside 349 372

3905 GCF_001910985.1_ASM191098v1 Escherichia coli Proteobacteria; Gammaproteobacteria; Enterobacterales; Enterobacteriaceae; Escherichia
MNNEETFYQAMRRQGVTRRSFLKYCSLAATSLGLGAGMAPKIAWALENKPRIPVVWIHGL WP_001058323.1
MULTISPECIES: hydrogenase-1 small chain [Proteobacteria] Length: 372\nNumber of predicted TMHs: 2\nExp number of AAs in TMHs: 37.2608499999999\nExp number, first 60 AAs: 19.52923\nTotal prob of N-in: 0.94351\nPOSSIBLE N-term signal sequence\ninside 1 19\nTMhelix 20 39\noutside 40 325\nTMhelix 326 348\ninside 349 372

3906 GCF_001910925.1_ASM191092v1 Escherichia coli Proteobacteria; Gammaproteobacteria; Enterobacterales; Enterobacteriaceae; Escherichia
MNNEETFYQAMRRQGVTRRSFLKYCSLAATSLGLGAGMAPKIAWALENKPRIPVVWIHGL WP_001058323.1
MULTISPECIES: hydrogenase-1 small chain [Proteobacteria] Length: 372\nNumber of predicted TMHs: 2\nExp number of AAs in TMHs: 37.2608499999999\nExp number, first 60 AAs: 19.52923\nTotal prob of N-in: 0.94351\nPOSSIBLE N-term signal sequence\ninside 1 19\nTMhelix 20 39\noutside 40 325\nTMhelix 326 348\ninside 349 372

3907 GCF_001910855.1_ASM191085v1 Escherichia coli Proteobacteria; Gammaproteobacteria; Enterobacterales; Enterobacteriaceae; Escherichia
MNNEETFYQAMRRQGVTRRSFLKYCSLAATSLGLGAGMAPKIAWALENKPRIPVVWIHGL WP_001058323.1
MULTISPECIES: hydrogenase-1 small chain [Proteobacteria] Length: 372\nNumber of predicted TMHs: 2\nExp number of AAs in TMHs: 37.2608499999999\nExp number, first 60 AAs: 19.52923\nTotal prob of N-in: 0.94351\nPOSSIBLE N-term signal sequence\ninside 1 19\nTMhelix 20 39\noutside 40 325\nTMhelix 326 348\ninside 349 372

3908 GCF_001911015.1_ASM191101v1 Escherichia coli Proteobacteria; Gammaproteobacteria; Enterobacterales; Enterobacteriaceae; Escherichia
MTGDNTLIHSHGINRRDFMKLCAALAATMGLSSKAAAEMAESVTNPQRPPVIWIGAQECT WP_000145410.1
MULTISPECIES: hydrogenase-2 small chain [Proteobacteria] Length: 372\nNumber of predicted TMHs: 1\nExp number of AAs in TMHs: 23.72226\nExp number, first 60 AAs: 0.22038\nTotal prob of N-in: 0.12994\noutside 1 331\nTMhelix 332 354\ninside 355 372

3909 GCF_001911075.1_ASM191107v1 Escherichia coli Proteobacteria; Gammaproteobacteria; Enterobacterales; Enterobacteriaceae; Escherichia
MTGDNTLIHSHGINRRDFMKLCAALAATMGLSSKAAAEMAESVTNPQRPPVIWIGAQECT WP_000145410.1
MULTISPECIES: hydrogenase-2 small chain [Proteobacteria] Length: 372\nNumber of predicted TMHs: 1\nExp number of AAs in TMHs: 23.72226\nExp number, first 60 AAs: 0.22038\nTotal prob of N-in: 0.12994\noutside 1 331\nTMhelix 332 354\ninside 355 372

3910 GCF_001911075.1_ASM191107v1 Escherichia coli Proteobacteria; Gammaproteobacteria; Enterobacterales; Enterobacteriaceae; Escherichia
MNNEETFYQAMRRQGVTRRSFLKYCSLAATSLGLGAGMAPKIAWALENKPRIPVVWIHGL WP_001058323.1
MULTISPECIES: hydrogenase-1 small chain [Proteobacteria] Length: 372\nNumber of predicted TMHs:

2\nExp number of AAs in TMHs: 37.2608499999999\nExp number, first 60 AAs: 19.52923\nTotal prob of N-in: 0.94351\nPOSSIBLE N-term signal sequence\ninside 1 19\nTMhelix 20 39\noutside 40 325\nTMhelix 326 348\ninside 349 372

3911 GCF_001911015.1_ASM191101v1 Escherichia coli Proteobacteria; Gammaproteobacteria; Enterobacterales; Enterobacteriaceae; Escherichia MNNEETFYQAMRRQGVTRRSFLKYCSLAATSLGLGAGMAPKIAWALENKPRIPVVWIHGL WP_001058323.1 MULTISPECIES: hydrogenase-1 small chain [Proteobacteria] Length: 372\nNumber of predicted TMHs: 2\nExp number of AAs in TMHs: 37.2608499999999\nExp number, first 60 AAs: 19.52923\nTotal prob of N-in: 0.94351\nPOSSIBLE N-term signal sequence\ninside 1 19\nTMhelix 20 39\noutside 40 325\nTMhelix 326 348\ninside 349 372

3912 GCF_001910985.1_ASM191098v1 Escherichia coli Proteobacteria; Gammaproteobacteria; Enterobacterales; Enterobacteriaceae; Escherichia MTGDNTLIHSHGINRRDFMKLCAALAATMGLSSKAAAEMAESVTNPQRPPVIWIGAQECT WP_001685818.1 MULTISPECIES: hydrogenase-2 small chain [Enterobacteriaceae] Length: 372\nNumber of predicted TMHs: 1\nExp number of AAs in TMHs: 23.82505\nExp number, first 60 AAs: 0.21974\nTotal prob of N-in: 0.13379\noutside 1 331\nTMhelix 332 354\ninside 355 372

3913 GCF_001911165.1_ASM191116v1 Escherichia coli Proteobacteria; Gammaproteobacteria; Enterobacterales; Enterobacteriaceae; Escherichia MTGDNTLIHSHGINRRDFMKLCAALAATMGLSSKAAAEMAESVTNPQRPPVIWIGAQECT WP_000145410.1 MULTISPECIES: hydrogenase-2 small chain [Proteobacteria] Length: 372\nNumber of predicted TMHs: 1\nExp number of AAs in TMHs: 23.72226\nExp number, first 60 AAs: 0.22038\nTotal prob of N-in: 0.12994\noutside 1 331\nTMhelix 332 354\ninside 355 372

3914 GCF_001911145.1_ASM191114v1 Escherichia coli Proteobacteria; Gammaproteobacteria; Enterobacterales; Enterobacteriaceae; Escherichia MTGDNTLIHSHGINRRDFMKLCAALAATMGLSSKAAAEMAESVTNPQRPPVIWIGAQECT WP_000145410.1 MULTISPECIES: hydrogenase-2 small chain [Proteobacteria] Length: 372\nNumber of predicted TMHs: 1\nExp number of AAs in TMHs: 23.72226\nExp number, first 60 AAs: 0.22038\nTotal prob of N-in: 0.12994\noutside 1 331\nTMhelix 332 354\ninside 355 372

3915 GCF_001911235.1_ASM191123v1 Escherichia coli Proteobacteria; Gammaproteobacteria; Enterobacterales; Enterobacteriaceae; Escherichia MTGDNTLIHSHGINRRDFMKLCAALAATMGLSSKAAAEMAESVTNPQRPPVIWIGAQECT WP_000145410.1 MULTISPECIES: hydrogenase-2 small chain [Proteobacteria] Length: 372\nNumber of predicted TMHs: 1\nExp number of AAs in TMHs: 23.72226\nExp number, first 60 AAs: 0.22038\nTotal prob of N-in: 0.12994\noutside 1 331\nTMhelix 332 354\ninside 355 372

3916 GCF_001911085.1_ASM191108v1 Escherichia coli Proteobacteria; Gammaproteobacteria; Enterobacterales; Enterobacteriaceae; Escherichia MNNEETFYQAMRRQGVTRRSFLKYCSLAATSLGLGAGMAPKIAWALENKPRIPVVWIHGL WP_001058323.1 MULTISPECIES: hydrogenase-1 small chain [Proteobacteria] Length: 372\nNumber of predicted TMHs: 2\nExp number of AAs in TMHs: 37.2608499999999\nExp number, first 60 AAs: 19.52923\nTotal prob of N-in: 0.94351\nPOSSIBLE N-term signal sequence\ninside 1 19\nTMhelix 20 39\noutside 40 325\nTMhelix 326 348\ninside 349 372

3917 GCF_001911505.1_ASM191150v1 Escherichia coli Proteobacteria; Gammaproteobacteria; Enterobacterales; Enterobacteriaceae; Escherichia MNNEETFYQAMRRQGVTRRSFLKYCSLAATSLGLGAGMAPKIAWALENKPRIPVVWIHGL WP_063501981.1 hydrogenase [Escherichia coli] Length: 372\nNumber of predicted TMHs: 2\nExp number of AAs in TMHs: 37.06322\nExp number, first 60 AAs: 19.52932\nTotal prob of N-in: 0.94340\nPOSSIBLE N-term signal sequence\ninside 1 19\nTMhelix 20 39\noutside 40 325\nTMhelix 326 348\ninside 349 372

3918 GCF_001911145.1_ASM191114v1 Escherichia coli Proteobacteria; Gammaproteobacteria; Enterobacterales; Enterobacteriaceae; Escherichia MNNEETFYQAMRRQGVTRRSFLKYCSLAATSLGLGAGMAPKIAWALENKPRIPVVWIHGL WP_074501023.1 hydrogenase [Escherichia coli] Length: 372\nNumber of predicted TMHs: 2\nExp number of AAs in TMHs: 37.2643699999999\nExp number, first 60 AAs: 19.52926\nTotal prob of N-in: 0.94351\nPOSSIBLE N-term signal sequence\ninside 1 19\nTMhelix 20 39\noutside 40 325\nTMhelix 326 348\ninside 349 372

3919 GCF_001911615.1_ASM191161v1 Escherichia coli Proteobacteria; Gammaproteobacteria; Enterobacterales; Enterobacteriaceae; Escherichia
MTGDNTLIHSHGINRRDFMKLCAALAATMGLSSKAAAEMAESVTNPQRPPVIWIGAQECT WP_000145410.1
MULTISPECIES: hydrogenase-2 small chain [Proteobacteria] Length: 372\nNumber of predicted TMHs: 1\nExp number of AAs in TMHs: 23.72226\nExp number, first 60 AAs: 0.22038\nTotal prob of N-in: 0.12994\noutside 1 331\nTMhelix 332 354\ninside 355 372

3920 GCF_001911675.1_ASM191167v1 Escherichia coli Proteobacteria; Gammaproteobacteria; Enterobacterales; Enterobacteriaceae; Escherichia
MTGDNTLIHSHGINRRDFMKLCAALAATMGLSSKAAAEMAESVTNPQRPPVIWIGAQECT WP_000145410.1
MULTISPECIES: hydrogenase-2 small chain [Proteobacteria] Length: 372\nNumber of predicted TMHs: 1\nExp number of AAs in TMHs: 23.72226\nExp number, first 60 AAs: 0.22038\nTotal prob of N-in: 0.12994\noutside 1 331\nTMhelix 332 354\ninside 355 372

3921 GCF_001911725.1_ASM191172v1 Escherichia coli Proteobacteria; Gammaproteobacteria; Enterobacterales; Enterobacteriaceae; Escherichia
MTGDNTLIHSHGINRRDFMKLCAALAATMGLSSKAAAEMAESVTNPQRPPVIWIGAQECT WP_000145410.1
MULTISPECIES: hydrogenase-2 small chain [Proteobacteria] Length: 372\nNumber of predicted TMHs: 1\nExp number of AAs in TMHs: 23.72226\nExp number, first 60 AAs: 0.22038\nTotal prob of N-in: 0.12994\noutside 1 331\nTMhelix 332 354\ninside 355 372

3922 GCF_001911775.1_ASM191177v1 Escherichia coli Proteobacteria; Gammaproteobacteria; Enterobacterales; Enterobacteriaceae; Escherichia
MTGDNTLIHSHGINRRDFMKLCAALAATMGLSSKAAAEMAESVTNPQRPPVIWIGAQECT WP_000145410.1
MULTISPECIES: hydrogenase-2 small chain [Proteobacteria] Length: 372\nNumber of predicted TMHs: 1\nExp number of AAs in TMHs: 23.72226\nExp number, first 60 AAs: 0.22038\nTotal prob of N-in: 0.12994\noutside 1 331\nTMhelix 332 354\ninside 355 372

3923 GCF_001911615.1_ASM191161v1 Escherichia coli Proteobacteria; Gammaproteobacteria; Enterobacterales; Enterobacteriaceae; Escherichia
MNNEETFYQAMRRQGVTRRSFLKYCSLAATSLGLGAGMAPKIAWALENKPRIPVVWIHGL WP_001058323.1
MULTISPECIES: hydrogenase-1 small chain [Proteobacteria] Length: 372\nNumber of predicted TMHs: 2\nExp number of AAs in TMHs: 37.2608499999999\nExp number, first 60 AAs: 19.52923\nTotal prob of N-in: 0.94351\nPOSSIBLE N-term signal sequence\ninside 1 19\nTMhelix 20 39\noutside 40 325\nTMhelix 326 348\ninside 349 372

3924 GCF_001911865.1_ASM191186v1 Escherichia coli Proteobacteria; Gammaproteobacteria; Enterobacterales; Enterobacteriaceae; Escherichia
MTGDNTLIHSHGINRRDFMKLCAALAATMGLSSKAAAEMAESVTNPQRPPVIWIGAQECT WP_000145410.1
MULTISPECIES: hydrogenase-2 small chain [Proteobacteria] Length: 372\nNumber of predicted TMHs: 1\nExp number of AAs in TMHs: 23.72226\nExp number, first 60 AAs: 0.22038\nTotal prob of N-in: 0.12994\noutside 1 331\nTMhelix 332 354\ninside 355 372

3925 GCF_001911825.1_ASM191182v1 Escherichia coli Proteobacteria; Gammaproteobacteria; Enterobacterales; Enterobacteriaceae; Escherichia
MNNEETFYQAMRRQGVTRRSFLKYCSLAATSLGLGAGMAPKIAWALENKPRIPVVWIHGL WP_001058323.1
MULTISPECIES: hydrogenase-1 small chain [Proteobacteria] Length: 372\nNumber of predicted TMHs: 2\nExp number of AAs in TMHs: 37.2608499999999\nExp number, first 60 AAs: 19.52923\nTotal prob of N-in: 0.94351\nPOSSIBLE N-term signal sequence\ninside 1 19\nTMhelix 20 39\noutside 40 325\nTMhelix 326 348\ninside 349 372

3926 GCF_001911925.1_ASM191192v1 Escherichia coli Proteobacteria; Gammaproteobacteria; Enterobacterales; Enterobacteriaceae; Escherichia
MTGDNTLIHSHGINRRDFMKLCAALAATMGLSSKAAAEMAESVTNPQRPPVIWIGAQECT WP_000145410.1
MULTISPECIES: hydrogenase-2 small chain [Proteobacteria] Length: 372\nNumber of predicted TMHs: 1\nExp number of AAs in TMHs: 23.72226\nExp number, first 60 AAs: 0.22038\nTotal prob of N-in: 0.12994\noutside 1 331\nTMhelix 332 354\ninside 355 372

3927 GCF_001911935.1_ASM191193v1 Escherichia coli Proteobacteria; Gammaproteobacteria; Enterobacterales; Enterobacteriaceae; Escherichia
MTGDNTLIHSHGINRRDFMKLCAALAATMGLSSKAAAEMAESVTNPQRPPVIWIGAQECT WP_000145410.1

MULTISPECIES: hydrogenase-2 small chain [Proteobacteria] Length: 372\nNumber of predicted TMHs: 1\nExp number of AAs in TMHs: 23.72226\nExp number, first 60 AAs: 0.22038\nTotal prob of N-in: 0.12994\noutside 1 331\nTMhelix 332 354\ninside 355 372

3928 GCF_001911905.1_ASM191190v1 Escherichia coli Proteobacteria; Gammaproteobacteria; Enterobacterales; Enterobacteriaceae; Escherichia
MNNEETFYQAMRRQGVTRRSFLKYCSLAATSLGLGAGMAPKIAWALENKPRIPVVWIHGL WP_001058323.1
MULTISPECIES: hydrogenase-1 small chain [Proteobacteria] Length: 372\nNumber of predicted TMHs: 2\nExp number of AAs in TMHs: 37.2608499999999\nExp number, first 60 AAs: 19.52923\nTotal prob of N-in: 0.94351\nPOSSIBLE N-term signal sequence\ninside 1 19\nTMhelix 20 39\noutside 40 325\nTMhelix 326 348\ninside 349 372

3929 GCF_001911865.1_ASM191186v1 Escherichia coli Proteobacteria; Gammaproteobacteria; Enterobacterales; Enterobacteriaceae; Escherichia
MNNEETFYQAMRRQGVTRRSFLKYCSLAATSLGLGAGMAPKIAWALENKPRIPVVWIHGL WP_001058323.1
MULTISPECIES: hydrogenase-1 small chain [Proteobacteria] Length: 372\nNumber of predicted TMHs: 2\nExp number of AAs in TMHs: 37.2608499999999\nExp number, first 60 AAs: 19.52923\nTotal prob of N-in: 0.94351\nPOSSIBLE N-term signal sequence\ninside 1 19\nTMhelix 20 39\noutside 40 325\nTMhelix 326 348\ninside 349 372

3930 GCF_001911845.1_ASM191184v1 Escherichia coli Proteobacteria; Gammaproteobacteria; Enterobacterales; Enterobacteriaceae; Escherichia
MNNEETFYQAMRRQGVTRRSFLKYCSLAATSLGLGAGMAPKIAWALENKPRIPVVWIHGL WP_001295940.1
MULTISPECIES: hydrogenase [Enterobacteriaceae] Length: 372\nNumber of predicted TMHs: 2\nExp number of AAs in TMHs: 37.2608899999999\nExp number, first 60 AAs: 19.52923\nTotal prob of N-in: 0.94351\nPOSSIBLE N-term signal sequence\ninside 1 19\nTMhelix 20 39\noutside 40 325\nTMhelix 326 348\ninside 349 372

3931 GCF_001912035.1_ASM191203v1 Escherichia coli Proteobacteria; Gammaproteobacteria; Enterobacterales; Enterobacteriaceae; Escherichia
MTGDNTLIHSHGINRRDFMKLCAALAATMGLSSKAAAEMAESVTNPQRPPVIWIGAQECT WP_000145410.1
MULTISPECIES: hydrogenase-2 small chain [Proteobacteria] Length: 372\nNumber of predicted TMHs: 1\nExp number of AAs in TMHs: 23.72226\nExp number, first 60 AAs: 0.22038\nTotal prob of N-in: 0.12994\noutside 1 331\nTMhelix 332 354\ninside 355 372

3932 GCF_001912055.1_ASM191205v1 Escherichia coli Proteobacteria; Gammaproteobacteria; Enterobacterales; Enterobacteriaceae; Escherichia
MTGDNTLIHSHGINRRDFMKLCAALAATMGLSSKAAAEMAESVTNPQRPPVIWIGAQECT WP_000145410.1
MULTISPECIES: hydrogenase-2 small chain [Proteobacteria] Length: 372\nNumber of predicted TMHs: 1\nExp number of AAs in TMHs: 23.72226\nExp number, first 60 AAs: 0.22038\nTotal prob of N-in: 0.12994\noutside 1 331\nTMhelix 332 354\ninside 355 372

3933 GCF_001912055.1_ASM191205v1 Escherichia coli Proteobacteria; Gammaproteobacteria; Enterobacterales; Enterobacteriaceae; Escherichia
MNNEETFYQAMRRQGVTRRSFLKYCSLAATSLGLGAGMAPKIAWALENKPRIPVVWIHGL WP_001058323.1
MULTISPECIES: hydrogenase-1 small chain [Proteobacteria] Length: 372\nNumber of predicted TMHs: 2\nExp number of AAs in TMHs: 37.2608499999999\nExp number, first 60 AAs: 19.52923\nTotal prob of N-in: 0.94351\nPOSSIBLE N-term signal sequence\ninside 1 19\nTMhelix 20 39\noutside 40 325\nTMhelix 326 348\ninside 349 372

3934 GCF_001912105.1_ASM191210v1 Escherichia coli Proteobacteria; Gammaproteobacteria; Enterobacterales; Enterobacteriaceae; Escherichia
MTGDNTLIHSHGINRRDFMKLCAALAATMGLSSKAAAEMAESVTNPQRPPVIWIGAQECT WP_000145410.1
MULTISPECIES: hydrogenase-2 small chain [Proteobacteria] Length: 372\nNumber of predicted TMHs: 1\nExp number of AAs in TMHs: 23.72226\nExp number, first 60 AAs: 0.22038\nTotal prob of N-in: 0.12994\noutside 1 331\nTMhelix 332 354\ninside 355 372

3935 GCF_001912085.1_ASM191208v1 Escherichia coli Proteobacteria; Gammaproteobacteria; Enterobacterales; Enterobacteriaceae; Escherichia
MTGDNTLIHSHGINRRDFMKLCAALAATMGLSSKAAAEMAESVTNPQRPPVIWIGAQECT WP_000145410.1
MULTISPECIES: hydrogenase-2 small chain [Proteobacteria] Length: 372\nNumber of predicted TMHs:

1\nExp number of AAs in TMHs: 23.72226\nExp number, first 60 AAs: 0.22038\nTotal prob of N-in: 0.12994\noutside 1 331\nTMhelix 332 354\ninside 355 372

3936 GCF_001912115.1_ASM191211v1 Escherichia coli Proteobacteria; Gammaproteobacteria; Enterobacterales; Enterobacteriaceae; Escherichia
MTGDNTLIHSHGINRRDFMKLCAALAATMGLSSKAAAEMAESVTNPQRPPVIWIGAQECT WP_000145410.1
MULTISPECIES: hydrogenase-2 small chain [Proteobacteria] Length: 372\nNumber of predicted TMHs: 1\nExp number of AAs in TMHs: 23.72226\nExp number, first 60 AAs: 0.22038\nTotal prob of N-in: 0.12994\noutside 1 331\nTMhelix 332 354\ninside 355 372

3937 GCF_001912035.1_ASM191203v1 Escherichia coli Proteobacteria; Gammaproteobacteria; Enterobacterales; Enterobacteriaceae; Escherichia
MNNEETFYQAMRRQGVTRRSFLKYCSLAATSLGLGAGMAPKIAWALENKPRIPVVWIHGL WP_001058323.1
MULTISPECIES: hydrogenase-1 small chain [Proteobacteria] Length: 372\nNumber of predicted TMHs: 2\nExp number of AAs in TMHs: 37.2608499999999\nExp number, first 60 AAs: 19.52923\nTotal prob of N-in: 0.94351\nPOSSIBLE N-term signal sequence\ninside 1 19\nTMhelix 20 39\noutside 40 325\nTMhelix 326 348\ninside 349 372

3938 GCF_001912115.1_ASM191211v1 Escherichia coli Proteobacteria; Gammaproteobacteria; Enterobacterales; Enterobacteriaceae; Escherichia
MNNEETFYQAMRRQGVTRRSFLKYCSLAATSLGLGAGMAPKIAWALENKPRIPVVWIHGL WP_001058323.1
MULTISPECIES: hydrogenase-1 small chain [Proteobacteria] Length: 372\nNumber of predicted TMHs: 2\nExp number of AAs in TMHs: 37.2608499999999\nExp number, first 60 AAs: 19.52923\nTotal prob of N-in: 0.94351\nPOSSIBLE N-term signal sequence\ninside 1 19\nTMhelix 20 39\noutside 40 325\nTMhelix 326 348\ninside 349 372

3939 GCF_001912105.1_ASM191210v1 Escherichia coli Proteobacteria; Gammaproteobacteria; Enterobacterales; Enterobacteriaceae; Escherichia
MNNEETFYQAMRRQGVTRRSFLKYCSLAATSLGLGAGMAPKIAWALENKPRIPVVWIHGL WP_001058323.1
MULTISPECIES: hydrogenase-1 small chain [Proteobacteria] Length: 372\nNumber of predicted TMHs: 2\nExp number of AAs in TMHs: 37.2608499999999\nExp number, first 60 AAs: 19.52923\nTotal prob of N-in: 0.94351\nPOSSIBLE N-term signal sequence\ninside 1 19\nTMhelix 20 39\noutside 40 325\nTMhelix 326 348\ninside 349 372

3940 GCF_001912005.1_ASM191200v1 Escherichia coli Proteobacteria; Gammaproteobacteria; Enterobacterales; Enterobacteriaceae; Escherichia
MTGDNTLIHSHGINRRDFMKLCAALAATMGLSSKAAAEMAESVTNPQRPPVIWIGAQECT WP_074534817.1
hydrogenase 2 small subunit [Escherichia coli] Length: 372\nNumber of predicted TMHs: 1\nExp number of AAs in TMHs: 23.7182\nExp number, first 60 AAs: 0.22037\nTotal prob of N-in: 0.12983\noutside 1 331\nTMhelix 332 354\ninside 355 372

3941 GCF_001912225.1_ASM191222v1 Escherichia coli Proteobacteria; Gammaproteobacteria; Enterobacterales; Enterobacteriaceae; Escherichia
MTGDNTLIHSHGINRRDFMKLCAALAATMGLSSKAAAEMAESVTNPQRPPVIWIGAQECT WP_000145410.1
MULTISPECIES: hydrogenase-2 small chain [Proteobacteria] Length: 372\nNumber of predicted TMHs: 1\nExp number of AAs in TMHs: 23.72226\nExp number, first 60 AAs: 0.22038\nTotal prob of N-in: 0.12994\noutside 1 331\nTMhelix 332 354\ninside 355 372

3942 GCF_001912235.1_ASM191223v1 Escherichia coli Proteobacteria; Gammaproteobacteria; Enterobacterales; Enterobacteriaceae; Escherichia
MTGDNTLIHSHGINRRDFMKLCAALAATMGLSSKAAAEMAESVTNPQRPPVIWIGAQECT WP_000145410.1
MULTISPECIES: hydrogenase-2 small chain [Proteobacteria] Length: 372\nNumber of predicted TMHs: 1\nExp number of AAs in TMHs: 23.72226\nExp number, first 60 AAs: 0.22038\nTotal prob of N-in: 0.12994\noutside 1 331\nTMhelix 332 354\ninside 355 372

3943 GCF_001912225.1_ASM191222v1 Escherichia coli Proteobacteria; Gammaproteobacteria; Enterobacterales; Enterobacteriaceae; Escherichia
MNNEETFYQAMRRQGVTRRSFLKYCSLAATSLGLGAGMAPKIAWALENKPRIPVVWIHGL WP_001058323.1
MULTISPECIES: hydrogenase-1 small chain [Proteobacteria] Length: 372\nNumber of predicted TMHs: 2\nExp number of AAs in TMHs: 37.2608499999999\nExp number, first 60 AAs: 19.52923\nTotal prob of N-in:

0.94351\nPOSSIBLE N-term signal sequence\ninside 1 19\nTMhelix 20 39\noutside 40 325\nTMhelix 326 348\ninside 349 372

3944 GCF_001912235.1_ASM191223v1 Escherichia coli Proteobacteria; Gammaproteobacteria; Enterobacterales; Enterobacteriaceae; Escherichia MNNEETFYQAMRRQGVTRRSFLKYCSLAATSLGLGAGMAPKIAWALENKPRIPVVWIHGL WP_001353311.1 MULTISPECIES: hydrogenase [Escherichia] Length: 372\nNumber of predicted TMHs: 2\nExp number of AAs in TMHs: 37.2638699999999\nExp number, first 60 AAs: 19.52924\nTotal prob of N-in: 0.94351\nPOSSIBLE N-term signal sequence\ninside 1 19\nTMhelix 20 39\noutside 40 325\nTMhelix 326 348\ninside 349 372

3945 GCF_001912345.1_ASM191234v1 Escherichia coli Proteobacteria; Gammaproteobacteria; Enterobacterales; Enterobacteriaceae; Escherichia MTGDNTLIHSHGINRRDFMKLCAALAATMGLSSKAAAEMAESVTNPQRPPVIWIGAQECT WP_000145410.1 MULTISPECIES: hydrogenase-2 small chain [Proteobacteria] Length: 372\nNumber of predicted TMHs: 1\nExp number of AAs in TMHs: 23.72226\nExp number, first 60 AAs: 0.22038\nTotal prob of N-in: 0.12994\noutside 1 331\nTMhelix 332 354\ninside 355 372

3946 GCF_001912275.1_ASM191227v1 Escherichia coli Proteobacteria; Gammaproteobacteria; Enterobacterales; Enterobacteriaceae; Escherichia MNNEETFYQAMRRQGVTRRSFLKYCSLAATSLGLGAGMAPKIAWALENKPRIPVVWIHGL WP_001058323.1 MULTISPECIES: hydrogenase-1 small chain [Proteobacteria] Length: 372\nNumber of predicted TMHs: 2\nExp number of AAs in TMHs: 37.2608499999999\nExp number, first 60 AAs: 19.52923\nTotal prob of N-in: 0.94351\nPOSSIBLE N-term signal sequence\ninside 1 19\nTMhelix 20 39\noutside 40 325\nTMhelix 326 348\ninside 349 372

3947 GCF_001912365.1_ASM191236v1 Escherichia coli Proteobacteria; Gammaproteobacteria; Enterobacterales; Enterobacteriaceae; Escherichia MTGDNTLIHSHGINRRDFMKLCAALAATMGLSSKAAAEMAESVTNPQRPPVIWIGAQECT WP_000145410.1 MULTISPECIES: hydrogenase-2 small chain [Proteobacteria] Length: 372\nNumber of predicted TMHs: 1\nExp number of AAs in TMHs: 23.72226\nExp number, first 60 AAs: 0.22038\nTotal prob of N-in: 0.12994\noutside 1 331\nTMhelix 332 354\ninside 355 372

3948 GCF_001912375.1_ASM191237v1 Escherichia coli Proteobacteria; Gammaproteobacteria; Enterobacterales; Enterobacteriaceae; Escherichia MTGDNTLIHSHGINRRDFMKLCAALAATMGLSSKAAAEMAESVTNPQRPPVIWIGAQECT WP_000145410.1 MULTISPECIES: hydrogenase-2 small chain [Proteobacteria] Length: 372\nNumber of predicted TMHs: 1\nExp number of AAs in TMHs: 23.72226\nExp number, first 60 AAs: 0.22038\nTotal prob of N-in: 0.12994\noutside 1 331\nTMhelix 332 354\ninside 355 372

3949 GCF_001912375.1_ASM191237v1 Escherichia coli Proteobacteria; Gammaproteobacteria; Enterobacterales; Enterobacteriaceae; Escherichia MNNEETFYQAMRRQGVTRRSFLKYCSLAATSLGLGAGMAPKIAWALENKPRIPVVWIHGL WP_001058323.1 MULTISPECIES: hydrogenase-1 small chain [Proteobacteria] Length: 372\nNumber of predicted TMHs: 2\nExp number of AAs in TMHs: 37.2608499999999\nExp number, first 60 AAs: 19.52923\nTotal prob of N-in: 0.94351\nPOSSIBLE N-term signal sequence\ninside 1 19\nTMhelix 20 39\noutside 40 325\nTMhelix 326 348\ninside 349 372

3950 GCF_001912365.1_ASM191236v1 Escherichia coli Proteobacteria; Gammaproteobacteria; Enterobacterales; Enterobacteriaceae; Escherichia MNNEETFYQAMRRQGVTRRSFLKYCSLAATSLGLGAGMAPKIAWALENKPRIPVVWIHGL WP_001058323.1 MULTISPECIES: hydrogenase-1 small chain [Proteobacteria] Length: 372\nNumber of predicted TMHs: 2\nExp number of AAs in TMHs: 37.2608499999999\nExp number, first 60 AAs: 19.52923\nTotal prob of N-in: 0.94351\nPOSSIBLE N-term signal sequence\ninside 1 19\nTMhelix 20 39\noutside 40 325\nTMhelix 326 348\ninside 349 372

3951 GCF_001912305.1_ASM191230v1 Escherichia coli Proteobacteria; Gammaproteobacteria; Enterobacterales; Enterobacteriaceae; Escherichia MNNEETFYQAMRRQGVTRRSFLKYCSLAATSLGLGAGMAPKIAWALENKPRIPVVWIHGL WP_001058320.1 hydrogenase [Escherichia coli] Length: 372\nNumber of predicted TMHs: 2\nExp number of AAs in TMHs: 37.26082\nExp number, first 60 AAs: 19.52923\nTotal prob of N-in: 0.94351\nPOSSIBLE N-term signal sequence\ninside 1 19\nTMhelix 20 39\noutside 40 325\nTMhelix 326 348\ninside 349 372

3952 GCF_001912445.1_ASM191244v1 Escherichia coli Proteobacteria; Gammaproteobacteria; Enterobacterales; Enterobacteriaceae; Escherichia
MNNEETFYQAMRRQGVTRRSFLKYCSLAATSLGLGAGMAPKIAWALENKPRIPVVWIHGL WP_001058323.1
MULTISPECIES: hydrogenase-1 small chain [Proteobacteria] Length: 372\nNumber of predicted TMHs: 2\nExp number of AAs in TMHs: 37.2608499999999\nExp number, first 60 AAs: 19.52923\nTotal prob of N-in: 0.94351\nPOSSIBLE N-term signal sequence\ninside 1 19\nTMhelix 20 39\noutside 40 325\nTMhelix 326 348\ninside 349 372

3953 GCF_001912345.1_ASM191234v1 Escherichia coli Proteobacteria; Gammaproteobacteria; Enterobacterales; Enterobacteriaceae; Escherichia
MNNEETFYQAMRRQGVTRRSFLKYCSLAATSLGLGAGMAPKIAWALENKPRIPVVWIHGL WP_001058323.1
MULTISPECIES: hydrogenase-1 small chain [Proteobacteria] Length: 372\nNumber of predicted TMHs: 2\nExp number of AAs in TMHs: 37.2608499999999\nExp number, first 60 AAs: 19.52923\nTotal prob of N-in: 0.94351\nPOSSIBLE N-term signal sequence\ninside 1 19\nTMhelix 20 39\noutside 40 325\nTMhelix 326 348\ninside 349 372

3954 GCF_001912455.1_ASM191245v1 Escherichia coli Proteobacteria; Gammaproteobacteria; Enterobacterales; Enterobacteriaceae; Escherichia
MTGDNTLIHSHGINRRDFMKLCAALAATMGLSSKAAAEMAESVTNPQRPPVIWIGAQECT WP_000145410.1
MULTISPECIES: hydrogenase-2 small chain [Proteobacteria] Length: 372\nNumber of predicted TMHs: 1\nExp number of AAs in TMHs: 23.72226\nExp number, first 60 AAs: 0.22038\nTotal prob of N-in: 0.12994\noutside 1 331\nTMhelix 332 354\ninside 355 372

3955 GCF_001912475.1_ASM191247v1 Escherichia coli Proteobacteria; Gammaproteobacteria; Enterobacterales; Enterobacteriaceae; Escherichia
MTGDNTLIHSHGINRRDFMKLCAALAATMGLSSKAAAEMAESVTNPQRPPVIWIGAQECT WP_000145410.1
MULTISPECIES: hydrogenase-2 small chain [Proteobacteria] Length: 372\nNumber of predicted TMHs: 1\nExp number of AAs in TMHs: 23.72226\nExp number, first 60 AAs: 0.22038\nTotal prob of N-in: 0.12994\noutside 1 331\nTMhelix 332 354\ninside 355 372

3956 GCF_001912455.1_ASM191245v1 Escherichia coli Proteobacteria; Gammaproteobacteria; Enterobacterales; Enterobacteriaceae; Escherichia
MNNEETFYQAMRRQGVTRRSFLKYCSLAATSLGLGAGMAPKIAWALENKPRIPVVWIHGL WP_001058323.1
MULTISPECIES: hydrogenase-1 small chain [Proteobacteria] Length: 372\nNumber of predicted TMHs: 2\nExp number of AAs in TMHs: 37.2608499999999\nExp number, first 60 AAs: 19.52923\nTotal prob of N-in: 0.94351\nPOSSIBLE N-term signal sequence\ninside 1 19\nTMhelix 20 39\noutside 40 325\nTMhelix 326 348\ninside 349 372

3957 GCF_001912525.1_ASM191252v1 Escherichia coli Proteobacteria; Gammaproteobacteria; Enterobacterales; Enterobacteriaceae; Escherichia
MTGDNTLIHSHGINRRDFMKLCAALAATMGLSSKAAAEMAESVTNPQRPPVIWIGAQECT WP_000145410.1
MULTISPECIES: hydrogenase-2 small chain [Proteobacteria] Length: 372\nNumber of predicted TMHs: 1\nExp number of AAs in TMHs: 23.72226\nExp number, first 60 AAs: 0.22038\nTotal prob of N-in: 0.12994\noutside 1 331\nTMhelix 332 354\ninside 355 372

3958 GCF_001912485.1_ASM191248v1 Escherichia coli Proteobacteria; Gammaproteobacteria; Enterobacterales; Enterobacteriaceae; Escherichia
MTGDNTLIHSHGINRRDFMKLCAALAATMGLSSKAAAEMAESVTNPQRPPVIWIGAQECT WP_000145410.1
MULTISPECIES: hydrogenase-2 small chain [Proteobacteria] Length: 372\nNumber of predicted TMHs: 1\nExp number of AAs in TMHs: 23.72226\nExp number, first 60 AAs: 0.22038\nTotal prob of N-in: 0.12994\noutside 1 331\nTMhelix 332 354\ninside 355 372

3959 GCF_001912565.1_ASM191256v1 Escherichia coli Proteobacteria; Gammaproteobacteria; Enterobacterales; Enterobacteriaceae; Escherichia
MNNEETFYQAMRRQGVTRRSFLKYCSLAATSLGLGAGMAPKIAWALENKPRIPVVWIHGL WP_001353311.1
MULTISPECIES: hydrogenase [Escherichia] Length: 372\nNumber of predicted TMHs: 2\nExp number of AAs in TMHs: 37.2638699999999\nExp number, first 60 AAs: 19.52924\nTotal prob of N-in: 0.94351\nPOSSIBLE N-term signal sequence\ninside 1 19\nTMhelix 20 39\noutside 40 325\nTMhelix 326 348\ninside 349 372

3960 GCF_001912485.1_ASM191248v1 Escherichia coli Proteobacteria; Gammaproteobacteria; Enterobacterales; Enterobacteriaceae; Escherichia

MNNEETFYQAMRRQGVTRRSFLKYCSLAATSLGLGAGMAPKIAWALENKPRIPVVWIHGL WP_001058323.1 MULTISPECIES: hydrogenase-1 small chain [Proteobacteria] Length: 372\nNumber of predicted TMHs: 2\nExp number of AAs in TMHs: 37.2608499999999\nExp number, first 60 AAs: 19.52923\nTotal prob of N-in: 0.94351\nPOSSIBLE N-term signal sequence\ninside 1 19\nTMhelix 20 39\noutside 40 325\nTMhelix 326 348\ninside 349 372

3961 GCF_001912625.1_ASM191262v1 Escherichia coli Proteobacteria; Gammaproteobacteria; Enterobacterales; Enterobacteriaceae; Escherichia MTGDNTLIHSHGINRRDFMKLCAALAATMGLSSKAAAEMAESVTNPQRPPVIWIGAQECT WP_000145410.1 MULTISPECIES: hydrogenase-2 small chain [Proteobacteria] Length: 372\nNumber of predicted TMHs: 1\nExp number of AAs in TMHs: 23.72226\nExp number, first 60 AAs: 0.22038\nTotal prob of N-in: 0.12994\noutside 1 331\nTMhelix 332 354\ninside 355 372

3962 GCF_001912625.1_ASM191262v1 Escherichia coli Proteobacteria; Gammaproteobacteria; Enterobacterales; Enterobacteriaceae; Escherichia MNNEETFYQAMRRQGVTRRSFLKYCSLAATSLGLGAGMAPKIAWALENKPRIPVVWIHGL WP_001058323.1 MULTISPECIES: hydrogenase-1 small chain [Proteobacteria] Length: 372\nNumber of predicted TMHs: 2\nExp number of AAs in TMHs: 37.2608499999999\nExp number, first 60 AAs: 19.52923\nTotal prob of N-in: 0.94351\nPOSSIBLE N-term signal sequence\ninside 1 19\nTMhelix 20 39\noutside 40 325\nTMhelix 326 348\ninside 349 372

3963 GCF_001912525.1_ASM191252v1 Escherichia coli Proteobacteria; Gammaproteobacteria; Enterobacterales; Enterobacteriaceae; Escherichia MNNEETFYQAMRRQGVTRRSFLKYCSLAATSLGLGAGMAPKIAWALENKPRIPVVWIHGL WP_001353311.1 MULTISPECIES: hydrogenase [Escherichia] Length: 372\nNumber of predicted TMHs: 2\nExp number of AAs in TMHs: 37.2638699999999\nExp number, first 60 AAs: 19.52924\nTotal prob of N-in: 0.94351\nPOSSIBLE N-term signal sequence\ninside 1 19\nTMhelix 20 39\noutside 40 325\nTMhelix 326 348\ninside 349 372

3964 GCF_001912665.1_ASM191266v1 Escherichia coli Proteobacteria; Gammaproteobacteria; Enterobacterales; Enterobacteriaceae; Escherichia MTGDNTLIHSHGINRRDFMKLCAALAATMGLSSKAAAEMAESVTNPQRPPVIWIGAQECT WP_000145410.1 MULTISPECIES: hydrogenase-2 small chain [Proteobacteria] Length: 372\nNumber of predicted TMHs: 1\nExp number of AAs in TMHs: 23.72226\nExp number, first 60 AAs: 0.22038\nTotal prob of N-in: 0.12994\noutside 1 331\nTMhelix 332 354\ninside 355 372

3965 GCF_001912675.1_ASM191267v1 Escherichia coli Proteobacteria; Gammaproteobacteria; Enterobacterales; Enterobacteriaceae; Escherichia MTGDNTLIHSHGINRRDFMKLCAALAATMGLSSKAAAEMAESVTNPQRPPVIWIGAQECT WP_000145410.1 MULTISPECIES: hydrogenase-2 small chain [Proteobacteria] Length: 372\nNumber of predicted TMHs: 1\nExp number of AAs in TMHs: 23.72226\nExp number, first 60 AAs: 0.22038\nTotal prob of N-in: 0.12994\noutside 1 331\nTMhelix 332 354\ninside 355 372

3966 GCF_001912555.1_ASM191255v1 Escherichia coli Proteobacteria; Gammaproteobacteria; Enterobacterales; Enterobacteriaceae; Escherichia MNNEETFYQAMRRQGVTRRSFLKYCSLAATSLGLGAGMAPKIAWALENKPRIPVVWIHGL WP_074507484.1 hydrogenase [Escherichia coli] Length: 372\nNumber of predicted TMHs: 2\nExp number of AAs in TMHs: 37.3347499999999\nExp number, first 60 AAs: 19.52915\nTotal prob of N-in: 0.94351\nPOSSIBLE N-term signal sequence\ninside 1 19\nTMhelix 20 39\noutside 40 325\nTMhelix 326 348\ninside 349 372

3967 GCF_001912705.1_ASM191270v1 Escherichia coli Proteobacteria; Gammaproteobacteria; Enterobacterales; Enterobacteriaceae; Escherichia MNNEETFYQAMRRQGVTRRSFLKYCSLAATSLGLGAGMAPKIAWALENKPRIPVVWIHGL WP_001058323.1 MULTISPECIES: hydrogenase-1 small chain [Proteobacteria] Length: 372\nNumber of predicted TMHs: 2\nExp number of AAs in TMHs: 37.2608499999999\nExp number, first 60 AAs: 19.52923\nTotal prob of N-in: 0.94351\nPOSSIBLE N-term signal sequence\ninside 1 19\nTMhelix 20 39\noutside 40 325\nTMhelix 326 348\ninside 349 372

3968 GCF_001932515.1_ASM193251v1 Escherichia coli Proteobacteria; Gammaproteobacteria; Enterobacterales; Enterobacteriaceae; Escherichia MTGDNTLIHSHGINRRDFMKLCAALAATMGLSSKAAAEMAESVTNPQRPPVIWIGAQECT WP_000145410.1 MULTISPECIES: hydrogenase-2 small chain [Proteobacteria] Length: 372\nNumber of predicted TMHs:

1\nExp number of AAs in TMHs: 23.72226\nExp number, first 60 AAs: 0.22038\nTotal prob of N-in: 0.12994\noutside 1 331\nTMhelix 332 354\ninside 355 372

3969 GCF_001936315.1_ASM193631v1 Escherichia coli Proteobacteria; Gammaproteobacteria; Enterobacterales; Enterobacteriaceae; Escherichia
MTGDNTLIHSHGINRRDFMKLCAALAATMGLSSKAAAEMAESVTNPQRPPVIWIGAQECT WP_000145410.1
MULTISPECIES: hydrogenase-2 small chain [Proteobacteria] Length: 372\nNumber of predicted TMHs: 1\nExp number of AAs in TMHs: 23.72226\nExp number, first 60 AAs: 0.22038\nTotal prob of N-in: 0.12994\noutside 1 331\nTMhelix 332 354\ninside 355 372

3970 GCF_001937255.1_ASM193725v1 Escherichia coli Proteobacteria; Gammaproteobacteria; Enterobacterales; Enterobacteriaceae; Escherichia
MTGDNTLIHSHGINRRDFMKLCAALAATMGLSSKAAAEMAESVTNPQRPPVIWIGAQECT WP_000145410.1
MULTISPECIES: hydrogenase-2 small chain [Proteobacteria] Length: 372\nNumber of predicted TMHs: 1\nExp number of AAs in TMHs: 23.72226\nExp number, first 60 AAs: 0.22038\nTotal prob of N-in: 0.12994\noutside 1 331\nTMhelix 332 354\ninside 355 372

3971 GCF_001937985.1_ASM193798v1 Escherichia coli Proteobacteria; Gammaproteobacteria; Enterobacterales; Enterobacteriaceae; Escherichia
MTGDNTLIHSHGINRRDFMKLCAALAATMGLSSKAAAEMAESVTNPQRPPVIWIGAQECT WP_000145410.1
MULTISPECIES: hydrogenase-2 small chain [Proteobacteria] Length: 372\nNumber of predicted TMHs: 1\nExp number of AAs in TMHs: 23.72226\nExp number, first 60 AAs: 0.22038\nTotal prob of N-in: 0.12994\noutside 1 331\nTMhelix 332 354\ninside 355 372

3972 GCF_001938625.2_ASM193862v2 Escherichia coli Proteobacteria; Gammaproteobacteria; Enterobacterales; Enterobacteriaceae; Escherichia
MTGDNTLIHSHGINRRDFMKLCAALAATMGLSSKAAAEMAESVTNPQRPPVIWIGAQECT WP_000145410.1
MULTISPECIES: hydrogenase-2 small chain [Proteobacteria] Length: 372\nNumber of predicted TMHs: 1\nExp number of AAs in TMHs: 23.72226\nExp number, first 60 AAs: 0.22038\nTotal prob of N-in: 0.12994\noutside 1 331\nTMhelix 332 354\ninside 355 372

3973 GCF_001940485.1_ASM194048v1 Escherichia coli Proteobacteria; Gammaproteobacteria; Enterobacterales; Enterobacteriaceae; Escherichia
MTGDNTLIHSHGINRRDFMKLCAALAATMGLSSKAAAEMAESVTNPQRPPVIWIGAQECT WP_000145410.1
MULTISPECIES: hydrogenase-2 small chain [Proteobacteria] Length: 372\nNumber of predicted TMHs: 1\nExp number of AAs in TMHs: 23.72226\nExp number, first 60 AAs: 0.22038\nTotal prob of N-in: 0.12994\noutside 1 331\nTMhelix 332 354\ninside 355 372

3974 GCF_001942065.1_ASM194206v1 Escherichia coli Proteobacteria; Gammaproteobacteria; Enterobacterales; Enterobacteriaceae; Escherichia
MTGDNTLIHSHGINRRDFMKLCAALAATMGLSSKAAAEMAESVTNPQRPPVIWIGAQECT WP_000145410.1
MULTISPECIES: hydrogenase-2 small chain [Proteobacteria] Length: 372\nNumber of predicted TMHs: 1\nExp number of AAs in TMHs: 23.72226\nExp number, first 60 AAs: 0.22038\nTotal prob of N-in: 0.12994\noutside 1 331\nTMhelix 332 354\ninside 355 372

3975 GCF_001942085.1_ASM194208v1 Escherichia coli Proteobacteria; Gammaproteobacteria; Enterobacterales; Enterobacteriaceae; Escherichia
MTGDNTLIHSHGINRRDFMKLCAALAATMGLSSKAAAEMAESVTNPQRPPVIWIGAQECT WP_000145410.1
MULTISPECIES: hydrogenase-2 small chain [Proteobacteria] Length: 372\nNumber of predicted TMHs: 1\nExp number of AAs in TMHs: 23.72226\nExp number, first 60 AAs: 0.22038\nTotal prob of N-in: 0.12994\noutside 1 331\nTMhelix 332 354\ninside 355 372

3976 GCF_001942095.1_ASM194209v1 Escherichia coli Proteobacteria; Gammaproteobacteria; Enterobacterales; Enterobacteriaceae; Escherichia
MTGDNTLIHSHGINRRDFMKLCAALAATMGLSSKAAAEMAESVTNPQRPPVIWIGAQECT WP_000145410.1
MULTISPECIES: hydrogenase-2 small chain [Proteobacteria] Length: 372\nNumber of predicted TMHs: 1\nExp number of AAs in TMHs: 23.72226\nExp number, first 60 AAs: 0.22038\nTotal prob of N-in: 0.12994\noutside 1 331\nTMhelix 332 354\ninside 355 372

3977 GCF_001942085.1_ASM194208v1 Escherichia coli Proteobacteria; Gammaproteobacteria; Enterobacterales; Enterobacteriaceae; Escherichia

MNNEETFYQAMRRQGVTRRSFLKYCSLAATSLGLGAGMAPKIAWALENKPRIPVVWIHGL WP_001058323.1 MULTISPECIES: hydrogenase-1 small chain [Proteobacteria] Length: 372\nNumber of predicted TMHs: 2\nExp number of AAs in TMHs: 37.2608499999999\nExp number, first 60 AAs: 19.52923\nTotal prob of N-in: 0.94351\nPOSSIBLE N-term signal sequence\ninside 1 19\nTMhelix 20 39\noutside 40 325\nTMhelix 326 348\ninside 349 372

3978 GCF_001942105.1_86J1 Escherichia coli Proteobacteria; Gammaproteobacteria; Enterobacterales; Enterobacteriaceae; Escherichia MTGDNTLIHSHGINRRDFMKLCAALAATMGLSSKAAAEMAESVTNPQRPPVIWIGAQECT WP_000145410.1 MULTISPECIES: hydrogenase-2 small chain [Proteobacteria] Length: 372\nNumber of predicted TMHs: 1\nExp number of AAs in TMHs: 23.72226\nExp number, first 60 AAs: 0.22038\nTotal prob of N-in: 0.12994\noutside 1 331\nTMhelix 332 354\ninside 355 372

3979 GCF_001942095.1_ASM194209v1 Escherichia coli Proteobacteria; Gammaproteobacteria; Enterobacterales; Enterobacteriaceae; Escherichia MNNEETFYQAMRRQGVTRRSFLKYCSLAATSLGLGAGMAPKIAWALENKPRIPVVWIHGL WP_001058323.1 MULTISPECIES: hydrogenase-1 small chain [Proteobacteria] Length: 372\nNumber of predicted TMHs: 2\nExp number of AAs in TMHs: 37.2608499999999\nExp number, first 60 AAs: 19.52923\nTotal prob of N-in: 0.94351\nPOSSIBLE N-term signal sequence\ninside 1 19\nTMhelix 20 39\noutside 40 325\nTMhelix 326 348\ninside 349 372

3980 GCF_001942175.1_ASM194217v1 Escherichia coli Proteobacteria; Gammaproteobacteria; Enterobacterales; Enterobacteriaceae; Escherichia MTGDNTLIHSHGINRRDFMKLCAALAATMGLSSKAAAEMAESVTNPQRPPVIWIGAQECT WP_000145410.1 MULTISPECIES: hydrogenase-2 small chain [Proteobacteria] Length: 372\nNumber of predicted TMHs: 1\nExp number of AAs in TMHs: 23.72226\nExp number, first 60 AAs: 0.22038\nTotal prob of N-in: 0.12994\noutside 1 331\nTMhelix 332 354\ninside 355 372

3981 GCF_001950695.1_ASM195069v1 Escherichia coli Proteobacteria; Gammaproteobacteria; Enterobacterales; Enterobacteriaceae; Escherichia MTGDNTLIHSHGINRRDFMKLCAALAATMGLSSKAAAEMAESVTNPQRPPVIWIGAQECT WP_000145410.1 MULTISPECIES: hydrogenase-2 small chain [Proteobacteria] Length: 372\nNumber of predicted TMHs: 1\nExp number of AAs in TMHs: 23.72226\nExp number, first 60 AAs: 0.22038\nTotal prob of N-in: 0.12994\noutside 1 331\nTMhelix 332 354\ninside 355 372

3982 GCF_001942175.1_ASM194217v1 Escherichia coli Proteobacteria; Gammaproteobacteria; Enterobacterales; Enterobacteriaceae; Escherichia MNNEETFYQAMRRQGVTRRSFLKYCSLAATSLGLGAGMAPKIAWALENKPRIPVVWIHGL WP_001058323.1 MULTISPECIES: hydrogenase-1 small chain [Proteobacteria] Length: 372\nNumber of predicted TMHs: 2\nExp number of AAs in TMHs: 37.2608499999999\nExp number, first 60 AAs: 19.52923\nTotal prob of N-in: 0.94351\nPOSSIBLE N-term signal sequence\ninside 1 19\nTMhelix 20 39\noutside 40 325\nTMhelix 326 348\ninside 349 372

3983 GCF_001950695.1_ASM195069v1 Escherichia coli Proteobacteria; Gammaproteobacteria; Enterobacterales; Enterobacteriaceae; Escherichia MNNEETFYQAMRRQGVTRRSFLKYCSLAATSLGLGAGMAPKIAWALENKPRIPVVWIHGL WP_001058323.1 MULTISPECIES: hydrogenase-1 small chain [Proteobacteria] Length: 372\nNumber of predicted TMHs: 2\nExp number of AAs in TMHs: 37.2608499999999\nExp number, first 60 AAs: 19.52923\nTotal prob of N-in: 0.94351\nPOSSIBLE N-term signal sequence\ninside 1 19\nTMhelix 20 39\noutside 40 325\nTMhelix 326 348\ninside 349 372

3984 GCF_001942105.1_86J1 Escherichia coli Proteobacteria; Gammaproteobacteria; Enterobacterales; Enterobacteriaceae; Escherichia MNNEETFYQAMRRQGVTRRSFLKYCSLAATSLGLGAGMAPKIAWALENKPRIPVVWIHGL WP_001058323.1 MULTISPECIES: hydrogenase-1 small chain [Proteobacteria] Length: 372\nNumber of predicted TMHs: 2\nExp number of AAs in TMHs: 37.2608499999999\nExp number, first 60 AAs: 19.52923\nTotal prob of N-in: 0.94351\nPOSSIBLE N-term signal sequence\ninside 1 19\nTMhelix 20 39\noutside 40 325\nTMhelix 326 348\ninside 349 372

3985 GCF_001950715.1_ASM195071v1 Escherichia coli Proteobacteria; Gammaproteobacteria; Enterobacterales; Enterobacteriaceae; Escherichia MTGDNTLIHSHGINRRDFMKLCAALAATMGLSSKAAAEMAESVTNPQRPPVIWIGAQECT WP_000145410.1 MULTISPECIES: hydrogenase-2 small chain [Proteobacteria] Length: 372\nNumber of predicted TMHs:

1\nExp number of AAs in TMHs: 23.72226\nExp number, first 60 AAs: 0.22038\nTotal prob of N-in: 0.12994\noutside 1 331\nTMhelix 332 354\ninside 355 372

3986 GCF_001957215.1_ASM195721v1 Escherichia coli Proteobacteria; Gammaproteobacteria; Enterobacterales; Enterobacteriaceae; Escherichia
MNNEETFYQAMRRQGVTRRSFLKYCSLAATSLGLGAGMAPKIAWALENKPRIPVVWIHGL WP_001058323.1
MULTISPECIES: hydrogenase-1 small chain [Proteobacteria] Length: 372\nNumber of predicted TMHs: 2\nExp number of AAs in TMHs: 37.2608499999999\nExp number, first 60 AAs: 19.52923\nTotal prob of N-in: 0.94351\nPOSSIBLE N-term signal sequence\ninside 1 19\nTMhelix 20 39\noutside 40 325\nTMhelix 326 348\ninside 349 372

3987 GCF_001950775.1_ASM195077v1 Escherichia coli Proteobacteria; Gammaproteobacteria; Enterobacterales; Enterobacteriaceae; Escherichia
MNNEETFYQAMRRQGVTRRSFLKYCSLAATSLGLGAGMAPKIAWALENKPRIPVVWIHGL WP_001058323.1
MULTISPECIES: hydrogenase-1 small chain [Proteobacteria] Length: 372\nNumber of predicted TMHs: 2\nExp number of AAs in TMHs: 37.2608499999999\nExp number, first 60 AAs: 19.52923\nTotal prob of N-in: 0.94351\nPOSSIBLE N-term signal sequence\ninside 1 19\nTMhelix 20 39\noutside 40 325\nTMhelix 326 348\ninside 349 372

3988 GCF_001957225.1_ASM195722v1 Escherichia coli Proteobacteria; Gammaproteobacteria; Enterobacterales; Enterobacteriaceae; Escherichia
MTGDNTLIHSHGINRRDFMKLCAALAATMGLSSKAAAEMAESVTNPQRPPVIWIGAQECT WP_000145410.1
MULTISPECIES: hydrogenase-2 small chain [Proteobacteria] Length: 372\nNumber of predicted TMHs: 1\nExp number of AAs in TMHs: 23.72226\nExp number, first 60 AAs: 0.22038\nTotal prob of N-in: 0.12994\noutside 1 331\nTMhelix 332 354\ninside 355 372

3989 GCF_001957225.1_ASM195722v1 Escherichia coli Proteobacteria; Gammaproteobacteria; Enterobacterales; Enterobacteriaceae; Escherichia
MNNEETFYQAMRRQGVTRRSFLKYCSLAATSLGLGAGMAPKIAWALENKPRIPVVWIHGL WP_001058323.1
MULTISPECIES: hydrogenase-1 small chain [Proteobacteria] Length: 372\nNumber of predicted TMHs: 2\nExp number of AAs in TMHs: 37.2608499999999\nExp number, first 60 AAs: 19.52923\nTotal prob of N-in: 0.94351\nPOSSIBLE N-term signal sequence\ninside 1 19\nTMhelix 20 39\noutside 40 325\nTMhelix 326 348\ninside 349 372

3990 GCF_001950735.1_ASM195073v1 Escherichia coli Proteobacteria; Gammaproteobacteria; Enterobacterales; Enterobacteriaceae; Escherichia
MNNEETFYQAMRRQGVTRRSFLKYCSLAATSLGLGAGMAPKIAWALENKPRIPVVWIHGL WP_001301613.1
hydrogenase [Escherichia coli] Length: 372\nNumber of predicted TMHs: 2\nExp number of AAs in TMHs: 37.46598\nExp number, first 60 AAs: 19.53139\nTotal prob of N-in: 0.94364\nPOSSIBLE N-term signal sequence\ninside 1 19\nTMhelix 20 39\noutside 40 325\nTMhelix 326 348\ninside 349 372

3991 GCF_001969285.1_ASM196928v1 Escherichia coli Proteobacteria; Gammaproteobacteria; Enterobacterales; Enterobacteriaceae; Escherichia
MTGDNTLIHSHGINRRDFMKLCAALAATMGLSSKAAAEMAESVTNPQRPPVIWIGAQECT WP_000145410.1
MULTISPECIES: hydrogenase-2 small chain [Proteobacteria] Length: 372\nNumber of predicted TMHs: 1\nExp number of AAs in TMHs: 23.72226\nExp number, first 60 AAs: 0.22038\nTotal prob of N-in: 0.12994\noutside 1 331\nTMhelix 332 354\ninside 355 372

3992 GCF_001957205.1_ASM195720v1 Escherichia coli Proteobacteria; Gammaproteobacteria; Enterobacterales; Enterobacteriaceae; Escherichia
MNNEETFYQAMRRQGVTRRSFLKYCSLAATSLGLGAGMAPKIAWALENKPRIPVVWIHGL WP_001058323.1
MULTISPECIES: hydrogenase-1 small chain [Proteobacteria] Length: 372\nNumber of predicted TMHs: 2\nExp number of AAs in TMHs: 37.2608499999999\nExp number, first 60 AAs: 19.52923\nTotal prob of N-in: 0.94351\nPOSSIBLE N-term signal sequence\ninside 1 19\nTMhelix 20 39\noutside 40 325\nTMhelix 326 348\ninside 349 372

3993 GCF_001981465.1_ASM198146v1 Escherichia coli Proteobacteria; Gammaproteobacteria; Enterobacterales; Enterobacteriaceae; Escherichia
MTGDNTLIHSHGINRRDFMKLCAALAATMGLSSKAAAEMAESVTNPQRPPVIWIGAQECT WP_000145410.1
MULTISPECIES: hydrogenase-2 small chain [Proteobacteria] Length: 372\nNumber of predicted TMHs:

1\nExp number of AAs in TMHs: 23.72226\nExp number, first 60 AAs: 0.22038\nTotal prob of N-in: 0.12994\noutside 1 331\nTMhelix 332 354\ninside 355 372

3994 GCF_001981465.1_ASM198146v1 Escherichia coli Proteobacteria; Gammaproteobacteria; Enterobacterales; Enterobacteriaceae; Escherichia MNNEETFYQAMRRQGVTRRSFLKYCSLAATSLGLGAGMAPKIAWALENKPRIPVVWIHGL WP_001058323.1 MULTISPECIES: hydrogenase-1 small chain [Proteobacteria] Length: 372\nNumber of predicted TMHs: 2\nExp number of AAs in TMHs: 37.2608499999999\nExp number, first 60 AAs: 19.52923\nTotal prob of N-in: 0.94351\nPOSSIBLE N-term signal sequence\ninside 1 19\nTMhelix 20 39\noutside 40 325\nTMhelix 326 348\ninside 349 372

3995 GCF_001984855.1_ASM198485v1 Escherichia coli Proteobacteria; Gammaproteobacteria; Enterobacterales; Enterobacteriaceae; Escherichia MTGDNTLIHSHGINRRDFMKLCAALAATMGLSSKAAAEMAESVTNPQRPPVIWIGAQECT WP_000145410.1 MULTISPECIES: hydrogenase-2 small chain [Proteobacteria] Length: 372\nNumber of predicted TMHs: 1\nExp number of AAs in TMHs: 23.72226\nExp number, first 60 AAs: 0.22038\nTotal prob of N-in: 0.12994\noutside 1 331\nTMhelix 332 354\ninside 355 372

3996 GCF_001990545.1_ASM199054v1 Escherichia coli Proteobacteria; Gammaproteobacteria; Enterobacterales; Enterobacteriaceae; Escherichia MTGDNTLIHSHGINRRDFMKLCAALAATMGLSSKAAAEMAESVTNPQRPPVIWIGAQECT WP_000145410.1 MULTISPECIES: hydrogenase-2 small chain [Proteobacteria] Length: 372\nNumber of predicted TMHs: 1\nExp number of AAs in TMHs: 23.72226\nExp number, first 60 AAs: 0.22038\nTotal prob of N-in: 0.12994\noutside 1 331\nTMhelix 332 354\ninside 355 372

3997 GCF_001984855.1_ASM198485v1 Escherichia coli Proteobacteria; Gammaproteobacteria; Enterobacterales; Enterobacteriaceae; Escherichia MNNEETFYQAMRRQGVTRRSFLKYCSLAATSLGLGAGMAPKIAWALENKPRIPVVWIHGL WP_001058323.1 MULTISPECIES: hydrogenase-1 small chain [Proteobacteria] Length: 372\nNumber of predicted TMHs: 2\nExp number of AAs in TMHs: 37.2608499999999\nExp number, first 60 AAs: 19.52923\nTotal prob of N-in: 0.94351\nPOSSIBLE N-term signal sequence\ninside 1 19\nTMhelix 20 39\noutside 40 325\nTMhelix 326 348\ninside 349 372

3998 GCF_001984845.1_ASM198484v1 Escherichia coli Proteobacteria; Gammaproteobacteria; Enterobacterales; Enterobacteriaceae; Escherichia MNNEETFYQAMRRQGVTRRSFLKYCSLAATSLGLGAGMAPKIAWALENKPRIPVVWIHGL WP_001058320.1 hydrogenase [Escherichia coli] Length: 372\nNumber of predicted TMHs: 2\nExp number of AAs in TMHs: 37.26082\nExp number, first 60 AAs: 19.52923\nTotal prob of N-in: 0.94351\nPOSSIBLE N-term signal sequence\ninside 1 19\nTMhelix 20 39\noutside 40 325\nTMhelix 326 348\ninside 349 372

3999 GCF_001990845.1_ASM199084v1 Escherichia coli Proteobacteria; Gammaproteobacteria; Enterobacterales; Enterobacteriaceae; Escherichia MTGDNTLIHSHGINRRDFMKLCAALAATMGLSSKAAAEMAESVTNPQRPPVIWIGAQECT WP_000145410.1 MULTISPECIES: hydrogenase-2 small chain [Proteobacteria] Length: 372\nNumber of predicted TMHs: 1\nExp number of AAs in TMHs: 23.72226\nExp number, first 60 AAs: 0.22038\nTotal prob of N-in: 0.12994\noutside 1 331\nTMhelix 332 354\ninside 355 372

4000 GCF_001990855.1_ASM199085v1 Escherichia coli Proteobacteria; Gammaproteobacteria; Enterobacterales; Enterobacteriaceae; Escherichia MTGDNTLIHSHGINRRDFMKLCAALAATMGLSSKAAAEMAESVTNPQRPPVIWIGAQECT WP_000145410.1 MULTISPECIES: hydrogenase-2 small chain [Proteobacteria] Length: 372\nNumber of predicted TMHs: 1\nExp number of AAs in TMHs: 23.72226\nExp number, first 60 AAs: 0.22038\nTotal prob of N-in: 0.12994\noutside 1 331\nTMhelix 332 354\ninside 355 372

4001 GCF_001990865.1_ASM199086v1 Escherichia coli Proteobacteria; Gammaproteobacteria; Enterobacterales; Enterobacteriaceae; Escherichia MTGDNTLIHSHGINRRDFMKLCAALAATMGLSSKAAAEMAESVTNPQRPPVIWIGAQECT WP_000145410.1 MULTISPECIES: hydrogenase-2 small chain [Proteobacteria] Length: 372\nNumber of predicted TMHs: 1\nExp number of AAs in TMHs: 23.72226\nExp number, first 60 AAs: 0.22038\nTotal prob of N-in: 0.12994\noutside 1 331\nTMhelix 332 354\ninside 355 372

4002 GCF_000782315.1_ASM78231v1 Escherichia coli Proteobacteria; Gammaproteobacteria; Enterobacterales; Enterobacteriaceae; Escherichia
MNNEETFYQAMRRQGVTRRSFLKYCSLAATSLGLGAGMAPKIAWALENKPRIPVVWIHGL WP_001295940.1
MULTISPECIES: hydrogenase [Enterobacteriaceae] Length: 372\nNumber of predicted TMHs: 2\nExp number of AAs in TMHs: 37.2608899999999\nExp number, first 60 AAs: 19.52923\nTotal prob of N-in: 0.94351\nPOSSIBLE N-term signal sequence\ninside 1 19\nTMhelix 20 39\noutside 40 325\nTMhelix 326 348\ninside 349 372

4003 GCF_001994975.1_ASM199497v1 Escherichia coli Proteobacteria; Gammaproteobacteria; Enterobacterales; Enterobacteriaceae; Escherichia
MTGDNTLIHSHGINRRDFMKLCAALAATMGLSSKAAAEMAESVTNPQRPPVIWIGAQECT WP_000145410.1
MULTISPECIES: hydrogenase-2 small chain [Proteobacteria] Length: 372\nNumber of predicted TMHs: 1\nExp number of AAs in TMHs: 23.72226\nExp number, first 60 AAs: 0.22038\nTotal prob of N-in: 0.12994\noutside 1 331\nTMhelix 332 354\ninside 355 372

4004 GCF_001990935.1_ASM199093v1 Escherichia coli Proteobacteria; Gammaproteobacteria; Enterobacterales; Enterobacteriaceae; Escherichia
MNNEETFYQAMRRQGVTRRSFLKYCSLAATSLGLGAGMAPKIAWALENKPRIPVVWIHGL WP_001058323.1
MULTISPECIES: hydrogenase-1 small chain [Proteobacteria] Length: 372\nNumber of predicted TMHs: 2\nExp number of AAs in TMHs: 37.2608499999999\nExp number, first 60 AAs: 19.52923\nTotal prob of N-in: 0.94351\nPOSSIBLE N-term signal sequence\ninside 1 19\nTMhelix 20 39\noutside 40 325\nTMhelix 326 348\ninside 349 372

4005 GCF_001994975.1_ASM199497v1 Escherichia coli Proteobacteria; Gammaproteobacteria; Enterobacterales; Enterobacteriaceae; Escherichia
MNNEETFYQAMRRQGVTRRSFLKYCSLAATSLGLGAGMAPKIAWALENKPRIPVVWIHGL WP_001058323.1
MULTISPECIES: hydrogenase-1 small chain [Proteobacteria] Length: 372\nNumber of predicted TMHs: 2\nExp number of AAs in TMHs: 37.2608499999999\nExp number, first 60 AAs: 19.52923\nTotal prob of N-in: 0.94351\nPOSSIBLE N-term signal sequence\ninside 1 19\nTMhelix 20 39\noutside 40 325\nTMhelix 326 348\ninside 349 372

4006 GCF_001990965.1_ASM199096v1 Escherichia coli Proteobacteria; Gammaproteobacteria; Enterobacterales; Enterobacteriaceae; Escherichia
MNNEETFYQAMRRQGVTRRSFLKYCSLAATSLGLGAGMAPKIAWALENKPRIPVVWIHGL WP_001058323.1
MULTISPECIES: hydrogenase-1 small chain [Proteobacteria] Length: 372\nNumber of predicted TMHs: 2\nExp number of AAs in TMHs: 37.2608499999999\nExp number, first 60 AAs: 19.52923\nTotal prob of N-in: 0.94351\nPOSSIBLE N-term signal sequence\ninside 1 19\nTMhelix 20 39\noutside 40 325\nTMhelix 326 348\ninside 349 372

4007 GCF_001990875.1_ASM199087v1 Escherichia coli Proteobacteria; Gammaproteobacteria; Enterobacterales; Enterobacteriaceae; Escherichia
MNNEETFYQAMRRQGVTRRSFLKYCSLAATSLGLGAGMAPKIAWALENKPRIPVVWIHGL WP_001058320.1
hydrogenase [Escherichia coli] Length: 372\nNumber of predicted TMHs: 2\nExp number of AAs in TMHs: 37.26082\nExp number, first 60 AAs: 19.52923\nTotal prob of N-in: 0.94351\nPOSSIBLE N-term signal sequence\ninside 1 19\nTMhelix 20 39\noutside 40 325\nTMhelix 326 348\ninside 349 372

4008 GCF_001994955.1_ASM199495v1 Escherichia coli Proteobacteria; Gammaproteobacteria; Enterobacterales; Enterobacteriaceae; Escherichia
MTGDNTLIHSHGINRRDFMKLCAALAATMGLSSKAAAEMAESVTNPQRPPVIWIGAQECT WP_000145410.1
MULTISPECIES: hydrogenase-2 small chain [Proteobacteria] Length: 372\nNumber of predicted TMHs: 1\nExp number of AAs in TMHs: 23.72226\nExp number, first 60 AAs: 0.22038\nTotal prob of N-in: 0.12994\noutside 1 331\nTMhelix 332 354\ninside 355 372

4009 GCF_001994955.1_ASM199495v1 Escherichia coli Proteobacteria; Gammaproteobacteria; Enterobacterales; Enterobacteriaceae; Escherichia
MNNEETFYQAMRRQGVTRRSFLKYCSLAATSLGLGAGMAPKIAWALENKPRIPVVWIHGL WP_001058323.1
MULTISPECIES: hydrogenase-1 small chain [Proteobacteria] Length: 372\nNumber of predicted TMHs: 2\nExp number of AAs in TMHs: 37.2608499999999\nExp number, first 60 AAs: 19.52923\nTotal prob of N-in: 0.94351\nPOSSIBLE N-term signal sequence\ninside 1 19\nTMhelix 20 39\noutside 40 325\nTMhelix 326 348\ninside 349 372

4010 GCF_001999185.1_ASM199918v1 Escherichia coli Proteobacteria; Gammaproteobacteria; Enterobacterales; Enterobacteriaceae; Escherichia
MTGDNTLIHSHGINRRDFMKLCAALAATMGLSSKAAAEMAESVTNPQRPPVIWIGAQECT WP_000145410.1
MULTISPECIES: hydrogenase-2 small chain [Proteobacteria] Length: 372\nNumber of predicted TMHs: 1\nExp number of AAs in TMHs: 23.72226\nExp number, first 60 AAs: 0.22038\nTotal prob of N-in: 0.12994\noutside 1 331\nTMhelix 332 354\ninside 355 372

4011 GCF_002001535.1_ASM200153v1 Escherichia coli Proteobacteria; Gammaproteobacteria; Enterobacterales; Enterobacteriaceae; Escherichia
MTGDNTLIHSHGINRRDFMKLCAALAATMGLSSKAAAEMAESVTNPQRPPVIWIGAQECT WP_000145410.1
MULTISPECIES: hydrogenase-2 small chain [Proteobacteria] Length: 372\nNumber of predicted TMHs: 1\nExp number of AAs in TMHs: 23.72226\nExp number, first 60 AAs: 0.22038\nTotal prob of N-in: 0.12994\noutside 1 331\nTMhelix 332 354\ninside 355 372

4012 GCF_002001585.1_ASM200158v1 Escherichia coli Proteobacteria; Gammaproteobacteria; Enterobacterales; Enterobacteriaceae; Escherichia
MTGDNTLIHSHGINRRDFMKLCAALAATMGLSSKAAAEMAESVTNPQRPPVIWIGAQECT WP_000145410.1
MULTISPECIES: hydrogenase-2 small chain [Proteobacteria] Length: 372\nNumber of predicted TMHs: 1\nExp number of AAs in TMHs: 23.72226\nExp number, first 60 AAs: 0.22038\nTotal prob of N-in: 0.12994\noutside 1 331\nTMhelix 332 354\ninside 355 372

4013 GCF_002001585.1_ASM200158v1 Escherichia coli Proteobacteria; Gammaproteobacteria; Enterobacterales; Enterobacteriaceae; Escherichia
MNNEETFYQAMRRQGVTRRSFLKYCSLAATSLGLGAGMAPKIAWALENKPRIPVVWIHGL WP_001058323.1
MULTISPECIES: hydrogenase-1 small chain [Proteobacteria] Length: 372\nNumber of predicted TMHs: 2\nExp number of AAs in TMHs: 37.2608499999999\nExp number, first 60 AAs: 19.52923\nTotal prob of N-in: 0.94351\nPOSSIBLE N-term signal sequence\ninside 1 19\nTMhelix 20 39\noutside 40 325\nTMhelix 326 348\ninside 349 372

4014 GCF_001997065.1_ASM199706v1 Escherichia coli Proteobacteria; Gammaproteobacteria; Enterobacterales; Enterobacteriaceae; Escherichia
MNNEETFYQAMRRQGVTRRSFLKYCSLAATSLGLGAGMAPKIAWALENKPRIPVVWIHGL WP_001058323.1
MULTISPECIES: hydrogenase-1 small chain [Proteobacteria] Length: 372\nNumber of predicted TMHs: 2\nExp number of AAs in TMHs: 37.2608499999999\nExp number, first 60 AAs: 19.52923\nTotal prob of N-in: 0.94351\nPOSSIBLE N-term signal sequence\ninside 1 19\nTMhelix 20 39\noutside 40 325\nTMhelix 326 348\ninside 349 372

4015 GCF_001997045.1_ASM199704v1 Escherichia coli Proteobacteria; Gammaproteobacteria; Enterobacterales; Enterobacteriaceae; Escherichia
MNNEETFYQAMRRQGVTRRSFLKYCSLAATSLGLGAGMAPKIAWALENKPRIPVVWIHGL WP_001301613.1
hydrogenase [Escherichia coli] Length: 372\nNumber of predicted TMHs: 2\nExp number of AAs in TMHs: 37.46598\nExp number, first 60 AAs: 19.53139\nTotal prob of N-in: 0.94364\nPOSSIBLE N-term signal sequence\ninside 1 19\nTMhelix 20 39\noutside 40 325\nTMhelix 326 348\ninside 349 372

4016 GCF_001997075.1_ASM199707v1 Escherichia coli Proteobacteria; Gammaproteobacteria; Enterobacterales; Enterobacteriaceae; Escherichia
MNNEETFYQAMRRQGVTRRSFLKYCSLAATSLGLGAGMAPKIAWALENKPRIPVVWIHGL WP_001058323.1
MULTISPECIES: hydrogenase-1 small chain [Proteobacteria] Length: 372\nNumber of predicted TMHs: 2\nExp number of AAs in TMHs: 37.2608499999999\nExp number, first 60 AAs: 19.52923\nTotal prob of N-in: 0.94351\nPOSSIBLE N-term signal sequence\ninside 1 19\nTMhelix 20 39\noutside 40 325\nTMhelix 326 348\ninside 349 372

4017 GCF_002001635.1_ASM200163v1 Escherichia coli Proteobacteria; Gammaproteobacteria; Enterobacterales; Enterobacteriaceae; Escherichia
MTGDNTLIHSHGINRRDFMKLCAALAATMGLSSKAAAEMAESVTNPQRPPVIWIGAQECT WP_000145410.1
MULTISPECIES: hydrogenase-2 small chain [Proteobacteria] Length: 372\nNumber of predicted TMHs: 1\nExp number of AAs in TMHs: 23.72226\nExp number, first 60 AAs: 0.22038\nTotal prob of N-in: 0.12994\noutside 1 331\nTMhelix 332 354\ninside 355 372

4018 GCF_002001525.1_ASM200152v1 Escherichia coli Proteobacteria; Gammaproteobacteria; Enterobacterales; Enterobacteriaceae; Escherichia

MNNEETFYQAMRRQGVTRRSFLKYCSLAATSLGLGAGMAPKIAWALENKPRIPVVWIHGL WP_001295940.1 MULTISPECIES: hydrogenase [Enterobacteriaceae] Length: 372\nNumber of predicted TMHs: 2\nExp number of AAs in TMHs: 37.2608899999999\nExp number, first 60 AAs: 19.52923\nTotal prob of N-in: 0.94351\nPOSSIBLE N-term signal sequence\ninside 1 19\nTMhelix 20 39\noutside 40 325\nTMhelix 326 348\ninside 349 372

4019 GCF_002001665.1_ASM200166v1 Escherichia coli Proteobacteria; Gammaproteobacteria; Enterobacterales; Enterobacteriaceae; Escherichia MTGDNTLIHSHGINRRDFMKLCAALAATMGLSSKAAAEMAESVTNPQRPPVIWIGAQECT WP_000145410.1 MULTISPECIES: hydrogenase-2 small chain [Proteobacteria] Length: 372\nNumber of predicted TMHs: 1\nExp number of AAs in TMHs: 23.72226\nExp number, first 60 AAs: 0.22038\nTotal prob of N-in: 0.12994\noutside 1 331\nTMhelix 332 354\ninside 355 372

4020 GCF_002001635.1_ASM200163v1 Escherichia coli Proteobacteria; Gammaproteobacteria; Enterobacterales; Enterobacteriaceae; Escherichia MNNEETFYQAMRRQGVTRRSFLKYCSLAATSLGLGAGMAPKIAWALENKPRIPVVWIHGL WP_001058323.1 MULTISPECIES: hydrogenase-1 small chain [Proteobacteria] Length: 372\nNumber of predicted TMHs: 2\nExp number of AAs in TMHs: 37.2608499999999\nExp number, first 60 AAs: 19.52923\nTotal prob of N-in: 0.94351\nPOSSIBLE N-term signal sequence\ninside 1 19\nTMhelix 20 39\noutside 40 325\nTMhelix 326 348\ninside 349 372

4021 GCF_002001525.1_ASM200152v1 Escherichia coli Proteobacteria; Gammaproteobacteria; Enterobacterales; Enterobacteriaceae; Escherichia MTGDNTLIHSHGINRRDFMKLCAALAATMGLSSKAAAEMAESVTNPQRPPVIWIGAQECT WP_077459392.1 hydrogenase 2 small subunit [Escherichia coli] Length: 372\nNumber of predicted TMHs: 1\nExp number of AAs in TMHs: 23.66671\nExp number, first 60 AAs: 0.22041\nTotal prob of N-in: 0.12079\noutside 1 331\nTMhelix 332 354\ninside 355 372

4022 GCF_002001685.1_ASM200168v1 Escherichia coli Proteobacteria; Gammaproteobacteria; Enterobacterales; Enterobacteriaceae; Escherichia MTGDNTLIHSHGINRRDFMKLCAALAATMGLSSKAAAEMAESVTNPQRPPVIWIGAQECT WP_000145410.1 MULTISPECIES: hydrogenase-2 small chain [Proteobacteria] Length: 372\nNumber of predicted TMHs: 1\nExp number of AAs in TMHs: 23.72226\nExp number, first 60 AAs: 0.22038\nTotal prob of N-in: 0.12994\noutside 1 331\nTMhelix 332 354\ninside 355 372

4023 GCF_002001705.1_ASM200170v1 Escherichia coli Proteobacteria; Gammaproteobacteria; Enterobacterales; Enterobacteriaceae; Escherichia MTGDNTLIHSHGINRRDFMKLCAALAATMGLSSKAAAEMAESVTNPQRPPVIWIGAQECT WP_000145410.1 MULTISPECIES: hydrogenase-2 small chain [Proteobacteria] Length: 372\nNumber of predicted TMHs: 1\nExp number of AAs in TMHs: 23.72226\nExp number, first 60 AAs: 0.22038\nTotal prob of N-in: 0.12994\noutside 1 331\nTMhelix 332 354\ninside 355 372

4024 GCF_002001845.1_ASM200184v1 Escherichia coli Proteobacteria; Gammaproteobacteria; Enterobacterales; Enterobacteriaceae; Escherichia MNNEETFYQAMRRQGVTRRSFLKYCSLAATSLGLGAGMAPKIAWALENKPRIPVVWIHGL WP_001058323.1 MULTISPECIES: hydrogenase-1 small chain [Proteobacteria] Length: 372\nNumber of predicted TMHs: 2\nExp number of AAs in TMHs: 37.2608499999999\nExp number, first 60 AAs: 19.52923\nTotal prob of N-in: 0.94351\nPOSSIBLE N-term signal sequence\ninside 1 19\nTMhelix 20 39\noutside 40 325\nTMhelix 326 348\ninside 349 372

4025 GCF_002001825.1_ASM200182v1 Escherichia coli Proteobacteria; Gammaproteobacteria; Enterobacterales; Enterobacteriaceae; Escherichia MNNEETFYQAMRRQGVTRRSFLKYCSLAATSLGLGAGMAPKIAWALENKPRIPVVWIHGL WP_001058323.1 MULTISPECIES: hydrogenase-1 small chain [Proteobacteria] Length: 372\nNumber of predicted TMHs: 2\nExp number of AAs in TMHs: 37.2608499999999\nExp number, first 60 AAs: 19.52923\nTotal prob of N-in: 0.94351\nPOSSIBLE N-term signal sequence\ninside 1 19\nTMhelix 20 39\noutside 40 325\nTMhelix 326 348\ninside 349 372

4026 GCF_002001715.1_ASM200171v1 Escherichia coli Proteobacteria; Gammaproteobacteria; Enterobacterales; Enterobacteriaceae; Escherichia MNNEETFYQAMRRQGVTRRSFLKYCSLAATSLGLGAGMAPKIAWALENKPRIPVVWIHGL WP_001058323.1

MULTISPECIES: hydrogenase-1 small chain [Proteobacteria] Length: 372\nNumber of predicted TMHs: 2\nExp number of AAs in TMHs: 37.2608499999999\nExp number, first 60 AAs: 19.52923\nTotal prob of N-in: 0.94351\nPOSSIBLE N-term signal sequence\ninside 1 19\nTMhelix 20 39\noutside 40 325\nTMhelix 326 348\ninside 349 372

4027 GCF_002001785.1_ASM200178v1 Escherichia coli Proteobacteria; Gammaproteobacteria; Enterobacterales; Enterobacteriaceae; Escherichia MNNEETFYQAMRRQGVTRRSFLKYCSLAATSLGLGAGMAPKIAWALENKPRIPVVWIHGL WP_001058323.1 MULTISPECIES: hydrogenase-1 small chain [Proteobacteria] Length: 372\nNumber of predicted TMHs: 2\nExp number of AAs in TMHs: 37.2608499999999\nExp number, first 60 AAs: 19.52923\nTotal prob of N-in: 0.94351\nPOSSIBLE N-term signal sequence\ninside 1 19\nTMhelix 20 39\noutside 40 325\nTMhelix 326 348\ninside 349 372

4028 GCF_002001855.1_ASM200185v1 Escherichia coli Proteobacteria; Gammaproteobacteria; Enterobacterales; Enterobacteriaceae; Escherichia MTGDNTLIHSHGINRRDFMKLCAALAATMGLSSKAAAEMAESVTNPQRPPVIWIGAQECT WP_000145410.1 MULTISPECIES: hydrogenase-2 small chain [Proteobacteria] Length: 372\nNumber of predicted TMHs: 1\nExp number of AAs in TMHs: 23.72226\nExp number, first 60 AAs: 0.22038\nTotal prob of N-in: 0.12994\noutside 1 331\nTMhelix 332 354\ninside 355 372

4029 GCF_002001855.1_ASM200185v1 Escherichia coli Proteobacteria; Gammaproteobacteria; Enterobacterales; Enterobacteriaceae; Escherichia MNNEETFYQAMRRQGVTRRSFLKYCSLAATSLGLGAGMAPKIAWALENKPRIPVVWIHGL WP_001058323.1 MULTISPECIES: hydrogenase-1 small chain [Proteobacteria] Length: 372\nNumber of predicted TMHs: 2\nExp number of AAs in TMHs: 37.2608499999999\nExp number, first 60 AAs: 19.52923\nTotal prob of N-in: 0.94351\nPOSSIBLE N-term signal sequence\ninside 1 19\nTMhelix 20 39\noutside 40 325\nTMhelix 326 348\ninside 349 372

4030 GCF_002001805.1_ASM200180v1 Escherichia coli Proteobacteria; Gammaproteobacteria; Enterobacterales; Enterobacteriaceae; Escherichia MNNEETFYQAMRRQGVTRRSFLKYCSLAATSLGLGAGMAPKIAWALENKPRIPVVWIHGL WP_001295940.1 MULTISPECIES: hydrogenase [Enterobacteriaceae] Length: 372\nNumber of predicted TMHs: 2\nExp number of AAs in TMHs: 37.2608899999999\nExp number, first 60 AAs: 19.52923\nTotal prob of N-in: 0.94351\nPOSSIBLE N-term signal sequence\ninside 1 19\nTMhelix 20 39\noutside 40 325\nTMhelix 326 348\ninside 349 372

4031 GCF_002001905.1_ASM200190v1 Escherichia coli Proteobacteria; Gammaproteobacteria; Enterobacterales; Enterobacteriaceae; Escherichia MTGDNTLIHSHGINRRDFMKLCAALAATMGLSSKAAAEMAESVTNPQRPPVIWIGAQECT WP_000145410.1 MULTISPECIES: hydrogenase-2 small chain [Proteobacteria] Length: 372\nNumber of predicted TMHs: 1\nExp number of AAs in TMHs: 23.72226\nExp number, first 60 AAs: 0.22038\nTotal prob of N-in: 0.12994\noutside 1 331\nTMhelix 332 354\ninside 355 372

4032 GCF_002001885.1_ASM200188v1 Escherichia coli Proteobacteria; Gammaproteobacteria; Enterobacterales; Enterobacteriaceae; Escherichia MTGDNTLIHSHGINRRDFMKLCAALAATMGLSSKAAAEMAESVTNPQRPPVIWIGAQECT WP_000145410.1 MULTISPECIES: hydrogenase-2 small chain [Proteobacteria] Length: 372\nNumber of predicted TMHs: 1\nExp number of AAs in TMHs: 23.72226\nExp number, first 60 AAs: 0.22038\nTotal prob of N-in: 0.12994\noutside 1 331\nTMhelix 332 354\ninside 355 372

4033 GCF_002001915.1_ASM200191v1 Escherichia coli Proteobacteria; Gammaproteobacteria; Enterobacterales; Enterobacteriaceae; Escherichia MTGDNTLIHSHGINRRDFMKLCAALAATMGLSSKAAAEMAESVTNPQRPPVIWIGAQECT WP_000145410.1 MULTISPECIES: hydrogenase-2 small chain [Proteobacteria] Length: 372\nNumber of predicted TMHs: 1\nExp number of AAs in TMHs: 23.72226\nExp number, first 60 AAs: 0.22038\nTotal prob of N-in: 0.12994\noutside 1 331\nTMhelix 332 354\ninside 355 372

4034 GCF_002001905.1_ASM200190v1 Escherichia coli Proteobacteria; Gammaproteobacteria; Enterobacterales; Enterobacteriaceae; Escherichia MNNEETFYQAMRRQGVTRRSFLKYCSLAATSLGLGAGMAPKIAWALENKPRIPVVWIHGL WP_001058323.1 MULTISPECIES: hydrogenase-1 small chain [Proteobacteria] Length: 372\nNumber of predicted TMHs:

2\nExp number of AAs in TMHs: 37.2608499999999\nExp number, first 60 AAs: 19.52923\nTotal prob of N-in: 0.94351\nPOSSIBLE N-term signal sequence\ninside 1 19\nTMhelix 20 39\noutside 40 325\nTMhelix 326 348\ninside 349 372

4035 GCF_002001915.1_ASM200191v1 Escherichia coli Proteobacteria; Gammaproteobacteria; Enterobacterales; Enterobacteriaceae; Escherichia
MNNEETFYQAMRRQGVTRRSFLKYCSLAATSLGLGAGMAPKIAWALENKPRIPVVWIHGL WP_001058323.1
MULTISPECIES: hydrogenase-1 small chain [Proteobacteria] Length: 372\nNumber of predicted TMHs: 2\nExp number of AAs in TMHs: 37.2608499999999\nExp number, first 60 AAs: 19.52923\nTotal prob of N-in: 0.94351\nPOSSIBLE N-term signal sequence\ninside 1 19\nTMhelix 20 39\noutside 40 325\nTMhelix 326 348\ninside 349 372

4036 GCF_002001885.1_ASM200188v1 Escherichia coli Proteobacteria; Gammaproteobacteria; Enterobacterales; Enterobacteriaceae; Escherichia
MNNEETFYQAMRRQGVTRRSFLKYCSLAATSLGLGAGMAPKIAWALENKPRIPVVWIHGL WP_021557704.1
hydrogenase-1 small chain [Escherichia coli] Length: 372\nNumber of predicted TMHs: 2\nExp number of AAs in TMHs: 37.2609699999999\nExp number, first 60 AAs: 19.52923\nTotal prob of N-in: 0.94351\nPOSSIBLE N-term signal sequence\ninside 1 19\nTMhelix 20 39\noutside 40 325\nTMhelix 326 348\ninside 349 372

4037 GCF_002001765.1_ASM200176v1 Escherichia coli Proteobacteria; Gammaproteobacteria; Enterobacterales; Enterobacteriaceae; Escherichia
MNNEETFYQAMRRQGVTRRSFLKYCSLAATSLGLGAGMAPKIAWALENKPRIPVVWIHGL WP_077575616.1
[Ni/Fe] hydrogenase small subunit [Escherichia coli] Length: 372\nNumber of predicted TMHs: 2\nExp number of AAs in TMHs: 42.1360199999999\nExp number, first 60 AAs: 19.52048\nTotal prob of N-in: 0.94624\nPOSSIBLE N-term signal sequence\ninside 1 19\nTMhelix 20 39\noutside 40 325\nTMhelix 326 348\ninside 349 372

4038 GCF_002001965.1_ASM200196v1 Escherichia coli Proteobacteria; Gammaproteobacteria; Enterobacterales; Enterobacteriaceae; Escherichia
MNNEETFYQAMRRQGVTRRSFLKYCSLAATSLGLGAGMAPKIAWALENKPRIPVVWIHGL WP_001058323.1
MULTISPECIES: hydrogenase-1 small chain [Proteobacteria] Length: 372\nNumber of predicted TMHs: 2\nExp number of AAs in TMHs: 37.2608499999999\nExp number, first 60 AAs: 19.52923\nTotal prob of N-in: 0.94351\nPOSSIBLE N-term signal sequence\ninside 1 19\nTMhelix 20 39\noutside 40 325\nTMhelix 326 348\ninside 349 372

4039 GCF_002002075.1_ASM200207v1 Escherichia coli Proteobacteria; Gammaproteobacteria; Enterobacterales; Enterobacteriaceae; Escherichia
MTGDNTLIHSHGINRRDFMKLCAALAATMGLSSKAAAEMAESVTNPQRPPVIWIGAQECT WP_000145410.1
MULTISPECIES: hydrogenase-2 small chain [Proteobacteria] Length: 372\nNumber of predicted TMHs: 1\nExp number of AAs in TMHs: 23.72226\nExp number, first 60 AAs: 0.22038\nTotal prob of N-in: 0.12994\noutside 1 331\nTMhelix 332 354\ninside 355 372

4040 GCF_002002075.1_ASM200207v1 Escherichia coli Proteobacteria; Gammaproteobacteria; Enterobacterales; Enterobacteriaceae; Escherichia
MNNEETFYQAMRRQGVTRRSFLKYCSLAATSLGLGAGMAPKIAWALENKPRIPVVWIHGL WP_021557704.1
hydrogenase-1 small chain [Escherichia coli] Length: 372\nNumber of predicted TMHs: 2\nExp number of AAs in TMHs: 37.2609699999999\nExp number, first 60 AAs: 19.52923\nTotal prob of N-in: 0.94351\nPOSSIBLE N-term signal sequence\ninside 1 19\nTMhelix 20 39\noutside 40 325\nTMhelix 326 348\ninside 349 372

4041 GCF_002002005.1_ASM200200v1 Escherichia coli Proteobacteria; Gammaproteobacteria; Enterobacterales; Enterobacteriaceae; Escherichia
MNNEETFYQAMRRQGVTRRSFLKYCSLAATSLGLGAGMAPKIAWALENKPRIPVVWIHGL WP_001058323.1
MULTISPECIES: hydrogenase-1 small chain [Proteobacteria] Length: 372\nNumber of predicted TMHs: 2\nExp number of AAs in TMHs: 37.2608499999999\nExp number, first 60 AAs: 19.52923\nTotal prob of N-in: 0.94351\nPOSSIBLE N-term signal sequence\ninside 1 19\nTMhelix 20 39\noutside 40 325\nTMhelix 326 348\ninside 349 372

4042 GCF_002002045.1_ASM200204v1 Escherichia coli Proteobacteria; Gammaproteobacteria; Enterobacterales; Enterobacteriaceae; Escherichia
MNNEETFYQAMRRQGVTRRSFLKYCSLAATSLGLGAGMAPKIAWALENKPRIPVVWIHGL WP_001058323.1
MULTISPECIES: hydrogenase-1 small chain [Proteobacteria] Length: 372\nNumber of predicted TMHs:

2\nExp number of AAs in TMHs: 37.2608499999999\nExp number, first 60 AAs: 19.52923\nTotal prob of N-in: 0.94351\nPOSSIBLE N-term signal sequence\ninside 1 19\nTMhelix 20 39\noutside 40 325\nTMhelix 326 348\ninside 349 372

4043 GCF_002002055.1_ASM200205v1 Escherichia coli Proteobacteria; Gammaproteobacteria; Enterobacterales; Enterobacteriaceae; Escherichia MNNEETFYQAMRRQGVTRRSFLKYCSLAATSLGLGAGMAPKIAWALENKPRIPVVWIHGL WP_001058323.1 MULTISPECIES: hydrogenase-1 small chain [Proteobacteria] Length: 372\nNumber of predicted TMHs: 2\nExp number of AAs in TMHs: 37.2608499999999\nExp number, first 60 AAs: 19.52923\nTotal prob of N-in: 0.94351\nPOSSIBLE N-term signal sequence\ninside 1 19\nTMhelix 20 39\noutside 40 325\nTMhelix 326 348\ninside 349 372

4044 GCF_002001995.1_ASM200199v1 Escherichia coli Proteobacteria; Gammaproteobacteria; Enterobacterales; Enterobacteriaceae; Escherichia MNNEETFYQAMRRQGVTRRSFLKYCSLAATSLGLGAGMAPKIAWALENKPRIPVVWIHGL WP_001295940.1 MULTISPECIES: hydrogenase [Enterobacteriaceae] Length: 372\nNumber of predicted TMHs: 2\nExp number of AAs in TMHs: 37.2608899999999\nExp number, first 60 AAs: 19.52923\nTotal prob of N-in: 0.94351\nPOSSIBLE N-term signal sequence\ninside 1 19\nTMhelix 20 39\noutside 40 325\nTMhelix 326 348\ninside 349 372

4045 GCF_002002175.1_ASM200217v1 Escherichia coli Proteobacteria; Gammaproteobacteria; Enterobacterales; Enterobacteriaceae; Escherichia MTGDNTLIHSHGINRRDFMKLCAALAATMGLSSKAAAEMAESVTNPQRPPVIWIGAQECT WP_000145410.1 MULTISPECIES: hydrogenase-2 small chain [Proteobacteria] Length: 372\nNumber of predicted TMHs: 1\nExp number of AAs in TMHs: 23.72226\nExp number, first 60 AAs: 0.22038\nTotal prob of N-in: 0.12994\noutside 1 331\nTMhelix 332 354\ninside 355 372

4046 GCF_002002195.1_ASM200219v1 Escherichia coli Proteobacteria; Gammaproteobacteria; Enterobacterales; Enterobacteriaceae; Escherichia MTGDNTLIHSHGINRRDFMKLCAALAATMGLSSKAAAEMAESVTNPQRPPVIWIGAQECT WP_000145410.1 MULTISPECIES: hydrogenase-2 small chain [Proteobacteria] Length: 372\nNumber of predicted TMHs: 1\nExp number of AAs in TMHs: 23.72226\nExp number, first 60 AAs: 0.22038\nTotal prob of N-in: 0.12994\noutside 1 331\nTMhelix 332 354\ninside 355 372

4047 GCF_002002125.1_ASM200212v1 Escherichia coli Proteobacteria; Gammaproteobacteria; Enterobacterales; Enterobacteriaceae; Escherichia MNNEETFYQAMRRQGVTRRSFLKYCSLAATSLGLGAGMAPKIAWALENKPRIPVVWIHGL WP_001058323.1 MULTISPECIES: hydrogenase-1 small chain [Proteobacteria] Length: 372\nNumber of predicted TMHs: 2\nExp number of AAs in TMHs: 37.2608499999999\nExp number, first 60 AAs: 19.52923\nTotal prob of N-in: 0.94351\nPOSSIBLE N-term signal sequence\ninside 1 19\nTMhelix 20 39\noutside 40 325\nTMhelix 326 348\ninside 349 372

4048 GCF_002002235.1_ASM200223v1 Escherichia coli Proteobacteria; Gammaproteobacteria; Enterobacterales; Enterobacteriaceae; Escherichia MTGDNTLIHSHGINRRDFMKLCAALAATMGLSSKAAAEMAESVTNPQRPPVIWIGAQECT WP_000145410.1 MULTISPECIES: hydrogenase-2 small chain [Proteobacteria] Length: 372\nNumber of predicted TMHs: 1\nExp number of AAs in TMHs: 23.72226\nExp number, first 60 AAs: 0.22038\nTotal prob of N-in: 0.12994\noutside 1 331\nTMhelix 332 354\ninside 355 372

4049 GCF_002002225.1_ASM200222v1 Escherichia coli Proteobacteria; Gammaproteobacteria; Enterobacterales; Enterobacteriaceae; Escherichia MTGDNTLIHSHGINRRDFMKLCAALAATMGLSSKAAAEMAESVTNPQRPPVIWIGAQECT WP_000145418.1 hydrogenase 2 small subunit [Escherichia coli] Length: 372\nNumber of predicted TMHs: 1\nExp number of AAs in TMHs: 23.72313\nExp number, first 60 AAs: 0.22034\nTotal prob of N-in: 0.12997\noutside 1 331\nTMhelix 332 354\ninside 355 372

4050 GCF_002002255.1_ASM200225v1 Escherichia coli Proteobacteria; Gammaproteobacteria; Enterobacterales; Enterobacteriaceae; Escherichia MTGDNTLIHSHGINRRDFMKLCAALAATMGLSSKAAAEMAESVTNPQRPPVIWIGAQECT WP_000145410.1 MULTISPECIES: hydrogenase-2 small chain [Proteobacteria] Length: 372\nNumber of predicted TMHs:

1\nExp number of AAs in TMHs: 23.72226\nExp number, first 60 AAs: 0.22038\nTotal prob of N-in: 0.12994\noutside 1 331\nTMhelix 332 354\ninside 355 372

4051 GCF_002002305.1_ASM200230v1 Escherichia coli Proteobacteria; Gammaproteobacteria; Enterobacterales; Enterobacteriaceae; Escherichia
MTGDNTLIHSHGINRRDFMKLCAALAATMGLSSKAAAEMAESVTNPQRPPVIWIGAQECT WP_000145410.1
MULTISPECIES: hydrogenase-2 small chain [Proteobacteria] Length: 372\nNumber of predicted TMHs: 1\nExp number of AAs in TMHs: 23.72226\nExp number, first 60 AAs: 0.22038\nTotal prob of N-in: 0.12994\noutside 1 331\nTMhelix 332 354\ninside 355 372

4052 GCF_002002285.1_ASM200228v1 Escherichia coli Proteobacteria; Gammaproteobacteria; Enterobacterales; Enterobacteriaceae; Escherichia
MTGDNTLIHSHGINRRDFMKLCAALAATMGLSSKAAAEMAESVTNPQRPPVIWIGAQECT WP_000145410.1
MULTISPECIES: hydrogenase-2 small chain [Proteobacteria] Length: 372\nNumber of predicted TMHs: 1\nExp number of AAs in TMHs: 23.72226\nExp number, first 60 AAs: 0.22038\nTotal prob of N-in: 0.12994\noutside 1 331\nTMhelix 332 354\ninside 355 372

4053 GCF_002002325.1_ASM200232v1 Escherichia coli Proteobacteria; Gammaproteobacteria; Enterobacterales; Enterobacteriaceae; Escherichia
MNNEETFYQAMRRQGVTRRSFLKYCSLAATSLGLGAGMAPKIAWALENKPRIPVVWIHGL WP_001058323.1
MULTISPECIES: hydrogenase-1 small chain [Proteobacteria] Length: 372\nNumber of predicted TMHs: 2\nExp number of AAs in TMHs: 37.2608499999999\nExp number, first 60 AAs: 19.52923\nTotal prob of N-in: 0.94351\nPOSSIBLE N-term signal sequence\ninside 1 19\nTMhelix 20 39\noutside 40 325\nTMhelix 326 348\ninside 349 372

4054 GCF_002002285.1_ASM200228v1 Escherichia coli Proteobacteria; Gammaproteobacteria; Enterobacterales; Enterobacteriaceae; Escherichia
MNNEETFYQAMRRQGVTRRSFLKYCSLAATSLGLGAGMAPKIAWALENKPRIPVVWIHGL WP_001058323.1
MULTISPECIES: hydrogenase-1 small chain [Proteobacteria] Length: 372\nNumber of predicted TMHs: 2\nExp number of AAs in TMHs: 37.2608499999999\nExp number, first 60 AAs: 19.52923\nTotal prob of N-in: 0.94351\nPOSSIBLE N-term signal sequence\ninside 1 19\nTMhelix 20 39\noutside 40 325\nTMhelix 326 348\ninside 349 372

4055 GCF_002002255.1_ASM200225v1 Escherichia coli Proteobacteria; Gammaproteobacteria; Enterobacterales; Enterobacteriaceae; Escherichia
MNNEETFYQAMRRQGVTRRSFLKYCSLAATSLGLGAGMAPKIAWALENKPRIPVVWIHGL WP_001058323.1
MULTISPECIES: hydrogenase-1 small chain [Proteobacteria] Length: 372\nNumber of predicted TMHs: 2\nExp number of AAs in TMHs: 37.2608499999999\nExp number, first 60 AAs: 19.52923\nTotal prob of N-in: 0.94351\nPOSSIBLE N-term signal sequence\ninside 1 19\nTMhelix 20 39\noutside 40 325\nTMhelix 326 348\ninside 349 372

4056 GCF_002002305.1_ASM200230v1 Escherichia coli Proteobacteria; Gammaproteobacteria; Enterobacterales; Enterobacteriaceae; Escherichia
MNNEETFYQAMRRQGVTRRSFLKYCSLAATSLGLGAGMAPKIAWALENKPRIPVVWIHGL WP_001058323.1
MULTISPECIES: hydrogenase-1 small chain [Proteobacteria] Length: 372\nNumber of predicted TMHs: 2\nExp number of AAs in TMHs: 37.2608499999999\nExp number, first 60 AAs: 19.52923\nTotal prob of N-in: 0.94351\nPOSSIBLE N-term signal sequence\ninside 1 19\nTMhelix 20 39\noutside 40 325\nTMhelix 326 348\ninside 349 372

4057 GCF_002002465.1_ASM200246v1 Escherichia coli Proteobacteria; Gammaproteobacteria; Enterobacterales; Enterobacteriaceae; Escherichia
MTGDNTLIHSHGINRRDFMKLCAALAATMGLSSKAAAEMAESVTNPQRPPVIWIGAQECT WP_000145410.1
MULTISPECIES: hydrogenase-2 small chain [Proteobacteria] Length: 372\nNumber of predicted TMHs: 1\nExp number of AAs in TMHs: 23.72226\nExp number, first 60 AAs: 0.22038\nTotal prob of N-in: 0.12994\noutside 1 331\nTMhelix 332 354\ninside 355 372

4058 GCF_002002545.1_ASM200254v1 Escherichia coli Proteobacteria; Gammaproteobacteria; Enterobacterales; Enterobacteriaceae; Escherichia
MTGDNTLIHSHGINRRDFMKLCAALAATMGLSSKAAAEMAESVTNPQRPPVIWIGAQECT WP_000145410.1
MULTISPECIES: hydrogenase-2 small chain [Proteobacteria] Length: 372\nNumber of predicted TMHs:

1\nExp number of AAs in TMHs: 23.72226\nExp number, first 60 AAs: 0.22038\nTotal prob of N-in: 0.12994\noutside 1 331\nTMhelix 332 354\ninside 355 372

4059 GCF_002002585.1_ASM200258v1 Escherichia coli Proteobacteria; Gammaproteobacteria; Enterobacterales; Enterobacteriaceae; Escherichia MTGDNTLIHSHGINRRDFMKLCAALAATMGLSSKAAAEMAESVTNPQRPPVIWIGAQECT WP_000145410.1 MULTISPECIES: hydrogenase-2 small chain [Proteobacteria] Length: 372\nNumber of predicted TMHs: 1\nExp number of AAs in TMHs: 23.72226\nExp number, first 60 AAs: 0.22038\nTotal prob of N-in: 0.12994\noutside 1 331\nTMhelix 332 354\ninside 355 372

4060 GCF_002002585.1_ASM200258v1 Escherichia coli Proteobacteria; Gammaproteobacteria; Enterobacterales; Enterobacteriaceae; Escherichia MNNEETFYQAMRRQGVTRRSFLKYCSLAATSLGLGAGMAPKIAWALENKPRIPVVWIHGL WP_001058323.1 MULTISPECIES: hydrogenase-1 small chain [Proteobacteria] Length: 372\nNumber of predicted TMHs: 2\nExp number of AAs in TMHs: 37.2608499999999\nExp number, first 60 AAs: 19.52923\nTotal prob of N-in: 0.94351\nPOSSIBLE N-term signal sequence\ninside 1 19\nTMhelix 20 39\noutside 40 325\nTMhelix 326 348\ninside 349 372

4061 GCF_002002845.1_ASM200284v1 Escherichia coli Proteobacteria; Gammaproteobacteria; Enterobacterales; Enterobacteriaceae; Escherichia MTGDNTLIHSHGINRRDFMKLCAALAATMGLSSKAAAEMAESVTNPQRPPVIWIGAQECT WP_000145410.1 MULTISPECIES: hydrogenase-2 small chain [Proteobacteria] Length: 372\nNumber of predicted TMHs: 1\nExp number of AAs in TMHs: 23.72226\nExp number, first 60 AAs: 0.22038\nTotal prob of N-in: 0.12994\noutside 1 331\nTMhelix 332 354\ninside 355 372

4062 GCF_002002775.1_ASM200277v1 Escherichia coli Proteobacteria; Gammaproteobacteria; Enterobacterales; Enterobacteriaceae; Escherichia MTGDNTLIHSHGINRRDFMKLCAALAATMGLSSKAAAEMAESVTNPQRPPVIWIGAQECT WP_000145410.1 MULTISPECIES: hydrogenase-2 small chain [Proteobacteria] Length: 372\nNumber of predicted TMHs: 1\nExp number of AAs in TMHs: 23.72226\nExp number, first 60 AAs: 0.22038\nTotal prob of N-in: 0.12994\noutside 1 331\nTMhelix 332 354\ninside 355 372

4063 GCF_002002845.1_ASM200284v1 Escherichia coli Proteobacteria; Gammaproteobacteria; Enterobacterales; Enterobacteriaceae; Escherichia MNNEETFYQAMRRQGVTRRSFLKYCSLAATSLGLGAGMAPKIAWALENKPRIPVVWIHGL WP_001058323.1 MULTISPECIES: hydrogenase-1 small chain [Proteobacteria] Length: 372\nNumber of predicted TMHs: 2\nExp number of AAs in TMHs: 37.2608499999999\nExp number, first 60 AAs: 19.52923\nTotal prob of N-in: 0.94351\nPOSSIBLE N-term signal sequence\ninside 1 19\nTMhelix 20 39\noutside 40 325\nTMhelix 326 348\ninside 349 372

4064 GCF_002002775.1_ASM200277v1 Escherichia coli Proteobacteria; Gammaproteobacteria; Enterobacterales; Enterobacteriaceae; Escherichia MNNEETFYQAMRRQGVTRRSFLKYCSLAATSLGLGAGMAPKIAWALENKPRIPVVWIHGL WP_001058323.1 MULTISPECIES: hydrogenase-1 small chain [Proteobacteria] Length: 372\nNumber of predicted TMHs: 2\nExp number of AAs in TMHs: 37.2608499999999\nExp number, first 60 AAs: 19.52923\nTotal prob of N-in: 0.94351\nPOSSIBLE N-term signal sequence\ninside 1 19\nTMhelix 20 39\noutside 40 325\nTMhelix 326 348\ninside 349 372

4065 GCF_002002595.1_ASM200259v1 Escherichia coli Proteobacteria; Gammaproteobacteria; Enterobacterales; Enterobacteriaceae; Escherichia MNNEETFYQAMRRQGVTRRSFLKYCSLAATSLGLGAGMAPKIAWALENKPRIPVVWIHGL WP_001058323.1 MULTISPECIES: hydrogenase-1 small chain [Proteobacteria] Length: 372\nNumber of predicted TMHs: 2\nExp number of AAs in TMHs: 37.2608499999999\nExp number, first 60 AAs: 19.52923\nTotal prob of N-in: 0.94351\nPOSSIBLE N-term signal sequence\ninside 1 19\nTMhelix 20 39\noutside 40 325\nTMhelix 326 348\ninside 349 372

4066 GCF_002006865.1_ASM200686v1 Escherichia coli Proteobacteria; Gammaproteobacteria; Enterobacterales; Enterobacteriaceae; Escherichia MNNEETFYQAMRRQGVTRRSFLKYCSLAATSLGLGAGMAPKIAWALENKPRIPVVWIHGL WP_001058323.1 MULTISPECIES: hydrogenase-1 small chain [Proteobacteria] Length: 372\nNumber of predicted TMHs: 2\nExp number of AAs in TMHs: 37.2608499999999\nExp number, first 60 AAs: 19.52923\nTotal prob of N-in:

0.94351\nPOSSIBLE N-term signal sequence\ninside 1 19\nTMhelix 20 39\noutside 40 325\nTMhelix 326 348\ninside 349 372

4067 GCF_002007705.1_ASM200770v1 Escherichia coli Proteobacteria; Gammaproteobacteria; Enterobacterales; Enterobacteriaceae; Escherichia MTGDNTLIHSHGINRRDFMKLCAALAATMGLSSKAAAEMAESVTNPQRPPVIWIGAQECT WP_000145410.1 MULTISPECIES: hydrogenase-2 small chain [Proteobacteria] Length: 372\nNumber of predicted TMHs: 1\nExp number of AAs in TMHs: 23.72226\nExp number, first 60 AAs: 0.22038\nTotal prob of N-in: 0.12994\noutside 1 331\nTMhelix 332 354\ninside 355 372

4068 GCF_002009315.1_ASM200931v1 Escherichia coli Proteobacteria; Gammaproteobacteria; Enterobacterales; Enterobacteriaceae; Escherichia MTGDNTLIHSHGINRRDFMKLCAALAATMGLSSKAAAEMAESVTNPQRPPVIWIGAQECT WP_000145410.1 MULTISPECIES: hydrogenase-2 small chain [Proteobacteria] Length: 372\nNumber of predicted TMHs: 1\nExp number of AAs in TMHs: 23.72226\nExp number, first 60 AAs: 0.22038\nTotal prob of N-in: 0.12994\noutside 1 331\nTMhelix 332 354\ninside 355 372

4069 GCF_002011945.1_ASM201194v1 Escherichia coli Proteobacteria; Gammaproteobacteria; Enterobacterales; Enterobacteriaceae; Escherichia MTGDNTLIHSHGINRRDFMKLCAALAATMGLSSKAAAEMAESVTNPQRPPVIWIGAQECT WP_000145410.1 MULTISPECIES: hydrogenase-2 small chain [Proteobacteria] Length: 372\nNumber of predicted TMHs: 1\nExp number of AAs in TMHs: 23.72226\nExp number, first 60 AAs: 0.22038\nTotal prob of N-in: 0.12994\noutside 1 331\nTMhelix 332 354\ninside 355 372

4070 GCF_002011985.1_ASM201198v1 Escherichia coli Proteobacteria; Gammaproteobacteria; Enterobacterales; Enterobacteriaceae; Escherichia MTGDNTLIHSHGINRRDFMKLCAALAATMGLSSKAAAEMAESVTNPQRPPVIWIGAQECT WP_000145410.1 MULTISPECIES: hydrogenase-2 small chain [Proteobacteria] Length: 372\nNumber of predicted TMHs: 1\nExp number of AAs in TMHs: 23.72226\nExp number, first 60 AAs: 0.22038\nTotal prob of N-in: 0.12994\noutside 1 331\nTMhelix 332 354\ninside 355 372

4071 GCF_002009315.1_ASM200931v1 Escherichia coli Proteobacteria; Gammaproteobacteria; Enterobacterales; Enterobacteriaceae; Escherichia MNNEETFYQAMRRQGVTRRSFLKYCSLAATSLGLGAGMAPKIAWALENKPRIPVVWIHGL WP_001058323.1 MULTISPECIES: hydrogenase-1 small chain [Proteobacteria] Length: 372\nNumber of predicted TMHs: 2\nExp number of AAs in TMHs: 37.2608499999999\nExp number, first 60 AAs: 19.52923\nTotal prob of N-in: 0.94351\nPOSSIBLE N-term signal sequence\ninside 1 19\nTMhelix 20 39\noutside 40 325\nTMhelix 326 348\ninside 349 372

4072 GCF_002011985.1_ASM201198v1 Escherichia coli Proteobacteria; Gammaproteobacteria; Enterobacterales; Enterobacteriaceae; Escherichia MNNEETFYQAMRRQGVTRRSFLKYCSLAATSLGLGAGMAPKIAWALENKPRIPVVWIHGL WP_001058323.1 MULTISPECIES: hydrogenase-1 small chain [Proteobacteria] Length: 372\nNumber of predicted TMHs: 2\nExp number of AAs in TMHs: 37.2608499999999\nExp number, first 60 AAs: 19.52923\nTotal prob of N-in: 0.94351\nPOSSIBLE N-term signal sequence\ninside 1 19\nTMhelix 20 39\noutside 40 325\nTMhelix 326 348\ninside 349 372

4073 GCF_002012045.1_ASM201204v1 Escherichia coli Proteobacteria; Gammaproteobacteria; Enterobacterales; Enterobacteriaceae; Escherichia MTGDNTLIHSHGINRRDFMKLCAALAATMGLSSKAAAEMAESVTNPQRPPVIWIGAQECT WP_000145410.1 MULTISPECIES: hydrogenase-2 small chain [Proteobacteria] Length: 372\nNumber of predicted TMHs: 1\nExp number of AAs in TMHs: 23.72226\nExp number, first 60 AAs: 0.22038\nTotal prob of N-in: 0.12994\noutside 1 331\nTMhelix 332 354\ninside 355 372

4074 GCF_002012065.1_ASM201206v1 Escherichia coli Proteobacteria; Gammaproteobacteria; Enterobacterales; Enterobacteriaceae; Escherichia MTGDNTLIHSHGINRRDFMKLCAALAATMGLSSKAAAEMVESVTNPQRPPVIWIGAQECT WP_000145423.1 hydrogenase 2 small subunit [Escherichia coli] Length: 372\nNumber of predicted TMHs: 1\nExp number of AAs in TMHs: 23.78201\nExp number, first 60 AAs: 0.28355\nTotal prob of N-in: 0.13257\noutside 1 331\nTMhelix 332 354\ninside 355 372

4075 GCF_002012085.1_ASM201208v1 Escherichia coli Proteobacteria; Gammaproteobacteria; Enterobacterales; Enterobacteriaceae; Escherichia
MTGDNTLIHSHGINRRDFMKLCAALAATMGLSSKAAAEMVESVTNPQRPPVIWIGAQECT WP_000145423.1
hydrogenase 2 small subunit [Escherichia coli] Length: 372\nNumber of predicted TMHs: 1\nExp number of AAs in TMHs: 23.78201\nExp number, first 60 AAs: 0.28355\nTotal prob of N-in: 0.13257\noutside 1 331\nTMhelix 332 354\ninside 355 372

4076 GCF_002028825.1_ASM202882v1 Escherichia coli Proteobacteria; Gammaproteobacteria; Enterobacterales; Enterobacteriaceae; Escherichia
MTGDNTLIHSHGINRRDFMKLCAALAATMGLSSKAAAEMAESVTNPQRPPVIWIGAQECT WP_000145410.1
MULTISPECIES: hydrogenase-2 small chain [Proteobacteria] Length: 372\nNumber of predicted TMHs: 1\nExp number of AAs in TMHs: 23.72226\nExp number, first 60 AAs: 0.22038\nTotal prob of N-in: 0.12994\noutside 1 331\nTMhelix 332 354\ninside 355 372

4077 GCF_002015195.1_ASM201519v1 Escherichia coli Proteobacteria; Gammaproteobacteria; Enterobacterales; Enterobacteriaceae; Escherichia
MTGDNTLIHSHGINRRDFMKLCAALAATMGLSSKAAAEMAESVTNPQRPPVIWIGAQECT WP_000145410.1
MULTISPECIES: hydrogenase-2 small chain [Proteobacteria] Length: 372\nNumber of predicted TMHs: 1\nExp number of AAs in TMHs: 23.72226\nExp number, first 60 AAs: 0.22038\nTotal prob of N-in: 0.12994\noutside 1 331\nTMhelix 332 354\ninside 355 372

4078 GCF_002012045.1_ASM201204v1 Escherichia coli Proteobacteria; Gammaproteobacteria; Enterobacterales; Enterobacteriaceae; Escherichia
MNNEETFYQAMRRQGVTRRSFLKYCSLAATSLGLGAGMAPKIAWALENKPRIPVVWIHGL WP_001058323.1
MULTISPECIES: hydrogenase-1 small chain [Proteobacteria] Length: 372\nNumber of predicted TMHs: 2\nExp number of AAs in TMHs: 37.2608499999999\nExp number, first 60 AAs: 19.52923\nTotal prob of N-in: 0.94351\nPOSSIBLE N-term signal sequence\ninside 1 19\nTMhelix 20 39\noutside 40 325\nTMhelix 326 348\ninside 349 372

4079 GCF_002012005.1_ASM201200v1 Escherichia coli Proteobacteria; Gammaproteobacteria; Enterobacterales; Enterobacteriaceae; Escherichia
MNNEETFYQAMRRQGVTRRSFLKYCSLAATSLGLGAGMAPKIAWALENKPRIPVVWIHGL WP_001058320.1
hydrogenase [Escherichia coli] Length: 372\nNumber of predicted TMHs: 2\nExp number of AAs in TMHs: 37.26082\nExp number, first 60 AAs: 19.52923\nTotal prob of N-in: 0.94351\nPOSSIBLE N-term signal sequence\ninside 1 19\nTMhelix 20 39\noutside 40 325\nTMhelix 326 348\ninside 349 372

4080 GCF_000782315.1_ASM78231v1 Escherichia coli Proteobacteria; Gammaproteobacteria; Enterobacterales; Enterobacteriaceae; Escherichia
MTGDNTLIHSHGINRRDFMKLCAALAATMGLSSKAAAEMAESVTNPQRPPVIWIGAQECT WP_000145410.1
MULTISPECIES: hydrogenase-2 small chain [Proteobacteria] Length: 372\nNumber of predicted TMHs: 1\nExp number of AAs in TMHs: 23.72226\nExp number, first 60 AAs: 0.22038\nTotal prob of N-in: 0.12994\noutside 1 331\nTMhelix 332 354\ninside 355 372

4081 GCF_002012145.1_ASM201214v1 Escherichia coli Proteobacteria; Gammaproteobacteria; Enterobacterales; Enterobacteriaceae; Escherichia
MNNEETFYQAMRRQGVTRRSFLKYCSLAATSLGLGAGMAPKIAWALENKPRIPVVWIHGL WP_001295940.1
MULTISPECIES: hydrogenase [Enterobacteriaceae] Length: 372\nNumber of predicted TMHs: 2\nExp number of AAs in TMHs: 37.2608899999999\nExp number, first 60 AAs: 19.52923\nTotal prob of N-in: 0.94351\nPOSSIBLE N-term signal sequence\ninside 1 19\nTMhelix 20 39\noutside 40 325\nTMhelix 326 348\ninside 349 372

4082 GCF_002012165.1_ASM201216v1 Escherichia coli Proteobacteria; Gammaproteobacteria; Enterobacterales; Enterobacteriaceae; Escherichia
MNNEETFYQAMRRQGVTRRSFLKYCSLAATSLGLGAGMAPKIAWALENKPRIPVVWIHGL WP_001295940.1
MULTISPECIES: hydrogenase [Enterobacteriaceae] Length: 372\nNumber of predicted TMHs: 2\nExp number of AAs in TMHs: 37.2608899999999\nExp number, first 60 AAs: 19.52923\nTotal prob of N-in: 0.94351\nPOSSIBLE N-term signal sequence\ninside 1 19\nTMhelix 20 39\noutside 40 325\nTMhelix 326 348\ninside 349 372

4083 GCF_002012065.1_ASM201206v1 Escherichia coli Proteobacteria; Gammaproteobacteria; Enterobacterales; Enterobacteriaceae; Escherichia

MNNEETFYQAMRRQGVTRRSFLKYCSLAATSLGLGAGMAPKIAWALENKPRIPVVWIHGL WP_001295940.1 MULTISPECIES: hydrogenase [Enterobacteriaceae] Length: 372\nNumber of predicted TMHs: 2\nExp number of AAs in TMHs: 37.2608899999999\nExp number, first 60 AAs: 19.52923\nTotal prob of N-in: 0.94351\nPOSSIBLE N-term signal sequence\ninside 1 19\nTMhelix 20 39\noutside 40 325\nTMhelix 326 348\ninside 349 372

4084 GCF_002012085.1_ASM201208v1 Escherichia coli Proteobacteria; Gammaproteobacteria; Enterobacterales; Enterobacteriaceae; Escherichia MNNEETFYQAMRRQGVTRRSFLKYCSLAATSLGLGAGMAPKIAWALENKPRIPVVWIHGL WP_001295940.1 MULTISPECIES: hydrogenase [Enterobacteriaceae] Length: 372\nNumber of predicted TMHs: 2\nExp number of AAs in TMHs: 37.2608899999999\nExp number, first 60 AAs: 19.52923\nTotal prob of N-in: 0.94351\nPOSSIBLE N-term signal sequence\ninside 1 19\nTMhelix 20 39\noutside 40 325\nTMhelix 326 348\ninside 349 372

4085 GCF_002012185.1_ASM201218v1 Escherichia coli Proteobacteria; Gammaproteobacteria; Enterobacterales; Enterobacteriaceae; Escherichia MTGDNTLIHSHGINRRDFMKLCAALAATMGLSSKAAAEMAESVTNPQRPPVIWIGAQECT WP_000145410.1 MULTISPECIES: hydrogenase-2 small chain [Proteobacteria] Length: 372\nNumber of predicted TMHs: 1\nExp number of AAs in TMHs: 23.72226\nExp number, first 60 AAs: 0.22038\nTotal prob of N-in: 0.12994\noutside 1 331\nTMhelix 332 354\ninside 355 372

4086 GCF_002012225.1_ASM201222v1 Escherichia coli Proteobacteria; Gammaproteobacteria; Enterobacterales; Enterobacteriaceae; Escherichia MTGDNTLIHSHGINRRDFMKLCAALAATMGLSSKAAAEMAESVTNPQRPPVIWIGAQECT WP_000145410.1 MULTISPECIES: hydrogenase-2 small chain [Proteobacteria] Length: 372\nNumber of predicted TMHs: 1\nExp number of AAs in TMHs: 23.72226\nExp number, first 60 AAs: 0.22038\nTotal prob of N-in: 0.12994\noutside 1 331\nTMhelix 332 354\ninside 355 372

4087 GCF_002012205.1_ASM201220v1 Escherichia coli Proteobacteria; Gammaproteobacteria; Enterobacterales; Enterobacteriaceae; Escherichia MTGDNTLIHSHGINRRDFMKLCAALAATMGLSSKAAAEMVESVTNPQRPPVIWIGAQECT WP_000145423.1 hydrogenase 2 small subunit [Escherichia coli] Length: 372\nNumber of predicted TMHs: 1\nExp number of AAs in TMHs: 23.78201\nExp number, first 60 AAs: 0.28355\nTotal prob of N-in: 0.13257\noutside 1 331\nTMhelix 332 354\ninside 355 372

4088 GCF_002015235.1_ASM201523v1 Escherichia coli Proteobacteria; Gammaproteobacteria; Enterobacterales; Enterobacteriaceae; Escherichia MTGDNTLIHSHGINRRDFMKLCAALAATMGLSSKAAAEMAESVTNPQRPPVIWIGAQECT WP_000145410.1 MULTISPECIES: hydrogenase-2 small chain [Proteobacteria] Length: 372\nNumber of predicted TMHs: 1\nExp number of AAs in TMHs: 23.72226\nExp number, first 60 AAs: 0.22038\nTotal prob of N-in: 0.12994\noutside 1 331\nTMhelix 332 354\ninside 355 372

4089 GCF_002015205.1_ASM201520v1 Escherichia coli Proteobacteria; Gammaproteobacteria; Enterobacterales; Enterobacteriaceae; Escherichia MNNEETFYQAMRRQGVTRRSFLKYCSLAATSLGLGAGMAPKIAWALENKPRIPVVWIHGL WP_024246676.1 hydrogenase-1 small chain [Escherichia coli] Length: 372\nNumber of predicted TMHs: 2\nExp number of AAs in TMHs: 37.8214899999999\nExp number, first 60 AAs: 19.52326\nTotal prob of N-in: 0.94395\nPOSSIBLE N-term signal sequence\ninside 1 19\nTMhelix 20 39\noutside 40 325\nTMhelix 326 348\ninside 349 372

4090 GCF_002015235.1_ASM201523v1 Escherichia coli Proteobacteria; Gammaproteobacteria; Enterobacterales; Enterobacteriaceae; Escherichia MNNEETFYQAMRRQGVTRRSFLKYCSLAATSLGLGAGMAPKIAWALENKPRIPVVWIHGL WP_024246676.1 hydrogenase-1 small chain [Escherichia coli] Length: 372\nNumber of predicted TMHs: 2\nExp number of AAs in TMHs: 37.8214899999999\nExp number, first 60 AAs: 19.52326\nTotal prob of N-in: 0.94395\nPOSSIBLE N-term signal sequence\ninside 1 19\nTMhelix 20 39\noutside 40 325\nTMhelix 326 348\ninside 349 372

4091 GCF_002012205.1_ASM201220v1 Escherichia coli Proteobacteria; Gammaproteobacteria; Enterobacterales; Enterobacteriaceae; Escherichia MNNEETFYQAMRRQGVTRRSFLKYCSLAATSLGLGAGMAPKIAWALENKPRIPVVWIHGL WP_001295940.1 MULTISPECIES: hydrogenase [Enterobacteriaceae] Length: 372\nNumber of predicted TMHs: 2\nExp number of AAs in TMHs: 37.2608899999999\nExp number, first 60 AAs: 19.52923\nTotal prob of N-in:

0.94351\nPOSSIBLE N-term signal sequence\ninside 1 19\nTMhelix 20 39\noutside 40 325\nTMhelix 326 348\ninside 349 372

4092 GCF_002012245.1_ASM201224v1 Escherichia coli Proteobacteria; Gammaproteobacteria; Enterobacterales; Enterobacteriaceae; Escherichia MNNEETFYQAMRRQGVTRRSFLKYCSLAATSLGLGAGMAPKIAWALENKPRIPVVWIHGL WP_001295940.1 MULTISPECIES: hydrogenase [Enterobacteriaceae] Length: 372\nNumber of predicted TMHs: 2\nExp number of AAs in TMHs: 37.2608899999999\nExp number, first 60 AAs: 19.52923\nTotal prob of N-in: 0.94351\nPOSSIBLE N-term signal sequence\ninside 1 19\nTMhelix 20 39\noutside 40 325\nTMhelix 326 348\ninside 349 372

4093 GCF_002015255.1_ASM201525v1 Escherichia coli Proteobacteria; Gammaproteobacteria; Enterobacterales; Enterobacteriaceae; Escherichia MTGDNTLIHSHGINRRDFMKLCAALAATMGLSSKAAAEMAESVTNPQRPPVIWIGAQECT WP_000145410.1 MULTISPECIES: hydrogenase-2 small chain [Proteobacteria] Length: 372\nNumber of predicted TMHs: 1\nExp number of AAs in TMHs: 23.72226\nExp number, first 60 AAs: 0.22038\nTotal prob of N-in: 0.12994\noutside 1 331\nTMhelix 332 354\ninside 355 372

4094 GCF_002015265.1_ASM201526v1 Escherichia coli Proteobacteria; Gammaproteobacteria; Enterobacterales; Enterobacteriaceae; Escherichia MTGDNTLIHSHGINRRDFMKLCAALAATMGLSSKAAAEMAESVTNPQRPPVIWIGAQECT WP_000145410.1 MULTISPECIES: hydrogenase-2 small chain [Proteobacteria] Length: 372\nNumber of predicted TMHs: 1\nExp number of AAs in TMHs: 23.72226\nExp number, first 60 AAs: 0.22038\nTotal prob of N-in: 0.12994\noutside 1 331\nTMhelix 332 354\ninside 355 372

4095 GCF_002015325.1_ASM201532v1 Escherichia coli Proteobacteria; Gammaproteobacteria; Enterobacterales; Enterobacteriaceae; Escherichia MTGDNTLIHSHGINRRDFMKLCAALAATMGLSSKAAAEMAESVTNPQRPPVIWIGAQECT WP_000145410.1 MULTISPECIES: hydrogenase-2 small chain [Proteobacteria] Length: 372\nNumber of predicted TMHs: 1\nExp number of AAs in TMHs: 23.72226\nExp number, first 60 AAs: 0.22038\nTotal prob of N-in: 0.12994\noutside 1 331\nTMhelix 332 354\ninside 355 372

4096 GCF_002015355.1_ASM201535v1 Escherichia coli Proteobacteria; Gammaproteobacteria; Enterobacterales; Enterobacteriaceae; Escherichia MTGDNTLIHSHGINRRDFMKLCAALAATMGLSSKAAAEMAESVTNPQRPPVIWIGAQECT WP_000145410.1 MULTISPECIES: hydrogenase-2 small chain [Proteobacteria] Length: 372\nNumber of predicted TMHs: 1\nExp number of AAs in TMHs: 23.72226\nExp number, first 60 AAs: 0.22038\nTotal prob of N-in: 0.12994\noutside 1 331\nTMhelix 332 354\ninside 355 372

4097 GCF_002015365.1_ASM201536v1 Escherichia coli Proteobacteria; Gammaproteobacteria; Enterobacterales; Enterobacteriaceae; Escherichia MTGDNTLIHSHGINRRDFMKLCAALAATMGLSSKAAAEMAESVTNPQRPPVIWIGAQECT WP_000145410.1 MULTISPECIES: hydrogenase-2 small chain [Proteobacteria] Length: 372\nNumber of predicted TMHs: 1\nExp number of AAs in TMHs: 23.72226\nExp number, first 60 AAs: 0.22038\nTotal prob of N-in: 0.12994\noutside 1 331\nTMhelix 332 354\ninside 355 372

4098 GCF_002015375.1_ASM201537v1 Escherichia coli Proteobacteria; Gammaproteobacteria; Enterobacterales; Enterobacteriaceae; Escherichia MTGDNTLIHSHGINRRDFMKLCAALAATMGLSSKAAAEMAESVTNPQRPPVIWIGAQECT WP_000145410.1 MULTISPECIES: hydrogenase-2 small chain [Proteobacteria] Length: 372\nNumber of predicted TMHs: 1\nExp number of AAs in TMHs: 23.72226\nExp number, first 60 AAs: 0.22038\nTotal prob of N-in: 0.12994\noutside 1 331\nTMhelix 332 354\ninside 355 372

4099 GCF_002015385.1_ASM201538v1 Escherichia coli Proteobacteria; Gammaproteobacteria; Enterobacterales; Enterobacteriaceae; Escherichia MTGDNTLIHSHGINRRDFMKLCAALAATMGLSSKAAAEMAESVTNPQRPPVIWIGAQECT WP_000145410.1 MULTISPECIES: hydrogenase-2 small chain [Proteobacteria] Length: 372\nNumber of predicted TMHs: 1\nExp number of AAs in TMHs: 23.72226\nExp number, first 60 AAs: 0.22038\nTotal prob of N-in: 0.12994\noutside 1 331\nTMhelix 332 354\ninside 355 372

4100 GCF_002015435.1_ASM201543v1 Escherichia coli Proteobacteria; Gammaproteobacteria; Enterobacterales; Enterobacteriaceae; Escherichia
MTGDNTLIHSHGINRRDFMKLCAALAATMGLSSKAAAEMAESVTNPQRPPVIWIGAQECT WP_000145410.1
MULTISPECIES: hydrogenase-2 small chain [Proteobacteria] Length: 372\nNumber of predicted TMHs: 1\nExp number of AAs in TMHs: 23.72226\nExp number, first 60 AAs: 0.22038\nTotal prob of N-in: 0.12994\noutside 1 331\nTMhelix 332 354\ninside 355 372

4101 GCF_002015455.1_ASM201545v1 Escherichia coli Proteobacteria; Gammaproteobacteria; Enterobacterales; Enterobacteriaceae; Escherichia
MTGDNTLIHSHGINRRDFMKLCAALAATMGLSSKAAAEMAESVTNPQRPPVIWIGAQECT WP_000145410.1
MULTISPECIES: hydrogenase-2 small chain [Proteobacteria] Length: 372\nNumber of predicted TMHs: 1\nExp number of AAs in TMHs: 23.72226\nExp number, first 60 AAs: 0.22038\nTotal prob of N-in: 0.12994\noutside 1 331\nTMhelix 332 354\ninside 355 372

4102 GCF_002015455.1_ASM201545v1 Escherichia coli Proteobacteria; Gammaproteobacteria; Enterobacterales; Enterobacteriaceae; Escherichia
MNNEETFYQAMRRQGVTRRSFLKYCSLAATSLGLGAGMAPKIAWALENKPRIPVVWIHGL WP_024246676.1
hydrogenase-1 small chain [Escherichia coli] Length: 372\nNumber of predicted TMHs: 2\nExp number of AAs in TMHs: 37.8214899999999\nExp number, first 60 AAs: 19.52326\nTotal prob of N-in: 0.94395\nPOSSIBLE N-term signal sequence\ninside 1 19\nTMhelix 20 39\noutside 40 325\nTMhelix 326 348\ninside 349 372

4103 GCF_002015545.1_ASM201554v1 Escherichia coli Proteobacteria; Gammaproteobacteria; Enterobacterales; Enterobacteriaceae; Escherichia
MTGDNTLIHSHGINRRDFMKLCAALAATMGLSSKAAAEMAESVTNPQRPPVIWIGAQECT WP_000145410.1
MULTISPECIES: hydrogenase-2 small chain [Proteobacteria] Length: 372\nNumber of predicted TMHs: 1\nExp number of AAs in TMHs: 23.72226\nExp number, first 60 AAs: 0.22038\nTotal prob of N-in: 0.12994\noutside 1 331\nTMhelix 332 354\ninside 355 372

4104 GCF_002015555.1_ASM201555v1 Escherichia coli Proteobacteria; Gammaproteobacteria; Enterobacterales; Enterobacteriaceae; Escherichia
MTGDNTLIHSHGINRRDFMKLCAALAATMGLSSKAAAEMAESVTNPQRPPVIWIGAQECT WP_000145410.1
MULTISPECIES: hydrogenase-2 small chain [Proteobacteria] Length: 372\nNumber of predicted TMHs: 1\nExp number of AAs in TMHs: 23.72226\nExp number, first 60 AAs: 0.22038\nTotal prob of N-in: 0.12994\noutside 1 331\nTMhelix 332 354\ninside 355 372

4105 GCF_002015595.1_ASM201559v1 Escherichia coli Proteobacteria; Gammaproteobacteria; Enterobacterales; Enterobacteriaceae; Escherichia
MTGDNTLIHSHGINRRDFMKLCAALAATMGLSSKAAAEMAESVTNPQRPPVIWIGAQECT WP_000145410.1
MULTISPECIES: hydrogenase-2 small chain [Proteobacteria] Length: 372\nNumber of predicted TMHs: 1\nExp number of AAs in TMHs: 23.72226\nExp number, first 60 AAs: 0.22038\nTotal prob of N-in: 0.12994\noutside 1 331\nTMhelix 332 354\ninside 355 372

4106 GCF_002015625.1_ASM201562v1 Escherichia coli Proteobacteria; Gammaproteobacteria; Enterobacterales; Enterobacteriaceae; Escherichia
MTGDNTLIHSHGINRRDFMKLCAALAATMGLSSKAAAEMAESVTNPQRPPVIWIGAQECT WP_000145410.1
MULTISPECIES: hydrogenase-2 small chain [Proteobacteria] Length: 372\nNumber of predicted TMHs: 1\nExp number of AAs in TMHs: 23.72226\nExp number, first 60 AAs: 0.22038\nTotal prob of N-in: 0.12994\noutside 1 331\nTMhelix 332 354\ninside 355 372

4107 GCF_002015615.1_ASM201561v1 Escherichia coli Proteobacteria; Gammaproteobacteria; Enterobacterales; Enterobacteriaceae; Escherichia
MTGDNTLIHSHGINRRDFMKLCAALAATMGLSSKAAAEMAESVTNPQRPPVIWIGAQECT WP_000145410.1
MULTISPECIES: hydrogenase-2 small chain [Proteobacteria] Length: 372\nNumber of predicted TMHs: 1\nExp number of AAs in TMHs: 23.72226\nExp number, first 60 AAs: 0.22038\nTotal prob of N-in: 0.12994\noutside 1 331\nTMhelix 332 354\ninside 355 372

4108 GCF_002015635.1_ASM201563v1 Escherichia coli Proteobacteria; Gammaproteobacteria; Enterobacterales; Enterobacteriaceae; Escherichia
MTGDNTLIHSHGINRRDFMKLCAALAATMGLSSKAAAEMAESVTNPQRPPVIWIGAQECT WP_000145410.1
MULTISPECIES: hydrogenase-2 small chain [Proteobacteria] Length: 372\nNumber of predicted TMHs:

1\nExp number of AAs in TMHs: 23.72226\nExp number, first 60 AAs: 0.22038\nTotal prob of N-in: 0.12994\noutside 1 331\nTMhelix 332 354\ninside 355 372

4109 GCF_002015675.1_ASM201567v1 Escherichia coli Proteobacteria; Gammaproteobacteria; Enterobacterales; Enterobacteriaceae; Escherichia
MTGDNTLIHSHGINRRDFMKLCAALAATMGLSSKAAAEMAESVTNPQRPPVIWIGAQECT WP_000145410.1
MULTISPECIES: hydrogenase-2 small chain [Proteobacteria] Length: 372\nNumber of predicted TMHs: 1\nExp number of AAs in TMHs: 23.72226\nExp number, first 60 AAs: 0.22038\nTotal prob of N-in: 0.12994\noutside 1 331\nTMhelix 332 354\ninside 355 372

4110 GCF_002015695.1_ASM201569v1 Escherichia coli Proteobacteria; Gammaproteobacteria; Enterobacterales; Enterobacteriaceae; Escherichia
MTGDNTLIHSHGINRRDFMKLCAALAATMGLSSKAAAEMAESVTNPQRPPVIWIGAQECT WP_000145410.1
MULTISPECIES: hydrogenase-2 small chain [Proteobacteria] Length: 372\nNumber of predicted TMHs: 1\nExp number of AAs in TMHs: 23.72226\nExp number, first 60 AAs: 0.22038\nTotal prob of N-in: 0.12994\noutside 1 331\nTMhelix 332 354\ninside 355 372

4111 GCF_002015705.1_ASM201570v1 Escherichia coli Proteobacteria; Gammaproteobacteria; Enterobacterales; Enterobacteriaceae; Escherichia
MNNEETFYQAMRRQGVTRRSFLKYCSLAATSLGLGAGMAPKIAWALENKPRIPVVWIHGL WP_001295940.1
MULTISPECIES: hydrogenase [Enterobacteriaceae] Length: 372\nNumber of predicted TMHs: 2\nExp number of AAs in TMHs: 37.2608899999999\nExp number, first 60 AAs: 19.52923\nTotal prob of N-in: 0.94351\nPOSSIBLE N-term signal sequence\ninside 1 19\nTMhelix 20 39\noutside 40 325\nTMhelix 326 348\ninside 349 372

4112 GCF_002015615.1_ASM201561v1 Escherichia coli Proteobacteria; Gammaproteobacteria; Enterobacterales; Enterobacteriaceae; Escherichia
MNNEETFYQAMRRQGVTRRSFLKYCSLAATSLGLGAGMAPKIAWALENKPRIPVVWIHGL WP_001295940.1
MULTISPECIES: hydrogenase [Enterobacteriaceae] Length: 372\nNumber of predicted TMHs: 2\nExp number of AAs in TMHs: 37.2608899999999\nExp number, first 60 AAs: 19.52923\nTotal prob of N-in: 0.94351\nPOSSIBLE N-term signal sequence\ninside 1 19\nTMhelix 20 39\noutside 40 325\nTMhelix 326 348\ninside 349 372

4113 GCF_002015595.1_ASM201559v1 Escherichia coli Proteobacteria; Gammaproteobacteria; Enterobacterales; Enterobacteriaceae; Escherichia
MNNEETFYQAMRRQGVTRRSFLKYCSLAATSLGLGAGMAPKIAWALENKPRIPVVWIHGL WP_001295940.1
MULTISPECIES: hydrogenase [Enterobacteriaceae] Length: 372\nNumber of predicted TMHs: 2\nExp number of AAs in TMHs: 37.2608899999999\nExp number, first 60 AAs: 19.52923\nTotal prob of N-in: 0.94351\nPOSSIBLE N-term signal sequence\ninside 1 19\nTMhelix 20 39\noutside 40 325\nTMhelix 326 348\ninside 349 372

4114 GCF_002015635.1_ASM201563v1 Escherichia coli Proteobacteria; Gammaproteobacteria; Enterobacterales; Enterobacteriaceae; Escherichia
MNNEETFYQAMRRQGVTRRSFLKYCSLAATSLGLGAGMAPKIAWALENKPRIPVVWIHGL WP_001295940.1
MULTISPECIES: hydrogenase [Enterobacteriaceae] Length: 372\nNumber of predicted TMHs: 2\nExp number of AAs in TMHs: 37.2608899999999\nExp number, first 60 AAs: 19.52923\nTotal prob of N-in: 0.94351\nPOSSIBLE N-term signal sequence\ninside 1 19\nTMhelix 20 39\noutside 40 325\nTMhelix 326 348\ninside 349 372

4115 GCF_002015765.1_ASM201576v1 Escherichia coli Proteobacteria; Gammaproteobacteria; Enterobacterales; Enterobacteriaceae; Escherichia
MTGDNTLIHSHGINRRDFMKLCAALAATMGLSSKAAAEMAESVTNPQRPPVIWIGAQECT WP_000145410.1
MULTISPECIES: hydrogenase-2 small chain [Proteobacteria] Length: 372\nNumber of predicted TMHs: 1\nExp number of AAs in TMHs: 23.72226\nExp number, first 60 AAs: 0.22038\nTotal prob of N-in: 0.12994\noutside 1 331\nTMhelix 332 354\ninside 355 372

4116 GCF_002015775.1_ASM201577v1 Escherichia coli Proteobacteria; Gammaproteobacteria; Enterobacterales; Enterobacteriaceae; Escherichia
MTGDNTLIHSHGINRRDFMKLCAALAATMGLSSKAAAEMAESVTNPQRPPVIWIGAQECT WP_000145410.1
MULTISPECIES: hydrogenase-2 small chain [Proteobacteria] Length: 372\nNumber of predicted TMHs:

1\nExp number of AAs in TMHs: 23.72226\nExp number, first 60 AAs: 0.22038\nTotal prob of N-in: 0.12994\noutside 1 331\nTMhelix 332 354\ninside 355 372

4117 GCF_002015815.1_ASM201581v1 Escherichia coli Proteobacteria; Gammaproteobacteria; Enterobacterales; Enterobacteriaceae; Escherichia
MTGDNTLIHSHGINRRDFMKLCAALAATMGLSSKAAAEMAESVTNPQRPPVIWIGAQECT WP_000145410.1
MULTISPECIES: hydrogenase-2 small chain [Proteobacteria] Length: 372\nNumber of predicted TMHs: 1\nExp number of AAs in TMHs: 23.72226\nExp number, first 60 AAs: 0.22038\nTotal prob of N-in: 0.12994\noutside 1 331\nTMhelix 332 354\ninside 355 372

4118 GCF_002015895.1_ASM201589v1 Escherichia coli Proteobacteria; Gammaproteobacteria; Enterobacterales; Enterobacteriaceae; Escherichia
MTGDNTLIHSHGINRRDFMKLCAALAATMGLSSKAAAEMAESVTNPQRPPVIWIGAQECT WP_000145410.1
MULTISPECIES: hydrogenase-2 small chain [Proteobacteria] Length: 372\nNumber of predicted TMHs: 1\nExp number of AAs in TMHs: 23.72226\nExp number, first 60 AAs: 0.22038\nTotal prob of N-in: 0.12994\noutside 1 331\nTMhelix 332 354\ninside 355 372

4119 GCF_002015935.1_ASM201593v1 Escherichia coli Proteobacteria; Gammaproteobacteria; Enterobacterales; Enterobacteriaceae; Escherichia
MTGDNTLIHSHGINRRDFMKLCAALAATMGLSSKAAAEMAESVTNPQRPPVIWIGAQECT WP_000145410.1
MULTISPECIES: hydrogenase-2 small chain [Proteobacteria] Length: 372\nNumber of predicted TMHs: 1\nExp number of AAs in TMHs: 23.72226\nExp number, first 60 AAs: 0.22038\nTotal prob of N-in: 0.12994\noutside 1 331\nTMhelix 332 354\ninside 355 372

4120 GCF_002015735.1_ASM201573v1 Escherichia coli Proteobacteria; Gammaproteobacteria; Enterobacterales; Enterobacteriaceae; Escherichia
MTGDNTLIHSHGINRRDFMKLCAALAATMGLSSKAAAEMAESVTNPQRPPVIWIGAQECT WP_078187771.1
hydrogenase 2 small subunit [Escherichia coli] Length: 372\nNumber of predicted TMHs: 1\nExp number of AAs in TMHs: 23.11869\nExp number, first 60 AAs: 0.22152\nTotal prob of N-in: 0.15789\noutside 1 331\nTMhelix 332 354\ninside 355 372

4121 GCF_002015755.1_ASM201575v1 Escherichia coli Proteobacteria; Gammaproteobacteria; Enterobacterales; Enterobacteriaceae; Escherichia
MTGDNTLIHSHGINRRDFMKLCAALAATMGLSSKAAAEMAESVTNPQRPPVIWIGAQECT WP_078187771.1
hydrogenase 2 small subunit [Escherichia coli] Length: 372\nNumber of predicted TMHs: 1\nExp number of AAs in TMHs: 23.11869\nExp number, first 60 AAs: 0.22152\nTotal prob of N-in: 0.15789\noutside 1 331\nTMhelix 332 354\ninside 355 372

4122 GCF_002015765.1_ASM201576v1 Escherichia coli Proteobacteria; Gammaproteobacteria; Enterobacterales; Enterobacteriaceae; Escherichia
MNNEETFYQAMRRQGVTRRSFLKYCSLAATSLGLGAGMAPKIAWALENKPRIPVVWIHGL WP_001295940.1
MULTISPECIES: hydrogenase [Enterobacteriaceae] Length: 372\nNumber of predicted TMHs: 2\nExp number of AAs in TMHs: 37.2608899999999\nExp number, first 60 AAs: 19.52923\nTotal prob of N-in: 0.94351\nPOSSIBLE N-term signal sequence\ninside 1 19\nTMhelix 20 39\noutside 40 325\nTMhelix 326 348\ninside 349 372

4123 GCF_002015775.1_ASM201577v1 Escherichia coli Proteobacteria; Gammaproteobacteria; Enterobacterales; Enterobacteriaceae; Escherichia
MNNEETFYQAMRRQGVTRRSFLKYCSLAATSLGLGAGMAPKIAWALENKPRIPVVWIHGL WP_001295940.1
MULTISPECIES: hydrogenase [Enterobacteriaceae] Length: 372\nNumber of predicted TMHs: 2\nExp number of AAs in TMHs: 37.2608899999999\nExp number, first 60 AAs: 19.52923\nTotal prob of N-in: 0.94351\nPOSSIBLE N-term signal sequence\ninside 1 19\nTMhelix 20 39\noutside 40 325\nTMhelix 326 348\ninside 349 372

4124 GCF_002015815.1_ASM201581v1 Escherichia coli Proteobacteria; Gammaproteobacteria; Enterobacterales; Enterobacteriaceae; Escherichia
MNNEETFYQAMRRQGVTRRSFLKYCSLAATSLGLGAGMAPKIAWALENKPRIPVVWIHGL WP_001295940.1
MULTISPECIES: hydrogenase [Enterobacteriaceae] Length: 372\nNumber of predicted TMHs: 2\nExp number of AAs in TMHs: 37.2608899999999\nExp number, first 60 AAs: 19.52923\nTotal prob of N-in: 0.94351\nPOSSIBLE N-term signal sequence\ninside 1 19\nTMhelix 20 39\noutside 40 325\nTMhelix 326 348\ninside 349 372

4125 GCF_002015885.1_ASM201588v1 Escherichia coli Proteobacteria; Gammaproteobacteria; Enterobacterales; Enterobacteriaceae; Escherichia
MNNEETFYQAMRRQGVTRRSFLKYCSLAATSLGLGAGMAPKIAWALENKPRIPVVWIHGL WP_001295940.1
MULTISPECIES: hydrogenase [Enterobacteriaceae] Length: 372\nNumber of predicted TMHs: 2\nExp number of AAs in TMHs: 37.2608899999999\nExp number, first 60 AAs: 19.52923\nTotal prob of N-in: 0.94351\nPOSSIBLE N-term signal sequence\ninside 1 19\nTMhelix 20 39\noutside 40 325\nTMhelix 326 348\ninside 349 372

4126 GCF_002015975.1_ASM201597v1 Escherichia coli Proteobacteria; Gammaproteobacteria; Enterobacterales; Enterobacteriaceae; Escherichia
MTGDNTLIHSHGINRRDFMKLCAALAATMGLSSKAAAEMAESVTNPQRPPVIWIGAQECT WP_000145410.1
MULTISPECIES: hydrogenase-2 small chain [Proteobacteria] Length: 372\nNumber of predicted TMHs: 1\nExp number of AAs in TMHs: 23.72226\nExp number, first 60 AAs: 0.22038\nTotal prob of N-in: 0.12994\noutside 1 331\nTMhelix 332 354\ninside 355 372

4127 GCF_002015975.1_ASM201597v1 Escherichia coli Proteobacteria; Gammaproteobacteria; Enterobacterales; Enterobacteriaceae; Escherichia
MNNEETFYQAMRRQGVTRRSFLKYCSLAATSLGLGAGMAPKIAWALENKPRIPVVWIHGL WP_001295940.1
MULTISPECIES: hydrogenase [Enterobacteriaceae] Length: 372\nNumber of predicted TMHs: 2\nExp number of AAs in TMHs: 37.2608899999999\nExp number, first 60 AAs: 19.52923\nTotal prob of N-in: 0.94351\nPOSSIBLE N-term signal sequence\ninside 1 19\nTMhelix 20 39\noutside 40 325\nTMhelix 326 348\ninside 349 372

4128 GCF_002015895.1_ASM201589v1 Escherichia coli Proteobacteria; Gammaproteobacteria; Enterobacterales; Enterobacteriaceae; Escherichia
MNNEETFYQAMRRQGVTRRSFLKYCSLAATSLGLGAGMAPKIAWALENKPRIPVVWIHGL WP_001295940.1
MULTISPECIES: hydrogenase [Enterobacteriaceae] Length: 372\nNumber of predicted TMHs: 2\nExp number of AAs in TMHs: 37.2608899999999\nExp number, first 60 AAs: 19.52923\nTotal prob of N-in: 0.94351\nPOSSIBLE N-term signal sequence\ninside 1 19\nTMhelix 20 39\noutside 40 325\nTMhelix 326 348\ninside 349 372

4129 GCF_002015935.1_ASM201593v1 Escherichia coli Proteobacteria; Gammaproteobacteria; Enterobacterales; Enterobacteriaceae; Escherichia
MNNEETFYQAMRRQGVTRRSFLKYCSLAATSLGLGAGMAPKIAWALENKPRIPVVWIHGL WP_001295940.1
MULTISPECIES: hydrogenase [Enterobacteriaceae] Length: 372\nNumber of predicted TMHs: 2\nExp number of AAs in TMHs: 37.2608899999999\nExp number, first 60 AAs: 19.52923\nTotal prob of N-in: 0.94351\nPOSSIBLE N-term signal sequence\ninside 1 19\nTMhelix 20 39\noutside 40 325\nTMhelix 326 348\ninside 349 372

4130 GCF_002016015.1_ASM201601v1 Escherichia coli Proteobacteria; Gammaproteobacteria; Enterobacterales; Enterobacteriaceae; Escherichia
MTGDNTLIHSHGINRRDFMKLCAALAATMGLSSKAAAEMAESVTNPQRPPVIWIGAQECT WP_000145410.1
MULTISPECIES: hydrogenase-2 small chain [Proteobacteria] Length: 372\nNumber of predicted TMHs: 1\nExp number of AAs in TMHs: 23.72226\nExp number, first 60 AAs: 0.22038\nTotal prob of N-in: 0.12994\noutside 1 331\nTMhelix 332 354\ninside 355 372

4131 GCF_002016055.1_ASM201605v1 Escherichia coli Proteobacteria; Gammaproteobacteria; Enterobacterales; Enterobacteriaceae; Escherichia
MTGDNTLIHSHGINRRDFMKLCAALAATMGLSSKAAAEMAESVTNPQRPPVIWIGAQECT WP_000145410.1
MULTISPECIES: hydrogenase-2 small chain [Proteobacteria] Length: 372\nNumber of predicted TMHs: 1\nExp number of AAs in TMHs: 23.72226\nExp number, first 60 AAs: 0.22038\nTotal prob of N-in: 0.12994\noutside 1 331\nTMhelix 332 354\ninside 355 372

4132 GCF_002016015.1_ASM201601v1 Escherichia coli Proteobacteria; Gammaproteobacteria; Enterobacterales; Enterobacteriaceae; Escherichia
MNNEETFYQAMRRQGVTRRSFLKYCSLAATSLGLGAGMAPKIAWALENKPRIPVVWIHGL WP_001295940.1
MULTISPECIES: hydrogenase [Enterobacteriaceae] Length: 372\nNumber of predicted TMHs: 2\nExp number of AAs in TMHs: 37.2608899999999\nExp number, first 60 AAs: 19.52923\nTotal prob of N-in: 0.94351\nPOSSIBLE N-term signal sequence\ninside 1 19\nTMhelix 20 39\noutside 40 325\nTMhelix 326 348\ninside 349 372

4133 GCF_002016055.1_ASM201605v1 Escherichia coli Proteobacteria; Gammaproteobacteria; Enterobacterales; Enterobacteriaceae; Escherichia
MNNEETFYQAMRRQGVTRRSFLKYCSLAATSLGLGAGMAPKIAWALENKPRIPVVWIHGL WP_001295940.1
MULTISPECIES: hydrogenase [Enterobacteriaceae] Length: 372\nNumber of predicted TMHs: 2\nExp number of AAs in TMHs: 37.2608899999999\nExp number, first 60 AAs: 19.52923\nTotal prob of N-in: 0.94351\nPOSSIBLE N-term signal sequence\ninside 1 19\nTMhelix 20 39\noutside 40 325\nTMhelix 326 348\ninside 349 372

4134 GCF_002016025.1_ASM201602v1 Escherichia coli Proteobacteria; Gammaproteobacteria; Enterobacterales; Enterobacteriaceae; Escherichia
MNNEETFYQAMRRQGVTRRSFLKYCSLAATSLGLGAGMAPKIAWALENKPRIPVVWIHGL WP_001295940.1
MULTISPECIES: hydrogenase [Enterobacteriaceae] Length: 372\nNumber of predicted TMHs: 2\nExp number of AAs in TMHs: 37.2608899999999\nExp number, first 60 AAs: 19.52923\nTotal prob of N-in: 0.94351\nPOSSIBLE N-term signal sequence\ninside 1 19\nTMhelix 20 39\noutside 40 325\nTMhelix 326 348\ninside 349 372

4135 GCF_002016065.1_ASM201606v1 Escherichia coli Proteobacteria; Gammaproteobacteria; Enterobacterales; Enterobacteriaceae; Escherichia
MNNEETFYQAMRRQGVTRRSFLKYCSLAATSLGLGAGMAPKIAWALENKPRIPVVWIHGL WP_001295940.1
MULTISPECIES: hydrogenase [Enterobacteriaceae] Length: 372\nNumber of predicted TMHs: 2\nExp number of AAs in TMHs: 37.2608899999999\nExp number, first 60 AAs: 19.52923\nTotal prob of N-in: 0.94351\nPOSSIBLE N-term signal sequence\ninside 1 19\nTMhelix 20 39\noutside 40 325\nTMhelix 326 348\ninside 349 372

4136 GCF_002016195.1_ASM201619v1 Escherichia coli Proteobacteria; Gammaproteobacteria; Enterobacterales; Enterobacteriaceae; Escherichia
MTGDNTLIHSHGINRRDFMKLCAALAATMGLSSKAAAEMAESVTNPQRPPVIWIGAQECT WP_000145410.1
MULTISPECIES: hydrogenase-2 small chain [Proteobacteria] Length: 372\nNumber of predicted TMHs: 1\nExp number of AAs in TMHs: 23.72226\nExp number, first 60 AAs: 0.22038\nTotal prob of N-in: 0.12994\noutside 1 331\nTMhelix 332 354\ninside 355 372

4137 GCF_002016195.1_ASM201619v1 Escherichia coli Proteobacteria; Gammaproteobacteria; Enterobacterales; Enterobacteriaceae; Escherichia
MNNEETFYQAMRRQGVTRRSFLKYCSLAATSLGLGAGMAPKIAWALENKPRIPVVWIHGL WP_001295940.1
MULTISPECIES: hydrogenase [Enterobacteriaceae] Length: 372\nNumber of predicted TMHs: 2\nExp number of AAs in TMHs: 37.2608899999999\nExp number, first 60 AAs: 19.52923\nTotal prob of N-in: 0.94351\nPOSSIBLE N-term signal sequence\ninside 1 19\nTMhelix 20 39\noutside 40 325\nTMhelix 326 348\ninside 349 372

4138 GCF_002016075.1_ASM201607v1 Escherichia coli Proteobacteria; Gammaproteobacteria; Enterobacterales; Enterobacteriaceae; Escherichia
MNNEETFYQAMRRQGVTRRSFLKYCSLAATSLGLGAGMAPKIAWALENKPRIPVVWIHGL WP_001295940.1
MULTISPECIES: hydrogenase [Enterobacteriaceae] Length: 372\nNumber of predicted TMHs: 2\nExp number of AAs in TMHs: 37.2608899999999\nExp number, first 60 AAs: 19.52923\nTotal prob of N-in: 0.94351\nPOSSIBLE N-term signal sequence\ninside 1 19\nTMhelix 20 39\noutside 40 325\nTMhelix 326 348\ninside 349 372

4139 GCF_002016295.1_ASM201629v1 Escherichia coli Proteobacteria; Gammaproteobacteria; Enterobacterales; Enterobacteriaceae; Escherichia
MTGDNTLIHSHGINRRDFMKLCAALAATMGLSSKAAAEMAESVTNPQRPPVIWIGAQECT WP_000145410.1
MULTISPECIES: hydrogenase-2 small chain [Proteobacteria] Length: 372\nNumber of predicted TMHs: 1\nExp number of AAs in TMHs: 23.72226\nExp number, first 60 AAs: 0.22038\nTotal prob of N-in: 0.12994\noutside 1 331\nTMhelix 332 354\ninside 355 372

4140 GCF_002016305.1_ASM201630v1 Escherichia coli Proteobacteria; Gammaproteobacteria; Enterobacterales; Enterobacteriaceae; Escherichia
MTGDNTLIHSHGINRRDFMKLCAALAATMGLSSKAAAEMAESVTNPQRPPVIWIGAQECT WP_000145410.1
MULTISPECIES: hydrogenase-2 small chain [Proteobacteria] Length: 372\nNumber of predicted TMHs: 1\nExp number of AAs in TMHs: 23.72226\nExp number, first 60 AAs: 0.22038\nTotal prob of N-in: 0.12994\noutside 1 331\nTMhelix 332 354\ninside 355 372

4141 GCF_002016355.1_ASM201635v1 Escherichia coli Proteobacteria; Gammaproteobacteria; Enterobacterales; Enterobacteriaceae; Escherichia
MTGDNTLIHSHGINRRDFMKLCAALAATMGLSSKAAAEMAESVTNPQRPPVIWIGAQECT WP_000145410.1
MULTISPECIES: hydrogenase-2 small chain [Proteobacteria] Length: 372\nNumber of predicted TMHs: 1\nExp number of AAs in TMHs: 23.72226\nExp number, first 60 AAs: 0.22038\nTotal prob of N-in: 0.12994\noutside 1 331\nTMhelix 332 354\ninside 355 372

4142 GCF_002016375.1_ASM201637v1 Escherichia coli Proteobacteria; Gammaproteobacteria; Enterobacterales; Enterobacteriaceae; Escherichia
MTGDNTLIHSHGINRRDFMKLCAALAATMGLSSKAAAEMAESVTNPQRPPVIWIGAQECT WP_000145410.1
MULTISPECIES: hydrogenase-2 small chain [Proteobacteria] Length: 372\nNumber of predicted TMHs: 1\nExp number of AAs in TMHs: 23.72226\nExp number, first 60 AAs: 0.22038\nTotal prob of N-in: 0.12994\noutside 1 331\nTMhelix 332 354\ninside 355 372

4143 GCF_002016405.1_ASM201640v1 Escherichia coli Proteobacteria; Gammaproteobacteria; Enterobacterales; Enterobacteriaceae; Escherichia
MTGDNTLIHSHGINRRDFMKLCAALAATMGLSSKAAAEMAESVTNPQRPPVIWIGAQECT WP_000145410.1
MULTISPECIES: hydrogenase-2 small chain [Proteobacteria] Length: 372\nNumber of predicted TMHs: 1\nExp number of AAs in TMHs: 23.72226\nExp number, first 60 AAs: 0.22038\nTotal prob of N-in: 0.12994\noutside 1 331\nTMhelix 332 354\ninside 355 372

4144 GCF_002016385.1_ASM201638v1 Escherichia coli Proteobacteria; Gammaproteobacteria; Enterobacterales; Enterobacteriaceae; Escherichia
MTGDNTLIHSHGINRRDFMKLCAALAATMGLSSKAAAEMAESVTNPQRPPVIWIGAQECT WP_000145410.1
MULTISPECIES: hydrogenase-2 small chain [Proteobacteria] Length: 372\nNumber of predicted TMHs: 1\nExp number of AAs in TMHs: 23.72226\nExp number, first 60 AAs: 0.22038\nTotal prob of N-in: 0.12994\noutside 1 331\nTMhelix 332 354\ninside 355 372

4145 GCF_002016355.1_ASM201635v1 Escherichia coli Proteobacteria; Gammaproteobacteria; Enterobacterales; Enterobacteriaceae; Escherichia
MNNEETFYQAMRRQGVTRRSFLKYCSLAATSLGLGAGMAPKIAWALENKPRIPVVWIHGL WP_001058323.1
MULTISPECIES: hydrogenase-1 small chain [Proteobacteria] Length: 372\nNumber of predicted TMHs: 2\nExp number of AAs in TMHs: 37.2608499999999\nExp number, first 60 AAs: 19.52923\nTotal prob of N-in: 0.94351\nPOSSIBLE N-term signal sequence\ninside 1 19\nTMhelix 20 39\noutside 40 325\nTMhelix 326 348\ninside 349 372

4146 GCF_002016495.1_ASM201649v1 Escherichia coli Proteobacteria; Gammaproteobacteria; Enterobacterales; Enterobacteriaceae; Escherichia
MTGDNTLIHSHGINRRDFMKLCAALAATMGLSSKAAAEMAESVTNPQRPPVIWIGAQECT WP_000145410.1
MULTISPECIES: hydrogenase-2 small chain [Proteobacteria] Length: 372\nNumber of predicted TMHs: 1\nExp number of AAs in TMHs: 23.72226\nExp number, first 60 AAs: 0.22038\nTotal prob of N-in: 0.12994\noutside 1 331\nTMhelix 332 354\ninside 355 372

4147 GCF_002016225.1_ASM201622v1 Escherichia coli Proteobacteria; Gammaproteobacteria; Enterobacterales; Enterobacteriaceae; Escherichia
MNNEETFYQAMRRQGVTRRSFLKYCSLAATSLGLGAGMAPKIAWALENKPRIPVVWIHGL WP_001295940.1
MULTISPECIES: hydrogenase [Enterobacteriaceae] Length: 372\nNumber of predicted TMHs: 2\nExp number of AAs in TMHs: 37.2608899999999\nExp number, first 60 AAs: 19.52923\nTotal prob of N-in: 0.94351\nPOSSIBLE N-term signal sequence\ninside 1 19\nTMhelix 20 39\noutside 40 325\nTMhelix 326 348\ninside 349 372

4148 GCF_002016295.1_ASM201629v1 Escherichia coli Proteobacteria; Gammaproteobacteria; Enterobacterales; Enterobacteriaceae; Escherichia
MNNEETFYQAMRRQGVTRRSFLKYCSLAATSLGLGAGMAPKIAWALENKPRIPVVWIHGL WP_001295940.1
MULTISPECIES: hydrogenase [Enterobacteriaceae] Length: 372\nNumber of predicted TMHs: 2\nExp number of AAs in TMHs: 37.2608899999999\nExp number, first 60 AAs: 19.52923\nTotal prob of N-in: 0.94351\nPOSSIBLE N-term signal sequence\ninside 1 19\nTMhelix 20 39\noutside 40 325\nTMhelix 326 348\ninside 349 372

4149 GCF_002016385.1_ASM201638v1 Escherichia coli Proteobacteria; Gammaproteobacteria; Enterobacterales; Enterobacteriaceae; Escherichia

MNNEETFYQAMRRQGVTRRSFLKYCSLAATSLGLGAGMAPKIAWALENKPRIPVVWIHGL WP_001058323.1 MULTISPECIES: hydrogenase-1 small chain [Proteobacteria] Length: 372\nNumber of predicted TMHs: 2\nExp number of AAs in TMHs: 37.2608499999999\nExp number, first 60 AAs: 19.52923\nTotal prob of N-in: 0.94351\nPOSSIBLE N-term signal sequence\ninside 1 19\nTMhelix 20 39\noutside 40 325\nTMhelix 326 348\ninside 349 372

4150 GCF_002016435.1_ASM201643v1 Escherichia coli Proteobacteria; Gammaproteobacteria; Enterobacterales; Enterobacteriaceae; Escherichia MTGDNTLIHSHGINRRDFMKLCAALAATMGLSSKAAAEMAESVTNPQRPPVIWIGAQECT WP_000145410.1 MULTISPECIES: hydrogenase-2 small chain [Proteobacteria] Length: 372\nNumber of predicted TMHs: 1\nExp number of AAs in TMHs: 23.72226\nExp number, first 60 AAs: 0.22038\nTotal prob of N-in: 0.12994\noutside 1 331\nTMhelix 332 354\ninside 355 372

4151 GCF_002016455.1_ASM201645v1 Escherichia coli Proteobacteria; Gammaproteobacteria; Enterobacterales; Enterobacteriaceae; Escherichia MTGDNTLIHSHGINRRDFMKLCAALAATMGLSSKAAAEMAESVTNPQRPPVIWIGAQECT WP_000145410.1 MULTISPECIES: hydrogenase-2 small chain [Proteobacteria] Length: 372\nNumber of predicted TMHs: 1\nExp number of AAs in TMHs: 23.72226\nExp number, first 60 AAs: 0.22038\nTotal prob of N-in: 0.12994\noutside 1 331\nTMhelix 332 354\ninside 355 372

4152 GCF_002016465.1_ASM201646v1 Escherichia coli Proteobacteria; Gammaproteobacteria; Enterobacterales; Enterobacteriaceae; Escherichia MTGDNTLIHSHGINRRDFMKLCAALAATMGLSSKAAAEMAESVTNPQRPPVIWIGAQECT WP_000145410.1 MULTISPECIES: hydrogenase-2 small chain [Proteobacteria] Length: 372\nNumber of predicted TMHs: 1\nExp number of AAs in TMHs: 23.72226\nExp number, first 60 AAs: 0.22038\nTotal prob of N-in: 0.12994\noutside 1 331\nTMhelix 332 354\ninside 355 372

4153 GCF_002016475.1_ASM201647v1 Escherichia coli Proteobacteria; Gammaproteobacteria; Enterobacterales; Enterobacteriaceae; Escherichia MTGDNTLIHSHGINRRDFMKLCAALAATMGLSSKAAAEMAESVTNPQRPPVIWIGAQECT WP_000145410.1 MULTISPECIES: hydrogenase-2 small chain [Proteobacteria] Length: 372\nNumber of predicted TMHs: 1\nExp number of AAs in TMHs: 23.72226\nExp number, first 60 AAs: 0.22038\nTotal prob of N-in: 0.12994\noutside 1 331\nTMhelix 332 354\ninside 355 372

4154 GCF_002018815.1_ASM201881v1 Escherichia coli Proteobacteria; Gammaproteobacteria; Enterobacterales; Enterobacteriaceae; Escherichia MNNEETFYQAMRRQGVTRRSFLKYCSLAATSLGLGAGMAPKIAWALENKPRIPVVWIHGL WP_001058323.1 MULTISPECIES: hydrogenase-1 small chain [Proteobacteria] Length: 372\nNumber of predicted TMHs: 2\nExp number of AAs in TMHs: 37.2608499999999\nExp number, first 60 AAs: 19.52923\nTotal prob of N-in: 0.94351\nPOSSIBLE N-term signal sequence\ninside 1 19\nTMhelix 20 39\noutside 40 325\nTMhelix 326 348\ninside 349 372

4155 GCF_002016455.1_ASM201645v1 Escherichia coli Proteobacteria; Gammaproteobacteria; Enterobacterales; Enterobacteriaceae; Escherichia MNNEETFYQAMRRQGVTRRSFLKYCSLAATSLGLGAGMAPKIAWALENKPRIPVVWIHGL WP_001058323.1 MULTISPECIES: hydrogenase-1 small chain [Proteobacteria] Length: 372\nNumber of predicted TMHs: 2\nExp number of AAs in TMHs: 37.2608499999999\nExp number, first 60 AAs: 19.52923\nTotal prob of N-in: 0.94351\nPOSSIBLE N-term signal sequence\ninside 1 19\nTMhelix 20 39\noutside 40 325\nTMhelix 326 348\ninside 349 372

4156 GCF_002016465.1_ASM201646v1 Escherichia coli Proteobacteria; Gammaproteobacteria; Enterobacterales; Enterobacteriaceae; Escherichia MNNEETFYQAMRRQGVTRRSFLKYCSLAATSLGLGAGMAPKIAWALENKPRIPVVWIHGL WP_001058323.1 MULTISPECIES: hydrogenase-1 small chain [Proteobacteria] Length: 372\nNumber of predicted TMHs: 2\nExp number of AAs in TMHs: 37.2608499999999\nExp number, first 60 AAs: 19.52923\nTotal prob of N-in: 0.94351\nPOSSIBLE N-term signal sequence\ninside 1 19\nTMhelix 20 39\noutside 40 325\nTMhelix 326 348\ninside 349 372

4157 GCF_002016475.1_ASM201647v1 Escherichia coli Proteobacteria; Gammaproteobacteria; Enterobacterales; Enterobacteriaceae; Escherichia MNNEETFYQAMRRQGVTRRSFLKYCSLAATSLGLGAGMAPKIAWALENKPRIPVVWIHGL WP_001058323.1

MULTISPECIES: hydrogenase-1 small chain [Proteobacteria] Length: 372\nNumber of predicted TMHs: 2\nExp number of AAs in TMHs: 37.2608499999999\nExp number, first 60 AAs: 19.52923\nTotal prob of N-in: 0.94351\nPOSSIBLE N-term signal sequence\ninside 1 19\nTMhelix 20 39\noutside 40 325\nTMhelix 326 348\ninside 349 372

4158 GCF_002016495.1_ASM201649v1 Escherichia coli Proteobacteria; Gammaproteobacteria; Enterobacterales; Enterobacteriaceae; Escherichia
MNNEETFYQAMRRQGVTRRSFLKYCSLAATSLGLGAGMAPKIAWALENKPRIPVVWIHGL WP_001058323.1
MULTISPECIES: hydrogenase-1 small chain [Proteobacteria] Length: 372\nNumber of predicted TMHs: 2\nExp number of AAs in TMHs: 37.2608499999999\nExp number, first 60 AAs: 19.52923\nTotal prob of N-in: 0.94351\nPOSSIBLE N-term signal sequence\ninside 1 19\nTMhelix 20 39\noutside 40 325\nTMhelix 326 348\ninside 349 372

4159 GCF_002028665.1_ASM202866v1 Escherichia coli Proteobacteria; Gammaproteobacteria; Enterobacterales; Enterobacteriaceae; Escherichia
MTGDNTLIHSHGINRRDFMKLCAALAATMGLSSKAAAEMAESVTNPQRPPVIWIGAQECT WP_000145410.1
MULTISPECIES: hydrogenase-2 small chain [Proteobacteria] Length: 372\nNumber of predicted TMHs: 1\nExp number of AAs in TMHs: 23.72226\nExp number, first 60 AAs: 0.22038\nTotal prob of N-in: 0.12994\noutside 1 331\nTMhelix 332 354\ninside 355 372

4160 GCF_002028705.1_ASM202870v1 Escherichia coli Proteobacteria; Gammaproteobacteria; Enterobacterales; Enterobacteriaceae; Escherichia
MTGDNTLIHSHGINRRDFMKLCAALAATMGLSSKAAAEMAESVTNPQRPPVIWIGAQECT WP_000145410.1
MULTISPECIES: hydrogenase-2 small chain [Proteobacteria] Length: 372\nNumber of predicted TMHs: 1\nExp number of AAs in TMHs: 23.72226\nExp number, first 60 AAs: 0.22038\nTotal prob of N-in: 0.12994\noutside 1 331\nTMhelix 332 354\ninside 355 372

4161 GCF_002028715.1_ASM202871v1 Escherichia coli Proteobacteria; Gammaproteobacteria; Enterobacterales; Enterobacteriaceae; Escherichia
MTGDNTLIHSHGINRRDFMKLCAALAATMGLSSKAAAEMAESVTNPQRPPVIWIGAQECT WP_000145410.1
MULTISPECIES: hydrogenase-2 small chain [Proteobacteria] Length: 372\nNumber of predicted TMHs: 1\nExp number of AAs in TMHs: 23.72226\nExp number, first 60 AAs: 0.22038\nTotal prob of N-in: 0.12994\noutside 1 331\nTMhelix 332 354\ninside 355 372

4162 GCF_002028745.1_ASM202874v1 Escherichia coli Proteobacteria; Gammaproteobacteria; Enterobacterales; Enterobacteriaceae; Escherichia
MTGDNTLIHSHGINRRDFMKLCAALAATMGLSSKAAAEMAESVTNPQRPPVIWIGAQECT WP_000145410.1
MULTISPECIES: hydrogenase-2 small chain [Proteobacteria] Length: 372\nNumber of predicted TMHs: 1\nExp number of AAs in TMHs: 23.72226\nExp number, first 60 AAs: 0.22038\nTotal prob of N-in: 0.12994\noutside 1 331\nTMhelix 332 354\ninside 355 372

4163 GCF_002028755.1_ASM202875v1 Escherichia coli Proteobacteria; Gammaproteobacteria; Enterobacterales; Enterobacteriaceae; Escherichia
MTGDNTLIHSHGINRRDFMKLCAALAATMGLSSKAAAEMAESVTNPQRPPVIWIGAQECT WP_000145410.1
MULTISPECIES: hydrogenase-2 small chain [Proteobacteria] Length: 372\nNumber of predicted TMHs: 1\nExp number of AAs in TMHs: 23.72226\nExp number, first 60 AAs: 0.22038\nTotal prob of N-in: 0.12994\noutside 1 331\nTMhelix 332 354\ninside 355 372

4164 GCF_002028785.1_ASM202878v1 Escherichia coli Proteobacteria; Gammaproteobacteria; Enterobacterales; Enterobacteriaceae; Escherichia
MTGDNTLIHSHGINRRDFMKLCAALAATMGLSSKAAAEMAESVTNPQRPPVIWIGAQECT WP_000145410.1
MULTISPECIES: hydrogenase-2 small chain [Proteobacteria] Length: 372\nNumber of predicted TMHs: 1\nExp number of AAs in TMHs: 23.72226\nExp number, first 60 AAs: 0.22038\nTotal prob of N-in: 0.12994\noutside 1 331\nTMhelix 332 354\ninside 355 372

4165 GCF_002028865.1_ASM202886v1 Escherichia coli Proteobacteria; Gammaproteobacteria; Enterobacterales; Enterobacteriaceae; Escherichia
MTGDNTLIHSHGINRRDFMKLCAALAATMGLSSKAAAEMAESVTNPQRPPVIWIGAQECT WP_000145410.1
MULTISPECIES: hydrogenase-2 small chain [Proteobacteria] Length: 372\nNumber of predicted TMHs: 1\nExp number of AAs in TMHs: 23.72226\nExp number, first 60 AAs: 0.22038\nTotal prob of N-in: 0.12994\noutside 1 331\nTMhelix 332 354\ninside 355 372

4166 GCF_002028905.1_ASM202890v1 Escherichia coli Proteobacteria; Gammaproteobacteria; Enterobacterales; Enterobacteriaceae; Escherichia
MTGDNTLIHSHGINRRDFMKLCAALAATMGLSSKAAAEMAESVTNPQRPPVIWIGAQECT WP_000145410.1
MULTISPECIES: hydrogenase-2 small chain [Proteobacteria] Length: 372\nNumber of predicted TMHs: 1\nExp number of AAs in TMHs: 23.72226\nExp number, first 60 AAs: 0.22038\nTotal prob of N-in: 0.12994\noutside 1 331\nTMhelix 332 354\ninside 355 372

4167 GCF_002028915.1_ASM202891v1 Escherichia coli Proteobacteria; Gammaproteobacteria; Enterobacterales; Enterobacteriaceae; Escherichia
MTGDNTLIHSHGINRRDFMKLCAALAATMGLSSKAAAEMAESVTNPQRPPVIWIGAQECT WP_000145410.1
MULTISPECIES: hydrogenase-2 small chain [Proteobacteria] Length: 372\nNumber of predicted TMHs: 1\nExp number of AAs in TMHs: 23.72226\nExp number, first 60 AAs: 0.22038\nTotal prob of N-in: 0.12994\noutside 1 331\nTMhelix 332 354\ninside 355 372

4168 GCF_002028945.1_ASM202894v1 Escherichia coli Proteobacteria; Gammaproteobacteria; Enterobacterales; Enterobacteriaceae; Escherichia
MTGDNTLIHSHGINRRDFMKLCAALAATMGLSSKAAAEMAESVTNPQRPPVIWIGAQECT WP_000145410.1
MULTISPECIES: hydrogenase-2 small chain [Proteobacteria] Length: 372\nNumber of predicted TMHs: 1\nExp number of AAs in TMHs: 23.72226\nExp number, first 60 AAs: 0.22038\nTotal prob of N-in: 0.12994\noutside 1 331\nTMhelix 332 354\ninside 355 372

4169 GCF_002028955.1_ASM202895v1 Escherichia coli Proteobacteria; Gammaproteobacteria; Enterobacterales; Enterobacteriaceae; Escherichia
MTGDNTLIHSHGINRRDFMKLCAALAATMGLSSKAAAEMAESVTNPQRPPVIWIGAQECT WP_000145410.1
MULTISPECIES: hydrogenase-2 small chain [Proteobacteria] Length: 372\nNumber of predicted TMHs: 1\nExp number of AAs in TMHs: 23.72226\nExp number, first 60 AAs: 0.22038\nTotal prob of N-in: 0.12994\noutside 1 331\nTMhelix 332 354\ninside 355 372

4170 GCF_002028965.1_ASM202896v1 Escherichia coli Proteobacteria; Gammaproteobacteria; Enterobacterales; Enterobacteriaceae; Escherichia
MTGDNTLIHSHGINRRDFMKLCAALAATMGLSSKAAAEMAESVTNPQRPPVIWIGAQECT WP_000145410.1
MULTISPECIES: hydrogenase-2 small chain [Proteobacteria] Length: 372\nNumber of predicted TMHs: 1\nExp number of AAs in TMHs: 23.72226\nExp number, first 60 AAs: 0.22038\nTotal prob of N-in: 0.12994\noutside 1 331\nTMhelix 332 354\ninside 355 372

4171 GCF_002029005.1_ASM202900v1 Escherichia coli Proteobacteria; Gammaproteobacteria; Enterobacterales; Enterobacteriaceae; Escherichia
MTGDNTLIHSHGINRRDFMKLCAALAATMGLSSKAAAEMAESVTNPQRPPVIWIGAQECT WP_000145410.1
MULTISPECIES: hydrogenase-2 small chain [Proteobacteria] Length: 372\nNumber of predicted TMHs: 1\nExp number of AAs in TMHs: 23.72226\nExp number, first 60 AAs: 0.22038\nTotal prob of N-in: 0.12994\noutside 1 331\nTMhelix 332 354\ninside 355 372

4172 GCF_002029015.1_ASM202901v1 Escherichia coli Proteobacteria; Gammaproteobacteria; Enterobacterales; Enterobacteriaceae; Escherichia
MTGDNTLIHSHGINRRDFMKLCAALAATMGLSSKAAAEMAESVTNPQRPPVIWIGAQECT WP_000145410.1
MULTISPECIES: hydrogenase-2 small chain [Proteobacteria] Length: 372\nNumber of predicted TMHs: 1\nExp number of AAs in TMHs: 23.72226\nExp number, first 60 AAs: 0.22038\nTotal prob of N-in: 0.12994\noutside 1 331\nTMhelix 332 354\ninside 355 372

4173 GCF_002056065.1_ASM205606v1 Escherichia coli Proteobacteria; Gammaproteobacteria; Enterobacterales; Enterobacteriaceae; Escherichia
MTGDNTLIHSHGINRRDFMKLCAALAATMGLSSKAAAEMAESVTNPQRPPVIWIGAQECT WP_000145410.1
MULTISPECIES: hydrogenase-2 small chain [Proteobacteria] Length: 372\nNumber of predicted TMHs: 1\nExp number of AAs in TMHs: 23.72226\nExp number, first 60 AAs: 0.22038\nTotal prob of N-in: 0.12994\noutside 1 331\nTMhelix 332 354\ninside 355 372

4174 GCF_002055635.1_ASM205563v1 Escherichia coli Proteobacteria; Gammaproteobacteria; Enterobacterales; Enterobacteriaceae; Escherichia
MTGDNTLIHSHGINRRDFMKLCAALAATMGLSSKAAAEMAESVTNPQRPPVIWIGAQECT WP_000145410.1
MULTISPECIES: hydrogenase-2 small chain [Proteobacteria] Length: 372\nNumber of predicted TMHs:

1\nExp number of AAs in TMHs: 23.72226\nExp number, first 60 AAs: 0.22038\nTotal prob of N-in: 0.12994\noutside 1 331\nTMhelix 332 354\ninside 355 372

4175 GCF_002057245.1_ASM205724v1 Escherichia coli Proteobacteria; Gammaproteobacteria; Enterobacterales; Enterobacteriaceae; Escherichia MTGDNTLIHSHGINRRDFMKLCAALAATMGLSSKAAAEMAESVTNPQRPPVIWIGAQECT WP_000145410.1 MULTISPECIES: hydrogenase-2 small chain [Proteobacteria] Length: 372\nNumber of predicted TMHs: 1\nExp number of AAs in TMHs: 23.72226\nExp number, first 60 AAs: 0.22038\nTotal prob of N-in: 0.12994\noutside 1 331\nTMhelix 332 354\ninside 355 372

4176 GCF_002056635.1_ASM205663v1 Escherichia coli Proteobacteria; Gammaproteobacteria; Enterobacterales; Enterobacteriaceae; Escherichia MTGDNTLIHSHGINRRDFMKLCAALAATMGLSSKAAAEMVESVTNPQRPPVIWIGAQECT WP_000145423.1 hydrogenase 2 small subunit [Escherichia coli] Length: 372\nNumber of predicted TMHs: 1\nExp number of AAs in TMHs: 23.78201\nExp number, first 60 AAs: 0.28355\nTotal prob of N-in: 0.13257\noutside 1 331\nTMhelix 332 354\ninside 355 372

4177 GCF_002056145.1_ASM205614v1 Escherichia coli Proteobacteria; Gammaproteobacteria; Enterobacterales; Enterobacteriaceae; Escherichia MTGDNTLIHSHGINRRDFMKLCAALAATMGLSSKAAAEMAESVTNPQRPPVIWIGAQECT WP_000145410.1 MULTISPECIES: hydrogenase-2 small chain [Proteobacteria] Length: 372\nNumber of predicted TMHs: 1\nExp number of AAs in TMHs: 23.72226\nExp number, first 60 AAs: 0.22038\nTotal prob of N-in: 0.12994\noutside 1 331\nTMhelix 332 354\ninside 355 372

4178 GCF_002056065.1_ASM205606v1 Escherichia coli Proteobacteria; Gammaproteobacteria; Enterobacterales; Enterobacteriaceae; Escherichia MNNEETFYQAMRRQGVTRRSFLKYCSLAATSLGLGAGMAPKIAWALENKPRIPVVWIHGL WP_001058323.1 MULTISPECIES: hydrogenase-1 small chain [Proteobacteria] Length: 372\nNumber of predicted TMHs: 2\nExp number of AAs in TMHs: 37.2608499999999\nExp number, first 60 AAs: 19.52923\nTotal prob of N-in: 0.94351\nPOSSIBLE N-term signal sequence\ninside 1 19\nTMhelix 20 39\noutside 40 325\nTMhelix 326 348\ninside 349 372

4179 GCF_002057355.1_ASM205735v1 Escherichia coli Proteobacteria; Gammaproteobacteria; Enterobacterales; Enterobacteriaceae; Escherichia MTGDNTLIHSHGINRRDFMKLCAALAATMGLSSKAAAEMAESVTNPQRPPVIWIGAQECT WP_000145410.1 MULTISPECIES: hydrogenase-2 small chain [Proteobacteria] Length: 372\nNumber of predicted TMHs: 1\nExp number of AAs in TMHs: 23.72226\nExp number, first 60 AAs: 0.22038\nTotal prob of N-in: 0.12994\noutside 1 331\nTMhelix 332 354\ninside 355 372

4180 GCF_002075205.1_102549 Escherichia coli Proteobacteria; Gammaproteobacteria; Enterobacterales; Enterobacteriaceae; Escherichia MTGDNTLIHSHGINRRDFMKLCAALAATMGLSSKAAAEMAESVTNPQRPPVIWIGAQECT WP_000145410.1 MULTISPECIES: hydrogenase-2 small chain [Proteobacteria] Length: 372\nNumber of predicted TMHs: 1\nExp number of AAs in TMHs: 23.72226\nExp number, first 60 AAs: 0.22038\nTotal prob of N-in: 0.12994\noutside 1 331\nTMhelix 332 354\ninside 355 372

4181 GCF_002056635.1_ASM205663v1 Escherichia coli Proteobacteria; Gammaproteobacteria; Enterobacterales; Enterobacteriaceae; Escherichia MNNEETFYQAMRRQGVTRRSFLKYCSLAATSLGLGAGMAPKIAWALENKPRIPVVWIHGL WP_001295940.1 MULTISPECIES: hydrogenase [Enterobacteriaceae] Length: 372\nNumber of predicted TMHs: 2\nExp number of AAs in TMHs: 37.2608899999999\nExp number, first 60 AAs: 19.52923\nTotal prob of N-in: 0.94351\nPOSSIBLE N-term signal sequence\ninside 1 19\nTMhelix 20 39\noutside 40 325\nTMhelix 326 348\ninside 349 372

4182 GCF_002075125.1_103334 Escherichia coli Proteobacteria; Gammaproteobacteria; Enterobacterales; Enterobacteriaceae; Escherichia MTGDNTLIHSHGINRRDFMKLCAALAATMGLSSKAAAEMAESVTNPQRPPVIWIGAQECT WP_000145410.1 MULTISPECIES: hydrogenase-2 small chain [Proteobacteria] Length: 372\nNumber of predicted TMHs: 1\nExp number of AAs in TMHs: 23.72226\nExp number, first 60 AAs: 0.22038\nTotal prob of N-in: 0.12994\noutside 1 331\nTMhelix 332 354\ninside 355 372

4183 GCF_002075135.1_300757 Escherichia coli Proteobacteria; Gammaproteobacteria; Enterobacterales; Enterobacteriaceae; Escherichia MTGDNTLIHSHGINRRDFMKLCAALAATMGLSSKAAAEMAESVTNPQRPPVIWIGAQECT

WP_000145410.1 MULTISPECIES: hydrogenase-2 small chain [Proteobacteria] Length: 372\nNumber of predicted TMHs: 1\nExp number of AAs in TMHs: 23.72226\nExp number, first 60 AAs: 0.22038\nTotal prob of N-in: 0.12994\noutside 1 331\nTMhelix 332 354\ninside 355 372

4184 GCF_002075175.1_102556 Escherichia coli Proteobacteria; Gammaproteobacteria; Enterobacterales; Enterobacteriaceae; Escherichia MTGDNTLIHSHGINRRDFMKLCAALAATMGLSSKAAAEMAESVTNPQRPPVIWIGAQECT WP_000145410.1 MULTISPECIES: hydrogenase-2 small chain [Proteobacteria] Length: 372\nNumber of predicted TMHs: 1\nExp number of AAs in TMHs: 23.72226\nExp number, first 60 AAs: 0.22038\nTotal prob of N-in: 0.12994\noutside 1 331\nTMhelix 332 354\ninside 355 372

4185 GCF_002075165.1_102667 Escherichia coli Proteobacteria; Gammaproteobacteria; Enterobacterales; Enterobacteriaceae; Escherichia MTGDNTLIHSHGINRRDFMKLCAALAATMGLSSKAAAEMAESVTNPQRPPVIWIGAQECT WP_000145410.1 MULTISPECIES: hydrogenase-2 small chain [Proteobacteria] Length: 372\nNumber of predicted TMHs: 1\nExp number of AAs in TMHs: 23.72226\nExp number, first 60 AAs: 0.22038\nTotal prob of N-in: 0.12994\noutside 1 331\nTMhelix 332 354\ninside 355 372

4186 GCF_002075135.1_300757 Escherichia coli Proteobacteria; Gammaproteobacteria; Enterobacterales; Enterobacteriaceae; Escherichia MNNEETFYQAMRRQGVTRRSFLKYCSLAATSLGLGAGMAPKIAWALENKPRIPVVWIHGL WP_001058323.1 MULTISPECIES: hydrogenase-1 small chain [Proteobacteria] Length: 372\nNumber of predicted TMHs: 2\nExp number of AAs in TMHs: 37.2608499999999\nExp number, first 60 AAs: 19.52923\nTotal prob of N-in: 0.94351\nPOSSIBLE N-term signal sequence\ninside 1 19\nTMhelix 20 39\noutside 40 325\nTMhelix 326 348\ninside 349 372

4187 GCF_002075175.1_102556 Escherichia coli Proteobacteria; Gammaproteobacteria; Enterobacterales; Enterobacteriaceae; Escherichia MNNEETFYQAMRRQGVTRRSFLKYCSLAATSLGLGAGMAPKIAWALENKPRIPVVWIHGL WP_001058323.1 MULTISPECIES: hydrogenase-1 small chain [Proteobacteria] Length: 372\nNumber of predicted TMHs: 2\nExp number of AAs in TMHs: 37.2608499999999\nExp number, first 60 AAs: 19.52923\nTotal prob of N-in: 0.94351\nPOSSIBLE N-term signal sequence\ninside 1 19\nTMhelix 20 39\noutside 40 325\nTMhelix 326 348\ninside 349 372

4188 GCF_002078275.1_ASM207827v1 Escherichia coli Proteobacteria; Gammaproteobacteria; Enterobacterales; Enterobacteriaceae; Escherichia MTGDNTLIHSHGINRRDFMKLCAALAATMGLSSKAAAEMAESVTNPQRPPVIWIGAQECT WP_000145410.1 MULTISPECIES: hydrogenase-2 small chain [Proteobacteria] Length: 372\nNumber of predicted TMHs: 1\nExp number of AAs in TMHs: 23.72226\nExp number, first 60 AAs: 0.22038\nTotal prob of N-in: 0.12994\noutside 1 331\nTMhelix 332 354\ninside 355 372

4189 GCF_002075225.1_102687 Escherichia coli Proteobacteria; Gammaproteobacteria; Enterobacterales; Enterobacteriaceae; Escherichia MNNEETFYQAMRRQGVTRRSFLKYCSLAATSLGLGAGMAPKIAWALENKPRIPVVWIHGL WP_001058323.1 MULTISPECIES: hydrogenase-1 small chain [Proteobacteria] Length: 372\nNumber of predicted TMHs: 2\nExp number of AAs in TMHs: 37.2608499999999\nExp number, first 60 AAs: 19.52923\nTotal prob of N-in: 0.94351\nPOSSIBLE N-term signal sequence\ninside 1 19\nTMhelix 20 39\noutside 40 325\nTMhelix 326 348\ninside 349 372

4190 GCF_002075295.1_103604 Escherichia coli Proteobacteria; Gammaproteobacteria; Enterobacterales; Enterobacteriaceae; Escherichia MNNEETFYQAMRRQGVTRRSFLKYCSLAATSLGLGAGMAPKIAWALENKPRIPVVWIHGL WP_001295940.1 MULTISPECIES: hydrogenase [Enterobacteriaceae] Length: 372\nNumber of predicted TMHs: 2\nExp number of AAs in TMHs: 37.2608899999999\nExp number, first 60 AAs: 19.52923\nTotal prob of N-in: 0.94351\nPOSSIBLE N-term signal sequence\ninside 1 19\nTMhelix 20 39\noutside 40 325\nTMhelix 326 348\ninside 349 372

4191 GCF_002078275.1_ASM207827v1 Escherichia coli Proteobacteria; Gammaproteobacteria; Enterobacterales; Enterobacteriaceae; Escherichia MNNEETFYQAMRRQGVTRRSFLKYCSLAATSLGLGAGMAPKIAWALENKPRIPVVWIHGL WP_001058323.1 MULTISPECIES: hydrogenase-1 small chain [Proteobacteria] Length: 372\nNumber of predicted TMHs: 2\nExp number of AAs in TMHs: 37.2608499999999\nExp number, first 60 AAs: 19.52923\nTotal prob of N-in: 0.94351\nPOSSIBLE N-term signal sequence\ninside 1 19\nTMhelix 20 39\noutside 40 325\nTMhelix 326 348\ninside 349 372

4192 GCF_002075165.1_102667 Escherichia coli Proteobacteria; Gammaproteobacteria; Enterobacterales; Enterobacteriaceae; Escherichia MNNEETFYQAMRRQGVTRRSFLKYCSLAATSLGLGAGMAPKIAWALENKPRIPVVWIHGL

WP_001295940.1 MULTISPECIES: hydrogenase [Enterobacteriaceae] Length: 372\nNumber of predicted TMHs: 2\nExp number of AAs in TMHs: 37.2608899999999\nExp number, first 60 AAs: 19.52923\nTotal prob of N-in: 0.94351\nPOSSIBLE N-term signal sequence\ninside 1 19\nTMhelix 20 39\noutside 40 325\nTMhelix 326 348\ninside 349 372

4193 GCF_002078295.1_ASM207829v1 Escherichia coli Proteobacteria; Gammaproteobacteria; Enterobacterales; Enterobacteriaceae; Escherichia MTGDNTLIHSHGINRRDFMKLCAALAATMGLSSKAAAEMAESVTNPQRPPVIWIGAQECT WP_000145410.1 MULTISPECIES: hydrogenase-2 small chain [Proteobacteria] Length: 372\nNumber of predicted TMHs: 1\nExp number of AAs in TMHs: 23.72226\nExp number, first 60 AAs: 0.22038\nTotal prob of N-in: 0.12994\noutside 1 331\nTMhelix 332 354\ninside 355 372

4194 GCF_002080555.1_ASM208055v1 Escherichia coli Proteobacteria; Gammaproteobacteria; Enterobacterales; Enterobacteriaceae; Escherichia MNNEETFYQAMRRQGVTRRSFLKYCSLAATSLGLGAGMAPKIAWALENKPRIPVVWIHGL WP_001058323.1 MULTISPECIES: hydrogenase-1 small chain [Proteobacteria] Length: 372\nNumber of predicted TMHs: 2\nExp number of AAs in TMHs: 37.2608499999999\nExp number, first 60 AAs: 19.52923\nTotal prob of N-in: 0.94351\nPOSSIBLE N-term signal sequence\ninside 1 19\nTMhelix 20 39\noutside 40 325\nTMhelix 326 348\ninside 349 372

4195 GCF_002080565.1_ASM208056v1 Escherichia coli Proteobacteria; Gammaproteobacteria; Enterobacterales; Enterobacteriaceae; Escherichia MNNEETFYQAMRRQGVTRRSFLKYCSLAATSLGLGAGMAPKIAWALENKPRIPVVWIHGL WP_001058323.1 MULTISPECIES: hydrogenase-1 small chain [Proteobacteria] Length: 372\nNumber of predicted TMHs: 2\nExp number of AAs in TMHs: 37.2608499999999\nExp number, first 60 AAs: 19.52923\nTotal prob of N-in: 0.94351\nPOSSIBLE N-term signal sequence\ninside 1 19\nTMhelix 20 39\noutside 40 325\nTMhelix 326 348\ninside 349 372

4196 GCF_002080595.1_ASM208059v1 Escherichia coli Proteobacteria; Gammaproteobacteria; Enterobacterales; Enterobacteriaceae; Escherichia MNNEETFYQAMRRQGVTRRSFLKYCSLAATSLGLGAGMAPKIAWALENKPRIPVVWIHGL WP_001058323.1 MULTISPECIES: hydrogenase-1 small chain [Proteobacteria] Length: 372\nNumber of predicted TMHs: 2\nExp number of AAs in TMHs: 37.2608499999999\nExp number, first 60 AAs: 19.52923\nTotal prob of N-in: 0.94351\nPOSSIBLE N-term signal sequence\ninside 1 19\nTMhelix 20 39\noutside 40 325\nTMhelix 326 348\ninside 349 372

4197 GCF_002080505.1_ASM208050v1 Escherichia coli Proteobacteria; Gammaproteobacteria; Enterobacterales; Enterobacteriaceae; Escherichia MNNEETFYQAMRRQGVTRRSFLKYCSLAATSLGLGAGMAPKIAWALENKPRIPVVWIHGL WP_001058323.1 MULTISPECIES: hydrogenase-1 small chain [Proteobacteria] Length: 372\nNumber of predicted TMHs: 2\nExp number of AAs in TMHs: 37.2608499999999\nExp number, first 60 AAs: 19.52923\nTotal prob of N-in: 0.94351\nPOSSIBLE N-term signal sequence\ninside 1 19\nTMhelix 20 39\noutside 40 325\nTMhelix 326 348\ninside 349 372

4198 GCF_002080605.1_ASM208060v1 Escherichia coli Proteobacteria; Gammaproteobacteria; Enterobacterales; Enterobacteriaceae; Escherichia MTGDNTLIHSHGINRRDFMKLCAALAATMGLSSKAAAEMAESVTNPQRPPVIWIGAQECT WP_000145410.1 MULTISPECIES: hydrogenase-2 small chain [Proteobacteria] Length: 372\nNumber of predicted TMHs: 1\nExp number of AAs in TMHs: 23.72226\nExp number, first 60 AAs: 0.22038\nTotal prob of N-in: 0.12994\noutside 1 331\nTMhelix 332 354\ninside 355 372

4199 GCF_002080635.1_ASM208063v1 Escherichia coli Proteobacteria; Gammaproteobacteria; Enterobacterales; Enterobacteriaceae; Escherichia MTGDNTLIHSHGINRRDFMKLCAALAATMGLSSKAAAEMAESVTNPQRPPVIWIGAQECT WP_000145410.1 MULTISPECIES: hydrogenase-2 small chain [Proteobacteria] Length: 372\nNumber of predicted TMHs: 1\nExp number of AAs in TMHs: 23.72226\nExp number, first 60 AAs: 0.22038\nTotal prob of N-in: 0.12994\noutside 1 331\nTMhelix 332 354\ninside 355 372

4200 GCF_002080495.1_ASM208049v1 Escherichia coli Proteobacteria; Gammaproteobacteria; Enterobacterales; Enterobacteriaceae; Escherichia MNNEETFYQAMRRQGVTRRSFLKYCSLAATSLGLGAGMAPKIAWALENKPRIPVVWIHGL WP_001058323.1

MULTISPECIES: hydrogenase-1 small chain [Proteobacteria] Length: 372\nNumber of predicted TMHs: 2\nExp number of AAs in TMHs: 37.2608499999999\nExp number, first 60 AAs: 19.52923\nTotal prob of N-in: 0.94351\nPOSSIBLE N-term signal sequence\ninside 1 19\nTMhelix 20 39\noutside 40 325\nTMhelix 326 348\ninside 349 372

4201 GCF_002080685.1_ASM208068v1 Escherichia coli Proteobacteria; Gammaproteobacteria; Enterobacterales; Enterobacteriaceae; Escherichia MTGDNTLIHSHGINRRDFMKLCAALAATMGLSSKAAAEMAESVTNPQRPPVIWIGAQECT WP_000145410.1 MULTISPECIES: hydrogenase-2 small chain [Proteobacteria] Length: 372\nNumber of predicted TMHs: 1\nExp number of AAs in TMHs: 23.72226\nExp number, first 60 AAs: 0.22038\nTotal prob of N-in: 0.12994\noutside 1 331\nTMhelix 332 354\ninside 355 372

4202 GCF_002080685.1_ASM208068v1 Escherichia coli Proteobacteria; Gammaproteobacteria; Enterobacterales; Enterobacteriaceae; Escherichia MNNEETFYQAMRRQGVTRRSFLKYCSLAATSLGLGAGMAPKIAWALENKPRIPVVWIHGL WP_001058323.1 MULTISPECIES: hydrogenase-1 small chain [Proteobacteria] Length: 372\nNumber of predicted TMHs: 2\nExp number of AAs in TMHs: 37.2608499999999\nExp number, first 60 AAs: 19.52923\nTotal prob of N-in: 0.94351\nPOSSIBLE N-term signal sequence\ninside 1 19\nTMhelix 20 39\noutside 40 325\nTMhelix 326 348\ninside 349 372

4203 GCF_002080725.1_ASM208072v1 Escherichia coli Proteobacteria; Gammaproteobacteria; Enterobacterales; Enterobacteriaceae; Escherichia MTGDNTLIHSHGINRRDFMKLCAALAATMGLSSKAAAEMAESVTNPQRPPVIWIGAQECT WP_000145410.1 MULTISPECIES: hydrogenase-2 small chain [Proteobacteria] Length: 372\nNumber of predicted TMHs: 1\nExp number of AAs in TMHs: 23.72226\nExp number, first 60 AAs: 0.22038\nTotal prob of N-in: 0.12994\noutside 1 331\nTMhelix 332 354\ninside 355 372

4204 GCF_002087455.1_ASM208745v1 Escherichia coli Proteobacteria; Gammaproteobacteria; Enterobacterales; Enterobacteriaceae; Escherichia MTGDNTLIHSHGINRRDFMKLCAALAATMGLSSKAAAEMVESVTNPQRPPVIWIGAQECT WP_000145423.1 hydrogenase 2 small subunit [Escherichia coli] Length: 372\nNumber of predicted TMHs: 1\nExp number of AAs in TMHs: 23.78201\nExp number, first 60 AAs: 0.28355\nTotal prob of N-in: 0.13257\noutside 1 331\nTMhelix 332 354\ninside 355 372

4205 GCF_002087465.1_ASM208746v1 Escherichia coli Proteobacteria; Gammaproteobacteria; Enterobacterales; Enterobacteriaceae; Escherichia MTGDNTLIHSHGINRRDFMKLCAALAATMGLSSKAAAEMVESVTNPQRPPVIWIGAQECT WP_000145423.1 hydrogenase 2 small subunit [Escherichia coli] Length: 372\nNumber of predicted TMHs: 1\nExp number of AAs in TMHs: 23.78201\nExp number, first 60 AAs: 0.28355\nTotal prob of N-in: 0.13257\noutside 1 331\nTMhelix 332 354\ninside 355 372

4206 GCF_002087505.1_Esch_coli_OS_09_001_TOC1_V5 Escherichia coli Proteobacteria; Gammaproteobacteria; Enterobacterales; Enterobacteriaceae; Escherichia MTGDNTLIHSHGINRRDFMKLCAALAATMGLSSKAAAEMAESVTNPQRPPVIWIGAQECT WP_000145410.1 MULTISPECIES: hydrogenase-2 small chain [Proteobacteria] Length: 372\nNumber of predicted TMHs: 1\nExp number of AAs in TMHs: 23.72226\nExp number, first 60 AAs: 0.22038\nTotal prob of N-in: 0.12994\noutside 1 331\nTMhelix 332 354\ninside 355 372

4207 GCF_002087635.1_ASM208763v1 Escherichia coli Proteobacteria; Gammaproteobacteria; Enterobacterales; Enterobacteriaceae; Escherichia MTGDNTLIHSHGINRRDFMKLCAALAATMGLSSKAAAEMAESVTNPQRPPVIWIGAQECT WP_000145410.1 MULTISPECIES: hydrogenase-2 small chain [Proteobacteria] Length: 372\nNumber of predicted TMHs: 1\nExp number of AAs in TMHs: 23.72226\nExp number, first 60 AAs: 0.22038\nTotal prob of N-in: 0.12994\noutside 1 331\nTMhelix 332 354\ninside 355 372

4208 GCF_002087695.1_ASM208769v1 Escherichia coli Proteobacteria; Gammaproteobacteria; Enterobacterales; Enterobacteriaceae; Escherichia MTGDNTLIHSHGINRRDFMKLCAALAATMGLSSKAAAEMAESVTNPQRPPVIWIGAQECT WP_000145410.1 MULTISPECIES: hydrogenase-2 small chain [Proteobacteria] Length: 372\nNumber of predicted TMHs: 1\nExp number of AAs in TMHs: 23.72226\nExp number, first 60 AAs: 0.22038\nTotal prob of N-in: 0.12994\noutside 1 331\nTMhelix 332 354\ninside 355 372

4209 GCF_002087635.1_ASM208763v1 Escherichia coli Proteobacteria; Gammaproteobacteria; Enterobacterales; Enterobacteriaceae; Escherichia
MNNEETFYQAMRRQGVTRRSFLKYCSLAATSLGLGAGMAPKIAWALENKPRIPVVWIHGL WP_001058323.1
MULTISPECIES: hydrogenase-1 small chain [Proteobacteria] Length: 372\nNumber of predicted TMHs: 2\nExp number of AAs in TMHs: 37.2608499999999\nExp number, first 60 AAs: 19.52923\nTotal prob of N-in: 0.94351\nPOSSIBLE N-term signal sequence\ninside 1 19\nTMhelix 20 39\noutside 40 325\nTMhelix 326 348\ninside 349 372

4210 GCF_002087695.1_ASM208769v1 Escherichia coli Proteobacteria; Gammaproteobacteria; Enterobacterales; Enterobacteriaceae; Escherichia
MNNEETFYQAMRRQGVTRRSFLKYCSLAATSLGLGAGMAPKIAWALENKPRIPVVWIHGL WP_001058323.1
MULTISPECIES: hydrogenase-1 small chain [Proteobacteria] Length: 372\nNumber of predicted TMHs: 2\nExp number of AAs in TMHs: 37.2608499999999\nExp number, first 60 AAs: 19.52923\nTotal prob of N-in: 0.94351\nPOSSIBLE N-term signal sequence\ninside 1 19\nTMhelix 20 39\noutside 40 325\nTMhelix 326 348\ninside 349 372

4211 GCF_002087505.1_Esch_coli_OS_09_001_TOC1_V5 Escherichia coli Proteobacteria; Gammaproteobacteria; Enterobacterales; Enterobacteriaceae; Escherichia
MNNEETFYQAMRRQGVTRRSFLKYCSLAATSLGLGAGMAPKIAWALENKPRIPVVWIHGL WP_001058323.1
MULTISPECIES: hydrogenase-1 small chain [Proteobacteria] Length: 372\nNumber of predicted TMHs: 2\nExp number of AAs in TMHs: 37.2608499999999\nExp number, first 60 AAs: 19.52923\nTotal prob of N-in: 0.94351\nPOSSIBLE N-term signal sequence\ninside 1 19\nTMhelix 20 39\noutside 40 325\nTMhelix 326 348\ninside 349 372

4212 GCF_002087725.1_ASM208772v1 Escherichia coli Proteobacteria; Gammaproteobacteria; Enterobacterales; Enterobacteriaceae; Escherichia
MTGDNTLIHSHGINRRDFMKLCAALAATMGLSSKAAAEMAESVTNPQRPPVIWIGAQECT WP_000145410.1
MULTISPECIES: hydrogenase-2 small chain [Proteobacteria] Length: 372\nNumber of predicted TMHs: 1\nExp number of AAs in TMHs: 23.72226\nExp number, first 60 AAs: 0.22038\nTotal prob of N-in: 0.12994\noutside 1 331\nTMhelix 332 354\ninside 355 372

4213 GCF_002087725.1_ASM208772v1 Escherichia coli Proteobacteria; Gammaproteobacteria; Enterobacterales; Enterobacteriaceae; Escherichia
MNNEETFYQAMRRQGVTRRSFLKYCSLAATSLGLGAGMAPKIAWALENKPRIPVVWIHGL WP_001058323.1
MULTISPECIES: hydrogenase-1 small chain [Proteobacteria] Length: 372\nNumber of predicted TMHs: 2\nExp number of AAs in TMHs: 37.2608499999999\nExp number, first 60 AAs: 19.52923\nTotal prob of N-in: 0.94351\nPOSSIBLE N-term signal sequence\ninside 1 19\nTMhelix 20 39\noutside 40 325\nTMhelix 326 348\ninside 349 372

4214 GCF_002087825.1_ASM208782v1 Escherichia coli Proteobacteria; Gammaproteobacteria; Enterobacterales; Enterobacteriaceae; Escherichia
MTGDNTLIHSHGINRRDFMKLCAALAATMGLSSKAAAEMAESVTNPQRPPVIWIGAQECT WP_000145410.1
MULTISPECIES: hydrogenase-2 small chain [Proteobacteria] Length: 372\nNumber of predicted TMHs: 1\nExp number of AAs in TMHs: 23.72226\nExp number, first 60 AAs: 0.22038\nTotal prob of N-in: 0.12994\noutside 1 331\nTMhelix 332 354\ninside 355 372

4215 GCF_002087815.1_ASM208781v1 Escherichia coli Proteobacteria; Gammaproteobacteria; Enterobacterales; Enterobacteriaceae; Escherichia
MNNEETFYQAMRRQGVTRRSFLKYCSLAATSLGLGAGMAPKIAWALENKPRIPVVWIHGL WP_001058323.1
MULTISPECIES: hydrogenase-1 small chain [Proteobacteria] Length: 372\nNumber of predicted TMHs: 2\nExp number of AAs in TMHs: 37.2608499999999\nExp number, first 60 AAs: 19.52923\nTotal prob of N-in: 0.94351\nPOSSIBLE N-term signal sequence\ninside 1 19\nTMhelix 20 39\noutside 40 325\nTMhelix 326 348\ninside 349 372

4216 GCF_002087945.1_ASM208794v1 Escherichia coli Proteobacteria; Gammaproteobacteria; Enterobacterales; Enterobacteriaceae; Escherichia
MTGDNTLIHSHGINRRDFMKLCAALAATMGLSSKAAAEMAESVTNPQRPPVIWIGAQECT WP_000145410.1
MULTISPECIES: hydrogenase-2 small chain [Proteobacteria] Length: 372\nNumber of predicted TMHs: 1\nExp number of AAs in TMHs: 23.72226\nExp number, first 60 AAs: 0.22038\nTotal prob of N-in: 0.12994\noutside 1 331\nTMhelix 332 354\ninside 355 372

4217 GCF_002087985.1_ASM208798v1 Escherichia coli Proteobacteria; Gammaproteobacteria; Enterobacterales; Enterobacteriaceae; Escherichia
MTGDNTLIHSHGINRRDFMKLCAALAATMGLSSKAAAEMAESVTNPQRPPVIWIGAQECT WP_000145410.1
MULTISPECIES: hydrogenase-2 small chain [Proteobacteria] Length: 372\nNumber of predicted TMHs: 1\nExp number of AAs in TMHs: 23.72226\nExp number, first 60 AAs: 0.22038\nTotal prob of N-in: 0.12994\noutside 1 331\nTMhelix 332 354\ninside 355 372

4218 GCF_002087775.1_ASM208777v1 Escherichia coli Proteobacteria; Gammaproteobacteria; Enterobacterales; Enterobacteriaceae; Escherichia
MNNEETFYQAMRRQGVTRRSFLKYCSLAATSLGLGAGMAPKIAWALENKPRIPVVWIHGL WP_001058323.1
MULTISPECIES: hydrogenase-1 small chain [Proteobacteria] Length: 372\nNumber of predicted TMHs: 2\nExp number of AAs in TMHs: 37.2608499999999\nExp number, first 60 AAs: 19.52923\nTotal prob of N-in: 0.94351\nPOSSIBLE N-term signal sequence\ninside 1 19\nTMhelix 20 39\noutside 40 325\nTMhelix 326 348\ninside 349 372

4219 GCF_002090355.1_ASM209035v1 Escherichia coli Proteobacteria; Gammaproteobacteria; Enterobacterales; Enterobacteriaceae; Escherichia
MTGDNTLIHSHGINRRDFMKLCAALAATMGLSSKAAAEMAESVTNPQRPPVIWIGAQECT WP_000145410.1
MULTISPECIES: hydrogenase-2 small chain [Proteobacteria] Length: 372\nNumber of predicted TMHs: 1\nExp number of AAs in TMHs: 23.72226\nExp number, first 60 AAs: 0.22038\nTotal prob of N-in: 0.12994\noutside 1 331\nTMhelix 332 354\ninside 355 372

4220 GCF_002087985.1_ASM208798v1 Escherichia coli Proteobacteria; Gammaproteobacteria; Enterobacterales; Enterobacteriaceae; Escherichia
MNNEETFYQAMRRQGVTRRSFLKYCSLAATSLGLGAGMAPKIAWALENKPRIPVVWIHGL WP_001058323.1
MULTISPECIES: hydrogenase-1 small chain [Proteobacteria] Length: 372\nNumber of predicted TMHs: 2\nExp number of AAs in TMHs: 37.2608499999999\nExp number, first 60 AAs: 19.52923\nTotal prob of N-in: 0.94351\nPOSSIBLE N-term signal sequence\ninside 1 19\nTMhelix 20 39\noutside 40 325\nTMhelix 326 348\ninside 349 372

4221 GCF_002093035.1_ASM209303v1 Escherichia coli Proteobacteria; Gammaproteobacteria; Enterobacterales; Enterobacteriaceae; Escherichia
MNNEETFYQAMRRQGVTRRSFLKYCSLAATSLGLGAGMAPKIAWALENKPRIPVVWIHGL WP_001058323.1
MULTISPECIES: hydrogenase-1 small chain [Proteobacteria] Length: 372\nNumber of predicted TMHs: 2\nExp number of AAs in TMHs: 37.2608499999999\nExp number, first 60 AAs: 19.52923\nTotal prob of N-in: 0.94351\nPOSSIBLE N-term signal sequence\ninside 1 19\nTMhelix 20 39\noutside 40 325\nTMhelix 326 348\ninside 349 372

4222 GCF_002087825.1_ASM208782v1 Escherichia coli Proteobacteria; Gammaproteobacteria; Enterobacterales; Enterobacteriaceae; Escherichia
MNNEETFYQAMRRQGVTRRSFLKYCSLAATSLGLGAGMAPKIAWALENKPRIPVVWIHGL WP_001058323.1
MULTISPECIES: hydrogenase-1 small chain [Proteobacteria] Length: 372\nNumber of predicted TMHs: 2\nExp number of AAs in TMHs: 37.2608499999999\nExp number, first 60 AAs: 19.52923\nTotal prob of N-in: 0.94351\nPOSSIBLE N-term signal sequence\ninside 1 19\nTMhelix 20 39\noutside 40 325\nTMhelix 326 348\ninside 349 372

4223 GCF_002099435.1_ASM209943v1 Escherichia coli Proteobacteria; Gammaproteobacteria; Enterobacterales; Enterobacteriaceae; Escherichia
MTGDNTLIHSHGINRRDFMKLCAALAATMGLSSKAAAEMAESVTNPQRPPVIWIGAQECT WP_000145410.1
MULTISPECIES: hydrogenase-2 small chain [Proteobacteria] Length: 372\nNumber of predicted TMHs: 1\nExp number of AAs in TMHs: 23.72226\nExp number, first 60 AAs: 0.22038\nTotal prob of N-in: 0.12994\noutside 1 331\nTMhelix 332 354\ninside 355 372

4224 GCF_002099435.1_ASM209943v1 Escherichia coli Proteobacteria; Gammaproteobacteria; Enterobacterales; Enterobacteriaceae; Escherichia
MNNEETFYQAMRRQGVTRRSFLKYCSLAATSLGLGAGMAPKIAWALENKPRIPVVWIHGL WP_001058323.1
MULTISPECIES: hydrogenase-1 small chain [Proteobacteria] Length: 372\nNumber of predicted TMHs: 2\nExp number of AAs in TMHs: 37.2608499999999\nExp number, first 60 AAs: 19.52923\nTotal prob of N-in: 0.94351\nPOSSIBLE N-term signal sequence\ninside 1 19\nTMhelix 20 39\noutside 40 325\nTMhelix 326 348\ninside 349 372

4225 GCF_002087945.1_ASM208794v1 Escherichia coli Proteobacteria; Gammaproteobacteria; Enterobacterales; Enterobacteriaceae; Escherichia
MNNEETFYQAMRRQGVTRRSFLKYCSLAATSLGLGAGMAPKIAWALENKPRIPVVWIHGL WP_001058323.1
MULTISPECIES: hydrogenase-1 small chain [Proteobacteria] Length: 372\nNumber of predicted TMHs: 2\nExp number of AAs in TMHs: 37.2608499999999\nExp number, first 60 AAs: 19.52923\nTotal prob of N-in: 0.94351\nPOSSIBLE N-term signal sequence\ninside 1 19\nTMhelix 20 39\noutside 40 325\nTMhelix 326 348\ninside 349 372

4226 GCF_002099455.1_ASM209945v1 Escherichia coli Proteobacteria; Gammaproteobacteria; Enterobacterales; Enterobacteriaceae; Escherichia
MTGDNTLIHSHGINRRDFMKLCAALAATMGLSSKAAAEMAESVTNPQRPPVIWIGAQECT WP_000145410.1
MULTISPECIES: hydrogenase-2 small chain [Proteobacteria] Length: 372\nNumber of predicted TMHs: 1\nExp number of AAs in TMHs: 23.72226\nExp number, first 60 AAs: 0.22038\nTotal prob of N-in: 0.12994\noutside 1 331\nTMhelix 332 354\ninside 355 372

4227 GCF_002099555.1_ASM209955v1 Escherichia coli Proteobacteria; Gammaproteobacteria; Enterobacterales; Enterobacteriaceae; Escherichia
MNNEETFYQAMRRQGVTRRSFLKYCSLAATSLGLGAGMAPKIAWALENKPRIPVVWIHGL WP_001058323.1
MULTISPECIES: hydrogenase-1 small chain [Proteobacteria] Length: 372\nNumber of predicted TMHs: 2\nExp number of AAs in TMHs: 37.2608499999999\nExp number, first 60 AAs: 19.52923\nTotal prob of N-in: 0.94351\nPOSSIBLE N-term signal sequence\ninside 1 19\nTMhelix 20 39\noutside 40 325\nTMhelix 326 348\ninside 349 372

4228 GCF_002099495.1_ASM209949v1 Escherichia coli Proteobacteria; Gammaproteobacteria; Enterobacterales; Enterobacteriaceae; Escherichia
MNNEETFYQAMRRQGVTRRSFLKYCSLAATSLGLGAGMAPKIAWALENKPRIPVVWIHGL WP_001058323.1
MULTISPECIES: hydrogenase-1 small chain [Proteobacteria] Length: 372\nNumber of predicted TMHs: 2\nExp number of AAs in TMHs: 37.2608499999999\nExp number, first 60 AAs: 19.52923\nTotal prob of N-in: 0.94351\nPOSSIBLE N-term signal sequence\ninside 1 19\nTMhelix 20 39\noutside 40 325\nTMhelix 326 348\ninside 349 372

4229 GCF_002099585.1_ASM209958v1 Escherichia coli Proteobacteria; Gammaproteobacteria; Enterobacterales; Enterobacteriaceae; Escherichia
MTGDNTLIHSHGINRRDFMKLCAALAATMGLSSKAAAEMAESVTNPQRPPVIWIGAQECT WP_000145410.1
MULTISPECIES: hydrogenase-2 small chain [Proteobacteria] Length: 372\nNumber of predicted TMHs: 1\nExp number of AAs in TMHs: 23.72226\nExp number, first 60 AAs: 0.22038\nTotal prob of N-in: 0.12994\noutside 1 331\nTMhelix 332 354\ninside 355 372

4230 GCF_002099615.1_ASM209961v1 Escherichia coli Proteobacteria; Gammaproteobacteria; Enterobacterales; Enterobacteriaceae; Escherichia
MTGDNTLIHSHGINRRDFMKLCAALAATMGLSSKAAAEMAESVTNPQRPPVIWIGAQECT WP_000145410.1
MULTISPECIES: hydrogenase-2 small chain [Proteobacteria] Length: 372\nNumber of predicted TMHs: 1\nExp number of AAs in TMHs: 23.72226\nExp number, first 60 AAs: 0.22038\nTotal prob of N-in: 0.12994\noutside 1 331\nTMhelix 332 354\ninside 355 372

4231 GCF_002099605.1_ASM209960v1 Escherichia coli Proteobacteria; Gammaproteobacteria; Enterobacterales; Enterobacteriaceae; Escherichia
MTGDNTLIHSHGINRRDFMKLCAALAATMGLSSKAAAEMAESVTNPQRPPVIWIGAQECT WP_000145410.1
MULTISPECIES: hydrogenase-2 small chain [Proteobacteria] Length: 372\nNumber of predicted TMHs: 1\nExp number of AAs in TMHs: 23.72226\nExp number, first 60 AAs: 0.22038\nTotal prob of N-in: 0.12994\noutside 1 331\nTMhelix 332 354\ninside 355 372

4232 GCF_002099585.1_ASM209958v1 Escherichia coli Proteobacteria; Gammaproteobacteria; Enterobacterales; Enterobacteriaceae; Escherichia
MNNEETFYQAMRRQGVTRRSFLKYCSLAATSLGLGAGMAPKIAWALENKPRIPVVWIHGL WP_001058323.1
MULTISPECIES: hydrogenase-1 small chain [Proteobacteria] Length: 372\nNumber of predicted TMHs: 2\nExp number of AAs in TMHs: 37.2608499999999\nExp number, first 60 AAs: 19.52923\nTotal prob of N-in: 0.94351\nPOSSIBLE N-term signal sequence\ninside 1 19\nTMhelix 20 39\noutside 40 325\nTMhelix 326 348\ninside 349 372

4233 GCF_002099615.1_ASM209961v1 Escherichia coli Proteobacteria; Gammaproteobacteria; Enterobacterales; Enterobacteriaceae; Escherichia
MNNEETFYQAMRRQGVTRRSFLKYCSLAATSLGLGAGMAPKIAWALENKPRIPVVWIHGL WP_001058323.1
MULTISPECIES: hydrogenase-1 small chain [Proteobacteria] Length: 372\nNumber of predicted TMHs: 2\nExp number of AAs in TMHs: 37.2608499999999\nExp number, first 60 AAs: 19.52923\nTotal prob of N-in: 0.94351\nPOSSIBLE N-term signal sequence\ninside 1 19\nTMhelix 20 39\noutside 40 325\nTMhelix 326 348\ninside 349 372

4234 GCF_002099665.1_ASM209966v1 Escherichia coli Proteobacteria; Gammaproteobacteria; Enterobacterales; Enterobacteriaceae; Escherichia
MNNEETFYQAMRRQGVTRRSFLKYCSLAATSLGLGAGMAPKIAWALENKPRIPVVWIHGL WP_001058323.1
MULTISPECIES: hydrogenase-1 small chain [Proteobacteria] Length: 372\nNumber of predicted TMHs: 2\nExp number of AAs in TMHs: 37.2608499999999\nExp number, first 60 AAs: 19.52923\nTotal prob of N-in: 0.94351\nPOSSIBLE N-term signal sequence\ninside 1 19\nTMhelix 20 39\noutside 40 325\nTMhelix 326 348\ninside 349 372

4235 GCF_002099535.1_ASM209953v1 Escherichia coli Proteobacteria; Gammaproteobacteria; Enterobacterales; Enterobacteriaceae; Escherichia
MNNEETFYQAMRRQGVTRRSFLKYCSLAATSLGLGAGMAPKIAWALENKPRIPVVWIHGL WP_001058323.1
MULTISPECIES: hydrogenase-1 small chain [Proteobacteria] Length: 372\nNumber of predicted TMHs: 2\nExp number of AAs in TMHs: 37.2608499999999\nExp number, first 60 AAs: 19.52923\nTotal prob of N-in: 0.94351\nPOSSIBLE N-term signal sequence\ninside 1 19\nTMhelix 20 39\noutside 40 325\nTMhelix 326 348\ninside 349 372

4236 GCF_002099695.1_ASM209969v1 Escherichia coli Proteobacteria; Gammaproteobacteria; Enterobacterales; Enterobacteriaceae; Escherichia
MTGDNTLIHSHGINRRDFMKLCAALAATMGLSSKAAAEMAESVTNPQRPPVIWIGAQECT WP_000145410.1
MULTISPECIES: hydrogenase-2 small chain [Proteobacteria] Length: 372\nNumber of predicted TMHs: 1\nExp number of AAs in TMHs: 23.72226\nExp number, first 60 AAs: 0.22038\nTotal prob of N-in: 0.12994\noutside 1 331\nTMhelix 332 354\ninside 355 372

4237 GCF_002099715.1_ASM209971v1 Escherichia coli Proteobacteria; Gammaproteobacteria; Enterobacterales; Enterobacteriaceae; Escherichia
MTGDNTLIHSHGINRRDFMKLCAALAATMGLSSKAAAEMAESVTNPQRPPVIWIGAQECT WP_000145410.1
MULTISPECIES: hydrogenase-2 small chain [Proteobacteria] Length: 372\nNumber of predicted TMHs: 1\nExp number of AAs in TMHs: 23.72226\nExp number, first 60 AAs: 0.22038\nTotal prob of N-in: 0.12994\noutside 1 331\nTMhelix 332 354\ninside 355 372

4238 GCF_002099735.1_ASM209973v1 Escherichia coli Proteobacteria; Gammaproteobacteria; Enterobacterales; Enterobacteriaceae; Escherichia
MTGDNTLIHSHGINRRDFMKLCAALAATMGLSSKAAAEMAESVTNPQRPPVIWIGAQECT WP_000145410.1
MULTISPECIES: hydrogenase-2 small chain [Proteobacteria] Length: 372\nNumber of predicted TMHs: 1\nExp number of AAs in TMHs: 23.72226\nExp number, first 60 AAs: 0.22038\nTotal prob of N-in: 0.12994\noutside 1 331\nTMhelix 332 354\ninside 355 372

4239 GCF_002099765.1_ASM209976v1 Escherichia coli Proteobacteria; Gammaproteobacteria; Enterobacterales; Enterobacteriaceae; Escherichia
MTGDNTLIHSHGINRRDFMKLCAALAATMGLSSKAAAEMAESVTNPQRPPVIWIGAQECT WP_000145410.1
MULTISPECIES: hydrogenase-2 small chain [Proteobacteria] Length: 372\nNumber of predicted TMHs: 1\nExp number of AAs in TMHs: 23.72226\nExp number, first 60 AAs: 0.22038\nTotal prob of N-in: 0.12994\noutside 1 331\nTMhelix 332 354\ninside 355 372

4240 GCF_002099785.1_ASM209978v1 Escherichia coli Proteobacteria; Gammaproteobacteria; Enterobacterales; Enterobacteriaceae; Escherichia
MTGDNTLIHSHGINRRDFMKLCAALAATMGLSSKAAAEMAESVTNPQRPPVIWIGAQECT WP_000145410.1
MULTISPECIES: hydrogenase-2 small chain [Proteobacteria] Length: 372\nNumber of predicted TMHs: 1\nExp number of AAs in TMHs: 23.72226\nExp number, first 60 AAs: 0.22038\nTotal prob of N-in: 0.12994\noutside 1 331\nTMhelix 332 354\ninside 355 372

4241 GCF_002099805.1_ASM209980v1 Escherichia coli Proteobacteria; Gammaproteobacteria; Enterobacterales; Enterobacteriaceae; Escherichia

MNNEETFYQAMRRQGVTRRSFLKYCSLAATSLGLGAGMAPKIAWALENKPRIPVVWIHGL WP_001058323.1 MULTISPECIES: hydrogenase-1 small chain [Proteobacteria] Length: 372\nNumber of predicted TMHs: 2\nExp number of AAs in TMHs: 37.2608499999999\nExp number, first 60 AAs: 19.52923\nTotal prob of N-in: 0.94351\nPOSSIBLE N-term signal sequence\ninside 1 19\nTMhelix 20 39\noutside 40 325\nTMhelix 326 348\ninside 349 372

4242 GCF_002099715.1_ASM209971v1 Escherichia coli Proteobacteria; Gammaproteobacteria; Enterobacterales; Enterobacteriaceae; Escherichia MNNEETFYQAMRRQGVTRRSFLKYCSLAATSLGLGAGMAPKIAWALENKPRIPVVWIHGL WP_001058323.1 MULTISPECIES: hydrogenase-1 small chain [Proteobacteria] Length: 372\nNumber of predicted TMHs: 2\nExp number of AAs in TMHs: 37.2608499999999\nExp number, first 60 AAs: 19.52923\nTotal prob of N-in: 0.94351\nPOSSIBLE N-term signal sequence\ninside 1 19\nTMhelix 20 39\noutside 40 325\nTMhelix 326 348\ninside 349 372

4243 GCF_002099735.1_ASM209973v1 Escherichia coli Proteobacteria; Gammaproteobacteria; Enterobacterales; Enterobacteriaceae; Escherichia MNNEETFYQAMRRQGVTRRSFLKYCSLAATSLGLGAGMAPKIAWALENKPRIPVVWIHGL WP_001058323.1 MULTISPECIES: hydrogenase-1 small chain [Proteobacteria] Length: 372\nNumber of predicted TMHs: 2\nExp number of AAs in TMHs: 37.2608499999999\nExp number, first 60 AAs: 19.52923\nTotal prob of N-in: 0.94351\nPOSSIBLE N-term signal sequence\ninside 1 19\nTMhelix 20 39\noutside 40 325\nTMhelix 326 348\ninside 349 372

4244 GCF_002099875.1_ASM209987v1 Escherichia coli Proteobacteria; Gammaproteobacteria; Enterobacterales; Enterobacteriaceae; Escherichia MTGDNTLIHSHGINRRDFMKLCAALAATMGLSSKAAAEMAESVTNPQRPPVIWIGAQECT WP_000145410.1 MULTISPECIES: hydrogenase-2 small chain [Proteobacteria] Length: 372\nNumber of predicted TMHs: 1\nExp number of AAs in TMHs: 23.72226\nExp number, first 60 AAs: 0.22038\nTotal prob of N-in: 0.12994\noutside 1 331\nTMhelix 332 354\ninside 355 372

4245 GCF_002099875.1_ASM209987v1 Escherichia coli Proteobacteria; Gammaproteobacteria; Enterobacterales; Enterobacteriaceae; Escherichia MNNEETFYQAMRRQGVTRRSFLKYCSLAATSLGLGAGMAPKIAWALENKPRIPVVWIHGL WP_001058323.1 MULTISPECIES: hydrogenase-1 small chain [Proteobacteria] Length: 372\nNumber of predicted TMHs: 2\nExp number of AAs in TMHs: 37.2608499999999\nExp number, first 60 AAs: 19.52923\nTotal prob of N-in: 0.94351\nPOSSIBLE N-term signal sequence\ninside 1 19\nTMhelix 20 39\noutside 40 325\nTMhelix 326 348\ninside 349 372

4246 GCF_002099765.1_ASM209976v1 Escherichia coli Proteobacteria; Gammaproteobacteria; Enterobacterales; Enterobacteriaceae; Escherichia MNNEETFYQAMRRQGVTRRSFLKYCSLAATSLGLGAGMAPKIAWALENKPRIPVVWIHGL WP_001058323.1 MULTISPECIES: hydrogenase-1 small chain [Proteobacteria] Length: 372\nNumber of predicted TMHs: 2\nExp number of AAs in TMHs: 37.2608499999999\nExp number, first 60 AAs: 19.52923\nTotal prob of N-in: 0.94351\nPOSSIBLE N-term signal sequence\ninside 1 19\nTMhelix 20 39\noutside 40 325\nTMhelix 326 348\ninside 349 372

4247 GCF_002099885.1_ASM209988v1 Escherichia coli Proteobacteria; Gammaproteobacteria; Enterobacterales; Enterobacteriaceae; Escherichia MTGDNTLIHSHGINRRDFMKLCAALAATMGLSSKAAAEMAESVTNPQRPPVIWIGAQECT WP_000145410.1 MULTISPECIES: hydrogenase-2 small chain [Proteobacteria] Length: 372\nNumber of predicted TMHs: 1\nExp number of AAs in TMHs: 23.72226\nExp number, first 60 AAs: 0.22038\nTotal prob of N-in: 0.12994\noutside 1 331\nTMhelix 332 354\ninside 355 372

4248 GCF_002099945.1_ASM209994v1 Escherichia coli Proteobacteria; Gammaproteobacteria; Enterobacterales; Enterobacteriaceae; Escherichia MTGDNTLIHSHGINRRDFMKLCAALAATMGLSSKAAAEMAESVTNPQRPPVIWIGAQECT WP_000145410.1 MULTISPECIES: hydrogenase-2 small chain [Proteobacteria] Length: 372\nNumber of predicted TMHs: 1\nExp number of AAs in TMHs: 23.72226\nExp number, first 60 AAs: 0.22038\nTotal prob of N-in: 0.12994\noutside 1 331\nTMhelix 332 354\ninside 355 372

4249 GCF_002100025.1_ASM210002v1 Escherichia coli Proteobacteria; Gammaproteobacteria; Enterobacterales; Enterobacteriaceae; Escherichia

MTGDNTLIHSHGINRRDFMKLCAALAATMGLSSKAAAEMAESVTNPQRPPVIWIGAQECT WP_000145410.1 MULTISPECIES: hydrogenase-2 small chain [Proteobacteria] Length: 372\nNumber of predicted TMHs: 1\nExp number of AAs in TMHs: 23.72226\nExp number, first 60 AAs: 0.22038\nTotal prob of N-in: 0.12994\noutside 1 331\nTMhelix 332 354\ninside 355 372

4250 GCF_002104125.1_ASM210412v1 Escherichia coli Proteobacteria; Gammaproteobacteria; Enterobacterales; Enterobacteriaceae; Escherichia MTGDNTLIHSHGINRRDFMKLCAALAATMGLSSKAAAEMAESVTNPQRPPVIWIGAQECT WP_000145410.1 MULTISPECIES: hydrogenase-2 small chain [Proteobacteria] Length: 372\nNumber of predicted TMHs: 1\nExp number of AAs in TMHs: 23.72226\nExp number, first 60 AAs: 0.22038\nTotal prob of N-in: 0.12994\noutside 1 331\nTMhelix 332 354\ninside 355 372

4251 GCF_002100055.1_ASM210005v1 Escherichia coli Proteobacteria; Gammaproteobacteria; Enterobacterales; Enterobacteriaceae; Escherichia MTGDNTLIHSHGINRRDFMKLCAALAATMGLSSKAAAEMAESVTNPQRPPVIWIGAQECT WP_000145410.1 MULTISPECIES: hydrogenase-2 small chain [Proteobacteria] Length: 372\nNumber of predicted TMHs: 1\nExp number of AAs in TMHs: 23.72226\nExp number, first 60 AAs: 0.22038\nTotal prob of N-in: 0.12994\noutside 1 331\nTMhelix 332 354\ninside 355 372

4252 GCF_002104405.1_ASM210440v1 Escherichia coli Proteobacteria; Gammaproteobacteria; Enterobacterales; Enterobacteriaceae; Escherichia MTGDNTLIHSHGINRRDFMKLCAALAATMGLSSKAAAEMAESVTNPQRPPVIWIGAQECT WP_000145410.1 MULTISPECIES: hydrogenase-2 small chain [Proteobacteria] Length: 372\nNumber of predicted TMHs: 1\nExp number of AAs in TMHs: 23.72226\nExp number, first 60 AAs: 0.22038\nTotal prob of N-in: 0.12994\noutside 1 331\nTMhelix 332 354\ninside 355 372

4253 GCF_002100045.1_ASM210004v1 Escherichia coli Proteobacteria; Gammaproteobacteria; Enterobacterales; Enterobacteriaceae; Escherichia MTGDNTLIHSHGINRRDFMKLCAALAATMGLSSKAAAEMAESVTNPQRPPVIWIGAQECT WP_000145410.1 MULTISPECIES: hydrogenase-2 small chain [Proteobacteria] Length: 372\nNumber of predicted TMHs: 1\nExp number of AAs in TMHs: 23.72226\nExp number, first 60 AAs: 0.22038\nTotal prob of N-in: 0.12994\noutside 1 331\nTMhelix 332 354\ninside 355 372

4254 GCF_002099955.1_ASM209995v1 Escherichia coli Proteobacteria; Gammaproteobacteria; Enterobacterales; Enterobacteriaceae; Escherichia MTGDNTLIHSHGINRRDFMKLCAALAATMGLSSKAAAEMAESVTNPQRPPVIWIGAQECT WP_000145410.1 MULTISPECIES: hydrogenase-2 small chain [Proteobacteria] Length: 372\nNumber of predicted TMHs: 1\nExp number of AAs in TMHs: 23.72226\nExp number, first 60 AAs: 0.22038\nTotal prob of N-in: 0.12994\noutside 1 331\nTMhelix 332 354\ninside 355 372

4255 GCF_002100105.1_ASM210010v1 Escherichia coli Proteobacteria; Gammaproteobacteria; Enterobacterales; Enterobacteriaceae; Escherichia MTGDNTLIHSHGINRRDFMKLCAALAATMGLSSKAAAEMAESVTNPQRPPVIWIGAQECT WP_000145410.1 MULTISPECIES: hydrogenase-2 small chain [Proteobacteria] Length: 372\nNumber of predicted TMHs: 1\nExp number of AAs in TMHs: 23.72226\nExp number, first 60 AAs: 0.22038\nTotal prob of N-in: 0.12994\noutside 1 331\nTMhelix 332 354\ninside 355 372

4256 GCF_002099945.1_ASM209994v1 Escherichia coli Proteobacteria; Gammaproteobacteria; Enterobacterales; Enterobacteriaceae; Escherichia MNNEETFYQAMRRQGVTRRSFLKYCSLAATSLGLGAGMAPKIAWALENKPRIPVVWIHGL WP_001058323.1 MULTISPECIES: hydrogenase-1 small chain [Proteobacteria] Length: 372\nNumber of predicted TMHs: 2\nExp number of AAs in TMHs: 37.2608499999999\nExp number, first 60 AAs: 19.52923\nTotal prob of N-in: 0.94351\nPOSSIBLE N-term signal sequence\ninside 1 19\nTMhelix 20 39\noutside 40 325\nTMhelix 326 348\ninside 349 372

4257 GCF_002100025.1_ASM210002v1 Escherichia coli Proteobacteria; Gammaproteobacteria; Enterobacterales; Enterobacteriaceae; Escherichia MNNEETFYQAMRRQGVTRRSFLKYCSLAATSLGLGAGMAPKIAWALENKPRIPVVWIHGL WP_001058323.1 MULTISPECIES: hydrogenase-1 small chain [Proteobacteria] Length: 372\nNumber of predicted TMHs: 2\nExp number of AAs in TMHs: 37.2608499999999\nExp number, first 60 AAs: 19.52923\nTotal prob of N-in:

0.94351\nPOSSIBLE N-term signal sequence\ninside 1 19\nTMhelix 20 39\noutside 40 325\nTMhelix 326 348\ninside 349 372

4258 GCF_002104125.1_ASM210412v1 Escherichia coli Proteobacteria; Gammaproteobacteria; Enterobacterales; Enterobacteriaceae; Escherichia
MNNEETFYQAMRRQGVTRRSFLKYCSLAATSLGLGAGMAPKIAWALENKPRIPVVWIHGL WP_001058323.1
MULTISPECIES: hydrogenase-1 small chain [Proteobacteria] Length: 372\nNumber of predicted TMHs: 2\nExp number of AAs in TMHs: 37.2608499999999\nExp number, first 60 AAs: 19.52923\nTotal prob of N-in: 0.94351\nPOSSIBLE N-term signal sequence\ninside 1 19\nTMhelix 20 39\noutside 40 325\nTMhelix 326 348\ninside 349 372

4259 GCF_002100055.1_ASM210005v1 Escherichia coli Proteobacteria; Gammaproteobacteria; Enterobacterales; Enterobacteriaceae; Escherichia
MNNEETFYQAMRRQGVTRRSFLKYCSLAATSLGLGAGMAPKIAWALENKPRIPVVWIHGL WP_001295940.1
MULTISPECIES: hydrogenase [Enterobacteriaceae] Length: 372\nNumber of predicted TMHs: 2\nExp number of AAs in TMHs: 37.2608899999999\nExp number, first 60 AAs: 19.52923\nTotal prob of N-in: 0.94351\nPOSSIBLE N-term signal sequence\ninside 1 19\nTMhelix 20 39\noutside 40 325\nTMhelix 326 348\ninside 349 372

4260 GCF_002100105.1_ASM210010v1 Escherichia coli Proteobacteria; Gammaproteobacteria; Enterobacterales; Enterobacteriaceae; Escherichia
MNNEETFYQAMRRQGVTRRSFLKYCSLAATSLGLGAGMAPKIAWALENKPRIPVVWIHGL WP_001295940.1
MULTISPECIES: hydrogenase [Enterobacteriaceae] Length: 372\nNumber of predicted TMHs: 2\nExp number of AAs in TMHs: 37.2608899999999\nExp number, first 60 AAs: 19.52923\nTotal prob of N-in: 0.94351\nPOSSIBLE N-term signal sequence\ninside 1 19\nTMhelix 20 39\noutside 40 325\nTMhelix 326 348\ninside 349 372

4261 GCF_002100035.1_ASM210003v1 Escherichia coli Proteobacteria; Gammaproteobacteria; Enterobacterales; Enterobacteriaceae; Escherichia
MNNEETFYQAMRRQGVTRRSFLKYCSLAATSLGLGAGMAPKIAWALENKPRIPVVWIHGL WP_001295940.1
MULTISPECIES: hydrogenase [Enterobacteriaceae] Length: 372\nNumber of predicted TMHs: 2\nExp number of AAs in TMHs: 37.2608899999999\nExp number, first 60 AAs: 19.52923\nTotal prob of N-in: 0.94351\nPOSSIBLE N-term signal sequence\ninside 1 19\nTMhelix 20 39\noutside 40 325\nTMhelix 326 348\ninside 349 372

4262 GCF_002104225.1_ASM210422v1 Escherichia coli Proteobacteria; Gammaproteobacteria; Enterobacterales; Enterobacteriaceae; Escherichia
MNNEETFYQAMRRQGVTRRSFLKYCSLAATSLGLGAGMAPKIAWALENKPRIPVVWIHGL WP_001295940.1
MULTISPECIES: hydrogenase [Enterobacteriaceae] Length: 372\nNumber of predicted TMHs: 2\nExp number of AAs in TMHs: 37.2608899999999\nExp number, first 60 AAs: 19.52923\nTotal prob of N-in: 0.94351\nPOSSIBLE N-term signal sequence\ninside 1 19\nTMhelix 20 39\noutside 40 325\nTMhelix 326 348\ninside 349 372

4263 GCF_002104485.1_ASM210448v1 Escherichia coli Proteobacteria; Gammaproteobacteria; Enterobacterales; Enterobacteriaceae; Escherichia
MNNEETFYQAMRRQGVTRRSFLKYCSLAATSLGLGAGMAPKIAWALENKPRIPVVWIHGL WP_001295940.1
MULTISPECIES: hydrogenase [Enterobacteriaceae] Length: 372\nNumber of predicted TMHs: 2\nExp number of AAs in TMHs: 37.2608899999999\nExp number, first 60 AAs: 19.52923\nTotal prob of N-in: 0.94351\nPOSSIBLE N-term signal sequence\ninside 1 19\nTMhelix 20 39\noutside 40 325\nTMhelix 326 348\ninside 349 372

4264 GCF_002114355.1_ASM211435v1 Escherichia coli Proteobacteria; Gammaproteobacteria; Enterobacterales; Enterobacteriaceae; Escherichia
MTGDNTLIHSHGINRRDFMKLCAALAATMGLSSKAAAEMAESVTNPQRPPVIWIGAQECT WP_000145410.1
MULTISPECIES: hydrogenase-2 small chain [Proteobacteria] Length: 372\nNumber of predicted TMHs: 1\nExp number of AAs in TMHs: 23.72226\nExp number, first 60 AAs: 0.22038\nTotal prob of N-in: 0.12994\noutside 1 331\nTMhelix 332 354\ninside 355 372

4265 GCF_002114355.1_ASM211435v1 Escherichia coli Proteobacteria; Gammaproteobacteria; Enterobacterales; Enterobacteriaceae; Escherichia
MNNEETFYQAMRRQGVTRRSFLKYCSLAATSLGLGAGMAPKIAWALENKPRIPVVWIHGL WP_001058323.1

MULTISPECIES: hydrogenase-1 small chain [Proteobacteria] Length: 372\nNumber of predicted TMHs: 2\nExp number of AAs in TMHs: 37.2608499999999\nExp number, first 60 AAs: 19.52923\nTotal prob of N-in: 0.94351\nPOSSIBLE N-term signal sequence\ninside 1 19\nTMhelix 20 39\noutside 40 325\nTMhelix 326 348\ninside 349 372

4266 GCF_002116465.1_ASM211646v1 Escherichia coli Proteobacteria; Gammaproteobacteria; Enterobacterales; Enterobacteriaceae; Escherichia
MTGDNTLIHSHGINRRDFMKLCAALAATMGLSSKAAAEMAESVTNPQRPPVIWIGAQECT WP_000145410.1
MULTISPECIES: hydrogenase-2 small chain [Proteobacteria] Length: 372\nNumber of predicted TMHs: 1\nExp number of AAs in TMHs: 23.72226\nExp number, first 60 AAs: 0.22038\nTotal prob of N-in: 0.12994\noutside 1 331\nTMhelix 332 354\ninside 355 372

4267 GCF_002116485.1_ASM211648v1 Escherichia coli Proteobacteria; Gammaproteobacteria; Enterobacterales; Enterobacteriaceae; Escherichia
MNNEETFYQAMRRQGVTRRSFLKYCSLAATSLGLGAGMAPKIAWALENKPRIPVVWIHGL WP_001058323.1
MULTISPECIES: hydrogenase-1 small chain [Proteobacteria] Length: 372\nNumber of predicted TMHs: 2\nExp number of AAs in TMHs: 37.2608499999999\nExp number, first 60 AAs: 19.52923\nTotal prob of N-in: 0.94351\nPOSSIBLE N-term signal sequence\ninside 1 19\nTMhelix 20 39\noutside 40 325\nTMhelix 326 348\ninside 349 372

4268 GCF_002116565.1_ASM211656v1 Escherichia coli Proteobacteria; Gammaproteobacteria; Enterobacterales; Enterobacteriaceae; Escherichia
MTGDNTLIHSHGINRRDFMKLCAALAATMGLSSKAAAEMAESVTNPQRPPVIWIGAQECT WP_000145410.1
MULTISPECIES: hydrogenase-2 small chain [Proteobacteria] Length: 372\nNumber of predicted TMHs: 1\nExp number of AAs in TMHs: 23.72226\nExp number, first 60 AAs: 0.22038\nTotal prob of N-in: 0.12994\noutside 1 331\nTMhelix 332 354\ninside 355 372

4269 GCF_002116555.1_ASM211655v1 Escherichia coli Proteobacteria; Gammaproteobacteria; Enterobacterales; Enterobacteriaceae; Escherichia
MTGDNTLIHSHGINRRDFMKLCAALAATMGLSSKAAAEMAESVTNPQRPPVIWIGAQECT WP_000145410.1
MULTISPECIES: hydrogenase-2 small chain [Proteobacteria] Length: 372\nNumber of predicted TMHs: 1\nExp number of AAs in TMHs: 23.72226\nExp number, first 60 AAs: 0.22038\nTotal prob of N-in: 0.12994\noutside 1 331\nTMhelix 332 354\ninside 355 372

4270 GCF_002116565.1_ASM211656v1 Escherichia coli Proteobacteria; Gammaproteobacteria; Enterobacterales; Enterobacteriaceae; Escherichia
MNNEETFYQAMRRQGVTRRSFLKYCSLAATSLGLGAGMAPKIAWALENKPRIPVVWIHGL WP_001058323.1
MULTISPECIES: hydrogenase-1 small chain [Proteobacteria] Length: 372\nNumber of predicted TMHs: 2\nExp number of AAs in TMHs: 37.2608499999999\nExp number, first 60 AAs: 19.52923\nTotal prob of N-in: 0.94351\nPOSSIBLE N-term signal sequence\ninside 1 19\nTMhelix 20 39\noutside 40 325\nTMhelix 326 348\ninside 349 372

4271 GCF_002116555.1_ASM211655v1 Escherichia coli Proteobacteria; Gammaproteobacteria; Enterobacterales; Enterobacteriaceae; Escherichia
MNNEETFYQAMRRQGVTRRSFLKYCSLAATSLGLGAGMAPKIAWALENKPRIPVVWIHGL WP_001058323.1
MULTISPECIES: hydrogenase-1 small chain [Proteobacteria] Length: 372\nNumber of predicted TMHs: 2\nExp number of AAs in TMHs: 37.2608499999999\nExp number, first 60 AAs: 19.52923\nTotal prob of N-in: 0.94351\nPOSSIBLE N-term signal sequence\ninside 1 19\nTMhelix 20 39\noutside 40 325\nTMhelix 326 348\ninside 349 372

4272 GCF_002116465.1_ASM211646v1 Escherichia coli Proteobacteria; Gammaproteobacteria; Enterobacterales; Enterobacteriaceae; Escherichia
MNNEETFYQAMRRQGVTRRSFLKYCSLAATSLGLGAGMAPKIAWALENKPRIPVVWIHGL WP_001058323.1
MULTISPECIES: hydrogenase-1 small chain [Proteobacteria] Length: 372\nNumber of predicted TMHs: 2\nExp number of AAs in TMHs: 37.2608499999999\nExp number, first 60 AAs: 19.52923\nTotal prob of N-in: 0.94351\nPOSSIBLE N-term signal sequence\ninside 1 19\nTMhelix 20 39\noutside 40 325\nTMhelix 326 348\ninside 349 372

4273 GCF_002116495.1_ASM211649v1 Escherichia coli Proteobacteria; Gammaproteobacteria; Enterobacterales; Enterobacteriaceae; Escherichia
MNNEETFYQAMRRQGVTRRSFLKYCSLAATSLGLGAGMAPKIAWALENKPRIPVVWIHGL WP_001058323.1

MULTISPECIES: hydrogenase-1 small chain [Proteobacteria] Length: 372\nNumber of predicted TMHs: 2\nExp number of AAs in TMHs: 37.2608499999999\nExp number, first 60 AAs: 19.52923\nTotal prob of N-in: 0.94351\nPOSSIBLE N-term signal sequence\ninside 1 19\nTMhelix 20 39\noutside 40 325\nTMhelix 326 348\ninside 349 372

4274 GCF_002133365.1_ASM213336v1 Escherichia coli Proteobacteria; Gammaproteobacteria; Enterobacterales; Enterobacteriaceae; Escherichia
MTGDNTLIHSHGINRRDFMKLCAALAATMGLSSKAAAEMAESVTNPQRPPVIWIGAQECT WP_000145410.1
MULTISPECIES: hydrogenase-2 small chain [Proteobacteria] Length: 372\nNumber of predicted TMHs: 1\nExp number of AAs in TMHs: 23.72226\nExp number, first 60 AAs: 0.22038\nTotal prob of N-in: 0.12994\noutside 1 331\nTMhelix 332 354\ninside 355 372

4275 GCF_002133385.1_ASM213338v1 Escherichia coli Proteobacteria; Gammaproteobacteria; Enterobacterales; Enterobacteriaceae; Escherichia
MTGDNTLIHSHGINRRDFMKLCAALAATMGLSSKAAAEMAESVTNPQRPPVIWIGAQECT WP_000145410.1
MULTISPECIES: hydrogenase-2 small chain [Proteobacteria] Length: 372\nNumber of predicted TMHs: 1\nExp number of AAs in TMHs: 23.72226\nExp number, first 60 AAs: 0.22038\nTotal prob of N-in: 0.12994\noutside 1 331\nTMhelix 332 354\ninside 355 372

4276 GCF_002120205.1_ASM212020v1 Escherichia coli Proteobacteria; Gammaproteobacteria; Enterobacterales; Enterobacteriaceae; Escherichia
MNNEETFYQAMRRQGVTRRSFLKYCSLAATSLGLGAGMAPKIAWALENKPRIPVVWIHGL WP_001295940.1
MULTISPECIES: hydrogenase [Enterobacteriaceae] Length: 372\nNumber of predicted TMHs: 2\nExp number of AAs in TMHs: 37.2608899999999\nExp number, first 60 AAs: 19.52923\nTotal prob of N-in: 0.94351\nPOSSIBLE N-term signal sequence\ninside 1 19\nTMhelix 20 39\noutside 40 325\nTMhelix 326 348\ninside 349 372

4277 GCF_002133405.1_ASM213340v1 Escherichia coli Proteobacteria; Gammaproteobacteria; Enterobacterales; Enterobacteriaceae; Escherichia
MTGDNTLIHSHGINRRDFMKLCAALAATMGLSSKAAAEMAESVTNPQRPPVIWIGAQECT WP_000145410.1
MULTISPECIES: hydrogenase-2 small chain [Proteobacteria] Length: 372\nNumber of predicted TMHs: 1\nExp number of AAs in TMHs: 23.72226\nExp number, first 60 AAs: 0.22038\nTotal prob of N-in: 0.12994\noutside 1 331\nTMhelix 332 354\ninside 355 372

4278 GCF_002133425.1_ASM213342v1 Escherichia coli Proteobacteria; Gammaproteobacteria; Enterobacterales; Enterobacteriaceae; Escherichia
MTGDNTLIHSHGINRRDFMKLCAALAATMGLSSKAAAEMAESVTNPQRPPVIWIGAQECT WP_000145410.1
MULTISPECIES: hydrogenase-2 small chain [Proteobacteria] Length: 372\nNumber of predicted TMHs: 1\nExp number of AAs in TMHs: 23.72226\nExp number, first 60 AAs: 0.22038\nTotal prob of N-in: 0.12994\noutside 1 331\nTMhelix 332 354\ninside 355 372

4279 GCF_002133435.1_ASM213343v1 Escherichia coli Proteobacteria; Gammaproteobacteria; Enterobacterales; Enterobacteriaceae; Escherichia
MTGDNTLIHSHGINRRDFMKLCAALAATMGLSSKAAAEMAESVTNPQRPPVIWIGAQECT WP_000145410.1
MULTISPECIES: hydrogenase-2 small chain [Proteobacteria] Length: 372\nNumber of predicted TMHs: 1\nExp number of AAs in TMHs: 23.72226\nExp number, first 60 AAs: 0.22038\nTotal prob of N-in: 0.12994\noutside 1 331\nTMhelix 332 354\ninside 355 372

4280 GCF_002133405.1_ASM213340v1 Escherichia coli Proteobacteria; Gammaproteobacteria; Enterobacterales; Enterobacteriaceae; Escherichia
MNNEETFYQAMRRQGVTRRSFLKYCSLAATSLGLGAGMAPKIAWALENKPRIPVVWIHGL WP_001058323.1
MULTISPECIES: hydrogenase-1 small chain [Proteobacteria] Length: 372\nNumber of predicted TMHs: 2\nExp number of AAs in TMHs: 37.2608499999999\nExp number, first 60 AAs: 19.52923\nTotal prob of N-in: 0.94351\nPOSSIBLE N-term signal sequence\ninside 1 19\nTMhelix 20 39\noutside 40 325\nTMhelix 326 348\ninside 349 372

4281 GCF_002133505.1_ASM213350v1 Escherichia coli Proteobacteria; Gammaproteobacteria; Enterobacterales; Enterobacteriaceae; Escherichia
MTGDNTLIHSHGINRRDFMKLCAALAATMGLSSKAAAEMAESVTNPQRPPVIWIGAQECT WP_000145410.1
MULTISPECIES: hydrogenase-2 small chain [Proteobacteria] Length: 372\nNumber of predicted TMHs:

1\nExp number of AAs in TMHs: 23.72226\nExp number, first 60 AAs: 0.22038\nTotal prob of N-in: 0.12994\noutside 1 331\nTMhelix 332 354\ninside 355 372

4282 GCF_002133515.1_ASM213351v1 Escherichia coli Proteobacteria; Gammaproteobacteria; Enterobacterales; Enterobacteriaceae; Escherichia MNNEETFYQAMRRQGVTRRSFLKYCSLAATSLGLGAGMAPKIAWALENKPRIPVVWIHGL WP_001058323.1 MULTISPECIES: hydrogenase-1 small chain [Proteobacteria] Length: 372\nNumber of predicted TMHs: 2\nExp number of AAs in TMHs: 37.2608499999999\nExp number, first 60 AAs: 19.52923\nTotal prob of N-in: 0.94351\nPOSSIBLE N-term signal sequence\ninside 1 19\nTMhelix 20 39\noutside 40 325\nTMhelix 326 348\ninside 349 372

4283 GCF_002133455.1_ASM213345v1 Escherichia coli Proteobacteria; Gammaproteobacteria; Enterobacterales; Enterobacteriaceae; Escherichia MNNEETFYQAMRRQGVTRRSFLKYCSLAATSLGLGAGMAPKIAWALENKPRIPVVWIHGL WP_001058323.1 MULTISPECIES: hydrogenase-1 small chain [Proteobacteria] Length: 372\nNumber of predicted TMHs: 2\nExp number of AAs in TMHs: 37.2608499999999\nExp number, first 60 AAs: 19.52923\nTotal prob of N-in: 0.94351\nPOSSIBLE N-term signal sequence\ninside 1 19\nTMhelix 20 39\noutside 40 325\nTMhelix 326 348\ninside 349 372

4284 GCF_002133385.1_ASM213338v1 Escherichia coli Proteobacteria; Gammaproteobacteria; Enterobacterales; Enterobacteriaceae; Escherichia MNNEETFYQAMRRQGVTRRSFLKYCSLAATSLGLGAGMAPKIAWALENKPRIPVVWIHGL WP_001058323.1 MULTISPECIES: hydrogenase-1 small chain [Proteobacteria] Length: 372\nNumber of predicted TMHs: 2\nExp number of AAs in TMHs: 37.2608499999999\nExp number, first 60 AAs: 19.52923\nTotal prob of N-in: 0.94351\nPOSSIBLE N-term signal sequence\ninside 1 19\nTMhelix 20 39\noutside 40 325\nTMhelix 326 348\ninside 349 372

4285 GCF_002133535.1_ASM213353v1 Escherichia coli Proteobacteria; Gammaproteobacteria; Enterobacterales; Enterobacteriaceae; Escherichia MTGDNTLIHSHGINRRDFMKLCAALAATMGLSSKAAAEMAESVTNPQRPPVIWIGAQECT WP_000145410.1 MULTISPECIES: hydrogenase-2 small chain [Proteobacteria] Length: 372\nNumber of predicted TMHs: 1\nExp number of AAs in TMHs: 23.72226\nExp number, first 60 AAs: 0.22038\nTotal prob of N-in: 0.12994\noutside 1 331\nTMhelix 332 354\ninside 355 372

4286 GCF_002133525.1_ASM213352v1 Escherichia coli Proteobacteria; Gammaproteobacteria; Enterobacterales; Enterobacteriaceae; Escherichia MTGDNTLIHSHGINRRDFMKLCAALAATMGLSSKAAAEMAESVTNPQRPPVIWIGAQECT WP_000145410.1 MULTISPECIES: hydrogenase-2 small chain [Proteobacteria] Length: 372\nNumber of predicted TMHs: 1\nExp number of AAs in TMHs: 23.72226\nExp number, first 60 AAs: 0.22038\nTotal prob of N-in: 0.12994\noutside 1 331\nTMhelix 332 354\ninside 355 372

4287 GCF_002133585.1_ASM213358v1 Escherichia coli Proteobacteria; Gammaproteobacteria; Enterobacterales; Enterobacteriaceae; Escherichia MTGDNTLIHSHGINRRDFMKLCAALAATMGLSSKAAAEMAESVTNPQRPPVIWIGAQECT WP_000145410.1 MULTISPECIES: hydrogenase-2 small chain [Proteobacteria] Length: 372\nNumber of predicted TMHs: 1\nExp number of AAs in TMHs: 23.72226\nExp number, first 60 AAs: 0.22038\nTotal prob of N-in: 0.12994\noutside 1 331\nTMhelix 332 354\ninside 355 372

4288 GCF_002133605.1_ASM213360v1 Escherichia coli Proteobacteria; Gammaproteobacteria; Enterobacterales; Enterobacteriaceae; Escherichia MTGDNTLIHSHGINRRDFMKLCAALAATMGLSSKAAAEMAESVTNPQRPPVIWIGAQECT WP_000145410.1 MULTISPECIES: hydrogenase-2 small chain [Proteobacteria] Length: 372\nNumber of predicted TMHs: 1\nExp number of AAs in TMHs: 23.72226\nExp number, first 60 AAs: 0.22038\nTotal prob of N-in: 0.12994\noutside 1 331\nTMhelix 332 354\ninside 355 372

4289 GCF_002133615.1_ASM213361v1 Escherichia coli Proteobacteria; Gammaproteobacteria; Enterobacterales; Enterobacteriaceae; Escherichia MTGDNTLIHSHGINRRDFMKLCAALAATMGLSSKAAAEMAESVTNPQRPPVIWIGAQECT WP_000145410.1 MULTISPECIES: hydrogenase-2 small chain [Proteobacteria] Length: 372\nNumber of predicted TMHs: 1\nExp number of AAs in TMHs: 23.72226\nExp number, first 60 AAs: 0.22038\nTotal prob of N-in: 0.12994\noutside 1 331\nTMhelix 332 354\ninside 355 372

4290 GCF_002133665.1_ASM213366v1 Escherichia coli Proteobacteria; Gammaproteobacteria; Enterobacterales; Enterobacteriaceae; Escherichia
MTGDNTLIHSHGINRRDFMKLCAALAATMGLSSKAAAEMAESVTNPQRPPVIWIGAQECT WP_000145410.1
MULTISPECIES: hydrogenase-2 small chain [Proteobacteria] Length: 372\nNumber of predicted TMHs: 1\nExp number of AAs in TMHs: 23.72226\nExp number, first 60 AAs: 0.22038\nTotal prob of N-in: 0.12994\noutside 1 331\nTMhelix 332 354\ninside 355 372

4291 GCF_002133675.1_ASM213367v1 Escherichia coli Proteobacteria; Gammaproteobacteria; Enterobacterales; Enterobacteriaceae; Escherichia
MTGDNTLIHSHGINRRDFMKLCAALAATMGLSSKAAAEMAESVTNPQRPPVIWIGAQECT WP_000145410.1
MULTISPECIES: hydrogenase-2 small chain [Proteobacteria] Length: 372\nNumber of predicted TMHs: 1\nExp number of AAs in TMHs: 23.72226\nExp number, first 60 AAs: 0.22038\nTotal prob of N-in: 0.12994\noutside 1 331\nTMhelix 332 354\ninside 355 372

4292 GCF_002133695.1_ASM213369v1 Escherichia coli Proteobacteria; Gammaproteobacteria; Enterobacterales; Enterobacteriaceae; Escherichia
MTGDNTLIHSHGINRRDFMKLCAALAATMGLSSKAAAEMAESVTNPQRPPVIWIGAQECT WP_000145410.1
MULTISPECIES: hydrogenase-2 small chain [Proteobacteria] Length: 372\nNumber of predicted TMHs: 1\nExp number of AAs in TMHs: 23.72226\nExp number, first 60 AAs: 0.22038\nTotal prob of N-in: 0.12994\noutside 1 331\nTMhelix 332 354\ninside 355 372

4293 GCF_002133685.1_ASM213368v1 Escherichia coli Proteobacteria; Gammaproteobacteria; Enterobacterales; Enterobacteriaceae; Escherichia
MTGDNTLIHSHGINRRDFMKLCAALAATMGLSSKAAAEMAESVTNPQRPPVIWIGAQECT WP_000145410.1
MULTISPECIES: hydrogenase-2 small chain [Proteobacteria] Length: 372\nNumber of predicted TMHs: 1\nExp number of AAs in TMHs: 23.72226\nExp number, first 60 AAs: 0.22038\nTotal prob of N-in: 0.12994\noutside 1 331\nTMhelix 332 354\ninside 355 372

4294 GCF_002133745.1_ASM213374v1 Escherichia coli Proteobacteria; Gammaproteobacteria; Enterobacterales; Enterobacteriaceae; Escherichia
MTGDNTLIHSHGINRRDFMKLCAALAATMGLSSKAAAEMAESVTNPQRPPVIWIGAQECT WP_000145410.1
MULTISPECIES: hydrogenase-2 small chain [Proteobacteria] Length: 372\nNumber of predicted TMHs: 1\nExp number of AAs in TMHs: 23.72226\nExp number, first 60 AAs: 0.22038\nTotal prob of N-in: 0.12994\noutside 1 331\nTMhelix 332 354\ninside 355 372

4295 GCF_002133605.1_ASM213360v1 Escherichia coli Proteobacteria; Gammaproteobacteria; Enterobacterales; Enterobacteriaceae; Escherichia
MNNEETFYQAMRRQGVTRRSFLKYCSLAATSLGLGAGMAPKIAWALENKPRIPVVWIHGL WP_001058323.1
MULTISPECIES: hydrogenase-1 small chain [Proteobacteria] Length: 372\nNumber of predicted TMHs: 2\nExp number of AAs in TMHs: 37.2608499999999\nExp number, first 60 AAs: 19.52923\nTotal prob of N-in: 0.94351\nPOSSIBLE N-term signal sequence\ninside 1 19\nTMhelix 20 39\noutside 40 325\nTMhelix 326 348\ninside 349 372

4296 GCF_002133615.1_ASM213361v1 Escherichia coli Proteobacteria; Gammaproteobacteria; Enterobacterales; Enterobacteriaceae; Escherichia
MNNEETFYQAMRRQGVTRRSFLKYCSLAATSLGLGAGMAPKIAWALENKPRIPVVWIHGL WP_001058323.1
MULTISPECIES: hydrogenase-1 small chain [Proteobacteria] Length: 372\nNumber of predicted TMHs: 2\nExp number of AAs in TMHs: 37.2608499999999\nExp number, first 60 AAs: 19.52923\nTotal prob of N-in: 0.94351\nPOSSIBLE N-term signal sequence\ninside 1 19\nTMhelix 20 39\noutside 40 325\nTMhelix 326 348\ninside 349 372

4297 GCF_002133825.1_ASM213382v1 Escherichia coli Proteobacteria; Gammaproteobacteria; Enterobacterales; Enterobacteriaceae; Escherichia
MTGDNTLIHSHGINRRDFMKLCAALAATMGLSSKAAAEMAESVTNPQRPPVIWIGAQECT WP_000145410.1
MULTISPECIES: hydrogenase-2 small chain [Proteobacteria] Length: 372\nNumber of predicted TMHs: 1\nExp number of AAs in TMHs: 23.72226\nExp number, first 60 AAs: 0.22038\nTotal prob of N-in: 0.12994\noutside 1 331\nTMhelix 332 354\ninside 355 372

4298 GCF_002133825.1_ASM213382v1 Escherichia coli Proteobacteria; Gammaproteobacteria; Enterobacterales; Enterobacteriaceae; Escherichia
MNNEETFYQAMRRQGVTRRSFLKYCSLAATSLGLGVGMAPKIAWALENKPRIPVVWIHGL WP_073714848.1

hydrogenase [Shigella boydii] Length: 372\nNumber of predicted TMHs: 2\nExp number of AAs in TMHs: 37.83336\nExp number, first 60 AAs: 20.10829\nTotal prob of N-in: 0.96228\nPOSSIBLE N-term signal sequence\ninside 1 20\nTMhelix 21 43\noutside 44 325\nTMhelix 326 348\ninside 349 372

4299 GCF_002133675.1_ASM213367v1 Escherichia coli Proteobacteria; Gammaproteobacteria; Enterobacterales; Enterobacteriaceae; Escherichia MNNEETFYQAMRRQGVTRRSFLKYCSLAATSLGLGAGMAPKIAWALENKPRIPVVWIHGL WP_001058323.1 MULTISPECIES: hydrogenase-1 small chain [Proteobacteria] Length: 372\nNumber of predicted TMHs: 2\nExp number of AAs in TMHs: 37.2608499999999\nExp number, first 60 AAs: 19.52923\nTotal prob of N-in: 0.94351\nPOSSIBLE N-term signal sequence\ninside 1 19\nTMhelix 20 39\noutside 40 325\nTMhelix 326 348\ninside 349 372

4300 GCF_002133695.1_ASM213369v1 Escherichia coli Proteobacteria; Gammaproteobacteria; Enterobacterales; Enterobacteriaceae; Escherichia MNNEETFYQAMRRQGVTRRSFLKYCSLAATSLGLGAGMAPKIAWALENKPRIPVVWIHGL WP_001058323.1 MULTISPECIES: hydrogenase-1 small chain [Proteobacteria] Length: 372\nNumber of predicted TMHs: 2\nExp number of AAs in TMHs: 37.2608499999999\nExp number, first 60 AAs: 19.52923\nTotal prob of N-in: 0.94351\nPOSSIBLE N-term signal sequence\ninside 1 19\nTMhelix 20 39\noutside 40 325\nTMhelix 326 348\ninside 349 372

4301 GCF_002133745.1_ASM213374v1 Escherichia coli Proteobacteria; Gammaproteobacteria; Enterobacterales; Enterobacteriaceae; Escherichia MNNEETFYQAMRRQGVTRRSFLKYCSLAATSLGLGAGMAPKIAWALENKPRIPVVWIHGL WP_001058323.1 MULTISPECIES: hydrogenase-1 small chain [Proteobacteria] Length: 372\nNumber of predicted TMHs: 2\nExp number of AAs in TMHs: 37.2608499999999\nExp number, first 60 AAs: 19.52923\nTotal prob of N-in: 0.94351\nPOSSIBLE N-term signal sequence\ninside 1 19\nTMhelix 20 39\noutside 40 325\nTMhelix 326 348\ninside 349 372

4302 GCF_002133835.1_ASM213383v1 Escherichia coli Proteobacteria; Gammaproteobacteria; Enterobacterales; Enterobacteriaceae; Escherichia MTGDNTLIHSHGINRRDFMKLCAALAATMGLSSKAAAEMAESVTNPQRPPVIWIGAQECT WP_000145410.1 MULTISPECIES: hydrogenase-2 small chain [Proteobacteria] Length: 372\nNumber of predicted TMHs: 1\nExp number of AAs in TMHs: 23.72226\nExp number, first 60 AAs: 0.22038\nTotal prob of N-in: 0.12994\noutside 1 331\nTMhelix 332 354\ninside 355 372

4303 GCF_002133835.1_ASM213383v1 Escherichia coli Proteobacteria; Gammaproteobacteria; Enterobacterales; Enterobacteriaceae; Escherichia MNNEETFYQAMRRQGVTRRSFLKYCSLAATSLGLGAGMAPKIAWALENKPRIPVVWIHGL WP_001058323.1 MULTISPECIES: hydrogenase-1 small chain [Proteobacteria] Length: 372\nNumber of predicted TMHs: 2\nExp number of AAs in TMHs: 37.2608499999999\nExp number, first 60 AAs: 19.52923\nTotal prob of N-in: 0.94351\nPOSSIBLE N-term signal sequence\ninside 1 19\nTMhelix 20 39\noutside 40 325\nTMhelix 326 348\ninside 349 372

4304 GCF_002134005.1_ASM213400v1 Escherichia coli Proteobacteria; Gammaproteobacteria; Enterobacterales; Enterobacteriaceae; Escherichia MTGDNTLIHSHGINRRDFMKLCAALAATMGLSSKAAAEMAESVTNPQRPPVIWIGAQECT WP_000145410.1 MULTISPECIES: hydrogenase-2 small chain [Proteobacteria] Length: 372\nNumber of predicted TMHs: 1\nExp number of AAs in TMHs: 23.72226\nExp number, first 60 AAs: 0.22038\nTotal prob of N-in: 0.12994\noutside 1 331\nTMhelix 332 354\ninside 355 372

4305 GCF_002133915.1_ASM213391v1 Escherichia coli Proteobacteria; Gammaproteobacteria; Enterobacterales; Enterobacteriaceae; Escherichia MTGDNTLIHSHGINRRDFMKLCAALAATMGLSSKAAAEMAESVTNPQRPPVIWIGAQECT WP_000145410.1 MULTISPECIES: hydrogenase-2 small chain [Proteobacteria] Length: 372\nNumber of predicted TMHs: 1\nExp number of AAs in TMHs: 23.72226\nExp number, first 60 AAs: 0.22038\nTotal prob of N-in: 0.12994\noutside 1 331\nTMhelix 332 354\ninside 355 372

4306 GCF_002133925.1_ASM213392v1 Escherichia coli Proteobacteria; Gammaproteobacteria; Enterobacterales; Enterobacteriaceae; Escherichia MTGDNTLIHSHGINRRDFMKLCAALAATMGLSSKAAAEMAESVTNPQRPPVIWIGAQECT WP_000145410.1 MULTISPECIES: hydrogenase-2 small chain [Proteobacteria] Length: 372\nNumber of predicted TMHs:

1\nExp number of AAs in TMHs: 23.72226\nExp number, first 60 AAs: 0.22038\nTotal prob of N-in: 0.12994\noutside 1 331\nTMhelix 332 354\ninside 355 372

4307 GCF_002133985.1_ASM213398v1 Escherichia coli Proteobacteria; Gammaproteobacteria; Enterobacterales; Enterobacteriaceae; Escherichia MTGDNTLIHSHGINRRDFMKLCAALAATMGLSSKAAAEMAESVTNPQRPPVIWIGAQECT WP_000145410.1 MULTISPECIES: hydrogenase-2 small chain [Proteobacteria] Length: 372\nNumber of predicted TMHs: 1\nExp number of AAs in TMHs: 23.72226\nExp number, first 60 AAs: 0.22038\nTotal prob of N-in: 0.12994\noutside 1 331\nTMhelix 332 354\ninside 355 372

4308 GCF_002134005.1_ASM213400v1 Escherichia coli Proteobacteria; Gammaproteobacteria; Enterobacterales; Enterobacteriaceae; Escherichia MNNEETFYQAMRRQGVTRRSFLKYCSLAATSLGLGAGMAPKIAWALENKPRIPVVWIHGL WP_001058323.1 MULTISPECIES: hydrogenase-1 small chain [Proteobacteria] Length: 372\nNumber of predicted TMHs: 2\nExp number of AAs in TMHs: 37.2608499999999\nExp number, first 60 AAs: 19.52923\nTotal prob of N-in: 0.94351\nPOSSIBLE N-term signal sequence\ninside 1 19\nTMhelix 20 39\noutside 40 325\nTMhelix 326 348\ninside 349 372

4309 GCF_002133915.1_ASM213391v1 Escherichia coli Proteobacteria; Gammaproteobacteria; Enterobacterales; Enterobacteriaceae; Escherichia MNNEETFYQAMRRQGVTRRSFLKYCSLAATSLGLGAGMAPKIAWALENKPRIPVVWIHGL WP_001058323.1 MULTISPECIES: hydrogenase-1 small chain [Proteobacteria] Length: 372\nNumber of predicted TMHs: 2\nExp number of AAs in TMHs: 37.2608499999999\nExp number, first 60 AAs: 19.52923\nTotal prob of N-in: 0.94351\nPOSSIBLE N-term signal sequence\ninside 1 19\nTMhelix 20 39\noutside 40 325\nTMhelix 326 348\ninside 349 372

4310 GCF_002134075.1_ASM213407v1 Escherichia coli Proteobacteria; Gammaproteobacteria; Enterobacterales; Enterobacteriaceae; Escherichia MTGDNTLIHSHGINRRDFMKLCAALAATMGLSSKAAAEMAESVTNPQRPPVIWIGAQECT WP_000145410.1 MULTISPECIES: hydrogenase-2 small chain [Proteobacteria] Length: 372\nNumber of predicted TMHs: 1\nExp number of AAs in TMHs: 23.72226\nExp number, first 60 AAs: 0.22038\nTotal prob of N-in: 0.12994\noutside 1 331\nTMhelix 332 354\ninside 355 372

4311 GCF_002134075.1_ASM213407v1 Escherichia coli Proteobacteria; Gammaproteobacteria; Enterobacterales; Enterobacteriaceae; Escherichia MNNEETFYQAMRRQGVTRRSFLKYCSLAATSLGLGAGMAPKIAWALENKPRIPVVWIHGL WP_001058323.1 MULTISPECIES: hydrogenase-1 small chain [Proteobacteria] Length: 372\nNumber of predicted TMHs: 2\nExp number of AAs in TMHs: 37.2608499999999\nExp number, first 60 AAs: 19.52923\nTotal prob of N-in: 0.94351\nPOSSIBLE N-term signal sequence\ninside 1 19\nTMhelix 20 39\noutside 40 325\nTMhelix 326 348\ninside 349 372

4312 GCF_002133985.1_ASM213398v1 Escherichia coli Proteobacteria; Gammaproteobacteria; Enterobacterales; Enterobacteriaceae; Escherichia MNNEETFYQAMRRQGVTRRSFLKYCSLAATSLGLGAGMAPKIAWALENKPRIPVVWIHGL WP_001295940.1 MULTISPECIES: hydrogenase [Enterobacteriaceae] Length: 372\nNumber of predicted TMHs: 2\nExp number of AAs in TMHs: 37.2608899999999\nExp number, first 60 AAs: 19.52923\nTotal prob of N-in: 0.94351\nPOSSIBLE N-term signal sequence\ninside 1 19\nTMhelix 20 39\noutside 40 325\nTMhelix 326 348\ninside 349 372

4313 GCF_002134085.1_ASM213408v1 Escherichia coli Proteobacteria; Gammaproteobacteria; Enterobacterales; Enterobacteriaceae; Escherichia MTGDNTLIHSHGINRRDFMKLCAALAATMGLSSKAAAEMAESVTNPQRPPVIWIGAQECT WP_000145410.1 MULTISPECIES: hydrogenase-2 small chain [Proteobacteria] Length: 372\nNumber of predicted TMHs: 1\nExp number of AAs in TMHs: 23.72226\nExp number, first 60 AAs: 0.22038\nTotal prob of N-in: 0.12994\noutside 1 331\nTMhelix 332 354\ninside 355 372

4314 GCF_002134105.1_ASM213410v1 Escherichia coli Proteobacteria; Gammaproteobacteria; Enterobacterales; Enterobacteriaceae; Escherichia MTGDNTLIHSHGINRRDFMKLCAALAATMGLSSKAAAEMAESVTNPQRPPVIWIGAQECT WP_000145410.1 MULTISPECIES: hydrogenase-2 small chain [Proteobacteria] Length: 372\nNumber of predicted TMHs:

1\nExp number of AAs in TMHs: 23.72226\nExp number, first 60 AAs: 0.22038\nTotal prob of N-in: 0.12994\noutside 1 331\nTMhelix 332 354\ninside 355 372

4315 GCF_002134085.1_ASM213408v1 Escherichia coli Proteobacteria; Gammaproteobacteria; Enterobacterales; Enterobacteriaceae; Escherichia
MNNEETFYQAMRRQGVTRRSFLKYCSLAATSLGLGAGMAPKIAWALENKPRIPVVWIHGL WP_001058323.1
MULTISPECIES: hydrogenase-1 small chain [Proteobacteria] Length: 372\nNumber of predicted TMHs: 2\nExp number of AAs in TMHs: 37.2608499999999\nExp number, first 60 AAs: 19.52923\nTotal prob of N-in: 0.94351\nPOSSIBLE N-term signal sequence\ninside 1 19\nTMhelix 20 39\noutside 40 325\nTMhelix 326 348\ninside 349 372

4316 GCF_002134105.1_ASM213410v1 Escherichia coli Proteobacteria; Gammaproteobacteria; Enterobacterales; Enterobacteriaceae; Escherichia
MNNEETFYQAMRRQGVTRRSFLKYCSLAATSLGLGAGMAPKIAWALENKPRIPVVWIHGL WP_001301613.1
hydrogenase [Escherichia coli] Length: 372\nNumber of predicted TMHs: 2\nExp number of AAs in TMHs: 37.46598\nExp number, first 60 AAs: 19.53139\nTotal prob of N-in: 0.94364\nPOSSIBLE N-term signal sequence\ninside 1 19\nTMhelix 20 39\noutside 40 325\nTMhelix 326 348\ninside 349 372

4317 GCF_002134155.1_ASM213415v1 Escherichia coli Proteobacteria; Gammaproteobacteria; Enterobacterales; Enterobacteriaceae; Escherichia
MNNEETFYQAMRRQGVTRRSFLKYCSLAATSLGLGAGMAPKIAWALENKPRIPVVWIHGL WP_001295940.1
MULTISPECIES: hydrogenase [Enterobacteriaceae] Length: 372\nNumber of predicted TMHs: 2\nExp number of AAs in TMHs: 37.2608899999999\nExp number, first 60 AAs: 19.52923\nTotal prob of N-in: 0.94351\nPOSSIBLE N-term signal sequence\ninside 1 19\nTMhelix 20 39\noutside 40 325\nTMhelix 326 348\ninside 349 372

4318 GCF_002134265.1_ASM213426v1 Escherichia coli Proteobacteria; Gammaproteobacteria; Enterobacterales; Enterobacteriaceae; Escherichia
MTGDNTLIHSHGINRRDFMKLCAALAATMGLSSKAAAEMAESVTNPQRPPVIWIGAQECT WP_000145410.1
MULTISPECIES: hydrogenase-2 small chain [Proteobacteria] Length: 372\nNumber of predicted TMHs: 1\nExp number of AAs in TMHs: 23.72226\nExp number, first 60 AAs: 0.22038\nTotal prob of N-in: 0.12994\noutside 1 331\nTMhelix 332 354\ninside 355 372

4319 GCF_002134265.1_ASM213426v1 Escherichia coli Proteobacteria; Gammaproteobacteria; Enterobacterales; Enterobacteriaceae; Escherichia
MNNEETFYQAMRRQGVTRRSFLKYCSLAATSLGLGAGMAPKIAWALENKPRIPVVWIHGL WP_014639883.1
hydrogenase [Escherichia coli] Length: 372\nNumber of predicted TMHs: 2\nExp number of AAs in TMHs: 37.26086\nExp number, first 60 AAs: 19.52923\nTotal prob of N-in: 0.94351\nPOSSIBLE N-term signal sequence\ninside 1 19\nTMhelix 20 39\noutside 40 325\nTMhelix 326 348\ninside 349 372

4320 GCF_002134145.1_ASM213414v1 Escherichia coli Proteobacteria; Gammaproteobacteria; Enterobacterales; Enterobacteriaceae; Escherichia
MNNEETFYQAMRRQGVTRRSFLKYCSLAATSLGLGAGMAPKIAWALENKPRIPVVWIHGL WP_001058323.1
MULTISPECIES: hydrogenase-1 small chain [Proteobacteria] Length: 372\nNumber of predicted TMHs: 2\nExp number of AAs in TMHs: 37.2608499999999\nExp number, first 60 AAs: 19.52923\nTotal prob of N-in: 0.94351\nPOSSIBLE N-term signal sequence\ninside 1 19\nTMhelix 20 39\noutside 40 325\nTMhelix 326 348\ninside 349 372

4321 GCF_002134285.1_ASM213428v1 Escherichia coli Proteobacteria; Gammaproteobacteria; Enterobacterales; Enterobacteriaceae; Escherichia
MTGDNTLIHSHGINRRDFMKLCAALAATMGLSSKAAAEMAESVTNPQRPPVIWIGAQECT WP_000145410.1
MULTISPECIES: hydrogenase-2 small chain [Proteobacteria] Length: 372\nNumber of predicted TMHs: 1\nExp number of AAs in TMHs: 23.72226\nExp number, first 60 AAs: 0.22038\nTotal prob of N-in: 0.12994\noutside 1 331\nTMhelix 332 354\ninside 355 372

4322 GCF_002134275.1_ASM213427v1 Escherichia coli Proteobacteria; Gammaproteobacteria; Enterobacterales; Enterobacteriaceae; Escherichia
MTGDNTLIHSHGINRRDFMKLCAALAATMGLSSKAAAEMAESVTNPQRPPVIWIGAQECT WP_000145410.1
MULTISPECIES: hydrogenase-2 small chain [Proteobacteria] Length: 372\nNumber of predicted TMHs: 1\nExp number of AAs in TMHs: 23.72226\nExp number, first 60 AAs: 0.22038\nTotal prob of N-in: 0.12994\noutside 1 331\nTMhelix 332 354\ninside 355 372

4323	GCF_002134275.1_ASM213427v1	Escherichia coli	Proteobacteria; Gammaproteobacteria; Enterobacterales; Enterobacteriaceae; Escherichia
	MNNEETFYQAMRRQGVTRRSFLKYCSLAATSLGLGAGMAPKIAWALENKPRIPVVWIHGL	WP_001295940.1
	MULTISPECIES: hydrogenase [Enterobacteriaceae]	Length: 372\nNumber of predicted TMHs: 2\nExp number of AAs in TMHs: 37.2608899999999\nExp number, first 60 AAs: 19.52923\nTotal prob of N-in: 0.94351\nPOSSIBLE N-term signal sequence\ninside 1 19\nTMhelix 20 39\noutside 40 325\nTMhelix 326 348\ninside 349 372

4324	GCF_002134215.1_ASM213421v1	Escherichia coli	Proteobacteria; Gammaproteobacteria; Enterobacterales; Enterobacteriaceae; Escherichia
	MNNEETFYQAMRRQGVTRRSFLKYCSLAATSLGLGAGMAPKIAWALENKPRIPVVWIHGL	WP_001353311.1
	MULTISPECIES: hydrogenase [Escherichia]	Length: 372\nNumber of predicted TMHs: 2\nExp number of AAs in TMHs: 37.2638699999999\nExp number, first 60 AAs: 19.52924\nTotal prob of N-in: 0.94351\nPOSSIBLE N-term signal sequence\ninside 1 19\nTMhelix 20 39\noutside 40 325\nTMhelix 326 348\ninside 349 372

4325	GCF_002134385.1_ASM213438v1	Escherichia coli	Proteobacteria; Gammaproteobacteria; Enterobacterales; Enterobacteriaceae; Escherichia
	MTGDNTLIHSHGINRRDFMKLCAALAATMGLSSKAAAEMAESVTNPQRPPVIWIGAQECT	WP_000145410.1
	MULTISPECIES: hydrogenase-2 small chain [Proteobacteria]	Length: 372\nNumber of predicted TMHs: 1\nExp number of AAs in TMHs: 23.72226\nExp number, first 60 AAs: 0.22038\nTotal prob of N-in: 0.12994\noutside 1 331\nTMhelix 332 354\ninside 355 372

4326	GCF_002134395.1_ASM213439v1	Escherichia coli	Proteobacteria; Gammaproteobacteria; Enterobacterales; Enterobacteriaceae; Escherichia
	MTGDNTLIHSHGINRRDFMKLCAALAATMGLSSKAAAEMAESVTNPQRPPVIWIGAQECT	WP_000145410.1
	MULTISPECIES: hydrogenase-2 small chain [Proteobacteria]	Length: 372\nNumber of predicted TMHs: 1\nExp number of AAs in TMHs: 23.72226\nExp number, first 60 AAs: 0.22038\nTotal prob of N-in: 0.12994\noutside 1 331\nTMhelix 332 354\ninside 355 372

4327	GCF_002134345.1_ASM213434v1	Escherichia coli	Proteobacteria; Gammaproteobacteria; Enterobacterales; Enterobacteriaceae; Escherichia
	MTGDNTLIHSHGINRRDFMKLCAALAATMGLSSKAAAEMAESVTNPQRPPVIWIGAQECT	WP_000145410.1
	MULTISPECIES: hydrogenase-2 small chain [Proteobacteria]	Length: 372\nNumber of predicted TMHs: 1\nExp number of AAs in TMHs: 23.72226\nExp number, first 60 AAs: 0.22038\nTotal prob of N-in: 0.12994\noutside 1 331\nTMhelix 332 354\ninside 355 372

4328	GCF_002134395.1_ASM213439v1	Escherichia coli	Proteobacteria; Gammaproteobacteria; Enterobacterales; Enterobacteriaceae; Escherichia
	MNNEETFYQAMRRQGVTRRSFLKYCSLAATSLGLGAGMAPKIAWALENKPRIPVVWIHGL	WP_001058323.1
	MULTISPECIES: hydrogenase-1 small chain [Proteobacteria]	Length: 372\nNumber of predicted TMHs: 2\nExp number of AAs in TMHs: 37.2608499999999\nExp number, first 60 AAs: 19.52923\nTotal prob of N-in: 0.94351\nPOSSIBLE N-term signal sequence\ninside 1 19\nTMhelix 20 39\noutside 40 325\nTMhelix 326 348\ninside 349 372

4329	GCF_002134455.1_ASM213445v1	Escherichia coli	Proteobacteria; Gammaproteobacteria; Enterobacterales; Enterobacteriaceae; Escherichia
	MTGDNTLIHSHGINRRDFMKLCAALAATMGLSSKAAAEMAESVTNPQRPPVIWIGAQECT	WP_000145410.1
	MULTISPECIES: hydrogenase-2 small chain [Proteobacteria]	Length: 372\nNumber of predicted TMHs: 1\nExp number of AAs in TMHs: 23.72226\nExp number, first 60 AAs: 0.22038\nTotal prob of N-in: 0.12994\noutside 1 331\nTMhelix 332 354\ninside 355 372

4330	GCF_002134405.1_ASM213440v1	Escherichia coli	Proteobacteria; Gammaproteobacteria; Enterobacterales; Enterobacteriaceae; Escherichia
	MTGDNTLIHSHGINRRDFMKLCAALAATMGLSSKAAAEMAESVTNPQRPPVIWIGAQECT	WP_000145410.1
	MULTISPECIES: hydrogenase-2 small chain [Proteobacteria]	Length: 372\nNumber of predicted TMHs: 1\nExp number of AAs in TMHs: 23.72226\nExp number, first 60 AAs: 0.22038\nTotal prob of N-in: 0.12994\noutside 1 331\nTMhelix 332 354\ninside 355 372

4331	GCF_002134505.1_ASM213450v1	Escherichia coli	Proteobacteria; Gammaproteobacteria; Enterobacterales; Enterobacteriaceae; Escherichia
	MNNEETFYQAMRRQGVTRRSFLKYCSLAATSLGLGAGMAPKIAWALENKPRIPVVWIHGL	WP_001301613.1

hydrogenase [Escherichia coli]	Length: 372\nNumber of predicted TMHs:  2\nExp number of AAs in TMHs: 37.46598\nExp number, first 60 AAs:  19.53139\nTotal prob of N-in:   0.94364\nPOSSIBLE N-term signal sequence\ninside   1   19\nTMhelix   20   39\noutside   40   325\nTMhelix   326   348\ninside   349   372

4332	GCF_002134525.1_ASM213452v1	Escherichia coli	Proteobacteria; Gammaproteobacteria; Enterobacterales; Enterobacteriaceae; Escherichia
MNNEETFYQAMRRQGVTRRSFLKYCSLAATSLGLGAGMAPKIAWALENKPRIPVVWIHGL	WP_001359719.1
hydrogenase [Escherichia coli]	Length: 372\nNumber of predicted TMHs:  2\nExp number of AAs in TMHs: 37.2176\nExp number, first 60 AAs:  19.53448\nTotal prob of N-in:   0.94360\nPOSSIBLE N-term signal sequence\ninside   1   19\nTMhelix   20   39\noutside   40   325\nTMhelix   326   348\ninside   349   372

4333	GCF_002134455.1_ASM213445v1	Escherichia coli	Proteobacteria; Gammaproteobacteria; Enterobacterales; Enterobacteriaceae; Escherichia
MNNEETFYQAMRRQGVTRRSFLKYCSLAATSLGLGAGMAPKIAWALENKPRIPVVWIHGL	WP_001058323.1
MULTISPECIES: hydrogenase-1 small chain [Proteobacteria]	Length: 372\nNumber of predicted TMHs: 2\nExp number of AAs in TMHs: 37.2608499999999\nExp number, first 60 AAs:  19.52923\nTotal prob of N-in: 0.94351\nPOSSIBLE N-term signal sequence\ninside   1   19\nTMhelix   20   39\noutside   40   325\nTMhelix   326 348\ninside   349   372

4334	GCF_002134385.1_ASM213438v1	Escherichia coli	Proteobacteria; Gammaproteobacteria; Enterobacterales; Enterobacteriaceae; Escherichia
MNNEETFYQAMRRQGVTRRSFLKYCSLAATSLGLGAGMAPKIAWALENKPRIPVVWIHGL	WP_001301613.1
hydrogenase [Escherichia coli]	Length: 372\nNumber of predicted TMHs:  2\nExp number of AAs in TMHs: 37.46598\nExp number, first 60 AAs:  19.53139\nTotal prob of N-in:   0.94364\nPOSSIBLE N-term signal sequence\ninside   1   19\nTMhelix   20   39\noutside   40   325\nTMhelix   326   348\ninside   349   372

4335	GCF_002134565.1_ASM213456v1	Escherichia coli	Proteobacteria; Gammaproteobacteria; Enterobacterales; Enterobacteriaceae; Escherichia
MTGDNTLIHSHGINRRDFMKLCAALAATMGLSSKAAAEMAESVTNPQRPPVIWIGAQECT	WP_000145410.1
MULTISPECIES: hydrogenase-2 small chain [Proteobacteria]	Length: 372\nNumber of predicted TMHs: 1\nExp number of AAs in TMHs: 23.72226\nExp number, first 60 AAs:  0.22038\nTotal prob of N-in:   0.12994\noutside   1 331\nTMhelix   332   354\ninside   355   372

4336	GCF_002134565.1_ASM213456v1	Escherichia coli	Proteobacteria; Gammaproteobacteria; Enterobacterales; Enterobacteriaceae; Escherichia
MNNEETFYQAMRRQGVTRRSFLKYCSLAATSLGLGAGMAPKIAWALENKPRIPVVWIHGL	WP_001058323.1
MULTISPECIES: hydrogenase-1 small chain [Proteobacteria]	Length: 372\nNumber of predicted TMHs: 2\nExp number of AAs in TMHs: 37.2608499999999\nExp number, first 60 AAs:  19.52923\nTotal prob of N-in: 0.94351\nPOSSIBLE N-term signal sequence\ninside   1   19\nTMhelix   20   39\noutside   40   325\nTMhelix   326 348\ninside   349   372

4337	GCF_002134405.1_ASM213440v1	Escherichia coli	Proteobacteria; Gammaproteobacteria; Enterobacterales; Enterobacteriaceae; Escherichia
MNNEETFYQAMRRQGVTRRSFLKYCSLAATSLGLGAGMAPKIAWALENKPRIPVVWIHGL	WP_001295940.1
MULTISPECIES: hydrogenase [Enterobacteriaceae]	Length: 372\nNumber of predicted TMHs:  2\nExp number of AAs in TMHs: 37.2608899999999\nExp number, first 60 AAs:  19.52923\nTotal prob of N-in: 0.94351\nPOSSIBLE N-term signal sequence\ninside   1   19\nTMhelix   20   39\noutside   40   325\nTMhelix   326 348\ninside   349   372

4338	GCF_002134725.1_ASM213472v1	Escherichia coli	Proteobacteria; Gammaproteobacteria; Enterobacterales; Enterobacteriaceae; Escherichia
MTGDNTLIHSHGINRRDFMKLCAALAATMGLSSKAAAEMAESVTNPQRPPVIWIGAQECT	WP_000145410.1
MULTISPECIES: hydrogenase-2 small chain [Proteobacteria]	Length: 372\nNumber of predicted TMHs: 1\nExp number of AAs in TMHs: 23.72226\nExp number, first 60 AAs:  0.22038\nTotal prob of N-in:   0.12994\noutside   1 331\nTMhelix   332   354\ninside   355   372

4339	GCF_002134695.1_ASM213469v1	Escherichia coli	Proteobacteria; Gammaproteobacteria; Enterobacterales; Enterobacteriaceae; Escherichia
MTGDNTLIHSHGINRRDFMKLCAALAATMGLSSKAAAEMAESVTNPQRPPVIWIGAQECT	WP_000145410.1
MULTISPECIES: hydrogenase-2 small chain [Proteobacteria]	Length: 372\nNumber of predicted TMHs:

1\nExp number of AAs in TMHs: 23.72226\nExp number, first 60 AAs: 0.22038\nTotal prob of N-in: 0.12994\noutside 1 331\nTMhelix 332 354\ninside 355 372

4340 GCF_002142695.1_ASM214269v1 Escherichia coli Proteobacteria; Gammaproteobacteria; Enterobacterales; Enterobacteriaceae; Escherichia
MTGDNTLIHSHGINRRDFMKLCAALATTMGLSSKAAAEMAESVTNPQRPPVIWIGAQECT WP_023149629.1
MULTISPECIES: hydrogenase-2 small chain [Enterobacteriaceae] Length: 372\nNumber of predicted TMHs: 1\nExp number of AAs in TMHs: 23.63603\nExp number, first 60 AAs: 0.12971\nTotal prob of N-in: 0.12608\noutside 1 331\nTMhelix 332 354\ninside 355 372

4341 GCF_002134625.1_ASM213462v1 Escherichia coli Proteobacteria; Gammaproteobacteria; Enterobacterales; Enterobacteriaceae; Escherichia
MNNEETFYQAMRRQGVTRRSFLKYCSLAATSLGLGAGMAPKIAWALENKPRIPVVWIHGL WP_001058323.1
MULTISPECIES: hydrogenase-1 small chain [Proteobacteria] Length: 372\nNumber of predicted TMHs: 2\nExp number of AAs in TMHs: 37.2608499999999\nExp number, first 60 AAs: 19.52923\nTotal prob of N-in: 0.94351\nPOSSIBLE N-term signal sequence\ninside 1 19\nTMhelix 20 39\noutside 40 325\nTMhelix 326 348\ninside 349 372

4342 GCF_002134685.1_ASM213468v1 Escherichia coli Proteobacteria; Gammaproteobacteria; Enterobacterales; Enterobacteriaceae; Escherichia
MTGDNTLIHSHGINRRDFMKLCAALAATMGLSSKAAAEMAESVTNPQRPPVIWIGAQECT WP_000145410.1
MULTISPECIES: hydrogenase-2 small chain [Proteobacteria] Length: 372\nNumber of predicted TMHs: 1\nExp number of AAs in TMHs: 23.72226\nExp number, first 60 AAs: 0.22038\nTotal prob of N-in: 0.12994\noutside 1 331\nTMhelix 332 354\ninside 355 372

4343 GCF_002134745.1_ASM213474v1 Escherichia coli Proteobacteria; Gammaproteobacteria; Enterobacterales; Enterobacteriaceae; Escherichia
MTGDNTLIHSHGINRRDFMKLCAALAATMGLSSKAAAEMAESVTNPQRPPVIWIGAQECT WP_000145410.1
MULTISPECIES: hydrogenase-2 small chain [Proteobacteria] Length: 372\nNumber of predicted TMHs: 1\nExp number of AAs in TMHs: 23.72226\nExp number, first 60 AAs: 0.22038\nTotal prob of N-in: 0.12994\noutside 1 331\nTMhelix 332 354\ninside 355 372

4344 GCF_002134745.1_ASM213474v1 Escherichia coli Proteobacteria; Gammaproteobacteria; Enterobacterales; Enterobacteriaceae; Escherichia
MNNEETFYQAMRRQGVTRRSFLKYCSLAATSLGLGAGMAPKIAWALENKPRIPVVWIHGL WP_001058323.1
MULTISPECIES: hydrogenase-1 small chain [Proteobacteria] Length: 372\nNumber of predicted TMHs: 2\nExp number of AAs in TMHs: 37.2608499999999\nExp number, first 60 AAs: 19.52923\nTotal prob of N-in: 0.94351\nPOSSIBLE N-term signal sequence\ninside 1 19\nTMhelix 20 39\noutside 40 325\nTMhelix 326 348\ninside 349 372

4345 GCF_002134635.1_ASM213463v1 Escherichia coli Proteobacteria; Gammaproteobacteria; Enterobacterales; Enterobacteriaceae; Escherichia
MNNEETFYQAMRRQGVTRRSFLKYCSLAATSLGLGAGMAPKIAWALENKPRIPVVWIHGL WP_001058323.1
MULTISPECIES: hydrogenase-1 small chain [Proteobacteria] Length: 372\nNumber of predicted TMHs: 2\nExp number of AAs in TMHs: 37.2608499999999\nExp number, first 60 AAs: 19.52923\nTotal prob of N-in: 0.94351\nPOSSIBLE N-term signal sequence\ninside 1 19\nTMhelix 20 39\noutside 40 325\nTMhelix 326 348\ninside 349 372

4346 GCF_002134585.1_ASM213458v1 Escherichia coli Proteobacteria; Gammaproteobacteria; Enterobacterales; Enterobacteriaceae; Escherichia
MNNEETFYQAMRRQGVTRRSFLKYCSLAATSLGLGAGMAPKIAWALENKPRIPVVWIHGL WP_001301613.1
hydrogenase [Escherichia coli] Length: 372\nNumber of predicted TMHs: 2\nExp number of AAs in TMHs: 37.46598\nExp number, first 60 AAs: 19.53139\nTotal prob of N-in: 0.94364\nPOSSIBLE N-term signal sequence\ninside 1 19\nTMhelix 20 39\noutside 40 325\nTMhelix 326 348\ninside 349 372

4347 GCF_002142355.1_ASM214235v1 Escherichia coli Proteobacteria; Gammaproteobacteria; Enterobacterales; Enterobacteriaceae; Escherichia
MTGDNTLIHSHGINRRDFMKLCAALAATMGLSSKAAAEMAESVTNPQRPPVIWIGAQECT WP_000145410.1
MULTISPECIES: hydrogenase-2 small chain [Proteobacteria] Length: 372\nNumber of predicted TMHs: 1\nExp number of AAs in TMHs: 23.72226\nExp number, first 60 AAs: 0.22038\nTotal prob of N-in: 0.12994\noutside 1 331\nTMhelix 332 354\ninside 355 372

4348 GCF_002142365.1_ASM214236v1 Escherichia coli Proteobacteria; Gammaproteobacteria; Enterobacterales; Enterobacteriaceae; Escherichia
MTGDNTLIHSHGINRRDFMKLCAALAATMGLSSKAAAEMAESVTNPQRPPVIWIGAQECT WP_000145410.1
MULTISPECIES: hydrogenase-2 small chain [Proteobacteria] Length: 372\nNumber of predicted TMHs: 1\nExp number of AAs in TMHs: 23.72226\nExp number, first 60 AAs: 0.22038\nTotal prob of N-in: 0.12994\noutside 1 331\nTMhelix 332 354\ninside 355 372

4349 GCF_002142415.1_ASM214241v1 Escherichia coli Proteobacteria; Gammaproteobacteria; Enterobacterales; Enterobacteriaceae; Escherichia
MTGDNTLIHSHGINRRDFMKLCAALAATMGLSSKAAAEMAESVTNPQRPPVIWIGAQECT WP_000145410.1
MULTISPECIES: hydrogenase-2 small chain [Proteobacteria] Length: 372\nNumber of predicted TMHs: 1\nExp number of AAs in TMHs: 23.72226\nExp number, first 60 AAs: 0.22038\nTotal prob of N-in: 0.12994\noutside 1 331\nTMhelix 332 354\ninside 355 372

4350 GCF_002142355.1_ASM214235v1 Escherichia coli Proteobacteria; Gammaproteobacteria; Enterobacterales; Enterobacteriaceae; Escherichia
MNNEETFYQAMRRQGVTRRSFLKYCSLAATSLGLGAGMAPKIAWALENKPRIPVVWIHGL WP_001058323.1
MULTISPECIES: hydrogenase-1 small chain [Proteobacteria] Length: 372\nNumber of predicted TMHs: 2\nExp number of AAs in TMHs: 37.2608499999999\nExp number, first 60 AAs: 19.52923\nTotal prob of N-in: 0.94351\nPOSSIBLE N-term signal sequence\ninside 1 19\nTMhelix 20 39\noutside 40 325\nTMhelix 326 348\ninside 349 372

4351 GCF_002142365.1_ASM214236v1 Escherichia coli Proteobacteria; Gammaproteobacteria; Enterobacterales; Enterobacteriaceae; Escherichia
MNNEETFYQAMRRQGVTRRSFLKYCSLAATSLGLGAGMAPKIAWALENKPRIPVVWIHGL WP_001058323.1
MULTISPECIES: hydrogenase-1 small chain [Proteobacteria] Length: 372\nNumber of predicted TMHs: 2\nExp number of AAs in TMHs: 37.2608499999999\nExp number, first 60 AAs: 19.52923\nTotal prob of N-in: 0.94351\nPOSSIBLE N-term signal sequence\ninside 1 19\nTMhelix 20 39\noutside 40 325\nTMhelix 326 348\ninside 349 372

4352 GCF_002144055.1_ASM214405v1 Escherichia coli Proteobacteria; Gammaproteobacteria; Enterobacterales; Enterobacteriaceae; Escherichia
MTGDNTLIHSHGINRRDFMKLCAALAATMGLSSKAAAEMAESVTNPQRPPVIWIGAQECT WP_000145410.1
MULTISPECIES: hydrogenase-2 small chain [Proteobacteria] Length: 372\nNumber of predicted TMHs: 1\nExp number of AAs in TMHs: 23.72226\nExp number, first 60 AAs: 0.22038\nTotal prob of N-in: 0.12994\noutside 1 331\nTMhelix 332 354\ninside 355 372

4353 GCF_002151245.1_ASM215124v1 Escherichia coli Proteobacteria; Gammaproteobacteria; Enterobacterales; Enterobacteriaceae; Escherichia
MTGDNTLIHSHGINRRDFMKLCAALATTMGLSSKAAAEMAESVTNPQRPPVIWIGAQECT WP_023149629.1
MULTISPECIES: hydrogenase-2 small chain [Enterobacteriaceae] Length: 372\nNumber of predicted TMHs: 1\nExp number of AAs in TMHs: 23.63603\nExp number, first 60 AAs: 0.12971\nTotal prob of N-in: 0.12608\noutside 1 331\nTMhelix 332 354\ninside 355 372

4354 GCF_002142435.1_ASM214243v1 Escherichia coli Proteobacteria; Gammaproteobacteria; Enterobacterales; Enterobacteriaceae; Escherichia
MNNEETFYQAMRRQGVTRRSFLKYCSLAATSLGLGAGMAPKIAWALENKPRIPVVWIHGL WP_001058323.1
MULTISPECIES: hydrogenase-1 small chain [Proteobacteria] Length: 372\nNumber of predicted TMHs: 2\nExp number of AAs in TMHs: 37.2608499999999\nExp number, first 60 AAs: 19.52923\nTotal prob of N-in: 0.94351\nPOSSIBLE N-term signal sequence\ninside 1 19\nTMhelix 20 39\noutside 40 325\nTMhelix 326 348\ninside 349 372

4355 GCF_002142675.1_ASM214267v1 Escherichia coli Proteobacteria; Gammaproteobacteria; Enterobacterales; Enterobacteriaceae; Escherichia
MNNEETFYQAMRRQGVTRRSFLKYCSLAATSLGLGAGMAPKIAWALENKPRIPVVWIHGL WP_001058323.1
MULTISPECIES: hydrogenase-1 small chain [Proteobacteria] Length: 372\nNumber of predicted TMHs: 2\nExp number of AAs in TMHs: 37.2608499999999\nExp number, first 60 AAs: 19.52923\nTotal prob of N-in: 0.94351\nPOSSIBLE N-term signal sequence\ninside 1 19\nTMhelix 20 39\noutside 40 325\nTMhelix 326 348\ninside 349 372

4356 GCF_002142445.1_ASM214244v1 Escherichia coli Proteobacteria; Gammaproteobacteria; Enterobacterales; Enterobacteriaceae; Escherichia
MNNEETFYQAMRRQGVTRRSFLKYCSLAATSLGLGAGMAPKIAWALENKPRIPVVWIHGL WP_001058323.1
MULTISPECIES: hydrogenase-1 small chain [Proteobacteria] Length: 372\nNumber of predicted TMHs: 2\nExp number of AAs in TMHs: 37.2608499999999\nExp number, first 60 AAs: 19.52923\nTotal prob of N-in: 0.94351\nPOSSIBLE N-term signal sequence\ninside 1 19\nTMhelix 20 39\noutside 40 325\nTMhelix 326 348\ninside 349 372

4357 GCF_002144095.1_ASM214409v1 Escherichia coli Proteobacteria; Gammaproteobacteria; Enterobacterales; Enterobacteriaceae; Escherichia
MTGDNTLIHSHGINRRDFMKLCAALAATMGLSSKAAAEMAESVTNPQRPPVIWIGAQECT WP_000145410.1
MULTISPECIES: hydrogenase-2 small chain [Proteobacteria] Length: 372\nNumber of predicted TMHs: 1\nExp number of AAs in TMHs: 23.72226\nExp number, first 60 AAs: 0.22038\nTotal prob of N-in: 0.12994\noutside 1 331\nTMhelix 332 354\ninside 355 372

4358 GCF_002144105.1_ASM214410v1 Escherichia coli Proteobacteria; Gammaproteobacteria; Enterobacterales; Enterobacteriaceae; Escherichia
MTGDNTLIHSHGINRRDFMKLCAALAATMGLSSKAAAEMAESVTNPQRPPVIWIGAQECT WP_000145410.1
MULTISPECIES: hydrogenase-2 small chain [Proteobacteria] Length: 372\nNumber of predicted TMHs: 1\nExp number of AAs in TMHs: 23.72226\nExp number, first 60 AAs: 0.22038\nTotal prob of N-in: 0.12994\noutside 1 331\nTMhelix 332 354\ninside 355 372

4359 GCF_002144055.1_ASM214405v1 Escherichia coli Proteobacteria; Gammaproteobacteria; Enterobacterales; Enterobacteriaceae; Escherichia
MNNEETFYQAMRRQGVTRRSFLKYCSLAATSLGLGAGMAPKIAWALENKPRIPVVWIHGL WP_001301613.1
hydrogenase [Escherichia coli] Length: 372\nNumber of predicted TMHs: 2\nExp number of AAs in TMHs: 37.46598\nExp number, first 60 AAs: 19.53139\nTotal prob of N-in: 0.94364\nPOSSIBLE N-term signal sequence\ninside 1 19\nTMhelix 20 39\noutside 40 325\nTMhelix 326 348\ninside 349 372

4360 GCF_002144185.1_ASM214418v1 Escherichia coli Proteobacteria; Gammaproteobacteria; Enterobacterales; Enterobacteriaceae; Escherichia
MTGDNTLIHSHGINRRDFMKLCAALAATMGLSSKAAAEMAESVTNPQRPPVIWIGAQECT WP_000145410.1
MULTISPECIES: hydrogenase-2 small chain [Proteobacteria] Length: 372\nNumber of predicted TMHs: 1\nExp number of AAs in TMHs: 23.72226\nExp number, first 60 AAs: 0.22038\nTotal prob of N-in: 0.12994\noutside 1 331\nTMhelix 332 354\ninside 355 372

4361 GCF_002144185.1_ASM214418v1 Escherichia coli Proteobacteria; Gammaproteobacteria; Enterobacterales; Enterobacteriaceae; Escherichia
MNNEETFYQAMRRQGVTRRSFLKYCSLAATSLGLGAGMAPKIAWALENKPRIPVVWIHGL WP_001058323.1
MULTISPECIES: hydrogenase-1 small chain [Proteobacteria] Length: 372\nNumber of predicted TMHs: 2\nExp number of AAs in TMHs: 37.2608499999999\nExp number, first 60 AAs: 19.52923\nTotal prob of N-in: 0.94351\nPOSSIBLE N-term signal sequence\ninside 1 19\nTMhelix 20 39\noutside 40 325\nTMhelix 326 348\ninside 349 372

4362 GCF_002144095.1_ASM214409v1 Escherichia coli Proteobacteria; Gammaproteobacteria; Enterobacterales; Enterobacteriaceae; Escherichia
MNNEETFYQAMRRQGVTRRSFLKYCSLAATSLGLGAGMAPKIAWALENKPRIPVVWIHGL WP_001301613.1
hydrogenase [Escherichia coli] Length: 372\nNumber of predicted TMHs: 2\nExp number of AAs in TMHs: 37.46598\nExp number, first 60 AAs: 19.53139\nTotal prob of N-in: 0.94364\nPOSSIBLE N-term signal sequence\ninside 1 19\nTMhelix 20 39\noutside 40 325\nTMhelix 326 348\ninside 349 372

4363 GCF_002144105.1_ASM214410v1 Escherichia coli Proteobacteria; Gammaproteobacteria; Enterobacterales; Enterobacteriaceae; Escherichia
MNNEETFYQAMRRQGVTRRSFLKYCSLAATSLGLGAGMAPKIAWALENKPRIPVVWIHGL WP_001301613.1
hydrogenase [Escherichia coli] Length: 372\nNumber of predicted TMHs: 2\nExp number of AAs in TMHs: 37.46598\nExp number, first 60 AAs: 19.53139\nTotal prob of N-in: 0.94364\nPOSSIBLE N-term signal sequence\ninside 1 19\nTMhelix 20 39\noutside 40 325\nTMhelix 326 348\ninside 349 372

4364 GCF_002152065.1_ASM215206v1 Escherichia coli Proteobacteria; Gammaproteobacteria; Enterobacterales; Enterobacteriaceae; Escherichia
MTGDNTLIHSHGINRRDFMKLCAALAATMGLSSKAAAEMAESVTNPQRPPVIWIGAQECT WP_000145410.1

MULTISPECIES: hydrogenase-2 small chain [Proteobacteria] Length: 372\nNumber of predicted TMHs: 1\nExp number of AAs in TMHs: 23.72226\nExp number, first 60 AAs: 0.22038\nTotal prob of N-in: 0.12994\noutside 1 331\nTMhelix 332 354\ninside 355 372

4365 GCF_002151245.1_ASM215124v1 Escherichia coli Proteobacteria; Gammaproteobacteria; Enterobacterales; Enterobacteriaceae; Escherichia MNNEETFYQAMRRQGVTRRSFLKYCSLAATSLGLGAGMAPKIAWALENKPRIPVVWIHGL WP_001058323.1 MULTISPECIES: hydrogenase-1 small chain [Proteobacteria] Length: 372\nNumber of predicted TMHs: 2\nExp number of AAs in TMHs: 37.2608499999999\nExp number, first 60 AAs: 19.52923\nTotal prob of N-in: 0.94351\nPOSSIBLE N-term signal sequence\ninside 1 19\nTMhelix 20 39\noutside 40 325\nTMhelix 326 348\ninside 349 372

4366 GCF_002152135.1_ASM215213v1 Escherichia coli Proteobacteria; Gammaproteobacteria; Enterobacterales; Enterobacteriaceae; Escherichia MTGDNTLIHSHGINRRDFMKLCAALAATMGLSSKAAAEMAESVTNPQRPPVIWIGAQECT WP_000145410.1 MULTISPECIES: hydrogenase-2 small chain [Proteobacteria] Length: 372\nNumber of predicted TMHs: 1\nExp number of AAs in TMHs: 23.72226\nExp number, first 60 AAs: 0.22038\nTotal prob of N-in: 0.12994\noutside 1 331\nTMhelix 332 354\ninside 355 372

4367 GCF_002152115.1_ASM215211v1 Escherichia coli Proteobacteria; Gammaproteobacteria; Enterobacterales; Enterobacteriaceae; Escherichia MTGDNTLIHSHGINRRDFMKLCAALAATMGLSSKAAAEMAESVTNPQRPPVIWIGAQECT WP_000145410.1 MULTISPECIES: hydrogenase-2 small chain [Proteobacteria] Length: 372\nNumber of predicted TMHs: 1\nExp number of AAs in TMHs: 23.72226\nExp number, first 60 AAs: 0.22038\nTotal prob of N-in: 0.12994\noutside 1 331\nTMhelix 332 354\ninside 355 372

4368 GCF_002152025.1_ASM215202v1 Escherichia coli Proteobacteria; Gammaproteobacteria; Enterobacterales; Enterobacteriaceae; Escherichia MNNEETFYQAMRRQGVTRRSFLKYCSLAATSLGLGAGMAPKIAWALENKPRIPVVWIHGL WP_001058323.1 MULTISPECIES: hydrogenase-1 small chain [Proteobacteria] Length: 372\nNumber of predicted TMHs: 2\nExp number of AAs in TMHs: 37.2608499999999\nExp number, first 60 AAs: 19.52923\nTotal prob of N-in: 0.94351\nPOSSIBLE N-term signal sequence\ninside 1 19\nTMhelix 20 39\noutside 40 325\nTMhelix 326 348\ninside 349 372

4369 GCF_002152135.1_ASM215213v1 Escherichia coli Proteobacteria; Gammaproteobacteria; Enterobacterales; Enterobacteriaceae; Escherichia MNNEETFYQAMRRQGVTRRSFLKYCSLAATSLGLGAGMAPKIAWALENKPRIPVVWIHGL WP_001058323.1 MULTISPECIES: hydrogenase-1 small chain [Proteobacteria] Length: 372\nNumber of predicted TMHs: 2\nExp number of AAs in TMHs: 37.2608499999999\nExp number, first 60 AAs: 19.52923\nTotal prob of N-in: 0.94351\nPOSSIBLE N-term signal sequence\ninside 1 19\nTMhelix 20 39\noutside 40 325\nTMhelix 326 348\ninside 349 372

4370 GCF_002152075.1_ASM215207v1 Escherichia coli Proteobacteria; Gammaproteobacteria; Enterobacterales; Enterobacteriaceae; Escherichia MNNEETFYQAMRRQGVTRRSFLKYCSLAATSLGLGAGMAPKIAWALENKPRIPVVWIHGL WP_001295940.1 MULTISPECIES: hydrogenase [Enterobacteriaceae] Length: 372\nNumber of predicted TMHs: 2\nExp number of AAs in TMHs: 37.2608899999999\nExp number, first 60 AAs: 19.52923\nTotal prob of N-in: 0.94351\nPOSSIBLE N-term signal sequence\ninside 1 19\nTMhelix 20 39\noutside 40 325\nTMhelix 326 348\ninside 349 372

4371 GCF_002152215.1_ASM215221v1 Escherichia coli Proteobacteria; Gammaproteobacteria; Enterobacterales; Enterobacteriaceae; Escherichia MTGDNTLIHSHGINRRDFMKLCAALAATMGLSSKAAAEMAESVTNPQRPPVIWIGAQECT WP_000145410.1 MULTISPECIES: hydrogenase-2 small chain [Proteobacteria] Length: 372\nNumber of predicted TMHs: 1\nExp number of AAs in TMHs: 23.72226\nExp number, first 60 AAs: 0.22038\nTotal prob of N-in: 0.12994\noutside 1 331\nTMhelix 332 354\ninside 355 372

4372 GCF_002152215.1_ASM215221v1 Escherichia coli Proteobacteria; Gammaproteobacteria; Enterobacterales; Enterobacteriaceae; Escherichia MNNEETFYQAMRRQGVTRRSFLKYCSLAATSLGLGAGMAPKIAWALENKPRIPVVWIHGL WP_001295940.1 MULTISPECIES: hydrogenase [Enterobacteriaceae] Length: 372\nNumber of predicted TMHs: 2\nExp

number of AAs in TMHs: 37.2608899999999\nExp number, first 60 AAs: 19.52923\nTotal prob of N-in: 0.94351\nPOSSIBLE N-term signal sequence\ninside 1 19\nTMhelix 20 39\noutside 40 325\nTMhelix 326 348\ninside 349 372

4373 GCF_002152115.1_ASM215211v1 Escherichia coli Proteobacteria; Gammaproteobacteria; Enterobacterales; Enterobacteriaceae; Escherichia
MNNEETFYQAMRRQGVTRRSFLKYCSLAATSLGLGAGMAPKIAWALENKPRIPVVWIHGL WP_001295940.1
MULTISPECIES: hydrogenase [Enterobacteriaceae] Length: 372\nNumber of predicted TMHs: 2\nExp number of AAs in TMHs: 37.2608899999999\nExp number, first 60 AAs: 19.52923\nTotal prob of N-in: 0.94351\nPOSSIBLE N-term signal sequence\ninside 1 19\nTMhelix 20 39\noutside 40 325\nTMhelix 326 348\ninside 349 372

4374 GCF_002152225.1_ASM215222v1 Escherichia coli Proteobacteria; Gammaproteobacteria; Enterobacterales; Enterobacteriaceae; Escherichia
MTGDNTLIHSHGINRRDFMKLCAALAATMGLSSKAAAEMVESVTNPQRPPVIWIGAQECT WP_000145423.1
hydrogenase 2 small subunit [Escherichia coli] Length: 372\nNumber of predicted TMHs: 1\nExp number of AAs in TMHs: 23.78201\nExp number, first 60 AAs: 0.28355\nTotal prob of N-in: 0.13257\noutside 1 331\nTMhelix 332 354\ninside 355 372

4375 GCF_002152255.1_ASM215225v1 Escherichia coli Proteobacteria; Gammaproteobacteria; Enterobacterales; Enterobacteriaceae; Escherichia
MTGDNTLIHSHGINRRDFMKLCAALAATMGLSSKAAAEMAESVTNPQRPPVIWIGAQECT WP_000145410.1
MULTISPECIES: hydrogenase-2 small chain [Proteobacteria] Length: 372\nNumber of predicted TMHs: 1\nExp number of AAs in TMHs: 23.72226\nExp number, first 60 AAs: 0.22038\nTotal prob of N-in: 0.12994\noutside 1 331\nTMhelix 332 354\ninside 355 372

4376 GCF_002152275.1_ASM215227v1 Escherichia coli Proteobacteria; Gammaproteobacteria; Enterobacterales; Enterobacteriaceae; Escherichia
MTGDNTLIHSHGINRRDFMKLCAALAATMGLSSKAAAEMAESVTNPQRPPVIWIGAQECT WP_000145410.1
MULTISPECIES: hydrogenase-2 small chain [Proteobacteria] Length: 372\nNumber of predicted TMHs: 1\nExp number of AAs in TMHs: 23.72226\nExp number, first 60 AAs: 0.22038\nTotal prob of N-in: 0.12994\noutside 1 331\nTMhelix 332 354\ninside 355 372

4377 GCF_002152275.1_ASM215227v1 Escherichia coli Proteobacteria; Gammaproteobacteria; Enterobacterales; Enterobacteriaceae; Escherichia
MNNEETFYQAMRRQGVTRRSFLKYCSLAATSLGLGAGMAPKIAWALENKPRIPVVWIHGL WP_001058323.1
MULTISPECIES: hydrogenase-1 small chain [Proteobacteria] Length: 372\nNumber of predicted TMHs: 2\nExp number of AAs in TMHs: 37.2608499999999\nExp number, first 60 AAs: 19.52923\nTotal prob of N-in: 0.94351\nPOSSIBLE N-term signal sequence\ninside 1 19\nTMhelix 20 39\noutside 40 325\nTMhelix 326 348\ninside 349 372

4378 GCF_002153985.1_ASM215398v1 Escherichia coli Proteobacteria; Gammaproteobacteria; Enterobacterales; Enterobacteriaceae; Escherichia
MTGDNTLIHSHGINRRDFMKLCAALAATMGLSSKAAAEMAESVTNPQRPPVIWIGAQECT WP_000145410.1
MULTISPECIES: hydrogenase-2 small chain [Proteobacteria] Length: 372\nNumber of predicted TMHs: 1\nExp number of AAs in TMHs: 23.72226\nExp number, first 60 AAs: 0.22038\nTotal prob of N-in: 0.12994\noutside 1 331\nTMhelix 332 354\ninside 355 372

4379 GCF_002153015.1_ASM215301v1 Escherichia coli Proteobacteria; Gammaproteobacteria; Enterobacterales; Enterobacteriaceae; Escherichia
MNNEETFYQAMRRQGVTRRSFLKYCSLAATSLGLGAGMAPKIAWALENKPRIPVVWIHGL WP_001058323.1
MULTISPECIES: hydrogenase-1 small chain [Proteobacteria] Length: 372\nNumber of predicted TMHs: 2\nExp number of AAs in TMHs: 37.2608499999999\nExp number, first 60 AAs: 19.52923\nTotal prob of N-in: 0.94351\nPOSSIBLE N-term signal sequence\ninside 1 19\nTMhelix 20 39\noutside 40 325\nTMhelix 326 348\ninside 349 372

4380 GCF_002154035.1_ASM215403v1 Escherichia coli Proteobacteria; Gammaproteobacteria; Enterobacterales; Enterobacteriaceae; Escherichia
MTGDNTLIHSHGINRRDFMKLCAALAATMGLSSKAAAEMAESVTNPQRPPVIWIGAQECT WP_000145410.1
MULTISPECIES: hydrogenase-2 small chain [Proteobacteria] Length: 372\nNumber of predicted TMHs:

1\nExp number of AAs in TMHs: 23.72226\nExp number, first 60 AAs: 0.22038\nTotal prob of N-in: 0.12994\noutside 1 331\nTMhelix 332 354\ninside 355 372

4381 GCF_002154075.1_ASM215407v1 Escherichia coli Proteobacteria; Gammaproteobacteria; Enterobacterales; Enterobacteriaceae; Escherichia
MTGDNTLIHSHGINRRDFMKLCAALAATMGLSSKAAAEMAESVTNPQRPPVIWIGAQECT WP_000145410.1
MULTISPECIES: hydrogenase-2 small chain [Proteobacteria] Length: 372\nNumber of predicted TMHs: 1\nExp number of AAs in TMHs: 23.72226\nExp number, first 60 AAs: 0.22038\nTotal prob of N-in: 0.12994\noutside 1 331\nTMhelix 332 354\ninside 355 372

4382 GCF_002153005.1_ASM215300v1 Escherichia coli Proteobacteria; Gammaproteobacteria; Enterobacterales; Enterobacteriaceae; Escherichia
MNNEETFYQAMRRQGVTRRSFLKYCSLAATSLGLGAGMAPKIAWALENKPRIPVVWIHGL WP_001295940.1
MULTISPECIES: hydrogenase [Enterobacteriaceae] Length: 372\nNumber of predicted TMHs: 2\nExp number of AAs in TMHs: 37.2608899999999\nExp number, first 60 AAs: 19.52923\nTotal prob of N-in: 0.94351\nPOSSIBLE N-term signal sequence\ninside 1 19\nTMhelix 20 39\noutside 40 325\nTMhelix 326 348\ninside 349 372

4383 GCF_002153985.1_ASM215398v1 Escherichia coli Proteobacteria; Gammaproteobacteria; Enterobacterales; Enterobacteriaceae; Escherichia
MNNEETFYQAMRRQGVTRRSFLKYCSLAATSLGLGAGMAPKIAWALENKPRIPVVWIHGL WP_001058323.1
MULTISPECIES: hydrogenase-1 small chain [Proteobacteria] Length: 372\nNumber of predicted TMHs: 2\nExp number of AAs in TMHs: 37.2608499999999\nExp number, first 60 AAs: 19.52923\nTotal prob of N-in: 0.94351\nPOSSIBLE N-term signal sequence\ninside 1 19\nTMhelix 20 39\noutside 40 325\nTMhelix 326 348\ninside 349 372

4384 GCF_002154185.1_ASM215418v1 Escherichia coli Proteobacteria; Gammaproteobacteria; Enterobacterales; Enterobacteriaceae; Escherichia
MTGDNTLIHSHGINRRDFMKLCAALAATMGLSSKAAAEMAESVTNPQRPPVIWIGAQECT WP_000145410.1
MULTISPECIES: hydrogenase-2 small chain [Proteobacteria] Length: 372\nNumber of predicted TMHs: 1\nExp number of AAs in TMHs: 23.72226\nExp number, first 60 AAs: 0.22038\nTotal prob of N-in: 0.12994\noutside 1 331\nTMhelix 332 354\ninside 355 372

4385 GCF_002154185.1_ASM215418v1 Escherichia coli Proteobacteria; Gammaproteobacteria; Enterobacterales; Enterobacteriaceae; Escherichia
MNNEETFYQAMRRQGVTRRSFLKYCSLAATSLGLGAGMAPKIAWALENKPRIPVVWIHGL WP_001058323.1
MULTISPECIES: hydrogenase-1 small chain [Proteobacteria] Length: 372\nNumber of predicted TMHs: 2\nExp number of AAs in TMHs: 37.2608499999999\nExp number, first 60 AAs: 19.52923\nTotal prob of N-in: 0.94351\nPOSSIBLE N-term signal sequence\ninside 1 19\nTMhelix 20 39\noutside 40 325\nTMhelix 326 348\ninside 349 372

4386 GCF_002154035.1_ASM215403v1 Escherichia coli Proteobacteria; Gammaproteobacteria; Enterobacterales; Enterobacteriaceae; Escherichia
MNNEETFYQAMRRQGVTRRSFLKYCSLAATSLGLGAGMAPKIAWALENKPRIPVVWIHGL WP_001058323.1
MULTISPECIES: hydrogenase-1 small chain [Proteobacteria] Length: 372\nNumber of predicted TMHs: 2\nExp number of AAs in TMHs: 37.2608499999999\nExp number, first 60 AAs: 19.52923\nTotal prob of N-in: 0.94351\nPOSSIBLE N-term signal sequence\ninside 1 19\nTMhelix 20 39\noutside 40 325\nTMhelix 326 348\ninside 349 372

4387 GCF_002154285.1_ASM215428v1 Escherichia coli Proteobacteria; Gammaproteobacteria; Enterobacterales; Enterobacteriaceae; Escherichia
MTGDNTLIHSHGINRRDFMKLCAALAATMGLSSKAAAEMAESVTNPQRPPVIWIGAQECT WP_000145410.1
MULTISPECIES: hydrogenase-2 small chain [Proteobacteria] Length: 372\nNumber of predicted TMHs: 1\nExp number of AAs in TMHs: 23.72226\nExp number, first 60 AAs: 0.22038\nTotal prob of N-in: 0.12994\noutside 1 331\nTMhelix 332 354\ninside 355 372

4388 GCF_002154295.1_ASM215429v1 Escherichia coli Proteobacteria; Gammaproteobacteria; Enterobacterales; Enterobacteriaceae; Escherichia
MTGDNTLIHSHGINRRDFMKLCAALAATMGLSSKAAAEMAESVTNPQRPPVIWIGAQECT WP_000145410.1
MULTISPECIES: hydrogenase-2 small chain [Proteobacteria] Length: 372\nNumber of predicted TMHs:

1\nExp number of AAs in TMHs: 23.72226\nExp number, first 60 AAs: 0.22038\nTotal prob of N-in: 0.12994\noutside 1 331\nTMhelix 332 354\ninside 355 372

4389 GCF_002154755.1_ASM215475v1 Escherichia coli Proteobacteria; Gammaproteobacteria; Enterobacterales; Enterobacteriaceae; Escherichia MTGDNTLIHSHGINRRDFMKLCAALAATMGLSSKAAAEMAESVTNPQRPPVIWIGAQECT WP_000145410.1 MULTISPECIES: hydrogenase-2 small chain [Proteobacteria] Length: 372\nNumber of predicted TMHs: 1\nExp number of AAs in TMHs: 23.72226\nExp number, first 60 AAs: 0.22038\nTotal prob of N-in: 0.12994\noutside 1 331\nTMhelix 332 354\ninside 355 372

4390 GCF_002154765.1_ASM215476v1 Escherichia coli Proteobacteria; Gammaproteobacteria; Enterobacterales; Enterobacteriaceae; Escherichia MTGDNTLIHSHGINRRDFMKLCAALAATMGLSSKAAAEMAESVTNPQRPPVIWIGAQECT WP_000145410.1 MULTISPECIES: hydrogenase-2 small chain [Proteobacteria] Length: 372\nNumber of predicted TMHs: 1\nExp number of AAs in TMHs: 23.72226\nExp number, first 60 AAs: 0.22038\nTotal prob of N-in: 0.12994\noutside 1 331\nTMhelix 332 354\ninside 355 372

4391 GCF_002155015.1_ASM215501v1 Escherichia coli Proteobacteria; Gammaproteobacteria; Enterobacterales; Enterobacteriaceae; Escherichia MTGDNTLIHSHGINRRDFMKLCAALAATMGLSSKAAAEMAESVTNPQRPPVIWIGAQECT WP_000145410.1 MULTISPECIES: hydrogenase-2 small chain [Proteobacteria] Length: 372\nNumber of predicted TMHs: 1\nExp number of AAs in TMHs: 23.72226\nExp number, first 60 AAs: 0.22038\nTotal prob of N-in: 0.12994\noutside 1 331\nTMhelix 332 354\ninside 355 372

4392 GCF_002155015.1_ASM215501v1 Escherichia coli Proteobacteria; Gammaproteobacteria; Enterobacterales; Enterobacteriaceae; Escherichia MNNEETFYQAMRRQGVTRRSFLKYCSLAATSLGLGAGMAPKIAWALENKPRIPVVWIHGL WP_001058323.1 MULTISPECIES: hydrogenase-1 small chain [Proteobacteria] Length: 372\nNumber of predicted TMHs: 2\nExp number of AAs in TMHs: 37.2608499999999\nExp number, first 60 AAs: 19.52923\nTotal prob of N-in: 0.94351\nPOSSIBLE N-term signal sequence\ninside 1 19\nTMhelix 20 39\noutside 40 325\nTMhelix 326 348\ninside 349 372

4393 GCF_002154925.1_ASM215492v1 Escherichia coli Proteobacteria; Gammaproteobacteria; Enterobacterales; Enterobacteriaceae; Escherichia MNNEETFYQAMRRQGVTRRSFLKYCSLAATSLGLGAGMAPKIAWALENKPRIPVVWIHGL WP_001058323.1 MULTISPECIES: hydrogenase-1 small chain [Proteobacteria] Length: 372\nNumber of predicted TMHs: 2\nExp number of AAs in TMHs: 37.2608499999999\nExp number, first 60 AAs: 19.52923\nTotal prob of N-in: 0.94351\nPOSSIBLE N-term signal sequence\ninside 1 19\nTMhelix 20 39\noutside 40 325\nTMhelix 326 348\ninside 349 372

4394 GCF_002154295.1_ASM215429v1 Escherichia coli Proteobacteria; Gammaproteobacteria; Enterobacterales; Enterobacteriaceae; Escherichia MNNEETFYQAMRRQGVTRRSFLKYCSLAATSLGLGAGMAPKIAWALENKPRIPVVWIHGL WP_001295940.1 MULTISPECIES: hydrogenase [Enterobacteriaceae] Length: 372\nNumber of predicted TMHs: 2\nExp number of AAs in TMHs: 37.2608899999999\nExp number, first 60 AAs: 19.52923\nTotal prob of N-in: 0.94351\nPOSSIBLE N-term signal sequence\ninside 1 19\nTMhelix 20 39\noutside 40 325\nTMhelix 326 348\ninside 349 372

4395 GCF_002155125.1_ASM215512v1 Escherichia coli Proteobacteria; Gammaproteobacteria; Enterobacterales; Enterobacteriaceae; Escherichia MTGDNTLIHSHGINRRDFMKLCAALAATMGLSSKAAAEMAESVTNPQRPPVIWIGAQECT WP_000145410.1 MULTISPECIES: hydrogenase-2 small chain [Proteobacteria] Length: 372\nNumber of predicted TMHs: 1\nExp number of AAs in TMHs: 23.72226\nExp number, first 60 AAs: 0.22038\nTotal prob of N-in: 0.12994\noutside 1 331\nTMhelix 332 354\ninside 355 372

4396 GCF_002155205.1_ASM215520v1 Escherichia coli Proteobacteria; Gammaproteobacteria; Enterobacterales; Enterobacteriaceae; Escherichia MTGDNTLIHSHGINRRDFMKLCAALAATMGLSSKAAAEMAESVTNPQRPPVIWIGAQECT WP_000145410.1 MULTISPECIES: hydrogenase-2 small chain [Proteobacteria] Length: 372\nNumber of predicted TMHs: 1\nExp number of AAs in TMHs: 23.72226\nExp number, first 60 AAs: 0.22038\nTotal prob of N-in: 0.12994\noutside 1 331\nTMhelix 332 354\ninside 355 372

4397 GCF_002155315.1_ASM215531v1 Escherichia coli Proteobacteria; Gammaproteobacteria; Enterobacterales; Enterobacteriaceae; Escherichia
MTGDNTLIHSHGINRRDFMKLCAALAATMGLSSKAAAEMAESVTNPQRPPVIWIGAQECT WP_000145410.1
MULTISPECIES: hydrogenase-2 small chain [Proteobacteria] Length: 372\nNumber of predicted TMHs: 1\nExp number of AAs in TMHs: 23.72226\nExp number, first 60 AAs: 0.22038\nTotal prob of N-in: 0.12994\noutside 1 331\nTMhelix 332 354\ninside 355 372

4398 GCF_002155445.1_ASM215544v1 Escherichia coli Proteobacteria; Gammaproteobacteria; Enterobacterales; Enterobacteriaceae; Escherichia
MNNEETFYQAMRRQGVTRRSFLKYCSLAATSLGLGAGMAPKIAWALENKPRIPVVWIHGL WP_001058323.1
MULTISPECIES: hydrogenase-1 small chain [Proteobacteria] Length: 372\nNumber of predicted TMHs: 2\nExp number of AAs in TMHs: 37.2608499999999\nExp number, first 60 AAs: 19.52923\nTotal prob of N-in: 0.94351\nPOSSIBLE N-term signal sequence\ninside 1 19\nTMhelix 20 39\noutside 40 325\nTMhelix 326 348\ninside 349 372

4399 GCF_002155315.1_ASM215531v1 Escherichia coli Proteobacteria; Gammaproteobacteria; Enterobacterales; Enterobacteriaceae; Escherichia
MNNEETFYQAMRRQGVTRRSFLKYCSLAATSLGLGAGMAPKIAWALENKPRIPVVWIHGL WP_001058323.1
MULTISPECIES: hydrogenase-1 small chain [Proteobacteria] Length: 372\nNumber of predicted TMHs: 2\nExp number of AAs in TMHs: 37.2608499999999\nExp number, first 60 AAs: 19.52923\nTotal prob of N-in: 0.94351\nPOSSIBLE N-term signal sequence\ninside 1 19\nTMhelix 20 39\noutside 40 325\nTMhelix 326 348\ninside 349 372

4400 GCF_002155455.1_ASM215545v1 Escherichia coli Proteobacteria; Gammaproteobacteria; Enterobacterales; Enterobacteriaceae; Escherichia
MTGDNTLIHSHGINRRDFMKLCAALAATMGLSSKAAAEMAESVTNPQRPPVIWIGAQECT WP_000145410.1
MULTISPECIES: hydrogenase-2 small chain [Proteobacteria] Length: 372\nNumber of predicted TMHs: 1\nExp number of AAs in TMHs: 23.72226\nExp number, first 60 AAs: 0.22038\nTotal prob of N-in: 0.12994\noutside 1 331\nTMhelix 332 354\ninside 355 372

4401 GCF_002155375.1_ASM215537v1 Escherichia coli Proteobacteria; Gammaproteobacteria; Enterobacterales; Enterobacteriaceae; Escherichia
MNNEETFYQAMRRQGVTRRSFLKYCSLAATSLGLGAGMAPKIAWALENKPRIPVVWIHGL WP_001058323.1
MULTISPECIES: hydrogenase-1 small chain [Proteobacteria] Length: 372\nNumber of predicted TMHs: 2\nExp number of AAs in TMHs: 37.2608499999999\nExp number, first 60 AAs: 19.52923\nTotal prob of N-in: 0.94351\nPOSSIBLE N-term signal sequence\ninside 1 19\nTMhelix 20 39\noutside 40 325\nTMhelix 326 348\ninside 349 372

4402 GCF_002155375.1_ASM215537v1 Escherichia coli Proteobacteria; Gammaproteobacteria; Enterobacterales; Enterobacteriaceae; Escherichia
MTGDNTLIHSHGINRRDFMKLCAALAATMGLSSKAAAEMAESVTNPQRPPVIWIGAQECT WP_032298447.1
hydrogenase 2 small subunit [Escherichia coli] Length: 372\nNumber of predicted TMHs: 1\nExp number of AAs in TMHs: 23.69807\nExp number, first 60 AAs: 0.221\nTotal prob of N-in: 0.12853\noutside 1 331\nTMhelix 332 354\ninside 355 372

4403 GCF_002155455.1_ASM215545v1 Escherichia coli Proteobacteria; Gammaproteobacteria; Enterobacterales; Enterobacteriaceae; Escherichia
MNNEETFYQAMRRQGVTRRSFLKYCSLAATSLGLGAGMAPKIAWALENKPRIPVVWIHGL WP_001058320.1
hydrogenase [Escherichia coli] Length: 372\nNumber of predicted TMHs: 2\nExp number of AAs in TMHs: 37.26082\nExp number, first 60 AAs: 19.52923\nTotal prob of N-in: 0.94351\nPOSSIBLE N-term signal sequence\ninside 1 19\nTMhelix 20 39\noutside 40 325\nTMhelix 326 348\ninside 349 372

4404 GCF_002155325.1_ASM215532v1 Escherichia coli Proteobacteria; Gammaproteobacteria; Enterobacterales; Enterobacteriaceae; Escherichia
MNNEETFYQAMRRQGVTRRSFLKYCSLAATSLGLGAGMAPKIAWALENKPRIPVVWIHGL WP_001058320.1
hydrogenase [Escherichia coli] Length: 372\nNumber of predicted TMHs: 2\nExp number of AAs in TMHs: 37.26082\nExp number, first 60 AAs: 19.52923\nTotal prob of N-in: 0.94351\nPOSSIBLE N-term signal sequence\ninside 1 19\nTMhelix 20 39\noutside 40 325\nTMhelix 326 348\ninside 349 372

4405 GCF_002155515.1_ASM215551v1 Escherichia coli Proteobacteria; Gammaproteobacteria; Enterobacterales; Enterobacteriaceae; Escherichia

MTGDNTLIHSHGINRRDFMKLCAALAATMGLSSKAAAEMAESVTNPQRPPVIWIGAQECT WP_000145410.1 MULTISPECIES: hydrogenase-2 small chain [Proteobacteria] Length: 372\nNumber of predicted TMHs: 1\nExp number of AAs in TMHs: 23.72226\nExp number, first 60 AAs: 0.22038\nTotal prob of N-in: 0.12994\noutside 1 331\nTMhelix 332 354\ninside 355 372

4406 GCF_002155535.1_ASM215553v1 Escherichia coli Proteobacteria; Gammaproteobacteria; Enterobacterales; Enterobacteriaceae; Escherichia MTGDNTLIHSHGINRRDFMKLCAALAATMGLSSKAAAEMAESVTNPQRPPVIWIGAQECT WP_000145410.1 MULTISPECIES: hydrogenase-2 small chain [Proteobacteria] Length: 372\nNumber of predicted TMHs: 1\nExp number of AAs in TMHs: 23.72226\nExp number, first 60 AAs: 0.22038\nTotal prob of N-in: 0.12994\noutside 1 331\nTMhelix 332 354\ninside 355 372

4407 GCF_002155515.1_ASM215551v1 Escherichia coli Proteobacteria; Gammaproteobacteria; Enterobacterales; Enterobacteriaceae; Escherichia MNNEETFYQAMRRQGVTRRSFLKYCSLAATSLGLGAGMAPKIAWALENKPRIPVVWIHGL WP_001058323.1 MULTISPECIES: hydrogenase-1 small chain [Proteobacteria] Length: 372\nNumber of predicted TMHs: 2\nExp number of AAs in TMHs: 37.2608499999999\nExp number, first 60 AAs: 19.52923\nTotal prob of N-in: 0.94351\nPOSSIBLE N-term signal sequence\ninside 1 19\nTMhelix 20 39\noutside 40 325\nTMhelix 326 348\ninside 349 372

4408 GCF_002155565.1_ASM215556v1 Escherichia coli Proteobacteria; Gammaproteobacteria; Enterobacterales; Enterobacteriaceae; Escherichia MTGDNTLIHSHGINRRDFMKLCAALAATMGLSSKAAAEMAESVTNPQRPPVIWIGAQECT WP_000145410.1 MULTISPECIES: hydrogenase-2 small chain [Proteobacteria] Length: 372\nNumber of predicted TMHs: 1\nExp number of AAs in TMHs: 23.72226\nExp number, first 60 AAs: 0.22038\nTotal prob of N-in: 0.12994\noutside 1 331\nTMhelix 332 354\ninside 355 372

4409 GCF_002155575.1_ASM215557v1 Escherichia coli Proteobacteria; Gammaproteobacteria; Enterobacterales; Enterobacteriaceae; Escherichia MTGDNTLIHSHGINRRDFMKLCAALAATMGLSSKAAAEMAESVTNPQRPPVIWIGAQECT WP_000145410.1 MULTISPECIES: hydrogenase-2 small chain [Proteobacteria] Length: 372\nNumber of predicted TMHs: 1\nExp number of AAs in TMHs: 23.72226\nExp number, first 60 AAs: 0.22038\nTotal prob of N-in: 0.12994\noutside 1 331\nTMhelix 332 354\ninside 355 372

4410 GCF_002155585.1_ASM215558v1 Escherichia coli Proteobacteria; Gammaproteobacteria; Enterobacterales; Enterobacteriaceae; Escherichia MTGDNTLIHSHGINRRDFMKLCAALAATMGLSSKAAAEMAESVTNPQRPPVIWIGAQECT WP_000145410.1 MULTISPECIES: hydrogenase-2 small chain [Proteobacteria] Length: 372\nNumber of predicted TMHs: 1\nExp number of AAs in TMHs: 23.72226\nExp number, first 60 AAs: 0.22038\nTotal prob of N-in: 0.12994\noutside 1 331\nTMhelix 332 354\ninside 355 372

4411 GCF_002155595.1_ASM215559v1 Escherichia coli Proteobacteria; Gammaproteobacteria; Enterobacterales; Enterobacteriaceae; Escherichia MTGDNTLIHSHGINRRDFMKLCAALAATMGLSSKAAAEMAESVTNPQRPPVIWIGAQECT WP_000145410.1 MULTISPECIES: hydrogenase-2 small chain [Proteobacteria] Length: 372\nNumber of predicted TMHs: 1\nExp number of AAs in TMHs: 23.72226\nExp number, first 60 AAs: 0.22038\nTotal prob of N-in: 0.12994\noutside 1 331\nTMhelix 332 354\ninside 355 372

4412 GCF_002155775.1_ASM215577v1 Escherichia coli Proteobacteria; Gammaproteobacteria; Enterobacterales; Enterobacteriaceae; Escherichia MNNEETFYQAMRRQGVTRRSFLKYCSLAATSLGLGAGMAPKIAWALENKPRIPVVWIHGL WP_001058320.1 hydrogenase [Escherichia coli] Length: 372\nNumber of predicted TMHs: 2\nExp number of AAs in TMHs: 37.26082\nExp number, first 60 AAs: 19.52923\nTotal prob of N-in: 0.94351\nPOSSIBLE N-term signal sequence\ninside 1 19\nTMhelix 20 39\noutside 40 325\nTMhelix 326 348\ninside 349 372

4413 GCF_002155565.1_ASM215556v1 Escherichia coli Proteobacteria; Gammaproteobacteria; Enterobacterales; Enterobacteriaceae; Escherichia MNNEETFYQAMRRQGVTRRSFLKYCSLAATSLGLGAGMAPKIAWALENKPRIPVVWIHGL WP_001058323.1 MULTISPECIES: hydrogenase-1 small chain [Proteobacteria] Length: 372\nNumber of predicted TMHs: 2\nExp number of AAs in TMHs: 37.2608499999999\nExp number, first 60 AAs: 19.52923\nTotal prob of N-in:

0.94351\nPOSSIBLE N-term signal sequence\ninside 1 19\nTMhelix 20 39\noutside 40 325\nTMhelix 326 348\ninside 349 372

4414 GCF_002155575.1_ASM215557v1 Escherichia coli Proteobacteria; Gammaproteobacteria; Enterobacterales; Enterobacteriaceae; Escherichia
MNNEETFYQAMRRQGVTRRSFLKYCSLAATSLGLGAGMAPKIAWALENKPRIPVVWIHGL WP_001058323.1
MULTISPECIES: hydrogenase-1 small chain [Proteobacteria] Length: 372\nNumber of predicted TMHs: 2\nExp number of AAs in TMHs: 37.2608499999999\nExp number, first 60 AAs: 19.52923\nTotal prob of N-in: 0.94351\nPOSSIBLE N-term signal sequence\ninside 1 19\nTMhelix 20 39\noutside 40 325\nTMhelix 326 348\ninside 349 372

4415 GCF_002156825.1_ASM215682v1 Escherichia coli Proteobacteria; Gammaproteobacteria; Enterobacterales; Enterobacteriaceae; Escherichia
MTGDNTLIHSHGINRRDFMKLCAALAATMGLSSKAAAEMAESVTNPQRPPVIWIGAQECT WP_000145410.1
MULTISPECIES: hydrogenase-2 small chain [Proteobacteria] Length: 372\nNumber of predicted TMHs: 1\nExp number of AAs in TMHs: 23.72226\nExp number, first 60 AAs: 0.22038\nTotal prob of N-in: 0.12994\noutside 1 331\nTMhelix 332 354\ninside 355 372

4416 GCF_002156825.1_ASM215682v1 Escherichia coli Proteobacteria; Gammaproteobacteria; Enterobacterales; Enterobacteriaceae; Escherichia
MNNEETFYQAMRRQGVTRRSFLKYCSLAATSLGLGAGMAPKIAWALENKPRIPVVWIHGL WP_001058323.1
MULTISPECIES: hydrogenase-1 small chain [Proteobacteria] Length: 372\nNumber of predicted TMHs: 2\nExp number of AAs in TMHs: 37.2608499999999\nExp number, first 60 AAs: 19.52923\nTotal prob of N-in: 0.94351\nPOSSIBLE N-term signal sequence\ninside 1 19\nTMhelix 20 39\noutside 40 325\nTMhelix 326 348\ninside 349 372

4417 GCF_002155585.1_ASM215558v1 Escherichia coli Proteobacteria; Gammaproteobacteria; Enterobacterales; Enterobacteriaceae; Escherichia
MNNEETFYQAMRRQGVTRRSFLKYCSLAATSLGLGAGMAPKIAWALENKPRIPVVWIHGL WP_001058323.1
MULTISPECIES: hydrogenase-1 small chain [Proteobacteria] Length: 372\nNumber of predicted TMHs: 2\nExp number of AAs in TMHs: 37.2608499999999\nExp number, first 60 AAs: 19.52923\nTotal prob of N-in: 0.94351\nPOSSIBLE N-term signal sequence\ninside 1 19\nTMhelix 20 39\noutside 40 325\nTMhelix 326 348\ninside 349 372

4418 GCF_002155825.1_ASM215582v1 Escherichia coli Proteobacteria; Gammaproteobacteria; Enterobacterales; Enterobacteriaceae; Escherichia
MNNEETFYQAMRRQGVTRRSFLKYCSLAATSLGLGAGMAPKIAWALENKPRIPVVWIHGL WP_001058323.1
MULTISPECIES: hydrogenase-1 small chain [Proteobacteria] Length: 372\nNumber of predicted TMHs: 2\nExp number of AAs in TMHs: 37.2608499999999\nExp number, first 60 AAs: 19.52923\nTotal prob of N-in: 0.94351\nPOSSIBLE N-term signal sequence\ninside 1 19\nTMhelix 20 39\noutside 40 325\nTMhelix 326 348\ninside 349 372

4419 GCF_002164115.1_ASM216411v1 Escherichia coli Proteobacteria; Gammaproteobacteria; Enterobacterales; Enterobacteriaceae; Escherichia
MTGDNTLIHSHGINRRDFMKLCAALAATMGLSSKAAAEMAESVTNPQRPPVIWIGAQECT WP_000145410.1
MULTISPECIES: hydrogenase-2 small chain [Proteobacteria] Length: 372\nNumber of predicted TMHs: 1\nExp number of AAs in TMHs: 23.72226\nExp number, first 60 AAs: 0.22038\nTotal prob of N-in: 0.12994\noutside 1 331\nTMhelix 332 354\ninside 355 372

4420 GCF_002164045.1_ASM216404v1 Escherichia coli Proteobacteria; Gammaproteobacteria; Enterobacterales; Enterobacteriaceae; Escherichia
MTGDNTLIHSHGINRRDFMKLCAALAATMGLSSKAAAEMAESVTNPQRPPVIWIGAQECT WP_000145410.1
MULTISPECIES: hydrogenase-2 small chain [Proteobacteria] Length: 372\nNumber of predicted TMHs: 1\nExp number of AAs in TMHs: 23.72226\nExp number, first 60 AAs: 0.22038\nTotal prob of N-in: 0.12994\noutside 1 331\nTMhelix 332 354\ninside 355 372

4421 GCF_002164175.1_ASM216417v1 Escherichia coli Proteobacteria; Gammaproteobacteria; Enterobacterales; Enterobacteriaceae; Escherichia
MTGDNTLIHSHGINRRDFMKLCAALAATMGLSSKAAAEMAESVTNPQRPPVIWIGAQECT WP_000145410.1
MULTISPECIES: hydrogenase-2 small chain [Proteobacteria] Length: 372\nNumber of predicted TMHs:

1\nExp number of AAs in TMHs: 23.72226\nExp number, first 60 AAs: 0.22038\nTotal prob of N-in: 0.12994\noutside 1 331\nTMhelix 332 354\ninside 355 372

4422 GCF_002164205.1_ASM216420v1 Escherichia coli Proteobacteria; Gammaproteobacteria; Enterobacterales; Enterobacteriaceae; Escherichia
MTGDNTLIHSHGINRRDFMKLCAALAATMGLSSKAAAEMAESVTNPQRPPVIWIGAQECT WP_000145410.1
MULTISPECIES: hydrogenase-2 small chain [Proteobacteria] Length: 372\nNumber of predicted TMHs: 1\nExp number of AAs in TMHs: 23.72226\nExp number, first 60 AAs: 0.22038\nTotal prob of N-in: 0.12994\noutside 1 331\nTMhelix 332 354\ninside 355 372

4423 GCF_002164365.1_ASM216436v1 Escherichia coli Proteobacteria; Gammaproteobacteria; Enterobacterales; Enterobacteriaceae; Escherichia
MTGDNTLIHSHGINRRDFMKLCAALAATMGLSSKAAAEMAESVTNPQRPPVIWIGAQECT WP_000145410.1
MULTISPECIES: hydrogenase-2 small chain [Proteobacteria] Length: 372\nNumber of predicted TMHs: 1\nExp number of AAs in TMHs: 23.72226\nExp number, first 60 AAs: 0.22038\nTotal prob of N-in: 0.12994\noutside 1 331\nTMhelix 332 354\ninside 355 372

4424 GCF_002164335.1_ASM216433v1 Escherichia coli Proteobacteria; Gammaproteobacteria; Enterobacterales; Enterobacteriaceae; Escherichia
MTGDNTLIHSHGINRRDFMKLCAALAATMGLSSKAAAEMAESVTNPQRPPVIWIGAQECT WP_000145410.1
MULTISPECIES: hydrogenase-2 small chain [Proteobacteria] Length: 372\nNumber of predicted TMHs: 1\nExp number of AAs in TMHs: 23.72226\nExp number, first 60 AAs: 0.22038\nTotal prob of N-in: 0.12994\noutside 1 331\nTMhelix 332 354\ninside 355 372

4425 GCF_002163655.1_ASM216365v1 Escherichia coli Proteobacteria; Gammaproteobacteria; Enterobacterales; Enterobacteriaceae; Escherichia
MNNEETFYQAMRRQGVTRRSFLKYCSLAATSLGLGAGMAPKIAWALENKPRIPVVWIHGL WP_001295940.1
MULTISPECIES: hydrogenase [Enterobacteriaceae] Length: 372\nNumber of predicted TMHs: 2\nExp number of AAs in TMHs: 37.2608899999999\nExp number, first 60 AAs: 19.52923\nTotal prob of N-in: 0.94351\nPOSSIBLE N-term signal sequence\ninside 1 19\nTMhelix 20 39\noutside 40 325\nTMhelix 326 348\ninside 349 372

4426 GCF_002164435.1_ASM216443v1 Escherichia coli Proteobacteria; Gammaproteobacteria; Enterobacterales; Enterobacteriaceae; Escherichia
MTGDNTLIHSHGINRRDFMKLCAALAATMGLSSKAAAEMAESVTNPQRPPVIWIGAQECT WP_000145410.1
MULTISPECIES: hydrogenase-2 small chain [Proteobacteria] Length: 372\nNumber of predicted TMHs: 1\nExp number of AAs in TMHs: 23.72226\nExp number, first 60 AAs: 0.22038\nTotal prob of N-in: 0.12994\noutside 1 331\nTMhelix 332 354\ninside 355 372

4427 GCF_002164205.1_ASM216420v1 Escherichia coli Proteobacteria; Gammaproteobacteria; Enterobacterales; Enterobacteriaceae; Escherichia
MNNEETFYQAMRRQGVTRRSFLKYCSLAATSLGLGAGMAPKIAWALENKPRIPVVWIHGL WP_001301613.1
hydrogenase [Escherichia coli] Length: 372\nNumber of predicted TMHs: 2\nExp number of AAs in TMHs: 37.46598\nExp number, first 60 AAs: 19.53139\nTotal prob of N-in: 0.94364\nPOSSIBLE N-term signal sequence\ninside 1 19\nTMhelix 20 39\noutside 40 325\nTMhelix 326 348\ninside 349 372

4428 GCF_002164365.1_ASM216436v1 Escherichia coli Proteobacteria; Gammaproteobacteria; Enterobacterales; Enterobacteriaceae; Escherichia
MNNEETFYQAMRRQGVTRRSFLKYCSLAATSLGLGAGMAPKIAWALENKPRIPVVWIHGL WP_001301613.1
hydrogenase [Escherichia coli] Length: 372\nNumber of predicted TMHs: 2\nExp number of AAs in TMHs: 37.46598\nExp number, first 60 AAs: 19.53139\nTotal prob of N-in: 0.94364\nPOSSIBLE N-term signal sequence\ninside 1 19\nTMhelix 20 39\noutside 40 325\nTMhelix 326 348\ninside 349 372

4429 GCF_002164335.1_ASM216433v1 Escherichia coli Proteobacteria; Gammaproteobacteria; Enterobacterales; Enterobacteriaceae; Escherichia
MNNEETFYQAMRRQGVTRRSFLKYCSLAATSLGLGAGMAPKIAWALENKPRIPVVWIHGL WP_001301613.1
hydrogenase [Escherichia coli] Length: 372\nNumber of predicted TMHs: 2\nExp number of AAs in TMHs: 37.46598\nExp number, first 60 AAs: 19.53139\nTotal prob of N-in: 0.94364\nPOSSIBLE N-term signal sequence\ninside 1 19\nTMhelix 20 39\noutside 40 325\nTMhelix 326 348\ninside 349 372

4430 GCF_002164645.1_ASM216464v1 Escherichia coli Proteobacteria; Gammaproteobacteria; Enterobacterales; Enterobacteriaceae; Escherichia
MTGDNTLIHSHGINRRDFMKLCAALAATMGLSSKAAAEMAESVTNPQRPPVIWIGAQECT WP_000145410.1
MULTISPECIES: hydrogenase-2 small chain [Proteobacteria] Length: 372\nNumber of predicted TMHs: 1\nExp number of AAs in TMHs: 23.72226\nExp number, first 60 AAs: 0.22038\nTotal prob of N-in: 0.12994\noutside 1 331\nTMhelix 332 354\ninside 355 372

4431 GCF_002164935.1_ASM216493v1 Escherichia coli Proteobacteria; Gammaproteobacteria; Enterobacterales; Enterobacteriaceae; Escherichia
MTGDNTLIHSHGINRRDFMKLCAALAATMGLSSKAAAEMAESVTNPQRPPVIWIGAQECT WP_000145410.1
MULTISPECIES: hydrogenase-2 small chain [Proteobacteria] Length: 372\nNumber of predicted TMHs: 1\nExp number of AAs in TMHs: 23.72226\nExp number, first 60 AAs: 0.22038\nTotal prob of N-in: 0.12994\noutside 1 331\nTMhelix 332 354\ninside 355 372

4432 GCF_002164975.1_ASM216497v1 Escherichia coli Proteobacteria; Gammaproteobacteria; Enterobacterales; Enterobacteriaceae; Escherichia
MTGDNTLIHSHGINRRDFMKLCAALAATMGLSSKAAAEMAESVTNPQRPPVIWIGAQECT WP_000145410.1
MULTISPECIES: hydrogenase-2 small chain [Proteobacteria] Length: 372\nNumber of predicted TMHs: 1\nExp number of AAs in TMHs: 23.72226\nExp number, first 60 AAs: 0.22038\nTotal prob of N-in: 0.12994\noutside 1 331\nTMhelix 332 354\ninside 355 372

4433 GCF_002164645.1_ASM216464v1 Escherichia coli Proteobacteria; Gammaproteobacteria; Enterobacterales; Enterobacteriaceae; Escherichia
MNNEETFYQAMRRQGVTRRSFLKYCSLAATSLGLGAGMAPKIAWALENKPRIPVVWIHGL WP_001058323.1
MULTISPECIES: hydrogenase-1 small chain [Proteobacteria] Length: 372\nNumber of predicted TMHs: 2\nExp number of AAs in TMHs: 37.2608499999999\nExp number, first 60 AAs: 19.52923\nTotal prob of N-in: 0.94351\nPOSSIBLE N-term signal sequence\ninside 1 19\nTMhelix 20 39\noutside 40 325\nTMhelix 326 348\ninside 349 372

4434 GCF_002165055.1_ASM216505v1 Escherichia coli Proteobacteria; Gammaproteobacteria; Enterobacterales; Enterobacteriaceae; Escherichia
MNNEETFYQAMRRQGVTRRSFLKYCSLAATSLGLGAGMAPKIAWALENKPRIPVVWIHGL WP_001058323.1
MULTISPECIES: hydrogenase-1 small chain [Proteobacteria] Length: 372\nNumber of predicted TMHs: 2\nExp number of AAs in TMHs: 37.2608499999999\nExp number, first 60 AAs: 19.52923\nTotal prob of N-in: 0.94351\nPOSSIBLE N-term signal sequence\ninside 1 19\nTMhelix 20 39\noutside 40 325\nTMhelix 326 348\ninside 349 372

4435 GCF_002165995.1_ASM216599v1 Escherichia coli Proteobacteria; Gammaproteobacteria; Enterobacterales; Enterobacteriaceae; Escherichia
MNNEETFYQAMRRQGVTRRSFLKYCSLAATSLGLGAGMAPKIAWALENKPRIPVVWIHGL WP_001058323.1
MULTISPECIES: hydrogenase-1 small chain [Proteobacteria] Length: 372\nNumber of predicted TMHs: 2\nExp number of AAs in TMHs: 37.2608499999999\nExp number, first 60 AAs: 19.52923\nTotal prob of N-in: 0.94351\nPOSSIBLE N-term signal sequence\ninside 1 19\nTMhelix 20 39\noutside 40 325\nTMhelix 326 348\ninside 349 372

4436 GCF_002164975.1_ASM216497v1 Escherichia coli Proteobacteria; Gammaproteobacteria; Enterobacterales; Enterobacteriaceae; Escherichia
MNNEETFYQAMRRQGVTRRSFLKYCSLAATSLGLGAGMAPKIAWALENKPRIPVVWIHGL WP_001058320.1
hydrogenase [Escherichia coli] Length: 372\nNumber of predicted TMHs: 2\nExp number of AAs in TMHs: 37.26082\nExp number, first 60 AAs: 19.52923\nTotal prob of N-in: 0.94351\nPOSSIBLE N-term signal sequence\ninside 1 19\nTMhelix 20 39\noutside 40 325\nTMhelix 326 348\ninside 349 372

4437 GCF_002165035.1_ASM216503v1 Escherichia coli Proteobacteria; Gammaproteobacteria; Enterobacterales; Enterobacteriaceae; Escherichia
MNNEETFYQAMRRQGVTRRSFLKYCSLAATSLGLGAGMAPKIAWALENKPRIPVVWIHGL WP_001058323.1
MULTISPECIES: hydrogenase-1 small chain [Proteobacteria] Length: 372\nNumber of predicted TMHs: 2\nExp number of AAs in TMHs: 37.2608499999999\nExp number, first 60 AAs: 19.52923\nTotal prob of N-in: 0.94351\nPOSSIBLE N-term signal sequence\ninside 1 19\nTMhelix 20 39\noutside 40 325\nTMhelix 326 348\ninside 349 372

4438 GCF_002166025.1_ASM216602v1 Escherichia coli Proteobacteria; Gammaproteobacteria; Enterobacterales; Enterobacteriaceae; Escherichia
MTGDNTLIHSHGINRRDFMKLCAALAATMGLSSKAAAEMAESVTNPQRPPVIWIGAQECT WP_000145410.1
MULTISPECIES: hydrogenase-2 small chain [Proteobacteria] Length: 372\nNumber of predicted TMHs: 1\nExp number of AAs in TMHs: 23.72226\nExp number, first 60 AAs: 0.22038\nTotal prob of N-in: 0.12994\noutside 1 331\nTMhelix 332 354\ninside 355 372

4439 GCF_002166095.1_ASM216609v1 Escherichia coli Proteobacteria; Gammaproteobacteria; Enterobacterales; Enterobacteriaceae; Escherichia
MTGDNTLIHSHGINRRDFMKLCAALAATMGLSSKAAAEMAESVTNPQRPPVIWIGAQECT WP_000145410.1
MULTISPECIES: hydrogenase-2 small chain [Proteobacteria] Length: 372\nNumber of predicted TMHs: 1\nExp number of AAs in TMHs: 23.72226\nExp number, first 60 AAs: 0.22038\nTotal prob of N-in: 0.12994\noutside 1 331\nTMhelix 332 354\ninside 355 372

4440 GCF_002166185.1_ASM216618v1 Escherichia coli Proteobacteria; Gammaproteobacteria; Enterobacterales; Enterobacteriaceae; Escherichia
MTGDNTLIHSHGINRRDFMKLCAALAATMGLSSKAAAEMAESVTNPQRPPVIWIGAQECT WP_000145410.1
MULTISPECIES: hydrogenase-2 small chain [Proteobacteria] Length: 372\nNumber of predicted TMHs: 1\nExp number of AAs in TMHs: 23.72226\nExp number, first 60 AAs: 0.22038\nTotal prob of N-in: 0.12994\noutside 1 331\nTMhelix 332 354\ninside 355 372

4441 GCF_002166125.1_ASM216612v1 Escherichia coli Proteobacteria; Gammaproteobacteria; Enterobacterales; Enterobacteriaceae; Escherichia
MTGDNTLIHSHGINRRDFMKLCAALAATMGLSSKAAAEMAESVTNPQRPPVIWIGAQECT WP_000145410.1
MULTISPECIES: hydrogenase-2 small chain [Proteobacteria] Length: 372\nNumber of predicted TMHs: 1\nExp number of AAs in TMHs: 23.72226\nExp number, first 60 AAs: 0.22038\nTotal prob of N-in: 0.12994\noutside 1 331\nTMhelix 332 354\ninside 355 372

4442 GCF_002166025.1_ASM216602v1 Escherichia coli Proteobacteria; Gammaproteobacteria; Enterobacterales; Enterobacteriaceae; Escherichia
MNNEETFYQAMRRQGVTRRSFLKYCSLAATSLGLGAGMAPKIAWALENKPRIPVVWIHGL WP_001058323.1
MULTISPECIES: hydrogenase-1 small chain [Proteobacteria] Length: 372\nNumber of predicted TMHs: 2\nExp number of AAs in TMHs: 37.2608499999999\nExp number, first 60 AAs: 19.52923\nTotal prob of N-in: 0.94351\nPOSSIBLE N-term signal sequence\ninside 1 19\nTMhelix 20 39\noutside 40 325\nTMhelix 326 348\ninside 349 372

4443 GCF_002166235.1_ASM216623v1 Escherichia coli Proteobacteria; Gammaproteobacteria; Enterobacterales; Enterobacteriaceae; Escherichia
MTGDNTLIHSHGINRRDFMKLCAALAATMGLSSKAAAEMAESVTNPQRPPVIWIGAQECT WP_000145410.1
MULTISPECIES: hydrogenase-2 small chain [Proteobacteria] Length: 372\nNumber of predicted TMHs: 1\nExp number of AAs in TMHs: 23.72226\nExp number, first 60 AAs: 0.22038\nTotal prob of N-in: 0.12994\noutside 1 331\nTMhelix 332 354\ninside 355 372

4444 GCF_002166255.1_ASM216625v1 Escherichia coli Proteobacteria; Gammaproteobacteria; Enterobacterales; Enterobacteriaceae; Escherichia
MTGDNTLIHSHGINRRDFMKLCAALAATMGLSSKAAAEMVESVTNPQRPPVIWIGAQECT WP_000145423.1
hydrogenase 2 small subunit [Escherichia coli] Length: 372\nNumber of predicted TMHs: 1\nExp number of AAs in TMHs: 23.78201\nExp number, first 60 AAs: 0.28355\nTotal prob of N-in: 0.13257\noutside 1 331\nTMhelix 332 354\ninside 355 372

4445 GCF_002166255.1_ASM216625v1 Escherichia coli Proteobacteria; Gammaproteobacteria; Enterobacterales; Enterobacteriaceae; Escherichia
MNNEETFYQAMRRQGVTRRSFLKYCSLAATSLGLGAGMAPKIAWALENKPRIPVVWIHGL WP_001058323.1
MULTISPECIES: hydrogenase-1 small chain [Proteobacteria] Length: 372\nNumber of predicted TMHs: 2\nExp number of AAs in TMHs: 37.2608499999999\nExp number, first 60 AAs: 19.52923\nTotal prob of N-in: 0.94351\nPOSSIBLE N-term signal sequence\ninside 1 19\nTMhelix 20 39\noutside 40 325\nTMhelix 326 348\ninside 349 372

4446 GCF_002165415.1_ASM216541v1 Escherichia coli Proteobacteria; Gammaproteobacteria; Enterobacterales; Enterobacteriaceae; Escherichia
MNNEETFYQAMRRQGVTRRSFLKYCSLAATSLGLGAGMAPKIAWALENKPRIPVVWIHGL WP_001301613.1

hydrogenase [Escherichia coli] Length: 372\nNumber of predicted TMHs: 2\nExp number of AAs in TMHs: 37.46598\nExp number, first 60 AAs: 19.53139\nTotal prob of N-in: 0.94364\nPOSSIBLE N-term signal sequence\ninside 1 19\nTMhelix 20 39\noutside 40 325\nTMhelix 326 348\ninside 349 372

4447 GCF_002166455.1_ASM216645v1 Escherichia coli Proteobacteria; Gammaproteobacteria; Enterobacterales; Enterobacteriaceae; Escherichia MTGDNTLIHSHGINRRDFMKLCAALAATMGLSSKAAAEMAESVTNPQRPPVIWIGAQECT WP_000145410.1 MULTISPECIES: hydrogenase-2 small chain [Proteobacteria] Length: 372\nNumber of predicted TMHs: 1\nExp number of AAs in TMHs: 23.72226\nExp number, first 60 AAs: 0.22038\nTotal prob of N-in: 0.12994\noutside 1 331\nTMhelix 332 354\ninside 355 372

4448 GCF_002166505.1_ASM216650v1 Escherichia coli Proteobacteria; Gammaproteobacteria; Enterobacterales; Enterobacteriaceae; Escherichia MTGDNTLIHSHGINRRDFMKLCAALAATMGLSSKAAAEMVESVTNPQRPPVIWIGAQECT WP_000145423.1 hydrogenase 2 small subunit [Escherichia coli] Length: 372\nNumber of predicted TMHs: 1\nExp number of AAs in TMHs: 23.78201\nExp number, first 60 AAs: 0.28355\nTotal prob of N-in: 0.13257\noutside 1 331\nTMhelix 332 354\ninside 355 372

4449 GCF_002166535.1_ASM216653v1 Escherichia coli Proteobacteria; Gammaproteobacteria; Enterobacterales; Enterobacteriaceae; Escherichia MTGDNTLIHSHGINRRDFMKLCAALAATMGLSSKAAAEMAESVTNPQRPPVIWIGAQECT WP_000145410.1 MULTISPECIES: hydrogenase-2 small chain [Proteobacteria] Length: 372\nNumber of predicted TMHs: 1\nExp number of AAs in TMHs: 23.72226\nExp number, first 60 AAs: 0.22038\nTotal prob of N-in: 0.12994\noutside 1 331\nTMhelix 332 354\ninside 355 372

4450 GCF_002166505.1_ASM216650v1 Escherichia coli Proteobacteria; Gammaproteobacteria; Enterobacterales; Enterobacteriaceae; Escherichia MNNEETFYQAMRRQGVTRRSFLKYCSLAATSLGLGAGMAPKIAWALENKPRIPVVWIHGL WP_001058323.1 MULTISPECIES: hydrogenase-1 small chain [Proteobacteria] Length: 372\nNumber of predicted TMHs: 2\nExp number of AAs in TMHs: 37.2608499999999\nExp number, first 60 AAs: 19.52923\nTotal prob of N-in: 0.94351\nPOSSIBLE N-term signal sequence\ninside 1 19\nTMhelix 20 39\noutside 40 325\nTMhelix 326 348\ninside 349 372

4451 GCF_002166455.1_ASM216645v1 Escherichia coli Proteobacteria; Gammaproteobacteria; Enterobacterales; Enterobacteriaceae; Escherichia MNNEETFYQAMRRQGVTRRSFLKYCSLAATSLGLGAGMAPKIAWALENKPRIPVVWIHGL WP_001058323.1 MULTISPECIES: hydrogenase-1 small chain [Proteobacteria] Length: 372\nNumber of predicted TMHs: 2\nExp number of AAs in TMHs: 37.2608499999999\nExp number, first 60 AAs: 19.52923\nTotal prob of N-in: 0.94351\nPOSSIBLE N-term signal sequence\ninside 1 19\nTMhelix 20 39\noutside 40 325\nTMhelix 326 348\ninside 349 372

4452 GCF_002166535.1_ASM216653v1 Escherichia coli Proteobacteria; Gammaproteobacteria; Enterobacterales; Enterobacteriaceae; Escherichia MNNEETFYQAMRRQGVTRRSFLKYCSLAATSLGLGAGMAPKIAWALENKPRIPVVWIHGL WP_001058320.1 hydrogenase [Escherichia coli] Length: 372\nNumber of predicted TMHs: 2\nExp number of AAs in TMHs: 37.26082\nExp number, first 60 AAs: 19.52923\nTotal prob of N-in: 0.94351\nPOSSIBLE N-term signal sequence\ninside 1 19\nTMhelix 20 39\noutside 40 325\nTMhelix 326 348\ninside 349 372

4453 GCF_002166235.1_ASM216623v1 Escherichia coli Proteobacteria; Gammaproteobacteria; Enterobacterales; Enterobacteriaceae; Escherichia MNNEETFYQAMRRQGVTRRSFLKYCSLAATSLGLGAGMAPKIAWALENKPRIPVVWIHGL WP_001058323.1 MULTISPECIES: hydrogenase-1 small chain [Proteobacteria] Length: 372\nNumber of predicted TMHs: 2\nExp number of AAs in TMHs: 37.2608499999999\nExp number, first 60 AAs: 19.52923\nTotal prob of N-in: 0.94351\nPOSSIBLE N-term signal sequence\ninside 1 19\nTMhelix 20 39\noutside 40 325\nTMhelix 326 348\ninside 349 372

4454 GCF_002173285.1_ASM217328v1 Escherichia coli Proteobacteria; Gammaproteobacteria; Enterobacterales; Enterobacteriaceae; Escherichia MNNEETFYQAMRRQGVTRRSFLKYCSLAATSLGLGAGMAPKIAWALENKPRIPVVWIHGL WP_001058323.1 MULTISPECIES: hydrogenase-1 small chain [Proteobacteria] Length: 372\nNumber of predicted TMHs: 2\nExp number of AAs in TMHs: 37.2608499999999\nExp number, first 60 AAs: 19.52923\nTotal prob of N-in:

0.94351\nPOSSIBLE N-term signal sequence\ninside 1 19\nTMhelix 20 39\noutside 40 325\nTMhelix 326 348\ninside 349 372

4455 GCF_002166645.1_ASM216664v1 Escherichia coli Proteobacteria; Gammaproteobacteria; Enterobacterales; Enterobacteriaceae; Escherichia MNNEETFYQAMRRQGVTRRSFLKYCSLAATSLGLGAGMAPKIAWALENKPRIPVVWIHGL WP_001058323.1 MULTISPECIES: hydrogenase-1 small chain [Proteobacteria] Length: 372\nNumber of predicted TMHs: 2\nExp number of AAs in TMHs: 37.2608499999999\nExp number, first 60 AAs: 19.52923\nTotal prob of N-in: 0.94351\nPOSSIBLE N-term signal sequence\ninside 1 19\nTMhelix 20 39\noutside 40 325\nTMhelix 326 348\ninside 349 372

4456 GCF_002166705.1_ASM216670v1 Escherichia coli Proteobacteria; Gammaproteobacteria; Enterobacterales; Enterobacteriaceae; Escherichia MNNEETFYQAMRRQGVTRRSFLKYCSLAATSLGLGAGMAPKIAWALENKPRIPVVWIHGL WP_001058323.1 MULTISPECIES: hydrogenase-1 small chain [Proteobacteria] Length: 372\nNumber of predicted TMHs: 2\nExp number of AAs in TMHs: 37.2608499999999\nExp number, first 60 AAs: 19.52923\nTotal prob of N-in: 0.94351\nPOSSIBLE N-term signal sequence\ninside 1 19\nTMhelix 20 39\noutside 40 325\nTMhelix 326 348\ninside 349 372

4457 GCF_002173355.1_ASM217335v1 Escherichia coli Proteobacteria; Gammaproteobacteria; Enterobacterales; Enterobacteriaceae; Escherichia MTGDNTLIHSHGINRRDFMKLCAALAATMGLSSKAAAEMAESVTNPQRPPVIWIGAQECT WP_000145410.1 MULTISPECIES: hydrogenase-2 small chain [Proteobacteria] Length: 372\nNumber of predicted TMHs: 1\nExp number of AAs in TMHs: 23.72226\nExp number, first 60 AAs: 0.22038\nTotal prob of N-in: 0.12994\noutside 1 331\nTMhelix 332 354\ninside 355 372

4458 GCF_002173355.1_ASM217335v1 Escherichia coli Proteobacteria; Gammaproteobacteria; Enterobacterales; Enterobacteriaceae; Escherichia MNNEETFYQAMRRQGVTRRSFLKYCSLAATSLGLGAGMAPKIAWALENKPRIPVVWIHGL WP_001058323.1 MULTISPECIES: hydrogenase-1 small chain [Proteobacteria] Length: 372\nNumber of predicted TMHs: 2\nExp number of AAs in TMHs: 37.2608499999999\nExp number, first 60 AAs: 19.52923\nTotal prob of N-in: 0.94351\nPOSSIBLE N-term signal sequence\ninside 1 19\nTMhelix 20 39\noutside 40 325\nTMhelix 326 348\ninside 349 372

4459 GCF_002166755.1_ASM216675v1 Escherichia coli Proteobacteria; Gammaproteobacteria; Enterobacterales; Enterobacteriaceae; Escherichia MNNEETFYQAMRRQGVTRRSFLKYCSLAATSLGLGAGMAPKIAWALENKPRIPVVWIHGL WP_001058323.1 MULTISPECIES: hydrogenase-1 small chain [Proteobacteria] Length: 372\nNumber of predicted TMHs: 2\nExp number of AAs in TMHs: 37.2608499999999\nExp number, first 60 AAs: 19.52923\nTotal prob of N-in: 0.94351\nPOSSIBLE N-term signal sequence\ninside 1 19\nTMhelix 20 39\noutside 40 325\nTMhelix 326 348\ninside 349 372

4460 GCF_002175295.1_ASM217529v1 Escherichia coli Proteobacteria; Gammaproteobacteria; Enterobacterales; Enterobacteriaceae; Escherichia MTGDNTLIHSHGINRRDFMKLCAALAATMGLSSKAAAEMAESVTNPQRPPVIWIGAQECT WP_000145410.1 MULTISPECIES: hydrogenase-2 small chain [Proteobacteria] Length: 372\nNumber of predicted TMHs: 1\nExp number of AAs in TMHs: 23.72226\nExp number, first 60 AAs: 0.22038\nTotal prob of N-in: 0.12994\noutside 1 331\nTMhelix 332 354\ninside 355 372

4461 GCF_002173265.1_ASM217326v1 Escherichia coli Proteobacteria; Gammaproteobacteria; Enterobacterales; Enterobacteriaceae; Escherichia MNNEETFYQAMRRQGVTRRSFLKYCSLAATSLGLGAGMAPKIAWALENKPRIPVVWIHGL WP_001058323.1 MULTISPECIES: hydrogenase-1 small chain [Proteobacteria] Length: 372\nNumber of predicted TMHs: 2\nExp number of AAs in TMHs: 37.2608499999999\nExp number, first 60 AAs: 19.52923\nTotal prob of N-in: 0.94351\nPOSSIBLE N-term signal sequence\ninside 1 19\nTMhelix 20 39\noutside 40 325\nTMhelix 326 348\ninside 349 372

4462 GCF_002175385.1_ASM217538v1 Escherichia coli Proteobacteria; Gammaproteobacteria; Enterobacterales; Enterobacteriaceae; Escherichia MTGDNTLIHSHGINRRDFMKLCAALAATMGLSSKAAAEMAESVTNPQRPPVIWIGAQECT WP_000145410.1 MULTISPECIES: hydrogenase-2 small chain [Proteobacteria] Length: 372\nNumber of predicted TMHs:

1\nExp number of AAs in TMHs: 23.72226\nExp number, first 60 AAs: 0.22038\nTotal prob of N-in: 0.12994\noutside 1 331\nTMhelix 332 354\ninside 355 372

4463 GCF_002175455.1_ASM217545v1 Escherichia coli Proteobacteria; Gammaproteobacteria; Enterobacterales; Enterobacteriaceae; Escherichia MTGDNTLIHSHGINRRDFMKLCAALAATMGLSSKAAAEMAESVTNPQRPPVIWIGAQECT WP_000145410.1 MULTISPECIES: hydrogenase-2 small chain [Proteobacteria] Length: 372\nNumber of predicted TMHs: 1\nExp number of AAs in TMHs: 23.72226\nExp number, first 60 AAs: 0.22038\nTotal prob of N-in: 0.12994\noutside 1 331\nTMhelix 332 354\ninside 355 372

4464 GCF_002175475.1_ASM217547v1 Escherichia coli Proteobacteria; Gammaproteobacteria; Enterobacterales; Enterobacteriaceae; Escherichia MTGDNTLIHSHGINRRDFMKLCAALAATMGLSSKAAAEMAESVTNPQRPPVIWIGAQECT WP_000145410.1 MULTISPECIES: hydrogenase-2 small chain [Proteobacteria] Length: 372\nNumber of predicted TMHs: 1\nExp number of AAs in TMHs: 23.72226\nExp number, first 60 AAs: 0.22038\nTotal prob of N-in: 0.12994\noutside 1 331\nTMhelix 332 354\ninside 355 372

4465 GCF_002175545.1_ASM217554v1 Escherichia coli Proteobacteria; Gammaproteobacteria; Enterobacterales; Enterobacteriaceae; Escherichia MTGDNTLIHSHGINRRDFMKLCAALAATMGLSSKAAAEMAESVTNPQRPPVIWIGAQECT WP_000145410.1 MULTISPECIES: hydrogenase-2 small chain [Proteobacteria] Length: 372\nNumber of predicted TMHs: 1\nExp number of AAs in TMHs: 23.72226\nExp number, first 60 AAs: 0.22038\nTotal prob of N-in: 0.12994\noutside 1 331\nTMhelix 332 354\ninside 355 372

4466 GCF_002175575.1_ASM217557v1 Escherichia coli Proteobacteria; Gammaproteobacteria; Enterobacterales; Enterobacteriaceae; Escherichia MTGDNTLIHSHGINRRDFMKLCAALAATMGLSSKAAAEMAESVTNPQRPPVIWIGAQECT WP_000145410.1 MULTISPECIES: hydrogenase-2 small chain [Proteobacteria] Length: 372\nNumber of predicted TMHs: 1\nExp number of AAs in TMHs: 23.72226\nExp number, first 60 AAs: 0.22038\nTotal prob of N-in: 0.12994\noutside 1 331\nTMhelix 332 354\ninside 355 372

4467 GCF_002175645.1_ASM217564v1 Escherichia coli Proteobacteria; Gammaproteobacteria; Enterobacterales; Enterobacteriaceae; Escherichia MTGDNTLIHSHGINRRDFMKLCAALAATMGLSSKAAAEMAESVTNPQRPPVIWIGAQECT WP_000145410.1 MULTISPECIES: hydrogenase-2 small chain [Proteobacteria] Length: 372\nNumber of predicted TMHs: 1\nExp number of AAs in TMHs: 23.72226\nExp number, first 60 AAs: 0.22038\nTotal prob of N-in: 0.12994\noutside 1 331\nTMhelix 332 354\ninside 355 372

4468 GCF_002175725.1_ASM217572v1 Escherichia coli Proteobacteria; Gammaproteobacteria; Enterobacterales; Enterobacteriaceae; Escherichia MTGDNTLIHSHGINRRDFMKLCAALAATMGLSSKAAAEMAESVTNPQRPPVIWIGAQECT WP_000145410.1 MULTISPECIES: hydrogenase-2 small chain [Proteobacteria] Length: 372\nNumber of predicted TMHs: 1\nExp number of AAs in TMHs: 23.72226\nExp number, first 60 AAs: 0.22038\nTotal prob of N-in: 0.12994\noutside 1 331\nTMhelix 332 354\ninside 355 372

4469 GCF_002175785.1_ASM217578v1 Escherichia coli Proteobacteria; Gammaproteobacteria; Enterobacterales; Enterobacteriaceae; Escherichia MNNEETFYQAMRRQGVTRRSFLKYCSLAATSLGLGAGMAPKIAWALENKPRIPVVWIHGL WP_001301613.1 hydrogenase [Escherichia coli] Length: 372\nNumber of predicted TMHs: 2\nExp number of AAs in TMHs: 37.46598\nExp number, first 60 AAs: 19.53139\nTotal prob of N-in: 0.94364\nPOSSIBLE N-term signal sequence\ninside 1 19\nTMhelix 20 39\noutside 40 325\nTMhelix 326 348\ninside 349 372

4470 GCF_002175855.1_ASM217585v1 Escherichia coli Proteobacteria; Gammaproteobacteria; Enterobacterales; Enterobacteriaceae; Escherichia MNNEETFYQAMRRQGVTRRSFLKYCSLAATSLGLGAGMAPKIAWALENKPRIPVVWIHGL WP_001301613.1 hydrogenase [Escherichia coli] Length: 372\nNumber of predicted TMHs: 2\nExp number of AAs in TMHs: 37.46598\nExp number, first 60 AAs: 19.53139\nTotal prob of N-in: 0.94364\nPOSSIBLE N-term signal sequence\ninside 1 19\nTMhelix 20 39\noutside 40 325\nTMhelix 326 348\ninside 349 372

4471 GCF_002175455.1_ASM217545v1 Escherichia coli Proteobacteria; Gammaproteobacteria; Enterobacterales; Enterobacteriaceae; Escherichia

MNNEETFYQAMRRQGVTRRSFLKYCSLAATSLGLGAGMAPKIAWALENKPRIPVVWIHGL WP_001301613.1 hydrogenase [Escherichia coli] Length: 372\nNumber of predicted TMHs: 2\nExp number of AAs in TMHs: 37.46598\nExp number, first 60 AAs: 19.53139\nTotal prob of N-in: 0.94364\nPOSSIBLE N-term signal sequence\ninside 1 19\nTMhelix 20 39\noutside 40 325\nTMhelix 326 348\ninside 349 372

4472 GCF_002175545.1_ASM217554v1 Escherichia coli Proteobacteria; Gammaproteobacteria; Enterobacterales; Enterobacteriaceae; Escherichia MNNEETFYQAMRRQGVTRRSFLKYCSLAATSLGLGAGMAPKIAWALENKPRIPVVWIHGL WP_001301613.1 hydrogenase [Escherichia coli] Length: 372\nNumber of predicted TMHs: 2\nExp number of AAs in TMHs: 37.46598\nExp number, first 60 AAs: 19.53139\nTotal prob of N-in: 0.94364\nPOSSIBLE N-term signal sequence\ninside 1 19\nTMhelix 20 39\noutside 40 325\nTMhelix 326 348\ninside 349 372

4473 GCF_002175475.1_ASM217547v1 Escherichia coli Proteobacteria; Gammaproteobacteria; Enterobacterales; Enterobacteriaceae; Escherichia MNNEETFYQAMRRQGVTRRSFLKYCSLAATSLGLGAGMAPKIAWALENKPRIPVVWIHGL WP_001301613.1 hydrogenase [Escherichia coli] Length: 372\nNumber of predicted TMHs: 2\nExp number of AAs in TMHs: 37.46598\nExp number, first 60 AAs: 19.53139\nTotal prob of N-in: 0.94364\nPOSSIBLE N-term signal sequence\ninside 1 19\nTMhelix 20 39\noutside 40 325\nTMhelix 326 348\ninside 349 372

4474 GCF_002175965.1_ASM217596v1 Escherichia coli Proteobacteria; Gammaproteobacteria; Enterobacterales; Enterobacteriaceae; Escherichia MTGDNTLIHSHGINRRDFMKLCAALAATMGLSSKAAAEMAESVTNPQRPPVIWIGAQECT WP_000145410.1 MULTISPECIES: hydrogenase-2 small chain [Proteobacteria] Length: 372\nNumber of predicted TMHs: 1\nExp number of AAs in TMHs: 23.72226\nExp number, first 60 AAs: 0.22038\nTotal prob of N-in: 0.12994\noutside 1 331\nTMhelix 332 354\ninside 355 372

4475 GCF_002176035.1_ASM217603v1 Escherichia coli Proteobacteria; Gammaproteobacteria; Enterobacterales; Enterobacteriaceae; Escherichia MNNEETFYQAMRRQGVTRRSFLKYCSLAATSLGLGAGMAPKIAWALENKPRIPVVWIHGL WP_001301613.1 hydrogenase [Escherichia coli] Length: 372\nNumber of predicted TMHs: 2\nExp number of AAs in TMHs: 37.46598\nExp number, first 60 AAs: 19.53139\nTotal prob of N-in: 0.94364\nPOSSIBLE N-term signal sequence\ninside 1 19\nTMhelix 20 39\noutside 40 325\nTMhelix 326 348\ninside 349 372

4476 GCF_002176095.1_ASM217609v1 Escherichia coli Proteobacteria; Gammaproteobacteria; Enterobacterales; Enterobacteriaceae; Escherichia MNNEETFYQAMRRQGVTRRSFLKYCSLAATSLGLGAGMAPKIAWALENKPRIPVVWIHGL WP_001301613.1 hydrogenase [Escherichia coli] Length: 372\nNumber of predicted TMHs: 2\nExp number of AAs in TMHs: 37.46598\nExp number, first 60 AAs: 19.53139\nTotal prob of N-in: 0.94364\nPOSSIBLE N-term signal sequence\ninside 1 19\nTMhelix 20 39\noutside 40 325\nTMhelix 326 348\ninside 349 372

4477 GCF_002176105.1_ASM217610v1 Escherichia coli Proteobacteria; Gammaproteobacteria; Enterobacterales; Enterobacteriaceae; Escherichia MNNEETFYQAMRRQGVTRRSFLKYCSLAATSLGLGAGMAPKIAWALENKPRIPVVWIHGL WP_001301613.1 hydrogenase [Escherichia coli] Length: 372\nNumber of predicted TMHs: 2\nExp number of AAs in TMHs: 37.46598\nExp number, first 60 AAs: 19.53139\nTotal prob of N-in: 0.94364\nPOSSIBLE N-term signal sequence\ninside 1 19\nTMhelix 20 39\noutside 40 325\nTMhelix 326 348\ninside 349 372

4478 GCF_002175875.1_ASM217587v1 Escherichia coli Proteobacteria; Gammaproteobacteria; Enterobacterales; Enterobacteriaceae; Escherichia MNNEETFYQAMRRQGVTRRSFLKYCSLAATSLGLGAGMAPKIAWALENKPRIPVVWIHGL WP_001301613.1 hydrogenase [Escherichia coli] Length: 372\nNumber of predicted TMHs: 2\nExp number of AAs in TMHs: 37.46598\nExp number, first 60 AAs: 19.53139\nTotal prob of N-in: 0.94364\nPOSSIBLE N-term signal sequence\ninside 1 19\nTMhelix 20 39\noutside 40 325\nTMhelix 326 348\ninside 349 372

4479 GCF_002175945.1_ASM217594v1 Escherichia coli Proteobacteria; Gammaproteobacteria; Enterobacterales; Enterobacteriaceae; Escherichia MNNEETFYQAMRRQGVTRRSFLKYCSLAATSLGLGAGMAPKIAWALENKPRIPVVWIHGL WP_001301613.1 hydrogenase [Escherichia coli] Length: 372\nNumber of predicted TMHs: 2\nExp number of AAs in TMHs: 37.46598\nExp number, first 60 AAs: 19.53139\nTotal prob of N-in: 0.94364\nPOSSIBLE N-term signal sequence\ninside 1 19\nTMhelix 20 39\noutside 40 325\nTMhelix 326 348\ninside 349 372

4480 GCF_002176135.1_ASM217613v1 Escherichia coli Proteobacteria; Gammaproteobacteria; Enterobacterales; Enterobacteriaceae; Escherichia
MTGDNTLIHSHGINRRDFMKLCAALAATMGLSSKAAAEMAESVTNPQRPPVIWIGAQECT WP_000145410.1
MULTISPECIES: hydrogenase-2 small chain [Proteobacteria] Length: 372\nNumber of predicted TMHs: 1\nExp number of AAs in TMHs: 23.72226\nExp number, first 60 AAs: 0.22038\nTotal prob of N-in: 0.12994\noutside 1 331\nTMhelix 332 354\ninside 355 372

4481 GCF_002176185.1_ASM217618v1 Escherichia coli Proteobacteria; Gammaproteobacteria; Enterobacterales; Enterobacteriaceae; Escherichia
MTGDNTLIHSHGINRRDFMKLCAALAATMGLSSKAAAEMAESVTNPQRPPVIWIGAQECT WP_000145410.1
MULTISPECIES: hydrogenase-2 small chain [Proteobacteria] Length: 372\nNumber of predicted TMHs: 1\nExp number of AAs in TMHs: 23.72226\nExp number, first 60 AAs: 0.22038\nTotal prob of N-in: 0.12994\noutside 1 331\nTMhelix 332 354\ninside 355 372

4482 GCF_002176635.1_ASM217663v1 Escherichia coli Proteobacteria; Gammaproteobacteria; Enterobacterales; Enterobacteriaceae; Escherichia
MNNEETFYQAMRRQGVTRRSFLKYCSLAATSLGLGAGMAPKIAWALENKPRIPVVWIHGL WP_001301613.1
hydrogenase [Escherichia coli] Length: 372\nNumber of predicted TMHs: 2\nExp number of AAs in TMHs: 37.46598\nExp number, first 60 AAs: 19.53139\nTotal prob of N-in: 0.94364\nPOSSIBLE N-term signal sequence\ninside 1 19\nTMhelix 20 39\noutside 40 325\nTMhelix 326 348\ninside 349 372

4483 GCF_002176255.1_ASM217625v1 Escherichia coli Proteobacteria; Gammaproteobacteria; Enterobacterales; Enterobacteriaceae; Escherichia
MNNEETFYQAMRRQGVTRRSFLKYCSLAATSLGLGAGMAPKIAWALENKPRIPVVWIHGL WP_001301613.1
hydrogenase [Escherichia coli] Length: 372\nNumber of predicted TMHs: 2\nExp number of AAs in TMHs: 37.46598\nExp number, first 60 AAs: 19.53139\nTotal prob of N-in: 0.94364\nPOSSIBLE N-term signal sequence\ninside 1 19\nTMhelix 20 39\noutside 40 325\nTMhelix 326 348\ninside 349 372

4484 GCF_002176135.1_ASM217613v1 Escherichia coli Proteobacteria; Gammaproteobacteria; Enterobacterales; Enterobacteriaceae; Escherichia
MNNEETFYQAMRRQGVTRRSFLKYCSLAATSLGLGAGMAPKIAWALENKPRIPVVWIHGL WP_001301613.1
hydrogenase [Escherichia coli] Length: 372\nNumber of predicted TMHs: 2\nExp number of AAs in TMHs: 37.46598\nExp number, first 60 AAs: 19.53139\nTotal prob of N-in: 0.94364\nPOSSIBLE N-term signal sequence\ninside 1 19\nTMhelix 20 39\noutside 40 325\nTMhelix 326 348\ninside 349 372

4485 GCF_002176185.1_ASM217618v1 Escherichia coli Proteobacteria; Gammaproteobacteria; Enterobacterales; Enterobacteriaceae; Escherichia
MNNEETFYQAMRRQGVTRRSFLKYCSLAATSLGLGAGMAPKIAWALENKPRIPVVWIHGL WP_001301613.1
hydrogenase [Escherichia coli] Length: 372\nNumber of predicted TMHs: 2\nExp number of AAs in TMHs: 37.46598\nExp number, first 60 AAs: 19.53139\nTotal prob of N-in: 0.94364\nPOSSIBLE N-term signal sequence\ninside 1 19\nTMhelix 20 39\noutside 40 325\nTMhelix 326 348\ninside 349 372

4486 GCF_002176275.1_ASM217627v1 Escherichia coli Proteobacteria; Gammaproteobacteria; Enterobacterales; Enterobacteriaceae; Escherichia
MNNEETFYQAMRRQGVTRRSFLKYCSLAATSLGLGAGMAPKIAWALENKPRIPVVWIHGL WP_001301613.1
hydrogenase [Escherichia coli] Length: 372\nNumber of predicted TMHs: 2\nExp number of AAs in TMHs: 37.46598\nExp number, first 60 AAs: 19.53139\nTotal prob of N-in: 0.94364\nPOSSIBLE N-term signal sequence\ninside 1 19\nTMhelix 20 39\noutside 40 325\nTMhelix 326 348\ninside 349 372

4487 GCF_002176795.1_ASM217679v1 Escherichia coli Proteobacteria; Gammaproteobacteria; Enterobacterales; Enterobacteriaceae; Escherichia
MTGDNTLIHSHGINRRDFMKLCAALAATMGLSSKAAAEMAESVTNPQRPPVIWIGAQECT WP_000145410.1
MULTISPECIES: hydrogenase-2 small chain [Proteobacteria] Length: 372\nNumber of predicted TMHs: 1\nExp number of AAs in TMHs: 23.72226\nExp number, first 60 AAs: 0.22038\nTotal prob of N-in: 0.12994\noutside 1 331\nTMhelix 332 354\ninside 355 372

4488 GCF_002176665.1_ASM217666v1 Escherichia coli Proteobacteria; Gammaproteobacteria; Enterobacterales; Enterobacteriaceae; Escherichia
MTGDNTLIHSHGINRRDFMKLCAALAATMGLSSKAAAEMAESVTNPQRPPVIWIGAQECT WP_000145410.1
MULTISPECIES: hydrogenase-2 small chain [Proteobacteria] Length: 372\nNumber of predicted TMHs:

1\nExp number of AAs in TMHs: 23.72226\nExp number, first 60 AAs: 0.22038\nTotal prob of N-in: 0.12994\noutside 1 331\nTMhelix 332 354\ninside 355 372

4489 GCF_002182025.1_ASM218202v1 Escherichia coli Proteobacteria; Gammaproteobacteria; Enterobacterales; Enterobacteriaceae; Escherichia
MTGDNTLIHSHGINRRDFMKLCAALAATMGLSSKAAAEMAESVTNPQRPPVIWIGAQECT WP_000145410.1
MULTISPECIES: hydrogenase-2 small chain [Proteobacteria] Length: 372\nNumber of predicted TMHs: 1\nExp number of AAs in TMHs: 23.72226\nExp number, first 60 AAs: 0.22038\nTotal prob of N-in: 0.12994\noutside 1 331\nTMhelix 332 354\ninside 355 372

4490 GCF_002189755.1_ASM218975v1 Escherichia coli Proteobacteria; Gammaproteobacteria; Enterobacterales; Enterobacteriaceae; Escherichia
MTGDNTLIHSHGINRRDFMKLCAALAATMGLSSKAAAEMAESVTNPQRPPVIWIGAQECT WP_000145410.1
MULTISPECIES: hydrogenase-2 small chain [Proteobacteria] Length: 372\nNumber of predicted TMHs: 1\nExp number of AAs in TMHs: 23.72226\nExp number, first 60 AAs: 0.22038\nTotal prob of N-in: 0.12994\noutside 1 331\nTMhelix 332 354\ninside 355 372

4491 GCF_002189815.1_ASM218981v1 Escherichia coli Proteobacteria; Gammaproteobacteria; Enterobacterales; Enterobacteriaceae; Escherichia
MTGDNTLIHSHGINRRDFMKLCAALAATMGLSSKAAAEMAESVTNPQRPPVIWIGAQECT WP_000145410.1
MULTISPECIES: hydrogenase-2 small chain [Proteobacteria] Length: 372\nNumber of predicted TMHs: 1\nExp number of AAs in TMHs: 23.72226\nExp number, first 60 AAs: 0.22038\nTotal prob of N-in: 0.12994\noutside 1 331\nTMhelix 332 354\ninside 355 372

4492 GCF_002189825.1_ASM218982v1 Escherichia coli Proteobacteria; Gammaproteobacteria; Enterobacterales; Enterobacteriaceae; Escherichia
MTGDNTLIHSHGINRRDFMKLCAALAATMGLSSKAAAEMAESVTNPQRPPVIWIGAQECT WP_000145410.1
MULTISPECIES: hydrogenase-2 small chain [Proteobacteria] Length: 372\nNumber of predicted TMHs: 1\nExp number of AAs in TMHs: 23.72226\nExp number, first 60 AAs: 0.22038\nTotal prob of N-in: 0.12994\noutside 1 331\nTMhelix 332 354\ninside 355 372

4493 GCF_002189875.1_ASM218987v1 Escherichia coli Proteobacteria; Gammaproteobacteria; Enterobacterales; Enterobacteriaceae; Escherichia
MTGDNTLIHSHGINRRDFMKLCAALAATMGLSSKAAAEMAESVTNPQRPPVIWIGAQECT WP_000145410.1
MULTISPECIES: hydrogenase-2 small chain [Proteobacteria] Length: 372\nNumber of predicted TMHs: 1\nExp number of AAs in TMHs: 23.72226\nExp number, first 60 AAs: 0.22038\nTotal prob of N-in: 0.12994\noutside 1 331\nTMhelix 332 354\ninside 355 372

4494 GCF_002189905.1_ASM218990v1 Escherichia coli Proteobacteria; Gammaproteobacteria; Enterobacterales; Enterobacteriaceae; Escherichia
MTGDNTLIHSHGINRRDFMKLCAALAATMGLSSKAAAEMAESVTNPQRPPVIWIGAQECT WP_000145410.1
MULTISPECIES: hydrogenase-2 small chain [Proteobacteria] Length: 372\nNumber of predicted TMHs: 1\nExp number of AAs in TMHs: 23.72226\nExp number, first 60 AAs: 0.22038\nTotal prob of N-in: 0.12994\noutside 1 331\nTMhelix 332 354\ninside 355 372

4495 GCF_002189905.1_ASM218990v1 Escherichia coli Proteobacteria; Gammaproteobacteria; Enterobacterales; Enterobacteriaceae; Escherichia
MNNEETFYQAMRRQGVTRRSFLKYCSLAATSLGLGAGMAPKIAWALENKPRIPVVWIHGL WP_001058323.1
MULTISPECIES: hydrogenase-1 small chain [Proteobacteria] Length: 372\nNumber of predicted TMHs: 2\nExp number of AAs in TMHs: 37.2608499999999\nExp number, first 60 AAs: 19.52923\nTotal prob of N-in: 0.94351\nPOSSIBLE N-term signal sequence\ninside 1 19\nTMhelix 20 39\noutside 40 325\nTMhelix 326 348\ninside 349 372

4496 GCF_002190375.1_ASM219037v1 Escherichia coli Proteobacteria; Gammaproteobacteria; Enterobacterales; Enterobacteriaceae; Escherichia
MTGDNTLIHSHGINRRDFMKLCAALAATMGLSSKAAAEMAESVTNPQRPPVIWIGAQECT WP_000145410.1
MULTISPECIES: hydrogenase-2 small chain [Proteobacteria] Length: 372\nNumber of predicted TMHs: 1\nExp number of AAs in TMHs: 23.72226\nExp number, first 60 AAs: 0.22038\nTotal prob of N-in: 0.12994\noutside 1 331\nTMhelix 332 354\ninside 355 372

4497 GCF_002190005.1_ASM219000v1 Escherichia coli Proteobacteria; Gammaproteobacteria; Enterobacterales; Enterobacteriaceae; Escherichia
MNNEETFYQAMRRQGVTRRSFLKYCSLAATSLGLGAGMAPKIAWALENKPRIPVVWIHGL WP_001058323.1
MULTISPECIES: hydrogenase-1 small chain [Proteobacteria] Length: 372\nNumber of predicted TMHs: 2\nExp number of AAs in TMHs: 37.2608499999999\nExp number, first 60 AAs: 19.52923\nTotal prob of N-in: 0.94351\nPOSSIBLE N-term signal sequence\ninside 1 19\nTMhelix 20 39\noutside 40 325\nTMhelix 326 348\ninside 349 372

4498 GCF_002189825.1_ASM218982v1 Escherichia coli Proteobacteria; Gammaproteobacteria; Enterobacterales; Enterobacteriaceae; Escherichia
MNNEETFYQAMRRQGVTRRSFLKYCSLAATSLGLGAGMAPKIAWALENKPRIPVVWIHGL WP_001058323.1
MULTISPECIES: hydrogenase-1 small chain [Proteobacteria] Length: 372\nNumber of predicted TMHs: 2\nExp number of AAs in TMHs: 37.2608499999999\nExp number, first 60 AAs: 19.52923\nTotal prob of N-in: 0.94351\nPOSSIBLE N-term signal sequence\ninside 1 19\nTMhelix 20 39\noutside 40 325\nTMhelix 326 348\ninside 349 372

4499 GCF_002179775.1_ASM217977v1 Escherichia coli Proteobacteria; Gammaproteobacteria; Enterobacterales; Enterobacteriaceae; Escherichia
MNNEETFYQAMRRQGVTRRSFLKYCSLAATSLGLGAGMAPKIAWALENKPRIPVVWIHGL WP_001301613.1
hydrogenase [Escherichia coli] Length: 372\nNumber of predicted TMHs: 2\nExp number of AAs in TMHs: 37.46598\nExp number, first 60 AAs: 19.53139\nTotal prob of N-in: 0.94364\nPOSSIBLE N-term signal sequence\ninside 1 19\nTMhelix 20 39\noutside 40 325\nTMhelix 326 348\ninside 349 372

4500 GCF_002190075.1_ASM219007v1 Escherichia coli Proteobacteria; Gammaproteobacteria; Enterobacterales; Enterobacteriaceae; Escherichia
MTGDNTLIHSHGINRRDFMKLCAALAATMGLSSKAAAEMAESVTNPQRPPVIWIGAQECT WP_000145410.1
MULTISPECIES: hydrogenase-2 small chain [Proteobacteria] Length: 372\nNumber of predicted TMHs: 1\nExp number of AAs in TMHs: 23.72226\nExp number, first 60 AAs: 0.22038\nTotal prob of N-in: 0.12994\noutside 1 331\nTMhelix 332 354\ninside 355 372

4501 GCF_002190105.1_ASM219010v1 Escherichia coli Proteobacteria; Gammaproteobacteria; Enterobacterales; Enterobacteriaceae; Escherichia
MTGDNTLIHSHGINRRDFMKLCAALAATMGLSSKAAAEMAESVTNPQRPPVIWIGAQECT WP_000145410.1
MULTISPECIES: hydrogenase-2 small chain [Proteobacteria] Length: 372\nNumber of predicted TMHs: 1\nExp number of AAs in TMHs: 23.72226\nExp number, first 60 AAs: 0.22038\nTotal prob of N-in: 0.12994\noutside 1 331\nTMhelix 332 354\ninside 355 372

4502 GCF_002190225.1_ASM219022v1 Escherichia coli Proteobacteria; Gammaproteobacteria; Enterobacterales; Enterobacteriaceae; Escherichia
MTGDNTLIHSHGINRRDFMKLCAALAATMGLSSKAAAEMAESVTNPQRPPVIWIGAQECT WP_000145410.1
MULTISPECIES: hydrogenase-2 small chain [Proteobacteria] Length: 372\nNumber of predicted TMHs: 1\nExp number of AAs in TMHs: 23.72226\nExp number, first 60 AAs: 0.22038\nTotal prob of N-in: 0.12994\noutside 1 331\nTMhelix 332 354\ninside 355 372

4503 GCF_002190165.1_ASM219016v1 Escherichia coli Proteobacteria; Gammaproteobacteria; Enterobacterales; Enterobacteriaceae; Escherichia
MTGDNTLIHSHGINRRDFMKLCAALAATMGLSSKAAAEMAESVTNPQRPPVIWIGAQECT WP_000145410.1
MULTISPECIES: hydrogenase-2 small chain [Proteobacteria] Length: 372\nNumber of predicted TMHs: 1\nExp number of AAs in TMHs: 23.72226\nExp number, first 60 AAs: 0.22038\nTotal prob of N-in: 0.12994\noutside 1 331\nTMhelix 332 354\ninside 355 372

4504 GCF_002190595.1_ASM219059v1 Escherichia coli Proteobacteria; Gammaproteobacteria; Enterobacterales; Enterobacteriaceae; Escherichia
MTGDNTLIHSHGINRRDFMKLCAALAATMGLSSKAAAEMAESVTNPQRPPVIWIGAQECT WP_000145410.1
MULTISPECIES: hydrogenase-2 small chain [Proteobacteria] Length: 372\nNumber of predicted TMHs: 1\nExp number of AAs in TMHs: 23.72226\nExp number, first 60 AAs: 0.22038\nTotal prob of N-in: 0.12994\noutside 1 331\nTMhelix 332 354\ninside 355 372

4505 GCF_002190595.1_ASM219059v1 Escherichia coli Proteobacteria; Gammaproteobacteria; Enterobacterales; Enterobacteriaceae; Escherichia
MNNEETFYQAMRRQGVTRRSFLKYCSLAATSLGLGAGMAPKIAWALENKPRIPVVWIHGL WP_001058323.1

MULTISPECIES: hydrogenase-1 small chain [Proteobacteria]	Length: 372\nNumber of predicted TMHs: 2\nExp number of AAs in TMHs: 37.2608499999999\nExp number, first 60 AAs: 19.52923\nTotal prob of N-in: 0.94351\nPOSSIBLE N-term signal sequence\ninside 1 19\nTMhelix 20 39\noutside 40 325\nTMhelix 326 348\ninside 349 372

4506	GCF_002190535.1_ASM219053v1	Escherichia coli	Proteobacteria; Gammaproteobacteria; Enterobacterales; Enterobacteriaceae; Escherichia
MNNEETFYQAMRRQGVTRRSFLKYCSLAATSLGLGAGMAPKIAWALENKPRIPVVWIHGL	WP_001058323.1
MULTISPECIES: hydrogenase-1 small chain [Proteobacteria]	Length: 372\nNumber of predicted TMHs: 2\nExp number of AAs in TMHs: 37.2608499999999\nExp number, first 60 AAs: 19.52923\nTotal prob of N-in: 0.94351\nPOSSIBLE N-term signal sequence\ninside 1 19\nTMhelix 20 39\noutside 40 325\nTMhelix 326 348\ninside 349 372

4507	GCF_002190475.1_ASM219047v1	Escherichia coli	Proteobacteria; Gammaproteobacteria; Enterobacterales; Enterobacteriaceae; Escherichia
MNNEETFYQAMRRQGVTRRSFLKYCSLAATSLGLGAGMAPKIAWALENKPRIPVVWIHGL	WP_001058323.1
MULTISPECIES: hydrogenase-1 small chain [Proteobacteria]	Length: 372\nNumber of predicted TMHs: 2\nExp number of AAs in TMHs: 37.2608499999999\nExp number, first 60 AAs: 19.52923\nTotal prob of N-in: 0.94351\nPOSSIBLE N-term signal sequence\ninside 1 19\nTMhelix 20 39\noutside 40 325\nTMhelix 326 348\ninside 349 372

4508	GCF_002190075.1_ASM219007v1	Escherichia coli	Proteobacteria; Gammaproteobacteria; Enterobacterales; Enterobacteriaceae; Escherichia
MNNEETFYQAMRRQGVTRRSFLKYCSLAATSLGLGAGMAPKIAWALENKPRIPVVWIHGL	WP_001058320.1
hydrogenase [Escherichia coli]	Length: 372\nNumber of predicted TMHs: 2\nExp number of AAs in TMHs: 37.26082\nExp number, first 60 AAs: 19.52923\nTotal prob of N-in: 0.94351\nPOSSIBLE N-term signal sequence\ninside 1 19\nTMhelix 20 39\noutside 40 325\nTMhelix 326 348\ninside 349 372

4509	GCF_002190165.1_ASM219016v1	Escherichia coli	Proteobacteria; Gammaproteobacteria; Enterobacterales; Enterobacteriaceae; Escherichia
MNNEETFYQAMRRQGVTRRSFLKYCSLAATSLGLGAGMAPKIAWALENKPRIPVVWIHGL	WP_001058320.1
hydrogenase [Escherichia coli]	Length: 372\nNumber of predicted TMHs: 2\nExp number of AAs in TMHs: 37.26082\nExp number, first 60 AAs: 19.52923\nTotal prob of N-in: 0.94351\nPOSSIBLE N-term signal sequence\ninside 1 19\nTMhelix 20 39\noutside 40 325\nTMhelix 326 348\ninside 349 372

4510	GCF_002190625.1_ASM219062v1	Escherichia coli	Proteobacteria; Gammaproteobacteria; Enterobacterales; Enterobacteriaceae; Escherichia
MNNEETFYQAMRRQGVTRRSFLKYCSLAATSLGLGAGMAPKIAWALENKPRIPVVWIHGL	WP_001295940.1
MULTISPECIES: hydrogenase [Enterobacteriaceae]	Length: 372\nNumber of predicted TMHs: 2\nExp number of AAs in TMHs: 37.2608899999999\nExp number, first 60 AAs: 19.52923\nTotal prob of N-in: 0.94351\nPOSSIBLE N-term signal sequence\ninside 1 19\nTMhelix 20 39\noutside 40 325\nTMhelix 326 348\ninside 349 372

4511	GCF_002190705.1_ASM219070v1	Escherichia coli	Proteobacteria; Gammaproteobacteria; Enterobacterales; Enterobacteriaceae; Escherichia
MTGDNTLIHSHGINRRDFMKLCAALAATMGLSSKAAAEMAESVTNPQRPPVIWIGAQECT	WP_000145410.1
MULTISPECIES: hydrogenase-2 small chain [Proteobacteria]	Length: 372\nNumber of predicted TMHs: 1\nExp number of AAs in TMHs: 23.72226\nExp number, first 60 AAs: 0.22038\nTotal prob of N-in: 0.12994\noutside 1 331\nTMhelix 332 354\ninside 355 372

4512	GCF_002190815.1_ASM219081v1	Escherichia coli	Proteobacteria; Gammaproteobacteria; Enterobacterales; Enterobacteriaceae; Escherichia
MTGDNTLIHSHGINRRDFMKLCAALAATMGLSSKAAAEMAESVTNPQRPPVIWIGAQECT	WP_000145410.1
MULTISPECIES: hydrogenase-2 small chain [Proteobacteria]	Length: 372\nNumber of predicted TMHs: 1\nExp number of AAs in TMHs: 23.72226\nExp number, first 60 AAs: 0.22038\nTotal prob of N-in: 0.12994\noutside 1 331\nTMhelix 332 354\ninside 355 372

4513	GCF_002190775.1_ASM219077v1	Escherichia coli	Proteobacteria; Gammaproteobacteria; Enterobacterales; Enterobacteriaceae; Escherichia
MTGDNTLIHSHGINRRDFMKLCAALAATMGLSSKAAAEMAESVTNPQRPPVIWIGAQECT	WP_000145410.1
MULTISPECIES: hydrogenase-2 small chain [Proteobacteria]	Length: 372\nNumber of predicted TMHs:

1\nExp number of AAs in TMHs: 23.72226\nExp number, first 60 AAs: 0.22038\nTotal prob of N-in: 0.12994\noutside 1 331\nTMhelix 332 354\ninside 355 372

4514 GCF_002190875.1_ASM219087v1 Escherichia coli Proteobacteria; Gammaproteobacteria; Enterobacterales; Enterobacteriaceae; Escherichia
MTGDNTLIHSHGINRRDFMKLCAALAATMGLSSKAAAEMAESVTNPQRPPVIWIGAQECT WP_000145410.1
MULTISPECIES: hydrogenase-2 small chain [Proteobacteria] Length: 372\nNumber of predicted TMHs: 1\nExp number of AAs in TMHs: 23.72226\nExp number, first 60 AAs: 0.22038\nTotal prob of N-in: 0.12994\noutside 1 331\nTMhelix 332 354\ninside 355 372

4515 GCF_002190935.1_ASM219093v1 Escherichia coli Proteobacteria; Gammaproteobacteria; Enterobacterales; Enterobacteriaceae; Escherichia
MTGDNTLIHSHGINRRDFMKLCAALAATMGLSSKAAAEMAESVTNPQRPPVIWIGAQECT WP_000145410.1
MULTISPECIES: hydrogenase-2 small chain [Proteobacteria] Length: 372\nNumber of predicted TMHs: 1\nExp number of AAs in TMHs: 23.72226\nExp number, first 60 AAs: 0.22038\nTotal prob of N-in: 0.12994\noutside 1 331\nTMhelix 332 354\ninside 355 372

4516 GCF_002190975.1_ASM219097v1 Escherichia coli Proteobacteria; Gammaproteobacteria; Enterobacterales; Enterobacteriaceae; Escherichia
MTGDNTLIHSHGINRRDFMKLCAALAATMGLSSKAAAEMAESVTNPQRPPVIWIGAQECT WP_000145410.1
MULTISPECIES: hydrogenase-2 small chain [Proteobacteria] Length: 372\nNumber of predicted TMHs: 1\nExp number of AAs in TMHs: 23.72226\nExp number, first 60 AAs: 0.22038\nTotal prob of N-in: 0.12994\noutside 1 331\nTMhelix 332 354\ninside 355 372

4517 GCF_002193165.1_ASM219316v1 Escherichia coli Proteobacteria; Gammaproteobacteria; Enterobacterales; Enterobacteriaceae; Escherichia
MTGDNTLIHSHGINRRDFMKLCAALAATMGLSSKAAAEMAESVTNPQRPPVIWIGAQECT WP_000145410.1
MULTISPECIES: hydrogenase-2 small chain [Proteobacteria] Length: 372\nNumber of predicted TMHs: 1\nExp number of AAs in TMHs: 23.72226\nExp number, first 60 AAs: 0.22038\nTotal prob of N-in: 0.12994\noutside 1 331\nTMhelix 332 354\ninside 355 372

4518 GCF_002193165.1_ASM219316v1 Escherichia coli Proteobacteria; Gammaproteobacteria; Enterobacterales; Enterobacteriaceae; Escherichia
MNNEETFYQAMRRQGVTRRSFLKYCSLAATSLGLGAGMAPKIAWALENKPRIPVVWIHGL WP_001058323.1
MULTISPECIES: hydrogenase-1 small chain [Proteobacteria] Length: 372\nNumber of predicted TMHs: 2\nExp number of AAs in TMHs: 37.2608499999999\nExp number, first 60 AAs: 19.52923\nTotal prob of N-in: 0.94351\nPOSSIBLE N-term signal sequence\ninside 1 19\nTMhelix 20 39\noutside 40 325\nTMhelix 326 348\ninside 349 372

4519 GCF_002190875.1_ASM219087v1 Escherichia coli Proteobacteria; Gammaproteobacteria; Enterobacterales; Enterobacteriaceae; Escherichia
MNNEETFYQAMRRQGVTRRSFLKYCSLAATSLGLGAGMAPKIAWALENKPRIPVVWIHGL WP_001295940.1
MULTISPECIES: hydrogenase [Enterobacteriaceae] Length: 372\nNumber of predicted TMHs: 2\nExp number of AAs in TMHs: 37.2608899999999\nExp number, first 60 AAs: 19.52923\nTotal prob of N-in: 0.94351\nPOSSIBLE N-term signal sequence\ninside 1 19\nTMhelix 20 39\noutside 40 325\nTMhelix 326 348\ninside 349 372

4520 GCF_002193205.1_ASM219320v1 Escherichia coli Proteobacteria; Gammaproteobacteria; Enterobacterales; Enterobacteriaceae; Escherichia
MTGDNTLIHSHGINRRDFMKLCAALAATMGLSSKAAAEMAESVTNPQRPPVIWIGAQECT WP_000145410.1
MULTISPECIES: hydrogenase-2 small chain [Proteobacteria] Length: 372\nNumber of predicted TMHs: 1\nExp number of AAs in TMHs: 23.72226\nExp number, first 60 AAs: 0.22038\nTotal prob of N-in: 0.12994\noutside 1 331\nTMhelix 332 354\ninside 355 372

4521 GCF_002193275.1_ASM219327v1 Escherichia coli Proteobacteria; Gammaproteobacteria; Enterobacterales; Enterobacteriaceae; Escherichia
MTGDNTLIHSHGINRRDFMKLCAALAATMGLSSKAAAEMAESVTNPQRPPVIWIGAQECT WP_000145410.1
MULTISPECIES: hydrogenase-2 small chain [Proteobacteria] Length: 372\nNumber of predicted TMHs: 1\nExp number of AAs in TMHs: 23.72226\nExp number, first 60 AAs: 0.22038\nTotal prob of N-in: 0.12994\noutside 1 331\nTMhelix 332 354\ninside 355 372

4522	GCF_002193315.1_ASM219331v1	Escherichia coli	Proteobacteria; Gammaproteobacteria; Enterobacterales; Enterobacteriaceae; Escherichia
	MTGDNTLIHSHGINRRDFMKLCAALAATMGLSSKAAAEMAESVTNPQRPPVIWIGAQECT	WP_000145410.1
	MULTISPECIES: hydrogenase-2 small chain [Proteobacteria]	Length: 372\nNumber of predicted TMHs: 1\nExp number of AAs in TMHs: 23.72226\nExp number, first 60 AAs: 0.22038\nTotal prob of N-in: 0.12994\noutside 1 331\nTMhelix 332 354\ninside 355 372

4523	GCF_002193475.1_ASM219347v1	Escherichia coli	Proteobacteria; Gammaproteobacteria; Enterobacterales; Enterobacteriaceae; Escherichia
	MTGDNTLIHSHGINRRDFMKLCAALAATMGLSSKAAAEMAESVTNPQRPPVIWIGAQECT	WP_000145410.1
	MULTISPECIES: hydrogenase-2 small chain [Proteobacteria]	Length: 372\nNumber of predicted TMHs: 1\nExp number of AAs in TMHs: 23.72226\nExp number, first 60 AAs: 0.22038\nTotal prob of N-in: 0.12994\noutside 1 331\nTMhelix 332 354\ninside 355 372

4524	GCF_002193315.1_ASM219331v1	Escherichia coli	Proteobacteria; Gammaproteobacteria; Enterobacterales; Enterobacteriaceae; Escherichia
	MNNEETFYQAMRRQGVTRRSFLKYCSLAATSLGLGAGMAPKIAWALENKPRIPVVWIHGL	WP_001058323.1
	MULTISPECIES: hydrogenase-1 small chain [Proteobacteria]	Length: 372\nNumber of predicted TMHs: 2\nExp number of AAs in TMHs: 37.2608499999999\nExp number, first 60 AAs: 19.52923\nTotal prob of N-in: 0.94351\nPOSSIBLE N-term signal sequence\ninside 1 19\nTMhelix 20 39\noutside 40 325\nTMhelix 326 348\ninside 349 372

4525	GCF_002193495.1_ASM219349v1	Escherichia coli	Proteobacteria; Gammaproteobacteria; Enterobacterales; Enterobacteriaceae; Escherichia
	MNNEETFYQAMRRQGVTRRSFLKYCSLAATSLGLGAGMAPKIAWALENKPRIPVVWIHGL	WP_001058323.1
	MULTISPECIES: hydrogenase-1 small chain [Proteobacteria]	Length: 372\nNumber of predicted TMHs: 2\nExp number of AAs in TMHs: 37.2608499999999\nExp number, first 60 AAs: 19.52923\nTotal prob of N-in: 0.94351\nPOSSIBLE N-term signal sequence\ninside 1 19\nTMhelix 20 39\noutside 40 325\nTMhelix 326 348\ninside 349 372

4526	GCF_002194645.1_ASM219464v1	Escherichia coli	Proteobacteria; Gammaproteobacteria; Enterobacterales; Enterobacteriaceae; Escherichia
	MTGDNTLIHSHGINRRDFMKLCAALAATMGLSSKAAAEMAESVTNPQRPPVIWIGAQECT	WP_000145410.1
	MULTISPECIES: hydrogenase-2 small chain [Proteobacteria]	Length: 372\nNumber of predicted TMHs: 1\nExp number of AAs in TMHs: 23.72226\nExp number, first 60 AAs: 0.22038\nTotal prob of N-in: 0.12994\noutside 1 331\nTMhelix 332 354\ninside 355 372

4527	GCF_002194645.1_ASM219464v1	Escherichia coli	Proteobacteria; Gammaproteobacteria; Enterobacterales; Enterobacteriaceae; Escherichia
	MNNEETFYQAMRRQGVTRRSFLKYCSLAATSLGLGAGMAPKIAWALENKPRIPVVWIHGL	WP_001301613.1
	hydrogenase [Escherichia coli]	Length: 372\nNumber of predicted TMHs: 2\nExp number of AAs in TMHs: 37.46598\nExp number, first 60 AAs: 19.53139\nTotal prob of N-in: 0.94364\nPOSSIBLE N-term signal sequence\ninside 1 19\nTMhelix 20 39\noutside 40 325\nTMhelix 326 348\ninside 349 372

4528	GCF_002193205.1_ASM219320v1	Escherichia coli	Proteobacteria; Gammaproteobacteria; Enterobacterales; Enterobacteriaceae; Escherichia
	MNNEETFYQAMRRQGVTRRSFLKYCSLAATSLGLGAGMAPKIAWALENKPRIPVVWIHGL	WP_001295940.1
	MULTISPECIES: hydrogenase [Enterobacteriaceae]	Length: 372\nNumber of predicted TMHs: 2\nExp number of AAs in TMHs: 37.2608899999999\nExp number, first 60 AAs: 19.52923\nTotal prob of N-in: 0.94351\nPOSSIBLE N-term signal sequence\ninside 1 19\nTMhelix 20 39\noutside 40 325\nTMhelix 326 348\ninside 349 372

4529	GCF_002194695.1_ASM219469v1	Escherichia coli	Proteobacteria; Gammaproteobacteria; Enterobacterales; Enterobacteriaceae; Escherichia
	MTGDNTLIHSHGINRRDFMKLCAALAATMGLSSKAAAEMAESVTNPQRPPVIWIGAQECT	WP_000145410.1
	MULTISPECIES: hydrogenase-2 small chain [Proteobacteria]	Length: 372\nNumber of predicted TMHs: 1\nExp number of AAs in TMHs: 23.72226\nExp number, first 60 AAs: 0.22038\nTotal prob of N-in: 0.12994\noutside 1 331\nTMhelix 332 354\ninside 355 372

4530	GCF_002194755.1_ASM219475v1	Escherichia coli	Proteobacteria; Gammaproteobacteria; Enterobacterales; Enterobacteriaceae; Escherichia

MTGDNTLIHSHGINRRDFMKLCAALAATMGLSSKAAAEMAESVTNPQRPPVIWIGAQECT WP_000145410.1 MULTISPECIES: hydrogenase-2 small chain [Proteobacteria] Length: 372\nNumber of predicted TMHs: 1\nExp number of AAs in TMHs: 23.72226\nExp number, first 60 AAs: 0.22038\nTotal prob of N-in: 0.12994\noutside 1 331\nTMhelix 332 354\ninside 355 372

4531 GCF_002194955.1_ASM219495v1 Escherichia coli Proteobacteria; Gammaproteobacteria; Enterobacterales; Enterobacteriaceae; Escherichia MTGDNTLIHSHGINRRDFMKLCAALAATMGLSSKAAAEMAESVTNPQRPPVIWIGAQECT WP_000145410.1 MULTISPECIES: hydrogenase-2 small chain [Proteobacteria] Length: 372\nNumber of predicted TMHs: 1\nExp number of AAs in TMHs: 23.72226\nExp number, first 60 AAs: 0.22038\nTotal prob of N-in: 0.12994\noutside 1 331\nTMhelix 332 354\ninside 355 372

4532 GCF_002195045.1_ASM219504v1 Escherichia coli Proteobacteria; Gammaproteobacteria; Enterobacterales; Enterobacteriaceae; Escherichia MTGDNTLIHSHGINRRDFMKLCAALAATMGLSSKAAAEMAESVTNPQRPPVIWIGAQECT WP_000145410.1 MULTISPECIES: hydrogenase-2 small chain [Proteobacteria] Length: 372\nNumber of predicted TMHs: 1\nExp number of AAs in TMHs: 23.72226\nExp number, first 60 AAs: 0.22038\nTotal prob of N-in: 0.12994\noutside 1 331\nTMhelix 332 354\ninside 355 372

4533 GCF_002195015.1_ASM219501v1 Escherichia coli Proteobacteria; Gammaproteobacteria; Enterobacterales; Enterobacteriaceae; Escherichia MTGDNTLIHSHGINRRDFMKLCAALAATMGLSSKAAAEMAESVTNPQRPPVIWIGAQECT WP_000145410.1 MULTISPECIES: hydrogenase-2 small chain [Proteobacteria] Length: 372\nNumber of predicted TMHs: 1\nExp number of AAs in TMHs: 23.72226\nExp number, first 60 AAs: 0.22038\nTotal prob of N-in: 0.12994\noutside 1 331\nTMhelix 332 354\ninside 355 372

4534 GCF_002195825.1_ASM219582v1 Escherichia coli Proteobacteria; Gammaproteobacteria; Enterobacterales; Enterobacteriaceae; Escherichia MTGDNTLIHSHGINRRDFMKLCAALAATMGLSSKAAAEMAESVTNPQRPPVIWIGAQECT WP_000145410.1 MULTISPECIES: hydrogenase-2 small chain [Proteobacteria] Length: 372\nNumber of predicted TMHs: 1\nExp number of AAs in TMHs: 23.72226\nExp number, first 60 AAs: 0.22038\nTotal prob of N-in: 0.12994\noutside 1 331\nTMhelix 332 354\ninside 355 372

4535 GCF_002194815.1_ASM219481v1 Escherichia coli Proteobacteria; Gammaproteobacteria; Enterobacterales; Enterobacteriaceae; Escherichia MNNEETFYQAMRRQGVTRRSFLKYCSLAATSLGLGAGMAPKIAWALENKPRIPVVWIHGL WP_001301613.1 hydrogenase [Escherichia coli] Length: 372\nNumber of predicted TMHs: 2\nExp number of AAs in TMHs: 37.46598\nExp number, first 60 AAs: 19.53139\nTotal prob of N-in: 0.94364\nPOSSIBLE N-term signal sequence\ninside 1 19\nTMhelix 20 39\noutside 40 325\nTMhelix 326 348\ninside 349 372

4536 GCF_002195115.1_ASM219511v1 Escherichia coli Proteobacteria; Gammaproteobacteria; Enterobacterales; Enterobacteriaceae; Escherichia MTGDNTLIHSHGINRRDFMKLCAALAATMGLSSKAAAEMAESVTNPQRPPVIWIGAQECT WP_000145410.1 MULTISPECIES: hydrogenase-2 small chain [Proteobacteria] Length: 372\nNumber of predicted TMHs: 1\nExp number of AAs in TMHs: 23.72226\nExp number, first 60 AAs: 0.22038\nTotal prob of N-in: 0.12994\noutside 1 331\nTMhelix 332 354\ninside 355 372

4537 GCF_002195135.1_ASM219513v1 Escherichia coli Proteobacteria; Gammaproteobacteria; Enterobacterales; Enterobacteriaceae; Escherichia MTGDNTLIHSHGINRRDFMKLCAALAATMGLSSKAAAEMAESVTNPQRPPVIWIGAQECT WP_000145410.1 MULTISPECIES: hydrogenase-2 small chain [Proteobacteria] Length: 372\nNumber of predicted TMHs: 1\nExp number of AAs in TMHs: 23.72226\nExp number, first 60 AAs: 0.22038\nTotal prob of N-in: 0.12994\noutside 1 331\nTMhelix 332 354\ninside 355 372

4538 GCF_002195255.1_ASM219525v1 Escherichia coli Proteobacteria; Gammaproteobacteria; Enterobacterales; Enterobacteriaceae; Escherichia MTGDNTLIHSHGINRRDFMKLCAALAATMGLSSKAAAEMAESVTNPQRPPVIWIGAQECT WP_000145410.1 MULTISPECIES: hydrogenase-2 small chain [Proteobacteria] Length: 372\nNumber of predicted TMHs: 1\nExp number of AAs in TMHs: 23.72226\nExp number, first 60 AAs: 0.22038\nTotal prob of N-in: 0.12994\noutside 1 331\nTMhelix 332 354\ninside 355 372

4539 GCF_002194915.1_ASM219491v1 Escherichia coli Proteobacteria; Gammaproteobacteria; Enterobacterales; Enterobacteriaceae; Escherichia
MNNEETFYQAMRRQGVTRRSFLKYCSLAATSLGLGAGMAPKIAWALENKPRIPVVWIHGL WP_001301613.1
hydrogenase [Escherichia coli] Length: 372\nNumber of predicted TMHs: 2\nExp number of AAs in TMHs: 37.46598\nExp number, first 60 AAs: 19.53139\nTotal prob of N-in: 0.94364\nPOSSIBLE N-term signal sequence\ninside 1 19\nTMhelix 20 39\noutside 40 325\nTMhelix 326 348\ninside 349 372

4540 GCF_900011015.1_ED680_contigs Escherichia coli Proteobacteria; Gammaproteobacteria; Enterobacterales; Enterobacteriaceae; Escherichia
MTGDNTLIHSHGINRRDFMKLCAALAATMGLSSKAAAEMAESVTNPQRPPVIWIGAQECT WP_000145410.1
MULTISPECIES: hydrogenase-2 small chain [Proteobacteria] Length: 372\nNumber of predicted TMHs: 1\nExp number of AAs in TMHs: 23.72226\nExp number, first 60 AAs: 0.22038\nTotal prob of N-in: 0.12994\noutside 1 331\nTMhelix 332 354\ninside 355 372

4541 GCF_900011015.1_ED680_contigs Escherichia coli Proteobacteria; Gammaproteobacteria; Enterobacterales; Enterobacteriaceae; Escherichia
MNNEETFYQAMRRQGVTRRSFLKYCSLAATSLGLGAGMAPKIAWALENKPRIPVVWIHGL WP_001301613.1
hydrogenase [Escherichia coli] Length: 372\nNumber of predicted TMHs: 2\nExp number of AAs in TMHs: 37.46598\nExp number, first 60 AAs: 19.53139\nTotal prob of N-in: 0.94364\nPOSSIBLE N-term signal sequence\ninside 1 19\nTMhelix 20 39\noutside 40 325\nTMhelix 326 348\ninside 349 372

4542 GCF_002194955.1_ASM219495v1 Escherichia coli Proteobacteria; Gammaproteobacteria; Enterobacterales; Enterobacteriaceae; Escherichia
MNNEETFYQAMRRQGVTRRSFLKYCSLAATSLGLGAGMAPKIAWALENKPRIPVVWIHGL WP_001301613.1
hydrogenase [Escherichia coli] Length: 372\nNumber of predicted TMHs: 2\nExp number of AAs in TMHs: 37.46598\nExp number, first 60 AAs: 19.53139\nTotal prob of N-in: 0.94364\nPOSSIBLE N-term signal sequence\ninside 1 19\nTMhelix 20 39\noutside 40 325\nTMhelix 326 348\ninside 349 372

4543 GCF_002195045.1_ASM219504v1 Escherichia coli Proteobacteria; Gammaproteobacteria; Enterobacterales; Enterobacteriaceae; Escherichia
MNNEETFYQAMRRQGVTRRSFLKYCSLAATSLGLGAGMAPKIAWALENKPRIPVVWIHGL WP_001301613.1
hydrogenase [Escherichia coli] Length: 372\nNumber of predicted TMHs: 2\nExp number of AAs in TMHs: 37.46598\nExp number, first 60 AAs: 19.53139\nTotal prob of N-in: 0.94364\nPOSSIBLE N-term signal sequence\ninside 1 19\nTMhelix 20 39\noutside 40 325\nTMhelix 326 348\ninside 349 372

4544 GCF_002195015.1_ASM219501v1 Escherichia coli Proteobacteria; Gammaproteobacteria; Enterobacterales; Enterobacteriaceae; Escherichia
MNNEETFYQAMRRQGVTRRSFLKYCSLAATSLGLGAGMAPKIAWALENKPRIPVVWIHGL WP_001301613.1
hydrogenase [Escherichia coli] Length: 372\nNumber of predicted TMHs: 2\nExp number of AAs in TMHs: 37.46598\nExp number, first 60 AAs: 19.53139\nTotal prob of N-in: 0.94364\nPOSSIBLE N-term signal sequence\ninside 1 19\nTMhelix 20 39\noutside 40 325\nTMhelix 326 348\ninside 349 372

4545 GCF_002195395.1_ASM219539v1 Escherichia coli Proteobacteria; Gammaproteobacteria; Enterobacterales; Enterobacteriaceae; Escherichia
MTGDNTLIHSHGINRRDFMKLCAALAATMGLSSKAAAEMAESVTNPQRPPVIWIGAQECT WP_000145410.1
MULTISPECIES: hydrogenase-2 small chain [Proteobacteria] Length: 372\nNumber of predicted TMHs: 1\nExp number of AAs in TMHs: 23.72226\nExp number, first 60 AAs: 0.22038\nTotal prob of N-in: 0.12994\noutside 1 331\nTMhelix 332 354\ninside 355 372

4546 GCF_002195485.1_ASM219548v1 Escherichia coli Proteobacteria; Gammaproteobacteria; Enterobacterales; Enterobacteriaceae; Escherichia
MTGDNTLIHSHGINRRDFMKLCAALAATMGLSSKAAAEMAESVTNPQRPPVIWIGAQECT WP_000145410.1
MULTISPECIES: hydrogenase-2 small chain [Proteobacteria] Length: 372\nNumber of predicted TMHs: 1\nExp number of AAs in TMHs: 23.72226\nExp number, first 60 AAs: 0.22038\nTotal prob of N-in: 0.12994\noutside 1 331\nTMhelix 332 354\ninside 355 372

4547 GCF_002195535.1_ASM219553v1 Escherichia coli Proteobacteria; Gammaproteobacteria; Enterobacterales; Enterobacteriaceae; Escherichia
MTGDNTLIHSHGINRRDFMKLCAALAATMGLSSKAAAEMAESVTNPQRPPVIWIGAQECT WP_000145410.1
MULTISPECIES: hydrogenase-2 small chain [Proteobacteria] Length: 372\nNumber of predicted TMHs:

1\nExp number of AAs in TMHs: 23.72226\nExp number, first 60 AAs: 0.22038\nTotal prob of N-in: 0.12994\noutside 1 331\nTMhelix 332 354\ninside 355 372

4548 GCF_002195595.1_ASM219559v1 Escherichia coli Proteobacteria; Gammaproteobacteria; Enterobacterales; Enterobacteriaceae; Escherichia
MTGDNTLIHSHGINRRDFMKLCAALAATMGLSSKAAAEMAESVTNPQRPPVIWIGAQECT WP_000145410.1
MULTISPECIES: hydrogenase-2 small chain [Proteobacteria] Length: 372\nNumber of predicted TMHs: 1\nExp number of AAs in TMHs: 23.72226\nExp number, first 60 AAs: 0.22038\nTotal prob of N-in: 0.12994\noutside 1 331\nTMhelix 332 354\ninside 355 372

4549 GCF_002195625.1_ASM219562v1 Escherichia coli Proteobacteria; Gammaproteobacteria; Enterobacterales; Enterobacteriaceae; Escherichia
MTGDNTLIHSHGINRRDFMKLCAALAATMGLSSKAAAEMAESVTNPQRPPVIWIGAQECT WP_000145410.1
MULTISPECIES: hydrogenase-2 small chain [Proteobacteria] Length: 372\nNumber of predicted TMHs: 1\nExp number of AAs in TMHs: 23.72226\nExp number, first 60 AAs: 0.22038\nTotal prob of N-in: 0.12994\noutside 1 331\nTMhelix 332 354\ninside 355 372

4550 GCF_002195695.1_ASM219569v1 Escherichia coli Proteobacteria; Gammaproteobacteria; Enterobacterales; Enterobacteriaceae; Escherichia
MTGDNTLIHSHGINRRDFMKLCAALAATMGLSSKAAAEMAESVTNPQRPPVIWIGAQECT WP_000145410.1
MULTISPECIES: hydrogenase-2 small chain [Proteobacteria] Length: 372\nNumber of predicted TMHs: 1\nExp number of AAs in TMHs: 23.72226\nExp number, first 60 AAs: 0.22038\nTotal prob of N-in: 0.12994\noutside 1 331\nTMhelix 332 354\ninside 355 372

4551 GCF_002195795.1_ASM219579v1 Escherichia coli Proteobacteria; Gammaproteobacteria; Enterobacterales; Enterobacteriaceae; Escherichia
MTGDNTLIHSHGINRRDFMKLCAALAATMGLSSKAAAEMAESVTNPQRPPVIWIGAQECT WP_000145410.1
MULTISPECIES: hydrogenase-2 small chain [Proteobacteria] Length: 372\nNumber of predicted TMHs: 1\nExp number of AAs in TMHs: 23.72226\nExp number, first 60 AAs: 0.22038\nTotal prob of N-in: 0.12994\noutside 1 331\nTMhelix 332 354\ninside 355 372

4552 GCF_002195745.1_ASM219574v1 Escherichia coli Proteobacteria; Gammaproteobacteria; Enterobacterales; Enterobacteriaceae; Escherichia
MTGDNTLIHSHGINRRDFMKLCAALAATMGLSSKAAAEMAESVTNPQRPPVIWIGAQECT WP_000145410.1
MULTISPECIES: hydrogenase-2 small chain [Proteobacteria] Length: 372\nNumber of predicted TMHs: 1\nExp number of AAs in TMHs: 23.72226\nExp number, first 60 AAs: 0.22038\nTotal prob of N-in: 0.12994\noutside 1 331\nTMhelix 332 354\ninside 355 372

4553 GCF_002195895.1_ASM219589v1 Escherichia coli Proteobacteria; Gammaproteobacteria; Enterobacterales; Enterobacteriaceae; Escherichia
MTGDNTLIHSHGINRRDFMKLCAALAATMGLSSKAAAEMAESVTNPQRPPVIWIGAQECT WP_000145410.1
MULTISPECIES: hydrogenase-2 small chain [Proteobacteria] Length: 372\nNumber of predicted TMHs: 1\nExp number of AAs in TMHs: 23.72226\nExp number, first 60 AAs: 0.22038\nTotal prob of N-in: 0.12994\noutside 1 331\nTMhelix 332 354\ninside 355 372

4554 GCF_002196625.1_ASM219662v1 Escherichia coli Proteobacteria; Gammaproteobacteria; Enterobacterales; Enterobacteriaceae; Escherichia
MNNEETFYQAMRRQGVTRRSFLKYCSLAATSLGLGAGMAPKIAWALENKPRIPVVWIHGL WP_001058323.1
MULTISPECIES: hydrogenase-1 small chain [Proteobacteria] Length: 372\nNumber of predicted TMHs: 2\nExp number of AAs in TMHs: 37.2608499999999\nExp number, first 60 AAs: 19.52923\nTotal prob of N-in: 0.94351\nPOSSIBLE N-term signal sequence\ninside 1 19\nTMhelix 20 39\noutside 40 325\nTMhelix 326 348\ninside 349 372

4555 GCF_002195625.1_ASM219562v1 Escherichia coli Proteobacteria; Gammaproteobacteria; Enterobacterales; Enterobacteriaceae; Escherichia
MNNEETFYQAMRRQGVTRRSFLKYCSLAATSLGLGAGMAPKIAWALENKPRIPVVWIHGL WP_001301613.1
hydrogenase [Escherichia coli] Length: 372\nNumber of predicted TMHs: 2\nExp number of AAs in TMHs: 37.46598\nExp number, first 60 AAs: 19.53139\nTotal prob of N-in: 0.94364\nPOSSIBLE N-term signal sequence\ninside 1 19\nTMhelix 20 39\noutside 40 325\nTMhelix 326 348\ninside 349 372

4556 GCF_002195745.1_ASM219574v1 Escherichia coli Proteobacteria; Gammaproteobacteria; Enterobacterales; Enterobacteriaceae; Escherichia
MNNEETFYQAMRRQGVTRRSFLKYCSLAATSLGLGAGMAPKIAWALENKPRIPVVWIHGL WP_001301613.1
hydrogenase [Escherichia coli] Length: 372\nNumber of predicted TMHs: 2\nExp number of AAs in TMHs: 37.46598\nExp number, first 60 AAs: 19.53139\nTotal prob of N-in: 0.94364\nPOSSIBLE N-term signal sequence\ninside 1 19\nTMhelix 20 39\noutside 40 325\nTMhelix 326 348\ninside 349 372

4557 GCF_002195695.1_ASM219569v1 Escherichia coli Proteobacteria; Gammaproteobacteria; Enterobacterales; Enterobacteriaceae; Escherichia
MNNEETFYQAMRRQGVTRRSFLKYCSLAATSLGLGAGMAPKIAWALENKPRIPVVWIHGL WP_001301613.1
hydrogenase [Escherichia coli] Length: 372\nNumber of predicted TMHs: 2\nExp number of AAs in TMHs: 37.46598\nExp number, first 60 AAs: 19.53139\nTotal prob of N-in: 0.94364\nPOSSIBLE N-term signal sequence\ninside 1 19\nTMhelix 20 39\noutside 40 325\nTMhelix 326 348\ninside 349 372

4558 GCF_002195795.1_ASM219579v1 Escherichia coli Proteobacteria; Gammaproteobacteria; Enterobacterales; Enterobacteriaceae; Escherichia
MNNEETFYQAMRRQGVTRRSFLKYCSLAATSLGLGAGMAPKIAWALENKPRIPVVWIHGL WP_001301613.1
hydrogenase [Escherichia coli] Length: 372\nNumber of predicted TMHs: 2\nExp number of AAs in TMHs: 37.46598\nExp number, first 60 AAs: 19.53139\nTotal prob of N-in: 0.94364\nPOSSIBLE N-term signal sequence\ninside 1 19\nTMhelix 20 39\noutside 40 325\nTMhelix 326 348\ninside 349 372

4559 GCF_002196645.1_ASM219664v1 Escherichia coli Proteobacteria; Gammaproteobacteria; Enterobacterales; Enterobacteriaceae; Escherichia
MTGDNTLIHSHGINRRDFMKLCAALAATMGLSSKAAAEMAESVTNPQRPPVIWIGAQECT WP_000145410.1
MULTISPECIES: hydrogenase-2 small chain [Proteobacteria] Length: 372\nNumber of predicted TMHs: 1\nExp number of AAs in TMHs: 23.72226\nExp number, first 60 AAs: 0.22038\nTotal prob of N-in: 0.12994\noutside 1 331\nTMhelix 332 354\ninside 355 372

4560 GCF_002196645.1_ASM219664v1 Escherichia coli Proteobacteria; Gammaproteobacteria; Enterobacterales; Enterobacteriaceae; Escherichia
MNNEETFYQAMRRQGVTRRSFLKYCSLAATSLGLGAGMAPKIAWALENKPRIPVVWIHGL WP_001058323.1
MULTISPECIES: hydrogenase-1 small chain [Proteobacteria] Length: 372\nNumber of predicted TMHs: 2\nExp number of AAs in TMHs: 37.2608499999999\nExp number, first 60 AAs: 19.52923\nTotal prob of N-in: 0.94351\nPOSSIBLE N-term signal sequence\ninside 1 19\nTMhelix 20 39\noutside 40 325\nTMhelix 326 348\ninside 349 372

4561 GCF_900008835.1_ED195_contigs Escherichia coli Proteobacteria; Gammaproteobacteria; Enterobacterales; Enterobacteriaceae; Escherichia
MTGDNTLIHSHGINRRDFMKLCAALAATMGLSSKAAAEMAESVTNPQRPPVIWIGAQECT WP_000145410.1
MULTISPECIES: hydrogenase-2 small chain [Proteobacteria] Length: 372\nNumber of predicted TMHs: 1\nExp number of AAs in TMHs: 23.72226\nExp number, first 60 AAs: 0.22038\nTotal prob of N-in: 0.12994\noutside 1 331\nTMhelix 332 354\ninside 355 372

4562 GCF_900008835.1_ED195_contigs Escherichia coli Proteobacteria; Gammaproteobacteria; Enterobacterales; Enterobacteriaceae; Escherichia
MNNEETFYQAMRRQGVTRRSFLKYCSLAATSLGLGAGMAPKIAWALENKPRIPVVWIHGL WP_001058323.1
MULTISPECIES: hydrogenase-1 small chain [Proteobacteria] Length: 372\nNumber of predicted TMHs: 2\nExp number of AAs in TMHs: 37.2608499999999\nExp number, first 60 AAs: 19.52923\nTotal prob of N-in: 0.94351\nPOSSIBLE N-term signal sequence\ninside 1 19\nTMhelix 20 39\noutside 40 325\nTMhelix 326 348\ninside 349 372

4563 GCF_002205095.1_ASM220509v1 Escherichia coli Proteobacteria; Gammaproteobacteria; Enterobacterales; Enterobacteriaceae; Escherichia
MNNEETFYQAMRRQGVTRRSFLKYCSLAATSLGLGAGMAPKIAWALENKPRIPVVWIHGL WP_001058323.1
MULTISPECIES: hydrogenase-1 small chain [Proteobacteria] Length: 372\nNumber of predicted TMHs: 2\nExp number of AAs in TMHs: 37.2608499999999\nExp number, first 60 AAs: 19.52923\nTotal prob of N-in: 0.94351\nPOSSIBLE N-term signal sequence\ninside 1 19\nTMhelix 20 39\noutside 40 325\nTMhelix 326 348\ninside 349 372

4564 GCF_002196745.1_ASM219674v1 Escherichia coli Proteobacteria; Gammaproteobacteria; Enterobacterales; Enterobacteriaceae; Escherichia

MNNEETFYQAMRRQGVTRRSFLKYCSLAATSLGLGAGMAPKIAWALENKPRIPVVWIHGL WP_001058323.1 MULTISPECIES: hydrogenase-1 small chain [Proteobacteria] Length: 372\nNumber of predicted TMHs: 2\nExp number of AAs in TMHs: 37.2608499999999\nExp number, first 60 AAs: 19.52923\nTotal prob of N-in: 0.94351\nPOSSIBLE N-term signal sequence\ninside 1 19\nTMhelix 20 39\noutside 40 325\nTMhelix 326 348\ninside 349 372

4565 GCF_002201835.1_ASM220183v1 Escherichia coli Proteobacteria; Gammaproteobacteria; Enterobacterales; Enterobacteriaceae; Escherichia MNNEETFYQAMRRQGVTRRSFLKYCSLAATSLGLGAGMAPKIAWALENKPRIPVVWIHGL WP_001058323.1 MULTISPECIES: hydrogenase-1 small chain [Proteobacteria] Length: 372\nNumber of predicted TMHs: 2\nExp number of AAs in TMHs: 37.2608499999999\nExp number, first 60 AAs: 19.52923\nTotal prob of N-in: 0.94351\nPOSSIBLE N-term signal sequence\ninside 1 19\nTMhelix 20 39\noutside 40 325\nTMhelix 326 348\ninside 349 372

4566 GCF_002201835.1_ASM220183v1 Escherichia coli Proteobacteria; Gammaproteobacteria; Enterobacterales; Enterobacteriaceae; Escherichia MTGDNTLIHSHGINRRDFMKLCAALATTMGLSSKAAAEMAESVTNPQRPPVIWIGAQECT WP_023149629.1 MULTISPECIES: hydrogenase-2 small chain [Enterobacteriaceae] Length: 372\nNumber of predicted TMHs: 1\nExp number of AAs in TMHs: 23.63603\nExp number, first 60 AAs: 0.12971\nTotal prob of N-in: 0.12608\noutside 1 331\nTMhelix 332 354\ninside 355 372

4567 GCF_002196825.1_ASM219682v1 Escherichia coli Proteobacteria; Gammaproteobacteria; Enterobacterales; Enterobacteriaceae; Escherichia MNNEETFYQAMRRQGVTRRSFLKYCSLAATSLGLGAGMAPKIAWALENKPRIPVVWIHGL WP_001058323.1 MULTISPECIES: hydrogenase-1 small chain [Proteobacteria] Length: 372\nNumber of predicted TMHs: 2\nExp number of AAs in TMHs: 37.2608499999999\nExp number, first 60 AAs: 19.52923\nTotal prob of N-in: 0.94351\nPOSSIBLE N-term signal sequence\ninside 1 19\nTMhelix 20 39\noutside 40 325\nTMhelix 326 348\ninside 349 372

4568 GCF_900015195.1_ED666_contigs Escherichia coli Proteobacteria; Gammaproteobacteria; Enterobacterales; Enterobacteriaceae; Escherichia MTGDNTLIHSHGINRRDFMKLCAALAATMGLSSKAAAEMAESVTNPQRPPVIWIGAQECT WP_000145410.1 MULTISPECIES: hydrogenase-2 small chain [Proteobacteria] Length: 372\nNumber of predicted TMHs: 1\nExp number of AAs in TMHs: 23.72226\nExp number, first 60 AAs: 0.22038\nTotal prob of N-in: 0.12994\noutside 1 331\nTMhelix 332 354\ninside 355 372

4569 GCF_900015725.1_ED604_contigs Escherichia coli Proteobacteria; Gammaproteobacteria; Enterobacterales; Enterobacteriaceae; Escherichia MTGDNTLIHSHGINRRDFMKLCAALAATMGLSSKAAAEMAESVTNPQRPPVIWIGAQECT WP_000145410.1 MULTISPECIES: hydrogenase-2 small chain [Proteobacteria] Length: 372\nNumber of predicted TMHs: 1\nExp number of AAs in TMHs: 23.72226\nExp number, first 60 AAs: 0.22038\nTotal prob of N-in: 0.12994\noutside 1 331\nTMhelix 332 354\ninside 355 372

4570 GCF_900015745.1_ED654_contigs Escherichia coli Proteobacteria; Gammaproteobacteria; Enterobacterales; Enterobacteriaceae; Escherichia MTGDNTLIHSHGINRRDFMKLCAALAATMGLSSKAAAEMAESVTNPQRPPVIWIGAQECT WP_000145410.1 MULTISPECIES: hydrogenase-2 small chain [Proteobacteria] Length: 372\nNumber of predicted TMHs: 1\nExp number of AAs in TMHs: 23.72226\nExp number, first 60 AAs: 0.22038\nTotal prob of N-in: 0.12994\noutside 1 331\nTMhelix 332 354\ninside 355 372

4571 GCF_900015795.1_ED075_contigs Escherichia coli Proteobacteria; Gammaproteobacteria; Enterobacterales; Enterobacteriaceae; Escherichia MTGDNTLIHSHGINRRDFMKLCAALAATMGLSSKAAAEMAESVTNPQRPPVIWIGAQECT WP_000145410.1 MULTISPECIES: hydrogenase-2 small chain [Proteobacteria] Length: 372\nNumber of predicted TMHs: 1\nExp number of AAs in TMHs: 23.72226\nExp number, first 60 AAs: 0.22038\nTotal prob of N-in: 0.12994\noutside 1 331\nTMhelix 332 354\ninside 355 372

4572 GCF_900015775.1_ED413_contigs Escherichia coli Proteobacteria; Gammaproteobacteria; Enterobacterales; Enterobacteriaceae; Escherichia MTGDNTLIHSHGINRRDFMKLCAALAATMGLSSKAAAEMAESVTNPQRPPVIWIGAQECT WP_000145410.1 MULTISPECIES: hydrogenase-2 small chain [Proteobacteria] Length: 372\nNumber of predicted TMHs:

1\nExp number of AAs in TMHs: 23.72226\nExp number, first 60 AAs: 0.22038\nTotal prob of N-in: 0.12994\noutside 1 331\nTMhelix 332 354\ninside 355 372

4573 GCF_900015875.1_ED766_contigs Escherichia coli Proteobacteria; Gammaproteobacteria; Enterobacterales; Enterobacteriaceae; Escherichia
MNNEETFYQAMRRQGVTRRSFLKYCSLAATSLGLGAGMAPKIAWALENKPRIPVVWIHGL WP_001058323.1
MULTISPECIES: hydrogenase-1 small chain [Proteobacteria] Length: 372\nNumber of predicted TMHs: 2\nExp number of AAs in TMHs: 37.2608499999999\nExp number, first 60 AAs: 19.52923\nTotal prob of N-in: 0.94351\nPOSSIBLE N-term signal sequence\ninside 1 19\nTMhelix 20 39\noutside 40 325\nTMhelix 326 348\ninside 349 372

4574 GCF_900015195.1_ED666_contigs Escherichia coli Proteobacteria; Gammaproteobacteria; Enterobacterales; Enterobacteriaceae; Escherichia
MNNEETFYQAMRRQGVTRRSFLKYCSLAATSLGLGAGMAPKIAWALENKPRIPVVWIHGL WP_001301613.1
hydrogenase [Escherichia coli] Length: 372\nNumber of predicted TMHs: 2\nExp number of AAs in TMHs: 37.46598\nExp number, first 60 AAs: 19.53139\nTotal prob of N-in: 0.94364\nPOSSIBLE N-term signal sequence\ninside 1 19\nTMhelix 20 39\noutside 40 325\nTMhelix 326 348\ninside 349 372

4575 GCF_900015795.1_ED075_contigs Escherichia coli Proteobacteria; Gammaproteobacteria; Enterobacterales; Enterobacteriaceae; Escherichia
MNNEETFYQAMRRQGVTRRSFLKYCSLAATSLGLGAGMAPKIAWALENKPRIPVVWIHGL WP_001058323.1
MULTISPECIES: hydrogenase-1 small chain [Proteobacteria] Length: 372\nNumber of predicted TMHs: 2\nExp number of AAs in TMHs: 37.2608499999999\nExp number, first 60 AAs: 19.52923\nTotal prob of N-in: 0.94351\nPOSSIBLE N-term signal sequence\ninside 1 19\nTMhelix 20 39\noutside 40 325\nTMhelix 326 348\ninside 349 372

4576 GCF_900015895.1_ED699_contigs Escherichia coli Proteobacteria; Gammaproteobacteria; Enterobacterales; Enterobacteriaceae; Escherichia
MTGDNTLIHSHGINRRDFMKLCAALAATMGLSSKAAAEMAESVTNPQRPPVIWIGAQECT WP_000145410.1
MULTISPECIES: hydrogenase-2 small chain [Proteobacteria] Length: 372\nNumber of predicted TMHs: 1\nExp number of AAs in TMHs: 23.72226\nExp number, first 60 AAs: 0.22038\nTotal prob of N-in: 0.12994\noutside 1 331\nTMhelix 332 354\ninside 355 372

4577 GCF_900015695.1_ED647_contigs Escherichia coli Proteobacteria; Gammaproteobacteria; Enterobacterales; Enterobacteriaceae; Escherichia
MNNEETFYQAMRRQGVTRRSFLKYCSLAATSLGLGAGMAPKIAWALENKPRIPVVWIHGL WP_001301613.1
hydrogenase [Escherichia coli] Length: 372\nNumber of predicted TMHs: 2\nExp number of AAs in TMHs: 37.46598\nExp number, first 60 AAs: 19.53139\nTotal prob of N-in: 0.94364\nPOSSIBLE N-term signal sequence\ninside 1 19\nTMhelix 20 39\noutside 40 325\nTMhelix 326 348\ninside 349 372

4578 GCF_900015695.1_ED647_contigs Escherichia coli Proteobacteria; Gammaproteobacteria; Enterobacterales; Enterobacteriaceae; Escherichia
MTGDNTLIHSHGINRRDFMKLCAALAATMGLSSKAAAEMAESVTNPQRPPVIWIGAQECT WP_069192915.1
hydrogenase 2 small subunit [Escherichia coli] Length: 372\nNumber of predicted TMHs: 1\nExp number of AAs in TMHs: 23.38409\nExp number, first 60 AAs: 0.22086\nTotal prob of N-in: 0.11608\noutside 1 331\nTMhelix 332 354\ninside 355 372

4579 GCF_900015895.1_ED699_contigs Escherichia coli Proteobacteria; Gammaproteobacteria; Enterobacterales; Enterobacteriaceae; Escherichia
MNNEETFYQAMRRQGVTRRSFLKYCSLAATSLGLGAGMAPKIAWALENKPRIPVVWIHGL WP_001301613.1
hydrogenase [Escherichia coli] Length: 372\nNumber of predicted TMHs: 2\nExp number of AAs in TMHs: 37.46598\nExp number, first 60 AAs: 19.53139\nTotal prob of N-in: 0.94364\nPOSSIBLE N-term signal sequence\ninside 1 19\nTMhelix 20 39\noutside 40 325\nTMhelix 326 348\ninside 349 372

4580 GCF_900016025.1_BCW5739_contigs Escherichia coli Proteobacteria; Gammaproteobacteria; Enterobacterales; Enterobacteriaceae; Escherichia
MTGDNTLIHSHGINRRDFMKLCAALAATMGLSSKAAAEMVESVTNPQRPPVIWIGAQECT WP_000145423.1
hydrogenase 2 small subunit [Escherichia coli] Length: 372\nNumber of predicted TMHs: 1\nExp number of AAs in TMHs: 23.78201\nExp number, first 60 AAs: 0.28355\nTotal prob of N-in: 0.13257\noutside 1 331\nTMhelix 332 354\ninside 355 372

4581 GCF_900016045.1_BCW5743_contigs Escherichia coli Proteobacteria; Gammaproteobacteria; Enterobacterales; Enterobacteriaceae; Escherichia MTGDNTLIHSHGINRRDFMKLCAALAATMGLSSKAAAEMVESVTNPQRPPVIWIGAQECT WP_000145423.1 hydrogenase 2 small subunit [Escherichia coli] Length: 372\nNumber of predicted TMHs: 1\nExp number of AAs in TMHs: 23.78201\nExp number, first 60 AAs: 0.28355\nTotal prob of N-in: 0.13257\noutside 1 331\nTMhelix 332 354\ninside 355 372

4582 GCF_900016075.1_ED361_contigs Escherichia coli Proteobacteria; Gammaproteobacteria; Enterobacterales; Enterobacteriaceae; Escherichia MTGDNTLIHSHGINRRDFMKLCAALAATMGLSSKAAAEMAESVTNPQRPPVIWIGAQECT WP_000145410.1 MULTISPECIES: hydrogenase-2 small chain [Proteobacteria] Length: 372\nNumber of predicted TMHs: 1\nExp number of AAs in TMHs: 23.72226\nExp number, first 60 AAs: 0.22038\nTotal prob of N-in: 0.12994\noutside 1 331\nTMhelix 332 354\ninside 355 372

4583 GCF_900016365.1_M884_contigs Escherichia coli Proteobacteria; Gammaproteobacteria; Enterobacterales; Enterobacteriaceae; Escherichia MTGDNTLIHSHGINRRDFMKLCAALAATMGLSSKAAAEMAESVTNPQRPPVIWIGAQECT WP_000145410.1 MULTISPECIES: hydrogenase-2 small chain [Proteobacteria] Length: 372\nNumber of predicted TMHs: 1\nExp number of AAs in TMHs: 23.72226\nExp number, first 60 AAs: 0.22038\nTotal prob of N-in: 0.12994\noutside 1 331\nTMhelix 332 354\ninside 355 372

4584 GCF_900016025.1_BCW5739_contigs Escherichia coli Proteobacteria; Gammaproteobacteria; Enterobacterales; Enterobacteriaceae; Escherichia MNNEETFYQAMRRQGVTRRSFLKYCSLAATSLGLGAGMAPKIAWALENKPRIPVVWIHGL WP_001295940.1 MULTISPECIES: hydrogenase [Enterobacteriaceae] Length: 372\nNumber of predicted TMHs: 2\nExp number of AAs in TMHs: 37.2608899999999\nExp number, first 60 AAs: 19.52923\nTotal prob of N-in: 0.94351\nPOSSIBLE N-term signal sequence\ninside 1 19\nTMhelix 20 39\noutside 40 325\nTMhelix 326 348\ninside 349 372

4585 GCF_900039545.1_F25OS1 Escherichia coli Proteobacteria; Gammaproteobacteria; Enterobacterales; Enterobacteriaceae; Escherichia MTGDNTLIHSHGINRRDFMKLCAALAATMGLSSKAAAEMAESVTNPQRPPVIWIGAQECT WP_000145410.1 MULTISPECIES: hydrogenase-2 small chain [Proteobacteria] Length: 372\nNumber of predicted TMHs: 1\nExp number of AAs in TMHs: 23.72226\nExp number, first 60 AAs: 0.22038\nTotal prob of N-in: 0.12994\noutside 1 331\nTMhelix 332 354\ninside 355 372

4586 GCF_900039545.1_F25OS1 Escherichia coli Proteobacteria; Gammaproteobacteria; Enterobacterales; Enterobacteriaceae; Escherichia MNNEETFYQAMRRQGVTRRSFLKYCSLAATSLGLGAGMAPKIAWALENKPRIPVVWIHGL WP_001058323.1 MULTISPECIES: hydrogenase-1 small chain [Proteobacteria] Length: 372\nNumber of predicted TMHs: 2\nExp number of AAs in TMHs: 37.2608499999999\nExp number, first 60 AAs: 19.52923\nTotal prob of N-in: 0.94351\nPOSSIBLE N-term signal sequence\ninside 1 19\nTMhelix 20 39\noutside 40 325\nTMhelix 326 348\ninside 349 372

4587 GCF_900087675.1_RL158 Escherichia coli Proteobacteria; Gammaproteobacteria; Enterobacterales; Enterobacteriaceae; Escherichia MTGDNTLIHSHGINRRDFMKLCAALAATMGLSSKAAAEMAESVTNPQRPPVIWIGAQECT WP_000145410.1 MULTISPECIES: hydrogenase-2 small chain [Proteobacteria] Length: 372\nNumber of predicted TMHs: 1\nExp number of AAs in TMHs: 23.72226\nExp number, first 60 AAs: 0.22038\nTotal prob of N-in: 0.12994\noutside 1 331\nTMhelix 332 354\ninside 355 372

4588 GCF_900044035.1_F25P5 Escherichia coli Proteobacteria; Gammaproteobacteria; Enterobacterales; Enterobacteriaceae; Escherichia MNNEETFYQAMRRQGVTRRSFLKYCSLAATSLGLGAGMAPKIAWALENKPRIPVVWIHGL WP_001058323.1 MULTISPECIES: hydrogenase-1 small chain [Proteobacteria] Length: 372\nNumber of predicted TMHs: 2\nExp number of AAs in TMHs: 37.2608499999999\nExp number, first 60 AAs: 19.52923\nTotal prob of N-in: 0.94351\nPOSSIBLE N-term signal sequence\ninside 1 19\nTMhelix 20 39\noutside 40 325\nTMhelix 326 348\ninside 349 372

4589 GCF_900087675.1_RL158 Escherichia coli Proteobacteria; Gammaproteobacteria; Enterobacterales; Enterobacteriaceae; Escherichia MNNEETFYQAMRRQGVTRRSFLKYCSLAATSLGLGAGMAPKIAWALENKPRIPVVWIHGL WP_001058323.1 MULTISPECIES: hydrogenase-1 small chain [Proteobacteria] Length: 372\nNumber of predicted TMHs: 2\nExp number of AAs in TMHs: 37.2608499999999\nExp number, first 60 AAs: 19.52923\nTotal prob of N-in: 0.94351\nPOSSIBLE N-term signal sequence\ninside 1 19\nTMhelix 20 39\noutside 40 325\nTMhelix 326 348\ninside 349 372

4590 GCF_900016095.1_ED377_contigs Escherichia coli Proteobacteria; Gammaproteobacteria; Enterobacterales; Enterobacteriaceae; Escherichia MNNEETFYQAMRRQGVTRRSFLKYCSLAATSLGLGAGMAPKIAWALENKPRIPVVWIHGL WP_001295940.1 MULTISPECIES: hydrogenase [Enterobacteriaceae] Length: 372\nNumber of predicted TMHs: 2\nExp number of AAs in TMHs: 37.2608899999999\nExp number, first 60 AAs: 19.52923\nTotal prob of N-in: 0.94351\nPOSSIBLE N-term signal sequence\ninside 1 19\nTMhelix 20 39\noutside 40 325\nTMhelix 326 348\ninside 349 372

4591 GCF_900087565.2_ASM90008756v2 Escherichia coli Proteobacteria; Gammaproteobacteria; Enterobacterales; Enterobacteriaceae; Escherichia MNNEETFYQAMRRQGVTRRSFLKYCSLAATSLGLGAGMAPKIAWALENKPRIPVVWIHGL WP_001058323.1 MULTISPECIES: hydrogenase-1 small chain [Proteobacteria] Length: 372\nNumber of predicted TMHs: 2\nExp number of AAs in TMHs: 37.2608499999999\nExp number, first 60 AAs: 19.52923\nTotal prob of N-in: 0.94351\nPOSSIBLE N-term signal sequence\ninside 1 19\nTMhelix 20 39\noutside 40 325\nTMhelix 326 348\ninside 349 372

4592 GCF_900016075.1_ED361_contigs Escherichia coli Proteobacteria; Gammaproteobacteria; Enterobacterales; Enterobacteriaceae; Escherichia MNNEETFYQAMRRQGVTRRSFLKYCSLAATSLGLGAGMAPKIAWALENKPRIPVVWIHGL WP_062898093.1 hydrogenase [Escherichia coli] Length: 372\nNumber of predicted TMHs: 2\nExp number of AAs in TMHs: 37.5090799999999\nExp number, first 60 AAs: 19.52755\nTotal prob of N-in: 0.94365\nPOSSIBLE N-term signal sequence\ninside 1 19\nTMhelix 20 39\noutside 40 325\nTMhelix 326 348\ninside 349 372

4593 GCF_900087745.1_RL138 Escherichia coli Proteobacteria; Gammaproteobacteria; Enterobacterales; Enterobacteriaceae; Escherichia MTGDNTLIHSHGINRRDFMKLCAALAATMGLSSKAAAEMAESVTNPQRPPVIWIGAQECT WP_000145410.1 MULTISPECIES: hydrogenase-2 small chain [Proteobacteria] Length: 372\nNumber of predicted TMHs: 1\nExp number of AAs in TMHs: 23.72226\nExp number, first 60 AAs: 0.22038\nTotal prob of N-in: 0.12994\noutside 1 331\nTMhelix 332 354\ninside 355 372

4594 GCF_900092615.1_PRJEB14041 Escherichia coli Proteobacteria; Gammaproteobacteria; Enterobacterales; Enterobacteriaceae; Escherichia MNNEETFYQAMRRQGVTRRSFLKYCSLAATSLGLGAGMAPKIAWALENKPRIPVVWIHGL WP_001058323.1 MULTISPECIES: hydrogenase-1 small chain [Proteobacteria] Length: 372\nNumber of predicted TMHs: 2\nExp number of AAs in TMHs: 37.2608499999999\nExp number, first 60 AAs: 19.52923\nTotal prob of N-in: 0.94351\nPOSSIBLE N-term signal sequence\ninside 1 19\nTMhelix 20 39\noutside 40 325\nTMhelix 326 348\ninside 349 372

4595 GCF_900093865.1_scaffolds_C318 Escherichia coli Proteobacteria; Gammaproteobacteria; Enterobacterales; Enterobacteriaceae; Escherichia MTGDNTLIHSHGINRRDFMKLCAALAATMGLSSKAAAEMAESVTNPQRPPVIWIGAQECT WP_000145410.1 MULTISPECIES: hydrogenase-2 small chain [Proteobacteria] Length: 372\nNumber of predicted TMHs: 1\nExp number of AAs in TMHs: 23.72226\nExp number, first 60 AAs: 0.22038\nTotal prob of N-in: 0.12994\noutside 1 331\nTMhelix 332 354\ninside 355 372

4596 GCF_900093845.1_scaffolds_20160103 Escherichia coli Proteobacteria; Gammaproteobacteria; Enterobacterales; Enterobacteriaceae; Escherichia MTGDNTLIHSHGINRRDFMKLCAALAATMGLSSKAAAEMAESVTNPQRPPVIWIGAQECT WP_000145410.1 MULTISPECIES: hydrogenase-2 small chain [Proteobacteria] Length: 372\nNumber of predicted TMHs: 1\nExp number of AAs in TMHs: 23.72226\nExp number, first 60 AAs: 0.22038\nTotal prob of N-in: 0.12994\noutside 1 331\nTMhelix 332 354\ninside 355 372

4597 GCF_900093895.1_scaffolds_C281 Escherichia coli Proteobacteria; Gammaproteobacteria; Enterobacterales; Enterobacteriaceae; Escherichia MTGDNTLIHSHGINRRDFMKLCAALAATMGLSSKAAAEMAESVTNPQRPPVIWIGAQECT WP_000145410.1 MULTISPECIES: hydrogenase-2 small chain [Proteobacteria] Length: 372\nNumber of predicted TMHs: 1\nExp number of AAs in TMHs: 23.72226\nExp number, first 60 AAs: 0.22038\nTotal prob of N-in: 0.12994\noutside 1 331\nTMhelix 332 354\ninside 355 372

4598 GCF_900093915.1_scaffolds_20160020 Escherichia coli Proteobacteria; Gammaproteobacteria; Enterobacterales; Enterobacteriaceae; Escherichia MTGDNTLIHSHGINRRDFMKLCAALAATMGLSSKAAAEMAESVTNPQRPPVIWIGAQECT WP_000145410.1

MULTISPECIES: hydrogenase-2 small chain [Proteobacteria] Length: 372\nNumber of predicted TMHs: 1\nExp number of AAs in TMHs: 23.72226\nExp number, first 60 AAs: 0.22038\nTotal prob of N-in: 0.12994\noutside 1 331\nTMhelix 332 354\ninside 355 372

4599 GCF_900093945.1_scaffolds_20151316 Escherichia coli Proteobacteria; Gammaproteobacteria; Enterobacterales; Enterobacteriaceae; Escherichia
MTGDNTLIHSHGINRRDFMKLCAALAATMGLSSKAAAEMAESVTNPQRPPVIWIGAQECT WP_000145410.1
MULTISPECIES: hydrogenase-2 small chain [Proteobacteria] Length: 372\nNumber of predicted TMHs: 1\nExp number of AAs in TMHs: 23.72226\nExp number, first 60 AAs: 0.22038\nTotal prob of N-in: 0.12994\noutside 1 331\nTMhelix 332 354\ninside 355 372

4600 GCF_900093975.1_scaffolds_20160018 Escherichia coli Proteobacteria; Gammaproteobacteria; Enterobacterales; Enterobacteriaceae; Escherichia
MTGDNTLIHSHGINRRDFMKLCAALAATMGLSSKAAAEMAESVTNPQRPPVIWIGAQECT WP_000145410.1
MULTISPECIES: hydrogenase-2 small chain [Proteobacteria] Length: 372\nNumber of predicted TMHs: 1\nExp number of AAs in TMHs: 23.72226\nExp number, first 60 AAs: 0.22038\nTotal prob of N-in: 0.12994\noutside 1 331\nTMhelix 332 354\ninside 355 372

4601 GCF_001700095.1_ASM170009v1 Escherichia coli Proteobacteria; Gammaproteobacteria; Enterobacterales; Enterobacteriaceae; Escherichia
MTGDNTLIHSHGINRRDFMKLCAALAATMGLSSKAAAEMAESVTNPQRPPVIWIGAQECT WP_000145410.1
MULTISPECIES: hydrogenase-2 small chain [Proteobacteria] Length: 372\nNumber of predicted TMHs: 1\nExp number of AAs in TMHs: 23.72226\nExp number, first 60 AAs: 0.22038\nTotal prob of N-in: 0.12994\noutside 1 331\nTMhelix 332 354\ninside 355 372

4602 GCF_900093975.1_scaffolds_20160018 Escherichia coli Proteobacteria; Gammaproteobacteria; Enterobacterales; Enterobacteriaceae; Escherichia
MNNEETFYQAMRRQGVTRRSFLKYCSLAATSLGLGAGMAPKIAWALENKPRIPVVWIHGL WP_001058323.1
MULTISPECIES: hydrogenase-1 small chain [Proteobacteria] Length: 372\nNumber of predicted TMHs: 2\nExp number of AAs in TMHs: 37.2608499999999\nExp number, first 60 AAs: 19.52923\nTotal prob of N-in: 0.94351\nPOSSIBLE N-term signal sequence\ninside 1 19\nTMhelix 20 39\noutside 40 325\nTMhelix 326 348\ninside 349 372

4603 GCF_900093845.1_scaffolds_20160103 Escherichia coli Proteobacteria; Gammaproteobacteria; Enterobacterales; Enterobacteriaceae; Escherichia
MNNEETFYQAMRRQGVTRRSFLKYCSLAATSLGLGAGMAPKIAWALENKPRIPVVWIHGL WP_001295940.1
MULTISPECIES: hydrogenase [Enterobacteriaceae] Length: 372\nNumber of predicted TMHs: 2\nExp number of AAs in TMHs: 37.2608899999999\nExp number, first 60 AAs: 19.52923\nTotal prob of N-in: 0.94351\nPOSSIBLE N-term signal sequence\ninside 1 19\nTMhelix 20 39\noutside 40 325\nTMhelix 326 348\ninside 349 372

4604 GCF_900096795.1_Ecoli_AG100_Sample3_Doxycycline_Assembly Escherichia coli Proteobacteria; Gammaproteobacteria; Enterobacterales; Enterobacteriaceae; Escherichia
MTGDNTLIHSHGINRRDFMKLCAALAATMGLSSKAAAEMAESVTNPQRPPVIWIGAQECT WP_000145410.1
MULTISPECIES: hydrogenase-2 small chain [Proteobacteria] Length: 372\nNumber of predicted TMHs: 1\nExp number of AAs in TMHs: 23.72226\nExp number, first 60 AAs: 0.22038\nTotal prob of N-in: 0.12994\noutside 1 331\nTMhelix 332 354\ninside 355 372

4605 GCF_900096825.1_Ecoli_AG100_Sample1_M9_Assembly Escherichia coli Proteobacteria; Gammaproteobacteria; Enterobacterales; Enterobacteriaceae; Escherichia
MTGDNTLIHSHGINRRDFMKLCAALAATMGLSSKAAAEMAESVTNPQRPPVIWIGAQECT WP_000145410.1
MULTISPECIES: hydrogenase-2 small chain [Proteobacteria] Length: 372\nNumber of predicted TMHs: 1\nExp number of AAs in TMHs: 23.72226\nExp number, first 60 AAs: 0.22038\nTotal prob of N-in: 0.12994\noutside 1 331\nTMhelix 332 354\ninside 355 372

4606 GCF_900096795.1_Ecoli_AG100_Sample3_Doxycycline_Assembly Escherichia coli Proteobacteria; Gammaproteobacteria; Enterobacterales; Enterobacteriaceae; Escherichia
MNNEETFYQAMRRQGVTRRSFLKYCSLAATSLGLGAGMAPKIAWALENKPRIPVVWIHGL WP_001058323.1
MULTISPECIES: hydrogenase-1 small chain [Proteobacteria] Length: 372\nNumber of predicted TMHs: 2\nExp number of AAs in TMHs: 37.2608499999999\nExp number, first 60 AAs: 19.52923\nTotal prob of N-in:

0.94351\nPOSSIBLE N-term signal sequence\ninside 1 19\nTMhelix 20 39\noutside 40 325\nTMhelix 326 348\ninside 349 372

4607 GCF_900096825.1_Ecoli_AG100_Sample1_M9_Assembly Escherichia coli Proteobacteria; Gammaproteobacteria; Enterobacterales; Enterobacteriaceae; Escherichia MNNEETFYQAMRRQGVTRRSFLKYCSLAATSLGLGAGMAPKIAWALENKPRIPVVWIHGL WP_001058323.1 MULTISPECIES: hydrogenase-1 small chain [Proteobacteria] Length: 372\nNumber of predicted TMHs: 2\nExp number of AAs in TMHs: 37.2608499999999\nExp number, first 60 AAs: 19.52923\nTotal prob of N-in: 0.94351\nPOSSIBLE N-term signal sequence\ninside 1 19\nTMhelix 20 39\noutside 40 325\nTMhelix 326 348\ninside 349 372

4608 GCF_900093945.1_scaffolds_20151316 Escherichia coli Proteobacteria; Gammaproteobacteria; Enterobacterales; Enterobacteriaceae; Escherichia MNNEETFYQAMRRQGVTRRSFLKYCSLAATSLGLGAGMAPKIAWALENKPRIPVVWIHGL WP_001295940.1 MULTISPECIES: hydrogenase [Enterobacteriaceae] Length: 372\nNumber of predicted TMHs: 2\nExp number of AAs in TMHs: 37.2608899999999\nExp number, first 60 AAs: 19.52923\nTotal prob of N-in: 0.94351\nPOSSIBLE N-term signal sequence\ninside 1 19\nTMhelix 20 39\noutside 40 325\nTMhelix 326 348\ninside 349 372

4609 GCF_900156555.1_RCEC007_PRJEB11777_wgs.embl.gz Escherichia coli Proteobacteria; Gammaproteobacteria; Enterobacterales; Enterobacteriaceae; Escherichia MTGDNTLIHSHGINRRDFMKLCAALAATMGLSSKAAAEMAESVTNPQRPPVIWIGAQECT WP_000145410.1 MULTISPECIES: hydrogenase-2 small chain [Proteobacteria] Length: 372\nNumber of predicted TMHs: 1\nExp number of AAs in TMHs: 23.72226\nExp number, first 60 AAs: 0.22038\nTotal prob of N-in: 0.12994\noutside 1 331\nTMhelix 332 354\ninside 355 372

4610 GCF_900166735.1_Isolate_11 Escherichia coli Proteobacteria; Gammaproteobacteria; Enterobacterales; Enterobacteriaceae; Escherichia MTGDNTLIHSHGINRRDFMKLCAALAATMGLSSKAAAEMVESVTNPQRPPVIWIGAQECT WP_000145423.1 hydrogenase 2 small subunit [Escherichia coli] Length: 372\nNumber of predicted TMHs: 1\nExp number of AAs in TMHs: 23.78201\nExp number, first 60 AAs: 0.28355\nTotal prob of N-in: 0.13257\noutside 1 331\nTMhelix 332 354\ninside 355 372

4611 GCF_900166765.1_Isolate_8 Escherichia coli Proteobacteria; Gammaproteobacteria; Enterobacterales; Enterobacteriaceae; Escherichia MTGDNTLIHSHGINRRDFMKLCAALAATMGLSSKAAAEMVESVTNPQRPPVIWIGAQECT WP_000145423.1 hydrogenase 2 small subunit [Escherichia coli] Length: 372\nNumber of predicted TMHs: 1\nExp number of AAs in TMHs: 23.78201\nExp number, first 60 AAs: 0.28355\nTotal prob of N-in: 0.13257\noutside 1 331\nTMhelix 332 354\ninside 355 372

4612 GCF_900166705.1_Isolate_13 Escherichia coli Proteobacteria; Gammaproteobacteria; Enterobacterales; Enterobacteriaceae; Escherichia MNNEETFYQAMRRQGVTRRSFLKYCSLAATSLGLGAGMAPKIAWALENKPRIPVVWIHGL WP_001295940.1 MULTISPECIES: hydrogenase [Enterobacteriaceae] Length: 372\nNumber of predicted TMHs: 2\nExp number of AAs in TMHs: 37.2608899999999\nExp number, first 60 AAs: 19.52923\nTotal prob of N-in: 0.94351\nPOSSIBLE N-term signal sequence\ninside 1 19\nTMhelix 20 39\noutside 40 325\nTMhelix 326 348\ninside 349 372

4613 GCF_900166735.1_Isolate_11 Escherichia coli Proteobacteria; Gammaproteobacteria; Enterobacterales; Enterobacteriaceae; Escherichia MNNEETFYQAMRRQGVTRRSFLKYCSLAATSLGLGAGMAPKIAWALENKPRIPVVWIHGL WP_001295940.1 MULTISPECIES: hydrogenase [Enterobacteriaceae] Length: 372\nNumber of predicted TMHs: 2\nExp number of AAs in TMHs: 37.2608899999999\nExp number, first 60 AAs: 19.52923\nTotal prob of N-in: 0.94351\nPOSSIBLE N-term signal sequence\ninside 1 19\nTMhelix 20 39\noutside 40 325\nTMhelix 326 348\ninside 349 372

4614 GCF_900166765.1_Isolate_8 Escherichia coli Proteobacteria; Gammaproteobacteria; Enterobacterales; Enterobacteriaceae; Escherichia MNNEETFYQAMRRQGVTRRSFLKYCSLAATSLGLGAGMAPKIAWALENKPRIPVVWIHGL WP_001295940.1 MULTISPECIES: hydrogenase [Enterobacteriaceae] Length: 372\nNumber of predicted TMHs: 2\nExp number of AAs in TMHs: 37.2608899999999\nExp number, first 60 AAs: 19.52923\nTotal prob of N-in: 0.94351\nPOSSIBLE N-term signal sequence\ninside 1 19\nTMhelix 20 39\noutside 40 325\nTMhelix 326 348\ninside 349 372

4615 GCF_900108745.1_PEI-B-P-19 Escherichia coli Proteobacteria; Gammaproteobacteria; Enterobacterales; Enterobacteriaceae; Escherichia
MNNEETFYQAMRRQGVTRRSFLKYCSLAATSLGLGAGMAPKIAWALENKPRIPVVWIHGL WP_001058323.1
MULTISPECIES: hydrogenase-1 small chain [Proteobacteria] Length: 372\nNumber of predicted TMHs: 2\nExp number of AAs in TMHs: 37.2608499999999\nExp number, first 60 AAs: 19.52923\nTotal prob of N-in: 0.94351\nPOSSIBLE N-term signal sequence\ninside 1 19\nTMhelix 20 39\noutside 40 325\nTMhelix 326 348\ninside 349 372

4616 GCF_900177355.1_IMG-taxon_2608642167_annotated_assembly Escherichia coli Proteobacteria; Gammaproteobacteria; Enterobacterales; Enterobacteriaceae; Escherichia
MTGDNTLIHSHGINRRDFMKLCAALAATMGLSSKAAAEMAESVTNPQRPPVIWIGAQECT WP_000145410.1
MULTISPECIES: hydrogenase-2 small chain [Proteobacteria] Length: 372\nNumber of predicted TMHs: 1\nExp number of AAs in TMHs: 23.72226\nExp number, first 60 AAs: 0.22038\nTotal prob of N-in: 0.12994\noutside 1 331\nTMhelix 332 354\ninside 355 372

4617 GCF_900166845.1_Isolate_21 Escherichia coli Proteobacteria; Gammaproteobacteria; Enterobacterales; Enterobacteriaceae; Escherichia
MNNEETFYQAMRRQGVTRRSFLKYCSLAATSLGLGAGMAPKIAWALENKPRIPVVWIHGL WP_001058320.1
hydrogenase [Escherichia coli] Length: 372\nNumber of predicted TMHs: 2\nExp number of AAs in TMHs: 37.26082\nExp number, first 60 AAs: 19.52923\nTotal prob of N-in: 0.94351\nPOSSIBLE N-term signal sequence\ninside 1 19\nTMhelix 20 39\noutside 40 325\nTMhelix 326 348\ninside 349 372

4618 GCF_900177355.1_IMG-taxon_2608642167_annotated_assembly Escherichia coli Proteobacteria; Gammaproteobacteria; Enterobacterales; Enterobacteriaceae; Escherichia
MNNEETFYQAMRRQGVTRRSFLKYCSLAATSLGLGAGMAPKIAWALENKPRIPVVWIHGL WP_001058323.1
MULTISPECIES: hydrogenase-1 small chain [Proteobacteria] Length: 372\nNumber of predicted TMHs: 2\nExp number of AAs in TMHs: 37.2608499999999\nExp number, first 60 AAs: 19.52923\nTotal prob of N-in: 0.94351\nPOSSIBLE N-term signal sequence\ninside 1 19\nTMhelix 20 39\noutside 40 325\nTMhelix 326 348\ninside 349 372

4619 GCF_001058915.1_ASM105891v1 Escherichia coli Proteobacteria; Gammaproteobacteria; Enterobacterales; Enterobacteriaceae; Escherichia
MTGDNTLIHSHGINRRDFMKLCAALAATMGLSSKAAAEMVESVTNPQRPPVIWIGAQECT WP_001556793.1
hydrogenase-2 small chain [Escherichia coli] Length: 372\nNumber of predicted TMHs: 1\nExp number of AAs in TMHs: 23.78201\nExp number, first 60 AAs: 0.28355\nTotal prob of N-in: 0.13257\noutside 1 331\nTMhelix 332 354\ninside 355 372

4620 GCF_001058915.1_ASM105891v1 Escherichia coli Proteobacteria; Gammaproteobacteria; Enterobacterales; Enterobacteriaceae; Escherichia
MNNEETFYQAMRRQGVTRRSFLKYCSLAATSLGLGAGMAPKIAWALENKPRIPVVWIHGL WP_001538532.1
hydrogenase-1 small chain [Escherichia coli] Length: 372\nNumber of predicted TMHs: 2\nExp number of AAs in TMHs: 37.27041\nExp number, first 60 AAs: 19.52927\nTotal prob of N-in: 0.94352\nPOSSIBLE N-term signal sequence\ninside 1 19\nTMhelix 20 39\noutside 40 325\nTMhelix 326 348\ninside 349 372

4621 GCF_000782255.1_ASM78225v1 Escherichia coli Proteobacteria; Gammaproteobacteria; Enterobacterales; Enterobacteriaceae; Escherichia
MNNEETFYQAMRRQGVTRRSFLKYCSLAATSLGLGAGMAPKIAWALENKPRIPVVWIHGL WP_001295940.1
MULTISPECIES: hydrogenase [Enterobacteriaceae] Length: 372\nNumber of predicted TMHs: 2\nExp number of AAs in TMHs: 37.2608899999999\nExp number, first 60 AAs: 19.52923\nTotal prob of N-in: 0.94351\nPOSSIBLE N-term signal sequence\ninside 1 19\nTMhelix 20 39\noutside 40 325\nTMhelix 326 348\ninside 349 372

4622 GCF_900093995.1_scaffolds_20160019 Escherichia coli Proteobacteria; Gammaproteobacteria; Enterobacterales; Enterobacteriaceae; Escherichia
MTGDNTLIHSHGINRRDFMKLCAALAATMGLSSKAAAEMAESVTNPQRPPVIWIGAQECT WP_000145410.1
MULTISPECIES: hydrogenase-2 small chain [Proteobacteria] Length: 372\nNumber of predicted TMHs: 1\nExp number of AAs in TMHs: 23.72226\nExp number, first 60 AAs: 0.22038\nTotal prob of N-in: 0.12994\noutside 1 331\nTMhelix 332 354\ninside 355 372

4623 GCF_001693635.1_ASM169363v1 Escherichia coli Proteobacteria; Gammaproteobacteria; Enterobacterales; Enterobacteriaceae; Escherichia

MNNEETFYQAMRRQGVTRRSFLKYCSLAATSLGLGAGMAPKIAWALENKPRIPVVWIHGL WP_001295940.1 MULTISPECIES: hydrogenase [Enterobacteriaceae] Length: 372\nNumber of predicted TMHs: 2\nExp number of AAs in TMHs: 37.2608899999999\nExp number, first 60 AAs: 19.52923\nTotal prob of N-in: 0.94351\nPOSSIBLE N-term signal sequence\ninside 1 19\nTMhelix 20 39\noutside 40 325\nTMhelix 326 348\ninside 349 372

4624 GCF_001700405.1_ASM170040v1 Escherichia coli Proteobacteria; Gammaproteobacteria; Enterobacterales; Enterobacteriaceae; Escherichia MTGDNTLIHSHGINRRDFMKLCAALAATMGLSSKAAAEMAESVTNPQRPPVIWIGAQECT WP_000145410.1 MULTISPECIES: hydrogenase-2 small chain [Proteobacteria] Length: 372\nNumber of predicted TMHs: 1\nExp number of AAs in TMHs: 23.72226\nExp number, first 60 AAs: 0.22038\nTotal prob of N-in: 0.12994\noutside 1 331\nTMhelix 332 354\ninside 355 372

4625 GCF_001709165.1_ASM170916v1 Escherichia coli Proteobacteria; Gammaproteobacteria; Enterobacterales; Enterobacteriaceae; Escherichia MTGDNTLIHSHGINRRDFMKLCAALAATMGLSSKAAAEMAESVTNPQRPPVIWIGAQECT WP_000145410.1 MULTISPECIES: hydrogenase-2 small chain [Proteobacteria] Length: 372\nNumber of predicted TMHs: 1\nExp number of AAs in TMHs: 23.72226\nExp number, first 60 AAs: 0.22038\nTotal prob of N-in: 0.12994\noutside 1 331\nTMhelix 332 354\ninside 355 372

4626 GCF_001696775.1_ASM169677v1 Escherichia coli Proteobacteria; Gammaproteobacteria; Enterobacterales; Enterobacteriaceae; Escherichia MNNEETFYQAMRRQGVTRRSFLKYCSLAATSLGLGAGMAPKIAWALENKPRIPVVWIHGL WP_001058323.1 MULTISPECIES: hydrogenase-1 small chain [Proteobacteria] Length: 372\nNumber of predicted TMHs: 2\nExp number of AAs in TMHs: 37.2608499999999\nExp number, first 60 AAs: 19.52923\nTotal prob of N-in: 0.94351\nPOSSIBLE N-term signal sequence\ninside 1 19\nTMhelix 20 39\noutside 40 325\nTMhelix 326 348\ninside 349 372

4627 GCF_001705405.1_ASM170540v1 Escherichia coli Proteobacteria; Gammaproteobacteria; Enterobacterales; Enterobacteriaceae; Escherichia MTGDNTLIHSHGINRRDFMKLCAALAATMGLSSKAAAEMAESVTNPQRPPVIWIGAQECT WP_000145410.1 MULTISPECIES: hydrogenase-2 small chain [Proteobacteria] Length: 372\nNumber of predicted TMHs: 1\nExp number of AAs in TMHs: 23.72226\nExp number, first 60 AAs: 0.22038\nTotal prob of N-in: 0.12994\noutside 1 331\nTMhelix 332 354\ninside 355 372

4628 GCF_001705405.1_ASM170540v1 Escherichia coli Proteobacteria; Gammaproteobacteria; Enterobacterales; Enterobacteriaceae; Escherichia MNNEETFYQAMRRQGVTRRSFLKYCSLAATSLGLGAGMAPKIAWALENKPRIPVVWIHGL WP_001301613.1 hydrogenase [Escherichia coli] Length: 372\nNumber of predicted TMHs: 2\nExp number of AAs in TMHs: 37.46598\nExp number, first 60 AAs: 19.53139\nTotal prob of N-in: 0.94364\nPOSSIBLE N-term signal sequence\ninside 1 19\nTMhelix 20 39\noutside 40 325\nTMhelix 326 348\ninside 349 372

4629 GCF_001742485.1_ASM174248v1 Escherichia coli Proteobacteria; Gammaproteobacteria; Enterobacterales; Enterobacteriaceae; Escherichia MTGDNTLIHSHGINRRDFMKLCAALAATMGLSSKAAAEMAESVTNPQRPPVIWIGAQECT WP_000145410.1 MULTISPECIES: hydrogenase-2 small chain [Proteobacteria] Length: 372\nNumber of predicted TMHs: 1\nExp number of AAs in TMHs: 23.72226\nExp number, first 60 AAs: 0.22038\nTotal prob of N-in: 0.12994\noutside 1 331\nTMhelix 332 354\ninside 355 372

4630 GCF_001742545.1_ASM174254v1 Escherichia coli Proteobacteria; Gammaproteobacteria; Enterobacterales; Enterobacteriaceae; Escherichia MTGDNTLIHSHGINRRDFMKLCAALAATMGLSSKAAAEMAESVTNPQRPPVIWIGAQECT WP_000145410.1 MULTISPECIES: hydrogenase-2 small chain [Proteobacteria] Length: 372\nNumber of predicted TMHs: 1\nExp number of AAs in TMHs: 23.72226\nExp number, first 60 AAs: 0.22038\nTotal prob of N-in: 0.12994\noutside 1 331\nTMhelix 332 354\ninside 355 372

4631 GCF_001742575.1_ASM174257v1 Escherichia coli Proteobacteria; Gammaproteobacteria; Enterobacterales; Enterobacteriaceae; Escherichia MTGDNTLIHSHGINRRDFMKLCAALAATMGLSSKAAAEMAESVTNPQRPPVIWIGAQECT WP_000145410.1 MULTISPECIES: hydrogenase-2 small chain [Proteobacteria] Length: 372\nNumber of predicted TMHs:

1\nExp number of AAs in TMHs: 23.72226\nExp number, first 60 AAs: 0.22038\nTotal prob of N-in: 0.12994\noutside 1 331\nTMhelix 332 354\ninside 355 372

4632 GCF_001742645.1_ASM174264v1 Escherichia coli Proteobacteria; Gammaproteobacteria; Enterobacterales; Enterobacteriaceae; Escherichia
MTGDNTLIHSHGINRRDFMKLCAALAATMGLSSKAAAEMAESVTNPQRPPVIWIGAQECT WP_000145406.1
hydrogenase-2 small chain [Escherichia coli] Length: 372\nNumber of predicted TMHs: 1\nExp number of AAs in TMHs: 23.72251\nExp number, first 60 AAs: 0.22027\nTotal prob of N-in: 0.13006\noutside 1 331\nTMhelix 332 354\ninside 355 372

4633 GCF_001742655.1_ASM174265v1 Escherichia coli Proteobacteria; Gammaproteobacteria; Enterobacterales; Enterobacteriaceae; Escherichia
MTGDNTLIHSHGINRRDFMKLCAALAATMGLSSKAAAEMAESVTNPQRPPVIWIGAQECT WP_000145406.1
hydrogenase-2 small chain [Escherichia coli] Length: 372\nNumber of predicted TMHs: 1\nExp number of AAs in TMHs: 23.72251\nExp number, first 60 AAs: 0.22027\nTotal prob of N-in: 0.13006\noutside 1 331\nTMhelix 332 354\ninside 355 372

4634 GCF_001742545.1_ASM174254v1 Escherichia coli Proteobacteria; Gammaproteobacteria; Enterobacterales; Enterobacteriaceae; Escherichia
MNNEETFYQAMRRQGVTRRSFLKYCSLAATSLGLGAGMAPKIAWALENKPRIPVVWIHGL WP_001058323.1
MULTISPECIES: hydrogenase-1 small chain [Proteobacteria] Length: 372\nNumber of predicted TMHs: 2\nExp number of AAs in TMHs: 37.2608499999999\nExp number, first 60 AAs: 19.52923\nTotal prob of N-in: 0.94351\nPOSSIBLE N-term signal sequence\ninside 1 19\nTMhelix 20 39\noutside 40 325\nTMhelix 326 348\ninside 349 372

4635 GCF_001742575.1_ASM174257v1 Escherichia coli Proteobacteria; Gammaproteobacteria; Enterobacterales; Enterobacteriaceae; Escherichia
MNNEETFYQAMRRQGVTRRSFLKYCSLAATSLGLGAGMAPKIAWALENKPRIPVVWIHGL WP_001058323.1
MULTISPECIES: hydrogenase-1 small chain [Proteobacteria] Length: 372\nNumber of predicted TMHs: 2\nExp number of AAs in TMHs: 37.2608499999999\nExp number, first 60 AAs: 19.52923\nTotal prob of N-in: 0.94351\nPOSSIBLE N-term signal sequence\ninside 1 19\nTMhelix 20 39\noutside 40 325\nTMhelix 326 348\ninside 349 372

4636 GCF_001748605.1_ASM174860v1 Escherichia coli Proteobacteria; Gammaproteobacteria; Enterobacterales; Enterobacteriaceae; Escherichia
MNNEETFYQAMRRQGVTRRSFLKYCSLAATSLGLGAGMAPKIAWALENKPRIPVVWIHGL WP_001058323.1
MULTISPECIES: hydrogenase-1 small chain [Proteobacteria] Length: 372\nNumber of predicted TMHs: 2\nExp number of AAs in TMHs: 37.2608499999999\nExp number, first 60 AAs: 19.52923\nTotal prob of N-in: 0.94351\nPOSSIBLE N-term signal sequence\ninside 1 19\nTMhelix 20 39\noutside 40 325\nTMhelix 326 348\ninside 349 372

4637 GCF_001742485.1_ASM174248v1 Escherichia coli Proteobacteria; Gammaproteobacteria; Enterobacterales; Enterobacteriaceae; Escherichia
MNNEETFYQAMRRQGVTRRSFLKYCSLAATSLGLGAGMAPKIAWALENKPRIPVVWIHGL WP_001058323.1
MULTISPECIES: hydrogenase-1 small chain [Proteobacteria] Length: 372\nNumber of predicted TMHs: 2\nExp number of AAs in TMHs: 37.2608499999999\nExp number, first 60 AAs: 19.52923\nTotal prob of N-in: 0.94351\nPOSSIBLE N-term signal sequence\ninside 1 19\nTMhelix 20 39\noutside 40 325\nTMhelix 326 348\ninside 349 372

4638 GCF_001748535.1_ASM174853v1 Escherichia coli Proteobacteria; Gammaproteobacteria; Enterobacterales; Enterobacteriaceae; Escherichia
MTGDNTLIHSHGINRRDFMKLCAALAATMGLSSKAAAEMAESVTNPQRPPVIWIGAQECT WP_000145410.1
MULTISPECIES: hydrogenase-2 small chain [Proteobacteria] Length: 372\nNumber of predicted TMHs: 1\nExp number of AAs in TMHs: 23.72226\nExp number, first 60 AAs: 0.22038\nTotal prob of N-in: 0.12994\noutside 1 331\nTMhelix 332 354\ninside 355 372

4639 GCF_001748535.1_ASM174853v1 Escherichia coli Proteobacteria; Gammaproteobacteria; Enterobacterales; Enterobacteriaceae; Escherichia
MNNEETFYQAMRRQGVTRRSFLKYCSLAATSLGLGAGMAPKIAWALENKPRIPVVWIHGL WP_001058323.1
MULTISPECIES: hydrogenase-1 small chain [Proteobacteria] Length: 372\nNumber of predicted TMHs: 2\nExp number of AAs in TMHs: 37.2608499999999\nExp number, first 60 AAs: 19.52923\nTotal prob of N-in:

0.94351\nPOSSIBLE N-term signal sequence\ninside 1 19\nTMhelix 20 39\noutside 40 325\nTMhelix 326 348\ninside 349 372

4640 GCF_001742655.1_ASM174265v1 Escherichia coli Proteobacteria; Gammaproteobacteria; Enterobacterales; Enterobacteriaceae; Escherichia MNNEETFYQAMRRQGVTRRSFLKYCSLAATSLGLGAGMAPKIAWALENKPRIPVVWIHGL WP_001058323.1 MULTISPECIES: hydrogenase-1 small chain [Proteobacteria] Length: 372\nNumber of predicted TMHs: 2\nExp number of AAs in TMHs: 37.2608499999999\nExp number, first 60 AAs: 19.52923\nTotal prob of N-in: 0.94351\nPOSSIBLE N-term signal sequence\ninside 1 19\nTMhelix 20 39\noutside 40 325\nTMhelix 326 348\ninside 349 372

4641 GCF_001748735.1_ASM174873v1 Escherichia coli Proteobacteria; Gammaproteobacteria; Enterobacterales; Enterobacteriaceae; Escherichia MTGDNTLIHSHGINRRDFMKLCAALAATMGLSSKAAAEMAESVTNPQRPPVIWIGAQECT WP_000145410.1 MULTISPECIES: hydrogenase-2 small chain [Proteobacteria] Length: 372\nNumber of predicted TMHs: 1\nExp number of AAs in TMHs: 23.72226\nExp number, first 60 AAs: 0.22038\nTotal prob of N-in: 0.12994\noutside 1 331\nTMhelix 332 354\ninside 355 372

4642 GCF_001748625.1_ASM174862v1 Escherichia coli Proteobacteria; Gammaproteobacteria; Enterobacterales; Enterobacteriaceae; Escherichia MTGDNTLIHSHGINRRDFMKLCAALAATMGLSSKAAAEMAESVTNPQRPPVIWIGAQECT WP_000145410.1 MULTISPECIES: hydrogenase-2 small chain [Proteobacteria] Length: 372\nNumber of predicted TMHs: 1\nExp number of AAs in TMHs: 23.72226\nExp number, first 60 AAs: 0.22038\nTotal prob of N-in: 0.12994\noutside 1 331\nTMhelix 332 354\ninside 355 372

4643 GCF_001748905.1_ASM174890v1 Escherichia coli Proteobacteria; Gammaproteobacteria; Enterobacterales; Enterobacteriaceae; Escherichia MNNEETFYQAMRRQGVTRRSFLKYCSLAATSLGLGAGMAPKIAWALENKPRIPVVWIHGL WP_001058323.1 MULTISPECIES: hydrogenase-1 small chain [Proteobacteria] Length: 372\nNumber of predicted TMHs: 2\nExp number of AAs in TMHs: 37.2608499999999\nExp number, first 60 AAs: 19.52923\nTotal prob of N-in: 0.94351\nPOSSIBLE N-term signal sequence\ninside 1 19\nTMhelix 20 39\noutside 40 325\nTMhelix 326 348\ninside 349 372

4644 GCF_001749055.1_ASM174905v1 Escherichia coli Proteobacteria; Gammaproteobacteria; Enterobacterales; Enterobacteriaceae; Escherichia MNNEETFYQAMRRQGVTRRSFLKYCSLAATSLGLGAGMAPKIAWALENKPRIPVVWIHGL WP_001058323.1 MULTISPECIES: hydrogenase-1 small chain [Proteobacteria] Length: 372\nNumber of predicted TMHs: 2\nExp number of AAs in TMHs: 37.2608499999999\nExp number, first 60 AAs: 19.52923\nTotal prob of N-in: 0.94351\nPOSSIBLE N-term signal sequence\ninside 1 19\nTMhelix 20 39\noutside 40 325\nTMhelix 326 348\ninside 349 372

4645 GCF_001749145.1_ASM174914v1 Escherichia coli Proteobacteria; Gammaproteobacteria; Enterobacterales; Enterobacteriaceae; Escherichia MTGDNTLIHSHGINRRDFMKLCAALAATMGLSSKAAAEMAESVTNPQRPPVIWIGAQECT WP_000145410.1 MULTISPECIES: hydrogenase-2 small chain [Proteobacteria] Length: 372\nNumber of predicted TMHs: 1\nExp number of AAs in TMHs: 23.72226\nExp number, first 60 AAs: 0.22038\nTotal prob of N-in: 0.12994\noutside 1 331\nTMhelix 332 354\ninside 355 372

4646 GCF_001749205.1_ASM174920v1 Escherichia coli Proteobacteria; Gammaproteobacteria; Enterobacterales; Enterobacteriaceae; Escherichia MTGDNTLIHSHGINRRDFMKLCAALAATMGLSSKAAAEMAESVTNPQRPPVIWIGAQECT WP_000145410.1 MULTISPECIES: hydrogenase-2 small chain [Proteobacteria] Length: 372\nNumber of predicted TMHs: 1\nExp number of AAs in TMHs: 23.72226\nExp number, first 60 AAs: 0.22038\nTotal prob of N-in: 0.12994\noutside 1 331\nTMhelix 332 354\ninside 355 372

4647 GCF_001749145.1_ASM174914v1 Escherichia coli Proteobacteria; Gammaproteobacteria; Enterobacterales; Enterobacteriaceae; Escherichia MNNEETFYQAMRRQGVTRRSFLKYCSLAATSLGLGAGMAPKIAWALENKPRIPVVWIHGL WP_001058323.1 MULTISPECIES: hydrogenase-1 small chain [Proteobacteria] Length: 372\nNumber of predicted TMHs: 2\nExp number of AAs in TMHs: 37.2608499999999\nExp number, first 60 AAs: 19.52923\nTotal prob of N-in:

0.94351\nPOSSIBLE N-term signal sequence\ninside 1 19\nTMhelix 20 39\noutside 40 325\nTMhelix 326 348\ninside 349 372

4648 GCF_001749205.1_ASM174920v1 Escherichia coli Proteobacteria; Gammaproteobacteria; Enterobacterales; Enterobacteriaceae; Escherichia MNNEETFYQAMRRQGVTRRSFLKYCSLAATSLGLGAGMAPKIAWALENKPRIPVVWIHGL WP_001058323.1 MULTISPECIES: hydrogenase-1 small chain [Proteobacteria] Length: 372\nNumber of predicted TMHs: 2\nExp number of AAs in TMHs: 37.2608499999999\nExp number, first 60 AAs: 19.52923\nTotal prob of N-in: 0.94351\nPOSSIBLE N-term signal sequence\ninside 1 19\nTMhelix 20 39\noutside 40 325\nTMhelix 326 348\ninside 349 372

4649 GCF_001749075.1_ASM174907v1 Escherichia coli Proteobacteria; Gammaproteobacteria; Enterobacterales; Enterobacteriaceae; Escherichia MNNEETFYQAMRRQGVTRRSFLKYCSLAATSLGLGAGMAPKIAWALENKPRIPVVWIHGL WP_001058323.1 MULTISPECIES: hydrogenase-1 small chain [Proteobacteria] Length: 372\nNumber of predicted TMHs: 2\nExp number of AAs in TMHs: 37.2608499999999\nExp number, first 60 AAs: 19.52923\nTotal prob of N-in: 0.94351\nPOSSIBLE N-term signal sequence\ninside 1 19\nTMhelix 20 39\noutside 40 325\nTMhelix 326 348\ninside 349 372

4650 GCF_001748835.1_ASM174883v1 Escherichia coli Proteobacteria; Gammaproteobacteria; Enterobacterales; Enterobacteriaceae; Escherichia MNNEETFYQAMRRQGVTRRSFLKYCSLAATSLGLGAGMAPKIAWALENKPRIPVVWIHGL WP_001295940.1 MULTISPECIES: hydrogenase [Enterobacteriaceae] Length: 372\nNumber of predicted TMHs: 2\nExp number of AAs in TMHs: 37.2608899999999\nExp number, first 60 AAs: 19.52923\nTotal prob of N-in: 0.94351\nPOSSIBLE N-term signal sequence\ninside 1 19\nTMhelix 20 39\noutside 40 325\nTMhelix 326 348\ninside 349 372

4651 GCF_001749385.1_ASM174938v1 Escherichia coli Proteobacteria; Gammaproteobacteria; Enterobacterales; Enterobacteriaceae; Escherichia MTGDNTLIHSHGINRRDFMKLCAALAATMGLSSKAAAEMAESVTNPQRPPVIWIGAQECT WP_000145410.1 MULTISPECIES: hydrogenase-2 small chain [Proteobacteria] Length: 372\nNumber of predicted TMHs: 1\nExp number of AAs in TMHs: 23.72226\nExp number, first 60 AAs: 0.22038\nTotal prob of N-in: 0.12994\noutside 1 331\nTMhelix 332 354\ninside 355 372

4652 GCF_001749535.1_ASM174953v1 Escherichia coli Proteobacteria; Gammaproteobacteria; Enterobacterales; Enterobacteriaceae; Escherichia MTGDNTLIHSHGINRRDFMKLCAALAATMGLSSKAAAEMAESVTNPQRPPVIWIGAQECT WP_000145410.1 MULTISPECIES: hydrogenase-2 small chain [Proteobacteria] Length: 372\nNumber of predicted TMHs: 1\nExp number of AAs in TMHs: 23.72226\nExp number, first 60 AAs: 0.22038\nTotal prob of N-in: 0.12994\noutside 1 331\nTMhelix 332 354\ninside 355 372

4653 GCF_001749505.1_ASM174950v1 Escherichia coli Proteobacteria; Gammaproteobacteria; Enterobacterales; Enterobacteriaceae; Escherichia MTGDNTLIHSHGINRRDFMKLCAALAATMGLSSKAAAEMAESVTNPQRPPVIWIGAQECT WP_000145410.1 MULTISPECIES: hydrogenase-2 small chain [Proteobacteria] Length: 372\nNumber of predicted TMHs: 1\nExp number of AAs in TMHs: 23.72226\nExp number, first 60 AAs: 0.22038\nTotal prob of N-in: 0.12994\noutside 1 331\nTMhelix 332 354\ninside 355 372

4654 GCF_001749435.1_ASM174943v1 Escherichia coli Proteobacteria; Gammaproteobacteria; Enterobacterales; Enterobacteriaceae; Escherichia MTGDNTLIHSHGINRRDFMKLCAALAATMGLSSKAAAEMAESVTNPQRPPVIWIGAQECT WP_000145410.1 MULTISPECIES: hydrogenase-2 small chain [Proteobacteria] Length: 372\nNumber of predicted TMHs: 1\nExp number of AAs in TMHs: 23.72226\nExp number, first 60 AAs: 0.22038\nTotal prob of N-in: 0.12994\noutside 1 331\nTMhelix 332 354\ninside 355 372

4655 GCF_001749595.1_ASM174959v1 Escherichia coli Proteobacteria; Gammaproteobacteria; Enterobacterales; Enterobacteriaceae; Escherichia MTGDNTLIHSHGINRRDFMKLCAALAATMGLSSKAAAEMAESVTNPQRPPVIWIGAQECT WP_000145410.1 MULTISPECIES: hydrogenase-2 small chain [Proteobacteria] Length: 372\nNumber of predicted TMHs: 1\nExp number of AAs in TMHs: 23.72226\nExp number, first 60 AAs: 0.22038\nTotal prob of N-in: 0.12994\noutside 1 331\nTMhelix 332 354\ninside 355 372

4656 GCF_001749635.1_ASM174963v1 Escherichia coli Proteobacteria; Gammaproteobacteria; Enterobacterales; Enterobacteriaceae; Escherichia
MTGDNTLIHSHGINRRDFMKLCAALAATMGLSSKAAAEMAESVTNPQRPPVIWIGAQECT WP_000145410.1
MULTISPECIES: hydrogenase-2 small chain [Proteobacteria] Length: 372\nNumber of predicted TMHs: 1\nExp number of AAs in TMHs: 23.72226\nExp number, first 60 AAs: 0.22038\nTotal prob of N-in: 0.12994\noutside 1 331\nTMhelix 332 354\ninside 355 372

4657 GCF_001749705.1_ASM174970v1 Escherichia coli Proteobacteria; Gammaproteobacteria; Enterobacterales; Enterobacteriaceae; Escherichia
MTGDNTLIHSHGINRRDFMKLCAALAATMGLSSKAAAEMAESVTNPQRPPVIWIGAQECT WP_000145410.1
MULTISPECIES: hydrogenase-2 small chain [Proteobacteria] Length: 372\nNumber of predicted TMHs: 1\nExp number of AAs in TMHs: 23.72226\nExp number, first 60 AAs: 0.22038\nTotal prob of N-in: 0.12994\noutside 1 331\nTMhelix 332 354\ninside 355 372

4658 GCF_001749435.1_ASM174943v1 Escherichia coli Proteobacteria; Gammaproteobacteria; Enterobacterales; Enterobacteriaceae; Escherichia
MNNEETFYQAMRRQGVTRRSFLKYCSLAATSLGLGAGMAPKIAWALENKPRIPVVWIHGL WP_001295940.1
MULTISPECIES: hydrogenase [Enterobacteriaceae] Length: 372\nNumber of predicted TMHs: 2\nExp number of AAs in TMHs: 37.2608899999999\nExp number, first 60 AAs: 19.52923\nTotal prob of N-in: 0.94351\nPOSSIBLE N-term signal sequence\ninside 1 19\nTMhelix 20 39\noutside 40 325\nTMhelix 326 348\ninside 349 372

4659 GCF_001749705.1_ASM174970v1 Escherichia coli Proteobacteria; Gammaproteobacteria; Enterobacterales; Enterobacteriaceae; Escherichia
MNNEETFYQAMRRQGVTRRSFLKYCSLAATSLGLGAGMAPKIAWALENKPRIPVVWIHGL WP_001058323.1
MULTISPECIES: hydrogenase-1 small chain [Proteobacteria] Length: 372\nNumber of predicted TMHs: 2\nExp number of AAs in TMHs: 37.2608499999999\nExp number, first 60 AAs: 19.52923\nTotal prob of N-in: 0.94351\nPOSSIBLE N-term signal sequence\ninside 1 19\nTMhelix 20 39\noutside 40 325\nTMhelix 326 348\ninside 349 372

4660 GCF_001806285.1_ASM180628v1 Escherichia coli Proteobacteria; Gammaproteobacteria; Enterobacterales; Enterobacteriaceae; Escherichia
MTGDNTLIHSHGINRRDFMKLCAALAATMGLSSKAAAEMAESVTNPQRPPVIWIGAQECT WP_000145410.1
MULTISPECIES: hydrogenase-2 small chain [Proteobacteria] Length: 372\nNumber of predicted TMHs: 1\nExp number of AAs in TMHs: 23.72226\nExp number, first 60 AAs: 0.22038\nTotal prob of N-in: 0.12994\noutside 1 331\nTMhelix 332 354\ninside 355 372

4661 GCF_001758245.1_ASM175824v1 Escherichia coli Proteobacteria; Gammaproteobacteria; Enterobacterales; Enterobacteriaceae; Escherichia
MTGDNTLIHSHGINRRDFMKLCAALAATMGLSSKAAAEMAESVTNPQRPPVIWIGAQECT WP_000145410.1
MULTISPECIES: hydrogenase-2 small chain [Proteobacteria] Length: 372\nNumber of predicted TMHs: 1\nExp number of AAs in TMHs: 23.72226\nExp number, first 60 AAs: 0.22038\nTotal prob of N-in: 0.12994\noutside 1 331\nTMhelix 332 354\ninside 355 372

4662 GCF_001865195.1_ASM186519v1 Escherichia coli Proteobacteria; Gammaproteobacteria; Enterobacterales; Enterobacteriaceae; Escherichia
MNNEETFYQAMRRQGVTRRSFLKYCSLAATSLGLGAGMAPKIAWALENKPRIPVVWIHGL WP_001058320.1
hydrogenase [Escherichia coli] Length: 372\nNumber of predicted TMHs: 2\nExp number of AAs in TMHs: 37.26082\nExp number, first 60 AAs: 19.52923\nTotal prob of N-in: 0.94351\nPOSSIBLE N-term signal sequence\ninside 1 19\nTMhelix 20 39\noutside 40 325\nTMhelix 326 348\ninside 349 372

4663 GCF_001865725.1_ASM186572v1 Escherichia coli Proteobacteria; Gammaproteobacteria; Enterobacterales; Enterobacteriaceae; Escherichia
MNNEETFYQAMRRQGVTRRSFLKYCSLAATSLGLGAGMAPKIAWALENKPRIPVVWIHGL WP_001058323.1
MULTISPECIES: hydrogenase-1 small chain [Proteobacteria] Length: 372\nNumber of predicted TMHs: 2\nExp number of AAs in TMHs: 37.2608499999999\nExp number, first 60 AAs: 19.52923\nTotal prob of N-in: 0.94351\nPOSSIBLE N-term signal sequence\ninside 1 19\nTMhelix 20 39\noutside 40 325\nTMhelix 326 348\ninside 349 372

4664 GCF_001865975.1_ASM186597v1 Escherichia coli Proteobacteria; Gammaproteobacteria; Enterobacterales; Enterobacteriaceae; Escherichia

MNNEETFYQAMRRQGVTRRSFLKYCSLAATSLGLGAGMAPKIAWALENKPRIPVVWIHGL WP_001058323.1 MULTISPECIES: hydrogenase-1 small chain [Proteobacteria] Length: 372\nNumber of predicted TMHs: 2\nExp number of AAs in TMHs: 37.2608499999999\nExp number, first 60 AAs: 19.52923\nTotal prob of N-in: 0.94351\nPOSSIBLE N-term signal sequence\ninside 1 19\nTMhelix 20 39\noutside 40 325\nTMhelix 326 348\ninside 349 372

4665 GCF_001806285.1_ASM180628v1 Escherichia coli Proteobacteria; Gammaproteobacteria; Enterobacterales; Enterobacteriaceae; Escherichia MNNEETFYQAMRRQGVTRRSFLKYCSLAATSLGLGAGMAPKIAWALENKPRIPVVWIHGL WP_001301613.1 hydrogenase [Escherichia coli] Length: 372\nNumber of predicted TMHs: 2\nExp number of AAs in TMHs: 37.46598\nExp number, first 60 AAs: 19.53139\nTotal prob of N-in: 0.94364\nPOSSIBLE N-term signal sequence\ninside 1 19\nTMhelix 20 39\noutside 40 325\nTMhelix 326 348\ninside 349 372

4666 GCF_001816315.1_ASM181631v1 Escherichia coli Proteobacteria; Gammaproteobacteria; Enterobacterales; Enterobacteriaceae; Escherichia MNNEETFYQAMRRQGVTRRSFLKYCSLAATSLGLGAGMAPKIAWALENKPRIPVVWIHGL WP_001058323.1 MULTISPECIES: hydrogenase-1 small chain [Proteobacteria] Length: 372\nNumber of predicted TMHs: 2\nExp number of AAs in TMHs: 37.2608499999999\nExp number, first 60 AAs: 19.52923\nTotal prob of N-in: 0.94351\nPOSSIBLE N-term signal sequence\ninside 1 19\nTMhelix 20 39\noutside 40 325\nTMhelix 326 348\ninside 349 372

4667 GCF_001865185.1_ASM186518v1 Escherichia coli Proteobacteria; Gammaproteobacteria; Enterobacterales; Enterobacteriaceae; Escherichia MNNEETFYQAMRRQGVTRRSFLKYCSLAATSLGLGAGMAPKIAWALENKPRIPVVWIHGL WP_001058323.1 MULTISPECIES: hydrogenase-1 small chain [Proteobacteria] Length: 372\nNumber of predicted TMHs: 2\nExp number of AAs in TMHs: 37.2608499999999\nExp number, first 60 AAs: 19.52923\nTotal prob of N-in: 0.94351\nPOSSIBLE N-term signal sequence\ninside 1 19\nTMhelix 20 39\noutside 40 325\nTMhelix 326 348\ninside 349 372

4668 GCF_001881315.1_ASM188131v1 Escherichia coli Proteobacteria; Gammaproteobacteria; Enterobacterales; Enterobacteriaceae; Escherichia MTGDNTLIHSHGINRRDFMKLCAALAATMGLSSKAAAEMAESVTNPQRPPVIWIGAQECT WP_000145410.1 MULTISPECIES: hydrogenase-2 small chain [Proteobacteria] Length: 372\nNumber of predicted TMHs: 1\nExp number of AAs in TMHs: 23.72226\nExp number, first 60 AAs: 0.22038\nTotal prob of N-in: 0.12994\noutside 1 331\nTMhelix 332 354\ninside 355 372

4669 GCF_001881275.1_ASM188127v1 Escherichia coli Proteobacteria; Gammaproteobacteria; Enterobacterales; Enterobacteriaceae; Escherichia MTGDNTLIHSHGINRRDFMKLCAALAATMGLSSKAAAEMAESVTNPQRPPVIWIGAQECT WP_000145410.1 MULTISPECIES: hydrogenase-2 small chain [Proteobacteria] Length: 372\nNumber of predicted TMHs: 1\nExp number of AAs in TMHs: 23.72226\nExp number, first 60 AAs: 0.22038\nTotal prob of N-in: 0.12994\noutside 1 331\nTMhelix 332 354\ninside 355 372

4670 GCF_001881385.1_ASM188138v1 Escherichia coli Proteobacteria; Gammaproteobacteria; Enterobacterales; Enterobacteriaceae; Escherichia MTGDNTLIHSHGINRRDFMKLCAALAATMGLSSKAAAEMAESVTNPQRPPVIWIGAQECT WP_000145410.1 MULTISPECIES: hydrogenase-2 small chain [Proteobacteria] Length: 372\nNumber of predicted TMHs: 1\nExp number of AAs in TMHs: 23.72226\nExp number, first 60 AAs: 0.22038\nTotal prob of N-in: 0.12994\noutside 1 331\nTMhelix 332 354\ninside 355 372

4671 GCF_001881425.1_ASM188142v1 Escherichia coli Proteobacteria; Gammaproteobacteria; Enterobacterales; Enterobacteriaceae; Escherichia MTGDNTLIHSHGINRRDFMKLCAALAATMGLSSKAAAEMAESVTNPQRPPVIWIGAQECT WP_000145410.1 MULTISPECIES: hydrogenase-2 small chain [Proteobacteria] Length: 372\nNumber of predicted TMHs: 1\nExp number of AAs in TMHs: 23.72226\nExp number, first 60 AAs: 0.22038\nTotal prob of N-in: 0.12994\noutside 1 331\nTMhelix 332 354\ninside 355 372

4672 GCF_001881505.1_ASM188150v1 Escherichia coli Proteobacteria; Gammaproteobacteria; Enterobacterales; Enterobacteriaceae; Escherichia MTGDNTLIHSHGINRRDFMKLCAALAATMGLSSKAAAEMAESVTNPQRPPVIWIGAQECT WP_000145410.1 MULTISPECIES: hydrogenase-2 small chain [Proteobacteria] Length: 372\nNumber of predicted TMHs:

1\nExp number of AAs in TMHs: 23.72226\nExp number, first 60 AAs: 0.22038\nTotal prob of N-in: 0.12994\noutside 1 331\nTMhelix 332 354\ninside 355 372

4673 GCF_001881595.3_ASM188159v3 Escherichia coli Proteobacteria; Gammaproteobacteria; Enterobacterales; Enterobacteriaceae; Escherichia
MTGDNTLIHSHGINRRDFMKLCAALAATMGLSSKAAAEMAESVTNPQRPPVIWIGAQECT WP_000145410.1
MULTISPECIES: hydrogenase-2 small chain [Proteobacteria] Length: 372\nNumber of predicted TMHs: 1\nExp number of AAs in TMHs: 23.72226\nExp number, first 60 AAs: 0.22038\nTotal prob of N-in: 0.12994\noutside 1 331\nTMhelix 332 354\ninside 355 372

4674 GCF_001881605.1_ASM188160v1 Escherichia coli Proteobacteria; Gammaproteobacteria; Enterobacterales; Enterobacteriaceae; Escherichia
MTGDNTLIHSHGINRRDFMKLCAALAATMGLSSKAAAEMAESVTNPQRPPVIWIGAQECT WP_000145410.1
MULTISPECIES: hydrogenase-2 small chain [Proteobacteria] Length: 372\nNumber of predicted TMHs: 1\nExp number of AAs in TMHs: 23.72226\nExp number, first 60 AAs: 0.22038\nTotal prob of N-in: 0.12994\noutside 1 331\nTMhelix 332 354\ninside 355 372

4675 GCF_001881645.1_ASM188164v1 Escherichia coli Proteobacteria; Gammaproteobacteria; Enterobacterales; Enterobacteriaceae; Escherichia
MTGDNTLIHSHGINRRDFMKLCAALAATMGLSSKAAAEMAESVTNPQRPPVIWIGAQECT WP_000145410.1
MULTISPECIES: hydrogenase-2 small chain [Proteobacteria] Length: 372\nNumber of predicted TMHs: 1\nExp number of AAs in TMHs: 23.72226\nExp number, first 60 AAs: 0.22038\nTotal prob of N-in: 0.12994\noutside 1 331\nTMhelix 332 354\ninside 355 372

4676 GCF_001886935.1_ASM188693v1 Escherichia coli Proteobacteria; Gammaproteobacteria; Enterobacterales; Enterobacteriaceae; Escherichia
MTGDNTLIHSHGINRRDFMKLCAALAATMGLSSKAAAEMAESVTNPQRPPVIWIGAQECT WP_000145410.1
MULTISPECIES: hydrogenase-2 small chain [Proteobacteria] Length: 372\nNumber of predicted TMHs: 1\nExp number of AAs in TMHs: 23.72226\nExp number, first 60 AAs: 0.22038\nTotal prob of N-in: 0.12994\noutside 1 331\nTMhelix 332 354\ninside 355 372

4677 GCF_001890225.1_ASM189022v1 Escherichia coli Proteobacteria; Gammaproteobacteria; Enterobacterales; Enterobacteriaceae; Escherichia
MTGDNTLIHSHGINRRDFMKLCAALAATMGLSSKAAAEMAESVTNPQRPPVIWIGAQECT WP_000145410.1
MULTISPECIES: hydrogenase-2 small chain [Proteobacteria] Length: 372\nNumber of predicted TMHs: 1\nExp number of AAs in TMHs: 23.72226\nExp number, first 60 AAs: 0.22038\nTotal prob of N-in: 0.12994\noutside 1 331\nTMhelix 332 354\ninside 355 372

4678 GCF_001884885.1_ASM188488v1 Escherichia coli Proteobacteria; Gammaproteobacteria; Enterobacterales; Enterobacteriaceae; Escherichia
MNNEETFYQAMRRQGVTRRSFLKYCSLAATSLGLGAGMAPKIAWALENKPRIPVVWIHGL WP_001301613.1
hydrogenase [Escherichia coli] Length: 372\nNumber of predicted TMHs: 2\nExp number of AAs in TMHs: 37.46598\nExp number, first 60 AAs: 19.53139\nTotal prob of N-in: 0.94364\nPOSSIBLE N-term signal sequence\ninside 1 19\nTMhelix 20 39\noutside 40 325\nTMhelix 326 348\ninside 349 372

4679 GCF_001890265.1_ASM189026v1 Escherichia coli Proteobacteria; Gammaproteobacteria; Enterobacterales; Enterobacteriaceae; Escherichia
MTGDNTLIHSHGINRRDFMKLCAALAATMGLSSKAAAEMAESVTNPQRPPVIWIGAQECT WP_000145410.1
MULTISPECIES: hydrogenase-2 small chain [Proteobacteria] Length: 372\nNumber of predicted TMHs: 1\nExp number of AAs in TMHs: 23.72226\nExp number, first 60 AAs: 0.22038\nTotal prob of N-in: 0.12994\noutside 1 331\nTMhelix 332 354\ninside 355 372

4680 GCF_001890325.1_ASM189032v1 Escherichia coli Proteobacteria; Gammaproteobacteria; Enterobacterales; Enterobacteriaceae; Escherichia
MTGDNTLIHSHGINRRDFMKLCAALAATMGLSSKAAAEMAESVTNPQRPPVIWIGAQECT WP_000145410.1
MULTISPECIES: hydrogenase-2 small chain [Proteobacteria] Length: 372\nNumber of predicted TMHs: 1\nExp number of AAs in TMHs: 23.72226\nExp number, first 60 AAs: 0.22038\nTotal prob of N-in: 0.12994\noutside 1 331\nTMhelix 332 354\ninside 355 372

4681 GCF_001884945.1_ASM188494v1 Escherichia coli Proteobacteria; Gammaproteobacteria; Enterobacterales; Enterobacteriaceae; Escherichia

MNNEETFYQAMRRQGVTRRSFLKYCSLAATSLGLGAGMAPKIAWALENKPRIPVVWIHGL WP_001301613.1 hydrogenase [Escherichia coli] Length: 372\nNumber of predicted TMHs: 2\nExp number of AAs in TMHs: 37.46598\nExp number, first 60 AAs: 19.53139\nTotal prob of N-in: 0.94364\nPOSSIBLE N-term signal sequence\ninside 1 19\nTMhelix 20 39\noutside 40 325\nTMhelix 326 348\ninside 349 372

4682 GCF_001884885.1_ASM188488v1 Escherichia coli Proteobacteria; Gammaproteobacteria; Enterobacterales; Enterobacteriaceae; Escherichia MTGDNTLIHSHGINRRDFMKLCAALAATMGLSSKAAAEMAESVTNPQRPPVIWIGAQECS WP_061069243.1 hydrogenase 2 small subunit [Escherichia coli] Length: 372\nNumber of predicted TMHs: 1\nExp number of AAs in TMHs: 23.72219\nExp number, first 60 AAs: 0.22034\nTotal prob of N-in: 0.12994\noutside 1 331\nTMhelix 332 354\ninside 355 372

4683 GCF_001890365.1_ASM189036v1 Escherichia coli Proteobacteria; Gammaproteobacteria; Enterobacterales; Enterobacteriaceae; Escherichia MNNEETFYQAMRRQGVTRRSFLKYCSLAATSLGLGAGMAPKIAWALENKPRIPVVWIHGL WP_001058323.1 MULTISPECIES: hydrogenase-1 small chain [Proteobacteria] Length: 372\nNumber of predicted TMHs: 2\nExp number of AAs in TMHs: 37.2608499999999\nExp number, first 60 AAs: 19.52923\nTotal prob of N-in: 0.94351\nPOSSIBLE N-term signal sequence\ninside 1 19\nTMhelix 20 39\noutside 40 325\nTMhelix 326 348\ninside 349 372

4684 GCF_001886935.1_ASM188693v1 Escherichia coli Proteobacteria; Gammaproteobacteria; Enterobacterales; Enterobacteriaceae; Escherichia MNNEETFYQAMRRQGVTRRSFLKYCSLAATSLGLGAGMAPKIAWALENKPRIPVVWIHGL WP_001295940.1 MULTISPECIES: hydrogenase [Enterobacteriaceae] Length: 372\nNumber of predicted TMHs: 2\nExp number of AAs in TMHs: 37.2608899999999\nExp number, first 60 AAs: 19.52923\nTotal prob of N-in: 0.94351\nPOSSIBLE N-term signal sequence\ninside 1 19\nTMhelix 20 39\noutside 40 325\nTMhelix 326 348\ninside 349 372

4685 GCF_001891235.1_ASM189123v1 Escherichia coli Proteobacteria; Gammaproteobacteria; Enterobacterales; Enterobacteriaceae; Escherichia MTGDNTLIHSHGINRRDFMKLCAALAATMGLSSKAAAEMAESVTNPQRPPVIWIGAQECT WP_000145410.1 MULTISPECIES: hydrogenase-2 small chain [Proteobacteria] Length: 372\nNumber of predicted TMHs: 1\nExp number of AAs in TMHs: 23.72226\nExp number, first 60 AAs: 0.22038\nTotal prob of N-in: 0.12994\noutside 1 331\nTMhelix 332 354\ninside 355 372

4686 GCF_001891365.1_ASM189136v1 Escherichia coli Proteobacteria; Gammaproteobacteria; Enterobacterales; Enterobacteriaceae; Escherichia MTGDNTLIHSHGINRRDFMKLCAALAATMGLSSKAAAEMAESVTNPQRPPVIWIGAQECT WP_000145410.1 MULTISPECIES: hydrogenase-2 small chain [Proteobacteria] Length: 372\nNumber of predicted TMHs: 1\nExp number of AAs in TMHs: 23.72226\nExp number, first 60 AAs: 0.22038\nTotal prob of N-in: 0.12994\noutside 1 331\nTMhelix 332 354\ninside 355 372

4687 GCF_001891295.1_ASM189129v1 Escherichia coli Proteobacteria; Gammaproteobacteria; Enterobacterales; Enterobacteriaceae; Escherichia MTGDNTLIHSHGINRRDFMKLCAALAATMGLSSKAAAEMAESVTNPQRPPVIWIGAQECT WP_000145410.1 MULTISPECIES: hydrogenase-2 small chain [Proteobacteria] Length: 372\nNumber of predicted TMHs: 1\nExp number of AAs in TMHs: 23.72226\nExp number, first 60 AAs: 0.22038\nTotal prob of N-in: 0.12994\noutside 1 331\nTMhelix 332 354\ninside 355 372

4688 GCF_001891385.1_ASM189138v1 Escherichia coli Proteobacteria; Gammaproteobacteria; Enterobacterales; Enterobacteriaceae; Escherichia MTGDNTLIHSHGINRRDFMKLCAALAATMGLSSKAAAEMAESVTNPQRPPVIWIGAQECT WP_000145410.1 MULTISPECIES: hydrogenase-2 small chain [Proteobacteria] Length: 372\nNumber of predicted TMHs: 1\nExp number of AAs in TMHs: 23.72226\nExp number, first 60 AAs: 0.22038\nTotal prob of N-in: 0.12994\noutside 1 331\nTMhelix 332 354\ninside 355 372

4689 GCF_001891235.1_ASM189123v1 Escherichia coli Proteobacteria; Gammaproteobacteria; Enterobacterales; Enterobacteriaceae; Escherichia MNNEETFYQAMRRQGVTRRSFLKYCSLAATSLGLGAGMAPKIAWALENKPRIPVVWIHGL WP_001295940.1 MULTISPECIES: hydrogenase [Enterobacteriaceae] Length: 372\nNumber of predicted TMHs: 2\nExp number of AAs in TMHs: 37.2608899999999\nExp number, first 60 AAs: 19.52923\nTotal prob of N-in:

0.94351\nPOSSIBLE N-term signal sequence\ninside 1 19\nTMhelix 20 39\noutside 40 325\nTMhelix 326 348\ninside 349 372

4690 GCF_001891555.1_ASM189155v1 Escherichia coli Proteobacteria; Gammaproteobacteria; Enterobacterales; Enterobacteriaceae; Escherichia MTGDNTLIHSHGINRRDFMKLCAALAATMGLSSKAAAEMAESVTNPQRPPVIWIGAQECT WP_000145410.1 MULTISPECIES: hydrogenase-2 small chain [Proteobacteria] Length: 372\nNumber of predicted TMHs: 1\nExp number of AAs in TMHs: 23.72226\nExp number, first 60 AAs: 0.22038\nTotal prob of N-in: 0.12994\noutside 1 331\nTMhelix 332 354\ninside 355 372

4691 GCF_001891535.1_ASM189153v1 Escherichia coli Proteobacteria; Gammaproteobacteria; Enterobacterales; Enterobacteriaceae; Escherichia MNNEETFYQAMRRQGVTRRSFLKYCSLAATSLGLGAGMAPKIAWALENKPRIPVVWIHGL WP_001058323.1 MULTISPECIES: hydrogenase-1 small chain [Proteobacteria] Length: 372\nNumber of predicted TMHs: 2\nExp number of AAs in TMHs: 37.2608499999999\nExp number, first 60 AAs: 19.52923\nTotal prob of N-in: 0.94351\nPOSSIBLE N-term signal sequence\ninside 1 19\nTMhelix 20 39\noutside 40 325\nTMhelix 326 348\ninside 349 372

4692 GCF_001891465.1_ASM189146v1 Escherichia coli Proteobacteria; Gammaproteobacteria; Enterobacterales; Enterobacteriaceae; Escherichia MNNEETFYQAMRRQGVTRRSFLKYCSLAATSLGLGAGMAPKIAWALENKPRIPVVWIHGL WP_001058323.1 MULTISPECIES: hydrogenase-1 small chain [Proteobacteria] Length: 372\nNumber of predicted TMHs: 2\nExp number of AAs in TMHs: 37.2608499999999\nExp number, first 60 AAs: 19.52923\nTotal prob of N-in: 0.94351\nPOSSIBLE N-term signal sequence\ninside 1 19\nTMhelix 20 39\noutside 40 325\nTMhelix 326 348\ninside 349 372

4693 GCF_000782215.1_ASM78221v1 Escherichia coli Proteobacteria; Gammaproteobacteria; Enterobacterales; Enterobacteriaceae; Escherichia MNNEETFYQAMRRQGVTRRSFLKYCSLAATSLGLGAGMAPKIAWALENKPRIPVVWIHGL WP_001295940.1 MULTISPECIES: hydrogenase [Enterobacteriaceae] Length: 372\nNumber of predicted TMHs: 2\nExp number of AAs in TMHs: 37.2608899999999\nExp number, first 60 AAs: 19.52923\nTotal prob of N-in: 0.94351\nPOSSIBLE N-term signal sequence\ninside 1 19\nTMhelix 20 39\noutside 40 325\nTMhelix 326 348\ninside 349 372

4694 GCF_000782115.1_ASM78211v1 Escherichia coli Proteobacteria; Gammaproteobacteria; Enterobacterales; Enterobacteriaceae; Escherichia MNNEETFYQAMRRQGVTRRSFLKYCSLAATSLGLGAGMAPKIAWALENKPRIPVVWIHGL WP_001058323.1 MULTISPECIES: hydrogenase-1 small chain [Proteobacteria] Length: 372\nNumber of predicted TMHs: 2\nExp number of AAs in TMHs: 37.2608499999999\nExp number, first 60 AAs: 19.52923\nTotal prob of N-in: 0.94351\nPOSSIBLE N-term signal sequence\ninside 1 19\nTMhelix 20 39\noutside 40 325\nTMhelix 326 348\ninside 349 372

4695 GCF_000782115.1_ASM78211v1 Escherichia coli Proteobacteria; Gammaproteobacteria; Enterobacterales; Enterobacteriaceae; Escherichia MTGDNTLIHSHGINRRDFMKLCAALAATMGLSSKAAAEMAESVTNPQRPPVIWIGAQECT WP_000145410.1 MULTISPECIES: hydrogenase-2 small chain [Proteobacteria] Length: 372\nNumber of predicted TMHs: 1\nExp number of AAs in TMHs: 23.72226\nExp number, first 60 AAs: 0.22038\nTotal prob of N-in: 0.12994\noutside 1 331\nTMhelix 332 354\ninside 355 372

4696 GCF_000782055.1_ASM78205v1 Escherichia coli Proteobacteria; Gammaproteobacteria; Enterobacterales; Enterobacteriaceae; Escherichia MTGDNTLIHSHGINRRDFMKLCAALAATMGLSSKAAAEMAESVTNPQRPPVIWIGAQECT WP_000145410.1 MULTISPECIES: hydrogenase-2 small chain [Proteobacteria] Length: 372\nNumber of predicted TMHs: 1\nExp number of AAs in TMHs: 23.72226\nExp number, first 60 AAs: 0.22038\nTotal prob of N-in: 0.12994\noutside 1 331\nTMhelix 332 354\ninside 355 372

4697 GCF_000782015.1_ASM78201v1 Escherichia coli Proteobacteria; Gammaproteobacteria; Enterobacterales; Enterobacteriaceae; Escherichia MNNEETFYQAMRRQGVTRRSFLKYCSLAATSLGLGAGMAPKIAWALENKPRIPVVWIHGL WP_001295940.1 MULTISPECIES: hydrogenase [Enterobacteriaceae] Length: 372\nNumber of predicted TMHs: 2\nExp number of AAs in TMHs: 37.2608899999999\nExp number, first 60 AAs: 19.52923\nTotal prob of N-in:

0.94351\nPOSSIBLE N-term signal sequence\ninside 1 19\nTMhelix 20 39\noutside 40 325\nTMhelix 326 348\ninside 349 372

4698 GCF_000782015.1_ASM78201v1 Escherichia coli Proteobacteria; Gammaproteobacteria; Enterobacterales; Enterobacteriaceae; Escherichia MTGDNTLIHSHGINRRDFMKLCAALAATMGLSSKAAAEMAESVTNPQRPPVIWIGAQECT WP_000145410.1 MULTISPECIES: hydrogenase-2 small chain [Proteobacteria] Length: 372\nNumber of predicted TMHs: 1\nExp number of AAs in TMHs: 23.72226\nExp number, first 60 AAs: 0.22038\nTotal prob of N-in: 0.12994\noutside 1 331\nTMhelix 332 354\ninside 355 372

4699 GCF_000781855.1_ASM78185v1 Escherichia coli Proteobacteria; Gammaproteobacteria; Enterobacterales; Enterobacteriaceae; Escherichia MTGDNTLIHSHGINRRDFMKLCAALAATMGLSSKAAAEMAESVTNPQRPPVIWIGAQECT WP_000145410.1 MULTISPECIES: hydrogenase-2 small chain [Proteobacteria] Length: 372\nNumber of predicted TMHs: 1\nExp number of AAs in TMHs: 23.72226\nExp number, first 60 AAs: 0.22038\nTotal prob of N-in: 0.12994\noutside 1 331\nTMhelix 332 354\ninside 355 372

4700 GCF_001058315.1_ASM105831v1 Escherichia coli Proteobacteria; Gammaproteobacteria; Enterobacterales; Enterobacteriaceae; Escherichia MNNEETFYQAMRRQGVTRRSFLKYCSLAATSLGLGAGMAPKIAWALENKPRIPVVWIHGL WP_001295940.1 MULTISPECIES: hydrogenase [Enterobacteriaceae] Length: 372\nNumber of predicted TMHs: 2\nExp number of AAs in TMHs: 37.2608899999999\nExp number, first 60 AAs: 19.52923\nTotal prob of N-in: 0.94351\nPOSSIBLE N-term signal sequence\ninside 1 19\nTMhelix 20 39\noutside 40 325\nTMhelix 326 348\ninside 349 372

4701 GCF_001058315.1_ASM105831v1 Escherichia coli Proteobacteria; Gammaproteobacteria; Enterobacterales; Enterobacteriaceae; Escherichia MTGDNTLIHSHGINRRDFMKLCAALAATMGLSSKAAAEMAESVTNPQRPPVIWIGAQECT WP_000145410.1 MULTISPECIES: hydrogenase-2 small chain [Proteobacteria] Length: 372\nNumber of predicted TMHs: 1\nExp number of AAs in TMHs: 23.72226\nExp number, first 60 AAs: 0.22038\nTotal prob of N-in: 0.12994\noutside 1 331\nTMhelix 332 354\ninside 355 372

4702 GCF_000781735.1_ASM78173v1 Escherichia coli Proteobacteria; Gammaproteobacteria; Enterobacterales; Enterobacteriaceae; Escherichia MNNEETFYQAMRRQGVTRRSFLKYCSLAATSLGLGAGMAPKIAWALENKPRIPVVWIHGL WP_001295940.1 MULTISPECIES: hydrogenase [Enterobacteriaceae] Length: 372\nNumber of predicted TMHs: 2\nExp number of AAs in TMHs: 37.2608899999999\nExp number, first 60 AAs: 19.52923\nTotal prob of N-in: 0.94351\nPOSSIBLE N-term signal sequence\ninside 1 19\nTMhelix 20 39\noutside 40 325\nTMhelix 326 348\ninside 349 372

4703 GCF_000781735.1_ASM78173v1 Escherichia coli Proteobacteria; Gammaproteobacteria; Enterobacterales; Enterobacteriaceae; Escherichia MTGDNTLIHSHGINRRDFMKLCAALAATMGLSSKAAAEMAESVTNPQRPPVIWIGAQECT WP_000145410.1 MULTISPECIES: hydrogenase-2 small chain [Proteobacteria] Length: 372\nNumber of predicted TMHs: 1\nExp number of AAs in TMHs: 23.72226\nExp number, first 60 AAs: 0.22038\nTotal prob of N-in: 0.12994\noutside 1 331\nTMhelix 332 354\ninside 355 372

4704 GCF_000781715.1_ASM78171v1 Escherichia coli Proteobacteria; Gammaproteobacteria; Enterobacterales; Enterobacteriaceae; Escherichia MNNEETFYQAMRRQGVTRRSFLKYCSLAATSLGLGAGMAPKIAWALENKPRIPVVWIHGL WP_001058323.1 MULTISPECIES: hydrogenase-1 small chain [Proteobacteria] Length: 372\nNumber of predicted TMHs: 2\nExp number of AAs in TMHs: 37.2608499999999\nExp number, first 60 AAs: 19.52923\nTotal prob of N-in: 0.94351\nPOSSIBLE N-term signal sequence\ninside 1 19\nTMhelix 20 39\noutside 40 325\nTMhelix 326 348\ninside 349 372

4705 GCF_000781715.1_ASM78171v1 Escherichia coli Proteobacteria; Gammaproteobacteria; Enterobacterales; Enterobacteriaceae; Escherichia MTGDNTLIHSHGINRRDFMKLCAALAATMGLSSKAAAEMAESVTNPQRPPVIWIGAQECT WP_000145410.1 MULTISPECIES: hydrogenase-2 small chain [Proteobacteria] Length: 372\nNumber of predicted TMHs: 1\nExp number of AAs in TMHs: 23.72226\nExp number, first 60 AAs: 0.22038\nTotal prob of N-in: 0.12994\noutside 1 331\nTMhelix 332 354\ninside 355 372

4706 GCF_001058225.1_ASM105822v1 Escherichia coli Proteobacteria; Gammaproteobacteria; Enterobacterales; Enterobacteriaceae; Escherichia
MNNEETFYQAMRRQGVTRRSFLKYCSLAATSLGLGAGMAPKIAWALENKPRIPVVWIHGL WP_001295940.1
MULTISPECIES: hydrogenase [Enterobacteriaceae] Length: 372\nNumber of predicted TMHs: 2\nExp number of AAs in TMHs: 37.2608899999999\nExp number, first 60 AAs: 19.52923\nTotal prob of N-in: 0.94351\nPOSSIBLE N-term signal sequence\ninside 1 19\nTMhelix 20 39\noutside 40 325\nTMhelix 326 348\ninside 349 372

4707 GCF_001058225.1_ASM105822v1 Escherichia coli Proteobacteria; Gammaproteobacteria; Enterobacterales; Enterobacteriaceae; Escherichia
MTGDNTLIHSHGINRRDFMKLCAALAATMGLSSKAAAEMAESVTNPQRPPVIWIGAQECT WP_000145410.1
MULTISPECIES: hydrogenase-2 small chain [Proteobacteria] Length: 372\nNumber of predicted TMHs: 1\nExp number of AAs in TMHs: 23.72226\nExp number, first 60 AAs: 0.22038\nTotal prob of N-in: 0.12994\noutside 1 331\nTMhelix 332 354\ninside 355 372

4708 GCF_000781485.1_ASM78148v1 Escherichia coli Proteobacteria; Gammaproteobacteria; Enterobacterales; Enterobacteriaceae; Escherichia
MNNEETFYQAMRRQGVTRRSFLKYCSLAATSLGLGAGMAPKIAWALENKPRIPVVWIHGL WP_001295940.1
MULTISPECIES: hydrogenase [Enterobacteriaceae] Length: 372\nNumber of predicted TMHs: 2\nExp number of AAs in TMHs: 37.2608899999999\nExp number, first 60 AAs: 19.52923\nTotal prob of N-in: 0.94351\nPOSSIBLE N-term signal sequence\ninside 1 19\nTMhelix 20 39\noutside 40 325\nTMhelix 326 348\ninside 349 372

4709 GCF_000781485.1_ASM78148v1 Escherichia coli Proteobacteria; Gammaproteobacteria; Enterobacterales; Enterobacteriaceae; Escherichia
MTGDNTLIHSHGINRRDFMKLCAALAATMGLSSKAAAEMVESVTNPQRPPVIWIGAQECT WP_000145423.1
hydrogenase 2 small subunit [Escherichia coli] Length: 372\nNumber of predicted TMHs: 1\nExp number of AAs in TMHs: 23.78201\nExp number, first 60 AAs: 0.28355\nTotal prob of N-in: 0.13257\noutside 1 331\nTMhelix 332 354\ninside 355 372

4710 GCF_001269325.1_YE6 Escherichia coli Proteobacteria; Gammaproteobacteria; Enterobacterales; Enterobacteriaceae; Escherichia MNNEETFYQAMRRQGVTRRSFLKYCSLAATSLGLGAGMAPKIAWALENKPRIPVVWIHGL
WP_001295940.1 MULTISPECIES: hydrogenase [Enterobacteriaceae] Length: 372\nNumber of predicted TMHs: 2\nExp number of AAs in TMHs: 37.2608899999999\nExp number, first 60 AAs: 19.52923\nTotal prob of N-in: 0.94351\nPOSSIBLE N-term signal sequence\ninside 1 19\nTMhelix 20 39\noutside 40 325\nTMhelix 326 348\ninside 349 372

4711 GCF_001277775.1_7790_1_60 Escherichia coli Proteobacteria; Gammaproteobacteria; Enterobacterales; Enterobacteriaceae; Escherichia
MNNEETFYQAMRRQGVTRRSFLKYCSLAATSLGLGAGMAPKIAWALENKPRIPVVWIHGL WP_001295940.1
MULTISPECIES: hydrogenase [Enterobacteriaceae] Length: 372\nNumber of predicted TMHs: 2\nExp number of AAs in TMHs: 37.2608899999999\nExp number, first 60 AAs: 19.52923\nTotal prob of N-in: 0.94351\nPOSSIBLE N-term signal sequence\ninside 1 19\nTMhelix 20 39\noutside 40 325\nTMhelix 326 348\ninside 349 372

4712 GCF_000798255.1_Escherichia_coli_CVM_N36093PS_v._1.0 Escherichia coli Proteobacteria; Gammaproteobacteria; Enterobacterales; Enterobacteriaceae; Escherichia
MNNEETFYQAMRRQGVTRRSFLKYCSLAATSLGLGAGMAPKIAWALENKPRIPVVWIHGL WP_001058323.1
MULTISPECIES: hydrogenase-1 small chain [Proteobacteria] Length: 372\nNumber of predicted TMHs: 2\nExp number of AAs in TMHs: 37.2608499999999\nExp number, first 60 AAs: 19.52923\nTotal prob of N-in: 0.94351\nPOSSIBLE N-term signal sequence\ninside 1 19\nTMhelix 20 39\noutside 40 325\nTMhelix 326 348\ninside 349 372

4713 GCF_001277575.1_7553_7_61 Escherichia coli Proteobacteria; Gammaproteobacteria; Enterobacterales; Enterobacteriaceae; Escherichia
MNNEETFYQAMRRQGVTRRSFLKYCSLAATSLGLGAGMAPKIAWALENKPRIPVVWIHGL WP_001295940.1
MULTISPECIES: hydrogenase [Enterobacteriaceae] Length: 372\nNumber of predicted TMHs: 2\nExp number of AAs in TMHs: 37.2608899999999\nExp number, first 60 AAs: 19.52923\nTotal prob of N-in: 0.94351\nPOSSIBLE N-term signal sequence\ninside 1 19\nTMhelix 20 39\noutside 40 325\nTMhelix 326 348\ninside 349 372

4714 GCF_000797995.1_Escherichia_coli_CVM_N36613PS_v._1.0 Escherichia coli Proteobacteria; Gammaproteobacteria; Enterobacterales; Enterobacteriaceae; Escherichia MTGDNTLIHSHGINRRDFMKLCAALAATMGLSSKAAAEMAESVTNPQRPPVIWIGAQECT WP_000145410.1 MULTISPECIES: hydrogenase-2 small chain [Proteobacteria] Length: 372\nNumber of predicted TMHs: 1\nExp number of AAs in TMHs: 23.72226\nExp number, first 60 AAs: 0.22038\nTotal prob of N-in: 0.12994\noutside 1 331\nTMhelix 332 354\ninside 355 372

4715 GCF_000797895.1_Escherichia_coli_CVM_N38834PS_v._1.0 Escherichia coli Proteobacteria; Gammaproteobacteria; Enterobacterales; Enterobacteriaceae; Escherichia MNNEETFYQAMRRQGVTRRSFLKYCSLAATSLGLGAGMAPKIAWALENKPRIPVVWIHGL WP_001058323.1 MULTISPECIES: hydrogenase-1 small chain [Proteobacteria] Length: 372\nNumber of predicted TMHs: 2\nExp number of AAs in TMHs: 37.2608499999999\nExp number, first 60 AAs: 19.52923\nTotal prob of N-in: 0.94351\nPOSSIBLE N-term signal sequence\ninside 1 19\nTMhelix 20 39\noutside 40 325\nTMhelix 326 348\ninside 349 372

4716 GCF_001284785.1_7748_7_28 Escherichia coli Proteobacteria; Gammaproteobacteria; Enterobacterales; Enterobacteriaceae; Escherichia MNNEETFYQAMRRQGVTRRSFLKYCSLAATSLGLGAGMAPKIAWALENKPRIPVVWIHGL WP_001359719.1 hydrogenase [Escherichia coli] Length: 372\nNumber of predicted TMHs: 2\nExp number of AAs in TMHs: 37.2176\nExp number, first 60 AAs: 19.53448\nTotal prob of N-in: 0.94360\nPOSSIBLE N-term signal sequence\ninside 1 19\nTMhelix 20 39\noutside 40 325\nTMhelix 326 348\ninside 349 372

4717 GCF_001284185.1_8205_3_69 Escherichia coli Proteobacteria; Gammaproteobacteria; Enterobacterales; Enterobacteriaceae; Escherichia MTGDNTLIHSHGINRRDFMKLCAALAATMGLSSKAAAEMAESVTNPQRPPVIWIGAQECT WP_000145410.1 MULTISPECIES: hydrogenase-2 small chain [Proteobacteria] Length: 372\nNumber of predicted TMHs: 1\nExp number of AAs in TMHs: 23.72226\nExp number, first 60 AAs: 0.22038\nTotal prob of N-in: 0.12994\noutside 1 331\nTMhelix 332 354\ninside 355 372

4718 GCF_001277535.1_7748_7_32 Escherichia coli Proteobacteria; Gammaproteobacteria; Enterobacterales; Enterobacteriaceae; Escherichia MNNEETFYQAMRRQGVTRRSFLKYCSLAATSLGLGAGMAPKIAWALENKPRIPVVWIHGL WP_001058323.1 MULTISPECIES: hydrogenase-1 small chain [Proteobacteria] Length: 372\nNumber of predicted TMHs: 2\nExp number of AAs in TMHs: 37.2608499999999\nExp number, first 60 AAs: 19.52923\nTotal prob of N-in: 0.94351\nPOSSIBLE N-term signal sequence\ninside 1 19\nTMhelix 20 39\noutside 40 325\nTMhelix 326 348\ninside 349 372

4719 GCF_001284605.1_7748_7_27 Escherichia coli Proteobacteria; Gammaproteobacteria; Enterobacterales; Enterobacteriaceae; Escherichia MNNEETFYQAMRRQGVTRRSFLKYCSLAATSLGLGAGMAPKIAWALENKPRIPVVWIHGL WP_001058323.1 MULTISPECIES: hydrogenase-1 small chain [Proteobacteria] Length: 372\nNumber of predicted TMHs: 2\nExp number of AAs in TMHs: 37.2608499999999\nExp number, first 60 AAs: 19.52923\nTotal prob of N-in: 0.94351\nPOSSIBLE N-term signal sequence\ninside 1 19\nTMhelix 20 39\noutside 40 325\nTMhelix 326 348\ninside 349 372

4720 GCF_000797995.1_Escherichia_coli_CVM_N36613PS_v._1.0 Escherichia coli Proteobacteria; Gammaproteobacteria; Enterobacterales; Enterobacteriaceae; Escherichia MNNEETFYQAMRRQGVTRRSFLKYCSLAATSLGLGAGMAPKIAWALENKPRIPVVWIHGL WP_001058323.1 MULTISPECIES: hydrogenase-1 small chain [Proteobacteria] Length: 372\nNumber of predicted TMHs: 2\nExp number of AAs in TMHs: 37.2608499999999\nExp number, first 60 AAs: 19.52923\nTotal prob of N-in: 0.94351\nPOSSIBLE N-term signal sequence\ninside 1 19\nTMhelix 20 39\noutside 40 325\nTMhelix 326 348\ninside 349 372

4721 GCF_001277735.1_7748_7_15 Escherichia coli Proteobacteria; Gammaproteobacteria; Enterobacterales; Enterobacteriaceae; Escherichia MNNEETFYQAMRRQGVTRRSFLKYCSLAATSLGLGAGMAPKIAWALENKPRIPVVWIHGL WP_001058323.1 MULTISPECIES: hydrogenase-1 small chain [Proteobacteria] Length: 372\nNumber of predicted TMHs: 2\nExp number of AAs in TMHs: 37.2608499999999\nExp number, first 60 AAs: 19.52923\nTotal prob of N-in: 0.94351\nPOSSIBLE N-term signal sequence\ninside 1 19\nTMhelix 20 39\noutside 40 325\nTMhelix 326 348\ninside 349 372

4722 GCF_000798095.1_Escherichia_coli_CVM_N36312PS_v._1.0 Escherichia coli Proteobacteria; Gammaproteobacteria; Enterobacterales; Enterobacteriaceae; Escherichia
MNNEETFYQAMRRQGVTRRSFLKYCSLAATSLGLGAGMAPKIAWALENKPRIPVVWIHGL WP_001058323.1
MULTISPECIES: hydrogenase-1 small chain [Proteobacteria] Length: 372\nNumber of predicted TMHs: 2\nExp number of AAs in TMHs: 37.2608499999999\nExp number, first 60 AAs: 19.52923\nTotal prob of N-in: 0.94351\nPOSSIBLE N-term signal sequence\ninside 1 19\nTMhelix 20 39\noutside 40 325\nTMhelix 326 348\ninside 349 372

4723 GCF_001284305.1_7748_7_43 Escherichia coli Proteobacteria; Gammaproteobacteria; Enterobacterales; Enterobacteriaceae; Escherichia
MNNEETFYQAMRRQGVTRRSFLKYCSLAATSLGLGAGMAPKIAWALENKPRIPVVWIHGL WP_001058323.1
MULTISPECIES: hydrogenase-1 small chain [Proteobacteria] Length: 372\nNumber of predicted TMHs: 2\nExp number of AAs in TMHs: 37.2608499999999\nExp number, first 60 AAs: 19.52923\nTotal prob of N-in: 0.94351\nPOSSIBLE N-term signal sequence\ninside 1 19\nTMhelix 20 39\noutside 40 325\nTMhelix 326 348\ninside 349 372

4724 GCF_001284385.1_8205_3_52 Escherichia coli Proteobacteria; Gammaproteobacteria; Enterobacterales; Enterobacteriaceae; Escherichia
MTGDNTLIHSHGINRRDFMKLCAALAATMGLSSKAAAEMAESVTNPQRPPVIWIGAQECT WP_000145410.1
MULTISPECIES: hydrogenase-2 small chain [Proteobacteria] Length: 372\nNumber of predicted TMHs: 1\nExp number of AAs in TMHs: 23.72226\nExp number, first 60 AAs: 0.22038\nTotal prob of N-in: 0.12994\noutside 1 331\nTMhelix 332 354\ninside 355 372

4725 GCF_001284385.1_8205_3_52 Escherichia coli Proteobacteria; Gammaproteobacteria; Enterobacterales; Enterobacteriaceae; Escherichia
MNNEETFYQAMRRQGVTRRSFLKYCSLAATSLGLGAGMAPKIAWALENKPRIPVVWIHGL WP_053876928.1
hydrogenase [Escherichia coli] Length: 372\nNumber of predicted TMHs: 2\nExp number of AAs in TMHs: 37.26524\nExp number, first 60 AAs: 19.52925\nTotal prob of N-in: 0.94351\nPOSSIBLE N-term signal sequence\ninside 1 19\nTMhelix 20 39\noutside 40 325\nTMhelix 326 348\ninside 349 372

4726 GCF_001269245.1_YE5 Escherichia coli Proteobacteria; Gammaproteobacteria; Enterobacterales; Enterobacteriaceae; Escherichia MNNEETFYQAMRRQGVTRRSFLKYCSLAATSLGLGAGMAPKIAWALENKPRIPVVWIHGL
WP_001058323.1 MULTISPECIES: hydrogenase-1 small chain [Proteobacteria] Length: 372\nNumber of predicted TMHs: 2\nExp number of AAs in TMHs: 37.2608499999999\nExp number, first 60 AAs: 19.52923\nTotal prob of N-in: 0.94351\nPOSSIBLE N-term signal sequence\ninside 1 19\nTMhelix 20 39\noutside 40 325\nTMhelix 326 348\ninside 349 372

4727 GCF_001269245.1_YE5 Escherichia coli Proteobacteria; Gammaproteobacteria; Enterobacterales; Enterobacteriaceae; Escherichia MTGDNTLIHSHGINRRDFMKLCAALAATMGLSSKAAAEMAESVTNPQRPPVIWIGAQECT
WP_000145410.1 MULTISPECIES: hydrogenase-2 small chain [Proteobacteria] Length: 372\nNumber of predicted TMHs: 1\nExp number of AAs in TMHs: 23.72226\nExp number, first 60 AAs: 0.22038\nTotal prob of N-in: 0.12994\noutside 1 331\nTMhelix 332 354\ninside 355 372

4728 GCF_000798215.1_Escherichia_coli_CVM_N36620PS_v._1.0 Escherichia coli Proteobacteria; Gammaproteobacteria; Enterobacterales; Enterobacteriaceae; Escherichia
MNNEETFYQAMRRQGVTRRSFLKYCSLAATSLGLGAGMAPKIAWALENKPRIPVVWIHGL WP_001058323.1
MULTISPECIES: hydrogenase-1 small chain [Proteobacteria] Length: 372\nNumber of predicted TMHs: 2\nExp number of AAs in TMHs: 37.2608499999999\nExp number, first 60 AAs: 19.52923\nTotal prob of N-in: 0.94351\nPOSSIBLE N-term signal sequence\ninside 1 19\nTMhelix 20 39\noutside 40 325\nTMhelix 326 348\ninside 349 372

4729 GCF_000798155.1_Escherichia_coli_CVM_N34077PS_v._1.0 Escherichia coli Proteobacteria; Gammaproteobacteria; Enterobacterales; Enterobacteriaceae; Escherichia
MTGDNTLIHSHGINRRDFMKLCAALAATMGLSSKAAAEMAESVTNPQRPPVIWIGAQECT WP_000145410.1
MULTISPECIES: hydrogenase-2 small chain [Proteobacteria] Length: 372\nNumber of predicted TMHs: 1\nExp number of AAs in TMHs: 23.72226\nExp number, first 60 AAs: 0.22038\nTotal prob of N-in: 0.12994\noutside 1 331\nTMhelix 332 354\ninside 355 372

4730 GCF_001284445.1_7790_1_76 Escherichia coli Proteobacteria; Gammaproteobacteria; Enterobacterales; Enterobacteriaceae; Escherichia
MNNEETFYQAMRRQGVTRRSFLKYCSLAATSLGLGAGMAPKIAWALENKPRIPVVWIHGL WP_001058323.1

MULTISPECIES: hydrogenase-1 small chain [Proteobacteria] Length: 372\nNumber of predicted TMHs: 2\nExp number of AAs in TMHs: 37.2608499999999\nExp number, first 60 AAs: 19.52923\nTotal prob of N-in: 0.94351\nPOSSIBLE N-term signal sequence\ninside 1 19\nTMhelix 20 39\noutside 40 325\nTMhelix 326 348\ninside 349 372

4731 GCF_001284505.1_7748_7_26 Escherichia coli Proteobacteria; Gammaproteobacteria; Enterobacterales; Enterobacteriaceae; Escherichia MNNEETFYQAMRRQGVTRRSFLKYCSLAATSLGLGAGMAPKIAWALENKPRIPVVWIHGL WP_024228890.1 hydrogenase [Escherichia coli] Length: 372\nNumber of predicted TMHs: 2\nExp number of AAs in TMHs: 38.3063\nExp number, first 60 AAs: 19.52635\nTotal prob of N-in: 0.94360\nPOSSIBLE N-term signal sequence\ninside 1 19\nTMhelix 20 39\noutside 40 325\nTMhelix 326 348\ninside 349 372

4732 GCF_001284345.1_8205_3_61 Escherichia coli Proteobacteria; Gammaproteobacteria; Enterobacterales; Enterobacteriaceae; Escherichia MNNEETFYQAMRRQGVTRRSFLKYCSLAATSLGLGAGMAPKIAWALENKPRIPVVWIHGL WP_001295940.1 MULTISPECIES: hydrogenase [Enterobacteriaceae] Length: 372\nNumber of predicted TMHs: 2\nExp number of AAs in TMHs: 37.2608899999999\nExp number, first 60 AAs: 19.52923\nTotal prob of N-in: 0.94351\nPOSSIBLE N-term signal sequence\ninside 1 19\nTMhelix 20 39\noutside 40 325\nTMhelix 326 348\ninside 349 372

4733 GCF_001284785.1_7748_7_28 Escherichia coli Proteobacteria; Gammaproteobacteria; Enterobacterales; Enterobacteriaceae; Escherichia MTGDNTLIHSHGINRRDFMKLCAALAATMGLSSKAAAEMAESVTNPQRPPVIWIGAQECT WP_000145410.1 MULTISPECIES: hydrogenase-2 small chain [Proteobacteria] Length: 372\nNumber of predicted TMHs: 1\nExp number of AAs in TMHs: 23.72226\nExp number, first 60 AAs: 0.22038\nTotal prob of N-in: 0.12994\noutside 1 331\nTMhelix 332 354\ninside 355 372

4734 GCF_001284005.1_7748_7_41 Escherichia coli Proteobacteria; Gammaproteobacteria; Enterobacterales; Enterobacteriaceae; Escherichia MTGDNTLIHSHGINRRDFMKLCAALAATMGLSSKAAAEMAESVTNPQRPPVIWIGAQECT WP_000145410.1 MULTISPECIES: hydrogenase-2 small chain [Proteobacteria] Length: 372\nNumber of predicted TMHs: 1\nExp number of AAs in TMHs: 23.72226\nExp number, first 60 AAs: 0.22038\nTotal prob of N-in: 0.12994\noutside 1 331\nTMhelix 332 354\ninside 355 372

4735 GCF_001269145.1_YE20 Escherichia coli Proteobacteria; Gammaproteobacteria; Enterobacterales; Enterobacteriaceae; Escherichia MNNEETFYQAMRRQGVTRRSFLKYCSLAATSLGLGAGMAPKIAWALENKPRIPVVWIHGL WP_001295940.1 MULTISPECIES: hydrogenase [Enterobacteriaceae] Length: 372\nNumber of predicted TMHs: 2\nExp number of AAs in TMHs: 37.2608899999999\nExp number, first 60 AAs: 19.52923\nTotal prob of N-in: 0.94351\nPOSSIBLE N-term signal sequence\ninside 1 19\nTMhelix 20 39\noutside 40 325\nTMhelix 326 348\ninside 349 372

4736 GCF_000798215.1_Escherichia_coli_CVM_N36620PS_v._1.0 Escherichia coli Proteobacteria; Gammaproteobacteria; Enterobacterales; Enterobacteriaceae; Escherichia MTGDNTLIHSHGINRRDFMKLCAALAATMGLSSKAAAEMAESVTNPQRPPVIWIGAQECT WP_000145410.1 MULTISPECIES: hydrogenase-2 small chain [Proteobacteria] Length: 372\nNumber of predicted TMHs: 1\nExp number of AAs in TMHs: 23.72226\nExp number, first 60 AAs: 0.22038\nTotal prob of N-in: 0.12994\noutside 1 331\nTMhelix 332 354\ninside 355 372

4737 GCF_001277535.1_7748_7_32 Escherichia coli Proteobacteria; Gammaproteobacteria; Enterobacterales; Enterobacteriaceae; Escherichia MTGDNTLIHSHGINRRDFMKLCAALAATMGLSSKAAAEMAESVTNPQRPPVIWIGAQECT WP_000145410.1 MULTISPECIES: hydrogenase-2 small chain [Proteobacteria] Length: 372\nNumber of predicted TMHs: 1\nExp number of AAs in TMHs: 23.72226\nExp number, first 60 AAs: 0.22038\nTotal prob of N-in: 0.12994\noutside 1 331\nTMhelix 332 354\ninside 355 372

4738 GCF_001283965.1_8205_3_5Escherichia coli Proteobacteria; Gammaproteobacteria; Enterobacterales; Enterobacteriaceae; Escherichia MTGDNTLIHSHGINRRDFMKLCAALAATMGLSSKAAAEMAESVTNPQRPPVIWIGAQECT WP_000145410.1 MULTISPECIES: hydrogenase-2 small chain [Proteobacteria] Length: 372\nNumber of predicted TMHs: 1\nExp number of AAs in TMHs: 23.72226\nExp number, first 60 AAs: 0.22038\nTotal prob of N-in: 0.12994\noutside 1 331\nTMhelix 332 354\ninside 355 372

4739	GCF_001284885.1_8205_3_19	Escherichia coli	Proteobacteria; Gammaproteobacteria; Enterobacterales; Enterobacteriaceae; Escherichia
MNNEETFYQAMRRQGVTRRSFLKYCSLAATSLGLGAGMAPKIAWALENKPRIPVVWIHGL	WP_001058323.1
MULTISPECIES: hydrogenase-1 small chain [Proteobacteria]	Length: 372\nNumber of predicted TMHs: 2\nExp number of AAs in TMHs: 37.2608499999999\nExp number, first 60 AAs: 19.52923\nTotal prob of N-in: 0.94351\nPOSSIBLE N-term signal sequence\ninside 1 19\nTMhelix 20 39\noutside 40 325\nTMhelix 326 348\ninside 349 372

4740	GCF_000798055.1_Escherichia_coli_CVM_N33922PS_v._1.0	Escherichia coli	Proteobacteria; Gammaproteobacteria; Enterobacterales; Enterobacteriaceae; Escherichia
MNNEETFYQAMRRQGVTRRSFLKYCSLAATSLGLGAGMAPKIAWALENKPRIPVVWIHGL	WP_001058323.1
MULTISPECIES: hydrogenase-1 small chain [Proteobacteria]	Length: 372\nNumber of predicted TMHs: 2\nExp number of AAs in TMHs: 37.2608499999999\nExp number, first 60 AAs: 19.52923\nTotal prob of N-in: 0.94351\nPOSSIBLE N-term signal sequence\ninside 1 19\nTMhelix 20 39\noutside 40 325\nTMhelix 326 348\ninside 349 372

4741	GCF_001268765.1_YE12	Escherichia coli	Proteobacteria; Gammaproteobacteria; Enterobacterales; Enterobacteriaceae; Escherichia	MNNEETFYQAMRRQGVTRRSFLKYCSLAATSLGLGAGMAPKIAWALENKPRIPVVWIHGL
WP_001058323.1	MULTISPECIES: hydrogenase-1 small chain [Proteobacteria]	Length: 372\nNumber of predicted TMHs: 2\nExp number of AAs in TMHs: 37.2608499999999\nExp number, first 60 AAs: 19.52923\nTotal prob of N-in: 0.94351\nPOSSIBLE N-term signal sequence\ninside 1 19\nTMhelix 20 39\noutside 40 325\nTMhelix 326 348\ninside 349 372

4742	GCF_001268805.1_YE22	Escherichia coli	Proteobacteria; Gammaproteobacteria; Enterobacterales; Enterobacteriaceae; Escherichia	MNNEETFYQAMRRQGVTRRSFLKYCSLAATSLGLGAGMAPKIAWALENKPRIPVVWIHGL
WP_001058323.1	MULTISPECIES: hydrogenase-1 small chain [Proteobacteria]	Length: 372\nNumber of predicted TMHs: 2\nExp number of AAs in TMHs: 37.2608499999999\nExp number, first 60 AAs: 19.52923\nTotal prob of N-in: 0.94351\nPOSSIBLE N-term signal sequence\ninside 1 19\nTMhelix 20 39\noutside 40 325\nTMhelix 326 348\ninside 349 372

4743	GCF_001268805.1_YE22	Escherichia coli	Proteobacteria; Gammaproteobacteria; Enterobacterales; Enterobacteriaceae; Escherichia	MTGDNTLIHSHGINRRDFMKLCAALAATMGLSSKAAAEMAESVTNPQRPPVIWIGAQECT
WP_032298447.1	hydrogenase 2 small subunit [Escherichia coli]	Length: 372\nNumber of predicted TMHs: 1\nExp number of AAs in TMHs: 23.69807\nExp number, first 60 AAs: 0.221\nTotal prob of N-in: 0.12853\noutside 1 331\nTMhelix 332 354\ninside 355 372

4744	GCF_001268845.1_YE3	Escherichia coli	Proteobacteria; Gammaproteobacteria; Enterobacterales; Enterobacteriaceae; Escherichia	MTGDNTLIHSHGINRRDFMKLCAALAATMGLSSKAAAEMAESVTNPQRPPVIWIGAQECT
WP_000145410.1	MULTISPECIES: hydrogenase-2 small chain [Proteobacteria]	Length: 372\nNumber of predicted TMHs: 1\nExp number of AAs in TMHs: 23.72226\nExp number, first 60 AAs: 0.22038\nTotal prob of N-in: 0.12994\noutside 1 331\nTMhelix 332 354\ninside 355 372

4745	GCF_001268845.1_YE3	Escherichia coli	Proteobacteria; Gammaproteobacteria; Enterobacterales; Enterobacteriaceae; Escherichia	MNNEETFYQAMRRQGVTRRSFLKYCSLAATSLGLGAGMAPKIAWALENKPRIPVVWIHGL
WP_001058323.1	MULTISPECIES: hydrogenase-1 small chain [Proteobacteria]	Length: 372\nNumber of predicted TMHs: 2\nExp number of AAs in TMHs: 37.2608499999999\nExp number, first 60 AAs: 19.52923\nTotal prob of N-in: 0.94351\nPOSSIBLE N-term signal sequence\ninside 1 19\nTMhelix 20 39\noutside 40 325\nTMhelix 326 348\ninside 349 372

4746	GCF_001268885.1_1.EC2731.1	Escherichia coli	Proteobacteria; Gammaproteobacteria; Enterobacterales; Enterobacteriaceae; Escherichia
MTGDNTLIHSHGINRRDFMKLCAALAATMGLSSKAAAEMAESVTNPQRPPVIWIGAQECT	WP_000145410.1
MULTISPECIES: hydrogenase-2 small chain [Proteobacteria]	Length: 372\nNumber of predicted TMHs: 1\nExp number of AAs in TMHs: 23.72226\nExp number, first 60 AAs: 0.22038\nTotal prob of N-in: 0.12994\noutside 1 331\nTMhelix 332 354\ninside 355 372

4747	GCF_001268885.1_1.EC2731.1	Escherichia coli	Proteobacteria; Gammaproteobacteria; Enterobacterales; Enterobacteriaceae; Escherichia
MNNEETFYQAMRRQGVTRRSFLKYCSLAATSLGLGAGMAPKIAWALENKPRIPVVWIHGL	WP_001295940.1
MULTISPECIES: hydrogenase [Enterobacteriaceae]	Length: 372\nNumber of predicted TMHs: 2\nExp number of AAs in TMHs: 37.2608899999999\nExp number, first 60 AAs: 19.52923\nTotal prob of N-in:

0.94351\nPOSSIBLE N-term signal sequence\ninside 1 19\nTMhelix 20 39\noutside 40 325\nTMhelix 326 348\ninside 349 372

4748 GCF_001268945.1_YE4 Escherichia coli Proteobacteria; Gammaproteobacteria; Enterobacterales; Enterobacteriaceae; Escherichia MTGDNTLIHSHGINRRDFMKLCAALAATMGLSSKAAAEMAESVTNPQRPPVIWIGAQECT WP_000145410.1 MULTISPECIES: hydrogenase-2 small chain [Proteobacteria] Length: 372\nNumber of predicted TMHs: 1\nExp number of AAs in TMHs: 23.72226\nExp number, first 60 AAs: 0.22038\nTotal prob of N-in: 0.12994\noutside 1 331\nTMhelix 332 354\ninside 355 372

4749 GCF_001268945.1_YE4 Escherichia coli Proteobacteria; Gammaproteobacteria; Enterobacterales; Enterobacteriaceae; Escherichia MNNEETFYQAMRRQGVTRRSFLKYCSLAATSLGLGAGMAPKIAWALENKPRIPVVWIHGL WP_001058323.1 MULTISPECIES: hydrogenase-1 small chain [Proteobacteria] Length: 372\nNumber of predicted TMHs: 2\nExp number of AAs in TMHs: 37.2608499999999\nExp number, first 60 AAs: 19.52923\nTotal prob of N-in: 0.94351\nPOSSIBLE N-term signal sequence\ninside 1 19\nTMhelix 20 39\noutside 40 325\nTMhelix 326 348\ninside 349 372

4750 GCF_001268965.1_1.ECD03.1 Escherichia coli Proteobacteria; Gammaproteobacteria; Enterobacterales; Enterobacteriaceae; Escherichia MTGDNTLIHSHGINRRDFMKLCAALAATMGLSSKAAAEMAESVTNPQRPPVIWIGAQECT WP_000145410.1 MULTISPECIES: hydrogenase-2 small chain [Proteobacteria] Length: 372\nNumber of predicted TMHs: 1\nExp number of AAs in TMHs: 23.72226\nExp number, first 60 AAs: 0.22038\nTotal prob of N-in: 0.12994\noutside 1 331\nTMhelix 332 354\ninside 355 372

4751 GCF_001268965.1_1.ECD03.1 Escherichia coli Proteobacteria; Gammaproteobacteria; Enterobacterales; Enterobacteriaceae; Escherichia MNNEETFYQAMRRQGVTRRSFLKYCSLAATSLGLGAGMAPKIAWALENKPRIPVVWIHGL WP_001295940.1 MULTISPECIES: hydrogenase [Enterobacteriaceae] Length: 372\nNumber of predicted TMHs: 2\nExp number of AAs in TMHs: 37.2608899999999\nExp number, first 60 AAs: 19.52923\nTotal prob of N-in: 0.94351\nPOSSIBLE N-term signal sequence\ninside 1 19\nTMhelix 20 39\noutside 40 325\nTMhelix 326 348\ninside 349 372

4752 GCF_001269005.1_YE18 Escherichia coli Proteobacteria; Gammaproteobacteria; Enterobacterales; Enterobacteriaceae; Escherichia MTGDNTLIHSHGINRRDFMKLCAALAATMGLSSKAAAEMAESVTNPQRPPVIWIGAQECT WP_000145410.1 MULTISPECIES: hydrogenase-2 small chain [Proteobacteria] Length: 372\nNumber of predicted TMHs: 1\nExp number of AAs in TMHs: 23.72226\nExp number, first 60 AAs: 0.22038\nTotal prob of N-in: 0.12994\noutside 1 331\nTMhelix 332 354\ninside 355 372

4753 GCF_001269005.1_YE18 Escherichia coli Proteobacteria; Gammaproteobacteria; Enterobacterales; Enterobacteriaceae; Escherichia MNNEETFYQAMRRQGVTRRSFLKYCSLAATSLGLGAGMAPKIAWALENKPRIPVVWIHGL WP_001058323.1 MULTISPECIES: hydrogenase-1 small chain [Proteobacteria] Length: 372\nNumber of predicted TMHs: 2\nExp number of AAs in TMHs: 37.2608499999999\nExp number, first 60 AAs: 19.52923\nTotal prob of N-in: 0.94351\nPOSSIBLE N-term signal sequence\ninside 1 19\nTMhelix 20 39\noutside 40 325\nTMhelix 326 348\ninside 349 372

4754 GCF_000797585.1_Escherichia_coli_CVM_N37059PS_v._1.0 Escherichia coli Proteobacteria; Gammaproteobacteria; Enterobacterales; Enterobacteriaceae; Escherichia MTGDNTLIHSHGINRRDFMKLCAALAATMGLSSKAAAEMAESVTNPQRPPVIWIGAQECT WP_000145410.1 MULTISPECIES: hydrogenase-2 small chain [Proteobacteria] Length: 372\nNumber of predicted TMHs: 1\nExp number of AAs in TMHs: 23.72226\nExp number, first 60 AAs: 0.22038\nTotal prob of N-in: 0.12994\noutside 1 331\nTMhelix 332 354\ninside 355 372

4755 GCF_000797585.1_Escherichia_coli_CVM_N37059PS_v._1.0 Escherichia coli Proteobacteria; Gammaproteobacteria; Enterobacterales; Enterobacteriaceae; Escherichia MNNEETFYQAMRRQGVTRRSFLKYCSLAATSLGLGAGMAPKIAWALENKPRIPVVWIHGL WP_001058323.1 MULTISPECIES: hydrogenase-1 small chain [Proteobacteria] Length: 372\nNumber of predicted TMHs: 2\nExp number of AAs in TMHs: 37.2608499999999\nExp number, first 60 AAs: 19.52923\nTotal prob of N-in: 0.94351\nPOSSIBLE N-term signal sequence\ninside 1 19\nTMhelix 20 39\noutside 40 325\nTMhelix 326 348\ninside 349 372

4756 GCF_000946315.1_Ec26A_L1_MIRA_assembly Escherichia coli Proteobacteria; Gammaproteobacteria; Enterobacterales; Enterobacteriaceae; Escherichia

MTGDNTLIHSHGINRRDFMKLCAALAATMGLSSKAAAEMAESVTNPQRPPVIWIGAQECT WP_000145410.1 MULTISPECIES: hydrogenase-2 small chain [Proteobacteria] Length: 372\nNumber of predicted TMHs: 1\nExp number of AAs in TMHs: 23.72226\nExp number, first 60 AAs: 0.22038\nTotal prob of N-in: 0.12994\noutside 1 331\nTMhelix 332 354\ninside 355 372

4757 GCF_000946275.1_FHI97 Escherichia coli Proteobacteria; Gammaproteobacteria; Enterobacterales; Enterobacteriaceae; Escherichia MTGDNTLIHSHGINRRDFMKLCAALAATMGLSSKAAAEMAESVTNPQRPPVIWIGAQECT WP_000145410.1 MULTISPECIES: hydrogenase-2 small chain [Proteobacteria] Length: 372\nNumber of predicted TMHs: 1\nExp number of AAs in TMHs: 23.72226\nExp number, first 60 AAs: 0.22038\nTotal prob of N-in: 0.12994\noutside 1 331\nTMhelix 332 354\ninside 355 372

4758 GCF_000942815.1_Ec138B_L2_MIRA_assembly Escherichia coli Proteobacteria; Gammaproteobacteria; Enterobacterales; Enterobacteriaceae; Escherichia MNNEETFYQAMRRQGVTRRSFLKYCSLAATSLGLGAGMAPKIAWALENKPRIPVVWIHGL WP_001058323.1 MULTISPECIES: hydrogenase-1 small chain [Proteobacteria] Length: 372\nNumber of predicted TMHs: 2\nExp number of AAs in TMHs: 37.2608499999999\nExp number, first 60 AAs: 19.52923\nTotal prob of N-in: 0.94351\nPOSSIBLE N-term signal sequence\ninside 1 19\nTMhelix 20 39\noutside 40 325\nTMhelix 326 348\ninside 349 372

4759 GCF_000943355.2_FHI42 Escherichia coli Proteobacteria; Gammaproteobacteria; Enterobacterales; Enterobacteriaceae; Escherichia MTGDNTLIHSHGINRRDFMKLCAALAATMGLSSKAAAEMAESVTNPQRPPVIWIGAQECT WP_000145410.1 MULTISPECIES: hydrogenase-2 small chain [Proteobacteria] Length: 372\nNumber of predicted TMHs: 1\nExp number of AAs in TMHs: 23.72226\nExp number, first 60 AAs: 0.22038\nTotal prob of N-in: 0.12994\noutside 1 331\nTMhelix 332 354\ninside 355 372

4760 GCF_000945615.2_FHI28 Escherichia coli Proteobacteria; Gammaproteobacteria; Enterobacterales; Enterobacteriaceae; Escherichia MTGDNTLIHSHGINRRDFMKLCAALAATMGLSSKAAAEMAESVTNPQRPPVIWIGAQECT WP_000145410.1 MULTISPECIES: hydrogenase-2 small chain [Proteobacteria] Length: 372\nNumber of predicted TMHs: 1\nExp number of AAs in TMHs: 23.72226\nExp number, first 60 AAs: 0.22038\nTotal prob of N-in: 0.12994\noutside 1 331\nTMhelix 332 354\ninside 355 372

4761 GCF_000941995.1_FHI59 Escherichia coli Proteobacteria; Gammaproteobacteria; Enterobacterales; Enterobacteriaceae; Escherichia MTGDNTLIHSHGINRRDFMKLCAALAATMGLSSKAAAEMAESVTNPQRPPVIWIGAQECT WP_000145410.1 MULTISPECIES: hydrogenase-2 small chain [Proteobacteria] Length: 372\nNumber of predicted TMHs: 1\nExp number of AAs in TMHs: 23.72226\nExp number, first 60 AAs: 0.22038\nTotal prob of N-in: 0.12994\noutside 1 331\nTMhelix 332 354\ninside 355 372

4762 GCF_000797605.1_Escherichia_coli_CVM_N36963PS_v._1.0 Escherichia coli Proteobacteria; Gammaproteobacteria; Enterobacterales; Enterobacteriaceae; Escherichia MTGDNTLIHSHGINRRDFMKLCAALAATMGLSSKAAAEMAESVTNPQRPPVIWIGAQECT WP_000145410.1 MULTISPECIES: hydrogenase-2 small chain [Proteobacteria] Length: 372\nNumber of predicted TMHs: 1\nExp number of AAs in TMHs: 23.72226\nExp number, first 60 AAs: 0.22038\nTotal prob of N-in: 0.12994\noutside 1 331\nTMhelix 332 354\ninside 355 372

4763 GCF_000797605.1_Escherichia_coli_CVM_N36963PS_v._1.0 Escherichia coli Proteobacteria; Gammaproteobacteria; Enterobacterales; Enterobacteriaceae; Escherichia MNNEETFYQAMRRQGVTRRSFLKYCSLAATSLGLGAGMAPKIAWALENKPRIPVVWIHGL WP_001058323.1 MULTISPECIES: hydrogenase-1 small chain [Proteobacteria] Length: 372\nNumber of predicted TMHs: 2\nExp number of AAs in TMHs: 37.2608499999999\nExp number, first 60 AAs: 19.52923\nTotal prob of N-in: 0.94351\nPOSSIBLE N-term signal sequence\ninside 1 19\nTMhelix 20 39\noutside 40 325\nTMhelix 326 348\ninside 349 372

4764 GCF_000797695.1_Escherichia_coli_CVM_N38733PS_v._1.0 Escherichia coli Proteobacteria; Gammaproteobacteria; Enterobacterales; Enterobacteriaceae; Escherichia MTGDNTLIHSHGINRRDFMKLCAALAATMGLSSKAAAEMAESVTNPQRPPVIWIGAQECT WP_000145410.1 MULTISPECIES: hydrogenase-2 small chain [Proteobacteria] Length: 372\nNumber of predicted TMHs: 1\nExp number of AAs in TMHs: 23.72226\nExp number, first 60 AAs: 0.22038\nTotal prob of N-in: 0.12994\noutside 1 331\nTMhelix 332 354\ninside 355 372

4765 GCF_000797695.1_Escherichia_coli_CVM_N38733PS_v._1.0 Escherichia coli Proteobacteria; Gammaproteobacteria; Enterobacterales; Enterobacteriaceae; Escherichia

MNNEETFYQAMRRQGVTRRSFLKYCSLAATSLGLGAGMAPKIAWALENKPRIPVVWIHGL WP_001295940.1 MULTISPECIES: hydrogenase [Enterobacteriaceae] Length: 372\nNumber of predicted TMHs: 2\nExp number of AAs in TMHs: 37.2608899999999\nExp number, first 60 AAs: 19.52923\nTotal prob of N-in: 0.94351\nPOSSIBLE N-term signal sequence\ninside 1 19\nTMhelix 20 39\noutside 40 325\nTMhelix 326 348\ninside 349 372

4766 GCF_000797755.1_Escherichia_coli_CVM_N33443PS_v._1.0 Escherichia coli Proteobacteria; Gammaproteobacteria; Enterobacterales; Enterobacteriaceae; Escherichia MTGDNTLIHSHGINRRDFMKLCAALAATMGLSSKAAAEMAESVTNPQRPPVIWIGAQECT WP_000145410.1 MULTISPECIES: hydrogenase-2 small chain [Proteobacteria] Length: 372\nNumber of predicted TMHs: 1\nExp number of AAs in TMHs: 23.72226\nExp number, first 60 AAs: 0.22038\nTotal prob of N-in: 0.12994\noutside 1 331\nTMhelix 332 354\ninside 355 372

4767 GCF_000797755.1_Escherichia_coli_CVM_N33443PS_v._1.0 Escherichia coli Proteobacteria; Gammaproteobacteria; Enterobacterales; Enterobacteriaceae; Escherichia MNNEETFYQAMRRQGVTRRSFLKYCSLAATSLGLGAGMAPKIAWALENKPRIPVVWIHGL WP_001058323.1 MULTISPECIES: hydrogenase-1 small chain [Proteobacteria] Length: 372\nNumber of predicted TMHs: 2\nExp number of AAs in TMHs: 37.2608499999999\nExp number, first 60 AAs: 19.52923\nTotal prob of N-in: 0.94351\nPOSSIBLE N-term signal sequence\ninside 1 19\nTMhelix 20 39\noutside 40 325\nTMhelix 326 348\ninside 349 372

4768 GCF_000797815.1_Escherichia_coli_CVM_N41556PS_v._1.0 Escherichia coli Proteobacteria; Gammaproteobacteria; Enterobacterales; Enterobacteriaceae; Escherichia MTGDNTLIHSHGINRRDFMKLCAALAATMGLSSKAAAEMAESVTNPQRPPVIWIGAQECT WP_000145410.1 MULTISPECIES: hydrogenase-2 small chain [Proteobacteria] Length: 372\nNumber of predicted TMHs: 1\nExp number of AAs in TMHs: 23.72226\nExp number, first 60 AAs: 0.22038\nTotal prob of N-in: 0.12994\noutside 1 331\nTMhelix 332 354\ninside 355 372

4769 GCF_000797815.1_Escherichia_coli_CVM_N41556PS_v._1.0 Escherichia coli Proteobacteria; Gammaproteobacteria; Enterobacterales; Enterobacteriaceae; Escherichia MNNEETFYQAMRRQGVTRRSFLKYCSLAATSLGLGAGMAPKIAWALENKPRIPVVWIHGL WP_001058323.1 MULTISPECIES: hydrogenase-1 small chain [Proteobacteria] Length: 372\nNumber of predicted TMHs: 2\nExp number of AAs in TMHs: 37.2608499999999\nExp number, first 60 AAs: 19.52923\nTotal prob of N-in: 0.94351\nPOSSIBLE N-term signal sequence\ninside 1 19\nTMhelix 20 39\noutside 40 325\nTMhelix 326 348\ninside 349 372

4770 GCF_000942815.1_Ec138B_L2_MIRA_assembly Escherichia coli Proteobacteria; Gammaproteobacteria; Enterobacterales; Enterobacteriaceae; Escherichia MTGDNTLIHSHGINRRDFMKLCAALAATMGLSSKAAAEMAESVTNPQRPPVIWIGAQECT WP_000145410.1 MULTISPECIES: hydrogenase-2 small chain [Proteobacteria] Length: 372\nNumber of predicted TMHs: 1\nExp number of AAs in TMHs: 23.72226\nExp number, first 60 AAs: 0.22038\nTotal prob of N-in: 0.12994\noutside 1 331\nTMhelix 332 354\ninside 355 372

4771 GCF_000940655.1_FHI70 Escherichia coli Proteobacteria; Gammaproteobacteria; Enterobacterales; Enterobacteriaceae; Escherichia MNNEETFYQAMRRQGVTRRSFLKYCSLAATSLGLGAGMAPKIAWALENKPRIPVVWIHGL WP_001058323.1 MULTISPECIES: hydrogenase-1 small chain [Proteobacteria] Length: 372\nNumber of predicted TMHs: 2\nExp number of AAs in TMHs: 37.2608499999999\nExp number, first 60 AAs: 19.52923\nTotal prob of N-in: 0.94351\nPOSSIBLE N-term signal sequence\ninside 1 19\nTMhelix 20 39\noutside 40 325\nTMhelix 326 348\ninside 349 372

4772 GCF_000940115.1_FHI20 Escherichia coli Proteobacteria; Gammaproteobacteria; Enterobacterales; Enterobacteriaceae; Escherichia MNNEETFYQAMRRQGVTRRSFLKYCSLAATSLGLGAGMAPKIAWALENKPRIPVVWIHGL WP_001058323.1 MULTISPECIES: hydrogenase-1 small chain [Proteobacteria] Length: 372\nNumber of predicted TMHs: 2\nExp number of AAs in TMHs: 37.2608499999999\nExp number, first 60 AAs: 19.52923\nTotal prob of N-in: 0.94351\nPOSSIBLE N-term signal sequence\ninside 1 19\nTMhelix 20 39\noutside 40 325\nTMhelix 326 348\ninside 349 372

4773 GCF_000940455.1_138B_L2_assembly Escherichia coli Proteobacteria; Gammaproteobacteria; Enterobacterales; Enterobacteriaceae; Escherichia MNNEETFYQAMRRQGVTRRSFLKYCSLAATSLGLGAGMAPKIAWALENKPRIPVVWIHGL WP_001058323.1 MULTISPECIES: hydrogenase-1 small chain [Proteobacteria] Length: 372\nNumber of predicted TMHs:

2\nExp number of AAs in TMHs: 37.2608499999999\nExp number, first 60 AAs: 19.52923\nTotal prob of N-in: 0.94351\nPOSSIBLE N-term signal sequence\ninside 1 19\nTMhelix 20 39\noutside 40 325\nTMhelix 326 348\ninside 349 372

4774 GCF_000940655.1_FHI70 Escherichia coli Proteobacteria; Gammaproteobacteria; Enterobacterales; Enterobacteriaceae; Escherichia MTGDNTLIHSHGINRRDFMKLCAALAATMGLSSKAAAEMAESVTNPQRPPVIWIGAQECT WP_000145410.1 MULTISPECIES: hydrogenase-2 small chain [Proteobacteria] Length: 372\nNumber of predicted TMHs: 1\nExp number of AAs in TMHs: 23.72226\nExp number, first 60 AAs: 0.22038\nTotal prob of N-in: 0.12994\noutside 1 331\nTMhelix 332 354\ninside 355 372

4775 GCF_000940455.1_138B_L2_assembly Escherichia coli Proteobacteria; Gammaproteobacteria; Enterobacterales; Enterobacteriaceae; Escherichia MTGDNTLIHSHGINRRDFMKLCAALAATMGLSSKAAAEMAESVTNPQRPPVIWIGAQECT WP_000145410.1 MULTISPECIES: hydrogenase-2 small chain [Proteobacteria] Length: 372\nNumber of predicted TMHs: 1\nExp number of AAs in TMHs: 23.72226\nExp number, first 60 AAs: 0.22038\nTotal prob of N-in: 0.12994\noutside 1 331\nTMhelix 332 354\ninside 355 372

4776 GCF_000940115.1_FHI20 Escherichia coli Proteobacteria; Gammaproteobacteria; Enterobacterales; Enterobacteriaceae; Escherichia MTGDNTLIHSHGINRRDFMKLCAALAATMGLSSKAAAEMAESVTNPQRPPVIWIGAQECT WP_000145410.1 MULTISPECIES: hydrogenase-2 small chain [Proteobacteria] Length: 372\nNumber of predicted TMHs: 1\nExp number of AAs in TMHs: 23.72226\nExp number, first 60 AAs: 0.22038\nTotal prob of N-in: 0.12994\noutside 1 331\nTMhelix 332 354\ninside 355 372

4777 GCF_000939195.1_FHI9 Escherichia coli Proteobacteria; Gammaproteobacteria; Enterobacterales; Enterobacteriaceae; Escherichia MNNEETFYQAMRRQGVTRRSFLKYCSLAATSLGLGAGMAPKIAWALENKPRIPVVWIHGL WP_001058323.1 MULTISPECIES: hydrogenase-1 small chain [Proteobacteria] Length: 372\nNumber of predicted TMHs: 2\nExp number of AAs in TMHs: 37.2608499999999\nExp number, first 60 AAs: 19.52923\nTotal prob of N-in: 0.94351\nPOSSIBLE N-term signal sequence\ninside 1 19\nTMhelix 20 39\noutside 40 325\nTMhelix 326 348\ninside 349 372

4778 GCF_000939195.1_FHI9 Escherichia coli Proteobacteria; Gammaproteobacteria; Enterobacterales; Enterobacteriaceae; Escherichia MTGDNTLIHSHGINRRDFMKLCAALAATMGLSSKAAAEMAESVTNPQRPPVIWIGAQECT WP_000145410.1 MULTISPECIES: hydrogenase-2 small chain [Proteobacteria] Length: 372\nNumber of predicted TMHs: 1\nExp number of AAs in TMHs: 23.72226\nExp number, first 60 AAs: 0.22038\nTotal prob of N-in: 0.12994\noutside 1 331\nTMhelix 332 354\ninside 355 372

4779 GCF_000939035.1_Ec26A_L2_MIRA_assembly Escherichia coli Proteobacteria; Gammaproteobacteria; Enterobacterales; Enterobacteriaceae; Escherichia MTGDNTLIHSHGINRRDFMKLCAALAATMGLSSKAAAEMAESVTNPQRPPVIWIGAQECT WP_000145410.1 MULTISPECIES: hydrogenase-2 small chain [Proteobacteria] Length: 372\nNumber of predicted TMHs: 1\nExp number of AAs in TMHs: 23.72226\nExp number, first 60 AAs: 0.22038\nTotal prob of N-in: 0.12994\noutside 1 331\nTMhelix 332 354\ninside 355 372

4780 GCF_000937855.1_FHI23 Escherichia coli Proteobacteria; Gammaproteobacteria; Enterobacterales; Enterobacteriaceae; Escherichia MNNEETFYQAMRRQGVTRRSFLKYCSLAATSLGLGAGMAPKIAWALENKPRIPVVWIHGL WP_001058323.1 MULTISPECIES: hydrogenase-1 small chain [Proteobacteria] Length: 372\nNumber of predicted TMHs: 2\nExp number of AAs in TMHs: 37.2608499999999\nExp number, first 60 AAs: 19.52923\nTotal prob of N-in: 0.94351\nPOSSIBLE N-term signal sequence\ninside 1 19\nTMhelix 20 39\noutside 40 325\nTMhelix 326 348\ninside 349 372

4781 GCF_000938695.1_FHI83 Escherichia coli Proteobacteria; Gammaproteobacteria; Enterobacterales; Enterobacteriaceae; Escherichia MTGDNTLIHSHGINRRDFMKLCAALAATMGLSSKAAAEMAESVTNPQRPPVIWIGAQECT WP_000145410.1 MULTISPECIES: hydrogenase-2 small chain [Proteobacteria] Length: 372\nNumber of predicted TMHs: 1\nExp number of AAs in TMHs: 23.72226\nExp number, first 60 AAs: 0.22038\nTotal prob of N-in: 0.12994\noutside 1 331\nTMhelix 332 354\ninside 355 372

4782 GCF_000937355.1_46A_L2_assembly Escherichia coli Proteobacteria; Gammaproteobacteria; Enterobacterales; Enterobacteriaceae; Escherichia MNNEETFYQAMRRQGVTRRSFLKYCSLAATSLGLGAGMAPKIAWALENKPRIPVVWIHGL WP_001058323.1 MULTISPECIES: hydrogenase-1 small chain [Proteobacteria] Length: 372\nNumber of predicted TMHs: 2\nExp number of AAs in TMHs: 37.2608499999999\nExp number, first 60 AAs: 19.52923\nTotal prob of N-in:

0.94351\nPOSSIBLE N-term signal sequence\ninside 1 19\nTMhelix 20 39\noutside 40 325\nTMhelix 326 348\ninside 349 372

4783 GCF_000937855.1_FHI23 Escherichia coli Proteobacteria; Gammaproteobacteria; Enterobacterales; Enterobacteriaceae; Escherichia MTGDNTLIHSHGINRRDFMKLCAALAATMGLSSKAAAEMAESVTNPQRPPVIWIGAQECT WP_000145410.1 MULTISPECIES: hydrogenase-2 small chain [Proteobacteria] Length: 372\nNumber of predicted TMHs: 1\nExp number of AAs in TMHs: 23.72226\nExp number, first 60 AAs: 0.22038\nTotal prob of N-in: 0.12994\noutside 1 331\nTMhelix 332 354\ninside 355 372

4784 GCF_000937415.1_FHI98 Escherichia coli Proteobacteria; Gammaproteobacteria; Enterobacterales; Enterobacteriaceae; Escherichia MTGDNTLIHSHGINRRDFMKLCAALAATMGLSSKAAAEMAESVTNPQRPPVIWIGAQECT WP_000145410.1 MULTISPECIES: hydrogenase-2 small chain [Proteobacteria] Length: 372\nNumber of predicted TMHs: 1\nExp number of AAs in TMHs: 23.72226\nExp number, first 60 AAs: 0.22038\nTotal prob of N-in: 0.12994\noutside 1 331\nTMhelix 332 354\ninside 355 372

4785 GCF_000937355.1_46A_L2_assembly Escherichia coli Proteobacteria; Gammaproteobacteria; Enterobacterales; Enterobacteriaceae; Escherichia MTGDNTLIHSHGINRRDFMKLCAALAATMGLSSKAAAEMAESVTNPQRPPVIWIGAQECT WP_000145410.1 MULTISPECIES: hydrogenase-2 small chain [Proteobacteria] Length: 372\nNumber of predicted TMHs: 1\nExp number of AAs in TMHs: 23.72226\nExp number, first 60 AAs: 0.22038\nTotal prob of N-in: 0.12994\noutside 1 331\nTMhelix 332 354\ninside 355 372

4786 GCF_001269045.1_1.EC2721.1 Escherichia coli Proteobacteria; Gammaproteobacteria; Enterobacterales; Enterobacteriaceae; Escherichia MTGDNTLIHSHGINRRDFMKLCAALAATMGLSSKAAAEMAESVTNPQRPPVIWIGAQECT WP_000145410.1 MULTISPECIES: hydrogenase-2 small chain [Proteobacteria] Length: 372\nNumber of predicted TMHs: 1\nExp number of AAs in TMHs: 23.72226\nExp number, first 60 AAs: 0.22038\nTotal prob of N-in: 0.12994\noutside 1 331\nTMhelix 332 354\ninside 355 372

4787 GCF_001269045.1_1.EC2721.1 Escherichia coli Proteobacteria; Gammaproteobacteria; Enterobacterales; Enterobacteriaceae; Escherichia MNNEETFYQAMRRQGVTRRSFLKYCSLAATSLGLGAGMAPKIAWALENKPRIPVVWIHGL WP_001058323.1 MULTISPECIES: hydrogenase-1 small chain [Proteobacteria] Length: 372\nNumber of predicted TMHs: 2\nExp number of AAs in TMHs: 37.2608499999999\nExp number, first 60 AAs: 19.52923\nTotal prob of N-in: 0.94351\nPOSSIBLE N-term signal sequence\ninside 1 19\nTMhelix 20 39\noutside 40 325\nTMhelix 326 348\ninside 349 372

4788 GCF_000936495.1_FHI102 Escherichia coli Proteobacteria; Gammaproteobacteria; Enterobacterales; Enterobacteriaceae; Escherichia MNNEETFYQAMRRQGVTRRSFLKYCSLAATSLGLGAGMAPKIAWALENKPRIPVVWIHGL WP_001058323.1 MULTISPECIES: hydrogenase-1 small chain [Proteobacteria] Length: 372\nNumber of predicted TMHs: 2\nExp number of AAs in TMHs: 37.2608499999999\nExp number, first 60 AAs: 19.52923\nTotal prob of N-in: 0.94351\nPOSSIBLE N-term signal sequence\ninside 1 19\nTMhelix 20 39\noutside 40 325\nTMhelix 326 348\ninside 349 372

4789 GCF_000782035.1_ASM78203v1 Escherichia coli Proteobacteria; Gammaproteobacteria; Enterobacterales; Enterobacteriaceae; Escherichia MNNEETFYQAMRRQGVTRRSFLKYCSLAATSLGLGAGMAPKIAWALENKPRIPVVWIHGL WP_001058323.1 MULTISPECIES: hydrogenase-1 small chain [Proteobacteria] Length: 372\nNumber of predicted TMHs: 2\nExp number of AAs in TMHs: 37.2608499999999\nExp number, first 60 AAs: 19.52923\nTotal prob of N-in: 0.94351\nPOSSIBLE N-term signal sequence\ninside 1 19\nTMhelix 20 39\noutside 40 325\nTMhelix 326 348\ninside 349 372

4790 GCF_000948745.1_ASM94874v1 Escherichia coli Proteobacteria; Gammaproteobacteria; Enterobacterales; Enterobacteriaceae; Escherichia MTGDNTLIHSHGINRRDFMKLCAALAATMGLSSKAAAEMAESVTNPQRPPVIWIGAQECT WP_000145410.1 MULTISPECIES: hydrogenase-2 small chain [Proteobacteria] Length: 372\nNumber of predicted TMHs: 1\nExp number of AAs in TMHs: 23.72226\nExp number, first 60 AAs: 0.22038\nTotal prob of N-in: 0.12994\noutside 1 331\nTMhelix 332 354\ninside 355 372

4791 GCF_000948715.1_ASM94871v1 Escherichia coli Proteobacteria; Gammaproteobacteria; Enterobacterales; Enterobacteriaceae; Escherichia

MTGDNTLIHSHGINRRDFMKLCAALAATMGLSSKAAAEMAESVTNPQRPPVIWIGAQECT WP_000145410.1 MULTISPECIES: hydrogenase-2 small chain [Proteobacteria] Length: 372\nNumber of predicted TMHs: 1\nExp number of AAs in TMHs: 23.72226\nExp number, first 60 AAs: 0.22038\nTotal prob of N-in: 0.12994\noutside 1 331\nTMhelix 332 354\ninside 355 372

4792 GCF_000948055.1_ASM94805v1 Escherichia coli Proteobacteria; Gammaproteobacteria; Enterobacterales; Enterobacteriaceae; Escherichia MNNEETFYQAMRRQGVTRRSFLKYCSLAATSLGLGAGMAPKIAWALENKPRIPVVWIHGL WP_001058323.1 MULTISPECIES: hydrogenase-1 small chain [Proteobacteria] Length: 372\nNumber of predicted TMHs: 2\nExp number of AAs in TMHs: 37.2608499999999\nExp number, first 60 AAs: 19.52923\nTotal prob of N-in: 0.94351\nPOSSIBLE N-term signal sequence\ninside 1 19\nTMhelix 20 39\noutside 40 325\nTMhelix 326 348\ninside 349 372

4793 GCF_000948625.1_ASM94862v1 Escherichia coli Proteobacteria; Gammaproteobacteria; Enterobacterales; Enterobacteriaceae; Escherichia MNNEETFYQAMRRQGVTRRSFLKYCSLAATSLGLGAGMAPKIAWALENKPRIPVVWIHGL WP_001295940.1 MULTISPECIES: hydrogenase [Enterobacteriaceae] Length: 372\nNumber of predicted TMHs: 2\nExp number of AAs in TMHs: 37.2608899999999\nExp number, first 60 AAs: 19.52923\nTotal prob of N-in: 0.94351\nPOSSIBLE N-term signal sequence\ninside 1 19\nTMhelix 20 39\noutside 40 325\nTMhelix 326 348\ninside 349 372

4794 GCF_000782415.1_ASM78241v1 Escherichia coli Proteobacteria; Gammaproteobacteria; Enterobacterales; Enterobacteriaceae; Escherichia MTGDNTLIHSHGINRRDFMKLCAALAATMGLSSKAAAEMAESVTNPQRPPVIWIGAQECT WP_000145410.1 MULTISPECIES: hydrogenase-2 small chain [Proteobacteria] Length: 372\nNumber of predicted TMHs: 1\nExp number of AAs in TMHs: 23.72226\nExp number, first 60 AAs: 0.22038\nTotal prob of N-in: 0.12994\noutside 1 331\nTMhelix 332 354\ninside 355 372

4795 GCF_000782415.1_ASM78241v1 Escherichia coli Proteobacteria; Gammaproteobacteria; Enterobacterales; Enterobacteriaceae; Escherichia MNNEETFYQAMRRQGVTRRSFLKYCSLAATSLGLGAGMAPKIAWALENKPRIPVVWIHGL WP_001058323.1 MULTISPECIES: hydrogenase-1 small chain [Proteobacteria] Length: 372\nNumber of predicted TMHs: 2\nExp number of AAs in TMHs: 37.2608499999999\nExp number, first 60 AAs: 19.52923\nTotal prob of N-in: 0.94351\nPOSSIBLE N-term signal sequence\ninside 1 19\nTMhelix 20 39\noutside 40 325\nTMhelix 326 348\ninside 349 372

4796 GCF_000782475.1_ASM78247v1 Escherichia coli Proteobacteria; Gammaproteobacteria; Enterobacterales; Enterobacteriaceae; Escherichia MTGDNTLIHSHGINRRDFMKLCAALAATMGLSSKAAAEMAESVTNPQRPPVIWIGAQECT WP_000145410.1 MULTISPECIES: hydrogenase-2 small chain [Proteobacteria] Length: 372\nNumber of predicted TMHs: 1\nExp number of AAs in TMHs: 23.72226\nExp number, first 60 AAs: 0.22038\nTotal prob of N-in: 0.12994\noutside 1 331\nTMhelix 332 354\ninside 355 372

4797 GCF_000782475.1_ASM78247v1 Escherichia coli Proteobacteria; Gammaproteobacteria; Enterobacterales; Enterobacteriaceae; Escherichia MNNEETFYQAMRRQGVTRRSFLKYCSLAATSLGLGAGMAPKIAWALENKPRIPVVWIHGL WP_001058323.1 MULTISPECIES: hydrogenase-1 small chain [Proteobacteria] Length: 372\nNumber of predicted TMHs: 2\nExp number of AAs in TMHs: 37.2608499999999\nExp number, first 60 AAs: 19.52923\nTotal prob of N-in: 0.94351\nPOSSIBLE N-term signal sequence\ninside 1 19\nTMhelix 20 39\noutside 40 325\nTMhelix 326 348\ninside 349 372

4798 GCF_000948605.1_ASM94860v1 Escherichia coli Proteobacteria; Gammaproteobacteria; Enterobacterales; Enterobacteriaceae; Escherichia MNNEETFYQAMRRQGVTRRSFLKYCSLAATSLGLGAGMAPKIAWALENKPRIPVVWIHGL WP_001058323.1 MULTISPECIES: hydrogenase-1 small chain [Proteobacteria] Length: 372\nNumber of predicted TMHs: 2\nExp number of AAs in TMHs: 37.2608499999999\nExp number, first 60 AAs: 19.52923\nTotal prob of N-in: 0.94351\nPOSSIBLE N-term signal sequence\ninside 1 19\nTMhelix 20 39\noutside 40 325\nTMhelix 326 348\ninside 349 372

4799 GCF_000948605.1_ASM94860v1 Escherichia coli Proteobacteria; Gammaproteobacteria; Enterobacterales; Enterobacteriaceae; Escherichia

MTGDNTLIHSHGINRRDFMKLCAALAATMGLSSKAAAEMAESVTNPQRPPVIWIGAQECT WP_000145410.1 MULTISPECIES: hydrogenase-2 small chain [Proteobacteria] Length: 372\nNumber of predicted TMHs: 1\nExp number of AAs in TMHs: 23.72226\nExp number, first 60 AAs: 0.22038\nTotal prob of N-in: 0.12994\noutside 1 331\nTMhelix 332 354\ninside 355 372

4800 GCF_001058965.1_ASM105896v1 Escherichia coli Proteobacteria; Gammaproteobacteria; Enterobacterales; Enterobacteriaceae; Escherichia MTGDNTLIHSHGINRRDFMKLCAALAATMGLSSKAAAEMVESVTNPQRPPVIWIGAQECT WP_000145423.1 hydrogenase 2 small subunit [Escherichia coli] Length: 372\nNumber of predicted TMHs: 1\nExp number of AAs in TMHs: 23.78201\nExp number, first 60 AAs: 0.28355\nTotal prob of N-in: 0.13257\noutside 1 331\nTMhelix 332 354\ninside 355 372

4801 GCF_001059075.1_ASM105907v1 Escherichia coli Proteobacteria; Gammaproteobacteria; Enterobacterales; Enterobacteriaceae; Escherichia MTGDNTLIHSHGINRRDFMKLCAALAATMGLSSKAAAEMAESVTNPQRPPVIWIGAQECT WP_000145410.1 MULTISPECIES: hydrogenase-2 small chain [Proteobacteria] Length: 372\nNumber of predicted TMHs: 1\nExp number of AAs in TMHs: 23.72226\nExp number, first 60 AAs: 0.22038\nTotal prob of N-in: 0.12994\noutside 1 331\nTMhelix 332 354\ninside 355 372

4802 GCF_001058965.1_ASM105896v1 Escherichia coli Proteobacteria; Gammaproteobacteria; Enterobacterales; Enterobacteriaceae; Escherichia MNNEETFYQAMRRQGVTRRSFLKYCSLAATSLGLGAGMAPKIAWALENKPRIPVVWIHGL WP_001295940.1 MULTISPECIES: hydrogenase [Enterobacteriaceae] Length: 372\nNumber of predicted TMHs: 2\nExp number of AAs in TMHs: 37.2608899999999\nExp number, first 60 AAs: 19.52923\nTotal prob of N-in: 0.94351\nPOSSIBLE N-term signal sequence\ninside 1 19\nTMhelix 20 39\noutside 40 325\nTMhelix 326 348\ninside 349 372

4803 GCF_001059075.1_ASM105907v1 Escherichia coli Proteobacteria; Gammaproteobacteria; Enterobacterales; Enterobacteriaceae; Escherichia MNNEETFYQAMRRQGVTRRSFLKYCSLAATSLGLGAGMAPKIAWALENKPRIPVVWIHGL WP_001295940.1 MULTISPECIES: hydrogenase [Enterobacteriaceae] Length: 372\nNumber of predicted TMHs: 2\nExp number of AAs in TMHs: 37.2608899999999\nExp number, first 60 AAs: 19.52923\nTotal prob of N-in: 0.94351\nPOSSIBLE N-term signal sequence\ninside 1 19\nTMhelix 20 39\noutside 40 325\nTMhelix 326 348\ninside 349 372

4804 GCF_000948435.1_ASM94843v1 Escherichia coli Proteobacteria; Gammaproteobacteria; Enterobacterales; Enterobacteriaceae; Escherichia MTGDNTLIHSHGINRRDFMKLCAALAATMGLSSKAAAEMAESVTNPQRPPVIWIGAQECT WP_000145410.1 MULTISPECIES: hydrogenase-2 small chain [Proteobacteria] Length: 372\nNumber of predicted TMHs: 1\nExp number of AAs in TMHs: 23.72226\nExp number, first 60 AAs: 0.22038\nTotal prob of N-in: 0.12994\noutside 1 331\nTMhelix 332 354\ninside 355 372

4805 GCF_000948055.1_ASM94805v1 Escherichia coli Proteobacteria; Gammaproteobacteria; Enterobacterales; Enterobacteriaceae; Escherichia MTGDNTLIHSHGINRRDFMKLCAALAATMGLSSKAAAEMAESVTNPQRPPVIWIGAQECT WP_000145410.1 MULTISPECIES: hydrogenase-2 small chain [Proteobacteria] Length: 372\nNumber of predicted TMHs: 1\nExp number of AAs in TMHs: 23.72226\nExp number, first 60 AAs: 0.22038\nTotal prob of N-in: 0.12994\noutside 1 331\nTMhelix 332 354\ninside 355 372

4806 GCF_000947995.1_ASM94799v1 Escherichia coli Proteobacteria; Gammaproteobacteria; Enterobacterales; Enterobacteriaceae; Escherichia MTGDNTLIHSHGINRRDFMKLCAALAATMGLSSKAAAEMAESVTNPQRPPVIWIGAQECT WP_000145410.1 MULTISPECIES: hydrogenase-2 small chain [Proteobacteria] Length: 372\nNumber of predicted TMHs: 1\nExp number of AAs in TMHs: 23.72226\nExp number, first 60 AAs: 0.22038\nTotal prob of N-in: 0.12994\noutside 1 331\nTMhelix 332 354\ninside 355 372

4807 GCF_000947975.1_ASM94797v1 Escherichia coli Proteobacteria; Gammaproteobacteria; Enterobacterales; Enterobacteriaceae; Escherichia MTGDNTLIHSHGINRRDFMKLCAALAATMGLSSKAAAEMAESVTNPQRPPVIWIGAQECT WP_000145410.1 MULTISPECIES: hydrogenase-2 small chain [Proteobacteria] Length: 372\nNumber of predicted TMHs:

1\nExp number of AAs in TMHs: 23.72226\nExp number, first 60 AAs: 0.22038\nTotal prob of N-in: 0.12994\noutside 1 331\nTMhelix 332 354\ninside 355 372

4808 GCF_000945615.2_FHI28 Escherichia coli Proteobacteria; Gammaproteobacteria; Enterobacterales; Enterobacteriaceae; Escherichia MNNEETFYQAMRRQGVTRRSFLKYCSLAATSLGLGAGMAPKIAWAPENKPRIPVVWIHGL WP_044804018.1 hydrogenase [Escherichia coli] Length: 372\nNumber of predicted TMHs: 2\nExp number of AAs in TMHs: 36.79941\nExp number, first 60 AAs: 18.91621\nTotal prob of N-in: 0.91758\nPOSSIBLE N-term signal sequence\ninside 1 19\nTMhelix 20 39\noutside 40 325\nTMhelix 326 348\ninside 349 372

4809 GCF_000782515.1_ASM78251v1 Escherichia coli Proteobacteria; Gammaproteobacteria; Enterobacterales; Enterobacteriaceae; Escherichia MTGDNTLIHSHGINRRDFMKLCAALAATMGLSSKAAAEMVESVTNPQRPPVIWIGAQECT WP_000145423.1 hydrogenase 2 small subunit [Escherichia coli] Length: 372\nNumber of predicted TMHs: 1\nExp number of AAs in TMHs: 23.78201\nExp number, first 60 AAs: 0.28355\nTotal prob of N-in: 0.13257\noutside 1 331\nTMhelix 332 354\ninside 355 372

4810 GCF_000782515.1_ASM78251v1 Escherichia coli Proteobacteria; Gammaproteobacteria; Enterobacterales; Enterobacteriaceae; Escherichia MNNEETFYQAMRRQGVTRRSFLKYCSLAATSLGLGAGMAPKIAWALENKPRIPVVWIHGL WP_001295940.1 MULTISPECIES: hydrogenase [Enterobacteriaceae] Length: 372\nNumber of predicted TMHs: 2\nExp number of AAs in TMHs: 37.2608899999999\nExp number, first 60 AAs: 19.52923\nTotal prob of N-in: 0.94351\nPOSSIBLE N-term signal sequence\ninside 1 19\nTMhelix 20 39\noutside 40 325\nTMhelix 326 348\ninside 349 372

4811 GCF_000782535.1_ASM78253v1 Escherichia coli Proteobacteria; Gammaproteobacteria; Enterobacterales; Enterobacteriaceae; Escherichia MTGDNTLIHSHGINRRDFMKLCAALAATMGLSSKAAAEMVESVTNPQRPPVIWIGAQECT WP_000145423.1 hydrogenase 2 small subunit [Escherichia coli] Length: 372\nNumber of predicted TMHs: 1\nExp number of AAs in TMHs: 23.78201\nExp number, first 60 AAs: 0.28355\nTotal prob of N-in: 0.13257\noutside 1 331\nTMhelix 332 354\ninside 355 372

4812 GCF_000782535.1_ASM78253v1 Escherichia coli Proteobacteria; Gammaproteobacteria; Enterobacterales; Enterobacteriaceae; Escherichia MNNEETFYQAMRRQGVTRRSFLKYCSLAATSLGLGAGMAPKIAWALENKPRIPVVWIHGL WP_001295940.1 MULTISPECIES: hydrogenase [Enterobacteriaceae] Length: 372\nNumber of predicted TMHs: 2\nExp number of AAs in TMHs: 37.2608899999999\nExp number, first 60 AAs: 19.52923\nTotal prob of N-in: 0.94351\nPOSSIBLE N-term signal sequence\ninside 1 19\nTMhelix 20 39\noutside 40 325\nTMhelix 326 348\ninside 349 372

4813 GCF_000782595.1_ASM78259v1 Escherichia coli Proteobacteria; Gammaproteobacteria; Enterobacterales; Enterobacteriaceae; Escherichia MTGDNTLIHSHGINRRDFMKLCAALAATMGLSSKAAAEMAESVTNPQRPPVIWIGAQECT WP_000145410.1 MULTISPECIES: hydrogenase-2 small chain [Proteobacteria] Length: 372\nNumber of predicted TMHs: 1\nExp number of AAs in TMHs: 23.72226\nExp number, first 60 AAs: 0.22038\nTotal prob of N-in: 0.12994\noutside 1 331\nTMhelix 332 354\ninside 355 372

4814 GCF_000782595.1_ASM78259v1 Escherichia coli Proteobacteria; Gammaproteobacteria; Enterobacterales; Enterobacteriaceae; Escherichia MNNEETFYQAMRRQGVTRRSFLKYCSLAATSLGLGAGMAPKIAWALENKPRIPVVWIHGL WP_001058320.1 hydrogenase [Escherichia coli] Length: 372\nNumber of predicted TMHs: 2\nExp number of AAs in TMHs: 37.26082\nExp number, first 60 AAs: 19.52923\nTotal prob of N-in: 0.94351\nPOSSIBLE N-term signal sequence\ninside 1 19\nTMhelix 20 39\noutside 40 325\nTMhelix 326 348\ninside 349 372

4815 GCF_001059105.1_ASM105910v1 Escherichia coli Proteobacteria; Gammaproteobacteria; Enterobacterales; Enterobacteriaceae; Escherichia MTGDNTLIHSHGINRRDFMKLCAALAATMGLSSKAAAEMAESVTNPQRPPVIWIGAQECT WP_000145410.1 MULTISPECIES: hydrogenase-2 small chain [Proteobacteria] Length: 372\nNumber of predicted TMHs: 1\nExp number of AAs in TMHs: 23.72226\nExp number, first 60 AAs: 0.22038\nTotal prob of N-in: 0.12994\noutside 1 331\nTMhelix 332 354\ninside 355 372

4816 GCF_001059105.1_ASM105910v1 Escherichia coli Proteobacteria; Gammaproteobacteria; Enterobacterales; Enterobacteriaceae; Escherichia
MNNEETFYQAMRRQGVTRRSFLKYCSLAATSLGLGAGMAPKIAWALENKPRIPVVWIHGL WP_001058323.1
MULTISPECIES: hydrogenase-1 small chain [Proteobacteria] Length: 372\nNumber of predicted TMHs: 2\nExp number of AAs in TMHs: 37.2608499999999\nExp number, first 60 AAs: 19.52923\nTotal prob of N-in: 0.94351\nPOSSIBLE N-term signal sequence\ninside 1 19\nTMhelix 20 39\noutside 40 325\nTMhelix 326 348\ninside 349 372

4817 GCF_000782655.1_ASM78265v1 Escherichia coli Proteobacteria; Gammaproteobacteria; Enterobacterales; Enterobacteriaceae; Escherichia
MTGDNTLIHSHGINRRDFMKLCAALAATMGLSSKAAAEMVESVTNPQRPPVIWIGAQECT WP_000145423.1
hydrogenase 2 small subunit [Escherichia coli] Length: 372\nNumber of predicted TMHs: 1\nExp number of AAs in TMHs: 23.78201\nExp number, first 60 AAs: 0.28355\nTotal prob of N-in: 0.13257\noutside 1 331\nTMhelix 332 354\ninside 355 372

4818 GCF_000782655.1_ASM78265v1 Escherichia coli Proteobacteria; Gammaproteobacteria; Enterobacterales; Enterobacteriaceae; Escherichia
MNNEETFYQAMRRQGVTRRSFLKYCSLAATSLGLGAGMAPKIAWALENKPRIPVVWIHGL WP_001538532.1
hydrogenase-1 small chain [Escherichia coli] Length: 372\nNumber of predicted TMHs: 2\nExp number of AAs in TMHs: 37.27041\nExp number, first 60 AAs: 19.52927\nTotal prob of N-in: 0.94352\nPOSSIBLE N-term signal sequence\ninside 1 19\nTMhelix 20 39\noutside 40 325\nTMhelix 326 348\ninside 349 372

4819 GCF_000782695.1_ASM78269v1 Escherichia coli Proteobacteria; Gammaproteobacteria; Enterobacterales; Enterobacteriaceae; Escherichia
MTGDNTLIHSHGINRRDFMKLCAALAATMGLSSKAAAEMAESVTNPQRPPVIWIGAQECT WP_000145410.1
MULTISPECIES: hydrogenase-2 small chain [Proteobacteria] Length: 372\nNumber of predicted TMHs: 1\nExp number of AAs in TMHs: 23.72226\nExp number, first 60 AAs: 0.22038\nTotal prob of N-in: 0.12994\noutside 1 331\nTMhelix 332 354\ninside 355 372

4820 GCF_000782755.1_ASM78275v1 Escherichia coli Proteobacteria; Gammaproteobacteria; Enterobacterales; Enterobacteriaceae; Escherichia
MTGDNTLIHSHGINRRDFMKLCAALAATMGLSSKAAAEMAESVTNPQRPPVIWIGAQECT WP_000145410.1
MULTISPECIES: hydrogenase-2 small chain [Proteobacteria] Length: 372\nNumber of predicted TMHs: 1\nExp number of AAs in TMHs: 23.72226\nExp number, first 60 AAs: 0.22038\nTotal prob of N-in: 0.12994\noutside 1 331\nTMhelix 332 354\ninside 355 372

4821 GCF_000782755.1_ASM78275v1 Escherichia coli Proteobacteria; Gammaproteobacteria; Enterobacterales; Enterobacteriaceae; Escherichia
MNNEETFYQAMRRQGVTRRSFLKYCSLAATSLGLGAGMAPKIAWALENKPRIPVVWIHGL WP_001058323.1
MULTISPECIES: hydrogenase-1 small chain [Proteobacteria] Length: 372\nNumber of predicted TMHs: 2\nExp number of AAs in TMHs: 37.2608499999999\nExp number, first 60 AAs: 19.52923\nTotal prob of N-in: 0.94351\nPOSSIBLE N-term signal sequence\ninside 1 19\nTMhelix 20 39\noutside 40 325\nTMhelix 326 348\ninside 349 372

4822 GCF_000782795.1_ASM78279v1 Escherichia coli Proteobacteria; Gammaproteobacteria; Enterobacterales; Enterobacteriaceae; Escherichia
MNNEETFYQAMRRQGVTRRSFLKYCSLAATSLGLGAGMAPKIAWALENKPRIPVVWIHGL WP_001058323.1
MULTISPECIES: hydrogenase-1 small chain [Proteobacteria] Length: 372\nNumber of predicted TMHs: 2\nExp number of AAs in TMHs: 37.2608499999999\nExp number, first 60 AAs: 19.52923\nTotal prob of N-in: 0.94351\nPOSSIBLE N-term signal sequence\ninside 1 19\nTMhelix 20 39\noutside 40 325\nTMhelix 326 348\ninside 349 372

4823 GCF_000782795.1_ASM78279v1 Escherichia coli Proteobacteria; Gammaproteobacteria; Enterobacterales; Enterobacteriaceae; Escherichia
MTGDNTLIHSHGINRRDFMKLCAALAATMGLSSKAAAEMAESVTNPQRPPVIWIGAQECT WP_032298447.1
hydrogenase 2 small subunit [Escherichia coli] Length: 372\nNumber of predicted TMHs: 1\nExp number of AAs in TMHs: 23.69807\nExp number, first 60 AAs: 0.221\nTotal prob of N-in: 0.12853\noutside 1 331\nTMhelix 332 354\ninside 355 372

4824 GCF_001059485.1_ASM105948v1 Escherichia coli Proteobacteria; Gammaproteobacteria; Enterobacterales; Enterobacteriaceae; Escherichia

MTGDNTLIHSHGINRRDFMKLCAALAATMGLSSKAAAEMVESVTNPQRPPVIWIGAQECT WP_000145423.1 hydrogenase 2 small subunit [Escherichia coli] Length: 372\nNumber of predicted TMHs: 1\nExp number of AAs in TMHs: 23.78201\nExp number, first 60 AAs: 0.28355\nTotal prob of N-in: 0.13257\noutside 1 331\nTMhelix 332 354\ninside 355 372

4825 GCF_001059485.1_ASM105948v1 Escherichia coli Proteobacteria; Gammaproteobacteria; Enterobacterales; Enterobacteriaceae; Escherichia MNNEETFYQAMRRQGVTRRSFLKYCSLAATSLGLGAGMAPKIAWALENKPRIPVVWIHGL WP_001295940.1 MULTISPECIES: hydrogenase [Enterobacteriaceae] Length: 372\nNumber of predicted TMHs: 2\nExp number of AAs in TMHs: 37.2608899999999\nExp number, first 60 AAs: 19.52923\nTotal prob of N-in: 0.94351\nPOSSIBLE N-term signal sequence\ninside 1 19\nTMhelix 20 39\noutside 40 325\nTMhelix 326 348\ninside 349 372

4826 GCF_001059655.1_ASM105965v1 Escherichia coli Proteobacteria; Gammaproteobacteria; Enterobacterales; Enterobacteriaceae; Escherichia MTGDNTLIHSHGINRRDFMKLCAALAATMGLSSKAAAEMVESVTNPQRPPVIWIGAQECT WP_000145423.1 hydrogenase 2 small subunit [Escherichia coli] Length: 372\nNumber of predicted TMHs: 1\nExp number of AAs in TMHs: 23.78201\nExp number, first 60 AAs: 0.28355\nTotal prob of N-in: 0.13257\noutside 1 331\nTMhelix 332 354\ninside 355 372

4827 GCF_001059795.1_ASM105979v1 Escherichia coli Proteobacteria; Gammaproteobacteria; Enterobacterales; Enterobacteriaceae; Escherichia MTGDNTLIHSHGINRRDFMKLCAALAATMGLSSKAAAEMAESVTNPQRPPVIWIGAQECT WP_000145410.1 MULTISPECIES: hydrogenase-2 small chain [Proteobacteria] Length: 372\nNumber of predicted TMHs: 1\nExp number of AAs in TMHs: 23.72226\nExp number, first 60 AAs: 0.22038\nTotal prob of N-in: 0.12994\noutside 1 331\nTMhelix 332 354\ninside 355 372

4828 GCF_001059655.1_ASM105965v1 Escherichia coli Proteobacteria; Gammaproteobacteria; Enterobacterales; Enterobacteriaceae; Escherichia MNNEETFYQAMRRQGVTRRSFLKYCSLAATSLGLGAGMAPKIAWALENKPRIPVVWIHGL WP_001295940.1 MULTISPECIES: hydrogenase [Enterobacteriaceae] Length: 372\nNumber of predicted TMHs: 2\nExp number of AAs in TMHs: 37.2608899999999\nExp number, first 60 AAs: 19.52923\nTotal prob of N-in: 0.94351\nPOSSIBLE N-term signal sequence\ninside 1 19\nTMhelix 20 39\noutside 40 325\nTMhelix 326 348\ninside 349 372

4829 GCF_001059795.1_ASM105979v1 Escherichia coli Proteobacteria; Gammaproteobacteria; Enterobacterales; Enterobacteriaceae; Escherichia MNNEETFYQAMRRQGVTRRSFLKYCSLAATSLGLGAGMAPKIAWALENKPRIPVVWIHGL WP_001058323.1 MULTISPECIES: hydrogenase-1 small chain [Proteobacteria] Length: 372\nNumber of predicted TMHs: 2\nExp number of AAs in TMHs: 37.2608499999999\nExp number, first 60 AAs: 19.52923\nTotal prob of N-in: 0.94351\nPOSSIBLE N-term signal sequence\ninside 1 19\nTMhelix 20 39\noutside 40 325\nTMhelix 326 348\ninside 349 372

4830 GCF_000947945.1_ASM94794v1 Escherichia coli Proteobacteria; Gammaproteobacteria; Enterobacterales; Enterobacteriaceae; Escherichia MNNEETFYQAMRRQGVTRRSFLKYCSLAATSLGLGAGMAPKIAWALENKPRIPVVWIHGL WP_001058323.1 MULTISPECIES: hydrogenase-1 small chain [Proteobacteria] Length: 372\nNumber of predicted TMHs: 2\nExp number of AAs in TMHs: 37.2608499999999\nExp number, first 60 AAs: 19.52923\nTotal prob of N-in: 0.94351\nPOSSIBLE N-term signal sequence\ninside 1 19\nTMhelix 20 39\noutside 40 325\nTMhelix 326 348\ninside 349 372

4831 GCF_000947945.1_ASM94794v1 Escherichia coli Proteobacteria; Gammaproteobacteria; Enterobacterales; Enterobacteriaceae; Escherichia MTGDNTLIHSHGINRRDFMKLCAALAATMGLSSKAAAEMAESVTNPQRPPVIWIGAQECT WP_000145410.1 MULTISPECIES: hydrogenase-2 small chain [Proteobacteria] Length: 372\nNumber of predicted TMHs: 1\nExp number of AAs in TMHs: 23.72226\nExp number, first 60 AAs: 0.22038\nTotal prob of N-in: 0.12994\noutside 1 331\nTMhelix 332 354\ninside 355 372

4832 GCF_001063395.1_ASM106339v1 Escherichia coli Proteobacteria; Gammaproteobacteria; Enterobacterales; Enterobacteriaceae; Escherichia MTGDNTLIHSHGINRRDFMKLCAALAATMGLSSKAAAEMVESVTNPQRPPVIWIGAQECT WP_000145423.1

hydrogenase 2 small subunit [Escherichia coli] Length: 372\nNumber of predicted TMHs: 1\nExp number of AAs in TMHs: 23.78201\nExp number, first 60 AAs: 0.28355\nTotal prob of N-in: 0.13257\noutside 1 331\nTMhelix 332 354\ninside 355 372

4833 GCF_001063395.1_ASM106339v1 Escherichia coli Proteobacteria; Gammaproteobacteria; Enterobacterales; Enterobacteriaceae; Escherichia MNNEETFYQAMRRQGVTRRSFLKYCSLAATSLGLGAGMAPKIAWALENKPRIPVVWIHGL WP_001295940.1 MULTISPECIES: hydrogenase [Enterobacteriaceae] Length: 372\nNumber of predicted TMHs: 2\nExp number of AAs in TMHs: 37.2608899999999\nExp number, first 60 AAs: 19.52923\nTotal prob of N-in: 0.94351\nPOSSIBLE N-term signal sequence\ninside 1 19\nTMhelix 20 39\noutside 40 325\nTMhelix 326 348\ninside 349 372

4834 GCF_001064885.1_ASM106488v1 Escherichia coli Proteobacteria; Gammaproteobacteria; Enterobacterales; Enterobacteriaceae; Escherichia MTGDNTLIHSHGINRRDFMKLCAALAATMGLSSKAAAEMVESVTNPQRPPVIWIGAQECT WP_000145423.1 hydrogenase 2 small subunit [Escherichia coli] Length: 372\nNumber of predicted TMHs: 1\nExp number of AAs in TMHs: 23.78201\nExp number, first 60 AAs: 0.28355\nTotal prob of N-in: 0.13257\noutside 1 331\nTMhelix 332 354\ninside 355 372

4835 GCF_001064885.1_ASM106488v1 Escherichia coli Proteobacteria; Gammaproteobacteria; Enterobacterales; Enterobacteriaceae; Escherichia MNNEETFYQAMRRQGVTRRSFLKYCSLAATSLGLGAGMAPKIAWALENKPRIPVVWIHGL WP_001295940.1 MULTISPECIES: hydrogenase [Enterobacteriaceae] Length: 372\nNumber of predicted TMHs: 2\nExp number of AAs in TMHs: 37.2608899999999\nExp number, first 60 AAs: 19.52923\nTotal prob of N-in: 0.94351\nPOSSIBLE N-term signal sequence\ninside 1 19\nTMhelix 20 39\noutside 40 325\nTMhelix 326 348\ninside 349 372

4836 GCF_001067225.1_ASM106722v1 Escherichia coli Proteobacteria; Gammaproteobacteria; Enterobacterales; Enterobacteriaceae; Escherichia MTGDNTLIHSHGINRRDFMKLCAALAATMGLSSKAAAEMAESVTNPQRPPVIWIGAQECT WP_000145410.1 MULTISPECIES: hydrogenase-2 small chain [Proteobacteria] Length: 372\nNumber of predicted TMHs: 1\nExp number of AAs in TMHs: 23.72226\nExp number, first 60 AAs: 0.22038\nTotal prob of N-in: 0.12994\noutside 1 331\nTMhelix 332 354\ninside 355 372

4837 GCF_001067225.1_ASM106722v1 Escherichia coli Proteobacteria; Gammaproteobacteria; Enterobacterales; Enterobacteriaceae; Escherichia MNNEETFYQAMRRQGVTRRSFLKYCSLAATSLGLGAGMAPKIAWALENKPRIPVVWIHGL WP_001058323.1 MULTISPECIES: hydrogenase-1 small chain [Proteobacteria] Length: 372\nNumber of predicted TMHs: 2\nExp number of AAs in TMHs: 37.2608499999999\nExp number, first 60 AAs: 19.52923\nTotal prob of N-in: 0.94351\nPOSSIBLE N-term signal sequence\ninside 1 19\nTMhelix 20 39\noutside 40 325\nTMhelix 326 348\ninside 349 372

4838 GCF_000784925.1_ASM78492v1 Escherichia coli Proteobacteria; Gammaproteobacteria; Enterobacterales; Enterobacteriaceae; Escherichia MTGDNTLIHSHGINRRDFMKLCAALAATMGLSSKAAAEMAESVTNPQRPPVIWIGAQECT WP_000145410.1 MULTISPECIES: hydrogenase-2 small chain [Proteobacteria] Length: 372\nNumber of predicted TMHs: 1\nExp number of AAs in TMHs: 23.72226\nExp number, first 60 AAs: 0.22038\nTotal prob of N-in: 0.12994\noutside 1 331\nTMhelix 332 354\ninside 355 372

4839 GCF_000784925.1_ASM78492v1 Escherichia coli Proteobacteria; Gammaproteobacteria; Enterobacterales; Enterobacteriaceae; Escherichia MNNEETFYQAMRRQGVTRRSFLKYCSLAATSLGLGAGMAPKIAWALENKPRIPVVWIHGL WP_001058323.1 MULTISPECIES: hydrogenase-1 small chain [Proteobacteria] Length: 372\nNumber of predicted TMHs: 2\nExp number of AAs in TMHs: 37.2608499999999\nExp number, first 60 AAs: 19.52923\nTotal prob of N-in: 0.94351\nPOSSIBLE N-term signal sequence\ninside 1 19\nTMhelix 20 39\noutside 40 325\nTMhelix 326 348\ninside 349 372

4840 GCF_000785255.1_ASM78525v1 Escherichia coli Proteobacteria; Gammaproteobacteria; Enterobacterales; Enterobacteriaceae; Escherichia MTGDNTLIHSHGINRRDFMKLCAALAATMGLSSKAAAEMAESVTNPQRPPVIWIGAQECT WP_000145410.1 MULTISPECIES: hydrogenase-2 small chain [Proteobacteria] Length: 372\nNumber of predicted TMHs:

1\nExp number of AAs in TMHs: 23.72226\nExp number, first 60 AAs: 0.22038\nTotal prob of N-in: 0.12994\noutside 1 331\nTMhelix 332 354\ninside 355 372

4841 GCF_000785255.1_ASM78525v1 Escherichia coli Proteobacteria; Gammaproteobacteria; Enterobacterales; Enterobacteriaceae; Escherichia MNNEETFYQAMRRQGVTRRSFLKYCSLAATSLGLGAGMAPKIAWALENKPRIPVVWIHGL WP_001058323.1 MULTISPECIES: hydrogenase-1 small chain [Proteobacteria] Length: 372\nNumber of predicted TMHs: 2\nExp number of AAs in TMHs: 37.2608499999999\nExp number, first 60 AAs: 19.52923\nTotal prob of N-in: 0.94351\nPOSSIBLE N-term signal sequence\ninside 1 19\nTMhelix 20 39\noutside 40 325\nTMhelix 326 348\ninside 349 372

4842 GCF_000785765.1_ASM78576v1 Escherichia coli Proteobacteria; Gammaproteobacteria; Enterobacterales; Enterobacteriaceae; Escherichia MTGDNTLIHSHGINRRDFMKLCAALAATMGLSSKAAAEMAESVTNPQRPPVIWIGAQECT WP_000145410.1 MULTISPECIES: hydrogenase-2 small chain [Proteobacteria] Length: 372\nNumber of predicted TMHs: 1\nExp number of AAs in TMHs: 23.72226\nExp number, first 60 AAs: 0.22038\nTotal prob of N-in: 0.12994\noutside 1 331\nTMhelix 332 354\ninside 355 372

4843 GCF_000785765.1_ASM78576v1 Escherichia coli Proteobacteria; Gammaproteobacteria; Enterobacterales; Enterobacteriaceae; Escherichia MNNEETFYQAMRRQGVTRRSFLKYCSLAATSLGLGAGMAPKIAWALENKPRIPVVWIHGL WP_001058323.1 MULTISPECIES: hydrogenase-1 small chain [Proteobacteria] Length: 372\nNumber of predicted TMHs: 2\nExp number of AAs in TMHs: 37.2608499999999\nExp number, first 60 AAs: 19.52923\nTotal prob of N-in: 0.94351\nPOSSIBLE N-term signal sequence\ninside 1 19\nTMhelix 20 39\noutside 40 325\nTMhelix 326 348\ninside 349 372

4844 GCF_000785795.1_ASM78579v1 Escherichia coli Proteobacteria; Gammaproteobacteria; Enterobacterales; Enterobacteriaceae; Escherichia MTGDNTLIHSHGINRRDFMKLCAALAATMGLSSKAAAEMAESVTNPQRPPVIWIGAQECT WP_000145410.1 MULTISPECIES: hydrogenase-2 small chain [Proteobacteria] Length: 372\nNumber of predicted TMHs: 1\nExp number of AAs in TMHs: 23.72226\nExp number, first 60 AAs: 0.22038\nTotal prob of N-in: 0.12994\noutside 1 331\nTMhelix 332 354\ninside 355 372

4845 GCF_000785795.1_ASM78579v1 Escherichia coli Proteobacteria; Gammaproteobacteria; Enterobacterales; Enterobacteriaceae; Escherichia MNNEETFYQAMRRQGVTRRSFLKYCSLAATSLGLGAGMAPKIAWALENKPRIPVVWIHGL WP_001058323.1 MULTISPECIES: hydrogenase-1 small chain [Proteobacteria] Length: 372\nNumber of predicted TMHs: 2\nExp number of AAs in TMHs: 37.2608499999999\nExp number, first 60 AAs: 19.52923\nTotal prob of N-in: 0.94351\nPOSSIBLE N-term signal sequence\ninside 1 19\nTMhelix 20 39\noutside 40 325\nTMhelix 326 348\ninside 349 372

4846 GCF_001268625.1_YE17 Escherichia coli Proteobacteria; Gammaproteobacteria; Enterobacterales; Enterobacteriaceae; Escherichia MTGDNTLIHSHGINRRDFMKLCAALAATMGLSSKAAAEMAESVTNPQRPPVIWIGAQECT WP_000145410.1 MULTISPECIES: hydrogenase-2 small chain [Proteobacteria] Length: 372\nNumber of predicted TMHs: 1\nExp number of AAs in TMHs: 23.72226\nExp number, first 60 AAs: 0.22038\nTotal prob of N-in: 0.12994\noutside 1 331\nTMhelix 332 354\ninside 355 372

4847 GCF_001268625.1_YE17 Escherichia coli Proteobacteria; Gammaproteobacteria; Enterobacterales; Enterobacteriaceae; Escherichia MNNEETFYQAMRRQGVTRRSFLKYCSLAATSLGLGAGMAPKIAWALENKPRIPVVWIHGL WP_001058323.1 MULTISPECIES: hydrogenase-1 small chain [Proteobacteria] Length: 372\nNumber of predicted TMHs: 2\nExp number of AAs in TMHs: 37.2608499999999\nExp number, first 60 AAs: 19.52923\nTotal prob of N-in: 0.94351\nPOSSIBLE N-term signal sequence\ninside 1 19\nTMhelix 20 39\noutside 40 325\nTMhelix 326 348\ninside 349 372

4848 GCF_001268665.1_1.ECAK83.1 Escherichia coli Proteobacteria; Gammaproteobacteria; Enterobacterales; Enterobacteriaceae; Escherichia MTGDNTLIHSHGINRRDFMKLCAALAATMGLSSKAAAEMAESVTNPQRPPVIWIGAQECT WP_000145410.1 MULTISPECIES: hydrogenase-2 small chain [Proteobacteria] Length: 372\nNumber of predicted TMHs: 1\nExp number of AAs in TMHs: 23.72226\nExp number, first 60 AAs: 0.22038\nTotal prob of N-in: 0.12994\noutside 1 331\nTMhelix 332 354\ninside 355 372

4849 GCF_001268665.1_1.ECAK83.1 Escherichia coli Proteobacteria; Gammaproteobacteria; Enterobacterales; Enterobacteriaceae; Escherichia
MNNEETFYQAMRRQGVTRRSFLKYCSLAATSLGLGAGMAPKIAWALENKPRIPVVWIHGL WP_001058323.1
MULTISPECIES: hydrogenase-1 small chain [Proteobacteria] Length: 372\nNumber of predicted TMHs: 2\nExp number of AAs in TMHs: 37.2608499999999\nExp number, first 60 AAs: 19.52923\nTotal prob of N-in: 0.94351\nPOSSIBLE N-term signal sequence\ninside 1 19\nTMhelix 20 39\noutside 40 325\nTMhelix 326 348\ninside 349 372

4850 GCF_001268705.1_YE21 Escherichia coli Proteobacteria; Gammaproteobacteria; Enterobacterales; Enterobacteriaceae; Escherichia MTGDNTLIHSHGINRRDFMKLCAALAATMGLSSKAAAEMAESVTNPQRPPVIWIGAQECT
WP_000145410.1 MULTISPECIES: hydrogenase-2 small chain [Proteobacteria] Length: 372\nNumber of predicted TMHs: 1\nExp number of AAs in TMHs: 23.72226\nExp number, first 60 AAs: 0.22038\nTotal prob of N-in: 0.12994\noutside 1 331\nTMhelix 332 354\ninside 355 372

4851 GCF_001268705.1_YE21 Escherichia coli Proteobacteria; Gammaproteobacteria; Enterobacterales; Enterobacteriaceae; Escherichia MNNEETFYQAMRRQGVTRRSFLKYCSLAATSLGLGAGMAPKIAWALENKPRIPVVWIHGL
WP_001295940.1 MULTISPECIES: hydrogenase [Enterobacteriaceae] Length: 372\nNumber of predicted TMHs: 2\nExp number of AAs in TMHs: 37.2608899999999\nExp number, first 60 AAs: 19.52923\nTotal prob of N-in: 0.94351\nPOSSIBLE N-term signal sequence\ninside 1 19\nTMhelix 20 39\noutside 40 325\nTMhelix 326 348\ninside 349 372

4852 GCF_001268765.1_YE12 Escherichia coli Proteobacteria; Gammaproteobacteria; Enterobacterales; Enterobacteriaceae; Escherichia MTGDNTLIHSHGINRRDFMKLCAALAATMGLSSKAAAEMAESVTNPQRPPVIWIGAQECT
WP_000145410.1 MULTISPECIES: hydrogenase-2 small chain [Proteobacteria] Length: 372\nNumber of predicted TMHs: 1\nExp number of AAs in TMHs: 23.72226\nExp number, first 60 AAs: 0.22038\nTotal prob of N-in: 0.12994\noutside 1 331\nTMhelix 332 354\ninside 355 372

4853 GCF_000936245.1_FHI7 Escherichia coli Proteobacteria; Gammaproteobacteria; Enterobacterales; Enterobacteriaceae; Escherichia MNNEETFYQAMRRQGVTRRSFLKYCSLAATSLGLGAGMAPKIAWALENKPRIPVVWIHGL
WP_001058323.1 MULTISPECIES: hydrogenase-1 small chain [Proteobacteria] Length: 372\nNumber of predicted TMHs: 2\nExp number of AAs in TMHs: 37.2608499999999\nExp number, first 60 AAs: 19.52923\nTotal prob of N-in: 0.94351\nPOSSIBLE N-term signal sequence\ninside 1 19\nTMhelix 20 39\noutside 40 325\nTMhelix 326 348\ninside 349 372

4854 GCF_000936635.1_FHI11 Escherichia coli Proteobacteria; Gammaproteobacteria; Enterobacterales; Enterobacteriaceae; Escherichia MNNEETFYQAMRRQGVTRRSFLKYCSLAATSLGLGAGMAPKIAWALENKPRIPVVWIHGL
WP_001058323.1 MULTISPECIES: hydrogenase-1 small chain [Proteobacteria] Length: 372\nNumber of predicted TMHs: 2\nExp number of AAs in TMHs: 37.2608499999999\nExp number, first 60 AAs: 19.52923\nTotal prob of N-in: 0.94351\nPOSSIBLE N-term signal sequence\ninside 1 19\nTMhelix 20 39\noutside 40 325\nTMhelix 326 348\ninside 349 372

4855 GCF_000937135.2_FHI74 Escherichia coli Proteobacteria; Gammaproteobacteria; Enterobacterales; Enterobacteriaceae; Escherichia MNNEETFYQAMRRQGVTRRSFLKYCSLAATSLGLGAGMAPKIAWALENKPRIPVVWIHGL
WP_001058323.1 MULTISPECIES: hydrogenase-1 small chain [Proteobacteria] Length: 372\nNumber of predicted TMHs: 2\nExp number of AAs in TMHs: 37.2608499999999\nExp number, first 60 AAs: 19.52923\nTotal prob of N-in: 0.94351\nPOSSIBLE N-term signal sequence\ninside 1 19\nTMhelix 20 39\noutside 40 325\nTMhelix 326 348\ninside 349 372

4856 GCF_000937135.2_FHI74 Escherichia coli Proteobacteria; Gammaproteobacteria; Enterobacterales; Enterobacteriaceae; Escherichia MTGDNTLIHSHGINRRDFMKLCAALAATMGLSSKAAAEMAESVTNPQRPPVIWIGAQECT
WP_000145410.1 MULTISPECIES: hydrogenase-2 small chain [Proteobacteria] Length: 372\nNumber of predicted TMHs: 1\nExp number of AAs in TMHs: 23.72226\nExp number, first 60 AAs: 0.22038\nTotal prob of N-in: 0.12994\noutside 1 331\nTMhelix 332 354\ninside 355 372

4857 GCF_000936635.1_FHI11 Escherichia coli Proteobacteria; Gammaproteobacteria; Enterobacterales; Enterobacteriaceae; Escherichia MTGDNTLIHSHGINRRDFMKLCAALAATMGLSSKAAAEMAESVTNPQRPPVIWIGAQECT
WP_000145410.1 MULTISPECIES: hydrogenase-2 small chain [Proteobacteria] Length: 372\nNumber of predicted TMHs: 1\nExp number of AAs in TMHs: 23.72226\nExp number, first 60 AAs: 0.22038\nTotal prob of N-in: 0.12994\noutside 1 331\nTMhelix 332 354\ninside 355 372

4858 GCF_000935515.1_ASM93551v1 Escherichia coli Proteobacteria; Gammaproteobacteria; Enterobacterales; Enterobacteriaceae; Escherichia MTGDNTLIHSHGINRRDFMKLCAALAATMGLSSKAAAEMAESVTNPQRPPVIWIGAQECT WP_000145410.1 MULTISPECIES: hydrogenase-2 small chain [Proteobacteria] Length: 372\nNumber of predicted TMHs: 1\nExp number of AAs in TMHs: 23.72226\nExp number, first 60 AAs: 0.22038\nTotal prob of N-in: 0.12994\noutside 1 331\nTMhelix 332 354\ninside 355 372

4859 GCF_000936105.1_Ec57A_A8_MIRA_assembly Escherichia coli Proteobacteria; Gammaproteobacteria; Enterobacterales; Enterobacteriaceae; Escherichia MTGDNTLIHSHGINRRDFMKLCAALAATMGLSSKAAAEMVESVTNPQRPPVIWIGAQECT WP_000145423.1 hydrogenase 2 small subunit [Escherichia coli] Length: 372\nNumber of predicted TMHs: 1\nExp number of AAs in TMHs: 23.78201\nExp number, first 60 AAs: 0.28355\nTotal prob of N-in: 0.13257\noutside 1 331\nTMhelix 332 354\ninside 355 372

4860 GCF_000934865.1_ASM93486v1 Escherichia coli Proteobacteria; Gammaproteobacteria; Enterobacterales; Enterobacteriaceae; Escherichia MNNEETFYQAMRRQGVTRRSFLKYCSLAATSLGLGAGMAPKIAWALENKPRIPVVWIHGL WP_001301613.1 hydrogenase [Escherichia coli] Length: 372\nNumber of predicted TMHs: 2\nExp number of AAs in TMHs: 37.46598\nExp number, first 60 AAs: 19.53139\nTotal prob of N-in: 0.94364\nPOSSIBLE N-term signal sequence\ninside 1 19\nTMhelix 20 39\noutside 40 325\nTMhelix 326 348\ninside 349 372

4861 GCF_000935445.1_ASM93544v1 Escherichia coli Proteobacteria; Gammaproteobacteria; Enterobacterales; Enterobacteriaceae; Escherichia MNNEETFYQAMRRQGVTRRSFLKYCSLAATSLGLGAGMAPKIAWALENKPRIPVVWIHGL WP_001058323.1 MULTISPECIES: hydrogenase-1 small chain [Proteobacteria] Length: 372\nNumber of predicted TMHs: 2\nExp number of AAs in TMHs: 37.2608499999999\nExp number, first 60 AAs: 19.52923\nTotal prob of N-in: 0.94351\nPOSSIBLE N-term signal sequence\ninside 1 19\nTMhelix 20 39\noutside 40 325\nTMhelix 326 348\ninside 349 372

4862 GCF_000935475.1_ASM93547v1 Escherichia coli Proteobacteria; Gammaproteobacteria; Enterobacterales; Enterobacteriaceae; Escherichia MNNEETFYQAMRRQGVTRRSFLKYCSLAATSLGLGAGMAPKIAWALENKPRIPVVWIHGL WP_001058323.1 MULTISPECIES: hydrogenase-1 small chain [Proteobacteria] Length: 372\nNumber of predicted TMHs: 2\nExp number of AAs in TMHs: 37.2608499999999\nExp number, first 60 AAs: 19.52923\nTotal prob of N-in: 0.94351\nPOSSIBLE N-term signal sequence\ninside 1 19\nTMhelix 20 39\noutside 40 325\nTMhelix 326 348\ninside 349 372

4863 GCF_000935905.1_FHI21 Escherichia coli Proteobacteria; Gammaproteobacteria; Enterobacterales; Enterobacteriaceae; Escherichia MTGDNTLIHSHGINRRDFMKLCAALAATMGLSSKAAAEMAESVTNPQRPPVIWIGAQECT WP_000145410.1 MULTISPECIES: hydrogenase-2 small chain [Proteobacteria] Length: 372\nNumber of predicted TMHs: 1\nExp number of AAs in TMHs: 23.72226\nExp number, first 60 AAs: 0.22038\nTotal prob of N-in: 0.12994\noutside 1 331\nTMhelix 332 354\ninside 355 372

4864 GCF_000935365.1_ASM93536v1 Escherichia coli Proteobacteria; Gammaproteobacteria; Enterobacterales; Enterobacteriaceae; Escherichia MTGDNTLIHSHGINRRDFMKLCAALAATMGLSSKAAAEMAESVTNPQRPPVIWIGAQECT WP_000145410.1 MULTISPECIES: hydrogenase-2 small chain [Proteobacteria] Length: 372\nNumber of predicted TMHs: 1\nExp number of AAs in TMHs: 23.72226\nExp number, first 60 AAs: 0.22038\nTotal prob of N-in: 0.12994\noutside 1 331\nTMhelix 332 354\ninside 355 372

4865 GCF_000935345.1_ASM93534v1 Escherichia coli Proteobacteria; Gammaproteobacteria; Enterobacterales; Enterobacteriaceae; Escherichia MNNEETFYQAMRRQGVTRRSFLKYCSLAATSLGLGAGMAPKIAWALENKPRIPVVWIHGL WP_001058323.1 MULTISPECIES: hydrogenase-1 small chain [Proteobacteria] Length: 372\nNumber of predicted TMHs: 2\nExp number of AAs in TMHs: 37.2608499999999\nExp number, first 60 AAs: 19.52923\nTotal prob of N-in: 0.94351\nPOSSIBLE N-term signal sequence\ninside 1 19\nTMhelix 20 39\noutside 40 325\nTMhelix 326 348\ninside 349 372

4866 GCF_000935345.1_ASM93534v1 Escherichia coli Proteobacteria; Gammaproteobacteria; Enterobacterales; Enterobacteriaceae; Escherichia MTGDNTLIHSHGINRRDFMKLCAALAATMGLSSKAAAEMAESVTNPQRPPVIWIGAQECT WP_000145410.1

MULTISPECIES: hydrogenase-2 small chain [Proteobacteria] Length: 372\nNumber of predicted TMHs: 1\nExp number of AAs in TMHs: 23.72226\nExp number, first 60 AAs: 0.22038\nTotal prob of N-in: 0.12994\noutside 1 331\nTMhelix 332 354\ninside 355 372

4867 GCF_000826805.1_ASM82680v1 Escherichia coli Proteobacteria; Gammaproteobacteria; Enterobacterales; Enterobacteriaceae; Escherichia MNNEETFYQAMRRQGVTRRSFLKYCSLAATSLGLGAGMAPKIAWALENKPRIPVVWIHGL WP_001295940.1 MULTISPECIES: hydrogenase [Enterobacteriaceae] Length: 372\nNumber of predicted TMHs: 2\nExp number of AAs in TMHs: 37.2608899999999\nExp number, first 60 AAs: 19.52923\nTotal prob of N-in: 0.94351\nPOSSIBLE N-term signal sequence\ninside 1 19\nTMhelix 20 39\noutside 40 325\nTMhelix 326 348\ninside 349 372

4868 GCF_000835055.1_734_3 Escherichia coli Proteobacteria; Gammaproteobacteria; Enterobacterales; Enterobacteriaceae; Escherichia MNNEETFYQAMRRQGVTRRSFLKYCSLAATSLGLGAGMAPKIAWALENKPRIPVVWIHGL WP_001058320.1 hydrogenase [Escherichia coli] Length: 372\nNumber of predicted TMHs: 2\nExp number of AAs in TMHs: 37.26082\nExp number, first 60 AAs: 19.52923\nTotal prob of N-in: 0.94351\nPOSSIBLE N-term signal sequence\ninside 1 19\nTMhelix 20 39\noutside 40 325\nTMhelix 326 348\ninside 349 372

4869 GCF_000833635.2_ASM83363v2 Escherichia coli Proteobacteria; Gammaproteobacteria; Enterobacterales; Enterobacteriaceae; Escherichia MNNEETFYQAMRRQGVTRRSFLKYCSLAATSLGLGAGMAPKIAWALENKPRIPVVWIHGL WP_001058323.1 MULTISPECIES: hydrogenase-1 small chain [Proteobacteria] Length: 372\nNumber of predicted TMHs: 2\nExp number of AAs in TMHs: 37.2608499999999\nExp number, first 60 AAs: 19.52923\nTotal prob of N-in: 0.94351\nPOSSIBLE N-term signal sequence\ninside 1 19\nTMhelix 20 39\noutside 40 325\nTMhelix 326 348\ninside 349 372

4870 GCF_000934865.1_ASM93486v1 Escherichia coli Proteobacteria; Gammaproteobacteria; Enterobacterales; Enterobacteriaceae; Escherichia MTGDNTLIHSHGINRRDFMKLCAALAATMGLSSKAAAEMAESVTNPQRPPVIWIGAQECT WP_000145410.1 MULTISPECIES: hydrogenase-2 small chain [Proteobacteria] Length: 372\nNumber of predicted TMHs: 1\nExp number of AAs in TMHs: 23.72226\nExp number, first 60 AAs: 0.22038\nTotal prob of N-in: 0.12994\noutside 1 331\nTMhelix 332 354\ninside 355 372

4871 GCF_000931565.1_ASM93156v1 Escherichia coli Proteobacteria; Gammaproteobacteria; Enterobacterales; Enterobacteriaceae; Escherichia MTGDNTLIHSHGINRRDFMKLCAALAATMGLSSKAAAEMVESVTNPQRPPVIWIGAQECT WP_000145423.1 hydrogenase 2 small subunit [Escherichia coli] Length: 372\nNumber of predicted TMHs: 1\nExp number of AAs in TMHs: 23.78201\nExp number, first 60 AAs: 0.28355\nTotal prob of N-in: 0.13257\noutside 1 331\nTMhelix 332 354\ninside 355 372

4872 GCF_000835055.1_734_3 Escherichia coli Proteobacteria; Gammaproteobacteria; Enterobacterales; Enterobacteriaceae; Escherichia MTGDNTLIHSHGINRRDFMKLCAALAATMGLSSKAAAEMAESVTNPQRPPVIWIGAQECT WP_000145410.1 MULTISPECIES: hydrogenase-2 small chain [Proteobacteria] Length: 372\nNumber of predicted TMHs: 1\nExp number of AAs in TMHs: 23.72226\nExp number, first 60 AAs: 0.22038\nTotal prob of N-in: 0.12994\noutside 1 331\nTMhelix 332 354\ninside 355 372

4873 GCF_000819345.1_Escherichia_coli_CVM_N33591PS_v1.0 Escherichia coli Proteobacteria; Gammaproteobacteria; Enterobacterales; Enterobacteriaceae; Escherichia MNNEETFYQAMRRQGVTRRSFLKYCSLAATSLGLGAGMAPKIAWALENKPRIPVVWIHGL WP_001058323.1 MULTISPECIES: hydrogenase-1 small chain [Proteobacteria] Length: 372\nNumber of predicted TMHs: 2\nExp number of AAs in TMHs: 37.2608499999999\nExp number, first 60 AAs: 19.52923\nTotal prob of N-in: 0.94351\nPOSSIBLE N-term signal sequence\ninside 1 19\nTMhelix 20 39\noutside 40 325\nTMhelix 326 348\ninside 349 372

4874 GCF_000819265.1_Escherichia_coli_CVM_N38428PS_v1.0 Escherichia coli Proteobacteria; Gammaproteobacteria; Enterobacterales; Enterobacteriaceae; Escherichia MTGDNTLIHSHGINRRDFMKLCAALAATMGLSSKAAAEMAESVTNPQRPPVIWIGAQECT WP_000145410.1 MULTISPECIES: hydrogenase-2 small chain [Proteobacteria] Length: 372\nNumber of predicted TMHs: 1\nExp number of AAs in TMHs: 23.72226\nExp number, first 60 AAs: 0.22038\nTotal prob of N-in: 0.12994\noutside 1 331\nTMhelix 332 354\ninside 355 372

4875 GCF_000819065.1_Escherichia_coli_CVM_N34228PS_v1.0 Escherichia coli Proteobacteria; Gammaproteobacteria; Enterobacterales; Enterobacteriaceae; Escherichia
MNNEETFYQAMRRQGVTRRSFLKYCSLAATSLGLGAGMAPKIAWALENKPRIPVVWIHGL WP_001058323.1
MULTISPECIES: hydrogenase-1 small chain [Proteobacteria] Length: 372\nNumber of predicted TMHs: 2\nExp number of AAs in TMHs: 37.2608499999999\nExp number, first 60 AAs: 19.52923\nTotal prob of N-in: 0.94351\nPOSSIBLE N-term signal sequence\ninside 1 19\nTMhelix 20 39\noutside 40 325\nTMhelix 326 348\ninside 349 372

4876 GCF_000818995.1_Escherichia_coli_CVM_N34152PS_v1.0 Escherichia coli Proteobacteria; Gammaproteobacteria; Enterobacterales; Enterobacteriaceae; Escherichia
MNNEETFYQAMRRQGVTRRSFLKYCSLAATSLGLGAGMAPKIAWALENKPRIPVVWIHGL WP_001058323.1
MULTISPECIES: hydrogenase-1 small chain [Proteobacteria] Length: 372\nNumber of predicted TMHs: 2\nExp number of AAs in TMHs: 37.2608499999999\nExp number, first 60 AAs: 19.52923\nTotal prob of N-in: 0.94351\nPOSSIBLE N-term signal sequence\ninside 1 19\nTMhelix 20 39\noutside 40 325\nTMhelix 326 348\ninside 349 372

4877 GCF_000819185.1_Escherichia_coli_CVM_N36113PS_v1.0 Escherichia coli Proteobacteria; Gammaproteobacteria; Enterobacterales; Enterobacteriaceae; Escherichia
MNNEETFYQAMRRQGVTRRSFLKYCSLAATSLGLGAGMAPKIAWALENKPRIPVVWIHGL WP_001058323.1
MULTISPECIES: hydrogenase-1 small chain [Proteobacteria] Length: 372\nNumber of predicted TMHs: 2\nExp number of AAs in TMHs: 37.2608499999999\nExp number, first 60 AAs: 19.52923\nTotal prob of N-in: 0.94351\nPOSSIBLE N-term signal sequence\ninside 1 19\nTMhelix 20 39\noutside 40 325\nTMhelix 326 348\ninside 349 372

4878 GCF_000819165.1_Escherichia_coli_CVM_N35912PS_v1.0 Escherichia coli Proteobacteria; Gammaproteobacteria; Enterobacterales; Enterobacteriaceae; Escherichia
MNNEETFYQAMRRQGVTRRSFLKYCSLAATSLGLGAGMAPKIAWALENKPRIPVVWIHGL WP_001058323.1
MULTISPECIES: hydrogenase-1 small chain [Proteobacteria] Length: 372\nNumber of predicted TMHs: 2\nExp number of AAs in TMHs: 37.2608499999999\nExp number, first 60 AAs: 19.52923\nTotal prob of N-in: 0.94351\nPOSSIBLE N-term signal sequence\ninside 1 19\nTMhelix 20 39\noutside 40 325\nTMhelix 326 348\ninside 349 372

4879 GCF_000819225.1_Escherichia_coli_CVM_N36601PS_v1.0 Escherichia coli Proteobacteria; Gammaproteobacteria; Enterobacterales; Enterobacteriaceae; Escherichia
MNNEETFYQAMRRQGVTRRSFLKYCSLAATSLGLGAGMAPKIAWALENKPRIPVVWIHGL WP_001058323.1
MULTISPECIES: hydrogenase-1 small chain [Proteobacteria] Length: 372\nNumber of predicted TMHs: 2\nExp number of AAs in TMHs: 37.2608499999999\nExp number, first 60 AAs: 19.52923\nTotal prob of N-in: 0.94351\nPOSSIBLE N-term signal sequence\ninside 1 19\nTMhelix 20 39\noutside 40 325\nTMhelix 326 348\ninside 349 372

4880 GCF_000819165.1_Escherichia_coli_CVM_N35912PS_v1.0 Escherichia coli Proteobacteria; Gammaproteobacteria; Enterobacterales; Enterobacteriaceae; Escherichia
MTGDNTLIHSHGINRRDFMKLCAALAATMGLSSKAAAEMAESVTNPQRPPVIWIGAQECT WP_000145410.1
MULTISPECIES: hydrogenase-2 small chain [Proteobacteria] Length: 372\nNumber of predicted TMHs: 1\nExp number of AAs in TMHs: 23.72226\nExp number, first 60 AAs: 0.22038\nTotal prob of N-in: 0.12994\noutside 1 331\nTMhelix 332 354\ninside 355 372

4881 GCF_000819125.1_Escherichia_coli_CVM_N34564PS_v1.0 Escherichia coli Proteobacteria; Gammaproteobacteria; Enterobacterales; Enterobacteriaceae; Escherichia
MTGDNTLIHSHGINRRDFMKLCAALAATMGLSSKAAAEMAESVTNPQRPPVIWIGAQECT WP_000145410.1
MULTISPECIES: hydrogenase-2 small chain [Proteobacteria] Length: 372\nNumber of predicted TMHs: 1\nExp number of AAs in TMHs: 23.72226\nExp number, first 60 AAs: 0.22038\nTotal prob of N-in: 0.12994\noutside 1 331\nTMhelix 332 354\ninside 355 372

4882 GCF_000819005.1_Escherichia_coli_CVM_N33825PS_v1.0 Escherichia coli Proteobacteria; Gammaproteobacteria; Enterobacterales; Enterobacteriaceae; Escherichia
MTGDNTLIHSHGINRRDFMKLCAALAATMGLSSKAAAEMAESVTNPQRPPVIWIGAQECT WP_000145410.1
MULTISPECIES: hydrogenase-2 small chain [Proteobacteria] Length: 372\nNumber of predicted TMHs: 1\nExp number of AAs in TMHs: 23.72226\nExp number, first 60 AAs: 0.22038\nTotal prob of N-in: 0.12994\noutside 1 331\nTMhelix 332 354\ninside 355 372

4883 GCF_000819065.1_Escherichia_coli_CVM_N34228PS_v1.0 Escherichia coli Proteobacteria; Gammaproteobacteria; Enterobacterales; Enterobacteriaceae; Escherichia
MTGDNTLIHSHGINRRDFMKLCAALAATMGLSSKAAAEMAESVTNPQRPPVIWIGAQECT WP_000145410.1
MULTISPECIES: hydrogenase-2 small chain [Proteobacteria] Length: 372\nNumber of predicted TMHs: 1\nExp number of AAs in TMHs: 23.72226\nExp number, first 60 AAs: 0.22038\nTotal prob of N-in: 0.12994\noutside 1 331\nTMhelix 332 354\ninside 355 372

4884 GCF_000817375.1_Iso_11_A5 Escherichia coli Proteobacteria; Gammaproteobacteria; Enterobacterales; Enterobacteriaceae; Escherichia
MNNEETFYQAMRRQGVTRRSFLKYCSLAATSLGLGAGMAPKIAWALENKPRIPVVWIHGL WP_001058323.1
MULTISPECIES: hydrogenase-1 small chain [Proteobacteria] Length: 372\nNumber of predicted TMHs: 2\nExp number of AAs in TMHs: 37.2608499999999\nExp number, first 60 AAs: 19.52923\nTotal prob of N-in: 0.94351\nPOSSIBLE N-term signal sequence\ninside 1 19\nTMhelix 20 39\noutside 40 325\nTMhelix 326 348\ninside 349 372

4885 GCF_000818995.1_Escherichia_coli_CVM_N34152PS_v1.0 Escherichia coli Proteobacteria; Gammaproteobacteria; Enterobacterales; Enterobacteriaceae; Escherichia
MTGDNTLIHSHGINRRDFMKLCAALAATMGLSSKAAAEMAESVTNPQRPPVIWIGAQECT WP_000145410.1
MULTISPECIES: hydrogenase-2 small chain [Proteobacteria] Length: 372\nNumber of predicted TMHs: 1\nExp number of AAs in TMHs: 23.72226\nExp number, first 60 AAs: 0.22038\nTotal prob of N-in: 0.12994\noutside 1 331\nTMhelix 332 354\ninside 355 372

4886 GCF_000817375.1_Iso_11_A5 Escherichia coli Proteobacteria; Gammaproteobacteria; Enterobacterales; Enterobacteriaceae; Escherichia
MTGDNTLIHSHGINRRDFMKLCAALAATMGLSSKAAAEMVESVTNPQRPPVIWIGAQECT WP_000145423.1
hydrogenase 2 small subunit [Escherichia coli] Length: 372\nNumber of predicted TMHs: 1\nExp number of AAs in TMHs: 23.78201\nExp number, first 60 AAs: 0.28355\nTotal prob of N-in: 0.13257\noutside 1 331\nTMhelix 332 354\ninside 355 372

4887 GCF_000812345.1_ASM81234v1 Escherichia coli Proteobacteria; Gammaproteobacteria; Enterobacterales; Enterobacteriaceae; Escherichia
MNNEETFYQAMRRQGVTRRSFLKYCSLAATSLGLGAGMAPKIAWALENKPRIPVVWIHGL WP_001058323.1
MULTISPECIES: hydrogenase-1 small chain [Proteobacteria] Length: 372\nNumber of predicted TMHs: 2\nExp number of AAs in TMHs: 37.2608499999999\nExp number, first 60 AAs: 19.52923\nTotal prob of N-in: 0.94351\nPOSSIBLE N-term signal sequence\ninside 1 19\nTMhelix 20 39\noutside 40 325\nTMhelix 326 348\ninside 349 372

4888 GCF_000812235.1_ASM81223v1 Escherichia coli Proteobacteria; Gammaproteobacteria; Enterobacterales; Enterobacteriaceae; Escherichia
MNNEETFYQAMRRQGVTRRSFLKYCSLAATSLGLGAGMAPKIAWALENKPRIPVVWIHGL WP_001058323.1
MULTISPECIES: hydrogenase-1 small chain [Proteobacteria] Length: 372\nNumber of predicted TMHs: 2\nExp number of AAs in TMHs: 37.2608499999999\nExp number, first 60 AAs: 19.52923\nTotal prob of N-in: 0.94351\nPOSSIBLE N-term signal sequence\ninside 1 19\nTMhelix 20 39\noutside 40 325\nTMhelix 326 348\ninside 349 372

4889 GCF_000812715.1_ASM81271v1 Escherichia coli Proteobacteria; Gammaproteobacteria; Enterobacterales; Enterobacteriaceae; Escherichia
MNNEETFYQAMRRQGVTRRSFLKYCSLAATSLGLGAGMAPKIAWALENKPRIPVVWIHGL WP_032161483.1
hydrogenase [Escherichia coli] Length: 372\nNumber of predicted TMHs: 2\nExp number of AAs in TMHs: 38.07604\nExp number, first 60 AAs: 19.52977\nTotal prob of N-in: 0.94353\nPOSSIBLE N-term signal sequence\ninside 1 19\nTMhelix 20 39\noutside 40 325\nTMhelix 326 348\ninside 349 372

4890 GCF_000812765.1_ASM81276v1 Escherichia coli Proteobacteria; Gammaproteobacteria; Enterobacterales; Enterobacteriaceae; Escherichia
MNNEETFYQAMRRQGVTRRSFLKYCSLAATSLGLGAGMAPKIAWALENKPRIPVVWIHGL WP_001353311.1
MULTISPECIES: hydrogenase [Escherichia] Length: 372\nNumber of predicted TMHs: 2\nExp number of AAs in TMHs: 37.2638699999999\nExp number, first 60 AAs: 19.52924\nTotal prob of N-in: 0.94351\nPOSSIBLE N-term signal sequence\ninside 1 19\nTMhelix 20 39\noutside 40 325\nTMhelix 326 348\ninside 349 372

4891 GCF_000812765.1_ASM81276v1 Escherichia coli Proteobacteria; Gammaproteobacteria; Enterobacterales; Enterobacteriaceae; Escherichia

MTGDNTLIHSHGINRRDFMKLCAALAATMGLSSKAAAEMAESVTNPQRPPVIWIGAQECT WP_000145410.1 MULTISPECIES: hydrogenase-2 small chain [Proteobacteria] Length: 372\nNumber of predicted TMHs: 1\nExp number of AAs in TMHs: 23.72226\nExp number, first 60 AAs: 0.22038\nTotal prob of N-in: 0.12994\noutside 1 331\nTMhelix 332 354\ninside 355 372

4892 GCF_000812715.1_ASM81271v1 Escherichia coli Proteobacteria; Gammaproteobacteria; Enterobacterales; Enterobacteriaceae; Escherichia MTGDNTLIHSHGINRRDFMKLCAALAATMGLSSKAAAEMAESVTNPQRPPVIWIGAQECT WP_000145410.1 MULTISPECIES: hydrogenase-2 small chain [Proteobacteria] Length: 372\nNumber of predicted TMHs: 1\nExp number of AAs in TMHs: 23.72226\nExp number, first 60 AAs: 0.22038\nTotal prob of N-in: 0.12994\noutside 1 331\nTMhelix 332 354\ninside 355 372

4893 GCF_000812645.1_ASM81264v1 Escherichia coli Proteobacteria; Gammaproteobacteria; Enterobacterales; Enterobacteriaceae; Escherichia MTGDNTLIHSHGINRRDFMKLCAALAATMGLSSKAAAEMAESVTNPQRPPVIWIGAQECT WP_000145410.1 MULTISPECIES: hydrogenase-2 small chain [Proteobacteria] Length: 372\nNumber of predicted TMHs: 1\nExp number of AAs in TMHs: 23.72226\nExp number, first 60 AAs: 0.22038\nTotal prob of N-in: 0.12994\noutside 1 331\nTMhelix 332 354\ninside 355 372

4894 GCF_000812305.1_ASM81230v1 Escherichia coli Proteobacteria; Gammaproteobacteria; Enterobacterales; Enterobacteriaceae; Escherichia MTGDNTLIHSHGINRRDFMKLCAALAATMGLSSKAAAEMAESVTNPQRPPVIWIGAQECT WP_000145410.1 MULTISPECIES: hydrogenase-2 small chain [Proteobacteria] Length: 372\nNumber of predicted TMHs: 1\nExp number of AAs in TMHs: 23.72226\nExp number, first 60 AAs: 0.22038\nTotal prob of N-in: 0.12994\noutside 1 331\nTMhelix 332 354\ninside 355 372

4895 GCF_000812365.1_ASM81236v1 Escherichia coli Proteobacteria; Gammaproteobacteria; Enterobacterales; Enterobacteriaceae; Escherichia MTGDNTLIHSHGINRRDFMKLCAALAATMGLSSKAAAEMAESVTNPQRPPVIWIGAQECT WP_000145410.1 MULTISPECIES: hydrogenase-2 small chain [Proteobacteria] Length: 372\nNumber of predicted TMHs: 1\nExp number of AAs in TMHs: 23.72226\nExp number, first 60 AAs: 0.22038\nTotal prob of N-in: 0.12994\noutside 1 331\nTMhelix 332 354\ninside 355 372

4896 GCF_000812345.1_ASM81234v1 Escherichia coli Proteobacteria; Gammaproteobacteria; Enterobacterales; Enterobacteriaceae; Escherichia MTGDNTLIHSHGINRRDFMKLCAALAATMGLSSKAAAEMAESVTNPQRPPVIWIGAQECT WP_000145410.1 MULTISPECIES: hydrogenase-2 small chain [Proteobacteria] Length: 372\nNumber of predicted TMHs: 1\nExp number of AAs in TMHs: 23.72226\nExp number, first 60 AAs: 0.22038\nTotal prob of N-in: 0.12994\noutside 1 331\nTMhelix 332 354\ninside 355 372

4897 GCF_000812235.1_ASM81223v1 Escherichia coli Proteobacteria; Gammaproteobacteria; Enterobacterales; Enterobacteriaceae; Escherichia MTGDNTLIHSHGINRRDFMKLCAALAATMGLSSKAAAEMAESVTNPQRPPVIWIGAQECT WP_000145410.1 MULTISPECIES: hydrogenase-2 small chain [Proteobacteria] Length: 372\nNumber of predicted TMHs: 1\nExp number of AAs in TMHs: 23.72226\nExp number, first 60 AAs: 0.22038\nTotal prob of N-in: 0.12994\noutside 1 331\nTMhelix 332 354\ninside 355 372

4898 GCF_000806195.1_ASM80619v1 Escherichia coli Proteobacteria; Gammaproteobacteria; Enterobacterales; Enterobacteriaceae; Escherichia MNNEETFYQAMRRQGVTRRSFLKYCSLAATSLGLGAGMAPKIAWALENKPRIPVVWIHGL WP_001295940.1 MULTISPECIES: hydrogenase [Enterobacteriaceae] Length: 372\nNumber of predicted TMHs: 2\nExp number of AAs in TMHs: 37.2608899999999\nExp number, first 60 AAs: 19.52923\nTotal prob of N-in: 0.94351\nPOSSIBLE N-term signal sequence\ninside 1 19\nTMhelix 20 39\noutside 40 325\nTMhelix 326 348\ninside 349 372

4899 GCF_000806195.1_ASM80619v1 Escherichia coli Proteobacteria; Gammaproteobacteria; Enterobacterales; Enterobacteriaceae; Escherichia MTGDNTLIHSHGINRRDFMKLCAALAATMGLSSKAAAEMAESVTNPQRPPVIWIGAQECT WP_000145410.1 MULTISPECIES: hydrogenase-2 small chain [Proteobacteria] Length: 372\nNumber of predicted TMHs: 1\nExp number of AAs in TMHs: 23.72226\nExp number, first 60 AAs: 0.22038\nTotal prob of N-in: 0.12994\noutside 1 331\nTMhelix 332 354\ninside 355 372

4900 GCF_000805835.1_ASM80583v1 Escherichia coli Proteobacteria; Gammaproteobacteria; Enterobacterales; Enterobacteriaceae; Escherichia
MNNEETFYQAMRRQGVTRRSFLKYCSLAATSLGLGAGMAPKIAWALENKPRIPVVWIHGL WP_001058323.1
MULTISPECIES: hydrogenase-1 small chain [Proteobacteria] Length: 372\nNumber of predicted TMHs: 2\nExp number of AAs in TMHs: 37.2608499999999\nExp number, first 60 AAs: 19.52923\nTotal prob of N-in: 0.94351\nPOSSIBLE N-term signal sequence\ninside 1 19\nTMhelix 20 39\noutside 40 325\nTMhelix 326 348\ninside 349 372

4901 GCF_000804365.1_ASM80436v1 Escherichia coli Proteobacteria; Gammaproteobacteria; Enterobacterales; Enterobacteriaceae; Escherichia
MTGDNTLIHSHGINRRDFMKLCAALAATMGLSSKAAAEMAESVTNPQRPPVIWIGAQECT WP_000145410.1
MULTISPECIES: hydrogenase-2 small chain [Proteobacteria] Length: 372\nNumber of predicted TMHs: 1\nExp number of AAs in TMHs: 23.72226\nExp number, first 60 AAs: 0.22038\nTotal prob of N-in: 0.12994\noutside 1 331\nTMhelix 332 354\ninside 355 372

4902 GCF_000804325.1_ASM80432v1 Escherichia coli Proteobacteria; Gammaproteobacteria; Enterobacterales; Enterobacteriaceae; Escherichia
MTGDNTLIHSHGINRRDFMKLCAALAATMGLSSKAAAEMAESVTNPQRPPVIWIGAQECT WP_000145410.1
MULTISPECIES: hydrogenase-2 small chain [Proteobacteria] Length: 372\nNumber of predicted TMHs: 1\nExp number of AAs in TMHs: 23.72226\nExp number, first 60 AAs: 0.22038\nTotal prob of N-in: 0.12994\noutside 1 331\nTMhelix 332 354\ninside 355 372

4903 GCF_000798515.1_Escherichia_coli_CVM_N33561PS_v._1.0 Escherichia coli Proteobacteria; Gammaproteobacteria; Enterobacterales; Enterobacteriaceae; Escherichia
MNNEETFYQAMRRQGVTRRSFLKYCSLAATSLGLGAGMAPKIAWALENKPRIPVVWIHGL WP_042973597.1
hydrogenase [Escherichia coli] Length: 372\nNumber of predicted TMHs: 2\nExp number of AAs in TMHs: 38.17092\nExp number, first 60 AAs: 19.52878\nTotal prob of N-in: 0.94355\nPOSSIBLE N-term signal sequence\ninside 1 19\nTMhelix 20 39\noutside 40 325\nTMhelix 326 348\ninside 349 372

4904 GCF_000801185.2_ASM80118v2 Escherichia coli Proteobacteria; Gammaproteobacteria; Enterobacterales; Enterobacteriaceae; Escherichia
MNNEETFYQAMRRQGVTRRSFLKYCSLAATSLGLGAGMAPKIAWALENKPRIPVVWIHGL WP_001295940.1
MULTISPECIES: hydrogenase [Enterobacteriaceae] Length: 372\nNumber of predicted TMHs: 2\nExp number of AAs in TMHs: 37.2608899999999\nExp number, first 60 AAs: 19.52923\nTotal prob of N-in: 0.94351\nPOSSIBLE N-term signal sequence\ninside 1 19\nTMhelix 20 39\noutside 40 325\nTMhelix 326 348\ninside 349 372

4905 GCF_000798355.1_Escherichia_coli_CVM_N33707PS_v._1.0 Escherichia coli Proteobacteria; Gammaproteobacteria; Enterobacterales; Enterobacteriaceae; Escherichia
MNNEETFYQAMRRQGVTRRSFLKYCSLAATSLGLGAGMAPKIAWALENKPRIPVVWIHGL WP_042965784.1
hydrogenase [Escherichia coli] Length: 372\nNumber of predicted TMHs: 2\nExp number of AAs in TMHs: 37.2608099999999\nExp number, first 60 AAs: 19.52923\nTotal prob of N-in: 0.94351\nPOSSIBLE N-term signal sequence\ninside 1 19\nTMhelix 20 39\noutside 40 325\nTMhelix 326 348\ninside 349 372

4906 GCF_000798395.1_Escherichia_coli_CVM_N34543PS_v._1.0 Escherichia coli Proteobacteria; Gammaproteobacteria; Enterobacterales; Enterobacteriaceae; Escherichia
MNNEETFYQAMRRQGVTRRSFLKYCSLAATSLGLGAGMAPKIAWALENKPRIPVVWIHGL WP_001295940.1
MULTISPECIES: hydrogenase [Enterobacteriaceae] Length: 372\nNumber of predicted TMHs: 2\nExp number of AAs in TMHs: 37.2608899999999\nExp number, first 60 AAs: 19.52923\nTotal prob of N-in: 0.94351\nPOSSIBLE N-term signal sequence\ninside 1 19\nTMhelix 20 39\noutside 40 325\nTMhelix 326 348\ninside 349 372

4907 GCF_000798495.1_Escherichia_coli_CVM_N36918PS_v._1.0 Escherichia coli Proteobacteria; Gammaproteobacteria; Enterobacterales; Enterobacteriaceae; Escherichia
MNNEETFYQAMRRQGVTRRSFLKYCSLAATSLGLGAGMAPKIAWALENKPRIPVVWIHGL WP_001058323.1
MULTISPECIES: hydrogenase-1 small chain [Proteobacteria] Length: 372\nNumber of predicted TMHs: 2\nExp number of AAs in TMHs: 37.2608499999999\nExp number, first 60 AAs: 19.52923\nTotal prob of N-in: 0.94351\nPOSSIBLE N-term signal sequence\ninside 1 19\nTMhelix 20 39\noutside 40 325\nTMhelix 326 348\ninside 349 372

4908 GCF_000800495.1_Escherichia_coli_CVM_N38823PS_v._1.0 Escherichia coli Proteobacteria; Gammaproteobacteria; Enterobacterales; Enterobacteriaceae; Escherichia MNNEETFYQAMRRQGVTRRSFLKYCSLAATSLGLGAGMAPKIAWALENKPRIPVVWIHGL WP_001295940.1 MULTISPECIES: hydrogenase [Enterobacteriaceae] Length: 372\nNumber of predicted TMHs: 2\nExp number of AAs in TMHs: 37.2608899999999\nExp number, first 60 AAs: 19.52923\nTotal prob of N-in: 0.94351\nPOSSIBLE N-term signal sequence\ninside 1 19\nTMhelix 20 39\noutside 40 325\nTMhelix 326 348\ninside 349 372

4909 GCF_000798595.1_Escherichia_coli_CVM_N33601PS_v._1.0 Escherichia coli Proteobacteria; Gammaproteobacteria; Enterobacterales; Enterobacteriaceae; Escherichia MNNEETFYQAMRRQGVTRRSFLKYCSLAATSLGLGAGMAPKIAWALENKPRIPVVWIHGL WP_001058323.1 MULTISPECIES: hydrogenase-1 small chain [Proteobacteria] Length: 372\nNumber of predicted TMHs: 2\nExp number of AAs in TMHs: 37.2608499999999\nExp number, first 60 AAs: 19.52923\nTotal prob of N-in: 0.94351\nPOSSIBLE N-term signal sequence\ninside 1 19\nTMhelix 20 39\noutside 40 325\nTMhelix 326 348\ninside 349 372

4910 GCF_000798595.1_Escherichia_coli_CVM_N33601PS_v._1.0 Escherichia coli Proteobacteria; Gammaproteobacteria; Enterobacterales; Enterobacteriaceae; Escherichia MTGDNTLIHSHGINRRDFMKLCAALAATMGLSSKAAAEMAESVTNPQRPPVIWIGAQECT WP_000145410.1 MULTISPECIES: hydrogenase-2 small chain [Proteobacteria] Length: 372\nNumber of predicted TMHs: 1\nExp number of AAs in TMHs: 23.72226\nExp number, first 60 AAs: 0.22038\nTotal prob of N-in: 0.12994\noutside 1 331\nTMhelix 332 354\ninside 355 372

4911 GCF_000800495.1_Escherichia_coli_CVM_N38823PS_v._1.0 Escherichia coli Proteobacteria; Gammaproteobacteria; Enterobacterales; Enterobacteriaceae; Escherichia MTGDNTLIHSHGINRRDFMKLCAALAATMGLSSKAAAEMAESVTNPQRPPVIWIGAQECT WP_000145410.1 MULTISPECIES: hydrogenase-2 small chain [Proteobacteria] Length: 372\nNumber of predicted TMHs: 1\nExp number of AAs in TMHs: 23.72226\nExp number, first 60 AAs: 0.22038\nTotal prob of N-in: 0.12994\noutside 1 331\nTMhelix 332 354\ninside 355 372

4912 GCF_000798295.1_Escherichia_coli_CVM_N33651PS_v._1.0 Escherichia coli Proteobacteria; Gammaproteobacteria; Enterobacterales; Enterobacteriaceae; Escherichia MNNEETFYQAMRRQGVTRRSFLKYCSLAATSLGLGAGMAPKIAWALENKPRIPVVWIHGL WP_001295940.1 MULTISPECIES: hydrogenase [Enterobacteriaceae] Length: 372\nNumber of predicted TMHs: 2\nExp number of AAs in TMHs: 37.2608899999999\nExp number, first 60 AAs: 19.52923\nTotal prob of N-in: 0.94351\nPOSSIBLE N-term signal sequence\ninside 1 19\nTMhelix 20 39\noutside 40 325\nTMhelix 326 348\ninside 349 372

4913 GCF_000798335.1_Escherichia_coli_CVM_N33687PS_v._1.0 Escherichia coli Proteobacteria; Gammaproteobacteria; Enterobacterales; Enterobacteriaceae; Escherichia MNNEETFYQAMRRQGVTRRSFLKYCSLAATSLGLGAGMAPKIAWALENKPRIPVVWIHGL WP_001058323.1 MULTISPECIES: hydrogenase-1 small chain [Proteobacteria] Length: 372\nNumber of predicted TMHs: 2\nExp number of AAs in TMHs: 37.2608499999999\nExp number, first 60 AAs: 19.52923\nTotal prob of N-in: 0.94351\nPOSSIBLE N-term signal sequence\ninside 1 19\nTMhelix 20 39\noutside 40 325\nTMhelix 326 348\ninside 349 372

4914 GCF_000798395.1_Escherichia_coli_CVM_N34543PS_v._1.0 Escherichia coli Proteobacteria; Gammaproteobacteria; Enterobacterales; Enterobacteriaceae; Escherichia MTGDNTLIHSHGINRRDFMKLCAALAATMGLSSKAAAEMAESVTNPQRPPVIWIGAQECT WP_000145410.1 MULTISPECIES: hydrogenase-2 small chain [Proteobacteria] Length: 372\nNumber of predicted TMHs: 1\nExp number of AAs in TMHs: 23.72226\nExp number, first 60 AAs: 0.22038\nTotal prob of N-in: 0.12994\noutside 1 331\nTMhelix 332 354\ninside 355 372

4915 GCF_000798355.1_Escherichia_coli_CVM_N33707PS_v._1.0 Escherichia coli Proteobacteria; Gammaproteobacteria; Enterobacterales; Enterobacteriaceae; Escherichia MTGDNTLIHSHGINRRDFMKLCAALAATMGLSSKAAAEMAESVTNPQRPPVIWIGAQECT WP_000145410.1 MULTISPECIES: hydrogenase-2 small chain [Proteobacteria] Length: 372\nNumber of predicted TMHs: 1\nExp number of AAs in TMHs: 23.72226\nExp number, first 60 AAs: 0.22038\nTotal prob of N-in: 0.12994\noutside 1 331\nTMhelix 332 354\ninside 355 372

4916 GCF_000798335.1_Escherichia_coli_CVM_N33687PS_v._1.0 Escherichia coli Proteobacteria; Gammaproteobacteria; Enterobacterales; Enterobacteriaceae; Escherichia
MTGDNTLIHSHGINRRDFMKLCAALAATMGLSSKAAAEMAESVTNPQRPPVIWIGAQECT WP_000145410.1
MULTISPECIES: hydrogenase-2 small chain [Proteobacteria] Length: 372\nNumber of predicted TMHs: 1\nExp number of AAs in TMHs: 23.72226\nExp number, first 60 AAs: 0.22038\nTotal prob of N-in: 0.12994\noutside 1 331\nTMhelix 332 354\ninside 355 372

4917 GCF_000798295.1_Escherichia_coli_CVM_N33651PS_v._1.0 Escherichia coli Proteobacteria; Gammaproteobacteria; Enterobacterales; Enterobacteriaceae; Escherichia
MTGDNTLIHSHGINRRDFMKLCAALAATMGLSSKAAAEMAESVTNPQRPPVIWIGAQECT WP_000145410.1
MULTISPECIES: hydrogenase-2 small chain [Proteobacteria] Length: 372\nNumber of predicted TMHs: 1\nExp number of AAs in TMHs: 23.72226\nExp number, first 60 AAs: 0.22038\nTotal prob of N-in: 0.12994\noutside 1 331\nTMhelix 332 354\ninside 355 372

4918 GCF_000798195.1_Escherichia_coli_CVM_N33742PS_v._1.0 Escherichia coli Proteobacteria; Gammaproteobacteria; Enterobacterales; Enterobacteriaceae; Escherichia
MNNEETFYQAMRRQGVTRRSFLKYCSLAATSLGLGAGMAPKIAWALENKPRIPVVWIHGL WP_001058323.1
MULTISPECIES: hydrogenase-1 small chain [Proteobacteria] Length: 372\nNumber of predicted TMHs: 2\nExp number of AAs in TMHs: 37.2608499999999\nExp number, first 60 AAs: 19.52923\nTotal prob of N-in: 0.94351\nPOSSIBLE N-term signal sequence\ninside 1 19\nTMhelix 20 39\noutside 40 325\nTMhelix 326 348\ninside 349 372

4919 GCF_000798275.1_Escherichia_coli_CVM_N33640PS_v._1.0 Escherichia coli Proteobacteria; Gammaproteobacteria; Enterobacterales; Enterobacteriaceae; Escherichia
MNNEETFYQAMRRQGVTRRSFLKYCSLAATSLGLGAGMAPKIAWALENKPRIPVVWIHGL WP_001058323.1
MULTISPECIES: hydrogenase-1 small chain [Proteobacteria] Length: 372\nNumber of predicted TMHs: 2\nExp number of AAs in TMHs: 37.2608499999999\nExp number, first 60 AAs: 19.52923\nTotal prob of N-in: 0.94351\nPOSSIBLE N-term signal sequence\ninside 1 19\nTMhelix 20 39\noutside 40 325\nTMhelix 326 348\ninside 349 372

4920 GCF_000798275.1_Escherichia_coli_CVM_N33640PS_v._1.0 Escherichia coli Proteobacteria; Gammaproteobacteria; Enterobacterales; Enterobacteriaceae; Escherichia
MTGDNTLIHSHGINRRDFMKLCAALAATMGLSSKAAAEMAESVTNPQRPPVIWIGAQECT WP_000145410.1
MULTISPECIES: hydrogenase-2 small chain [Proteobacteria] Length: 372\nNumber of predicted TMHs: 1\nExp number of AAs in TMHs: 23.72226\nExp number, first 60 AAs: 0.22038\nTotal prob of N-in: 0.12994\noutside 1 331\nTMhelix 332 354\ninside 355 372

4921 GCF_000798175.1_Escherichia_coli_CVM_N41586PS_v._1.0 Escherichia coli Proteobacteria; Gammaproteobacteria; Enterobacterales; Enterobacteriaceae; Escherichia
MNNEETFYQAMRRQGVTRRSFLKYCSLAATSLGLGAGMAPKIAWALENKPRIPVVWIHGL WP_001058320.1
hydrogenase [Escherichia coli] Length: 372\nNumber of predicted TMHs: 2\nExp number of AAs in TMHs: 37.26082\nExp number, first 60 AAs: 19.52923\nTotal prob of N-in: 0.94351\nPOSSIBLE N-term signal sequence\ninside 1 19\nTMhelix 20 39\noutside 40 325\nTMhelix 326 348\ninside 349 372

4922 GCF_000798135.1_Escherichia_coli_CVM_N34054PS_v._1.0 Escherichia coli Proteobacteria; Gammaproteobacteria; Enterobacterales; Enterobacteriaceae; Escherichia
MNNEETFYQAMRRQGVTRRSFLKYCSLAATSLGLGAGMAPKIAWALENKPRIPVVWIHGL WP_001058323.1
MULTISPECIES: hydrogenase-1 small chain [Proteobacteria] Length: 372\nNumber of predicted TMHs: 2\nExp number of AAs in TMHs: 37.2608499999999\nExp number, first 60 AAs: 19.52923\nTotal prob of N-in: 0.94351\nPOSSIBLE N-term signal sequence\ninside 1 19\nTMhelix 20 39\noutside 40 325\nTMhelix 326 348\ninside 349 372

4923 GCF_000798235.1_Escherichia_coli_CVM_N36834PS_v._1.0 Escherichia coli Proteobacteria; Gammaproteobacteria; Enterobacterales; Enterobacteriaceae; Escherichia
MTGDNTLIHSHGINRRDFMKLCAALAATMGLSSKAAAEMAESVTNPQRPPVIWIGAQECT WP_000145410.1
MULTISPECIES: hydrogenase-2 small chain [Proteobacteria] Length: 372\nNumber of predicted TMHs: 1\nExp number of AAs in TMHs: 23.72226\nExp number, first 60 AAs: 0.22038\nTotal prob of N-in: 0.12994\noutside 1 331\nTMhelix 332 354\ninside 355 372

4924 GCF_000798195.1_Escherichia_coli_CVM_N33742PS_v._1.0 Escherichia coli Proteobacteria; Gammaproteobacteria; Enterobacterales; Enterobacteriaceae; Escherichia

MTGDNTLIHSHGINRRDFMKLCAALAATMGLSSKAAAEMAESVTNPQRPPVIWIGAQECT WP_000145410.1 MULTISPECIES: hydrogenase-2 small chain [Proteobacteria] Length: 372\nNumber of predicted TMHs: 1\nExp number of AAs in TMHs: 23.72226\nExp number, first 60 AAs: 0.22038\nTotal prob of N-in: 0.12994\noutside 1 331\nTMhelix 332 354\ninside 355 372

4925 GCF_000798175.1_Escherichia_coli_CVM_N41586PS_v._1.0 Escherichia coli Proteobacteria; Gammaproteobacteria; Enterobacterales; Enterobacteriaceae; Escherichia MTGDNTLIHSHGINRRDFMKLCAALAATMGLSSKAAAEMAESVTNPQRPPVIWIGAQECT WP_000145410.1 MULTISPECIES: hydrogenase-2 small chain [Proteobacteria] Length: 372\nNumber of predicted TMHs: 1\nExp number of AAs in TMHs: 23.72226\nExp number, first 60 AAs: 0.22038\nTotal prob of N-in: 0.12994\noutside 1 331\nTMhelix 332 354\ninside 355 372

4926 GCF_000797955.1_Escherichia_coli_CVM_N36400PS_v._1.0 Escherichia coli Proteobacteria; Gammaproteobacteria; Enterobacterales; Enterobacteriaceae; Escherichia MTGDNTLIHSHGINRRDFMKLCAALAATMGLSSKAAAEMAESVTNPQRPPVIWIGAQECT WP_000145410.1 MULTISPECIES: hydrogenase-2 small chain [Proteobacteria] Length: 372\nNumber of predicted TMHs: 1\nExp number of AAs in TMHs: 23.72226\nExp number, first 60 AAs: 0.22038\nTotal prob of N-in: 0.12994\noutside 1 331\nTMhelix 332 354\ninside 355 372

4927 GCF_000797975.1_Escherichia_coli_CVM_N36609PS_v._1.0 Escherichia coli Proteobacteria; Gammaproteobacteria; Enterobacterales; Enterobacteriaceae; Escherichia MTGDNTLIHSHGINRRDFMKLCAALAATMGLSSKAAAEMAESVTNPQRPPVIWIGAQECT WP_000145410.1 MULTISPECIES: hydrogenase-2 small chain [Proteobacteria] Length: 372\nNumber of predicted TMHs: 1\nExp number of AAs in TMHs: 23.72226\nExp number, first 60 AAs: 0.22038\nTotal prob of N-in: 0.12994\noutside 1 331\nTMhelix 332 354\ninside 355 372

4928 GCF_000797915.1_Escherichia_coli_CVM_N41498PS_v._1.0 Escherichia coli Proteobacteria; Gammaproteobacteria; Enterobacterales; Enterobacteriaceae; Escherichia MTGDNTLIHSHGINRRDFMKLCAALAATMGLSSKAAAEMAESVTNPQRPPVIWIGAQECT WP_000145410.1 MULTISPECIES: hydrogenase-2 small chain [Proteobacteria] Length: 372\nNumber of predicted TMHs: 1\nExp number of AAs in TMHs: 23.72226\nExp number, first 60 AAs: 0.22038\nTotal prob of N-in: 0.12994\noutside 1 331\nTMhelix 332 354\ninside 355 372

4929 GCF_000797795.1_Escherichia_coli_CVM_N34562PS_v._1.0 Escherichia coli Proteobacteria; Gammaproteobacteria; Enterobacterales; Enterobacteriaceae; Escherichia MNNEETFYQAMRRQGVTRRSFLKYCSLAATSLGLGAGMAPKIAWALENKPRIPVVWIHGL WP_001058323.1 MULTISPECIES: hydrogenase-1 small chain [Proteobacteria] Length: 372\nNumber of predicted TMHs: 2\nExp number of AAs in TMHs: 37.2608499999999\nExp number, first 60 AAs: 19.52923\nTotal prob of N-in: 0.94351\nPOSSIBLE N-term signal sequence\ninside 1 19\nTMhelix 20 39\noutside 40 325\nTMhelix 326 348\ninside 349 372

4930 GCF_000797875.1_Escherichia_coli_CVM_N38833PS_v._1.0 Escherichia coli Proteobacteria; Gammaproteobacteria; Enterobacterales; Enterobacteriaceae; Escherichia MTGDNTLIHSHGINRRDFMKLCAALAATMGLSSKAAAEMAESVTNPQRPPVIWIGAQECT WP_000145410.1 MULTISPECIES: hydrogenase-2 small chain [Proteobacteria] Length: 372\nNumber of predicted TMHs: 1\nExp number of AAs in TMHs: 23.72226\nExp number, first 60 AAs: 0.22038\nTotal prob of N-in: 0.12994\noutside 1 331\nTMhelix 332 354\ninside 355 372

4931 GCF_000797855.1_Escherichia_coli_CVM_N33806PS_v._1.0 Escherichia coli Proteobacteria; Gammaproteobacteria; Enterobacterales; Enterobacteriaceae; Escherichia MTGDNTLIHSHGINRRDFMKLCAALAATMGLSSKAAAEMAESVTNPQRPPVIWIGAQECT WP_000145410.1 MULTISPECIES: hydrogenase-2 small chain [Proteobacteria] Length: 372\nNumber of predicted TMHs: 1\nExp number of AAs in TMHs: 23.72226\nExp number, first 60 AAs: 0.22038\nTotal prob of N-in: 0.12994\noutside 1 331\nTMhelix 332 354\ninside 355 372

4932 GCF_000797795.1_Escherichia_coli_CVM_N34562PS_v._1.0 Escherichia coli Proteobacteria; Gammaproteobacteria; Enterobacterales; Enterobacteriaceae; Escherichia MTGDNTLIHSHGINRRDFMKLCAALAATMGLSSKAAAEMAESVTNPQRPPVIWIGAQECT WP_000145410.1 MULTISPECIES: hydrogenase-2 small chain [Proteobacteria] Length: 372\nNumber of predicted TMHs: 1\nExp number of AAs in TMHs: 23.72226\nExp number, first 60 AAs: 0.22038\nTotal prob of N-in: 0.12994\noutside 1 331\nTMhelix 332 354\ninside 355 372

4933 GCF_000797595.1_Escherichia_coli_CVM_N36225PS_v._1.0 Escherichia coli Proteobacteria; Gammaproteobacteria; Enterobacterales; Enterobacteriaceae; Escherichia
MNNEETFYQAMRRQGVTRRSFLKYCSLAATSLGLGAGMAPKIAWALENKPRIPVVWIHGL WP_001058323.1
MULTISPECIES: hydrogenase-1 small chain [Proteobacteria] Length: 372\nNumber of predicted TMHs: 2\nExp number of AAs in TMHs: 37.2608499999999\nExp number, first 60 AAs: 19.52923\nTotal prob of N-in: 0.94351\nPOSSIBLE N-term signal sequence\ninside 1 19\nTMhelix 20 39\noutside 40 325\nTMhelix 326 348\ninside 349 372

4934 GCF_000797655.1_Escherichia_coli_CVM_N36401PS_v._1.0 Escherichia coli Proteobacteria; Gammaproteobacteria; Enterobacterales; Enterobacteriaceae; Escherichia
MNNEETFYQAMRRQGVTRRSFLKYCSLAATSLGLGAGMAPKIAWALENKPRIPVVWIHGL WP_001058323.1
MULTISPECIES: hydrogenase-1 small chain [Proteobacteria] Length: 372\nNumber of predicted TMHs: 2\nExp number of AAs in TMHs: 37.2608499999999\nExp number, first 60 AAs: 19.52923\nTotal prob of N-in: 0.94351\nPOSSIBLE N-term signal sequence\ninside 1 19\nTMhelix 20 39\noutside 40 325\nTMhelix 326 348\ninside 349 372

4935 GCF_000797775.1_Escherichia_coli_CVM_N34322PS_v._1.0 Escherichia coli Proteobacteria; Gammaproteobacteria; Enterobacterales; Enterobacteriaceae; Escherichia
MNNEETFYQAMRRQGVTRRSFLKYCSLAATSLGLGAGMAPKIAWALENKPRIPVVWIHGL WP_001295940.1
MULTISPECIES: hydrogenase [Enterobacteriaceae] Length: 372\nNumber of predicted TMHs: 2\nExp number of AAs in TMHs: 37.2608899999999\nExp number, first 60 AAs: 19.52923\nTotal prob of N-in: 0.94351\nPOSSIBLE N-term signal sequence\ninside 1 19\nTMhelix 20 39\noutside 40 325\nTMhelix 326 348\ninside 349 372

4936 GCF_000797775.1_Escherichia_coli_CVM_N34322PS_v._1.0 Escherichia coli Proteobacteria; Gammaproteobacteria; Enterobacterales; Enterobacteriaceae; Escherichia
MTGDNTLIHSHGINRRDFMKLCAALAATMGLSSKAAAEMAESVTNPQRPPVIWIGAQECT WP_000145410.1
MULTISPECIES: hydrogenase-2 small chain [Proteobacteria] Length: 372\nNumber of predicted TMHs: 1\nExp number of AAs in TMHs: 23.72226\nExp number, first 60 AAs: 0.22038\nTotal prob of N-in: 0.12994\noutside 1 331\nTMhelix 332 354\ninside 355 372

4937 GCF_000785865.1_ASM78586v1 Escherichia coli Proteobacteria; Gammaproteobacteria; Enterobacterales; Enterobacteriaceae; Escherichia
MNNEETFYQAMRRQGVTRRSFLKYCSLAATSLGLGAGMAPKIAWALENKPRIPVVWIHGL WP_001058323.1
MULTISPECIES: hydrogenase-1 small chain [Proteobacteria] Length: 372\nNumber of predicted TMHs: 2\nExp number of AAs in TMHs: 37.2608499999999\nExp number, first 60 AAs: 19.52923\nTotal prob of N-in: 0.94351\nPOSSIBLE N-term signal sequence\ninside 1 19\nTMhelix 20 39\noutside 40 325\nTMhelix 326 348\ninside 349 372

4938 GCF_000797575.1_Escherichia_coli_CVM_N37067PS_v._1.0 Escherichia coli Proteobacteria; Gammaproteobacteria; Enterobacterales; Enterobacteriaceae; Escherichia
MNNEETFYQAMRRQGVTRRSFLKYCSLAATSLGLGAGMAPKIAWALENKPRIPVVWIHGL WP_001058323.1
MULTISPECIES: hydrogenase-1 small chain [Proteobacteria] Length: 372\nNumber of predicted TMHs: 2\nExp number of AAs in TMHs: 37.2608499999999\nExp number, first 60 AAs: 19.52923\nTotal prob of N-in: 0.94351\nPOSSIBLE N-term signal sequence\ninside 1 19\nTMhelix 20 39\noutside 40 325\nTMhelix 326 348\ninside 349 372

4939 GCF_000797675.1_Escherichia_coli_CVM_N38663PS_v._1.0 Escherichia coli Proteobacteria; Gammaproteobacteria; Enterobacterales; Enterobacteriaceae; Escherichia
MNNEETFYQAMRRQGVTRRSFLKYCSLAATSLGLGAGMAPKIAWALENKPRIPVVWIHGL WP_001058323.1
MULTISPECIES: hydrogenase-1 small chain [Proteobacteria] Length: 372\nNumber of predicted TMHs: 2\nExp number of AAs in TMHs: 37.2608499999999\nExp number, first 60 AAs: 19.52923\nTotal prob of N-in: 0.94351\nPOSSIBLE N-term signal sequence\ninside 1 19\nTMhelix 20 39\noutside 40 325\nTMhelix 326 348\ninside 349 372

4940 GCF_000785355.1_ASM78535v1 Escherichia coli Proteobacteria; Gammaproteobacteria; Enterobacterales; Enterobacteriaceae; Escherichia
MNNEETFYQAMRRQGVTRRSFLKYCSLAATSLGLGAGMAPKIAWALENKPRIPVVWIHGL WP_001058323.1
MULTISPECIES: hydrogenase-1 small chain [Proteobacteria] Length: 372\nNumber of predicted TMHs: 2\nExp number of AAs in TMHs: 37.2608499999999\nExp number, first 60 AAs: 19.52923\nTotal prob of N-in:

0.94351\nPOSSIBLE N-term signal sequence\ninside 1 19\nTMhelix 20 39\noutside 40 325\nTMhelix 326 348\ninside 349 372

4941 GCF_000785355.1_ASM78535v1 Escherichia coli Proteobacteria; Gammaproteobacteria; Enterobacterales; Enterobacteriaceae; Escherichia
MTGDNTLIHSHGINRRDFMKLCAALAATMGLSSKAAAEMAESVTNPQRPPVIWIGAQECT WP_000145410.1
MULTISPECIES: hydrogenase-2 small chain [Proteobacteria] Length: 372\nNumber of predicted TMHs: 1\nExp number of AAs in TMHs: 23.72226\nExp number, first 60 AAs: 0.22038\nTotal prob of N-in: 0.12994\noutside 1 331\nTMhelix 332 354\ninside 355 372

4942 GCF_000782815.1_ASM78281v1 Escherichia coli Proteobacteria; Gammaproteobacteria; Enterobacterales; Enterobacteriaceae; Escherichia
MNNEETFYQAMRRQGVTRRSFLKYCSLAATSLGLGAGMAPKIAWALENKPRIPVVWIHGL WP_001058323.1
MULTISPECIES: hydrogenase-1 small chain [Proteobacteria] Length: 372\nNumber of predicted TMHs: 2\nExp number of AAs in TMHs: 37.2608499999999\nExp number, first 60 AAs: 19.52923\nTotal prob of N-in: 0.94351\nPOSSIBLE N-term signal sequence\ninside 1 19\nTMhelix 20 39\noutside 40 325\nTMhelix 326 348\ninside 349 372

4943 GCF_000785325.1_ASM78532v1 Escherichia coli Proteobacteria; Gammaproteobacteria; Enterobacterales; Enterobacteriaceae; Escherichia
MNNEETFYQAMRRQGVTRRSFLKYCSLAATSLGLGAGMAPKIAWALENKPRIPVVWIHGL WP_001058323.1
MULTISPECIES: hydrogenase-1 small chain [Proteobacteria] Length: 372\nNumber of predicted TMHs: 2\nExp number of AAs in TMHs: 37.2608499999999\nExp number, first 60 AAs: 19.52923\nTotal prob of N-in: 0.94351\nPOSSIBLE N-term signal sequence\ninside 1 19\nTMhelix 20 39\noutside 40 325\nTMhelix 326 348\ninside 349 372

4944 GCF_000785325.1_ASM78532v1 Escherichia coli Proteobacteria; Gammaproteobacteria; Enterobacterales; Enterobacteriaceae; Escherichia
MTGDNTLIHSHGINRRDFMKLCAALAATMGLSSKAAAEMAESVTNPQRPPVIWIGAQECT WP_000145410.1
MULTISPECIES: hydrogenase-2 small chain [Proteobacteria] Length: 372\nNumber of predicted TMHs: 1\nExp number of AAs in TMHs: 23.72226\nExp number, first 60 AAs: 0.22038\nTotal prob of N-in: 0.12994\noutside 1 331\nTMhelix 332 354\ninside 355 372

4945 GCF_000782615.1_ASM78261v1 Escherichia coli Proteobacteria; Gammaproteobacteria; Enterobacterales; Enterobacteriaceae; Escherichia
MNNEETFYQAMRRQGVTRRSFLKYCSLAATSLGLGAGMAPKIAWALENKPRIPVVWIHGL WP_001295940.1
MULTISPECIES: hydrogenase [Enterobacteriaceae] Length: 372\nNumber of predicted TMHs: 2\nExp number of AAs in TMHs: 37.2608899999999\nExp number, first 60 AAs: 19.52923\nTotal prob of N-in: 0.94351\nPOSSIBLE N-term signal sequence\ninside 1 19\nTMhelix 20 39\noutside 40 325\nTMhelix 326 348\ninside 349 372

4946 GCF_000782775.1_ASM78277v1 Escherichia coli Proteobacteria; Gammaproteobacteria; Enterobacterales; Enterobacteriaceae; Escherichia
MNNEETFYQAMRRQGVTRRSFLKYCSLAATSLGLGAGMAPKIAWALENKPRIPVVWIHGL WP_001058323.1
MULTISPECIES: hydrogenase-1 small chain [Proteobacteria] Length: 372\nNumber of predicted TMHs: 2\nExp number of AAs in TMHs: 37.2608499999999\nExp number, first 60 AAs: 19.52923\nTotal prob of N-in: 0.94351\nPOSSIBLE N-term signal sequence\ninside 1 19\nTMhelix 20 39\noutside 40 325\nTMhelix 326 348\ninside 349 372

4947 GCF_000785235.1_ASM78523v1 Escherichia coli Proteobacteria; Gammaproteobacteria; Enterobacterales; Enterobacteriaceae; Escherichia
MNNEETFYQAMRRQGVTRRSFLKYCSLAATSLGLGAGMAPKIAWALENKPRIPVVWIHGL WP_001058323.1
MULTISPECIES: hydrogenase-1 small chain [Proteobacteria] Length: 372\nNumber of predicted TMHs: 2\nExp number of AAs in TMHs: 37.2608499999999\nExp number, first 60 AAs: 19.52923\nTotal prob of N-in: 0.94351\nPOSSIBLE N-term signal sequence\ninside 1 19\nTMhelix 20 39\noutside 40 325\nTMhelix 326 348\ninside 349 372

4948 GCF_000785235.1_ASM78523v1 Escherichia coli Proteobacteria; Gammaproteobacteria; Enterobacterales; Enterobacteriaceae; Escherichia
MTGDNTLIHSHGINRRDFMKLCAALAATMGLSSKAAAEMAESVTNPQRPPVIWIGAQECT WP_000145410.1
MULTISPECIES: hydrogenase-2 small chain [Proteobacteria] Length: 372\nNumber of predicted TMHs:

1\nExp number of AAs in TMHs: 23.72226\nExp number, first 60 AAs:  0.22038\nTotal prob of N-in:   0.12994\noutside   1   331\nTMhelix   332   354\ninside   355   372

4949 GCF_000782635.1_ASM78263v1 Escherichia coli Proteobacteria; Gammaproteobacteria; Enterobacterales; Enterobacteriaceae; Escherichia MNNEETFYQAMRRQGVTRRSFLKYCSLAATSLGLGAGMAPKIAWALENKPRIPVVWIHGL WP_001538532.1 hydrogenase-1 small chain [Escherichia coli] Length: 372\nNumber of predicted TMHs:  2\nExp number of AAs in TMHs: 37.27041\nExp number, first 60 AAs:  19.52927\nTotal prob of N-in:   0.94352\nPOSSIBLE N-term signal sequence\ninside   1   19\nTMhelix   20   39\noutside   40   325\nTMhelix   326   348\ninside   349   372

4950 GCF_000783335.1_ASM78333v1 Escherichia coli Proteobacteria; Gammaproteobacteria; Enterobacterales; Enterobacteriaceae; Escherichia MNNEETFYQAMRRQGVTRRSFLKYCSLAATSLGLGAGMAPKIAWALENKPRIPVVWIHGL WP_001058323.1 MULTISPECIES: hydrogenase-1 small chain [Proteobacteria] Length: 372\nNumber of predicted TMHs: 2\nExp number of AAs in TMHs: 37.2608499999999\nExp number, first 60 AAs:  19.52923\nTotal prob of N-in: 0.94351\nPOSSIBLE N-term signal sequence\ninside   1   19\nTMhelix   20   39\noutside   40   325\nTMhelix   326   348\ninside   349   372

4951 GCF_000783335.1_ASM78333v1 Escherichia coli Proteobacteria; Gammaproteobacteria; Enterobacterales; Enterobacteriaceae; Escherichia MTGDNTLIHSHGINRRDFMKLCAALAATMGLSSKAAAEMAESVTNPQRPPVIWIGAQECT WP_000145410.1 MULTISPECIES: hydrogenase-2 small chain [Proteobacteria] Length: 372\nNumber of predicted TMHs: 1\nExp number of AAs in TMHs: 23.72226\nExp number, first 60 AAs:  0.22038\nTotal prob of N-in:   0.12994\noutside   1   331\nTMhelix   332   354\ninside   355   372

4952 GCF_000782815.1_ASM78281v1 Escherichia coli Proteobacteria; Gammaproteobacteria; Enterobacterales; Enterobacteriaceae; Escherichia MTGDNTLIHSHGINRRDFMKLCAALAATMGLSSKAAAEMAESVTNPQRPPVIWIGAQECT WP_000145410.1 MULTISPECIES: hydrogenase-2 small chain [Proteobacteria] Length: 372\nNumber of predicted TMHs: 1\nExp number of AAs in TMHs: 23.72226\nExp number, first 60 AAs:  0.22038\nTotal prob of N-in:   0.12994\noutside   1   331\nTMhelix   332   354\ninside   355   372

4953 GCF_000782775.1_ASM78277v1 Escherichia coli Proteobacteria; Gammaproteobacteria; Enterobacterales; Enterobacteriaceae; Escherichia MTGDNTLIHSHGINRRDFMKLCAALAATMGLSSKAAAEMAESVTNPQRPPVIWIGAQECT WP_000145410.1 MULTISPECIES: hydrogenase-2 small chain [Proteobacteria] Length: 372\nNumber of predicted TMHs: 1\nExp number of AAs in TMHs: 23.72226\nExp number, first 60 AAs:  0.22038\nTotal prob of N-in:   0.12994\noutside   1   331\nTMhelix   332   354\ninside   355   372

4954 GCF_000782495.1_ASM78249v1 Escherichia coli Proteobacteria; Gammaproteobacteria; Enterobacterales; Enterobacteriaceae; Escherichia MNNEETFYQAMRRQGVTRRSFLKYCSLAATSLGLGAGMAPKIAWALENKPRIPVVWIHGL WP_001295940.1 MULTISPECIES: hydrogenase [Enterobacteriaceae] Length: 372\nNumber of predicted TMHs:  2\nExp number of AAs in TMHs: 37.2608899999999\nExp number, first 60 AAs:  19.52923\nTotal prob of N-in: 0.94351\nPOSSIBLE N-term signal sequence\ninside   1   19\nTMhelix   20   39\noutside   40   325\nTMhelix   326   348\ninside   349   372

4955 GCF_000782555.1_ASM78255v1 Escherichia coli Proteobacteria; Gammaproteobacteria; Enterobacterales; Enterobacteriaceae; Escherichia MNNEETFYQAMRRQGVTRRSFLKYCSLAATSLGLGAGMAPKIAWALENKPRIPVVWIHGL WP_001058320.1 hydrogenase [Escherichia coli] Length: 372\nNumber of predicted TMHs:  2\nExp number of AAs in TMHs: 37.26082\nExp number, first 60 AAs:  19.52923\nTotal prob of N-in:   0.94351\nPOSSIBLE N-term signal sequence\ninside   1   19\nTMhelix   20   39\noutside   40   325\nTMhelix   326   348\ninside   349   372

4956 GCF_000782575.1_ASM78257v1 Escherichia coli Proteobacteria; Gammaproteobacteria; Enterobacterales; Enterobacteriaceae; Escherichia MNNEETFYQAMRRQGVTRRSFLKYCSLAATSLGLGAGMAPKIAWALENKPRIPVVWIHGL WP_001058323.1 MULTISPECIES: hydrogenase-1 small chain [Proteobacteria] Length: 372\nNumber of predicted TMHs: 2\nExp number of AAs in TMHs: 37.2608499999999\nExp number, first 60 AAs:  19.52923\nTotal prob of N-in: 0.94351\nPOSSIBLE N-term signal sequence\ninside   1   19\nTMhelix   20   39\noutside   40   325\nTMhelix   326   348\ninside   349   372

4957 GCF_000782575.1_ASM78257v1 Escherichia coli Proteobacteria; Gammaproteobacteria; Enterobacterales; Enterobacteriaceae; Escherichia
MTGDNTLIHSHGINRRDFMKLCAALAATMGLSSKAAAEMAESVTNPQRPPVIWIGAQECT WP_000145410.1
MULTISPECIES: hydrogenase-2 small chain [Proteobacteria] Length: 372\nNumber of predicted TMHs: 1\nExp number of AAs in TMHs: 23.72226\nExp number, first 60 AAs: 0.22038\nTotal prob of N-in: 0.12994\noutside 1 331\nTMhelix 332 354\ninside 355 372

4958 GCF_000782555.1_ASM78255v1 Escherichia coli Proteobacteria; Gammaproteobacteria; Enterobacterales; Enterobacteriaceae; Escherichia
MTGDNTLIHSHGINRRDFMKLCAALAATMGLSSKAAAEMAESVTNPQRPPVIWIGAQECT WP_000145410.1
MULTISPECIES: hydrogenase-2 small chain [Proteobacteria] Length: 372\nNumber of predicted TMHs: 1\nExp number of AAs in TMHs: 23.72226\nExp number, first 60 AAs: 0.22038\nTotal prob of N-in: 0.12994\noutside 1 331\nTMhelix 332 354\ninside 355 372

4959 GCF_000782495.1_ASM78249v1 Escherichia coli Proteobacteria; Gammaproteobacteria; Enterobacterales; Enterobacteriaceae; Escherichia
MTGDNTLIHSHGINRRDFMKLCAALAATMGLSSKAAAEMVESVTNPQRPPVIWIGAQECT WP_000145423.1
hydrogenase 2 small subunit [Escherichia coli] Length: 372\nNumber of predicted TMHs: 1\nExp number of AAs in TMHs: 23.78201\nExp number, first 60 AAs: 0.28355\nTotal prob of N-in: 0.13257\noutside 1 331\nTMhelix 332 354\ninside 355 372

4960 GCF_000782195.1_ASM78219v1 Escherichia coli Proteobacteria; Gammaproteobacteria; Enterobacterales; Enterobacteriaceae; Escherichia
MNNEETFYQAMRRQGVTRRSFLKYCSLAATSLGLGAGMAPKIAWALENKPRIPVVWIHGL WP_001295940.1
MULTISPECIES: hydrogenase [Enterobacteriaceae] Length: 372\nNumber of predicted TMHs: 2\nExp number of AAs in TMHs: 37.2608899999999\nExp number, first 60 AAs: 19.52923\nTotal prob of N-in: 0.94351\nPOSSIBLE N-term signal sequence\ninside 1 19\nTMhelix 20 39\noutside 40 325\nTMhelix 326 348\ninside 349 372

4961 GCF_000782095.1_ASM78209v1 Escherichia coli Proteobacteria; Gammaproteobacteria; Enterobacterales; Enterobacteriaceae; Escherichia
MTGDNTLIHSHGINRRDFMKLCAALATTMGLSSKAAAEMAESVTNPQRPPVIWIGAQECT WP_023149629.1
MULTISPECIES: hydrogenase-2 small chain [Enterobacteriaceae] Length: 372\nNumber of predicted TMHs: 1\nExp number of AAs in TMHs: 23.63603\nExp number, first 60 AAs: 0.12971\nTotal prob of N-in: 0.12608\noutside 1 331\nTMhelix 332 354\ninside 355 372

4962 GCF_000782295.1_ASM78229v1 Escherichia coli Proteobacteria; Gammaproteobacteria; Enterobacterales; Enterobacteriaceae; Escherichia
MNNEETFYQAMRRQGVTRRSFLKYCSLAATSLGLGAGMAPKIAWALENKPRIPVVWIHGL WP_001295940.1
MULTISPECIES: hydrogenase [Enterobacteriaceae] Length: 372\nNumber of predicted TMHs: 2\nExp number of AAs in TMHs: 37.2608899999999\nExp number, first 60 AAs: 19.52923\nTotal prob of N-in: 0.94351\nPOSSIBLE N-term signal sequence\ninside 1 19\nTMhelix 20 39\noutside 40 325\nTMhelix 326 348\ninside 349 372

4963 GCF_000782395.1_ASM78239v1 Escherichia coli Proteobacteria; Gammaproteobacteria; Enterobacterales; Enterobacteriaceae; Escherichia
MNNEETFYQAMRRQGVTRRSFLKYCSLAATSLGLGAGMAPKIAWALENKPRIPVVWIHGL WP_001058323.1
MULTISPECIES: hydrogenase-1 small chain [Proteobacteria] Length: 372\nNumber of predicted TMHs: 2\nExp number of AAs in TMHs: 37.2608499999999\nExp number, first 60 AAs: 19.52923\nTotal prob of N-in: 0.94351\nPOSSIBLE N-term signal sequence\ninside 1 19\nTMhelix 20 39\noutside 40 325\nTMhelix 326 348\ninside 349 372

4964 GCF_000782275.1_ASM78227v1 Escherichia coli Proteobacteria; Gammaproteobacteria; Enterobacterales; Enterobacteriaceae; Escherichia
MNNEETFYQAMRRQGVTRRSFLKYCSLAATSLGLGAGMAPKIAWALENKPRIPVVWIHGL WP_001058323.1
MULTISPECIES: hydrogenase-1 small chain [Proteobacteria] Length: 372\nNumber of predicted TMHs: 2\nExp number of AAs in TMHs: 37.2608499999999\nExp number, first 60 AAs: 19.52923\nTotal prob of N-in: 0.94351\nPOSSIBLE N-term signal sequence\ninside 1 19\nTMhelix 20 39\noutside 40 325\nTMhelix 326 348\ninside 349 372

4965 GCF_000782395.1_ASM78239v1 Escherichia coli Proteobacteria; Gammaproteobacteria; Enterobacterales; Enterobacteriaceae; Escherichia
MTGDNTLIHSHGINRRDFMKLCAALAATMGLSSKAAAEMAESVTNPQRPPVIWIGAQECT WP_000145410.1
MULTISPECIES: hydrogenase-2 small chain [Proteobacteria] Length: 372\nNumber of predicted TMHs: 1\nExp number of AAs in TMHs: 23.72226\nExp number, first 60 AAs: 0.22038\nTotal prob of N-in: 0.12994\noutside 1 331\nTMhelix 332 354\ninside 355 372

4966 GCF_000782355.1_ASM78235v1 Escherichia coli Proteobacteria; Gammaproteobacteria; Enterobacterales; Enterobacteriaceae; Escherichia
MTGDNTLIHSHGINRRDFMKLCAALAATMGLSSKAAAEMAESVTNPQRPPVIWIGAQECT WP_000145410.1
MULTISPECIES: hydrogenase-2 small chain [Proteobacteria] Length: 372\nNumber of predicted TMHs: 1\nExp number of AAs in TMHs: 23.72226\nExp number, first 60 AAs: 0.22038\nTotal prob of N-in: 0.12994\noutside 1 331\nTMhelix 332 354\ninside 355 372

4967 GCF_000782275.1_ASM78227v1 Escherichia coli Proteobacteria; Gammaproteobacteria; Enterobacterales; Enterobacteriaceae; Escherichia
MTGDNTLIHSHGINRRDFMKLCAALAATMGLSSKAAAEMAESVTNPQRPPVIWIGAQECT WP_000145410.1
MULTISPECIES: hydrogenase-2 small chain [Proteobacteria] Length: 372\nNumber of predicted TMHs: 1\nExp number of AAs in TMHs: 23.72226\nExp number, first 60 AAs: 0.22038\nTotal prob of N-in: 0.12994\noutside 1 331\nTMhelix 332 354\ninside 355 372

4968 GCF_000781965.1_ASM78196v1 Escherichia coli Proteobacteria; Gammaproteobacteria; Enterobacterales; Enterobacteriaceae; Escherichia
MNNEETFYQAMRRQGVTRRSFLKYCSLAATSLGLGAGMAPKIAWALENKPRIPVVWIHGL WP_001058323.1
MULTISPECIES: hydrogenase-1 small chain [Proteobacteria] Length: 372\nNumber of predicted TMHs: 2\nExp number of AAs in TMHs: 37.2608499999999\nExp number, first 60 AAs: 19.52923\nTotal prob of N-in: 0.94351\nPOSSIBLE N-term signal sequence\ninside 1 19\nTMhelix 20 39\noutside 40 325\nTMhelix 326 348\ninside 349 372

4969 GCF_000782075.1_ASM78207v1 Escherichia coli Proteobacteria; Gammaproteobacteria; Enterobacterales; Enterobacteriaceae; Escherichia
MNNEETFYQAMRRQGVTRRSFLKYCSLAATSLGLGAGMAPKIAWALENKPRIPVVWIHGL WP_001058323.1
MULTISPECIES: hydrogenase-1 small chain [Proteobacteria] Length: 372\nNumber of predicted TMHs: 2\nExp number of AAs in TMHs: 37.2608499999999\nExp number, first 60 AAs: 19.52923\nTotal prob of N-in: 0.94351\nPOSSIBLE N-term signal sequence\ninside 1 19\nTMhelix 20 39\noutside 40 325\nTMhelix 326 348\ninside 349 372

4970 GCF_000782075.1_ASM78207v1 Escherichia coli Proteobacteria; Gammaproteobacteria; Enterobacterales; Enterobacteriaceae; Escherichia
MTGDNTLIHSHGINRRDFMKLCAALAATMGLSSKAAAEMAESVTNPQRPPVIWIGAQECT WP_000145410.1
MULTISPECIES: hydrogenase-2 small chain [Proteobacteria] Length: 372\nNumber of predicted TMHs: 1\nExp number of AAs in TMHs: 23.72226\nExp number, first 60 AAs: 0.22038\nTotal prob of N-in: 0.12994\noutside 1 331\nTMhelix 332 354\ninside 355 372

4971 GCF_000782035.1_ASM78203v1 Escherichia coli Proteobacteria; Gammaproteobacteria; Enterobacterales; Enterobacteriaceae; Escherichia
MTGDNTLIHSHGINRRDFMKLCAALAATMGLSSKAAAEMAESVTNPQRPPVIWIGAQECT WP_000145410.1
MULTISPECIES: hydrogenase-2 small chain [Proteobacteria] Length: 372\nNumber of predicted TMHs: 1\nExp number of AAs in TMHs: 23.72226\nExp number, first 60 AAs: 0.22038\nTotal prob of N-in: 0.12994\noutside 1 331\nTMhelix 332 354\ninside 355 372

4972 GCF_000781895.1_ASM78189v1 Escherichia coli Proteobacteria; Gammaproteobacteria; Enterobacterales; Enterobacteriaceae; Escherichia
MNNEETFYQAMRRQGVTRRSFLKYCSLAATSLGLGAGMAPKIAWALENKPRIPVVWIHGL WP_001058323.1
MULTISPECIES: hydrogenase-1 small chain [Proteobacteria] Length: 372\nNumber of predicted TMHs: 2\nExp number of AAs in TMHs: 37.2608499999999\nExp number, first 60 AAs: 19.52923\nTotal prob of N-in: 0.94351\nPOSSIBLE N-term signal sequence\ninside 1 19\nTMhelix 20 39\noutside 40 325\nTMhelix 326 348\ninside 349 372

4973 GCF_000781995.1_ASM78199v1 Escherichia coli Proteobacteria; Gammaproteobacteria; Enterobacterales; Enterobacteriaceae; Escherichia

MNNEETFYQAMRRQGVTRRSFLKYCSLAATSLGLGAGMAPKIAWALENKPRIPVVWIHGL WP_001058323.1 MULTISPECIES: hydrogenase-1 small chain [Proteobacteria] Length: 372\nNumber of predicted TMHs: 2\nExp number of AAs in TMHs: 37.2608499999999\nExp number, first 60 AAs: 19.52923\nTotal prob of N-in: 0.94351\nPOSSIBLE N-term signal sequence\ninside 1 19\nTMhelix 20 39\noutside 40 325\nTMhelix 326 348\ninside 349 372

4974 GCF_000781805.1_ASM78180v1 Escherichia coli Proteobacteria; Gammaproteobacteria; Enterobacterales; Enterobacteriaceae; Escherichia MNNEETFYQAMRRQGVTRRSFLKYCSLAATSLGLGAGMAPKIAWALENKPRIPVVWIHGL WP_001058323.1 MULTISPECIES: hydrogenase-1 small chain [Proteobacteria] Length: 372\nNumber of predicted TMHs: 2\nExp number of AAs in TMHs: 37.2608499999999\nExp number, first 60 AAs: 19.52923\nTotal prob of N-in: 0.94351\nPOSSIBLE N-term signal sequence\ninside 1 19\nTMhelix 20 39\noutside 40 325\nTMhelix 326 348\ninside 349 372

4975 GCF_000781835.1_ASM78183v1 Escherichia coli Proteobacteria; Gammaproteobacteria; Enterobacterales; Enterobacteriaceae; Escherichia MNNEETFYQAMRRQGVTRRSFLKYCSLAATSLGLGAGMAPKIAWALENKPRIPVVWIHGL WP_001538532.1 hydrogenase-1 small chain [Escherichia coli] Length: 372\nNumber of predicted TMHs: 2\nExp number of AAs in TMHs: 37.27041\nExp number, first 60 AAs: 19.52927\nTotal prob of N-in: 0.94352\nPOSSIBLE N-term signal sequence\ninside 1 19\nTMhelix 20 39\noutside 40 325\nTMhelix 326 348\ninside 349 372

4976 GCF_000781835.1_ASM78183v1 Escherichia coli Proteobacteria; Gammaproteobacteria; Enterobacterales; Enterobacteriaceae; Escherichia MTGDNTLIHSHGINRRDFMKLCAALAATMGLSSKAAAEMVESVTNPQRPPVIWIGAQECT WP_000145423.1 hydrogenase 2 small subunit [Escherichia coli] Length: 372\nNumber of predicted TMHs: 1\nExp number of AAs in TMHs: 23.78201\nExp number, first 60 AAs: 0.28355\nTotal prob of N-in: 0.13257\noutside 1 331\nTMhelix 332 354\ninside 355 372

4977 GCF_000781605.1_ASM78160v1 Escherichia coli Proteobacteria; Gammaproteobacteria; Enterobacterales; Enterobacteriaceae; Escherichia MNNEETFYQAMRRQGVTRRSFLKYCSLAATSLGLGAGMAPKIAWALENKPRIPVVWIHGL WP_001058323.1 MULTISPECIES: hydrogenase-1 small chain [Proteobacteria] Length: 372\nNumber of predicted TMHs: 2\nExp number of AAs in TMHs: 37.2608499999999\nExp number, first 60 AAs: 19.52923\nTotal prob of N-in: 0.94351\nPOSSIBLE N-term signal sequence\ninside 1 19\nTMhelix 20 39\noutside 40 325\nTMhelix 326 348\ninside 349 372

4978 GCF_000781675.1_ASM78167v1 Escherichia coli Proteobacteria; Gammaproteobacteria; Enterobacterales; Enterobacteriaceae; Escherichia MNNEETFYQAMRRQGVTRRSFLKYCSLAATSLGLGAGMAPKIAWALENKPRIPVVWIHGL WP_001058323.1 MULTISPECIES: hydrogenase-1 small chain [Proteobacteria] Length: 372\nNumber of predicted TMHs: 2\nExp number of AAs in TMHs: 37.2608499999999\nExp number, first 60 AAs: 19.52923\nTotal prob of N-in: 0.94351\nPOSSIBLE N-term signal sequence\ninside 1 19\nTMhelix 20 39\noutside 40 325\nTMhelix 326 348\ninside 349 372

4979 GCF_000781655.1_ASM78165v1 Escherichia coli Proteobacteria; Gammaproteobacteria; Enterobacterales; Enterobacteriaceae; Escherichia MNNEETFYQAMRRQGVTRRSFLKYCSLAATSLGLGAGMAPKIAWALENKPRIPVVWIHGL WP_001058323.1 MULTISPECIES: hydrogenase-1 small chain [Proteobacteria] Length: 372\nNumber of predicted TMHs: 2\nExp number of AAs in TMHs: 37.2608499999999\nExp number, first 60 AAs: 19.52923\nTotal prob of N-in: 0.94351\nPOSSIBLE N-term signal sequence\ninside 1 19\nTMhelix 20 39\noutside 40 325\nTMhelix 326 348\ninside 349 372

4980 GCF_000781435.1_ASM78143v1 Escherichia coli Proteobacteria; Gammaproteobacteria; Enterobacterales; Enterobacteriaceae; Escherichia MNNEETFYQAMRRQGVTRRSFLKYCSLAATSLGLGAGMAPKIAWALENKPRIPVVWIHGL WP_001058323.1 MULTISPECIES: hydrogenase-1 small chain [Proteobacteria] Length: 372\nNumber of predicted TMHs: 2\nExp number of AAs in TMHs: 37.2608499999999\nExp number, first 60 AAs: 19.52923\nTotal prob of N-in: 0.94351\nPOSSIBLE N-term signal sequence\ninside 1 19\nTMhelix 20 39\noutside 40 325\nTMhelix 326 348\ninside 349 372

4981 GCF_000781355.1_ASM78135v1 Escherichia coli Proteobacteria; Gammaproteobacteria; Enterobacterales; Enterobacteriaceae; Escherichia
MNNEETFYQAMRRQGVTRRSFLKYCSLAATSLGLGAGMAPKIAWALENKPRIPVVWIHGL WP_001058323.1
MULTISPECIES: hydrogenase-1 small chain [Proteobacteria] Length: 372\nNumber of predicted TMHs: 2\nExp number of AAs in TMHs: 37.2608499999999\nExp number, first 60 AAs: 19.52923\nTotal prob of N-in: 0.94351\nPOSSIBLE N-term signal sequence\ninside 1 19\nTMhelix 20 39\noutside 40 325\nTMhelix 326 348\ninside 349 372

4982 GCF_000781505.1_ASM78150v1 Escherichia coli Proteobacteria; Gammaproteobacteria; Enterobacterales; Enterobacteriaceae; Escherichia
MNNEETFYQAMRRQGVTRRSFLKYCSLAATSLGLGAGMAPKIAWALENKPRIPVVWIHGL WP_001295940.1
MULTISPECIES: hydrogenase [Enterobacteriaceae] Length: 372\nNumber of predicted TMHs: 2\nExp number of AAs in TMHs: 37.2608899999999\nExp number, first 60 AAs: 19.52923\nTotal prob of N-in: 0.94351\nPOSSIBLE N-term signal sequence\ninside 1 19\nTMhelix 20 39\noutside 40 325\nTMhelix 326 348\ninside 349 372

4983 GCF_000781555.1_ASM78155v1 Escherichia coli Proteobacteria; Gammaproteobacteria; Enterobacterales; Enterobacteriaceae; Escherichia
MTGDNTLIHSHGINRRDFMKLCAALAATMGLSSKAAAEMAESVTNPQRPPVIWIGAQECT WP_000145410.1
MULTISPECIES: hydrogenase-2 small chain [Proteobacteria] Length: 372\nNumber of predicted TMHs: 1\nExp number of AAs in TMHs: 23.72226\nExp number, first 60 AAs: 0.22038\nTotal prob of N-in: 0.12994\noutside 1 331\nTMhelix 332 354\ninside 355 372

4984 GCF_000781505.1_ASM78150v1 Escherichia coli Proteobacteria; Gammaproteobacteria; Enterobacterales; Enterobacteriaceae; Escherichia
MTGDNTLIHSHGINRRDFMKLCAALAATMGLSSKAAAEMVESVTNPQRPPVIWIGAQECT WP_000145423.1
hydrogenase 2 small subunit [Escherichia coli] Length: 372\nNumber of predicted TMHs: 1\nExp number of AAs in TMHs: 23.78201\nExp number, first 60 AAs: 0.28355\nTotal prob of N-in: 0.13257\noutside 1 331\nTMhelix 332 354\ninside 355 372

4985 GCF_000781455.1_ASM78145v1 Escherichia coli Proteobacteria; Gammaproteobacteria; Enterobacterales; Enterobacteriaceae; Escherichia
MNNEETFYQAMRRQGVTRRSFLKYCSLAATSLGLGAGMAPKIAWALENKPRIPVVWIHGL WP_001058323.1
MULTISPECIES: hydrogenase-1 small chain [Proteobacteria] Length: 372\nNumber of predicted TMHs: 2\nExp number of AAs in TMHs: 37.2608499999999\nExp number, first 60 AAs: 19.52923\nTotal prob of N-in: 0.94351\nPOSSIBLE N-term signal sequence\ninside 1 19\nTMhelix 20 39\noutside 40 325\nTMhelix 326 348\ninside 349 372

4986 GCF_000781255.1_ASM78125v1 Escherichia coli Proteobacteria; Gammaproteobacteria; Enterobacterales; Enterobacteriaceae; Escherichia
MNNEETFYQAMRRQGVTRRSFLKYCSLAATSLGLGAGMAPKIAWALENKPRIPVVWIHGL WP_001058323.1
MULTISPECIES: hydrogenase-1 small chain [Proteobacteria] Length: 372\nNumber of predicted TMHs: 2\nExp number of AAs in TMHs: 37.2608499999999\nExp number, first 60 AAs: 19.52923\nTotal prob of N-in: 0.94351\nPOSSIBLE N-term signal sequence\ninside 1 19\nTMhelix 20 39\noutside 40 325\nTMhelix 326 348\ninside 349 372

4987 GCF_000781295.1_ASM78129v1 Escherichia coli Proteobacteria; Gammaproteobacteria; Enterobacterales; Enterobacteriaceae; Escherichia
MNNEETFYQAMRRQGVTRRSFLKYCSLAATSLGLGAGMAPKIAWALENKPRIPVVWIHGL WP_001058323.1
MULTISPECIES: hydrogenase-1 small chain [Proteobacteria] Length: 372\nNumber of predicted TMHs: 2\nExp number of AAs in TMHs: 37.2608499999999\nExp number, first 60 AAs: 19.52923\nTotal prob of N-in: 0.94351\nPOSSIBLE N-term signal sequence\ninside 1 19\nTMhelix 20 39\noutside 40 325\nTMhelix 326 348\ninside 349 372

4988 GCF_000781295.1_ASM78129v1 Escherichia coli Proteobacteria; Gammaproteobacteria; Enterobacterales; Enterobacteriaceae; Escherichia
MTGDNTLIHSHGINRRDFMKLCAALAATMGLSSKAAAEMAESVTNPQRPPVIWIGAQECT WP_000145418.1
hydrogenase 2 small subunit [Escherichia coli] Length: 372\nNumber of predicted TMHs: 1\nExp number of AAs in TMHs: 23.72313\nExp number, first 60 AAs: 0.22034\nTotal prob of N-in: 0.12997\noutside 1 331\nTMhelix 332 354\ninside 355 372

4989 GCF_000781235.1_ASM78123v1 Escherichia coli Proteobacteria; Gammaproteobacteria; Enterobacterales; Enterobacteriaceae; Escherichia MTGDNTLIHSHGINRRDFMKLCAALAATMGLSSKAAAEMAESVTNPQRPPVIWIGAQECT WP_000145410.1 MULTISPECIES: hydrogenase-2 small chain [Proteobacteria] Length: 372\nNumber of predicted TMHs: 1\nExp number of AAs in TMHs: 23.72226\nExp number, first 60 AAs: 0.22038\nTotal prob of N-in: 0.12994\noutside 1 331\nTMhelix 332 354\ninside 355 372

4990 GCF_000781255.1_ASM78125v1 Escherichia coli Proteobacteria; Gammaproteobacteria; Enterobacterales; Enterobacteriaceae; Escherichia MTGDNTLIHSHGINRRDFMKLCAALAATMGLSSKAAAEMAESVTNPQRPPVIWIGAQECT WP_000145410.1 MULTISPECIES: hydrogenase-2 small chain [Proteobacteria] Length: 372\nNumber of predicted TMHs: 1\nExp number of AAs in TMHs: 23.72226\nExp number, first 60 AAs: 0.22038\nTotal prob of N-in: 0.12994\noutside 1 331\nTMhelix 332 354\ninside 355 372

4991 GCF_000781075.1_ASM78107v1 Escherichia coli Proteobacteria; Gammaproteobacteria; Enterobacterales; Enterobacteriaceae; Escherichia MNNEETFYQAMRRQGVTRRSFLKYCSLAATSLGLGAGMAPKIAWALENKPRIPVVWIHGL WP_001058323.1 MULTISPECIES: hydrogenase-1 small chain [Proteobacteria] Length: 372\nNumber of predicted TMHs: 2\nExp number of AAs in TMHs: 37.2608499999999\nExp number, first 60 AAs: 19.52923\nTotal prob of N-in: 0.94351\nPOSSIBLE N-term signal sequence\ninside 1 19\nTMhelix 20 39\noutside 40 325\nTMhelix 326 348\ninside 349 372

4992 GCF_000781195.1_ASM78119v1 Escherichia coli Proteobacteria; Gammaproteobacteria; Enterobacterales; Enterobacteriaceae; Escherichia MNNEETFYQAMRRQGVTRRSFLKYCSLAATSLGLGAGMAPKIAWALENKPRIPVVWIHGL WP_001058323.1 MULTISPECIES: hydrogenase-1 small chain [Proteobacteria] Length: 372\nNumber of predicted TMHs: 2\nExp number of AAs in TMHs: 37.2608499999999\nExp number, first 60 AAs: 19.52923\nTotal prob of N-in: 0.94351\nPOSSIBLE N-term signal sequence\ninside 1 19\nTMhelix 20 39\noutside 40 325\nTMhelix 326 348\ninside 349 372

4993 GCF_000781145.1_ASM78114v1 Escherichia coli Proteobacteria; Gammaproteobacteria; Enterobacterales; Enterobacteriaceae; Escherichia MNNEETFYQAMRRQGVTRRSFLKYCSLAATSLGLGAGMAPKIAWALENKPRIPVVWIHGL WP_001295940.1 MULTISPECIES: hydrogenase [Enterobacteriaceae] Length: 372\nNumber of predicted TMHs: 2\nExp number of AAs in TMHs: 37.2608899999999\nExp number, first 60 AAs: 19.52923\nTotal prob of N-in: 0.94351\nPOSSIBLE N-term signal sequence\ninside 1 19\nTMhelix 20 39\noutside 40 325\nTMhelix 326 348\ninside 349 372

4994 GCF_000781195.1_ASM78119v1 Escherichia coli Proteobacteria; Gammaproteobacteria; Enterobacterales; Enterobacteriaceae; Escherichia MTGDNTLIHSHGINRRDFMKLCAALAATMGLSSKAAAEMAESVTNPQRPPVIWIGAQECT WP_000145410.1 MULTISPECIES: hydrogenase-2 small chain [Proteobacteria] Length: 372\nNumber of predicted TMHs: 1\nExp number of AAs in TMHs: 23.72226\nExp number, first 60 AAs: 0.22038\nTotal prob of N-in: 0.12994\noutside 1 331\nTMhelix 332 354\ninside 355 372

4995 GCF_000781145.1_ASM78114v1 Escherichia coli Proteobacteria; Gammaproteobacteria; Enterobacterales; Enterobacteriaceae; Escherichia MTGDNTLIHSHGINRRDFMKLCAALAATMGLSSKAAAEMAESVTNPQRPPVIWIGAQECT WP_000145410.1 MULTISPECIES: hydrogenase-2 small chain [Proteobacteria] Length: 372\nNumber of predicted TMHs: 1\nExp number of AAs in TMHs: 23.72226\nExp number, first 60 AAs: 0.22038\nTotal prob of N-in: 0.12994\noutside 1 331\nTMhelix 332 354\ninside 355 372

4996 GCF_000780895.1_ASM78089v1 Escherichia coli Proteobacteria; Gammaproteobacteria; Enterobacterales; Enterobacteriaceae; Escherichia MNNEETFYQAMRRQGVTRRSFLKYCSLAATSLGLGAGMAPKIAWALENKPRIPVVWIHGL WP_001295940.1 MULTISPECIES: hydrogenase [Enterobacteriaceae] Length: 372\nNumber of predicted TMHs: 2\nExp number of AAs in TMHs: 37.2608899999999\nExp number, first 60 AAs: 19.52923\nTotal prob of N-in: 0.94351\nPOSSIBLE N-term signal sequence\ninside 1 19\nTMhelix 20 39\noutside 40 325\nTMhelix 326 348\ninside 349 372

4997 GCF_000780965.1_ASM78096v1 Escherichia coli Proteobacteria; Gammaproteobacteria; Enterobacterales; Enterobacteriaceae; Escherichia
MNNEETFYQAMRRQGVTRRSFLKYCSLAATSLGLGAGMAPKIAWALENKPRIPVVWIHGL WP_001058323.1
MULTISPECIES: hydrogenase-1 small chain [Proteobacteria] Length: 372\nNumber of predicted TMHs: 2\nExp number of AAs in TMHs: 37.2608499999999\nExp number, first 60 AAs: 19.52923\nTotal prob of N-in: 0.94351\nPOSSIBLE N-term signal sequence\ninside 1 19\nTMhelix 20 39\noutside 40 325\nTMhelix 326 348\ninside 349 372

4998 GCF_000781115.1_ASM78111v1 Escherichia coli Proteobacteria; Gammaproteobacteria; Enterobacterales; Enterobacteriaceae; Escherichia
MNNEETFYQAMRRQGVTRRSFLKYCSLAATSLGLGAGMAPKIAWALENKPRIPVVWIHGL WP_001295940.1
MULTISPECIES: hydrogenase [Enterobacteriaceae] Length: 372\nNumber of predicted TMHs: 2\nExp number of AAs in TMHs: 37.2608899999999\nExp number, first 60 AAs: 19.52923\nTotal prob of N-in: 0.94351\nPOSSIBLE N-term signal sequence\ninside 1 19\nTMhelix 20 39\noutside 40 325\nTMhelix 326 348\ninside 349 372

4999 GCF_000781115.1_ASM78111v1 Escherichia coli Proteobacteria; Gammaproteobacteria; Enterobacterales; Enterobacteriaceae; Escherichia
MTGDNTLIHSHGINRRDFMKLCAALAATMGLSSKAAAEMAESVTNPQRPPVIWIGAQECT WP_000145410.1
MULTISPECIES: hydrogenase-2 small chain [Proteobacteria] Length: 372\nNumber of predicted TMHs: 1\nExp number of AAs in TMHs: 23.72226\nExp number, first 60 AAs: 0.22038\nTotal prob of N-in: 0.12994\noutside 1 331\nTMhelix 332 354\ninside 355 372

5000 GCF_000780815.1_ASM78081v1 Escherichia coli Proteobacteria; Gammaproteobacteria; Enterobacterales; Enterobacteriaceae; Escherichia
MNNEETFYQAMRRQGVTRRSFLKYCSLAATSLGLGAGMAPKIAWALENKPRIPVVWIHGL WP_001058323.1
MULTISPECIES: hydrogenase-1 small chain [Proteobacteria] Length: 372\nNumber of predicted TMHs: 2\nExp number of AAs in TMHs: 37.2608499999999\nExp number, first 60 AAs: 19.52923\nTotal prob of N-in: 0.94351\nPOSSIBLE N-term signal sequence\ninside 1 19\nTMhelix 20 39\noutside 40 325\nTMhelix 326 348\ninside 349 372

5001 GCF_000780815.1_ASM78081v1 Escherichia coli Proteobacteria; Gammaproteobacteria; Enterobacterales; Enterobacteriaceae; Escherichia
MTGDNTLIHSHGINRRDFMKLCAALAATMGLSSKAAAEMAESVTNPQRPPVIWIGAQECT WP_000145410.1
MULTISPECIES: hydrogenase-2 small chain [Proteobacteria] Length: 372\nNumber of predicted TMHs: 1\nExp number of AAs in TMHs: 23.72226\nExp number, first 60 AAs: 0.22038\nTotal prob of N-in: 0.12994\noutside 1 331\nTMhelix 332 354\ninside 355 372

5002 GCF_000780655.1_ASM78065v1 Escherichia coli Proteobacteria; Gammaproteobacteria; Enterobacterales; Enterobacteriaceae; Escherichia
MNNEETFYQAMRRQGVTRRSFLKYCSLAATSLGLGAGMAPKIAWALENKPRIPVVWIHGL WP_001058323.1
MULTISPECIES: hydrogenase-1 small chain [Proteobacteria] Length: 372\nNumber of predicted TMHs: 2\nExp number of AAs in TMHs: 37.2608499999999\nExp number, first 60 AAs: 19.52923\nTotal prob of N-in: 0.94351\nPOSSIBLE N-term signal sequence\ninside 1 19\nTMhelix 20 39\noutside 40 325\nTMhelix 326 348\ninside 349 372

5003 GCF_000780795.1_ASM78079v1 Escherichia coli Proteobacteria; Gammaproteobacteria; Enterobacterales; Enterobacteriaceae; Escherichia
MNNEETFYQAMRRQGVTRRSFLKYCSLAATSLGLGAGMAPKIAWALENKPRIPVVWIHGL WP_001058323.1
MULTISPECIES: hydrogenase-1 small chain [Proteobacteria] Length: 372\nNumber of predicted TMHs: 2\nExp number of AAs in TMHs: 37.2608499999999\nExp number, first 60 AAs: 19.52923\nTotal prob of N-in: 0.94351\nPOSSIBLE N-term signal sequence\ninside 1 19\nTMhelix 20 39\noutside 40 325\nTMhelix 326 348\ninside 349 372

5004 GCF_000780795.1_ASM78079v1 Escherichia coli Proteobacteria; Gammaproteobacteria; Enterobacterales; Enterobacteriaceae; Escherichia
MTGDNTLIHSHGINRRDFMKLCAALAATMGLSSKAAAEMAESVTNPQRPPVIWIGAQECT WP_000145410.1
MULTISPECIES: hydrogenase-2 small chain [Proteobacteria] Length: 372\nNumber of predicted TMHs: 1\nExp number of AAs in TMHs: 23.72226\nExp number, first 60 AAs: 0.22038\nTotal prob of N-in: 0.12994\noutside 1 331\nTMhelix 332 354\ninside 355 372

5005 GCF_000780755.1_ASM78075v1 Escherichia coli Proteobacteria; Gammaproteobacteria; Enterobacterales; Enterobacteriaceae; Escherichia
MTGDNTLIHSHGINRRDFMKLCAALAATMGLSSKAAAEMAESVTNPQRPPVIWIGAQECT WP_000145410.1
MULTISPECIES: hydrogenase-2 small chain [Proteobacteria] Length: 372\nNumber of predicted TMHs: 1\nExp number of AAs in TMHs: 23.72226\nExp number, first 60 AAs: 0.22038\nTotal prob of N-in: 0.12994\noutside 1 331\nTMhelix 332 354\ninside 355 372

5006 GCF_000781075.1_ASM78107v1 Escherichia coli Proteobacteria; Gammaproteobacteria; Enterobacterales; Enterobacteriaceae; Escherichia
MTGDNTLIHSHGINRRDFMKLCAALAATMGLSSKAAAEMAESVTNPQRPPVIWIGAQECT WP_000145410.1
MULTISPECIES: hydrogenase-2 small chain [Proteobacteria] Length: 372\nNumber of predicted TMHs: 1\nExp number of AAs in TMHs: 23.72226\nExp number, first 60 AAs: 0.22038\nTotal prob of N-in: 0.12994\noutside 1 331\nTMhelix 332 354\ninside 355 372

5007 GCF_000780655.1_ASM78065v1 Escherichia coli Proteobacteria; Gammaproteobacteria; Enterobacterales; Enterobacteriaceae; Escherichia
MTGDNTLIHSHGINRRDFMKLCAALAATMGLSSKAAAEMAESVTNPQRPPVIWIGAQECT WP_000145410.1
MULTISPECIES: hydrogenase-2 small chain [Proteobacteria] Length: 372\nNumber of predicted TMHs: 1\nExp number of AAs in TMHs: 23.72226\nExp number, first 60 AAs: 0.22038\nTotal prob of N-in: 0.12994\noutside 1 331\nTMhelix 332 354\ninside 355 372

5008 GCF_000780615.1_ASM78061v1 Escherichia coli Proteobacteria; Gammaproteobacteria; Enterobacterales; Enterobacteriaceae; Escherichia
MTGDNTLIHSHGINRRDFMKLCAALAATMGLSSKAAAEMVESVTNPQRPPVIWIGAQECT WP_000145423.1
hydrogenase 2 small subunit [Escherichia coli] Length: 372\nNumber of predicted TMHs: 1\nExp number of AAs in TMHs: 23.78201\nExp number, first 60 AAs: 0.28355\nTotal prob of N-in: 0.13257\noutside 1 331\nTMhelix 332 354\ninside 355 372

5009 GCF_000780595.1_ASM78059v1 Escherichia coli Proteobacteria; Gammaproteobacteria; Enterobacterales; Enterobacteriaceae; Escherichia
MTGDNTLIHSHGINRRDFMKLCAALAATMGLSSKAAAEMAESVTNPQRPPVIWIGAQECT WP_000145410.1
MULTISPECIES: hydrogenase-2 small chain [Proteobacteria] Length: 372\nNumber of predicted TMHs: 1\nExp number of AAs in TMHs: 23.72226\nExp number, first 60 AAs: 0.22038\nTotal prob of N-in: 0.12994\noutside 1 331\nTMhelix 332 354\ninside 355 372

5010 GCF_000780555.1_ASM78055v1 Escherichia coli Proteobacteria; Gammaproteobacteria; Enterobacterales; Enterobacteriaceae; Escherichia
MTGDNTLIHSHGINRRDFMKLCAALAATMGLSSKAAAEMAESVTNPQRPPVIWIGAQECT WP_000145410.1
MULTISPECIES: hydrogenase-2 small chain [Proteobacteria] Length: 372\nNumber of predicted TMHs: 1\nExp number of AAs in TMHs: 23.72226\nExp number, first 60 AAs: 0.22038\nTotal prob of N-in: 0.12994\noutside 1 331\nTMhelix 332 354\ninside 355 372

5011 GCF_000780495.1_ASM78049v1 Escherichia coli Proteobacteria; Gammaproteobacteria; Enterobacterales; Enterobacteriaceae; Escherichia
MTGDNTLIHSHGINRRDFMKLCAALAATMGLSSKAAAEMVESVTNPQRPPVIWIGAQECT WP_000145423.1
hydrogenase 2 small subunit [Escherichia coli] Length: 372\nNumber of predicted TMHs: 1\nExp number of AAs in TMHs: 23.78201\nExp number, first 60 AAs: 0.28355\nTotal prob of N-in: 0.13257\noutside 1 331\nTMhelix 332 354\ninside 355 372

5012 GCF_000780475.1_ASM78047v1 Escherichia coli Proteobacteria; Gammaproteobacteria; Enterobacterales; Enterobacteriaceae; Escherichia
MTGDNTLIHSHGINRRDFMKLCAALAATMGLSSKAAAEMVESVTNPQRPPVIWIGAQECT WP_000145423.1
hydrogenase 2 small subunit [Escherichia coli] Length: 372\nNumber of predicted TMHs: 1\nExp number of AAs in TMHs: 23.78201\nExp number, first 60 AAs: 0.28355\nTotal prob of N-in: 0.13257\noutside 1 331\nTMhelix 332 354\ninside 355 372

5013 GCF_000780335.1_ASM78033v1 Escherichia coli Proteobacteria; Gammaproteobacteria; Enterobacterales; Enterobacteriaceae; Escherichia
MNNEETFYQAMRRQGVTRRSFLKYCSLAATSLGLGAGMAPKIAWALENKPRIPVVWIHGL WP_001058323.1
MULTISPECIES: hydrogenase-1 small chain [Proteobacteria] Length: 372\nNumber of predicted TMHs: 2\nExp number of AAs in TMHs: 37.2608499999999\nExp number, first 60 AAs: 19.52923\nTotal prob of N-in:

0.94351\nPOSSIBLE N-term signal sequence\ninside 1 19\nTMhelix 20 39\noutside 40 325\nTMhelix 326 348\ninside 349 372

5014 GCF_000780395.1_ASM78039v1 Escherichia coli Proteobacteria; Gammaproteobacteria; Enterobacterales; Enterobacteriaceae; Escherichia MTGDNTLIHSHGINRRDFMKLCAALAATMGLSSKAAAEMVESVTNPQRPPVIWIGAQECT WP_000145423.1 hydrogenase 2 small subunit [Escherichia coli] Length: 372\nNumber of predicted TMHs: 1\nExp number of AAs in TMHs: 23.78201\nExp number, first 60 AAs: 0.28355\nTotal prob of N-in: 0.13257\noutside 1 331\nTMhelix 332 354\ninside 355 372

5015 GCF_000780375.1_ASM78037v1 Escherichia coli Proteobacteria; Gammaproteobacteria; Enterobacterales; Enterobacteriaceae; Escherichia MTGDNTLIHSHGINRRDFMKLCAALAATMGLSSKAAAEMVESVTNPQRPPVIWIGAQECT WP_000145423.1 hydrogenase 2 small subunit [Escherichia coli] Length: 372\nNumber of predicted TMHs: 1\nExp number of AAs in TMHs: 23.78201\nExp number, first 60 AAs: 0.28355\nTotal prob of N-in: 0.13257\noutside 1 331\nTMhelix 332 354\ninside 355 372

5016 GCF_000780335.1_ASM78033v1 Escherichia coli Proteobacteria; Gammaproteobacteria; Enterobacterales; Enterobacteriaceae; Escherichia MTGDNTLIHSHGINRRDFMKLCAALAATMGLSSKAAAEMAESVTNPQRPPVIWIGAQECT WP_000145410.1 MULTISPECIES: hydrogenase-2 small chain [Proteobacteria] Length: 372\nNumber of predicted TMHs: 1\nExp number of AAs in TMHs: 23.72226\nExp number, first 60 AAs: 0.22038\nTotal prob of N-in: 0.12994\noutside 1 331\nTMhelix 332 354\ninside 355 372

5017 GCF_000780195.1_ASM78019v1 Escherichia coli Proteobacteria; Gammaproteobacteria; Enterobacterales; Enterobacteriaceae; Escherichia MNNEETFYQAMRRQGVTRRSFLKYCSLAATSLGLGAGMAPKIAWALENKPRIPVVWIHGL WP_001058323.1 MULTISPECIES: hydrogenase-1 small chain [Proteobacteria] Length: 372\nNumber of predicted TMHs: 2\nExp number of AAs in TMHs: 37.2608499999999\nExp number, first 60 AAs: 19.52923\nTotal prob of N-in: 0.94351\nPOSSIBLE N-term signal sequence\ninside 1 19\nTMhelix 20 39\noutside 40 325\nTMhelix 326 348\ninside 349 372

5018 GCF_000780295.1_ASM78029v1 Escherichia coli Proteobacteria; Gammaproteobacteria; Enterobacterales; Enterobacteriaceae; Escherichia MTGDNTLIHSHGINRRDFMKLCAALAATMGLSSKAAAEMAESVTNPQRPPVIWIGAQECT WP_000145410.1 MULTISPECIES: hydrogenase-2 small chain [Proteobacteria] Length: 372\nNumber of predicted TMHs: 1\nExp number of AAs in TMHs: 23.72226\nExp number, first 60 AAs: 0.22038\nTotal prob of N-in: 0.12994\noutside 1 331\nTMhelix 332 354\ninside 355 372

5019 GCF_000780275.1_ASM78027v1 Escherichia coli Proteobacteria; Gammaproteobacteria; Enterobacterales; Enterobacteriaceae; Escherichia MTGDNTLIHSHGINRRDFMKLCAALAATMGLSSKAAAEMAESVTNPQRPPVIWIGAQECT WP_000145410.1 MULTISPECIES: hydrogenase-2 small chain [Proteobacteria] Length: 372\nNumber of predicted TMHs: 1\nExp number of AAs in TMHs: 23.72226\nExp number, first 60 AAs: 0.22038\nTotal prob of N-in: 0.12994\noutside 1 331\nTMhelix 332 354\ninside 355 372

5020 GCF_000780235.1_ASM78023v1 Escherichia coli Proteobacteria; Gammaproteobacteria; Enterobacterales; Enterobacteriaceae; Escherichia MTGDNTLIHSHGINRRDFMKLCAALAATMGLSSKAAAEMAESVTNPQRPPVIWIGAQECT WP_000145410.1 MULTISPECIES: hydrogenase-2 small chain [Proteobacteria] Length: 372\nNumber of predicted TMHs: 1\nExp number of AAs in TMHs: 23.72226\nExp number, first 60 AAs: 0.22038\nTotal prob of N-in: 0.12994\noutside 1 331\nTMhelix 332 354\ninside 355 372

5021 GCF_000780195.1_ASM78019v1 Escherichia coli Proteobacteria; Gammaproteobacteria; Enterobacterales; Enterobacteriaceae; Escherichia MTGDNTLIHSHGINRRDFMKLCAALAATMGLSSKAAAEMAESVTNPQRPPVIWIGAQECT WP_000145410.1 MULTISPECIES: hydrogenase-2 small chain [Proteobacteria] Length: 372\nNumber of predicted TMHs: 1\nExp number of AAs in TMHs: 23.72226\nExp number, first 60 AAs: 0.22038\nTotal prob of N-in: 0.12994\noutside 1 331\nTMhelix 332 354\ninside 355 372

5022 GCF_000779875.1_ASM77987v1 Escherichia coli Proteobacteria; Gammaproteobacteria; Enterobacterales; Enterobacteriaceae; Escherichia
MNNEETFYQAMRRQGVTRRSFLKYCSLAATSLGLGAGMAPKIAWALENKPRIPVVWIHGL WP_001295940.1
MULTISPECIES: hydrogenase [Enterobacteriaceae] Length: 372\nNumber of predicted TMHs: 2\nExp number of AAs in TMHs: 37.2608899999999\nExp number, first 60 AAs: 19.52923\nTotal prob of N-in: 0.94351\nPOSSIBLE N-term signal sequence\ninside 1 19\nTMhelix 20 39\noutside 40 325\nTMhelix 326 348\ninside 349 372

5023 GCF_000780135.1_ASM78013v1 Escherichia coli Proteobacteria; Gammaproteobacteria; Enterobacterales; Enterobacteriaceae; Escherichia
MNNEETFYQAMRRQGVTRRSFLKYCSLAATSLGLGAGMAPKIAWALENKPRIPVVWIHGL WP_001295940.1
MULTISPECIES: hydrogenase [Enterobacteriaceae] Length: 372\nNumber of predicted TMHs: 2\nExp number of AAs in TMHs: 37.2608899999999\nExp number, first 60 AAs: 19.52923\nTotal prob of N-in: 0.94351\nPOSSIBLE N-term signal sequence\ninside 1 19\nTMhelix 20 39\noutside 40 325\nTMhelix 326 348\ninside 349 372

5024 GCF_000780075.1_ASM78007v1 Escherichia coli Proteobacteria; Gammaproteobacteria; Enterobacterales; Enterobacteriaceae; Escherichia
MNNEETFYQAMRRQGVTRRSFLKYCSLAATSLGLGAGMAPKIAWALENKPRIPVVWIHGL WP_001295940.1
MULTISPECIES: hydrogenase [Enterobacteriaceae] Length: 372\nNumber of predicted TMHs: 2\nExp number of AAs in TMHs: 37.2608899999999\nExp number, first 60 AAs: 19.52923\nTotal prob of N-in: 0.94351\nPOSSIBLE N-term signal sequence\ninside 1 19\nTMhelix 20 39\noutside 40 325\nTMhelix 326 348\ninside 349 372

5025 GCF_000780135.1_ASM78013v1 Escherichia coli Proteobacteria; Gammaproteobacteria; Enterobacterales; Enterobacteriaceae; Escherichia
MTGDNTLIHSHGINRRDFMKLCAALAATMGLSSKAAAEMVESVTNPQRPPVIWIGAQECT WP_000145423.1
hydrogenase 2 small subunit [Escherichia coli] Length: 372\nNumber of predicted TMHs: 1\nExp number of AAs in TMHs: 23.78201\nExp number, first 60 AAs: 0.28355\nTotal prob of N-in: 0.13257\noutside 1 331\nTMhelix 332 354\ninside 355 372

5026 GCF_000780175.1_ASM78017v1 Escherichia coli Proteobacteria; Gammaproteobacteria; Enterobacterales; Enterobacteriaceae; Escherichia
MTGDNTLIHSHGINRRDFMKLCAALAATMGLSSKAAAEMVESVTNPQRPPVIWIGAQECT WP_000145423.1
hydrogenase 2 small subunit [Escherichia coli] Length: 372\nNumber of predicted TMHs: 1\nExp number of AAs in TMHs: 23.78201\nExp number, first 60 AAs: 0.28355\nTotal prob of N-in: 0.13257\noutside 1 331\nTMhelix 332 354\ninside 355 372

5027 GCF_000780075.1_ASM78007v1 Escherichia coli Proteobacteria; Gammaproteobacteria; Enterobacterales; Enterobacteriaceae; Escherichia
MTGDNTLIHSHGINRRDFMKLCAALAATMGLSSKAAAEMAESVTNPQRPPVIWIGAQECT WP_000145410.1
MULTISPECIES: hydrogenase-2 small chain [Proteobacteria] Length: 372\nNumber of predicted TMHs: 1\nExp number of AAs in TMHs: 23.72226\nExp number, first 60 AAs: 0.22038\nTotal prob of N-in: 0.12994\noutside 1 331\nTMhelix 332 354\ninside 355 372

5028 GCF_000779975.1_ASM77997v1 Escherichia coli Proteobacteria; Gammaproteobacteria; Enterobacterales; Enterobacteriaceae; Escherichia
MTGDNTLIHSHGINRRDFMKLCAALAATMGLSSKAAAEMAESVTNPQRPPVIWIGAQECT WP_000145410.1
MULTISPECIES: hydrogenase-2 small chain [Proteobacteria] Length: 372\nNumber of predicted TMHs: 1\nExp number of AAs in TMHs: 23.72226\nExp number, first 60 AAs: 0.22038\nTotal prob of N-in: 0.12994\noutside 1 331\nTMhelix 332 354\ninside 355 372

5029 GCF_000779915.1_ASM77991v1 Escherichia coli Proteobacteria; Gammaproteobacteria; Enterobacterales; Enterobacteriaceae; Escherichia
MTGDNTLIHSHGINRRDFMKLCAALAATMGLSSKAAAEMAESVTNPQRPPVIWIGAQECT WP_000145410.1
MULTISPECIES: hydrogenase-2 small chain [Proteobacteria] Length: 372\nNumber of predicted TMHs: 1\nExp number of AAs in TMHs: 23.72226\nExp number, first 60 AAs: 0.22038\nTotal prob of N-in: 0.12994\noutside 1 331\nTMhelix 332 354\ninside 355 372

5030 GCF_000779795.1_ASM77979v1 Escherichia coli Proteobacteria; Gammaproteobacteria; Enterobacterales; Enterobacteriaceae; Escherichia

MNNEETFYQAMRRQGVTRRSFLKYCSLAATSLGLGAGMAPKIAWALENKPRIPVVWIHGL WP_001058323.1 MULTISPECIES: hydrogenase-1 small chain [Proteobacteria] Length: 372\nNumber of predicted TMHs: 2\nExp number of AAs in TMHs: 37.2608499999999\nExp number, first 60 AAs: 19.52923\nTotal prob of N-in: 0.94351\nPOSSIBLE N-term signal sequence\ninside 1 19\nTMhelix 20 39\noutside 40 325\nTMhelix 326 348\ninside 349 372

5031 GCF_000779815.1_ASM77981v1 Escherichia coli Proteobacteria; Gammaproteobacteria; Enterobacterales; Enterobacteriaceae; Escherichia MNNEETFYQAMRRQGVTRRSFLKYCSLAATSLGLGAGMAPKIAWALENKPRIPVVWIHGL WP_001058323.1 MULTISPECIES: hydrogenase-1 small chain [Proteobacteria] Length: 372\nNumber of predicted TMHs: 2\nExp number of AAs in TMHs: 37.2608499999999\nExp number, first 60 AAs: 19.52923\nTotal prob of N-in: 0.94351\nPOSSIBLE N-term signal sequence\ninside 1 19\nTMhelix 20 39\noutside 40 325\nTMhelix 326 348\ninside 349 372

5032 GCF_000779875.1_ASM77987v1 Escherichia coli Proteobacteria; Gammaproteobacteria; Enterobacterales; Enterobacteriaceae; Escherichia MTGDNTLIHSHGINRRDFMKLCAALAATMGLSSKAAAEMVESVTNPQRPPVIWIGAQECT WP_000145423.1 hydrogenase 2 small subunit [Escherichia coli] Length: 372\nNumber of predicted TMHs: 1\nExp number of AAs in TMHs: 23.78201\nExp number, first 60 AAs: 0.28355\nTotal prob of N-in: 0.13257\noutside 1 331\nTMhelix 332 354\ninside 355 372

5033 GCF_000779815.1_ASM77981v1 Escherichia coli Proteobacteria; Gammaproteobacteria; Enterobacterales; Enterobacteriaceae; Escherichia MTGDNTLIHSHGINRRDFMKLCAALAATMGLSSKAAAEMAESVTNPQRPPVIWIGAQECT WP_000145410.1 MULTISPECIES: hydrogenase-2 small chain [Proteobacteria] Length: 372\nNumber of predicted TMHs: 1\nExp number of AAs in TMHs: 23.72226\nExp number, first 60 AAs: 0.22038\nTotal prob of N-in: 0.12994\noutside 1 331\nTMhelix 332 354\ninside 355 372

5034 GCF_000779795.1_ASM77979v1 Escherichia coli Proteobacteria; Gammaproteobacteria; Enterobacterales; Enterobacteriaceae; Escherichia MTGDNTLIHSHGINRRDFMKLCAALAATMGLSSKAAAEMAESVTNPQRPPVIWIGAQECT WP_000145410.1 MULTISPECIES: hydrogenase-2 small chain [Proteobacteria] Length: 372\nNumber of predicted TMHs: 1\nExp number of AAs in TMHs: 23.72226\nExp number, first 60 AAs: 0.22038\nTotal prob of N-in: 0.12994\noutside 1 331\nTMhelix 332 354\ninside 355 372

5035 GCF_000779515.1_ASM77951v1 Escherichia coli Proteobacteria; Gammaproteobacteria; Enterobacterales; Enterobacteriaceae; Escherichia MNNEETFYQAMRRQGVTRRSFLKYCSLAATSLGLGAGMAPKIAWALENKPRIPVVWIHGL WP_001538532.1 hydrogenase-1 small chain [Escherichia coli] Length: 372\nNumber of predicted TMHs: 2\nExp number of AAs in TMHs: 37.27041\nExp number, first 60 AAs: 19.52927\nTotal prob of N-in: 0.94352\nPOSSIBLE N-term signal sequence\ninside 1 19\nTMhelix 20 39\noutside 40 325\nTMhelix 326 348\ninside 349 372

5036 GCF_000779635.1_ASM77963v1 Escherichia coli Proteobacteria; Gammaproteobacteria; Enterobacterales; Enterobacteriaceae; Escherichia MNNEETFYQAMRRQGVTRRSFLKYCSLAATSLGLGAGMAPKIAWALENKPRIPVVWIHGL WP_001058323.1 MULTISPECIES: hydrogenase-1 small chain [Proteobacteria] Length: 372\nNumber of predicted TMHs: 2\nExp number of AAs in TMHs: 37.2608499999999\nExp number, first 60 AAs: 19.52923\nTotal prob of N-in: 0.94351\nPOSSIBLE N-term signal sequence\ninside 1 19\nTMhelix 20 39\noutside 40 325\nTMhelix 326 348\ninside 349 372

5037 GCF_000779635.1_ASM77963v1 Escherichia coli Proteobacteria; Gammaproteobacteria; Enterobacterales; Enterobacteriaceae; Escherichia MTGDNTLIHSHGINRRDFMKLCAALAATMGLSSKAAAEMVESVTNPQRPPVIWIGAQECT WP_000145423.1 hydrogenase 2 small subunit [Escherichia coli] Length: 372\nNumber of predicted TMHs: 1\nExp number of AAs in TMHs: 23.78201\nExp number, first 60 AAs: 0.28355\nTotal prob of N-in: 0.13257\noutside 1 331\nTMhelix 332 354\ninside 355 372

5038 GCF_000779475.1_ASM77947v1 Escherichia coli Proteobacteria; Gammaproteobacteria; Enterobacterales; Enterobacteriaceae; Escherichia MNNEETFYQAMRRQGVTRRSFLKYCSLAATSLGLGAGMAPKIAWALENKPRIPVVWIHGL WP_001058323.1 MULTISPECIES: hydrogenase-1 small chain [Proteobacteria] Length: 372\nNumber of predicted TMHs:

2\nExp number of AAs in TMHs: 37.2608499999999\nExp number, first 60 AAs: 19.52923\nTotal prob of N-in: 0.94351\nPOSSIBLE N-term signal sequence\ninside 1 19\nTMhelix 20 39\noutside 40 325\nTMhelix 326 348\ninside 349 372

5039 GCF_000779545.1_ASM77954v1 Escherichia coli Proteobacteria; Gammaproteobacteria; Enterobacterales; Enterobacteriaceae; Escherichia MTGDNTLIHSHGINRRDFMKLCAALAATMGLSSKAAAEMAESVTNPQRPPVIWIGAQECT WP_000145410.1 MULTISPECIES: hydrogenase-2 small chain [Proteobacteria] Length: 372\nNumber of predicted TMHs: 1\nExp number of AAs in TMHs: 23.72226\nExp number, first 60 AAs: 0.22038\nTotal prob of N-in: 0.12994\noutside 1 331\nTMhelix 332 354\ninside 355 372

5040 GCF_000779515.1_ASM77951v1 Escherichia coli Proteobacteria; Gammaproteobacteria; Enterobacterales; Enterobacteriaceae; Escherichia MTGDNTLIHSHGINRRDFMKLCAALAATMGLSSKAAAEMVESVTNPQRPPVIWIGAQECT WP_000145423.1 hydrogenase 2 small subunit [Escherichia coli] Length: 372\nNumber of predicted TMHs: 1\nExp number of AAs in TMHs: 23.78201\nExp number, first 60 AAs: 0.28355\nTotal prob of N-in: 0.13257\noutside 1 331\nTMhelix 332 354\ninside 355 372

5041 GCF_000779275.1_ASM77927v1 Escherichia coli Proteobacteria; Gammaproteobacteria; Enterobacterales; Enterobacteriaceae; Escherichia MNNEETFYQAMRRQGVTRRSFLKYCSLAATSLGLGAGMAPKIAWALENKPRIPVVWIHGL WP_001058323.1 MULTISPECIES: hydrogenase-1 small chain [Proteobacteria] Length: 372\nNumber of predicted TMHs: 2\nExp number of AAs in TMHs: 37.2608499999999\nExp number, first 60 AAs: 19.52923\nTotal prob of N-in: 0.94351\nPOSSIBLE N-term signal sequence\ninside 1 19\nTMhelix 20 39\noutside 40 325\nTMhelix 326 348\ninside 349 372

5042 GCF_000779335.1_ASM77933v1 Escherichia coli Proteobacteria; Gammaproteobacteria; Enterobacterales; Enterobacteriaceae; Escherichia MTGDNTLIHSHGINRRDFMKLCAALAATMGLSSKAAAEMAESVTNPQRPPVIWIGAQECT WP_000145410.1 MULTISPECIES: hydrogenase-2 small chain [Proteobacteria] Length: 372\nNumber of predicted TMHs: 1\nExp number of AAs in TMHs: 23.72226\nExp number, first 60 AAs: 0.22038\nTotal prob of N-in: 0.12994\noutside 1 331\nTMhelix 332 354\ninside 355 372

5043 GCF_000779275.1_ASM77927v1 Escherichia coli Proteobacteria; Gammaproteobacteria; Enterobacterales; Enterobacteriaceae; Escherichia MTGDNTLIHSHGINRRDFMKLCAALAATMGLSSKAAAEMAESVTNPQRPPVIWIGAQECT WP_000145410.1 MULTISPECIES: hydrogenase-2 small chain [Proteobacteria] Length: 372\nNumber of predicted TMHs: 1\nExp number of AAs in TMHs: 23.72226\nExp number, first 60 AAs: 0.22038\nTotal prob of N-in: 0.12994\noutside 1 331\nTMhelix 332 354\ninside 355 372

5044 GCF_000779265.1_ASM77926v1 Escherichia coli Proteobacteria; Gammaproteobacteria; Enterobacterales; Enterobacteriaceae; Escherichia MTGDNTLIHSHGINRRDFMKLCAALAATMGLSSKAAAEMVESVTNPQRPPVIWIGAQECT WP_000145423.1 hydrogenase 2 small subunit [Escherichia coli] Length: 372\nNumber of predicted TMHs: 1\nExp number of AAs in TMHs: 23.78201\nExp number, first 60 AAs: 0.28355\nTotal prob of N-in: 0.13257\noutside 1 331\nTMhelix 332 354\ninside 355 372

5045 GCF_000779025.1_ASM77902v1 Escherichia coli Proteobacteria; Gammaproteobacteria; Enterobacterales; Enterobacteriaceae; Escherichia MNNEETFYQAMRRQGVTRRSFLKYCSLAATSLGLGAGMAPKIAWALENKPRIPVVWIHGL WP_001058323.1 MULTISPECIES: hydrogenase-1 small chain [Proteobacteria] Length: 372\nNumber of predicted TMHs: 2\nExp number of AAs in TMHs: 37.2608499999999\nExp number, first 60 AAs: 19.52923\nTotal prob of N-in: 0.94351\nPOSSIBLE N-term signal sequence\ninside 1 19\nTMhelix 20 39\noutside 40 325\nTMhelix 326 348\ninside 349 372

5046 GCF_000779215.1_ASM77921v1 Escherichia coli Proteobacteria; Gammaproteobacteria; Enterobacterales; Enterobacteriaceae; Escherichia MNNEETFYQAMRRQGVTRRSFLKYCSLAATSLGLGAGMAPKIAWALENKPRIPVVWIHGL WP_021520630.1 hydrogenase-1 small chain [Escherichia coli] Length: 372\nNumber of predicted TMHs: 2\nExp number of AAs in TMHs: 37.8197599999999\nExp number, first 60 AAs: 19.53295\nTotal prob of N-in: 0.94396\nPOSSIBLE N-term signal sequence\ninside 1 19\nTMhelix 20 39\noutside 40 325\nTMhelix 326 348\ninside 349 372

5047 GCF_000779155.1_ASM77915v1 Escherichia coli Proteobacteria; Gammaproteobacteria; Enterobacterales; Enterobacteriaceae; Escherichia
MNNEETFYQAMRRQGVTRRSFLKYCSLAATSLGLGAGMAPKIAWALENKPRIPVVWIHGL WP_001058323.1
MULTISPECIES: hydrogenase-1 small chain [Proteobacteria] Length: 372\nNumber of predicted TMHs: 2\nExp number of AAs in TMHs: 37.2608499999999\nExp number, first 60 AAs: 19.52923\nTotal prob of N-in: 0.94351\nPOSSIBLE N-term signal sequence\ninside 1 19\nTMhelix 20 39\noutside 40 325\nTMhelix 326 348\ninside 349 372

5048 GCF_000778955.1_ASM77895v1 Escherichia coli Proteobacteria; Gammaproteobacteria; Enterobacterales; Enterobacteriaceae; Escherichia
MTGDNTLIHSHGINRRDFMKLCAALAATMGLSSKAAAEMAESVTNPQRPPVIWIGAQECT WP_000145410.1
MULTISPECIES: hydrogenase-2 small chain [Proteobacteria] Length: 372\nNumber of predicted TMHs: 1\nExp number of AAs in TMHs: 23.72226\nExp number, first 60 AAs: 0.22038\nTotal prob of N-in: 0.12994\noutside 1 331\nTMhelix 332 354\ninside 355 372

5049 GCF_000778685.1_ASM77868v1 Escherichia coli Proteobacteria; Gammaproteobacteria; Enterobacterales; Enterobacteriaceae; Escherichia
MNNEETFYQAMRRQGVTRRSFLKYCSLAATSLGLGAGMAPKIAWALENKPRIPVVWIHGL WP_001058323.1
MULTISPECIES: hydrogenase-1 small chain [Proteobacteria] Length: 372\nNumber of predicted TMHs: 2\nExp number of AAs in TMHs: 37.2608499999999\nExp number, first 60 AAs: 19.52923\nTotal prob of N-in: 0.94351\nPOSSIBLE N-term signal sequence\ninside 1 19\nTMhelix 20 39\noutside 40 325\nTMhelix 326 348\ninside 349 372

5050 GCF_000778915.1_ASM77891v1 Escherichia coli Proteobacteria; Gammaproteobacteria; Enterobacterales; Enterobacteriaceae; Escherichia
MNNEETFYQAMRRQGVTRRSFLKYCSLAATSLGLGAGMAPKIAWALENKPRIPVVWIHGL WP_001058323.1
MULTISPECIES: hydrogenase-1 small chain [Proteobacteria] Length: 372\nNumber of predicted TMHs: 2\nExp number of AAs in TMHs: 37.2608499999999\nExp number, first 60 AAs: 19.52923\nTotal prob of N-in: 0.94351\nPOSSIBLE N-term signal sequence\ninside 1 19\nTMhelix 20 39\noutside 40 325\nTMhelix 326 348\ninside 349 372

5051 GCF_000778915.1_ASM77891v1 Escherichia coli Proteobacteria; Gammaproteobacteria; Enterobacterales; Enterobacteriaceae; Escherichia
MTGDNTLIHSHGINRRDFMKLCAALAATMGLSSKAAAEMAESVTNPQRPPVIWIGAQECT WP_000145410.1
MULTISPECIES: hydrogenase-2 small chain [Proteobacteria] Length: 372\nNumber of predicted TMHs: 1\nExp number of AAs in TMHs: 23.72226\nExp number, first 60 AAs: 0.22038\nTotal prob of N-in: 0.12994\noutside 1 331\nTMhelix 332 354\ninside 355 372

5052 GCF_000778745.1_ASM77874v1 Escherichia coli Proteobacteria; Gammaproteobacteria; Enterobacterales; Enterobacteriaceae; Escherichia
MNNEETFYQAMRRQGVTRRSFLKYCSLAATSLGLGAGMAPKIAWALENKPRIPVVWIHGL WP_001058323.1
MULTISPECIES: hydrogenase-1 small chain [Proteobacteria] Length: 372\nNumber of predicted TMHs: 2\nExp number of AAs in TMHs: 37.2608499999999\nExp number, first 60 AAs: 19.52923\nTotal prob of N-in: 0.94351\nPOSSIBLE N-term signal sequence\ninside 1 19\nTMhelix 20 39\noutside 40 325\nTMhelix 326 348\ninside 349 372

5053 GCF_000778815.1_ASM77881v1 Escherichia coli Proteobacteria; Gammaproteobacteria; Enterobacterales; Enterobacteriaceae; Escherichia
MNNEETFYQAMRRQGVTRRSFLKYCSLAATSLGLGAGMAPKIAWALENKPRIPVVWIHGL WP_001058323.1
MULTISPECIES: hydrogenase-1 small chain [Proteobacteria] Length: 372\nNumber of predicted TMHs: 2\nExp number of AAs in TMHs: 37.2608499999999\nExp number, first 60 AAs: 19.52923\nTotal prob of N-in: 0.94351\nPOSSIBLE N-term signal sequence\ninside 1 19\nTMhelix 20 39\noutside 40 325\nTMhelix 326 348\ninside 349 372

5054 GCF_000778815.1_ASM77881v1 Escherichia coli Proteobacteria; Gammaproteobacteria; Enterobacterales; Enterobacteriaceae; Escherichia
MTGDNTLIHSHGINRRDFMKLCAALAATMGLSSKAAAEMAESVTNPQRPPVIWIGAQECT WP_000145410.1
MULTISPECIES: hydrogenase-2 small chain [Proteobacteria] Length: 372\nNumber of predicted TMHs: 1\nExp number of AAs in TMHs: 23.72226\nExp number, first 60 AAs: 0.22038\nTotal prob of N-in: 0.12994\noutside 1 331\nTMhelix 332 354\ninside 355 372

5055 GCF_000778725.1_ASM77872v1 Escherichia coli Proteobacteria; Gammaproteobacteria; Enterobacterales; Enterobacteriaceae; Escherichia
MNNEETFYQAMRRQGVTRRSFLKYCSLAATSLGLGAGMAPKIAWALENKPRIPVVWIHGL WP_001058323.1
MULTISPECIES: hydrogenase-1 small chain [Proteobacteria] Length: 372\nNumber of predicted TMHs: 2\nExp number of AAs in TMHs: 37.2608499999999\nExp number, first 60 AAs: 19.52923\nTotal prob of N-in: 0.94351\nPOSSIBLE N-term signal sequence\ninside 1 19\nTMhelix 20 39\noutside 40 325\nTMhelix 326 348\ninside 349 372

5056 GCF_000778785.1_ASM77878v1 Escherichia coli Proteobacteria; Gammaproteobacteria; Enterobacterales; Enterobacteriaceae; Escherichia
MNNEETFYQAMRRQGVTRRSFLKYCSLAATSLGLGAGMAPKIAWALENKPRIPVVWIHGL WP_001058323.1
MULTISPECIES: hydrogenase-1 small chain [Proteobacteria] Length: 372\nNumber of predicted TMHs: 2\nExp number of AAs in TMHs: 37.2608499999999\nExp number, first 60 AAs: 19.52923\nTotal prob of N-in: 0.94351\nPOSSIBLE N-term signal sequence\ninside 1 19\nTMhelix 20 39\noutside 40 325\nTMhelix 326 348\ninside 349 372

5057 GCF_000778585.1_ASM77858v1 Escherichia coli Proteobacteria; Gammaproteobacteria; Enterobacterales; Enterobacteriaceae; Escherichia
MNNEETFYQAMRRQGVTRRSFLKYCSLAATSLGLGAGMAPKIAWALENKPRIPVVWIHGL WP_001058323.1
MULTISPECIES: hydrogenase-1 small chain [Proteobacteria] Length: 372\nNumber of predicted TMHs: 2\nExp number of AAs in TMHs: 37.2608499999999\nExp number, first 60 AAs: 19.52923\nTotal prob of N-in: 0.94351\nPOSSIBLE N-term signal sequence\ninside 1 19\nTMhelix 20 39\noutside 40 325\nTMhelix 326 348\ninside 349 372

5058 GCF_000778785.1_ASM77878v1 Escherichia coli Proteobacteria; Gammaproteobacteria; Enterobacterales; Enterobacteriaceae; Escherichia
MTGDNTLIHSHGINRRDFMKLCAALAATMGLSSKAAAEMAESVTNPQRPPVIWIGAQECT WP_000145410.1
MULTISPECIES: hydrogenase-2 small chain [Proteobacteria] Length: 372\nNumber of predicted TMHs: 1\nExp number of AAs in TMHs: 23.72226\nExp number, first 60 AAs: 0.22038\nTotal prob of N-in: 0.12994\noutside 1 331\nTMhelix 332 354\ninside 355 372

5059 GCF_000778745.1_ASM77874v1 Escherichia coli Proteobacteria; Gammaproteobacteria; Enterobacterales; Enterobacteriaceae; Escherichia
MTGDNTLIHSHGINRRDFMKLCAALAATMGLSSKAAAEMAESVTNPQRPPVIWIGAQECT WP_000145410.1
MULTISPECIES: hydrogenase-2 small chain [Proteobacteria] Length: 372\nNumber of predicted TMHs: 1\nExp number of AAs in TMHs: 23.72226\nExp number, first 60 AAs: 0.22038\nTotal prob of N-in: 0.12994\noutside 1 331\nTMhelix 332 354\ninside 355 372

5060 GCF_000778335.1_ASM77833v1 Escherichia coli Proteobacteria; Gammaproteobacteria; Enterobacterales; Enterobacteriaceae; Escherichia
MNNEETFYQAMRRQGVTRRSFLKYCSLAATSLGLGAGMAPKIAWALENKPRIPVVWIHGL WP_001058323.1
MULTISPECIES: hydrogenase-1 small chain [Proteobacteria] Length: 372\nNumber of predicted TMHs: 2\nExp number of AAs in TMHs: 37.2608499999999\nExp number, first 60 AAs: 19.52923\nTotal prob of N-in: 0.94351\nPOSSIBLE N-term signal sequence\ninside 1 19\nTMhelix 20 39\noutside 40 325\nTMhelix 326 348\ninside 349 372

5061 GCF_000779585.1_ASM77958v1 Escherichia coli Proteobacteria; Gammaproteobacteria; Enterobacterales; Enterobacteriaceae; Escherichia
MTGDNTLIHSHGINRRDFMKLCAALAATMGLSSKAAAEMAESVTNPQRPPVIWIGAQECT WP_000145410.1
MULTISPECIES: hydrogenase-2 small chain [Proteobacteria] Length: 372\nNumber of predicted TMHs: 1\nExp number of AAs in TMHs: 23.72226\nExp number, first 60 AAs: 0.22038\nTotal prob of N-in: 0.12994\noutside 1 331\nTMhelix 332 354\ninside 355 372

5062 GCF_000778465.1_ASM77846v1 Escherichia coli Proteobacteria; Gammaproteobacteria; Enterobacterales; Enterobacteriaceae; Escherichia
MTGDNTLIHSHGINRRDFMKLCAALAATMGLSSKAAAEMAESVTNPQRPPVIWIGAQECT WP_000145410.1
MULTISPECIES: hydrogenase-2 small chain [Proteobacteria] Length: 372\nNumber of predicted TMHs: 1\nExp number of AAs in TMHs: 23.72226\nExp number, first 60 AAs: 0.22038\nTotal prob of N-in: 0.12994\noutside 1 331\nTMhelix 332 354\ninside 355 372

5063 GCF_000778485.1_ASM77848v1 Escherichia coli Proteobacteria; Gammaproteobacteria; Enterobacterales; Enterobacteriaceae; Escherichia
MTGDNTLIHSHGINRRDFMKLCAALAATMGLSSKAAAEMVESVTNPQRPPVIWIGAQECT WP_000145423.1
hydrogenase 2 small subunit [Escherichia coli] Length: 372\nNumber of predicted TMHs: 1\nExp number of AAs in TMHs: 23.78201\nExp number, first 60 AAs: 0.28355\nTotal prob of N-in: 0.13257\noutside 1 331\nTMhelix 332 354\ninside 355 372

5064 GCF_000778435.1_ASM77843v1 Escherichia coli Proteobacteria; Gammaproteobacteria; Enterobacterales; Enterobacteriaceae; Escherichia
MTGDNTLIHSHGINRRDFMKLCAALAATMGLSSKAAAEMAESVTNPQRPPVIWIGAQECT WP_000145410.1
MULTISPECIES: hydrogenase-2 small chain [Proteobacteria] Length: 372\nNumber of predicted TMHs: 1\nExp number of AAs in TMHs: 23.72226\nExp number, first 60 AAs: 0.22038\nTotal prob of N-in: 0.12994\noutside 1 331\nTMhelix 332 354\ninside 355 372

5065 GCF_000778355.1_ASM77835v1 Escherichia coli Proteobacteria; Gammaproteobacteria; Enterobacterales; Enterobacteriaceae; Escherichia
MTGDNTLIHSHGINRRDFMKLCAALAATMGLSSKAAAEMAESVTNPQRPPVIWIGAQECT WP_000145410.1
MULTISPECIES: hydrogenase-2 small chain [Proteobacteria] Length: 372\nNumber of predicted TMHs: 1\nExp number of AAs in TMHs: 23.72226\nExp number, first 60 AAs: 0.22038\nTotal prob of N-in: 0.12994\noutside 1 331\nTMhelix 332 354\ninside 355 372

5066 GCF_000778395.1_ASM77839v1 Escherichia coli Proteobacteria; Gammaproteobacteria; Enterobacterales; Enterobacteriaceae; Escherichia
MTGDNTLIHSHGINRRDFMKLCAALAATMGLSSKAAAEMAESVTNPQRPPVIWIGAQECT WP_000145410.1
MULTISPECIES: hydrogenase-2 small chain [Proteobacteria] Length: 372\nNumber of predicted TMHs: 1\nExp number of AAs in TMHs: 23.72226\nExp number, first 60 AAs: 0.22038\nTotal prob of N-in: 0.12994\noutside 1 331\nTMhelix 332 354\ninside 355 372

5067 GCF_000778335.1_ASM77833v1 Escherichia coli Proteobacteria; Gammaproteobacteria; Enterobacterales; Enterobacteriaceae; Escherichia
MTGDNTLIHSHGINRRDFMKLCAALAATMGLSSKAAAEMAESVTNPQRPPVIWIGAQECT WP_000145410.1
MULTISPECIES: hydrogenase-2 small chain [Proteobacteria] Length: 372\nNumber of predicted TMHs: 1\nExp number of AAs in TMHs: 23.72226\nExp number, first 60 AAs: 0.22038\nTotal prob of N-in: 0.12994\noutside 1 331\nTMhelix 332 354\ninside 355 372

5068 GCF_000777605.1_ASM77760v1 Escherichia coli Proteobacteria; Gammaproteobacteria; Enterobacterales; Enterobacteriaceae; Escherichia
MNNEETFYQAMRRQGVTRRSFLKYCSLAATSLGLGAGMAPKIAWALENKPRIPVVWIHGL WP_001058323.1
MULTISPECIES: hydrogenase-1 small chain [Proteobacteria] Length: 372\nNumber of predicted TMHs: 2\nExp number of AAs in TMHs: 37.2608499999999\nExp number, first 60 AAs: 19.52923\nTotal prob of N-in: 0.94351\nPOSSIBLE N-term signal sequence\ninside 1 19\nTMhelix 20 39\noutside 40 325\nTMhelix 326 348\ninside 349 372

5069 GCF_000777895.1_ASM77789v1 Escherichia coli Proteobacteria; Gammaproteobacteria; Enterobacterales; Enterobacteriaceae; Escherichia
MNNEETFYQAMRRQGVTRRSFLKYCSLAATSLGLGAGMAPKIAWALENKPRIPVVWIHGL WP_001058323.1
MULTISPECIES: hydrogenase-1 small chain [Proteobacteria] Length: 372\nNumber of predicted TMHs: 2\nExp number of AAs in TMHs: 37.2608499999999\nExp number, first 60 AAs: 19.52923\nTotal prob of N-in: 0.94351\nPOSSIBLE N-term signal sequence\ninside 1 19\nTMhelix 20 39\noutside 40 325\nTMhelix 326 348\ninside 349 372

5070 GCF_000777835.1_ASM77783v1 Escherichia coli Proteobacteria; Gammaproteobacteria; Enterobacterales; Enterobacteriaceae; Escherichia
MNNEETFYQAMRRQGVTRRSFLKYCSLAATSLGLGAGMAPKIAWALENKPRIPVVWIHGL WP_001058323.1
MULTISPECIES: hydrogenase-1 small chain [Proteobacteria] Length: 372\nNumber of predicted TMHs: 2\nExp number of AAs in TMHs: 37.2608499999999\nExp number, first 60 AAs: 19.52923\nTotal prob of N-in: 0.94351\nPOSSIBLE N-term signal sequence\ninside 1 19\nTMhelix 20 39\noutside 40 325\nTMhelix 326 348\ninside 349 372

5071 GCF_000777895.1_ASM77789v1 Escherichia coli Proteobacteria; Gammaproteobacteria; Enterobacterales; Enterobacteriaceae; Escherichia

MTGDNTLIHSHGINRRDFMKLCAALAATMGLSSKAAAEMAESVTNPQRPPVIWIGAQECT WP_000145410.1 MULTISPECIES: hydrogenase-2 small chain [Proteobacteria] Length: 372\nNumber of predicted TMHs: 1\nExp number of AAs in TMHs: 23.72226\nExp number, first 60 AAs: 0.22038\nTotal prob of N-in: 0.12994\noutside 1 331\nTMhelix 332 354\ninside 355 372

5072 GCF_000777835.1_ASM77783v1 Escherichia coli Proteobacteria; Gammaproteobacteria; Enterobacterales; Enterobacteriaceae; Escherichia MTGDNTLIHSHGINRRDFMKLCAALAATMGLSSKAAAEMAESVTNPQRPPVIWIGAQECT WP_000145410.1 MULTISPECIES: hydrogenase-2 small chain [Proteobacteria] Length: 372\nNumber of predicted TMHs: 1\nExp number of AAs in TMHs: 23.72226\nExp number, first 60 AAs: 0.22038\nTotal prob of N-in: 0.12994\noutside 1 331\nTMhelix 332 354\ninside 355 372

5073 GCF_000777655.1_ASM77765v1 Escherichia coli Proteobacteria; Gammaproteobacteria; Enterobacterales; Enterobacteriaceae; Escherichia MNNEETFYQAMRRQGVTRRSFLKYCSLAATSLGLGAGMAPKIAWALENKPRIPVVWIHGL WP_001058323.1 MULTISPECIES: hydrogenase-1 small chain [Proteobacteria] Length: 372\nNumber of predicted TMHs: 2\nExp number of AAs in TMHs: 37.2608499999999\nExp number, first 60 AAs: 19.52923\nTotal prob of N-in: 0.94351\nPOSSIBLE N-term signal sequence\ninside 1 19\nTMhelix 20 39\noutside 40 325\nTMhelix 326 348\ninside 349 372

5074 GCF_000777735.1_ASM77773v1 Escherichia coli Proteobacteria; Gammaproteobacteria; Enterobacterales; Enterobacteriaceae; Escherichia MNNEETFYQAMRRQGVTRRSFLKYCSLAATSLGLGAGMAPKIAWALENKPRIPVVWIHGL WP_001058323.1 MULTISPECIES: hydrogenase-1 small chain [Proteobacteria] Length: 372\nNumber of predicted TMHs: 2\nExp number of AAs in TMHs: 37.2608499999999\nExp number, first 60 AAs: 19.52923\nTotal prob of N-in: 0.94351\nPOSSIBLE N-term signal sequence\ninside 1 19\nTMhelix 20 39\noutside 40 325\nTMhelix 326 348\ninside 349 372

5075 GCF_000777735.1_ASM77773v1 Escherichia coli Proteobacteria; Gammaproteobacteria; Enterobacterales; Enterobacteriaceae; Escherichia MTGDNTLIHSHGINRRDFMKLCAALAATMGLSSKAAAEMAESVTNPQRPPVIWIGAQECT WP_000145410.1 MULTISPECIES: hydrogenase-2 small chain [Proteobacteria] Length: 372\nNumber of predicted TMHs: 1\nExp number of AAs in TMHs: 23.72226\nExp number, first 60 AAs: 0.22038\nTotal prob of N-in: 0.12994\noutside 1 331\nTMhelix 332 354\ninside 355 372

5076 GCF_000777435.1_ASM77743v1 Escherichia coli Proteobacteria; Gammaproteobacteria; Enterobacterales; Enterobacteriaceae; Escherichia MTGDNTLIHSHGINRRDFMKLCAALAATMGLSSKAAAEMAESVTNPQRPPVIWIGAQECT WP_050009873.1 hydrogenase 2 small subunit [Escherichia coli] Length: 372\nNumber of predicted TMHs: 1\nExp number of AAs in TMHs: 23.71371\nExp number, first 60 AAs: 0.2205\nTotal prob of N-in: 0.12969\noutside 1 331\nTMhelix 332 354\ninside 355 372

5077 GCF_000777535.1_ASM77753v1 Escherichia coli Proteobacteria; Gammaproteobacteria; Enterobacterales; Enterobacteriaceae; Escherichia MNNEETFYQAMRRQGVTRRSFLKYCSLAATSLGLGAGMAPKIAWALENKPRIPVVWIHGL WP_001058323.1 MULTISPECIES: hydrogenase-1 small chain [Proteobacteria] Length: 372\nNumber of predicted TMHs: 2\nExp number of AAs in TMHs: 37.2608499999999\nExp number, first 60 AAs: 19.52923\nTotal prob of N-in: 0.94351\nPOSSIBLE N-term signal sequence\ninside 1 19\nTMhelix 20 39\noutside 40 325\nTMhelix 326 348\ninside 349 372

5078 GCF_000777695.1_ASM77769v1 Escherichia coli Proteobacteria; Gammaproteobacteria; Enterobacterales; Enterobacteriaceae; Escherichia MNNEETFYQAMRRQGVTRRSFLKYCSLAATSLGLGAGMAPKIAWALENKPRIPVVWIHGL WP_001058323.1 MULTISPECIES: hydrogenase-1 small chain [Proteobacteria] Length: 372\nNumber of predicted TMHs: 2\nExp number of AAs in TMHs: 37.2608499999999\nExp number, first 60 AAs: 19.52923\nTotal prob of N-in: 0.94351\nPOSSIBLE N-term signal sequence\ninside 1 19\nTMhelix 20 39\noutside 40 325\nTMhelix 326 348\ninside 349 372

5079 GCF_000777695.1_ASM77769v1 Escherichia coli Proteobacteria; Gammaproteobacteria; Enterobacterales; Enterobacteriaceae; Escherichia MTGDNTLIHSHGINRRDFMKLCAALAATMGLSSKAAAEMAESVTNPQRPPVIWIGAQECT WP_000145410.1

MULTISPECIES: hydrogenase-2 small chain [Proteobacteria] Length: 372\nNumber of predicted TMHs: 1\nExp number of AAs in TMHs: 23.72226\nExp number, first 60 AAs: 0.22038\nTotal prob of N-in: 0.12994\noutside 1 331\nTMhelix 332 354\ninside 355 372

5080 GCF_000777495.1_ASM77749v1 Escherichia coli Proteobacteria; Gammaproteobacteria; Enterobacterales; Enterobacteriaceae; Escherichia
MNNEETFYQAMRRQGVTRRSFLKYCSLAATSLGLGAGMAPKIAWALENKPRIPVVWIHGL WP_001058323.1
MULTISPECIES: hydrogenase-1 small chain [Proteobacteria] Length: 372\nNumber of predicted TMHs: 2\nExp number of AAs in TMHs: 37.2608499999999\nExp number, first 60 AAs: 19.52923\nTotal prob of N-in: 0.94351\nPOSSIBLE N-term signal sequence\ninside 1 19\nTMhelix 20 39\noutside 40 325\nTMhelix 326 348\ninside 349 372

5081 GCF_000777535.1_ASM77753v1 Escherichia coli Proteobacteria; Gammaproteobacteria; Enterobacterales; Enterobacteriaceae; Escherichia
MTGDNTLIHSHGINRRDFMKLCAALAATMGLSSKAAAEMAESVTNPQRPPVIWIGAQECT WP_000145410.1
MULTISPECIES: hydrogenase-2 small chain [Proteobacteria] Length: 372\nNumber of predicted TMHs: 1\nExp number of AAs in TMHs: 23.72226\nExp number, first 60 AAs: 0.22038\nTotal prob of N-in: 0.12994\noutside 1 331\nTMhelix 332 354\ninside 355 372

5082 GCF_000777435.1_ASM77743v1 Escherichia coli Proteobacteria; Gammaproteobacteria; Enterobacterales; Enterobacteriaceae; Escherichia
MNNEETFYQAMRRQGVTRRSFLKYCSLAATSLGLGAGMAPKIAWALENKPRIPVVWIHGL WP_001058323.1
MULTISPECIES: hydrogenase-1 small chain [Proteobacteria] Length: 372\nNumber of predicted TMHs: 2\nExp number of AAs in TMHs: 37.2608499999999\nExp number, first 60 AAs: 19.52923\nTotal prob of N-in: 0.94351\nPOSSIBLE N-term signal sequence\ninside 1 19\nTMhelix 20 39\noutside 40 325\nTMhelix 326 348\ninside 349 372

5083 GCF_000777195.1_ASM77719v1 Escherichia coli Proteobacteria; Gammaproteobacteria; Enterobacterales; Enterobacteriaceae; Escherichia
MNNEETFYQAMRRQGVTRRSFLKYCSLAATSLGLGAGMAPKIAWALENKPRIPVVWIHGL WP_001295940.1
MULTISPECIES: hydrogenase [Enterobacteriaceae] Length: 372\nNumber of predicted TMHs: 2\nExp number of AAs in TMHs: 37.2608899999999\nExp number, first 60 AAs: 19.52923\nTotal prob of N-in: 0.94351\nPOSSIBLE N-term signal sequence\ninside 1 19\nTMhelix 20 39\noutside 40 325\nTMhelix 326 348\ninside 349 372

5084 GCF_000777475.1_ASM77747v1 Escherichia coli Proteobacteria; Gammaproteobacteria; Enterobacterales; Enterobacteriaceae; Escherichia
MNNEETFYQAMRRQGVTRRSFLKYCSLAATSLGLGAGMAPKIAWALENKPRIPVVWIHGL WP_001295940.1
MULTISPECIES: hydrogenase [Enterobacteriaceae] Length: 372\nNumber of predicted TMHs: 2\nExp number of AAs in TMHs: 37.2608899999999\nExp number, first 60 AAs: 19.52923\nTotal prob of N-in: 0.94351\nPOSSIBLE N-term signal sequence\ninside 1 19\nTMhelix 20 39\noutside 40 325\nTMhelix 326 348\ninside 349 372

5085 GCF_000777475.1_ASM77747v1 Escherichia coli Proteobacteria; Gammaproteobacteria; Enterobacterales; Enterobacteriaceae; Escherichia
MTGDNTLIHSHGINRRDFMKLCAALAATMGLSSKAAAEMAESVTNPQRPPVIWIGAQECT WP_000145410.1
MULTISPECIES: hydrogenase-2 small chain [Proteobacteria] Length: 372\nNumber of predicted TMHs: 1\nExp number of AAs in TMHs: 23.72226\nExp number, first 60 AAs: 0.22038\nTotal prob of N-in: 0.12994\noutside 1 331\nTMhelix 332 354\ninside 355 372

5086 GCF_000777235.1_ASM77723v1 Escherichia coli Proteobacteria; Gammaproteobacteria; Enterobacterales; Enterobacteriaceae; Escherichia
MNNEETFYQAMRRQGVTRRSFLKYCSLAATSLGLGAGMAPKIAWALENKPRIPVVWIHGL WP_001538532.1
hydrogenase-1 small chain [Escherichia coli] Length: 372\nNumber of predicted TMHs: 2\nExp number of AAs in TMHs: 37.27041\nExp number, first 60 AAs: 19.52927\nTotal prob of N-in: 0.94352\nPOSSIBLE N-term signal sequence\ninside 1 19\nTMhelix 20 39\noutside 40 325\nTMhelix 326 348\ninside 349 372

5087 GCF_000777155.1_ASM77715v1 Escherichia coli Proteobacteria; Gammaproteobacteria; Enterobacterales; Enterobacteriaceae; Escherichia
MNNEETFYQAMRRQGVTRRSFLKYCSLAATSLGLGAGMAPKIAWALENKPRIPVVWIHGL WP_001295940.1
MULTISPECIES: hydrogenase [Enterobacteriaceae] Length: 372\nNumber of predicted TMHs: 2\nExp

number of AAs in TMHs: 37.2608899999999\nExp number, first 60 AAs: 19.52923\nTotal prob of N-in: 0.94351\nPOSSIBLE N-term signal sequence\ninside 1 19\nTMhelix 20 39\noutside 40 325\nTMhelix 326 348\ninside 349 372

5088 GCF_000777095.1_ASM77709v1 Escherichia coli Proteobacteria; Gammaproteobacteria; Enterobacterales; Enterobacteriaceae; Escherichia
MTGDNTLIHSHGINRRDFMKLCAALAATMGLSSKAAAEMAESVTNPQRPPVIWIGAQECT WP_042097182.1
hydrogenase 2 small subunit [Escherichia coli] Length: 372\nNumber of predicted TMHs: 1\nExp number of AAs in TMHs: 23.26313\nExp number, first 60 AAs: 0.22087\nTotal prob of N-in: 0.13038\noutside 1 331\nTMhelix 332 354\ninside 355 372

5089 GCF_000308975.1_ASM30897v2 Escherichia coli Proteobacteria; Gammaproteobacteria; Enterobacterales; Enterobacteriaceae; Escherichia
MTGDNTLIHSHGINRRDFMKLCAALAATMGLSSKAAAEMAESVTNPQRPPVIWIGAQECT WP_000145410.1
MULTISPECIES: hydrogenase-2 small chain [Proteobacteria] Length: 372\nNumber of predicted TMHs: 1\nExp number of AAs in TMHs: 23.72226\nExp number, first 60 AAs: 0.22038\nTotal prob of N-in: 0.12994\noutside 1 331\nTMhelix 332 354\ninside 355 372

5090 GCF_000308975.1_ASM30897v2 Escherichia coli Proteobacteria; Gammaproteobacteria; Enterobacterales; Enterobacteriaceae; Escherichia
MNNEETFYQAMRRQGVTRRSFLKYCSLAATSLGLGAGMAPKIAWALENKPRIPVVWIHGL WP_001058320.1
hydrogenase [Escherichia coli] Length: 372\nNumber of predicted TMHs: 2\nExp number of AAs in TMHs: 37.26082\nExp number, first 60 AAs: 19.52923\nTotal prob of N-in: 0.94351\nPOSSIBLE N-term signal sequence\ninside 1 19\nTMhelix 20 39\noutside 40 325\nTMhelix 326 348\ninside 349 372

5091 GCF_000777135.1_ASM77713v1 Escherichia coli Proteobacteria; Gammaproteobacteria; Enterobacterales; Enterobacteriaceae; Escherichia
MNNEETFYQAMRRQGVTRRSFLKYCSLAATSLGLGAGMAPKIAWALENKPRIPVVWIHGL WP_001058323.1
MULTISPECIES: hydrogenase-1 small chain [Proteobacteria] Length: 372\nNumber of predicted TMHs: 2\nExp number of AAs in TMHs: 37.2608499999999\nExp number, first 60 AAs: 19.52923\nTotal prob of N-in: 0.94351\nPOSSIBLE N-term signal sequence\ninside 1 19\nTMhelix 20 39\noutside 40 325\nTMhelix 326 348\ninside 349 372

5092 GCF_000777195.1_ASM77719v1 Escherichia coli Proteobacteria; Gammaproteobacteria; Enterobacterales; Enterobacteriaceae; Escherichia
MTGDNTLIHSHGINRRDFMKLCAALAATMGLSSKAAAEMAESVTNPQRPPVIWIGAQECT WP_000145410.1
MULTISPECIES: hydrogenase-2 small chain [Proteobacteria] Length: 372\nNumber of predicted TMHs: 1\nExp number of AAs in TMHs: 23.72226\nExp number, first 60 AAs: 0.22038\nTotal prob of N-in: 0.12994\noutside 1 331\nTMhelix 332 354\ninside 355 372

5093 GCF_000777095.1_ASM77709v1 Escherichia coli Proteobacteria; Gammaproteobacteria; Enterobacterales; Enterobacteriaceae; Escherichia
MNNEETFYQAMRRQGVTRRSFLKYCSLAATSLGLGAGMAPKIAWALENKPRIPVVWIHGL WP_001058323.1
MULTISPECIES: hydrogenase-1 small chain [Proteobacteria] Length: 372\nNumber of predicted TMHs: 2\nExp number of AAs in TMHs: 37.2608499999999\nExp number, first 60 AAs: 19.52923\nTotal prob of N-in: 0.94351\nPOSSIBLE N-term signal sequence\ninside 1 19\nTMhelix 20 39\noutside 40 325\nTMhelix 326 348\ninside 349 372

5094 GCF_000777155.1_ASM77715v1 Escherichia coli Proteobacteria; Gammaproteobacteria; Enterobacterales; Enterobacteriaceae; Escherichia
MTGDNTLIHSHGINRRDFMKLCAALAATMGLSSKAAAEMVESVTNPQRPPVIWIGAQECT WP_000145423.1
hydrogenase 2 small subunit [Escherichia coli] Length: 372\nNumber of predicted TMHs: 1\nExp number of AAs in TMHs: 23.78201\nExp number, first 60 AAs: 0.28355\nTotal prob of N-in: 0.13257\noutside 1 331\nTMhelix 332 354\ninside 355 372

5095 GCF_000777135.1_ASM77713v1 Escherichia coli Proteobacteria; Gammaproteobacteria; Enterobacterales; Enterobacteriaceae; Escherichia
MTGDNTLIHSHGINRRDFMKLCAALAATMGLSSKAAAEMAESVTNPQRPPVIWIGAQECT WP_000145410.1
MULTISPECIES: hydrogenase-2 small chain [Proteobacteria] Length: 372\nNumber of predicted TMHs: 1\nExp number of AAs in TMHs: 23.72226\nExp number, first 60 AAs: 0.22038\nTotal prob of N-in: 0.12994\noutside 1 331\nTMhelix 332 354\ninside 355 372

5096 GCF_000776745.1_ASM77674v1 Escherichia coli Proteobacteria; Gammaproteobacteria; Enterobacterales; Enterobacteriaceae; Escherichia
MNNEETFYQAMRRQGVTRRSFLKYCSLAATSLGLGAGMAPKIAWALENKPRIPVVWIHGL WP_021520630.1
hydrogenase-1 small chain [Escherichia coli] Length: 372\nNumber of predicted TMHs: 2\nExp number of AAs in TMHs: 37.8197599999999\nExp number, first 60 AAs: 19.53295\nTotal prob of N-in: 0.94396\nPOSSIBLE N-term signal sequence\ninside 1 19\nTMhelix 20 39\noutside 40 325\nTMhelix 326 348\ninside 349 372

5097 GCF_000776935.1_ASM77693v1 Escherichia coli Proteobacteria; Gammaproteobacteria; Enterobacterales; Enterobacteriaceae; Escherichia
MTGDNTLIHSHGINRRDFMKLCAALAATMGLSSKAAAEMAESVTNPQRPPVIWIGAQECT WP_000145410.1
MULTISPECIES: hydrogenase-2 small chain [Proteobacteria] Length: 372\nNumber of predicted TMHs: 1\nExp number of AAs in TMHs: 23.72226\nExp number, first 60 AAs: 0.22038\nTotal prob of N-in: 0.12994\noutside 1 331\nTMhelix 332 354\ninside 355 372

5098 GCF_000776915.1_ASM77691v1 Escherichia coli Proteobacteria; Gammaproteobacteria; Enterobacterales; Enterobacteriaceae; Escherichia
MTGDNTLIHSHGINRRDFMKLCAALAATMGLSSKAAAEMAESVTNPQRPPVIWIGAQECT WP_000145410.1
MULTISPECIES: hydrogenase-2 small chain [Proteobacteria] Length: 372\nNumber of predicted TMHs: 1\nExp number of AAs in TMHs: 23.72226\nExp number, first 60 AAs: 0.22038\nTotal prob of N-in: 0.12994\noutside 1 331\nTMhelix 332 354\ninside 355 372

5099 GCF_000776815.1_ASM77681v1 Escherichia coli Proteobacteria; Gammaproteobacteria; Enterobacterales; Enterobacteriaceae; Escherichia
MTGDNTLIHSHGINRRDFMKLCAALAATMGLSSKAAAEMAESVTNPQRPPVIWIGAQECT WP_000145410.1
MULTISPECIES: hydrogenase-2 small chain [Proteobacteria] Length: 372\nNumber of predicted TMHs: 1\nExp number of AAs in TMHs: 23.72226\nExp number, first 60 AAs: 0.22038\nTotal prob of N-in: 0.12994\noutside 1 331\nTMhelix 332 354\ninside 355 372

5100 GCF_000776855.1_ASM77685v1 Escherichia coli Proteobacteria; Gammaproteobacteria; Enterobacterales; Enterobacteriaceae; Escherichia
MTGDNTLIHSHGINRRDFMKLCAALAATMGLSSKAAAEMAESVTNPQRPPVIWIGAQECT WP_000145410.1
MULTISPECIES: hydrogenase-2 small chain [Proteobacteria] Length: 372\nNumber of predicted TMHs: 1\nExp number of AAs in TMHs: 23.72226\nExp number, first 60 AAs: 0.22038\nTotal prob of N-in: 0.12994\noutside 1 331\nTMhelix 332 354\ninside 355 372

5101 GCF_000776775.1_ASM77677v1 Escherichia coli Proteobacteria; Gammaproteobacteria; Enterobacterales; Enterobacteriaceae; Escherichia
MTGDNTLIHSHGINRRDFMKLCAALAATMGLSSKAAAEMAESVTNPQRPPVIWIGAQECT WP_000145410.1
MULTISPECIES: hydrogenase-2 small chain [Proteobacteria] Length: 372\nNumber of predicted TMHs: 1\nExp number of AAs in TMHs: 23.72226\nExp number, first 60 AAs: 0.22038\nTotal prob of N-in: 0.12994\noutside 1 331\nTMhelix 332 354\ninside 355 372

5102 GCF_000776675.1_ASM77667v1 Escherichia coli Proteobacteria; Gammaproteobacteria; Enterobacterales; Enterobacteriaceae; Escherichia
MNNEETFYQAMRRQGVTRRSFLKYCSLAATSLGLGAGMAPKIAWALENKPRIPVVWIHGL WP_001058323.1
MULTISPECIES: hydrogenase-1 small chain [Proteobacteria] Length: 372\nNumber of predicted TMHs: 2\nExp number of AAs in TMHs: 37.2608499999999\nExp number, first 60 AAs: 19.52923\nTotal prob of N-in: 0.94351\nPOSSIBLE N-term signal sequence\ninside 1 19\nTMhelix 20 39\noutside 40 325\nTMhelix 326 348\ninside 349 372

5103 GCF_000776565.1_ASM77656v1 Escherichia coli Proteobacteria; Gammaproteobacteria; Enterobacterales; Enterobacteriaceae; Escherichia
MTGDNTLIHSHGINRRDFMKLCAALAATMGLSSKAAAEMAESVTNPQRPPVIWIGAQECT WP_000145410.1
MULTISPECIES: hydrogenase-2 small chain [Proteobacteria] Length: 372\nNumber of predicted TMHs: 1\nExp number of AAs in TMHs: 23.72226\nExp number, first 60 AAs: 0.22038\nTotal prob of N-in: 0.12994\noutside 1 331\nTMhelix 332 354\ninside 355 372

5104 GCF_000776435.1_ASM77643v1 Escherichia coli Proteobacteria; Gammaproteobacteria; Enterobacterales; Enterobacteriaceae; Escherichia
MNNEETFYQAMRRQGVTRRSFLKYCSLAATSLGLGAGMAPKIAWALENKPRIPVVWIHGL WP_001058323.1
MULTISPECIES: hydrogenase-1 small chain [Proteobacteria] Length: 372\nNumber of predicted TMHs:

2\nExp number of AAs in TMHs: 37.2608499999999\nExp number, first 60 AAs: 19.52923\nTotal prob of N-in: 0.94351\nPOSSIBLE N-term signal sequence\ninside 1 19\nTMhelix 20 39\noutside 40 325\nTMhelix 326 348\ninside 349 372

5105 GCF_000776415.1_ASM77641v1 Escherichia coli Proteobacteria; Gammaproteobacteria; Enterobacterales; Enterobacteriaceae; Escherichia MTGDNTLIHSHGINRRDFMKLCAALAATMGLSSKAAAEMAESVTNPQRPPVIWIGAQECT WP_021519494.1 hydrogenase-2 small chain [Escherichia coli] Length: 372\nNumber of predicted TMHs: 1\nExp number of AAs in TMHs: 23.72331\nExp number, first 60 AAs: 0.22039\nTotal prob of N-in: 0.12995\noutside 1 331\nTMhelix 332 354\ninside 355 372

5106 GCF_000776535.1_ASM77653v1 Escherichia coli Proteobacteria; Gammaproteobacteria; Enterobacterales; Enterobacteriaceae; Escherichia MNNEETFYQAMRRQGVTRRSFLKYCSLAATSLGLGAGMAPKIAWALENKPRIPVVWIHGL WP_001058323.1 MULTISPECIES: hydrogenase-1 small chain [Proteobacteria] Length: 372\nNumber of predicted TMHs: 2\nExp number of AAs in TMHs: 37.2608499999999\nExp number, first 60 AAs: 19.52923\nTotal prob of N-in: 0.94351\nPOSSIBLE N-term signal sequence\ninside 1 19\nTMhelix 20 39\noutside 40 325\nTMhelix 326 348\ninside 349 372

5107 GCF_000776415.1_ASM77641v1 Escherichia coli Proteobacteria; Gammaproteobacteria; Enterobacterales; Enterobacteriaceae; Escherichia MNNEETFYQAMRRQGVTRRSFLKYCSLAATSLGLGAGMAPKIAWALENKPRIPVVWIHGL WP_001058323.1 MULTISPECIES: hydrogenase-1 small chain [Proteobacteria] Length: 372\nNumber of predicted TMHs: 2\nExp number of AAs in TMHs: 37.2608499999999\nExp number, first 60 AAs: 19.52923\nTotal prob of N-in: 0.94351\nPOSSIBLE N-term signal sequence\ninside 1 19\nTMhelix 20 39\noutside 40 325\nTMhelix 326 348\ninside 349 372

5108 GCF_000776535.1_ASM77653v1 Escherichia coli Proteobacteria; Gammaproteobacteria; Enterobacterales; Enterobacteriaceae; Escherichia MTGDNTLIHSHGINRRDFMKLCAALAATMGLSSKAAAEMAESVTNPQRPPVIWIGAQECT WP_000145410.1 MULTISPECIES: hydrogenase-2 small chain [Proteobacteria] Length: 372\nNumber of predicted TMHs: 1\nExp number of AAs in TMHs: 23.72226\nExp number, first 60 AAs: 0.22038\nTotal prob of N-in: 0.12994\noutside 1 331\nTMhelix 332 354\ninside 355 372

5109 GCF_000776475.1_ASM77647v1 Escherichia coli Proteobacteria; Gammaproteobacteria; Enterobacterales; Enterobacteriaceae; Escherichia MTGDNTLIHSHGINRRDFMKLCAALAATMGLSSKAAAEMAESVTNPQRPPVIWIGAQECT WP_000145410.1 MULTISPECIES: hydrogenase-2 small chain [Proteobacteria] Length: 372\nNumber of predicted TMHs: 1\nExp number of AAs in TMHs: 23.72226\nExp number, first 60 AAs: 0.22038\nTotal prob of N-in: 0.12994\noutside 1 331\nTMhelix 332 354\ninside 355 372

5110 GCF_000776215.1_ASM77621v1 Escherichia coli Proteobacteria; Gammaproteobacteria; Enterobacterales; Enterobacteriaceae; Escherichia MTGDNTLIHSHGINRRDFMKLCAALAATMGLSSKAAAEMAESVTNPQRPPVIWIGAQECT WP_000145410.1 MULTISPECIES: hydrogenase-2 small chain [Proteobacteria] Length: 372\nNumber of predicted TMHs: 1\nExp number of AAs in TMHs: 23.72226\nExp number, first 60 AAs: 0.22038\nTotal prob of N-in: 0.12994\noutside 1 331\nTMhelix 332 354\ninside 355 372

5111 GCF_000776175.1_ASM77617v1 Escherichia coli Proteobacteria; Gammaproteobacteria; Enterobacterales; Enterobacteriaceae; Escherichia MTGDNTLIHSHGINRRDFMKLCAALAATMGLSSKAAAEMAESVTNPQRPPVIWIGAQECT WP_000145410.1 MULTISPECIES: hydrogenase-2 small chain [Proteobacteria] Length: 372\nNumber of predicted TMHs: 1\nExp number of AAs in TMHs: 23.72226\nExp number, first 60 AAs: 0.22038\nTotal prob of N-in: 0.12994\noutside 1 331\nTMhelix 332 354\ninside 355 372

5112 GCF_000776135.1_ASM77613v1 Escherichia coli Proteobacteria; Gammaproteobacteria; Enterobacterales; Enterobacteriaceae; Escherichia MTGDNTLIHSHGINRRDFMKLCAALAATMGLSSKAAAEMAESVTNPQRPPVIWIGAQECT WP_000145410.1 MULTISPECIES: hydrogenase-2 small chain [Proteobacteria] Length: 372\nNumber of predicted TMHs: 1\nExp number of AAs in TMHs: 23.72226\nExp number, first 60 AAs: 0.22038\nTotal prob of N-in: 0.12994\noutside 1 331\nTMhelix 332 354\ninside 355 372

5113 GCF_000770275.1_ASM77027v1 Escherichia coli Proteobacteria; Gammaproteobacteria; Enterobacterales; Enterobacteriaceae; Escherichia MNNEETFYQAMRRQGVTRRSFLKYCSLAATSLGLGAGMAPKIAWALENKPRIPVVWIHGL WP_001295940.1 MULTISPECIES: hydrogenase [Enterobacteriaceae] Length: 372\nNumber of predicted TMHs: 2\nExp number of AAs in TMHs: 37.2608899999999\nExp number, first 60 AAs: 19.52923\nTotal prob of N-in: 0.94351\nPOSSIBLE N-term signal sequence\ninside 1 19\nTMhelix 20 39\noutside 40 325\nTMhelix 326 348\ninside 349 372

5114 GCF_000773595.1_ASM77359v1 Escherichia coli Proteobacteria; Gammaproteobacteria; Enterobacterales; Enterobacteriaceae; Escherichia MNNEETFYQAMRRQGVTRRSFLKYCSLAATSLGLGAGMAPKIAWALENKPRIPVVWIHGL WP_001058323.1 MULTISPECIES: hydrogenase-1 small chain [Proteobacteria] Length: 372\nNumber of predicted TMHs: 2\nExp number of AAs in TMHs: 37.2608499999999\nExp number, first 60 AAs: 19.52923\nTotal prob of N-in: 0.94351\nPOSSIBLE N-term signal sequence\ninside 1 19\nTMhelix 20 39\noutside 40 325\nTMhelix 326 348\ninside 349 372

5115 GCF_000773475.1_ASM77347v1 Escherichia coli Proteobacteria; Gammaproteobacteria; Enterobacterales; Enterobacteriaceae; Escherichia MNNEETFYQAMRRQGVTRRSFLKYCSLAATSLGLGAGMAPKIAWALENKPRIPVVWIHGL WP_001058323.1 MULTISPECIES: hydrogenase-1 small chain [Proteobacteria] Length: 372\nNumber of predicted TMHs: 2\nExp number of AAs in TMHs: 37.2608499999999\nExp number, first 60 AAs: 19.52923\nTotal prob of N-in: 0.94351\nPOSSIBLE N-term signal sequence\ninside 1 19\nTMhelix 20 39\noutside 40 325\nTMhelix 326 348\ninside 349 372

5116 GCF_000770055.1_ASM77005v1 Escherichia coli Proteobacteria; Gammaproteobacteria; Enterobacterales; Enterobacteriaceae; Escherichia MNNEETFYQAMRRQGVTRRSFLKYCSLAATSLGLGAGMAPKIAWALENKPRIPVVWIHGL WP_001058323.1 MULTISPECIES: hydrogenase-1 small chain [Proteobacteria] Length: 372\nNumber of predicted TMHs: 2\nExp number of AAs in TMHs: 37.2608499999999\nExp number, first 60 AAs: 19.52923\nTotal prob of N-in: 0.94351\nPOSSIBLE N-term signal sequence\ninside 1 19\nTMhelix 20 39\noutside 40 325\nTMhelix 326 348\ninside 349 372

5117 GCF_000776435.1_ASM77643v1 Escherichia coli Proteobacteria; Gammaproteobacteria; Enterobacterales; Enterobacteriaceae; Escherichia MTGDNTLIHSHGINRRDFMKLCAALAATMGLSSKAAAEMVESVTNPQRPPVIWIGAQECT WP_000145423.1 hydrogenase 2 small subunit [Escherichia coli] Length: 372\nNumber of predicted TMHs: 1\nExp number of AAs in TMHs: 23.78201\nExp number, first 60 AAs: 0.28355\nTotal prob of N-in: 0.13257\noutside 1 331\nTMhelix 332 354\ninside 355 372

5118 GCF_000776035.1_ASM77603v1 Escherichia coli Proteobacteria; Gammaproteobacteria; Enterobacterales; Enterobacteriaceae; Escherichia MNNEETFYQAMRRQGVTRRSFLKYCSLAATSLGLGAGMAPKIAWALENKPRIPVVWIHGL WP_001058323.1 MULTISPECIES: hydrogenase-1 small chain [Proteobacteria] Length: 372\nNumber of predicted TMHs: 2\nExp number of AAs in TMHs: 37.2608499999999\nExp number, first 60 AAs: 19.52923\nTotal prob of N-in: 0.94351\nPOSSIBLE N-term signal sequence\ninside 1 19\nTMhelix 20 39\noutside 40 325\nTMhelix 326 348\ninside 349 372

5119 GCF_000773555.1_ASM77355v1 Escherichia coli Proteobacteria; Gammaproteobacteria; Enterobacterales; Enterobacteriaceae; Escherichia MNNEETFYQAMRRQGVTRRSFLKYCSLAATSLGLGAGMAPKIAWALENKPRIPVVWIHGL WP_001058323.1 MULTISPECIES: hydrogenase-1 small chain [Proteobacteria] Length: 372\nNumber of predicted TMHs: 2\nExp number of AAs in TMHs: 37.2608499999999\nExp number, first 60 AAs: 19.52923\nTotal prob of N-in: 0.94351\nPOSSIBLE N-term signal sequence\ninside 1 19\nTMhelix 20 39\noutside 40 325\nTMhelix 326 348\ninside 349 372

5120 GCF_000776055.1_ASM77605v1 Escherichia coli Proteobacteria; Gammaproteobacteria; Enterobacterales; Enterobacteriaceae; Escherichia MTGDNTLIHSHGINRRDFMKLCAALAATMGLSSKAAAEMAESVTNPQRPPVIWIGAQECT WP_000145410.1 MULTISPECIES: hydrogenase-2 small chain [Proteobacteria] Length: 372\nNumber of predicted TMHs: 1\nExp number of AAs in TMHs: 23.72226\nExp number, first 60 AAs: 0.22038\nTotal prob of N-in: 0.12994\noutside 1 331\nTMhelix 332 354\ninside 355 372

5121 GCF_000776035.1_ASM77603v1 Escherichia coli Proteobacteria; Gammaproteobacteria; Enterobacterales; Enterobacteriaceae; Escherichia
MTGDNTLIHSHGINRRDFMKLCAALAATMGLSSKAAAEMAESVTNPQRPPVIWIGAQECT WP_000145410.1
MULTISPECIES: hydrogenase-2 small chain [Proteobacteria] Length: 372\nNumber of predicted TMHs: 1\nExp number of AAs in TMHs: 23.72226\nExp number, first 60 AAs: 0.22038\nTotal prob of N-in: 0.12994\noutside 1 331\nTMhelix 332 354\ninside 355 372

5122 GCF_000773595.1_ASM77359v1 Escherichia coli Proteobacteria; Gammaproteobacteria; Enterobacterales; Enterobacteriaceae; Escherichia
MTGDNTLIHSHGINRRDFMKLCAALAATMGLSSKAAAEMAESVTNPQRPPVIWIGAQECT WP_000145410.1
MULTISPECIES: hydrogenase-2 small chain [Proteobacteria] Length: 372\nNumber of predicted TMHs: 1\nExp number of AAs in TMHs: 23.72226\nExp number, first 60 AAs: 0.22038\nTotal prob of N-in: 0.12994\noutside 1 331\nTMhelix 332 354\ninside 355 372

5123 GCF_000762385.1_ASM76238v1 Escherichia coli Proteobacteria; Gammaproteobacteria; Enterobacterales; Enterobacteriaceae; Escherichia
MNNEETFYQAMRRQGVTRRSFLKYCSLAATSLGLGAGMAPKIAWALENKPRIPVVWIHGL WP_001349947.1
hydrogenase [Escherichia coli] Length: 372\nNumber of predicted TMHs: 2\nExp number of AAs in TMHs: 37.2615299999999\nExp number, first 60 AAs: 19.52923\nTotal prob of N-in: 0.94351\nPOSSIBLE N-term signal sequence\ninside 1 19\nTMhelix 20 39\noutside 40 325\nTMhelix 326 348\ninside 349 372

5124 GCF_000768505.1_ASM76850v1 Escherichia coli Proteobacteria; Gammaproteobacteria; Enterobacterales; Enterobacteriaceae; Escherichia
MNNEETFYQAMRRQGVTRRSFLKYCSLAATSLGLGAGMAPKIAWALENKPRIPVVWIHGL WP_001058323.1
MULTISPECIES: hydrogenase-1 small chain [Proteobacteria] Length: 372\nNumber of predicted TMHs: 2\nExp number of AAs in TMHs: 37.2608499999999\nExp number, first 60 AAs: 19.52923\nTotal prob of N-in: 0.94351\nPOSSIBLE N-term signal sequence\ninside 1 19\nTMhelix 20 39\noutside 40 325\nTMhelix 326 348\ninside 349 372

5125 GCF_000768505.1_ASM76850v1 Escherichia coli Proteobacteria; Gammaproteobacteria; Enterobacterales; Enterobacteriaceae; Escherichia
MTGDNTLIHSHGINRRDFMKLCAALAATMGLSSKAAAEMAESVTNPQRPPVIWIGAQECT WP_000145410.1
MULTISPECIES: hydrogenase-2 small chain [Proteobacteria] Length: 372\nNumber of predicted TMHs: 1\nExp number of AAs in TMHs: 23.72226\nExp number, first 60 AAs: 0.22038\nTotal prob of N-in: 0.12994\noutside 1 331\nTMhelix 332 354\ninside 355 372

5126 GCF_000753535.1_FHI54 Escherichia coli Proteobacteria; Gammaproteobacteria; Enterobacterales; Enterobacteriaceae; Escherichia MNNEETFYQAMRRQGVTRRSFLKYCSLAATSLGLGAGMAPKIAWALENKPRIPVVWIHGL
WP_001717971.1 hydrogenase-1 small chain [Escherichia coli] Length: 372\nNumber of predicted TMHs: 2\nExp number of AAs in TMHs: 38.00708\nExp number, first 60 AAs: 19.5286\nTotal prob of N-in: 0.94354\nPOSSIBLE N-term signal sequence\ninside 1 19\nTMhelix 20 39\noutside 40 325\nTMhelix 326 348\ninside 349 372

5127 GCF_000768465.1_ASM76846v1 Escherichia coli Proteobacteria; Gammaproteobacteria; Enterobacterales; Enterobacteriaceae; Escherichia
MNNEETFYQAMRRQGVTRRSFLKYCSLAATSLGLGAGMAPKIAWALENKPRIPVVWIHGL WP_001058323.1
MULTISPECIES: hydrogenase-1 small chain [Proteobacteria] Length: 372\nNumber of predicted TMHs: 2\nExp number of AAs in TMHs: 37.2608499999999\nExp number, first 60 AAs: 19.52923\nTotal prob of N-in: 0.94351\nPOSSIBLE N-term signal sequence\ninside 1 19\nTMhelix 20 39\noutside 40 325\nTMhelix 326 348\ninside 349 372

5128 GCF_000768465.1_ASM76846v1 Escherichia coli Proteobacteria; Gammaproteobacteria; Enterobacterales; Enterobacteriaceae; Escherichia
MTGDNTLIHSHGINRRDFMKLCAALAATMGLSSKAAAEMAESVTNPQRPPVIWIGAQECT WP_000145410.1
MULTISPECIES: hydrogenase-2 small chain [Proteobacteria] Length: 372\nNumber of predicted TMHs: 1\nExp number of AAs in TMHs: 23.72226\nExp number, first 60 AAs: 0.22038\nTotal prob of N-in: 0.12994\noutside 1 331\nTMhelix 332 354\ninside 355 372

5129 GCF_000768415.1_ASM76841v1 Escherichia coli Proteobacteria; Gammaproteobacteria; Enterobacterales; Enterobacteriaceae; Escherichia
MNNEETFYQAMRRQGVTRRSFLKYCSLAATSLGLGAGMAPKIAWALENKPRIPVVWIHGL WP_001058323.1
MULTISPECIES: hydrogenase-1 small chain [Proteobacteria] Length: 372\nNumber of predicted TMHs:

2\nExp number of AAs in TMHs: 37.2608499999999\nExp number, first 60 AAs: 19.52923\nTotal prob of N-in: 0.94351\nPOSSIBLE N-term signal sequence\ninside 1 19\nTMhelix 20 39\noutside 40 325\nTMhelix 326 348\ninside 349 372

5130 GCF_000768415.1_ASM76841v1 Escherichia coli Proteobacteria; Gammaproteobacteria; Enterobacterales; Enterobacteriaceae; Escherichia MTGDNTLIHSHGINRRDFMKLCAALAATMGLSSKAAAEMAESVTNPQRPPVIWIGAQECT WP_000145410.1 MULTISPECIES: hydrogenase-2 small chain [Proteobacteria] Length: 372\nNumber of predicted TMHs: 1\nExp number of AAs in TMHs: 23.72226\nExp number, first 60 AAs: 0.22038\nTotal prob of N-in: 0.12994\noutside 1 331\nTMhelix 332 354\ninside 355 372

5131 GCF_000762385.1_ASM76238v1 Escherichia coli Proteobacteria; Gammaproteobacteria; Enterobacterales; Enterobacteriaceae; Escherichia MTGDNTLIHSHGINRRDFMKLCAALAATMGLSSKAAAEMAESVTNPQRPPVIWIGAQECT WP_000145410.1 MULTISPECIES: hydrogenase-2 small chain [Proteobacteria] Length: 372\nNumber of predicted TMHs: 1\nExp number of AAs in TMHs: 23.72226\nExp number, first 60 AAs: 0.22038\nTotal prob of N-in: 0.12994\noutside 1 331\nTMhelix 332 354\ninside 355 372

5132 GCF_000754855.1_ASM75485v1 Escherichia coli Proteobacteria; Gammaproteobacteria; Enterobacterales; Enterobacteriaceae; Escherichia MTGDNTLIHSHGINRRDFMKLCAALAATMGLSSKAAAEMAESVTNPQRPPVIWIGAQECT WP_000145410.1 MULTISPECIES: hydrogenase-2 small chain [Proteobacteria] Length: 372\nNumber of predicted TMHs: 1\nExp number of AAs in TMHs: 23.72226\nExp number, first 60 AAs: 0.22038\nTotal prob of N-in: 0.12994\noutside 1 331\nTMhelix 332 354\ninside 355 372

5133 GCF_000755445.1_ASM75544v1 Escherichia coli Proteobacteria; Gammaproteobacteria; Enterobacterales; Enterobacteriaceae; Escherichia MTGDNTLIHSHGINRRDFMKLCAALAATMGLSSKAAAEMAESVTNPQRPPVIWIGAQECT WP_000145410.1 MULTISPECIES: hydrogenase-2 small chain [Proteobacteria] Length: 372\nNumber of predicted TMHs: 1\nExp number of AAs in TMHs: 23.72226\nExp number, first 60 AAs: 0.22038\nTotal prob of N-in: 0.12994\noutside 1 331\nTMhelix 332 354\ninside 355 372

5134 GCF_000753535.1_FHI54 Escherichia coli Proteobacteria; Gammaproteobacteria; Enterobacterales; Enterobacteriaceae; Escherichia MTGDNTLIHSHGINRRDFMKLCAALAATMGLSSKAAAEMAESVTNPQRPPVIWIGAQECT WP_000145410.1 MULTISPECIES: hydrogenase-2 small chain [Proteobacteria] Length: 372\nNumber of predicted TMHs: 1\nExp number of AAs in TMHs: 23.72226\nExp number, first 60 AAs: 0.22038\nTotal prob of N-in: 0.12994\noutside 1 331\nTMhelix 332 354\ninside 355 372

5135 GCF_000753515.1_FHI13 Escherichia coli Proteobacteria; Gammaproteobacteria; Enterobacterales; Enterobacteriaceae; Escherichia MTGDNTLIHSHGINRRDFMKLCAALAATMGLSSKAAAEMAESVTNPQRPPVIWIGAQECT WP_000145410.1 MULTISPECIES: hydrogenase-2 small chain [Proteobacteria] Length: 372\nNumber of predicted TMHs: 1\nExp number of AAs in TMHs: 23.72226\nExp number, first 60 AAs: 0.22038\nTotal prob of N-in: 0.12994\noutside 1 331\nTMhelix 332 354\ninside 355 372

5136 GCF_000753315.1_FHI35 Escherichia coli Proteobacteria; Gammaproteobacteria; Enterobacterales; Enterobacteriaceae; Escherichia MTGDNTLIHSHGINRRDFMKLCAALAATMGLSSKAAAEMAESVTNPQRPPVIWIGAQECT WP_000145410.1 MULTISPECIES: hydrogenase-2 small chain [Proteobacteria] Length: 372\nNumber of predicted TMHs: 1\nExp number of AAs in TMHs: 23.72226\nExp number, first 60 AAs: 0.22038\nTotal prob of N-in: 0.12994\noutside 1 331\nTMhelix 332 354\ninside 355 372

5137 GCF_000753155.1_FHI88 Escherichia coli Proteobacteria; Gammaproteobacteria; Enterobacterales; Enterobacteriaceae; Escherichia MNNEETFYQAMRRQGVTRRSFLKYCSLAATSLGLGAGMAPKIAWALENKPRIPVVWIHGL WP_001058320.1 hydrogenase [Escherichia coli] Length: 372\nNumber of predicted TMHs: 2\nExp number of AAs in TMHs: 37.26082\nExp number, first 60 AAs: 19.52923\nTotal prob of N-in: 0.94351\nPOSSIBLE N-term signal sequence\ninside 1 19\nTMhelix 20 39\noutside 40 325\nTMhelix 326 348\ninside 349 372

5138 GCF_000753155.1_FHI88 Escherichia coli Proteobacteria; Gammaproteobacteria; Enterobacterales; Enterobacteriaceae; Escherichia MTGDNTLIHSHGINRRDFMKLCAALAATMGLSSKAAAEMAESVTNPQRPPVIWIGAQECT WP_000145410.1 MULTISPECIES: hydrogenase-2 small chain [Proteobacteria] Length: 372\nNumber of predicted TMHs: 1\nExp number of AAs in TMHs: 23.72226\nExp number, first 60 AAs: 0.22038\nTotal prob of N-in: 0.12994\noutside 1 331\nTMhelix 332 354\ninside 355 372

5139 GCF_000752915.1_FHI77 Escherichia coli Proteobacteria; Gammaproteobacteria; Enterobacterales; Enterobacteriaceae; Escherichia MNNEETFYQAMRRQGVTRRSFLKYCSLAATSLGLGAGMAPKIAWALENKPRIPVVWIHGL WP_001058323.1 MULTISPECIES: hydrogenase-1 small chain [Proteobacteria] Length: 372\nNumber of predicted TMHs: 2\nExp number of AAs in TMHs: 37.2608499999999\nExp number, first 60 AAs: 19.52923\nTotal prob of N-in: 0.94351\nPOSSIBLE N-term signal sequence\ninside 1 19\nTMhelix 20 39\noutside 40 325\nTMhelix 326 348\ninside 349 372

5140 GCF_000752875.1_FHI39 Escherichia coli Proteobacteria; Gammaproteobacteria; Enterobacterales; Enterobacteriaceae; Escherichia MNNEETFYQAMRRQGVTRRSFLKYCSLAATSLGLGAGMAPKIAWALENKPRIPVVWIHGL WP_001058323.1 MULTISPECIES: hydrogenase-1 small chain [Proteobacteria] Length: 372\nNumber of predicted TMHs: 2\nExp number of AAs in TMHs: 37.2608499999999\nExp number, first 60 AAs: 19.52923\nTotal prob of N-in: 0.94351\nPOSSIBLE N-term signal sequence\ninside 1 19\nTMhelix 20 39\noutside 40 325\nTMhelix 326 348\ninside 349 372

5141 GCF_000752955.1_FHI91 Escherichia coli Proteobacteria; Gammaproteobacteria; Enterobacterales; Enterobacteriaceae; Escherichia MNNEETFYQAMRRQGVTRRSFLKYCSLAATSLGLGAGMAPKIAWALENKPRIPVVWIHGL WP_001058323.1 MULTISPECIES: hydrogenase-1 small chain [Proteobacteria] Length: 372\nNumber of predicted TMHs: 2\nExp number of AAs in TMHs: 37.2608499999999\nExp number, first 60 AAs: 19.52923\nTotal prob of N-in: 0.94351\nPOSSIBLE N-term signal sequence\ninside 1 19\nTMhelix 20 39\noutside 40 325\nTMhelix 326 348\ninside 349 372

5142 GCF_000752735.1_FHI56 Escherichia coli Proteobacteria; Gammaproteobacteria; Enterobacterales; Enterobacteriaceae; Escherichia MNNEETFYQAMRRQGVTRRSFLKYCSLAATSLGLGAGMAPKIAWALENKPRIPVVWIHGL WP_001295940.1 MULTISPECIES: hydrogenase [Enterobacteriaceae] Length: 372\nNumber of predicted TMHs: 2\nExp number of AAs in TMHs: 37.2608899999999\nExp number, first 60 AAs: 19.52923\nTotal prob of N-in: 0.94351\nPOSSIBLE N-term signal sequence\ninside 1 19\nTMhelix 20 39\noutside 40 325\nTMhelix 326 348\ninside 349 372

5143 GCF_000752835.1_FHI53 Escherichia coli Proteobacteria; Gammaproteobacteria; Enterobacterales; Enterobacteriaceae; Escherichia MNNEETFYQAMRRQGVTRRSFLKYCSLAATSLGLGAGMAPKIAWALENKPRIPVVWIHGL WP_001058323.1 MULTISPECIES: hydrogenase-1 small chain [Proteobacteria] Length: 372\nNumber of predicted TMHs: 2\nExp number of AAs in TMHs: 37.2608499999999\nExp number, first 60 AAs: 19.52923\nTotal prob of N-in: 0.94351\nPOSSIBLE N-term signal sequence\ninside 1 19\nTMhelix 20 39\noutside 40 325\nTMhelix 326 348\ninside 349 372

5144 GCF_000752835.1_FHI53 Escherichia coli Proteobacteria; Gammaproteobacteria; Enterobacterales; Enterobacteriaceae; Escherichia MTGDNTLIHSHGINRRDFMKLCAALAATMGLSSKAAAEMAESVTNPQRPPVIWIGAQECT WP_000145410.1 MULTISPECIES: hydrogenase-2 small chain [Proteobacteria] Length: 372\nNumber of predicted TMHs: 1\nExp number of AAs in TMHs: 23.72226\nExp number, first 60 AAs: 0.22038\nTotal prob of N-in: 0.12994\noutside 1 331\nTMhelix 332 354\ninside 355 372

5145 GCF_000752635.1_FHI84 Escherichia coli Proteobacteria; Gammaproteobacteria; Enterobacterales; Enterobacteriaceae; Escherichia MNNEETFYQAMRRQGVTRRSFLKYCSLAATSLGLGAGMAPKIAWALENKPRIPVVWIHGL WP_001058323.1 MULTISPECIES: hydrogenase-1 small chain [Proteobacteria] Length: 372\nNumber of predicted TMHs: 2\nExp number of AAs in TMHs: 37.2608499999999\nExp number, first 60 AAs: 19.52923\nTotal prob of N-in: 0.94351\nPOSSIBLE N-term signal sequence\ninside 1 19\nTMhelix 20 39\noutside 40 325\nTMhelix 326 348\ninside 349 372

5146 GCF_000752775.1_FHI5 Escherichia coli Proteobacteria; Gammaproteobacteria; Enterobacterales; Enterobacteriaceae; Escherichia MTGDNTLIHSHGINRRDFMKLCAALAATMGLSSKAAAEMAESVTNPQRPPVIWIGAQECT WP_000145410.1 MULTISPECIES: hydrogenase-2 small chain [Proteobacteria] Length: 372\nNumber of predicted TMHs: 1\nExp number of AAs in TMHs: 23.72226\nExp number, first 60 AAs: 0.22038\nTotal prob of N-in: 0.12994\noutside 1 331\nTMhelix 332 354\ninside 355 372

5147 GCF_000752735.1_FHI56 Escherichia coli Proteobacteria; Gammaproteobacteria; Enterobacterales; Enterobacteriaceae; Escherichia MTGDNTLIHSHGINRRDFMKLCAALAATMGLSSKAAAEMAESVTNPQRPPVIWIGAQECT WP_000145410.1 MULTISPECIES: hydrogenase-2 small chain [Proteobacteria] Length: 372\nNumber of predicted TMHs: 1\nExp number of AAs in TMHs: 23.72226\nExp number, first 60 AAs: 0.22038\nTotal prob of N-in: 0.12994\noutside 1 331\nTMhelix 332 354\ninside 355 372

5148 GCF_000752655.1_FHI47 Escherichia coli Proteobacteria; Gammaproteobacteria; Enterobacterales; Enterobacteriaceae; Escherichia MTGDNTLIHSHGINRRDFMKLCAALAATMGLSSKAAAEMAESVTNPQRPPVIWIGAQECT WP_000145410.1 MULTISPECIES: hydrogenase-2 small chain [Proteobacteria] Length: 372\nNumber of predicted TMHs: 1\nExp number of AAs in TMHs: 23.72226\nExp number, first 60 AAs: 0.22038\nTotal prob of N-in: 0.12994\noutside 1 331\nTMhelix 332 354\ninside 355 372

5149 GCF_000752635.1_FHI84 Escherichia coli Proteobacteria; Gammaproteobacteria; Enterobacterales; Enterobacteriaceae; Escherichia MTGDNTLIHSHGINRRDFMKLCAALAATMGLSSKAAAEMAESVTNPQRPPVIWIGAQECT WP_000145410.1 MULTISPECIES: hydrogenase-2 small chain [Proteobacteria] Length: 372\nNumber of predicted TMHs: 1\nExp number of AAs in TMHs: 23.72226\nExp number, first 60 AAs: 0.22038\nTotal prob of N-in: 0.12994\noutside 1 331\nTMhelix 332 354\ninside 355 372

5150 GCF_000752495.1_FHI2 Escherichia coli Proteobacteria; Gammaproteobacteria; Enterobacterales; Enterobacteriaceae; Escherichia MNNEETFYQAMRRQGVTRRSFLKYCSLAATSLGLGAGMAPKIAWALENKPRIPVVWIHGL WP_001058323.1 MULTISPECIES: hydrogenase-1 small chain [Proteobacteria] Length: 372\nNumber of predicted TMHs: 2\nExp number of AAs in TMHs: 37.2608499999999\nExp number, first 60 AAs: 19.52923\nTotal prob of N-in: 0.94351\nPOSSIBLE N-term signal sequence\ninside 1 19\nTMhelix 20 39\noutside 40 325\nTMhelix 326 348\ninside 349 372

5151 GCF_000752555.1_FHI76 Escherichia coli Proteobacteria; Gammaproteobacteria; Enterobacterales; Enterobacteriaceae; Escherichia MTGDNTLIHSHGINRRDFMKLCAALAATMGLSSKAAAEMVESVTNPQRPPVIWIGAQECT WP_000145423.1 hydrogenase 2 small subunit [Escherichia coli] Length: 372\nNumber of predicted TMHs: 1\nExp number of AAs in TMHs: 23.78201\nExp number, first 60 AAs: 0.28355\nTotal prob of N-in: 0.13257\noutside 1 331\nTMhelix 332 354\ninside 355 372

5152 GCF_000752415.1_FHI68 Escherichia coli Proteobacteria; Gammaproteobacteria; Enterobacterales; Enterobacteriaceae; Escherichia MNNEETFYQAMRRQGVTRRSFLKYCSLAATSLGLGAGMAPKIAWALENKPRIPVVWIHGL WP_001058323.1 MULTISPECIES: hydrogenase-1 small chain [Proteobacteria] Length: 372\nNumber of predicted TMHs: 2\nExp number of AAs in TMHs: 37.2608499999999\nExp number, first 60 AAs: 19.52923\nTotal prob of N-in: 0.94351\nPOSSIBLE N-term signal sequence\ninside 1 19\nTMhelix 20 39\noutside 40 325\nTMhelix 326 348\ninside 349 372

5153 GCF_000752495.1_FHI2 Escherichia coli Proteobacteria; Gammaproteobacteria; Enterobacterales; Enterobacteriaceae; Escherichia MTGDNTLIHSHGINRRDFMKLCAALAATMGLSSKAAAEMAESVTNPQRPPVIWIGAQECT WP_000145410.1 MULTISPECIES: hydrogenase-2 small chain [Proteobacteria] Length: 372\nNumber of predicted TMHs: 1\nExp number of AAs in TMHs: 23.72226\nExp number, first 60 AAs: 0.22038\nTotal prob of N-in: 0.12994\noutside 1 331\nTMhelix 332 354\ninside 355 372

5154 GCF_000752435.1_FHI1 Escherichia coli Proteobacteria; Gammaproteobacteria; Enterobacterales; Enterobacteriaceae; Escherichia MTGDNTLIHSHGINRRDFMKLCAALAATMGLSSKAAAEMAESVTNPQRPPVIWIGAQECT WP_000145410.1 MULTISPECIES: hydrogenase-2 small chain [Proteobacteria] Length: 372\nNumber of predicted TMHs: 1\nExp number of AAs in TMHs: 23.72226\nExp number, first 60 AAs: 0.22038\nTotal prob of N-in: 0.12994\noutside 1 331\nTMhelix 332 354\ninside 355 372

5155 GCF_000752315.1_FHI32 Escherichia coli Proteobacteria; Gammaproteobacteria; Enterobacterales; Enterobacteriaceae; Escherichia MTGDNTLIHSHGINRRDFMKLCAALAATMGLSSKAAAEMAESVTNPQRPPVIWIGAQECT WP_000145410.1 MULTISPECIES: hydrogenase-2 small chain [Proteobacteria] Length: 372\nNumber of predicted TMHs: 1\nExp number of AAs in TMHs: 23.72226\nExp number, first 60 AAs: 0.22038\nTotal prob of N-in: 0.12994\noutside 1 331\nTMhelix 332 354\ninside 355 372

5156 GCF_000752415.1_FHI68 Escherichia coli Proteobacteria; Gammaproteobacteria; Enterobacterales; Enterobacteriaceae; Escherichia MTGDNTLIHSHGINRRDFMKLCAALAATMGLSSKAAAEMVESVTNPQRPPVIWIGAQECT WP_000145423.1 hydrogenase 2 small subunit [Escherichia coli] Length: 372\nNumber of predicted TMHs: 1\nExp number of AAs in TMHs: 23.78201\nExp number, first 60 AAs: 0.28355\nTotal prob of N-in: 0.13257\noutside 1 331\nTMhelix 332 354\ninside 355 372

5157 GCF_000752135.1_FHI51 Escherichia coli Proteobacteria; Gammaproteobacteria; Enterobacterales; Enterobacteriaceae; Escherichia MNNEETFYQAMRRQGVTRRSFLKYCSLAATSLGLGAGMAPKIAWALENKPRIPVVWIHGL WP_024246676.1 hydrogenase-1 small chain [Escherichia coli] Length: 372\nNumber of predicted TMHs: 2\nExp number of AAs in TMHs: 37.8214899999999\nExp number, first 60 AAs: 19.52326\nTotal prob of N-in:

0.94395\nPOSSIBLE N-term signal sequence\ninside 1 19\nTMhelix 20 39\noutside 40 325\nTMhelix 326 348\ninside 349 372

5158 GCF_000752135.1_FHI51 Escherichia coli Proteobacteria; Gammaproteobacteria; Enterobacterales; Enterobacteriaceae; Escherichia MTGDNTLIHSHGINRRDFMKLCAALAATMGLSSKAAAEMAESVTNPQRPPVIWIGAQECT WP_000145410.1 MULTISPECIES: hydrogenase-2 small chain [Proteobacteria] Length: 372\nNumber of predicted TMHs: 1\nExp number of AAs in TMHs: 23.72226\nExp number, first 60 AAs: 0.22038\nTotal prob of N-in: 0.12994\noutside 1 331\nTMhelix 332 354\ninside 355 372

5159 GCF_000751315.1_assembly1 Escherichia coli Proteobacteria; Gammaproteobacteria; Enterobacterales; Enterobacteriaceae; Escherichia MNNEETFYQAMRRQGVTRRSFLKYCSLAATSLGLGAGMAPKIAWALENKPRIPVVWIHGL WP_001058323.1 MULTISPECIES: hydrogenase-1 small chain [Proteobacteria] Length: 372\nNumber of predicted TMHs: 2\nExp number of AAs in TMHs: 37.2608499999999\nExp number, first 60 AAs: 19.52923\nTotal prob of N-in: 0.94351\nPOSSIBLE N-term signal sequence\ninside 1 19\nTMhelix 20 39\noutside 40 325\nTMhelix 326 348\ninside 349 372

5160 GCF_000751315.1_assembly1 Escherichia coli Proteobacteria; Gammaproteobacteria; Enterobacterales; Enterobacteriaceae; Escherichia MTGDNTLIHSHGINRRDFMKLCAALAATMGLSSKAAAEMAESVTNPQRPPVIWIGAQECT WP_000145410.1 MULTISPECIES: hydrogenase-2 small chain [Proteobacteria] Length: 372\nNumber of predicted TMHs: 1\nExp number of AAs in TMHs: 23.72226\nExp number, first 60 AAs: 0.22038\nTotal prob of N-in: 0.12994\noutside 1 331\nTMhelix 332 354\ninside 355 372

5161 GCF_000736735.1_ASM73673v1 Escherichia coli Proteobacteria; Gammaproteobacteria; Enterobacterales; Enterobacteriaceae; Escherichia MNNEETFYQAMRRQGVTRRSFLKYCSLAATSLGLGAGMAPKIAWALENKPRIPVVWIHGL WP_001058323.1 MULTISPECIES: hydrogenase-1 small chain [Proteobacteria] Length: 372\nNumber of predicted TMHs: 2\nExp number of AAs in TMHs: 37.2608499999999\nExp number, first 60 AAs: 19.52923\nTotal prob of N-in: 0.94351\nPOSSIBLE N-term signal sequence\ninside 1 19\nTMhelix 20 39\noutside 40 325\nTMhelix 326 348\ninside 349 372

5162 GCF_000731455.1_V1 Escherichia coli Proteobacteria; Gammaproteobacteria; Enterobacterales; Enterobacteriaceae; Escherichia MNNEETFYQAMRRQGVTRRSFLKYCSLAATSLGLGAGMAPKIAWALENKPRIPVVWIHGL WP_001058323.1 MULTISPECIES: hydrogenase-1 small chain [Proteobacteria] Length: 372\nNumber of predicted TMHs: 2\nExp number of AAs in TMHs: 37.2608499999999\nExp number, first 60 AAs: 19.52923\nTotal prob of N-in: 0.94351\nPOSSIBLE N-term signal sequence\ninside 1 19\nTMhelix 20 39\noutside 40 325\nTMhelix 326 348\ninside 349 372

5163 GCF_000715035.1_UCD_JA23_pb Escherichia coli Proteobacteria; Gammaproteobacteria; Enterobacterales; Enterobacteriaceae; Escherichia MNNEETFYQAMRRQGVTRRSFLKYCSLAATSLGLGAGMAPKIAWALENKPRIPVVWIHGL WP_001058323.1 MULTISPECIES: hydrogenase-1 small chain [Proteobacteria] Length: 372\nNumber of predicted TMHs: 2\nExp number of AAs in TMHs: 37.2608499999999\nExp number, first 60 AAs: 19.52923\nTotal prob of N-in: 0.94351\nPOSSIBLE N-term signal sequence\ninside 1 19\nTMhelix 20 39\noutside 40 325\nTMhelix 326 348\ninside 349 372

5164 GCF_000708165.1_E_coli_EC2.contigs Escherichia coli Proteobacteria; Gammaproteobacteria; Enterobacterales; Enterobacteriaceae; Escherichia MNNEETFYQAMRRQGVTRRSFLKYCSLAATSLGLGAGMAPKIAWALENKPRIPVVWIHGL WP_001058323.1 MULTISPECIES: hydrogenase-1 small chain [Proteobacteria] Length: 372\nNumber of predicted TMHs: 2\nExp number of AAs in TMHs: 37.2608499999999\nExp number, first 60 AAs: 19.52923\nTotal prob of N-in: 0.94351\nPOSSIBLE N-term signal sequence\ninside 1 19\nTMhelix 20 39\noutside 40 325\nTMhelix 326 348\ninside 349 372

5165 GCF_000696835.1_ASM69683v1 Escherichia coli Proteobacteria; Gammaproteobacteria; Enterobacterales; Enterobacteriaceae; Escherichia MNNEETFYQAMRRQGVTRRSFLKYCSLAATSLGLGAGMAPKIAWALENKPRIPVVWIHGL WP_001058323.1 MULTISPECIES: hydrogenase-1 small chain [Proteobacteria] Length: 372\nNumber of predicted TMHs: 2\nExp number of AAs in TMHs: 37.2608499999999\nExp number, first 60 AAs: 19.52923\nTotal prob of N-in:

0.94351\nPOSSIBLE N-term signal sequence\ninside 1 19\nTMhelix 20 39\noutside 40 325\nTMhelix 326 348\ninside 349 372

5166 GCF_000696835.1_ASM69683v1 Escherichia coli Proteobacteria; Gammaproteobacteria; Enterobacterales; Enterobacteriaceae; Escherichia MTGDNTLIHSHGINRRDFMKLCAALAATMGLSSKAAAEMAESVTNPQRPPVIWIGAQECT WP_000145410.1 MULTISPECIES: hydrogenase-2 small chain [Proteobacteria] Length: 372\nNumber of predicted TMHs: 1\nExp number of AAs in TMHs: 23.72226\nExp number, first 60 AAs: 0.22038\nTotal prob of N-in: 0.12994\noutside 1 331\nTMhelix 332 354\ninside 355 372

5167 GCF_000695505.1_SCB34 Escherichia coli Proteobacteria; Gammaproteobacteria; Enterobacterales; Enterobacteriaceae; Escherichia MNNEETFYQAMRRQGVTRRSFLKYCSLAATSLGLGAGMAPKIAWALENKPRIPVVWIHGL WP_001295940.1 MULTISPECIES: hydrogenase [Enterobacteriaceae] Length: 372\nNumber of predicted TMHs: 2\nExp number of AAs in TMHs: 37.2608899999999\nExp number, first 60 AAs: 19.52923\nTotal prob of N-in: 0.94351\nPOSSIBLE N-term signal sequence\ninside 1 19\nTMhelix 20 39\noutside 40 325\nTMhelix 326 348\ninside 349 372

5168 GCF_000695135.1_Ecoli97-3250v1 Escherichia coli Proteobacteria; Gammaproteobacteria; Enterobacterales; Enterobacteriaceae; Escherichia MNNEETFYQAMRRQGVTRRSFLKYCSLAATSLGLGAGMAPKIAWALENKPRIPVVWIHGL WP_001295940.1 MULTISPECIES: hydrogenase [Enterobacteriaceae] Length: 372\nNumber of predicted TMHs: 2\nExp number of AAs in TMHs: 37.2608899999999\nExp number, first 60 AAs: 19.52923\nTotal prob of N-in: 0.94351\nPOSSIBLE N-term signal sequence\ninside 1 19\nTMhelix 20 39\noutside 40 325\nTMhelix 326 348\ninside 349 372

5169 GCF_000695175.1_EcoliMI10-01v1 Escherichia coli Proteobacteria; Gammaproteobacteria; Enterobacterales; Enterobacteriaceae; Escherichia MNNEETFYQAMRRQGVTRRSFLKYCSLAATSLGLGAGMAPKIAWALENKPRIPVVWIHGL WP_001058323.1 MULTISPECIES: hydrogenase-1 small chain [Proteobacteria] Length: 372\nNumber of predicted TMHs: 2\nExp number of AAs in TMHs: 37.2608499999999\nExp number, first 60 AAs: 19.52923\nTotal prob of N-in: 0.94351\nPOSSIBLE N-term signal sequence\ninside 1 19\nTMhelix 20 39\noutside 40 325\nTMhelix 326 348\ninside 349 372

5170 GCF_000691025.1_S6662 Escherichia coli Proteobacteria; Gammaproteobacteria; Enterobacterales; Enterobacteriaceae; Escherichia MNNEETFYQAMRRQGVTRRSFLKYCSLAATSLGLGAGMAPKIAWALENKPRIPVVWIHGL WP_001295940.1 MULTISPECIES: hydrogenase [Enterobacteriaceae] Length: 372\nNumber of predicted TMHs: 2\nExp number of AAs in TMHs: 37.2608899999999\nExp number, first 60 AAs: 19.52923\nTotal prob of N-in: 0.94351\nPOSSIBLE N-term signal sequence\ninside 1 19\nTMhelix 20 39\noutside 40 325\nTMhelix 326 348\ninside 349 372

5171 GCF_000690985.1_S7005 Escherichia coli Proteobacteria; Gammaproteobacteria; Enterobacterales; Enterobacteriaceae; Escherichia MNNEETFYQAMRRQGVTRRSFLKYCSLAATSLGLGAGMAPKIAWALENKPRIPVVWIHGL WP_001295940.1 MULTISPECIES: hydrogenase [Enterobacteriaceae] Length: 372\nNumber of predicted TMHs: 2\nExp number of AAs in TMHs: 37.2608899999999\nExp number, first 60 AAs: 19.52923\nTotal prob of N-in: 0.94351\nPOSSIBLE N-term signal sequence\ninside 1 19\nTMhelix 20 39\noutside 40 325\nTMhelix 326 348\ninside 349 372

5172 GCF_000691085.1_S6995 Escherichia coli Proteobacteria; Gammaproteobacteria; Enterobacterales; Enterobacteriaceae; Escherichia MTGDNTLIHSHGINRRDFMKLCAALAATMGLSSKAAAEMAESVTNPQRPPVIWIGAQECT WP_000145410.1 MULTISPECIES: hydrogenase-2 small chain [Proteobacteria] Length: 372\nNumber of predicted TMHs: 1\nExp number of AAs in TMHs: 23.72226\nExp number, first 60 AAs: 0.22038\nTotal prob of N-in: 0.12994\noutside 1 331\nTMhelix 332 354\ninside 355 372

5173 GCF_000691025.1_S6662 Escherichia coli Proteobacteria; Gammaproteobacteria; Enterobacterales; Enterobacteriaceae; Escherichia MTGDNTLIHSHGINRRDFMKLCAALAATMGLSSKAAAEMAESVTNPQRPPVIWIGAQECT WP_000145410.1 MULTISPECIES: hydrogenase-2 small chain [Proteobacteria] Length: 372\nNumber of predicted TMHs: 1\nExp number of AAs in TMHs: 23.72226\nExp number, first 60 AAs: 0.22038\nTotal prob of N-in: 0.12994\noutside 1 331\nTMhelix 332 354\ninside 355 372

5174 GCF_000691065.1_S6966 Escherichia coli Proteobacteria; Gammaproteobacteria; Enterobacterales; Enterobacteriaceae; Escherichia MTGDNTLIHSHGINRRDFMKLCAALAATMGLSSKAAAEMAESVTNPQRPPVIWIGAQECT

WP_000145410.1 MULTISPECIES: hydrogenase-2 small chain [Proteobacteria] Length: 372\nNumber of predicted TMHs: 1\nExp number of AAs in TMHs: 23.72226\nExp number, first 60 AAs: 0.22038\nTotal prob of N-in: 0.12994\noutside 1 331\nTMhelix 332 354\ninside 355 372

5175 GCF_000690985.1_S7005 Escherichia coli Proteobacteria; Gammaproteobacteria; Enterobacterales; Enterobacteriaceae; Escherichia MTGDNTLIHSHGINRRDFMKLCAALAATMGLSSKAAAEMAESVTNPQHPPVIWIGAQECT WP_000145403.1 hydrogenase 2 small subunit [Escherichia coli] Length: 372\nNumber of predicted TMHs: 1\nExp number of AAs in TMHs: 23.66365\nExp number, first 60 AAs: 0.1537\nTotal prob of N-in: 0.12709\noutside 1 331\nTMhelix 332 354\ninside 355 372

5176 GCF_000681435.1_IEH71520_1.0 Escherichia coli Proteobacteria; Gammaproteobacteria; Enterobacterales; Enterobacteriaceae; Escherichia MNNEETFYQAMRRQGVTRRSFLKYCSLAATSLGLGAGMAPKIAWALENKPRIPVVWIHGL WP_001295940.1 MULTISPECIES: hydrogenase [Enterobacteriaceae] Length: 372\nNumber of predicted TMHs: 2\nExp number of AAs in TMHs: 37.2608899999999\nExp number, first 60 AAs: 19.52923\nTotal prob of N-in: 0.94351\nPOSSIBLE N-term signal sequence\ninside 1 19\nTMhelix 20 39\noutside 40 325\nTMhelix 326 348\ninside 349 372

5177 GCF_000647795.1_ASM64779v2 Escherichia coli Proteobacteria; Gammaproteobacteria; Enterobacterales; Enterobacteriaceae; Escherichia MNNEETFYQAMRRQGVTRRSFLKYCSLAATSLGLGAGMAPKIAWALENKPRIPVVWIHGL WP_032161483.1 hydrogenase [Escherichia coli] Length: 372\nNumber of predicted TMHs: 2\nExp number of AAs in TMHs: 38.07604\nExp number, first 60 AAs: 19.52977\nTotal prob of N-in: 0.94353\nPOSSIBLE N-term signal sequence\ninside 1 19\nTMhelix 20 39\noutside 40 325\nTMhelix 326 348\ninside 349 372

5178 GCF_000599765.2_a5_miseq_linux_20140113 Escherichia coli Proteobacteria; Gammaproteobacteria; Enterobacterales; Enterobacteriaceae; Escherichia MTGDNTLIHSHGINRRDFMKLCAALAATMGLSSKAAAEMAESVTNPQRPPVIWIGAQECT WP_000145410.1 MULTISPECIES: hydrogenase-2 small chain [Proteobacteria] Length: 372\nNumber of predicted TMHs: 1\nExp number of AAs in TMHs: 23.72226\nExp number, first 60 AAs: 0.22038\nTotal prob of N-in: 0.12994\noutside 1 331\nTMhelix 332 354\ninside 355 372

5179 GCF_000599665.1_ASM59966v1 Escherichia coli Proteobacteria; Gammaproteobacteria; Enterobacterales; Enterobacteriaceae; Escherichia MNNEETFYQAMRRQGVTRRSFLKYCSLAATSLGLGAGMAPKIAWALENKPRIPVVWIHGL WP_001058323.1 MULTISPECIES: hydrogenase-1 small chain [Proteobacteria] Length: 372\nNumber of predicted TMHs: 2\nExp number of AAs in TMHs: 37.2608499999999\nExp number, first 60 AAs: 19.52923\nTotal prob of N-in: 0.94351\nPOSSIBLE N-term signal sequence\ninside 1 19\nTMhelix 20 39\noutside 40 325\nTMhelix 326 348\ninside 349 372

5180 GCF_000599685.1_ASM59968v1 Escherichia coli Proteobacteria; Gammaproteobacteria; Enterobacterales; Enterobacteriaceae; Escherichia MNNEETFYQAMRRQGVTRRSFLKYCSLAATSLGLGAGMAPKIAWALENKPRIPVVWIHGL WP_001058323.1 MULTISPECIES: hydrogenase-1 small chain [Proteobacteria] Length: 372\nNumber of predicted TMHs: 2\nExp number of AAs in TMHs: 37.2608499999999\nExp number, first 60 AAs: 19.52923\nTotal prob of N-in: 0.94351\nPOSSIBLE N-term signal sequence\ninside 1 19\nTMhelix 20 39\noutside 40 325\nTMhelix 326 348\ninside 349 372

5181 GCF_000599685.1_ASM59968v1 Escherichia coli Proteobacteria; Gammaproteobacteria; Enterobacterales; Enterobacteriaceae; Escherichia MTGDNTLIHSHGINRRDFMKLCAALAATMGLSSKAAAEMAESVTNPQRPPVIWIGAQECT WP_000145410.1 MULTISPECIES: hydrogenase-2 small chain [Proteobacteria] Length: 372\nNumber of predicted TMHs: 1\nExp number of AAs in TMHs: 23.72226\nExp number, first 60 AAs: 0.22038\nTotal prob of N-in: 0.12994\noutside 1 331\nTMhelix 332 354\ninside 355 372

5182 GCF_000599625.1_ASM59962v1 Escherichia coli Proteobacteria; Gammaproteobacteria; Enterobacterales; Enterobacteriaceae; Escherichia MNNEETFYQAMRRQGVTRRSFLKYCSLAATSLGLGAGMAPKIAWALENKPRIPVVWIHGL WP_001058323.1 MULTISPECIES: hydrogenase-1 small chain [Proteobacteria] Length: 372\nNumber of predicted TMHs: 2\nExp number of AAs in TMHs: 37.2608499999999\nExp number, first 60 AAs: 19.52923\nTotal prob of N-in:

0.94351\nPOSSIBLE N-term signal sequence\ninside 1 19\nTMhelix 20 39\noutside 40 325\nTMhelix 326 348\ninside 349 372

5183 GCF_000597845.1_ASM59784v1 Escherichia coli Proteobacteria; Gammaproteobacteria; Enterobacterales; Enterobacteriaceae; Escherichia MNNEETFYQAMRRQGVTRRSFLKYCSLAATSLGLGAGMAPKIAWALENKPRIPVVWIHGL WP_001058323.1 MULTISPECIES: hydrogenase-1 small chain [Proteobacteria] Length: 372\nNumber of predicted TMHs: 2\nExp number of AAs in TMHs: 37.2608499999999\nExp number, first 60 AAs: 19.52923\nTotal prob of N-in: 0.94351\nPOSSIBLE N-term signal sequence\ninside 1 19\nTMhelix 20 39\noutside 40 325\nTMhelix 326 348\ninside 349 372

5184 GCF_000599665.1_ASM59966v1 Escherichia coli Proteobacteria; Gammaproteobacteria; Enterobacterales; Enterobacteriaceae; Escherichia MTGDNTLIHSHGINRRDFMKLCAALAATMGLSSKAAAEMAESVTNPQRPPVIWIGAQECT WP_000145410.1 MULTISPECIES: hydrogenase-2 small chain [Proteobacteria] Length: 372\nNumber of predicted TMHs: 1\nExp number of AAs in TMHs: 23.72226\nExp number, first 60 AAs: 0.22038\nTotal prob of N-in: 0.12994\noutside 1 331\nTMhelix 332 354\ninside 355 372

5185 GCF_000599625.1_ASM59962v1 Escherichia coli Proteobacteria; Gammaproteobacteria; Enterobacterales; Enterobacteriaceae; Escherichia MTGDNTLIHSHGINRRDFMKLCAALAATMGLSSKAAAEMAESVTNPQRPPVIWIGAQECT WP_000145410.1 MULTISPECIES: hydrogenase-2 small chain [Proteobacteria] Length: 372\nNumber of predicted TMHs: 1\nExp number of AAs in TMHs: 23.72226\nExp number, first 60 AAs: 0.22038\nTotal prob of N-in: 0.12994\noutside 1 331\nTMhelix 332 354\ninside 355 372

5186 GCF_000597845.1_ASM59784v1 Escherichia coli Proteobacteria; Gammaproteobacteria; Enterobacterales; Enterobacteriaceae; Escherichia MTGDNTLIHSHGINRRDFMKLCAALAATMGLSSKAAAEMAESVTNPQRPPVIWIGAQECT WP_000145410.1 MULTISPECIES: hydrogenase-2 small chain [Proteobacteria] Length: 372\nNumber of predicted TMHs: 1\nExp number of AAs in TMHs: 23.72226\nExp number, first 60 AAs: 0.22038\nTotal prob of N-in: 0.12994\noutside 1 331\nTMhelix 332 354\ninside 355 372

5187 GCF_000948715.1_ASM94871v1 Escherichia coli Proteobacteria; Gammaproteobacteria; Enterobacterales; Enterobacteriaceae; Escherichia MNNEETFYQAMRRQGVTRRSFLKYCSLAATSLGLGAGMAPKIAWALENKPRIPVVWIHGL WP_001058323.1 MULTISPECIES: hydrogenase-1 small chain [Proteobacteria] Length: 372\nNumber of predicted TMHs: 2\nExp number of AAs in TMHs: 37.2608499999999\nExp number, first 60 AAs: 19.52923\nTotal prob of N-in: 0.94351\nPOSSIBLE N-term signal sequence\ninside 1 19\nTMhelix 20 39\noutside 40 325\nTMhelix 326 348\ninside 349 372

5188 GCF_000941935.1_FHI82 Escherichia coli Proteobacteria; Gammaproteobacteria; Enterobacterales; Enterobacteriaceae; Escherichia MTGDNTLIHSHGINRRDFMKLCAALAATMGLSSKAAAEMAESVTNPQRPPVIWIGAQECT WP_000145410.1 MULTISPECIES: hydrogenase-2 small chain [Proteobacteria] Length: 372\nNumber of predicted TMHs: 1\nExp number of AAs in TMHs: 23.72226\nExp number, first 60 AAs: 0.22038\nTotal prob of N-in: 0.12994\noutside 1 331\nTMhelix 332 354\ninside 355 372

5189 GCF_000948615.1_ASM94861v1 Escherichia coli Proteobacteria; Gammaproteobacteria; Enterobacterales; Enterobacteriaceae; Escherichia MNNEETFYQAMRRQGVTRRSFLKYCSLAATSLGLGAGMAPKIAWALENKPRIPVVWIHGL WP_001301613.1 hydrogenase [Escherichia coli] Length: 372\nNumber of predicted TMHs: 2\nExp number of AAs in TMHs: 37.46598\nExp number, first 60 AAs: 19.53139\nTotal prob of N-in: 0.94364\nPOSSIBLE N-term signal sequence\ninside 1 19\nTMhelix 20 39\noutside 40 325\nTMhelix 326 348\ninside 349 372

5190 GCF_001285325.1_7748_7_23 Escherichia coli Proteobacteria; Gammaproteobacteria; Enterobacterales; Enterobacteriaceae; Escherichia MTGDNTLIHSHGINRRDFMKLCAALAATMGLSSKAAAEMAESVTNPQRPPVIWIGAQECT WP_000145410.1 MULTISPECIES: hydrogenase-2 small chain [Proteobacteria] Length: 372\nNumber of predicted TMHs: 1\nExp number of AAs in TMHs: 23.72226\nExp number, first 60 AAs: 0.22038\nTotal prob of N-in: 0.12994\noutside 1 331\nTMhelix 332 354\ninside 355 372

5191 GCF_000948535.1_ASM94853v1 Escherichia coli Proteobacteria; Gammaproteobacteria; Enterobacterales; Enterobacteriaceae; Escherichia
MNNEETFYQAMRRQGVTRRSFLKYCSLAATSLGLGAGMAPKIAWALENKPRIPVVWIHGL WP_001058323.1
MULTISPECIES: hydrogenase-1 small chain [Proteobacteria] Length: 372\nNumber of predicted TMHs: 2\nExp number of AAs in TMHs: 37.2608499999999\nExp number, first 60 AAs: 19.52923\nTotal prob of N-in: 0.94351\nPOSSIBLE N-term signal sequence\ninside 1 19\nTMhelix 20 39\noutside 40 325\nTMhelix 326 348\ninside 349 372

5192 GCF_000948765.1_ASM94876v1 Escherichia coli Proteobacteria; Gammaproteobacteria; Enterobacterales; Enterobacteriaceae; Escherichia
MTGDNTLIHSHGINRRDFMKLCAALAATMGLSSKAAAEMAESVTNPQRPPVIWIGAQECT WP_000145410.1
MULTISPECIES: hydrogenase-2 small chain [Proteobacteria] Length: 372\nNumber of predicted TMHs: 1\nExp number of AAs in TMHs: 23.72226\nExp number, first 60 AAs: 0.22038\nTotal prob of N-in: 0.12994\noutside 1 331\nTMhelix 332 354\ninside 355 372

5193 GCF_000691105.1_S6685 Escherichia coli Proteobacteria; Gammaproteobacteria; Enterobacterales; Enterobacteriaceae; Escherichia MTGDNTLIHSHGINRRDFMKLCAALAATMGLSSKAAAEMAESVTNPQRPPVIWIGAQECT
WP_000145410.1 MULTISPECIES: hydrogenase-2 small chain [Proteobacteria] Length: 372\nNumber of predicted TMHs: 1\nExp number of AAs in TMHs: 23.72226\nExp number, first 60 AAs: 0.22038\nTotal prob of N-in: 0.12994\noutside 1 331\nTMhelix 332 354\ninside 355 372

5194 GCF_000691045.1_S7380 Escherichia coli Proteobacteria; Gammaproteobacteria; Enterobacterales; Enterobacteriaceae; Escherichia MNNEETFYQAMRRQGVTRRSFLKYCSLAATSLGLGAGMAPKIAWALENKPRIPVVWIHGL
WP_001295940.1 MULTISPECIES: hydrogenase [Enterobacteriaceae] Length: 372\nNumber of predicted TMHs: 2\nExp number of AAs in TMHs: 37.2608899999999\nExp number, first 60 AAs: 19.52923\nTotal prob of N-in: 0.94351\nPOSSIBLE N-term signal sequence\ninside 1 19\nTMhelix 20 39\noutside 40 325\nTMhelix 326 348\ninside 349 372

5195 GCF_000948815.1_ASM94881v1 Escherichia coli Proteobacteria; Gammaproteobacteria; Enterobacterales; Enterobacteriaceae; Escherichia
MTGDNTLIHSHGINRRDFMKLCAALAATMGLSSKAAAEMAESVTNPQRPPVIWIGAQECT WP_000145410.1
MULTISPECIES: hydrogenase-2 small chain [Proteobacteria] Length: 372\nNumber of predicted TMHs: 1\nExp number of AAs in TMHs: 23.72226\nExp number, first 60 AAs: 0.22038\nTotal prob of N-in: 0.12994\noutside 1 331\nTMhelix 332 354\ninside 355 372

5196 GCF_000691005.1_S5274 Escherichia coli Proteobacteria; Gammaproteobacteria; Enterobacterales; Enterobacteriaceae; Escherichia MNNEETFYQAMRRQGVTRRSFLKYCSLAATSLGLGAGMAPKIAWALENKPRIPVVWIHGL
WP_001295940.1 MULTISPECIES: hydrogenase [Enterobacteriaceae] Length: 372\nNumber of predicted TMHs: 2\nExp number of AAs in TMHs: 37.2608899999999\nExp number, first 60 AAs: 19.52923\nTotal prob of N-in: 0.94351\nPOSSIBLE N-term signal sequence\ninside 1 19\nTMhelix 20 39\noutside 40 325\nTMhelix 326 348\ninside 349 372

5197 GCF_000948815.1_ASM94881v1 Escherichia coli Proteobacteria; Gammaproteobacteria; Enterobacterales; Enterobacteriaceae; Escherichia
MNNEETFYQAMRRQGVTRRSFLKYCSLAATSLGLGAGMAPKIAWALENKPRIPVVWIHGL WP_001058323.1
MULTISPECIES: hydrogenase-1 small chain [Proteobacteria] Length: 372\nNumber of predicted TMHs: 2\nExp number of AAs in TMHs: 37.2608499999999\nExp number, first 60 AAs: 19.52923\nTotal prob of N-in: 0.94351\nPOSSIBLE N-term signal sequence\ninside 1 19\nTMhelix 20 39\noutside 40 325\nTMhelix 326 348\ninside 349 372

5198 GCF_000691005.1_S5274 Escherichia coli Proteobacteria; Gammaproteobacteria; Enterobacterales; Enterobacteriaceae; Escherichia MTGDNTLIHSHGINRRDFMKLCAALAATMGLSSKAAAEMAESVTNPQRPPVIWIGAQECT
WP_000145410.1 MULTISPECIES: hydrogenase-2 small chain [Proteobacteria] Length: 372\nNumber of predicted TMHs: 1\nExp number of AAs in TMHs: 23.72226\nExp number, first 60 AAs: 0.22038\nTotal prob of N-in: 0.12994\noutside 1 331\nTMhelix 332 354\ninside 355 372

5199 GCF_000695115.1_Ecoli97-3250v1 Escherichia coli Proteobacteria; Gammaproteobacteria; Enterobacterales; Enterobacteriaceae; Escherichia
MTGDNTLIHSHGINRRDFMKLCAALAATMGLSSKAAAEMAESVTNPQRPPVIWIGAQECT WP_000145410.1
MULTISPECIES: hydrogenase-2 small chain [Proteobacteria] Length: 372\nNumber of predicted TMHs:

1\nExp number of AAs in TMHs: 23.72226\nExp number, first 60 AAs: 0.22038\nTotal prob of N-in: 0.12994\noutside 1 331\nTMhelix 332 354\ninside 355 372

5200 GCF_000695115.1_Ecoli97-3250v1 Escherichia coli Proteobacteria; Gammaproteobacteria; Enterobacterales; Enterobacteriaceae; Escherichia MNNEETFYQAMRRQGVTRRSFLKYCSLAATSLGLGAGMAPKIAWALENKPRIPVVWIHGL WP_001058323.1 MULTISPECIES: hydrogenase-1 small chain [Proteobacteria] Length: 372\nNumber of predicted TMHs: 2\nExp number of AAs in TMHs: 37.2608499999999\nExp number, first 60 AAs: 19.52923\nTotal prob of N-in: 0.94351\nPOSSIBLE N-term signal sequence\ninside 1 19\nTMhelix 20 39\noutside 40 325\nTMhelix 326 348\ninside 349 372

5201 GCF_001285405.1_7790_1_74 Escherichia coli Proteobacteria; Gammaproteobacteria; Enterobacterales; Enterobacteriaceae; Escherichia MTGDNTLIHSHGINRRDFMKLCAALAATMGLSSKAAAEMAESVTNPQRPPVIWIGAQECT WP_000145410.1 MULTISPECIES: hydrogenase-2 small chain [Proteobacteria] Length: 372\nNumber of predicted TMHs: 1\nExp number of AAs in TMHs: 23.72226\nExp number, first 60 AAs: 0.22038\nTotal prob of N-in: 0.12994\noutside 1 331\nTMhelix 332 354\ninside 355 372

5202 GCF_000951735.1_ASM95173v1 Escherichia coli Proteobacteria; Gammaproteobacteria; Enterobacterales; Enterobacteriaceae; Escherichia MTGDNTLIHSHGINRRDFMKLCAALAATMGLSSKAAAEMAESVTNPQRPPVIWIGAQECT WP_000145410.1 MULTISPECIES: hydrogenase-2 small chain [Proteobacteria] Length: 372\nNumber of predicted TMHs: 1\nExp number of AAs in TMHs: 23.72226\nExp number, first 60 AAs: 0.22038\nTotal prob of N-in: 0.12994\noutside 1 331\nTMhelix 332 354\ninside 355 372

5203 GCF_000951735.1_ASM95173v1 Escherichia coli Proteobacteria; Gammaproteobacteria; Enterobacterales; Enterobacteriaceae; Escherichia MNNEETFYQAMRRQGVTRRSFLKYCSLAATSLGLGAGMAPKIAWALENKPRIPVVWIHGL WP_001058323.1 MULTISPECIES: hydrogenase-1 small chain [Proteobacteria] Length: 372\nNumber of predicted TMHs: 2\nExp number of AAs in TMHs: 37.2608499999999\nExp number, first 60 AAs: 19.52923\nTotal prob of N-in: 0.94351\nPOSSIBLE N-term signal sequence\ninside 1 19\nTMhelix 20 39\noutside 40 325\nTMhelix 326 348\ninside 349 372

5204 GCF_001285185.1_7748_7_6Escherichia coli Proteobacteria; Gammaproteobacteria; Enterobacterales; Enterobacteriaceae; Escherichia MNNEETFYQAMRRQGVTRRSFLKYCSLAATSLGLGAGMAPKIAWALENKPRIPVVWIHGL WP_001295940.1 MULTISPECIES: hydrogenase [Enterobacteriaceae] Length: 372\nNumber of predicted TMHs: 2\nExp number of AAs in TMHs: 37.2608899999999\nExp number, first 60 AAs: 19.52923\nTotal prob of N-in: 0.94351\nPOSSIBLE N-term signal sequence\ninside 1 19\nTMhelix 20 39\noutside 40 325\nTMhelix 326 348\ninside 349 372

5205 GCF_001285165.1_8205_8_82 Escherichia coli Proteobacteria; Gammaproteobacteria; Enterobacterales; Enterobacteriaceae; Escherichia MTGDNTLIHSHGINRRDFMKLCAALAATMGLSSKAAAEMAESVTNPQRPPVIWIGAQECT WP_000145410.1 MULTISPECIES: hydrogenase-2 small chain [Proteobacteria] Length: 372\nNumber of predicted TMHs: 1\nExp number of AAs in TMHs: 23.72226\nExp number, first 60 AAs: 0.22038\nTotal prob of N-in: 0.12994\noutside 1 331\nTMhelix 332 354\ninside 355 372

5206 GCF_000951755.1_ASM95175v1 Escherichia coli Proteobacteria; Gammaproteobacteria; Enterobacterales; Enterobacteriaceae; Escherichia MTGDNTLIHSHGINRRDFMKLCAALAATMGLSSKAAAEMAESVTNPQRPPVIWIGAQECT WP_000145410.1 MULTISPECIES: hydrogenase-2 small chain [Proteobacteria] Length: 372\nNumber of predicted TMHs: 1\nExp number of AAs in TMHs: 23.72226\nExp number, first 60 AAs: 0.22038\nTotal prob of N-in: 0.12994\noutside 1 331\nTMhelix 332 354\ninside 355 372

5207 GCF_000647815.1_ASM64781v3 Escherichia coli Proteobacteria; Gammaproteobacteria; Enterobacterales; Enterobacteriaceae; Escherichia MTGDNTLIHSHGINRRDFMKLCAALAATMGLSSKAAAEMAESVTNPQRPPVIWIGAQECT WP_000145410.1 MULTISPECIES: hydrogenase-2 small chain [Proteobacteria] Length: 372\nNumber of predicted TMHs: 1\nExp number of AAs in TMHs: 23.72226\nExp number, first 60 AAs: 0.22038\nTotal prob of N-in: 0.12994\noutside 1 331\nTMhelix 332 354\ninside 355 372

5208 GCF_001285445.1_8205_3_65 Escherichia coli Proteobacteria; Gammaproteobacteria; Enterobacterales; Enterobacteriaceae; Escherichia MTGDNTLIHSHGINRRDFMKLCAALAATMGLSSKAAAEMAESVTNPQRPPVIWIGAQECT WP_000145410.1 MULTISPECIES: hydrogenase-2 small chain [Proteobacteria] Length: 372\nNumber of predicted TMHs: 1\nExp number of AAs in TMHs: 23.72226\nExp number, first 60 AAs: 0.22038\nTotal prob of N-in: 0.12994\noutside 1 331\nTMhelix 332 354\ninside 355 372

5209 GCF_000951895.1_FHI30 Escherichia coli Proteobacteria; Gammaproteobacteria; Enterobacterales; Enterobacteriaceae; Escherichia MTGDNTLIHSHGINRRDFMKLCAALAATMGLSSKAAAEMAESVTNPQRPPVIWIGAQECT WP_045148915.1 hydrogenase 2 small subunit [Escherichia coli] Length: 372\nNumber of predicted TMHs: 1\nExp number of AAs in TMHs: 23.7345\nExp number, first 60 AAs: 0.22038\nTotal prob of N-in: 0.13029\noutside 1 331\nTMhelix 332 354\ninside 355 372

5210 GCF_001265575.1_702324 Escherichia coli Proteobacteria; Gammaproteobacteria; Enterobacterales; Enterobacteriaceae; Escherichia MTGDNTLIHSHGINRRDFMKLCAALAATMGLSSKAAAEMAESVTNPQRPPVIWIGAQECT WP_000145410.1 MULTISPECIES: hydrogenase-2 small chain [Proteobacteria] Length: 372\nNumber of predicted TMHs: 1\nExp number of AAs in TMHs: 23.72226\nExp number, first 60 AAs: 0.22038\nTotal prob of N-in: 0.12994\noutside 1 331\nTMhelix 332 354\ninside 355 372

5211 GCF_000947985.1_ASM94798v1 Escherichia coli Proteobacteria; Gammaproteobacteria; Enterobacterales; Enterobacteriaceae; Escherichia MTGDNTLIHSHGINRRDFMKLCAALAATMGLSSKAAAEMAESVTNPQRPPVIWIGAQECT WP_000145410.1 MULTISPECIES: hydrogenase-2 small chain [Proteobacteria] Length: 372\nNumber of predicted TMHs: 1\nExp number of AAs in TMHs: 23.72226\nExp number, first 60 AAs: 0.22038\nTotal prob of N-in: 0.12994\noutside 1 331\nTMhelix 332 354\ninside 355 372

5212 GCF_001265575.1_702324 Escherichia coli Proteobacteria; Gammaproteobacteria; Enterobacterales; Enterobacteriaceae; Escherichia MNNEETFYQAMRRQGVTRRSFLKYCSLAATSLGLGAGMAPKIAWALENKPRIPVVWIHGL WP_001295940.1 MULTISPECIES: hydrogenase [Enterobacteriaceae] Length: 372\nNumber of predicted TMHs: 2\nExp number of AAs in TMHs: 37.2608899999999\nExp number, first 60 AAs: 19.52923\nTotal prob of N-in: 0.94351\nPOSSIBLE N-term signal sequence\ninside 1 19\nTMhelix 20 39\noutside 40 325\nTMhelix 326 348\ninside 349 372

5213 GCF_000945955.1_FHI85 Escherichia coli Proteobacteria; Gammaproteobacteria; Enterobacterales; Enterobacteriaceae; Escherichia MNNEETFYQAMRRQGVTRRSFLKYCSLAATSLGLGAGMAPKIAWALENKPRIPVVWIHGL WP_001058323.1 MULTISPECIES: hydrogenase-1 small chain [Proteobacteria] Length: 372\nNumber of predicted TMHs: 2\nExp number of AAs in TMHs: 37.2608499999999\nExp number, first 60 AAs: 19.52923\nTotal prob of N-in: 0.94351\nPOSSIBLE N-term signal sequence\ninside 1 19\nTMhelix 20 39\noutside 40 325\nTMhelix 326 348\ninside 349 372

5214 GCF_001268425.1_1.EC2993.1 Escherichia coli Proteobacteria; Gammaproteobacteria; Enterobacterales; Enterobacteriaceae; Escherichia MNNEETFYQAMRRQGVTRRSFLKYCSLAATSLGLGAGMAPKIAWALENKPRIPVVWIHGL WP_001058323.1 MULTISPECIES: hydrogenase-1 small chain [Proteobacteria] Length: 372\nNumber of predicted TMHs: 2\nExp number of AAs in TMHs: 37.2608499999999\nExp number, first 60 AAs: 19.52923\nTotal prob of N-in: 0.94351\nPOSSIBLE N-term signal sequence\ninside 1 19\nTMhelix 20 39\noutside 40 325\nTMhelix 326 348\ninside 349 372

5215 GCF_000952955.1_EcRV308Chr Escherichia coli Proteobacteria; Gammaproteobacteria; Enterobacterales; Enterobacteriaceae; Escherichia MTGDNTLIHSHGINRRDFMKLCAALAATMGLSSKAAAEMAESVTNPQRPPVIWIGAQECT WP_000145410.1 MULTISPECIES: hydrogenase-2 small chain [Proteobacteria] Length: 372\nNumber of predicted TMHs: 1\nExp number of AAs in TMHs: 23.72226\nExp number, first 60 AAs: 0.22038\nTotal prob of N-in: 0.12994\noutside 1 331\nTMhelix 332 354\ninside 355 372

5216 GCF_000952955.1_EcRV308Chr Escherichia coli Proteobacteria; Gammaproteobacteria; Enterobacterales; Enterobacteriaceae; Escherichia MNNEETFYQAMRRQGVTRRSFLKYCSLAATSLGLGAGMAPKIAWALENKPRIPVVWIHGL WP_001058323.1 MULTISPECIES: hydrogenase-1 small chain [Proteobacteria] Length: 372\nNumber of predicted TMHs: 2\nExp number of AAs in TMHs: 37.2608499999999\nExp number, first 60 AAs: 19.52923\nTotal prob of N-in:

0.94351\nPOSSIBLE N-term signal sequence\ninside 1 19\nTMhelix 20 39\noutside 40 325\nTMhelix 326 348\ninside 349 372

5217 GCF_000941935.1_FHI82 Escherichia coli Proteobacteria; Gammaproteobacteria; Enterobacterales; Enterobacteriaceae; Escherichia MNNEETFYQAMRRQGVTRRSFLKYCSLAATSLGLGAGMAPKIAWALENKPRIPVVWIHGL WP_001058323.1 MULTISPECIES: hydrogenase-1 small chain [Proteobacteria] Length: 372\nNumber of predicted TMHs: 2\nExp number of AAs in TMHs: 37.2608499999999\nExp number, first 60 AAs: 19.52923\nTotal prob of N-in: 0.94351\nPOSSIBLE N-term signal sequence\ninside 1 19\nTMhelix 20 39\noutside 40 325\nTMhelix 326 348\ninside 349 372

5218 GCF_000965545.1_ASM96554v1 Escherichia coli Proteobacteria; Gammaproteobacteria; Enterobacterales; Enterobacteriaceae; Escherichia MTGDNTLIHSHGINRRDFMKLCAALAATMGLSSKAAAEMAESVTNPQRPPVIWIGAQECT WP_000145410.1 MULTISPECIES: hydrogenase-2 small chain [Proteobacteria] Length: 372\nNumber of predicted TMHs: 1\nExp number of AAs in TMHs: 23.72226\nExp number, first 60 AAs: 0.22038\nTotal prob of N-in: 0.12994\noutside 1 331\nTMhelix 332 354\ninside 355 372

5219 GCF_000940035.2_FHI89 Escherichia coli Proteobacteria; Gammaproteobacteria; Enterobacterales; Enterobacteriaceae; Escherichia MTGDNTLIHSHGINRRDFMKLCAALAATMGLSSKAAAEMAESVTNPQRPPVIWIGAQECT WP_000145410.1 MULTISPECIES: hydrogenase-2 small chain [Proteobacteria] Length: 372\nNumber of predicted TMHs: 1\nExp number of AAs in TMHs: 23.72226\nExp number, first 60 AAs: 0.22038\nTotal prob of N-in: 0.12994\noutside 1 331\nTMhelix 332 354\ninside 355 372

5220 GCF_000939695.1_4d11B_assembly Escherichia coli Proteobacteria; Gammaproteobacteria; Enterobacterales; Enterobacteriaceae; Escherichia MNNEETFYQAMRRQGVTRRSFLKYCSLAATSLGLGAGMAPKIAWALENKPRIPVVWIHGL WP_001058323.1 MULTISPECIES: hydrogenase-1 small chain [Proteobacteria] Length: 372\nNumber of predicted TMHs: 2\nExp number of AAs in TMHs: 37.2608499999999\nExp number, first 60 AAs: 19.52923\nTotal prob of N-in: 0.94351\nPOSSIBLE N-term signal sequence\ninside 1 19\nTMhelix 20 39\noutside 40 325\nTMhelix 326 348\ninside 349 372

5221 GCF_000939695.1_4d11B_assembly Escherichia coli Proteobacteria; Gammaproteobacteria; Enterobacterales; Enterobacteriaceae; Escherichia MTGDNTLIHSHGINRRDFMKLCAALAATMGLSSKAAAEMAESVTNPQRPPVIWIGAQECT WP_000145410.1 MULTISPECIES: hydrogenase-2 small chain [Proteobacteria] Length: 372\nNumber of predicted TMHs: 1\nExp number of AAs in TMHs: 23.72226\nExp number, first 60 AAs: 0.22038\nTotal prob of N-in: 0.12994\noutside 1 331\nTMhelix 332 354\ninside 355 372

5222 GCF_000939135.1_46A_L1_assembly Escherichia coli Proteobacteria; Gammaproteobacteria; Enterobacterales; Enterobacteriaceae; Escherichia MNNEETFYQAMRRQGVTRRSFLKYCSLAATSLGLGAGMAPKIAWALENKPRIPVVWIHGL WP_001058323.1 MULTISPECIES: hydrogenase-1 small chain [Proteobacteria] Length: 372\nNumber of predicted TMHs: 2\nExp number of AAs in TMHs: 37.2608499999999\nExp number, first 60 AAs: 19.52923\nTotal prob of N-in: 0.94351\nPOSSIBLE N-term signal sequence\ninside 1 19\nTMhelix 20 39\noutside 40 325\nTMhelix 326 348\ninside 349 372

5223 GCF_000939135.1_46A_L1_assembly Escherichia coli Proteobacteria; Gammaproteobacteria; Enterobacterales; Enterobacteriaceae; Escherichia MTGDNTLIHSHGINRRDFMKLCAALAATMGLSSKAAAEMAESVTNPQRPPVIWIGAQECT WP_000145410.1 MULTISPECIES: hydrogenase-2 small chain [Proteobacteria] Length: 372\nNumber of predicted TMHs: 1\nExp number of AAs in TMHs: 23.72226\nExp number, first 60 AAs: 0.22038\nTotal prob of N-in: 0.12994\noutside 1 331\nTMhelix 332 354\ninside 355 372

5224 GCF_001285645.1_8205_3_23 Escherichia coli Proteobacteria; Gammaproteobacteria; Enterobacterales; Enterobacteriaceae; Escherichia MNNEETFYQAMRRQGVTRRSFLKYCSLAATSLGLGAGMAPKIAWALENKPRIPVVWIHGL WP_001058323.1 MULTISPECIES: hydrogenase-1 small chain [Proteobacteria] Length: 372\nNumber of predicted TMHs: 2\nExp number of AAs in TMHs: 37.2608499999999\nExp number, first 60 AAs: 19.52923\nTotal prob of N-in: 0.94351\nPOSSIBLE N-term signal sequence\ninside 1 19\nTMhelix 20 39\noutside 40 325\nTMhelix 326 348\ninside 349 372

5225 GCF_000965655.1_ASM96565v1 Escherichia coli Proteobacteria; Gammaproteobacteria; Enterobacterales; Enterobacteriaceae; Escherichia MNNEETFYQAMRRQGVTRRSFLKYCSLAATSLGLGAGMAPKIAWALENKPRIPVVWIHGL WP_001058323.1 MULTISPECIES: hydrogenase-1 small chain [Proteobacteria] Length: 372\nNumber of predicted TMHs: 2\nExp number of AAs in TMHs: 37.2608499999999\nExp number, first 60 AAs: 19.52923\nTotal prob of N-in: 0.94351\nPOSSIBLE N-term signal sequence\ninside 1 19\nTMhelix 20 39\noutside 40 325\nTMhelix 326 348\ninside 349 372

5226 GCF_001285685.1_8205_3_75 Escherichia coli Proteobacteria; Gammaproteobacteria; Enterobacterales; Enterobacteriaceae; Escherichia MTGDNTLIHSHGINRRDFMKLCAALAATMGLSSKAAAEMAESVTNPQRPPVIWIGAQECT WP_000145410.1 MULTISPECIES: hydrogenase-2 small chain [Proteobacteria] Length: 372\nNumber of predicted TMHs: 1\nExp number of AAs in TMHs: 23.72226\nExp number, first 60 AAs: 0.22038\nTotal prob of N-in: 0.12994\noutside 1 331\nTMhelix 332 354\ninside 355 372

5227 GCF_000965655.1_ASM96565v1 Escherichia coli Proteobacteria; Gammaproteobacteria; Enterobacterales; Enterobacteriaceae; Escherichia MTGDNTLIHSHGINRRDFMKLCAALAATMGLSSKAAAEMAESVTNPQRPPVIWIGAQECT WP_045904146.1 hydrogenase 2 small subunit [Escherichia coli] Length: 372\nNumber of predicted TMHs: 1\nExp number of AAs in TMHs: 22.83694\nExp number, first 60 AAs: 0.22071\nTotal prob of N-in: 0.12690\noutside 1 331\nTMhelix 332 354\ninside 355 372

5228 GCF_001285685.1_8205_3_75 Escherichia coli Proteobacteria; Gammaproteobacteria; Enterobacterales; Enterobacteriaceae; Escherichia MNNEETFYQAMRRQGVTRRSFLKYCSLAATSLGLGAGMAPKIAWALENKPRIPVVWIHGL WP_001295940.1 MULTISPECIES: hydrogenase [Enterobacteriaceae] Length: 372\nNumber of predicted TMHs: 2\nExp number of AAs in TMHs: 37.2608899999999\nExp number, first 60 AAs: 19.52923\nTotal prob of N-in: 0.94351\nPOSSIBLE N-term signal sequence\ninside 1 19\nTMhelix 20 39\noutside 40 325\nTMhelix 326 348\ninside 349 372

5229 GCF_000937715.2_FHI65 Escherichia coli Proteobacteria; Gammaproteobacteria; Enterobacterales; Enterobacteriaceae; Escherichia MNNEETFYQAMRRQGVTRRSFLKYCSLAATSLGLGAGMAPKIAWALENKPRIPVVWIHGL WP_001295940.1 MULTISPECIES: hydrogenase [Enterobacteriaceae] Length: 372\nNumber of predicted TMHs: 2\nExp number of AAs in TMHs: 37.2608899999999\nExp number, first 60 AAs: 19.52923\nTotal prob of N-in: 0.94351\nPOSSIBLE N-term signal sequence\ninside 1 19\nTMhelix 20 39\noutside 40 325\nTMhelix 326 348\ninside 349 372

5230 GCF_001285745.1_7748_7_35 Escherichia coli Proteobacteria; Gammaproteobacteria; Enterobacterales; Enterobacteriaceae; Escherichia MNNEETFYQAMRRQGVTRRSFLKYCSLAATSLGLGAGMAPKIAWALENKPRIPVVWIHGL WP_001295940.1 MULTISPECIES: hydrogenase [Enterobacteriaceae] Length: 372\nNumber of predicted TMHs: 2\nExp number of AAs in TMHs: 37.2608899999999\nExp number, first 60 AAs: 19.52923\nTotal prob of N-in: 0.94351\nPOSSIBLE N-term signal sequence\ninside 1 19\nTMhelix 20 39\noutside 40 325\nTMhelix 326 348\ninside 349 372

5231 GCF_001285785.1_8205_3_49 Escherichia coli Proteobacteria; Gammaproteobacteria; Enterobacterales; Enterobacteriaceae; Escherichia MTGDNTLIHSHGINRRDFMKLCAALAATMGLSSKAAAEMAESVTNPQRPPVIWIGAQECT WP_000145410.1 MULTISPECIES: hydrogenase-2 small chain [Proteobacteria] Length: 372\nNumber of predicted TMHs: 1\nExp number of AAs in TMHs: 23.72226\nExp number, first 60 AAs: 0.22038\nTotal prob of N-in: 0.12994\noutside 1 331\nTMhelix 332 354\ninside 355 372

5232 GCF_001285785.1_8205_3_49 Escherichia coli Proteobacteria; Gammaproteobacteria; Enterobacterales; Enterobacteriaceae; Escherichia MNNEETFYQAMRRQGVTRRSFLKYCSLAATSLGLGAGMAPKIAWALENKPRIPVVWIHGL WP_001058323.1 MULTISPECIES: hydrogenase-1 small chain [Proteobacteria] Length: 372\nNumber of predicted TMHs: 2\nExp number of AAs in TMHs: 37.2608499999999\nExp number, first 60 AAs: 19.52923\nTotal prob of N-in: 0.94351\nPOSSIBLE N-term signal sequence\ninside 1 19\nTMhelix 20 39\noutside 40 325\nTMhelix 326 348\ninside 349 372

5233 GCF_000798555.1_Escherichia_coli_CVM_N36903PS_v._1.0 Escherichia coli Proteobacteria; Gammaproteobacteria; Enterobacterales; Enterobacteriaceae; Escherichia
MTGDNTLIHSHGINRRDFMKLCAALAATMGLSSKAAAEMAESVTNPQRPPVIWIGAQECT WP_000145410.1
MULTISPECIES: hydrogenase-2 small chain [Proteobacteria] Length: 372\nNumber of predicted TMHs: 1\nExp number of AAs in TMHs: 23.72226\nExp number, first 60 AAs: 0.22038\nTotal prob of N-in: 0.12994\noutside 1 331\nTMhelix 332 354\ninside 355 372

5234 GCF_000798555.1_Escherichia_coli_CVM_N36903PS_v._1.0 Escherichia coli Proteobacteria; Gammaproteobacteria; Enterobacterales; Enterobacteriaceae; Escherichia
MNNEETFYQAMRRQGVTRRSFLKYCSLAATSLGLGAGMAPKIAWALENKPRIPVVWIHGL WP_001295940.1
MULTISPECIES: hydrogenase [Enterobacteriaceae] Length: 372\nNumber of predicted TMHs: 2\nExp number of AAs in TMHs: 37.2608899999999\nExp number, first 60 AAs: 19.52923\nTotal prob of N-in: 0.94351\nPOSSIBLE N-term signal sequence\ninside 1 19\nTMhelix 20 39\noutside 40 325\nTMhelix 326 348\ninside 349 372

5235 GCF_001077875.1_ASM107787v1 Escherichia coli Proteobacteria; Gammaproteobacteria; Enterobacterales; Enterobacteriaceae; Escherichia
MTGDNTLIHSHGINRRDFMKLCAALAATMGLSSKAAAEMAESVTNPQRPPVIWIGAQECT WP_000145410.1
MULTISPECIES: hydrogenase-2 small chain [Proteobacteria] Length: 372\nNumber of predicted TMHs: 1\nExp number of AAs in TMHs: 23.72226\nExp number, first 60 AAs: 0.22038\nTotal prob of N-in: 0.12994\noutside 1 331\nTMhelix 332 354\ninside 355 372

5236 GCF_001077875.1_ASM107787v1 Escherichia coli Proteobacteria; Gammaproteobacteria; Enterobacterales; Enterobacteriaceae; Escherichia
MNNEETFYQAMRRQGVTRRSFLKYCSLAATSLGLGAGMAPKIAWALENKPRIPVVWIHGL WP_001058323.1
MULTISPECIES: hydrogenase-1 small chain [Proteobacteria] Length: 372\nNumber of predicted TMHs: 2\nExp number of AAs in TMHs: 37.2608499999999\nExp number, first 60 AAs: 19.52923\nTotal prob of N-in: 0.94351\nPOSSIBLE N-term signal sequence\ninside 1 19\nTMhelix 20 39\noutside 40 325\nTMhelix 326 348\ninside 349 372

5237 GCF_001077955.1_ASM107795v1 Escherichia coli Proteobacteria; Gammaproteobacteria; Enterobacterales; Enterobacteriaceae; Escherichia
MNNEETFYQAMRRQGVTRRSFLKYCSLAATSLGLGAGMAPKIAWALENKSRIPVVWIHGL WP_049595080.1
hydrogenase [Escherichia coli] Length: 372\nNumber of predicted TMHs: 2\nExp number of AAs in TMHs: 37.1898399999999\nExp number, first 60 AAs: 19.44607\nTotal prob of N-in: 0.94159\nPOSSIBLE N-term signal sequence\ninside 1 19\nTMhelix 20 39\noutside 40 325\nTMhelix 326 348\ninside 349 372

5238 GCF_001182805.1_ASM118280v1 Escherichia coli Proteobacteria; Gammaproteobacteria; Enterobacterales; Enterobacteriaceae; Escherichia
MTGDNTLIHSHGINRRDFMKLCAALAATMGLSSKAAAEMAESVTNPQRPPVIWIGAQECT WP_000145410.1
MULTISPECIES: hydrogenase-2 small chain [Proteobacteria] Length: 372\nNumber of predicted TMHs: 1\nExp number of AAs in TMHs: 23.72226\nExp number, first 60 AAs: 0.22038\nTotal prob of N-in: 0.12994\noutside 1 331\nTMhelix 332 354\ninside 355 372

5239 GCF_001182805.1_ASM118280v1 Escherichia coli Proteobacteria; Gammaproteobacteria; Enterobacterales; Enterobacteriaceae; Escherichia
MNNEETFYQAMRRQGVTRRSFLKYCSLAATSLGLGAGMAPKIAWALENKPRIPVVWIHGL WP_001295940.1
MULTISPECIES: hydrogenase [Enterobacteriaceae] Length: 372\nNumber of predicted TMHs: 2\nExp number of AAs in TMHs: 37.2608899999999\nExp number, first 60 AAs: 19.52923\nTotal prob of N-in: 0.94351\nPOSSIBLE N-term signal sequence\ninside 1 19\nTMhelix 20 39\noutside 40 325\nTMhelix 326 348\ninside 349 372

5240 GCF_000800675.1_ASM80067v1 Escherichia coli Proteobacteria; Gammaproteobacteria; Enterobacterales; Enterobacteriaceae; Escherichia
MNNEETFYQAMRRQGVTRRSFLKYCSLAATSLGLGAGMAPKIAWALENKPRIPVVWIHGL WP_001058323.1
MULTISPECIES: hydrogenase-1 small chain [Proteobacteria] Length: 372\nNumber of predicted TMHs: 2\nExp number of AAs in TMHs: 37.2608499999999\nExp number, first 60 AAs: 19.52923\nTotal prob of N-in: 0.94351\nPOSSIBLE N-term signal sequence\ninside 1 19\nTMhelix 20 39\noutside 40 325\nTMhelix 326 348\ninside 349 372

5241 GCF_001285045.1_7790_1_67 Escherichia coli Proteobacteria; Gammaproteobacteria; Enterobacterales; Enterobacteriaceae; Escherichia MNNEETFYQAMRRQGVTRRSFLKYCSLAATSLGLGAGMAPKIAWALENKPRIPVVWIHGL WP_001058323.1 MULTISPECIES: hydrogenase-1 small chain [Proteobacteria] Length: 372\nNumber of predicted TMHs: 2\nExp number of AAs in TMHs: 37.2608499999999\nExp number, first 60 AAs: 19.52923\nTotal prob of N-in: 0.94351\nPOSSIBLE N-term signal sequence\ninside 1 19\nTMhelix 20 39\noutside 40 325\nTMhelix 326 348\ninside 349 372

5242 GCF_001285045.1_7790_1_67 Escherichia coli Proteobacteria; Gammaproteobacteria; Enterobacterales; Enterobacteriaceae; Escherichia MTGDNTLIHSHGINRRDFMKLCAALAATMGLSSKAAAEMAESVTNPQRPPVIWIGAQECT WP_000145410.1 MULTISPECIES: hydrogenase-2 small chain [Proteobacteria] Length: 372\nNumber of predicted TMHs: 1\nExp number of AAs in TMHs: 23.72226\nExp number, first 60 AAs: 0.22038\nTotal prob of N-in: 0.12994\noutside 1 331\nTMhelix 332 354\ninside 355 372

5243 GCF_001286105.1_8205_3_6Escherichia coli Proteobacteria; Gammaproteobacteria; Enterobacterales; Enterobacteriaceae; Escherichia MTGDNTLIHSHGINRRDFMKLCAALAATMGLSSKAAAEMAESVTNPQRPPVIWIGAQECT WP_000145410.1 MULTISPECIES: hydrogenase-2 small chain [Proteobacteria] Length: 372\nNumber of predicted TMHs: 1\nExp number of AAs in TMHs: 23.72226\nExp number, first 60 AAs: 0.22038\nTotal prob of N-in: 0.12994\noutside 1 331\nTMhelix 332 354\ninside 355 372

5244 GCF_001286105.1_8205_3_6Escherichia coli Proteobacteria; Gammaproteobacteria; Enterobacterales; Enterobacteriaceae; Escherichia MNNEETFYQAMRRQGVTRRSFLKYCSLAATSLGLGAGMAPKIAWALENKPRIPVVWIHGL WP_001295940.1 MULTISPECIES: hydrogenase [Enterobacteriaceae] Length: 372\nNumber of predicted TMHs: 2\nExp number of AAs in TMHs: 37.2608899999999\nExp number, first 60 AAs: 19.52923\nTotal prob of N-in: 0.94351\nPOSSIBLE N-term signal sequence\ninside 1 19\nTMhelix 20 39\noutside 40 325\nTMhelix 326 348\ninside 349 372

5245 GCF_000969495.1_ASM96949v1 Escherichia coli Proteobacteria; Gammaproteobacteria; Enterobacterales; Enterobacteriaceae; Escherichia MNNEETFYQAMRRQGVTRRSFLKYCSLAATSLGLGAGMAPKIAWALENKPRIPVVWIHGL WP_001058323.1 MULTISPECIES: hydrogenase-1 small chain [Proteobacteria] Length: 372\nNumber of predicted TMHs: 2\nExp number of AAs in TMHs: 37.2608499999999\nExp number, first 60 AAs: 19.52923\nTotal prob of N-in: 0.94351\nPOSSIBLE N-term signal sequence\ninside 1 19\nTMhelix 20 39\noutside 40 325\nTMhelix 326 348\ninside 349 372

5246 GCF_000819355.1_Escherichia_coli_CVM_N33574PS_v1.0 Escherichia coli Proteobacteria; Gammaproteobacteria; Enterobacterales; Enterobacteriaceae; Escherichia MTGDNTLIHSHGINRRDFMKLCAALAATMGLSSKAAAEMAESVTNPQRPPVIWIGAQECT WP_000145410.1 MULTISPECIES: hydrogenase-2 small chain [Proteobacteria] Length: 372\nNumber of predicted TMHs: 1\nExp number of AAs in TMHs: 23.72226\nExp number, first 60 AAs: 0.22038\nTotal prob of N-in: 0.12994\noutside 1 331\nTMhelix 332 354\ninside 355 372

5247 GCF_000819355.1_Escherichia_coli_CVM_N33574PS_v1.0 Escherichia coli Proteobacteria; Gammaproteobacteria; Enterobacterales; Enterobacteriaceae; Escherichia MNNEETFYQAMRRQGVTRRSFLKYCSLAATSLGLGAGMAPKIAWALENKPRIPVVWIHGL WP_001058323.1 MULTISPECIES: hydrogenase-1 small chain [Proteobacteria] Length: 372\nNumber of predicted TMHs: 2\nExp number of AAs in TMHs: 37.2608499999999\nExp number, first 60 AAs: 19.52923\nTotal prob of N-in: 0.94351\nPOSSIBLE N-term signal sequence\ninside 1 19\nTMhelix 20 39\noutside 40 325\nTMhelix 326 348\ninside 349 372

5248 GCF_001268465.1_YE28 Escherichia coli Proteobacteria; Gammaproteobacteria; Enterobacterales; Enterobacteriaceae; Escherichia MTGDNTLIHSHGINRRDFMKLCAALAATMGLSSKAAAEMAESVTNPQRPPVIWIGAQECT WP_000145410.1 MULTISPECIES: hydrogenase-2 small chain [Proteobacteria] Length: 372\nNumber of predicted TMHs: 1\nExp number of AAs in TMHs: 23.72226\nExp number, first 60 AAs: 0.22038\nTotal prob of N-in: 0.12994\noutside 1 331\nTMhelix 332 354\ninside 355 372

5249 GCF_001268465.1_YE28 Escherichia coli Proteobacteria; Gammaproteobacteria; Enterobacterales; Enterobacteriaceae; Escherichia MNNEETFYQAMRRQGVTRRSFLKYCSLAATSLGLGAGMAPKIAWALENKPRIPVVWIHGL WP_001058323.1 MULTISPECIES: hydrogenase-1 small chain [Proteobacteria] Length: 372\nNumber of predicted TMHs: 2\nExp number of AAs in TMHs: 37.2608499999999\nExp number, first 60 AAs: 19.52923\nTotal prob of

N-in: 0.94351\nPOSSIBLE N-term signal sequence\ninside 1 19\nTMhelix 20 39\noutside 40 325\nTMhelix 326 348\ninside 349 372

5250 GCF_001286245.1_8205_8_71 Escherichia coli Proteobacteria; Gammaproteobacteria; Enterobacterales; Enterobacteriaceae; Escherichia
MTGDNTLIHSHGINRRDFMKLCAALAATMGLSSKAAAEMAESVTNPQRPPVIWIGAQECT WP_000145410.1
MULTISPECIES: hydrogenase-2 small chain [Proteobacteria] Length: 372\nNumber of predicted TMHs: 1\nExp number of AAs in TMHs: 23.72226\nExp number, first 60 AAs: 0.22038\nTotal prob of N-in: 0.12994\noutside 1 331\nTMhelix 332 354\ninside 355 372

5251 GCF_001183685.1_ASM118368v1 Escherichia coli Proteobacteria; Gammaproteobacteria; Enterobacterales; Enterobacteriaceae; Escherichia
MNNEETFYQAMRRQGVTRRSFLKYCSLAATSLGLGAGMAPKIAWALENKPRIPVVWIHGL WP_001058323.1
MULTISPECIES: hydrogenase-1 small chain [Proteobacteria] Length: 372\nNumber of predicted TMHs: 2\nExp number of AAs in TMHs: 37.2608499999999\nExp number, first 60 AAs: 19.52923\nTotal prob of N-in: 0.94351\nPOSSIBLE N-term signal sequence\ninside 1 19\nTMhelix 20 39\noutside 40 325\nTMhelix 326 348\ninside 349 372

5252 GCF_001268565.1_1.EC2923.1 Escherichia coli Proteobacteria; Gammaproteobacteria; Enterobacterales; Enterobacteriaceae; Escherichia
MTGDNTLIHSHGINRRDFMKLCAALAATMGLSSKAAAEMAESVTNPQRPPVIWIGAQECT WP_000145410.1
MULTISPECIES: hydrogenase-2 small chain [Proteobacteria] Length: 372\nNumber of predicted TMHs: 1\nExp number of AAs in TMHs: 23.72226\nExp number, first 60 AAs: 0.22038\nTotal prob of N-in: 0.12994\noutside 1 331\nTMhelix 332 354\ninside 355 372

5253 GCF_001056205.1_ASM105620v1 Escherichia coli Proteobacteria; Gammaproteobacteria; Enterobacterales; Enterobacteriaceae; Escherichia
MTGDNTLIHSHGINRRDFMKLCAALAATMGLSSKAAAEMVESVTNPQRPPVIWIGAQECT WP_000145423.1
hydrogenase 2 small subunit [Escherichia coli] Length: 372\nNumber of predicted TMHs: 1\nExp number of AAs in TMHs: 23.78201\nExp number, first 60 AAs: 0.28355\nTotal prob of N-in: 0.13257\noutside 1 331\nTMhelix 332 354\ninside 355 372

5254 GCF_001056205.1_ASM105620v1 Escherichia coli Proteobacteria; Gammaproteobacteria; Enterobacterales; Enterobacteriaceae; Escherichia
MNNEETFYQAMRRQGVTRRSFLKYCSLAATSLGLGAGMAPKIAWALENKPRIPVVWIHGL WP_001295940.1
MULTISPECIES: hydrogenase [Enterobacteriaceae] Length: 372\nNumber of predicted TMHs: 2\nExp number of AAs in TMHs: 37.2608899999999\nExp number, first 60 AAs: 19.52923\nTotal prob of N-in: 0.94351\nPOSSIBLE N-term signal sequence\ninside 1 19\nTMhelix 20 39\noutside 40 325\nTMhelix 326 348\ninside 349 372

5255 GCF_001285005.1_8205_3_22 Escherichia coli Proteobacteria; Gammaproteobacteria; Enterobacterales; Enterobacteriaceae; Escherichia
MNNEETFYQAMRRQGVTRRSFLKYCSLAATSLGLGAGMAPKIAWALENKPRIPVVWIHGL WP_001295940.1
MULTISPECIES: hydrogenase [Enterobacteriaceae] Length: 372\nNumber of predicted TMHs: 2\nExp number of AAs in TMHs: 37.2608899999999\nExp number, first 60 AAs: 19.52923\nTotal prob of N-in: 0.94351\nPOSSIBLE N-term signal sequence\ninside 1 19\nTMhelix 20 39\noutside 40 325\nTMhelix 326 348\ninside 349 372

5256 GCF_001058145.1_ASM105814v1 Escherichia coli Proteobacteria; Gammaproteobacteria; Enterobacterales; Enterobacteriaceae; Escherichia
MTGDNTLIHSHGINRRDFMKLCAALAATMGLSSKAAAEMAESVTNPQRPPVIWIGAQECT WP_000145410.1
MULTISPECIES: hydrogenase-2 small chain [Proteobacteria] Length: 372\nNumber of predicted TMHs: 1\nExp number of AAs in TMHs: 23.72226\nExp number, first 60 AAs: 0.22038\nTotal prob of N-in: 0.12994\noutside 1 331\nTMhelix 332 354\ninside 355 372

5257 GCF_001058175.1_ASM105817v1 Escherichia coli Proteobacteria; Gammaproteobacteria; Enterobacterales; Enterobacteriaceae; Escherichia
MTGDNTLIHSHGINRRDFMKLCAALAATMGLSSKAAAEMVESVTNPQRPPVIWIGAQECT WP_000145423.1
hydrogenase 2 small subunit [Escherichia coli] Length: 372\nNumber of predicted TMHs: 1\nExp number of AAs in TMHs: 23.78201\nExp number, first 60 AAs: 0.28355\nTotal prob of N-in: 0.13257\noutside 1 331\nTMhelix 332 354\ninside 355 372

5258 GCF_000714915.1_UCD_JA17_pb Escherichia coli Proteobacteria; Gammaproteobacteria; Enterobacterales; Enterobacteriaceae; Escherichia MNNEETFYQAMRRQGVTRRSFLKYCSLAATSLGLGAGMAPKIAWALENKPRIPVVWIHGL WP_001058323.1 MULTISPECIES: hydrogenase-1 small chain [Proteobacteria] Length: 372\nNumber of predicted TMHs: 2\nExp number of AAs in TMHs: 37.2608499999999\nExp number, first 60 AAs: 19.52923\nTotal prob of N-in: 0.94351\nPOSSIBLE N-term signal sequence\ninside 1 19\nTMhelix 20 39\noutside 40 325\nTMhelix 326 348\ninside 349 372

5259 GCF_000752115.1_FHI46 Escherichia coli Proteobacteria; Gammaproteobacteria; Enterobacterales; Enterobacteriaceae; Escherichia MTGDNTLIHSHGINRRDFMKLCAALAATMGLSSKAAAEMAESVTNPQRPPVIWIGAQECT WP_000145410.1 MULTISPECIES: hydrogenase-2 small chain [Proteobacteria] Length: 372\nNumber of predicted TMHs: 1\nExp number of AAs in TMHs: 23.72226\nExp number, first 60 AAs: 0.22038\nTotal prob of N-in: 0.12994\noutside 1 331\nTMhelix 332 354\ninside 355 372

5260 GCF_000752155.1_FHI52 Escherichia coli Proteobacteria; Gammaproteobacteria; Enterobacterales; Enterobacteriaceae; Escherichia MTGDNTLIHSHGINRRDFMKLCAALAATMGLSSKAAAEMAESVTNPQRPPVIWIGAQECT WP_000145410.1 MULTISPECIES: hydrogenase-2 small chain [Proteobacteria] Length: 372\nNumber of predicted TMHs: 1\nExp number of AAs in TMHs: 23.72226\nExp number, first 60 AAs: 0.22038\nTotal prob of N-in: 0.12994\noutside 1 331\nTMhelix 332 354\ninside 355 372

5261 GCF_000752195.1_FHI3 Escherichia coli Proteobacteria; Gammaproteobacteria; Enterobacterales; Enterobacteriaceae; Escherichia MTGDNTLIHSHGINRRDFMKLCAALAATMGLSSKAAAEMAESVTNPQRPPVIWIGAQECT WP_000145410.1 MULTISPECIES: hydrogenase-2 small chain [Proteobacteria] Length: 372\nNumber of predicted TMHs: 1\nExp number of AAs in TMHs: 23.72226\nExp number, first 60 AAs: 0.22038\nTotal prob of N-in: 0.12994\noutside 1 331\nTMhelix 332 354\ninside 355 372

5262 GCF_000749545.1_ASM74954v1 Escherichia coli Proteobacteria; Gammaproteobacteria; Enterobacterales; Enterobacteriaceae; Escherichia MTGDNTLIHSHGINRRDFMKLCAALAATMGLSSKAAAEMAESVTNPQRPPVIWIGAQECT WP_000145410.1 MULTISPECIES: hydrogenase-2 small chain [Proteobacteria] Length: 372\nNumber of predicted TMHs: 1\nExp number of AAs in TMHs: 23.72226\nExp number, first 60 AAs: 0.22038\nTotal prob of N-in: 0.12994\noutside 1 331\nTMhelix 332 354\ninside 355 372

5263 GCF_000749575.1_ASM74957v1 Escherichia coli Proteobacteria; Gammaproteobacteria; Enterobacterales; Enterobacteriaceae; Escherichia MTGDNTLIHSHGINRRDFMKLCAALAATMGLSSKAAAEMAESVTNPQRPPVIWIGAQECT WP_000145410.1 MULTISPECIES: hydrogenase-2 small chain [Proteobacteria] Length: 372\nNumber of predicted TMHs: 1\nExp number of AAs in TMHs: 23.72226\nExp number, first 60 AAs: 0.22038\nTotal prob of N-in: 0.12994\noutside 1 331\nTMhelix 332 354\ninside 355 372

5264 GCF_000781095.1_ASM78109v1 Escherichia coli Proteobacteria; Gammaproteobacteria; Enterobacterales; Enterobacteriaceae; Escherichia MTGDNTLIHSHGINRRDFMKLCAALAATMGLSSKAAAEMAESVTNPQRPPVIWIGAQECT WP_000145410.1 MULTISPECIES: hydrogenase-2 small chain [Proteobacteria] Length: 372\nNumber of predicted TMHs: 1\nExp number of AAs in TMHs: 23.72226\nExp number, first 60 AAs: 0.22038\nTotal prob of N-in: 0.12994\noutside 1 331\nTMhelix 332 354\ninside 355 372

5265 GCF_000752615.1_StOlav143 Escherichia coli Proteobacteria; Gammaproteobacteria; Enterobacterales; Enterobacteriaceae; Escherichia MTGDNTLIHSHGINRRDFMKLCAALAATMGLSSKAAAEMAESVTNPQRPPVIWIGAQECT WP_000145410.1 MULTISPECIES: hydrogenase-2 small chain [Proteobacteria] Length: 372\nNumber of predicted TMHs: 1\nExp number of AAs in TMHs: 23.72226\nExp number, first 60 AAs: 0.22038\nTotal prob of N-in: 0.12994\noutside 1 331\nTMhelix 332 354\ninside 355 372

5266 GCF_000752695.1_FHI96 Escherichia coli Proteobacteria; Gammaproteobacteria; Enterobacterales; Enterobacteriaceae; Escherichia MTGDNTLIHSHGINRRDFMKLCAALAATMGLSSKAAAEMAESVTNPQRPPVIWIGAQECT WP_000145410.1 MULTISPECIES: hydrogenase-2 small chain [Proteobacteria] Length: 372\nNumber of predicted TMHs: 1\nExp number of AAs in TMHs: 23.72226\nExp number, first 60 AAs: 0.22038\nTotal prob of N-in: 0.12994\noutside 1 331\nTMhelix 332 354\ninside 355 372

5267 GCF_000781175.1_ASM78117v1 Escherichia coli Proteobacteria; Gammaproteobacteria; Enterobacterales; Enterobacteriaceae; Escherichia MTGDNTLIHSHGINRRDFMKLCAALAATMGLSSKAAAEMAESVTNPQRPPVIWIGAQECT WP_000145410.1 MULTISPECIES: hydrogenase-2 small chain [Proteobacteria] Length: 372\nNumber of predicted TMHs: 1\nExp number of AAs in TMHs: 23.72226\nExp number, first 60 AAs: 0.22038\nTotal prob of N-in: 0.12994\noutside 1 331\nTMhelix 332 354\ninside 355 372

5268 GCF_000781215.1_ASM78121v1 Escherichia coli Proteobacteria; Gammaproteobacteria; Enterobacterales; Enterobacteriaceae; Escherichia MTGDNTLIHSHGINRRDFMKLCAALAATMGLSSKAAAEMAESVTNPQRPPVIWIGAQECT WP_000145410.1 MULTISPECIES: hydrogenase-2 small chain [Proteobacteria] Length: 372\nNumber of predicted TMHs: 1\nExp number of AAs in TMHs: 23.72226\nExp number, first 60 AAs: 0.22038\nTotal prob of N-in: 0.12994\noutside 1 331\nTMhelix 332 354\ninside 355 372

5269 GCF_000752895.1_FHI101 Escherichia coli Proteobacteria; Gammaproteobacteria; Enterobacterales; Enterobacteriaceae; Escherichia MTGDNTLIHSHGINRRDFMKLCAALAATMGLSSKAAAEMAESVTNPQRPPVIWIGAQECT WP_000145410.1 MULTISPECIES: hydrogenase-2 small chain [Proteobacteria] Length: 372\nNumber of predicted TMHs: 1\nExp number of AAs in TMHs: 23.72226\nExp number, first 60 AAs: 0.22038\nTotal prob of N-in: 0.12994\noutside 1 331\nTMhelix 332 354\ninside 355 372

5270 GCF_000752695.1_FHI96 Escherichia coli Proteobacteria; Gammaproteobacteria; Enterobacterales; Enterobacteriaceae; Escherichia MNNEETFYQAMRRQGVTRRSFLKYCSLAATSLGLGAGMAPKIAWALENKPRIPVVWIHGL WP_001058323.1 MULTISPECIES: hydrogenase-1 small chain [Proteobacteria] Length: 372\nNumber of predicted TMHs: 2\nExp number of AAs in TMHs: 37.2608499999999\nExp number, first 60 AAs: 19.52923\nTotal prob of N-in: 0.94351\nPOSSIBLE N-term signal sequence\ninside 1 19\nTMhelix 20 39\noutside 40 325\nTMhelix 326 348\ninside 349 372

5271 GCF_000752615.1_StOlav143 Escherichia coli Proteobacteria; Gammaproteobacteria; Enterobacterales; Enterobacteriaceae; Escherichia MNNEETFYQAMRRQGVTRRSFLKYCSLAATSLGLGAGMAPKIAWALENKPRIPVVWIHGL WP_001295940.1 MULTISPECIES: hydrogenase [Enterobacteriaceae] Length: 372\nNumber of predicted TMHs: 2\nExp number of AAs in TMHs: 37.2608899999999\nExp number, first 60 AAs: 19.52923\nTotal prob of N-in: 0.94351\nPOSSIBLE N-term signal sequence\ninside 1 19\nTMhelix 20 39\noutside 40 325\nTMhelix 326 348\ninside 349 372

5272 GCF_000752815.1_FHI37 Escherichia coli Proteobacteria; Gammaproteobacteria; Enterobacterales; Enterobacteriaceae; Escherichia MTGDNTLIHSHGINRRDFMKLCAALAATMGLSSKAAAEMAESVTNPQRPPVIWIGAQECT WP_000145410.1 MULTISPECIES: hydrogenase-2 small chain [Proteobacteria] Length: 372\nNumber of predicted TMHs: 1\nExp number of AAs in TMHs: 23.72226\nExp number, first 60 AAs: 0.22038\nTotal prob of N-in: 0.12994\noutside 1 331\nTMhelix 332 354\ninside 355 372

5273 GCF_000752755.1_FHI22 Escherichia coli Proteobacteria; Gammaproteobacteria; Enterobacterales; Enterobacteriaceae; Escherichia MTGDNTLIHSHGINRRDFMKLCAALAATMGLSSKAAAEMAESVTNPQRPPVIWIGAQECT WP_000145410.1 MULTISPECIES: hydrogenase-2 small chain [Proteobacteria] Length: 372\nNumber of predicted TMHs: 1\nExp number of AAs in TMHs: 23.72226\nExp number, first 60 AAs: 0.22038\nTotal prob of N-in: 0.12994\noutside 1 331\nTMhelix 332 354\ninside 355 372

5274 GCF_000754845.1_ASM75484v1 Escherichia coli Proteobacteria; Gammaproteobacteria; Enterobacterales; Enterobacteriaceae; Escherichia MNNEETFYQAMRRQGVTRRSFLKYCSLAATSLGLGAGMAPKIAWALENKPRIPVVWIHGL WP_001058323.1 MULTISPECIES: hydrogenase-1 small chain [Proteobacteria] Length: 372\nNumber of predicted TMHs: 2\nExp number of AAs in TMHs: 37.2608499999999\nExp number, first 60 AAs: 19.52923\nTotal prob of N-in: 0.94351\nPOSSIBLE N-term signal sequence\ninside 1 19\nTMhelix 20 39\noutside 40 325\nTMhelix 326 348\ninside 349 372

5275 GCF_000753195.1_FHI17 Escherichia coli Proteobacteria; Gammaproteobacteria; Enterobacterales; Enterobacteriaceae; Escherichia MNNEETFYQAMRRQGVTRRSFLKYCSLAATSLGLGAGMAPKIAWALENKPRIPVVWIHGL WP_001058323.1 MULTISPECIES: hydrogenase-1 small chain [Proteobacteria] Length: 372\nNumber of predicted TMHs: 2\nExp number of AAs in TMHs: 37.2608499999999\nExp number, first 60 AAs: 19.52923\nTotal prob of N-in: 0.94351\nPOSSIBLE N-term signal sequence\ninside 1 19\nTMhelix 20 39\noutside 40 325\nTMhelix 326 348\ninside 349 372

5276 GCF_000752935.1_FHI49 Escherichia coli Proteobacteria; Gammaproteobacteria; Enterobacterales; Enterobacteriaceae; Escherichia MNNEETFYQAMRRQGVTRRSFLKYCSLAATSLGLGAGMAPKIAWALENKPRIPVVWIHGL WP_001295940.1 MULTISPECIES: hydrogenase [Enterobacteriaceae] Length: 372\nNumber of predicted TMHs: 2\nExp number of AAs in TMHs: 37.2608899999999\nExp number, first 60 AAs: 19.52923\nTotal prob of N-in: 0.94351\nPOSSIBLE N-term signal sequence\ninside 1 19\nTMhelix 20 39\noutside 40 325\nTMhelix 326 348\ninside 349 372

5277 GCF_000753115.1_FHI45 Escherichia coli Proteobacteria; Gammaproteobacteria; Enterobacterales; Enterobacteriaceae; Escherichia MNNEETFYQAMRRQGVTRRSFLKYCSLAATSLGLGAGMAPKIAWALENKPRIPVVWIHGL WP_001295940.1 MULTISPECIES: hydrogenase [Enterobacteriaceae] Length: 372\nNumber of predicted TMHs: 2\nExp number of AAs in TMHs: 37.2608899999999\nExp number, first 60 AAs: 19.52923\nTotal prob of N-in: 0.94351\nPOSSIBLE N-term signal sequence\ninside 1 19\nTMhelix 20 39\noutside 40 325\nTMhelix 326 348\ninside 349 372

5278 GCF_000754865.1_ASM75486v1 Escherichia coli Proteobacteria; Gammaproteobacteria; Enterobacterales; Enterobacteriaceae; Escherichia MNNEETFYQAMRRQGVTRRSFLKYCSLAATSLGLGAGMAPKIAWALENKPRIPVVWIHGL WP_001058323.1 MULTISPECIES: hydrogenase-1 small chain [Proteobacteria] Length: 372\nNumber of predicted TMHs: 2\nExp number of AAs in TMHs: 37.2608499999999\nExp number, first 60 AAs: 19.52923\nTotal prob of N-in: 0.94351\nPOSSIBLE N-term signal sequence\ninside 1 19\nTMhelix 20 39\noutside 40 325\nTMhelix 326 348\ninside 349 372

5279 GCF_000768125.1_ASM76812v1 Escherichia coli Proteobacteria; Gammaproteobacteria; Enterobacterales; Enterobacteriaceae; Escherichia MTGDNTLIHSHGINRRDFMKLCAALAATMGLSSKAAAEMAESVTNPQRPPVIWIGAQECT WP_000145410.1 MULTISPECIES: hydrogenase-2 small chain [Proteobacteria] Length: 372\nNumber of predicted TMHs: 1\nExp number of AAs in TMHs: 23.72226\nExp number, first 60 AAs: 0.22038\nTotal prob of N-in: 0.12994\noutside 1 331\nTMhelix 332 354\ninside 355 372

5280 GCF_000776065.1_ASM77606v1 Escherichia coli Proteobacteria; Gammaproteobacteria; Enterobacterales; Enterobacteriaceae; Escherichia MTGDNTLIHSHGINRRDFMKLCAALAATMGLSSKAAAEMVESVTNPQRPPVIWIGAQECT WP_000145423.1 hydrogenase 2 small subunit [Escherichia coli] Length: 372\nNumber of predicted TMHs: 1\nExp number of AAs in TMHs: 23.78201\nExp number, first 60 AAs: 0.28355\nTotal prob of N-in: 0.13257\noutside 1 331\nTMhelix 332 354\ninside 355 372

5281 GCF_000776115.1_ASM77611v1 Escherichia coli Proteobacteria; Gammaproteobacteria; Enterobacterales; Enterobacteriaceae; Escherichia MTGDNTLIHSHGINRRDFMKLCAALAATMGLSSKAAAEMAESVTNPQRPPVIWIGAQECT WP_000145410.1 MULTISPECIES: hydrogenase-2 small chain [Proteobacteria] Length: 372\nNumber of predicted TMHs: 1\nExp number of AAs in TMHs: 23.72226\nExp number, first 60 AAs: 0.22038\nTotal prob of N-in: 0.12994\noutside 1 331\nTMhelix 332 354\ninside 355 372

5282 GCF_000776065.1_ASM77606v1 Escherichia coli Proteobacteria; Gammaproteobacteria; Enterobacterales; Enterobacteriaceae; Escherichia MNNEETFYQAMRRQGVTRRSFLKYCSLAATSLGLGAGMAPKIAWALENKPRIPVVWIHGL WP_001058323.1 MULTISPECIES: hydrogenase-1 small chain [Proteobacteria] Length: 372\nNumber of predicted TMHs: 2\nExp number of AAs in TMHs: 37.2608499999999\nExp number, first 60 AAs: 19.52923\nTotal prob of N-in: 0.94351\nPOSSIBLE N-term signal sequence\ninside 1 19\nTMhelix 20 39\noutside 40 325\nTMhelix 326 348\ninside 349 372

5283 GCF_000776115.1_ASM77611v1 Escherichia coli Proteobacteria; Gammaproteobacteria; Enterobacterales; Enterobacteriaceae; Escherichia MNNEETFYQAMRRQGVTRRSFLKYCSLAATSLGLGAGMAPKIAWALENKPRIPVVWIHGL WP_001295940.1 MULTISPECIES: hydrogenase [Enterobacteriaceae] Length: 372\nNumber of predicted TMHs: 2\nExp number of AAs in TMHs: 37.2608899999999\nExp number, first 60 AAs: 19.52923\nTotal prob of N-in: 0.94351\nPOSSIBLE N-term signal sequence\ninside 1 19\nTMhelix 20 39\noutside 40 325\nTMhelix 326 348\ninside 349 372

5284 GCF_000770285.1_ASM77028v1 Escherichia coli Proteobacteria; Gammaproteobacteria; Enterobacterales; Enterobacteriaceae; Escherichia

MNNEETFYQAMRRQGVTRRSFLKYCSLAATSLGLGAGMAPKIAWALENKPRIPVVWIHGL WP_001295940.1 MULTISPECIES: hydrogenase [Enterobacteriaceae] Length: 372\nNumber of predicted TMHs: 2\nExp number of AAs in TMHs: 37.2608899999999\nExp number, first 60 AAs: 19.52923\nTotal prob of N-in: 0.94351\nPOSSIBLE N-term signal sequence\ninside 1 19\nTMhelix 20 39\noutside 40 325\nTMhelix 326 348\ninside 349 372

5285 GCF_000770255.1_ASM77025v1 Escherichia coli Proteobacteria; Gammaproteobacteria; Enterobacterales; Enterobacteriaceae; Escherichia MNNEETFYQAMRRQGVTRRSFLKYCSLAATSLGLGAGMAPKIAWALENKPRIPVVWIHGL WP_047603468.1 hydrogenase [Escherichia coli] Length: 372\nNumber of predicted TMHs: 2\nExp number of AAs in TMHs: 39.28018\nExp number, first 60 AAs: 19.5238\nTotal prob of N-in: 0.94367\nPOSSIBLE N-term signal sequence\ninside 1 19\nTMhelix 20 39\noutside 40 326\nTMhelix 327 349\ninside 350 372

5286 GCF_000776155.1_ASM77615v1 Escherichia coli Proteobacteria; Gammaproteobacteria; Enterobacterales; Enterobacteriaceae; Escherichia MNNEETFYQAMRRQGVTRRSFLKYCSLAATSLGLGAGMAPKIAWALENKPRIPVVWIHGL WP_001538532.1 hydrogenase-1 small chain [Escherichia coli] Length: 372\nNumber of predicted TMHs: 2\nExp number of AAs in TMHs: 37.27041\nExp number, first 60 AAs: 19.52927\nTotal prob of N-in: 0.94352\nPOSSIBLE N-term signal sequence\ninside 1 19\nTMhelix 20 39\noutside 40 325\nTMhelix 326 348\ninside 349 372

5287 GCF_000776455.1_ASM77645v1 Escherichia coli Proteobacteria; Gammaproteobacteria; Enterobacterales; Enterobacteriaceae; Escherichia MNNEETFYQAMRRQGVTRRSFLKYCSLAATSLGLGAGMAPKIAWALENKPRIPVVWIHGL WP_001058323.1 MULTISPECIES: hydrogenase-1 small chain [Proteobacteria] Length: 372\nNumber of predicted TMHs: 2\nExp number of AAs in TMHs: 37.2608499999999\nExp number, first 60 AAs: 19.52923\nTotal prob of N-in: 0.94351\nPOSSIBLE N-term signal sequence\ninside 1 19\nTMhelix 20 39\noutside 40 325\nTMhelix 326 348\ninside 349 372

5288 GCF_000776505.1_ASM77650v1 Escherichia coli Proteobacteria; Gammaproteobacteria; Enterobacterales; Enterobacteriaceae; Escherichia MNNEETFYQAMRRQGVTRRSFLKYCSLAATSLGLGAGMAPKIAWALENKPRIPVVWIHGL WP_001058323.1 MULTISPECIES: hydrogenase-1 small chain [Proteobacteria] Length: 372\nNumber of predicted TMHs: 2\nExp number of AAs in TMHs: 37.2608499999999\nExp number, first 60 AAs: 19.52923\nTotal prob of N-in: 0.94351\nPOSSIBLE N-term signal sequence\ninside 1 19\nTMhelix 20 39\noutside 40 325\nTMhelix 326 348\ninside 349 372

5289 GCF_000776695.1_ASM77669v1 Escherichia coli Proteobacteria; Gammaproteobacteria; Enterobacterales; Enterobacteriaceae; Escherichia MTGDNTLIHSHGINRRDFMKLCAALAATMGLSSKAAAEMAESVTNPQRPPVIWIGAQECT WP_000145410.1 MULTISPECIES: hydrogenase-2 small chain [Proteobacteria] Length: 372\nNumber of predicted TMHs: 1\nExp number of AAs in TMHs: 23.72226\nExp number, first 60 AAs: 0.22038\nTotal prob of N-in: 0.12994\noutside 1 331\nTMhelix 332 354\ninside 355 372

5290 GCF_000776735.1_ASM77673v1 Escherichia coli Proteobacteria; Gammaproteobacteria; Enterobacterales; Enterobacteriaceae; Escherichia MTGDNTLIHSHGINRRDFMKLCAALAATMGLSSKAAAEMAESVTNPQRPPVIWIGAQECT WP_000145410.1 MULTISPECIES: hydrogenase-2 small chain [Proteobacteria] Length: 372\nNumber of predicted TMHs: 1\nExp number of AAs in TMHs: 23.72226\nExp number, first 60 AAs: 0.22038\nTotal prob of N-in: 0.12994\noutside 1 331\nTMhelix 332 354\ninside 355 372

5291 GCF_000776695.1_ASM77669v1 Escherichia coli Proteobacteria; Gammaproteobacteria; Enterobacterales; Enterobacteriaceae; Escherichia MNNEETFYQAMRRQGVTRRSFLKYCSLAATSLGLGAGMAPKIAWALENKPRIPVVWIHGL WP_001058323.1 MULTISPECIES: hydrogenase-1 small chain [Proteobacteria] Length: 372\nNumber of predicted TMHs: 2\nExp number of AAs in TMHs: 37.2608499999999\nExp number, first 60 AAs: 19.52923\nTotal prob of N-in: 0.94351\nPOSSIBLE N-term signal sequence\ninside 1 19\nTMhelix 20 39\noutside 40 325\nTMhelix 326 348\ninside 349 372

5292 GCF_000776395.1_ASM77639v1 Escherichia coli Proteobacteria; Gammaproteobacteria; Enterobacterales; Enterobacteriaceae; Escherichia MNNEETFYQAMRRQGVTRRSFLKYCSLAATSLGLGAGMAPKIAWALENKPRIPVVWIHGL WP_001295940.1

MULTISPECIES: hydrogenase [Enterobacteriaceae] Length: 372\nNumber of predicted TMHs: 2\nExp number of AAs in TMHs: 37.2608899999999\nExp number, first 60 AAs: 19.52923\nTotal prob of N-in: 0.94351\nPOSSIBLE N-term signal sequence\ninside 1 19\nTMhelix 20 39\noutside 40 325\nTMhelix 326 348\ninside 349 372

5293 GCF_000777105.1_ASM77710v1 Escherichia coli Proteobacteria; Gammaproteobacteria; Enterobacterales; Enterobacteriaceae; Escherichia
MNNEETFYQAMRRQGVTRRSFLKYCSLAATSLGLGAGMAPKIAWALENKPRIPVVWIHGL WP_001538532.1
hydrogenase-1 small chain [Escherichia coli] Length: 372\nNumber of predicted TMHs: 2\nExp number of AAs in TMHs: 37.27041\nExp number, first 60 AAs: 19.52927\nTotal prob of N-in: 0.94352\nPOSSIBLE N-term signal sequence\ninside 1 19\nTMhelix 20 39\noutside 40 325\nTMhelix 326 348\ninside 349 372

5294 GCF_000777355.1_ASM77735v1 Escherichia coli Proteobacteria; Gammaproteobacteria; Enterobacterales; Enterobacteriaceae; Escherichia
MTGDNTLIHSHGINRRDFMKLCAALAATMGLSSKAAAEMAESVTNPQRPPVIWIGAQECT WP_000145410.1
MULTISPECIES: hydrogenase-2 small chain [Proteobacteria] Length: 372\nNumber of predicted TMHs: 1\nExp number of AAs in TMHs: 23.72226\nExp number, first 60 AAs: 0.22038\nTotal prob of N-in: 0.12994\noutside 1 331\nTMhelix 332 354\ninside 355 372

5295 GCF_000777355.1_ASM77735v1 Escherichia coli Proteobacteria; Gammaproteobacteria; Enterobacterales; Enterobacteriaceae; Escherichia
MNNEETFYQAMRRQGVTRRSFLKYCSLAATSLGLGAGMAPKIAWALENKPRIPVVWIHGL WP_001058323.1
MULTISPECIES: hydrogenase-1 small chain [Proteobacteria] Length: 372\nNumber of predicted TMHs: 2\nExp number of AAs in TMHs: 37.2608499999999\nExp number, first 60 AAs: 19.52923\nTotal prob of N-in: 0.94351\nPOSSIBLE N-term signal sequence\ninside 1 19\nTMhelix 20 39\noutside 40 325\nTMhelix 326 348\ninside 349 372

5296 GCF_000776835.1_ASM77683v1 Escherichia coli Proteobacteria; Gammaproteobacteria; Enterobacterales; Enterobacteriaceae; Escherichia
MNNEETFYQAMRRQGVTRRSFLKYCSLAATSLGLGAGMAPKIAWALENKPRIPVVWIHGL WP_001295940.1
MULTISPECIES: hydrogenase [Enterobacteriaceae] Length: 372\nNumber of predicted TMHs: 2\nExp number of AAs in TMHs: 37.2608899999999\nExp number, first 60 AAs: 19.52923\nTotal prob of N-in: 0.94351\nPOSSIBLE N-term signal sequence\ninside 1 19\nTMhelix 20 39\noutside 40 325\nTMhelix 326 348\ninside 349 372

5297 GCF_000776925.1_ASM77692v1 Escherichia coli Proteobacteria; Gammaproteobacteria; Enterobacterales; Enterobacteriaceae; Escherichia
MNNEETFYQAMRRQGVTRRSFLKYCSLAATSLGLGAGMAPKIAWALENKPRIPVVWIHGL WP_001295940.1
MULTISPECIES: hydrogenase [Enterobacteriaceae] Length: 372\nNumber of predicted TMHs: 2\nExp number of AAs in TMHs: 37.2608899999999\nExp number, first 60 AAs: 19.52923\nTotal prob of N-in: 0.94351\nPOSSIBLE N-term signal sequence\ninside 1 19\nTMhelix 20 39\noutside 40 325\nTMhelix 326 348\ninside 349 372

5298 GCF_000776995.1_ASM77699v1 Escherichia coli Proteobacteria; Gammaproteobacteria; Enterobacterales; Enterobacteriaceae; Escherichia
MNNEETFYQAMRRQGVTRRSFLKYCSLAATSLGLGAGMAPKIAWALENKPRIPVVWIHGL WP_001295940.1
MULTISPECIES: hydrogenase [Enterobacteriaceae] Length: 372\nNumber of predicted TMHs: 2\nExp number of AAs in TMHs: 37.2608899999999\nExp number, first 60 AAs: 19.52923\nTotal prob of N-in: 0.94351\nPOSSIBLE N-term signal sequence\ninside 1 19\nTMhelix 20 39\noutside 40 325\nTMhelix 326 348\ninside 349 372

5299 GCF_000777415.1_ASM77741v1 Escherichia coli Proteobacteria; Gammaproteobacteria; Enterobacterales; Enterobacteriaceae; Escherichia
MTGDNTLIHSHGINRRDFMKLCAALAATMGLSSKAAAEMAESVTNPQRPPVIWIGAQECT WP_000145410.1
MULTISPECIES: hydrogenase-2 small chain [Proteobacteria] Length: 372\nNumber of predicted TMHs: 1\nExp number of AAs in TMHs: 23.72226\nExp number, first 60 AAs: 0.22038\nTotal prob of N-in: 0.12994\noutside 1 331\nTMhelix 332 354\ninside 355 372

5300 GCF_000777675.1_ASM77767v1 Escherichia coli Proteobacteria; Gammaproteobacteria; Enterobacterales; Enterobacteriaceae; Escherichia
MNNEETFYQAMRRQGVTRRSFLKYCSLAATSLGLGAGMAPKIAWALENKPRIPVVWIHGL WP_001058323.1

MULTISPECIES: hydrogenase-1 small chain [Proteobacteria] Length: 372\nNumber of predicted TMHs: 2\nExp number of AAs in TMHs: 37.2608499999999\nExp number, first 60 AAs: 19.52923\nTotal prob of N-in: 0.94351\nPOSSIBLE N-term signal sequence\ninside 1 19\nTMhelix 20 39\noutside 40 325\nTMhelix 326 348\ninside 349 372

5301 GCF_000777415.1_ASM77741v1 Escherichia coli Proteobacteria; Gammaproteobacteria; Enterobacterales; Enterobacteriaceae; Escherichia MNNEETFYQAMRRQGVTRRSFLKYCSLAATSLGLGAGMAPKIAWALENKPRIPVVWIHGL WP_042092595.1 hydrogenase [Escherichia coli] Length: 372\nNumber of predicted TMHs: 2\nExp number of AAs in TMHs: 37.2507099999999\nExp number, first 60 AAs: 19.52643\nTotal prob of N-in: 0.94349\nPOSSIBLE N-term signal sequence\ninside 1 19\nTMhelix 20 39\noutside 40 325\nTMhelix 326 348\ninside 349 372

5302 GCF_000937095.2_FHI87 Escherichia coli Proteobacteria; Gammaproteobacteria; Enterobacterales; Enterobacteriaceae; Escherichia MTGDNTLIHSHGINRRDFMKLCAALAATMGLSSKAAAEMAESVTNPQRPPVIWIGAQECT WP_000145410.1 MULTISPECIES: hydrogenase-2 small chain [Proteobacteria] Length: 372\nNumber of predicted TMHs: 1\nExp number of AAs in TMHs: 23.72226\nExp number, first 60 AAs: 0.22038\nTotal prob of N-in: 0.12994\noutside 1 331\nTMhelix 332 354\ninside 355 372

5303 GCF_001187545.1_ASM118754v1 Escherichia coli Proteobacteria; Gammaproteobacteria; Enterobacterales; Enterobacteriaceae; Escherichia MTGDNTLIHSHGINRRDFMKLCAALAATMGLSSKAAAEMAESVTNPQRPPVIWIGAQECT WP_000145410.1 MULTISPECIES: hydrogenase-2 small chain [Proteobacteria] Length: 372\nNumber of predicted TMHs: 1\nExp number of AAs in TMHs: 23.72226\nExp number, first 60 AAs: 0.22038\nTotal prob of N-in: 0.12994\noutside 1 331\nTMhelix 332 354\ninside 355 372

5304 GCF_000777555.1_ASM77755v1 Escherichia coli Proteobacteria; Gammaproteobacteria; Enterobacterales; Enterobacteriaceae; Escherichia MNNEETFYQAMRRQGVTRRSFLKYCSLAATSLGLGAGMAPKIAWALENKPRIPVVWIHGL WP_001295940.1 MULTISPECIES: hydrogenase [Enterobacteriaceae] Length: 372\nNumber of predicted TMHs: 2\nExp number of AAs in TMHs: 37.2608899999999\nExp number, first 60 AAs: 19.52923\nTotal prob of N-in: 0.94351\nPOSSIBLE N-term signal sequence\ninside 1 19\nTMhelix 20 39\noutside 40 325\nTMhelix 326 348\ninside 349 372

5305 GCF_000778055.1_ASM77805v1 Escherichia coli Proteobacteria; Gammaproteobacteria; Enterobacterales; Enterobacteriaceae; Escherichia MTGDNTLIHSHGINRRDFMKLCAALAATMGLSSKAAAEMAESVTNPQRPPVIWIGAQECT WP_000145410.1 MULTISPECIES: hydrogenase-2 small chain [Proteobacteria] Length: 372\nNumber of predicted TMHs: 1\nExp number of AAs in TMHs: 23.72226\nExp number, first 60 AAs: 0.22038\nTotal prob of N-in: 0.12994\noutside 1 331\nTMhelix 332 354\ninside 355 372

5306 GCF_000778055.1_ASM77805v1 Escherichia coli Proteobacteria; Gammaproteobacteria; Enterobacterales; Enterobacteriaceae; Escherichia MNNEETFYQAMRRQGVTRRSFLKYCSLAATSLGLGAGMAPKIAWALENKPRIPVVWIHGL WP_001058323.1 MULTISPECIES: hydrogenase-1 small chain [Proteobacteria] Length: 372\nNumber of predicted TMHs: 2\nExp number of AAs in TMHs: 37.2608499999999\nExp number, first 60 AAs: 19.52923\nTotal prob of N-in: 0.94351\nPOSSIBLE N-term signal sequence\ninside 1 19\nTMhelix 20 39\noutside 40 325\nTMhelix 326 348\ninside 349 372

5307 GCF_000777515.1_ASM77751v1 Escherichia coli Proteobacteria; Gammaproteobacteria; Enterobacterales; Enterobacteriaceae; Escherichia MNNEETFYQAMRRQGVTRRSFLKYCSLAATSLGLGAGMAPKIAWALENKPRIPVVWIHGL WP_042110493.1 hydrogenase [Escherichia coli] Length: 372\nNumber of predicted TMHs: 2\nExp number of AAs in TMHs: 42.1359999999999\nExp number, first 60 AAs: 19.5205\nTotal prob of N-in: 0.94624\nPOSSIBLE N-term signal sequence\ninside 1 19\nTMhelix 20 39\noutside 40 325\nTMhelix 326 348\ninside 349 372

5308 GCF_000778105.1_ASM77810v1 Escherichia coli Proteobacteria; Gammaproteobacteria; Enterobacterales; Enterobacteriaceae; Escherichia MTGDNTLIHSHGINRRDFMKLCAALAATMGLSSKAAAEMAESVTNPQRPPVIWIGAQECT WP_000145410.1 MULTISPECIES: hydrogenase-2 small chain [Proteobacteria] Length: 372\nNumber of predicted TMHs: 1\nExp number of AAs in TMHs: 23.72226\nExp number, first 60 AAs: 0.22038\nTotal prob of N-in: 0.12994\noutside 1 331\nTMhelix 332 354\ninside 355 372

5309 GCF_000778275.1_ASM77827v1 Escherichia coli Proteobacteria; Gammaproteobacteria; Enterobacterales; Enterobacteriaceae; Escherichia
MNNEETFYQAMRRQGVTRRSFLKYCSLAATSLGLGAGMAPKIAWALENKPRIPVVWIHGL WP_021520630.1
hydrogenase-1 small chain [Escherichia coli] Length: 372\nNumber of predicted TMHs: 2\nExp number of AAs in TMHs: 37.8197599999999\nExp number, first 60 AAs: 19.53295\nTotal prob of N-in: 0.94396\nPOSSIBLE N-term signal sequence\ninside 1 19\nTMhelix 20 39\noutside 40 325\nTMhelix 326 348\ninside 349 372

5310 GCF_000778895.1_ASM77889v1 Escherichia coli Proteobacteria; Gammaproteobacteria; Enterobacterales; Enterobacteriaceae; Escherichia
MTGDNTLIHSHGINRRDFMKLCAALAATMGLSSKAAAEMAESVTNPQRPPVIWIGAQECT WP_000145410.1
MULTISPECIES: hydrogenase-2 small chain [Proteobacteria] Length: 372\nNumber of predicted TMHs: 1\nExp number of AAs in TMHs: 23.72226\nExp number, first 60 AAs: 0.22038\nTotal prob of N-in: 0.12994\noutside 1 331\nTMhelix 332 354\ninside 355 372

5311 GCF_000778895.1_ASM77889v1 Escherichia coli Proteobacteria; Gammaproteobacteria; Enterobacterales; Enterobacteriaceae; Escherichia
MNNEETFYQAMRRQGVTRRSFLKYCSLAATSLGLGAGMAPKIAWALENKPRIPVVWIHGL WP_001058323.1
MULTISPECIES: hydrogenase-1 small chain [Proteobacteria] Length: 372\nNumber of predicted TMHs: 2\nExp number of AAs in TMHs: 37.2608499999999\nExp number, first 60 AAs: 19.52923\nTotal prob of N-in: 0.94351\nPOSSIBLE N-term signal sequence\ninside 1 19\nTMhelix 20 39\noutside 40 325\nTMhelix 326 348\ninside 349 372

5312 GCF_000778475.1_ASM77847v1 Escherichia coli Proteobacteria; Gammaproteobacteria; Enterobacterales; Enterobacteriaceae; Escherichia
MNNEETFYQAMRRQGVTRRSFLKYCSLAATSLGLGAGMAPKIAWALENKPRIPVVWIHGL WP_001295940.1
MULTISPECIES: hydrogenase [Enterobacteriaceae] Length: 372\nNumber of predicted TMHs: 2\nExp number of AAs in TMHs: 37.2608899999999\nExp number, first 60 AAs: 19.52923\nTotal prob of N-in: 0.94351\nPOSSIBLE N-term signal sequence\ninside 1 19\nTMhelix 20 39\noutside 40 325\nTMhelix 326 348\ninside 349 372

5313 GCF_000778935.1_ASM77893v1 Escherichia coli Proteobacteria; Gammaproteobacteria; Enterobacterales; Enterobacteriaceae; Escherichia
MTGDNTLIHSHGINRRDFMKLCAALAATMGLSSKAAAEMVESVTNPQRPPVIWIGAQECT WP_000145423.1
hydrogenase 2 small subunit [Escherichia coli] Length: 372\nNumber of predicted TMHs: 1\nExp number of AAs in TMHs: 23.78201\nExp number, first 60 AAs: 0.28355\nTotal prob of N-in: 0.13257\noutside 1 331\nTMhelix 332 354\ninside 355 372

5314 GCF_000779055.1_ASM77905v1 Escherichia coli Proteobacteria; Gammaproteobacteria; Enterobacterales; Enterobacteriaceae; Escherichia
MTGDNTLIHSHGINRRDFMKLCAALAATMGLSSKAAAEMAESVTNPQRPPVIWIGAQECT WP_000145410.1
MULTISPECIES: hydrogenase-2 small chain [Proteobacteria] Length: 372\nNumber of predicted TMHs: 1\nExp number of AAs in TMHs: 23.72226\nExp number, first 60 AAs: 0.22038\nTotal prob of N-in: 0.12994\noutside 1 331\nTMhelix 332 354\ninside 355 372

5315 GCF_000778965.1_ASM77896v1 Escherichia coli Proteobacteria; Gammaproteobacteria; Enterobacterales; Enterobacteriaceae; Escherichia
MTGDNTLIHSHGINRRDFMKLCAALAATMGLSSKAAAEMAESVTNPQRPPVIWIGAQECT WP_000145410.1
MULTISPECIES: hydrogenase-2 small chain [Proteobacteria] Length: 372\nNumber of predicted TMHs: 1\nExp number of AAs in TMHs: 23.72226\nExp number, first 60 AAs: 0.22038\nTotal prob of N-in: 0.12994\noutside 1 331\nTMhelix 332 354\ninside 355 372

5316 GCF_000779255.1_ASM77925v1 Escherichia coli Proteobacteria; Gammaproteobacteria; Enterobacterales; Enterobacteriaceae; Escherichia
MNNEETFYQAMRRQGVTRRSFLKYCSLAATSLGLGAGMAPKIAWALENKPRIPVVWIHGL WP_001058323.1
MULTISPECIES: hydrogenase-1 small chain [Proteobacteria] Length: 372\nNumber of predicted TMHs: 2\nExp number of AAs in TMHs: 37.2608499999999\nExp number, first 60 AAs: 19.52923\nTotal prob of N-in: 0.94351\nPOSSIBLE N-term signal sequence\ninside 1 19\nTMhelix 20 39\noutside 40 325\nTMhelix 326 348\ninside 349 372

5317 GCF_000778935.1_ASM77893v1 Escherichia coli Proteobacteria; Gammaproteobacteria; Enterobacterales; Enterobacteriaceae; Escherichia

MNNEETFYQAMRRQGVTRRSFLKYCSLAATSLGLGAGMAPKIAWALENKPRIPVVWIHGL WP_001538532.1 hydrogenase-1 small chain [Escherichia coli] Length: 372\nNumber of predicted TMHs: 2\nExp number of AAs in TMHs: 37.27041\nExp number, first 60 AAs: 19.52927\nTotal prob of N-in: 0.94352\nPOSSIBLE N-term signal sequence\ninside 1 19\nTMhelix 20 39\noutside 40 325\nTMhelix 326 348\ninside 349 372

5318 GCF_001284925.1_7790_1_88 Escherichia coli Proteobacteria; Gammaproteobacteria; Enterobacterales; Enterobacteriaceae; Escherichia MNNEETFYQAMRRQGVTRRSFLKYCSLAATSLGLGAGMAPKIAWALENKPRIPVVWIHGL WP_001058323.1 MULTISPECIES: hydrogenase-1 small chain [Proteobacteria] Length: 372\nNumber of predicted TMHs: 2\nExp number of AAs in TMHs: 37.2608499999999\nExp number, first 60 AAs: 19.52923\nTotal prob of N-in: 0.94351\nPOSSIBLE N-term signal sequence\ninside 1 19\nTMhelix 20 39\noutside 40 325\nTMhelix 326 348\ninside 349 372

5319 GCF_000779095.1_ASM77909v1 Escherichia coli Proteobacteria; Gammaproteobacteria; Enterobacterales; Enterobacteriaceae; Escherichia MNNEETFYQAMRRQGVTRRSFLKYCSLAATSLGLGAGMAPKIAWALENKPRIPVVWIHGL WP_001058323.1 MULTISPECIES: hydrogenase-1 small chain [Proteobacteria] Length: 372\nNumber of predicted TMHs: 2\nExp number of AAs in TMHs: 37.2608499999999\nExp number, first 60 AAs: 19.52923\nTotal prob of N-in: 0.94351\nPOSSIBLE N-term signal sequence\ninside 1 19\nTMhelix 20 39\noutside 40 325\nTMhelix 326 348\ninside 349 372

5320 GCF_000779295.1_ASM77929v1 Escherichia coli Proteobacteria; Gammaproteobacteria; Enterobacterales; Enterobacteriaceae; Escherichia MNNEETFYQAMRRQGVTRRSFLKYCSLAATSLGLGAGMAPKIAWALENKPRIPVVWIHGL WP_001058323.1 MULTISPECIES: hydrogenase-1 small chain [Proteobacteria] Length: 372\nNumber of predicted TMHs: 2\nExp number of AAs in TMHs: 37.2608499999999\nExp number, first 60 AAs: 19.52923\nTotal prob of N-in: 0.94351\nPOSSIBLE N-term signal sequence\ninside 1 19\nTMhelix 20 39\noutside 40 325\nTMhelix 326 348\ninside 349 372

5321 GCF_000778965.1_ASM77896v1 Escherichia coli Proteobacteria; Gammaproteobacteria; Enterobacterales; Enterobacteriaceae; Escherichia MNNEETFYQAMRRQGVTRRSFLKYCSLAATSLGLGAGMAPKIAWALENKPRIPVVWIHGL WP_001058323.1 MULTISPECIES: hydrogenase-1 small chain [Proteobacteria] Length: 372\nNumber of predicted TMHs: 2\nExp number of AAs in TMHs: 37.2608499999999\nExp number, first 60 AAs: 19.52923\nTotal prob of N-in: 0.94351\nPOSSIBLE N-term signal sequence\ninside 1 19\nTMhelix 20 39\noutside 40 325\nTMhelix 326 348\ninside 349 372

5322 GCF_001191045.1_CFSAN026843_01-NGE.clc Escherichia coli Proteobacteria; Gammaproteobacteria; Enterobacterales; Enterobacteriaceae; Escherichia MTGDNTLIHSHGINRRDFMKLCAALAATMGLSSKAAAEMAESVTNPQRPPVIWIGAQECT WP_000145410.1 MULTISPECIES: hydrogenase-2 small chain [Proteobacteria] Length: 372\nNumber of predicted TMHs: 1\nExp number of AAs in TMHs: 23.72226\nExp number, first 60 AAs: 0.22038\nTotal prob of N-in: 0.12994\noutside 1 331\nTMhelix 332 354\ninside 355 372

5323 GCF_000779055.1_ASM77905v1 Escherichia coli Proteobacteria; Gammaproteobacteria; Enterobacterales; Enterobacteriaceae; Escherichia MNNEETFYQAMRRQGVTRRSFLKYCSLAATSLGLGAGMAPKIAWALENKPRIPVVWIHGL WP_001058323.1 MULTISPECIES: hydrogenase-1 small chain [Proteobacteria] Length: 372\nNumber of predicted TMHs: 2\nExp number of AAs in TMHs: 37.2608499999999\nExp number, first 60 AAs: 19.52923\nTotal prob of N-in: 0.94351\nPOSSIBLE N-term signal sequence\ninside 1 19\nTMhelix 20 39\noutside 40 325\nTMhelix 326 348\ninside 349 372

5324 GCF_000779455.1_ASM77945v1 Escherichia coli Proteobacteria; Gammaproteobacteria; Enterobacterales; Enterobacteriaceae; Escherichia MTGDNTLIHSHGINRRDFMKLCAALAATMGLSSKAAAEMAESVTNPQRPPVIWIGAQECT WP_000145410.1 MULTISPECIES: hydrogenase-2 small chain [Proteobacteria] Length: 372\nNumber of predicted TMHs: 1\nExp number of AAs in TMHs: 23.72226\nExp number, first 60 AAs: 0.22038\nTotal prob of N-in: 0.12994\noutside 1 331\nTMhelix 332 354\ninside 355 372

5325 GCF_000779935.1_ASM77993v1 Escherichia coli Proteobacteria; Gammaproteobacteria; Enterobacterales; Enterobacteriaceae; Escherichia

MNNEETFYQAMRRQGVTRRSFLKYCSLAATSLGLGAGMAPKIAWALENKPRIPVVWIHGL WP_001295940.1 MULTISPECIES: hydrogenase [Enterobacteriaceae] Length: 372\nNumber of predicted TMHs: 2\nExp number of AAs in TMHs: 37.2608899999999\nExp number, first 60 AAs: 19.52923\nTotal prob of N-in: 0.94351\nPOSSIBLE N-term signal sequence\ninside 1 19\nTMhelix 20 39\noutside 40 325\nTMhelix 326 348\ninside 349 372

5326 GCF_000936085.2_FHI90 Escherichia coli Proteobacteria; Gammaproteobacteria; Enterobacterales; Enterobacteriaceae; Escherichia MNNEETFYQAMRRQGVTRRSFLKYCSLAATSLGLGAGMAPKIAWALENKPRIPVVWIHGL WP_001058323.1 MULTISPECIES: hydrogenase-1 small chain [Proteobacteria] Length: 372\nNumber of predicted TMHs: 2\nExp number of AAs in TMHs: 37.2608499999999\nExp number, first 60 AAs: 19.52923\nTotal prob of N-in: 0.94351\nPOSSIBLE N-term signal sequence\ninside 1 19\nTMhelix 20 39\noutside 40 325\nTMhelix 326 348\ninside 349 372

5327 GCF_000779675.1_ASM77967v1 Escherichia coli Proteobacteria; Gammaproteobacteria; Enterobacterales; Enterobacteriaceae; Escherichia MNNEETFYQAMRRQGVTRRSFLKYCSLAATSLGLGAGMAPKIAWALENKPRIPVVWIHGL WP_001058323.1 MULTISPECIES: hydrogenase-1 small chain [Proteobacteria] Length: 372\nNumber of predicted TMHs: 2\nExp number of AAs in TMHs: 37.2608499999999\nExp number, first 60 AAs: 19.52923\nTotal prob of N-in: 0.94351\nPOSSIBLE N-term signal sequence\ninside 1 19\nTMhelix 20 39\noutside 40 325\nTMhelix 326 348\ninside 349 372

5328 GCF_000779895.1_ASM77989v1 Escherichia coli Proteobacteria; Gammaproteobacteria; Enterobacterales; Enterobacteriaceae; Escherichia MTGDNTLIHSHGINRRDFMKLCAALAATMGLSSKAAAEMVESVTNPQRPPVIWIGAQECT WP_000145423.1 hydrogenase 2 small subunit [Escherichia coli] Length: 372\nNumber of predicted TMHs: 1\nExp number of AAs in TMHs: 23.78201\nExp number, first 60 AAs: 0.28355\nTotal prob of N-in: 0.13257\noutside 1 331\nTMhelix 332 354\ninside 355 372

5329 GCF_000780035.1_ASM78003v1 Escherichia coli Proteobacteria; Gammaproteobacteria; Enterobacterales; Enterobacteriaceae; Escherichia MTGDNTLIHSHGINRRDFMKLCAALAATMGLSSKAAAEMAESVTNPQRPPVIWIGAQECT WP_000145410.1 MULTISPECIES: hydrogenase-2 small chain [Proteobacteria] Length: 372\nNumber of predicted TMHs: 1\nExp number of AAs in TMHs: 23.72226\nExp number, first 60 AAs: 0.22038\nTotal prob of N-in: 0.12994\noutside 1 331\nTMhelix 332 354\ninside 355 372

5330 GCF_000780015.1_ASM78001v1 Escherichia coli Proteobacteria; Gammaproteobacteria; Enterobacterales; Enterobacteriaceae; Escherichia MTGDNTLIHSHGINRRDFMKLCAALAATMGLSSKAAAEMVESVTNPQRPPVIWIGAQECT WP_000145423.1 hydrogenase 2 small subunit [Escherichia coli] Length: 372\nNumber of predicted TMHs: 1\nExp number of AAs in TMHs: 23.78201\nExp number, first 60 AAs: 0.28355\nTotal prob of N-in: 0.13257\noutside 1 331\nTMhelix 332 354\ninside 355 372

5331 GCF_000780035.1_ASM78003v1 Escherichia coli Proteobacteria; Gammaproteobacteria; Enterobacterales; Enterobacteriaceae; Escherichia MNNEETFYQAMRRQGVTRRSFLKYCSLAATSLGLGAGMAPKIAWALENKPRIPVVWIHGL WP_001058323.1 MULTISPECIES: hydrogenase-1 small chain [Proteobacteria] Length: 372\nNumber of predicted TMHs: 2\nExp number of AAs in TMHs: 37.2608499999999\nExp number, first 60 AAs: 19.52923\nTotal prob of N-in: 0.94351\nPOSSIBLE N-term signal sequence\ninside 1 19\nTMhelix 20 39\noutside 40 325\nTMhelix 326 348\ninside 349 372

5332 GCF_000936085.2_FHI90 Escherichia coli Proteobacteria; Gammaproteobacteria; Enterobacterales; Enterobacteriaceae; Escherichia MTGDNTLIHSHGINRRDFMKLCAALAATMGLSSKAAAEMAESVTNPQRPPVIWIGAQECT WP_000145410.1 MULTISPECIES: hydrogenase-2 small chain [Proteobacteria] Length: 372\nNumber of predicted TMHs: 1\nExp number of AAs in TMHs: 23.72226\nExp number, first 60 AAs: 0.22038\nTotal prob of N-in: 0.12994\noutside 1 331\nTMhelix 332 354\ninside 355 372

5333 GCF_000779895.1_ASM77989v1 Escherichia coli Proteobacteria; Gammaproteobacteria; Enterobacterales; Enterobacteriaceae; Escherichia MNNEETFYQAMRRQGVTRRSFLKYCSLAATSLGLGAGMAPKIAWALENKPRIPVVWIHGL WP_001295940.1 MULTISPECIES: hydrogenase [Enterobacteriaceae] Length: 372\nNumber of predicted TMHs: 2\nExp number of AAs in TMHs: 37.2608899999999\nExp number, first 60 AAs: 19.52923\nTotal prob of N-in:

0.94351\nPOSSIBLE N-term signal sequence\ninside 1 19\nTMhelix 20 39\noutside 40 325\nTMhelix 326 348\ninside 349 372

5334 GCF_000780455.1_ASM78045v1 Escherichia coli Proteobacteria; Gammaproteobacteria; Enterobacterales; Enterobacteriaceae; Escherichia MTGDNTLIHSHGINRRDFMKLCAALAATMGLSSKAAAEMVESVTNPQRPPVIWIGAQECT WP_000145423.1 hydrogenase 2 small subunit [Escherichia coli] Length: 372\nNumber of predicted TMHs: 1\nExp number of AAs in TMHs: 23.78201\nExp number, first 60 AAs: 0.28355\nTotal prob of N-in: 0.13257\noutside 1 331\nTMhelix 332 354\ninside 355 372

5335 GCF_000830035.1_ASM83003v1 Escherichia coli Proteobacteria; Gammaproteobacteria; Enterobacterales; Enterobacteriaceae; Escherichia MTGDNTLIHSHGINRRDFMKLCAALAATMGLSSKAAAEMAESVTNPQRPPVIWIGAQECT WP_000145410.1 MULTISPECIES: hydrogenase-2 small chain [Proteobacteria] Length: 372\nNumber of predicted TMHs: 1\nExp number of AAs in TMHs: 23.72226\nExp number, first 60 AAs: 0.22038\nTotal prob of N-in: 0.12994\noutside 1 331\nTMhelix 332 354\ninside 355 372

5336 GCF_000830035.1_ASM83003v1 Escherichia coli Proteobacteria; Gammaproteobacteria; Enterobacterales; Enterobacteriaceae; Escherichia MNNEETFYQAMRRQGVTRRSFLKYCSLAATSLGLGAGMAPKIAWALENKPRIPVVWIHGL WP_001058323.1 MULTISPECIES: hydrogenase-1 small chain [Proteobacteria] Length: 372\nNumber of predicted TMHs: 2\nExp number of AAs in TMHs: 37.2608499999999\nExp number, first 60 AAs: 19.52923\nTotal prob of N-in: 0.94351\nPOSSIBLE N-term signal sequence\ninside 1 19\nTMhelix 20 39\noutside 40 325\nTMhelix 326 348\ninside 349 372

5337 GCF_000780535.1_ASM78053v1 Escherichia coli Proteobacteria; Gammaproteobacteria; Enterobacterales; Enterobacteriaceae; Escherichia MTGDNTLIHSHGINRRDFMKLCAALAATMGLSSKAAAEMAESVTNPQRPPVIWIGAQECT WP_000145410.1 MULTISPECIES: hydrogenase-2 small chain [Proteobacteria] Length: 372\nNumber of predicted TMHs: 1\nExp number of AAs in TMHs: 23.72226\nExp number, first 60 AAs: 0.22038\nTotal prob of N-in: 0.12994\noutside 1 331\nTMhelix 332 354\ninside 355 372

5338 GCF_000831335.1_Escherichia_coli_CVM_N34086PS_v1.0 Escherichia coli Proteobacteria; Gammaproteobacteria; Enterobacterales; Enterobacteriaceae; Escherichia MTGDNTLIHSHGINRRDFMKLCAALAATMGLSSKAAAEMAESVTNPQRPPVIWIGAQECT WP_000145410.1 MULTISPECIES: hydrogenase-2 small chain [Proteobacteria] Length: 372\nNumber of predicted TMHs: 1\nExp number of AAs in TMHs: 23.72226\nExp number, first 60 AAs: 0.22038\nTotal prob of N-in: 0.12994\noutside 1 331\nTMhelix 332 354\ninside 355 372

5339 GCF_000831335.1_Escherichia_coli_CVM_N34086PS_v1.0 Escherichia coli Proteobacteria; Gammaproteobacteria; Enterobacterales; Enterobacteriaceae; Escherichia MNNEETFYQAMRRQGVTRRSFLKYCSLAATSLGLGAGMAPKIAWALENKPRIPVVWIHGL WP_001058323.1 MULTISPECIES: hydrogenase-1 small chain [Proteobacteria] Length: 372\nNumber of predicted TMHs: 2\nExp number of AAs in TMHs: 37.2608499999999\nExp number, first 60 AAs: 19.52923\nTotal prob of N-in: 0.94351\nPOSSIBLE N-term signal sequence\ninside 1 19\nTMhelix 20 39\noutside 40 325\nTMhelix 326 348\ninside 349 372

5340 GCF_000780575.1_ASM78057v1 Escherichia coli Proteobacteria; Gammaproteobacteria; Enterobacterales; Enterobacteriaceae; Escherichia MTGDNTLIHSHGINRRDFMKLCAALAATMGLSSKAAAEMAESVTNPQRPPVIWIGAQECT WP_000145410.1 MULTISPECIES: hydrogenase-2 small chain [Proteobacteria] Length: 372\nNumber of predicted TMHs: 1\nExp number of AAs in TMHs: 23.72226\nExp number, first 60 AAs: 0.22038\nTotal prob of N-in: 0.12994\noutside 1 331\nTMhelix 332 354\ninside 355 372

5341 GCF_000780635.1_ASM78063v1 Escherichia coli Proteobacteria; Gammaproteobacteria; Enterobacterales; Enterobacteriaceae; Escherichia MTGDNTLIHSHGINRRDFMKLCAALAATMGLSSKAAAEMVESVTNPQRPPVIWIGAQECT WP_000145423.1 hydrogenase 2 small subunit [Escherichia coli] Length: 372\nNumber of predicted TMHs: 1\nExp number of AAs in TMHs: 23.78201\nExp number, first 60 AAs: 0.28355\nTotal prob of N-in: 0.13257\noutside 1 331\nTMhelix 332 354\ninside 355 372

5342 GCF_000780675.1_ASM78067v1 Escherichia coli Proteobacteria; Gammaproteobacteria; Enterobacterales; Enterobacteriaceae; Escherichia
MTGDNTLIHSHGINRRDFMKLCAALAATMGLSSKAAAEMAESVTNPQRPPVIWIGAQECT WP_000145410.1
MULTISPECIES: hydrogenase-2 small chain [Proteobacteria] Length: 372\nNumber of predicted TMHs: 1\nExp number of AAs in TMHs: 23.72226\nExp number, first 60 AAs: 0.22038\nTotal prob of N-in: 0.12994\noutside 1 331\nTMhelix 332 354\ninside 355 372

5343 GCF_000780575.1_ASM78057v1 Escherichia coli Proteobacteria; Gammaproteobacteria; Enterobacterales; Enterobacteriaceae; Escherichia
MNNEETFYQAMRRQGVTRRSFLKYCSLAATSLGLGAGMAPKIAWALENKPRIPVVWIHGL WP_001058323.1
MULTISPECIES: hydrogenase-1 small chain [Proteobacteria] Length: 372\nNumber of predicted TMHs: 2\nExp number of AAs in TMHs: 37.2608499999999\nExp number, first 60 AAs: 19.52923\nTotal prob of N-in: 0.94351\nPOSSIBLE N-term signal sequence\ninside 1 19\nTMhelix 20 39\noutside 40 325\nTMhelix 326 348\ninside 349 372

5344 GCF_000780535.1_ASM78053v1 Escherichia coli Proteobacteria; Gammaproteobacteria; Enterobacterales; Enterobacteriaceae; Escherichia
MNNEETFYQAMRRQGVTRRSFLKYCSLAATSLGLGAGMAPKIAWALENKPRIPVVWIHGL WP_001058320.1
hydrogenase [Escherichia coli] Length: 372\nNumber of predicted TMHs: 2\nExp number of AAs in TMHs: 37.26082\nExp number, first 60 AAs: 19.52923\nTotal prob of N-in: 0.94351\nPOSSIBLE N-term signal sequence\ninside 1 19\nTMhelix 20 39\noutside 40 325\nTMhelix 326 348\ninside 349 372

5345 GCF_000780925.1_ASM78092v1 Escherichia coli Proteobacteria; Gammaproteobacteria; Enterobacterales; Enterobacteriaceae; Escherichia
MTGDNTLIHSHGINRRDFMKLCAALAATMGLSSKAAAEMAESVTNPQRPPVIWIGAQECT WP_000145410.1
MULTISPECIES: hydrogenase-2 small chain [Proteobacteria] Length: 372\nNumber of predicted TMHs: 1\nExp number of AAs in TMHs: 23.72226\nExp number, first 60 AAs: 0.22038\nTotal prob of N-in: 0.12994\noutside 1 331\nTMhelix 332 354\ninside 355 372

5346 GCF_000780995.1_ASM78099v1 Escherichia coli Proteobacteria; Gammaproteobacteria; Enterobacterales; Enterobacteriaceae; Escherichia
MTGDNTLIHSHGINRRDFMKLCAALAATMGLSSKAAAEMAESVTNPQRPPVIWIGAQECT WP_000145410.1
MULTISPECIES: hydrogenase-2 small chain [Proteobacteria] Length: 372\nNumber of predicted TMHs: 1\nExp number of AAs in TMHs: 23.72226\nExp number, first 60 AAs: 0.22038\nTotal prob of N-in: 0.12994\noutside 1 331\nTMhelix 332 354\ninside 355 372

5347 GCF_000781045.1_ASM78104v1 Escherichia coli Proteobacteria; Gammaproteobacteria; Enterobacterales; Enterobacteriaceae; Escherichia
MTGDNTLIHSHGINRRDFMKLCAALAATMGLSSKAAAEMAESVTNPQRPPVIWIGAQECT WP_000145410.1
MULTISPECIES: hydrogenase-2 small chain [Proteobacteria] Length: 372\nNumber of predicted TMHs: 1\nExp number of AAs in TMHs: 23.72226\nExp number, first 60 AAs: 0.22038\nTotal prob of N-in: 0.12994\noutside 1 331\nTMhelix 332 354\ninside 355 372

5348 GCF_000935435.1_ASM93543v1 Escherichia coli Proteobacteria; Gammaproteobacteria; Enterobacterales; Enterobacteriaceae; Escherichia
MNNEETFYQAMRRQGVTRRSFLKYCSLAATSLGLGAGMAPKIAWALENKPRIPVVWIHGL WP_001058323.1
MULTISPECIES: hydrogenase-1 small chain [Proteobacteria] Length: 372\nNumber of predicted TMHs: 2\nExp number of AAs in TMHs: 37.2608499999999\nExp number, first 60 AAs: 19.52923\nTotal prob of N-in: 0.94351\nPOSSIBLE N-term signal sequence\ninside 1 19\nTMhelix 20 39\noutside 40 325\nTMhelix 326 348\ninside 349 372

5349 GCF_000780845.1_ASM78084v1 Escherichia coli Proteobacteria; Gammaproteobacteria; Enterobacterales; Enterobacteriaceae; Escherichia
MNNEETFYQAMRRQGVTRRSFLKYCSLAATSLGLGAGMAPKIAWALENKPRIPVVWIHGL WP_001058323.1
MULTISPECIES: hydrogenase-1 small chain [Proteobacteria] Length: 372\nNumber of predicted TMHs: 2\nExp number of AAs in TMHs: 37.2608499999999\nExp number, first 60 AAs: 19.52923\nTotal prob of N-in: 0.94351\nPOSSIBLE N-term signal sequence\ninside 1 19\nTMhelix 20 39\noutside 40 325\nTMhelix 326 348\ninside 349 372

5350 GCF_001286705.1_8205_8_76 Escherichia coli Proteobacteria; Gammaproteobacteria; Enterobacterales; Enterobacteriaceae; Escherichia

MTGDNTLIHSHGINRRDFMKLCAALAATMGLSSKAAAEMAESVTNPQRPPVIWIGAQECT WP_000145410.1 MULTISPECIES: hydrogenase-2 small chain [Proteobacteria] Length: 372\nNumber of predicted TMHs: 1\nExp number of AAs in TMHs: 23.72226\nExp number, first 60 AAs: 0.22038\nTotal prob of N-in: 0.12994\noutside 1 331\nTMhelix 332 354\ninside 355 372

5351 GCF_001297985.1_ASM129798v1 Escherichia coli Proteobacteria; Gammaproteobacteria; Enterobacterales; Enterobacteriaceae; Escherichia MTGDNTLIHSHGINRRDFMKLCAALAATMGLSSKAAAEMAESVTNPQRPPVIWIGAQECT WP_000145410.1 MULTISPECIES: hydrogenase-2 small chain [Proteobacteria] Length: 372\nNumber of predicted TMHs: 1\nExp number of AAs in TMHs: 23.72226\nExp number, first 60 AAs: 0.22038\nTotal prob of N-in: 0.12994\noutside 1 331\nTMhelix 332 354\ninside 355 372

5352 GCF_000935375.1_ASM93537v1 Escherichia coli Proteobacteria; Gammaproteobacteria; Enterobacterales; Enterobacteriaceae; Escherichia MNNEETFYQAMRRQGVTRRSFLKYCSLAATSLGLGAGMAPKIAWALENKPRIPVVWIHGL WP_001058323.1 MULTISPECIES: hydrogenase-1 small chain [Proteobacteria] Length: 372\nNumber of predicted TMHs: 2\nExp number of AAs in TMHs: 37.2608499999999\nExp number, first 60 AAs: 19.52923\nTotal prob of N-in: 0.94351\nPOSSIBLE N-term signal sequence\ninside 1 19\nTMhelix 20 39\noutside 40 325\nTMhelix 326 348\ninside 349 372

5353 GCF_001306685.1_ASM130668v1 Escherichia coli Proteobacteria; Gammaproteobacteria; Enterobacterales; Enterobacteriaceae; Escherichia MNNEETFYQAMRRQGVTRRSFLKYCSLAATSLGLGAGMAPKIAWALENKPRIPVVWIHGL WP_001058323.1 MULTISPECIES: hydrogenase-1 small chain [Proteobacteria] Length: 372\nNumber of predicted TMHs: 2\nExp number of AAs in TMHs: 37.2608499999999\nExp number, first 60 AAs: 19.52923\nTotal prob of N-in: 0.94351\nPOSSIBLE N-term signal sequence\ninside 1 19\nTMhelix 20 39\noutside 40 325\nTMhelix 326 348\ninside 349 372

5354 GCF_001306685.1_ASM130668v1 Escherichia coli Proteobacteria; Gammaproteobacteria; Enterobacterales; Enterobacteriaceae; Escherichia MTGDNTLIHSHGINRRDFMKLCAALAATMGLSSKAAAEMAESVTNPQRPPVIWIGAQECT WP_000145410.1 MULTISPECIES: hydrogenase-2 small chain [Proteobacteria] Length: 372\nNumber of predicted TMHs: 1\nExp number of AAs in TMHs: 23.72226\nExp number, first 60 AAs: 0.22038\nTotal prob of N-in: 0.12994\noutside 1 331\nTMhelix 332 354\ninside 355 372

5355 GCF_001306635.1_ASM130663v1 Escherichia coli Proteobacteria; Gammaproteobacteria; Enterobacterales; Enterobacteriaceae; Escherichia MTGDNTLIHSHGINRRDFMKLCAALAATMGLSSKAAAEMAESVTNPQRPPVIWIGAQECT WP_000145410.1 MULTISPECIES: hydrogenase-2 small chain [Proteobacteria] Length: 372\nNumber of predicted TMHs: 1\nExp number of AAs in TMHs: 23.72226\nExp number, first 60 AAs: 0.22038\nTotal prob of N-in: 0.12994\noutside 1 331\nTMhelix 332 354\ninside 355 372

5356 GCF_001306585.1_ASM130658v1 Escherichia coli Proteobacteria; Gammaproteobacteria; Enterobacterales; Enterobacteriaceae; Escherichia MNNEETFYQAMRRQGVTRRSFLKYCSLAATSLGLGAGMAPKIAWALENKPRIPVVWIHGL WP_001058323.1 MULTISPECIES: hydrogenase-1 small chain [Proteobacteria] Length: 372\nNumber of predicted TMHs: 2\nExp number of AAs in TMHs: 37.2608499999999\nExp number, first 60 AAs: 19.52923\nTotal prob of N-in: 0.94351\nPOSSIBLE N-term signal sequence\ninside 1 19\nTMhelix 20 39\noutside 40 325\nTMhelix 326 348\ninside 349 372

5357 GCF_000833145.1_ASM83314v1 Escherichia coli Proteobacteria; Gammaproteobacteria; Enterobacterales; Enterobacteriaceae; Escherichia MTGDNTLIHSHGINRRDFMKLCAALAATMGLSSKAAAEMAESVTNPQRPPVIWIGAQECT WP_000145410.1 MULTISPECIES: hydrogenase-2 small chain [Proteobacteria] Length: 372\nNumber of predicted TMHs: 1\nExp number of AAs in TMHs: 23.72226\nExp number, first 60 AAs: 0.22038\nTotal prob of N-in: 0.12994\noutside 1 331\nTMhelix 332 354\ninside 355 372

5358 GCF_000835045.1_ASM83504v1 Escherichia coli Proteobacteria; Gammaproteobacteria; Enterobacterales; Enterobacteriaceae; Escherichia MNNEETFYQAMRRQGVTRRSFLKYCSLAATSLGLGAGMAPKIAWALENKPRIPVVWIHGL WP_001058320.1 hydrogenase [Escherichia coli] Length: 372\nNumber of predicted TMHs: 2\nExp number of AAs in

TMHs: 37.26082\nExp number, first 60 AAs: 19.52923\nTotal prob of N-in: 0.94351\nPOSSIBLE N-term signal sequence\ninside 1 19\nTMhelix 20 39\noutside 40 325\nTMhelix 326 348\ninside 349 372

5359 GCF_000835045.1_ASM83504v1 Escherichia coli Proteobacteria; Gammaproteobacteria; Enterobacterales; Enterobacteriaceae; Escherichia MTGDNTLIHSHGINRRDFMKLCAALAATMGLSSKAAAEMAESVTNPQRPPVIWIGAQECT WP_000145410.1 MULTISPECIES: hydrogenase-2 small chain [Proteobacteria] Length: 372\nNumber of predicted TMHs: 1\nExp number of AAs in TMHs: 23.72226\nExp number, first 60 AAs: 0.22038\nTotal prob of N-in: 0.12994\noutside 1 331\nTMhelix 332 354\ninside 355 372

5360 GCF_001283845.1_7790_1_73 Escherichia coli Proteobacteria; Gammaproteobacteria; Enterobacterales; Enterobacteriaceae; Escherichia MNNEETFYQAMRRQGVTRRSFLKYCSLAATSLGLGAGMAPKIAWALENKPRIPVVWIHGL WP_001058323.1 MULTISPECIES: hydrogenase-1 small chain [Proteobacteria] Length: 372\nNumber of predicted TMHs: 2\nExp number of AAs in TMHs: 37.2608499999999\nExp number, first 60 AAs: 19.52923\nTotal prob of N-in: 0.94351\nPOSSIBLE N-term signal sequence\ninside 1 19\nTMhelix 20 39\noutside 40 325\nTMhelix 326 348\ninside 349 372

5361 GCF_001283885.1_7748_7_18 Escherichia coli Proteobacteria; Gammaproteobacteria; Enterobacterales; Enterobacteriaceae; Escherichia MNNEETFYQAMRRQGVTRRSFLKYCSLAATSLGLGAGMAPKIAWALENKPRIPVVWIHGL WP_001295940.1 MULTISPECIES: hydrogenase [Enterobacteriaceae] Length: 372\nNumber of predicted TMHs: 2\nExp number of AAs in TMHs: 37.2608899999999\nExp number, first 60 AAs: 19.52923\nTotal prob of N-in: 0.94351\nPOSSIBLE N-term signal sequence\ninside 1 19\nTMhelix 20 39\noutside 40 325\nTMhelix 326 348\ninside 349 372

5362 GCF_001269145.1_YE20 Escherichia coli Proteobacteria; Gammaproteobacteria; Enterobacterales; Enterobacteriaceae; Escherichia MTGDNTLIHSHGINRRDFMKLCAALAATMGLSSKAAAEMAESVTNPQRPPVIWIGAQECT WP_000145410.1 MULTISPECIES: hydrogenase-2 small chain [Proteobacteria] Length: 372\nNumber of predicted TMHs: 1\nExp number of AAs in TMHs: 23.72226\nExp number, first 60 AAs: 0.22038\nTotal prob of N-in: 0.12994\noutside 1 331\nTMhelix 332 354\ninside 355 372

5363 GCF_001283965.1_8205_3_5Escherichia coli Proteobacteria; Gammaproteobacteria; Enterobacterales; Enterobacteriaceae; Escherichia MNNEETFYQAMRRQGVTRRSFLKYCSLAATSLGLGAGMAPKIAWALENKPRIPVVWIHGL WP_001295940.1 MULTISPECIES: hydrogenase [Enterobacteriaceae] Length: 372\nNumber of predicted TMHs: 2\nExp number of AAs in TMHs: 37.2608899999999\nExp number, first 60 AAs: 19.52923\nTotal prob of N-in: 0.94351\nPOSSIBLE N-term signal sequence\ninside 1 19\nTMhelix 20 39\noutside 40 325\nTMhelix 326 348\ninside 349 372

5364 GCF_001284065.1_8205_3_57 Escherichia coli Proteobacteria; Gammaproteobacteria; Enterobacterales; Enterobacteriaceae; Escherichia MTGDNTLIHSHGINRRDFMKLCAALAATMGLSSKAAAEMAESVTNPQRPPVIWIGAQECT WP_000145410.1 MULTISPECIES: hydrogenase-2 small chain [Proteobacteria] Length: 372\nNumber of predicted TMHs: 1\nExp number of AAs in TMHs: 23.72226\nExp number, first 60 AAs: 0.22038\nTotal prob of N-in: 0.12994\noutside 1 331\nTMhelix 332 354\ninside 355 372

5365 GCF_001284065.1_8205_3_57 Escherichia coli Proteobacteria; Gammaproteobacteria; Enterobacterales; Enterobacteriaceae; Escherichia MNNEETFYQAMRRQGVTRRSFLKYCSLAATSLGLGAGMAPKIAWALENKPRIPVVWIHGL WP_001058323.1 MULTISPECIES: hydrogenase-1 small chain [Proteobacteria] Length: 372\nNumber of predicted TMHs: 2\nExp number of AAs in TMHs: 37.2608499999999\nExp number, first 60 AAs: 19.52923\nTotal prob of N-in: 0.94351\nPOSSIBLE N-term signal sequence\ninside 1 19\nTMhelix 20 39\noutside 40 325\nTMhelix 326 348\ninside 349 372

5366 GCF_001284285.1_8205_8_81 Escherichia coli Proteobacteria; Gammaproteobacteria; Enterobacterales; Enterobacteriaceae; Escherichia MTGDNTLIHSHGINRRDFMKLCAALAATMGLSSKAAAEMAESVTNPQRPPVIWIGAQECT WP_000145410.1 MULTISPECIES: hydrogenase-2 small chain [Proteobacteria] Length: 372\nNumber of predicted TMHs: 1\nExp number of AAs in TMHs: 23.72226\nExp number, first 60 AAs: 0.22038\nTotal prob of N-in: 0.12994\noutside 1 331\nTMhelix 332 354\ninside 355 372

5367 GCF_000798155.1_Escherichia_coli_CVM_N34077PS_v._1.0 Escherichia coli Proteobacteria; Gammaproteobacteria; Enterobacterales; Enterobacteriaceae; Escherichia MNNEETFYQAMRRQGVTRRSFLKYCSLAATSLGLGAGMAPKIAWALENKPRIPVVWIHGL WP_001295940.1 MULTISPECIES: hydrogenase [Enterobacteriaceae] Length: 372\nNumber of predicted TMHs: 2\nExp number of AAs in TMHs: 37.2608899999999\nExp number, first 60 AAs: 19.52923\nTotal prob of N-in: 0.94351\nPOSSIBLE N-term signal sequence\ninside 1 19\nTMhelix 20 39\noutside 40 325\nTMhelix 326 348\ninside 349 372

5368 GCF_001284725.1_7748_7_36 Escherichia coli Proteobacteria; Gammaproteobacteria; Enterobacterales; Enterobacteriaceae; Escherichia MNNEETFYQAMRRQGVTRRSFLKYCSLAATSLGLGAGMAPKIAWALENKPRIPVVWIHGL WP_001058323.1 MULTISPECIES: hydrogenase-1 small chain [Proteobacteria] Length: 372\nNumber of predicted TMHs: 2\nExp number of AAs in TMHs: 37.2608499999999\nExp number, first 60 AAs: 19.52923\nTotal prob of N-in: 0.94351\nPOSSIBLE N-term signal sequence\ninside 1 19\nTMhelix 20 39\noutside 40 325\nTMhelix 326 348\ninside 349 372

5369 GCF_001284725.1_7748_7_36 Escherichia coli Proteobacteria; Gammaproteobacteria; Enterobacterales; Enterobacteriaceae; Escherichia MTGDNTLIHSHGINRRDFMKLCAALAATMGLSSKAAAEMAESVTNPQRPPVIWIGAQECT WP_000145410.1 MULTISPECIES: hydrogenase-2 small chain [Proteobacteria] Length: 372\nNumber of predicted TMHs: 1\nExp number of AAs in TMHs: 23.72226\nExp number, first 60 AAs: 0.22038\nTotal prob of N-in: 0.12994\noutside 1 331\nTMhelix 332 354\ninside 355 372

5370 GCF_001277435.1_7748_7_2Escherichia coli Proteobacteria; Gammaproteobacteria; Enterobacterales; Enterobacteriaceae; Escherichia MNNEETFYQAMRRQGVTRRSFLKYCSLAATSLGLGAGMAPKIAWALENKPRIPVVWIHGL WP_001359719.1 hydrogenase [Escherichia coli] Length: 372\nNumber of predicted TMHs: 2\nExp number of AAs in TMHs: 37.2176\nExp number, first 60 AAs: 19.53448\nTotal prob of N-in: 0.94360\nPOSSIBLE N-term signal sequence\ninside 1 19\nTMhelix 20 39\noutside 40 325\nTMhelix 326 348\ninside 349 372

5371 GCF_000797835.1_Escherichia_coli_CVM_N33804PS_v._1.0 Escherichia coli Proteobacteria; Gammaproteobacteria; Enterobacterales; Enterobacteriaceae; Escherichia MNNEETFYQAMRRQGVTRRSFLKYCSLAATSLGLGAGMAPKIAWALENKPRIPVVWIHGL WP_001295940.1 MULTISPECIES: hydrogenase [Enterobacteriaceae] Length: 372\nNumber of predicted TMHs: 2\nExp number of AAs in TMHs: 37.2608899999999\nExp number, first 60 AAs: 19.52923\nTotal prob of N-in: 0.94351\nPOSSIBLE N-term signal sequence\ninside 1 19\nTMhelix 20 39\noutside 40 325\nTMhelix 326 348\ninside 349 372

5372 GCF_001284105.1_8205_3_63 Escherichia coli Proteobacteria; Gammaproteobacteria; Enterobacterales; Enterobacteriaceae; Escherichia MTGDNTLIHSHGINRRDFMKLCAALAATMGLSSKAAAEMAESVTNPQRPPVIWIGAQECT WP_000145410.1 MULTISPECIES: hydrogenase-2 small chain [Proteobacteria] Length: 372\nNumber of predicted TMHs: 1\nExp number of AAs in TMHs: 23.72226\nExp number, first 60 AAs: 0.22038\nTotal prob of N-in: 0.12994\noutside 1 331\nTMhelix 332 354\ninside 355 372

5373 GCF_001276585.2_ASM127658v2 Escherichia coli Proteobacteria; Gammaproteobacteria; Enterobacterales; Enterobacteriaceae; Escherichia MTGDNTLIHSHGINRRDFMKLCAALAATMGLSSKAAAEMAESVTNPQRPPVIWIGAQECT WP_000145410.1 MULTISPECIES: hydrogenase-2 small chain [Proteobacteria] Length: 372\nNumber of predicted TMHs: 1\nExp number of AAs in TMHs: 23.72226\nExp number, first 60 AAs: 0.22038\nTotal prob of N-in: 0.12994\noutside 1 331\nTMhelix 332 354\ninside 355 372

5374 GCF_001283885.1_7748_7_18 Escherichia coli Proteobacteria; Gammaproteobacteria; Enterobacterales; Enterobacteriaceae; Escherichia MTGDNTLIHSHGINRRDFMKLCAALAATMGLSSKAAAEMAESVTNPQRPPVIWIGAQECT WP_000145410.1 MULTISPECIES: hydrogenase-2 small chain [Proteobacteria] Length: 372\nNumber of predicted TMHs: 1\nExp number of AAs in TMHs: 23.72226\nExp number, first 60 AAs: 0.22038\nTotal prob of N-in: 0.12994\noutside 1 331\nTMhelix 332 354\ninside 355 372

5375 GCF_001269185.1_YE13 Escherichia coli Proteobacteria; Gammaproteobacteria; Enterobacterales; Enterobacteriaceae; Escherichia MNNEETFYQAMRRQGVTRRSFLKYCSLAATSLGLGAGMAPKIAWALENKPRIPVVWIHGL WP_001295940.1 MULTISPECIES: hydrogenase [Enterobacteriaceae] Length: 372\nNumber of

predicted TMHs: 2\nExp number of AAs in TMHs: 37.2608899999999\nExp number, first 60 AAs: 19.52923\nTotal prob of N-in: 0.94351\nPOSSIBLE N-term signal sequence\ninside 1 19\nTMhelix 20 39\noutside 40 325\nTMhelix 326 348\ninside 349 372

5376 GCF_001284825.1_7790_1_50 Escherichia coli Proteobacteria; Gammaproteobacteria; Enterobacterales; Enterobacteriaceae; Escherichia MTGDNTLIHSHGINRRDFMKLCAALAATMGLSSKAAAEMAESVTNPQRPPVIWIGAQECT WP_000145410.1 MULTISPECIES: hydrogenase-2 small chain [Proteobacteria] Length: 372\nNumber of predicted TMHs: 1\nExp number of AAs in TMHs: 23.72226\nExp number, first 60 AAs: 0.22038\nTotal prob of N-in: 0.12994\noutside 1 331\nTMhelix 332 354\ninside 355 372

5377 GCF_001284665.1_7748_7_34 Escherichia coli Proteobacteria; Gammaproteobacteria; Enterobacterales; Enterobacteriaceae; Escherichia MTGDNTLIHSHGINRRDFMKLCAALAATMGLSSKAAAEMAESVTNPQRPPVIWIGAQECT WP_000145410.1 MULTISPECIES: hydrogenase-2 small chain [Proteobacteria] Length: 372\nNumber of predicted TMHs: 1\nExp number of AAs in TMHs: 23.72226\nExp number, first 60 AAs: 0.22038\nTotal prob of N-in: 0.12994\noutside 1 331\nTMhelix 332 354\ninside 355 372

5378 GCF_000797935.1_Escherichia_coli_CVM_N36396PS_v._1.0 Escherichia coli Proteobacteria; Gammaproteobacteria; Enterobacterales; Enterobacteriaceae; Escherichia MTGDNTLIHSHGINRRDFMKLCAALAATMGLSSKAAAEMAESVTNPQRPPVIWIGAQECT WP_000145410.1 MULTISPECIES: hydrogenase-2 small chain [Proteobacteria] Length: 372\nNumber of predicted TMHs: 1\nExp number of AAs in TMHs: 23.72226\nExp number, first 60 AAs: 0.22038\nTotal prob of N-in: 0.12994\noutside 1 331\nTMhelix 332 354\ninside 355 372

5379 GCF_001277475.1_7790_1_89 Escherichia coli Proteobacteria; Gammaproteobacteria; Enterobacterales; Enterobacteriaceae; Escherichia MTGDNTLIHSHGINRRDFMKLCAALAATMGLSSKAAAEMAESVTNPQRPPVIWIGAQECT WP_000145410.1 MULTISPECIES: hydrogenase-2 small chain [Proteobacteria] Length: 372\nNumber of predicted TMHs: 1\nExp number of AAs in TMHs: 23.72226\nExp number, first 60 AAs: 0.22038\nTotal prob of N-in: 0.12994\noutside 1 331\nTMhelix 332 354\ninside 355 372

5380 GCF_001277475.1_7790_1_89 Escherichia coli Proteobacteria; Gammaproteobacteria; Enterobacterales; Enterobacteriaceae; Escherichia MNNEETFYQAMRRQGVTRRSFLKYCSLAATSLGLGAGMAPKIAWALENKPRIPVVWIHGL WP_001295940.1 MULTISPECIES: hydrogenase [Enterobacteriaceae] Length: 372\nNumber of predicted TMHs: 2\nExp number of AAs in TMHs: 37.2608899999999\nExp number, first 60 AAs: 19.52923\nTotal prob of N-in: 0.94351\nPOSSIBLE N-term signal sequence\ninside 1 19\nTMhelix 20 39\noutside 40 325\nTMhelix 326 348\ninside 349 372

5381 GCF_001284545.1_7790_1_53 Escherichia coli Proteobacteria; Gammaproteobacteria; Enterobacterales; Enterobacteriaceae; Escherichia MNNEETFYQAMRRQGVTRRSFLKYCSLAATSLGLGAGMAPKIAWALENKPRIPVVWIHGL WP_053884447.1 hydrogenase [Escherichia coli] Length: 372\nNumber of predicted TMHs: 2\nExp number of AAs in TMHs: 37.6111999999999\nExp number, first 60 AAs: 19.5124\nTotal prob of N-in: 0.94377\nPOSSIBLE N-term signal sequence\ninside 1 19\nTMhelix 20 39\noutside 40 325\nTMhelix 326 348\ninside 349 372

5382 GCF_001277575.1_7553_7_61 Escherichia coli Proteobacteria; Gammaproteobacteria; Enterobacterales; Enterobacteriaceae; Escherichia MTGDNTLIHSHGINRRDFMKLCAALAATMGLSSKAAAEMAESVTNPQRPPVIWIGAQECT WP_000145410.1 MULTISPECIES: hydrogenase-2 small chain [Proteobacteria] Length: 372\nNumber of predicted TMHs: 1\nExp number of AAs in TMHs: 23.72226\nExp number, first 60 AAs: 0.22038\nTotal prob of N-in: 0.12994\noutside 1 331\nTMhelix 332 354\ninside 355 372

5383 GCF_001277635.1_7790_1_49 Escherichia coli Proteobacteria; Gammaproteobacteria; Enterobacterales; Enterobacteriaceae; Escherichia MNNEETFYQAMRRQGVTRRSFLKYCSLAATSLGLGAGMAPKIAWALENKPRIPVVWIHGL WP_001058323.1 MULTISPECIES: hydrogenase-1 small chain [Proteobacteria] Length: 372\nNumber of predicted TMHs: 2\nExp number of AAs in TMHs: 37.2608499999999\nExp number, first 60 AAs: 19.52923\nTotal prob of N-in: 0.94351\nPOSSIBLE N-term signal sequence\ninside 1 19\nTMhelix 20 39\noutside 40 325\nTMhelix 326 348\ninside 349 372

5384 GCF_001283785.1_8205_3_70 Escherichia coli Proteobacteria; Gammaproteobacteria; Enterobacterales; Enterobacteriaceae; Escherichia
MTGDNTLIHSHGINRRDFMKLCAALAATMGLSSKAAAEMAESVTNPQRPPVIWIGAQECT WP_000145410.1
MULTISPECIES: hydrogenase-2 small chain [Proteobacteria] Length: 372\nNumber of predicted TMHs: 1\nExp number of AAs in TMHs: 23.72226\nExp number, first 60 AAs: 0.22038\nTotal prob of N-in: 0.12994\noutside 1 331\nTMhelix 332 354\ninside 355 372

5385 GCF_001283845.1_7790_1_73 Escherichia coli Proteobacteria; Gammaproteobacteria; Enterobacterales; Enterobacteriaceae; Escherichia
MTGDNTLIHSHGINRRDFMKLCAALAATMGLSSKAAAEMAESVTNPQRPPVIWIGAQECT WP_000145410.1
MULTISPECIES: hydrogenase-2 small chain [Proteobacteria] Length: 372\nNumber of predicted TMHs: 1\nExp number of AAs in TMHs: 23.72226\nExp number, first 60 AAs: 0.22038\nTotal prob of N-in: 0.12994\noutside 1 331\nTMhelix 332 354\ninside 355 372

5386 GCF_001269105.1_1.ECAD27.1 Escherichia coli Proteobacteria; Gammaproteobacteria; Enterobacterales; Enterobacteriaceae; Escherichia
MTGDNTLIHSHGINRRDFMKLCAALAATMGLSSKAAAEMAESVTNPQRPPVIWIGAQECT WP_000145410.1
MULTISPECIES: hydrogenase-2 small chain [Proteobacteria] Length: 372\nNumber of predicted TMHs: 1\nExp number of AAs in TMHs: 23.72226\nExp number, first 60 AAs: 0.22038\nTotal prob of N-in: 0.12994\noutside 1 331\nTMhelix 332 354\ninside 355 372

5387 GCF_001269105.1_1.ECAD27.1 Escherichia coli Proteobacteria; Gammaproteobacteria; Enterobacterales; Enterobacteriaceae; Escherichia
MNNEETFYQAMRRQGVTRRSFLKYCSLAATSLGLGAGMAPKIAWALENKPRIPVVWIHGL WP_053266014.1
hydrogenase [Escherichia coli] Length: 372\nNumber of predicted TMHs: 2\nExp number of AAs in TMHs: 37.25792\nExp number, first 60 AAs: 19.52926\nTotal prob of N-in: 0.94351\nPOSSIBLE N-term signal sequence\ninside 1 19\nTMhelix 20 39\noutside 40 325\nTMhelix 326 348\ninside 349 372

5388 GCF_000798255.1_Escherichia_coli_CVM_N36093PS_v._1.0 Escherichia coli Proteobacteria; Gammaproteobacteria; Enterobacterales; Enterobacteriaceae; Escherichia
MTGDNTLIHSHGINRRDFMKLCAALAATMGLSSKAAAEMAESVTNPQRPPVIWIGAQECT WP_000145410.1
MULTISPECIES: hydrogenase-2 small chain [Proteobacteria] Length: 372\nNumber of predicted TMHs: 1\nExp number of AAs in TMHs: 23.72226\nExp number, first 60 AAs: 0.22038\nTotal prob of N-in: 0.12994\noutside 1 331\nTMhelix 332 354\ninside 355 372

5389 GCF_001284305.1_7748_7_43 Escherichia coli Proteobacteria; Gammaproteobacteria; Enterobacterales; Enterobacteriaceae; Escherichia
MTGDNTLIHSHGINRRDFMKLCAALAATMGLSSKAAAEMAESVTNPQRPPVIWIGAQECT WP_000145410.1
MULTISPECIES: hydrogenase-2 small chain [Proteobacteria] Length: 372\nNumber of predicted TMHs: 1\nExp number of AAs in TMHs: 23.72226\nExp number, first 60 AAs: 0.22038\nTotal prob of N-in: 0.12994\noutside 1 331\nTMhelix 332 354\ninside 355 372

5390 GCF_001284285.1_8205_8_81 Escherichia coli Proteobacteria; Gammaproteobacteria; Enterobacterales; Enterobacteriaceae; Escherichia
MNNEETFYQAMRRQGVTRRSFLKYCSLAATSLGLGAGMAPKIAWALENKPRIPVVWIHGL WP_001058323.1
MULTISPECIES: hydrogenase-1 small chain [Proteobacteria] Length: 372\nNumber of predicted TMHs: 2\nExp number of AAs in TMHs: 37.2608499999999\nExp number, first 60 AAs: 19.52923\nTotal prob of N-in: 0.94351\nPOSSIBLE N-term signal sequence\ninside 1 19\nTMhelix 20 39\noutside 40 325\nTMhelix 326 348\ninside 349 372

5391 GCF_001283765.1_7790_1_80 Escherichia coli Proteobacteria; Gammaproteobacteria; Enterobacterales; Enterobacteriaceae; Escherichia
MNNEETFYQAMRRQGVTRRSFLKYCSLAATSLGLGAGMAPKIAWALENKPRIPVVWIHGL WP_001058323.1
MULTISPECIES: hydrogenase-1 small chain [Proteobacteria] Length: 372\nNumber of predicted TMHs: 2\nExp number of AAs in TMHs: 37.2608499999999\nExp number, first 60 AAs: 19.52923\nTotal prob of N-in: 0.94351\nPOSSIBLE N-term signal sequence\ninside 1 19\nTMhelix 20 39\noutside 40 325\nTMhelix 326 348\ninside 349 372

5392 GCF_001660165.1_ASM166016v1 Escherichia coli Proteobacteria; Gammaproteobacteria; Enterobacterales; Enterobacteriaceae; Escherichia
MTGDNTLIHSHGINRRDFMKLCAALAATMGLSSKAAAEMAESVTNPQRPPVIWIGAQECT WP_000145410.1

MULTISPECIES: hydrogenase-2 small chain [Proteobacteria] Length: 372\nNumber of predicted TMHs: 1\nExp number of AAs in TMHs: 23.72226\nExp number, first 60 AAs: 0.22038\nTotal prob of N-in: 0.12994\noutside 1 331\nTMhelix 332 354\ninside 355 372

5393 GCF_001660085.1_ASM166008v1 Escherichia coli Proteobacteria; Gammaproteobacteria; Enterobacterales; Enterobacteriaceae; Escherichia
MNNEETFYQAMRRQGVTRRSFLKYCSLAATSLGLGAGMAPKIAWALENKPRIPVVWIHGL WP_001058323.1
MULTISPECIES: hydrogenase-1 small chain [Proteobacteria] Length: 372\nNumber of predicted TMHs: 2\nExp number of AAs in TMHs: 37.2608499999999\nExp number, first 60 AAs: 19.52923\nTotal prob of N-in: 0.94351\nPOSSIBLE N-term signal sequence\ninside 1 19\nTMhelix 20 39\noutside 40 325\nTMhelix 326 348\ninside 349 372

5394 GCF_001660165.1_ASM166016v1 Escherichia coli Proteobacteria; Gammaproteobacteria; Enterobacterales; Enterobacteriaceae; Escherichia
MNNEETFYQAMRRQGVTRRSFLKYCSLAATSLGLGAGMAPKIAWALENKPRIPVVWIHGL WP_001058323.1
MULTISPECIES: hydrogenase-1 small chain [Proteobacteria] Length: 372\nNumber of predicted TMHs: 2\nExp number of AAs in TMHs: 37.2608499999999\nExp number, first 60 AAs: 19.52923\nTotal prob of N-in: 0.94351\nPOSSIBLE N-term signal sequence\ninside 1 19\nTMhelix 20 39\noutside 40 325\nTMhelix 326 348\ninside 349 372

5395 GCF_001659805.1_ASM165980v1 Escherichia coli Proteobacteria; Gammaproteobacteria; Enterobacterales; Enterobacteriaceae; Escherichia
MNNEETFYQAMRRQGVTRRSFLKYCSLAATSLGLGAGMAPKIAWALENKPRIPVVWIHGL WP_001295940.1
MULTISPECIES: hydrogenase [Enterobacteriaceae] Length: 372\nNumber of predicted TMHs: 2\nExp number of AAs in TMHs: 37.2608899999999\nExp number, first 60 AAs: 19.52923\nTotal prob of N-in: 0.94351\nPOSSIBLE N-term signal sequence\ninside 1 19\nTMhelix 20 39\noutside 40 325\nTMhelix 326 348\ninside 349 372

5396 GCF_001660255.1_ASM166025v1 Escherichia coli Proteobacteria; Gammaproteobacteria; Enterobacterales; Enterobacteriaceae; Escherichia
MTGDNTLIHSHGINRRDFMKLCAALAATMGLSSKAAAEMAESVTNPQRPPVIWIGAQECT WP_000145410.1
MULTISPECIES: hydrogenase-2 small chain [Proteobacteria] Length: 372\nNumber of predicted TMHs: 1\nExp number of AAs in TMHs: 23.72226\nExp number, first 60 AAs: 0.22038\nTotal prob of N-in: 0.12994\noutside 1 331\nTMhelix 332 354\ninside 355 372

5397 GCF_001660255.1_ASM166025v1 Escherichia coli Proteobacteria; Gammaproteobacteria; Enterobacterales; Enterobacteriaceae; Escherichia
MNNEETFYQAMRRQGVTRRSFLKYCSLAATSLGLGAGMAPKIAWALENKPRIPVVWIHGL WP_001058323.1
MULTISPECIES: hydrogenase-1 small chain [Proteobacteria] Length: 372\nNumber of predicted TMHs: 2\nExp number of AAs in TMHs: 37.2608499999999\nExp number, first 60 AAs: 19.52923\nTotal prob of N-in: 0.94351\nPOSSIBLE N-term signal sequence\ninside 1 19\nTMhelix 20 39\noutside 40 325\nTMhelix 326 348\ninside 349 372

5398 GCF_001660105.1_ASM166010v1 Escherichia coli Proteobacteria; Gammaproteobacteria; Enterobacterales; Enterobacteriaceae; Escherichia
MNNEETFYQAMRRQGVTRRSFLKYCSLAATSLGLGAGMAPKIAWALENKPRIPVVWIHGL WP_001295940.1
MULTISPECIES: hydrogenase [Enterobacteriaceae] Length: 372\nNumber of predicted TMHs: 2\nExp number of AAs in TMHs: 37.2608899999999\nExp number, first 60 AAs: 19.52923\nTotal prob of N-in: 0.94351\nPOSSIBLE N-term signal sequence\ninside 1 19\nTMhelix 20 39\noutside 40 325\nTMhelix 326 348\ninside 349 372

5399 GCF_001660275.1_ASM166027v1 Escherichia coli Proteobacteria; Gammaproteobacteria; Enterobacterales; Enterobacteriaceae; Escherichia
MTGDNTLIHSHGINRRDFMKLCAALAATMGLSSKAAAEMAESVTNPQRPPVIWIGAQECT WP_000145410.1
MULTISPECIES: hydrogenase-2 small chain [Proteobacteria] Length: 372\nNumber of predicted TMHs: 1\nExp number of AAs in TMHs: 23.72226\nExp number, first 60 AAs: 0.22038\nTotal prob of N-in: 0.12994\noutside 1 331\nTMhelix 332 354\ninside 355 372

5400 GCF_001660325.1_ASM166032v1 Escherichia coli Proteobacteria; Gammaproteobacteria; Enterobacterales; Enterobacteriaceae; Escherichia
MTGDNTLIHSHGINRRDFMKLCAALAATMGLSSKAAAEMAESVTNPQRPPVIWIGAQECT WP_000145410.1

MULTISPECIES: hydrogenase-2 small chain [Proteobacteria] Length: 372\nNumber of predicted TMHs: 1\nExp number of AAs in TMHs: 23.72226\nExp number, first 60 AAs: 0.22038\nTotal prob of N-in: 0.12994\noutside 1 331\nTMhelix 332 354\ninside 355 372

5401 GCF_001660585.1_ASM166058v1 Escherichia coli Proteobacteria; Gammaproteobacteria; Enterobacterales; Enterobacteriaceae; Escherichia
MTGDNTLIHSHGINRRDFMKLCAALAATMGLSSKAAAEMVESVTNPQRPPVIWIGAQECT WP_000145423.1
hydrogenase 2 small subunit [Escherichia coli] Length: 372\nNumber of predicted TMHs: 1\nExp number of AAs in TMHs: 23.78201\nExp number, first 60 AAs: 0.28355\nTotal prob of N-in: 0.13257\noutside 1 331\nTMhelix 332 354\ninside 355 372

5402 GCF_001660545.2_ASM166054v3 Escherichia coli Proteobacteria; Gammaproteobacteria; Enterobacterales; Enterobacteriaceae; Escherichia
MTGDNTLIHSHGINRRDFMKLCAALAATMGLSSKAAAEMAESVTNPQRPPVIWIGAQECT WP_000145410.1
MULTISPECIES: hydrogenase-2 small chain [Proteobacteria] Length: 372\nNumber of predicted TMHs: 1\nExp number of AAs in TMHs: 23.72226\nExp number, first 60 AAs: 0.22038\nTotal prob of N-in: 0.12994\noutside 1 331\nTMhelix 332 354\ninside 355 372

5403 GCF_001660345.1_ASM166034v1 Escherichia coli Proteobacteria; Gammaproteobacteria; Enterobacterales; Enterobacteriaceae; Escherichia
MTGDNTLIHSHGINRRDFMKLCAALAATMGLSSKAAAEMAESVTNPQRPPVIWIGAQECT WP_000145410.1
MULTISPECIES: hydrogenase-2 small chain [Proteobacteria] Length: 372\nNumber of predicted TMHs: 1\nExp number of AAs in TMHs: 23.72226\nExp number, first 60 AAs: 0.22038\nTotal prob of N-in: 0.12994\noutside 1 331\nTMhelix 332 354\ninside 355 372

5404 GCF_001663075.1_ASM166307v1 Escherichia coli Proteobacteria; Gammaproteobacteria; Enterobacterales; Enterobacteriaceae; Escherichia
MTGDNTLIHSHGINRRDFMKLCAALAATMGLSSKAAAEMAESVTNPQRPPVIWIGAQECT WP_000145410.1
MULTISPECIES: hydrogenase-2 small chain [Proteobacteria] Length: 372\nNumber of predicted TMHs: 1\nExp number of AAs in TMHs: 23.72226\nExp number, first 60 AAs: 0.22038\nTotal prob of N-in: 0.12994\noutside 1 331\nTMhelix 332 354\ninside 355 372

5405 GCF_001675145.1_ASM167514v1 Escherichia coli Proteobacteria; Gammaproteobacteria; Enterobacterales; Enterobacteriaceae; Escherichia
MTGDNTLIHSHGINRRDFMKLCAALAATMGLSSKAAAEMAESVTNPQRPPVIWIGAQECT WP_000145410.1
MULTISPECIES: hydrogenase-2 small chain [Proteobacteria] Length: 372\nNumber of predicted TMHs: 1\nExp number of AAs in TMHs: 23.72226\nExp number, first 60 AAs: 0.22038\nTotal prob of N-in: 0.12994\noutside 1 331\nTMhelix 332 354\ninside 355 372

5406 GCF_001663475.1_ASM166347v1 Escherichia coli Proteobacteria; Gammaproteobacteria; Enterobacterales; Enterobacteriaceae; Escherichia
MTGDNTLIHSHGINRRDFMKLCAALAATMGLSSKAAAEMAESVTNPQRPPVIWIGAQECT WP_000145410.1
MULTISPECIES: hydrogenase-2 small chain [Proteobacteria] Length: 372\nNumber of predicted TMHs: 1\nExp number of AAs in TMHs: 23.72226\nExp number, first 60 AAs: 0.22038\nTotal prob of N-in: 0.12994\noutside 1 331\nTMhelix 332 354\ninside 355 372

5407 GCF_001663475.1_ASM166347v1 Escherichia coli Proteobacteria; Gammaproteobacteria; Enterobacterales; Enterobacteriaceae; Escherichia
MNNEETFYQAMRRQGVTRRSFLKYCSLAATSLGLGAGMAPKIAWALENKPRIPVVWIHGL WP_001058323.1
MULTISPECIES: hydrogenase-1 small chain [Proteobacteria] Length: 372\nNumber of predicted TMHs: 2\nExp number of AAs in TMHs: 37.2608499999999\nExp number, first 60 AAs: 19.52923\nTotal prob of N-in: 0.94351\nPOSSIBLE N-term signal sequence\ninside 1 19\nTMhelix 20 39\noutside 40 325\nTMhelix 326 348\ninside 349 372

5408 GCF_001660275.1_ASM166027v1 Escherichia coli Proteobacteria; Gammaproteobacteria; Enterobacterales; Enterobacteriaceae; Escherichia
MNNEETFYQAMRRQGVTRRSFLKYCSLAATSLGLGAGMAPKIAWALENKPRIPVVWIHGL WP_001058323.1
MULTISPECIES: hydrogenase-1 small chain [Proteobacteria] Length: 372\nNumber of predicted TMHs: 2\nExp number of AAs in TMHs: 37.2608499999999\nExp number, first 60 AAs: 19.52923\nTotal prob of N-in: 0.94351\nPOSSIBLE N-term signal sequence\ninside 1 19\nTMhelix 20 39\noutside 40 325\nTMhelix 326 348\ninside 349 372

5409 GCF_001676975.1_ASM167697v1 Escherichia coli Proteobacteria; Gammaproteobacteria; Enterobacterales; Enterobacteriaceae; Escherichia
MTGDNTLIHSHGINRRDFMKLCAALAATMGLSSKAAAEMAESVTNPQRPPVIWIGAQECT WP_000145410.1
MULTISPECIES: hydrogenase-2 small chain [Proteobacteria] Length: 372\nNumber of predicted TMHs: 1\nExp number of AAs in TMHs: 23.72226\nExp number, first 60 AAs: 0.22038\nTotal prob of N-in: 0.12994\noutside 1 331\nTMhelix 332 354\ninside 355 372

5410 GCF_001676995.1_ASM167699v1 Escherichia coli Proteobacteria; Gammaproteobacteria; Enterobacterales; Enterobacteriaceae; Escherichia
MTGDNTLIHSHGINRRDFMKLCAALAATMGLSSKAAAEMAESVTNPQRPPVIWIGAQECT WP_000145410.1
MULTISPECIES: hydrogenase-2 small chain [Proteobacteria] Length: 372\nNumber of predicted TMHs: 1\nExp number of AAs in TMHs: 23.72226\nExp number, first 60 AAs: 0.22038\nTotal prob of N-in: 0.12994\noutside 1 331\nTMhelix 332 354\ninside 355 372

5411 GCF_001676975.1_ASM167697v1 Escherichia coli Proteobacteria; Gammaproteobacteria; Enterobacterales; Enterobacteriaceae; Escherichia
MNNEETFYQAMRRQGVTRRSFLKYCSLAATSLGLGAGMAPKIAWALENKPRIPVVWIHGL WP_001058323.1
MULTISPECIES: hydrogenase-1 small chain [Proteobacteria] Length: 372\nNumber of predicted TMHs: 2\nExp number of AAs in TMHs: 37.2608499999999\nExp number, first 60 AAs: 19.52923\nTotal prob of N-in: 0.94351\nPOSSIBLE N-term signal sequence\ninside 1 19\nTMhelix 20 39\noutside 40 325\nTMhelix 326 348\ninside 349 372

5412 GCF_001676995.1_ASM167699v1 Escherichia coli Proteobacteria; Gammaproteobacteria; Enterobacterales; Enterobacteriaceae; Escherichia
MNNEETFYQAMRRQGVTRRSFLKYCSLAATSLGLGAGMAPKIAWALENKPRIPVVWIHGL WP_001295940.1
MULTISPECIES: hydrogenase [Enterobacteriaceae] Length: 372\nNumber of predicted TMHs: 2\nExp number of AAs in TMHs: 37.2608899999999\nExp number, first 60 AAs: 19.52923\nTotal prob of N-in: 0.94351\nPOSSIBLE N-term signal sequence\ninside 1 19\nTMhelix 20 39\noutside 40 325\nTMhelix 326 348\ninside 349 372

5413 GCF_001660325.1_ASM166032v1 Escherichia coli Proteobacteria; Gammaproteobacteria; Enterobacterales; Enterobacteriaceae; Escherichia
MNNEETFYQAMRRQGVTRRSFLKYCSLAATSLGLGAGMAPKIAWALENKPRIPVVWIHGL WP_001058323.1
MULTISPECIES: hydrogenase-1 small chain [Proteobacteria] Length: 372\nNumber of predicted TMHs: 2\nExp number of AAs in TMHs: 37.2608499999999\nExp number, first 60 AAs: 19.52923\nTotal prob of N-in: 0.94351\nPOSSIBLE N-term signal sequence\ninside 1 19\nTMhelix 20 39\noutside 40 325\nTMhelix 326 348\ninside 349 372

5414 GCF_001660585.1_ASM166058v1 Escherichia coli Proteobacteria; Gammaproteobacteria; Enterobacterales; Enterobacteriaceae; Escherichia
MNNEETFYQAMRRQGVTRRSFLKYCSLAATSLGLGAGMAPKIAWALENKPRIPVVWIHGL WP_001058323.1
MULTISPECIES: hydrogenase-1 small chain [Proteobacteria] Length: 372\nNumber of predicted TMHs: 2\nExp number of AAs in TMHs: 37.2608499999999\nExp number, first 60 AAs: 19.52923\nTotal prob of N-in: 0.94351\nPOSSIBLE N-term signal sequence\ninside 1 19\nTMhelix 20 39\noutside 40 325\nTMhelix 326 348\ninside 349 372

5415 GCF_001677615.2_ASM167761v2 Escherichia coli Proteobacteria; Gammaproteobacteria; Enterobacterales; Enterobacteriaceae; Escherichia
MTGDNTLIHSHGINRRDFMKLCAALAATMGLSSKAAAEMAESVTNPQRPPVIWIGAQECT WP_000145410.1
MULTISPECIES: hydrogenase-2 small chain [Proteobacteria] Length: 372\nNumber of predicted TMHs: 1\nExp number of AAs in TMHs: 23.72226\nExp number, first 60 AAs: 0.22038\nTotal prob of N-in: 0.12994\noutside 1 331\nTMhelix 332 354\ninside 355 372

5416 GCF_001677625.2_ASM167762v2 Escherichia coli Proteobacteria; Gammaproteobacteria; Enterobacterales; Enterobacteriaceae; Escherichia
MTGDNTLIHSHGINRRDFMKLCAALAATMGLSSKAAAEMAESVTNPQRPPVIWIGAQECT WP_000145410.1
MULTISPECIES: hydrogenase-2 small chain [Proteobacteria] Length: 372\nNumber of predicted TMHs: 1\nExp number of AAs in TMHs: 23.72226\nExp number, first 60 AAs: 0.22038\nTotal prob of N-in: 0.12994\noutside 1 331\nTMhelix 332 354\ninside 355 372

5417 GCF_001677645.2_ASM167764v2 Escherichia coli Proteobacteria; Gammaproteobacteria; Enterobacterales; Enterobacteriaceae; Escherichia
MTGDNTLIHSHGINRRDFMKLCAALAATMGLSSKAAAEMAESVTNPQRPPVIWIGAQECT WP_000145410.1
MULTISPECIES: hydrogenase-2 small chain [Proteobacteria] Length: 372\nNumber of predicted TMHs: 1\nExp number of AAs in TMHs: 23.72226\nExp number, first 60 AAs: 0.22038\nTotal prob of N-in: 0.12994\noutside 1 331\nTMhelix 332 354\ninside 355 372

5418 GCF_001677705.2_ASM167770v2 Escherichia coli Proteobacteria; Gammaproteobacteria; Enterobacterales; Enterobacteriaceae; Escherichia
MTGDNTLIHSHGINRRDFMKLCAALAATMGLSSKAAAEMAESVTNPQRPPVIWIGAQECT WP_000145410.1
MULTISPECIES: hydrogenase-2 small chain [Proteobacteria] Length: 372\nNumber of predicted TMHs: 1\nExp number of AAs in TMHs: 23.72226\nExp number, first 60 AAs: 0.22038\nTotal prob of N-in: 0.12994\noutside 1 331\nTMhelix 332 354\ninside 355 372

5419 GCF_001677515.1_ASM167751v1 Escherichia coli Proteobacteria; Gammaproteobacteria; Enterobacterales; Enterobacteriaceae; Escherichia
MNNEETFYQAMRRQGVTRRSFLKYCSLAATSLGLGAGMAPKIAWALENKPRIPVVWIHGL WP_001058323.1
MULTISPECIES: hydrogenase-1 small chain [Proteobacteria] Length: 372\nNumber of predicted TMHs: 2\nExp number of AAs in TMHs: 37.2608499999999\nExp number, first 60 AAs: 19.52923\nTotal prob of N-in: 0.94351\nPOSSIBLE N-term signal sequence\ninside 1 19\nTMhelix 20 39\noutside 40 325\nTMhelix 326 348\ninside 349 372

5420 GCF_001677775.2_ASM167777v2 Escherichia coli Proteobacteria; Gammaproteobacteria; Enterobacterales; Enterobacteriaceae; Escherichia
MTGDNTLIHSHGINRRDFMKLCAALAATMGLSSKAAAEMAESVTNPQRPPVIWIGAQECT WP_000145410.1
MULTISPECIES: hydrogenase-2 small chain [Proteobacteria] Length: 372\nNumber of predicted TMHs: 1\nExp number of AAs in TMHs: 23.72226\nExp number, first 60 AAs: 0.22038\nTotal prob of N-in: 0.12994\noutside 1 331\nTMhelix 332 354\ninside 355 372

5421 GCF_001677715.2_ASM167771v2 Escherichia coli Proteobacteria; Gammaproteobacteria; Enterobacterales; Enterobacteriaceae; Escherichia
MTGDNTLIHSHGINRRDFMKLCAALAATMGLSSKAAAEMAESVTNPQRPPVIWIGAQECT WP_000145410.1
MULTISPECIES: hydrogenase-2 small chain [Proteobacteria] Length: 372\nNumber of predicted TMHs: 1\nExp number of AAs in TMHs: 23.72226\nExp number, first 60 AAs: 0.22038\nTotal prob of N-in: 0.12994\noutside 1 331\nTMhelix 332 354\ninside 355 372

5422 GCF_001677795.2_ASM167779v2 Escherichia coli Proteobacteria; Gammaproteobacteria; Enterobacterales; Enterobacteriaceae; Escherichia
MTGDNTLIHSHGINRRDFMKLCAALAATMGLSSKAAAEMAESVTNPQRPPVIWIGAQECT WP_000145410.1
MULTISPECIES: hydrogenase-2 small chain [Proteobacteria] Length: 372\nNumber of predicted TMHs: 1\nExp number of AAs in TMHs: 23.72226\nExp number, first 60 AAs: 0.22038\nTotal prob of N-in: 0.12994\noutside 1 331\nTMhelix 332 354\ninside 355 372

5423 GCF_001677805.2_ASM167780v2 Escherichia coli Proteobacteria; Gammaproteobacteria; Enterobacterales; Enterobacteriaceae; Escherichia
MTGDNTLIHSHGINRRDFMKLCAALAATMGLSSKAAAEMAESVTNPQRPPVIWIGAQECT WP_000145410.1
MULTISPECIES: hydrogenase-2 small chain [Proteobacteria] Length: 372\nNumber of predicted TMHs: 1\nExp number of AAs in TMHs: 23.72226\nExp number, first 60 AAs: 0.22038\nTotal prob of N-in: 0.12994\noutside 1 331\nTMhelix 332 354\ninside 355 372

5424 GCF_001677865.2_ASM167786v2 Escherichia coli Proteobacteria; Gammaproteobacteria; Enterobacterales; Enterobacteriaceae; Escherichia
MTGDNTLIHSHGINRRDFMKLCAALAATMGLSSKAAAEMAESVTNPQRPPVIWIGAQECT WP_000145410.1
MULTISPECIES: hydrogenase-2 small chain [Proteobacteria] Length: 372\nNumber of predicted TMHs: 1\nExp number of AAs in TMHs: 23.72226\nExp number, first 60 AAs: 0.22038\nTotal prob of N-in: 0.12994\noutside 1 331\nTMhelix 332 354\ninside 355 372

5425 GCF_001677715.2_ASM167771v2 Escherichia coli Proteobacteria; Gammaproteobacteria; Enterobacterales; Enterobacteriaceae; Escherichia
MNNEETFYQAMRRQGVTRRSFLKYCSLAATSLGLGAGMAPKIAWALENKPRIPVVWIHGL WP_001301613.1
hydrogenase [Escherichia coli] Length: 372\nNumber of predicted TMHs: 2\nExp number of AAs in

TMHs: 37.46598\nExp number, first 60 AAs: 19.53139\nTotal prob of N-in: 0.94364\nPOSSIBLE N-term signal sequence\ninside 1 19\nTMhelix 20 39\noutside 40 325\nTMhelix 326 348\ninside 349 372

5426 GCF_001677535.2_ASM167753v2 Escherichia coli Proteobacteria; Gammaproteobacteria; Enterobacterales; Enterobacteriaceae; Escherichia MNNEETFYQAMRRQGVTRRSFLKYCSLAATSLGLGAGMAPKIAWALENKPRIPVVWIHGL WP_001301613.1 hydrogenase [Escherichia coli] Length: 372\nNumber of predicted TMHs: 2\nExp number of AAs in TMHs: 37.46598\nExp number, first 60 AAs: 19.53139\nTotal prob of N-in: 0.94364\nPOSSIBLE N-term signal sequence\ninside 1 19\nTMhelix 20 39\noutside 40 325\nTMhelix 326 348\ninside 349 372

5427 GCF_001677625.2_ASM167762v2 Escherichia coli Proteobacteria; Gammaproteobacteria; Enterobacterales; Enterobacteriaceae; Escherichia MNNEETFYQAMRRQGVTRRSFLKYCSLAATSLGLGAGMAPKIAWALENKPRIPVVWIHGL WP_001301613.1 hydrogenase [Escherichia coli] Length: 372\nNumber of predicted TMHs: 2\nExp number of AAs in TMHs: 37.46598\nExp number, first 60 AAs: 19.53139\nTotal prob of N-in: 0.94364\nPOSSIBLE N-term signal sequence\ninside 1 19\nTMhelix 20 39\noutside 40 325\nTMhelix 326 348\ninside 349 372

5428 GCF_001677555.2_ASM167755v2 Escherichia coli Proteobacteria; Gammaproteobacteria; Enterobacterales; Enterobacteriaceae; Escherichia MNNEETFYQAMRRQGVTRRSFLKYCSLAATSLGLGAGMAPKIAWALENKPRIPVVWIHGL WP_001301613.1 hydrogenase [Escherichia coli] Length: 372\nNumber of predicted TMHs: 2\nExp number of AAs in TMHs: 37.46598\nExp number, first 60 AAs: 19.53139\nTotal prob of N-in: 0.94364\nPOSSIBLE N-term signal sequence\ninside 1 19\nTMhelix 20 39\noutside 40 325\nTMhelix 326 348\ninside 349 372

5429 GCF_001678965.1_ASM167896v1 Escherichia coli Proteobacteria; Gammaproteobacteria; Enterobacterales; Enterobacteriaceae; Escherichia MTGDNTLIHSHGINRRDFMKLCAALAATMGLSSKAAAEMAESVTNPQRPPVIWIGAQECT WP_000145410.1 MULTISPECIES: hydrogenase-2 small chain [Proteobacteria] Length: 372\nNumber of predicted TMHs: 1\nExp number of AAs in TMHs: 23.72226\nExp number, first 60 AAs: 0.22038\nTotal prob of N-in: 0.12994\noutside 1 331\nTMhelix 332 354\ninside 355 372

5430 GCF_001683575.1_NGF4 Escherichia coli Proteobacteria; Gammaproteobacteria; Enterobacterales; Enterobacteriaceae; Escherichia MTGDNTLIHSHGINRRDFMKLCAALAATMGLSSKAAAEMVESVTNPQRPPVIWIGAQECT WP_000145423.1 hydrogenase 2 small subunit [Escherichia coli] Length: 372\nNumber of predicted TMHs: 1\nExp number of AAs in TMHs: 23.78201\nExp number, first 60 AAs: 0.28355\nTotal prob of N-in: 0.13257\noutside 1 331\nTMhelix 332 354\ninside 355 372

5431 GCF_001683585.1_NGF3 Escherichia coli Proteobacteria; Gammaproteobacteria; Enterobacterales; Enterobacteriaceae; Escherichia MTGDNTLIHSHGINRRDFMKLCAALAATMGLSSKAAAEMVESVTNPQRPPVIWIGAQECT WP_000145423.1 hydrogenase 2 small subunit [Escherichia coli] Length: 372\nNumber of predicted TMHs: 1\nExp number of AAs in TMHs: 23.78201\nExp number, first 60 AAs: 0.28355\nTotal prob of N-in: 0.13257\noutside 1 331\nTMhelix 332 354\ninside 355 372

5432 GCF_001683575.1_NGF4 Escherichia coli Proteobacteria; Gammaproteobacteria; Enterobacterales; Enterobacteriaceae; Escherichia MNNEETFYQAMRRQGVTRRSFLKYCSLAATSLGLGAGMAPKIAWALENKPRIPVVWIHGL WP_001058323.1 MULTISPECIES: hydrogenase-1 small chain [Proteobacteria] Length: 372\nNumber of predicted TMHs: 2\nExp number of AAs in TMHs: 37.2608499999999\nExp number, first 60 AAs: 19.52923\nTotal prob of N-in: 0.94351\nPOSSIBLE N-term signal sequence\ninside 1 19\nTMhelix 20 39\noutside 40 325\nTMhelix 326 348\ninside 349 372

5433 GCF_001683585.1_NGF3 Escherichia coli Proteobacteria; Gammaproteobacteria; Enterobacterales; Enterobacteriaceae; Escherichia MNNEETFYQAMRRQGVTRRSFLKYCSLAATSLGLGAGMAPKIAWALENKPRIPVVWIHGL WP_001058323.1 MULTISPECIES: hydrogenase-1 small chain [Proteobacteria] Length: 372\nNumber of predicted TMHs: 2\nExp number of AAs in TMHs: 37.2608499999999\nExp number, first 60 AAs: 19.52923\nTotal prob of N-in: 0.94351\nPOSSIBLE N-term signal sequence\ninside 1 19\nTMhelix 20 39\noutside 40 325\nTMhelix 326 348\ninside 349 372

5434 GCF_001678965.1_ASM167896v1 Escherichia coli Proteobacteria; Gammaproteobacteria; Enterobacterales; Enterobacteriaceae; Escherichia MNNEETFYQAMRRQGVTRRSFLKYCSLAATSLGLGAGMAPKIAWALENKPRIPVVWIHGL WP_001295940.1 MULTISPECIES: hydrogenase [Enterobacteriaceae] Length: 372\nNumber of predicted TMHs: 2\nExp

number of AAs in TMHs: 37.2608899999999\nExp number, first 60 AAs:  19.52923\nTotal prob of N-in:  0.94351\nPOSSIBLE N-term signal sequence\ninside   1   19\nTMhelix   20   39\noutside   40   325\nTMhelix   326   348\ninside   349   372

5435 GCF_001677805.2_ASM167780v2 Escherichia coli Proteobacteria; Gammaproteobacteria; Enterobacterales; Enterobacteriaceae; Escherichia MNNEETFYQAMRRQGVTRRSFLKYCSLAATSLGLGAGMAPKIAWALENKPRIPVVWIHGL WP_001301613.1 hydrogenase [Escherichia coli] Length: 372\nNumber of predicted TMHs:  2\nExp number of AAs in TMHs: 37.46598\nExp number, first 60 AAs:  19.53139\nTotal prob of N-in:  0.94364\nPOSSIBLE N-term signal sequence\ninside   1   19\nTMhelix   20   39\noutside   40   325\nTMhelix   326   348\ninside   349   372

5436 GCF_001682305.2_ASM168230v2 Escherichia coli Proteobacteria; Gammaproteobacteria; Enterobacterales; Enterobacteriaceae; Escherichia MTGDNTLIHSHGINRRDFMKLCAALAATMGLSSKAAAEMAESVTNPQRPPVIWIGAQECT WP_000145410.1 MULTISPECIES: hydrogenase-2 small chain [Proteobacteria] Length: 372\nNumber of predicted TMHs:  1\nExp number of AAs in TMHs: 23.72226\nExp number, first 60 AAs:  0.22038\nTotal prob of N-in:  0.12994\noutside   1   331\nTMhelix   332   354\ninside   355   372

5437 GCF_001677865.2_ASM167786v2 Escherichia coli Proteobacteria; Gammaproteobacteria; Enterobacterales; Enterobacteriaceae; Escherichia MNNEETFYQAMRRQGVTRRSFLKYCSLAATSLGLGAGMAPKIAWALENKPRIPVVWIHGL WP_001301613.1 hydrogenase [Escherichia coli] Length: 372\nNumber of predicted TMHs:  2\nExp number of AAs in TMHs: 37.46598\nExp number, first 60 AAs:  19.53139\nTotal prob of N-in:  0.94364\nPOSSIBLE N-term signal sequence\ninside   1   19\nTMhelix   20   39\noutside   40   325\nTMhelix   326   348\ninside   349   372

5438 GCF_001692795.1_ASM169279v1 Escherichia coli Proteobacteria; Gammaproteobacteria; Enterobacterales; Enterobacteriaceae; Escherichia MTGDNTLIHSHGINRRDFMKLCAALAATMGLSSKAAAEMVESVTNPQRPPVIWIGAQECT WP_000145423.1 hydrogenase 2 small subunit [Escherichia coli] Length: 372\nNumber of predicted TMHs:  1\nExp number of AAs in TMHs: 23.78201\nExp number, first 60 AAs:  0.28355\nTotal prob of N-in:  0.13257\noutside   1   331\nTMhelix   332   354\ninside   355   372

5439 GCF_001692775.1_ASM169277v1 Escherichia coli Proteobacteria; Gammaproteobacteria; Enterobacterales; Enterobacteriaceae; Escherichia MTGDNTLIHSHGINRRDFMKLCAALAATMGLSSKAAAEMVESVTNPQRPPVIWIGAQECT WP_000145423.1 hydrogenase 2 small subunit [Escherichia coli] Length: 372\nNumber of predicted TMHs:  1\nExp number of AAs in TMHs: 23.78201\nExp number, first 60 AAs:  0.28355\nTotal prob of N-in:  0.13257\noutside   1   331\nTMhelix   332   354\ninside   355   372

5440 GCF_001692805.1_ASM169280v1 Escherichia coli Proteobacteria; Gammaproteobacteria; Enterobacterales; Enterobacteriaceae; Escherichia MTGDNTLIHSHGINRRDFMKLCAALAATMGLSSKAAAEMVESVTNPQRPPVIWIGAQECT WP_000145423.1 hydrogenase 2 small subunit [Escherichia coli] Length: 372\nNumber of predicted TMHs:  1\nExp number of AAs in TMHs: 23.78201\nExp number, first 60 AAs:  0.28355\nTotal prob of N-in:  0.13257\noutside   1   331\nTMhelix   332   354\ninside   355   372

5441 GCF_001693315.1_ASM169331v1 Escherichia coli Proteobacteria; Gammaproteobacteria; Enterobacterales; Enterobacteriaceae; Escherichia MTGDNTLIHSHGINRRDFMKLCAALAATMGLSSKAAAEMAESVTNPQRPPVIWIGAQECT WP_000145410.1 MULTISPECIES: hydrogenase-2 small chain [Proteobacteria] Length: 372\nNumber of predicted TMHs:  1\nExp number of AAs in TMHs: 23.72226\nExp number, first 60 AAs:  0.22038\nTotal prob of N-in:  0.12994\noutside   1   331\nTMhelix   332   354\ninside   355   372

5442 GCF_001693555.1_ASM169355v1 Escherichia coli Proteobacteria; Gammaproteobacteria; Enterobacterales; Enterobacteriaceae; Escherichia MTGDNTLIHSHGINRRDFMKLCAALAATMGLSSKAAAEMAESVTNPQRPPVIWIGAQECT WP_000145410.1 MULTISPECIES: hydrogenase-2 small chain [Proteobacteria] Length: 372\nNumber of predicted TMHs:  1\nExp number of AAs in TMHs: 23.72226\nExp number, first 60 AAs:  0.22038\nTotal prob of N-in:  0.12994\noutside   1   331\nTMhelix   332   354\ninside   355   372

5443 GCF_001703435.1_ASM170343v1 Escherichia coli Proteobacteria; Gammaproteobacteria; Enterobacterales; Enterobacteriaceae; Escherichia
MTGDNTLIHSHGINRRDFMKLCAALAATMGLSSKAAAEMAESVTNPQRPPVIWIGAQECT WP_000145410.1
MULTISPECIES: hydrogenase-2 small chain [Proteobacteria] Length: 372\nNumber of predicted TMHs: 1\nExp number of AAs in TMHs: 23.72226\nExp number, first 60 AAs: 0.22038\nTotal prob of N-in: 0.12994\noutside 1 331\nTMhelix 332 354\ninside 355 372

5444 GCF_001692805.1_ASM169280v1 Escherichia coli Proteobacteria; Gammaproteobacteria; Enterobacterales; Enterobacteriaceae; Escherichia
MNNEETFYQAMRRQGVTRRSFLKYCSLAATSLGLGAGMAPKIAWALENKPRIPVVWIHGL WP_001058323.1
MULTISPECIES: hydrogenase-1 small chain [Proteobacteria] Length: 372\nNumber of predicted TMHs: 2\nExp number of AAs in TMHs: 37.2608499999999\nExp number, first 60 AAs: 19.52923\nTotal prob of N-in: 0.94351\nPOSSIBLE N-term signal sequence\ninside 1 19\nTMhelix 20 39\noutside 40 325\nTMhelix 326 348\ninside 349 372

5445 GCF_001693315.1_ASM169331v1 Escherichia coli Proteobacteria; Gammaproteobacteria; Enterobacterales; Enterobacteriaceae; Escherichia
MNNEETFYQAMRRQGVTRRSFLKYCSLAATSLGLGAGMAPKIAWALENKPRIPVVWIHGL WP_001058323.1
MULTISPECIES: hydrogenase-1 small chain [Proteobacteria] Length: 372\nNumber of predicted TMHs: 2\nExp number of AAs in TMHs: 37.2608499999999\nExp number, first 60 AAs: 19.52923\nTotal prob of N-in: 0.94351\nPOSSIBLE N-term signal sequence\ninside 1 19\nTMhelix 20 39\noutside 40 325\nTMhelix 326 348\ninside 349 372

5446 GCF_001693555.1_ASM169355v1 Escherichia coli Proteobacteria; Gammaproteobacteria; Enterobacterales; Enterobacteriaceae; Escherichia
MNNEETFYQAMRRQGVTRRSFLKYCSLAATSLGLGAGMAPKIAWALENKPRIPVVWIHGL WP_001058323.1
MULTISPECIES: hydrogenase-1 small chain [Proteobacteria] Length: 372\nNumber of predicted TMHs: 2\nExp number of AAs in TMHs: 37.2608499999999\nExp number, first 60 AAs: 19.52923\nTotal prob of N-in: 0.94351\nPOSSIBLE N-term signal sequence\ninside 1 19\nTMhelix 20 39\noutside 40 325\nTMhelix 326 348\ninside 349 372

5447 GCF_001696345.1_ASM169634v1 Escherichia coli Proteobacteria; Gammaproteobacteria; Enterobacterales; Enterobacteriaceae; Escherichia
MTGDNTLIHSHGINRRDFMKLCAALAATMGLSSKAAAEMAESVTNPQRPPVIWIGAQECT WP_000145410.1
MULTISPECIES: hydrogenase-2 small chain [Proteobacteria] Length: 372\nNumber of predicted TMHs: 1\nExp number of AAs in TMHs: 23.72226\nExp number, first 60 AAs: 0.22038\nTotal prob of N-in: 0.12994\noutside 1 331\nTMhelix 332 354\ninside 355 372

5448 GCF_001700015.1_ASM170001v1 Escherichia coli Proteobacteria; Gammaproteobacteria; Enterobacterales; Enterobacteriaceae; Escherichia
MTGDNTLIHSHGINRRDFMKLCAALAATMGLSSKAAAEMAESVTNPQRPPVIWIGAQECT WP_000145410.1
MULTISPECIES: hydrogenase-2 small chain [Proteobacteria] Length: 372\nNumber of predicted TMHs: 1\nExp number of AAs in TMHs: 23.72226\nExp number, first 60 AAs: 0.22038\nTotal prob of N-in: 0.12994\noutside 1 331\nTMhelix 332 354\ninside 355 372

5449 GCF_001700045.1_ASM170004v1 Escherichia coli Proteobacteria; Gammaproteobacteria; Enterobacterales; Enterobacteriaceae; Escherichia
MTGDNTLIHSHGINRRDFMKLCAALAATMGLSSKAAAEMAESVTNPQRPPVIWIGAQECT WP_000145410.1
MULTISPECIES: hydrogenase-2 small chain [Proteobacteria] Length: 372\nNumber of predicted TMHs: 1\nExp number of AAs in TMHs: 23.72226\nExp number, first 60 AAs: 0.22038\nTotal prob of N-in: 0.12994\noutside 1 331\nTMhelix 332 354\ninside 355 372

5450 GCF_001700035.1_ASM170003v1 Escherichia coli Proteobacteria; Gammaproteobacteria; Enterobacterales; Enterobacteriaceae; Escherichia
MTGDNTLIHSHGINRRDFMKLCAALAATMGLSSKAAAEMAESVTNPQRPPVIWIGAQECT WP_000145410.1
MULTISPECIES: hydrogenase-2 small chain [Proteobacteria] Length: 372\nNumber of predicted TMHs: 1\nExp number of AAs in TMHs: 23.72226\nExp number, first 60 AAs: 0.22038\nTotal prob of N-in: 0.12994\noutside 1 331\nTMhelix 332 354\ninside 355 372

5451 GCF_001704035.1_ASM170403v1 Escherichia coli Proteobacteria; Gammaproteobacteria; Enterobacterales; Enterobacteriaceae; Escherichia

MTGDNTLIHSHGINRRDFMKLCAALAATMGLSSKAAAEMAESVTNPQRPPVIWIGAQECT WP_000145410.1 MULTISPECIES: hydrogenase-2 small chain [Proteobacteria] Length: 372\nNumber of predicted TMHs: 1\nExp number of AAs in TMHs: 23.72226\nExp number, first 60 AAs: 0.22038\nTotal prob of N-in: 0.12994\noutside 1 331\nTMhelix 332 354\ninside 355 372

5452 GCF_001700105.1_ASM170010v1 Escherichia coli Proteobacteria; Gammaproteobacteria; Enterobacterales; Enterobacteriaceae; Escherichia MTGDNTLIHSHGINRRDFMKLCAALAATMGLSSKAAAEMAESVTNPQRPPVIWIGAQECT WP_000145410.1 MULTISPECIES: hydrogenase-2 small chain [Proteobacteria] Length: 372\nNumber of predicted TMHs: 1\nExp number of AAs in TMHs: 23.72226\nExp number, first 60 AAs: 0.22038\nTotal prob of N-in: 0.12994\noutside 1 331\nTMhelix 332 354\ninside 355 372

5453 GCF_001704835.1_ASM170483v1 Escherichia coli Proteobacteria; Gammaproteobacteria; Enterobacterales; Enterobacteriaceae; Escherichia MTGDNTLIHSHGINRRDFMKLCAALAATMGLSSKAAAEMAESVTNPQRPPVIWIGAQECT WP_000145410.1 MULTISPECIES: hydrogenase-2 small chain [Proteobacteria] Length: 372\nNumber of predicted TMHs: 1\nExp number of AAs in TMHs: 23.72226\nExp number, first 60 AAs: 0.22038\nTotal prob of N-in: 0.12994\noutside 1 331\nTMhelix 332 354\ninside 355 372

5454 GCF_001704835.1_ASM170483v1 Escherichia coli Proteobacteria; Gammaproteobacteria; Enterobacterales; Enterobacteriaceae; Escherichia MNNEETFYQAMRRQGVTRRSFLKYCSLAATSLGLGAGMAPKIAWALENKPRIPVVWIHGL WP_001058323.1 MULTISPECIES: hydrogenase-1 small chain [Proteobacteria] Length: 372\nNumber of predicted TMHs: 2\nExp number of AAs in TMHs: 37.2608499999999\nExp number, first 60 AAs: 19.52923\nTotal prob of N-in: 0.94351\nPOSSIBLE N-term signal sequence\ninside 1 19\nTMhelix 20 39\noutside 40 325\nTMhelix 326 348\ninside 349 372

5455 GCF_001696345.1_ASM169634v1 Escherichia coli Proteobacteria; Gammaproteobacteria; Enterobacterales; Enterobacteriaceae; Escherichia MNNEETFYQAMRRQGVTRRSFLKYCSLAATSLGLGAGMAPKIAWALENKPRIPVVWIHGL WP_001058323.1 MULTISPECIES: hydrogenase-1 small chain [Proteobacteria] Length: 372\nNumber of predicted TMHs: 2\nExp number of AAs in TMHs: 37.2608499999999\nExp number, first 60 AAs: 19.52923\nTotal prob of N-in: 0.94351\nPOSSIBLE N-term signal sequence\ninside 1 19\nTMhelix 20 39\noutside 40 325\nTMhelix 326 348\ninside 349 372

5456 GCF_001708145.1_ASM170814v1 Escherichia coli Proteobacteria; Gammaproteobacteria; Enterobacterales; Enterobacteriaceae; Escherichia MTGDNTLIHSHGINRRDFMKLCAALAATMGLSSKAAAEMAESVTNPQRPPVIWIGAQECT WP_000145410.1 MULTISPECIES: hydrogenase-2 small chain [Proteobacteria] Length: 372\nNumber of predicted TMHs: 1\nExp number of AAs in TMHs: 23.72226\nExp number, first 60 AAs: 0.22038\nTotal prob of N-in: 0.12994\noutside 1 331\nTMhelix 332 354\ninside 355 372

5457 GCF_001708145.1_ASM170814v1 Escherichia coli Proteobacteria; Gammaproteobacteria; Enterobacterales; Enterobacteriaceae; Escherichia MNNEETFYQAMRRQGVTRRSFLKYCSLAATSLGLGAGMAPKIAWALENKPRIPVVWIHGL WP_001058320.1 hydrogenase [Escherichia coli] Length: 372\nNumber of predicted TMHs: 2\nExp number of AAs in TMHs: 37.26082\nExp number, first 60 AAs: 19.52923\nTotal prob of N-in: 0.94351\nPOSSIBLE N-term signal sequence\ninside 1 19\nTMhelix 20 39\noutside 40 325\nTMhelix 326 348\ninside 349 372

5458 GCF_001713545.1_ASM171354v1 Escherichia coli Proteobacteria; Gammaproteobacteria; Enterobacterales; Enterobacteriaceae; Escherichia MNNEETFYQAMRRQGVTRRSFLKYCSLAATSLGLGAGMAPKIAWALENKPRIPVVWIHGL WP_001058323.1 MULTISPECIES: hydrogenase-1 small chain [Proteobacteria] Length: 372\nNumber of predicted TMHs: 2\nExp number of AAs in TMHs: 37.2608499999999\nExp number, first 60 AAs: 19.52923\nTotal prob of N-in: 0.94351\nPOSSIBLE N-term signal sequence\ninside 1 19\nTMhelix 20 39\noutside 40 325\nTMhelix 326 348\ninside 349 372

5459 GCF_001713575.1_ASM171357v1 Escherichia coli Proteobacteria; Gammaproteobacteria; Enterobacterales; Enterobacteriaceae; Escherichia MNNEETFYQAMRRQGVTRRSFLKYCSLAATSLGLGAGMAPKIAWALENKPRIPVVWIHGL WP_069067259.1 hydrogenase [Escherichia coli] Length: 372\nNumber of predicted TMHs: 2\nExp number of AAs in

TMHs: 37.25908\nExp number, first 60 AAs: 19.52926\nTotal prob of N-in: 0.94351\nPOSSIBLE N-term signal sequence\ninside 1 19\nTMhelix 20 39\noutside 40 325\nTMhelix 326 348\ninside 349 372

5460 GCF_001703435.1_ASM170343v1 Escherichia coli Proteobacteria; Gammaproteobacteria; Enterobacterales; Enterobacteriaceae; Escherichia
MNNEETFYQAMRRQGVTRRSFLKYCSLAATSLGLGAGMAPKIAWALENKPRIPVVWIHGL WP_047603468.1
hydrogenase [Escherichia coli] Length: 372\nNumber of predicted TMHs: 2\nExp number of AAs in TMHs: 39.28018\nExp number, first 60 AAs: 19.5238\nTotal prob of N-in: 0.94367\nPOSSIBLE N-term signal sequence\ninside 1 19\nTMhelix 20 39\noutside 40 326\nTMhelix 327 349\ninside 350 372

5461 GCF_001721125.1_ASM172112v1 Escherichia coli Proteobacteria; Gammaproteobacteria; Enterobacterales; Enterobacteriaceae; Escherichia
MTGDNTLIHSHGINRRDFMKLCAALAATMGLSSKAAAEMAESVTNPQRPPVIWIGAQECT WP_000145410.1
MULTISPECIES: hydrogenase-2 small chain [Proteobacteria] Length: 372\nNumber of predicted TMHs: 1\nExp number of AAs in TMHs: 23.72226\nExp number, first 60 AAs: 0.22038\nTotal prob of N-in: 0.12994\noutside 1 331\nTMhelix 332 354\ninside 355 372

5462 GCF_001713585.1_ASM171358v1 Escherichia coli Proteobacteria; Gammaproteobacteria; Enterobacterales; Enterobacteriaceae; Escherichia
MTGDNTLIHSHGINRRDFMKLCAALAATMGLSSKAAAEMAESVTNPQRPPVIWIGAQECT WP_000145410.1
MULTISPECIES: hydrogenase-2 small chain [Proteobacteria] Length: 372\nNumber of predicted TMHs: 1\nExp number of AAs in TMHs: 23.72226\nExp number, first 60 AAs: 0.22038\nTotal prob of N-in: 0.12994\noutside 1 331\nTMhelix 332 354\ninside 355 372

5463 GCF_001721525.1_ASM172152v1 Escherichia coli Proteobacteria; Gammaproteobacteria; Enterobacterales; Enterobacteriaceae; Escherichia
MTGDNTLIHSHGINRRDFMKLCAALAATMGLSSKAAAEMAESVTNPQRPPVIWIGAQECT WP_000145410.1
MULTISPECIES: hydrogenase-2 small chain [Proteobacteria] Length: 372\nNumber of predicted TMHs: 1\nExp number of AAs in TMHs: 23.72226\nExp number, first 60 AAs: 0.22038\nTotal prob of N-in: 0.12994\noutside 1 331\nTMhelix 332 354\ninside 355 372

5464 GCF_001728785.1_ASM172878v1 Escherichia coli Proteobacteria; Gammaproteobacteria; Enterobacterales; Enterobacteriaceae; Escherichia
MTGDNTLIHSHGINRRDFMKLCAALAATMGLSSKAAAEMAESVTNPQRPPVIWIGAQECT WP_000145410.1
MULTISPECIES: hydrogenase-2 small chain [Proteobacteria] Length: 372\nNumber of predicted TMHs: 1\nExp number of AAs in TMHs: 23.72226\nExp number, first 60 AAs: 0.22038\nTotal prob of N-in: 0.12994\noutside 1 331\nTMhelix 332 354\ninside 355 372

5465 GCF_001730745.1_ASM173074v1 Escherichia coli Proteobacteria; Gammaproteobacteria; Enterobacterales; Enterobacteriaceae; Escherichia
MTGDNTLIHSHGINRRDFMKLCAALAATMGLSSKAAAEMAESVTNPQRPPVIWIGAQECT WP_000145410.1
MULTISPECIES: hydrogenase-2 small chain [Proteobacteria] Length: 372\nNumber of predicted TMHs: 1\nExp number of AAs in TMHs: 23.72226\nExp number, first 60 AAs: 0.22038\nTotal prob of N-in: 0.12994\noutside 1 331\nTMhelix 332 354\ninside 355 372

5466 GCF_001721525.1_ASM172152v1 Escherichia coli Proteobacteria; Gammaproteobacteria; Enterobacterales; Enterobacteriaceae; Escherichia
MNNEETFYQAMRRQGVTRRSFLKYCSLAATSLGLGAGMAPKIAWALENKPRIPVVWIHGL WP_001058323.1
MULTISPECIES: hydrogenase-1 small chain [Proteobacteria] Length: 372\nNumber of predicted TMHs: 2\nExp number of AAs in TMHs: 37.2608499999999\nExp number, first 60 AAs: 19.52923\nTotal prob of N-in: 0.94351\nPOSSIBLE N-term signal sequence\ninside 1 19\nTMhelix 20 39\noutside 40 325\nTMhelix 326 348\ninside 349 372

5467 GCF_001742465.1_ASM174246v1 Escherichia coli Proteobacteria; Gammaproteobacteria; Enterobacterales; Enterobacteriaceae; Escherichia
MTGDNTLIHSHGINRRDFMKLCAALAATMGLSSKAAAEMAESVTNPQRPPVIWIGAQECT WP_000145410.1
MULTISPECIES: hydrogenase-2 small chain [Proteobacteria] Length: 372\nNumber of predicted TMHs: 1\nExp number of AAs in TMHs: 23.72226\nExp number, first 60 AAs: 0.22038\nTotal prob of N-in: 0.12994\noutside 1 331\nTMhelix 332 354\ninside 355 372

5468 GCF_001742475.1_ASM174247v1 Escherichia coli Proteobacteria; Gammaproteobacteria; Enterobacterales; Enterobacteriaceae; Escherichia
MTGDNTLIHSHGINRRDFMKLCAALAATMGLSSKAAAEMAESVTNPQRPPVIWIGAQECT WP_000145410.1
MULTISPECIES: hydrogenase-2 small chain [Proteobacteria] Length: 372\nNumber of predicted TMHs: 1\nExp number of AAs in TMHs: 23.72226\nExp number, first 60 AAs: 0.22038\nTotal prob of N-in: 0.12994\noutside 1 331\nTMhelix 332 354\ninside 355 372

5469 GCF_001742495.1_ASM174249v1 Escherichia coli Proteobacteria; Gammaproteobacteria; Enterobacterales; Enterobacteriaceae; Escherichia
MTGDNTLIHSHGINRRDFMKLCAALAATMGLSSKAAAEMAESVTNPQRPPVIWIGAQECT WP_000145410.1
MULTISPECIES: hydrogenase-2 small chain [Proteobacteria] Length: 372\nNumber of predicted TMHs: 1\nExp number of AAs in TMHs: 23.72226\nExp number, first 60 AAs: 0.22038\nTotal prob of N-in: 0.12994\noutside 1 331\nTMhelix 332 354\ninside 355 372

5470 GCF_001742465.1_ASM174246v1 Escherichia coli Proteobacteria; Gammaproteobacteria; Enterobacterales; Enterobacteriaceae; Escherichia
MNNEETFYQAMRRQGVTRRSFLKYCSLAATSLGLGAGMAPKIAWALENKPRIPVVWIHGL WP_001058323.1
MULTISPECIES: hydrogenase-1 small chain [Proteobacteria] Length: 372\nNumber of predicted TMHs: 2\nExp number of AAs in TMHs: 37.2608499999999\nExp number, first 60 AAs: 19.52923\nTotal prob of N-in: 0.94351\nPOSSIBLE N-term signal sequence\ninside 1 19\nTMhelix 20 39\noutside 40 325\nTMhelix 326 348\ninside 349 372

5471 GCF_001748525.1_ASM174852v1 Escherichia coli Proteobacteria; Gammaproteobacteria; Enterobacterales; Enterobacteriaceae; Escherichia
MTGDNTLIHSHGINRRDFMKLCAALAATMGLSSKAAAEMAESVTNPQRPPVIWIGAQECT WP_000145410.1
MULTISPECIES: hydrogenase-2 small chain [Proteobacteria] Length: 372\nNumber of predicted TMHs: 1\nExp number of AAs in TMHs: 23.72226\nExp number, first 60 AAs: 0.22038\nTotal prob of N-in: 0.12994\noutside 1 331\nTMhelix 332 354\ninside 355 372

5472 GCF_001742475.1_ASM174247v1 Escherichia coli Proteobacteria; Gammaproteobacteria; Enterobacterales; Enterobacteriaceae; Escherichia
MNNEETFYQAMRRQGVTRRSFLKYCSLAATSLGLGAGMAPKIAWALENKPRIPVVWIHGL WP_001058323.1
MULTISPECIES: hydrogenase-1 small chain [Proteobacteria] Length: 372\nNumber of predicted TMHs: 2\nExp number of AAs in TMHs: 37.2608499999999\nExp number, first 60 AAs: 19.52923\nTotal prob of N-in: 0.94351\nPOSSIBLE N-term signal sequence\ninside 1 19\nTMhelix 20 39\noutside 40 325\nTMhelix 326 348\ninside 349 372

5473 GCF_001742495.1_ASM174249v1 Escherichia coli Proteobacteria; Gammaproteobacteria; Enterobacterales; Enterobacteriaceae; Escherichia
MNNEETFYQAMRRQGVTRRSFLKYCSLAATSLGLGAGMAPKIAWALENKPRIPVVWIHGL WP_001058323.1
MULTISPECIES: hydrogenase-1 small chain [Proteobacteria] Length: 372\nNumber of predicted TMHs: 2\nExp number of AAs in TMHs: 37.2608499999999\nExp number, first 60 AAs: 19.52923\nTotal prob of N-in: 0.94351\nPOSSIBLE N-term signal sequence\ninside 1 19\nTMhelix 20 39\noutside 40 325\nTMhelix 326 348\ninside 349 372

5474 GCF_001721125.1_ASM172112v1 Escherichia coli Proteobacteria; Gammaproteobacteria; Enterobacterales; Enterobacteriaceae; Escherichia
MNNEETFYQAMRRQGVTRRSFLKYCSLAATSLGLGAGMAPKIAWALENKPRIPVVWIHGL WP_001058323.1
MULTISPECIES: hydrogenase-1 small chain [Proteobacteria] Length: 372\nNumber of predicted TMHs: 2\nExp number of AAs in TMHs: 37.2608499999999\nExp number, first 60 AAs: 19.52923\nTotal prob of N-in: 0.94351\nPOSSIBLE N-term signal sequence\ninside 1 19\nTMhelix 20 39\noutside 40 325\nTMhelix 326 348\ninside 349 372

5475 GCF_001742555.1_ASM174255v1 Escherichia coli Proteobacteria; Gammaproteobacteria; Enterobacterales; Enterobacteriaceae; Escherichia
MTGDNTLIHSHGINRRDFMKLCAALAATMGLSSKAAAEMAESVTNPQRPPVIWIGAQECT WP_000145410.1
MULTISPECIES: hydrogenase-2 small chain [Proteobacteria] Length: 372\nNumber of predicted TMHs: 1\nExp number of AAs in TMHs: 23.72226\nExp number, first 60 AAs: 0.22038\nTotal prob of N-in: 0.12994\noutside 1 331\nTMhelix 332 354\ninside 355 372

5476 GCF_001742565.1_ASM174256v1 Escherichia coli Proteobacteria; Gammaproteobacteria; Enterobacterales; Enterobacteriaceae; Escherichia
MTGDNTLIHSHGINRRDFMKLCAALAATMGLSSKAAAEMAESVTNPQRPPVIWIGAQECT WP_000145410.1
MULTISPECIES: hydrogenase-2 small chain [Proteobacteria] Length: 372\nNumber of predicted TMHs: 1\nExp number of AAs in TMHs: 23.72226\nExp number, first 60 AAs: 0.22038\nTotal prob of N-in: 0.12994\noutside 1 331\nTMhelix 332 354\ninside 355 372

5477 GCF_001742625.1_ASM174262v1 Escherichia coli Proteobacteria; Gammaproteobacteria; Enterobacterales; Enterobacteriaceae; Escherichia
MTGDNTLIHSHGINRRDFMKLCAALAATMGLSSKAAAEMAESVTNPQRPPVIWIGAQECT WP_000145406.1
hydrogenase-2 small chain [Escherichia coli] Length: 372\nNumber of predicted TMHs: 1\nExp number of AAs in TMHs: 23.72251\nExp number, first 60 AAs: 0.22027\nTotal prob of N-in: 0.13006\noutside 1 331\nTMhelix 332 354\ninside 355 372

5478 GCF_001742565.1_ASM174256v1 Escherichia coli Proteobacteria; Gammaproteobacteria; Enterobacterales; Enterobacteriaceae; Escherichia
MNNEETFYQAMRRQGVTRRSFLKYCSLAATSLGLGAGMAPKIAWALENKPRIPVVWIHGL WP_001058323.1
MULTISPECIES: hydrogenase-1 small chain [Proteobacteria] Length: 372\nNumber of predicted TMHs: 2\nExp number of AAs in TMHs: 37.2608499999999\nExp number, first 60 AAs: 19.52923\nTotal prob of N-in: 0.94351\nPOSSIBLE N-term signal sequence\ninside 1 19\nTMhelix 20 39\noutside 40 325\nTMhelix 326 348\ninside 349 372

5479 GCF_001748555.1_ASM174855v1 Escherichia coli Proteobacteria; Gammaproteobacteria; Enterobacterales; Enterobacteriaceae; Escherichia
MTGDNTLIHSHGINRRDFMKLCAALAATMGLSSKAAAEMAESVTNPQRPPVIWIGAQECT WP_000145410.1
MULTISPECIES: hydrogenase-2 small chain [Proteobacteria] Length: 372\nNumber of predicted TMHs: 1\nExp number of AAs in TMHs: 23.72226\nExp number, first 60 AAs: 0.22038\nTotal prob of N-in: 0.12994\noutside 1 331\nTMhelix 332 354\ninside 355 372

5480 GCF_001748545.1_ASM174854v1 Escherichia coli Proteobacteria; Gammaproteobacteria; Enterobacterales; Enterobacteriaceae; Escherichia
MTGDNTLIHSHGINRRDFMKLCAALAATMGLSSKAAAEMAESVTNPQRPPVIWIGAQECT WP_000145410.1
MULTISPECIES: hydrogenase-2 small chain [Proteobacteria] Length: 372\nNumber of predicted TMHs: 1\nExp number of AAs in TMHs: 23.72226\nExp number, first 60 AAs: 0.22038\nTotal prob of N-in: 0.12994\noutside 1 331\nTMhelix 332 354\ninside 355 372

5481 GCF_001748615.1_ASM174861v1 Escherichia coli Proteobacteria; Gammaproteobacteria; Enterobacterales; Enterobacteriaceae; Escherichia
MTGDNTLIHSHGINRRDFMKLCAALAATMGLSSKAAAEMAESVTNPQRPPVIWIGAQECT WP_000145410.1
MULTISPECIES: hydrogenase-2 small chain [Proteobacteria] Length: 372\nNumber of predicted TMHs: 1\nExp number of AAs in TMHs: 23.72226\nExp number, first 60 AAs: 0.22038\nTotal prob of N-in: 0.12994\noutside 1 331\nTMhelix 332 354\ninside 355 372

5482 GCF_001742725.1_ASM174272v1 Escherichia coli Proteobacteria; Gammaproteobacteria; Enterobacterales; Enterobacteriaceae; Escherichia
MNNEETFYQAMRRQGVTRRSFLKYCSLAATSLGLGAGMAPKIAWALENKPRIPVVWIHGL WP_001058323.1
MULTISPECIES: hydrogenase-1 small chain [Proteobacteria] Length: 372\nNumber of predicted TMHs: 2\nExp number of AAs in TMHs: 37.2608499999999\nExp number, first 60 AAs: 19.52923\nTotal prob of N-in: 0.94351\nPOSSIBLE N-term signal sequence\ninside 1 19\nTMhelix 20 39\noutside 40 325\nTMhelix 326 348\ninside 349 372

5483 GCF_001748635.1_ASM174863v1 Escherichia coli Proteobacteria; Gammaproteobacteria; Enterobacterales; Enterobacteriaceae; Escherichia
MTGDNTLIHSHGINRRDFMKLCAALAATMGLSSKAAAEMAESVTNPQRPPVIWIGAQECT WP_000145410.1
MULTISPECIES: hydrogenase-2 small chain [Proteobacteria] Length: 372\nNumber of predicted TMHs: 1\nExp number of AAs in TMHs: 23.72226\nExp number, first 60 AAs: 0.22038\nTotal prob of N-in: 0.12994\noutside 1 331\nTMhelix 332 354\ninside 355 372

5484 GCF_001748635.1_ASM174863v1 Escherichia coli Proteobacteria; Gammaproteobacteria; Enterobacterales; Enterobacteriaceae; Escherichia
MNNEETFYQAMRRQGVTRRSFLKYCSLAATSLGLGAGMAPKIAWALENKPRIPVVWIHGL WP_001295940.1

MULTISPECIES: hydrogenase [Enterobacteriaceae] Length: 372\nNumber of predicted TMHs: 2\nExp number of AAs in TMHs: 37.2608899999999\nExp number, first 60 AAs: 19.52923\nTotal prob of N-in: 0.94351\nPOSSIBLE N-term signal sequence\ninside 1 19\nTMhelix 20 39\noutside 40 325\nTMhelix 326 348\ninside 349 372

5485 GCF_001742705.1_ASM174270v1 Escherichia coli Proteobacteria; Gammaproteobacteria; Enterobacterales; Enterobacteriaceae; Escherichia
MTGDNTLIHSHGINRRDFMKLCAALAATMGLSSKAAAEMAESVTNPQRPPVIWIGAQECT WP_069723081.1
hydrogenase 2 small subunit [Escherichia coli] Length: 372\nNumber of predicted TMHs: 1\nExp number of AAs in TMHs: 23.03253\nExp number, first 60 AAs: 0.22057\nTotal prob of N-in: 0.12305\noutside 1 331\nTMhelix 332 354\ninside 355 372

5486 GCF_001748685.1_ASM174868v1 Escherichia coli Proteobacteria; Gammaproteobacteria; Enterobacterales; Enterobacteriaceae; Escherichia
MTGDNTLIHSHGINRRDFMKLCAALAATMGLSSKAAAEMAESVTNPQRPPVIWIGAQECT WP_000145410.1
MULTISPECIES: hydrogenase-2 small chain [Proteobacteria] Length: 372\nNumber of predicted TMHs: 1\nExp number of AAs in TMHs: 23.72226\nExp number, first 60 AAs: 0.22038\nTotal prob of N-in: 0.12994\noutside 1 331\nTMhelix 332 354\ninside 355 372

5487 GCF_001748685.1_ASM174868v1 Escherichia coli Proteobacteria; Gammaproteobacteria; Enterobacterales; Enterobacteriaceae; Escherichia
MNNEETFYQAMRRQGVTRRSFLKYCSLAATSLGLGAGMAPKIAWALENKPRIPVVWIHGL WP_001058323.1
MULTISPECIES: hydrogenase-1 small chain [Proteobacteria] Length: 372\nNumber of predicted TMHs: 2\nExp number of AAs in TMHs: 37.2608499999999\nExp number, first 60 AAs: 19.52923\nTotal prob of N-in: 0.94351\nPOSSIBLE N-term signal sequence\ninside 1 19\nTMhelix 20 39\noutside 40 325\nTMhelix 326 348\ninside 349 372

5488 GCF_001748555.1_ASM174855v1 Escherichia coli Proteobacteria; Gammaproteobacteria; Enterobacterales; Enterobacteriaceae; Escherichia
MNNEETFYQAMRRQGVTRRSFLKYCSLAATSLGLGAGMAPKIAWALENKPRIPVVWIHGL WP_001058323.1
MULTISPECIES: hydrogenase-1 small chain [Proteobacteria] Length: 372\nNumber of predicted TMHs: 2\nExp number of AAs in TMHs: 37.2608499999999\nExp number, first 60 AAs: 19.52923\nTotal prob of N-in: 0.94351\nPOSSIBLE N-term signal sequence\ninside 1 19\nTMhelix 20 39\noutside 40 325\nTMhelix 326 348\ninside 349 372

5489 GCF_001748765.1_ASM174876v1 Escherichia coli Proteobacteria; Gammaproteobacteria; Enterobacterales; Enterobacteriaceae; Escherichia
MTGDNTLIHSHGINRRDFMKLCAALAATMGLSSKAAAEMAESVTNPQRPPVIWIGAQECT WP_000145410.1
MULTISPECIES: hydrogenase-2 small chain [Proteobacteria] Length: 372\nNumber of predicted TMHs: 1\nExp number of AAs in TMHs: 23.72226\nExp number, first 60 AAs: 0.22038\nTotal prob of N-in: 0.12994\noutside 1 331\nTMhelix 332 354\ninside 355 372

5490 GCF_001748765.1_ASM174876v1 Escherichia coli Proteobacteria; Gammaproteobacteria; Enterobacterales; Enterobacteriaceae; Escherichia
MNNEETFYQAMRRQGVTRRSFLKYCSLAATSLGLGAGMAPKIAWALENKPRIPVVWIHGL WP_069912832.1
hydrogenase [Escherichia coli] Length: 372\nNumber of predicted TMHs: 2\nExp number of AAs in TMHs: 37.3694099999999\nExp number, first 60 AAs: 19.52905\nTotal prob of N-in: 0.94357\nPOSSIBLE N-term signal sequence\ninside 1 19\nTMhelix 20 39\noutside 40 325\nTMhelix 326 348\ninside 349 372

5491 GCF_001748545.1_ASM174854v1 Escherichia coli Proteobacteria; Gammaproteobacteria; Enterobacterales; Enterobacteriaceae; Escherichia
MNNEETFYQAMRRQGVTRRSFLKYCSLAATSLGLGAGMAPKIAWALENKPRIPVVWIHGL WP_001058323.1
MULTISPECIES: hydrogenase-1 small chain [Proteobacteria] Length: 372\nNumber of predicted TMHs: 2\nExp number of AAs in TMHs: 37.2608499999999\nExp number, first 60 AAs: 19.52923\nTotal prob of N-in: 0.94351\nPOSSIBLE N-term signal sequence\ninside 1 19\nTMhelix 20 39\noutside 40 325\nTMhelix 326 348\ninside 349 372

5492 GCF_001748805.1_ASM174880v1 Escherichia coli Proteobacteria; Gammaproteobacteria; Enterobacterales; Enterobacteriaceae; Escherichia
MTGDNTLIHSHGINRRDFMKLCAALAATMGLSSKAAAEMAESVTNPQRPPVIWIGAQECT WP_000145410.1
MULTISPECIES: hydrogenase-2 small chain [Proteobacteria] Length: 372\nNumber of predicted TMHs:

1\nExp number of AAs in TMHs: 23.72226\nExp number, first 60 AAs: 0.22038\nTotal prob of N-in: 0.12994\noutside 1 331\nTMhelix 332 354\ninside 355 372

5493 GCF_001748885.1_ASM174888v1 Escherichia coli Proteobacteria; Gammaproteobacteria; Enterobacterales; Enterobacteriaceae; Escherichia
MNNEETFYQAMRRQGVTRRSFLKYCSLAATSLGLGAGMAPKIAWALENKPRIPVVWIHGL WP_001058323.1
MULTISPECIES: hydrogenase-1 small chain [Proteobacteria] Length: 372\nNumber of predicted TMHs: 2\nExp number of AAs in TMHs: 37.2608499999999\nExp number, first 60 AAs: 19.52923\nTotal prob of N-in: 0.94351\nPOSSIBLE N-term signal sequence\ninside 1 19\nTMhelix 20 39\noutside 40 325\nTMhelix 326 348\ninside 349 372

5494 GCF_001748615.1_ASM174861v1 Escherichia coli Proteobacteria; Gammaproteobacteria; Enterobacterales; Enterobacteriaceae; Escherichia
MNNEETFYQAMRRQGVTRRSFLKYCSLAATSLGLGAGMAPKIAWALENKPRIPVVWIHGL WP_001295940.1
MULTISPECIES: hydrogenase [Enterobacteriaceae] Length: 372\nNumber of predicted TMHs: 2\nExp number of AAs in TMHs: 37.2608899999999\nExp number, first 60 AAs: 19.52923\nTotal prob of N-in: 0.94351\nPOSSIBLE N-term signal sequence\ninside 1 19\nTMhelix 20 39\noutside 40 325\nTMhelix 326 348\ninside 349 372

5495 GCF_001748895.1_ASM174889v1 Escherichia coli Proteobacteria; Gammaproteobacteria; Enterobacterales; Enterobacteriaceae; Escherichia
MTGDNTLIHSHGINRRDFMKLCAALAATMGLSSKAAAEMAESVTNPQRPPVIWIGAQECT WP_000145410.1
MULTISPECIES: hydrogenase-2 small chain [Proteobacteria] Length: 372\nNumber of predicted TMHs: 1\nExp number of AAs in TMHs: 23.72226\nExp number, first 60 AAs: 0.22038\nTotal prob of N-in: 0.12994\noutside 1 331\nTMhelix 332 354\ninside 355 372

5496 GCF_001749045.1_ASM174904v1 Escherichia coli Proteobacteria; Gammaproteobacteria; Enterobacterales; Enterobacteriaceae; Escherichia
MTGDNTLIHSHGINRRDFMKLCAALAATMGLSSKAAAEMAESVTNPQRPPVIWIGAQECT WP_000145410.1
MULTISPECIES: hydrogenase-2 small chain [Proteobacteria] Length: 372\nNumber of predicted TMHs: 1\nExp number of AAs in TMHs: 23.72226\nExp number, first 60 AAs: 0.22038\nTotal prob of N-in: 0.12994\noutside 1 331\nTMhelix 332 354\ninside 355 372

5497 GCF_001748915.1_ASM174891v1 Escherichia coli Proteobacteria; Gammaproteobacteria; Enterobacterales; Enterobacteriaceae; Escherichia
MTGDNTLIHSHGINRRDFMKLCAALAATMGLSSKAAAEMAESVTNPQRPPVIWIGAQECT WP_000145410.1
MULTISPECIES: hydrogenase-2 small chain [Proteobacteria] Length: 372\nNumber of predicted TMHs: 1\nExp number of AAs in TMHs: 23.72226\nExp number, first 60 AAs: 0.22038\nTotal prob of N-in: 0.12994\noutside 1 331\nTMhelix 332 354\ninside 355 372

5498 GCF_001748985.1_ASM174898v1 Escherichia coli Proteobacteria; Gammaproteobacteria; Enterobacterales; Enterobacteriaceae; Escherichia
MTGDNTLIHSHGINRRDFMKLCAALAATMGLSSKAAAEMAESVTNPQRPPVIWIGAQECT WP_000145410.1
MULTISPECIES: hydrogenase-2 small chain [Proteobacteria] Length: 372\nNumber of predicted TMHs: 1\nExp number of AAs in TMHs: 23.72226\nExp number, first 60 AAs: 0.22038\nTotal prob of N-in: 0.12994\noutside 1 331\nTMhelix 332 354\ninside 355 372

5499 GCF_001748975.1_ASM174897v1 Escherichia coli Proteobacteria; Gammaproteobacteria; Enterobacterales; Enterobacteriaceae; Escherichia
MTGDNTLIHSHGINRRDFMKLCAALAATMGLSSKAAAEMAESVTNPQRPPVIWIGAQECT WP_000145410.1
MULTISPECIES: hydrogenase-2 small chain [Proteobacteria] Length: 372\nNumber of predicted TMHs: 1\nExp number of AAs in TMHs: 23.72226\nExp number, first 60 AAs: 0.22038\nTotal prob of N-in: 0.12994\noutside 1 331\nTMhelix 332 354\ninside 355 372

5500 GCF_001749125.1_ASM174912v1 Escherichia coli Proteobacteria; Gammaproteobacteria; Enterobacterales; Enterobacteriaceae; Escherichia
MTGDNTLIHSHGINRRDFMKLCAALAATMGLSSKAAAEMAESVTNPQRPPVIWIGAQECT WP_000145410.1
MULTISPECIES: hydrogenase-2 small chain [Proteobacteria] Length: 372\nNumber of predicted TMHs: 1\nExp number of AAs in TMHs: 23.72226\nExp number, first 60 AAs: 0.22038\nTotal prob of N-in: 0.12994\noutside 1 331\nTMhelix 332 354\ninside 355 372

5501 GCF_001749065.1_ASM174906v1 Escherichia coli Proteobacteria; Gammaproteobacteria; Enterobacterales; Enterobacteriaceae; Escherichia
MTGDNTLIHSHGINRRDFMKLCAALAATMGLSSKAAAEMAESVTNPQRPPVIWIGAQECT WP_000145410.1
MULTISPECIES: hydrogenase-2 small chain [Proteobacteria] Length: 372\nNumber of predicted TMHs: 1\nExp number of AAs in TMHs: 23.72226\nExp number, first 60 AAs: 0.22038\nTotal prob of N-in: 0.12994\noutside 1 331\nTMhelix 332 354\ninside 355 372

5502 GCF_001749045.1_ASM174904v1 Escherichia coli Proteobacteria; Gammaproteobacteria; Enterobacterales; Enterobacteriaceae; Escherichia
MNNEETFYQAMRRQGVTRRSFLKYCSLAATSLGLGAGMAPKIAWALENKPRIPVVWIHGL WP_001058323.1
MULTISPECIES: hydrogenase-1 small chain [Proteobacteria] Length: 372\nNumber of predicted TMHs: 2\nExp number of AAs in TMHs: 37.2608499999999\nExp number, first 60 AAs: 19.52923\nTotal prob of N-in: 0.94351\nPOSSIBLE N-term signal sequence\ninside 1 19\nTMhelix 20 39\noutside 40 325\nTMhelix 326 348\ninside 349 372

5503 GCF_001749135.1_ASM174913v1 Escherichia coli Proteobacteria; Gammaproteobacteria; Enterobacterales; Enterobacteriaceae; Escherichia
MTGDNTLIHSHGINRRDFMKLCAALAATMGLSSKAAAEMAESVTNPQRPPVIWIGAQECT WP_000145410.1
MULTISPECIES: hydrogenase-2 small chain [Proteobacteria] Length: 372\nNumber of predicted TMHs: 1\nExp number of AAs in TMHs: 23.72226\nExp number, first 60 AAs: 0.22038\nTotal prob of N-in: 0.12994\noutside 1 331\nTMhelix 332 354\ninside 355 372

5504 GCF_001749155.1_ASM174915v1 Escherichia coli Proteobacteria; Gammaproteobacteria; Enterobacterales; Enterobacteriaceae; Escherichia
MTGDNTLIHSHGINRRDFMKLCAALAATMGLSSKAAAEMAESVTNPQRPPVIWIGAQECT WP_000145410.1
MULTISPECIES: hydrogenase-2 small chain [Proteobacteria] Length: 372\nNumber of predicted TMHs: 1\nExp number of AAs in TMHs: 23.72226\nExp number, first 60 AAs: 0.22038\nTotal prob of N-in: 0.12994\noutside 1 331\nTMhelix 332 354\ninside 355 372

5505 GCF_001749155.1_ASM174915v1 Escherichia coli Proteobacteria; Gammaproteobacteria; Enterobacterales; Enterobacteriaceae; Escherichia
MNNEETFYQAMRRQGVTRRSFLKYCSLAATSLGLGAGMAPKIAWALENKPRIPVVWIHGL WP_001058323.1
MULTISPECIES: hydrogenase-1 small chain [Proteobacteria] Length: 372\nNumber of predicted TMHs: 2\nExp number of AAs in TMHs: 37.2608499999999\nExp number, first 60 AAs: 19.52923\nTotal prob of N-in: 0.94351\nPOSSIBLE N-term signal sequence\ninside 1 19\nTMhelix 20 39\noutside 40 325\nTMhelix 326 348\ninside 349 372

5506 GCF_001749135.1_ASM174913v1 Escherichia coli Proteobacteria; Gammaproteobacteria; Enterobacterales; Enterobacteriaceae; Escherichia
MNNEETFYQAMRRQGVTRRSFLKYCSLAATSLGLGAGMAPKIAWALENKPRIPVVWIHGL WP_001295940.1
MULTISPECIES: hydrogenase [Enterobacteriaceae] Length: 372\nNumber of predicted TMHs: 2\nExp number of AAs in TMHs: 37.2608899999999\nExp number, first 60 AAs: 19.52923\nTotal prob of N-in: 0.94351\nPOSSIBLE N-term signal sequence\ninside 1 19\nTMhelix 20 39\noutside 40 325\nTMhelix 326 348\ninside 349 372

5507 GCF_001748985.1_ASM174898v1 Escherichia coli Proteobacteria; Gammaproteobacteria; Enterobacterales; Enterobacteriaceae; Escherichia
MNNEETFYQAMRRQGVTRRSFLKYCSLAATSLGLGAGMAPKIAWALENKPRIPVVWIHGL WP_001295940.1
MULTISPECIES: hydrogenase [Enterobacteriaceae] Length: 372\nNumber of predicted TMHs: 2\nExp number of AAs in TMHs: 37.2608899999999\nExp number, first 60 AAs: 19.52923\nTotal prob of N-in: 0.94351\nPOSSIBLE N-term signal sequence\ninside 1 19\nTMhelix 20 39\noutside 40 325\nTMhelix 326 348\ninside 349 372

5508 GCF_001749285.1_ASM174928v1 Escherichia coli Proteobacteria; Gammaproteobacteria; Enterobacterales; Enterobacteriaceae; Escherichia
MTGDNTLIHSHGINRRDFMKLCAALAATMGLSSKAAAEMAESVTNPQRPPVIWIGAQECT WP_000145410.1
MULTISPECIES: hydrogenase-2 small chain [Proteobacteria] Length: 372\nNumber of predicted TMHs: 1\nExp number of AAs in TMHs: 23.72226\nExp number, first 60 AAs: 0.22038\nTotal prob of N-in: 0.12994\noutside 1 331\nTMhelix 332 354\ninside 355 372

5509 GCF_001749285.1_ASM174928v1 Escherichia coli Proteobacteria; Gammaproteobacteria; Enterobacterales; Enterobacteriaceae; Escherichia
MNNEETFYQAMRRQGVTRRSFLKYCSLAATSLGLGAGMAPKIAWALENKPRIPVVWIHGL WP_001058323.1
MULTISPECIES: hydrogenase-1 small chain [Proteobacteria] Length: 372\nNumber of predicted TMHs: 2\nExp number of AAs in TMHs: 37.2608499999999\nExp number, first 60 AAs: 19.52923\nTotal prob of N-in: 0.94351\nPOSSIBLE N-term signal sequence\ninside 1 19\nTMhelix 20 39\noutside 40 325\nTMhelix 326 348\ninside 349 372

5510 GCF_001749065.1_ASM174906v1 Escherichia coli Proteobacteria; Gammaproteobacteria; Enterobacterales; Enterobacteriaceae; Escherichia
MNNEETFYQAMRRQGVTRRSFLKYCSLAATSLGLGAGMAPKIAWALENKPRIPVVWIHGL WP_001058323.1
MULTISPECIES: hydrogenase-1 small chain [Proteobacteria] Length: 372\nNumber of predicted TMHs: 2\nExp number of AAs in TMHs: 37.2608499999999\nExp number, first 60 AAs: 19.52923\nTotal prob of N-in: 0.94351\nPOSSIBLE N-term signal sequence\ninside 1 19\nTMhelix 20 39\noutside 40 325\nTMhelix 326 348\ninside 349 372

5511 GCF_001748895.1_ASM174889v1 Escherichia coli Proteobacteria; Gammaproteobacteria; Enterobacterales; Enterobacteriaceae; Escherichia
MNNEETFYQAMRRQGVTRRSFLKYCSLAATSLGLGAGMAPKIAWALENKPRIPVVWIHGL WP_052934366.1
hydrogenase [Escherichia coli] Length: 372\nNumber of predicted TMHs: 2\nExp number of AAs in TMHs: 36.76923\nExp number, first 60 AAs: 19.5319\nTotal prob of N-in: 0.94313\nPOSSIBLE N-term signal sequence\ninside 1 19\nTMhelix 20 39\noutside 40 325\nTMhelix 326 348\ninside 349 372

5512 GCF_001749315.1_ASM174931v1 Escherichia coli Proteobacteria; Gammaproteobacteria; Enterobacterales; Enterobacteriaceae; Escherichia
MTGDNTLIHSHGINRRDFMKLCAALAATMGLSSKAAAEMAESVTNPQRPPVIWIGAQECT WP_000145410.1
MULTISPECIES: hydrogenase-2 small chain [Proteobacteria] Length: 372\nNumber of predicted TMHs: 1\nExp number of AAs in TMHs: 23.72226\nExp number, first 60 AAs: 0.22038\nTotal prob of N-in: 0.12994\noutside 1 331\nTMhelix 332 354\ninside 355 372

5513 GCF_001749375.1_ASM174937v1 Escherichia coli Proteobacteria; Gammaproteobacteria; Enterobacterales; Enterobacteriaceae; Escherichia
MTGDNTLIHSHGINRRDFMKLCAALAATMGLSSKAAAEMAESVTNPQRPPVIWIGAQECT WP_000145410.1
MULTISPECIES: hydrogenase-2 small chain [Proteobacteria] Length: 372\nNumber of predicted TMHs: 1\nExp number of AAs in TMHs: 23.72226\nExp number, first 60 AAs: 0.22038\nTotal prob of N-in: 0.12994\noutside 1 331\nTMhelix 332 354\ninside 355 372

5514 GCF_001749415.1_ASM174941v1 Escherichia coli Proteobacteria; Gammaproteobacteria; Enterobacterales; Enterobacteriaceae; Escherichia
MTGDNTLIHSHGINRRDFMKLCAALAATMGLSSKAAAEMAESVTNPQRPPVIWIGAQECT WP_000145410.1
MULTISPECIES: hydrogenase-2 small chain [Proteobacteria] Length: 372\nNumber of predicted TMHs: 1\nExp number of AAs in TMHs: 23.72226\nExp number, first 60 AAs: 0.22038\nTotal prob of N-in: 0.12994\noutside 1 331\nTMhelix 332 354\ninside 355 372

5515 GCF_001749455.1_ASM174945v1 Escherichia coli Proteobacteria; Gammaproteobacteria; Enterobacterales; Enterobacteriaceae; Escherichia
MTGDNTLIHSHGINRRDFMKLCAALAATMGLSSKAAAEMAESVTNPQRPPVIWIGAQECT WP_000145410.1
MULTISPECIES: hydrogenase-2 small chain [Proteobacteria] Length: 372\nNumber of predicted TMHs: 1\nExp number of AAs in TMHs: 23.72226\nExp number, first 60 AAs: 0.22038\nTotal prob of N-in: 0.12994\noutside 1 331\nTMhelix 332 354\ninside 355 372

5516 GCF_001749315.1_ASM174931v1 Escherichia coli Proteobacteria; Gammaproteobacteria; Enterobacterales; Enterobacteriaceae; Escherichia
MNNEETFYQAMRRQGVTRRSFLKYCSLAATSLGLGAGMAPKIAWALENKPRIPVVWIHGL WP_001058323.1
MULTISPECIES: hydrogenase-1 small chain [Proteobacteria] Length: 372\nNumber of predicted TMHs: 2\nExp number of AAs in TMHs: 37.2608499999999\nExp number, first 60 AAs: 19.52923\nTotal prob of N-in: 0.94351\nPOSSIBLE N-term signal sequence\ninside 1 19\nTMhelix 20 39\noutside 40 325\nTMhelix 326 348\ninside 349 372

5517 GCF_001749525.1_ASM174952v1 Escherichia coli Proteobacteria; Gammaproteobacteria; Enterobacterales; Enterobacteriaceae; Escherichia

MTGDNTLIHSHGINRRDFMKLCAALAATMGLSSKAAAEMAESVTNPQRPPVIWIGAQECT WP_000145410.1 MULTISPECIES: hydrogenase-2 small chain [Proteobacteria] Length: 372\nNumber of predicted TMHs: 1\nExp number of AAs in TMHs: 23.72226\nExp number, first 60 AAs: 0.22038\nTotal prob of N-in: 0.12994\noutside 1 331\nTMhelix 332 354\ninside 355 372

5518 GCF_001749565.1_ASM174956v1 Escherichia coli Proteobacteria; Gammaproteobacteria; Enterobacterales; Enterobacteriaceae; Escherichia MTGDNTLIHSHGINRRDFMKLCAALAATMGLSSKAAAEMAESVTNPQRPPVIWIGAQECT WP_000145410.1 MULTISPECIES: hydrogenase-2 small chain [Proteobacteria] Length: 372\nNumber of predicted TMHs: 1\nExp number of AAs in TMHs: 23.72226\nExp number, first 60 AAs: 0.22038\nTotal prob of N-in: 0.12994\noutside 1 331\nTMhelix 332 354\ninside 355 372

5519 GCF_001749565.1_ASM174956v1 Escherichia coli Proteobacteria; Gammaproteobacteria; Enterobacterales; Enterobacteriaceae; Escherichia MNNEETFYQAMRRQGVTRRSFLKYCSLAATSLGLGAGMAPKIAWALENKPRIPVVWIHGL WP_001058323.1 MULTISPECIES: hydrogenase-1 small chain [Proteobacteria] Length: 372\nNumber of predicted TMHs: 2\nExp number of AAs in TMHs: 37.2608499999999\nExp number, first 60 AAs: 19.52923\nTotal prob of N-in: 0.94351\nPOSSIBLE N-term signal sequence\ninside 1 19\nTMhelix 20 39\noutside 40 325\nTMhelix 326 348\ninside 349 372

5520 GCF_001749675.1_ASM174967v1 Escherichia coli Proteobacteria; Gammaproteobacteria; Enterobacterales; Enterobacteriaceae; Escherichia MTGDNTLIHSHGINRRDFMKLCAALAATMGLSSKAAAEMAESVTNPQRPPVIWIGAQECT WP_000145410.1 MULTISPECIES: hydrogenase-2 small chain [Proteobacteria] Length: 372\nNumber of predicted TMHs: 1\nExp number of AAs in TMHs: 23.72226\nExp number, first 60 AAs: 0.22038\nTotal prob of N-in: 0.12994\noutside 1 331\nTMhelix 332 354\ninside 355 372

5521 GCF_001749585.1_ASM174958v1 Escherichia coli Proteobacteria; Gammaproteobacteria; Enterobacterales; Enterobacteriaceae; Escherichia MNNEETFYQAMRRQGVTRRSFLKYCSLAATSLGLGAGMAPKIAWALENKPRIPVVWIHGL WP_001295940.1 MULTISPECIES: hydrogenase [Enterobacteriaceae] Length: 372\nNumber of predicted TMHs: 2\nExp number of AAs in TMHs: 37.2608899999999\nExp number, first 60 AAs: 19.52923\nTotal prob of N-in: 0.94351\nPOSSIBLE N-term signal sequence\ninside 1 19\nTMhelix 20 39\noutside 40 325\nTMhelix 326 348\ninside 349 372

5522 GCF_001749455.1_ASM174945v1 Escherichia coli Proteobacteria; Gammaproteobacteria; Enterobacterales; Enterobacteriaceae; Escherichia MNNEETFYQAMRRQGVTRRSFLKYCSLAATSLGLGAGMAPKIAWALENKPRIPVVWIHGL WP_001058323.1 MULTISPECIES: hydrogenase-1 small chain [Proteobacteria] Length: 372\nNumber of predicted TMHs: 2\nExp number of AAs in TMHs: 37.2608499999999\nExp number, first 60 AAs: 19.52923\nTotal prob of N-in: 0.94351\nPOSSIBLE N-term signal sequence\ninside 1 19\nTMhelix 20 39\noutside 40 325\nTMhelix 326 348\ninside 349 372

5523 GCF_001749625.1_ASM174962v1 Escherichia coli Proteobacteria; Gammaproteobacteria; Enterobacterales; Enterobacteriaceae; Escherichia MTGDNTLIHSHGINRRDFMKLCAALAATMGLSSKAAAEMAESVTNPQRPPVIWIGAQECT WP_000145410.1 MULTISPECIES: hydrogenase-2 small chain [Proteobacteria] Length: 372\nNumber of predicted TMHs: 1\nExp number of AAs in TMHs: 23.72226\nExp number, first 60 AAs: 0.22038\nTotal prob of N-in: 0.12994\noutside 1 331\nTMhelix 332 354\ninside 355 372

5524 GCF_001749665.1_ASM174966v1 Escherichia coli Proteobacteria; Gammaproteobacteria; Enterobacterales; Enterobacteriaceae; Escherichia MTGDNTLIHSHGINRRDFMKLCAALAATMGLSSKAAAEMAESVTNPQRPPVIWIGAQECT WP_000145410.1 MULTISPECIES: hydrogenase-2 small chain [Proteobacteria] Length: 372\nNumber of predicted TMHs: 1\nExp number of AAs in TMHs: 23.72226\nExp number, first 60 AAs: 0.22038\nTotal prob of N-in: 0.12994\noutside 1 331\nTMhelix 332 354\ninside 355 372

5525 GCF_001749625.1_ASM174962v1 Escherichia coli Proteobacteria; Gammaproteobacteria; Enterobacterales; Enterobacteriaceae; Escherichia MNNEETFYQAMRRQGVTRRSFLKYCSLAATSLGLGAGMAPKIAWALENKPRIPVVWIHGL WP_001295940.1 MULTISPECIES: hydrogenase [Enterobacteriaceae] Length: 372\nNumber of predicted TMHs: 2\nExp

number of AAs in TMHs: 37.2608899999999\nExp number, first 60 AAs:  19.52923\nTotal prob of N-in:  0.94351\nPOSSIBLE N-term signal sequence\ninside   1   19\nTMhelix   20   39\noutside   40   325\nTMhelix   326   348\ninside   349   372

5526 GCF_001749665.1_ASM174966v1 Escherichia coli Proteobacteria; Gammaproteobacteria; Enterobacterales; Enterobacteriaceae; Escherichia
MNNEETFYQAMRRQGVTRRSFLKYCSLAATSLGLGAGMAPKIAWALENKPRIPVVWIHGL WP_001295940.1
MULTISPECIES: hydrogenase [Enterobacteriaceae] Length: 372\nNumber of predicted TMHs:  2\nExp number of AAs in TMHs: 37.2608899999999\nExp number, first 60 AAs:  19.52923\nTotal prob of N-in:  0.94351\nPOSSIBLE N-term signal sequence\ninside   1   19\nTMhelix   20   39\noutside   40   325\nTMhelix   326   348\ninside   349   372

5527 GCF_001749525.1_ASM174952v1 Escherichia coli Proteobacteria; Gammaproteobacteria; Enterobacterales; Enterobacteriaceae; Escherichia
MNNEETFYQAMRRQGVTRRSFLKYCSLAATSLGLGAGMAPKIAWALENKPRIPVVWIHGL WP_001058323.1
MULTISPECIES: hydrogenase-1 small chain [Proteobacteria] Length: 372\nNumber of predicted TMHs:  2\nExp number of AAs in TMHs: 37.2608499999999\nExp number, first 60 AAs:  19.52923\nTotal prob of N-in:  0.94351\nPOSSIBLE N-term signal sequence\ninside   1   19\nTMhelix   20   39\noutside   40   325\nTMhelix   326   348\ninside   349   372

5528 GCF_001750845.1_ASM175084v1 Escherichia coli Proteobacteria; Gammaproteobacteria; Enterobacterales; Enterobacteriaceae; Escherichia
MTGDNTLIHSHGINRRDFMKLCAALAATMGLSSKAAAEMAESVTNPQRPPVIWIGAQECT WP_000145410.1
MULTISPECIES: hydrogenase-2 small chain [Proteobacteria] Length: 372\nNumber of predicted TMHs:  1\nExp number of AAs in TMHs: 23.72226\nExp number, first 60 AAs:  0.22038\nTotal prob of N-in:   0.12994\noutside   1   331\nTMhelix   332   354\ninside   355   372

5529 GCF_001749675.1_ASM174967v1 Escherichia coli Proteobacteria; Gammaproteobacteria; Enterobacterales; Enterobacteriaceae; Escherichia
MNNEETFYQAMRRQGVTRRSFLKYCSLAATSLGLGAGMAPKIAWALENKPRIPVVWIHGL WP_001295940.1
MULTISPECIES: hydrogenase [Enterobacteriaceae] Length: 372\nNumber of predicted TMHs:  2\nExp number of AAs in TMHs: 37.2608899999999\nExp number, first 60 AAs:  19.52923\nTotal prob of N-in:  0.94351\nPOSSIBLE N-term signal sequence\ninside   1   19\nTMhelix   20   39\noutside   40   325\nTMhelix   326   348\ninside   349   372

5530 GCF_001806265.1_ASM180626v1 Escherichia coli Proteobacteria; Gammaproteobacteria; Enterobacterales; Enterobacteriaceae; Escherichia
MTGDNTLIHSHGINRRDFMKLCAALAATMGLSSKAAAEMAESVTNPQRPPVIWIGAQECT WP_000145410.1
MULTISPECIES: hydrogenase-2 small chain [Proteobacteria] Length: 372\nNumber of predicted TMHs:  1\nExp number of AAs in TMHs: 23.72226\nExp number, first 60 AAs:  0.22038\nTotal prob of N-in:   0.12994\noutside   1   331\nTMhelix   332   354\ninside   355   372

5531 GCF_001860505.1_ASM186050v1 Escherichia coli Proteobacteria; Gammaproteobacteria; Enterobacterales; Enterobacteriaceae; Escherichia
MTGDNTLIHSHGINRRDFMKLCAALAATMGLSSKAAAEMAESVTNPQRPPVIWIGAQECT WP_000145410.1
MULTISPECIES: hydrogenase-2 small chain [Proteobacteria] Length: 372\nNumber of predicted TMHs:  1\nExp number of AAs in TMHs: 23.72226\nExp number, first 60 AAs:  0.22038\nTotal prob of N-in:   0.12994\noutside   1   331\nTMhelix   332   354\ninside   355   372

5532 GCF_001865295.1_ASM186529v1 Escherichia coli Proteobacteria; Gammaproteobacteria; Enterobacterales; Enterobacteriaceae; Escherichia
MTGDNTLIHSHGINRRDFMKLCAALAATMGLSSKAAAEMAESVTNPQRPPVIWIGAQECT WP_000145410.1
MULTISPECIES: hydrogenase-2 small chain [Proteobacteria] Length: 372\nNumber of predicted TMHs:  1\nExp number of AAs in TMHs: 23.72226\nExp number, first 60 AAs:  0.22038\nTotal prob of N-in:   0.12994\noutside   1   331\nTMhelix   332   354\ninside   355   372

5533 GCF_001860505.1_ASM186050v1 Escherichia coli Proteobacteria; Gammaproteobacteria; Enterobacterales; Enterobacteriaceae; Escherichia
MNNEETFYQAMRRQGVTRRSFLKYCSLAATSLGLGAGMAPKIAWALENKPRIPVVWIHGL WP_001058323.1
MULTISPECIES: hydrogenase-1 small chain [Proteobacteria] Length: 372\nNumber of predicted TMHs:  2\nExp number of AAs in TMHs: 37.2608499999999\nExp number, first 60 AAs:  19.52923\nTotal prob of N-in:

0.94351\nPOSSIBLE N-term signal sequence\ninside 1 19\nTMhelix 20 39\noutside 40 325\nTMhelix 326 348\ninside 349 372

5534 GCF_001865295.1_ASM186529v1 Escherichia coli Proteobacteria; Gammaproteobacteria; Enterobacterales; Enterobacteriaceae; Escherichia MNNEETFYQAMRRQGVTRRSFLKYCSLAATSLGLGAGMAPKIAWALENKPRIPVVWIHGL WP_001301613.1 hydrogenase [Escherichia coli] Length: 372\nNumber of predicted TMHs: 2\nExp number of AAs in TMHs: 37.46598\nExp number, first 60 AAs: 19.53139\nTotal prob of N-in: 0.94364\nPOSSIBLE N-term signal sequence\ninside 1 19\nTMhelix 20 39\noutside 40 325\nTMhelix 326 348\ninside 349 372

5535 GCF_001806265.1_ASM180626v1 Escherichia coli Proteobacteria; Gammaproteobacteria; Enterobacterales; Enterobacteriaceae; Escherichia MNNEETFYQAMRRQGVTRRSFLKYCSLAATSLGLGAGMAPKIAWALENKPRIPVVWIHGL WP_001301613.1 hydrogenase [Escherichia coli] Length: 372\nNumber of predicted TMHs: 2\nExp number of AAs in TMHs: 37.46598\nExp number, first 60 AAs: 19.53139\nTotal prob of N-in: 0.94364\nPOSSIBLE N-term signal sequence\ninside 1 19\nTMhelix 20 39\noutside 40 325\nTMhelix 326 348\ninside 349 372

5536 GCF_001865895.1_ASM186589v1 Escherichia coli Proteobacteria; Gammaproteobacteria; Enterobacterales; Enterobacteriaceae; Escherichia MTGDNTLIHSHGINRRDFMKLCAALAATMGLSSKAAAEMAESVTNPQRPPVIWIGAQECT WP_000145410.1 MULTISPECIES: hydrogenase-2 small chain [Proteobacteria] Length: 372\nNumber of predicted TMHs: 1\nExp number of AAs in TMHs: 23.72226\nExp number, first 60 AAs: 0.22038\nTotal prob of N-in: 0.12994\noutside 1 331\nTMhelix 332 354\ninside 355 372

5537 GCF_001865925.1_ASM186592v1 Escherichia coli Proteobacteria; Gammaproteobacteria; Enterobacterales; Enterobacteriaceae; Escherichia MTGDNTLIHSHGINRRDFMKLCAALAATMGLSSKAAAEMAESVTNPQRPPVIWIGAQECT WP_000145410.1 MULTISPECIES: hydrogenase-2 small chain [Proteobacteria] Length: 372\nNumber of predicted TMHs: 1\nExp number of AAs in TMHs: 23.72226\nExp number, first 60 AAs: 0.22038\nTotal prob of N-in: 0.12994\noutside 1 331\nTMhelix 332 354\ninside 355 372

5538 GCF_001865915.1_ASM186591v1 Escherichia coli Proteobacteria; Gammaproteobacteria; Enterobacterales; Enterobacteriaceae; Escherichia MTGDNTLIHSHGINRRDFMKLCAALAATMGLSSKAAAEMAESVTNPQRPPVIWIGAQECT WP_000145410.1 MULTISPECIES: hydrogenase-2 small chain [Proteobacteria] Length: 372\nNumber of predicted TMHs: 1\nExp number of AAs in TMHs: 23.72226\nExp number, first 60 AAs: 0.22038\nTotal prob of N-in: 0.12994\noutside 1 331\nTMhelix 332 354\ninside 355 372

5539 GCF_001865925.1_ASM186592v1 Escherichia coli Proteobacteria; Gammaproteobacteria; Enterobacterales; Enterobacteriaceae; Escherichia MNNEETFYQAMRRQGVTRRSFLKYCSLAATSLGLGAGMAPKIAWALENKPRIPVVWIHGL WP_001058323.1 MULTISPECIES: hydrogenase-1 small chain [Proteobacteria] Length: 372\nNumber of predicted TMHs: 2\nExp number of AAs in TMHs: 37.2608499999999\nExp number, first 60 AAs: 19.52923\nTotal prob of N-in: 0.94351\nPOSSIBLE N-term signal sequence\ninside 1 19\nTMhelix 20 39\noutside 40 325\nTMhelix 326 348\ninside 349 372

5540 GCF_001881105.1_ASM188110v1 Escherichia coli Proteobacteria; Gammaproteobacteria; Enterobacterales; Enterobacteriaceae; Escherichia MTGDNTLIHSHGINRRDFMKLCAALAATMGLSSKAAAEMAESVTNPQRPPVIWIGAQECT WP_000145410.1 MULTISPECIES: hydrogenase-2 small chain [Proteobacteria] Length: 372\nNumber of predicted TMHs: 1\nExp number of AAs in TMHs: 23.72226\nExp number, first 60 AAs: 0.22038\nTotal prob of N-in: 0.12994\noutside 1 331\nTMhelix 332 354\ninside 355 372

5541 GCF_001865985.1_ASM186598v1 Escherichia coli Proteobacteria; Gammaproteobacteria; Enterobacterales; Enterobacteriaceae; Escherichia MNNEETFYQAMRRQGVTRRSFLKYCSLAATSLGLGAGMAPKIAWALENKPRIPVVWIHGL WP_001058323.1 MULTISPECIES: hydrogenase-1 small chain [Proteobacteria] Length: 372\nNumber of predicted TMHs: 2\nExp number of AAs in TMHs: 37.2608499999999\nExp number, first 60 AAs: 19.52923\nTotal prob of N-in: 0.94351\nPOSSIBLE N-term signal sequence\ninside 1 19\nTMhelix 20 39\noutside 40 325\nTMhelix 326 348\ninside 349 372

5542 GCF_001865915.1_ASM186591v1 Escherichia coli Proteobacteria; Gammaproteobacteria; Enterobacterales; Enterobacteriaceae; Escherichia
MNNEETFYQAMRRQGVTRRSFLKYCSLAATSLGLGAGMAPKIAWALENKPRIPVVWIHGL WP_001058323.1
MULTISPECIES: hydrogenase-1 small chain [Proteobacteria] Length: 372\nNumber of predicted TMHs: 2\nExp number of AAs in TMHs: 37.2608499999999\nExp number, first 60 AAs: 19.52923\nTotal prob of N-in: 0.94351\nPOSSIBLE N-term signal sequence\ninside 1 19\nTMhelix 20 39\noutside 40 325\nTMhelix 326 348\ninside 349 372

5543 GCF_001881145.1_ASM188114v1 Escherichia coli Proteobacteria; Gammaproteobacteria; Enterobacterales; Enterobacteriaceae; Escherichia
MTGDNTLIHSHGINRRDFMKLCAALAATMGLSSKAAAEMAESVTNPQRPPVIWIGAQECT WP_000145410.1
MULTISPECIES: hydrogenase-2 small chain [Proteobacteria] Length: 372\nNumber of predicted TMHs: 1\nExp number of AAs in TMHs: 23.72226\nExp number, first 60 AAs: 0.22038\nTotal prob of N-in: 0.12994\noutside 1 331\nTMhelix 332 354\ninside 355 372

5544 GCF_001874785.1_ASM187478v1 Escherichia coli Proteobacteria; Gammaproteobacteria; Enterobacterales; Enterobacteriaceae; Escherichia
MNNEETFYQAMRRQGVTRRSFLKYCSLAATSLGLGAGMAPKIAWALENKPRIPVVWIHGL WP_001295940.1
MULTISPECIES: hydrogenase [Enterobacteriaceae] Length: 372\nNumber of predicted TMHs: 2\nExp number of AAs in TMHs: 37.2608899999999\nExp number, first 60 AAs: 19.52923\nTotal prob of N-in: 0.94351\nPOSSIBLE N-term signal sequence\ninside 1 19\nTMhelix 20 39\noutside 40 325\nTMhelix 326 348\ninside 349 372

5545 GCF_001881155.1_ASM188115v1 Escherichia coli Proteobacteria; Gammaproteobacteria; Enterobacterales; Enterobacteriaceae; Escherichia
MTGDNTLIHSHGINRRDFMKLCAALAATMGLSSKAAAEMAESVTNPQRPPVIWIGAQECT WP_000145410.1
MULTISPECIES: hydrogenase-2 small chain [Proteobacteria] Length: 372\nNumber of predicted TMHs: 1\nExp number of AAs in TMHs: 23.72226\nExp number, first 60 AAs: 0.22038\nTotal prob of N-in: 0.12994\noutside 1 331\nTMhelix 332 354\ninside 355 372

5546 GCF_001881205.1_ASM188120v1 Escherichia coli Proteobacteria; Gammaproteobacteria; Enterobacterales; Enterobacteriaceae; Escherichia
MTGDNTLIHSHGINRRDFMKLCAALAATMGLSSKAAAEMAESVTNPQRPPVIWIGAQECT WP_000145410.1
MULTISPECIES: hydrogenase-2 small chain [Proteobacteria] Length: 372\nNumber of predicted TMHs: 1\nExp number of AAs in TMHs: 23.72226\nExp number, first 60 AAs: 0.22038\nTotal prob of N-in: 0.12994\noutside 1 331\nTMhelix 332 354\ninside 355 372

5547 GCF_001881235.1_ASM188123v1 Escherichia coli Proteobacteria; Gammaproteobacteria; Enterobacterales; Enterobacteriaceae; Escherichia
MTGDNTLIHSHGINRRDFMKLCAALAATMGLSSKAAAEMAESVTNPQRPPVIWIGAQECT WP_000145410.1
MULTISPECIES: hydrogenase-2 small chain [Proteobacteria] Length: 372\nNumber of predicted TMHs: 1\nExp number of AAs in TMHs: 23.72226\nExp number, first 60 AAs: 0.22038\nTotal prob of N-in: 0.12994\noutside 1 331\nTMhelix 332 354\ninside 355 372

5548 GCF_001881245.1_ASM188124v1 Escherichia coli Proteobacteria; Gammaproteobacteria; Enterobacterales; Enterobacteriaceae; Escherichia
MTGDNTLIHSHGINRRDFMKLCAALAATMGLSSKAAAEMAESVTNPQRPPVIWIGAQECT WP_000145410.1
MULTISPECIES: hydrogenase-2 small chain [Proteobacteria] Length: 372\nNumber of predicted TMHs: 1\nExp number of AAs in TMHs: 23.72226\nExp number, first 60 AAs: 0.22038\nTotal prob of N-in: 0.12994\noutside 1 331\nTMhelix 332 354\ninside 355 372

5549 GCF_001881305.1_ASM188130v1 Escherichia coli Proteobacteria; Gammaproteobacteria; Enterobacterales; Enterobacteriaceae; Escherichia
MTGDNTLIHSHGINRRDFMKLCAALAATMGLSSKAAAEMAESVTNPQRPPVIWIGAQECT WP_000145410.1
MULTISPECIES: hydrogenase-2 small chain [Proteobacteria] Length: 372\nNumber of predicted TMHs: 1\nExp number of AAs in TMHs: 23.72226\nExp number, first 60 AAs: 0.22038\nTotal prob of N-in: 0.12994\noutside 1 331\nTMhelix 332 354\ninside 355 372

5550 GCF_001881345.1_ASM188134v1 Escherichia coli Proteobacteria; Gammaproteobacteria; Enterobacterales; Enterobacteriaceae; Escherichia
MTGDNTLIHSHGINRRDFMKLCAALAATMGLSSKAAAEMAESVTNPQRPPVIWIGAQECT WP_000145410.1

MULTISPECIES: hydrogenase-2 small chain [Proteobacteria] Length: 372\nNumber of predicted TMHs: 1\nExp number of AAs in TMHs: 23.72226\nExp number, first 60 AAs: 0.22038\nTotal prob of N-in: 0.12994\noutside 1 331\nTMhelix 332 354\ninside 355 372

5551 GCF_001881355.1_ASM188135v1 Escherichia coli Proteobacteria; Gammaproteobacteria; Enterobacterales; Enterobacteriaceae; Escherichia
MTGDNTLIHSHGINRRDFMKLCAALAATMGLSSKAAAEMAESVTNPQRPPVIWIGAQECT WP_000145410.1
MULTISPECIES: hydrogenase-2 small chain [Proteobacteria] Length: 372\nNumber of predicted TMHs: 1\nExp number of AAs in TMHs: 23.72226\nExp number, first 60 AAs: 0.22038\nTotal prob of N-in: 0.12994\noutside 1 331\nTMhelix 332 354\ninside 355 372

5552 GCF_001881435.1_ASM188143v1 Escherichia coli Proteobacteria; Gammaproteobacteria; Enterobacterales; Enterobacteriaceae; Escherichia
MTGDNTLIHSHGINRRDFMKLCAALAATMGLSSKAAAEMAESVTNPQRPPVIWIGAQECT WP_000145410.1
MULTISPECIES: hydrogenase-2 small chain [Proteobacteria] Length: 372\nNumber of predicted TMHs: 1\nExp number of AAs in TMHs: 23.72226\nExp number, first 60 AAs: 0.22038\nTotal prob of N-in: 0.12994\noutside 1 331\nTMhelix 332 354\ninside 355 372

5553 GCF_001881395.1_ASM188139v1 Escherichia coli Proteobacteria; Gammaproteobacteria; Enterobacterales; Enterobacteriaceae; Escherichia
MTGDNTLIHSHGINRRDFMKLCAALAATMGLSSKAAAEMAESVTNPQRPPVIWIGAQECT WP_000145410.1
MULTISPECIES: hydrogenase-2 small chain [Proteobacteria] Length: 372\nNumber of predicted TMHs: 1\nExp number of AAs in TMHs: 23.72226\nExp number, first 60 AAs: 0.22038\nTotal prob of N-in: 0.12994\noutside 1 331\nTMhelix 332 354\ninside 355 372

5554 GCF_001884865.1_ASM188486v1 Escherichia coli Proteobacteria; Gammaproteobacteria; Enterobacterales; Enterobacteriaceae; Escherichia
MTGDNTLIHSHGINRRDFMKLCAALAATMGLSSKAAAEMAESVTNPQRPPVIWIGAQECS WP_061069243.1
hydrogenase 2 small subunit [Escherichia coli] Length: 372\nNumber of predicted TMHs: 1\nExp number of AAs in TMHs: 23.72219\nExp number, first 60 AAs: 0.22034\nTotal prob of N-in: 0.12994\noutside 1 331\nTMhelix 332 354\ninside 355 372

5555 GCF_001890245.1_ASM189024v1 Escherichia coli Proteobacteria; Gammaproteobacteria; Enterobacterales; Enterobacteriaceae; Escherichia
MTGDNTLIHSHGINRRDFMKLCAALAATMGLSSKAAAEMAESVTNPQRPPVIWIGAQECT WP_000145410.1
MULTISPECIES: hydrogenase-2 small chain [Proteobacteria] Length: 372\nNumber of predicted TMHs: 1\nExp number of AAs in TMHs: 23.72226\nExp number, first 60 AAs: 0.22038\nTotal prob of N-in: 0.12994\noutside 1 331\nTMhelix 332 354\ninside 355 372

5556 GCF_001884895.1_ASM188489v1 Escherichia coli Proteobacteria; Gammaproteobacteria; Enterobacterales; Enterobacteriaceae; Escherichia
MNNEETFYQAMRRQGVTRRSFLKYCSLAATSLGLGAGMAPKIAWALENKPRIPVVWIHGL WP_001301613.1
hydrogenase [Escherichia coli] Length: 372\nNumber of predicted TMHs: 2\nExp number of AAs in TMHs: 37.46598\nExp number, first 60 AAs: 19.53139\nTotal prob of N-in: 0.94364\nPOSSIBLE N-term signal sequence\ninside 1 19\nTMhelix 20 39\noutside 40 325\nTMhelix 326 348\ninside 349 372

5557 GCF_001890205.1_ASM189020v1 Escherichia coli Proteobacteria; Gammaproteobacteria; Enterobacterales; Enterobacteriaceae; Escherichia
MTGDNTLIHSHGINRRDFMKLCAALAATMGLSSKAAAEMAESVTNPQRPPVIWIGAQECT WP_000145410.1
MULTISPECIES: hydrogenase-2 small chain [Proteobacteria] Length: 372\nNumber of predicted TMHs: 1\nExp number of AAs in TMHs: 23.72226\nExp number, first 60 AAs: 0.22038\nTotal prob of N-in: 0.12994\noutside 1 331\nTMhelix 332 354\ninside 355 372

5558 GCF_001886535.1_ASM188653v1 Escherichia coli Proteobacteria; Gammaproteobacteria; Enterobacterales; Enterobacteriaceae; Escherichia
MNNEETFYQAMRRQGVTRRSFLKYCSLAATSLGLGAGMAPKIAWALENKPRIPVVWIHGL WP_001058323.1
MULTISPECIES: hydrogenase-1 small chain [Proteobacteria] Length: 372\nNumber of predicted TMHs: 2\nExp number of AAs in TMHs: 37.2608499999999\nExp number, first 60 AAs: 19.52923\nTotal prob of N-in: 0.94351\nPOSSIBLE N-term signal sequence\ninside 1 19\nTMhelix 20 39\noutside 40 325\nTMhelix 326 348\ninside 349 372

5559 GCF_001886555.1_ASM188655v1 Escherichia coli Proteobacteria; Gammaproteobacteria; Enterobacterales; Enterobacteriaceae; Escherichia
MNNEETFYQAMRRQGVTRRSFLKYCSLAATSLGLGAGMAPKIAWALENKPRIPVVWIHGL WP_001058323.1
MULTISPECIES: hydrogenase-1 small chain [Proteobacteria] Length: 372\nNumber of predicted TMHs: 2\nExp number of AAs in TMHs: 37.2608499999999\nExp number, first 60 AAs: 19.52923\nTotal prob of N-in: 0.94351\nPOSSIBLE N-term signal sequence\ninside 1 19\nTMhelix 20 39\noutside 40 325\nTMhelix 326 348\ninside 349 372

5560 GCF_001886755.1_ASM188675v1 Escherichia coli Proteobacteria; Gammaproteobacteria; Enterobacterales; Enterobacteriaceae; Escherichia
MNNEETFYQAMRRQGVTRRSFLKYCSLAATSLGLGAGMAPKIAWALENKPRIPVVWIHGL WP_001058323.1
MULTISPECIES: hydrogenase-1 small chain [Proteobacteria] Length: 372\nNumber of predicted TMHs: 2\nExp number of AAs in TMHs: 37.2608499999999\nExp number, first 60 AAs: 19.52923\nTotal prob of N-in: 0.94351\nPOSSIBLE N-term signal sequence\ninside 1 19\nTMhelix 20 39\noutside 40 325\nTMhelix 326 348\ninside 349 372

5561 GCF_001884875.1_ASM188487v1 Escherichia coli Proteobacteria; Gammaproteobacteria; Enterobacterales; Enterobacteriaceae; Escherichia
MTGDNTLIHSHGINRRDFMKLCAALAATMGLSSKAAAEMAESVTNPQRPPVIWIGAQECS WP_061069243.1
hydrogenase 2 small subunit [Escherichia coli] Length: 372\nNumber of predicted TMHs: 1\nExp number of AAs in TMHs: 23.72219\nExp number, first 60 AAs: 0.22034\nTotal prob of N-in: 0.12994\noutside 1 331\nTMhelix 332 354\ninside 355 372

5562 GCF_001884875.1_ASM188487v1 Escherichia coli Proteobacteria; Gammaproteobacteria; Enterobacterales; Enterobacteriaceae; Escherichia
MNNEETFYQAMRRQGVTRRSFLKYCSLAATSLGLGAGMAPKIAWALENKPRIPVVWIHGL WP_071813750.1
hydrogenase [Escherichia coli] Length: 372\nNumber of predicted TMHs: 2\nExp number of AAs in TMHs: 37.58713\nExp number, first 60 AAs: 19.53582\nTotal prob of N-in: 0.94353\nPOSSIBLE N-term signal sequence\ninside 1 19\nTMhelix 20 39\noutside 40 325\nTMhelix 326 348\ninside 349 372

5563 GCF_001886755.1_ASM188675v1 Escherichia coli Proteobacteria; Gammaproteobacteria; Enterobacterales; Enterobacteriaceae; Escherichia
MTGDNTLIHSHGINRRDFMKLCAALATTMGLSSKAAAEMAESVTNPQRPPVIWIGAQECT WP_023149629.1
MULTISPECIES: hydrogenase-2 small chain [Enterobacteriaceae] Length: 372\nNumber of predicted TMHs: 1\nExp number of AAs in TMHs: 23.63603\nExp number, first 60 AAs: 0.12971\nTotal prob of N-in: 0.12608\noutside 1 331\nTMhelix 332 354\ninside 355 372

5564 GCF_001884895.1_ASM188489v1 Escherichia coli Proteobacteria; Gammaproteobacteria; Enterobacterales; Enterobacteriaceae; Escherichia
MTGDNTLIHSHGINRRDFMKLCAALAATMGLSSKAAAEMAESVTNPQRPPVIWIGAQECS WP_061069243.1
hydrogenase 2 small subunit [Escherichia coli] Length: 372\nNumber of predicted TMHs: 1\nExp number of AAs in TMHs: 23.72219\nExp number, first 60 AAs: 0.22034\nTotal prob of N-in: 0.12994\noutside 1 331\nTMhelix 332 354\ninside 355 372

5565 GCF_001884955.1_ASM188495v1 Escherichia coli Proteobacteria; Gammaproteobacteria; Enterobacterales; Enterobacteriaceae; Escherichia
MNNEETFYQAMRRQGVTRRSFLKYCSLAATSLGLGAGMAPKIAWALENKPRIPVVWIHGL WP_001301613.1
hydrogenase [Escherichia coli] Length: 372\nNumber of predicted TMHs: 2\nExp number of AAs in TMHs: 37.46598\nExp number, first 60 AAs: 19.53139\nTotal prob of N-in: 0.94364\nPOSSIBLE N-term signal sequence\ninside 1 19\nTMhelix 20 39\noutside 40 325\nTMhelix 326 348\ninside 349 372

5566 GCF_001884965.1_ASM188496v1 Escherichia coli Proteobacteria; Gammaproteobacteria; Enterobacterales; Enterobacteriaceae; Escherichia
MNNEETFYQAMRRQGVTRRSFLKYCSLAATSLGLGAGMAPKIAWALENKPRIPVVWIHGL WP_001301613.1
hydrogenase [Escherichia coli] Length: 372\nNumber of predicted TMHs: 2\nExp number of AAs in TMHs: 37.46598\nExp number, first 60 AAs: 19.53139\nTotal prob of N-in: 0.94364\nPOSSIBLE N-term signal sequence\ninside 1 19\nTMhelix 20 39\noutside 40 325\nTMhelix 326 348\ninside 349 372

5567 GCF_001885025.1_ASM188502v1 Escherichia coli Proteobacteria; Gammaproteobacteria; Enterobacterales; Enterobacteriaceae; Escherichia
MNNEETFYQAMRRQGVTRRSFLKYCSLAATSLGLGAGMAPKIAWALENKPRIPVVWIHGL WP_001301613.1

hydrogenase [Escherichia coli] Length: 372\nNumber of predicted TMHs: 2\nExp number of AAs in TMHs: 37.46598\nExp number, first 60 AAs: 19.53139\nTotal prob of N-in: 0.94364\nPOSSIBLE N-term signal sequence\ninside 1 19\nTMhelix 20 39\noutside 40 325\nTMhelix 326 348\ninside 349 372

5568 GCF_001890345.1_ASM189034v1 Escherichia coli Proteobacteria; Gammaproteobacteria; Enterobacterales; Enterobacteriaceae; Escherichia
MTGDNTLIHSHGINRRDFMKLCAALAATMGLSSKAAAEMAESVTNPQRPPVIWIGAQECT WP_000145410.1
MULTISPECIES: hydrogenase-2 small chain [Proteobacteria] Length: 372\nNumber of predicted TMHs: 1\nExp number of AAs in TMHs: 23.72226\nExp number, first 60 AAs: 0.22038\nTotal prob of N-in: 0.12994\noutside 1 331\nTMhelix 332 354\ninside 355 372

5569 GCF_001888075.1_ASM188807v1 Escherichia coli Proteobacteria; Gammaproteobacteria; Enterobacterales; Enterobacteriaceae; Escherichia
MNNEETFYQAMRRQGVTRRSFLKYCSLAATSLGLGAGMAPKIAWAQENKPRIPVVWIHGL WP_072327948.1
hydrogenase [Escherichia coli] Length: 372\nNumber of predicted TMHs: 2\nExp number of AAs in TMHs: 36.59361\nExp number, first 60 AAs: 18.81285\nTotal prob of N-in: 0.91312\nPOSSIBLE N-term signal sequence\ninside 1 19\nTMhelix 20 39\noutside 40 325\nTMhelix 326 348\ninside 349 372

5570 GCF_001891215.1_ASM189121v1 Escherichia coli Proteobacteria; Gammaproteobacteria; Enterobacterales; Enterobacteriaceae; Escherichia
MTGDNTLIHSHGINRRDFMKLCAALAATMGLSSKAAAEMAESVTNPQRPPVIWIGAQECT WP_000145410.1
MULTISPECIES: hydrogenase-2 small chain [Proteobacteria] Length: 372\nNumber of predicted TMHs: 1\nExp number of AAs in TMHs: 23.72226\nExp number, first 60 AAs: 0.22038\nTotal prob of N-in: 0.12994\noutside 1 331\nTMhelix 332 354\ninside 355 372

5571 GCF_001890305.1_ASM189030v1 Escherichia coli Proteobacteria; Gammaproteobacteria; Enterobacterales; Enterobacteriaceae; Escherichia
MNNEETFYQAMRRQGVTRRSFLKYCSLAATSLGLGAGMAPKIAWALENKPRIPVVWIHGL WP_001301613.1
hydrogenase [Escherichia coli] Length: 372\nNumber of predicted TMHs: 2\nExp number of AAs in TMHs: 37.46598\nExp number, first 60 AAs: 19.53139\nTotal prob of N-in: 0.94364\nPOSSIBLE N-term signal sequence\ninside 1 19\nTMhelix 20 39\noutside 40 325\nTMhelix 326 348\ninside 349 372

5572 GCF_001891215.1_ASM189121v1 Escherichia coli Proteobacteria; Gammaproteobacteria; Enterobacterales; Enterobacteriaceae; Escherichia
MNNEETFYQAMRRQGVTRRSFLKYCSLAATSLGLGAGMAPKIAWALENKPRIPVVWIHGL WP_001295940.1
MULTISPECIES: hydrogenase [Enterobacteriaceae] Length: 372\nNumber of predicted TMHs: 2\nExp number of AAs in TMHs: 37.2608899999999\nExp number, first 60 AAs: 19.52923\nTotal prob of N-in: 0.94351\nPOSSIBLE N-term signal sequence\ninside 1 19\nTMhelix 20 39\noutside 40 325\nTMhelix 326 348\ninside 349 372

5573 GCF_001890245.1_ASM189024v1 Escherichia coli Proteobacteria; Gammaproteobacteria; Enterobacterales; Enterobacteriaceae; Escherichia
MNNEETFYQAMRRQGVTRRSFLKYCSLAATSLGLGAGMAPKIAWALENKPRIPVVWIHGL WP_001301613.1
hydrogenase [Escherichia coli] Length: 372\nNumber of predicted TMHs: 2\nExp number of AAs in TMHs: 37.46598\nExp number, first 60 AAs: 19.53139\nTotal prob of N-in: 0.94364\nPOSSIBLE N-term signal sequence\ninside 1 19\nTMhelix 20 39\noutside 40 325\nTMhelix 326 348\ninside 349 372

5574 GCF_001890205.1_ASM189020v1 Escherichia coli Proteobacteria; Gammaproteobacteria; Enterobacterales; Enterobacteriaceae; Escherichia
MNNEETFYQAMRRQGVTRRSFLKYCSLAATSLGLGAGMAPKIAWALENKPRIPVVWIHGL WP_001301613.1
hydrogenase [Escherichia coli] Length: 372\nNumber of predicted TMHs: 2\nExp number of AAs in TMHs: 37.46598\nExp number, first 60 AAs: 19.53139\nTotal prob of N-in: 0.94364\nPOSSIBLE N-term signal sequence\ninside 1 19\nTMhelix 20 39\noutside 40 325\nTMhelix 326 348\ninside 349 372

5575 GCF_001891225.1_ASM189122v1 Escherichia coli Proteobacteria; Gammaproteobacteria; Enterobacterales; Enterobacteriaceae; Escherichia
MTGDNTLIHSHGINRRDFMKLCAALAATMGLSSKAAAEMAESVTNPQRPPVIWIGAQECT WP_000145410.1
MULTISPECIES: hydrogenase-2 small chain [Proteobacteria] Length: 372\nNumber of predicted TMHs: 1\nExp number of AAs in TMHs: 23.72226\nExp number, first 60 AAs: 0.22038\nTotal prob of N-in: 0.12994\noutside 1 331\nTMhelix 332 354\ninside 355 372

5576 GCF_001891285.1_ASM189128v1 Escherichia coli Proteobacteria; Gammaproteobacteria; Enterobacterales; Enterobacteriaceae; Escherichia
MTGDNTLIHSHGINRRDFMKLCAALAATMGLSSKAAAEMAESVTNPQRPPVIWIGAQECT WP_000145410.1
MULTISPECIES: hydrogenase-2 small chain [Proteobacteria] Length: 372\nNumber of predicted TMHs: 1\nExp number of AAs in TMHs: 23.72226\nExp number, first 60 AAs: 0.22038\nTotal prob of N-in: 0.12994\noutside 1 331\nTMhelix 332 354\ninside 355 372

5577 GCF_001891305.1_ASM189130v1 Escherichia coli Proteobacteria; Gammaproteobacteria; Enterobacterales; Enterobacteriaceae; Escherichia
MTGDNTLIHSHGINRRDFMKLCAALAATMGLSSKAAAEMAESVTNPQRPPVIWIGAQECT WP_000145410.1
MULTISPECIES: hydrogenase-2 small chain [Proteobacteria] Length: 372\nNumber of predicted TMHs: 1\nExp number of AAs in TMHs: 23.72226\nExp number, first 60 AAs: 0.22038\nTotal prob of N-in: 0.12994\noutside 1 331\nTMhelix 332 354\ninside 355 372

5578 GCF_001891315.1_ASM189131v1 Escherichia coli Proteobacteria; Gammaproteobacteria; Enterobacterales; Enterobacteriaceae; Escherichia
MTGDNTLIHSHGINRRDFMKLCAALAATMGLSSKAAAEMAESVTNPQRPPVIWIGAQECT WP_000145410.1
MULTISPECIES: hydrogenase-2 small chain [Proteobacteria] Length: 372\nNumber of predicted TMHs: 1\nExp number of AAs in TMHs: 23.72226\nExp number, first 60 AAs: 0.22038\nTotal prob of N-in: 0.12994\noutside 1 331\nTMhelix 332 354\ninside 355 372

5579 GCF_001891375.1_ASM189137v1 Escherichia coli Proteobacteria; Gammaproteobacteria; Enterobacterales; Enterobacteriaceae; Escherichia
MTGDNTLIHSHGINRRDFMKLCAALAATMGLSSKAAAEMAESVTNPQRPPVIWIGAQECT WP_000145410.1
MULTISPECIES: hydrogenase-2 small chain [Proteobacteria] Length: 372\nNumber of predicted TMHs: 1\nExp number of AAs in TMHs: 23.72226\nExp number, first 60 AAs: 0.22038\nTotal prob of N-in: 0.12994\noutside 1 331\nTMhelix 332 354\ninside 355 372

5580 GCF_001891395.1_ASM189139v1 Escherichia coli Proteobacteria; Gammaproteobacteria; Enterobacterales; Enterobacteriaceae; Escherichia
MTGDNTLIHSHGINRRDFMKLCAALAATMGLSSKAAAEMAESVTNPQRPPVIWIGAQECT WP_000145410.1
MULTISPECIES: hydrogenase-2 small chain [Proteobacteria] Length: 372\nNumber of predicted TMHs: 1\nExp number of AAs in TMHs: 23.72226\nExp number, first 60 AAs: 0.22038\nTotal prob of N-in: 0.12994\noutside 1 331\nTMhelix 332 354\ninside 355 372

5581 GCF_001891765.1_ASM189176v1 Escherichia coli Proteobacteria; Gammaproteobacteria; Enterobacterales; Enterobacteriaceae; Escherichia
MTGDNTLIHSHGINRRDFMKLCAALAATMGLSSKAAAEMAESVTNPQRPPVIWIGAQECT WP_000145410.1
MULTISPECIES: hydrogenase-2 small chain [Proteobacteria] Length: 372\nNumber of predicted TMHs: 1\nExp number of AAs in TMHs: 23.72226\nExp number, first 60 AAs: 0.22038\nTotal prob of N-in: 0.12994\noutside 1 331\nTMhelix 332 354\ninside 355 372

5582 GCF_001891305.1_ASM189130v1 Escherichia coli Proteobacteria; Gammaproteobacteria; Enterobacterales; Enterobacteriaceae; Escherichia
MNNEETFYQAMRRQGVTRRSFLKYCSLAATSLGLGAGMAPKIAWALENKPRIPVVWIHGL WP_001295940.1
MULTISPECIES: hydrogenase [Enterobacteriaceae] Length: 372\nNumber of predicted TMHs: 2\nExp number of AAs in TMHs: 37.2608899999999\nExp number, first 60 AAs: 19.52923\nTotal prob of N-in: 0.94351\nPOSSIBLE N-term signal sequence\ninside 1 19\nTMhelix 20 39\noutside 40 325\nTMhelix 326 348\ninside 349 372

5583 GCF_001891285.1_ASM189128v1 Escherichia coli Proteobacteria; Gammaproteobacteria; Enterobacterales; Enterobacteriaceae; Escherichia
MNNEETFYQAMRRQGVTRRSFLKYCSLAATSLGLGAGMAPKIAWALENKPRIPVVWIHGL WP_001295940.1
MULTISPECIES: hydrogenase [Enterobacteriaceae] Length: 372\nNumber of predicted TMHs: 2\nExp number of AAs in TMHs: 37.2608899999999\nExp number, first 60 AAs: 19.52923\nTotal prob of N-in: 0.94351\nPOSSIBLE N-term signal sequence\ninside 1 19\nTMhelix 20 39\noutside 40 325\nTMhelix 326 348\ninside 349 372

5584 GCF_001891315.1_ASM189131v1 Escherichia coli Proteobacteria; Gammaproteobacteria; Enterobacterales; Enterobacteriaceae; Escherichia
MNNEETFYQAMRRQGVTRRSFLKYCSLAATSLGLGAGMAPKIAWALENKPRIPVVWIHGL WP_001295940.1

MULTISPECIES: hydrogenase [Enterobacteriaceae] Length: 372\nNumber of predicted TMHs: 2\nExp number of AAs in TMHs: 37.2608899999999\nExp number, first 60 AAs: 19.52923\nTotal prob of N-in: 0.94351\nPOSSIBLE N-term signal sequence\ninside 1 19\nTMhelix 20 39\noutside 40 325\nTMhelix 326 348\ninside 349 372

5585 GCF_001891375.1_ASM189137v1 Escherichia coli Proteobacteria; Gammaproteobacteria; Enterobacterales; Enterobacteriaceae; Escherichia
MNNEETFYQAMRRQGVTRRSFLKYCSLAATSLGLGAGMAPKIAWALENKPRIPVVWIHGL WP_001295940.1
MULTISPECIES: hydrogenase [Enterobacteriaceae] Length: 372\nNumber of predicted TMHs: 2\nExp number of AAs in TMHs: 37.2608899999999\nExp number, first 60 AAs: 19.52923\nTotal prob of N-in: 0.94351\nPOSSIBLE N-term signal sequence\ninside 1 19\nTMhelix 20 39\noutside 40 325\nTMhelix 326 348\ninside 349 372

5586 GCF_001891525.1_ASM189152v1 Escherichia coli Proteobacteria; Gammaproteobacteria; Enterobacterales; Enterobacteriaceae; Escherichia
MTGDNTLIHSHGINRRDFMKLCAALAATMGLSSKAAAEMAESVTNPQRPPVIWIGAQECT WP_000145410.1
MULTISPECIES: hydrogenase-2 small chain [Proteobacteria] Length: 372\nNumber of predicted TMHs: 1\nExp number of AAs in TMHs: 23.72226\nExp number, first 60 AAs: 0.22038\nTotal prob of N-in: 0.12994\noutside 1 331\nTMhelix 332 354\ninside 355 372

5587 GCF_001891525.1_ASM189152v1 Escherichia coli Proteobacteria; Gammaproteobacteria; Enterobacterales; Enterobacteriaceae; Escherichia
MNNEETFYQAMRRQGVTRRSFLKYCSLAATSLGLGAGMAPKIAWALENKPRIPVVWIHGL WP_001295940.1
MULTISPECIES: hydrogenase [Enterobacteriaceae] Length: 372\nNumber of predicted TMHs: 2\nExp number of AAs in TMHs: 37.2608899999999\nExp number, first 60 AAs: 19.52923\nTotal prob of N-in: 0.94351\nPOSSIBLE N-term signal sequence\ninside 1 19\nTMhelix 20 39\noutside 40 325\nTMhelix 326 348\ninside 349 372

5588 GCF_001891395.1_ASM189139v1 Escherichia coli Proteobacteria; Gammaproteobacteria; Enterobacterales; Enterobacteriaceae; Escherichia
MNNEETFYQAMRRQGVTRRSFLKYCSLAATSLGLGAGMAPKIAWALENKPRIPVVWIHGL WP_001295940.1
MULTISPECIES: hydrogenase [Enterobacteriaceae] Length: 372\nNumber of predicted TMHs: 2\nExp number of AAs in TMHs: 37.2608899999999\nExp number, first 60 AAs: 19.52923\nTotal prob of N-in: 0.94351\nPOSSIBLE N-term signal sequence\ninside 1 19\nTMhelix 20 39\noutside 40 325\nTMhelix 326 348\ninside 349 372

5589 GCF_001891545.1_ASM189154v1 Escherichia coli Proteobacteria; Gammaproteobacteria; Enterobacterales; Enterobacteriaceae; Escherichia
MTGDNTLIHSHGINRRDFMKLCAALAATMGLSSKAAAEMAESVTNPQRPPVIWIGAQECT WP_000145410.1
MULTISPECIES: hydrogenase-2 small chain [Proteobacteria] Length: 372\nNumber of predicted TMHs: 1\nExp number of AAs in TMHs: 23.72226\nExp number, first 60 AAs: 0.22038\nTotal prob of N-in: 0.12994\noutside 1 331\nTMhelix 332 354\ninside 355 372

5590 GCF_001891625.1_ASM189162v1 Escherichia coli Proteobacteria; Gammaproteobacteria; Enterobacterales; Enterobacteriaceae; Escherichia
MTGDNTLIHSHGINRRDFMKLCAALAATMGLSSKAAAEMAESVTNPQRPPVIWIGAQECT WP_000145410.1
MULTISPECIES: hydrogenase-2 small chain [Proteobacteria] Length: 372\nNumber of predicted TMHs: 1\nExp number of AAs in TMHs: 23.72226\nExp number, first 60 AAs: 0.22038\nTotal prob of N-in: 0.12994\noutside 1 331\nTMhelix 332 354\ninside 355 372

5591 GCF_001891605.1_ASM189160v1 Escherichia coli Proteobacteria; Gammaproteobacteria; Enterobacterales; Enterobacteriaceae; Escherichia
MTGDNTLIHSHGINRRDFMKLCAALAATMGLSSKAAAEMAESVTNPQRPPVIWIGAQECT WP_000145410.1
MULTISPECIES: hydrogenase-2 small chain [Proteobacteria] Length: 372\nNumber of predicted TMHs: 1\nExp number of AAs in TMHs: 23.72226\nExp number, first 60 AAs: 0.22038\nTotal prob of N-in: 0.12994\noutside 1 331\nTMhelix 332 354\ninside 355 372

5592 GCF_001891655.1_ASM189165v1 Escherichia coli Proteobacteria; Gammaproteobacteria; Enterobacterales; Enterobacteriaceae; Escherichia
MTGDNTLIHSHGINRRDFMKLCAALAATMGLSSKAAAEMAESVTNPQRPPVIWIGAQECT WP_000145410.1
MULTISPECIES: hydrogenase-2 small chain [Proteobacteria] Length: 372\nNumber of predicted TMHs:

1\nExp number of AAs in TMHs: 23.72226\nExp number, first 60 AAs: 0.22038\nTotal prob of N-in: 0.12994\noutside 1 331\nTMhelix 332 354\ninside 355 372

5593 GCF_001891725.1_ASM189172v1 Escherichia coli Proteobacteria; Gammaproteobacteria; Enterobacterales; Enterobacteriaceae; Escherichia
MTGDNTLIHSHGINRRDFMKLCAALAATMGLSSKAAAEMAESVTNPQRPPVIWIGAQECT WP_000145410.1
MULTISPECIES: hydrogenase-2 small chain [Proteobacteria] Length: 372\nNumber of predicted TMHs: 1\nExp number of AAs in TMHs: 23.72226\nExp number, first 60 AAs: 0.22038\nTotal prob of N-in: 0.12994\noutside 1 331\nTMhelix 332 354\ninside 355 372

5594 GCF_001891795.1_ASM189179v1 Escherichia coli Proteobacteria; Gammaproteobacteria; Enterobacterales; Enterobacteriaceae; Escherichia
MTGDNTLIHSHGINRRDFMKLCAALAATMGLSSKAAAEMAESVTNPQRPPVIWIGAQECT WP_000145410.1
MULTISPECIES: hydrogenase-2 small chain [Proteobacteria] Length: 372\nNumber of predicted TMHs: 1\nExp number of AAs in TMHs: 23.72226\nExp number, first 60 AAs: 0.22038\nTotal prob of N-in: 0.12994\noutside 1 331\nTMhelix 332 354\ninside 355 372

5595 GCF_001891695.1_ASM189169v1 Escherichia coli Proteobacteria; Gammaproteobacteria; Enterobacterales; Enterobacteriaceae; Escherichia
MTGDNTLIHSHGINRRDFMKLCAALAATMGLSSKAAAEMAESVTNPQRPPVIWIGAQECT WP_000145410.1
MULTISPECIES: hydrogenase-2 small chain [Proteobacteria] Length: 372\nNumber of predicted TMHs: 1\nExp number of AAs in TMHs: 23.72226\nExp number, first 60 AAs: 0.22038\nTotal prob of N-in: 0.12994\noutside 1 331\nTMhelix 332 354\ninside 355 372

5596 GCF_001891835.1_ASM189183v1 Escherichia coli Proteobacteria; Gammaproteobacteria; Enterobacterales; Enterobacteriaceae; Escherichia
MTGDNTLIHSHGINRRDFMKLCAALAATMGLSSKAAAEMAESVTNPQRPPVIWIGAQECT WP_000145410.1
MULTISPECIES: hydrogenase-2 small chain [Proteobacteria] Length: 372\nNumber of predicted TMHs: 1\nExp number of AAs in TMHs: 23.72226\nExp number, first 60 AAs: 0.22038\nTotal prob of N-in: 0.12994\noutside 1 331\nTMhelix 332 354\ninside 355 372

5597 GCF_001891885.1_ASM189188v1 Escherichia coli Proteobacteria; Gammaproteobacteria; Enterobacterales; Enterobacteriaceae; Escherichia
MNNEETFYQAMRRQGVTRRSFLKYCSLAATSLGLGAGMAPKIAWALENKPRIPVVWIHGL WP_001058323.1
MULTISPECIES: hydrogenase-1 small chain [Proteobacteria] Length: 372\nNumber of predicted TMHs: 2\nExp number of AAs in TMHs: 37.2608499999999\nExp number, first 60 AAs: 19.52923\nTotal prob of N-in: 0.94351\nPOSSIBLE N-term signal sequence\ninside 1 19\nTMhelix 20 39\noutside 40 325\nTMhelix 326 348\ninside 349 372

5598 GCF_001891605.1_ASM189160v1 Escherichia coli Proteobacteria; Gammaproteobacteria; Enterobacterales; Enterobacteriaceae; Escherichia
MNNEETFYQAMRRQGVTRRSFLKYCSLAATSLGLGAGMAPKIAWALENKPRIPVVWIHGL WP_001295940.1
MULTISPECIES: hydrogenase [Enterobacteriaceae] Length: 372\nNumber of predicted TMHs: 2\nExp number of AAs in TMHs: 37.2608899999999\nExp number, first 60 AAs: 19.52923\nTotal prob of N-in: 0.94351\nPOSSIBLE N-term signal sequence\ninside 1 19\nTMhelix 20 39\noutside 40 325\nTMhelix 326 348\ninside 349 372

5599 GCF_001891795.1_ASM189179v1 Escherichia coli Proteobacteria; Gammaproteobacteria; Enterobacterales; Enterobacteriaceae; Escherichia
MNNEETFYQAMRRQGVTRRSFLKYCSLAATSLGLGAGMAPKIAWALENKPRIPVVWIHGL WP_001295940.1
MULTISPECIES: hydrogenase [Enterobacteriaceae] Length: 372\nNumber of predicted TMHs: 2\nExp number of AAs in TMHs: 37.2608899999999\nExp number, first 60 AAs: 19.52923\nTotal prob of N-in: 0.94351\nPOSSIBLE N-term signal sequence\ninside 1 19\nTMhelix 20 39\noutside 40 325\nTMhelix 326 348\ninside 349 372

5600 GCF_001891625.1_ASM189162v1 Escherichia coli Proteobacteria; Gammaproteobacteria; Enterobacterales; Enterobacteriaceae; Escherichia
MNNEETFYQAMRRQGVTRRSFLKYCSLAATSLGLGAGMAPKIAWALENKPRIPVVWIHGL WP_001295940.1
MULTISPECIES: hydrogenase [Enterobacteriaceae] Length: 372\nNumber of predicted TMHs: 2\nExp number of AAs in TMHs: 37.2608899999999\nExp number, first 60 AAs: 19.52923\nTotal prob of N-in:

0.94351\nPOSSIBLE N-term signal sequence\ninside 1 19\nTMhelix 20 39\noutside 40 325\nTMhelix 326 348\ninside 349 372

5601 GCF_001891655.1_ASM189165v1 Escherichia coli Proteobacteria; Gammaproteobacteria; Enterobacterales; Enterobacteriaceae; Escherichia
MNNEETFYQAMRRQGVTRRSFLKYCSLAATSLGLGAGMAPKIAWALENKPRIPVVWIHGL WP_001295940.1
MULTISPECIES: hydrogenase [Enterobacteriaceae] Length: 372\nNumber of predicted TMHs: 2\nExp number of AAs in TMHs: 37.2608899999999\nExp number, first 60 AAs: 19.52923\nTotal prob of N-in: 0.94351\nPOSSIBLE N-term signal sequence\ninside 1 19\nTMhelix 20 39\noutside 40 325\nTMhelix 326 348\ninside 349 372

5602 GCF_001891945.1_ASM189194v1 Escherichia coli Proteobacteria; Gammaproteobacteria; Enterobacterales; Enterobacteriaceae; Escherichia
MTGDNTLIHSHGINRRDFMKLCAALAATMGLSSKAAAEMAESVTNPQRPPVIWIGAQECT WP_000145410.1
MULTISPECIES: hydrogenase-2 small chain [Proteobacteria] Length: 372\nNumber of predicted TMHs: 1\nExp number of AAs in TMHs: 23.72226\nExp number, first 60 AAs: 0.22038\nTotal prob of N-in: 0.12994\noutside 1 331\nTMhelix 332 354\ninside 355 372

5603 GCF_001891945.1_ASM189194v1 Escherichia coli Proteobacteria; Gammaproteobacteria; Enterobacterales; Enterobacteriaceae; Escherichia
MNNEETFYQAMRRQGVTRRSFLKYCSLAATSLGLGAGMAPKIAWALENKPRIPVVWIHGL WP_001058323.1
MULTISPECIES: hydrogenase-1 small chain [Proteobacteria] Length: 372\nNumber of predicted TMHs: 2\nExp number of AAs in TMHs: 37.2608499999999\nExp number, first 60 AAs: 19.52923\nTotal prob of N-in: 0.94351\nPOSSIBLE N-term signal sequence\ninside 1 19\nTMhelix 20 39\noutside 40 325\nTMhelix 326 348\ninside 349 372

5604 GCF_001891725.1_ASM189172v1 Escherichia coli Proteobacteria; Gammaproteobacteria; Enterobacterales; Enterobacteriaceae; Escherichia
MNNEETFYQAMRRQGVTRRSFLKYCSLAATSLGLGAGMAPKIAWALENKPRIPVVWIHGL WP_001295940.1
MULTISPECIES: hydrogenase [Enterobacteriaceae] Length: 372\nNumber of predicted TMHs: 2\nExp number of AAs in TMHs: 37.2608899999999\nExp number, first 60 AAs: 19.52923\nTotal prob of N-in: 0.94351\nPOSSIBLE N-term signal sequence\ninside 1 19\nTMhelix 20 39\noutside 40 325\nTMhelix 326 348\ninside 349 372

5605 GCF_001891965.1_ASM189196v1 Escherichia coli Proteobacteria; Gammaproteobacteria; Enterobacterales; Enterobacteriaceae; Escherichia
MTGDNTLIHSHGINRRDFMKLCAALAATMGLSSKAAAEMAESVTNPQRPPVIWIGAQECT WP_000145410.1
MULTISPECIES: hydrogenase-2 small chain [Proteobacteria] Length: 372\nNumber of predicted TMHs: 1\nExp number of AAs in TMHs: 23.72226\nExp number, first 60 AAs: 0.22038\nTotal prob of N-in: 0.12994\noutside 1 331\nTMhelix 332 354\ninside 355 372

5606 GCF_001891995.1_ASM189199v1 Escherichia coli Proteobacteria; Gammaproteobacteria; Enterobacterales; Enterobacteriaceae; Escherichia
MTGDNTLIHSHGINRRDFMKLCAALAATMGLSSKAAAEMAESVTNPQRPPVIWIGAQECT WP_000145410.1
MULTISPECIES: hydrogenase-2 small chain [Proteobacteria] Length: 372\nNumber of predicted TMHs: 1\nExp number of AAs in TMHs: 23.72226\nExp number, first 60 AAs: 0.22038\nTotal prob of N-in: 0.12994\noutside 1 331\nTMhelix 332 354\ninside 355 372

5607 GCF_001892045.1_ASM189204v1 Escherichia coli Proteobacteria; Gammaproteobacteria; Enterobacterales; Enterobacteriaceae; Escherichia
MTGDNTLIHSHGINRRDFMKLCAALAATMGLSSKAAAEMAESVTNPQRPPVIWIGAQECT WP_000145410.1
MULTISPECIES: hydrogenase-2 small chain [Proteobacteria] Length: 372\nNumber of predicted TMHs: 1\nExp number of AAs in TMHs: 23.72226\nExp number, first 60 AAs: 0.22038\nTotal prob of N-in: 0.12994\noutside 1 331\nTMhelix 332 354\ninside 355 372

5608 GCF_001892065.1_ASM189206v1 Escherichia coli Proteobacteria; Gammaproteobacteria; Enterobacterales; Enterobacteriaceae; Escherichia
MTGDNTLIHSHGINRRDFMKLCAALAATMGLSSKAAAEMAESVTNPQRPPVIWIGAQECT WP_000145410.1
MULTISPECIES: hydrogenase-2 small chain [Proteobacteria] Length: 372\nNumber of predicted TMHs: 1\nExp number of AAs in TMHs: 23.72226\nExp number, first 60 AAs: 0.22038\nTotal prob of N-in: 0.12994\noutside 1 331\nTMhelix 332 354\ninside 355 372

5609 GCF_001892105.1_ASM189210v1 Escherichia coli Proteobacteria; Gammaproteobacteria; Enterobacterales; Enterobacteriaceae; Escherichia
MTGDNTLIHSHGINRRDFMKLCAALAATMGLSSKAAAEMAESVTNPQRPPVIWIGAQECT WP_000145410.1
MULTISPECIES: hydrogenase-2 small chain [Proteobacteria] Length: 372\nNumber of predicted TMHs: 1\nExp number of AAs in TMHs: 23.72226\nExp number, first 60 AAs: 0.22038\nTotal prob of N-in: 0.12994\noutside 1 331\nTMhelix 332 354\ninside 355 372

5610 GCF_001892135.1_ASM189213v1 Escherichia coli Proteobacteria; Gammaproteobacteria; Enterobacterales; Enterobacteriaceae; Escherichia
MTGDNTLIHSHGINRRDFMKLCAALAATMGLSSKAAAEMAESVTNPQRPPVIWIGAQECT WP_000145410.1
MULTISPECIES: hydrogenase-2 small chain [Proteobacteria] Length: 372\nNumber of predicted TMHs: 1\nExp number of AAs in TMHs: 23.72226\nExp number, first 60 AAs: 0.22038\nTotal prob of N-in: 0.12994\noutside 1 331\nTMhelix 332 354\ninside 355 372

5611 GCF_001892185.1_ASM189218v1 Escherichia coli Proteobacteria; Gammaproteobacteria; Enterobacterales; Enterobacteriaceae; Escherichia
MTGDNTLIHSHGINRRDFMKLCAALAATMGLSSKAAAEMAESVTNPQRPPVIWIGAQECT WP_000145410.1
MULTISPECIES: hydrogenase-2 small chain [Proteobacteria] Length: 372\nNumber of predicted TMHs: 1\nExp number of AAs in TMHs: 23.72226\nExp number, first 60 AAs: 0.22038\nTotal prob of N-in: 0.12994\noutside 1 331\nTMhelix 332 354\ninside 355 372

5612 GCF_001892205.1_ASM189220v1 Escherichia coli Proteobacteria; Gammaproteobacteria; Enterobacterales; Enterobacteriaceae; Escherichia
MTGDNTLIHSHGINRRDFMKLCAALAATMGLSSKAAAEMAESVTNPQRPPVIWIGAQECT WP_000145410.1
MULTISPECIES: hydrogenase-2 small chain [Proteobacteria] Length: 372\nNumber of predicted TMHs: 1\nExp number of AAs in TMHs: 23.72226\nExp number, first 60 AAs: 0.22038\nTotal prob of N-in: 0.12994\noutside 1 331\nTMhelix 332 354\ninside 355 372

5613 GCF_001892205.1_ASM189220v1 Escherichia coli Proteobacteria; Gammaproteobacteria; Enterobacterales; Enterobacteriaceae; Escherichia
MNNEETFYQAMRRQGVTRRSFLKYCSLAATSLGLGAGMAPKIAWALENKPRIPVVWIHGL WP_001058323.1
MULTISPECIES: hydrogenase-1 small chain [Proteobacteria] Length: 372\nNumber of predicted TMHs: 2\nExp number of AAs in TMHs: 37.2608499999999\nExp number, first 60 AAs: 19.52923\nTotal prob of N-in: 0.94351\nPOSSIBLE N-term signal sequence\ninside 1 19\nTMhelix 20 39\noutside 40 325\nTMhelix 326 348\ninside 349 372

5614 GCF_001891965.1_ASM189196v1 Escherichia coli Proteobacteria; Gammaproteobacteria; Enterobacterales; Enterobacteriaceae; Escherichia
MNNEETFYQAMRRQGVTRRSFLKYCSLAATSLGLGAGMAPKIAWALENKPRIPVVWIHGL WP_001058323.1
MULTISPECIES: hydrogenase-1 small chain [Proteobacteria] Length: 372\nNumber of predicted TMHs: 2\nExp number of AAs in TMHs: 37.2608499999999\nExp number, first 60 AAs: 19.52923\nTotal prob of N-in: 0.94351\nPOSSIBLE N-term signal sequence\ninside 1 19\nTMhelix 20 39\noutside 40 325\nTMhelix 326 348\ninside 349 372

5615 GCF_001892245.1_ASM189224v1 Escherichia coli Proteobacteria; Gammaproteobacteria; Enterobacterales; Enterobacteriaceae; Escherichia
MTGDNTLIHSHGINRRDFMKLCAALAATMGLSSKAAAEMAESVTNPQRPPVIWIGAQECT WP_000145410.1
MULTISPECIES: hydrogenase-2 small chain [Proteobacteria] Length: 372\nNumber of predicted TMHs: 1\nExp number of AAs in TMHs: 23.72226\nExp number, first 60 AAs: 0.22038\nTotal prob of N-in: 0.12994\noutside 1 331\nTMhelix 332 354\ninside 355 372

5616 GCF_001892275.1_ASM189227v1 Escherichia coli Proteobacteria; Gammaproteobacteria; Enterobacterales; Enterobacteriaceae; Escherichia
MTGDNTLIHSHGINRRDFMKLCAALAATMGLSSKAAAEMAESVTNPQRPPVIWIGAQECT WP_000145410.1
MULTISPECIES: hydrogenase-2 small chain [Proteobacteria] Length: 372\nNumber of predicted TMHs: 1\nExp number of AAs in TMHs: 23.72226\nExp number, first 60 AAs: 0.22038\nTotal prob of N-in: 0.12994\noutside 1 331\nTMhelix 332 354\ninside 355 372

5617 GCF_001892245.1_ASM189224v1 Escherichia coli Proteobacteria; Gammaproteobacteria; Enterobacterales; Enterobacteriaceae; Escherichia
MNNEETFYQAMRRQGVTRRSFLKYCSLAATSLGLGAGMAPKIAWALENKPRIPVVWIHGL WP_001295940.1

MULTISPECIES: hydrogenase [Enterobacteriaceae]	Length: 372\nNumber of predicted TMHs:  2\nExp number of AAs in TMHs: 37.2608899999999\nExp number, first 60 AAs:  19.52923\nTotal prob of N-in: 0.94351\nPOSSIBLE N-term signal sequence\ninside   1   19\nTMhelix   20   39\noutside   40   325\nTMhelix   326   348\ninside   349   372

5618	GCF_001892045.1_ASM189204v1	Escherichia coli	Proteobacteria; Gammaproteobacteria; Enterobacterales; Enterobacteriaceae; Escherichia
MNNEETFYQAMRRQGVTRRSFLKYCSLAATSLGLGAGMAPKIAWALENKPRIPVVWIHGL	WP_001295940.1
MULTISPECIES: hydrogenase [Enterobacteriaceae]	Length: 372\nNumber of predicted TMHs:  2\nExp number of AAs in TMHs: 37.2608899999999\nExp number, first 60 AAs:  19.52923\nTotal prob of N-in: 0.94351\nPOSSIBLE N-term signal sequence\ninside   1   19\nTMhelix   20   39\noutside   40   325\nTMhelix   326   348\ninside   349   372

5619	GCF_001892405.1_ASM189240v1	Escherichia coli	Proteobacteria; Gammaproteobacteria; Enterobacterales; Enterobacteriaceae; Escherichia
MTGDNTLIHSHGINRRDFMKLCAALAATMGLSSKAAAEMAESVTNPQRPPVIWIGAQECT	WP_000145410.1
MULTISPECIES: hydrogenase-2 small chain [Proteobacteria]	Length: 372\nNumber of predicted TMHs: 1\nExp number of AAs in TMHs: 23.72226\nExp number, first 60 AAs:  0.22038\nTotal prob of N-in:   0.12994\noutside   1   331\nTMhelix   332   354\ninside   355   372

5620	GCF_001892435.1_ASM189243v1	Escherichia coli	Proteobacteria; Gammaproteobacteria; Enterobacterales; Enterobacteriaceae; Escherichia
MTGDNTLIHSHGINRRDFMKLCAALAATMGLSSKAAAEMAESVTNPQRPPVIWIGAQECT	WP_000145410.1
MULTISPECIES: hydrogenase-2 small chain [Proteobacteria]	Length: 372\nNumber of predicted TMHs: 1\nExp number of AAs in TMHs: 23.72226\nExp number, first 60 AAs:  0.22038\nTotal prob of N-in:   0.12994\noutside   1   331\nTMhelix   332   354\ninside   355   372

5621	GCF_001892275.1_ASM189227v1	Escherichia coli	Proteobacteria; Gammaproteobacteria; Enterobacterales; Enterobacteriaceae; Escherichia
MNNEETFYQAMRRQGVTRRSFLKYCSLAATSLGLGAGMAPKIAWALENKPRIPVVWIHGL	WP_001295940.1
MULTISPECIES: hydrogenase [Enterobacteriaceae]	Length: 372\nNumber of predicted TMHs:  2\nExp number of AAs in TMHs: 37.2608899999999\nExp number, first 60 AAs:  19.52923\nTotal prob of N-in: 0.94351\nPOSSIBLE N-term signal sequence\ninside   1   19\nTMhelix   20   39\noutside   40   325\nTMhelix   326   348\ninside   349   372

5622	GCF_001892505.1_ASM189250v1	Escherichia coli	Proteobacteria; Gammaproteobacteria; Enterobacterales; Enterobacteriaceae; Escherichia
MTGDNTLIHSHGINRRDFMKLCAALAATMGLSSKAAAEMAESVTNPQRPPVIWIGAQECT	WP_000145410.1
MULTISPECIES: hydrogenase-2 small chain [Proteobacteria]	Length: 372\nNumber of predicted TMHs: 1\nExp number of AAs in TMHs: 23.72226\nExp number, first 60 AAs:  0.22038\nTotal prob of N-in:   0.12994\noutside   1   331\nTMhelix   332   354\ninside   355   372

5623	GCF_001892595.1_ASM189259v1	Escherichia coli	Proteobacteria; Gammaproteobacteria; Enterobacterales; Enterobacteriaceae; Escherichia
MTGDNTLIHSHGINRRDFMKLCAALAATMGLSSKAAAEMAESVTNPQRPPVIWIGAQECT	WP_000145410.1
MULTISPECIES: hydrogenase-2 small chain [Proteobacteria]	Length: 372\nNumber of predicted TMHs: 1\nExp number of AAs in TMHs: 23.72226\nExp number, first 60 AAs:  0.22038\nTotal prob of N-in:   0.12994\noutside   1   331\nTMhelix   332   354\ninside   355   372

5624	GCF_001892485.1_ASM189248v1	Escherichia coli	Proteobacteria; Gammaproteobacteria; Enterobacterales; Enterobacteriaceae; Escherichia
MTGDNTLIHSHGINRRDFMKLCAALAATMGLSSKAAAEMAESVTNPQRPPVIWIGAQECT	WP_000145410.1
MULTISPECIES: hydrogenase-2 small chain [Proteobacteria]	Length: 372\nNumber of predicted TMHs: 1\nExp number of AAs in TMHs: 23.72226\nExp number, first 60 AAs:  0.22038\nTotal prob of N-in:   0.12994\noutside   1   331\nTMhelix   332   354\ninside   355   372

5625	GCF_001892585.1_ASM189258v1	Escherichia coli	Proteobacteria; Gammaproteobacteria; Enterobacterales; Enterobacteriaceae; Escherichia
MTGDNTLIHSHGINRRDFMKLCAALAATMGLSSKAAAEMAESVTNPQRPPVIWIGAQECT	WP_000145410.1
MULTISPECIES: hydrogenase-2 small chain [Proteobacteria]	Length: 372\nNumber of predicted TMHs:

1\nExp number of AAs in TMHs: 23.72226\nExp number, first 60 AAs: 0.22038\nTotal prob of N-in: 0.12994\noutside 1 331\nTMhelix 332 354\ninside 355 372

5626 GCF_001892525.1_ASM189252v1 Escherichia coli Proteobacteria; Gammaproteobacteria; Enterobacterales; Enterobacteriaceae; Escherichia
MTGDNTLIHSHGINRRDFMKLCAALAATMGLSSKAAAEMAESVTNPQRPPVIWIGAQECT WP_000145410.1
MULTISPECIES: hydrogenase-2 small chain [Proteobacteria] Length: 372\nNumber of predicted TMHs: 1\nExp number of AAs in TMHs: 23.72226\nExp number, first 60 AAs: 0.22038\nTotal prob of N-in: 0.12994\noutside 1 331\nTMhelix 332 354\ninside 355 372

5627 GCF_001892335.1_ASM189233v1 Escherichia coli Proteobacteria; Gammaproteobacteria; Enterobacterales; Enterobacteriaceae; Escherichia
MTGDNTLIHSHGINRRDFMKLCAALATTMGLSSKAAAEMAESVTNPQRPPVIWIGAQECT WP_023149629.1
MULTISPECIES: hydrogenase-2 small chain [Enterobacteriaceae] Length: 372\nNumber of predicted TMHs: 1\nExp number of AAs in TMHs: 23.63603\nExp number, first 60 AAs: 0.12971\nTotal prob of N-in: 0.12608\noutside 1 331\nTMhelix 332 354\ninside 355 372

5628 GCF_001892645.1_ASM189264v1 Escherichia coli Proteobacteria; Gammaproteobacteria; Enterobacterales; Enterobacteriaceae; Escherichia
MTGDNTLIHSHGINRRDFMKLCAALAATMGLSSKAAAEMAESVTNPQRPPVIWIGAQECT WP_000145410.1
MULTISPECIES: hydrogenase-2 small chain [Proteobacteria] Length: 372\nNumber of predicted TMHs: 1\nExp number of AAs in TMHs: 23.72226\nExp number, first 60 AAs: 0.22038\nTotal prob of N-in: 0.12994\noutside 1 331\nTMhelix 332 354\ninside 355 372

5629 GCF_001892355.1_ASM189235v1 Escherichia coli Proteobacteria; Gammaproteobacteria; Enterobacterales; Enterobacteriaceae; Escherichia
MNNEETFYQAMRRQGVTRRSFLKYCSLAATSLGLGAGMAPKIAWALENKPRIPVVWIHGL WP_001295940.1
MULTISPECIES: hydrogenase [Enterobacteriaceae] Length: 372\nNumber of predicted TMHs: 2\nExp number of AAs in TMHs: 37.2608899999999\nExp number, first 60 AAs: 19.52923\nTotal prob of N-in: 0.94351\nPOSSIBLE N-term signal sequence\ninside 1 19\nTMhelix 20 39\noutside 40 325\nTMhelix 326 348\ninside 349 372

5630 GCF_001892675.1_ASM189267v1 Escherichia coli Proteobacteria; Gammaproteobacteria; Enterobacterales; Enterobacteriaceae; Escherichia
MTGDNTLIHSHGINRRDFMKLCAALAATMGLSSKAAAEMAESVTNPQRPPVIWIGAQECT WP_000145410.1
MULTISPECIES: hydrogenase-2 small chain [Proteobacteria] Length: 372\nNumber of predicted TMHs: 1\nExp number of AAs in TMHs: 23.72226\nExp number, first 60 AAs: 0.22038\nTotal prob of N-in: 0.12994\noutside 1 331\nTMhelix 332 354\ninside 355 372

5631 GCF_001892595.1_ASM189259v1 Escherichia coli Proteobacteria; Gammaproteobacteria; Enterobacterales; Enterobacteriaceae; Escherichia
MNNEETFYQAMRRQGVTRRSFLKYCSLAATSLGLGAGMAPKIAWALENKPRIPVVWIHGL WP_001058323.1
MULTISPECIES: hydrogenase-1 small chain [Proteobacteria] Length: 372\nNumber of predicted TMHs: 2\nExp number of AAs in TMHs: 37.2608499999999\nExp number, first 60 AAs: 19.52923\nTotal prob of N-in: 0.94351\nPOSSIBLE N-term signal sequence\ninside 1 19\nTMhelix 20 39\noutside 40 325\nTMhelix 326 348\ninside 349 372

5632 GCF_001892405.1_ASM189240v1 Escherichia coli Proteobacteria; Gammaproteobacteria; Enterobacterales; Enterobacteriaceae; Escherichia
MNNEETFYQAMRRQGVTRRSFLKYCSLAATSLGLGAGMAPKIAWALENKPRIPVVWIHGL WP_001058323.1
MULTISPECIES: hydrogenase-1 small chain [Proteobacteria] Length: 372\nNumber of predicted TMHs: 2\nExp number of AAs in TMHs: 37.2608499999999\nExp number, first 60 AAs: 19.52923\nTotal prob of N-in: 0.94351\nPOSSIBLE N-term signal sequence\ninside 1 19\nTMhelix 20 39\noutside 40 325\nTMhelix 326 348\ninside 349 372

5633 GCF_001892585.1_ASM189258v1 Escherichia coli Proteobacteria; Gammaproteobacteria; Enterobacterales; Enterobacteriaceae; Escherichia
MNNEETFYQAMRRQGVTRRSFLKYCSLAATSLGLGAGMAPKIAWALENKPRIPVVWIHGL WP_001058323.1
MULTISPECIES: hydrogenase-1 small chain [Proteobacteria] Length: 372\nNumber of predicted TMHs: 2\nExp number of AAs in TMHs: 37.2608499999999\nExp number, first 60 AAs: 19.52923\nTotal prob of N-in:

0.94351\nPOSSIBLE N-term signal sequence\ninside 1 19\nTMhelix 20 39\noutside 40 325\nTMhelix 326 348\ninside 349 372

5634 GCF_001892525.1_ASM189252v1 Escherichia coli Proteobacteria; Gammaproteobacteria; Enterobacterales; Enterobacteriaceae; Escherichia
MNNEETFYQAMRRQGVTRRSFLKYCSLAATSLGLGAGMAPKIAWALENKPRIPVVWIHGL WP_001058323.1
MULTISPECIES: hydrogenase-1 small chain [Proteobacteria] Length: 372\nNumber of predicted TMHs: 2\nExp number of AAs in TMHs: 37.2608499999999\nExp number, first 60 AAs: 19.52923\nTotal prob of N-in: 0.94351\nPOSSIBLE N-term signal sequence\ninside 1 19\nTMhelix 20 39\noutside 40 325\nTMhelix 326 348\ninside 349 372

5635 GCF_001892505.1_ASM189250v1 Escherichia coli Proteobacteria; Gammaproteobacteria; Enterobacterales; Enterobacteriaceae; Escherichia
MNNEETFYQAMRRQGVTRRSFLKYCSLAATSLGLGAGMAPKIAWALENKPRIPVVWIHGL WP_001058323.1
MULTISPECIES: hydrogenase-1 small chain [Proteobacteria] Length: 372\nNumber of predicted TMHs: 2\nExp number of AAs in TMHs: 37.2608499999999\nExp number, first 60 AAs: 19.52923\nTotal prob of N-in: 0.94351\nPOSSIBLE N-term signal sequence\ninside 1 19\nTMhelix 20 39\noutside 40 325\nTMhelix 326 348\ninside 349 372

5636 GCF_001892685.1_ASM189268v1 Escherichia coli Proteobacteria; Gammaproteobacteria; Enterobacterales; Enterobacteriaceae; Escherichia
MTGDNTLIHSHGINRRDFMKLCAALAATMGLSSKAAAEMAESVTNPQRPPVIWIGAQECT WP_000145410.1
MULTISPECIES: hydrogenase-2 small chain [Proteobacteria] Length: 372\nNumber of predicted TMHs: 1\nExp number of AAs in TMHs: 23.72226\nExp number, first 60 AAs: 0.22038\nTotal prob of N-in: 0.12994\noutside 1 331\nTMhelix 332 354\ninside 355 372

5637 GCF_001892685.1_ASM189268v1 Escherichia coli Proteobacteria; Gammaproteobacteria; Enterobacterales; Enterobacteriaceae; Escherichia
MNNEETFYQAMRRQGVTRRSFLKYCSLAATSLGLGAGMAPKIAWALENKPRIPVVWIHGL WP_001295940.1
MULTISPECIES: hydrogenase [Enterobacteriaceae] Length: 372\nNumber of predicted TMHs: 2\nExp number of AAs in TMHs: 37.2608899999999\nExp number, first 60 AAs: 19.52923\nTotal prob of N-in: 0.94351\nPOSSIBLE N-term signal sequence\ninside 1 19\nTMhelix 20 39\noutside 40 325\nTMhelix 326 348\ninside 349 372

5638 GCF_001892435.1_ASM189243v1 Escherichia coli Proteobacteria; Gammaproteobacteria; Enterobacterales; Enterobacteriaceae; Escherichia
MNNEETFYQAMRRQGVTRRSFLKYCSLAATSLGLGAGMAPKIAWALENKPRIPVVWIHGL WP_001295940.1
MULTISPECIES: hydrogenase [Enterobacteriaceae] Length: 372\nNumber of predicted TMHs: 2\nExp number of AAs in TMHs: 37.2608899999999\nExp number, first 60 AAs: 19.52923\nTotal prob of N-in: 0.94351\nPOSSIBLE N-term signal sequence\ninside 1 19\nTMhelix 20 39\noutside 40 325\nTMhelix 326 348\ninside 349 372

5639 GCF_001892485.1_ASM189248v1 Escherichia coli Proteobacteria; Gammaproteobacteria; Enterobacterales; Enterobacteriaceae; Escherichia
MNNEETFYQAMRRQGVTRRSFLKYCSLAATSLGLGAGMAPKIAWALENKPRIPVVWIHGL WP_001295940.1
MULTISPECIES: hydrogenase [Enterobacteriaceae] Length: 372\nNumber of predicted TMHs: 2\nExp number of AAs in TMHs: 37.2608899999999\nExp number, first 60 AAs: 19.52923\nTotal prob of N-in: 0.94351\nPOSSIBLE N-term signal sequence\ninside 1 19\nTMhelix 20 39\noutside 40 325\nTMhelix 326 348\ninside 349 372

5640 GCF_001892935.1_ASM189293v1 Escherichia coli Proteobacteria; Gammaproteobacteria; Enterobacterales; Enterobacteriaceae; Escherichia
MTGDNTLIHSHGINRRDFMKLCAALAATMGLSSKAAAEMAESVTNPQRPPVIWIGAQECT WP_000145410.1
MULTISPECIES: hydrogenase-2 small chain [Proteobacteria] Length: 372\nNumber of predicted TMHs: 1\nExp number of AAs in TMHs: 23.72226\nExp number, first 60 AAs: 0.22038\nTotal prob of N-in: 0.12994\noutside 1 331\nTMhelix 332 354\ninside 355 372

5641 GCF_001893015.1_ASM189301v1 Escherichia coli Proteobacteria; Gammaproteobacteria; Enterobacterales; Enterobacteriaceae; Escherichia
MTGDNTLIHSHGINRRDFMKLCAALAATMGLSSKAAAEMAESVTNPQRPPVIWIGAQECT WP_000145410.1
MULTISPECIES: hydrogenase-2 small chain [Proteobacteria] Length: 372\nNumber of predicted TMHs:

1\nExp number of AAs in TMHs: 23.72226\nExp number, first 60 AAs: 0.22038\nTotal prob of N-in: 0.12994\noutside 1 331\nTMhelix 332 354\ninside 355 372

5642 GCF_001892955.1_ASM189295v1 Escherichia coli Proteobacteria; Gammaproteobacteria; Enterobacterales; Enterobacteriaceae; Escherichia
MTGDNTLIHSHGINRRDFMKLCAALAATMGLSSKAAAEMAESVTNPQRPPVIWIGAQECT WP_000145410.1
MULTISPECIES: hydrogenase-2 small chain [Proteobacteria] Length: 372\nNumber of predicted TMHs: 1\nExp number of AAs in TMHs: 23.72226\nExp number, first 60 AAs: 0.22038\nTotal prob of N-in: 0.12994\noutside 1 331\nTMhelix 332 354\ninside 355 372

5643 GCF_001893015.1_ASM189301v1 Escherichia coli Proteobacteria; Gammaproteobacteria; Enterobacterales; Enterobacteriaceae; Escherichia
MNNEETFYQAMRRQGVTRRSFLKYCSLAATSLGLGAGMAPKIAWALENKSRIPVVWIHGL WP_049595080.1
hydrogenase [Escherichia coli] Length: 372\nNumber of predicted TMHs: 2\nExp number of AAs in TMHs: 37.1898399999999\nExp number, first 60 AAs: 19.44607\nTotal prob of N-in: 0.94159\nPOSSIBLE N-term signal sequence\ninside 1 19\nTMhelix 20 39\noutside 40 325\nTMhelix 326 348\ninside 349 372

5644 GCF_001892955.1_ASM189295v1 Escherichia coli Proteobacteria; Gammaproteobacteria; Enterobacterales; Enterobacteriaceae; Escherichia
MNNEETFYQAMRRQGVTRRSFLKYCSLAATSLGLGAGMAPKIAWALENKPRIPVVWIHGL WP_001295940.1
MULTISPECIES: hydrogenase [Enterobacteriaceae] Length: 372\nNumber of predicted TMHs: 2\nExp number of AAs in TMHs: 37.2608899999999\nExp number, first 60 AAs: 19.52923\nTotal prob of N-in: 0.94351\nPOSSIBLE N-term signal sequence\ninside 1 19\nTMhelix 20 39\noutside 40 325\nTMhelix 326 348\ninside 349 372

5645 GCF_001892745.1_ASM189274v1 Escherichia coli Proteobacteria; Gammaproteobacteria; Enterobacterales; Enterobacteriaceae; Escherichia
MTGDNTLIHSHGINRRDFMKLCAALATTMGLSSKAAAEMAESVTNPQRPPVIWIGAQECT WP_023149629.1
MULTISPECIES: hydrogenase-2 small chain [Enterobacteriaceae] Length: 372\nNumber of predicted TMHs: 1\nExp number of AAs in TMHs: 23.63603\nExp number, first 60 AAs: 0.12971\nTotal prob of N-in: 0.12608\noutside 1 331\nTMhelix 332 354\ninside 355 372

5646 GCF_001893085.1_ASM189308v1 Escherichia coli Proteobacteria; Gammaproteobacteria; Enterobacterales; Enterobacteriaceae; Escherichia
MTGDNTLIHSHGINRRDFMKLCAALAATMGLSSKAAAEMAESVTNPQRPPVIWIGAQECT WP_000145410.1
MULTISPECIES: hydrogenase-2 small chain [Proteobacteria] Length: 372\nNumber of predicted TMHs: 1\nExp number of AAs in TMHs: 23.72226\nExp number, first 60 AAs: 0.22038\nTotal prob of N-in: 0.12994\noutside 1 331\nTMhelix 332 354\ninside 355 372

5647 GCF_001893035.1_ASM189303v1 Escherichia coli Proteobacteria; Gammaproteobacteria; Enterobacterales; Enterobacteriaceae; Escherichia
MTGDNTLIHSHGINRRDFMKLCAALAATMGLSSKAAAEMAESVTNPQRPPVIWIGAQECT WP_000145410.1
MULTISPECIES: hydrogenase-2 small chain [Proteobacteria] Length: 372\nNumber of predicted TMHs: 1\nExp number of AAs in TMHs: 23.72226\nExp number, first 60 AAs: 0.22038\nTotal prob of N-in: 0.12994\noutside 1 331\nTMhelix 332 354\ninside 355 372

5648 GCF_001893085.1_ASM189308v1 Escherichia coli Proteobacteria; Gammaproteobacteria; Enterobacterales; Enterobacteriaceae; Escherichia
MNNEETFYQAMRRQGVTRRSFLKYCSLAATSLGLGAGMAPKIAWALENKPRIPVVWIHGL WP_001058323.1
MULTISPECIES: hydrogenase-1 small chain [Proteobacteria] Length: 372\nNumber of predicted TMHs: 2\nExp number of AAs in TMHs: 37.2608499999999\nExp number, first 60 AAs: 19.52923\nTotal prob of N-in: 0.94351\nPOSSIBLE N-term signal sequence\ninside 1 19\nTMhelix 20 39\noutside 40 325\nTMhelix 326 348\ninside 349 372

5649 GCF_001893035.1_ASM189303v1 Escherichia coli Proteobacteria; Gammaproteobacteria; Enterobacterales; Enterobacteriaceae; Escherichia
MNNEETFYQAMRRQGVTRRSFLKYCSLAATSLGLGAGMAPKIAWALENKPRIPVVWIHGL WP_001058323.1
MULTISPECIES: hydrogenase-1 small chain [Proteobacteria] Length: 372\nNumber of predicted TMHs: 2\nExp number of AAs in TMHs: 37.2608499999999\nExp number, first 60 AAs: 19.52923\nTotal prob of N-in: 0.94351\nPOSSIBLE N-term signal sequence\ninside 1 19\nTMhelix 20 39\noutside 40 325\nTMhelix 326 348\ninside 349 372

5650 GCF_001892825.1_ASM189282v1 Escherichia coli Proteobacteria; Gammaproteobacteria; Enterobacterales; Enterobacteriaceae; Escherichia
MNNEETFYQAMRRQGVTRRSFLKYCSLAATSLGLGAGMAPKIAWALENKPRIPVVWIHGL WP_001058323.1
MULTISPECIES: hydrogenase-1 small chain [Proteobacteria] Length: 372\nNumber of predicted TMHs: 2\nExp number of AAs in TMHs: 37.2608499999999\nExp number, first 60 AAs: 19.52923\nTotal prob of N-in: 0.94351\nPOSSIBLE N-term signal sequence\ninside 1 19\nTMhelix 20 39\noutside 40 325\nTMhelix 326 348\ninside 349 372

5651 GCF_001892765.1_ASM189276v1 Escherichia coli Proteobacteria; Gammaproteobacteria; Enterobacterales; Enterobacteriaceae; Escherichia
MNNEETFYQAMRRQGVTRRSFLKYCSLAATSLGLGAGMAPKIAWALENKPRIPVVWIHGL WP_001058323.1
MULTISPECIES: hydrogenase-1 small chain [Proteobacteria] Length: 372\nNumber of predicted TMHs: 2\nExp number of AAs in TMHs: 37.2608499999999\nExp number, first 60 AAs: 19.52923\nTotal prob of N-in: 0.94351\nPOSSIBLE N-term signal sequence\ninside 1 19\nTMhelix 20 39\noutside 40 325\nTMhelix 326 348\ninside 349 372

5652 GCF_001893125.1_ASM189312v1 Escherichia coli Proteobacteria; Gammaproteobacteria; Enterobacterales; Enterobacteriaceae; Escherichia
MTGDNTLIHSHGINRRDFMKLCAALAATMGLSSKAAAEMAESVTNPQRPPVIWIGAQECT WP_000145410.1
MULTISPECIES: hydrogenase-2 small chain [Proteobacteria] Length: 372\nNumber of predicted TMHs: 1\nExp number of AAs in TMHs: 23.72226\nExp number, first 60 AAs: 0.22038\nTotal prob of N-in: 0.12994\noutside 1 331\nTMhelix 332 354\ninside 355 372

5653 GCF_001893125.1_ASM189312v1 Escherichia coli Proteobacteria; Gammaproteobacteria; Enterobacterales; Enterobacteriaceae; Escherichia
MNNEETFYQAMRRQGVTRRSFLKYCSLAATSLGLGAGMAPKIAWALENKPRIPVVWIHGL WP_001058323.1
MULTISPECIES: hydrogenase-1 small chain [Proteobacteria] Length: 372\nNumber of predicted TMHs: 2\nExp number of AAs in TMHs: 37.2608499999999\nExp number, first 60 AAs: 19.52923\nTotal prob of N-in: 0.94351\nPOSSIBLE N-term signal sequence\ninside 1 19\nTMhelix 20 39\noutside 40 325\nTMhelix 326 348\ninside 349 372

5654 GCF_001893285.1_ASM189328v1 Escherichia coli Proteobacteria; Gammaproteobacteria; Enterobacterales; Enterobacteriaceae; Escherichia
MTGDNTLIHSHGINRRDFMKLCAALAATMGLSSKAAAEMAESVTNPQRPPVIWIGAQECT WP_000145410.1
MULTISPECIES: hydrogenase-2 small chain [Proteobacteria] Length: 372\nNumber of predicted TMHs: 1\nExp number of AAs in TMHs: 23.72226\nExp number, first 60 AAs: 0.22038\nTotal prob of N-in: 0.12994\noutside 1 331\nTMhelix 332 354\ninside 355 372

5655 GCF_001893315.1_ASM189331v1 Escherichia coli Proteobacteria; Gammaproteobacteria; Enterobacterales; Enterobacteriaceae; Escherichia
MTGDNTLIHSHGINRRDFMKLCAALAATMGLSSKAAAEMAESVTNPQRPPVIWIGAQECT WP_000145410.1
MULTISPECIES: hydrogenase-2 small chain [Proteobacteria] Length: 372\nNumber of predicted TMHs: 1\nExp number of AAs in TMHs: 23.72226\nExp number, first 60 AAs: 0.22038\nTotal prob of N-in: 0.12994\noutside 1 331\nTMhelix 332 354\ninside 355 372

5656 GCF_001893215.1_ASM189321v1 Escherichia coli Proteobacteria; Gammaproteobacteria; Enterobacterales; Enterobacteriaceae; Escherichia
MNNEETFYQAMRRQGVTRRSFLKYCSLAATSLGLGAGMAPKIAWALENKPRIPVVWIHGL WP_001058323.1
MULTISPECIES: hydrogenase-1 small chain [Proteobacteria] Length: 372\nNumber of predicted TMHs: 2\nExp number of AAs in TMHs: 37.2608499999999\nExp number, first 60 AAs: 19.52923\nTotal prob of N-in: 0.94351\nPOSSIBLE N-term signal sequence\ninside 1 19\nTMhelix 20 39\noutside 40 325\nTMhelix 326 348\ninside 349 372

5657 GCF_001893365.1_ASM189336v1 Escherichia coli Proteobacteria; Gammaproteobacteria; Enterobacterales; Enterobacteriaceae; Escherichia
MTGDNTLIHSHGINRRDFMKLCAALAATMGLSSKAAAEMAESVTNPQRPPVIWIGAQECT WP_000145410.1
MULTISPECIES: hydrogenase-2 small chain [Proteobacteria] Length: 372\nNumber of predicted TMHs: 1\nExp number of AAs in TMHs: 23.72226\nExp number, first 60 AAs: 0.22038\nTotal prob of N-in: 0.12994\noutside 1 331\nTMhelix 332 354\ninside 355 372

5658 GCF_001893395.1_ASM189339v1 Escherichia coli Proteobacteria; Gammaproteobacteria; Enterobacterales; Enterobacteriaceae; Escherichia MTGDNTLIHSHGINRRDFMKLCAALAATMGLSSKAAAEMAESVTNPQRPPVIWIGAQECT WP_000145410.1 MULTISPECIES: hydrogenase-2 small chain [Proteobacteria] Length: 372\nNumber of predicted TMHs: 1\nExp number of AAs in TMHs: 23.72226\nExp number, first 60 AAs: 0.22038\nTotal prob of N-in: 0.12994\noutside 1 331\nTMhelix 332 354\ninside 355 372

5659 GCF_001893375.1_ASM189337v1 Escherichia coli Proteobacteria; Gammaproteobacteria; Enterobacterales; Enterobacteriaceae; Escherichia MTGDNTLIHSHGINRRDFMKLCAALAATMGLSSKAAAEMAESVTNPQRPPVIWIGAQECT WP_000145410.1 MULTISPECIES: hydrogenase-2 small chain [Proteobacteria] Length: 372\nNumber of predicted TMHs: 1\nExp number of AAs in TMHs: 23.72226\nExp number, first 60 AAs: 0.22038\nTotal prob of N-in: 0.12994\noutside 1 331\nTMhelix 332 354\ninside 355 372

5660 GCF_001893465.1_ASM189346v1 Escherichia coli Proteobacteria; Gammaproteobacteria; Enterobacterales; Enterobacteriaceae; Escherichia MTGDNTLIHSHGINRRDFMKLCAALAATMGLSSKAAAEMAESVTNPQRPPVIWIGAQECT WP_000145410.1 MULTISPECIES: hydrogenase-2 small chain [Proteobacteria] Length: 372\nNumber of predicted TMHs: 1\nExp number of AAs in TMHs: 23.72226\nExp number, first 60 AAs: 0.22038\nTotal prob of N-in: 0.12994\noutside 1 331\nTMhelix 332 354\ninside 355 372

5661 GCF_001893455.1_ASM189345v1 Escherichia coli Proteobacteria; Gammaproteobacteria; Enterobacterales; Enterobacteriaceae; Escherichia MTGDNTLIHSHGINRRDFMKLCAALAATMGLSSKAAAEMAESVTNPQRPPVIWIGAQECT WP_000145410.1 MULTISPECIES: hydrogenase-2 small chain [Proteobacteria] Length: 372\nNumber of predicted TMHs: 1\nExp number of AAs in TMHs: 23.72226\nExp number, first 60 AAs: 0.22038\nTotal prob of N-in: 0.12994\noutside 1 331\nTMhelix 332 354\ninside 355 372

5662 GCF_001893395.1_ASM189339v1 Escherichia coli Proteobacteria; Gammaproteobacteria; Enterobacterales; Enterobacteriaceae; Escherichia MNNEETFYQAMRRQGVTRRSFLKYCSLAATSLGLGAGMAPKIAWALENKPRIPVVWIHGL WP_001058323.1 MULTISPECIES: hydrogenase-1 small chain [Proteobacteria] Length: 372\nNumber of predicted TMHs: 2\nExp number of AAs in TMHs: 37.2608499999999\nExp number, first 60 AAs: 19.52923\nTotal prob of N-in: 0.94351\nPOSSIBLE N-term signal sequence\ninside 1 19\nTMhelix 20 39\noutside 40 325\nTMhelix 326 348\ninside 349 372

5663 GCF_001893465.1_ASM189346v1 Escherichia coli Proteobacteria; Gammaproteobacteria; Enterobacterales; Enterobacteriaceae; Escherichia MNNEETFYQAMRRQGVTRRSFLKYCSLAATSLGLGAGMAPKIAWALENKPRIPVVWIHGL WP_001058323.1 MULTISPECIES: hydrogenase-1 small chain [Proteobacteria] Length: 372\nNumber of predicted TMHs: 2\nExp number of AAs in TMHs: 37.2608499999999\nExp number, first 60 AAs: 19.52923\nTotal prob of N-in: 0.94351\nPOSSIBLE N-term signal sequence\ninside 1 19\nTMhelix 20 39\noutside 40 325\nTMhelix 326 348\ninside 349 372

5664 GCF_001893375.1_ASM189337v1 Escherichia coli Proteobacteria; Gammaproteobacteria; Enterobacterales; Enterobacteriaceae; Escherichia MNNEETFYQAMRRQGVTRRSFLKYCSLAATSLGLGAGMAPKIAWALENKPRIPVVWIHGL WP_001058323.1 MULTISPECIES: hydrogenase-1 small chain [Proteobacteria] Length: 372\nNumber of predicted TMHs: 2\nExp number of AAs in TMHs: 37.2608499999999\nExp number, first 60 AAs: 19.52923\nTotal prob of N-in: 0.94351\nPOSSIBLE N-term signal sequence\ninside 1 19\nTMhelix 20 39\noutside 40 325\nTMhelix 326 348\ninside 349 372

5665 GCF_001893285.1_ASM189328v1 Escherichia coli Proteobacteria; Gammaproteobacteria; Enterobacterales; Enterobacteriaceae; Escherichia MNNEETFYQAMRRQGVTRRSFLKYCSLAATSLGLGAGMAPKIAWALENKPRIPVVWIHGL WP_001058323.1 MULTISPECIES: hydrogenase-1 small chain [Proteobacteria] Length: 372\nNumber of predicted TMHs: 2\nExp number of AAs in TMHs: 37.2608499999999\nExp number, first 60 AAs: 19.52923\nTotal prob of N-in: 0.94351\nPOSSIBLE N-term signal sequence\ninside 1 19\nTMhelix 20 39\noutside 40 325\nTMhelix 326 348\ninside 349 372

5666 GCF_001893545.1_ASM189354v1 Escherichia coli Proteobacteria; Gammaproteobacteria; Enterobacterales; Enterobacteriaceae; Escherichia
MTGDNTLIHSHGINRRDFMKLCAALAATMGLSSKAAAEMAESVTNPQRPPVIWIGAQECT WP_000145410.1
MULTISPECIES: hydrogenase-2 small chain [Proteobacteria] Length: 372\nNumber of predicted TMHs: 1\nExp number of AAs in TMHs: 23.72226\nExp number, first 60 AAs: 0.22038\nTotal prob of N-in: 0.12994\noutside 1 331\nTMhelix 332 354\ninside 355 372

5667 GCF_001893555.1_ASM189355v1 Escherichia coli Proteobacteria; Gammaproteobacteria; Enterobacterales; Enterobacteriaceae; Escherichia
MNNEETFYQAMRRQGVTRRSFLKYCSLAATSLGLGAGMAPKIAWALENKPRIPVVWIHGL WP_001058323.1
MULTISPECIES: hydrogenase-1 small chain [Proteobacteria] Length: 372\nNumber of predicted TMHs: 2\nExp number of AAs in TMHs: 37.2608499999999\nExp number, first 60 AAs: 19.52923\nTotal prob of N-in: 0.94351\nPOSSIBLE N-term signal sequence\ninside 1 19\nTMhelix 20 39\noutside 40 325\nTMhelix 326 348\ninside 349 372

5668 GCF_001893615.1_ASM189361v1 Escherichia coli Proteobacteria; Gammaproteobacteria; Enterobacterales; Enterobacteriaceae; Escherichia
MNNEETFYQAMRRQGVTRRSFLKYCSLAATSLGLGAGMAPKIAWALENKPRIPVVWIHGL WP_001058323.1
MULTISPECIES: hydrogenase-1 small chain [Proteobacteria] Length: 372\nNumber of predicted TMHs: 2\nExp number of AAs in TMHs: 37.2608499999999\nExp number, first 60 AAs: 19.52923\nTotal prob of N-in: 0.94351\nPOSSIBLE N-term signal sequence\ninside 1 19\nTMhelix 20 39\noutside 40 325\nTMhelix 326 348\ninside 349 372

5669 GCF_001893365.1_ASM189336v1 Escherichia coli Proteobacteria; Gammaproteobacteria; Enterobacterales; Enterobacteriaceae; Escherichia
MNNEETFYQAMRRQGVTRRSFLKYCSLAATSLGLGAGMAPKIAWALENKPRIPVVWIHGL WP_063501981.1
hydrogenase [Escherichia coli] Length: 372\nNumber of predicted TMHs: 2\nExp number of AAs in TMHs: 37.06322\nExp number, first 60 AAs: 19.52932\nTotal prob of N-in: 0.94340\nPOSSIBLE N-term signal sequence\ninside 1 19\nTMhelix 20 39\noutside 40 325\nTMhelix 326 348\ninside 349 372

5670 GCF_001893455.1_ASM189345v1 Escherichia coli Proteobacteria; Gammaproteobacteria; Enterobacterales; Enterobacteriaceae; Escherichia
MNNEETFYQAMRRQGVTRRSFLKYCSLAATSLGLGAGMAPKIAWALENKPRIPVVWIHGL WP_001295940.1
MULTISPECIES: hydrogenase [Enterobacteriaceae] Length: 372\nNumber of predicted TMHs: 2\nExp number of AAs in TMHs: 37.2608899999999\nExp number, first 60 AAs: 19.52923\nTotal prob of N-in: 0.94351\nPOSSIBLE N-term signal sequence\ninside 1 19\nTMhelix 20 39\noutside 40 325\nTMhelix 326 348\ninside 349 372

5671 GCF_001893635.1_ASM189363v1 Escherichia coli Proteobacteria; Gammaproteobacteria; Enterobacterales; Enterobacteriaceae; Escherichia
MTGDNTLIHSHGINRRDFMKLCAALAATMGLSSKAAAEMAESVTNPQRPPVIWIGAQECT WP_000145410.1
MULTISPECIES: hydrogenase-2 small chain [Proteobacteria] Length: 372\nNumber of predicted TMHs: 1\nExp number of AAs in TMHs: 23.72226\nExp number, first 60 AAs: 0.22038\nTotal prob of N-in: 0.12994\noutside 1 331\nTMhelix 332 354\ninside 355 372

5672 GCF_001893685.1_ASM189368v1 Escherichia coli Proteobacteria; Gammaproteobacteria; Enterobacterales; Enterobacteriaceae; Escherichia
MTGDNTLIHSHGINRRDFMKLCAALAATMGLSSKAAAEMAESVTNPQRPPVIWIGAQECT WP_000145410.1
MULTISPECIES: hydrogenase-2 small chain [Proteobacteria] Length: 372\nNumber of predicted TMHs: 1\nExp number of AAs in TMHs: 23.72226\nExp number, first 60 AAs: 0.22038\nTotal prob of N-in: 0.12994\noutside 1 331\nTMhelix 332 354\ninside 355 372

5673 GCF_001893705.1_ASM189370v1 Escherichia coli Proteobacteria; Gammaproteobacteria; Enterobacterales; Enterobacteriaceae; Escherichia
MTGDNTLIHSHGINRRDFMKLCAALAATMGLSSKAAAEMAESVTNPQRPPVIWIGAQECT WP_000145410.1
MULTISPECIES: hydrogenase-2 small chain [Proteobacteria] Length: 372\nNumber of predicted TMHs: 1\nExp number of AAs in TMHs: 23.72226\nExp number, first 60 AAs: 0.22038\nTotal prob of N-in: 0.12994\noutside 1 331\nTMhelix 332 354\ninside 355 372

5674 GCF_001893635.1_ASM189363v1 Escherichia coli Proteobacteria; Gammaproteobacteria; Enterobacterales; Enterobacteriaceae; Escherichia

MNNEETFYQAMRRQGVTRRSFLKYCSLAATSLGLGAGMAPKIAWALENKPRIPVVWIHGL WP_001058323.1 MULTISPECIES: hydrogenase-1 small chain [Proteobacteria] Length: 372\nNumber of predicted TMHs: 2\nExp number of AAs in TMHs: 37.2608499999999\nExp number, first 60 AAs: 19.52923\nTotal prob of N-in: 0.94351\nPOSSIBLE N-term signal sequence\ninside 1 19\nTMhelix 20 39\noutside 40 325\nTMhelix 326 348\ninside 349 372

5675 GCF_001893765.1_ASM189376v1 Escherichia coli Proteobacteria; Gammaproteobacteria; Enterobacterales; Enterobacteriaceae; Escherichia MTGDNTLIHSHGINRRDFMKLCAALAATMGLSSKAAAEMAESVTNPQRPPVIWIGAQECT WP_000145410.1 MULTISPECIES: hydrogenase-2 small chain [Proteobacteria] Length: 372\nNumber of predicted TMHs: 1\nExp number of AAs in TMHs: 23.72226\nExp number, first 60 AAs: 0.22038\nTotal prob of N-in: 0.12994\noutside 1 331\nTMhelix 332 354\ninside 355 372

5676 GCF_001893875.1_ASM189387v1 Escherichia coli Proteobacteria; Gammaproteobacteria; Enterobacterales; Enterobacteriaceae; Escherichia MTGDNTLIHSHGINRRDFMKLCAALAATMGLSSKAAAEMAESVTNPQRPPVIWIGAQECT WP_000145410.1 MULTISPECIES: hydrogenase-2 small chain [Proteobacteria] Length: 372\nNumber of predicted TMHs: 1\nExp number of AAs in TMHs: 23.72226\nExp number, first 60 AAs: 0.22038\nTotal prob of N-in: 0.12994\noutside 1 331\nTMhelix 332 354\ninside 355 372

5677 GCF_001893785.1_ASM189378v1 Escherichia coli Proteobacteria; Gammaproteobacteria; Enterobacterales; Enterobacteriaceae; Escherichia MTGDNTLIHSHGINRRDFMKLCAALAATMGLSSKAAAEMAESVTNPQRPPVIWIGAQECT WP_000145410.1 MULTISPECIES: hydrogenase-2 small chain [Proteobacteria] Length: 372\nNumber of predicted TMHs: 1\nExp number of AAs in TMHs: 23.72226\nExp number, first 60 AAs: 0.22038\nTotal prob of N-in: 0.12994\noutside 1 331\nTMhelix 332 354\ninside 355 372

5678 GCF_001893845.1_ASM189384v1 Escherichia coli Proteobacteria; Gammaproteobacteria; Enterobacterales; Enterobacteriaceae; Escherichia MTGDNTLIHSHGINRRDFMKLCAALAATMGLSSKAAAEMAESVTNPQRPPVIWIGAQECT WP_000145410.1 MULTISPECIES: hydrogenase-2 small chain [Proteobacteria] Length: 372\nNumber of predicted TMHs: 1\nExp number of AAs in TMHs: 23.72226\nExp number, first 60 AAs: 0.22038\nTotal prob of N-in: 0.12994\noutside 1 331\nTMhelix 332 354\ninside 355 372

5679 GCF_001893935.1_ASM189393v1 Escherichia coli Proteobacteria; Gammaproteobacteria; Enterobacterales; Enterobacteriaceae; Escherichia MTGDNTLIHSHGINRRDFMKLCAALAATMGLSSKAAAEMAESVTNPQRPPVIWIGAQECT WP_000145410.1 MULTISPECIES: hydrogenase-2 small chain [Proteobacteria] Length: 372\nNumber of predicted TMHs: 1\nExp number of AAs in TMHs: 23.72226\nExp number, first 60 AAs: 0.22038\nTotal prob of N-in: 0.12994\noutside 1 331\nTMhelix 332 354\ninside 355 372

5680 GCF_001893785.1_ASM189378v1 Escherichia coli Proteobacteria; Gammaproteobacteria; Enterobacterales; Enterobacteriaceae; Escherichia MNNEETFYQAMRRQGVTRRSFLKYCSLAATSLGLGAGMAPKIAWALENKPRIPVVWIHGL WP_001058323.1 MULTISPECIES: hydrogenase-1 small chain [Proteobacteria] Length: 372\nNumber of predicted TMHs: 2\nExp number of AAs in TMHs: 37.2608499999999\nExp number, first 60 AAs: 19.52923\nTotal prob of N-in: 0.94351\nPOSSIBLE N-term signal sequence\ninside 1 19\nTMhelix 20 39\noutside 40 325\nTMhelix 326 348\ninside 349 372

5681 GCF_001893915.1_ASM189391v1 Escherichia coli Proteobacteria; Gammaproteobacteria; Enterobacterales; Enterobacteriaceae; Escherichia MNNEETFYQAMRRQGVTRRSFLKYCSLAATSLGLGAGMAPKIAWALENKPRIPVVWIHGL WP_001295940.1 MULTISPECIES: hydrogenase [Enterobacteriaceae] Length: 372\nNumber of predicted TMHs: 2\nExp number of AAs in TMHs: 37.2608899999999\nExp number, first 60 AAs: 19.52923\nTotal prob of N-in: 0.94351\nPOSSIBLE N-term signal sequence\ninside 1 19\nTMhelix 20 39\noutside 40 325\nTMhelix 326 348\ninside 349 372

5682 GCF_001893765.1_ASM189376v1 Escherichia coli Proteobacteria; Gammaproteobacteria; Enterobacterales; Enterobacteriaceae; Escherichia MNNEETFYQAMRRQGVTRRSFLKYCSLAATSLGLGAGMAPKIAWALENKPRIPVVWIHGL WP_001058323.1 MULTISPECIES: hydrogenase-1 small chain [Proteobacteria] Length: 372\nNumber of predicted TMHs:

2\nExp number of AAs in TMHs: 37.2608499999999\nExp number, first 60 AAs: 19.52923\nTotal prob of N-in: 0.94351\nPOSSIBLE N-term signal sequence\ninside 1 19\nTMhelix 20 39\noutside 40 325\nTMhelix 326 348\ninside 349 372

5683 GCF_001893875.1_ASM189387v1 Escherichia coli Proteobacteria; Gammaproteobacteria; Enterobacterales; Enterobacteriaceae; Escherichia MNNEETFYQAMRRQGVTRRSFLKYCSLAATSLGLGAGMAPKIAWALENKPRIPVVWIHGL WP_001058323.1 MULTISPECIES: hydrogenase-1 small chain [Proteobacteria] Length: 372\nNumber of predicted TMHs: 2\nExp number of AAs in TMHs: 37.2608499999999\nExp number, first 60 AAs: 19.52923\nTotal prob of N-in: 0.94351\nPOSSIBLE N-term signal sequence\ninside 1 19\nTMhelix 20 39\noutside 40 325\nTMhelix 326 348\ninside 349 372

5684 GCF_001894025.1_ASM189402v1 Escherichia coli Proteobacteria; Gammaproteobacteria; Enterobacterales; Enterobacteriaceae; Escherichia MTGDNTLIHSHGINRRDFMKLCAALAATMGLSSKAAAEMAESVTNPQRPPVIWIGAQECT WP_000145410.1 MULTISPECIES: hydrogenase-2 small chain [Proteobacteria] Length: 372\nNumber of predicted TMHs: 1\nExp number of AAs in TMHs: 23.72226\nExp number, first 60 AAs: 0.22038\nTotal prob of N-in: 0.12994\noutside 1 331\nTMhelix 332 354\ninside 355 372

5685 GCF_001894075.1_ASM189407v1 Escherichia coli Proteobacteria; Gammaproteobacteria; Enterobacterales; Enterobacteriaceae; Escherichia MTGDNTLIHSHGINRRDFMKLCAALAATMGLSSKAAAEMAESVTNPQRPPVIWIGAQECT WP_000145410.1 MULTISPECIES: hydrogenase-2 small chain [Proteobacteria] Length: 372\nNumber of predicted TMHs: 1\nExp number of AAs in TMHs: 23.72226\nExp number, first 60 AAs: 0.22038\nTotal prob of N-in: 0.12994\noutside 1 331\nTMhelix 332 354\ninside 355 372

5686 GCF_001894025.1_ASM189402v1 Escherichia coli Proteobacteria; Gammaproteobacteria; Enterobacterales; Enterobacteriaceae; Escherichia MNNEETFYQAMRRQGVTRRSFLKYCSLAATSLGLGAGMAPKIAWALENKPRIPVVWIHGL WP_001058323.1 MULTISPECIES: hydrogenase-1 small chain [Proteobacteria] Length: 372\nNumber of predicted TMHs: 2\nExp number of AAs in TMHs: 37.2608499999999\nExp number, first 60 AAs: 19.52923\nTotal prob of N-in: 0.94351\nPOSSIBLE N-term signal sequence\ninside 1 19\nTMhelix 20 39\noutside 40 325\nTMhelix 326 348\ninside 349 372

5687 GCF_001894125.1_ASM189412v1 Escherichia coli Proteobacteria; Gammaproteobacteria; Enterobacterales; Enterobacteriaceae; Escherichia MTGDNTLIHSHGINRRDFMKLCAALAATMGLSSKAAAEMAESVTNPQRPPVIWIGAQECT WP_000145410.1 MULTISPECIES: hydrogenase-2 small chain [Proteobacteria] Length: 372\nNumber of predicted TMHs: 1\nExp number of AAs in TMHs: 23.72226\nExp number, first 60 AAs: 0.22038\nTotal prob of N-in: 0.12994\noutside 1 331\nTMhelix 332 354\ninside 355 372

5688 GCF_001894225.1_ASM189422v1 Escherichia coli Proteobacteria; Gammaproteobacteria; Enterobacterales; Enterobacteriaceae; Escherichia MTGDNTLIHSHGINRRDFMKLCAALAATMGLSSKAAAEMAESVTNPQRPPVIWIGAQECT WP_000145410.1 MULTISPECIES: hydrogenase-2 small chain [Proteobacteria] Length: 372\nNumber of predicted TMHs: 1\nExp number of AAs in TMHs: 23.72226\nExp number, first 60 AAs: 0.22038\nTotal prob of N-in: 0.12994\noutside 1 331\nTMhelix 332 354\ninside 355 372

5689 GCF_001894235.1_ASM189423v1 Escherichia coli Proteobacteria; Gammaproteobacteria; Enterobacterales; Enterobacteriaceae; Escherichia MTGDNTLIHSHGINRRDFMKLCAALAATMGLSSKAAAEMAESVTNPQRPPVIWIGAQECT WP_000145410.1 MULTISPECIES: hydrogenase-2 small chain [Proteobacteria] Length: 372\nNumber of predicted TMHs: 1\nExp number of AAs in TMHs: 23.72226\nExp number, first 60 AAs: 0.22038\nTotal prob of N-in: 0.12994\noutside 1 331\nTMhelix 332 354\ninside 355 372

5690 GCF_001894165.1_ASM189416v1 Escherichia coli Proteobacteria; Gammaproteobacteria; Enterobacterales; Enterobacteriaceae; Escherichia MTGDNTLIHSHGINRRDFMKLCAALAATMGLSSKAAAEMAESVTNPQRPPVIWIGAQECT WP_000145410.1 MULTISPECIES: hydrogenase-2 small chain [Proteobacteria] Length: 372\nNumber of predicted TMHs: 1\nExp number of AAs in TMHs: 23.72226\nExp number, first 60 AAs: 0.22038\nTotal prob of N-in: 0.12994\noutside 1 331\nTMhelix 332 354\ninside 355 372

5691 GCF_001894255.1_ASM189425v1 Escherichia coli Proteobacteria; Gammaproteobacteria; Enterobacterales; Enterobacteriaceae; Escherichia
MTGDNTLIHSHGINRRDFMKLCAALAATMGLSSKAAAEMAESVTNPQRPPVIWIGAQECT WP_000145410.1
MULTISPECIES: hydrogenase-2 small chain [Proteobacteria] Length: 372\nNumber of predicted TMHs: 1\nExp number of AAs in TMHs: 23.72226\nExp number, first 60 AAs: 0.22038\nTotal prob of N-in: 0.12994\noutside 1 331\nTMhelix 332 354\ninside 355 372

5692 GCF_001894165.1_ASM189416v1 Escherichia coli Proteobacteria; Gammaproteobacteria; Enterobacterales; Enterobacteriaceae; Escherichia
MNNEETFYQAMRRQGVTRRSFLKYCSLAATSLGLGAGMAPKIAWALENKPRIPVVWIHGL WP_001058323.1
MULTISPECIES: hydrogenase-1 small chain [Proteobacteria] Length: 372\nNumber of predicted TMHs: 2\nExp number of AAs in TMHs: 37.2608499999999\nExp number, first 60 AAs: 19.52923\nTotal prob of N-in: 0.94351\nPOSSIBLE N-term signal sequence\ninside 1 19\nTMhelix 20 39\noutside 40 325\nTMhelix 326 348\ninside 349 372

5693 GCF_001894255.1_ASM189425v1 Escherichia coli Proteobacteria; Gammaproteobacteria; Enterobacterales; Enterobacteriaceae; Escherichia
MNNEETFYQAMRRQGVTRRSFLKYCSLAATSLGLGAGMAPKIAWALENKPRIPVVWIHGL WP_001295940.1
MULTISPECIES: hydrogenase [Enterobacteriaceae] Length: 372\nNumber of predicted TMHs: 2\nExp number of AAs in TMHs: 37.2608899999999\nExp number, first 60 AAs: 19.52923\nTotal prob of N-in: 0.94351\nPOSSIBLE N-term signal sequence\ninside 1 19\nTMhelix 20 39\noutside 40 325\nTMhelix 326 348\ninside 349 372

5694 GCF_001894465.1_ASM189446v1 Escherichia coli Proteobacteria; Gammaproteobacteria; Enterobacterales; Enterobacteriaceae; Escherichia
MTGDNTLIHSHGINRRDFMKLCAALAATMGLSSKAAAEMAESVTNPQRPPVIWIGAQECT WP_000145410.1
MULTISPECIES: hydrogenase-2 small chain [Proteobacteria] Length: 372\nNumber of predicted TMHs: 1\nExp number of AAs in TMHs: 23.72226\nExp number, first 60 AAs: 0.22038\nTotal prob of N-in: 0.12994\noutside 1 331\nTMhelix 332 354\ninside 355 372

5695 GCF_001894125.1_ASM189412v1 Escherichia coli Proteobacteria; Gammaproteobacteria; Enterobacterales; Enterobacteriaceae; Escherichia
MNNEETFYQAMRRQGVTRRSFLKYCSLAATSLGLGAGMAPKIAWALENKPRIPVVWIHGL WP_001058320.1
hydrogenase [Escherichia coli] Length: 372\nNumber of predicted TMHs: 2\nExp number of AAs in TMHs: 37.26082\nExp number, first 60 AAs: 19.52923\nTotal prob of N-in: 0.94351\nPOSSIBLE N-term signal sequence\ninside 1 19\nTMhelix 20 39\noutside 40 325\nTMhelix 326 348\ninside 349 372

5696 GCF_001894415.1_ASM189441v1 Escherichia coli Proteobacteria; Gammaproteobacteria; Enterobacterales; Enterobacteriaceae; Escherichia
MTGDNTLIHSHGINRRDFMKLCAALAATMGLSSKAAAEMAESVTNPQRPPVIWIGAQECT WP_000145410.1
MULTISPECIES: hydrogenase-2 small chain [Proteobacteria] Length: 372\nNumber of predicted TMHs: 1\nExp number of AAs in TMHs: 23.72226\nExp number, first 60 AAs: 0.22038\nTotal prob of N-in: 0.12994\noutside 1 331\nTMhelix 332 354\ninside 355 372

5697 GCF_001894145.1_ASM189414v1 Escherichia coli Proteobacteria; Gammaproteobacteria; Enterobacterales; Enterobacteriaceae; Escherichia
MNNEETFYQAMRRQGVTRRSFLKYCSLAATSLGLGAGMAPKIAWALENKPRIPVVWIHGL WP_001058323.1
MULTISPECIES: hydrogenase-1 small chain [Proteobacteria] Length: 372\nNumber of predicted TMHs: 2\nExp number of AAs in TMHs: 37.2608499999999\nExp number, first 60 AAs: 19.52923\nTotal prob of N-in: 0.94351\nPOSSIBLE N-term signal sequence\ninside 1 19\nTMhelix 20 39\noutside 40 325\nTMhelix 326 348\ninside 349 372

5698 GCF_001894415.1_ASM189441v1 Escherichia coli Proteobacteria; Gammaproteobacteria; Enterobacterales; Enterobacteriaceae; Escherichia
MNNEETFYQAMRRQGVTRRSFLKYCSLAATSLGLGAGMAPKIAWALENKPRIPVVWIHGL WP_001058323.1
MULTISPECIES: hydrogenase-1 small chain [Proteobacteria] Length: 372\nNumber of predicted TMHs: 2\nExp number of AAs in TMHs: 37.2608499999999\nExp number, first 60 AAs: 19.52923\nTotal prob of N-in: 0.94351\nPOSSIBLE N-term signal sequence\ninside 1 19\nTMhelix 20 39\noutside 40 325\nTMhelix 326 348\ninside 349 372

5699	GCF_001894145.1_ASM189414v1	Escherichia coli	Proteobacteria; Gammaproteobacteria; Enterobacterales; Enterobacteriaceae; Escherichia
MTGDNTLIHSHGINRRDFMKLCAALATTMGLSSKAAAEMAESVTNPQRPPVIWIGAQECT	WP_023149629.1
MULTISPECIES: hydrogenase-2 small chain [Enterobacteriaceae]	Length: 372\nNumber of predicted TMHs: 1\nExp number of AAs in TMHs: 23.63603\nExp number, first 60 AAs: 0.12971\nTotal prob of N-in: 0.12608\noutside 1 331\nTMhelix 332 354\ninside 355 372

5700	GCF_001894225.1_ASM189422v1	Escherichia coli	Proteobacteria; Gammaproteobacteria; Enterobacterales; Enterobacteriaceae; Escherichia
MNNEETFYQAMRRQGVTRRSFLKYCSLAATSLGLGAGMAPKIAWALENKPRIPVVWIHGL	WP_001058323.1
MULTISPECIES: hydrogenase-1 small chain [Proteobacteria]	Length: 372\nNumber of predicted TMHs: 2\nExp number of AAs in TMHs: 37.2608499999999\nExp number, first 60 AAs: 19.52923\nTotal prob of N-in: 0.94351\nPOSSIBLE N-term signal sequence\ninside 1 19\nTMhelix 20 39\noutside 40 325\nTMhelix 326 348\ninside 349 372

5701	GCF_001894235.1_ASM189423v1	Escherichia coli	Proteobacteria; Gammaproteobacteria; Enterobacterales; Enterobacteriaceae; Escherichia
MNNEETFYQAMRRQGVTRRSFLKYCSLAATSLGLGAGMAPKIAWALENKPRIPVVWIHGL	WP_001058323.1
MULTISPECIES: hydrogenase-1 small chain [Proteobacteria]	Length: 372\nNumber of predicted TMHs: 2\nExp number of AAs in TMHs: 37.2608499999999\nExp number, first 60 AAs: 19.52923\nTotal prob of N-in: 0.94351\nPOSSIBLE N-term signal sequence\ninside 1 19\nTMhelix 20 39\noutside 40 325\nTMhelix 326 348\ninside 349 372

5702	GCF_001894475.1_ASM189447v1	Escherichia coli	Proteobacteria; Gammaproteobacteria; Enterobacterales; Enterobacteriaceae; Escherichia
MTGDNTLIHSHGINRRDFMKLCAALAATMGLSSKAAAEMAESVTNPQRPPVIWIGAQECT	WP_000145410.1
MULTISPECIES: hydrogenase-2 small chain [Proteobacteria]	Length: 372\nNumber of predicted TMHs: 1\nExp number of AAs in TMHs: 23.72226\nExp number, first 60 AAs: 0.22038\nTotal prob of N-in: 0.12994\noutside 1 331\nTMhelix 332 354\ninside 355 372

5703	GCF_001894465.1_ASM189446v1	Escherichia coli	Proteobacteria; Gammaproteobacteria; Enterobacterales; Enterobacteriaceae; Escherichia
MNNEETFYQAMRRQGVTRRSFLKYCSLAATSLGLGAGMAPKIAWALENKPRIPVVWIHGL	WP_001058323.1
MULTISPECIES: hydrogenase-1 small chain [Proteobacteria]	Length: 372\nNumber of predicted TMHs: 2\nExp number of AAs in TMHs: 37.2608499999999\nExp number, first 60 AAs: 19.52923\nTotal prob of N-in: 0.94351\nPOSSIBLE N-term signal sequence\ninside 1 19\nTMhelix 20 39\noutside 40 325\nTMhelix 326 348\ninside 349 372

5704	GCF_001894555.1_ASM189455v1	Escherichia coli	Proteobacteria; Gammaproteobacteria; Enterobacterales; Enterobacteriaceae; Escherichia
MTGDNTLIHSHGINRRDFMKLCAALAATMGLSSKAAAEMAESVTNPQRPPVIWIGAQECT	WP_000145410.1
MULTISPECIES: hydrogenase-2 small chain [Proteobacteria]	Length: 372\nNumber of predicted TMHs: 1\nExp number of AAs in TMHs: 23.72226\nExp number, first 60 AAs: 0.22038\nTotal prob of N-in: 0.12994\noutside 1 331\nTMhelix 332 354\ninside 355 372

5705	GCF_001894505.1_ASM189450v1	Escherichia coli	Proteobacteria; Gammaproteobacteria; Enterobacterales; Enterobacteriaceae; Escherichia
MTGDNTLIHSHGINRRDFMKLCAALAATMGLSSKAAAEMAESVTNPQRPPVIWIGAQECT	WP_000145410.1
MULTISPECIES: hydrogenase-2 small chain [Proteobacteria]	Length: 372\nNumber of predicted TMHs: 1\nExp number of AAs in TMHs: 23.72226\nExp number, first 60 AAs: 0.22038\nTotal prob of N-in: 0.12994\noutside 1 331\nTMhelix 332 354\ninside 355 372

5706	GCF_001894625.1_ASM189462v1	Escherichia coli	Proteobacteria; Gammaproteobacteria; Enterobacterales; Enterobacteriaceae; Escherichia
MTGDNTLIHSHGINRRDFMKLCAALAATMGLSSKAAAEMAESVTNPQRPPVIWIGAQECT	WP_000145410.1
MULTISPECIES: hydrogenase-2 small chain [Proteobacteria]	Length: 372\nNumber of predicted TMHs: 1\nExp number of AAs in TMHs: 23.72226\nExp number, first 60 AAs: 0.22038\nTotal prob of N-in: 0.12994\noutside 1 331\nTMhelix 332 354\ninside 355 372

5707	GCF_001894565.1_ASM189456v1	Escherichia coli	Proteobacteria; Gammaproteobacteria; Enterobacterales; Enterobacteriaceae; Escherichia

MTGDNTLIHSHGINRRDFMKLCAALAATMGLSSKAAAEMAESVTNPQRPPVIWIGAQECT WP_000145410.1 MULTISPECIES: hydrogenase-2 small chain [Proteobacteria] Length: 372\nNumber of predicted TMHs: 1\nExp number of AAs in TMHs: 23.72226\nExp number, first 60 AAs: 0.22038\nTotal prob of N-in: 0.12994\noutside 1 331\nTMhelix 332 354\ninside 355 372

5708 GCF_001894645.1_ASM189464v1 Escherichia coli Proteobacteria; Gammaproteobacteria; Enterobacterales; Enterobacteriaceae; Escherichia MTGDNTLIHSHGINRRDFMKLCAALAATMGLSSKAAAEMAESVTNPQRPPVIWIGAQECT WP_000145410.1 MULTISPECIES: hydrogenase-2 small chain [Proteobacteria] Length: 372\nNumber of predicted TMHs: 1\nExp number of AAs in TMHs: 23.72226\nExp number, first 60 AAs: 0.22038\nTotal prob of N-in: 0.12994\noutside 1 331\nTMhelix 332 354\ninside 355 372

5709 GCF_001894685.1_ASM189468v1 Escherichia coli Proteobacteria; Gammaproteobacteria; Enterobacterales; Enterobacteriaceae; Escherichia MTGDNTLIHSHGINRRDFMKLCAALAATMGLSSKAAAEMAESVTNPQRPPVIWIGAQECT WP_000145410.1 MULTISPECIES: hydrogenase-2 small chain [Proteobacteria] Length: 372\nNumber of predicted TMHs: 1\nExp number of AAs in TMHs: 23.72226\nExp number, first 60 AAs: 0.22038\nTotal prob of N-in: 0.12994\noutside 1 331\nTMhelix 332 354\ninside 355 372

5710 GCF_001894645.1_ASM189464v1 Escherichia coli Proteobacteria; Gammaproteobacteria; Enterobacterales; Enterobacteriaceae; Escherichia MNNEETFYQAMRRQGVTRRSFLKYCSLAATSLGLGAGMAPKIAWALENKPRIPVVWIHGL WP_001058323.1 MULTISPECIES: hydrogenase-1 small chain [Proteobacteria] Length: 372\nNumber of predicted TMHs: 2\nExp number of AAs in TMHs: 37.2608499999999\nExp number, first 60 AAs: 19.52923\nTotal prob of N-in: 0.94351\nPOSSIBLE N-term signal sequence\ninside 1 19\nTMhelix 20 39\noutside 40 325\nTMhelix 326 348\ninside 349 372

5711 GCF_001894565.1_ASM189456v1 Escherichia coli Proteobacteria; Gammaproteobacteria; Enterobacterales; Enterobacteriaceae; Escherichia MNNEETFYQAMRRQGVTRRSFLKYCSLAATSLGLGAGMAPKIAWALENKPRIPVVWIHGL WP_001058323.1 MULTISPECIES: hydrogenase-1 small chain [Proteobacteria] Length: 372\nNumber of predicted TMHs: 2\nExp number of AAs in TMHs: 37.2608499999999\nExp number, first 60 AAs: 19.52923\nTotal prob of N-in: 0.94351\nPOSSIBLE N-term signal sequence\ninside 1 19\nTMhelix 20 39\noutside 40 325\nTMhelix 326 348\ninside 349 372

5712 GCF_001894685.1_ASM189468v1 Escherichia coli Proteobacteria; Gammaproteobacteria; Enterobacterales; Enterobacteriaceae; Escherichia MNNEETFYQAMRRQGVTRRSFLKYCSLAATSLGLGAGMAPKIAWALENKPRIPVVWIHGL WP_001058323.1 MULTISPECIES: hydrogenase-1 small chain [Proteobacteria] Length: 372\nNumber of predicted TMHs: 2\nExp number of AAs in TMHs: 37.2608499999999\nExp number, first 60 AAs: 19.52923\nTotal prob of N-in: 0.94351\nPOSSIBLE N-term signal sequence\ninside 1 19\nTMhelix 20 39\noutside 40 325\nTMhelix 326 348\ninside 349 372

5713 GCF_001894475.1_ASM189447v1 Escherichia coli Proteobacteria; Gammaproteobacteria; Enterobacterales; Enterobacteriaceae; Escherichia MNNEETFYQAMRRQGVTRRSFLKYCSLAATSLGLGAGMAPKIAWALENKPRIPVVWIHGL WP_001295940.1 MULTISPECIES: hydrogenase [Enterobacteriaceae] Length: 372\nNumber of predicted TMHs: 2\nExp number of AAs in TMHs: 37.2608899999999\nExp number, first 60 AAs: 19.52923\nTotal prob of N-in: 0.94351\nPOSSIBLE N-term signal sequence\ninside 1 19\nTMhelix 20 39\noutside 40 325\nTMhelix 326 348\ninside 349 372

5714 GCF_001894715.1_ASM189471v1 Escherichia coli Proteobacteria; Gammaproteobacteria; Enterobacterales; Enterobacteriaceae; Escherichia MTGDNTLIHSHGINRRDFMKLCAALAATMGLSSKAAAEMAESVTNPQRPPVIWIGAQECT WP_000145410.1 MULTISPECIES: hydrogenase-2 small chain [Proteobacteria] Length: 372\nNumber of predicted TMHs: 1\nExp number of AAs in TMHs: 23.72226\nExp number, first 60 AAs: 0.22038\nTotal prob of N-in: 0.12994\noutside 1 331\nTMhelix 332 354\ninside 355 372

5715 GCF_001900295.1_ASM190029v1 Escherichia coli Proteobacteria; Gammaproteobacteria; Enterobacterales; Enterobacteriaceae; Escherichia MTGDNTLIHSHGINRRDFMKLCAALAATMGLSSKAAAEMAESVTNPQRPPVIWIGAQECT WP_000145410.1

MULTISPECIES: hydrogenase-2 small chain [Proteobacteria] Length: 372\nNumber of predicted TMHs: 1\nExp number of AAs in TMHs: 23.72226\nExp number, first 60 AAs: 0.22038\nTotal prob of N-in: 0.12994\noutside 1 331\nTMhelix 332 354\ninside 355 372

5716 GCF_001900395.1_ASM190039v1 Escherichia coli Proteobacteria; Gammaproteobacteria; Enterobacterales; Enterobacteriaceae; Escherichia
MNNEETFYQAMRRQGVTRRSFLKYCSLAATSLGLGAGMAPKIAWALENKPRIPVVWIHGL WP_001058323.1
MULTISPECIES: hydrogenase-1 small chain [Proteobacteria] Length: 372\nNumber of predicted TMHs: 2\nExp number of AAs in TMHs: 37.2608499999999\nExp number, first 60 AAs: 19.52923\nTotal prob of N-in: 0.94351\nPOSSIBLE N-term signal sequence\ninside 1 19\nTMhelix 20 39\noutside 40 325\nTMhelix 326 348\ninside 349 372

5717 GCF_001894555.1_ASM189455v1 Escherichia coli Proteobacteria; Gammaproteobacteria; Enterobacterales; Enterobacteriaceae; Escherichia
MNNEETFYQAMRRQGVTRRSFLKYCSLAATSLGLGAGMAPKIAWALENKPRIPVVWIHGL WP_001295940.1
MULTISPECIES: hydrogenase [Enterobacteriaceae] Length: 372\nNumber of predicted TMHs: 2\nExp number of AAs in TMHs: 37.2608899999999\nExp number, first 60 AAs: 19.52923\nTotal prob of N-in: 0.94351\nPOSSIBLE N-term signal sequence\ninside 1 19\nTMhelix 20 39\noutside 40 325\nTMhelix 326 348\ninside 349 372

5718 GCF_001894625.1_ASM189462v1 Escherichia coli Proteobacteria; Gammaproteobacteria; Enterobacterales; Enterobacteriaceae; Escherichia
MNNEETFYQAMRRQGVTRRSFLKYCSLAATSLGLGAGMAPKIAWALENKPRIPVVWIHGL WP_001295940.1
MULTISPECIES: hydrogenase [Enterobacteriaceae] Length: 372\nNumber of predicted TMHs: 2\nExp number of AAs in TMHs: 37.2608899999999\nExp number, first 60 AAs: 19.52923\nTotal prob of N-in: 0.94351\nPOSSIBLE N-term signal sequence\ninside 1 19\nTMhelix 20 39\noutside 40 325\nTMhelix 326 348\ninside 349 372

5719 GCF_001900335.1_ASM190033v1 Escherichia coli Proteobacteria; Gammaproteobacteria; Enterobacterales; Enterobacteriaceae; Escherichia
MNNEETFYQAMRRQGVTRRSFLKYCSLAATSLGLGAGMAPKIAWALENKPRIPVVWIHGL WP_001058323.1
MULTISPECIES: hydrogenase-1 small chain [Proteobacteria] Length: 372\nNumber of predicted TMHs: 2\nExp number of AAs in TMHs: 37.2608499999999\nExp number, first 60 AAs: 19.52923\nTotal prob of N-in: 0.94351\nPOSSIBLE N-term signal sequence\ninside 1 19\nTMhelix 20 39\noutside 40 325\nTMhelix 326 348\ninside 349 372

5720 GCF_001894715.1_ASM189471v1 Escherichia coli Proteobacteria; Gammaproteobacteria; Enterobacterales; Enterobacteriaceae; Escherichia
MNNEETFYQAMRRQGVTRRSFLKYCSLAATSLGLGAGMAPKIAWALENKPRIPVVWIHGL WP_001295940.1
MULTISPECIES: hydrogenase [Enterobacteriaceae] Length: 372\nNumber of predicted TMHs: 2\nExp number of AAs in TMHs: 37.2608899999999\nExp number, first 60 AAs: 19.52923\nTotal prob of N-in: 0.94351\nPOSSIBLE N-term signal sequence\ninside 1 19\nTMhelix 20 39\noutside 40 325\nTMhelix 326 348\ninside 349 372

5721 GCF_001900435.1_ASM190043v1 Escherichia coli Proteobacteria; Gammaproteobacteria; Enterobacterales; Enterobacteriaceae; Escherichia
MTGDNTLIHSHGINRRDFMKLCAALAATMGLSSKAAAEMAESVTNPQRPPVIWIGAQECT WP_000145410.1
MULTISPECIES: hydrogenase-2 small chain [Proteobacteria] Length: 372\nNumber of predicted TMHs: 1\nExp number of AAs in TMHs: 23.72226\nExp number, first 60 AAs: 0.22038\nTotal prob of N-in: 0.12994\noutside 1 331\nTMhelix 332 354\ninside 355 372

5722 GCF_001900475.1_ASM190047v1 Escherichia coli Proteobacteria; Gammaproteobacteria; Enterobacterales; Enterobacteriaceae; Escherichia
MTGDNTLIHSHGINRRDFMKLCAALAATMGLSSKAAAEMAESVTNPQRPPVIWIGAQECT WP_000145410.1
MULTISPECIES: hydrogenase-2 small chain [Proteobacteria] Length: 372\nNumber of predicted TMHs: 1\nExp number of AAs in TMHs: 23.72226\nExp number, first 60 AAs: 0.22038\nTotal prob of N-in: 0.12994\noutside 1 331\nTMhelix 332 354\ninside 355 372

5723 GCF_001900575.1_ASM190057v1 Escherichia coli Proteobacteria; Gammaproteobacteria; Enterobacterales; Enterobacteriaceae; Escherichia
MTGDNTLIHSHGINRRDFMKLCAALAATMGLSSKAAAEMAESVTNPQRPPVIWIGAQECT WP_000145410.1

MULTISPECIES: hydrogenase-2 small chain [Proteobacteria] Length: 372\nNumber of predicted TMHs: 1\nExp number of AAs in TMHs: 23.72226\nExp number, first 60 AAs: 0.22038\nTotal prob of N-in: 0.12994\noutside 1 331\nTMhelix 332 354\ninside 355 372

5724 GCF_001900515.1_ASM190051v1 Escherichia coli Proteobacteria; Gammaproteobacteria; Enterobacterales; Enterobacteriaceae; Escherichia MTGDNTLIHSHGINRRDFMKLCAALAATMGLSSKAAAEMAESVTNPQRPPVIWIGAQECT WP_000145410.1 MULTISPECIES: hydrogenase-2 small chain [Proteobacteria] Length: 372\nNumber of predicted TMHs: 1\nExp number of AAs in TMHs: 23.72226\nExp number, first 60 AAs: 0.22038\nTotal prob of N-in: 0.12994\noutside 1 331\nTMhelix 332 354\ninside 355 372

5725 GCF_001900535.1_ASM190053v1 Escherichia coli Proteobacteria; Gammaproteobacteria; Enterobacterales; Enterobacteriaceae; Escherichia MTGDNTLIHSHGINRRDFMKLCAALAATMGLSSKAAAEMAESVTNPQRPPVIWIGAQECT WP_000145410.1 MULTISPECIES: hydrogenase-2 small chain [Proteobacteria] Length: 372\nNumber of predicted TMHs: 1\nExp number of AAs in TMHs: 23.72226\nExp number, first 60 AAs: 0.22038\nTotal prob of N-in: 0.12994\noutside 1 331\nTMhelix 332 354\ninside 355 372

5726 GCF_001900615.1_ASM190061v1 Escherichia coli Proteobacteria; Gammaproteobacteria; Enterobacterales; Enterobacteriaceae; Escherichia MTGDNTLIHSHGINRRDFMKLCAALAATMGLSSKAAAEMAESVTNPQRPPVIWIGAQECT WP_000145410.1 MULTISPECIES: hydrogenase-2 small chain [Proteobacteria] Length: 372\nNumber of predicted TMHs: 1\nExp number of AAs in TMHs: 23.72226\nExp number, first 60 AAs: 0.22038\nTotal prob of N-in: 0.12994\noutside 1 331\nTMhelix 332 354\ninside 355 372

5727 GCF_001900675.1_ASM190067v1 Escherichia coli Proteobacteria; Gammaproteobacteria; Enterobacterales; Enterobacteriaceae; Escherichia MTGDNTLIHSHGINRRDFMKLCAALAATMGLSSKAAAEMAESVTNPQRPPVIWIGAQECT WP_000145410.1 MULTISPECIES: hydrogenase-2 small chain [Proteobacteria] Length: 372\nNumber of predicted TMHs: 1\nExp number of AAs in TMHs: 23.72226\nExp number, first 60 AAs: 0.22038\nTotal prob of N-in: 0.12994\noutside 1 331\nTMhelix 332 354\ninside 355 372

5728 GCF_001900635.1_ASM190063v1 Escherichia coli Proteobacteria; Gammaproteobacteria; Enterobacterales; Enterobacteriaceae; Escherichia MTGDNTLIHSHGINRRDFMKLCAALAATMGLSSKAAAEMAESVTNPQRPPVIWIGAQECT WP_000145410.1 MULTISPECIES: hydrogenase-2 small chain [Proteobacteria] Length: 372\nNumber of predicted TMHs: 1\nExp number of AAs in TMHs: 23.72226\nExp number, first 60 AAs: 0.22038\nTotal prob of N-in: 0.12994\noutside 1 331\nTMhelix 332 354\ninside 355 372

5729 GCF_001900635.1_ASM190063v1 Escherichia coli Proteobacteria; Gammaproteobacteria; Enterobacterales; Enterobacteriaceae; Escherichia MNNEETFYQAMRRQGVTRRSFLKYCSLAATSLGLGAGMAPKIAWALENKPRIPVVWIHGL WP_001058320.1 hydrogenase [Escherichia coli] Length: 372\nNumber of predicted TMHs: 2\nExp number of AAs in TMHs: 37.26082\nExp number, first 60 AAs: 19.52923\nTotal prob of N-in: 0.94351\nPOSSIBLE N-term signal sequence\ninside 1 19\nTMhelix 20 39\noutside 40 325\nTMhelix 326 348\ninside 349 372

5730 GCF_001900475.1_ASM190047v1 Escherichia coli Proteobacteria; Gammaproteobacteria; Enterobacterales; Enterobacteriaceae; Escherichia MNNEETFYQAMRRQGVTRRSFLKYCSLAATSLGLGAGMAPKIAWALENKPRIPVVWIHGL WP_001058323.1 MULTISPECIES: hydrogenase-1 small chain [Proteobacteria] Length: 372\nNumber of predicted TMHs: 2\nExp number of AAs in TMHs: 37.2608499999999\nExp number, first 60 AAs: 19.52923\nTotal prob of N-in: 0.94351\nPOSSIBLE N-term signal sequence\ninside 1 19\nTMhelix 20 39\noutside 40 325\nTMhelix 326 348\ninside 349 372

5731 GCF_001900675.1_ASM190067v1 Escherichia coli Proteobacteria; Gammaproteobacteria; Enterobacterales; Enterobacteriaceae; Escherichia MNNEETFYQAMRRQGVTRRSFLKYCSLAATSLGLGAGMAPKIAWALENKPRIPVVWIHGL WP_001058323.1 MULTISPECIES: hydrogenase-1 small chain [Proteobacteria] Length: 372\nNumber of predicted TMHs: 2\nExp number of AAs in TMHs: 37.2608499999999\nExp number, first 60 AAs: 19.52923\nTotal prob of N-in: 0.94351\nPOSSIBLE N-term signal sequence\ninside 1 19\nTMhelix 20 39\noutside 40 325\nTMhelix 326 348\ninside 349 372

5732 GCF_001900515.1_ASM190051v1 Escherichia coli Proteobacteria; Gammaproteobacteria; Enterobacterales; Enterobacteriaceae; Escherichia
MNNEETFYQAMRRQGVTRRSFLKYCSLAATSLGLGAGMAPKIAWALENKPRIPVVWIHGL WP_001295940.1
MULTISPECIES: hydrogenase [Enterobacteriaceae] Length: 372\nNumber of predicted TMHs: 2\nExp number of AAs in TMHs: 37.2608899999999\nExp number, first 60 AAs: 19.52923\nTotal prob of N-in: 0.94351\nPOSSIBLE N-term signal sequence\ninside 1 19\nTMhelix 20 39\noutside 40 325\nTMhelix 326 348\ninside 349 372

5733 GCF_001900615.1_ASM190061v1 Escherichia coli Proteobacteria; Gammaproteobacteria; Enterobacterales; Enterobacteriaceae; Escherichia
MNNEETFYQAMRRQGVTRRSFLKYCSLAATSLGLGAGMAPKIAWALENKPRIPVVWIHGL WP_073544303.1
hydrogenase [Escherichia coli] Length: 372\nNumber of predicted TMHs: 2\nExp number of AAs in TMHs: 37.2612399999999\nExp number, first 60 AAs: 19.52921\nTotal prob of N-in: 0.94351\nPOSSIBLE N-term signal sequence\ninside 1 19\nTMhelix 20 39\noutside 40 325\nTMhelix 326 348\ninside 349 372

5734 GCF_001900835.1_ASM190083v1 Escherichia coli Proteobacteria; Gammaproteobacteria; Enterobacterales; Enterobacteriaceae; Escherichia
MNNEETFYQAMRRQGVTRRSFLKYCSLAATSLGLGAGMAPKIAWALENKPRIPVVWIHGL WP_001058323.1
MULTISPECIES: hydrogenase-1 small chain [Proteobacteria] Length: 372\nNumber of predicted TMHs: 2\nExp number of AAs in TMHs: 37.2608499999999\nExp number, first 60 AAs: 19.52923\nTotal prob of N-in: 0.94351\nPOSSIBLE N-term signal sequence\ninside 1 19\nTMhelix 20 39\noutside 40 325\nTMhelix 326 348\ninside 349 372

5735 GCF_001900885.1_ASM190088v1 Escherichia coli Proteobacteria; Gammaproteobacteria; Enterobacterales; Enterobacteriaceae; Escherichia
MTGDNTLIHSHGINRRDFMKLCAALAATMGLSSKAAAEMAESVTNPQRPPVIWIGAQECT WP_000145410.1
MULTISPECIES: hydrogenase-2 small chain [Proteobacteria] Length: 372\nNumber of predicted TMHs: 1\nExp number of AAs in TMHs: 23.72226\nExp number, first 60 AAs: 0.22038\nTotal prob of N-in: 0.12994\noutside 1 331\nTMhelix 332 354\ninside 355 372

5736 GCF_001900925.1_ASM190092v1 Escherichia coli Proteobacteria; Gammaproteobacteria; Enterobacterales; Enterobacteriaceae; Escherichia
MTGDNTLIHSHGINRRDFMKLCAALAATMGLSSKAAAEMAESVTNPQRPPVIWIGAQECT WP_000145410.1
MULTISPECIES: hydrogenase-2 small chain [Proteobacteria] Length: 372\nNumber of predicted TMHs: 1\nExp number of AAs in TMHs: 23.72226\nExp number, first 60 AAs: 0.22038\nTotal prob of N-in: 0.12994\noutside 1 331\nTMhelix 332 354\ninside 355 372

5737 GCF_001900795.1_ASM190079v1 Escherichia coli Proteobacteria; Gammaproteobacteria; Enterobacterales; Enterobacteriaceae; Escherichia
MTGDNTLIHSHGINRRDFMKLCAALAATMGLSSKAAAEMAESVTNPQRPPVIWIGAQECT WP_032298447.1
hydrogenase 2 small subunit [Escherichia coli] Length: 372\nNumber of predicted TMHs: 1\nExp number of AAs in TMHs: 23.69807\nExp number, first 60 AAs: 0.221\nTotal prob of N-in: 0.12853\noutside 1 331\nTMhelix 332 354\ninside 355 372

5738 GCF_001900985.1_ASM190098v1 Escherichia coli Proteobacteria; Gammaproteobacteria; Enterobacterales; Enterobacteriaceae; Escherichia
MTGDNTLIHSHGINRRDFMKLCAALAATMGLSSKAAAEMAESVTNPQRPPVIWIGAQECT WP_000145410.1
MULTISPECIES: hydrogenase-2 small chain [Proteobacteria] Length: 372\nNumber of predicted TMHs: 1\nExp number of AAs in TMHs: 23.72226\nExp number, first 60 AAs: 0.22038\nTotal prob of N-in: 0.12994\noutside 1 331\nTMhelix 332 354\ninside 355 372

5739 GCF_001900965.1_ASM190096v1 Escherichia coli Proteobacteria; Gammaproteobacteria; Enterobacterales; Enterobacteriaceae; Escherichia
MTGDNTLIHSHGINRRDFMKLCAALAATMGLSSKAAAEMAESVTNPQRPPVIWIGAQECT WP_000145410.1
MULTISPECIES: hydrogenase-2 small chain [Proteobacteria] Length: 372\nNumber of predicted TMHs: 1\nExp number of AAs in TMHs: 23.72226\nExp number, first 60 AAs: 0.22038\nTotal prob of N-in: 0.12994\noutside 1 331\nTMhelix 332 354\ninside 355 372

5740 GCF_001901025.1_ASM190102v1 Escherichia coli Proteobacteria; Gammaproteobacteria; Enterobacterales; Enterobacteriaceae; Escherichia
MTGDNTLIHSHGINRRDFMKLCAALAATMGLSSKAAAEMAESVTNPQRPPVIWIGAQECT WP_000145410.1

MULTISPECIES: hydrogenase-2 small chain [Proteobacteria] Length: 372\nNumber of predicted TMHs: 1\nExp number of AAs in TMHs: 23.72226\nExp number, first 60 AAs: 0.22038\nTotal prob of N-in: 0.12994\noutside 1 331\nTMhelix 332 354\ninside 355 372

5741 GCF_001901065.1_ASM190106v1 Escherichia coli Proteobacteria; Gammaproteobacteria; Enterobacterales; Enterobacteriaceae; Escherichia
MTGDNTLIHSHGINRRDFMKLCAALAATMGLSSKAAAEMAESVTNPQRPPVIWIGAQECT WP_000145410.1
MULTISPECIES: hydrogenase-2 small chain [Proteobacteria] Length: 372\nNumber of predicted TMHs: 1\nExp number of AAs in TMHs: 23.72226\nExp number, first 60 AAs: 0.22038\nTotal prob of N-in: 0.12994\noutside 1 331\nTMhelix 332 354\ninside 355 372

5742 GCF_001901065.1_ASM190106v1 Escherichia coli Proteobacteria; Gammaproteobacteria; Enterobacterales; Enterobacteriaceae; Escherichia
MNNEETFYQAMRRQGVTRRSFLKYCSLAATSLGLGAGMAPKIAWALENKPRIPVVWIHGL WP_001058323.1
MULTISPECIES: hydrogenase-1 small chain [Proteobacteria] Length: 372\nNumber of predicted TMHs: 2\nExp number of AAs in TMHs: 37.2608499999999\nExp number, first 60 AAs: 19.52923\nTotal prob of N-in: 0.94351\nPOSSIBLE N-term signal sequence\ninside 1 19\nTMhelix 20 39\noutside 40 325\nTMhelix 326 348\ninside 349 372

5743 GCF_001901165.1_ASM190116v1 Escherichia coli Proteobacteria; Gammaproteobacteria; Enterobacterales; Enterobacteriaceae; Escherichia
MNNEETFYQAMRRQGVTRRSFLKYCSLAATSLGLGAGMAPKIAWALENKPRIPVVWIHGL WP_001058323.1
MULTISPECIES: hydrogenase-1 small chain [Proteobacteria] Length: 372\nNumber of predicted TMHs: 2\nExp number of AAs in TMHs: 37.2608499999999\nExp number, first 60 AAs: 19.52923\nTotal prob of N-in: 0.94351\nPOSSIBLE N-term signal sequence\ninside 1 19\nTMhelix 20 39\noutside 40 325\nTMhelix 326 348\ninside 349 372

5744 GCF_001900835.1_ASM190083v1 Escherichia coli Proteobacteria; Gammaproteobacteria; Enterobacterales; Enterobacteriaceae; Escherichia
MTGDNTLIHSHGINRRDFMKLCAALATTMGLSSKAAAEMAESVTNPQRPPVIWIGAQECT WP_023149629.1
MULTISPECIES: hydrogenase-2 small chain [Enterobacteriaceae] Length: 372\nNumber of predicted TMHs: 1\nExp number of AAs in TMHs: 23.63603\nExp number, first 60 AAs: 0.12971\nTotal prob of N-in: 0.12608\noutside 1 331\nTMhelix 332 354\ninside 355 372

5745 GCF_001900985.1_ASM190098v1 Escherichia coli Proteobacteria; Gammaproteobacteria; Enterobacterales; Enterobacteriaceae; Escherichia
MNNEETFYQAMRRQGVTRRSFLKYCSLAATSLGLGAGMAPKIAWALENKPRIPVVWIHGL WP_001058323.1
MULTISPECIES: hydrogenase-1 small chain [Proteobacteria] Length: 372\nNumber of predicted TMHs: 2\nExp number of AAs in TMHs: 37.2608499999999\nExp number, first 60 AAs: 19.52923\nTotal prob of N-in: 0.94351\nPOSSIBLE N-term signal sequence\ninside 1 19\nTMhelix 20 39\noutside 40 325\nTMhelix 326 348\ninside 349 372

5746 GCF_001901185.1_ASM190118v1 Escherichia coli Proteobacteria; Gammaproteobacteria; Enterobacterales; Enterobacteriaceae; Escherichia
MTGDNTLIHSHGINRRDFMKLCAALAATMGLSSKAAAEMAESVTNPQRPPVIWIGAQECT WP_000145410.1
MULTISPECIES: hydrogenase-2 small chain [Proteobacteria] Length: 372\nNumber of predicted TMHs: 1\nExp number of AAs in TMHs: 23.72226\nExp number, first 60 AAs: 0.22038\nTotal prob of N-in: 0.12994\noutside 1 331\nTMhelix 332 354\ninside 355 372

5747 GCF_001901315.1_ASM190131v1 Escherichia coli Proteobacteria; Gammaproteobacteria; Enterobacterales; Enterobacteriaceae; Escherichia
MTGDNTLIHSHGINRRDFMKLCAALAATMGLSSKAAAEMAESVTNPQRPPVIWIGAQECT WP_000145410.1
MULTISPECIES: hydrogenase-2 small chain [Proteobacteria] Length: 372\nNumber of predicted TMHs: 1\nExp number of AAs in TMHs: 23.72226\nExp number, first 60 AAs: 0.22038\nTotal prob of N-in: 0.12994\noutside 1 331\nTMhelix 332 354\ninside 355 372

5748 GCF_001901315.1_ASM190131v1 Escherichia coli Proteobacteria; Gammaproteobacteria; Enterobacterales; Enterobacteriaceae; Escherichia
MNNEETFYQAMRRQGVTRRSFLKYCSLAATSLGLGAGMAPKIAWALENKPRIPVVWIHGL WP_001058323.1
MULTISPECIES: hydrogenase-1 small chain [Proteobacteria] Length: 372\nNumber of predicted TMHs: 2\nExp number of AAs in TMHs: 37.2608499999999\nExp number, first 60 AAs: 19.52923\nTotal prob of N-in:

0.94351\nPOSSIBLE N-term signal sequence\ninside 1 19\nTMhelix 20 39\noutside 40 325\nTMhelix 326 348\ninside 349 372

5749 GCF_001901185.1_ASM190118v1 Escherichia coli Proteobacteria; Gammaproteobacteria; Enterobacterales; Enterobacteriaceae; Escherichia
MNNEETFYQAMRRQGVTRRSFLKYCSLAATSLGLGAGMAPKIAWALENKPRIPVVWIHGL WP_001058323.1
MULTISPECIES: hydrogenase-1 small chain [Proteobacteria] Length: 372\nNumber of predicted TMHs: 2\nExp number of AAs in TMHs: 37.2608499999999\nExp number, first 60 AAs: 19.52923\nTotal prob of N-in: 0.94351\nPOSSIBLE N-term signal sequence\ninside 1 19\nTMhelix 20 39\noutside 40 325\nTMhelix 326 348\ninside 349 372

5750 GCF_001900925.1_ASM190092v1 Escherichia coli Proteobacteria; Gammaproteobacteria; Enterobacterales; Enterobacteriaceae; Escherichia
MNNEETFYQAMRRQGVTRRSFLKYCSLAATSLGLGAGMAPKIAWALENKPRIPVVWIHGL WP_001295940.1
MULTISPECIES: hydrogenase [Enterobacteriaceae] Length: 372\nNumber of predicted TMHs: 2\nExp number of AAs in TMHs: 37.2608899999999\nExp number, first 60 AAs: 19.52923\nTotal prob of N-in: 0.94351\nPOSSIBLE N-term signal sequence\ninside 1 19\nTMhelix 20 39\noutside 40 325\nTMhelix 326 348\ninside 349 372

5751 GCF_001900885.1_ASM190088v1 Escherichia coli Proteobacteria; Gammaproteobacteria; Enterobacterales; Enterobacteriaceae; Escherichia
MNNEETFYQAMRRQGVTRRSFLKYCSLAATSLGLGAGMAPKIAWALENKPRIPVVWIHGL WP_001295940.1
MULTISPECIES: hydrogenase [Enterobacteriaceae] Length: 372\nNumber of predicted TMHs: 2\nExp number of AAs in TMHs: 37.2608899999999\nExp number, first 60 AAs: 19.52923\nTotal prob of N-in: 0.94351\nPOSSIBLE N-term signal sequence\ninside 1 19\nTMhelix 20 39\noutside 40 325\nTMhelix 326 348\ninside 349 372

5752 GCF_001900965.1_ASM190096v1 Escherichia coli Proteobacteria; Gammaproteobacteria; Enterobacterales; Enterobacteriaceae; Escherichia
MNNEETFYQAMRRQGVTRRSFLKYCSLAATSLGLGAGMAPKIAWALENKPRIPVVWIHGL WP_001295940.1
MULTISPECIES: hydrogenase [Enterobacteriaceae] Length: 372\nNumber of predicted TMHs: 2\nExp number of AAs in TMHs: 37.2608899999999\nExp number, first 60 AAs: 19.52923\nTotal prob of N-in: 0.94351\nPOSSIBLE N-term signal sequence\ninside 1 19\nTMhelix 20 39\noutside 40 325\nTMhelix 326 348\ninside 349 372

5753 GCF_001902655.1_ASM190265v1 Escherichia coli Proteobacteria; Gammaproteobacteria; Enterobacterales; Enterobacteriaceae; Escherichia
MTGDNTLIHSHGINRRDFMKLCAALAATMGLSSKAAAEMAESVTNPQRPPVIWIGAQECT WP_000145410.1
MULTISPECIES: hydrogenase-2 small chain [Proteobacteria] Length: 372\nNumber of predicted TMHs: 1\nExp number of AAs in TMHs: 23.72226\nExp number, first 60 AAs: 0.22038\nTotal prob of N-in: 0.12994\noutside 1 331\nTMhelix 332 354\ninside 355 372

5754 GCF_001901425.1_ASM190142v1 Escherichia coli Proteobacteria; Gammaproteobacteria; Enterobacterales; Enterobacteriaceae; Escherichia
MTGDNTLIHSHGINRRDFMKLCAALAATMGLSSKAAAEMAESVTNPQRPPVIWIGAQECT WP_000145410.1
MULTISPECIES: hydrogenase-2 small chain [Proteobacteria] Length: 372\nNumber of predicted TMHs: 1\nExp number of AAs in TMHs: 23.72226\nExp number, first 60 AAs: 0.22038\nTotal prob of N-in: 0.12994\noutside 1 331\nTMhelix 332 354\ninside 355 372

5755 GCF_001901465.1_ASM190146v1 Escherichia coli Proteobacteria; Gammaproteobacteria; Enterobacterales; Enterobacteriaceae; Escherichia
MTGDNTLIHSHGINRRDFMKLCAALAATMGLSSKAAAEMAESVTNPQRPPVIWIGAQECT WP_073535658.1
hydrogenase 2 small subunit [Escherichia coli] Length: 372\nNumber of predicted TMHs: 1\nExp number of AAs in TMHs: 23.78776\nExp number, first 60 AAs: 0.2204\nTotal prob of N-in: 0.13104\noutside 1 331\nTMhelix 332 354\ninside 355 372

5756 GCF_001907385.1_ASM190738v1 Escherichia coli Proteobacteria; Gammaproteobacteria; Enterobacterales; Enterobacteriaceae; Escherichia
MTGDNTLIHSHGINRRDFMKLCAALAATMGLSSKAAAEMAESVTNPQRPPVIWIGAQECT WP_038339805.1
hydrogenase 2 small subunit [Escherichia coli] Length: 372\nNumber of predicted TMHs: 1\nExp number of

AAs in TMHs: 23.86829\nExp number, first 60 AAs:  0.22037\nTotal prob of N-in:   0.12535\noutside   1   332\nTMhelix   333   355\ninside   356   372

5757 GCF_001907335.1_ASM190733v1 Escherichia coli Proteobacteria; Gammaproteobacteria; Enterobacterales; Enterobacteriaceae; Escherichia MNNEETFYQAMRRQGVTRRSFLKYCSLAATSLGLGAGMAPKIAWALENKPRIPVVWIHGL WP_001058323.1 MULTISPECIES: hydrogenase-1 small chain [Proteobacteria] Length: 372\nNumber of predicted TMHs: 2\nExp number of AAs in TMHs: 37.2608499999999\nExp number, first 60 AAs:  19.52923\nTotal prob of N-in: 0.94351\nPOSSIBLE N-term signal sequence\ninside   1   19\nTMhelix   20   39\noutside   40   325\nTMhelix   326   348\ninside   349   372

5758 GCF_001907455.1_ASM190745v1 Escherichia coli Proteobacteria; Gammaproteobacteria; Enterobacterales; Enterobacteriaceae; Escherichia MTGDNTLIHSHGINRRDFMKLCAALAATMGLSSKAAAEMAESVTNPQRPPVIWIGAQECT WP_000145410.1 MULTISPECIES: hydrogenase-2 small chain [Proteobacteria] Length: 372\nNumber of predicted TMHs: 1\nExp number of AAs in TMHs: 23.72226\nExp number, first 60 AAs:  0.22038\nTotal prob of N-in:   0.12994\noutside   1   331\nTMhelix   332   354\ninside   355   372

5759 GCF_001907455.1_ASM190745v1 Escherichia coli Proteobacteria; Gammaproteobacteria; Enterobacterales; Enterobacteriaceae; Escherichia MNNEETFYQAMRRQGVTRRSFLKYCSLAATSLGLGAGMAPKIAWALENKPRIPVVWIHGL WP_001058323.1 MULTISPECIES: hydrogenase-1 small chain [Proteobacteria] Length: 372\nNumber of predicted TMHs: 2\nExp number of AAs in TMHs: 37.2608499999999\nExp number, first 60 AAs:  19.52923\nTotal prob of N-in: 0.94351\nPOSSIBLE N-term signal sequence\ninside   1   19\nTMhelix   20   39\noutside   40   325\nTMhelix   326   348\ninside   349   372

5760 GCF_001907325.1_ASM190732v1 Escherichia coli Proteobacteria; Gammaproteobacteria; Enterobacterales; Enterobacteriaceae; Escherichia MNNEETFYQAMRRQGVTRRSFLKYCSLAATSLGLGAGMAPKIAWALENKPRIPVVWIHGL WP_001058323.1 MULTISPECIES: hydrogenase-1 small chain [Proteobacteria] Length: 372\nNumber of predicted TMHs: 2\nExp number of AAs in TMHs: 37.2608499999999\nExp number, first 60 AAs:  19.52923\nTotal prob of N-in: 0.94351\nPOSSIBLE N-term signal sequence\ninside   1   19\nTMhelix   20   39\noutside   40   325\nTMhelix   326   348\ninside   349   372

5761 GCF_001902735.1_ASM190273v1 Escherichia coli Proteobacteria; Gammaproteobacteria; Enterobacterales; Enterobacteriaceae; Escherichia MNNEETFYQAMRRQGVTRRSFLKYCSLAATSLGLGAGMAPKIAWALENKPRIPVVWIHGL WP_001058323.1 MULTISPECIES: hydrogenase-1 small chain [Proteobacteria] Length: 372\nNumber of predicted TMHs: 2\nExp number of AAs in TMHs: 37.2608499999999\nExp number, first 60 AAs:  19.52923\nTotal prob of N-in: 0.94351\nPOSSIBLE N-term signal sequence\ninside   1   19\nTMhelix   20   39\noutside   40   325\nTMhelix   326   348\ninside   349   372

5762 GCF_001902675.1_ASM190267v1 Escherichia coli Proteobacteria; Gammaproteobacteria; Enterobacterales; Enterobacteriaceae; Escherichia MNNEETFYQAMRRQGVTRRSFLKYCSLAATSLGLGAGMAPKIAWALENKPRIPVVWIHGL WP_001058323.1 MULTISPECIES: hydrogenase-1 small chain [Proteobacteria] Length: 372\nNumber of predicted TMHs: 2\nExp number of AAs in TMHs: 37.2608499999999\nExp number, first 60 AAs:  19.52923\nTotal prob of N-in: 0.94351\nPOSSIBLE N-term signal sequence\ninside   1   19\nTMhelix   20   39\noutside   40   325\nTMhelix   326   348\ninside   349   372

5763 GCF_001907495.1_ASM190749v1 Escherichia coli Proteobacteria; Gammaproteobacteria; Enterobacterales; Enterobacteriaceae; Escherichia MTGDNTLIHSHGINRRDFMKLCAALAATMGLSSKAAAEMAESVTNPQRPPVIWIGAQECT WP_000145410.1 MULTISPECIES: hydrogenase-2 small chain [Proteobacteria] Length: 372\nNumber of predicted TMHs: 1\nExp number of AAs in TMHs: 23.72226\nExp number, first 60 AAs:  0.22038\nTotal prob of N-in:   0.12994\noutside   1   331\nTMhelix   332   354\ninside   355   372

5764 GCF_001907495.1_ASM190749v1 Escherichia coli Proteobacteria; Gammaproteobacteria; Enterobacterales; Enterobacteriaceae; Escherichia MNNEETFYQAMRRQGVTRRSFLKYCSLAATSLGLGAGMAPKIAWALENKPRIPVVWIHGL WP_001058323.1 MULTISPECIES: hydrogenase-1 small chain [Proteobacteria] Length: 372\nNumber of predicted TMHs:

2\nExp number of AAs in TMHs: 37.2608499999999\nExp number, first 60 AAs: 19.52923\nTotal prob of N-in: 0.94351\nPOSSIBLE N-term signal sequence\ninside 1 19\nTMhelix 20 39\noutside 40 325\nTMhelix 326 348\ninside 349 372

5765 GCF_001907435.1_ASM190743v1 Escherichia coli Proteobacteria; Gammaproteobacteria; Enterobacterales; Enterobacteriaceae; Escherichia MNNEETFYQAMRRQGVTRRSFLKYCSLAATSLGLGAGMAPKIAWALENKPRIPVVWIHGL WP_001058323.1 MULTISPECIES: hydrogenase-1 small chain [Proteobacteria] Length: 372\nNumber of predicted TMHs: 2\nExp number of AAs in TMHs: 37.2608499999999\nExp number, first 60 AAs: 19.52923\nTotal prob of N-in: 0.94351\nPOSSIBLE N-term signal sequence\ninside 1 19\nTMhelix 20 39\noutside 40 325\nTMhelix 326 348\ninside 349 372

5766 GCF_001907535.1_ASM190753v1 Escherichia coli Proteobacteria; Gammaproteobacteria; Enterobacterales; Enterobacteriaceae; Escherichia MNNEETFYQAMRRQGVTRRSFLKYCSLAATSLGLGAGMAPKIAWALENKPRIPVVWIHGL WP_001058323.1 MULTISPECIES: hydrogenase-1 small chain [Proteobacteria] Length: 372\nNumber of predicted TMHs: 2\nExp number of AAs in TMHs: 37.2608499999999\nExp number, first 60 AAs: 19.52923\nTotal prob of N-in: 0.94351\nPOSSIBLE N-term signal sequence\ninside 1 19\nTMhelix 20 39\noutside 40 325\nTMhelix 326 348\ninside 349 372

5767 GCF_001907535.1_ASM190753v1 Escherichia coli Proteobacteria; Gammaproteobacteria; Enterobacterales; Enterobacteriaceae; Escherichia MTGDNTLIHSHGINRRDFMKLCAALAATMGLSSKAAAEMAESVTNPQRPPVIWIGAQECT WP_038339805.1 hydrogenase 2 small subunit [Escherichia coli] Length: 372\nNumber of predicted TMHs: 1\nExp number of AAs in TMHs: 23.86829\nExp number, first 60 AAs: 0.22037\nTotal prob of N-in: 0.12535\noutside 1 332\nTMhelix 333 355\ninside 356 372

5768 GCF_001910475.1_ASM191047v1 Escherichia coli Proteobacteria; Gammaproteobacteria; Enterobacterales; Enterobacteriaceae; Escherichia MNNEETFYQAMRRQGVTRRSFLKYCSLAATSLGLGAGMAPKIAWALENKPRIPVVWIHGL WP_001058323.1 MULTISPECIES: hydrogenase-1 small chain [Proteobacteria] Length: 372\nNumber of predicted TMHs: 2\nExp number of AAs in TMHs: 37.2608499999999\nExp number, first 60 AAs: 19.52923\nTotal prob of N-in: 0.94351\nPOSSIBLE N-term signal sequence\ninside 1 19\nTMhelix 20 39\noutside 40 325\nTMhelix 326 348\ninside 349 372

5769 GCF_001910905.1_ASM191090v1 Escherichia coli Proteobacteria; Gammaproteobacteria; Enterobacterales; Enterobacteriaceae; Escherichia MTGDNTLIHSHGINRRDFMKLCAALAATMGLSSKAAAEMAESVTNPQRPPVIWIGAQECT WP_000145410.1 MULTISPECIES: hydrogenase-2 small chain [Proteobacteria] Length: 372\nNumber of predicted TMHs: 1\nExp number of AAs in TMHs: 23.72226\nExp number, first 60 AAs: 0.22038\nTotal prob of N-in: 0.12994\noutside 1 331\nTMhelix 332 354\ninside 355 372

5770 GCF_001910995.1_ASM191099v1 Escherichia coli Proteobacteria; Gammaproteobacteria; Enterobacterales; Enterobacteriaceae; Escherichia MTGDNTLIHSHGINRRDFMKLCAALAATMGLSSKAAAEMAESVTNPQRPPVIWIGAQECT WP_000145410.1 MULTISPECIES: hydrogenase-2 small chain [Proteobacteria] Length: 372\nNumber of predicted TMHs: 1\nExp number of AAs in TMHs: 23.72226\nExp number, first 60 AAs: 0.22038\nTotal prob of N-in: 0.12994\noutside 1 331\nTMhelix 332 354\ninside 355 372

5771 GCF_001910935.1_ASM191093v1 Escherichia coli Proteobacteria; Gammaproteobacteria; Enterobacterales; Enterobacteriaceae; Escherichia MTGDNTLIHSHGINRRDFMKLCAALAATMGLSSKAAAEMAESVTNPQRPPVIWIGAQECT WP_000145410.1 MULTISPECIES: hydrogenase-2 small chain [Proteobacteria] Length: 372\nNumber of predicted TMHs: 1\nExp number of AAs in TMHs: 23.72226\nExp number, first 60 AAs: 0.22038\nTotal prob of N-in: 0.12994\noutside 1 331\nTMhelix 332 354\ninside 355 372

5772 GCF_001910915.1_ASM191091v1 Escherichia coli Proteobacteria; Gammaproteobacteria; Enterobacterales; Enterobacteriaceae; Escherichia MTGDNTLIHSHGINRRDFMKLCAALAATMGLSSKAAAEMAESVTNPQRPPVIWIGAQECT WP_000145410.1 MULTISPECIES: hydrogenase-2 small chain [Proteobacteria] Length: 372\nNumber of predicted TMHs:

1\nExp number of AAs in TMHs: 23.72226\nExp number, first 60 AAs: 0.22038\nTotal prob of N-in: 0.12994\noutside 1 331\nTMhelix 332 354\ninside 355 372

5773 GCF_001911005.1_ASM191100v1 Escherichia coli Proteobacteria; Gammaproteobacteria; Enterobacterales; Enterobacteriaceae; Escherichia
MTGDNTLIHSHGINRRDFMKLCAALAATMGLSSKAAAEMAESVTNPQRPPVIWIGAQECT WP_000145410.1
MULTISPECIES: hydrogenase-2 small chain [Proteobacteria] Length: 372\nNumber of predicted TMHs: 1\nExp number of AAs in TMHs: 23.72226\nExp number, first 60 AAs: 0.22038\nTotal prob of N-in: 0.12994\noutside 1 331\nTMhelix 332 354\ninside 355 372

5774 GCF_001910935.1_ASM191093v1 Escherichia coli Proteobacteria; Gammaproteobacteria; Enterobacterales; Enterobacteriaceae; Escherichia
MNNEETFYQAMRRQGVTRRSFLKYCSLAATSLGLGAGMAPKIAWALENKPRIPVVWIHGL WP_001058323.1
MULTISPECIES: hydrogenase-1 small chain [Proteobacteria] Length: 372\nNumber of predicted TMHs: 2\nExp number of AAs in TMHs: 37.2608499999999\nExp number, first 60 AAs: 19.52923\nTotal prob of N-in: 0.94351\nPOSSIBLE N-term signal sequence\ninside 1 19\nTMhelix 20 39\noutside 40 325\nTMhelix 326 348\ninside 349 372

5775 GCF_001910905.1_ASM191090v1 Escherichia coli Proteobacteria; Gammaproteobacteria; Enterobacterales; Enterobacteriaceae; Escherichia
MNNEETFYQAMRRQGVTRRSFLKYCSLAATSLGLGAGMAPKIAWALENKPRIPVVWIHGL WP_001058320.1
hydrogenase [Escherichia coli] Length: 372\nNumber of predicted TMHs: 2\nExp number of AAs in TMHs: 37.26082\nExp number, first 60 AAs: 19.52923\nTotal prob of N-in: 0.94351\nPOSSIBLE N-term signal sequence\ninside 1 19\nTMhelix 20 39\noutside 40 325\nTMhelix 326 348\ninside 349 372

5776 GCF_001910845.1_ASM191084v1 Escherichia coli Proteobacteria; Gammaproteobacteria; Enterobacterales; Enterobacteriaceae; Escherichia
MNNEETFYQAMRRQGVTRRSFLKYCSLAATSLGLGAGMAPKIAWALENKPRIPVVWIHGL WP_001058323.1
MULTISPECIES: hydrogenase-1 small chain [Proteobacteria] Length: 372\nNumber of predicted TMHs: 2\nExp number of AAs in TMHs: 37.2608499999999\nExp number, first 60 AAs: 19.52923\nTotal prob of N-in: 0.94351\nPOSSIBLE N-term signal sequence\ninside 1 19\nTMhelix 20 39\noutside 40 325\nTMhelix 326 348\ninside 349 372

5777 GCF_001907545.1_ASM190754v1 Escherichia coli Proteobacteria; Gammaproteobacteria; Enterobacterales; Enterobacteriaceae; Escherichia
MTGDNTLIHSHGINRRDFMKLCAALAATMGLSSKAAAEMAESVTNPQRPPVIWIGAQECT WP_038339805.1
hydrogenase 2 small subunit [Escherichia coli] Length: 372\nNumber of predicted TMHs: 1\nExp number of AAs in TMHs: 23.86829\nExp number, first 60 AAs: 0.22037\nTotal prob of N-in: 0.12535\noutside 1 332\nTMhelix 333 355\ninside 356 372

5778 GCF_001910825.1_ASM191082v1 Escherichia coli Proteobacteria; Gammaproteobacteria; Enterobacterales; Enterobacteriaceae; Escherichia
MNNEETFYQAMRRQGVTRRSFLKYCSLAATSLGLGAGMAPKIAWALENKPRIPVVWIHGL WP_001058323.1
MULTISPECIES: hydrogenase-1 small chain [Proteobacteria] Length: 372\nNumber of predicted TMHs: 2\nExp number of AAs in TMHs: 37.2608499999999\nExp number, first 60 AAs: 19.52923\nTotal prob of N-in: 0.94351\nPOSSIBLE N-term signal sequence\ninside 1 19\nTMhelix 20 39\noutside 40 325\nTMhelix 326 348\ninside 349 372

5779 GCF_001911065.1_ASM191106v1 Escherichia coli Proteobacteria; Gammaproteobacteria; Enterobacterales; Enterobacteriaceae; Escherichia
MTGDNTLIHSHGINRRDFMKLCAALAATMGLSSKAAAEMAESVTNPQRPPVIWIGAQECT WP_000145410.1
MULTISPECIES: hydrogenase-2 small chain [Proteobacteria] Length: 372\nNumber of predicted TMHs: 1\nExp number of AAs in TMHs: 23.72226\nExp number, first 60 AAs: 0.22038\nTotal prob of N-in: 0.12994\noutside 1 331\nTMhelix 332 354\ninside 355 372

5780 GCF_001911065.1_ASM191106v1 Escherichia coli Proteobacteria; Gammaproteobacteria; Enterobacterales; Enterobacteriaceae; Escherichia
MNNEETFYQAMRRQGVTRRSFLKYCSLAATSLGLGAGMAPKIAWALENKPRIPVVWIHGL WP_001353311.1
MULTISPECIES: hydrogenase [Escherichia] Length: 372\nNumber of predicted TMHs: 2\nExp number of AAs in TMHs: 37.2638699999999\nExp number, first 60 AAs: 19.52924\nTotal prob of N-in: 0.94351\nPOSSIBLE N-term signal sequence\ninside 1 19\nTMhelix 20 39\noutside 40 325\nTMhelix 326 348\ninside 349 372

5781 GCF_001910915.1_ASM191091v1 Escherichia coli Proteobacteria; Gammaproteobacteria; Enterobacterales; Enterobacteriaceae; Escherichia
MNNEETFYQAMRRQGVTRRSFLKYCSLAATSLGLGAGMAPKIAWALENKPRIPVVWIHGL WP_001058323.1
MULTISPECIES: hydrogenase-1 small chain [Proteobacteria] Length: 372\nNumber of predicted TMHs: 2\nExp number of AAs in TMHs: 37.2608499999999\nExp number, first 60 AAs: 19.52923\nTotal prob of N-in: 0.94351\nPOSSIBLE N-term signal sequence\ninside 1 19\nTMhelix 20 39\noutside 40 325\nTMhelix 326 348\ninside 349 372

5782 GCF_001911005.1_ASM191100v1 Escherichia coli Proteobacteria; Gammaproteobacteria; Enterobacterales; Enterobacteriaceae; Escherichia
MNNEETFYQAMRRQGVTRRSFLKYCSLAATSLGLGAGMAPKIAWALENKPRIPVVWIHGL WP_001058323.1
MULTISPECIES: hydrogenase-1 small chain [Proteobacteria] Length: 372\nNumber of predicted TMHs: 2\nExp number of AAs in TMHs: 37.2608499999999\nExp number, first 60 AAs: 19.52923\nTotal prob of N-in: 0.94351\nPOSSIBLE N-term signal sequence\ninside 1 19\nTMhelix 20 39\noutside 40 325\nTMhelix 326 348\ninside 349 372

5783 GCF_001907935.1_ASM190793v1 Escherichia coli Proteobacteria; Gammaproteobacteria; Enterobacterales; Enterobacteriaceae; Escherichia
MNNEETFYQAMRRQGVTRRSFLKYCSLAATSLGLGAGMAPKIAWALENKPRIPVVWIHGL WP_001295940.1
MULTISPECIES: hydrogenase [Enterobacteriaceae] Length: 372\nNumber of predicted TMHs: 2\nExp number of AAs in TMHs: 37.2608899999999\nExp number, first 60 AAs: 19.52923\nTotal prob of N-in: 0.94351\nPOSSIBLE N-term signal sequence\ninside 1 19\nTMhelix 20 39\noutside 40 325\nTMhelix 326 348\ninside 349 372

5784 GCF_001910995.1_ASM191099v1 Escherichia coli Proteobacteria; Gammaproteobacteria; Enterobacterales; Enterobacteriaceae; Escherichia
MNNEETFYQAMRRQGVTRRSFLKYCSLAATSLGLGAGMAPKIAWALENKPRIPVVWIHGL WP_001353311.1
MULTISPECIES: hydrogenase [Escherichia] Length: 372\nNumber of predicted TMHs: 2\nExp number of AAs in TMHs: 37.2638699999999\nExp number, first 60 AAs: 19.52924\nTotal prob of N-in: 0.94351\nPOSSIBLE N-term signal sequence\ninside 1 19\nTMhelix 20 39\noutside 40 325\nTMhelix 326 348\ninside 349 372

5785 GCF_001911095.1_ASM191109v1 Escherichia coli Proteobacteria; Gammaproteobacteria; Enterobacterales; Enterobacteriaceae; Escherichia
MTGDNTLIHSHGINRRDFMKLCAALAATMGLSSKAAAEMAESVTNPQRPPVIWIGAQECT WP_000145410.1
MULTISPECIES: hydrogenase-2 small chain [Proteobacteria] Length: 372\nNumber of predicted TMHs: 1\nExp number of AAs in TMHs: 23.72226\nExp number, first 60 AAs: 0.22038\nTotal prob of N-in: 0.12994\noutside 1 331\nTMhelix 332 354\ninside 355 372

5786 GCF_001911225.1_ASM191122v1 Escherichia coli Proteobacteria; Gammaproteobacteria; Enterobacterales; Enterobacteriaceae; Escherichia
MTGDNTLIHSHGINRRDFMKLCAALAATMGLSSKAAAEMAESVTNPQRPPVIWIGAQECT WP_000145410.1
MULTISPECIES: hydrogenase-2 small chain [Proteobacteria] Length: 372\nNumber of predicted TMHs: 1\nExp number of AAs in TMHs: 23.72226\nExp number, first 60 AAs: 0.22038\nTotal prob of N-in: 0.12994\noutside 1 331\nTMhelix 332 354\ninside 355 372

5787 GCF_001911175.1_ASM191117v1 Escherichia coli Proteobacteria; Gammaproteobacteria; Enterobacterales; Enterobacteriaceae; Escherichia
MTGDNTLIHSHGINRRDFMKLCAALAATMGLSSKAAAEMAESVTNPQRPPVIWIGAQECT WP_000145410.1
MULTISPECIES: hydrogenase-2 small chain [Proteobacteria] Length: 372\nNumber of predicted TMHs: 1\nExp number of AAs in TMHs: 23.72226\nExp number, first 60 AAs: 0.22038\nTotal prob of N-in: 0.12994\noutside 1 331\nTMhelix 332 354\ninside 355 372

5788 GCF_001911155.1_ASM191115v1 Escherichia coli Proteobacteria; Gammaproteobacteria; Enterobacterales; Enterobacteriaceae; Escherichia
MTGDNTLIHSHGINRRDFMKLCAALAATMGLSSKAAAEMAESVTNPQRPPVIWIGAQECT WP_000145410.1
MULTISPECIES: hydrogenase-2 small chain [Proteobacteria] Length: 372\nNumber of predicted TMHs: 1\nExp number of AAs in TMHs: 23.72226\nExp number, first 60 AAs: 0.22038\nTotal prob of N-in: 0.12994\noutside 1 331\nTMhelix 332 354\ninside 355 372

5789 GCF_001911315.1_ASM191131v1 Escherichia coli Proteobacteria; Gammaproteobacteria; Enterobacterales; Enterobacteriaceae; Escherichia

MTGDNTLIHSHGINRRDFMKLCAALAATMGLSSKAAAEMAESVTNPQRPPVIWIGAQECT WP_000145410.1 MULTISPECIES: hydrogenase-2 small chain [Proteobacteria] Length: 372\nNumber of predicted TMHs: 1\nExp number of AAs in TMHs: 23.72226\nExp number, first 60 AAs: 0.22038\nTotal prob of N-in: 0.12994\noutside 1 331\nTMhelix 332 354\ninside 355 372

5790 GCF_001911245.1_ASM191124v1 Escherichia coli Proteobacteria; Gammaproteobacteria; Enterobacterales; Enterobacteriaceae; Escherichia MTGDNTLIHSHGINRRDFMKLCAALAATMGLSSKAAAEMAESVTNPQRPPVIWIGAQECT WP_000145410.1 MULTISPECIES: hydrogenase-2 small chain [Proteobacteria] Length: 372\nNumber of predicted TMHs: 1\nExp number of AAs in TMHs: 23.72226\nExp number, first 60 AAs: 0.22038\nTotal prob of N-in: 0.12994\noutside 1 331\nTMhelix 332 354\ninside 355 372

5791 GCF_001911305.1_ASM191130v1 Escherichia coli Proteobacteria; Gammaproteobacteria; Enterobacterales; Enterobacteriaceae; Escherichia MTGDNTLIHSHGINRRDFMKLCAALAATMGLSSKAAAEMAESVTNPQRPPVIWIGAQECT WP_000145410.1 MULTISPECIES: hydrogenase-2 small chain [Proteobacteria] Length: 372\nNumber of predicted TMHs: 1\nExp number of AAs in TMHs: 23.72226\nExp number, first 60 AAs: 0.22038\nTotal prob of N-in: 0.12994\noutside 1 331\nTMhelix 332 354\ninside 355 372

5792 GCF_001911315.1_ASM191131v1 Escherichia coli Proteobacteria; Gammaproteobacteria; Enterobacterales; Enterobacteriaceae; Escherichia MNNEETFYQAMRRQGVTRRSFLKYCSLAATSLGLGAGMAPKIAWALENKPRIPVVWIHGL WP_001058323.1 MULTISPECIES: hydrogenase-1 small chain [Proteobacteria] Length: 372\nNumber of predicted TMHs: 2\nExp number of AAs in TMHs: 37.2608499999999\nExp number, first 60 AAs: 19.52923\nTotal prob of N-in: 0.94351\nPOSSIBLE N-term signal sequence\ninside 1 19\nTMhelix 20 39\noutside 40 325\nTMhelix 326 348\ninside 349 372

5793 GCF_001911465.1_ASM191146v1 Escherichia coli Proteobacteria; Gammaproteobacteria; Enterobacterales; Enterobacteriaceae; Escherichia MTGDNTLIHSHGINRRDFMKLCAALAATMGLSSKAAAEMAESVTNPQRPPVIWIGAQECT WP_000145410.1 MULTISPECIES: hydrogenase-2 small chain [Proteobacteria] Length: 372\nNumber of predicted TMHs: 1\nExp number of AAs in TMHs: 23.72226\nExp number, first 60 AAs: 0.22038\nTotal prob of N-in: 0.12994\noutside 1 331\nTMhelix 332 354\ninside 355 372

5794 GCF_001911465.1_ASM191146v1 Escherichia coli Proteobacteria; Gammaproteobacteria; Enterobacterales; Enterobacteriaceae; Escherichia MNNEETFYQAMRRQGVTRRSFLKYCSLAATSLGLGAGMAPKIAWALENKPRIPVVWIHGL WP_001058323.1 MULTISPECIES: hydrogenase-1 small chain [Proteobacteria] Length: 372\nNumber of predicted TMHs: 2\nExp number of AAs in TMHs: 37.2608499999999\nExp number, first 60 AAs: 19.52923\nTotal prob of N-in: 0.94351\nPOSSIBLE N-term signal sequence\ninside 1 19\nTMhelix 20 39\noutside 40 325\nTMhelix 326 348\ninside 349 372

5795 GCF_001911155.1_ASM191115v1 Escherichia coli Proteobacteria; Gammaproteobacteria; Enterobacterales; Enterobacteriaceae; Escherichia MNNEETFYQAMRRQGVTRRSFLKYCSLAATSLGLGAGMAPKIAWALENKPRIPVVWIHGL WP_001058323.1 MULTISPECIES: hydrogenase-1 small chain [Proteobacteria] Length: 372\nNumber of predicted TMHs: 2\nExp number of AAs in TMHs: 37.2608499999999\nExp number, first 60 AAs: 19.52923\nTotal prob of N-in: 0.94351\nPOSSIBLE N-term signal sequence\ninside 1 19\nTMhelix 20 39\noutside 40 325\nTMhelix 326 348\ninside 349 372

5796 GCF_001911335.1_ASM191133v1 Escherichia coli Proteobacteria; Gammaproteobacteria; Enterobacterales; Enterobacteriaceae; Escherichia MNNEETFYQAMRRQGVTRRSFLKYCSLAATSLGLGAGMAPKIAWALENKPRIPVVWIHGL WP_001295940.1 MULTISPECIES: hydrogenase [Enterobacteriaceae] Length: 372\nNumber of predicted TMHs: 2\nExp number of AAs in TMHs: 37.2608899999999\nExp number, first 60 AAs: 19.52923\nTotal prob of N-in: 0.94351\nPOSSIBLE N-term signal sequence\ninside 1 19\nTMhelix 20 39\noutside 40 325\nTMhelix 326 348\ninside 349 372

5797 GCF_001911545.1_ASM191154v1 Escherichia coli Proteobacteria; Gammaproteobacteria; Enterobacterales; Enterobacteriaceae; Escherichia MTGDNTLIHSHGINRRDFMKLCAALAATMGLSSKAAAEMAESVTNPQRPPVIWIGAQECT WP_000145410.1

MULTISPECIES: hydrogenase-2 small chain [Proteobacteria] Length: 372\nNumber of predicted TMHs: 1\nExp number of AAs in TMHs: 23.72226\nExp number, first 60 AAs: 0.22038\nTotal prob of N-in: 0.12994\noutside 1 331\nTMhelix 332 354\ninside 355 372

5798 GCF_001911495.1_ASM191149v1 Escherichia coli Proteobacteria; Gammaproteobacteria; Enterobacterales; Enterobacteriaceae; Escherichia
MTGDNTLIHSHGINRRDFMKLCAALAATMGLSSKAAAEMAESVTNPQRPPVIWIGAQECT WP_000145410.1
MULTISPECIES: hydrogenase-2 small chain [Proteobacteria] Length: 372\nNumber of predicted TMHs: 1\nExp number of AAs in TMHs: 23.72226\nExp number, first 60 AAs: 0.22038\nTotal prob of N-in: 0.12994\noutside 1 331\nTMhelix 332 354\ninside 355 372

5799 GCF_001911595.1_ASM191159v1 Escherichia coli Proteobacteria; Gammaproteobacteria; Enterobacterales; Enterobacteriaceae; Escherichia
MTGDNTLIHSHGINRRDFMKLCAALAATMGLSSKAAAEMAESVTNPQRPPVIWIGAQECT WP_000145410.1
MULTISPECIES: hydrogenase-2 small chain [Proteobacteria] Length: 372\nNumber of predicted TMHs: 1\nExp number of AAs in TMHs: 23.72226\nExp number, first 60 AAs: 0.22038\nTotal prob of N-in: 0.12994\noutside 1 331\nTMhelix 332 354\ninside 355 372

5800 GCF_001911555.1_ASM191155v1 Escherichia coli Proteobacteria; Gammaproteobacteria; Enterobacterales; Enterobacteriaceae; Escherichia
MTGDNTLIHSHGINRRDFMKLCAALAATMGLSSKAAAEMAESVTNPQRPPVIWIGAQECT WP_000145410.1
MULTISPECIES: hydrogenase-2 small chain [Proteobacteria] Length: 372\nNumber of predicted TMHs: 1\nExp number of AAs in TMHs: 23.72226\nExp number, first 60 AAs: 0.22038\nTotal prob of N-in: 0.12994\noutside 1 331\nTMhelix 332 354\ninside 355 372

5801 GCF_001911545.1_ASM191154v1 Escherichia coli Proteobacteria; Gammaproteobacteria; Enterobacterales; Enterobacteriaceae; Escherichia
MNNEETFYQAMRRQGVTRRSFLKYCSLAATSLGLGAGMAPKIAWALENKPRIPVVWIHGL WP_001058323.1
MULTISPECIES: hydrogenase-1 small chain [Proteobacteria] Length: 372\nNumber of predicted TMHs: 2\nExp number of AAs in TMHs: 37.2608499999999\nExp number, first 60 AAs: 19.52923\nTotal prob of N-in: 0.94351\nPOSSIBLE N-term signal sequence\ninside 1 19\nTMhelix 20 39\noutside 40 325\nTMhelix 326 348\ninside 349 372

5802 GCF_001911665.1_ASM191166v1 Escherichia coli Proteobacteria; Gammaproteobacteria; Enterobacterales; Enterobacteriaceae; Escherichia
MTGDNTLIHSHGINRRDFMKLCAALAATMGLSSKAAAEMAESVTNPQRPPVIWIGAQECT WP_000145410.1
MULTISPECIES: hydrogenase-2 small chain [Proteobacteria] Length: 372\nNumber of predicted TMHs: 1\nExp number of AAs in TMHs: 23.72226\nExp number, first 60 AAs: 0.22038\nTotal prob of N-in: 0.12994\noutside 1 331\nTMhelix 332 354\ninside 355 372

5803 GCF_001911635.1_ASM191163v1 Escherichia coli Proteobacteria; Gammaproteobacteria; Enterobacterales; Enterobacteriaceae; Escherichia
MTGDNTLIHSHGINRRDFMKLCAALAATMGLSSKAAAEMAESVTNPQRPPVIWIGAQECT WP_000145410.1
MULTISPECIES: hydrogenase-2 small chain [Proteobacteria] Length: 372\nNumber of predicted TMHs: 1\nExp number of AAs in TMHs: 23.72226\nExp number, first 60 AAs: 0.22038\nTotal prob of N-in: 0.12994\noutside 1 331\nTMhelix 332 354\ninside 355 372

5804 GCF_001911695.1_ASM191169v1 Escherichia coli Proteobacteria; Gammaproteobacteria; Enterobacterales; Enterobacteriaceae; Escherichia
MTGDNTLIHSHGINRRDFMKLCAALAATMGLSSKAAAEMAESVTNPQRPPVIWIGAQECT WP_000145410.1
MULTISPECIES: hydrogenase-2 small chain [Proteobacteria] Length: 372\nNumber of predicted TMHs: 1\nExp number of AAs in TMHs: 23.72226\nExp number, first 60 AAs: 0.22038\nTotal prob of N-in: 0.12994\noutside 1 331\nTMhelix 332 354\ninside 355 372

5805 GCF_001911755.1_ASM191175v1 Escherichia coli Proteobacteria; Gammaproteobacteria; Enterobacterales; Enterobacteriaceae; Escherichia
MTGDNTLIHSHGINRRDFMKLCAALAATMGLSSKAAAEMAESVTNPQRPPVIWIGAQECT WP_000145410.1
MULTISPECIES: hydrogenase-2 small chain [Proteobacteria] Length: 372\nNumber of predicted TMHs: 1\nExp number of AAs in TMHs: 23.72226\nExp number, first 60 AAs: 0.22038\nTotal prob of N-in: 0.12994\noutside 1 331\nTMhelix 332 354\ninside 355 372

5806 GCF_001911555.1_ASM191155v1 Escherichia coli Proteobacteria; Gammaproteobacteria; Enterobacterales; Enterobacteriaceae; Escherichia
MNNEETFYQAMRRQGVTRRSFLKYCSLAATSLGLGAGMAPKIAWALENKPRIPVVWIHGL WP_001058323.1
MULTISPECIES: hydrogenase-1 small chain [Proteobacteria] Length: 372\nNumber of predicted TMHs: 2\nExp number of AAs in TMHs: 37.2608499999999\nExp number, first 60 AAs: 19.52923\nTotal prob of N-in: 0.94351\nPOSSIBLE N-term signal sequence\ninside 1 19\nTMhelix 20 39\noutside 40 325\nTMhelix 326 348\ninside 349 372

5807 GCF_001911595.1_ASM191159v1 Escherichia coli Proteobacteria; Gammaproteobacteria; Enterobacterales; Enterobacteriaceae; Escherichia
MNNEETFYQAMRRQGVTRRSFLKYCSLAATSLGLGAGMAPKIAWALENKPRIPVVWIHGL WP_001058323.1
MULTISPECIES: hydrogenase-1 small chain [Proteobacteria] Length: 372\nNumber of predicted TMHs: 2\nExp number of AAs in TMHs: 37.2608499999999\nExp number, first 60 AAs: 19.52923\nTotal prob of N-in: 0.94351\nPOSSIBLE N-term signal sequence\ninside 1 19\nTMhelix 20 39\noutside 40 325\nTMhelix 326 348\ninside 349 372

5808 GCF_001911745.1_ASM191174v1 Escherichia coli Proteobacteria; Gammaproteobacteria; Enterobacterales; Enterobacteriaceae; Escherichia
MNNEETFYQAMRRQGVTRRSFLKYCSLAATSLGLGAGMAPKIAWALENKPRIPVVWIHGL WP_001058323.1
MULTISPECIES: hydrogenase-1 small chain [Proteobacteria] Length: 372\nNumber of predicted TMHs: 2\nExp number of AAs in TMHs: 37.2608499999999\nExp number, first 60 AAs: 19.52923\nTotal prob of N-in: 0.94351\nPOSSIBLE N-term signal sequence\ninside 1 19\nTMhelix 20 39\noutside 40 325\nTMhelix 326 348\ninside 349 372

5809 GCF_001911745.1_ASM191174v1 Escherichia coli Proteobacteria; Gammaproteobacteria; Enterobacterales; Enterobacteriaceae; Escherichia
MTGDNTLIHSHGINRRDFMKLCAALAATMGLSSKAAAEMAESVTNPQRPPVIWIGAQECT WP_074534817.1
hydrogenase 2 small subunit [Escherichia coli] Length: 372\nNumber of predicted TMHs: 1\nExp number of AAs in TMHs: 23.7182\nExp number, first 60 AAs: 0.22037\nTotal prob of N-in: 0.12983\noutside 1 331\nTMhelix 332 354\ninside 355 372

5810 GCF_001911695.1_ASM191169v1 Escherichia coli Proteobacteria; Gammaproteobacteria; Enterobacterales; Enterobacteriaceae; Escherichia
MNNEETFYQAMRRQGVTRRSFLKYCSLAATSLGLGAGMAPKIAWALENKPRIPVVWIHGL WP_001058323.1
MULTISPECIES: hydrogenase-1 small chain [Proteobacteria] Length: 372\nNumber of predicted TMHs: 2\nExp number of AAs in TMHs: 37.2608499999999\nExp number, first 60 AAs: 19.52923\nTotal prob of N-in: 0.94351\nPOSSIBLE N-term signal sequence\ninside 1 19\nTMhelix 20 39\noutside 40 325\nTMhelix 326 348\ninside 349 372

5811 GCF_001911635.1_ASM191163v1 Escherichia coli Proteobacteria; Gammaproteobacteria; Enterobacterales; Enterobacteriaceae; Escherichia
MNNEETFYQAMRRQGVTRRSFLKYCSLAATSLGLGAGMAPKIAWALENKPRIPVVWIHGL WP_001295940.1
MULTISPECIES: hydrogenase [Enterobacteriaceae] Length: 372\nNumber of predicted TMHs: 2\nExp number of AAs in TMHs: 37.2608899999999\nExp number, first 60 AAs: 19.52923\nTotal prob of N-in: 0.94351\nPOSSIBLE N-term signal sequence\ninside 1 19\nTMhelix 20 39\noutside 40 325\nTMhelix 326 348\ninside 349 372

5812 GCF_001911805.1_ASM191180v1 Escherichia coli Proteobacteria; Gammaproteobacteria; Enterobacterales; Enterobacteriaceae; Escherichia
MNNEETFYQAMRRQGVTRRSFLKYCSLAATSLGLGAGMAPKIAWALENKPRIPVVWIHGL WP_001058323.1
MULTISPECIES: hydrogenase-1 small chain [Proteobacteria] Length: 372\nNumber of predicted TMHs: 2\nExp number of AAs in TMHs: 37.2608499999999\nExp number, first 60 AAs: 19.52923\nTotal prob of N-in: 0.94351\nPOSSIBLE N-term signal sequence\ninside 1 19\nTMhelix 20 39\noutside 40 325\nTMhelix 326 348\ninside 349 372

5813 GCF_002155845.1_ASM215584v1 Escherichia coli Proteobacteria; Gammaproteobacteria; Enterobacterales; Enterobacteriaceae; Escherichia
MTGDNTLIHSHGINRRDFMKLCAALAATMGLSSKAAAEMVESVTNPQRPPVIWIGAQECT WP_000145423.1
hydrogenase 2 small subunit [Escherichia coli] Length: 372\nNumber of predicted TMHs: 1\nExp number of AAs in TMHs: 23.78201\nExp number, first 60 AAs: 0.28355\nTotal prob of N-in: 0.13257\noutside 1 331\nTMhelix 332 354\ninside 355 372

5814 GCF_002155835.1_ASM215583v1 Escherichia coli Proteobacteria; Gammaproteobacteria; Enterobacterales; Enterobacteriaceae; Escherichia
MTGDNTLIHSHGINRRDFMKLCAALAATMGLSSKAAAEMAESVTNPQRPPVIWIGAQECT WP_000145410.1
MULTISPECIES: hydrogenase-2 small chain [Proteobacteria] Length: 372\nNumber of predicted TMHs: 1\nExp number of AAs in TMHs: 23.72226\nExp number, first 60 AAs: 0.22038\nTotal prob of N-in: 0.12994\noutside 1 331\nTMhelix 332 354\ninside 355 372

5815 GCF_002163275.1_ASM216327v1 Escherichia coli Proteobacteria; Gammaproteobacteria; Enterobacterales; Enterobacteriaceae; Escherichia
MTGDNTLIHSHGINRRDFMKLCAALAATMGLSSKAAAEMAESVTNPQRPPVIWIGAQECT WP_000145410.1
MULTISPECIES: hydrogenase-2 small chain [Proteobacteria] Length: 372\nNumber of predicted TMHs: 1\nExp number of AAs in TMHs: 23.72226\nExp number, first 60 AAs: 0.22038\nTotal prob of N-in: 0.12994\noutside 1 331\nTMhelix 332 354\ninside 355 372

5816 GCF_002160185.1_ASM216018v1 Escherichia coli Proteobacteria; Gammaproteobacteria; Enterobacterales; Enterobacteriaceae; Escherichia
MNNEETFYQAMRRQGVTRRSFLKYCSLAATSLGLGAGMAPKIAWALENKPRIPVVWIHGL WP_001058323.1
MULTISPECIES: hydrogenase-1 small chain [Proteobacteria] Length: 372\nNumber of predicted TMHs: 2\nExp number of AAs in TMHs: 37.2608499999999\nExp number, first 60 AAs: 19.52923\nTotal prob of N-in: 0.94351\nPOSSIBLE N-term signal sequence\ninside 1 19\nTMhelix 20 39\noutside 40 325\nTMhelix 326 348\ninside 349 372

5817 GCF_002163275.1_ASM216327v1 Escherichia coli Proteobacteria; Gammaproteobacteria; Enterobacterales; Enterobacteriaceae; Escherichia
MNNEETFYQAMRRQGVTRRSFLKYCSLAATSLGLGAGMAPKIAWALENKPRIPVVWIHGL WP_001058323.1
MULTISPECIES: hydrogenase-1 small chain [Proteobacteria] Length: 372\nNumber of predicted TMHs: 2\nExp number of AAs in TMHs: 37.2608499999999\nExp number, first 60 AAs: 19.52923\nTotal prob of N-in: 0.94351\nPOSSIBLE N-term signal sequence\ninside 1 19\nTMhelix 20 39\noutside 40 325\nTMhelix 326 348\ninside 349 372

5818 GCF_002163295.1_ASM216329v1 Escherichia coli Proteobacteria; Gammaproteobacteria; Enterobacterales; Enterobacteriaceae; Escherichia
MTGDNTLIHSHGINRRDFMKLCAALAATMGLSSKAAAEMVESVTNPQRPPVIWIGAQECT WP_000145423.1
hydrogenase 2 small subunit [Escherichia coli] Length: 372\nNumber of predicted TMHs: 1\nExp number of AAs in TMHs: 23.78201\nExp number, first 60 AAs: 0.28355\nTotal prob of N-in: 0.13257\noutside 1 331\nTMhelix 332 354\ninside 355 372

5819 GCF_002163695.1_ASM216369v1 Escherichia coli Proteobacteria; Gammaproteobacteria; Enterobacterales; Enterobacteriaceae; Escherichia
MTGDNTLIHSHGINRRDFMKLCAALAATMGLSSKAAAEMAESVTNPQRPPVIWIGAQECT WP_000145410.1
MULTISPECIES: hydrogenase-2 small chain [Proteobacteria] Length: 372\nNumber of predicted TMHs: 1\nExp number of AAs in TMHs: 23.72226\nExp number, first 60 AAs: 0.22038\nTotal prob of N-in: 0.12994\noutside 1 331\nTMhelix 332 354\ninside 355 372

5820 GCF_002163955.1_ASM216395v1 Escherichia coli Proteobacteria; Gammaproteobacteria; Enterobacterales; Enterobacteriaceae; Escherichia
MTGDNTLIHSHGINRRDFMKLCAALAATMGLSSKAAAEMAESVTNPQRPPVIWIGAQECT WP_000145410.1
MULTISPECIES: hydrogenase-2 small chain [Proteobacteria] Length: 372\nNumber of predicted TMHs: 1\nExp number of AAs in TMHs: 23.72226\nExp number, first 60 AAs: 0.22038\nTotal prob of N-in: 0.12994\noutside 1 331\nTMhelix 332 354\ninside 355 372

5821 GCF_002164015.1_ASM216401v1 Escherichia coli Proteobacteria; Gammaproteobacteria; Enterobacterales; Enterobacteriaceae; Escherichia
MTGDNTLIHSHGINRRDFMKLCAALAATMGLSSKAAAEMAESVTNPQRPPVIWIGAQECT WP_000145410.1
MULTISPECIES: hydrogenase-2 small chain [Proteobacteria] Length: 372\nNumber of predicted TMHs: 1\nExp number of AAs in TMHs: 23.72226\nExp number, first 60 AAs: 0.22038\nTotal prob of N-in: 0.12994\noutside 1 331\nTMhelix 332 354\ninside 355 372

5822 GCF_002164035.1_ASM216403v1 Escherichia coli Proteobacteria; Gammaproteobacteria; Enterobacterales; Enterobacteriaceae; Escherichia
MTGDNTLIHSHGINRRDFMKLCAALAATMGLSSKAAAEMAESVTNPQRPPVIWIGAQECT WP_000145410.1

MULTISPECIES: hydrogenase-2 small chain [Proteobacteria] Length: 372\nNumber of predicted TMHs: 1\nExp number of AAs in TMHs: 23.72226\nExp number, first 60 AAs: 0.22038\nTotal prob of N-in: 0.12994\noutside 1 331\nTMhelix 332 354\ninside 355 372

5823 GCF_002163955.1_ASM216395v1 Escherichia coli Proteobacteria; Gammaproteobacteria; Enterobacterales; Enterobacteriaceae; Escherichia
MNNEETFYQAMRRQGVTRRSFLKYCSLAATSLGLGAGMAPKIAWALENKPRIPVVWIHGL WP_001058323.1
MULTISPECIES: hydrogenase-1 small chain [Proteobacteria] Length: 372\nNumber of predicted TMHs: 2\nExp number of AAs in TMHs: 37.2608499999999\nExp number, first 60 AAs: 19.52923\nTotal prob of N-in: 0.94351\nPOSSIBLE N-term signal sequence\ninside 1 19\nTMhelix 20 39\noutside 40 325\nTMhelix 326 348\ninside 349 372

5824 GCF_002163695.1_ASM216369v1 Escherichia coli Proteobacteria; Gammaproteobacteria; Enterobacterales; Enterobacteriaceae; Escherichia
MNNEETFYQAMRRQGVTRRSFLKYCSLAATSLGLGAGMAPKIAWALENKPRIPVVWIHGL WP_001058323.1
MULTISPECIES: hydrogenase-1 small chain [Proteobacteria] Length: 372\nNumber of predicted TMHs: 2\nExp number of AAs in TMHs: 37.2608499999999\nExp number, first 60 AAs: 19.52923\nTotal prob of N-in: 0.94351\nPOSSIBLE N-term signal sequence\ninside 1 19\nTMhelix 20 39\noutside 40 325\nTMhelix 326 348\ninside 349 372

5825 GCF_002163295.1_ASM216329v1 Escherichia coli Proteobacteria; Gammaproteobacteria; Enterobacterales; Enterobacteriaceae; Escherichia
MNNEETFYQAMRRQGVTRRSFLKYCSLAATSLGLGAGMAPKIAWALENKPRIPVVWIHGL WP_001295940.1
MULTISPECIES: hydrogenase [Enterobacteriaceae] Length: 372\nNumber of predicted TMHs: 2\nExp number of AAs in TMHs: 37.2608899999999\nExp number, first 60 AAs: 19.52923\nTotal prob of N-in: 0.94351\nPOSSIBLE N-term signal sequence\ninside 1 19\nTMhelix 20 39\noutside 40 325\nTMhelix 326 348\ninside 349 372

5826 GCF_002164255.1_ASM216425v1 Escherichia coli Proteobacteria; Gammaproteobacteria; Enterobacterales; Enterobacteriaceae; Escherichia
MTGDNTLIHSHGINRRDFMKLCAALAATMGLSSKAAAEMAESVTNPQRPPVIWIGAQECT WP_000145410.1
MULTISPECIES: hydrogenase-2 small chain [Proteobacteria] Length: 372\nNumber of predicted TMHs: 1\nExp number of AAs in TMHs: 23.72226\nExp number, first 60 AAs: 0.22038\nTotal prob of N-in: 0.12994\noutside 1 331\nTMhelix 332 354\ninside 355 372

5827 GCF_002164285.1_ASM216428v1 Escherichia coli Proteobacteria; Gammaproteobacteria; Enterobacterales; Enterobacteriaceae; Escherichia
MTGDNTLIHSHGINRRDFMKLCAALAATMGLSSKAAAEMAESVTNPQRPPVIWIGAQECT WP_000145410.1
MULTISPECIES: hydrogenase-2 small chain [Proteobacteria] Length: 372\nNumber of predicted TMHs: 1\nExp number of AAs in TMHs: 23.72226\nExp number, first 60 AAs: 0.22038\nTotal prob of N-in: 0.12994\noutside 1 331\nTMhelix 332 354\ninside 355 372

5828 GCF_002164255.1_ASM216425v1 Escherichia coli Proteobacteria; Gammaproteobacteria; Enterobacterales; Enterobacteriaceae; Escherichia
MNNEETFYQAMRRQGVTRRSFLKYCSLAATSLGLGAGMAPKIAWALENKPRIPVVWIHGL WP_001301613.1
hydrogenase [Escherichia coli] Length: 372\nNumber of predicted TMHs: 2\nExp number of AAs in TMHs: 37.46598\nExp number, first 60 AAs: 19.53139\nTotal prob of N-in: 0.94364\nPOSSIBLE N-term signal sequence\ninside 1 19\nTMhelix 20 39\noutside 40 325\nTMhelix 326 348\ninside 349 372

5829 GCF_002164285.1_ASM216428v1 Escherichia coli Proteobacteria; Gammaproteobacteria; Enterobacterales; Enterobacteriaceae; Escherichia
MNNEETFYQAMRRQGVTRRSFLKYCSLAATSLGLGAGMAPKIAWALENKPRIPVVWIHGL WP_001301613.1
hydrogenase [Escherichia coli] Length: 372\nNumber of predicted TMHs: 2\nExp number of AAs in TMHs: 37.46598\nExp number, first 60 AAs: 19.53139\nTotal prob of N-in: 0.94364\nPOSSIBLE N-term signal sequence\ninside 1 19\nTMhelix 20 39\noutside 40 325\nTMhelix 326 348\ninside 349 372

5830 GCF_002164015.1_ASM216401v1 Escherichia coli Proteobacteria; Gammaproteobacteria; Enterobacterales; Enterobacteriaceae; Escherichia
MNNEETFYQAMRRQGVTRRSFLKYCSLAATSLGLGAGMAPKIAWALENKPRIPVVWIHGL WP_001301613.1
hydrogenase [Escherichia coli] Length: 372\nNumber of predicted TMHs: 2\nExp number of AAs in TMHs: 37.46598\nExp number, first 60 AAs: 19.53139\nTotal prob of N-in: 0.94364\nPOSSIBLE N-term signal sequence\ninside 1 19\nTMhelix 20 39\noutside 40 325\nTMhelix 326 348\ninside 349 372

TMHs: 37.46598\nExp number, first 60 AAs: 19.53139\nTotal prob of N-in: 0.94364\nPOSSIBLE N-term signal sequence\ninside 1 19\nTMhelix 20 39\noutside 40 325\nTMhelix 326 348\ninside 349 372

5831 GCF_002164035.1_ASM216403v1 Escherichia coli Proteobacteria; Gammaproteobacteria; Enterobacterales; Enterobacteriaceae; Escherichia
MNNEETFYQAMRRQGVTRRSFLKYCSLAATSLGLGAGMAPKIAWALENKPRIPVVWIHGL WP_001301613.1
hydrogenase [Escherichia coli] Length: 372\nNumber of predicted TMHs: 2\nExp number of AAs in TMHs: 37.46598\nExp number, first 60 AAs: 19.53139\nTotal prob of N-in: 0.94364\nPOSSIBLE N-term signal sequence\ninside 1 19\nTMhelix 20 39\noutside 40 325\nTMhelix 326 348\ninside 349 372

5832 GCF_002164295.1_ASM216429v1 Escherichia coli Proteobacteria; Gammaproteobacteria; Enterobacterales; Enterobacteriaceae; Escherichia
MTGDNTLIHSHGINRRDFMKLCAALAATMGLSSKAAAEMAESVTNPQRPPVIWIGAQECT WP_000145410.1
MULTISPECIES: hydrogenase-2 small chain [Proteobacteria] Length: 372\nNumber of predicted TMHs: 1\nExp number of AAs in TMHs: 23.72226\nExp number, first 60 AAs: 0.22038\nTotal prob of N-in: 0.12994\noutside 1 331\nTMhelix 332 354\ninside 355 372

5833 GCF_002164355.1_ASM216435v1 Escherichia coli Proteobacteria; Gammaproteobacteria; Enterobacterales; Enterobacteriaceae; Escherichia
MTGDNTLIHSHGINRRDFMKLCAALAATMGLSSKAAAEMAESVTNPQRPPVIWIGAQECT WP_000145410.1
MULTISPECIES: hydrogenase-2 small chain [Proteobacteria] Length: 372\nNumber of predicted TMHs: 1\nExp number of AAs in TMHs: 23.72226\nExp number, first 60 AAs: 0.22038\nTotal prob of N-in: 0.12994\noutside 1 331\nTMhelix 332 354\ninside 355 372

5834 GCF_002164295.1_ASM216429v1 Escherichia coli Proteobacteria; Gammaproteobacteria; Enterobacterales; Enterobacteriaceae; Escherichia
MNNEETFYQAMRRQGVTRRSFLKYCSLAATSLGLGAGMAPKIAWALENKPRIPVVWIHGL WP_001301613.1
hydrogenase [Escherichia coli] Length: 372\nNumber of predicted TMHs: 2\nExp number of AAs in TMHs: 37.46598\nExp number, first 60 AAs: 19.53139\nTotal prob of N-in: 0.94364\nPOSSIBLE N-term signal sequence\ninside 1 19\nTMhelix 20 39\noutside 40 325\nTMhelix 326 348\ninside 349 372

5835 GCF_002164355.1_ASM216435v1 Escherichia coli Proteobacteria; Gammaproteobacteria; Enterobacterales; Enterobacteriaceae; Escherichia
MNNEETFYQAMRRQGVTRRSFLKYCSLAATSLGLGAGMAPKIAWALENKPRIPVVWIHGL WP_001301613.1
hydrogenase [Escherichia coli] Length: 372\nNumber of predicted TMHs: 2\nExp number of AAs in TMHs: 37.46598\nExp number, first 60 AAs: 19.53139\nTotal prob of N-in: 0.94364\nPOSSIBLE N-term signal sequence\ninside 1 19\nTMhelix 20 39\noutside 40 325\nTMhelix 326 348\ninside 349 372

5836 GCF_002164185.1_ASM216418v1 Escherichia coli Proteobacteria; Gammaproteobacteria; Enterobacterales; Enterobacteriaceae; Escherichia
MNNEETFYQAMRRQGVTRRSFLKYCSLAATSLGLGAGMAPKIAWALENKPRIPVVWIHGL WP_001301613.1
hydrogenase [Escherichia coli] Length: 372\nNumber of predicted TMHs: 2\nExp number of AAs in TMHs: 37.46598\nExp number, first 60 AAs: 19.53139\nTotal prob of N-in: 0.94364\nPOSSIBLE N-term signal sequence\ninside 1 19\nTMhelix 20 39\noutside 40 325\nTMhelix 326 348\ninside 349 372

5837 GCF_002164105.1_ASM216410v1 Escherichia coli Proteobacteria; Gammaproteobacteria; Enterobacterales; Enterobacteriaceae; Escherichia
MNNEETFYQAMRRQGVTRRSFLKYCSLAATSLGLGAGMAPKIAWALENKPRIPVVWIHGL WP_001301613.1
hydrogenase [Escherichia coli] Length: 372\nNumber of predicted TMHs: 2\nExp number of AAs in TMHs: 37.46598\nExp number, first 60 AAs: 19.53139\nTotal prob of N-in: 0.94364\nPOSSIBLE N-term signal sequence\ninside 1 19\nTMhelix 20 39\noutside 40 325\nTMhelix 326 348\ninside 349 372

5838 GCF_002164415.1_ASM216441v1 Escherichia coli Proteobacteria; Gammaproteobacteria; Enterobacterales; Enterobacteriaceae; Escherichia
MTGDNTLIHSHGINRRDFMKLCAALAATMGLSSKAAAEMAESVTNPQRPPVIWIGAQECT WP_000145410.1
MULTISPECIES: hydrogenase-2 small chain [Proteobacteria] Length: 372\nNumber of predicted TMHs: 1\nExp number of AAs in TMHs: 23.72226\nExp number, first 60 AAs: 0.22038\nTotal prob of N-in: 0.12994\noutside 1 331\nTMhelix 332 354\ninside 355 372

5839 GCF_002164535.1_ASM216453v1 Escherichia coli Proteobacteria; Gammaproteobacteria; Enterobacterales; Enterobacteriaceae; Escherichia

MTGDNTLIHSHGINRRDFMKLCAALAATMGLSSKAAAEMAESVTNPQRPPVIWIGAQECT WP_000145410.1 MULTISPECIES: hydrogenase-2 small chain [Proteobacteria] Length: 372\nNumber of predicted TMHs: 1\nExp number of AAs in TMHs: 23.72226\nExp number, first 60 AAs: 0.22038\nTotal prob of N-in: 0.12994\noutside 1 331\nTMhelix 332 354\ninside 355 372

5840 GCF_002164635.1_ASM216463v1 Escherichia coli Proteobacteria; Gammaproteobacteria; Enterobacterales; Enterobacteriaceae; Escherichia MNNEETFYQAMRRQGVTRRSFLKYCSLAATSLGLGAGMAPKIAWALENKPRIPVVWIHGL WP_001058323.1 MULTISPECIES: hydrogenase-1 small chain [Proteobacteria] Length: 372\nNumber of predicted TMHs: 2\nExp number of AAs in TMHs: 37.2608499999999\nExp number, first 60 AAs: 19.52923\nTotal prob of N-in: 0.94351\nPOSSIBLE N-term signal sequence\ninside 1 19\nTMhelix 20 39\noutside 40 325\nTMhelix 326 348\ninside 349 372

5841 GCF_002164915.1_ASM216491v1 Escherichia coli Proteobacteria; Gammaproteobacteria; Enterobacterales; Enterobacteriaceae; Escherichia MTGDNTLIHSHGINRRDFMKLCAALAATMGLSSKAAAEMAESVTNPQRPPVIWIGAQECT WP_000145410.1 MULTISPECIES: hydrogenase-2 small chain [Proteobacteria] Length: 372\nNumber of predicted TMHs: 1\nExp number of AAs in TMHs: 23.72226\nExp number, first 60 AAs: 0.22038\nTotal prob of N-in: 0.12994\noutside 1 331\nTMhelix 332 354\ninside 355 372

5842 GCF_002164865.1_ASM216486v1 Escherichia coli Proteobacteria; Gammaproteobacteria; Enterobacterales; Enterobacteriaceae; Escherichia MTGDNTLIHSHGINRRDFMKLCAALAATMGLSSKAAAEMAESVTNPQRPPVIWIGAQECT WP_000145410.1 MULTISPECIES: hydrogenase-2 small chain [Proteobacteria] Length: 372\nNumber of predicted TMHs: 1\nExp number of AAs in TMHs: 23.72226\nExp number, first 60 AAs: 0.22038\nTotal prob of N-in: 0.12994\noutside 1 331\nTMhelix 332 354\ninside 355 372

5843 GCF_002164865.1_ASM216486v1 Escherichia coli Proteobacteria; Gammaproteobacteria; Enterobacterales; Enterobacteriaceae; Escherichia MNNEETFYQAMRRQGVTRRSFLKYCSLAATSLGLGAGMAPKIAWALENKPRIPVVWIHGL WP_001058320.1 hydrogenase [Escherichia coli] Length: 372\nNumber of predicted TMHs: 2\nExp number of AAs in TMHs: 37.26082\nExp number, first 60 AAs: 19.52923\nTotal prob of N-in: 0.94351\nPOSSIBLE N-term signal sequence\ninside 1 19\nTMhelix 20 39\noutside 40 325\nTMhelix 326 348\ninside 349 372

5844 GCF_002164915.1_ASM216491v1 Escherichia coli Proteobacteria; Gammaproteobacteria; Enterobacterales; Enterobacteriaceae; Escherichia MNNEETFYQAMRRQGVTRRSFLKYCSLAATSLGLGAGMAPKIAWALENKPRIPVVWIHGL WP_001295940.1 MULTISPECIES: hydrogenase [Enterobacteriaceae] Length: 372\nNumber of predicted TMHs: 2\nExp number of AAs in TMHs: 37.2608899999999\nExp number, first 60 AAs: 19.52923\nTotal prob of N-in: 0.94351\nPOSSIBLE N-term signal sequence\ninside 1 19\nTMhelix 20 39\noutside 40 325\nTMhelix 326 348\ninside 349 372

5845 GCF_002164375.1_ASM216437v1 Escherichia coli Proteobacteria; Gammaproteobacteria; Enterobacterales; Enterobacteriaceae; Escherichia MNNEETFYQAMRRQGVTRRSFLKYCSLAATSLGLGAGMAPKIAWALENKPRIPVVWIHGL WP_001301613.1 hydrogenase [Escherichia coli] Length: 372\nNumber of predicted TMHs: 2\nExp number of AAs in TMHs: 37.46598\nExp number, first 60 AAs: 19.53139\nTotal prob of N-in: 0.94364\nPOSSIBLE N-term signal sequence\ninside 1 19\nTMhelix 20 39\noutside 40 325\nTMhelix 326 348\ninside 349 372

5846 GCF_002164955.1_ASM216495v1 Escherichia coli Proteobacteria; Gammaproteobacteria; Enterobacterales; Enterobacteriaceae; Escherichia MTGDNTLIHSHGINRRDFMKLCAALAATMGLSSKAAAEMAESVTNPQRPPVIWIGAQECT WP_000145410.1 MULTISPECIES: hydrogenase-2 small chain [Proteobacteria] Length: 372\nNumber of predicted TMHs: 1\nExp number of AAs in TMHs: 23.72226\nExp number, first 60 AAs: 0.22038\nTotal prob of N-in: 0.12994\noutside 1 331\nTMhelix 332 354\ninside 355 372

5847 GCF_002164985.1_ASM216498v1 Escherichia coli Proteobacteria; Gammaproteobacteria; Enterobacterales; Enterobacteriaceae; Escherichia MTGDNTLIHSHGINRRDFMKLCAALAATMGLSSKAAAEMAESVTNPQRPPVIWIGAQECT WP_000145410.1 MULTISPECIES: hydrogenase-2 small chain [Proteobacteria] Length: 372\nNumber of predicted TMHs:

1\nExp number of AAs in TMHs: 23.72226\nExp number, first 60 AAs: 0.22038\nTotal prob of N-in: 0.12994\noutside 1 331\nTMhelix 332 354\ninside 355 372

5848 GCF_002164955.1_ASM216495v1 Escherichia coli Proteobacteria; Gammaproteobacteria; Enterobacterales; Enterobacteriaceae; Escherichia
MNNEETFYQAMRRQGVTRRSFLKYCSLAATSLGLGAGMAPKIAWALENKPRIPVVWIHGL WP_001058323.1
MULTISPECIES: hydrogenase-1 small chain [Proteobacteria] Length: 372\nNumber of predicted TMHs: 2\nExp number of AAs in TMHs: 37.2608499999999\nExp number, first 60 AAs: 19.52923\nTotal prob of N-in: 0.94351\nPOSSIBLE N-term signal sequence\ninside 1 19\nTMhelix 20 39\noutside 40 325\nTMhelix 326 348\ninside 349 372

5849 GCF_002164985.1_ASM216498v1 Escherichia coli Proteobacteria; Gammaproteobacteria; Enterobacterales; Enterobacteriaceae; Escherichia
MNNEETFYQAMRRQGVTRRSFLKYCSLAATSLGLGAGMAPKIAWALENKPRIPVVWIHGL WP_001058323.1
MULTISPECIES: hydrogenase-1 small chain [Proteobacteria] Length: 372\nNumber of predicted TMHs: 2\nExp number of AAs in TMHs: 37.2608499999999\nExp number, first 60 AAs: 19.52923\nTotal prob of N-in: 0.94351\nPOSSIBLE N-term signal sequence\ninside 1 19\nTMhelix 20 39\noutside 40 325\nTMhelix 326 348\ninside 349 372

5850 GCF_002164455.1_ASM216445v1 Escherichia coli Proteobacteria; Gammaproteobacteria; Enterobacterales; Enterobacteriaceae; Escherichia
MNNEETFYQAMRRQGVTRRSFLKYCSLAATSLGLGAGMAPKIAWALENKPRIPVVWIHGL WP_001301613.1
hydrogenase [Escherichia coli] Length: 372\nNumber of predicted TMHs: 2\nExp number of AAs in TMHs: 37.46598\nExp number, first 60 AAs: 19.53139\nTotal prob of N-in: 0.94364\nPOSSIBLE N-term signal sequence\ninside 1 19\nTMhelix 20 39\noutside 40 325\nTMhelix 326 348\ninside 349 372

5851 GCF_002165045.1_ASM216504v1 Escherichia coli Proteobacteria; Gammaproteobacteria; Enterobacterales; Enterobacteriaceae; Escherichia
MTGDNTLIHSHGINRRDFMKLCAALAATMGLSSKAAAEMAESVTNPQRPPVIWIGAQECT WP_000145410.1
MULTISPECIES: hydrogenase-2 small chain [Proteobacteria] Length: 372\nNumber of predicted TMHs: 1\nExp number of AAs in TMHs: 23.72226\nExp number, first 60 AAs: 0.22038\nTotal prob of N-in: 0.12994\noutside 1 331\nTMhelix 332 354\ninside 355 372

5852 GCF_002165045.1_ASM216504v1 Escherichia coli Proteobacteria; Gammaproteobacteria; Enterobacterales; Enterobacteriaceae; Escherichia
MNNEETFYQAMRRQGVTRRSFLKYCSLAATSLGLGAGMAPKIAWALENKPRIPVVWIHGL WP_001058323.1
MULTISPECIES: hydrogenase-1 small chain [Proteobacteria] Length: 372\nNumber of predicted TMHs: 2\nExp number of AAs in TMHs: 37.2608499999999\nExp number, first 60 AAs: 19.52923\nTotal prob of N-in: 0.94351\nPOSSIBLE N-term signal sequence\ninside 1 19\nTMhelix 20 39\noutside 40 325\nTMhelix 326 348\ninside 349 372

5853 GCF_002165965.1_ASM216596v1 Escherichia coli Proteobacteria; Gammaproteobacteria; Enterobacterales; Enterobacteriaceae; Escherichia
MTGDNTLIHSHGINRRDFMKLCAALAATMGLSSKAAAEMAESVTNPQRPPVIWIGAQECT WP_000145410.1
MULTISPECIES: hydrogenase-2 small chain [Proteobacteria] Length: 372\nNumber of predicted TMHs: 1\nExp number of AAs in TMHs: 23.72226\nExp number, first 60 AAs: 0.22038\nTotal prob of N-in: 0.12994\noutside 1 331\nTMhelix 332 354\ninside 355 372

5854 GCF_002165115.1_ASM216511v1 Escherichia coli Proteobacteria; Gammaproteobacteria; Enterobacterales; Enterobacteriaceae; Escherichia
MTGDNTLIHSHGINRRDFMKLCAALAATMGLSSKAAAEMAESVTNPQRPPVIWIGAQECT WP_000145410.1
MULTISPECIES: hydrogenase-2 small chain [Proteobacteria] Length: 372\nNumber of predicted TMHs: 1\nExp number of AAs in TMHs: 23.72226\nExp number, first 60 AAs: 0.22038\nTotal prob of N-in: 0.12994\noutside 1 331\nTMhelix 332 354\ninside 355 372

5855 GCF_002165955.1_ASM216595v1 Escherichia coli Proteobacteria; Gammaproteobacteria; Enterobacterales; Enterobacteriaceae; Escherichia
MTGDNTLIHSHGINRRDFMKLCAALAATMGLSSKAAAEMAESVTNPQRPPVIWIGAQECT WP_000145410.1
MULTISPECIES: hydrogenase-2 small chain [Proteobacteria] Length: 372\nNumber of predicted TMHs: 1\nExp number of AAs in TMHs: 23.72226\nExp number, first 60 AAs: 0.22038\nTotal prob of N-in: 0.12994\noutside 1 331\nTMhelix 332 354\ninside 355 372

5856 GCF_002166005.1_ASM216600v1 Escherichia coli Proteobacteria; Gammaproteobacteria; Enterobacterales; Enterobacteriaceae; Escherichia
MTGDNTLIHSHGINRRDFMKLCAALAATMGLSSKAAAEMAESVTNPQRPPVIWIGAQECT WP_000145410.1
MULTISPECIES: hydrogenase-2 small chain [Proteobacteria] Length: 372\nNumber of predicted TMHs: 1\nExp number of AAs in TMHs: 23.72226\nExp number, first 60 AAs: 0.22038\nTotal prob of N-in: 0.12994\noutside 1 331\nTMhelix 332 354\ninside 355 372

5857 GCF_002166075.1_ASM216607v1 Escherichia coli Proteobacteria; Gammaproteobacteria; Enterobacterales; Enterobacteriaceae; Escherichia
MTGDNTLIHSHGINRRDFMKLCAALAATMGLSSKAAAEMAESVTNPQRPPVIWIGAQECT WP_000145410.1
MULTISPECIES: hydrogenase-2 small chain [Proteobacteria] Length: 372\nNumber of predicted TMHs: 1\nExp number of AAs in TMHs: 23.72226\nExp number, first 60 AAs: 0.22038\nTotal prob of N-in: 0.12994\noutside 1 331\nTMhelix 332 354\ninside 355 372

5858 GCF_002166035.1_ASM216603v1 Escherichia coli Proteobacteria; Gammaproteobacteria; Enterobacterales; Enterobacteriaceae; Escherichia
MTGDNTLIHSHGINRRDFMKLCAALAATMGLSSKAAAEMAESVTNPQRPPVIWIGAQECT WP_000145410.1
MULTISPECIES: hydrogenase-2 small chain [Proteobacteria] Length: 372\nNumber of predicted TMHs: 1\nExp number of AAs in TMHs: 23.72226\nExp number, first 60 AAs: 0.22038\nTotal prob of N-in: 0.12994\noutside 1 331\nTMhelix 332 354\ninside 355 372

5859 GCF_002166035.1_ASM216603v1 Escherichia coli Proteobacteria; Gammaproteobacteria; Enterobacterales; Enterobacteriaceae; Escherichia
MNNEETFYQAMRRQGVTRRSFLKYCSLAATSLGLGAGMAPKIAWALENKPRIPVVWIHGL WP_001058323.1
MULTISPECIES: hydrogenase-1 small chain [Proteobacteria] Length: 372\nNumber of predicted TMHs: 2\nExp number of AAs in TMHs: 37.2608499999999\nExp number, first 60 AAs: 19.52923\nTotal prob of N-in: 0.94351\nPOSSIBLE N-term signal sequence\ninside 1 19\nTMhelix 20 39\noutside 40 325\nTMhelix 326 348\ninside 349 372

5860 GCF_002166375.1_ASM216637v1 Escherichia coli Proteobacteria; Gammaproteobacteria; Enterobacterales; Enterobacteriaceae; Escherichia
MTGDNTLIHSHGINRRDFMKLCAALAATMGLSSKAAAEMAESVTNPQRPPVIWIGAQECT WP_000145410.1
MULTISPECIES: hydrogenase-2 small chain [Proteobacteria] Length: 372\nNumber of predicted TMHs: 1\nExp number of AAs in TMHs: 23.72226\nExp number, first 60 AAs: 0.22038\nTotal prob of N-in: 0.12994\noutside 1 331\nTMhelix 332 354\ninside 355 372

5861 GCF_002165915.1_ASM216591v1 Escherichia coli Proteobacteria; Gammaproteobacteria; Enterobacterales; Enterobacteriaceae; Escherichia
MNNEETFYQAMRRQGVTRRSFLKYCSLAATSLGLGAGMAPKIAWALENKPRIPVVWIHGL WP_001058323.1
MULTISPECIES: hydrogenase-1 small chain [Proteobacteria] Length: 372\nNumber of predicted TMHs: 2\nExp number of AAs in TMHs: 37.2608499999999\nExp number, first 60 AAs: 19.52923\nTotal prob of N-in: 0.94351\nPOSSIBLE N-term signal sequence\ninside 1 19\nTMhelix 20 39\noutside 40 325\nTMhelix 326 348\ninside 349 372

5862 GCF_002165095.1_ASM216509v1 Escherichia coli Proteobacteria; Gammaproteobacteria; Enterobacterales; Enterobacteriaceae; Escherichia
MNNEETFYQAMRRQGVTRRSFLKYCSLAATSLGLGAGMAPKIAWALENKPRIPVVWIHGL WP_001295940.1
MULTISPECIES: hydrogenase [Enterobacteriaceae] Length: 372\nNumber of predicted TMHs: 2\nExp number of AAs in TMHs: 37.2608899999999\nExp number, first 60 AAs: 19.52923\nTotal prob of N-in: 0.94351\nPOSSIBLE N-term signal sequence\ninside 1 19\nTMhelix 20 39\noutside 40 325\nTMhelix 326 348\ninside 349 372

5863 GCF_002165115.1_ASM216511v1 Escherichia coli Proteobacteria; Gammaproteobacteria; Enterobacterales; Enterobacteriaceae; Escherichia
MNNEETFYQAMRRQGVTRRSFLKYCSLAATSLGLGAGMAPKIAWALENKPRIPVVWIHGL WP_001058320.1
hydrogenase [Escherichia coli] Length: 372\nNumber of predicted TMHs: 2\nExp number of AAs in TMHs: 37.26082\nExp number, first 60 AAs: 19.52923\nTotal prob of N-in: 0.94351\nPOSSIBLE N-term signal sequence\ninside 1 19\nTMhelix 20 39\noutside 40 325\nTMhelix 326 348\ninside 349 372

5864 GCF_002165965.1_ASM216596v1 Escherichia coli Proteobacteria; Gammaproteobacteria; Enterobacterales; Enterobacteriaceae; Escherichia

MNNEETFYQAMRRQGVTRRSFLKYCSLAATSLGLGAGMAPKIAWALENKPRIPVVWIHGL WP_001058323.1 MULTISPECIES: hydrogenase-1 small chain [Proteobacteria] Length: 372\nNumber of predicted TMHs: 2\nExp number of AAs in TMHs: 37.2608499999999\nExp number, first 60 AAs: 19.52923\nTotal prob of N-in: 0.94351\nPOSSIBLE N-term signal sequence\ninside 1 19\nTMhelix 20 39\noutside 40 325\nTMhelix 326 348\ninside 349 372

5865 GCF_002165955.1_ASM216595v1 Escherichia coli Proteobacteria; Gammaproteobacteria; Enterobacterales; Enterobacteriaceae; Escherichia MNNEETFYQAMRRQGVTRRSFLKYCSLAATSLGLGAGMAPKIAWALENKPRIPVVWIHGL WP_001058323.1 MULTISPECIES: hydrogenase-1 small chain [Proteobacteria] Length: 372\nNumber of predicted TMHs: 2\nExp number of AAs in TMHs: 37.2608499999999\nExp number, first 60 AAs: 19.52923\nTotal prob of N-in: 0.94351\nPOSSIBLE N-term signal sequence\ninside 1 19\nTMhelix 20 39\noutside 40 325\nTMhelix 326 348\ninside 349 372

5866 GCF_002166005.1_ASM216600v1 Escherichia coli Proteobacteria; Gammaproteobacteria; Enterobacterales; Enterobacteriaceae; Escherichia MNNEETFYQAMRRQGVTRRSFLKYCSLAATSLGLGAGMAPKIAWALENKPRIPVVWIHGL WP_001058323.1 MULTISPECIES: hydrogenase-1 small chain [Proteobacteria] Length: 372\nNumber of predicted TMHs: 2\nExp number of AAs in TMHs: 37.2608499999999\nExp number, first 60 AAs: 19.52923\nTotal prob of N-in: 0.94351\nPOSSIBLE N-term signal sequence\ninside 1 19\nTMhelix 20 39\noutside 40 325\nTMhelix 326 348\ninside 349 372

5867 GCF_002166115.1_ASM216611v1 Escherichia coli Proteobacteria; Gammaproteobacteria; Enterobacterales; Enterobacteriaceae; Escherichia MTGDNTLIHSHGINRRDFMKLCAALAATMGLSSKAAAEMAESVTNPQRPPVIWIGAQECT WP_000145410.1 MULTISPECIES: hydrogenase-2 small chain [Proteobacteria] Length: 372\nNumber of predicted TMHs: 1\nExp number of AAs in TMHs: 23.72226\nExp number, first 60 AAs: 0.22038\nTotal prob of N-in: 0.12994\noutside 1 331\nTMhelix 332 354\ninside 355 372

5868 GCF_002166195.1_ASM216619v1 Escherichia coli Proteobacteria; Gammaproteobacteria; Enterobacterales; Enterobacteriaceae; Escherichia MNNEETFYQAMRRQGVTRRSFLKYCSLAATSLGLGAGMAPKIAWALENKPRIPVVWIHGL WP_001058323.1 MULTISPECIES: hydrogenase-1 small chain [Proteobacteria] Length: 372\nNumber of predicted TMHs: 2\nExp number of AAs in TMHs: 37.2608499999999\nExp number, first 60 AAs: 19.52923\nTotal prob of N-in: 0.94351\nPOSSIBLE N-term signal sequence\ninside 1 19\nTMhelix 20 39\noutside 40 325\nTMhelix 326 348\ninside 349 372

5869 GCF_002166415.1_ASM216641v1 Escherichia coli Proteobacteria; Gammaproteobacteria; Enterobacterales; Enterobacteriaceae; Escherichia MTGDNTLIHSHGINRRDFMKLCAALAATMGLSSKAAAEMAESVTNPQRPPVIWIGAQECT WP_000145410.1 MULTISPECIES: hydrogenase-2 small chain [Proteobacteria] Length: 372\nNumber of predicted TMHs: 1\nExp number of AAs in TMHs: 23.72226\nExp number, first 60 AAs: 0.22038\nTotal prob of N-in: 0.12994\noutside 1 331\nTMhelix 332 354\ninside 355 372

5870 GCF_002166265.1_ASM216626v1 Escherichia coli Proteobacteria; Gammaproteobacteria; Enterobacterales; Enterobacteriaceae; Escherichia MTGDNTLIHSHGINRRDFMKLCAALAATMGLSSKAAAEMAESVTNPQRPPVIWIGAQECT WP_000145410.1 MULTISPECIES: hydrogenase-2 small chain [Proteobacteria] Length: 372\nNumber of predicted TMHs: 1\nExp number of AAs in TMHs: 23.72226\nExp number, first 60 AAs: 0.22038\nTotal prob of N-in: 0.12994\noutside 1 331\nTMhelix 332 354\ninside 355 372

5871 GCF_002166265.1_ASM216626v1 Escherichia coli Proteobacteria; Gammaproteobacteria; Enterobacterales; Enterobacteriaceae; Escherichia MNNEETFYQAMRRQGVTRRSFLKYCSLAATSLGLGAGMAPKIAWALENKPRIPVVWIHGL WP_001058323.1 MULTISPECIES: hydrogenase-1 small chain [Proteobacteria] Length: 372\nNumber of predicted TMHs: 2\nExp number of AAs in TMHs: 37.2608499999999\nExp number, first 60 AAs: 19.52923\nTotal prob of N-in: 0.94351\nPOSSIBLE N-term signal sequence\ninside 1 19\nTMhelix 20 39\noutside 40 325\nTMhelix 326 348\ninside 349 372

5872 GCF_002166415.1_ASM216641v1 Escherichia coli Proteobacteria; Gammaproteobacteria; Enterobacterales; Enterobacteriaceae; Escherichia

MNNEETFYQAMRRQGVTRRSFLKYCSLAATSLGLGAGMAPKIAWALENKPRIPVVWIHGL WP_001058323.1 MULTISPECIES: hydrogenase-1 small chain [Proteobacteria] Length: 372\nNumber of predicted TMHs: 2\nExp number of AAs in TMHs: 37.2608499999999\nExp number, first 60 AAs: 19.52923\nTotal prob of N-in: 0.94351\nPOSSIBLE N-term signal sequence\ninside 1 19\nTMhelix 20 39\noutside 40 325\nTMhelix 326 348\ninside 349 372

5873 GCF_002166135.1_ASM216613v1 Escherichia coli Proteobacteria; Gammaproteobacteria; Enterobacterales; Enterobacteriaceae; Escherichia MNNEETFYQAMRRQGVTRRSFLKYCSLAATSLGLGAGMAPKIAWALENKPRIPVVWIHGL WP_001058323.1 MULTISPECIES: hydrogenase-1 small chain [Proteobacteria] Length: 372\nNumber of predicted TMHs: 2\nExp number of AAs in TMHs: 37.2608499999999\nExp number, first 60 AAs: 19.52923\nTotal prob of N-in: 0.94351\nPOSSIBLE N-term signal sequence\ninside 1 19\nTMhelix 20 39\noutside 40 325\nTMhelix 326 348\ninside 349 372

5874 GCF_002166345.1_ASM216634v1 Escherichia coli Proteobacteria; Gammaproteobacteria; Enterobacterales; Enterobacteriaceae; Escherichia MTGDNTLIHSHGINRRDFMKLCAALAATMGLSSKAAAEMAESVTNPQRPPVIWIGAQECT WP_000145410.1 MULTISPECIES: hydrogenase-2 small chain [Proteobacteria] Length: 372\nNumber of predicted TMHs: 1\nExp number of AAs in TMHs: 23.72226\nExp number, first 60 AAs: 0.22038\nTotal prob of N-in: 0.12994\noutside 1 331\nTMhelix 332 354\ninside 355 372

5875 GCF_002166375.1_ASM216637v1 Escherichia coli Proteobacteria; Gammaproteobacteria; Enterobacterales; Enterobacteriaceae; Escherichia MNNEETFYQAMRRQGVTRRSFLKYCSLAATSLGLGAGMAPKIAWALENKPRIPVVWIHGL WP_001058323.1 MULTISPECIES: hydrogenase-1 small chain [Proteobacteria] Length: 372\nNumber of predicted TMHs: 2\nExp number of AAs in TMHs: 37.2608499999999\nExp number, first 60 AAs: 19.52923\nTotal prob of N-in: 0.94351\nPOSSIBLE N-term signal sequence\ninside 1 19\nTMhelix 20 39\noutside 40 325\nTMhelix 326 348\ninside 349 372

5876 GCF_002166175.1_ASM216617v1 Escherichia coli Proteobacteria; Gammaproteobacteria; Enterobacterales; Enterobacteriaceae; Escherichia MNNEETFYQAMRRQGVTRRSFLKYCSLAATSLGLGAGMAPKIAWALENKPRIPVVWIHGL WP_001058323.1 MULTISPECIES: hydrogenase-1 small chain [Proteobacteria] Length: 372\nNumber of predicted TMHs: 2\nExp number of AAs in TMHs: 37.2608499999999\nExp number, first 60 AAs: 19.52923\nTotal prob of N-in: 0.94351\nPOSSIBLE N-term signal sequence\ninside 1 19\nTMhelix 20 39\noutside 40 325\nTMhelix 326 348\ninside 349 372

5877 GCF_002166465.1_ASM216646v1 Escherichia coli Proteobacteria; Gammaproteobacteria; Enterobacterales; Enterobacteriaceae; Escherichia MTGDNTLIHSHGINRRDFMKLCAALAATMGLSSKAAAEMVESVTNPQRPPVIWIGAQECT WP_000145423.1 hydrogenase 2 small subunit [Escherichia coli] Length: 372\nNumber of predicted TMHs: 1\nExp number of AAs in TMHs: 23.78201\nExp number, first 60 AAs: 0.28355\nTotal prob of N-in: 0.13257\noutside 1 331\nTMhelix 332 354\ninside 355 372

5878 GCF_002166495.1_ASM216649v1 Escherichia coli Proteobacteria; Gammaproteobacteria; Enterobacterales; Enterobacteriaceae; Escherichia MTGDNTLIHSHGINRRDFMKLCAALAATMGLSSKAAAEMAESVTNPQRPPVIWIGAQECT WP_000145410.1 MULTISPECIES: hydrogenase-2 small chain [Proteobacteria] Length: 372\nNumber of predicted TMHs: 1\nExp number of AAs in TMHs: 23.72226\nExp number, first 60 AAs: 0.22038\nTotal prob of N-in: 0.12994\noutside 1 331\nTMhelix 332 354\ninside 355 372

5879 GCF_002166465.1_ASM216646v1 Escherichia coli Proteobacteria; Gammaproteobacteria; Enterobacterales; Enterobacteriaceae; Escherichia MNNEETFYQAMRRQGVTRRSFLKYCSLAATSLGLGAGMAPKIAWALENKPRIPVVWIHGL WP_001058323.1 MULTISPECIES: hydrogenase-1 small chain [Proteobacteria] Length: 372\nNumber of predicted TMHs: 2\nExp number of AAs in TMHs: 37.2608499999999\nExp number, first 60 AAs: 19.52923\nTotal prob of N-in: 0.94351\nPOSSIBLE N-term signal sequence\ninside 1 19\nTMhelix 20 39\noutside 40 325\nTMhelix 326 348\ninside 349 372

5880 GCF_002166495.1_ASM216649v1 Escherichia coli Proteobacteria; Gammaproteobacteria; Enterobacterales; Enterobacteriaceae; Escherichia

MNNEETFYQAMRRQGVTRRSFLKYCSLAATSLGLGAGMAPKIAWALENKPRIPVVWIHGL WP_001058323.1 MULTISPECIES: hydrogenase-1 small chain [Proteobacteria] Length: 372\nNumber of predicted TMHs: 2\nExp number of AAs in TMHs: 37.2608499999999\nExp number, first 60 AAs: 19.52923\nTotal prob of N-in: 0.94351\nPOSSIBLE N-term signal sequence\ninside 1 19\nTMhelix 20 39\noutside 40 325\nTMhelix 326 348\ninside 349 372

5881 GCF_002166595.1_ASM216659v1 Escherichia coli Proteobacteria; Gammaproteobacteria; Enterobacterales; Enterobacteriaceae; Escherichia MNNEETFYQAMRRQGVTRRSFLKYCSLAATSLGLGAGMAPKIAWALENKPRIPVVWIHGL WP_001058323.1 MULTISPECIES: hydrogenase-1 small chain [Proteobacteria] Length: 372\nNumber of predicted TMHs: 2\nExp number of AAs in TMHs: 37.2608499999999\nExp number, first 60 AAs: 19.52923\nTotal prob of N-in: 0.94351\nPOSSIBLE N-term signal sequence\ninside 1 19\nTMhelix 20 39\noutside 40 325\nTMhelix 326 348\ninside 349 372

5882 GCF_002166565.1_ASM216656v1 Escherichia coli Proteobacteria; Gammaproteobacteria; Enterobacterales; Enterobacteriaceae; Escherichia MNNEETFYQAMRRQGVTRRSFLKYCSLAATSLGLGAGMAPKIAWALENKPRIPVVWIHGL WP_001058320.1 hydrogenase [Escherichia coli] Length: 372\nNumber of predicted TMHs: 2\nExp number of AAs in TMHs: 37.26082\nExp number, first 60 AAs: 19.52923\nTotal prob of N-in: 0.94351\nPOSSIBLE N-term signal sequence\ninside 1 19\nTMhelix 20 39\noutside 40 325\nTMhelix 326 348\ninside 349 372

5883 GCF_002166715.1_ASM216671v1 Escherichia coli Proteobacteria; Gammaproteobacteria; Enterobacterales; Enterobacteriaceae; Escherichia MTGDNTLIHSHGINRRDFMKLCAALAATMGLSSKAAAEMAESVTNPQRPPVIWIGAQECT WP_000145410.1 MULTISPECIES: hydrogenase-2 small chain [Proteobacteria] Length: 372\nNumber of predicted TMHs: 1\nExp number of AAs in TMHs: 23.72226\nExp number, first 60 AAs: 0.22038\nTotal prob of N-in: 0.12994\noutside 1 331\nTMhelix 332 354\ninside 355 372

5884 GCF_002166685.1_ASM216668v1 Escherichia coli Proteobacteria; Gammaproteobacteria; Enterobacterales; Enterobacteriaceae; Escherichia MTGDNTLIHSHGINRRDFMKLCAALAATMGLSSKAAAEMAESVTNPQRPPVIWIGAQECT WP_000145410.1 MULTISPECIES: hydrogenase-2 small chain [Proteobacteria] Length: 372\nNumber of predicted TMHs: 1\nExp number of AAs in TMHs: 23.72226\nExp number, first 60 AAs: 0.22038\nTotal prob of N-in: 0.12994\noutside 1 331\nTMhelix 332 354\ninside 355 372

5885 GCF_002173125.1_ASM217312v1 Escherichia coli Proteobacteria; Gammaproteobacteria; Enterobacterales; Enterobacteriaceae; Escherichia MTGDNTLIHSHGINRRDFMKLCAALAATMGLSSKAAAEMAESVTNPQRPPVIWIGAQECT WP_000145410.1 MULTISPECIES: hydrogenase-2 small chain [Proteobacteria] Length: 372\nNumber of predicted TMHs: 1\nExp number of AAs in TMHs: 23.72226\nExp number, first 60 AAs: 0.22038\nTotal prob of N-in: 0.12994\noutside 1 331\nTMhelix 332 354\ninside 355 372

5886 GCF_002173195.1_ASM217319v1 Escherichia coli Proteobacteria; Gammaproteobacteria; Enterobacterales; Enterobacteriaceae; Escherichia MTGDNTLIHSHGINRRDFMKLCAALAATMGLSSKAAAEMAESVTNPQRPPVIWIGAQECT WP_000145410.1 MULTISPECIES: hydrogenase-2 small chain [Proteobacteria] Length: 372\nNumber of predicted TMHs: 1\nExp number of AAs in TMHs: 23.72226\nExp number, first 60 AAs: 0.22038\nTotal prob of N-in: 0.12994\noutside 1 331\nTMhelix 332 354\ninside 355 372

5887 GCF_002173145.1_ASM217314v1 Escherichia coli Proteobacteria; Gammaproteobacteria; Enterobacterales; Enterobacteriaceae; Escherichia MTGDNTLIHSHGINRRDFMKLCAALAATMGLSSKAAAEMAESVTNPQRPPVIWIGAQECT WP_000145410.1 MULTISPECIES: hydrogenase-2 small chain [Proteobacteria] Length: 372\nNumber of predicted TMHs: 1\nExp number of AAs in TMHs: 23.72226\nExp number, first 60 AAs: 0.22038\nTotal prob of N-in: 0.12994\noutside 1 331\nTMhelix 332 354\ninside 355 372

5888 GCF_002173215.1_ASM217321v1 Escherichia coli Proteobacteria; Gammaproteobacteria; Enterobacterales; Enterobacteriaceae; Escherichia MTGDNTLIHSHGINRRDFMKLCAALAATMGLSSKAAAEMAESVTNPQRPPVIWIGAQECT WP_000145410.1 MULTISPECIES: hydrogenase-2 small chain [Proteobacteria] Length: 372\nNumber of predicted TMHs:

1\nExp number of AAs in TMHs: 23.72226\nExp number, first 60 AAs: 0.22038\nTotal prob of N-in: 0.12994\noutside 1 331\nTMhelix 332 354\ninside 355 372

5889 GCF_002173255.1_ASM217325v1 Escherichia coli Proteobacteria; Gammaproteobacteria; Enterobacterales; Enterobacteriaceae; Escherichia
MTGDNTLIHSHGINRRDFMKLCAALAATMGLSSKAAAEMAESVTNPQRPPVIWIGAQECT WP_000145410.1
MULTISPECIES: hydrogenase-2 small chain [Proteobacteria] Length: 372\nNumber of predicted TMHs: 1\nExp number of AAs in TMHs: 23.72226\nExp number, first 60 AAs: 0.22038\nTotal prob of N-in: 0.12994\noutside 1 331\nTMhelix 332 354\ninside 355 372

5890 GCF_002166675.1_ASM216667v1 Escherichia coli Proteobacteria; Gammaproteobacteria; Enterobacterales; Enterobacteriaceae; Escherichia
MNNEETFYQAMRRQGVTRRSFLKYCSLAATSLGLGAGMAPKIAWALENKPRIPVVWIHGL WP_001058323.1
MULTISPECIES: hydrogenase-1 small chain [Proteobacteria] Length: 372\nNumber of predicted TMHs: 2\nExp number of AAs in TMHs: 37.2608499999999\nExp number, first 60 AAs: 19.52923\nTotal prob of N-in: 0.94351\nPOSSIBLE N-term signal sequence\ninside 1 19\nTMhelix 20 39\noutside 40 325\nTMhelix 326 348\ninside 349 372

5891 GCF_002166715.1_ASM216671v1 Escherichia coli Proteobacteria; Gammaproteobacteria; Enterobacterales; Enterobacteriaceae; Escherichia
MNNEETFYQAMRRQGVTRRSFLKYCSLAATSLGLGAGMAPKIAWALENKPRIPVVWIHGL WP_001058323.1
MULTISPECIES: hydrogenase-1 small chain [Proteobacteria] Length: 372\nNumber of predicted TMHs: 2\nExp number of AAs in TMHs: 37.2608499999999\nExp number, first 60 AAs: 19.52923\nTotal prob of N-in: 0.94351\nPOSSIBLE N-term signal sequence\ninside 1 19\nTMhelix 20 39\noutside 40 325\nTMhelix 326 348\ninside 349 372

5892 GCF_002166685.1_ASM216668v1 Escherichia coli Proteobacteria; Gammaproteobacteria; Enterobacterales; Enterobacteriaceae; Escherichia
MNNEETFYQAMRRQGVTRRSFLKYCSLAATSLGLGAGMAPKIAWALENKPRIPVVWIHGL WP_001058323.1
MULTISPECIES: hydrogenase-1 small chain [Proteobacteria] Length: 372\nNumber of predicted TMHs: 2\nExp number of AAs in TMHs: 37.2608499999999\nExp number, first 60 AAs: 19.52923\nTotal prob of N-in: 0.94351\nPOSSIBLE N-term signal sequence\ninside 1 19\nTMhelix 20 39\noutside 40 325\nTMhelix 326 348\ninside 349 372

5893 GCF_002173125.1_ASM217312v1 Escherichia coli Proteobacteria; Gammaproteobacteria; Enterobacterales; Enterobacteriaceae; Escherichia
MNNEETFYQAMRRQGVTRRSFLKYCSLAATSLGLGAGMAPKIAWALENKPRIPVVWIHGL WP_001058323.1
MULTISPECIES: hydrogenase-1 small chain [Proteobacteria] Length: 372\nNumber of predicted TMHs: 2\nExp number of AAs in TMHs: 37.2608499999999\nExp number, first 60 AAs: 19.52923\nTotal prob of N-in: 0.94351\nPOSSIBLE N-term signal sequence\ninside 1 19\nTMhelix 20 39\noutside 40 325\nTMhelix 326 348\ninside 349 372

5894 GCF_002173195.1_ASM217319v1 Escherichia coli Proteobacteria; Gammaproteobacteria; Enterobacterales; Enterobacteriaceae; Escherichia
MNNEETFYQAMRRQGVTRRSFLKYCSLAATSLGLGAGMAPKIAWALENKPRIPVVWIHGL WP_001058323.1
MULTISPECIES: hydrogenase-1 small chain [Proteobacteria] Length: 372\nNumber of predicted TMHs: 2\nExp number of AAs in TMHs: 37.2608499999999\nExp number, first 60 AAs: 19.52923\nTotal prob of N-in: 0.94351\nPOSSIBLE N-term signal sequence\ninside 1 19\nTMhelix 20 39\noutside 40 325\nTMhelix 326 348\ninside 349 372

5895 GCF_002173365.1_ASM217336v1 Escherichia coli Proteobacteria; Gammaproteobacteria; Enterobacterales; Enterobacteriaceae; Escherichia
MTGDNTLIHSHGINRRDFMKLCAALAATMGLSSKAAAEMAESVTNPQRPPVIWIGAQECT WP_000145410.1
MULTISPECIES: hydrogenase-2 small chain [Proteobacteria] Length: 372\nNumber of predicted TMHs: 1\nExp number of AAs in TMHs: 23.72226\nExp number, first 60 AAs: 0.22038\nTotal prob of N-in: 0.12994\noutside 1 331\nTMhelix 332 354\ninside 355 372

5896 GCF_002173345.1_ASM217334v1 Escherichia coli Proteobacteria; Gammaproteobacteria; Enterobacterales; Enterobacteriaceae; Escherichia
MNNEETFYQAMRRQGVTRRSFLKYCSLAATSLGLGAGMAPKIAWALENKPRIPVVWIHGL WP_001058323.1
MULTISPECIES: hydrogenase-1 small chain [Proteobacteria] Length: 372\nNumber of predicted TMHs:

2\nExp number of AAs in TMHs: 37.2608499999999\nExp number, first 60 AAs: 19.52923\nTotal prob of N-in: 0.94351\nPOSSIBLE N-term signal sequence\ninside 1 19\nTMhelix 20 39\noutside 40 325\nTMhelix 326 348\ninside 349 372

5897 GCF_002173335.1_ASM217333v1 Escherichia coli Proteobacteria; Gammaproteobacteria; Enterobacterales; Enterobacteriaceae; Escherichia MNNEETFYQAMRRQGVTRRSFLKYCSLAATSLGLGAGMAPKIAWALENKPRIPVVWIHGL WP_001058323.1 MULTISPECIES: hydrogenase-1 small chain [Proteobacteria] Length: 372\nNumber of predicted TMHs: 2\nExp number of AAs in TMHs: 37.2608499999999\nExp number, first 60 AAs: 19.52923\nTotal prob of N-in: 0.94351\nPOSSIBLE N-term signal sequence\ninside 1 19\nTMhelix 20 39\noutside 40 325\nTMhelix 326 348\ninside 349 372

5898 GCF_002175315.1_ASM217531v1 Escherichia coli Proteobacteria; Gammaproteobacteria; Enterobacterales; Enterobacteriaceae; Escherichia MTGDNTLIHSHGINRRDFMKLCAALAATMGLSSKAAAEMAESVTNPQRPPVIWIGAQECT WP_000145410.1 MULTISPECIES: hydrogenase-2 small chain [Proteobacteria] Length: 372\nNumber of predicted TMHs: 1\nExp number of AAs in TMHs: 23.72226\nExp number, first 60 AAs: 0.22038\nTotal prob of N-in: 0.12994\noutside 1 331\nTMhelix 332 354\ninside 355 372

5899 GCF_002175375.1_ASM217537v1 Escherichia coli Proteobacteria; Gammaproteobacteria; Enterobacterales; Enterobacteriaceae; Escherichia MTGDNTLIHSHGINRRDFMKLCAALAATMGLSSKAAAEMAESVTNPQRPPVIWIGAQECT WP_000145410.1 MULTISPECIES: hydrogenase-2 small chain [Proteobacteria] Length: 372\nNumber of predicted TMHs: 1\nExp number of AAs in TMHs: 23.72226\nExp number, first 60 AAs: 0.22038\nTotal prob of N-in: 0.12994\noutside 1 331\nTMhelix 332 354\ninside 355 372

5900 GCF_002175315.1_ASM217531v1 Escherichia coli Proteobacteria; Gammaproteobacteria; Enterobacterales; Enterobacteriaceae; Escherichia MNNEETFYQAMRRQGVTRRSFLKYCSLAATSLGLGAGMAPKIAWALENKPRIPVVWIHGL WP_001301613.1 hydrogenase [Escherichia coli] Length: 372\nNumber of predicted TMHs: 2\nExp number of AAs in TMHs: 37.46598\nExp number, first 60 AAs: 19.53139\nTotal prob of N-in: 0.94364\nPOSSIBLE N-term signal sequence\ninside 1 19\nTMhelix 20 39\noutside 40 325\nTMhelix 326 348\ninside 349 372

5901 GCF_002175375.1_ASM217537v1 Escherichia coli Proteobacteria; Gammaproteobacteria; Enterobacterales; Enterobacteriaceae; Escherichia MNNEETFYQAMRRQGVTRRSFLKYCSLAATSLGLGAGMAPKIAWALENKPRIPVVWIHGL WP_001301613.1 hydrogenase [Escherichia coli] Length: 372\nNumber of predicted TMHs: 2\nExp number of AAs in TMHs: 37.46598\nExp number, first 60 AAs: 19.53139\nTotal prob of N-in: 0.94364\nPOSSIBLE N-term signal sequence\ninside 1 19\nTMhelix 20 39\noutside 40 325\nTMhelix 326 348\ninside 349 372

5902 GCF_002173365.1_ASM217336v1 Escherichia coli Proteobacteria; Gammaproteobacteria; Enterobacterales; Enterobacteriaceae; Escherichia MNNEETFYQAMRRQGVTRRSFLKYCSLAATSLGLGAGMAPKIAWALENKPRIPVVWIHGL WP_001058323.1 MULTISPECIES: hydrogenase-1 small chain [Proteobacteria] Length: 372\nNumber of predicted TMHs: 2\nExp number of AAs in TMHs: 37.2608499999999\nExp number, first 60 AAs: 19.52923\nTotal prob of N-in: 0.94351\nPOSSIBLE N-term signal sequence\ninside 1 19\nTMhelix 20 39\noutside 40 325\nTMhelix 326 348\ninside 349 372

5903 GCF_002175405.1_ASM217540v1 Escherichia coli Proteobacteria; Gammaproteobacteria; Enterobacterales; Enterobacteriaceae; Escherichia MTGDNTLIHSHGINRRDFMKLCAALAATMGLSSKAAAEMAESVTNPQRPPVIWIGAQECT WP_000145410.1 MULTISPECIES: hydrogenase-2 small chain [Proteobacteria] Length: 372\nNumber of predicted TMHs: 1\nExp number of AAs in TMHs: 23.72226\nExp number, first 60 AAs: 0.22038\nTotal prob of N-in: 0.12994\noutside 1 331\nTMhelix 332 354\ninside 355 372

5904 GCF_002175395.1_ASM217539v1 Escherichia coli Proteobacteria; Gammaproteobacteria; Enterobacterales; Enterobacteriaceae; Escherichia MTGDNTLIHSHGINRRDFMKLCAALAATMGLSSKAAAEMAESVTNPQRPPVIWIGAQECT WP_000145410.1 MULTISPECIES: hydrogenase-2 small chain [Proteobacteria] Length: 372\nNumber of predicted TMHs: 1\nExp number of AAs in TMHs: 23.72226\nExp number, first 60 AAs: 0.22038\nTotal prob of N-in: 0.12994\noutside 1 331\nTMhelix 332 354\ninside 355 372

5905 GCF_002175465.1_ASM217546v1 Escherichia coli Proteobacteria; Gammaproteobacteria; Enterobacterales; Enterobacteriaceae; Escherichia
MTGDNTLIHSHGINRRDFMKLCAALAATMGLSSKAAAEMAESVTNPQRPPVIWIGAQECT WP_000145410.1
MULTISPECIES: hydrogenase-2 small chain [Proteobacteria] Length: 372\nNumber of predicted TMHs: 1\nExp number of AAs in TMHs: 23.72226\nExp number, first 60 AAs: 0.22038\nTotal prob of N-in: 0.12994\noutside 1 331\nTMhelix 332 354\ninside 355 372

5906 GCF_002175485.1_ASM217548v1 Escherichia coli Proteobacteria; Gammaproteobacteria; Enterobacterales; Enterobacteriaceae; Escherichia
MTGDNTLIHSHGINRRDFMKLCAALAATMGLSSKAAAEMAESVTNPQRPPVIWIGAQECT WP_000145410.1
MULTISPECIES: hydrogenase-2 small chain [Proteobacteria] Length: 372\nNumber of predicted TMHs: 1\nExp number of AAs in TMHs: 23.72226\nExp number, first 60 AAs: 0.22038\nTotal prob of N-in: 0.12994\noutside 1 331\nTMhelix 332 354\ninside 355 372

5907 GCF_002175535.1_ASM217553v1 Escherichia coli Proteobacteria; Gammaproteobacteria; Enterobacterales; Enterobacteriaceae; Escherichia
MTGDNTLIHSHGINRRDFMKLCAALAATMGLSSKAAAEMAESVTNPQRPPVIWIGAQECT WP_000145410.1
MULTISPECIES: hydrogenase-2 small chain [Proteobacteria] Length: 372\nNumber of predicted TMHs: 1\nExp number of AAs in TMHs: 23.72226\nExp number, first 60 AAs: 0.22038\nTotal prob of N-in: 0.12994\noutside 1 331\nTMhelix 332 354\ninside 355 372

5908 GCF_002175625.1_ASM217562v1 Escherichia coli Proteobacteria; Gammaproteobacteria; Enterobacterales; Enterobacteriaceae; Escherichia
MNNEETFYQAMRRQGVTRRSFLKYCSLAATSLGLGAGMAPKIAWALENKPRIPVVWIHGL WP_001301613.1
hydrogenase [Escherichia coli] Length: 372\nNumber of predicted TMHs: 2\nExp number of AAs in TMHs: 37.46598\nExp number, first 60 AAs: 19.53139\nTotal prob of N-in: 0.94364\nPOSSIBLE N-term signal sequence\ninside 1 19\nTMhelix 20 39\noutside 40 325\nTMhelix 326 348\ninside 349 372

5909 GCF_002175465.1_ASM217546v1 Escherichia coli Proteobacteria; Gammaproteobacteria; Enterobacterales; Enterobacteriaceae; Escherichia
MNNEETFYQAMRRQGVTRRSFLKYCSLAATSLGLGAGMAPKIAWALENKPRIPVVWIHGL WP_001301613.1
hydrogenase [Escherichia coli] Length: 372\nNumber of predicted TMHs: 2\nExp number of AAs in TMHs: 37.46598\nExp number, first 60 AAs: 19.53139\nTotal prob of N-in: 0.94364\nPOSSIBLE N-term signal sequence\ninside 1 19\nTMhelix 20 39\noutside 40 325\nTMhelix 326 348\ninside 349 372

5910 GCF_002175705.1_ASM217570v1 Escherichia coli Proteobacteria; Gammaproteobacteria; Enterobacterales; Enterobacteriaceae; Escherichia
MTGDNTLIHSHGINRRDFMKLCAALAATMGLSSKAAAEMAESVTNPQRPPVIWIGAQECT WP_000145410.1
MULTISPECIES: hydrogenase-2 small chain [Proteobacteria] Length: 372\nNumber of predicted TMHs: 1\nExp number of AAs in TMHs: 23.72226\nExp number, first 60 AAs: 0.22038\nTotal prob of N-in: 0.12994\noutside 1 331\nTMhelix 332 354\ninside 355 372

5911 GCF_002175705.1_ASM217570v1 Escherichia coli Proteobacteria; Gammaproteobacteria; Enterobacterales; Enterobacteriaceae; Escherichia
MNNEETFYQAMRRQGVTRRSFLKYCSLAATSLGLGAGMAPKIAWALENKPRIPVVWIHGL WP_001301613.1
hydrogenase [Escherichia coli] Length: 372\nNumber of predicted TMHs: 2\nExp number of AAs in TMHs: 37.46598\nExp number, first 60 AAs: 19.53139\nTotal prob of N-in: 0.94364\nPOSSIBLE N-term signal sequence\ninside 1 19\nTMhelix 20 39\noutside 40 325\nTMhelix 326 348\ninside 349 372

5912 GCF_002175485.1_ASM217548v1 Escherichia coli Proteobacteria; Gammaproteobacteria; Enterobacterales; Enterobacteriaceae; Escherichia
MNNEETFYQAMRRQGVTRRSFLKYCSLAATSLGLGAGMAPKIAWALENKPRIPVVWIHGL WP_001301613.1
hydrogenase [Escherichia coli] Length: 372\nNumber of predicted TMHs: 2\nExp number of AAs in TMHs: 37.46598\nExp number, first 60 AAs: 19.53139\nTotal prob of N-in: 0.94364\nPOSSIBLE N-term signal sequence\ninside 1 19\nTMhelix 20 39\noutside 40 325\nTMhelix 326 348\ninside 349 372

5913 GCF_002175535.1_ASM217553v1 Escherichia coli Proteobacteria; Gammaproteobacteria; Enterobacterales; Enterobacteriaceae; Escherichia
MNNEETFYQAMRRQGVTRRSFLKYCSLAATSLGLGAGMAPKIAWALENKPRIPVVWIHGL WP_001301613.1
hydrogenase [Escherichia coli] Length: 372\nNumber of predicted TMHs: 2\nExp number of AAs in

TMHs: 37.46598\nExp number, first 60 AAs: 19.53139\nTotal prob of N-in: 0.94364\nPOSSIBLE N-term signal sequence\ninside 1 19\nTMhelix 20 39\noutside 40 325\nTMhelix 326 348\ninside 349 372

5914 GCF_002175715.1_ASM217571v1 Escherichia coli Proteobacteria; Gammaproteobacteria; Enterobacterales; Enterobacteriaceae; Escherichia MTGDNTLIHSHGINRRDFMKLCAALAATMGLSSKAAAEMAESVTNPQRPPVIWIGAQECT WP_000145410.1 MULTISPECIES: hydrogenase-2 small chain [Proteobacteria] Length: 372\nNumber of predicted TMHs: 1\nExp number of AAs in TMHs: 23.72226\nExp number, first 60 AAs: 0.22038\nTotal prob of N-in: 0.12994\noutside 1 331\nTMhelix 332 354\ninside 355 372

5915 GCF_002175775.1_ASM217577v1 Escherichia coli Proteobacteria; Gammaproteobacteria; Enterobacterales; Enterobacteriaceae; Escherichia MTGDNTLIHSHGINRRDFMKLCAALAATMGLSSKAAAEMAESVTNPQRPPVIWIGAQECT WP_000145410.1 MULTISPECIES: hydrogenase-2 small chain [Proteobacteria] Length: 372\nNumber of predicted TMHs: 1\nExp number of AAs in TMHs: 23.72226\nExp number, first 60 AAs: 0.22038\nTotal prob of N-in: 0.12994\noutside 1 331\nTMhelix 332 354\ninside 355 372

5916 GCF_002175715.1_ASM217571v1 Escherichia coli Proteobacteria; Gammaproteobacteria; Enterobacterales; Enterobacteriaceae; Escherichia MNNEETFYQAMRRQGVTRRSFLKYCSLAATSLGLGAGMAPKIAWALENKPRIPVVWIHGL WP_001301613.1 hydrogenase [Escherichia coli] Length: 372\nNumber of predicted TMHs: 2\nExp number of AAs in TMHs: 37.46598\nExp number, first 60 AAs: 19.53139\nTotal prob of N-in: 0.94364\nPOSSIBLE N-term signal sequence\ninside 1 19\nTMhelix 20 39\noutside 40 325\nTMhelix 326 348\ninside 349 372

5917 GCF_002175775.1_ASM217577v1 Escherichia coli Proteobacteria; Gammaproteobacteria; Enterobacterales; Enterobacteriaceae; Escherichia MNNEETFYQAMRRQGVTRRSFLKYCSLAATSLGLGAGMAPKIAWALENKPRIPVVWIHGL WP_001301613.1 hydrogenase [Escherichia coli] Length: 372\nNumber of predicted TMHs: 2\nExp number of AAs in TMHs: 37.46598\nExp number, first 60 AAs: 19.53139\nTotal prob of N-in: 0.94364\nPOSSIBLE N-term signal sequence\ninside 1 19\nTMhelix 20 39\noutside 40 325\nTMhelix 326 348\ninside 349 372

5918 GCF_002175555.1_ASM217555v1 Escherichia coli Proteobacteria; Gammaproteobacteria; Enterobacterales; Enterobacteriaceae; Escherichia MNNEETFYQAMRRQGVTRRSFLKYCSLAATSLGLGAGMAPKIAWALENKPRIPVVWIHGL WP_001301613.1 hydrogenase [Escherichia coli] Length: 372\nNumber of predicted TMHs: 2\nExp number of AAs in TMHs: 37.46598\nExp number, first 60 AAs: 19.53139\nTotal prob of N-in: 0.94364\nPOSSIBLE N-term signal sequence\ninside 1 19\nTMhelix 20 39\noutside 40 325\nTMhelix 326 348\ninside 349 372

5919 GCF_002175795.1_ASM217579v1 Escherichia coli Proteobacteria; Gammaproteobacteria; Enterobacterales; Enterobacteriaceae; Escherichia MTGDNTLIHSHGINRRDFMKLCAALAATMGLSSKAAAEMAESVTNPQRPPVIWIGAQECT WP_000145410.1 MULTISPECIES: hydrogenase-2 small chain [Proteobacteria] Length: 372\nNumber of predicted TMHs: 1\nExp number of AAs in TMHs: 23.72226\nExp number, first 60 AAs: 0.22038\nTotal prob of N-in: 0.12994\noutside 1 331\nTMhelix 332 354\ninside 355 372

5920 GCF_002175805.1_ASM217580v1 Escherichia coli Proteobacteria; Gammaproteobacteria; Enterobacterales; Enterobacteriaceae; Escherichia MTGDNTLIHSHGINRRDFMKLCAALAATMGLSSKAAAEMAESVTNPQRPPVIWIGAQECT WP_000145410.1 MULTISPECIES: hydrogenase-2 small chain [Proteobacteria] Length: 372\nNumber of predicted TMHs: 1\nExp number of AAs in TMHs: 23.72226\nExp number, first 60 AAs: 0.22038\nTotal prob of N-in: 0.12994\noutside 1 331\nTMhelix 332 354\ninside 355 372

5921 GCF_002175865.1_ASM217586v1 Escherichia coli Proteobacteria; Gammaproteobacteria; Enterobacterales; Enterobacteriaceae; Escherichia MTGDNTLIHSHGINRRDFMKLCAALAATMGLSSKAAAEMAESVTNPQRPPVIWIGAQECT WP_000145410.1 MULTISPECIES: hydrogenase-2 small chain [Proteobacteria] Length: 372\nNumber of predicted TMHs: 1\nExp number of AAs in TMHs: 23.72226\nExp number, first 60 AAs: 0.22038\nTotal prob of N-in: 0.12994\noutside 1 331\nTMhelix 332 354\ninside 355 372

5922 GCF_002175955.1_ASM217595v1 Escherichia coli Proteobacteria; Gammaproteobacteria; Enterobacterales; Enterobacteriaceae; Escherichia

MNNEETFYQAMRRQGVTRRSFLKYCSLAATSLGLGAGMAPKIAWALENKPRIPVVWIHGL WP_001301613.1 hydrogenase [Escherichia coli] Length: 372\nNumber of predicted TMHs: 2\nExp number of AAs in TMHs: 37.46598\nExp number, first 60 AAs: 19.53139\nTotal prob of N-in: 0.94364\nPOSSIBLE N-term signal sequence\ninside 1 19\nTMhelix 20 39\noutside 40 325\nTMhelix 326 348\ninside 349 372

5923 GCF_002175795.1_ASM217579v1 Escherichia coli Proteobacteria; Gammaproteobacteria; Enterobacterales; Enterobacteriaceae; Escherichia MNNEETFYQAMRRQGVTRRSFLKYCSLAATSLGLGAGMAPKIAWALENKPRIPVVWIHGL WP_001301613.1 hydrogenase [Escherichia coli] Length: 372\nNumber of predicted TMHs: 2\nExp number of AAs in TMHs: 37.46598\nExp number, first 60 AAs: 19.53139\nTotal prob of N-in: 0.94364\nPOSSIBLE N-term signal sequence\ninside 1 19\nTMhelix 20 39\noutside 40 325\nTMhelix 326 348\ninside 349 372

5924 GCF_002176025.1_ASM217602v1 Escherichia coli Proteobacteria; Gammaproteobacteria; Enterobacterales; Enterobacteriaceae; Escherichia MTGDNTLIHSHGINRRDFMKLCAALAATMGLSSKAAAEMAESVTNPQRPPVIWIGAQECT WP_000145410.1 MULTISPECIES: hydrogenase-2 small chain [Proteobacteria] Length: 372\nNumber of predicted TMHs: 1\nExp number of AAs in TMHs: 23.72226\nExp number, first 60 AAs: 0.22038\nTotal prob of N-in: 0.12994\noutside 1 331\nTMhelix 332 354\ninside 355 372

5925 GCF_002175865.1_ASM217586v1 Escherichia coli Proteobacteria; Gammaproteobacteria; Enterobacterales; Enterobacteriaceae; Escherichia MNNEETFYQAMRRQGVTRRSFLKYCSLAATSLGLGAGMAPKIAWALENKPRIPVVWIHGL WP_001301613.1 hydrogenase [Escherichia coli] Length: 372\nNumber of predicted TMHs: 2\nExp number of AAs in TMHs: 37.46598\nExp number, first 60 AAs: 19.53139\nTotal prob of N-in: 0.94364\nPOSSIBLE N-term signal sequence\ninside 1 19\nTMhelix 20 39\noutside 40 325\nTMhelix 326 348\ninside 349 372

5926 GCF_002176045.1_ASM217604v1 Escherichia coli Proteobacteria; Gammaproteobacteria; Enterobacterales; Enterobacteriaceae; Escherichia MTGDNTLIHSHGINRRDFMKLCAALAATMGLSSKAAAEMAESVTNPQRPPVIWIGAQECT WP_000145410.1 MULTISPECIES: hydrogenase-2 small chain [Proteobacteria] Length: 372\nNumber of predicted TMHs: 1\nExp number of AAs in TMHs: 23.72226\nExp number, first 60 AAs: 0.22038\nTotal prob of N-in: 0.12994\noutside 1 331\nTMhelix 332 354\ninside 355 372

5927 GCF_002175885.1_ASM217588v1 Escherichia coli Proteobacteria; Gammaproteobacteria; Enterobacterales; Enterobacteriaceae; Escherichia MNNEETFYQAMRRQGVTRRSFLKYCSLAATSLGLGAGMAPKIAWALENKPRIPVVWIHGL WP_001301613.1 hydrogenase [Escherichia coli] Length: 372\nNumber of predicted TMHs: 2\nExp number of AAs in TMHs: 37.46598\nExp number, first 60 AAs: 19.53139\nTotal prob of N-in: 0.94364\nPOSSIBLE N-term signal sequence\ninside 1 19\nTMhelix 20 39\noutside 40 325\nTMhelix 326 348\ninside 349 372

5928 GCF_002175935.1_ASM217593v1 Escherichia coli Proteobacteria; Gammaproteobacteria; Enterobacterales; Enterobacteriaceae; Escherichia MNNEETFYQAMRRQGVTRRSFLKYCSLAATSLGLGAGMAPKIAWALENKPRIPVVWIHGL WP_001301613.1 hydrogenase [Escherichia coli] Length: 372\nNumber of predicted TMHs: 2\nExp number of AAs in TMHs: 37.46598\nExp number, first 60 AAs: 19.53139\nTotal prob of N-in: 0.94364\nPOSSIBLE N-term signal sequence\ninside 1 19\nTMhelix 20 39\noutside 40 325\nTMhelix 326 348\ninside 349 372

5929 GCF_002176115.1_ASM217611v1 Escherichia coli Proteobacteria; Gammaproteobacteria; Enterobacterales; Enterobacteriaceae; Escherichia MTGDNTLIHSHGINRRDFMKLCAALAATMGLSSKAAAEMAESVTNPQRPPVIWIGAQECT WP_000145410.1 MULTISPECIES: hydrogenase-2 small chain [Proteobacteria] Length: 372\nNumber of predicted TMHs: 1\nExp number of AAs in TMHs: 23.72226\nExp number, first 60 AAs: 0.22038\nTotal prob of N-in: 0.12994\noutside 1 331\nTMhelix 332 354\ninside 355 372

5930 GCF_002176175.1_ASM217617v1 Escherichia coli Proteobacteria; Gammaproteobacteria; Enterobacterales; Enterobacteriaceae; Escherichia MTGDNTLIHSHGINRRDFMKLCAALAATMGLSSKAAAEMAESVTNPQRPPVIWIGAQECT WP_000145410.1 MULTISPECIES: hydrogenase-2 small chain [Proteobacteria] Length: 372\nNumber of predicted TMHs: 1\nExp number of AAs in TMHs: 23.72226\nExp number, first 60 AAs: 0.22038\nTotal prob of N-in: 0.12994\noutside 1 331\nTMhelix 332 354\ninside 355 372

5931 GCF_002176215.1_ASM217621v1 Escherichia coli Proteobacteria; Gammaproteobacteria; Enterobacterales; Enterobacteriaceae; Escherichia
MTGDNTLIHSHGINRRDFMKLCAALAATMGLSSKAAAEMAESVTNPQRPPVIWIGAQECT WP_000145410.1
MULTISPECIES: hydrogenase-2 small chain [Proteobacteria] Length: 372\nNumber of predicted TMHs: 1\nExp number of AAs in TMHs: 23.72226\nExp number, first 60 AAs: 0.22038\nTotal prob of N-in: 0.12994\noutside 1 331\nTMhelix 332 354\ninside 355 372

5932 GCF_002176195.1_ASM217619v1 Escherichia coli Proteobacteria; Gammaproteobacteria; Enterobacterales; Enterobacteriaceae; Escherichia
MTGDNTLIHSHGINRRDFMKLCAALAATMGLSSKAAAEMAESVTNPQRPPVIWIGAQECT WP_000145410.1
MULTISPECIES: hydrogenase-2 small chain [Proteobacteria] Length: 372\nNumber of predicted TMHs: 1\nExp number of AAs in TMHs: 23.72226\nExp number, first 60 AAs: 0.22038\nTotal prob of N-in: 0.12994\noutside 1 331\nTMhelix 332 354\ninside 355 372

5933 GCF_002176265.1_ASM217626v1 Escherichia coli Proteobacteria; Gammaproteobacteria; Enterobacterales; Enterobacteriaceae; Escherichia
MTGDNTLIHSHGINRRDFMKLCAALAATMGLSSKAAAEMAESVTNPQRPPVIWIGAQECT WP_000145410.1
MULTISPECIES: hydrogenase-2 small chain [Proteobacteria] Length: 372\nNumber of predicted TMHs: 1\nExp number of AAs in TMHs: 23.72226\nExp number, first 60 AAs: 0.22038\nTotal prob of N-in: 0.12994\noutside 1 331\nTMhelix 332 354\ninside 355 372

5934 GCF_002176295.1_ASM217629v1 Escherichia coli Proteobacteria; Gammaproteobacteria; Enterobacterales; Enterobacteriaceae; Escherichia
MTGDNTLIHSHGINRRDFMKLCAALAATMGLSSKAAAEMAESVTNPQRPPVIWIGAQECT WP_000145410.1
MULTISPECIES: hydrogenase-2 small chain [Proteobacteria] Length: 372\nNumber of predicted TMHs: 1\nExp number of AAs in TMHs: 23.72226\nExp number, first 60 AAs: 0.22038\nTotal prob of N-in: 0.12994\noutside 1 331\nTMhelix 332 354\ninside 355 372

5935 GCF_002176335.1_ASM217633v1 Escherichia coli Proteobacteria; Gammaproteobacteria; Enterobacterales; Enterobacteriaceae; Escherichia
MTGDNTLIHSHGINRRDFMKLCAALAATMGLSSKAAAEMAESVTNPQRPPVIWIGAQECT WP_000145410.1
MULTISPECIES: hydrogenase-2 small chain [Proteobacteria] Length: 372\nNumber of predicted TMHs: 1\nExp number of AAs in TMHs: 23.72226\nExp number, first 60 AAs: 0.22038\nTotal prob of N-in: 0.12994\noutside 1 331\nTMhelix 332 354\ninside 355 372

5936 GCF_002176015.1_ASM217601v1 Escherichia coli Proteobacteria; Gammaproteobacteria; Enterobacterales; Enterobacteriaceae; Escherichia
MNNEETFYQAMRRQGVTRRSFLKYCSLAATSLGLGAGMAPKIAWALENKPRIPVVWIHGL WP_001301613.1
hydrogenase [Escherichia coli] Length: 372\nNumber of predicted TMHs: 2\nExp number of AAs in TMHs: 37.46598\nExp number, first 60 AAs: 19.53139\nTotal prob of N-in: 0.94364\nPOSSIBLE N-term signal sequence\ninside 1 19\nTMhelix 20 39\noutside 40 325\nTMhelix 326 348\ninside 349 372

5937 GCF_002176475.1_ASM217647v1 Escherichia coli Proteobacteria; Gammaproteobacteria; Enterobacterales; Enterobacteriaceae; Escherichia
MTGDNTLIHSHGINRRDFMKLCAALAATMGLSSKAAAEMAESVTNPQRPPVIWIGAQECT WP_000145410.1
MULTISPECIES: hydrogenase-2 small chain [Proteobacteria] Length: 372\nNumber of predicted TMHs: 1\nExp number of AAs in TMHs: 23.72226\nExp number, first 60 AAs: 0.22038\nTotal prob of N-in: 0.12994\noutside 1 331\nTMhelix 332 354\ninside 355 372

5938 GCF_002176175.1_ASM217617v1 Escherichia coli Proteobacteria; Gammaproteobacteria; Enterobacterales; Enterobacteriaceae; Escherichia
MNNEETFYQAMRRQGVTRRSFLKYCSLAATSLGLGAGMAPKIAWALENKPRIPVVWIHGL WP_001301613.1
hydrogenase [Escherichia coli] Length: 372\nNumber of predicted TMHs: 2\nExp number of AAs in TMHs: 37.46598\nExp number, first 60 AAs: 19.53139\nTotal prob of N-in: 0.94364\nPOSSIBLE N-term signal sequence\ninside 1 19\nTMhelix 20 39\noutside 40 325\nTMhelix 326 348\ninside 349 372

5939 GCF_002176505.1_ASM217650v1 Escherichia coli Proteobacteria; Gammaproteobacteria; Enterobacterales; Enterobacteriaceae; Escherichia
MTGDNTLIHSHGINRRDFMKLCAALAATMGLSSKAAAEMAESVTNPQRPPVIWIGAQECT WP_000145410.1
MULTISPECIES: hydrogenase-2 small chain [Proteobacteria] Length: 372\nNumber of predicted TMHs:

1\nExp number of AAs in TMHs: 23.72226\nExp number, first 60 AAs: 0.22038\nTotal prob of N-in: 0.12994\noutside 1 331\nTMhelix 332 354\ninside 355 372

5940 GCF_002176295.1_ASM217629v1 Escherichia coli Proteobacteria; Gammaproteobacteria; Enterobacterales; Enterobacteriaceae; Escherichia
MNNEETFYQAMRRQGVTRRSFLKYCSLAATSLGLGAGMAPKIAWALENKPRIPVVWIHGL WP_001301613.1
hydrogenase [Escherichia coli] Length: 372\nNumber of predicted TMHs: 2\nExp number of AAs in TMHs: 37.46598\nExp number, first 60 AAs: 19.53139\nTotal prob of N-in: 0.94364\nPOSSIBLE N-term signal sequence\ninside 1 19\nTMhelix 20 39\noutside 40 325\nTMhelix 326 348\ninside 349 372

5941 GCF_002176265.1_ASM217626v1 Escherichia coli Proteobacteria; Gammaproteobacteria; Enterobacterales; Enterobacteriaceae; Escherichia
MNNEETFYQAMRRQGVTRRSFLKYCSLAATSLGLGAGMAPKIAWALENKPRIPVVWIHGL WP_001301613.1
hydrogenase [Escherichia coli] Length: 372\nNumber of predicted TMHs: 2\nExp number of AAs in TMHs: 37.46598\nExp number, first 60 AAs: 19.53139\nTotal prob of N-in: 0.94364\nPOSSIBLE N-term signal sequence\ninside 1 19\nTMhelix 20 39\noutside 40 325\nTMhelix 326 348\ninside 349 372

5942 GCF_002176215.1_ASM217621v1 Escherichia coli Proteobacteria; Gammaproteobacteria; Enterobacterales; Enterobacteriaceae; Escherichia
MNNEETFYQAMRRQGVTRRSFLKYCSLAATSLGLGAGMAPKIAWALENKPRIPVVWIHGL WP_001301613.1
hydrogenase [Escherichia coli] Length: 372\nNumber of predicted TMHs: 2\nExp number of AAs in TMHs: 37.46598\nExp number, first 60 AAs: 19.53139\nTotal prob of N-in: 0.94364\nPOSSIBLE N-term signal sequence\ninside 1 19\nTMhelix 20 39\noutside 40 325\nTMhelix 326 348\ninside 349 372

5943 GCF_002176745.1_ASM217674v1 Escherichia coli Proteobacteria; Gammaproteobacteria; Enterobacterales; Enterobacteriaceae; Escherichia
MTGDNTLIHSHGINRRDFMKLCAALAATMGLSSKAAAEMAESVTNPQRPPVIWIGAQECT WP_000145410.1
MULTISPECIES: hydrogenase-2 small chain [Proteobacteria] Length: 372\nNumber of predicted TMHs: 1\nExp number of AAs in TMHs: 23.72226\nExp number, first 60 AAs: 0.22038\nTotal prob of N-in: 0.12994\noutside 1 331\nTMhelix 332 354\ninside 355 372

5944 GCF_002176195.1_ASM217619v1 Escherichia coli Proteobacteria; Gammaproteobacteria; Enterobacterales; Enterobacteriaceae; Escherichia
MNNEETFYQAMRRQGVTRRSFLKYCSLAATSLGLGAGMAPKIAWALENKPRIPVVWIHGL WP_001301613.1
hydrogenase [Escherichia coli] Length: 372\nNumber of predicted TMHs: 2\nExp number of AAs in TMHs: 37.46598\nExp number, first 60 AAs: 19.53139\nTotal prob of N-in: 0.94364\nPOSSIBLE N-term signal sequence\ninside 1 19\nTMhelix 20 39\noutside 40 325\nTMhelix 326 348\ninside 349 372

5945 GCF_002176335.1_ASM217633v1 Escherichia coli Proteobacteria; Gammaproteobacteria; Enterobacterales; Enterobacteriaceae; Escherichia
MNNEETFYQAMRRQGVTRRSFLKYCSLAATSLGLGAGMAPKIAWALENKPRIPVVWIHGL WP_001301613.1
hydrogenase [Escherichia coli] Length: 372\nNumber of predicted TMHs: 2\nExp number of AAs in TMHs: 37.46598\nExp number, first 60 AAs: 19.53139\nTotal prob of N-in: 0.94364\nPOSSIBLE N-term signal sequence\ninside 1 19\nTMhelix 20 39\noutside 40 325\nTMhelix 326 348\ninside 349 372

5946 GCF_002176515.1_ASM217651v1 Escherichia coli Proteobacteria; Gammaproteobacteria; Enterobacterales; Enterobacteriaceae; Escherichia
MTGDNTLIHSHGINRRDFMKLCAALAATMGLSSKAAAEMAESVTNPQRPPVIWIGAQECT WP_000145410.1
MULTISPECIES: hydrogenase-2 small chain [Proteobacteria] Length: 372\nNumber of predicted TMHs: 1\nExp number of AAs in TMHs: 23.72226\nExp number, first 60 AAs: 0.22038\nTotal prob of N-in: 0.12994\noutside 1 331\nTMhelix 332 354\ninside 355 372

5947 GCF_002176515.1_ASM217651v1 Escherichia coli Proteobacteria; Gammaproteobacteria; Enterobacterales; Enterobacteriaceae; Escherichia
MNNEETFYQAMRRQGVTRRSFLKYCSLAATSLGLGAGMAPKIAWALENKPRIPVVWIHGL WP_001301613.1
hydrogenase [Escherichia coli] Length: 372\nNumber of predicted TMHs: 2\nExp number of AAs in TMHs: 37.46598\nExp number, first 60 AAs: 19.53139\nTotal prob of N-in: 0.94364\nPOSSIBLE N-term signal sequence\ninside 1 19\nTMhelix 20 39\noutside 40 325\nTMhelix 326 348\ninside 349 372

5948 GCF_002176355.1_ASM217635v1 Escherichia coli Proteobacteria; Gammaproteobacteria; Enterobacterales; Enterobacteriaceae; Escherichia

MNNEETFYQAMRRQGVTRRSFLKYCSLAATSLGLGAGMAPKIAWALENKPRIPVVWIHGL WP_001301613.1 hydrogenase [Escherichia coli] Length: 372\nNumber of predicted TMHs: 2\nExp number of AAs in TMHs: 37.46598\nExp number, first 60 AAs: 19.53139\nTotal prob of N-in: 0.94364\nPOSSIBLE N-term signal sequence\ninside 1 19\nTMhelix 20 39\noutside 40 325\nTMhelix 326 348\ninside 349 372

5949 GCF_002176645.1_ASM217664v1 Escherichia coli Proteobacteria; Gammaproteobacteria; Enterobacterales; Enterobacteriaceae; Escherichia MTGDNTLIHSHGINRRDFMKLCAALAATMGLSSKAAAEMAESVTNPQRPPVIWIGAQECT WP_000145410.1 MULTISPECIES: hydrogenase-2 small chain [Proteobacteria] Length: 372\nNumber of predicted TMHs: 1\nExp number of AAs in TMHs: 23.72226\nExp number, first 60 AAs: 0.22038\nTotal prob of N-in: 0.12994\noutside 1 331\nTMhelix 332 354\ninside 355 372

5950 GCF_002176735.1_ASM217673v1 Escherichia coli Proteobacteria; Gammaproteobacteria; Enterobacterales; Enterobacteriaceae; Escherichia MTGDNTLIHSHGINRRDFMKLCAALAATMGLSSKAAAEMAESVTNPQRPPVIWIGAQECT WP_000145410.1 MULTISPECIES: hydrogenase-2 small chain [Proteobacteria] Length: 372\nNumber of predicted TMHs: 1\nExp number of AAs in TMHs: 23.72226\nExp number, first 60 AAs: 0.22038\nTotal prob of N-in: 0.12994\noutside 1 331\nTMhelix 332 354\ninside 355 372

5951 GCF_002176505.1_ASM217650v1 Escherichia coli Proteobacteria; Gammaproteobacteria; Enterobacterales; Enterobacteriaceae; Escherichia MNNEETFYQAMRRQGVTRRSFLKYCSLAATSLGLGAGMAPKIAWALENKPRIPVVWIHGL WP_001301613.1 hydrogenase [Escherichia coli] Length: 372\nNumber of predicted TMHs: 2\nExp number of AAs in TMHs: 37.46598\nExp number, first 60 AAs: 19.53139\nTotal prob of N-in: 0.94364\nPOSSIBLE N-term signal sequence\ninside 1 19\nTMhelix 20 39\noutside 40 325\nTMhelix 326 348\ninside 349 372

5952 GCF_002188315.1_ASM218831v1 Escherichia coli Proteobacteria; Gammaproteobacteria; Enterobacterales; Enterobacteriaceae; Escherichia MTGDNTLIHSHGINRRDFMKLCAALAATMGLSSKAAAEMAESVTNPQRPPVIWIGAQECT WP_000145410.1 MULTISPECIES: hydrogenase-2 small chain [Proteobacteria] Length: 372\nNumber of predicted TMHs: 1\nExp number of AAs in TMHs: 23.72226\nExp number, first 60 AAs: 0.22038\nTotal prob of N-in: 0.12994\noutside 1 331\nTMhelix 332 354\ninside 355 372

5953 GCF_002188315.1_ASM218831v1 Escherichia coli Proteobacteria; Gammaproteobacteria; Enterobacterales; Enterobacteriaceae; Escherichia MNNEETFYQAMRRQGVTRRSFLKYCSLAATSLGLGAGMAPKIAWALENKPRIPVVWIHGL WP_001058323.1 MULTISPECIES: hydrogenase-1 small chain [Proteobacteria] Length: 372\nNumber of predicted TMHs: 2\nExp number of AAs in TMHs: 37.2608499999999\nExp number, first 60 AAs: 19.52923\nTotal prob of N-in: 0.94351\nPOSSIBLE N-term signal sequence\ninside 1 19\nTMhelix 20 39\noutside 40 325\nTMhelix 326 348\ninside 349 372

5954 GCF_002180815.1_ASM218081v1 Escherichia coli Proteobacteria; Gammaproteobacteria; Enterobacterales; Enterobacteriaceae; Escherichia MNNEETFYQAMRRQGVTRRSFLKYCSLAATSLGLGAGMAPKIAWALENKPRIPVVWIHGL WP_001058323.1 MULTISPECIES: hydrogenase-1 small chain [Proteobacteria] Length: 372\nNumber of predicted TMHs: 2\nExp number of AAs in TMHs: 37.2608499999999\nExp number, first 60 AAs: 19.52923\nTotal prob of N-in: 0.94351\nPOSSIBLE N-term signal sequence\ninside 1 19\nTMhelix 20 39\noutside 40 325\nTMhelix 326 348\ninside 349 372

5955 GCF_002189745.1_ASM218974v1 Escherichia coli Proteobacteria; Gammaproteobacteria; Enterobacterales; Enterobacteriaceae; Escherichia MTGDNTLIHSHGINRRDFMKLCAALAATMGLSSKAAAEMAESVTNPQRPPVIWIGAQECT WP_000145410.1 MULTISPECIES: hydrogenase-2 small chain [Proteobacteria] Length: 372\nNumber of predicted TMHs: 1\nExp number of AAs in TMHs: 23.72226\nExp number, first 60 AAs: 0.22038\nTotal prob of N-in: 0.12994\noutside 1 331\nTMhelix 332 354\ninside 355 372

5956 GCF_002189735.1_ASM218973v1 Escherichia coli Proteobacteria; Gammaproteobacteria; Enterobacterales; Enterobacteriaceae; Escherichia MTGDNTLIHSHGINRRDFMKLCAALAATMGLSSKAAAEMAESVTNPQRPPVIWIGAQECT WP_000145410.1 MULTISPECIES: hydrogenase-2 small chain [Proteobacteria] Length: 372\nNumber of predicted TMHs:

1\nExp number of AAs in TMHs: 23.72226\nExp number, first 60 AAs: 0.22038\nTotal prob of N-in: 0.12994\noutside 1 331\nTMhelix 332 354\ninside 355 372

5957 GCF_002189765.1_ASM218976v1 Escherichia coli Proteobacteria; Gammaproteobacteria; Enterobacterales; Enterobacteriaceae; Escherichia
MTGDNTLIHSHGINRRDFMKLCAALAATMGLSSKAAAEMAESVTNPQRPPVIWIGAQECT WP_000145410.1
MULTISPECIES: hydrogenase-2 small chain [Proteobacteria] Length: 372\nNumber of predicted TMHs: 1\nExp number of AAs in TMHs: 23.72226\nExp number, first 60 AAs: 0.22038\nTotal prob of N-in: 0.12994\noutside 1 331\nTMhelix 332 354\ninside 355 372

5958 GCF_002189745.1_ASM218974v1 Escherichia coli Proteobacteria; Gammaproteobacteria; Enterobacterales; Enterobacteriaceae; Escherichia
MNNEETFYQAMRRQGVTRRSFLKYCSLAATSLGLGAGMAPKIAWALENKPRIPVVWIHGL WP_001058323.1
MULTISPECIES: hydrogenase-1 small chain [Proteobacteria] Length: 372\nNumber of predicted TMHs: 2\nExp number of AAs in TMHs: 37.2608499999999\nExp number, first 60 AAs: 19.52923\nTotal prob of N-in: 0.94351\nPOSSIBLE N-term signal sequence\ninside 1 19\nTMhelix 20 39\noutside 40 325\nTMhelix 326 348\ninside 349 372

5959 GCF_002189735.1_ASM218973v1 Escherichia coli Proteobacteria; Gammaproteobacteria; Enterobacterales; Enterobacteriaceae; Escherichia
MNNEETFYQAMRRQGVTRRSFLKYCSLAATSLGLGAGMAPKIAWALENKPRIPVVWIHGL WP_001058323.1
MULTISPECIES: hydrogenase-1 small chain [Proteobacteria] Length: 372\nNumber of predicted TMHs: 2\nExp number of AAs in TMHs: 37.2608499999999\nExp number, first 60 AAs: 19.52923\nTotal prob of N-in: 0.94351\nPOSSIBLE N-term signal sequence\ninside 1 19\nTMhelix 20 39\noutside 40 325\nTMhelix 326 348\ninside 349 372

5960 GCF_002176735.1_ASM217673v1 Escherichia coli Proteobacteria; Gammaproteobacteria; Enterobacterales; Enterobacteriaceae; Escherichia
MNNEETFYQAMRRQGVTRRSFLKYCSLAATSLGLGAGMAPKIAWALENKPRIPVVWIHGL WP_001301613.1
hydrogenase [Escherichia coli] Length: 372\nNumber of predicted TMHs: 2\nExp number of AAs in TMHs: 37.46598\nExp number, first 60 AAs: 19.53139\nTotal prob of N-in: 0.94364\nPOSSIBLE N-term signal sequence\ninside 1 19\nTMhelix 20 39\noutside 40 325\nTMhelix 326 348\ninside 349 372

5961 GCF_002176645.1_ASM217664v1 Escherichia coli Proteobacteria; Gammaproteobacteria; Enterobacterales; Enterobacteriaceae; Escherichia
MNNEETFYQAMRRQGVTRRSFLKYCSLAATSLGLGAGMAPKIAWALENKPRIPVVWIHGL WP_001301613.1
hydrogenase [Escherichia coli] Length: 372\nNumber of predicted TMHs: 2\nExp number of AAs in TMHs: 37.46598\nExp number, first 60 AAs: 19.53139\nTotal prob of N-in: 0.94364\nPOSSIBLE N-term signal sequence\ninside 1 19\nTMhelix 20 39\noutside 40 325\nTMhelix 326 348\ninside 349 372

5962 GCF_002181625.1_ASM218162v1 Escherichia coli Proteobacteria; Gammaproteobacteria; Enterobacterales; Enterobacteriaceae; Escherichia
MNNEETFYQAMRRQGVTRRSFLKYCSLAATSLGLGAGMAPKIAWALENKPRIPVVWIHGL WP_001295940.1
MULTISPECIES: hydrogenase [Enterobacteriaceae] Length: 372\nNumber of predicted TMHs: 2\nExp number of AAs in TMHs: 37.2608899999999\nExp number, first 60 AAs: 19.52923\nTotal prob of N-in: 0.94351\nPOSSIBLE N-term signal sequence\ninside 1 19\nTMhelix 20 39\noutside 40 325\nTMhelix 326 348\ninside 349 372

5963 GCF_002176745.1_ASM217674v1 Escherichia coli Proteobacteria; Gammaproteobacteria; Enterobacterales; Enterobacteriaceae; Escherichia
MNNEETFYQAMRRQGVTRRSFLKYCSLAATSLGLGAGMAPKIAWALENKPRIPVVWIHGL WP_001301613.1
hydrogenase [Escherichia coli] Length: 372\nNumber of predicted TMHs: 2\nExp number of AAs in TMHs: 37.46598\nExp number, first 60 AAs: 19.53139\nTotal prob of N-in: 0.94364\nPOSSIBLE N-term signal sequence\ninside 1 19\nTMhelix 20 39\noutside 40 325\nTMhelix 326 348\ninside 349 372

5964 GCF_002179755.1_ASM217975v1 Escherichia coli Proteobacteria; Gammaproteobacteria; Enterobacterales; Enterobacteriaceae; Escherichia
MNNEETFYQAMRRQGVTRRSFLKYCSLAATSLGLGAGMAPKIAWALENKPRIPVVWIHGL WP_001301613.1
hydrogenase [Escherichia coli] Length: 372\nNumber of predicted TMHs: 2\nExp number of AAs in TMHs: 37.46598\nExp number, first 60 AAs: 19.53139\nTotal prob of N-in: 0.94364\nPOSSIBLE N-term signal sequence\ninside 1 19\nTMhelix 20 39\noutside 40 325\nTMhelix 326 348\ninside 349 372

5965 GCF_002189835.1_ASM218983v1 Escherichia coli Proteobacteria; Gammaproteobacteria; Enterobacterales; Enterobacteriaceae; Escherichia
MTGDNTLIHSHGINRRDFMKLCAALAATMGLSSKAAAEMAESVTNPQRPPVIWIGAQECT WP_000145410.1
MULTISPECIES: hydrogenase-2 small chain [Proteobacteria] Length: 372\nNumber of predicted TMHs: 1\nExp number of AAs in TMHs: 23.72226\nExp number, first 60 AAs: 0.22038\nTotal prob of N-in: 0.12994\noutside 1 331\nTMhelix 332 354\ninside 355 372

5966 GCF_002189835.1_ASM218983v1 Escherichia coli Proteobacteria; Gammaproteobacteria; Enterobacterales; Enterobacteriaceae; Escherichia
MNNEETFYQAMRRQGVTRRSFLKYCSLAATSLGLGAGMAPKIAWALENKPRIPVVWIHGL WP_001058323.1
MULTISPECIES: hydrogenase-1 small chain [Proteobacteria] Length: 372\nNumber of predicted TMHs: 2\nExp number of AAs in TMHs: 37.2608499999999\nExp number, first 60 AAs: 19.52923\nTotal prob of N-in: 0.94351\nPOSSIBLE N-term signal sequence\ninside 1 19\nTMhelix 20 39\noutside 40 325\nTMhelix 326 348\ninside 349 372

5967 GCF_002189765.1_ASM218976v1 Escherichia coli Proteobacteria; Gammaproteobacteria; Enterobacterales; Enterobacteriaceae; Escherichia
MNNEETFYQAMRRQGVTRRSFLKYCSLAATSLGLGAGMAPKIAWALENKPRIPVVWIHGL WP_001058323.1
MULTISPECIES: hydrogenase-1 small chain [Proteobacteria] Length: 372\nNumber of predicted TMHs: 2\nExp number of AAs in TMHs: 37.2608499999999\nExp number, first 60 AAs: 19.52923\nTotal prob of N-in: 0.94351\nPOSSIBLE N-term signal sequence\ninside 1 19\nTMhelix 20 39\noutside 40 325\nTMhelix 326 348\ninside 349 372

5968 GCF_002189895.1_ASM218989v1 Escherichia coli Proteobacteria; Gammaproteobacteria; Enterobacterales; Enterobacteriaceae; Escherichia
MTGDNTLIHSHGINRRDFMKLCAALAATMGLSSKAAAEMAESVTNPQRPPVIWIGAQECT WP_000145410.1
MULTISPECIES: hydrogenase-2 small chain [Proteobacteria] Length: 372\nNumber of predicted TMHs: 1\nExp number of AAs in TMHs: 23.72226\nExp number, first 60 AAs: 0.22038\nTotal prob of N-in: 0.12994\noutside 1 331\nTMhelix 332 354\ninside 355 372

5969 GCF_002189945.1_ASM218994v1 Escherichia coli Proteobacteria; Gammaproteobacteria; Enterobacterales; Enterobacteriaceae; Escherichia
MTGDNTLIHSHGINRRDFMKLCAALAATMGLSSKAAAEMVESVTNPQRPPVIWIGAQECT WP_000145423.1
hydrogenase 2 small subunit [Escherichia coli] Length: 372\nNumber of predicted TMHs: 1\nExp number of AAs in TMHs: 23.78201\nExp number, first 60 AAs: 0.28355\nTotal prob of N-in: 0.13257\noutside 1 331\nTMhelix 332 354\ninside 355 372

5970 GCF_002189985.1_ASM218998v1 Escherichia coli Proteobacteria; Gammaproteobacteria; Enterobacterales; Enterobacteriaceae; Escherichia
MNNEETFYQAMRRQGVTRRSFLKYCSLAATSLGLGAGMAPKIAWALENKPRIPVVWIHGL WP_001295940.1
MULTISPECIES: hydrogenase [Enterobacteriaceae] Length: 372\nNumber of predicted TMHs: 2\nExp number of AAs in TMHs: 37.2608899999999\nExp number, first 60 AAs: 19.52923\nTotal prob of N-in: 0.94351\nPOSSIBLE N-term signal sequence\ninside 1 19\nTMhelix 20 39\noutside 40 325\nTMhelix 326 348\ninside 349 372

5971 GCF_002189945.1_ASM218994v1 Escherichia coli Proteobacteria; Gammaproteobacteria; Enterobacterales; Enterobacteriaceae; Escherichia
MNNEETFYQAMRRQGVTRRSFLKYCSLAATSLGLGAGMAPKIAWALENKPRIPVVWIHGL WP_001538532.1
hydrogenase-1 small chain [Escherichia coli] Length: 372\nNumber of predicted TMHs: 2\nExp number of AAs in TMHs: 37.27041\nExp number, first 60 AAs: 19.52927\nTotal prob of N-in: 0.94352\nPOSSIBLE N-term signal sequence\ninside 1 19\nTMhelix 20 39\noutside 40 325\nTMhelix 326 348\ninside 349 372

5972 GCF_002190035.1_ASM219003v1 Escherichia coli Proteobacteria; Gammaproteobacteria; Enterobacterales; Enterobacteriaceae; Escherichia
MNNEETFYQAMRRQGVTRRSFLKYCSLAATSLGLGAGMAPKIAWALENKPRIPVVWIHGL WP_001058323.1
MULTISPECIES: hydrogenase-1 small chain [Proteobacteria] Length: 372\nNumber of predicted TMHs: 2\nExp number of AAs in TMHs: 37.2608499999999\nExp number, first 60 AAs: 19.52923\nTotal prob of N-in: 0.94351\nPOSSIBLE N-term signal sequence\ninside 1 19\nTMhelix 20 39\noutside 40 325\nTMhelix 326 348\ninside 349 372

5973 GCF_002190145.1_ASM219014v1 Escherichia coli Proteobacteria; Gammaproteobacteria; Enterobacterales; Enterobacteriaceae; Escherichia
MTGDNTLIHSHGINRRDFMKLCAALAATMGLSSKAAAEMAESVTNPQRPPVIWIGAQECT WP_000145410.1
MULTISPECIES: hydrogenase-2 small chain [Proteobacteria] Length: 372\nNumber of predicted TMHs: 1\nExp number of AAs in TMHs: 23.72226\nExp number, first 60 AAs: 0.22038\nTotal prob of N-in: 0.12994\noutside 1 331\nTMhelix 332 354\ninside 355 372

5974 GCF_002190155.1_ASM219015v1 Escherichia coli Proteobacteria; Gammaproteobacteria; Enterobacterales; Enterobacteriaceae; Escherichia
MTGDNTLIHSHGINRRDFMKLCAALAATMGLSSKAAAEMAESVTNPQRPPVIWIGAQECT WP_000145410.1
MULTISPECIES: hydrogenase-2 small chain [Proteobacteria] Length: 372\nNumber of predicted TMHs: 1\nExp number of AAs in TMHs: 23.72226\nExp number, first 60 AAs: 0.22038\nTotal prob of N-in: 0.12994\noutside 1 331\nTMhelix 332 354\ninside 355 372

5975 GCF_002190245.1_ASM219024v1 Escherichia coli Proteobacteria; Gammaproteobacteria; Enterobacterales; Enterobacteriaceae; Escherichia
MTGDNTLIHSHGINRRDFMKLCAALAATMGLSSKAAAEMAESVTNPQRPPVIWIGAQECT WP_000145410.1
MULTISPECIES: hydrogenase-2 small chain [Proteobacteria] Length: 372\nNumber of predicted TMHs: 1\nExp number of AAs in TMHs: 23.72226\nExp number, first 60 AAs: 0.22038\nTotal prob of N-in: 0.12994\noutside 1 331\nTMhelix 332 354\ninside 355 372

5976 GCF_002190205.1_ASM219020v1 Escherichia coli Proteobacteria; Gammaproteobacteria; Enterobacterales; Enterobacteriaceae; Escherichia
MTGDNTLIHSHGINRRDFMKLCAALAATMGLSSKAAAEMAESVTNPQRPPVIWIGAQECT WP_000145410.1
MULTISPECIES: hydrogenase-2 small chain [Proteobacteria] Length: 372\nNumber of predicted TMHs: 1\nExp number of AAs in TMHs: 23.72226\nExp number, first 60 AAs: 0.22038\nTotal prob of N-in: 0.12994\noutside 1 331\nTMhelix 332 354\ninside 355 372

5977 GCF_002190145.1_ASM219014v1 Escherichia coli Proteobacteria; Gammaproteobacteria; Enterobacterales; Enterobacteriaceae; Escherichia
MNNEETFYQAMRRQGVTRRSFLKYCSLAATSLGLGAGMAPKIAWALENKPRIPVVWIHGL WP_001058323.1
MULTISPECIES: hydrogenase-1 small chain [Proteobacteria] Length: 372\nNumber of predicted TMHs: 2\nExp number of AAs in TMHs: 37.2608499999999\nExp number, first 60 AAs: 19.52923\nTotal prob of N-in: 0.94351\nPOSSIBLE N-term signal sequence\ninside 1 19\nTMhelix 20 39\noutside 40 325\nTMhelix 326 348\ninside 349 372

5978 GCF_002190245.1_ASM219024v1 Escherichia coli Proteobacteria; Gammaproteobacteria; Enterobacterales; Enterobacteriaceae; Escherichia
MNNEETFYQAMRRQGVTRRSFLKYCSLAATSLGLGAGMAPKIAWALENKPRIPVVWIHGL WP_001058323.1
MULTISPECIES: hydrogenase-1 small chain [Proteobacteria] Length: 372\nNumber of predicted TMHs: 2\nExp number of AAs in TMHs: 37.2608499999999\nExp number, first 60 AAs: 19.52923\nTotal prob of N-in: 0.94351\nPOSSIBLE N-term signal sequence\ninside 1 19\nTMhelix 20 39\noutside 40 325\nTMhelix 326 348\ninside 349 372

5979 GCF_002190085.1_ASM219008v1 Escherichia coli Proteobacteria; Gammaproteobacteria; Enterobacterales; Enterobacteriaceae; Escherichia
MNNEETFYQAMRRQGVTRRSFLKYCSLAATSLGLGAGMAPKIAWALENKPRIPVVWIHGL WP_001058323.1
MULTISPECIES: hydrogenase-1 small chain [Proteobacteria] Length: 372\nNumber of predicted TMHs: 2\nExp number of AAs in TMHs: 37.2608499999999\nExp number, first 60 AAs: 19.52923\nTotal prob of N-in: 0.94351\nPOSSIBLE N-term signal sequence\ninside 1 19\nTMhelix 20 39\noutside 40 325\nTMhelix 326 348\ninside 349 372

5980 GCF_002190275.1_ASM219027v1 Escherichia coli Proteobacteria; Gammaproteobacteria; Enterobacterales; Enterobacteriaceae; Escherichia
MTGDNTLIHSHGINRRDFMKLCAALAATMGLSSKAAAEMAESVTNPQRPPVIWIGAQECT WP_000145410.1
MULTISPECIES: hydrogenase-2 small chain [Proteobacteria] Length: 372\nNumber of predicted TMHs: 1\nExp number of AAs in TMHs: 23.72226\nExp number, first 60 AAs: 0.22038\nTotal prob of N-in: 0.12994\noutside 1 331\nTMhelix 332 354\ninside 355 372

5981 GCF_002190275.1_ASM219027v1 Escherichia coli Proteobacteria; Gammaproteobacteria; Enterobacterales; Enterobacteriaceae; Escherichia

MNNEETFYQAMRRQGVTRRSFLKYCSLAATSLGLGAGMAPKIAWALENKPRIPVVWIHGL WP_001058323.1 MULTISPECIES: hydrogenase-1 small chain [Proteobacteria] Length: 372\nNumber of predicted TMHs: 2\nExp number of AAs in TMHs: 37.2608499999999\nExp number, first 60 AAs: 19.52923\nTotal prob of N-in: 0.94351\nPOSSIBLE N-term signal sequence\ninside 1 19\nTMhelix 20 39\noutside 40 325\nTMhelix 326 348\ninside 349 372

5982 GCF_002190155.1_ASM219015v1 Escherichia coli Proteobacteria; Gammaproteobacteria; Enterobacterales; Enterobacteriaceae; Escherichia MNNEETFYQAMRRQGVTRRSFLKYCSLAATSLGLGAGMAPKIAWALENKPRIPVVWIHGL WP_001058323.1 MULTISPECIES: hydrogenase-1 small chain [Proteobacteria] Length: 372\nNumber of predicted TMHs: 2\nExp number of AAs in TMHs: 37.2608499999999\nExp number, first 60 AAs: 19.52923\nTotal prob of N-in: 0.94351\nPOSSIBLE N-term signal sequence\ninside 1 19\nTMhelix 20 39\noutside 40 325\nTMhelix 326 348\ninside 349 372

5983 GCF_002190295.1_ASM219029v1 Escherichia coli Proteobacteria; Gammaproteobacteria; Enterobacterales; Enterobacteriaceae; Escherichia MNNEETFYQAMRRQGVTRRSFLKYCSLAATSLGLGAGMAPKIAWALENKPRIPVVWIHGL WP_001058323.1 MULTISPECIES: hydrogenase-1 small chain [Proteobacteria] Length: 372\nNumber of predicted TMHs: 2\nExp number of AAs in TMHs: 37.2608499999999\nExp number, first 60 AAs: 19.52923\nTotal prob of N-in: 0.94351\nPOSSIBLE N-term signal sequence\ninside 1 19\nTMhelix 20 39\noutside 40 325\nTMhelix 326 348\ninside 349 372

5984 GCF_002190515.1_ASM219051v1 Escherichia coli Proteobacteria; Gammaproteobacteria; Enterobacterales; Enterobacteriaceae; Escherichia MTGDNTLIHSHGINRRDFMKLCAALAATMGLSSKAAAEMAESVTNPQRPPVIWIGAQECT WP_000145410.1 MULTISPECIES: hydrogenase-2 small chain [Proteobacteria] Length: 372\nNumber of predicted TMHs: 1\nExp number of AAs in TMHs: 23.72226\nExp number, first 60 AAs: 0.22038\nTotal prob of N-in: 0.12994\noutside 1 331\nTMhelix 332 354\ninside 355 372

5985 GCF_002190515.1_ASM219051v1 Escherichia coli Proteobacteria; Gammaproteobacteria; Enterobacterales; Enterobacteriaceae; Escherichia MNNEETFYQAMRRQGVTRRSFLKYCSLAATSLGLGAGMAPKIAWALENKPRIPVVWIHGL WP_001058323.1 MULTISPECIES: hydrogenase-1 small chain [Proteobacteria] Length: 372\nNumber of predicted TMHs: 2\nExp number of AAs in TMHs: 37.2608499999999\nExp number, first 60 AAs: 19.52923\nTotal prob of N-in: 0.94351\nPOSSIBLE N-term signal sequence\ninside 1 19\nTMhelix 20 39\noutside 40 325\nTMhelix 326 348\ninside 349 372

5986 GCF_002190365.1_ASM219036v1 Escherichia coli Proteobacteria; Gammaproteobacteria; Enterobacterales; Enterobacteriaceae; Escherichia MNNEETFYQAMRRQGVTRRSFLKYCSLAATSLGLGAGMAPKIAWALENKPRIPVVWIHGL WP_001058323.1 MULTISPECIES: hydrogenase-1 small chain [Proteobacteria] Length: 372\nNumber of predicted TMHs: 2\nExp number of AAs in TMHs: 37.2608499999999\nExp number, first 60 AAs: 19.52923\nTotal prob of N-in: 0.94351\nPOSSIBLE N-term signal sequence\ninside 1 19\nTMhelix 20 39\noutside 40 325\nTMhelix 326 348\ninside 349 372

5987 GCF_002190395.1_ASM219039v1 Escherichia coli Proteobacteria; Gammaproteobacteria; Enterobacterales; Enterobacteriaceae; Escherichia MNNEETFYQAMRRQGVTRRSFLKYCSLAATSLGLGAGMAPKIAWALENKPRIPVVWIHGL WP_001058323.1 MULTISPECIES: hydrogenase-1 small chain [Proteobacteria] Length: 372\nNumber of predicted TMHs: 2\nExp number of AAs in TMHs: 37.2608499999999\nExp number, first 60 AAs: 19.52923\nTotal prob of N-in: 0.94351\nPOSSIBLE N-term signal sequence\ninside 1 19\nTMhelix 20 39\noutside 40 325\nTMhelix 326 348\ninside 349 372

5988 GCF_002190455.1_ASM219045v1 Escherichia coli Proteobacteria; Gammaproteobacteria; Enterobacterales; Enterobacteriaceae; Escherichia MNNEETFYQAMRRQGVTRRSFLKYCSLAATSLGLGAGMAPKIAWALENKPRIPVVWIHGL WP_001058323.1 MULTISPECIES: hydrogenase-1 small chain [Proteobacteria] Length: 372\nNumber of predicted TMHs: 2\nExp number of AAs in TMHs: 37.2608499999999\nExp number, first 60 AAs: 19.52923\nTotal prob of N-in: 0.94351\nPOSSIBLE N-term signal sequence\ninside 1 19\nTMhelix 20 39\noutside 40 325\nTMhelix 326 348\ninside 349 372

5989 GCF_002190355.1_ASM219035v1 Escherichia coli Proteobacteria; Gammaproteobacteria; Enterobacterales; Enterobacteriaceae; Escherichia
MNNEETFYQAMRRQGVTRRSFLKYCSLAATSLGLGAGMAPKIAWALENKPRIPVVWIHGL WP_001058323.1
MULTISPECIES: hydrogenase-1 small chain [Proteobacteria] Length: 372\nNumber of predicted TMHs: 2\nExp number of AAs in TMHs: 37.2608499999999\nExp number, first 60 AAs: 19.52923\nTotal prob of N-in: 0.94351\nPOSSIBLE N-term signal sequence\ninside 1 19\nTMhelix 20 39\noutside 40 325\nTMhelix 326 348\ninside 349 372

5990 GCF_002190465.1_ASM219046v1 Escherichia coli Proteobacteria; Gammaproteobacteria; Enterobacterales; Enterobacteriaceae; Escherichia
MNNEETFYQAMRRQGVTRRSFLKYCSLAATSLGLGAGMAPKIAWALENKPRIPVVWIHGL WP_001058323.1
MULTISPECIES: hydrogenase-1 small chain [Proteobacteria] Length: 372\nNumber of predicted TMHs: 2\nExp number of AAs in TMHs: 37.2608499999999\nExp number, first 60 AAs: 19.52923\nTotal prob of N-in: 0.94351\nPOSSIBLE N-term signal sequence\ninside 1 19\nTMhelix 20 39\noutside 40 325\nTMhelix 326 348\ninside 349 372

5991 GCF_002190545.1_ASM219054v1 Escherichia coli Proteobacteria; Gammaproteobacteria; Enterobacterales; Enterobacteriaceae; Escherichia
MTGDNTLIHSHGINRRDFMKLCAALAATMGLSSKAAAEMAESVTNPQRPPVIWIGAQECT WP_000145410.1
MULTISPECIES: hydrogenase-2 small chain [Proteobacteria] Length: 372\nNumber of predicted TMHs: 1\nExp number of AAs in TMHs: 23.72226\nExp number, first 60 AAs: 0.22038\nTotal prob of N-in: 0.12994\noutside 1 331\nTMhelix 332 354\ninside 355 372

5992 GCF_002190605.1_ASM219060v1 Escherichia coli Proteobacteria; Gammaproteobacteria; Enterobacterales; Enterobacteriaceae; Escherichia
MTGDNTLIHSHGINRRDFMKLCAALAATMGLSSKAAAEMAESVTNPQRPPVIWIGAQECT WP_000145410.1
MULTISPECIES: hydrogenase-2 small chain [Proteobacteria] Length: 372\nNumber of predicted TMHs: 1\nExp number of AAs in TMHs: 23.72226\nExp number, first 60 AAs: 0.22038\nTotal prob of N-in: 0.12994\noutside 1 331\nTMhelix 332 354\ninside 355 372

5993 GCF_002190555.1_ASM219055v1 Escherichia coli Proteobacteria; Gammaproteobacteria; Enterobacterales; Enterobacteriaceae; Escherichia
MTGDNTLIHSHGINRRDFMKLCAALAATMGLSSKAAAEMAESVTNPQRPPVIWIGAQECT WP_000145410.1
MULTISPECIES: hydrogenase-2 small chain [Proteobacteria] Length: 372\nNumber of predicted TMHs: 1\nExp number of AAs in TMHs: 23.72226\nExp number, first 60 AAs: 0.22038\nTotal prob of N-in: 0.12994\noutside 1 331\nTMhelix 332 354\ninside 355 372

5994 GCF_002190685.1_ASM219068v1 Escherichia coli Proteobacteria; Gammaproteobacteria; Enterobacterales; Enterobacteriaceae; Escherichia
MTGDNTLIHSHGINRRDFMKLCAALAATMGLSSKAAAEMAESVTNPQRPPVIWIGAQECT WP_000145410.1
MULTISPECIES: hydrogenase-2 small chain [Proteobacteria] Length: 372\nNumber of predicted TMHs: 1\nExp number of AAs in TMHs: 23.72226\nExp number, first 60 AAs: 0.22038\nTotal prob of N-in: 0.12994\noutside 1 331\nTMhelix 332 354\ninside 355 372

5995 GCF_002190545.1_ASM219054v1 Escherichia coli Proteobacteria; Gammaproteobacteria; Enterobacterales; Enterobacteriaceae; Escherichia
MNNEETFYQAMRRQGVTRRSFLKYCSLAATSLGLGAGMAPKIAWALENKPRIPVVWIHGL WP_001058323.1
MULTISPECIES: hydrogenase-1 small chain [Proteobacteria] Length: 372\nNumber of predicted TMHs: 2\nExp number of AAs in TMHs: 37.2608499999999\nExp number, first 60 AAs: 19.52923\nTotal prob of N-in: 0.94351\nPOSSIBLE N-term signal sequence\ninside 1 19\nTMhelix 20 39\noutside 40 325\nTMhelix 326 348\ninside 349 372

5996 GCF_002190715.1_ASM219071v1 Escherichia coli Proteobacteria; Gammaproteobacteria; Enterobacterales; Enterobacteriaceae; Escherichia
MTGDNTLIHSHGINRRDFMKLCAALAATMGLSSKAAAEMAESVTNPQRPPVIWIGAQECT WP_000145410.1
MULTISPECIES: hydrogenase-2 small chain [Proteobacteria] Length: 372\nNumber of predicted TMHs: 1\nExp number of AAs in TMHs: 23.72226\nExp number, first 60 AAs: 0.22038\nTotal prob of N-in: 0.12994\noutside 1 331\nTMhelix 332 354\ninside 355 372

5997 GCF_002190555.1_ASM219055v1 Escherichia coli Proteobacteria; Gammaproteobacteria; Enterobacterales; Enterobacteriaceae; Escherichia

MNNEETFYQAMRRQGVTRRSFLKYCSLAATSLGLGAGMAPKIAWALENKPRIPVVWIHGL WP_001058323.1 MULTISPECIES: hydrogenase-1 small chain [Proteobacteria] Length: 372\nNumber of predicted TMHs: 2\nExp number of AAs in TMHs: 37.2608499999999\nExp number, first 60 AAs: 19.52923\nTotal prob of N-in: 0.94351\nPOSSIBLE N-term signal sequence\ninside 1 19\nTMhelix 20 39\noutside 40 325\nTMhelix 326 348\ninside 349 372

5998 GCF_002190605.1_ASM219060v1 Escherichia coli Proteobacteria; Gammaproteobacteria; Enterobacterales; Enterobacteriaceae; Escherichia MNNEETFYQAMRRQGVTRRSFLKYCSLAATSLGLGAGMAPKIAWALENKPRIPVVWIHGL WP_001058323.1 MULTISPECIES: hydrogenase-1 small chain [Proteobacteria] Length: 372\nNumber of predicted TMHs: 2\nExp number of AAs in TMHs: 37.2608499999999\nExp number, first 60 AAs: 19.52923\nTotal prob of N-in: 0.94351\nPOSSIBLE N-term signal sequence\ninside 1 19\nTMhelix 20 39\noutside 40 325\nTMhelix 326 348\ninside 349 372

5999 GCF_002190745.1_ASM219074v1 Escherichia coli Proteobacteria; Gammaproteobacteria; Enterobacterales; Enterobacteriaceae; Escherichia MTGDNTLIHSHGINRRDFMKLCAALAATMGLSSKAAAEMAESVTNPQRPPVIWIGAQECT WP_000145410.1 MULTISPECIES: hydrogenase-2 small chain [Proteobacteria] Length: 372\nNumber of predicted TMHs: 1\nExp number of AAs in TMHs: 23.72226\nExp number, first 60 AAs: 0.22038\nTotal prob of N-in: 0.12994\noutside 1 331\nTMhelix 332 354\ninside 355 372

6000 GCF_002190745.1_ASM219074v1 Escherichia coli Proteobacteria; Gammaproteobacteria; Enterobacterales; Enterobacteriaceae; Escherichia MNNEETFYQAMRRQGVTRRSFLKYCSLAATSLGLGAGMAPKIAWALENKPRIPVVWIHGL WP_001058323.1 MULTISPECIES: hydrogenase-1 small chain [Proteobacteria] Length: 372\nNumber of predicted TMHs: 2\nExp number of AAs in TMHs: 37.2608499999999\nExp number, first 60 AAs: 19.52923\nTotal prob of N-in: 0.94351\nPOSSIBLE N-term signal sequence\ninside 1 19\nTMhelix 20 39\noutside 40 325\nTMhelix 326 348\ninside 349 372

6001 GCF_002190685.1_ASM219068v1 Escherichia coli Proteobacteria; Gammaproteobacteria; Enterobacterales; Enterobacteriaceae; Escherichia MNNEETFYQAMRRQGVTRRSFLKYCSLAATSLGLGAGMAPKIAWALENKPRIPVVWIHGL WP_001058323.1 MULTISPECIES: hydrogenase-1 small chain [Proteobacteria] Length: 372\nNumber of predicted TMHs: 2\nExp number of AAs in TMHs: 37.2608499999999\nExp number, first 60 AAs: 19.52923\nTotal prob of N-in: 0.94351\nPOSSIBLE N-term signal sequence\ninside 1 19\nTMhelix 20 39\noutside 40 325\nTMhelix 326 348\ninside 349 372

6002 GCF_002190785.1_ASM219078v1 Escherichia coli Proteobacteria; Gammaproteobacteria; Enterobacterales; Enterobacteriaceae; Escherichia MTGDNTLIHSHGINRRDFMKLCAALAATMGLSSKAAAEMAESVTNPQRPPVIWIGAQECT WP_000145410.1 MULTISPECIES: hydrogenase-2 small chain [Proteobacteria] Length: 372\nNumber of predicted TMHs: 1\nExp number of AAs in TMHs: 23.72226\nExp number, first 60 AAs: 0.22038\nTotal prob of N-in: 0.12994\noutside 1 331\nTMhelix 332 354\ninside 355 372

6003 GCF_002190865.1_ASM219086v1 Escherichia coli Proteobacteria; Gammaproteobacteria; Enterobacterales; Enterobacteriaceae; Escherichia MTGDNTLIHSHGINRRDFMKLCAALAATMGLSSKAAAEMAESVTNPQRPPVIWIGAQECT WP_000145410.1 MULTISPECIES: hydrogenase-2 small chain [Proteobacteria] Length: 372\nNumber of predicted TMHs: 1\nExp number of AAs in TMHs: 23.72226\nExp number, first 60 AAs: 0.22038\nTotal prob of N-in: 0.12994\noutside 1 331\nTMhelix 332 354\ninside 355 372

6004 GCF_002190955.1_ASM219095v1 Escherichia coli Proteobacteria; Gammaproteobacteria; Enterobacterales; Enterobacteriaceae; Escherichia MTGDNTLIHSHGINRRDFMKLCAALAATMGLSSKAAAEMAESVTNPQRPPVIWIGAQECT WP_000145410.1 MULTISPECIES: hydrogenase-2 small chain [Proteobacteria] Length: 372\nNumber of predicted TMHs: 1\nExp number of AAs in TMHs: 23.72226\nExp number, first 60 AAs: 0.22038\nTotal prob of N-in: 0.12994\noutside 1 331\nTMhelix 332 354\ninside 355 372

6005 GCF_002191025.1_ASM219102v1 Escherichia coli Proteobacteria; Gammaproteobacteria; Enterobacterales; Enterobacteriaceae; Escherichia MTGDNTLIHSHGINRRDFMKLCAALAATMGLSSKAAAEMAESVTNPQRPPVIWIGAQECT WP_000145410.1

MULTISPECIES: hydrogenase-2 small chain [Proteobacteria] Length: 372\nNumber of predicted TMHs: 1\nExp number of AAs in TMHs: 23.72226\nExp number, first 60 AAs: 0.22038\nTotal prob of N-in: 0.12994\noutside 1 331\nTMhelix 332 354\ninside 355 372

6006 GCF_002190955.1_ASM219095v1 Escherichia coli Proteobacteria; Gammaproteobacteria; Enterobacterales; Enterobacteriaceae; Escherichia MNNEETFYQAMRRQGVTRRSFLKYCSLAATSLGLGAGMAPKIAWALENKPRIPVVWIHGL WP_001058323.1 MULTISPECIES: hydrogenase-1 small chain [Proteobacteria] Length: 372\nNumber of predicted TMHs: 2\nExp number of AAs in TMHs: 37.2608499999999\nExp number, first 60 AAs: 19.52923\nTotal prob of N-in: 0.94351\nPOSSIBLE N-term signal sequence\ninside 1 19\nTMhelix 20 39\noutside 40 325\nTMhelix 326 348\ninside 349 372

6007 GCF_002190945.1_ASM219094v1 Escherichia coli Proteobacteria; Gammaproteobacteria; Enterobacterales; Enterobacteriaceae; Escherichia MNNEETFYQAMRRQGVTRRSFLKYCSLAATSLGLGAGMAPKIAWALENKPRIPVVWIHGL WP_001058323.1 MULTISPECIES: hydrogenase-1 small chain [Proteobacteria] Length: 372\nNumber of predicted TMHs: 2\nExp number of AAs in TMHs: 37.2608499999999\nExp number, first 60 AAs: 19.52923\nTotal prob of N-in: 0.94351\nPOSSIBLE N-term signal sequence\ninside 1 19\nTMhelix 20 39\noutside 40 325\nTMhelix 326 348\ninside 349 372

6008 GCF_002190885.1_ASM219088v1 Escherichia coli Proteobacteria; Gammaproteobacteria; Enterobacterales; Enterobacteriaceae; Escherichia MNNEETFYQAMRRQGVTRRSFLKYCSLAATSLGLGAGMAPKIAWALENKPRIPVVWIHGL WP_001295940.1 MULTISPECIES: hydrogenase [Enterobacteriaceae] Length: 372\nNumber of predicted TMHs: 2\nExp number of AAs in TMHs: 37.2608899999999\nExp number, first 60 AAs: 19.52923\nTotal prob of N-in: 0.94351\nPOSSIBLE N-term signal sequence\ninside 1 19\nTMhelix 20 39\noutside 40 325\nTMhelix 326 348\ninside 349 372

6009 GCF_002191025.1_ASM219102v1 Escherichia coli Proteobacteria; Gammaproteobacteria; Enterobacterales; Enterobacteriaceae; Escherichia MNNEETFYQAMRRQGVTRRSFLKYCSLAATSLGLGAGMAPKIAWALENKPRIPVVWIHGL WP_001058323.1 MULTISPECIES: hydrogenase-1 small chain [Proteobacteria] Length: 372\nNumber of predicted TMHs: 2\nExp number of AAs in TMHs: 37.2608499999999\nExp number, first 60 AAs: 19.52923\nTotal prob of N-in: 0.94351\nPOSSIBLE N-term signal sequence\ninside 1 19\nTMhelix 20 39\noutside 40 325\nTMhelix 326 348\ninside 349 372

6010 GCF_002191075.1_ASM219107v1 Escherichia coli Proteobacteria; Gammaproteobacteria; Enterobacterales; Enterobacteriaceae; Escherichia MNNEETFYQAMRRQGVTRRSFLKYCSLAATSLGLGAGMAPKIAWALENKPRIPVVWIHGL WP_001058323.1 MULTISPECIES: hydrogenase-1 small chain [Proteobacteria] Length: 372\nNumber of predicted TMHs: 2\nExp number of AAs in TMHs: 37.2608499999999\nExp number, first 60 AAs: 19.52923\nTotal prob of N-in: 0.94351\nPOSSIBLE N-term signal sequence\ninside 1 19\nTMhelix 20 39\noutside 40 325\nTMhelix 326 348\ninside 349 372

6011 GCF_002192275.1_ASM219227v1 Escherichia coli Proteobacteria; Gammaproteobacteria; Enterobacterales; Enterobacteriaceae; Escherichia MNNEETFYQAMRRQGVTRRSFLKYCSLAATSLGLGAGMAPKIAWALENKPRIPVVWIHGL WP_001058323.1 MULTISPECIES: hydrogenase-1 small chain [Proteobacteria] Length: 372\nNumber of predicted TMHs: 2\nExp number of AAs in TMHs: 37.2608499999999\nExp number, first 60 AAs: 19.52923\nTotal prob of N-in: 0.94351\nPOSSIBLE N-term signal sequence\ninside 1 19\nTMhelix 20 39\noutside 40 325\nTMhelix 326 348\ninside 349 372

6012 GCF_002193195.1_ASM219319v1 Escherichia coli Proteobacteria; Gammaproteobacteria; Enterobacterales; Enterobacteriaceae; Escherichia MTGDNTLIHSHGINRRDFMKLCAALAATMGLSSKAAAEMAESVTNPQRPPVIWIGAQECT WP_000145410.1 MULTISPECIES: hydrogenase-2 small chain [Proteobacteria] Length: 372\nNumber of predicted TMHs: 1\nExp number of AAs in TMHs: 23.72226\nExp number, first 60 AAs: 0.22038\nTotal prob of N-in: 0.12994\noutside 1 331\nTMhelix 332 354\ninside 355 372

6013 GCF_002193095.1_ASM219309v1 Escherichia coli Proteobacteria; Gammaproteobacteria; Enterobacterales; Enterobacteriaceae; Escherichia

MTGDNTLIHSHGINRRDFMKLCAALAATMGLSSKAAAEMVESVTNPQRPPVIWIGAQECT WP_000145423.1 hydrogenase 2 small subunit [Escherichia coli] Length: 372\nNumber of predicted TMHs: 1\nExp number of AAs in TMHs: 23.78201\nExp number, first 60 AAs: 0.28355\nTotal prob of N-in: 0.13257\noutside 1 331\nTMhelix 332 354\ninside 355 372

6014 GCF_002193245.1_ASM219324v1 Escherichia coli Proteobacteria; Gammaproteobacteria; Enterobacterales; Enterobacteriaceae; Escherichia MTGDNTLIHSHGINRRDFMKLCAALAATMGLSSKAAAEMAESVTNPQRPPVIWIGAQECT WP_000145410.1 MULTISPECIES: hydrogenase-2 small chain [Proteobacteria] Length: 372\nNumber of predicted TMHs: 1\nExp number of AAs in TMHs: 23.72226\nExp number, first 60 AAs: 0.22038\nTotal prob of N-in: 0.12994\noutside 1 331\nTMhelix 332 354\ninside 355 372

6015 GCF_002193235.1_ASM219323v1 Escherichia coli Proteobacteria; Gammaproteobacteria; Enterobacterales; Enterobacteriaceae; Escherichia MTGDNTLIHSHGINRRDFMKLCAALAATMGLSSKAAAEMAESVTNPQRPPVIWIGAQECT WP_000145410.1 MULTISPECIES: hydrogenase-2 small chain [Proteobacteria] Length: 372\nNumber of predicted TMHs: 1\nExp number of AAs in TMHs: 23.72226\nExp number, first 60 AAs: 0.22038\nTotal prob of N-in: 0.12994\noutside 1 331\nTMhelix 332 354\ninside 355 372

6016 GCF_002193195.1_ASM219319v1 Escherichia coli Proteobacteria; Gammaproteobacteria; Enterobacterales; Enterobacteriaceae; Escherichia MNNEETFYQAMRRQGVTRRSFLKYCSLAATSLGLGAGMAPKIAWALENKPRIPVVWIHGL WP_001058323.1 MULTISPECIES: hydrogenase-1 small chain [Proteobacteria] Length: 372\nNumber of predicted TMHs: 2\nExp number of AAs in TMHs: 37.2608499999999\nExp number, first 60 AAs: 19.52923\nTotal prob of N-in: 0.94351\nPOSSIBLE N-term signal sequence\ninside 1 19\nTMhelix 20 39\noutside 40 325\nTMhelix 326 348\ninside 349 372

6017 GCF_002193235.1_ASM219323v1 Escherichia coli Proteobacteria; Gammaproteobacteria; Enterobacterales; Enterobacteriaceae; Escherichia MNNEETFYQAMRRQGVTRRSFLKYCSLAATSLGLGAGMAPKIAWALENKPRIPVVWIHGL WP_001058323.1 MULTISPECIES: hydrogenase-1 small chain [Proteobacteria] Length: 372\nNumber of predicted TMHs: 2\nExp number of AAs in TMHs: 37.2608499999999\nExp number, first 60 AAs: 19.52923\nTotal prob of N-in: 0.94351\nPOSSIBLE N-term signal sequence\ninside 1 19\nTMhelix 20 39\noutside 40 325\nTMhelix 326 348\ninside 349 372

6018 GCF_002193385.1_ASM219338v1 Escherichia coli Proteobacteria; Gammaproteobacteria; Enterobacterales; Enterobacteriaceae; Escherichia MTGDNTLIHSHGINRRDFMKLCAALAATMGLSSKAAAEMAESVTNPQRPPVIWIGAQECT WP_000145410.1 MULTISPECIES: hydrogenase-2 small chain [Proteobacteria] Length: 372\nNumber of predicted TMHs: 1\nExp number of AAs in TMHs: 23.72226\nExp number, first 60 AAs: 0.22038\nTotal prob of N-in: 0.12994\noutside 1 331\nTMhelix 332 354\ninside 355 372

6019 GCF_002193325.1_ASM219332v1 Escherichia coli Proteobacteria; Gammaproteobacteria; Enterobacterales; Enterobacteriaceae; Escherichia MNNEETFYQAMRRQGVTRRSFLKYCSLAATSLGLGAGMAPKIAWALENKPRIPVVWIHGL WP_001058323.1 MULTISPECIES: hydrogenase-1 small chain [Proteobacteria] Length: 372\nNumber of predicted TMHs: 2\nExp number of AAs in TMHs: 37.2608499999999\nExp number, first 60 AAs: 19.52923\nTotal prob of N-in: 0.94351\nPOSSIBLE N-term signal sequence\ninside 1 19\nTMhelix 20 39\noutside 40 325\nTMhelix 326 348\ninside 349 372

6020 GCF_002193445.1_ASM219344v1 Escherichia coli Proteobacteria; Gammaproteobacteria; Enterobacterales; Enterobacteriaceae; Escherichia MTGDNTLIHSHGINRRDFMKLCAALAATMGLSSKAAAEMAESVTNPQRPPVIWIGAQECT WP_000145410.1 MULTISPECIES: hydrogenase-2 small chain [Proteobacteria] Length: 372\nNumber of predicted TMHs: 1\nExp number of AAs in TMHs: 23.72226\nExp number, first 60 AAs: 0.22038\nTotal prob of N-in: 0.12994\noutside 1 331\nTMhelix 332 354\ninside 355 372

6021 GCF_002193435.1_ASM219343v1 Escherichia coli Proteobacteria; Gammaproteobacteria; Enterobacterales; Enterobacteriaceae; Escherichia MTGDNTLIHSHGINRRDFMKLCAALAATMGLSSKAAAEMAESVTNPQRPPVIWIGAQECT WP_000145410.1 MULTISPECIES: hydrogenase-2 small chain [Proteobacteria] Length: 372\nNumber of predicted TMHs:

1\nExp number of AAs in TMHs: 23.72226\nExp number, first 60 AAs: 0.22038\nTotal prob of N-in: 0.12994\noutside 1 331\nTMhelix 332 354\ninside 355 372

6022 GCF_002193445.1_ASM219344v1 Escherichia coli Proteobacteria; Gammaproteobacteria; Enterobacterales; Enterobacteriaceae; Escherichia MNNEETFYQAMRRQGVTRRSFLKYCSLAATSLGLGAGMAPKIAWALENKPRIPVVWIHGL WP_001058323.1 MULTISPECIES: hydrogenase-1 small chain [Proteobacteria] Length: 372\nNumber of predicted TMHs: 2\nExp number of AAs in TMHs: 37.2608499999999\nExp number, first 60 AAs: 19.52923\nTotal prob of N-in: 0.94351\nPOSSIBLE N-term signal sequence\ninside 1 19\nTMhelix 20 39\noutside 40 325\nTMhelix 326 348\ninside 349 372

6023 GCF_002193385.1_ASM219338v1 Escherichia coli Proteobacteria; Gammaproteobacteria; Enterobacterales; Enterobacteriaceae; Escherichia MNNEETFYQAMRRQGVTRRSFLKYCSLAATSLGLGAGMAPKIAWALENKPRIPVVWIHGL WP_001058323.1 MULTISPECIES: hydrogenase-1 small chain [Proteobacteria] Length: 372\nNumber of predicted TMHs: 2\nExp number of AAs in TMHs: 37.2608499999999\nExp number, first 60 AAs: 19.52923\nTotal prob of N-in: 0.94351\nPOSSIBLE N-term signal sequence\ninside 1 19\nTMhelix 20 39\noutside 40 325\nTMhelix 326 348\ninside 349 372

6024 GCF_002193095.1_ASM219309v1 Escherichia coli Proteobacteria; Gammaproteobacteria; Enterobacterales; Enterobacteriaceae; Escherichia MNNEETFYQAMRRQGVTRRSFLKYCSLAATSLGLGAGMAPKIAWALENKPRIPVVWIHGL WP_001295940.1 MULTISPECIES: hydrogenase [Enterobacteriaceae] Length: 372\nNumber of predicted TMHs: 2\nExp number of AAs in TMHs: 37.2608899999999\nExp number, first 60 AAs: 19.52923\nTotal prob of N-in: 0.94351\nPOSSIBLE N-term signal sequence\ninside 1 19\nTMhelix 20 39\noutside 40 325\nTMhelix 326 348\ninside 349 372

6025 GCF_002193155.1_ASM219315v1 Escherichia coli Proteobacteria; Gammaproteobacteria; Enterobacterales; Enterobacteriaceae; Escherichia MTGDNTLIHSHGINRRDFMKLCAALAATMGLSSKAAAEMAESVTNPQRPPVIWIGAQECT WP_088144389.1 hydrogenase 2 small subunit [Escherichia coli] Length: 372\nNumber of predicted TMHs: 1\nExp number of AAs in TMHs: 23.7257\nExp number, first 60 AAs: 0.22034\nTotal prob of N-in: 0.13001\noutside 1 331\nTMhelix 332 354\ninside 355 372

6026 GCF_002194555.1_ASM219455v1 Escherichia coli Proteobacteria; Gammaproteobacteria; Enterobacterales; Enterobacteriaceae; Escherichia MTGDNTLIHSHGINRRDFMKLCAALAATMGLSSKAAAEMAESVTNPQRPPVIWIGAQECT WP_000145410.1 MULTISPECIES: hydrogenase-2 small chain [Proteobacteria] Length: 372\nNumber of predicted TMHs: 1\nExp number of AAs in TMHs: 23.72226\nExp number, first 60 AAs: 0.22038\nTotal prob of N-in: 0.12994\noutside 1 331\nTMhelix 332 354\ninside 355 372

6027 GCF_002193155.1_ASM219315v1 Escherichia coli Proteobacteria; Gammaproteobacteria; Enterobacterales; Enterobacteriaceae; Escherichia MNNEETFYQTMRRQGVTRRSFLKYCSLAATSLGLGAGMAPKIAWALENKPRIPVVWIHGL WP_088144456.1 [Ni/Fe] hydrogenase small subunit [Escherichia coli] Length: 372\nNumber of predicted TMHs: 2\nExp number of AAs in TMHs: 37.5080899999999\nExp number, first 60 AAs: 19.77085\nTotal prob of N-in: 0.95522\nPOSSIBLE N-term signal sequence\ninside 1 19\nTMhelix 20 39\noutside 40 325\nTMhelix 326 348\ninside 349 372

6028 GCF_002194555.1_ASM219455v1 Escherichia coli Proteobacteria; Gammaproteobacteria; Enterobacterales; Enterobacteriaceae; Escherichia MNNEETFYQAMRRQGVTRRSFLKYCSLAATSLGLGAGMAPKIAWALENKPRIPVVWIHGL WP_001301613.1 hydrogenase [Escherichia coli] Length: 372\nNumber of predicted TMHs: 2\nExp number of AAs in TMHs: 37.46598\nExp number, first 60 AAs: 19.53139\nTotal prob of N-in: 0.94364\nPOSSIBLE N-term signal sequence\ninside 1 19\nTMhelix 20 39\noutside 40 325\nTMhelix 326 348\ninside 349 372

6029 GCF_002193435.1_ASM219343v1 Escherichia coli Proteobacteria; Gammaproteobacteria; Enterobacterales; Enterobacteriaceae; Escherichia MNNEETFYQAMRRQGVTRRSFLKYCSLAATSLGLGAGMAPKIAWALENKPRIPVVWIHGL WP_001058323.1 MULTISPECIES: hydrogenase-1 small chain [Proteobacteria] Length: 372\nNumber of predicted TMHs: 2\nExp number of AAs in TMHs: 37.2608499999999\nExp number, first 60 AAs: 19.52923\nTotal prob of N-in:

0.94351\nPOSSIBLE N-term signal sequence\ninside 1 19\nTMhelix 20 39\noutside 40 325\nTMhelix 326 348\ninside 349 372

6030 GCF_002193325.1_ASM219332v1 Escherichia coli Proteobacteria; Gammaproteobacteria; Enterobacterales; Enterobacteriaceae; Escherichia MTGDNTLIHSHGINRRDFMKLCAALATTMGLSSKAAAEMAESVTNPQRPPVIWIGAQECT WP_023149629.1 MULTISPECIES: hydrogenase-2 small chain [Enterobacteriaceae] Length: 372\nNumber of predicted TMHs: 1\nExp number of AAs in TMHs: 23.63603\nExp number, first 60 AAs: 0.12971\nTotal prob of N-in: 0.12608\noutside 1 331\nTMhelix 332 354\ninside 355 372

6031 GCF_002194865.1_ASM219486v1 Escherichia coli Proteobacteria; Gammaproteobacteria; Enterobacterales; Enterobacteriaceae; Escherichia MTGDNTLIHSHGINRRDFMKLCAALAATMGLSSKAAAEMAESVTNPQRPPVIWIGAQECT WP_000145410.1 MULTISPECIES: hydrogenase-2 small chain [Proteobacteria] Length: 372\nNumber of predicted TMHs: 1\nExp number of AAs in TMHs: 23.72226\nExp number, first 60 AAs: 0.22038\nTotal prob of N-in: 0.12994\noutside 1 331\nTMhelix 332 354\ninside 355 372

6032 GCF_002194875.1_ASM219487v1 Escherichia coli Proteobacteria; Gammaproteobacteria; Enterobacterales; Enterobacteriaceae; Escherichia MTGDNTLIHSHGINRRDFMKLCAALAATMGLSSKAAAEMAESVTNPQRPPVIWIGAQECT WP_000145410.1 MULTISPECIES: hydrogenase-2 small chain [Proteobacteria] Length: 372\nNumber of predicted TMHs: 1\nExp number of AAs in TMHs: 23.72226\nExp number, first 60 AAs: 0.22038\nTotal prob of N-in: 0.12994\noutside 1 331\nTMhelix 332 354\ninside 355 372

6033 GCF_002194935.1_ASM219493v1 Escherichia coli Proteobacteria; Gammaproteobacteria; Enterobacterales; Enterobacteriaceae; Escherichia MTGDNTLIHSHGINRRDFMKLCAALAATMGLSSKAAAEMAESVTNPQRPPVIWIGAQECT WP_000145410.1 MULTISPECIES: hydrogenase-2 small chain [Proteobacteria] Length: 372\nNumber of predicted TMHs: 1\nExp number of AAs in TMHs: 23.72226\nExp number, first 60 AAs: 0.22038\nTotal prob of N-in: 0.12994\noutside 1 331\nTMhelix 332 354\ninside 355 372

6034 GCF_002194835.1_ASM219483v1 Escherichia coli Proteobacteria; Gammaproteobacteria; Enterobacterales; Enterobacteriaceae; Escherichia MNNEETFYQAMRRQGVTRRSFLKYCSLAATSLGLGAGMAPKIAWALENKPRIPVVWIHGL WP_001301613.1 hydrogenase [Escherichia coli] Length: 372\nNumber of predicted TMHs: 2\nExp number of AAs in TMHs: 37.46598\nExp number, first 60 AAs: 19.53139\nTotal prob of N-in: 0.94364\nPOSSIBLE N-term signal sequence\ninside 1 19\nTMhelix 20 39\noutside 40 325\nTMhelix 326 348\ninside 349 372

6035 GCF_002194865.1_ASM219486v1 Escherichia coli Proteobacteria; Gammaproteobacteria; Enterobacterales; Enterobacteriaceae; Escherichia MNNEETFYQAMRRQGVTRRSFLKYCSLAATSLGLGAGMAPKIAWALENKPRIPVVWIHGL WP_001301613.1 hydrogenase [Escherichia coli] Length: 372\nNumber of predicted TMHs: 2\nExp number of AAs in TMHs: 37.46598\nExp number, first 60 AAs: 19.53139\nTotal prob of N-in: 0.94364\nPOSSIBLE N-term signal sequence\ninside 1 19\nTMhelix 20 39\noutside 40 325\nTMhelix 326 348\ninside 349 372

6036 GCF_002194875.1_ASM219487v1 Escherichia coli Proteobacteria; Gammaproteobacteria; Enterobacterales; Enterobacteriaceae; Escherichia MNNEETFYQAMRRQGVTRRSFLKYCSLAATSLGLGAGMAPKIAWALENKPRIPVVWIHGL WP_001301613.1 hydrogenase [Escherichia coli] Length: 372\nNumber of predicted TMHs: 2\nExp number of AAs in TMHs: 37.46598\nExp number, first 60 AAs: 19.53139\nTotal prob of N-in: 0.94364\nPOSSIBLE N-term signal sequence\ninside 1 19\nTMhelix 20 39\noutside 40 325\nTMhelix 326 348\ninside 349 372

6037 GCF_002195125.1_ASM219512v1 Escherichia coli Proteobacteria; Gammaproteobacteria; Enterobacterales; Enterobacteriaceae; Escherichia MTGDNTLIHSHGINRRDFMKLCAALAATMGLSSKAAAEMAESVTNPQRPPVIWIGAQECT WP_000145410.1 MULTISPECIES: hydrogenase-2 small chain [Proteobacteria] Length: 372\nNumber of predicted TMHs: 1\nExp number of AAs in TMHs: 23.72226\nExp number, first 60 AAs: 0.22038\nTotal prob of N-in: 0.12994\noutside 1 331\nTMhelix 332 354\ninside 355 372

6038 GCF_002194605.1_ASM219460v1 Escherichia coli Proteobacteria; Gammaproteobacteria; Enterobacterales; Enterobacteriaceae; Escherichia

MNNEETFYQAMRRQGVTRRSFLKYCSLAATSLGLGAGMAPKIAWALENKPRIPVVWIHGL WP_001301613.1 hydrogenase [Escherichia coli] Length: 372\nNumber of predicted TMHs: 2\nExp number of AAs in TMHs: 37.46598\nExp number, first 60 AAs: 19.53139\nTotal prob of N-in: 0.94364\nPOSSIBLE N-term signal sequence\ninside 1 19\nTMhelix 20 39\noutside 40 325\nTMhelix 326 348\ninside 349 372

6039 GCF_002194625.1_ASM219462v1 Escherichia coli Proteobacteria; Gammaproteobacteria; Enterobacterales; Enterobacteriaceae; Escherichia MNNEETFYQAMRRQGVTRRSFLKYCSLAATSLGLGAGMAPKIAWALENKPRIPVVWIHGL WP_001301613.1 hydrogenase [Escherichia coli] Length: 372\nNumber of predicted TMHs: 2\nExp number of AAs in TMHs: 37.46598\nExp number, first 60 AAs: 19.53139\nTotal prob of N-in: 0.94364\nPOSSIBLE N-term signal sequence\ninside 1 19\nTMhelix 20 39\noutside 40 325\nTMhelix 326 348\ninside 349 372

6040 GCF_002194965.1_ASM219496v1 Escherichia coli Proteobacteria; Gammaproteobacteria; Enterobacterales; Enterobacteriaceae; Escherichia MTGDNTLIHSHGINRRDFMKLCAALAATMGLSSKAAAEMAESVTNPQRPPVIWIGAQECT WP_000145410.1 MULTISPECIES: hydrogenase-2 small chain [Proteobacteria] Length: 372\nNumber of predicted TMHs: 1\nExp number of AAs in TMHs: 23.72226\nExp number, first 60 AAs: 0.22038\nTotal prob of N-in: 0.12994\noutside 1 331\nTMhelix 332 354\ninside 355 372

6041 GCF_002194965.1_ASM219496v1 Escherichia coli Proteobacteria; Gammaproteobacteria; Enterobacterales; Enterobacteriaceae; Escherichia MNNEETFYQAMRRQGVTRRSFLKYCSLAATSLGLGAGMAPKIAWALENKPRIPVVWIHGL WP_001301613.1 hydrogenase [Escherichia coli] Length: 372\nNumber of predicted TMHs: 2\nExp number of AAs in TMHs: 37.46598\nExp number, first 60 AAs: 19.53139\nTotal prob of N-in: 0.94364\nPOSSIBLE N-term signal sequence\ninside 1 19\nTMhelix 20 39\noutside 40 325\nTMhelix 326 348\ninside 349 372

6042 GCF_002194675.1_ASM219467v1 Escherichia coli Proteobacteria; Gammaproteobacteria; Enterobacterales; Enterobacteriaceae; Escherichia MNNEETFYQAMRRQGVTRRSFLKYCSLAATSLGLGAGMAPKIAWALENKPRIPVVWIHGL WP_001301613.1 hydrogenase [Escherichia coli] Length: 372\nNumber of predicted TMHs: 2\nExp number of AAs in TMHs: 37.46598\nExp number, first 60 AAs: 19.53139\nTotal prob of N-in: 0.94364\nPOSSIBLE N-term signal sequence\ninside 1 19\nTMhelix 20 39\noutside 40 325\nTMhelix 326 348\ninside 349 372

6043 GCF_002194715.1_ASM219471v1 Escherichia coli Proteobacteria; Gammaproteobacteria; Enterobacterales; Enterobacteriaceae; Escherichia MNNEETFYQAMRRQGVTRRSFLKYCSLAATSLGLGAGMAPKIAWALENKPRIPVVWIHGL WP_001301613.1 hydrogenase [Escherichia coli] Length: 372\nNumber of predicted TMHs: 2\nExp number of AAs in TMHs: 37.46598\nExp number, first 60 AAs: 19.53139\nTotal prob of N-in: 0.94364\nPOSSIBLE N-term signal sequence\ninside 1 19\nTMhelix 20 39\noutside 40 325\nTMhelix 326 348\ninside 349 372

6044 GCF_002194765.1_ASM219476v1 Escherichia coli Proteobacteria; Gammaproteobacteria; Enterobacterales; Enterobacteriaceae; Escherichia MNNEETFYQAMRRQGVTRRSFLKYCSLAATSLGLGAGMAPKIAWALENKPRIPVVWIHGL WP_001301613.1 hydrogenase [Escherichia coli] Length: 372\nNumber of predicted TMHs: 2\nExp number of AAs in TMHs: 37.46598\nExp number, first 60 AAs: 19.53139\nTotal prob of N-in: 0.94364\nPOSSIBLE N-term signal sequence\ninside 1 19\nTMhelix 20 39\noutside 40 325\nTMhelix 326 348\ninside 349 372

6045 GCF_002194705.1_ASM219470v1 Escherichia coli Proteobacteria; Gammaproteobacteria; Enterobacterales; Enterobacteriaceae; Escherichia MNNEETFYQAMRRQGVTRRSFLKYCSLAATSLGLGAGMAPKIAWALENKPRIPVVWIHGL WP_001301613.1 hydrogenase [Escherichia coli] Length: 372\nNumber of predicted TMHs: 2\nExp number of AAs in TMHs: 37.46598\nExp number, first 60 AAs: 19.53139\nTotal prob of N-in: 0.94364\nPOSSIBLE N-term signal sequence\ninside 1 19\nTMhelix 20 39\noutside 40 325\nTMhelix 326 348\ninside 349 372

6046 GCF_002194775.1_ASM219477v1 Escherichia coli Proteobacteria; Gammaproteobacteria; Enterobacterales; Enterobacteriaceae; Escherichia MNNEETFYQAMRRQGVTRRSFLKYCSLAATSLGLGAGMAPKIAWALENKPRIPVVWIHGL WP_001301613.1 hydrogenase [Escherichia coli] Length: 372\nNumber of predicted TMHs: 2\nExp number of AAs in TMHs: 37.46598\nExp number, first 60 AAs: 19.53139\nTotal prob of N-in: 0.94364\nPOSSIBLE N-term signal sequence\ninside 1 19\nTMhelix 20 39\noutside 40 325\nTMhelix 326 348\ninside 349 372

6047 GCF_002195035.1_ASM219503v1 Escherichia coli Proteobacteria; Gammaproteobacteria; Enterobacterales; Enterobacteriaceae; Escherichia
MTGDNTLIHSHGINRRDFMKLCAALAATMGLSSKAAAEMAESVTNPQRPPVIWIGAQECT WP_000145410.1
MULTISPECIES: hydrogenase-2 small chain [Proteobacteria] Length: 372\nNumber of predicted TMHs: 1\nExp number of AAs in TMHs: 23.72226\nExp number, first 60 AAs: 0.22038\nTotal prob of N-in: 0.12994\noutside 1 331\nTMhelix 332 354\ninside 355 372

6048 GCF_002194975.1_ASM219497v1 Escherichia coli Proteobacteria; Gammaproteobacteria; Enterobacterales; Enterobacteriaceae; Escherichia
MTGDNTLIHSHGINRRDFMKLCAALAATMGLSSKAAAEMAESVTNPQRPPVIWIGAQECT WP_000145410.1
MULTISPECIES: hydrogenase-2 small chain [Proteobacteria] Length: 372\nNumber of predicted TMHs: 1\nExp number of AAs in TMHs: 23.72226\nExp number, first 60 AAs: 0.22038\nTotal prob of N-in: 0.12994\noutside 1 331\nTMhelix 332 354\ninside 355 372

6049 GCF_002195055.1_ASM219505v1 Escherichia coli Proteobacteria; Gammaproteobacteria; Enterobacterales; Enterobacteriaceae; Escherichia
MTGDNTLIHSHGINRRDFMKLCAALAATMGLSSKAAAEMAESVTNPQRPPVIWIGAQECT WP_000145410.1
MULTISPECIES: hydrogenase-2 small chain [Proteobacteria] Length: 372\nNumber of predicted TMHs: 1\nExp number of AAs in TMHs: 23.72226\nExp number, first 60 AAs: 0.22038\nTotal prob of N-in: 0.12994\noutside 1 331\nTMhelix 332 354\ninside 355 372

6050 GCF_002195075.1_ASM219507v1 Escherichia coli Proteobacteria; Gammaproteobacteria; Enterobacterales; Enterobacteriaceae; Escherichia
MTGDNTLIHSHGINRRDFMKLCAALAATMGLSSKAAAEMAESVTNPQRPPVIWIGAQECT WP_000145410.1
MULTISPECIES: hydrogenase-2 small chain [Proteobacteria] Length: 372\nNumber of predicted TMHs: 1\nExp number of AAs in TMHs: 23.72226\nExp number, first 60 AAs: 0.22038\nTotal prob of N-in: 0.12994\noutside 1 331\nTMhelix 332 354\ninside 355 372

6051 GCF_002195175.1_ASM219517v1 Escherichia coli Proteobacteria; Gammaproteobacteria; Enterobacterales; Enterobacteriaceae; Escherichia
MTGDNTLIHSHGINRRDFMKLCAALAATMGLSSKAAAEMAESVTNPQRPPVIWIGAQECT WP_000145410.1
MULTISPECIES: hydrogenase-2 small chain [Proteobacteria] Length: 372\nNumber of predicted TMHs: 1\nExp number of AAs in TMHs: 23.72226\nExp number, first 60 AAs: 0.22038\nTotal prob of N-in: 0.12994\noutside 1 331\nTMhelix 332 354\ninside 355 372

6052 GCF_002195245.1_ASM219524v1 Escherichia coli Proteobacteria; Gammaproteobacteria; Enterobacterales; Enterobacteriaceae; Escherichia
MNNEETFYQAMRRQGVTRRSFLKYCSLAATSLGLGAGMAPKIAWALENKPRIPVVWIHGL WP_001301613.1
hydrogenase [Escherichia coli] Length: 372\nNumber of predicted TMHs: 2\nExp number of AAs in TMHs: 37.46598\nExp number, first 60 AAs: 19.53139\nTotal prob of N-in: 0.94364\nPOSSIBLE N-term signal sequence\ninside 1 19\nTMhelix 20 39\noutside 40 325\nTMhelix 326 348\ninside 349 372

6053 GCF_002195265.1_ASM219526v1 Escherichia coli Proteobacteria; Gammaproteobacteria; Enterobacterales; Enterobacteriaceae; Escherichia
MNNEETFYQAMRRQGVTRRSFLKYCSLAATSLGLGAGMAPKIAWALENKPRIPVVWIHGL WP_001301613.1
hydrogenase [Escherichia coli] Length: 372\nNumber of predicted TMHs: 2\nExp number of AAs in TMHs: 37.46598\nExp number, first 60 AAs: 19.53139\nTotal prob of N-in: 0.94364\nPOSSIBLE N-term signal sequence\ninside 1 19\nTMhelix 20 39\noutside 40 325\nTMhelix 326 348\ninside 349 372

6054 GCF_002195335.1_ASM219533v1 Escherichia coli Proteobacteria; Gammaproteobacteria; Enterobacterales; Enterobacteriaceae; Escherichia
MNNEETFYQAMRRQGVTRRSFLKYCSLAATSLGLGAGMAPKIAWALENKPRIPVVWIHGL WP_001301613.1
hydrogenase [Escherichia coli] Length: 372\nNumber of predicted TMHs: 2\nExp number of AAs in TMHs: 37.46598\nExp number, first 60 AAs: 19.53139\nTotal prob of N-in: 0.94364\nPOSSIBLE N-term signal sequence\ninside 1 19\nTMhelix 20 39\noutside 40 325\nTMhelix 326 348\ninside 349 372

6055 GCF_002195035.1_ASM219503v1 Escherichia coli Proteobacteria; Gammaproteobacteria; Enterobacterales; Enterobacteriaceae; Escherichia
MNNEETFYQAMRRQGVTRRSFLKYCSLAATSLGLGAGMAPKIAWALENKPRIPVVWIHGL WP_001301613.1
hydrogenase [Escherichia coli] Length: 372\nNumber of predicted TMHs: 2\nExp number of AAs in

TMHs: 37.46598\nExp number, first 60 AAs: 19.53139\nTotal prob of N-in: 0.94364\nPOSSIBLE N-term signal sequence\ninside 1 19\nTMhelix 20 39\noutside 40 325\nTMhelix 326 348\ninside 349 372

6056 GCF_002194975.1_ASM219497v1 Escherichia coli Proteobacteria; Gammaproteobacteria; Enterobacterales; Enterobacteriaceae; Escherichia
MNNEETFYQAMRRQGVTRRSFLKYCSLAATSLGLGAGMAPKIAWALENKPRIPVVWIHGL WP_001301613.1
hydrogenase [Escherichia coli] Length: 372\nNumber of predicted TMHs: 2\nExp number of AAs in TMHs: 37.46598\nExp number, first 60 AAs: 19.53139\nTotal prob of N-in: 0.94364\nPOSSIBLE N-term signal sequence\ninside 1 19\nTMhelix 20 39\noutside 40 325\nTMhelix 326 348\ninside 349 372

6057 GCF_002195355.1_ASM219535v1 Escherichia coli Proteobacteria; Gammaproteobacteria; Enterobacterales; Enterobacteriaceae; Escherichia
MTGDNTLIHSHGINRRDFMKLCAALAATMGLSSKAAAEMAESVTNPQRPPVIWIGAQECT WP_000145410.1
MULTISPECIES: hydrogenase-2 small chain [Proteobacteria] Length: 372\nNumber of predicted TMHs: 1\nExp number of AAs in TMHs: 23.72226\nExp number, first 60 AAs: 0.22038\nTotal prob of N-in: 0.12994\noutside 1 331\nTMhelix 332 354\ninside 355 372

6058 GCF_002195355.1_ASM219535v1 Escherichia coli Proteobacteria; Gammaproteobacteria; Enterobacterales; Enterobacteriaceae; Escherichia
MNNEETFYQAMRRQGVTRRSFLKYCSLAATSLGLGAGMAPKIAWALENKPRIPVVWIHGL WP_001301613.1
hydrogenase [Escherichia coli] Length: 372\nNumber of predicted TMHs: 2\nExp number of AAs in TMHs: 37.46598\nExp number, first 60 AAs: 19.53139\nTotal prob of N-in: 0.94364\nPOSSIBLE N-term signal sequence\ninside 1 19\nTMhelix 20 39\noutside 40 325\nTMhelix 326 348\ninside 349 372

6059 GCF_002195055.1_ASM219505v1 Escherichia coli Proteobacteria; Gammaproteobacteria; Enterobacterales; Enterobacteriaceae; Escherichia
MNNEETFYQAMRRQGVTRRSFLKYCSLAATSLGLGAGMAPKIAWALENKPRIPVVWIHGL WP_001301613.1
hydrogenase [Escherichia coli] Length: 372\nNumber of predicted TMHs: 2\nExp number of AAs in TMHs: 37.46598\nExp number, first 60 AAs: 19.53139\nTotal prob of N-in: 0.94364\nPOSSIBLE N-term signal sequence\ninside 1 19\nTMhelix 20 39\noutside 40 325\nTMhelix 326 348\ninside 349 372

6060 GCF_002195075.1_ASM219507v1 Escherichia coli Proteobacteria; Gammaproteobacteria; Enterobacterales; Enterobacteriaceae; Escherichia
MNNEETFYQAMRRQGVTRRSFLKYCSLAATSLGLGAGMAPKIAWALENKPRIPVVWIHGL WP_001301613.1
hydrogenase [Escherichia coli] Length: 372\nNumber of predicted TMHs: 2\nExp number of AAs in TMHs: 37.46598\nExp number, first 60 AAs: 19.53139\nTotal prob of N-in: 0.94364\nPOSSIBLE N-term signal sequence\ninside 1 19\nTMhelix 20 39\noutside 40 325\nTMhelix 326 348\ninside 349 372

6061 GCF_002195125.1_ASM219512v1 Escherichia coli Proteobacteria; Gammaproteobacteria; Enterobacterales; Enterobacteriaceae; Escherichia
MNNEETFYQAMRRQGVTRRSFLKYCSLAATSLGLGAGMAPKIAWALENKPRIPVVWIHGL WP_001301613.1
hydrogenase [Escherichia coli] Length: 372\nNumber of predicted TMHs: 2\nExp number of AAs in TMHs: 37.46598\nExp number, first 60 AAs: 19.53139\nTotal prob of N-in: 0.94364\nPOSSIBLE N-term signal sequence\ninside 1 19\nTMhelix 20 39\noutside 40 325\nTMhelix 326 348\ninside 349 372

6062 GCF_002195175.1_ASM219517v1 Escherichia coli Proteobacteria; Gammaproteobacteria; Enterobacterales; Enterobacteriaceae; Escherichia
MNNEETFYQAMRRQGVTRRSFLKYCSLAATSLGLGAGMAPKIAWALENKPRIPVVWIHGL WP_001301613.1
hydrogenase [Escherichia coli] Length: 372\nNumber of predicted TMHs: 2\nExp number of AAs in TMHs: 37.46598\nExp number, first 60 AAs: 19.53139\nTotal prob of N-in: 0.94364\nPOSSIBLE N-term signal sequence\ninside 1 19\nTMhelix 20 39\noutside 40 325\nTMhelix 326 348\ninside 349 372

6063 GCF_002195185.1_ASM219518v1 Escherichia coli Proteobacteria; Gammaproteobacteria; Enterobacterales; Enterobacteriaceae; Escherichia
MNNEETFYQAMRRQGVTRRSFLKYCSLAATSLGLGAGMAPKIAWALENKPRIPVVWIHGL WP_001301613.1
hydrogenase [Escherichia coli] Length: 372\nNumber of predicted TMHs: 2\nExp number of AAs in TMHs: 37.46598\nExp number, first 60 AAs: 19.53139\nTotal prob of N-in: 0.94364\nPOSSIBLE N-term signal sequence\ninside 1 19\nTMhelix 20 39\noutside 40 325\nTMhelix 326 348\ninside 349 372

6064 GCF_002195405.1_ASM219540v1 Escherichia coli Proteobacteria; Gammaproteobacteria; Enterobacterales; Enterobacteriaceae; Escherichia

MTGDNTLIHSHGINRRDFMKLCAALAATMGLSSKAAAEMAESVTNPQRPPVIWIGAQECT WP_000145410.1 MULTISPECIES: hydrogenase-2 small chain [Proteobacteria] Length: 372\nNumber of predicted TMHs: 1\nExp number of AAs in TMHs: 23.72226\nExp number, first 60 AAs: 0.22038\nTotal prob of N-in: 0.12994\noutside 1 331\nTMhelix 332 354\ninside 355 372

6065 GCF_002195445.1_ASM219544v1 Escherichia coli Proteobacteria; Gammaproteobacteria; Enterobacterales; Enterobacteriaceae; Escherichia MTGDNTLIHSHGINRRDFMKLCAALAATMGLSSKAAAEMAESVTNPQRPPVIWIGAQECT WP_000145410.1 MULTISPECIES: hydrogenase-2 small chain [Proteobacteria] Length: 372\nNumber of predicted TMHs: 1\nExp number of AAs in TMHs: 23.72226\nExp number, first 60 AAs: 0.22038\nTotal prob of N-in: 0.12994\noutside 1 331\nTMhelix 332 354\ninside 355 372

6066 GCF_002195475.1_ASM219547v1 Escherichia coli Proteobacteria; Gammaproteobacteria; Enterobacterales; Enterobacteriaceae; Escherichia MTGDNTLIHSHGINRRDFMKLCAALAATMGLSSKAAAEMAESVTNPQRPPVIWIGAQECT WP_000145410.1 MULTISPECIES: hydrogenase-2 small chain [Proteobacteria] Length: 372\nNumber of predicted TMHs: 1\nExp number of AAs in TMHs: 23.72226\nExp number, first 60 AAs: 0.22038\nTotal prob of N-in: 0.12994\noutside 1 331\nTMhelix 332 354\ninside 355 372

6067 GCF_002195515.1_ASM219551v1 Escherichia coli Proteobacteria; Gammaproteobacteria; Enterobacterales; Enterobacteriaceae; Escherichia MTGDNTLIHSHGINRRDFMKLCAALAATMGLSSKAAAEMAESVTNPQRPPVIWIGAQECT WP_000145410.1 MULTISPECIES: hydrogenase-2 small chain [Proteobacteria] Length: 372\nNumber of predicted TMHs: 1\nExp number of AAs in TMHs: 23.72226\nExp number, first 60 AAs: 0.22038\nTotal prob of N-in: 0.12994\noutside 1 331\nTMhelix 332 354\ninside 355 372

6068 GCF_002195525.1_ASM219552v1 Escherichia coli Proteobacteria; Gammaproteobacteria; Enterobacterales; Enterobacteriaceae; Escherichia MTGDNTLIHSHGINRRDFMKLCAALAATMGLSSKAAAEMAESVTNPQRPPVIWIGAQECT WP_000145410.1 MULTISPECIES: hydrogenase-2 small chain [Proteobacteria] Length: 372\nNumber of predicted TMHs: 1\nExp number of AAs in TMHs: 23.72226\nExp number, first 60 AAs: 0.22038\nTotal prob of N-in: 0.12994\noutside 1 331\nTMhelix 332 354\ninside 355 372

6069 GCF_002195565.1_ASM219556v1 Escherichia coli Proteobacteria; Gammaproteobacteria; Enterobacterales; Enterobacteriaceae; Escherichia MTGDNTLIHSHGINRRDFMKLCAALAATMGLSSKAAAEMAESVTNPQRPPVIWIGAQECT WP_000145410.1 MULTISPECIES: hydrogenase-2 small chain [Proteobacteria] Length: 372\nNumber of predicted TMHs: 1\nExp number of AAs in TMHs: 23.72226\nExp number, first 60 AAs: 0.22038\nTotal prob of N-in: 0.12994\noutside 1 331\nTMhelix 332 354\ninside 355 372

6070 GCF_002195605.1_ASM219560v1 Escherichia coli Proteobacteria; Gammaproteobacteria; Enterobacterales; Enterobacteriaceae; Escherichia MTGDNTLIHSHGINRRDFMKLCAALAATMGLSSKAAAEMAESVTNPQRPPVIWIGAQECT WP_000145410.1 MULTISPECIES: hydrogenase-2 small chain [Proteobacteria] Length: 372\nNumber of predicted TMHs: 1\nExp number of AAs in TMHs: 23.72226\nExp number, first 60 AAs: 0.22038\nTotal prob of N-in: 0.12994\noutside 1 331\nTMhelix 332 354\ninside 355 372

6071 GCF_002195615.1_ASM219561v1 Escherichia coli Proteobacteria; Gammaproteobacteria; Enterobacterales; Enterobacteriaceae; Escherichia MTGDNTLIHSHGINRRDFMKLCAALAATMGLSSKAAAEMAESVTNPQRPPVIWIGAQECT WP_000145410.1 MULTISPECIES: hydrogenase-2 small chain [Proteobacteria] Length: 372\nNumber of predicted TMHs: 1\nExp number of AAs in TMHs: 23.72226\nExp number, first 60 AAs: 0.22038\nTotal prob of N-in: 0.12994\noutside 1 331\nTMhelix 332 354\ninside 355 372

6072 GCF_002195675.1_ASM219567v1 Escherichia coli Proteobacteria; Gammaproteobacteria; Enterobacterales; Enterobacteriaceae; Escherichia MTGDNTLIHSHGINRRDFMKLCAALAATMGLSSKAAAEMAESVTNPQRPPVIWIGAQECT WP_000145410.1 MULTISPECIES: hydrogenase-2 small chain [Proteobacteria] Length: 372\nNumber of predicted TMHs: 1\nExp number of AAs in TMHs: 23.72226\nExp number, first 60 AAs: 0.22038\nTotal prob of N-in: 0.12994\noutside 1 331\nTMhelix 332 354\ninside 355 372

6073 GCF_002195705.1_ASM219570v1 Escherichia coli Proteobacteria; Gammaproteobacteria; Enterobacterales; Enterobacteriaceae; Escherichia
MTGDNTLIHSHGINRRDFMKLCAALAATMGLSSKAAAEMAESVTNPQRPPVIWIGAQECT WP_000145410.1
MULTISPECIES: hydrogenase-2 small chain [Proteobacteria] Length: 372\nNumber of predicted TMHs: 1\nExp number of AAs in TMHs: 23.72226\nExp number, first 60 AAs: 0.22038\nTotal prob of N-in: 0.12994\noutside 1 331\nTMhelix 332 354\ninside 355 372

6074 GCF_002195775.1_ASM219577v1 Escherichia coli Proteobacteria; Gammaproteobacteria; Enterobacterales; Enterobacteriaceae; Escherichia
MNNEETFYQAMRRQGVTRRSFLKYCSLAATSLGLGAGMAPKIAWALENKPRIPVVWIHGL WP_001301613.1
hydrogenase [Escherichia coli] Length: 372\nNumber of predicted TMHs: 2\nExp number of AAs in TMHs: 37.46598\nExp number, first 60 AAs: 19.53139\nTotal prob of N-in: 0.94364\nPOSSIBLE N-term signal sequence\ninside 1 19\nTMhelix 20 39\noutside 40 325\nTMhelix 326 348\ninside 349 372

6075 GCF_002195445.1_ASM219544v1 Escherichia coli Proteobacteria; Gammaproteobacteria; Enterobacterales; Enterobacteriaceae; Escherichia
MNNEETFYQAMRRQGVTRRSFLKYCSLAATSLGLGAGMAPKIAWALENKPRIPVVWIHGL WP_001301613.1
hydrogenase [Escherichia coli] Length: 372\nNumber of predicted TMHs: 2\nExp number of AAs in TMHs: 37.46598\nExp number, first 60 AAs: 19.53139\nTotal prob of N-in: 0.94364\nPOSSIBLE N-term signal sequence\ninside 1 19\nTMhelix 20 39\noutside 40 325\nTMhelix 326 348\ninside 349 372

6076 GCF_002195475.1_ASM219547v1 Escherichia coli Proteobacteria; Gammaproteobacteria; Enterobacterales; Enterobacteriaceae; Escherichia
MNNEETFYQAMRRQGVTRRSFLKYCSLAATSLGLGAGMAPKIAWALENKPRIPVVWIHGL WP_001301613.1
hydrogenase [Escherichia coli] Length: 372\nNumber of predicted TMHs: 2\nExp number of AAs in TMHs: 37.46598\nExp number, first 60 AAs: 19.53139\nTotal prob of N-in: 0.94364\nPOSSIBLE N-term signal sequence\ninside 1 19\nTMhelix 20 39\noutside 40 325\nTMhelix 326 348\ninside 349 372

6077 GCF_002195515.1_ASM219551v1 Escherichia coli Proteobacteria; Gammaproteobacteria; Enterobacterales; Enterobacteriaceae; Escherichia
MNNEETFYQAMRRQGVTRRSFLKYCSLAATSLGLGAGMAPKIAWALENKPRIPVVWIHGL WP_001301613.1
hydrogenase [Escherichia coli] Length: 372\nNumber of predicted TMHs: 2\nExp number of AAs in TMHs: 37.46598\nExp number, first 60 AAs: 19.53139\nTotal prob of N-in: 0.94364\nPOSSIBLE N-term signal sequence\ninside 1 19\nTMhelix 20 39\noutside 40 325\nTMhelix 326 348\ninside 349 372

6078 GCF_002195565.1_ASM219556v1 Escherichia coli Proteobacteria; Gammaproteobacteria; Enterobacterales; Enterobacteriaceae; Escherichia
MNNEETFYQAMRRQGVTRRSFLKYCSLAATSLGLGAGMAPKIAWALENKPRIPVVWIHGL WP_001301613.1
hydrogenase [Escherichia coli] Length: 372\nNumber of predicted TMHs: 2\nExp number of AAs in TMHs: 37.46598\nExp number, first 60 AAs: 19.53139\nTotal prob of N-in: 0.94364\nPOSSIBLE N-term signal sequence\ninside 1 19\nTMhelix 20 39\noutside 40 325\nTMhelix 326 348\ninside 349 372

6079 GCF_002195525.1_ASM219552v1 Escherichia coli Proteobacteria; Gammaproteobacteria; Enterobacterales; Enterobacteriaceae; Escherichia
MNNEETFYQAMRRQGVTRRSFLKYCSLAATSLGLGAGMAPKIAWALENKPRIPVVWIHGL WP_001301613.1
hydrogenase [Escherichia coli] Length: 372\nNumber of predicted TMHs: 2\nExp number of AAs in TMHs: 37.46598\nExp number, first 60 AAs: 19.53139\nTotal prob of N-in: 0.94364\nPOSSIBLE N-term signal sequence\ninside 1 19\nTMhelix 20 39\noutside 40 325\nTMhelix 326 348\ninside 349 372

6080 GCF_002195605.1_ASM219560v1 Escherichia coli Proteobacteria; Gammaproteobacteria; Enterobacterales; Enterobacteriaceae; Escherichia
MNNEETFYQAMRRQGVTRRSFLKYCSLAATSLGLGAGMAPKIAWALENKPRIPVVWIHGL WP_001301613.1
hydrogenase [Escherichia coli] Length: 372\nNumber of predicted TMHs: 2\nExp number of AAs in TMHs: 37.46598\nExp number, first 60 AAs: 19.53139\nTotal prob of N-in: 0.94364\nPOSSIBLE N-term signal sequence\ninside 1 19\nTMhelix 20 39\noutside 40 325\nTMhelix 326 348\ninside 349 372

6081 GCF_002195805.1_ASM219580v1 Escherichia coli Proteobacteria; Gammaproteobacteria; Enterobacterales; Enterobacteriaceae; Escherichia
MTGDNTLIHSHGINRRDFMKLCAALAATMGLSSKAAAEMAESVTNPQRPPVIWIGAQECT WP_000145410.1
MULTISPECIES: hydrogenase-2 small chain [Proteobacteria] Length: 372\nNumber of predicted TMHs:

1\nExp number of AAs in TMHs: 23.72226\nExp number, first 60 AAs: 0.22038\nTotal prob of N-in: 0.12994\noutside 1 331\nTMhelix 332 354\ninside 355 372

6082 GCF_002195815.1_ASM219581v1 Escherichia coli Proteobacteria; Gammaproteobacteria; Enterobacterales; Enterobacteriaceae; Escherichia
MTGDNTLIHSHGINRRDFMKLCAALAATMGLSSKAAAEMAESVTNPQRPPVIWIGAQECT WP_000145410.1
MULTISPECIES: hydrogenase-2 small chain [Proteobacteria] Length: 372\nNumber of predicted TMHs: 1\nExp number of AAs in TMHs: 23.72226\nExp number, first 60 AAs: 0.22038\nTotal prob of N-in: 0.12994\noutside 1 331\nTMhelix 332 354\ninside 355 372

6083 GCF_002196495.1_ASM219649v1 Escherichia coli Proteobacteria; Gammaproteobacteria; Enterobacterales; Enterobacteriaceae; Escherichia
MTGDNTLIHSHGINRRDFMKLCAALAATMGLSSKAAAEMAESVTNPQRPPVIWIGAQECT WP_000145410.1
MULTISPECIES: hydrogenase-2 small chain [Proteobacteria] Length: 372\nNumber of predicted TMHs: 1\nExp number of AAs in TMHs: 23.72226\nExp number, first 60 AAs: 0.22038\nTotal prob of N-in: 0.12994\noutside 1 331\nTMhelix 332 354\ninside 355 372

6084 GCF_002195875.1_ASM219587v1 Escherichia coli Proteobacteria; Gammaproteobacteria; Enterobacterales; Enterobacteriaceae; Escherichia
MTGDNTLIHSHGINRRDFMKLCAALAATMGLSSKAAAEMAESVTNPQRPPVIWIGAQECT WP_000145410.1
MULTISPECIES: hydrogenase-2 small chain [Proteobacteria] Length: 372\nNumber of predicted TMHs: 1\nExp number of AAs in TMHs: 23.72226\nExp number, first 60 AAs: 0.22038\nTotal prob of N-in: 0.12994\noutside 1 331\nTMhelix 332 354\ninside 355 372

6085 GCF_002195905.1_ASM219590v1 Escherichia coli Proteobacteria; Gammaproteobacteria; Enterobacterales; Enterobacteriaceae; Escherichia
MTGDNTLIHSHGINRRDFMKLCAALAATMGLSSKAAAEMAESVTNPQRPPVIWIGAQECT WP_000145410.1
MULTISPECIES: hydrogenase-2 small chain [Proteobacteria] Length: 372\nNumber of predicted TMHs: 1\nExp number of AAs in TMHs: 23.72226\nExp number, first 60 AAs: 0.22038\nTotal prob of N-in: 0.12994\noutside 1 331\nTMhelix 332 354\ninside 355 372

6086 GCF_002195885.1_ASM219588v1 Escherichia coli Proteobacteria; Gammaproteobacteria; Enterobacterales; Enterobacteriaceae; Escherichia
MTGDNTLIHSHGINRRDFMKLCAALAATMGLSSKAAAEMAESVTNPQRPPVIWIGAQECT WP_000145410.1
MULTISPECIES: hydrogenase-2 small chain [Proteobacteria] Length: 372\nNumber of predicted TMHs: 1\nExp number of AAs in TMHs: 23.72226\nExp number, first 60 AAs: 0.22038\nTotal prob of N-in: 0.12994\noutside 1 331\nTMhelix 332 354\ninside 355 372

6087 GCF_002196615.1_ASM219661v1 Escherichia coli Proteobacteria; Gammaproteobacteria; Enterobacterales; Enterobacteriaceae; Escherichia
MTGDNTLIHSHGINRRDFMKLCAALAATMGLSSKAAAEMAESVTNPQRPPVIWIGAQECT WP_000145410.1
MULTISPECIES: hydrogenase-2 small chain [Proteobacteria] Length: 372\nNumber of predicted TMHs: 1\nExp number of AAs in TMHs: 23.72226\nExp number, first 60 AAs: 0.22038\nTotal prob of N-in: 0.12994\noutside 1 331\nTMhelix 332 354\ninside 355 372

6088 GCF_002196635.1_ASM219663v1 Escherichia coli Proteobacteria; Gammaproteobacteria; Enterobacterales; Enterobacteriaceae; Escherichia
MTGDNTLIHSHGINRRDFMKLCAALAATMGLSSKAAAEMAESVTNPQRPPVIWIGAQECT WP_000145410.1
MULTISPECIES: hydrogenase-2 small chain [Proteobacteria] Length: 372\nNumber of predicted TMHs: 1\nExp number of AAs in TMHs: 23.72226\nExp number, first 60 AAs: 0.22038\nTotal prob of N-in: 0.12994\noutside 1 331\nTMhelix 332 354\ninside 355 372

6089 GCF_002195805.1_ASM219580v1 Escherichia coli Proteobacteria; Gammaproteobacteria; Enterobacterales; Enterobacteriaceae; Escherichia
MNNEETFYQAMRRQGVTRRSFLKYCSLAATSLGLGAGMAPKIAWALENKPRIPVVWIHGL WP_001301613.1
hydrogenase [Escherichia coli] Length: 372\nNumber of predicted TMHs: 2\nExp number of AAs in TMHs: 37.46598\nExp number, first 60 AAs: 19.53139\nTotal prob of N-in: 0.94364\nPOSSIBLE N-term signal sequence\ninside 1 19\nTMhelix 20 39\noutside 40 325\nTMhelix 326 348\ninside 349 372

6090 GCF_002195815.1_ASM219581v1 Escherichia coli Proteobacteria; Gammaproteobacteria; Enterobacterales; Enterobacteriaceae; Escherichia

MNNEETFYQAMRRQGVTRRSFLKYCSLAATSLGLGAGMAPKIAWALENKPRIPVVWIHGL WP_001301613.1 hydrogenase [Escherichia coli] Length: 372\nNumber of predicted TMHs: 2\nExp number of AAs in TMHs: 37.46598\nExp number, first 60 AAs: 19.53139\nTotal prob of N-in: 0.94364\nPOSSIBLE N-term signal sequence\ninside 1 19\nTMhelix 20 39\noutside 40 325\nTMhelix 326 348\ninside 349 372

6091 GCF_002198005.1_ASM219800v1 Escherichia coli Proteobacteria; Gammaproteobacteria; Enterobacterales; Enterobacteriaceae; Escherichia MTGDNTLIHSHGINRRDFMKLCAALAATMGLSSKAAAEMAESVTNPQRPPVIWIGAQECT WP_000145410.1 MULTISPECIES: hydrogenase-2 small chain [Proteobacteria] Length: 372\nNumber of predicted TMHs: 1\nExp number of AAs in TMHs: 23.72226\nExp number, first 60 AAs: 0.22038\nTotal prob of N-in: 0.12994\noutside 1 331\nTMhelix 332 354\ninside 355 372

6092 GCF_002198005.1_ASM219800v1 Escherichia coli Proteobacteria; Gammaproteobacteria; Enterobacterales; Enterobacteriaceae; Escherichia MNNEETFYQAMRRQGVTRRSFLKYCSLAATSLGLGAGMAPKIAWALENKPRIPVVWIHGL WP_001301613.1 hydrogenase [Escherichia coli] Length: 372\nNumber of predicted TMHs: 2\nExp number of AAs in TMHs: 37.46598\nExp number, first 60 AAs: 19.53139\nTotal prob of N-in: 0.94364\nPOSSIBLE N-term signal sequence\ninside 1 19\nTMhelix 20 39\noutside 40 325\nTMhelix 326 348\ninside 349 372

6093 GCF_002196615.1_ASM219661v1 Escherichia coli Proteobacteria; Gammaproteobacteria; Enterobacterales; Enterobacteriaceae; Escherichia MNNEETFYQAMRRQGVTRRSFLKYCSLAATSLGLGAGMAPKIAWALENKPRIPVVWIHGL WP_001295940.1 MULTISPECIES: hydrogenase [Enterobacteriaceae] Length: 372\nNumber of predicted TMHs: 2\nExp number of AAs in TMHs: 37.2608899999999\nExp number, first 60 AAs: 19.52923\nTotal prob of N-in: 0.94351\nPOSSIBLE N-term signal sequence\ninside 1 19\nTMhelix 20 39\noutside 40 325\nTMhelix 326 348\ninside 349 372

6094 GCF_002195905.1_ASM219590v1 Escherichia coli Proteobacteria; Gammaproteobacteria; Enterobacterales; Enterobacteriaceae; Escherichia MNNEETFYQAMRRQGVTRRSFLKYCSLAATSLGLGAGMAPKIAWALENKPRIPVVWIHGL WP_001301613.1 hydrogenase [Escherichia coli] Length: 372\nNumber of predicted TMHs: 2\nExp number of AAs in TMHs: 37.46598\nExp number, first 60 AAs: 19.53139\nTotal prob of N-in: 0.94364\nPOSSIBLE N-term signal sequence\ninside 1 19\nTMhelix 20 39\noutside 40 325\nTMhelix 326 348\ninside 349 372

6095 GCF_002195885.1_ASM219588v1 Escherichia coli Proteobacteria; Gammaproteobacteria; Enterobacterales; Enterobacteriaceae; Escherichia MNNEETFYQAMRRQGVTRRSFLKYCSLAATSLGLGAGMAPKIAWALENKPRIPVVWIHGL WP_001301613.1 hydrogenase [Escherichia coli] Length: 372\nNumber of predicted TMHs: 2\nExp number of AAs in TMHs: 37.46598\nExp number, first 60 AAs: 19.53139\nTotal prob of N-in: 0.94364\nPOSSIBLE N-term signal sequence\ninside 1 19\nTMhelix 20 39\noutside 40 325\nTMhelix 326 348\ninside 349 372

6096 GCF_002195875.1_ASM219587v1 Escherichia coli Proteobacteria; Gammaproteobacteria; Enterobacterales; Enterobacteriaceae; Escherichia MNNEETFYQAMRRQGVTRRSFLKYCSLAATSLGLGAGMAPKIAWALENKPRIPVVWIHGL WP_001301613.1 hydrogenase [Escherichia coli] Length: 372\nNumber of predicted TMHs: 2\nExp number of AAs in TMHs: 37.46598\nExp number, first 60 AAs: 19.53139\nTotal prob of N-in: 0.94364\nPOSSIBLE N-term signal sequence\ninside 1 19\nTMhelix 20 39\noutside 40 325\nTMhelix 326 348\ninside 349 372

6097 GCF_002205945.1_ASM220594v1 Escherichia coli Proteobacteria; Gammaproteobacteria; Enterobacterales; Enterobacteriaceae; Escherichia MTGDNTLIHSHGINRRDFMKLCAALAATMGLSSKAAAEMAESVTNPQRPPVIWIGAQECT WP_000145410.1 MULTISPECIES: hydrogenase-2 small chain [Proteobacteria] Length: 372\nNumber of predicted TMHs: 1\nExp number of AAs in TMHs: 23.72226\nExp number, first 60 AAs: 0.22038\nTotal prob of N-in: 0.12994\noutside 1 331\nTMhelix 332 354\ninside 355 372

6098 GCF_002202175.1_ASM220217v1 Escherichia coli Proteobacteria; Gammaproteobacteria; Enterobacterales; Enterobacteriaceae; Escherichia MTGDNTLIHSHGINRRDFMKLCAALAATMGLSSKAAAEMVESVTNPQRPPVIWIGAQECT WP_000145423.1 hydrogenase 2 small subunit [Escherichia coli] Length: 372\nNumber of predicted TMHs: 1\nExp number of AAs in TMHs: 23.78201\nExp number, first 60 AAs: 0.28355\nTotal prob of N-in: 0.13257\noutside 1 331\nTMhelix 332 354\ninside 355 372

6099 GCF_002207985.1_ASM220798v1 Escherichia coli Proteobacteria; Gammaproteobacteria; Enterobacterales; Enterobacteriaceae; Escherichia
MTGDNTLIHSHGINRRDFMKLCAALAATMGLSSKAAAEMAESVTNPQRPPVIWIGAQECT WP_000145410.1
MULTISPECIES: hydrogenase-2 small chain [Proteobacteria] Length: 372\nNumber of predicted TMHs: 1\nExp number of AAs in TMHs: 23.72226\nExp number, first 60 AAs: 0.22038\nTotal prob of N-in: 0.12994\noutside 1 331\nTMhelix 332 354\ninside 355 372

6100 GCF_002208025.1_ASM220802v1 Escherichia coli Proteobacteria; Gammaproteobacteria; Enterobacterales; Enterobacteriaceae; Escherichia
MTGDNTLIHSHGINRRDFMKLCAALAATMGLSSKAAAEMAESVTNPQRPPVIWIGAQECT WP_000145410.1
MULTISPECIES: hydrogenase-2 small chain [Proteobacteria] Length: 372\nNumber of predicted TMHs: 1\nExp number of AAs in TMHs: 23.72226\nExp number, first 60 AAs: 0.22038\nTotal prob of N-in: 0.12994\noutside 1 331\nTMhelix 332 354\ninside 355 372

6101 GCF_002205945.1_ASM220594v1 Escherichia coli Proteobacteria; Gammaproteobacteria; Enterobacterales; Enterobacteriaceae; Escherichia
MNNEETFYQAMRRQGVTRRSFLKYCSLAATSLGLGAGMAPKIAWALENKPRIPVVWIHGL WP_001058323.1
MULTISPECIES: hydrogenase-1 small chain [Proteobacteria] Length: 372\nNumber of predicted TMHs: 2\nExp number of AAs in TMHs: 37.2608499999999\nExp number, first 60 AAs: 19.52923\nTotal prob of N-in: 0.94351\nPOSSIBLE N-term signal sequence\ninside 1 19\nTMhelix 20 39\noutside 40 325\nTMhelix 326 348\ninside 349 372

6102 GCF_900008845.1_ED423_contigs Escherichia coli Proteobacteria; Gammaproteobacteria; Enterobacterales; Enterobacteriaceae; Escherichia
MTGDNTLIHSHGINRRDFMKLCAALAATMGLSSKAAAEMAESVTNPQRPPVIWIGAQECT WP_000145410.1
MULTISPECIES: hydrogenase-2 small chain [Proteobacteria] Length: 372\nNumber of predicted TMHs: 1\nExp number of AAs in TMHs: 23.72226\nExp number, first 60 AAs: 0.22038\nTotal prob of N-in: 0.12994\noutside 1 331\nTMhelix 332 354\ninside 355 372

6103 GCF_900008845.1_ED423_contigs Escherichia coli Proteobacteria; Gammaproteobacteria; Enterobacterales; Enterobacteriaceae; Escherichia
MNNEETFYQAMRRQGVTRRSFLKYCSLAATSLGLGAGMAPKIAWALENKPRIPVVWIHGL WP_001058323.1
MULTISPECIES: hydrogenase-1 small chain [Proteobacteria] Length: 372\nNumber of predicted TMHs: 2\nExp number of AAs in TMHs: 37.2608499999999\nExp number, first 60 AAs: 19.52923\nTotal prob of N-in: 0.94351\nPOSSIBLE N-term signal sequence\ninside 1 19\nTMhelix 20 39\noutside 40 325\nTMhelix 326 348\ninside 349 372

6104 GCF_002208025.1_ASM220802v1 Escherichia coli Proteobacteria; Gammaproteobacteria; Enterobacterales; Enterobacteriaceae; Escherichia
MNNEETFYQAMRRQGVTRRSFLKYCSLAATSLGLGAGMAPKIAWALENKPRIPVVWIHGL WP_001058320.1
hydrogenase [Escherichia coli] Length: 372\nNumber of predicted TMHs: 2\nExp number of AAs in TMHs: 37.26082\nExp number, first 60 AAs: 19.52923\nTotal prob of N-in: 0.94351\nPOSSIBLE N-term signal sequence\ninside 1 19\nTMhelix 20 39\noutside 40 325\nTMhelix 326 348\ninside 349 372

6105 GCF_002202175.1_ASM220217v1 Escherichia coli Proteobacteria; Gammaproteobacteria; Enterobacterales; Enterobacteriaceae; Escherichia
MNNEETFYQAMRRQGVTRRSFLKYCSLAATSLGLGAGMAPKIAWALENKPRIPVVWIHGL WP_001295940.1
MULTISPECIES: hydrogenase [Enterobacteriaceae] Length: 372\nNumber of predicted TMHs: 2\nExp number of AAs in TMHs: 37.2608899999999\nExp number, first 60 AAs: 19.52923\nTotal prob of N-in: 0.94351\nPOSSIBLE N-term signal sequence\ninside 1 19\nTMhelix 20 39\noutside 40 325\nTMhelix 326 348\ninside 349 372

6106 GCF_900010045.1_ED082_contigs Escherichia coli Proteobacteria; Gammaproteobacteria; Enterobacterales; Enterobacteriaceae; Escherichia
MTGDNTLIHSHGINRRDFMKLCAALAATMGLSSKAAAEMAESVTNPQRPPVIWIGAQECT WP_000145410.1
MULTISPECIES: hydrogenase-2 small chain [Proteobacteria] Length: 372\nNumber of predicted TMHs: 1\nExp number of AAs in TMHs: 23.72226\nExp number, first 60 AAs: 0.22038\nTotal prob of N-in: 0.12994\noutside 1 331\nTMhelix 332 354\ninside 355 372

6107 GCF_900010045.1_ED082_contigs Escherichia coli Proteobacteria; Gammaproteobacteria; Enterobacterales; Enterobacteriaceae; Escherichia

MNNEETFYQAMRRQGVTRRSFLKYCSLAATSLGLGAGMAPKIAWALENKPRIPVVWIHGL WP_001058323.1 MULTISPECIES: hydrogenase-1 small chain [Proteobacteria] Length: 372\nNumber of predicted TMHs: 2\nExp number of AAs in TMHs: 37.2608499999999\nExp number, first 60 AAs: 19.52923\nTotal prob of N-in: 0.94351\nPOSSIBLE N-term signal sequence\ninside 1 19\nTMhelix 20 39\noutside 40 325\nTMhelix 326 348\ninside 349 372

6108 GCF_900010055.1_ED010_contigs Escherichia coli Proteobacteria; Gammaproteobacteria; Enterobacterales; Enterobacteriaceae; Escherichia MTGDNTLIHSHGINRRDFMKLCAALAATMGLSSKAAAEMAESVTNPQRPPVIWIGAQECT WP_000145410.1 MULTISPECIES: hydrogenase-2 small chain [Proteobacteria] Length: 372\nNumber of predicted TMHs: 1\nExp number of AAs in TMHs: 23.72226\nExp number, first 60 AAs: 0.22038\nTotal prob of N-in: 0.12994\noutside 1 331\nTMhelix 332 354\ninside 355 372

6109 GCF_900013265.1_ED664_contigs Escherichia coli Proteobacteria; Gammaproteobacteria; Enterobacterales; Enterobacteriaceae; Escherichia MTGDNTLIHSHGINRRDFMKLCAALAATMGLSSKAAAEMAESVTNPQRPPVIWIGAQECT WP_000145410.1 MULTISPECIES: hydrogenase-2 small chain [Proteobacteria] Length: 372\nNumber of predicted TMHs: 1\nExp number of AAs in TMHs: 23.72226\nExp number, first 60 AAs: 0.22038\nTotal prob of N-in: 0.12994\noutside 1 331\nTMhelix 332 354\ninside 355 372

6110 GCF_900013275.1_ED017_contigs Escherichia coli Proteobacteria; Gammaproteobacteria; Enterobacterales; Enterobacteriaceae; Escherichia MTGDNTLIHSHGINRRDFMKLCAALAATMGLSSKAAAEMAESVTNPQRPPVIWIGAQECT WP_000145410.1 MULTISPECIES: hydrogenase-2 small chain [Proteobacteria] Length: 372\nNumber of predicted TMHs: 1\nExp number of AAs in TMHs: 23.72226\nExp number, first 60 AAs: 0.22038\nTotal prob of N-in: 0.12994\noutside 1 331\nTMhelix 332 354\ninside 355 372

6111 GCF_900013275.1_ED017_contigs Escherichia coli Proteobacteria; Gammaproteobacteria; Enterobacterales; Enterobacteriaceae; Escherichia MNNEETFYQAMRRQGVTRRSFLKYCSLAATSLGLGAGMAPKIAWALENKPRIPVVWIHGL WP_001058323.1 MULTISPECIES: hydrogenase-1 small chain [Proteobacteria] Length: 372\nNumber of predicted TMHs: 2\nExp number of AAs in TMHs: 37.2608499999999\nExp number, first 60 AAs: 19.52923\nTotal prob of N-in: 0.94351\nPOSSIBLE N-term signal sequence\ninside 1 19\nTMhelix 20 39\noutside 40 325\nTMhelix 326 348\ninside 349 372

6112 GCF_900015375.1_ED668_contigs Escherichia coli Proteobacteria; Gammaproteobacteria; Enterobacterales; Enterobacteriaceae; Escherichia MTGDNTLIHSHGINRRDFMKLCAALAATMGLSSKAAAEMAESVTNPQRPPVIWIGAQECT WP_000145410.1 MULTISPECIES: hydrogenase-2 small chain [Proteobacteria] Length: 372\nNumber of predicted TMHs: 1\nExp number of AAs in TMHs: 23.72226\nExp number, first 60 AAs: 0.22038\nTotal prob of N-in: 0.12994\noutside 1 331\nTMhelix 332 354\ninside 355 372

6113 GCF_900014325.1_ED657_contigs Escherichia coli Proteobacteria; Gammaproteobacteria; Enterobacterales; Enterobacteriaceae; Escherichia MTGDNTLIHSHGINRRDFMKLCAALAATMGLSSKAAAEMAESVTNPQRPPVIWIGAQECT WP_000145410.1 MULTISPECIES: hydrogenase-2 small chain [Proteobacteria] Length: 372\nNumber of predicted TMHs: 1\nExp number of AAs in TMHs: 23.72226\nExp number, first 60 AAs: 0.22038\nTotal prob of N-in: 0.12994\noutside 1 331\nTMhelix 332 354\ninside 355 372

6114 GCF_900015385.1_ED676_contigs Escherichia coli Proteobacteria; Gammaproteobacteria; Enterobacterales; Enterobacteriaceae; Escherichia MTGDNTLIHSHGINRRDFMKLCAALAATMGLSSKAAAEMAESVTNPQRPPVIWIGAQECT WP_000145410.1 MULTISPECIES: hydrogenase-2 small chain [Proteobacteria] Length: 372\nNumber of predicted TMHs: 1\nExp number of AAs in TMHs: 23.72226\nExp number, first 60 AAs: 0.22038\nTotal prob of N-in: 0.12994\noutside 1 331\nTMhelix 332 354\ninside 355 372

6115 GCF_900011005.1_ED728_contigs Escherichia coli Proteobacteria; Gammaproteobacteria; Enterobacterales; Enterobacteriaceae; Escherichia MNNEETFYQAMRRQGVTRRSFLKYCSLAATSLGLGAGMAPKIAWALENKPRIPVVWIHGL WP_001295940.1 MULTISPECIES: hydrogenase [Enterobacteriaceae] Length: 372\nNumber of predicted TMHs: 2\nExp number of AAs in TMHs: 37.2608899999999\nExp number, first 60 AAs: 19.52923\nTotal prob of N-in:

0.94351\nPOSSIBLE N-term signal sequence\ninside 1 19\nTMhelix 20 39\noutside 40 325\nTMhelix 326 348\ninside 349 372

6116 GCF_900015705.1_ED644_contigs Escherichia coli Proteobacteria; Gammaproteobacteria; Enterobacterales; Enterobacteriaceae; Escherichia
MTGDNTLIHSHGINRRDFMKLCAALAATMGLSSKAAAEMAESVTNPQRPPVIWIGAQECT WP_000145410.1
MULTISPECIES: hydrogenase-2 small chain [Proteobacteria] Length: 372\nNumber of predicted TMHs: 1\nExp number of AAs in TMHs: 23.72226\nExp number, first 60 AAs: 0.22038\nTotal prob of N-in: 0.12994\noutside 1 331\nTMhelix 332 354\ninside 355 372

6117 GCF_900015385.1_ED676_contigs Escherichia coli Proteobacteria; Gammaproteobacteria; Enterobacterales; Enterobacteriaceae; Escherichia
MNNEETFYQAMRRQGVTRRSFLKYCSLAATSLGLGAGMAPKIAWALENKPRIPVVWIHGL WP_001058323.1
MULTISPECIES: hydrogenase-1 small chain [Proteobacteria] Length: 372\nNumber of predicted TMHs: 2\nExp number of AAs in TMHs: 37.2608499999999\nExp number, first 60 AAs: 19.52923\nTotal prob of N-in: 0.94351\nPOSSIBLE N-term signal sequence\ninside 1 19\nTMhelix 20 39\noutside 40 325\nTMhelix 326 348\ninside 349 372

6118 GCF_900015705.1_ED644_contigs Escherichia coli Proteobacteria; Gammaproteobacteria; Enterobacterales; Enterobacteriaceae; Escherichia
MNNEETFYQAMRRQGVTRRSFLKYCSLAATSLGLGAGMAPKIAWALENKPRIPVVWIHGL WP_001301613.1
hydrogenase [Escherichia coli] Length: 372\nNumber of predicted TMHs: 2\nExp number of AAs in TMHs: 37.46598\nExp number, first 60 AAs: 19.53139\nTotal prob of N-in: 0.94364\nPOSSIBLE N-term signal sequence\ninside 1 19\nTMhelix 20 39\noutside 40 325\nTMhelix 326 348\ninside 349 372

6119 GCF_900013265.1_ED664_contigs Escherichia coli Proteobacteria; Gammaproteobacteria; Enterobacterales; Enterobacteriaceae; Escherichia
MNNEETFYQAMRRQGVTRRSFLKYCSLAATSLGLGAGMAPKIAWALENKPRIPVVWIHGL WP_001058323.1
MULTISPECIES: hydrogenase-1 small chain [Proteobacteria] Length: 372\nNumber of predicted TMHs: 2\nExp number of AAs in TMHs: 37.2608499999999\nExp number, first 60 AAs: 19.52923\nTotal prob of N-in: 0.94351\nPOSSIBLE N-term signal sequence\ninside 1 19\nTMhelix 20 39\noutside 40 325\nTMhelix 326 348\ninside 349 372

6120 GCF_900010055.1_ED010_contigs Escherichia coli Proteobacteria; Gammaproteobacteria; Enterobacterales; Enterobacteriaceae; Escherichia
MNNEETFYQAMRRQGVTRRSFLKYCSLAATSLGLGAGMAPKIAWALENKPRIPVVWIHGL WP_001301613.1
hydrogenase [Escherichia coli] Length: 372\nNumber of predicted TMHs: 2\nExp number of AAs in TMHs: 37.46598\nExp number, first 60 AAs: 19.53139\nTotal prob of N-in: 0.94364\nPOSSIBLE N-term signal sequence\ninside 1 19\nTMhelix 20 39\noutside 40 325\nTMhelix 326 348\ninside 349 372

6121 GCF_900015715.1_ED449_contigs Escherichia coli Proteobacteria; Gammaproteobacteria; Enterobacterales; Enterobacteriaceae; Escherichia
MTGDNTLIHSHGINRRDFMKLCAALAATMGLSSKAAAEMAESVTNPQRPPVIWIGAQECT WP_000145410.1
MULTISPECIES: hydrogenase-2 small chain [Proteobacteria] Length: 372\nNumber of predicted TMHs: 1\nExp number of AAs in TMHs: 23.72226\nExp number, first 60 AAs: 0.22038\nTotal prob of N-in: 0.12994\noutside 1 331\nTMhelix 332 354\ninside 355 372

6122 GCF_900015785.1_ED073_contigs Escherichia coli Proteobacteria; Gammaproteobacteria; Enterobacterales; Enterobacteriaceae; Escherichia
MNNEETFYQAMRRQGVTRRSFLKYCSLAATSLGLGAGMAPKIAWALENKPRIPVVWIHGL WP_001058323.1
MULTISPECIES: hydrogenase-1 small chain [Proteobacteria] Length: 372\nNumber of predicted TMHs: 2\nExp number of AAs in TMHs: 37.2608499999999\nExp number, first 60 AAs: 19.52923\nTotal prob of N-in: 0.94351\nPOSSIBLE N-term signal sequence\ninside 1 19\nTMhelix 20 39\noutside 40 325\nTMhelix 326 348\ninside 349 372

6123 GCF_900015765.1_ED414_contigs Escherichia coli Proteobacteria; Gammaproteobacteria; Enterobacterales; Enterobacteriaceae; Escherichia
MNNEETFYQAMRRQGVTRRSFLKYCSLAATSLGLGAGMAPKIAWALENKPRIPVVWIHGL WP_001301613.1
hydrogenase [Escherichia coli] Length: 372\nNumber of predicted TMHs: 2\nExp number of AAs in TMHs: 37.46598\nExp number, first 60 AAs: 19.53139\nTotal prob of N-in: 0.94364\nPOSSIBLE N-term signal sequence\ninside 1 19\nTMhelix 20 39\noutside 40 325\nTMhelix 326 348\ninside 349 372

6124 GCF_900015805.1_ED287_contigs Escherichia coli Proteobacteria; Gammaproteobacteria; Enterobacterales; Enterobacteriaceae; Escherichia
MNNEETFYQAMRRQGVTRRSFLKYCSLAATSLGLGAGMAPKIAWALENKPRIPVVWIHGL WP_001295940.1
MULTISPECIES: hydrogenase [Enterobacteriaceae] Length: 372\nNumber of predicted TMHs: 2\nExp number of AAs in TMHs: 37.2608899999999\nExp number, first 60 AAs: 19.52923\nTotal prob of N-in: 0.94351\nPOSSIBLE N-term signal sequence\ninside 1 19\nTMhelix 20 39\noutside 40 325\nTMhelix 326 348\ninside 349 372

6125 GCF_900015375.1_ED668_contigs Escherichia coli Proteobacteria; Gammaproteobacteria; Enterobacterales; Enterobacteriaceae; Escherichia
MNNEETFYQAMRRQGVTRRSFLKYCSLAATSLGLGAGMAPKIAWALENKPRIPVVWIHGL WP_001301613.1
hydrogenase [Escherichia coli] Length: 372\nNumber of predicted TMHs: 2\nExp number of AAs in TMHs: 37.46598\nExp number, first 60 AAs: 19.53139\nTotal prob of N-in: 0.94364\nPOSSIBLE N-term signal sequence\ninside 1 19\nTMhelix 20 39\noutside 40 325\nTMhelix 326 348\ninside 349 372

6126 GCF_900015405.1_E._coli_102701 Escherichia coli Proteobacteria; Gammaproteobacteria; Enterobacterales; Enterobacteriaceae; Escherichia
MTGDNTLIHSHGINRRDFMKLCAALATTMGLSSKAAAEMAESVTNPQRPPVIWIGAQECT WP_023149629.1
MULTISPECIES: hydrogenase-2 small chain [Enterobacteriaceae] Length: 372\nNumber of predicted TMHs: 1\nExp number of AAs in TMHs: 23.63603\nExp number, first 60 AAs: 0.12971\nTotal prob of N-in: 0.12608\noutside 1 331\nTMhelix 332 354\ninside 355 372

6127 GCF_900015405.1_E._coli_102701 Escherichia coli Proteobacteria; Gammaproteobacteria; Enterobacterales; Enterobacteriaceae; Escherichia
MNNEETFYQAMRRQGVTRRSFLKYCSLAATSLGLGAGMAPKIAWALENKPRIPVVWIHGL WP_073961310.1
hydrogenase [Escherichia coli] Length: 372\nNumber of predicted TMHs: 2\nExp number of AAs in TMHs: 37.29723\nExp number, first 60 AAs: 19.53583\nTotal prob of N-in: 0.94357\nPOSSIBLE N-term signal sequence\ninside 1 19\nTMhelix 20 39\noutside 40 325\nTMhelix 326 348\ninside 349 372

6128 GCF_900015835.1_ED142_contigs Escherichia coli Proteobacteria; Gammaproteobacteria; Enterobacterales; Enterobacteriaceae; Escherichia
MTGDNTLIHSHGINRRDFMKLCAALAATMGLSSKAAAEMAESVTNPQRPPVIWIGAQECT WP_000145410.1
MULTISPECIES: hydrogenase-2 small chain [Proteobacteria] Length: 372\nNumber of predicted TMHs: 1\nExp number of AAs in TMHs: 23.72226\nExp number, first 60 AAs: 0.22038\nTotal prob of N-in: 0.12994\noutside 1 331\nTMhelix 332 354\ninside 355 372

6129 GCF_900015885.1_ED729_contigs Escherichia coli Proteobacteria; Gammaproteobacteria; Enterobacterales; Enterobacteriaceae; Escherichia
MNNEETFYQAMRRQGVTRRSFLKYCSLAATSLGLGAGMAPKIAWALENKPRIPVVWIHGL WP_001058323.1
MULTISPECIES: hydrogenase-1 small chain [Proteobacteria] Length: 372\nNumber of predicted TMHs: 2\nExp number of AAs in TMHs: 37.2608499999999\nExp number, first 60 AAs: 19.52923\nTotal prob of N-in: 0.94351\nPOSSIBLE N-term signal sequence\ninside 1 19\nTMhelix 20 39\noutside 40 325\nTMhelix 326 348\ninside 349 372

6130 GCF_900015985.1_M856_contigs Escherichia coli Proteobacteria; Gammaproteobacteria; Enterobacterales; Enterobacteriaceae; Escherichia
MNNEETFYQAMRRQGVTRRSFLKYCSLAATSLGLGAGMAPKIAWALENKPRIPVVWIHGL WP_001295940.1
MULTISPECIES: hydrogenase [Enterobacteriaceae] Length: 372\nNumber of predicted TMHs: 2\nExp number of AAs in TMHs: 37.2608899999999\nExp number, first 60 AAs: 19.52923\nTotal prob of N-in: 0.94351\nPOSSIBLE N-term signal sequence\ninside 1 19\nTMhelix 20 39\noutside 40 325\nTMhelix 326 348\ninside 349 372

6131 GCF_900015905.1_ED679_contigs Escherichia coli Proteobacteria; Gammaproteobacteria; Enterobacterales; Enterobacteriaceae; Escherichia
MNNEETFYQAMRRQGVTRRSFLKYCSLAATSLGLGAGMAPKIAWALENKPRIPVVWIHGL WP_001301613.1
hydrogenase [Escherichia coli] Length: 372\nNumber of predicted TMHs: 2\nExp number of AAs in TMHs: 37.46598\nExp number, first 60 AAs: 19.53139\nTotal prob of N-in: 0.94364\nPOSSIBLE N-term signal sequence\ninside 1 19\nTMhelix 20 39\noutside 40 325\nTMhelix 326 348\ninside 349 372

6132 GCF_900015835.1_ED142_contigs Escherichia coli Proteobacteria; Gammaproteobacteria; Enterobacterales; Enterobacteriaceae; Escherichia

MNNEETFYQAMRRQGVTRRSFLKYCSLAATSLGLGAGMAPKIAWALENKPRIPVVWIHGL WP_001058323.1 MULTISPECIES: hydrogenase-1 small chain [Proteobacteria] Length: 372\nNumber of predicted TMHs: 2\nExp number of AAs in TMHs: 37.2608499999999\nExp number, first 60 AAs: 19.52923\nTotal prob of N-in: 0.94351\nPOSSIBLE N-term signal sequence\ninside 1 19\nTMhelix 20 39\noutside 40 325\nTMhelix 326 348\ninside 349 372

6133 GCF_900016005.1_M859_contigs Escherichia coli Proteobacteria; Gammaproteobacteria; Enterobacterales; Enterobacteriaceae; Escherichia MNNEETFYQAMRRQGVTRRSFLKYCSLAATSLGLGAGMAPKIAWALENKPRIPVVWIHGL WP_001058323.1 MULTISPECIES: hydrogenase-1 small chain [Proteobacteria] Length: 372\nNumber of predicted TMHs: 2\nExp number of AAs in TMHs: 37.2608499999999\nExp number, first 60 AAs: 19.52923\nTotal prob of N-in: 0.94351\nPOSSIBLE N-term signal sequence\ninside 1 19\nTMhelix 20 39\noutside 40 325\nTMhelix 326 348\ninside 349 372

6134 GCF_900016005.1_M859_contigs Escherichia coli Proteobacteria; Gammaproteobacteria; Enterobacterales; Enterobacteriaceae; Escherichia MTGDNTLIHSHGINRRDFMKLCAALAATMGLSSKAAAEMAESVTNPQRPPVIWIGAQECT WP_024245965.1 hydrogenase 2 small subunit [Escherichia coli] Length: 372\nNumber of predicted TMHs: 1\nExp number of AAs in TMHs: 24.15412\nExp number, first 60 AAs: 0.21967\nTotal prob of N-in: 0.14289\noutside 1 331\nTMhelix 332 354\ninside 355 372

6135 GCF_900015855.1_ED180_contigs Escherichia coli Proteobacteria; Gammaproteobacteria; Enterobacterales; Enterobacteriaceae; Escherichia MNNEETFYQAMRRQGVTRRSFLKYCSLAATSLGLGAGMAPKIAWALENKPRIPVVWIHGL WP_001058323.1 MULTISPECIES: hydrogenase-1 small chain [Proteobacteria] Length: 372\nNumber of predicted TMHs: 2\nExp number of AAs in TMHs: 37.2608499999999\nExp number, first 60 AAs: 19.52923\nTotal prob of N-in: 0.94351\nPOSSIBLE N-term signal sequence\ninside 1 19\nTMhelix 20 39\noutside 40 325\nTMhelix 326 348\ninside 349 372

6136 GCF_900015865.1_ED411_contigs Escherichia coli Proteobacteria; Gammaproteobacteria; Enterobacterales; Enterobacteriaceae; Escherichia MNNEETFYQAMRRQGVTRRSFLKYCSLAATSLGLGAGMAPKIAWALENKPRIPVVWIHGL WP_001058323.1 MULTISPECIES: hydrogenase-1 small chain [Proteobacteria] Length: 372\nNumber of predicted TMHs: 2\nExp number of AAs in TMHs: 37.2608499999999\nExp number, first 60 AAs: 19.52923\nTotal prob of N-in: 0.94351\nPOSSIBLE N-term signal sequence\ninside 1 19\nTMhelix 20 39\noutside 40 325\nTMhelix 326 348\ninside 349 372

6137 GCF_900016015.1_BCW5746_contigs Escherichia coli Proteobacteria; Gammaproteobacteria; Enterobacterales; Enterobacteriaceae; Escherichia MTGDNTLIHSHGINRRDFMKLCAALAATMGLSSKAAAEMVESVTNPQRPPVIWIGAQECT WP_000145423.1 hydrogenase 2 small subunit [Escherichia coli] Length: 372\nNumber of predicted TMHs: 1\nExp number of AAs in TMHs: 23.78201\nExp number, first 60 AAs: 0.28355\nTotal prob of N-in: 0.13257\noutside 1 331\nTMhelix 332 354\ninside 355 372

6138 GCF_900016035.1_BCW5711_contigs Escherichia coli Proteobacteria; Gammaproteobacteria; Enterobacterales; Enterobacteriaceae; Escherichia MTGDNTLIHSHGINRRDFMKLCAALAATMGLSSKAAAEMVESVTNPQRPPVIWIGAQECT WP_000145423.1 hydrogenase 2 small subunit [Escherichia coli] Length: 372\nNumber of predicted TMHs: 1\nExp number of AAs in TMHs: 23.78201\nExp number, first 60 AAs: 0.28355\nTotal prob of N-in: 0.13257\noutside 1 331\nTMhelix 332 354\ninside 355 372

6139 GCF_900016055.1_BCW5717_contigs Escherichia coli Proteobacteria; Gammaproteobacteria; Enterobacterales; Enterobacteriaceae; Escherichia MTGDNTLIHSHGINRRDFMKLCAALAATMGLSSKAAAEMVESVTNPQRPPVIWIGAQECT WP_000145423.1 hydrogenase 2 small subunit [Escherichia coli] Length: 372\nNumber of predicted TMHs: 1\nExp number of AAs in TMHs: 23.78201\nExp number, first 60 AAs: 0.28355\nTotal prob of N-in: 0.13257\noutside 1 331\nTMhelix 332 354\ninside 355 372

6140 GCF_900016015.1_BCW5746_contigs Escherichia coli Proteobacteria; Gammaproteobacteria; Enterobacterales; Enterobacteriaceae; Escherichia MNNEETFYQAMRRQGVTRRSFLKYCSLAATSLGLGAGMAPKIAWALENKPRIPVVWIHGL WP_001295940.1

MULTISPECIES: hydrogenase [Enterobacteriaceae] Length: 372\nNumber of predicted TMHs: 2\nExp number of AAs in TMHs: 37.2608899999999\nExp number, first 60 AAs: 19.52923\nTotal prob of N-in: 0.94351\nPOSSIBLE N-term signal sequence\ninside 1 19\nTMhelix 20 39\noutside 40 325\nTMhelix 326 348\ninside 349 372

6141 GCF_900016035.1_BCW5711_contigs Escherichia coli Proteobacteria; Gammaproteobacteria; Enterobacterales; Enterobacteriaceae; Escherichia MNNEETFYQAMRRQGVTRRSFLKYCSLAATSLGLGAGMAPKIAWALENKPRIPVVWIHGL WP_001295940.1 MULTISPECIES: hydrogenase [Enterobacteriaceae] Length: 372\nNumber of predicted TMHs: 2\nExp number of AAs in TMHs: 37.2608899999999\nExp number, first 60 AAs: 19.52923\nTotal prob of N-in: 0.94351\nPOSSIBLE N-term signal sequence\ninside 1 19\nTMhelix 20 39\noutside 40 325\nTMhelix 326 348\ninside 349 372

6142 GCF_900041715.1_NG-7574_Bell_ERRESI Escherichia coli Proteobacteria; Gammaproteobacteria; Enterobacterales; Enterobacteriaceae; Escherichia MTGDNTLIHSHGINRRDFMKLCAALAATMGLSSKAAAEMAESVTNPQRPPVIWIGAQECT WP_000145410.1 MULTISPECIES: hydrogenase-2 small chain [Proteobacteria] Length: 372\nNumber of predicted TMHs: 1\nExp number of AAs in TMHs: 23.72226\nExp number, first 60 AAs: 0.22038\nTotal prob of N-in: 0.12994\noutside 1 331\nTMhelix 332 354\ninside 355 372

6143 GCF_900042795.1_F1L3 Escherichia coli Proteobacteria; Gammaproteobacteria; Enterobacterales; Enterobacteriaceae; Escherichia MTGDNTLIHSHGINRRDFMKLCAALAATMGLSSKAAAEMAESVTNPQRPPVIWIGAQECT WP_000145410.1 MULTISPECIES: hydrogenase-2 small chain [Proteobacteria] Length: 372\nNumber of predicted TMHs: 1\nExp number of AAs in TMHs: 23.72226\nExp number, first 60 AAs: 0.22038\nTotal prob of N-in: 0.12994\noutside 1 331\nTMhelix 332 354\ninside 355 372

6144 GCF_900016085.1_ED366_contigs Escherichia coli Proteobacteria; Gammaproteobacteria; Enterobacterales; Enterobacteriaceae; Escherichia MNNEETFYQAMRRQGVTRRSFLKYCSLAATSLGLGAGMAPKIAWALENKPRIPVVWIHGL WP_001295940.1 MULTISPECIES: hydrogenase [Enterobacteriaceae] Length: 372\nNumber of predicted TMHs: 2\nExp number of AAs in TMHs: 37.2608899999999\nExp number, first 60 AAs: 19.52923\nTotal prob of N-in: 0.94351\nPOSSIBLE N-term signal sequence\ninside 1 19\nTMhelix 20 39\noutside 40 325\nTMhelix 326 348\ninside 349 372

6145 GCF_900044045.1_F13P4 Escherichia coli Proteobacteria; Gammaproteobacteria; Enterobacterales; Enterobacteriaceae; Escherichia MTGDNTLIHSHGINRRDFMKLCAALAATMGLSSKAAAEMAESVTNPQRPPVIWIGAQECT WP_000145410.1 MULTISPECIES: hydrogenase-2 small chain [Proteobacteria] Length: 372\nNumber of predicted TMHs: 1\nExp number of AAs in TMHs: 23.72226\nExp number, first 60 AAs: 0.22038\nTotal prob of N-in: 0.12994\noutside 1 331\nTMhelix 332 354\ninside 355 372

6146 GCF_900016065.1_ED363_contigs Escherichia coli Proteobacteria; Gammaproteobacteria; Enterobacterales; Enterobacteriaceae; Escherichia MNNEETFYQAMRRQGVTRRSFLKYCSLAATSLGLGAGMAPKIAWALENKPRIPVVWIHGL WP_001295940.1 MULTISPECIES: hydrogenase [Enterobacteriaceae] Length: 372\nNumber of predicted TMHs: 2\nExp number of AAs in TMHs: 37.2608899999999\nExp number, first 60 AAs: 19.52923\nTotal prob of N-in: 0.94351\nPOSSIBLE N-term signal sequence\ninside 1 19\nTMhelix 20 39\noutside 40 325\nTMhelix 326 348\ninside 349 372

6147 GCF_900016065.1_ED363_contigs Escherichia coli Proteobacteria; Gammaproteobacteria; Enterobacterales; Enterobacteriaceae; Escherichia MTGDNTLIHSHGINRRDFMKLCAALAATMGLSSKAAAEMAESVTNPQRPPVIWIGAQECT WP_074459508.1 hydrogenase 2 small subunit [Escherichia coli] Length: 372\nNumber of predicted TMHs: 1\nExp number of AAs in TMHs: 23.72079\nExp number, first 60 AAs: 0.22038\nTotal prob of N-in: 0.12990\noutside 1 331\nTMhelix 332 354\ninside 355 372

6148 GCF_900044045.1_F13P4 Escherichia coli Proteobacteria; Gammaproteobacteria; Enterobacterales; Enterobacteriaceae; Escherichia MNNEETFYQAMRRQGVTRRSFLKYCSLAATSLGLGAGMAPKIAWALENKPRIPVVWIHGL WP_001058323.1 MULTISPECIES: hydrogenase-1 small chain [Proteobacteria] Length: 372\nNumber of predicted TMHs: 2\nExp number of AAs in TMHs: 37.2608499999999\nExp number, first 60 AAs: 19.52923\nTotal prob of N-in: 0.94351\nPOSSIBLE N-term signal sequence\ninside 1 19\nTMhelix 20 39\noutside 40 325\nTMhelix 326 348\ninside 349 372

6149 GCF_900042775.1_F1B2 Escherichia coli Proteobacteria; Gammaproteobacteria; Enterobacterales; Enterobacteriaceae; Escherichia MNNEETFYQAMRRQGVTRRSFLKYCSLAATSLGLGAGMAPKIAWALENKPRIPVVWIHGL WP_001058323.1 MULTISPECIES: hydrogenase-1 small chain [Proteobacteria] Length: 372\nNumber of predicted TMHs: 2\nExp number of AAs in TMHs: 37.2608499999999\nExp number, first 60 AAs: 19.52923\nTotal prob of N-in: 0.94351\nPOSSIBLE N-term signal sequence\ninside 1 19\nTMhelix 20 39\noutside 40 325\nTMhelix 326 348\ninside 349 372

6150 GCF_900087705.1_RL465 Escherichia coli Proteobacteria; Gammaproteobacteria; Enterobacterales; Enterobacteriaceae; Escherichia MTGDNTLIHSHGINRRDFMKLCAALAATMGLSSKAAAEMAESVTNPQRPPVIWIGAQECT WP_000145410.1 MULTISPECIES: hydrogenase-2 small chain [Proteobacteria] Length: 372\nNumber of predicted TMHs: 1\nExp number of AAs in TMHs: 23.72226\nExp number, first 60 AAs: 0.22038\nTotal prob of N-in: 0.12994\noutside 1 331\nTMhelix 332 354\ninside 355 372

6151 GCF_900087705.1_RL465 Escherichia coli Proteobacteria; Gammaproteobacteria; Enterobacterales; Enterobacteriaceae; Escherichia MNNEETFYQAMRRQGVTRRSFLKYCSLAATSLGLGAGMAPKIAWALENKPRIPVVWIHGL WP_001058320.1 hydrogenase [Escherichia coli] Length: 372\nNumber of predicted TMHs: 2\nExp number of AAs in TMHs: 37.26082\nExp number, first 60 AAs: 19.52923\nTotal prob of N-in: 0.94351\nPOSSIBLE N-term signal sequence\ninside 1 19\nTMhelix 20 39\noutside 40 325\nTMhelix 326 348\ninside 349 372

6152 GCF_900039535.1_F13P5 Escherichia coli Proteobacteria; Gammaproteobacteria; Enterobacterales; Enterobacteriaceae; Escherichia MNNEETFYQAMRRQGVTRRSFLKYCSLAATSLGLGAGMAPKIAWALENKPRIPVVWIHGL WP_001058323.1 MULTISPECIES: hydrogenase-1 small chain [Proteobacteria] Length: 372\nNumber of predicted TMHs: 2\nExp number of AAs in TMHs: 37.2608499999999\nExp number, first 60 AAs: 19.52923\nTotal prob of N-in: 0.94351\nPOSSIBLE N-term signal sequence\ninside 1 19\nTMhelix 20 39\noutside 40 325\nTMhelix 326 348\ninside 349 372

6153 GCF_900088915.1_ASM90008891v1 Escherichia coli Proteobacteria; Gammaproteobacteria; Enterobacterales; Enterobacteriaceae; Escherichia MTGDNTLIHSHGINRRDFMKLCAALAATMGLSSKAAAEMAESVTNPQRPPVIWIGAQECT WP_000145410.1 MULTISPECIES: hydrogenase-2 small chain [Proteobacteria] Length: 372\nNumber of predicted TMHs: 1\nExp number of AAs in TMHs: 23.72226\nExp number, first 60 AAs: 0.22038\nTotal prob of N-in: 0.12994\noutside 1 331\nTMhelix 332 354\ninside 355 372

6154 GCF_900087775.1_RL145 Escherichia coli Proteobacteria; Gammaproteobacteria; Enterobacterales; Enterobacteriaceae; Escherichia MTGDNTLIHSHGINRRDFMKLCAALAATMGLSSKAAAEMAESVTNPQRPPVIWIGAQECT WP_000145410.1 MULTISPECIES: hydrogenase-2 small chain [Proteobacteria] Length: 372\nNumber of predicted TMHs: 1\nExp number of AAs in TMHs: 23.72226\nExp number, first 60 AAs: 0.22038\nTotal prob of N-in: 0.12994\noutside 1 331\nTMhelix 332 354\ninside 355 372

6155 GCF_900088915.1_ASM90008891v1 Escherichia coli Proteobacteria; Gammaproteobacteria; Enterobacterales; Enterobacteriaceae; Escherichia MNNEETFYQAMRRQGVTRRSFLKYCSLAATSLGLGAGMAPKIAWALENKPRIPVVWIHGL WP_001058323.1 MULTISPECIES: hydrogenase-1 small chain [Proteobacteria] Length: 372\nNumber of predicted TMHs: 2\nExp number of AAs in TMHs: 37.2608499999999\nExp number, first 60 AAs: 19.52923\nTotal prob of N-in: 0.94351\nPOSSIBLE N-term signal sequence\ninside 1 19\nTMhelix 20 39\noutside 40 325\nTMhelix 326 348\ninside 349 372

6156 GCF_900087775.1_RL145 Escherichia coli Proteobacteria; Gammaproteobacteria; Enterobacterales; Enterobacteriaceae; Escherichia MNNEETFYQAMRRQGVTRRSFLKYCSLAATSLGLGAGMAPKIAWALENKPRIPVVWIHGL WP_001058320.1 hydrogenase [Escherichia coli] Length: 372\nNumber of predicted TMHs: 2\nExp number of AAs in TMHs: 37.26082\nExp number, first 60 AAs: 19.52923\nTotal prob of N-in: 0.94351\nPOSSIBLE N-term signal sequence\ninside 1 19\nTMhelix 20 39\noutside 40 325\nTMhelix 326 348\ninside 349 372

6157 GCF_900042795.1_F1L3 Escherichia coli Proteobacteria; Gammaproteobacteria; Enterobacterales; Enterobacteriaceae; Escherichia MNNEETFYQAMRRQGVTRRSFLKYCSLAATSLGLGAGMAPKIAWALENKPRIPVVWIHGL WP_001058320.1 hydrogenase [Escherichia coli] Length: 372\nNumber of predicted TMHs: 2\nExp number of AAs in TMHs: 37.26082\nExp number, first 60 AAs: 19.52923\nTotal prob of N-in: 0.94351\nPOSSIBLE N-term signal sequence\ninside 1 19\nTMhelix 20 39\noutside 40 325\nTMhelix 326 348\ninside 349 372

6158 GCF_900016115.1_ED430_contigs Escherichia coli Proteobacteria; Gammaproteobacteria; Enterobacterales; Enterobacteriaceae; Escherichia

MNNEETFYQAMRRQGVTRRSFLKYCSLAATSLGLGAGMAPKIAWALENKPRIPVVWIHGL WP_001295940.1 MULTISPECIES: hydrogenase [Enterobacteriaceae] Length: 372\nNumber of predicted TMHs: 2\nExp number of AAs in TMHs: 37.2608899999999\nExp number, first 60 AAs: 19.52923\nTotal prob of N-in: 0.94351\nPOSSIBLE N-term signal sequence\ninside 1 19\nTMhelix 20 39\noutside 40 325\nTMhelix 326 348\ninside 349 372

6159 GCF_900016135.1_ED444_contigs Escherichia coli Proteobacteria; Gammaproteobacteria; Enterobacterales; Enterobacteriaceae; Escherichia MNNEETFYQAMRRQGVTRRSFLKYCSLAATSLGLGAGMAPKIAWALENKPRIPVVWIHGL WP_001295940.1 MULTISPECIES: hydrogenase [Enterobacteriaceae] Length: 372\nNumber of predicted TMHs: 2\nExp number of AAs in TMHs: 37.2608899999999\nExp number, first 60 AAs: 19.52923\nTotal prob of N-in: 0.94351\nPOSSIBLE N-term signal sequence\ninside 1 19\nTMhelix 20 39\noutside 40 325\nTMhelix 326 348\ninside 349 372

6160 GCF_900041715.1_NG-7574_Bell_ERRESI Escherichia coli Proteobacteria; Gammaproteobacteria; Enterobacterales; Enterobacteriaceae; Escherichia MNNEETFYQAMRRQGVTRRSFLKYCSLAATSLGLGAGMAPKIAWALENKPRIPVVWIHGL WP_001295940.1 MULTISPECIES: hydrogenase [Enterobacteriaceae] Length: 372\nNumber of predicted TMHs: 2\nExp number of AAs in TMHs: 37.2608899999999\nExp number, first 60 AAs: 19.52923\nTotal prob of N-in: 0.94351\nPOSSIBLE N-term signal sequence\ninside 1 19\nTMhelix 20 39\noutside 40 325\nTMhelix 326 348\ninside 349 372

6161 GCF_900088965.1_ASM90008896v1 Escherichia coli Proteobacteria; Gammaproteobacteria; Enterobacterales; Enterobacteriaceae; Escherichia MTGDNTLIHSHGINRRDFMKLCAALAATMGLSSKAAAEMAESVTNPQRPPVIWIGAQECT WP_000145410.1 MULTISPECIES: hydrogenase-2 small chain [Proteobacteria] Length: 372\nNumber of predicted TMHs: 1\nExp number of AAs in TMHs: 23.72226\nExp number, first 60 AAs: 0.22038\nTotal prob of N-in: 0.12994\noutside 1 331\nTMhelix 332 354\ninside 355 372

6162 GCF_900089795.1_CQ09 Escherichia coli Proteobacteria; Gammaproteobacteria; Enterobacterales; Enterobacteriaceae; Escherichia MNNEETFYQAMRRQGVTRRSFLKYCSLAATSLGLGAGMAPKIAWALENKPRIPVVWIHGL WP_001058320.1 hydrogenase [Escherichia coli] Length: 372\nNumber of predicted TMHs: 2\nExp number of AAs in TMHs: 37.26082\nExp number, first 60 AAs: 19.52923\nTotal prob of N-in: 0.94351\nPOSSIBLE N-term signal sequence\ninside 1 19\nTMhelix 20 39\noutside 40 325\nTMhelix 326 348\ninside 349 372

6163 GCF_900093885.1_scaffolds_20151359 Escherichia coli Proteobacteria; Gammaproteobacteria; Enterobacterales; Enterobacteriaceae; Escherichia MTGDNTLIHSHGINRRDFMKLCAALAATMGLSSKAAAEMAESVTNPQRPPVIWIGAQECT WP_000145410.1 MULTISPECIES: hydrogenase-2 small chain [Proteobacteria] Length: 372\nNumber of predicted TMHs: 1\nExp number of AAs in TMHs: 23.72226\nExp number, first 60 AAs: 0.22038\nTotal prob of N-in: 0.12994\noutside 1 331\nTMhelix 332 354\ninside 355 372

6164 GCF_900093905.1_scaffolds_C322 Escherichia coli Proteobacteria; Gammaproteobacteria; Enterobacterales; Enterobacteriaceae; Escherichia MTGDNTLIHSHGINRRDFMKLCAALAATMGLSSKAAAEMAESVTNPQRPPVIWIGAQECT WP_000145410.1 MULTISPECIES: hydrogenase-2 small chain [Proteobacteria] Length: 372\nNumber of predicted TMHs: 1\nExp number of AAs in TMHs: 23.72226\nExp number, first 60 AAs: 0.22038\nTotal prob of N-in: 0.12994\noutside 1 331\nTMhelix 332 354\ninside 355 372

6165 GCF_900093925.1_scaffolds_20151473 Escherichia coli Proteobacteria; Gammaproteobacteria; Enterobacterales; Enterobacteriaceae; Escherichia MTGDNTLIHSHGINRRDFMKLCAALAATMGLSSKAAAEMAESVTNPQRPPVIWIGAQECT WP_000145410.1 MULTISPECIES: hydrogenase-2 small chain [Proteobacteria] Length: 372\nNumber of predicted TMHs: 1\nExp number of AAs in TMHs: 23.72226\nExp number, first 60 AAs: 0.22038\nTotal prob of N-in: 0.12994\noutside 1 331\nTMhelix 332 354\ninside 355 372

6166 GCF_900093905.1_scaffolds_C322 Escherichia coli Proteobacteria; Gammaproteobacteria; Enterobacterales; Enterobacteriaceae; Escherichia MNNEETFYQAMRRQGVTRRSFLKYCSLAATSLGLGAGMAPKIAWALENKPRIPVVWIHGL WP_001058323.1 MULTISPECIES: hydrogenase-1 small chain [Proteobacteria] Length: 372\nNumber of predicted TMHs: 2\nExp number of AAs in TMHs: 37.2608499999999\nExp number, first 60 AAs: 19.52923\nTotal prob of N-in:

0.94351\nPOSSIBLE N-term signal sequence\ninside 1 19\nTMhelix 20 39\noutside 40 325\nTMhelix 326 348\ninside 349 372

6167 GCF_900093885.1_scaffolds_20151359 Escherichia coli Proteobacteria; Gammaproteobacteria; Enterobacterales; Enterobacteriaceae; Escherichia
MNNEETFYQAMRRQGVTRRSFLKYCSLAATSLGLGAGMAPKIAWALENKPRIPVVWIHGL WP_001295940.1
MULTISPECIES: hydrogenase [Enterobacteriaceae] Length: 372\nNumber of predicted TMHs: 2\nExp number of AAs in TMHs: 37.2608899999999\nExp number, first 60 AAs: 19.52923\nTotal prob of N-in: 0.94351\nPOSSIBLE N-term signal sequence\ninside 1 19\nTMhelix 20 39\noutside 40 325\nTMhelix 326 348\ninside 349 372

6168 GCF_900093925.1_scaffolds_20151473 Escherichia coli Proteobacteria; Gammaproteobacteria; Enterobacterales; Enterobacteriaceae; Escherichia
MNNEETFYQAMRRQGVTRRSFLKYCSLAATSLGLGAGMAPKIAWALENKPRIPVVWIHGL WP_001295940.1
MULTISPECIES: hydrogenase [Enterobacteriaceae] Length: 372\nNumber of predicted TMHs: 2\nExp number of AAs in TMHs: 37.2608899999999\nExp number, first 60 AAs: 19.52923\nTotal prob of N-in: 0.94351\nPOSSIBLE N-term signal sequence\ninside 1 19\nTMhelix 20 39\noutside 40 325\nTMhelix 326 348\ninside 349 372

6169 GCF_900093875.1_scaffolds_C333 Escherichia coli Proteobacteria; Gammaproteobacteria; Enterobacterales; Enterobacteriaceae; Escherichia
MNNEETFYQAMRRQGVTRRSFLKYCSLAATSLGLGAGMAPKIAWALENKPRIPVVWIHGL WP_001058323.1
MULTISPECIES: hydrogenase-1 small chain [Proteobacteria] Length: 372\nNumber of predicted TMHs: 2\nExp number of AAs in TMHs: 37.2608499999999\nExp number, first 60 AAs: 19.52923\nTotal prob of N-in: 0.94351\nPOSSIBLE N-term signal sequence\ninside 1 19\nTMhelix 20 39\noutside 40 325\nTMhelix 326 348\ninside 349 372

6170 GCF_900093825.1_scaffolds_20160015 Escherichia coli Proteobacteria; Gammaproteobacteria; Enterobacterales; Enterobacteriaceae; Escherichia
MNNEETFYQAMRRQGVTRRSFLKYCSLAATSLGLGAGMAPKIAWALENKPRIPVVWIHGL WP_001058323.1
MULTISPECIES: hydrogenase-1 small chain [Proteobacteria] Length: 372\nNumber of predicted TMHs: 2\nExp number of AAs in TMHs: 37.2608499999999\nExp number, first 60 AAs: 19.52923\nTotal prob of N-in: 0.94351\nPOSSIBLE N-term signal sequence\ninside 1 19\nTMhelix 20 39\noutside 40 325\nTMhelix 326 348\ninside 349 372

6171 GCF_900093965.1_scaffolds_20160014 Escherichia coli Proteobacteria; Gammaproteobacteria; Enterobacterales; Enterobacteriaceae; Escherichia
MTGDNTLIHSHGINRRDFMKLCAALAATMGLSSKAAAEMAESVTNPQRPPVIWIGAQECT WP_000145410.1
MULTISPECIES: hydrogenase-2 small chain [Proteobacteria] Length: 372\nNumber of predicted TMHs: 1\nExp number of AAs in TMHs: 23.72226\nExp number, first 60 AAs: 0.22038\nTotal prob of N-in: 0.12994\noutside 1 331\nTMhelix 332 354\ninside 355 372

6172 GCF_900093955.1_scaffolds_20160016 Escherichia coli Proteobacteria; Gammaproteobacteria; Enterobacterales; Enterobacteriaceae; Escherichia
MTGDNTLIHSHGINRRDFMKLCAALAATMGLSSKAAAEMAESVTNPQRPPVIWIGAQECT WP_000145410.1
MULTISPECIES: hydrogenase-2 small chain [Proteobacteria] Length: 372\nNumber of predicted TMHs: 1\nExp number of AAs in TMHs: 23.72226\nExp number, first 60 AAs: 0.22038\nTotal prob of N-in: 0.12994\noutside 1 331\nTMhelix 332 354\ninside 355 372

6173 GCF_900093965.1_scaffolds_20160014 Escherichia coli Proteobacteria; Gammaproteobacteria; Enterobacterales; Enterobacteriaceae; Escherichia
MNNEETFYQAMRRQGVTRRSFLKYCSLAATSLGLGAGMAPKIAWALENKPRIPVVWIHGL WP_001058323.1
MULTISPECIES: hydrogenase-1 small chain [Proteobacteria] Length: 372\nNumber of predicted TMHs: 2\nExp number of AAs in TMHs: 37.2608499999999\nExp number, first 60 AAs: 19.52923\nTotal prob of N-in: 0.94351\nPOSSIBLE N-term signal sequence\ninside 1 19\nTMhelix 20 39\noutside 40 325\nTMhelix 326 348\ninside 349 372

6174 GCF_900093955.1_scaffolds_20160016 Escherichia coli Proteobacteria; Gammaproteobacteria; Enterobacterales; Enterobacteriaceae; Escherichia
MNNEETFYQAMRRQGVTRRSFLKYCSLAATSLGLGAGMAPKIAWALENKPRIPVVWIHGL WP_001295940.1
MULTISPECIES: hydrogenase [Enterobacteriaceae] Length: 372\nNumber of predicted TMHs: 2\nExp

number of AAs in TMHs: 37.2608899999999\nExp number, first 60 AAs: 19.52923\nTotal prob of N-in: 0.94351\nPOSSIBLE N-term signal sequence\ninside 1 19\nTMhelix 20 39\noutside 40 325\nTMhelix 326 348\ninside 349 372

6175 GCF_900093855.1_scaffolds_20160007 Escherichia coli Proteobacteria; Gammaproteobacteria; Enterobacterales; Enterobacteriaceae; Escherichia MNNEETFYQAMRRQGVTRRSFLKYCSLAATSLGLGAGMAPKIAWALENKPRIPVVWIHGL WP_001058323.1 MULTISPECIES: hydrogenase-1 small chain [Proteobacteria] Length: 372\nNumber of predicted TMHs: 2\nExp number of AAs in TMHs: 37.2608499999999\nExp number, first 60 AAs: 19.52923\nTotal prob of N-in: 0.94351\nPOSSIBLE N-term signal sequence\ninside 1 19\nTMhelix 20 39\noutside 40 325\nTMhelix 326 348\ninside 349 372

6176 GCF_900096835.1_Ecoli_AG100_Sample3_Wildtype_Assembly Escherichia coli Proteobacteria; Gammaproteobacteria; Enterobacterales; Enterobacteriaceae; Escherichia MTGDNTLIHSHGINRRDFMKLCAALAATMGLSSKAAAEMAESVTNPQRPPVIWIGAQECT WP_000145410.1 MULTISPECIES: hydrogenase-2 small chain [Proteobacteria] Length: 372\nNumber of predicted TMHs: 1\nExp number of AAs in TMHs: 23.72226\nExp number, first 60 AAs: 0.22038\nTotal prob of N-in: 0.12994\noutside 1 331\nTMhelix 332 354\ninside 355 372

6177 GCF_900096815.1_Ecoli_AG100_Sample2_M9_Assembly Escherichia coli Proteobacteria; Gammaproteobacteria; Enterobacterales; Enterobacteriaceae; Escherichia MNNEETFYQAMRRQGVTRRSFLKYCSLAATSLGLGAGMAPKIAWALENKPRIPVVWIHGL WP_001058323.1 MULTISPECIES: hydrogenase-1 small chain [Proteobacteria] Length: 372\nNumber of predicted TMHs: 2\nExp number of AAs in TMHs: 37.2608499999999\nExp number, first 60 AAs: 19.52923\nTotal prob of N-in: 0.94351\nPOSSIBLE N-term signal sequence\ninside 1 19\nTMhelix 20 39\noutside 40 325\nTMhelix 326 348\ninside 349 372

6178 GCF_900096835.1_Ecoli_AG100_Sample3_Wildtype_Assembly Escherichia coli Proteobacteria; Gammaproteobacteria; Enterobacterales; Enterobacteriaceae; Escherichia MNNEETFYQAMRRQGVTRRSFLKYCSLAATSLGLGAGMAPKIAWALENKPRIPVVWIHGL WP_001058323.1 MULTISPECIES: hydrogenase-1 small chain [Proteobacteria] Length: 372\nNumber of predicted TMHs: 2\nExp number of AAs in TMHs: 37.2608499999999\nExp number, first 60 AAs: 19.52923\nTotal prob of N-in: 0.94351\nPOSSIBLE N-term signal sequence\ninside 1 19\nTMhelix 20 39\noutside 40 325\nTMhelix 326 348\ninside 349 372

6179 GCF_900094015.1_scaffolds_20151012 Escherichia coli Proteobacteria; Gammaproteobacteria; Enterobacterales; Enterobacteriaceae; Escherichia MNNEETFYQAMRRQGVTRRSFLKYCSLAATSLGLGAGMAPKIAWALENKPRIPVVWIHGL WP_001058323.1 MULTISPECIES: hydrogenase-1 small chain [Proteobacteria] Length: 372\nNumber of predicted TMHs: 2\nExp number of AAs in TMHs: 37.2608499999999\nExp number, first 60 AAs: 19.52923\nTotal prob of N-in: 0.94351\nPOSSIBLE N-term signal sequence\ninside 1 19\nTMhelix 20 39\noutside 40 325\nTMhelix 326 348\ninside 349 372

6180 GCF_900096855.1_Ecoli_AG100_Sample2_Wildtype_Assembly Escherichia coli Proteobacteria; Gammaproteobacteria; Enterobacterales; Enterobacteriaceae; Escherichia MTGDNTLIHSHGINRRDFMKLCAALAATMGLSSKAAAEMAESVTNPQRPPVIWIGAQECT WP_000145410.1 MULTISPECIES: hydrogenase-2 small chain [Proteobacteria] Length: 372\nNumber of predicted TMHs: 1\nExp number of AAs in TMHs: 23.72226\nExp number, first 60 AAs: 0.22038\nTotal prob of N-in: 0.12994\noutside 1 331\nTMhelix 332 354\ninside 355 372

6181 GCF_900096865.1_Ecoli_AG100_Sample1_Doxycycline_Assembly Escherichia coli Proteobacteria; Gammaproteobacteria; Enterobacterales; Enterobacteriaceae; Escherichia MTGDNTLIHSHGINRRDFMKLCAALAATMGLSSKAAAEMAESVTNPQRPPVIWIGAQECT WP_000145410.1 MULTISPECIES: hydrogenase-2 small chain [Proteobacteria] Length: 372\nNumber of predicted TMHs: 1\nExp number of AAs in TMHs: 23.72226\nExp number, first 60 AAs: 0.22038\nTotal prob of N-in: 0.12994\noutside 1 331\nTMhelix 332 354\ninside 355 372

6182 GCF_900096855.1_Ecoli_AG100_Sample2_Wildtype_Assembly Escherichia coli Proteobacteria; Gammaproteobacteria; Enterobacterales; Enterobacteriaceae; Escherichia MNNEETFYQAMRRQGVTRRSFLKYCSLAATSLGLGAGMAPKIAWALENKPRIPVVWIHGL WP_001058323.1 MULTISPECIES: hydrogenase-1 small chain [Proteobacteria] Length: 372\nNumber of predicted TMHs:

2\nExp number of AAs in TMHs: 37.2608499999999\nExp number, first 60 AAs: 19.52923\nTotal prob of N-in: 0.94351\nPOSSIBLE N-term signal sequence\ninside 1 19\nTMhelix 20 39\noutside 40 325\nTMhelix 326 348\ninside 349 372

6183 GCF_900096865.1_Ecoli_AG100_Sample1_Doxycycline_Assembly Escherichia coli Proteobacteria; Gammaproteobacteria; Enterobacterales; Enterobacteriaceae; Escherichia MNNEETFYQAMRRQGVTRRSFLKYCSLAATSLGLGAGMAPKIAWALENKPRIPVVWIHGL WP_001058323.1 MULTISPECIES: hydrogenase-1 small chain [Proteobacteria] Length: 372\nNumber of predicted TMHs: 2\nExp number of AAs in TMHs: 37.2608499999999\nExp number, first 60 AAs: 19.52923\nTotal prob of N-in: 0.94351\nPOSSIBLE N-term signal sequence\ninside 1 19\nTMhelix 20 39\noutside 40 325\nTMhelix 326 348\ninside 349 372

6184 GCF_900094005.1_scaffolds_20160006 Escherichia coli Proteobacteria; Gammaproteobacteria; Enterobacterales; Enterobacteriaceae; Escherichia MNNEETFYQAMRRQGVTRRSFLKYCSLAATSLGLGAGMAPKIAWALENKPRIPVVWIHGL WP_001295940.1 MULTISPECIES: hydrogenase [Enterobacteriaceae] Length: 372\nNumber of predicted TMHs: 2\nExp number of AAs in TMHs: 37.2608899999999\nExp number, first 60 AAs: 19.52923\nTotal prob of N-in: 0.94351\nPOSSIBLE N-term signal sequence\ninside 1 19\nTMhelix 20 39\noutside 40 325\nTMhelix 326 348\ninside 349 372

6185 GCF_900096785.1_Ecoli_AG100_Sample2_Doxycycline_Assembly Escherichia coli Proteobacteria; Gammaproteobacteria; Enterobacterales; Enterobacteriaceae; Escherichia MNNEETFYQAMRRQGVTRRSFLKYCSLAATSLGLGAGMAPKIAWALENKPRIPVVWIHGL WP_001058323.1 MULTISPECIES: hydrogenase-1 small chain [Proteobacteria] Length: 372\nNumber of predicted TMHs: 2\nExp number of AAs in TMHs: 37.2608499999999\nExp number, first 60 AAs: 19.52923\nTotal prob of N-in: 0.94351\nPOSSIBLE N-term signal sequence\ninside 1 19\nTMhelix 20 39\noutside 40 325\nTMhelix 326 348\ninside 349 372

6186 GCF_900110665.1_IMG-taxon_2608642106_annotated_assembly Escherichia coli Proteobacteria; Gammaproteobacteria; Enterobacterales; Enterobacteriaceae; Escherichia MTGDNTLIHSHGINRRDFMKLCAALAATMGLSSKAAAEMAESVTNPQRPPVIWIGAQECT WP_000145410.1 MULTISPECIES: hydrogenase-2 small chain [Proteobacteria] Length: 372\nNumber of predicted TMHs: 1\nExp number of AAs in TMHs: 23.72226\nExp number, first 60 AAs: 0.22038\nTotal prob of N-in: 0.12994\noutside 1 331\nTMhelix 332 354\ninside 355 372

6187 GCF_900128765.1_ASM90012876v1 Escherichia coli Proteobacteria; Gammaproteobacteria; Enterobacterales; Enterobacteriaceae; Escherichia MTGDNTLIHSHGINRRDFMKLCAALAATMGLSSKAAAEMAESVTNPQRPPVIWIGAQECT WP_000145410.1 MULTISPECIES: hydrogenase-2 small chain [Proteobacteria] Length: 372\nNumber of predicted TMHs: 1\nExp number of AAs in TMHs: 23.72226\nExp number, first 60 AAs: 0.22038\nTotal prob of N-in: 0.12994\noutside 1 331\nTMhelix 332 354\ninside 355 372

6188 GCF_900166685.1_Isolate_2 Escherichia coli Proteobacteria; Gammaproteobacteria; Enterobacterales; Enterobacteriaceae; Escherichia MTGDNTLIHSHGINRRDFMKLCAALAATMGLSSKAAAEMVESVTNPQRPPVIWIGAQECT WP_000145423.1 hydrogenase 2 small subunit [Escherichia coli] Length: 372\nNumber of predicted TMHs: 1\nExp number of AAs in TMHs: 23.78201\nExp number, first 60 AAs: 0.28355\nTotal prob of N-in: 0.13257\noutside 1 331\nTMhelix 332 354\ninside 355 372

6189 GCF_900110665.1_IMG-taxon_2608642106_annotated_assembly Escherichia coli Proteobacteria; Gammaproteobacteria; Enterobacterales; Enterobacteriaceae; Escherichia MNNEETFYQAMRRQGVTRRSFLKYCSLAATSLGLGAGMAPKIAWALENKPRIPVVWIHGL WP_001058323.1 MULTISPECIES: hydrogenase-1 small chain [Proteobacteria] Length: 372\nNumber of predicted TMHs: 2\nExp number of AAs in TMHs: 37.2608499999999\nExp number, first 60 AAs: 19.52923\nTotal prob of N-in: 0.94351\nPOSSIBLE N-term signal sequence\ninside 1 19\nTMhelix 20 39\noutside 40 325\nTMhelix 326 348\ninside 349 372

6190 GCF_900166785.1_Isolate_4 Escherichia coli Proteobacteria; Gammaproteobacteria; Enterobacterales; Enterobacteriaceae; Escherichia MTGDNTLIHSHGINRRDFMKLCAALAATMGLSSKAAAEMVESVTNPQRPPVIWIGAQECT WP_000145423.1 hydrogenase 2 small subunit [Escherichia coli] Length: 372\nNumber of predicted TMHs: 1\nExp number of AAs in TMHs: 23.78201\nExp number, first 60 AAs: 0.28355\nTotal prob of N-in: 0.13257\noutside 1 331\nTMhelix 332 354\ninside 355 372

6191 GCF_900166805.1_Isolate_7 Escherichia coli Proteobacteria; Gammaproteobacteria; Enterobacterales; Enterobacteriaceae; Escherichia MTGDNTLIHSHGINRRDFMKLCAALAATMGLSSKAAAEMVESVTNPQRPPVIWIGAQECT WP_000145423.1 hydrogenase 2 small subunit [Escherichia coli] Length: 372\nNumber of predicted TMHs: 1\nExp number of AAs in TMHs: 23.78201\nExp number, first 60 AAs: 0.28355\nTotal prob of N-in: 0.13257\noutside 1 331\nTMhelix 332 354\ninside 355 372

6192 GCF_900166755.1_Isolate_17 Escherichia coli Proteobacteria; Gammaproteobacteria; Enterobacterales; Enterobacteriaceae; Escherichia MNNEETFYQAMRRQGVTRRSFLKYCSLAATSLGLGAGMAPKIAWALENKPRIPVVWIHGL WP_001058323.1 MULTISPECIES: hydrogenase-1 small chain [Proteobacteria] Length: 372\nNumber of predicted TMHs: 2\nExp number of AAs in TMHs: 37.2608499999999\nExp number, first 60 AAs: 19.52923\nTotal prob of N-in: 0.94351\nPOSSIBLE N-term signal sequence\ninside 1 19\nTMhelix 20 39\noutside 40 325\nTMhelix 326 348\ninside 349 372

6193 GCF_900166815.1_Isolate_1 Escherichia coli Proteobacteria; Gammaproteobacteria; Enterobacterales; Enterobacteriaceae; Escherichia MTGDNTLIHSHGINRRDFMKLCAALAATMGLSSKAAAEMVESVTNPQRPPVIWIGAQECT WP_000145423.1 hydrogenase 2 small subunit [Escherichia coli] Length: 372\nNumber of predicted TMHs: 1\nExp number of AAs in TMHs: 23.78201\nExp number, first 60 AAs: 0.28355\nTotal prob of N-in: 0.13257\noutside 1 331\nTMhelix 332 354\ninside 355 372

6194 GCF_900166835.1_Isolate_6 Escherichia coli Proteobacteria; Gammaproteobacteria; Enterobacterales; Enterobacteriaceae; Escherichia MTGDNTLIHSHGINRRDFMKLCAALAATMGLSSKAAAEMVESVTNPQRPPVIWIGAQECT WP_000145423.1 hydrogenase 2 small subunit [Escherichia coli] Length: 372\nNumber of predicted TMHs: 1\nExp number of AAs in TMHs: 23.78201\nExp number, first 60 AAs: 0.28355\nTotal prob of N-in: 0.13257\noutside 1 331\nTMhelix 332 354\ninside 355 372

6195 GCF_900174625.1_WI1-reUpload Escherichia coli Proteobacteria; Gammaproteobacteria; Enterobacterales; Enterobacteriaceae; Escherichia MTGDNTLIHSHGINRRDFMKLCAALAATMGLSSKAAAEMAESVTNPQRPPVIWIGAQECT WP_000145410.1 MULTISPECIES: hydrogenase-2 small chain [Proteobacteria] Length: 372\nNumber of predicted TMHs: 1\nExp number of AAs in TMHs: 23.72226\nExp number, first 60 AAs: 0.22038\nTotal prob of N-in: 0.12994\noutside 1 331\nTMhelix 332 354\ninside 355 372

6196 GCF_900166855.1_Isolate_5 Escherichia coli Proteobacteria; Gammaproteobacteria; Enterobacterales; Enterobacteriaceae; Escherichia MTGDNTLIHSHGINRRDFMKLCAALAATMGLSSKAAAEMVESVTNPQRPPVIWIGAQECT WP_000145423.1 hydrogenase 2 small subunit [Escherichia coli] Length: 372\nNumber of predicted TMHs: 1\nExp number of AAs in TMHs: 23.78201\nExp number, first 60 AAs: 0.28355\nTotal prob of N-in: 0.13257\noutside 1 331\nTMhelix 332 354\ninside 355 372

6197 GCF_900174625.1_WI1-reUpload Escherichia coli Proteobacteria; Gammaproteobacteria; Enterobacterales; Enterobacteriaceae; Escherichia MNNEETFYQAMRRQGVTRRSFLKYCSLAATSLGLGAGMAPKIAWALENKPRIPVVWIHGL WP_001058323.1 MULTISPECIES: hydrogenase-1 small chain [Proteobacteria] Length: 372\nNumber of predicted TMHs: 2\nExp number of AAs in TMHs: 37.2608499999999\nExp number, first 60 AAs: 19.52923\nTotal prob of N-in: 0.94351\nPOSSIBLE N-term signal sequence\ninside 1 19\nTMhelix 20 39\noutside 40 325\nTMhelix 326 348\ninside 349 372

6198 GCF_900166815.1_Isolate_1 Escherichia coli Proteobacteria; Gammaproteobacteria; Enterobacterales; Enterobacteriaceae; Escherichia MNNEETFYQAMRRQGVTRRSFLKYCSLAATSLGLGAGMAPKIAWALENKPRIPVVWIHGL WP_001295940.1 MULTISPECIES: hydrogenase [Enterobacteriaceae] Length: 372\nNumber of predicted TMHs: 2\nExp number of AAs in TMHs: 37.2608899999999\nExp number, first 60 AAs: 19.52923\nTotal prob of N-in: 0.94351\nPOSSIBLE N-term signal sequence\ninside 1 19\nTMhelix 20 39\noutside 40 325\nTMhelix 326 348\ninside 349 372

6199 GCF_900166695.1_Isolate_3 Escherichia coli Proteobacteria; Gammaproteobacteria; Enterobacterales; Enterobacteriaceae; Escherichia MNNEETFYQAMRRQGVTRRSFLKYCSLAATSLGLGAGMAPKIAWALENKPRIPVVWIHGL WP_001295940.1 MULTISPECIES: hydrogenase [Enterobacteriaceae] Length: 372\nNumber of predicted TMHs: 2\nExp number of AAs in TMHs: 37.2608899999999\nExp number, first 60 AAs: 19.52923\nTotal prob of N-in: 0.94351\nPOSSIBLE N-term signal sequence\ninside 1 19\nTMhelix 20 39\noutside 40 325\nTMhelix 326 348\ninside 349 372

6200 GCF_900166715.1_Isolate_16 Escherichia coli Proteobacteria; Gammaproteobacteria; Enterobacterales; Enterobacteriaceae; Escherichia MNNEETFYQAMRRQGVTRRSFLKYCSLAATSLGLGAGMAPKIAWALENKPRIPVVWIHGL WP_001295940.1 MULTISPECIES: hydrogenase [Enterobacteriaceae] Length: 372\nNumber of predicted TMHs: 2\nExp number of AAs in TMHs: 37.2608899999999\nExp number, first 60 AAs: 19.52923\nTotal prob of N-in: 0.94351\nPOSSIBLE N-term signal sequence\ninside 1 19\nTMhelix 20 39\noutside 40 325\nTMhelix 326 348\ninside 349 372

6201 GCF_900166835.1_Isolate_6 Escherichia coli Proteobacteria; Gammaproteobacteria; Enterobacterales; Enterobacteriaceae; Escherichia MNNEETFYQAMRRQGVTRRSFLKYCSLAATSLGLGAGMAPKIAWALENKPRIPVVWIHGL WP_001295940.1 MULTISPECIES: hydrogenase [Enterobacteriaceae] Length: 372\nNumber of predicted TMHs: 2\nExp number of AAs in TMHs: 37.2608899999999\nExp number, first 60 AAs: 19.52923\nTotal prob of N-in: 0.94351\nPOSSIBLE N-term signal sequence\ninside 1 19\nTMhelix 20 39\noutside 40 325\nTMhelix 326 348\ninside 349 372

6202 GCF_900166855.1_Isolate_5 Escherichia coli Proteobacteria; Gammaproteobacteria; Enterobacterales; Enterobacteriaceae; Escherichia MNNEETFYQAMRRQGVTRRSFLKYCSLAATSLGLGAGMAPKIAWALENKPRIPVVWIHGL WP_001295940.1 MULTISPECIES: hydrogenase [Enterobacteriaceae] Length: 372\nNumber of predicted TMHs: 2\nExp number of AAs in TMHs: 37.2608899999999\nExp number, first 60 AAs: 19.52923\nTotal prob of N-in: 0.94351\nPOSSIBLE N-term signal sequence\ninside 1 19\nTMhelix 20 39\noutside 40 325\nTMhelix 326 348\ninside 349 372

6203 GCF_000599825.1_a5_miseq_linux_20140113 Escherichia coli Proteobacteria; Gammaproteobacteria; Enterobacterales; Enterobacteriaceae; Escherichia MNNEETFYQAMRRQGVTRRSFLKYCSLAATSLGLGAGMAPKIAWALENKPRIPVVWIHGL WP_001058323.1 MULTISPECIES: hydrogenase-1 small chain [Proteobacteria] Length: 372\nNumber of predicted TMHs: 2\nExp number of AAs in TMHs: 37.2608499999999\nExp number, first 60 AAs: 19.52923\nTotal prob of N-in: 0.94351\nPOSSIBLE N-term signal sequence\ninside 1 19\nTMhelix 20 39\noutside 40 325\nTMhelix 326 348\ninside 349 372

6204 GCF_000599785.2_a5_miseq_linux_20140113 Escherichia coli Proteobacteria; Gammaproteobacteria; Enterobacterales; Enterobacteriaceae; Escherichia MNNEETFYQAMRRQGVTRRSFLKYCSLAATSLGLGAGMAPKIAWALENKPRIPVVWIHGL WP_001295940.1 MULTISPECIES: hydrogenase [Enterobacteriaceae] Length: 372\nNumber of predicted TMHs: 2\nExp number of AAs in TMHs: 37.2608899999999\nExp number, first 60 AAs: 19.52923\nTotal prob of N-in: 0.94351\nPOSSIBLE N-term signal sequence\ninside 1 19\nTMhelix 20 39\noutside 40 325\nTMhelix 326 348\ninside 349 372

6205 GCF_000752535.1_FHI67 Escherichia coli Proteobacteria; Gammaproteobacteria; Enterobacterales; Enterobacteriaceae; Escherichia MTGDNTLIHSHGINRRDFMKLCAALAATMGLSSKAAAEMAESVTNPQRPPVIWIGAQECT WP_000145410.1 MULTISPECIES: hydrogenase-2 small chain [Proteobacteria] Length: 372\nNumber of predicted TMHs: 1\nExp number of AAs in TMHs: 23.72226\nExp number, first 60 AAs: 0.22038\nTotal prob of N-in: 0.12994\noutside 1 331\nTMhelix 332 354\ninside 355 372

6206 GCF_000647795.1_ASM64779v2 Escherichia coli Proteobacteria; Gammaproteobacteria; Enterobacterales; Enterobacteriaceae; Escherichia MTGDNTLIHSHGINRRDFMKLCAALAATMGLSSKAAAEMAESVTNPQRPPVIWIGAQECT WP_000145410.1 MULTISPECIES: hydrogenase-2 small chain [Proteobacteria] Length: 372\nNumber of predicted TMHs: 1\nExp number of AAs in TMHs: 23.72226\nExp number, first 60 AAs: 0.22038\nTotal prob of N-in: 0.12994\noutside 1 331\nTMhelix 332 354\ninside 355 372

6207 GCF_000681435.1_IEH71520_1.0 Escherichia coli Proteobacteria; Gammaproteobacteria; Enterobacterales; Enterobacteriaceae; Escherichia MTGDNTLIHSHGINRRDFMKLCAALAATMGLSSKAAAEMVESVTNPQRPPVIWIGAQECT WP_000145423.1 hydrogenase 2 small subunit [Escherichia coli] Length: 372\nNumber of predicted TMHs: 1\nExp number of AAs in TMHs: 23.78201\nExp number, first 60 AAs: 0.28355\nTotal prob of N-in: 0.13257\noutside 1 331\nTMhelix 332 354\ninside 355 372

6208 GCF_000691065.1_S6966 Escherichia coli Proteobacteria; Gammaproteobacteria; Enterobacterales; Enterobacteriaceae; Escherichia MNNEETFYQAMRRQGVTRRSFLKYCSLAATSLGLGAGMAPKIAWALENKPRIPVVWIHGL WP_001058323.1 MULTISPECIES: hydrogenase-1 small chain [Proteobacteria] Length: 372\nNumber of

predicted TMHs: 2\nExp number of AAs in TMHs: 37.2608499999999\nExp number, first 60 AAs: 19.52923\nTotal prob of N-in: 0.94351\nPOSSIBLE N-term signal sequence\ninside 1 19\nTMhelix 20 39\noutside 40 325\nTMhelix 326 348\ninside 349 372

6209 GCF_000691085.1_S6995 Escherichia coli Proteobacteria; Gammaproteobacteria; Enterobacterales; Enterobacteriaceae; Escherichia MNNEETFYQAMRRQGVTRRSFLKYCSLAATSLGLGAGMAPKIAWALENKPRIPVVWIHGL WP_001058323.1 MULTISPECIES: hydrogenase-1 small chain [Proteobacteria] Length: 372\nNumber of predicted TMHs: 2\nExp number of AAs in TMHs: 37.2608499999999\nExp number, first 60 AAs: 19.52923\nTotal prob of N-in: 0.94351\nPOSSIBLE N-term signal sequence\ninside 1 19\nTMhelix 20 39\noutside 40 325\nTMhelix 326 348\ninside 349 372

6210 GCF_000695135.1_Ecoli97-3250v1 Escherichia coli Proteobacteria; Gammaproteobacteria; Enterobacterales; Enterobacteriaceae; Escherichia MTGDNTLIHSHGINRRDFMKLCAALAATMGLSSKAAAEMAESVTNPQRPPVIWIGAQECT WP_000145410.1 MULTISPECIES: hydrogenase-2 small chain [Proteobacteria] Length: 372\nNumber of predicted TMHs: 1\nExp number of AAs in TMHs: 23.72226\nExp number, first 60 AAs: 0.22038\nTotal prob of N-in: 0.12994\noutside 1 331\nTMhelix 332 354\ninside 355 372

6211 GCF_000695175.1_EcoliMI10-01v1 Escherichia coli Proteobacteria; Gammaproteobacteria; Enterobacterales; Enterobacteriaceae; Escherichia MTGDNTLIHSHGINRRDFMKLCAALAATMGLSSKAAAEMAESVTNPQRPPVIWIGAQECT WP_000145410.1 MULTISPECIES: hydrogenase-2 small chain [Proteobacteria] Length: 372\nNumber of predicted TMHs: 1\nExp number of AAs in TMHs: 23.72226\nExp number, first 60 AAs: 0.22038\nTotal prob of N-in: 0.12994\noutside 1 331\nTMhelix 332 354\ninside 355 372

6212 GCF_000695505.1_SCB34 Escherichia coli Proteobacteria; Gammaproteobacteria; Enterobacterales; Enterobacteriaceae; Escherichia MTGDNTLIHSHGINRRDFMKLCAALAATMGLSSKAAAEMVESVTNPQRPPVIWIGAQECT WP_000145423.1 hydrogenase 2 small subunit [Escherichia coli] Length: 372\nNumber of predicted TMHs: 1\nExp number of AAs in TMHs: 23.78201\nExp number, first 60 AAs: 0.28355\nTotal prob of N-in: 0.13257\noutside 1 331\nTMhelix 332 354\ninside 355 372

6213 GCF_000715035.1_UCD_JA23_pb Escherichia coli Proteobacteria; Gammaproteobacteria; Enterobacterales; Enterobacteriaceae; Escherichia MTGDNTLIHSHGINRRDFMKLCAALAATMGLSSKAAAEMAESVTNPQRPPVIWIGAQECT WP_000145410.1 MULTISPECIES: hydrogenase-2 small chain [Proteobacteria] Length: 372\nNumber of predicted TMHs: 1\nExp number of AAs in TMHs: 23.72226\nExp number, first 60 AAs: 0.22038\nTotal prob of N-in: 0.12994\noutside 1 331\nTMhelix 332 354\ninside 355 372

6214 GCF_000731455.1_V1 Escherichia coli Proteobacteria; Gammaproteobacteria; Enterobacterales; Enterobacteriaceae; Escherichia MTGDNTLIHSHGINRRDFMKLCAALAATMGLSSKAAAEMAESVTNPQRPPVIWIGAQECT WP_000145410.1 MULTISPECIES: hydrogenase-2 small chain [Proteobacteria] Length: 372\nNumber of predicted TMHs: 1\nExp number of AAs in TMHs: 23.72226\nExp number, first 60 AAs: 0.22038\nTotal prob of N-in: 0.12994\noutside 1 331\nTMhelix 332 354\ninside 355 372

6215 GCF_000752535.1_FHI67 Escherichia coli Proteobacteria; Gammaproteobacteria; Enterobacterales; Enterobacteriaceae; Escherichia MNNEETFYQAMRRQGVTRRSFLKYCSLAATSLGLGAGMAPKIAWALENKPRIPVVWIHGL WP_001058323.1 MULTISPECIES: hydrogenase-1 small chain [Proteobacteria] Length: 372\nNumber of predicted TMHs: 2\nExp number of AAs in TMHs: 37.2608499999999\nExp number, first 60 AAs: 19.52923\nTotal prob of N-in: 0.94351\nPOSSIBLE N-term signal sequence\ninside 1 19\nTMhelix 20 39\noutside 40 325\nTMhelix 326 348\ninside 349 372

6216 GCF_000752255.1_FHI60 Escherichia coli Proteobacteria; Gammaproteobacteria; Enterobacterales; Enterobacteriaceae; Escherichia MNNEETFYQAMRRQGVTRRSFLKYCSLAATSLGLGAGMAPKIAWALENKPRIPVVWIHGL WP_001058323.1 MULTISPECIES: hydrogenase-1 small chain [Proteobacteria] Length: 372\nNumber of predicted TMHs: 2\nExp number of AAs in TMHs: 37.2608499999999\nExp number, first 60 AAs: 19.52923\nTotal prob of N-in: 0.94351\nPOSSIBLE N-term signal sequence\ninside 1 19\nTMhelix 20 39\noutside 40 325\nTMhelix 326 348\ninside 349 372

6217 GCF_000752555.1_FHI76 Escherichia coli Proteobacteria; Gammaproteobacteria; Enterobacterales; Enterobacteriaceae; Escherichia MNNEETFYQAMRRQGVTRRSFLKYCSLAATSLGLGAGMAPKIAWALENKPRIPVVWIHGL WP_001058323.1 MULTISPECIES: hydrogenase-1 small chain [Proteobacteria] Length: 372\nNumber of

predicted TMHs:  2\nExp number of AAs in TMHs: 37.2608499999999\nExp number, first 60 AAs:  19.52923\nTotal prob of N-in:  0.94351\nPOSSIBLE N-term signal sequence\ninside   1   19\nTMhelix   20   39\noutside   40   325\nTMhelix   326   348\ninside   349   372

6218 GCF_000752315.1_FHI32 Escherichia coli Proteobacteria; Gammaproteobacteria; Enterobacterales; Enterobacteriaceae; Escherichia MNNEETFYQAMRRQGVTRRSFLKYCSLAATSLGLGAGMAPKIAWALENKPRIPVVWIHGL WP_001058323.1 MULTISPECIES: hydrogenase-1 small chain [Proteobacteria] Length: 372\nNumber of predicted TMHs:  2\nExp number of AAs in TMHs: 37.2608499999999\nExp number, first 60 AAs:  19.52923\nTotal prob of N-in:  0.94351\nPOSSIBLE N-term signal sequence\ninside   1   19\nTMhelix   20   39\noutside   40   325\nTMhelix   326   348\ninside   349   372

6219 GCF_000752595.1_FHI16 Escherichia coli Proteobacteria; Gammaproteobacteria; Enterobacterales; Enterobacteriaceae; Escherichia MTGDNTLIHSHGINRRDFMKLCAALAATMGLSSKAAAEMAESVTNPQRPPVIWIGAQECT WP_000145410.1 MULTISPECIES: hydrogenase-2 small chain [Proteobacteria] Length: 372\nNumber of predicted TMHs:  1\nExp number of AAs in TMHs: 23.72226\nExp number, first 60 AAs:  0.22038\nTotal prob of N-in:  0.12994\noutside   1   331\nTMhelix   332   354\ninside   355   372

6220 GCF_000752595.1_FHI16 Escherichia coli Proteobacteria; Gammaproteobacteria; Enterobacterales; Enterobacteriaceae; Escherichia MNNEETFYQAMRRQGVTRRSFLKYCSLAATSLGLGAGMAPKIAWALENKPRIPVVWIHGL WP_001058323.1 MULTISPECIES: hydrogenase-1 small chain [Proteobacteria] Length: 372\nNumber of predicted TMHs:  2\nExp number of AAs in TMHs: 37.2608499999999\nExp number, first 60 AAs:  19.52923\nTotal prob of N-in:  0.94351\nPOSSIBLE N-term signal sequence\ninside   1   19\nTMhelix   20   39\noutside   40   325\nTMhelix   326   348\ninside   349   372

6221 GCF_000752435.1_FHI1 Escherichia coli Proteobacteria; Gammaproteobacteria; Enterobacterales; Enterobacteriaceae; Escherichia MNNEETFYQAMRRQGVTRRSFLKYCSLAATSLGLGAGMAPKIAWALENKPRIPVVWIHGL WP_001058323.1 MULTISPECIES: hydrogenase-1 small chain [Proteobacteria] Length: 372\nNumber of predicted TMHs:  2\nExp number of AAs in TMHs: 37.2608499999999\nExp number, first 60 AAs:  19.52923\nTotal prob of N-in:  0.94351\nPOSSIBLE N-term signal sequence\ninside   1   19\nTMhelix   20   39\noutside   40   325\nTMhelix   326   348\ninside   349   372

6222 GCF_000753075.1_FHI73 Escherichia coli Proteobacteria; Gammaproteobacteria; Enterobacterales; Enterobacteriaceae; Escherichia MNNEETFYQAMRRQGVTRRSFLKYCSLAATSLGLGAGMAPKIAWALENKPRIPVVWIHGL WP_001058323.1 MULTISPECIES: hydrogenase-1 small chain [Proteobacteria] Length: 372\nNumber of predicted TMHs:  2\nExp number of AAs in TMHs: 37.2608499999999\nExp number, first 60 AAs:  19.52923\nTotal prob of N-in:  0.94351\nPOSSIBLE N-term signal sequence\ninside   1   19\nTMhelix   20   39\noutside   40   325\nTMhelix   326   348\ninside   349   372

6223 GCF_000752775.1_FHI5 Escherichia coli Proteobacteria; Gammaproteobacteria; Enterobacterales; Enterobacteriaceae; Escherichia MNNEETFYQAMRRQGVTRRSFLKYCSLAATSLGLGAGMAPKIAWALENKPRIPVVWIHGL WP_001301613.1 hydrogenase [Escherichia coli] Length: 372\nNumber of predicted TMHs:  2\nExp number of AAs in TMHs: 37.46598\nExp number, first 60 AAs:  19.53139\nTotal prob of N-in:   0.94364\nPOSSIBLE N-term signal sequence\ninside   1   19\nTMhelix   20   39\noutside   40   325\nTMhelix   326   348\ninside   349   372

6224 GCF_000753215.1_FHI36 Escherichia coli Proteobacteria; Gammaproteobacteria; Enterobacterales; Enterobacteriaceae; Escherichia MTGDNTLIHSHGINRRDFMKLCAALAATMGLSSKAAAEMAESVTNPQRPPVIWIGAQECT WP_000145410.1 MULTISPECIES: hydrogenase-2 small chain [Proteobacteria] Length: 372\nNumber of predicted TMHs:  1\nExp number of AAs in TMHs: 23.72226\nExp number, first 60 AAs:  0.22038\nTotal prob of N-in:  0.12994\noutside   1   331\nTMhelix   332   354\ninside   355   372

6225 GCF_000753255.1_FHI86 Escherichia coli Proteobacteria; Gammaproteobacteria; Enterobacterales; Enterobacteriaceae; Escherichia MTGDNTLIHSHGINRRDFMKLCAALAATMGLSSKAAAEMAESVTNPQRPPVIWIGAQECT WP_000145410.1 MULTISPECIES: hydrogenase-2 small chain [Proteobacteria] Length: 372\nNumber of predicted TMHs:  1\nExp number of AAs in TMHs: 23.72226\nExp number, first 60 AAs:  0.22038\nTotal prob of N-in:  0.12994\noutside   1   331\nTMhelix   332   354\ninside   355   372

6226 GCF_000753215.1_FHI36 Escherichia coli Proteobacteria; Gammaproteobacteria; Enterobacterales; Enterobacteriaceae; Escherichia MNNEETFYQAMRRQGVTRRSFLKYCSLAATSLGLGAGMAPKIAWALENKPRIPVVWIHGL WP_001058323.1 MULTISPECIES: hydrogenase-1 small chain [Proteobacteria] Length: 372\nNumber of predicted TMHs:  2\nExp number of AAs in TMHs: 37.2608499999999\nExp number, first 60 AAs:  19.52923\nTotal prob of

N-in: 0.94351\nPOSSIBLE N-term signal sequence\ninside 1 19\nTMhelix 20 39\noutside 40 325\nTMhelix 326 348\ninside 349 372

6227 GCF_000753255.1_FHI86 Escherichia coli Proteobacteria; Gammaproteobacteria; Enterobacterales; Enterobacteriaceae; Escherichia MNNEETFYQAMRRQGVTRRSFLKYCSLAATSLGLGAGMAPKIAWALENKPRIPVVWIHGL WP_001058320.1 hydrogenase [Escherichia coli] Length: 372\nNumber of predicted TMHs: 2\nExp number of AAs in TMHs: 37.26082\nExp number, first 60 AAs: 19.52923\nTotal prob of N-in: 0.94351\nPOSSIBLE N-term signal sequence\ninside 1 19\nTMhelix 20 39\noutside 40 325\nTMhelix 326 348\ninside 349 372

6228 GCF_000752975.1_StOlav172 Escherichia coli Proteobacteria; Gammaproteobacteria; Enterobacterales; Enterobacteriaceae; Escherichia MNNEETFYQAMRRQGVTRRSFLKYCSLAATSLGLGAGMAPKIAWALENKPRIPVVWIHGL WP_001295940.1 MULTISPECIES: hydrogenase [Enterobacteriaceae] Length: 372\nNumber of predicted TMHs: 2\nExp number of AAs in TMHs: 37.2608899999999\nExp number, first 60 AAs: 19.52923\nTotal prob of N-in: 0.94351\nPOSSIBLE N-term signal sequence\ninside 1 19\nTMhelix 20 39\noutside 40 325\nTMhelix 326 348\ninside 349 372

6229 GCF_000770045.1_ASM77004v1 Escherichia coli Proteobacteria; Gammaproteobacteria; Enterobacterales; Enterobacteriaceae; Escherichia MTGDNTLIHSHGINRRDFMKLCAALAATMGLSSKAAAEMAESVTNPQRPPVIWIGAQECT WP_000145410.1 MULTISPECIES: hydrogenase-2 small chain [Proteobacteria] Length: 372\nNumber of predicted TMHs: 1\nExp number of AAs in TMHs: 23.72226\nExp number, first 60 AAs: 0.22038\nTotal prob of N-in: 0.12994\noutside 1 331\nTMhelix 332 354\ninside 355 372

6230 GCF_000770055.1_ASM77005v1 Escherichia coli Proteobacteria; Gammaproteobacteria; Enterobacterales; Enterobacteriaceae; Escherichia MTGDNTLIHSHGINRRDFMKLCAALAATMGLSSKAAAEMAESVTNPQRPPVIWIGAQECT WP_000145410.1 MULTISPECIES: hydrogenase-2 small chain [Proteobacteria] Length: 372\nNumber of predicted TMHs: 1\nExp number of AAs in TMHs: 23.72226\nExp number, first 60 AAs: 0.22038\nTotal prob of N-in: 0.12994\noutside 1 331\nTMhelix 332 354\ninside 355 372

6231 GCF_000770275.1_ASM77027v1 Escherichia coli Proteobacteria; Gammaproteobacteria; Enterobacterales; Enterobacteriaceae; Escherichia MTGDNTLIHSHGINRRDFMKLCAALAATMGLSSKAAAEMVESVTNPQRPPVIWIGAQECT WP_000145423.1 hydrogenase 2 small subunit [Escherichia coli] Length: 372\nNumber of predicted TMHs: 1\nExp number of AAs in TMHs: 23.78201\nExp number, first 60 AAs: 0.28355\nTotal prob of N-in: 0.13257\noutside 1 331\nTMhelix 332 354\ninside 355 372

6232 GCF_000770045.1_ASM77004v1 Escherichia coli Proteobacteria; Gammaproteobacteria; Enterobacterales; Enterobacteriaceae; Escherichia MNNEETFYQAMRRQGVTRRSFLKYCSLAATSLGLGAGMAPKIAWALENKPRIPVVWIHGL WP_001058323.1 MULTISPECIES: hydrogenase-1 small chain [Proteobacteria] Length: 372\nNumber of predicted TMHs: 2\nExp number of AAs in TMHs: 37.2608499999999\nExp number, first 60 AAs: 19.52923\nTotal prob of N-in: 0.94351\nPOSSIBLE N-term signal sequence\ninside 1 19\nTMhelix 20 39\noutside 40 325\nTMhelix 326 348\ninside 349 372

6233 GCF_000773475.1_ASM77347v1 Escherichia coli Proteobacteria; Gammaproteobacteria; Enterobacterales; Enterobacteriaceae; Escherichia MTGDNTLIHSHGINRRDFMKLCAALAATMGLSSKAAAEMAESVTNPQRPPVIWIGAQECT WP_000145410.1 MULTISPECIES: hydrogenase-2 small chain [Proteobacteria] Length: 372\nNumber of predicted TMHs: 1\nExp number of AAs in TMHs: 23.72226\nExp number, first 60 AAs: 0.22038\nTotal prob of N-in: 0.12994\noutside 1 331\nTMhelix 332 354\ninside 355 372

6234 GCF_000773435.1_ASM77343v1 Escherichia coli Proteobacteria; Gammaproteobacteria; Enterobacterales; Enterobacteriaceae; Escherichia MTGDNTLIHSHGINRRDFMKLCAALAATMGLSSKAAAEMAESVTNPQRPPVIWIGAQECT WP_000145410.1 MULTISPECIES: hydrogenase-2 small chain [Proteobacteria] Length: 372\nNumber of predicted TMHs: 1\nExp number of AAs in TMHs: 23.72226\nExp number, first 60 AAs: 0.22038\nTotal prob of N-in: 0.12994\noutside 1 331\nTMhelix 332 354\ninside 355 372

6235 GCF_000773555.1_ASM77355v1 Escherichia coli Proteobacteria; Gammaproteobacteria; Enterobacterales; Enterobacteriaceae; Escherichia
MTGDNTLIHSHGINRRDFMKLCAALAATMGLSSKAAAEMAESVTNPQRPPVIWIGAQECT WP_000145410.1
MULTISPECIES: hydrogenase-2 small chain [Proteobacteria] Length: 372\nNumber of predicted TMHs: 1\nExp number of AAs in TMHs: 23.72226\nExp number, first 60 AAs: 0.22038\nTotal prob of N-in: 0.12994\noutside 1 331\nTMhelix 332 354\ninside 355 372

6236 GCF_000776335.1_ASM77633v1 Escherichia coli Proteobacteria; Gammaproteobacteria; Enterobacterales; Enterobacteriaceae; Escherichia
MNNEETFYQAMRRQGVTRRSFLKYCSLAATSLGLGAGMAPKIAWALENKPRIPVVWIHGL WP_001058323.1
MULTISPECIES: hydrogenase-1 small chain [Proteobacteria] Length: 372\nNumber of predicted TMHs: 2\nExp number of AAs in TMHs: 37.2608499999999\nExp number, first 60 AAs: 19.52923\nTotal prob of N-in: 0.94351\nPOSSIBLE N-term signal sequence\ninside 1 19\nTMhelix 20 39\noutside 40 325\nTMhelix 326 348\ninside 349 372

6237 GCF_000776615.1_ASM77661v1 Escherichia coli Proteobacteria; Gammaproteobacteria; Enterobacterales; Enterobacteriaceae; Escherichia
MTGDNTLIHSHGINRRDFMKLCAALAATMGLSSKAAAEMAESVTNPQRPPVIWIGAQECT WP_000145410.1
MULTISPECIES: hydrogenase-2 small chain [Proteobacteria] Length: 372\nNumber of predicted TMHs: 1\nExp number of AAs in TMHs: 23.72226\nExp number, first 60 AAs: 0.22038\nTotal prob of N-in: 0.12994\noutside 1 331\nTMhelix 332 354\ninside 355 372

6238 GCF_000776655.1_ASM77665v1 Escherichia coli Proteobacteria; Gammaproteobacteria; Enterobacterales; Enterobacteriaceae; Escherichia
MTGDNTLIHSHGINRRDFMKLCAALAATMGLSSKAAAEMAESVTNPQRPPVIWIGAQECT WP_000145410.1
MULTISPECIES: hydrogenase-2 small chain [Proteobacteria] Length: 372\nNumber of predicted TMHs: 1\nExp number of AAs in TMHs: 23.72226\nExp number, first 60 AAs: 0.22038\nTotal prob of N-in: 0.12994\noutside 1 331\nTMhelix 332 354\ninside 355 372

6239 GCF_000776675.1_ASM77667v1 Escherichia coli Proteobacteria; Gammaproteobacteria; Enterobacterales; Enterobacteriaceae; Escherichia
MTGDNTLIHSHGINRRDFMKLCAALAATMGLSSKAAAEMAESVTNPQRPPVIWIGAQECT WP_000145410.1
MULTISPECIES: hydrogenase-2 small chain [Proteobacteria] Length: 372\nNumber of predicted TMHs: 1\nExp number of AAs in TMHs: 23.72226\nExp number, first 60 AAs: 0.22038\nTotal prob of N-in: 0.12994\noutside 1 331\nTMhelix 332 354\ninside 355 372

6240 GCF_000776615.1_ASM77661v1 Escherichia coli Proteobacteria; Gammaproteobacteria; Enterobacterales; Enterobacteriaceae; Escherichia
MNNEETFYQAMRRQGVTRRSFLKYCSLAATSLGLGAGMAPKIAWALENKPRIPVVWIHGL WP_001058323.1
MULTISPECIES: hydrogenase-1 small chain [Proteobacteria] Length: 372\nNumber of predicted TMHs: 2\nExp number of AAs in TMHs: 37.2608499999999\nExp number, first 60 AAs: 19.52923\nTotal prob of N-in: 0.94351\nPOSSIBLE N-term signal sequence\ninside 1 19\nTMhelix 20 39\noutside 40 325\nTMhelix 326 348\ninside 349 372

6241 GCF_000776715.1_ASM77671v1 Escherichia coli Proteobacteria; Gammaproteobacteria; Enterobacterales; Enterobacteriaceae; Escherichia
MTGDNTLIHSHGINRRDFMKLCAALAATMGLSSKAAAEMAESVTNPQRPPVIWIGAQECT WP_000145410.1
MULTISPECIES: hydrogenase-2 small chain [Proteobacteria] Length: 372\nNumber of predicted TMHs: 1\nExp number of AAs in TMHs: 23.72226\nExp number, first 60 AAs: 0.22038\nTotal prob of N-in: 0.12994\noutside 1 331\nTMhelix 332 354\ninside 355 372

6242 GCF_000776745.1_ASM77674v1 Escherichia coli Proteobacteria; Gammaproteobacteria; Enterobacterales; Enterobacteriaceae; Escherichia
MTGDNTLIHSHGINRRDFMKLCAALAATMGLSSKAAAEMAESVTNPQRPPVIWIGAQECT WP_000145418.1
hydrogenase 2 small subunit [Escherichia coli] Length: 372\nNumber of predicted TMHs: 1\nExp number of AAs in TMHs: 23.72313\nExp number, first 60 AAs: 0.22034\nTotal prob of N-in: 0.12997\noutside 1 331\nTMhelix 332 354\ninside 355 372

6243 GCF_000777265.1_ASM77726v1 Escherichia coli Proteobacteria; Gammaproteobacteria; Enterobacterales; Enterobacteriaceae; Escherichia
MTGDNTLIHSHGINRRDFMKLCAALAATMGLSSKAAAEMAESVTNPQRPPVIWIGAQECT WP_000145410.1

MULTISPECIES: hydrogenase-2 small chain [Proteobacteria] Length: 372\nNumber of predicted TMHs: 1\nExp number of AAs in TMHs: 23.72226\nExp number, first 60 AAs: 0.22038\nTotal prob of N-in: 0.12994\noutside 1 331\nTMhelix 332 354\ninside 355 372

6244 GCF_000777395.1_ASM77739v1 Escherichia coli Proteobacteria; Gammaproteobacteria; Enterobacterales; Enterobacteriaceae; Escherichia
MTGDNTLIHSHGINRRDFMKLCAALAATMGLSSKAAAEMAESVTNPQRPPVIWIGAQECT WP_000145410.1
MULTISPECIES: hydrogenase-2 small chain [Proteobacteria] Length: 372\nNumber of predicted TMHs: 1\nExp number of AAs in TMHs: 23.72226\nExp number, first 60 AAs: 0.22038\nTotal prob of N-in: 0.12994\noutside 1 331\nTMhelix 332 354\ninside 355 372

6245 GCF_000777375.1_ASM77737v1 Escherichia coli Proteobacteria; Gammaproteobacteria; Enterobacterales; Enterobacteriaceae; Escherichia
MTGDNTLIHSHGINRRDFMKLCAALAATMGLSSKAAAEMAESVTNPQRPPVIWIGAQECT WP_000145410.1
MULTISPECIES: hydrogenase-2 small chain [Proteobacteria] Length: 372\nNumber of predicted TMHs: 1\nExp number of AAs in TMHs: 23.72226\nExp number, first 60 AAs: 0.22038\nTotal prob of N-in: 0.12994\noutside 1 331\nTMhelix 332 354\ninside 355 372

6246 GCF_000777325.1_ASM77732v1 Escherichia coli Proteobacteria; Gammaproteobacteria; Enterobacterales; Enterobacteriaceae; Escherichia
MTGDNTLIHSHGINRRDFMKLCAALAATMGLSSKAAAEMAESVTNPQRPPVIWIGAQECT WP_000145410.1
MULTISPECIES: hydrogenase-2 small chain [Proteobacteria] Length: 372\nNumber of predicted TMHs: 1\nExp number of AAs in TMHs: 23.72226\nExp number, first 60 AAs: 0.22038\nTotal prob of N-in: 0.12994\noutside 1 331\nTMhelix 332 354\ninside 355 372

6247 GCF_000777265.1_ASM77726v1 Escherichia coli Proteobacteria; Gammaproteobacteria; Enterobacterales; Enterobacteriaceae; Escherichia
MNNEETFYQAMRRQGVTRRSFLKYCSLAATSLGLGAGMAPKIAWALENKPRIPVVWIHGL WP_001058323.1
MULTISPECIES: hydrogenase-1 small chain [Proteobacteria] Length: 372\nNumber of predicted TMHs: 2\nExp number of AAs in TMHs: 37.2608499999999\nExp number, first 60 AAs: 19.52923\nTotal prob of N-in: 0.94351\nPOSSIBLE N-term signal sequence\ninside 1 19\nTMhelix 20 39\noutside 40 325\nTMhelix 326 348\ninside 349 372

6248 GCF_000777395.1_ASM77739v1 Escherichia coli Proteobacteria; Gammaproteobacteria; Enterobacterales; Enterobacteriaceae; Escherichia
MNNEETFYQAMRRQGVTRRSFLKYCSLAATSLGLGAGMAPKIAWALENKPRIPVVWIHGL WP_001058323.1
MULTISPECIES: hydrogenase-1 small chain [Proteobacteria] Length: 372\nNumber of predicted TMHs: 2\nExp number of AAs in TMHs: 37.2608499999999\nExp number, first 60 AAs: 19.52923\nTotal prob of N-in: 0.94351\nPOSSIBLE N-term signal sequence\ninside 1 19\nTMhelix 20 39\noutside 40 325\nTMhelix 326 348\ninside 349 372

6249 GCF_000777325.1_ASM77732v1 Escherichia coli Proteobacteria; Gammaproteobacteria; Enterobacterales; Enterobacteriaceae; Escherichia
MNNEETFYQAMRRQGVTRRSFLKYCSLAATSLGLGAGMAPKIAWALENKPRIPVVWIHGL WP_001058323.1
MULTISPECIES: hydrogenase-1 small chain [Proteobacteria] Length: 372\nNumber of predicted TMHs: 2\nExp number of AAs in TMHs: 37.2608499999999\nExp number, first 60 AAs: 19.52923\nTotal prob of N-in: 0.94351\nPOSSIBLE N-term signal sequence\ninside 1 19\nTMhelix 20 39\noutside 40 325\nTMhelix 326 348\ninside 349 372

6250 GCF_000777995.1_ASM77799v1 Escherichia coli Proteobacteria; Gammaproteobacteria; Enterobacterales; Enterobacteriaceae; Escherichia
MTGDNTLIHSHGINRRDFMKLCAALAATMGLSSKAAAEMAESVTNPQRPPVIWIGAQECT WP_000145410.1
MULTISPECIES: hydrogenase-2 small chain [Proteobacteria] Length: 372\nNumber of predicted TMHs: 1\nExp number of AAs in TMHs: 23.72226\nExp number, first 60 AAs: 0.22038\nTotal prob of N-in: 0.12994\noutside 1 331\nTMhelix 332 354\ninside 355 372

6251 GCF_000778075.1_ASM77807v1 Escherichia coli Proteobacteria; Gammaproteobacteria; Enterobacterales; Enterobacteriaceae; Escherichia
MTGDNTLIHSHGINRRDFMKLCAALAATMGLSSKAAAEMAESVTNPQRPPVIWIGAQECT WP_000145410.1
MULTISPECIES: hydrogenase-2 small chain [Proteobacteria] Length: 372\nNumber of predicted TMHs:

1\nExp number of AAs in TMHs: 23.72226\nExp number, first 60 AAs: 0.22038\nTotal prob of N-in: 0.12994\noutside 1 331\nTMhelix 332 354\ninside 355 372

6252 GCF_000778095.1_ASM77809v1 Escherichia coli Proteobacteria; Gammaproteobacteria; Enterobacterales; Enterobacteriaceae; Escherichia
MTGDNTLIHSHGINRRDFMKLCAALAATMGLSSKAAAEMAESVTNPQRPPVIWIGAQECT WP_000145410.1
MULTISPECIES: hydrogenase-2 small chain [Proteobacteria] Length: 372\nNumber of predicted TMHs: 1\nExp number of AAs in TMHs: 23.72226\nExp number, first 60 AAs: 0.22038\nTotal prob of N-in: 0.12994\noutside 1 331\nTMhelix 332 354\ninside 355 372

6253 GCF_000777995.1_ASM77799v1 Escherichia coli Proteobacteria; Gammaproteobacteria; Enterobacterales; Enterobacteriaceae; Escherichia
MNNEETFYQAMRRQGVTRRSFLKYCSLAATSLGLGAGMAPKIAWALENKPRIPVVWIHGL WP_001058323.1
MULTISPECIES: hydrogenase-1 small chain [Proteobacteria] Length: 372\nNumber of predicted TMHs: 2\nExp number of AAs in TMHs: 37.2608499999999\nExp number, first 60 AAs: 19.52923\nTotal prob of N-in: 0.94351\nPOSSIBLE N-term signal sequence\ninside 1 19\nTMhelix 20 39\noutside 40 325\nTMhelix 326 348\ninside 349 372

6254 GCF_000778135.1_ASM77813v1 Escherichia coli Proteobacteria; Gammaproteobacteria; Enterobacterales; Enterobacteriaceae; Escherichia
MTGDNTLIHSHGINRRDFMKLCAALAATMGLSSKAAAEMAESVTNPQRPPVIWIGAQECT WP_000145410.1
MULTISPECIES: hydrogenase-2 small chain [Proteobacteria] Length: 372\nNumber of predicted TMHs: 1\nExp number of AAs in TMHs: 23.72226\nExp number, first 60 AAs: 0.22038\nTotal prob of N-in: 0.12994\noutside 1 331\nTMhelix 332 354\ninside 355 372

6255 GCF_000778215.1_ASM77821v1 Escherichia coli Proteobacteria; Gammaproteobacteria; Enterobacterales; Enterobacteriaceae; Escherichia
MTGDNTLIHSHGINRRDFMKLCAALAATMGLSSKAAAEMAESVTNPQRPPVIWIGAQECT WP_000145410.1
MULTISPECIES: hydrogenase-2 small chain [Proteobacteria] Length: 372\nNumber of predicted TMHs: 1\nExp number of AAs in TMHs: 23.72226\nExp number, first 60 AAs: 0.22038\nTotal prob of N-in: 0.12994\noutside 1 331\nTMhelix 332 354\ninside 355 372

6256 GCF_000778235.1_ASM77823v1 Escherichia coli Proteobacteria; Gammaproteobacteria; Enterobacterales; Enterobacteriaceae; Escherichia
MTGDNTLIHSHGINRRDFMKLCAALAATMGLSSKAAAEMAESVTNPQRPPVIWIGAQECT WP_000145410.1
MULTISPECIES: hydrogenase-2 small chain [Proteobacteria] Length: 372\nNumber of predicted TMHs: 1\nExp number of AAs in TMHs: 23.72226\nExp number, first 60 AAs: 0.22038\nTotal prob of N-in: 0.12994\noutside 1 331\nTMhelix 332 354\ninside 355 372

6257 GCF_000778565.1_ASM77856v1 Escherichia coli Proteobacteria; Gammaproteobacteria; Enterobacterales; Enterobacteriaceae; Escherichia
MTGDNTLIHSHGINRRDFMKLCAALAATMGLSSKAAAEMAESVTNPQRPPVIWIGAQECT WP_000145410.1
MULTISPECIES: hydrogenase-2 small chain [Proteobacteria] Length: 372\nNumber of predicted TMHs: 1\nExp number of AAs in TMHs: 23.72226\nExp number, first 60 AAs: 0.22038\nTotal prob of N-in: 0.12994\noutside 1 331\nTMhelix 332 354\ninside 355 372

6258 GCF_000778565.1_ASM77856v1 Escherichia coli Proteobacteria; Gammaproteobacteria; Enterobacterales; Enterobacteriaceae; Escherichia
MNNEETFYQAMRRQGVTRRSFLKYCSLAATSLGLGAGMAPKIAWALENKPRIPVVWIHGL WP_001058323.1
MULTISPECIES: hydrogenase-1 small chain [Proteobacteria] Length: 372\nNumber of predicted TMHs: 2\nExp number of AAs in TMHs: 37.2608499999999\nExp number, first 60 AAs: 19.52923\nTotal prob of N-in: 0.94351\nPOSSIBLE N-term signal sequence\ninside 1 19\nTMhelix 20 39\noutside 40 325\nTMhelix 326 348\ninside 349 372

6259 GCF_000778435.1_ASM77843v1 Escherichia coli Proteobacteria; Gammaproteobacteria; Enterobacterales; Enterobacteriaceae; Escherichia
MNNEETFYQAMRRQGVTRRSFLKYCSLAATSLGLGAGMAPKIAWALENKPRIPVVWIHGL WP_001058323.1
MULTISPECIES: hydrogenase-1 small chain [Proteobacteria] Length: 372\nNumber of predicted TMHs: 2\nExp number of AAs in TMHs: 37.2608499999999\nExp number, first 60 AAs: 19.52923\nTotal prob of N-in: 0.94351\nPOSSIBLE N-term signal sequence\ninside 1 19\nTMhelix 20 39\noutside 40 325\nTMhelix 326 348\ninside 349 372

6260 GCF_000778485.1_ASM77848v1 Escherichia coli Proteobacteria; Gammaproteobacteria; Enterobacterales; Enterobacteriaceae; Escherichia
MNNEETFYQAMRRQGVTRRSFLKYCSLAATSLGLGAGMAPKIAWALENKPRIPVVWIHGL WP_001058323.1
MULTISPECIES: hydrogenase-1 small chain [Proteobacteria] Length: 372\nNumber of predicted TMHs: 2\nExp number of AAs in TMHs: 37.2608499999999\nExp number, first 60 AAs: 19.52923\nTotal prob of N-in: 0.94351\nPOSSIBLE N-term signal sequence\ninside 1 19\nTMhelix 20 39\noutside 40 325\nTMhelix 326 348\ninside 349 372

6261 GCF_000778585.1_ASM77858v1 Escherichia coli Proteobacteria; Gammaproteobacteria; Enterobacterales; Enterobacteriaceae; Escherichia
MTGDNTLIHSHGINRRDFMKLCAALAATMGLSSKAAAEMAESVTNPQRPPVIWIGAQECT WP_000145410.1
MULTISPECIES: hydrogenase-2 small chain [Proteobacteria] Length: 372\nNumber of predicted TMHs: 1\nExp number of AAs in TMHs: 23.72226\nExp number, first 60 AAs: 0.22038\nTotal prob of N-in: 0.12994\noutside 1 331\nTMhelix 332 354\ninside 355 372

6262 GCF_000778725.1_ASM77872v1 Escherichia coli Proteobacteria; Gammaproteobacteria; Enterobacterales; Enterobacteriaceae; Escherichia
MTGDNTLIHSHGINRRDFMKLCAALAATMGLSSKAAAEMAESVTNPQRPPVIWIGAQECT WP_000145410.1
MULTISPECIES: hydrogenase-2 small chain [Proteobacteria] Length: 372\nNumber of predicted TMHs: 1\nExp number of AAs in TMHs: 23.72226\nExp number, first 60 AAs: 0.22038\nTotal prob of N-in: 0.12994\noutside 1 331\nTMhelix 332 354\ninside 355 372

6263 GCF_000778685.1_ASM77868v1 Escherichia coli Proteobacteria; Gammaproteobacteria; Enterobacterales; Enterobacteriaceae; Escherichia
MTGDNTLIHSHGINRRDFMKLCAALAATMGLSSKAAAEMAESVTNPQRPPVIWIGAQECT WP_000145410.1
MULTISPECIES: hydrogenase-2 small chain [Proteobacteria] Length: 372\nNumber of predicted TMHs: 1\nExp number of AAs in TMHs: 23.72226\nExp number, first 60 AAs: 0.22038\nTotal prob of N-in: 0.12994\noutside 1 331\nTMhelix 332 354\ninside 355 372

6264 GCF_000779335.1_ASM77933v1 Escherichia coli Proteobacteria; Gammaproteobacteria; Enterobacterales; Enterobacteriaceae; Escherichia
MNNEETFYQAMRRQGVTRRSFLKYCSLAATSLGLGAGMAPKIAWALENKPRIPVVWIHGL WP_001058323.1
MULTISPECIES: hydrogenase-1 small chain [Proteobacteria] Length: 372\nNumber of predicted TMHs: 2\nExp number of AAs in TMHs: 37.2608499999999\nExp number, first 60 AAs: 19.52923\nTotal prob of N-in: 0.94351\nPOSSIBLE N-term signal sequence\ninside 1 19\nTMhelix 20 39\noutside 40 325\nTMhelix 326 348\ninside 349 372

6265 GCF_000779265.1_ASM77926v1 Escherichia coli Proteobacteria; Gammaproteobacteria; Enterobacterales; Enterobacteriaceae; Escherichia
MNNEETFYQAMRRQGVTRRSFLKYCSLAATSLGLGAGMAPKIAWALENKPRIPVVWIHGL WP_001295940.1
MULTISPECIES: hydrogenase [Enterobacteriaceae] Length: 372\nNumber of predicted TMHs: 2\nExp number of AAs in TMHs: 37.2608899999999\nExp number, first 60 AAs: 19.52923\nTotal prob of N-in: 0.94351\nPOSSIBLE N-term signal sequence\ninside 1 19\nTMhelix 20 39\noutside 40 325\nTMhelix 326 348\ninside 349 372

6266 GCF_000779125.1_ASM77912v1 Escherichia coli Proteobacteria; Gammaproteobacteria; Enterobacterales; Enterobacteriaceae; Escherichia
MNNEETFYQAMRRQGVTRRSFLKYCSLAATSLGLGAGMAPKIAWALENKPRIPVVWIHGL WP_001058323.1
MULTISPECIES: hydrogenase-1 small chain [Proteobacteria] Length: 372\nNumber of predicted TMHs: 2\nExp number of AAs in TMHs: 37.2608499999999\nExp number, first 60 AAs: 19.52923\nTotal prob of N-in: 0.94351\nPOSSIBLE N-term signal sequence\ninside 1 19\nTMhelix 20 39\noutside 40 325\nTMhelix 326 348\ninside 349 372

6267 GCF_000779395.1_ASM77939v1 Escherichia coli Proteobacteria; Gammaproteobacteria; Enterobacterales; Enterobacteriaceae; Escherichia
MTGDNTLIHSHGINRRDFMKLCAALAATMGLSSKAAAEMAESVTNPQRPPVIWIGAQECT WP_000145410.1
MULTISPECIES: hydrogenase-2 small chain [Proteobacteria] Length: 372\nNumber of predicted TMHs: 1\nExp number of AAs in TMHs: 23.72226\nExp number, first 60 AAs: 0.22038\nTotal prob of N-in: 0.12994\noutside 1 331\nTMhelix 332 354\ninside 355 372

6268 GCF_000779395.1_ASM77939v1 Escherichia coli Proteobacteria; Gammaproteobacteria; Enterobacterales; Enterobacteriaceae; Escherichia
MNNEETFYQAMRRQGVTRRSFLKYCSLAATSLGLGAGMAPKIAWALENKPRIPVVWIHGL WP_001058323.1
MULTISPECIES: hydrogenase-1 small chain [Proteobacteria] Length: 372\nNumber of predicted TMHs: 2\nExp number of AAs in TMHs: 37.2608499999999\nExp number, first 60 AAs: 19.52923\nTotal prob of N-in: 0.94351\nPOSSIBLE N-term signal sequence\ninside 1 19\nTMhelix 20 39\noutside 40 325\nTMhelix 326 348\ninside 349 372

6269 GCF_000779425.1_ASM77942v1 Escherichia coli Proteobacteria; Gammaproteobacteria; Enterobacterales; Enterobacteriaceae; Escherichia
MTGDNTLIHSHGINRRDFMKLCAALAATMGLSSKAAAEMAESVTNPQRPPVIWIGAQECT WP_000145410.1
MULTISPECIES: hydrogenase-2 small chain [Proteobacteria] Length: 372\nNumber of predicted TMHs: 1\nExp number of AAs in TMHs: 23.72226\nExp number, first 60 AAs: 0.22038\nTotal prob of N-in: 0.12994\noutside 1 331\nTMhelix 332 354\ninside 355 372

6270 GCF_000779475.1_ASM77947v1 Escherichia coli Proteobacteria; Gammaproteobacteria; Enterobacterales; Enterobacteriaceae; Escherichia
MTGDNTLIHSHGINRRDFMKLCAALAATMGLSSKAAAEMAESVTNPQRPPVIWIGAQECT WP_000145410.1
MULTISPECIES: hydrogenase-2 small chain [Proteobacteria] Length: 372\nNumber of predicted TMHs: 1\nExp number of AAs in TMHs: 23.72226\nExp number, first 60 AAs: 0.22038\nTotal prob of N-in: 0.12994\noutside 1 331\nTMhelix 332 354\ninside 355 372

6271 GCF_000779955.1_ASM77995v1 Escherichia coli Proteobacteria; Gammaproteobacteria; Enterobacterales; Enterobacteriaceae; Escherichia
MTGDNTLIHSHGINRRDFMKLCAALAATMGLSSKAAAEMAESVTNPQRPPVIWIGAQECT WP_000145410.1
MULTISPECIES: hydrogenase-2 small chain [Proteobacteria] Length: 372\nNumber of predicted TMHs: 1\nExp number of AAs in TMHs: 23.72226\nExp number, first 60 AAs: 0.22038\nTotal prob of N-in: 0.12994\noutside 1 331\nTMhelix 332 354\ninside 355 372

6272 GCF_000779975.1_ASM77997v1 Escherichia coli Proteobacteria; Gammaproteobacteria; Enterobacterales; Enterobacteriaceae; Escherichia
MNNEETFYQAMRRQGVTRRSFLKYCSLAATSLGLGAGMAPKIAWALENKPRIPVVWIHGL WP_001058323.1
MULTISPECIES: hydrogenase-1 small chain [Proteobacteria] Length: 372\nNumber of predicted TMHs: 2\nExp number of AAs in TMHs: 37.2608499999999\nExp number, first 60 AAs: 19.52923\nTotal prob of N-in: 0.94351\nPOSSIBLE N-term signal sequence\ninside 1 19\nTMhelix 20 39\noutside 40 325\nTMhelix 326 348\ninside 349 372

6273 GCF_000779915.1_ASM77991v1 Escherichia coli Proteobacteria; Gammaproteobacteria; Enterobacterales; Enterobacteriaceae; Escherichia
MNNEETFYQAMRRQGVTRRSFLKYCSLAATSLGLGAGMAPKIAWALENKPRIPVVWIHGL WP_001058323.1
MULTISPECIES: hydrogenase-1 small chain [Proteobacteria] Length: 372\nNumber of predicted TMHs: 2\nExp number of AAs in TMHs: 37.2608499999999\nExp number, first 60 AAs: 19.52923\nTotal prob of N-in: 0.94351\nPOSSIBLE N-term signal sequence\ninside 1 19\nTMhelix 20 39\noutside 40 325\nTMhelix 326 348\ninside 349 372

6274 GCF_000779955.1_ASM77995v1 Escherichia coli Proteobacteria; Gammaproteobacteria; Enterobacterales; Enterobacteriaceae; Escherichia
MNNEETFYQAMRRQGVTRRSFLKYCSLAATSLGLGAGMAPKIAWALENKPRIPVVWIHGL WP_001058323.1
MULTISPECIES: hydrogenase-1 small chain [Proteobacteria] Length: 372\nNumber of predicted TMHs: 2\nExp number of AAs in TMHs: 37.2608499999999\nExp number, first 60 AAs: 19.52923\nTotal prob of N-in: 0.94351\nPOSSIBLE N-term signal sequence\ninside 1 19\nTMhelix 20 39\noutside 40 325\nTMhelix 326 348\ninside 349 372

6275 GCF_000779745.1_ASM77974v1 Escherichia coli Proteobacteria; Gammaproteobacteria; Enterobacterales; Enterobacteriaceae; Escherichia
MNNEETFYQAMRRQGVTRRSFLKYCSLAATSLGLGAGMAPKIAWALENKPRIPVVWIHGL WP_001295940.1
MULTISPECIES: hydrogenase [Enterobacteriaceae] Length: 372\nNumber of predicted TMHs: 2\nExp number of AAs in TMHs: 37.2608899999999\nExp number, first 60 AAs: 19.52923\nTotal prob of N-in: 0.94351\nPOSSIBLE N-term signal sequence\ninside 1 19\nTMhelix 20 39\noutside 40 325\nTMhelix 326 348\ninside 349 372

6276 GCF_000780095.1_ASM78009v1 Escherichia coli Proteobacteria; Gammaproteobacteria; Enterobacterales; Enterobacteriaceae; Escherichia
MTGDNTLIHSHGINRRDFMKLCAALAATMGLSSKAAAEMAESVTNPQRPPVIWIGAQECT WP_000145410.1
MULTISPECIES: hydrogenase-2 small chain [Proteobacteria] Length: 372\nNumber of predicted TMHs: 1\nExp number of AAs in TMHs: 23.72226\nExp number, first 60 AAs: 0.22038\nTotal prob of N-in: 0.12994\noutside 1 331\nTMhelix 332 354\ninside 355 372

6277 GCF_000780475.1_ASM78047v1 Escherichia coli Proteobacteria; Gammaproteobacteria; Enterobacterales; Enterobacteriaceae; Escherichia
MNNEETFYQAMRRQGVTRRSFLKYCSLAATSLGLGAGMAPKIAWALENKPRIPVVWIHGL WP_001538532.1
hydrogenase-1 small chain [Escherichia coli] Length: 372\nNumber of predicted TMHs: 2\nExp number of AAs in TMHs: 37.27041\nExp number, first 60 AAs: 19.52927\nTotal prob of N-in: 0.94352\nPOSSIBLE N-term signal sequence\ninside 1 19\nTMhelix 20 39\noutside 40 325\nTMhelix 326 348\ninside 349 372

6278 GCF_000780715.1_ASM78071v1 Escherichia coli Proteobacteria; Gammaproteobacteria; Enterobacterales; Enterobacteriaceae; Escherichia
MTGDNTLIHSHGINRRDFMKLCAALAATMGLSSKAAAEMAESVTNPQRPPVIWIGAQECT WP_000145410.1
MULTISPECIES: hydrogenase-2 small chain [Proteobacteria] Length: 372\nNumber of predicted TMHs: 1\nExp number of AAs in TMHs: 23.72226\nExp number, first 60 AAs: 0.22038\nTotal prob of N-in: 0.12994\noutside 1 331\nTMhelix 332 354\ninside 355 372

6279 GCF_000780695.1_ASM78069v1 Escherichia coli Proteobacteria; Gammaproteobacteria; Enterobacterales; Enterobacteriaceae; Escherichia
MTGDNTLIHSHGINRRDFMKLCAALAATMGLSSKAAAEMAESVTNPQRPPVIWIGAQECT WP_000145410.1
MULTISPECIES: hydrogenase-2 small chain [Proteobacteria] Length: 372\nNumber of predicted TMHs: 1\nExp number of AAs in TMHs: 23.72226\nExp number, first 60 AAs: 0.22038\nTotal prob of N-in: 0.12994\noutside 1 331\nTMhelix 332 354\ninside 355 372

6280 GCF_000780715.1_ASM78071v1 Escherichia coli Proteobacteria; Gammaproteobacteria; Enterobacterales; Enterobacteriaceae; Escherichia
MNNEETFYQAMRRQGVTRRSFLKYCSLAATSLGLGAGMAPKIAWALENKPRIPVVWIHGL WP_001058323.1
MULTISPECIES: hydrogenase-1 small chain [Proteobacteria] Length: 372\nNumber of predicted TMHs: 2\nExp number of AAs in TMHs: 37.2608499999999\nExp number, first 60 AAs: 19.52923\nTotal prob of N-in: 0.94351\nPOSSIBLE N-term signal sequence\ninside 1 19\nTMhelix 20 39\noutside 40 325\nTMhelix 326 348\ninside 349 372

6281 GCF_000780715.1_ASM78071v1 Escherichia coli Proteobacteria; Gammaproteobacteria; Enterobacterales; Enterobacteriaceae; Escherichia
MKGVLEKCTVEDFNYISATLDSYVSFTNDKRRKTLLSAYQNNPALHAELISLIDTQIKYF WP_001515206.1 MULTISPECIES: hypothetical protein [Enterobacteriaceae] Length: 249\nNumber of predicted TMHs: 2\nExp number of AAs in TMHs: 39.68696\nExp number, first 60 AAs: 0.01961\nTotal prob of N-in: 0.09053\noutside 1 157\nTMhelix 158 180\ninside 181 200\nTMhelix 201 220\noutside 221 249

6282 GCF_000780555.1_ASM78055v1 Escherichia coli Proteobacteria; Gammaproteobacteria; Enterobacterales; Enterobacteriaceae; Escherichia
MNNEETFYQAMRRQGVTRRSFLKYCSLAATSLGLGAGMAPKIAWALENKPRIPVVWIHGL WP_001058320.1
hydrogenase [Escherichia coli] Length: 372\nNumber of predicted TMHs: 2\nExp number of AAs in TMHs: 37.26082\nExp number, first 60 AAs: 19.52923\nTotal prob of N-in: 0.94351\nPOSSIBLE N-term signal sequence\ninside 1 19\nTMhelix 20 39\noutside 40 325\nTMhelix 326 348\ninside 349 372

6283 GCF_000780495.1_ASM78049v1 Escherichia coli Proteobacteria; Gammaproteobacteria; Enterobacterales; Enterobacteriaceae; Escherichia
MNNEETFYQAMRRQGVTRRSFLKYCSLAATSLGLGAGMAPKIAWALENKPRIPVVWIHGL WP_001295940.1
MULTISPECIES: hydrogenase [Enterobacteriaceae] Length: 372\nNumber of predicted TMHs: 2\nExp number of AAs in TMHs: 37.2608899999999\nExp number, first 60 AAs: 19.52923\nTotal prob of N-in: 0.94351\nPOSSIBLE N-term signal sequence\ninside 1 19\nTMhelix 20 39\noutside 40 325\nTMhelix 326 348\ninside 349 372

6284 GCF_000781235.1_ASM78123v1 Escherichia coli Proteobacteria; Gammaproteobacteria; Enterobacterales; Enterobacteriaceae; Escherichia
MNNEETFYQAMRRQGVTRRSFLKYCSLAATSLGLGAGMAPKIAWALENKPRIPVVWIHGL WP_001058323.1

MULTISPECIES: hydrogenase-1 small chain [Proteobacteria] Length: 372\nNumber of predicted TMHs: 2\nExp number of AAs in TMHs: 37.2608499999999\nExp number, first 60 AAs: 19.52923\nTotal prob of N-in: 0.94351\nPOSSIBLE N-term signal sequence\ninside 1 19\nTMhelix 20 39\noutside 40 325\nTMhelix 326 348\ninside 349 372

6285 GCF_000781005.1_ASM78100v1 Escherichia coli Proteobacteria; Gammaproteobacteria; Enterobacterales; Enterobacteriaceae; Escherichia MNNEETFYQAMRRQGVTRRSFLKYCSLAATSLGLGAGMAPKIAWALENKPRIPVVWIHGL WP_001295940.1 MULTISPECIES: hydrogenase [Enterobacteriaceae] Length: 372\nNumber of predicted TMHs: 2\nExp number of AAs in TMHs: 37.2608899999999\nExp number, first 60 AAs: 19.52923\nTotal prob of N-in: 0.94351\nPOSSIBLE N-term signal sequence\ninside 1 19\nTMhelix 20 39\noutside 40 325\nTMhelix 326 348\ninside 349 372

6286 GCF_000781335.1_ASM78133v1 Escherichia coli Proteobacteria; Gammaproteobacteria; Enterobacterales; Enterobacteriaceae; Escherichia MTGDNTLIHSHGINRRDFMKLCAALAATMGLSSKAAAEMVESVTNPQRPPVIWIGAQECT WP_000145423.1 hydrogenase 2 small subunit [Escherichia coli] Length: 372\nNumber of predicted TMHs: 1\nExp number of AAs in TMHs: 23.78201\nExp number, first 60 AAs: 0.28355\nTotal prob of N-in: 0.13257\noutside 1 331\nTMhelix 332 354\ninside 355 372

6287 GCF_000781355.1_ASM78135v1 Escherichia coli Proteobacteria; Gammaproteobacteria; Enterobacterales; Enterobacteriaceae; Escherichia MTGDNTLIHSHGINRRDFMKLCAALAATMGLSSKAAAEMAESVTNPQRPPVIWIGAQECT WP_000145410.1 MULTISPECIES: hydrogenase-2 small chain [Proteobacteria] Length: 372\nNumber of predicted TMHs: 1\nExp number of AAs in TMHs: 23.72226\nExp number, first 60 AAs: 0.22038\nTotal prob of N-in: 0.12994\noutside 1 331\nTMhelix 332 354\ninside 355 372

6288 GCF_000781385.1_ASM78138v1 Escherichia coli Proteobacteria; Gammaproteobacteria; Enterobacterales; Enterobacteriaceae; Escherichia MTGDNTLIHSHGINRRDFMKLCAALAATMGLSSKAAAEMAESVTNPQRPPVIWIGAQECT WP_000145410.1 MULTISPECIES: hydrogenase-2 small chain [Proteobacteria] Length: 372\nNumber of predicted TMHs: 1\nExp number of AAs in TMHs: 23.72226\nExp number, first 60 AAs: 0.22038\nTotal prob of N-in: 0.12994\noutside 1 331\nTMhelix 332 354\ninside 355 372

6289 GCF_000781455.1_ASM78145v1 Escherichia coli Proteobacteria; Gammaproteobacteria; Enterobacterales; Enterobacteriaceae; Escherichia MTGDNTLIHSHGINRRDFMKLCAALAATMGLSSKAAAEMAESVTNPQRPPVIWIGAQECT WP_000145410.1 MULTISPECIES: hydrogenase-2 small chain [Proteobacteria] Length: 372\nNumber of predicted TMHs: 1\nExp number of AAs in TMHs: 23.72226\nExp number, first 60 AAs: 0.22038\nTotal prob of N-in: 0.12994\noutside 1 331\nTMhelix 332 354\ninside 355 372

6290 GCF_000781435.1_ASM78143v1 Escherichia coli Proteobacteria; Gammaproteobacteria; Enterobacterales; Enterobacteriaceae; Escherichia MTGDNTLIHSHGINRRDFMKLCAALAATMGLSSKAAAEMAESVTNPQRPPVIWIGAQECT WP_000145410.1 MULTISPECIES: hydrogenase-2 small chain [Proteobacteria] Length: 372\nNumber of predicted TMHs: 1\nExp number of AAs in TMHs: 23.72226\nExp number, first 60 AAs: 0.22038\nTotal prob of N-in: 0.12994\noutside 1 331\nTMhelix 332 354\ninside 355 372

6291 GCF_000781875.1_ASM78187v1 Escherichia coli Proteobacteria; Gammaproteobacteria; Enterobacterales; Enterobacteriaceae; Escherichia MTGDNTLIHSHGINRRDFMKLCAALAATMGLSSKAAAEMAESVTNPQRPPVIWIGAQECT WP_000145410.1 MULTISPECIES: hydrogenase-2 small chain [Proteobacteria] Length: 372\nNumber of predicted TMHs: 1\nExp number of AAs in TMHs: 23.72226\nExp number, first 60 AAs: 0.22038\nTotal prob of N-in: 0.12994\noutside 1 331\nTMhelix 332 354\ninside 355 372

6292 GCF_000781895.1_ASM78189v1 Escherichia coli Proteobacteria; Gammaproteobacteria; Enterobacterales; Enterobacteriaceae; Escherichia MTGDNTLIHSHGINRRDFMKLCAALAATMGLSSKAAAEMAESVTNPQRPPVIWIGAQECT WP_000145410.1 MULTISPECIES: hydrogenase-2 small chain [Proteobacteria] Length: 372\nNumber of predicted TMHs: 1\nExp number of AAs in TMHs: 23.72226\nExp number, first 60 AAs: 0.22038\nTotal prob of N-in: 0.12994\noutside 1 331\nTMhelix 332 354\ninside 355 372

6293 GCF_000781965.1_ASM78196v1 Escherichia coli Proteobacteria; Gammaproteobacteria; Enterobacterales; Enterobacteriaceae; Escherichia
MTGDNTLIHSHGINRRDFMKLCAALAATMGLSSKAAAEMVESVTNPQRPPVIWIGAQECT WP_000145423.1
hydrogenase 2 small subunit [Escherichia coli] Length: 372\nNumber of predicted TMHs: 1\nExp number of AAs in TMHs: 23.78201\nExp number, first 60 AAs: 0.28355\nTotal prob of N-in: 0.13257\noutside 1 331\nTMhelix 332 354\ninside 355 372

6294 GCF_000781935.1_ASM78193v1 Escherichia coli Proteobacteria; Gammaproteobacteria; Enterobacterales; Enterobacteriaceae; Escherichia
MNNEETFYQAMRRQGVTRRSFLKYCSLAATSLGLGAGMAPKIAWALENKPRIPVVWIHGL WP_001058323.1
MULTISPECIES: hydrogenase-1 small chain [Proteobacteria] Length: 372\nNumber of predicted TMHs: 2\nExp number of AAs in TMHs: 37.2608499999999\nExp number, first 60 AAs: 19.52923\nTotal prob of N-in: 0.94351\nPOSSIBLE N-term signal sequence\ninside 1 19\nTMhelix 20 39\noutside 40 325\nTMhelix 326 348\ninside 349 372

6295 GCF_000781875.1_ASM78187v1 Escherichia coli Proteobacteria; Gammaproteobacteria; Enterobacterales; Enterobacteriaceae; Escherichia
MNNEETFYQAMRRQGVTRRSFLKYCSLAATSLGLGAGMAPKIAWALENKPRIPVVWIHGL WP_001058323.1
MULTISPECIES: hydrogenase-1 small chain [Proteobacteria] Length: 372\nNumber of predicted TMHs: 2\nExp number of AAs in TMHs: 37.2608499999999\nExp number, first 60 AAs: 19.52923\nTotal prob of N-in: 0.94351\nPOSSIBLE N-term signal sequence\ninside 1 19\nTMhelix 20 39\noutside 40 325\nTMhelix 326 348\ninside 349 372

6296 GCF_000781755.1_ASM78175v1 Escherichia coli Proteobacteria; Gammaproteobacteria; Enterobacterales; Enterobacteriaceae; Escherichia
MNNEETFYQAMRRQGVTRRSFLKYCSLAATSLGLGAGMAPKIAWALENKPRIPVVWIHGL WP_001295940.1
MULTISPECIES: hydrogenase [Enterobacteriaceae] Length: 372\nNumber of predicted TMHs: 2\nExp number of AAs in TMHs: 37.2608899999999\nExp number, first 60 AAs: 19.52923\nTotal prob of N-in: 0.94351\nPOSSIBLE N-term signal sequence\ninside 1 19\nTMhelix 20 39\noutside 40 325\nTMhelix 326 348\ninside 349 372

6297 GCF_000781995.1_ASM78199v1 Escherichia coli Proteobacteria; Gammaproteobacteria; Enterobacterales; Enterobacteriaceae; Escherichia
MTGDNTLIHSHGINRRDFMKLCAALAATMGLSSKAAAEMVESVTNPQRPPVIWIGAQECT WP_000145423.1
hydrogenase 2 small subunit [Escherichia coli] Length: 372\nNumber of predicted TMHs: 1\nExp number of AAs in TMHs: 23.78201\nExp number, first 60 AAs: 0.28355\nTotal prob of N-in: 0.13257\noutside 1 331\nTMhelix 332 354\ninside 355 372

6298 GCF_000782135.1_ASM78213v1 Escherichia coli Proteobacteria; Gammaproteobacteria; Enterobacterales; Enterobacteriaceae; Escherichia
MNNEETFYQAMRRQGVTRRSFLKYCSLAATSLGLGAGMAPKIAWALENKPRIPVVWIHGL WP_001295940.1
MULTISPECIES: hydrogenase [Enterobacteriaceae] Length: 372\nNumber of predicted TMHs: 2\nExp number of AAs in TMHs: 37.2608899999999\nExp number, first 60 AAs: 19.52923\nTotal prob of N-in: 0.94351\nPOSSIBLE N-term signal sequence\ninside 1 19\nTMhelix 20 39\noutside 40 325\nTMhelix 326 348\ninside 349 372

6299 GCF_000782235.1_ASM78223v1 Escherichia coli Proteobacteria; Gammaproteobacteria; Enterobacterales; Enterobacteriaceae; Escherichia
MNNEETFYQAMRRQGVTRRSFLKYCSLAATSLGLGAGMAPKIAWALENKPRIPVVWIHGL WP_001295940.1
MULTISPECIES: hydrogenase [Enterobacteriaceae] Length: 372\nNumber of predicted TMHs: 2\nExp number of AAs in TMHs: 37.2608899999999\nExp number, first 60 AAs: 19.52923\nTotal prob of N-in: 0.94351\nPOSSIBLE N-term signal sequence\ninside 1 19\nTMhelix 20 39\noutside 40 325\nTMhelix 326 348\ninside 349 372

6300 GCF_000782435.1_ASM78243v1 Escherichia coli Proteobacteria; Gammaproteobacteria; Enterobacterales; Enterobacteriaceae; Escherichia
MTGDNTLIHSHGINRRDFMKLCAALAATMGLSSKAAAEMAESVTNPQRPPVIWIGAQECT WP_000145410.1
MULTISPECIES: hydrogenase-2 small chain [Proteobacteria] Length: 372\nNumber of predicted TMHs: 1\nExp number of AAs in TMHs: 23.72226\nExp number, first 60 AAs: 0.22038\nTotal prob of N-in: 0.12994\noutside 1 331\nTMhelix 332 354\ninside 355 372

6301 GCF_000782435.1_ASM78243v1 Escherichia coli Proteobacteria; Gammaproteobacteria; Enterobacterales; Enterobacteriaceae; Escherichia
MNNEETFYQAMRRQGVTRRSFLKYCSLAATSLGLGAGMAPKIAWALENKPRIPVVWIHGL WP_001058323.1
MULTISPECIES: hydrogenase-1 small chain [Proteobacteria] Length: 372\nNumber of predicted TMHs: 2\nExp number of AAs in TMHs: 37.2608499999999\nExp number, first 60 AAs: 19.52923\nTotal prob of N-in: 0.94351\nPOSSIBLE N-term signal sequence\ninside 1 19\nTMhelix 20 39\noutside 40 325\nTMhelix 326 348\ninside 349 372

6302 GCF_000782175.1_ASM78217v1 Escherichia coli Proteobacteria; Gammaproteobacteria; Enterobacterales; Enterobacteriaceae; Escherichia
MNNEETFYQAMRRQGVTRRSFLKYCSLAATSLGLGAGMAPKIAWALENKPRIPVVWIHGL WP_001295940.1
MULTISPECIES: hydrogenase [Enterobacteriaceae] Length: 372\nNumber of predicted TMHs: 2\nExp number of AAs in TMHs: 37.2608899999999\nExp number, first 60 AAs: 19.52923\nTotal prob of N-in: 0.94351\nPOSSIBLE N-term signal sequence\ninside 1 19\nTMhelix 20 39\noutside 40 325\nTMhelix 326 348\ninside 349 372

6303 GCF_000782455.1_ASM78245v1 Escherichia coli Proteobacteria; Gammaproteobacteria; Enterobacterales; Enterobacteriaceae; Escherichia
MTGDNTLIHSHGINRRDFMKLCAALAATMGLSSKAAAEMAESVTNPQRPPVIWIGAQECT WP_000145410.1
MULTISPECIES: hydrogenase-2 small chain [Proteobacteria] Length: 372\nNumber of predicted TMHs: 1\nExp number of AAs in TMHs: 23.72226\nExp number, first 60 AAs: 0.22038\nTotal prob of N-in: 0.12994\noutside 1 331\nTMhelix 332 354\ninside 355 372

6304 GCF_000782455.1_ASM78245v1 Escherichia coli Proteobacteria; Gammaproteobacteria; Enterobacterales; Enterobacteriaceae; Escherichia
MNNEETFYQAMRRQGVTRRSFLKYCSLAATSLGLGAGMAPKIAWALENKPRIPVVWIHGL WP_001058323.1
MULTISPECIES: hydrogenase-1 small chain [Proteobacteria] Length: 372\nNumber of predicted TMHs: 2\nExp number of AAs in TMHs: 37.2608499999999\nExp number, first 60 AAs: 19.52923\nTotal prob of N-in: 0.94351\nPOSSIBLE N-term signal sequence\ninside 1 19\nTMhelix 20 39\noutside 40 325\nTMhelix 326 348\ninside 349 372

6305 GCF_000782675.1_ASM78267v1 Escherichia coli Proteobacteria; Gammaproteobacteria; Enterobacterales; Enterobacteriaceae; Escherichia
MNNEETFYQAMRRQGVTRRSFLKYCSLAATSLGLGAGMAPKIAWALENKPRIPVVWIHGL WP_001295940.1
MULTISPECIES: hydrogenase [Enterobacteriaceae] Length: 372\nNumber of predicted TMHs: 2\nExp number of AAs in TMHs: 37.2608899999999\nExp number, first 60 AAs: 19.52923\nTotal prob of N-in: 0.94351\nPOSSIBLE N-term signal sequence\ninside 1 19\nTMhelix 20 39\noutside 40 325\nTMhelix 326 348\ninside 349 372

6306 GCF_000782715.1_ASM78271v1 Escherichia coli Proteobacteria; Gammaproteobacteria; Enterobacterales; Enterobacteriaceae; Escherichia
MNNEETFYQAMRRQGVTRRSFLKYCSLAATSLGLGAGMAPKIAWALENKPRIPVVWIHGL WP_001295940.1
MULTISPECIES: hydrogenase [Enterobacteriaceae] Length: 372\nNumber of predicted TMHs: 2\nExp number of AAs in TMHs: 37.2608899999999\nExp number, first 60 AAs: 19.52923\nTotal prob of N-in: 0.94351\nPOSSIBLE N-term signal sequence\ninside 1 19\nTMhelix 20 39\noutside 40 325\nTMhelix 326 348\ninside 349 372

6307 GCF_000785775.1_ASM78577v1 Escherichia coli Proteobacteria; Gammaproteobacteria; Enterobacterales; Enterobacteriaceae; Escherichia
MTGDNTLIHSHGINRRDFMKLCAALAATMGLSSKAAAEMAESVTNPQRPPVIWIGAQECT WP_000145410.1
MULTISPECIES: hydrogenase-2 small chain [Proteobacteria] Length: 372\nNumber of predicted TMHs: 1\nExp number of AAs in TMHs: 23.72226\nExp number, first 60 AAs: 0.22038\nTotal prob of N-in: 0.12994\noutside 1 331\nTMhelix 332 354\ninside 355 372

6308 GCF_000797575.1_Escherichia_coli_CVM_N37067PS_v._1.0 Escherichia coli Proteobacteria; Gammaproteobacteria; Enterobacterales; Enterobacteriaceae; Escherichia
MTGDNTLIHSHGINRRDFMKLCAALAATMGLSSKAAAEMAESVTNPQRPPVIWIGAQECT WP_000145410.1
MULTISPECIES: hydrogenase-2 small chain [Proteobacteria] Length: 372\nNumber of predicted TMHs: 1\nExp number of AAs in TMHs: 23.72226\nExp number, first 60 AAs: 0.22038\nTotal prob of N-in: 0.12994\noutside 1 331\nTMhelix 332 354\ninside 355 372

6309 GCF_000785865.1_ASM78586v1 Escherichia coli Proteobacteria; Gammaproteobacteria; Enterobacterales; Enterobacteriaceae; Escherichia
MTGDNTLIHSHGINRRDFMKLCAALAATMGLSSKAAAEMAESVTNPQRPPVIWIGAQECT WP_000145410.1
MULTISPECIES: hydrogenase-2 small chain [Proteobacteria] Length: 372\nNumber of predicted TMHs: 1\nExp number of AAs in TMHs: 23.72226\nExp number, first 60 AAs: 0.22038\nTotal prob of N-in: 0.12994\noutside 1 331\nTMhelix 332 354\ninside 355 372

6310 GCF_000785775.1_ASM78577v1 Escherichia coli Proteobacteria; Gammaproteobacteria; Enterobacterales; Enterobacteriaceae; Escherichia
MNNEETFYQAMRRQGVTRRSFLKYCSLAATSLGLGAGMAPKIAWALENKPRIPVVWIHGL WP_001058323.1
MULTISPECIES: hydrogenase-1 small chain [Proteobacteria] Length: 372\nNumber of predicted TMHs: 2\nExp number of AAs in TMHs: 37.2608499999999\nExp number, first 60 AAs: 19.52923\nTotal prob of N-in: 0.94351\nPOSSIBLE N-term signal sequence\ninside 1 19\nTMhelix 20 39\noutside 40 325\nTMhelix 326 348\ninside 349 372

6311 GCF_000797595.1_Escherichia_coli_CVM_N36225PS_v._1.0 Escherichia coli Proteobacteria; Gammaproteobacteria; Enterobacterales; Enterobacteriaceae; Escherichia
MTGDNTLIHSHGINRRDFMKLCAALAATMGLSSKAAAEMAESVTNPQRPPVIWIGAQECT WP_000145410.1
MULTISPECIES: hydrogenase-2 small chain [Proteobacteria] Length: 372\nNumber of predicted TMHs: 1\nExp number of AAs in TMHs: 23.72226\nExp number, first 60 AAs: 0.22038\nTotal prob of N-in: 0.12994\noutside 1 331\nTMhelix 332 354\ninside 355 372

6312 GCF_000798115.1_Escherichia_coli_CVM_N36393PS_v._1.0 Escherichia coli Proteobacteria; Gammaproteobacteria; Enterobacterales; Enterobacteriaceae; Escherichia
MNNEETFYQAMRRQGVTRRSFLKYCSLAATSLGLGAGMAPKIAWALENKPRIPVVWIHGL WP_001058323.1
MULTISPECIES: hydrogenase-1 small chain [Proteobacteria] Length: 372\nNumber of predicted TMHs: 2\nExp number of AAs in TMHs: 37.2608499999999\nExp number, first 60 AAs: 19.52923\nTotal prob of N-in: 0.94351\nPOSSIBLE N-term signal sequence\ninside 1 19\nTMhelix 20 39\noutside 40 325\nTMhelix 326 348\ninside 349 372

6313 GCF_000797875.1_Escherichia_coli_CVM_N38833PS_v._1.0 Escherichia coli Proteobacteria; Gammaproteobacteria; Enterobacterales; Enterobacteriaceae; Escherichia
MNNEETFYQAMRRQGVTRRSFLKYCSLAATSLGLGAGMAPKIAWALENKPRIPVVWIHGL WP_001295940.1
MULTISPECIES: hydrogenase [Enterobacteriaceae] Length: 372\nNumber of predicted TMHs: 2\nExp number of AAs in TMHs: 37.2608899999999\nExp number, first 60 AAs: 19.52923\nTotal prob of N-in: 0.94351\nPOSSIBLE N-term signal sequence\ninside 1 19\nTMhelix 20 39\noutside 40 325\nTMhelix 326 348\ninside 349 372

6314 GCF_000797915.1_Escherichia_coli_CVM_N41498PS_v._1.0 Escherichia coli Proteobacteria; Gammaproteobacteria; Enterobacterales; Enterobacteriaceae; Escherichia
MNNEETFYQAMRRQGVTRRSFLKYCSLAATSLGLGAGMAPKIAWALENKPRIPVVWIHGL WP_001058323.1
MULTISPECIES: hydrogenase-1 small chain [Proteobacteria] Length: 372\nNumber of predicted TMHs: 2\nExp number of AAs in TMHs: 37.2608499999999\nExp number, first 60 AAs: 19.52923\nTotal prob of N-in: 0.94351\nPOSSIBLE N-term signal sequence\ninside 1 19\nTMhelix 20 39\noutside 40 325\nTMhelix 326 348\ninside 349 372

6315 GCF_000798075.1_Escherichia_coli_CVM_N33849PS_v._1.0 Escherichia coli Proteobacteria; Gammaproteobacteria; Enterobacterales; Enterobacteriaceae; Escherichia
MTGDNTLIHSHGINRRDFMKLCAALAATMGLSSKAAAEMAESVTNPQRPPVIWIGAQECT WP_000145410.1
MULTISPECIES: hydrogenase-2 small chain [Proteobacteria] Length: 372\nNumber of predicted TMHs: 1\nExp number of AAs in TMHs: 23.72226\nExp number, first 60 AAs: 0.22038\nTotal prob of N-in: 0.12994\noutside 1 331\nTMhelix 332 354\ninside 355 372

6316 GCF_000798075.1_Escherichia_coli_CVM_N33849PS_v._1.0 Escherichia coli Proteobacteria; Gammaproteobacteria; Enterobacterales; Enterobacteriaceae; Escherichia
MNNEETFYQAMRRQGVTRRSFLKYCSLAATSLGLGAGMAPKIAWALENKPRIPVVWIHGL WP_001058323.1
MULTISPECIES: hydrogenase-1 small chain [Proteobacteria] Length: 372\nNumber of predicted TMHs: 2\nExp number of AAs in TMHs: 37.2608499999999\nExp number, first 60 AAs: 19.52923\nTotal prob of N-in: 0.94351\nPOSSIBLE N-term signal sequence\ninside 1 19\nTMhelix 20 39\noutside 40 325\nTMhelix 326 348\ninside 349 372

6317 GCF_000797975.1_Escherichia_coli_CVM_N36609PS_v._1.0 Escherichia coli Proteobacteria; Gammaproteobacteria; Enterobacterales; Enterobacteriaceae; Escherichia
MNNEETFYQAMRRQGVTRRSFLKYCSLAATSLGLGAGMAPKIAWALENKPRIPVVWIHGL WP_001058323.1
MULTISPECIES: hydrogenase-1 small chain [Proteobacteria] Length: 372\nNumber of predicted TMHs: 2\nExp number of AAs in TMHs: 37.2608499999999\nExp number, first 60 AAs: 19.52923\nTotal prob of N-in: 0.94351\nPOSSIBLE N-term signal sequence\ninside 1 19\nTMhelix 20 39\noutside 40 325\nTMhelix 326 348\ninside 349 372

6318 GCF_000798135.1_Escherichia_coli_CVM_N34054PS_v._1.0 Escherichia coli Proteobacteria; Gammaproteobacteria; Enterobacterales; Enterobacteriaceae; Escherichia
MTGDNTLIHSHGINRRDFMKLCAALAATMGLSSKAAAEMAESVTNPQRPPVIWIGAQECT WP_000145410.1
MULTISPECIES: hydrogenase-2 small chain [Proteobacteria] Length: 372\nNumber of predicted TMHs: 1\nExp number of AAs in TMHs: 23.72226\nExp number, first 60 AAs: 0.22038\nTotal prob of N-in: 0.12994\noutside 1 331\nTMhelix 332 354\ninside 355 372

6319 GCF_000798575.1_Escherichia_coli_CVM_N36404PS_v._1.0 Escherichia coli Proteobacteria; Gammaproteobacteria; Enterobacterales; Enterobacteriaceae; Escherichia
MNNEETFYQAMRRQGVTRRSFLKYCSLAATSLGLGAGMAPKIAWALENKPRIPVVWIHGL WP_001058323.1
MULTISPECIES: hydrogenase-1 small chain [Proteobacteria] Length: 372\nNumber of predicted TMHs: 2\nExp number of AAs in TMHs: 37.2608499999999\nExp number, first 60 AAs: 19.52923\nTotal prob of N-in: 0.94351\nPOSSIBLE N-term signal sequence\ninside 1 19\nTMhelix 20 39\noutside 40 325\nTMhelix 326 348\ninside 349 372

6320 GCF_000798455.1_Escherichia_coli_CVM_N36158PS_v._1.0 Escherichia coli Proteobacteria; Gammaproteobacteria; Enterobacterales; Enterobacteriaceae; Escherichia
MNNEETFYQAMRRQGVTRRSFLKYCSLAATSLGLGAGMAPKIAWALENKPRIPVVWIHGL WP_001295940.1
MULTISPECIES: hydrogenase [Enterobacteriaceae] Length: 372\nNumber of predicted TMHs: 2\nExp number of AAs in TMHs: 37.2608899999999\nExp number, first 60 AAs: 19.52923\nTotal prob of N-in: 0.94351\nPOSSIBLE N-term signal sequence\ninside 1 19\nTMhelix 20 39\noutside 40 325\nTMhelix 326 348\ninside 349 372

6321 GCF_000801165.1_ASM80116v1 Escherichia coli Proteobacteria; Gammaproteobacteria; Enterobacterales; Enterobacteriaceae; Escherichia
MTGDNTLIHSHGINRRDFMKLCAALAATMGLSSKAAAEMAESVTNPQRPPVIWIGAQECT WP_000145410.1
MULTISPECIES: hydrogenase-2 small chain [Proteobacteria] Length: 372\nNumber of predicted TMHs: 1\nExp number of AAs in TMHs: 23.72226\nExp number, first 60 AAs: 0.22038\nTotal prob of N-in: 0.12994\noutside 1 331\nTMhelix 332 354\ninside 355 372

6322 GCF_000801185.2_ASM80118v2 Escherichia coli Proteobacteria; Gammaproteobacteria; Enterobacterales; Enterobacteriaceae; Escherichia
MTGDNTLIHSHGINRRDFMKLCAALAATMGLSSKAAAEMAESVTNPQRPPVIWIGAQECT WP_000145410.1
MULTISPECIES: hydrogenase-2 small chain [Proteobacteria] Length: 372\nNumber of predicted TMHs: 1\nExp number of AAs in TMHs: 23.72226\nExp number, first 60 AAs: 0.22038\nTotal prob of N-in: 0.12994\noutside 1 331\nTMhelix 332 354\ninside 355 372

6323 GCF_000801165.1_ASM80116v1 Escherichia coli Proteobacteria; Gammaproteobacteria; Enterobacterales; Enterobacteriaceae; Escherichia
MNNEETFYQAMRRQGVTRRSFLKYCSLAATSLGLGAGMAPKIAWALENKPRIPVVWIHGL WP_001058323.1
MULTISPECIES: hydrogenase-1 small chain [Proteobacteria] Length: 372\nNumber of predicted TMHs: 2\nExp number of AAs in TMHs: 37.2608499999999\nExp number, first 60 AAs: 19.52923\nTotal prob of N-in: 0.94351\nPOSSIBLE N-term signal sequence\ninside 1 19\nTMhelix 20 39\noutside 40 325\nTMhelix 326 348\ninside 349 372

6324 GCF_000804365.1_ASM80436v1 Escherichia coli Proteobacteria; Gammaproteobacteria; Enterobacterales; Enterobacteriaceae; Escherichia
MNNEETFYQAMRRQGVTRRSFLKYCSLAATSLGLGAGMAPKIAWALENKPRIPVVWIHGL WP_001058323.1
MULTISPECIES: hydrogenase-1 small chain [Proteobacteria] Length: 372\nNumber of predicted TMHs: 2\nExp number of AAs in TMHs: 37.2608499999999\nExp number, first 60 AAs: 19.52923\nTotal prob of N-in: 0.94351\nPOSSIBLE N-term signal sequence\ninside 1 19\nTMhelix 20 39\noutside 40 325\nTMhelix 326 348\ninside 349 372

6325 GCF_000806245.1_ASM80624v1 Escherichia coli Proteobacteria; Gammaproteobacteria; Enterobacterales; Enterobacteriaceae; Escherichia
MTGDNTLIHSHGINRRDFMKLCAALAATMGLSSKAAAEMAESVTNPQRPPVIWIGAQECT WP_000145410.1
MULTISPECIES: hydrogenase-2 small chain [Proteobacteria] Length: 372\nNumber of predicted TMHs: 1\nExp number of AAs in TMHs: 23.72226\nExp number, first 60 AAs: 0.22038\nTotal prob of N-in: 0.12994\noutside 1 331\nTMhelix 332 354\ninside 355 372

6326 GCF_000806265.1_ASM80626v1 Escherichia coli Proteobacteria; Gammaproteobacteria; Enterobacterales; Enterobacteriaceae; Escherichia
MTGDNTLIHSHGINRRDFMKLCAALAATMGLSSKAAAEMAESVTNPQRPPVIWIGAQECT WP_000145410.1
MULTISPECIES: hydrogenase-2 small chain [Proteobacteria] Length: 372\nNumber of predicted TMHs: 1\nExp number of AAs in TMHs: 23.72226\nExp number, first 60 AAs: 0.22038\nTotal prob of N-in: 0.12994\noutside 1 331\nTMhelix 332 354\ninside 355 372

6327 GCF_000807555.1_ASM80755v1 Escherichia coli Proteobacteria; Gammaproteobacteria; Enterobacterales; Enterobacteriaceae; Escherichia
MTGDNTLIHSHGINRRDFMKLCAALAATMGLSSKAAAEMAESVTNPQRPPVIWIGAQECT WP_000145410.1
MULTISPECIES: hydrogenase-2 small chain [Proteobacteria] Length: 372\nNumber of predicted TMHs: 1\nExp number of AAs in TMHs: 23.72226\nExp number, first 60 AAs: 0.22038\nTotal prob of N-in: 0.12994\noutside 1 331\nTMhelix 332 354\ninside 355 372

6328 GCF_000806245.1_ASM80624v1 Escherichia coli Proteobacteria; Gammaproteobacteria; Enterobacterales; Enterobacteriaceae; Escherichia
MNNEETFYQAMRRQGVTRRSFLKYCSLAATSLGLGAGMAPKIAWALENKPRIPVVWIHGL WP_001058323.1
MULTISPECIES: hydrogenase-1 small chain [Proteobacteria] Length: 372\nNumber of predicted TMHs: 2\nExp number of AAs in TMHs: 37.2608499999999\nExp number, first 60 AAs: 19.52923\nTotal prob of N-in: 0.94351\nPOSSIBLE N-term signal sequence\ninside 1 19\nTMhelix 20 39\noutside 40 325\nTMhelix 326 348\ninside 349 372

6329 GCF_000807555.1_ASM80755v1 Escherichia coli Proteobacteria; Gammaproteobacteria; Enterobacterales; Enterobacteriaceae; Escherichia
MNNEETFYQAMRRQGVTRRSFLKYCSLAATSLGLGAGMAPKIAWALENKPRIPVVWIHGL WP_001058323.1
MULTISPECIES: hydrogenase-1 small chain [Proteobacteria] Length: 372\nNumber of predicted TMHs: 2\nExp number of AAs in TMHs: 37.2608499999999\nExp number, first 60 AAs: 19.52923\nTotal prob of N-in: 0.94351\nPOSSIBLE N-term signal sequence\ninside 1 19\nTMhelix 20 39\noutside 40 325\nTMhelix 326 348\ninside 349 372

6330 GCF_000806265.1_ASM80626v1 Escherichia coli Proteobacteria; Gammaproteobacteria; Enterobacterales; Enterobacteriaceae; Escherichia
MNNEETFYQAMRRQGVTRRSFLKYCSLAATSLGLGAGMAPKIVWALENKPRIPVVWIHGL WP_001398876.1
hydrogenase-1 small chain [Escherichia coli] Length: 372\nNumber of predicted TMHs: 2\nExp number of AAs in TMHs: 37.60268\nExp number, first 60 AAs: 19.84677\nTotal prob of N-in: 0.94621\nPOSSIBLE N-term signal sequence\ninside 1 20\nTMhelix 21 43\noutside 44 325\nTMhelix 326 348\ninside 349 372

6331 GCF_000812365.1_ASM81236v1 Escherichia coli Proteobacteria; Gammaproteobacteria; Enterobacterales; Enterobacteriaceae; Escherichia
MNNEETFYQAMRRQGVTRRSFLKYCSLAATSLGLGAGMAPKIAWALENKPRIPVVWIHGL WP_001058323.1
MULTISPECIES: hydrogenase-1 small chain [Proteobacteria] Length: 372\nNumber of predicted TMHs: 2\nExp number of AAs in TMHs: 37.2608499999999\nExp number, first 60 AAs: 19.52923\nTotal prob of N-in: 0.94351\nPOSSIBLE N-term signal sequence\ninside 1 19\nTMhelix 20 39\noutside 40 325\nTMhelix 326 348\ninside 349 372

6332 GCF_000814525.1_ASM81452v1 Escherichia coli Proteobacteria; Gammaproteobacteria; Enterobacterales; Enterobacteriaceae; Escherichia
MTGDNTLIHSHGINRRDFMKLCAALAATMGLSSKAAAEMAESVTNPQRPPVIWIGAQECT WP_000145410.1
MULTISPECIES: hydrogenase-2 small chain [Proteobacteria] Length: 372\nNumber of predicted TMHs: 1\nExp number of AAs in TMHs: 23.72226\nExp number, first 60 AAs: 0.22038\nTotal prob of N-in: 0.12994\noutside 1 331\nTMhelix 332 354\ninside 355 372

6333 GCF_000814525.1_ASM81452v1 Escherichia coli Proteobacteria; Gammaproteobacteria; Enterobacterales; Enterobacteriaceae; Escherichia

MNNEETFYQAMRRQGVTRRSFLKYCSLAATSLGLGAGMAPKIAWALENKPRIPVVWIHGL WP_001058323.1 MULTISPECIES: hydrogenase-1 small chain [Proteobacteria] Length: 372\nNumber of predicted TMHs: 2\nExp number of AAs in TMHs: 37.2608499999999\nExp number, first 60 AAs: 19.52923\nTotal prob of N-in: 0.94351\nPOSSIBLE N-term signal sequence\ninside 1 19\nTMhelix 20 39\noutside 40 325\nTMhelix 326 348\ninside 349 372

6334 GCF_000812645.1_ASM81264v1 Escherichia coli Proteobacteria; Gammaproteobacteria; Enterobacterales; Enterobacteriaceae; Escherichia MNNEETFYQAMRRQGVTRRSFLKYCSLAATSLGLGAGMAPKIAWALENKPRIPVVWIHGL WP_001058323.1 MULTISPECIES: hydrogenase-1 small chain [Proteobacteria] Length: 372\nNumber of predicted TMHs: 2\nExp number of AAs in TMHs: 37.2608499999999\nExp number, first 60 AAs: 19.52923\nTotal prob of N-in: 0.94351\nPOSSIBLE N-term signal sequence\ninside 1 19\nTMhelix 20 39\noutside 40 325\nTMhelix 326 348\ninside 349 372

6335 GCF_000812305.1_ASM81230v1 Escherichia coli Proteobacteria; Gammaproteobacteria; Enterobacterales; Enterobacteriaceae; Escherichia MNNEETFYQAMRRQGVTRRSFLKYCSLAATSLGLGAGMAPKIAWALENKPRIPVVWIHGL WP_001295940.1 MULTISPECIES: hydrogenase [Enterobacteriaceae] Length: 372\nNumber of predicted TMHs: 2\nExp number of AAs in TMHs: 37.2608899999999\nExp number, first 60 AAs: 19.52923\nTotal prob of N-in: 0.94351\nPOSSIBLE N-term signal sequence\ninside 1 19\nTMhelix 20 39\noutside 40 325\nTMhelix 326 348\ninside 349 372

6336 GCF_000817355.1_SCB12 Escherichia coli Proteobacteria; Gammaproteobacteria; Enterobacterales; Enterobacteriaceae; Escherichia MTGDNTLIHSHGINRRDFMKLCAALAATMGLSSKAAAEMAESVTNPQRPPVIWIGAQECT WP_000145410.1 MULTISPECIES: hydrogenase-2 small chain [Proteobacteria] Length: 372\nNumber of predicted TMHs: 1\nExp number of AAs in TMHs: 23.72226\nExp number, first 60 AAs: 0.22038\nTotal prob of N-in: 0.12994\noutside 1 331\nTMhelix 332 354\ninside 355 372

6337 GCF_000819285.1_Escherichia_coli_CVM_N38796PS_v1.0 Escherichia coli Proteobacteria; Gammaproteobacteria; Enterobacterales; Enterobacteriaceae; Escherichia MTGDNTLIHSHGINRRDFMKLCAALAATMGLSSKAAAEMAESVTNPQRPPVIWIGAQECT WP_000145410.1 MULTISPECIES: hydrogenase-2 small chain [Proteobacteria] Length: 372\nNumber of predicted TMHs: 1\nExp number of AAs in TMHs: 23.72226\nExp number, first 60 AAs: 0.22038\nTotal prob of N-in: 0.12994\noutside 1 331\nTMhelix 332 354\ninside 355 372

6338 GCF_000819325.1_Escherichia_coli_CVM_N38381PS_v1.0 Escherichia coli Proteobacteria; Gammaproteobacteria; Enterobacterales; Enterobacteriaceae; Escherichia MTGDNTLIHSHGINRRDFMKLCAALAATMGLSSKAAAEMAESVTNPQRPPVIWIGAQECT WP_000145410.1 MULTISPECIES: hydrogenase-2 small chain [Proteobacteria] Length: 372\nNumber of predicted TMHs: 1\nExp number of AAs in TMHs: 23.72226\nExp number, first 60 AAs: 0.22038\nTotal prob of N-in: 0.12994\noutside 1 331\nTMhelix 332 354\ninside 355 372

6339 GCF_000819265.1_Escherichia_coli_CVM_N38428PS_v1.0 Escherichia coli Proteobacteria; Gammaproteobacteria; Enterobacterales; Enterobacteriaceae; Escherichia MNNEETFYQAMRRQGVTRRSFLKYCSLAATSLGLGAGMAPKIAWALENKPRIPVVWIHGL WP_001058323.1 MULTISPECIES: hydrogenase-1 small chain [Proteobacteria] Length: 372\nNumber of predicted TMHs: 2\nExp number of AAs in TMHs: 37.2608499999999\nExp number, first 60 AAs: 19.52923\nTotal prob of N-in: 0.94351\nPOSSIBLE N-term signal sequence\ninside 1 19\nTMhelix 20 39\noutside 40 325\nTMhelix 326 348\ninside 349 372

6340 GCF_000819345.1_Escherichia_coli_CVM_N33591PS_v1.0 Escherichia coli Proteobacteria; Gammaproteobacteria; Enterobacterales; Enterobacteriaceae; Escherichia MTGDNTLIHSHGINRRDFMKLCAALAATMGLSSKAAAEMAESVTNPQRPPVIWIGAQECT WP_000145410.1 MULTISPECIES: hydrogenase-2 small chain [Proteobacteria] Length: 372\nNumber of predicted TMHs: 1\nExp number of AAs in TMHs: 23.72226\nExp number, first 60 AAs: 0.22038\nTotal prob of N-in: 0.12994\noutside 1 331\nTMhelix 332 354\ninside 355 372

6341 GCF_000819285.1_Escherichia_coli_CVM_N38796PS_v1.0 Escherichia coli Proteobacteria; Gammaproteobacteria; Enterobacterales; Enterobacteriaceae; Escherichia MNNEETFYQAMRRQGVTRRSFLKYCSLAATSLGLGAGMAPKIAWALENKPRIPVVWIHGL WP_001058323.1 MULTISPECIES: hydrogenase-1 small chain [Proteobacteria] Length: 372\nNumber of predicted TMHs:

2\nExp number of AAs in TMHs: 37.2608499999999\nExp number, first 60 AAs: 19.52923\nTotal prob of N-in: 0.94351\nPOSSIBLE N-term signal sequence\ninside 1 19\nTMhelix 20 39\noutside 40 325\nTMhelix 326 348\ninside 349 372

6342 GCF_000819365.1_Escherichia_coli_CVM_N33653PS_v1.0 Escherichia coli Proteobacteria; Gammaproteobacteria; Enterobacterales; Enterobacteriaceae; Escherichia MTGDNTLIHSHGINRRDFMKLCAALAATMGLSSKAAAEMAESVTNPQRPPVIWIGAQECT WP_000145410.1 MULTISPECIES: hydrogenase-2 small chain [Proteobacteria] Length: 372\nNumber of predicted TMHs: 1\nExp number of AAs in TMHs: 23.72226\nExp number, first 60 AAs: 0.22038\nTotal prob of N-in: 0.12994\noutside 1 331\nTMhelix 332 354\ninside 355 372

6343 GCF_000819365.1_Escherichia_coli_CVM_N33653PS_v1.0 Escherichia coli Proteobacteria; Gammaproteobacteria; Enterobacterales; Enterobacteriaceae; Escherichia MNNEETFYQAMRRQGVTRRSFLKYCSLAATSLGLGAGMAPKIAWALENKPRIPVVWIHGL WP_001058323.1 MULTISPECIES: hydrogenase-1 small chain [Proteobacteria] Length: 372\nNumber of predicted TMHs: 2\nExp number of AAs in TMHs: 37.2608499999999\nExp number, first 60 AAs: 19.52923\nTotal prob of N-in: 0.94351\nPOSSIBLE N-term signal sequence\ninside 1 19\nTMhelix 20 39\noutside 40 325\nTMhelix 326 348\ninside 349 372

6344 GCF_000935425.1_ASM93542v1 Escherichia coli Proteobacteria; Gammaproteobacteria; Enterobacterales; Enterobacteriaceae; Escherichia MTGDNTLIHSHGINRRDFMKLCAALAATMGLSSKAAAEMAESVTNPQRPPVIWIGAQECT WP_000145410.1 MULTISPECIES: hydrogenase-2 small chain [Proteobacteria] Length: 372\nNumber of predicted TMHs: 1\nExp number of AAs in TMHs: 23.72226\nExp number, first 60 AAs: 0.22038\nTotal prob of N-in: 0.12994\noutside 1 331\nTMhelix 332 354\ninside 355 372

6345 GCF_000935365.1_ASM93536v1 Escherichia coli Proteobacteria; Gammaproteobacteria; Enterobacterales; Enterobacteriaceae; Escherichia MNNEETFYQAMRRQGVTRRSFLKYCSLAATSLGLGAGMAPKIAWALENKPRIPVVWIHGL WP_001058323.1 MULTISPECIES: hydrogenase-1 small chain [Proteobacteria] Length: 372\nNumber of predicted TMHs: 2\nExp number of AAs in TMHs: 37.2608499999999\nExp number, first 60 AAs: 19.52923\nTotal prob of N-in: 0.94351\nPOSSIBLE N-term signal sequence\ninside 1 19\nTMhelix 20 39\noutside 40 325\nTMhelix 326 348\ninside 349 372

6346 GCF_000935425.1_ASM93542v1 Escherichia coli Proteobacteria; Gammaproteobacteria; Enterobacterales; Enterobacteriaceae; Escherichia MNNEETFYQAMRRQGVTRRSFLKYCSLAATSLGLGAGMAPKIAWALENKPRIPVVWIHGL WP_001058323.1 MULTISPECIES: hydrogenase-1 small chain [Proteobacteria] Length: 372\nNumber of predicted TMHs: 2\nExp number of AAs in TMHs: 37.2608499999999\nExp number, first 60 AAs: 19.52923\nTotal prob of N-in: 0.94351\nPOSSIBLE N-term signal sequence\ninside 1 19\nTMhelix 20 39\noutside 40 325\nTMhelix 326 348\ninside 349 372

6347 GCF_000931565.1_ASM93156v1 Escherichia coli Proteobacteria; Gammaproteobacteria; Enterobacterales; Enterobacteriaceae; Escherichia MNNEETFYQAMRRQGVTRRSFLKYCSLAATSLGLGAGMAPKIAWALENKPRIPVVWIHGL WP_001295940.1 MULTISPECIES: hydrogenase [Enterobacteriaceae] Length: 372\nNumber of predicted TMHs: 2\nExp number of AAs in TMHs: 37.2608899999999\nExp number, first 60 AAs: 19.52923\nTotal prob of N-in: 0.94351\nPOSSIBLE N-term signal sequence\ninside 1 19\nTMhelix 20 39\noutside 40 325\nTMhelix 326 348\ninside 349 372

6348 GCF_000935475.1_ASM93547v1 Escherichia coli Proteobacteria; Gammaproteobacteria; Enterobacterales; Enterobacteriaceae; Escherichia MTGDNTLIHSHGINRRDFMKLCAALAATMGLSSKAAAEMAESVTNPQRPPVIWIGAQECT WP_000145410.1 MULTISPECIES: hydrogenase-2 small chain [Proteobacteria] Length: 372\nNumber of predicted TMHs: 1\nExp number of AAs in TMHs: 23.72226\nExp number, first 60 AAs: 0.22038\nTotal prob of N-in: 0.12994\noutside 1 331\nTMhelix 332 354\ninside 355 372

6349 GCF_000935445.1_ASM93544v1 Escherichia coli Proteobacteria; Gammaproteobacteria; Enterobacterales; Enterobacteriaceae; Escherichia MTGDNTLIHSHGINRRDFMKLCAALAATMGLSSKAAAEMAESVTNPQRPPVIWIGAQECT WP_000145410.1 MULTISPECIES: hydrogenase-2 small chain [Proteobacteria] Length: 372\nNumber of predicted TMHs:

1\nExp number of AAs in TMHs: 23.72226\nExp number, first 60 AAs: 0.22038\nTotal prob of N-in: 0.12994\noutside 1 331\nTMhelix 332 354\ninside 355 372

6350 GCF_000935555.1_ASM93555v1 Escherichia coli Proteobacteria; Gammaproteobacteria; Enterobacterales; Enterobacteriaceae; Escherichia MTGDNTLIHSHGINRRDFMKLCAALAATMGLSSKAAAEMAESVTNPQRPPVIWIGAQECT WP_000145410.1 MULTISPECIES: hydrogenase-2 small chain [Proteobacteria] Length: 372\nNumber of predicted TMHs: 1\nExp number of AAs in TMHs: 23.72226\nExp number, first 60 AAs: 0.22038\nTotal prob of N-in: 0.12994\noutside 1 331\nTMhelix 332 354\ninside 355 372

6351 GCF_000939955.1_FHI79 Escherichia coli Proteobacteria; Gammaproteobacteria; Enterobacterales; Enterobacteriaceae; Escherichia MNNEETFYQAMRRQGVTRRSFLKYCSLAATSLGLGAGMAPKIAWALENKPRIPVVWIHGL WP_001058323.1 MULTISPECIES: hydrogenase-1 small chain [Proteobacteria] Length: 372\nNumber of predicted TMHs: 2\nExp number of AAs in TMHs: 37.2608499999999\nExp number, first 60 AAs: 19.52923\nTotal prob of N-in: 0.94351\nPOSSIBLE N-term signal sequence\ninside 1 19\nTMhelix 20 39\noutside 40 325\nTMhelix 326 348\ninside 349 372

6352 GCF_000941395.1_FHI8 Escherichia coli Proteobacteria; Gammaproteobacteria; Enterobacterales; Enterobacteriaceae; Escherichia MTGDNTLIHSHGINRRDFMKLCAALAATMGLSSKAAAEMAESVTNPQRPPVIWIGAQECT WP_000145410.1 MULTISPECIES: hydrogenase-2 small chain [Proteobacteria] Length: 372\nNumber of predicted TMHs: 1\nExp number of AAs in TMHs: 23.72226\nExp number, first 60 AAs: 0.22038\nTotal prob of N-in: 0.12994\noutside 1 331\nTMhelix 332 354\ninside 355 372

6353 GCF_000941535.1_2d8A_assembly Escherichia coli Proteobacteria; Gammaproteobacteria; Enterobacterales; Enterobacteriaceae; Escherichia MTGDNTLIHSHGINRRDFMKLCAALAATMGLSSKAAAEMVESVTNPQRPPVIWIGAQECT WP_000145423.1 hydrogenase 2 small subunit [Escherichia coli] Length: 372\nNumber of predicted TMHs: 1\nExp number of AAs in TMHs: 23.78201\nExp number, first 60 AAs: 0.28355\nTotal prob of N-in: 0.13257\noutside 1 331\nTMhelix 332 354\ninside 355 372

6354 GCF_000941535.1_2d8A_assembly Escherichia coli Proteobacteria; Gammaproteobacteria; Enterobacterales; Enterobacteriaceae; Escherichia MNNEETFYQAMRRQGVTRRSFLKYCSLAATSLGLGAGMAPKIAWALENKPRIPVVWIHGL WP_001058323.1 MULTISPECIES: hydrogenase-1 small chain [Proteobacteria] Length: 372\nNumber of predicted TMHs: 2\nExp number of AAs in TMHs: 37.2608499999999\nExp number, first 60 AAs: 19.52923\nTotal prob of N-in: 0.94351\nPOSSIBLE N-term signal sequence\ninside 1 19\nTMhelix 20 39\noutside 40 325\nTMhelix 326 348\ninside 349 372

6355 GCF_000941395.1_FHI8 Escherichia coli Proteobacteria; Gammaproteobacteria; Enterobacterales; Enterobacteriaceae; Escherichia MNNEETFYQAMRRQGVTRRSFLKYCSLAATSLGLGAGMAPKIAWALENKPRIPVVWIHGL WP_044781592.1 hydrogenase [Escherichia coli] Length: 372\nNumber of predicted TMHs: 2\nExp number of AAs in TMHs: 36.91858\nExp number, first 60 AAs: 19.53522\nTotal prob of N-in: 0.94324\nPOSSIBLE N-term signal sequence\ninside 1 19\nTMhelix 20 39\noutside 40 325\nTMhelix 326 348\ninside 349 372

6356 GCF_000939755.1_FHI43 Escherichia coli Proteobacteria; Gammaproteobacteria; Enterobacterales; Enterobacteriaceae; Escherichia MNNEETFYQAMRRQGVTRRSFLKYCSLAATSLGLGAGMAPKIAWALENKPRIPVVWIHGL WP_044723380.1 hydrogenase [Escherichia coli] Length: 372\nNumber of predicted TMHs: 2\nExp number of AAs in TMHs: 37.2612299999999\nExp number, first 60 AAs: 19.52923\nTotal prob of N-in: 0.94351\nPOSSIBLE N-term signal sequence\ninside 1 19\nTMhelix 20 39\noutside 40 325\nTMhelix 326 348\ninside 349 372

6357 GCF_000941895.1_FHI6 Escherichia coli Proteobacteria; Gammaproteobacteria; Enterobacterales; Enterobacteriaceae; Escherichia MTGDNTLIHSHGINRRDFMKLCAALAATMGLSSKAAAEMAESVTNPQRPPVIWIGAQECT WP_000145410.1 MULTISPECIES: hydrogenase-2 small chain [Proteobacteria] Length: 372\nNumber of predicted TMHs: 1\nExp number of AAs in TMHs: 23.72226\nExp number, first 60 AAs: 0.22038\nTotal prob of N-in: 0.12994\noutside 1 331\nTMhelix 332 354\ninside 355 372

6358 GCF_000948465.1_ASM94846v1 Escherichia coli Proteobacteria; Gammaproteobacteria; Enterobacterales; Enterobacteriaceae; Escherichia MTGDNTLIHSHGINRRDFMKLCAALAATMGLSSKAAAEMAESVTNPQRPPVIWIGAQECT WP_000145410.1 MULTISPECIES: hydrogenase-2 small chain [Proteobacteria] Length: 372\nNumber of predicted TMHs:

1\nExp number of AAs in TMHs: 23.72226\nExp number, first 60 AAs: 0.22038\nTotal prob of N-in: 0.12994\noutside 1 331\nTMhelix 332 354\ninside 355 372

6359 GCF_000948555.1_ASM94855v1 Escherichia coli Proteobacteria; Gammaproteobacteria; Enterobacterales; Enterobacteriaceae; Escherichia MTGDNTLIHSHGINRRDFMKLCAALAATMGLSSKAAAEMAESVTNPQRPPVIWIGAQECT WP_000145410.1 MULTISPECIES: hydrogenase-2 small chain [Proteobacteria] Length: 372\nNumber of predicted TMHs: 1\nExp number of AAs in TMHs: 23.72226\nExp number, first 60 AAs: 0.22038\nTotal prob of N-in: 0.12994\noutside 1 331\nTMhelix 332 354\ninside 355 372

6360 GCF_000948595.1_ASM94859v1 Escherichia coli Proteobacteria; Gammaproteobacteria; Enterobacterales; Enterobacteriaceae; Escherichia MTGDNTLIHSHGINRRDFMKLCAALAATMGLSSKAAAEMAESVTNPQRPPVIWIGAQECT WP_000145410.1 MULTISPECIES: hydrogenase-2 small chain [Proteobacteria] Length: 372\nNumber of predicted TMHs: 1\nExp number of AAs in TMHs: 23.72226\nExp number, first 60 AAs: 0.22038\nTotal prob of N-in: 0.12994\noutside 1 331\nTMhelix 332 354\ninside 355 372

6361 GCF_000947975.1_ASM94797v1 Escherichia coli Proteobacteria; Gammaproteobacteria; Enterobacterales; Enterobacteriaceae; Escherichia MNNEETFYQAMRRQGVTRRSFLKYCSLAATSLGLGAGMAPKIAWALENKPRIPVVWIHGL WP_001058323.1 MULTISPECIES: hydrogenase-1 small chain [Proteobacteria] Length: 372\nNumber of predicted TMHs: 2\nExp number of AAs in TMHs: 37.2608499999999\nExp number, first 60 AAs: 19.52923\nTotal prob of N-in: 0.94351\nPOSSIBLE N-term signal sequence\ninside 1 19\nTMhelix 20 39\noutside 40 325\nTMhelix 326 348\ninside 349 372

6362 GCF_000948465.1_ASM94846v1 Escherichia coli Proteobacteria; Gammaproteobacteria; Enterobacterales; Enterobacteriaceae; Escherichia MNNEETFYQAMRRQGVTRRSFLKYCSLAATSLGLGAGMAPKIAWALENKPRIPVVWIHGL WP_001058323.1 MULTISPECIES: hydrogenase-1 small chain [Proteobacteria] Length: 372\nNumber of predicted TMHs: 2\nExp number of AAs in TMHs: 37.2608499999999\nExp number, first 60 AAs: 19.52923\nTotal prob of N-in: 0.94351\nPOSSIBLE N-term signal sequence\ninside 1 19\nTMhelix 20 39\noutside 40 325\nTMhelix 326 348\ninside 349 372

6363 GCF_000947995.1_ASM94799v1 Escherichia coli Proteobacteria; Gammaproteobacteria; Enterobacterales; Enterobacteriaceae; Escherichia MNNEETFYQAMRRQGVTRRSFLKYCSLAATSLGLGAGMAPKIAWALENKPRIPVVWIHGL WP_001058323.1 MULTISPECIES: hydrogenase-1 small chain [Proteobacteria] Length: 372\nNumber of predicted TMHs: 2\nExp number of AAs in TMHs: 37.2608499999999\nExp number, first 60 AAs: 19.52923\nTotal prob of N-in: 0.94351\nPOSSIBLE N-term signal sequence\ninside 1 19\nTMhelix 20 39\noutside 40 325\nTMhelix 326 348\ninside 349 372

6364 GCF_000948625.1_ASM94862v1 Escherichia coli Proteobacteria; Gammaproteobacteria; Enterobacterales; Enterobacteriaceae; Escherichia MTGDNTLIHSHGINRRDFMKLCAALAATMGLSSKAAAEMAESVTNPQRPPVIWIGAQECT WP_000145410.1 MULTISPECIES: hydrogenase-2 small chain [Proteobacteria] Length: 372\nNumber of predicted TMHs: 1\nExp number of AAs in TMHs: 23.72226\nExp number, first 60 AAs: 0.22038\nTotal prob of N-in: 0.12994\noutside 1 331\nTMhelix 332 354\ninside 355 372

6365 GCF_000948825.1_ASM94882v1 Escherichia coli Proteobacteria; Gammaproteobacteria; Enterobacterales; Enterobacteriaceae; Escherichia MTGDNTLIHSHGINRRDFMKLCAALAATMGLSSKAAAEMAESVTNPQRPPVIWIGAQECT WP_000145410.1 MULTISPECIES: hydrogenase-2 small chain [Proteobacteria] Length: 372\nNumber of predicted TMHs: 1\nExp number of AAs in TMHs: 23.72226\nExp number, first 60 AAs: 0.22038\nTotal prob of N-in: 0.12994\noutside 1 331\nTMhelix 332 354\ninside 355 372

6366 GCF_000948885.1_ASM94888v1 Escherichia coli Proteobacteria; Gammaproteobacteria; Enterobacterales; Enterobacteriaceae; Escherichia MTGDNTLIHSHGINRRDFMKLCAALAATMGLSSKAAAEMAESVTNPQRPPVIWIGAQECT WP_000145410.1 MULTISPECIES: hydrogenase-2 small chain [Proteobacteria] Length: 372\nNumber of predicted TMHs: 1\nExp number of AAs in TMHs: 23.72226\nExp number, first 60 AAs: 0.22038\nTotal prob of N-in: 0.12994\noutside 1 331\nTMhelix 332 354\ninside 355 372

6367 GCF_000948905.1_ASM94890v1 Escherichia coli Proteobacteria; Gammaproteobacteria; Enterobacterales; Enterobacteriaceae; Escherichia
MTGDNTLIHSHGINRRDFMKLCAALAATMGLSSKAAAEMAESVTNPQRPPVIWIGAQECT WP_000145410.1
MULTISPECIES: hydrogenase-2 small chain [Proteobacteria] Length: 372\nNumber of predicted TMHs: 1\nExp number of AAs in TMHs: 23.72226\nExp number, first 60 AAs: 0.22038\nTotal prob of N-in: 0.12994\noutside 1 331\nTMhelix 332 354\ninside 355 372

6368 GCF_000948955.1_ASM94895v1 Escherichia coli Proteobacteria; Gammaproteobacteria; Enterobacterales; Enterobacteriaceae; Escherichia
MTGDNTLIHSHGINRRDFMKLCAALAATMGLSSKAAAEMAESVTNPQRPPVIWIGAQECT WP_000145410.1
MULTISPECIES: hydrogenase-2 small chain [Proteobacteria] Length: 372\nNumber of predicted TMHs: 1\nExp number of AAs in TMHs: 23.72226\nExp number, first 60 AAs: 0.22038\nTotal prob of N-in: 0.12994\noutside 1 331\nTMhelix 332 354\ninside 355 372

6369 GCF_000948835.1_ASM94883v1 Escherichia coli Proteobacteria; Gammaproteobacteria; Enterobacterales; Enterobacteriaceae; Escherichia
MNNEETFYQAMRRQGVTRRSFLKYCSLAATSLGLGAGMAPKIAWALENKPRIPVVWIHGL WP_001058323.1
MULTISPECIES: hydrogenase-1 small chain [Proteobacteria] Length: 372\nNumber of predicted TMHs: 2\nExp number of AAs in TMHs: 37.2608499999999\nExp number, first 60 AAs: 19.52923\nTotal prob of N-in: 0.94351\nPOSSIBLE N-term signal sequence\ninside 1 19\nTMhelix 20 39\noutside 40 325\nTMhelix 326 348\ninside 349 372

6370 GCF_000951745.1_ASM95174v1 Escherichia coli Proteobacteria; Gammaproteobacteria; Enterobacterales; Enterobacteriaceae; Escherichia
MTGDNTLIHSHGINRRDFMKLCAALAATMGLSSKAAAEMAESVTNPQRPPVIWIGAQECT WP_000145410.1
MULTISPECIES: hydrogenase-2 small chain [Proteobacteria] Length: 372\nNumber of predicted TMHs: 1\nExp number of AAs in TMHs: 23.72226\nExp number, first 60 AAs: 0.22038\nTotal prob of N-in: 0.12994\noutside 1 331\nTMhelix 332 354\ninside 355 372

6371 GCF_000951855.1_FHI14 Escherichia coli Proteobacteria; Gammaproteobacteria; Enterobacterales; Enterobacteriaceae; Escherichia MTGDNTLIHSHGINRRDFMKLCAALAATMGLSSKAAAEMAESVTNPQRPPVIWIGAQECT
WP_000145410.1 MULTISPECIES: hydrogenase-2 small chain [Proteobacteria] Length: 372\nNumber of predicted TMHs: 1\nExp number of AAs in TMHs: 23.72226\nExp number, first 60 AAs: 0.22038\nTotal prob of N-in: 0.12994\noutside 1 331\nTMhelix 332 354\ninside 355 372

6372 GCF_000952305.1_ASM95230v1 Escherichia coli Proteobacteria; Gammaproteobacteria; Enterobacterales; Enterobacteriaceae; Escherichia
MNNEETFYQAMRRQGVTRRSFLKYCSLAATSLGLGAGMAPKIAWALENKPRIPVVWIHGL WP_001295940.1
MULTISPECIES: hydrogenase [Enterobacteriaceae] Length: 372\nNumber of predicted TMHs: 2\nExp number of AAs in TMHs: 37.2608899999999\nExp number, first 60 AAs: 19.52923\nTotal prob of N-in: 0.94351\nPOSSIBLE N-term signal sequence\ninside 1 19\nTMhelix 20 39\noutside 40 325\nTMhelix 326 348\ninside 349 372

6373 GCF_000948825.1_ASM94882v1 Escherichia coli Proteobacteria; Gammaproteobacteria; Enterobacterales; Enterobacteriaceae; Escherichia
MNNEETFYQAMRRQGVTRRSFLKYCSLAATSLGLGAGMAPKIAWALENKPRIPVVWIHGL WP_032204866.1
hydrogenase [Escherichia coli] Length: 372\nNumber of predicted TMHs: 2\nExp number of AAs in TMHs: 37.2621299999999\nExp number, first 60 AAs: 19.52923\nTotal prob of N-in: 0.94351\nPOSSIBLE N-term signal sequence\ninside 1 19\nTMhelix 20 39\noutside 40 325\nTMhelix 326 348\ninside 349 372

6374 GCF_000953515.1_EcHMS174Chr Escherichia coli Proteobacteria; Gammaproteobacteria; Enterobacterales; Enterobacteriaceae; Escherichia
MTGDNTLIHSHGINRRDFMKLCAALAATMGLSSKAAAEMAESVTNPQRPPVIWIGAQECT WP_000145410.1
MULTISPECIES: hydrogenase-2 small chain [Proteobacteria] Length: 372\nNumber of predicted TMHs: 1\nExp number of AAs in TMHs: 23.72226\nExp number, first 60 AAs: 0.22038\nTotal prob of N-in: 0.12994\noutside 1 331\nTMhelix 332 354\ninside 355 372

6375 GCF_000953515.1_EcHMS174Chr Escherichia coli Proteobacteria; Gammaproteobacteria; Enterobacterales; Enterobacteriaceae; Escherichia
MNNEETFYQAMRRQGVTRRSFLKYCSLAATSLGLGAGMAPKIAWALENKPRIPVVWIHGL WP_001058323.1
MULTISPECIES: hydrogenase-1 small chain [Proteobacteria] Length: 372\nNumber of predicted TMHs:

2\nExp number of AAs in TMHs: 37.2608499999999\nExp number, first 60 AAs:  19.52923\nTotal prob of N-in: 0.94351\nPOSSIBLE N-term signal sequence\ninside   1   19\nTMhelix   20   39\noutside   40   325\nTMhelix   326 348\ninside   349   372

6376 GCF_000948885.1_ASM94888v1 Escherichia coli Proteobacteria; Gammaproteobacteria; Enterobacterales; Enterobacteriaceae; Escherichia MNNEETFYQAMRRQGVTRRSFLKYCSLAATSLGLGAGMAPKIAWALENKPRIPVVWIHGL WP_001295940.1 MULTISPECIES: hydrogenase [Enterobacteriaceae] Length: 372\nNumber of predicted TMHs:  2\nExp number of AAs in TMHs: 37.2608899999999\nExp number, first 60 AAs:  19.52923\nTotal prob of N-in: 0.94351\nPOSSIBLE N-term signal sequence\ninside   1   19\nTMhelix   20   39\noutside   40   325\nTMhelix   326 348\ninside   349   372

6377 GCF_000948955.1_ASM94895v1 Escherichia coli Proteobacteria; Gammaproteobacteria; Enterobacterales; Enterobacteriaceae; Escherichia MNNEETFYQAMRRQGVTRRSFLKYCSLAATSLGLGAGMAPKIAWALENKPRIPVVWIHGL WP_001301613.1 hydrogenase [Escherichia coli] Length: 372\nNumber of predicted TMHs:  2\nExp number of AAs in TMHs: 37.46598\nExp number, first 60 AAs:  19.53139\nTotal prob of N-in:   0.94364\nPOSSIBLE N-term signal sequence\ninside   1   19\nTMhelix   20   39\noutside   40   325\nTMhelix   326   348\ninside   349   372

6378 GCF_000954035.1_ASM95403v1 Escherichia coli Proteobacteria; Gammaproteobacteria; Enterobacterales; Enterobacteriaceae; Escherichia MTGDNTLIHSHGINRRDFMKLCAALAATMGLSSKAAAEMAESVTNPQRPPVIWIGAQECT WP_000145410.1 MULTISPECIES: hydrogenase-2 small chain [Proteobacteria] Length: 372\nNumber of predicted TMHs: 1\nExp number of AAs in TMHs: 23.72226\nExp number, first 60 AAs:  0.22038\nTotal prob of N-in:   0.12994\noutside   1 331\nTMhelix   332   354\ninside   355   372

6379 GCF_000965715.1_ASM96571v1 Escherichia coli Proteobacteria; Gammaproteobacteria; Enterobacterales; Enterobacteriaceae; Escherichia MTGDNTLIHSHGINRRDFMKLCAALAATMGLSSKAAAEMAESVTNPQRPPVIWIGAQECT WP_000145410.1 MULTISPECIES: hydrogenase-2 small chain [Proteobacteria] Length: 372\nNumber of predicted TMHs: 1\nExp number of AAs in TMHs: 23.72226\nExp number, first 60 AAs:  0.22038\nTotal prob of N-in:   0.12994\noutside   1 331\nTMhelix   332   354\ninside   355   372

6380 GCF_000965555.1_ASM96555v1 Escherichia coli Proteobacteria; Gammaproteobacteria; Enterobacterales; Enterobacteriaceae; Escherichia MNNEETFYQAMRRQGVTRRSFLKYCSLAATSLGLGAGMAPKIAWALENKPRIPVVWIHGL WP_001058323.1 MULTISPECIES: hydrogenase-1 small chain [Proteobacteria] Length: 372\nNumber of predicted TMHs: 2\nExp number of AAs in TMHs: 37.2608499999999\nExp number, first 60 AAs:  19.52923\nTotal prob of N-in: 0.94351\nPOSSIBLE N-term signal sequence\ninside   1   19\nTMhelix   20   39\noutside   40   325\nTMhelix   326 348\ninside   349   372

6381 GCF_000966935.1_ASM96693v1 Escherichia coli Proteobacteria; Gammaproteobacteria; Enterobacterales; Enterobacteriaceae; Escherichia MTGDNTLIHSHGINRRDFMKLCAALAATMGLSSKAAAEMAESVTNPQRPPVIWIGAQECT WP_000145410.1 MULTISPECIES: hydrogenase-2 small chain [Proteobacteria] Length: 372\nNumber of predicted TMHs: 1\nExp number of AAs in TMHs: 23.72226\nExp number, first 60 AAs:  0.22038\nTotal prob of N-in:   0.12994\noutside   1 331\nTMhelix   332   354\ninside   355   372

6382 GCF_000954035.1_ASM95403v1 Escherichia coli Proteobacteria; Gammaproteobacteria; Enterobacterales; Enterobacteriaceae; Escherichia MNNEETFYQAMRRQGVTRRSFLKYCSLAATSLGLGAGMAPKIAWALENKPRIPVVWIHGL WP_001295940.1 MULTISPECIES: hydrogenase [Enterobacteriaceae] Length: 372\nNumber of predicted TMHs:  2\nExp number of AAs in TMHs: 37.2608899999999\nExp number, first 60 AAs:  19.52923\nTotal prob of N-in: 0.94351\nPOSSIBLE N-term signal sequence\ninside   1   19\nTMhelix   20   39\noutside   40   325\nTMhelix   326 348\ninside   349   372

6383 GCF_000965715.1_ASM96571v1 Escherichia coli Proteobacteria; Gammaproteobacteria; Enterobacterales; Enterobacteriaceae; Escherichia MNNEETFYQAMRRQGVTRRSFLKYCSLAATSLGLGAGMAPKIAWALENKPRIPVVWIHGL WP_001058323.1 MULTISPECIES: hydrogenase-1 small chain [Proteobacteria] Length: 372\nNumber of predicted TMHs: 2\nExp number of AAs in TMHs: 37.2608499999999\nExp number, first 60 AAs:  19.52923\nTotal prob of N-in:

0.94351\nPOSSIBLE N-term signal sequence\ninside 1 19\nTMhelix 20 39\noutside 40 325\nTMhelix 326 348\ninside 349 372

6384 GCF_000987525.1_ASM98752v1 Escherichia coli Proteobacteria; Gammaproteobacteria; Enterobacterales; Enterobacteriaceae; Escherichia MTGDNTLIHSHGINRRDFMKLCAALAATMGLSSKAAAEMAESVTNPQRPPVIWIGAQECT WP_000145410.1 MULTISPECIES: hydrogenase-2 small chain [Proteobacteria] Length: 372\nNumber of predicted TMHs: 1\nExp number of AAs in TMHs: 23.72226\nExp number, first 60 AAs: 0.22038\nTotal prob of N-in: 0.12994\noutside 1 331\nTMhelix 332 354\ninside 355 372

6385 GCF_000966935.1_ASM96693v1 Escherichia coli Proteobacteria; Gammaproteobacteria; Enterobacterales; Enterobacteriaceae; Escherichia MNNEETFYQAMRRQGVTRRSFLKYCSLAATSLGLGAGMAPKIAWALENKPRIPVVWIHGL WP_044723380.1 hydrogenase [Escherichia coli] Length: 372\nNumber of predicted TMHs: 2\nExp number of AAs in TMHs: 37.2612299999999\nExp number, first 60 AAs: 19.52923\nTotal prob of N-in: 0.94351\nPOSSIBLE N-term signal sequence\ninside 1 19\nTMhelix 20 39\noutside 40 325\nTMhelix 326 348\ninside 349 372

6386 GCF_000981065.1_ASM98106v1 Escherichia coli Proteobacteria; Gammaproteobacteria; Enterobacterales; Enterobacteriaceae; Escherichia MTGDNTLIHSHGINRRDFMKLCAALAATMGLSSKAAAEMAESVTNPQRPPVIWIGAQECT WP_000145410.1 MULTISPECIES: hydrogenase-2 small chain [Proteobacteria] Length: 372\nNumber of predicted TMHs: 1\nExp number of AAs in TMHs: 23.72226\nExp number, first 60 AAs: 0.22038\nTotal prob of N-in: 0.12994\noutside 1 331\nTMhelix 332 354\ninside 355 372

6387 GCF_000986635.1_ASM98663v1 Escherichia coli Proteobacteria; Gammaproteobacteria; Enterobacterales; Enterobacteriaceae; Escherichia MTGDNTLIHSHGINRRDFMKLCAALAATMGLSSKAAAEMAESVTNPQRPPVIWIGAQECT WP_000145410.1 MULTISPECIES: hydrogenase-2 small chain [Proteobacteria] Length: 372\nNumber of predicted TMHs: 1\nExp number of AAs in TMHs: 23.72226\nExp number, first 60 AAs: 0.22038\nTotal prob of N-in: 0.12994\noutside 1 331\nTMhelix 332 354\ninside 355 372

6388 GCF_000987555.1_ASM98755v1 Escherichia coli Proteobacteria; Gammaproteobacteria; Enterobacterales; Enterobacteriaceae; Escherichia MTGDNTLIHSHGINRRDFMKLCAALAATMGLSSKAAAEMAESVTNPQRPPVIWIGAQECT WP_000145410.1 MULTISPECIES: hydrogenase-2 small chain [Proteobacteria] Length: 372\nNumber of predicted TMHs: 1\nExp number of AAs in TMHs: 23.72226\nExp number, first 60 AAs: 0.22038\nTotal prob of N-in: 0.12994\noutside 1 331\nTMhelix 332 354\ninside 355 372

6389 GCF_000988355.1_ASM98835v1 Escherichia coli Proteobacteria; Gammaproteobacteria; Enterobacterales; Enterobacteriaceae; Escherichia MTGDNTLIHSHGINRRDFMKLCAALAATMGLSSKAAAEMAESVTNPQRPPVIWIGAQECT WP_000145410.1 MULTISPECIES: hydrogenase-2 small chain [Proteobacteria] Length: 372\nNumber of predicted TMHs: 1\nExp number of AAs in TMHs: 23.72226\nExp number, first 60 AAs: 0.22038\nTotal prob of N-in: 0.12994\noutside 1 331\nTMhelix 332 354\ninside 355 372

6390 GCF_000987555.1_ASM98755v1 Escherichia coli Proteobacteria; Gammaproteobacteria; Enterobacterales; Enterobacteriaceae; Escherichia MNNEETFYQAMRRQGVTRRSFLKYCSLAATSLGLGAGMAPKIAWALENKPRIPVVWIHGL WP_001058320.1 hydrogenase [Escherichia coli] Length: 372\nNumber of predicted TMHs: 2\nExp number of AAs in TMHs: 37.26082\nExp number, first 60 AAs: 19.52923\nTotal prob of N-in: 0.94351\nPOSSIBLE N-term signal sequence\ninside 1 19\nTMhelix 20 39\noutside 40 325\nTMhelix 326 348\ninside 349 372

6391 GCF_000988355.1_ASM98835v1 Escherichia coli Proteobacteria; Gammaproteobacteria; Enterobacterales; Enterobacteriaceae; Escherichia MNNEETFYQAMRRQGVTRRSFLKYCSLAATSLGLGAGMAPKIAWALENKPRIPVVWIHGL WP_001058323.1 MULTISPECIES: hydrogenase-1 small chain [Proteobacteria] Length: 372\nNumber of predicted TMHs: 2\nExp number of AAs in TMHs: 37.2608499999999\nExp number, first 60 AAs: 19.52923\nTotal prob of N-in: 0.94351\nPOSSIBLE N-term signal sequence\ninside 1 19\nTMhelix 20 39\noutside 40 325\nTMhelix 326 348\ninside 349 372

6392 GCF_000988385.1_ASM98838v1 Escherichia coli Proteobacteria; Gammaproteobacteria; Enterobacterales; Enterobacteriaceae; Escherichia
MTGDNTLIHSHGINRRDFMKLCAALAATMGLSSKAAAEMAESVTNPQRPPVIWIGAQECT WP_000145410.1
MULTISPECIES: hydrogenase-2 small chain [Proteobacteria] Length: 372\nNumber of predicted TMHs: 1\nExp number of AAs in TMHs: 23.72226\nExp number, first 60 AAs: 0.22038\nTotal prob of N-in: 0.12994\noutside 1 331\nTMhelix 332 354\ninside 355 372

6393 GCF_000988385.1_ASM98838v1 Escherichia coli Proteobacteria; Gammaproteobacteria; Enterobacterales; Enterobacteriaceae; Escherichia
MNNEETFYQAMRRQGVTRRSFLKYCSLAATSLGLGAGMAPKIAWALENKPRIPVVWIHGL WP_001058323.1
MULTISPECIES: hydrogenase-1 small chain [Proteobacteria] Length: 372\nNumber of predicted TMHs: 2\nExp number of AAs in TMHs: 37.2608499999999\nExp number, first 60 AAs: 19.52923\nTotal prob of N-in: 0.94351\nPOSSIBLE N-term signal sequence\ninside 1 19\nTMhelix 20 39\noutside 40 325\nTMhelix 326 348\ninside 349 372

6394 GCF_000986635.1_ASM98663v1 Escherichia coli Proteobacteria; Gammaproteobacteria; Enterobacterales; Enterobacteriaceae; Escherichia
MNNEETFYQAMRRQGVTRRSFLKYCSLAATSLGLGAGMAPKIAWALENKPRIPVVWIHGL WP_001301613.1
hydrogenase [Escherichia coli] Length: 372\nNumber of predicted TMHs: 2\nExp number of AAs in TMHs: 37.46598\nExp number, first 60 AAs: 19.53139\nTotal prob of N-in: 0.94364\nPOSSIBLE N-term signal sequence\ninside 1 19\nTMhelix 20 39\noutside 40 325\nTMhelix 326 348\ninside 349 372

6395 GCF_001005605.1_ASM100560v1 Escherichia coli Proteobacteria; Gammaproteobacteria; Enterobacterales; Enterobacteriaceae; Escherichia
MTGDNTLIHSHGINRRDFMKLCAALAATMGLSSKAAAEMAESVTNPQRPPVIWIGAQECT WP_000145410.1
MULTISPECIES: hydrogenase-2 small chain [Proteobacteria] Length: 372\nNumber of predicted TMHs: 1\nExp number of AAs in TMHs: 23.72226\nExp number, first 60 AAs: 0.22038\nTotal prob of N-in: 0.12994\noutside 1 331\nTMhelix 332 354\ninside 355 372

6396 GCF_001005685.1_ASM100568v1 Escherichia coli Proteobacteria; Gammaproteobacteria; Enterobacterales; Enterobacteriaceae; Escherichia
MTGDNTLIHSHGINRRDFMKLCAALAATMGLSSKAAAEMAESVTNPQRPPVIWIGAQECT WP_000145410.1
MULTISPECIES: hydrogenase-2 small chain [Proteobacteria] Length: 372\nNumber of predicted TMHs: 1\nExp number of AAs in TMHs: 23.72226\nExp number, first 60 AAs: 0.22038\nTotal prob of N-in: 0.12994\noutside 1 331\nTMhelix 332 354\ninside 355 372

6397 GCF_001007915.1_ASM100791v1 Escherichia coli Proteobacteria; Gammaproteobacteria; Enterobacterales; Enterobacteriaceae; Escherichia
MTGDNTLIHSHGINRRDFMKLCAALAATMGLSSKAAAEMAESVTNPQRPPVIWIGAQECT WP_000145410.1
MULTISPECIES: hydrogenase-2 small chain [Proteobacteria] Length: 372\nNumber of predicted TMHs: 1\nExp number of AAs in TMHs: 23.72226\nExp number, first 60 AAs: 0.22038\nTotal prob of N-in: 0.12994\noutside 1 331\nTMhelix 332 354\ninside 355 372

6398 GCF_001011995.1_CFSAN026838_01.0Escherichia coli Proteobacteria; Gammaproteobacteria; Enterobacterales; Enterobacteriaceae; Escherichia
MTGDNTLIHSHGINRRDFMKLCAALAATMGLSSKAAAEMAESVTNPQRPPVIWIGAQECT WP_000145410.1
MULTISPECIES: hydrogenase-2 small chain [Proteobacteria] Length: 372\nNumber of predicted TMHs: 1\nExp number of AAs in TMHs: 23.72226\nExp number, first 60 AAs: 0.22038\nTotal prob of N-in: 0.12994\noutside 1 331\nTMhelix 332 354\ninside 355 372

6399 GCF_001012015.1_CFSAN026799_01.0Escherichia coli Proteobacteria; Gammaproteobacteria; Enterobacterales; Enterobacteriaceae; Escherichia
MTGDNTLIHSHGINRRDFMKLCAALAATMGLSSKAAAEMAESVTNPQRPPVIWIGAQECT WP_000145410.1
MULTISPECIES: hydrogenase-2 small chain [Proteobacteria] Length: 372\nNumber of predicted TMHs: 1\nExp number of AAs in TMHs: 23.72226\nExp number, first 60 AAs: 0.22038\nTotal prob of N-in: 0.12994\noutside 1 331\nTMhelix 332 354\ninside 355 372

6400 GCF_001012025.1_CFSAN026796_01.0Escherichia coli Proteobacteria; Gammaproteobacteria; Enterobacterales; Enterobacteriaceae; Escherichia
MTGDNTLIHSHGINRRDFMKLCAALAATMGLSSKAAAEMAESVTNPQRPPVIWIGAQECT WP_000145410.1
MULTISPECIES: hydrogenase-2 small chain [Proteobacteria] Length: 372\nNumber of predicted TMHs:

1\nExp number of AAs in TMHs: 23.72226\nExp number, first 60 AAs: 0.22038\nTotal prob of N-in: 0.12994\noutside 1 331\nTMhelix 332 354\ninside 355 372

6401 GCF_001012345.1_CFSAN026809_01.0 Escherichia coli Proteobacteria; Gammaproteobacteria; Enterobacterales; Enterobacteriaceae; Escherichia MTGDNTLIHSHGINRRDFMKLCAALAATMGLSSKAAAEMAESVTNPQRPPVIWIGAQECT WP_000145410.1 MULTISPECIES: hydrogenase-2 small chain [Proteobacteria] Length: 372\nNumber of predicted TMHs: 1\nExp number of AAs in TMHs: 23.72226\nExp number, first 60 AAs: 0.22038\nTotal prob of N-in: 0.12994\noutside 1 331\nTMhelix 332 354\ninside 355 372

6402 GCF_001005685.1_ASM100568v1 Escherichia coli Proteobacteria; Gammaproteobacteria; Enterobacterales; Enterobacteriaceae; Escherichia MNNEETFYQAMRRQGVTRRSFLKYCSLAATSLGLGAGMAPKIAWALENKPRIPVVWIHGL WP_001058323.1 MULTISPECIES: hydrogenase-1 small chain [Proteobacteria] Length: 372\nNumber of predicted TMHs: 2\nExp number of AAs in TMHs: 37.2608499999999\nExp number, first 60 AAs: 19.52923\nTotal prob of N-in: 0.94351\nPOSSIBLE N-term signal sequence\ninside 1 19\nTMhelix 20 39\noutside 40 325\nTMhelix 326 348\ninside 349 372

6403 GCF_001007915.1_ASM100791v1 Escherichia coli Proteobacteria; Gammaproteobacteria; Enterobacterales; Enterobacteriaceae; Escherichia MNNEETFYQAMRRQGVTRRSFLKYCSLAATSLGLGAGMAPKIAWALENKPRIPVVWIHGL WP_001058323.1 MULTISPECIES: hydrogenase-1 small chain [Proteobacteria] Length: 372\nNumber of predicted TMHs: 2\nExp number of AAs in TMHs: 37.2608499999999\nExp number, first 60 AAs: 19.52923\nTotal prob of N-in: 0.94351\nPOSSIBLE N-term signal sequence\ninside 1 19\nTMhelix 20 39\noutside 40 325\nTMhelix 326 348\ninside 349 372

6404 GCF_001012025.1_CFSAN026796_01.0 Escherichia coli Proteobacteria; Gammaproteobacteria; Enterobacterales; Enterobacteriaceae; Escherichia MNNEETFYQAMRRQGVTRRSFLKYCSLAATSLGLGAGMAPKIAWALENKPRIPVVWIHGL WP_001058323.1 MULTISPECIES: hydrogenase-1 small chain [Proteobacteria] Length: 372\nNumber of predicted TMHs: 2\nExp number of AAs in TMHs: 37.2608499999999\nExp number, first 60 AAs: 19.52923\nTotal prob of N-in: 0.94351\nPOSSIBLE N-term signal sequence\ninside 1 19\nTMhelix 20 39\noutside 40 325\nTMhelix 326 348\ninside 349 372

6405 GCF_001011995.1_CFSAN026838_01.0 Escherichia coli Proteobacteria; Gammaproteobacteria; Enterobacterales; Enterobacteriaceae; Escherichia MNNEETFYQAMRRQGVTRRSFLKYCSLAATSLGLGAGMAPKIAWALENKPRIPVVWIHGL WP_001058323.1 MULTISPECIES: hydrogenase-1 small chain [Proteobacteria] Length: 372\nNumber of predicted TMHs: 2\nExp number of AAs in TMHs: 37.2608499999999\nExp number, first 60 AAs: 19.52923\nTotal prob of N-in: 0.94351\nPOSSIBLE N-term signal sequence\ninside 1 19\nTMhelix 20 39\noutside 40 325\nTMhelix 326 348\ninside 349 372

6406 GCF_001012175.1_CFSAN026787_02.0 Escherichia coli Proteobacteria; Gammaproteobacteria; Enterobacterales; Enterobacteriaceae; Escherichia MTGDNTLIHSHGINRRDFMKLCAALAATMGLSSKAAAEMAESVTNPQRPPVIWIGAQECT WP_000145410.1 MULTISPECIES: hydrogenase-2 small chain [Proteobacteria] Length: 372\nNumber of predicted TMHs: 1\nExp number of AAs in TMHs: 23.72226\nExp number, first 60 AAs: 0.22038\nTotal prob of N-in: 0.12994\noutside 1 331\nTMhelix 332 354\ninside 355 372

6407 GCF_001012195.1_CFSAN026788_02.0 Escherichia coli Proteobacteria; Gammaproteobacteria; Enterobacterales; Enterobacteriaceae; Escherichia MTGDNTLIHSHGINRRDFMKLCAALAATMGLSSKAAAEMAESVTNPQRPPVIWIGAQECT WP_000145410.1 MULTISPECIES: hydrogenase-2 small chain [Proteobacteria] Length: 372\nNumber of predicted TMHs: 1\nExp number of AAs in TMHs: 23.72226\nExp number, first 60 AAs: 0.22038\nTotal prob of N-in: 0.12994\noutside 1 331\nTMhelix 332 354\ninside 355 372

6408 GCF_001012255.1_CFSAN026790_01.0 Escherichia coli Proteobacteria; Gammaproteobacteria; Enterobacterales; Enterobacteriaceae; Escherichia MTGDNTLIHSHGINRRDFMKLCAALAATMGLSSKAAAEMAESVTNPQRPPVIWIGAQECT WP_000145410.1 MULTISPECIES: hydrogenase-2 small chain [Proteobacteria] Length: 372\nNumber of predicted TMHs:

1\nExp number of AAs in TMHs: 23.72226\nExp number, first 60 AAs: 0.22038\nTotal prob of N-in: 0.12994\noutside 1 331\nTMhelix 332 354\ninside 355 372

6409 GCF_001012315.1_CFSAN026801_01.0 Escherichia coli Proteobacteria; Gammaproteobacteria; Enterobacterales; Enterobacteriaceae; Escherichia MNNEETFYQAMRRQGVTRRSFLKYCSLAATSLGLGAGMAPKIAWALENKPRIPVVWIHGL WP_001058323.1 MULTISPECIES: hydrogenase-1 small chain [Proteobacteria] Length: 372\nNumber of predicted TMHs: 2\nExp number of AAs in TMHs: 37.2608499999999\nExp number, first 60 AAs: 19.52923\nTotal prob of N-in: 0.94351\nPOSSIBLE N-term signal sequence\ninside 1 19\nTMhelix 20 39\noutside 40 325\nTMhelix 326 348\ninside 349 372

6410 GCF_001012445.1_CFSAN026826_02.0 Escherichia coli Proteobacteria; Gammaproteobacteria; Enterobacterales; Enterobacteriaceae; Escherichia MNNEETFYQAMRRQGVTRRSFLKYCSLAATSLGLGAGMAPKIAWALENKPRIPVVWIHGL WP_001295940.1 MULTISPECIES: hydrogenase [Enterobacteriaceae] Length: 372\nNumber of predicted TMHs: 2\nExp number of AAs in TMHs: 37.2608899999999\nExp number, first 60 AAs: 19.52923\nTotal prob of N-in: 0.94351\nPOSSIBLE N-term signal sequence\ninside 1 19\nTMhelix 20 39\noutside 40 325\nTMhelix 326 348\ninside 349 372

6411 GCF_001012405.1_CFSAN026815_01.0 Escherichia coli Proteobacteria; Gammaproteobacteria; Enterobacterales; Enterobacteriaceae; Escherichia MNNEETFYQAMRRQGVTRRSFLKYCSLAATSLGLGAGIAPKIAWALENKPRIPVVWIHGL WP_023982790.1 hydrogenase [Escherichia coli] Length: 372\nNumber of predicted TMHs: 2\nExp number of AAs in TMHs: 37.8310099999999\nExp number, first 60 AAs: 20.15722\nTotal prob of N-in: 0.95896\nPOSSIBLE N-term signal sequence\ninside 1 20\nTMhelix 21 43\noutside 44 325\nTMhelix 326 348\ninside 349 372

6412 GCF_001012255.1_CFSAN026790_01.0 Escherichia coli Proteobacteria; Gammaproteobacteria; Enterobacterales; Enterobacteriaceae; Escherichia MNNEETFYQAMRRQGVTRRSFLKYCSLAATSLGLGAGMAPKIAWALENKPRIPVVWIHGL WP_001058323.1 MULTISPECIES: hydrogenase-1 small chain [Proteobacteria] Length: 372\nNumber of predicted TMHs: 2\nExp number of AAs in TMHs: 37.2608499999999\nExp number, first 60 AAs: 19.52923\nTotal prob of N-in: 0.94351\nPOSSIBLE N-term signal sequence\ninside 1 19\nTMhelix 20 39\noutside 40 325\nTMhelix 326 348\ninside 349 372

6413 GCF_001012395.1_CFSAN026813_01.0 Escherichia coli Proteobacteria; Gammaproteobacteria; Enterobacterales; Enterobacteriaceae; Escherichia MNNEETFYQAMRRQGVTRRSFLKYCSLAATSLGLGAGMAPKIAWALENKPRIPVVWIHGL WP_001058323.1 MULTISPECIES: hydrogenase-1 small chain [Proteobacteria] Length: 372\nNumber of predicted TMHs: 2\nExp number of AAs in TMHs: 37.2608499999999\nExp number, first 60 AAs: 19.52923\nTotal prob of N-in: 0.94351\nPOSSIBLE N-term signal sequence\ninside 1 19\nTMhelix 20 39\noutside 40 325\nTMhelix 326 348\ninside 349 372

6414 GCF_001012275.1_CFSAN026791_01.0 Escherichia coli Proteobacteria; Gammaproteobacteria; Enterobacterales; Enterobacteriaceae; Escherichia MNNEETFYQAMRRQGVTRRSFLKYCSLAATSLGLGAGMAPKIAWALENKPRIPVVWIHGL WP_001058323.1 MULTISPECIES: hydrogenase-1 small chain [Proteobacteria] Length: 372\nNumber of predicted TMHs: 2\nExp number of AAs in TMHs: 37.2608499999999\nExp number, first 60 AAs: 19.52923\nTotal prob of N-in: 0.94351\nPOSSIBLE N-term signal sequence\ninside 1 19\nTMhelix 20 39\noutside 40 325\nTMhelix 326 348\ninside 349 372

6415 GCF_001012495.1_CFSAN026835_01.0 Escherichia coli Proteobacteria; Gammaproteobacteria; Enterobacterales; Enterobacteriaceae; Escherichia MTGDNTLIHSHGINRRDFMKLCAALAATMGLSSKAAAEMAESVTNPQRPPVIWIGAQECT WP_000145410.1 MULTISPECIES: hydrogenase-2 small chain [Proteobacteria] Length: 372\nNumber of predicted TMHs: 1\nExp number of AAs in TMHs: 23.72226\nExp number, first 60 AAs: 0.22038\nTotal prob of N-in: 0.12994\noutside 1 331\nTMhelix 332 354\ninside 355 372

6416 GCF_001021595.1_ASM102159v1 Escherichia coli Proteobacteria; Gammaproteobacteria; Enterobacterales; Enterobacteriaceae; Escherichia MTGDNTLIHSHGINRRDFMKLCAALAATMGLSSKAAAEMAESVTNPQRPPVIWIGAQECT WP_000145410.1 MULTISPECIES: hydrogenase-2 small chain [Proteobacteria] Length: 372\nNumber of predicted TMHs:

1\nExp number of AAs in TMHs: 23.72226\nExp number, first 60 AAs: 0.22038\nTotal prob of N-in: 0.12994\noutside 1 331\nTMhelix 332 354\ninside 355 372

6417 GCF_001029415.1_ASM102941v1 Escherichia coli Proteobacteria; Gammaproteobacteria; Enterobacterales; Enterobacteriaceae; Escherichia
MTGDNTLIHSHGINRRDFMKLCAALAATMGLSSKAAAEMVESVTNPQRPPVIWIGAQECT WP_000145423.1
hydrogenase 2 small subunit [Escherichia coli] Length: 372\nNumber of predicted TMHs: 1\nExp number of AAs in TMHs: 23.78201\nExp number, first 60 AAs: 0.28355\nTotal prob of N-in: 0.13257\noutside 1 331\nTMhelix 332 354\ninside 355 372

6418 GCF_001014795.1_ASM101479v1 Escherichia coli Proteobacteria; Gammaproteobacteria; Enterobacterales; Enterobacteriaceae; Escherichia
MNNEETFYQAMRRQGVTRRSFLKYCSLAATSLGLGAGMAPKIAWALENKPRIPVVWIHGL WP_001058323.1
MULTISPECIES: hydrogenase-1 small chain [Proteobacteria] Length: 372\nNumber of predicted TMHs: 2\nExp number of AAs in TMHs: 37.2608499999999\nExp number, first 60 AAs: 19.52923\nTotal prob of N-in: 0.94351\nPOSSIBLE N-term signal sequence\ninside 1 19\nTMhelix 20 39\noutside 40 325\nTMhelix 326 348\ninside 349 372

6419 GCF_001030285.1_Esch_coli_BWH59_V1 Escherichia coli Proteobacteria; Gammaproteobacteria; Enterobacterales; Enterobacteriaceae; Escherichia
MTGDNTLIHSHGINRRDFMKLCAALAATMGLSSKAAAEMAESVTNPQRPPVIWIGAQECT WP_000145410.1
MULTISPECIES: hydrogenase-2 small chain [Proteobacteria] Length: 372\nNumber of predicted TMHs: 1\nExp number of AAs in TMHs: 23.72226\nExp number, first 60 AAs: 0.22038\nTotal prob of N-in: 0.12994\noutside 1 331\nTMhelix 332 354\ninside 355 372

6420 GCF_001030325.1_Esch_coli_BWH61_V1 Escherichia coli Proteobacteria; Gammaproteobacteria; Enterobacterales; Enterobacteriaceae; Escherichia
MTGDNTLIHSHGINRRDFMKLCAALAATMGLSSKAAAEMVESVTNPQRPPVIWIGAQECT WP_000145423.1
hydrogenase 2 small subunit [Escherichia coli] Length: 372\nNumber of predicted TMHs: 1\nExp number of AAs in TMHs: 23.78201\nExp number, first 60 AAs: 0.28355\nTotal prob of N-in: 0.13257\noutside 1 331\nTMhelix 332 354\ninside 355 372

6421 GCF_001030285.1_Esch_coli_BWH59_V1 Escherichia coli Proteobacteria; Gammaproteobacteria; Enterobacterales; Enterobacteriaceae; Escherichia
MNNEETFYQAMRRQGVTRRSFLKYCSLAATSLGLGAGMAPKIAWALENKPRIPVVWIHGL WP_001058323.1
MULTISPECIES: hydrogenase-1 small chain [Proteobacteria] Length: 372\nNumber of predicted TMHs: 2\nExp number of AAs in TMHs: 37.2608499999999\nExp number, first 60 AAs: 19.52923\nTotal prob of N-in: 0.94351\nPOSSIBLE N-term signal sequence\ninside 1 19\nTMhelix 20 39\noutside 40 325\nTMhelix 326 348\ninside 349 372

6422 GCF_001014795.1_ASM101479v1 Escherichia coli Proteobacteria; Gammaproteobacteria; Enterobacterales; Enterobacteriaceae; Escherichia
MTGDNTLIHSHGINRRDFMKLCAALATTMGLSSKAAAEMAESVTNPQRPPVIWIGAQECT WP_023149629.1
MULTISPECIES: hydrogenase-2 small chain [Enterobacteriaceae] Length: 372\nNumber of predicted TMHs: 1\nExp number of AAs in TMHs: 23.63603\nExp number, first 60 AAs: 0.12971\nTotal prob of N-in: 0.12608\noutside 1 331\nTMhelix 332 354\ninside 355 372

6423 GCF_001030425.1_Esch_coli_MGH121_V1 Escherichia coli Proteobacteria; Gammaproteobacteria; Enterobacterales; Enterobacteriaceae; Escherichia
MTGDNTLIHSHGINRRDFMKLCAALAATMGLSSKAAAEMAESVTNPQRPPVIWIGAQECT WP_000145410.1
MULTISPECIES: hydrogenase-2 small chain [Proteobacteria] Length: 372\nNumber of predicted TMHs: 1\nExp number of AAs in TMHs: 23.72226\nExp number, first 60 AAs: 0.22038\nTotal prob of N-in: 0.12994\noutside 1 331\nTMhelix 332 354\ninside 355 372

6424 GCF_001030425.1_Esch_coli_MGH121_V1 Escherichia coli Proteobacteria; Gammaproteobacteria; Enterobacterales; Enterobacteriaceae; Escherichia
MNNEETFYQAMRRQGVTRRSFLKYCSLAATSLGLGAGMAPKIAWALENKPRIPVVWIHGL WP_001058323.1
MULTISPECIES: hydrogenase-1 small chain [Proteobacteria] Length: 372\nNumber of predicted TMHs: 2\nExp number of AAs in TMHs: 37.2608499999999\nExp number, first 60 AAs: 19.52923\nTotal prob of N-in: 0.94351\nPOSSIBLE N-term signal sequence\ninside 1 19\nTMhelix 20 39\noutside 40 325\nTMhelix 326 348\ninside 349 372

6425 GCF_001029415.1_ASM102941v1 Escherichia coli Proteobacteria; Gammaproteobacteria; Enterobacterales; Enterobacteriaceae; Escherichia
MNNEETFYQAMRRQGVTRRSFLKYCSLAATSLGLGAGMAPKIAWALENKPRIPVVWIHGL WP_001295940.1
MULTISPECIES: hydrogenase [Enterobacteriaceae] Length: 372\nNumber of predicted TMHs: 2\nExp number of AAs in TMHs: 37.2608899999999\nExp number, first 60 AAs: 19.52923\nTotal prob of N-in: 0.94351\nPOSSIBLE N-term signal sequence\ninside 1 19\nTMhelix 20 39\noutside 40 325\nTMhelix 326 348\ninside 349 372

6426 GCF_001012615.1_CFSAN026816_01.0Escherichia coli Proteobacteria; Gammaproteobacteria; Enterobacterales; Enterobacteriaceae; Escherichia
MTGDNTLIHSHGINRRDFMKLCAALAATMGLSSKAAAEMAESVTNPQRPPVIWIGAQECT WP_000145410.1
MULTISPECIES: hydrogenase-2 small chain [Proteobacteria] Length: 372\nNumber of predicted TMHs: 1\nExp number of AAs in TMHs: 23.72226\nExp number, first 60 AAs: 0.22038\nTotal prob of N-in: 0.12994\noutside 1 331\nTMhelix 332 354\ninside 355 372

6427 GCF_001030435.1_Esch_coli_MGH122_V1 Escherichia coli Proteobacteria; Gammaproteobacteria; Enterobacterales; Enterobacteriaceae; Escherichia
MTGDNTLIHSHGINRRDFMKLCAALAATMGLSSKAAAEMAESVTNPQRPPVIWIGAQECT WP_000145410.1
MULTISPECIES: hydrogenase-2 small chain [Proteobacteria] Length: 372\nNumber of predicted TMHs: 1\nExp number of AAs in TMHs: 23.72226\nExp number, first 60 AAs: 0.22038\nTotal prob of N-in: 0.12994\noutside 1 331\nTMhelix 332 354\ninside 355 372

6428 GCF_001030505.1_Esch_coli_BIDMC101_V1 Escherichia coli Proteobacteria; Gammaproteobacteria; Enterobacterales; Enterobacteriaceae; Escherichia
MTGDNTLIHSHGINRRDFMKLCAALAATMGLSSKAAAEMAESVTNPQRPPVIWIGAQECT WP_000145410.1
MULTISPECIES: hydrogenase-2 small chain [Proteobacteria] Length: 372\nNumber of predicted TMHs: 1\nExp number of AAs in TMHs: 23.72226\nExp number, first 60 AAs: 0.22038\nTotal prob of N-in: 0.12994\noutside 1 331\nTMhelix 332 354\ninside 355 372

6429 GCF_001030545.1_Esch_coli_BIDMC103_V1 Escherichia coli Proteobacteria; Gammaproteobacteria; Enterobacterales; Enterobacteriaceae; Escherichia
MNNEETFYQAMRRQGVTRRSFLKYCSLAATSLGLGAGMAPKIAWALENKPRIPVVWIHGL WP_001058323.1
MULTISPECIES: hydrogenase-1 small chain [Proteobacteria] Length: 372\nNumber of predicted TMHs: 2\nExp number of AAs in TMHs: 37.2608499999999\nExp number, first 60 AAs: 19.52923\nTotal prob of N-in: 0.94351\nPOSSIBLE N-term signal sequence\ninside 1 19\nTMhelix 20 39\noutside 40 325\nTMhelix 326 348\ninside 349 372

6430 GCF_001032735.1_Esch_coli_CHS199_V1 Escherichia coli Proteobacteria; Gammaproteobacteria; Enterobacterales; Enterobacteriaceae; Escherichia
MTGDNTLIHSHGINRRDFMKLCAALAATMGLSSKAAAEMAESVTNPQRPPVIWIGAQECT WP_000145410.1
MULTISPECIES: hydrogenase-2 small chain [Proteobacteria] Length: 372\nNumber of predicted TMHs: 1\nExp number of AAs in TMHs: 23.72226\nExp number, first 60 AAs: 0.22038\nTotal prob of N-in: 0.12994\noutside 1 331\nTMhelix 332 354\ninside 355 372

6431 GCF_001012575.1_CFSAN026845_01.0Escherichia coli Proteobacteria; Gammaproteobacteria; Enterobacterales; Enterobacteriaceae; Escherichia
MTGDNTLIHSHGINRRDFMKLCAALAATMGLSSKAAAEMAESVTNPQRPPVIWIGAQECT WP_000145410.1
MULTISPECIES: hydrogenase-2 small chain [Proteobacteria] Length: 372\nNumber of predicted TMHs: 1\nExp number of AAs in TMHs: 23.72226\nExp number, first 60 AAs: 0.22038\nTotal prob of N-in: 0.12994\noutside 1 331\nTMhelix 332 354\ninside 355 372

6432 GCF_001032735.1_Esch_coli_CHS199_V1 Escherichia coli Proteobacteria; Gammaproteobacteria; Enterobacterales; Enterobacteriaceae; Escherichia
MNNEETFYQAMRRQGVTRRSFLKYCSLAATSLGLGAGMAPKIAWALENKPRIPVVWIHGL WP_001058323.1
MULTISPECIES: hydrogenase-1 small chain [Proteobacteria] Length: 372\nNumber of predicted TMHs: 2\nExp number of AAs in TMHs: 37.2608499999999\nExp number, first 60 AAs: 19.52923\nTotal prob of N-in: 0.94351\nPOSSIBLE N-term signal sequence\ninside 1 19\nTMhelix 20 39\noutside 40 325\nTMhelix 326 348\ninside 349 372

6433 GCF_001012575.1_CFSAN026845_01.0Escherichia coli Proteobacteria; Gammaproteobacteria; Enterobacterales; Enterobacteriaceae; Escherichia

MNNEETFYQAMRRQGVTRRSFLKYCSLAATSLGLGAGMAPKIAWALENKPRIPVVWIHGL WP_001058323.1 MULTISPECIES: hydrogenase-1 small chain [Proteobacteria] Length: 372\nNumber of predicted TMHs: 2\nExp number of AAs in TMHs: 37.2608499999999\nExp number, first 60 AAs: 19.52923\nTotal prob of N-in: 0.94351\nPOSSIBLE N-term signal sequence\ninside 1 19\nTMhelix 20 39\noutside 40 325\nTMhelix 326 348\ninside 349 372

6434 GCF_001030585.1_Esch_coli_BIDMC105_V1 Escherichia coli Proteobacteria; Gammaproteobacteria; Enterobacterales; Enterobacteriaceae; Escherichia MNNEETFYQAMRRQGVTRRSFLKYCSLAATSLGLGAGMAPKIAWALENKPRIPVVWIHGL WP_001538532.1 hydrogenase-1 small chain [Escherichia coli] Length: 372\nNumber of predicted TMHs: 2\nExp number of AAs in TMHs: 37.27041\nExp number, first 60 AAs: 19.52927\nTotal prob of N-in: 0.94352\nPOSSIBLE N-term signal sequence\ninside 1 19\nTMhelix 20 39\noutside 40 325\nTMhelix 326 348\ninside 349 372

6435 GCF_001039145.1_ASM103914v1 Escherichia coli Proteobacteria; Gammaproteobacteria; Enterobacterales; Enterobacteriaceae; Escherichia MTGDNTLIHSHGINRRDFMKLCAALAATMGLSSKAAAEMAESVTNPQRPPVIWIGAQECT WP_000145410.1 MULTISPECIES: hydrogenase-2 small chain [Proteobacteria] Length: 372\nNumber of predicted TMHs: 1\nExp number of AAs in TMHs: 23.72226\nExp number, first 60 AAs: 0.22038\nTotal prob of N-in: 0.12994\noutside 1 331\nTMhelix 332 354\ninside 355 372

6436 GCF_001052705.1_ASM105270v1 Escherichia coli Proteobacteria; Gammaproteobacteria; Enterobacterales; Enterobacteriaceae; Escherichia MTGDNTLIHSHGINRRDFMKLCAALAATMGLSSKAAAEMAESVTNPQRPPVIWIGAQECT WP_000145410.1 MULTISPECIES: hydrogenase-2 small chain [Proteobacteria] Length: 372\nNumber of predicted TMHs: 1\nExp number of AAs in TMHs: 23.72226\nExp number, first 60 AAs: 0.22038\nTotal prob of N-in: 0.12994\noutside 1 331\nTMhelix 332 354\ninside 355 372

6437 GCF_001052705.1_ASM105270v1 Escherichia coli Proteobacteria; Gammaproteobacteria; Enterobacterales; Enterobacteriaceae; Escherichia MNNEETFYQAMRRQGVTRRSFLKYCSLAATSLGLGAGMAPKIAWALENKPRIPVVWIHGL WP_001058323.1 MULTISPECIES: hydrogenase-1 small chain [Proteobacteria] Length: 372\nNumber of predicted TMHs: 2\nExp number of AAs in TMHs: 37.2608499999999\nExp number, first 60 AAs: 19.52923\nTotal prob of N-in: 0.94351\nPOSSIBLE N-term signal sequence\ninside 1 19\nTMhelix 20 39\noutside 40 325\nTMhelix 326 348\ninside 349 372

6438 GCF_001052125.1_ASM105212v1 Escherichia coli Proteobacteria; Gammaproteobacteria; Enterobacterales; Enterobacteriaceae; Escherichia MNNEETFYQAMRRQGVTRRSFLKYCSLAATSLGLGAGMAPKIAWALENKPRIPVVWIHGL WP_001058323.1 MULTISPECIES: hydrogenase-1 small chain [Proteobacteria] Length: 372\nNumber of predicted TMHs: 2\nExp number of AAs in TMHs: 37.2608499999999\nExp number, first 60 AAs: 19.52923\nTotal prob of N-in: 0.94351\nPOSSIBLE N-term signal sequence\ninside 1 19\nTMhelix 20 39\noutside 40 325\nTMhelix 326 348\ninside 349 372

6439 GCF_001039145.1_ASM103914v1 Escherichia coli Proteobacteria; Gammaproteobacteria; Enterobacterales; Enterobacteriaceae; Escherichia MNNEETFYQAMRRQGVTRRSFLKYCSLAATSLGLGAGMAPKIAWALENKPRIPVVWIHGL WP_001058323.1 MULTISPECIES: hydrogenase-1 small chain [Proteobacteria] Length: 372\nNumber of predicted TMHs: 2\nExp number of AAs in TMHs: 37.2608499999999\nExp number, first 60 AAs: 19.52923\nTotal prob of N-in: 0.94351\nPOSSIBLE N-term signal sequence\ninside 1 19\nTMhelix 20 39\noutside 40 325\nTMhelix 326 348\ninside 349 372

6440 GCF_001039155.1_ASM103915v1 Escherichia coli Proteobacteria; Gammaproteobacteria; Enterobacterales; Enterobacteriaceae; Escherichia MNNEETFYQAMRRQGVTRRSFLKYCSLAATSLGLGAGMAPKIAWALENKPRIPVVWIHGL WP_001058323.1 MULTISPECIES: hydrogenase-1 small chain [Proteobacteria] Length: 372\nNumber of predicted TMHs: 2\nExp number of AAs in TMHs: 37.2608499999999\nExp number, first 60 AAs: 19.52923\nTotal prob of N-in: 0.94351\nPOSSIBLE N-term signal sequence\ninside 1 19\nTMhelix 20 39\noutside 40 325\nTMhelix 326 348\ninside 349 372

6441 GCF_001052855.1_ASM105285v1 Escherichia coli Proteobacteria; Gammaproteobacteria; Enterobacterales; Enterobacteriaceae; Escherichia

MTGDNTLIHSHGINRRDFMKLCAALAATMGLSSKAAAEMAESVTNPQRPPVIWIGAQECT WP_000145410.1 MULTISPECIES: hydrogenase-2 small chain [Proteobacteria] Length: 372\nNumber of predicted TMHs: 1\nExp number of AAs in TMHs: 23.72226\nExp number, first 60 AAs: 0.22038\nTotal prob of N-in: 0.12994\noutside 1 331\nTMhelix 332 354\ninside 355 372

6442 GCF_001053245.1_ASM105324v1 Escherichia coli Proteobacteria; Gammaproteobacteria; Enterobacterales; Enterobacteriaceae; Escherichia MNNEETFYQAMRRQGVTRRSFLKYCSLAATSLGLGAGMAPKIAWALENKPRIPVVWIHGL WP_001058323.1 MULTISPECIES: hydrogenase-1 small chain [Proteobacteria] Length: 372\nNumber of predicted TMHs: 2\nExp number of AAs in TMHs: 37.2608499999999\nExp number, first 60 AAs: 19.52923\nTotal prob of N-in: 0.94351\nPOSSIBLE N-term signal sequence\ninside 1 19\nTMhelix 20 39\noutside 40 325\nTMhelix 326 348\ninside 349 372

6443 GCF_001053065.1_ASM105306v1 Escherichia coli Proteobacteria; Gammaproteobacteria; Enterobacterales; Enterobacteriaceae; Escherichia MNNEETFYQAMRRQGVTRRSFLKYCSLAATSLGLGAGMAPKIAWALENKPRIPVVWIHGL WP_001058323.1 MULTISPECIES: hydrogenase-1 small chain [Proteobacteria] Length: 372\nNumber of predicted TMHs: 2\nExp number of AAs in TMHs: 37.2608499999999\nExp number, first 60 AAs: 19.52923\nTotal prob of N-in: 0.94351\nPOSSIBLE N-term signal sequence\ninside 1 19\nTMhelix 20 39\noutside 40 325\nTMhelix 326 348\ninside 349 372

6444 GCF_001054615.1_ASM105461v1 Escherichia coli Proteobacteria; Gammaproteobacteria; Enterobacterales; Enterobacteriaceae; Escherichia MTGDNTLIHSHGINRRDFMKLCAALAATMGLSSKAAAEMVESVTNPQRPPVIWIGAQECT WP_000145423.1 hydrogenase 2 small subunit [Escherichia coli] Length: 372\nNumber of predicted TMHs: 1\nExp number of AAs in TMHs: 23.78201\nExp number, first 60 AAs: 0.28355\nTotal prob of N-in: 0.13257\noutside 1 331\nTMhelix 332 354\ninside 355 372

6445 GCF_001054815.1_ASM105481v1 Escherichia coli Proteobacteria; Gammaproteobacteria; Enterobacterales; Enterobacteriaceae; Escherichia MTGDNTLIHSHGINRRDFMKLCAALAATMGLSSKAAAEMAESVTNPQRPPVIWIGAQECT WP_000145410.1 MULTISPECIES: hydrogenase-2 small chain [Proteobacteria] Length: 372\nNumber of predicted TMHs: 1\nExp number of AAs in TMHs: 23.72226\nExp number, first 60 AAs: 0.22038\nTotal prob of N-in: 0.12994\noutside 1 331\nTMhelix 332 354\ninside 355 372

6446 GCF_001055685.1_ASM105568v1 Escherichia coli Proteobacteria; Gammaproteobacteria; Enterobacterales; Enterobacteriaceae; Escherichia MTGDNTLIHSHGINRRDFMKLCAALAATMGLSSKAAAEMVESVTNPQRPPVIWIGAQECT WP_000145423.1 hydrogenase 2 small subunit [Escherichia coli] Length: 372\nNumber of predicted TMHs: 1\nExp number of AAs in TMHs: 23.78201\nExp number, first 60 AAs: 0.28355\nTotal prob of N-in: 0.13257\noutside 1 331\nTMhelix 332 354\ninside 355 372

6447 GCF_001054845.1_ASM105484v1 Escherichia coli Proteobacteria; Gammaproteobacteria; Enterobacterales; Enterobacteriaceae; Escherichia MTGDNTLIHSHGINRRDFMKLCAALAATMGLSSKAAAEMAESVTNPQRPPVIWIGAQECT WP_000145410.1 MULTISPECIES: hydrogenase-2 small chain [Proteobacteria] Length: 372\nNumber of predicted TMHs: 1\nExp number of AAs in TMHs: 23.72226\nExp number, first 60 AAs: 0.22038\nTotal prob of N-in: 0.12994\noutside 1 331\nTMhelix 332 354\ninside 355 372

6448 GCF_001056025.1_ASM105602v1 Escherichia coli Proteobacteria; Gammaproteobacteria; Enterobacterales; Enterobacteriaceae; Escherichia MTGDNTLIHSHGINRRDFMKLCAALAATMGLSSKAAAEMVESVTNPQRPPVIWIGAQECT WP_000145423.1 hydrogenase 2 small subunit [Escherichia coli] Length: 372\nNumber of predicted TMHs: 1\nExp number of AAs in TMHs: 23.78201\nExp number, first 60 AAs: 0.28355\nTotal prob of N-in: 0.13257\noutside 1 331\nTMhelix 332 354\ninside 355 372

6449 GCF_001054175.1_ASM105417v1 Escherichia coli Proteobacteria; Gammaproteobacteria; Enterobacterales; Enterobacteriaceae; Escherichia MNNEETFYQAMRRQGVTRRSFLKYCSLAATSLGLGAGMAPKIAWALENKPRIPVVWIHGL WP_001058323.1 MULTISPECIES: hydrogenase-1 small chain [Proteobacteria] Length: 372\nNumber of predicted TMHs: 2\nExp number of AAs in TMHs: 37.2608499999999\nExp number, first 60 AAs: 19.52923\nTotal prob of N-in:

0.94351\nPOSSIBLE N-term signal sequence\ninside 1 19\nTMhelix 20 39\noutside 40 325\nTMhelix 326 348\ninside 349 372

6450 GCF_001056215.1_ASM105621v1 Escherichia coli Proteobacteria; Gammaproteobacteria; Enterobacterales; Enterobacteriaceae; Escherichia MTGDNTLIHSHGINRRDFMKLCAALAATMGLSSKAAAEMAESVTNPQRPPVIWIGAQECT WP_000145410.1 MULTISPECIES: hydrogenase-2 small chain [Proteobacteria] Length: 372\nNumber of predicted TMHs: 1\nExp number of AAs in TMHs: 23.72226\nExp number, first 60 AAs: 0.22038\nTotal prob of N-in: 0.12994\noutside 1 331\nTMhelix 332 354\ninside 355 372

6451 GCF_001056845.1_ASM105684v1 Escherichia coli Proteobacteria; Gammaproteobacteria; Enterobacterales; Enterobacteriaceae; Escherichia MTGDNTLIHSHGINRRDFMKLCAALAATMGLSSKAAAEMAESVTNPQRPPVIWIGAQECT WP_000145410.1 MULTISPECIES: hydrogenase-2 small chain [Proteobacteria] Length: 372\nNumber of predicted TMHs: 1\nExp number of AAs in TMHs: 23.72226\nExp number, first 60 AAs: 0.22038\nTotal prob of N-in: 0.12994\noutside 1 331\nTMhelix 332 354\ninside 355 372

6452 GCF_001056895.1_ASM105689v1 Escherichia coli Proteobacteria; Gammaproteobacteria; Enterobacterales; Enterobacteriaceae; Escherichia MTGDNTLIHSHGINRRDFMKLCAALAATMGLSSKAAAEMAESVTNPQRPPVIWIGAQECT WP_000145410.1 MULTISPECIES: hydrogenase-2 small chain [Proteobacteria] Length: 372\nNumber of predicted TMHs: 1\nExp number of AAs in TMHs: 23.72226\nExp number, first 60 AAs: 0.22038\nTotal prob of N-in: 0.12994\noutside 1 331\nTMhelix 332 354\ninside 355 372

6453 GCF_001056815.1_ASM105681v1 Escherichia coli Proteobacteria; Gammaproteobacteria; Enterobacterales; Enterobacteriaceae; Escherichia MTGDNTLIHSHGINRRDFMKLCAALAATMGLSSKAAAEMAESVTNPQRPPVIWIGAQECT WP_000145410.1 MULTISPECIES: hydrogenase-2 small chain [Proteobacteria] Length: 372\nNumber of predicted TMHs: 1\nExp number of AAs in TMHs: 23.72226\nExp number, first 60 AAs: 0.22038\nTotal prob of N-in: 0.12994\noutside 1 331\nTMhelix 332 354\ninside 355 372

6454 GCF_001056895.1_ASM105689v1 Escherichia coli Proteobacteria; Gammaproteobacteria; Enterobacterales; Enterobacteriaceae; Escherichia MNNEETFYQAMRRQGVTRRSFLKYCSLAATSLGLGAGMAPKIAWALENKPRIPVVWIHGL WP_001058323.1 MULTISPECIES: hydrogenase-1 small chain [Proteobacteria] Length: 372\nNumber of predicted TMHs: 2\nExp number of AAs in TMHs: 37.2608499999999\nExp number, first 60 AAs: 19.52923\nTotal prob of N-in: 0.94351\nPOSSIBLE N-term signal sequence\ninside 1 19\nTMhelix 20 39\noutside 40 325\nTMhelix 326 348\ninside 349 372

6455 GCF_001056215.1_ASM105621v1 Escherichia coli Proteobacteria; Gammaproteobacteria; Enterobacterales; Enterobacteriaceae; Escherichia MNNEETFYQAMRRQGVTRRSFLKYCSLAATSLGLGAGMAPKIAWALENKPRIPVVWIHGL WP_001058323.1 MULTISPECIES: hydrogenase-1 small chain [Proteobacteria] Length: 372\nNumber of predicted TMHs: 2\nExp number of AAs in TMHs: 37.2608499999999\nExp number, first 60 AAs: 19.52923\nTotal prob of N-in: 0.94351\nPOSSIBLE N-term signal sequence\ninside 1 19\nTMhelix 20 39\noutside 40 325\nTMhelix 326 348\ninside 349 372

6456 GCF_001057225.1_ASM105722v1 Escherichia coli Proteobacteria; Gammaproteobacteria; Enterobacterales; Enterobacteriaceae; Escherichia MTGDNTLIHSHGINRRDFMKLCAALAATMGLSSKAAAEMVESVTNPQRPPVIWIGAQECT WP_000145423.1 hydrogenase 2 small subunit [Escherichia coli] Length: 372\nNumber of predicted TMHs: 1\nExp number of AAs in TMHs: 23.78201\nExp number, first 60 AAs: 0.28355\nTotal prob of N-in: 0.13257\noutside 1 331\nTMhelix 332 354\ninside 355 372

6457 GCF_001057065.1_ASM105706v1 Escherichia coli Proteobacteria; Gammaproteobacteria; Enterobacterales; Enterobacteriaceae; Escherichia MTGDNTLIHSHGINRRDFMKLCAALAATMGLSSKAAAEMAESVTNPQRPPVIWIGAQECT WP_000145410.1 MULTISPECIES: hydrogenase-2 small chain [Proteobacteria] Length: 372\nNumber of predicted TMHs: 1\nExp number of AAs in TMHs: 23.72226\nExp number, first 60 AAs: 0.22038\nTotal prob of N-in: 0.12994\noutside 1 331\nTMhelix 332 354\ninside 355 372

6458 GCF_001057445.1_ASM105744v1 Escherichia coli Proteobacteria; Gammaproteobacteria; Enterobacterales; Enterobacteriaceae; Escherichia
MTGDNTLIHSHGINRRDFMKLCAALAATMGLSSKAAAEMAESVTNPQRPPVIWIGAQECT WP_000145410.1
MULTISPECIES: hydrogenase-2 small chain [Proteobacteria] Length: 372\nNumber of predicted TMHs: 1\nExp number of AAs in TMHs: 23.72226\nExp number, first 60 AAs: 0.22038\nTotal prob of N-in: 0.12994\noutside 1 331\nTMhelix 332 354\ninside 355 372

6459 GCF_001057815.1_ASM105781v1 Escherichia coli Proteobacteria; Gammaproteobacteria; Enterobacterales; Enterobacteriaceae; Escherichia
MTGDNTLIHSHGINRRDFMKLCAALAATMGLSSKAAAEMVESVTNPQRPPVIWIGAQECT WP_000145423.1
hydrogenase 2 small subunit [Escherichia coli] Length: 372\nNumber of predicted TMHs: 1\nExp number of AAs in TMHs: 23.78201\nExp number, first 60 AAs: 0.28355\nTotal prob of N-in: 0.13257\noutside 1 331\nTMhelix 332 354\ninside 355 372

6460 GCF_001057975.1_ASM105797v1 Escherichia coli Proteobacteria; Gammaproteobacteria; Enterobacterales; Enterobacteriaceae; Escherichia
MTGDNTLIHSHGINRRDFMKLCAALAATMGLSSKAAAEMAESVTNPQRPPVIWIGAQECT WP_000145410.1
MULTISPECIES: hydrogenase-2 small chain [Proteobacteria] Length: 372\nNumber of predicted TMHs: 1\nExp number of AAs in TMHs: 23.72226\nExp number, first 60 AAs: 0.22038\nTotal prob of N-in: 0.12994\noutside 1 331\nTMhelix 332 354\ninside 355 372

6461 GCF_001058205.1_ASM105820v1 Escherichia coli Proteobacteria; Gammaproteobacteria; Enterobacterales; Enterobacteriaceae; Escherichia
MTGDNTLIHSHGINRRDFMKLCAALAATMGLSSKAAAEMVESVTNPQRPPVIWIGAQECT WP_000145423.1
hydrogenase 2 small subunit [Escherichia coli] Length: 372\nNumber of predicted TMHs: 1\nExp number of AAs in TMHs: 23.78201\nExp number, first 60 AAs: 0.28355\nTotal prob of N-in: 0.13257\noutside 1 331\nTMhelix 332 354\ninside 355 372

6462 GCF_001058255.1_ASM105825v1 Escherichia coli Proteobacteria; Gammaproteobacteria; Enterobacterales; Enterobacteriaceae; Escherichia
MTGDNTLIHSHGINRRDFMKLCAALAATMGLSSKAAAEMVESVTNPQRPPVIWIGAQECT WP_000145423.1
hydrogenase 2 small subunit [Escherichia coli] Length: 372\nNumber of predicted TMHs: 1\nExp number of AAs in TMHs: 23.78201\nExp number, first 60 AAs: 0.28355\nTotal prob of N-in: 0.13257\noutside 1 331\nTMhelix 332 354\ninside 355 372

6463 GCF_001057445.1_ASM105744v1 Escherichia coli Proteobacteria; Gammaproteobacteria; Enterobacterales; Enterobacteriaceae; Escherichia
MNNEETFYQAMRRQGVTRRSFLKYCSLAATSLGLGAGMAPKIAWALENKPRIPVVWIHGL WP_001058323.1
MULTISPECIES: hydrogenase-1 small chain [Proteobacteria] Length: 372\nNumber of predicted TMHs: 2\nExp number of AAs in TMHs: 37.2608499999999\nExp number, first 60 AAs: 19.52923\nTotal prob of N-in: 0.94351\nPOSSIBLE N-term signal sequence\ninside 1 19\nTMhelix 20 39\noutside 40 325\nTMhelix 326 348\ninside 349 372

6464 GCF_001058895.1_ASM105889v1 Escherichia coli Proteobacteria; Gammaproteobacteria; Enterobacterales; Enterobacteriaceae; Escherichia
MTGDNTLIHSHGINRRDFMKLCAALAATMGLSSKAAAEMAESVTNPQRPPVIWIGAQECT WP_000145410.1
MULTISPECIES: hydrogenase-2 small chain [Proteobacteria] Length: 372\nNumber of predicted TMHs: 1\nExp number of AAs in TMHs: 23.72226\nExp number, first 60 AAs: 0.22038\nTotal prob of N-in: 0.12994\noutside 1 331\nTMhelix 332 354\ninside 355 372

6465 GCF_001058795.1_ASM105879v1 Escherichia coli Proteobacteria; Gammaproteobacteria; Enterobacterales; Enterobacteriaceae; Escherichia
MNNEETFYQAMRRQGVTRRSFLKYCSLAATSLGLGAGMAPKIAWALENKPRIPVVWIHGL WP_001058323.1
MULTISPECIES: hydrogenase-1 small chain [Proteobacteria] Length: 372\nNumber of predicted TMHs: 2\nExp number of AAs in TMHs: 37.2608499999999\nExp number, first 60 AAs: 19.52923\nTotal prob of N-in: 0.94351\nPOSSIBLE N-term signal sequence\ninside 1 19\nTMhelix 20 39\noutside 40 325\nTMhelix 326 348\ninside 349 372

6466 GCF_001058895.1_ASM105889v1 Escherichia coli Proteobacteria; Gammaproteobacteria; Enterobacterales; Enterobacteriaceae; Escherichia
MNNEETFYQAMRRQGVTRRSFLKYCSLAATSLGLGAGMAPKIAWALENKPRIPVVWIHGL WP_001058323.1

MULTISPECIES: hydrogenase-1 small chain [Proteobacteria] Length: 372\nNumber of predicted TMHs: 2\nExp number of AAs in TMHs: 37.2608499999999\nExp number, first 60 AAs: 19.52923\nTotal prob of N-in: 0.94351\nPOSSIBLE N-term signal sequence\ninside 1 19\nTMhelix 20 39\noutside 40 325\nTMhelix 326 348\ninside 349 372

6467 GCF_001057815.1_ASM105781v1 Escherichia coli Proteobacteria; Gammaproteobacteria; Enterobacterales; Enterobacteriaceae; Escherichia MNNEETFYQAMRRQGVTRRSFLKYCSLAATSLGLGAGMAPKIAWALENKPRIPVVWIHGL WP_001295940.1 MULTISPECIES: hydrogenase [Enterobacteriaceae] Length: 372\nNumber of predicted TMHs: 2\nExp number of AAs in TMHs: 37.2608899999999\nExp number, first 60 AAs: 19.52923\nTotal prob of N-in: 0.94351\nPOSSIBLE N-term signal sequence\ninside 1 19\nTMhelix 20 39\noutside 40 325\nTMhelix 326 348\ninside 349 372

6468 GCF_001058205.1_ASM105820v1 Escherichia coli Proteobacteria; Gammaproteobacteria; Enterobacterales; Enterobacteriaceae; Escherichia MNNEETFYQAMRRQGVTRRSFLKYCSLAATSLGLGAGMAPKIAWALENKPRIPVVWIHGL WP_001295940.1 MULTISPECIES: hydrogenase [Enterobacteriaceae] Length: 372\nNumber of predicted TMHs: 2\nExp number of AAs in TMHs: 37.2608899999999\nExp number, first 60 AAs: 19.52923\nTotal prob of N-in: 0.94351\nPOSSIBLE N-term signal sequence\ninside 1 19\nTMhelix 20 39\noutside 40 325\nTMhelix 326 348\ninside 349 372

6469 GCF_001059575.1_ASM105957v1 Escherichia coli Proteobacteria; Gammaproteobacteria; Enterobacterales; Enterobacteriaceae; Escherichia MTGDNTLIHSHGINRRDFMKLCAALAATMGLSSKAAAEMAESVTNPQRPPVIWIGAQECT WP_000145410.1 MULTISPECIES: hydrogenase-2 small chain [Proteobacteria] Length: 372\nNumber of predicted TMHs: 1\nExp number of AAs in TMHs: 23.72226\nExp number, first 60 AAs: 0.22038\nTotal prob of N-in: 0.12994\noutside 1 331\nTMhelix 332 354\ninside 355 372

6470 GCF_001059455.1_ASM105945v1 Escherichia coli Proteobacteria; Gammaproteobacteria; Enterobacterales; Enterobacteriaceae; Escherichia MTGDNTLIHSHGINRRDFMKLCAALAATMGLSSKAAAEMAESVTNPQRPPVIWIGAQECT WP_000145410.1 MULTISPECIES: hydrogenase-2 small chain [Proteobacteria] Length: 372\nNumber of predicted TMHs: 1\nExp number of AAs in TMHs: 23.72226\nExp number, first 60 AAs: 0.22038\nTotal prob of N-in: 0.12994\noutside 1 331\nTMhelix 332 354\ninside 355 372

6471 GCF_001059715.1_ASM105971v1 Escherichia coli Proteobacteria; Gammaproteobacteria; Enterobacterales; Enterobacteriaceae; Escherichia MTGDNTLIHSHGINRRDFMKLCAALAATMGLSSKAAAEMAESVTNPQRPPVIWIGAQECT WP_000145410.1 MULTISPECIES: hydrogenase-2 small chain [Proteobacteria] Length: 372\nNumber of predicted TMHs: 1\nExp number of AAs in TMHs: 23.72226\nExp number, first 60 AAs: 0.22038\nTotal prob of N-in: 0.12994\noutside 1 331\nTMhelix 332 354\ninside 355 372

6472 GCF_001058375.1_ASM105837v1 Escherichia coli Proteobacteria; Gammaproteobacteria; Enterobacterales; Enterobacteriaceae; Escherichia MNNEETFYQAMRRQGVTRRSFLKYCSLAATSLGLGAGMAPKIAWALENKPRIPVVWIHGL WP_001295940.1 MULTISPECIES: hydrogenase [Enterobacteriaceae] Length: 372\nNumber of predicted TMHs: 2\nExp number of AAs in TMHs: 37.2608899999999\nExp number, first 60 AAs: 19.52923\nTotal prob of N-in: 0.94351\nPOSSIBLE N-term signal sequence\ninside 1 19\nTMhelix 20 39\noutside 40 325\nTMhelix 326 348\ninside 349 372

6473 GCF_001059715.1_ASM105971v1 Escherichia coli Proteobacteria; Gammaproteobacteria; Enterobacterales; Enterobacteriaceae; Escherichia MNNEETFYQAMRRQGVTRRSFLKYCSLAATSLGLGAGMAPKIAWALENKPRIPVVWIHGL WP_001058323.1 MULTISPECIES: hydrogenase-1 small chain [Proteobacteria] Length: 372\nNumber of predicted TMHs: 2\nExp number of AAs in TMHs: 37.2608499999999\nExp number, first 60 AAs: 19.52923\nTotal prob of N-in: 0.94351\nPOSSIBLE N-term signal sequence\ninside 1 19\nTMhelix 20 39\noutside 40 325\nTMhelix 326 348\ninside 349 372

6474 GCF_001059015.1_ASM105901v1 Escherichia coli Proteobacteria; Gammaproteobacteria; Enterobacterales; Enterobacteriaceae; Escherichia MNNEETFYQAMRRQGVTRRSFLKYCSLAATSLGLGAGMAPKIAWALENKPRIPVVWIHGL WP_001058323.1

MULTISPECIES: hydrogenase-1 small chain [Proteobacteria] Length: 372\nNumber of predicted TMHs: 2\nExp number of AAs in TMHs: 37.2608499999999\nExp number, first 60 AAs: 19.52923\nTotal prob of N-in: 0.94351\nPOSSIBLE N-term signal sequence\ninside 1 19\nTMhelix 20 39\noutside 40 325\nTMhelix 326 348\ninside 349 372

6475 GCF_001059735.1_ASM105973v1 Escherichia coli Proteobacteria; Gammaproteobacteria; Enterobacterales; Enterobacteriaceae; Escherichia
MTGDNTLIHSHGINRRDFMKLCAALAATMGLSSKAAAEMVESVTNPQRPPVIWIGAQECT WP_000145423.1
hydrogenase 2 small subunit [Escherichia coli] Length: 372\nNumber of predicted TMHs: 1\nExp number of AAs in TMHs: 23.78201\nExp number, first 60 AAs: 0.28355\nTotal prob of N-in: 0.13257\noutside 1 331\nTMhelix 332 354\ninside 355 372

6476 GCF_001063835.1_ASM106383v1 Escherichia coli Proteobacteria; Gammaproteobacteria; Enterobacterales; Enterobacteriaceae; Escherichia
MTGDNTLIHSHGINRRDFMKLCAALAATMGLSSKAAAEMAESVTNPQRPPVIWIGAQECT WP_000145410.1
MULTISPECIES: hydrogenase-2 small chain [Proteobacteria] Length: 372\nNumber of predicted TMHs: 1\nExp number of AAs in TMHs: 23.72226\nExp number, first 60 AAs: 0.22038\nTotal prob of N-in: 0.12994\noutside 1 331\nTMhelix 332 354\ninside 355 372

6477 GCF_001063835.1_ASM106383v1 Escherichia coli Proteobacteria; Gammaproteobacteria; Enterobacterales; Enterobacteriaceae; Escherichia
MNNEETFYQAMRRQGVTRRSFLKYCSLAATSLGLGAGMAPKIAWALENKPRIPVVWIHGL WP_001058323.1
MULTISPECIES: hydrogenase-1 small chain [Proteobacteria] Length: 372\nNumber of predicted TMHs: 2\nExp number of AAs in TMHs: 37.2608499999999\nExp number, first 60 AAs: 19.52923\nTotal prob of N-in: 0.94351\nPOSSIBLE N-term signal sequence\ninside 1 19\nTMhelix 20 39\noutside 40 325\nTMhelix 326 348\ninside 349 372

6478 GCF_001062415.1_ASM106241v1 Escherichia coli Proteobacteria; Gammaproteobacteria; Enterobacterales; Enterobacteriaceae; Escherichia
MNNEETFYQAMRRQGVTRRSFLKYCSLAATSLGLGAGMAPKIAWALENKPRIPVVWIHGL WP_001295940.1
MULTISPECIES: hydrogenase [Enterobacteriaceae] Length: 372\nNumber of predicted TMHs: 2\nExp number of AAs in TMHs: 37.2608899999999\nExp number, first 60 AAs: 19.52923\nTotal prob of N-in: 0.94351\nPOSSIBLE N-term signal sequence\ninside 1 19\nTMhelix 20 39\noutside 40 325\nTMhelix 326 348\ninside 349 372

6479 GCF_001063685.1_ASM106368v1 Escherichia coli Proteobacteria; Gammaproteobacteria; Enterobacterales; Enterobacteriaceae; Escherichia
MNNEETFYQAMRRQGVTRRSFLKYCSLAATSLGLGAGMAPKIAWALENKPRIPVVWIHGL WP_001295940.1
MULTISPECIES: hydrogenase [Enterobacteriaceae] Length: 372\nNumber of predicted TMHs: 2\nExp number of AAs in TMHs: 37.2608899999999\nExp number, first 60 AAs: 19.52923\nTotal prob of N-in: 0.94351\nPOSSIBLE N-term signal sequence\ninside 1 19\nTMhelix 20 39\noutside 40 325\nTMhelix 326 348\ninside 349 372

6480 GCF_001059095.1_ASM105909v1 Escherichia coli Proteobacteria; Gammaproteobacteria; Enterobacterales; Enterobacteriaceae; Escherichia
MNNEETFYQAMRRQGVTRRSFLKYCSLAATSLGLGAGMAPKIAWALENKPRIPVVWIHGL WP_001295940.1
MULTISPECIES: hydrogenase [Enterobacteriaceae] Length: 372\nNumber of predicted TMHs: 2\nExp number of AAs in TMHs: 37.2608899999999\nExp number, first 60 AAs: 19.52923\nTotal prob of N-in: 0.94351\nPOSSIBLE N-term signal sequence\ninside 1 19\nTMhelix 20 39\noutside 40 325\nTMhelix 326 348\ninside 349 372

6481 GCF_001065165.1_ASM106516v1 Escherichia coli Proteobacteria; Gammaproteobacteria; Enterobacterales; Enterobacteriaceae; Escherichia
MTGDNTLIHSHGINRRDFMKLCAALAATMGLSSKAAAEMAESVTNPQRPPVIWIGAQECT WP_000145410.1
MULTISPECIES: hydrogenase-2 small chain [Proteobacteria] Length: 372\nNumber of predicted TMHs: 1\nExp number of AAs in TMHs: 23.72226\nExp number, first 60 AAs: 0.22038\nTotal prob of N-in: 0.12994\noutside 1 331\nTMhelix 332 354\ninside 355 372

6482 GCF_001076105.1_ASM107610v1 Escherichia coli Proteobacteria; Gammaproteobacteria; Enterobacterales; Enterobacteriaceae; Escherichia
MTGDNTLIHSHGINRRDFMKLCAALAATMGLSSKAAAEMAESVTNPQRPPVIWIGAQECT WP_000145410.1

MULTISPECIES: hydrogenase-2 small chain [Proteobacteria] Length: 372\nNumber of predicted TMHs: 1\nExp number of AAs in TMHs: 23.72226\nExp number, first 60 AAs: 0.22038\nTotal prob of N-in: 0.12994\noutside 1 331\nTMhelix 332 354\ninside 355 372

6483 GCF_001067425.1_ASM106742v1 Escherichia coli Proteobacteria; Gammaproteobacteria; Enterobacterales; Enterobacteriaceae; Escherichia
MNNEETFYQAMRRQGVTRRSFLKYCSLAATSLGLGAGMAPKIAWALENKPRIPVVWIHGL WP_001058323.1
MULTISPECIES: hydrogenase-1 small chain [Proteobacteria] Length: 372\nNumber of predicted TMHs: 2\nExp number of AAs in TMHs: 37.2608499999999\nExp number, first 60 AAs: 19.52923\nTotal prob of N-in: 0.94351\nPOSSIBLE N-term signal sequence\ninside 1 19\nTMhelix 20 39\noutside 40 325\nTMhelix 326 348\ninside 349 372

6484 GCF_001182885.1_ASM118288v1 Escherichia coli Proteobacteria; Gammaproteobacteria; Enterobacterales; Enterobacteriaceae; Escherichia
MTGDNTLIHSHGINRRDFMKLCAALAATMGLSSKAAAEMAESVTNPQRPPVIWIGAQECT WP_000145410.1
MULTISPECIES: hydrogenase-2 small chain [Proteobacteria] Length: 372\nNumber of predicted TMHs: 1\nExp number of AAs in TMHs: 23.72226\nExp number, first 60 AAs: 0.22038\nTotal prob of N-in: 0.12994\noutside 1 331\nTMhelix 332 354\ninside 355 372

6485 GCF_001182815.1_ASM118281v1 Escherichia coli Proteobacteria; Gammaproteobacteria; Enterobacterales; Enterobacteriaceae; Escherichia
MNNEETFYQAMRRQGVTRRSFLKYCSLAATSLGLGAGMAPKIAWALENKPRIPVVWIHGL WP_001058323.1
MULTISPECIES: hydrogenase-1 small chain [Proteobacteria] Length: 372\nNumber of predicted TMHs: 2\nExp number of AAs in TMHs: 37.2608499999999\nExp number, first 60 AAs: 19.52923\nTotal prob of N-in: 0.94351\nPOSSIBLE N-term signal sequence\ninside 1 19\nTMhelix 20 39\noutside 40 325\nTMhelix 326 348\ninside 349 372

6486 GCF_001182885.1_ASM118288v1 Escherichia coli Proteobacteria; Gammaproteobacteria; Enterobacterales; Enterobacteriaceae; Escherichia
MNNEETFYQAMRRQGVTRRSFLKYCSLAATSLGLGAGMAPKIAWALENKPRIPVVWIHGL WP_001058323.1
MULTISPECIES: hydrogenase-1 small chain [Proteobacteria] Length: 372\nNumber of predicted TMHs: 2\nExp number of AAs in TMHs: 37.2608499999999\nExp number, first 60 AAs: 19.52923\nTotal prob of N-in: 0.94351\nPOSSIBLE N-term signal sequence\ninside 1 19\nTMhelix 20 39\noutside 40 325\nTMhelix 326 348\ninside 349 372

6487 GCF_001076105.1_ASM107610v1 Escherichia coli Proteobacteria; Gammaproteobacteria; Enterobacterales; Enterobacteriaceae; Escherichia
MNNEETFYQAMRRQGVTRRSFLKYCSLAATSLGLGAGMAPKIAWALENKPRIPVVWIHGL WP_001058323.1
MULTISPECIES: hydrogenase-1 small chain [Proteobacteria] Length: 372\nNumber of predicted TMHs: 2\nExp number of AAs in TMHs: 37.2608499999999\nExp number, first 60 AAs: 19.52923\nTotal prob of N-in: 0.94351\nPOSSIBLE N-term signal sequence\ninside 1 19\nTMhelix 20 39\noutside 40 325\nTMhelix 326 348\ninside 349 372

6488 GCF_001077865.1_ASM107786v1 Escherichia coli Proteobacteria; Gammaproteobacteria; Enterobacterales; Enterobacteriaceae; Escherichia
MNNEETFYQAMRRQGVTRRSFLKYCSLAATSLGLGAGMAPKIAWALENKPRIPVVWIHGL WP_001058323.1
MULTISPECIES: hydrogenase-1 small chain [Proteobacteria] Length: 372\nNumber of predicted TMHs: 2\nExp number of AAs in TMHs: 37.2608499999999\nExp number, first 60 AAs: 19.52923\nTotal prob of N-in: 0.94351\nPOSSIBLE N-term signal sequence\ninside 1 19\nTMhelix 20 39\noutside 40 325\nTMhelix 326 348\ninside 349 372

6489 GCF_001077935.1_ASM107793v1 Escherichia coli Proteobacteria; Gammaproteobacteria; Enterobacterales; Enterobacteriaceae; Escherichia
MNNEETFYQAMRRQGVTRRSFLKYCSLAATSLGLGAGMAPKIAWALENKPRIPVVWIHGL WP_001058323.1
MULTISPECIES: hydrogenase-1 small chain [Proteobacteria] Length: 372\nNumber of predicted TMHs: 2\nExp number of AAs in TMHs: 37.2608499999999\nExp number, first 60 AAs: 19.52923\nTotal prob of N-in: 0.94351\nPOSSIBLE N-term signal sequence\ninside 1 19\nTMhelix 20 39\noutside 40 325\nTMhelix 326 348\ninside 349 372

6490 GCF_001191055.1_CFSAN026776_02-NGE.clc Escherichia coli Proteobacteria; Gammaproteobacteria; Enterobacterales; Enterobacteriaceae; Escherichia

MTGDNTLIHSHGINRRDFMKLCAALAATMGLSSKAAAEMAESVTNPQRPPVIWIGAQECT WP_000145410.1 MULTISPECIES: hydrogenase-2 small chain [Proteobacteria] Length: 372\nNumber of predicted TMHs: 1\nExp number of AAs in TMHs: 23.72226\nExp number, first 60 AAs: 0.22038\nTotal prob of N-in: 0.12994\noutside 1 331\nTMhelix 332 354\ninside 355 372

6491 GCF_001191115.1_CFSAN026783_02-NGE.clc Escherichia coli Proteobacteria; Gammaproteobacteria; Enterobacterales; Enterobacteriaceae; Escherichia MTGDNTLIHSHGINRRDFMKLCAALAATMGLSSKAAAEMAESVTNPQRPPVIWIGAQECT WP_000145410.1 MULTISPECIES: hydrogenase-2 small chain [Proteobacteria] Length: 372\nNumber of predicted TMHs: 1\nExp number of AAs in TMHs: 23.72226\nExp number, first 60 AAs: 0.22038\nTotal prob of N-in: 0.12994\noutside 1 331\nTMhelix 332 354\ninside 355 372

6492 GCF_001191125.1_CFSAN026804_01-NGE.clc Escherichia coli Proteobacteria; Gammaproteobacteria; Enterobacterales; Enterobacteriaceae; Escherichia MTGDNTLIHSHGINRRDFMKLCAALAATMGLSSKAAAEMAESVTNPQRPPVIWIGAQECT WP_000145410.1 MULTISPECIES: hydrogenase-2 small chain [Proteobacteria] Length: 372\nNumber of predicted TMHs: 1\nExp number of AAs in TMHs: 23.72226\nExp number, first 60 AAs: 0.22038\nTotal prob of N-in: 0.12994\noutside 1 331\nTMhelix 332 354\ninside 355 372

6493 GCF_001191185.1_CFSAN026847_01-NGE.clc Escherichia coli Proteobacteria; Gammaproteobacteria; Enterobacterales; Enterobacteriaceae; Escherichia MTGDNTLIHSHGINRRDFMKLCAALAATMGLSSKAAAEMAESVTNPQRPPVIWIGAQECT WP_000145410.1 MULTISPECIES: hydrogenase-2 small chain [Proteobacteria] Length: 372\nNumber of predicted TMHs: 1\nExp number of AAs in TMHs: 23.72226\nExp number, first 60 AAs: 0.22038\nTotal prob of N-in: 0.12994\noutside 1 331\nTMhelix 332 354\ninside 355 372

6494 GCF_001191025.1_CFSAN026792_01-NGE.clc Escherichia coli Proteobacteria; Gammaproteobacteria; Enterobacterales; Enterobacteriaceae; Escherichia MNNEETFYQAMRRQGVTRRSFLKYCSLAATSLGLGAGMAPKIAWALENKPRIPVVWIHGL WP_001058323.1 MULTISPECIES: hydrogenase-1 small chain [Proteobacteria] Length: 372\nNumber of predicted TMHs: 2\nExp number of AAs in TMHs: 37.2608499999999\nExp number, first 60 AAs: 19.52923\nTotal prob of N-in: 0.94351\nPOSSIBLE N-term signal sequence\ninside 1 19\nTMhelix 20 39\noutside 40 325\nTMhelix 326 348\ninside 349 372

6495 GCF_001191195.1_CFSAN026846_01-NGE.clc Escherichia coli Proteobacteria; Gammaproteobacteria; Enterobacterales; Enterobacteriaceae; Escherichia MTGDNTLIHSHGINRRDFMKLCAALAATMGLSSKAAAEMAESVTNPQRPPVIWIGAQECT WP_000145410.1 MULTISPECIES: hydrogenase-2 small chain [Proteobacteria] Length: 372\nNumber of predicted TMHs: 1\nExp number of AAs in TMHs: 23.72226\nExp number, first 60 AAs: 0.22038\nTotal prob of N-in: 0.12994\noutside 1 331\nTMhelix 332 354\ninside 355 372

6496 GCF_001191215.1_CFSAN026817_01-NGE.clc Escherichia coli Proteobacteria; Gammaproteobacteria; Enterobacterales; Enterobacteriaceae; Escherichia MTGDNTLIHSHGINRRDFMKLCAALAATMGLSSKAAAEMAESVTNPQRPPVIWIGAQECT WP_000145410.1 MULTISPECIES: hydrogenase-2 small chain [Proteobacteria] Length: 372\nNumber of predicted TMHs: 1\nExp number of AAs in TMHs: 23.72226\nExp number, first 60 AAs: 0.22038\nTotal prob of N-in: 0.12994\noutside 1 331\nTMhelix 332 354\ninside 355 372

6497 GCF_001191055.1_CFSAN026776_02-NGE.clc Escherichia coli Proteobacteria; Gammaproteobacteria; Enterobacterales; Enterobacteriaceae; Escherichia MNNEETFYQAMRRQGVTRRSFLKYCSLAATSLGLGAGMAPKIAWALENKPRIPVVWIHGL WP_001295940.1 MULTISPECIES: hydrogenase [Enterobacteriaceae] Length: 372\nNumber of predicted TMHs: 2\nExp number of AAs in TMHs: 37.2608899999999\nExp number, first 60 AAs: 19.52923\nTotal prob of N-in: 0.94351\nPOSSIBLE N-term signal sequence\ninside 1 19\nTMhelix 20 39\noutside 40 325\nTMhelix 326 348\ninside 349 372

6498 GCF_001191035.1_CFSAN026821_01-NGE.clc Escherichia coli Proteobacteria; Gammaproteobacteria; Enterobacterales; Enterobacteriaceae; Escherichia MNNEETFYQAMRRQGVTRRSFLKYCSLAATSLGLGAGMAPKIAWALENKPRIPVVWIHGL WP_001295940.1 MULTISPECIES: hydrogenase [Enterobacteriaceae] Length: 372\nNumber of predicted TMHs: 2\nExp number of AAs in TMHs: 37.2608899999999\nExp number, first 60 AAs: 19.52923\nTotal prob of N-in:

0.94351\nPOSSIBLE N-term signal sequence\ninside 1 19\nTMhelix 20 39\noutside 40 325\nTMhelix 326 348\ninside 349 372

6499 GCF_001191275.1_CFSAN026786_02-NGE.clc Escherichia coli Proteobacteria; Gammaproteobacteria; Enterobacterales; Enterobacteriaceae; Escherichia MNNEETFYQAMRRQGVTRRSFLKYCSLAATSLGLGAGMAPKIAWALENKPRIPVVWIHGL WP_001058323.1 MULTISPECIES: hydrogenase-1 small chain [Proteobacteria] Length: 372\nNumber of predicted TMHs: 2\nExp number of AAs in TMHs: 37.2608499999999\nExp number, first 60 AAs: 19.52923\nTotal prob of N-in: 0.94351\nPOSSIBLE N-term signal sequence\ninside 1 19\nTMhelix 20 39\noutside 40 325\nTMhelix 326 348\ninside 349 372

6500 GCF_001191345.1_CFSAN026795_01-NGE.clc Escherichia coli Proteobacteria; Gammaproteobacteria; Enterobacterales; Enterobacteriaceae; Escherichia MTGDNTLIHSHGINRRDFMKLCAALAATMGLSSKAAAEMAESVTNPQRPPVIWIGAQECT WP_000145410.1 MULTISPECIES: hydrogenase-2 small chain [Proteobacteria] Length: 372\nNumber of predicted TMHs: 1\nExp number of AAs in TMHs: 23.72226\nExp number, first 60 AAs: 0.22038\nTotal prob of N-in: 0.12994\noutside 1 331\nTMhelix 332 354\ninside 355 372

6501 GCF_001191355.1_CFSAN026832_01-NGE.clc Escherichia coli Proteobacteria; Gammaproteobacteria; Enterobacterales; Enterobacteriaceae; Escherichia MTGDNTLIHSHGINRRDFMKLCAALAATMGLSSKAAAEMAESVTNPQRPPVIWIGAQECT WP_000145410.1 MULTISPECIES: hydrogenase-2 small chain [Proteobacteria] Length: 372\nNumber of predicted TMHs: 1\nExp number of AAs in TMHs: 23.72226\nExp number, first 60 AAs: 0.22038\nTotal prob of N-in: 0.12994\noutside 1 331\nTMhelix 332 354\ninside 355 372

6502 GCF_001191355.1_CFSAN026832_01-NGE.clc Escherichia coli Proteobacteria; Gammaproteobacteria; Enterobacterales; Enterobacteriaceae; Escherichia MNNEETFYQAMRRQGVTRRSFLKYCSLAATSLGLGAGMAPKIAWALENKPRIPVVWIHGL WP_001058323.1 MULTISPECIES: hydrogenase-1 small chain [Proteobacteria] Length: 372\nNumber of predicted TMHs: 2\nExp number of AAs in TMHs: 37.2608499999999\nExp number, first 60 AAs: 19.52923\nTotal prob of N-in: 0.94351\nPOSSIBLE N-term signal sequence\ninside 1 19\nTMhelix 20 39\noutside 40 325\nTMhelix 326 348\ninside 349 372

6503 GCF_001191345.1_CFSAN026795_01-NGE.clc Escherichia coli Proteobacteria; Gammaproteobacteria; Enterobacterales; Enterobacteriaceae; Escherichia MNNEETFYQAMRRQGVTRRSFLKYCSLAATSLGLGAGMAPKIAWALENKPRIPVVWIHGL WP_001058323.1 MULTISPECIES: hydrogenase-1 small chain [Proteobacteria] Length: 372\nNumber of predicted TMHs: 2\nExp number of AAs in TMHs: 37.2608499999999\nExp number, first 60 AAs: 19.52923\nTotal prob of N-in: 0.94351\nPOSSIBLE N-term signal sequence\ninside 1 19\nTMhelix 20 39\noutside 40 325\nTMhelix 326 348\ninside 349 372

6504 GCF_001262775.1_2089-2 Escherichia coli Proteobacteria; Gammaproteobacteria; Enterobacterales; Enterobacteriaceae; Escherichia MTGDNTLIHSHGINRRDFMKLCAALAATMGLSSKAAAEMAESVTNPQRPPVIWIGAQECT WP_000145410.1 MULTISPECIES: hydrogenase-2 small chain [Proteobacteria] Length: 372\nNumber of predicted TMHs: 1\nExp number of AAs in TMHs: 23.72226\nExp number, first 60 AAs: 0.22038\nTotal prob of N-in: 0.12994\noutside 1 331\nTMhelix 332 354\ninside 355 372

6505 GCF_001262775.1_2089-2 Escherichia coli Proteobacteria; Gammaproteobacteria; Enterobacterales; Enterobacteriaceae; Escherichia MNNEETFYQAMRRQGVTRRSFLKYCSLAATSLGLGAGMAPKIAWALENKPRIPVVWIHGL WP_001295940.1 MULTISPECIES: hydrogenase [Enterobacteriaceae] Length: 372\nNumber of predicted TMHs: 2\nExp number of AAs in TMHs: 37.2608899999999\nExp number, first 60 AAs: 19.52923\nTotal prob of N-in: 0.94351\nPOSSIBLE N-term signal sequence\ninside 1 19\nTMhelix 20 39\noutside 40 325\nTMhelix 326 348\ninside 349 372

6506 GCF_001191395.1_CFSAN026824_01-NGE.clc Escherichia coli Proteobacteria; Gammaproteobacteria; Enterobacterales; Enterobacteriaceae; Escherichia MNNEETFYQAMRRQGVTRRSFLKYCSLAATSLGLGAGMAPKIAWALENKPRIPVVWIHGL WP_001295940.1 MULTISPECIES: hydrogenase [Enterobacteriaceae] Length: 372\nNumber of predicted TMHs: 2\nExp number of AAs in TMHs: 37.2608899999999\nExp number, first 60 AAs: 19.52923\nTotal prob of N-in: 0.94351\nPOSSIBLE N-term signal sequence\ninside 1 19\nTMhelix 20 39\noutside 40 325\nTMhelix 326 348\ninside 349 372

6507 GCF_001191455.1_CFSAN026831_01-NGE.clc Escherichia coli Proteobacteria; Gammaproteobacteria; Enterobacterales; Enterobacteriaceae; Escherichia MNNEETFYQAMRRQGVTRRSFLKYCSLAATSLGLGAGMAPKIAWALENKPRIPVVWIHGL WP_001058323.1 MULTISPECIES: hydrogenase-1 small chain [Proteobacteria] Length: 372\nNumber of predicted TMHs: 2\nExp number of AAs in TMHs: 37.2608499999999\nExp number, first 60 AAs: 19.52923\nTotal prob of N-in: 0.94351\nPOSSIBLE N-term signal sequence\ninside 1 19\nTMhelix 20 39\noutside 40 325\nTMhelix 326 348\ninside 349 372

6508 GCF_001262795.1_7712-3 Escherichia coli Proteobacteria; Gammaproteobacteria; Enterobacterales; Enterobacteriaceae; Escherichia MTGDNTLIHSHGINRRDFMKLCAALAATMGLSSKAAAEMAESVTNPQRPPVIWIGAQECT WP_000145410.1 MULTISPECIES: hydrogenase-2 small chain [Proteobacteria] Length: 372\nNumber of predicted TMHs: 1\nExp number of AAs in TMHs: 23.72226\nExp number, first 60 AAs: 0.22038\nTotal prob of N-in: 0.12994\noutside 1 331\nTMhelix 332 354\ninside 355 372

6509 GCF_001262855.1_ASM126285v1 Escherichia coli Proteobacteria; Gammaproteobacteria; Enterobacterales; Enterobacteriaceae; Escherichia MTGDNTLIHSHGINRRDFMKLCAALAATMGLSSKAAAEMAESVTNPQRPPVIWIGAQECT WP_000145410.1 MULTISPECIES: hydrogenase-2 small chain [Proteobacteria] Length: 372\nNumber of predicted TMHs: 1\nExp number of AAs in TMHs: 23.72226\nExp number, first 60 AAs: 0.22038\nTotal prob of N-in: 0.12994\noutside 1 331\nTMhelix 332 354\ninside 355 372

6510 GCF_001262795.1_7712-3 Escherichia coli Proteobacteria; Gammaproteobacteria; Enterobacterales; Enterobacteriaceae; Escherichia MNNEETFYQAMRRQGVTRRSFLKYCSLAATSLGLGAGMAPKIAWALENKPRIPVVWIHGL WP_001295940.1 MULTISPECIES: hydrogenase [Enterobacteriaceae] Length: 372\nNumber of predicted TMHs: 2\nExp number of AAs in TMHs: 37.2608899999999\nExp number, first 60 AAs: 19.52923\nTotal prob of N-in: 0.94351\nPOSSIBLE N-term signal sequence\ninside 1 19\nTMhelix 20 39\noutside 40 325\nTMhelix 326 348\ninside 349 372

6511 GCF_001262955.1_96-1156 Escherichia coli Proteobacteria; Gammaproteobacteria; Enterobacterales; Enterobacteriaceae; Escherichia MTGDNTLIHSHGINRRDFMKLCAALAATMGLSSKAAAEMAESVTNPQRPPVIWIGAQECT WP_000145410.1 MULTISPECIES: hydrogenase-2 small chain [Proteobacteria] Length: 372\nNumber of predicted TMHs: 1\nExp number of AAs in TMHs: 23.72226\nExp number, first 60 AAs: 0.22038\nTotal prob of N-in: 0.12994\noutside 1 331\nTMhelix 332 354\ninside 355 372

6512 GCF_001262965.1_1223-4 Escherichia coli Proteobacteria; Gammaproteobacteria; Enterobacterales; Enterobacteriaceae; Escherichia MTGDNTLIHSHGINRRDFMKLCAALAATMGLSSKAAAEMAESVTNPQRPPVIWIGAQECT WP_000145410.1 MULTISPECIES: hydrogenase-2 small chain [Proteobacteria] Length: 372\nNumber of predicted TMHs: 1\nExp number of AAs in TMHs: 23.72226\nExp number, first 60 AAs: 0.22038\nTotal prob of N-in: 0.12994\noutside 1 331\nTMhelix 332 354\ninside 355 372

6513 GCF_001263015.1_23-second Escherichia coli Proteobacteria; Gammaproteobacteria; Enterobacterales; Enterobacteriaceae; Escherichia MTGDNTLIHSHGINRRDFMKLCAALAATMGLSSKAAAEMAESVTNPQRPPVIWIGAQECT WP_000145410.1 MULTISPECIES: hydrogenase-2 small chain [Proteobacteria] Length: 372\nNumber of predicted TMHs: 1\nExp number of AAs in TMHs: 23.72226\nExp number, first 60 AAs: 0.22038\nTotal prob of N-in: 0.12994\noutside 1 331\nTMhelix 332 354\ninside 355 372

6514 GCF_001262955.1_96-1156 Escherichia coli Proteobacteria; Gammaproteobacteria; Enterobacterales; Enterobacteriaceae; Escherichia MNNEETFYQAMRRQGVTRRSFLKYCSLAATSLGLGAGMAPKIAWALENKPRIPVVWIHGL WP_001058323.1 MULTISPECIES: hydrogenase-1 small chain [Proteobacteria] Length: 372\nNumber of predicted TMHs: 2\nExp number of AAs in TMHs: 37.2608499999999\nExp number, first 60 AAs: 19.52923\nTotal prob of N-in: 0.94351\nPOSSIBLE N-term signal sequence\ninside 1 19\nTMhelix 20 39\noutside 40 325\nTMhelix 326 348\ninside 349 372

6515 GCF_001263735.1_ASM126373v1 Escherichia coli Proteobacteria; Gammaproteobacteria; Enterobacterales; Enterobacteriaceae; Escherichia MTGDNTLIHSHGINRRDFMKLCAALAATMGLSSKAAAEMAESVTNPQRPPVIWIGAQECT WP_000145410.1 MULTISPECIES: hydrogenase-2 small chain [Proteobacteria] Length: 372\nNumber of predicted TMHs: 1\nExp number of AAs in TMHs: 23.72226\nExp number, first 60 AAs: 0.22038\nTotal prob of N-in: 0.12994\noutside 1 331\nTMhelix 332 354\ninside 355 372

6516 GCF_001264195.1_ASM126419v1 Escherichia coli Proteobacteria; Gammaproteobacteria; Enterobacterales; Enterobacteriaceae; Escherichia
MTGDNTLIHSHGINRRDFMKLCAALAATMGLSSKAAAEMAESVTNPQRPPVIWIGAQECT WP_000145410.1
MULTISPECIES: hydrogenase-2 small chain [Proteobacteria] Length: 372\nNumber of predicted TMHs: 1\nExp number of AAs in TMHs: 23.72226\nExp number, first 60 AAs: 0.22038\nTotal prob of N-in: 0.12994\noutside 1 331\nTMhelix 332 354\ninside 355 372

6517 GCF_001263065.1_16118-2 Escherichia coli Proteobacteria; Gammaproteobacteria; Enterobacterales; Enterobacteriaceae; Escherichia MTGDNTLIHSHGINRRDFMKLCAALAATMGLSSKAAAEMAESVTNPQRPPVIWIGAQECT
WP_000145410.1 MULTISPECIES: hydrogenase-2 small chain [Proteobacteria] Length: 372\nNumber of predicted TMHs: 1\nExp number of AAs in TMHs: 23.72226\nExp number, first 60 AAs: 0.22038\nTotal prob of N-in: 0.12994\noutside 1 331\nTMhelix 332 354\ninside 355 372

6518 GCF_001265185.1_102550 Escherichia coli Proteobacteria; Gammaproteobacteria; Enterobacterales; Enterobacteriaceae; Escherichia MTGDNTLIHSHGINRRDFMKLCAALAATMGLSSKAAAEMAESVTNPQRPPVIWIGAQECT
WP_000145410.1 MULTISPECIES: hydrogenase-2 small chain [Proteobacteria] Length: 372\nNumber of predicted TMHs: 1\nExp number of AAs in TMHs: 23.72226\nExp number, first 60 AAs: 0.22038\nTotal prob of N-in: 0.12994\noutside 1 331\nTMhelix 332 354\ninside 355 372

6519 GCF_001265185.1_102550 Escherichia coli Proteobacteria; Gammaproteobacteria; Enterobacterales; Enterobacteriaceae; Escherichia MNNEETFYQAMRRQGVTRRSFLKYCSLAATSLGLGAGMAPKIAWALENKPRIPVVWIHGL
WP_001058323.1 MULTISPECIES: hydrogenase-1 small chain [Proteobacteria] Length: 372\nNumber of predicted TMHs: 2\nExp number of AAs in TMHs: 37.2608499999999\nExp number, first 60 AAs: 19.52923\nTotal prob of N-in: 0.94351\nPOSSIBLE N-term signal sequence\ninside 1 19\nTMhelix 20 39\noutside 40 325\nTMhelix 326 348\ninside 349 372

6520 GCF_001263015.1_23-second Escherichia coli Proteobacteria; Gammaproteobacteria; Enterobacterales; Enterobacteriaceae; Escherichia
MNNEETFYQAMRRQGVTRRSFLKYCSLAATSLGLGAGMAPKIAWALENKPRIPVVWIHGL WP_001295940.1
MULTISPECIES: hydrogenase [Enterobacteriaceae] Length: 372\nNumber of predicted TMHs: 2\nExp number of AAs in TMHs: 37.2608899999999\nExp number, first 60 AAs: 19.52923\nTotal prob of N-in: 0.94351\nPOSSIBLE N-term signal sequence\ninside 1 19\nTMhelix 20 39\noutside 40 325\nTMhelix 326 348\ninside 349 372

6521 GCF_001265495.1_401150 Escherichia coli Proteobacteria; Gammaproteobacteria; Enterobacterales; Enterobacteriaceae; Escherichia MTGDNTLIHSHGINRRDFMKLCAALAATMGLSSKAAAEMAESVTNPQRPPVIWIGAQECT
WP_000145410.1 MULTISPECIES: hydrogenase-2 small chain [Proteobacteria] Length: 372\nNumber of predicted TMHs: 1\nExp number of AAs in TMHs: 23.72226\nExp number, first 60 AAs: 0.22038\nTotal prob of N-in: 0.12994\noutside 1 331\nTMhelix 332 354\ninside 355 372

6522 GCF_001265535.1_401954 Escherichia coli Proteobacteria; Gammaproteobacteria; Enterobacterales; Enterobacteriaceae; Escherichia MTGDNTLIHSHGINRRDFMKLCAALAATMGLSSKAAAEMAESVTNPQRPPVIWIGAQECT
WP_000145410.1 MULTISPECIES: hydrogenase-2 small chain [Proteobacteria] Length: 372\nNumber of predicted TMHs: 1\nExp number of AAs in TMHs: 23.72226\nExp number, first 60 AAs: 0.22038\nTotal prob of N-in: 0.12994\noutside 1 331\nTMhelix 332 354\ninside 355 372

6523 GCF_001265595.1_703533 Escherichia coli Proteobacteria; Gammaproteobacteria; Enterobacterales; Enterobacteriaceae; Escherichia MTGDNTLIHSHGINRRDFMKLCAALAATMGLSSKAAAEMAESVTNPQHPPVIWIGAQECT
WP_000145403.1 hydrogenase 2 small subunit [Escherichia coli] Length: 372\nNumber of predicted TMHs: 1\nExp number of AAs in TMHs: 23.66365\nExp number, first 60 AAs: 0.1537\nTotal prob of N-in: 0.12709\noutside 1 331\nTMhelix 332 354\ninside 355 372

6524 GCF_001265625.1_100854 Escherichia coli Proteobacteria; Gammaproteobacteria; Enterobacterales; Enterobacteriaceae; Escherichia MTGDNTLIHSHGINRRDFMKLCAALAATMGLSSKAAAEMAESVTNPQRPPVIWIGAQECT
WP_000145410.1 MULTISPECIES: hydrogenase-2 small chain [Proteobacteria] Length: 372\nNumber of predicted TMHs: 1\nExp number of AAs in TMHs: 23.72226\nExp number, first 60 AAs: 0.22038\nTotal prob of N-in: 0.12994\noutside 1 331\nTMhelix 332 354\ninside 355 372

6525 GCF_001265495.1_401150 Escherichia coli Proteobacteria; Gammaproteobacteria; Enterobacterales; Enterobacteriaceae; Escherichia MNNEETFYQAMRRQGVTRRSFLKYCSLAATSLGLGAGMAPKIAWALENKPRIPVVWIHGL
WP_001058323.1 MULTISPECIES: hydrogenase-1 small chain [Proteobacteria] Length: 372\nNumber of

predicted TMHs:  2\nExp number of AAs in TMHs: 37.2608499999999\nExp number, first 60 AAs:  19.52923\nTotal prob of N-in:  0.94351\nPOSSIBLE N-term signal sequence\ninside  1  19\nTMhelix  20  39\noutside  40  325\nTMhelix  326  348\ninside  349  372

6526 GCF_001265625.1_100854 Escherichia coli Proteobacteria; Gammaproteobacteria; Enterobacterales; Enterobacteriaceae; Escherichia MNNEETFYQAMRRQGVTRRSFLKYCSLAATSLGLGAGMAPKIAWALENKPRIPVVWIHGL WP_001058323.1 MULTISPECIES: hydrogenase-1 small chain [Proteobacteria] Length: 372\nNumber of predicted TMHs:  2\nExp number of AAs in TMHs: 37.2608499999999\nExp number, first 60 AAs:  19.52923\nTotal prob of N-in:  0.94351\nPOSSIBLE N-term signal sequence\ninside  1  19\nTMhelix  20  39\noutside  40  325\nTMhelix  326  348\ninside  349  372

6527 GCF_001265535.1_401954 Escherichia coli Proteobacteria; Gammaproteobacteria; Enterobacterales; Enterobacteriaceae; Escherichia MNNEETFYQAMRRQGVTRRSFLKYCSLAATSLGLGAGMAPKIAWALENKPRIPVVWIHGL WP_001295940.1 MULTISPECIES: hydrogenase [Enterobacteriaceae] Length: 372\nNumber of predicted TMHs:  2\nExp number of AAs in TMHs: 37.2608899999999\nExp number, first 60 AAs:  19.52923\nTotal prob of N-in:  0.94351\nPOSSIBLE N-term signal sequence\ninside  1  19\nTMhelix  20  39\noutside  40  325\nTMhelix  326  348\ninside  349  372

6528 GCF_001265765.1_300214 Escherichia coli Proteobacteria; Gammaproteobacteria; Enterobacterales; Enterobacteriaceae; Escherichia MNNEETFYQAMRRQGVTRRSFLKYCSLAATSLGLGAGMAPKIAWALENKPRIPVVWIHGL WP_001058323.1 MULTISPECIES: hydrogenase-1 small chain [Proteobacteria] Length: 372\nNumber of predicted TMHs:  2\nExp number of AAs in TMHs: 37.2608499999999\nExp number, first 60 AAs:  19.52923\nTotal prob of N-in:  0.94351\nPOSSIBLE N-term signal sequence\ninside  1  19\nTMhelix  20  39\noutside  40  325\nTMhelix  326  348\ninside  349  372

6529 GCF_001265795.1_300231 Escherichia coli Proteobacteria; Gammaproteobacteria; Enterobacterales; Enterobacteriaceae; Escherichia MTGDNTLIHSHGINRRDFMKLCAALAATMGLSSKAAAEMAESVTNPQRPPVIWIGAQECT WP_000145410.1 MULTISPECIES: hydrogenase-2 small chain [Proteobacteria] Length: 372\nNumber of predicted TMHs:  1\nExp number of AAs in TMHs: 23.72226\nExp number, first 60 AAs:  0.22038\nTotal prob of N-in:  0.12994\noutside  1  331\nTMhelix  332  354\ninside  355  372

6530 GCF_001265795.1_300231 Escherichia coli Proteobacteria; Gammaproteobacteria; Enterobacterales; Enterobacteriaceae; Escherichia MNNEETFYQAMRRQGVTRRSFLKYCSLAATSLGLGAGMAPKIAWALENKPRIPVVWIHGL WP_001058323.1 MULTISPECIES: hydrogenase-1 small chain [Proteobacteria] Length: 372\nNumber of predicted TMHs:  2\nExp number of AAs in TMHs: 37.2608499999999\nExp number, first 60 AAs:  19.52923\nTotal prob of N-in:  0.94351\nPOSSIBLE N-term signal sequence\ninside  1  19\nTMhelix  20  39\noutside  40  325\nTMhelix  326  348\ninside  349  372

6531 GCF_001265595.1_703533 Escherichia coli Proteobacteria; Gammaproteobacteria; Enterobacterales; Enterobacteriaceae; Escherichia MNNEETFYQAMRRQGVTRRSFLKYCSLAATSLGLGAGMAPKIAWALENKPRIPVVWIHGL WP_001295940.1 MULTISPECIES: hydrogenase [Enterobacteriaceae] Length: 372\nNumber of predicted TMHs:  2\nExp number of AAs in TMHs: 37.2608899999999\nExp number, first 60 AAs:  19.52923\nTotal prob of N-in:  0.94351\nPOSSIBLE N-term signal sequence\ninside  1  19\nTMhelix  20  39\noutside  40  325\nTMhelix  326  348\ninside  349  372

6532 GCF_001265835.1_302687 Escherichia coli Proteobacteria; Gammaproteobacteria; Enterobacterales; Enterobacteriaceae; Escherichia MTGDNTLIHSHGINRRDFMKLCAALAATMGLSSKAAAEMAESVTNPQRPPVIWIGAQECT WP_000145410.1 MULTISPECIES: hydrogenase-2 small chain [Proteobacteria] Length: 372\nNumber of predicted TMHs:  1\nExp number of AAs in TMHs: 23.72226\nExp number, first 60 AAs:  0.22038\nTotal prob of N-in:  0.12994\noutside  1  331\nTMhelix  332  354\ninside  355  372

6533 GCF_001265875.1_303145 Escherichia coli Proteobacteria; Gammaproteobacteria; Enterobacterales; Enterobacteriaceae; Escherichia MTGDNTLIHSHGINRRDFMKLCAALAATMGLSSKAAAEMAESVTNPQRPPVIWIGAQECT WP_000145410.1 MULTISPECIES: hydrogenase-2 small chain [Proteobacteria] Length: 372\nNumber of predicted TMHs:  1\nExp number of AAs in TMHs: 23.72226\nExp number, first 60 AAs:  0.22038\nTotal prob of N-in:  0.12994\noutside  1  331\nTMhelix  332  354\ninside  355  372

6534 GCF_001266015.1_100100 Escherichia coli Proteobacteria; Gammaproteobacteria; Enterobacterales; Enterobacteriaceae; Escherichia MTGDNTLIHSHGINRRDFMKLCAALAATMGLSSKAAAEMAESVTNPQRPPVIWIGAQECT WP_000145410.1 MULTISPECIES: hydrogenase-2 small chain [Proteobacteria] Length: 372\nNumber of

predicted TMHs: 1\nExp number of AAs in TMHs: 23.72226\nExp number, first 60 AAs: 0.22038\nTotal prob of N-in: 0.12994\noutside 1 331\nTMhelix 332 354\ninside 355 372

6535 GCF_001265975.1_402310 Escherichia coli Proteobacteria; Gammaproteobacteria; Enterobacterales; Enterobacteriaceae; Escherichia MNNEETFYQAMRRQGVTRRSFLKYCSLAATSLGLGAGMAPKIAWALENKPRIPVVWIHGL WP_001058323.1 MULTISPECIES: hydrogenase-1 small chain [Proteobacteria] Length: 372\nNumber of predicted TMHs: 2\nExp number of AAs in TMHs: 37.2608499999999\nExp number, first 60 AAs: 19.52923\nTotal prob of N-in: 0.94351\nPOSSIBLE N-term signal sequence\ninside 1 19\nTMhelix 20 39\noutside 40 325\nTMhelix 326 348\ninside 349 372

6536 GCF_001266005.1_702423 Escherichia coli Proteobacteria; Gammaproteobacteria; Enterobacterales; Enterobacteriaceae; Escherichia MTGDNTLIHSHGINRRDFMKLCAALAATMGLSSKAAAEMAESVTNPQHPPVIWIGAQECT WP_000145403.1 hydrogenase 2 small subunit [Escherichia coli] Length: 372\nNumber of predicted TMHs: 1\nExp number of AAs in TMHs: 23.66365\nExp number, first 60 AAs: 0.1537\nTotal prob of N-in: 0.12709\noutside 1 331\nTMhelix 332 354\ninside 355 372

6537 GCF_001266005.1_702423 Escherichia coli Proteobacteria; Gammaproteobacteria; Enterobacterales; Enterobacteriaceae; Escherichia MNNEETFYQAMRRQGVTRRSFLKYCSLAATSLGLGAGMAPKIAWALENKPRIPVVWIHGL WP_001295940.1 MULTISPECIES: hydrogenase [Enterobacteriaceae] Length: 372\nNumber of predicted TMHs: 2\nExp number of AAs in TMHs: 37.2608899999999\nExp number, first 60 AAs: 19.52923\nTotal prob of N-in: 0.94351\nPOSSIBLE N-term signal sequence\ninside 1 19\nTMhelix 20 39\noutside 40 325\nTMhelix 326 348\ninside 349 372

6538 GCF_001265735.1_200146 Escherichia coli Proteobacteria; Gammaproteobacteria; Enterobacterales; Enterobacteriaceae; Escherichia MNNEETFYQAMRRQGVTRRSFLKYCSLAATSLGLGAGMAPKIAWALENKPRIPVVWIHGL WP_052923131.1 hydrogenase [Escherichia coli] Length: 372\nNumber of predicted TMHs: 2\nExp number of AAs in TMHs: 36.3624399999999\nExp number, first 60 AAs: 19.53053\nTotal prob of N-in: 0.94346\nPOSSIBLE N-term signal sequence\ninside 1 19\nTMhelix 20 39\noutside 40 325\nTMhelix 326 348\ninside 349 372

6539 GCF_001265915.1_401210 Escherichia coli Proteobacteria; Gammaproteobacteria; Enterobacterales; Enterobacteriaceae; Escherichia MNNEETFYQAMRRQGVTRRSFLKYCSLAATSLGLGAGMAPKIAWALENKPRIPVVWIHGL WP_001058323.1 MULTISPECIES: hydrogenase-1 small chain [Proteobacteria] Length: 372\nNumber of predicted TMHs: 2\nExp number of AAs in TMHs: 37.2608499999999\nExp number, first 60 AAs: 19.52923\nTotal prob of N-in: 0.94351\nPOSSIBLE N-term signal sequence\ninside 1 19\nTMhelix 20 39\noutside 40 325\nTMhelix 326 348\ninside 349 372

6540 GCF_001266115.1_102535 Escherichia coli Proteobacteria; Gammaproteobacteria; Enterobacterales; Enterobacteriaceae; Escherichia MTGDNTLIHSHGINRRDFMKLCAALAATMGLSSKAAAEMAESVTNPQRPPVIWIGAQECT WP_000145410.1 MULTISPECIES: hydrogenase-2 small chain [Proteobacteria] Length: 372\nNumber of predicted TMHs: 1\nExp number of AAs in TMHs: 23.72226\nExp number, first 60 AAs: 0.22038\nTotal prob of N-in: 0.12994\noutside 1 331\nTMhelix 332 354\ninside 355 372

6541 GCF_001266125.1_103338 Escherichia coli Proteobacteria; Gammaproteobacteria; Enterobacterales; Enterobacteriaceae; Escherichia MNNEETFYQAMRRQGVTRRSFLKYCSLAATSLGLGAGMAPKIAWALENKPRIPVVWIHGL WP_001058323.1 MULTISPECIES: hydrogenase-1 small chain [Proteobacteria] Length: 372\nNumber of predicted TMHs: 2\nExp number of AAs in TMHs: 37.2608499999999\nExp number, first 60 AAs: 19.52923\nTotal prob of N-in: 0.94351\nPOSSIBLE N-term signal sequence\ninside 1 19\nTMhelix 20 39\noutside 40 325\nTMhelix 326 348\ninside 349 372

6542 GCF_001266225.1_302137 Escherichia coli Proteobacteria; Gammaproteobacteria; Enterobacterales; Enterobacteriaceae; Escherichia MTGDNTLIHSHGINRRDFMKLCAALAATMGLSSKAAAEMAESVTNPQRPPVIWIGAQECT WP_000145410.1 MULTISPECIES: hydrogenase-2 small chain [Proteobacteria] Length: 372\nNumber of predicted TMHs: 1\nExp number of AAs in TMHs: 23.72226\nExp number, first 60 AAs: 0.22038\nTotal prob of N-in: 0.12994\noutside 1 331\nTMhelix 332 354\ninside 355 372

6543 GCF_001266305.1_303301 Escherichia coli Proteobacteria; Gammaproteobacteria; Enterobacterales; Enterobacteriaceae; Escherichia MTGDNTLIHSHGINRRDFMKLCAALAATMGLSSKAAAEMAESVTNPQRPPVIWIGAQECT WP_000145410.1 MULTISPECIES: hydrogenase-2 small chain [Proteobacteria] Length: 372\nNumber of predicted TMHs: 1\nExp number of AAs in TMHs: 23.72226\nExp number, first 60 AAs: 0.22038\nTotal prob of N-in: 0.12994\noutside 1 331\nTMhelix 332 354\ninside 355 372

6544 GCF_001266215.1_300847 Escherichia coli Proteobacteria; Gammaproteobacteria; Enterobacterales; Enterobacteriaceae; Escherichia MNNEETFYQAMRRQGVTRRSFLKYCSLAATSLGLGAGMAPKIAWALENKPRIPVVWIHGL WP_001295940.1 MULTISPECIES: hydrogenase [Enterobacteriaceae] Length: 372\nNumber of predicted TMHs: 2\nExp number of AAs in TMHs: 37.2608899999999\nExp number, first 60 AAs: 19.52923\nTotal prob of N-in: 0.94351\nPOSSIBLE N-term signal sequence\ninside 1 19\nTMhelix 20 39\noutside 40 325\nTMhelix 326 348\ninside 349 372

6545 GCF_001266275.1_303139 Escherichia coli Proteobacteria; Gammaproteobacteria; Enterobacterales; Enterobacteriaceae; Escherichia MTGDNTLIHSHGINRRDFMKLCAALAATMGLSSKAAAEMAESVTNPQRPPVIWIGAQECT WP_000145410.1 MULTISPECIES: hydrogenase-2 small chain [Proteobacteria] Length: 372\nNumber of predicted TMHs: 1\nExp number of AAs in TMHs: 23.72226\nExp number, first 60 AAs: 0.22038\nTotal prob of N-in: 0.12994\noutside 1 331\nTMhelix 332 354\ninside 355 372

6546 GCF_001266225.1_302137 Escherichia coli Proteobacteria; Gammaproteobacteria; Enterobacterales; Enterobacteriaceae; Escherichia MNNEETFYQAMRRQGVTRRSFLKYCSLAATSLGLGAGMAPKIAWALENKPRIPVVWIHGL WP_001058323.1 MULTISPECIES: hydrogenase-1 small chain [Proteobacteria] Length: 372\nNumber of predicted TMHs: 2\nExp number of AAs in TMHs: 37.2608499999999\nExp number, first 60 AAs: 19.52923\nTotal prob of N-in: 0.94351\nPOSSIBLE N-term signal sequence\ninside 1 19\nTMhelix 20 39\noutside 40 325\nTMhelix 326 348\ninside 349 372

6547 GCF_001266375.1_401675 Escherichia coli Proteobacteria; Gammaproteobacteria; Enterobacterales; Enterobacteriaceae; Escherichia MTGDNTLIHSHGINRRDFMKLCAALAATMGLSSKAAAEMAESVTNPQRPPVIWIGAQECT WP_000145410.1 MULTISPECIES: hydrogenase-2 small chain [Proteobacteria] Length: 372\nNumber of predicted TMHs: 1\nExp number of AAs in TMHs: 23.72226\nExp number, first 60 AAs: 0.22038\nTotal prob of N-in: 0.12994\noutside 1 331\nTMhelix 332 354\ninside 355 372

6548 GCF_001267335.1_403116 Escherichia coli Proteobacteria; Gammaproteobacteria; Enterobacterales; Enterobacteriaceae; Escherichia MNNEETFYQAMRRQGVTRRSFLKYCSLAATSLGLGAGMAPKIAWALENKPRIPVVWIHGL WP_001058323.1 MULTISPECIES: hydrogenase-1 small chain [Proteobacteria] Length: 372\nNumber of predicted TMHs: 2\nExp number of AAs in TMHs: 37.2608499999999\nExp number, first 60 AAs: 19.52923\nTotal prob of N-in: 0.94351\nPOSSIBLE N-term signal sequence\ninside 1 19\nTMhelix 20 39\noutside 40 325\nTMhelix 326 348\ninside 349 372

6549 GCF_001266305.1_303301 Escherichia coli Proteobacteria; Gammaproteobacteria; Enterobacterales; Enterobacteriaceae; Escherichia MNNEETFYQAMRRQGVTRRSFLKYCSLAATSLGLGAGMAPKIAWALENKPRIPVVWIHGL WP_001295940.1 MULTISPECIES: hydrogenase [Enterobacteriaceae] Length: 372\nNumber of predicted TMHs: 2\nExp number of AAs in TMHs: 37.2608899999999\nExp number, first 60 AAs: 19.52923\nTotal prob of N-in: 0.94351\nPOSSIBLE N-term signal sequence\ninside 1 19\nTMhelix 20 39\noutside 40 325\nTMhelix 326 348\ninside 349 372

6550 GCF_001267355.1_401091 Escherichia coli Proteobacteria; Gammaproteobacteria; Enterobacterales; Enterobacteriaceae; Escherichia MTGDNTLIHSHGINRRDFMKLCAALAATMGLSSKAAAEMAESVTNPQRPPVIWIGAQECT WP_000145410.1 MULTISPECIES: hydrogenase-2 small chain [Proteobacteria] Length: 372\nNumber of predicted TMHs: 1\nExp number of AAs in TMHs: 23.72226\nExp number, first 60 AAs: 0.22038\nTotal prob of N-in: 0.12994\noutside 1 331\nTMhelix 332 354\ninside 355 372

6551 GCF_001268205.1_1.EC2733.1 Escherichia coli Proteobacteria; Gammaproteobacteria; Enterobacterales; Enterobacteriaceae; Escherichia MTGDNTLIHSHGINRRDFMKLCAALAATMGLSSKAAAEMAESVTNPQRPPVIWIGAQECT WP_000145410.1 MULTISPECIES: hydrogenase-2 small chain [Proteobacteria] Length: 372\nNumber of predicted TMHs: 1\nExp number of AAs in TMHs: 23.72226\nExp number, first 60 AAs: 0.22038\nTotal prob of N-in: 0.12994\noutside 1 331\nTMhelix 332 354\ninside 355 372

6552 GCF_001268225.1_1.EC2990.1 Escherichia coli Proteobacteria; Gammaproteobacteria; Enterobacterales; Enterobacteriaceae; Escherichia MTGDNTLIHSHGINRRDFMKLCAALAATMGLSSKAAAEMAESVTNPQRPPVIWIGAQECT WP_000145410.1 MULTISPECIES: hydrogenase-2 small chain [Proteobacteria] Length: 372\nNumber of predicted TMHs: 1\nExp number of AAs in TMHs: 23.72226\nExp number, first 60 AAs: 0.22038\nTotal prob of N-in: 0.12994\noutside 1 331\nTMhelix 332 354\ninside 355 372

6553 GCF_001268265.1_1.EC2996.1 Escherichia coli Proteobacteria; Gammaproteobacteria; Enterobacterales; Enterobacteriaceae; Escherichia MTGDNTLIHSHGINRRDFMKLCAALAATMGLSSKAAAEMAESVTNPQRPPVIWIGAQECT WP_000145410.1 MULTISPECIES: hydrogenase-2 small chain [Proteobacteria] Length: 372\nNumber of predicted TMHs: 1\nExp number of AAs in TMHs: 23.72226\nExp number, first 60 AAs: 0.22038\nTotal prob of N-in: 0.12994\noutside 1 331\nTMhelix 332 354\ninside 355 372

6554 GCF_001268305.1_YE15 Escherichia coli Proteobacteria; Gammaproteobacteria; Enterobacterales; Enterobacteriaceae; Escherichia MTGDNTLIHSHGINRRDFMKLCAALAATMGLSSKAAAEMAESVTNPQRPPVIWIGAQECT WP_000145410.1 MULTISPECIES: hydrogenase-2 small chain [Proteobacteria] Length: 372\nNumber of predicted TMHs: 1\nExp number of AAs in TMHs: 23.72226\nExp number, first 60 AAs: 0.22038\nTotal prob of N-in: 0.12994\noutside 1 331\nTMhelix 332 354\ninside 355 372

6555 GCF_001268405.1_1.ECD11.1 Escherichia coli Proteobacteria; Gammaproteobacteria; Enterobacterales; Enterobacteriaceae; Escherichia MTGDNTLIHSHGINRRDFMKLCAALAATMGLSSKAAAEMAESVTNPQRPPVIWIGAQECT WP_000145410.1 MULTISPECIES: hydrogenase-2 small chain [Proteobacteria] Length: 372\nNumber of predicted TMHs: 1\nExp number of AAs in TMHs: 23.72226\nExp number, first 60 AAs: 0.22038\nTotal prob of N-in: 0.12994\noutside 1 331\nTMhelix 332 354\ninside 355 372

6556 GCF_001268365.1_YE30 Escherichia coli Proteobacteria; Gammaproteobacteria; Enterobacterales; Enterobacteriaceae; Escherichia MNNEETFYQAMRRQGVTRRSFLKYCSLAATSLGLGAGMAPKIAWALENKPRIPVVWIHGL WP_001058323.1 MULTISPECIES: hydrogenase-1 small chain [Proteobacteria] Length: 372\nNumber of predicted TMHs: 2\nExp number of AAs in TMHs: 37.2608499999999\nExp number, first 60 AAs: 19.52923\nTotal prob of N-in: 0.94351\nPOSSIBLE N-term signal sequence\ninside 1 19\nTMhelix 20 39\noutside 40 325\nTMhelix 326 348\ninside 349 372

6557 GCF_001268405.1_1.ECD11.1 Escherichia coli Proteobacteria; Gammaproteobacteria; Enterobacterales; Enterobacteriaceae; Escherichia MNNEETFYQAMRRQGVTRRSFLKYCSLAATSLGLGAGMAPKIAWALENKPRIPVVWIHGL WP_001058323.1 MULTISPECIES: hydrogenase-1 small chain [Proteobacteria] Length: 372\nNumber of predicted TMHs: 2\nExp number of AAs in TMHs: 37.2608499999999\nExp number, first 60 AAs: 19.52923\nTotal prob of N-in: 0.94351\nPOSSIBLE N-term signal sequence\ninside 1 19\nTMhelix 20 39\noutside 40 325\nTMhelix 326 348\ninside 349 372

6558 GCF_001268345.1_YE1 Escherichia coli Proteobacteria; Gammaproteobacteria; Enterobacterales; Enterobacteriaceae; Escherichia MNNEETFYQAMRRQGVTRRSFLKYCSLAATSLGLGAGMAPKIAWALENKPRIPVVWIHGL WP_001058323.1 MULTISPECIES: hydrogenase-1 small chain [Proteobacteria] Length: 372\nNumber of predicted TMHs: 2\nExp number of AAs in TMHs: 37.2608499999999\nExp number, first 60 AAs: 19.52923\nTotal prob of N-in: 0.94351\nPOSSIBLE N-term signal sequence\ninside 1 19\nTMhelix 20 39\noutside 40 325\nTMhelix 326 348\ninside 349 372

6559 GCF_001268445.1_1.EC2997.1 Escherichia coli Proteobacteria; Gammaproteobacteria; Enterobacterales; Enterobacteriaceae; Escherichia MTGDNTLIHSHGINRRDFMKLCAALAATMGLSSKAAAEMAESVTNPQRPPVIWIGAQECT WP_000145410.1 MULTISPECIES: hydrogenase-2 small chain [Proteobacteria] Length: 372\nNumber of predicted TMHs: 1\nExp number of AAs in TMHs: 23.72226\nExp number, first 60 AAs: 0.22038\nTotal prob of N-in: 0.12994\noutside 1 331\nTMhelix 332 354\ninside 355 372

6560 GCF_001268485.1_YE24 Escherichia coli Proteobacteria; Gammaproteobacteria; Enterobacterales; Enterobacteriaceae; Escherichia MTGDNTLIHSHGINRRDFMKLCAALAATMGLSSKAAAEMAESVTNPQRPPVIWIGAQECT WP_000145410.1 MULTISPECIES: hydrogenase-2 small chain [Proteobacteria] Length: 372\nNumber of predicted TMHs: 1\nExp number of AAs in TMHs: 23.72226\nExp number, first 60 AAs: 0.22038\nTotal prob of N-in: 0.12994\noutside 1 331\nTMhelix 332 354\ninside 355 372

6561 GCF_001268445.1_1.EC2997.1 Escherichia coli Proteobacteria; Gammaproteobacteria; Enterobacterales; Enterobacteriaceae; Escherichia MNNEETFYQAMRRQGVTRRSFLKYCSLAATSLGLGAGMAPKIAWALENKPRIPVVWIHGL WP_001058320.1 hydrogenase [Escherichia coli] Length: 372\nNumber of predicted TMHs: 2\nExp number of AAs in TMHs: 37.26082\nExp number, first 60 AAs: 19.52923\nTotal prob of N-in: 0.94351\nPOSSIBLE N-term signal sequence\ninside 1 19\nTMhelix 20 39\noutside 40 325\nTMhelix 326 348\ninside 349 372

6562 GCF_001268725.1_YE7 Escherichia coli Proteobacteria; Gammaproteobacteria; Enterobacterales; Enterobacteriaceae; Escherichia MTGDNTLIHSHGINRRDFMKLCAALAATMGLSSKAAAEMAESVTNPQRPPVIWIGAQECT WP_000145410.1 MULTISPECIES: hydrogenase-2 small chain [Proteobacteria] Length: 372\nNumber of predicted TMHs: 1\nExp number of AAs in TMHs: 23.72226\nExp number, first 60 AAs: 0.22038\nTotal prob of N-in: 0.12994\noutside 1 331\nTMhelix 332 354\ninside 355 372

6563 GCF_001268505.1_YE16 Escherichia coli Proteobacteria; Gammaproteobacteria; Enterobacterales; Enterobacteriaceae; Escherichia MTGDNTLIHSHGINRRDFMKLCAALAATMGLSSKAAAEMAESVTNPQRPPVIWIGAQECT WP_053263529.1 hydrogenase 2 small subunit [Escherichia coli] Length: 372\nNumber of predicted TMHs: 1\nExp number of AAs in TMHs: 23.79108\nExp number, first 60 AAs: 0.22018\nTotal prob of N-in: 0.12753\noutside 1 331\nTMhelix 332 354\ninside 355 372

6564 GCF_001268545.1_YE11 Escherichia coli Proteobacteria; Gammaproteobacteria; Enterobacterales; Enterobacteriaceae; Escherichia MNNEETFYQAMRRQGVTRRSFLKYCSLAATSLGLGAGMAPKIAWALENKPRIPVVWIHGL WP_001295940.1 MULTISPECIES: hydrogenase [Enterobacteriaceae] Length: 372\nNumber of predicted TMHs: 2\nExp number of AAs in TMHs: 37.2608899999999\nExp number, first 60 AAs: 19.52923\nTotal prob of N-in: 0.94351\nPOSSIBLE N-term signal sequence\ninside 1 19\nTMhelix 20 39\noutside 40 325\nTMhelix 326 348\ninside 349 372

6565 GCF_001268585.1_YE14 Escherichia coli Proteobacteria; Gammaproteobacteria; Enterobacterales; Enterobacteriaceae; Escherichia MNNEETFYQAMRRQGVTRRSFLKYCSLAATSLGLGAGMAPKIAWALENKPRIPVVWIHGL WP_001058323.1 MULTISPECIES: hydrogenase-1 small chain [Proteobacteria] Length: 372\nNumber of predicted TMHs: 2\nExp number of AAs in TMHs: 37.2608499999999\nExp number, first 60 AAs: 19.52923\nTotal prob of N-in: 0.94351\nPOSSIBLE N-term signal sequence\ninside 1 19\nTMhelix 20 39\noutside 40 325\nTMhelix 326 348\ninside 349 372

6566 GCF_001268605.1_YE29 Escherichia coli Proteobacteria; Gammaproteobacteria; Enterobacterales; Enterobacteriaceae; Escherichia MNNEETFYQAMRRQGVTRRSFLKYCSLAATSLGLGAGMAPKIAWALENKPRIPVVWIHGL WP_001058323.1 MULTISPECIES: hydrogenase-1 small chain [Proteobacteria] Length: 372\nNumber of predicted TMHs: 2\nExp number of AAs in TMHs: 37.2608499999999\nExp number, first 60 AAs: 19.52923\nTotal prob of N-in: 0.94351\nPOSSIBLE N-term signal sequence\ninside 1 19\nTMhelix 20 39\noutside 40 325\nTMhelix 326 348\ninside 349 372

6567 GCF_001268645.1_YE2 Escherichia coli Proteobacteria; Gammaproteobacteria; Enterobacterales; Enterobacteriaceae; Escherichia MNNEETFYQAMRRQGVTRRSFLKYCSLAATSLGLGAGMAPKIAWALENKPRIPVVWIHGL WP_001058323.1 MULTISPECIES: hydrogenase-1 small chain [Proteobacteria] Length: 372\nNumber of predicted TMHs: 2\nExp number of AAs in TMHs: 37.2608499999999\nExp number, first 60 AAs: 19.52923\nTotal prob of N-in: 0.94351\nPOSSIBLE N-term signal sequence\ninside 1 19\nTMhelix 20 39\noutside 40 325\nTMhelix 326 348\ninside 349 372

6568 GCF_001268685.1_YE19 Escherichia coli Proteobacteria; Gammaproteobacteria; Enterobacterales; Enterobacteriaceae; Escherichia MNNEETFYQAMRRQGVTRRSFLKYCSLAATSLGLGAGMAPKIAWALENKPRIPVVWIHGL WP_001058323.1 MULTISPECIES: hydrogenase-1 small chain [Proteobacteria] Length: 372\nNumber of predicted TMHs: 2\nExp number of AAs in TMHs: 37.2608499999999\nExp number, first 60 AAs: 19.52923\nTotal prob of N-in: 0.94351\nPOSSIBLE N-term signal sequence\ninside 1 19\nTMhelix 20 39\noutside 40 325\nTMhelix 326 348\ninside 349 372

6569 GCF_001268905.1_YE10 Escherichia coli Proteobacteria; Gammaproteobacteria; Enterobacterales; Enterobacteriaceae; Escherichia MTGDNTLIHSHGINRRDFMKLCAALAATMGLSSKAAAEMAESVTNPQRPPVIWIGAQECT WP_000145410.1 MULTISPECIES: hydrogenase-2 small chain [Proteobacteria] Length: 372\nNumber of predicted TMHs: 1\nExp number of AAs in TMHs: 23.72226\nExp number, first 60 AAs: 0.22038\nTotal prob of N-in: 0.12994\noutside 1 331\nTMhelix 332 354\ninside 355 372

6570 GCF_001268925.1_1.EC2732.1 Escherichia coli Proteobacteria; Gammaproteobacteria; Enterobacterales; Enterobacteriaceae; Escherichia MTGDNTLIHSHGINRRDFMKLCAALAATMGLSSKAAAEMAESVTNPQRPPVIWIGAQECT WP_000145410.1 MULTISPECIES: hydrogenase-2 small chain [Proteobacteria] Length: 372\nNumber of predicted TMHs: 1\nExp number of AAs in TMHs: 23.72226\nExp number, first 60 AAs: 0.22038\nTotal prob of N-in: 0.12994\noutside 1 331\nTMhelix 332 354\ninside 355 372

6571 GCF_001268985.1_1.ECB31.1 Escherichia coli Proteobacteria; Gammaproteobacteria; Enterobacterales; Enterobacteriaceae; Escherichia
MTGDNTLIHSHGINRRDFMKLCAALAATMGLSSKAAAEMAESVTNPQRPPVIWIGAQECT WP_000145410.1
MULTISPECIES: hydrogenase-2 small chain [Proteobacteria] Length: 372\nNumber of predicted TMHs: 1\nExp number of AAs in TMHs: 23.72226\nExp number, first 60 AAs: 0.22038\nTotal prob of N-in: 0.12994\noutside 1 331\nTMhelix 332 354\ninside 355 372

6572 GCF_001269025.1_1.EC2989.1 Escherichia coli Proteobacteria; Gammaproteobacteria; Enterobacterales; Enterobacteriaceae; Escherichia
MTGDNTLIHSHGINRRDFMKLCAALAATMGLSSKAAAEMAESVTNPQRPPVIWIGAQECT WP_000145410.1
MULTISPECIES: hydrogenase-2 small chain [Proteobacteria] Length: 372\nNumber of predicted TMHs: 1\nExp number of AAs in TMHs: 23.72226\nExp number, first 60 AAs: 0.22038\nTotal prob of N-in: 0.12994\noutside 1 331\nTMhelix 332 354\ninside 355 372

6573 GCF_001268865.1_1.EC2739.1 Escherichia coli Proteobacteria; Gammaproteobacteria; Enterobacterales; Enterobacteriaceae; Escherichia
MNNEETFYQAMRRQGVTRRSFLKYCSLAATSLGLGAGMAPKIAWALENKPRIPVVWIHGL WP_001058323.1
MULTISPECIES: hydrogenase-1 small chain [Proteobacteria] Length: 372\nNumber of predicted TMHs: 2\nExp number of AAs in TMHs: 37.2608499999999\nExp number, first 60 AAs: 19.52923\nTotal prob of N-in: 0.94351\nPOSSIBLE N-term signal sequence\ninside 1 19\nTMhelix 20 39\noutside 40 325\nTMhelix 326 348\ninside 349 372

6574 GCF_001268785.1_YE25 Escherichia coli Proteobacteria; Gammaproteobacteria; Enterobacterales; Enterobacteriaceae; Escherichia MNNEETFYQAMRRQGVTRRSFLKYCSLAATSLGLGAGMAPKIAWALENKPRIPVVWIHGL
WP_001295940.1 MULTISPECIES: hydrogenase [Enterobacteriaceae] Length: 372\nNumber of predicted TMHs: 2\nExp number of AAs in TMHs: 37.2608899999999\nExp number, first 60 AAs: 19.52923\nTotal prob of N-in: 0.94351\nPOSSIBLE N-term signal sequence\ninside 1 19\nTMhelix 20 39\noutside 40 325\nTMhelix 326 348\ninside 349 372

6575 GCF_001269065.1_1.EC2729.1 Escherichia coli Proteobacteria; Gammaproteobacteria; Enterobacterales; Enterobacteriaceae; Escherichia
MTGDNTLIHSHGINRRDFMKLCAALAATMGLSSKAAAEMAESVTNPQRPPVIWIGAQECT WP_000145410.1
MULTISPECIES: hydrogenase-2 small chain [Proteobacteria] Length: 372\nNumber of predicted TMHs: 1\nExp number of AAs in TMHs: 23.72226\nExp number, first 60 AAs: 0.22038\nTotal prob of N-in: 0.12994\noutside 1 331\nTMhelix 332 354\ninside 355 372

6576 GCF_001268925.1_1.EC2732.1 Escherichia coli Proteobacteria; Gammaproteobacteria; Enterobacterales; Enterobacteriaceae; Escherichia
MNNEETFYQAMRRQGVTRRSFLKYCSLAATSLGLGAGMAPKIAWALENKPRIPVVWIHGL WP_001058323.1
MULTISPECIES: hydrogenase-1 small chain [Proteobacteria] Length: 372\nNumber of predicted TMHs: 2\nExp number of AAs in TMHs: 37.2608499999999\nExp number, first 60 AAs: 19.52923\nTotal prob of N-in: 0.94351\nPOSSIBLE N-term signal sequence\ninside 1 19\nTMhelix 20 39\noutside 40 325\nTMhelix 326 348\ninside 349 372

6577 GCF_001269205.1_1.EC2995.1 Escherichia coli Proteobacteria; Gammaproteobacteria; Enterobacterales; Enterobacteriaceae; Escherichia
MTGDNTLIHSHGINRRDFMKLCAALAATMGLSSKAAAEMAESVTNPQRPPVIWIGAQECT WP_000145410.1
MULTISPECIES: hydrogenase-2 small chain [Proteobacteria] Length: 372\nNumber of predicted TMHs: 1\nExp number of AAs in TMHs: 23.72226\nExp number, first 60 AAs: 0.22038\nTotal prob of N-in: 0.12994\noutside 1 331\nTMhelix 332 354\ninside 355 372

6578 GCF_001269205.1_1.EC2995.1 Escherichia coli Proteobacteria; Gammaproteobacteria; Enterobacterales; Enterobacteriaceae; Escherichia
MNNEETFYQAMRRQGVTRRSFLKYCSLAATSLGLGAGMAPKIAWALENKPRIPVVWIHGL WP_053264900.1
hydrogenase [Escherichia coli] Length: 372\nNumber of predicted TMHs: 2\nExp number of AAs in TMHs: 37.24714\nExp number, first 60 AAs: 19.52921\nTotal prob of N-in: 0.94351\nPOSSIBLE N-term signal sequence\ninside 1 19\nTMhelix 20 39\noutside 40 325\nTMhelix 326 348\ninside 349 372

6579 GCF_001269025.1_1.EC2989.1 Escherichia coli Proteobacteria; Gammaproteobacteria; Enterobacterales; Enterobacteriaceae; Escherichia
MNNEETFYQAMRRQGVTRRSFLKYCSLAATSLGLGAGMAPKIAWALENKPRIPVVWIHGL WP_001295940.1

MULTISPECIES: hydrogenase [Enterobacteriaceae] Length: 372\nNumber of predicted TMHs: 2\nExp number of AAs in TMHs: 37.2608899999999\nExp number, first 60 AAs: 19.52923\nTotal prob of N-in: 0.94351\nPOSSIBLE N-term signal sequence\ninside 1 19\nTMhelix 20 39\noutside 40 325\nTMhelix 326 348\ninside 349 372

6580 GCF_001269125.1_YE26 Escherichia coli Proteobacteria; Gammaproteobacteria; Enterobacterales; Enterobacteriaceae; Escherichia MNNEETFYQAMRRQGVTRRSFLKYCSLAATSLGLGAGMAPKIAWALENKPRIPVVWIHGL WP_001058323.1 MULTISPECIES: hydrogenase-1 small chain [Proteobacteria] Length: 372\nNumber of predicted TMHs: 2\nExp number of AAs in TMHs: 37.2608499999999\nExp number, first 60 AAs: 19.52923\nTotal prob of N-in: 0.94351\nPOSSIBLE N-term signal sequence\ninside 1 19\nTMhelix 20 39\noutside 40 325\nTMhelix 326 348\ninside 349 372

6581 GCF_001269225.1_1.EC2987.1 Escherichia coli Proteobacteria; Gammaproteobacteria; Enterobacterales; Enterobacteriaceae; Escherichia MTGDNTLIHSHGINRRDFMKLCAALAATMGLSSKAAAEMAESVTNPQRPPVIWIGAQECT WP_000145410.1 MULTISPECIES: hydrogenase-2 small chain [Proteobacteria] Length: 372\nNumber of predicted TMHs: 1\nExp number of AAs in TMHs: 23.72226\nExp number, first 60 AAs: 0.22038\nTotal prob of N-in: 0.12994\noutside 1 331\nTMhelix 332 354\ninside 355 372

6582 GCF_001269265.1_1.EC2723.1 Escherichia coli Proteobacteria; Gammaproteobacteria; Enterobacterales; Enterobacteriaceae; Escherichia MTGDNTLIHSHGINRRDFMKLCAALAATMGLSSKAAAEMAESVTNPQRPPVIWIGAQECT WP_000145410.1 MULTISPECIES: hydrogenase-2 small chain [Proteobacteria] Length: 372\nNumber of predicted TMHs: 1\nExp number of AAs in TMHs: 23.72226\nExp number, first 60 AAs: 0.22038\nTotal prob of N-in: 0.12994\noutside 1 331\nTMhelix 332 354\ninside 355 372

6583 GCF_001277455.1_7553_7_63 Escherichia coli Proteobacteria; Gammaproteobacteria; Enterobacterales; Enterobacteriaceae; Escherichia MTGDNTLIHSHGINRRDFMKLCAALAATMGLSSKAAAEMAESVTNPQRPPVIWIGAQECT WP_000145410.1 MULTISPECIES: hydrogenase-2 small chain [Proteobacteria] Length: 372\nNumber of predicted TMHs: 1\nExp number of AAs in TMHs: 23.72226\nExp number, first 60 AAs: 0.22038\nTotal prob of N-in: 0.12994\noutside 1 331\nTMhelix 332 354\ninside 355 372

6584 GCF_001277455.1_7553_7_63 Escherichia coli Proteobacteria; Gammaproteobacteria; Enterobacterales; Enterobacteriaceae; Escherichia MNNEETFYQAMRRQGVTRRSFLKYCSLAATSLGLGAGMAPKIAWALENKPRIPVVWIHGL WP_001058323.1 MULTISPECIES: hydrogenase-1 small chain [Proteobacteria] Length: 372\nNumber of predicted TMHs: 2\nExp number of AAs in TMHs: 37.2608499999999\nExp number, first 60 AAs: 19.52923\nTotal prob of N-in: 0.94351\nPOSSIBLE N-term signal sequence\ninside 1 19\nTMhelix 20 39\noutside 40 325\nTMhelix 326 348\ninside 349 372

6585 GCF_001269225.1_1.EC2987.1 Escherichia coli Proteobacteria; Gammaproteobacteria; Enterobacterales; Enterobacteriaceae; Escherichia MNNEETFYQAMRRQGVTRRSFLKYCSLAATSLGLGAGMAPKIAWALENKPRIPVVWIHGL WP_001295940.1 MULTISPECIES: hydrogenase [Enterobacteriaceae] Length: 372\nNumber of predicted TMHs: 2\nExp number of AAs in TMHs: 37.2608899999999\nExp number, first 60 AAs: 19.52923\nTotal prob of N-in: 0.94351\nPOSSIBLE N-term signal sequence\ninside 1 19\nTMhelix 20 39\noutside 40 325\nTMhelix 326 348\ninside 349 372

6586 GCF_001277495.1_7790_1_79 Escherichia coli Proteobacteria; Gammaproteobacteria; Enterobacterales; Enterobacteriaceae; Escherichia MTGDNTLIHSHGINRRDFMKLCAALAATMGLSSKAAAEMAESVTNPQRPPVIWIGAQECT WP_000145410.1 MULTISPECIES: hydrogenase-2 small chain [Proteobacteria] Length: 372\nNumber of predicted TMHs: 1\nExp number of AAs in TMHs: 23.72226\nExp number, first 60 AAs: 0.22038\nTotal prob of N-in: 0.12994\noutside 1 331\nTMhelix 332 354\ninside 355 372

6587 GCF_001277495.1_7790_1_79 Escherichia coli Proteobacteria; Gammaproteobacteria; Enterobacterales; Enterobacteriaceae; Escherichia MNNEETFYQAMRRQGVTRRSFLKYCSLAATSLGLGAGMAPKIAWALENKPRIPVVWIHGL WP_001058320.1 hydrogenase [Escherichia coli] Length: 372\nNumber of predicted TMHs: 2\nExp number of AAs in

TMHs: 37.26082\nExp number, first 60 AAs: 19.52923\nTotal prob of N-in: 0.94351\nPOSSIBLE N-term signal sequence\ninside 1 19\nTMhelix 20 39\noutside 40 325\nTMhelix 326 348\ninside 349 372

6588 GCF_001276265.1_ASM127626v1 Escherichia coli Proteobacteria; Gammaproteobacteria; Enterobacterales; Enterobacteriaceae; Escherichia
MNNEETFYQAMRRQGVTRRSFLKYCSLAATSLGLGAGMAPKIAWALENKPRIPVVWIHGL WP_001058320.1
hydrogenase [Escherichia coli] Length: 372\nNumber of predicted TMHs: 2\nExp number of AAs in TMHs: 37.26082\nExp number, first 60 AAs: 19.52923\nTotal prob of N-in: 0.94351\nPOSSIBLE N-term signal sequence\ninside 1 19\nTMhelix 20 39\noutside 40 325\nTMhelix 326 348\ninside 349 372

6589 GCF_001277515.1_7790_1_59 Escherichia coli Proteobacteria; Gammaproteobacteria; Enterobacterales; Enterobacteriaceae; Escherichia
MTGDNTLIHSHGINRRDFMKLCAALAATMGLSSKAAAEMAESVTNPQRPPVIWIGAQECT WP_000145410.1
MULTISPECIES: hydrogenase-2 small chain [Proteobacteria] Length: 372\nNumber of predicted TMHs: 1\nExp number of AAs in TMHs: 23.72226\nExp number, first 60 AAs: 0.22038\nTotal prob of N-in: 0.12994\noutside 1 331\nTMhelix 332 354\ninside 355 372

6590 GCF_001277615.1_7790_1_70 Escherichia coli Proteobacteria; Gammaproteobacteria; Enterobacterales; Enterobacteriaceae; Escherichia
MNNEETFYQAMRRQGVTRRSFLKYCSLAATSLGLGAGMAPKIAWALENKPRIPVVWIHGL WP_001295940.1
MULTISPECIES: hydrogenase [Enterobacteriaceae] Length: 372\nNumber of predicted TMHs: 2\nExp number of AAs in TMHs: 37.2608899999999\nExp number, first 60 AAs: 19.52923\nTotal prob of N-in: 0.94351\nPOSSIBLE N-term signal sequence\ninside 1 19\nTMhelix 20 39\noutside 40 325\nTMhelix 326 348\ninside 349 372

6591 GCF_001277395.1_7790_1_92 Escherichia coli Proteobacteria; Gammaproteobacteria; Enterobacterales; Enterobacteriaceae; Escherichia
MNNEETFYQAMRRQGVTRRSFLKYCSLAATSLGLGAGMAPKIAWALENKPRIPVVWIHGL WP_001295940.1
MULTISPECIES: hydrogenase [Enterobacteriaceae] Length: 372\nNumber of predicted TMHs: 2\nExp number of AAs in TMHs: 37.2608899999999\nExp number, first 60 AAs: 19.52923\nTotal prob of N-in: 0.94351\nPOSSIBLE N-term signal sequence\ninside 1 19\nTMhelix 20 39\noutside 40 325\nTMhelix 326 348\ninside 349 372

6592 GCF_001277695.1_7748_7_33 Escherichia coli Proteobacteria; Gammaproteobacteria; Enterobacterales; Enterobacteriaceae; Escherichia
MTGDNTLIHSHGINRRDFMKLCAALAATMGLSSKAAAEMAESVTNPQRPPVIWIGAQECT WP_000145410.1
MULTISPECIES: hydrogenase-2 small chain [Proteobacteria] Length: 372\nNumber of predicted TMHs: 1\nExp number of AAs in TMHs: 23.72226\nExp number, first 60 AAs: 0.22038\nTotal prob of N-in: 0.12994\noutside 1 331\nTMhelix 332 354\ninside 355 372

6593 GCF_001277655.1_7553_7_71 Escherichia coli Proteobacteria; Gammaproteobacteria; Enterobacterales; Enterobacteriaceae; Escherichia
MTGDNTLIHSHGINRRDFMKLCAALAATMGLSSKAAAEMAESVTNPQRPPVIWIGAQECT WP_000145410.1
MULTISPECIES: hydrogenase-2 small chain [Proteobacteria] Length: 372\nNumber of predicted TMHs: 1\nExp number of AAs in TMHs: 23.72226\nExp number, first 60 AAs: 0.22038\nTotal prob of N-in: 0.12994\noutside 1 331\nTMhelix 332 354\ninside 355 372

6594 GCF_001277715.1_7790_1_58 Escherichia coli Proteobacteria; Gammaproteobacteria; Enterobacterales; Enterobacteriaceae; Escherichia
MTGDNTLIHSHGINRRDFMKLCAALAATMGLSSKAAAEMAESVTNPQRPPVIWIGAQECT WP_000145410.1
MULTISPECIES: hydrogenase-2 small chain [Proteobacteria] Length: 372\nNumber of predicted TMHs: 1\nExp number of AAs in TMHs: 23.72226\nExp number, first 60 AAs: 0.22038\nTotal prob of N-in: 0.12994\noutside 1 331\nTMhelix 332 354\ninside 355 372

6595 GCF_001277715.1_7790_1_58 Escherichia coli Proteobacteria; Gammaproteobacteria; Enterobacterales; Enterobacteriaceae; Escherichia
MNNEETFYQAMRRQGVTRRSFLKYCSLAATSLGLGAGMAPKIAWALENKPRIPVVWIHGL WP_001058323.1
MULTISPECIES: hydrogenase-1 small chain [Proteobacteria] Length: 372\nNumber of predicted TMHs: 2\nExp number of AAs in TMHs: 37.2608499999999\nExp number, first 60 AAs: 19.52923\nTotal prob of N-in: 0.94351\nPOSSIBLE N-term signal sequence\ninside 1 19\nTMhelix 20 39\noutside 40 325\nTMhelix 326 348\ninside 349 372

6596 GCF_001281775.1_ASM128177v1 Escherichia coli Proteobacteria; Gammaproteobacteria; Enterobacterales; Enterobacteriaceae; Escherichia
MNNEETFYQAMRRQGVTRRSFLKYCSLAATSLGLGAGMAPKIAWALENKPRIPVVWIHGL WP_001301613.1
hydrogenase [Escherichia coli] Length: 372\nNumber of predicted TMHs: 2\nExp number of AAs in TMHs: 37.46598\nExp number, first 60 AAs: 19.53139\nTotal prob of N-in: 0.94364\nPOSSIBLE N-term signal sequence\ninside 1 19\nTMhelix 20 39\noutside 40 325\nTMhelix 326 348\ninside 349 372

6597 GCF_001283705.1_7748_7_16 Escherichia coli Proteobacteria; Gammaproteobacteria; Enterobacterales; Enterobacteriaceae; Escherichia
MTGDNTLIHSHGINRRDFMKLCAALAATMGLSSKAAAEMAESVTNPQRPPVIWIGAQECT WP_000145410.1
MULTISPECIES: hydrogenase-2 small chain [Proteobacteria] Length: 372\nNumber of predicted TMHs: 1\nExp number of AAs in TMHs: 23.72226\nExp number, first 60 AAs: 0.22038\nTotal prob of N-in: 0.12994\noutside 1 331\nTMhelix 332 354\ninside 355 372

6598 GCF_001281685.1_ASM128168v1 Escherichia coli Proteobacteria; Gammaproteobacteria; Enterobacterales; Enterobacteriaceae; Escherichia
MNNEETFYQAMRRQGVTRRSFLKYCSLAATSLGLGAGMAPKIAWALENKPRIPVVWIHGL WP_001301613.1
hydrogenase [Escherichia coli] Length: 372\nNumber of predicted TMHs: 2\nExp number of AAs in TMHs: 37.46598\nExp number, first 60 AAs: 19.53139\nTotal prob of N-in: 0.94364\nPOSSIBLE N-term signal sequence\ninside 1 19\nTMhelix 20 39\noutside 40 325\nTMhelix 326 348\ninside 349 372

6599 GCF_001281985.1_ASM128198v1 Escherichia coli Proteobacteria; Gammaproteobacteria; Enterobacterales; Enterobacteriaceae; Escherichia
MTGDNTLIHSHGINRRDFMKLCAALAATMGLSSKAAAEMAESVTNPQRPPVIWIGAQECT WP_000145410.1
MULTISPECIES: hydrogenase-2 small chain [Proteobacteria] Length: 372\nNumber of predicted TMHs: 1\nExp number of AAs in TMHs: 23.72226\nExp number, first 60 AAs: 0.22038\nTotal prob of N-in: 0.12994\noutside 1 331\nTMhelix 332 354\ninside 355 372

6600 GCF_001281985.1_ASM128198v1 Escherichia coli Proteobacteria; Gammaproteobacteria; Enterobacterales; Enterobacteriaceae; Escherichia
MNNEETFYQAMRRQGVTRRSFLKYCSLAATSLGLGAGMAPKIAWALENKPRIPVVWIHGL WP_001301613.1
hydrogenase [Escherichia coli] Length: 372\nNumber of predicted TMHs: 2\nExp number of AAs in TMHs: 37.46598\nExp number, first 60 AAs: 19.53139\nTotal prob of N-in: 0.94364\nPOSSIBLE N-term signal sequence\ninside 1 19\nTMhelix 20 39\noutside 40 325\nTMhelix 326 348\ninside 349 372

6601 GCF_001281755.1_ASM128175v1 Escherichia coli Proteobacteria; Gammaproteobacteria; Enterobacterales; Enterobacteriaceae; Escherichia
MNNEETFYQAMRRQGVTRRSFLKYCSLAATSLGLGAGMAPKIAWALENKPRIPVVWIHGL WP_001301613.1
hydrogenase [Escherichia coli] Length: 372\nNumber of predicted TMHs: 2\nExp number of AAs in TMHs: 37.46598\nExp number, first 60 AAs: 19.53139\nTotal prob of N-in: 0.94364\nPOSSIBLE N-term signal sequence\ninside 1 19\nTMhelix 20 39\noutside 40 325\nTMhelix 326 348\ninside 349 372

6602 GCF_001282025.1_ASM128202v1 Escherichia coli Proteobacteria; Gammaproteobacteria; Enterobacterales; Enterobacteriaceae; Escherichia
MTGDNTLIHSHGINRRDFMKLCAALAATMGLSSKAAAEMAESVTNPQRPPVIWIGAQECT WP_000145410.1
MULTISPECIES: hydrogenase-2 small chain [Proteobacteria] Length: 372\nNumber of predicted TMHs: 1\nExp number of AAs in TMHs: 23.72226\nExp number, first 60 AAs: 0.22038\nTotal prob of N-in: 0.12994\noutside 1 331\nTMhelix 332 354\ninside 355 372

6603 GCF_001282215.1_ASM128221v1 Escherichia coli Proteobacteria; Gammaproteobacteria; Enterobacterales; Enterobacteriaceae; Escherichia
MTGDNTLIHSHGINRRDFMKLCAALAATMGLSSKAAAEMAESVTNPQRPPVIWIGAQECT WP_000145410.1
MULTISPECIES: hydrogenase-2 small chain [Proteobacteria] Length: 372\nNumber of predicted TMHs: 1\nExp number of AAs in TMHs: 23.72226\nExp number, first 60 AAs: 0.22038\nTotal prob of N-in: 0.12994\noutside 1 331\nTMhelix 332 354\ninside 355 372

6604 GCF_001282235.1_ASM128223v1 Escherichia coli Proteobacteria; Gammaproteobacteria; Enterobacterales; Enterobacteriaceae; Escherichia
MTGDNTLIHSHGINRRDFMKLCAALAATMGLSSKAAAEMAESVTNPQRPPVIWIGAQECT WP_000145410.1
MULTISPECIES: hydrogenase-2 small chain [Proteobacteria] Length: 372\nNumber of predicted TMHs:

1\nExp number of AAs in TMHs: 23.72226\nExp number, first 60 AAs: 0.22038\nTotal prob of N-in: 0.12994\noutside 1 331\nTMhelix 332 354\ninside 355 372

6605 GCF_001282215.1_ASM128221v1 Escherichia coli Proteobacteria; Gammaproteobacteria; Enterobacterales; Enterobacteriaceae; Escherichia
MNNEETFYQAMRRQGVTRRSFLKYCSLAATSLGLGAGMAPKIAWALENKPRIPVVWIHGL WP_001058320.1
hydrogenase [Escherichia coli] Length: 372\nNumber of predicted TMHs: 2\nExp number of AAs in TMHs: 37.26082\nExp number, first 60 AAs: 19.52923\nTotal prob of N-in: 0.94351\nPOSSIBLE N-term signal sequence\ninside 1 19\nTMhelix 20 39\noutside 40 325\nTMhelix 326 348\ninside 349 372

6606 GCF_001282235.1_ASM128223v1 Escherichia coli Proteobacteria; Gammaproteobacteria; Enterobacterales; Enterobacteriaceae; Escherichia
MNNEETFYQAMRRQGVTRRSFLKYCSLAATSLGLGAGMAPKIAWALENKPRIPVVWIHGL WP_001058320.1
hydrogenase [Escherichia coli] Length: 372\nNumber of predicted TMHs: 2\nExp number of AAs in TMHs: 37.26082\nExp number, first 60 AAs: 19.52923\nTotal prob of N-in: 0.94351\nPOSSIBLE N-term signal sequence\ninside 1 19\nTMhelix 20 39\noutside 40 325\nTMhelix 326 348\ninside 349 372

6607 GCF_001281885.1_ASM128188v1 Escherichia coli Proteobacteria; Gammaproteobacteria; Enterobacterales; Enterobacteriaceae; Escherichia
MNNEETFYQAMRRQGVTRRSFLKYCSLAATSLGLGAGMAPKIAWALENKPRIPVVWIHGL WP_001301613.1
hydrogenase [Escherichia coli] Length: 372\nNumber of predicted TMHs: 2\nExp number of AAs in TMHs: 37.46598\nExp number, first 60 AAs: 19.53139\nTotal prob of N-in: 0.94364\nPOSSIBLE N-term signal sequence\ninside 1 19\nTMhelix 20 39\noutside 40 325\nTMhelix 326 348\ninside 349 372

6608 GCF_001281915.1_ASM128191v1 Escherichia coli Proteobacteria; Gammaproteobacteria; Enterobacterales; Enterobacteriaceae; Escherichia
MNNEETFYQAMRRQGVTRRSFLKYCSLAATSLGLGAGMAPKIAWALENKPRIPVVWIHGL WP_001301613.1
hydrogenase [Escherichia coli] Length: 372\nNumber of predicted TMHs: 2\nExp number of AAs in TMHs: 37.46598\nExp number, first 60 AAs: 19.53139\nTotal prob of N-in: 0.94364\nPOSSIBLE N-term signal sequence\ninside 1 19\nTMhelix 20 39\noutside 40 325\nTMhelix 326 348\ninside 349 372

6609 GCF_001281965.1_ASM128196v1 Escherichia coli Proteobacteria; Gammaproteobacteria; Enterobacterales; Enterobacteriaceae; Escherichia
MNNEETFYQAMRRQGVTRRSFLKYCSLAATSLGLGAGMAPKIAWALENKPRIPVVWIHGL WP_001301613.1
hydrogenase [Escherichia coli] Length: 372\nNumber of predicted TMHs: 2\nExp number of AAs in TMHs: 37.46598\nExp number, first 60 AAs: 19.53139\nTotal prob of N-in: 0.94364\nPOSSIBLE N-term signal sequence\ninside 1 19\nTMhelix 20 39\noutside 40 325\nTMhelix 326 348\ninside 349 372

6610 GCF_001282375.1_ASM128237v1 Escherichia coli Proteobacteria; Gammaproteobacteria; Enterobacterales; Enterobacteriaceae; Escherichia
MNNEETFYQAMRRQGVTRRSFLKYCSLAATSLGLGAGMAPKIAWALENKPRIPVVWIHGL WP_001058320.1
hydrogenase [Escherichia coli] Length: 372\nNumber of predicted TMHs: 2\nExp number of AAs in TMHs: 37.26082\nExp number, first 60 AAs: 19.52923\nTotal prob of N-in: 0.94351\nPOSSIBLE N-term signal sequence\ninside 1 19\nTMhelix 20 39\noutside 40 325\nTMhelix 326 348\ninside 349 372

6611 GCF_001282335.1_ASM128233v1 Escherichia coli Proteobacteria; Gammaproteobacteria; Enterobacterales; Enterobacteriaceae; Escherichia
MNNEETFYQAMRRQGVTRRSFLKYCSLAATSLGLGAGMAPKIAWALENKPRIPVVWIHGL WP_001058320.1
hydrogenase [Escherichia coli] Length: 372\nNumber of predicted TMHs: 2\nExp number of AAs in TMHs: 37.26082\nExp number, first 60 AAs: 19.52923\nTotal prob of N-in: 0.94351\nPOSSIBLE N-term signal sequence\ninside 1 19\nTMhelix 20 39\noutside 40 325\nTMhelix 326 348\ninside 349 372

6612 GCF_001283445.1_8205_3_67 Escherichia coli Proteobacteria; Gammaproteobacteria; Enterobacterales; Enterobacteriaceae; Escherichia
MTGDNTLIHSHGINRRDFMKLCAALAATMGLSSKAAAEMAESVTNPQRPPVIWIGAQECT WP_000145410.1
MULTISPECIES: hydrogenase-2 small chain [Proteobacteria] Length: 372\nNumber of predicted TMHs: 1\nExp number of AAs in TMHs: 23.72226\nExp number, first 60 AAs: 0.22038\nTotal prob of N-in: 0.12994\noutside 1 331\nTMhelix 332 354\ninside 355 372

6613 GCF_001283305.1_8205_3_48 Escherichia coli Proteobacteria; Gammaproteobacteria; Enterobacterales; Enterobacteriaceae; Escherichia

MTGDNTLIHSHGINRRDFMKLCAALAATMGLSSKAAAEMAESVTNPQRPPVIWIGAQECT WP_000145410.1 MULTISPECIES: hydrogenase-2 small chain [Proteobacteria] Length: 372\nNumber of predicted TMHs: 1\nExp number of AAs in TMHs: 23.72226\nExp number, first 60 AAs: 0.22038\nTotal prob of N-in: 0.12994\noutside 1 331\nTMhelix 332 354\ninside 355 372

6614 GCF_001283485.1_7790_1_83 Escherichia coli Proteobacteria; Gammaproteobacteria; Enterobacterales; Enterobacteriaceae; Escherichia MTGDNTLIHSHGINRRDFMKLCAALAATMGLSSKAAAEMAESVTNPQRPPVIWIGAQECT WP_000145410.1 MULTISPECIES: hydrogenase-2 small chain [Proteobacteria] Length: 372\nNumber of predicted TMHs: 1\nExp number of AAs in TMHs: 23.72226\nExp number, first 60 AAs: 0.22038\nTotal prob of N-in: 0.12994\noutside 1 331\nTMhelix 332 354\ninside 355 372

6615 GCF_001283445.1_8205_3_67 Escherichia coli Proteobacteria; Gammaproteobacteria; Enterobacterales; Enterobacteriaceae; Escherichia MNNEETFYQAMRRQGVTRRSFLKYCSLAATSLGLGAGMAPKIAWALENKPRIPVVWIHGL WP_001058323.1 MULTISPECIES: hydrogenase-1 small chain [Proteobacteria] Length: 372\nNumber of predicted TMHs: 2\nExp number of AAs in TMHs: 37.2608499999999\nExp number, first 60 AAs: 19.52923\nTotal prob of N-in: 0.94351\nPOSSIBLE N-term signal sequence\ninside 1 19\nTMhelix 20 39\noutside 40 325\nTMhelix 326 348\ninside 349 372

6616 GCF_001283485.1_7790_1_83 Escherichia coli Proteobacteria; Gammaproteobacteria; Enterobacterales; Enterobacteriaceae; Escherichia MNNEETFYQAMRRQGVTRRSFLKYCSLAATSLGLGAGMAPKIAWALENKPRIPVVWIHGL WP_001058323.1 MULTISPECIES: hydrogenase-1 small chain [Proteobacteria] Length: 372\nNumber of predicted TMHs: 2\nExp number of AAs in TMHs: 37.2608499999999\nExp number, first 60 AAs: 19.52923\nTotal prob of N-in: 0.94351\nPOSSIBLE N-term signal sequence\ninside 1 19\nTMhelix 20 39\noutside 40 325\nTMhelix 326 348\ninside 349 372

6617 GCF_001283365.1_7790_1_96 Escherichia coli Proteobacteria; Gammaproteobacteria; Enterobacterales; Enterobacteriaceae; Escherichia MNNEETFYQAMRRQGVTRRSFLKYCSLAATSLGLGAGMAPKIAWALENKPRIPVVWIHGL WP_001058323.1 MULTISPECIES: hydrogenase-1 small chain [Proteobacteria] Length: 372\nNumber of predicted TMHs: 2\nExp number of AAs in TMHs: 37.2608499999999\nExp number, first 60 AAs: 19.52923\nTotal prob of N-in: 0.94351\nPOSSIBLE N-term signal sequence\ninside 1 19\nTMhelix 20 39\noutside 40 325\nTMhelix 326 348\ninside 349 372

6618 GCF_001283585.1_8205_3_77 Escherichia coli Proteobacteria; Gammaproteobacteria; Enterobacterales; Enterobacteriaceae; Escherichia MTGDNTLIHSHGINRRDFMKLCAALAATMGLSSKAAAEMAESVTNPQRPPVIWIGAQECT WP_000145410.1 MULTISPECIES: hydrogenase-2 small chain [Proteobacteria] Length: 372\nNumber of predicted TMHs: 1\nExp number of AAs in TMHs: 23.72226\nExp number, first 60 AAs: 0.22038\nTotal prob of N-in: 0.12994\noutside 1 331\nTMhelix 332 354\ninside 355 372

6619 GCF_001283605.1_8205_3_25 Escherichia coli Proteobacteria; Gammaproteobacteria; Enterobacterales; Enterobacteriaceae; Escherichia MTGDNTLIHSHGINRRDFMKLCAALAATMGLSSKAAAEMAESVTNPQRPPVIWIGAQECT WP_000145410.1 MULTISPECIES: hydrogenase-2 small chain [Proteobacteria] Length: 372\nNumber of predicted TMHs: 1\nExp number of AAs in TMHs: 23.72226\nExp number, first 60 AAs: 0.22038\nTotal prob of N-in: 0.12994\noutside 1 331\nTMhelix 332 354\ninside 355 372

6620 GCF_001283665.1_7790_1_75 Escherichia coli Proteobacteria; Gammaproteobacteria; Enterobacterales; Enterobacteriaceae; Escherichia MTGDNTLIHSHGINRRDFMKLCAALAATMGLSSKAAAEMAESVTNPQRPPVIWIGAQECT WP_000145410.1 MULTISPECIES: hydrogenase-2 small chain [Proteobacteria] Length: 372\nNumber of predicted TMHs: 1\nExp number of AAs in TMHs: 23.72226\nExp number, first 60 AAs: 0.22038\nTotal prob of N-in: 0.12994\noutside 1 331\nTMhelix 332 354\ninside 355 372

6621 GCF_001283545.1_7790_1_82 Escherichia coli Proteobacteria; Gammaproteobacteria; Enterobacterales; Enterobacteriaceae; Escherichia MNNEETFYQAMRRQGVTRRSFLKYCSLAATSLGLGAGMAPKIAWALENKPRIPVVWIHGL WP_001058323.1 MULTISPECIES: hydrogenase-1 small chain [Proteobacteria] Length: 372\nNumber of predicted TMHs:

2\nExp number of AAs in TMHs: 37.2608499999999\nExp number, first 60 AAs: 19.52923\nTotal prob of N-in: 0.94351\nPOSSIBLE N-term signal sequence\ninside 1 19\nTMhelix 20 39\noutside 40 325\nTMhelix 326 348\ninside 349 372

6622 GCF_001283725.1_7748_7_9Escherichia coli Proteobacteria; Gammaproteobacteria; Enterobacterales; Enterobacteriaceae; Escherichia MTGDNTLIHSHGINRRDFMKLCAALAATMGLSSKAAAEMAESVTNPQRPPVIWIGAQECT WP_000145410.1 MULTISPECIES: hydrogenase-2 small chain [Proteobacteria] Length: 372\nNumber of predicted TMHs: 1\nExp number of AAs in TMHs: 23.72226\nExp number, first 60 AAs: 0.22038\nTotal prob of N-in: 0.12994\noutside 1 331\nTMhelix 332 354\ninside 355 372

6623 GCF_001283725.1_7748_7_9Escherichia coli Proteobacteria; Gammaproteobacteria; Enterobacterales; Enterobacteriaceae; Escherichia MNNEETFYQAMRRQGVTRRSFLKYCSLAATSLGLGAGMAPKIAWALENKPRIPVVWIHGL WP_001058323.1 MULTISPECIES: hydrogenase-1 small chain [Proteobacteria] Length: 372\nNumber of predicted TMHs: 2\nExp number of AAs in TMHs: 37.2608499999999\nExp number, first 60 AAs: 19.52923\nTotal prob of N-in: 0.94351\nPOSSIBLE N-term signal sequence\ninside 1 19\nTMhelix 20 39\noutside 40 325\nTMhelix 326 348\ninside 349 372

6624 GCF_001283865.1_8205_3_32 Escherichia coli Proteobacteria; Gammaproteobacteria; Enterobacterales; Enterobacteriaceae; Escherichia MNNEETFYQAMRRQGVTRRSFLKYCSLAATSLGLGAGMAPKIAWALENKPRIPVVWIHGL WP_053876928.1 hydrogenase [Escherichia coli] Length: 372\nNumber of predicted TMHs: 2\nExp number of AAs in TMHs: 37.26524\nExp number, first 60 AAs: 19.52925\nTotal prob of N-in: 0.94351\nPOSSIBLE N-term signal sequence\ninside 1 19\nTMhelix 20 39\noutside 40 325\nTMhelix 326 348\ninside 349 372

6625 GCF_001283585.1_8205_3_77 Escherichia coli Proteobacteria; Gammaproteobacteria; Enterobacterales; Enterobacteriaceae; Escherichia MNNEETFYQAMRRQGVTRRSFLKYCSLAATSLGLGAGMAPKIAWALENKPRIPVVWIHGL WP_001295940.1 MULTISPECIES: hydrogenase [Enterobacteriaceae] Length: 372\nNumber of predicted TMHs: 2\nExp number of AAs in TMHs: 37.2608899999999\nExp number, first 60 AAs: 19.52923\nTotal prob of N-in: 0.94351\nPOSSIBLE N-term signal sequence\ninside 1 19\nTMhelix 20 39\noutside 40 325\nTMhelix 326 348\ninside 349 372

6626 GCF_001283605.1_8205_3_25 Escherichia coli Proteobacteria; Gammaproteobacteria; Enterobacterales; Enterobacteriaceae; Escherichia MNNEETFYQAMRRQGVTRRSFLKYCSLAATSLGLGAGMAPKIAWALENKPRIPVVWIHGL WP_001295940.1 MULTISPECIES: hydrogenase [Enterobacteriaceae] Length: 372\nNumber of predicted TMHs: 2\nExp number of AAs in TMHs: 37.2608899999999\nExp number, first 60 AAs: 19.52923\nTotal prob of N-in: 0.94351\nPOSSIBLE N-term signal sequence\ninside 1 19\nTMhelix 20 39\noutside 40 325\nTMhelix 326 348\ninside 349 372

6627 GCF_001283925.1_7748_7_31 Escherichia coli Proteobacteria; Gammaproteobacteria; Enterobacterales; Enterobacteriaceae; Escherichia MTGDNTLIHSHGINRRDFMKLCAALAATMGLSSKAAAEMAESVTNPQRPPVIWIGAQECT WP_000145410.1 MULTISPECIES: hydrogenase-2 small chain [Proteobacteria] Length: 372\nNumber of predicted TMHs: 1\nExp number of AAs in TMHs: 23.72226\nExp number, first 60 AAs: 0.22038\nTotal prob of N-in: 0.12994\noutside 1 331\nTMhelix 332 354\ninside 355 372

6628 GCF_001283945.1_7790_1_87 Escherichia coli Proteobacteria; Gammaproteobacteria; Enterobacterales; Enterobacteriaceae; Escherichia MTGDNTLIHSHGINRRDFMKLCAALAATMGLSSKAAAEMAESVTNPQRPPVIWIGAQECT WP_000145410.1 MULTISPECIES: hydrogenase-2 small chain [Proteobacteria] Length: 372\nNumber of predicted TMHs: 1\nExp number of AAs in TMHs: 23.72226\nExp number, first 60 AAs: 0.22038\nTotal prob of N-in: 0.12994\noutside 1 331\nTMhelix 332 354\ninside 355 372

6629 GCF_001283985.1_7553_7_67 Escherichia coli Proteobacteria; Gammaproteobacteria; Enterobacterales; Enterobacteriaceae; Escherichia MTGDNTLIHSHGINRRDFMKLCAALAATMGLSSKAAAEMAESVTNPQRPPVIWIGAQECT WP_000145406.1 hydrogenase-2 small chain [Escherichia coli] Length: 372\nNumber of predicted TMHs: 1\nExp number of AAs in TMHs: 23.72251\nExp number, first 60 AAs: 0.22027\nTotal prob of N-in: 0.13006\noutside 1 331\nTMhelix 332 354\ninside 355 372

6630 GCF_001284125.1_8205_3_28 Escherichia coli Proteobacteria; Gammaproteobacteria; Enterobacterales; Enterobacteriaceae; Escherichia
MTGDNTLIHSHGINRRDFMKLCAALAATMGLSSKAAAEMAESVTNPQRPPVIWIGAQECT WP_000145410.1
MULTISPECIES: hydrogenase-2 small chain [Proteobacteria] Length: 372\nNumber of predicted TMHs: 1\nExp number of AAs in TMHs: 23.72226\nExp number, first 60 AAs: 0.22038\nTotal prob of N-in: 0.12994\noutside 1 331\nTMhelix 332 354\ninside 355 372

6631 GCF_001284025.1_7790_1_52 Escherichia coli Proteobacteria; Gammaproteobacteria; Enterobacterales; Enterobacteriaceae; Escherichia
MNNEETFYQAMRRQGVTRRSFLKYCSLAATSLGLGAGMAPKIAWALENKPRIPVVWIHGL WP_001058323.1
MULTISPECIES: hydrogenase-1 small chain [Proteobacteria] Length: 372\nNumber of predicted TMHs: 2\nExp number of AAs in TMHs: 37.2608499999999\nExp number, first 60 AAs: 19.52923\nTotal prob of N-in: 0.94351\nPOSSIBLE N-term signal sequence\ninside 1 19\nTMhelix 20 39\noutside 40 325\nTMhelix 326 348\ninside 349 372

6632 GCF_001284165.1_8205_3_33 Escherichia coli Proteobacteria; Gammaproteobacteria; Enterobacterales; Enterobacteriaceae; Escherichia
MTGDNTLIHSHGINRRDFMKLCAALAATMGLSSKAAAEMAESVTNPQRPPVIWIGAQECT WP_000145410.1
MULTISPECIES: hydrogenase-2 small chain [Proteobacteria] Length: 372\nNumber of predicted TMHs: 1\nExp number of AAs in TMHs: 23.72226\nExp number, first 60 AAs: 0.22038\nTotal prob of N-in: 0.12994\noutside 1 331\nTMhelix 332 354\ninside 355 372

6633 GCF_001284045.1_8205_3_14 Escherichia coli Proteobacteria; Gammaproteobacteria; Enterobacterales; Enterobacteriaceae; Escherichia
MNNEETFYQAMRRQGVTRRSFLKYCSLAATSLGLGAGMAPKIAWALENKPRIPVVWIHGL WP_001058323.1
MULTISPECIES: hydrogenase-1 small chain [Proteobacteria] Length: 372\nNumber of predicted TMHs: 2\nExp number of AAs in TMHs: 37.2608499999999\nExp number, first 60 AAs: 19.52923\nTotal prob of N-in: 0.94351\nPOSSIBLE N-term signal sequence\ninside 1 19\nTMhelix 20 39\noutside 40 325\nTMhelix 326 348\ninside 349 372

6634 GCF_001284205.1_7748_7_12 Escherichia coli Proteobacteria; Gammaproteobacteria; Enterobacterales; Enterobacteriaceae; Escherichia
MTGDNTLIHSHGINRRDFMKLCAALAATMGLSSKAAAEMAESVTNPQRPPVIWIGAQECT WP_000145410.1
MULTISPECIES: hydrogenase-2 small chain [Proteobacteria] Length: 372\nNumber of predicted TMHs: 1\nExp number of AAs in TMHs: 23.72226\nExp number, first 60 AAs: 0.22038\nTotal prob of N-in: 0.12994\noutside 1 331\nTMhelix 332 354\ninside 355 372

6635 GCF_001284265.1_7790_1_57 Escherichia coli Proteobacteria; Gammaproteobacteria; Enterobacterales; Enterobacteriaceae; Escherichia
MTGDNTLIHSHGINRRDFMKLCAALAATMGLSSKAAAEMAESVTNPQRPPVIWIGAQECT WP_000145410.1
MULTISPECIES: hydrogenase-2 small chain [Proteobacteria] Length: 372\nNumber of predicted TMHs: 1\nExp number of AAs in TMHs: 23.72226\nExp number, first 60 AAs: 0.22038\nTotal prob of N-in: 0.12994\noutside 1 331\nTMhelix 332 354\ninside 355 372

6636 GCF_001284325.1_8205_8_80 Escherichia coli Proteobacteria; Gammaproteobacteria; Enterobacterales; Enterobacteriaceae; Escherichia
MNNEETFYQAMRRQGVTRRSFLKYCSLAATSLGLGAGMAPKIAWALENKPRIPVVWIHGL WP_001295940.1
MULTISPECIES: hydrogenase [Enterobacteriaceae] Length: 372\nNumber of predicted TMHs: 2\nExp number of AAs in TMHs: 37.2608899999999\nExp number, first 60 AAs: 19.52923\nTotal prob of N-in: 0.94351\nPOSSIBLE N-term signal sequence\ninside 1 19\nTMhelix 20 39\noutside 40 325\nTMhelix 326 348\ninside 349 372

6637 GCF_001284125.1_8205_3_28 Escherichia coli Proteobacteria; Gammaproteobacteria; Enterobacterales; Enterobacteriaceae; Escherichia
MNNEETFYQAMRRQGVTRRSFLKYCSLAATSLGLGAGMAPKIAWALENKPRIPVVWIHGL WP_001058323.1
MULTISPECIES: hydrogenase-1 small chain [Proteobacteria] Length: 372\nNumber of predicted TMHs: 2\nExp number of AAs in TMHs: 37.2608499999999\nExp number, first 60 AAs: 19.52923\nTotal prob of N-in: 0.94351\nPOSSIBLE N-term signal sequence\ninside 1 19\nTMhelix 20 39\noutside 40 325\nTMhelix 326 348\ninside 349 372

6638 GCF_001284165.1_8205_3_33 Escherichia coli Proteobacteria; Gammaproteobacteria; Enterobacterales; Enterobacteriaceae; Escherichia
MNNEETFYQAMRRQGVTRRSFLKYCSLAATSLGLGAGMAPKIAWALENKPRIPVVWIHGL WP_001058323.1
MULTISPECIES: hydrogenase-1 small chain [Proteobacteria] Length: 372\nNumber of predicted TMHs: 2\nExp number of AAs in TMHs: 37.2608499999999\nExp number, first 60 AAs: 19.52923\nTotal prob of N-in: 0.94351\nPOSSIBLE N-term signal sequence\ninside 1 19\nTMhelix 20 39\noutside 40 325\nTMhelix 326 348\ninside 349 372

6639 GCF_001284365.1_8205_8_72 Escherichia coli Proteobacteria; Gammaproteobacteria; Enterobacterales; Enterobacteriaceae; Escherichia
MTGDNTLIHSHGINRRDFMKLCAALAATMGLSSKAAAEMAESVTNPQRPPVIWIGAQECT WP_000145410.1
MULTISPECIES: hydrogenase-2 small chain [Proteobacteria] Length: 372\nNumber of predicted TMHs: 1\nExp number of AAs in TMHs: 23.72226\nExp number, first 60 AAs: 0.22038\nTotal prob of N-in: 0.12994\noutside 1 331\nTMhelix 332 354\ninside 355 372

6640 GCF_001284405.1_8205_3_59 Escherichia coli Proteobacteria; Gammaproteobacteria; Enterobacterales; Enterobacteriaceae; Escherichia
MTGDNTLIHSHGINRRDFMKLCAALAATMGLSSKAAAEMAESVTNPQRPPVIWIGAQECT WP_000145410.1
MULTISPECIES: hydrogenase-2 small chain [Proteobacteria] Length: 372\nNumber of predicted TMHs: 1\nExp number of AAs in TMHs: 23.72226\nExp number, first 60 AAs: 0.22038\nTotal prob of N-in: 0.12994\noutside 1 331\nTMhelix 332 354\ninside 355 372

6641 GCF_001284425.1_8205_8_85 Escherichia coli Proteobacteria; Gammaproteobacteria; Enterobacterales; Enterobacteriaceae; Escherichia
MTGDNTLIHSHGINRRDFMKLCAALAATMGLSSKAAAEMAESVTNPQRPPVIWIGAQECT WP_000145410.1
MULTISPECIES: hydrogenase-2 small chain [Proteobacteria] Length: 372\nNumber of predicted TMHs: 1\nExp number of AAs in TMHs: 23.72226\nExp number, first 60 AAs: 0.22038\nTotal prob of N-in: 0.12994\noutside 1 331\nTMhelix 332 354\ninside 355 372

6642 GCF_001284465.1_7790_1_51 Escherichia coli Proteobacteria; Gammaproteobacteria; Enterobacterales; Enterobacteriaceae; Escherichia
MTGDNTLIHSHGINRRDFMKLCAALAATMGLSSKAAAEMAESVTNPQRPPVIWIGAQECT WP_000145410.1
MULTISPECIES: hydrogenase-2 small chain [Proteobacteria] Length: 372\nNumber of predicted TMHs: 1\nExp number of AAs in TMHs: 23.72226\nExp number, first 60 AAs: 0.22038\nTotal prob of N-in: 0.12994\noutside 1 331\nTMhelix 332 354\ninside 355 372

6643 GCF_001284465.1_7790_1_51 Escherichia coli Proteobacteria; Gammaproteobacteria; Enterobacterales; Enterobacteriaceae; Escherichia
MNNEETFYQAMRRQGVTRRSFLKYCSLAATSLGLGAGMAPKIAWALENKPRIPVVWIHGL WP_001058323.1
MULTISPECIES: hydrogenase-1 small chain [Proteobacteria] Length: 372\nNumber of predicted TMHs: 2\nExp number of AAs in TMHs: 37.2608499999999\nExp number, first 60 AAs: 19.52923\nTotal prob of N-in: 0.94351\nPOSSIBLE N-term signal sequence\ninside 1 19\nTMhelix 20 39\noutside 40 325\nTMhelix 326 348\ninside 349 372

6644 GCF_001284425.1_8205_8_85 Escherichia coli Proteobacteria; Gammaproteobacteria; Enterobacterales; Enterobacteriaceae; Escherichia
MNNEETFYQAMRRQGVTRRSFLKYCSLAATSLGLGAGMAPKIAWALENKPRIPVVWIHGL WP_001058323.1
MULTISPECIES: hydrogenase-1 small chain [Proteobacteria] Length: 372\nNumber of predicted TMHs: 2\nExp number of AAs in TMHs: 37.2608499999999\nExp number, first 60 AAs: 19.52923\nTotal prob of N-in: 0.94351\nPOSSIBLE N-term signal sequence\ninside 1 19\nTMhelix 20 39\noutside 40 325\nTMhelix 326 348\ninside 349 372

6645 GCF_001284625.1_8205_3_62 Escherichia coli Proteobacteria; Gammaproteobacteria; Enterobacterales; Enterobacteriaceae; Escherichia
MTGDNTLIHSHGINRRDFMKLCAALAATMGLSSKAAAEMAESVTNPQRPPVIWIGAQECT WP_000145410.1
MULTISPECIES: hydrogenase-2 small chain [Proteobacteria] Length: 372\nNumber of predicted TMHs: 1\nExp number of AAs in TMHs: 23.72226\nExp number, first 60 AAs: 0.22038\nTotal prob of N-in: 0.12994\noutside 1 331\nTMhelix 332 354\ninside 355 372

6646 GCF_001284625.1_8205_3_62 Escherichia coli Proteobacteria; Gammaproteobacteria; Enterobacterales; Enterobacteriaceae; Escherichia

MNNEETFYQAMRRQGVTRRSFLKYCSLAATSLGLGAGMAPKIAWALENKPRIPVVWIHGL WP_001058323.1 MULTISPECIES: hydrogenase-1 small chain [Proteobacteria] Length: 372\nNumber of predicted TMHs: 2\nExp number of AAs in TMHs: 37.2608499999999\nExp number, first 60 AAs: 19.52923\nTotal prob of N-in: 0.94351\nPOSSIBLE N-term signal sequence\ninside 1 19\nTMhelix 20 39\noutside 40 325\nTMhelix 326 348\ninside 349 372

6647 GCF_001284405.1_8205_3_59 Escherichia coli Proteobacteria; Gammaproteobacteria; Enterobacterales; Enterobacteriaceae; Escherichia MNNEETFYQAMRRQGVTRRSFLKYCSLAATSLGLGAGMAPKIAWALENKPRIPVVWIHGL WP_024237484.1 hydrogenase-1 small chain [Escherichia coli] Length: 372\nNumber of predicted TMHs: 2\nExp number of AAs in TMHs: 37.2691999999999\nExp number, first 60 AAs: 19.53117\nTotal prob of N-in: 0.94352\nPOSSIBLE N-term signal sequence\ninside 1 19\nTMhelix 20 39\noutside 40 325\nTMhelix 326 348\ninside 349 372

6648 GCF_001284745.1_7790_1_91 Escherichia coli Proteobacteria; Gammaproteobacteria; Enterobacterales; Enterobacteriaceae; Escherichia MTGDNTLIHSHGINRRDFMKLCAALAATMGLSSKAAAEMAESVTNPQRPPVIWIGAQECT WP_000145410.1 MULTISPECIES: hydrogenase-2 small chain [Proteobacteria] Length: 372\nNumber of predicted TMHs: 1\nExp number of AAs in TMHs: 23.72226\nExp number, first 60 AAs: 0.22038\nTotal prob of N-in: 0.12994\noutside 1 331\nTMhelix 332 354\ninside 355 372

6649 GCF_001284705.1_8205_3_17 Escherichia coli Proteobacteria; Gammaproteobacteria; Enterobacterales; Enterobacteriaceae; Escherichia MTGDNTLIHSHGINRRDFMKLCAALAATMGLSSKAAAEMAESVTNPQRPPVIWIGAQECT WP_000145410.1 MULTISPECIES: hydrogenase-2 small chain [Proteobacteria] Length: 372\nNumber of predicted TMHs: 1\nExp number of AAs in TMHs: 23.72226\nExp number, first 60 AAs: 0.22038\nTotal prob of N-in: 0.12994\noutside 1 331\nTMhelix 332 354\ninside 355 372

6650 GCF_001284845.1_8205_8_83 Escherichia coli Proteobacteria; Gammaproteobacteria; Enterobacterales; Enterobacteriaceae; Escherichia MTGDNTLIHSHGINRRDFMKLCAALAATMGLSSKAAAEMAESVTNPQRPPVIWIGAQECT WP_000145410.1 MULTISPECIES: hydrogenase-2 small chain [Proteobacteria] Length: 372\nNumber of predicted TMHs: 1\nExp number of AAs in TMHs: 23.72226\nExp number, first 60 AAs: 0.22038\nTotal prob of N-in: 0.12994\noutside 1 331\nTMhelix 332 354\ninside 355 372

6651 GCF_001284485.1_8205_3_8Escherichia coli Proteobacteria; Gammaproteobacteria; Enterobacterales; Enterobacteriaceae; Escherichia MTGDNTLIHSHGISRRDFMKLCAALAATMGLSSKAAAEMAESVTNPQRPPVIWIGAQECT WP_053879127.1 hydrogenase 2 small subunit [Escherichia coli] Length: 372\nNumber of predicted TMHs: 1\nExp number of AAs in TMHs: 23.66488\nExp number, first 60 AAs: 0.15896\nTotal prob of N-in: 0.12726\noutside 1 331\nTMhelix 332 354\ninside 355 372

6652 GCF_001284865.1_8205_3_72 Escherichia coli Proteobacteria; Gammaproteobacteria; Enterobacterales; Enterobacteriaceae; Escherichia MTGDNTLIHSHGINRRDFMKLCAALAATMGLSSKAAAEMAESVTNPQRPPVIWIGAQECT WP_000145410.1 MULTISPECIES: hydrogenase-2 small chain [Proteobacteria] Length: 372\nNumber of predicted TMHs: 1\nExp number of AAs in TMHs: 23.72226\nExp number, first 60 AAs: 0.22038\nTotal prob of N-in: 0.12994\noutside 1 331\nTMhelix 332 354\ninside 355 372

6653 GCF_001284905.1_8205_3_31 Escherichia coli Proteobacteria; Gammaproteobacteria; Enterobacterales; Enterobacteriaceae; Escherichia MTGDNTLIHSHGINRRDFMKLCAALAATMGLSSKAAAEMAESVTNPQRPPVIWIGAQECT WP_000145410.1 MULTISPECIES: hydrogenase-2 small chain [Proteobacteria] Length: 372\nNumber of predicted TMHs: 1\nExp number of AAs in TMHs: 23.72226\nExp number, first 60 AAs: 0.22038\nTotal prob of N-in: 0.12994\noutside 1 331\nTMhelix 332 354\ninside 355 372

6654 GCF_001284945.1_8205_3_21 Escherichia coli Proteobacteria; Gammaproteobacteria; Enterobacterales; Enterobacteriaceae; Escherichia MTGDNTLIHSHGINRRDFMKLCAALAATMGLSSKAAAEMAESVTNPQRPPVIWIGAQECT WP_000145410.1 MULTISPECIES: hydrogenase-2 small chain [Proteobacteria] Length: 372\nNumber of predicted TMHs: 1\nExp number of AAs in TMHs: 23.72226\nExp number, first 60 AAs: 0.22038\nTotal prob of N-in: 0.12994\noutside 1 331\nTMhelix 332 354\ninside 355 372

6655 GCF_001284945.1_8205_3_21 Escherichia coli Proteobacteria; Gammaproteobacteria; Enterobacterales; Enterobacteriaceae; Escherichia
MNNEETFYQAMRRQGVTRRSFLKYCSLAATSLGLGAGMAPKIAWALENKPRIPVVWIHGL WP_001295940.1
MULTISPECIES: hydrogenase [Enterobacteriaceae] Length: 372\nNumber of predicted TMHs: 2\nExp number of AAs in TMHs: 37.2608899999999\nExp number, first 60 AAs: 19.52923\nTotal prob of N-in: 0.94351\nPOSSIBLE N-term signal sequence\ninside 1 19\nTMhelix 20 39\noutside 40 325\nTMhelix 326 348\ninside 349 372

6656 GCF_001284845.1_8205_8_83 Escherichia coli Proteobacteria; Gammaproteobacteria; Enterobacterales; Enterobacteriaceae; Escherichia
MNNEETFYQAMRRQGVTRRSFLKYCSLAATSLGLGAGMAPKIAWALENKPRIPVVWIHGL WP_001058323.1
MULTISPECIES: hydrogenase-1 small chain [Proteobacteria] Length: 372\nNumber of predicted TMHs: 2\nExp number of AAs in TMHs: 37.2608499999999\nExp number, first 60 AAs: 19.52923\nTotal prob of N-in: 0.94351\nPOSSIBLE N-term signal sequence\ninside 1 19\nTMhelix 20 39\noutside 40 325\nTMhelix 326 348\ninside 349 372

6657 GCF_001285085.1_8205_3_12 Escherichia coli Proteobacteria; Gammaproteobacteria; Enterobacterales; Enterobacteriaceae; Escherichia
MNNEETFYQAMRRQGVTRRSFLKYCSLAATSLGLGAGMAPKIAWALENKPRIPVVWIHGL WP_001058323.1
MULTISPECIES: hydrogenase-1 small chain [Proteobacteria] Length: 372\nNumber of predicted TMHs: 2\nExp number of AAs in TMHs: 37.2608499999999\nExp number, first 60 AAs: 19.52923\nTotal prob of N-in: 0.94351\nPOSSIBLE N-term signal sequence\ninside 1 19\nTMhelix 20 39\noutside 40 325\nTMhelix 326 348\ninside 349 372

6658 GCF_001285065.1_8205_3_27 Escherichia coli Proteobacteria; Gammaproteobacteria; Enterobacterales; Enterobacteriaceae; Escherichia
MNNEETFYQAMRRQGVTRRSFLKYCSLAATSLGLGAGMAPKIAWALENKPRIPVVWIHGL WP_001058320.1
hydrogenase [Escherichia coli] Length: 372\nNumber of predicted TMHs: 2\nExp number of AAs in TMHs: 37.26082\nExp number, first 60 AAs: 19.52923\nTotal prob of N-in: 0.94351\nPOSSIBLE N-term signal sequence\ninside 1 19\nTMhelix 20 39\noutside 40 325\nTMhelix 326 348\ninside 349 372

6659 GCF_001285025.1_7748_7_46 Escherichia coli Proteobacteria; Gammaproteobacteria; Enterobacterales; Enterobacteriaceae; Escherichia
MNNEETFYQAMRRQGVTRRSFLKYCSLAATSLGLGAGMAPKIAWALENKPRIPVVWIHGL WP_001295940.1
MULTISPECIES: hydrogenase [Enterobacteriaceae] Length: 372\nNumber of predicted TMHs: 2\nExp number of AAs in TMHs: 37.2608899999999\nExp number, first 60 AAs: 19.52923\nTotal prob of N-in: 0.94351\nPOSSIBLE N-term signal sequence\ninside 1 19\nTMhelix 20 39\noutside 40 325\nTMhelix 326 348\ninside 349 372

6660 GCF_001284805.1_7748_7_14 Escherichia coli Proteobacteria; Gammaproteobacteria; Enterobacterales; Enterobacteriaceae; Escherichia
MNNEETFYQAMRRQGVTRRSFLKYCSLAATSLGLGAGMAPKIAWALENKPRIPVVWIHGL WP_052935581.1
hydrogenase [Escherichia coli] Length: 372\nNumber of predicted TMHs: 2\nExp number of AAs in TMHs: 37.2610899999999\nExp number, first 60 AAs: 19.52923\nTotal prob of N-in: 0.94351\nPOSSIBLE N-term signal sequence\ninside 1 19\nTMhelix 20 39\noutside 40 325\nTMhelix 326 348\ninside 349 372

6661 GCF_001285125.1_7553_7_72 Escherichia coli Proteobacteria; Gammaproteobacteria; Enterobacterales; Enterobacteriaceae; Escherichia
MTGDNTLIHSHGINRRDFMKLCAALAATMGLSSKAAAEMAESVTNPQRPPVIWIGAQECT WP_000145410.1
MULTISPECIES: hydrogenase-2 small chain [Proteobacteria] Length: 372\nNumber of predicted TMHs: 1\nExp number of AAs in TMHs: 23.72226\nExp number, first 60 AAs: 0.22038\nTotal prob of N-in: 0.12994\noutside 1 331\nTMhelix 332 354\ninside 355 372

6662 GCF_001285245.1_8205_3_18 Escherichia coli Proteobacteria; Gammaproteobacteria; Enterobacterales; Enterobacteriaceae; Escherichia
MTGDNTLIHSHGINRRDFMKLCAALAATMGLSSKAAAEMAESVTNPQRPPVIWIGAQECT WP_000145410.1
MULTISPECIES: hydrogenase-2 small chain [Proteobacteria] Length: 372\nNumber of predicted TMHs: 1\nExp number of AAs in TMHs: 23.72226\nExp number, first 60 AAs: 0.22038\nTotal prob of N-in: 0.12994\noutside 1 331\nTMhelix 332 354\ninside 355 372

6663 GCF_001285205.1_8205_3_78 Escherichia coli Proteobacteria; Gammaproteobacteria; Enterobacterales; Enterobacteriaceae; Escherichia
MTGDNTLIHSHGINRRDFMKLCAALAATMGLSSKAAAEMAESVTNPQRPPVIWIGAQECT WP_000145410.1
MULTISPECIES: hydrogenase-2 small chain [Proteobacteria] Length: 372\nNumber of predicted TMHs: 1\nExp number of AAs in TMHs: 23.72226\nExp number, first 60 AAs: 0.22038\nTotal prob of N-in: 0.12994\noutside 1 331\nTMhelix 332 354\ninside 355 372

6664 GCF_001285425.1_7790_1_85 Escherichia coli Proteobacteria; Gammaproteobacteria; Enterobacterales; Enterobacteriaceae; Escherichia
MNNEETFYQAMRRQGVTRRSFLKYCSLAATSLGLGAGMAPKIAWALENKPRIPVVWIHGL WP_001058323.1
MULTISPECIES: hydrogenase-1 small chain [Proteobacteria] Length: 372\nNumber of predicted TMHs: 2\nExp number of AAs in TMHs: 37.2608499999999\nExp number, first 60 AAs: 19.52923\nTotal prob of N-in: 0.94351\nPOSSIBLE N-term signal sequence\ninside 1 19\nTMhelix 20 39\noutside 40 325\nTMhelix 326 348\ninside 349 372

6665 GCF_001285625.1_7790_1_84 Escherichia coli Proteobacteria; Gammaproteobacteria; Enterobacterales; Enterobacteriaceae; Escherichia
MTGDNTLIHSHGINRRDFMKLCAALAATMGLSSKAAAEMAESVTNPQRPPVIWIGAQECT WP_000145410.1
MULTISPECIES: hydrogenase-2 small chain [Proteobacteria] Length: 372\nNumber of predicted TMHs: 1\nExp number of AAs in TMHs: 23.72226\nExp number, first 60 AAs: 0.22038\nTotal prob of N-in: 0.12994\noutside 1 331\nTMhelix 332 354\ninside 355 372

6666 GCF_001285305.1_8205_3_50 Escherichia coli Proteobacteria; Gammaproteobacteria; Enterobacterales; Enterobacteriaceae; Escherichia
MNNEETFYQAMRRQGVTRRSFLKYCSLAATSLGLGAGMAPKIAWALENKPRIPVVWIHGL WP_001295940.1
MULTISPECIES: hydrogenase [Enterobacteriaceae] Length: 372\nNumber of predicted TMHs: 2\nExp number of AAs in TMHs: 37.2608899999999\nExp number, first 60 AAs: 19.52923\nTotal prob of N-in: 0.94351\nPOSSIBLE N-term signal sequence\ninside 1 19\nTMhelix 20 39\noutside 40 325\nTMhelix 326 348\ninside 349 372

6667 GCF_001285365.1_8205_3_7Escherichia coli Proteobacteria; Gammaproteobacteria; Enterobacterales; Enterobacteriaceae; Escherichia MNNEETFYQAMRRQGVTRRSFLKYCSLAATSLGLGAGMAPKIAWALENKPRIPVVWIHGL
WP_001058320.1 hydrogenase [Escherichia coli] Length: 372\nNumber of predicted TMHs: 2\nExp number of AAs in TMHs: 37.26082\nExp number, first 60 AAs: 19.52923\nTotal prob of N-in: 0.94351\nPOSSIBLE N-term signal sequence\ninside 1 19\nTMhelix 20 39\noutside 40 325\nTMhelix 326 348\ninside 349 372

6668 GCF_001285705.1_8205_3_56 Escherichia coli Proteobacteria; Gammaproteobacteria; Enterobacterales; Enterobacteriaceae; Escherichia
MTGDNTLIHSHGINRRDFMKLCAALAATMGLSSKAAAEMAESVTNPQRPPVIWIGAQECT WP_000145410.1
MULTISPECIES: hydrogenase-2 small chain [Proteobacteria] Length: 372\nNumber of predicted TMHs: 1\nExp number of AAs in TMHs: 23.72226\nExp number, first 60 AAs: 0.22038\nTotal prob of N-in: 0.12994\noutside 1 331\nTMhelix 332 354\ninside 355 372

6669 GCF_001285605.1_8205_8_74 Escherichia coli Proteobacteria; Gammaproteobacteria; Enterobacterales; Enterobacteriaceae; Escherichia
MNNEETFYQAMRRQGVTRRSFLKYCSLAATSLGLGAGMAPKIAWALENKPRIPVVWIHGL WP_001058323.1
MULTISPECIES: hydrogenase-1 small chain [Proteobacteria] Length: 372\nNumber of predicted TMHs: 2\nExp number of AAs in TMHs: 37.2608499999999\nExp number, first 60 AAs: 19.52923\nTotal prob of N-in: 0.94351\nPOSSIBLE N-term signal sequence\ninside 1 19\nTMhelix 20 39\noutside 40 325\nTMhelix 326 348\ninside 349 372

6670 GCF_001285465.1_7790_1_63 Escherichia coli Proteobacteria; Gammaproteobacteria; Enterobacterales; Enterobacteriaceae; Escherichia
MNNEETFYQAMRRQGVTRRSFLKYCSLAATSLGLGAGMAPKIAWALENKPRIPVVWIHGL WP_001058323.1
MULTISPECIES: hydrogenase-1 small chain [Proteobacteria] Length: 372\nNumber of predicted TMHs: 2\nExp number of AAs in TMHs: 37.2608499999999\nExp number, first 60 AAs: 19.52923\nTotal prob of N-in: 0.94351\nPOSSIBLE N-term signal sequence\ninside 1 19\nTMhelix 20 39\noutside 40 325\nTMhelix 326 348\ninside 349 372

6671 GCF_001285765.1_7790_1_66 Escherichia coli Proteobacteria; Gammaproteobacteria; Enterobacterales; Enterobacteriaceae; Escherichia

MTGDNTLIHSHGINRRDFMKLCAALAATMGLSSKAAAEMAESVTNPQRPPVIWIGAQECT WP_000145410.1 MULTISPECIES: hydrogenase-2 small chain [Proteobacteria] Length: 372\nNumber of predicted TMHs: 1\nExp number of AAs in TMHs: 23.72226\nExp number, first 60 AAs: 0.22038\nTotal prob of N-in: 0.12994\noutside 1 331\nTMhelix 332 354\ninside 355 372

6672 GCF_001285805.1_7790_1_56 Escherichia coli Proteobacteria; Gammaproteobacteria; Enterobacterales; Enterobacteriaceae; Escherichia MTGDNTLIHSHGINRRDFMKLCAALAATMGLSSKAAAEMAESVTNPQRPPVIWIGAQECT WP_000145410.1 MULTISPECIES: hydrogenase-2 small chain [Proteobacteria] Length: 372\nNumber of predicted TMHs: 1\nExp number of AAs in TMHs: 23.72226\nExp number, first 60 AAs: 0.22038\nTotal prob of N-in: 0.12994\noutside 1 331\nTMhelix 332 354\ninside 355 372

6673 GCF_001285765.1_7790_1_66 Escherichia coli Proteobacteria; Gammaproteobacteria; Enterobacterales; Enterobacteriaceae; Escherichia MNNEETFYQAMRRQGVTRRSFLKYCSLAATSLGLGAGMAPKIAWALENKPRIPVVWIHGL WP_001295940.1 MULTISPECIES: hydrogenase [Enterobacteriaceae] Length: 372\nNumber of predicted TMHs: 2\nExp number of AAs in TMHs: 37.2608899999999\nExp number, first 60 AAs: 19.52923\nTotal prob of N-in: 0.94351\nPOSSIBLE N-term signal sequence\ninside 1 19\nTMhelix 20 39\noutside 40 325\nTMhelix 326 348\ninside 349 372

6674 GCF_001285805.1_7790_1_56 Escherichia coli Proteobacteria; Gammaproteobacteria; Enterobacterales; Enterobacteriaceae; Escherichia MNNEETFYQAMRRQGVTRRSFLKYCSLAATSLGLGAGMAPKIAWALENKPRIPVVWIHGL WP_052935581.1 hydrogenase [Escherichia coli] Length: 372\nNumber of predicted TMHs: 2\nExp number of AAs in TMHs: 37.2610899999999\nExp number, first 60 AAs: 19.52923\nTotal prob of N-in: 0.94351\nPOSSIBLE N-term signal sequence\ninside 1 19\nTMhelix 20 39\noutside 40 325\nTMhelix 326 348\ninside 349 372

6675 GCF_001285525.1_7748_7_4Escherichia coli Proteobacteria; Gammaproteobacteria; Enterobacterales; Enterobacteriaceae; Escherichia MNNEETFYQAMRRQGVTRRSFLKYCSLAATSLGLGAGMAPKIAWALENKPRIPVVWIHGL WP_001295940.1 MULTISPECIES: hydrogenase [Enterobacteriaceae] Length: 372\nNumber of predicted TMHs: 2\nExp number of AAs in TMHs: 37.2608899999999\nExp number, first 60 AAs: 19.52923\nTotal prob of N-in: 0.94351\nPOSSIBLE N-term signal sequence\ninside 1 19\nTMhelix 20 39\noutside 40 325\nTMhelix 326 348\ninside 349 372

6676 GCF_001285825.1_7748_7_20 Escherichia coli Proteobacteria; Gammaproteobacteria; Enterobacterales; Enterobacteriaceae; Escherichia MTGDNTLIHSHGINRRDFMKLCAALAATMGLSSKAAAEMVESVTNPQRPPVIWIGAQECT WP_000145423.1 hydrogenase 2 small subunit [Escherichia coli] Length: 372\nNumber of predicted TMHs: 1\nExp number of AAs in TMHs: 23.78201\nExp number, first 60 AAs: 0.28355\nTotal prob of N-in: 0.13257\noutside 1 331\nTMhelix 332 354\ninside 355 372

6677 GCF_001285865.1_7553_7_64 Escherichia coli Proteobacteria; Gammaproteobacteria; Enterobacterales; Enterobacteriaceae; Escherichia MTGDNTLIHSHGINRRDFMKLCAALAATMGLSSKAAAEMAESVTNPQRPPVIWIGAQECT WP_000145410.1 MULTISPECIES: hydrogenase-2 small chain [Proteobacteria] Length: 372\nNumber of predicted TMHs: 1\nExp number of AAs in TMHs: 23.72226\nExp number, first 60 AAs: 0.22038\nTotal prob of N-in: 0.12994\noutside 1 331\nTMhelix 332 354\ninside 355 372

6678 GCF_001286025.1_8205_3_11 Escherichia coli Proteobacteria; Gammaproteobacteria; Enterobacterales; Enterobacteriaceae; Escherichia MNNEETFYQAMRRQGVTRRSFLKYCSLAATSLGLGAGMAPKIAWALENKPRIPVVWIHGL WP_001058323.1 MULTISPECIES: hydrogenase-1 small chain [Proteobacteria] Length: 372\nNumber of predicted TMHs: 2\nExp number of AAs in TMHs: 37.2608499999999\nExp number, first 60 AAs: 19.52923\nTotal prob of N-in: 0.94351\nPOSSIBLE N-term signal sequence\ninside 1 19\nTMhelix 20 39\noutside 40 325\nTMhelix 326 348\ninside 349 372

6679 GCF_001286125.1_8205_3_71 Escherichia coli Proteobacteria; Gammaproteobacteria; Enterobacterales; Enterobacteriaceae; Escherichia MTGDNTLIHSHGINRRDFMKLCAALAATMGLSSKAAAEMAESVTNPQRPPVIWIGAQECT WP_000145410.1 MULTISPECIES: hydrogenase-2 small chain [Proteobacteria] Length: 372\nNumber of predicted TMHs:

1\nExp number of AAs in TMHs: 23.72226\nExp number, first 60 AAs: 0.22038\nTotal prob of N-in: 0.12994\noutside 1 331\nTMhelix 332 354\ninside 355 372

6680 GCF_001286165.1_7748_7_44 Escherichia coli Proteobacteria; Gammaproteobacteria; Enterobacterales; Enterobacteriaceae; Escherichia MTGDNTLIHSHGINRRDFMKLCAALAATMGLSSKAAAEMAESVTNPQHPPVIWIGAQECT WP_000145403.1 hydrogenase 2 small subunit [Escherichia coli] Length: 372\nNumber of predicted TMHs: 1\nExp number of AAs in TMHs: 23.66365\nExp number, first 60 AAs: 0.1537\nTotal prob of N-in: 0.12709\noutside 1 331\nTMhelix 332 354\ninside 355 372

6681 GCF_001286045.1_7790_1_68 Escherichia coli Proteobacteria; Gammaproteobacteria; Enterobacterales; Enterobacteriaceae; Escherichia MNNEETFYQAMRRQGVTRRSFLKYCSLAATSLGLGAGMAPKIAWALENKPRIPVVWIHGL WP_001295940.1 MULTISPECIES: hydrogenase [Enterobacteriaceae] Length: 372\nNumber of predicted TMHs: 2\nExp number of AAs in TMHs: 37.2608899999999\nExp number, first 60 AAs: 19.52923\nTotal prob of N-in: 0.94351\nPOSSIBLE N-term signal sequence\ninside 1 19\nTMhelix 20 39\noutside 40 325\nTMhelix 326 348\ninside 349 372

6682 GCF_001286025.1_8205_3_11 Escherichia coli Proteobacteria; Gammaproteobacteria; Enterobacterales; Enterobacteriaceae; Escherichia MTGDNTLIHSHGINRRDFMKLCAALAATMGLSSKAAAEMAESVTNPQRPPVIWIGAQECT WP_053889560.1 hydrogenase 2 small subunit [Escherichia coli] Length: 372\nNumber of predicted TMHs: 1\nExp number of AAs in TMHs: 23.96182\nExp number, first 60 AAs: 0.2204\nTotal prob of N-in: 0.13316\noutside 1 332\nTMhelix 333 355\ninside 356 372

6683 GCF_001285945.1_8205_3_9Escherichia coli Proteobacteria; Gammaproteobacteria; Enterobacterales; Enterobacteriaceae; Escherichia MNNEETFYQAMRRQGVTRRSFLKYCSLAATSLGLGAGMAPKIAWALENKPRIPVVWIHGL WP_001058323.1 MULTISPECIES: hydrogenase-1 small chain [Proteobacteria] Length: 372\nNumber of predicted TMHs: 2\nExp number of AAs in TMHs: 37.2608499999999\nExp number, first 60 AAs: 19.52923\nTotal prob of N-in: 0.94351\nPOSSIBLE N-term signal sequence\ninside 1 19\nTMhelix 20 39\noutside 40 325\nTMhelix 326 348\ninside 349 372

6684 GCF_001286185.1_7790_1_77 Escherichia coli Proteobacteria; Gammaproteobacteria; Enterobacterales; Enterobacteriaceae; Escherichia MTGDNTLIHSHGINRRDFMKLCAALAATMGLSSKAAAEMAESVTNPQRPPVIWIGAQECT WP_000145410.1 MULTISPECIES: hydrogenase-2 small chain [Proteobacteria] Length: 372\nNumber of predicted TMHs: 1\nExp number of AAs in TMHs: 23.72226\nExp number, first 60 AAs: 0.22038\nTotal prob of N-in: 0.12994\noutside 1 331\nTMhelix 332 354\ninside 355 372

6685 GCF_001286305.1_8205_3_81 Escherichia coli Proteobacteria; Gammaproteobacteria; Enterobacterales; Enterobacteriaceae; Escherichia MNNEETFYQAMRRQGVTRRSFLKYCSLAATSLGLGAGMAPKIAWALENKPRIPVVWIHGL WP_001058323.1 MULTISPECIES: hydrogenase-1 small chain [Proteobacteria] Length: 372\nNumber of predicted TMHs: 2\nExp number of AAs in TMHs: 37.2608499999999\nExp number, first 60 AAs: 19.52923\nTotal prob of N-in: 0.94351\nPOSSIBLE N-term signal sequence\ninside 1 19\nTMhelix 20 39\noutside 40 325\nTMhelix 326 348\ninside 349 372

6686 GCF_001286125.1_8205_3_71 Escherichia coli Proteobacteria; Gammaproteobacteria; Enterobacterales; Enterobacteriaceae; Escherichia MNNEETFYQAMRRQGVTRRSFLKYCSLAATSLGLGAGMAPKIAWALENKPRIPVVWIHGL WP_001295940.1 MULTISPECIES: hydrogenase [Enterobacteriaceae] Length: 372\nNumber of predicted TMHs: 2\nExp number of AAs in TMHs: 37.2608899999999\nExp number, first 60 AAs: 19.52923\nTotal prob of N-in: 0.94351\nPOSSIBLE N-term signal sequence\ninside 1 19\nTMhelix 20 39\noutside 40 325\nTMhelix 326 348\ninside 349 372

6687 GCF_001286365.1_7748_7_22 Escherichia coli Proteobacteria; Gammaproteobacteria; Enterobacterales; Enterobacteriaceae; Escherichia MTGDNTLIHSHGINRRDFMKLCAALAATMGLSSKAAAEMAESVTNPQRPPVIWIGAQECT WP_000145410.1 MULTISPECIES: hydrogenase-2 small chain [Proteobacteria] Length: 372\nNumber of predicted TMHs: 1\nExp number of AAs in TMHs: 23.72226\nExp number, first 60 AAs: 0.22038\nTotal prob of N-in: 0.12994\noutside 1 331\nTMhelix 332 354\ninside 355 372

6688 GCF_001286405.1_7748_7_17 Escherichia coli Proteobacteria; Gammaproteobacteria; Enterobacterales; Enterobacteriaceae; Escherichia
MTGDNTLIHSHGINRRDFMKLCAALAATMGLSSKAAAEMAESVTNPQRPPVIWIGAQECT WP_000145410.1
MULTISPECIES: hydrogenase-2 small chain [Proteobacteria] Length: 372\nNumber of predicted TMHs: 1\nExp number of AAs in TMHs: 23.72226\nExp number, first 60 AAs: 0.22038\nTotal prob of N-in: 0.12994\noutside 1 331\nTMhelix 332 354\ninside 355 372

6689 GCF_001286405.1_7748_7_17 Escherichia coli Proteobacteria; Gammaproteobacteria; Enterobacterales; Enterobacteriaceae; Escherichia
MNNEETFYQAMRRQGVTRRSFLKYCSLAATSLGLGAGMAPKIAWALENKPRIPVVWIHGL WP_001058323.1
MULTISPECIES: hydrogenase-1 small chain [Proteobacteria] Length: 372\nNumber of predicted TMHs: 2\nExp number of AAs in TMHs: 37.2608499999999\nExp number, first 60 AAs: 19.52923\nTotal prob of N-in: 0.94351\nPOSSIBLE N-term signal sequence\ninside 1 19\nTMhelix 20 39\noutside 40 325\nTMhelix 326 348\ninside 349 372

6690 GCF_001286365.1_7748_7_22 Escherichia coli Proteobacteria; Gammaproteobacteria; Enterobacterales; Enterobacteriaceae; Escherichia
MNNEETFYQAMRRQGVTRRSFLKYCSLAATSLGLGAGMAPKIAWALENKPRIPVVWIHGL WP_001058323.1
MULTISPECIES: hydrogenase-1 small chain [Proteobacteria] Length: 372\nNumber of predicted TMHs: 2\nExp number of AAs in TMHs: 37.2608499999999\nExp number, first 60 AAs: 19.52923\nTotal prob of N-in: 0.94351\nPOSSIBLE N-term signal sequence\ninside 1 19\nTMhelix 20 39\noutside 40 325\nTMhelix 326 348\ninside 349 372

6691 GCF_001286165.1_7748_7_44 Escherichia coli Proteobacteria; Gammaproteobacteria; Enterobacterales; Enterobacteriaceae; Escherichia
MNNEETFYQAMRRQGVTRRSFLKYCSLAATSLGLGAGMAPKIAWALENKPRIPVVWIHGL WP_001295940.1
MULTISPECIES: hydrogenase [Enterobacteriaceae] Length: 372\nNumber of predicted TMHs: 2\nExp number of AAs in TMHs: 37.2608899999999\nExp number, first 60 AAs: 19.52923\nTotal prob of N-in: 0.94351\nPOSSIBLE N-term signal sequence\ninside 1 19\nTMhelix 20 39\noutside 40 325\nTMhelix 326 348\ninside 349 372

6692 GCF_001286465.1_7748_7_3Escherichia coli Proteobacteria; Gammaproteobacteria; Enterobacterales; Enterobacteriaceae; Escherichia MTGDNTLIHSHGINRRDFMKLCAALAATMGLSSKAAAEMAESVTNPQRPPVIWIGAQECT
WP_024245965.1 hydrogenase 2 small subunit [Escherichia coli] Length: 372\nNumber of predicted TMHs: 1\nExp number of AAs in TMHs: 24.15412\nExp number, first 60 AAs: 0.21967\nTotal prob of N-in: 0.14289\noutside 1 331\nTMhelix 332 354\ninside 355 372

6693 GCF_001286645.1_8205_3_66 Escherichia coli Proteobacteria; Gammaproteobacteria; Enterobacterales; Enterobacteriaceae; Escherichia
MTGDNTLIHSHGINRRDFMKLCAALAATMGLSSKAAAEMAESVTNPQRPPVIWIGAQECT WP_000145410.1
MULTISPECIES: hydrogenase-2 small chain [Proteobacteria] Length: 372\nNumber of predicted TMHs: 1\nExp number of AAs in TMHs: 23.72226\nExp number, first 60 AAs: 0.22038\nTotal prob of N-in: 0.12994\noutside 1 331\nTMhelix 332 354\ninside 355 372

6694 GCF_001286505.1_8205_3_64 Escherichia coli Proteobacteria; Gammaproteobacteria; Enterobacterales; Enterobacteriaceae; Escherichia
MNNEETFYQAMRRQGVTRRSFLKYCSLAATSLGLGAGMAPKIAWALENKPRIPVVWIHGL WP_024237484.1
hydrogenase-1 small chain [Escherichia coli] Length: 372\nNumber of predicted TMHs: 2\nExp number of AAs in TMHs: 37.2691999999999\nExp number, first 60 AAs: 19.53117\nTotal prob of N-in: 0.94352\nPOSSIBLE N-term signal sequence\ninside 1 19\nTMhelix 20 39\noutside 40 325\nTMhelix 326 348\ninside 349 372

6695 GCF_001286685.1_8205_8_79 Escherichia coli Proteobacteria; Gammaproteobacteria; Enterobacterales; Enterobacteriaceae; Escherichia
MTGDNTLIHSHGINRRDFMKLCAALAATMGLSSKAAAEMAESVTNPQRPPVIWIGAQECT WP_000145410.1
MULTISPECIES: hydrogenase-2 small chain [Proteobacteria] Length: 372\nNumber of predicted TMHs: 1\nExp number of AAs in TMHs: 23.72226\nExp number, first 60 AAs: 0.22038\nTotal prob of N-in: 0.12994\noutside 1 331\nTMhelix 332 354\ninside 355 372

6696 GCF_001286505.1_8205_3_64 Escherichia coli Proteobacteria; Gammaproteobacteria; Enterobacterales; Enterobacteriaceae; Escherichia
MTGDNTLIHSHGINRRDFMKLCAALAATMGLSSKAAAEMAESVTNPQRPPVIWIGAQECT WP_053897311.1

hydrogenase 2 small subunit [Escherichia coli] Length: 372\nNumber of predicted TMHs: 1\nExp number of AAs in TMHs: 23.7222\nExp number, first 60 AAs: 0.22035\nTotal prob of N-in: 0.12994\noutside 1 331\nTMhelix 332 354\ninside 355 372

6697 GCF_001286685.1_8205_8_79 Escherichia coli Proteobacteria; Gammaproteobacteria; Enterobacterales; Enterobacteriaceae; Escherichia MNNEETFYQAMRRQGVTRRSFLKYCSLAATSLGLGAGMAPKIAWALENKPRIPVVWIHGL WP_001058320.1 hydrogenase [Escherichia coli] Length: 372\nNumber of predicted TMHs: 2\nExp number of AAs in TMHs: 37.26082\nExp number, first 60 AAs: 19.52923\nTotal prob of N-in: 0.94351\nPOSSIBLE N-term signal sequence\ninside 1 19\nTMhelix 20 39\noutside 40 325\nTMhelix 326 348\ninside 349 372

6698 GCF_001286645.1_8205_3_66 Escherichia coli Proteobacteria; Gammaproteobacteria; Enterobacterales; Enterobacteriaceae; Escherichia MNNEETFYQAMRRQGVTRRSFLKYCSLAATSLGLGAGMAPKIAWALENKPRIPVVWIHGL WP_001295940.1 MULTISPECIES: hydrogenase [Enterobacteriaceae] Length: 372\nNumber of predicted TMHs: 2\nExp number of AAs in TMHs: 37.2608899999999\nExp number, first 60 AAs: 19.52923\nTotal prob of N-in: 0.94351\nPOSSIBLE N-term signal sequence\ninside 1 19\nTMhelix 20 39\noutside 40 325\nTMhelix 326 348\ninside 349 372

6699 GCF_001309465.1_ASM130946v1 Escherichia coli Proteobacteria; Gammaproteobacteria; Enterobacterales; Enterobacteriaceae; Escherichia MTGDNTLIHSHGINRRDFMKLCAALAATMGLSSKAAAEMAESVTNPQRPPVIWIGAQECT WP_000145410.1 MULTISPECIES: hydrogenase-2 small chain [Proteobacteria] Length: 372\nNumber of predicted TMHs: 1\nExp number of AAs in TMHs: 23.72226\nExp number, first 60 AAs: 0.22038\nTotal prob of N-in: 0.12994\noutside 1 331\nTMhelix 332 354\ninside 355 372

6700 GCF_001306605.1_ASM130660v1 Escherichia coli Proteobacteria; Gammaproteobacteria; Enterobacterales; Enterobacteriaceae; Escherichia MTGDNTLIHSHGINRRDFMKLCAALAATMGLSSKAAAEMVESVTNPQRPPVIWIGAQECT WP_000145423.1 hydrogenase 2 small subunit [Escherichia coli] Length: 372\nNumber of predicted TMHs: 1\nExp number of AAs in TMHs: 23.78201\nExp number, first 60 AAs: 0.28355\nTotal prob of N-in: 0.13257\noutside 1 331\nTMhelix 332 354\ninside 355 372

6701 GCF_001306675.1_JA18 Escherichia coli Proteobacteria; Gammaproteobacteria; Enterobacterales; Enterobacteriaceae; Escherichia MTGDNTLIHSHGINRRDFMKLCAALAATMGLSSKAAAEMAESVTNPQRPPVIWIGAQECT WP_000145410.1 MULTISPECIES: hydrogenase-2 small chain [Proteobacteria] Length: 372\nNumber of predicted TMHs: 1\nExp number of AAs in TMHs: 23.72226\nExp number, first 60 AAs: 0.22038\nTotal prob of N-in: 0.12994\noutside 1 331\nTMhelix 332 354\ninside 355 372

6702 GCF_001306575.1_JA09 Escherichia coli Proteobacteria; Gammaproteobacteria; Enterobacterales; Enterobacteriaceae; Escherichia MTGDNTLIHSHGINRRDFMKLCAALAATMGLSSKAAAEMAESVTNPQRPPVIWIGAQECT WP_000145410.1 MULTISPECIES: hydrogenase-2 small chain [Proteobacteria] Length: 372\nNumber of predicted TMHs: 1\nExp number of AAs in TMHs: 23.72226\nExp number, first 60 AAs: 0.22038\nTotal prob of N-in: 0.12994\noutside 1 331\nTMhelix 332 354\ninside 355 372

6703 GCF_001306655.1_JA16 Escherichia coli Proteobacteria; Gammaproteobacteria; Enterobacterales; Enterobacteriaceae; Escherichia MTGDNTLIHSHGINRRDFMKLCAALAATMGLSSKAAAEMVESVTNPQRPPVIWIGAQECT WP_000145423.1 hydrogenase 2 small subunit [Escherichia coli] Length: 372\nNumber of predicted TMHs: 1\nExp number of AAs in TMHs: 23.78201\nExp number, first 60 AAs: 0.28355\nTotal prob of N-in: 0.13257\noutside 1 331\nTMhelix 332 354\ninside 355 372

6704 GCF_001297995.1_ASM129799v1 Escherichia coli Proteobacteria; Gammaproteobacteria; Enterobacterales; Enterobacteriaceae; Escherichia MTGDNTLIHSHGINRRDFMKLCAALAATMGLSSKAAAEMAESVTNPQRPPVIWIGAQECT WP_000145410.1 MULTISPECIES: hydrogenase-2 small chain [Proteobacteria] Length: 372\nNumber of predicted TMHs: 1\nExp number of AAs in TMHs: 23.72226\nExp number, first 60 AAs: 0.22038\nTotal prob of N-in: 0.12994\noutside 1 331\nTMhelix 332 354\ninside 355 372

6705 GCF_001308125.1_ASM130812v1 Escherichia coli Proteobacteria; Gammaproteobacteria; Enterobacterales; Enterobacteriaceae; Escherichia MTGDNTLIHSHGINRRDFMKLCAALAATMGLSSKAAAEMAESVTNPQRPPVIWIGAQECT WP_000145410.1

MULTISPECIES: hydrogenase-2 small chain [Proteobacteria] Length: 372\nNumber of predicted TMHs: 1\nExp number of AAs in TMHs: 23.72226\nExp number, first 60 AAs: 0.22038\nTotal prob of N-in: 0.12994\noutside 1 331\nTMhelix 332 354\ninside 355 372

6706 GCF_001306575.1_JA09 Escherichia coli Proteobacteria; Gammaproteobacteria; Enterobacterales; Enterobacteriaceae; Escherichia MNNEETFYQAMRRQGVTRRSFLKYCSLAATSLGLGAGMAPKIAWALENKPRIPVVWIHGL WP_001058323.1 MULTISPECIES: hydrogenase-1 small chain [Proteobacteria] Length: 372\nNumber of predicted TMHs: 2\nExp number of AAs in TMHs: 37.2608499999999\nExp number, first 60 AAs: 19.52923\nTotal prob of N-in: 0.94351\nPOSSIBLE N-term signal sequence\ninside 1 19\nTMhelix 20 39\noutside 40 325\nTMhelix 326 348\ninside 349 372

6707 GCF_001306675.1_JA18 Escherichia coli Proteobacteria; Gammaproteobacteria; Enterobacterales; Enterobacteriaceae; Escherichia MNNEETFYQAMRRQGVTRRSFLKYCSLAATSLGLGAGMAPKIAWALENKPRIPVVWIHGL WP_001058323.1 MULTISPECIES: hydrogenase-1 small chain [Proteobacteria] Length: 372\nNumber of predicted TMHs: 2\nExp number of AAs in TMHs: 37.2608499999999\nExp number, first 60 AAs: 19.52923\nTotal prob of N-in: 0.94351\nPOSSIBLE N-term signal sequence\ninside 1 19\nTMhelix 20 39\noutside 40 325\nTMhelix 326 348\ninside 349 372

6708 GCF_001308125.1_ASM130812v1 Escherichia coli Proteobacteria; Gammaproteobacteria; Enterobacterales; Enterobacteriaceae; Escherichia MNNEETFYQAMRRQGVTRRSFLKYCSLAATSLGLGAGMAPKIAWALENKPRIPVVWIHGL WP_001058323.1 MULTISPECIES: hydrogenase-1 small chain [Proteobacteria] Length: 372\nNumber of predicted TMHs: 2\nExp number of AAs in TMHs: 37.2608499999999\nExp number, first 60 AAs: 19.52923\nTotal prob of N-in: 0.94351\nPOSSIBLE N-term signal sequence\ninside 1 19\nTMhelix 20 39\noutside 40 325\nTMhelix 326 348\ninside 349 372

6709 GCF_001309465.1_ASM130946v1 Escherichia coli Proteobacteria; Gammaproteobacteria; Enterobacterales; Enterobacteriaceae; Escherichia MNNEETFYQAMRRQGVTRRSFLKYCSLAATSLGLGAGMAPKIAWALENKPRIPVVWIHGL WP_001058323.1 MULTISPECIES: hydrogenase-1 small chain [Proteobacteria] Length: 372\nNumber of predicted TMHs: 2\nExp number of AAs in TMHs: 37.2608499999999\nExp number, first 60 AAs: 19.52923\nTotal prob of N-in: 0.94351\nPOSSIBLE N-term signal sequence\ninside 1 19\nTMhelix 20 39\noutside 40 325\nTMhelix 326 348\ninside 349 372

6710 GCF_001308165.1_ASM130816v1 Escherichia coli Proteobacteria; Gammaproteobacteria; Enterobacterales; Enterobacteriaceae; Escherichia MNNEETFYQAMRRQGVTRRSFLKYCSLAATSLGLGAGMAPKIAWALENKPRIPVVWIHGL WP_001058323.1 MULTISPECIES: hydrogenase-1 small chain [Proteobacteria] Length: 372\nNumber of predicted TMHs: 2\nExp number of AAs in TMHs: 37.2608499999999\nExp number, first 60 AAs: 19.52923\nTotal prob of N-in: 0.94351\nPOSSIBLE N-term signal sequence\ninside 1 19\nTMhelix 20 39\noutside 40 325\nTMhelix 326 348\ninside 349 372

6711 GCF_001306605.1_ASM130660v1 Escherichia coli Proteobacteria; Gammaproteobacteria; Enterobacterales; Enterobacteriaceae; Escherichia MNNEETFYQAMRRQGVTRRSFLKYCSLAATSLGLGAGMAPKIAWALENKPRIPVVWIHGL WP_001295940.1 MULTISPECIES: hydrogenase [Enterobacteriaceae] Length: 372\nNumber of predicted TMHs: 2\nExp number of AAs in TMHs: 37.2608899999999\nExp number, first 60 AAs: 19.52923\nTotal prob of N-in: 0.94351\nPOSSIBLE N-term signal sequence\ninside 1 19\nTMhelix 20 39\noutside 40 325\nTMhelix 326 348\ninside 349 372

6712 GCF_001309565.1_ASM130956v1 Escherichia coli Proteobacteria; Gammaproteobacteria; Enterobacterales; Enterobacteriaceae; Escherichia MNNEETFYQAMRRQGVTRRSFLKYCSLAATSLGLGAGMAPKIAWALENKPRIPVVWIHGL WP_001058323.1 MULTISPECIES: hydrogenase-1 small chain [Proteobacteria] Length: 372\nNumber of predicted TMHs: 2\nExp number of AAs in TMHs: 37.2608499999999\nExp number, first 60 AAs: 19.52923\nTotal prob of N-in: 0.94351\nPOSSIBLE N-term signal sequence\ninside 1 19\nTMhelix 20 39\noutside 40 325\nTMhelix 326 348\ninside 349 372

6713 GCF_001309735.1_ASM130973v1 Escherichia coli Proteobacteria; Gammaproteobacteria; Enterobacterales; Enterobacteriaceae; Escherichia MTGDNTLIHSHGINRRDFMKLCAALAATMGLSSKAAAEMAESVTNPQRPPVIWIGAQECT WP_000145410.1

MULTISPECIES: hydrogenase-2 small chain [Proteobacteria] Length: 372\nNumber of predicted TMHs: 1\nExp number of AAs in TMHs: 23.72226\nExp number, first 60 AAs: 0.22038\nTotal prob of N-in: 0.12994\noutside 1 331\nTMhelix 332 354\ninside 355 372

6714 GCF_001309805.1_ASM130980v1 Escherichia coli Proteobacteria; Gammaproteobacteria; Enterobacterales; Enterobacteriaceae; Escherichia MTGDNTLIHSHGINRRDFMKLCAALAATMGLSSKAAAEMAESVTNPQRPPVIWIGAQECT WP_000145410.1 MULTISPECIES: hydrogenase-2 small chain [Proteobacteria] Length: 372\nNumber of predicted TMHs: 1\nExp number of AAs in TMHs: 23.72226\nExp number, first 60 AAs: 0.22038\nTotal prob of N-in: 0.12994\noutside 1 331\nTMhelix 332 354\ninside 355 372

6715 GCF_001309825.1_ASM130982v1 Escherichia coli Proteobacteria; Gammaproteobacteria; Enterobacterales; Enterobacteriaceae; Escherichia MTGDNTLIHSHGINRRDFMKLCAALAATMGLSSKAAAEMAESVTNPQRPPVIWIGAQECT WP_000145410.1 MULTISPECIES: hydrogenase-2 small chain [Proteobacteria] Length: 372\nNumber of predicted TMHs: 1\nExp number of AAs in TMHs: 23.72226\nExp number, first 60 AAs: 0.22038\nTotal prob of N-in: 0.12994\noutside 1 331\nTMhelix 332 354\ninside 355 372

6716 GCF_001309885.1_ASM130988v1 Escherichia coli Proteobacteria; Gammaproteobacteria; Enterobacterales; Enterobacteriaceae; Escherichia MTGDNTLIHSHGINRRDFMKLCAALAATMGLSSKAAAEMAESVTNPQRPPVIWIGAQECT WP_000145410.1 MULTISPECIES: hydrogenase-2 small chain [Proteobacteria] Length: 372\nNumber of predicted TMHs: 1\nExp number of AAs in TMHs: 23.72226\nExp number, first 60 AAs: 0.22038\nTotal prob of N-in: 0.12994\noutside 1 331\nTMhelix 332 354\ninside 355 372

6717 GCF_001309895.1_ASM130989v1 Escherichia coli Proteobacteria; Gammaproteobacteria; Enterobacterales; Enterobacteriaceae; Escherichia MTGDNTLIHSHGINRRDFMKLCAALAATMGLSSKAAAEMAESVTNPQRPPVIWIGAQECT WP_000145410.1 MULTISPECIES: hydrogenase-2 small chain [Proteobacteria] Length: 372\nNumber of predicted TMHs: 1\nExp number of AAs in TMHs: 23.72226\nExp number, first 60 AAs: 0.22038\nTotal prob of N-in: 0.12994\noutside 1 331\nTMhelix 332 354\ninside 355 372

6718 GCF_001309925.1_ASM130992v1 Escherichia coli Proteobacteria; Gammaproteobacteria; Enterobacterales; Enterobacteriaceae; Escherichia MTGDNTLIHSHGINRRDFMKLCAALAATMGLSSKAAAEMAESVTNPQRPPVIWIGAQECT WP_000145410.1 MULTISPECIES: hydrogenase-2 small chain [Proteobacteria] Length: 372\nNumber of predicted TMHs: 1\nExp number of AAs in TMHs: 23.72226\nExp number, first 60 AAs: 0.22038\nTotal prob of N-in: 0.12994\noutside 1 331\nTMhelix 332 354\ninside 355 372

6719 GCF_001309985.1_ASM130998v1 Escherichia coli Proteobacteria; Gammaproteobacteria; Enterobacterales; Enterobacteriaceae; Escherichia MTGDNTLIHSHGINRRDFMKLCAALAATMGLSSKAAAEMAESVTNPQRPPVIWIGAQECT WP_000145410.1 MULTISPECIES: hydrogenase-2 small chain [Proteobacteria] Length: 372\nNumber of predicted TMHs: 1\nExp number of AAs in TMHs: 23.72226\nExp number, first 60 AAs: 0.22038\nTotal prob of N-in: 0.12994\noutside 1 331\nTMhelix 332 354\ninside 355 372

6720 GCF_001309595.1_ASM130959v1 Escherichia coli Proteobacteria; Gammaproteobacteria; Enterobacterales; Enterobacteriaceae; Escherichia MNNEETFYQAMRRQGVTRRSFLKYCSLAATSLGLGAGMAPKIAWALENKPRIPVVWIHGL WP_001058323.1 MULTISPECIES: hydrogenase-1 small chain [Proteobacteria] Length: 372\nNumber of predicted TMHs: 2\nExp number of AAs in TMHs: 37.2608499999999\nExp number, first 60 AAs: 19.52923\nTotal prob of N-in: 0.94351\nPOSSIBLE N-term signal sequence\ninside 1 19\nTMhelix 20 39\noutside 40 325\nTMhelix 326 348\ninside 349 372

6721 GCF_001309635.1_ASM130963v1 Escherichia coli Proteobacteria; Gammaproteobacteria; Enterobacterales; Enterobacteriaceae; Escherichia MNNEETFYQAMRRQGVTRRSFLKYCSLAATSLGLGAGIAPKIAWALENKPRIPVVWIHGL WP_023982790.1 hydrogenase [Escherichia coli] Length: 372\nNumber of predicted TMHs: 2\nExp number of AAs in TMHs: 37.8310099999999\nExp number, first 60 AAs: 20.15722\nTotal prob of N-in: 0.95896\nPOSSIBLE N-term signal sequence\ninside 1 20\nTMhelix 21 43\noutside 44 325\nTMhelix 326 348\ninside 349 372

6722 GCF_001309925.1_ASM130992v1 Escherichia coli Proteobacteria; Gammaproteobacteria; Enterobacterales; Enterobacteriaceae; Escherichia
MNNEETFYQAMRRQGVTRRSFLKYCSLAATSLGLGAGMAPKIAWALENKPRIPVVWIHGL WP_001301613.1
hydrogenase [Escherichia coli] Length: 372\nNumber of predicted TMHs: 2\nExp number of AAs in TMHs: 37.46598\nExp number, first 60 AAs: 19.53139\nTotal prob of N-in: 0.94364\nPOSSIBLE N-term signal sequence\ninside 1 19\nTMhelix 20 39\noutside 40 325\nTMhelix 326 348\ninside 349 372

6723 GCF_001309895.1_ASM130989v1 Escherichia coli Proteobacteria; Gammaproteobacteria; Enterobacterales; Enterobacteriaceae; Escherichia
MNNEETFYQAMRRQGVTRRSFLKYCSLAATSLGLGAGMAPKIAWALENKPRIPVVWIHGL WP_001301613.1
hydrogenase [Escherichia coli] Length: 372\nNumber of predicted TMHs: 2\nExp number of AAs in TMHs: 37.46598\nExp number, first 60 AAs: 19.53139\nTotal prob of N-in: 0.94364\nPOSSIBLE N-term signal sequence\ninside 1 19\nTMhelix 20 39\noutside 40 325\nTMhelix 326 348\ninside 349 372

6724 GCF_001309985.1_ASM130998v1 Escherichia coli Proteobacteria; Gammaproteobacteria; Enterobacterales; Enterobacteriaceae; Escherichia
MNNEETFYQAMRRQGVTRRSFLKYCSLAATSLGLGAGMAPKIAWALENKPRIPVVWIHGL WP_001301613.1
hydrogenase [Escherichia coli] Length: 372\nNumber of predicted TMHs: 2\nExp number of AAs in TMHs: 37.46598\nExp number, first 60 AAs: 19.53139\nTotal prob of N-in: 0.94364\nPOSSIBLE N-term signal sequence\ninside 1 19\nTMhelix 20 39\noutside 40 325\nTMhelix 326 348\ninside 349 372

6725 GCF_001316875.2_v02 Escherichia coli Proteobacteria; Gammaproteobacteria; Enterobacterales; Enterobacteriaceae; Escherichia MTGDNTLIHSHGINRRDFMKLCAALAATMGLSSKAAAEMAESVTNPQRPPVIWIGAQECT
WP_000145410.1 MULTISPECIES: hydrogenase-2 small chain [Proteobacteria] Length: 372\nNumber of predicted TMHs: 1\nExp number of AAs in TMHs: 23.72226\nExp number, first 60 AAs: 0.22038\nTotal prob of N-in: 0.12994\noutside 1 331\nTMhelix 332 354\ninside 355 372

6726 GCF_001412895.1_ASM141289v1 Escherichia coli Proteobacteria; Gammaproteobacteria; Enterobacterales; Enterobacteriaceae; Escherichia
MTGDNTLIHSHGINRRDFMKLCAALAATMGLSSKAAAEMAESVTNPQRPPVIWIGAQECT WP_000145410.1
MULTISPECIES: hydrogenase-2 small chain [Proteobacteria] Length: 372\nNumber of predicted TMHs: 1\nExp number of AAs in TMHs: 23.72226\nExp number, first 60 AAs: 0.22038\nTotal prob of N-in: 0.12994\noutside 1 331\nTMhelix 332 354\ninside 355 372

6727 GCF_001309995.1_ASM130999v1 Escherichia coli Proteobacteria; Gammaproteobacteria; Enterobacterales; Enterobacteriaceae; Escherichia
MTGDNTLIHSHGINRRDFMKLCAALAATMGLSSKAAAEMAESVTNPQRPPVIWIGAQECT WP_000145410.1
MULTISPECIES: hydrogenase-2 small chain [Proteobacteria] Length: 372\nNumber of predicted TMHs: 1\nExp number of AAs in TMHs: 23.72226\nExp number, first 60 AAs: 0.22038\nTotal prob of N-in: 0.12994\noutside 1 331\nTMhelix 332 354\ninside 355 372

6728 GCF_001413425.1_ASM141342v1 Escherichia coli Proteobacteria; Gammaproteobacteria; Enterobacterales; Enterobacteriaceae; Escherichia
MTGDNTLIHSHGINRRDFMKLCAALAATMGLSSKAAAEMAESVTNPQRPPVIWIGAQECT WP_000145410.1
MULTISPECIES: hydrogenase-2 small chain [Proteobacteria] Length: 372\nNumber of predicted TMHs: 1\nExp number of AAs in TMHs: 23.72226\nExp number, first 60 AAs: 0.22038\nTotal prob of N-in: 0.12994\noutside 1 331\nTMhelix 332 354\ninside 355 372

6729 GCF_001413455.1_ASM141345v1 Escherichia coli Proteobacteria; Gammaproteobacteria; Enterobacterales; Enterobacteriaceae; Escherichia
MTGDNTLIHSHGINRRDFMKLCAALAATMGLSSKAAAEMAESVTNPQRPPVIWIGAQECT WP_000145410.1
MULTISPECIES: hydrogenase-2 small chain [Proteobacteria] Length: 372\nNumber of predicted TMHs: 1\nExp number of AAs in TMHs: 23.72226\nExp number, first 60 AAs: 0.22038\nTotal prob of N-in: 0.12994\noutside 1 331\nTMhelix 332 354\ninside 355 372

6730 GCF_001413425.1_ASM141342v1 Escherichia coli Proteobacteria; Gammaproteobacteria; Enterobacterales; Enterobacteriaceae; Escherichia
MNNEETFYQAMRRQGVTRRSFLKYCSLAATSLGLGAGMAPKIAWALENKPRIPVVWIHGL WP_001058323.1
MULTISPECIES: hydrogenase-1 small chain [Proteobacteria] Length: 372\nNumber of predicted TMHs: 2\nExp number of AAs in TMHs: 37.2608499999999\nExp number, first 60 AAs: 19.52923\nTotal prob of N-in:

0.94351\nPOSSIBLE N-term signal sequence\ninside 1 19\nTMhelix 20 39\noutside 40 325\nTMhelix 326 348\ninside 349 372

6731 GCF_001413455.1_ASM141345v1 Escherichia coli Proteobacteria; Gammaproteobacteria; Enterobacterales; Enterobacteriaceae; Escherichia MNNEETFYQAMRRQGVTRRSFLKYCSLAATSLGLGAGMAPKIAWALENKPRIPVVWIHGL WP_001058323.1 MULTISPECIES: hydrogenase-1 small chain [Proteobacteria] Length: 372\nNumber of predicted TMHs: 2\nExp number of AAs in TMHs: 37.2608499999999\nExp number, first 60 AAs: 19.52923\nTotal prob of N-in: 0.94351\nPOSSIBLE N-term signal sequence\ninside 1 19\nTMhelix 20 39\noutside 40 325\nTMhelix 326 348\ninside 349 372

6732 GCF_001412975.1_ASM141297v1 Escherichia coli Proteobacteria; Gammaproteobacteria; Enterobacterales; Enterobacteriaceae; Escherichia MNNEETFYQAMRRQGVTRRSFLKYCSLAATSLGLGAGMAPKIAWALENKPRIPVVWIHGL WP_001058323.1 MULTISPECIES: hydrogenase-1 small chain [Proteobacteria] Length: 372\nNumber of predicted TMHs: 2\nExp number of AAs in TMHs: 37.2608499999999\nExp number, first 60 AAs: 19.52923\nTotal prob of N-in: 0.94351\nPOSSIBLE N-term signal sequence\ninside 1 19\nTMhelix 20 39\noutside 40 325\nTMhelix 326 348\ninside 349 372

6733 GCF_001316875.2_v02 Escherichia coli Proteobacteria; Gammaproteobacteria; Enterobacterales; Enterobacteriaceae; Escherichia MNNEETFYQAMRRQGVTRRSFLKYCSLAATSLGLGAGMAPKIAWALENKPRIPVVWIHGL WP_001058323.1 MULTISPECIES: hydrogenase-1 small chain [Proteobacteria] Length: 372\nNumber of predicted TMHs: 2\nExp number of AAs in TMHs: 37.2608499999999\nExp number, first 60 AAs: 19.52923\nTotal prob of N-in: 0.94351\nPOSSIBLE N-term signal sequence\ninside 1 19\nTMhelix 20 39\noutside 40 325\nTMhelix 326 348\ninside 349 372

6734 GCF_001413485.1_ASM141348v1 Escherichia coli Proteobacteria; Gammaproteobacteria; Enterobacterales; Enterobacteriaceae; Escherichia MTGDNTLIHSHGINRRDFMKLCAALAATMGLSSKAAAEMAESVTNPQRPPVIWIGAQECT WP_000145410.1 MULTISPECIES: hydrogenase-2 small chain [Proteobacteria] Length: 372\nNumber of predicted TMHs: 1\nExp number of AAs in TMHs: 23.72226\nExp number, first 60 AAs: 0.22038\nTotal prob of N-in: 0.12994\noutside 1 331\nTMhelix 332 354\ninside 355 372

6735 GCF_001413555.1_ASM141355v1 Escherichia coli Proteobacteria; Gammaproteobacteria; Enterobacterales; Enterobacteriaceae; Escherichia MTGDNTLIHSHGINRRDFMKLCAALAATMGLSSKAAAEMAESVTNPQRPPVIWIGAQECT WP_000145410.1 MULTISPECIES: hydrogenase-2 small chain [Proteobacteria] Length: 372\nNumber of predicted TMHs: 1\nExp number of AAs in TMHs: 23.72226\nExp number, first 60 AAs: 0.22038\nTotal prob of N-in: 0.12994\noutside 1 331\nTMhelix 332 354\ninside 355 372

6736 GCF_001413595.1_ASM141359v1 Escherichia coli Proteobacteria; Gammaproteobacteria; Enterobacterales; Enterobacteriaceae; Escherichia MTGDNTLIHSHGINRRDFMKLCAALAATMGLSSKAAAEMAESVTNPQRPPVIWIGAQECT WP_000145410.1 MULTISPECIES: hydrogenase-2 small chain [Proteobacteria] Length: 372\nNumber of predicted TMHs: 1\nExp number of AAs in TMHs: 23.72226\nExp number, first 60 AAs: 0.22038\nTotal prob of N-in: 0.12994\noutside 1 331\nTMhelix 332 354\ninside 355 372

6737 GCF_001413555.1_ASM141355v1 Escherichia coli Proteobacteria; Gammaproteobacteria; Enterobacterales; Enterobacteriaceae; Escherichia MNNEETFYQAMRRQGVTRRSFLKYCSLAATSLGLGAGMAPKIAWALENKPRIPVVWIHGL WP_001058323.1 MULTISPECIES: hydrogenase-1 small chain [Proteobacteria] Length: 372\nNumber of predicted TMHs: 2\nExp number of AAs in TMHs: 37.2608499999999\nExp number, first 60 AAs: 19.52923\nTotal prob of N-in: 0.94351\nPOSSIBLE N-term signal sequence\ninside 1 19\nTMhelix 20 39\noutside 40 325\nTMhelix 326 348\ninside 349 372

6738 GCF_001413635.1_ASM141363v1 Escherichia coli Proteobacteria; Gammaproteobacteria; Enterobacterales; Enterobacteriaceae; Escherichia MTGDNTLIHSHGINRRDFMKLCAALAATMGLSSKAAAEMAESVTNPQRPPVIWIGAQECT WP_000145410.1 MULTISPECIES: hydrogenase-2 small chain [Proteobacteria] Length: 372\nNumber of predicted TMHs: 1\nExp number of AAs in TMHs: 23.72226\nExp number, first 60 AAs: 0.22038\nTotal prob of N-in: 0.12994\noutside 1 331\nTMhelix 332 354\ninside 355 372

6739 GCF_001413645.1_ASM141364v1 Escherichia coli Proteobacteria; Gammaproteobacteria; Enterobacterales; Enterobacteriaceae; Escherichia
MTGDNTLIHSHGINRRDFMKLCAALAATMGLSSKAAAEMAESVTNPQRPPVIWIGAQECT WP_000145410.1
MULTISPECIES: hydrogenase-2 small chain [Proteobacteria] Length: 372\nNumber of predicted TMHs: 1\nExp number of AAs in TMHs: 23.72226\nExp number, first 60 AAs: 0.22038\nTotal prob of N-in: 0.12994\noutside 1 331\nTMhelix 332 354\ninside 355 372

6740 GCF_001413685.1_ASM141368v1 Escherichia coli Proteobacteria; Gammaproteobacteria; Enterobacterales; Enterobacteriaceae; Escherichia
MTGDNTLIHSHGINRRDFMKLCAALAATMGLSSKAAAEMAESVTNPQRPPVIWIGAQECT WP_000145410.1
MULTISPECIES: hydrogenase-2 small chain [Proteobacteria] Length: 372\nNumber of predicted TMHs: 1\nExp number of AAs in TMHs: 23.72226\nExp number, first 60 AAs: 0.22038\nTotal prob of N-in: 0.12994\noutside 1 331\nTMhelix 332 354\ninside 355 372

6741 GCF_001413735.1_ASM141373v1 Escherichia coli Proteobacteria; Gammaproteobacteria; Enterobacterales; Enterobacteriaceae; Escherichia
MTGDNTLIHSHGINRRDFMKLCAALAATMGLSSKAAAEMAESVTNPQRPPVIWIGAQECT WP_000145410.1
MULTISPECIES: hydrogenase-2 small chain [Proteobacteria] Length: 372\nNumber of predicted TMHs: 1\nExp number of AAs in TMHs: 23.72226\nExp number, first 60 AAs: 0.22038\nTotal prob of N-in: 0.12994\noutside 1 331\nTMhelix 332 354\ninside 355 372

6742 GCF_001413825.1_ASM141382v1 Escherichia coli Proteobacteria; Gammaproteobacteria; Enterobacterales; Enterobacteriaceae; Escherichia
MTGDNTLIHSHGINRRDFMKLCAALAATMGLSSKAAAEMAESVTNPQRPPVIWIGAQECT WP_000145410.1
MULTISPECIES: hydrogenase-2 small chain [Proteobacteria] Length: 372\nNumber of predicted TMHs: 1\nExp number of AAs in TMHs: 23.72226\nExp number, first 60 AAs: 0.22038\nTotal prob of N-in: 0.12994\noutside 1 331\nTMhelix 332 354\ninside 355 372

6743 GCF_001413825.1_ASM141382v1 Escherichia coli Proteobacteria; Gammaproteobacteria; Enterobacterales; Enterobacteriaceae; Escherichia
MNNEETFYQAMRRQGVTRRSFLKYCSLAATSLGLGAGMAPKIAWALENKPRIPVVWIHGL WP_001058323.1
MULTISPECIES: hydrogenase-1 small chain [Proteobacteria] Length: 372\nNumber of predicted TMHs: 2\nExp number of AAs in TMHs: 37.2608499999999\nExp number, first 60 AAs: 19.52923\nTotal prob of N-in: 0.94351\nPOSSIBLE N-term signal sequence\ninside 1 19\nTMhelix 20 39\noutside 40 325\nTMhelix 326 348\ninside 349 372

6744 GCF_001413685.1_ASM141368v1 Escherichia coli Proteobacteria; Gammaproteobacteria; Enterobacterales; Enterobacteriaceae; Escherichia
MNNEETFYQAMRRQGVTRRSFLKYCSLAATSLGLGAGMAPKIAWALENKPRIPVVWIHGL WP_001058323.1
MULTISPECIES: hydrogenase-1 small chain [Proteobacteria] Length: 372\nNumber of predicted TMHs: 2\nExp number of AAs in TMHs: 37.2608499999999\nExp number, first 60 AAs: 19.52923\nTotal prob of N-in: 0.94351\nPOSSIBLE N-term signal sequence\ninside 1 19\nTMhelix 20 39\noutside 40 325\nTMhelix 326 348\ninside 349 372

6745 GCF_001413645.1_ASM141364v1 Escherichia coli Proteobacteria; Gammaproteobacteria; Enterobacterales; Enterobacteriaceae; Escherichia
MNNEETFYQAMRRQGVTRRSFLKYCSLAATSLGLGAGMAPKIAWALENKPRIPVVWIHGL WP_001058323.1
MULTISPECIES: hydrogenase-1 small chain [Proteobacteria] Length: 372\nNumber of predicted TMHs: 2\nExp number of AAs in TMHs: 37.2608499999999\nExp number, first 60 AAs: 19.52923\nTotal prob of N-in: 0.94351\nPOSSIBLE N-term signal sequence\ninside 1 19\nTMhelix 20 39\noutside 40 325\nTMhelix 326 348\ninside 349 372

6746 GCF_001413875.1_ASM141387v1 Escherichia coli Proteobacteria; Gammaproteobacteria; Enterobacterales; Enterobacteriaceae; Escherichia
MTGDNTLIHSHGINRRDFMKLCAALAATMGLSSKAAAEMAESVTNPQRPPVIWIGAQECT WP_000145410.1
MULTISPECIES: hydrogenase-2 small chain [Proteobacteria] Length: 372\nNumber of predicted TMHs: 1\nExp number of AAs in TMHs: 23.72226\nExp number, first 60 AAs: 0.22038\nTotal prob of N-in: 0.12994\noutside 1 331\nTMhelix 332 354\ninside 355 372

6747 GCF_001413905.1_ASM141390v1 Escherichia coli Proteobacteria; Gammaproteobacteria; Enterobacterales; Enterobacteriaceae; Escherichia

MTGDNTLIHSHGINRRDFMKLCAALAATMGLSSKAAAEMAESVTNPQRPPVIWIGAQECT WP_000145410.1 MULTISPECIES: hydrogenase-2 small chain [Proteobacteria] Length: 372\nNumber of predicted TMHs: 1\nExp number of AAs in TMHs: 23.72226\nExp number, first 60 AAs: 0.22038\nTotal prob of N-in: 0.12994\noutside 1 331\nTMhelix 332 354\ninside 355 372

6748 GCF_001419785.1_ASM141978v1 Escherichia coli Proteobacteria; Gammaproteobacteria; Enterobacterales; Enterobacteriaceae; Escherichia MTGDNTLIHSHGINRRDFMKLCAALAATMGLSSKAAAEMAESVTNPQRPPVIWIGAQECT WP_000145410.1 MULTISPECIES: hydrogenase-2 small chain [Proteobacteria] Length: 372\nNumber of predicted TMHs: 1\nExp number of AAs in TMHs: 23.72226\nExp number, first 60 AAs: 0.22038\nTotal prob of N-in: 0.12994\noutside 1 331\nTMhelix 332 354\ninside 355 372

6749 GCF_001420045.1_ASM142004v1 Escherichia coli Proteobacteria; Gammaproteobacteria; Enterobacterales; Enterobacteriaceae; Escherichia MTGDNTLIHSHGINRRDFMKLCAALAATMGLSSKAAAEMAESVTNPQRPPVIWIGAQECT WP_000145410.1 MULTISPECIES: hydrogenase-2 small chain [Proteobacteria] Length: 372\nNumber of predicted TMHs: 1\nExp number of AAs in TMHs: 23.72226\nExp number, first 60 AAs: 0.22038\nTotal prob of N-in: 0.12994\noutside 1 331\nTMhelix 332 354\ninside 355 372

6750 GCF_001420135.1_ASM142013v1 Escherichia coli Proteobacteria; Gammaproteobacteria; Enterobacterales; Enterobacteriaceae; Escherichia MTGDNTLIHSHGINRRDFMKLCAALAATMGLSSKAAAEMAESVTNPQRPPVIWIGAQECT WP_000145410.1 MULTISPECIES: hydrogenase-2 small chain [Proteobacteria] Length: 372\nNumber of predicted TMHs: 1\nExp number of AAs in TMHs: 23.72226\nExp number, first 60 AAs: 0.22038\nTotal prob of N-in: 0.12994\noutside 1 331\nTMhelix 332 354\ninside 355 372

6751 GCF_001420075.1_ASM142007v1 Escherichia coli Proteobacteria; Gammaproteobacteria; Enterobacterales; Enterobacteriaceae; Escherichia MTGDNTLIHSHGINRRDFMKLCAALAATMGLSSKAAAEMAESVTNPQRPPVIWIGAQECT WP_000145410.1 MULTISPECIES: hydrogenase-2 small chain [Proteobacteria] Length: 372\nNumber of predicted TMHs: 1\nExp number of AAs in TMHs: 23.72226\nExp number, first 60 AAs: 0.22038\nTotal prob of N-in: 0.12994\noutside 1 331\nTMhelix 332 354\ninside 355 372

6752 GCF_001420065.1_ASM142006v1 Escherichia coli Proteobacteria; Gammaproteobacteria; Enterobacterales; Enterobacteriaceae; Escherichia MTGDNTLIHSHGINRRDFMKLCAALAATMGLSSKAAAEMAESVTNPQRPPVIWIGAQECT WP_000145410.1 MULTISPECIES: hydrogenase-2 small chain [Proteobacteria] Length: 372\nNumber of predicted TMHs: 1\nExp number of AAs in TMHs: 23.72226\nExp number, first 60 AAs: 0.22038\nTotal prob of N-in: 0.12994\noutside 1 331\nTMhelix 332 354\ninside 355 372

6753 GCF_001420935.1_ASM142093v1 Escherichia coli Proteobacteria; Gammaproteobacteria; Enterobacterales; Enterobacteriaceae; Escherichia MTGDNTLIHSHGINRRDFMKLCAALAATMGLSSKAAAEMAESVTNPQRPPVIWIGAQECT WP_000145410.1 MULTISPECIES: hydrogenase-2 small chain [Proteobacteria] Length: 372\nNumber of predicted TMHs: 1\nExp number of AAs in TMHs: 23.72226\nExp number, first 60 AAs: 0.22038\nTotal prob of N-in: 0.12994\noutside 1 331\nTMhelix 332 354\ninside 355 372

6754 GCF_001420935.1_ASM142093v1 Escherichia coli Proteobacteria; Gammaproteobacteria; Enterobacterales; Enterobacteriaceae; Escherichia MNNEETFYQAMRRQGVTRRSFLKYCSLAATSLGLGAGMAPKIAWALENKPRIPVVWIHGL WP_001058323.1 MULTISPECIES: hydrogenase-1 small chain [Proteobacteria] Length: 372\nNumber of predicted TMHs: 2\nExp number of AAs in TMHs: 37.2608499999999\nExp number, first 60 AAs: 19.52923\nTotal prob of N-in: 0.94351\nPOSSIBLE N-term signal sequence\ninside 1 19\nTMhelix 20 39\noutside 40 325\nTMhelix 326 348\ninside 349 372

6755 GCF_001419895.1_ASM141989v1 Escherichia coli Proteobacteria; Gammaproteobacteria; Enterobacterales; Enterobacteriaceae; Escherichia MNNEETFYQAMRRQGVTRRSFLKYCSLAATSLGLGAGMAPKIAWALENKPRIPVVWIHGL WP_001058323.1 MULTISPECIES: hydrogenase-1 small chain [Proteobacteria] Length: 372\nNumber of predicted TMHs: 2\nExp number of AAs in TMHs: 37.2608499999999\nExp number, first 60 AAs: 19.52923\nTotal prob of N-in:

0.94351\nPOSSIBLE N-term signal sequence\ninside 1 19\nTMhelix 20 39\noutside 40 325\nTMhelix 326 348\ninside 349 372

6756 GCF_001420135.1_ASM142013v1 Escherichia coli Proteobacteria; Gammaproteobacteria; Enterobacterales; Enterobacteriaceae; Escherichia
MNNEETFYQAMRRQGVTRRSFLKYCSLAATSLGLGAGMAPKIAWALENKPRIPVVWIHGL WP_001058323.1
MULTISPECIES: hydrogenase-1 small chain [Proteobacteria] Length: 372\nNumber of predicted TMHs: 2\nExp number of AAs in TMHs: 37.2608499999999\nExp number, first 60 AAs: 19.52923\nTotal prob of N-in: 0.94351\nPOSSIBLE N-term signal sequence\ninside 1 19\nTMhelix 20 39\noutside 40 325\nTMhelix 326 348\ninside 349 372

6757 GCF_001420065.1_ASM142006v1 Escherichia coli Proteobacteria; Gammaproteobacteria; Enterobacterales; Enterobacteriaceae; Escherichia
MNNEETFYQAMRRQGVTRRSFLKYCSLAATSLGLGAGMAPKIAWALENKPRIPVVWIHGL WP_001058323.1
MULTISPECIES: hydrogenase-1 small chain [Proteobacteria] Length: 372\nNumber of predicted TMHs: 2\nExp number of AAs in TMHs: 37.2608499999999\nExp number, first 60 AAs: 19.52923\nTotal prob of N-in: 0.94351\nPOSSIBLE N-term signal sequence\ninside 1 19\nTMhelix 20 39\noutside 40 325\nTMhelix 326 348\ninside 349 372

6758 GCF_001420075.1_ASM142007v1 Escherichia coli Proteobacteria; Gammaproteobacteria; Enterobacterales; Enterobacteriaceae; Escherichia
MNNEETFYQAMRRQGVTRRSFLKYCSLAATSLGLGAGMAPKIAWALENKPRIPVVWIHGL WP_001058323.1
MULTISPECIES: hydrogenase-1 small chain [Proteobacteria] Length: 372\nNumber of predicted TMHs: 2\nExp number of AAs in TMHs: 37.2608499999999\nExp number, first 60 AAs: 19.52923\nTotal prob of N-in: 0.94351\nPOSSIBLE N-term signal sequence\ninside 1 19\nTMhelix 20 39\noutside 40 325\nTMhelix 326 348\ninside 349 372

6759 GCF_001440495.1_ASM144049v1 Escherichia coli Proteobacteria; Gammaproteobacteria; Enterobacterales; Enterobacteriaceae; Escherichia
MTGDNTLIHSHGINRRDFMKLCAALAATMGLSSKAAAEMAESVTNPQRPPVIWIGAQECT WP_000145410.1
MULTISPECIES: hydrogenase-2 small chain [Proteobacteria] Length: 372\nNumber of predicted TMHs: 1\nExp number of AAs in TMHs: 23.72226\nExp number, first 60 AAs: 0.22038\nTotal prob of N-in: 0.12994\noutside 1 331\nTMhelix 332 354\ninside 355 372

6760 GCF_001440575.1_ASM144057v1 Escherichia coli Proteobacteria; Gammaproteobacteria; Enterobacterales; Enterobacteriaceae; Escherichia
MTGDNTLIHSHGINRRDFMKLCAALAATMGLSSKAAAEMAESVTNPQRPPVIWIGAQECT WP_000145410.1
MULTISPECIES: hydrogenase-2 small chain [Proteobacteria] Length: 372\nNumber of predicted TMHs: 1\nExp number of AAs in TMHs: 23.72226\nExp number, first 60 AAs: 0.22038\nTotal prob of N-in: 0.12994\noutside 1 331\nTMhelix 332 354\ninside 355 372

6761 GCF_001440645.1_ASM144064v1 Escherichia coli Proteobacteria; Gammaproteobacteria; Enterobacterales; Enterobacteriaceae; Escherichia
MNNEETFYQAMRRQGVTRRSFLKSCSLAATSLGLGAGMAPKIAWALENKPRIPVVWIHGL WP_001058318.1
hydrogenase [Escherichia coli] Length: 372\nNumber of predicted TMHs: 1\nExp number of AAs in TMHs: 33.59637\nExp number, first 60 AAs: 15.81976\nTotal prob of N-in: 0.77394\nPOSSIBLE N-term signal sequence\noutside 1 325\nTMhelix 326 348\ninside 349 372

6762 GCF_001440615.1_ASM144061v1 Escherichia coli Proteobacteria; Gammaproteobacteria; Enterobacterales; Enterobacteriaceae; Escherichia
MNNEETFYQAMRRQGVTRRSFLKSCSLAATSLGLGAGMAPKIAWALENKPRIPVVWIHGL WP_001058318.1
hydrogenase [Escherichia coli] Length: 372\nNumber of predicted TMHs: 1\nExp number of AAs in TMHs: 33.59637\nExp number, first 60 AAs: 15.81976\nTotal prob of N-in: 0.77394\nPOSSIBLE N-term signal sequence\noutside 1 325\nTMhelix 326 348\ninside 349 372

6763 GCF_001440665.1_ASM144066v1 Escherichia coli Proteobacteria; Gammaproteobacteria; Enterobacterales; Enterobacteriaceae; Escherichia
MNNEETFYQAMRRQGVTRRSFLKSCSLAATSLGLGAGMAPKIAWALENKPRIPVVWIHGL WP_001058318.1
hydrogenase [Escherichia coli] Length: 372\nNumber of predicted TMHs: 1\nExp number of AAs in TMHs: 33.59637\nExp number, first 60 AAs: 15.81976\nTotal prob of N-in: 0.77394\nPOSSIBLE N-term signal sequence\noutside 1 325\nTMhelix 326 348\ninside 349 372

6764 GCF_001440735.1_ASM144073v1 Escherichia coli Proteobacteria; Gammaproteobacteria; Enterobacterales; Enterobacteriaceae; Escherichia
MNNEETFYQAMRRQGVTRRSFLKSCSLAATSLGLGAGMAPKIAWALENKPRIPVVWIHGL WP_001058318.1
hydrogenase [Escherichia coli] Length: 372\nNumber of predicted TMHs: 1\nExp number of AAs in TMHs: 33.59637\nExp number, first 60 AAs: 15.81976\nTotal prob of N-in: 0.77394\nPOSSIBLE N-term signal sequence\noutside 1 325\nTMhelix 326 348\ninside 349 372

6765 GCF_001442495.1_ASM144249v1 Escherichia coli Proteobacteria; Gammaproteobacteria; Enterobacterales; Enterobacteriaceae; Escherichia
MTGDNTLIHSHGINRRDFMKLCAALAATMGLSSKAAAEMAESVTNPQRPPVIWIGAQECT WP_000145410.1
MULTISPECIES: hydrogenase-2 small chain [Proteobacteria] Length: 372\nNumber of predicted TMHs: 1\nExp number of AAs in TMHs: 23.72226\nExp number, first 60 AAs: 0.22038\nTotal prob of N-in: 0.12994\noutside 1 331\nTMhelix 332 354\ninside 355 372

6766 GCF_001442495.1_ASM144249v1 Escherichia coli Proteobacteria; Gammaproteobacteria; Enterobacterales; Enterobacteriaceae; Escherichia
MNNEETFYQAMRRQGVTRRSFLKYCSLAATSLGLGAGMAPKIAWALENKPRIPVVWIHGL WP_001058320.1
hydrogenase [Escherichia coli] Length: 372\nNumber of predicted TMHs: 2\nExp number of AAs in TMHs: 37.26082\nExp number, first 60 AAs: 19.52923\nTotal prob of N-in: 0.94351\nPOSSIBLE N-term signal sequence\ninside 1 19\nTMhelix 20 39\noutside 40 325\nTMhelix 326 348\ninside 349 372

6767 GCF_001440495.1_ASM144049v1 Escherichia coli Proteobacteria; Gammaproteobacteria; Enterobacterales; Enterobacteriaceae; Escherichia
MNNEETFYQAMRRQGVTRRSFLKYCSLAATSLGLGAGMAPKIAWALENKPRIPVVWXHGL WP_057980725.1
hydrogenase [Escherichia coli] Length: 372\nNumber of predicted TMHs: 2\nExp number of AAs in TMHs: 37.2484799999999\nExp number, first 60 AAs: 19.52547\nTotal prob of N-in: 0.94349\nPOSSIBLE N-term signal sequence\ninside 1 19\nTMhelix 20 39\noutside 40 325\nTMhelix 326 348\ninside 349 372

6768 GCF_001463115.1_ASM146311v1 Escherichia coli Proteobacteria; Gammaproteobacteria; Enterobacterales; Enterobacteriaceae; Escherichia
MTGDNTLIHSHGINRRDFMKLCAALAATMGLSSKAAAEMAESVTNPQRPPVIWIGAQECT WP_000145410.1
MULTISPECIES: hydrogenase-2 small chain [Proteobacteria] Length: 372\nNumber of predicted TMHs: 1\nExp number of AAs in TMHs: 23.72226\nExp number, first 60 AAs: 0.22038\nTotal prob of N-in: 0.12994\noutside 1 331\nTMhelix 332 354\ninside 355 372

6769 GCF_001463205.1_ASM146320v1 Escherichia coli Proteobacteria; Gammaproteobacteria; Enterobacterales; Enterobacteriaceae; Escherichia
MTGDNTLIHSHGINRRDFMKLCAALAATMGLSSKAAAEMAESVTNPQRPPVIWIGAQECT WP_000145410.1
MULTISPECIES: hydrogenase-2 small chain [Proteobacteria] Length: 372\nNumber of predicted TMHs: 1\nExp number of AAs in TMHs: 23.72226\nExp number, first 60 AAs: 0.22038\nTotal prob of N-in: 0.12994\noutside 1 331\nTMhelix 332 354\ninside 355 372

6770 GCF_001455385.1_ASM145538v1 Escherichia coli Proteobacteria; Gammaproteobacteria; Enterobacterales; Enterobacteriaceae; Escherichia
MNNEETFYQAMRRQGVTRRSFLKYCSLAATSLGLGAGMAPKIAWALENKPRIPVVWIHGL WP_001058323.1
MULTISPECIES: hydrogenase-1 small chain [Proteobacteria] Length: 372\nNumber of predicted TMHs: 2\nExp number of AAs in TMHs: 37.2608499999999\nExp number, first 60 AAs: 19.52923\nTotal prob of N-in: 0.94351\nPOSSIBLE N-term signal sequence\ninside 1 19\nTMhelix 20 39\noutside 40 325\nTMhelix 326 348\ninside 349 372

6771 GCF_001463115.1_ASM146311v1 Escherichia coli Proteobacteria; Gammaproteobacteria; Enterobacterales; Enterobacteriaceae; Escherichia
MNNEETFYQAMRRQGVTRRSFLKYCSLAATSLGLGAGMAPKIAWALENKPRIPVVWIHGL WP_001058323.1
MULTISPECIES: hydrogenase-1 small chain [Proteobacteria] Length: 372\nNumber of predicted TMHs: 2\nExp number of AAs in TMHs: 37.2608499999999\nExp number, first 60 AAs: 19.52923\nTotal prob of N-in: 0.94351\nPOSSIBLE N-term signal sequence\ninside 1 19\nTMhelix 20 39\noutside 40 325\nTMhelix 326 348\ninside 349 372

6772 GCF_001463205.1_ASM146320v1 Escherichia coli Proteobacteria; Gammaproteobacteria; Enterobacterales; Enterobacteriaceae; Escherichia
MNNEETFYQAMRRQGVTRRSFLKYCSLAATSLGLGAGMAPKIAWALENKPRIPVVWIHGL WP_001058323.1

MULTISPECIES: hydrogenase-1 small chain [Proteobacteria] Length: 372\nNumber of predicted TMHs: 2\nExp number of AAs in TMHs: 37.2608499999999\nExp number, first 60 AAs: 19.52923\nTotal prob of N-in: 0.94351\nPOSSIBLE N-term signal sequence\ninside 1 19\nTMhelix 20 39\noutside 40 325\nTMhelix 326 348\ninside 349 372

6773 GCF_001448025.1_ASM144802v1 Escherichia coli Proteobacteria; Gammaproteobacteria; Enterobacterales; Enterobacteriaceae; Escherichia
MNNEETFYQAMRRQGVTRRSFLKYCSLAATSLGLGAGMAPKIAWALENKPRIPVVWIHGL WP_001058323.1
MULTISPECIES: hydrogenase-1 small chain [Proteobacteria] Length: 372\nNumber of predicted TMHs: 2\nExp number of AAs in TMHs: 37.2608499999999\nExp number, first 60 AAs: 19.52923\nTotal prob of N-in: 0.94351\nPOSSIBLE N-term signal sequence\ninside 1 19\nTMhelix 20 39\noutside 40 325\nTMhelix 326 348\ninside 349 372

6774 GCF_001462975.1_ASM146297v1 Escherichia coli Proteobacteria; Gammaproteobacteria; Enterobacterales; Enterobacteriaceae; Escherichia
MNNEETFYQAMRRQGVTRRSFLKYCSLAATSLGLGAGMAPKIAWALENKPRIPVVWIHGL WP_001058323.1
MULTISPECIES: hydrogenase-1 small chain [Proteobacteria] Length: 372\nNumber of predicted TMHs: 2\nExp number of AAs in TMHs: 37.2608499999999\nExp number, first 60 AAs: 19.52923\nTotal prob of N-in: 0.94351\nPOSSIBLE N-term signal sequence\ninside 1 19\nTMhelix 20 39\noutside 40 325\nTMhelix 326 348\ninside 349 372

6775 GCF_001463585.1_ASM146358v1 Escherichia coli Proteobacteria; Gammaproteobacteria; Enterobacterales; Enterobacteriaceae; Escherichia
MTGDNTLIHSHGINRRDFMKLCAALAATMGLSSKAAAEMAESVTNPQRPPVIWIGAQECT WP_000145410.1
MULTISPECIES: hydrogenase-2 small chain [Proteobacteria] Length: 372\nNumber of predicted TMHs: 1\nExp number of AAs in TMHs: 23.72226\nExp number, first 60 AAs: 0.22038\nTotal prob of N-in: 0.12994\noutside 1 331\nTMhelix 332 354\ninside 355 372

6776 GCF_001463455.1_ASM146345v1 Escherichia coli Proteobacteria; Gammaproteobacteria; Enterobacterales; Enterobacteriaceae; Escherichia
MTGDNTLIHSHGINRRDFMKLCAALAATMGLSSKAAAEMAESVTNPQRPPVIWIGAQECT WP_000145410.1
MULTISPECIES: hydrogenase-2 small chain [Proteobacteria] Length: 372\nNumber of predicted TMHs: 1\nExp number of AAs in TMHs: 23.72226\nExp number, first 60 AAs: 0.22038\nTotal prob of N-in: 0.12994\noutside 1 331\nTMhelix 332 354\ninside 355 372

6777 GCF_001463645.1_ASM146364v1 Escherichia coli Proteobacteria; Gammaproteobacteria; Enterobacterales; Enterobacteriaceae; Escherichia
MTGDNTLIHSHGINRRDFMKLCAALAATMGLSSKAAAEMAESVTNPQRPPVIWIGAQECT WP_000145410.1
MULTISPECIES: hydrogenase-2 small chain [Proteobacteria] Length: 372\nNumber of predicted TMHs: 1\nExp number of AAs in TMHs: 23.72226\nExp number, first 60 AAs: 0.22038\nTotal prob of N-in: 0.12994\noutside 1 331\nTMhelix 332 354\ninside 355 372

6778 GCF_001484085.1_ASM148408v1 Escherichia coli Proteobacteria; Gammaproteobacteria; Enterobacterales; Enterobacteriaceae; Escherichia
MTGDNTLIHSHGINRRDFMKLCAALAATMGLSSKAAAEMAESVTNPQRPPVIWIGAQECT WP_000145410.1
MULTISPECIES: hydrogenase-2 small chain [Proteobacteria] Length: 372\nNumber of predicted TMHs: 1\nExp number of AAs in TMHs: 23.72226\nExp number, first 60 AAs: 0.22038\nTotal prob of N-in: 0.12994\noutside 1 331\nTMhelix 332 354\ninside 355 372

6779 GCF_001469815.1_ASM146981v1 Escherichia coli Proteobacteria; Gammaproteobacteria; Enterobacterales; Enterobacteriaceae; Escherichia
MTGDNTLIHSHGINRRDFMKLCAALAATMGLSSKAAAEMVESVTNPQRPPVIWIGAQECT WP_000145423.1
hydrogenase 2 small subunit [Escherichia coli] Length: 372\nNumber of predicted TMHs: 1\nExp number of AAs in TMHs: 23.78201\nExp number, first 60 AAs: 0.28355\nTotal prob of N-in: 0.13257\noutside 1 331\nTMhelix 332 354\ninside 355 372

6780 GCF_001519245.1_ASM151924v1 Escherichia coli Proteobacteria; Gammaproteobacteria; Enterobacterales; Enterobacteriaceae; Escherichia
MTGDNTLIHSHGINRRDFMKLCAALAATMGLSSKAAAEMAESVTNPQRPPVIWIGAQECT WP_000145410.1
MULTISPECIES: hydrogenase-2 small chain [Proteobacteria] Length: 372\nNumber of predicted TMHs:

1\nExp number of AAs in TMHs: 23.72226\nExp number, first 60 AAs: 0.22038\nTotal prob of N-in: 0.12994\noutside 1 331\nTMhelix 332 354\ninside 355 372

6781 GCF_001463585.1_ASM146358v1 Escherichia coli Proteobacteria; Gammaproteobacteria; Enterobacterales; Enterobacteriaceae; Escherichia
MNNEETFYQAMRRQGVTRRSFLKYCSLAATSLGLGAGMAPKIAWALENKPRIPVVWIHGL WP_001058323.1
MULTISPECIES: hydrogenase-1 small chain [Proteobacteria] Length: 372\nNumber of predicted TMHs: 2\nExp number of AAs in TMHs: 37.2608499999999\nExp number, first 60 AAs: 19.52923\nTotal prob of N-in: 0.94351\nPOSSIBLE N-term signal sequence\ninside 1 19\nTMhelix 20 39\noutside 40 325\nTMhelix 326 348\ninside 349 372

6782 GCF_001463645.1_ASM146364v1 Escherichia coli Proteobacteria; Gammaproteobacteria; Enterobacterales; Enterobacteriaceae; Escherichia
MNNEETFYQAMRRQGVTRRSFLKYCSLAATSLGLGAGMAPKIAWALENKPRIPVVWIHGL WP_001058320.1
hydrogenase [Escherichia coli] Length: 372\nNumber of predicted TMHs: 2\nExp number of AAs in TMHs: 37.26082\nExp number, first 60 AAs: 19.52923\nTotal prob of N-in: 0.94351\nPOSSIBLE N-term signal sequence\ninside 1 19\nTMhelix 20 39\noutside 40 325\nTMhelix 326 348\ninside 349 372

6783 GCF_001484355.1_ASM148435v1 Escherichia coli Proteobacteria; Gammaproteobacteria; Enterobacterales; Enterobacteriaceae; Escherichia
MTGDNTLIHSHGINRRDFMKLCAALAATMGLSSKAAAEMAESVTNPQRPPVIWIGAQECT WP_000145410.1
MULTISPECIES: hydrogenase-2 small chain [Proteobacteria] Length: 372\nNumber of predicted TMHs: 1\nExp number of AAs in TMHs: 23.72226\nExp number, first 60 AAs: 0.22038\nTotal prob of N-in: 0.12994\noutside 1 331\nTMhelix 332 354\ninside 355 372

6784 GCF_001484355.1_ASM148435v1 Escherichia coli Proteobacteria; Gammaproteobacteria; Enterobacterales; Enterobacteriaceae; Escherichia
MNNEETFYQAMRRQGVTRRSFLKYCSLAATSLGLGAGMAPKIAWALENKPRIPVVWIHGL WP_001058323.1
MULTISPECIES: hydrogenase-1 small chain [Proteobacteria] Length: 372\nNumber of predicted TMHs: 2\nExp number of AAs in TMHs: 37.2608499999999\nExp number, first 60 AAs: 19.52923\nTotal prob of N-in: 0.94351\nPOSSIBLE N-term signal sequence\ninside 1 19\nTMhelix 20 39\noutside 40 325\nTMhelix 326 348\ninside 349 372

6785 GCF_001484085.1_ASM148408v1 Escherichia coli Proteobacteria; Gammaproteobacteria; Enterobacterales; Enterobacteriaceae; Escherichia
MNNEETFYQAMRRQGVTRRSFLKYCSLAATSLGLGAGMAPKIAWALENKPRIPVVWIHGL WP_001717971.1
hydrogenase-1 small chain [Escherichia coli] Length: 372\nNumber of predicted TMHs: 2\nExp number of AAs in TMHs: 38.00708\nExp number, first 60 AAs: 19.5286\nTotal prob of N-in: 0.94354\nPOSSIBLE N-term signal sequence\ninside 1 19\nTMhelix 20 39\noutside 40 325\nTMhelix 326 348\ninside 349 372

6786 GCF_001484405.1_ASM148440v1 Escherichia coli Proteobacteria; Gammaproteobacteria; Enterobacterales; Enterobacteriaceae; Escherichia
MTGDNTLIHSHGINRRDFMKLCAALAATMGLSSKAAAEMAESVTNPQRPPVIWIGAQECT WP_000145410.1
MULTISPECIES: hydrogenase-2 small chain [Proteobacteria] Length: 372\nNumber of predicted TMHs: 1\nExp number of AAs in TMHs: 23.72226\nExp number, first 60 AAs: 0.22038\nTotal prob of N-in: 0.12994\noutside 1 331\nTMhelix 332 354\ninside 355 372

6787 GCF_001484395.1_ASM148439v1 Escherichia coli Proteobacteria; Gammaproteobacteria; Enterobacterales; Enterobacteriaceae; Escherichia
MTGDNTLIHSHGINRRDFMKLCAALAATMGLSSKAAAEMAESVTNPQRPPVIWIGAQECT WP_000145410.1
MULTISPECIES: hydrogenase-2 small chain [Proteobacteria] Length: 372\nNumber of predicted TMHs: 1\nExp number of AAs in TMHs: 23.72226\nExp number, first 60 AAs: 0.22038\nTotal prob of N-in: 0.12994\noutside 1 331\nTMhelix 332 354\ninside 355 372

6788 GCF_001509675.1_ASM150967v1 Escherichia coli Proteobacteria; Gammaproteobacteria; Enterobacterales; Enterobacteriaceae; Escherichia
MTGDNTLIHSHGINRRDFMKLCAALAATMGLSSKAAAEMAESVTNPQRPPVIWIGAQECT WP_000145410.1
MULTISPECIES: hydrogenase-2 small chain [Proteobacteria] Length: 372\nNumber of predicted TMHs: 1\nExp number of AAs in TMHs: 23.72226\nExp number, first 60 AAs: 0.22038\nTotal prob of N-in: 0.12994\noutside 1 331\nTMhelix 332 354\ninside 355 372

6789 GCF_001509745.1_ASM150974v1 Escherichia coli Proteobacteria; Gammaproteobacteria; Enterobacterales; Enterobacteriaceae; Escherichia
MTGDNTLIHSHGINRRDFMKLCAALAATMGLSSKAAAEMAESVTNPQRPPVIWIGAQECT WP_000145410.1
MULTISPECIES: hydrogenase-2 small chain [Proteobacteria] Length: 372\nNumber of predicted TMHs: 1\nExp number of AAs in TMHs: 23.72226\nExp number, first 60 AAs: 0.22038\nTotal prob of N-in: 0.12994\noutside 1 331\nTMhelix 332 354\ninside 355 372

6790 GCF_001519215.1_ASM151921v1 Escherichia coli Proteobacteria; Gammaproteobacteria; Enterobacterales; Enterobacteriaceae; Escherichia
MTGDNTLIHSHGINRRDFMKLCAALAATMGLSSKAAAEMAESVTNPQRPPVIWIGAQECT WP_000145410.1
MULTISPECIES: hydrogenase-2 small chain [Proteobacteria] Length: 372\nNumber of predicted TMHs: 1\nExp number of AAs in TMHs: 23.72226\nExp number, first 60 AAs: 0.22038\nTotal prob of N-in: 0.12994\noutside 1 331\nTMhelix 332 354\ninside 355 372

6791 GCF_001484455.1_ASM148445v1 Escherichia coli Proteobacteria; Gammaproteobacteria; Enterobacterales; Enterobacteriaceae; Escherichia
MNNEETFYQAMRRQGVTRRSFLKYCSLAATSLGLGAGMAPKIAWALENKPRIPVVWIHGL WP_001058323.1
MULTISPECIES: hydrogenase-1 small chain [Proteobacteria] Length: 372\nNumber of predicted TMHs: 2\nExp number of AAs in TMHs: 37.2608499999999\nExp number, first 60 AAs: 19.52923\nTotal prob of N-in: 0.94351\nPOSSIBLE N-term signal sequence\ninside 1 19\nTMhelix 20 39\noutside 40 325\nTMhelix 326 348\ninside 349 372

6792 GCF_001484485.1_ASM148448v1 Escherichia coli Proteobacteria; Gammaproteobacteria; Enterobacterales; Enterobacteriaceae; Escherichia
MNNEETFYQAMRRQGVTRRSFLKYCSLAATSLGLGAGMAPKIAWALENKPRIPVVWIHGL WP_001058323.1
MULTISPECIES: hydrogenase-1 small chain [Proteobacteria] Length: 372\nNumber of predicted TMHs: 2\nExp number of AAs in TMHs: 37.2608499999999\nExp number, first 60 AAs: 19.52923\nTotal prob of N-in: 0.94351\nPOSSIBLE N-term signal sequence\ninside 1 19\nTMhelix 20 39\noutside 40 325\nTMhelix 326 348\ninside 349 372

6793 GCF_001513635.1_ASM151363v1 Escherichia coli Proteobacteria; Gammaproteobacteria; Enterobacterales; Enterobacteriaceae; Escherichia
MTGDNTLIHSHGINRRDFMKLCAALAATMGLSSKAAAEMVESVTNPQRPPVIWIGAQECT WP_000145423.1
hydrogenase 2 small subunit [Escherichia coli] Length: 372\nNumber of predicted TMHs: 1\nExp number of AAs in TMHs: 23.78201\nExp number, first 60 AAs: 0.28355\nTotal prob of N-in: 0.13257\noutside 1 331\nTMhelix 332 354\ninside 355 372

6794 GCF_001517685.1_NG-7574_Bell_ERRESI Escherichia coli Proteobacteria; Gammaproteobacteria; Enterobacterales; Enterobacteriaceae; Escherichia
MTGDNTLIHSHGINRRDFMKLCAALAATMGLSSKAAAEMAESVTNPQRPPVIWIGAQECT WP_000145410.1
MULTISPECIES: hydrogenase-2 small chain [Proteobacteria] Length: 372\nNumber of predicted TMHs: 1\nExp number of AAs in TMHs: 23.72226\nExp number, first 60 AAs: 0.22038\nTotal prob of N-in: 0.12994\noutside 1 331\nTMhelix 332 354\ninside 355 372

6795 GCF_001509745.1_ASM150974v1 Escherichia coli Proteobacteria; Gammaproteobacteria; Enterobacterales; Enterobacteriaceae; Escherichia
MNNEETFYQAMRRQGVTRRSFLKYCSLAATSLGLGAGMAPKIAWALENKPRIPVVWIHGL WP_001058323.1
MULTISPECIES: hydrogenase-1 small chain [Proteobacteria] Length: 372\nNumber of predicted TMHs: 2\nExp number of AAs in TMHs: 37.2608499999999\nExp number, first 60 AAs: 19.52923\nTotal prob of N-in: 0.94351\nPOSSIBLE N-term signal sequence\ninside 1 19\nTMhelix 20 39\noutside 40 325\nTMhelix 326 348\ninside 349 372

6796 GCF_001518365.1_ASM151836v1 Escherichia coli Proteobacteria; Gammaproteobacteria; Enterobacterales; Enterobacteriaceae; Escherichia
MTGDNTLIHSHGINRRDFMKLCAALAATMGLSSKAAAEMAESVTNPQRPPVIWIGAQECT WP_000145410.1
MULTISPECIES: hydrogenase-2 small chain [Proteobacteria] Length: 372\nNumber of predicted TMHs: 1\nExp number of AAs in TMHs: 23.72226\nExp number, first 60 AAs: 0.22038\nTotal prob of N-in: 0.12994\noutside 1 331\nTMhelix 332 354\ninside 355 372

6797 GCF_001519115.1_ASM151911v1 Escherichia coli Proteobacteria; Gammaproteobacteria; Enterobacterales; Enterobacteriaceae; Escherichia

MTGDNTLIHSHGINRRDFMKLCAALAATMGLSSKAAAEMAESVTNPQRPPVIWIGAQECT WP_000145410.1 MULTISPECIES: hydrogenase-2 small chain [Proteobacteria] Length: 372\nNumber of predicted TMHs: 1\nExp number of AAs in TMHs: 23.72226\nExp number, first 60 AAs: 0.22038\nTotal prob of N-in: 0.12994\noutside 1 331\nTMhelix 332 354\ninside 355 372

6798 GCF_001519135.1_ASM151913v1 Escherichia coli Proteobacteria; Gammaproteobacteria; Enterobacterales; Enterobacteriaceae; Escherichia MTGDNTLIHSHGINRRDFMKLCAALAATMGLSSKAAAEMAESVTNPQRPPVIWIGAQECT WP_000145410.1 MULTISPECIES: hydrogenase-2 small chain [Proteobacteria] Length: 372\nNumber of predicted TMHs: 1\nExp number of AAs in TMHs: 23.72226\nExp number, first 60 AAs: 0.22038\nTotal prob of N-in: 0.12994\noutside 1 331\nTMhelix 332 354\ninside 355 372

6799 GCF_001518365.1_ASM151836v1 Escherichia coli Proteobacteria; Gammaproteobacteria; Enterobacterales; Enterobacteriaceae; Escherichia MNNEETFYQAMRRQGVTRRSFLKYCSLAATSLGLGAGMAPKIAWALENKPRIPVVWIHGL WP_001058323.1 MULTISPECIES: hydrogenase-1 small chain [Proteobacteria] Length: 372\nNumber of predicted TMHs: 2\nExp number of AAs in TMHs: 37.2608499999999\nExp number, first 60 AAs: 19.52923\nTotal prob of N-in: 0.94351\nPOSSIBLE N-term signal sequence\ninside 1 19\nTMhelix 20 39\noutside 40 325\nTMhelix 326 348\ninside 349 372

6800 GCF_001519115.1_ASM151911v1 Escherichia coli Proteobacteria; Gammaproteobacteria; Enterobacterales; Enterobacteriaceae; Escherichia MNNEETFYQAMRRQGVTRRSFLKYCSLAATSLGLGAGMAPKIAWALENKPRIPVVWIHGL WP_001058323.1 MULTISPECIES: hydrogenase-1 small chain [Proteobacteria] Length: 372\nNumber of predicted TMHs: 2\nExp number of AAs in TMHs: 37.2608499999999\nExp number, first 60 AAs: 19.52923\nTotal prob of N-in: 0.94351\nPOSSIBLE N-term signal sequence\ninside 1 19\nTMhelix 20 39\noutside 40 325\nTMhelix 326 348\ninside 349 372

6801 GCF_001519195.1_ASM151919v1 Escherichia coli Proteobacteria; Gammaproteobacteria; Enterobacterales; Enterobacteriaceae; Escherichia MTGDNTLIHSHGINRRDFMKLCAALAATMGLSSKAAAEMAESVTNPQRPPVIWIGAQECT WP_000145410.1 MULTISPECIES: hydrogenase-2 small chain [Proteobacteria] Length: 372\nNumber of predicted TMHs: 1\nExp number of AAs in TMHs: 23.72226\nExp number, first 60 AAs: 0.22038\nTotal prob of N-in: 0.12994\noutside 1 331\nTMhelix 332 354\ninside 355 372

6802 GCF_001519395.1_ASM151939v1 Escherichia coli Proteobacteria; Gammaproteobacteria; Enterobacterales; Enterobacteriaceae; Escherichia MTGDNTLIHSHGINRRDFMKLCAALAATMGLSSKAAAEMAESVTNPQRPPVIWIGAQECT WP_000145410.1 MULTISPECIES: hydrogenase-2 small chain [Proteobacteria] Length: 372\nNumber of predicted TMHs: 1\nExp number of AAs in TMHs: 23.72226\nExp number, first 60 AAs: 0.22038\nTotal prob of N-in: 0.12994\noutside 1 331\nTMhelix 332 354\ninside 355 372

6803 GCF_001519405.1_ASM151940v1 Escherichia coli Proteobacteria; Gammaproteobacteria; Enterobacterales; Enterobacteriaceae; Escherichia MTGDNTLIHSHGINRRDFMKLCAALAATMGLSSKAAAEMAESVTNPQRPPVIWIGAQECT WP_000145410.1 MULTISPECIES: hydrogenase-2 small chain [Proteobacteria] Length: 372\nNumber of predicted TMHs: 1\nExp number of AAs in TMHs: 23.72226\nExp number, first 60 AAs: 0.22038\nTotal prob of N-in: 0.12994\noutside 1 331\nTMhelix 332 354\ninside 355 372

6804 GCF_001519395.1_ASM151939v1 Escherichia coli Proteobacteria; Gammaproteobacteria; Enterobacterales; Enterobacteriaceae; Escherichia MNNEETFYQAMRRQGVTRRSFLKYCSLAATSLGLGAGMAPKIAWALENKPRIPVVWIHGL WP_001058323.1 MULTISPECIES: hydrogenase-1 small chain [Proteobacteria] Length: 372\nNumber of predicted TMHs: 2\nExp number of AAs in TMHs: 37.2608499999999\nExp number, first 60 AAs: 19.52923\nTotal prob of N-in: 0.94351\nPOSSIBLE N-term signal sequence\ninside 1 19\nTMhelix 20 39\noutside 40 325\nTMhelix 326 348\ninside 349 372

6805 GCF_001519405.1_ASM151940v1 Escherichia coli Proteobacteria; Gammaproteobacteria; Enterobacterales; Enterobacteriaceae; Escherichia MNNEETFYQAMRRQGVTRRSFLKYCSLAATSLGLGAGMAPKIAWALENKPRIPVVWIHGL WP_001295940.1 MULTISPECIES: hydrogenase [Enterobacteriaceae] Length: 372\nNumber of predicted TMHs: 2\nExp

number of AAs in TMHs: 37.2608899999999\nExp number, first 60 AAs: 19.52923\nTotal prob of N-in: 0.94351\nPOSSIBLE N-term signal sequence\ninside 1 19\nTMhelix 20 39\noutside 40 325\nTMhelix 326 348\ninside 349 372

6806 GCF_001519215.1_ASM151921v1 Escherichia coli Proteobacteria; Gammaproteobacteria; Enterobacterales; Enterobacteriaceae; Escherichia MNNEETFYQAMRRQGVTRRSFLKYCSLAATSLGLGAGMAPKIAWALENKPRIPVVWIHGL WP_001058323.1 MULTISPECIES: hydrogenase-1 small chain [Proteobacteria] Length: 372\nNumber of predicted TMHs: 2\nExp number of AAs in TMHs: 37.2608499999999\nExp number, first 60 AAs: 19.52923\nTotal prob of N-in: 0.94351\nPOSSIBLE N-term signal sequence\ninside 1 19\nTMhelix 20 39\noutside 40 325\nTMhelix 326 348\ninside 349 372

6807 GCF_001519445.1_ASM151944v1 Escherichia coli Proteobacteria; Gammaproteobacteria; Enterobacterales; Enterobacteriaceae; Escherichia MTGDNTLIHSHGINRRDFMKLCAALAATMGLSSKAAAEMAESVTNPQRPPVIWIGAQECT WP_000145410.1 MULTISPECIES: hydrogenase-2 small chain [Proteobacteria] Length: 372\nNumber of predicted TMHs: 1\nExp number of AAs in TMHs: 23.72226\nExp number, first 60 AAs: 0.22038\nTotal prob of N-in: 0.12994\noutside 1 331\nTMhelix 332 354\ninside 355 372

6808 GCF_001519485.1_ASM151948v1 Escherichia coli Proteobacteria; Gammaproteobacteria; Enterobacterales; Enterobacteriaceae; Escherichia MTGDNTLIHSHGINRRDFMKLCAALAATMGLSSKAAAEMAESVTNPQRPPVIWIGAQECT WP_000145410.1 MULTISPECIES: hydrogenase-2 small chain [Proteobacteria] Length: 372\nNumber of predicted TMHs: 1\nExp number of AAs in TMHs: 23.72226\nExp number, first 60 AAs: 0.22038\nTotal prob of N-in: 0.12994\noutside 1 331\nTMhelix 332 354\ninside 355 372

6809 GCF_001519615.1_ASM151961v1 Escherichia coli Proteobacteria; Gammaproteobacteria; Enterobacterales; Enterobacteriaceae; Escherichia MTGDNTLIHSHGINRRDFMKLCAALAATMGLSSKAAAEMAESVTNPQRPPVIWIGAQECT WP_000145410.1 MULTISPECIES: hydrogenase-2 small chain [Proteobacteria] Length: 372\nNumber of predicted TMHs: 1\nExp number of AAs in TMHs: 23.72226\nExp number, first 60 AAs: 0.22038\nTotal prob of N-in: 0.12994\noutside 1 331\nTMhelix 332 354\ninside 355 372

6810 GCF_001519645.1_ASM151964v1 Escherichia coli Proteobacteria; Gammaproteobacteria; Enterobacterales; Enterobacteriaceae; Escherichia MTGDNTLIHSHGINRRDFMKLCAALAATMGLSSKAAAEMAESVTNPQRPPVIWIGAQECT WP_000145410.1 MULTISPECIES: hydrogenase-2 small chain [Proteobacteria] Length: 372\nNumber of predicted TMHs: 1\nExp number of AAs in TMHs: 23.72226\nExp number, first 60 AAs: 0.22038\nTotal prob of N-in: 0.12994\noutside 1 331\nTMhelix 332 354\ninside 355 372

6811 GCF_001519685.1_ASM151968v1 Escherichia coli Proteobacteria; Gammaproteobacteria; Enterobacterales; Enterobacteriaceae; Escherichia MTGDNTLIHSHGINRRDFMKLCAALAATMGLSSKAAAEMAESVTNPQRPPVIWIGAQECT WP_000145410.1 MULTISPECIES: hydrogenase-2 small chain [Proteobacteria] Length: 372\nNumber of predicted TMHs: 1\nExp number of AAs in TMHs: 23.72226\nExp number, first 60 AAs: 0.22038\nTotal prob of N-in: 0.12994\noutside 1 331\nTMhelix 332 354\ninside 355 372

6812 GCF_001519735.1_ASM151973v1 Escherichia coli Proteobacteria; Gammaproteobacteria; Enterobacterales; Enterobacteriaceae; Escherichia MTGDNTLIHSHGINRRDFMKLCAALAATMGLSSKAAAEMAESVTNPQRPPVIWIGAQECT WP_000145410.1 MULTISPECIES: hydrogenase-2 small chain [Proteobacteria] Length: 372\nNumber of predicted TMHs: 1\nExp number of AAs in TMHs: 23.72226\nExp number, first 60 AAs: 0.22038\nTotal prob of N-in: 0.12994\noutside 1 331\nTMhelix 332 354\ninside 355 372

6813 GCF_001519615.1_ASM151961v1 Escherichia coli Proteobacteria; Gammaproteobacteria; Enterobacterales; Enterobacteriaceae; Escherichia MNNEETFYQAMRRQGVTRRSFLKYCSLAATSLGLGAGMAPKIAWALENKPRIPVVWIHGL WP_001058323.1 MULTISPECIES: hydrogenase-1 small chain [Proteobacteria] Length: 372\nNumber of predicted TMHs: 2\nExp number of AAs in TMHs: 37.2608499999999\nExp number, first 60 AAs: 19.52923\nTotal prob of N-in: 0.94351\nPOSSIBLE N-term signal sequence\ninside 1 19\nTMhelix 20 39\noutside 40 325\nTMhelix 326 348\ninside 349 372

6814 GCF_001519595.1_ASM151959v1 Escherichia coli Proteobacteria; Gammaproteobacteria; Enterobacterales; Enterobacteriaceae; Escherichia
MNNEETFYQAMRRQGVTRRSFLKYCSLAATSLGLGAGMAPKIAWALENKPRIPVVWIHGL WP_001058323.1
MULTISPECIES: hydrogenase-1 small chain [Proteobacteria] Length: 372\nNumber of predicted TMHs: 2\nExp number of AAs in TMHs: 37.2608499999999\nExp number, first 60 AAs: 19.52923\nTotal prob of N-in: 0.94351\nPOSSIBLE N-term signal sequence\ninside 1 19\nTMhelix 20 39\noutside 40 325\nTMhelix 326 348\ninside 349 372

6815 GCF_001519755.1_ASM151975v1 Escherichia coli Proteobacteria; Gammaproteobacteria; Enterobacterales; Enterobacteriaceae; Escherichia
MTGDNTLIHSHGINRRDFMKLCAALAATMGLSSKAAAEMAESVTNPQRPPVIWIGAQECT WP_000145410.1
MULTISPECIES: hydrogenase-2 small chain [Proteobacteria] Length: 372\nNumber of predicted TMHs: 1\nExp number of AAs in TMHs: 23.72226\nExp number, first 60 AAs: 0.22038\nTotal prob of N-in: 0.12994\noutside 1 331\nTMhelix 332 354\ninside 355 372

6816 GCF_001519685.1_ASM151968v1 Escherichia coli Proteobacteria; Gammaproteobacteria; Enterobacterales; Enterobacteriaceae; Escherichia
MNNEETFYQAMRRQGVTRRSFLKYCSLAATSLGLGAGMAPKIAWALENKPRIPVVWIHGL WP_001058323.1
MULTISPECIES: hydrogenase-1 small chain [Proteobacteria] Length: 372\nNumber of predicted TMHs: 2\nExp number of AAs in TMHs: 37.2608499999999\nExp number, first 60 AAs: 19.52923\nTotal prob of N-in: 0.94351\nPOSSIBLE N-term signal sequence\ninside 1 19\nTMhelix 20 39\noutside 40 325\nTMhelix 326 348\ninside 349 372

6817 GCF_001519895.1_ASM151989v1 Escherichia coli Proteobacteria; Gammaproteobacteria; Enterobacterales; Enterobacteriaceae; Escherichia
MTGDNTLIHSHGINRRDFMKLCAALAATMGLSSKAAAEMAESVTNPQRPPVIWIGAQECT WP_000145410.1
MULTISPECIES: hydrogenase-2 small chain [Proteobacteria] Length: 372\nNumber of predicted TMHs: 1\nExp number of AAs in TMHs: 23.72226\nExp number, first 60 AAs: 0.22038\nTotal prob of N-in: 0.12994\noutside 1 331\nTMhelix 332 354\ninside 355 372

6818 GCF_001519935.1_ASM151993v1 Escherichia coli Proteobacteria; Gammaproteobacteria; Enterobacterales; Enterobacteriaceae; Escherichia
MTGDNTLIHSHGINRRDFMKLCAALAATMGLSSKAAAEMVESVTNPQRPPVIWIGAQECT WP_000145423.1
hydrogenase 2 small subunit [Escherichia coli] Length: 372\nNumber of predicted TMHs: 1\nExp number of AAs in TMHs: 23.78201\nExp number, first 60 AAs: 0.28355\nTotal prob of N-in: 0.13257\noutside 1 331\nTMhelix 332 354\ninside 355 372

6819 GCF_001519945.1_ASM151994v1 Escherichia coli Proteobacteria; Gammaproteobacteria; Enterobacterales; Enterobacteriaceae; Escherichia
MTGDNTLIHSHGINRRDFMKLCAALAATMGLSSKAAAEMAESVTNPQRPPVIWIGAQECT WP_000145410.1
MULTISPECIES: hydrogenase-2 small chain [Proteobacteria] Length: 372\nNumber of predicted TMHs: 1\nExp number of AAs in TMHs: 23.72226\nExp number, first 60 AAs: 0.22038\nTotal prob of N-in: 0.12994\noutside 1 331\nTMhelix 332 354\ninside 355 372

6820 GCF_001520055.1_ASM152005v1 Escherichia coli Proteobacteria; Gammaproteobacteria; Enterobacterales; Enterobacteriaceae; Escherichia
MTGDNTLIHSHGINRRDFMKLCAALAATMGLSSKAAAEMAESVTNPQRPPVIWIGAQECT WP_000145410.1
MULTISPECIES: hydrogenase-2 small chain [Proteobacteria] Length: 372\nNumber of predicted TMHs: 1\nExp number of AAs in TMHs: 23.72226\nExp number, first 60 AAs: 0.22038\nTotal prob of N-in: 0.12994\noutside 1 331\nTMhelix 332 354\ninside 355 372

6821 GCF_001519995.1_ASM151999v1 Escherichia coli Proteobacteria; Gammaproteobacteria; Enterobacterales; Enterobacteriaceae; Escherichia
MTGDNTLIHSHGINRRDFMKLCAALAATMGLSSKAAAEMVESVTNPQRPPVIWIGAQECT WP_000145423.1
hydrogenase 2 small subunit [Escherichia coli] Length: 372\nNumber of predicted TMHs: 1\nExp number of AAs in TMHs: 23.78201\nExp number, first 60 AAs: 0.28355\nTotal prob of N-in: 0.13257\noutside 1 331\nTMhelix 332 354\ninside 355 372

6822 GCF_001520035.1_ASM152003v1 Escherichia coli Proteobacteria; Gammaproteobacteria; Enterobacterales; Enterobacteriaceae; Escherichia
MTGDNTLIHSHGINRRDFMKLCAALAATMGLSSKAAAEMVESVTNPQRPPVIWIGAQECT WP_000145423.1

hydrogenase 2 small subunit [Escherichia coli] Length: 372\nNumber of predicted TMHs: 1\nExp number of AAs in TMHs: 23.78201\nExp number, first 60 AAs: 0.28355\nTotal prob of N-in: 0.13257\noutside 1 331\nTMhelix 332 354\ninside 355 372

6823 GCF_001520595.1_ASM152059v1 Escherichia coli Proteobacteria; Gammaproteobacteria; Enterobacterales; Enterobacteriaceae; Escherichia MTGDNTLIHSHGINRRDFMKLCAALAATMGLSSKAAAEMAESVTNPQRPPVIWIGAQECT WP_000145410.1 MULTISPECIES: hydrogenase-2 small chain [Proteobacteria] Length: 372\nNumber of predicted TMHs: 1\nExp number of AAs in TMHs: 23.72226\nExp number, first 60 AAs: 0.22038\nTotal prob of N-in: 0.12994\noutside 1 331\nTMhelix 332 354\ninside 355 372

6824 GCF_001520145.1_ASM152014v1 Escherichia coli Proteobacteria; Gammaproteobacteria; Enterobacterales; Enterobacteriaceae; Escherichia MTGDNTLIHSHGINRRDFMKLCAALAATMGLSSKAAAEMAESVTNPQRPPVIWIGAQECT WP_000145410.1 MULTISPECIES: hydrogenase-2 small chain [Proteobacteria] Length: 372\nNumber of predicted TMHs: 1\nExp number of AAs in TMHs: 23.72226\nExp number, first 60 AAs: 0.22038\nTotal prob of N-in: 0.12994\noutside 1 331\nTMhelix 332 354\ninside 355 372

6825 GCF_001520195.1_ASM152019v1 Escherichia coli Proteobacteria; Gammaproteobacteria; Enterobacterales; Enterobacteriaceae; Escherichia MTGDNTLIHSHGINRRDFMKLCAALAATMGLSSKAAAEMAESVTNPQRPPVIWIGAQECT WP_000145410.1 MULTISPECIES: hydrogenase-2 small chain [Proteobacteria] Length: 372\nNumber of predicted TMHs: 1\nExp number of AAs in TMHs: 23.72226\nExp number, first 60 AAs: 0.22038\nTotal prob of N-in: 0.12994\noutside 1 331\nTMhelix 332 354\ninside 355 372

6826 GCF_001520235.1_ASM152023v1 Escherichia coli Proteobacteria; Gammaproteobacteria; Enterobacterales; Enterobacteriaceae; Escherichia MTGDNTLIHSHGINRRDFMKLCAALAATMGLSSKAAAEMAESVTNPQRPPVIWIGAQECT WP_000145410.1 MULTISPECIES: hydrogenase-2 small chain [Proteobacteria] Length: 372\nNumber of predicted TMHs: 1\nExp number of AAs in TMHs: 23.72226\nExp number, first 60 AAs: 0.22038\nTotal prob of N-in: 0.12994\noutside 1 331\nTMhelix 332 354\ninside 355 372

6827 GCF_001520255.1_ASM152025v1 Escherichia coli Proteobacteria; Gammaproteobacteria; Enterobacterales; Enterobacteriaceae; Escherichia MTGDNTLIHSHGINRRDFMKLCAALAATMGLSSKAAAEMAESVTNPQRPPVIWIGAQECT WP_000145410.1 MULTISPECIES: hydrogenase-2 small chain [Proteobacteria] Length: 372\nNumber of predicted TMHs: 1\nExp number of AAs in TMHs: 23.72226\nExp number, first 60 AAs: 0.22038\nTotal prob of N-in: 0.12994\noutside 1 331\nTMhelix 332 354\ninside 355 372

6828 GCF_001520335.1_ASM152033v1 Escherichia coli Proteobacteria; Gammaproteobacteria; Enterobacterales; Enterobacteriaceae; Escherichia MTGDNTLIHSHGINRRDFMKLCAALAATMGLSSKAAAEMAESVTNPQRPPVIWIGAQECT WP_000145410.1 MULTISPECIES: hydrogenase-2 small chain [Proteobacteria] Length: 372\nNumber of predicted TMHs: 1\nExp number of AAs in TMHs: 23.72226\nExp number, first 60 AAs: 0.22038\nTotal prob of N-in: 0.12994\noutside 1 331\nTMhelix 332 354\ninside 355 372

6829 GCF_001520235.1_ASM152023v1 Escherichia coli Proteobacteria; Gammaproteobacteria; Enterobacterales; Enterobacteriaceae; Escherichia MNNEETFYQAMRRQGVTRRSFLKYCSLAATSLGLGAGMAPKIAWALENKPRIPVVWIHGL WP_001058323.1 MULTISPECIES: hydrogenase-1 small chain [Proteobacteria] Length: 372\nNumber of predicted TMHs: 2\nExp number of AAs in TMHs: 37.2608499999999\nExp number, first 60 AAs: 19.52923\nTotal prob of N-in: 0.94351\nPOSSIBLE N-term signal sequence\ninside 1 19\nTMhelix 20 39\noutside 40 325\nTMhelix 326 348\ninside 349 372

6830 GCF_001520145.1_ASM152014v1 Escherichia coli Proteobacteria; Gammaproteobacteria; Enterobacterales; Enterobacteriaceae; Escherichia MNNEETFYQAMRRQGVTRRSFLKYCSLAATSLGLGAGMAPKIAWALENKPRIPVVWIHGL WP_001058323.1 MULTISPECIES: hydrogenase-1 small chain [Proteobacteria] Length: 372\nNumber of predicted TMHs: 2\nExp number of AAs in TMHs: 37.2608499999999\nExp number, first 60 AAs: 19.52923\nTotal prob of N-in: 0.94351\nPOSSIBLE N-term signal sequence\ninside 1 19\nTMhelix 20 39\noutside 40 325\nTMhelix 326 348\ninside 349 372

6831 GCF_001520355.1_ASM152035v1 Escherichia coli Proteobacteria; Gammaproteobacteria; Enterobacterales; Enterobacteriaceae; Escherichia
MTGDNTLIHSHGINRRDFMKLCAALAATMGLSSKAAAEMVESVTNPQRPPVIWIGAQECT WP_000145423.1
hydrogenase 2 small subunit [Escherichia coli] Length: 372\nNumber of predicted TMHs: 1\nExp number of AAs in TMHs: 23.78201\nExp number, first 60 AAs: 0.28355\nTotal prob of N-in: 0.13257\noutside 1 331\nTMhelix 332 354\ninside 355 372

6832 GCF_001520395.1_ASM152039v1 Escherichia coli Proteobacteria; Gammaproteobacteria; Enterobacterales; Enterobacteriaceae; Escherichia
MTGDNTLIHSHGINRRDFMKLCAALAATMGLSSKAAAEMAESVTNPQRPPVIWIGAQECT WP_000145410.1
MULTISPECIES: hydrogenase-2 small chain [Proteobacteria] Length: 372\nNumber of predicted TMHs: 1\nExp number of AAs in TMHs: 23.72226\nExp number, first 60 AAs: 0.22038\nTotal prob of N-in: 0.12994\noutside 1 331\nTMhelix 332 354\ninside 355 372

6833 GCF_001520455.1_ASM152045v1 Escherichia coli Proteobacteria; Gammaproteobacteria; Enterobacterales; Enterobacteriaceae; Escherichia
MTGDNTLIHSHGINRRDFMKLCAALAATMGLSSKAAAEMAESVTNPQRPPVIWIGAQECT WP_000145410.1
MULTISPECIES: hydrogenase-2 small chain [Proteobacteria] Length: 372\nNumber of predicted TMHs: 1\nExp number of AAs in TMHs: 23.72226\nExp number, first 60 AAs: 0.22038\nTotal prob of N-in: 0.12994\noutside 1 331\nTMhelix 332 354\ninside 355 372

6834 GCF_001520495.1_ASM152049v1 Escherichia coli Proteobacteria; Gammaproteobacteria; Enterobacterales; Enterobacteriaceae; Escherichia
MTGDNTLIHSHGINRRDFMKLCAALAATMGLSSKAAAEMAESVTNPQRPPVIWIGAQECT WP_000145410.1
MULTISPECIES: hydrogenase-2 small chain [Proteobacteria] Length: 372\nNumber of predicted TMHs: 1\nExp number of AAs in TMHs: 23.72226\nExp number, first 60 AAs: 0.22038\nTotal prob of N-in: 0.12994\noutside 1 331\nTMhelix 332 354\ninside 355 372

6835 GCF_001520535.1_ASM152053v1 Escherichia coli Proteobacteria; Gammaproteobacteria; Enterobacterales; Enterobacteriaceae; Escherichia
MTGDNTLIHSHGINRRDFMKLCAALAATMGLSSKAAAEMVESVTNPQRPPVIWIGAQECT WP_000145423.1
hydrogenase 2 small subunit [Escherichia coli] Length: 372\nNumber of predicted TMHs: 1\nExp number of AAs in TMHs: 23.78201\nExp number, first 60 AAs: 0.28355\nTotal prob of N-in: 0.13257\noutside 1 331\nTMhelix 332 354\ninside 355 372

6836 GCF_001520555.1_ASM152055v1 Escherichia coli Proteobacteria; Gammaproteobacteria; Enterobacterales; Enterobacteriaceae; Escherichia
MTGDNTLIHSHGINRRDFMKLCAALAATMGLSSKAAAEMAESVTNPQRPPVIWIGAQECT WP_000145410.1
MULTISPECIES: hydrogenase-2 small chain [Proteobacteria] Length: 372\nNumber of predicted TMHs: 1\nExp number of AAs in TMHs: 23.72226\nExp number, first 60 AAs: 0.22038\nTotal prob of N-in: 0.12994\noutside 1 331\nTMhelix 332 354\ninside 355 372

6837 GCF_001520635.1_ASM152063v1 Escherichia coli Proteobacteria; Gammaproteobacteria; Enterobacterales; Enterobacteriaceae; Escherichia
MTGDNTLIHSHGINRRDFMKLCAALAATMGLSSKAAAEMVESVTNPQRPPVIWIGAQECT WP_000145423.1
hydrogenase 2 small subunit [Escherichia coli] Length: 372\nNumber of predicted TMHs: 1\nExp number of AAs in TMHs: 23.78201\nExp number, first 60 AAs: 0.28355\nTotal prob of N-in: 0.13257\noutside 1 331\nTMhelix 332 354\ninside 355 372

6838 GCF_001520635.1_ASM152063v1 Escherichia coli Proteobacteria; Gammaproteobacteria; Enterobacterales; Enterobacteriaceae; Escherichia
MNNEETFYQAMRRQGVTRRSFLKYCSLAATSLGLGAGMAPKIAWALENKPRIPVVWIHGL WP_001295940.1
MULTISPECIES: hydrogenase [Enterobacteriaceae] Length: 372\nNumber of predicted TMHs: 2\nExp number of AAs in TMHs: 37.2608899999999\nExp number, first 60 AAs: 19.52923\nTotal prob of N-in: 0.94351\nPOSSIBLE N-term signal sequence\ninside 1 19\nTMhelix 20 39\noutside 40 325\nTMhelix 326 348\ninside 349 372

6839 GCF_001520735.1_ASM152073v1 Escherichia coli Proteobacteria; Gammaproteobacteria; Enterobacterales; Enterobacteriaceae; Escherichia
MTGDNTLIHSHGINRRDFMKLCAALAATMGLSSKAAAEMAESVTNPQRPPVIWIGAQECT WP_000145418.1
hydrogenase 2 small subunit [Escherichia coli] Length: 372\nNumber of predicted TMHs: 1\nExp number of

AAs in TMHs: 23.72313\nExp number, first 60 AAs:  0.22034\nTotal prob of N-in:   0.12997\noutside   1   331\nTMhelix 332   354\ninside   355   372

6840 GCF_001520735.1_ASM152073v1 Escherichia coli Proteobacteria; Gammaproteobacteria; Enterobacterales; Enterobacteriaceae; Escherichia
MNNEETFYQAMRRQGVTRRSFLKYCSLAATSLGLGAGMAPKIAWALENKPRIPVVWIHGL WP_001058323.1
MULTISPECIES: hydrogenase-1 small chain [Proteobacteria] Length: 372\nNumber of predicted TMHs: 2\nExp number of AAs in TMHs: 37.2608499999999\nExp number, first 60 AAs:  19.52923\nTotal prob of N-in: 0.94351\nPOSSIBLE N-term signal sequence\ninside   1   19\nTMhelix   20   39\noutside   40   325\nTMhelix   326 348\ninside   349   372

6841 GCF_001520695.1_ASM152069v1 Escherichia coli Proteobacteria; Gammaproteobacteria; Enterobacterales; Enterobacteriaceae; Escherichia
MNNEETFYQAMRRQGVTRRSFLKYCSLAATSLGLGAGMAPKIAWALENKPRIPVVWIHGL WP_001295940.1
MULTISPECIES: hydrogenase [Enterobacteriaceae] Length: 372\nNumber of predicted TMHs:  2\nExp number of AAs in TMHs: 37.2608899999999\nExp number, first 60 AAs:  19.52923\nTotal prob of N-in: 0.94351\nPOSSIBLE N-term signal sequence\ninside   1   19\nTMhelix   20   39\noutside   40   325\nTMhelix   326 348\ninside   349   372

6842 GCF_001520555.1_ASM152055v1 Escherichia coli Proteobacteria; Gammaproteobacteria; Enterobacterales; Enterobacteriaceae; Escherichia
MNNEETFYQAMRRQGVTRRSFLKYCSLAATSLGLGAGMAPKIAWALENKPRIPVVWIHGL WP_001058323.1
MULTISPECIES: hydrogenase-1 small chain [Proteobacteria] Length: 372\nNumber of predicted TMHs: 2\nExp number of AAs in TMHs: 37.2608499999999\nExp number, first 60 AAs:  19.52923\nTotal prob of N-in: 0.94351\nPOSSIBLE N-term signal sequence\ninside   1   19\nTMhelix   20   39\noutside   40   325\nTMhelix   326 348\ninside   349   372

6843 GCF_001520535.1_ASM152053v1 Escherichia coli Proteobacteria; Gammaproteobacteria; Enterobacterales; Enterobacteriaceae; Escherichia
MNNEETFYQAMRRQGVTRRSFLKYCSLAATSLGLGAGMAPKIAWALENKPRIPVVWIHGL WP_001295940.1
MULTISPECIES: hydrogenase [Enterobacteriaceae] Length: 372\nNumber of predicted TMHs:  2\nExp number of AAs in TMHs: 37.2608899999999\nExp number, first 60 AAs:  19.52923\nTotal prob of N-in: 0.94351\nPOSSIBLE N-term signal sequence\ninside   1   19\nTMhelix   20   39\noutside   40   325\nTMhelix   326 348\ninside   349   372

6844 GCF_001520755.1_ASM152075v1 Escherichia coli Proteobacteria; Gammaproteobacteria; Enterobacterales; Enterobacteriaceae; Escherichia
MTGDNTLIHSHGINRRDFMKLCAALAATMGLSSKAAAEMVESVTNPQRPPVIWIGAQECT WP_000145423.1
hydrogenase 2 small subunit [Escherichia coli] Length: 372\nNumber of predicted TMHs:  1\nExp number of AAs in TMHs: 23.78201\nExp number, first 60 AAs:  0.28355\nTotal prob of N-in:   0.13257\noutside   1   331\nTMhelix 332   354\ninside   355   372

6845 GCF_001520795.1_ASM152079v1 Escherichia coli Proteobacteria; Gammaproteobacteria; Enterobacterales; Enterobacteriaceae; Escherichia
MTGDNTLIHSHGINRRDFMKLCAALAATMGLSSKAAAEMVESVTNPQRPPVIWIGAQECT WP_000145423.1
hydrogenase 2 small subunit [Escherichia coli] Length: 372\nNumber of predicted TMHs:  1\nExp number of AAs in TMHs: 23.78201\nExp number, first 60 AAs:  0.28355\nTotal prob of N-in:   0.13257\noutside   1   331\nTMhelix 332   354\ninside   355   372

6846 GCF_001520955.1_ASM152095v1 Escherichia coli Proteobacteria; Gammaproteobacteria; Enterobacterales; Enterobacteriaceae; Escherichia
MTGDNTLIHSHGINRRDFMKLCAALAATMGLSSKAAAEMAESVTNPQRPPVIWIGAQECT WP_000145410.1
MULTISPECIES: hydrogenase-2 small chain [Proteobacteria] Length: 372\nNumber of predicted TMHs: 1\nExp number of AAs in TMHs: 23.72226\nExp number, first 60 AAs:  0.22038\nTotal prob of N-in:   0.12994\noutside   1 331\nTMhelix   332   354\ninside   355   372

6847 GCF_001520995.1_ASM152099v1 Escherichia coli Proteobacteria; Gammaproteobacteria; Enterobacterales; Enterobacteriaceae; Escherichia
MTGDNTLIHSHGINRRDFMKLCAALAATMGLSSKAAAEMAESVTNPQRPPVIWIGAQECT WP_000145410.1
MULTISPECIES: hydrogenase-2 small chain [Proteobacteria] Length: 372\nNumber of predicted TMHs:

1\nExp number of AAs in TMHs: 23.72226\nExp number, first 60 AAs: 0.22038\nTotal prob of N-in: 0.12994\noutside 1 331\nTMhelix 332 354\ninside 355 372

6848 GCF_001521025.1_ASM152102v1 Escherichia coli Proteobacteria; Gammaproteobacteria; Enterobacterales; Enterobacteriaceae; Escherichia
MTGDNTLIHSHGINRRDFMKLCAALAATMGLSSKAAAEMAESVTNPQRPPVIWIGAQECT WP_000145410.1
MULTISPECIES: hydrogenase-2 small chain [Proteobacteria] Length: 372\nNumber of predicted TMHs: 1\nExp number of AAs in TMHs: 23.72226\nExp number, first 60 AAs: 0.22038\nTotal prob of N-in: 0.12994\noutside 1 331\nTMhelix 332 354\ninside 355 372

6849 GCF_001520895.1_ASM152089v1 Escherichia coli Proteobacteria; Gammaproteobacteria; Enterobacterales; Enterobacteriaceae; Escherichia
MNNEETFYQAMRRQGVTRRSFLKYCSLAATSLGLGAGMAPKIAWALENKPRIPVVWIHGL WP_001058323.1
MULTISPECIES: hydrogenase-1 small chain [Proteobacteria] Length: 372\nNumber of predicted TMHs: 2\nExp number of AAs in TMHs: 37.2608499999999\nExp number, first 60 AAs: 19.52923\nTotal prob of N-in: 0.94351\nPOSSIBLE N-term signal sequence\ninside 1 19\nTMhelix 20 39\noutside 40 325\nTMhelix 326 348\ninside 349 372

6850 GCF_001520935.1_ASM152093v1 Escherichia coli Proteobacteria; Gammaproteobacteria; Enterobacterales; Enterobacteriaceae; Escherichia
MNNEETFYQAMRRQGVTRRSFLKYCSLAATSLGLGAGMAPKIAWALENKPRIPVVWIHGL WP_001058323.1
MULTISPECIES: hydrogenase-1 small chain [Proteobacteria] Length: 372\nNumber of predicted TMHs: 2\nExp number of AAs in TMHs: 37.2608499999999\nExp number, first 60 AAs: 19.52923\nTotal prob of N-in: 0.94351\nPOSSIBLE N-term signal sequence\ninside 1 19\nTMhelix 20 39\noutside 40 325\nTMhelix 326 348\ninside 349 372

6851 GCF_001520755.1_ASM152075v1 Escherichia coli Proteobacteria; Gammaproteobacteria; Enterobacterales; Enterobacteriaceae; Escherichia
MNNEETFYQAMRRQGVTRRSFLKYCSLAATSLGLGAGMAPKIAWALENKPRIPVVWIHGL WP_001295940.1
MULTISPECIES: hydrogenase [Enterobacteriaceae] Length: 372\nNumber of predicted TMHs: 2\nExp number of AAs in TMHs: 37.2608899999999\nExp number, first 60 AAs: 19.52923\nTotal prob of N-in: 0.94351\nPOSSIBLE N-term signal sequence\ninside 1 19\nTMhelix 20 39\noutside 40 325\nTMhelix 326 348\ninside 349 372

6852 GCF_001521135.1_ASM152113v1 Escherichia coli Proteobacteria; Gammaproteobacteria; Enterobacterales; Enterobacteriaceae; Escherichia
MTGDNTLIHSHGINRRDFMKLCAALAATMGLSSKAAAEMVESVTNPQRPPVIWIGAQECT WP_000145423.1
hydrogenase 2 small subunit [Escherichia coli] Length: 372\nNumber of predicted TMHs: 1\nExp number of AAs in TMHs: 23.78201\nExp number, first 60 AAs: 0.28355\nTotal prob of N-in: 0.13257\noutside 1 331\nTMhelix 332 354\ninside 355 372

6853 GCF_001521135.1_ASM152113v1 Escherichia coli Proteobacteria; Gammaproteobacteria; Enterobacterales; Enterobacteriaceae; Escherichia
MNNEETFYQAMRRQGVTRRSFLKYCSLAATSLGLGAGMAPKIAWALENKPRIPVVWIHGL WP_001295940.1
MULTISPECIES: hydrogenase [Enterobacteriaceae] Length: 372\nNumber of predicted TMHs: 2\nExp number of AAs in TMHs: 37.2608899999999\nExp number, first 60 AAs: 19.52923\nTotal prob of N-in: 0.94351\nPOSSIBLE N-term signal sequence\ninside 1 19\nTMhelix 20 39\noutside 40 325\nTMhelix 326 348\ninside 349 372

6854 GCF_001520995.1_ASM152099v1 Escherichia coli Proteobacteria; Gammaproteobacteria; Enterobacterales; Enterobacteriaceae; Escherichia
MNNEETFYQAMRRQGVTRRSFLKYCSLAATSLGLGAGMAPKIAWALENKPRIPVVWIHGL WP_001058323.1
MULTISPECIES: hydrogenase-1 small chain [Proteobacteria] Length: 372\nNumber of predicted TMHs: 2\nExp number of AAs in TMHs: 37.2608499999999\nExp number, first 60 AAs: 19.52923\nTotal prob of N-in: 0.94351\nPOSSIBLE N-term signal sequence\ninside 1 19\nTMhelix 20 39\noutside 40 325\nTMhelix 326 348\ninside 349 372

6855 GCF_001521195.1_ASM152119v1 Escherichia coli Proteobacteria; Gammaproteobacteria; Enterobacterales; Enterobacteriaceae; Escherichia
MTGDNTLIHSHGINRRDFMKLCAALAATMGLSSKAAAEMAESVTNPQRPPVIWIGAQECT WP_000145418.1
hydrogenase 2 small subunit [Escherichia coli] Length: 372\nNumber of predicted TMHs: 1\nExp number of

AAs in TMHs: 23.72313\nExp number, first 60 AAs: 0.22034\nTotal prob of N-in: 0.12997\noutside 1 331\nTMhelix 332 354\ninside 355 372

6856 GCF_001521155.1_ASM152115v1 Escherichia coli Proteobacteria; Gammaproteobacteria; Enterobacterales; Enterobacteriaceae; Escherichia MTGDNTLIHSHGINRRDFMKLCAALAATMGLSSKAAAEMAESVTNPQRPPVIWIGAQECT WP_000145410.1 MULTISPECIES: hydrogenase-2 small chain [Proteobacteria] Length: 372\nNumber of predicted TMHs: 1\nExp number of AAs in TMHs: 23.72226\nExp number, first 60 AAs: 0.22038\nTotal prob of N-in: 0.12994\noutside 1 331\nTMhelix 332 354\ninside 355 372

6857 GCF_001521275.1_ASM152127v1 Escherichia coli Proteobacteria; Gammaproteobacteria; Enterobacterales; Enterobacteriaceae; Escherichia MTGDNTLIHSHGINRRDFMKLCAALAATMGLSSKAAAEMAESVTNPQRPPVIWIGAQECT WP_000145410.1 MULTISPECIES: hydrogenase-2 small chain [Proteobacteria] Length: 372\nNumber of predicted TMHs: 1\nExp number of AAs in TMHs: 23.72226\nExp number, first 60 AAs: 0.22038\nTotal prob of N-in: 0.12994\noutside 1 331\nTMhelix 332 354\ninside 355 372

6858 GCF_001521155.1_ASM152115v1 Escherichia coli Proteobacteria; Gammaproteobacteria; Enterobacterales; Enterobacteriaceae; Escherichia MNNEETFYQAMRRQGVTRRSFLKYCSLAATSLGLGAGMAPKIAWALENKPRIPVVWIHGL WP_001058323.1 MULTISPECIES: hydrogenase-1 small chain [Proteobacteria] Length: 372\nNumber of predicted TMHs: 2\nExp number of AAs in TMHs: 37.2608499999999\nExp number, first 60 AAs: 19.52923\nTotal prob of N-in: 0.94351\nPOSSIBLE N-term signal sequence\ninside 1 19\nTMhelix 20 39\noutside 40 325\nTMhelix 326 348\ninside 349 372

6859 GCF_001521395.1_ASM152139v1 Escherichia coli Proteobacteria; Gammaproteobacteria; Enterobacterales; Enterobacteriaceae; Escherichia MTGDNTLIHSHGINRRDFMKLCAALAATMGLSSKAAAEMVESVTNPQRPPVIWIGAQECT WP_000145423.1 hydrogenase 2 small subunit [Escherichia coli] Length: 372\nNumber of predicted TMHs: 1\nExp number of AAs in TMHs: 23.78201\nExp number, first 60 AAs: 0.28355\nTotal prob of N-in: 0.13257\noutside 1 331\nTMhelix 332 354\ninside 355 372

6860 GCF_001521435.1_ASM152143v1 Escherichia coli Proteobacteria; Gammaproteobacteria; Enterobacterales; Enterobacteriaceae; Escherichia MNNEETFYQAMRRQGVTRRSFLKYCSLAATSLGLGAGMAPKIAWALENKPRIPVVWIHGL WP_001058323.1 MULTISPECIES: hydrogenase-1 small chain [Proteobacteria] Length: 372\nNumber of predicted TMHs: 2\nExp number of AAs in TMHs: 37.2608499999999\nExp number, first 60 AAs: 19.52923\nTotal prob of N-in: 0.94351\nPOSSIBLE N-term signal sequence\ninside 1 19\nTMhelix 20 39\noutside 40 325\nTMhelix 326 348\ninside 349 372

6861 GCF_001521285.1_ASM152128v1 Escherichia coli Proteobacteria; Gammaproteobacteria; Enterobacterales; Enterobacteriaceae; Escherichia MNNEETFYQAMRRQGVTRRSFLKYCSLAATSLGLGAGMAPKIAWALENKPRIPVVWIHGL WP_001058323.1 MULTISPECIES: hydrogenase-1 small chain [Proteobacteria] Length: 372\nNumber of predicted TMHs: 2\nExp number of AAs in TMHs: 37.2608499999999\nExp number, first 60 AAs: 19.52923\nTotal prob of N-in: 0.94351\nPOSSIBLE N-term signal sequence\ninside 1 19\nTMhelix 20 39\noutside 40 325\nTMhelix 326 348\ninside 349 372

6862 GCF_001521475.1_ASM152147v1 Escherichia coli Proteobacteria; Gammaproteobacteria; Enterobacterales; Enterobacteriaceae; Escherichia MNNEETFYQAMRRQGVTRRSFLKYCSLAATSLGLGAGMAPKIAWALENKPRIPVVWIHGL WP_001058323.1 MULTISPECIES: hydrogenase-1 small chain [Proteobacteria] Length: 372\nNumber of predicted TMHs: 2\nExp number of AAs in TMHs: 37.2608499999999\nExp number, first 60 AAs: 19.52923\nTotal prob of N-in: 0.94351\nPOSSIBLE N-term signal sequence\ninside 1 19\nTMhelix 20 39\noutside 40 325\nTMhelix 326 348\ninside 349 372

6863 GCF_001521495.1_ASM152149v1 Escherichia coli Proteobacteria; Gammaproteobacteria; Enterobacterales; Enterobacteriaceae; Escherichia MTGDNTLIHSHGINRRDFMKLCAALAATMGLSSKAAAEMVESVTNPQRPPVIWIGAQECT WP_000145423.1 hydrogenase 2 small subunit [Escherichia coli] Length: 372\nNumber of predicted TMHs: 1\nExp number of

AAs in TMHs: 23.78201\nExp number, first 60 AAs: 0.28355\nTotal prob of N-in: 0.13257\noutside 1 331\nTMhelix 332 354\ninside 355 372

6864 GCF_001521535.1_ASM152153v1 Escherichia coli Proteobacteria; Gammaproteobacteria; Enterobacterales; Enterobacteriaceae; Escherichia MTGDNTLIHSHGINRRDFMKLCAALAATMGLSSKAAAEMVESVTNPQRPPVIWIGAQECT WP_000145423.1 hydrogenase 2 small subunit [Escherichia coli] Length: 372\nNumber of predicted TMHs: 1\nExp number of AAs in TMHs: 23.78201\nExp number, first 60 AAs: 0.28355\nTotal prob of N-in: 0.13257\noutside 1 331\nTMhelix 332 354\ninside 355 372

6865 GCF_001521635.1_ASM152163v1 Escherichia coli Proteobacteria; Gammaproteobacteria; Enterobacterales; Enterobacteriaceae; Escherichia MNNEETFYQAMRRQGVTRRSFLKYCSLAATSLGLGAGMAPKIAWALENKPRIPVVWIHGL WP_001058323.1 MULTISPECIES: hydrogenase-1 small chain [Proteobacteria] Length: 372\nNumber of predicted TMHs: 2\nExp number of AAs in TMHs: 37.2608499999999\nExp number, first 60 AAs: 19.52923\nTotal prob of N-in: 0.94351\nPOSSIBLE N-term signal sequence\ninside 1 19\nTMhelix 20 39\noutside 40 325\nTMhelix 326 348\ninside 349 372

6866 GCF_001521595.1_ASM152159v1 Escherichia coli Proteobacteria; Gammaproteobacteria; Enterobacterales; Enterobacteriaceae; Escherichia MNNEETFYQAMRRQGVTRRSFLKYCSLAATSLGLGAGMAPKIAWALENKPRIPVVWIHGL WP_001058323.1 MULTISPECIES: hydrogenase-1 small chain [Proteobacteria] Length: 372\nNumber of predicted TMHs: 2\nExp number of AAs in TMHs: 37.2608499999999\nExp number, first 60 AAs: 19.52923\nTotal prob of N-in: 0.94351\nPOSSIBLE N-term signal sequence\ninside 1 19\nTMhelix 20 39\noutside 40 325\nTMhelix 326 348\ninside 349 372

6867 GCF_001521395.1_ASM152139v1 Escherichia coli Proteobacteria; Gammaproteobacteria; Enterobacterales; Enterobacteriaceae; Escherichia MNNEETFYQAMRRQGVTRRSFLKYCSLAATSLGLGAGMAPKIAWALENKPRIPVVWIHGL WP_001295940.1 MULTISPECIES: hydrogenase [Enterobacteriaceae] Length: 372\nNumber of predicted TMHs: 2\nExp number of AAs in TMHs: 37.2608899999999\nExp number, first 60 AAs: 19.52923\nTotal prob of N-in: 0.94351\nPOSSIBLE N-term signal sequence\ninside 1 19\nTMhelix 20 39\noutside 40 325\nTMhelix 326 348\ninside 349 372

6868 GCF_001521675.1_ASM152167v1 Escherichia coli Proteobacteria; Gammaproteobacteria; Enterobacterales; Enterobacteriaceae; Escherichia MTGDNTLIHSHGINRRDFMKLCAALAATMGLSSKAAAEMVESVTNPQRPPVIWIGAQECT WP_000145423.1 hydrogenase 2 small subunit [Escherichia coli] Length: 372\nNumber of predicted TMHs: 1\nExp number of AAs in TMHs: 23.78201\nExp number, first 60 AAs: 0.28355\nTotal prob of N-in: 0.13257\noutside 1 331\nTMhelix 332 354\ninside 355 372

6869 GCF_001521695.1_ASM152169v1 Escherichia coli Proteobacteria; Gammaproteobacteria; Enterobacterales; Enterobacteriaceae; Escherichia MTGDNTLIHSHGINRRDFMKLCAALAATMGLSSKAAAEMAESVTNPQRPPVIWIGAQECT WP_000145410.1 MULTISPECIES: hydrogenase-2 small chain [Proteobacteria] Length: 372\nNumber of predicted TMHs: 1\nExp number of AAs in TMHs: 23.72226\nExp number, first 60 AAs: 0.22038\nTotal prob of N-in: 0.12994\noutside 1 331\nTMhelix 332 354\ninside 355 372

6870 GCF_001524895.1_ASM152489v1 Escherichia coli Proteobacteria; Gammaproteobacteria; Enterobacterales; Enterobacteriaceae; Escherichia MTGDNTLIHSHGINRRDFMKLCAALAATMGLSSKAAAEMVESVTNPQRPPVIWIGAQECT WP_000145423.1 hydrogenase 2 small subunit [Escherichia coli] Length: 372\nNumber of predicted TMHs: 1\nExp number of AAs in TMHs: 23.78201\nExp number, first 60 AAs: 0.28355\nTotal prob of N-in: 0.13257\noutside 1 331\nTMhelix 332 354\ninside 355 372

6871 GCF_001542545.1_ASM154254v1 Escherichia coli Proteobacteria; Gammaproteobacteria; Enterobacterales; Enterobacteriaceae; Escherichia MTGDNTLIHSHGINRRDFMKLCAALAATMGLSSKAAAEMAESVTNPQRPPVIWIGAQECT WP_000145410.1 MULTISPECIES: hydrogenase-2 small chain [Proteobacteria] Length: 372\nNumber of predicted TMHs: 1\nExp number of AAs in TMHs: 23.72226\nExp number, first 60 AAs: 0.22038\nTotal prob of N-in: 0.12994\noutside 1 331\nTMhelix 332 354\ninside 355 372

6872 GCF_001550005.1_ASM155000v1 Escherichia coli Proteobacteria; Gammaproteobacteria; Enterobacterales; Enterobacteriaceae; Escherichia
MNNEETFYQAMRRQGVTRRSFLKYCSLAATSLGLGAGMAPKIAWALENKPRIPVVWIHGL WP_001058323.1
MULTISPECIES: hydrogenase-1 small chain [Proteobacteria] Length: 372\nNumber of predicted TMHs: 2\nExp number of AAs in TMHs: 37.2608499999999\nExp number, first 60 AAs: 19.52923\nTotal prob of N-in: 0.94351\nPOSSIBLE N-term signal sequence\ninside 1 19\nTMhelix 20 39\noutside 40 325\nTMhelix 326 348\ninside 349 372

6873 GCF_001554295.1_ASM155429v1 Escherichia coli Proteobacteria; Gammaproteobacteria; Enterobacterales; Enterobacteriaceae; Escherichia
MNNEETFYQAMRRQGVTRRSFLKYCSLAATSLGLGAGMAPKIAWALENKPRIPVVWIHGL WP_001295940.1
MULTISPECIES: hydrogenase [Enterobacteriaceae] Length: 372\nNumber of predicted TMHs: 2\nExp number of AAs in TMHs: 37.2608899999999\nExp number, first 60 AAs: 19.52923\nTotal prob of N-in: 0.94351\nPOSSIBLE N-term signal sequence\ninside 1 19\nTMhelix 20 39\noutside 40 325\nTMhelix 326 348\ninside 349 372

6874 GCF_001555495.1_ASM155549v1 Escherichia coli Proteobacteria; Gammaproteobacteria; Enterobacterales; Enterobacteriaceae; Escherichia
MNNEETFYQAMRRQGVTRRSFLKYCSLAATSLGLGAGMAPKIAWALENKPRIPVVWIHGL WP_001295940.1
MULTISPECIES: hydrogenase [Enterobacteriaceae] Length: 372\nNumber of predicted TMHs: 2\nExp number of AAs in TMHs: 37.2608899999999\nExp number, first 60 AAs: 19.52923\nTotal prob of N-in: 0.94351\nPOSSIBLE N-term signal sequence\ninside 1 19\nTMhelix 20 39\noutside 40 325\nTMhelix 326 348\ninside 349 372

6875 GCF_001521675.1_ASM152167v1 Escherichia coli Proteobacteria; Gammaproteobacteria; Enterobacterales; Enterobacteriaceae; Escherichia
MNNEETFYQAMRRQGVTRRSFLKYCSLAATSLGLGAGMAPKIAWALENKPRIPVVWIHGL WP_001295940.1
MULTISPECIES: hydrogenase [Enterobacteriaceae] Length: 372\nNumber of predicted TMHs: 2\nExp number of AAs in TMHs: 37.2608899999999\nExp number, first 60 AAs: 19.52923\nTotal prob of N-in: 0.94351\nPOSSIBLE N-term signal sequence\ninside 1 19\nTMhelix 20 39\noutside 40 325\nTMhelix 326 348\ninside 349 372

6876 GCF_001524895.1_ASM152489v1 Escherichia coli Proteobacteria; Gammaproteobacteria; Enterobacterales; Enterobacteriaceae; Escherichia
MNNEETFYQAMRRQGVTRRSFLKYCSLAATSLGLGAGMAPKIAWALENKPRIPVVWIHGL WP_001295940.1
MULTISPECIES: hydrogenase [Enterobacteriaceae] Length: 372\nNumber of predicted TMHs: 2\nExp number of AAs in TMHs: 37.2608899999999\nExp number, first 60 AAs: 19.52923\nTotal prob of N-in: 0.94351\nPOSSIBLE N-term signal sequence\ninside 1 19\nTMhelix 20 39\noutside 40 325\nTMhelix 326 348\ninside 349 372

6877 GCF_001561255.1_ASM156125v1 Escherichia coli Proteobacteria; Gammaproteobacteria; Enterobacterales; Enterobacteriaceae; Escherichia
MTGDNTLIHSHGINRRDFMKLCAALAATMGLSSKAAAEMVESVTNPQRPPVIWIGAQECT WP_000145423.1
hydrogenase 2 small subunit [Escherichia coli] Length: 372\nNumber of predicted TMHs: 1\nExp number of AAs in TMHs: 23.78201\nExp number, first 60 AAs: 0.28355\nTotal prob of N-in: 0.13257\noutside 1 331\nTMhelix 332 354\ninside 355 372

6878 GCF_001561285.1_ASM156128v1 Escherichia coli Proteobacteria; Gammaproteobacteria; Enterobacterales; Enterobacteriaceae; Escherichia
MTGDNTLIHSHGINRRDFMKLCAALAATMGLSSKAAAEMVESVTNPQRPPVIWIGAQECT WP_000145423.1
hydrogenase 2 small subunit [Escherichia coli] Length: 372\nNumber of predicted TMHs: 1\nExp number of AAs in TMHs: 23.78201\nExp number, first 60 AAs: 0.28355\nTotal prob of N-in: 0.13257\noutside 1 331\nTMhelix 332 354\ninside 355 372

6879 GCF_001561195.1_ASM156119v1 Escherichia coli Proteobacteria; Gammaproteobacteria; Enterobacterales; Enterobacteriaceae; Escherichia
MNNEETFYQAMRRQGVTRRSFLKYCSLAATSLGLGAGMAPKIAWALENKPRIPVVWIHGL WP_001295940.1
MULTISPECIES: hydrogenase [Enterobacteriaceae] Length: 372\nNumber of predicted TMHs: 2\nExp number of AAs in TMHs: 37.2608899999999\nExp number, first 60 AAs: 19.52923\nTotal prob of N-in: 0.94351\nPOSSIBLE N-term signal sequence\ninside 1 19\nTMhelix 20 39\noutside 40 325\nTMhelix 326 348\ninside 349 372

6880 GCF_001561015.1_ASM156101v1 Escherichia coli Proteobacteria; Gammaproteobacteria; Enterobacterales; Enterobacteriaceae; Escherichia
MNNEETFYQAMRRQGVTRRSFLKYCSLAATSLGLGAGMAPKIAWALENKPRIPVVWIHGL WP_001295940.1
MULTISPECIES: hydrogenase [Enterobacteriaceae] Length: 372\nNumber of predicted TMHs: 2\nExp number of AAs in TMHs: 37.2608899999999\nExp number, first 60 AAs: 19.52923\nTotal prob of N-in: 0.94351\nPOSSIBLE N-term signal sequence\ninside 1 19\nTMhelix 20 39\noutside 40 325\nTMhelix 326 348\ninside 349 372

6881 GCF_001561415.1_ASM156141v1 Escherichia coli Proteobacteria; Gammaproteobacteria; Enterobacterales; Enterobacteriaceae; Escherichia
MTGDNTLIHSHGINRRDFMKLCAALAATMGLSSKAAAEMVESVTNPQRPPVIWIGAQECT WP_000145423.1
hydrogenase 2 small subunit [Escherichia coli] Length: 372\nNumber of predicted TMHs: 1\nExp number of AAs in TMHs: 23.78201\nExp number, first 60 AAs: 0.28355\nTotal prob of N-in: 0.13257\noutside 1 331\nTMhelix 332 354\ninside 355 372

6882 GCF_001561355.1_ASM156135v1 Escherichia coli Proteobacteria; Gammaproteobacteria; Enterobacterales; Enterobacteriaceae; Escherichia
MNNEETFYQAMRRQGVTRRSFLKYCSLAATSLGLGAGMAPKIAWALENKPRIPVVWIHGL WP_001058323.1
MULTISPECIES: hydrogenase-1 small chain [Proteobacteria] Length: 372\nNumber of predicted TMHs: 2\nExp number of AAs in TMHs: 37.2608499999999\nExp number, first 60 AAs: 19.52923\nTotal prob of N-in: 0.94351\nPOSSIBLE N-term signal sequence\ninside 1 19\nTMhelix 20 39\noutside 40 325\nTMhelix 326 348\ninside 349 372

6883 GCF_001561375.1_ASM156137v1 Escherichia coli Proteobacteria; Gammaproteobacteria; Enterobacterales; Enterobacteriaceae; Escherichia
MNNEETFYQAMRRQGVTRRSFLKYCSLAATSLGLGAGMAPKIAWALENKPRIPVVWIHGL WP_001295940.1
MULTISPECIES: hydrogenase [Enterobacteriaceae] Length: 372\nNumber of predicted TMHs: 2\nExp number of AAs in TMHs: 37.2608899999999\nExp number, first 60 AAs: 19.52923\nTotal prob of N-in: 0.94351\nPOSSIBLE N-term signal sequence\ninside 1 19\nTMhelix 20 39\noutside 40 325\nTMhelix 326 348\ninside 349 372

6884 GCF_001561415.1_ASM156141v1 Escherichia coli Proteobacteria; Gammaproteobacteria; Enterobacterales; Enterobacteriaceae; Escherichia
MNNEETFYQAMRRQGVTRRSFLKYCSLAATSLGLGAGMAPKIAWALENKPRIPVVWIHGL WP_001295940.1
MULTISPECIES: hydrogenase [Enterobacteriaceae] Length: 372\nNumber of predicted TMHs: 2\nExp number of AAs in TMHs: 37.2608899999999\nExp number, first 60 AAs: 19.52923\nTotal prob of N-in: 0.94351\nPOSSIBLE N-term signal sequence\ninside 1 19\nTMhelix 20 39\noutside 40 325\nTMhelix 326 348\ninside 349 372

6885 GCF_001561205.1_ASM156120v1 Escherichia coli Proteobacteria; Gammaproteobacteria; Enterobacterales; Enterobacteriaceae; Escherichia
MNNEETFYQAMRRQGVTRRSFLKYCSLAATSLGLGAGMAPKIAWALENKPRIPVVWIHGL WP_001295940.1
MULTISPECIES: hydrogenase [Enterobacteriaceae] Length: 372\nNumber of predicted TMHs: 2\nExp number of AAs in TMHs: 37.2608899999999\nExp number, first 60 AAs: 19.52923\nTotal prob of N-in: 0.94351\nPOSSIBLE N-term signal sequence\ninside 1 19\nTMhelix 20 39\noutside 40 325\nTMhelix 326 348\ninside 349 372

6886 GCF_001561445.1_ASM156144v1 Escherichia coli Proteobacteria; Gammaproteobacteria; Enterobacterales; Enterobacteriaceae; Escherichia
MTGDNTLIHSHGINRRDFMKLCAALAATMGLSSKAAAEMAESVTNPQRPPVIWIGAQECT WP_000145410.1
MULTISPECIES: hydrogenase-2 small chain [Proteobacteria] Length: 372\nNumber of predicted TMHs: 1\nExp number of AAs in TMHs: 23.72226\nExp number, first 60 AAs: 0.22038\nTotal prob of N-in: 0.12994\noutside 1 331\nTMhelix 332 354\ninside 355 372

6887 GCF_001561495.1_ASM156149v1 Escherichia coli Proteobacteria; Gammaproteobacteria; Enterobacterales; Enterobacteriaceae; Escherichia
MTGDNTLIHSHGINRRDFMKLCAALAATMGLSSKAAAEMVESVTNPQRPPVIWIGAQECT WP_000145423.1
hydrogenase 2 small subunit [Escherichia coli] Length: 372\nNumber of predicted TMHs: 1\nExp number of AAs in TMHs: 23.78201\nExp number, first 60 AAs: 0.28355\nTotal prob of N-in: 0.13257\noutside 1 331\nTMhelix 332 354\ninside 355 372

6888 GCF_001561595.1_ASM156159v1 Escherichia coli Proteobacteria; Gammaproteobacteria; Enterobacterales; Enterobacteriaceae; Escherichia
MTGDNTLIHSHGINRRDFMKLCAALAATMGLSSKAAAEMVESVTNPQRPPVIWIGAQECT WP_000145423.1
hydrogenase 2 small subunit [Escherichia coli] Length: 372\nNumber of predicted TMHs: 1\nExp number of AAs in TMHs: 23.78201\nExp number, first 60 AAs: 0.28355\nTotal prob of N-in: 0.13257\noutside 1 331\nTMhelix 332 354\ninside 355 372

6889 GCF_001561655.1_ASM156165v1 Escherichia coli Proteobacteria; Gammaproteobacteria; Enterobacterales; Enterobacteriaceae; Escherichia
MTGDNTLIHSHGINRRDFMKLCAALAATMGLSSKAAAEMVESVTNPQRPPVIWIGAQECT WP_000145423.1
hydrogenase 2 small subunit [Escherichia coli] Length: 372\nNumber of predicted TMHs: 1\nExp number of AAs in TMHs: 23.78201\nExp number, first 60 AAs: 0.28355\nTotal prob of N-in: 0.13257\noutside 1 331\nTMhelix 332 354\ninside 355 372

6890 GCF_001561635.1_ASM156163v1 Escherichia coli Proteobacteria; Gammaproteobacteria; Enterobacterales; Enterobacteriaceae; Escherichia
MTGDNTLIHSHGINRRDFMKLCAALAATMGLSSKAAAEMAESVTNPQRPPVIWIGAQECT WP_000145410.1
MULTISPECIES: hydrogenase-2 small chain [Proteobacteria] Length: 372\nNumber of predicted TMHs: 1\nExp number of AAs in TMHs: 23.72226\nExp number, first 60 AAs: 0.22038\nTotal prob of N-in: 0.12994\noutside 1 331\nTMhelix 332 354\ninside 355 372

6891 GCF_001561685.1_ASM156168v1 Escherichia coli Proteobacteria; Gammaproteobacteria; Enterobacterales; Enterobacteriaceae; Escherichia
MTGDNTLIHSHGINRRDFMKLCAALAATMGLSSKAAAEMVESVTNPQRPPVIWIGAQECT WP_000145423.1
hydrogenase 2 small subunit [Escherichia coli] Length: 372\nNumber of predicted TMHs: 1\nExp number of AAs in TMHs: 23.78201\nExp number, first 60 AAs: 0.28355\nTotal prob of N-in: 0.13257\noutside 1 331\nTMhelix 332 354\ninside 355 372

6892 GCF_001561735.1_ASM156173v1 Escherichia coli Proteobacteria; Gammaproteobacteria; Enterobacterales; Enterobacteriaceae; Escherichia
MTGDNTLIHSHGINRRDFMKLCAALAATMGLSSKAAAEMAESVTNPQRPPVIWIGAQECT WP_000145410.1
MULTISPECIES: hydrogenase-2 small chain [Proteobacteria] Length: 372\nNumber of predicted TMHs: 1\nExp number of AAs in TMHs: 23.72226\nExp number, first 60 AAs: 0.22038\nTotal prob of N-in: 0.12994\noutside 1 331\nTMhelix 332 354\ninside 355 372

6893 GCF_001562335.1_ASM156233v1 Escherichia coli Proteobacteria; Gammaproteobacteria; Enterobacterales; Enterobacteriaceae; Escherichia
MTGDNTLIHSHGINRRDFMKLCAALATTMGLSSKAAAEMAESVTNPQRPPVIWIGAQECT WP_023149629.1
MULTISPECIES: hydrogenase-2 small chain [Enterobacteriaceae] Length: 372\nNumber of predicted TMHs: 1\nExp number of AAs in TMHs: 23.63603\nExp number, first 60 AAs: 0.12971\nTotal prob of N-in: 0.12608\noutside 1 331\nTMhelix 332 354\ninside 355 372

6894 GCF_001561635.1_ASM156163v1 Escherichia coli Proteobacteria; Gammaproteobacteria; Enterobacterales; Enterobacteriaceae; Escherichia
MNNEETFYQAMRRQGVTRRSFLKYCSLAATSLGLGAGMAPKIAWALENKPRIPVVWIHGL WP_001058323.1
MULTISPECIES: hydrogenase-1 small chain [Proteobacteria] Length: 372\nNumber of predicted TMHs: 2\nExp number of AAs in TMHs: 37.2608499999999\nExp number, first 60 AAs: 19.52923\nTotal prob of N-in: 0.94351\nPOSSIBLE N-term signal sequence\ninside 1 19\nTMhelix 20 39\noutside 40 325\nTMhelix 326 348\ninside 349 372

6895 GCF_001561755.1_ASM156175v1 Escherichia coli Proteobacteria; Gammaproteobacteria; Enterobacterales; Enterobacteriaceae; Escherichia
MTGDNTLIHSHGINRRDFMKLCAALAATMGLSSKAAAEMVESVTNPQRPPVIWIGAQECT WP_000145423.1
hydrogenase 2 small subunit [Escherichia coli] Length: 372\nNumber of predicted TMHs: 1\nExp number of AAs in TMHs: 23.78201\nExp number, first 60 AAs: 0.28355\nTotal prob of N-in: 0.13257\noutside 1 331\nTMhelix 332 354\ninside 355 372

6896 GCF_001561735.1_ASM156173v1 Escherichia coli Proteobacteria; Gammaproteobacteria; Enterobacterales; Enterobacteriaceae; Escherichia
MNNEETFYQAMRRQGVTRRSFLKYCSLAATSLGLGAGMAPKIAWALENKPRIPVVWIHGL WP_001058323.1
MULTISPECIES: hydrogenase-1 small chain [Proteobacteria] Length: 372\nNumber of predicted TMHs:

2\nExp number of AAs in TMHs: 37.2608499999999\nExp number, first 60 AAs: 19.52923\nTotal prob of N-in: 0.94351\nPOSSIBLE N-term signal sequence\ninside 1 19\nTMhelix 20 39\noutside 40 325\nTMhelix 326 348\ninside 349 372

6897 GCF_001561595.1_ASM156159v1 Escherichia coli Proteobacteria; Gammaproteobacteria; Enterobacterales; Enterobacteriaceae; Escherichia MNNEETFYQAMRRQGVTRRSFLKYCSLAATSLGLGAGMAPKIAWALENKPRIPVVWIHGL WP_001295940.1 MULTISPECIES: hydrogenase [Enterobacteriaceae] Length: 372\nNumber of predicted TMHs: 2\nExp number of AAs in TMHs: 37.2608899999999\nExp number, first 60 AAs: 19.52923\nTotal prob of N-in: 0.94351\nPOSSIBLE N-term signal sequence\ninside 1 19\nTMhelix 20 39\noutside 40 325\nTMhelix 326 348\ninside 349 372

6898 GCF_001561845.1_ASM156184v1 Escherichia coli Proteobacteria; Gammaproteobacteria; Enterobacterales; Enterobacteriaceae; Escherichia MTGDNTLIHSHGINRRDFMKLCAALAATMGLSSKAAAEMVESVTNPQRPPVIWIGAQECT WP_000145423.1 hydrogenase 2 small subunit [Escherichia coli] Length: 372\nNumber of predicted TMHs: 1\nExp number of AAs in TMHs: 23.78201\nExp number, first 60 AAs: 0.28355\nTotal prob of N-in: 0.13257\noutside 1 331\nTMhelix 332 354\ninside 355 372

6899 GCF_001561845.1_ASM156184v1 Escherichia coli Proteobacteria; Gammaproteobacteria; Enterobacterales; Enterobacteriaceae; Escherichia MNNEETFYQAMRRQGVTRRSFLKYCSLAATSLGLGAGMAPKIAWALENKPRIPVVWIHGL WP_001295940.1 MULTISPECIES: hydrogenase [Enterobacteriaceae] Length: 372\nNumber of predicted TMHs: 2\nExp number of AAs in TMHs: 37.2608899999999\nExp number, first 60 AAs: 19.52923\nTotal prob of N-in: 0.94351\nPOSSIBLE N-term signal sequence\ninside 1 19\nTMhelix 20 39\noutside 40 325\nTMhelix 326 348\ninside 349 372

6900 GCF_001561655.1_ASM156165v1 Escherichia coli Proteobacteria; Gammaproteobacteria; Enterobacterales; Enterobacteriaceae; Escherichia MNNEETFYQAMRRQGVTRRSFLKYCSLAATSLGLGAGMAPKIAWALENKPRIPVVWIHGL WP_001295940.1 MULTISPECIES: hydrogenase [Enterobacteriaceae] Length: 372\nNumber of predicted TMHs: 2\nExp number of AAs in TMHs: 37.2608899999999\nExp number, first 60 AAs: 19.52923\nTotal prob of N-in: 0.94351\nPOSSIBLE N-term signal sequence\ninside 1 19\nTMhelix 20 39\noutside 40 325\nTMhelix 326 348\ninside 349 372

6901 GCF_001561685.1_ASM156168v1 Escherichia coli Proteobacteria; Gammaproteobacteria; Enterobacterales; Enterobacteriaceae; Escherichia MNNEETFYQAMRRQGVTRRSFLKYCSLAATSLGLGAGMAPKIAWALENKPRIPVVWIHGL WP_001295940.1 MULTISPECIES: hydrogenase [Enterobacteriaceae] Length: 372\nNumber of predicted TMHs: 2\nExp number of AAs in TMHs: 37.2608899999999\nExp number, first 60 AAs: 19.52923\nTotal prob of N-in: 0.94351\nPOSSIBLE N-term signal sequence\ninside 1 19\nTMhelix 20 39\noutside 40 325\nTMhelix 326 348\ninside 349 372

6902 GCF_001562375.1_ASM156237v1 Escherichia coli Proteobacteria; Gammaproteobacteria; Enterobacterales; Enterobacteriaceae; Escherichia MTGDNTLIHSHGINRRDFMKLCAALAATMGLSSKAAAEMAESVTNPQRPPVIWIGAQECT WP_000145410.1 MULTISPECIES: hydrogenase-2 small chain [Proteobacteria] Length: 372\nNumber of predicted TMHs: 1\nExp number of AAs in TMHs: 23.72226\nExp number, first 60 AAs: 0.22038\nTotal prob of N-in: 0.12994\noutside 1 331\nTMhelix 332 354\ninside 355 372

6903 GCF_001562855.1_ASM156285v1 Escherichia coli Proteobacteria; Gammaproteobacteria; Enterobacterales; Enterobacteriaceae; Escherichia MNNEETFYQAMRRQGVTRRSFLKYCSLAATSLGLGAGMAPKIAWALENKPRIPVVWIHGL WP_001058323.1 MULTISPECIES: hydrogenase-1 small chain [Proteobacteria] Length: 372\nNumber of predicted TMHs: 2\nExp number of AAs in TMHs: 37.2608499999999\nExp number, first 60 AAs: 19.52923\nTotal prob of N-in: 0.94351\nPOSSIBLE N-term signal sequence\ninside 1 19\nTMhelix 20 39\noutside 40 325\nTMhelix 326 348\ninside 349 372

6904 GCF_001562715.1_ASM156271v1 Escherichia coli Proteobacteria; Gammaproteobacteria; Enterobacterales; Enterobacteriaceae; Escherichia MNNEETFYQAMRRQGVTRRSFLKYCSLAATSLGLGAGMAPKIAWALENKPRIPVVWIHGL WP_001058323.1

MULTISPECIES: hydrogenase-1 small chain [Proteobacteria] Length: 372\nNumber of predicted TMHs: 2\nExp number of AAs in TMHs: 37.2608499999999\nExp number, first 60 AAs: 19.52923\nTotal prob of N-in: 0.94351\nPOSSIBLE N-term signal sequence\ninside 1 19\nTMhelix 20 39\noutside 40 325\nTMhelix 326 348\ninside 349 372

6905 GCF_001563385.1_ASM156338v1 Escherichia coli Proteobacteria; Gammaproteobacteria; Enterobacterales; Enterobacteriaceae; Escherichia
MTGDNTLIHSHGINRRDFMKLCAALAATMGLSSKAAAEMAESVTNPQRPPVIWIGAQECT WP_000145410.1
MULTISPECIES: hydrogenase-2 small chain [Proteobacteria] Length: 372\nNumber of predicted TMHs: 1\nExp number of AAs in TMHs: 23.72226\nExp number, first 60 AAs: 0.22038\nTotal prob of N-in: 0.12994\noutside 1 331\nTMhelix 332 354\ninside 355 372

6906 GCF_001563405.1_ASM156340v1 Escherichia coli Proteobacteria; Gammaproteobacteria; Enterobacterales; Enterobacteriaceae; Escherichia
MTGDNTLIHSHGINRRDFMKLCAALAATMGLSSKAAAEMAESVTNPQRPPVIWIGAQECT WP_000145410.1
MULTISPECIES: hydrogenase-2 small chain [Proteobacteria] Length: 372\nNumber of predicted TMHs: 1\nExp number of AAs in TMHs: 23.72226\nExp number, first 60 AAs: 0.22038\nTotal prob of N-in: 0.12994\noutside 1 331\nTMhelix 332 354\ninside 355 372

6907 GCF_001563715.1_ASM156371v1 Escherichia coli Proteobacteria; Gammaproteobacteria; Enterobacterales; Enterobacteriaceae; Escherichia
MTGDNTLIHSHGINRRDFMKLCAALAATMGLSSKAAAEMAESVTNPQRPPVIWIGAQECT WP_000145410.1
MULTISPECIES: hydrogenase-2 small chain [Proteobacteria] Length: 372\nNumber of predicted TMHs: 1\nExp number of AAs in TMHs: 23.72226\nExp number, first 60 AAs: 0.22038\nTotal prob of N-in: 0.12994\noutside 1 331\nTMhelix 332 354\ninside 355 372

6908 GCF_001563385.1_ASM156338v1 Escherichia coli Proteobacteria; Gammaproteobacteria; Enterobacterales; Enterobacteriaceae; Escherichia
MNNEETFYQAMRRQGVTRRSFLKYCSLAATSLGLGAGMAPKIAWALENKPRIPVVWIHGL WP_001058323.1
MULTISPECIES: hydrogenase-1 small chain [Proteobacteria] Length: 372\nNumber of predicted TMHs: 2\nExp number of AAs in TMHs: 37.2608499999999\nExp number, first 60 AAs: 19.52923\nTotal prob of N-in: 0.94351\nPOSSIBLE N-term signal sequence\ninside 1 19\nTMhelix 20 39\noutside 40 325\nTMhelix 326 348\ninside 349 372

6909 GCF_001566615.1_ASM156661v1 Escherichia coli Proteobacteria; Gammaproteobacteria; Enterobacterales; Enterobacteriaceae; Escherichia
MTGDNTLIHSHGINRRDFMKLCAALAATMGLSSKAAAEMAESVTNPQRPPVIWIGAQECT WP_000145410.1
MULTISPECIES: hydrogenase-2 small chain [Proteobacteria] Length: 372\nNumber of predicted TMHs: 1\nExp number of AAs in TMHs: 23.72226\nExp number, first 60 AAs: 0.22038\nTotal prob of N-in: 0.12994\noutside 1 331\nTMhelix 332 354\ninside 355 372

6910 GCF_001571595.1_ASM157159v1 Escherichia coli Proteobacteria; Gammaproteobacteria; Enterobacterales; Enterobacteriaceae; Escherichia
MTGDNTLIHSHGINRRDFMKLCAALAATMGLSSKAAAEMAESVTNPQRPPVIWIGAQECT WP_000145410.1
MULTISPECIES: hydrogenase-2 small chain [Proteobacteria] Length: 372\nNumber of predicted TMHs: 1\nExp number of AAs in TMHs: 23.72226\nExp number, first 60 AAs: 0.22038\nTotal prob of N-in: 0.12994\noutside 1 331\nTMhelix 332 354\ninside 355 372

6911 GCF_001571645.1_ASM157164v1 Escherichia coli Proteobacteria; Gammaproteobacteria; Enterobacterales; Enterobacteriaceae; Escherichia
MTGDNTLIHSHGINRRDFMKLCAALAATMGLSSKAAAEMAESVTNPQRPPVIWIGAQECT WP_000145418.1
hydrogenase 2 small subunit [Escherichia coli] Length: 372\nNumber of predicted TMHs: 1\nExp number of AAs in TMHs: 23.72313\nExp number, first 60 AAs: 0.22034\nTotal prob of N-in: 0.12997\noutside 1 331\nTMhelix 332 354\ninside 355 372

6912 GCF_001571495.1_ASM157149v1 Escherichia coli Proteobacteria; Gammaproteobacteria; Enterobacterales; Enterobacteriaceae; Escherichia
MNNEETFYQAMRRQGVTRRSFLKYCSLAATSLGLGAGMAPKIAWALENKPRIPVVWIHGL WP_001058323.1
MULTISPECIES: hydrogenase-1 small chain [Proteobacteria] Length: 372\nNumber of predicted TMHs: 2\nExp number of AAs in TMHs: 37.2608499999999\nExp number, first 60 AAs: 19.52923\nTotal prob of N-in:

0.94351\nPOSSIBLE N-term signal sequence\ninside 1 19\nTMhelix 20 39\noutside 40 325\nTMhelix 326 348\ninside 349 372

6913 GCF_001566615.1_ASM156661v1 Escherichia coli Proteobacteria; Gammaproteobacteria; Enterobacterales; Enterobacteriaceae; Escherichia
MNNEETFYQAMRRQGVTRRSFLKYCSLAATSLGLGAGMAPKIAWALENKPRIPVVWIHGL WP_001295940.1
MULTISPECIES: hydrogenase [Enterobacteriaceae] Length: 372\nNumber of predicted TMHs: 2\nExp number of AAs in TMHs: 37.2608899999999\nExp number, first 60 AAs: 19.52923\nTotal prob of N-in: 0.94351\nPOSSIBLE N-term signal sequence\ninside 1 19\nTMhelix 20 39\noutside 40 325\nTMhelix 326 348\ninside 349 372

6914 GCF_001571725.1_ASM157172v1 Escherichia coli Proteobacteria; Gammaproteobacteria; Enterobacterales; Enterobacteriaceae; Escherichia
MTGDNTLIHSHGINRRDFMKLCAALAATMGLSSKAAAEMAESVTNPQRPPVIWIGAQECT WP_000145410.1
MULTISPECIES: hydrogenase-2 small chain [Proteobacteria] Length: 372\nNumber of predicted TMHs: 1\nExp number of AAs in TMHs: 23.72226\nExp number, first 60 AAs: 0.22038\nTotal prob of N-in: 0.12994\noutside 1 331\nTMhelix 332 354\ninside 355 372

6915 GCF_001571745.1_ASM157174v1 Escherichia coli Proteobacteria; Gammaproteobacteria; Enterobacterales; Enterobacteriaceae; Escherichia
MTGDNTLIHSHGINRRDFMKLCAALAATMGLSSKAAAEMVESVTNPQRPPVIWIGAQECT WP_000145423.1
hydrogenase 2 small subunit [Escherichia coli] Length: 372\nNumber of predicted TMHs: 1\nExp number of AAs in TMHs: 23.78201\nExp number, first 60 AAs: 0.28355\nTotal prob of N-in: 0.13257\noutside 1 331\nTMhelix 332 354\ninside 355 372

6916 GCF_001571595.1_ASM157159v1 Escherichia coli Proteobacteria; Gammaproteobacteria; Enterobacterales; Enterobacteriaceae; Escherichia
MNNEETFYQAMRRQGVTRRSFLKYCSLAATSLGLGAGMAPKIAWALENKPRIPVVWIHGL WP_001058323.1
MULTISPECIES: hydrogenase-1 small chain [Proteobacteria] Length: 372\nNumber of predicted TMHs: 2\nExp number of AAs in TMHs: 37.2608499999999\nExp number, first 60 AAs: 19.52923\nTotal prob of N-in: 0.94351\nPOSSIBLE N-term signal sequence\ninside 1 19\nTMhelix 20 39\noutside 40 325\nTMhelix 326 348\ninside 349 372

6917 GCF_001571645.1_ASM157164v1 Escherichia coli Proteobacteria; Gammaproteobacteria; Enterobacterales; Enterobacteriaceae; Escherichia
MNNEETFYQAMRRQGVTRRSFLKYCSLAATSLGLGAGMAPKIAWALENKPRIPVVWIHGL WP_001058323.1
MULTISPECIES: hydrogenase-1 small chain [Proteobacteria] Length: 372\nNumber of predicted TMHs: 2\nExp number of AAs in TMHs: 37.2608499999999\nExp number, first 60 AAs: 19.52923\nTotal prob of N-in: 0.94351\nPOSSIBLE N-term signal sequence\ninside 1 19\nTMhelix 20 39\noutside 40 325\nTMhelix 326 348\ninside 349 372

6918 GCF_001571845.1_ASM157184v1 Escherichia coli Proteobacteria; Gammaproteobacteria; Enterobacterales; Enterobacteriaceae; Escherichia
MTGDNTLIHSHGINRRDFMKLCAALAATMGLSSKAAAEMAESVTNPQRPPVIWIGAQECT WP_000145410.1
MULTISPECIES: hydrogenase-2 small chain [Proteobacteria] Length: 372\nNumber of predicted TMHs: 1\nExp number of AAs in TMHs: 23.72226\nExp number, first 60 AAs: 0.22038\nTotal prob of N-in: 0.12994\noutside 1 331\nTMhelix 332 354\ninside 355 372

6919 GCF_001571905.1_ASM157190v1 Escherichia coli Proteobacteria; Gammaproteobacteria; Enterobacterales; Enterobacteriaceae; Escherichia
MTGDNTLIHSHGINRRDFMKLCAALAATMGLSSKAAAEMAESVTNPQRPPVIWIGAQECT WP_000145410.1
MULTISPECIES: hydrogenase-2 small chain [Proteobacteria] Length: 372\nNumber of predicted TMHs: 1\nExp number of AAs in TMHs: 23.72226\nExp number, first 60 AAs: 0.22038\nTotal prob of N-in: 0.12994\noutside 1 331\nTMhelix 332 354\ninside 355 372

6920 GCF_001571845.1_ASM157184v1 Escherichia coli Proteobacteria; Gammaproteobacteria; Enterobacterales; Enterobacteriaceae; Escherichia
MNNEETFYQAMRRQGVTRRSFLKYCSLAATSLGLGAGMAPKIAWALENKPRIPVVWIHGL WP_001058323.1
MULTISPECIES: hydrogenase-1 small chain [Proteobacteria] Length: 372\nNumber of predicted TMHs: 2\nExp number of AAs in TMHs: 37.2608499999999\nExp number, first 60 AAs: 19.52923\nTotal prob of N-in:

0.94351\nPOSSIBLE N-term signal sequence\ninside 1 19\nTMhelix 20 39\noutside 40 325\nTMhelix 326 348\ninside 349 372

6921 GCF_001571905.1_ASM157190v1 Escherichia coli Proteobacteria; Gammaproteobacteria; Enterobacterales; Enterobacteriaceae; Escherichia
MNNEETFYQAMRRQGVTRRSFLKYCSLAATSLGLGAGMAPKIAWALENKPRIPVVWIHGL WP_001058323.1
MULTISPECIES: hydrogenase-1 small chain [Proteobacteria] Length: 372\nNumber of predicted TMHs: 2\nExp number of AAs in TMHs: 37.2608499999999\nExp number, first 60 AAs: 19.52923\nTotal prob of N-in: 0.94351\nPOSSIBLE N-term signal sequence\ninside 1 19\nTMhelix 20 39\noutside 40 325\nTMhelix 326 348\ninside 349 372

6922 GCF_001571745.1_ASM157174v1 Escherichia coli Proteobacteria; Gammaproteobacteria; Enterobacterales; Enterobacteriaceae; Escherichia
MNNEETFYQAMRRQGVTRRSFLKYCSLAATSLGLGAGMAPKIAWALENKPRIPVVWIHGL WP_001538532.1
hydrogenase-1 small chain [Escherichia coli] Length: 372\nNumber of predicted TMHs: 2\nExp number of AAs in TMHs: 37.27041\nExp number, first 60 AAs: 19.52927\nTotal prob of N-in: 0.94352\nPOSSIBLE N-term signal sequence\ninside 1 19\nTMhelix 20 39\noutside 40 325\nTMhelix 326 348\ninside 349 372

6923 GCF_001571925.1_ASM157192v1 Escherichia coli Proteobacteria; Gammaproteobacteria; Enterobacterales; Enterobacteriaceae; Escherichia
MTGDNTLIHSHGINRRDFMKLCAALAATMGLSSKAAAEMAESVTNPQRPPVIWIGAQECT WP_000145410.1
MULTISPECIES: hydrogenase-2 small chain [Proteobacteria] Length: 372\nNumber of predicted TMHs: 1\nExp number of AAs in TMHs: 23.72226\nExp number, first 60 AAs: 0.22038\nTotal prob of N-in: 0.12994\noutside 1 331\nTMhelix 332 354\ninside 355 372

6924 GCF_001571965.1_ASM157196v1 Escherichia coli Proteobacteria; Gammaproteobacteria; Enterobacterales; Enterobacteriaceae; Escherichia
MTGDNTLIHSHGINRRDFMKLCAALAATMGLSSKAAAEMAESVTNPQRPPVIWIGAQECT WP_000145410.1
MULTISPECIES: hydrogenase-2 small chain [Proteobacteria] Length: 372\nNumber of predicted TMHs: 1\nExp number of AAs in TMHs: 23.72226\nExp number, first 60 AAs: 0.22038\nTotal prob of N-in: 0.12994\noutside 1 331\nTMhelix 332 354\ninside 355 372

6925 GCF_001571995.1_ASM157199v1 Escherichia coli Proteobacteria; Gammaproteobacteria; Enterobacterales; Enterobacteriaceae; Escherichia
MTGDNTLIHSHGINRRDFMKLCAALAATMGLSSKAAAEMAESVTNPQRPPVIWIGAQECT WP_000145410.1
MULTISPECIES: hydrogenase-2 small chain [Proteobacteria] Length: 372\nNumber of predicted TMHs: 1\nExp number of AAs in TMHs: 23.72226\nExp number, first 60 AAs: 0.22038\nTotal prob of N-in: 0.12994\noutside 1 331\nTMhelix 332 354\ninside 355 372

6926 GCF_001572045.1_ASM157204v1 Escherichia coli Proteobacteria; Gammaproteobacteria; Enterobacterales; Enterobacteriaceae; Escherichia
MTGDNTLIHSHGINRRDFMKLCAALAATMGLSSKAAAEMAESVTNPQRPPVIWIGAQECT WP_000145410.1
MULTISPECIES: hydrogenase-2 small chain [Proteobacteria] Length: 372\nNumber of predicted TMHs: 1\nExp number of AAs in TMHs: 23.72226\nExp number, first 60 AAs: 0.22038\nTotal prob of N-in: 0.12994\noutside 1 331\nTMhelix 332 354\ninside 355 372

6927 GCF_001572135.1_ASM157213v1 Escherichia coli Proteobacteria; Gammaproteobacteria; Enterobacterales; Enterobacteriaceae; Escherichia
MTGDNTLIHSHGINRRDFMKLCAALAATMGLSSKAAAEMAESVTNPQRPPVIWIGAQECT WP_000145410.1
MULTISPECIES: hydrogenase-2 small chain [Proteobacteria] Length: 372\nNumber of predicted TMHs: 1\nExp number of AAs in TMHs: 23.72226\nExp number, first 60 AAs: 0.22038\nTotal prob of N-in: 0.12994\noutside 1 331\nTMhelix 332 354\ninside 355 372

6928 GCF_001572185.1_ASM157218v1 Escherichia coli Proteobacteria; Gammaproteobacteria; Enterobacterales; Enterobacteriaceae; Escherichia
MTGDNTLIHSHGINRRDFMKLCAALAATMGLSSKAAAEMAESVTNPQRPPVIWIGAQECT WP_000145410.1
MULTISPECIES: hydrogenase-2 small chain [Proteobacteria] Length: 372\nNumber of predicted TMHs: 1\nExp number of AAs in TMHs: 23.72226\nExp number, first 60 AAs: 0.22038\nTotal prob of N-in: 0.12994\noutside 1 331\nTMhelix 332 354\ninside 355 372

6929 GCF_001572535.1_ASM157253v1 Escherichia coli Proteobacteria; Gammaproteobacteria; Enterobacterales; Enterobacteriaceae; Escherichia
MTGDNTLIHSHGINRRDFMKLCAALAATMGLSSKAAAEMAESVTNPQRPPVIWIGAQECT WP_000145410.1
MULTISPECIES: hydrogenase-2 small chain [Proteobacteria] Length: 372\nNumber of predicted TMHs: 1\nExp number of AAs in TMHs: 23.72226\nExp number, first 60 AAs: 0.22038\nTotal prob of N-in: 0.12994\noutside 1 331\nTMhelix 332 354\ninside 355 372

6930 GCF_001572135.1_ASM157213v1 Escherichia coli Proteobacteria; Gammaproteobacteria; Enterobacterales; Enterobacteriaceae; Escherichia
MNNEETFYQAMRRQGVTRRSFLKYCSLAATSLGLGAGMAPKIAWALENKPRIPVVWIHGL WP_001058323.1
MULTISPECIES: hydrogenase-1 small chain [Proteobacteria] Length: 372\nNumber of predicted TMHs: 2\nExp number of AAs in TMHs: 37.2608499999999\nExp number, first 60 AAs: 19.52923\nTotal prob of N-in: 0.94351\nPOSSIBLE N-term signal sequence\ninside 1 19\nTMhelix 20 39\noutside 40 325\nTMhelix 326 348\ninside 349 372

6931 GCF_001572185.1_ASM157218v1 Escherichia coli Proteobacteria; Gammaproteobacteria; Enterobacterales; Enterobacteriaceae; Escherichia
MNNEETFYQAMRRQGVTRRSFLKYCSLAATSLGLGAGMAPKIAWALENKPRIPVVWIHGL WP_001058323.1
MULTISPECIES: hydrogenase-1 small chain [Proteobacteria] Length: 372\nNumber of predicted TMHs: 2\nExp number of AAs in TMHs: 37.2608499999999\nExp number, first 60 AAs: 19.52923\nTotal prob of N-in: 0.94351\nPOSSIBLE N-term signal sequence\ninside 1 19\nTMhelix 20 39\noutside 40 325\nTMhelix 326 348\ninside 349 372

6932 GCF_001572375.1_ASM157237v1 Escherichia coli Proteobacteria; Gammaproteobacteria; Enterobacterales; Enterobacteriaceae; Escherichia
MTGDNTLIHSHGINRRDFMKLCAALAATMGLSSKAAAEMAESVTNPQRPPVIWIGAQECT WP_000145410.1
MULTISPECIES: hydrogenase-2 small chain [Proteobacteria] Length: 372\nNumber of predicted TMHs: 1\nExp number of AAs in TMHs: 23.72226\nExp number, first 60 AAs: 0.22038\nTotal prob of N-in: 0.12994\noutside 1 331\nTMhelix 332 354\ninside 355 372

6933 GCF_001572425.1_ASM157242v1 Escherichia coli Proteobacteria; Gammaproteobacteria; Enterobacterales; Enterobacteriaceae; Escherichia
MTGDNTLIHSHGINRRDFMKLCAALAATMGLSSKAAAEMAESVTNPQRPPVIWIGAQECT WP_000145410.1
MULTISPECIES: hydrogenase-2 small chain [Proteobacteria] Length: 372\nNumber of predicted TMHs: 1\nExp number of AAs in TMHs: 23.72226\nExp number, first 60 AAs: 0.22038\nTotal prob of N-in: 0.12994\noutside 1 331\nTMhelix 332 354\ninside 355 372

6934 GCF_001572445.1_ASM157244v1 Escherichia coli Proteobacteria; Gammaproteobacteria; Enterobacterales; Enterobacteriaceae; Escherichia
MTGDNTLIHSHGINRRDFMKLCAALAATMGLSSKAAAEMAESVTNPQRPPVIWIGAQECT WP_000145410.1
MULTISPECIES: hydrogenase-2 small chain [Proteobacteria] Length: 372\nNumber of predicted TMHs: 1\nExp number of AAs in TMHs: 23.72226\nExp number, first 60 AAs: 0.22038\nTotal prob of N-in: 0.12994\noutside 1 331\nTMhelix 332 354\ninside 355 372

6935 GCF_001572485.1_ASM157248v1 Escherichia coli Proteobacteria; Gammaproteobacteria; Enterobacterales; Enterobacteriaceae; Escherichia
MTGDNTLIHSHGINRRDFMKLCAALAATMGLSSKAAAEMAESVTNPQRPPVIWIGAQECT WP_000145410.1
MULTISPECIES: hydrogenase-2 small chain [Proteobacteria] Length: 372\nNumber of predicted TMHs: 1\nExp number of AAs in TMHs: 23.72226\nExp number, first 60 AAs: 0.22038\nTotal prob of N-in: 0.12994\noutside 1 331\nTMhelix 332 354\ninside 355 372

6936 GCF_001572375.1_ASM157237v1 Escherichia coli Proteobacteria; Gammaproteobacteria; Enterobacterales; Enterobacteriaceae; Escherichia
MNNEETFYQAMRRQGVTRRSFLKYCSLAATSLGLGAGMAPKIAWALENKPRIPVVWIHGL WP_001058323.1
MULTISPECIES: hydrogenase-1 small chain [Proteobacteria] Length: 372\nNumber of predicted TMHs: 2\nExp number of AAs in TMHs: 37.2608499999999\nExp number, first 60 AAs: 19.52923\nTotal prob of N-in: 0.94351\nPOSSIBLE N-term signal sequence\ninside 1 19\nTMhelix 20 39\noutside 40 325\nTMhelix 326 348\ninside 349 372

6937 GCF_001572635.1_ASM157263v1 Escherichia coli Proteobacteria; Gammaproteobacteria; Enterobacterales; Enterobacteriaceae; Escherichia

MNNEETFYQAMRRQGVTRRSFLKYCSLAATSLGLGAGMAPKIAWALENKPRIPVVWIHGL WP_001058323.1 MULTISPECIES: hydrogenase-1 small chain [Proteobacteria] Length: 372\nNumber of predicted TMHs: 2\nExp number of AAs in TMHs: 37.2608499999999\nExp number, first 60 AAs: 19.52923\nTotal prob of N-in: 0.94351\nPOSSIBLE N-term signal sequence\ninside 1 19\nTMhelix 20 39\noutside 40 325\nTMhelix 326 348\ninside 349 372

6938 GCF_001572445.1_ASM157244v1 Escherichia coli Proteobacteria; Gammaproteobacteria; Enterobacterales; Enterobacteriaceae; Escherichia MNNEETFYQAMRRQGVTRRSFLKYCSLAATSLGLGAGMAPKIAWALENKPRIPVVWIHGL WP_001058323.1 MULTISPECIES: hydrogenase-1 small chain [Proteobacteria] Length: 372\nNumber of predicted TMHs: 2\nExp number of AAs in TMHs: 37.2608499999999\nExp number, first 60 AAs: 19.52923\nTotal prob of N-in: 0.94351\nPOSSIBLE N-term signal sequence\ninside 1 19\nTMhelix 20 39\noutside 40 325\nTMhelix 326 348\ninside 349 372

6939 GCF_001575285.1_ASM157528v1 Escherichia coli Proteobacteria; Gammaproteobacteria; Enterobacterales; Enterobacteriaceae; Escherichia MTGDNTLIHSHGINRRDFMKLCAALAATMGLSSKAAAEMAESVTNPQRPPVIWIGAQECT WP_000145410.1 MULTISPECIES: hydrogenase-2 small chain [Proteobacteria] Length: 372\nNumber of predicted TMHs: 1\nExp number of AAs in TMHs: 23.72226\nExp number, first 60 AAs: 0.22038\nTotal prob of N-in: 0.12994\noutside 1 331\nTMhelix 332 354\ninside 355 372

6940 GCF_001572615.1_ASM157261v1 Escherichia coli Proteobacteria; Gammaproteobacteria; Enterobacterales; Enterobacteriaceae; Escherichia MTGDNTLIHSHGINRRDFMKLCAALAATMGLSSKAAAEMAESVTNPQRPPVIWIGAQECT WP_000145410.1 MULTISPECIES: hydrogenase-2 small chain [Proteobacteria] Length: 372\nNumber of predicted TMHs: 1\nExp number of AAs in TMHs: 23.72226\nExp number, first 60 AAs: 0.22038\nTotal prob of N-in: 0.12994\noutside 1 331\nTMhelix 332 354\ninside 355 372

6941 GCF_001572615.1_ASM157261v1 Escherichia coli Proteobacteria; Gammaproteobacteria; Enterobacterales; Enterobacteriaceae; Escherichia MNNEETFYQAMRRQGVTRRSFLKYCSLAATSLGLGAGMAPKIAWALENKPRIPVVWIHGL WP_001058323.1 MULTISPECIES: hydrogenase-1 small chain [Proteobacteria] Length: 372\nNumber of predicted TMHs: 2\nExp number of AAs in TMHs: 37.2608499999999\nExp number, first 60 AAs: 19.52923\nTotal prob of N-in: 0.94351\nPOSSIBLE N-term signal sequence\ninside 1 19\nTMhelix 20 39\noutside 40 325\nTMhelix 326 348\ninside 349 372

6942 GCF_001575285.1_ASM157528v1 Escherichia coli Proteobacteria; Gammaproteobacteria; Enterobacterales; Enterobacteriaceae; Escherichia MNNEETFYQAMRRQGVTRRSFLKYCSLAATSLGLGAGMAPKIAWALENKPRIPVVWIHGL WP_001058323.1 MULTISPECIES: hydrogenase-1 small chain [Proteobacteria] Length: 372\nNumber of predicted TMHs: 2\nExp number of AAs in TMHs: 37.2608499999999\nExp number, first 60 AAs: 19.52923\nTotal prob of N-in: 0.94351\nPOSSIBLE N-term signal sequence\ninside 1 19\nTMhelix 20 39\noutside 40 325\nTMhelix 326 348\ninside 349 372

6943 GCF_001575395.1_ASM157539v1 Escherichia coli Proteobacteria; Gammaproteobacteria; Enterobacterales; Enterobacteriaceae; Escherichia MNNEETFYQAMRRQGVTRRSFLKYCSLAATSLGLGAGMAPKIAWALENKPRIPVVWIHGL WP_001058323.1 MULTISPECIES: hydrogenase-1 small chain [Proteobacteria] Length: 372\nNumber of predicted TMHs: 2\nExp number of AAs in TMHs: 37.2608499999999\nExp number, first 60 AAs: 19.52923\nTotal prob of N-in: 0.94351\nPOSSIBLE N-term signal sequence\ninside 1 19\nTMhelix 20 39\noutside 40 325\nTMhelix 326 348\ninside 349 372

6944 GCF_001575375.1_ASM157537v1 Escherichia coli Proteobacteria; Gammaproteobacteria; Enterobacterales; Enterobacteriaceae; Escherichia MNNEETFYQAMRRQGVTRRSFLKYCSLAATSLGLGAGMAPKIAWALENKPRIPVVWIHGL WP_001058323.1 MULTISPECIES: hydrogenase-1 small chain [Proteobacteria] Length: 372\nNumber of predicted TMHs: 2\nExp number of AAs in TMHs: 37.2608499999999\nExp number, first 60 AAs: 19.52923\nTotal prob of N-in: 0.94351\nPOSSIBLE N-term signal sequence\ninside 1 19\nTMhelix 20 39\noutside 40 325\nTMhelix 326 348\ninside 349 372

6945 GCF_001575455.1_ASM157545v1 Escherichia coli Proteobacteria; Gammaproteobacteria; Enterobacterales; Enterobacteriaceae; Escherichia
MNNEETFYQAMRRQGVTRRSFLKYCSLAATSLGLGAGMAPKIAWALENKPRIPVVWIHGL WP_001058323.1
MULTISPECIES: hydrogenase-1 small chain [Proteobacteria] Length: 372\nNumber of predicted TMHs: 2\nExp number of AAs in TMHs: 37.2608499999999\nExp number, first 60 AAs: 19.52923\nTotal prob of N-in: 0.94351\nPOSSIBLE N-term signal sequence\ninside 1 19\nTMhelix 20 39\noutside 40 325\nTMhelix 326 348\ninside 349 372

6946 GCF_001575545.1_ASM157554v1 Escherichia coli Proteobacteria; Gammaproteobacteria; Enterobacterales; Enterobacteriaceae; Escherichia
MTGDNTLIHSHGINRRDFMKLCAALAATMGLSSKAAAEMAESVTNPQRPPVIWIGAQECT WP_000145410.1
MULTISPECIES: hydrogenase-2 small chain [Proteobacteria] Length: 372\nNumber of predicted TMHs: 1\nExp number of AAs in TMHs: 23.72226\nExp number, first 60 AAs: 0.22038\nTotal prob of N-in: 0.12994\noutside 1 331\nTMhelix 332 354\ninside 355 372

6947 GCF_001575555.1_ASM157555v1 Escherichia coli Proteobacteria; Gammaproteobacteria; Enterobacterales; Enterobacteriaceae; Escherichia
MTGDNTLIHSHGINRRDFMKLCAALAATMGLSSKAAAEMAESVTNPQRPPVIWIGAQECT WP_000145410.1
MULTISPECIES: hydrogenase-2 small chain [Proteobacteria] Length: 372\nNumber of predicted TMHs: 1\nExp number of AAs in TMHs: 23.72226\nExp number, first 60 AAs: 0.22038\nTotal prob of N-in: 0.12994\noutside 1 331\nTMhelix 332 354\ninside 355 372

6948 GCF_001575675.1_ASM157567v1 Escherichia coli Proteobacteria; Gammaproteobacteria; Enterobacterales; Enterobacteriaceae; Escherichia
MTGDNTLIHSHGINRRDFMKLCAALAATMGLSSKAAAEMAESVTNPQRPPVIWIGAQECT WP_000145410.1
MULTISPECIES: hydrogenase-2 small chain [Proteobacteria] Length: 372\nNumber of predicted TMHs: 1\nExp number of AAs in TMHs: 23.72226\nExp number, first 60 AAs: 0.22038\nTotal prob of N-in: 0.12994\noutside 1 331\nTMhelix 332 354\ninside 355 372

6949 GCF_001575615.1_ASM157561v1 Escherichia coli Proteobacteria; Gammaproteobacteria; Enterobacterales; Enterobacteriaceae; Escherichia
MTGDNTLIHSHGINRRDFMKLCAALAATMGLSSKAAAEMAESVTNPQRPPVIWIGAQECT WP_000145410.1
MULTISPECIES: hydrogenase-2 small chain [Proteobacteria] Length: 372\nNumber of predicted TMHs: 1\nExp number of AAs in TMHs: 23.72226\nExp number, first 60 AAs: 0.22038\nTotal prob of N-in: 0.12994\noutside 1 331\nTMhelix 332 354\ninside 355 372

6950 GCF_001575635.1_ASM157563v1 Escherichia coli Proteobacteria; Gammaproteobacteria; Enterobacterales; Enterobacteriaceae; Escherichia
MNNEETFYQAMRRQGVTRRSFLKYCSLAATSLGLGAGMAPKIAWALENKPRIPVVWIHGL WP_001058323.1
MULTISPECIES: hydrogenase-1 small chain [Proteobacteria] Length: 372\nNumber of predicted TMHs: 2\nExp number of AAs in TMHs: 37.2608499999999\nExp number, first 60 AAs: 19.52923\nTotal prob of N-in: 0.94351\nPOSSIBLE N-term signal sequence\ninside 1 19\nTMhelix 20 39\noutside 40 325\nTMhelix 326 348\ninside 349 372

6951 GCF_001575555.1_ASM157555v1 Escherichia coli Proteobacteria; Gammaproteobacteria; Enterobacterales; Enterobacteriaceae; Escherichia
MNNEETFYQAMRRQGVTRRSFLKYCSLAATSLGLGAGMAPKIAWALENKPRIPVVWIHGL WP_001058323.1
MULTISPECIES: hydrogenase-1 small chain [Proteobacteria] Length: 372\nNumber of predicted TMHs: 2\nExp number of AAs in TMHs: 37.2608499999999\nExp number, first 60 AAs: 19.52923\nTotal prob of N-in: 0.94351\nPOSSIBLE N-term signal sequence\ninside 1 19\nTMhelix 20 39\noutside 40 325\nTMhelix 326 348\ninside 349 372

6952 GCF_001575775.1_ASM157577v1 Escherichia coli Proteobacteria; Gammaproteobacteria; Enterobacterales; Enterobacteriaceae; Escherichia
MTGDNTLIHSHGINRRDFMKLCAALAATMGLSSKAAAEMAESVTNPQRPPVIWIGAQECT WP_000145410.1
MULTISPECIES: hydrogenase-2 small chain [Proteobacteria] Length: 372\nNumber of predicted TMHs: 1\nExp number of AAs in TMHs: 23.72226\nExp number, first 60 AAs: 0.22038\nTotal prob of N-in: 0.12994\noutside 1 331\nTMhelix 332 354\ninside 355 372

6953 GCF_001575755.1_ASM157575v1 Escherichia coli Proteobacteria; Gammaproteobacteria; Enterobacterales; Enterobacteriaceae; Escherichia

MNNEETFYQAMRRQGVTRRSFLKYCSLAATSLGLGAGMAPKIAWALENKPRIPVVWIHGL WP_001058323.1 MULTISPECIES: hydrogenase-1 small chain [Proteobacteria] Length: 372\nNumber of predicted TMHs: 2\nExp number of AAs in TMHs: 37.2608499999999\nExp number, first 60 AAs: 19.52923\nTotal prob of N-in: 0.94351\nPOSSIBLE N-term signal sequence\ninside 1 19\nTMhelix 20 39\noutside 40 325\nTMhelix 326 348\ninside 349 372

6954 GCF_001575705.1_ASM157570v1 Escherichia coli Proteobacteria; Gammaproteobacteria; Enterobacterales; Enterobacteriaceae; Escherichia MNNEETFYQAMRRQGVTRRSFLKYCSLAATSLGLGAGMAPKIAWALENKPRIPVVWIHGL WP_001058323.1 MULTISPECIES: hydrogenase-1 small chain [Proteobacteria] Length: 372\nNumber of predicted TMHs: 2\nExp number of AAs in TMHs: 37.2608499999999\nExp number, first 60 AAs: 19.52923\nTotal prob of N-in: 0.94351\nPOSSIBLE N-term signal sequence\ninside 1 19\nTMhelix 20 39\noutside 40 325\nTMhelix 326 348\ninside 349 372

6955 GCF_001575795.1_ASM157579v1 Escherichia coli Proteobacteria; Gammaproteobacteria; Enterobacterales; Enterobacteriaceae; Escherichia MTGDNTLIHSHGINRRDFMKLCAALAATMGLSSKAAAEMAESVTNPQRPPVIWIGAQECT WP_000145410.1 MULTISPECIES: hydrogenase-2 small chain [Proteobacteria] Length: 372\nNumber of predicted TMHs: 1\nExp number of AAs in TMHs: 23.72226\nExp number, first 60 AAs: 0.22038\nTotal prob of N-in: 0.12994\noutside 1 331\nTMhelix 332 354\ninside 355 372

6956 GCF_001575855.1_ASM157585v1 Escherichia coli Proteobacteria; Gammaproteobacteria; Enterobacterales; Enterobacteriaceae; Escherichia MTGDNTLIHSHGINRRDFMKLCAALAATMGLSSKAAAEMAESVTNPQRPPVIWIGAQECT WP_000145410.1 MULTISPECIES: hydrogenase-2 small chain [Proteobacteria] Length: 372\nNumber of predicted TMHs: 1\nExp number of AAs in TMHs: 23.72226\nExp number, first 60 AAs: 0.22038\nTotal prob of N-in: 0.12994\noutside 1 331\nTMhelix 332 354\ninside 355 372

6957 GCF_001575945.1_ASM157594v1 Escherichia coli Proteobacteria; Gammaproteobacteria; Enterobacterales; Enterobacteriaceae; Escherichia MTGDNTLIHSHGINRRDFMKLCAALAATMGLSSKAAAEMAESVTNPQRPPVIWIGAQECT WP_000145410.1 MULTISPECIES: hydrogenase-2 small chain [Proteobacteria] Length: 372\nNumber of predicted TMHs: 1\nExp number of AAs in TMHs: 23.72226\nExp number, first 60 AAs: 0.22038\nTotal prob of N-in: 0.12994\noutside 1 331\nTMhelix 332 354\ninside 355 372

6958 GCF_001576035.1_ASM157603v1 Escherichia coli Proteobacteria; Gammaproteobacteria; Enterobacterales; Enterobacteriaceae; Escherichia MTGDNTLIHSHGINRRDFMKLCAALAATMGLSSKAAAEMAESVTNPQRPPVIWIGAQECT WP_000145410.1 MULTISPECIES: hydrogenase-2 small chain [Proteobacteria] Length: 372\nNumber of predicted TMHs: 1\nExp number of AAs in TMHs: 23.72226\nExp number, first 60 AAs: 0.22038\nTotal prob of N-in: 0.12994\noutside 1 331\nTMhelix 332 354\ninside 355 372

6959 GCF_001576015.1_ASM157601v1 Escherichia coli Proteobacteria; Gammaproteobacteria; Enterobacterales; Enterobacteriaceae; Escherichia MTGDNTLIHSHGINRRDFMKLCAALAATMGLSSKAAAEMAESVTNPQRPPVIWIGAQECT WP_000145410.1 MULTISPECIES: hydrogenase-2 small chain [Proteobacteria] Length: 372\nNumber of predicted TMHs: 1\nExp number of AAs in TMHs: 23.72226\nExp number, first 60 AAs: 0.22038\nTotal prob of N-in: 0.12994\noutside 1 331\nTMhelix 332 354\ninside 355 372

6960 GCF_001576015.1_ASM157601v1 Escherichia coli Proteobacteria; Gammaproteobacteria; Enterobacterales; Enterobacteriaceae; Escherichia MNNEETFYQAMRRQGVTRRSFLKYCSLAATSLGLGAGMAPKIAWALENKPRIPVVWIHGL WP_001058323.1 MULTISPECIES: hydrogenase-1 small chain [Proteobacteria] Length: 372\nNumber of predicted TMHs: 2\nExp number of AAs in TMHs: 37.2608499999999\nExp number, first 60 AAs: 19.52923\nTotal prob of N-in: 0.94351\nPOSSIBLE N-term signal sequence\ninside 1 19\nTMhelix 20 39\noutside 40 325\nTMhelix 326 348\ninside 349 372

6961 GCF_001575915.1_ASM157591v1 Escherichia coli Proteobacteria; Gammaproteobacteria; Enterobacterales; Enterobacteriaceae; Escherichia MNNEETFYQAMRRQGVTRRSFLKYCSLAATSLGLGAGMAPKIAWALENKPRIPVVWIHGL WP_001058323.1 MULTISPECIES: hydrogenase-1 small chain [Proteobacteria] Length: 372\nNumber of predicted TMHs:

2\nExp number of AAs in TMHs: 37.2608499999999\nExp number, first 60 AAs: 19.52923\nTotal prob of N-in: 0.94351\nPOSSIBLE N-term signal sequence\ninside 1 19\nTMhelix 20 39\noutside 40 325\nTMhelix 326 348\ninside 349 372

6962 GCF_001576105.1_ASM157610v1 Escherichia coli Proteobacteria; Gammaproteobacteria; Enterobacterales; Enterobacteriaceae; Escherichia MTGDNTLIHSHGINRRDFMKLCAALAATMGLSSKAAAEMAESVTNPQRPPVIWIGAQECT WP_000145410.1 MULTISPECIES: hydrogenase-2 small chain [Proteobacteria] Length: 372\nNumber of predicted TMHs: 1\nExp number of AAs in TMHs: 23.72226\nExp number, first 60 AAs: 0.22038\nTotal prob of N-in: 0.12994\noutside 1 331\nTMhelix 332 354\ninside 355 372

6963 GCF_001576195.1_ASM157619v1 Escherichia coli Proteobacteria; Gammaproteobacteria; Enterobacterales; Enterobacteriaceae; Escherichia MTGDNTLIHSHGINRRDFMKLCAALAATMGLSSKAAAEMAESVTNPQRPPVIWIGAQECT WP_000145410.1 MULTISPECIES: hydrogenase-2 small chain [Proteobacteria] Length: 372\nNumber of predicted TMHs: 1\nExp number of AAs in TMHs: 23.72226\nExp number, first 60 AAs: 0.22038\nTotal prob of N-in: 0.12994\noutside 1 331\nTMhelix 332 354\ninside 355 372

6964 GCF_001576295.1_ASM157629v1 Escherichia coli Proteobacteria; Gammaproteobacteria; Enterobacterales; Enterobacteriaceae; Escherichia MTGDNTLIHSHGINRRDFMKLCAALAATMGLSSKAAAEMAESVTNPQRPPVIWIGAQECT WP_000145410.1 MULTISPECIES: hydrogenase-2 small chain [Proteobacteria] Length: 372\nNumber of predicted TMHs: 1\nExp number of AAs in TMHs: 23.72226\nExp number, first 60 AAs: 0.22038\nTotal prob of N-in: 0.12994\noutside 1 331\nTMhelix 332 354\ninside 355 372

6965 GCF_001576195.1_ASM157619v1 Escherichia coli Proteobacteria; Gammaproteobacteria; Enterobacterales; Enterobacteriaceae; Escherichia MNNEETFYQAMRRQGVTRRSFLKYCSLAATSLGLGAGMAPKIAWALENKPRIPVVWIHGL WP_001058323.1 MULTISPECIES: hydrogenase-1 small chain [Proteobacteria] Length: 372\nNumber of predicted TMHs: 2\nExp number of AAs in TMHs: 37.2608499999999\nExp number, first 60 AAs: 19.52923\nTotal prob of N-in: 0.94351\nPOSSIBLE N-term signal sequence\ninside 1 19\nTMhelix 20 39\noutside 40 325\nTMhelix 326 348\ninside 349 372

6966 GCF_001576375.1_ASM157637v1 Escherichia coli Proteobacteria; Gammaproteobacteria; Enterobacterales; Enterobacteriaceae; Escherichia MTGDNTLIHSHGINRRDFMKLCAALAATMGLSSKAAAEMAESVTNPQRPPVIWIGAQECT WP_000145410.1 MULTISPECIES: hydrogenase-2 small chain [Proteobacteria] Length: 372\nNumber of predicted TMHs: 1\nExp number of AAs in TMHs: 23.72226\nExp number, first 60 AAs: 0.22038\nTotal prob of N-in: 0.12994\noutside 1 331\nTMhelix 332 354\ninside 355 372

6967 GCF_001576315.1_ASM157631v1 Escherichia coli Proteobacteria; Gammaproteobacteria; Enterobacterales; Enterobacteriaceae; Escherichia MTGDNTLIHSHGINRRDFMKLCAALAATMGLSSKAAAEMAESVTNPQRPPVIWIGAQECT WP_000145410.1 MULTISPECIES: hydrogenase-2 small chain [Proteobacteria] Length: 372\nNumber of predicted TMHs: 1\nExp number of AAs in TMHs: 23.72226\nExp number, first 60 AAs: 0.22038\nTotal prob of N-in: 0.12994\noutside 1 331\nTMhelix 332 354\ninside 355 372

6968 GCF_001576235.1_ASM157623v1 Escherichia coli Proteobacteria; Gammaproteobacteria; Enterobacterales; Enterobacteriaceae; Escherichia MNNEETFYQAMRRQGVTRRSFLKYCSLAATSLGLGAGMAPKIAWALENKPRIPVVWIHGL WP_001058323.1 MULTISPECIES: hydrogenase-1 small chain [Proteobacteria] Length: 372\nNumber of predicted TMHs: 2\nExp number of AAs in TMHs: 37.2608499999999\nExp number, first 60 AAs: 19.52923\nTotal prob of N-in: 0.94351\nPOSSIBLE N-term signal sequence\ninside 1 19\nTMhelix 20 39\noutside 40 325\nTMhelix 326 348\ninside 349 372

6969 GCF_001576465.1_ASM157646v1 Escherichia coli Proteobacteria; Gammaproteobacteria; Enterobacterales; Enterobacteriaceae; Escherichia MTGDNTLIHSHGINRRDFMKLCAALAATMGLSSKAAAEMAESVTNPQRPPVIWIGAQECT WP_000145410.1 MULTISPECIES: hydrogenase-2 small chain [Proteobacteria] Length: 372\nNumber of predicted TMHs: 1\nExp number of AAs in TMHs: 23.72226\nExp number, first 60 AAs: 0.22038\nTotal prob of N-in: 0.12994\noutside 1 331\nTMhelix 332 354\ninside 355 372

6970 GCF_001576385.1_ASM157638v1 Escherichia coli Proteobacteria; Gammaproteobacteria; Enterobacterales; Enterobacteriaceae; Escherichia
MTGDNTLIHSHGINRRDFMKLCAALAATMGLSSKAAAEMAESVTNPQRPPVIWIGAQECT WP_000145410.1
MULTISPECIES: hydrogenase-2 small chain [Proteobacteria] Length: 372\nNumber of predicted TMHs: 1\nExp number of AAs in TMHs: 23.72226\nExp number, first 60 AAs: 0.22038\nTotal prob of N-in: 0.12994\noutside 1 331\nTMhelix 332 354\ninside 355 372

6971 GCF_001576435.1_ASM157643v1 Escherichia coli Proteobacteria; Gammaproteobacteria; Enterobacterales; Enterobacteriaceae; Escherichia
MTGDNTLIHSHGINRRDFMKLCAALAATMGLSSKAAAEMAESVTNPQRPPVIWIGAQECT WP_032298447.1
hydrogenase 2 small subunit [Escherichia coli] Length: 372\nNumber of predicted TMHs: 1\nExp number of AAs in TMHs: 23.69807\nExp number, first 60 AAs: 0.221\nTotal prob of N-in: 0.12853\noutside 1 331\nTMhelix 332 354\ninside 355 372

6972 GCF_001576295.1_ASM157629v1 Escherichia coli Proteobacteria; Gammaproteobacteria; Enterobacterales; Enterobacteriaceae; Escherichia
MNNEETFYQAMRRQGVTRRSFLKYCSLAATSLGLGAGMAPKIAWALENKPRIPVVWIHGL WP_061348854.1
hydrogenase [Escherichia coli] Length: 372\nNumber of predicted TMHs: 2\nExp number of AAs in TMHs: 37.2608499999999\nExp number, first 60 AAs: 19.52923\nTotal prob of N-in: 0.94351\nPOSSIBLE N-term signal sequence\ninside 1 19\nTMhelix 20 39\noutside 40 325\nTMhelix 326 348\ninside 349 372

6973 GCF_001576495.1_ASM157649v1 Escherichia coli Proteobacteria; Gammaproteobacteria; Enterobacterales; Enterobacteriaceae; Escherichia
MNNEETFYQAMRRQGVTRRSFLKYCSLAATSLGLGAGMAPKIAWALENKPRIPVVWIHGL WP_001058323.1
MULTISPECIES: hydrogenase-1 small chain [Proteobacteria] Length: 372\nNumber of predicted TMHs: 2\nExp number of AAs in TMHs: 37.2608499999999\nExp number, first 60 AAs: 19.52923\nTotal prob of N-in: 0.94351\nPOSSIBLE N-term signal sequence\ninside 1 19\nTMhelix 20 39\noutside 40 325\nTMhelix 326 348\ninside 349 372

6974 GCF_001596865.1_ASM159686v1 Escherichia coli Proteobacteria; Gammaproteobacteria; Enterobacterales; Enterobacteriaceae; Escherichia
MTGDNTLIHSHGINRRDFMKLCAALAATMGLSSKAAAEMAESVTNPQRPPVIWIGAQECT WP_000145410.1
MULTISPECIES: hydrogenase-2 small chain [Proteobacteria] Length: 372\nNumber of predicted TMHs: 1\nExp number of AAs in TMHs: 23.72226\nExp number, first 60 AAs: 0.22038\nTotal prob of N-in: 0.12994\noutside 1 331\nTMhelix 332 354\ninside 355 372

6975 GCF_001606435.1_ASM160643v1 Escherichia coli Proteobacteria; Gammaproteobacteria; Enterobacterales; Enterobacteriaceae; Escherichia
MNNEETFYQAMRRQGVTRRSFLKYCSLAATSLGLGAGMAPKIAWALENKPRIPVVWIHGL WP_001058323.1
MULTISPECIES: hydrogenase-1 small chain [Proteobacteria] Length: 372\nNumber of predicted TMHs: 2\nExp number of AAs in TMHs: 37.2608499999999\nExp number, first 60 AAs: 19.52923\nTotal prob of N-in: 0.94351\nPOSSIBLE N-term signal sequence\ninside 1 19\nTMhelix 20 39\noutside 40 325\nTMhelix 326 348\ninside 349 372

6976 GCF_001606375.1_ASM160637v1 Escherichia coli Proteobacteria; Gammaproteobacteria; Enterobacterales; Enterobacteriaceae; Escherichia
MNNEETFYQAMRRQGVTRRSFLKYCSLAATSLGLGAGMAPKIAWALENKPRIPVVWIHGL WP_001058323.1
MULTISPECIES: hydrogenase-1 small chain [Proteobacteria] Length: 372\nNumber of predicted TMHs: 2\nExp number of AAs in TMHs: 37.2608499999999\nExp number, first 60 AAs: 19.52923\nTotal prob of N-in: 0.94351\nPOSSIBLE N-term signal sequence\ninside 1 19\nTMhelix 20 39\noutside 40 325\nTMhelix 326 348\ninside 349 372

6977 GCF_001596865.1_ASM159686v1 Escherichia coli Proteobacteria; Gammaproteobacteria; Enterobacterales; Enterobacteriaceae; Escherichia
MNNEETFYQAMRRQGVTRRSFLKYCSLAATSLGLGAGMAPKIAWALENKPRIPVVWIHGL WP_001295940.1
MULTISPECIES: hydrogenase [Enterobacteriaceae] Length: 372\nNumber of predicted TMHs: 2\nExp number of AAs in TMHs: 37.2608899999999\nExp number, first 60 AAs: 19.52923\nTotal prob of N-in: 0.94351\nPOSSIBLE N-term signal sequence\ninside 1 19\nTMhelix 20 39\noutside 40 325\nTMhelix 326 348\ninside 349 372

6978 GCF_001606595.1_ASM160659v1 Escherichia coli Proteobacteria; Gammaproteobacteria; Enterobacterales; Enterobacteriaceae; Escherichia
MTGDNTLIHSHGINRRDFMKLCAALAATMGLSSKAAAEMAESVTNPQRPPVIWIGAQECT WP_000145410.1
MULTISPECIES: hydrogenase-2 small chain [Proteobacteria] Length: 372\nNumber of predicted TMHs: 1\nExp number of AAs in TMHs: 23.72226\nExp number, first 60 AAs: 0.22038\nTotal prob of N-in: 0.12994\noutside 1 331\nTMhelix 332 354\ninside 355 372

6979 GCF_001606525.1_ASM160652v1 Escherichia coli Proteobacteria; Gammaproteobacteria; Enterobacterales; Enterobacteriaceae; Escherichia
MNNEETFYQAMRRQGVTRRSFLKYCSLAATSLGLGAGMAPKIAWALENKPRIPVVWIHGL WP_001058323.1
MULTISPECIES: hydrogenase-1 small chain [Proteobacteria] Length: 372\nNumber of predicted TMHs: 2\nExp number of AAs in TMHs: 37.2608499999999\nExp number, first 60 AAs: 19.52923\nTotal prob of N-in: 0.94351\nPOSSIBLE N-term signal sequence\ninside 1 19\nTMhelix 20 39\noutside 40 325\nTMhelix 326 348\ninside 349 372

6980 GCF_001606675.1_ASM160667v1 Escherichia coli Proteobacteria; Gammaproteobacteria; Enterobacterales; Enterobacteriaceae; Escherichia
MNNEETFYQAMRRQGVTRRSFLKYCSLAATSLGLGAGMAPKIAWALENKPRIPVVWIHGL WP_001058323.1
MULTISPECIES: hydrogenase-1 small chain [Proteobacteria] Length: 372\nNumber of predicted TMHs: 2\nExp number of AAs in TMHs: 37.2608499999999\nExp number, first 60 AAs: 19.52923\nTotal prob of N-in: 0.94351\nPOSSIBLE N-term signal sequence\ninside 1 19\nTMhelix 20 39\noutside 40 325\nTMhelix 326 348\ninside 349 372

6981 GCF_001606625.1_ASM160662v1 Escherichia coli Proteobacteria; Gammaproteobacteria; Enterobacterales; Enterobacteriaceae; Escherichia
MNNEETFYQAMRRQGVTRRSFLKYCSLAATSLGLGAGMAPKIAWALENKPRIPVVWIHGL WP_001295940.1
MULTISPECIES: hydrogenase [Enterobacteriaceae] Length: 372\nNumber of predicted TMHs: 2\nExp number of AAs in TMHs: 37.2608899999999\nExp number, first 60 AAs: 19.52923\nTotal prob of N-in: 0.94351\nPOSSIBLE N-term signal sequence\ninside 1 19\nTMhelix 20 39\noutside 40 325\nTMhelix 326 348\ninside 349 372

6982 GCF_001606475.1_ASM160647v1 Escherichia coli Proteobacteria; Gammaproteobacteria; Enterobacterales; Enterobacteriaceae; Escherichia
MTGDNTLIHSHGINRRDFMKLCAALAATMGLSSKAAAEMAESVTNPQRPPVIWIGAQECT WP_024245965.1
hydrogenase 2 small subunit [Escherichia coli] Length: 372\nNumber of predicted TMHs: 1\nExp number of AAs in TMHs: 24.15412\nExp number, first 60 AAs: 0.21967\nTotal prob of N-in: 0.14289\noutside 1 331\nTMhelix 332 354\ninside 355 372

6983 GCF_001606695.1_ASM160669v1 Escherichia coli Proteobacteria; Gammaproteobacteria; Enterobacterales; Enterobacteriaceae; Escherichia
MTGDNTLIHSHGINRRDFMKLCAALAATMGLSSKAAAEMAESVTNPQRPPVIWIGAQECT WP_000145410.1
MULTISPECIES: hydrogenase-2 small chain [Proteobacteria] Length: 372\nNumber of predicted TMHs: 1\nExp number of AAs in TMHs: 23.72226\nExp number, first 60 AAs: 0.22038\nTotal prob of N-in: 0.12994\noutside 1 331\nTMhelix 332 354\ninside 355 372

6984 GCF_001606775.1_ASM160677v1 Escherichia coli Proteobacteria; Gammaproteobacteria; Enterobacterales; Enterobacteriaceae; Escherichia
MTGDNTLIHSHGINRRDFMKLCAALAATMGLSSKAAAEMAESVTNPQRPPVIWIGAQECT WP_000145410.1
MULTISPECIES: hydrogenase-2 small chain [Proteobacteria] Length: 372\nNumber of predicted TMHs: 1\nExp number of AAs in TMHs: 23.72226\nExp number, first 60 AAs: 0.22038\nTotal prob of N-in: 0.12994\noutside 1 331\nTMhelix 332 354\ninside 355 372

6985 GCF_001606725.1_ASM160672v1 Escherichia coli Proteobacteria; Gammaproteobacteria; Enterobacterales; Enterobacteriaceae; Escherichia
MTGDNTLIHSHGINRRDFMKLCAALAATMGLSSKAAAEMAESVTNPQRPPVIWIGAQECT WP_000145410.1
MULTISPECIES: hydrogenase-2 small chain [Proteobacteria] Length: 372\nNumber of predicted TMHs: 1\nExp number of AAs in TMHs: 23.72226\nExp number, first 60 AAs: 0.22038\nTotal prob of N-in: 0.12994\noutside 1 331\nTMhelix 332 354\ninside 355 372

6986 GCF_001606895.1_ASM160689v1 Escherichia coli Proteobacteria; Gammaproteobacteria; Enterobacterales; Enterobacteriaceae; Escherichia

MTGDNTLIHSHGINRRDFMKLCAALAATMGLSSKAAAEMAESVTNPQRPPVIWIGAQECT WP_000145410.1 MULTISPECIES: hydrogenase-2 small chain [Proteobacteria] Length: 372\nNumber of predicted TMHs: 1\nExp number of AAs in TMHs: 23.72226\nExp number, first 60 AAs: 0.22038\nTotal prob of N-in: 0.12994\noutside 1 331\nTMhelix 332 354\ninside 355 372

6987 GCF_001606865.1_ASM160686v1 Escherichia coli Proteobacteria; Gammaproteobacteria; Enterobacterales; Enterobacteriaceae; Escherichia MTGDNTLIHSHGINRRDFMKLCAALAATMGLSSKAAAEMAESVTNPQRPPVIWIGAQECT WP_000145410.1 MULTISPECIES: hydrogenase-2 small chain [Proteobacteria] Length: 372\nNumber of predicted TMHs: 1\nExp number of AAs in TMHs: 23.72226\nExp number, first 60 AAs: 0.22038\nTotal prob of N-in: 0.12994\noutside 1 331\nTMhelix 332 354\ninside 355 372

6988 GCF_001606935.1_ASM160693v1 Escherichia coli Proteobacteria; Gammaproteobacteria; Enterobacterales; Enterobacteriaceae; Escherichia MTGDNTLIHSHGINRRDFMKLCAALAATMGLSSKAAAEMAESVTNPQRPPVIWIGAQECT WP_000145410.1 MULTISPECIES: hydrogenase-2 small chain [Proteobacteria] Length: 372\nNumber of predicted TMHs: 1\nExp number of AAs in TMHs: 23.72226\nExp number, first 60 AAs: 0.22038\nTotal prob of N-in: 0.12994\noutside 1 331\nTMhelix 332 354\ninside 355 372

6989 GCF_001606785.1_ASM160678v1 Escherichia coli Proteobacteria; Gammaproteobacteria; Enterobacterales; Enterobacteriaceae; Escherichia MTGDNTLIHSHGINRRDFMKLCAALAATMGLSSKAAAEMAESVTNPQRPPVIWIGAQECT WP_000145410.1 MULTISPECIES: hydrogenase-2 small chain [Proteobacteria] Length: 372\nNumber of predicted TMHs: 1\nExp number of AAs in TMHs: 23.72226\nExp number, first 60 AAs: 0.22038\nTotal prob of N-in: 0.12994\noutside 1 331\nTMhelix 332 354\ninside 355 372

6990 GCF_001606835.1_ASM160683v1 Escherichia coli Proteobacteria; Gammaproteobacteria; Enterobacterales; Enterobacteriaceae; Escherichia MNNEETFYQAMRRQGVTRRSFLKYCSLAATSLGLGAGMAPKIAWALENKPRIPVVWIHGL WP_001058323.1 MULTISPECIES: hydrogenase-1 small chain [Proteobacteria] Length: 372\nNumber of predicted TMHs: 2\nExp number of AAs in TMHs: 37.2608499999999\nExp number, first 60 AAs: 19.52923\nTotal prob of N-in: 0.94351\nPOSSIBLE N-term signal sequence\ninside 1 19\nTMhelix 20 39\noutside 40 325\nTMhelix 326 348\ninside 349 372

6991 GCF_001606835.1_ASM160683v1 Escherichia coli Proteobacteria; Gammaproteobacteria; Enterobacterales; Enterobacteriaceae; Escherichia MTGDNTLIHSHGINRRDFMKLCAALAATMGLSSKAAAEMAESVTNPQRPPVIWIGAQECT WP_024245965.1 hydrogenase 2 small subunit [Escherichia coli] Length: 372\nNumber of predicted TMHs: 1\nExp number of AAs in TMHs: 24.15412\nExp number, first 60 AAs: 0.21967\nTotal prob of N-in: 0.14289\noutside 1 331\nTMhelix 332 354\ninside 355 372

6992 GCF_001606775.1_ASM160677v1 Escherichia coli Proteobacteria; Gammaproteobacteria; Enterobacterales; Enterobacteriaceae; Escherichia MNNEETFYQAMRRQGVTRRSFLKYCSLAATSLGLGAGMAPKIAWALENKPRIPVVWIHGL WP_001058323.1 MULTISPECIES: hydrogenase-1 small chain [Proteobacteria] Length: 372\nNumber of predicted TMHs: 2\nExp number of AAs in TMHs: 37.2608499999999\nExp number, first 60 AAs: 19.52923\nTotal prob of N-in: 0.94351\nPOSSIBLE N-term signal sequence\ninside 1 19\nTMhelix 20 39\noutside 40 325\nTMhelix 326 348\ninside 349 372

6993 GCF_001606975.1_ASM160697v1 Escherichia coli Proteobacteria; Gammaproteobacteria; Enterobacterales; Enterobacteriaceae; Escherichia MTGDNTLIHSHGINRRDFMKLCAALAATMGLSSKAAAEMAESVTNPQRPPVIWIGAQECT WP_000145410.1 MULTISPECIES: hydrogenase-2 small chain [Proteobacteria] Length: 372\nNumber of predicted TMHs: 1\nExp number of AAs in TMHs: 23.72226\nExp number, first 60 AAs: 0.22038\nTotal prob of N-in: 0.12994\noutside 1 331\nTMhelix 332 354\ninside 355 372

6994 GCF_001606935.1_ASM160693v1 Escherichia coli Proteobacteria; Gammaproteobacteria; Enterobacterales; Enterobacteriaceae; Escherichia MNNEETFYQAMRRQGVTRRSFLKYCSLAATSLGLGAGMAPKIAWALENKPRIPVVWIHGL WP_001058323.1 MULTISPECIES: hydrogenase-1 small chain [Proteobacteria] Length: 372\nNumber of predicted TMHs: 2\nExp number of AAs in TMHs: 37.2608499999999\nExp number, first 60 AAs: 19.52923\nTotal prob of N-in:

0.94351\nPOSSIBLE N-term signal sequence\ninside 1 19\nTMhelix 20 39\noutside 40 325\nTMhelix 326 348\ninside 349 372

6995 GCF_001607025.1_ASM160702v1 Escherichia coli Proteobacteria; Gammaproteobacteria; Enterobacterales; Enterobacteriaceae; Escherichia
MTGDNTLIHSHGINRRDFMKLCAALAATMGLSSKAAAEMAESVTNPQRPPVIWIGAQECT WP_000145410.1
MULTISPECIES: hydrogenase-2 small chain [Proteobacteria] Length: 372\nNumber of predicted TMHs: 1\nExp number of AAs in TMHs: 23.72226\nExp number, first 60 AAs: 0.22038\nTotal prob of N-in: 0.12994\noutside 1 331\nTMhelix 332 354\ninside 355 372

6996 GCF_001607025.1_ASM160702v1 Escherichia coli Proteobacteria; Gammaproteobacteria; Enterobacterales; Enterobacteriaceae; Escherichia
MNNEETFYQAMRRQGVTRRSFLKYCSLAATSLGLGAGMAPKIAWALENKPRIPVVWIHGL WP_001058323.1
MULTISPECIES: hydrogenase-1 small chain [Proteobacteria] Length: 372\nNumber of predicted TMHs: 2\nExp number of AAs in TMHs: 37.2608499999999\nExp number, first 60 AAs: 19.52923\nTotal prob of N-in: 0.94351\nPOSSIBLE N-term signal sequence\ninside 1 19\nTMhelix 20 39\noutside 40 325\nTMhelix 326 348\ninside 349 372

6997 GCF_001606725.1_ASM160672v1 Escherichia coli Proteobacteria; Gammaproteobacteria; Enterobacterales; Enterobacteriaceae; Escherichia
MNNEETFYQAMRRQGVTRRSFLKYCSLAATSLGLGAGMAPKIAWALENKPRIPVVWIHGL WP_001295940.1
MULTISPECIES: hydrogenase [Enterobacteriaceae] Length: 372\nNumber of predicted TMHs: 2\nExp number of AAs in TMHs: 37.2608899999999\nExp number, first 60 AAs: 19.52923\nTotal prob of N-in: 0.94351\nPOSSIBLE N-term signal sequence\ninside 1 19\nTMhelix 20 39\noutside 40 325\nTMhelix 326 348\ninside 349 372

6998 GCF_001607075.1_ASM160707v1 Escherichia coli Proteobacteria; Gammaproteobacteria; Enterobacterales; Enterobacteriaceae; Escherichia
MTGDNTLIHSHGINRRDFMKLCAALAATMGLSSKAAAEMAESVTNPQRPPVIWIGAQECT WP_000145410.1
MULTISPECIES: hydrogenase-2 small chain [Proteobacteria] Length: 372\nNumber of predicted TMHs: 1\nExp number of AAs in TMHs: 23.72226\nExp number, first 60 AAs: 0.22038\nTotal prob of N-in: 0.12994\noutside 1 331\nTMhelix 332 354\ninside 355 372

6999 GCF_001607095.1_ASM160709v1 Escherichia coli Proteobacteria; Gammaproteobacteria; Enterobacterales; Enterobacteriaceae; Escherichia
MTGDNTLIHSHGINRRDFMKLCAALAATMGLSSKAAAEMAESVTNPQRPPVIWIGAQECT WP_000145410.1
MULTISPECIES: hydrogenase-2 small chain [Proteobacteria] Length: 372\nNumber of predicted TMHs: 1\nExp number of AAs in TMHs: 23.72226\nExp number, first 60 AAs: 0.22038\nTotal prob of N-in: 0.12994\noutside 1 331\nTMhelix 332 354\ninside 355 372

7000 GCF_001607125.1_ASM160712v1 Escherichia coli Proteobacteria; Gammaproteobacteria; Enterobacterales; Enterobacteriaceae; Escherichia
MTGDNTLIHSHGINRRDFMKLCAALAATMGLSSKAAAEMAESVTNPQRPPVIWIGAQECT WP_000145410.1
MULTISPECIES: hydrogenase-2 small chain [Proteobacteria] Length: 372\nNumber of predicted TMHs: 1\nExp number of AAs in TMHs: 23.72226\nExp number, first 60 AAs: 0.22038\nTotal prob of N-in: 0.12994\noutside 1 331\nTMhelix 332 354\ninside 355 372

7001 GCF_001607285.1_ASM160728v1 Escherichia coli Proteobacteria; Gammaproteobacteria; Enterobacterales; Enterobacteriaceae; Escherichia
MNNEETFYQAMRRQGVTRRSFLKYCSLAATSLGLGAGMAPKIAWALENKPRIPVVWIHGL WP_001058323.1
MULTISPECIES: hydrogenase-1 small chain [Proteobacteria] Length: 372\nNumber of predicted TMHs: 2\nExp number of AAs in TMHs: 37.2608499999999\nExp number, first 60 AAs: 19.52923\nTotal prob of N-in: 0.94351\nPOSSIBLE N-term signal sequence\ninside 1 19\nTMhelix 20 39\noutside 40 325\nTMhelix 326 348\ninside 349 372

7002 GCF_001607265.1_ASM160726v1 Escherichia coli Proteobacteria; Gammaproteobacteria; Enterobacterales; Enterobacteriaceae; Escherichia
MNNEETFYQAMRRQGVTRRSFLKSCSLAATSLGLGAGMAPKIAWALENKPRIPVVWIHGL WP_001058318.1
hydrogenase [Escherichia coli] Length: 372\nNumber of predicted TMHs: 1\nExp number of AAs in TMHs: 33.59637\nExp number, first 60 AAs: 15.81976\nTotal prob of N-in: 0.77394\nPOSSIBLE N-term signal sequence\noutside 1 325\nTMhelix 326 348\ninside 349 372

7003	GCF_001607095.1_ASM160709v1	Escherichia coli	Proteobacteria; Gammaproteobacteria; Enterobacterales; Enterobacteriaceae; Escherichia
	MNNEETFYQAMRRQGVTRRSFLKYCSLAATSLGLGAGMAPKIAWALENKPRIPVVWIHGL	WP_001058323.1
	MULTISPECIES: hydrogenase-1 small chain [Proteobacteria]	Length: 372\nNumber of predicted TMHs: 2\nExp number of AAs in TMHs: 37.2608499999999\nExp number, first 60 AAs: 19.52923\nTotal prob of N-in: 0.94351\nPOSSIBLE N-term signal sequence\ninside 1 19\nTMhelix 20 39\noutside 40 325\nTMhelix 326 348\ninside 349 372

7004	GCF_001607335.1_ASM160733v1	Escherichia coli	Proteobacteria; Gammaproteobacteria; Enterobacterales; Enterobacteriaceae; Escherichia
	MTGDNTLIHSHGINRRDFMKLCAALAATMGLSSKAAAEMAESVTNPQRPPVIWIGAQECT	WP_000145410.1
	MULTISPECIES: hydrogenase-2 small chain [Proteobacteria]	Length: 372\nNumber of predicted TMHs: 1\nExp number of AAs in TMHs: 23.72226\nExp number, first 60 AAs: 0.22038\nTotal prob of N-in: 0.12994\noutside 1 331\nTMhelix 332 354\ninside 355 372

7005	GCF_001607335.1_ASM160733v1	Escherichia coli	Proteobacteria; Gammaproteobacteria; Enterobacterales; Enterobacteriaceae; Escherichia
	MNNEETFYQAMRRQGVTRRSFLKYCSLAATSLGLGAGMAPKIAWALENKPRIPVVWIHGL	WP_001058323.1
	MULTISPECIES: hydrogenase-1 small chain [Proteobacteria]	Length: 372\nNumber of predicted TMHs: 2\nExp number of AAs in TMHs: 37.2608499999999\nExp number, first 60 AAs: 19.52923\nTotal prob of N-in: 0.94351\nPOSSIBLE N-term signal sequence\ninside 1 19\nTMhelix 20 39\noutside 40 325\nTMhelix 326 348\ninside 349 372

7006	GCF_001607225.1_ASM160722v1	Escherichia coli	Proteobacteria; Gammaproteobacteria; Enterobacterales; Enterobacteriaceae; Escherichia
	MNNEETFYQAMRRQGVTRRSFLKYCSLAATSLGLGAGMAPKIAWALENKPRIPVVWIHGL	WP_001295940.1
	MULTISPECIES: hydrogenase [Enterobacteriaceae]	Length: 372\nNumber of predicted TMHs: 2\nExp number of AAs in TMHs: 37.2608899999999\nExp number, first 60 AAs: 19.52923\nTotal prob of N-in: 0.94351\nPOSSIBLE N-term signal sequence\ninside 1 19\nTMhelix 20 39\noutside 40 325\nTMhelix 326 348\ninside 349 372

7007	GCF_001607455.1_ASM160745v1	Escherichia coli	Proteobacteria; Gammaproteobacteria; Enterobacterales; Enterobacteriaceae; Escherichia
	MTGDNTLIHSHGINRRDFMKLCAALAATMGLSSKAAAEMAESVTNPQRPPVIWIGAQECT	WP_000145410.1
	MULTISPECIES: hydrogenase-2 small chain [Proteobacteria]	Length: 372\nNumber of predicted TMHs: 1\nExp number of AAs in TMHs: 23.72226\nExp number, first 60 AAs: 0.22038\nTotal prob of N-in: 0.12994\noutside 1 331\nTMhelix 332 354\ninside 355 372

7008	GCF_001607495.1_ASM160749v1	Escherichia coli	Proteobacteria; Gammaproteobacteria; Enterobacterales; Enterobacteriaceae; Escherichia
	MTGDNTLIHSHGINRRDFMKLCAALAATMGLSSKAAAEMAESVTNPQRPPVIWIGAQECT	WP_000145410.1
	MULTISPECIES: hydrogenase-2 small chain [Proteobacteria]	Length: 372\nNumber of predicted TMHs: 1\nExp number of AAs in TMHs: 23.72226\nExp number, first 60 AAs: 0.22038\nTotal prob of N-in: 0.12994\noutside 1 331\nTMhelix 332 354\ninside 355 372

7009	GCF_001607525.1_ASM160752v1	Escherichia coli	Proteobacteria; Gammaproteobacteria; Enterobacterales; Enterobacteriaceae; Escherichia
	MTGDNTLIHSHGINRRDFMKLCAALAATMGLSSKAAAEMVESVTNPQRPPVIWIGAQECT	WP_000145423.1
	hydrogenase 2 small subunit [Escherichia coli]	Length: 372\nNumber of predicted TMHs: 1\nExp number of AAs in TMHs: 23.78201\nExp number, first 60 AAs: 0.28355\nTotal prob of N-in: 0.13257\noutside 1 331\nTMhelix 332 354\ninside 355 372

7010	GCF_001607575.1_ASM160757v1	Escherichia coli	Proteobacteria; Gammaproteobacteria; Enterobacterales; Enterobacteriaceae; Escherichia
	MTGDNTLIHSHGINRRDFMKLCAALAATMGLSSKAAAEMAESVTNPQRPPVIWIGAQECT	WP_000145410.1
	MULTISPECIES: hydrogenase-2 small chain [Proteobacteria]	Length: 372\nNumber of predicted TMHs: 1\nExp number of AAs in TMHs: 23.72226\nExp number, first 60 AAs: 0.22038\nTotal prob of N-in: 0.12994\noutside 1 331\nTMhelix 332 354\ninside 355 372

7011	GCF_001607595.1_ASM160759v1	Escherichia coli	Proteobacteria; Gammaproteobacteria; Enterobacterales; Enterobacteriaceae; Escherichia

MTGDNTLIHSHGINRRDFMKLCAALAATMGLSSKAAAEMAESVTNPQRPPVIWIGAQECT WP_000145410.1 MULTISPECIES: hydrogenase-2 small chain [Proteobacteria] Length: 372\nNumber of predicted TMHs: 1\nExp number of AAs in TMHs: 23.72226\nExp number, first 60 AAs: 0.22038\nTotal prob of N-in: 0.12994\noutside 1 331\nTMhelix 332 354\ninside 355 372

7012 GCF_001607635.1_ASM160763v1 Escherichia coli Proteobacteria; Gammaproteobacteria; Enterobacterales; Enterobacteriaceae; Escherichia MTGDNTLIHSHGINRRDFMKLCAALAATMGLSSKAAAEMAESVTNPQRPPVIWIGAQECT WP_000145410.1 MULTISPECIES: hydrogenase-2 small chain [Proteobacteria] Length: 372\nNumber of predicted TMHs: 1\nExp number of AAs in TMHs: 23.72226\nExp number, first 60 AAs: 0.22038\nTotal prob of N-in: 0.12994\noutside 1 331\nTMhelix 332 354\ninside 355 372

7013 GCF_001607685.1_ASM160768v1 Escherichia coli Proteobacteria; Gammaproteobacteria; Enterobacterales; Enterobacteriaceae; Escherichia MTGDNTLIHSHGINRRDFMKLCAALAATMGLSSKAAAEMAESVTNPQRPPVIWIGAQECT WP_000145410.1 MULTISPECIES: hydrogenase-2 small chain [Proteobacteria] Length: 372\nNumber of predicted TMHs: 1\nExp number of AAs in TMHs: 23.72226\nExp number, first 60 AAs: 0.22038\nTotal prob of N-in: 0.12994\noutside 1 331\nTMhelix 332 354\ninside 355 372

7014 GCF_001607525.1_ASM160752v1 Escherichia coli Proteobacteria; Gammaproteobacteria; Enterobacterales; Enterobacteriaceae; Escherichia MNNEETFYQAMRRQGVTRRSFLKYCSLAATSLGLGAGMAPKIAWALENKPRIPVVWIHGL WP_001295940.1 MULTISPECIES: hydrogenase [Enterobacteriaceae] Length: 372\nNumber of predicted TMHs: 2\nExp number of AAs in TMHs: 37.2608899999999\nExp number, first 60 AAs: 19.52923\nTotal prob of N-in: 0.94351\nPOSSIBLE N-term signal sequence\ninside 1 19\nTMhelix 20 39\noutside 40 325\nTMhelix 326 348\ninside 349 372

7015 GCF_001607455.1_ASM160745v1 Escherichia coli Proteobacteria; Gammaproteobacteria; Enterobacterales; Enterobacteriaceae; Escherichia MNNEETFYQAMRRQGVTRRSFLKYCSLAATSLGLGAGMAPKIAWALENKPRIPVVWIHGL WP_001295940.1 MULTISPECIES: hydrogenase [Enterobacteriaceae] Length: 372\nNumber of predicted TMHs: 2\nExp number of AAs in TMHs: 37.2608899999999\nExp number, first 60 AAs: 19.52923\nTotal prob of N-in: 0.94351\nPOSSIBLE N-term signal sequence\ninside 1 19\nTMhelix 20 39\noutside 40 325\nTMhelix 326 348\ninside 349 372

7016 GCF_001607575.1_ASM160757v1 Escherichia coli Proteobacteria; Gammaproteobacteria; Enterobacterales; Enterobacteriaceae; Escherichia MNNEETFYQAMRRQGVTRRSFLKYCSLAATSLGLGAGMAPKIAWALENKPRIPVVWIHGL WP_001058323.1 MULTISPECIES: hydrogenase-1 small chain [Proteobacteria] Length: 372\nNumber of predicted TMHs: 2\nExp number of AAs in TMHs: 37.2608499999999\nExp number, first 60 AAs: 19.52923\nTotal prob of N-in: 0.94351\nPOSSIBLE N-term signal sequence\ninside 1 19\nTMhelix 20 39\noutside 40 325\nTMhelix 326 348\ninside 349 372

7017 GCF_001607635.1_ASM160763v1 Escherichia coli Proteobacteria; Gammaproteobacteria; Enterobacterales; Enterobacteriaceae; Escherichia MNNEETFYQAMRRQGVTRRSFLKYCSLAATSLGLGAGMAPKIAWALENKPRIPVVWIHGL WP_001058323.1 MULTISPECIES: hydrogenase-1 small chain [Proteobacteria] Length: 372\nNumber of predicted TMHs: 2\nExp number of AAs in TMHs: 37.2608499999999\nExp number, first 60 AAs: 19.52923\nTotal prob of N-in: 0.94351\nPOSSIBLE N-term signal sequence\ninside 1 19\nTMhelix 20 39\noutside 40 325\nTMhelix 326 348\ninside 349 372

7018 GCF_001607595.1_ASM160759v1 Escherichia coli Proteobacteria; Gammaproteobacteria; Enterobacterales; Enterobacteriaceae; Escherichia MNNEETFYQAMRRQGVTRRSFLKYCSLAATSLGLGAGMAPKIAWALENKPRIPVVWIHGL WP_001058323.1 MULTISPECIES: hydrogenase-1 small chain [Proteobacteria] Length: 372\nNumber of predicted TMHs: 2\nExp number of AAs in TMHs: 37.2608499999999\nExp number, first 60 AAs: 19.52923\nTotal prob of N-in: 0.94351\nPOSSIBLE N-term signal sequence\ninside 1 19\nTMhelix 20 39\noutside 40 325\nTMhelix 326 348\ninside 349 372

7019 GCF_001607765.1_ASM160776v1 Escherichia coli Proteobacteria; Gammaproteobacteria; Enterobacterales; Enterobacteriaceae; Escherichia

MNNEETFYQAMRRQGVTRRSFLKYCSLAATSLGLGAGMAPKIAWALENKPRIPVVWIHGL WP_001058323.1 MULTISPECIES: hydrogenase-1 small chain [Proteobacteria] Length: 372\nNumber of predicted TMHs: 2\nExp number of AAs in TMHs: 37.2608499999999\nExp number, first 60 AAs: 19.52923\nTotal prob of N-in: 0.94351\nPOSSIBLE N-term signal sequence\ninside 1 19\nTMhelix 20 39\noutside 40 325\nTMhelix 326 348\ninside 349 372

7020 GCF_001607685.1_ASM160768v1 Escherichia coli Proteobacteria; Gammaproteobacteria; Enterobacterales; Enterobacteriaceae; Escherichia MNNEETFYQAMRRQGVTRRSFLKYCSLAATSLGLGAGMAPKIAWALENKPRIPVVWIHGL WP_001058323.1 MULTISPECIES: hydrogenase-1 small chain [Proteobacteria] Length: 372\nNumber of predicted TMHs: 2\nExp number of AAs in TMHs: 37.2608499999999\nExp number, first 60 AAs: 19.52923\nTotal prob of N-in: 0.94351\nPOSSIBLE N-term signal sequence\ninside 1 19\nTMhelix 20 39\noutside 40 325\nTMhelix 326 348\ninside 349 372

7021 GCF_001607835.1_ASM160783v1 Escherichia coli Proteobacteria; Gammaproteobacteria; Enterobacterales; Enterobacteriaceae; Escherichia MTGDNTLIHSHGINRRDFMKLCAALAATMGLSSKAAAEMAESVTNPQRPPVIWIGAQECT WP_000145410.1 MULTISPECIES: hydrogenase-2 small chain [Proteobacteria] Length: 372\nNumber of predicted TMHs: 1\nExp number of AAs in TMHs: 23.72226\nExp number, first 60 AAs: 0.22038\nTotal prob of N-in: 0.12994\noutside 1 331\nTMhelix 332 354\ninside 355 372

7022 GCF_001608085.1_ASM160808v1 Escherichia coli Proteobacteria; Gammaproteobacteria; Enterobacterales; Enterobacteriaceae; Escherichia MTGDNTLIHSHGINRRDFMKLCAALAATMGLSSKAAAEMAESVTNPQRPPVIWIGAQECT WP_000145410.1 MULTISPECIES: hydrogenase-2 small chain [Proteobacteria] Length: 372\nNumber of predicted TMHs: 1\nExp number of AAs in TMHs: 23.72226\nExp number, first 60 AAs: 0.22038\nTotal prob of N-in: 0.12994\noutside 1 331\nTMhelix 332 354\ninside 355 372

7023 GCF_001608015.1_ASM160801v1 Escherichia coli Proteobacteria; Gammaproteobacteria; Enterobacterales; Enterobacteriaceae; Escherichia MNNEETFYQAMRRQGVTRRSFLKYCSLAATSLGLGAGMAPKIAWALENKPRIPVVWIHGL WP_001058323.1 MULTISPECIES: hydrogenase-1 small chain [Proteobacteria] Length: 372\nNumber of predicted TMHs: 2\nExp number of AAs in TMHs: 37.2608499999999\nExp number, first 60 AAs: 19.52923\nTotal prob of N-in: 0.94351\nPOSSIBLE N-term signal sequence\ninside 1 19\nTMhelix 20 39\noutside 40 325\nTMhelix 326 348\ninside 349 372

7024 GCF_001608085.1_ASM160808v1 Escherichia coli Proteobacteria; Gammaproteobacteria; Enterobacterales; Enterobacteriaceae; Escherichia MNNEETFYQAMRRQGVTRRSFLKYCSLAATSLGLGAGMAPKIAWALENKPRIPVVWIHGL WP_001058323.1 MULTISPECIES: hydrogenase-1 small chain [Proteobacteria] Length: 372\nNumber of predicted TMHs: 2\nExp number of AAs in TMHs: 37.2608499999999\nExp number, first 60 AAs: 19.52923\nTotal prob of N-in: 0.94351\nPOSSIBLE N-term signal sequence\ninside 1 19\nTMhelix 20 39\noutside 40 325\nTMhelix 326 348\ninside 349 372

7025 GCF_001607995.1_ASM160799v1 Escherichia coli Proteobacteria; Gammaproteobacteria; Enterobacterales; Enterobacteriaceae; Escherichia MNNEETFYQAMRRQGVTRRSFLKSCSLAATSLGLGAGMAPKIAWALENKPRIPVVWIHGL WP_001058318.1 hydrogenase [Escherichia coli] Length: 372\nNumber of predicted TMHs: 1\nExp number of AAs in TMHs: 33.59637\nExp number, first 60 AAs: 15.81976\nTotal prob of N-in: 0.77394\nPOSSIBLE N-term signal sequence\noutside 1 325\nTMhelix 326 348\ninside 349 372

7026 GCF_001607875.1_ASM160787v1 Escherichia coli Proteobacteria; Gammaproteobacteria; Enterobacterales; Enterobacteriaceae; Escherichia MNNEETFYQAMRRQGVTRRSFLKYCSLAATSLGLGAGMAPKIAWALENKPRIPVVWIHGL WP_001058323.1 MULTISPECIES: hydrogenase-1 small chain [Proteobacteria] Length: 372\nNumber of predicted TMHs: 2\nExp number of AAs in TMHs: 37.2608499999999\nExp number, first 60 AAs: 19.52923\nTotal prob of N-in: 0.94351\nPOSSIBLE N-term signal sequence\ninside 1 19\nTMhelix 20 39\noutside 40 325\nTMhelix 326 348\ninside 349 372

7027 GCF_001607925.1_ASM160792v1 Escherichia coli Proteobacteria; Gammaproteobacteria; Enterobacterales; Enterobacteriaceae; Escherichia

MNNEETFYQAMRRQGVTRRSFLKYCSLAATSLGLGAGMAPKIAWALENKPRIPVVWIHGL WP_001058323.1 MULTISPECIES: hydrogenase-1 small chain [Proteobacteria] Length: 372\nNumber of predicted TMHs: 2\nExp number of AAs in TMHs: 37.2608499999999\nExp number, first 60 AAs: 19.52923\nTotal prob of N-in: 0.94351\nPOSSIBLE N-term signal sequence\ninside 1 19\nTMhelix 20 39\noutside 40 325\nTMhelix 326 348\ninside 349 372

7028 GCF_001609255.1_ASM160925v1 Escherichia coli Proteobacteria; Gammaproteobacteria; Enterobacterales; Enterobacteriaceae; Escherichia MTGDNTLIHSHGINRRDFMKLCAALAATMGLSSKAAAEMAESVTNPQRPPVIWIGAQECT WP_000145410.1 MULTISPECIES: hydrogenase-2 small chain [Proteobacteria] Length: 372\nNumber of predicted TMHs: 1\nExp number of AAs in TMHs: 23.72226\nExp number, first 60 AAs: 0.22038\nTotal prob of N-in: 0.12994\noutside 1 331\nTMhelix 332 354\ninside 355 372

7029 GCF_001609215.1_ASM160921v1 Escherichia coli Proteobacteria; Gammaproteobacteria; Enterobacterales; Enterobacteriaceae; Escherichia MNNEETFYQAMRRQGVTRRSFLKYCSLAATSLGLGAGMAPKIAWALENKPRIPVVWIHGL WP_001058323.1 MULTISPECIES: hydrogenase-1 small chain [Proteobacteria] Length: 372\nNumber of predicted TMHs: 2\nExp number of AAs in TMHs: 37.2608499999999\nExp number, first 60 AAs: 19.52923\nTotal prob of N-in: 0.94351\nPOSSIBLE N-term signal sequence\ninside 1 19\nTMhelix 20 39\noutside 40 325\nTMhelix 326 348\ninside 349 372

7030 GCF_001609255.1_ASM160925v1 Escherichia coli Proteobacteria; Gammaproteobacteria; Enterobacterales; Enterobacteriaceae; Escherichia MNNEETFYQAMRRQGVTRRSFLKYCSLAATSLGLGAGMAPKIAWALENKPRIPVVWIHGL WP_001058323.1 MULTISPECIES: hydrogenase-1 small chain [Proteobacteria] Length: 372\nNumber of predicted TMHs: 2\nExp number of AAs in TMHs: 37.2608499999999\nExp number, first 60 AAs: 19.52923\nTotal prob of N-in: 0.94351\nPOSSIBLE N-term signal sequence\ninside 1 19\nTMhelix 20 39\noutside 40 325\nTMhelix 326 348\ninside 349 372

7031 GCF_001608265.1_ASM160826v1 Escherichia coli Proteobacteria; Gammaproteobacteria; Enterobacterales; Enterobacteriaceae; Escherichia MNNEETFYQAMRRQGVTRRSFLKYCSLAATSLGLGAGMAPKIAWALENKPRIPVVWIHGL WP_001058323.1 MULTISPECIES: hydrogenase-1 small chain [Proteobacteria] Length: 372\nNumber of predicted TMHs: 2\nExp number of AAs in TMHs: 37.2608499999999\nExp number, first 60 AAs: 19.52923\nTotal prob of N-in: 0.94351\nPOSSIBLE N-term signal sequence\ninside 1 19\nTMhelix 20 39\noutside 40 325\nTMhelix 326 348\ninside 349 372

7032 GCF_001609265.1_ASM160926v1 Escherichia coli Proteobacteria; Gammaproteobacteria; Enterobacterales; Enterobacteriaceae; Escherichia MTGDNTLIHSHGINRRDFMKLCAALAATMGLSSKAAAEMAESVTNPQRPPVIWIGAQECT WP_000145410.1 MULTISPECIES: hydrogenase-2 small chain [Proteobacteria] Length: 372\nNumber of predicted TMHs: 1\nExp number of AAs in TMHs: 23.72226\nExp number, first 60 AAs: 0.22038\nTotal prob of N-in: 0.12994\noutside 1 331\nTMhelix 332 354\ninside 355 372

7033 GCF_001609315.1_ASM160931v1 Escherichia coli Proteobacteria; Gammaproteobacteria; Enterobacterales; Enterobacteriaceae; Escherichia MTGDNTLIHSHGINRRDFMKLCAALAATMGLSSKAAAEMAESVTNPQRPPVIWIGAQECT WP_000145410.1 MULTISPECIES: hydrogenase-2 small chain [Proteobacteria] Length: 372\nNumber of predicted TMHs: 1\nExp number of AAs in TMHs: 23.72226\nExp number, first 60 AAs: 0.22038\nTotal prob of N-in: 0.12994\noutside 1 331\nTMhelix 332 354\ninside 355 372

7034 GCF_001609735.1_ASM160973v1 Escherichia coli Proteobacteria; Gammaproteobacteria; Enterobacterales; Enterobacteriaceae; Escherichia MTGDNTLIHSHGINRRDFMKLCAALAATMGLSSKAAAEMAESVTNPQRPPVIWIGAQECT WP_000145410.1 MULTISPECIES: hydrogenase-2 small chain [Proteobacteria] Length: 372\nNumber of predicted TMHs: 1\nExp number of AAs in TMHs: 23.72226\nExp number, first 60 AAs: 0.22038\nTotal prob of N-in: 0.12994\noutside 1 331\nTMhelix 332 354\ninside 355 372

7035 GCF_001609855.1_ASM160985v1 Escherichia coli Proteobacteria; Gammaproteobacteria; Enterobacterales; Enterobacteriaceae; Escherichia MTGDNTLIHSHGINRRDFMKLCAALAATMGLSSKAAAEMAESVTNPQRPPVIWIGAQECT WP_000145410.1

MULTISPECIES: hydrogenase-2 small chain [Proteobacteria] Length: 372\nNumber of predicted TMHs: 1\nExp number of AAs in TMHs: 23.72226\nExp number, first 60 AAs: 0.22038\nTotal prob of N-in: 0.12994\noutside 1 331\nTMhelix 332 354\ninside 355 372

7036 GCF_001614325.1_ASM161432v1 Escherichia coli Proteobacteria; Gammaproteobacteria; Enterobacterales; Enterobacteriaceae; Escherichia MTGDNTLIHSHGINRRDFMKLCAALAATMGLSSKAAAEMAESVTNPQRPPVIWIGAQECT WP_000145410.1 MULTISPECIES: hydrogenase-2 small chain [Proteobacteria] Length: 372\nNumber of predicted TMHs: 1\nExp number of AAs in TMHs: 23.72226\nExp number, first 60 AAs: 0.22038\nTotal prob of N-in: 0.12994\noutside 1 331\nTMhelix 332 354\ninside 355 372

7037 GCF_001609835.1_ASM160983v1 Escherichia coli Proteobacteria; Gammaproteobacteria; Enterobacterales; Enterobacteriaceae; Escherichia MNNEETFYQAMRRQGVTRRSFLKYCSLAATSLGLGAGMAPKIAWALENKPRIPVVWIHGL WP_001058323.1 MULTISPECIES: hydrogenase-1 small chain [Proteobacteria] Length: 372\nNumber of predicted TMHs: 2\nExp number of AAs in TMHs: 37.2608499999999\nExp number, first 60 AAs: 19.52923\nTotal prob of N-in: 0.94351\nPOSSIBLE N-term signal sequence\ninside 1 19\nTMhelix 20 39\noutside 40 325\nTMhelix 326 348\ninside 349 372

7038 GCF_001609745.1_ASM160974v1 Escherichia coli Proteobacteria; Gammaproteobacteria; Enterobacterales; Enterobacteriaceae; Escherichia MNNEETFYQAMRRQGVTRRSFLKYCSLAATSLGLGAGMAPKIAWALENKPRIPVVWIHGL WP_001058323.1 MULTISPECIES: hydrogenase-1 small chain [Proteobacteria] Length: 372\nNumber of predicted TMHs: 2\nExp number of AAs in TMHs: 37.2608499999999\nExp number, first 60 AAs: 19.52923\nTotal prob of N-in: 0.94351\nPOSSIBLE N-term signal sequence\ninside 1 19\nTMhelix 20 39\noutside 40 325\nTMhelix 326 348\ninside 349 372

7039 GCF_001609855.1_ASM160985v1 Escherichia coli Proteobacteria; Gammaproteobacteria; Enterobacterales; Enterobacteriaceae; Escherichia MNNEETFYQAMRRQGVTRRSFLKYCSLAATSLGLGAGMAPKIAWALENKPRIPVVWIHGL WP_001058323.1 MULTISPECIES: hydrogenase-1 small chain [Proteobacteria] Length: 372\nNumber of predicted TMHs: 2\nExp number of AAs in TMHs: 37.2608499999999\nExp number, first 60 AAs: 19.52923\nTotal prob of N-in: 0.94351\nPOSSIBLE N-term signal sequence\ninside 1 19\nTMhelix 20 39\noutside 40 325\nTMhelix 326 348\ninside 349 372

7040 GCF_001614395.1_ASM161439v1 Escherichia coli Proteobacteria; Gammaproteobacteria; Enterobacterales; Enterobacteriaceae; Escherichia MTGDNTLIHSHGINRRDFMKLCAALAATMGLSSKAAAEMAESVTNPQRPPVIWIGAQECT WP_000145410.1 MULTISPECIES: hydrogenase-2 small chain [Proteobacteria] Length: 372\nNumber of predicted TMHs: 1\nExp number of AAs in TMHs: 23.72226\nExp number, first 60 AAs: 0.22038\nTotal prob of N-in: 0.12994\noutside 1 331\nTMhelix 332 354\ninside 355 372

7041 GCF_001614495.1_ASM161449v1 Escherichia coli Proteobacteria; Gammaproteobacteria; Enterobacterales; Enterobacteriaceae; Escherichia MTGDNTLIHSHGINRRDFMKLCAALAATMGLSSKAAAEMAESVTNPQRPPVIWIGAQECT WP_000145410.1 MULTISPECIES: hydrogenase-2 small chain [Proteobacteria] Length: 372\nNumber of predicted TMHs: 1\nExp number of AAs in TMHs: 23.72226\nExp number, first 60 AAs: 0.22038\nTotal prob of N-in: 0.12994\noutside 1 331\nTMhelix 332 354\ninside 355 372

7042 GCF_001614555.1_ASM161455v1 Escherichia coli Proteobacteria; Gammaproteobacteria; Enterobacterales; Enterobacteriaceae; Escherichia MTGDNTLIHSHGINRRDFMKLCAALAATMGLSSKAAAEMAESVTNPQRPPVIWIGAQECT WP_000145410.1 MULTISPECIES: hydrogenase-2 small chain [Proteobacteria] Length: 372\nNumber of predicted TMHs: 1\nExp number of AAs in TMHs: 23.72226\nExp number, first 60 AAs: 0.22038\nTotal prob of N-in: 0.12994\noutside 1 331\nTMhelix 332 354\ninside 355 372

7043 GCF_001614615.1_ASM161461v1 Escherichia coli Proteobacteria; Gammaproteobacteria; Enterobacterales; Enterobacteriaceae; Escherichia MTGDNTLIHSHGINRRDFMKLCAALAATMGLSSKAAAEMAESVTNPQRPPVIWIGAQECT WP_000145410.1 MULTISPECIES: hydrogenase-2 small chain [Proteobacteria] Length: 372\nNumber of predicted TMHs:

1\nExp number of AAs in TMHs: 23.72226\nExp number, first 60 AAs: 0.22038\nTotal prob of N-in: 0.12994\noutside 1 331\nTMhelix 332 354\ninside 355 372

7044 GCF_001614375.1_ASM161437v1 Escherichia coli Proteobacteria; Gammaproteobacteria; Enterobacterales; Enterobacteriaceae; Escherichia
MNNEETFYQAMRRQGVTRRSFLKYCSLAATSLGLGAGMAPKIAWALENKPRIPVVWIHGL WP_001058323.1
MULTISPECIES: hydrogenase-1 small chain [Proteobacteria] Length: 372\nNumber of predicted TMHs: 2\nExp number of AAs in TMHs: 37.2608499999999\nExp number, first 60 AAs: 19.52923\nTotal prob of N-in: 0.94351\nPOSSIBLE N-term signal sequence\ninside 1 19\nTMhelix 20 39\noutside 40 325\nTMhelix 326 348\ninside 349 372

7045 GCF_001614565.1_ASM161456v1 Escherichia coli Proteobacteria; Gammaproteobacteria; Enterobacterales; Enterobacteriaceae; Escherichia
MTGDNTLIHSHGINRRDFMKLCAALAATMGLSSKAAAEMAESVTNPQRPPVIWIGAQECT WP_000145410.1
MULTISPECIES: hydrogenase-2 small chain [Proteobacteria] Length: 372\nNumber of predicted TMHs: 1\nExp number of AAs in TMHs: 23.72226\nExp number, first 60 AAs: 0.22038\nTotal prob of N-in: 0.12994\noutside 1 331\nTMhelix 332 354\ninside 355 372

7046 GCF_001614665.1_ASM161466v1 Escherichia coli Proteobacteria; Gammaproteobacteria; Enterobacterales; Enterobacteriaceae; Escherichia
MTGDNTLIHSHGINRRDFMKLCAALAATMGLSSKAAAEMAESVTNPQRPPVIWIGAQECT WP_000145410.1
MULTISPECIES: hydrogenase-2 small chain [Proteobacteria] Length: 372\nNumber of predicted TMHs: 1\nExp number of AAs in TMHs: 23.72226\nExp number, first 60 AAs: 0.22038\nTotal prob of N-in: 0.12994\noutside 1 331\nTMhelix 332 354\ninside 355 372

7047 GCF_001614555.1_ASM161455v1 Escherichia coli Proteobacteria; Gammaproteobacteria; Enterobacterales; Enterobacteriaceae; Escherichia
MNNEETFYQAMRRQGVTRRSFLKYCSLAATSLGLGAGMAPKIAWALENKPRIPVVWIHGL WP_001058323.1
MULTISPECIES: hydrogenase-1 small chain [Proteobacteria] Length: 372\nNumber of predicted TMHs: 2\nExp number of AAs in TMHs: 37.2608499999999\nExp number, first 60 AAs: 19.52923\nTotal prob of N-in: 0.94351\nPOSSIBLE N-term signal sequence\ninside 1 19\nTMhelix 20 39\noutside 40 325\nTMhelix 326 348\ninside 349 372

7048 GCF_001614715.1_ASM161471v1 Escherichia coli Proteobacteria; Gammaproteobacteria; Enterobacterales; Enterobacteriaceae; Escherichia
MNNEETFYQAMRRQGVTRRSFLKYCSLAATSLGLGAGMAPKIAWALENKPRIPVVWIHGL WP_001058323.1
MULTISPECIES: hydrogenase-1 small chain [Proteobacteria] Length: 372\nNumber of predicted TMHs: 2\nExp number of AAs in TMHs: 37.2608499999999\nExp number, first 60 AAs: 19.52923\nTotal prob of N-in: 0.94351\nPOSSIBLE N-term signal sequence\ninside 1 19\nTMhelix 20 39\noutside 40 325\nTMhelix 326 348\ninside 349 372

7049 GCF_001614775.1_ASM161477v1 Escherichia coli Proteobacteria; Gammaproteobacteria; Enterobacterales; Enterobacteriaceae; Escherichia
MTGDNTLIHSHGINRRDFMKLCAALAATMGLSSKAAAEMAESVTNPQRPPVIWIGAQECT WP_000145410.1
MULTISPECIES: hydrogenase-2 small chain [Proteobacteria] Length: 372\nNumber of predicted TMHs: 1\nExp number of AAs in TMHs: 23.72226\nExp number, first 60 AAs: 0.22038\nTotal prob of N-in: 0.12994\noutside 1 331\nTMhelix 332 354\ninside 355 372

7050 GCF_001614805.1_ASM161480v1 Escherichia coli Proteobacteria; Gammaproteobacteria; Enterobacterales; Enterobacteriaceae; Escherichia
MTGDNTLIHSHGINRRDFMKLCAALAATMGLSSKAAAEMAESVTNPQRPPVIWIGAQECT WP_000145410.1
MULTISPECIES: hydrogenase-2 small chain [Proteobacteria] Length: 372\nNumber of predicted TMHs: 1\nExp number of AAs in TMHs: 23.72226\nExp number, first 60 AAs: 0.22038\nTotal prob of N-in: 0.12994\noutside 1 331\nTMhelix 332 354\ninside 355 372

7051 GCF_001614875.1_ASM161487v1 Escherichia coli Proteobacteria; Gammaproteobacteria; Enterobacterales; Enterobacteriaceae; Escherichia
MTGDNTLIHSHGINRRDFMKLCAALAATMGLSSKAAAEMAESVTNPQRPPVIWIGAQECT WP_000145410.1
MULTISPECIES: hydrogenase-2 small chain [Proteobacteria] Length: 372\nNumber of predicted TMHs: 1\nExp number of AAs in TMHs: 23.72226\nExp number, first 60 AAs: 0.22038\nTotal prob of N-in: 0.12994\noutside 1 331\nTMhelix 332 354\ninside 355 372

7052 GCF_001614855.1_ASM161485v1 Escherichia coli Proteobacteria; Gammaproteobacteria; Enterobacterales; Enterobacteriaceae; Escherichia
MTGDNTLIHSHGINRRDFMKLCAALAATMGLSSKAAAEMAESVTNPQRPPVIWIGAQECT WP_000145410.1
MULTISPECIES: hydrogenase-2 small chain [Proteobacteria] Length: 372\nNumber of predicted TMHs: 1\nExp number of AAs in TMHs: 23.72226\nExp number, first 60 AAs: 0.22038\nTotal prob of N-in: 0.12994\noutside 1 331\nTMhelix 332 354\ninside 355 372

7053 GCF_001614965.1_ASM161496v1 Escherichia coli Proteobacteria; Gammaproteobacteria; Enterobacterales; Enterobacteriaceae; Escherichia
MTGDNTLIHSHGINRRDFMKLCAALAATMGLSSKAAAEMAESVTNPQRPPVIWIGAQECT WP_000145410.1
MULTISPECIES: hydrogenase-2 small chain [Proteobacteria] Length: 372\nNumber of predicted TMHs: 1\nExp number of AAs in TMHs: 23.72226\nExp number, first 60 AAs: 0.22038\nTotal prob of N-in: 0.12994\noutside 1 331\nTMhelix 332 354\ninside 355 372

7054 GCF_001615035.1_ASM161503v1 Escherichia coli Proteobacteria; Gammaproteobacteria; Enterobacterales; Enterobacteriaceae; Escherichia
MTGDNTLIHSHGINRRDFMKLCAALAATMGLSSKAAAEMAESVTNPQRPPVIWIGAQECT WP_000145410.1
MULTISPECIES: hydrogenase-2 small chain [Proteobacteria] Length: 372\nNumber of predicted TMHs: 1\nExp number of AAs in TMHs: 23.72226\nExp number, first 60 AAs: 0.22038\nTotal prob of N-in: 0.12994\noutside 1 331\nTMhelix 332 354\ninside 355 372

7055 GCF_001615035.1_ASM161503v1 Escherichia coli Proteobacteria; Gammaproteobacteria; Enterobacterales; Enterobacteriaceae; Escherichia
MNNEETFYQAMRRQGVTRRSFLKYCSLAATSLGLGAGMAPKIAWALENKPRIPVVWIHGL WP_001058320.1
hydrogenase [Escherichia coli] Length: 372\nNumber of predicted TMHs: 2\nExp number of AAs in TMHs: 37.26082\nExp number, first 60 AAs: 19.52923\nTotal prob of N-in: 0.94351\nPOSSIBLE N-term signal sequence\ninside 1 19\nTMhelix 20 39\noutside 40 325\nTMhelix 326 348\ninside 349 372

7056 GCF_001615065.1_ASM161506v1 Escherichia coli Proteobacteria; Gammaproteobacteria; Enterobacterales; Enterobacteriaceae; Escherichia
MNNEETFYQAMRRQGVTRRSFLKYCSLAATSLGLGAGMAPKIAWALENKPRIPVVWIHGL WP_001058323.1
MULTISPECIES: hydrogenase-1 small chain [Proteobacteria] Length: 372\nNumber of predicted TMHs: 2\nExp number of AAs in TMHs: 37.2608499999999\nExp number, first 60 AAs: 19.52923\nTotal prob of N-in: 0.94351\nPOSSIBLE N-term signal sequence\ninside 1 19\nTMhelix 20 39\noutside 40 325\nTMhelix 326 348\ninside 349 372

7057 GCF_001614855.1_ASM161485v1 Escherichia coli Proteobacteria; Gammaproteobacteria; Enterobacterales; Enterobacteriaceae; Escherichia
MNNEETFYQAMRRQGVTRRSFLKYCSLAATSLGLGAGMAPKIAWALENKPRIPVVWIHGL WP_001058323.1
MULTISPECIES: hydrogenase-1 small chain [Proteobacteria] Length: 372\nNumber of predicted TMHs: 2\nExp number of AAs in TMHs: 37.2608499999999\nExp number, first 60 AAs: 19.52923\nTotal prob of N-in: 0.94351\nPOSSIBLE N-term signal sequence\ninside 1 19\nTMhelix 20 39\noutside 40 325\nTMhelix 326 348\ninside 349 372

7058 GCF_001615095.1_ASM161509v1 Escherichia coli Proteobacteria; Gammaproteobacteria; Enterobacterales; Enterobacteriaceae; Escherichia
MTGDNTLIHSHGINRRDFMKLCAALAATMGLSSKAAAEMAESVTNPQRPPVIWIGAQECT WP_000145410.1
MULTISPECIES: hydrogenase-2 small chain [Proteobacteria] Length: 372\nNumber of predicted TMHs: 1\nExp number of AAs in TMHs: 23.72226\nExp number, first 60 AAs: 0.22038\nTotal prob of N-in: 0.12994\noutside 1 331\nTMhelix 332 354\ninside 355 372

7059 GCF_001614965.1_ASM161496v1 Escherichia coli Proteobacteria; Gammaproteobacteria; Enterobacterales; Enterobacteriaceae; Escherichia
MNNEETFYQAMRRQGVTRRSFLKYCSLAATSLGLGAGMAPKIAWALENKPRIPVVWIHGL WP_001058323.1
MULTISPECIES: hydrogenase-1 small chain [Proteobacteria] Length: 372\nNumber of predicted TMHs: 2\nExp number of AAs in TMHs: 37.2608499999999\nExp number, first 60 AAs: 19.52923\nTotal prob of N-in: 0.94351\nPOSSIBLE N-term signal sequence\ninside 1 19\nTMhelix 20 39\noutside 40 325\nTMhelix 326 348\ninside 349 372

7060 GCF_001615125.1_ASM161512v1 Escherichia coli Proteobacteria; Gammaproteobacteria; Enterobacterales; Enterobacteriaceae; Escherichia

MTGDNTLIHSHGINRRDFMKLCAALAATMGLSSKAAAEMAESVTNPQRPPVIWIGAQECT WP_000145410.1 MULTISPECIES: hydrogenase-2 small chain [Proteobacteria] Length: 372\nNumber of predicted TMHs: 1\nExp number of AAs in TMHs: 23.72226\nExp number, first 60 AAs: 0.22038\nTotal prob of N-in: 0.12994\noutside 1 331\nTMhelix 332 354\ninside 355 372

7061 GCF_001615295.1_ASM161529v1 Escherichia coli Proteobacteria; Gammaproteobacteria; Enterobacterales; Enterobacteriaceae; Escherichia MNNEETFYQAMRRQGVTRRSFLKYCSLAATSLGLGAGMAPKIAWALENKPRIPVVWIHGL WP_001058323.1 MULTISPECIES: hydrogenase-1 small chain [Proteobacteria] Length: 372\nNumber of predicted TMHs: 2\nExp number of AAs in TMHs: 37.2608499999999\nExp number, first 60 AAs: 19.52923\nTotal prob of N-in: 0.94351\nPOSSIBLE N-term signal sequence\ninside 1 19\nTMhelix 20 39\noutside 40 325\nTMhelix 326 348\ninside 349 372

7062 GCF_001615375.1_ASM161537v1 Escherichia coli Proteobacteria; Gammaproteobacteria; Enterobacterales; Enterobacteriaceae; Escherichia MTGDNTLIHSHGINRRDFMKLCAALAATMGLSSKAAAEMAESVTNPQRPPVIWIGAQECT WP_000145410.1 MULTISPECIES: hydrogenase-2 small chain [Proteobacteria] Length: 372\nNumber of predicted TMHs: 1\nExp number of AAs in TMHs: 23.72226\nExp number, first 60 AAs: 0.22038\nTotal prob of N-in: 0.12994\noutside 1 331\nTMhelix 332 354\ninside 355 372

7063 GCF_001615325.1_ASM161532v1 Escherichia coli Proteobacteria; Gammaproteobacteria; Enterobacterales; Enterobacteriaceae; Escherichia MTGDNTLIHSHGINRRDFMKLCAALAATMGLSSKAAAEMAESVTNPQRPPVIWIGAQECT WP_000145410.1 MULTISPECIES: hydrogenase-2 small chain [Proteobacteria] Length: 372\nNumber of predicted TMHs: 1\nExp number of AAs in TMHs: 23.72226\nExp number, first 60 AAs: 0.22038\nTotal prob of N-in: 0.12994\noutside 1 331\nTMhelix 332 354\ninside 355 372

7064 GCF_001615415.1_ASM161541v1 Escherichia coli Proteobacteria; Gammaproteobacteria; Enterobacterales; Enterobacteriaceae; Escherichia MTGDNTLIHSHGINRRDFMKLCAALAATMGLSSKAAAEMAESVTNPQRPPVIWIGAQECT WP_000145410.1 MULTISPECIES: hydrogenase-2 small chain [Proteobacteria] Length: 372\nNumber of predicted TMHs: 1\nExp number of AAs in TMHs: 23.72226\nExp number, first 60 AAs: 0.22038\nTotal prob of N-in: 0.12994\noutside 1 331\nTMhelix 332 354\ninside 355 372

7065 GCF_001615395.1_ASM161539v1 Escherichia coli Proteobacteria; Gammaproteobacteria; Enterobacterales; Enterobacteriaceae; Escherichia MTGDNTLIHSHGINRRDFMKLCAALAATMGLSSKAAAEMAESVTNPQRPPVIWIGAQECT WP_000145410.1 MULTISPECIES: hydrogenase-2 small chain [Proteobacteria] Length: 372\nNumber of predicted TMHs: 1\nExp number of AAs in TMHs: 23.72226\nExp number, first 60 AAs: 0.22038\nTotal prob of N-in: 0.12994\noutside 1 331\nTMhelix 332 354\ninside 355 372

7066 GCF_001615415.1_ASM161541v1 Escherichia coli Proteobacteria; Gammaproteobacteria; Enterobacterales; Enterobacteriaceae; Escherichia MNNEETFYQAMRRQGVTRRSFLKYCSLAATSLGLGAGMAPKIAWALENKPRIPVVWIHGL WP_001058323.1 MULTISPECIES: hydrogenase-1 small chain [Proteobacteria] Length: 372\nNumber of predicted TMHs: 2\nExp number of AAs in TMHs: 37.2608499999999\nExp number, first 60 AAs: 19.52923\nTotal prob of N-in: 0.94351\nPOSSIBLE N-term signal sequence\ninside 1 19\nTMhelix 20 39\noutside 40 325\nTMhelix 326 348\ninside 349 372

7067 GCF_001615275.1_ASM161527v1 Escherichia coli Proteobacteria; Gammaproteobacteria; Enterobacterales; Enterobacteriaceae; Escherichia MNNEETFYQAMRRQGVTRRSFLKYCSLAATSLGLGAGMAPKIAWALENKPRIPVVWIHGL WP_001295940.1 MULTISPECIES: hydrogenase [Enterobacteriaceae] Length: 372\nNumber of predicted TMHs: 2\nExp number of AAs in TMHs: 37.2608899999999\nExp number, first 60 AAs: 19.52923\nTotal prob of N-in: 0.94351\nPOSSIBLE N-term signal sequence\ninside 1 19\nTMhelix 20 39\noutside 40 325\nTMhelix 326 348\ninside 349 372

7068 GCF_001615395.1_ASM161539v1 Escherichia coli Proteobacteria; Gammaproteobacteria; Enterobacterales; Enterobacteriaceae; Escherichia MNNEETFYQAMRRQGVTRRSFLKYCSLAATSLGLGAGMAPKIAWALENKPRIPVVWIHGL WP_001058323.1 MULTISPECIES: hydrogenase-1 small chain [Proteobacteria] Length: 372\nNumber of predicted TMHs:

2\nExp number of AAs in TMHs: 37.2608499999999\nExp number, first 60 AAs: 19.52923\nTotal prob of N-in: 0.94351\nPOSSIBLE N-term signal sequence\ninside 1 19\nTMhelix 20 39\noutside 40 325\nTMhelix 326 348\ninside 349 372

7069 GCF_001615375.1_ASM161537v1 Escherichia coli Proteobacteria; Gammaproteobacteria; Enterobacterales; Enterobacteriaceae; Escherichia
MNNEETFYQAMRRQGVTRRSFLKYCSLAATSLGLGAGMAPKIAWALENKPRIPVVWIHGL WP_001295940.1
MULTISPECIES: hydrogenase [Enterobacteriaceae] Length: 372\nNumber of predicted TMHs: 2\nExp number of AAs in TMHs: 37.2608899999999\nExp number, first 60 AAs: 19.52923\nTotal prob of N-in: 0.94351\nPOSSIBLE N-term signal sequence\ninside 1 19\nTMhelix 20 39\noutside 40 325\nTMhelix 326 348\ninside 349 372

7070 GCF_001615595.1_ASM161559v1 Escherichia coli Proteobacteria; Gammaproteobacteria; Enterobacterales; Enterobacteriaceae; Escherichia
MTGDNTLIHSHGINRRDFMKLCAALAATMGLSSKAAAEMAESVTNPQRPPVIWIGAQECT WP_000145410.1
MULTISPECIES: hydrogenase-2 small chain [Proteobacteria] Length: 372\nNumber of predicted TMHs: 1\nExp number of AAs in TMHs: 23.72226\nExp number, first 60 AAs: 0.22038\nTotal prob of N-in: 0.12994\noutside 1 331\nTMhelix 332 354\ninside 355 372

7071 GCF_001615625.1_ASM161562v1 Escherichia coli Proteobacteria; Gammaproteobacteria; Enterobacterales; Enterobacteriaceae; Escherichia
MTGDNTLIHSHGINRRDFMKLCAALAATMGLSSKAAAEMAESVTNPQRPPVIWIGAQECT WP_000145410.1
MULTISPECIES: hydrogenase-2 small chain [Proteobacteria] Length: 372\nNumber of predicted TMHs: 1\nExp number of AAs in TMHs: 23.72226\nExp number, first 60 AAs: 0.22038\nTotal prob of N-in: 0.12994\noutside 1 331\nTMhelix 332 354\ninside 355 372

7072 GCF_001615675.1_ASM161567v1 Escherichia coli Proteobacteria; Gammaproteobacteria; Enterobacterales; Enterobacteriaceae; Escherichia
MTGDNTLIHSHGINRRDFMKLCAALAATMGLSSKAAAEMAESVTNPQRPPVIWIGAQECT WP_000145410.1
MULTISPECIES: hydrogenase-2 small chain [Proteobacteria] Length: 372\nNumber of predicted TMHs: 1\nExp number of AAs in TMHs: 23.72226\nExp number, first 60 AAs: 0.22038\nTotal prob of N-in: 0.12994\noutside 1 331\nTMhelix 332 354\ninside 355 372

7073 GCF_001615695.1_ASM161569v1 Escherichia coli Proteobacteria; Gammaproteobacteria; Enterobacterales; Enterobacteriaceae; Escherichia
MTGDNTLIHSHGINRRDFMKLCAALAATMGLSSKAAAEMAESVTNPQRPPVIWIGAQECT WP_000145410.1
MULTISPECIES: hydrogenase-2 small chain [Proteobacteria] Length: 372\nNumber of predicted TMHs: 1\nExp number of AAs in TMHs: 23.72226\nExp number, first 60 AAs: 0.22038\nTotal prob of N-in: 0.12994\noutside 1 331\nTMhelix 332 354\ninside 355 372

7074 GCF_001615735.1_ASM161573v1 Escherichia coli Proteobacteria; Gammaproteobacteria; Enterobacterales; Enterobacteriaceae; Escherichia
MTGDNTLIHSHGINRRDFMKLCAALAATMGLSSKAAAEMAESVTNPQRPPVIWIGAQECT WP_000145410.1
MULTISPECIES: hydrogenase-2 small chain [Proteobacteria] Length: 372\nNumber of predicted TMHs: 1\nExp number of AAs in TMHs: 23.72226\nExp number, first 60 AAs: 0.22038\nTotal prob of N-in: 0.12994\noutside 1 331\nTMhelix 332 354\ninside 355 372

7075 GCF_001615775.1_ASM161577v1 Escherichia coli Proteobacteria; Gammaproteobacteria; Enterobacterales; Enterobacteriaceae; Escherichia
MTGDNTLIHSHGINRRDFMKLCAALAATMGLSSKAAAEMAESVTNPQRPPVIWIGAQECT WP_000145410.1
MULTISPECIES: hydrogenase-2 small chain [Proteobacteria] Length: 372\nNumber of predicted TMHs: 1\nExp number of AAs in TMHs: 23.72226\nExp number, first 60 AAs: 0.22038\nTotal prob of N-in: 0.12994\noutside 1 331\nTMhelix 332 354\ninside 355 372

7076 GCF_001615835.1_ASM161583v1 Escherichia coli Proteobacteria; Gammaproteobacteria; Enterobacterales; Enterobacteriaceae; Escherichia
MTGDNTLIHSHGINRRDFMKLCAALAATMGLSSKAAAEMAESVTNPQRPPVIWIGAQECT WP_000145410.1
MULTISPECIES: hydrogenase-2 small chain [Proteobacteria] Length: 372\nNumber of predicted TMHs: 1\nExp number of AAs in TMHs: 23.72226\nExp number, first 60 AAs: 0.22038\nTotal prob of N-in: 0.12994\noutside 1 331\nTMhelix 332 354\ninside 355 372

7077 GCF_001615865.1_ASM161586v1 Escherichia coli Proteobacteria; Gammaproteobacteria; Enterobacterales; Enterobacteriaceae; Escherichia
MTGDNTLIHSHGINRRDFMKLCAALAATMGLSSKAAAEMAESVTNPQRPPVIWIGAQECT WP_000145410.1
MULTISPECIES: hydrogenase-2 small chain [Proteobacteria] Length: 372\nNumber of predicted TMHs: 1\nExp number of AAs in TMHs: 23.72226\nExp number, first 60 AAs: 0.22038\nTotal prob of N-in: 0.12994\noutside 1 331\nTMhelix 332 354\ninside 355 372

7078 GCF_001615785.1_ASM161578v1 Escherichia coli Proteobacteria; Gammaproteobacteria; Enterobacterales; Enterobacteriaceae; Escherichia
MTGDNTLIHSHGINRRDFMKLCAALAATMGLSSKAAAEMAESVTNPQRPPVIWIGAQECT WP_000145410.1
MULTISPECIES: hydrogenase-2 small chain [Proteobacteria] Length: 372\nNumber of predicted TMHs: 1\nExp number of AAs in TMHs: 23.72226\nExp number, first 60 AAs: 0.22038\nTotal prob of N-in: 0.12994\noutside 1 331\nTMhelix 332 354\ninside 355 372

7079 GCF_001615865.1_ASM161586v1 Escherichia coli Proteobacteria; Gammaproteobacteria; Enterobacterales; Enterobacteriaceae; Escherichia
MNNEETFYQAMRRQGVTRRSFLKYCSLAATSLGLGAGMAPKIAWALENKPRIPVVWIHGL WP_001058323.1
MULTISPECIES: hydrogenase-1 small chain [Proteobacteria] Length: 372\nNumber of predicted TMHs: 2\nExp number of AAs in TMHs: 37.2608499999999\nExp number, first 60 AAs: 19.52923\nTotal prob of N-in: 0.94351\nPOSSIBLE N-term signal sequence\ninside 1 19\nTMhelix 20 39\noutside 40 325\nTMhelix 326 348\ninside 349 372

7080 GCF_001615785.1_ASM161578v1 Escherichia coli Proteobacteria; Gammaproteobacteria; Enterobacterales; Enterobacteriaceae; Escherichia
MNNEETFYQAMRRQGVTRRSFLKYCSLAATSLGLGAGMAPKIAWALENKPRIPVVWIHGL WP_001058323.1
MULTISPECIES: hydrogenase-1 small chain [Proteobacteria] Length: 372\nNumber of predicted TMHs: 2\nExp number of AAs in TMHs: 37.2608499999999\nExp number, first 60 AAs: 19.52923\nTotal prob of N-in: 0.94351\nPOSSIBLE N-term signal sequence\ninside 1 19\nTMhelix 20 39\noutside 40 325\nTMhelix 326 348\ninside 349 372

7081 GCF_001615625.1_ASM161562v1 Escherichia coli Proteobacteria; Gammaproteobacteria; Enterobacterales; Enterobacteriaceae; Escherichia
MNNEETFYQAMRRQGVTRRSFLKYCSLAATSLGLGAGMAPKIAWALENKPRIPVVWIHGL WP_001058323.1
MULTISPECIES: hydrogenase-1 small chain [Proteobacteria] Length: 372\nNumber of predicted TMHs: 2\nExp number of AAs in TMHs: 37.2608499999999\nExp number, first 60 AAs: 19.52923\nTotal prob of N-in: 0.94351\nPOSSIBLE N-term signal sequence\ninside 1 19\nTMhelix 20 39\noutside 40 325\nTMhelix 326 348\ninside 349 372

7082 GCF_001615895.1_ASM161589v1 Escherichia coli Proteobacteria; Gammaproteobacteria; Enterobacterales; Enterobacteriaceae; Escherichia
MTGDNTLIHSHGINRRDFMKLCAALAATMGLSSKAAAEMAESVTNPQRPPVIWIGAQECT WP_000145410.1
MULTISPECIES: hydrogenase-2 small chain [Proteobacteria] Length: 372\nNumber of predicted TMHs: 1\nExp number of AAs in TMHs: 23.72226\nExp number, first 60 AAs: 0.22038\nTotal prob of N-in: 0.12994\noutside 1 331\nTMhelix 332 354\ninside 355 372

7083 GCF_001615895.1_ASM161589v1 Escherichia coli Proteobacteria; Gammaproteobacteria; Enterobacterales; Enterobacteriaceae; Escherichia
MNNEETFYQAMRRQGVTRRSFLKYCSLAATSLGLGAGMAPKIAWALENKPRIPVVWIHGL WP_001058323.1
MULTISPECIES: hydrogenase-1 small chain [Proteobacteria] Length: 372\nNumber of predicted TMHs: 2\nExp number of AAs in TMHs: 37.2608499999999\nExp number, first 60 AAs: 19.52923\nTotal prob of N-in: 0.94351\nPOSSIBLE N-term signal sequence\ninside 1 19\nTMhelix 20 39\noutside 40 325\nTMhelix 326 348\ninside 349 372

7084 GCF_001615835.1_ASM161583v1 Escherichia coli Proteobacteria; Gammaproteobacteria; Enterobacterales; Enterobacteriaceae; Escherichia
MNNEETFYQAMRRQGVTRRSFLKYCSLAATSLGLGAGMAPKIAWALENKPRIPVVWIHGL WP_001058323.1
MULTISPECIES: hydrogenase-1 small chain [Proteobacteria] Length: 372\nNumber of predicted TMHs: 2\nExp number of AAs in TMHs: 37.2608499999999\nExp number, first 60 AAs: 19.52923\nTotal prob of N-in: 0.94351\nPOSSIBLE N-term signal sequence\ninside 1 19\nTMhelix 20 39\noutside 40 325\nTMhelix 326 348\ninside 349 372

7085 GCF_001615675.1_ASM161567v1 Escherichia coli Proteobacteria; Gammaproteobacteria; Enterobacterales; Enterobacteriaceae; Escherichia
MNNEETFYQAMRRQGVTRRSFLKYCSLAATSLGLGAGMAPKIAWALENKPRIPVVWIHGL WP_001058320.1
hydrogenase [Escherichia coli] Length: 372\nNumber of predicted TMHs: 2\nExp number of AAs in TMHs: 37.26082\nExp number, first 60 AAs: 19.52923\nTotal prob of N-in: 0.94351\nPOSSIBLE N-term signal sequence\ninside 1 19\nTMhelix 20 39\noutside 40 325\nTMhelix 326 348\ninside 349 372

7086 GCF_001615775.1_ASM161577v1 Escherichia coli Proteobacteria; Gammaproteobacteria; Enterobacterales; Enterobacteriaceae; Escherichia
MNNEETFYQAMRRQGVTRRSFLKYCSLAATSLGLGAGMAPKIAWALENKPRIPVVWIHGL WP_001058323.1
MULTISPECIES: hydrogenase-1 small chain [Proteobacteria] Length: 372\nNumber of predicted TMHs: 2\nExp number of AAs in TMHs: 37.2608499999999\nExp number, first 60 AAs: 19.52923\nTotal prob of N-in: 0.94351\nPOSSIBLE N-term signal sequence\ninside 1 19\nTMhelix 20 39\noutside 40 325\nTMhelix 326 348\ninside 349 372

7087 GCF_001615735.1_ASM161573v1 Escherichia coli Proteobacteria; Gammaproteobacteria; Enterobacterales; Enterobacteriaceae; Escherichia
MNNEETFYQAMRRQGVTRRSFLKYCSLAATSLGLGAGMAPKIAWALENKPRIPVVWIHGL WP_001058323.1
MULTISPECIES: hydrogenase-1 small chain [Proteobacteria] Length: 372\nNumber of predicted TMHs: 2\nExp number of AAs in TMHs: 37.2608499999999\nExp number, first 60 AAs: 19.52923\nTotal prob of N-in: 0.94351\nPOSSIBLE N-term signal sequence\ninside 1 19\nTMhelix 20 39\noutside 40 325\nTMhelix 326 348\ninside 349 372

7088 GCF_001615565.1_ASM161556v1 Escherichia coli Proteobacteria; Gammaproteobacteria; Enterobacterales; Enterobacteriaceae; Escherichia
MNNEETFYQAMRRQGVTRRSFLKYCSLAATSLGLGAGMAPKIAWALENKPRIPVVWIHGL WP_001058323.1
MULTISPECIES: hydrogenase-1 small chain [Proteobacteria] Length: 372\nNumber of predicted TMHs: 2\nExp number of AAs in TMHs: 37.2608499999999\nExp number, first 60 AAs: 19.52923\nTotal prob of N-in: 0.94351\nPOSSIBLE N-term signal sequence\ninside 1 19\nTMhelix 20 39\noutside 40 325\nTMhelix 326 348\ninside 349 372

7089 GCF_001615975.1_ASM161597v1 Escherichia coli Proteobacteria; Gammaproteobacteria; Enterobacterales; Enterobacteriaceae; Escherichia
MNNEETFYQAMRRQGVTRRSFLKYCSLAATSLGLGAGMAPKIAWALENKPRIPVVWIHGL WP_001058323.1
MULTISPECIES: hydrogenase-1 small chain [Proteobacteria] Length: 372\nNumber of predicted TMHs: 2\nExp number of AAs in TMHs: 37.2608499999999\nExp number, first 60 AAs: 19.52923\nTotal prob of N-in: 0.94351\nPOSSIBLE N-term signal sequence\ninside 1 19\nTMhelix 20 39\noutside 40 325\nTMhelix 326 348\ninside 349 372

7090 GCF_001616195.1_ASM161619v1 Escherichia coli Proteobacteria; Gammaproteobacteria; Enterobacterales; Enterobacteriaceae; Escherichia
MTGDNTLIHSHGINRRDFMKLCAALAATMGLSSKAAAEMAESVTNPQRPPVIWIGAQECT WP_000145410.1
MULTISPECIES: hydrogenase-2 small chain [Proteobacteria] Length: 372\nNumber of predicted TMHs: 1\nExp number of AAs in TMHs: 23.72226\nExp number, first 60 AAs: 0.22038\nTotal prob of N-in: 0.12994\noutside 1 331\nTMhelix 332 354\ninside 355 372

7091 GCF_001616175.1_ASM161617v1 Escherichia coli Proteobacteria; Gammaproteobacteria; Enterobacterales; Enterobacteriaceae; Escherichia
MTGDNTLIHSHGINRRDFMKLCAALAATMGLSSKAAAEMAESVTNPQRPPVIWIGAQECT WP_000145410.1
MULTISPECIES: hydrogenase-2 small chain [Proteobacteria] Length: 372\nNumber of predicted TMHs: 1\nExp number of AAs in TMHs: 23.72226\nExp number, first 60 AAs: 0.22038\nTotal prob of N-in: 0.12994\noutside 1 331\nTMhelix 332 354\ninside 355 372

7092 GCF_001616135.1_ASM161613v1 Escherichia coli Proteobacteria; Gammaproteobacteria; Enterobacterales; Enterobacteriaceae; Escherichia
MTGDNTLIHSHGINRRDFMKLCAALAATMGLSSKAAAEMAESVTNPQRPPVIWIGAQECT WP_000145410.1
MULTISPECIES: hydrogenase-2 small chain [Proteobacteria] Length: 372\nNumber of predicted TMHs: 1\nExp number of AAs in TMHs: 23.72226\nExp number, first 60 AAs: 0.22038\nTotal prob of N-in: 0.12994\noutside 1 331\nTMhelix 332 354\ninside 355 372

7093 GCF_001616135.1_ASM161613v1 Escherichia coli Proteobacteria; Gammaproteobacteria; Enterobacterales; Enterobacteriaceae; Escherichia
MNNEETFYQAMRRQGVTRRSFLKYCSLAATSLGLGAGMAPKIAWALENKPRIPVVWIHGL WP_001058323.1
MULTISPECIES: hydrogenase-1 small chain [Proteobacteria] Length: 372\nNumber of predicted TMHs: 2\nExp number of AAs in TMHs: 37.2608499999999\nExp number, first 60 AAs: 19.52923\nTotal prob of N-in: 0.94351\nPOSSIBLE N-term signal sequence\ninside 1 19\nTMhelix 20 39\noutside 40 325\nTMhelix 326 348\ninside 349 372

7094 GCF_001616175.1_ASM161617v1 Escherichia coli Proteobacteria; Gammaproteobacteria; Enterobacterales; Enterobacteriaceae; Escherichia
MNNEETFYQAMRRQGVTRRSFLKYCSLAATSLGLGAGMAPKIAWALENKPRIPVVWIHGL WP_001058323.1
MULTISPECIES: hydrogenase-1 small chain [Proteobacteria] Length: 372\nNumber of predicted TMHs: 2\nExp number of AAs in TMHs: 37.2608499999999\nExp number, first 60 AAs: 19.52923\nTotal prob of N-in: 0.94351\nPOSSIBLE N-term signal sequence\ninside 1 19\nTMhelix 20 39\noutside 40 325\nTMhelix 326 348\ninside 349 372

7095 GCF_001615925.1_ASM161592v1 Escherichia coli Proteobacteria; Gammaproteobacteria; Enterobacterales; Enterobacteriaceae; Escherichia
MNNEETFYQAMRRQGVTRRSFLKYCSLAATSLGLGAGMAPKIAWALENKPRIPVVWIHGL WP_001058323.1
MULTISPECIES: hydrogenase-1 small chain [Proteobacteria] Length: 372\nNumber of predicted TMHs: 2\nExp number of AAs in TMHs: 37.2608499999999\nExp number, first 60 AAs: 19.52923\nTotal prob of N-in: 0.94351\nPOSSIBLE N-term signal sequence\ninside 1 19\nTMhelix 20 39\noutside 40 325\nTMhelix 326 348\ninside 349 372

7096 GCF_001616035.1_ASM161603v1 Escherichia coli Proteobacteria; Gammaproteobacteria; Enterobacterales; Enterobacteriaceae; Escherichia
MNNEETFYQAMRRQGVTRRSFLKYCSLAATSLGLGAGMAPKIAWALENKPRIPVVWIHGL WP_001058323.1
MULTISPECIES: hydrogenase-1 small chain [Proteobacteria] Length: 372\nNumber of predicted TMHs: 2\nExp number of AAs in TMHs: 37.2608499999999\nExp number, first 60 AAs: 19.52923\nTotal prob of N-in: 0.94351\nPOSSIBLE N-term signal sequence\ninside 1 19\nTMhelix 20 39\noutside 40 325\nTMhelix 326 348\ninside 349 372

7097 GCF_001616215.1_ASM161621v1 Escherichia coli Proteobacteria; Gammaproteobacteria; Enterobacterales; Enterobacteriaceae; Escherichia
MTGDNTLIHSHGINRRDFMKLCAALAATMGLSSKAAAEMAESVTNPQRPPVIWIGAQECT WP_000145410.1
MULTISPECIES: hydrogenase-2 small chain [Proteobacteria] Length: 372\nNumber of predicted TMHs: 1\nExp number of AAs in TMHs: 23.72226\nExp number, first 60 AAs: 0.22038\nTotal prob of N-in: 0.12994\noutside 1 331\nTMhelix 332 354\ninside 355 372

7098 GCF_001616215.1_ASM161621v1 Escherichia coli Proteobacteria; Gammaproteobacteria; Enterobacterales; Enterobacteriaceae; Escherichia
MNNEETFYQAMRRQGVTRRSFLKYCSLAATSLGLGAGMAPKIAWALENKPRIPVVWIHGL WP_001058323.1
MULTISPECIES: hydrogenase-1 small chain [Proteobacteria] Length: 372\nNumber of predicted TMHs: 2\nExp number of AAs in TMHs: 37.2608499999999\nExp number, first 60 AAs: 19.52923\nTotal prob of N-in: 0.94351\nPOSSIBLE N-term signal sequence\ninside 1 19\nTMhelix 20 39\noutside 40 325\nTMhelix 326 348\ninside 349 372

7099 GCF_001615985.1_ASM161598v1 Escherichia coli Proteobacteria; Gammaproteobacteria; Enterobacterales; Enterobacteriaceae; Escherichia
MNNEETFYQAMRRQGVTRRSFLKYCSLAATSLGLGAGMAPKIAWALENKPRIPVVWIHGL WP_001058323.1
MULTISPECIES: hydrogenase-1 small chain [Proteobacteria] Length: 372\nNumber of predicted TMHs: 2\nExp number of AAs in TMHs: 37.2608499999999\nExp number, first 60 AAs: 19.52923\nTotal prob of N-in: 0.94351\nPOSSIBLE N-term signal sequence\ninside 1 19\nTMhelix 20 39\noutside 40 325\nTMhelix 326 348\ninside 349 372

7100 GCF_001615535.1_ASM161553v1 Escherichia coli Proteobacteria; Gammaproteobacteria; Enterobacterales; Enterobacteriaceae; Escherichia
MTGDNTLIHSHGINRRDFMKLCAALAATMGLSSKAAAEMAESVTNPQRPPVIWIGAQECT WP_000145410.1
MULTISPECIES: hydrogenase-2 small chain [Proteobacteria] Length: 372\nNumber of predicted TMHs: 1\nExp number of AAs in TMHs: 23.72226\nExp number, first 60 AAs: 0.22038\nTotal prob of N-in: 0.12994\noutside 1 331\nTMhelix 332 354\ninside 355 372

7101 GCF_001616475.1_ASM161647v1 Escherichia coli Proteobacteria; Gammaproteobacteria; Enterobacterales; Enterobacteriaceae; Escherichia
MTGDNTLIHSHGINRRDFMKLCAALAATMGLSSKAAAEMAESVTNPQRPPVIWIGAQECT WP_000145410.1
MULTISPECIES: hydrogenase-2 small chain [Proteobacteria] Length: 372\nNumber of predicted TMHs: 1\nExp number of AAs in TMHs: 23.72226\nExp number, first 60 AAs: 0.22038\nTotal prob of N-in: 0.12994\noutside 1 331\nTMhelix 332 354\ninside 355 372

7102 GCF_001616435.1_ASM161643v1 Escherichia coli Proteobacteria; Gammaproteobacteria; Enterobacterales; Enterobacteriaceae; Escherichia
MTGDNTLIHSHGINRRDFMKLCAALAATMGLSSKAAAEMAESVTNPQRPPVIWIGAQECT WP_000145410.1
MULTISPECIES: hydrogenase-2 small chain [Proteobacteria] Length: 372\nNumber of predicted TMHs: 1\nExp number of AAs in TMHs: 23.72226\nExp number, first 60 AAs: 0.22038\nTotal prob of N-in: 0.12994\noutside 1 331\nTMhelix 332 354\ninside 355 372

7103 GCF_001616585.1_ASM161658v1 Escherichia coli Proteobacteria; Gammaproteobacteria; Enterobacterales; Enterobacteriaceae; Escherichia
MTGDNTLIHSHGINRRDFMKLCAALAATMGLSSKAAAEMAESVTNPQRPPVIWIGAQECT WP_000145410.1
MULTISPECIES: hydrogenase-2 small chain [Proteobacteria] Length: 372\nNumber of predicted TMHs: 1\nExp number of AAs in TMHs: 23.72226\nExp number, first 60 AAs: 0.22038\nTotal prob of N-in: 0.12994\noutside 1 331\nTMhelix 332 354\ninside 355 372

7104 GCF_001616495.1_ASM161649v1 Escherichia coli Proteobacteria; Gammaproteobacteria; Enterobacterales; Enterobacteriaceae; Escherichia
MTGDNTLIHSHGINRRDFMKLCAALAATMGLSSKAAAEMAESVTNPQRPPVIWIGAQECT WP_000145410.1
MULTISPECIES: hydrogenase-2 small chain [Proteobacteria] Length: 372\nNumber of predicted TMHs: 1\nExp number of AAs in TMHs: 23.72226\nExp number, first 60 AAs: 0.22038\nTotal prob of N-in: 0.12994\noutside 1 331\nTMhelix 332 354\ninside 355 372

7105 GCF_001616575.1_ASM161657v1 Escherichia coli Proteobacteria; Gammaproteobacteria; Enterobacterales; Enterobacteriaceae; Escherichia
MTGDNTLIHSHGINRRDFMKLCAALAATMGLSSKAAAEMAESVTNPQRPPVIWIGAQECT WP_000145410.1
MULTISPECIES: hydrogenase-2 small chain [Proteobacteria] Length: 372\nNumber of predicted TMHs: 1\nExp number of AAs in TMHs: 23.72226\nExp number, first 60 AAs: 0.22038\nTotal prob of N-in: 0.12994\noutside 1 331\nTMhelix 332 354\ninside 355 372

7106 GCF_001616535.1_ASM161653v1 Escherichia coli Proteobacteria; Gammaproteobacteria; Enterobacterales; Enterobacteriaceae; Escherichia
MTGDNTLIHSHGINRRDFMKLCAALAATMGLSSKAAAEMAESVTNPQRPPVIWIGAQECT WP_000145410.1
MULTISPECIES: hydrogenase-2 small chain [Proteobacteria] Length: 372\nNumber of predicted TMHs: 1\nExp number of AAs in TMHs: 23.72226\nExp number, first 60 AAs: 0.22038\nTotal prob of N-in: 0.12994\noutside 1 331\nTMhelix 332 354\ninside 355 372

7107 GCF_001616475.1_ASM161647v1 Escherichia coli Proteobacteria; Gammaproteobacteria; Enterobacterales; Enterobacteriaceae; Escherichia
MNNEETFYQAMRRQGVTRRSFLKYCSLAATSLGLGAGMAPKIAWALENKPRIPVVWIHGL WP_001058323.1
MULTISPECIES: hydrogenase-1 small chain [Proteobacteria] Length: 372\nNumber of predicted TMHs: 2\nExp number of AAs in TMHs: 37.2608499999999\nExp number, first 60 AAs: 19.52923\nTotal prob of N-in: 0.94351\nPOSSIBLE N-term signal sequence\ninside 1 19\nTMhelix 20 39\noutside 40 325\nTMhelix 326 348\ninside 349 372

7108 GCF_001616825.1_ASM161682v1 Escherichia coli Proteobacteria; Gammaproteobacteria; Enterobacterales; Enterobacteriaceae; Escherichia
MTGDNTLIHSHGINRRDFMKLCAALAATMGLSSKAAAEMAESVTNPQRPPVIWIGAQECT WP_000145410.1
MULTISPECIES: hydrogenase-2 small chain [Proteobacteria] Length: 372\nNumber of predicted TMHs: 1\nExp number of AAs in TMHs: 23.72226\nExp number, first 60 AAs: 0.22038\nTotal prob of N-in: 0.12994\noutside 1 331\nTMhelix 332 354\ninside 355 372

7109 GCF_001616435.1_ASM161643v1 Escherichia coli Proteobacteria; Gammaproteobacteria; Enterobacterales; Enterobacteriaceae; Escherichia
MNNEETFYQAMRRQGVTRRSFLKYCSLAATSLGLGAGMAPKIAWALENKPRIPVVWIHGL WP_001058323.1
MULTISPECIES: hydrogenase-1 small chain [Proteobacteria] Length: 372\nNumber of predicted TMHs:

2\nExp number of AAs in TMHs: 37.2608499999999\nExp number, first 60 AAs: 19.52923\nTotal prob of N-in: 0.94351\nPOSSIBLE N-term signal sequence\ninside 1 19\nTMhelix 20 39\noutside 40 325\nTMhelix 326 348\ninside 349 372

7110 GCF_001616535.1_ASM161653v1 Escherichia coli Proteobacteria; Gammaproteobacteria; Enterobacterales; Enterobacteriaceae; Escherichia MNNEETFYQAMRRQGVTRRSFLKYCSLAATSLGLGAGMAPKIAWALENKPRIPVVWIHGL WP_001058323.1 MULTISPECIES: hydrogenase-1 small chain [Proteobacteria] Length: 372\nNumber of predicted TMHs: 2\nExp number of AAs in TMHs: 37.2608499999999\nExp number, first 60 AAs: 19.52923\nTotal prob of N-in: 0.94351\nPOSSIBLE N-term signal sequence\ninside 1 19\nTMhelix 20 39\noutside 40 325\nTMhelix 326 348\ninside 349 372

7111 GCF_001616585.1_ASM161658v1 Escherichia coli Proteobacteria; Gammaproteobacteria; Enterobacterales; Enterobacteriaceae; Escherichia MNNEETFYQAMRRQGVTRRSFLKYCSLAATSLGLGAGMAPKIAWALENKPRIPVVWIHGL WP_001058323.1 MULTISPECIES: hydrogenase-1 small chain [Proteobacteria] Length: 372\nNumber of predicted TMHs: 2\nExp number of AAs in TMHs: 37.2608499999999\nExp number, first 60 AAs: 19.52923\nTotal prob of N-in: 0.94351\nPOSSIBLE N-term signal sequence\ninside 1 19\nTMhelix 20 39\noutside 40 325\nTMhelix 326 348\ninside 349 372

7112 GCF_001616665.1_ASM161666v1 Escherichia coli Proteobacteria; Gammaproteobacteria; Enterobacterales; Enterobacteriaceae; Escherichia MTGDNTLIHSHGINRRDFMKLCAALAATMGLSSKAAAEMAESVTNPQRPPVIWIGAQECT WP_000145410.1 MULTISPECIES: hydrogenase-2 small chain [Proteobacteria] Length: 372\nNumber of predicted TMHs: 1\nExp number of AAs in TMHs: 23.72226\nExp number, first 60 AAs: 0.22038\nTotal prob of N-in: 0.12994\noutside 1 331\nTMhelix 332 354\ninside 355 372

7113 GCF_001616695.1_ASM161669v1 Escherichia coli Proteobacteria; Gammaproteobacteria; Enterobacterales; Enterobacteriaceae; Escherichia MTGDNTLIHSHGINRRDFMKLCAALAATMGLSSKAAAEMAESVTNPQRPPVIWIGAQECT WP_000145410.1 MULTISPECIES: hydrogenase-2 small chain [Proteobacteria] Length: 372\nNumber of predicted TMHs: 1\nExp number of AAs in TMHs: 23.72226\nExp number, first 60 AAs: 0.22038\nTotal prob of N-in: 0.12994\noutside 1 331\nTMhelix 332 354\ninside 355 372

7114 GCF_001616735.1_ASM161673v1 Escherichia coli Proteobacteria; Gammaproteobacteria; Enterobacterales; Enterobacteriaceae; Escherichia MTGDNTLIHSHGINRRDFMKLCAALAATMGLSSKAAAEMAESVTNPQRPPVIWIGAQECT WP_000145410.1 MULTISPECIES: hydrogenase-2 small chain [Proteobacteria] Length: 372\nNumber of predicted TMHs: 1\nExp number of AAs in TMHs: 23.72226\nExp number, first 60 AAs: 0.22038\nTotal prob of N-in: 0.12994\noutside 1 331\nTMhelix 332 354\ninside 355 372

7115 GCF_001616775.1_ASM161677v1 Escherichia coli Proteobacteria; Gammaproteobacteria; Enterobacterales; Enterobacteriaceae; Escherichia MTGDNTLIHSHGINRRDFMKLCAALAATMGLSSKAAAEMAESVTNPQRPPVIWIGAQECT WP_000145410.1 MULTISPECIES: hydrogenase-2 small chain [Proteobacteria] Length: 372\nNumber of predicted TMHs: 1\nExp number of AAs in TMHs: 23.72226\nExp number, first 60 AAs: 0.22038\nTotal prob of N-in: 0.12994\noutside 1 331\nTMhelix 332 354\ninside 355 372

7116 GCF_001616785.1_ASM161678v1 Escherichia coli Proteobacteria; Gammaproteobacteria; Enterobacterales; Enterobacteriaceae; Escherichia MTGDNTLIHSHGINRRDFMKLCAALAATMGLSSKAAAEMAESVTNPQRPPVIWIGAQECT WP_000145410.1 MULTISPECIES: hydrogenase-2 small chain [Proteobacteria] Length: 372\nNumber of predicted TMHs: 1\nExp number of AAs in TMHs: 23.72226\nExp number, first 60 AAs: 0.22038\nTotal prob of N-in: 0.12994\noutside 1 331\nTMhelix 332 354\ninside 355 372

7117 GCF_001616875.1_ASM161687v1 Escherichia coli Proteobacteria; Gammaproteobacteria; Enterobacterales; Enterobacteriaceae; Escherichia MTGDNTLIHSHGINRRDFMKLCAALAATMGLSSKAAAEMAESVTNPQRPPVIWIGAQECT WP_000145410.1 MULTISPECIES: hydrogenase-2 small chain [Proteobacteria] Length: 372\nNumber of predicted TMHs: 1\nExp number of AAs in TMHs: 23.72226\nExp number, first 60 AAs: 0.22038\nTotal prob of N-in: 0.12994\noutside 1 331\nTMhelix 332 354\ninside 355 372

7118 GCF_001616895.1_ASM161689v1 Escherichia coli Proteobacteria; Gammaproteobacteria; Enterobacterales; Enterobacteriaceae; Escherichia
MTGDNTLIHSHGINRRDFMKLCAALAATMGLSSKAAAEMAESVTNPQRPPVIWIGAQECT WP_000145410.1
MULTISPECIES: hydrogenase-2 small chain [Proteobacteria] Length: 372\nNumber of predicted TMHs: 1\nExp number of AAs in TMHs: 23.72226\nExp number, first 60 AAs: 0.22038\nTotal prob of N-in: 0.12994\noutside 1 331\nTMhelix 332 354\ninside 355 372

7119 GCF_001616695.1_ASM161669v1 Escherichia coli Proteobacteria; Gammaproteobacteria; Enterobacterales; Enterobacteriaceae; Escherichia
MNNEETFYQAMRRQGVTRRSFLKYCSLAATSLGLGAGMAPKIAWALENKPRIPVVWIHGL WP_001058323.1
MULTISPECIES: hydrogenase-1 small chain [Proteobacteria] Length: 372\nNumber of predicted TMHs: 2\nExp number of AAs in TMHs: 37.2608499999999\nExp number, first 60 AAs: 19.52923\nTotal prob of N-in: 0.94351\nPOSSIBLE N-term signal sequence\ninside 1 19\nTMhelix 20 39\noutside 40 325\nTMhelix 326 348\ninside 349 372

7120 GCF_001616985.1_ASM161698v1 Escherichia coli Proteobacteria; Gammaproteobacteria; Enterobacterales; Enterobacteriaceae; Escherichia
MTGDNTLIHSHGINRRDFMKLCAALAATMGLSSKAAAEMAESVTNPQRPPVIWIGAQECT WP_000145410.1
MULTISPECIES: hydrogenase-2 small chain [Proteobacteria] Length: 372\nNumber of predicted TMHs: 1\nExp number of AAs in TMHs: 23.72226\nExp number, first 60 AAs: 0.22038\nTotal prob of N-in: 0.12994\noutside 1 331\nTMhelix 332 354\ninside 355 372

7121 GCF_001616985.1_ASM161698v1 Escherichia coli Proteobacteria; Gammaproteobacteria; Enterobacterales; Enterobacteriaceae; Escherichia
MNNEETFYQAMRRQGVTRRSFLKYCSLAATSLGLGAGMAPKIAWALENKPRIPVVWIHGL WP_001058323.1
MULTISPECIES: hydrogenase-1 small chain [Proteobacteria] Length: 372\nNumber of predicted TMHs: 2\nExp number of AAs in TMHs: 37.2608499999999\nExp number, first 60 AAs: 19.52923\nTotal prob of N-in: 0.94351\nPOSSIBLE N-term signal sequence\ninside 1 19\nTMhelix 20 39\noutside 40 325\nTMhelix 326 348\ninside 349 372

7122 GCF_001616735.1_ASM161673v1 Escherichia coli Proteobacteria; Gammaproteobacteria; Enterobacterales; Enterobacteriaceae; Escherichia
MNNEETFYQAMRRQGVTRRSFLKYCSLAATSLGLGAGMAPKIAWALENKPRIPVVWIHGL WP_001058323.1
MULTISPECIES: hydrogenase-1 small chain [Proteobacteria] Length: 372\nNumber of predicted TMHs: 2\nExp number of AAs in TMHs: 37.2608499999999\nExp number, first 60 AAs: 19.52923\nTotal prob of N-in: 0.94351\nPOSSIBLE N-term signal sequence\ninside 1 19\nTMhelix 20 39\noutside 40 325\nTMhelix 326 348\ninside 349 372

7123 GCF_001616935.1_ASM161693v1 Escherichia coli Proteobacteria; Gammaproteobacteria; Enterobacterales; Enterobacteriaceae; Escherichia
MTGDNTLIHSHGINRRDFMKLCAALAATMGLSSKAAAEMAESVTNPQRPPVIWIGAQECT WP_000145410.1
MULTISPECIES: hydrogenase-2 small chain [Proteobacteria] Length: 372\nNumber of predicted TMHs: 1\nExp number of AAs in TMHs: 23.72226\nExp number, first 60 AAs: 0.22038\nTotal prob of N-in: 0.12994\noutside 1 331\nTMhelix 332 354\ninside 355 372

7124 GCF_001616975.1_ASM161697v1 Escherichia coli Proteobacteria; Gammaproteobacteria; Enterobacterales; Enterobacteriaceae; Escherichia
MTGDNTLIHSHGINRRDFMKLCAALAATMGLSSKAAAEMAESVTNPQRPPVIWIGAQECT WP_000145410.1
MULTISPECIES: hydrogenase-2 small chain [Proteobacteria] Length: 372\nNumber of predicted TMHs: 1\nExp number of AAs in TMHs: 23.72226\nExp number, first 60 AAs: 0.22038\nTotal prob of N-in: 0.12994\noutside 1 331\nTMhelix 332 354\ninside 355 372

7125 GCF_001617025.1_ASM161702v1 Escherichia coli Proteobacteria; Gammaproteobacteria; Enterobacterales; Enterobacteriaceae; Escherichia
MTGDNTLIHSHGINRRDFMKLCAALAATMGLSSKAAAEMAESVTNPQRPPVIWIGAQECT WP_000145410.1
MULTISPECIES: hydrogenase-2 small chain [Proteobacteria] Length: 372\nNumber of predicted TMHs: 1\nExp number of AAs in TMHs: 23.72226\nExp number, first 60 AAs: 0.22038\nTotal prob of N-in: 0.12994\noutside 1 331\nTMhelix 332 354\ninside 355 372

7126 GCF_001616975.1_ASM161697v1 Escherichia coli Proteobacteria; Gammaproteobacteria; Enterobacterales; Enterobacteriaceae; Escherichia

MNNEETFYQAMRRQGVTRRSFLKYCSLAATSLGLGAGMAPKIAWALENKPRIPVVWIHGL WP_001058323.1 MULTISPECIES: hydrogenase-1 small chain [Proteobacteria] Length: 372\nNumber of predicted TMHs: 2\nExp number of AAs in TMHs: 37.2608499999999\nExp number, first 60 AAs: 19.52923\nTotal prob of N-in: 0.94351\nPOSSIBLE N-term signal sequence\ninside 1 19\nTMhelix 20 39\noutside 40 325\nTMhelix 326 348\ninside 349 372

7127 GCF_001616935.1_ASM161693v1 Escherichia coli Proteobacteria; Gammaproteobacteria; Enterobacterales; Enterobacteriaceae; Escherichia MNNEETFYQAMRRQGVTRRSFLKYCSLAATSLGLGAGMAPKIAWALENKPRIPVVWIHGL WP_001058323.1 MULTISPECIES: hydrogenase-1 small chain [Proteobacteria] Length: 372\nNumber of predicted TMHs: 2\nExp number of AAs in TMHs: 37.2608499999999\nExp number, first 60 AAs: 19.52923\nTotal prob of N-in: 0.94351\nPOSSIBLE N-term signal sequence\ninside 1 19\nTMhelix 20 39\noutside 40 325\nTMhelix 326 348\ninside 349 372

7128 GCF_001617025.1_ASM161702v1 Escherichia coli Proteobacteria; Gammaproteobacteria; Enterobacterales; Enterobacteriaceae; Escherichia MNNEETFYQAMRRQGVTRRSFLKYCSLAATSLGLGAGMAPKIAWALENKPRIPVVWIHGL WP_001058323.1 MULTISPECIES: hydrogenase-1 small chain [Proteobacteria] Length: 372\nNumber of predicted TMHs: 2\nExp number of AAs in TMHs: 37.2608499999999\nExp number, first 60 AAs: 19.52923\nTotal prob of N-in: 0.94351\nPOSSIBLE N-term signal sequence\ninside 1 19\nTMhelix 20 39\noutside 40 325\nTMhelix 326 348\ninside 349 372

7129 GCF_001616875.1_ASM161687v1 Escherichia coli Proteobacteria; Gammaproteobacteria; Enterobacterales; Enterobacteriaceae; Escherichia MNNEETFYQAMRRQGVTRRSFLKYCSLAATSLGLGAGMAPKIAWALENKPRIPVVWIHGL WP_001058323.1 MULTISPECIES: hydrogenase-1 small chain [Proteobacteria] Length: 372\nNumber of predicted TMHs: 2\nExp number of AAs in TMHs: 37.2608499999999\nExp number, first 60 AAs: 19.52923\nTotal prob of N-in: 0.94351\nPOSSIBLE N-term signal sequence\ninside 1 19\nTMhelix 20 39\noutside 40 325\nTMhelix 326 348\ninside 349 372

7130 GCF_001617215.1_ASM161721v1 Escherichia coli Proteobacteria; Gammaproteobacteria; Enterobacterales; Enterobacteriaceae; Escherichia MTGDNTLIHSHGINRRDFMKLCAALAATMGLSSKAAAEMAESVTNPQRPPVIWIGAQECT WP_000145410.1 MULTISPECIES: hydrogenase-2 small chain [Proteobacteria] Length: 372\nNumber of predicted TMHs: 1\nExp number of AAs in TMHs: 23.72226\nExp number, first 60 AAs: 0.22038\nTotal prob of N-in: 0.12994\noutside 1 331\nTMhelix 332 354\ninside 355 372

7131 GCF_001617265.1_ASM161726v1 Escherichia coli Proteobacteria; Gammaproteobacteria; Enterobacterales; Enterobacteriaceae; Escherichia MTGDNTLIHSHGINRRDFMKLCAALAATMGLSSKAAAEMAESVTNPQRPPVIWIGAQECT WP_000145410.1 MULTISPECIES: hydrogenase-2 small chain [Proteobacteria] Length: 372\nNumber of predicted TMHs: 1\nExp number of AAs in TMHs: 23.72226\nExp number, first 60 AAs: 0.22038\nTotal prob of N-in: 0.12994\noutside 1 331\nTMhelix 332 354\ninside 355 372

7132 GCF_001617115.1_ASM161711v1 Escherichia coli Proteobacteria; Gammaproteobacteria; Enterobacterales; Enterobacteriaceae; Escherichia MNNEETFYQAMRRQGVTRRSFLKYCSLAATSLGLGAGMAPKIAWALENKPRIPVVWIHGL WP_001058323.1 MULTISPECIES: hydrogenase-1 small chain [Proteobacteria] Length: 372\nNumber of predicted TMHs: 2\nExp number of AAs in TMHs: 37.2608499999999\nExp number, first 60 AAs: 19.52923\nTotal prob of N-in: 0.94351\nPOSSIBLE N-term signal sequence\ninside 1 19\nTMhelix 20 39\noutside 40 325\nTMhelix 326 348\ninside 349 372

7133 GCF_001618325.1_ASM161832v1 Escherichia coli Proteobacteria; Gammaproteobacteria; Enterobacterales; Enterobacteriaceae; Escherichia MTGDNTLIHSHGINRRDFMKLCAALAATMGLSSKAAAEMAESVTNPQRPPVIWIGAQECT WP_000145410.1 MULTISPECIES: hydrogenase-2 small chain [Proteobacteria] Length: 372\nNumber of predicted TMHs: 1\nExp number of AAs in TMHs: 23.72226\nExp number, first 60 AAs: 0.22038\nTotal prob of N-in: 0.12994\noutside 1 331\nTMhelix 332 354\ninside 355 372

7134 GCF_001618345.2_ASM161834v2 Escherichia coli Proteobacteria; Gammaproteobacteria; Enterobacterales; Enterobacteriaceae; Escherichia

MTGDNTLIHSHGINRRDFMKLCAALAATMGLSSKAAAEMAESVTNPQRPPVIWIGAQECT WP_000145410.1 MULTISPECIES: hydrogenase-2 small chain [Proteobacteria] Length: 372\nNumber of predicted TMHs: 1\nExp number of AAs in TMHs: 23.72226\nExp number, first 60 AAs: 0.22038\nTotal prob of N-in: 0.12994\noutside 1 331\nTMhelix 332 354\ninside 355 372

7135 GCF_001617225.1_ASM161722v1 Escherichia coli Proteobacteria; Gammaproteobacteria; Enterobacterales; Enterobacteriaceae; Escherichia MTGDNTLIHSHGINRRDFMKLCAALAATMGLSSKAAAEMAESVTNPQRPPVIWIGAQECT WP_000145410.1 MULTISPECIES: hydrogenase-2 small chain [Proteobacteria] Length: 372\nNumber of predicted TMHs: 1\nExp number of AAs in TMHs: 23.72226\nExp number, first 60 AAs: 0.22038\nTotal prob of N-in: 0.12994\noutside 1 331\nTMhelix 332 354\ninside 355 372

7136 GCF_001618325.1_ASM161832v1 Escherichia coli Proteobacteria; Gammaproteobacteria; Enterobacterales; Enterobacteriaceae; Escherichia MNNEETFYQAMRRQGVTRRSFLKYCSLAATSLGLGAGMAPKIAWALENKPRIPVVWIHGL WP_001058323.1 MULTISPECIES: hydrogenase-1 small chain [Proteobacteria] Length: 372\nNumber of predicted TMHs: 2\nExp number of AAs in TMHs: 37.2608499999999\nExp number, first 60 AAs: 19.52923\nTotal prob of N-in: 0.94351\nPOSSIBLE N-term signal sequence\ninside 1 19\nTMhelix 20 39\noutside 40 325\nTMhelix 326 348\ninside 349 372

7137 GCF_001618345.2_ASM161834v2 Escherichia coli Proteobacteria; Gammaproteobacteria; Enterobacterales; Enterobacteriaceae; Escherichia MNNEETFYQAMRRQGVTRRSFLKYCSLAATSLGLGAGMAPKIAWALENKPRIPVVWIHGL WP_001058323.1 MULTISPECIES: hydrogenase-1 small chain [Proteobacteria] Length: 372\nNumber of predicted TMHs: 2\nExp number of AAs in TMHs: 37.2608499999999\nExp number, first 60 AAs: 19.52923\nTotal prob of N-in: 0.94351\nPOSSIBLE N-term signal sequence\ninside 1 19\nTMhelix 20 39\noutside 40 325\nTMhelix 326 348\ninside 349 372

7138 GCF_001617225.1_ASM161722v1 Escherichia coli Proteobacteria; Gammaproteobacteria; Enterobacterales; Enterobacteriaceae; Escherichia MNNEETFYQAMRRQGVTRRSFLKYCSLAATSLGLGAGMAPKIAWALENKPRIPVVWIHGL WP_001058320.1 hydrogenase [Escherichia coli] Length: 372\nNumber of predicted TMHs: 2\nExp number of AAs in TMHs: 37.26082\nExp number, first 60 AAs: 19.52923\nTotal prob of N-in: 0.94351\nPOSSIBLE N-term signal sequence\ninside 1 19\nTMhelix 20 39\noutside 40 325\nTMhelix 326 348\ninside 349 372

7139 GCF_001617135.1_ASM161713v1 Escherichia coli Proteobacteria; Gammaproteobacteria; Enterobacterales; Enterobacteriaceae; Escherichia MNNEETFYQAMRRQGVTRRSFLKYCSLAATSLGLGAGMAPKIAWALENKPRIPVVWIHGL WP_001058323.1 MULTISPECIES: hydrogenase-1 small chain [Proteobacteria] Length: 372\nNumber of predicted TMHs: 2\nExp number of AAs in TMHs: 37.2608499999999\nExp number, first 60 AAs: 19.52923\nTotal prob of N-in: 0.94351\nPOSSIBLE N-term signal sequence\ninside 1 19\nTMhelix 20 39\noutside 40 325\nTMhelix 326 348\ninside 349 372

7140 GCF_001619085.1_ASM161908v1 Escherichia coli Proteobacteria; Gammaproteobacteria; Enterobacterales; Enterobacteriaceae; Escherichia MTGDNTLIHSHGINRRDFMKLCAALAATMGLSSKAAAEMAESVTNPQRPPVIWIGAQECT WP_000145410.1 MULTISPECIES: hydrogenase-2 small chain [Proteobacteria] Length: 372\nNumber of predicted TMHs: 1\nExp number of AAs in TMHs: 23.72226\nExp number, first 60 AAs: 0.22038\nTotal prob of N-in: 0.12994\noutside 1 331\nTMhelix 332 354\ninside 355 372

7141 GCF_001619065.1_ASM161906v1 Escherichia coli Proteobacteria; Gammaproteobacteria; Enterobacterales; Enterobacteriaceae; Escherichia MTGDNTLIHSHGINRRDFMKLCAALAATMGLSSKAAAEMAESVTNPQRPPVIWIGAQECT WP_000145410.1 MULTISPECIES: hydrogenase-2 small chain [Proteobacteria] Length: 372\nNumber of predicted TMHs: 1\nExp number of AAs in TMHs: 23.72226\nExp number, first 60 AAs: 0.22038\nTotal prob of N-in: 0.12994\noutside 1 331\nTMhelix 332 354\ninside 355 372

7142 GCF_001619085.1_ASM161908v1 Escherichia coli Proteobacteria; Gammaproteobacteria; Enterobacterales; Enterobacteriaceae; Escherichia MNNEETFYQAMRRQGVTRRSFLKYCSLAATSLGLGAGMAPKIAWALENKPRIPVVWIHGL WP_001301613.1 hydrogenase [Escherichia coli] Length: 372\nNumber of predicted TMHs: 2\nExp number of AAs in

TMHs: 37.46598\nExp number, first 60 AAs:  19.53139\nTotal prob of N-in:   0.94364\nPOSSIBLE N-term signal sequence\ninside   1   19\nTMhelix   20   39\noutside   40   325\nTMhelix   326   348\ninside   349   372

7143 GCF_001619065.1_ASM161906v1 Escherichia coli Proteobacteria; Gammaproteobacteria; Enterobacterales; Enterobacteriaceae; Escherichia
MNNEETFYQAMRRQGVTRRSFLKYCSLAATSLGLGAGMAPKIAWALENKPRIPVVWIHGL WP_001301613.1
hydrogenase [Escherichia coli] Length: 372\nNumber of predicted TMHs:  2\nExp number of AAs in TMHs: 37.46598\nExp number, first 60 AAs:  19.53139\nTotal prob of N-in:   0.94364\nPOSSIBLE N-term signal sequence\ninside   1   19\nTMhelix   20   39\noutside   40   325\nTMhelix   326   348\ninside   349   372

7144 GCF_001619205.1_ASM161920v1 Escherichia coli Proteobacteria; Gammaproteobacteria; Enterobacterales; Enterobacteriaceae; Escherichia
MTGDNTLIHSHGINRRDFMKLCAALAATMGLSSKAAAEMAESVTNPQRPPVIWIGAQECT WP_000145410.1
MULTISPECIES: hydrogenase-2 small chain [Proteobacteria] Length: 372\nNumber of predicted TMHs: 1\nExp number of AAs in TMHs: 23.72226\nExp number, first 60 AAs:  0.22038\nTotal prob of N-in:   0.12994\noutside   1   331\nTMhelix   332   354\ninside   355   372

7145 GCF_001620915.1_ASM162091v1 Escherichia coli Proteobacteria; Gammaproteobacteria; Enterobacterales; Enterobacteriaceae; Escherichia
MTGDNTLIHSHGINRRDFMKLCAALAATMGLSSKAAAEMAESVTNPQRPPVIWIGAQECT WP_000145410.1
MULTISPECIES: hydrogenase-2 small chain [Proteobacteria] Length: 372\nNumber of predicted TMHs: 1\nExp number of AAs in TMHs: 23.72226\nExp number, first 60 AAs:  0.22038\nTotal prob of N-in:   0.12994\noutside   1   331\nTMhelix   332   354\ninside   355   372

7146 GCF_001620985.1_ASM162098v1 Escherichia coli Proteobacteria; Gammaproteobacteria; Enterobacterales; Enterobacteriaceae; Escherichia
MTGDNTLIHSHGINRRDFMKLCAALAATMGLSSKAAAEMAESVTNPQRPPVIWIGAQECT WP_000145410.1
MULTISPECIES: hydrogenase-2 small chain [Proteobacteria] Length: 372\nNumber of predicted TMHs: 1\nExp number of AAs in TMHs: 23.72226\nExp number, first 60 AAs:  0.22038\nTotal prob of N-in:   0.12994\noutside   1   331\nTMhelix   332   354\ninside   355   372

7147 GCF_001621005.1_ASM162100v1 Escherichia coli Proteobacteria; Gammaproteobacteria; Enterobacterales; Enterobacteriaceae; Escherichia
MTGDNTLIHSHGINRRDFMKLCAALAATMGLSSKAAAEMAESVTNPQRPPVIWIGAQECT WP_000145410.1
MULTISPECIES: hydrogenase-2 small chain [Proteobacteria] Length: 372\nNumber of predicted TMHs: 1\nExp number of AAs in TMHs: 23.72226\nExp number, first 60 AAs:  0.22038\nTotal prob of N-in:   0.12994\noutside   1   331\nTMhelix   332   354\ninside   355   372

7148 GCF_001620925.1_ASM162092v1 Escherichia coli Proteobacteria; Gammaproteobacteria; Enterobacterales; Enterobacteriaceae; Escherichia
MTGDNTLIHSHGINRRDFMKLCAALAATMGLSSKAAAEMVESVTNPQRPPVIWIGAQECT WP_000145423.1
hydrogenase 2 small subunit [Escherichia coli] Length: 372\nNumber of predicted TMHs:  1\nExp number of AAs in TMHs: 23.78201\nExp number, first 60 AAs:  0.28355\nTotal prob of N-in:   0.13257\noutside   1   331\nTMhelix   332   354\ninside   355   372

7149 GCF_001621005.1_ASM162100v1 Escherichia coli Proteobacteria; Gammaproteobacteria; Enterobacterales; Enterobacteriaceae; Escherichia
MNNEETFYQAMRRQGVTRRSFLKYCSLAATSLGLGAGMAPKIAWALENKPRIPVVWIHGL WP_001058323.1
MULTISPECIES: hydrogenase-1 small chain [Proteobacteria] Length: 372\nNumber of predicted TMHs: 2\nExp number of AAs in TMHs: 37.2608499999999\nExp number, first 60 AAs:  19.52923\nTotal prob of N-in: 0.94351\nPOSSIBLE N-term signal sequence\ninside   1   19\nTMhelix   20   39\noutside   40   325\nTMhelix   326   348\ninside   349   372

7150 GCF_001620925.1_ASM162092v1 Escherichia coli Proteobacteria; Gammaproteobacteria; Enterobacterales; Enterobacteriaceae; Escherichia
MNNEETFYQAMRRQGVTRRSFLKYCSLAATSLGLGAGMAPKIAWALENKPRIPVVWIHGL WP_001295940.1
MULTISPECIES: hydrogenase [Enterobacteriaceae] Length: 372\nNumber of predicted TMHs:  2\nExp number of AAs in TMHs: 37.2608899999999\nExp number, first 60 AAs:  19.52923\nTotal prob of N-in: 0.94351\nPOSSIBLE N-term signal sequence\ninside   1   19\nTMhelix   20   39\noutside   40   325\nTMhelix   326   348\ninside   349   372

7151 GCF_001620925.1_ASM162092v1 Escherichia coli Proteobacteria; Gammaproteobacteria; Enterobacterales; Enterobacteriaceae; Escherichia
MNNNDLFQASRRRFLAQLGGLTVAGMLGPSLLTPRRATAAQAATEAVISKEGILTGSHWG WP_074158664.1
trimethylamine-N-oxide reductase [Escherichia coli] Length: 410\nNumber of predicted TMHs: 1\nExp number of AAs in TMHs: 32.52697\nExp number, first 60 AAs: 11.25077\nTotal prob of N-in: 0.29419\nPOSSIBLE N-term signal sequence\noutside 1 384\nTMhelix 385 407\ninside 408 410

7152 GCF_001620915.1_ASM162091v1 Escherichia coli Proteobacteria; Gammaproteobacteria; Enterobacterales; Enterobacteriaceae; Escherichia
MNNEETFYQAMRRQGVTRRSFLKYCSLAATSLGLGAGMAPKIAWALENKPRIPVVWIHGL WP_001058323.1
MULTISPECIES: hydrogenase-1 small chain [Proteobacteria] Length: 372\nNumber of predicted TMHs: 2\nExp number of AAs in TMHs: 37.2608499999999\nExp number, first 60 AAs: 19.52923\nTotal prob of N-in: 0.94351\nPOSSIBLE N-term signal sequence\ninside 1 19\nTMhelix 20 39\noutside 40 325\nTMhelix 326 348\ninside 349 372

7153 GCF_001621035.1_ASM162103v1 Escherichia coli Proteobacteria; Gammaproteobacteria; Enterobacterales; Enterobacteriaceae; Escherichia
MTGDNTLIHSHGINRRDFMKLCAALAATMGLSSKAAAEMVESVTNPQRPPVIWIGAQECT WP_000145423.1
hydrogenase 2 small subunit [Escherichia coli] Length: 372\nNumber of predicted TMHs: 1\nExp number of AAs in TMHs: 23.78201\nExp number, first 60 AAs: 0.28355\nTotal prob of N-in: 0.13257\noutside 1 331\nTMhelix 332 354\ninside 355 372

7154 GCF_001621075.1_ASM162107v1 Escherichia coli Proteobacteria; Gammaproteobacteria; Enterobacterales; Enterobacteriaceae; Escherichia
MTGDNTLIHSHGINRRDFMKLCAALAATMGLSSKAAAEMAESVTNPQRPPVIWIGAQECT WP_000145410.1
MULTISPECIES: hydrogenase-2 small chain [Proteobacteria] Length: 372\nNumber of predicted TMHs: 1\nExp number of AAs in TMHs: 23.72226\nExp number, first 60 AAs: 0.22038\nTotal prob of N-in: 0.12994\noutside 1 331\nTMhelix 332 354\ninside 355 372

7155 GCF_001621125.1_ASM162112v1 Escherichia coli Proteobacteria; Gammaproteobacteria; Enterobacterales; Enterobacteriaceae; Escherichia
MTGDNTLIHSHGINRRDFMKLCAALAATMGLSSKAAAEMAESVTNPQRPPVIWIGAQECT WP_000145410.1
MULTISPECIES: hydrogenase-2 small chain [Proteobacteria] Length: 372\nNumber of predicted TMHs: 1\nExp number of AAs in TMHs: 23.72226\nExp number, first 60 AAs: 0.22038\nTotal prob of N-in: 0.12994\noutside 1 331\nTMhelix 332 354\ninside 355 372

7156 GCF_001621075.1_ASM162107v1 Escherichia coli Proteobacteria; Gammaproteobacteria; Enterobacterales; Enterobacteriaceae; Escherichia
MNNEETFYQAMRRQGVTRRSFLKYCSLAATSLGLGAGMAPKIAWALENKPRIPVVWIHGL WP_001058323.1
MULTISPECIES: hydrogenase-1 small chain [Proteobacteria] Length: 372\nNumber of predicted TMHs: 2\nExp number of AAs in TMHs: 37.2608499999999\nExp number, first 60 AAs: 19.52923\nTotal prob of N-in: 0.94351\nPOSSIBLE N-term signal sequence\ninside 1 19\nTMhelix 20 39\noutside 40 325\nTMhelix 326 348\ninside 349 372

7157 GCF_001621205.1_ASM162120v1 Escherichia coli Proteobacteria; Gammaproteobacteria; Enterobacterales; Enterobacteriaceae; Escherichia
MTGDNTLIHSHGINRRDFMKLCAALAATMGLSSKAAAEMVESVTNPQRPPVIWIGAQECT WP_000145423.1
hydrogenase 2 small subunit [Escherichia coli] Length: 372\nNumber of predicted TMHs: 1\nExp number of AAs in TMHs: 23.78201\nExp number, first 60 AAs: 0.28355\nTotal prob of N-in: 0.13257\noutside 1 331\nTMhelix 332 354\ninside 355 372

7158 GCF_001621245.1_ASM162124v1 Escherichia coli Proteobacteria; Gammaproteobacteria; Enterobacterales; Enterobacteriaceae; Escherichia
MTGDNTLIHSHGINRRDFMKLCAALAATMGLSSKAAAEMVESVTNPQRPPVIWIGAQECT WP_000145423.1
hydrogenase 2 small subunit [Escherichia coli] Length: 372\nNumber of predicted TMHs: 1\nExp number of AAs in TMHs: 23.78201\nExp number, first 60 AAs: 0.28355\nTotal prob of N-in: 0.13257\noutside 1 331\nTMhelix 332 354\ninside 355 372

7159 GCF_001621275.1_ASM162127v1 Escherichia coli Proteobacteria; Gammaproteobacteria; Enterobacterales; Enterobacteriaceae; Escherichia
MTGDNTLIHSHGINRRDFMKLCAALAATMGLSSKAAAEMVESVTNPQRPPVIWIGAQECT WP_000145423.1

hydrogenase 2 small subunit [Escherichia coli]	Length: 372\nNumber of predicted TMHs: 1\nExp number of AAs in TMHs: 23.78201\nExp number, first 60 AAs: 0.28355\nTotal prob of N-in: 0.13257\noutside 1 331\nTMhelix 332 354\ninside 355 372

7160	GCF_001621345.1_ASM162134v1	Escherichia coli	Proteobacteria; Gammaproteobacteria; Enterobacterales; Enterobacteriaceae; Escherichia
MTGDNTLIHSHGINRRDFMKLCAALAATMGLSSKAAAEMAESVTNPQRPPVIWIGAQECT	WP_000145410.1
MULTISPECIES: hydrogenase-2 small chain [Proteobacteria]	Length: 372\nNumber of predicted TMHs: 1\nExp number of AAs in TMHs: 23.72226\nExp number, first 60 AAs: 0.22038\nTotal prob of N-in: 0.12994\noutside 1 331\nTMhelix 332 354\ninside 355 372

7161	GCF_001621385.1_ASM162138v1	Escherichia coli	Proteobacteria; Gammaproteobacteria; Enterobacterales; Enterobacteriaceae; Escherichia
MTGDNTLIHSHGINRRDFMKLCAALAATMGLSSKAAAEMVESVTNPQRPPVIWIGAQECT	WP_000145423.1
hydrogenase 2 small subunit [Escherichia coli]	Length: 372\nNumber of predicted TMHs: 1\nExp number of AAs in TMHs: 23.78201\nExp number, first 60 AAs: 0.28355\nTotal prob of N-in: 0.13257\noutside 1 331\nTMhelix 332 354\ninside 355 372

7162	GCF_001621305.1_ASM162130v1	Escherichia coli	Proteobacteria; Gammaproteobacteria; Enterobacterales; Enterobacteriaceae; Escherichia
MTGDNTLIHSHGINRRDFMKLCAALAATMGLSSKAAAEMVESVTNPQRPPVIWIGAQECT	WP_000145423.1
hydrogenase 2 small subunit [Escherichia coli]	Length: 372\nNumber of predicted TMHs: 1\nExp number of AAs in TMHs: 23.78201\nExp number, first 60 AAs: 0.28355\nTotal prob of N-in: 0.13257\noutside 1 331\nTMhelix 332 354\ninside 355 372

7163	GCF_001621405.1_ASM162140v1	Escherichia coli	Proteobacteria; Gammaproteobacteria; Enterobacterales; Enterobacteriaceae; Escherichia
MTGDNTLIHSHGINRRDFMKLCAALAATMGLSSKAAAEMAESVTNPQRPPVIWIGAQECT	WP_000145410.1
MULTISPECIES: hydrogenase-2 small chain [Proteobacteria]	Length: 372\nNumber of predicted TMHs: 1\nExp number of AAs in TMHs: 23.72226\nExp number, first 60 AAs: 0.22038\nTotal prob of N-in: 0.12994\noutside 1 331\nTMhelix 332 354\ninside 355 372

7164	GCF_001621435.1_ASM162143v1	Escherichia coli	Proteobacteria; Gammaproteobacteria; Enterobacterales; Enterobacteriaceae; Escherichia
MTGDNTLIHSHGINRRDFMKLCAALAATMGLSSKAAAEMAESVTNPQRPPVIWIGAQECT	WP_000145410.1
MULTISPECIES: hydrogenase-2 small chain [Proteobacteria]	Length: 372\nNumber of predicted TMHs: 1\nExp number of AAs in TMHs: 23.72226\nExp number, first 60 AAs: 0.22038\nTotal prob of N-in: 0.12994\noutside 1 331\nTMhelix 332 354\ninside 355 372

7165	GCF_001621485.1_ASM162148v1	Escherichia coli	Proteobacteria; Gammaproteobacteria; Enterobacterales; Enterobacteriaceae; Escherichia
MTGDNTLIHSHGINRRDFMKLCAALAATMGLSSKAAAEMVESVTNPQRPPVIWIGAQECT	WP_000145423.1
hydrogenase 2 small subunit [Escherichia coli]	Length: 372\nNumber of predicted TMHs: 1\nExp number of AAs in TMHs: 23.78201\nExp number, first 60 AAs: 0.28355\nTotal prob of N-in: 0.13257\noutside 1 331\nTMhelix 332 354\ninside 355 372

7166	GCF_001621505.1_ASM162150v1	Escherichia coli	Proteobacteria; Gammaproteobacteria; Enterobacterales; Enterobacteriaceae; Escherichia
MTGDNTLIHSHGINRRDFMKLCAALAATMGLSSKAAAEMVESVTNPQRPPVIWIGAQECT	WP_000145423.1
hydrogenase 2 small subunit [Escherichia coli]	Length: 372\nNumber of predicted TMHs: 1\nExp number of AAs in TMHs: 23.78201\nExp number, first 60 AAs: 0.28355\nTotal prob of N-in: 0.13257\noutside 1 331\nTMhelix 332 354\ninside 355 372

7167	GCF_001621435.1_ASM162143v1	Escherichia coli	Proteobacteria; Gammaproteobacteria; Enterobacterales; Enterobacteriaceae; Escherichia
MNNEETFYQAMRRQGVTRRSFLKYCSLAATSLGLGAGMAPKIAWALENKPRIPVVWIHGL	WP_001058320.1
hydrogenase [Escherichia coli]	Length: 372\nNumber of predicted TMHs: 2\nExp number of AAs in TMHs: 37.26082\nExp number, first 60 AAs: 19.52923\nTotal prob of N-in: 0.94351\nPOSSIBLE N-term signal sequence\ninside 1 19\nTMhelix 20 39\noutside 40 325\nTMhelix 326 348\ninside 349 372

7168	GCF_001621505.1_ASM162150v1	Escherichia coli	Proteobacteria; Gammaproteobacteria; Enterobacterales; Enterobacteriaceae; Escherichia
	MNNEETFYQAMRRQGVTRRSFLKYCSLAATSLGLGAGMAPKIAWALENKPRIPVVWIHGL	WP_001538532.1
	hydrogenase-1 small chain [Escherichia coli]	Length: 372\nNumber of predicted TMHs:  2\nExp number of AAs in TMHs: 37.27041\nExp number, first 60 AAs:  19.52927\nTotal prob of N-in:   0.94352\nPOSSIBLE N-term signal sequence\ninside   1   19\nTMhelix   20   39\noutside   40   325\nTMhelix   326   348\ninside   349   372

7169	GCF_001621485.1_ASM162148v1	Escherichia coli	Proteobacteria; Gammaproteobacteria; Enterobacterales; Enterobacteriaceae; Escherichia
	MNNEETFYQAMRRQGVTRRSFLKYCSLAATSLGLGAGMAPKIAWALENKPRIPVVWIHGL	WP_001295940.1
	MULTISPECIES: hydrogenase [Enterobacteriaceae]	Length: 372\nNumber of predicted TMHs:  2\nExp number of AAs in TMHs: 37.2608899999999\nExp number, first 60 AAs:  19.52923\nTotal prob of N-in: 0.94351\nPOSSIBLE N-term signal sequence\ninside   1   19\nTMhelix   20   39\noutside   40   325\nTMhelix   326 348\ninside   349   372

7170	GCF_001621345.1_ASM162134v1	Escherichia coli	Proteobacteria; Gammaproteobacteria; Enterobacterales; Enterobacteriaceae; Escherichia
	MNNEETFYQAMRRQGVTRRSFLKYCSLAATSLGLGAGMAPKIAWALENKPRIPVVWIHGL	WP_001058323.1
	MULTISPECIES: hydrogenase-1 small chain [Proteobacteria]	Length: 372\nNumber of predicted TMHs: 2\nExp number of AAs in TMHs: 37.2608499999999\nExp number, first 60 AAs:  19.52923\nTotal prob of N-in: 0.94351\nPOSSIBLE N-term signal sequence\ninside   1   19\nTMhelix   20   39\noutside   40   325\nTMhelix   326 348\ninside   349   372

7171	GCF_001621545.1_ASM162154v1	Escherichia coli	Proteobacteria; Gammaproteobacteria; Enterobacterales; Enterobacteriaceae; Escherichia
	MTGDNTLIHSHGINRRDFMKLCAALAATMGLSSKAAAEMAESVTNPQRPPVIWIGAQECT	WP_000145410.1
	MULTISPECIES: hydrogenase-2 small chain [Proteobacteria]	Length: 372\nNumber of predicted TMHs: 1\nExp number of AAs in TMHs: 23.72226\nExp number, first 60 AAs:  0.22038\nTotal prob of N-in:   0.12994\noutside   1 331\nTMhelix   332   354\ninside   355   372

7172	GCF_001621545.1_ASM162154v1	Escherichia coli	Proteobacteria; Gammaproteobacteria; Enterobacterales; Enterobacteriaceae; Escherichia
	MNNEETFYQAMRRQGVTRRSFLKYCSLAATSLGLGAGMAPKIAWALENKPRIPVVWIHGL	WP_001058323.1
	MULTISPECIES: hydrogenase-1 small chain [Proteobacteria]	Length: 372\nNumber of predicted TMHs: 2\nExp number of AAs in TMHs: 37.2608499999999\nExp number, first 60 AAs:  19.52923\nTotal prob of N-in: 0.94351\nPOSSIBLE N-term signal sequence\ninside   1   19\nTMhelix   20   39\noutside   40   325\nTMhelix   326 348\ninside   349   372

7173	GCF_001621405.1_ASM162140v1	Escherichia coli	Proteobacteria; Gammaproteobacteria; Enterobacterales; Enterobacteriaceae; Escherichia
	MNNEETFYQAMRRQGVTRRSFLKYCSLAATSLGLGAGMAPKIAWALENKPRIPVVWIHGL	WP_001058323.1
	MULTISPECIES: hydrogenase-1 small chain [Proteobacteria]	Length: 372\nNumber of predicted TMHs: 2\nExp number of AAs in TMHs: 37.2608499999999\nExp number, first 60 AAs:  19.52923\nTotal prob of N-in: 0.94351\nPOSSIBLE N-term signal sequence\ninside   1   19\nTMhelix   20   39\noutside   40   325\nTMhelix   326 348\ninside   349   372

7174	GCF_001621385.1_ASM162138v1	Escherichia coli	Proteobacteria; Gammaproteobacteria; Enterobacterales; Enterobacteriaceae; Escherichia
	MNNEETFYQAMRRQGVTRRSFLKYCSLAATSLGLGAGMAPKIAWALENKPRIPVVWIHGL	WP_001538532.1
	hydrogenase-1 small chain [Escherichia coli]	Length: 372\nNumber of predicted TMHs:  2\nExp number of AAs in TMHs: 37.27041\nExp number, first 60 AAs:  19.52927\nTotal prob of N-in:   0.94352\nPOSSIBLE N-term signal sequence\ninside   1   19\nTMhelix   20   39\noutside   40   325\nTMhelix   326   348\ninside   349   372

7175	GCF_001621205.1_ASM162120v1	Escherichia coli	Proteobacteria; Gammaproteobacteria; Enterobacterales; Enterobacteriaceae; Escherichia
	MNNEETFYQAMRRQGVTRRSFLKYCSLAATSLGLGAGMAPKIAWALENKPRIPVVWIHGL	WP_001295940.1
	MULTISPECIES: hydrogenase [Enterobacteriaceae]	Length: 372\nNumber of predicted TMHs:  2\nExp number of AAs in TMHs: 37.2608899999999\nExp number, first 60 AAs:  19.52923\nTotal prob of N-in: 0.94351\nPOSSIBLE N-term signal sequence\ninside   1   19\nTMhelix   20   39\noutside   40   325\nTMhelix   326 348\ninside   349   372

7176 GCF_001621245.1_ASM162124v1 Escherichia coli Proteobacteria; Gammaproteobacteria; Enterobacterales; Enterobacteriaceae; Escherichia
MNNEETFYQAMRRQGVTRRSFLKYCSLAATSLGLGAGMAPKIAWALENKPRIPVVWIHGL WP_001295940.1
MULTISPECIES: hydrogenase [Enterobacteriaceae] Length: 372\nNumber of predicted TMHs: 2\nExp number of AAs in TMHs: 37.2608899999999\nExp number, first 60 AAs: 19.52923\nTotal prob of N-in: 0.94351\nPOSSIBLE N-term signal sequence\ninside 1 19\nTMhelix 20 39\noutside 40 325\nTMhelix 326 348\ninside 349 372

7177 GCF_001621275.1_ASM162127v1 Escherichia coli Proteobacteria; Gammaproteobacteria; Enterobacterales; Enterobacteriaceae; Escherichia
MNNEETFYQAMRRQGVTRRSFLKYCSLAATSLGLGAGMAPKIAWALENKPRIPVVWIHGL WP_001295940.1
MULTISPECIES: hydrogenase [Enterobacteriaceae] Length: 372\nNumber of predicted TMHs: 2\nExp number of AAs in TMHs: 37.2608899999999\nExp number, first 60 AAs: 19.52923\nTotal prob of N-in: 0.94351\nPOSSIBLE N-term signal sequence\ninside 1 19\nTMhelix 20 39\noutside 40 325\nTMhelix 326 348\ninside 349 372

7178 GCF_001621765.1_ASM162176v1 Escherichia coli Proteobacteria; Gammaproteobacteria; Enterobacterales; Enterobacteriaceae; Escherichia
MTGDNTLIHSHGINRRDFMKLCAALAATMGLSSKAAAEMAESVTNPQRPPVIWIGAQECT WP_000145410.1
MULTISPECIES: hydrogenase-2 small chain [Proteobacteria] Length: 372\nNumber of predicted TMHs: 1\nExp number of AAs in TMHs: 23.72226\nExp number, first 60 AAs: 0.22038\nTotal prob of N-in: 0.12994\noutside 1 331\nTMhelix 332 354\ninside 355 372

7179 GCF_001621835.1_ASM162183v1 Escherichia coli Proteobacteria; Gammaproteobacteria; Enterobacterales; Enterobacteriaceae; Escherichia
MTGDNTLIHSHGINRRDFMKLCAALAATMGLSSKAAAEMVESVTNPQRPPVIWIGAQECT WP_000145423.1
hydrogenase 2 small subunit [Escherichia coli] Length: 372\nNumber of predicted TMHs: 1\nExp number of AAs in TMHs: 23.78201\nExp number, first 60 AAs: 0.28355\nTotal prob of N-in: 0.13257\noutside 1 331\nTMhelix 332 354\ninside 355 372

7180 GCF_001621885.1_ASM162188v1 Escherichia coli Proteobacteria; Gammaproteobacteria; Enterobacterales; Enterobacteriaceae; Escherichia
MTGDNTLIHSHGINRRDFMKLCAALAATMGLSSKAAAEMAESVTNPQRPPVIWIGAQECT WP_000145410.1
MULTISPECIES: hydrogenase-2 small chain [Proteobacteria] Length: 372\nNumber of predicted TMHs: 1\nExp number of AAs in TMHs: 23.72226\nExp number, first 60 AAs: 0.22038\nTotal prob of N-in: 0.12994\noutside 1 331\nTMhelix 332 354\ninside 355 372

7181 GCF_001621905.1_ASM162190v1 Escherichia coli Proteobacteria; Gammaproteobacteria; Enterobacterales; Enterobacteriaceae; Escherichia
MTGDNTLIHSHGINRRDFMKLCAALAATMGLSSKAAAEMAESVTNPQRPPVIWIGAQECT WP_000145410.1
MULTISPECIES: hydrogenase-2 small chain [Proteobacteria] Length: 372\nNumber of predicted TMHs: 1\nExp number of AAs in TMHs: 23.72226\nExp number, first 60 AAs: 0.22038\nTotal prob of N-in: 0.12994\noutside 1 331\nTMhelix 332 354\ninside 355 372

7182 GCF_001621905.1_ASM162190v1 Escherichia coli Proteobacteria; Gammaproteobacteria; Enterobacterales; Enterobacteriaceae; Escherichia
MNNEETFYQAMRRQGVTRRSFLKYCSLAATSLGLGAGMAPKIAWALENKPRIPVVWIHGL WP_001058323.1
MULTISPECIES: hydrogenase-1 small chain [Proteobacteria] Length: 372\nNumber of predicted TMHs: 2\nExp number of AAs in TMHs: 37.2608499999999\nExp number, first 60 AAs: 19.52923\nTotal prob of N-in: 0.94351\nPOSSIBLE N-term signal sequence\ninside 1 19\nTMhelix 20 39\noutside 40 325\nTMhelix 326 348\ninside 349 372

7183 GCF_001621885.1_ASM162188v1 Escherichia coli Proteobacteria; Gammaproteobacteria; Enterobacterales; Enterobacteriaceae; Escherichia
MNNEETFYQAMRRQGVTRRSFLKYCSLAATSLGLGAGMAPKIAWALENKPRIPVVWIHGL WP_001058323.1
MULTISPECIES: hydrogenase-1 small chain [Proteobacteria] Length: 372\nNumber of predicted TMHs: 2\nExp number of AAs in TMHs: 37.2608499999999\nExp number, first 60 AAs: 19.52923\nTotal prob of N-in: 0.94351\nPOSSIBLE N-term signal sequence\ninside 1 19\nTMhelix 20 39\noutside 40 325\nTMhelix 326 348\ninside 349 372

7184 GCF_001621765.1_ASM162176v1 Escherichia coli Proteobacteria; Gammaproteobacteria; Enterobacterales; Enterobacteriaceae; Escherichia
MNNEETFYQAMRRQGVTRRSFLKYCSLAATSLGLGAGMAPKIAWALENKPRIPVVWIHGL WP_001295940.1
MULTISPECIES: hydrogenase [Enterobacteriaceae] Length: 372\nNumber of predicted TMHs: 2\nExp number of AAs in TMHs: 37.2608899999999\nExp number, first 60 AAs: 19.52923\nTotal prob of N-in: 0.94351\nPOSSIBLE N-term signal sequence\ninside 1 19\nTMhelix 20 39\noutside 40 325\nTMhelix 326 348\ninside 349 372

7185 GCF_001621675.1_ASM162167v1 Escherichia coli Proteobacteria; Gammaproteobacteria; Enterobacterales; Enterobacteriaceae; Escherichia
MNNEETFYQAMRRQGVTRRSFLKYCSLAATSLGLGAGMAPKIAWALENKPRIPVVWIHGL WP_001058323.1
MULTISPECIES: hydrogenase-1 small chain [Proteobacteria] Length: 372\nNumber of predicted TMHs: 2\nExp number of AAs in TMHs: 37.2608499999999\nExp number, first 60 AAs: 19.52923\nTotal prob of N-in: 0.94351\nPOSSIBLE N-term signal sequence\ninside 1 19\nTMhelix 20 39\noutside 40 325\nTMhelix 326 348\ninside 349 372

7186 GCF_001621575.1_ASM162157v1 Escherichia coli Proteobacteria; Gammaproteobacteria; Enterobacterales; Enterobacteriaceae; Escherichia
MNNEETFYQAMRRQGVTRRSFLKYCSLAATSLGLGAGMAPKIAWALENKPRIPVVWIHGL WP_001295940.1
MULTISPECIES: hydrogenase [Enterobacteriaceae] Length: 372\nNumber of predicted TMHs: 2\nExp number of AAs in TMHs: 37.2608899999999\nExp number, first 60 AAs: 19.52923\nTotal prob of N-in: 0.94351\nPOSSIBLE N-term signal sequence\ninside 1 19\nTMhelix 20 39\noutside 40 325\nTMhelix 326 348\ninside 349 372

7187 GCF_001621585.1_ASM162158v1 Escherichia coli Proteobacteria; Gammaproteobacteria; Enterobacterales; Enterobacteriaceae; Escherichia
MNNEETFYQAMRRQGVTRRSFLKYCSLAATSLGLGAGMAPKIAWALENKPRIPVVWIHGL WP_001058323.1
MULTISPECIES: hydrogenase-1 small chain [Proteobacteria] Length: 372\nNumber of predicted TMHs: 2\nExp number of AAs in TMHs: 37.2608499999999\nExp number, first 60 AAs: 19.52923\nTotal prob of N-in: 0.94351\nPOSSIBLE N-term signal sequence\ninside 1 19\nTMhelix 20 39\noutside 40 325\nTMhelix 326 348\ninside 349 372

7188 GCF_001621945.1_ASM162194v1 Escherichia coli Proteobacteria; Gammaproteobacteria; Enterobacterales; Enterobacteriaceae; Escherichia
MTGDNTLIHSHGINRRDFMKLCAALAATMGLSSKAAAEMVESVTNPQRPPVIWIGAQECT WP_000145423.1
hydrogenase 2 small subunit [Escherichia coli] Length: 372\nNumber of predicted TMHs: 1\nExp number of AAs in TMHs: 23.78201\nExp number, first 60 AAs: 0.28355\nTotal prob of N-in: 0.13257\noutside 1 331\nTMhelix 332 354\ninside 355 372

7189 GCF_001621945.1_ASM162194v1 Escherichia coli Proteobacteria; Gammaproteobacteria; Enterobacterales; Enterobacteriaceae; Escherichia
MNNEETFYQAMRRQGVTRRSFLKYCSLAATSLGLGAGMAPKIAWALENKPRIPVVWIHGL WP_001295940.1
MULTISPECIES: hydrogenase [Enterobacteriaceae] Length: 372\nNumber of predicted TMHs: 2\nExp number of AAs in TMHs: 37.2608899999999\nExp number, first 60 AAs: 19.52923\nTotal prob of N-in: 0.94351\nPOSSIBLE N-term signal sequence\ninside 1 19\nTMhelix 20 39\noutside 40 325\nTMhelix 326 348\ninside 349 372

7190 GCF_001621705.1_ASM162170v1 Escherichia coli Proteobacteria; Gammaproteobacteria; Enterobacterales; Enterobacteriaceae; Escherichia
MNNEETFYQAMRRQGVTRRSFLKYCSLAATSLGLGAGMAPKIAWALENKPRIPVVWIHGL WP_001295940.1
MULTISPECIES: hydrogenase [Enterobacteriaceae] Length: 372\nNumber of predicted TMHs: 2\nExp number of AAs in TMHs: 37.2608899999999\nExp number, first 60 AAs: 19.52923\nTotal prob of N-in: 0.94351\nPOSSIBLE N-term signal sequence\ninside 1 19\nTMhelix 20 39\noutside 40 325\nTMhelix 326 348\ninside 349 372

7191 GCF_001621745.1_ASM162174v1 Escherichia coli Proteobacteria; Gammaproteobacteria; Enterobacterales; Enterobacteriaceae; Escherichia
MNNEETFYQAMRRQGVTRRSFLKYCSLAATSLGLGAGMAPKIAWALENKPRIPVVWIHGL WP_001295940.1
MULTISPECIES: hydrogenase [Enterobacteriaceae] Length: 372\nNumber of predicted TMHs: 2\nExp number of AAs in TMHs: 37.2608899999999\nExp number, first 60 AAs: 19.52923\nTotal prob of N-in:

0.94351\nPOSSIBLE N-term signal sequence\ninside 1 19\nTMhelix 20 39\noutside 40 325\nTMhelix 326 348\ninside 349 372

7192 GCF_001637625.1_ASM163762v1 Escherichia coli Proteobacteria; Gammaproteobacteria; Enterobacterales; Enterobacteriaceae; Escherichia
MTGDNTLIHSHGINRRDFMKLCAALAATMGLSSKAAAEMAESVTNPQRPPVIWIGAQECT WP_000145410.1
MULTISPECIES: hydrogenase-2 small chain [Proteobacteria] Length: 372\nNumber of predicted TMHs: 1\nExp number of AAs in TMHs: 23.72226\nExp number, first 60 AAs: 0.22038\nTotal prob of N-in: 0.12994\noutside 1 331\nTMhelix 332 354\ninside 355 372

7193 GCF_001622055.1_ASM162205v1 Escherichia coli Proteobacteria; Gammaproteobacteria; Enterobacterales; Enterobacteriaceae; Escherichia
MNNEETFYQAMRRQGVTRRSFLKYCSLAATSLGLGAGMAPKIAWALENKPRIPVVWIHGL WP_001058323.1
MULTISPECIES: hydrogenase-1 small chain [Proteobacteria] Length: 372\nNumber of predicted TMHs: 2\nExp number of AAs in TMHs: 37.2608499999999\nExp number, first 60 AAs: 19.52923\nTotal prob of N-in: 0.94351\nPOSSIBLE N-term signal sequence\ninside 1 19\nTMhelix 20 39\noutside 40 325\nTMhelix 326 348\ninside 349 372

7194 GCF_001637635.1_ASM163763v1 Escherichia coli Proteobacteria; Gammaproteobacteria; Enterobacterales; Enterobacteriaceae; Escherichia
MTGDNTLIHSHGINRRDFMKLCAALAATMGLSSKAAAEMAESVTNPQRPPVIWIGAQECT WP_000145410.1
MULTISPECIES: hydrogenase-2 small chain [Proteobacteria] Length: 372\nNumber of predicted TMHs: 1\nExp number of AAs in TMHs: 23.72226\nExp number, first 60 AAs: 0.22038\nTotal prob of N-in: 0.12994\noutside 1 331\nTMhelix 332 354\ninside 355 372

7195 GCF_001637785.1_ASM163778v1 Escherichia coli Proteobacteria; Gammaproteobacteria; Enterobacterales; Enterobacteriaceae; Escherichia
MTGDNTLIHSHGINRRDFMKLCAALAATMGLSSKAAAEMAESVTNPQRPPVIWIGAQECT WP_000145410.1
MULTISPECIES: hydrogenase-2 small chain [Proteobacteria] Length: 372\nNumber of predicted TMHs: 1\nExp number of AAs in TMHs: 23.72226\nExp number, first 60 AAs: 0.22038\nTotal prob of N-in: 0.12994\noutside 1 331\nTMhelix 332 354\ninside 355 372

7196 GCF_001637715.1_ASM163771v1 Escherichia coli Proteobacteria; Gammaproteobacteria; Enterobacterales; Enterobacteriaceae; Escherichia
MTGDNTLIHSHGINRRDFMKLCAALAATMGLSSKAAAEMAESVTNPQRPPVIWIGAQECT WP_000145410.1
MULTISPECIES: hydrogenase-2 small chain [Proteobacteria] Length: 372\nNumber of predicted TMHs: 1\nExp number of AAs in TMHs: 23.72226\nExp number, first 60 AAs: 0.22038\nTotal prob of N-in: 0.12994\noutside 1 331\nTMhelix 332 354\ninside 355 372

7197 GCF_001637655.1_ASM163765v1 Escherichia coli Proteobacteria; Gammaproteobacteria; Enterobacterales; Enterobacteriaceae; Escherichia
MTGDNTLIHSHGINRRDFMKLCAALAATMGLSSKAAAEMAESVTNPQRPPVIWIGAQECT WP_000145410.1
MULTISPECIES: hydrogenase-2 small chain [Proteobacteria] Length: 372\nNumber of predicted TMHs: 1\nExp number of AAs in TMHs: 23.72226\nExp number, first 60 AAs: 0.22038\nTotal prob of N-in: 0.12994\noutside 1 331\nTMhelix 332 354\ninside 355 372

7198 GCF_001637735.1_ASM163773v1 Escherichia coli Proteobacteria; Gammaproteobacteria; Enterobacterales; Enterobacteriaceae; Escherichia
MTGDNTLIHSHGINRRDFMKLCAALAATMGLSSKAAAEMAESVTNPQRPPVIWIGAQECT WP_000145410.1
MULTISPECIES: hydrogenase-2 small chain [Proteobacteria] Length: 372\nNumber of predicted TMHs: 1\nExp number of AAs in TMHs: 23.72226\nExp number, first 60 AAs: 0.22038\nTotal prob of N-in: 0.12994\noutside 1 331\nTMhelix 332 354\ninside 355 372

7199 GCF_001637825.1_ASM163782v1 Escherichia coli Proteobacteria; Gammaproteobacteria; Enterobacterales; Enterobacteriaceae; Escherichia
MTGDNTLIHSHGINRRDFMKLCAALAATMGLSSKAAAEMVESVTNPQRPPVIWIGAQECT WP_000145423.1
hydrogenase 2 small subunit [Escherichia coli] Length: 372\nNumber of predicted TMHs: 1\nExp number of AAs in TMHs: 23.78201\nExp number, first 60 AAs: 0.28355\nTotal prob of N-in: 0.13257\noutside 1 331\nTMhelix 332 354\ninside 355 372

7200 GCF_001637785.1_ASM163778v1 Escherichia coli Proteobacteria; Gammaproteobacteria; Enterobacterales; Enterobacteriaceae; Escherichia
MNNEETFYQAMRRQGVTRRSFLKYCSLAATSLGLGAGMAPKIAWALENKPRIPVVWIHGL WP_001058320.1
hydrogenase [Escherichia coli] Length: 372\nNumber of predicted TMHs: 2\nExp number of AAs in TMHs: 37.26082\nExp number, first 60 AAs: 19.52923\nTotal prob of N-in: 0.94351\nPOSSIBLE N-term signal sequence\ninside 1 19\nTMhelix 20 39\noutside 40 325\nTMhelix 326 348\ninside 349 372

7201 GCF_001637655.1_ASM163765v1 Escherichia coli Proteobacteria; Gammaproteobacteria; Enterobacterales; Enterobacteriaceae; Escherichia
MNNEETFYQAMRRQGVTRRSFLKYCSLAATSLGLGAGMAPKIAWALENKPRIPVVWIHGL WP_001058320.1
hydrogenase [Escherichia coli] Length: 372\nNumber of predicted TMHs: 2\nExp number of AAs in TMHs: 37.26082\nExp number, first 60 AAs: 19.52923\nTotal prob of N-in: 0.94351\nPOSSIBLE N-term signal sequence\ninside 1 19\nTMhelix 20 39\noutside 40 325\nTMhelix 326 348\ninside 349 372

7202 GCF_001622115.1_ASM162211v1 Escherichia coli Proteobacteria; Gammaproteobacteria; Enterobacterales; Enterobacteriaceae; Escherichia
MNNEETFYQAMRRQGVTRRSFLKYCSLAATSLGLGAGMAPKIAWALENKPRIPVVWIHGL WP_001538532.1
hydrogenase-1 small chain [Escherichia coli] Length: 372\nNumber of predicted TMHs: 2\nExp number of AAs in TMHs: 37.27041\nExp number, first 60 AAs: 19.52927\nTotal prob of N-in: 0.94352\nPOSSIBLE N-term signal sequence\ninside 1 19\nTMhelix 20 39\noutside 40 325\nTMhelix 326 348\ninside 349 372

7203 GCF_001622075.1_ASM162207v1 Escherichia coli Proteobacteria; Gammaproteobacteria; Enterobacterales; Enterobacteriaceae; Escherichia
MNNEETFYQAMRRQGVTRRSFLKYCSLAATSLGLGAGMAPKIAWALENKPRIPVVWIHGL WP_001058323.1
MULTISPECIES: hydrogenase-1 small chain [Proteobacteria] Length: 372\nNumber of predicted TMHs: 2\nExp number of AAs in TMHs: 37.2608499999999\nExp number, first 60 AAs: 19.52923\nTotal prob of N-in: 0.94351\nPOSSIBLE N-term signal sequence\ninside 1 19\nTMhelix 20 39\noutside 40 325\nTMhelix 326 348\ninside 349 372

7204 GCF_001637845.1_ASM163784v1 Escherichia coli Proteobacteria; Gammaproteobacteria; Enterobacterales; Enterobacteriaceae; Escherichia
MTGDNTLIHSHGINRRDFMKLCAALAATMGLSSKAAAEMAESVTNPQRPPVIWIGAQECT WP_000145410.1
MULTISPECIES: hydrogenase-2 small chain [Proteobacteria] Length: 372\nNumber of predicted TMHs: 1\nExp number of AAs in TMHs: 23.72226\nExp number, first 60 AAs: 0.22038\nTotal prob of N-in: 0.12994\noutside 1 331\nTMhelix 332 354\ninside 355 372

7205 GCF_001637825.1_ASM163782v1 Escherichia coli Proteobacteria; Gammaproteobacteria; Enterobacterales; Enterobacteriaceae; Escherichia
MNNEETFYQAMRRQGVTRRSFLKYCSLAATSLGLGAGMAPKIAWALENKPRIPVVWIHGL WP_001295940.1
MULTISPECIES: hydrogenase [Enterobacteriaceae] Length: 372\nNumber of predicted TMHs: 2\nExp number of AAs in TMHs: 37.2608899999999\nExp number, first 60 AAs: 19.52923\nTotal prob of N-in: 0.94351\nPOSSIBLE N-term signal sequence\ninside 1 19\nTMhelix 20 39\noutside 40 325\nTMhelix 326 348\ninside 349 372

7206 GCF_001637945.1_ASM163794v1 Escherichia coli Proteobacteria; Gammaproteobacteria; Enterobacterales; Enterobacteriaceae; Escherichia
MTGDNTLIHSHGINRRDFMKLCAALAATMGLSSKAAAEMVESVTNPQRPPVIWIGAQECT WP_000145423.1
hydrogenase 2 small subunit [Escherichia coli] Length: 372\nNumber of predicted TMHs: 1\nExp number of AAs in TMHs: 23.78201\nExp number, first 60 AAs: 0.28355\nTotal prob of N-in: 0.13257\noutside 1 331\nTMhelix 332 354\ninside 355 372

7207 GCF_001637985.1_ASM163798v1 Escherichia coli Proteobacteria; Gammaproteobacteria; Enterobacterales; Enterobacteriaceae; Escherichia
MTGDNTLIHSHGINRRDFMKLCAALAATMGLSSKAAAEMVESVTNPQRPPVIWIGAQECT WP_000145423.1
hydrogenase 2 small subunit [Escherichia coli] Length: 372\nNumber of predicted TMHs: 1\nExp number of AAs in TMHs: 23.78201\nExp number, first 60 AAs: 0.28355\nTotal prob of N-in: 0.13257\noutside 1 331\nTMhelix 332 354\ninside 355 372

7208 GCF_001637905.1_ASM163790v1 Escherichia coli Proteobacteria; Gammaproteobacteria; Enterobacterales; Enterobacteriaceae; Escherichia
MTGDNTLIHSHGINRRDFMKLCAALAATMGLSSKAAAEMVESVTNPQRPPVIWIGAQECT WP_000145423.1

hydrogenase 2 small subunit [Escherichia coli] Length: 372\nNumber of predicted TMHs: 1\nExp number of AAs in TMHs: 23.78201\nExp number, first 60 AAs: 0.28355\nTotal prob of N-in: 0.13257\noutside 1 331\nTMhelix 332 354\ninside 355 372

7209 GCF_001638025.1_ASM163802v1 Escherichia coli Proteobacteria; Gammaproteobacteria; Enterobacterales; Enterobacteriaceae; Escherichia MTGDNTLIHSHGINRRDFMKLCAALAATMGLSSKAAAEMAESVTNPQRPPVIWIGAQECT WP_000145410.1 MULTISPECIES: hydrogenase-2 small chain [Proteobacteria] Length: 372\nNumber of predicted TMHs: 1\nExp number of AAs in TMHs: 23.72226\nExp number, first 60 AAs: 0.22038\nTotal prob of N-in: 0.12994\noutside 1 331\nTMhelix 332 354\ninside 355 372

7210 GCF_001638045.1_ASM163804v1 Escherichia coli Proteobacteria; Gammaproteobacteria; Enterobacterales; Enterobacteriaceae; Escherichia MTGDNTLIHSHGINRRDFMKLCAALAATMGLSSKAAAEMVESVTNPQRPPVIWIGAQECT WP_000145423.1 hydrogenase 2 small subunit [Escherichia coli] Length: 372\nNumber of predicted TMHs: 1\nExp number of AAs in TMHs: 23.78201\nExp number, first 60 AAs: 0.28355\nTotal prob of N-in: 0.13257\noutside 1 331\nTMhelix 332 354\ninside 355 372

7211 GCF_001638065.1_ASM163806v1 Escherichia coli Proteobacteria; Gammaproteobacteria; Enterobacterales; Enterobacteriaceae; Escherichia MTGDNTLIHSHGINRRDFMKLCAALAATMGLSSKAAAEMVESVTNPQRPPVIWIGAQECT WP_000145423.1 hydrogenase 2 small subunit [Escherichia coli] Length: 372\nNumber of predicted TMHs: 1\nExp number of AAs in TMHs: 23.78201\nExp number, first 60 AAs: 0.28355\nTotal prob of N-in: 0.13257\noutside 1 331\nTMhelix 332 354\ninside 355 372

7212 GCF_001638125.1_ASM163812v1 Escherichia coli Proteobacteria; Gammaproteobacteria; Enterobacterales; Enterobacteriaceae; Escherichia MTGDNTLIHSHGINRRDFMKLCAALAATMGLSSKAAAEMVESVTNPQRPPVIWIGAQECT WP_000145423.1 hydrogenase 2 small subunit [Escherichia coli] Length: 372\nNumber of predicted TMHs: 1\nExp number of AAs in TMHs: 23.78201\nExp number, first 60 AAs: 0.28355\nTotal prob of N-in: 0.13257\noutside 1 331\nTMhelix 332 354\ninside 355 372

7213 GCF_001638135.1_ASM163813v1 Escherichia coli Proteobacteria; Gammaproteobacteria; Enterobacterales; Enterobacteriaceae; Escherichia MTGDNTLIHSHGINRRDFMKLCAALAATMGLSSKAAAEMAESVTNPQRPPVIWIGAQECT WP_000145410.1 MULTISPECIES: hydrogenase-2 small chain [Proteobacteria] Length: 372\nNumber of predicted TMHs: 1\nExp number of AAs in TMHs: 23.72226\nExp number, first 60 AAs: 0.22038\nTotal prob of N-in: 0.12994\noutside 1 331\nTMhelix 332 354\ninside 355 372

7214 GCF_001638165.1_ASM163816v1 Escherichia coli Proteobacteria; Gammaproteobacteria; Enterobacterales; Enterobacteriaceae; Escherichia MTGDNTLIHSHGINRRDFMKLCAALAATMGLSSKAAAEMVESVTNPQRPPVIWIGAQECT WP_000145423.1 hydrogenase 2 small subunit [Escherichia coli] Length: 372\nNumber of predicted TMHs: 1\nExp number of AAs in TMHs: 23.78201\nExp number, first 60 AAs: 0.28355\nTotal prob of N-in: 0.13257\noutside 1 331\nTMhelix 332 354\ninside 355 372

7215 GCF_001638225.1_ASM163822v1 Escherichia coli Proteobacteria; Gammaproteobacteria; Enterobacterales; Enterobacteriaceae; Escherichia MTGDNTLIHSHGINRRDFMKLCAALAATMGLSSKAAAEMVESVTNPQRPPVIWIGAQECT WP_000145423.1 hydrogenase 2 small subunit [Escherichia coli] Length: 372\nNumber of predicted TMHs: 1\nExp number of AAs in TMHs: 23.78201\nExp number, first 60 AAs: 0.28355\nTotal prob of N-in: 0.13257\noutside 1 331\nTMhelix 332 354\ninside 355 372

7216 GCF_001640145.1_ASM164014v1 Escherichia coli Proteobacteria; Gammaproteobacteria; Enterobacterales; Enterobacteriaceae; Escherichia MTGDNTLIHSHGINRRDFMKLCAALAATMGLSSKAAAEMAESVTNPQRPPVIWIGAQECT WP_000145411.1 hydrogenase-2 small chain [Escherichia coli] Length: 372\nNumber of predicted TMHs: 1\nExp number of AAs in TMHs: 23.70577\nExp number, first 60 AAs: 0.22039\nTotal prob of N-in: 0.12981\noutside 1 331\nTMhelix 332 354\ninside 355 372

7217 GCF_001638135.1_ASM163813v1 Escherichia coli Proteobacteria; Gammaproteobacteria; Enterobacterales; Enterobacteriaceae; Escherichia
MNNEETFYQAMRRQGVTRRSFLKYCSLAATSLGLGAGMAPKIAWALENKPRIPVVWIHGL WP_001058323.1
MULTISPECIES: hydrogenase-1 small chain [Proteobacteria] Length: 372\nNumber of predicted TMHs: 2\nExp number of AAs in TMHs: 37.2608499999999\nExp number, first 60 AAs: 19.52923\nTotal prob of N-in: 0.94351\nPOSSIBLE N-term signal sequence\ninside 1 19\nTMhelix 20 39\noutside 40 325\nTMhelix 326 348\ninside 349 372

7218 GCF_001640145.1_ASM164014v1 Escherichia coli Proteobacteria; Gammaproteobacteria; Enterobacterales; Enterobacteriaceae; Escherichia
MNNEETFYQAMRRQGVTRRSFLKYCSLAATSLGLGAGMAPKIAWALENKPRIPVVWIHGL WP_001058323.1
MULTISPECIES: hydrogenase-1 small chain [Proteobacteria] Length: 372\nNumber of predicted TMHs: 2\nExp number of AAs in TMHs: 37.2608499999999\nExp number, first 60 AAs: 19.52923\nTotal prob of N-in: 0.94351\nPOSSIBLE N-term signal sequence\ninside 1 19\nTMhelix 20 39\noutside 40 325\nTMhelix 326 348\ninside 349 372

7219 GCF_001638165.1_ASM163816v1 Escherichia coli Proteobacteria; Gammaproteobacteria; Enterobacterales; Enterobacteriaceae; Escherichia
MNNEETFYQAMRRQGVTRRSFLKYCSLAATSLGLGAGMAPKIAWALENKPRIPVVWIHGL WP_001295940.1
MULTISPECIES: hydrogenase [Enterobacteriaceae] Length: 372\nNumber of predicted TMHs: 2\nExp number of AAs in TMHs: 37.2608899999999\nExp number, first 60 AAs: 19.52923\nTotal prob of N-in: 0.94351\nPOSSIBLE N-term signal sequence\ninside 1 19\nTMhelix 20 39\noutside 40 325\nTMhelix 326 348\ninside 349 372

7220 GCF_001638065.1_ASM163806v1 Escherichia coli Proteobacteria; Gammaproteobacteria; Enterobacterales; Enterobacteriaceae; Escherichia
MNNEETFYQAMRRQGVTRRSFLKYCSLAATSLGLGAGMAPKIAWALENKPRIPVVWIHGL WP_001295940.1
MULTISPECIES: hydrogenase [Enterobacteriaceae] Length: 372\nNumber of predicted TMHs: 2\nExp number of AAs in TMHs: 37.2608899999999\nExp number, first 60 AAs: 19.52923\nTotal prob of N-in: 0.94351\nPOSSIBLE N-term signal sequence\ninside 1 19\nTMhelix 20 39\noutside 40 325\nTMhelix 326 348\ninside 349 372

7221 GCF_001638125.1_ASM163812v1 Escherichia coli Proteobacteria; Gammaproteobacteria; Enterobacterales; Enterobacteriaceae; Escherichia
MNNEETFYQAMRRQGVTRRSFLKYCSLAATSLGLGAGMAPKIAWALENKPRIPVVWIHGL WP_001295940.1
MULTISPECIES: hydrogenase [Enterobacteriaceae] Length: 372\nNumber of predicted TMHs: 2\nExp number of AAs in TMHs: 37.2608899999999\nExp number, first 60 AAs: 19.52923\nTotal prob of N-in: 0.94351\nPOSSIBLE N-term signal sequence\ninside 1 19\nTMhelix 20 39\noutside 40 325\nTMhelix 326 348\ninside 349 372

7222 GCF_001638225.1_ASM163822v1 Escherichia coli Proteobacteria; Gammaproteobacteria; Enterobacterales; Enterobacteriaceae; Escherichia
MNNEETFYQAMRRQGVTRRSFLKYCSLAATSLGLGAGMAPKIAWALENKPRIPVVWIHGL WP_001295940.1
MULTISPECIES: hydrogenase [Enterobacteriaceae] Length: 372\nNumber of predicted TMHs: 2\nExp number of AAs in TMHs: 37.2608899999999\nExp number, first 60 AAs: 19.52923\nTotal prob of N-in: 0.94351\nPOSSIBLE N-term signal sequence\ninside 1 19\nTMhelix 20 39\noutside 40 325\nTMhelix 326 348\ninside 349 372

7223 GCF_001641435.1_ASM164143v1 Escherichia coli Proteobacteria; Gammaproteobacteria; Enterobacterales; Enterobacteriaceae; Escherichia
MTGDNTLIHSHGINRRDFMKLCAALAATMGLSSKAAAEMAESVTNPQRPPVIWIGAQECT WP_000145410.1
MULTISPECIES: hydrogenase-2 small chain [Proteobacteria] Length: 372\nNumber of predicted TMHs: 1\nExp number of AAs in TMHs: 23.72226\nExp number, first 60 AAs: 0.22038\nTotal prob of N-in: 0.12994\noutside 1 331\nTMhelix 332 354\ninside 355 372

7224 GCF_001641865.1_ASM164186v1 Escherichia coli Proteobacteria; Gammaproteobacteria; Enterobacterales; Enterobacteriaceae; Escherichia
MTGDNTLIHSHGINRRDFMKLCAALAATMGLSSKAAAEMAESVTNPQRPPVIWIGAQECT WP_000145410.1
MULTISPECIES: hydrogenase-2 small chain [Proteobacteria] Length: 372\nNumber of predicted TMHs: 1\nExp number of AAs in TMHs: 23.72226\nExp number, first 60 AAs: 0.22038\nTotal prob of N-in: 0.12994\noutside 1 331\nTMhelix 332 354\ninside 355 372

7225 GCF_001641845.1_ASM164184v1 Escherichia coli Proteobacteria; Gammaproteobacteria; Enterobacterales; Enterobacteriaceae; Escherichia
MTGDNTLIHSHGINRRDFMKLCAALAATMGLSSKAAAEMAESVTNPQRPPVIWIGAQECT WP_000145410.1
MULTISPECIES: hydrogenase-2 small chain [Proteobacteria] Length: 372\nNumber of predicted TMHs: 1\nExp number of AAs in TMHs: 23.72226\nExp number, first 60 AAs: 0.22038\nTotal prob of N-in: 0.12994\noutside 1 331\nTMhelix 332 354\ninside 355 372

7226 GCF_001641925.1_ASM164192v1 Escherichia coli Proteobacteria; Gammaproteobacteria; Enterobacterales; Enterobacteriaceae; Escherichia
MTGDNTLIHSHGINRRDFMKLCAALAATMGLSSKAAAEMAESVTNPQRPPVIWIGAQECT WP_000145410.1
MULTISPECIES: hydrogenase-2 small chain [Proteobacteria] Length: 372\nNumber of predicted TMHs: 1\nExp number of AAs in TMHs: 23.72226\nExp number, first 60 AAs: 0.22038\nTotal prob of N-in: 0.12994\noutside 1 331\nTMhelix 332 354\ninside 355 372

7227 GCF_001641955.1_ASM164195v1 Escherichia coli Proteobacteria; Gammaproteobacteria; Enterobacterales; Enterobacteriaceae; Escherichia
MTGDNTLIHSHGINRRDFMKLCAALAATMGLSSKAAAEMAESVTNPQRPPVIWIGAQECT WP_000145410.1
MULTISPECIES: hydrogenase-2 small chain [Proteobacteria] Length: 372\nNumber of predicted TMHs: 1\nExp number of AAs in TMHs: 23.72226\nExp number, first 60 AAs: 0.22038\nTotal prob of N-in: 0.12994\noutside 1 331\nTMhelix 332 354\ninside 355 372

7228 GCF_001644175.1_ASM164417v1 Escherichia coli Proteobacteria; Gammaproteobacteria; Enterobacterales; Enterobacteriaceae; Escherichia
MTGDNTLIHSHGINRRDFMKLCAALAATMGLSSKAAAEMAESVTNPQRPPVIWIGAQECT WP_000145410.1
MULTISPECIES: hydrogenase-2 small chain [Proteobacteria] Length: 372\nNumber of predicted TMHs: 1\nExp number of AAs in TMHs: 23.72226\nExp number, first 60 AAs: 0.22038\nTotal prob of N-in: 0.12994\noutside 1 331\nTMhelix 332 354\ninside 355 372

7229 GCF_001644745.1_ASM164474v1 Escherichia coli Proteobacteria; Gammaproteobacteria; Enterobacterales; Enterobacteriaceae; Escherichia
MTGDNTLIHSHGINRRDFMKLCAALAATMGLSSKAAAEMAESVTNPQRPPVIWIGAQECT WP_000145410.1
MULTISPECIES: hydrogenase-2 small chain [Proteobacteria] Length: 372\nNumber of predicted TMHs: 1\nExp number of AAs in TMHs: 23.72226\nExp number, first 60 AAs: 0.22038\nTotal prob of N-in: 0.12994\noutside 1 331\nTMhelix 332 354\ninside 355 372

7230 GCF_001644745.1_ASM164474v1 Escherichia coli Proteobacteria; Gammaproteobacteria; Enterobacterales; Enterobacteriaceae; Escherichia
MNNEETFYQAMRRQGVTRRSFLKYCSLAATSLGLGAGMAPKIAWALENKPRIPVVWIHGL WP_001359719.1
hydrogenase [Escherichia coli] Length: 372\nNumber of predicted TMHs: 2\nExp number of AAs in TMHs: 37.2176\nExp number, first 60 AAs: 19.53448\nTotal prob of N-in: 0.94360\nPOSSIBLE N-term signal sequence\ninside 1 19\nTMhelix 20 39\noutside 40 325\nTMhelix 326 348\ninside 349 372

7231 GCF_001641435.1_ASM164143v1 Escherichia coli Proteobacteria; Gammaproteobacteria; Enterobacterales; Enterobacteriaceae; Escherichia
MNNEETFYQAMRRQGVTRRSFLKYCSLAATSLGLGAGMAPKIAWALENKPRIPVVWIHGL WP_001058323.1
MULTISPECIES: hydrogenase-1 small chain [Proteobacteria] Length: 372\nNumber of predicted TMHs: 2\nExp number of AAs in TMHs: 37.2608499999999\nExp number, first 60 AAs: 19.52923\nTotal prob of N-in: 0.94351\nPOSSIBLE N-term signal sequence\ninside 1 19\nTMhelix 20 39\noutside 40 325\nTMhelix 326 348\ninside 349 372

7232 GCF_001645245.1_ASM164524v1 Escherichia coli Proteobacteria; Gammaproteobacteria; Enterobacterales; Enterobacteriaceae; Escherichia
MTGDNTLIHSHGINRRDFMKLCAALAATMGLSSKAAAEMAESVTNPQRPPVIWIGAQECT WP_000145410.1
MULTISPECIES: hydrogenase-2 small chain [Proteobacteria] Length: 372\nNumber of predicted TMHs: 1\nExp number of AAs in TMHs: 23.72226\nExp number, first 60 AAs: 0.22038\nTotal prob of N-in: 0.12994\noutside 1 331\nTMhelix 332 354\ninside 355 372

7233 GCF_001645245.1_ASM164524v1 Escherichia coli Proteobacteria; Gammaproteobacteria; Enterobacterales; Enterobacteriaceae; Escherichia
MNNEETFYQAMRRQGVTRRSFLKYCSLAATSLGLGAGMAPKIAWALENKPRIPVVWIHGL WP_001058323.1
MULTISPECIES: hydrogenase-1 small chain [Proteobacteria] Length: 372\nNumber of predicted TMHs:

2\nExp number of AAs in TMHs: 37.2608499999999\nExp number, first 60 AAs: 19.52923\nTotal prob of N-in: 0.94351\nPOSSIBLE N-term signal sequence\ninside 1 19\nTMhelix 20 39\noutside 40 325\nTMhelix 326 348\ninside 349 372

7234 GCF_001641925.1_ASM164192v1 Escherichia coli Proteobacteria; Gammaproteobacteria; Enterobacterales; Enterobacteriaceae; Escherichia MNNEETFYQAMRRQGVTRRSFLKYCSLAATSLGLGAGMAPKIAWALENKPRIPVVWIHGL WP_001058320.1 hydrogenase [Escherichia coli] Length: 372\nNumber of predicted TMHs: 2\nExp number of AAs in TMHs: 37.26082\nExp number, first 60 AAs: 19.52923\nTotal prob of N-in: 0.94351\nPOSSIBLE N-term signal sequence\ninside 1 19\nTMhelix 20 39\noutside 40 325\nTMhelix 326 348\ninside 349 372

7235 GCF_001641865.1_ASM164186v1 Escherichia coli Proteobacteria; Gammaproteobacteria; Enterobacterales; Enterobacteriaceae; Escherichia MNNEETFYQAMRRQGVTRRSFLKYCSLAATSLGLGAGMAPKIAWALENKPRIPVVWIHGL WP_001058323.1 MULTISPECIES: hydrogenase-1 small chain [Proteobacteria] Length: 372\nNumber of predicted TMHs: 2\nExp number of AAs in TMHs: 37.2608499999999\nExp number, first 60 AAs: 19.52923\nTotal prob of N-in: 0.94351\nPOSSIBLE N-term signal sequence\ninside 1 19\nTMhelix 20 39\noutside 40 325\nTMhelix 326 348\ninside 349 372

7236 GCF_001641845.1_ASM164184v1 Escherichia coli Proteobacteria; Gammaproteobacteria; Enterobacterales; Enterobacteriaceae; Escherichia MNNEETFYQAMRRQGVTRRSFLKYCSLAATSLGLGAGMAPKIAWALENKPRIPVVWIHGL WP_001058323.1 MULTISPECIES: hydrogenase-1 small chain [Proteobacteria] Length: 372\nNumber of predicted TMHs: 2\nExp number of AAs in TMHs: 37.2608499999999\nExp number, first 60 AAs: 19.52923\nTotal prob of N-in: 0.94351\nPOSSIBLE N-term signal sequence\ninside 1 19\nTMhelix 20 39\noutside 40 325\nTMhelix 326 348\ninside 349 372

7237 GCF_001645235.1_ASM164523v1 Escherichia coli Proteobacteria; Gammaproteobacteria; Enterobacterales; Enterobacteriaceae; Escherichia MTGDNTLIHSHGINRRDFMKLCAALAATMGLSSKAAAEMAESVTNPQRPPVIWIGAQECT WP_000145410.1 MULTISPECIES: hydrogenase-2 small chain [Proteobacteria] Length: 372\nNumber of predicted TMHs: 1\nExp number of AAs in TMHs: 23.72226\nExp number, first 60 AAs: 0.22038\nTotal prob of N-in: 0.12994\noutside 1 331\nTMhelix 332 354\ninside 355 372

7238 GCF_001645235.1_ASM164523v1 Escherichia coli Proteobacteria; Gammaproteobacteria; Enterobacterales; Enterobacteriaceae; Escherichia MNNEETFYQAMRRQGVTRRSFLKYCSLAATSLGLGAGMAPKIAWALENKPRIPVVWIHGL WP_001295940.1 MULTISPECIES: hydrogenase [Enterobacteriaceae] Length: 372\nNumber of predicted TMHs: 2\nExp number of AAs in TMHs: 37.2608899999999\nExp number, first 60 AAs: 19.52923\nTotal prob of N-in: 0.94351\nPOSSIBLE N-term signal sequence\ninside 1 19\nTMhelix 20 39\noutside 40 325\nTMhelix 326 348\ninside 349 372

7239 GCF_001648715.1_ASM164871v1 Escherichia coli Proteobacteria; Gammaproteobacteria; Enterobacterales; Enterobacteriaceae; Escherichia MTGDNTLIHSHGINRRDFMKLCAALAATMGLSSKAAAEMAESVTNPQRPPVIWIGAQECT WP_000145410.1 MULTISPECIES: hydrogenase-2 small chain [Proteobacteria] Length: 372\nNumber of predicted TMHs: 1\nExp number of AAs in TMHs: 23.72226\nExp number, first 60 AAs: 0.22038\nTotal prob of N-in: 0.12994\noutside 1 331\nTMhelix 332 354\ninside 355 372

7240 GCF_001650505.1_ASM165050v1 Escherichia coli Proteobacteria; Gammaproteobacteria; Enterobacterales; Enterobacteriaceae; Escherichia MTGDNTLIHSHGINRRDFMKLCAALAATMGLSSKAAAEMAESVTNPQRPPVIWIGAQECT WP_000145410.1 MULTISPECIES: hydrogenase-2 small chain [Proteobacteria] Length: 372\nNumber of predicted TMHs: 1\nExp number of AAs in TMHs: 23.72226\nExp number, first 60 AAs: 0.22038\nTotal prob of N-in: 0.12994\noutside 1 331\nTMhelix 332 354\ninside 355 372

7241 GCF_001650515.1_ASM165051v1 Escherichia coli Proteobacteria; Gammaproteobacteria; Enterobacterales; Enterobacteriaceae; Escherichia MTGDNTLIHSHGINRRDFMKLCAALAATMGLSSKAAAEMAESVTNPQRPPVIWIGAQECT WP_000145410.1 MULTISPECIES: hydrogenase-2 small chain [Proteobacteria] Length: 372\nNumber of predicted TMHs:

1\nExp number of AAs in TMHs: 23.72226\nExp number, first 60 AAs: 0.22038\nTotal prob of N-in: 0.12994\noutside 1 331\nTMhelix 332 354\ninside 355 372

7242 GCF_001650565.1_ASM165056v1 Escherichia coli Proteobacteria; Gammaproteobacteria; Enterobacterales; Enterobacteriaceae; Escherichia MTGDNTLIHSHGINRRDFMKLCAALAATMGLSSKAAAEMAESVTNPQRPPVIWIGAQECT WP_000145410.1 MULTISPECIES: hydrogenase-2 small chain [Proteobacteria] Length: 372\nNumber of predicted TMHs: 1\nExp number of AAs in TMHs: 23.72226\nExp number, first 60 AAs: 0.22038\nTotal prob of N-in: 0.12994\noutside 1 331\nTMhelix 332 354\ninside 355 372

7243 GCF_001651605.1_ASM165160v1 Escherichia coli Proteobacteria; Gammaproteobacteria; Enterobacterales; Enterobacteriaceae; Escherichia MTGDNTLIHSHGINRRDFMKLCAALAATMGLSSKAAAEMAESVTNPQRPPVIWIGAQECT WP_000145410.1 MULTISPECIES: hydrogenase-2 small chain [Proteobacteria] Length: 372\nNumber of predicted TMHs: 1\nExp number of AAs in TMHs: 23.72226\nExp number, first 60 AAs: 0.22038\nTotal prob of N-in: 0.12994\noutside 1 331\nTMhelix 332 354\ninside 355 372

7244 GCF_001650585.1_ASM165058v1 Escherichia coli Proteobacteria; Gammaproteobacteria; Enterobacterales; Enterobacteriaceae; Escherichia MTGDNTLIHSHGINRRDFMKLCAALAATMGLSSKAAAEMAESVTNPQRPPVIWIGAQECT WP_000145410.1 MULTISPECIES: hydrogenase-2 small chain [Proteobacteria] Length: 372\nNumber of predicted TMHs: 1\nExp number of AAs in TMHs: 23.72226\nExp number, first 60 AAs: 0.22038\nTotal prob of N-in: 0.12994\noutside 1 331\nTMhelix 332 354\ninside 355 372

7245 GCF_001651625.1_ASM165162v1 Escherichia coli Proteobacteria; Gammaproteobacteria; Enterobacterales; Enterobacteriaceae; Escherichia MTGDNTLIHSHGINRRDFMKLCAALAATMGLSSKAAAEMAESVTNPQRPPVIWIGAQECT WP_000145410.1 MULTISPECIES: hydrogenase-2 small chain [Proteobacteria] Length: 372\nNumber of predicted TMHs: 1\nExp number of AAs in TMHs: 23.72226\nExp number, first 60 AAs: 0.22038\nTotal prob of N-in: 0.12994\noutside 1 331\nTMhelix 332 354\ninside 355 372

7246 GCF_001651635.1_ASM165163v1 Escherichia coli Proteobacteria; Gammaproteobacteria; Enterobacterales; Enterobacteriaceae; Escherichia MTGDNTLIHSHGINRRDFMKLCAALAATMGLSSKAAAEMVESVTNPQRPPVIWIGAQECT WP_000145423.1 hydrogenase 2 small subunit [Escherichia coli] Length: 372\nNumber of predicted TMHs: 1\nExp number of AAs in TMHs: 23.78201\nExp number, first 60 AAs: 0.28355\nTotal prob of N-in: 0.13257\noutside 1 331\nTMhelix 332 354\ninside 355 372

7247 GCF_001650585.1_ASM165058v1 Escherichia coli Proteobacteria; Gammaproteobacteria; Enterobacterales; Enterobacteriaceae; Escherichia MNNEETFYQAMRRQGVTRRSFLKYCSLAATSLGLGAGMAPKIAWALENKPRIPVVWIHGL WP_001058323.1 MULTISPECIES: hydrogenase-1 small chain [Proteobacteria] Length: 372\nNumber of predicted TMHs: 2\nExp number of AAs in TMHs: 37.2608499999999\nExp number, first 60 AAs: 19.52923\nTotal prob of N-in: 0.94351\nPOSSIBLE N-term signal sequence\ninside 1 19\nTMhelix 20 39\noutside 40 325\nTMhelix 326 348\ninside 349 372

7248 GCF_001651625.1_ASM165162v1 Escherichia coli Proteobacteria; Gammaproteobacteria; Enterobacterales; Enterobacteriaceae; Escherichia MNNEETFYQAMRRQGVTRRSFLKYCSLAATSLGLGAGMAPKIAWALENKPRIPVVWIHGL WP_001058323.1 MULTISPECIES: hydrogenase-1 small chain [Proteobacteria] Length: 372\nNumber of predicted TMHs: 2\nExp number of AAs in TMHs: 37.2608499999999\nExp number, first 60 AAs: 19.52923\nTotal prob of N-in: 0.94351\nPOSSIBLE N-term signal sequence\ninside 1 19\nTMhelix 20 39\noutside 40 325\nTMhelix 326 348\ninside 349 372

7249 GCF_001651635.1_ASM165163v1 Escherichia coli Proteobacteria; Gammaproteobacteria; Enterobacterales; Enterobacteriaceae; Escherichia MNNEETFYQAMRRQGVTRRSFLKYCSLAATSLGLGAGMAPKIAWALENKPRIPVVWIHGL WP_001058323.1 MULTISPECIES: hydrogenase-1 small chain [Proteobacteria] Length: 372\nNumber of predicted TMHs: 2\nExp number of AAs in TMHs: 37.2608499999999\nExp number, first 60 AAs: 19.52923\nTotal prob of N-in: 0.94351\nPOSSIBLE N-term signal sequence\ninside 1 19\nTMhelix 20 39\noutside 40 325\nTMhelix 326 348\ninside 349 372

7250 GCF_001650505.1_ASM165050v1 Escherichia coli Proteobacteria; Gammaproteobacteria; Enterobacterales; Enterobacteriaceae; Escherichia
MNNEETFYQAMRRQGVTRRSFLKYCSLAATSLGLGAGMAPKIAWALENKPRIPVVWIHGL WP_001058323.1
MULTISPECIES: hydrogenase-1 small chain [Proteobacteria] Length: 372\nNumber of predicted TMHs: 2\nExp number of AAs in TMHs: 37.2608499999999\nExp number, first 60 AAs: 19.52923\nTotal prob of N-in: 0.94351\nPOSSIBLE N-term signal sequence\ninside 1 19\nTMhelix 20 39\noutside 40 325\nTMhelix 326 348\ninside 349 372

7251 GCF_001650515.1_ASM165051v1 Escherichia coli Proteobacteria; Gammaproteobacteria; Enterobacterales; Enterobacteriaceae; Escherichia
MNNEETFYQAMRRQGVTRRSFLKYCSLAATSLGLGAGMAPKIAWALENKPRIPVVWIHGL WP_001058323.1
MULTISPECIES: hydrogenase-1 small chain [Proteobacteria] Length: 372\nNumber of predicted TMHs: 2\nExp number of AAs in TMHs: 37.2608499999999\nExp number, first 60 AAs: 19.52923\nTotal prob of N-in: 0.94351\nPOSSIBLE N-term signal sequence\ninside 1 19\nTMhelix 20 39\noutside 40 325\nTMhelix 326 348\ninside 349 372

7252 GCF_001651605.1_ASM165160v1 Escherichia coli Proteobacteria; Gammaproteobacteria; Enterobacterales; Enterobacteriaceae; Escherichia
MNNEETFYQAMRRQGVTRRSFLKYCSLAATSLGLGAGMAPKIAWALENKPRIPVVWIHGL WP_001058323.1
MULTISPECIES: hydrogenase-1 small chain [Proteobacteria] Length: 372\nNumber of predicted TMHs: 2\nExp number of AAs in TMHs: 37.2608499999999\nExp number, first 60 AAs: 19.52923\nTotal prob of N-in: 0.94351\nPOSSIBLE N-term signal sequence\ninside 1 19\nTMhelix 20 39\noutside 40 325\nTMhelix 326 348\ninside 349 372

7253 GCF_001652805.1_ASM165280v1 Escherichia coli Proteobacteria; Gammaproteobacteria; Enterobacterales; Enterobacteriaceae; Escherichia
MTGDNTLIHSHGINRRDFMKLCAALAATMGLSSKAAAEMVESVTNPQRPPVIWIGAQECT WP_000145423.1
hydrogenase 2 small subunit [Escherichia coli] Length: 372\nNumber of predicted TMHs: 1\nExp number of AAs in TMHs: 23.78201\nExp number, first 60 AAs: 0.28355\nTotal prob of N-in: 0.13257\noutside 1 331\nTMhelix 332 354\ninside 355 372

7254 GCF_001653555.1_ASM165355v1 Escherichia coli Proteobacteria; Gammaproteobacteria; Enterobacterales; Enterobacteriaceae; Escherichia
MTGDNTLIHSHGINRRDFMKLCAALAATMGLSSKAAAEMAESVTNPQRPPVIWIGAQECT WP_000145410.1
MULTISPECIES: hydrogenase-2 small chain [Proteobacteria] Length: 372\nNumber of predicted TMHs: 1\nExp number of AAs in TMHs: 23.72226\nExp number, first 60 AAs: 0.22038\nTotal prob of N-in: 0.12994\noutside 1 331\nTMhelix 332 354\ninside 355 372

7255 GCF_001657975.1_ASM165797v1 Escherichia coli Proteobacteria; Gammaproteobacteria; Enterobacterales; Enterobacteriaceae; Escherichia
MTGDNTLIHSHGINRRDFMKLCAALAATMGLSSKAAAEMAESVTNPQRPPVIWIGAQECT WP_000145410.1
MULTISPECIES: hydrogenase-2 small chain [Proteobacteria] Length: 372\nNumber of predicted TMHs: 1\nExp number of AAs in TMHs: 23.72226\nExp number, first 60 AAs: 0.22038\nTotal prob of N-in: 0.12994\noutside 1 331\nTMhelix 332 354\ninside 355 372

7256 GCF_001654955.1_ASM165495v1 Escherichia coli Proteobacteria; Gammaproteobacteria; Enterobacterales; Enterobacteriaceae; Escherichia
MTGDNTLIHSHGINRRDFMKLCAALAATMGLSSKAAAEMAESVTNPQRPPVIWIGAQECT WP_000145410.1
MULTISPECIES: hydrogenase-2 small chain [Proteobacteria] Length: 372\nNumber of predicted TMHs: 1\nExp number of AAs in TMHs: 23.72226\nExp number, first 60 AAs: 0.22038\nTotal prob of N-in: 0.12994\noutside 1 331\nTMhelix 332 354\ninside 355 372

7257 GCF_001653635.1_ASM165363v1 Escherichia coli Proteobacteria; Gammaproteobacteria; Enterobacterales; Enterobacteriaceae; Escherichia
MTGDNTLIHSHGINRRDFMKLCAALAATMGLSSKAAAEMAESVTNPQRPPVIWIGAQECT WP_000145410.1
MULTISPECIES: hydrogenase-2 small chain [Proteobacteria] Length: 372\nNumber of predicted TMHs: 1\nExp number of AAs in TMHs: 23.72226\nExp number, first 60 AAs: 0.22038\nTotal prob of N-in: 0.12994\noutside 1 331\nTMhelix 332 354\ninside 355 372

7258 GCF_001659805.1_ASM165980v1 Escherichia coli Proteobacteria; Gammaproteobacteria; Enterobacterales; Enterobacteriaceae; Escherichia

MTGDNTLIHSHGINRRDFMKLCAALAATMGLSSKAAAEMAESVTNPQRPPVIWIGAQECT WP_000145410.1 MULTISPECIES: hydrogenase-2 small chain [Proteobacteria] Length: 372\nNumber of predicted TMHs: 1\nExp number of AAs in TMHs: 23.72226\nExp number, first 60 AAs: 0.22038\nTotal prob of N-in: 0.12994\noutside 1 331\nTMhelix 332 354\ninside 355 372

7259 GCF_001660085.1_ASM166008v1 Escherichia coli Proteobacteria; Gammaproteobacteria; Enterobacterales; Enterobacteriaceae; Escherichia MTGDNTLIHSHGINRRDFMKLCAALAATMGLSSKAAAEMAESVTNPQRPPVIWIGAQECT WP_000145410.1 MULTISPECIES: hydrogenase-2 small chain [Proteobacteria] Length: 372\nNumber of predicted TMHs: 1\nExp number of AAs in TMHs: 23.72226\nExp number, first 60 AAs: 0.22038\nTotal prob of N-in: 0.12994\noutside 1 331\nTMhelix 332 354\ninside 355 372

7260 GCF_001660105.1_ASM166010v1 Escherichia coli Proteobacteria; Gammaproteobacteria; Enterobacterales; Enterobacteriaceae; Escherichia MTGDNTLIHSHGINRRDFMKLCAALAATMGLSSKAAAEMAESVTNPQRPPVIWIGAQECT WP_000145410.1 MULTISPECIES: hydrogenase-2 small chain [Proteobacteria] Length: 372\nNumber of predicted TMHs: 1\nExp number of AAs in TMHs: 23.72226\nExp number, first 60 AAs: 0.22038\nTotal prob of N-in: 0.12994\noutside 1 331\nTMhelix 332 354\ninside 355 372

7261 GCF_002018825.1_ASM201882v1 Escherichia coli Proteobacteria; Gammaproteobacteria; Enterobacterales; Enterobacteriaceae; Escherichia MTGDNTLIHSHGINRRDFMKLCAALATTMGLSSKAAAEMAESVTNPQRPPVIWIGAQECT WP_023149629.1 MULTISPECIES: hydrogenase-2 small chain [Enterobacteriaceae] Length: 372\nNumber of predicted TMHs: 1\nExp number of AAs in TMHs: 23.63603\nExp number, first 60 AAs: 0.12971\nTotal prob of N-in: 0.12608\noutside 1 331\nTMhelix 332 354\ninside 355 372

7262 GCF_002024905.1_ASM202490v1 Escherichia coli Proteobacteria; Gammaproteobacteria; Enterobacterales; Enterobacteriaceae; Escherichia MNNEETFYQAMRRQGVTRRSFLKYCSLAATSLGLGAGMAPKIAWALENKPRIPVVWIHGL WP_001058323.1 MULTISPECIES: hydrogenase-1 small chain [Proteobacteria] Length: 372\nNumber of predicted TMHs: 2\nExp number of AAs in TMHs: 37.2608499999999\nExp number, first 60 AAs: 19.52923\nTotal prob of N-in: 0.94351\nPOSSIBLE N-term signal sequence\ninside 1 19\nTMhelix 20 39\noutside 40 325\nTMhelix 326 348\ninside 349 372

7263 GCF_002028795.1_ASM202879v1 Escherichia coli Proteobacteria; Gammaproteobacteria; Enterobacterales; Enterobacteriaceae; Escherichia MTGDNTLIHSHGINRRDFMKLCAALAATMGLSSKAAAEMAESVTNPQRPPVIWIGAQECT WP_000145410.1 MULTISPECIES: hydrogenase-2 small chain [Proteobacteria] Length: 372\nNumber of predicted TMHs: 1\nExp number of AAs in TMHs: 23.72226\nExp number, first 60 AAs: 0.22038\nTotal prob of N-in: 0.12994\noutside 1 331\nTMhelix 332 354\ninside 355 372

7264 GCF_002028835.1_ASM202883v1 Escherichia coli Proteobacteria; Gammaproteobacteria; Enterobacterales; Enterobacteriaceae; Escherichia MTGDNTLIHSHGINRRDFMKLCAALAATMGLSSKAAAEMAESVTNPQRPPVIWIGAQECT WP_000145410.1 MULTISPECIES: hydrogenase-2 small chain [Proteobacteria] Length: 372\nNumber of predicted TMHs: 1\nExp number of AAs in TMHs: 23.72226\nExp number, first 60 AAs: 0.22038\nTotal prob of N-in: 0.12994\noutside 1 331\nTMhelix 332 354\ninside 355 372

7265 GCF_002028875.1_ASM202887v1 Escherichia coli Proteobacteria; Gammaproteobacteria; Enterobacterales; Enterobacteriaceae; Escherichia MTGDNTLIHSHGINRRDFMKLCAALAATMGLSSKAAAEMAESVTNPQRPPVIWIGAQECT WP_000145410.1 MULTISPECIES: hydrogenase-2 small chain [Proteobacteria] Length: 372\nNumber of predicted TMHs: 1\nExp number of AAs in TMHs: 23.72226\nExp number, first 60 AAs: 0.22038\nTotal prob of N-in: 0.12994\noutside 1 331\nTMhelix 332 354\ninside 355 372

7266 GCF_002029025.1_ASM202902v1 Escherichia coli Proteobacteria; Gammaproteobacteria; Enterobacterales; Enterobacteriaceae; Escherichia MTGDNTLIHSHGINRRDFMKLCAALAATMGLSSKAAAEMAESVTNPQRPPVIWIGAQECT WP_000145410.1 MULTISPECIES: hydrogenase-2 small chain [Proteobacteria] Length: 372\nNumber of predicted TMHs: 1\nExp number of AAs in TMHs: 23.72226\nExp number, first 60 AAs: 0.22038\nTotal prob of N-in: 0.12994\noutside 1 331\nTMhelix 332 354\ninside 355 372

7267 GCF_002029035.1_ASM202903v1 Escherichia coli Proteobacteria; Gammaproteobacteria; Enterobacterales; Enterobacteriaceae; Escherichia
MTGDNTLIHSHGINRRDFMKLCAALAATMGLSSKAAAEMAESVTNPQRPPVIWIGAQECT WP_000145410.1
MULTISPECIES: hydrogenase-2 small chain [Proteobacteria] Length: 372\nNumber of predicted TMHs: 1\nExp number of AAs in TMHs: 23.72226\nExp number, first 60 AAs: 0.22038\nTotal prob of N-in: 0.12994\noutside 1 331\nTMhelix 332 354\ninside 355 372

7268 GCF_002029085.1_ASM202908v1 Escherichia coli Proteobacteria; Gammaproteobacteria; Enterobacterales; Enterobacteriaceae; Escherichia
MTGDNTLIHSHGINRRDFMKLCAALAATMGLSSKAAAEMAESVTNPQRPPVIWIGAQECT WP_000145410.1
MULTISPECIES: hydrogenase-2 small chain [Proteobacteria] Length: 372\nNumber of predicted TMHs: 1\nExp number of AAs in TMHs: 23.72226\nExp number, first 60 AAs: 0.22038\nTotal prob of N-in: 0.12994\noutside 1 331\nTMhelix 332 354\ninside 355 372

7269 GCF_002029095.1_ASM202909v1 Escherichia coli Proteobacteria; Gammaproteobacteria; Enterobacterales; Enterobacteriaceae; Escherichia
MTGDNTLIHSHGINRRDFMKLCAALAATMGLSSKAAAEMAESVTNPQRPPVIWIGAQECT WP_000145410.1
MULTISPECIES: hydrogenase-2 small chain [Proteobacteria] Length: 372\nNumber of predicted TMHs: 1\nExp number of AAs in TMHs: 23.72226\nExp number, first 60 AAs: 0.22038\nTotal prob of N-in: 0.12994\noutside 1 331\nTMhelix 332 354\ninside 355 372

7270 GCF_002029115.1_ASM202911v1 Escherichia coli Proteobacteria; Gammaproteobacteria; Enterobacterales; Enterobacteriaceae; Escherichia
MTGDNTLIHSHGINRRDFMKLCAALAATMGLSSKAAAEMAESVTNPQRPPVIWIGAQECT WP_000145410.1
MULTISPECIES: hydrogenase-2 small chain [Proteobacteria] Length: 372\nNumber of predicted TMHs: 1\nExp number of AAs in TMHs: 23.72226\nExp number, first 60 AAs: 0.22038\nTotal prob of N-in: 0.12994\noutside 1 331\nTMhelix 332 354\ninside 355 372

7271 GCF_002029145.1_ASM202914v1 Escherichia coli Proteobacteria; Gammaproteobacteria; Enterobacterales; Enterobacteriaceae; Escherichia
MTGDNTLIHSHGINRRDFMKLCAALAATMGLSSKAAAEMAESVTNPQRPPVIWIGAQECT WP_000145410.1
MULTISPECIES: hydrogenase-2 small chain [Proteobacteria] Length: 372\nNumber of predicted TMHs: 1\nExp number of AAs in TMHs: 23.72226\nExp number, first 60 AAs: 0.22038\nTotal prob of N-in: 0.12994\noutside 1 331\nTMhelix 332 354\ninside 355 372

7272 GCF_002029165.1_ASM202916v1 Escherichia coli Proteobacteria; Gammaproteobacteria; Enterobacterales; Enterobacteriaceae; Escherichia
MTGDNTLIHSHGINRRDFMKLCAALAATMGLSSKAAAEMAESVTNPQRPPVIWIGAQECT WP_000145410.1
MULTISPECIES: hydrogenase-2 small chain [Proteobacteria] Length: 372\nNumber of predicted TMHs: 1\nExp number of AAs in TMHs: 23.72226\nExp number, first 60 AAs: 0.22038\nTotal prob of N-in: 0.12994\noutside 1 331\nTMhelix 332 354\ninside 355 372

7273 GCF_002055605.1_ASM205560v1 Escherichia coli Proteobacteria; Gammaproteobacteria; Enterobacterales; Enterobacteriaceae; Escherichia
MTGDNTLIHSHGINRRDFMKLCAALAATMGLSSKAAAEMAESVTNPQRPPVIWIGAQECT WP_000145410.1
MULTISPECIES: hydrogenase-2 small chain [Proteobacteria] Length: 372\nNumber of predicted TMHs: 1\nExp number of AAs in TMHs: 23.72226\nExp number, first 60 AAs: 0.22038\nTotal prob of N-in: 0.12994\noutside 1 331\nTMhelix 332 354\ninside 355 372

7274 GCF_002057355.1_ASM205735v1 Escherichia coli Proteobacteria; Gammaproteobacteria; Enterobacterales; Enterobacteriaceae; Escherichia
MNNEETFYQAMRRQGVTRRSFLKYCSLAATSLGLGAGMAPKIAWALENKPRIPVVWIHGL WP_001058323.1
MULTISPECIES: hydrogenase-1 small chain [Proteobacteria] Length: 372\nNumber of predicted TMHs: 2\nExp number of AAs in TMHs: 37.2608499999999\nExp number, first 60 AAs: 19.52923\nTotal prob of N-in: 0.94351\nPOSSIBLE N-term signal sequence\ninside 1 19\nTMhelix 20 39\noutside 40 325\nTMhelix 326 348\ninside 349 372

7275 GCF_002055605.1_ASM205560v1 Escherichia coli Proteobacteria; Gammaproteobacteria; Enterobacterales; Enterobacteriaceae; Escherichia
MNNEETFYQAMRRQGVTRRSFLKYCSLAATSLGLGAGMAPKIAWALENKPRIPVVWIHGL WP_001058323.1
MULTISPECIES: hydrogenase-1 small chain [Proteobacteria] Length: 372\nNumber of predicted TMHs:

2\nExp number of AAs in TMHs: 37.2608499999999\nExp number, first 60 AAs: 19.52923\nTotal prob of N-in: 0.94351\nPOSSIBLE N-term signal sequence\ninside 1 19\nTMhelix 20 39\noutside 40 325\nTMhelix 326 348\ninside 349 372

7276 GCF_002057245.1_ASM205724v1 Escherichia coli Proteobacteria; Gammaproteobacteria; Enterobacterales; Enterobacteriaceae; Escherichia MNNEETFYQAMRRQGVTRRSFLKYCSLAATSLGLGAGMAPKIAWALENKPRIPVVWIHGL WP_001058323.1 MULTISPECIES: hydrogenase-1 small chain [Proteobacteria] Length: 372\nNumber of predicted TMHs: 2\nExp number of AAs in TMHs: 37.2608499999999\nExp number, first 60 AAs: 19.52923\nTotal prob of N-in: 0.94351\nPOSSIBLE N-term signal sequence\ninside 1 19\nTMhelix 20 39\noutside 40 325\nTMhelix 326 348\ninside 349 372

7277 GCF_002056145.1_ASM205614v1 Escherichia coli Proteobacteria; Gammaproteobacteria; Enterobacterales; Enterobacteriaceae; Escherichia MNNEETFYQAMRRQGVTRRSFLKYCSLAATSLGLGAGMAPKIAWALENKPRIPVVWIHGL WP_001058320.1 hydrogenase [Escherichia coli] Length: 372\nNumber of predicted TMHs: 2\nExp number of AAs in TMHs: 37.26082\nExp number, first 60 AAs: 19.52923\nTotal prob of N-in: 0.94351\nPOSSIBLE N-term signal sequence\ninside 1 19\nTMhelix 20 39\noutside 40 325\nTMhelix 326 348\ninside 349 372

7278 GCF_002055635.1_ASM205563v1 Escherichia coli Proteobacteria; Gammaproteobacteria; Enterobacterales; Enterobacteriaceae; Escherichia MNNEETFYQAMRRQGVTRRSFLKYCSLAATSLGLGAGMAPKIAWALENKPRIPVVWIHGL WP_001295940.1 MULTISPECIES: hydrogenase [Enterobacteriaceae] Length: 372\nNumber of predicted TMHs: 2\nExp number of AAs in TMHs: 37.2608899999999\nExp number, first 60 AAs: 19.52923\nTotal prob of N-in: 0.94351\nPOSSIBLE N-term signal sequence\ninside 1 19\nTMhelix 20 39\noutside 40 325\nTMhelix 326 348\ninside 349 372

7279 GCF_002058765.1_ASM205876v1 Escherichia coli Proteobacteria; Gammaproteobacteria; Enterobacterales; Enterobacteriaceae; Escherichia MTGDNTLIHSHGINRRDFMKLCAALAATMGLSSKAAAEMAESVTNPQRPPVIWIGAQECT WP_000145410.1 MULTISPECIES: hydrogenase-2 small chain [Proteobacteria] Length: 372\nNumber of predicted TMHs: 1\nExp number of AAs in TMHs: 23.72226\nExp number, first 60 AAs: 0.22038\nTotal prob of N-in: 0.12994\noutside 1 331\nTMhelix 332 354\ninside 355 372

7280 GCF_002058765.1_ASM205876v1 Escherichia coli Proteobacteria; Gammaproteobacteria; Enterobacterales; Enterobacteriaceae; Escherichia MNNEETFYQAMRRQGVTRRSFLKYCSLAATSLGLGAGMAPKIAWALENKPRIPVVWIHGL WP_001058323.1 MULTISPECIES: hydrogenase-1 small chain [Proteobacteria] Length: 372\nNumber of predicted TMHs: 2\nExp number of AAs in TMHs: 37.2608499999999\nExp number, first 60 AAs: 19.52923\nTotal prob of N-in: 0.94351\nPOSSIBLE N-term signal sequence\ninside 1 19\nTMhelix 20 39\noutside 40 325\nTMhelix 326 348\ninside 349 372

7281 GCF_002075235.1_103414 Escherichia coli Proteobacteria; Gammaproteobacteria; Enterobacterales; Enterobacteriaceae; Escherichia MTGDNTLIHSHGINRRDFMKLCAALAATMGLSSKAAAEMAESVTNPQRPPVIWIGAQECT WP_000145410.1 MULTISPECIES: hydrogenase-2 small chain [Proteobacteria] Length: 372\nNumber of predicted TMHs: 1\nExp number of AAs in TMHs: 23.72226\nExp number, first 60 AAs: 0.22038\nTotal prob of N-in: 0.12994\noutside 1 331\nTMhelix 332 354\ninside 355 372

7282 GCF_002075265.1_102713 Escherichia coli Proteobacteria; Gammaproteobacteria; Enterobacterales; Enterobacteriaceae; Escherichia MTGDNTLIHSHGINRRDFMKLCAALAATMGLSSKAAAEMAESVTNPQRPPVIWIGAQECT WP_000145410.1 MULTISPECIES: hydrogenase-2 small chain [Proteobacteria] Length: 372\nNumber of predicted TMHs: 1\nExp number of AAs in TMHs: 23.72226\nExp number, first 60 AAs: 0.22038\nTotal prob of N-in: 0.12994\noutside 1 331\nTMhelix 332 354\ninside 355 372

7283 GCF_002075205.1_102549 Escherichia coli Proteobacteria; Gammaproteobacteria; Enterobacterales; Enterobacteriaceae; Escherichia MNNEETFYQAMRRQGVTRRSFLKYCSLAATSLGLGAGMAPKIAWALENKPRIPVVWIHGL WP_001295940.1 MULTISPECIES: hydrogenase [Enterobacteriaceae] Length: 372\nNumber of predicted TMHs: 2\nExp number of AAs in TMHs: 37.2608899999999\nExp number, first 60 AAs: 19.52923\nTotal prob of N-in: 0.94351\nPOSSIBLE N-term signal sequence\ninside 1 19\nTMhelix 20 39\noutside 40 325\nTMhelix 326 348\ninside 349 372

7284 GCF_002075235.1_103414 Escherichia coli Proteobacteria; Gammaproteobacteria; Enterobacterales; Enterobacteriaceae; Escherichia MNNEETFYQAMRRQGVTRRSFLKYCSLAATSLGLGAGMAPKIAWALENKPRIPVVWIHGL WP_001058323.1 MULTISPECIES: hydrogenase-1 small chain [Proteobacteria] Length: 372\nNumber of predicted TMHs: 2\nExp number of AAs in TMHs: 37.2608499999999\nExp number, first 60 AAs: 19.52923\nTotal prob of N-in: 0.94351\nPOSSIBLE N-term signal sequence\ninside 1 19\nTMhelix 20 39\noutside 40 325\nTMhelix 326 348\ninside 349 372

7285 GCF_002075125.1_103334 Escherichia coli Proteobacteria; Gammaproteobacteria; Enterobacterales; Enterobacteriaceae; Escherichia MNNEETFYQAMRRQGVTRRSFLKYCSLAATSLGLGAGMAPKIAWALENKPRIPVVWIHGL WP_001058323.1 MULTISPECIES: hydrogenase-1 small chain [Proteobacteria] Length: 372\nNumber of predicted TMHs: 2\nExp number of AAs in TMHs: 37.2608499999999\nExp number, first 60 AAs: 19.52923\nTotal prob of N-in: 0.94351\nPOSSIBLE N-term signal sequence\ninside 1 19\nTMhelix 20 39\noutside 40 325\nTMhelix 326 348\ninside 349 372

7286 GCF_002075295.1_103604 Escherichia coli Proteobacteria; Gammaproteobacteria; Enterobacterales; Enterobacteriaceae; Escherichia MTGDNTLIHSHGINRRDFMKLCAALAATMGLSSKAAAEMAESVTNPQRPPVIWIGAQECT WP_000145410.1 MULTISPECIES: hydrogenase-2 small chain [Proteobacteria] Length: 372\nNumber of predicted TMHs: 1\nExp number of AAs in TMHs: 23.72226\nExp number, first 60 AAs: 0.22038\nTotal prob of N-in: 0.12994\noutside 1 331\nTMhelix 332 354\ninside 355 372

7287 GCF_002075225.1_102687 Escherichia coli Proteobacteria; Gammaproteobacteria; Enterobacterales; Enterobacteriaceae; Escherichia MTGDNTLIHSHGINRRDFMKLCAALAATMGLSSKAAAEMAESVTNPQRPPVIWIGAQECT WP_000145410.1 MULTISPECIES: hydrogenase-2 small chain [Proteobacteria] Length: 372\nNumber of predicted TMHs: 1\nExp number of AAs in TMHs: 23.72226\nExp number, first 60 AAs: 0.22038\nTotal prob of N-in: 0.12994\noutside 1 331\nTMhelix 332 354\ninside 355 372

7288 GCF_002078295.1_ASM207829v1 Escherichia coli Proteobacteria; Gammaproteobacteria; Enterobacterales; Enterobacteriaceae; Escherichia MNNEETFYQAMRRQGVTRRSFLKYCSLAATSLGLGAGMAPKIAWALENKPRIPVVWIHGL WP_001058323.1 MULTISPECIES: hydrogenase-1 small chain [Proteobacteria] Length: 372\nNumber of predicted TMHs: 2\nExp number of AAs in TMHs: 37.2608499999999\nExp number, first 60 AAs: 19.52923\nTotal prob of N-in: 0.94351\nPOSSIBLE N-term signal sequence\ninside 1 19\nTMhelix 20 39\noutside 40 325\nTMhelix 326 348\ninside 349 372

7289 GCF_002080495.1_ASM208049v1 Escherichia coli Proteobacteria; Gammaproteobacteria; Enterobacterales; Enterobacteriaceae; Escherichia MTGDNTLIHSHGINRRDFMKLCAALAATMGLSSKAAAEMAESVTNPQRPPVIWIGAQECT WP_000145410.1 MULTISPECIES: hydrogenase-2 small chain [Proteobacteria] Length: 372\nNumber of predicted TMHs: 1\nExp number of AAs in TMHs: 23.72226\nExp number, first 60 AAs: 0.22038\nTotal prob of N-in: 0.12994\noutside 1 331\nTMhelix 332 354\ninside 355 372

7290 GCF_002080535.1_ASM208053v1 Escherichia coli Proteobacteria; Gammaproteobacteria; Enterobacterales; Enterobacteriaceae; Escherichia MTGDNTLIHSHGINRRDFMKLCAALAATMGLSSKAAAEMAESVTNPQRPPVIWIGAQECT WP_000145410.1 MULTISPECIES: hydrogenase-2 small chain [Proteobacteria] Length: 372\nNumber of predicted TMHs: 1\nExp number of AAs in TMHs: 23.72226\nExp number, first 60 AAs: 0.22038\nTotal prob of N-in: 0.12994\noutside 1 331\nTMhelix 332 354\ninside 355 372

7291 GCF_002080505.1_ASM208050v1 Escherichia coli Proteobacteria; Gammaproteobacteria; Enterobacterales; Enterobacteriaceae; Escherichia MTGDNTLIHSHGINRRDFMKLCAALAATMGLSSKAAAEMAESVTNPQRPPVIWIGAQECT WP_000145410.1 MULTISPECIES: hydrogenase-2 small chain [Proteobacteria] Length: 372\nNumber of predicted TMHs: 1\nExp number of AAs in TMHs: 23.72226\nExp number, first 60 AAs: 0.22038\nTotal prob of N-in: 0.12994\noutside 1 331\nTMhelix 332 354\ninside 355 372

7292 GCF_002080555.1_ASM208055v1 Escherichia coli Proteobacteria; Gammaproteobacteria; Enterobacterales; Enterobacteriaceae; Escherichia MTGDNTLIHSHGINRRDFMKLCAALAATMGLSSKAAAEMAESVTNPQRPPVIWIGAQECT WP_000145410.1 MULTISPECIES: hydrogenase-2 small chain [Proteobacteria] Length: 372\nNumber of predicted TMHs: 1\nExp number of AAs in TMHs: 23.72226\nExp number, first 60 AAs: 0.22038\nTotal prob of N-in: 0.12994\noutside 1 331\nTMhelix 332 354\ninside 355 372

7293 GCF_002080565.1_ASM208056v1 Escherichia coli Proteobacteria; Gammaproteobacteria; Enterobacterales; Enterobacteriaceae; Escherichia
MTGDNTLIHSHGINRRDFMKLCAALAATMGLSSKAAAEMAESVTNPQRPPVIWIGAQECT WP_000145410.1
MULTISPECIES: hydrogenase-2 small chain [Proteobacteria] Length: 372\nNumber of predicted TMHs: 1\nExp number of AAs in TMHs: 23.72226\nExp number, first 60 AAs: 0.22038\nTotal prob of N-in: 0.12994\noutside 1 331\nTMhelix 332 354\ninside 355 372

7294 GCF_002080595.1_ASM208059v1 Escherichia coli Proteobacteria; Gammaproteobacteria; Enterobacterales; Enterobacteriaceae; Escherichia
MTGDNTLIHSHGINRRDFMKLCAALAATMGLSSKAAAEMAESVTNPQRPPVIWIGAQECT WP_000145410.1
MULTISPECIES: hydrogenase-2 small chain [Proteobacteria] Length: 372\nNumber of predicted TMHs: 1\nExp number of AAs in TMHs: 23.72226\nExp number, first 60 AAs: 0.22038\nTotal prob of N-in: 0.12994\noutside 1 331\nTMhelix 332 354\ninside 355 372

7295 GCF_002087535.1_ASM208753v1 Escherichia coli Proteobacteria; Gammaproteobacteria; Enterobacterales; Enterobacteriaceae; Escherichia
MTGDNTLIHSHGINRRDFMKLCAALAATMGLSSKAAAEMAESVTNPQRPPVIWIGAQECT WP_000145410.1
MULTISPECIES: hydrogenase-2 small chain [Proteobacteria] Length: 372\nNumber of predicted TMHs: 1\nExp number of AAs in TMHs: 23.72226\nExp number, first 60 AAs: 0.22038\nTotal prob of N-in: 0.12994\noutside 1 331\nTMhelix 332 354\ninside 355 372

7296 GCF_002080635.1_ASM208063v1 Escherichia coli Proteobacteria; Gammaproteobacteria; Enterobacterales; Enterobacteriaceae; Escherichia
MNNEETFYQAMRRQGVTRRSFLKYCSLAATSLGLGAGMAPKIAWALENKPRIPVVWIHGL WP_001058323.1
MULTISPECIES: hydrogenase-1 small chain [Proteobacteria] Length: 372\nNumber of predicted TMHs: 2\nExp number of AAs in TMHs: 37.2608499999999\nExp number, first 60 AAs: 19.52923\nTotal prob of N-in: 0.94351\nPOSSIBLE N-term signal sequence\ninside 1 19\nTMhelix 20 39\noutside 40 325\nTMhelix 326 348\ninside 349 372

7297 GCF_002080605.1_ASM208060v1 Escherichia coli Proteobacteria; Gammaproteobacteria; Enterobacterales; Enterobacteriaceae; Escherichia
MNNEETFYQAMRRQGVTRRSFLKYCSLAATSLGLGAGMAPKIAWALENKPRIPVVWIHGL WP_001058323.1
MULTISPECIES: hydrogenase-1 small chain [Proteobacteria] Length: 372\nNumber of predicted TMHs: 2\nExp number of AAs in TMHs: 37.2608499999999\nExp number, first 60 AAs: 19.52923\nTotal prob of N-in: 0.94351\nPOSSIBLE N-term signal sequence\ninside 1 19\nTMhelix 20 39\noutside 40 325\nTMhelix 326 348\ninside 349 372

7298 GCF_002080535.1_ASM208053v1 Escherichia coli Proteobacteria; Gammaproteobacteria; Enterobacterales; Enterobacteriaceae; Escherichia
MNNEETFYQAMRRQGVTRRSFLKYCSLAATSLGLGAGMAPKIAWALENKPRIPVVWIHGL WP_001058323.1
MULTISPECIES: hydrogenase-1 small chain [Proteobacteria] Length: 372\nNumber of predicted TMHs: 2\nExp number of AAs in TMHs: 37.2608499999999\nExp number, first 60 AAs: 19.52923\nTotal prob of N-in: 0.94351\nPOSSIBLE N-term signal sequence\ninside 1 19\nTMhelix 20 39\noutside 40 325\nTMhelix 326 348\ninside 349 372

7299 GCF_002080645.1_ASM208064v1 Escherichia coli Proteobacteria; Gammaproteobacteria; Enterobacterales; Enterobacteriaceae; Escherichia
MTGDNTLIHSHGINRRDFMKLCAALAATMGLSSKAAAEMAESVTNPQRPPVIWIGAQECT WP_000145410.1
MULTISPECIES: hydrogenase-2 small chain [Proteobacteria] Length: 372\nNumber of predicted TMHs: 1\nExp number of AAs in TMHs: 23.72226\nExp number, first 60 AAs: 0.22038\nTotal prob of N-in: 0.12994\noutside 1 331\nTMhelix 332 354\ninside 355 372

7300 GCF_002080675.1_ASM208067v1 Escherichia coli Proteobacteria; Gammaproteobacteria; Enterobacterales; Enterobacteriaceae; Escherichia
MTGDNTLIHSHGINRRDFMKLCAALAATMGLSSKAAAEMAESVTNPQRPPVIWIGAQECT WP_000145410.1
MULTISPECIES: hydrogenase-2 small chain [Proteobacteria] Length: 372\nNumber of predicted TMHs: 1\nExp number of AAs in TMHs: 23.72226\nExp number, first 60 AAs: 0.22038\nTotal prob of N-in: 0.12994\noutside 1 331\nTMhelix 332 354\ninside 355 372

7301 GCF_002080645.1_ASM208064v1 Escherichia coli Proteobacteria; Gammaproteobacteria; Enterobacterales; Enterobacteriaceae; Escherichia

MNNEETFYQAMRRQGVTRRSFLKYCSLAATSLGLGAGMAPKIAWALENKPRIPVVWIHGL WP_001058323.1 MULTISPECIES: hydrogenase-1 small chain [Proteobacteria] Length: 372\nNumber of predicted TMHs: 2\nExp number of AAs in TMHs: 37.2608499999999\nExp number, first 60 AAs: 19.52923\nTotal prob of N-in: 0.94351\nPOSSIBLE N-term signal sequence\ninside 1 19\nTMhelix 20 39\noutside 40 325\nTMhelix 326 348\ninside 349 372

7302 GCF_002080675.1_ASM208067v1 Escherichia coli Proteobacteria; Gammaproteobacteria; Enterobacterales; Enterobacteriaceae; Escherichia MNNEETFYQAMRRQGVTRRSFLKYCSLAATSLGLGAGMAPKIAWALENKPRIPVVWIHGL WP_001058323.1 MULTISPECIES: hydrogenase-1 small chain [Proteobacteria] Length: 372\nNumber of predicted TMHs: 2\nExp number of AAs in TMHs: 37.2608499999999\nExp number, first 60 AAs: 19.52923\nTotal prob of N-in: 0.94351\nPOSSIBLE N-term signal sequence\ninside 1 19\nTMhelix 20 39\noutside 40 325\nTMhelix 326 348\ninside 349 372

7303 GCF_002087555.1_ASM208755v1 Escherichia coli Proteobacteria; Gammaproteobacteria; Enterobacterales; Enterobacteriaceae; Escherichia MTGDNTLIHSHGINRRDFMKLCAALAATMGLSSKAAAEMAESVTNPQRPPVIWIGAQECT WP_000145410.1 MULTISPECIES: hydrogenase-2 small chain [Proteobacteria] Length: 372\nNumber of predicted TMHs: 1\nExp number of AAs in TMHs: 23.72226\nExp number, first 60 AAs: 0.22038\nTotal prob of N-in: 0.12994\noutside 1 331\nTMhelix 332 354\ninside 355 372

7304 GCF_002087565.1_ASM208756v1 Escherichia coli Proteobacteria; Gammaproteobacteria; Enterobacterales; Enterobacteriaceae; Escherichia MTGDNTLIHSHGINRRDFMKLCAALAATMGLSSKAAAEMAESVTNPQRPPVIWIGAQECT WP_000145410.1 MULTISPECIES: hydrogenase-2 small chain [Proteobacteria] Length: 372\nNumber of predicted TMHs: 1\nExp number of AAs in TMHs: 23.72226\nExp number, first 60 AAs: 0.22038\nTotal prob of N-in: 0.12994\noutside 1 331\nTMhelix 332 354\ninside 355 372

7305 GCF_002087555.1_ASM208755v1 Escherichia coli Proteobacteria; Gammaproteobacteria; Enterobacterales; Enterobacteriaceae; Escherichia MNNEETFYQAMRRQGVTRRSFLKYCSLAATSLGLGAGMAPKIAWALENKPRIPVVWIHGL WP_001058323.1 MULTISPECIES: hydrogenase-1 small chain [Proteobacteria] Length: 372\nNumber of predicted TMHs: 2\nExp number of AAs in TMHs: 37.2608499999999\nExp number, first 60 AAs: 19.52923\nTotal prob of N-in: 0.94351\nPOSSIBLE N-term signal sequence\ninside 1 19\nTMhelix 20 39\noutside 40 325\nTMhelix 326 348\ninside 349 372

7306 GCF_002087565.1_ASM208756v1 Escherichia coli Proteobacteria; Gammaproteobacteria; Enterobacterales; Enterobacteriaceae; Escherichia MNNEETFYQAMRRQGVTRRSFLKYCSLAATSLGLGAGMAPKIAWALENKPRIPVVWIHGL WP_001058323.1 MULTISPECIES: hydrogenase-1 small chain [Proteobacteria] Length: 372\nNumber of predicted TMHs: 2\nExp number of AAs in TMHs: 37.2608499999999\nExp number, first 60 AAs: 19.52923\nTotal prob of N-in: 0.94351\nPOSSIBLE N-term signal sequence\ninside 1 19\nTMhelix 20 39\noutside 40 325\nTMhelix 326 348\ninside 349 372

7307 GCF_002080725.1_ASM208072v1 Escherichia coli Proteobacteria; Gammaproteobacteria; Enterobacterales; Enterobacteriaceae; Escherichia MNNEETFYQAMRRQGVTRRSFLKYCSLAATSLGLGAGMAPKIAWALENKPRIPVVWIHGL WP_001058323.1 MULTISPECIES: hydrogenase-1 small chain [Proteobacteria] Length: 372\nNumber of predicted TMHs: 2\nExp number of AAs in TMHs: 37.2608499999999\nExp number, first 60 AAs: 19.52923\nTotal prob of N-in: 0.94351\nPOSSIBLE N-term signal sequence\ninside 1 19\nTMhelix 20 39\noutside 40 325\nTMhelix 326 348\ninside 349 372

7308 GCF_002087615.1_ASM208761v1 Escherichia coli Proteobacteria; Gammaproteobacteria; Enterobacterales; Enterobacteriaceae; Escherichia MTGDNTLIHSHGINRRDFMKLCAALAATMGLSSKAAAEMAESVTNPQRPPVIWIGAQECT WP_000145410.1 MULTISPECIES: hydrogenase-2 small chain [Proteobacteria] Length: 372\nNumber of predicted TMHs: 1\nExp number of AAs in TMHs: 23.72226\nExp number, first 60 AAs: 0.22038\nTotal prob of N-in: 0.12994\noutside 1 331\nTMhelix 332 354\ninside 355 372

7309 GCF_002087575.1_ASM208757v1 Escherichia coli Proteobacteria; Gammaproteobacteria; Enterobacterales; Enterobacteriaceae; Escherichia

MTGDNTLIHSHGINRRDFMKLCAALAATMGLSSKAAAEMAESVTNPQRPPVIWIGAQECT WP_000145410.1 MULTISPECIES: hydrogenase-2 small chain [Proteobacteria] Length: 372\nNumber of predicted TMHs: 1\nExp number of AAs in TMHs: 23.72226\nExp number, first 60 AAs: 0.22038\nTotal prob of N-in: 0.12994\noutside 1 331\nTMhelix 332 354\ninside 355 372

7310 GCF_002087455.1_ASM208745v1 Escherichia coli Proteobacteria; Gammaproteobacteria; Enterobacterales; Enterobacteriaceae; Escherichia MNNEETFYQAMRRQGVTRRSFLKYCSLAATSLGLGAGMAPKIAWALENKPRIPVVWIHGL WP_001295940.1 MULTISPECIES: hydrogenase [Enterobacteriaceae] Length: 372\nNumber of predicted TMHs: 2\nExp number of AAs in TMHs: 37.2608899999999\nExp number, first 60 AAs: 19.52923\nTotal prob of N-in: 0.94351\nPOSSIBLE N-term signal sequence\ninside 1 19\nTMhelix 20 39\noutside 40 325\nTMhelix 326 348\ninside 349 372

7311 GCF_002087615.1_ASM208761v1 Escherichia coli Proteobacteria; Gammaproteobacteria; Enterobacterales; Enterobacteriaceae; Escherichia MNNEETFYQAMRRQGVTRRSFLKYCSLAATSLGLGAGMAPKIAWALENKPRIPVVWIHGL WP_001058323.1 MULTISPECIES: hydrogenase-1 small chain [Proteobacteria] Length: 372\nNumber of predicted TMHs: 2\nExp number of AAs in TMHs: 37.2608499999999\nExp number, first 60 AAs: 19.52923\nTotal prob of N-in: 0.94351\nPOSSIBLE N-term signal sequence\ninside 1 19\nTMhelix 20 39\noutside 40 325\nTMhelix 326 348\ninside 349 372

7312 GCF_002087465.1_ASM208746v1 Escherichia coli Proteobacteria; Gammaproteobacteria; Enterobacterales; Enterobacteriaceae; Escherichia MNNEETFYQAMRRQGVTRRSFLKYCSLAATSLGLGAGMAPKIAWALENKPRIPVVWIHGL WP_001295940.1 MULTISPECIES: hydrogenase [Enterobacteriaceae] Length: 372\nNumber of predicted TMHs: 2\nExp number of AAs in TMHs: 37.2608899999999\nExp number, first 60 AAs: 19.52923\nTotal prob of N-in: 0.94351\nPOSSIBLE N-term signal sequence\ninside 1 19\nTMhelix 20 39\noutside 40 325\nTMhelix 326 348\ninside 349 372

7313 GCF_002087575.1_ASM208757v1 Escherichia coli Proteobacteria; Gammaproteobacteria; Enterobacterales; Enterobacteriaceae; Escherichia MNNEETFYQAMRRQGVTRRSFLKYCSLAATSLGLGAGMAPKIAWALENKPRIPVVWIHGL WP_001058323.1 MULTISPECIES: hydrogenase-1 small chain [Proteobacteria] Length: 372\nNumber of predicted TMHs: 2\nExp number of AAs in TMHs: 37.2608499999999\nExp number, first 60 AAs: 19.52923\nTotal prob of N-in: 0.94351\nPOSSIBLE N-term signal sequence\ninside 1 19\nTMhelix 20 39\noutside 40 325\nTMhelix 326 348\ninside 349 372

7314 GCF_002087735.1_ASM208773v1 Escherichia coli Proteobacteria; Gammaproteobacteria; Enterobacterales; Enterobacteriaceae; Escherichia MTGDNTLIHSHGINRRDFMKLCAALAATMGLSSKAAAEMAESVTNPQRPPVIWIGAQECT WP_000145410.1 MULTISPECIES: hydrogenase-2 small chain [Proteobacteria] Length: 372\nNumber of predicted TMHs: 1\nExp number of AAs in TMHs: 23.72226\nExp number, first 60 AAs: 0.22038\nTotal prob of N-in: 0.12994\noutside 1 331\nTMhelix 332 354\ninside 355 372

7315 GCF_002087815.1_ASM208781v1 Escherichia coli Proteobacteria; Gammaproteobacteria; Enterobacterales; Enterobacteriaceae; Escherichia MTGDNTLIHSHGINRRDFMKLCAALAATMGLSSKAAAEMAESVTNPQRPPVIWIGAQECT WP_000145410.1 MULTISPECIES: hydrogenase-2 small chain [Proteobacteria] Length: 372\nNumber of predicted TMHs: 1\nExp number of AAs in TMHs: 23.72226\nExp number, first 60 AAs: 0.22038\nTotal prob of N-in: 0.12994\noutside 1 331\nTMhelix 332 354\ninside 355 372

7316 GCF_002087785.1_ASM208778v1 Escherichia coli Proteobacteria; Gammaproteobacteria; Enterobacterales; Enterobacteriaceae; Escherichia MTGDNTLIHSHGINRRDFMKLCAALAATMGLSSKAAAEMAESVTNPQRPPVIWIGAQECT WP_000145410.1 MULTISPECIES: hydrogenase-2 small chain [Proteobacteria] Length: 372\nNumber of predicted TMHs: 1\nExp number of AAs in TMHs: 23.72226\nExp number, first 60 AAs: 0.22038\nTotal prob of N-in: 0.12994\noutside 1 331\nTMhelix 332 354\ninside 355 372

7317 GCF_002090355.1_ASM209035v1 Escherichia coli Proteobacteria; Gammaproteobacteria; Enterobacterales; Enterobacteriaceae; Escherichia MNNEETFYQAMRRQGVTRRSFLKYCSLAATSLGLGAGMAPKIAWALENKPRIPVVWIHGL WP_001058323.1

MULTISPECIES: hydrogenase-1 small chain [Proteobacteria] Length: 372\nNumber of predicted TMHs: 2\nExp number of AAs in TMHs: 37.2608499999999\nExp number, first 60 AAs: 19.52923\nTotal prob of N-in: 0.94351\nPOSSIBLE N-term signal sequence\ninside 1 19\nTMhelix 20 39\noutside 40 325\nTMhelix 326 348\ninside 349 372

7318 GCF_002087775.1_ASM208777v1 Escherichia coli Proteobacteria; Gammaproteobacteria; Enterobacterales; Enterobacteriaceae; Escherichia
MTGDNTLIHSHGINRRDFMKLCAALAVTMGLSSKAAAEMAESVTNPQRPPVIWIGAQECT WP_083577138.1
hydrogenase 2 small subunit [Escherichia coli] Length: 372\nNumber of predicted TMHs: 1\nExp number of AAs in TMHs: 23.87436\nExp number, first 60 AAs: 0.38018\nTotal prob of N-in: 0.13675\noutside 1 331\nTMhelix 332 354\ninside 355 372

7319 GCF_002087785.1_ASM208778v1 Escherichia coli Proteobacteria; Gammaproteobacteria; Enterobacterales; Enterobacteriaceae; Escherichia
MNNEETFYQAMRRQGVTRRSFLKYCSLAATSLGLGAGMAPKIAWALENKPRIPVVWIHGL WP_001058323.1
MULTISPECIES: hydrogenase-1 small chain [Proteobacteria] Length: 372\nNumber of predicted TMHs: 2\nExp number of AAs in TMHs: 37.2608499999999\nExp number, first 60 AAs: 19.52923\nTotal prob of N-in: 0.94351\nPOSSIBLE N-term signal sequence\ninside 1 19\nTMhelix 20 39\noutside 40 325\nTMhelix 326 348\ninside 349 372

7320 GCF_002099405.1_ASM209940v1 Escherichia coli Proteobacteria; Gammaproteobacteria; Enterobacterales; Enterobacteriaceae; Escherichia
MTGDNTLIHSHGINRRDFMKLCAALAATMGLSSKAAAEMAESVTNPQRPPVIWIGAQECT WP_000145410.1
MULTISPECIES: hydrogenase-2 small chain [Proteobacteria] Length: 372\nNumber of predicted TMHs: 1\nExp number of AAs in TMHs: 23.72226\nExp number, first 60 AAs: 0.22038\nTotal prob of N-in: 0.12994\noutside 1 331\nTMhelix 332 354\ninside 355 372

7321 GCF_002093035.1_ASM209303v1 Escherichia coli Proteobacteria; Gammaproteobacteria; Enterobacterales; Enterobacteriaceae; Escherichia
MTGDNTLIHSHGINRRDFMKLCAALAATMGLSSKAAAEMAESVTNPQRPPVIWIGAQECT WP_000145410.1
MULTISPECIES: hydrogenase-2 small chain [Proteobacteria] Length: 372\nNumber of predicted TMHs: 1\nExp number of AAs in TMHs: 23.72226\nExp number, first 60 AAs: 0.22038\nTotal prob of N-in: 0.12994\noutside 1 331\nTMhelix 332 354\ninside 355 372

7322 GCF_002099405.1_ASM209940v1 Escherichia coli Proteobacteria; Gammaproteobacteria; Enterobacterales; Enterobacteriaceae; Escherichia
MNNEETFYQAMRRQGVTRRSFLKYCSLAATSLGLGAGMAPKIAWALENKPRIPVVWIHGL WP_001058323.1
MULTISPECIES: hydrogenase-1 small chain [Proteobacteria] Length: 372\nNumber of predicted TMHs: 2\nExp number of AAs in TMHs: 37.2608499999999\nExp number, first 60 AAs: 19.52923\nTotal prob of N-in: 0.94351\nPOSSIBLE N-term signal sequence\ninside 1 19\nTMhelix 20 39\noutside 40 325\nTMhelix 326 348\ninside 349 372

7323 GCF_002099465.1_ASM209946v1 Escherichia coli Proteobacteria; Gammaproteobacteria; Enterobacterales; Enterobacteriaceae; Escherichia
MTGDNTLIHSHGINRRDFMKLCAALAATMGLSSKAAAEMAESVTNPQRPPVIWIGAQECT WP_000145410.1
MULTISPECIES: hydrogenase-2 small chain [Proteobacteria] Length: 372\nNumber of predicted TMHs: 1\nExp number of AAs in TMHs: 23.72226\nExp number, first 60 AAs: 0.22038\nTotal prob of N-in: 0.12994\noutside 1 331\nTMhelix 332 354\ninside 355 372

7324 GCF_002099495.1_ASM209949v1 Escherichia coli Proteobacteria; Gammaproteobacteria; Enterobacterales; Enterobacteriaceae; Escherichia
MTGDNTLIHSHGINRRDFMKLCAALAATMGLSSKAAAEMAESVTNPQRPPVIWIGAQECT WP_000145410.1
MULTISPECIES: hydrogenase-2 small chain [Proteobacteria] Length: 372\nNumber of predicted TMHs: 1\nExp number of AAs in TMHs: 23.72226\nExp number, first 60 AAs: 0.22038\nTotal prob of N-in: 0.12994\noutside 1 331\nTMhelix 332 354\ninside 355 372

7325 GCF_002099525.1_ASM209952v1 Escherichia coli Proteobacteria; Gammaproteobacteria; Enterobacterales; Enterobacteriaceae; Escherichia
MTGDNTLIHSHGINRRDFMKLCAALAATMGLSSKAAAEMAESVTNPQRPPVIWIGAQECT WP_000145410.1
MULTISPECIES: hydrogenase-2 small chain [Proteobacteria] Length: 372\nNumber of predicted TMHs:

1\nExp number of AAs in TMHs: 23.72226\nExp number, first 60 AAs: 0.22038\nTotal prob of N-in: 0.12994\noutside 1 331\nTMhelix 332 354\ninside 355 372

7326 GCF_002099535.1_ASM209953v1 Escherichia coli Proteobacteria; Gammaproteobacteria; Enterobacterales; Enterobacteriaceae; Escherichia
MTGDNTLIHSHGINRRDFMKLCAALAATMGLSSKAAAEMAESVTNPQRPPVIWIGAQECT WP_000145410.1
MULTISPECIES: hydrogenase-2 small chain [Proteobacteria] Length: 372\nNumber of predicted TMHs: 1\nExp number of AAs in TMHs: 23.72226\nExp number, first 60 AAs: 0.22038\nTotal prob of N-in: 0.12994\noutside 1 331\nTMhelix 332 354\ninside 355 372

7327 GCF_002099455.1_ASM209945v1 Escherichia coli Proteobacteria; Gammaproteobacteria; Enterobacterales; Enterobacteriaceae; Escherichia
MNNEETFYQAMRRQGVTRRSFLKYCSLAATSLGLGAGMAPKIAWALENKPRIPVVWIHGL WP_001058323.1
MULTISPECIES: hydrogenase-1 small chain [Proteobacteria] Length: 372\nNumber of predicted TMHs: 2\nExp number of AAs in TMHs: 37.2608499999999\nExp number, first 60 AAs: 19.52923\nTotal prob of N-in: 0.94351\nPOSSIBLE N-term signal sequence\ninside 1 19\nTMhelix 20 39\noutside 40 325\nTMhelix 326 348\ninside 349 372

7328 GCF_002099465.1_ASM209946v1 Escherichia coli Proteobacteria; Gammaproteobacteria; Enterobacterales; Enterobacteriaceae; Escherichia
MNNEETFYQAMRRQGVTRRSFLKYCSLAATSLGLGAGMAPKIAWALENKPRIPVVWIHGL WP_001058323.1
MULTISPECIES: hydrogenase-1 small chain [Proteobacteria] Length: 372\nNumber of predicted TMHs: 2\nExp number of AAs in TMHs: 37.2608499999999\nExp number, first 60 AAs: 19.52923\nTotal prob of N-in: 0.94351\nPOSSIBLE N-term signal sequence\ninside 1 19\nTMhelix 20 39\noutside 40 325\nTMhelix 326 348\ninside 349 372

7329 GCF_002099555.1_ASM209955v1 Escherichia coli Proteobacteria; Gammaproteobacteria; Enterobacterales; Enterobacteriaceae; Escherichia
MTGDNTLIHSHGINRRDFMKLCAALAATMGLSSKAAAEMAESVTNPQRPPVIWIGAQECT WP_000145410.1
MULTISPECIES: hydrogenase-2 small chain [Proteobacteria] Length: 372\nNumber of predicted TMHs: 1\nExp number of AAs in TMHs: 23.72226\nExp number, first 60 AAs: 0.22038\nTotal prob of N-in: 0.12994\noutside 1 331\nTMhelix 332 354\ninside 355 372

7330 GCF_002099605.1_ASM209960v1 Escherichia coli Proteobacteria; Gammaproteobacteria; Enterobacterales; Enterobacteriaceae; Escherichia
MNNEETFYQAMRRQGVTRRSFLKYCSLAATSLGLGAGMAPKIAWALENKPRIPVVWIHGL WP_001058323.1
MULTISPECIES: hydrogenase-1 small chain [Proteobacteria] Length: 372\nNumber of predicted TMHs: 2\nExp number of AAs in TMHs: 37.2608499999999\nExp number, first 60 AAs: 19.52923\nTotal prob of N-in: 0.94351\nPOSSIBLE N-term signal sequence\ninside 1 19\nTMhelix 20 39\noutside 40 325\nTMhelix 326 348\ninside 349 372

7331 GCF_002099525.1_ASM209952v1 Escherichia coli Proteobacteria; Gammaproteobacteria; Enterobacterales; Enterobacteriaceae; Escherichia
MNNEETFYQAMRRQGVTRRSFLKYCSLAATSLGLGAGMAPKIAWALENKPRIPVVWIHGL WP_001058323.1
MULTISPECIES: hydrogenase-1 small chain [Proteobacteria] Length: 372\nNumber of predicted TMHs: 2\nExp number of AAs in TMHs: 37.2608499999999\nExp number, first 60 AAs: 19.52923\nTotal prob of N-in: 0.94351\nPOSSIBLE N-term signal sequence\ninside 1 19\nTMhelix 20 39\noutside 40 325\nTMhelix 326 348\ninside 349 372

7332 GCF_002099625.1_ASM209962v1 Escherichia coli Proteobacteria; Gammaproteobacteria; Enterobacterales; Enterobacteriaceae; Escherichia
MTGDNTLIHSHGINRRDFMKLCAALAATMGLSSKAAAEMAESVTNPQRPPVIWIGAQECT WP_000145410.1
MULTISPECIES: hydrogenase-2 small chain [Proteobacteria] Length: 372\nNumber of predicted TMHs: 1\nExp number of AAs in TMHs: 23.72226\nExp number, first 60 AAs: 0.22038\nTotal prob of N-in: 0.12994\noutside 1 331\nTMhelix 332 354\ninside 355 372

7333 GCF_002099665.1_ASM209966v1 Escherichia coli Proteobacteria; Gammaproteobacteria; Enterobacterales; Enterobacteriaceae; Escherichia
MTGDNTLIHSHGINRRDFMKLCAALAATMGLSSKAAAEMAESVTNPQRPPVIWIGAQECT WP_000145410.1
MULTISPECIES: hydrogenase-2 small chain [Proteobacteria] Length: 372\nNumber of predicted TMHs:

1\nExp number of AAs in TMHs: 23.72226\nExp number, first 60 AAs: 0.22038\nTotal prob of N-in: 0.12994\noutside 1 331\nTMhelix 332 354\ninside 355 372

7334 GCF_002099685.1_ASM209968v1 Escherichia coli Proteobacteria; Gammaproteobacteria; Enterobacterales; Enterobacteriaceae; Escherichia MTGDNTLIHSHGINRRDFMKLCAALAATMGLSSKAAAEMAESVTNPQRPPVIWIGAQECT WP_000145410.1 MULTISPECIES: hydrogenase-2 small chain [Proteobacteria] Length: 372\nNumber of predicted TMHs: 1\nExp number of AAs in TMHs: 23.72226\nExp number, first 60 AAs: 0.22038\nTotal prob of N-in: 0.12994\noutside 1 331\nTMhelix 332 354\ninside 355 372

7335 GCF_002099625.1_ASM209962v1 Escherichia coli Proteobacteria; Gammaproteobacteria; Enterobacterales; Enterobacteriaceae; Escherichia MNNEETFYQAMRRQGVTRRSFLKYCSLAATSLGLGAGMAPKIAWALENKPRIPVVWIHGL WP_001058323.1 MULTISPECIES: hydrogenase-1 small chain [Proteobacteria] Length: 372\nNumber of predicted TMHs: 2\nExp number of AAs in TMHs: 37.2608499999999\nExp number, first 60 AAs: 19.52923\nTotal prob of N-in: 0.94351\nPOSSIBLE N-term signal sequence\ninside 1 19\nTMhelix 20 39\noutside 40 325\nTMhelix 326 348\ninside 349 372

7336 GCF_002099685.1_ASM209968v1 Escherichia coli Proteobacteria; Gammaproteobacteria; Enterobacterales; Enterobacteriaceae; Escherichia MNNEETFYQAMRRQGVTRRSFLKYCSLAATSLGLGAGMAPKIAWALENKPRIPVVWIHGL WP_001058323.1 MULTISPECIES: hydrogenase-1 small chain [Proteobacteria] Length: 372\nNumber of predicted TMHs: 2\nExp number of AAs in TMHs: 37.2608499999999\nExp number, first 60 AAs: 19.52923\nTotal prob of N-in: 0.94351\nPOSSIBLE N-term signal sequence\ninside 1 19\nTMhelix 20 39\noutside 40 325\nTMhelix 326 348\ninside 349 372

7337 GCF_002099695.1_ASM209969v1 Escherichia coli Proteobacteria; Gammaproteobacteria; Enterobacterales; Enterobacteriaceae; Escherichia MNNEETFYQAMRRQGVTRRSFLKYCSLAATSLGLGAGMAPKIAWALENKPRIPVVWIHGL WP_001058323.1 MULTISPECIES: hydrogenase-1 small chain [Proteobacteria] Length: 372\nNumber of predicted TMHs: 2\nExp number of AAs in TMHs: 37.2608499999999\nExp number, first 60 AAs: 19.52923\nTotal prob of N-in: 0.94351\nPOSSIBLE N-term signal sequence\ninside 1 19\nTMhelix 20 39\noutside 40 325\nTMhelix 326 348\ninside 349 372

7338 GCF_002099795.1_ASM209979v1 Escherichia coli Proteobacteria; Gammaproteobacteria; Enterobacterales; Enterobacteriaceae; Escherichia MTGDNTLIHSHGINRRDFMKLCAALAATMGLSSKAAAEMAESVTNPQRPPVIWIGAQECT WP_000145410.1 MULTISPECIES: hydrogenase-2 small chain [Proteobacteria] Length: 372\nNumber of predicted TMHs: 1\nExp number of AAs in TMHs: 23.72226\nExp number, first 60 AAs: 0.22038\nTotal prob of N-in: 0.12994\noutside 1 331\nTMhelix 332 354\ninside 355 372

7339 GCF_002099815.1_ASM209981v1 Escherichia coli Proteobacteria; Gammaproteobacteria; Enterobacterales; Enterobacteriaceae; Escherichia MTGDNTLIHSHGINRRDFMKLCAALAATMGLSSKAAAEMAESVTNPQRPPVIWIGAQECT WP_000145410.1 MULTISPECIES: hydrogenase-2 small chain [Proteobacteria] Length: 372\nNumber of predicted TMHs: 1\nExp number of AAs in TMHs: 23.72226\nExp number, first 60 AAs: 0.22038\nTotal prob of N-in: 0.12994\noutside 1 331\nTMhelix 332 354\ninside 355 372

7340 GCF_002099865.1_ASM209986v1 Escherichia coli Proteobacteria; Gammaproteobacteria; Enterobacterales; Enterobacteriaceae; Escherichia MTGDNTLIHSHGINRRDFMKLCAALAATMGLSSKAAAEMAESVTNPQRPPVIWIGAQECT WP_000145410.1 MULTISPECIES: hydrogenase-2 small chain [Proteobacteria] Length: 372\nNumber of predicted TMHs: 1\nExp number of AAs in TMHs: 23.72226\nExp number, first 60 AAs: 0.22038\nTotal prob of N-in: 0.12994\noutside 1 331\nTMhelix 332 354\ninside 355 372

7341 GCF_002099805.1_ASM209980v1 Escherichia coli Proteobacteria; Gammaproteobacteria; Enterobacterales; Enterobacteriaceae; Escherichia MTGDNTLIHSHGINRRDFMKLCAALAATMGLSSKAAAEMAESVTNPQRPPVIWIGAQECT WP_000145410.1 MULTISPECIES: hydrogenase-2 small chain [Proteobacteria] Length: 372\nNumber of predicted TMHs: 1\nExp number of AAs in TMHs: 23.72226\nExp number, first 60 AAs: 0.22038\nTotal prob of N-in: 0.12994\noutside 1 331\nTMhelix 332 354\ninside 355 372

7342 GCF_002099795.1_ASM209979v1 Escherichia coli Proteobacteria; Gammaproteobacteria; Enterobacterales; Enterobacteriaceae; Escherichia
MNNEETFYQAMRRQGVTRRSFLKYCSLAATSLGLGAGMAPKIAWALENKPRIPVVWIHGL WP_001058323.1
MULTISPECIES: hydrogenase-1 small chain [Proteobacteria] Length: 372\nNumber of predicted TMHs: 2\nExp number of AAs in TMHs: 37.2608499999999\nExp number, first 60 AAs: 19.52923\nTotal prob of N-in: 0.94351\nPOSSIBLE N-term signal sequence\ninside 1 19\nTMhelix 20 39\noutside 40 325\nTMhelix 326 348\ninside 349 372

7343 GCF_002099815.1_ASM209981v1 Escherichia coli Proteobacteria; Gammaproteobacteria; Enterobacterales; Enterobacteriaceae; Escherichia
MNNEETFYQAMRRQGVTRRSFLKYCSLAATSLGLGAGMAPKIAWALENKPRIPVVWIHGL WP_001058323.1
MULTISPECIES: hydrogenase-1 small chain [Proteobacteria] Length: 372\nNumber of predicted TMHs: 2\nExp number of AAs in TMHs: 37.2608499999999\nExp number, first 60 AAs: 19.52923\nTotal prob of N-in: 0.94351\nPOSSIBLE N-term signal sequence\ninside 1 19\nTMhelix 20 39\noutside 40 325\nTMhelix 326 348\ninside 349 372

7344 GCF_002099885.1_ASM209988v1 Escherichia coli Proteobacteria; Gammaproteobacteria; Enterobacterales; Enterobacteriaceae; Escherichia
MNNEETFYQAMRRQGVTRRSFLKYCSLAATSLGLGAGMAPKIAWALENKPRIPVVWIHGL WP_001058323.1
MULTISPECIES: hydrogenase-1 small chain [Proteobacteria] Length: 372\nNumber of predicted TMHs: 2\nExp number of AAs in TMHs: 37.2608499999999\nExp number, first 60 AAs: 19.52923\nTotal prob of N-in: 0.94351\nPOSSIBLE N-term signal sequence\ninside 1 19\nTMhelix 20 39\noutside 40 325\nTMhelix 326 348\ninside 349 372

7345 GCF_002099785.1_ASM209978v1 Escherichia coli Proteobacteria; Gammaproteobacteria; Enterobacterales; Enterobacteriaceae; Escherichia
MNNEETFYQAMRRQGVTRRSFLKYCSLAATSLGLGAGMAPKIAWALENKPRIPVVWIHGL WP_001058323.1
MULTISPECIES: hydrogenase-1 small chain [Proteobacteria] Length: 372\nNumber of predicted TMHs: 2\nExp number of AAs in TMHs: 37.2608499999999\nExp number, first 60 AAs: 19.52923\nTotal prob of N-in: 0.94351\nPOSSIBLE N-term signal sequence\ninside 1 19\nTMhelix 20 39\noutside 40 325\nTMhelix 326 348\ninside 349 372

7346 GCF_002099895.1_ASM209989v1 Escherichia coli Proteobacteria; Gammaproteobacteria; Enterobacterales; Enterobacteriaceae; Escherichia
MTGDNTLIHSHGINRRDFMKLCAALAATMGLSSKAAAEMAESVTNPQRPPVIWIGAQECT WP_000145410.1
MULTISPECIES: hydrogenase-2 small chain [Proteobacteria] Length: 372\nNumber of predicted TMHs: 1\nExp number of AAs in TMHs: 23.72226\nExp number, first 60 AAs: 0.22038\nTotal prob of N-in: 0.12994\noutside 1 331\nTMhelix 332 354\ninside 355 372

7347 GCF_002099895.1_ASM209989v1 Escherichia coli Proteobacteria; Gammaproteobacteria; Enterobacterales; Enterobacteriaceae; Escherichia
MNNEETFYQAMRRQGVTRRSFLKYCSLAATSLGLGAGMAPKIAWALENKPRIPVVWIHGL WP_001058323.1
MULTISPECIES: hydrogenase-1 small chain [Proteobacteria] Length: 372\nNumber of predicted TMHs: 2\nExp number of AAs in TMHs: 37.2608499999999\nExp number, first 60 AAs: 19.52923\nTotal prob of N-in: 0.94351\nPOSSIBLE N-term signal sequence\ninside 1 19\nTMhelix 20 39\noutside 40 325\nTMhelix 326 348\ninside 349 372

7348 GCF_002099865.1_ASM209986v1 Escherichia coli Proteobacteria; Gammaproteobacteria; Enterobacterales; Enterobacteriaceae; Escherichia
MNNEETFYQAMRRQGVTRRSFLKYCSLAATSLGLGAGMAPKIAWALENKPRIPVVWIHGL WP_001058323.1
MULTISPECIES: hydrogenase-1 small chain [Proteobacteria] Length: 372\nNumber of predicted TMHs: 2\nExp number of AAs in TMHs: 37.2608499999999\nExp number, first 60 AAs: 19.52923\nTotal prob of N-in: 0.94351\nPOSSIBLE N-term signal sequence\ninside 1 19\nTMhelix 20 39\noutside 40 325\nTMhelix 326 348\ninside 349 372

7349 GCF_002099975.1_ASM209997v1 Escherichia coli Proteobacteria; Gammaproteobacteria; Enterobacterales; Enterobacteriaceae; Escherichia
MTGDNTLIHSHGINRRDFMKLCAALAATMGLSSKAAAEMAESVTNPQRPPVIWIGAQECT WP_000145410.1
MULTISPECIES: hydrogenase-2 small chain [Proteobacteria] Length: 372\nNumber of predicted TMHs: 1\nExp number of AAs in TMHs: 23.72226\nExp number, first 60 AAs: 0.22038\nTotal prob of N-in: 0.12994\noutside 1 331\nTMhelix 332 354\ninside 355 372

7350 GCF_002099965.1_ASM209996v1 Escherichia coli Proteobacteria; Gammaproteobacteria; Enterobacterales; Enterobacteriaceae; Escherichia
MTGDNTLIHSHGINRRDFMKLCAALAATMGLSSKAAAEMAESVTNPQRPPVIWIGAQECT WP_000145410.1
MULTISPECIES: hydrogenase-2 small chain [Proteobacteria] Length: 372\nNumber of predicted TMHs: 1\nExp number of AAs in TMHs: 23.72226\nExp number, first 60 AAs: 0.22038\nTotal prob of N-in: 0.12994\noutside 1 331\nTMhelix 332 354\ninside 355 372

7351 GCF_002100035.1_ASM210003v1 Escherichia coli Proteobacteria; Gammaproteobacteria; Enterobacterales; Enterobacteriaceae; Escherichia
MTGDNTLIHSHGINRRDFMKLCAALAATMGLSSKAAAEMAESVTNPQRPPVIWIGAQECT WP_000145410.1
MULTISPECIES: hydrogenase-2 small chain [Proteobacteria] Length: 372\nNumber of predicted TMHs: 1\nExp number of AAs in TMHs: 23.72226\nExp number, first 60 AAs: 0.22038\nTotal prob of N-in: 0.12994\noutside 1 331\nTMhelix 332 354\ninside 355 372

7352 GCF_002104225.1_ASM210422v1 Escherichia coli Proteobacteria; Gammaproteobacteria; Enterobacterales; Enterobacteriaceae; Escherichia
MTGDNTLIHSHGINRRDFMKLCAALAATMGLSSKAAAEMAESVTNPQRPPVIWIGAQECT WP_000145410.1
MULTISPECIES: hydrogenase-2 small chain [Proteobacteria] Length: 372\nNumber of predicted TMHs: 1\nExp number of AAs in TMHs: 23.72226\nExp number, first 60 AAs: 0.22038\nTotal prob of N-in: 0.12994\noutside 1 331\nTMhelix 332 354\ninside 355 372

7353 GCF_002099955.1_ASM209995v1 Escherichia coli Proteobacteria; Gammaproteobacteria; Enterobacterales; Enterobacteriaceae; Escherichia
MNNEETFYQAMRRQGVTRRSFLKYCSLAATSLGLGAGMAPKIAWALENKPRIPVVWIHGL WP_001058323.1
MULTISPECIES: hydrogenase-1 small chain [Proteobacteria] Length: 372\nNumber of predicted TMHs: 2\nExp number of AAs in TMHs: 37.2608499999999\nExp number, first 60 AAs: 19.52923\nTotal prob of N-in: 0.94351\nPOSSIBLE N-term signal sequence\ninside 1 19\nTMhelix 20 39\noutside 40 325\nTMhelix 326 348\ninside 349 372

7354 GCF_002100045.1_ASM210004v1 Escherichia coli Proteobacteria; Gammaproteobacteria; Enterobacterales; Enterobacteriaceae; Escherichia
MNNEETFYQAMRRQGVTRRSFLKYCSLAATSLGLGAGMAPKIAWALENKPRIPVVWIHGL WP_001295940.1
MULTISPECIES: hydrogenase [Enterobacteriaceae] Length: 372\nNumber of predicted TMHs: 2\nExp number of AAs in TMHs: 37.2608899999999\nExp number, first 60 AAs: 19.52923\nTotal prob of N-in: 0.94351\nPOSSIBLE N-term signal sequence\ninside 1 19\nTMhelix 20 39\noutside 40 325\nTMhelix 326 348\ninside 349 372

7355 GCF_002099965.1_ASM209996v1 Escherichia coli Proteobacteria; Gammaproteobacteria; Enterobacterales; Enterobacteriaceae; Escherichia
MNNEETFYQAMRRQGVTRRSFLKYCSLAATSLGLGAGMAPKIAWALENKPRIPVVWIHGL WP_001058323.1
MULTISPECIES: hydrogenase-1 small chain [Proteobacteria] Length: 372\nNumber of predicted TMHs: 2\nExp number of AAs in TMHs: 37.2608499999999\nExp number, first 60 AAs: 19.52923\nTotal prob of N-in: 0.94351\nPOSSIBLE N-term signal sequence\ninside 1 19\nTMhelix 20 39\noutside 40 325\nTMhelix 326 348\ninside 349 372

7356 GCF_002104405.1_ASM210440v1 Escherichia coli Proteobacteria; Gammaproteobacteria; Enterobacterales; Enterobacteriaceae; Escherichia
MNNEETFYQAMRRQGVTRRSFLKYCSLAATSLGLGAGMAPKIAWALENKPRIPVVWIHGL WP_001058323.1
MULTISPECIES: hydrogenase-1 small chain [Proteobacteria] Length: 372\nNumber of predicted TMHs: 2\nExp number of AAs in TMHs: 37.2608499999999\nExp number, first 60 AAs: 19.52923\nTotal prob of N-in: 0.94351\nPOSSIBLE N-term signal sequence\ninside 1 19\nTMhelix 20 39\noutside 40 325\nTMhelix 326 348\ninside 349 372

7357 GCF_002099975.1_ASM209997v1 Escherichia coli Proteobacteria; Gammaproteobacteria; Enterobacterales; Enterobacteriaceae; Escherichia
MNNEETFYQAMRRQGVTRRSFLKYCSLAATSLGLGAGMAPKIAWALENKPRIPVVWIHGL WP_001058323.1
MULTISPECIES: hydrogenase-1 small chain [Proteobacteria] Length: 372\nNumber of predicted TMHs: 2\nExp number of AAs in TMHs: 37.2608499999999\nExp number, first 60 AAs: 19.52923\nTotal prob of N-in: 0.94351\nPOSSIBLE N-term signal sequence\ninside 1 19\nTMhelix 20 39\noutside 40 325\nTMhelix 326 348\ninside 349 372

7358 GCF_002105735.1_ASM210573v1 Escherichia coli Proteobacteria; Gammaproteobacteria; Enterobacterales; Enterobacteriaceae; Escherichia
MTGDNTLIHSHGINRRDFMKLCAALAATMGLSSKAAAEMAESVTNPQRPPVIWIGAQECT WP_000145410.1
MULTISPECIES: hydrogenase-2 small chain [Proteobacteria] Length: 372\nNumber of predicted TMHs: 1\nExp number of AAs in TMHs: 23.72226\nExp number, first 60 AAs: 0.22038\nTotal prob of N-in: 0.12994\noutside 1 331\nTMhelix 332 354\ninside 355 372

7359 GCF_002109485.1_ASM210948v1 Escherichia coli Proteobacteria; Gammaproteobacteria; Enterobacterales; Enterobacteriaceae; Escherichia
MTGDNTLIHSHGINRRDFMKLCAALAATMGLSSKAAAEMAESVTNPQRPPVIWIGAQECT WP_000145410.1
MULTISPECIES: hydrogenase-2 small chain [Proteobacteria] Length: 372\nNumber of predicted TMHs: 1\nExp number of AAs in TMHs: 23.72226\nExp number, first 60 AAs: 0.22038\nTotal prob of N-in: 0.12994\noutside 1 331\nTMhelix 332 354\ninside 355 372

7360 GCF_002104645.1_ASM210464v1 Escherichia coli Proteobacteria; Gammaproteobacteria; Enterobacterales; Enterobacteriaceae; Escherichia
MTGDNTLIHSHGINRRDFMKLCAALAATMGLSSKAAAEMAESVTNPQRPPVIWIGAQECT WP_000145410.1
MULTISPECIES: hydrogenase-2 small chain [Proteobacteria] Length: 372\nNumber of predicted TMHs: 1\nExp number of AAs in TMHs: 23.72226\nExp number, first 60 AAs: 0.22038\nTotal prob of N-in: 0.12994\noutside 1 331\nTMhelix 332 354\ninside 355 372

7361 GCF_002104485.1_ASM210448v1 Escherichia coli Proteobacteria; Gammaproteobacteria; Enterobacterales; Enterobacteriaceae; Escherichia
MTGDNTLIHSHGINRRDFMKLCAALAATMGLSSKAAAEMVESVTNPQRPPVIWIGAQECT WP_000145423.1
hydrogenase 2 small subunit [Escherichia coli] Length: 372\nNumber of predicted TMHs: 1\nExp number of AAs in TMHs: 23.78201\nExp number, first 60 AAs: 0.28355\nTotal prob of N-in: 0.13257\noutside 1 331\nTMhelix 332 354\ninside 355 372

7362 GCF_002109445.1_ASM210944v1 Escherichia coli Proteobacteria; Gammaproteobacteria; Enterobacterales; Enterobacteriaceae; Escherichia
MTGDNTLIHSHGINRRDFMKLCAALAATMGLSSKAAAEMAESVTNPQRPPVIWIGAQECT WP_000145410.1
MULTISPECIES: hydrogenase-2 small chain [Proteobacteria] Length: 372\nNumber of predicted TMHs: 1\nExp number of AAs in TMHs: 23.72226\nExp number, first 60 AAs: 0.22038\nTotal prob of N-in: 0.12994\noutside 1 331\nTMhelix 332 354\ninside 355 372

7363 GCF_002105735.1_ASM210573v1 Escherichia coli Proteobacteria; Gammaproteobacteria; Enterobacterales; Enterobacteriaceae; Escherichia
MNNEETFYQAMRRQGVTRRSFLKYCSLAATSLGLGAGMAPKIAWALENKPRIPVVWIHGL WP_001058323.1
MULTISPECIES: hydrogenase-1 small chain [Proteobacteria] Length: 372\nNumber of predicted TMHs: 2\nExp number of AAs in TMHs: 37.2608499999999\nExp number, first 60 AAs: 19.52923\nTotal prob of N-in: 0.94351\nPOSSIBLE N-term signal sequence\ninside 1 19\nTMhelix 20 39\noutside 40 325\nTMhelix 326 348\ninside 349 372

7364 GCF_002109485.1_ASM210948v1 Escherichia coli Proteobacteria; Gammaproteobacteria; Enterobacterales; Enterobacteriaceae; Escherichia
MNNEETFYQAMRRQGVTRRSFLKYCSLAATSLGLGAGMAPKIAWALENKPRIPVVWIHGL WP_001058323.1
MULTISPECIES: hydrogenase-1 small chain [Proteobacteria] Length: 372\nNumber of predicted TMHs: 2\nExp number of AAs in TMHs: 37.2608499999999\nExp number, first 60 AAs: 19.52923\nTotal prob of N-in: 0.94351\nPOSSIBLE N-term signal sequence\ninside 1 19\nTMhelix 20 39\noutside 40 325\nTMhelix 326 348\ninside 349 372

7365 GCF_002104645.1_ASM210464v1 Escherichia coli Proteobacteria; Gammaproteobacteria; Enterobacterales; Enterobacteriaceae; Escherichia
MNNEETFYQAMRRQGVTRRSFLKYCSLAATSLGLGAGMAPKIAWALENKPRIPVVWIHGL WP_001058323.1
MULTISPECIES: hydrogenase-1 small chain [Proteobacteria] Length: 372\nNumber of predicted TMHs: 2\nExp number of AAs in TMHs: 37.2608499999999\nExp number, first 60 AAs: 19.52923\nTotal prob of N-in: 0.94351\nPOSSIBLE N-term signal sequence\ninside 1 19\nTMhelix 20 39\noutside 40 325\nTMhelix 326 348\ninside 349 372

7366 GCF_002109445.1_ASM210944v1 Escherichia coli Proteobacteria; Gammaproteobacteria; Enterobacterales; Enterobacteriaceae; Escherichia

MNNEETFYQAMRRQGVTRRSFLKYCSLAATSLGLGAGMAPKIAWALENKPRIPVVWIHGL WP_001058323.1 MULTISPECIES: hydrogenase-1 small chain [Proteobacteria] Length: 372\nNumber of predicted TMHs: 2\nExp number of AAs in TMHs: 37.2608499999999\nExp number, first 60 AAs: 19.52923\nTotal prob of N-in: 0.94351\nPOSSIBLE N-term signal sequence\ninside 1 19\nTMhelix 20 39\noutside 40 325\nTMhelix 326 348\ninside 349 372

7367 GCF_002116485.1_ASM211648v1 Escherichia coli Proteobacteria; Gammaproteobacteria; Enterobacterales; Enterobacteriaceae; Escherichia MTGDNTLIHSHGINRRDFMKLCAALAATMGLSSKAAAEMAESVTNPQRPPVIWIGAQECT WP_000145410.1 MULTISPECIES: hydrogenase-2 small chain [Proteobacteria] Length: 372\nNumber of predicted TMHs: 1\nExp number of AAs in TMHs: 23.72226\nExp number, first 60 AAs: 0.22038\nTotal prob of N-in: 0.12994\noutside 1 331\nTMhelix 332 354\ninside 355 372

7368 GCF_002116475.1_ASM211647v1 Escherichia coli Proteobacteria; Gammaproteobacteria; Enterobacterales; Enterobacteriaceae; Escherichia MTGDNTLIHSHGINRRDFMKLCAALAATMGLSSKAAAEMAESVTNPQRPPVIWIGAQECT WP_000145410.1 MULTISPECIES: hydrogenase-2 small chain [Proteobacteria] Length: 372\nNumber of predicted TMHs: 1\nExp number of AAs in TMHs: 23.72226\nExp number, first 60 AAs: 0.22038\nTotal prob of N-in: 0.12994\noutside 1 331\nTMhelix 332 354\ninside 355 372

7369 GCF_002116495.1_ASM211649v1 Escherichia coli Proteobacteria; Gammaproteobacteria; Enterobacterales; Enterobacteriaceae; Escherichia MTGDNTLIHSHGINRRDFMKLCAALAATMGLSSKAAAEMAESVTNPQRPPVIWIGAQECT WP_000145410.1 MULTISPECIES: hydrogenase-2 small chain [Proteobacteria] Length: 372\nNumber of predicted TMHs: 1\nExp number of AAs in TMHs: 23.72226\nExp number, first 60 AAs: 0.22038\nTotal prob of N-in: 0.12994\noutside 1 331\nTMhelix 332 354\ninside 355 372

7370 GCF_002115565.1_ASM211556v1 Escherichia coli Proteobacteria; Gammaproteobacteria; Enterobacterales; Enterobacteriaceae; Escherichia MNNEETFYQAMRRQGVTRRSFLKYCSLAATSLGLGAGMAPKIAWALENKPRIPVVWIHGL WP_001058323.1 MULTISPECIES: hydrogenase-1 small chain [Proteobacteria] Length: 372\nNumber of predicted TMHs: 2\nExp number of AAs in TMHs: 37.2608499999999\nExp number, first 60 AAs: 19.52923\nTotal prob of N-in: 0.94351\nPOSSIBLE N-term signal sequence\ninside 1 19\nTMhelix 20 39\noutside 40 325\nTMhelix 326 348\ninside 349 372

7371 GCF_002116545.1_ASM211654v1 Escherichia coli Proteobacteria; Gammaproteobacteria; Enterobacterales; Enterobacteriaceae; Escherichia MTGDNTLIHSHGINRRDFMKLCAALAATMGLSSKAAAEMAESVTNPQRPPVIWIGAQECT WP_000145410.1 MULTISPECIES: hydrogenase-2 small chain [Proteobacteria] Length: 372\nNumber of predicted TMHs: 1\nExp number of AAs in TMHs: 23.72226\nExp number, first 60 AAs: 0.22038\nTotal prob of N-in: 0.12994\noutside 1 331\nTMhelix 332 354\ninside 355 372

7372 GCF_002115565.1_ASM211556v1 Escherichia coli Proteobacteria; Gammaproteobacteria; Enterobacterales; Enterobacteriaceae; Escherichia MTGDNTLIHSHGINRRDFMKLCAALAATMGLSSKAAAEMAESVTNPQRPPVIWIGAQECT WP_032298447.1 hydrogenase 2 small subunit [Escherichia coli] Length: 372\nNumber of predicted TMHs: 1\nExp number of AAs in TMHs: 23.69807\nExp number, first 60 AAs: 0.221\nTotal prob of N-in: 0.12853\noutside 1 331\nTMhelix 332 354\ninside 355 372

7373 GCF_002116545.1_ASM211654v1 Escherichia coli Proteobacteria; Gammaproteobacteria; Enterobacterales; Enterobacteriaceae; Escherichia MNNEETFYQAMRRQGVTRRSFLKYCSLAATSLGLGAGMAPKIAWALENKPRIPVVWIHGL WP_085696533.1 [Ni/Fe] hydrogenase small subunit [Escherichia coli] Length: 372\nNumber of predicted TMHs: 2\nExp number of AAs in TMHs: 37.2608499999999\nExp number, first 60 AAs: 19.52923\nTotal prob of N-in: 0.94351\nPOSSIBLE N-term signal sequence\ninside 1 19\nTMhelix 20 39\noutside 40 325\nTMhelix 326 348\ninside 349 372

7374 GCF_002116715.1_ASM211671v1 Escherichia coli Proteobacteria; Gammaproteobacteria; Enterobacterales; Enterobacteriaceae; Escherichia MTGDNTLIHSHGINRRDFMKLCAALAATMGLSSKAAAEMAESVTNPQRPPVIWIGAQECT WP_000145410.1 MULTISPECIES: hydrogenase-2 small chain [Proteobacteria] Length: 372\nNumber of predicted TMHs:

1\nExp number of AAs in TMHs: 23.72226\nExp number, first 60 AAs: 0.22038\nTotal prob of N-in: 0.12994\noutside 1 331\nTMhelix 332 354\ninside 355 372

7375 GCF_002116715.1_ASM211671v1 Escherichia coli Proteobacteria; Gammaproteobacteria; Enterobacterales; Enterobacteriaceae; Escherichia MNNEETFYQAMRRQGVTRRSFLKYCSLAATSLGLGAGMAPKIAWALENKPRIPVVWIHGL WP_001295940.1 MULTISPECIES: hydrogenase [Enterobacteriaceae] Length: 372\nNumber of predicted TMHs: 2\nExp number of AAs in TMHs: 37.2608899999999\nExp number, first 60 AAs: 19.52923\nTotal prob of N-in: 0.94351\nPOSSIBLE N-term signal sequence\ninside 1 19\nTMhelix 20 39\noutside 40 325\nTMhelix 326 348\ninside 349 372

7376 GCF_002116475.1_ASM211647v1 Escherichia coli Proteobacteria; Gammaproteobacteria; Enterobacterales; Enterobacteriaceae; Escherichia MNNEETFYQAMRRQGVTRRSFLKYCSLAATSLGLGAGMAPKIAWALENKPRIPVVWIHGL WP_001058323.1 MULTISPECIES: hydrogenase-1 small chain [Proteobacteria] Length: 372\nNumber of predicted TMHs: 2\nExp number of AAs in TMHs: 37.2608499999999\nExp number, first 60 AAs: 19.52923\nTotal prob of N-in: 0.94351\nPOSSIBLE N-term signal sequence\ninside 1 19\nTMhelix 20 39\noutside 40 325\nTMhelix 326 348\ninside 349 372

7377 GCF_002118095.1_ASM211809v1 Escherichia coli Proteobacteria; Gammaproteobacteria; Enterobacterales; Enterobacteriaceae; Escherichia MTGDNTLIHSHGINRRDFMKLCAALAATMGLSSKAAAEMAESVTNPQRPPVIWIGAQECT WP_000145410.1 MULTISPECIES: hydrogenase-2 small chain [Proteobacteria] Length: 372\nNumber of predicted TMHs: 1\nExp number of AAs in TMHs: 23.72226\nExp number, first 60 AAs: 0.22038\nTotal prob of N-in: 0.12994\noutside 1 331\nTMhelix 332 354\ninside 355 372

7378 GCF_002120205.1_ASM212020v1 Escherichia coli Proteobacteria; Gammaproteobacteria; Enterobacterales; Enterobacteriaceae; Escherichia MTGDNTLIHSHGINRRDFMKLCAALAATMGLSSKAAAEMAESVTNPQRPPVIWIGAQECT WP_000145410.1 MULTISPECIES: hydrogenase-2 small chain [Proteobacteria] Length: 372\nNumber of predicted TMHs: 1\nExp number of AAs in TMHs: 23.72226\nExp number, first 60 AAs: 0.22038\nTotal prob of N-in: 0.12994\noutside 1 331\nTMhelix 332 354\ninside 355 372

7379 GCF_002118095.1_ASM211809v1 Escherichia coli Proteobacteria; Gammaproteobacteria; Enterobacterales; Enterobacteriaceae; Escherichia MNNEETFYQAMRRQGVTRRSFLKYCSLAATSLGLGAGMAPKIAWALENKPRIPVVWIHGL WP_001058323.1 MULTISPECIES: hydrogenase-1 small chain [Proteobacteria] Length: 372\nNumber of predicted TMHs: 2\nExp number of AAs in TMHs: 37.2608499999999\nExp number, first 60 AAs: 19.52923\nTotal prob of N-in: 0.94351\nPOSSIBLE N-term signal sequence\ninside 1 19\nTMhelix 20 39\noutside 40 325\nTMhelix 326 348\ninside 349 372

7380 GCF_002125925.1_ASM212592v1 Escherichia coli Proteobacteria; Gammaproteobacteria; Enterobacterales; Enterobacteriaceae; Escherichia MNNEETFYQAMRRQGVTRRSFLKYCSLAATSLGLGAGMAPKIAWALENKPRIPVVWIHGL WP_001058323.1 MULTISPECIES: hydrogenase-1 small chain [Proteobacteria] Length: 372\nNumber of predicted TMHs: 2\nExp number of AAs in TMHs: 37.2608499999999\nExp number, first 60 AAs: 19.52923\nTotal prob of N-in: 0.94351\nPOSSIBLE N-term signal sequence\ninside 1 19\nTMhelix 20 39\noutside 40 325\nTMhelix 326 348\ninside 349 372

7381 GCF_002133425.1_ASM213342v1 Escherichia coli Proteobacteria; Gammaproteobacteria; Enterobacterales; Enterobacteriaceae; Escherichia MNNEETFYQAMRRQGVTRRSFLKYCSLAATSLGLGAGMAPKIAWALENKPRIPVVWIHGL WP_001353311.1 MULTISPECIES: hydrogenase [Escherichia] Length: 372\nNumber of predicted TMHs: 2\nExp number of AAs in TMHs: 37.2638699999999\nExp number, first 60 AAs: 19.52924\nTotal prob of N-in: 0.94351\nPOSSIBLE N-term signal sequence\ninside 1 19\nTMhelix 20 39\noutside 40 325\nTMhelix 326 348\ninside 349 372

7382 GCF_002133435.1_ASM213343v1 Escherichia coli Proteobacteria; Gammaproteobacteria; Enterobacterales; Enterobacteriaceae; Escherichia MNNEETFYQAMRRQGVTRRSFLKYCSLAATSLGLGAGMAPKIAWALENKPRIPVVWIHGL WP_001301613.1 hydrogenase [Escherichia coli] Length: 372\nNumber of predicted TMHs: 2\nExp number of AAs in TMHs: 2\nExp number of AAs in

TMHs: 37.46598\nExp number, first 60 AAs: 19.53139\nTotal prob of N-in: 0.94364\nPOSSIBLE N-term signal sequence\ninside 1 19\nTMhelix 20 39\noutside 40 325\nTMhelix 326 348\ninside 349 372

7383 GCF_002125925.1_ASM212592v1 Escherichia coli Proteobacteria; Gammaproteobacteria; Enterobacterales; Enterobacteriaceae; Escherichia
MTGDNTLIHSHGINRRDFMKLCAALATTMGLSSKAAAEMAESVTNPQRPPVIWIGAQECT WP_023149629.1
MULTISPECIES: hydrogenase-2 small chain [Enterobacteriaceae] Length: 372\nNumber of predicted TMHs: 1\nExp number of AAs in TMHs: 23.63603\nExp number, first 60 AAs: 0.12971\nTotal prob of N-in: 0.12608\noutside 1 331\nTMhelix 332 354\ninside 355 372

7384 GCF_002133365.1_ASM213336v1 Escherichia coli Proteobacteria; Gammaproteobacteria; Enterobacterales; Enterobacteriaceae; Escherichia
MNNEETFYQAMRRQGVTRRSFLKYCSLAATSLGLGAGMAPKIAWALENKPRIPVVWIHGL WP_001058323.1
MULTISPECIES: hydrogenase-1 small chain [Proteobacteria] Length: 372\nNumber of predicted TMHs: 2\nExp number of AAs in TMHs: 37.2608499999999\nExp number, first 60 AAs: 19.52923\nTotal prob of N-in: 0.94351\nPOSSIBLE N-term signal sequence\ninside 1 19\nTMhelix 20 39\noutside 40 325\nTMhelix 326 348\ninside 349 372

7385 GCF_002133515.1_ASM213351v1 Escherichia coli Proteobacteria; Gammaproteobacteria; Enterobacterales; Enterobacteriaceae; Escherichia
MTGDNTLIHSHGINRRDFMKLCAALAATMGLSSKAAAEMAESVTNPQRPPVIWIGAQECT WP_000145410.1
MULTISPECIES: hydrogenase-2 small chain [Proteobacteria] Length: 372\nNumber of predicted TMHs: 1\nExp number of AAs in TMHs: 23.72226\nExp number, first 60 AAs: 0.22038\nTotal prob of N-in: 0.12994\noutside 1 331\nTMhelix 332 354\ninside 355 372

7386 GCF_002133455.1_ASM213345v1 Escherichia coli Proteobacteria; Gammaproteobacteria; Enterobacterales; Enterobacteriaceae; Escherichia
MTGDNTLIHSHGINRRDFMKLCAALAATMGLSSKAAAEMAESVTNPQRPPVIWIGAQECT WP_000145410.1
MULTISPECIES: hydrogenase-2 small chain [Proteobacteria] Length: 372\nNumber of predicted TMHs: 1\nExp number of AAs in TMHs: 23.72226\nExp number, first 60 AAs: 0.22038\nTotal prob of N-in: 0.12994\noutside 1 331\nTMhelix 332 354\ninside 355 372

7387 GCF_002133445.1_ASM213344v1 Escherichia coli Proteobacteria; Gammaproteobacteria; Enterobacterales; Enterobacteriaceae; Escherichia
MTGDNTLIHSHGINRRDFMKLCAALAATMGLSSKAAAEMAESVTNPQRPPVIWIGAQECT WP_000145410.1
MULTISPECIES: hydrogenase-2 small chain [Proteobacteria] Length: 372\nNumber of predicted TMHs: 1\nExp number of AAs in TMHs: 23.72226\nExp number, first 60 AAs: 0.22038\nTotal prob of N-in: 0.12994\noutside 1 331\nTMhelix 332 354\ninside 355 372

7388 GCF_002133525.1_ASM213352v1 Escherichia coli Proteobacteria; Gammaproteobacteria; Enterobacterales; Enterobacteriaceae; Escherichia
MNNEETFYQAMRRQGVTRRSFLKYCSLAATSLGLGAGMAPKIAWALENKPRIPVVWIHGL WP_001058323.1
MULTISPECIES: hydrogenase-1 small chain [Proteobacteria] Length: 372\nNumber of predicted TMHs: 2\nExp number of AAs in TMHs: 37.2608499999999\nExp number, first 60 AAs: 19.52923\nTotal prob of N-in: 0.94351\nPOSSIBLE N-term signal sequence\ninside 1 19\nTMhelix 20 39\noutside 40 325\nTMhelix 326 348\ninside 349 372

7389 GCF_002133585.1_ASM213358v1 Escherichia coli Proteobacteria; Gammaproteobacteria; Enterobacterales; Enterobacteriaceae; Escherichia
MNNEETFYQAMRRQGVTRRSFLKYCSLAATSLGLGAGMAPKIAWALENKPRIPVVWIHGL WP_001058323.1
MULTISPECIES: hydrogenase-1 small chain [Proteobacteria] Length: 372\nNumber of predicted TMHs: 2\nExp number of AAs in TMHs: 37.2608499999999\nExp number, first 60 AAs: 19.52923\nTotal prob of N-in: 0.94351\nPOSSIBLE N-term signal sequence\ninside 1 19\nTMhelix 20 39\noutside 40 325\nTMhelix 326 348\ninside 349 372

7390 GCF_002133535.1_ASM213353v1 Escherichia coli Proteobacteria; Gammaproteobacteria; Enterobacterales; Enterobacteriaceae; Escherichia
MNNEETFYQAMRRQGVTRRSFLKYCSLAATSLGLGAGMAPKIAWALENKPRIPVVWIHGL WP_001295940.1
MULTISPECIES: hydrogenase [Enterobacteriaceae] Length: 372\nNumber of predicted TMHs: 2\nExp number of AAs in TMHs: 37.2608899999999\nExp number, first 60 AAs: 19.52923\nTotal prob of N-in:

0.94351\nPOSSIBLE N-term signal sequence\ninside 1 19\nTMhelix 20 39\noutside 40 325\nTMhelix 326 348\ninside 349 372

7391 GCF_002133445.1_ASM213344v1 Escherichia coli Proteobacteria; Gammaproteobacteria; Enterobacterales; Enterobacteriaceae; Escherichia
MNNEETFYQAMRRQGVTRRSFLKYCSLAATSLGLGAGMAPKIAWALENKPRIPVVWIHGL WP_001058323.1
MULTISPECIES: hydrogenase-1 small chain [Proteobacteria] Length: 372\nNumber of predicted TMHs: 2\nExp number of AAs in TMHs: 37.2608499999999\nExp number, first 60 AAs: 19.52923\nTotal prob of N-in: 0.94351\nPOSSIBLE N-term signal sequence\ninside 1 19\nTMhelix 20 39\noutside 40 325\nTMhelix 326 348\ninside 349 372

7392 GCF_002133595.1_ASM213359v1 Escherichia coli Proteobacteria; Gammaproteobacteria; Enterobacterales; Enterobacteriaceae; Escherichia
MTGDNTLIHSHGINRRDFMKLCAALAATMGLSSKAAAEMAESVTNPQRPPVIWIGAQECT WP_000145410.1
MULTISPECIES: hydrogenase-2 small chain [Proteobacteria] Length: 372\nNumber of predicted TMHs: 1\nExp number of AAs in TMHs: 23.72226\nExp number, first 60 AAs: 0.22038\nTotal prob of N-in: 0.12994\noutside 1 331\nTMhelix 332 354\ninside 355 372

7393 GCF_002133595.1_ASM213359v1 Escherichia coli Proteobacteria; Gammaproteobacteria; Enterobacterales; Enterobacteriaceae; Escherichia
MNNEETFYQAMRRQGVTRRSFLKYCSLAATSLGLGAGMAPKIAWALENKPRIPVVWIHGL WP_001058323.1
MULTISPECIES: hydrogenase-1 small chain [Proteobacteria] Length: 372\nNumber of predicted TMHs: 2\nExp number of AAs in TMHs: 37.2608499999999\nExp number, first 60 AAs: 19.52923\nTotal prob of N-in: 0.94351\nPOSSIBLE N-term signal sequence\ninside 1 19\nTMhelix 20 39\noutside 40 325\nTMhelix 326 348\ninside 349 372

7394 GCF_002133505.1_ASM213350v1 Escherichia coli Proteobacteria; Gammaproteobacteria; Enterobacterales; Enterobacteriaceae; Escherichia
MNNEETFYQAMRRQGVTRRSFLKYCSLAATSLGLGAGMAPKIAWALENKPRIPVVWIHGL WP_001295940.1
MULTISPECIES: hydrogenase [Enterobacteriaceae] Length: 372\nNumber of predicted TMHs: 2\nExp number of AAs in TMHs: 37.2608899999999\nExp number, first 60 AAs: 19.52923\nTotal prob of N-in: 0.94351\nPOSSIBLE N-term signal sequence\ninside 1 19\nTMhelix 20 39\noutside 40 325\nTMhelix 326 348\ninside 349 372

7395 GCF_002133765.1_ASM213376v1 Escherichia coli Proteobacteria; Gammaproteobacteria; Enterobacterales; Enterobacteriaceae; Escherichia
MTGDNTLIHSHGINRRDFMKLCAALAATMGLSSKAAAEMAESVTNPQRPPVIWIGAQECT WP_000145410.1
MULTISPECIES: hydrogenase-2 small chain [Proteobacteria] Length: 372\nNumber of predicted TMHs: 1\nExp number of AAs in TMHs: 23.72226\nExp number, first 60 AAs: 0.22038\nTotal prob of N-in: 0.12994\noutside 1 331\nTMhelix 332 354\ninside 355 372

7396 GCF_002133755.1_ASM213375v1 Escherichia coli Proteobacteria; Gammaproteobacteria; Enterobacterales; Enterobacteriaceae; Escherichia
MTGDNTLIHSHGINRRDFMKLCAALAATMGLSSKAAAEMAESVTNPQRPPVIWIGAQECT WP_000145410.1
MULTISPECIES: hydrogenase-2 small chain [Proteobacteria] Length: 372\nNumber of predicted TMHs: 1\nExp number of AAs in TMHs: 23.72226\nExp number, first 60 AAs: 0.22038\nTotal prob of N-in: 0.12994\noutside 1 331\nTMhelix 332 354\ninside 355 372

7397 GCF_002133775.1_ASM213377v1 Escherichia coli Proteobacteria; Gammaproteobacteria; Enterobacterales; Enterobacteriaceae; Escherichia
MTGDNTLIHSHGINRRDFMKLCAALAATMGLSSKAAAEMAESVTNPQRPPVIWIGAQECT WP_000145410.1
MULTISPECIES: hydrogenase-2 small chain [Proteobacteria] Length: 372\nNumber of predicted TMHs: 1\nExp number of AAs in TMHs: 23.72226\nExp number, first 60 AAs: 0.22038\nTotal prob of N-in: 0.12994\noutside 1 331\nTMhelix 332 354\ninside 355 372

7398 GCF_002133765.1_ASM213376v1 Escherichia coli Proteobacteria; Gammaproteobacteria; Enterobacterales; Enterobacteriaceae; Escherichia
MNNEETFYQAMRRQGVTRRSFLKYCSLAATSLGLGAGMAPKIAWALENKPRIPVVWIHGL WP_001058323.1
MULTISPECIES: hydrogenase-1 small chain [Proteobacteria] Length: 372\nNumber of predicted TMHs: 2\nExp number of AAs in TMHs: 37.2608499999999\nExp number, first 60 AAs: 19.52923\nTotal prob of N-in:

0.94351\nPOSSIBLE N-term signal sequence\ninside 1 19\nTMhelix 20 39\noutside 40 325\nTMhelix 326 348\ninside 349 372

7399 GCF_002133755.1_ASM213375v1 Escherichia coli Proteobacteria; Gammaproteobacteria; Enterobacterales; Enterobacteriaceae; Escherichia MNNEETFYQAMRRQGVTRRSFLKYCSLAATSLGLGAGMAPKIAWALENKPRIPVVWIHGL WP_001295940.1 MULTISPECIES: hydrogenase [Enterobacteriaceae] Length: 372\nNumber of predicted TMHs: 2\nExp number of AAs in TMHs: 37.2608899999999\nExp number, first 60 AAs: 19.52923\nTotal prob of N-in: 0.94351\nPOSSIBLE N-term signal sequence\ninside 1 19\nTMhelix 20 39\noutside 40 325\nTMhelix 326 348\ninside 349 372

7400 GCF_002133775.1_ASM213377v1 Escherichia coli Proteobacteria; Gammaproteobacteria; Enterobacterales; Enterobacteriaceae; Escherichia MNNEETFYQAMRRQGVTRRSFLKYCSLAATSLGLGAGMAPKIAWALENKPRIPVVWIHGL WP_001301613.1 hydrogenase [Escherichia coli] Length: 372\nNumber of predicted TMHs: 2\nExp number of AAs in TMHs: 37.46598\nExp number, first 60 AAs: 19.53139\nTotal prob of N-in: 0.94364\nPOSSIBLE N-term signal sequence\ninside 1 19\nTMhelix 20 39\noutside 40 325\nTMhelix 326 348\ninside 349 372

7401 GCF_002133665.1_ASM213366v1 Escherichia coli Proteobacteria; Gammaproteobacteria; Enterobacterales; Enterobacteriaceae; Escherichia MNNEETFYQAMRRQGVTRRSFLKYCSLAATSLGLGAGMAPKIAWALENKPRIPVVWIHGL WP_001058323.1 MULTISPECIES: hydrogenase-1 small chain [Proteobacteria] Length: 372\nNumber of predicted TMHs: 2\nExp number of AAs in TMHs: 37.2608499999999\nExp number, first 60 AAs: 19.52923\nTotal prob of N-in: 0.94351\nPOSSIBLE N-term signal sequence\ninside 1 19\nTMhelix 20 39\noutside 40 325\nTMhelix 326 348\ninside 349 372

7402 GCF_002133845.1_ASM213384v1 Escherichia coli Proteobacteria; Gammaproteobacteria; Enterobacterales; Enterobacteriaceae; Escherichia MTGDNTLIHSHGINRRDFMKLCAALAATMGLSSKAAAEMAESVTNPQRPPVIWIGAQECT WP_000145410.1 MULTISPECIES: hydrogenase-2 small chain [Proteobacteria] Length: 372\nNumber of predicted TMHs: 1\nExp number of AAs in TMHs: 23.72226\nExp number, first 60 AAs: 0.22038\nTotal prob of N-in: 0.12994\noutside 1 331\nTMhelix 332 354\ninside 355 372

7403 GCF_002133905.1_ASM213390v1 Escherichia coli Proteobacteria; Gammaproteobacteria; Enterobacterales; Enterobacteriaceae; Escherichia MTGDNTLIHSHGINRRDFMKLCAALAATMGLSSKAAAEMAESVTNPQRPPVIWIGAQECT WP_000145410.1 MULTISPECIES: hydrogenase-2 small chain [Proteobacteria] Length: 372\nNumber of predicted TMHs: 1\nExp number of AAs in TMHs: 23.72226\nExp number, first 60 AAs: 0.22038\nTotal prob of N-in: 0.12994\noutside 1 331\nTMhelix 332 354\ninside 355 372

7404 GCF_002133855.1_ASM213385v1 Escherichia coli Proteobacteria; Gammaproteobacteria; Enterobacterales; Enterobacteriaceae; Escherichia MTGDNTLIHSHGINRRDFMKLCAALAATMGLSSKAAAEMAESVTNPQRPPVIWIGAQECT WP_000145410.1 MULTISPECIES: hydrogenase-2 small chain [Proteobacteria] Length: 372\nNumber of predicted TMHs: 1\nExp number of AAs in TMHs: 23.72226\nExp number, first 60 AAs: 0.22038\nTotal prob of N-in: 0.12994\noutside 1 331\nTMhelix 332 354\ninside 355 372

7405 GCF_002133905.1_ASM213390v1 Escherichia coli Proteobacteria; Gammaproteobacteria; Enterobacterales; Enterobacteriaceae; Escherichia MNNEETFYQAMRRQGVTRRSFLKYCSLAATSLGLGAGMAPKIAWALENKPRIPVVWIHGL WP_001058323.1 MULTISPECIES: hydrogenase-1 small chain [Proteobacteria] Length: 372\nNumber of predicted TMHs: 2\nExp number of AAs in TMHs: 37.2608499999999\nExp number, first 60 AAs: 19.52923\nTotal prob of N-in: 0.94351\nPOSSIBLE N-term signal sequence\ninside 1 19\nTMhelix 20 39\noutside 40 325\nTMhelix 326 348\ninside 349 372

7406 GCF_002133935.1_ASM213393v1 Escherichia coli Proteobacteria; Gammaproteobacteria; Enterobacterales; Enterobacteriaceae; Escherichia MTGDNTLIHSHGINRRDFMKLCAALAATMGLSSKAAAEMAESVTNPQRPPVIWIGAQECT WP_000145410.1 MULTISPECIES: hydrogenase-2 small chain [Proteobacteria] Length: 372\nNumber of predicted TMHs: 1\nExp number of AAs in TMHs: 23.72226\nExp number, first 60 AAs: 0.22038\nTotal prob of N-in: 0.12994\noutside 1 331\nTMhelix 332 354\ninside 355 372

7407 GCF_002133995.1_ASM213399v1 Escherichia coli Proteobacteria; Gammaproteobacteria; Enterobacterales; Enterobacteriaceae; Escherichia
MTGDNTLIHSHGINRRDFMKLCAALAATMGLSSKAAAEMAESVTNPQRPPVIWIGAQECT WP_000145410.1
MULTISPECIES: hydrogenase-2 small chain [Proteobacteria] Length: 372\nNumber of predicted TMHs: 1\nExp number of AAs in TMHs: 23.72226\nExp number, first 60 AAs: 0.22038\nTotal prob of N-in: 0.12994\noutside 1 331\nTMhelix 332 354\ninside 355 372

7408 GCF_002133995.1_ASM213399v1 Escherichia coli Proteobacteria; Gammaproteobacteria; Enterobacterales; Enterobacteriaceae; Escherichia
MNNEETFYQAMRRQGVTRRSFLKYCSLAATSLGLGAGMAPKIAWALENKPRIPVVWIHGL WP_001353311.1
MULTISPECIES: hydrogenase [Escherichia] Length: 372\nNumber of predicted TMHs: 2\nExp number of AAs in TMHs: 37.2638699999999\nExp number, first 60 AAs: 19.52924\nTotal prob of N-in: 0.94351\nPOSSIBLE N-term signal sequence\ninside 1 19\nTMhelix 20 39\noutside 40 325\nTMhelix 326 348\ninside 349 372

7409 GCF_002133855.1_ASM213385v1 Escherichia coli Proteobacteria; Gammaproteobacteria; Enterobacterales; Enterobacteriaceae; Escherichia
MNNEETFYQAMRRQGVTRRSFLKYCSLAATSLGLGAGMAPKIAWALENKPRIPVVWIHGL WP_001058323.1
MULTISPECIES: hydrogenase-1 small chain [Proteobacteria] Length: 372\nNumber of predicted TMHs: 2\nExp number of AAs in TMHs: 37.2608499999999\nExp number, first 60 AAs: 19.52923\nTotal prob of N-in: 0.94351\nPOSSIBLE N-term signal sequence\ninside 1 19\nTMhelix 20 39\noutside 40 325\nTMhelix 326 348\ninside 349 372

7410 GCF_002133935.1_ASM213393v1 Escherichia coli Proteobacteria; Gammaproteobacteria; Enterobacterales; Enterobacteriaceae; Escherichia
MNNEETFYQAMRRQGVTRRSFLKYCSLAATSLGLGAGMAPKIAWALENKPRIPVVWIHGL WP_001058323.1
MULTISPECIES: hydrogenase-1 small chain [Proteobacteria] Length: 372\nNumber of predicted TMHs: 2\nExp number of AAs in TMHs: 37.2608499999999\nExp number, first 60 AAs: 19.52923\nTotal prob of N-in: 0.94351\nPOSSIBLE N-term signal sequence\ninside 1 19\nTMhelix 20 39\noutside 40 325\nTMhelix 326 348\ninside 349 372

7411 GCF_002134035.1_ASM213403v1 Escherichia coli Proteobacteria; Gammaproteobacteria; Enterobacterales; Enterobacteriaceae; Escherichia
MTGDNTLIHSHGINRRDFMKLCAALAATMGLSSKAAAEMAESVTNPQRPPVIWIGAQECT WP_000145410.1
MULTISPECIES: hydrogenase-2 small chain [Proteobacteria] Length: 372\nNumber of predicted TMHs: 1\nExp number of AAs in TMHs: 23.72226\nExp number, first 60 AAs: 0.22038\nTotal prob of N-in: 0.12994\noutside 1 331\nTMhelix 332 354\ninside 355 372

7412 GCF_002134065.1_ASM213406v1 Escherichia coli Proteobacteria; Gammaproteobacteria; Enterobacterales; Enterobacteriaceae; Escherichia
MTGDNTLIHSHGINRRDFMKLCAALAATMGLSSKAAAEMAESVTNPQRPPVIWIGAQECT WP_000145410.1
MULTISPECIES: hydrogenase-2 small chain [Proteobacteria] Length: 372\nNumber of predicted TMHs: 1\nExp number of AAs in TMHs: 23.72226\nExp number, first 60 AAs: 0.22038\nTotal prob of N-in: 0.12994\noutside 1 331\nTMhelix 332 354\ninside 355 372

7413 GCF_002134035.1_ASM213403v1 Escherichia coli Proteobacteria; Gammaproteobacteria; Enterobacterales; Enterobacteriaceae; Escherichia
MNNEETFYQAMRRQGVTRRSFLKYCSLAATSLGLGAGMAPKIAWALENKPRIPVVWIHGL WP_001058323.1
MULTISPECIES: hydrogenase-1 small chain [Proteobacteria] Length: 372\nNumber of predicted TMHs: 2\nExp number of AAs in TMHs: 37.2608499999999\nExp number, first 60 AAs: 19.52923\nTotal prob of N-in: 0.94351\nPOSSIBLE N-term signal sequence\ninside 1 19\nTMhelix 20 39\noutside 40 325\nTMhelix 326 348\ninside 349 372

7414 GCF_002134065.1_ASM213406v1 Escherichia coli Proteobacteria; Gammaproteobacteria; Enterobacterales; Enterobacteriaceae; Escherichia
MNNEETFYQAMRRQGVTRRSFLKYCSLAATSLGLGAGMAPKIAWALENKPRIPVVWIHGL WP_001301613.1
hydrogenase [Escherichia coli] Length: 372\nNumber of predicted TMHs: 2\nExp number of AAs in TMHs: 37.46598\nExp number, first 60 AAs: 19.53139\nTotal prob of N-in: 0.94364\nPOSSIBLE N-term signal sequence\ninside 1 19\nTMhelix 20 39\noutside 40 325\nTMhelix 326 348\ninside 349 372

7415 GCF_002133845.1_ASM213384v1 Escherichia coli Proteobacteria; Gammaproteobacteria; Enterobacterales; Enterobacteriaceae; Escherichia

MNNEETFYQAMRRQGVTRRSFLKYCSLAATSLGLGAGMAPKIAWALENKPRIPVVWIHGL WP_001295940.1 MULTISPECIES: hydrogenase [Enterobacteriaceae] Length: 372\nNumber of predicted TMHs: 2\nExp number of AAs in TMHs: 37.2608899999999\nExp number, first 60 AAs: 19.52923\nTotal prob of N-in: 0.94351\nPOSSIBLE N-term signal sequence\ninside 1 19\nTMhelix 20 39\noutside 40 325\nTMhelix 326 348\ninside 349 372

7416 GCF_002133925.1_ASM213392v1 Escherichia coli Proteobacteria; Gammaproteobacteria; Enterobacterales; Enterobacteriaceae; Escherichia MNNEETFYQAMRRQGVTRRSFLKYCSLAATSLGLGVGMAPKIAWALENKPRIPVVWIHGL WP_073714848.1 hydrogenase [Shigella boydii] Length: 372\nNumber of predicted TMHs: 2\nExp number of AAs in TMHs: 37.83336\nExp number, first 60 AAs: 20.10829\nTotal prob of N-in: 0.96228\nPOSSIBLE N-term signal sequence\ninside 1 20\nTMhelix 21 43\noutside 44 325\nTMhelix 326 348\ninside 349 372

7417 GCF_002134145.1_ASM213414v1 Escherichia coli Proteobacteria; Gammaproteobacteria; Enterobacterales; Enterobacteriaceae; Escherichia MTGDNTLIHSHGINRRDFMKLCAALAATMGLSSKAAAEMAESVTNPQRPPVIWIGAQECT WP_000145410.1 MULTISPECIES: hydrogenase-2 small chain [Proteobacteria] Length: 372\nNumber of predicted TMHs: 1\nExp number of AAs in TMHs: 23.72226\nExp number, first 60 AAs: 0.22038\nTotal prob of N-in: 0.12994\noutside 1 331\nTMhelix 332 354\ninside 355 372

7418 GCF_002134155.1_ASM213415v1 Escherichia coli Proteobacteria; Gammaproteobacteria; Enterobacterales; Enterobacteriaceae; Escherichia MTGDNTLIHSHGINRRDFMKLCAALAATMGLSSKAAAEMVESVTNPQRPPVIWIGAQECT WP_000145423.1 hydrogenase 2 small subunit [Escherichia coli] Length: 372\nNumber of predicted TMHs: 1\nExp number of AAs in TMHs: 23.78201\nExp number, first 60 AAs: 0.28355\nTotal prob of N-in: 0.13257\noutside 1 331\nTMhelix 332 354\ninside 355 372

7419 GCF_002134175.1_ASM213417v1 Escherichia coli Proteobacteria; Gammaproteobacteria; Enterobacterales; Enterobacteriaceae; Escherichia MTGDNTLIHSHGINRRDFMKLCAALAATMGLSSKAAAEMAESVTNPQRPPVIWIGAQECT WP_000145410.1 MULTISPECIES: hydrogenase-2 small chain [Proteobacteria] Length: 372\nNumber of predicted TMHs: 1\nExp number of AAs in TMHs: 23.72226\nExp number, first 60 AAs: 0.22038\nTotal prob of N-in: 0.12994\noutside 1 331\nTMhelix 332 354\ninside 355 372

7420 GCF_002134215.1_ASM213421v1 Escherichia coli Proteobacteria; Gammaproteobacteria; Enterobacterales; Enterobacteriaceae; Escherichia MTGDNTLIHSHGINRRDFMKLCAALAATMGLSSKAAAEMAESVTNPQRPPVIWIGAQECT WP_000145410.1 MULTISPECIES: hydrogenase-2 small chain [Proteobacteria] Length: 372\nNumber of predicted TMHs: 1\nExp number of AAs in TMHs: 23.72226\nExp number, first 60 AAs: 0.22038\nTotal prob of N-in: 0.12994\noutside 1 331\nTMhelix 332 354\ninside 355 372

7421 GCF_002134205.1_ASM213420v1 Escherichia coli Proteobacteria; Gammaproteobacteria; Enterobacterales; Enterobacteriaceae; Escherichia MTGDNTLIHSHGINRRDFMKLCAALAATMGLSSKAAAEMAESVTNPQRPPVIWIGAQECT WP_000145410.1 MULTISPECIES: hydrogenase-2 small chain [Proteobacteria] Length: 372\nNumber of predicted TMHs: 1\nExp number of AAs in TMHs: 23.72226\nExp number, first 60 AAs: 0.22038\nTotal prob of N-in: 0.12994\noutside 1 331\nTMhelix 332 354\ninside 355 372

7422 GCF_002134235.1_ASM213423v1 Escherichia coli Proteobacteria; Gammaproteobacteria; Enterobacterales; Enterobacteriaceae; Escherichia MTGDNTLIHSHGINRRDFMKLCAALAATMGLSSKAAAEMAESVTNPQRPPVIWIGAQECT WP_000145410.1 MULTISPECIES: hydrogenase-2 small chain [Proteobacteria] Length: 372\nNumber of predicted TMHs: 1\nExp number of AAs in TMHs: 23.72226\nExp number, first 60 AAs: 0.22038\nTotal prob of N-in: 0.12994\noutside 1 331\nTMhelix 332 354\ninside 355 372

7423 GCF_002134445.1_ASM213444v1 Escherichia coli Proteobacteria; Gammaproteobacteria; Enterobacterales; Enterobacteriaceae; Escherichia MTGDNTLIHSHGINRRDFMKLCAALAATMGLSSKAAAEMAESVTNPQRPPVIWIGAQECT WP_000145410.1 MULTISPECIES: hydrogenase-2 small chain [Proteobacteria] Length: 372\nNumber of predicted TMHs: 1\nExp number of AAs in TMHs: 23.72226\nExp number, first 60 AAs: 0.22038\nTotal prob of N-in: 0.12994\noutside 1 331\nTMhelix 332 354\ninside 355 372

7424 GCF_002134285.1_ASM213428v1 Escherichia coli Proteobacteria; Gammaproteobacteria; Enterobacterales; Enterobacteriaceae; Escherichia
MNNEETFYQAMRRQGVTRRSFLKYCSLAATSLGLGAGMAPKIAWALENKPRIPVVWIHGL WP_001301613.1
hydrogenase [Escherichia coli] Length: 372\nNumber of predicted TMHs: 2\nExp number of AAs in TMHs: 37.46598\nExp number, first 60 AAs: 19.53139\nTotal prob of N-in: 0.94364\nPOSSIBLE N-term signal sequence\ninside 1 19\nTMhelix 20 39\noutside 40 325\nTMhelix 326 348\ninside 349 372

7425 GCF_002134205.1_ASM213420v1 Escherichia coli Proteobacteria; Gammaproteobacteria; Enterobacterales; Enterobacteriaceae; Escherichia
MNNEETFYQAMRRQGVTRRSFLKYCSLAATSLGLGAGMAPKIAWALENKPRIPVVWIHGL WP_001058323.1
MULTISPECIES: hydrogenase-1 small chain [Proteobacteria] Length: 372\nNumber of predicted TMHs: 2\nExp number of AAs in TMHs: 37.2608499999999\nExp number, first 60 AAs: 19.52923\nTotal prob of N-in: 0.94351\nPOSSIBLE N-term signal sequence\ninside 1 19\nTMhelix 20 39\noutside 40 325\nTMhelix 326 348\ninside 349 372

7426 GCF_002134325.1_ASM213432v1 Escherichia coli Proteobacteria; Gammaproteobacteria; Enterobacterales; Enterobacteriaceae; Escherichia
MTGDNTLIHSHGINRRDFMKLCAALAATMGLSSKAAAEMAESVTNPQRPPVIWIGAQECT WP_000145410.1
MULTISPECIES: hydrogenase-2 small chain [Proteobacteria] Length: 372\nNumber of predicted TMHs: 1\nExp number of AAs in TMHs: 23.72226\nExp number, first 60 AAs: 0.22038\nTotal prob of N-in: 0.12994\noutside 1 331\nTMhelix 332 354\ninside 355 372

7427 GCF_002134335.1_ASM213433v1 Escherichia coli Proteobacteria; Gammaproteobacteria; Enterobacterales; Enterobacteriaceae; Escherichia
MTGDNTLIHSHGINRRDFMKLCAALAATMGLSSKAAAEMAESVTNPQRPPVIWIGAQECT WP_000145410.1
MULTISPECIES: hydrogenase-2 small chain [Proteobacteria] Length: 372\nNumber of predicted TMHs: 1\nExp number of AAs in TMHs: 23.72226\nExp number, first 60 AAs: 0.22038\nTotal prob of N-in: 0.12994\noutside 1 331\nTMhelix 332 354\ninside 355 372

7428 GCF_002134325.1_ASM213432v1 Escherichia coli Proteobacteria; Gammaproteobacteria; Enterobacterales; Enterobacteriaceae; Escherichia
MNNEETFYQAMRRQGVTRRSFLKYCSLAATSLGLGAGMAPKIAWALENKPRIPVVWIHGL WP_001058323.1
MULTISPECIES: hydrogenase-1 small chain [Proteobacteria] Length: 372\nNumber of predicted TMHs: 2\nExp number of AAs in TMHs: 37.2608499999999\nExp number, first 60 AAs: 19.52923\nTotal prob of N-in: 0.94351\nPOSSIBLE N-term signal sequence\ninside 1 19\nTMhelix 20 39\noutside 40 325\nTMhelix 326 348\ninside 349 372

7429 GCF_002134335.1_ASM213433v1 Escherichia coli Proteobacteria; Gammaproteobacteria; Enterobacterales; Enterobacteriaceae; Escherichia
MNNEETFYQAMRRQGVTRRSFLKYCSLAATSLGLGAGMAPKIAWALENKPRIPVVWIHGL WP_086258182.1
[Ni/Fe] hydrogenase small subunit [Escherichia coli] Length: 372\nNumber of predicted TMHs: 2\nExp number of AAs in TMHs: 37.30752\nExp number, first 60 AAs: 19.54005\nTotal prob of N-in: 0.94357\nPOSSIBLE N-term signal sequence\ninside 1 19\nTMhelix 20 39\noutside 40 325\nTMhelix 326 348\ninside 349 372

7430 GCF_002134235.1_ASM213423v1 Escherichia coli Proteobacteria; Gammaproteobacteria; Enterobacterales; Enterobacteriaceae; Escherichia
MNNEETFYQAMRRQGVTRRSFLKYCSLAATSLGLGAGMAPKIAWALENKPRIPVVWIHGL WP_001295940.1
MULTISPECIES: hydrogenase [Enterobacteriaceae] Length: 372\nNumber of predicted TMHs: 2\nExp number of AAs in TMHs: 37.2608899999999\nExp number, first 60 AAs: 19.52923\nTotal prob of N-in: 0.94351\nPOSSIBLE N-term signal sequence\ninside 1 19\nTMhelix 20 39\noutside 40 325\nTMhelix 326 348\ninside 349 372

7431 GCF_002134465.1_ASM213446v1 Escherichia coli Proteobacteria; Gammaproteobacteria; Enterobacterales; Enterobacteriaceae; Escherichia
MTGDNTLIHSHGINRRDFMKLCAALAATMGLSSKAAAEMAESVTNPQRPPVIWIGAQECT WP_000145410.1
MULTISPECIES: hydrogenase-2 small chain [Proteobacteria] Length: 372\nNumber of predicted TMHs: 1\nExp number of AAs in TMHs: 23.72226\nExp number, first 60 AAs: 0.22038\nTotal prob of N-in: 0.12994\noutside 1 331\nTMhelix 332 354\ninside 355 372

7432 GCF_002134465.1_ASM213446v1 Escherichia coli Proteobacteria; Gammaproteobacteria; Enterobacterales; Enterobacteriaceae; Escherichia

MNNEETFYQAMRRQGVTRRSFLKYCSLAATSLGLGAGMAPKIAWALENKPRIPVVWIHGL WP_001058323.1 MULTISPECIES: hydrogenase-1 small chain [Proteobacteria] Length: 372\nNumber of predicted TMHs: 2\nExp number of AAs in TMHs: 37.2608499999999\nExp number, first 60 AAs: 19.52923\nTotal prob of N-in: 0.94351\nPOSSIBLE N-term signal sequence\ninside 1 19\nTMhelix 20 39\noutside 40 325\nTMhelix 326 348\ninside 349 372

7433 GCF_002134345.1_ASM213434v1 Escherichia coli Proteobacteria; Gammaproteobacteria; Enterobacterales; Enterobacteriaceae; Escherichia MNNEETFYQAMRRQGVTRRSFLKYCSLAATSLGLGAGMAPKIAWALENKPRIPVVWIHGL WP_001058323.1 MULTISPECIES: hydrogenase-1 small chain [Proteobacteria] Length: 372\nNumber of predicted TMHs: 2\nExp number of AAs in TMHs: 37.2608499999999\nExp number, first 60 AAs: 19.52923\nTotal prob of N-in: 0.94351\nPOSSIBLE N-term signal sequence\ninside 1 19\nTMhelix 20 39\noutside 40 325\nTMhelix 326 348\ninside 349 372

7434 GCF_002134505.1_ASM213450v1 Escherichia coli Proteobacteria; Gammaproteobacteria; Enterobacterales; Enterobacteriaceae; Escherichia MTGDNTLIHSHGINRRDFMKLCAALAATMGLSSKAAAEMAESVTNPQRPPVIWIGAQECT WP_000145410.1 MULTISPECIES: hydrogenase-2 small chain [Proteobacteria] Length: 372\nNumber of predicted TMHs: 1\nExp number of AAs in TMHs: 23.72226\nExp number, first 60 AAs: 0.22038\nTotal prob of N-in: 0.12994\noutside 1 331\nTMhelix 332 354\ninside 355 372

7435 GCF_002134515.1_ASM213451v1 Escherichia coli Proteobacteria; Gammaproteobacteria; Enterobacterales; Enterobacteriaceae; Escherichia MTGDNTLIHSHGINRRDFMKLCAALAATMGLSSKAAAEMAESVTNPQRPPVIWIGAQECT WP_000145410.1 MULTISPECIES: hydrogenase-2 small chain [Proteobacteria] Length: 372\nNumber of predicted TMHs: 1\nExp number of AAs in TMHs: 23.72226\nExp number, first 60 AAs: 0.22038\nTotal prob of N-in: 0.12994\noutside 1 331\nTMhelix 332 354\ninside 355 372

7436 GCF_002134525.1_ASM213452v1 Escherichia coli Proteobacteria; Gammaproteobacteria; Enterobacterales; Enterobacteriaceae; Escherichia MTGDNTLIHSHGINRRDFMKLCAALAATMGLSSKAAAEMAESVTNPQRPPVIWIGAQECT WP_000145410.1 MULTISPECIES: hydrogenase-2 small chain [Proteobacteria] Length: 372\nNumber of predicted TMHs: 1\nExp number of AAs in TMHs: 23.72226\nExp number, first 60 AAs: 0.22038\nTotal prob of N-in: 0.12994\noutside 1 331\nTMhelix 332 354\ninside 355 372

7437 GCF_002134515.1_ASM213451v1 Escherichia coli Proteobacteria; Gammaproteobacteria; Enterobacterales; Enterobacteriaceae; Escherichia MNNEETFYQAMRRQGVTRRSFLKYCSLAATSLGLGAGMAPKIAWALENKPRIPVVWIHGL WP_001058323.1 MULTISPECIES: hydrogenase-1 small chain [Proteobacteria] Length: 372\nNumber of predicted TMHs: 2\nExp number of AAs in TMHs: 37.2608499999999\nExp number, first 60 AAs: 19.52923\nTotal prob of N-in: 0.94351\nPOSSIBLE N-term signal sequence\ninside 1 19\nTMhelix 20 39\noutside 40 325\nTMhelix 326 348\ninside 349 372

7438 GCF_002134575.1_ASM213457v1 Escherichia coli Proteobacteria; Gammaproteobacteria; Enterobacterales; Enterobacteriaceae; Escherichia MTGDNTLIHSHGINRRDFMKLCAALAATMGLSSKAAAEMAESVTNPQRPPVIWIGAQECT WP_000145410.1 MULTISPECIES: hydrogenase-2 small chain [Proteobacteria] Length: 372\nNumber of predicted TMHs: 1\nExp number of AAs in TMHs: 23.72226\nExp number, first 60 AAs: 0.22038\nTotal prob of N-in: 0.12994\noutside 1 331\nTMhelix 332 354\ninside 355 372

7439 GCF_002134575.1_ASM213457v1 Escherichia coli Proteobacteria; Gammaproteobacteria; Enterobacterales; Enterobacteriaceae; Escherichia MNNEETFYQAMRRQGVTRRSFLKYCSLAATSLGLGAGMAPKIAWALENKPRIPVVWIHGL WP_001301613.1 hydrogenase [Escherichia coli] Length: 372\nNumber of predicted TMHs: 2\nExp number of AAs in TMHs: 37.46598\nExp number, first 60 AAs: 19.53139\nTotal prob of N-in: 0.94364\nPOSSIBLE N-term signal sequence\ninside 1 19\nTMhelix 20 39\noutside 40 325\nTMhelix 326 348\ninside 349 372

7440 GCF_002134445.1_ASM213444v1 Escherichia coli Proteobacteria; Gammaproteobacteria; Enterobacterales; Enterobacteriaceae; Escherichia MNNEETFYQAMRRQGVTRRSFLKYCSLAATSLGLGAGMAPKIAWALENKPRIPVVWIHGL WP_001301613.1 hydrogenase [Escherichia coli] Length: 372\nNumber of predicted TMHs: 2\nExp number of AAs in

TMHs: 37.46598\nExp number, first 60 AAs:  19.53139\nTotal prob of N-in:   0.94364\nPOSSIBLE N-term signal sequence\ninside   1   19\nTMhelix   20   39\noutside   40   325\nTMhelix   326   348\ninside   349   372

7441 GCF_002134625.1_ASM213462v1 Escherichia coli Proteobacteria; Gammaproteobacteria; Enterobacterales; Enterobacteriaceae; Escherichia MTGDNTLIHSHGINRRDFMKLCAALAATMGLSSKAAAEMAESVTNPQRPPVIWIGAQECT WP_000145410.1 MULTISPECIES: hydrogenase-2 small chain [Proteobacteria] Length: 372\nNumber of predicted TMHs: 1\nExp number of AAs in TMHs: 23.72226\nExp number, first 60 AAs:  0.22038\nTotal prob of N-in:   0.12994\noutside   1   331\nTMhelix   332   354\ninside   355   372

7442 GCF_002134585.1_ASM213458v1 Escherichia coli Proteobacteria; Gammaproteobacteria; Enterobacterales; Enterobacteriaceae; Escherichia MTGDNTLIHSHGINRRDFMKLCAALAATMGLSSKAAAEMAESVTNPQRPPVIWIGAQECT WP_000145410.1 MULTISPECIES: hydrogenase-2 small chain [Proteobacteria] Length: 372\nNumber of predicted TMHs: 1\nExp number of AAs in TMHs: 23.72226\nExp number, first 60 AAs:  0.22038\nTotal prob of N-in:   0.12994\noutside   1   331\nTMhelix   332   354\ninside   355   372

7443 GCF_002134635.1_ASM213463v1 Escherichia coli Proteobacteria; Gammaproteobacteria; Enterobacterales; Enterobacteriaceae; Escherichia MTGDNTLIHSHGINRRDFMKLCAALAATMGLSSKAAAEMAESVTNPQRPPVIWIGAQECT WP_000145410.1 MULTISPECIES: hydrogenase-2 small chain [Proteobacteria] Length: 372\nNumber of predicted TMHs: 1\nExp number of AAs in TMHs: 23.72226\nExp number, first 60 AAs:  0.22038\nTotal prob of N-in:   0.12994\noutside   1   331\nTMhelix   332   354\ninside   355   372

7444 GCF_002134645.1_ASM213464v1 Escherichia coli Proteobacteria; Gammaproteobacteria; Enterobacterales; Enterobacteriaceae; Escherichia MTGDNTLIHSHGINRRDFMKLCAALAATMGLSSKAAAEMAESVTNPQRPPVIWIGAQECT WP_000145410.1 MULTISPECIES: hydrogenase-2 small chain [Proteobacteria] Length: 372\nNumber of predicted TMHs: 1\nExp number of AAs in TMHs: 23.72226\nExp number, first 60 AAs:  0.22038\nTotal prob of N-in:   0.12994\noutside   1   331\nTMhelix   332   354\ninside   355   372

7445 GCF_002134685.1_ASM213468v1 Escherichia coli Proteobacteria; Gammaproteobacteria; Enterobacterales; Enterobacteriaceae; Escherichia MNNEETFYQAMRRQGVTRRSFLKYCSLAATSLGLGAGMAPKIAWALENKPRIPVVWIHGL WP_044723380.1 hydrogenase [Escherichia coli] Length: 372\nNumber of predicted TMHs:  2\nExp number of AAs in TMHs: 37.2612299999999\nExp number, first 60 AAs:  19.52923\nTotal prob of N-in:   0.94351\nPOSSIBLE N-term signal sequence\ninside   1   19\nTMhelix   20   39\noutside   40   325\nTMhelix   326   348\ninside   349   372

7446 GCF_002134645.1_ASM213464v1 Escherichia coli Proteobacteria; Gammaproteobacteria; Enterobacterales; Enterobacteriaceae; Escherichia MNNEETFYQAMRRQGVTRRSFLKYCSLAATSLGLGAGMAPKIAWALENKPRIPVVWIHGL WP_001058323.1 MULTISPECIES: hydrogenase-1 small chain [Proteobacteria] Length: 372\nNumber of predicted TMHs: 2\nExp number of AAs in TMHs: 37.2608499999999\nExp number, first 60 AAs:  19.52923\nTotal prob of N-in:   0.94351\nPOSSIBLE N-term signal sequence\ninside   1   19\nTMhelix   20   39\noutside   40   325\nTMhelix   326   348\ninside   349   372

7447 GCF_002134765.1_ASM213476v1 Escherichia coli Proteobacteria; Gammaproteobacteria; Enterobacterales; Enterobacteriaceae; Escherichia MTGDNTLIHSHGINRRDFMKLCAALAATMGLSSKAAAEMAESVTNPQRPPVIWIGAQECT WP_000145410.1 MULTISPECIES: hydrogenase-2 small chain [Proteobacteria] Length: 372\nNumber of predicted TMHs: 1\nExp number of AAs in TMHs: 23.72226\nExp number, first 60 AAs:  0.22038\nTotal prob of N-in:   0.12994\noutside   1   331\nTMhelix   332   354\ninside   355   372

7448 GCF_002134765.1_ASM213476v1 Escherichia coli Proteobacteria; Gammaproteobacteria; Enterobacterales; Enterobacteriaceae; Escherichia MNNEETFYQAMRRQGVTRRSFLKYCSLAATSLGLGAGMAPKIAWALENKPRIPVVWIHGL WP_001058320.1 hydrogenase [Escherichia coli] Length: 372\nNumber of predicted TMHs:  2\nExp number of AAs in TMHs: 37.26082\nExp number, first 60 AAs:  19.52923\nTotal prob of N-in:   0.94351\nPOSSIBLE N-term signal sequence\ninside   1   19\nTMhelix   20   39\noutside   40   325\nTMhelix   326   348\ninside   349   372

7449 GCF_002134725.1_ASM213472v1 Escherichia coli Proteobacteria; Gammaproteobacteria; Enterobacterales; Enterobacteriaceae; Escherichia
MNNEETFYQAMRRQGVTRRSFLKYCSLAATSLGLGAGMAPKIAWALENKPRIPVVWIHGL WP_001058323.1
MULTISPECIES: hydrogenase-1 small chain [Proteobacteria] Length: 372\nNumber of predicted TMHs: 2\nExp number of AAs in TMHs: 37.2608499999999\nExp number, first 60 AAs: 19.52923\nTotal prob of N-in: 0.94351\nPOSSIBLE N-term signal sequence\ninside 1 19\nTMhelix 20 39\noutside 40 325\nTMhelix 326 348\ninside 349 372

7450 GCF_002142335.1_ASM214233v1 Escherichia coli Proteobacteria; Gammaproteobacteria; Enterobacterales; Enterobacteriaceae; Escherichia
MTGDNTLIHSHGINRRDFMKLCAALAATMGLSSKAAAEMAESVTNPQRPPVIWIGAQECT WP_000145410.1
MULTISPECIES: hydrogenase-2 small chain [Proteobacteria] Length: 372\nNumber of predicted TMHs: 1\nExp number of AAs in TMHs: 23.72226\nExp number, first 60 AAs: 0.22038\nTotal prob of N-in: 0.12994\noutside 1 331\nTMhelix 332 354\ninside 355 372

7451 GCF_002142335.1_ASM214233v1 Escherichia coli Proteobacteria; Gammaproteobacteria; Enterobacterales; Enterobacteriaceae; Escherichia
MNNEETFYQAMRRQGVTRRSFLKYCSLAATSLGLGAGMAPKIAWALENKPRIPVVWIHGL WP_001058323.1
MULTISPECIES: hydrogenase-1 small chain [Proteobacteria] Length: 372\nNumber of predicted TMHs: 2\nExp number of AAs in TMHs: 37.2608499999999\nExp number, first 60 AAs: 19.52923\nTotal prob of N-in: 0.94351\nPOSSIBLE N-term signal sequence\ninside 1 19\nTMhelix 20 39\noutside 40 325\nTMhelix 326 348\ninside 349 372

7452 GCF_002142415.1_ASM214241v1 Escherichia coli Proteobacteria; Gammaproteobacteria; Enterobacterales; Enterobacteriaceae; Escherichia
MNNEETFYQAMRRQGVTRRSFLKYCSLAATSLGLGAGMAPKIAWALENKPRIPVVWIHGL WP_001058323.1
MULTISPECIES: hydrogenase-1 small chain [Proteobacteria] Length: 372\nNumber of predicted TMHs: 2\nExp number of AAs in TMHs: 37.2608499999999\nExp number, first 60 AAs: 19.52923\nTotal prob of N-in: 0.94351\nPOSSIBLE N-term signal sequence\ninside 1 19\nTMhelix 20 39\noutside 40 325\nTMhelix 326 348\ninside 349 372

7453 GCF_002134695.1_ASM213469v1 Escherichia coli Proteobacteria; Gammaproteobacteria; Enterobacterales; Enterobacteriaceae; Escherichia
MNNEETFYQAMRRQGVTRRSFLKYCSLAATSLGLGAGMAPKIAWALENKPRIPVVWIHGL WP_001301613.1
hydrogenase [Escherichia coli] Length: 372\nNumber of predicted TMHs: 2\nExp number of AAs in TMHs: 37.46598\nExp number, first 60 AAs: 19.53139\nTotal prob of N-in: 0.94364\nPOSSIBLE N-term signal sequence\ninside 1 19\nTMhelix 20 39\noutside 40 325\nTMhelix 326 348\ninside 349 372

7454 GCF_002142435.1_ASM214243v1 Escherichia coli Proteobacteria; Gammaproteobacteria; Enterobacterales; Enterobacteriaceae; Escherichia
MTGDNTLIHSHGINRRDFMKLCAALAATMGLSSKAAAEMAESVTNPQRPPVIWIGAQECT WP_000145410.1
MULTISPECIES: hydrogenase-2 small chain [Proteobacteria] Length: 372\nNumber of predicted TMHs: 1\nExp number of AAs in TMHs: 23.72226\nExp number, first 60 AAs: 0.22038\nTotal prob of N-in: 0.12994\noutside 1 331\nTMhelix 332 354\ninside 355 372

7455 GCF_002142445.1_ASM214244v1 Escherichia coli Proteobacteria; Gammaproteobacteria; Enterobacterales; Enterobacteriaceae; Escherichia
MTGDNTLIHSHGINRRDFMKLCAALAATMGLSSKAAAEMAESVTNPQRPPVIWIGAQECT WP_000145410.1
MULTISPECIES: hydrogenase-2 small chain [Proteobacteria] Length: 372\nNumber of predicted TMHs: 1\nExp number of AAs in TMHs: 23.72226\nExp number, first 60 AAs: 0.22038\nTotal prob of N-in: 0.12994\noutside 1 331\nTMhelix 332 354\ninside 355 372

7456 GCF_002142675.1_ASM214267v1 Escherichia coli Proteobacteria; Gammaproteobacteria; Enterobacterales; Enterobacteriaceae; Escherichia
MTGDNTLIHSHGINRRDFMKLCAALAATMGLSSKAAAEMAESVTNPQRPPVIWIGAQECT WP_000145410.1
MULTISPECIES: hydrogenase-2 small chain [Proteobacteria] Length: 372\nNumber of predicted TMHs: 1\nExp number of AAs in TMHs: 23.72226\nExp number, first 60 AAs: 0.22038\nTotal prob of N-in: 0.12994\noutside 1 331\nTMhelix 332 354\ninside 355 372

7457 GCF_002142715.1_ASM214271v1 Escherichia coli Proteobacteria; Gammaproteobacteria; Enterobacterales; Enterobacteriaceae; Escherichia

MTGDNTLIHSHGINRRDFMKLCAALAATMGLSSKAAAEMAESVTNPQRPPVIWIGAQECT WP_000145410.1 MULTISPECIES: hydrogenase-2 small chain [Proteobacteria] Length: 372\nNumber of predicted TMHs: 1\nExp number of AAs in TMHs: 23.72226\nExp number, first 60 AAs: 0.22038\nTotal prob of N-in: 0.12994\noutside 1 331\nTMhelix 332 354\ninside 355 372

7458 GCF_002144065.1_ASM214406v1 Escherichia coli Proteobacteria; Gammaproteobacteria; Enterobacterales; Enterobacteriaceae; Escherichia MTGDNTLIHSHGINRRDFMKLCAALAATMGLSSKAAAEMAESVTNPQRPPVIWIGAQECT WP_000145410.1 MULTISPECIES: hydrogenase-2 small chain [Proteobacteria] Length: 372\nNumber of predicted TMHs: 1\nExp number of AAs in TMHs: 23.72226\nExp number, first 60 AAs: 0.22038\nTotal prob of N-in: 0.12994\noutside 1 331\nTMhelix 332 354\ninside 355 372

7459 GCF_002142695.1_ASM214269v1 Escherichia coli Proteobacteria; Gammaproteobacteria; Enterobacterales; Enterobacteriaceae; Escherichia MNNEETFYQAMRRQGVTRRSFLKYCSLAATSLGLGAGMAPKIAWALENKPRIPVVWIHGL WP_001058323.1 MULTISPECIES: hydrogenase-1 small chain [Proteobacteria] Length: 372\nNumber of predicted TMHs: 2\nExp number of AAs in TMHs: 37.2608499999999\nExp number, first 60 AAs: 19.52923\nTotal prob of N-in: 0.94351\nPOSSIBLE N-term signal sequence\ninside 1 19\nTMhelix 20 39\noutside 40 325\nTMhelix 326 348\ninside 349 372

7460 GCF_002144135.1_ASM214413v1 Escherichia coli Proteobacteria; Gammaproteobacteria; Enterobacterales; Enterobacteriaceae; Escherichia MTGDNTLIHSHGINRRDFMKLCAALAATMGLSSKAAAEMAESVTNPQRPPVIWIGAQECT WP_000145410.1 MULTISPECIES: hydrogenase-2 small chain [Proteobacteria] Length: 372\nNumber of predicted TMHs: 1\nExp number of AAs in TMHs: 23.72226\nExp number, first 60 AAs: 0.22038\nTotal prob of N-in: 0.12994\noutside 1 331\nTMhelix 332 354\ninside 355 372

7461 GCF_002144165.1_ASM214416v1 Escherichia coli Proteobacteria; Gammaproteobacteria; Enterobacterales; Enterobacteriaceae; Escherichia MTGDNTLIHSHGINRRDFMKLCAALAATMGLSSKAAAEMAESVTNPQRPPVIWIGAQECT WP_000145410.1 MULTISPECIES: hydrogenase-2 small chain [Proteobacteria] Length: 372\nNumber of predicted TMHs: 1\nExp number of AAs in TMHs: 23.72226\nExp number, first 60 AAs: 0.22038\nTotal prob of N-in: 0.12994\noutside 1 331\nTMhelix 332 354\ninside 355 372

7462 GCF_002144145.1_ASM214414v1 Escherichia coli Proteobacteria; Gammaproteobacteria; Enterobacterales; Enterobacteriaceae; Escherichia MTGDNTLIHSHGINRRDFMKLCAALAATMGLSSKAAAEMAESVTNPQRPPVIWIGAQECT WP_000145410.1 MULTISPECIES: hydrogenase-2 small chain [Proteobacteria] Length: 372\nNumber of predicted TMHs: 1\nExp number of AAs in TMHs: 23.72226\nExp number, first 60 AAs: 0.22038\nTotal prob of N-in: 0.12994\noutside 1 331\nTMhelix 332 354\ninside 355 372

7463 GCF_002144065.1_ASM214406v1 Escherichia coli Proteobacteria; Gammaproteobacteria; Enterobacterales; Enterobacteriaceae; Escherichia MNNEETFYQAMRRQGVTRRSFLKYCSLAATSLGLGAGMAPKIAWALENKPRIPVVWIHGL WP_001301613.1 hydrogenase [Escherichia coli] Length: 372\nNumber of predicted TMHs: 2\nExp number of AAs in TMHs: 37.46598\nExp number, first 60 AAs: 19.53139\nTotal prob of N-in: 0.94364\nPOSSIBLE N-term signal sequence\ninside 1 19\nTMhelix 20 39\noutside 40 325\nTMhelix 326 348\ninside 349 372

7464 GCF_002142715.1_ASM214271v1 Escherichia coli Proteobacteria; Gammaproteobacteria; Enterobacterales; Enterobacteriaceae; Escherichia MNNEETFYQAMRRQGVTRRSFLKYCSLAATSLGLGAGMAPKIAWALENKPRIPVVWIHGL WP_001295940.1 MULTISPECIES: hydrogenase [Enterobacteriaceae] Length: 372\nNumber of predicted TMHs: 2\nExp number of AAs in TMHs: 37.2608899999999\nExp number, first 60 AAs: 19.52923\nTotal prob of N-in: 0.94351\nPOSSIBLE N-term signal sequence\ninside 1 19\nTMhelix 20 39\noutside 40 325\nTMhelix 326 348\ninside 349 372

7465 GCF_002144135.1_ASM214413v1 Escherichia coli Proteobacteria; Gammaproteobacteria; Enterobacterales; Enterobacteriaceae; Escherichia MNNEETFYQAMRRQGVTRRSFLKYCSLAATSLGLGAGMAPKIAWALENKPRIPVVWIHGL WP_001058323.1 MULTISPECIES: hydrogenase-1 small chain [Proteobacteria] Length: 372\nNumber of predicted TMHs: 2\nExp number of AAs in TMHs: 37.2608499999999\nExp number, first 60 AAs: 19.52923\nTotal prob of N-in:

0.94351\nPOSSIBLE N-term signal sequence\ninside 1 19\nTMhelix 20 39\noutside 40 325\nTMhelix 326 348\ninside 349 372

7466 GCF_002144165.1_ASM214416v1 Escherichia coli Proteobacteria; Gammaproteobacteria; Enterobacterales; Enterobacteriaceae; Escherichia
MNNEETFYQAMRRQGVTRRSFLKYCSLAATSLGLGAGMAPKIAWALENKPRIPVVWIHGL WP_001058323.1
MULTISPECIES: hydrogenase-1 small chain [Proteobacteria] Length: 372\nNumber of predicted TMHs: 2\nExp number of AAs in TMHs: 37.2608499999999\nExp number, first 60 AAs: 19.52923\nTotal prob of N-in: 0.94351\nPOSSIBLE N-term signal sequence\ninside 1 19\nTMhelix 20 39\noutside 40 325\nTMhelix 326 348\ninside 349 372

7467 GCF_002152015.1_ASM215201v1 Escherichia coli Proteobacteria; Gammaproteobacteria; Enterobacterales; Enterobacteriaceae; Escherichia
MTGDNTLIHSHGINRRDFMKLCAALAATMGLSSKAAAEMAESVTNPQRPPVIWIGAQECT WP_000145410.1
MULTISPECIES: hydrogenase-2 small chain [Proteobacteria] Length: 372\nNumber of predicted TMHs: 1\nExp number of AAs in TMHs: 23.72226\nExp number, first 60 AAs: 0.22038\nTotal prob of N-in: 0.12994\noutside 1 331\nTMhelix 332 354\ninside 355 372

7468 GCF_002152015.1_ASM215201v1 Escherichia coli Proteobacteria; Gammaproteobacteria; Enterobacterales; Enterobacteriaceae; Escherichia
MNNEETFYQAMRRQGVTRRSFLKYCSLAATSLGLGAGMAPKIAWALENKPRIPVVWIHGL WP_001058323.1
MULTISPECIES: hydrogenase-1 small chain [Proteobacteria] Length: 372\nNumber of predicted TMHs: 2\nExp number of AAs in TMHs: 37.2608499999999\nExp number, first 60 AAs: 19.52923\nTotal prob of N-in: 0.94351\nPOSSIBLE N-term signal sequence\ninside 1 19\nTMhelix 20 39\noutside 40 325\nTMhelix 326 348\ninside 349 372

7469 GCF_002144145.1_ASM214414v1 Escherichia coli Proteobacteria; Gammaproteobacteria; Enterobacterales; Enterobacteriaceae; Escherichia
MNNEETFYQAMRRQGVTRRSFLKYCSLAATSLGLGAGMAPKIAWALENKPRIPVVWIHGL WP_001058323.1
MULTISPECIES: hydrogenase-1 small chain [Proteobacteria] Length: 372\nNumber of predicted TMHs: 2\nExp number of AAs in TMHs: 37.2608499999999\nExp number, first 60 AAs: 19.52923\nTotal prob of N-in: 0.94351\nPOSSIBLE N-term signal sequence\ninside 1 19\nTMhelix 20 39\noutside 40 325\nTMhelix 326 348\ninside 349 372

7470 GCF_002152025.1_ASM215202v1 Escherichia coli Proteobacteria; Gammaproteobacteria; Enterobacterales; Enterobacteriaceae; Escherichia
MTGDNTLIHSHGINRRDFMKLCAALAATMGLSSKAAAEMAESVTNPQRPPVIWIGAQECT WP_000145410.1
MULTISPECIES: hydrogenase-2 small chain [Proteobacteria] Length: 372\nNumber of predicted TMHs: 1\nExp number of AAs in TMHs: 23.72226\nExp number, first 60 AAs: 0.22038\nTotal prob of N-in: 0.12994\noutside 1 331\nTMhelix 332 354\ninside 355 372

7471 GCF_002152075.1_ASM215207v1 Escherichia coli Proteobacteria; Gammaproteobacteria; Enterobacterales; Enterobacteriaceae; Escherichia
MTGDNTLIHSHGINRRDFMKLCAALAATMGLSSKAAAEMVESVTNPQRPPVIWIGAQECT WP_000145423.1
hydrogenase 2 small subunit [Escherichia coli] Length: 372\nNumber of predicted TMHs: 1\nExp number of AAs in TMHs: 23.78201\nExp number, first 60 AAs: 0.28355\nTotal prob of N-in: 0.13257\noutside 1 331\nTMhelix 332 354\ninside 355 372

7472 GCF_002152065.1_ASM215206v1 Escherichia coli Proteobacteria; Gammaproteobacteria; Enterobacterales; Enterobacteriaceae; Escherichia
MNNEETFYQAMRRQGVTRRSFLKYCSLAATSLGLGAGMAPKIAWALENKPRIPVVWIHGL WP_001058323.1
MULTISPECIES: hydrogenase-1 small chain [Proteobacteria] Length: 372\nNumber of predicted TMHs: 2\nExp number of AAs in TMHs: 37.2608499999999\nExp number, first 60 AAs: 19.52923\nTotal prob of N-in: 0.94351\nPOSSIBLE N-term signal sequence\ninside 1 19\nTMhelix 20 39\noutside 40 325\nTMhelix 326 348\ninside 349 372

7473 GCF_002152145.1_ASM215214v1 Escherichia coli Proteobacteria; Gammaproteobacteria; Enterobacterales; Enterobacteriaceae; Escherichia
MTGDNTLIHSHGINRRDFMKLCAALAATMGLSSKAAAEMVESVTNPQRPPVIWIGAQECT WP_000145423.1
hydrogenase 2 small subunit [Escherichia coli] Length: 372\nNumber of predicted TMHs: 1\nExp number of

AAs in TMHs: 23.78201\nExp number, first 60 AAs:  0.28355\nTotal prob of N-in:   0.13257\noutside   1   331\nTMhelix   332   354\ninside   355   372

7474 GCF_002152175.1_ASM215217v1 Escherichia coli Proteobacteria; Gammaproteobacteria; Enterobacterales; Enterobacteriaceae; Escherichia
MTGDNTLIHSHGINRRDFMKLCAALAATMGLSSKAAAEMVESVTNPQRPPVIWIGAQECT WP_000145423.1
hydrogenase 2 small subunit [Escherichia coli] Length: 372\nNumber of predicted TMHs:  1\nExp number of AAs in TMHs: 23.78201\nExp number, first 60 AAs:  0.28355\nTotal prob of N-in:   0.13257\noutside   1   331\nTMhelix   332   354\ninside   355   372

7475 GCF_002152185.1_ASM215218v1 Escherichia coli Proteobacteria; Gammaproteobacteria; Enterobacterales; Enterobacteriaceae; Escherichia
MTGDNTLIHSHGINRRDFMKLCAALAATMGLSSKAAAEMAESVTNPQRPPVIWIGAQECT WP_000145410.1
MULTISPECIES: hydrogenase-2 small chain [Proteobacteria] Length: 372\nNumber of predicted TMHs: 1\nExp number of AAs in TMHs: 23.72226\nExp number, first 60 AAs:  0.22038\nTotal prob of N-in:   0.12994\noutside   1   331\nTMhelix   332   354\ninside   355   372

7476 GCF_002152145.1_ASM215214v1 Escherichia coli Proteobacteria; Gammaproteobacteria; Enterobacterales; Enterobacteriaceae; Escherichia
MNNEETFYQAMRRQGVTRRSFLKYCSLAATSLGLGAGMAPKIAWALENKPRIPVVWIHGL WP_001538532.1
hydrogenase-1 small chain [Escherichia coli] Length: 372\nNumber of predicted TMHs:  2\nExp number of AAs in TMHs: 37.27041\nExp number, first 60 AAs:  19.52927\nTotal prob of N-in:   0.94352\nPOSSIBLE N-term signal sequence\ninside   1   19\nTMhelix   20   39\noutside   40   325\nTMhelix   326   348\ninside   349   372

7477 GCF_002152185.1_ASM215218v1 Escherichia coli Proteobacteria; Gammaproteobacteria; Enterobacterales; Enterobacteriaceae; Escherichia
MNNEETFYQAMRRQGVTRRSFLKYCSLAATSLGLGAGMAPKIAWALENKPRIPVVWIHGL WP_001058323.1
MULTISPECIES: hydrogenase-1 small chain [Proteobacteria] Length: 372\nNumber of predicted TMHs: 2\nExp number of AAs in TMHs: 37.2608499999999\nExp number, first 60 AAs:  19.52923\nTotal prob of N-in: 0.94351\nPOSSIBLE N-term signal sequence\ninside   1   19\nTMhelix   20   39\noutside   40   325\nTMhelix   326 348\ninside   349   372

7478 GCF_002152175.1_ASM215217v1 Escherichia coli Proteobacteria; Gammaproteobacteria; Enterobacterales; Enterobacteriaceae; Escherichia
MNNEETFYQAMRRQGVTRRSFLKYCSLAATSLGLGAGMAPKIAWALENKPRIPVVWIHGL WP_001295940.1
MULTISPECIES: hydrogenase [Enterobacteriaceae] Length: 372\nNumber of predicted TMHs:  2\nExp number of AAs in TMHs: 37.2608899999999\nExp number, first 60 AAs:  19.52923\nTotal prob of N-in: 0.94351\nPOSSIBLE N-term signal sequence\ninside   1   19\nTMhelix   20   39\noutside   40   325\nTMhelix   326 348\ninside   349   372

7479 GCF_002152995.1_ASM215299v1 Escherichia coli Proteobacteria; Gammaproteobacteria; Enterobacterales; Enterobacteriaceae; Escherichia
MTGDNTLIHSHGINRRDFMKLCAALAATMGLSSKAAAEMAESVTNPQRPPVIWIGAQECT WP_000145410.1
MULTISPECIES: hydrogenase-2 small chain [Proteobacteria] Length: 372\nNumber of predicted TMHs: 1\nExp number of AAs in TMHs: 23.72226\nExp number, first 60 AAs:  0.22038\nTotal prob of N-in:   0.12994\noutside   1   331\nTMhelix   332   354\ninside   355   372

7480 GCF_002152255.1_ASM215225v1 Escherichia coli Proteobacteria; Gammaproteobacteria; Enterobacterales; Enterobacteriaceae; Escherichia
MNNEETFYQAMRRQGVTRRSFLKYCSLAATSLGLGAGMAPKIAWALENKPRIPVVWIHGL WP_001058323.1
MULTISPECIES: hydrogenase-1 small chain [Proteobacteria] Length: 372\nNumber of predicted TMHs: 2\nExp number of AAs in TMHs: 37.2608499999999\nExp number, first 60 AAs:  19.52923\nTotal prob of N-in: 0.94351\nPOSSIBLE N-term signal sequence\ninside   1   19\nTMhelix   20   39\noutside   40   325\nTMhelix   326 348\ninside   349   372

7481 GCF_002152225.1_ASM215222v1 Escherichia coli Proteobacteria; Gammaproteobacteria; Enterobacterales; Enterobacteriaceae; Escherichia
MNNEETFYQAMRRQGVTRRSFLKYCSLAATSLGLGAGMAPKIAWALENKPRIPVVWIHGL WP_001058323.1
MULTISPECIES: hydrogenase-1 small chain [Proteobacteria] Length: 372\nNumber of predicted TMHs: 2\nExp number of AAs in TMHs: 37.2608499999999\nExp number, first 60 AAs:  19.52923\nTotal prob of N-in:

0.94351\nPOSSIBLE N-term signal sequence\ninside 1 19\nTMhelix 20 39\noutside 40 325\nTMhelix 326 348\ninside 349 372

7482 GCF_002152995.1_ASM215299v1 Escherichia coli Proteobacteria; Gammaproteobacteria; Enterobacterales; Enterobacteriaceae; Escherichia MNNEETFYQAMRRQGVTRRSFLKYCSLAATSLGLGAGMAPKIAWALENKPRIPVVWIHGL WP_001058323.1 MULTISPECIES: hydrogenase-1 small chain [Proteobacteria] Length: 372\nNumber of predicted TMHs: 2\nExp number of AAs in TMHs: 37.2608499999999\nExp number, first 60 AAs: 19.52923\nTotal prob of N-in: 0.94351\nPOSSIBLE N-term signal sequence\ninside 1 19\nTMhelix 20 39\noutside 40 325\nTMhelix 326 348\ninside 349 372

7483 GCF_002153055.1_ASM215305v1 Escherichia coli Proteobacteria; Gammaproteobacteria; Enterobacterales; Enterobacteriaceae; Escherichia MTGDNTLIHSHGINRRDFMKLCAALAATMGLSSKAAAEMAESVTNPQRPPVIWIGAQECT WP_000145410.1 MULTISPECIES: hydrogenase-2 small chain [Proteobacteria] Length: 372\nNumber of predicted TMHs: 1\nExp number of AAs in TMHs: 23.72226\nExp number, first 60 AAs: 0.22038\nTotal prob of N-in: 0.12994\noutside 1 331\nTMhelix 332 354\ninside 355 372

7484 GCF_002153055.1_ASM215305v1 Escherichia coli Proteobacteria; Gammaproteobacteria; Enterobacterales; Enterobacteriaceae; Escherichia MNNEETFYQAMRRQGVTRRSFLKYCSLAATSLGLGAGMAPKIAWALENKPRIPVVWIHGL WP_001058320.1 hydrogenase [Escherichia coli] Length: 372\nNumber of predicted TMHs: 2\nExp number of AAs in TMHs: 37.26082\nExp number, first 60 AAs: 19.52923\nTotal prob of N-in: 0.94351\nPOSSIBLE N-term signal sequence\ninside 1 19\nTMhelix 20 39\noutside 40 325\nTMhelix 326 348\ninside 349 372

7485 GCF_002154095.1_ASM215409v1 Escherichia coli Proteobacteria; Gammaproteobacteria; Enterobacterales; Enterobacteriaceae; Escherichia MTGDNTLIHSHGINRRDFMKLCAALAATMGLSSKAAAEMAESVTNPQRPPVIWIGAQECT WP_000145410.1 MULTISPECIES: hydrogenase-2 small chain [Proteobacteria] Length: 372\nNumber of predicted TMHs: 1\nExp number of AAs in TMHs: 23.72226\nExp number, first 60 AAs: 0.22038\nTotal prob of N-in: 0.12994\noutside 1 331\nTMhelix 332 354\ninside 355 372

7486 GCF_002154135.1_ASM215413v1 Escherichia coli Proteobacteria; Gammaproteobacteria; Enterobacterales; Enterobacteriaceae; Escherichia MTGDNTLIHSHGINRRDFMKLCAALAATMGLSSKAAAEMAESVTNPQRPPVIWIGAQECT WP_000145410.1 MULTISPECIES: hydrogenase-2 small chain [Proteobacteria] Length: 372\nNumber of predicted TMHs: 1\nExp number of AAs in TMHs: 23.72226\nExp number, first 60 AAs: 0.22038\nTotal prob of N-in: 0.12994\noutside 1 331\nTMhelix 332 354\ninside 355 372

7487 GCF_002154135.1_ASM215413v1 Escherichia coli Proteobacteria; Gammaproteobacteria; Enterobacterales; Enterobacteriaceae; Escherichia MNNEETFYQAMRRQGVTRRSFLKYCSLAATSLGLGAGMAPKIAWALENKPRIPVVWIHGL WP_001058323.1 MULTISPECIES: hydrogenase-1 small chain [Proteobacteria] Length: 372\nNumber of predicted TMHs: 2\nExp number of AAs in TMHs: 37.2608499999999\nExp number, first 60 AAs: 19.52923\nTotal prob of N-in: 0.94351\nPOSSIBLE N-term signal sequence\ninside 1 19\nTMhelix 20 39\noutside 40 325\nTMhelix 326 348\ninside 349 372

7488 GCF_002154095.1_ASM215409v1 Escherichia coli Proteobacteria; Gammaproteobacteria; Enterobacterales; Enterobacteriaceae; Escherichia MNNEETFYQAMRRQGVTRRSFLKYCSLAATSLGLGAGMAPKIAWALENKPRIPVVWIHGL WP_001058323.1 MULTISPECIES: hydrogenase-1 small chain [Proteobacteria] Length: 372\nNumber of predicted TMHs: 2\nExp number of AAs in TMHs: 37.2608499999999\nExp number, first 60 AAs: 19.52923\nTotal prob of N-in: 0.94351\nPOSSIBLE N-term signal sequence\ninside 1 19\nTMhelix 20 39\noutside 40 325\nTMhelix 326 348\ninside 349 372

7489 GCF_002153015.1_ASM215301v1 Escherichia coli Proteobacteria; Gammaproteobacteria; Enterobacterales; Enterobacteriaceae; Escherichia MTGDNTLIHSHGINRRDFMKLCAALATTMGLSSKAAAEMAESVTNPQRPPVIWIGAQECT WP_023149629.1 MULTISPECIES: hydrogenase-2 small chain [Enterobacteriaceae] Length: 372\nNumber of predicted TMHs: 1\nExp number of AAs in TMHs: 23.63603\nExp number, first 60 AAs: 0.12971\nTotal prob of N-in: 0.12608\noutside 1 331\nTMhelix 332 354\ninside 355 372

7490 GCF_002153005.1_ASM215300v1 Escherichia coli Proteobacteria; Gammaproteobacteria; Enterobacterales; Enterobacteriaceae; Escherichia
MTGDNTLIHSHGINRRDFMKLCAALAATMGLSSKAAAEMVESVTNPQRPPVIWIGAQECT WP_086557857.1
hydrogenase 2 small subunit [Escherichia coli] Length: 372\nNumber of predicted TMHs: 1\nExp number of AAs in TMHs: 23.78729\nExp number, first 60 AAs: 0.28355\nTotal prob of N-in: 0.13271\noutside 1 331\nTMhelix 332 354\ninside 355 372

7491 GCF_002154075.1_ASM215407v1 Escherichia coli Proteobacteria; Gammaproteobacteria; Enterobacterales; Enterobacteriaceae; Escherichia
MNNEETFYQAMRRQGVTRRSFLKYCSLAATSLGLGAGMAPKIAWALENKPRIPVVWIHGL WP_001058323.1
MULTISPECIES: hydrogenase-1 small chain [Proteobacteria] Length: 372\nNumber of predicted TMHs: 2\nExp number of AAs in TMHs: 37.2608499999999\nExp number, first 60 AAs: 19.52923\nTotal prob of N-in: 0.94351\nPOSSIBLE N-term signal sequence\ninside 1 19\nTMhelix 20 39\noutside 40 325\nTMhelix 326 348\ninside 349 372

7492 GCF_002154805.1_ASM215480v1 Escherichia coli Proteobacteria; Gammaproteobacteria; Enterobacterales; Enterobacteriaceae; Escherichia
MTGDNTLIHSHGINRRDFMKLCAALAATMGLSSKAAAEMAESVTNPQRPPVIWIGAQECT WP_000145410.1
MULTISPECIES: hydrogenase-2 small chain [Proteobacteria] Length: 372\nNumber of predicted TMHs: 1\nExp number of AAs in TMHs: 23.72226\nExp number, first 60 AAs: 0.22038\nTotal prob of N-in: 0.12994\noutside 1 331\nTMhelix 332 354\ninside 355 372

7493 GCF_002154925.1_ASM215492v1 Escherichia coli Proteobacteria; Gammaproteobacteria; Enterobacterales; Enterobacteriaceae; Escherichia
MTGDNTLIHSHGINRRDFMKLCAALAATMGLSSKAAAEMAESVTNPQRPPVIWIGAQECT WP_000145410.1
MULTISPECIES: hydrogenase-2 small chain [Proteobacteria] Length: 372\nNumber of predicted TMHs: 1\nExp number of AAs in TMHs: 23.72226\nExp number, first 60 AAs: 0.22038\nTotal prob of N-in: 0.12994\noutside 1 331\nTMhelix 332 354\ninside 355 372

7494 GCF_002155365.1_ASM215536v1 Escherichia coli Proteobacteria; Gammaproteobacteria; Enterobacterales; Enterobacteriaceae; Escherichia
MTGDNTLIHSHGINRRDFMKLCAALAATMGLSSKAAAEMAESVTNPQRPPVIWIGAQECT WP_000145410.1
MULTISPECIES: hydrogenase-2 small chain [Proteobacteria] Length: 372\nNumber of predicted TMHs: 1\nExp number of AAs in TMHs: 23.72226\nExp number, first 60 AAs: 0.22038\nTotal prob of N-in: 0.12994\noutside 1 331\nTMhelix 332 354\ninside 355 372

7495 GCF_002154285.1_ASM215428v1 Escherichia coli Proteobacteria; Gammaproteobacteria; Enterobacterales; Enterobacteriaceae; Escherichia
MNNEETFYQAMRRQGVTRRSFLKYCSLAATSLGLGAGMAPKIAWALENKPRIPVVWIHGL WP_001058323.1
MULTISPECIES: hydrogenase-1 small chain [Proteobacteria] Length: 372\nNumber of predicted TMHs: 2\nExp number of AAs in TMHs: 37.2608499999999\nExp number, first 60 AAs: 19.52923\nTotal prob of N-in: 0.94351\nPOSSIBLE N-term signal sequence\ninside 1 19\nTMhelix 20 39\noutside 40 325\nTMhelix 326 348\ninside 349 372

7496 GCF_002154755.1_ASM215475v1 Escherichia coli Proteobacteria; Gammaproteobacteria; Enterobacterales; Enterobacteriaceae; Escherichia
MNNEETFYQAMRRQGVTRRSFLKYCSLAATSLGLGAGMAPKIAWALENKPRIPVVWIHGL WP_001058323.1
MULTISPECIES: hydrogenase-1 small chain [Proteobacteria] Length: 372\nNumber of predicted TMHs: 2\nExp number of AAs in TMHs: 37.2608499999999\nExp number, first 60 AAs: 19.52923\nTotal prob of N-in: 0.94351\nPOSSIBLE N-term signal sequence\ninside 1 19\nTMhelix 20 39\noutside 40 325\nTMhelix 326 348\ninside 349 372

7497 GCF_002154765.1_ASM215476v1 Escherichia coli Proteobacteria; Gammaproteobacteria; Enterobacterales; Enterobacteriaceae; Escherichia
MNNEETFYQAMRRQGVTRRSFLKYCSLAATSLGLGAGMAPKIAWALENKPRIPVVWIHGL WP_001058323.1
MULTISPECIES: hydrogenase-1 small chain [Proteobacteria] Length: 372\nNumber of predicted TMHs: 2\nExp number of AAs in TMHs: 37.2608499999999\nExp number, first 60 AAs: 19.52923\nTotal prob of N-in: 0.94351\nPOSSIBLE N-term signal sequence\ninside 1 19\nTMhelix 20 39\noutside 40 325\nTMhelix 326 348\ninside 349 372

7498 GCF_002154805.1_ASM215480v1 Escherichia coli Proteobacteria; Gammaproteobacteria; Enterobacterales; Enterobacteriaceae; Escherichia
MNNEETFYQAMRRQGVTRRSFLKYCSLAATSLGLGAGMAPKIAWALENKPRIPVVWIHGL WP_001058323.1
MULTISPECIES: hydrogenase-1 small chain [Proteobacteria] Length: 372\nNumber of predicted TMHs: 2\nExp number of AAs in TMHs: 37.2608499999999\nExp number, first 60 AAs: 19.52923\nTotal prob of N-in: 0.94351\nPOSSIBLE N-term signal sequence\ninside 1 19\nTMhelix 20 39\noutside 40 325\nTMhelix 326 348\ninside 349 372

7499 GCF_002155325.1_ASM215532v1 Escherichia coli Proteobacteria; Gammaproteobacteria; Enterobacterales; Enterobacteriaceae; Escherichia
MTGDNTLIHSHGINRRDFMKLCAALAATMGLSSKAAAEMAESVTNPQRPPVIWIGAQECT WP_000145410.1
MULTISPECIES: hydrogenase-2 small chain [Proteobacteria] Length: 372\nNumber of predicted TMHs: 1\nExp number of AAs in TMHs: 23.72226\nExp number, first 60 AAs: 0.22038\nTotal prob of N-in: 0.12994\noutside 1 331\nTMhelix 332 354\ninside 355 372

7500 GCF_002155195.1_ASM215519v1 Escherichia coli Proteobacteria; Gammaproteobacteria; Enterobacterales; Enterobacteriaceae; Escherichia
MNNEETFYQAMRRQGVTRRSFLKYCSLAATSLGLGAGMAPKIAWALENKPRIPVVWIHGL WP_001058323.1
MULTISPECIES: hydrogenase-1 small chain [Proteobacteria] Length: 372\nNumber of predicted TMHs: 2\nExp number of AAs in TMHs: 37.2608499999999\nExp number, first 60 AAs: 19.52923\nTotal prob of N-in: 0.94351\nPOSSIBLE N-term signal sequence\ninside 1 19\nTMhelix 20 39\noutside 40 325\nTMhelix 326 348\ninside 349 372

7501 GCF_002155125.1_ASM215512v1 Escherichia coli Proteobacteria; Gammaproteobacteria; Enterobacterales; Enterobacteriaceae; Escherichia
MNNEETFYQAMRRQGVTRRSFLKYCSLAATSLGLGAGMAPKIAWALENKPRIPVVWIHGL WP_001058323.1
MULTISPECIES: hydrogenase-1 small chain [Proteobacteria] Length: 372\nNumber of predicted TMHs: 2\nExp number of AAs in TMHs: 37.2608499999999\nExp number, first 60 AAs: 19.52923\nTotal prob of N-in: 0.94351\nPOSSIBLE N-term signal sequence\ninside 1 19\nTMhelix 20 39\noutside 40 325\nTMhelix 326 348\ninside 349 372

7502 GCF_002155385.1_ASM215538v1 Escherichia coli Proteobacteria; Gammaproteobacteria; Enterobacterales; Enterobacteriaceae; Escherichia
MTGDNTLIHSHGINRRDFMKLCAALAATMGLSSKAAAEMAESVTNPQRPPVIWIGAQECT WP_000145410.1
MULTISPECIES: hydrogenase-2 small chain [Proteobacteria] Length: 372\nNumber of predicted TMHs: 1\nExp number of AAs in TMHs: 23.72226\nExp number, first 60 AAs: 0.22038\nTotal prob of N-in: 0.12994\noutside 1 331\nTMhelix 332 354\ninside 355 372

7503 GCF_002155445.1_ASM215544v1 Escherichia coli Proteobacteria; Gammaproteobacteria; Enterobacterales; Enterobacteriaceae; Escherichia
MTGDNTLIHSHGINRRDFMKLCAALAATMGLSSKAAAEMAESVTNPQRPPVIWIGAQECT WP_000145410.1
MULTISPECIES: hydrogenase-2 small chain [Proteobacteria] Length: 372\nNumber of predicted TMHs: 1\nExp number of AAs in TMHs: 23.72226\nExp number, first 60 AAs: 0.22038\nTotal prob of N-in: 0.12994\noutside 1 331\nTMhelix 332 354\ninside 355 372

7504 GCF_002155385.1_ASM215538v1 Escherichia coli Proteobacteria; Gammaproteobacteria; Enterobacterales; Enterobacteriaceae; Escherichia
MNNEETFYQAMRRQGVTRRSFLKYCSLAATSLGLGAGMAPKIAWALENKPRIPVVWIHGL WP_001058323.1
MULTISPECIES: hydrogenase-1 small chain [Proteobacteria] Length: 372\nNumber of predicted TMHs: 2\nExp number of AAs in TMHs: 37.2608499999999\nExp number, first 60 AAs: 19.52923\nTotal prob of N-in: 0.94351\nPOSSIBLE N-term signal sequence\ninside 1 19\nTMhelix 20 39\noutside 40 325\nTMhelix 326 348\ninside 349 372

7505 GCF_002155465.1_ASM215546v1 Escherichia coli Proteobacteria; Gammaproteobacteria; Enterobacterales; Enterobacteriaceae; Escherichia
MTGDNTLIHSHGINRRDFMKLCAALAATMGLSSKAAAEMAESVTNPQRPPVIWIGAQECT WP_000145410.1
MULTISPECIES: hydrogenase-2 small chain [Proteobacteria] Length: 372\nNumber of predicted TMHs: 1\nExp number of AAs in TMHs: 23.72226\nExp number, first 60 AAs: 0.22038\nTotal prob of N-in: 0.12994\noutside 1 331\nTMhelix 332 354\ninside 355 372

7506 GCF_002155465.1_ASM215546v1 Escherichia coli Proteobacteria; Gammaproteobacteria; Enterobacterales; Enterobacteriaceae; Escherichia
MNNEETFYQAMRRQGVTRRSFLKYCSLAATSLGLGAGMAPKIAWALENKPRIPVVWIHGL WP_001295940.1
MULTISPECIES: hydrogenase [Enterobacteriaceae] Length: 372\nNumber of predicted TMHs: 2\nExp number of AAs in TMHs: 37.2608899999999\nExp number, first 60 AAs: 19.52923\nTotal prob of N-in: 0.94351\nPOSSIBLE N-term signal sequence\ninside 1 19\nTMhelix 20 39\noutside 40 325\nTMhelix 326 348\ninside 349 372

7507 GCF_002155195.1_ASM215519v1 Escherichia coli Proteobacteria; Gammaproteobacteria; Enterobacterales; Enterobacteriaceae; Escherichia
MTGDNTLIHSHGINRRDFMKLCAALAATMGLSSKAAAEMAESVTNPQRPPVIWIGAQECT WP_086623878.1
hydrogenase 2 small subunit [Escherichia coli] Length: 372\nNumber of predicted TMHs: 1\nExp number of AAs in TMHs: 23.72025\nExp number, first 60 AAs: 0.22032\nTotal prob of N-in: 0.12990\noutside 1 331\nTMhelix 332 354\ninside 355 372

7508 GCF_002155365.1_ASM215536v1 Escherichia coli Proteobacteria; Gammaproteobacteria; Enterobacterales; Enterobacteriaceae; Escherichia
MNNEETFYQAMRRQGVTRRSFLKYCSLAATSLGLGAGMAPKIAWALENKPRIPVVWIHGL WP_001058323.1
MULTISPECIES: hydrogenase-1 small chain [Proteobacteria] Length: 372\nNumber of predicted TMHs: 2\nExp number of AAs in TMHs: 37.2608499999999\nExp number, first 60 AAs: 19.52923\nTotal prob of N-in: 0.94351\nPOSSIBLE N-term signal sequence\ninside 1 19\nTMhelix 20 39\noutside 40 325\nTMhelix 326 348\ninside 349 372

7509 GCF_002155505.1_ASM215550v1 Escherichia coli Proteobacteria; Gammaproteobacteria; Enterobacterales; Enterobacteriaceae; Escherichia
MTGDNTLIHSHGINRRDFMKLCAALAATMGLSSKAAAEMAESVTNPQRPPVIWIGAQECT WP_000145410.1
MULTISPECIES: hydrogenase-2 small chain [Proteobacteria] Length: 372\nNumber of predicted TMHs: 1\nExp number of AAs in TMHs: 23.72226\nExp number, first 60 AAs: 0.22038\nTotal prob of N-in: 0.12994\noutside 1 331\nTMhelix 332 354\ninside 355 372

7510 GCF_002155505.1_ASM215550v1 Escherichia coli Proteobacteria; Gammaproteobacteria; Enterobacterales; Enterobacteriaceae; Escherichia
MNNEETFYQAMRRQGVTRRSFLKYCSLAATSLGLGAGMAPKIAWALENKPRIPVVWIHGL WP_001058323.1
MULTISPECIES: hydrogenase-1 small chain [Proteobacteria] Length: 372\nNumber of predicted TMHs: 2\nExp number of AAs in TMHs: 37.2608499999999\nExp number, first 60 AAs: 19.52923\nTotal prob of N-in: 0.94351\nPOSSIBLE N-term signal sequence\ninside 1 19\nTMhelix 20 39\noutside 40 325\nTMhelix 326 348\ninside 349 372

7511 GCF_002155205.1_ASM215520v1 Escherichia coli Proteobacteria; Gammaproteobacteria; Enterobacterales; Enterobacteriaceae; Escherichia
MNNEETFYQAMRRQGVTRRSFLKYCSLAATSLGLGAGMAPKIAWALENKPRIPVVWIHGL WP_086619139.1
[Ni/Fe] hydrogenase small subunit [Escherichia coli] Length: 372\nNumber of predicted TMHs: 2\nExp number of AAs in TMHs: 37.26077\nExp number, first 60 AAs: 19.52923\nTotal prob of N-in: 0.94351\nPOSSIBLE N-term signal sequence\ninside 1 19\nTMhelix 20 39\noutside 40 325\nTMhelix 326 348\ninside 349 372

7512 GCF_002155645.1_ASM215564v1 Escherichia coli Proteobacteria; Gammaproteobacteria; Enterobacterales; Enterobacteriaceae; Escherichia
MTGDNTLIHSHGINRRDFMKLCAALAATMGLSSKAAAEMAESVTNPQRPPVIWIGAQECT WP_000145410.1
MULTISPECIES: hydrogenase-2 small chain [Proteobacteria] Length: 372\nNumber of predicted TMHs: 1\nExp number of AAs in TMHs: 23.72226\nExp number, first 60 AAs: 0.22038\nTotal prob of N-in: 0.12994\noutside 1 331\nTMhelix 332 354\ninside 355 372

7513 GCF_002155755.1_ASM215575v1 Escherichia coli Proteobacteria; Gammaproteobacteria; Enterobacterales; Enterobacteriaceae; Escherichia
MTGDNTLIHSHGINRRDFMKLCAALAATMGLSSKAAAEMAESVTNPQRPPVIWIGAQECT WP_000145410.1
MULTISPECIES: hydrogenase-2 small chain [Proteobacteria] Length: 372\nNumber of predicted TMHs: 1\nExp number of AAs in TMHs: 23.72226\nExp number, first 60 AAs: 0.22038\nTotal prob of N-in: 0.12994\noutside 1 331\nTMhelix 332 354\ninside 355 372

7514 GCF_002155755.1_ASM215575v1 Escherichia coli Proteobacteria; Gammaproteobacteria; Enterobacterales; Enterobacteriaceae; Escherichia

MNNEETFYQAMRRQGVTRRSFLKYCSLAATSLGLGAGMAPKIAWALENKPRIPVVWIHGL WP_001058323.1 MULTISPECIES: hydrogenase-1 small chain [Proteobacteria] Length: 372\nNumber of predicted TMHs: 2\nExp number of AAs in TMHs: 37.2608499999999\nExp number, first 60 AAs: 19.52923\nTotal prob of N-in: 0.94351\nPOSSIBLE N-term signal sequence\ninside 1 19\nTMhelix 20 39\noutside 40 325\nTMhelix 326 348\ninside 349 372

7515 GCF_002155595.1_ASM215559v1 Escherichia coli Proteobacteria; Gammaproteobacteria; Enterobacterales; Enterobacteriaceae; Escherichia MNNEETFYQAMRRQGVTRRSFLKYCSLAATSLGLGAGMAPKIAWALENKPRIPVVWIHGL WP_001058320.1 hydrogenase [Escherichia coli] Length: 372\nNumber of predicted TMHs: 2\nExp number of AAs in TMHs: 37.26082\nExp number, first 60 AAs: 19.52923\nTotal prob of N-in: 0.94351\nPOSSIBLE N-term signal sequence\ninside 1 19\nTMhelix 20 39\noutside 40 325\nTMhelix 326 348\ninside 349 372

7516 GCF_002155645.1_ASM215564v1 Escherichia coli Proteobacteria; Gammaproteobacteria; Enterobacterales; Enterobacteriaceae; Escherichia MNNEETFYQAMRRQGVTRRSFLKYCSLAATSLGLGAGMAPKIAWALENKPRIPVVWIHGL WP_001295940.1 MULTISPECIES: hydrogenase [Enterobacteriaceae] Length: 372\nNumber of predicted TMHs: 2\nExp number of AAs in TMHs: 37.2608899999999\nExp number, first 60 AAs: 19.52923\nTotal prob of N-in: 0.94351\nPOSSIBLE N-term signal sequence\ninside 1 19\nTMhelix 20 39\noutside 40 325\nTMhelix 326 348\ninside 349 372

7517 GCF_002155535.1_ASM215553v1 Escherichia coli Proteobacteria; Gammaproteobacteria; Enterobacterales; Enterobacteriaceae; Escherichia MNNEETFYQAMRRQGVTRRSFLKYCSLAATSLGLGAGMAPKIAWALENKPRIPVVWIHGL WP_001058320.1 hydrogenase [Escherichia coli] Length: 372\nNumber of predicted TMHs: 2\nExp number of AAs in TMHs: 37.26082\nExp number, first 60 AAs: 19.52923\nTotal prob of N-in: 0.94351\nPOSSIBLE N-term signal sequence\ninside 1 19\nTMhelix 20 39\noutside 40 325\nTMhelix 326 348\ninside 349 372

7518 GCF_002155775.1_ASM215577v1 Escherichia coli Proteobacteria; Gammaproteobacteria; Enterobacterales; Enterobacteriaceae; Escherichia MTGDNTLIHSHGINRRDFMKLCAALAATMGLSSKAAAEMAESVTNPQRPPVIWIGAQECT WP_000145410.1 MULTISPECIES: hydrogenase-2 small chain [Proteobacteria] Length: 372\nNumber of predicted TMHs: 1\nExp number of AAs in TMHs: 23.72226\nExp number, first 60 AAs: 0.22038\nTotal prob of N-in: 0.12994\noutside 1 331\nTMhelix 332 354\ninside 355 372

7519 GCF_002157815.1_ASM215781v1 Escherichia coli Proteobacteria; Gammaproteobacteria; Enterobacterales; Enterobacteriaceae; Escherichia MTGDNTLIHSHGINRRDFMKLCAALAATMGLSSKAAAEMAESVTNPQRPPVIWIGAQECT WP_000145410.1 MULTISPECIES: hydrogenase-2 small chain [Proteobacteria] Length: 372\nNumber of predicted TMHs: 1\nExp number of AAs in TMHs: 23.72226\nExp number, first 60 AAs: 0.22038\nTotal prob of N-in: 0.12994\noutside 1 331\nTMhelix 332 354\ninside 355 372

7520 GCF_002163195.1_ASM216319v1 Escherichia coli Proteobacteria; Gammaproteobacteria; Enterobacterales; Enterobacteriaceae; Escherichia MTGDNTLIHSHGINRRDFMKLCAALAATMGLSSKAAAEMAESVTNPQRPPVIWIGAQECT WP_000145410.1 MULTISPECIES: hydrogenase-2 small chain [Proteobacteria] Length: 372\nNumber of predicted TMHs: 1\nExp number of AAs in TMHs: 23.72226\nExp number, first 60 AAs: 0.22038\nTotal prob of N-in: 0.12994\noutside 1 331\nTMhelix 332 354\ninside 355 372

7521 GCF_002163655.1_ASM216365v1 Escherichia coli Proteobacteria; Gammaproteobacteria; Enterobacterales; Enterobacteriaceae; Escherichia MTGDNTLIHSHGINRRDFMKLCAALAATMGLSSKAAAEMAESVTNPQRPPVIWIGAQECT WP_000145410.1 MULTISPECIES: hydrogenase-2 small chain [Proteobacteria] Length: 372\nNumber of predicted TMHs: 1\nExp number of AAs in TMHs: 23.72226\nExp number, first 60 AAs: 0.22038\nTotal prob of N-in: 0.12994\noutside 1 331\nTMhelix 332 354\ninside 355 372

7522 GCF_002157815.1_ASM215781v1 Escherichia coli Proteobacteria; Gammaproteobacteria; Enterobacterales; Enterobacteriaceae; Escherichia MNNEETFYQAMRRQGVTRRSFLKYCSLAATSLGLGAGMAPKIAWALENKPRIPVVWIHGL WP_001058323.1 MULTISPECIES: hydrogenase-1 small chain [Proteobacteria] Length: 372\nNumber of predicted TMHs: 2\nExp number of AAs in TMHs: 37.2608499999999\nExp number, first 60 AAs: 19.52923\nTotal prob of N-in:

0.94351\nPOSSIBLE N-term signal sequence\ninside 1 19\nTMhelix 20 39\noutside 40 325\nTMhelix 326 348\ninside 349 372

7523 GCF_002163935.1_ASM216393v1 Escherichia coli Proteobacteria; Gammaproteobacteria; Enterobacterales; Enterobacteriaceae; Escherichia MTGDNTLIHSHGINRRDFMKLCAALAATMGLSSKAAAEMAESVTNPQRPPVIWIGAQECT WP_000145410.1 MULTISPECIES: hydrogenase-2 small chain [Proteobacteria] Length: 372\nNumber of predicted TMHs: 1\nExp number of AAs in TMHs: 23.72226\nExp number, first 60 AAs: 0.22038\nTotal prob of N-in: 0.12994\noutside 1 331\nTMhelix 332 354\ninside 355 372

7524 GCF_002155825.1_ASM215582v1 Escherichia coli Proteobacteria; Gammaproteobacteria; Enterobacterales; Enterobacteriaceae; Escherichia MTGDNTLIHSHGINRRDFMKLCAALATTMGLSSKAAAEMAESVTNPQRPPVIWIGAQECT WP_023149629.1 MULTISPECIES: hydrogenase-2 small chain [Enterobacteriaceae] Length: 372\nNumber of predicted TMHs: 1\nExp number of AAs in TMHs: 23.63603\nExp number, first 60 AAs: 0.12971\nTotal prob of N-in: 0.12608\noutside 1 331\nTMhelix 332 354\ninside 355 372

7525 GCF_002164025.1_ASM216402v1 Escherichia coli Proteobacteria; Gammaproteobacteria; Enterobacterales; Enterobacteriaceae; Escherichia MTGDNTLIHSHGINRRDFMKLCAALAATMGLSSKAAAEMAESVTNPQRPPVIWIGAQECT WP_000145410.1 MULTISPECIES: hydrogenase-2 small chain [Proteobacteria] Length: 372\nNumber of predicted TMHs: 1\nExp number of AAs in TMHs: 23.72226\nExp number, first 60 AAs: 0.22038\nTotal prob of N-in: 0.12994\noutside 1 331\nTMhelix 332 354\ninside 355 372

7526 GCF_002164435.1_ASM216443v1 Escherichia coli Proteobacteria; Gammaproteobacteria; Enterobacterales; Enterobacteriaceae; Escherichia MNNEETFYQAMRRQGVTRRSFLKYCSLAATSLGLGAGMAPKIAWALENKPRIPVVWIHGL WP_001301613.1 hydrogenase [Escherichia coli] Length: 372\nNumber of predicted TMHs: 2\nExp number of AAs in TMHs: 37.46598\nExp number, first 60 AAs: 19.53139\nTotal prob of N-in: 0.94364\nPOSSIBLE N-term signal sequence\ninside 1 19\nTMhelix 20 39\noutside 40 325\nTMhelix 326 348\ninside 349 372

7527 GCF_002163935.1_ASM216393v1 Escherichia coli Proteobacteria; Gammaproteobacteria; Enterobacterales; Enterobacteriaceae; Escherichia MNNEETFYQAMRRQGVTRRSFLKYCSLAATSLGLGAGMAPKIAWALENKPRIPVVWIHGL WP_001295940.1 MULTISPECIES: hydrogenase [Enterobacteriaceae] Length: 372\nNumber of predicted TMHs: 2\nExp number of AAs in TMHs: 37.2608899999999\nExp number, first 60 AAs: 19.52923\nTotal prob of N-in: 0.94351\nPOSSIBLE N-term signal sequence\ninside 1 19\nTMhelix 20 39\noutside 40 325\nTMhelix 326 348\ninside 349 372

7528 GCF_002164445.1_ASM216444v1 Escherichia coli Proteobacteria; Gammaproteobacteria; Enterobacterales; Enterobacteriaceae; Escherichia MTGDNTLIHSHGINRRDFMKLCAALAATMGLSSKAAAEMAESVTNPQRPPVIWIGAQECT WP_000145410.1 MULTISPECIES: hydrogenase-2 small chain [Proteobacteria] Length: 372\nNumber of predicted TMHs: 1\nExp number of AAs in TMHs: 23.72226\nExp number, first 60 AAs: 0.22038\nTotal prob of N-in: 0.12994\noutside 1 331\nTMhelix 332 354\ninside 355 372

7529 GCF_002164445.1_ASM216444v1 Escherichia coli Proteobacteria; Gammaproteobacteria; Enterobacterales; Enterobacteriaceae; Escherichia MNNEETFYQAMRRQGVTRRSFLKYCSLAATSLGLGAGMAPKIAWALENKPRIPVVWIHGL WP_001301613.1 hydrogenase [Escherichia coli] Length: 372\nNumber of predicted TMHs: 2\nExp number of AAs in TMHs: 37.46598\nExp number, first 60 AAs: 19.53139\nTotal prob of N-in: 0.94364\nPOSSIBLE N-term signal sequence\ninside 1 19\nTMhelix 20 39\noutside 40 325\nTMhelix 326 348\ninside 349 372

7530 GCF_002164025.1_ASM216402v1 Escherichia coli Proteobacteria; Gammaproteobacteria; Enterobacterales; Enterobacteriaceae; Escherichia MNNEETFYQAMRRQGVTRRSFLKYCSLAATSLGLGAGMAPKIAWALENKPRIPVVWIHGL WP_001301613.1 hydrogenase [Escherichia coli] Length: 372\nNumber of predicted TMHs: 2\nExp number of AAs in TMHs: 37.46598\nExp number, first 60 AAs: 19.53139\nTotal prob of N-in: 0.94364\nPOSSIBLE N-term signal sequence\ninside 1 19\nTMhelix 20 39\noutside 40 325\nTMhelix 326 348\ninside 349 372

7531 GCF_002164115.1_ASM216411v1 Escherichia coli Proteobacteria; Gammaproteobacteria; Enterobacterales; Enterobacteriaceae; Escherichia
MNNEETFYQAMRRQGVTRRSFLKYCSLAATSLGLGAGMAPKIAWALENKPRIPVVWIHGL WP_001301613.1
hydrogenase [Escherichia coli] Length: 372\nNumber of predicted TMHs: 2\nExp number of AAs in TMHs: 37.46598\nExp number, first 60 AAs: 19.53139\nTotal prob of N-in: 0.94364\nPOSSIBLE N-term signal sequence\ninside 1 19\nTMhelix 20 39\noutside 40 325\nTMhelix 326 348\ninside 349 372

7532 GCF_002164045.1_ASM216404v1 Escherichia coli Proteobacteria; Gammaproteobacteria; Enterobacterales; Enterobacteriaceae; Escherichia
MNNEETFYQAMRRQGVTRRSFLKYCSLAATSLGLGAGMAPKIAWALENKPRIPVVWIHGL WP_001301613.1
hydrogenase [Escherichia coli] Length: 372\nNumber of predicted TMHs: 2\nExp number of AAs in TMHs: 37.46598\nExp number, first 60 AAs: 19.53139\nTotal prob of N-in: 0.94364\nPOSSIBLE N-term signal sequence\ninside 1 19\nTMhelix 20 39\noutside 40 325\nTMhelix 326 348\ninside 349 372

7533 GCF_002164175.1_ASM216417v1 Escherichia coli Proteobacteria; Gammaproteobacteria; Enterobacterales; Enterobacteriaceae; Escherichia
MNNEETFYQAMRRQGVTRRSFLKYCSLAATSLGLGAGMAPKIAWALENKPRIPVVWIHGL WP_001301613.1
hydrogenase [Escherichia coli] Length: 372\nNumber of predicted TMHs: 2\nExp number of AAs in TMHs: 37.46598\nExp number, first 60 AAs: 19.53139\nTotal prob of N-in: 0.94364\nPOSSIBLE N-term signal sequence\ninside 1 19\nTMhelix 20 39\noutside 40 325\nTMhelix 326 348\ninside 349 372

7534 GCF_002164465.1_ASM216446v1 Escherichia coli Proteobacteria; Gammaproteobacteria; Enterobacterales; Enterobacteriaceae; Escherichia
MTGDNTLIHSHGINRRDFMKLCAALAATMGLSSKAAAEMAESVTNPQRPPVIWIGAQECT WP_000145410.1
MULTISPECIES: hydrogenase-2 small chain [Proteobacteria] Length: 372\nNumber of predicted TMHs: 1\nExp number of AAs in TMHs: 23.72226\nExp number, first 60 AAs: 0.22038\nTotal prob of N-in: 0.12994\noutside 1 331\nTMhelix 332 354\ninside 355 372

7535 GCF_002164525.1_ASM216452v1 Escherichia coli Proteobacteria; Gammaproteobacteria; Enterobacterales; Enterobacteriaceae; Escherichia
MTGDNTLIHSHGINRRDFMKLCAALAATMGLSSKAAAEMAESVTNPQRPPVIWIGAQECT WP_000145410.1
MULTISPECIES: hydrogenase-2 small chain [Proteobacteria] Length: 372\nNumber of predicted TMHs: 1\nExp number of AAs in TMHs: 23.72226\nExp number, first 60 AAs: 0.22038\nTotal prob of N-in: 0.12994\noutside 1 331\nTMhelix 332 354\ninside 355 372

7536 GCF_002164595.1_ASM216459v1 Escherichia coli Proteobacteria; Gammaproteobacteria; Enterobacterales; Enterobacteriaceae; Escherichia
MTGDNTLIHSHGINRRDFMKLCAALAATMGLSSKAAAEMAESVTNPQRPPVIWIGAQECT WP_000145410.1
MULTISPECIES: hydrogenase-2 small chain [Proteobacteria] Length: 372\nNumber of predicted TMHs: 1\nExp number of AAs in TMHs: 23.72226\nExp number, first 60 AAs: 0.22038\nTotal prob of N-in: 0.12994\noutside 1 331\nTMhelix 332 354\ninside 355 372

7537 GCF_002164595.1_ASM216459v1 Escherichia coli Proteobacteria; Gammaproteobacteria; Enterobacterales; Enterobacteriaceae; Escherichia
MNNEETFYQAMRRQGVTRRSFLKYCSLAATSLGLGAGMAPKIAWALENKPRIPVVWIHGL WP_001058323.1
MULTISPECIES: hydrogenase-1 small chain [Proteobacteria] Length: 372\nNumber of predicted TMHs: 2\nExp number of AAs in TMHs: 37.2608499999999\nExp number, first 60 AAs: 19.52923\nTotal prob of N-in: 0.94351\nPOSSIBLE N-term signal sequence\ninside 1 19\nTMhelix 20 39\noutside 40 325\nTMhelix 326 348\ninside 349 372

7538 GCF_002164465.1_ASM216446v1 Escherichia coli Proteobacteria; Gammaproteobacteria; Enterobacterales; Enterobacteriaceae; Escherichia
MNNEETFYQAMRRQGVTRRSFLKYCSLAATSLGLGAGMAPKIAWALENKPRIPVVWIHGL WP_001301613.1
hydrogenase [Escherichia coli] Length: 372\nNumber of predicted TMHs: 2\nExp number of AAs in TMHs: 37.46598\nExp number, first 60 AAs: 19.53139\nTotal prob of N-in: 0.94364\nPOSSIBLE N-term signal sequence\ninside 1 19\nTMhelix 20 39\noutside 40 325\nTMhelix 326 348\ninside 349 372

7539 GCF_002164525.1_ASM216452v1 Escherichia coli Proteobacteria; Gammaproteobacteria; Enterobacterales; Enterobacteriaceae; Escherichia
MNNEETFYQAMRRQGVTRRSFLKYCSLAATSLGLGAGMAPKIAWALENKPRIPVVWIHGL WP_001301613.1
hydrogenase [Escherichia coli] Length: 372\nNumber of predicted TMHs: 2\nExp number of AAs in

TMHs: 37.46598\nExp number, first 60 AAs: 19.53139\nTotal prob of N-in: 0.94364\nPOSSIBLE N-term signal sequence\ninside 1 19\nTMhelix 20 39\noutside 40 325\nTMhelix 326 348\ninside 349 372

7540 GCF_002165035.1_ASM216503v1 Escherichia coli Proteobacteria; Gammaproteobacteria; Enterobacterales; Enterobacteriaceae; Escherichia MTGDNTLIHSHGINRRDFMKLCAALAATMGLSSKAAAEMAESVTNPQRPPVIWIGAQECT WP_000145410.1 MULTISPECIES: hydrogenase-2 small chain [Proteobacteria] Length: 372\nNumber of predicted TMHs: 1\nExp number of AAs in TMHs: 23.72226\nExp number, first 60 AAs: 0.22038\nTotal prob of N-in: 0.12994\noutside 1 331\nTMhelix 332 354\ninside 355 372

7541 GCF_002164935.1_ASM216493v1 Escherichia coli Proteobacteria; Gammaproteobacteria; Enterobacterales; Enterobacteriaceae; Escherichia MNNEETFYQAMRRQGVTRRSFLKYCSLAATSLGLGAGMAPKIAWALENKPRIPVVWIHGL WP_001058323.1 MULTISPECIES: hydrogenase-1 small chain [Proteobacteria] Length: 372\nNumber of predicted TMHs: 2\nExp number of AAs in TMHs: 37.2608499999999\nExp number, first 60 AAs: 19.52923\nTotal prob of N-in: 0.94351\nPOSSIBLE N-term signal sequence\ninside 1 19\nTMhelix 20 39\noutside 40 325\nTMhelix 326 348\ninside 349 372

7542 GCF_002165925.1_ASM216592v1 Escherichia coli Proteobacteria; Gammaproteobacteria; Enterobacterales; Enterobacteriaceae; Escherichia MTGDNTLIHSHGINRRDFMKLCAALAATMGLSSKAAAEMAESVTNPQRPPVIWIGAQECT WP_000145410.1 MULTISPECIES: hydrogenase-2 small chain [Proteobacteria] Length: 372\nNumber of predicted TMHs: 1\nExp number of AAs in TMHs: 23.72226\nExp number, first 60 AAs: 0.22038\nTotal prob of N-in: 0.12994\noutside 1 331\nTMhelix 332 354\ninside 355 372

7543 GCF_002165055.1_ASM216505v1 Escherichia coli Proteobacteria; Gammaproteobacteria; Enterobacterales; Enterobacteriaceae; Escherichia MTGDNTLIHSHGINRRDFMKLCAALAATMGLSSKAAAEMAESVTNPQRPPVIWIGAQECT WP_000145410.1 MULTISPECIES: hydrogenase-2 small chain [Proteobacteria] Length: 372\nNumber of predicted TMHs: 1\nExp number of AAs in TMHs: 23.72226\nExp number, first 60 AAs: 0.22038\nTotal prob of N-in: 0.12994\noutside 1 331\nTMhelix 332 354\ninside 355 372

7544 GCF_002165415.1_ASM216541v1 Escherichia coli Proteobacteria; Gammaproteobacteria; Enterobacterales; Enterobacteriaceae; Escherichia MTGDNTLIHSHGINRRDFMKLCAALAATMGLSSKAAAEMAESVTNPQRPPVIWIGAQECT WP_000145410.1 MULTISPECIES: hydrogenase-2 small chain [Proteobacteria] Length: 372\nNumber of predicted TMHs: 1\nExp number of AAs in TMHs: 23.72226\nExp number, first 60 AAs: 0.22038\nTotal prob of N-in: 0.12994\noutside 1 331\nTMhelix 332 354\ninside 355 372

7545 GCF_002165995.1_ASM216599v1 Escherichia coli Proteobacteria; Gammaproteobacteria; Enterobacterales; Enterobacteriaceae; Escherichia MTGDNTLIHSHGINRRDFMKLCAALAATMGLSSKAAAEMAESVTNPQRPPVIWIGAQECT WP_000145410.1 MULTISPECIES: hydrogenase-2 small chain [Proteobacteria] Length: 372\nNumber of predicted TMHs: 1\nExp number of AAs in TMHs: 23.72226\nExp number, first 60 AAs: 0.22038\nTotal prob of N-in: 0.12994\noutside 1 331\nTMhelix 332 354\ninside 355 372

7546 GCF_002165925.1_ASM216592v1 Escherichia coli Proteobacteria; Gammaproteobacteria; Enterobacterales; Enterobacteriaceae; Escherichia MNNEETFYQAMRRQGVTRRSFLKYCSLAATSLGLGAGMAPKIAWALENKPRIPVVWIHGL WP_001058323.1 MULTISPECIES: hydrogenase-1 small chain [Proteobacteria] Length: 372\nNumber of predicted TMHs: 2\nExp number of AAs in TMHs: 37.2608499999999\nExp number, first 60 AAs: 19.52923\nTotal prob of N-in: 0.94351\nPOSSIBLE N-term signal sequence\ninside 1 19\nTMhelix 20 39\noutside 40 325\nTMhelix 326 348\ninside 349 372

7547 GCF_002166275.1_ASM216627v1 Escherichia coli Proteobacteria; Gammaproteobacteria; Enterobacterales; Enterobacteriaceae; Escherichia MTGDNTLIHSHGINRRDFMKLCAALAATMGLSSKAAAEMVESVTNPQRPPVIWIGAQECT WP_000145423.1 hydrogenase 2 small subunit [Escherichia coli] Length: 372\nNumber of predicted TMHs: 1\nExp number of AAs in TMHs: 23.78201\nExp number, first 60 AAs: 0.28355\nTotal prob of N-in: 0.13257\noutside 1 331\nTMhelix 332 354\ninside 355 372

7548 GCF_002166275.1_ASM216627v1 Escherichia coli Proteobacteria; Gammaproteobacteria; Enterobacterales; Enterobacteriaceae; Escherichia
MNNEETFYQAMRRQGVTRRSFLKYCSLAATSLGLGAGMAPKIAWALENKPRIPVVWIHGL WP_001058323.1
MULTISPECIES: hydrogenase-1 small chain [Proteobacteria] Length: 372\nNumber of predicted TMHs: 2\nExp number of AAs in TMHs: 37.2608499999999\nExp number, first 60 AAs: 19.52923\nTotal prob of N-in: 0.94351\nPOSSIBLE N-term signal sequence\ninside 1 19\nTMhelix 20 39\noutside 40 325\nTMhelix 326 348\ninside 349 372

7549 GCF_002166095.1_ASM216609v1 Escherichia coli Proteobacteria; Gammaproteobacteria; Enterobacterales; Enterobacteriaceae; Escherichia
MNNEETFYQAMRRQGVTRRSFLKYCSLAATSLGLGAGMAPKIAWALENKPRIPVVWIHGL WP_001058323.1
MULTISPECIES: hydrogenase-1 small chain [Proteobacteria] Length: 372\nNumber of predicted TMHs: 2\nExp number of AAs in TMHs: 37.2608499999999\nExp number, first 60 AAs: 19.52923\nTotal prob of N-in: 0.94351\nPOSSIBLE N-term signal sequence\ninside 1 19\nTMhelix 20 39\noutside 40 325\nTMhelix 326 348\ninside 349 372

7550 GCF_002166335.1_ASM216633v1 Escherichia coli Proteobacteria; Gammaproteobacteria; Enterobacterales; Enterobacteriaceae; Escherichia
MTGDNTLIHSHGINRRDFMKLCAALAATMGLSSKAAAEMAESVTNPQRPPVIWIGAQECT WP_000145410.1
MULTISPECIES: hydrogenase-2 small chain [Proteobacteria] Length: 372\nNumber of predicted TMHs: 1\nExp number of AAs in TMHs: 23.72226\nExp number, first 60 AAs: 0.22038\nTotal prob of N-in: 0.12994\noutside 1 331\nTMhelix 332 354\ninside 355 372

7551 GCF_002166125.1_ASM216612v1 Escherichia coli Proteobacteria; Gammaproteobacteria; Enterobacterales; Enterobacteriaceae; Escherichia
MNNEETFYQAMRRQGVTRRSFLKYCSLAATSLGLGAGMAPKIAWALENKPRIPVVWIHGL WP_001058323.1
MULTISPECIES: hydrogenase-1 small chain [Proteobacteria] Length: 372\nNumber of predicted TMHs: 2\nExp number of AAs in TMHs: 37.2608499999999\nExp number, first 60 AAs: 19.52923\nTotal prob of N-in: 0.94351\nPOSSIBLE N-term signal sequence\ninside 1 19\nTMhelix 20 39\noutside 40 325\nTMhelix 326 348\ninside 349 372

7552 GCF_002166405.1_ASM216640v1 Escherichia coli Proteobacteria; Gammaproteobacteria; Enterobacterales; Enterobacteriaceae; Escherichia
MTGDNTLIHSHGINRRDFMKLCAALAATMGLSSKAAAEMAESVTNPQRPPVIWIGAQECT WP_000145410.1
MULTISPECIES: hydrogenase-2 small chain [Proteobacteria] Length: 372\nNumber of predicted TMHs: 1\nExp number of AAs in TMHs: 23.72226\nExp number, first 60 AAs: 0.22038\nTotal prob of N-in: 0.12994\noutside 1 331\nTMhelix 332 354\ninside 355 372

7553 GCF_002166185.1_ASM216618v1 Escherichia coli Proteobacteria; Gammaproteobacteria; Enterobacterales; Enterobacteriaceae; Escherichia
MNNEETFYQAMRRQGVTRRSFLKYCSLAATSLGLGAGMAPKIAWALENKPRIPVVWIHGL WP_001058323.1
MULTISPECIES: hydrogenase-1 small chain [Proteobacteria] Length: 372\nNumber of predicted TMHs: 2\nExp number of AAs in TMHs: 37.2608499999999\nExp number, first 60 AAs: 19.52923\nTotal prob of N-in: 0.94351\nPOSSIBLE N-term signal sequence\ninside 1 19\nTMhelix 20 39\noutside 40 325\nTMhelix 326 348\ninside 349 372

7554 GCF_002166615.1_ASM216661v1 Escherichia coli Proteobacteria; Gammaproteobacteria; Enterobacterales; Enterobacteriaceae; Escherichia
MTGDNTLIHSHGINRRDFMKLCAALAATMGLSSKAAAEMAESVTNPQRPPVIWIGAQECT WP_000145410.1
MULTISPECIES: hydrogenase-2 small chain [Proteobacteria] Length: 372\nNumber of predicted TMHs: 1\nExp number of AAs in TMHs: 23.72226\nExp number, first 60 AAs: 0.22038\nTotal prob of N-in: 0.12994\noutside 1 331\nTMhelix 332 354\ninside 355 372

7555 GCF_002166705.1_ASM216670v1 Escherichia coli Proteobacteria; Gammaproteobacteria; Enterobacterales; Enterobacteriaceae; Escherichia
MTGDNTLIHSHGINRRDFMKLCAALAATMGLSSKAAAEMAESVTNPQRPPVIWIGAQECT WP_000145410.1
MULTISPECIES: hydrogenase-2 small chain [Proteobacteria] Length: 372\nNumber of predicted TMHs: 1\nExp number of AAs in TMHs: 23.72226\nExp number, first 60 AAs: 0.22038\nTotal prob of N-in: 0.12994\noutside 1 331\nTMhelix 332 354\ninside 355 372

7556 GCF_002166645.1_ASM216664v1 Escherichia coli Proteobacteria; Gammaproteobacteria; Enterobacterales; Enterobacteriaceae; Escherichia
MTGDNTLIHSHGINRRDFMKLCAALAATMGLSSKAAAEMAESVTNPQRPPVIWIGAQECT WP_000145410.1
MULTISPECIES: hydrogenase-2 small chain [Proteobacteria] Length: 372\nNumber of predicted TMHs: 1\nExp number of AAs in TMHs: 23.72226\nExp number, first 60 AAs: 0.22038\nTotal prob of N-in: 0.12994\noutside 1 331\nTMhelix 332 354\ninside 355 372

7557 GCF_002166755.1_ASM216675v1 Escherichia coli Proteobacteria; Gammaproteobacteria; Enterobacterales; Enterobacteriaceae; Escherichia
MTGDNTLIHSHGINRRDFMKLCAALAATMGLSSKAAAEMAESVTNPQRPPVIWIGAQECT WP_000145410.1
MULTISPECIES: hydrogenase-2 small chain [Proteobacteria] Length: 372\nNumber of predicted TMHs: 1\nExp number of AAs in TMHs: 23.72226\nExp number, first 60 AAs: 0.22038\nTotal prob of N-in: 0.12994\noutside 1 331\nTMhelix 332 354\ninside 355 372

7558 GCF_002173155.1_ASM217315v1 Escherichia coli Proteobacteria; Gammaproteobacteria; Enterobacterales; Enterobacteriaceae; Escherichia
MTGDNTLIHSHGINRRDFMKLCAALAATMGLSSKAAAEMAESVTNPQRPPVIWIGAQECT WP_000145410.1
MULTISPECIES: hydrogenase-2 small chain [Proteobacteria] Length: 372\nNumber of predicted TMHs: 1\nExp number of AAs in TMHs: 23.72226\nExp number, first 60 AAs: 0.22038\nTotal prob of N-in: 0.12994\noutside 1 331\nTMhelix 332 354\ninside 355 372

7559 GCF_002173205.1_ASM217320v1 Escherichia coli Proteobacteria; Gammaproteobacteria; Enterobacterales; Enterobacteriaceae; Escherichia
MTGDNTLIHSHGINRRDFMKLCAALAATMGLSSKAAAEMAESVTNPQRPPVIWIGAQECT WP_000145410.1
MULTISPECIES: hydrogenase-2 small chain [Proteobacteria] Length: 372\nNumber of predicted TMHs: 1\nExp number of AAs in TMHs: 23.72226\nExp number, first 60 AAs: 0.22038\nTotal prob of N-in: 0.12994\noutside 1 331\nTMhelix 332 354\ninside 355 372

7560 GCF_002173265.1_ASM217326v1 Escherichia coli Proteobacteria; Gammaproteobacteria; Enterobacterales; Enterobacteriaceae; Escherichia
MTGDNTLIHSHGINRRDFMKLCAALAATMGLSSKAAAEMAESVTNPQRPPVIWIGAQECT WP_000145410.1
MULTISPECIES: hydrogenase-2 small chain [Proteobacteria] Length: 372\nNumber of predicted TMHs: 1\nExp number of AAs in TMHs: 23.72226\nExp number, first 60 AAs: 0.22038\nTotal prob of N-in: 0.12994\noutside 1 331\nTMhelix 332 354\ninside 355 372

7561 GCF_002173285.1_ASM217328v1 Escherichia coli Proteobacteria; Gammaproteobacteria; Enterobacterales; Enterobacteriaceae; Escherichia
MTGDNTLIHSHGINRRDFMKLCAALAATMGLSSKAAAEMAESVTNPQRPPVIWIGAQECT WP_000145410.1
MULTISPECIES: hydrogenase-2 small chain [Proteobacteria] Length: 372\nNumber of predicted TMHs: 1\nExp number of AAs in TMHs: 23.72226\nExp number, first 60 AAs: 0.22038\nTotal prob of N-in: 0.12994\noutside 1 331\nTMhelix 332 354\ninside 355 372

7562 GCF_002175295.1_ASM217529v1 Escherichia coli Proteobacteria; Gammaproteobacteria; Enterobacterales; Enterobacteriaceae; Escherichia
MNNEETFYQAMRRQGVTRRSFLKYCSLAATSLGLGAGMAPKIAWALENKPRIPVVWIHGL WP_001301613.1
hydrogenase [Escherichia coli] Length: 372\nNumber of predicted TMHs: 2\nExp number of AAs in TMHs: 37.46598\nExp number, first 60 AAs: 19.53139\nTotal prob of N-in: 0.94364\nPOSSIBLE N-term signal sequence\ninside 1 19\nTMhelix 20 39\noutside 40 325\nTMhelix 326 348\ninside 349 372

7563 GCF_002173155.1_ASM217315v1 Escherichia coli Proteobacteria; Gammaproteobacteria; Enterobacterales; Enterobacteriaceae; Escherichia
MNNEETFYQAMRRQGVTRRSFLKYCSLAATSLGLGAGMAPKIAWALENKPRIPVVWIHGL WP_001058323.1
MULTISPECIES: hydrogenase-1 small chain [Proteobacteria] Length: 372\nNumber of predicted TMHs: 2\nExp number of AAs in TMHs: 37.2608499999999\nExp number, first 60 AAs: 19.52923\nTotal prob of N-in: 0.94351\nPOSSIBLE N-term signal sequence\ninside 1 19\nTMhelix 20 39\noutside 40 325\nTMhelix 326 348\ninside 349 372

7564 GCF_002175305.1_ASM217530v1 Escherichia coli Proteobacteria; Gammaproteobacteria; Enterobacterales; Enterobacteriaceae; Escherichia
MTGDNTLIHSHGINRRDFMKLCAALAATMGLSSKAAAEMAESVTNPQRPPVIWIGAQECT WP_000145410.1
MULTISPECIES: hydrogenase-2 small chain [Proteobacteria] Length: 372\nNumber of predicted TMHs:

1\nExp number of AAs in TMHs: 23.72226\nExp number, first 60 AAs: 0.22038\nTotal prob of N-in: 0.12994\noutside 1 331\nTMhelix 332 354\ninside 355 372

7565 GCF_002175305.1_ASM217530v1 Escherichia coli Proteobacteria; Gammaproteobacteria; Enterobacterales; Enterobacteriaceae; Escherichia
MNNEETFYQAMRRQGVTRRSFLKYCSLAATSLGLGAGMAPKIAWALENKPRIPVVWIHGL WP_001301613.1
hydrogenase [Escherichia coli] Length: 372\nNumber of predicted TMHs: 2\nExp number of AAs in TMHs: 37.46598\nExp number, first 60 AAs: 19.53139\nTotal prob of N-in: 0.94364\nPOSSIBLE N-term signal sequence\ninside 1 19\nTMhelix 20 39\noutside 40 325\nTMhelix 326 348\ninside 349 372

7566 GCF_002173205.1_ASM217320v1 Escherichia coli Proteobacteria; Gammaproteobacteria; Enterobacterales; Enterobacteriaceae; Escherichia
MNNEETFYQAMRRQGVTRRSFLKYCSLAATSLGLGAGMAPKIAWALENKPRIPVVWIHGL WP_001058323.1
MULTISPECIES: hydrogenase-1 small chain [Proteobacteria] Length: 372\nNumber of predicted TMHs: 2\nExp number of AAs in TMHs: 37.2608499999999\nExp number, first 60 AAs: 19.52923\nTotal prob of N-in: 0.94351\nPOSSIBLE N-term signal sequence\ninside 1 19\nTMhelix 20 39\noutside 40 325\nTMhelix 326 348\ninside 349 372

7567 GCF_002175325.1_ASM217532v1 Escherichia coli Proteobacteria; Gammaproteobacteria; Enterobacterales; Enterobacteriaceae; Escherichia
MTGDNTLIHSHGINRRDFMKLCAALAATMGLSSKAAAEMAESVTNPQRPPVIWIGAQECT WP_000145410.1
MULTISPECIES: hydrogenase-2 small chain [Proteobacteria] Length: 372\nNumber of predicted TMHs: 1\nExp number of AAs in TMHs: 23.72226\nExp number, first 60 AAs: 0.22038\nTotal prob of N-in: 0.12994\noutside 1 331\nTMhelix 332 354\ninside 355 372

7568 GCF_002175325.1_ASM217532v1 Escherichia coli Proteobacteria; Gammaproteobacteria; Enterobacterales; Enterobacteriaceae; Escherichia
MNNEETFYQAMRRQGVTRRSFLKYCSLAATSLGLGAGMAPKIAWALENKPRIPVVWIHGL WP_001301613.1
hydrogenase [Escherichia coli] Length: 372\nNumber of predicted TMHs: 2\nExp number of AAs in TMHs: 37.46598\nExp number, first 60 AAs: 19.53139\nTotal prob of N-in: 0.94364\nPOSSIBLE N-term signal sequence\ninside 1 19\nTMhelix 20 39\noutside 40 325\nTMhelix 326 348\ninside 349 372

7569 GCF_002175695.1_ASM217569v1 Escherichia coli Proteobacteria; Gammaproteobacteria; Enterobacterales; Enterobacteriaceae; Escherichia
MTGDNTLIHSHGINRRDFMKLCAALAATMGLSSKAAAEMAESVTNPQRPPVIWIGAQECT WP_000145410.1
MULTISPECIES: hydrogenase-2 small chain [Proteobacteria] Length: 372\nNumber of predicted TMHs: 1\nExp number of AAs in TMHs: 23.72226\nExp number, first 60 AAs: 0.22038\nTotal prob of N-in: 0.12994\noutside 1 331\nTMhelix 332 354\ninside 355 372

7570 GCF_002175695.1_ASM217569v1 Escherichia coli Proteobacteria; Gammaproteobacteria; Enterobacterales; Enterobacteriaceae; Escherichia
MNNEETFYQAMRRQGVTRRSFLKYCSLAATSLGLGAGMAPKIAWALENKPRIPVVWIHGL WP_001301613.1
hydrogenase [Escherichia coli] Length: 372\nNumber of predicted TMHs: 2\nExp number of AAs in TMHs: 37.46598\nExp number, first 60 AAs: 19.53139\nTotal prob of N-in: 0.94364\nPOSSIBLE N-term signal sequence\ninside 1 19\nTMhelix 20 39\noutside 40 325\nTMhelix 326 348\ninside 349 372

7571 GCF_002175385.1_ASM217538v1 Escherichia coli Proteobacteria; Gammaproteobacteria; Enterobacterales; Enterobacteriaceae; Escherichia
MNNEETFYQAMRRQGVTRRSFLKYCSLAATSLGLGAGMAPKIAWALENKPRIPVVWIHGL WP_001301613.1
hydrogenase [Escherichia coli] Length: 372\nNumber of predicted TMHs: 2\nExp number of AAs in TMHs: 37.46598\nExp number, first 60 AAs: 19.53139\nTotal prob of N-in: 0.94364\nPOSSIBLE N-term signal sequence\ninside 1 19\nTMhelix 20 39\noutside 40 325\nTMhelix 326 348\ninside 349 372

7572 GCF_002175785.1_ASM217578v1 Escherichia coli Proteobacteria; Gammaproteobacteria; Enterobacterales; Enterobacteriaceae; Escherichia
MTGDNTLIHSHGINRRDFMKLCAALAATMGLSSKAAAEMAESVTNPQRPPVIWIGAQECT WP_000145410.1
MULTISPECIES: hydrogenase-2 small chain [Proteobacteria] Length: 372\nNumber of predicted TMHs: 1\nExp number of AAs in TMHs: 23.72226\nExp number, first 60 AAs: 0.22038\nTotal prob of N-in: 0.12994\noutside 1 331\nTMhelix 332 354\ninside 355 372

7573 GCF_002175855.1_ASM217585v1 Escherichia coli Proteobacteria; Gammaproteobacteria; Enterobacterales; Enterobacteriaceae; Escherichia
MTGDNTLIHSHGINRRDFMKLCAALAATMGLSSKAAAEMAESVTNPQRPPVIWIGAQECT WP_000145410.1
MULTISPECIES: hydrogenase-2 small chain [Proteobacteria] Length: 372\nNumber of predicted TMHs: 1\nExp number of AAs in TMHs: 23.72226\nExp number, first 60 AAs: 0.22038\nTotal prob of N-in: 0.12994\noutside 1 331\nTMhelix 332 354\ninside 355 372

7574 GCF_002175875.1_ASM217587v1 Escherichia coli Proteobacteria; Gammaproteobacteria; Enterobacterales; Enterobacteriaceae; Escherichia
MTGDNTLIHSHGINRRDFMKLCAALAATMGLSSKAAAEMAESVTNPQRPPVIWIGAQECT WP_000145410.1
MULTISPECIES: hydrogenase-2 small chain [Proteobacteria] Length: 372\nNumber of predicted TMHs: 1\nExp number of AAs in TMHs: 23.72226\nExp number, first 60 AAs: 0.22038\nTotal prob of N-in: 0.12994\noutside 1 331\nTMhelix 332 354\ninside 355 372

7575 GCF_002175945.1_ASM217594v1 Escherichia coli Proteobacteria; Gammaproteobacteria; Enterobacterales; Enterobacteriaceae; Escherichia
MTGDNTLIHSHGINRRDFMKLCAALAATMGLSSKAAAEMAESVTNPQRPPVIWIGAQECT WP_000145410.1
MULTISPECIES: hydrogenase-2 small chain [Proteobacteria] Length: 372\nNumber of predicted TMHs: 1\nExp number of AAs in TMHs: 23.72226\nExp number, first 60 AAs: 0.22038\nTotal prob of N-in: 0.12994\noutside 1 331\nTMhelix 332 354\ninside 355 372

7576 GCF_002175965.1_ASM217596v1 Escherichia coli Proteobacteria; Gammaproteobacteria; Enterobacterales; Enterobacteriaceae; Escherichia
MNNEETFYQAMRRQGVTRRSFLKYCSLAATSLGLGAGMAPKIAWALENKPRIPVVWIHGL WP_001301613.1
hydrogenase [Escherichia coli] Length: 372\nNumber of predicted TMHs: 2\nExp number of AAs in TMHs: 37.46598\nExp number, first 60 AAs: 19.53139\nTotal prob of N-in: 0.94364\nPOSSIBLE N-term signal sequence\ninside 1 19\nTMhelix 20 39\noutside 40 325\nTMhelix 326 348\ninside 349 372

7577 GCF_002175575.1_ASM217557v1 Escherichia coli Proteobacteria; Gammaproteobacteria; Enterobacterales; Enterobacteriaceae; Escherichia
MNNEETFYQAMRRQGVTRRSFLKYCSLAATSLGLGAGMAPKIAWALENKPRIPVVWIHGL WP_001301613.1
hydrogenase [Escherichia coli] Length: 372\nNumber of predicted TMHs: 2\nExp number of AAs in TMHs: 37.46598\nExp number, first 60 AAs: 19.53139\nTotal prob of N-in: 0.94364\nPOSSIBLE N-term signal sequence\ninside 1 19\nTMhelix 20 39\noutside 40 325\nTMhelix 326 348\ninside 349 372

7578 GCF_002175645.1_ASM217564v1 Escherichia coli Proteobacteria; Gammaproteobacteria; Enterobacterales; Enterobacteriaceae; Escherichia
MNNEETFYQAMRRQGVTRRSFLKYCSLAATSLGLGAGMAPKIAWALENKPRIPVVWIHGL WP_001301613.1
hydrogenase [Escherichia coli] Length: 372\nNumber of predicted TMHs: 2\nExp number of AAs in TMHs: 37.46598\nExp number, first 60 AAs: 19.53139\nTotal prob of N-in: 0.94364\nPOSSIBLE N-term signal sequence\ninside 1 19\nTMhelix 20 39\noutside 40 325\nTMhelix 326 348\ninside 349 372

7579 GCF_002175725.1_ASM217572v1 Escherichia coli Proteobacteria; Gammaproteobacteria; Enterobacterales; Enterobacteriaceae; Escherichia
MNNEETFYQAMRRQGVTRRSFLKYCSLAATSLGLGAGMAPKIAWALENKPRIPVVWIHGL WP_001301613.1
hydrogenase [Escherichia coli] Length: 372\nNumber of predicted TMHs: 2\nExp number of AAs in TMHs: 37.46598\nExp number, first 60 AAs: 19.53139\nTotal prob of N-in: 0.94364\nPOSSIBLE N-term signal sequence\ninside 1 19\nTMhelix 20 39\noutside 40 325\nTMhelix 326 348\ninside 349 372

7580 GCF_002176035.1_ASM217603v1 Escherichia coli Proteobacteria; Gammaproteobacteria; Enterobacterales; Enterobacteriaceae; Escherichia
MTGDNTLIHSHGINRRDFMKLCAALAATMGLSSKAAAEMAESVTNPQRPPVIWIGAQECT WP_000145410.1
MULTISPECIES: hydrogenase-2 small chain [Proteobacteria] Length: 372\nNumber of predicted TMHs: 1\nExp number of AAs in TMHs: 23.72226\nExp number, first 60 AAs: 0.22038\nTotal prob of N-in: 0.12994\noutside 1 331\nTMhelix 332 354\ninside 355 372

7581 GCF_002176095.1_ASM217609v1 Escherichia coli Proteobacteria; Gammaproteobacteria; Enterobacterales; Enterobacteriaceae; Escherichia
MTGDNTLIHSHGINRRDFMKLCAALAATMGLSSKAAAEMAESVTNPQRPPVIWIGAQECT WP_000145410.1
MULTISPECIES: hydrogenase-2 small chain [Proteobacteria] Length: 372\nNumber of predicted TMHs:

1\nExp number of AAs in TMHs: 23.72226\nExp number, first 60 AAs: 0.22038\nTotal prob of N-in: 0.12994\noutside 1 331\nTMhelix 332 354\ninside 355 372

7582 GCF_002176105.1_ASM217610v1 Escherichia coli Proteobacteria; Gammaproteobacteria; Enterobacterales; Enterobacteriaceae; Escherichia
MTGDNTLIHSHGINRRDFMKLCAALAATMGLSSKAAAEMAESVTNPQRPPVIWIGAQECT WP_000145410.1
MULTISPECIES: hydrogenase-2 small chain [Proteobacteria] Length: 372\nNumber of predicted TMHs: 1\nExp number of AAs in TMHs: 23.72226\nExp number, first 60 AAs: 0.22038\nTotal prob of N-in: 0.12994\noutside 1 331\nTMhelix 332 354\ninside 355 372

7583 GCF_002176255.1_ASM217625v1 Escherichia coli Proteobacteria; Gammaproteobacteria; Enterobacterales; Enterobacteriaceae; Escherichia
MTGDNTLIHSHGINRRDFMKLCAALAATMGLSSKAAAEMAESVTNPQRPPVIWIGAQECT WP_000145410.1
MULTISPECIES: hydrogenase-2 small chain [Proteobacteria] Length: 372\nNumber of predicted TMHs: 1\nExp number of AAs in TMHs: 23.72226\nExp number, first 60 AAs: 0.22038\nTotal prob of N-in: 0.12994\noutside 1 331\nTMhelix 332 354\ninside 355 372

7584 GCF_002176275.1_ASM217627v1 Escherichia coli Proteobacteria; Gammaproteobacteria; Enterobacterales; Enterobacteriaceae; Escherichia
MTGDNTLIHSHGINRRDFMKLCAALAATMGLSSKAAAEMAESVTNPQRPPVIWIGAQECT WP_000145410.1
MULTISPECIES: hydrogenase-2 small chain [Proteobacteria] Length: 372\nNumber of predicted TMHs: 1\nExp number of AAs in TMHs: 23.72226\nExp number, first 60 AAs: 0.22038\nTotal prob of N-in: 0.12994\noutside 1 331\nTMhelix 332 354\ninside 355 372

7585 GCF_002176345.1_ASM217634v1 Escherichia coli Proteobacteria; Gammaproteobacteria; Enterobacterales; Enterobacteriaceae; Escherichia
MTGDNTLIHSHGINRRDFMKLCAALAATMGLSSKAAAEMAESVTNPQRPPVIWIGAQECT WP_000145410.1
MULTISPECIES: hydrogenase-2 small chain [Proteobacteria] Length: 372\nNumber of predicted TMHs: 1\nExp number of AAs in TMHs: 23.72226\nExp number, first 60 AAs: 0.22038\nTotal prob of N-in: 0.12994\noutside 1 331\nTMhelix 332 354\ninside 355 372

7586 GCF_002176385.1_ASM217638v1 Escherichia coli Proteobacteria; Gammaproteobacteria; Enterobacterales; Enterobacteriaceae; Escherichia
MTGDNTLIHSHGINRRDFMKLCAALAATMGLSSKAAAEMAESVTNPQRPPVIWIGAQECT WP_000145410.1
MULTISPECIES: hydrogenase-2 small chain [Proteobacteria] Length: 372\nNumber of predicted TMHs: 1\nExp number of AAs in TMHs: 23.72226\nExp number, first 60 AAs: 0.22038\nTotal prob of N-in: 0.12994\noutside 1 331\nTMhelix 332 354\ninside 355 372

7587 GCF_002176435.1_ASM217643v1 Escherichia coli Proteobacteria; Gammaproteobacteria; Enterobacterales; Enterobacteriaceae; Escherichia
MTGDNTLIHSHGINRRDFMKLCAALAATMGLSSKAAAEMAESVTNPQRPPVIWIGAQECT WP_000145410.1
MULTISPECIES: hydrogenase-2 small chain [Proteobacteria] Length: 372\nNumber of predicted TMHs: 1\nExp number of AAs in TMHs: 23.72226\nExp number, first 60 AAs: 0.22038\nTotal prob of N-in: 0.12994\noutside 1 331\nTMhelix 332 354\ninside 355 372

7588 GCF_002176495.1_ASM217649v1 Escherichia coli Proteobacteria; Gammaproteobacteria; Enterobacterales; Enterobacteriaceae; Escherichia
MTGDNTLIHSHGINRRDFMKLCAALAATMGLSSKAAAEMAESVTNPQRPPVIWIGAQECT WP_000145410.1
MULTISPECIES: hydrogenase-2 small chain [Proteobacteria] Length: 372\nNumber of predicted TMHs: 1\nExp number of AAs in TMHs: 23.72226\nExp number, first 60 AAs: 0.22038\nTotal prob of N-in: 0.12994\noutside 1 331\nTMhelix 332 354\ninside 355 372

7589 GCF_002176635.1_ASM217663v1 Escherichia coli Proteobacteria; Gammaproteobacteria; Enterobacterales; Enterobacteriaceae; Escherichia
MTGDNTLIHSHGINRRDFMKLCAALAATMGLSSKAAAEMAESVTNPQRPPVIWIGAQECT WP_000145410.1
MULTISPECIES: hydrogenase-2 small chain [Proteobacteria] Length: 372\nNumber of predicted TMHs: 1\nExp number of AAs in TMHs: 23.72226\nExp number, first 60 AAs: 0.22038\nTotal prob of N-in: 0.12994\noutside 1 331\nTMhelix 332 354\ninside 355 372

7590 GCF_002176495.1_ASM217649v1 Escherichia coli Proteobacteria; Gammaproteobacteria; Enterobacterales; Enterobacteriaceae; Escherichia

MNNEETFYQAMRRQGVTRRSFLKYCSLAATSLGLGAGMAPKIAWALENKPRIPVVWIHGL WP_001301613.1 hydrogenase [Escherichia coli] Length: 372\nNumber of predicted TMHs: 2\nExp number of AAs in TMHs: 37.46598\nExp number, first 60 AAs: 19.53139\nTotal prob of N-in: 0.94364\nPOSSIBLE N-term signal sequence\ninside 1 19\nTMhelix 20 39\noutside 40 325\nTMhelix 326 348\ninside 349 372

7591 GCF_002176665.1_ASM217666v1 Escherichia coli Proteobacteria; Gammaproteobacteria; Enterobacterales; Enterobacteriaceae; Escherichia MNNEETFYQAMRRQGVTRRSFLKYCSLAATSLGLGAGMAPKIAWALENKPRIPVVWIHGL WP_001301613.1 hydrogenase [Escherichia coli] Length: 372\nNumber of predicted TMHs: 2\nExp number of AAs in TMHs: 37.46598\nExp number, first 60 AAs: 19.53139\nTotal prob of N-in: 0.94364\nPOSSIBLE N-term signal sequence\ninside 1 19\nTMhelix 20 39\noutside 40 325\nTMhelix 326 348\ninside 349 372

7592 GCF_002176795.1_ASM217679v1 Escherichia coli Proteobacteria; Gammaproteobacteria; Enterobacterales; Enterobacteriaceae; Escherichia MNNEETFYQAMRRQGVTRRSFLKYCSLAATSLGLGAGMAPKIAWALENKPRIPVVWIHGL WP_001301613.1 hydrogenase [Escherichia coli] Length: 372\nNumber of predicted TMHs: 2\nExp number of AAs in TMHs: 37.46598\nExp number, first 60 AAs: 19.53139\nTotal prob of N-in: 0.94364\nPOSSIBLE N-term signal sequence\ninside 1 19\nTMhelix 20 39\noutside 40 325\nTMhelix 326 348\ninside 349 372

7593 GCF_002176345.1_ASM217634v1 Escherichia coli Proteobacteria; Gammaproteobacteria; Enterobacterales; Enterobacteriaceae; Escherichia MNNEETFYQAMRRQGVTRRSFLKYCSLAATSLGLGAGMAPKIAWALENKPRIPVVWIHGL WP_001301613.1 hydrogenase [Escherichia coli] Length: 372\nNumber of predicted TMHs: 2\nExp number of AAs in TMHs: 37.46598\nExp number, first 60 AAs: 19.53139\nTotal prob of N-in: 0.94364\nPOSSIBLE N-term signal sequence\ninside 1 19\nTMhelix 20 39\noutside 40 325\nTMhelix 326 348\ninside 349 372

7594 GCF_002176385.1_ASM217638v1 Escherichia coli Proteobacteria; Gammaproteobacteria; Enterobacterales; Enterobacteriaceae; Escherichia MNNEETFYQAMRRQGVTRRSFLKYCSLAATSLGLGAGMAPKIAWALENKPRIPVVWIHGL WP_001301613.1 hydrogenase [Escherichia coli] Length: 372\nNumber of predicted TMHs: 2\nExp number of AAs in TMHs: 37.46598\nExp number, first 60 AAs: 19.53139\nTotal prob of N-in: 0.94364\nPOSSIBLE N-term signal sequence\ninside 1 19\nTMhelix 20 39\noutside 40 325\nTMhelix 326 348\ninside 349 372

7595 GCF_002176435.1_ASM217643v1 Escherichia coli Proteobacteria; Gammaproteobacteria; Enterobacterales; Enterobacteriaceae; Escherichia MNNEETFYQAMRRQGVTRRSFLKYCSLAATSLGLGAGMAPKIAWALENKPRIPVVWIHGL WP_001301613.1 hydrogenase [Escherichia coli] Length: 372\nNumber of predicted TMHs: 2\nExp number of AAs in TMHs: 37.46598\nExp number, first 60 AAs: 19.53139\nTotal prob of N-in: 0.94364\nPOSSIBLE N-term signal sequence\ninside 1 19\nTMhelix 20 39\noutside 40 325\nTMhelix 326 348\ninside 349 372

7596 GCF_002179775.1_ASM217977v1 Escherichia coli Proteobacteria; Gammaproteobacteria; Enterobacterales; Enterobacteriaceae; Escherichia MTGDNTLIHSHGINRRDFMKLCAALAATMGLSSKAAAEMAESVTNPQRPPVIWIGAQECT WP_000145410.1 MULTISPECIES: hydrogenase-2 small chain [Proteobacteria] Length: 372\nNumber of predicted TMHs: 1\nExp number of AAs in TMHs: 23.72226\nExp number, first 60 AAs: 0.22038\nTotal prob of N-in: 0.12994\noutside 1 331\nTMhelix 332 354\ninside 355 372

7597 GCF_002189715.1_ASM218971v1 Escherichia coli Proteobacteria; Gammaproteobacteria; Enterobacterales; Enterobacteriaceae; Escherichia MTGDNTLIHSHGINRRDFMKLCAALAATMGLSSKAAAEMAESVTNPQRPPVIWIGAQECT WP_000145410.1 MULTISPECIES: hydrogenase-2 small chain [Proteobacteria] Length: 372\nNumber of predicted TMHs: 1\nExp number of AAs in TMHs: 23.72226\nExp number, first 60 AAs: 0.22038\nTotal prob of N-in: 0.12994\noutside 1 331\nTMhelix 332 354\ninside 355 372

7598 GCF_002189715.1_ASM218971v1 Escherichia coli Proteobacteria; Gammaproteobacteria; Enterobacterales; Enterobacteriaceae; Escherichia MNNEETFYQAMRRQGVTRRSFLKYCSLAATSLGLGAGMAPKIAWALENKPRIPVVWIHGL WP_001058323.1 MULTISPECIES: hydrogenase-1 small chain [Proteobacteria] Length: 372\nNumber of predicted TMHs: 2\nExp number of AAs in TMHs: 37.2608499999999\nExp number, first 60 AAs: 19.52923\nTotal prob of N-in: 0.94351\nPOSSIBLE N-term signal sequence\ninside 1 19\nTMhelix 20 39\noutside 40 325\nTMhelix 326 348\ninside 349 372

7599 GCF_002189955.1_ASM218995v1 Escherichia coli Proteobacteria; Gammaproteobacteria; Enterobacterales; Enterobacteriaceae; Escherichia
MTGDNTLIHSHGINRRDFMKLCAALAATMGLSSKAAAEMAESVTNPQRPPVIWIGAQECT WP_000145410.1
MULTISPECIES: hydrogenase-2 small chain [Proteobacteria] Length: 372\nNumber of predicted TMHs: 1\nExp number of AAs in TMHs: 23.72226\nExp number, first 60 AAs: 0.22038\nTotal prob of N-in: 0.12994\noutside 1 331\nTMhelix 332 354\ninside 355 372

7600 GCF_002189955.1_ASM218995v1 Escherichia coli Proteobacteria; Gammaproteobacteria; Enterobacterales; Enterobacteriaceae; Escherichia
MNNEETFYQAMRRQGVTRRSFLKYCSLAATSLGLGAGMAPKIAWALENKPRIPVVWIHGL WP_001058323.1
MULTISPECIES: hydrogenase-1 small chain [Proteobacteria] Length: 372\nNumber of predicted TMHs: 2\nExp number of AAs in TMHs: 37.2608499999999\nExp number, first 60 AAs: 19.52923\nTotal prob of N-in: 0.94351\nPOSSIBLE N-term signal sequence\ninside 1 19\nTMhelix 20 39\noutside 40 325\nTMhelix 326 348\ninside 349 372

7601 GCF_002182025.1_ASM218202v1 Escherichia coli Proteobacteria; Gammaproteobacteria; Enterobacterales; Enterobacteriaceae; Escherichia
MNNEETFYQAMRRQGVTRRSFLKYCSLAATSLGLGAGMAPKIAWALENKPRIPVVWIHGL WP_001058323.1
MULTISPECIES: hydrogenase-1 small chain [Proteobacteria] Length: 372\nNumber of predicted TMHs: 2\nExp number of AAs in TMHs: 37.2608499999999\nExp number, first 60 AAs: 19.52923\nTotal prob of N-in: 0.94351\nPOSSIBLE N-term signal sequence\ninside 1 19\nTMhelix 20 39\noutside 40 325\nTMhelix 326 348\ninside 349 372

7602 GCF_002189755.1_ASM218975v1 Escherichia coli Proteobacteria; Gammaproteobacteria; Enterobacterales; Enterobacteriaceae; Escherichia
MNNEETFYQAMRRQGVTRRSFLKYCSLAATSLGLGAGMAPKIAWALENKPRIPVVWIHGL WP_001058323.1
MULTISPECIES: hydrogenase-1 small chain [Proteobacteria] Length: 372\nNumber of predicted TMHs: 2\nExp number of AAs in TMHs: 37.2608499999999\nExp number, first 60 AAs: 19.52923\nTotal prob of N-in: 0.94351\nPOSSIBLE N-term signal sequence\ninside 1 19\nTMhelix 20 39\noutside 40 325\nTMhelix 326 348\ninside 349 372

7603 GCF_002189815.1_ASM218981v1 Escherichia coli Proteobacteria; Gammaproteobacteria; Enterobacterales; Enterobacteriaceae; Escherichia
MNNEETFYQAMRRQGVTRRSFLKYCSLAATSLGLGAGMAPKIAWALENKPRIPVVWIHGL WP_001058323.1
MULTISPECIES: hydrogenase-1 small chain [Proteobacteria] Length: 372\nNumber of predicted TMHs: 2\nExp number of AAs in TMHs: 37.2608499999999\nExp number, first 60 AAs: 19.52923\nTotal prob of N-in: 0.94351\nPOSSIBLE N-term signal sequence\ninside 1 19\nTMhelix 20 39\noutside 40 325\nTMhelix 326 348\ninside 349 372

7604 GCF_002190005.1_ASM219000v1 Escherichia coli Proteobacteria; Gammaproteobacteria; Enterobacterales; Enterobacteriaceae; Escherichia
MTGDNTLIHSHGINRRDFMKLCAALAATMGLSSKAAAEMAESVTNPQRPPVIWIGAQECT WP_000145410.1
MULTISPECIES: hydrogenase-2 small chain [Proteobacteria] Length: 372\nNumber of predicted TMHs: 1\nExp number of AAs in TMHs: 23.72226\nExp number, first 60 AAs: 0.22038\nTotal prob of N-in: 0.12994\noutside 1 331\nTMhelix 332 354\ninside 355 372

7605 GCF_002190285.1_ASM219028v1 Escherichia coli Proteobacteria; Gammaproteobacteria; Enterobacterales; Enterobacteriaceae; Escherichia
MTGDNTLIHSHGINRRDFMKLCAALAATMGLSSKAAAEMAESVTNPQRPPVIWIGAQECT WP_000145410.1
MULTISPECIES: hydrogenase-2 small chain [Proteobacteria] Length: 372\nNumber of predicted TMHs: 1\nExp number of AAs in TMHs: 23.72226\nExp number, first 60 AAs: 0.22038\nTotal prob of N-in: 0.12994\noutside 1 331\nTMhelix 332 354\ninside 355 372

7606 GCF_002190305.1_ASM219030v1 Escherichia coli Proteobacteria; Gammaproteobacteria; Enterobacterales; Enterobacteriaceae; Escherichia
MTGDNTLIHSHGINRRDFMKLCAALAATMGLSSKAAAEMAESVTNPQRPPVIWIGAQECT WP_000145410.1
MULTISPECIES: hydrogenase-2 small chain [Proteobacteria] Length: 372\nNumber of predicted TMHs: 1\nExp number of AAs in TMHs: 23.72226\nExp number, first 60 AAs: 0.22038\nTotal prob of N-in: 0.12994\noutside 1 331\nTMhelix 332 354\ninside 355 372

7607 GCF_002190435.1_ASM219043v1 Escherichia coli Proteobacteria; Gammaproteobacteria; Enterobacterales; Enterobacteriaceae; Escherichia
MTGDNTLIHSHGINRRDFMKLCAALAATMGLSSKAAAEMAESVTNPQRPPVIWIGAQECT WP_000145410.1
MULTISPECIES: hydrogenase-2 small chain [Proteobacteria] Length: 372\nNumber of predicted TMHs: 1\nExp number of AAs in TMHs: 23.72226\nExp number, first 60 AAs: 0.22038\nTotal prob of N-in: 0.12994\noutside 1 331\nTMhelix 332 354\ninside 355 372

7608 GCF_002190435.1_ASM219043v1 Escherichia coli Proteobacteria; Gammaproteobacteria; Enterobacterales; Enterobacteriaceae; Escherichia
MNNEETFYQAMRRQGVTRRSFLKYCSLAATSLGLGAGMAPKIAWALENKPRIPVVWIHGL WP_001058323.1
MULTISPECIES: hydrogenase-1 small chain [Proteobacteria] Length: 372\nNumber of predicted TMHs: 2\nExp number of AAs in TMHs: 37.2608499999999\nExp number, first 60 AAs: 19.52923\nTotal prob of N-in: 0.94351\nPOSSIBLE N-term signal sequence\ninside 1 19\nTMhelix 20 39\noutside 40 325\nTMhelix 326 348\ninside 349 372

7609 GCF_002190105.1_ASM219010v1 Escherichia coli Proteobacteria; Gammaproteobacteria; Enterobacterales; Enterobacteriaceae; Escherichia
MNNEETFYQAMRRQGVTRRSFLKYCSLAATSLGLGAGMAPKIAWALENKPRIPVVWIHGL WP_001058323.1
MULTISPECIES: hydrogenase-1 small chain [Proteobacteria] Length: 372\nNumber of predicted TMHs: 2\nExp number of AAs in TMHs: 37.2608499999999\nExp number, first 60 AAs: 19.52923\nTotal prob of N-in: 0.94351\nPOSSIBLE N-term signal sequence\ninside 1 19\nTMhelix 20 39\noutside 40 325\nTMhelix 326 348\ninside 349 372

7610 GCF_002190475.1_ASM219047v1 Escherichia coli Proteobacteria; Gammaproteobacteria; Enterobacterales; Enterobacteriaceae; Escherichia
MTGDNTLIHSHGINRRDFMKLCAALAATMGLSSKAAAEMAESVTNPQRPPVIWIGAQECT WP_000145410.1
MULTISPECIES: hydrogenase-2 small chain [Proteobacteria] Length: 372\nNumber of predicted TMHs: 1\nExp number of AAs in TMHs: 23.72226\nExp number, first 60 AAs: 0.22038\nTotal prob of N-in: 0.12994\noutside 1 331\nTMhelix 332 354\ninside 355 372

7611 GCF_002190535.1_ASM219053v1 Escherichia coli Proteobacteria; Gammaproteobacteria; Enterobacterales; Enterobacteriaceae; Escherichia
MTGDNTLIHSHGINRRDFMKLCAALAATMGLSSKAAAEMAESVTNPQRPPVIWIGAQECT WP_000145410.1
MULTISPECIES: hydrogenase-2 small chain [Proteobacteria] Length: 372\nNumber of predicted TMHs: 1\nExp number of AAs in TMHs: 23.72226\nExp number, first 60 AAs: 0.22038\nTotal prob of N-in: 0.12994\noutside 1 331\nTMhelix 332 354\ninside 355 372

7612 GCF_002190225.1_ASM219022v1 Escherichia coli Proteobacteria; Gammaproteobacteria; Enterobacterales; Enterobacteriaceae; Escherichia
MNNEETFYQAMRRQGVTRRSFLKYCSLAATSLGLGAGMAPKIAWALENKPRIPVVWIHGL WP_001058323.1
MULTISPECIES: hydrogenase-1 small chain [Proteobacteria] Length: 372\nNumber of predicted TMHs: 2\nExp number of AAs in TMHs: 37.2608499999999\nExp number, first 60 AAs: 19.52923\nTotal prob of N-in: 0.94351\nPOSSIBLE N-term signal sequence\ninside 1 19\nTMhelix 20 39\noutside 40 325\nTMhelix 326 348\ninside 349 372

7613 GCF_002190285.1_ASM219028v1 Escherichia coli Proteobacteria; Gammaproteobacteria; Enterobacterales; Enterobacteriaceae; Escherichia
MNNEETFYQAMRRQGVTRRSFLKYCSLAATSLGLGAGMAPKIAWALENKPRIPVVWIHGL WP_001058323.1
MULTISPECIES: hydrogenase-1 small chain [Proteobacteria] Length: 372\nNumber of predicted TMHs: 2\nExp number of AAs in TMHs: 37.2608499999999\nExp number, first 60 AAs: 19.52923\nTotal prob of N-in: 0.94351\nPOSSIBLE N-term signal sequence\ninside 1 19\nTMhelix 20 39\noutside 40 325\nTMhelix 326 348\ninside 349 372

7614 GCF_002190375.1_ASM219037v1 Escherichia coli Proteobacteria; Gammaproteobacteria; Enterobacterales; Enterobacteriaceae; Escherichia
MNNEETFYQAMRRQGVTRRSFLKYCSLAATSLGLGAGMAPKIAWALENKPRIPVVWIHGL WP_001058323.1
MULTISPECIES: hydrogenase-1 small chain [Proteobacteria] Length: 372\nNumber of predicted TMHs: 2\nExp number of AAs in TMHs: 37.2608499999999\nExp number, first 60 AAs: 19.52923\nTotal prob of N-in: 0.94351\nPOSSIBLE N-term signal sequence\ninside 1 19\nTMhelix 20 39\noutside 40 325\nTMhelix 326 348\ninside 349 372

7615 GCF_002190305.1_ASM219030v1 Escherichia coli Proteobacteria; Gammaproteobacteria; Enterobacterales; Enterobacteriaceae; Escherichia
MNNEETFYQAMRRQGVTRRSFLKYCSLAATSLGLGAGMAPKIAWALENKPRIPVVWIHGL WP_001058323.1
MULTISPECIES: hydrogenase-1 small chain [Proteobacteria] Length: 372\nNumber of predicted TMHs: 2\nExp number of AAs in TMHs: 37.2608499999999\nExp number, first 60 AAs: 19.52923\nTotal prob of N-in: 0.94351\nPOSSIBLE N-term signal sequence\ninside 1 19\nTMhelix 20 39\noutside 40 325\nTMhelix 326 348\ninside 349 372

7616 GCF_002190625.1_ASM219062v1 Escherichia coli Proteobacteria; Gammaproteobacteria; Enterobacterales; Enterobacteriaceae; Escherichia
MTGDNTLIHSHGINRRDFMKLCAALAATMGLSSKAAAEMAESVTNPQRPPVIWIGAQECT WP_000145410.1
MULTISPECIES: hydrogenase-2 small chain [Proteobacteria] Length: 372\nNumber of predicted TMHs: 1\nExp number of AAs in TMHs: 23.72226\nExp number, first 60 AAs: 0.22038\nTotal prob of N-in: 0.12994\noutside 1 331\nTMhelix 332 354\ninside 355 372

7617 GCF_002190645.1_ASM219064v1 Escherichia coli Proteobacteria; Gammaproteobacteria; Enterobacterales; Enterobacteriaceae; Escherichia
MTGDNTLIHSHGINRRDFMKLCAALAATMGLSSKAAAEMAESVTNPQRPPVIWIGAQECT WP_000145410.1
MULTISPECIES: hydrogenase-2 small chain [Proteobacteria] Length: 372\nNumber of predicted TMHs: 1\nExp number of AAs in TMHs: 23.72226\nExp number, first 60 AAs: 0.22038\nTotal prob of N-in: 0.12994\noutside 1 331\nTMhelix 332 354\ninside 355 372

7618 GCF_002190675.1_ASM219067v1 Escherichia coli Proteobacteria; Gammaproteobacteria; Enterobacterales; Enterobacteriaceae; Escherichia
MTGDNTLIHSHGINRRDFMKLCAALAATMGLSSKAAAEMAESVTNPQRPPVIWIGAQECT WP_000145410.1
MULTISPECIES: hydrogenase-2 small chain [Proteobacteria] Length: 372\nNumber of predicted TMHs: 1\nExp number of AAs in TMHs: 23.72226\nExp number, first 60 AAs: 0.22038\nTotal prob of N-in: 0.12994\noutside 1 331\nTMhelix 332 354\ninside 355 372

7619 GCF_002191035.1_ASM219103v1 Escherichia coli Proteobacteria; Gammaproteobacteria; Enterobacterales; Enterobacteriaceae; Escherichia
MTGDNTLIHSHGINRRDFMKLCAALAATMGLSSKAAAEMAESVTNPQRPPVIWIGAQECT WP_000145410.1
MULTISPECIES: hydrogenase-2 small chain [Proteobacteria] Length: 372\nNumber of predicted TMHs: 1\nExp number of AAs in TMHs: 23.72226\nExp number, first 60 AAs: 0.22038\nTotal prob of N-in: 0.12994\noutside 1 331\nTMhelix 332 354\ninside 355 372

7620 GCF_002190975.1_ASM219097v1 Escherichia coli Proteobacteria; Gammaproteobacteria; Enterobacterales; Enterobacteriaceae; Escherichia
MNNEETFYQAMRRQGVTRRSFLKYCSLAATSLGLGAGMAPKIAWALENKPRIPVVWIHGL WP_001058323.1
MULTISPECIES: hydrogenase-1 small chain [Proteobacteria] Length: 372\nNumber of predicted TMHs: 2\nExp number of AAs in TMHs: 37.2608499999999\nExp number, first 60 AAs: 19.52923\nTotal prob of N-in: 0.94351\nPOSSIBLE N-term signal sequence\ninside 1 19\nTMhelix 20 39\noutside 40 325\nTMhelix 326 348\ninside 349 372

7621 GCF_002190935.1_ASM219093v1 Escherichia coli Proteobacteria; Gammaproteobacteria; Enterobacterales; Enterobacteriaceae; Escherichia
MNNEETFYQAMRRQGVTRRSFLKYCSLAATSLGLGAGMAPKIAWALENKPRIPVVWIHGL WP_001058323.1
MULTISPECIES: hydrogenase-1 small chain [Proteobacteria] Length: 372\nNumber of predicted TMHs: 2\nExp number of AAs in TMHs: 37.2608499999999\nExp number, first 60 AAs: 19.52923\nTotal prob of N-in: 0.94351\nPOSSIBLE N-term signal sequence\ninside 1 19\nTMhelix 20 39\noutside 40 325\nTMhelix 326 348\ninside 349 372

7622 GCF_002190675.1_ASM219067v1 Escherichia coli Proteobacteria; Gammaproteobacteria; Enterobacterales; Enterobacteriaceae; Escherichia
MNNEETFYQAMRRQGVTRRSFLKYCSLAATSLGLGAGMAPKIAWALENKPRIPVVWIHGL WP_001058323.1
MULTISPECIES: hydrogenase-1 small chain [Proteobacteria] Length: 372\nNumber of predicted TMHs: 2\nExp number of AAs in TMHs: 37.2608499999999\nExp number, first 60 AAs: 19.52923\nTotal prob of N-in: 0.94351\nPOSSIBLE N-term signal sequence\ninside 1 19\nTMhelix 20 39\noutside 40 325\nTMhelix 326 348\ninside 349 372

7623 GCF_002191105.1_ASM219110v1 Escherichia coli Proteobacteria; Gammaproteobacteria; Enterobacterales; Enterobacteriaceae; Escherichia
MTGDNTLIHSHGINRRDFMKLCAALAATMGLSSKAAAEMAESVTNPQRPPVIWIGAQECT WP_000145410.1
MULTISPECIES: hydrogenase-2 small chain [Proteobacteria] Length: 372\nNumber of predicted TMHs: 1\nExp number of AAs in TMHs: 23.72226\nExp number, first 60 AAs: 0.22038\nTotal prob of N-in: 0.12994\noutside 1 331\nTMhelix 332 354\ninside 355 372

7624 GCF_002191105.1_ASM219110v1 Escherichia coli Proteobacteria; Gammaproteobacteria; Enterobacterales; Enterobacteriaceae; Escherichia
MNNEETFYQAMRRQGVTRRSFLKYCSLAATSLGLGAGMAPKIAWALENKPRIPVVWIHGL WP_001058323.1
MULTISPECIES: hydrogenase-1 small chain [Proteobacteria] Length: 372\nNumber of predicted TMHs: 2\nExp number of AAs in TMHs: 37.2608499999999\nExp number, first 60 AAs: 19.52923\nTotal prob of N-in: 0.94351\nPOSSIBLE N-term signal sequence\ninside 1 19\nTMhelix 20 39\noutside 40 325\nTMhelix 326 348\ninside 349 372

7625 GCF_002190645.1_ASM219064v1 Escherichia coli Proteobacteria; Gammaproteobacteria; Enterobacterales; Enterobacteriaceae; Escherichia
MNNEETFYQAMRRQGVTRRSFLKYCSLAATSLGLGAGMAPKIAWALENKPRIPVVWIHGL WP_001295940.1
MULTISPECIES: hydrogenase [Enterobacteriaceae] Length: 372\nNumber of predicted TMHs: 2\nExp number of AAs in TMHs: 37.2608899999999\nExp number, first 60 AAs: 19.52923\nTotal prob of N-in: 0.94351\nPOSSIBLE N-term signal sequence\ninside 1 19\nTMhelix 20 39\noutside 40 325\nTMhelix 326 348\ninside 349 372

7626 GCF_002192295.1_ASM219229v1 Escherichia coli Proteobacteria; Gammaproteobacteria; Enterobacterales; Enterobacteriaceae; Escherichia
MTGDNTLIHSHGINRRDFMKLCAALAATMGLSSKAAAEMAESVTNPQRPPVIWIGAQECT WP_000145410.1
MULTISPECIES: hydrogenase-2 small chain [Proteobacteria] Length: 372\nNumber of predicted TMHs: 1\nExp number of AAs in TMHs: 23.72226\nExp number, first 60 AAs: 0.22038\nTotal prob of N-in: 0.12994\noutside 1 331\nTMhelix 332 354\ninside 355 372

7627 GCF_002192295.1_ASM219229v1 Escherichia coli Proteobacteria; Gammaproteobacteria; Enterobacterales; Enterobacteriaceae; Escherichia
MNNEETFYQAMRRQGVTRRSFLKYCSLAATSLGLGAGMAPKIAWALENKPRIPVVWIHGL WP_001058323.1
MULTISPECIES: hydrogenase-1 small chain [Proteobacteria] Length: 372\nNumber of predicted TMHs: 2\nExp number of AAs in TMHs: 37.2608499999999\nExp number, first 60 AAs: 19.52923\nTotal prob of N-in: 0.94351\nPOSSIBLE N-term signal sequence\ninside 1 19\nTMhelix 20 39\noutside 40 325\nTMhelix 326 348\ninside 349 372

7628 GCF_002190815.1_ASM219081v1 Escherichia coli Proteobacteria; Gammaproteobacteria; Enterobacterales; Enterobacteriaceae; Escherichia
MNNEETFYQAMRRQGVTRRSFLKYCSLAATSLGLGAGMAPKIAWALENKPRIPVVWIHGL WP_001058323.1
MULTISPECIES: hydrogenase-1 small chain [Proteobacteria] Length: 372\nNumber of predicted TMHs: 2\nExp number of AAs in TMHs: 37.2608499999999\nExp number, first 60 AAs: 19.52923\nTotal prob of N-in: 0.94351\nPOSSIBLE N-term signal sequence\ninside 1 19\nTMhelix 20 39\noutside 40 325\nTMhelix 326 348\ninside 349 372

7629 GCF_002191035.1_ASM219103v1 Escherichia coli Proteobacteria; Gammaproteobacteria; Enterobacterales; Enterobacteriaceae; Escherichia
MNNEETFYQAMRRQGVTRRSFLKYCSLAATSLGLGAGMAPKIAWALENKPRIPVVWIHGL WP_001058323.1
MULTISPECIES: hydrogenase-1 small chain [Proteobacteria] Length: 372\nNumber of predicted TMHs: 2\nExp number of AAs in TMHs: 37.2608499999999\nExp number, first 60 AAs: 19.52923\nTotal prob of N-in: 0.94351\nPOSSIBLE N-term signal sequence\ninside 1 19\nTMhelix 20 39\noutside 40 325\nTMhelix 326 348\ninside 349 372

7630 GCF_002190775.1_ASM219077v1 Escherichia coli Proteobacteria; Gammaproteobacteria; Enterobacterales; Enterobacteriaceae; Escherichia
MNNEETFYQAMRRQGVTRRSFLKYCSLAATSLGLGAGMAPKIAWALENKPRIPVVWIHGL WP_001058323.1
MULTISPECIES: hydrogenase-1 small chain [Proteobacteria] Length: 372\nNumber of predicted TMHs: 2\nExp number of AAs in TMHs: 37.2608499999999\nExp number, first 60 AAs: 19.52923\nTotal prob of N-in: 0.94351\nPOSSIBLE N-term signal sequence\ninside 1 19\nTMhelix 20 39\noutside 40 325\nTMhelix 326 348\ninside 349 372

7631 GCF_002190705.1_ASM219070v1 Escherichia coli Proteobacteria; Gammaproteobacteria; Enterobacterales; Enterobacteriaceae; Escherichia
MNNEETFYQAMRRQGVTRRSFLKYCSLAATSLGLGAGMAPKIAWALENKPRIPVVWIHGL WP_001058323.1
MULTISPECIES: hydrogenase-1 small chain [Proteobacteria] Length: 372\nNumber of predicted TMHs: 2\nExp number of AAs in TMHs: 37.2608499999999\nExp number, first 60 AAs: 19.52923\nTotal prob of N-in: 0.94351\nPOSSIBLE N-term signal sequence\ninside 1 19\nTMhelix 20 39\noutside 40 325\nTMhelix 326 348\ninside 349 372

7632 GCF_002193415.1_ASM219341v1 Escherichia coli Proteobacteria; Gammaproteobacteria; Enterobacterales; Enterobacteriaceae; Escherichia
MTGDNTLIHSHGINRRDFMKLCAALAATMGLSSKAAAEMAESVTNPQRPPVIWIGAQECT WP_000145410.1
MULTISPECIES: hydrogenase-2 small chain [Proteobacteria] Length: 372\nNumber of predicted TMHs: 1\nExp number of AAs in TMHs: 23.72226\nExp number, first 60 AAs: 0.22038\nTotal prob of N-in: 0.12994\noutside 1 331\nTMhelix 332 354\ninside 355 372

7633 GCF_002193495.1_ASM219349v1 Escherichia coli Proteobacteria; Gammaproteobacteria; Enterobacterales; Enterobacteriaceae; Escherichia
MTGDNTLIHSHGINRRDFMKLCAALAATMGLSSKAAAEMAESVTNPQRPPVIWIGAQECT WP_000145410.1
MULTISPECIES: hydrogenase-2 small chain [Proteobacteria] Length: 372\nNumber of predicted TMHs: 1\nExp number of AAs in TMHs: 23.72226\nExp number, first 60 AAs: 0.22038\nTotal prob of N-in: 0.12994\noutside 1 331\nTMhelix 332 354\ninside 355 372

7634 GCF_002193375.1_ASM219337v1 Escherichia coli Proteobacteria; Gammaproteobacteria; Enterobacterales; Enterobacteriaceae; Escherichia
MTGDNTLIHSHGINRRDFMKLCAALAATMGLSSKAAAEMAESVTNPQRPPVIWIGAQECT WP_000145410.1
MULTISPECIES: hydrogenase-2 small chain [Proteobacteria] Length: 372\nNumber of predicted TMHs: 1\nExp number of AAs in TMHs: 23.72226\nExp number, first 60 AAs: 0.22038\nTotal prob of N-in: 0.12994\noutside 1 331\nTMhelix 332 354\ninside 355 372

7635 GCF_002193275.1_ASM219327v1 Escherichia coli Proteobacteria; Gammaproteobacteria; Enterobacterales; Enterobacteriaceae; Escherichia
MNNEETFYQAMRRQGVTRRSFLKYCSLAATSLGLGAGMAPKIAWALENKPRIPVVWIHGL WP_001058323.1
MULTISPECIES: hydrogenase-1 small chain [Proteobacteria] Length: 372\nNumber of predicted TMHs: 2\nExp number of AAs in TMHs: 37.2608499999999\nExp number, first 60 AAs: 19.52923\nTotal prob of N-in: 0.94351\nPOSSIBLE N-term signal sequence\ninside 1 19\nTMhelix 20 39\noutside 40 325\nTMhelix 326 348\ninside 349 372

7636 GCF_002194595.1_ASM219459v1 Escherichia coli Proteobacteria; Gammaproteobacteria; Enterobacterales; Enterobacteriaceae; Escherichia
MTGDNTLIHSHGINRRDFMKLCAALAATMGLSSKAAAEMAESVTNPQRPPVIWIGAQECT WP_000145410.1
MULTISPECIES: hydrogenase-2 small chain [Proteobacteria] Length: 372\nNumber of predicted TMHs: 1\nExp number of AAs in TMHs: 23.72226\nExp number, first 60 AAs: 0.22038\nTotal prob of N-in: 0.12994\noutside 1 331\nTMhelix 332 354\ninside 355 372

7637 GCF_002194595.1_ASM219459v1 Escherichia coli Proteobacteria; Gammaproteobacteria; Enterobacterales; Enterobacteriaceae; Escherichia
MNNEETFYQAMRRQGVTRRSFLKYCSLAATSLGLGAGMAPKIAWALENKPRIPVVWIHGL WP_001301613.1
hydrogenase [Escherichia coli] Length: 372\nNumber of predicted TMHs: 2\nExp number of AAs in TMHs: 37.46598\nExp number, first 60 AAs: 19.53139\nTotal prob of N-in: 0.94364\nPOSSIBLE N-term signal sequence\ninside 1 19\nTMhelix 20 39\noutside 40 325\nTMhelix 326 348\ninside 349 372

7638 GCF_002193475.1_ASM219347v1 Escherichia coli Proteobacteria; Gammaproteobacteria; Enterobacterales; Enterobacteriaceae; Escherichia
MNNEETFYQAMRRQGVTRRSFLKYCSLAATSLGLGAGMAPKIAWALENKPRIPVVWIHGL WP_001058323.1
MULTISPECIES: hydrogenase-1 small chain [Proteobacteria] Length: 372\nNumber of predicted TMHs: 2\nExp number of AAs in TMHs: 37.2608499999999\nExp number, first 60 AAs: 19.52923\nTotal prob of N-in: 0.94351\nPOSSIBLE N-term signal sequence\ninside 1 19\nTMhelix 20 39\noutside 40 325\nTMhelix 326 348\ninside 349 372

7639 GCF_002194695.1_ASM219469v1 Escherichia coli Proteobacteria; Gammaproteobacteria; Enterobacterales; Enterobacteriaceae; Escherichia

MNNEETFYQAMRRQGVTRRSFLKYCSLAATSLGLGAGMAPKIAWALENKPRIPVVWIHGL WP_001301613.1 hydrogenase [Escherichia coli] Length: 372\nNumber of predicted TMHs: 2\nExp number of AAs in TMHs: 37.46598\nExp number, first 60 AAs: 19.53139\nTotal prob of N-in: 0.94364\nPOSSIBLE N-term signal sequence\ninside 1 19\nTMhelix 20 39\noutside 40 325\nTMhelix 326 348\ninside 349 372

7640 GCF_002194755.1_ASM219475v1 Escherichia coli Proteobacteria; Gammaproteobacteria; Enterobacterales; Enterobacteriaceae; Escherichia MNNEETFYQAMRRQGVTRRSFLKYCSLAATSLGLGAGMAPKIAWALENKPRIPVVWIHGL WP_001301613.1 hydrogenase [Escherichia coli] Length: 372\nNumber of predicted TMHs: 2\nExp number of AAs in TMHs: 37.46598\nExp number, first 60 AAs: 19.53139\nTotal prob of N-in: 0.94364\nPOSSIBLE N-term signal sequence\ninside 1 19\nTMhelix 20 39\noutside 40 325\nTMhelix 326 348\ninside 349 372

7641 GCF_002193415.1_ASM219341v1 Escherichia coli Proteobacteria; Gammaproteobacteria; Enterobacterales; Enterobacteriaceae; Escherichia MNNEETFYQAMRRQGVTRRSFLKYCSLAATSLGLGAGMAPKIAWALENKPRIPVVWIHGL WP_001058323.1 MULTISPECIES: hydrogenase-1 small chain [Proteobacteria] Length: 372\nNumber of predicted TMHs: 2\nExp number of AAs in TMHs: 37.2608499999999\nExp number, first 60 AAs: 19.52923\nTotal prob of N-in: 0.94351\nPOSSIBLE N-term signal sequence\ninside 1 19\nTMhelix 20 39\noutside 40 325\nTMhelix 326 348\ninside 349 372

7642 GCF_002193375.1_ASM219337v1 Escherichia coli Proteobacteria; Gammaproteobacteria; Enterobacterales; Enterobacteriaceae; Escherichia MNNEETFYQAMRRQGVTRRSFLKYCSLAATSLGLGAGMAPKIAWALENKPRIPVVWIHGL WP_001058323.1 MULTISPECIES: hydrogenase-1 small chain [Proteobacteria] Length: 372\nNumber of predicted TMHs: 2\nExp number of AAs in TMHs: 37.2608499999999\nExp number, first 60 AAs: 19.52923\nTotal prob of N-in: 0.94351\nPOSSIBLE N-term signal sequence\ninside 1 19\nTMhelix 20 39\noutside 40 325\nTMhelix 326 348\ninside 349 372

7643 GCF_002194815.1_ASM219481v1 Escherichia coli Proteobacteria; Gammaproteobacteria; Enterobacterales; Enterobacteriaceae; Escherichia MTGDNTLIHSHGINRRDFMKLCAALAATMGLSSKAAAEMAESVTNPQRPPVIWIGAQECT WP_000145410.1 MULTISPECIES: hydrogenase-2 small chain [Proteobacteria] Length: 372\nNumber of predicted TMHs: 1\nExp number of AAs in TMHs: 23.72226\nExp number, first 60 AAs: 0.22038\nTotal prob of N-in: 0.12994\noutside 1 331\nTMhelix 332 354\ninside 355 372

7644 GCF_002194855.1_ASM219485v1 Escherichia coli Proteobacteria; Gammaproteobacteria; Enterobacterales; Enterobacteriaceae; Escherichia MTGDNTLIHSHGINRRDFMKLCAALAATMGLSSKAAAEMAESVTNPQRPPVIWIGAQECT WP_000145410.1 MULTISPECIES: hydrogenase-2 small chain [Proteobacteria] Length: 372\nNumber of predicted TMHs: 1\nExp number of AAs in TMHs: 23.72226\nExp number, first 60 AAs: 0.22038\nTotal prob of N-in: 0.12994\noutside 1 331\nTMhelix 332 354\ninside 355 372

7645 GCF_002194915.1_ASM219491v1 Escherichia coli Proteobacteria; Gammaproteobacteria; Enterobacterales; Enterobacteriaceae; Escherichia MTGDNTLIHSHGINRRDFMKLCAALAATMGLSSKAAAEMAESVTNPQRPPVIWIGAQECT WP_000145410.1 MULTISPECIES: hydrogenase-2 small chain [Proteobacteria] Length: 372\nNumber of predicted TMHs: 1\nExp number of AAs in TMHs: 23.72226\nExp number, first 60 AAs: 0.22038\nTotal prob of N-in: 0.12994\noutside 1 331\nTMhelix 332 354\ninside 355 372

7646 GCF_002195195.1_ASM219519v1 Escherichia coli Proteobacteria; Gammaproteobacteria; Enterobacterales; Enterobacteriaceae; Escherichia MTGDNTLIHSHGINRRDFMKLCAALAATMGLSSKAAAEMAESVTNPQRPPVIWIGAQECT WP_000145410.1 MULTISPECIES: hydrogenase-2 small chain [Proteobacteria] Length: 372\nNumber of predicted TMHs: 1\nExp number of AAs in TMHs: 23.72226\nExp number, first 60 AAs: 0.22038\nTotal prob of N-in: 0.12994\noutside 1 331\nTMhelix 332 354\ninside 355 372

7647 GCF_002195315.1_ASM219531v1 Escherichia coli Proteobacteria; Gammaproteobacteria; Enterobacterales; Enterobacteriaceae; Escherichia MTGDNTLIHSHGINRRDFMKLCAALAATMGLSSKAAAEMAESVTNPQRPPVIWIGAQECT WP_000145410.1 MULTISPECIES: hydrogenase-2 small chain [Proteobacteria] Length: 372\nNumber of predicted TMHs:

1\nExp number of AAs in TMHs: 23.72226\nExp number, first 60 AAs: 0.22038\nTotal prob of N-in: 0.12994\noutside 1 331\nTMhelix 332 354\ninside 355 372

7648 GCF_002195135.1_ASM219513v1 Escherichia coli Proteobacteria; Gammaproteobacteria; Enterobacterales; Enterobacteriaceae; Escherichia
MNNEETFYQAMRRQGVTRRSFLKYCSLAATSLGLGAGMAPKIAWALENKPRIPVVWIHGL WP_001301613.1
hydrogenase [Escherichia coli] Length: 372\nNumber of predicted TMHs: 2\nExp number of AAs in TMHs: 37.46598\nExp number, first 60 AAs: 19.53139\nTotal prob of N-in: 0.94364\nPOSSIBLE N-term signal sequence\ninside 1 19\nTMhelix 20 39\noutside 40 325\nTMhelix 326 348\ninside 349 372

7649 GCF_002194855.1_ASM219485v1 Escherichia coli Proteobacteria; Gammaproteobacteria; Enterobacterales; Enterobacteriaceae; Escherichia
MNNEETFYQAMRRQGVTRRSFLKYCSLAATSLGLGAGMAPKIAWALENKPRIPVVWIHGL WP_001301613.1
hydrogenase [Escherichia coli] Length: 372\nNumber of predicted TMHs: 2\nExp number of AAs in TMHs: 37.46598\nExp number, first 60 AAs: 19.53139\nTotal prob of N-in: 0.94364\nPOSSIBLE N-term signal sequence\ninside 1 19\nTMhelix 20 39\noutside 40 325\nTMhelix 326 348\ninside 349 372

7650 GCF_002195325.1_ASM219532v1 Escherichia coli Proteobacteria; Gammaproteobacteria; Enterobacterales; Enterobacteriaceae; Escherichia
MTGDNTLIHSHGINRRDFMKLCAALAATMGLSSKAAAEMAESVTNPQRPPVIWIGAQECT WP_000145410.1
MULTISPECIES: hydrogenase-2 small chain [Proteobacteria] Length: 372\nNumber of predicted TMHs: 1\nExp number of AAs in TMHs: 23.72226\nExp number, first 60 AAs: 0.22038\nTotal prob of N-in: 0.12994\noutside 1 331\nTMhelix 332 354\ninside 355 372

7651 GCF_002195325.1_ASM219532v1 Escherichia coli Proteobacteria; Gammaproteobacteria; Enterobacterales; Enterobacteriaceae; Escherichia
MNNEETFYQAMRRQGVTRRSFLKYCSLAATSLGLGAGMAPKIAWALENKPRIPVVWIHGL WP_001301613.1
hydrogenase [Escherichia coli] Length: 372\nNumber of predicted TMHs: 2\nExp number of AAs in TMHs: 37.46598\nExp number, first 60 AAs: 19.53139\nTotal prob of N-in: 0.94364\nPOSSIBLE N-term signal sequence\ninside 1 19\nTMhelix 20 39\noutside 40 325\nTMhelix 326 348\ninside 349 372

7652 GCF_002195395.1_ASM219539v1 Escherichia coli Proteobacteria; Gammaproteobacteria; Enterobacterales; Enterobacteriaceae; Escherichia
MNNEETFYQAMRRQGVTRRSFLKYCSLAATSLGLGAGMAPKIAWALENKPRIPVVWIHGL WP_001301613.1
hydrogenase [Escherichia coli] Length: 372\nNumber of predicted TMHs: 2\nExp number of AAs in TMHs: 37.46598\nExp number, first 60 AAs: 19.53139\nTotal prob of N-in: 0.94364\nPOSSIBLE N-term signal sequence\ninside 1 19\nTMhelix 20 39\noutside 40 325\nTMhelix 326 348\ninside 349 372

7653 GCF_002195115.1_ASM219511v1 Escherichia coli Proteobacteria; Gammaproteobacteria; Enterobacterales; Enterobacteriaceae; Escherichia
MNNEETFYQAMRRQGVTRRSFLKYCSLAATSLGLGAGMAPKIAWALENKPRIPVVWIHGL WP_001301613.1
hydrogenase [Escherichia coli] Length: 372\nNumber of predicted TMHs: 2\nExp number of AAs in TMHs: 37.46598\nExp number, first 60 AAs: 19.53139\nTotal prob of N-in: 0.94364\nPOSSIBLE N-term signal sequence\ninside 1 19\nTMhelix 20 39\noutside 40 325\nTMhelix 326 348\ninside 349 372

7654 GCF_002195195.1_ASM219519v1 Escherichia coli Proteobacteria; Gammaproteobacteria; Enterobacterales; Enterobacteriaceae; Escherichia
MNNEETFYQAMRRQGVTRRSFLKYCSLAATSLGLGAGMAPKIAWALENKPRIPVVWIHGL WP_001301613.1
hydrogenase [Escherichia coli] Length: 372\nNumber of predicted TMHs: 2\nExp number of AAs in TMHs: 37.46598\nExp number, first 60 AAs: 19.53139\nTotal prob of N-in: 0.94364\nPOSSIBLE N-term signal sequence\ninside 1 19\nTMhelix 20 39\noutside 40 325\nTMhelix 326 348\ninside 349 372

7655 GCF_002195435.1_ASM219543v1 Escherichia coli Proteobacteria; Gammaproteobacteria; Enterobacterales; Enterobacteriaceae; Escherichia
MTGDNTLIHSHGINRRDFMKLCAALAATMGLSSKAAAEMAESVTNPQRPPVIWIGAQECT WP_000145410.1
MULTISPECIES: hydrogenase-2 small chain [Proteobacteria] Length: 372\nNumber of predicted TMHs: 1\nExp number of AAs in TMHs: 23.72226\nExp number, first 60 AAs: 0.22038\nTotal prob of N-in: 0.12994\noutside 1 331\nTMhelix 332 354\ninside 355 372

7656 GCF_002195435.1_ASM219543v1 Escherichia coli Proteobacteria; Gammaproteobacteria; Enterobacterales; Enterobacteriaceae; Escherichia

MNNEETFYQAMRRQGVTRRSFLKYCSLAATSLGLGAGMAPKIAWALENKPRIPVVWIHGL WP_001301613.1 hydrogenase [Escherichia coli] Length: 372\nNumber of predicted TMHs: 2\nExp number of AAs in TMHs: 37.46598\nExp number, first 60 AAs: 19.53139\nTotal prob of N-in: 0.94364\nPOSSIBLE N-term signal sequence\ninside 1 19\nTMhelix 20 39\noutside 40 325\nTMhelix 326 348\ninside 349 372

7657 GCF_002195255.1_ASM219525v1 Escherichia coli Proteobacteria; Gammaproteobacteria; Enterobacterales; Enterobacteriaceae; Escherichia MNNEETFYQAMRRQGVTRRSFLKYCSLAATSLGLGAGMAPKIAWALENKPRIPVVWIHGL WP_001301613.1 hydrogenase [Escherichia coli] Length: 372\nNumber of predicted TMHs: 2\nExp number of AAs in TMHs: 37.46598\nExp number, first 60 AAs: 19.53139\nTotal prob of N-in: 0.94364\nPOSSIBLE N-term signal sequence\ninside 1 19\nTMhelix 20 39\noutside 40 325\nTMhelix 326 348\ninside 349 372

7658 GCF_002195315.1_ASM219531v1 Escherichia coli Proteobacteria; Gammaproteobacteria; Enterobacterales; Enterobacteriaceae; Escherichia MNNEETFYQAMRRQGVTRRSFLKYCSLAATSLGLGAGMAPKIAWALENKPRIPVVWIHGL WP_001301613.1 hydrogenase [Escherichia coli] Length: 372\nNumber of predicted TMHs: 2\nExp number of AAs in TMHs: 37.46598\nExp number, first 60 AAs: 19.53139\nTotal prob of N-in: 0.94364\nPOSSIBLE N-term signal sequence\ninside 1 19\nTMhelix 20 39\noutside 40 325\nTMhelix 326 348\ninside 349 372

7659 GCF_002196475.1_ASM219647v1 Escherichia coli Proteobacteria; Gammaproteobacteria; Enterobacterales; Enterobacteriaceae; Escherichia MTGDNTLIHSHGINRRDFMKLCAALAATMGLSSKAAAEMAESVTNPQRPPVIWIGAQECT WP_000145410.1 MULTISPECIES: hydrogenase-2 small chain [Proteobacteria] Length: 372\nNumber of predicted TMHs: 1\nExp number of AAs in TMHs: 23.72226\nExp number, first 60 AAs: 0.22038\nTotal prob of N-in: 0.12994\noutside 1 331\nTMhelix 332 354\ninside 355 372

7660 GCF_002196475.1_ASM219647v1 Escherichia coli Proteobacteria; Gammaproteobacteria; Enterobacterales; Enterobacteriaceae; Escherichia MNNEETFYQAMRRQGVTRRSFLKYCSLAATSLGLGAGMAPKIAWALENKPRIPVVWIHGL WP_001058323.1 MULTISPECIES: hydrogenase-1 small chain [Proteobacteria] Length: 372\nNumber of predicted TMHs: 2\nExp number of AAs in TMHs: 37.2608499999999\nExp number, first 60 AAs: 19.52923\nTotal prob of N-in: 0.94351\nPOSSIBLE N-term signal sequence\ninside 1 19\nTMhelix 20 39\noutside 40 325\nTMhelix 326 348\ninside 349 372

7661 GCF_002195895.1_ASM219589v1 Escherichia coli Proteobacteria; Gammaproteobacteria; Enterobacterales; Enterobacteriaceae; Escherichia MNNEETFYQAMRRQGVTRRSFLKYCSLAATSLGLGAGMAPKIAWALENKPRIPVVWIHGL WP_001301613.1 hydrogenase [Escherichia coli] Length: 372\nNumber of predicted TMHs: 2\nExp number of AAs in TMHs: 37.46598\nExp number, first 60 AAs: 19.53139\nTotal prob of N-in: 0.94364\nPOSSIBLE N-term signal sequence\ninside 1 19\nTMhelix 20 39\noutside 40 325\nTMhelix 326 348\ninside 349 372

7662 GCF_002195595.1_ASM219559v1 Escherichia coli Proteobacteria; Gammaproteobacteria; Enterobacterales; Enterobacteriaceae; Escherichia MNNEETFYQAMRRQGVTRRSFLKYCSLAATSLGLGAGMAPKIAWALENKPRIPVVWIHGL WP_001301613.1 hydrogenase [Escherichia coli] Length: 372\nNumber of predicted TMHs: 2\nExp number of AAs in TMHs: 37.46598\nExp number, first 60 AAs: 19.53139\nTotal prob of N-in: 0.94364\nPOSSIBLE N-term signal sequence\ninside 1 19\nTMhelix 20 39\noutside 40 325\nTMhelix 326 348\ninside 349 372

7663 GCF_002195535.1_ASM219553v1 Escherichia coli Proteobacteria; Gammaproteobacteria; Enterobacterales; Enterobacteriaceae; Escherichia MNNEETFYQAMRRQGVTRRSFLKYCSLAATSLGLGAGMAPKIAWALENKPRIPVVWIHGL WP_001301613.1 hydrogenase [Escherichia coli] Length: 372\nNumber of predicted TMHs: 2\nExp number of AAs in TMHs: 37.46598\nExp number, first 60 AAs: 19.53139\nTotal prob of N-in: 0.94364\nPOSSIBLE N-term signal sequence\ninside 1 19\nTMhelix 20 39\noutside 40 325\nTMhelix 326 348\ninside 349 372

7664 GCF_002195485.1_ASM219548v1 Escherichia coli Proteobacteria; Gammaproteobacteria; Enterobacterales; Enterobacteriaceae; Escherichia MNNEETFYQAMRRQGVTRRSFLKYCSLAATSLGLGAGMAPKIAWALENKPRIPVVWIHGL WP_001301613.1 hydrogenase [Escherichia coli] Length: 372\nNumber of predicted TMHs: 2\nExp number of AAs in TMHs: 37.46598\nExp number, first 60 AAs: 19.53139\nTotal prob of N-in: 0.94364\nPOSSIBLE N-term signal sequence\ninside 1 19\nTMhelix 20 39\noutside 40 325\nTMhelix 326 348\ninside 349 372

7665 GCF_002196625.1_ASM219662v1 Escherichia coli Proteobacteria; Gammaproteobacteria; Enterobacterales; Enterobacteriaceae; Escherichia
MTGDNTLIHSHGINRRDFMKLCAALAATMGLSSKAAAEMAESVTNPQRPPVIWIGAQECT WP_000145410.1
MULTISPECIES: hydrogenase-2 small chain [Proteobacteria] Length: 372\nNumber of predicted TMHs: 1\nExp number of AAs in TMHs: 23.72226\nExp number, first 60 AAs: 0.22038\nTotal prob of N-in: 0.12994\noutside 1 331\nTMhelix 332 354\ninside 355 372

7666 GCF_002195825.1_ASM219582v1 Escherichia coli Proteobacteria; Gammaproteobacteria; Enterobacterales; Enterobacteriaceae; Escherichia
MNNEETFYQAMRRQGVTRRSFLKYCSLAATSLGLGAGMAPKIAWALENKPRIPVVWIHGL WP_001301613.1
hydrogenase [Escherichia coli] Length: 372\nNumber of predicted TMHs: 2\nExp number of AAs in TMHs: 37.46598\nExp number, first 60 AAs: 19.53139\nTotal prob of N-in: 0.94364\nPOSSIBLE N-term signal sequence\ninside 1 19\nTMhelix 20 39\noutside 40 325\nTMhelix 326 348\ninside 349 372

7667 GCF_002196745.1_ASM219674v1 Escherichia coli Proteobacteria; Gammaproteobacteria; Enterobacterales; Enterobacteriaceae; Escherichia
MTGDNTLIHSHGINRRDFMKLCAALAATMGLSSKAAAEMAESVTNPQRPPVIWIGAQECT WP_000145410.1
MULTISPECIES: hydrogenase-2 small chain [Proteobacteria] Length: 372\nNumber of predicted TMHs: 1\nExp number of AAs in TMHs: 23.72226\nExp number, first 60 AAs: 0.22038\nTotal prob of N-in: 0.12994\noutside 1 331\nTMhelix 332 354\ninside 355 372

7668 GCF_002196825.1_ASM219682v1 Escherichia coli Proteobacteria; Gammaproteobacteria; Enterobacterales; Enterobacteriaceae; Escherichia
MTGDNTLIHSHGINRRDFMKLCAALAATMGLSSKAAAEMAESVTNPQRPPVIWIGAQECT WP_000145410.1
MULTISPECIES: hydrogenase-2 small chain [Proteobacteria] Length: 372\nNumber of predicted TMHs: 1\nExp number of AAs in TMHs: 23.72226\nExp number, first 60 AAs: 0.22038\nTotal prob of N-in: 0.12994\noutside 1 331\nTMhelix 332 354\ninside 355 372

7669 GCF_002205095.1_ASM220509v1 Escherichia coli Proteobacteria; Gammaproteobacteria; Enterobacterales; Enterobacteriaceae; Escherichia
MTGDNTLIHSHGINRRDFMKLCAALAATMGLSSKAAAEMAESVTNPQRPPVIWIGAQECT WP_000145410.1
MULTISPECIES: hydrogenase-2 small chain [Proteobacteria] Length: 372\nNumber of predicted TMHs: 1\nExp number of AAs in TMHs: 23.72226\nExp number, first 60 AAs: 0.22038\nTotal prob of N-in: 0.12994\noutside 1 331\nTMhelix 332 354\ninside 355 372

7670 GCF_900010065.1_ED392_contigs Escherichia coli Proteobacteria; Gammaproteobacteria; Enterobacterales; Enterobacteriaceae; Escherichia
MTGDNTLIHSHGINRRDFMKLCAALAATMGLSSKAAAEMAESVTNPQRPPVIWIGAQECT WP_000145410.1
MULTISPECIES: hydrogenase-2 small chain [Proteobacteria] Length: 372\nNumber of predicted TMHs: 1\nExp number of AAs in TMHs: 23.72226\nExp number, first 60 AAs: 0.22038\nTotal prob of N-in: 0.12994\noutside 1 331\nTMhelix 332 354\ninside 355 372

7671 GCF_900009485.1_ED608_contigs Escherichia coli Proteobacteria; Gammaproteobacteria; Enterobacterales; Enterobacteriaceae; Escherichia
MNNEETFYQAMRRQGVTRRSFLKYCSLAATSLGLGAGMAPKIAWALENKPRIPVVWIHGL WP_001058323.1
MULTISPECIES: hydrogenase-1 small chain [Proteobacteria] Length: 372\nNumber of predicted TMHs: 2\nExp number of AAs in TMHs: 37.2608499999999\nExp number, first 60 AAs: 19.52923\nTotal prob of N-in: 0.94351\nPOSSIBLE N-term signal sequence\ninside 1 19\nTMhelix 20 39\noutside 40 325\nTMhelix 326 348\ninside 349 372

7672 GCF_900014215.1_ED669_contigs Escherichia coli Proteobacteria; Gammaproteobacteria; Enterobacterales; Enterobacteriaceae; Escherichia
MTGDNTLIHSHGINRRDFMKLCAALAATMGLSSKAAAEMAESVTNPQRPPVIWIGAQECT WP_000145410.1
MULTISPECIES: hydrogenase-2 small chain [Proteobacteria] Length: 372\nNumber of predicted TMHs: 1\nExp number of AAs in TMHs: 23.72226\nExp number, first 60 AAs: 0.22038\nTotal prob of N-in: 0.12994\noutside 1 331\nTMhelix 332 354\ninside 355 372

7673 GCF_900009485.1_ED608_contigs Escherichia coli Proteobacteria; Gammaproteobacteria; Enterobacterales; Enterobacteriaceae; Escherichia
MTGDNTLIHSHGINRRDFMKLCAALAATMGLSGKAAAEMAESVTNPQRPPVIWIGAQECT WP_009448969.1
hydrogenase 2 small subunit [Escherichia coli] Length: 372\nNumber of predicted TMHs: 1\nExp number of

AAs in TMHs: 23.92168\nExp number, first 60 AAs: 0.43108\nTotal prob of N-in: 0.13898\noutside 1 331\nTMhelix 332 354\ninside 355 372

7674 GCF_900010065.1_ED392_contigs Escherichia coli Proteobacteria; Gammaproteobacteria; Enterobacterales; Enterobacteriaceae; Escherichia
MNNEETFYQAMRRQGVTRRSFLKYCSLAATSLGLGAGMAPKIAWALENKPRIPVVWIHGL WP_001058323.1
MULTISPECIES: hydrogenase-1 small chain [Proteobacteria] Length: 372\nNumber of predicted TMHs: 2\nExp number of AAs in TMHs: 37.2608499999999\nExp number, first 60 AAs: 19.52923\nTotal prob of N-in: 0.94351\nPOSSIBLE N-term signal sequence\ninside 1 19\nTMhelix 20 39\noutside 40 325\nTMhelix 326 348\ninside 349 372

7675 GCF_900014215.1_ED669_contigs Escherichia coli Proteobacteria; Gammaproteobacteria; Enterobacterales; Enterobacteriaceae; Escherichia
MNNEETFYQAMRRQGVTRRSFLKYCSLAATSLGLGAGMAPKIAWALENKPRIPVVWIHGL WP_001058323.1
MULTISPECIES: hydrogenase-1 small chain [Proteobacteria] Length: 372\nNumber of predicted TMHs: 2\nExp number of AAs in TMHs: 37.2608499999999\nExp number, first 60 AAs: 19.52923\nTotal prob of N-in: 0.94351\nPOSSIBLE N-term signal sequence\ninside 1 19\nTMhelix 20 39\noutside 40 325\nTMhelix 326 348\ninside 349 372

7676 GCF_900015845.1_ED653_contigs Escherichia coli Proteobacteria; Gammaproteobacteria; Enterobacterales; Enterobacteriaceae; Escherichia
MTGDNTLIHSHGINRRDFMKLCAALAATMGLSSKAAAEMAESVTNPQRPPVIWIGAQECT WP_000145410.1
MULTISPECIES: hydrogenase-2 small chain [Proteobacteria] Length: 372\nNumber of predicted TMHs: 1\nExp number of AAs in TMHs: 23.72226\nExp number, first 60 AAs: 0.22038\nTotal prob of N-in: 0.12994\noutside 1 331\nTMhelix 332 354\ninside 355 372

7677 GCF_900015825.1_ED415_contigs Escherichia coli Proteobacteria; Gammaproteobacteria; Enterobacterales; Enterobacteriaceae; Escherichia
MTGDNTLIHSHGINRRDFMKLCAALAATMGLSSKAAAEMAESVTNPQRPPVIWIGAQECT WP_000145410.1
MULTISPECIES: hydrogenase-2 small chain [Proteobacteria] Length: 372\nNumber of predicted TMHs: 1\nExp number of AAs in TMHs: 23.72226\nExp number, first 60 AAs: 0.22038\nTotal prob of N-in: 0.12994\noutside 1 331\nTMhelix 332 354\ninside 355 372

7678 GCF_900015395.1_ED672_contigs Escherichia coli Proteobacteria; Gammaproteobacteria; Enterobacterales; Enterobacteriaceae; Escherichia
MNNEETFYQAMRRQGVTRRSFLKYCSLAATSLGLGAGMAPKIAWALENKPRIPVVWIHGL WP_001058323.1
MULTISPECIES: hydrogenase-1 small chain [Proteobacteria] Length: 372\nNumber of predicted TMHs: 2\nExp number of AAs in TMHs: 37.2608499999999\nExp number, first 60 AAs: 19.52923\nTotal prob of N-in: 0.94351\nPOSSIBLE N-term signal sequence\ninside 1 19\nTMhelix 20 39\noutside 40 325\nTMhelix 326 348\ninside 349 372

7679 GCF_900015755.1_ED040_contigs Escherichia coli Proteobacteria; Gammaproteobacteria; Enterobacterales; Enterobacteriaceae; Escherichia
MIGDNTLIHSHGINRRDFMKLCAALAATMGLSSKAAAEMAESVTNPQRPPVIWIGAQECT WP_000578724.1
hydrogenase 2 small subunit [Escherichia coli] Length: 372\nNumber of predicted TMHs: 1\nExp number of AAs in TMHs: 23.05601\nExp number, first 60 AAs: 0.15431\nTotal prob of N-in: 0.09116\noutside 1 331\nTMhelix 332 354\ninside 355 372

7680 GCF_900015755.1_ED040_contigs Escherichia coli Proteobacteria; Gammaproteobacteria; Enterobacterales; Enterobacteriaceae; Escherichia
MNNEETFYQAMRRQGVTRRSFLKYCSLAATSLGLGAGMAPKIAWALENKPRIPVVWIHGL WP_001301613.1
hydrogenase [Escherichia coli] Length: 372\nNumber of predicted TMHs: 2\nExp number of AAs in TMHs: 37.46598\nExp number, first 60 AAs: 19.53139\nTotal prob of N-in: 0.94364\nPOSSIBLE N-term signal sequence\ninside 1 19\nTMhelix 20 39\noutside 40 325\nTMhelix 326 348\ninside 349 372

7681 GCF_900015745.1_ED654_contigs Escherichia coli Proteobacteria; Gammaproteobacteria; Enterobacterales; Enterobacteriaceae; Escherichia
MNNEETFYQAMRRQGVTRRSFLKYCSLAATSLGLGAGMAPKIAWALENKPRIPVVWIHGL WP_001058323.1
MULTISPECIES: hydrogenase-1 small chain [Proteobacteria] Length: 372\nNumber of predicted TMHs: 2\nExp number of AAs in TMHs: 37.2608499999999\nExp number, first 60 AAs: 19.52923\nTotal prob of N-in:

0.94351\nPOSSIBLE N-term signal sequence\ninside 1 19\nTMhelix 20 39\noutside 40 325\nTMhelix 326 348\ninside 349 372

7682 GCF_900015875.1_ED766_contigs Escherichia coli Proteobacteria; Gammaproteobacteria; Enterobacterales; Enterobacteriaceae; Escherichia MTGDNTLIHSHGINRRDFMKLCAALAATMGLSSKAAAEMAESVTNPQRPPVIWIGAQECT WP_000145410.1 MULTISPECIES: hydrogenase-2 small chain [Proteobacteria] Length: 372\nNumber of predicted TMHs: 1\nExp number of AAs in TMHs: 23.72226\nExp number, first 60 AAs: 0.22038\nTotal prob of N-in: 0.12994\noutside 1 331\nTMhelix 332 354\ninside 355 372

7683 GCF_900015725.1_ED604_contigs Escherichia coli Proteobacteria; Gammaproteobacteria; Enterobacterales; Enterobacteriaceae; Escherichia MNNEETFYQAMRRQGVTRRSFLKYCSLAATSLGLGAGMAPKIAWALENKPRIPVVWIHGL WP_001301613.1 hydrogenase [Escherichia coli] Length: 372\nNumber of predicted TMHs: 2\nExp number of AAs in TMHs: 37.46598\nExp number, first 60 AAs: 19.53139\nTotal prob of N-in: 0.94364\nPOSSIBLE N-term signal sequence\ninside 1 19\nTMhelix 20 39\noutside 40 325\nTMhelix 326 348\ninside 349 372

7684 GCF_900015825.1_ED415_contigs Escherichia coli Proteobacteria; Gammaproteobacteria; Enterobacterales; Enterobacteriaceae; Escherichia MNNEETFYQAMRRQGVTRRSFLKYCSLAATSLGLGAGMAPKIAWALENKPRIPVVWIHGL WP_001301613.1 hydrogenase [Escherichia coli] Length: 372\nNumber of predicted TMHs: 2\nExp number of AAs in TMHs: 37.46598\nExp number, first 60 AAs: 19.53139\nTotal prob of N-in: 0.94364\nPOSSIBLE N-term signal sequence\ninside 1 19\nTMhelix 20 39\noutside 40 325\nTMhelix 326 348\ninside 349 372

7685 GCF_900015775.1_ED413_contigs Escherichia coli Proteobacteria; Gammaproteobacteria; Enterobacterales; Enterobacteriaceae; Escherichia MNNEETFYQAMRRQGVTRRSFLKYCSLAATSLGLGAGMAPKIAWALENKPRIPVVWIHGL WP_001301613.1 hydrogenase [Escherichia coli] Length: 372\nNumber of predicted TMHs: 2\nExp number of AAs in TMHs: 37.46598\nExp number, first 60 AAs: 19.53139\nTotal prob of N-in: 0.94364\nPOSSIBLE N-term signal sequence\ninside 1 19\nTMhelix 20 39\noutside 40 325\nTMhelix 326 348\ninside 349 372

7686 GCF_900015995.1_M858_contigs Escherichia coli Proteobacteria; Gammaproteobacteria; Enterobacterales; Enterobacteriaceae; Escherichia MTGDNTLIHSHGINRRDFMKLCAALAATMGLSSKAAAEMVESVTNPQRPPVIWIGAQECT WP_000145423.1 hydrogenase 2 small subunit [Escherichia coli] Length: 372\nNumber of predicted TMHs: 1\nExp number of AAs in TMHs: 23.78201\nExp number, first 60 AAs: 0.28355\nTotal prob of N-in: 0.13257\noutside 1 331\nTMhelix 332 354\ninside 355 372

7687 GCF_900015995.1_M858_contigs Escherichia coli Proteobacteria; Gammaproteobacteria; Enterobacterales; Enterobacteriaceae; Escherichia MNNEETFYQAMRRQGVTRRSFLKYCSLAATSLGLGAGMAPKIAWALENKPRIPVVWIHGL WP_001295940.1 MULTISPECIES: hydrogenase [Enterobacteriaceae] Length: 372\nNumber of predicted TMHs: 2\nExp number of AAs in TMHs: 37.2608899999999\nExp number, first 60 AAs: 19.52923\nTotal prob of N-in: 0.94351\nPOSSIBLE N-term signal sequence\ninside 1 19\nTMhelix 20 39\noutside 40 325\nTMhelix 326 348\ninside 349 372

7688 GCF_900015845.1_ED653_contigs Escherichia coli Proteobacteria; Gammaproteobacteria; Enterobacterales; Enterobacteriaceae; Escherichia MNNEETFYQAMRRQGVTRRSFLKYCSLAATSLGLGAGMAPKIAWALENKPRIPVVWIHGL WP_001301613.1 hydrogenase [Escherichia coli] Length: 372\nNumber of predicted TMHs: 2\nExp number of AAs in TMHs: 37.46598\nExp number, first 60 AAs: 19.53139\nTotal prob of N-in: 0.94364\nPOSSIBLE N-term signal sequence\ninside 1 19\nTMhelix 20 39\noutside 40 325\nTMhelix 326 348\ninside 349 372

7689 GCF_900015955.1_JF733 Escherichia coli Proteobacteria; Gammaproteobacteria; Enterobacterales; Enterobacteriaceae; Escherichia MTGDNTLIHSHGINRRDFMKLCAALAATMGLSSKAAAEMAESVTNPQRPPVIWIGAQECT WP_000145410.1 MULTISPECIES: hydrogenase-2 small chain [Proteobacteria] Length: 372\nNumber of predicted TMHs: 1\nExp number of AAs in TMHs: 23.72226\nExp number, first 60 AAs: 0.22038\nTotal prob of N-in: 0.12994\noutside 1 331\nTMhelix 332 354\ninside 355 372

7690 GCF_900016045.1_BCW5743_contigs Escherichia coli Proteobacteria; Gammaproteobacteria; Enterobacterales; Enterobacteriaceae; Escherichia

MNNEETFYQAMRRQGVTRRSFLKYCSLAATSLGLGAGMAPKIAWALENKPRIPVVWIHGL WP_001295940.1 MULTISPECIES: hydrogenase [Enterobacteriaceae] Length: 372\nNumber of predicted TMHs: 2\nExp number of AAs in TMHs: 37.2608899999999\nExp number, first 60 AAs: 19.52923\nTotal prob of N-in: 0.94351\nPOSSIBLE N-term signal sequence\ninside 1 19\nTMhelix 20 39\noutside 40 325\nTMhelix 326 348\ninside 349 372

7691 GCF_900042785.1_F1L4 Escherichia coli Proteobacteria; Gammaproteobacteria; Enterobacterales; Enterobacteriaceae; Escherichia MTGDNTLIHSHGINRRDFMKLCAALAATMGLSSKAAAEMAESVTNPQRPPVIWIGAQECT WP_000145410.1 MULTISPECIES: hydrogenase-2 small chain [Proteobacteria] Length: 372\nNumber of predicted TMHs: 1\nExp number of AAs in TMHs: 23.72226\nExp number, first 60 AAs: 0.22038\nTotal prob of N-in: 0.12994\noutside 1 331\nTMhelix 332 354\ninside 355 372

7692 GCF_900042785.1_F1L4 Escherichia coli Proteobacteria; Gammaproteobacteria; Enterobacterales; Enterobacteriaceae; Escherichia MNNEETFYQAMRRQGVTRRSFLKYCSLAATSLGLGAGMAPKIAWALENKPRIPVVWIHGL WP_001058320.1 hydrogenase [Escherichia coli] Length: 372\nNumber of predicted TMHs: 2\nExp number of AAs in TMHs: 37.26082\nExp number, first 60 AAs: 19.52923\nTotal prob of N-in: 0.94351\nPOSSIBLE N-term signal sequence\ninside 1 19\nTMhelix 20 39\noutside 40 325\nTMhelix 326 348\ninside 349 372

7693 GCF_900016365.1_M884_contigs Escherichia coli Proteobacteria; Gammaproteobacteria; Enterobacterales; Enterobacteriaceae; Escherichia MNNEETFYQAMRRQGVTRRSFLKYCSLAATSLGLGAGMAPKIAWALENKPRIPVVWIHGL WP_001058323.1 MULTISPECIES: hydrogenase-1 small chain [Proteobacteria] Length: 372\nNumber of predicted TMHs: 2\nExp number of AAs in TMHs: 37.2608499999999\nExp number, first 60 AAs: 19.52923\nTotal prob of N-in: 0.94351\nPOSSIBLE N-term signal sequence\ninside 1 19\nTMhelix 20 39\noutside 40 325\nTMhelix 326 348\ninside 349 372

7694 GCF_900015955.1_JF733 Escherichia coli Proteobacteria; Gammaproteobacteria; Enterobacterales; Enterobacteriaceae; Escherichia MNNEETFYQAMRRQGVTRRSFLKYCSLAATSLGLGAGMAPKIAWALENKPRIPVVWIHGL WP_001058323.1 MULTISPECIES: hydrogenase-1 small chain [Proteobacteria] Length: 372\nNumber of predicted TMHs: 2\nExp number of AAs in TMHs: 37.2608499999999\nExp number, first 60 AAs: 19.52923\nTotal prob of N-in: 0.94351\nPOSSIBLE N-term signal sequence\ninside 1 19\nTMhelix 20 39\noutside 40 325\nTMhelix 326 348\ninside 349 372

7695 GCF_900087565.2_ASM90008756v2 Escherichia coli Proteobacteria; Gammaproteobacteria; Enterobacterales; Enterobacteriaceae; Escherichia MTGDNTLIHSHGINRRDFMKLCAALAATMGLSSKAAAEMAESVTNPQRPPVIWIGAQECT WP_000145410.1 MULTISPECIES: hydrogenase-2 small chain [Proteobacteria] Length: 372\nNumber of predicted TMHs: 1\nExp number of AAs in TMHs: 23.72226\nExp number, first 60 AAs: 0.22038\nTotal prob of N-in: 0.12994\noutside 1 331\nTMhelix 332 354\ninside 355 372

7696 GCF_900088905.1_ASM90008890v1 Escherichia coli Proteobacteria; Gammaproteobacteria; Enterobacterales; Enterobacteriaceae; Escherichia MTGDNTLIHSHGINRRDFMKLCAALAATMGLSSKAAAEMAESVTNPQRPPVIWIGAQECT WP_000145410.1 MULTISPECIES: hydrogenase-2 small chain [Proteobacteria] Length: 372\nNumber of predicted TMHs: 1\nExp number of AAs in TMHs: 23.72226\nExp number, first 60 AAs: 0.22038\nTotal prob of N-in: 0.12994\noutside 1 331\nTMhelix 332 354\ninside 355 372

7697 GCF_900089055.1_ASM90008905v1 Escherichia coli Proteobacteria; Gammaproteobacteria; Enterobacterales; Enterobacteriaceae; Escherichia MTGDNTLIHSHGINRRDFMKLCAALAATMGLSSKAAAEMAESVTNPQRPPVIWIGAQECT WP_000145410.1 MULTISPECIES: hydrogenase-2 small chain [Proteobacteria] Length: 372\nNumber of predicted TMHs: 1\nExp number of AAs in TMHs: 23.72226\nExp number, first 60 AAs: 0.22038\nTotal prob of N-in: 0.12994\noutside 1 331\nTMhelix 332 354\ninside 355 372

7698 GCF_900087745.1_RL138 Escherichia coli Proteobacteria; Gammaproteobacteria; Enterobacterales; Enterobacteriaceae; Escherichia MNNEETFYQAMRRQGVTRRSFLKYCSLAATSLGLGAGMAPKIAWALENKPRIPVVWIHGL WP_001295940.1 MULTISPECIES: hydrogenase [Enterobacteriaceae] Length: 372\nNumber of predicted TMHs: 2\nExp number of AAs in TMHs: 37.2608899999999\nExp number, first 60 AAs: 19.52923\nTotal prob of N-in: 0.94351\nPOSSIBLE N-term signal sequence\ninside 1 19\nTMhelix 20 39\noutside 40 325\nTMhelix 326 348\ninside 349 372

7699 GCF_900088905.1_ASM90008890v1 Escherichia coli Proteobacteria; Gammaproteobacteria; Enterobacterales; Enterobacteriaceae; Escherichia MNNEETFYQAMRRQGVTRRSFLKYCSLAATSLGLGAGMAPKIAWALENKPRIPVVWIHGL WP_001295940.1 MULTISPECIES: hydrogenase [Enterobacteriaceae] Length: 372\nNumber of predicted TMHs: 2\nExp number of AAs in TMHs: 37.2608899999999\nExp number, first 60 AAs: 19.52923\nTotal prob of N-in: 0.94351\nPOSSIBLE N-term signal sequence\ninside 1 19\nTMhelix 20 39\noutside 40 325\nTMhelix 326 348\ninside 349 372

7700 GCF_900089055.1_ASM90008905v1 Escherichia coli Proteobacteria; Gammaproteobacteria; Enterobacterales; Enterobacteriaceae; Escherichia MNNEETFYQAMRRQGVTRRSFLKYCSLAATSLGLGAGMAPKIAWALENKPRIPVVWIHGL WP_001295940.1 MULTISPECIES: hydrogenase [Enterobacteriaceae] Length: 372\nNumber of predicted TMHs: 2\nExp number of AAs in TMHs: 37.2608899999999\nExp number, first 60 AAs: 19.52923\nTotal prob of N-in: 0.94351\nPOSSIBLE N-term signal sequence\ninside 1 19\nTMhelix 20 39\noutside 40 325\nTMhelix 326 348\ninside 349 372

7701 GCF_900016095.1_ED377_contigs Escherichia coli Proteobacteria; Gammaproteobacteria; Enterobacterales; Enterobacteriaceae; Escherichia MTGDNTLIHSHGINRRDFMKLCAALAATMGLSSKAAAEMAESVTNPQRPPVIWIGAQECT WP_074459508.1 hydrogenase 2 small subunit [Escherichia coli] Length: 372\nNumber of predicted TMHs: 1\nExp number of AAs in TMHs: 23.72079\nExp number, first 60 AAs: 0.22038\nTotal prob of N-in: 0.12990\noutside 1 331\nTMhelix 332 354\ninside 355 372

7702 GCF_900044035.1_F25P5 Escherichia coli Proteobacteria; Gammaproteobacteria; Enterobacterales; Enterobacteriaceae; Escherichia MTGDNTLIHSHGINRRDFMKLCAALAATMGLSSKAAAEMAESVTNPQRPPVIWIGAQECT WP_064674092.1 hydrogenase 2 small subunit [Escherichia coli] Length: 372\nNumber of predicted TMHs: 1\nExp number of AAs in TMHs: 22.99701\nExp number, first 60 AAs: 0.22059\nTotal prob of N-in: 0.12347\noutside 1 331\nTMhelix 332 354\ninside 355 372

7703 GCF_900093995.1_scaffolds_20160019 Escherichia coli Proteobacteria; Gammaproteobacteria; Enterobacterales; Enterobacteriaceae; Escherichia MNNEETFYQAMRRQGVTRRSFLKYCSLAATSLGLGAGMAPKIAWALENKPRIPVVWIHGL WP_001058323.1 MULTISPECIES: hydrogenase-1 small chain [Proteobacteria] Length: 372\nNumber of predicted TMHs: 2\nExp number of AAs in TMHs: 37.2608499999999\nExp number, first 60 AAs: 19.52923\nTotal prob of N-in: 0.94351\nPOSSIBLE N-term signal sequence\ninside 1 19\nTMhelix 20 39\noutside 40 325\nTMhelix 326 348\ninside 349 372

7704 GCF_900093865.1_scaffolds_C318 Escherichia coli Proteobacteria; Gammaproteobacteria; Enterobacterales; Enterobacteriaceae; Escherichia MNNEETFYQAMRRQGVTRRSFLKYCSLAATSLGLGAGMAPKIAWALENKPRIPVVWIHGL WP_001058323.1 MULTISPECIES: hydrogenase-1 small chain [Proteobacteria] Length: 372\nNumber of predicted TMHs: 2\nExp number of AAs in TMHs: 37.2608499999999\nExp number, first 60 AAs: 19.52923\nTotal prob of N-in: 0.94351\nPOSSIBLE N-term signal sequence\ninside 1 19\nTMhelix 20 39\noutside 40 325\nTMhelix 326 348\ninside 349 372

7705 GCF_900093915.1_scaffolds_20160020 Escherichia coli Proteobacteria; Gammaproteobacteria; Enterobacterales; Enterobacteriaceae; Escherichia MNNEETFYQAMRRQGVTRRSFLKYCSLAATSLGLGAGMAPKIAWALENKPRIPVVWIHGL WP_001058323.1 MULTISPECIES: hydrogenase-1 small chain [Proteobacteria] Length: 372\nNumber of predicted TMHs: 2\nExp number of AAs in TMHs: 37.2608499999999\nExp number, first 60 AAs: 19.52923\nTotal prob of N-in: 0.94351\nPOSSIBLE N-term signal sequence\ninside 1 19\nTMhelix 20 39\noutside 40 325\nTMhelix 326 348\ninside 349 372

7706 GCF_900093895.1_scaffolds_C281 Escherichia coli Proteobacteria; Gammaproteobacteria; Enterobacterales; Enterobacteriaceae; Escherichia MNNEETFYQAMRRQGVTRRSFLKYCSLAATSLGLGAGMAPKIAWALENKPRIPVVWIHGL WP_001058323.1 MULTISPECIES: hydrogenase-1 small chain [Proteobacteria] Length: 372\nNumber of predicted TMHs: 2\nExp number of AAs in TMHs: 37.2608499999999\nExp number, first 60 AAs: 19.52923\nTotal prob of N-in: 0.94351\nPOSSIBLE N-term signal sequence\ninside 1 19\nTMhelix 20 39\noutside 40 325\nTMhelix 326 348\ninside 349 372

7707 GCF_900092615.1_PRJEB14041 Escherichia coli Proteobacteria; Gammaproteobacteria; Enterobacterales; Enterobacteriaceae; Escherichia MTGDNTLIHSHGINRRDFMKLCAALAATMGLSSKAAAEMAESVTNPQRPPVIWIGAQECT WP_066009678.1 hydrogenase 2 small subunit [Escherichia coli] Length: 372\nNumber of predicted TMHs: 1\nExp number of AAs in TMHs: 23.72258\nExp number, first 60 AAs: 0.22053\nTotal prob of N-in: 0.12995\noutside 1 331\nTMhelix 332 354\ninside 355 372

7708 GCF_900094025.1_scaffolds_C413 Escherichia coli Proteobacteria; Gammaproteobacteria; Enterobacterales; Enterobacteriaceae; Escherichia MTGDNTLIHSHGINRRDFMKLCAALAATMGLSSKAAAEMAESVTNPQRPPVIWIGAQECT WP_000145410.1 MULTISPECIES: hydrogenase-2 small chain [Proteobacteria] Length: 372\nNumber of predicted TMHs: 1\nExp number of AAs in TMHs: 23.72226\nExp number, first 60 AAs: 0.22038\nTotal prob of N-in: 0.12994\noutside 1 331\nTMhelix 332 354\ninside 355 372

7709 GCF_900094025.1_scaffolds_C413 Escherichia coli Proteobacteria; Gammaproteobacteria; Enterobacterales; Enterobacteriaceae; Escherichia MNNEETFYQAMRRQGVTRRSFLKYCSLAATSLGLGAGMAPKIAWALENKPRIPVVWIHGL WP_001058323.1 MULTISPECIES: hydrogenase-1 small chain [Proteobacteria] Length: 372\nNumber of predicted TMHs: 2\nExp number of AAs in TMHs: 37.2608499999999\nExp number, first 60 AAs: 19.52923\nTotal prob of N-in: 0.94351\nPOSSIBLE N-term signal sequence\ninside 1 19\nTMhelix 20 39\noutside 40 325\nTMhelix 326 348\ninside 349 372

7710 GCF_900096845.1_Ecoli_AG100_Sample3_M9_Assembly Escherichia coli Proteobacteria; Gammaproteobacteria; Enterobacterales; Enterobacteriaceae; Escherichia MTGDNTLIHSHGINRRDFMKLCAALAATMGLSSKAAAEMAESVTNPQRPPVIWIGAQECT WP_000145410.1 MULTISPECIES: hydrogenase-2 small chain [Proteobacteria] Length: 372\nNumber of predicted TMHs: 1\nExp number of AAs in TMHs: 23.72226\nExp number, first 60 AAs: 0.22038\nTotal prob of N-in: 0.12994\noutside 1 331\nTMhelix 332 354\ninside 355 372

7711 GCF_900097005.1_K802 Escherichia coli Proteobacteria; Gammaproteobacteria; Enterobacterales; Enterobacteriaceae; Escherichia MTGDNTLIHSHGINRRDFMKLCAALAATMGLSSKAAAEMAESVTNPQRPPVIWIGAQECT WP_000145410.1 MULTISPECIES: hydrogenase-2 small chain [Proteobacteria] Length: 372\nNumber of predicted TMHs: 1\nExp number of AAs in TMHs: 23.72226\nExp number, first 60 AAs: 0.22038\nTotal prob of N-in: 0.12994\noutside 1 331\nTMhelix 332 354\ninside 355 372

7712 GCF_900108745.1_PEI-B-P-19 Escherichia coli Proteobacteria; Gammaproteobacteria; Enterobacterales; Enterobacteriaceae; Escherichia MTGDNTLIHSHGINRRDFMKLCAALAATMGLSSKAAAEMAESVTNPQRPPVIWIGAQECT WP_000145410.1 MULTISPECIES: hydrogenase-2 small chain [Proteobacteria] Length: 372\nNumber of predicted TMHs: 1\nExp number of AAs in TMHs: 23.72226\nExp number, first 60 AAs: 0.22038\nTotal prob of N-in: 0.12994\noutside 1 331\nTMhelix 332 354\ninside 355 372

7713 GCF_900119685.1_26578_F01 Escherichia coli Proteobacteria; Gammaproteobacteria; Enterobacterales; Enterobacteriaceae; Escherichia MTGDNTLIHSHGINRRDFMKLCAALAATMGLSSKAAAEMVESVTNPQRPPVIWIGAQECT WP_000145423.1 hydrogenase 2 small subunit [Escherichia coli] Length: 372\nNumber of predicted TMHs: 1\nExp number of AAs in TMHs: 23.78201\nExp number, first 60 AAs: 0.28355\nTotal prob of N-in: 0.13257\noutside 1 331\nTMhelix 332 354\ninside 355 372

7714 GCF_900097005.1_K802 Escherichia coli Proteobacteria; Gammaproteobacteria; Enterobacterales; Enterobacteriaceae; Escherichia MNNEETFYQAMRRQGVTRRSFLKYCSLAATSLGLGAGMAPKIAWALENKPRIPVVWIHGL WP_001058323.1 MULTISPECIES: hydrogenase-1 small chain [Proteobacteria] Length: 372\nNumber of predicted TMHs: 2\nExp number of AAs in TMHs: 37.2608499999999\nExp number, first 60 AAs: 19.52923\nTotal prob of N-in: 0.94351\nPOSSIBLE N-term signal sequence\ninside 1 19\nTMhelix 20 39\noutside 40 325\nTMhelix 326 348\ninside 349 372

7715 GCF_900096845.1_Ecoli_AG100_Sample3_M9_Assembly Escherichia coli Proteobacteria; Gammaproteobacteria; Enterobacterales; Enterobacteriaceae; Escherichia MNNEETFYQAMRRQGVTRRSFLKYCSLAATSLGLGAGMAPKIAWALENKPRIPVVWIHGL WP_001058323.1 MULTISPECIES: hydrogenase-1 small chain [Proteobacteria] Length: 372\nNumber of predicted TMHs: 2\nExp number of AAs in TMHs: 37.2608499999999\nExp number, first 60 AAs: 19.52923\nTotal prob of N-in:

0.94351\nPOSSIBLE N-term signal sequence\ninside 1 19\nTMhelix 20 39\noutside 40 325\nTMhelix 326 348\ninside 349 372

7716 GCF_900166705.1_Isolate_13 Escherichia coli Proteobacteria; Gammaproteobacteria; Enterobacterales; Enterobacteriaceae; Escherichia MTGDNTLIHSHGINRRDFMKLCAALAATMGLSSKAAAEMVESVTNPQRPPVIWIGAQECT WP_000145423.1 hydrogenase 2 small subunit [Escherichia coli] Length: 372\nNumber of predicted TMHs: 1\nExp number of AAs in TMHs: 23.78201\nExp number, first 60 AAs: 0.28355\nTotal prob of N-in: 0.13257\noutside 1 331\nTMhelix 332 354\ninside 355 372

7717 GCF_900166795.1_Isolate_12 Escherichia coli Proteobacteria; Gammaproteobacteria; Enterobacterales; Enterobacteriaceae; Escherichia MTGDNTLIHSHGINRRDFMKLCAALAATMGLSSKAAAEMVESVTNPQRPPVIWIGAQECT WP_000145423.1 hydrogenase 2 small subunit [Escherichia coli] Length: 372\nNumber of predicted TMHs: 1\nExp number of AAs in TMHs: 23.78201\nExp number, first 60 AAs: 0.28355\nTotal prob of N-in: 0.13257\noutside 1 331\nTMhelix 332 354\ninside 355 372

7718 GCF_900166825.1_Isolate_18 Escherichia coli Proteobacteria; Gammaproteobacteria; Enterobacterales; Enterobacteriaceae; Escherichia MTGDNTLIHSHGINRRDFMKLCAALAATMGLSSKAAAEMAESVTNPQRPPVIWIGAQECT WP_000145410.1 MULTISPECIES: hydrogenase-2 small chain [Proteobacteria] Length: 372\nNumber of predicted TMHs: 1\nExp number of AAs in TMHs: 23.72226\nExp number, first 60 AAs: 0.22038\nTotal prob of N-in: 0.12994\noutside 1 331\nTMhelix 332 354\ninside 355 372

7719 GCF_900166795.1_Isolate_12 Escherichia coli Proteobacteria; Gammaproteobacteria; Enterobacterales; Enterobacteriaceae; Escherichia MNNEETFYQAMRRQGVTRRSFLKYCSLAATSLGLGAGMAPKIAWALENKPRIPVVWIHGL WP_001295940.1 MULTISPECIES: hydrogenase [Enterobacteriaceae] Length: 372\nNumber of predicted TMHs: 2\nExp number of AAs in TMHs: 37.2608899999999\nExp number, first 60 AAs: 19.52923\nTotal prob of N-in: 0.94351\nPOSSIBLE N-term signal sequence\ninside 1 19\nTMhelix 20 39\noutside 40 325\nTMhelix 326 348\ninside 349 372

7720 GCF_900166825.1_Isolate_18 Escherichia coli Proteobacteria; Gammaproteobacteria; Enterobacterales; Enterobacteriaceae; Escherichia MNNEETFYQAMRRQGVTRRSFLKYCSLAATSLGLGAGMAPKIAWALENKPRIPVVWIHGL WP_001058323.1 MULTISPECIES: hydrogenase-1 small chain [Proteobacteria] Length: 372\nNumber of predicted TMHs: 2\nExp number of AAs in TMHs: 37.2608499999999\nExp number, first 60 AAs: 19.52923\nTotal prob of N-in: 0.94351\nPOSSIBLE N-term signal sequence\ninside 1 19\nTMhelix 20 39\noutside 40 325\nTMhelix 326 348\ninside 349 372

7721 GCF_900119685.1_26578_F01 Escherichia coli Proteobacteria; Gammaproteobacteria; Enterobacterales; Enterobacteriaceae; Escherichia MNNEETFYQAMRRQGVTRRSFLKYCSLAATSLGLGAGMAPKIAWALENKPRIPVVWIHGL WP_001295940.1 MULTISPECIES: hydrogenase [Enterobacteriaceae] Length: 372\nNumber of predicted TMHs: 2\nExp number of AAs in TMHs: 37.2608899999999\nExp number, first 60 AAs: 19.52923\nTotal prob of N-in: 0.94351\nPOSSIBLE N-term signal sequence\ninside 1 19\nTMhelix 20 39\noutside 40 325\nTMhelix 326 348\ninside 349 372

7722 GCF_900166865.1_Isolate_15 Escherichia coli Proteobacteria; Gammaproteobacteria; Enterobacterales; Enterobacteriaceae; Escherichia MTGDNTLIHSHGINRRDFMKLCAALAATMGLSSKAAAEMVESVTNPQRPPVIWIGAQECT WP_000145423.1 hydrogenase 2 small subunit [Escherichia coli] Length: 372\nNumber of predicted TMHs: 1\nExp number of AAs in TMHs: 23.78201\nExp number, first 60 AAs: 0.28355\nTotal prob of N-in: 0.13257\noutside 1 331\nTMhelix 332 354\ninside 355 372

7723 GCF_900166845.1_Isolate_21 Escherichia coli Proteobacteria; Gammaproteobacteria; Enterobacterales; Enterobacteriaceae; Escherichia MTGDNTLIHSHGINRRDFMKLCAALAATMGLSSKAAAEMAESVTNPQRPPVIWIGAQECT WP_000145410.1 MULTISPECIES: hydrogenase-2 small chain [Proteobacteria] Length: 372\nNumber of predicted TMHs: 1\nExp number of AAs in TMHs: 23.72226\nExp number, first 60 AAs: 0.22038\nTotal prob of N-in: 0.12994\noutside 1 331\nTMhelix 332 354\ninside 355 372

7724 GCF_900166865.1_Isolate_15 Escherichia coli Proteobacteria; Gammaproteobacteria; Enterobacterales; Enterobacteriaceae; Escherichia
MNNEETFYQAMRRQGVTRRSFLKYCSLAATSLGLGAGMAPKIAWALENKPRIPVVWIHGL WP_001295940.1
MULTISPECIES: hydrogenase [Enterobacteriaceae] Length: 372\nNumber of predicted TMHs: 2\nExp number of AAs in TMHs: 37.2608899999999\nExp number, first 60 AAs: 19.52923\nTotal prob of N-in: 0.94351\nPOSSIBLE N-term signal sequence\ninside 1 19\nTMhelix 20 39\noutside 40 325\nTMhelix 326 348\ninside 349 372

7725 GCF_000806175.1_Assembly_535 Escherichia coli Proteobacteria; Gammaproteobacteria; Enterobacterales; Enterobacteriaceae; Escherichia
MTGDNTLIHSHGINRRDFMKLCAALAATMGLSSKAAAEMAESVTNPQRPPVIWIGAQECT WP_000145410.1
MULTISPECIES: hydrogenase-2 small chain [Proteobacteria] Length: 372\nNumber of predicted TMHs: 1\nExp number of AAs in TMHs: 23.72226\nExp number, first 60 AAs: 0.22038\nTotal prob of N-in: 0.12994\noutside 1 331\nTMhelix 332 354\ninside 355 372

7726 GCF_000804345.1_ASM80434v1 Escherichia coli Proteobacteria; Gammaproteobacteria; Enterobacterales; Enterobacteriaceae; Escherichia
MTGDNTLIHSHGINRRDFMKLCAALAATMGLSSKAAAEMAESVTNPQRPPVIWIGAQECT WP_000145410.1
MULTISPECIES: hydrogenase-2 small chain [Proteobacteria] Length: 372\nNumber of predicted TMHs: 1\nExp number of AAs in TMHs: 23.72226\nExp number, first 60 AAs: 0.22038\nTotal prob of N-in: 0.12994\noutside 1 331\nTMhelix 332 354\ninside 355 372

7727 GCF_000805815.1_ASM80581v1 Escherichia coli Proteobacteria; Gammaproteobacteria; Enterobacterales; Enterobacteriaceae; Escherichia
MTGDNTLIHSHGINRRDFMKLCAALAATMGLSSKAAAEMAESVTNPQRPPVIWIGAQECT WP_000145410.1
MULTISPECIES: hydrogenase-2 small chain [Proteobacteria] Length: 372\nNumber of predicted TMHs: 1\nExp number of AAs in TMHs: 23.72226\nExp number, first 60 AAs: 0.22038\nTotal prob of N-in: 0.12994\noutside 1 331\nTMhelix 332 354\ninside 355 372

7728 GCF_000806275.1_ASM80627v1 Escherichia coli Proteobacteria; Gammaproteobacteria; Enterobacterales; Enterobacteriaceae; Escherichia
MTGDNTLIHSHGINRRDFMKLCAALAATMGLSSKAAAEMAESVTNPQRPPVIWIGAQECT WP_000145410.1
MULTISPECIES: hydrogenase-2 small chain [Proteobacteria] Length: 372\nNumber of predicted TMHs: 1\nExp number of AAs in TMHs: 23.72226\nExp number, first 60 AAs: 0.22038\nTotal prob of N-in: 0.12994\noutside 1 331\nTMhelix 332 354\ninside 355 372

7729 GCF_000807575.1_ASM80757v1 Escherichia coli Proteobacteria; Gammaproteobacteria; Enterobacterales; Enterobacteriaceae; Escherichia
MTGDNTLIHSHGINRRDFMKLCAALAATMGLSSKAAAEMAESVTNPQRPPVIWIGAQECT WP_000145410.1
MULTISPECIES: hydrogenase-2 small chain [Proteobacteria] Length: 372\nNumber of predicted TMHs: 1\nExp number of AAs in TMHs: 23.72226\nExp number, first 60 AAs: 0.22038\nTotal prob of N-in: 0.12994\noutside 1 331\nTMhelix 332 354\ninside 355 372

7730 GCF_000807575.1_ASM80757v1 Escherichia coli Proteobacteria; Gammaproteobacteria; Enterobacterales; Enterobacteriaceae; Escherichia
MNNEETFYQAMRRQGVTRRSFLKYCSLAATSLGLGAGMAPKIAWALENKPRIPVVWIHGL WP_001058323.1
MULTISPECIES: hydrogenase-1 small chain [Proteobacteria] Length: 372\nNumber of predicted TMHs: 2\nExp number of AAs in TMHs: 37.2608499999999\nExp number, first 60 AAs: 19.52923\nTotal prob of N-in: 0.94351\nPOSSIBLE N-term signal sequence\ninside 1 19\nTMhelix 20 39\noutside 40 325\nTMhelix 326 348\ninside 349 372

7731 GCF_000805815.1_ASM80581v1 Escherichia coli Proteobacteria; Gammaproteobacteria; Enterobacterales; Enterobacteriaceae; Escherichia
MNNEETFYQAMRRQGVTRRSFLKYCSLAATSLGLGAGMAPKIAWALENKPRIPVVWIHGL WP_001058323.1
MULTISPECIES: hydrogenase-1 small chain [Proteobacteria] Length: 372\nNumber of predicted TMHs: 2\nExp number of AAs in TMHs: 37.2608499999999\nExp number, first 60 AAs: 19.52923\nTotal prob of N-in: 0.94351\nPOSSIBLE N-term signal sequence\ninside 1 19\nTMhelix 20 39\noutside 40 325\nTMhelix 326 348\ninside 349 372

7732 GCF_000806175.1_Assembly_535 Escherichia coli Proteobacteria; Gammaproteobacteria; Enterobacterales; Enterobacteriaceae; Escherichia

MNNEETFYQAMRRQGVTRRSFLKYCSLAATSLGLGAGMAPKIAWALENKPRIPVVWIHGL WP_001058323.1 MULTISPECIES: hydrogenase-1 small chain [Proteobacteria] Length: 372\nNumber of predicted TMHs: 2\nExp number of AAs in TMHs: 37.2608499999999\nExp number, first 60 AAs: 19.52923\nTotal prob of N-in: 0.94351\nPOSSIBLE N-term signal sequence\ninside 1 19\nTMhelix 20 39\noutside 40 325\nTMhelix 326 348\ninside 349 372

7733 GCF_000804345.1_ASM80434v1 Escherichia coli Proteobacteria; Gammaproteobacteria; Enterobacterales; Enterobacteriaceae; Escherichia MNNEETFYQAMRRQGVTRRSFLKYCSLAATSLGLGAGMAPKIAWALENKPRIPVVWIHGL WP_001058320.1 hydrogenase [Escherichia coli] Length: 372\nNumber of predicted TMHs: 2\nExp number of AAs in TMHs: 37.26082\nExp number, first 60 AAs: 19.52923\nTotal prob of N-in: 0.94351\nPOSSIBLE N-term signal sequence\ninside 1 19\nTMhelix 20 39\noutside 40 325\nTMhelix 326 348\ninside 349 372

7734 GCF_000807655.1_ASM80765v1 Escherichia coli Proteobacteria; Gammaproteobacteria; Enterobacterales; Enterobacteriaceae; Escherichia MTGDNTLIHSHGINRRDFMKLCAALAATMGLSSKAAAEMAESVTNPQRPPVIWIGAQECT WP_000145410.1 MULTISPECIES: hydrogenase-2 small chain [Proteobacteria] Length: 372\nNumber of predicted TMHs: 1\nExp number of AAs in TMHs: 23.72226\nExp number, first 60 AAs: 0.22038\nTotal prob of N-in: 0.12994\noutside 1 331\nTMhelix 332 354\ninside 355 372

7735 GCF_000807655.1_ASM80765v1 Escherichia coli Proteobacteria; Gammaproteobacteria; Enterobacterales; Enterobacteriaceae; Escherichia MNNEETFYQAMRRQGVTRRSFLKYCSLAATSLGLGAGMAPKIAWALENKPRIPVVWIHGL WP_001058323.1 MULTISPECIES: hydrogenase-1 small chain [Proteobacteria] Length: 372\nNumber of predicted TMHs: 2\nExp number of AAs in TMHs: 37.2608499999999\nExp number, first 60 AAs: 19.52923\nTotal prob of N-in: 0.94351\nPOSSIBLE N-term signal sequence\ninside 1 19\nTMhelix 20 39\noutside 40 325\nTMhelix 326 348\ninside 349 372

7736 GCF_000806275.1_ASM80627v1 Escherichia coli Proteobacteria; Gammaproteobacteria; Enterobacterales; Enterobacteriaceae; Escherichia MNNEETFYQAMRRQGVTRRSFLKYCSLAATSLGLGAGMAPKIAWALENKPRIPVVWIHGL WP_001058323.1 MULTISPECIES: hydrogenase-1 small chain [Proteobacteria] Length: 372\nNumber of predicted TMHs: 2\nExp number of AAs in TMHs: 37.2608499999999\nExp number, first 60 AAs: 19.52923\nTotal prob of N-in: 0.94351\nPOSSIBLE N-term signal sequence\ninside 1 19\nTMhelix 20 39\noutside 40 325\nTMhelix 326 348\ninside 349 372

7737 GCF_000812255.1_ASM81225v1 Escherichia coli Proteobacteria; Gammaproteobacteria; Enterobacterales; Enterobacteriaceae; Escherichia MTGDNTLIHSHGINRRDFMKLCAALAATMGLSSKAAAEMAESVTNPQRPPVIWIGAQECT WP_000145410.1 MULTISPECIES: hydrogenase-2 small chain [Proteobacteria] Length: 372\nNumber of predicted TMHs: 1\nExp number of AAs in TMHs: 23.72226\nExp number, first 60 AAs: 0.22038\nTotal prob of N-in: 0.12994\noutside 1 331\nTMhelix 332 354\ninside 355 372

7738 GCF_000812325.1_ASM81232v1 Escherichia coli Proteobacteria; Gammaproteobacteria; Enterobacterales; Enterobacteriaceae; Escherichia MTGDNTLIHSHGINRRDFMKLCAALAATMGLSSKAAAEMAESVTNPQRPPVIWIGAQECT WP_000145410.1 MULTISPECIES: hydrogenase-2 small chain [Proteobacteria] Length: 372\nNumber of predicted TMHs: 1\nExp number of AAs in TMHs: 23.72226\nExp number, first 60 AAs: 0.22038\nTotal prob of N-in: 0.12994\noutside 1 331\nTMhelix 332 354\ninside 355 372

7739 GCF_000812385.1_ASM81238v1 Escherichia coli Proteobacteria; Gammaproteobacteria; Enterobacterales; Enterobacteriaceae; Escherichia MTGDNTLIHSHGINRRDFMKLCAALAATMGLSSKAAAEMAESVTNPQRPPVIWIGAQECT WP_000145410.1 MULTISPECIES: hydrogenase-2 small chain [Proteobacteria] Length: 372\nNumber of predicted TMHs: 1\nExp number of AAs in TMHs: 23.72226\nExp number, first 60 AAs: 0.22038\nTotal prob of N-in: 0.12994\noutside 1 331\nTMhelix 332 354\ninside 355 372

7740 GCF_000812705.1_ASM81270v1 Escherichia coli Proteobacteria; Gammaproteobacteria; Enterobacterales; Enterobacteriaceae; Escherichia MTGDNTLIHSHGINRRDFMKLCAALAATMGLSSKAAAEMAESVTNPQRPPVIWIGAQECT WP_000145410.1 MULTISPECIES: hydrogenase-2 small chain [Proteobacteria] Length: 372\nNumber of predicted TMHs:

1\nExp number of AAs in TMHs: 23.72226\nExp number, first 60 AAs: 0.22038\nTotal prob of N-in: 0.12994\noutside 1 331\nTMhelix 332 354\ninside 355 372

7741 GCF_000806255.1_ASM80625v1 Escherichia coli Proteobacteria; Gammaproteobacteria; Enterobacterales; Enterobacteriaceae; Escherichia MNNEETFYQAMRRQGVTRRSFLKYCSLAATSLGLGAGMAPKIVWALENKPRIPVVWIHGL WP_001398876.1 hydrogenase-1 small chain [Escherichia coli] Length: 372\nNumber of predicted TMHs: 2\nExp number of AAs in TMHs: 37.60268\nExp number, first 60 AAs: 19.84677\nTotal prob of N-in: 0.94621\nPOSSIBLE N-term signal sequence\ninside 1 20\nTMhelix 21 43\noutside 44 325\nTMhelix 326 348\ninside 349 372

7742 GCF_000806255.1_ASM80625v1 Escherichia coli Proteobacteria; Gammaproteobacteria; Enterobacterales; Enterobacteriaceae; Escherichia MTGDNTLIHSHGINRRDFMKLCAALAATMGLSSKAAAEMAESVTNPQRPPVIWIGAQECT WP_032190500.1 hydrogenase 2 small subunit [Escherichia coli] Length: 405\nNumber of predicted TMHs: 1\nExp number of AAs in TMHs: 34.53127\nExp number, first 60 AAs: 0.3257\nTotal prob of N-in: 0.21863\noutside 1 364\nTMhelix 365 387\ninside 388 405

7743 GCF_000812255.1_ASM81225v1 Escherichia coli Proteobacteria; Gammaproteobacteria; Enterobacterales; Enterobacteriaceae; Escherichia MNNEETFYQAMRRQGVTRRSFLKYCSLAATSLGLGAGMAPKIAWALENKPRIPVVWIHGL WP_001058323.1 MULTISPECIES: hydrogenase-1 small chain [Proteobacteria] Length: 372\nNumber of predicted TMHs: 2\nExp number of AAs in TMHs: 37.2608499999999\nExp number, first 60 AAs: 19.52923\nTotal prob of N-in: 0.94351\nPOSSIBLE N-term signal sequence\ninside 1 19\nTMhelix 20 39\noutside 40 325\nTMhelix 326 348\ninside 349 372

7744 GCF_000812725.1_ASM81272v1 Escherichia coli Proteobacteria; Gammaproteobacteria; Enterobacterales; Enterobacteriaceae; Escherichia MTGDNTLIHSHGINRRDFMKLCAALAATMGLSSKAAAEMAESVTNPQRPPVIWIGAQECT WP_000145410.1 MULTISPECIES: hydrogenase-2 small chain [Proteobacteria] Length: 372\nNumber of predicted TMHs: 1\nExp number of AAs in TMHs: 23.72226\nExp number, first 60 AAs: 0.22038\nTotal prob of N-in: 0.12994\noutside 1 331\nTMhelix 332 354\ninside 355 372

7745 GCF_000812725.1_ASM81272v1 Escherichia coli Proteobacteria; Gammaproteobacteria; Enterobacterales; Enterobacteriaceae; Escherichia MNNEETFYQAMRRQGVTRRSFLKYCSLAATSLGLGAGMAPKIAWALENKPRIPVVWIHGL WP_001058323.1 MULTISPECIES: hydrogenase-1 small chain [Proteobacteria] Length: 372\nNumber of predicted TMHs: 2\nExp number of AAs in TMHs: 37.2608499999999\nExp number, first 60 AAs: 19.52923\nTotal prob of N-in: 0.94351\nPOSSIBLE N-term signal sequence\ninside 1 19\nTMhelix 20 39\noutside 40 325\nTMhelix 326 348\ninside 349 372

7746 GCF_000812325.1_ASM81232v1 Escherichia coli Proteobacteria; Gammaproteobacteria; Enterobacterales; Enterobacteriaceae; Escherichia MNNEETFYQAMRRQGVTRRSFLKYCSLAATSLGLGAGMAPKIAWALENKPRIPVVWIHGL WP_001058323.1 MULTISPECIES: hydrogenase-1 small chain [Proteobacteria] Length: 372\nNumber of predicted TMHs: 2\nExp number of AAs in TMHs: 37.2608499999999\nExp number, first 60 AAs: 19.52923\nTotal prob of N-in: 0.94351\nPOSSIBLE N-term signal sequence\ninside 1 19\nTMhelix 20 39\noutside 40 325\nTMhelix 326 348\ninside 349 372

7747 GCF_000812785.1_ASM81278v1 Escherichia coli Proteobacteria; Gammaproteobacteria; Enterobacterales; Enterobacteriaceae; Escherichia MTGDNTLIHSHGINRRDFMKLCAALAATMGLSSKAAAEMAESVTNPQRPPVIWIGAQECT WP_000145410.1 MULTISPECIES: hydrogenase-2 small chain [Proteobacteria] Length: 372\nNumber of predicted TMHs: 1\nExp number of AAs in TMHs: 23.72226\nExp number, first 60 AAs: 0.22038\nTotal prob of N-in: 0.12994\noutside 1 331\nTMhelix 332 354\ninside 355 372

7748 GCF_000812785.1_ASM81278v1 Escherichia coli Proteobacteria; Gammaproteobacteria; Enterobacterales; Enterobacteriaceae; Escherichia MNNEETFYQAMRRQGVTRRSFLKYCSLAATSLGLGAGMAPKIAWALENKPRIPVVWIHGL WP_038994518.1 hydrogenase-1 small chain [Escherichia coli] Length: 372\nNumber of predicted TMHs: 2\nExp number of AAs in TMHs: 37.2608999999999\nExp number, first 60 AAs: 19.52922\nTotal prob of N-in: 0.94351\nPOSSIBLE N-term signal sequence\ninside 1 19\nTMhelix 20 39\noutside 40 325\nTMhelix 326 348\ninside 349 372

7749 GCF_000812705.1_ASM81270v1 Escherichia coli Proteobacteria; Gammaproteobacteria; Enterobacterales; Enterobacteriaceae; Escherichia
MNNEETFYQAMRRQGVTRRSFLKYCSLAATSLGLGAGMAPKIAWALENKPRIPVVWIHGL WP_001058323.1
MULTISPECIES: hydrogenase-1 small chain [Proteobacteria] Length: 372\nNumber of predicted TMHs: 2\nExp number of AAs in TMHs: 37.2608499999999\nExp number, first 60 AAs: 19.52923\nTotal prob of N-in: 0.94351\nPOSSIBLE N-term signal sequence\ninside 1 19\nTMhelix 20 39\noutside 40 325\nTMhelix 326 348\ninside 349 372

7750 GCF_000817425.1_Iso_15_A5 Escherichia coli Proteobacteria; Gammaproteobacteria; Enterobacterales; Enterobacteriaceae; Escherichia
MTGDNTLIHSHGINRRDFMKLCAALAATMGLSSKAAAEMAESVTNPQRPPVIWIGAQECT WP_000145410.1
MULTISPECIES: hydrogenase-2 small chain [Proteobacteria] Length: 372\nNumber of predicted TMHs: 1\nExp number of AAs in TMHs: 23.72226\nExp number, first 60 AAs: 0.22038\nTotal prob of N-in: 0.12994\noutside 1 331\nTMhelix 332 354\ninside 355 372

7751 GCF_000814145.2_ASM81414v2 Escherichia coli Proteobacteria; Gammaproteobacteria; Enterobacterales; Enterobacteriaceae; Escherichia
MTGDNTLIHSHGINRRDFMKLCAALAATMGLSSKAAAEMAESVTNPQRPPVIWIGAQECT WP_000145410.1
MULTISPECIES: hydrogenase-2 small chain [Proteobacteria] Length: 372\nNumber of predicted TMHs: 1\nExp number of AAs in TMHs: 23.72226\nExp number, first 60 AAs: 0.22038\nTotal prob of N-in: 0.12994\noutside 1 331\nTMhelix 332 354\ninside 355 372

7752 GCF_000817425.1_Iso_15_A5 Escherichia coli Proteobacteria; Gammaproteobacteria; Enterobacterales; Enterobacteriaceae; Escherichia
MNNEETFYQAMRRQGVTRRSFLKYCSLAATSLGLGAGMAPKIAWALENKPRIPVVWIHGL WP_001058323.1
MULTISPECIES: hydrogenase-1 small chain [Proteobacteria] Length: 372\nNumber of predicted TMHs: 2\nExp number of AAs in TMHs: 37.2608499999999\nExp number, first 60 AAs: 19.52923\nTotal prob of N-in: 0.94351\nPOSSIBLE N-term signal sequence\ninside 1 19\nTMhelix 20 39\noutside 40 325\nTMhelix 326 348\ninside 349 372

7753 GCF_000814145.2_ASM81414v2 Escherichia coli Proteobacteria; Gammaproteobacteria; Enterobacterales; Enterobacteriaceae; Escherichia
MNNEETFYQAMRRQGVTRRSFLKYCSLAATSLGLGAGMAPKIAWALENKPRIPVVWIHGL WP_001058323.1
MULTISPECIES: hydrogenase-1 small chain [Proteobacteria] Length: 372\nNumber of predicted TMHs: 2\nExp number of AAs in TMHs: 37.2608499999999\nExp number, first 60 AAs: 19.52923\nTotal prob of N-in: 0.94351\nPOSSIBLE N-term signal sequence\ninside 1 19\nTMhelix 20 39\noutside 40 325\nTMhelix 326 348\ninside 349 372

7754 GCF_000812385.1_ASM81238v1 Escherichia coli Proteobacteria; Gammaproteobacteria; Enterobacterales; Enterobacteriaceae; Escherichia
MNNEETFYQAMRRQGVTRRSFLKYCSLAATSLGLGAGMAPKIAWALENKPRIPVVWIHGL WP_001295940.1
MULTISPECIES: hydrogenase [Enterobacteriaceae] Length: 372\nNumber of predicted TMHs: 2\nExp number of AAs in TMHs: 37.2608899999999\nExp number, first 60 AAs: 19.52923\nTotal prob of N-in: 0.94351\nPOSSIBLE N-term signal sequence\ninside 1 19\nTMhelix 20 39\noutside 40 325\nTMhelix 326 348\ninside 349 372

7755 GCF_000819015.1_Escherichia_coli_CVM_N33978PS_v1.0 Escherichia coli Proteobacteria; Gammaproteobacteria; Enterobacterales; Enterobacteriaceae; Escherichia
MTGDNTLIHSHGINRRDFMKLCAALAATMGLSSKAAAEMAESVTNPQRPPVIWIGAQECT WP_000145410.1
MULTISPECIES: hydrogenase-2 small chain [Proteobacteria] Length: 372\nNumber of predicted TMHs: 1\nExp number of AAs in TMHs: 23.72226\nExp number, first 60 AAs: 0.22038\nTotal prob of N-in: 0.12994\noutside 1 331\nTMhelix 332 354\ninside 355 372

7756 GCF_000818985.1_Escherichia_coli_CVM_N33857PS_v1.0 Escherichia coli Proteobacteria; Gammaproteobacteria; Enterobacterales; Enterobacteriaceae; Escherichia
MTGDNTLIHSHGINRRDFMKLCAALAATMGLSSKAAAEMAESVTNPQRPPVIWIGAQECT WP_000145410.1
MULTISPECIES: hydrogenase-2 small chain [Proteobacteria] Length: 372\nNumber of predicted TMHs: 1\nExp number of AAs in TMHs: 23.72226\nExp number, first 60 AAs: 0.22038\nTotal prob of N-in: 0.12994\noutside 1 331\nTMhelix 332 354\ninside 355 372

7757 GCF_000818985.1_Escherichia_coli_CVM_N33857PS_v1.0 Escherichia coli Proteobacteria; Gammaproteobacteria; Enterobacterales; Enterobacteriaceae; Escherichia
MNNEETFYQAMRRQGVTRRSFLKYCSLAATSLGLGAGMAPKIAWALENKPRIPVVWIHGL WP_001058323.1
MULTISPECIES: hydrogenase-1 small chain [Proteobacteria] Length: 372\nNumber of predicted TMHs: 2\nExp number of AAs in TMHs: 37.2608499999999\nExp number, first 60 AAs: 19.52923\nTotal prob of N-in: 0.94351\nPOSSIBLE N-term signal sequence\ninside 1 19\nTMhelix 20 39\noutside 40 325\nTMhelix 326 348\ninside 349 372

7758 GCF_000819085.1_Escherichia_coli_CVM_N33624PS_v1.0 Escherichia coli Proteobacteria; Gammaproteobacteria; Enterobacterales; Enterobacteriaceae; Escherichia
MTGDNTLIHSHGINRRDFMKLCAALAATMGLSSKAAAEMAESVTNPQRPPVIWIGAQECT WP_000145410.1
MULTISPECIES: hydrogenase-2 small chain [Proteobacteria] Length: 372\nNumber of predicted TMHs: 1\nExp number of AAs in TMHs: 23.72226\nExp number, first 60 AAs: 0.22038\nTotal prob of N-in: 0.12994\noutside 1 331\nTMhelix 332 354\ninside 355 372

7759 GCF_000819145.1_Escherichia_coli_CVM_N34577PS_v1.0 Escherichia coli Proteobacteria; Gammaproteobacteria; Enterobacterales; Enterobacteriaceae; Escherichia
MTGDNTLIHSHGINRRDFMKLCAALAATMGLSSKAAAEMAESVTNPQRPPVIWIGAQECT WP_000145410.1
MULTISPECIES: hydrogenase-2 small chain [Proteobacteria] Length: 372\nNumber of predicted TMHs: 1\nExp number of AAs in TMHs: 23.72226\nExp number, first 60 AAs: 0.22038\nTotal prob of N-in: 0.12994\noutside 1 331\nTMhelix 332 354\ninside 355 372

7760 GCF_000819015.1_Escherichia_coli_CVM_N33978PS_v1.0 Escherichia coli Proteobacteria; Gammaproteobacteria; Enterobacterales; Enterobacteriaceae; Escherichia
MNNEETFYQAMRRQGVTRRSFLKYCSLAATSLGLGAGMAPKIAWALENKPRIPVVWIHGL WP_001058323.1
MULTISPECIES: hydrogenase-1 small chain [Proteobacteria] Length: 372\nNumber of predicted TMHs: 2\nExp number of AAs in TMHs: 37.2608499999999\nExp number, first 60 AAs: 19.52923\nTotal prob of N-in: 0.94351\nPOSSIBLE N-term signal sequence\ninside 1 19\nTMhelix 20 39\noutside 40 325\nTMhelix 326 348\ninside 349 372

7761 GCF_000819205.1_Escherichia_coli_CVM_N36210PS_v1.0 Escherichia coli Proteobacteria; Gammaproteobacteria; Enterobacterales; Enterobacteriaceae; Escherichia
MTGDNTLIHSHGINRRDFMKLCAALAATMGLSSKAAAEMAESVTNPQRPPVIWIGAQECT WP_000145410.1
MULTISPECIES: hydrogenase-2 small chain [Proteobacteria] Length: 372\nNumber of predicted TMHs: 1\nExp number of AAs in TMHs: 23.72226\nExp number, first 60 AAs: 0.22038\nTotal prob of N-in: 0.12994\noutside 1 331\nTMhelix 332 354\ninside 355 372

7762 GCF_000819245.1_Escherichia_coli_CVM_N37069PS_v1.0 Escherichia coli Proteobacteria; Gammaproteobacteria; Enterobacterales; Enterobacteriaceae; Escherichia
MTGDNTLIHSHGINRRDFMKLCAALAATMGLSSKAAAEMAESVTNPQRPPVIWIGAQECT WP_000145410.1
MULTISPECIES: hydrogenase-2 small chain [Proteobacteria] Length: 372\nNumber of predicted TMHs: 1\nExp number of AAs in TMHs: 23.72226\nExp number, first 60 AAs: 0.22038\nTotal prob of N-in: 0.12994\noutside 1 331\nTMhelix 332 354\ninside 355 372

7763 GCF_000819305.1_Escherichia_coli_CVM_N36099PS_v1.0 Escherichia coli Proteobacteria; Gammaproteobacteria; Enterobacterales; Enterobacteriaceae; Escherichia
MTGDNTLIHSHGINRRDFMKLCAALAATMGLSSKAAAEMAESVTNPQRPPVIWIGAQECT WP_000145410.1
MULTISPECIES: hydrogenase-2 small chain [Proteobacteria] Length: 372\nNumber of predicted TMHs: 1\nExp number of AAs in TMHs: 23.72226\nExp number, first 60 AAs: 0.22038\nTotal prob of N-in: 0.12994\noutside 1 331\nTMhelix 332 354\ninside 355 372

7764 GCF_000819205.1_Escherichia_coli_CVM_N36210PS_v1.0 Escherichia coli Proteobacteria; Gammaproteobacteria; Enterobacterales; Enterobacteriaceae; Escherichia
MNNEETFYQAMRRQGVTRRSFLKYCSLAATSLGLGAGMAPKIAWALENKPRIPVVWIHGL WP_001058323.1
MULTISPECIES: hydrogenase-1 small chain [Proteobacteria] Length: 372\nNumber of predicted TMHs: 2\nExp number of AAs in TMHs: 37.2608499999999\nExp number, first 60 AAs: 19.52923\nTotal prob of N-in: 0.94351\nPOSSIBLE N-term signal sequence\ninside 1 19\nTMhelix 20 39\noutside 40 325\nTMhelix 326 348\ninside 349 372

7765 GCF_000819245.1_Escherichia_coli_CVM_N37069PS_v1.0 Escherichia coli Proteobacteria; Gammaproteobacteria; Enterobacterales; Enterobacteriaceae; Escherichia

MNNEETFYQAMRRQGVTRRSFLKYCSLAATSLGLGAGMAPKIAWALENKPRIPVVWIHGL WP_001058323.1 MULTISPECIES: hydrogenase-1 small chain [Proteobacteria] Length: 372\nNumber of predicted TMHs: 2\nExp number of AAs in TMHs: 37.2608499999999\nExp number, first 60 AAs: 19.52923\nTotal prob of N-in: 0.94351\nPOSSIBLE N-term signal sequence\ninside 1 19\nTMhelix 20 39\noutside 40 325\nTMhelix 326 348\ninside 349 372

7766 GCF_000819305.1_Escherichia_coli_CVM_N36099PS_v1.0 Escherichia coli Proteobacteria; Gammaproteobacteria; Enterobacterales; Enterobacteriaceae; Escherichia MNNEETFYQAMRRQGVTRRSFLKYCSLAATSLGLGAGMAPKIAWALENKPRIPVVWIHGL WP_001058323.1 MULTISPECIES: hydrogenase-1 small chain [Proteobacteria] Length: 372\nNumber of predicted TMHs: 2\nExp number of AAs in TMHs: 37.2608499999999\nExp number, first 60 AAs: 19.52923\nTotal prob of N-in: 0.94351\nPOSSIBLE N-term signal sequence\ninside 1 19\nTMhelix 20 39\noutside 40 325\nTMhelix 326 348\ninside 349 372

7767 GCF_000819085.1_Escherichia_coli_CVM_N33624PS_v1.0 Escherichia coli Proteobacteria; Gammaproteobacteria; Enterobacterales; Enterobacteriaceae; Escherichia MNNEETFYQAMRRQGVTRRSFLKYCSLAATSLGLGAGMAPKIAWALENKPRIPVVWIHGL WP_001058323.1 MULTISPECIES: hydrogenase-1 small chain [Proteobacteria] Length: 372\nNumber of predicted TMHs: 2\nExp number of AAs in TMHs: 37.2608499999999\nExp number, first 60 AAs: 19.52923\nTotal prob of N-in: 0.94351\nPOSSIBLE N-term signal sequence\ninside 1 19\nTMhelix 20 39\noutside 40 325\nTMhelix 326 348\ninside 349 372

7768 GCF_000819145.1_Escherichia_coli_CVM_N34577PS_v1.0 Escherichia coli Proteobacteria; Gammaproteobacteria; Enterobacterales; Enterobacteriaceae; Escherichia MNNEETFYQAMRRQGVTRRSFLKYCSLAATSLGLGAGMAPKIAWALENKPRIPVVWIHGL WP_042965784.1 hydrogenase [Escherichia coli] Length: 372\nNumber of predicted TMHs: 2\nExp number of AAs in TMHs: 37.2608099999999\nExp number, first 60 AAs: 19.52923\nTotal prob of N-in: 0.94351\nPOSSIBLE N-term signal sequence\ninside 1 19\nTMhelix 20 39\noutside 40 325\nTMhelix 326 348\ninside 349 372

7769 GCF_000987545.1_ASM98754v1 Escherichia coli Proteobacteria; Gammaproteobacteria; Enterobacterales; Enterobacteriaceae; Escherichia MTGDNTLIHSHGINRRDFMKLCAALAATMGLSSKAAAEMAESVTNPQRPPVIWIGAQECT WP_000145410.1 MULTISPECIES: hydrogenase-2 small chain [Proteobacteria] Length: 372\nNumber of predicted TMHs: 1\nExp number of AAs in TMHs: 23.72226\nExp number, first 60 AAs: 0.22038\nTotal prob of N-in: 0.12994\noutside 1 331\nTMhelix 332 354\ninside 355 372

7770 GCF_000987875.1_ASM98787v1 Escherichia coli Proteobacteria; Gammaproteobacteria; Enterobacterales; Enterobacteriaceae; Escherichia MTGDNTLIHSHGINRRDFMKLCAALAATMGLSSKAAAEMAESVTNPQRPPVIWIGAQECT WP_000145410.1 MULTISPECIES: hydrogenase-2 small chain [Proteobacteria] Length: 372\nNumber of predicted TMHs: 1\nExp number of AAs in TMHs: 23.72226\nExp number, first 60 AAs: 0.22038\nTotal prob of N-in: 0.12994\noutside 1 331\nTMhelix 332 354\ninside 355 372

7771 GCF_000988465.1_ASM98846v1 Escherichia coli Proteobacteria; Gammaproteobacteria; Enterobacterales; Enterobacteriaceae; Escherichia MTGDNTLIHSHGINRRDFMKLCAALAATMGLSSKAAAEMAESVTNPQRPPVIWIGAQECT WP_000145410.1 MULTISPECIES: hydrogenase-2 small chain [Proteobacteria] Length: 372\nNumber of predicted TMHs: 1\nExp number of AAs in TMHs: 23.72226\nExp number, first 60 AAs: 0.22038\nTotal prob of N-in: 0.12994\noutside 1 331\nTMhelix 332 354\ninside 355 372

7772 GCF_000987545.1_ASM98754v1 Escherichia coli Proteobacteria; Gammaproteobacteria; Enterobacterales; Enterobacteriaceae; Escherichia MNNEETFYQAMRRQGVTRRSFLKYCSLAATSLGLGAGMAPKIAWALENKPRIPVVWIHGL WP_001058320.1 hydrogenase [Escherichia coli] Length: 372\nNumber of predicted TMHs: 2\nExp number of AAs in TMHs: 37.26082\nExp number, first 60 AAs: 19.52923\nTotal prob of N-in: 0.94351\nPOSSIBLE N-term signal sequence\ninside 1 19\nTMhelix 20 39\noutside 40 325\nTMhelix 326 348\ninside 349 372

7773 GCF_001010165.1_ASM101016v1 Escherichia coli Proteobacteria; Gammaproteobacteria; Enterobacterales; Enterobacteriaceae; Escherichia MTGDNTLIHSHGINRRDFMKLCAALAATMGLSSKAAAEMAESVTNPQRPPVIWIGAQECT WP_000145410.1 MULTISPECIES: hydrogenase-2 small chain [Proteobacteria] Length: 372\nNumber of predicted TMHs:

1\nExp number of AAs in TMHs: 23.72226\nExp number, first 60 AAs: 0.22038\nTotal prob of N-in: 0.12994\noutside 1 331\nTMhelix 332 354\ninside 355 372

7774 GCF_001010195.1_ASM101019v1 Escherichia coli Proteobacteria; Gammaproteobacteria; Enterobacterales; Enterobacteriaceae; Escherichia
MTGDNTLIHSHGINRRDFMKLCAALAATMGLSSKAAAEMVESVTNPQRPPVIWIGAQECT WP_000145423.1
hydrogenase 2 small subunit [Escherichia coli] Length: 372\nNumber of predicted TMHs: 1\nExp number of AAs in TMHs: 23.78201\nExp number, first 60 AAs: 0.28355\nTotal prob of N-in: 0.13257\noutside 1 331\nTMhelix 332 354\ninside 355 372

7775 GCF_001012005.1_CFSAN026773_02.0Escherichia coli Proteobacteria; Gammaproteobacteria; Enterobacterales; Enterobacteriaceae; Escherichia
MTGDNTLIHSHGINRRDFMKLCAALAATMGLSSKAAAEMAESVTNPQRPPVIWIGAQECT WP_000145410.1
MULTISPECIES: hydrogenase-2 small chain [Proteobacteria] Length: 372\nNumber of predicted TMHs: 1\nExp number of AAs in TMHs: 23.72226\nExp number, first 60 AAs: 0.22038\nTotal prob of N-in: 0.12994\noutside 1 331\nTMhelix 332 354\ninside 355 372

7776 GCF_001010195.1_ASM101019v1 Escherichia coli Proteobacteria; Gammaproteobacteria; Enterobacterales; Enterobacteriaceae; Escherichia
MNNEETFYQAMRRQGVTRRSFLKYCSLAATSLGLGAGMAPKIAWALENKPRIPVVWIHGL WP_001058323.1
MULTISPECIES: hydrogenase-1 small chain [Proteobacteria] Length: 372\nNumber of predicted TMHs: 2\nExp number of AAs in TMHs: 37.2608499999999\nExp number, first 60 AAs: 19.52923\nTotal prob of N-in: 0.94351\nPOSSIBLE N-term signal sequence\ninside 1 19\nTMhelix 20 39\noutside 40 325\nTMhelix 326 348\ninside 349 372

7777 GCF_001012005.1_CFSAN026773_02.0Escherichia coli Proteobacteria; Gammaproteobacteria; Enterobacterales; Enterobacteriaceae; Escherichia
MNNEETFYQAMRRQGVTRRSFLKYCSLAATSLGLGAGMAPKIAWALENKPRIPVVWIHGL WP_001498899.1
hydrogenase-1 small chain [Escherichia coli] Length: 372\nNumber of predicted TMHs: 2\nExp number of AAs in TMHs: 39.34459\nExp number, first 60 AAs: 19.53328\nTotal prob of N-in: 0.94350\nPOSSIBLE N-term signal sequence\ninside 1 19\nTMhelix 20 39\noutside 40 325\nTMhelix 326 348\ninside 349 372

7778 GCF_000988465.1_ASM98846v1 Escherichia coli Proteobacteria; Gammaproteobacteria; Enterobacterales; Enterobacteriaceae; Escherichia
MNNEETFYQAMRRQGVTRRSFLKYCSLAATSLGLGAGMAPKIAWALENKPRIPVVWIHGL WP_001058323.1
MULTISPECIES: hydrogenase-1 small chain [Proteobacteria] Length: 372\nNumber of predicted TMHs: 2\nExp number of AAs in TMHs: 37.2608499999999\nExp number, first 60 AAs: 19.52923\nTotal prob of N-in: 0.94351\nPOSSIBLE N-term signal sequence\ninside 1 19\nTMhelix 20 39\noutside 40 325\nTMhelix 326 348\ninside 349 372

7779 GCF_001012075.1_CFSAN026805_01.0Escherichia coli Proteobacteria; Gammaproteobacteria; Enterobacterales; Enterobacteriaceae; Escherichia
MTGDNTLIHSHGINRRDFMKLCAALAATMGLSSKAAAEMAESVTNPQRPPVIWIGAQECT WP_000145410.1
MULTISPECIES: hydrogenase-2 small chain [Proteobacteria] Length: 372\nNumber of predicted TMHs: 1\nExp number of AAs in TMHs: 23.72226\nExp number, first 60 AAs: 0.22038\nTotal prob of N-in: 0.12994\noutside 1 331\nTMhelix 332 354\ninside 355 372

7780 GCF_001012075.1_CFSAN026805_01.0Escherichia coli Proteobacteria; Gammaproteobacteria; Enterobacterales; Enterobacteriaceae; Escherichia
MNNEETFYQAMRRQGVTRRSFLKYCSLAATSLGLGAGMAPKIAWALENKPRIPVVWIHGL WP_001058323.1
MULTISPECIES: hydrogenase-1 small chain [Proteobacteria] Length: 372\nNumber of predicted TMHs: 2\nExp number of AAs in TMHs: 37.2608499999999\nExp number, first 60 AAs: 19.52923\nTotal prob of N-in: 0.94351\nPOSSIBLE N-term signal sequence\ninside 1 19\nTMhelix 20 39\noutside 40 325\nTMhelix 326 348\ninside 349 372

7781 GCF_000987875.1_ASM98787v1 Escherichia coli Proteobacteria; Gammaproteobacteria; Enterobacterales; Enterobacteriaceae; Escherichia
MNNEETFYQAMRRQGVTRRSFLKYCSLAATSLGLGAGMAPKIAWALENKPRIPVVWIHGL WP_014639883.1
hydrogenase [Escherichia coli] Length: 372\nNumber of predicted TMHs: 2\nExp number of AAs in TMHs: 37.26086\nExp number, first 60 AAs: 19.52923\nTotal prob of N-in: 0.94351\nPOSSIBLE N-term signal sequence\ninside 1 19\nTMhelix 20 39\noutside 40 325\nTMhelix 326 348\ninside 349 372

7782 GCF_001012095.1_CFSAN026810_01.0 Escherichia coli Proteobacteria; Gammaproteobacteria; Enterobacterales; Enterobacteriaceae; Escherichia
MTGDNTLIHSHGINRRDFMKLCAALAATMGLSSKAAAEMAESVTNPQRPPVIWIGAQECT WP_000145410.1
MULTISPECIES: hydrogenase-2 small chain [Proteobacteria] Length: 372\nNumber of predicted TMHs: 1\nExp number of AAs in TMHs: 23.72226\nExp number, first 60 AAs: 0.22038\nTotal prob of N-in: 0.12994\noutside 1 331\nTMhelix 332 354\ninside 355 372

7783 GCF_001012095.1_CFSAN026810_01.0 Escherichia coli Proteobacteria; Gammaproteobacteria; Enterobacterales; Enterobacteriaceae; Escherichia
MNNEETFYQAMRRQGVTRRSFLKYCSLAATSLGLGAGMAPKIAWALENKPRIPVVWIHGL WP_001295940.1
MULTISPECIES: hydrogenase [Enterobacteriaceae] Length: 372\nNumber of predicted TMHs: 2\nExp number of AAs in TMHs: 37.2608899999999\nExp number, first 60 AAs: 19.52923\nTotal prob of N-in: 0.94351\nPOSSIBLE N-term signal sequence\ninside 1 19\nTMhelix 20 39\noutside 40 325\nTMhelix 326 348\ninside 349 372

7784 GCF_001012165.1_CFSAN026785_02.0 Escherichia coli Proteobacteria; Gammaproteobacteria; Enterobacterales; Enterobacteriaceae; Escherichia
MTGDNTLIHSHGINRRDFMKLCAALAATMGLSSKAAAEMAESVTNPQRPPVIWIGAQECT WP_000145410.1
MULTISPECIES: hydrogenase-2 small chain [Proteobacteria] Length: 372\nNumber of predicted TMHs: 1\nExp number of AAs in TMHs: 23.72226\nExp number, first 60 AAs: 0.22038\nTotal prob of N-in: 0.12994\noutside 1 331\nTMhelix 332 354\ninside 355 372

7785 GCF_001012235.1_CFSAN026789_01.0 Escherichia coli Proteobacteria; Gammaproteobacteria; Enterobacterales; Enterobacteriaceae; Escherichia
MTGDNTLIHSHGINRRDFMKLCAALAATMGLSSKAAAEMAESVTNPQRPPVIWIGAQECT WP_000145410.1
MULTISPECIES: hydrogenase-2 small chain [Proteobacteria] Length: 372\nNumber of predicted TMHs: 1\nExp number of AAs in TMHs: 23.72226\nExp number, first 60 AAs: 0.22038\nTotal prob of N-in: 0.12994\noutside 1 331\nTMhelix 332 354\ninside 355 372

7786 GCF_001012265.1_CFSAN026839_01.0 Escherichia coli Proteobacteria; Gammaproteobacteria; Enterobacterales; Enterobacteriaceae; Escherichia
MTGDNTLIHSHGINRRDFMKLCAALAATMGLSSKAAAEMAESVTNPQRPPVIWIGAQECT WP_000145410.1
MULTISPECIES: hydrogenase-2 small chain [Proteobacteria] Length: 372\nNumber of predicted TMHs: 1\nExp number of AAs in TMHs: 23.72226\nExp number, first 60 AAs: 0.22038\nTotal prob of N-in: 0.12994\noutside 1 331\nTMhelix 332 354\ninside 355 372

7787 GCF_001012355.1_CFSAN026812_01.0 Escherichia coli Proteobacteria; Gammaproteobacteria; Enterobacterales; Enterobacteriaceae; Escherichia
MTGDNTLIHSHGINRRDFMKLCAALAATMGLSSKAAAEMAESVTNPQRPPVIWIGAQECT WP_000145410.1
MULTISPECIES: hydrogenase-2 small chain [Proteobacteria] Length: 372\nNumber of predicted TMHs: 1\nExp number of AAs in TMHs: 23.72226\nExp number, first 60 AAs: 0.22038\nTotal prob of N-in: 0.12994\noutside 1 331\nTMhelix 332 354\ninside 355 372

7788 GCF_001012435.1_CFSAN026828_01.0 Escherichia coli Proteobacteria; Gammaproteobacteria; Enterobacterales; Enterobacteriaceae; Escherichia
MTGDNTLIHSHGINRRDFMKLCAALAATMGLSSKAAAEMAESVTNPQRPPVIWIGAQECT WP_000145410.1
MULTISPECIES: hydrogenase-2 small chain [Proteobacteria] Length: 372\nNumber of predicted TMHs: 1\nExp number of AAs in TMHs: 23.72226\nExp number, first 60 AAs: 0.22038\nTotal prob of N-in: 0.12994\noutside 1 331\nTMhelix 332 354\ninside 355 372

7789 GCF_001012475.1_CFSAN026830_01.0 Escherichia coli Proteobacteria; Gammaproteobacteria; Enterobacterales; Enterobacteriaceae; Escherichia
MTGDNTLIHSHGINRRDFMKLCAALAATMGLSSKAAAEMAESVTNPQRPPVIWIGAQECT WP_000145410.1
MULTISPECIES: hydrogenase-2 small chain [Proteobacteria] Length: 372\nNumber of predicted TMHs: 1\nExp number of AAs in TMHs: 23.72226\nExp number, first 60 AAs: 0.22038\nTotal prob of N-in: 0.12994\noutside 1 331\nTMhelix 332 354\ninside 355 372

7790 GCF_001052755.1_ASM105275v1 Escherichia coli Proteobacteria; Gammaproteobacteria; Enterobacterales; Enterobacteriaceae; Escherichia
MTGDNTLIHSHGINRRDFMKLCAALAATMGLSSKAAAEMAESVTNPQRPPVIWIGAQECT WP_000145410.1
MULTISPECIES: hydrogenase-2 small chain [Proteobacteria] Length: 372\nNumber of predicted TMHs:

1\nExp number of AAs in TMHs: 23.72226\nExp number, first 60 AAs: 0.22038\nTotal prob of N-in: 0.12994\noutside 1 331\nTMhelix 332 354\ninside 355 372

7791 GCF_001012235.1_CFSAN026789_01.0 Escherichia coli Proteobacteria; Gammaproteobacteria; Enterobacterales; Enterobacteriaceae; Escherichia
MNNEETFYQAMRRQGVTRRSFLKYCSLAATSLGLGAGMAPKIAWALENKPRIPVVWIHGL WP_001058323.1
MULTISPECIES: hydrogenase-1 small chain [Proteobacteria] Length: 372\nNumber of predicted TMHs: 2\nExp number of AAs in TMHs: 37.2608499999999\nExp number, first 60 AAs: 19.52923\nTotal prob of N-in: 0.94351\nPOSSIBLE N-term signal sequence\ninside 1 19\nTMhelix 20 39\noutside 40 325\nTMhelix 326 348\ninside 349 372

7792 GCF_001012355.1_CFSAN026812_01.0 Escherichia coli Proteobacteria; Gammaproteobacteria; Enterobacterales; Enterobacteriaceae; Escherichia
MNNEETFYQAMRRQGVTRRSFLKYCSLAATSLGLGAGMAPKIAWALENKPRIPVVWIHGL WP_001058323.1
MULTISPECIES: hydrogenase-1 small chain [Proteobacteria] Length: 372\nNumber of predicted TMHs: 2\nExp number of AAs in TMHs: 37.2608499999999\nExp number, first 60 AAs: 19.52923\nTotal prob of N-in: 0.94351\nPOSSIBLE N-term signal sequence\ninside 1 19\nTMhelix 20 39\noutside 40 325\nTMhelix 326 348\ninside 349 372

7793 GCF_001012165.1_CFSAN026785_02.0 Escherichia coli Proteobacteria; Gammaproteobacteria; Enterobacterales; Enterobacteriaceae; Escherichia
MNNEETFYQAMRRQGVTRRSFLKYCSLAATSLGLGAGMAPKIAWALENKPRIPVVWIHGL WP_001295940.1
MULTISPECIES: hydrogenase [Enterobacteriaceae] Length: 372\nNumber of predicted TMHs: 2\nExp number of AAs in TMHs: 37.2608899999999\nExp number, first 60 AAs: 19.52923\nTotal prob of N-in: 0.94351\nPOSSIBLE N-term signal sequence\ninside 1 19\nTMhelix 20 39\noutside 40 325\nTMhelix 326 348\ninside 349 372

7794 GCF_001012535.1_CFSAN026842_02.0 Escherichia coli Proteobacteria; Gammaproteobacteria; Enterobacterales; Enterobacteriaceae; Escherichia
MTGDNTLIHSHGINRRDFMKLCAALAATMGLSSKAAAEMAESVTNPQRPPVIWIGAQECT WP_000145410.1
MULTISPECIES: hydrogenase-2 small chain [Proteobacteria] Length: 372\nNumber of predicted TMHs: 1\nExp number of AAs in TMHs: 23.72226\nExp number, first 60 AAs: 0.22038\nTotal prob of N-in: 0.12994\noutside 1 331\nTMhelix 332 354\ninside 355 372

7795 GCF_001012335.1_CFSAN026802_01.0 Escherichia coli Proteobacteria; Gammaproteobacteria; Enterobacterales; Enterobacteriaceae; Escherichia
MNNEETFYQAMRRQGVTRRSFLKYCSLAATSLGLGAGMAPKIAWALENKPRIPVVWIHGL WP_001058323.1
MULTISPECIES: hydrogenase-1 small chain [Proteobacteria] Length: 372\nNumber of predicted TMHs: 2\nExp number of AAs in TMHs: 37.2608499999999\nExp number, first 60 AAs: 19.52923\nTotal prob of N-in: 0.94351\nPOSSIBLE N-term signal sequence\ninside 1 19\nTMhelix 20 39\noutside 40 325\nTMhelix 326 348\ninside 349 372

7796 GCF_001012535.1_CFSAN026842_02.0 Escherichia coli Proteobacteria; Gammaproteobacteria; Enterobacterales; Enterobacteriaceae; Escherichia
MNNEETFYQAMRRQGVTRRSFLKYCSLAATSLGLGAGMAPKIAWALENKPRIPVVWIHGL WP_001058323.1
MULTISPECIES: hydrogenase-1 small chain [Proteobacteria] Length: 372\nNumber of predicted TMHs: 2\nExp number of AAs in TMHs: 37.2608499999999\nExp number, first 60 AAs: 19.52923\nTotal prob of N-in: 0.94351\nPOSSIBLE N-term signal sequence\ninside 1 19\nTMhelix 20 39\noutside 40 325\nTMhelix 326 348\ninside 349 372

7797 GCF_001012335.1_CFSAN026802_01.0 Escherichia coli Proteobacteria; Gammaproteobacteria; Enterobacterales; Enterobacteriaceae; Escherichia
MTGDNTLIHSHGINRRDFMKLCAALAATMGLSSKAAAEMAESVTNPQRPPVIWIGAQECT WP_047086748.1
hydrogenase 2 small subunit [Escherichia coli] Length: 372\nNumber of predicted TMHs: 1\nExp number of AAs in TMHs: 24.06887\nExp number, first 60 AAs: 0.21829\nTotal prob of N-in: 0.14344\noutside 1 331\nTMhelix 332 354\ninside 355 372

7798 GCF_001012265.1_CFSAN026839_01.0 Escherichia coli Proteobacteria; Gammaproteobacteria; Enterobacterales; Enterobacteriaceae; Escherichia
MNNEETFYQAMRRQGVTRRSFLKYCSLAATSLGLGAGMAPKIAWALENKPRIPVVWIHGL WP_001058323.1
MULTISPECIES: hydrogenase-1 small chain [Proteobacteria] Length: 372\nNumber of predicted TMHs:

2\nExp number of AAs in TMHs: 37.2608499999999\nExp number, first 60 AAs: 19.52923\nTotal prob of N-in: 0.94351\nPOSSIBLE N-term signal sequence\ninside 1 19\nTMhelix 20 39\noutside 40 325\nTMhelix 326 348\ninside 349 372

7799 GCF_001012635.1_CFSAN026818_01.0 Escherichia coli Proteobacteria; Gammaproteobacteria; Enterobacterales; Enterobacteriaceae; Escherichia
MTGDNTLIHSHGINRRDFMKLCAALAATMGLSSKAAAEMAESVTNPQRPPVIWIGAQECT WP_000145410.1
MULTISPECIES: hydrogenase-2 small chain [Proteobacteria] Length: 372\nNumber of predicted TMHs: 1\nExp number of AAs in TMHs: 23.72226\nExp number, first 60 AAs: 0.22038\nTotal prob of N-in: 0.12994\noutside 1 331\nTMhelix 332 354\ninside 355 372

7800 GCF_001012585.1_CFSAN026774_02.0 Escherichia coli Proteobacteria; Gammaproteobacteria; Enterobacterales; Enterobacteriaceae; Escherichia
MTGDNTLIHSHGINRRDFMKLCAALAATMGLSSKAAAEMAESVTNPQRPPVIWIGAQECT WP_000145410.1
MULTISPECIES: hydrogenase-2 small chain [Proteobacteria] Length: 372\nNumber of predicted TMHs: 1\nExp number of AAs in TMHs: 23.72226\nExp number, first 60 AAs: 0.22038\nTotal prob of N-in: 0.12994\noutside 1 331\nTMhelix 332 354\ninside 355 372

7801 GCF_001012585.1_CFSAN026774_02.0 Escherichia coli Proteobacteria; Gammaproteobacteria; Enterobacterales; Enterobacteriaceae; Escherichia
MNNEETFYQAMRRQGVTRRSFLKYCSLAATSLGLGAGMAPKIAWALENKPRIPVVWIHGL WP_001498899.1
hydrogenase-1 small chain [Escherichia coli] Length: 372\nNumber of predicted TMHs: 2\nExp number of AAs in TMHs: 39.34459\nExp number, first 60 AAs: 19.53328\nTotal prob of N-in: 0.94350\nPOSSIBLE N-term signal sequence\ninside 1 19\nTMhelix 20 39\noutside 40 325\nTMhelix 326 348\ninside 349 372

7802 GCF_001012635.1_CFSAN026818_01.0 Escherichia coli Proteobacteria; Gammaproteobacteria; Enterobacterales; Enterobacteriaceae; Escherichia
MNNEETFYQAMRRQGVTRRSFLKYCSLAATSLGLGAGMAPKIAWALENKPRIPVVWIHGL WP_024246676.1
hydrogenase-1 small chain [Escherichia coli] Length: 372\nNumber of predicted TMHs: 2\nExp number of AAs in TMHs: 37.8214899999999\nExp number, first 60 AAs: 19.52326\nTotal prob of N-in: 0.94395\nPOSSIBLE N-term signal sequence\ninside 1 19\nTMhelix 20 39\noutside 40 325\nTMhelix 326 348\ninside 349 372

7803 GCF_001012435.1_CFSAN026828_01.0 Escherichia coli Proteobacteria; Gammaproteobacteria; Enterobacterales; Enterobacteriaceae; Escherichia
MNNEETFYQAMRRQGVTRRSFLKYCSLAATSLGLGAGMAPKIAWALENKPRIPVVWIHGL WP_001295940.1
MULTISPECIES: hydrogenase [Enterobacteriaceae] Length: 372\nNumber of predicted TMHs: 2\nExp number of AAs in TMHs: 37.2608899999999\nExp number, first 60 AAs: 19.52923\nTotal prob of N-in: 0.94351\nPOSSIBLE N-term signal sequence\ninside 1 19\nTMhelix 20 39\noutside 40 325\nTMhelix 326 348\ninside 349 372

7804 GCF_001012475.1_CFSAN026830_01.0 Escherichia coli Proteobacteria; Gammaproteobacteria; Enterobacterales; Enterobacteriaceae; Escherichia
MNNEETFYQAMRRQGVTRRSFLKYCSLAATSLGLGAGMAPKIAWALENKPRIPVVWIHGL WP_001058323.1
MULTISPECIES: hydrogenase-1 small chain [Proteobacteria] Length: 372\nNumber of predicted TMHs: 2\nExp number of AAs in TMHs: 37.2608499999999\nExp number, first 60 AAs: 19.52923\nTotal prob of N-in: 0.94351\nPOSSIBLE N-term signal sequence\ninside 1 19\nTMhelix 20 39\noutside 40 325\nTMhelix 326 348\ninside 349 372

7805 GCF_001030305.1_Esch_coli_BWH60_V1 Escherichia coli Proteobacteria; Gammaproteobacteria; Enterobacterales; Enterobacteriaceae; Escherichia
MTGDNTLIHSHGINRRDFMKLCAALAATMGLSSKAAAEMVESVTNPQRPPVIWIGAQECT WP_000145423.1
hydrogenase 2 small subunit [Escherichia coli] Length: 372\nNumber of predicted TMHs: 1\nExp number of AAs in TMHs: 23.78201\nExp number, first 60 AAs: 0.28355\nTotal prob of N-in: 0.13257\noutside 1 331\nTMhelix 332 354\ninside 355 372

7806 GCF_001030245.1_Esch_coli_BWH49_V1 Escherichia coli Proteobacteria; Gammaproteobacteria; Enterobacterales; Enterobacteriaceae; Escherichia
MTGDNTLIHSHGINRRDFMKLCAALAATMGLSSKAAAEMAESVTNPQRPPVIWIGAQECT WP_000145410.1
MULTISPECIES: hydrogenase-2 small chain [Proteobacteria] Length: 372\nNumber of predicted TMHs: 1\nExp number of AAs in TMHs: 23.72226\nExp number, first 60 AAs: 0.22038\nTotal prob of N-in: 0.12994\noutside 1 331\nTMhelix 332 354\ninside 355 372

7807	GCF_001030405.1_Esch_coli_MGH108_V1	Escherichia coli	Proteobacteria; Gammaproteobacteria; Enterobacterales; Enterobacteriaceae; Escherichia
	MTGDNTLIHSHGINRRDFMKLCAALAATMGLSSKAAAEMAESVTNPQRPPVIWIGAQECT	WP_000145410.1
	MULTISPECIES: hydrogenase-2 small chain [Proteobacteria]	Length: 372\nNumber of predicted TMHs: 1\nExp number of AAs in TMHs: 23.72226\nExp number, first 60 AAs: 0.22038\nTotal prob of N-in: 0.12994\noutside 1 331\nTMhelix 332 354\ninside 355 372

7808	GCF_001030355.1_Esch_coli_BWH50_V1	Escherichia coli	Proteobacteria; Gammaproteobacteria; Enterobacterales; Enterobacteriaceae; Escherichia
	MTGDNTLIHSHGINRRDFMKLCAALAATMGLSSKAAAEMAESVTNPQRPPVIWIGAQECT	WP_000145410.1
	MULTISPECIES: hydrogenase-2 small chain [Proteobacteria]	Length: 372\nNumber of predicted TMHs: 1\nExp number of AAs in TMHs: 23.72226\nExp number, first 60 AAs: 0.22038\nTotal prob of N-in: 0.12994\noutside 1 331\nTMhelix 332 354\ninside 355 372

7809	GCF_001030445.1_Esch_coli_BIDMC97_V1	Escherichia coli	Proteobacteria; Gammaproteobacteria; Enterobacterales; Enterobacteriaceae; Escherichia
	MTGDNTLIHSHGINRRDFMKLCAALAATMGLSSKAAAEMAESVTNPQRPPVIWIGAQECT	WP_000145410.1
	MULTISPECIES: hydrogenase-2 small chain [Proteobacteria]	Length: 372\nNumber of predicted TMHs: 1\nExp number of AAs in TMHs: 23.72226\nExp number, first 60 AAs: 0.22038\nTotal prob of N-in: 0.12994\noutside 1 331\nTMhelix 332 354\ninside 355 372

7810	GCF_001030515.1_Esch_coli_BIDMC102_V1	Escherichia coli	Proteobacteria; Gammaproteobacteria; Enterobacterales; Enterobacteriaceae; Escherichia
	MTGDNTLIHSHGINRRDFMKLCAALAATMGLSSKAAAEMAESVTNPQRPPVIWIGAQECT	WP_000145410.1
	MULTISPECIES: hydrogenase-2 small chain [Proteobacteria]	Length: 372\nNumber of predicted TMHs: 1\nExp number of AAs in TMHs: 23.72226\nExp number, first 60 AAs: 0.22038\nTotal prob of N-in: 0.12994\noutside 1 331\nTMhelix 332 354\ninside 355 372

7811	GCF_001030515.1_Esch_coli_BIDMC102_V1	Escherichia coli	Proteobacteria; Gammaproteobacteria; Enterobacterales; Enterobacteriaceae; Escherichia
	MNNEETFYQAMRRQGVTRRSFLKYCSLAATSLGLGAGMAPKIAWALENKPRIPVVWIHGL	WP_001058323.1
	MULTISPECIES: hydrogenase-1 small chain [Proteobacteria]	Length: 372\nNumber of predicted TMHs: 2\nExp number of AAs in TMHs: 37.2608499999999\nExp number, first 60 AAs: 19.52923\nTotal prob of N-in: 0.94351\nPOSSIBLE N-term signal sequence\ninside 1 19\nTMhelix 20 39\noutside 40 325\nTMhelix 326 348\ninside 349 372

7812	GCF_001030355.1_Esch_coli_BWH50_V1	Escherichia coli	Proteobacteria; Gammaproteobacteria; Enterobacterales; Enterobacteriaceae; Escherichia
	MNNEETFYQAMRRQGVTRRSFLKYCSLAATSLGLGAGMAPKIAWALENKPRIPVVWIHGL	WP_001058323.1
	MULTISPECIES: hydrogenase-1 small chain [Proteobacteria]	Length: 372\nNumber of predicted TMHs: 2\nExp number of AAs in TMHs: 37.2608499999999\nExp number, first 60 AAs: 19.52923\nTotal prob of N-in: 0.94351\nPOSSIBLE N-term signal sequence\ninside 1 19\nTMhelix 20 39\noutside 40 325\nTMhelix 326 348\ninside 349 372

7813	GCF_001030445.1_Esch_coli_BIDMC97_V1	Escherichia coli	Proteobacteria; Gammaproteobacteria; Enterobacterales; Enterobacteriaceae; Escherichia
	MNNEETFYQAMRRQGVTRRSFLKYCSLAATSLGLGAGMAPKIAWALENKPRIPVVWIHGL	WP_001058323.1
	MULTISPECIES: hydrogenase-1 small chain [Proteobacteria]	Length: 372\nNumber of predicted TMHs: 2\nExp number of AAs in TMHs: 37.2608499999999\nExp number, first 60 AAs: 19.52923\nTotal prob of N-in: 0.94351\nPOSSIBLE N-term signal sequence\ninside 1 19\nTMhelix 20 39\noutside 40 325\nTMhelix 326 348\ninside 349 372

7814	GCF_001030405.1_Esch_coli_MGH108_V1	Escherichia coli	Proteobacteria; Gammaproteobacteria; Enterobacterales; Enterobacteriaceae; Escherichia
	MNNEETFYQAMRRQGVTRRSFLKYCSLAATSLGLGAGMAPKIAWALENKPRIPVVWIHGL	WP_001058323.1
	MULTISPECIES: hydrogenase-1 small chain [Proteobacteria]	Length: 372\nNumber of predicted TMHs: 2\nExp number of AAs in TMHs: 37.2608499999999\nExp number, first 60 AAs: 19.52923\nTotal prob of N-in: 0.94351\nPOSSIBLE N-term signal sequence\ninside 1 19\nTMhelix 20 39\noutside 40 325\nTMhelix 326 348\ninside 349 372

7815 GCF_001030245.1_Esch_coli_BWH49_V1 Escherichia coli Proteobacteria; Gammaproteobacteria; Enterobacterales; Enterobacteriaceae; Escherichia
MNNEETFYQAMRRQGVTRRSFLKYCSLAATSLGLGAGMAPKIAWALENKPRIPVVWIHGL WP_001058323.1
MULTISPECIES: hydrogenase-1 small chain [Proteobacteria] Length: 372\nNumber of predicted TMHs: 2\nExp number of AAs in TMHs: 37.2608499999999\nExp number, first 60 AAs: 19.52923\nTotal prob of N-in: 0.94351\nPOSSIBLE N-term signal sequence\ninside 1 19\nTMhelix 20 39\noutside 40 325\nTMhelix 326 348\ninside 349 372

7816 GCF_001030555.1_Esch_coli_BIDMC104_V1 Escherichia coli Proteobacteria; Gammaproteobacteria; Enterobacterales; Enterobacteriaceae; Escherichia
MTGDNTLIHSHGINRRDFMKLCAALAATMGLSSKAAAEMAESVTNPQRPPVIWIGAQECT WP_000145410.1
MULTISPECIES: hydrogenase-2 small chain [Proteobacteria] Length: 372\nNumber of predicted TMHs: 1\nExp number of AAs in TMHs: 23.72226\nExp number, first 60 AAs: 0.22038\nTotal prob of N-in: 0.12994\noutside 1 331\nTMhelix 332 354\ninside 355 372

7817 GCF_001030625.1_Esch_coli_BIDMC111_V1 Escherichia coli Proteobacteria; Gammaproteobacteria; Enterobacterales; Enterobacteriaceae; Escherichia
MTGDNTLIHSHGINRRDFMKLCAALAATMGLSSKAAAEMVESVTNPQRPPVIWIGAQECT WP_000145423.1
hydrogenase 2 small subunit [Escherichia coli] Length: 372\nNumber of predicted TMHs: 1\nExp number of AAs in TMHs: 23.78201\nExp number, first 60 AAs: 0.28355\nTotal prob of N-in: 0.13257\noutside 1 331\nTMhelix 332 354\ninside 355 372

7818 GCF_001030665.1_Esch_coli_BIDMC114_V1 Escherichia coli Proteobacteria; Gammaproteobacteria; Enterobacterales; Enterobacteriaceae; Escherichia
MTGDNTLIHSHGINRRDFMKLCAALAATMGLSSKAAAEMAESVTNPQRPPVIWIGAQECT WP_000145410.1
MULTISPECIES: hydrogenase-2 small chain [Proteobacteria] Length: 372\nNumber of predicted TMHs: 1\nExp number of AAs in TMHs: 23.72226\nExp number, first 60 AAs: 0.22038\nTotal prob of N-in: 0.12994\noutside 1 331\nTMhelix 332 354\ninside 355 372

7819 GCF_001030555.1_Esch_coli_BIDMC104_V1 Escherichia coli Proteobacteria; Gammaproteobacteria; Enterobacterales; Enterobacteriaceae; Escherichia
MNNEETFYQAMRRQGVTRRSFLKYCSLAATSLGLGAGMAPKIAWALENKPRIPVVWIHGL WP_001058323.1
MULTISPECIES: hydrogenase-1 small chain [Proteobacteria] Length: 372\nNumber of predicted TMHs: 2\nExp number of AAs in TMHs: 37.2608499999999\nExp number, first 60 AAs: 19.52923\nTotal prob of N-in: 0.94351\nPOSSIBLE N-term signal sequence\ninside 1 19\nTMhelix 20 39\noutside 40 325\nTMhelix 326 348\ninside 349 372

7820 GCF_001030625.1_Esch_coli_BIDMC111_V1 Escherichia coli Proteobacteria; Gammaproteobacteria; Enterobacterales; Enterobacteriaceae; Escherichia
MNNEETFYQAMRRQGVTRRSFLKYCSLAATSLGLGAGMAPKIAWALENKPRIPVVWIHGL WP_001295940.1
MULTISPECIES: hydrogenase [Enterobacteriaceae] Length: 372\nNumber of predicted TMHs: 2\nExp number of AAs in TMHs: 37.2608899999999\nExp number, first 60 AAs: 19.52923\nTotal prob of N-in: 0.94351\nPOSSIBLE N-term signal sequence\ninside 1 19\nTMhelix 20 39\noutside 40 325\nTMhelix 326 348\ninside 349 372

7821 GCF_001030305.1_Esch_coli_BWH60_V1 Escherichia coli Proteobacteria; Gammaproteobacteria; Enterobacterales; Enterobacteriaceae; Escherichia
MNNEETFYQAMRRQGVTRRSFLKYCSLAATSLGLGAGMAPKIAWALENKPRIPVVWIHGL WP_001295940.1
MULTISPECIES: hydrogenase [Enterobacteriaceae] Length: 372\nNumber of predicted TMHs: 2\nExp number of AAs in TMHs: 37.2608899999999\nExp number, first 60 AAs: 19.52923\nTotal prob of N-in: 0.94351\nPOSSIBLE N-term signal sequence\ninside 1 19\nTMhelix 20 39\noutside 40 325\nTMhelix 326 348\ninside 349 372

7822 GCF_001039075.1_ASM103907v1 Escherichia coli Proteobacteria; Gammaproteobacteria; Enterobacterales; Enterobacteriaceae; Escherichia
MTGDNTLIHSHGINRRDFMKLCAALAATMGLSSKAAAEMAESVTNPQRPPVIWIGAQECT WP_000145410.1
MULTISPECIES: hydrogenase-2 small chain [Proteobacteria] Length: 372\nNumber of predicted TMHs: 1\nExp number of AAs in TMHs: 23.72226\nExp number, first 60 AAs: 0.22038\nTotal prob of N-in: 0.12994\noutside 1 331\nTMhelix 332 354\ninside 355 372

7823 GCF_001039135.1_ASM103913v1 Escherichia coli Proteobacteria; Gammaproteobacteria; Enterobacterales; Enterobacteriaceae; Escherichia
MTGDNTLIHSHGINRRDFMKLCAALAATMGLSSKAAAEMAESVTNPQRPPVIWIGAQECT WP_000145410.1
MULTISPECIES: hydrogenase-2 small chain [Proteobacteria] Length: 372\nNumber of predicted TMHs: 1\nExp number of AAs in TMHs: 23.72226\nExp number, first 60 AAs: 0.22038\nTotal prob of N-in: 0.12994\noutside 1 331\nTMhelix 332 354\ninside 355 372

7824 GCF_001039205.1_ASM103920v1 Escherichia coli Proteobacteria; Gammaproteobacteria; Enterobacterales; Enterobacteriaceae; Escherichia
MTGDNTLIHSHGINRRDFMKLCAALAATMGLSSKAAAEMAESVTNPQRPPVIWIGAQECT WP_000145410.1
MULTISPECIES: hydrogenase-2 small chain [Proteobacteria] Length: 372\nNumber of predicted TMHs: 1\nExp number of AAs in TMHs: 23.72226\nExp number, first 60 AAs: 0.22038\nTotal prob of N-in: 0.12994\noutside 1 331\nTMhelix 332 354\ninside 355 372

7825 GCF_001039225.1_ASM103922v1 Escherichia coli Proteobacteria; Gammaproteobacteria; Enterobacterales; Enterobacteriaceae; Escherichia
MTGDNTLIHSHGINRRDFMKLCAALAATMGLSSKAAAEMAESVTNPQRPPVIWIGAQECT WP_000145410.1
MULTISPECIES: hydrogenase-2 small chain [Proteobacteria] Length: 372\nNumber of predicted TMHs: 1\nExp number of AAs in TMHs: 23.72226\nExp number, first 60 AAs: 0.22038\nTotal prob of N-in: 0.12994\noutside 1 331\nTMhelix 332 354\ninside 355 372

7826 GCF_001039075.1_ASM103907v1 Escherichia coli Proteobacteria; Gammaproteobacteria; Enterobacterales; Enterobacteriaceae; Escherichia
MNNEETFYQAMRRQGVTRRSFLKYCSLAATSLGLGAGMAPKIAWALENKPRIPVVWIHGL WP_001058323.1
MULTISPECIES: hydrogenase-1 small chain [Proteobacteria] Length: 372\nNumber of predicted TMHs: 2\nExp number of AAs in TMHs: 37.2608499999999\nExp number, first 60 AAs: 19.52923\nTotal prob of N-in: 0.94351\nPOSSIBLE N-term signal sequence\ninside 1 19\nTMhelix 20 39\noutside 40 325\nTMhelix 326 348\ninside 349 372

7827 GCF_001039135.1_ASM103913v1 Escherichia coli Proteobacteria; Gammaproteobacteria; Enterobacterales; Enterobacteriaceae; Escherichia
MNNEETFYQAMRRQGVTRRSFLKYCSLAATSLGLGAGMAPKIAWALENKPRIPVVWIHGL WP_001058323.1
MULTISPECIES: hydrogenase-1 small chain [Proteobacteria] Length: 372\nNumber of predicted TMHs: 2\nExp number of AAs in TMHs: 37.2608499999999\nExp number, first 60 AAs: 19.52923\nTotal prob of N-in: 0.94351\nPOSSIBLE N-term signal sequence\ninside 1 19\nTMhelix 20 39\noutside 40 325\nTMhelix 326 348\ninside 349 372

7828 GCF_001052935.1_ASM105293v1 Escherichia coli Proteobacteria; Gammaproteobacteria; Enterobacterales; Enterobacteriaceae; Escherichia
MTGDNTLIHSHGINRRDFMKLCAALAATMGLSSKAAAEMAESVTNPQRPPVIWIGAQECT WP_000145418.1
hydrogenase 2 small subunit [Escherichia coli] Length: 372\nNumber of predicted TMHs: 1\nExp number of AAs in TMHs: 23.72313\nExp number, first 60 AAs: 0.22034\nTotal prob of N-in: 0.12997\noutside 1 331\nTMhelix 332 354\ninside 355 372

7829 GCF_001052935.1_ASM105293v1 Escherichia coli Proteobacteria; Gammaproteobacteria; Enterobacterales; Enterobacteriaceae; Escherichia
MNNEETFYQAMRRQGVTRRSFLKYCSLAATSLGLGAGMAPKIAWALENKPRIPVVWIHGL WP_001058323.1
MULTISPECIES: hydrogenase-1 small chain [Proteobacteria] Length: 372\nNumber of predicted TMHs: 2\nExp number of AAs in TMHs: 37.2608499999999\nExp number, first 60 AAs: 19.52923\nTotal prob of N-in: 0.94351\nPOSSIBLE N-term signal sequence\ninside 1 19\nTMhelix 20 39\noutside 40 325\nTMhelix 326 348\ninside 349 372

7830 GCF_001039205.1_ASM103920v1 Escherichia coli Proteobacteria; Gammaproteobacteria; Enterobacterales; Enterobacteriaceae; Escherichia
MNNEETFYQAMRRQGVTRRSFLKYCSLAATSLGLGAGMAPKIAWALENKPRIPVVWIHGL WP_001058323.1
MULTISPECIES: hydrogenase-1 small chain [Proteobacteria] Length: 372\nNumber of predicted TMHs: 2\nExp number of AAs in TMHs: 37.2608499999999\nExp number, first 60 AAs: 19.52923\nTotal prob of N-in: 0.94351\nPOSSIBLE N-term signal sequence\ninside 1 19\nTMhelix 20 39\noutside 40 325\nTMhelix 326 348\ninside 349 372

7831 GCF_001053295.1_ASM105329v1 Escherichia coli Proteobacteria; Gammaproteobacteria; Enterobacterales; Enterobacteriaceae; Escherichia
MTGDNTLIHSHGINRRDFMKLCAALAATMGLSSKAAAEMVESVTNPQRPPVIWIGAQECT WP_000145423.1
hydrogenase 2 small subunit [Escherichia coli] Length: 372\nNumber of predicted TMHs: 1\nExp number of AAs in TMHs: 23.78201\nExp number, first 60 AAs: 0.28355\nTotal prob of N-in: 0.13257\noutside 1 331\nTMhelix 332 354\ninside 355 372

7832 GCF_001053295.1_ASM105329v1 Escherichia coli Proteobacteria; Gammaproteobacteria; Enterobacterales; Enterobacteriaceae; Escherichia
MNNEETFYQAMRRQGVTRRSFLKYCSLAATSLGLGAGMAPKIAWALENKPRIPVVWIHGL WP_001295940.1
MULTISPECIES: hydrogenase [Enterobacteriaceae] Length: 372\nNumber of predicted TMHs: 2\nExp number of AAs in TMHs: 37.2608899999999\nExp number, first 60 AAs: 19.52923\nTotal prob of N-in: 0.94351\nPOSSIBLE N-term signal sequence\ninside 1 19\nTMhelix 20 39\noutside 40 325\nTMhelix 326 348\ninside 349 372

7833 GCF_001039225.1_ASM103922v1 Escherichia coli Proteobacteria; Gammaproteobacteria; Enterobacterales; Enterobacteriaceae; Escherichia
MNNEETFYQAMRRQGVTRRSFLKYCSLAATSLGLGAGMAPKIAWALENKPRIPVVWIHGL WP_001058323.1
MULTISPECIES: hydrogenase-1 small chain [Proteobacteria] Length: 372\nNumber of predicted TMHs: 2\nExp number of AAs in TMHs: 37.2608499999999\nExp number, first 60 AAs: 19.52923\nTotal prob of N-in: 0.94351\nPOSSIBLE N-term signal sequence\ninside 1 19\nTMhelix 20 39\noutside 40 325\nTMhelix 326 348\ninside 349 372

7834 GCF_001054095.1_ASM105409v1 Escherichia coli Proteobacteria; Gammaproteobacteria; Enterobacterales; Enterobacteriaceae; Escherichia
MTGDNTLIHSHGINRRDFMKLCAALAATMGLSSKAAAEMAESVTNPQRPPVIWIGAQECT WP_000145410.1
MULTISPECIES: hydrogenase-2 small chain [Proteobacteria] Length: 372\nNumber of predicted TMHs: 1\nExp number of AAs in TMHs: 23.72226\nExp number, first 60 AAs: 0.22038\nTotal prob of N-in: 0.12994\noutside 1 331\nTMhelix 332 354\ninside 355 372

7835 GCF_001054245.1_ASM105424v1 Escherichia coli Proteobacteria; Gammaproteobacteria; Enterobacterales; Enterobacteriaceae; Escherichia
MTGDNTLIHSHGINRRDFMKLCAALAATMGLSSKAAAEMAESVTNPQRPPVIWIGAQECT WP_000145410.1
MULTISPECIES: hydrogenase-2 small chain [Proteobacteria] Length: 372\nNumber of predicted TMHs: 1\nExp number of AAs in TMHs: 23.72226\nExp number, first 60 AAs: 0.22038\nTotal prob of N-in: 0.12994\noutside 1 331\nTMhelix 332 354\ninside 355 372

7836 GCF_001054245.1_ASM105424v1 Escherichia coli Proteobacteria; Gammaproteobacteria; Enterobacterales; Enterobacteriaceae; Escherichia
MNNEETFYQAMRRQGVTRRSFLKYCSLAATSLGLGAGMAPKIAWALENKPRIPVVWIHGL WP_001295940.1
MULTISPECIES: hydrogenase [Enterobacteriaceae] Length: 372\nNumber of predicted TMHs: 2\nExp number of AAs in TMHs: 37.2608899999999\nExp number, first 60 AAs: 19.52923\nTotal prob of N-in: 0.94351\nPOSSIBLE N-term signal sequence\ninside 1 19\nTMhelix 20 39\noutside 40 325\nTMhelix 326 348\ninside 349 372

7837 GCF_001052365.1_ASM105236v1 Escherichia coli Proteobacteria; Gammaproteobacteria; Enterobacterales; Enterobacteriaceae; Escherichia
MNNEETFYQAMRRQGVTRRSFLKYCSLAATSLGLGAGMAPKIAWALENKPRIPVVWIHGL WP_001058323.1
MULTISPECIES: hydrogenase-1 small chain [Proteobacteria] Length: 372\nNumber of predicted TMHs: 2\nExp number of AAs in TMHs: 37.2608499999999\nExp number, first 60 AAs: 19.52923\nTotal prob of N-in: 0.94351\nPOSSIBLE N-term signal sequence\ninside 1 19\nTMhelix 20 39\noutside 40 325\nTMhelix 326 348\ninside 349 372

7838 GCF_001055155.1_ASM105515v1 Escherichia coli Proteobacteria; Gammaproteobacteria; Enterobacterales; Enterobacteriaceae; Escherichia
MTGDNTLIHSHGINRRDFMKLCAALAATMGLSSKAAAEMAESVTNPQRPPVIWIGAQECT WP_000145410.1
MULTISPECIES: hydrogenase-2 small chain [Proteobacteria] Length: 372\nNumber of predicted TMHs: 1\nExp number of AAs in TMHs: 23.72226\nExp number, first 60 AAs: 0.22038\nTotal prob of N-in: 0.12994\noutside 1 331\nTMhelix 332 354\ninside 355 372

7839 GCF_001055155.1_ASM105515v1 Escherichia coli Proteobacteria; Gammaproteobacteria; Enterobacterales; Enterobacteriaceae; Escherichia
MNNEETFYQAMRRQGVTRRSFLKYCSLAATSLGLGAGMAPKIAWALENKPRIPVVWIHGL WP_001058323.1
MULTISPECIES: hydrogenase-1 small chain [Proteobacteria] Length: 372\nNumber of predicted TMHs: 2\nExp number of AAs in TMHs: 37.2608499999999\nExp number, first 60 AAs: 19.52923\nTotal prob of N-in: 0.94351\nPOSSIBLE N-term signal sequence\ninside 1 19\nTMhelix 20 39\noutside 40 325\nTMhelix 326 348\ninside 349 372

7840 GCF_001056015.1_ASM105601v1 Escherichia coli Proteobacteria; Gammaproteobacteria; Enterobacterales; Enterobacteriaceae; Escherichia
MTGDNTLIHSHGINRRDFMKLCAALAATMGLSSKAAAEMVESVTNPQRPPVIWIGAQECT WP_000145423.1
hydrogenase 2 small subunit [Escherichia coli] Length: 372\nNumber of predicted TMHs: 1\nExp number of AAs in TMHs: 23.78201\nExp number, first 60 AAs: 0.28355\nTotal prob of N-in: 0.13257\noutside 1 331\nTMhelix 332 354\ninside 355 372

7841 GCF_001052365.1_ASM105236v1 Escherichia coli Proteobacteria; Gammaproteobacteria; Enterobacterales; Enterobacteriaceae; Escherichia
MTGDNTLIHSHGINRRDFMKLCAALATTMGLSSKAAAEMAESVTNPQRPPVIWIGAQECT WP_023149629.1
MULTISPECIES: hydrogenase-2 small chain [Enterobacteriaceae] Length: 372\nNumber of predicted TMHs: 1\nExp number of AAs in TMHs: 23.63603\nExp number, first 60 AAs: 0.12971\nTotal prob of N-in: 0.12608\noutside 1 331\nTMhelix 332 354\ninside 355 372

7842 GCF_001191265.1_CFSAN026840_01-NGE.clc Escherichia coli Proteobacteria; Gammaproteobacteria; Enterobacterales; Enterobacteriaceae; Escherichia
MTGDNTLIHSHGINRRDFMKLCAALAATMGLSSKAAAEMAESVTNPQRPPVIWIGAQECT WP_000145410.1
MULTISPECIES: hydrogenase-2 small chain [Proteobacteria] Length: 372\nNumber of predicted TMHs: 1\nExp number of AAs in TMHs: 23.72226\nExp number, first 60 AAs: 0.22038\nTotal prob of N-in: 0.12994\noutside 1 331\nTMhelix 332 354\ninside 355 372

7843 GCF_001191315.1_CFSAN026781_02-NGE.clc Escherichia coli Proteobacteria; Gammaproteobacteria; Enterobacterales; Enterobacteriaceae; Escherichia
MTGDNTLIHSHGINRRDFMKLCAALAATMGLSSKAAAEMAESVTNPQRPPVIWIGAQECT WP_000145410.1
MULTISPECIES: hydrogenase-2 small chain [Proteobacteria] Length: 372\nNumber of predicted TMHs: 1\nExp number of AAs in TMHs: 23.72226\nExp number, first 60 AAs: 0.22038\nTotal prob of N-in: 0.12994\noutside 1 331\nTMhelix 332 354\ninside 355 372

7844 GCF_001191375.1_CFSAN026777_02-NGE.clc Escherichia coli Proteobacteria; Gammaproteobacteria; Enterobacterales; Enterobacteriaceae; Escherichia
MTGDNTLIHSHGINRRDFMKLCAALAATMGLSSKAAAEMAESVTNPQRPPVIWIGAQECT WP_000145410.1
MULTISPECIES: hydrogenase-2 small chain [Proteobacteria] Length: 372\nNumber of predicted TMHs: 1\nExp number of AAs in TMHs: 23.72226\nExp number, first 60 AAs: 0.22038\nTotal prob of N-in: 0.12994\noutside 1 331\nTMhelix 332 354\ninside 355 372

7845 GCF_001191425.1_CFSAN026798_01-NGE.clc Escherichia coli Proteobacteria; Gammaproteobacteria; Enterobacterales; Enterobacteriaceae; Escherichia
MTGDNTLIHSHGINRRDFMKLCAALAATMGLSSKAAAEMAESVTNPQRPPVIWIGAQECT WP_000145410.1
MULTISPECIES: hydrogenase-2 small chain [Proteobacteria] Length: 372\nNumber of predicted TMHs: 1\nExp number of AAs in TMHs: 23.72226\nExp number, first 60 AAs: 0.22038\nTotal prob of N-in: 0.12994\noutside 1 331\nTMhelix 332 354\ninside 355 372

7846 GCF_001191465.1_CFSAN026825_01-NGE.clc Escherichia coli Proteobacteria; Gammaproteobacteria; Enterobacterales; Enterobacteriaceae; Escherichia
MTGDNTLIHSHGINRRDFMKLCAALAATMGLSSKAAAEMAESVTNPQRPPVIWIGAQECT WP_000145410.1
MULTISPECIES: hydrogenase-2 small chain [Proteobacteria] Length: 372\nNumber of predicted TMHs: 1\nExp number of AAs in TMHs: 23.72226\nExp number, first 60 AAs: 0.22038\nTotal prob of N-in: 0.12994\noutside 1 331\nTMhelix 332 354\ninside 355 372

7847 GCF_001262455.1_ASM126245v1 Escherichia coli Proteobacteria; Gammaproteobacteria; Enterobacterales; Enterobacteriaceae; Escherichia
MTGDNTLIHSHGINRRDFMKLCAALAATMGLSSKAAAEMAESVTNPQRPPVIWIGAQECT WP_000145410.1
MULTISPECIES: hydrogenase-2 small chain [Proteobacteria] Length: 372\nNumber of predicted TMHs:

1\nExp number of AAs in TMHs: 23.72226\nExp number, first 60 AAs: 0.22038\nTotal prob of N-in: 0.12994\noutside 1 331\nTMhelix 332 354\ninside 355 372

7848 GCF_001191525.1_CFSAN026848_01-NGE.clc Escherichia coli Proteobacteria; Gammaproteobacteria; Enterobacterales; Enterobacteriaceae; Escherichia MTGDNTLIHSHGINRRDFMKLCAALAATMGLSSKAAAEMAESVTNPQRPPVIWIGAQECT WP_000145410.1 MULTISPECIES: hydrogenase-2 small chain [Proteobacteria] Length: 372\nNumber of predicted TMHs: 1\nExp number of AAs in TMHs: 23.72226\nExp number, first 60 AAs: 0.22038\nTotal prob of N-in: 0.12994\noutside 1 331\nTMhelix 332 354\ninside 355 372

7849 GCF_001191375.1_CFSAN026777_02-NGE.clc Escherichia coli Proteobacteria; Gammaproteobacteria; Enterobacterales; Enterobacteriaceae; Escherichia MNNEETFYQAMRRQGVTRRSFLKYCSLAATSLGLGAGMAPKIAWALENKPRIPVVWIHGL WP_001058323.1 MULTISPECIES: hydrogenase-1 small chain [Proteobacteria] Length: 372\nNumber of predicted TMHs: 2\nExp number of AAs in TMHs: 37.2608499999999\nExp number, first 60 AAs: 19.52923\nTotal prob of N-in: 0.94351\nPOSSIBLE N-term signal sequence\ninside 1 19\nTMhelix 20 39\noutside 40 325\nTMhelix 326 348\ninside 349 372

7850 GCF_001191265.1_CFSAN026840_01-NGE.clc Escherichia coli Proteobacteria; Gammaproteobacteria; Enterobacterales; Enterobacteriaceae; Escherichia MNNEETFYQAMRRQGVTRRSFLKYCSLAATSLGLGAGMAPKIAWALENKPRIPVVWIHGL WP_001058323.1 MULTISPECIES: hydrogenase-1 small chain [Proteobacteria] Length: 372\nNumber of predicted TMHs: 2\nExp number of AAs in TMHs: 37.2608499999999\nExp number, first 60 AAs: 19.52923\nTotal prob of N-in: 0.94351\nPOSSIBLE N-term signal sequence\ninside 1 19\nTMhelix 20 39\noutside 40 325\nTMhelix 326 348\ninside 349 372

7851 GCF_001056015.1_ASM105601v1 Escherichia coli Proteobacteria; Gammaproteobacteria; Enterobacterales; Enterobacteriaceae; Escherichia MNNEETFYQAMRRQGVTRRSFLKYCSLAATSLGLGAGMAPKIAWALENKPRIPVVWIHGL WP_001295940.1 MULTISPECIES: hydrogenase [Enterobacteriaceae] Length: 372\nNumber of predicted TMHs: 2\nExp number of AAs in TMHs: 37.2608899999999\nExp number, first 60 AAs: 19.52923\nTotal prob of N-in: 0.94351\nPOSSIBLE N-term signal sequence\ninside 1 19\nTMhelix 20 39\noutside 40 325\nTMhelix 326 348\ninside 349 372

7852 GCF_001262785.1_1027-4 Escherichia coli Proteobacteria; Gammaproteobacteria; Enterobacterales; Enterobacteriaceae; Escherichia MTGDNTLIHSHGINRRDFMKLCAALAATMGLSSKAAAEMAESVTNPQRPPVIWIGAQECT WP_000145410.1 MULTISPECIES: hydrogenase-2 small chain [Proteobacteria] Length: 372\nNumber of predicted TMHs: 1\nExp number of AAs in TMHs: 23.72226\nExp number, first 60 AAs: 0.22038\nTotal prob of N-in: 0.12994\noutside 1 331\nTMhelix 332 354\ninside 355 372

7853 GCF_001262805.1_15484-1 Escherichia coli Proteobacteria; Gammaproteobacteria; Enterobacterales; Enterobacteriaceae; Escherichia MTGDNTLIHSHGINRRDFMKLCAALAATMGLSSKAAAEMAESVTNPQRPPVIWIGAQECT WP_000145410.1 MULTISPECIES: hydrogenase-2 small chain [Proteobacteria] Length: 372\nNumber of predicted TMHs: 1\nExp number of AAs in TMHs: 23.72226\nExp number, first 60 AAs: 0.22038\nTotal prob of N-in: 0.12994\noutside 1 331\nTMhelix 332 354\ninside 355 372

7854 GCF_001262895.1_9916-1 Escherichia coli Proteobacteria; Gammaproteobacteria; Enterobacterales; Enterobacteriaceae; Escherichia MTGDNTLIHSHGINRRDFMKLCAALAATMGLSSKAAAEMAESVTNPQRPPVIWIGAQECT WP_000145410.1 MULTISPECIES: hydrogenase-2 small chain [Proteobacteria] Length: 372\nNumber of predicted TMHs: 1\nExp number of AAs in TMHs: 23.72226\nExp number, first 60 AAs: 0.22038\nTotal prob of N-in: 0.12994\noutside 1 331\nTMhelix 332 354\ninside 355 372

7855 GCF_001262785.1_1027-4 Escherichia coli Proteobacteria; Gammaproteobacteria; Enterobacterales; Enterobacteriaceae; Escherichia MNNEETFYQAMRRQGVTRRSFLKYCSLAATSLGLGAGMAPKIAWALENKPRIPVVWIHGL WP_001058323.1 MULTISPECIES: hydrogenase-1 small chain [Proteobacteria] Length: 372\nNumber of predicted TMHs: 2\nExp number of AAs in TMHs: 37.2608499999999\nExp number, first 60 AAs: 19.52923\nTotal prob of N-in: 0.94351\nPOSSIBLE N-term signal sequence\ninside 1 19\nTMhelix 20 39\noutside 40 325\nTMhelix 326 348\ninside 349 372

7856 GCF_001262895.1_9916-1 Escherichia coli Proteobacteria; Gammaproteobacteria; Enterobacterales; Enterobacteriaceae; Escherichia MNNEETFYQAMRRQGVTRRSFLKYCSLAATSLGLGAGMAPKIAWALENKPRIPVVWIHGL

WP_001058323.1 MULTISPECIES: hydrogenase-1 small chain [Proteobacteria] Length: 372\nNumber of predicted TMHs: 2\nExp number of AAs in TMHs: 37.2608499999999\nExp number, first 60 AAs: 19.52923\nTotal prob of N-in: 0.94351\nPOSSIBLE N-term signal sequence\ninside 1 19\nTMhelix 20 39\noutside 40 325\nTMhelix 326 348\ninside 349 372

7857 GCF_001262805.1_15484-1 Escherichia coli Proteobacteria; Gammaproteobacteria; Enterobacterales; Enterobacteriaceae; Escherichia MNNEETFYQAMRRQGVTRRSFLKYCSLAATSLGLGAGMAPKIAWALENKPRIPVVWIHGL WP_001295940.1 MULTISPECIES: hydrogenase [Enterobacteriaceae] Length: 372\nNumber of predicted TMHs: 2\nExp number of AAs in TMHs: 37.2608899999999\nExp number, first 60 AAs: 19.52923\nTotal prob of N-in: 0.94351\nPOSSIBLE N-term signal sequence\ninside 1 19\nTMhelix 20 39\noutside 40 325\nTMhelix 326 348\ninside 349 372

7858 GCF_001191315.1_CFSAN026781_02-NGE.clc Escherichia coli Proteobacteria; Gammaproteobacteria; Enterobacterales; Enterobacteriaceae; Escherichia MNNEETFYQAMRRQGVTRRSFLKYCSLAATSLGLGAGMAPKIAWALENKPRIPVVWIHGL WP_001295940.1 MULTISPECIES: hydrogenase [Enterobacteriaceae] Length: 372\nNumber of predicted TMHs: 2\nExp number of AAs in TMHs: 37.2608899999999\nExp number, first 60 AAs: 19.52923\nTotal prob of N-in: 0.94351\nPOSSIBLE N-term signal sequence\ninside 1 19\nTMhelix 20 39\noutside 40 325\nTMhelix 326 348\ninside 349 372

7859 GCF_001191525.1_CFSAN026848_01-NGE.clc Escherichia coli Proteobacteria; Gammaproteobacteria; Enterobacterales; Enterobacteriaceae; Escherichia MNNEETFYQAMRRQGVTRRSFLKYCSLAATSLGLGAGMAPKIAWALENKPRIPVVWIHGL WP_001058323.1 MULTISPECIES: hydrogenase-1 small chain [Proteobacteria] Length: 372\nNumber of predicted TMHs: 2\nExp number of AAs in TMHs: 37.2608499999999\nExp number, first 60 AAs: 19.52923\nTotal prob of N-in: 0.94351\nPOSSIBLE N-term signal sequence\ninside 1 19\nTMhelix 20 39\noutside 40 325\nTMhelix 326 348\ninside 349 372

7860 GCF_001191465.1_CFSAN026825_01-NGE.clc Escherichia coli Proteobacteria; Gammaproteobacteria; Enterobacterales; Enterobacteriaceae; Escherichia MNNEETFYQAMRRQGVTRRSFLKYCSLAATSLGLGAGMAPKIAWALENKPRIPVVWIHGL WP_001295940.1 MULTISPECIES: hydrogenase [Enterobacteriaceae] Length: 372\nNumber of predicted TMHs: 2\nExp number of AAs in TMHs: 37.2608899999999\nExp number, first 60 AAs: 19.52923\nTotal prob of N-in: 0.94351\nPOSSIBLE N-term signal sequence\ninside 1 19\nTMhelix 20 39\noutside 40 325\nTMhelix 326 348\ninside 349 372

7861 GCF_001262975.1_10674-1 Escherichia coli Proteobacteria; Gammaproteobacteria; Enterobacterales; Enterobacteriaceae; Escherichia MTGDNTLIHSHGINRRDFMKLCAALAATMGLSSKAAAEMAESVTNPQRPPVIWIGAQECT WP_000145410.1 MULTISPECIES: hydrogenase-2 small chain [Proteobacteria] Length: 372\nNumber of predicted TMHs: 1\nExp number of AAs in TMHs: 23.72226\nExp number, first 60 AAs: 0.22038\nTotal prob of N-in: 0.12994\noutside 1 331\nTMhelix 332 354\ninside 355 372

7862 GCF_001262935.1_1932 Escherichia coli Proteobacteria; Gammaproteobacteria; Enterobacterales; Enterobacteriaceae; Escherichia MTGDNTLIHSHGINRRDFMKLCAALAATMGLSSKAAAEMAESVTNPQRPPVIWIGAQECT WP_000145410.1 MULTISPECIES: hydrogenase-2 small chain [Proteobacteria] Length: 372\nNumber of predicted TMHs: 1\nExp number of AAs in TMHs: 23.72226\nExp number, first 60 AAs: 0.22038\nTotal prob of N-in: 0.12994\noutside 1 331\nTMhelix 332 354\ninside 355 372

7863 GCF_001263035.1_14 Escherichia coli Proteobacteria; Gammaproteobacteria; Enterobacterales; Enterobacteriaceae; Escherichia MTGDNTLIHSHGINRRDFMKLCAALAATMGLSSKAAAEMAESVTNPQRPPVIWIGAQECT WP_000145410.1 MULTISPECIES: hydrogenase-2 small chain [Proteobacteria] Length: 372\nNumber of predicted TMHs: 1\nExp number of AAs in TMHs: 23.72226\nExp number, first 60 AAs: 0.22038\nTotal prob of N-in: 0.12994\noutside 1 331\nTMhelix 332 354\ninside 355 372

7864 GCF_001264215.1_ASM126421v1 Escherichia coli Proteobacteria; Gammaproteobacteria; Enterobacterales; Enterobacteriaceae; Escherichia MTGDNTLIHSHGINRRDFMKLCAALAATMGLSSKAAAEMAESVTNPQRPPVIWIGAQECT WP_000145410.1 MULTISPECIES: hydrogenase-2 small chain [Proteobacteria] Length: 372\nNumber of predicted TMHs: 1\nExp number of AAs in TMHs: 23.72226\nExp number, first 60 AAs: 0.22038\nTotal prob of N-in: 0.12994\noutside 1 331\nTMhelix 332 354\ninside 355 372

7865 GCF_001264175.1_ASM126417v1 Escherichia coli Proteobacteria; Gammaproteobacteria; Enterobacterales; Enterobacteriaceae; Escherichia MTGDNTLIHSHGINRRDFMKLCAALAATMGLSSKAAAEMAESVTNPQRPPVIWIGAQECT WP_000145410.1 MULTISPECIES: hydrogenase-2 small chain [Proteobacteria] Length: 372\nNumber of predicted TMHs: 1\nExp number of AAs in TMHs: 23.72226\nExp number, first 60 AAs: 0.22038\nTotal prob of N-in: 0.12994\noutside 1 331\nTMhelix 332 354\ninside 355 372

7866 GCF_001262975.1_10674-1 Escherichia coli Proteobacteria; Gammaproteobacteria; Enterobacterales; Enterobacteriaceae; Escherichia MNNEETFYQAMRRQGVTRRSFLKYCSLAATSLGLGAGMAPKIAWALENKPRIPVVWIHGL WP_001058323.1 MULTISPECIES: hydrogenase-1 small chain [Proteobacteria] Length: 372\nNumber of predicted TMHs: 2\nExp number of AAs in TMHs: 37.2608499999999\nExp number, first 60 AAs: 19.52923\nTotal prob of N-in: 0.94351\nPOSSIBLE N-term signal sequence\ninside 1 19\nTMhelix 20 39\noutside 40 325\nTMhelix 326 348\ninside 349 372

7867 GCF_001262935.1_1932 Escherichia coli Proteobacteria; Gammaproteobacteria; Enterobacterales; Enterobacteriaceae; Escherichia MNNEETFYQAMRRQGVTRRSFLKYCSLAATSLGLGAGMAPKIAWALENKPRIPVVWIHGL WP_001058323.1 MULTISPECIES: hydrogenase-1 small chain [Proteobacteria] Length: 372\nNumber of predicted TMHs: 2\nExp number of AAs in TMHs: 37.2608499999999\nExp number, first 60 AAs: 19.52923\nTotal prob of N-in: 0.94351\nPOSSIBLE N-term signal sequence\ninside 1 19\nTMhelix 20 39\noutside 40 325\nTMhelix 326 348\ninside 349 372

7868 GCF_001265175.1_100343 Escherichia coli Proteobacteria; Gammaproteobacteria; Enterobacterales; Enterobacteriaceae; Escherichia MTGDNTLIHSHGINRRDFMKLCAALAATMGLSSKAAAEMAESVTNPQRPPVIWIGAQECT WP_000145410.1 MULTISPECIES: hydrogenase-2 small chain [Proteobacteria] Length: 372\nNumber of predicted TMHs: 1\nExp number of AAs in TMHs: 23.72226\nExp number, first 60 AAs: 0.22038\nTotal prob of N-in: 0.12994\noutside 1 331\nTMhelix 332 354\ninside 355 372

7869 GCF_001265175.1_100343 Escherichia coli Proteobacteria; Gammaproteobacteria; Enterobacterales; Enterobacteriaceae; Escherichia MNNEETFYQAMRRQGVTRRSFLKYCSLAATSLGLGAGMAPKIAWALENKPRIPVVWIHGL WP_001058323.1 MULTISPECIES: hydrogenase-1 small chain [Proteobacteria] Length: 372\nNumber of predicted TMHs: 2\nExp number of AAs in TMHs: 37.2608499999999\nExp number, first 60 AAs: 19.52923\nTotal prob of N-in: 0.94351\nPOSSIBLE N-term signal sequence\ninside 1 19\nTMhelix 20 39\noutside 40 325\nTMhelix 326 348\ninside 349 372

7870 GCF_001263035.1_14 Escherichia coli Proteobacteria; Gammaproteobacteria; Enterobacterales; Enterobacteriaceae; Escherichia MNNEETFYQAMRRQGVTRRSFLKYCSLAATSLGLGAGMAPKIAWALENKPRIPVVWIHGL WP_001058323.1 MULTISPECIES: hydrogenase-1 small chain [Proteobacteria] Length: 372\nNumber of predicted TMHs: 2\nExp number of AAs in TMHs: 37.2608499999999\nExp number, first 60 AAs: 19.52923\nTotal prob of N-in: 0.94351\nPOSSIBLE N-term signal sequence\ninside 1 19\nTMhelix 20 39\noutside 40 325\nTMhelix 326 348\ninside 349 372

7871 GCF_001265265.1_203741 Escherichia coli Proteobacteria; Gammaproteobacteria; Enterobacterales; Enterobacteriaceae; Escherichia MTGDNTLIHSHGINRRDFMKLCAALAATMGLSSKAAAEMAESVTNPQRPPVIWIGAQECT WP_000145410.1 MULTISPECIES: hydrogenase-2 small chain [Proteobacteria] Length: 372\nNumber of predicted TMHs: 1\nExp number of AAs in TMHs: 23.72226\nExp number, first 60 AAs: 0.22038\nTotal prob of N-in: 0.12994\noutside 1 331\nTMhelix 332 354\ninside 355 372

7872 GCF_001265335.1_302014 Escherichia coli Proteobacteria; Gammaproteobacteria; Enterobacterales; Enterobacteriaceae; Escherichia MTGDNTLIHSHGINRRDFMKLCAALAATMGLSSKAAAEMAESVTNPQRPPVIWIGAQECT WP_000145410.1 MULTISPECIES: hydrogenase-2 small chain [Proteobacteria] Length: 372\nNumber of predicted TMHs: 1\nExp number of AAs in TMHs: 23.72226\nExp number, first 60 AAs: 0.22038\nTotal prob of N-in: 0.12994\noutside 1 331\nTMhelix 332 354\ninside 355 372

7873 GCF_001265265.1_203741 Escherichia coli Proteobacteria; Gammaproteobacteria; Enterobacterales; Enterobacteriaceae; Escherichia MNNEETFYQAMRRQGVTRRSFLKYCSLAATSLGLGAGMAPKIAWALENKPRIPVVWIHGL WP_001058323.1 MULTISPECIES: hydrogenase-1 small chain [Proteobacteria] Length: 372\nNumber of predicted TMHs: 2\nExp number of AAs in TMHs: 37.2608499999999\nExp number, first 60 AAs: 19.52923\nTotal prob of N-in: 0.94351\nPOSSIBLE N-term signal sequence\ninside 1 19\nTMhelix 20 39\noutside 40 325\nTMhelix 326 348\ninside 349 372

7874 GCF_001265355.1_302048 Escherichia coli Proteobacteria; Gammaproteobacteria; Enterobacterales; Enterobacteriaceae; Escherichia MTGDNTLIHSHGINRRDFMKLCAALAATMGLSSKAAAEMAESVTNPQRPPVIWIGAQECT WP_000145410.1 MULTISPECIES: hydrogenase-2 small chain [Proteobacteria] Length: 372\nNumber of predicted TMHs: 1\nExp number of AAs in TMHs: 23.72226\nExp number, first 60 AAs: 0.22038\nTotal prob of N-in: 0.12994\noutside 1 331\nTMhelix 332 354\ninside 355 372

7875 GCF_001265355.1_302048 Escherichia coli Proteobacteria; Gammaproteobacteria; Enterobacterales; Enterobacteriaceae; Escherichia MNNEETFYQAMRRQGVTRRSFLKYCSLAATSLGLGAGMAPKIAWALENKPRIPVVWIHGL WP_001058323.1 MULTISPECIES: hydrogenase-1 small chain [Proteobacteria] Length: 372\nNumber of predicted TMHs: 2\nExp number of AAs in TMHs: 37.2608499999999\nExp number, first 60 AAs: 19.52923\nTotal prob of N-in: 0.94351\nPOSSIBLE N-term signal sequence\ninside 1 19\nTMhelix 20 39\noutside 40 325\nTMhelix 326 348\ninside 349 372

7876 GCF_001265335.1_302014 Escherichia coli Proteobacteria; Gammaproteobacteria; Enterobacterales; Enterobacteriaceae; Escherichia MNNEETFYQAMRRQGVTRRSFLKYCSLAATSLGLGAGMAPKIAWALENKPRIPVVWIHGL WP_001631259.1 hydrogenase-1 small chain [Escherichia coli] Length: 372\nNumber of predicted TMHs: 2\nExp number of AAs in TMHs: 37.14002\nExp number, first 60 AAs: 19.53586\nTotal prob of N-in: 0.94339\nPOSSIBLE N-term signal sequence\ninside 1 19\nTMhelix 20 39\noutside 40 325\nTMhelix 326 348\ninside 349 372

7877 GCF_001264175.1_ASM126417v1 Escherichia coli Proteobacteria; Gammaproteobacteria; Enterobacterales; Enterobacteriaceae; Escherichia MNNEETFYQAMRRQGVTRRSFLKYCSLAATSLGLGAGMAPKIAWALENKPRIPVVWIHGL WP_001058323.1 MULTISPECIES: hydrogenase-1 small chain [Proteobacteria] Length: 372\nNumber of predicted TMHs: 2\nExp number of AAs in TMHs: 37.2608499999999\nExp number, first 60 AAs: 19.52923\nTotal prob of N-in: 0.94351\nPOSSIBLE N-term signal sequence\ninside 1 19\nTMhelix 20 39\noutside 40 325\nTMhelix 326 348\ninside 349 372

7878 GCF_001265415.1_401031 Escherichia coli Proteobacteria; Gammaproteobacteria; Enterobacterales; Enterobacteriaceae; Escherichia MTGDNTLIHSHGINRRDFMKLCAALAATMGLSSKAAAEMAESVTNPQRPPVIWIGAQECT WP_000145408.1 hydrogenase 2 small subunit [Escherichia coli] Length: 372\nNumber of predicted TMHs: 1\nExp number of AAs in TMHs: 23.72255\nExp number, first 60 AAs: 0.22038\nTotal prob of N-in: 0.12995\noutside 1 331\nTMhelix 332 354\ninside 355 372

7879 GCF_001265515.1_401264 Escherichia coli Proteobacteria; Gammaproteobacteria; Enterobacterales; Enterobacteriaceae; Escherichia MTGDNTLIHSHGINRRDFMKLCAALAATMGLSSKAAAEMAESVTNPQRPPVIWIGAQECT WP_000145410.1 MULTISPECIES: hydrogenase-2 small chain [Proteobacteria] Length: 372\nNumber of predicted TMHs: 1\nExp number of AAs in TMHs: 23.72226\nExp number, first 60 AAs: 0.22038\nTotal prob of N-in: 0.12994\noutside 1 331\nTMhelix 332 354\ninside 355 372

7880 GCF_001265515.1_401264 Escherichia coli Proteobacteria; Gammaproteobacteria; Enterobacterales; Enterobacteriaceae; Escherichia MNNEETFYQAMRRQGVTRRSFLKYCSLAATSLGLGAGMAPKIAWALENKPRIPVVWIHGL WP_001058323.1 MULTISPECIES: hydrogenase-1 small chain [Proteobacteria] Length: 372\nNumber of predicted TMHs: 2\nExp number of AAs in TMHs: 37.2608499999999\nExp number, first 60 AAs: 19.52923\nTotal prob of N-in: 0.94351\nPOSSIBLE N-term signal sequence\ninside 1 19\nTMhelix 20 39\noutside 40 325\nTMhelix 326 348\ninside 349 372

7881 GCF_001265415.1_401031 Escherichia coli Proteobacteria; Gammaproteobacteria; Enterobacterales; Enterobacteriaceae; Escherichia MNNEETFYQAMRRQGVTRRSFLKYCSLAATSLGLGAGMAPKIAWALENKPRIPVVWIHGL WP_052931011.1 hydrogenase [Escherichia coli] Length: 372\nNumber of predicted TMHs: 2\nExp number of AAs in TMHs: 37.25877\nExp number, first 60 AAs: 19.52924\nTotal prob of N-in: 0.94351\nPOSSIBLE N-term signal sequence\ninside 1 19\nTMhelix 20 39\noutside 40 325\nTMhelix 326 348\ninside 349 372

7882 GCF_001264215.1_ASM126421v1 Escherichia coli Proteobacteria; Gammaproteobacteria; Enterobacterales; Enterobacteriaceae; Escherichia MNNEETFYQAMRRQGVTRRSFLKYCSLAATSLGLGAGMAPKIAWALENKPRIPVVWIHGL WP_052934366.1 hydrogenase [Escherichia coli] Length: 372\nNumber of predicted TMHs: 2\nExp number of AAs in TMHs: 36.76923\nExp number, first 60 AAs: 19.5319\nTotal prob of N-in: 0.94313\nPOSSIBLE N-term signal sequence\ninside 1 19\nTMhelix 20 39\noutside 40 325\nTMhelix 326 348\ninside 349 372

7883 GCF_001265655.1_102536 Escherichia coli Proteobacteria; Gammaproteobacteria; Enterobacterales; Enterobacteriaceae; Escherichia MTGDNTLIHSHGINRRDFMKLCAALAATMGLSSKAAAEMAESVTNPQRPPVIWIGAQECT

WP_000145410.1 MULTISPECIES: hydrogenase-2 small chain [Proteobacteria] Length: 372\nNumber of predicted TMHs: 1\nExp number of AAs in TMHs: 23.72226\nExp number, first 60 AAs: 0.22038\nTotal prob of N-in: 0.12994\noutside 1 331\nTMhelix 332 354\ninside 355 372

7884 GCF_001265755.1_300075 Escherichia coli Proteobacteria; Gammaproteobacteria; Enterobacterales; Enterobacteriaceae; Escherichia MTGDNTLIHSHGINRRDFMKLCAALAATMGLSSKAAAEMAESVTNPQRPPVIWIGAQECT WP_000145410.1 MULTISPECIES: hydrogenase-2 small chain [Proteobacteria] Length: 372\nNumber of predicted TMHs: 1\nExp number of AAs in TMHs: 23.72226\nExp number, first 60 AAs: 0.22038\nTotal prob of N-in: 0.12994\noutside 1 331\nTMhelix 332 354\ninside 355 372

7885 GCF_001265715.1_103578 Escherichia coli Proteobacteria; Gammaproteobacteria; Enterobacterales; Enterobacteriaceae; Escherichia MTGDNTLIHSHGINRRDFMKLCAALAATMGLSSKAAAEMAESVTNPQRPPVIWIGAQECT WP_000145410.1 MULTISPECIES: hydrogenase-2 small chain [Proteobacteria] Length: 372\nNumber of predicted TMHs: 1\nExp number of AAs in TMHs: 23.72226\nExp number, first 60 AAs: 0.22038\nTotal prob of N-in: 0.12994\noutside 1 331\nTMhelix 332 354\ninside 355 372

7886 GCF_001265655.1_102536 Escherichia coli Proteobacteria; Gammaproteobacteria; Enterobacterales; Enterobacteriaceae; Escherichia MNNEETFYQAMRRQGVTRRSFLKYCSLAATSLGLGAGMAPKIAWALENKPRIPVVWIHGL WP_001058323.1 MULTISPECIES: hydrogenase-1 small chain [Proteobacteria] Length: 372\nNumber of predicted TMHs: 2\nExp number of AAs in TMHs: 37.2608499999999\nExp number, first 60 AAs: 19.52923\nTotal prob of N-in: 0.94351\nPOSSIBLE N-term signal sequence\ninside 1 19\nTMhelix 20 39\noutside 40 325\nTMhelix 326 348\ninside 349 372

7887 GCF_001265845.1_302909 Escherichia coli Proteobacteria; Gammaproteobacteria; Enterobacterales; Enterobacteriaceae; Escherichia MTGDNTLIHSHGINRRDFMKLCAALAATMGLSSKAAAEMAESVTNPQRPPVIWIGAQECT WP_000145410.1 MULTISPECIES: hydrogenase-2 small chain [Proteobacteria] Length: 372\nNumber of predicted TMHs: 1\nExp number of AAs in TMHs: 23.72226\nExp number, first 60 AAs: 0.22038\nTotal prob of N-in: 0.12994\noutside 1 331\nTMhelix 332 354\ninside 355 372

7888 GCF_001265815.1_302150 Escherichia coli Proteobacteria; Gammaproteobacteria; Enterobacterales; Enterobacteriaceae; Escherichia MTGDNTLIHSHGINRRDFMKLCAALAATMGLSSKAAAEMAESVTNPQRPPVIWIGAQECT WP_000145410.1 MULTISPECIES: hydrogenase-2 small chain [Proteobacteria] Length: 372\nNumber of predicted TMHs: 1\nExp number of AAs in TMHs: 23.72226\nExp number, first 60 AAs: 0.22038\nTotal prob of N-in: 0.12994\noutside 1 331\nTMhelix 332 354\ninside 355 372

7889 GCF_001265885.1_400738 Escherichia coli Proteobacteria; Gammaproteobacteria; Enterobacterales; Enterobacteriaceae; Escherichia MTGDNTLIHSHGINRRDFMKLCAALAATMGLSSKAAAEMAESVTNPQRPPVIWIGAQECT WP_000145410.1 MULTISPECIES: hydrogenase-2 small chain [Proteobacteria] Length: 372\nNumber of predicted TMHs: 1\nExp number of AAs in TMHs: 23.72226\nExp number, first 60 AAs: 0.22038\nTotal prob of N-in: 0.12994\noutside 1 331\nTMhelix 332 354\ninside 355 372

7890 GCF_001265815.1_302150 Escherichia coli Proteobacteria; Gammaproteobacteria; Enterobacterales; Enterobacteriaceae; Escherichia MNNEETFYQAMRRQGVTRRSFLKYCSLAATSLGLGAGMAPKIAWALENKPRIPVVWIHGL WP_001058323.1 MULTISPECIES: hydrogenase-1 small chain [Proteobacteria] Length: 372\nNumber of predicted TMHs: 2\nExp number of AAs in TMHs: 37.2608499999999\nExp number, first 60 AAs: 19.52923\nTotal prob of N-in: 0.94351\nPOSSIBLE N-term signal sequence\ninside 1 19\nTMhelix 20 39\noutside 40 325\nTMhelix 326 348\ninside 349 372

7891 GCF_001265845.1_302909 Escherichia coli Proteobacteria; Gammaproteobacteria; Enterobacterales; Enterobacteriaceae; Escherichia MNNEETFYQAMRRQGVTRRSFLKYCSLAATSLGLGAGMAPKIAWALENKPRIPVVWIHGL WP_001058323.1 MULTISPECIES: hydrogenase-1 small chain [Proteobacteria] Length: 372\nNumber of predicted TMHs: 2\nExp number of AAs in TMHs: 37.2608499999999\nExp number, first 60 AAs: 19.52923\nTotal prob of N-in: 0.94351\nPOSSIBLE N-term signal sequence\ninside 1 19\nTMhelix 20 39\noutside 40 325\nTMhelix 326 348\ninside 349 372

7892 GCF_001265885.1_400738 Escherichia coli Proteobacteria; Gammaproteobacteria; Enterobacterales; Enterobacteriaceae; Escherichia MNNEETFYQAMRRQGVTRRSFLKYCSLAATSLGLGAGMAPKIAWALENKPRIPVVWIHGL WP_001058323.1 MULTISPECIES: hydrogenase-1 small chain [Proteobacteria] Length: 372\nNumber of predicted TMHs: 2\nExp number of AAs in TMHs: 37.2608499999999\nExp number, first 60 AAs: 19.52923\nTotal prob of N-in: 0.94351\nPOSSIBLE N-term signal sequence\ninside 1 19\nTMhelix 20 39\noutside 40 325\nTMhelix 326 348\ninside 349 372

7893 GCF_001265935.1_401817 Escherichia coli Proteobacteria; Gammaproteobacteria; Enterobacterales; Enterobacteriaceae; Escherichia MTGDNTLIHSHGINRRDFMKLCAALAATMGLSSKAAAEMAESVTNPQRPPVIWIGAQECT WP_000145410.1 MULTISPECIES: hydrogenase-2 small chain [Proteobacteria] Length: 372\nNumber of predicted TMHs: 1\nExp number of AAs in TMHs: 23.72226\nExp number, first 60 AAs: 0.22038\nTotal prob of N-in: 0.12994\noutside 1 331\nTMhelix 332 354\ninside 355 372

7894 GCF_001265935.1_401817 Escherichia coli Proteobacteria; Gammaproteobacteria; Enterobacterales; Enterobacteriaceae; Escherichia MNNEETFYQAMRRQGVTRRSFLKYCSLAATSLGLGAGMAPKIAWALENKPRIPVVWIHGL WP_001058323.1 MULTISPECIES: hydrogenase-1 small chain [Proteobacteria] Length: 372\nNumber of predicted TMHs: 2\nExp number of AAs in TMHs: 37.2608499999999\nExp number, first 60 AAs: 19.52923\nTotal prob of N-in: 0.94351\nPOSSIBLE N-term signal sequence\ninside 1 19\nTMhelix 20 39\noutside 40 325\nTMhelix 326 348\ninside 349 372

7895 GCF_001265755.1_300075 Escherichia coli Proteobacteria; Gammaproteobacteria; Enterobacterales; Enterobacteriaceae; Escherichia MNNEETFYQAMRRQGVTRRSFLKYCSLAATSLGLGAGMAPKIAWALENKPRIPVVWIHGL WP_001295940.1 MULTISPECIES: hydrogenase [Enterobacteriaceae] Length: 372\nNumber of predicted TMHs: 2\nExp number of AAs in TMHs: 37.2608899999999\nExp number, first 60 AAs: 19.52923\nTotal prob of N-in: 0.94351\nPOSSIBLE N-term signal sequence\ninside 1 19\nTMhelix 20 39\noutside 40 325\nTMhelix 326 348\ninside 349 372

7896 GCF_001266025.1_702626 Escherichia coli Proteobacteria; Gammaproteobacteria; Enterobacterales; Enterobacteriaceae; Escherichia MTGDNTLIHSHGINRRDFMKLCAALAATMGLSSKAAAEMAESVTNPQRPPVIWIGAQECT WP_000145410.1 MULTISPECIES: hydrogenase-2 small chain [Proteobacteria] Length: 372\nNumber of predicted TMHs: 1\nExp number of AAs in TMHs: 23.72226\nExp number, first 60 AAs: 0.22038\nTotal prob of N-in: 0.12994\noutside 1 331\nTMhelix 332 354\ninside 355 372

7897 GCF_001265995.1_402804 Escherichia coli Proteobacteria; Gammaproteobacteria; Enterobacterales; Enterobacteriaceae; Escherichia MTGDNTLIHSHGINRRDFMKLCAALAATMGLSSKAAAEMAESVTNPQRPPVIWIGAQECT WP_000145410.1 MULTISPECIES: hydrogenase-2 small chain [Proteobacteria] Length: 372\nNumber of predicted TMHs: 1\nExp number of AAs in TMHs: 23.72226\nExp number, first 60 AAs: 0.22038\nTotal prob of N-in: 0.12994\noutside 1 331\nTMhelix 332 354\ninside 355 372

7898 GCF_001266025.1_702626 Escherichia coli Proteobacteria; Gammaproteobacteria; Enterobacterales; Enterobacteriaceae; Escherichia MNNEETFYQAMRRQGVTRRSFLKYCSLAATSLGLGAGMAPKIAWALENKPRIPVVWIHGL WP_001295940.1 MULTISPECIES: hydrogenase [Enterobacteriaceae] Length: 372\nNumber of predicted TMHs: 2\nExp number of AAs in TMHs: 37.2608899999999\nExp number, first 60 AAs: 19.52923\nTotal prob of N-in: 0.94351\nPOSSIBLE N-term signal sequence\ninside 1 19\nTMhelix 20 39\noutside 40 325\nTMhelix 326 348\ninside 349 372

7899 GCF_001265995.1_402804 Escherichia coli Proteobacteria; Gammaproteobacteria; Enterobacterales; Enterobacteriaceae; Escherichia MNNEETFYQAMRRQGVTRRSFLKYCSLAATSLGLGAGMAPKIAWALENKPRIPVVWIHGL WP_001295940.1 MULTISPECIES: hydrogenase [Enterobacteriaceae] Length: 372\nNumber of predicted TMHs: 2\nExp number of AAs in TMHs: 37.2608899999999\nExp number, first 60 AAs: 19.52923\nTotal prob of N-in: 0.94351\nPOSSIBLE N-term signal sequence\ninside 1 19\nTMhelix 20 39\noutside 40 325\nTMhelix 326 348\ninside 349 372

7900 GCF_001266095.1_102132 Escherichia coli Proteobacteria; Gammaproteobacteria; Enterobacterales; Enterobacteriaceae; Escherichia MTGDNTLIHSHGINRRDFMKLCAALAATMGLSSKAAAEMAESVTNPQRPPVIWIGAQECT WP_000145410.1 MULTISPECIES: hydrogenase-2 small chain [Proteobacteria] Length: 372\nNumber of predicted TMHs: 1\nExp number of AAs in TMHs: 23.72226\nExp number, first 60 AAs: 0.22038\nTotal prob of N-in: 0.12994\noutside 1 331\nTMhelix 332 354\ninside 355 372

7901 GCF_001266095.1_102132 Escherichia coli Proteobacteria; Gammaproteobacteria; Enterobacterales; Enterobacteriaceae; Escherichia MNNEETFYQAMRRQGVTRRSFLKYCSLAATSLGLGAGMAPKIAWALENKPRIPVVWIHGL WP_001058323.1 MULTISPECIES: hydrogenase-1 small chain [Proteobacteria] Length: 372\nNumber of predicted TMHs: 2\nExp number of AAs in TMHs: 37.2608499999999\nExp number, first 60 AAs: 19.52923\nTotal prob of N-in: 0.94351\nPOSSIBLE N-term signal sequence\ninside 1 19\nTMhelix 20 39\noutside 40 325\nTMhelix 326 348\ninside 349 372

7902 GCF_001265715.1_103578 Escherichia coli Proteobacteria; Gammaproteobacteria; Enterobacterales; Enterobacteriaceae; Escherichia MNNEETFYQAMRRQGVTRRSFLKYCSLAATSLGLGAGMAPKIAWALENKPRIPVVWIHGL

WP_001295940.1 MULTISPECIES: hydrogenase [Enterobacteriaceae] Length: 372\nNumber of predicted TMHs: 2\nExp number of AAs in TMHs: 37.2608899999999\nExp number, first 60 AAs: 19.52923\nTotal prob of N-in: 0.94351\nPOSSIBLE N-term signal sequence\ninside 1 19\nTMhelix 20 39\noutside 40 325\nTMhelix 326 348\ninside 349 372

7903 GCF_001266155.1_103447 Escherichia coli Proteobacteria; Gammaproteobacteria; Enterobacterales; Enterobacteriaceae; Escherichia MTGDNTLIHSHGINRRDFMKLCAALAATMGLSSKAAAEMAESVTNPQRPPVIWIGAQECT WP_000145410.1 MULTISPECIES: hydrogenase-2 small chain [Proteobacteria] Length: 372\nNumber of predicted TMHs: 1\nExp number of AAs in TMHs: 23.72226\nExp number, first 60 AAs: 0.22038\nTotal prob of N-in: 0.12994\noutside 1 331\nTMhelix 332 354\ninside 355 372

7904 GCF_001266195.1_300469 Escherichia coli Proteobacteria; Gammaproteobacteria; Enterobacterales; Enterobacteriaceae; Escherichia MTGDNTLIHSHGINRRDFMKLCAALAATMGLSSKAAAEMAESVTNPQRPPVIWIGAQECT WP_000145410.1 MULTISPECIES: hydrogenase-2 small chain [Proteobacteria] Length: 372\nNumber of predicted TMHs: 1\nExp number of AAs in TMHs: 23.72226\nExp number, first 60 AAs: 0.22038\nTotal prob of N-in: 0.12994\noutside 1 331\nTMhelix 332 354\ninside 355 372

7905 GCF_001266255.1_302312 Escherichia coli Proteobacteria; Gammaproteobacteria; Enterobacterales; Enterobacteriaceae; Escherichia MTGDNTLIHSHGINRRDFMKLCAALAATMGLSSKAAAEMAESVTNPQRPPVIWIGAQECT WP_000145410.1 MULTISPECIES: hydrogenase-2 small chain [Proteobacteria] Length: 372\nNumber of predicted TMHs: 1\nExp number of AAs in TMHs: 23.72226\nExp number, first 60 AAs: 0.22038\nTotal prob of N-in: 0.12994\noutside 1 331\nTMhelix 332 354\ninside 355 372

7906 GCF_001266335.1_400929 Escherichia coli Proteobacteria; Gammaproteobacteria; Enterobacterales; Enterobacteriaceae; Escherichia MTGDNTLIHSHGINRRDFMKLCAALAATMGLSSKAAAEMAESVTNPQRPPVIWIGAQECT WP_000145410.1 MULTISPECIES: hydrogenase-2 small chain [Proteobacteria] Length: 372\nNumber of predicted TMHs: 1\nExp number of AAs in TMHs: 23.72226\nExp number, first 60 AAs: 0.22038\nTotal prob of N-in: 0.12994\noutside 1 331\nTMhelix 332 354\ninside 355 372

7907 GCF_001266385.1_402559 Escherichia coli Proteobacteria; Gammaproteobacteria; Enterobacterales; Enterobacteriaceae; Escherichia MTGDNTLIHSHGINRRDFMKLCAALAATMGLSSKAAAEMAESVTNPQRPPVIWIGAQECT WP_000145410.1 MULTISPECIES: hydrogenase-2 small chain [Proteobacteria] Length: 372\nNumber of predicted TMHs: 1\nExp number of AAs in TMHs: 23.72226\nExp number, first 60 AAs: 0.22038\nTotal prob of N-in: 0.12994\noutside 1 331\nTMhelix 332 354\ninside 355 372

7908 GCF_001266155.1_103447 Escherichia coli Proteobacteria; Gammaproteobacteria; Enterobacterales; Enterobacteriaceae; Escherichia MNNEETFYQAMRRQGVTRRSFLKYCSLAATSLGLGAGMAPKIAWALENKPRIPVVWIHGL WP_001058323.1 MULTISPECIES: hydrogenase-1 small chain [Proteobacteria] Length: 372\nNumber of predicted TMHs: 2\nExp number of AAs in TMHs: 37.2608499999999\nExp number, first 60 AAs: 19.52923\nTotal prob of N-in: 0.94351\nPOSSIBLE N-term signal sequence\ninside 1 19\nTMhelix 20 39\noutside 40 325\nTMhelix 326 348\ninside 349 372

7909 GCF_001266295.1_303341 Escherichia coli Proteobacteria; Gammaproteobacteria; Enterobacterales; Enterobacteriaceae; Escherichia MTGDNTLIHSHGINRRDFMKLCAALAATMGLSSKAAAEMAESVTNPQRPPVIWIGAQECT WP_000145410.1 MULTISPECIES: hydrogenase-2 small chain [Proteobacteria] Length: 372\nNumber of predicted TMHs: 1\nExp number of AAs in TMHs: 23.72226\nExp number, first 60 AAs: 0.22038\nTotal prob of N-in: 0.12994\noutside 1 331\nTMhelix 332 354\ninside 355 372

7910 GCF_001266295.1_303341 Escherichia coli Proteobacteria; Gammaproteobacteria; Enterobacterales; Enterobacteriaceae; Escherichia MNNEETFYQAMRRQGVTRRSFLKYCSLAATSLGLGAGMAPKIAWALENKPRIPVVWIHGL WP_001058323.1 MULTISPECIES: hydrogenase-1 small chain [Proteobacteria] Length: 372\nNumber of predicted TMHs: 2\nExp number of AAs in TMHs: 37.2608499999999\nExp number, first 60 AAs: 19.52923\nTotal prob of N-in: 0.94351\nPOSSIBLE N-term signal sequence\ninside 1 19\nTMhelix 20 39\noutside 40 325\nTMhelix 326 348\ninside 349 372

7911 GCF_001266385.1_402559 Escherichia coli Proteobacteria; Gammaproteobacteria; Enterobacterales; Enterobacteriaceae; Escherichia MNNEETFYQAMRRQGVTRRSFLKYCSLAATSLGLGAGMAPKIAWALENKPRIPVVWIHGL WP_001058323.1 MULTISPECIES: hydrogenase-1 small chain [Proteobacteria] Length: 372\nNumber of predicted TMHs: 2\nExp number of AAs in TMHs: 37.2608499999999\nExp number, first 60 AAs: 19.52923\nTotal prob of N-in: 0.94351\nPOSSIBLE N-term signal sequence\ninside 1 19\nTMhelix 20 39\noutside 40 325\nTMhelix 326 348\ninside 349 372

7912 GCF_001266405.1_402981 Escherichia coli Proteobacteria; Gammaproteobacteria; Enterobacterales; Enterobacteriaceae; Escherichia MTGDNTLIHSHGINRRDFMKLCAALAATMGLSSKAAAEMAESVTNPQRPPVIWIGAQECT WP_000145410.1 MULTISPECIES: hydrogenase-2 small chain [Proteobacteria] Length: 372\nNumber of predicted TMHs: 1\nExp number of AAs in TMHs: 23.72226\nExp number, first 60 AAs: 0.22038\nTotal prob of N-in: 0.12994\noutside 1 331\nTMhelix 332 354\ninside 355 372

7913 GCF_001266405.1_402981 Escherichia coli Proteobacteria; Gammaproteobacteria; Enterobacterales; Enterobacteriaceae; Escherichia MNNEETFYQAMRRQGVTRRSFLKYCSLAATSLGLGAGMAPKIAWALENKPRIPVVWIHGL WP_001058323.1 MULTISPECIES: hydrogenase-1 small chain [Proteobacteria] Length: 372\nNumber of predicted TMHs: 2\nExp number of AAs in TMHs: 37.2608499999999\nExp number, first 60 AAs: 19.52923\nTotal prob of N-in: 0.94351\nPOSSIBLE N-term signal sequence\ninside 1 19\nTMhelix 20 39\noutside 40 325\nTMhelix 326 348\ninside 349 372

7914 GCF_001266195.1_300469 Escherichia coli Proteobacteria; Gammaproteobacteria; Enterobacterales; Enterobacteriaceae; Escherichia MNNEETFYQAMRRQGVTRRSFLKYCSLAATSLGLGAGMAPKIAWALENKPRIPVVWIHGL WP_001295940.1 MULTISPECIES: hydrogenase [Enterobacteriaceae] Length: 372\nNumber of predicted TMHs: 2\nExp number of AAs in TMHs: 37.2608899999999\nExp number, first 60 AAs: 19.52923\nTotal prob of N-in: 0.94351\nPOSSIBLE N-term signal sequence\ninside 1 19\nTMhelix 20 39\noutside 40 325\nTMhelix 326 348\ninside 349 372

7915 GCF_001266255.1_302312 Escherichia coli Proteobacteria; Gammaproteobacteria; Enterobacterales; Enterobacteriaceae; Escherichia MNNEETFYQAMRRQGVTRRSFLKYCSLAATSLGLGAGMAPKIAWALENKPRIPVVWIHGL WP_001295940.1 MULTISPECIES: hydrogenase [Enterobacteriaceae] Length: 372\nNumber of predicted TMHs: 2\nExp number of AAs in TMHs: 37.2608899999999\nExp number, first 60 AAs: 19.52923\nTotal prob of N-in: 0.94351\nPOSSIBLE N-term signal sequence\ninside 1 19\nTMhelix 20 39\noutside 40 325\nTMhelix 326 348\ninside 349 372

7916 GCF_001266435.1_403341 Escherichia coli Proteobacteria; Gammaproteobacteria; Enterobacterales; Enterobacteriaceae; Escherichia MTGDNTLIHSHGINRRDFMKLCAALAATMGLSSKAAAEMAESVTNPQRPPVIWIGAQECT WP_000145410.1 MULTISPECIES: hydrogenase-2 small chain [Proteobacteria] Length: 372\nNumber of predicted TMHs: 1\nExp number of AAs in TMHs: 23.72226\nExp number, first 60 AAs: 0.22038\nTotal prob of N-in: 0.12994\noutside 1 331\nTMhelix 332 354\ninside 355 372

7917 GCF_001267345.1_303289 Escherichia coli Proteobacteria; Gammaproteobacteria; Enterobacterales; Enterobacteriaceae; Escherichia MTGDNTLIHSHGINRRDFMKLCAALAATMGLSSKAAAEMAESVTNPQRPPVIWIGAQECT WP_000145410.1 MULTISPECIES: hydrogenase-2 small chain [Proteobacteria] Length: 372\nNumber of predicted TMHs: 1\nExp number of AAs in TMHs: 23.72226\nExp number, first 60 AAs: 0.22038\nTotal prob of N-in: 0.12994\noutside 1 331\nTMhelix 332 354\ninside 355 372

7918 GCF_001267345.1_303289 Escherichia coli Proteobacteria; Gammaproteobacteria; Enterobacterales; Enterobacteriaceae; Escherichia MNNEETFYQAMRRQGVTRRSFLKYCSLAATSLGLGAGMAPKIAWALENKPRIPVVWIHGL WP_001058323.1 MULTISPECIES: hydrogenase-1 small chain [Proteobacteria] Length: 372\nNumber of predicted TMHs: 2\nExp number of AAs in TMHs: 37.2608499999999\nExp number, first 60 AAs: 19.52923\nTotal prob of N-in: 0.94351\nPOSSIBLE N-term signal sequence\ninside 1 19\nTMhelix 20 39\noutside 40 325\nTMhelix 326 348\ninside 349 372

7919 GCF_001266435.1_403341 Escherichia coli Proteobacteria; Gammaproteobacteria; Enterobacterales; Enterobacteriaceae; Escherichia MNNEETFYQAMRRQGVTRRSFLKYCSLAATSLGLGAGMAPKIAWALENKPRIPVVWIHGL WP_052923131.1 hydrogenase [Escherichia coli] Length: 372\nNumber of predicted TMHs: 2\nExp number of AAs in TMHs: 36.3624399999999\nExp number, first 60 AAs: 19.53053\nTotal prob of N-in: 0.94346\nPOSSIBLE N-term signal sequence\ninside 1 19\nTMhelix 20 39\noutside 40 325\nTMhelix 326 348\ninside 349 372

7920 GCF_001266335.1_400929 Escherichia coli Proteobacteria; Gammaproteobacteria; Enterobacterales; Enterobacteriaceae; Escherichia MNNEETFYQAMRRQGVTRRSFLKYCSLAATSLGLGAGMAPKIAWALENKPRIPVVWIHGL WP_001058323.1 MULTISPECIES: hydrogenase-1 small chain [Proteobacteria] Length: 372\nNumber of predicted TMHs: 2\nExp number of AAs in TMHs: 37.2608499999999\nExp number, first 60 AAs: 19.52923\nTotal prob of N-in: 0.94351\nPOSSIBLE N-term signal sequence\ninside 1 19\nTMhelix 20 39\noutside 40 325\nTMhelix 326 348\ninside 349 372

7921 GCF_001268185.1_1.EC2994.1 Escherichia coli Proteobacteria; Gammaproteobacteria; Enterobacterales; Enterobacteriaceae; Escherichia
MTGDNTLIHSHGINRRDFMKLCAALAATMGLSSKAAAEMAESVTNPQRPPVIWIGAQECT WP_000145410.1
MULTISPECIES: hydrogenase-2 small chain [Proteobacteria] Length: 372\nNumber of predicted TMHs: 1\nExp number of AAs in TMHs: 23.72226\nExp number, first 60 AAs: 0.22038\nTotal prob of N-in: 0.12994\noutside 1 331\nTMhelix 332 354\ninside 355 372

7922 GCF_001268285.1_1.ECAJ28.1 Escherichia coli Proteobacteria; Gammaproteobacteria; Enterobacterales; Enterobacteriaceae; Escherichia
MTGDNTLIHSHGINRRDFMKLCAALAATMGLSSKAAAEMAESVTNPQRPPVIWIGAQECT WP_000145410.1
MULTISPECIES: hydrogenase-2 small chain [Proteobacteria] Length: 372\nNumber of predicted TMHs: 1\nExp number of AAs in TMHs: 23.72226\nExp number, first 60 AAs: 0.22038\nTotal prob of N-in: 0.12994\noutside 1 331\nTMhelix 332 354\ninside 355 372

7923 GCF_001268245.1_1.EC2722.1 Escherichia coli Proteobacteria; Gammaproteobacteria; Enterobacterales; Enterobacteriaceae; Escherichia
MTGDNTLIHSHGINRRDFMKLCAALAATMGLSSKAAAEMAESVTNPQRPPVIWIGAQECT WP_000145410.1
MULTISPECIES: hydrogenase-2 small chain [Proteobacteria] Length: 372\nNumber of predicted TMHs: 1\nExp number of AAs in TMHs: 23.72226\nExp number, first 60 AAs: 0.22038\nTotal prob of N-in: 0.12994\noutside 1 331\nTMhelix 332 354\ninside 355 372

7924 GCF_001268185.1_1.EC2994.1 Escherichia coli Proteobacteria; Gammaproteobacteria; Enterobacterales; Enterobacteriaceae; Escherichia
MNNEETFYQAMRRQGVTRRSFLKYCSLAATSLGLGAGMAPKIAWALENKPRIPVVWIHGL WP_001058323.1
MULTISPECIES: hydrogenase-1 small chain [Proteobacteria] Length: 372\nNumber of predicted TMHs: 2\nExp number of AAs in TMHs: 37.2608499999999\nExp number, first 60 AAs: 19.52923\nTotal prob of N-in: 0.94351\nPOSSIBLE N-term signal sequence\ninside 1 19\nTMhelix 20 39\noutside 40 325\nTMhelix 326 348\ninside 349 372

7925 GCF_001268285.1_1.ECAJ28.1 Escherichia coli Proteobacteria; Gammaproteobacteria; Enterobacterales; Enterobacteriaceae; Escherichia
MNNEETFYQAMRRQGVTRRSFLKYCSLAATSLGLGAGMAPKIAWALENKPRIPVVWIHGL WP_001058323.1
MULTISPECIES: hydrogenase-1 small chain [Proteobacteria] Length: 372\nNumber of predicted TMHs: 2\nExp number of AAs in TMHs: 37.2608499999999\nExp number, first 60 AAs: 19.52923\nTotal prob of N-in: 0.94351\nPOSSIBLE N-term signal sequence\ninside 1 19\nTMhelix 20 39\noutside 40 325\nTMhelix 326 348\ninside 349 372

7926 GCF_001268325.1_YE23 Escherichia coli Proteobacteria; Gammaproteobacteria; Enterobacterales; Enterobacteriaceae; Escherichia MTGDNTLIHSHGINRRDFMKLCAALAATMGLSSKAAAEMAESVTNPQRPPVIWIGAQECT
WP_000145410.1 MULTISPECIES: hydrogenase-2 small chain [Proteobacteria] Length: 372\nNumber of predicted TMHs: 1\nExp number of AAs in TMHs: 23.72226\nExp number, first 60 AAs: 0.22038\nTotal prob of N-in: 0.12994\noutside 1 331\nTMhelix 332 354\ninside 355 372

7927 GCF_001268385.1_1.EC2719.1 Escherichia coli Proteobacteria; Gammaproteobacteria; Enterobacterales; Enterobacteriaceae; Escherichia
MTGDNTLIHSHGINRRDFMKLCAALAATMGLSSKAAAEMAESVTNPQRPPVIWIGAQECT WP_000145410.1
MULTISPECIES: hydrogenase-2 small chain [Proteobacteria] Length: 372\nNumber of predicted TMHs: 1\nExp number of AAs in TMHs: 23.72226\nExp number, first 60 AAs: 0.22038\nTotal prob of N-in: 0.12994\noutside 1 331\nTMhelix 332 354\ninside 355 372

7928 GCF_001268385.1_1.EC2719.1 Escherichia coli Proteobacteria; Gammaproteobacteria; Enterobacterales; Enterobacteriaceae; Escherichia
MNNEETFYQAMRRQGVTRRSFLKYCSLAATSLGLGAGMAPKIAWALENKPRIPVVWIHGL WP_001058323.1
MULTISPECIES: hydrogenase-1 small chain [Proteobacteria] Length: 372\nNumber of predicted TMHs: 2\nExp number of AAs in TMHs: 37.2608499999999\nExp number, first 60 AAs: 19.52923\nTotal prob of N-in: 0.94351\nPOSSIBLE N-term signal sequence\ninside 1 19\nTMhelix 20 39\noutside 40 325\nTMhelix 326 348\ninside 349 372

7929 GCF_001268245.1_1.EC2722.1 Escherichia coli Proteobacteria; Gammaproteobacteria; Enterobacterales; Enterobacteriaceae; Escherichia
MNNEETFYQAMRRQGVTRRSFLKYCSLAATSLGLGAGMAPKIAWALENKPRIPVVWIHGL WP_001058323.1

MULTISPECIES: hydrogenase-1 small chain [Proteobacteria] Length: 372\nNumber of predicted TMHs: 2\nExp number of AAs in TMHs: 37.2608499999999\nExp number, first 60 AAs: 19.52923\nTotal prob of N-in: 0.94351\nPOSSIBLE N-term signal sequence\ninside 1 19\nTMhelix 20 39\noutside 40 325\nTMhelix 326 348\ninside 349 372

7930 GCF_001280345.1_ASM128034v1 Escherichia coli Proteobacteria; Gammaproteobacteria; Enterobacterales; Enterobacteriaceae; Escherichia MTGDNTLIHSHGINRRDFMKLCAALAATMGLSSKAAAEMAESVTNPQRPPVIWIGAQECT WP_000145410.1 MULTISPECIES: hydrogenase-2 small chain [Proteobacteria] Length: 372\nNumber of predicted TMHs: 1\nExp number of AAs in TMHs: 23.72226\nExp number, first 60 AAs: 0.22038\nTotal prob of N-in: 0.12994\noutside 1 331\nTMhelix 332 354\ninside 355 372

7931 GCF_001280405.1_ASM128040v1 Escherichia coli Proteobacteria; Gammaproteobacteria; Enterobacterales; Enterobacteriaceae; Escherichia MTGDNTLIHSHGINRRDFMKLCAALAATMGLSSKAAAEMAESVTNPQRPPVIWIGAQECT WP_000145410.1 MULTISPECIES: hydrogenase-2 small chain [Proteobacteria] Length: 372\nNumber of predicted TMHs: 1\nExp number of AAs in TMHs: 23.72226\nExp number, first 60 AAs: 0.22038\nTotal prob of N-in: 0.12994\noutside 1 331\nTMhelix 332 354\ninside 355 372

7932 GCF_001268325.1_YE23 Escherichia coli Proteobacteria; Gammaproteobacteria; Enterobacterales; Enterobacteriaceae; Escherichia MNNEETFYQAMRRQGVTRRSFLKYCSLAATSLGLGAGMAPKIAWALENKPRIPVVWIHGL WP_001058323.1 MULTISPECIES: hydrogenase-1 small chain [Proteobacteria] Length: 372\nNumber of predicted TMHs: 2\nExp number of AAs in TMHs: 37.2608499999999\nExp number, first 60 AAs: 19.52923\nTotal prob of N-in: 0.94351\nPOSSIBLE N-term signal sequence\ninside 1 19\nTMhelix 20 39\noutside 40 325\nTMhelix 326 348\ninside 349 372

7933 GCF_001281725.1_ASM128172v1 Escherichia coli Proteobacteria; Gammaproteobacteria; Enterobacterales; Enterobacteriaceae; Escherichia MTGDNTLIHSHGINRRDFMKLCAALAATMGLSSKAAAEMAESVTNPQRPPVIWIGAQECT WP_000145410.1 MULTISPECIES: hydrogenase-2 small chain [Proteobacteria] Length: 372\nNumber of predicted TMHs: 1\nExp number of AAs in TMHs: 23.72226\nExp number, first 60 AAs: 0.22038\nTotal prob of N-in: 0.12994\noutside 1 331\nTMhelix 332 354\ninside 355 372

7934 GCF_001281845.1_ASM128184v1 Escherichia coli Proteobacteria; Gammaproteobacteria; Enterobacterales; Enterobacteriaceae; Escherichia MTGDNTLIHSHGINRRDFMKLCAALAATMGLSSKAAAEMAESVTNPQRPPVIWIGAQECT WP_000145410.1 MULTISPECIES: hydrogenase-2 small chain [Proteobacteria] Length: 372\nNumber of predicted TMHs: 1\nExp number of AAs in TMHs: 23.72226\nExp number, first 60 AAs: 0.22038\nTotal prob of N-in: 0.12994\noutside 1 331\nTMhelix 332 354\ninside 355 372

7935 GCF_001281795.1_ASM128179v1 Escherichia coli Proteobacteria; Gammaproteobacteria; Enterobacterales; Enterobacteriaceae; Escherichia MTGDNTLIHSHGINRRDFMKLCAALAATMGLSSKAAAEMAESVTNPQRPPVIWIGAQECT WP_000145410.1 MULTISPECIES: hydrogenase-2 small chain [Proteobacteria] Length: 372\nNumber of predicted TMHs: 1\nExp number of AAs in TMHs: 23.72226\nExp number, first 60 AAs: 0.22038\nTotal prob of N-in: 0.12994\noutside 1 331\nTMhelix 332 354\ninside 355 372

7936 GCF_001281905.1_ASM128190v1 Escherichia coli Proteobacteria; Gammaproteobacteria; Enterobacterales; Enterobacteriaceae; Escherichia MTGDNTLIHSHGINRRDFMKLCAALAATMGLSSKAAAEMAESVTNPQRPPVIWIGAQECT WP_000145410.1 MULTISPECIES: hydrogenase-2 small chain [Proteobacteria] Length: 372\nNumber of predicted TMHs: 1\nExp number of AAs in TMHs: 23.72226\nExp number, first 60 AAs: 0.22038\nTotal prob of N-in: 0.12994\noutside 1 331\nTMhelix 332 354\ninside 355 372

7937 GCF_001281925.1_ASM128192v1 Escherichia coli Proteobacteria; Gammaproteobacteria; Enterobacterales; Enterobacteriaceae; Escherichia MTGDNTLIHSHGINRRDFMKLCAALAATMGLSSKAAAEMAESVTNPQRPPVIWIGAQECT WP_000145410.1 MULTISPECIES: hydrogenase-2 small chain [Proteobacteria] Length: 372\nNumber of predicted TMHs: 1\nExp number of AAs in TMHs: 23.72226\nExp number, first 60 AAs: 0.22038\nTotal prob of N-in: 0.12994\noutside 1 331\nTMhelix 332 354\ninside 355 372

7938 GCF_001281995.1_ASM128199v1 Escherichia coli Proteobacteria; Gammaproteobacteria; Enterobacterales; Enterobacteriaceae; Escherichia
MTGDNTLIHSHGINRRDFMKLCAALAATMGLSSKAAAEMAESVTNPQRPPVIWIGAQECT WP_000145410.1
MULTISPECIES: hydrogenase-2 small chain [Proteobacteria] Length: 372\nNumber of predicted TMHs: 1\nExp number of AAs in TMHs: 23.72226\nExp number, first 60 AAs: 0.22038\nTotal prob of N-in: 0.12994\noutside 1 331\nTMhelix 332 354\ninside 355 372

7939 GCF_001282035.1_ASM128203v1 Escherichia coli Proteobacteria; Gammaproteobacteria; Enterobacterales; Enterobacteriaceae; Escherichia
MTGDNTLIHSHGINRRDFMKLCAALAATMGLSSKAAAEMAESVTNPQRPPVIWIGAQECT WP_000145410.1
MULTISPECIES: hydrogenase-2 small chain [Proteobacteria] Length: 372\nNumber of predicted TMHs: 1\nExp number of AAs in TMHs: 23.72226\nExp number, first 60 AAs: 0.22038\nTotal prob of N-in: 0.12994\noutside 1 331\nTMhelix 332 354\ninside 355 372

7940 GCF_001282195.1_ASM128219v1 Escherichia coli Proteobacteria; Gammaproteobacteria; Enterobacterales; Enterobacteriaceae; Escherichia
MTGDNTLIHSHGINRRDFMKLCAALAATMGLSSKAAAEMAESVTNPQRPPVIWIGAQECT WP_000145410.1
MULTISPECIES: hydrogenase-2 small chain [Proteobacteria] Length: 372\nNumber of predicted TMHs: 1\nExp number of AAs in TMHs: 23.72226\nExp number, first 60 AAs: 0.22038\nTotal prob of N-in: 0.12994\noutside 1 331\nTMhelix 332 354\ninside 355 372

7941 GCF_001282195.1_ASM128219v1 Escherichia coli Proteobacteria; Gammaproteobacteria; Enterobacterales; Enterobacteriaceae; Escherichia
MNNEETFYQAMRRQGVTRRSFLKYCSLAATSLGLGAGMAPKIAWALENKPRIPVVWIHGL WP_001058320.1
hydrogenase [Escherichia coli] Length: 372\nNumber of predicted TMHs: 2\nExp number of AAs in TMHs: 37.26082\nExp number, first 60 AAs: 19.52923\nTotal prob of N-in: 0.94351\nPOSSIBLE N-term signal sequence\ninside 1 19\nTMhelix 20 39\noutside 40 325\nTMhelix 326 348\ninside 349 372

7942 GCF_001281845.1_ASM128184v1 Escherichia coli Proteobacteria; Gammaproteobacteria; Enterobacterales; Enterobacteriaceae; Escherichia
MNNEETFYQAMRRQGVTRRSFLKYCSLAATSLGLGAGMAPKIAWALENKPRIPVVWIHGL WP_001301613.1
hydrogenase [Escherichia coli] Length: 372\nNumber of predicted TMHs: 2\nExp number of AAs in TMHs: 37.46598\nExp number, first 60 AAs: 19.53139\nTotal prob of N-in: 0.94364\nPOSSIBLE N-term signal sequence\ninside 1 19\nTMhelix 20 39\noutside 40 325\nTMhelix 326 348\ninside 349 372

7943 GCF_001281725.1_ASM128172v1 Escherichia coli Proteobacteria; Gammaproteobacteria; Enterobacterales; Enterobacteriaceae; Escherichia
MNNEETFYQAMRRQGVTRRSFLKYCSLAATSLGLGAGMAPKIAWALENKPRIPVVWIHGL WP_001301613.1
hydrogenase [Escherichia coli] Length: 372\nNumber of predicted TMHs: 2\nExp number of AAs in TMHs: 37.46598\nExp number, first 60 AAs: 19.53139\nTotal prob of N-in: 0.94364\nPOSSIBLE N-term signal sequence\ninside 1 19\nTMhelix 20 39\noutside 40 325\nTMhelix 326 348\ninside 349 372

7944 GCF_001281795.1_ASM128179v1 Escherichia coli Proteobacteria; Gammaproteobacteria; Enterobacterales; Enterobacteriaceae; Escherichia
MNNEETFYQAMRRQGVTRRSFLKYCSLAATSLGLGAGMAPKIAWALENKPRIPVVWIHGL WP_001301613.1
hydrogenase [Escherichia coli] Length: 372\nNumber of predicted TMHs: 2\nExp number of AAs in TMHs: 37.46598\nExp number, first 60 AAs: 19.53139\nTotal prob of N-in: 0.94364\nPOSSIBLE N-term signal sequence\ninside 1 19\nTMhelix 20 39\noutside 40 325\nTMhelix 326 348\ninside 349 372

7945 GCF_001282295.1_ASM128229v1 Escherichia coli Proteobacteria; Gammaproteobacteria; Enterobacterales; Enterobacteriaceae; Escherichia
MTGDNTLIHSHGINRRDFMKLCAALAATMGLSSKAAAEMAESVTNPQRPPVIWIGAQECT WP_000145410.1
MULTISPECIES: hydrogenase-2 small chain [Proteobacteria] Length: 372\nNumber of predicted TMHs: 1\nExp number of AAs in TMHs: 23.72226\nExp number, first 60 AAs: 0.22038\nTotal prob of N-in: 0.12994\noutside 1 331\nTMhelix 332 354\ninside 355 372

7946 GCF_001282225.1_ASM128222v1 Escherichia coli Proteobacteria; Gammaproteobacteria; Enterobacterales; Enterobacteriaceae; Escherichia
MTGDNTLIHSHGINRRDFMKLCAALAATMGLSSKAAAEMAESVTNPQRPPVIWIGAQECT WP_000145410.1
MULTISPECIES: hydrogenase-2 small chain [Proteobacteria] Length: 372\nNumber of predicted TMHs:

1\nExp number of AAs in TMHs: 23.72226\nExp number, first 60 AAs: 0.22038\nTotal prob of N-in: 0.12994\noutside 1 331\nTMhelix 332 354\ninside 355 372

7947 GCF_001282295.1_ASM128229v1 Escherichia coli Proteobacteria; Gammaproteobacteria; Enterobacterales; Enterobacteriaceae; Escherichia
MNNEETFYQAMRRQGVTRRSFLKYCSLAATSLGLGAGMAPKIAWALENKPRIPVVWIHGL WP_001058320.1
hydrogenase [Escherichia coli] Length: 372\nNumber of predicted TMHs: 2\nExp number of AAs in TMHs: 37.26082\nExp number, first 60 AAs: 19.52923\nTotal prob of N-in: 0.94351\nPOSSIBLE N-term signal sequence\ninside 1 19\nTMhelix 20 39\noutside 40 325\nTMhelix 326 348\ninside 349 372

7948 GCF_001282225.1_ASM128222v1 Escherichia coli Proteobacteria; Gammaproteobacteria; Enterobacterales; Enterobacteriaceae; Escherichia
MNNEETFYQAMRRQGVTRRSFLKYCSLAATSLGLGAGMAPKIAWALENKPRIPVVWIHGL WP_001058320.1
hydrogenase [Escherichia coli] Length: 372\nNumber of predicted TMHs: 2\nExp number of AAs in TMHs: 37.26082\nExp number, first 60 AAs: 19.52923\nTotal prob of N-in: 0.94351\nPOSSIBLE N-term signal sequence\ninside 1 19\nTMhelix 20 39\noutside 40 325\nTMhelix 326 348\ninside 349 372

7949 GCF_001281905.1_ASM128190v1 Escherichia coli Proteobacteria; Gammaproteobacteria; Enterobacterales; Enterobacteriaceae; Escherichia
MNNEETFYQAMRRQGVTRRSFLKYCSLAATSLGLGAGMAPKIAWALENKPRIPVVWIHGL WP_001301613.1
hydrogenase [Escherichia coli] Length: 372\nNumber of predicted TMHs: 2\nExp number of AAs in TMHs: 37.46598\nExp number, first 60 AAs: 19.53139\nTotal prob of N-in: 0.94364\nPOSSIBLE N-term signal sequence\ninside 1 19\nTMhelix 20 39\noutside 40 325\nTMhelix 326 348\ninside 349 372

7950 GCF_001282345.1_ASM128234v1 Escherichia coli Proteobacteria; Gammaproteobacteria; Enterobacterales; Enterobacteriaceae; Escherichia
MTGDNTLIHSHGINRRDFMKLCAALAATMGLSSKAAAEMAESVTNPQRPPVIWIGAQECT WP_000145410.1
MULTISPECIES: hydrogenase-2 small chain [Proteobacteria] Length: 372\nNumber of predicted TMHs: 1\nExp number of AAs in TMHs: 23.72226\nExp number, first 60 AAs: 0.22038\nTotal prob of N-in: 0.12994\noutside 1 331\nTMhelix 332 354\ninside 355 372

7951 GCF_001282345.1_ASM128234v1 Escherichia coli Proteobacteria; Gammaproteobacteria; Enterobacterales; Enterobacteriaceae; Escherichia
MNNEETFYQAMRRQGVTRRSFLKYCSLAATSLGLGAGMAPKIAWALENKPRIPVVWIHGL WP_001058320.1
hydrogenase [Escherichia coli] Length: 372\nNumber of predicted TMHs: 2\nExp number of AAs in TMHs: 37.26082\nExp number, first 60 AAs: 19.52923\nTotal prob of N-in: 0.94351\nPOSSIBLE N-term signal sequence\ninside 1 19\nTMhelix 20 39\noutside 40 325\nTMhelix 326 348\ninside 349 372

7952 GCF_001281925.1_ASM128192v1 Escherichia coli Proteobacteria; Gammaproteobacteria; Enterobacterales; Enterobacteriaceae; Escherichia
MNNEETFYQAMRRQGVTRRSFLKYCSLAATSLGLGAGMAPKIAWALENKPRIPVVWIHGL WP_001301613.1
hydrogenase [Escherichia coli] Length: 372\nNumber of predicted TMHs: 2\nExp number of AAs in TMHs: 37.46598\nExp number, first 60 AAs: 19.53139\nTotal prob of N-in: 0.94364\nPOSSIBLE N-term signal sequence\ninside 1 19\nTMhelix 20 39\noutside 40 325\nTMhelix 326 348\ninside 349 372

7953 GCF_001283165.1_7553_7_66 Escherichia coli Proteobacteria; Gammaproteobacteria; Enterobacterales; Enterobacteriaceae; Escherichia
MTGDNTLIHSHGINRRDFMKLCAALAATMGLSSKAAAEMAESVTNPQRPPVIWIGAQECT WP_000145410.1
MULTISPECIES: hydrogenase-2 small chain [Proteobacteria] Length: 372\nNumber of predicted TMHs: 1\nExp number of AAs in TMHs: 23.72226\nExp number, first 60 AAs: 0.22038\nTotal prob of N-in: 0.12994\noutside 1 331\nTMhelix 332 354\ninside 355 372

7954 GCF_001283325.1_8205_8_75 Escherichia coli Proteobacteria; Gammaproteobacteria; Enterobacterales; Enterobacteriaceae; Escherichia
MTGDNTLIHSHGINRRDFMKLCAALAATMGLSSKAAAEMAESVTNPQRPPVIWIGAQECT WP_000145410.1
MULTISPECIES: hydrogenase-2 small chain [Proteobacteria] Length: 372\nNumber of predicted TMHs: 1\nExp number of AAs in TMHs: 23.72226\nExp number, first 60 AAs: 0.22038\nTotal prob of N-in: 0.12994\noutside 1 331\nTMhelix 332 354\ninside 355 372

7955 GCF_001283265.1_7748_7_47 Escherichia coli Proteobacteria; Gammaproteobacteria; Enterobacterales; Enterobacteriaceae; Escherichia

MTGDNTLIHSHGINRRDFMKLCAALAATMGLSSKAAAEMAESVTNPQRPPVIWIGAQECT WP_000145410.1 MULTISPECIES: hydrogenase-2 small chain [Proteobacteria] Length: 372\nNumber of predicted TMHs: 1\nExp number of AAs in TMHs: 23.72226\nExp number, first 60 AAs: 0.22038\nTotal prob of N-in: 0.12994\noutside 1 331\nTMhelix 332 354\ninside 355 372

7956 GCF_001283385.1_7748_7_30 Escherichia coli Proteobacteria; Gammaproteobacteria; Enterobacterales; Enterobacteriaceae; Escherichia MTGDNTLIHSHGINRRDFMKLCAALAATMGLSSKAAAEMAESVTNPQRPPVIWIGAQECT WP_000145410.1 MULTISPECIES: hydrogenase-2 small chain [Proteobacteria] Length: 372\nNumber of predicted TMHs: 1\nExp number of AAs in TMHs: 23.72226\nExp number, first 60 AAs: 0.22038\nTotal prob of N-in: 0.12994\noutside 1 331\nTMhelix 332 354\ninside 355 372

7957 GCF_001283165.1_7553_7_66 Escherichia coli Proteobacteria; Gammaproteobacteria; Enterobacterales; Enterobacteriaceae; Escherichia MNNEETFYQAMRRQGVTRRSFLKYCSLAATSLGLGAGMAPKIAWALENKPRIPVVWIHGL WP_001058323.1 MULTISPECIES: hydrogenase-1 small chain [Proteobacteria] Length: 372\nNumber of predicted TMHs: 2\nExp number of AAs in TMHs: 37.2608499999999\nExp number, first 60 AAs: 19.52923\nTotal prob of N-in: 0.94351\nPOSSIBLE N-term signal sequence\ninside 1 19\nTMhelix 20 39\noutside 40 325\nTMhelix 326 348\ninside 349 372

7958 GCF_001283325.1_8205_8_75 Escherichia coli Proteobacteria; Gammaproteobacteria; Enterobacterales; Enterobacteriaceae; Escherichia MNNEETFYQAMRRQGVTRRSFLKYCSLAATSLGLGAGMAPKIAWALENKPRIPVVWIHGL WP_001058323.1 MULTISPECIES: hydrogenase-1 small chain [Proteobacteria] Length: 372\nNumber of predicted TMHs: 2\nExp number of AAs in TMHs: 37.2608499999999\nExp number, first 60 AAs: 19.52923\nTotal prob of N-in: 0.94351\nPOSSIBLE N-term signal sequence\ninside 1 19\nTMhelix 20 39\noutside 40 325\nTMhelix 326 348\ninside 349 372

7959 GCF_001281995.1_ASM128199v1 Escherichia coli Proteobacteria; Gammaproteobacteria; Enterobacterales; Enterobacteriaceae; Escherichia MNNEETFYQAMRRQGVTRRSFLKYCSLAATSLGLGAGMAPKIAWALENKPRIPVVWIHGL WP_053920217.1 hydrogenase [Escherichia coli] Length: 372\nNumber of predicted TMHs: 2\nExp number of AAs in TMHs: 37.45884\nExp number, first 60 AAs: 19.5295\nTotal prob of N-in: 0.94363\nPOSSIBLE N-term signal sequence\ninside 1 19\nTMhelix 20 39\noutside 40 325\nTMhelix 326 348\ninside 349 372

7960 GCF_001282035.1_ASM128203v1 Escherichia coli Proteobacteria; Gammaproteobacteria; Enterobacterales; Enterobacteriaceae; Escherichia MNNEETFYQAMRRQGVTRRSFLKYCSLAATSLGLGAGMAPKIAWALENKPRIPVVWIHGL WP_053920217.1 hydrogenase [Escherichia coli] Length: 372\nNumber of predicted TMHs: 2\nExp number of AAs in TMHs: 37.45884\nExp number, first 60 AAs: 19.5295\nTotal prob of N-in: 0.94363\nPOSSIBLE N-term signal sequence\ninside 1 19\nTMhelix 20 39\noutside 40 325\nTMhelix 326 348\ninside 349 372

7961 GCF_001283405.1_7790_1_65 Escherichia coli Proteobacteria; Gammaproteobacteria; Enterobacterales; Enterobacteriaceae; Escherichia MTGDNTLIHSHGINRRDFMKLCAALAATMGLSSKAAAEMAESVTNPQRPPVIWIGAQECT WP_000145410.1 MULTISPECIES: hydrogenase-2 small chain [Proteobacteria] Length: 372\nNumber of predicted TMHs: 1\nExp number of AAs in TMHs: 23.72226\nExp number, first 60 AAs: 0.22038\nTotal prob of N-in: 0.12994\noutside 1 331\nTMhelix 332 354\ninside 355 372

7962 GCF_001283425.1_7790_1_95 Escherichia coli Proteobacteria; Gammaproteobacteria; Enterobacterales; Enterobacteriaceae; Escherichia MTGDNTLIHSHGINRRDFMKLCAALAATMGLSSKAAAEMAESVTNPQRPPVIWIGAQECT WP_000145410.1 MULTISPECIES: hydrogenase-2 small chain [Proteobacteria] Length: 372\nNumber of predicted TMHs: 1\nExp number of AAs in TMHs: 23.72226\nExp number, first 60 AAs: 0.22038\nTotal prob of N-in: 0.12994\noutside 1 331\nTMhelix 332 354\ninside 355 372

7963 GCF_001283565.1_8205_3_26 Escherichia coli Proteobacteria; Gammaproteobacteria; Enterobacterales; Enterobacteriaceae; Escherichia MTGDNTLIHSHGINRRDFMKLCAALAATMGLSSKAAAEMAESVTNPQRPPVIWIGAQECT WP_000145410.1 MULTISPECIES: hydrogenase-2 small chain [Proteobacteria] Length: 372\nNumber of predicted TMHs:

1\nExp number of AAs in TMHs: 23.72226\nExp number, first 60 AAs: 0.22038\nTotal prob of N-in: 0.12994\noutside 1 331\nTMhelix 332 354\ninside 355 372

7964 GCF_001283465.1_8205_8_73 Escherichia coli Proteobacteria; Gammaproteobacteria; Enterobacterales; Enterobacteriaceae; Escherichia MTGDNTLIHSHGINRRDFMKLCAALAATMGLSSKAAAEMAESVTNPQRPPVIWIGAQECT WP_000145410.1 MULTISPECIES: hydrogenase-2 small chain [Proteobacteria] Length: 372\nNumber of predicted TMHs: 1\nExp number of AAs in TMHs: 23.72226\nExp number, first 60 AAs: 0.22038\nTotal prob of N-in: 0.12994\noutside 1 331\nTMhelix 332 354\ninside 355 372

7965 GCF_001283525.1_7553_7_69 Escherichia coli Proteobacteria; Gammaproteobacteria; Enterobacterales; Enterobacteriaceae; Escherichia MTGDNTLIHSHGINRRDFMKLCAALAATMGLSSKAAAEMAESVTNPQRPPVIWIGAQECT WP_000145410.1 MULTISPECIES: hydrogenase-2 small chain [Proteobacteria] Length: 372\nNumber of predicted TMHs: 1\nExp number of AAs in TMHs: 23.72226\nExp number, first 60 AAs: 0.22038\nTotal prob of N-in: 0.12994\noutside 1 331\nTMhelix 332 354\ninside 355 372

7966 GCF_001283565.1_8205_3_26 Escherichia coli Proteobacteria; Gammaproteobacteria; Enterobacterales; Enterobacteriaceae; Escherichia MNNEETFYQAMRRQGVTRRSFLKYCSLAATSLGLGAGMAPKIAWALENKPRIPVVWIHGL WP_001058323.1 MULTISPECIES: hydrogenase-1 small chain [Proteobacteria] Length: 372\nNumber of predicted TMHs: 2\nExp number of AAs in TMHs: 37.2608499999999\nExp number, first 60 AAs: 19.52923\nTotal prob of N-in: 0.94351\nPOSSIBLE N-term signal sequence\ninside 1 19\nTMhelix 20 39\noutside 40 325\nTMhelix 326 348\ninside 349 372

7967 GCF_001283385.1_7748_7_30 Escherichia coli Proteobacteria; Gammaproteobacteria; Enterobacterales; Enterobacteriaceae; Escherichia MNNEETFYQAMRRQGVTRRSFLKYCSLAATSLGLGAGMAPKIAWALENKPRIPVVWIHGL WP_001295940.1 MULTISPECIES: hydrogenase [Enterobacteriaceae] Length: 372\nNumber of predicted TMHs: 2\nExp number of AAs in TMHs: 37.2608899999999\nExp number, first 60 AAs: 19.52923\nTotal prob of N-in: 0.94351\nPOSSIBLE N-term signal sequence\ninside 1 19\nTMhelix 20 39\noutside 40 325\nTMhelix 326 348\ninside 349 372

7968 GCF_001283265.1_7748_7_47 Escherichia coli Proteobacteria; Gammaproteobacteria; Enterobacterales; Enterobacteriaceae; Escherichia MNNEETFYQAMRRQGVTRRSFLKYCSLAATSLGLGAGMAPKIAWALENKPRIPVVWIHGL WP_052923131.1 hydrogenase [Escherichia coli] Length: 372\nNumber of predicted TMHs: 2\nExp number of AAs in TMHs: 36.3624399999999\nExp number, first 60 AAs: 19.53053\nTotal prob of N-in: 0.94346\nPOSSIBLE N-term signal sequence\ninside 1 19\nTMhelix 20 39\noutside 40 325\nTMhelix 326 348\ninside 349 372

7969 GCF_001283645.1_7748_7_10 Escherichia coli Proteobacteria; Gammaproteobacteria; Enterobacterales; Enterobacteriaceae; Escherichia MTGDNTLIHSHGINRRDFMKLCAALAATMGLSSKAAAEMAESVTNPQRPPVIWIGAQECT WP_000145410.1 MULTISPECIES: hydrogenase-2 small chain [Proteobacteria] Length: 372\nNumber of predicted TMHs: 1\nExp number of AAs in TMHs: 23.72226\nExp number, first 60 AAs: 0.22038\nTotal prob of N-in: 0.12994\noutside 1 331\nTMhelix 332 354\ninside 355 372

7970 GCF_001283645.1_7748_7_10 Escherichia coli Proteobacteria; Gammaproteobacteria; Enterobacterales; Enterobacteriaceae; Escherichia MNNEETFYQAMRRQGVTRRSFLKYCSLAATSLGLGAGMAPKIAWALENKPRIPVVWIHGL WP_024237484.1 hydrogenase-1 small chain [Escherichia coli] Length: 372\nNumber of predicted TMHs: 2\nExp number of AAs in TMHs: 37.2691999999999\nExp number, first 60 AAs: 19.53117\nTotal prob of N-in: 0.94352\nPOSSIBLE N-term signal sequence\ninside 1 19\nTMhelix 20 39\noutside 40 325\nTMhelix 326 348\ninside 349 372

7971 GCF_001283465.1_8205_8_73 Escherichia coli Proteobacteria; Gammaproteobacteria; Enterobacterales; Enterobacteriaceae; Escherichia MNNEETFYQAMRRQGVTRRSFLKYCSLAATSLGLGAGMAPKIAWALENKPRIPVVWIHGL WP_001058320.1 hydrogenase [Escherichia coli] Length: 372\nNumber of predicted TMHs: 2\nExp number of AAs in TMHs: 37.26082\nExp number, first 60 AAs: 19.52923\nTotal prob of N-in: 0.94351\nPOSSIBLE N-term signal sequence\ninside 1 19\nTMhelix 20 39\noutside 40 325\nTMhelix 326 348\ninside 349 372

7972 GCF_001283745.1_8205_3_76 Escherichia coli Proteobacteria; Gammaproteobacteria; Enterobacterales; Enterobacteriaceae; Escherichia
MTGDNTLIHSHGINRRDFMKLCAALAATMGLSSKAAAEMAESVTNPQRPPVIWIGAQECT WP_000145410.1
MULTISPECIES: hydrogenase-2 small chain [Proteobacteria] Length: 372\nNumber of predicted TMHs: 1\nExp number of AAs in TMHs: 23.72226\nExp number, first 60 AAs: 0.22038\nTotal prob of N-in: 0.12994\noutside 1 331\nTMhelix 332 354\ninside 355 372

7973 GCF_001283685.1_7790_1_61 Escherichia coli Proteobacteria; Gammaproteobacteria; Enterobacterales; Enterobacteriaceae; Escherichia
MTGDNTLIHSHGINRRDFMKLCAALAATMGLSSKAAAEMAESVTNPQRPPVIWIGAQECT WP_000145410.1
MULTISPECIES: hydrogenase-2 small chain [Proteobacteria] Length: 372\nNumber of predicted TMHs: 1\nExp number of AAs in TMHs: 23.72226\nExp number, first 60 AAs: 0.22038\nTotal prob of N-in: 0.12994\noutside 1 331\nTMhelix 332 354\ninside 355 372

7974 GCF_001283745.1_8205_3_76 Escherichia coli Proteobacteria; Gammaproteobacteria; Enterobacterales; Enterobacteriaceae; Escherichia
MNNEETFYQAMRRQGVTRRSFLKYCSLAATSLGLGAGMAPKIAWALENKPRIPVVWIHGL WP_001058323.1
MULTISPECIES: hydrogenase-1 small chain [Proteobacteria] Length: 372\nNumber of predicted TMHs: 2\nExp number of AAs in TMHs: 37.2608499999999\nExp number, first 60 AAs: 19.52923\nTotal prob of N-in: 0.94351\nPOSSIBLE N-term signal sequence\ninside 1 19\nTMhelix 20 39\noutside 40 325\nTMhelix 326 348\ninside 349 372

7975 GCF_001283685.1_7790_1_61 Escherichia coli Proteobacteria; Gammaproteobacteria; Enterobacterales; Enterobacteriaceae; Escherichia
MNNEETFYQAMRRQGVTRRSFLKYCSLAATSLGLGAGMAPKIAWALENKPRIPVVWIHGL WP_001058320.1
hydrogenase [Escherichia coli] Length: 372\nNumber of predicted TMHs: 2\nExp number of AAs in TMHs: 37.26082\nExp number, first 60 AAs: 19.52923\nTotal prob of N-in: 0.94351\nPOSSIBLE N-term signal sequence\ninside 1 19\nTMhelix 20 39\noutside 40 325\nTMhelix 326 348\ninside 349 372

7976 GCF_001283405.1_7790_1_65 Escherichia coli Proteobacteria; Gammaproteobacteria; Enterobacterales; Enterobacteriaceae; Escherichia
MNNEETFYQAMRRQGVTRRSFLKYCSLAATSLGLGAGMAPKIAWALENKPRIPVVWIHGL WP_001295940.1
MULTISPECIES: hydrogenase [Enterobacteriaceae] Length: 372\nNumber of predicted TMHs: 2\nExp number of AAs in TMHs: 37.2608899999999\nExp number, first 60 AAs: 19.52923\nTotal prob of N-in: 0.94351\nPOSSIBLE N-term signal sequence\ninside 1 19\nTMhelix 20 39\noutside 40 325\nTMhelix 326 348\ninside 349 372

7977 GCF_001283425.1_7790_1_95 Escherichia coli Proteobacteria; Gammaproteobacteria; Enterobacterales; Enterobacteriaceae; Escherichia
MNNEETFYQAMRRQGVTRRSFLKYCSLAATSLGLGAGMAPKIAWALENKPRIPVVWIHGL WP_001295940.1
MULTISPECIES: hydrogenase [Enterobacteriaceae] Length: 372\nNumber of predicted TMHs: 2\nExp number of AAs in TMHs: 37.2608899999999\nExp number, first 60 AAs: 19.52923\nTotal prob of N-in: 0.94351\nPOSSIBLE N-term signal sequence\ninside 1 19\nTMhelix 20 39\noutside 40 325\nTMhelix 326 348\ninside 349 372

7978 GCF_001306715.1_ASM130671v1 Escherichia coli Proteobacteria; Gammaproteobacteria; Enterobacterales; Enterobacteriaceae; Escherichia
MTGDNTLIHSHGINRRDFMKLCAALAATMGLSSKAAAEMAESVTNPQRPPVIWIGAQECT WP_000145410.1
MULTISPECIES: hydrogenase-2 small chain [Proteobacteria] Length: 372\nNumber of predicted TMHs: 1\nExp number of AAs in TMHs: 23.72226\nExp number, first 60 AAs: 0.22038\nTotal prob of N-in: 0.12994\noutside 1 331\nTMhelix 332 354\ninside 355 372

7979 GCF_001309455.1_ASM130945v1 Escherichia coli Proteobacteria; Gammaproteobacteria; Enterobacterales; Enterobacteriaceae; Escherichia
MTGDNTLIHSHGINRRDFMKLCAALAATMGLSSKAAAEMAESVTNPQRPPVIWIGAQECT WP_000145410.1
MULTISPECIES: hydrogenase-2 small chain [Proteobacteria] Length: 372\nNumber of predicted TMHs: 1\nExp number of AAs in TMHs: 23.72226\nExp number, first 60 AAs: 0.22038\nTotal prob of N-in: 0.12994\noutside 1 331\nTMhelix 332 354\ninside 355 372

7980 GCF_001309475.1_ASM130947v1 Escherichia coli Proteobacteria; Gammaproteobacteria; Enterobacterales; Enterobacteriaceae; Escherichia

MTGDNTLIHSHGINRRDFMKLCAALAATMGLSSKAAAEMAESVTNPQRPPVIWIGAQECT WP_000145410.1 MULTISPECIES: hydrogenase-2 small chain [Proteobacteria] Length: 372\nNumber of predicted TMHs: 1\nExp number of AAs in TMHs: 23.72226\nExp number, first 60 AAs: 0.22038\nTotal prob of N-in: 0.12994\noutside 1 331\nTMhelix 332 354\ninside 355 372

7981 GCF_001306715.1_ASM130671v1 Escherichia coli Proteobacteria; Gammaproteobacteria; Enterobacterales; Enterobacteriaceae; Escherichia MNNEETFYQAMRRQGVTRRSFLKYCSLAATSLGLGAGMAPKIAWALENKPRIPVVWIHGL WP_001058323.1 MULTISPECIES: hydrogenase-1 small chain [Proteobacteria] Length: 372\nNumber of predicted TMHs: 2\nExp number of AAs in TMHs: 37.2608499999999\nExp number, first 60 AAs: 19.52923\nTotal prob of N-in: 0.94351\nPOSSIBLE N-term signal sequence\ninside 1 19\nTMhelix 20 39\noutside 40 325\nTMhelix 326 348\ninside 349 372

7982 GCF_001309455.1_ASM130945v1 Escherichia coli Proteobacteria; Gammaproteobacteria; Enterobacterales; Enterobacteriaceae; Escherichia MNNEETFYQAMRRQGVTRRSFLKYCSLAATSLGLGAGMAPKIAWALENKPRIPVVWIHGL WP_001058323.1 MULTISPECIES: hydrogenase-1 small chain [Proteobacteria] Length: 372\nNumber of predicted TMHs: 2\nExp number of AAs in TMHs: 37.2608499999999\nExp number, first 60 AAs: 19.52923\nTotal prob of N-in: 0.94351\nPOSSIBLE N-term signal sequence\ninside 1 19\nTMhelix 20 39\noutside 40 325\nTMhelix 326 348\ninside 349 372

7983 GCF_001283525.1_7553_7_69 Escherichia coli Proteobacteria; Gammaproteobacteria; Enterobacterales; Enterobacteriaceae; Escherichia MNNEETFYQAMRRQGVTRRSFLKYCSLAATSLGLGAGMAPKIAWALENKPRIPVVWIHGL WP_052923131.1 hydrogenase [Escherichia coli] Length: 372\nNumber of predicted TMHs: 2\nExp number of AAs in TMHs: 36.3624399999999\nExp number, first 60 AAs: 19.53053\nTotal prob of N-in: 0.94346\nPOSSIBLE N-term signal sequence\ninside 1 19\nTMhelix 20 39\noutside 40 325\nTMhelix 326 348\ninside 349 372

7984 GCF_001309555.1_ASM130955v1 Escherichia coli Proteobacteria; Gammaproteobacteria; Enterobacterales; Enterobacteriaceae; Escherichia MTGDNTLIHSHGINRRDFMKLCAALAATMGLSSKAAAEMAESVTNPQRPPVIWIGAQECT WP_000145410.1 MULTISPECIES: hydrogenase-2 small chain [Proteobacteria] Length: 372\nNumber of predicted TMHs: 1\nExp number of AAs in TMHs: 23.72226\nExp number, first 60 AAs: 0.22038\nTotal prob of N-in: 0.12994\noutside 1 331\nTMhelix 332 354\ninside 355 372

7985 GCF_001309615.1_ASM130961v1 Escherichia coli Proteobacteria; Gammaproteobacteria; Enterobacterales; Enterobacteriaceae; Escherichia MTGDNTLIHSHGINRRDFMKLCAALAATMGLSSKAAAEMAESVTNPQRPPVIWIGAQECT WP_000145410.1 MULTISPECIES: hydrogenase-2 small chain [Proteobacteria] Length: 372\nNumber of predicted TMHs: 1\nExp number of AAs in TMHs: 23.72226\nExp number, first 60 AAs: 0.22038\nTotal prob of N-in: 0.12994\noutside 1 331\nTMhelix 332 354\ninside 355 372

7986 GCF_001309715.1_ASM130971v1 Escherichia coli Proteobacteria; Gammaproteobacteria; Enterobacterales; Enterobacteriaceae; Escherichia MTGDNTLIHSHGINRRDFMKLCAALAATMGLSSKAAAEMAESVTNPQRPPVIWIGAQECT WP_000145410.1 MULTISPECIES: hydrogenase-2 small chain [Proteobacteria] Length: 372\nNumber of predicted TMHs: 1\nExp number of AAs in TMHs: 23.72226\nExp number, first 60 AAs: 0.22038\nTotal prob of N-in: 0.12994\noutside 1 331\nTMhelix 332 354\ninside 355 372

7987 GCF_001309555.1_ASM130955v1 Escherichia coli Proteobacteria; Gammaproteobacteria; Enterobacterales; Enterobacteriaceae; Escherichia MNNEETFYQAMRRQGVTRRSFLKYCSLAATSLGLGAGMAPKIAWALENKPRIPVVWIHGL WP_001058323.1 MULTISPECIES: hydrogenase-1 small chain [Proteobacteria] Length: 372\nNumber of predicted TMHs: 2\nExp number of AAs in TMHs: 37.2608499999999\nExp number, first 60 AAs: 19.52923\nTotal prob of N-in: 0.94351\nPOSSIBLE N-term signal sequence\ninside 1 19\nTMhelix 20 39\noutside 40 325\nTMhelix 326 348\ninside 349 372

7988 GCF_001309615.1_ASM130961v1 Escherichia coli Proteobacteria; Gammaproteobacteria; Enterobacterales; Enterobacteriaceae; Escherichia MNNEETFYQAMRRQGVTRRSFLKYCSLAATSLGLGAGMAPKIAWALENKPRIPVVWIHGL WP_001058323.1 MULTISPECIES: hydrogenase-1 small chain [Proteobacteria] Length: 372\nNumber of predicted TMHs:

2\nExp number of AAs in TMHs: 37.2608499999999\nExp number, first 60 AAs:  19.52923\nTotal prob of N-in:   0.94351\nPOSSIBLE N-term signal sequence\ninside   1   19\nTMhelix   20   39\noutside   40   325\nTMhelix   326   348\ninside   349   372

7989 GCF_001309715.1_ASM130971v1 Escherichia coli Proteobacteria; Gammaproteobacteria; Enterobacterales; Enterobacteriaceae; Escherichia MNNEETFYQAMRRQGVTRRSFLKYCSLAATSLGLGAGMAPKIAWALENKPRIPVVWIHGL WP_001301613.1 hydrogenase [Escherichia coli] Length: 372\nNumber of predicted TMHs:  2\nExp number of AAs in TMHs: 37.46598\nExp number, first 60 AAs:  19.53139\nTotal prob of N-in:   0.94364\nPOSSIBLE N-term signal sequence\ninside   1   19\nTMhelix   20   39\noutside   40   325\nTMhelix   326   348\ninside   349   372

7990 GCF_001309475.1_ASM130947v1 Escherichia coli Proteobacteria; Gammaproteobacteria; Enterobacterales; Enterobacteriaceae; Escherichia MNNEETFYQAMRRQGVTRRSFLKYCSLAATSLGLGAGMAPKIAWALENKPRIPVVWIHGL WP_001058323.1 MULTISPECIES: hydrogenase-1 small chain [Proteobacteria] Length: 372\nNumber of predicted TMHs:  2\nExp number of AAs in TMHs: 37.2608499999999\nExp number, first 60 AAs:  19.52923\nTotal prob of N-in:   0.94351\nPOSSIBLE N-term signal sequence\ninside   1   19\nTMhelix   20   39\noutside   40   325\nTMhelix   326   348\ninside   349   372

7991 GCF_001309645.1_ASM130964v1 Escherichia coli Proteobacteria; Gammaproteobacteria; Enterobacterales; Enterobacteriaceae; Escherichia MTGDNTLIHSHGINRRDFMKLCAALAATMGLSSKAAAEMAESVTNPQRPPVIWIGAQECT WP_000145410.1 MULTISPECIES: hydrogenase-2 small chain [Proteobacteria] Length: 372\nNumber of predicted TMHs:  1\nExp number of AAs in TMHs: 23.72226\nExp number, first 60 AAs:  0.22038\nTotal prob of N-in:   0.12994\noutside   1   331\nTMhelix   332   354\ninside   355   372

7992 GCF_001309645.1_ASM130964v1 Escherichia coli Proteobacteria; Gammaproteobacteria; Enterobacterales; Enterobacteriaceae; Escherichia MNNEETFYQAMRRQGVTRRSFLKYCSLAATSLGLGAGMAPKIAWALENKPRIPVVWIHGL WP_001058323.1 MULTISPECIES: hydrogenase-1 small chain [Proteobacteria] Length: 372\nNumber of predicted TMHs:  2\nExp number of AAs in TMHs: 37.2608499999999\nExp number, first 60 AAs:  19.52923\nTotal prob of N-in:   0.94351\nPOSSIBLE N-term signal sequence\ninside   1   19\nTMhelix   20   39\noutside   40   325\nTMhelix   326   348\ninside   349   372

7993 GCF_001309755.1_ASM130975v1 Escherichia coli Proteobacteria; Gammaproteobacteria; Enterobacterales; Enterobacteriaceae; Escherichia MTGDNTLIHSHGINRRDFMKLCAALAATMGLSSKAAAEMAESVTNPQRPPVIWIGAQECT WP_000145410.1 MULTISPECIES: hydrogenase-2 small chain [Proteobacteria] Length: 372\nNumber of predicted TMHs:  1\nExp number of AAs in TMHs: 23.72226\nExp number, first 60 AAs:  0.22038\nTotal prob of N-in:   0.12994\noutside   1   331\nTMhelix   332   354\ninside   355   372

7994 GCF_001309815.1_ASM130981v1 Escherichia coli Proteobacteria; Gammaproteobacteria; Enterobacterales; Enterobacteriaceae; Escherichia MTGDNTLIHSHGINRRDFMKLCAALAATMGLSSKAAAEMAESVTNPQRPPVIWIGAQECT WP_000145410.1 MULTISPECIES: hydrogenase-2 small chain [Proteobacteria] Length: 372\nNumber of predicted TMHs:  1\nExp number of AAs in TMHs: 23.72226\nExp number, first 60 AAs:  0.22038\nTotal prob of N-in:   0.12994\noutside   1   331\nTMhelix   332   354\ninside   355   372

7995 GCF_001309835.1_ASM130983v1 Escherichia coli Proteobacteria; Gammaproteobacteria; Enterobacterales; Enterobacteriaceae; Escherichia MTGDNTLIHSHGINRRDFMKLCAALAATMGLSSKAAAEMAESVTNPQRPPVIWIGAQECT WP_000145410.1 MULTISPECIES: hydrogenase-2 small chain [Proteobacteria] Length: 372\nNumber of predicted TMHs:  1\nExp number of AAs in TMHs: 23.72226\nExp number, first 60 AAs:  0.22038\nTotal prob of N-in:   0.12994\noutside   1   331\nTMhelix   332   354\ninside   355   372

7996 GCF_001309755.1_ASM130975v1 Escherichia coli Proteobacteria; Gammaproteobacteria; Enterobacterales; Enterobacteriaceae; Escherichia MNNEETFYQAMRRQGVTRRSFLKYCSLAATSLGLGAGMAPKIAWALENKPRIPVVWIHGL WP_001058323.1 MULTISPECIES: hydrogenase-1 small chain [Proteobacteria] Length: 372\nNumber of predicted TMHs:  2\nExp number of AAs in TMHs: 37.2608499999999\nExp number, first 60 AAs:  19.52923\nTotal prob of N-in:

0.94351\nPOSSIBLE N-term signal sequence\ninside 1 19\nTMhelix 20 39\noutside 40 325\nTMhelix 326 348\ninside 349 372

7997 GCF_001309905.1_ASM130990v1 Escherichia coli Proteobacteria; Gammaproteobacteria; Enterobacterales; Enterobacteriaceae; Escherichia MTGDNTLIHSHGINRRDFMKLCAALAATMGLSSKAAAEMAESVTNPQRPPVIWIGAQECT WP_000145410.1 MULTISPECIES: hydrogenase-2 small chain [Proteobacteria] Length: 372\nNumber of predicted TMHs: 1\nExp number of AAs in TMHs: 23.72226\nExp number, first 60 AAs: 0.22038\nTotal prob of N-in: 0.12994\noutside 1 331\nTMhelix 332 354\ninside 355 372

7998 GCF_001309965.1_ASM130996v1 Escherichia coli Proteobacteria; Gammaproteobacteria; Enterobacterales; Enterobacteriaceae; Escherichia MTGDNTLIHSHGINRRDFMKLCAALAATMGLSSKAAAEMAESVTNPQRPPVIWIGAQECT WP_000145410.1 MULTISPECIES: hydrogenase-2 small chain [Proteobacteria] Length: 372\nNumber of predicted TMHs: 1\nExp number of AAs in TMHs: 23.72226\nExp number, first 60 AAs: 0.22038\nTotal prob of N-in: 0.12994\noutside 1 331\nTMhelix 332 354\ninside 355 372

7999 GCF_001310005.1_ASM131000v1 Escherichia coli Proteobacteria; Gammaproteobacteria; Enterobacterales; Enterobacteriaceae; Escherichia MTGDNTLIHSHGINRRDFMKLCAALAATMGLSSKAAAEMAESVTNPQRPPVIWIGAQECT WP_000145410.1 MULTISPECIES: hydrogenase-2 small chain [Proteobacteria] Length: 372\nNumber of predicted TMHs: 1\nExp number of AAs in TMHs: 23.72226\nExp number, first 60 AAs: 0.22038\nTotal prob of N-in: 0.12994\noutside 1 331\nTMhelix 332 354\ninside 355 372

8000 GCF_001373235.1_26A_L2_assembly Escherichia coli Proteobacteria; Gammaproteobacteria; Enterobacterales; Enterobacteriaceae; Escherichia MTGDNTLIHSHGINRRDFMKLCAALAATMGLSSKAAAEMAESVTNPQRPPVIWIGAQECT WP_000145410.1 MULTISPECIES: hydrogenase-2 small chain [Proteobacteria] Length: 372\nNumber of predicted TMHs: 1\nExp number of AAs in TMHs: 23.72226\nExp number, first 60 AAs: 0.22038\nTotal prob of N-in: 0.12994\noutside 1 331\nTMhelix 332 354\ninside 355 372

8001 GCF_001412955.1_ASM141295v1 Escherichia coli Proteobacteria; Gammaproteobacteria; Enterobacterales; Enterobacteriaceae; Escherichia MTGDNTLIHSHGINRRDFMKLCAALAATMGLSSKAAAEMAESVTNPQRPPVIWIGAQECT WP_000145410.1 MULTISPECIES: hydrogenase-2 small chain [Proteobacteria] Length: 372\nNumber of predicted TMHs: 1\nExp number of AAs in TMHs: 23.72226\nExp number, first 60 AAs: 0.22038\nTotal prob of N-in: 0.12994\noutside 1 331\nTMhelix 332 354\ninside 355 372

8002 GCF_001373235.1_26A_L2_assembly Escherichia coli Proteobacteria; Gammaproteobacteria; Enterobacterales; Enterobacteriaceae; Escherichia MNNEETFYQAMRRQGVTRRSFLKYCSLAATSLGLGAGMAPKIAWALENKPRIPVVWIHGL WP_001058320.1 hydrogenase [Escherichia coli] Length: 372\nNumber of predicted TMHs: 2\nExp number of AAs in TMHs: 37.26082\nExp number, first 60 AAs: 19.52923\nTotal prob of N-in: 0.94351\nPOSSIBLE N-term signal sequence\ninside 1 19\nTMhelix 20 39\noutside 40 325\nTMhelix 326 348\ninside 349 372

8003 GCF_001310005.1_ASM131000v1 Escherichia coli Proteobacteria; Gammaproteobacteria; Enterobacterales; Enterobacteriaceae; Escherichia MNNEETFYQAMRRQGVTRRSFLKYCSLAATSLGLGAGMAPKIAWALENKPRIPVVWIHGL WP_001301613.1 hydrogenase [Escherichia coli] Length: 372\nNumber of predicted TMHs: 2\nExp number of AAs in TMHs: 37.46598\nExp number, first 60 AAs: 19.53139\nTotal prob of N-in: 0.94364\nPOSSIBLE N-term signal sequence\ninside 1 19\nTMhelix 20 39\noutside 40 325\nTMhelix 326 348\ninside 349 372

8004 GCF_001309815.1_ASM130981v1 Escherichia coli Proteobacteria; Gammaproteobacteria; Enterobacterales; Enterobacteriaceae; Escherichia MNNEETFYQAMRRQGVTRRSFLKYCSLAATSLGLGAGMAPKIAWALENKPRIPVVWIHGL WP_001058323.1 MULTISPECIES: hydrogenase-1 small chain [Proteobacteria] Length: 372\nNumber of predicted TMHs: 2\nExp number of AAs in TMHs: 37.2608499999999\nExp number, first 60 AAs: 19.52923\nTotal prob of N-in: 0.94351\nPOSSIBLE N-term signal sequence\ninside 1 19\nTMhelix 20 39\noutside 40 325\nTMhelix 326 348\ninside 349 372

8005 GCF_001412915.1_ASM141291v1 Escherichia coli Proteobacteria; Gammaproteobacteria; Enterobacterales; Enterobacteriaceae; Escherichia MTGDNTLIHSHGINRRDFMKLCAALAATMGLSSKAAAEMAESVTNPQRPPVIWIGAQECT WP_000145410.1 MULTISPECIES: hydrogenase-2 small chain [Proteobacteria] Length: 372\nNumber of predicted TMHs: 1\nExp number of AAs in TMHs: 23.72226\nExp number, first 60 AAs: 0.22038\nTotal prob of N-in: 0.12994\noutside 1 331\nTMhelix 332 354\ninside 355 372

8006 GCF_001412915.1_ASM141291v1 Escherichia coli Proteobacteria; Gammaproteobacteria; Enterobacterales; Enterobacteriaceae; Escherichia MNNEETFYQAMRRQGVTRRSFLKYCSLAATSLGLGAGMAPKIAWALENKPRIPVVWIHGL WP_001058323.1 MULTISPECIES: hydrogenase-1 small chain [Proteobacteria] Length: 372\nNumber of predicted TMHs: 2\nExp number of AAs in TMHs: 37.2608499999999\nExp number, first 60 AAs: 19.52923\nTotal prob of N-in: 0.94351\nPOSSIBLE N-term signal sequence\ninside 1 19\nTMhelix 20 39\noutside 40 325\nTMhelix 326 348\ninside 349 372

8007 GCF_001309835.1_ASM130983v1 Escherichia coli Proteobacteria; Gammaproteobacteria; Enterobacterales; Enterobacteriaceae; Escherichia MNNEETFYQAMRRQGVTRRSFLKYCSLAATSLGLGAGMAPKIAWALENKPRIPVVWIHGL WP_001058323.1 MULTISPECIES: hydrogenase-1 small chain [Proteobacteria] Length: 372\nNumber of predicted TMHs: 2\nExp number of AAs in TMHs: 37.2608499999999\nExp number, first 60 AAs: 19.52923\nTotal prob of N-in: 0.94351\nPOSSIBLE N-term signal sequence\ninside 1 19\nTMhelix 20 39\noutside 40 325\nTMhelix 326 348\ninside 349 372

8008 GCF_001412925.1_ASM141292v1 Escherichia coli Proteobacteria; Gammaproteobacteria; Enterobacterales; Enterobacteriaceae; Escherichia MTGDNTLIHSHGINRRDFMKLCAALAATMGLSSKAAAEMAESVTNPQRPPVIWIGAQECT WP_000145410.1 MULTISPECIES: hydrogenase-2 small chain [Proteobacteria] Length: 372\nNumber of predicted TMHs: 1\nExp number of AAs in TMHs: 23.72226\nExp number, first 60 AAs: 0.22038\nTotal prob of N-in: 0.12994\noutside 1 331\nTMhelix 332 354\ninside 355 372

8009 GCF_001412995.1_ASM141299v1 Escherichia coli Proteobacteria; Gammaproteobacteria; Enterobacterales; Enterobacteriaceae; Escherichia MTGDNTLIHSHGINRRDFMKLCAALAATMGLSSKAAAEMAESVTNPQRPPVIWIGAQECT WP_000145410.1 MULTISPECIES: hydrogenase-2 small chain [Proteobacteria] Length: 372\nNumber of predicted TMHs: 1\nExp number of AAs in TMHs: 23.72226\nExp number, first 60 AAs: 0.22038\nTotal prob of N-in: 0.12994\noutside 1 331\nTMhelix 332 354\ninside 355 372

8010 GCF_001412925.1_ASM141292v1 Escherichia coli Proteobacteria; Gammaproteobacteria; Enterobacterales; Enterobacteriaceae; Escherichia MNNEETFYQAMRRQGVTRRSFLKYCSLAATSLGLGAGMAPKIAWALENKPRIPVVWIHGL WP_001058323.1 MULTISPECIES: hydrogenase-1 small chain [Proteobacteria] Length: 372\nNumber of predicted TMHs: 2\nExp number of AAs in TMHs: 37.2608499999999\nExp number, first 60 AAs: 19.52923\nTotal prob of N-in: 0.94351\nPOSSIBLE N-term signal sequence\ninside 1 19\nTMhelix 20 39\noutside 40 325\nTMhelix 326 348\ninside 349 372

8011 GCF_001412955.1_ASM141295v1 Escherichia coli Proteobacteria; Gammaproteobacteria; Enterobacterales; Enterobacteriaceae; Escherichia MNNEETFYQAMRRQGVTRRSFLKYCSLAATSLGLGAGMAPKIAWALENKPRIPVVWIHGL WP_001058323.1 MULTISPECIES: hydrogenase-1 small chain [Proteobacteria] Length: 372\nNumber of predicted TMHs: 2\nExp number of AAs in TMHs: 37.2608499999999\nExp number, first 60 AAs: 19.52923\nTotal prob of N-in: 0.94351\nPOSSIBLE N-term signal sequence\ninside 1 19\nTMhelix 20 39\noutside 40 325\nTMhelix 326 348\ninside 349 372

8012 GCF_001412995.1_ASM141299v1 Escherichia coli Proteobacteria; Gammaproteobacteria; Enterobacterales; Enterobacteriaceae; Escherichia MNNEETFYQAMRRQGVTRRSFLKYCSLAATSLGLGAGMAPKIAWALENKPRIPVVWIHGL WP_001058323.1 MULTISPECIES: hydrogenase-1 small chain [Proteobacteria] Length: 372\nNumber of predicted TMHs: 2\nExp number of AAs in TMHs: 37.2608499999999\nExp number, first 60 AAs: 19.52923\nTotal prob of N-in: 0.94351\nPOSSIBLE N-term signal sequence\ninside 1 19\nTMhelix 20 39\noutside 40 325\nTMhelix 326 348\ninside 349 372

8013	GCF_001309905.1_ASM130990v1	Escherichia coli	Proteobacteria; Gammaproteobacteria; Enterobacterales; Enterobacteriaceae; Escherichia
	MNNEETFYQAMRRQGVTRRSFLKYCSLAATSLGLGAGMAPKIAWALENKPRIPVVWIHGL	WP_001301613.1
	hydrogenase [Escherichia coli]	Length: 372\nNumber of predicted TMHs: 2\nExp number of AAs in TMHs: 37.46598\nExp number, first 60 AAs: 19.53139\nTotal prob of N-in: 0.94364\nPOSSIBLE N-term signal sequence\ninside 1 19\nTMhelix 20 39\noutside 40 325\nTMhelix 326 348\ninside 349 372

8014	GCF_001413375.1_ASM141337v1	Escherichia coli	Proteobacteria; Gammaproteobacteria; Enterobacterales; Enterobacteriaceae; Escherichia
	MTGDNTLIHSHGINRRDFMKLCAALAATMGLSSKAAAEMAESVTNPQRPPVIWIGAQECT	WP_000145410.1
	MULTISPECIES: hydrogenase-2 small chain [Proteobacteria]	Length: 372\nNumber of predicted TMHs: 1\nExp number of AAs in TMHs: 23.72226\nExp number, first 60 AAs: 0.22038\nTotal prob of N-in: 0.12994\noutside 1 331\nTMhelix 332 354\ninside 355 372

8015	GCF_001413415.1_ASM141341v1	Escherichia coli	Proteobacteria; Gammaproteobacteria; Enterobacterales; Enterobacteriaceae; Escherichia
	MTGDNTLIHSHGINRRDFMKLCAALAATMGLSSKAAAEMAESVTNPQRPPVIWIGAQECT	WP_000145410.1
	MULTISPECIES: hydrogenase-2 small chain [Proteobacteria]	Length: 372\nNumber of predicted TMHs: 1\nExp number of AAs in TMHs: 23.72226\nExp number, first 60 AAs: 0.22038\nTotal prob of N-in: 0.12994\noutside 1 331\nTMhelix 332 354\ninside 355 372

8016	GCF_001413375.1_ASM141337v1	Escherichia coli	Proteobacteria; Gammaproteobacteria; Enterobacterales; Enterobacteriaceae; Escherichia
	MNNEETFYQAMRRQGVTRRSFLKYCSLAATSLGLGAGMAPKIAWALENKPRIPVVWIHGL	WP_001058323.1
	MULTISPECIES: hydrogenase-1 small chain [Proteobacteria]	Length: 372\nNumber of predicted TMHs: 2\nExp number of AAs in TMHs: 37.2608499999999\nExp number, first 60 AAs: 19.52923\nTotal prob of N-in: 0.94351\nPOSSIBLE N-term signal sequence\ninside 1 19\nTMhelix 20 39\noutside 40 325\nTMhelix 326 348\ninside 349 372

8017	GCF_001413415.1_ASM141341v1	Escherichia coli	Proteobacteria; Gammaproteobacteria; Enterobacterales; Enterobacteriaceae; Escherichia
	MNNEETFYQAMRRQGVTRRSFLKYCSLAATSLGLGAGMAPKIAWALENKPRIPVVWIHGL	WP_001058323.1
	MULTISPECIES: hydrogenase-1 small chain [Proteobacteria]	Length: 372\nNumber of predicted TMHs: 2\nExp number of AAs in TMHs: 37.2608499999999\nExp number, first 60 AAs: 19.52923\nTotal prob of N-in: 0.94351\nPOSSIBLE N-term signal sequence\ninside 1 19\nTMhelix 20 39\noutside 40 325\nTMhelix 326 348\ninside 349 372

8018	GCF_001309965.1_ASM130996v1	Escherichia coli	Proteobacteria; Gammaproteobacteria; Enterobacterales; Enterobacteriaceae; Escherichia
	MNNEETFYQAMRRQGVTRRSFLKYCSLAATSLGLGAGMAPKIAWALENKPRIPVVWIHGL	WP_001301613.1
	hydrogenase [Escherichia coli]	Length: 372\nNumber of predicted TMHs: 2\nExp number of AAs in TMHs: 37.46598\nExp number, first 60 AAs: 19.53139\nTotal prob of N-in: 0.94364\nPOSSIBLE N-term signal sequence\ninside 1 19\nTMhelix 20 39\noutside 40 325\nTMhelix 326 348\ninside 349 372

8019	GCF_001413475.1_ASM141347v1	Escherichia coli	Proteobacteria; Gammaproteobacteria; Enterobacterales; Enterobacteriaceae; Escherichia
	MTGDNTLIHSHGINRRDFMKLCAALAATMGLSSKAAAEMAESVTNPQRPPVIWIGAQECT	WP_000145410.1
	MULTISPECIES: hydrogenase-2 small chain [Proteobacteria]	Length: 372\nNumber of predicted TMHs: 1\nExp number of AAs in TMHs: 23.72226\nExp number, first 60 AAs: 0.22038\nTotal prob of N-in: 0.12994\noutside 1 331\nTMhelix 332 354\ninside 355 372

8020	GCF_001413515.1_ASM141351v1	Escherichia coli	Proteobacteria; Gammaproteobacteria; Enterobacterales; Enterobacteriaceae; Escherichia
	MTGDNTLIHSHGINRRDFMKLCAALAATMGLSSKAAAEMAESVTNPQRPPVIWIGAQECT	WP_000145410.1
	MULTISPECIES: hydrogenase-2 small chain [Proteobacteria]	Length: 372\nNumber of predicted TMHs: 1\nExp number of AAs in TMHs: 23.72226\nExp number, first 60 AAs: 0.22038\nTotal prob of N-in: 0.12994\noutside 1 331\nTMhelix 332 354\ninside 355 372

8021	GCF_001413475.1_ASM141347v1	Escherichia coli	Proteobacteria; Gammaproteobacteria; Enterobacterales; Enterobacteriaceae; Escherichia
	MNNEETFYQAMRRQGVTRRSFLKYCSLAATSLGLGAGMAPKIAWALENKPRIPVVWIHGL	WP_001058323.1

MULTISPECIES: hydrogenase-1 small chain [Proteobacteria] Length: 372\nNumber of predicted TMHs: 2\nExp number of AAs in TMHs: 37.2608499999999\nExp number, first 60 AAs: 19.52923\nTotal prob of N-in: 0.94351\nPOSSIBLE N-term signal sequence\ninside 1 19\nTMhelix 20 39\noutside 40 325\nTMhelix 326 348\ninside 349 372

8022 GCF_001413565.1_ASM141356v1 Escherichia coli Proteobacteria; Gammaproteobacteria; Enterobacterales; Enterobacteriaceae; Escherichia
MTGDNTLIHSHGINRRDFMKLCAALAATMGLSSKAAAEMAESVTNPQRPPVIWIGAQECT WP_000145410.1
MULTISPECIES: hydrogenase-2 small chain [Proteobacteria] Length: 372\nNumber of predicted TMHs: 1\nExp number of AAs in TMHs: 23.72226\nExp number, first 60 AAs: 0.22038\nTotal prob of N-in: 0.12994\noutside 1 331\nTMhelix 332 354\ninside 355 372

8023 GCF_001413515.1_ASM141351v1 Escherichia coli Proteobacteria; Gammaproteobacteria; Enterobacterales; Enterobacteriaceae; Escherichia
MNNEETFYQAMRRQGVTRRSFLKYCSLAATSLGLGAGMAPKIAWALENKPRIPVVWIHGL WP_001058323.1
MULTISPECIES: hydrogenase-1 small chain [Proteobacteria] Length: 372\nNumber of predicted TMHs: 2\nExp number of AAs in TMHs: 37.2608499999999\nExp number, first 60 AAs: 19.52923\nTotal prob of N-in: 0.94351\nPOSSIBLE N-term signal sequence\ninside 1 19\nTMhelix 20 39\noutside 40 325\nTMhelix 326 348\ninside 349 372

8024 GCF_001413605.1_ASM141360v1 Escherichia coli Proteobacteria; Gammaproteobacteria; Enterobacterales; Enterobacteriaceae; Escherichia
MTGDNTLIHSHGINRRDFMKLCAALAATMGLSSKAAAEMAESVTNPQRPPVIWIGAQECT WP_000145410.1
MULTISPECIES: hydrogenase-2 small chain [Proteobacteria] Length: 372\nNumber of predicted TMHs: 1\nExp number of AAs in TMHs: 23.72226\nExp number, first 60 AAs: 0.22038\nTotal prob of N-in: 0.12994\noutside 1 331\nTMhelix 332 354\ninside 355 372

8025 GCF_001413715.1_ASM141371v1 Escherichia coli Proteobacteria; Gammaproteobacteria; Enterobacterales; Enterobacteriaceae; Escherichia
MTGDNTLIHSHGINRRDFMKLCAALAATMGLSSKAAAEMAESVTNPQRPPVIWIGAQECT WP_000145410.1
MULTISPECIES: hydrogenase-2 small chain [Proteobacteria] Length: 372\nNumber of predicted TMHs: 1\nExp number of AAs in TMHs: 23.72226\nExp number, first 60 AAs: 0.22038\nTotal prob of N-in: 0.12994\noutside 1 331\nTMhelix 332 354\ninside 355 372

8026 GCF_001413675.1_ASM141367v1 Escherichia coli Proteobacteria; Gammaproteobacteria; Enterobacterales; Enterobacteriaceae; Escherichia
MTGDNTLIHSHGINRRDFMKLCAALAATMGLSSKAAAEMAESVTNPQRPPVIWIGAQECT WP_000145410.1
MULTISPECIES: hydrogenase-2 small chain [Proteobacteria] Length: 372\nNumber of predicted TMHs: 1\nExp number of AAs in TMHs: 23.72226\nExp number, first 60 AAs: 0.22038\nTotal prob of N-in: 0.12994\noutside 1 331\nTMhelix 332 354\ninside 355 372

8027 GCF_001413715.1_ASM141371v1 Escherichia coli Proteobacteria; Gammaproteobacteria; Enterobacterales; Enterobacteriaceae; Escherichia
MNNEETFYQAMRRQGVTRRSFLKYCSLAATSLGLGAGMAPKIAWALENKPRIPVVWIHGL WP_001058323.1
MULTISPECIES: hydrogenase-1 small chain [Proteobacteria] Length: 372\nNumber of predicted TMHs: 2\nExp number of AAs in TMHs: 37.2608499999999\nExp number, first 60 AAs: 19.52923\nTotal prob of N-in: 0.94351\nPOSSIBLE N-term signal sequence\ninside 1 19\nTMhelix 20 39\noutside 40 325\nTMhelix 326 348\ninside 349 372

8028 GCF_001413675.1_ASM141367v1 Escherichia coli Proteobacteria; Gammaproteobacteria; Enterobacterales; Enterobacteriaceae; Escherichia
MNNEETFYQAMRRQGVTRRSFLKYCSLAATSLGLGAGMAPKIAWALENKPRIPVVWIHGL WP_001058323.1
MULTISPECIES: hydrogenase-1 small chain [Proteobacteria] Length: 372\nNumber of predicted TMHs: 2\nExp number of AAs in TMHs: 37.2608499999999\nExp number, first 60 AAs: 19.52923\nTotal prob of N-in: 0.94351\nPOSSIBLE N-term signal sequence\ninside 1 19\nTMhelix 20 39\noutside 40 325\nTMhelix 326 348\ninside 349 372

8029 GCF_001413565.1_ASM141356v1 Escherichia coli Proteobacteria; Gammaproteobacteria; Enterobacterales; Enterobacteriaceae; Escherichia
MNNEETFYQAMRRQGVTRRSFLKYCSLAATSLGLGAGMAPKIAWALENKPRIPVVWIHGL WP_001058323.1
MULTISPECIES: hydrogenase-1 small chain [Proteobacteria] Length: 372\nNumber of predicted TMHs:

2\nExp number of AAs in TMHs: 37.2608499999999\nExp number, first 60 AAs: 19.52923\nTotal prob of N-in: 0.94351\nPOSSIBLE N-term signal sequence\ninside 1 19\nTMhelix 20 39\noutside 40 325\nTMhelix 326 348\ninside 349 372

8030 GCF_001413755.1_ASM141375v1 Escherichia coli Proteobacteria; Gammaproteobacteria; Enterobacterales; Enterobacteriaceae; Escherichia MTGDNTLIHSHGINRRDFMKLCAALAATMGLSSKAAAEMAESVTNPQRPPVIWIGAQECT WP_000145410.1 MULTISPECIES: hydrogenase-2 small chain [Proteobacteria] Length: 372\nNumber of predicted TMHs: 1\nExp number of AAs in TMHs: 23.72226\nExp number, first 60 AAs: 0.22038\nTotal prob of N-in: 0.12994\noutside 1 331\nTMhelix 332 354\ninside 355 372

8031 GCF_001413755.1_ASM141375v1 Escherichia coli Proteobacteria; Gammaproteobacteria; Enterobacterales; Enterobacteriaceae; Escherichia MNNEETFYQAMRRQGVTRRSFLKYCSLAATSLGLGAGMAPKIAWALENKPRIPVVWIHGL WP_001058323.1 MULTISPECIES: hydrogenase-1 small chain [Proteobacteria] Length: 372\nNumber of predicted TMHs: 2\nExp number of AAs in TMHs: 37.2608499999999\nExp number, first 60 AAs: 19.52923\nTotal prob of N-in: 0.94351\nPOSSIBLE N-term signal sequence\ninside 1 19\nTMhelix 20 39\noutside 40 325\nTMhelix 326 348\ninside 349 372

8032 GCF_001413605.1_ASM141360v1 Escherichia coli Proteobacteria; Gammaproteobacteria; Enterobacterales; Enterobacteriaceae; Escherichia MNNEETFYQAMRRQGVTRRSFLKYCSLAATSLGLGAGMAPKIAWALENKPRIPVVWIHGL WP_001058323.1 MULTISPECIES: hydrogenase-1 small chain [Proteobacteria] Length: 372\nNumber of predicted TMHs: 2\nExp number of AAs in TMHs: 37.2608499999999\nExp number, first 60 AAs: 19.52923\nTotal prob of N-in: 0.94351\nPOSSIBLE N-term signal sequence\ninside 1 19\nTMhelix 20 39\noutside 40 325\nTMhelix 326 348\ninside 349 372

8033 GCF_001413815.1_ASM141381v1 Escherichia coli Proteobacteria; Gammaproteobacteria; Enterobacterales; Enterobacteriaceae; Escherichia MTGDNTLIHSHGINRRDFMKLCAALAATMGLSSKAAAEMAESVTNPQRPPVIWIGAQECT WP_000145410.1 MULTISPECIES: hydrogenase-2 small chain [Proteobacteria] Length: 372\nNumber of predicted TMHs: 1\nExp number of AAs in TMHs: 23.72226\nExp number, first 60 AAs: 0.22038\nTotal prob of N-in: 0.12994\noutside 1 331\nTMhelix 332 354\ninside 355 372

8034 GCF_001413885.1_ASM141388v1 Escherichia coli Proteobacteria; Gammaproteobacteria; Enterobacterales; Enterobacteriaceae; Escherichia MTGDNTLIHSHGINRRDFMKLCAALAATMGLSSKAAAEMAESVTNPQRPPVIWIGAQECT WP_000145410.1 MULTISPECIES: hydrogenase-2 small chain [Proteobacteria] Length: 372\nNumber of predicted TMHs: 1\nExp number of AAs in TMHs: 23.72226\nExp number, first 60 AAs: 0.22038\nTotal prob of N-in: 0.12994\noutside 1 331\nTMhelix 332 354\ninside 355 372

8035 GCF_001419885.1_ASM141988v1 Escherichia coli Proteobacteria; Gammaproteobacteria; Enterobacterales; Enterobacteriaceae; Escherichia MTGDNTLIHSHGINRRDFMKLCAALAATMGLSSKAAAEMAESVTNPQRPPVIWIGAQECT WP_000145418.1 hydrogenase 2 small subunit [Escherichia coli] Length: 372\nNumber of predicted TMHs: 1\nExp number of AAs in TMHs: 23.72313\nExp number, first 60 AAs: 0.22034\nTotal prob of N-in: 0.12997\noutside 1 331\nTMhelix 332 354\ninside 355 372

8036 GCF_001413995.1_ASM141399v1 Escherichia coli Proteobacteria; Gammaproteobacteria; Enterobacterales; Enterobacteriaceae; Escherichia MTGDNTLIHSHGINRRDFMKLCAALAATMGLSSKAAAEMAESVTNPQRPPVIWIGAQECT WP_000145410.1 MULTISPECIES: hydrogenase-2 small chain [Proteobacteria] Length: 372\nNumber of predicted TMHs: 1\nExp number of AAs in TMHs: 23.72226\nExp number, first 60 AAs: 0.22038\nTotal prob of N-in: 0.12994\noutside 1 331\nTMhelix 332 354\ninside 355 372

8037 GCF_001413815.1_ASM141381v1 Escherichia coli Proteobacteria; Gammaproteobacteria; Enterobacterales; Enterobacteriaceae; Escherichia MNNEETFYQAMRRQGVTRRSFLKYCSLAATSLGLGAGMAPKIAWALENKPRIPVVWIHGL WP_001058323.1 MULTISPECIES: hydrogenase-1 small chain [Proteobacteria] Length: 372\nNumber of predicted TMHs: 2\nExp number of AAs in TMHs: 37.2608499999999\nExp number, first 60 AAs: 19.52923\nTotal prob of N-in:

0.94351\nPOSSIBLE N-term signal sequence\ninside 1 19\nTMhelix 20 39\noutside 40 325\nTMhelix 326 348\ninside 349 372

8038 GCF_001419845.1_ASM141984v1 Escherichia coli Proteobacteria; Gammaproteobacteria; Enterobacterales; Enterobacteriaceae; Escherichia MTGDNTLIHSHGINRRDFMKLCAALAATMGLSSKAAAEMAESVTNPQRPPVIWIGAQECT WP_000145410.1 MULTISPECIES: hydrogenase-2 small chain [Proteobacteria] Length: 372\nNumber of predicted TMHs: 1\nExp number of AAs in TMHs: 23.72226\nExp number, first 60 AAs: 0.22038\nTotal prob of N-in: 0.12994\noutside 1 331\nTMhelix 332 354\ninside 355 372

8039 GCF_001413885.1_ASM141388v1 Escherichia coli Proteobacteria; Gammaproteobacteria; Enterobacterales; Enterobacteriaceae; Escherichia MNNEETFYQAMRRQGVTRRSFLKYCSLAATSLGLGAGMAPKIAWALENKPRIPVVWIHGL WP_001058323.1 MULTISPECIES: hydrogenase-1 small chain [Proteobacteria] Length: 372\nNumber of predicted TMHs: 2\nExp number of AAs in TMHs: 37.2608499999999\nExp number, first 60 AAs: 19.52923\nTotal prob of N-in: 0.94351\nPOSSIBLE N-term signal sequence\ninside 1 19\nTMhelix 20 39\noutside 40 325\nTMhelix 326 348\ninside 349 372

8040 GCF_001419945.1_ASM141994v1 Escherichia coli Proteobacteria; Gammaproteobacteria; Enterobacterales; Enterobacteriaceae; Escherichia MTGDNTLIHSHGINRRDFMKLCAALAATMGLSSKAAAEMAESVTNPQRPPVIWIGAQECT WP_000145410.1 MULTISPECIES: hydrogenase-2 small chain [Proteobacteria] Length: 372\nNumber of predicted TMHs: 1\nExp number of AAs in TMHs: 23.72226\nExp number, first 60 AAs: 0.22038\nTotal prob of N-in: 0.12994\noutside 1 331\nTMhelix 332 354\ninside 355 372

8041 GCF_001419945.1_ASM141994v1 Escherichia coli Proteobacteria; Gammaproteobacteria; Enterobacterales; Enterobacteriaceae; Escherichia MNNEETFYQAMRRQGVTRRSFLKYCSLAATSLGLGAGMAPKIAWALENKPRIPVVWIHGL WP_057108217.1 hydrogenase [Escherichia coli] Length: 372\nNumber of predicted TMHs: 2\nExp number of AAs in TMHs: 38.98478\nExp number, first 60 AAs: 19.52916\nTotal prob of N-in: 0.94354\nPOSSIBLE N-term signal sequence\ninside 1 19\nTMhelix 20 39\noutside 40 325\nTMhelix 326 348\ninside 349 372

8042 GCF_001419845.1_ASM141984v1 Escherichia coli Proteobacteria; Gammaproteobacteria; Enterobacterales; Enterobacteriaceae; Escherichia MNNEETFYQAMRRQGVTRRSFLKYCSLAATSLGLGAGMAPKIAWALENKPRIPVVWIHGL WP_001058323.1 MULTISPECIES: hydrogenase-1 small chain [Proteobacteria] Length: 372\nNumber of predicted TMHs: 2\nExp number of AAs in TMHs: 37.2608499999999\nExp number, first 60 AAs: 19.52923\nTotal prob of N-in: 0.94351\nPOSSIBLE N-term signal sequence\ninside 1 19\nTMhelix 20 39\noutside 40 325\nTMhelix 326 348\ninside 349 372

8043 GCF_001419965.1_ASM141996v1 Escherichia coli Proteobacteria; Gammaproteobacteria; Enterobacterales; Enterobacteriaceae; Escherichia MTGDNTLIHSHGINRRDFMKLCAALAATMGLSSKAAAEMAESVTNPQRPPVIWIGAQECT WP_000145410.1 MULTISPECIES: hydrogenase-2 small chain [Proteobacteria] Length: 372\nNumber of predicted TMHs: 1\nExp number of AAs in TMHs: 23.72226\nExp number, first 60 AAs: 0.22038\nTotal prob of N-in: 0.12994\noutside 1 331\nTMhelix 332 354\ninside 355 372

8044 GCF_001419965.1_ASM141996v1 Escherichia coli Proteobacteria; Gammaproteobacteria; Enterobacterales; Enterobacteriaceae; Escherichia MNNEETFYQAMRRQGVTRRSFLKYCSLAATSLGLGAGMAPKIAWALENKPRIPVVWIHGL WP_001058323.1 MULTISPECIES: hydrogenase-1 small chain [Proteobacteria] Length: 372\nNumber of predicted TMHs: 2\nExp number of AAs in TMHs: 37.2608499999999\nExp number, first 60 AAs: 19.52923\nTotal prob of N-in: 0.94351\nPOSSIBLE N-term signal sequence\ninside 1 19\nTMhelix 20 39\noutside 40 325\nTMhelix 326 348\ninside 349 372

8045 GCF_001413995.1_ASM141399v1 Escherichia coli Proteobacteria; Gammaproteobacteria; Enterobacterales; Enterobacteriaceae; Escherichia MNNEETFYQAMRRQGVTRRSFLKYCSLAATSLGLGAGMAPKIAWALENKPRIPVVWIHGL WP_001295940.1 MULTISPECIES: hydrogenase [Enterobacteriaceae] Length: 372\nNumber of predicted TMHs: 2\nExp number of AAs in TMHs: 37.2608899999999\nExp number, first 60 AAs: 19.52923\nTotal prob of N-in:

0.94351\nPOSSIBLE N-term signal sequence\ninside 1 19\nTMhelix 20 39\noutside 40 325\nTMhelix 326 348\ninside 349 372

8046 GCF_001419885.1_ASM141988v1 Escherichia coli Proteobacteria; Gammaproteobacteria; Enterobacterales; Enterobacteriaceae; Escherichia
MNNEETFYQAMRRQGVTRRSFLKYCSLAATSLGLGAGMAPKIAWALENKPRIPVVWIHGL WP_001295940.1
MULTISPECIES: hydrogenase [Enterobacteriaceae] Length: 372\nNumber of predicted TMHs: 2\nExp number of AAs in TMHs: 37.2608899999999\nExp number, first 60 AAs: 19.52923\nTotal prob of N-in: 0.94351\nPOSSIBLE N-term signal sequence\ninside 1 19\nTMhelix 20 39\noutside 40 325\nTMhelix 326 348\ninside 349 372

8047 GCF_001420125.1_ASM142012v1 Escherichia coli Proteobacteria; Gammaproteobacteria; Enterobacterales; Enterobacteriaceae; Escherichia
MTGDNTLIHSHGINRRDFMKLCAALAATMGLSSKAAAEMAESVTNPQRPPVIWIGAQECT WP_000145410.1
MULTISPECIES: hydrogenase-2 small chain [Proteobacteria] Length: 372\nNumber of predicted TMHs: 1\nExp number of AAs in TMHs: 23.72226\nExp number, first 60 AAs: 0.22038\nTotal prob of N-in: 0.12994\noutside 1 331\nTMhelix 332 354\ninside 355 372

8048 GCF_001420955.1_ASM142095v1 Escherichia coli Proteobacteria; Gammaproteobacteria; Enterobacterales; Enterobacteriaceae; Escherichia
MTGDNTLIHSHGINRRDFMKLCAALAATMGLSSKAAAEMAESVTNPQRPPVIWIGAQECT WP_000145410.1
MULTISPECIES: hydrogenase-2 small chain [Proteobacteria] Length: 372\nNumber of predicted TMHs: 1\nExp number of AAs in TMHs: 23.72226\nExp number, first 60 AAs: 0.22038\nTotal prob of N-in: 0.12994\noutside 1 331\nTMhelix 332 354\ninside 355 372

8049 GCF_001420165.1_ASM142016v1 Escherichia coli Proteobacteria; Gammaproteobacteria; Enterobacterales; Enterobacteriaceae; Escherichia
MTGDNTLIHSHGINRRDFMKLCAALAATMGLSSKAAAEMAESVTNPQRPPVIWIGAQECT WP_000145410.1
MULTISPECIES: hydrogenase-2 small chain [Proteobacteria] Length: 372\nNumber of predicted TMHs: 1\nExp number of AAs in TMHs: 23.72226\nExp number, first 60 AAs: 0.22038\nTotal prob of N-in: 0.12994\noutside 1 331\nTMhelix 332 354\ninside 355 372

8050 GCF_001440355.1_ECO5_020 Escherichia coli Proteobacteria; Gammaproteobacteria; Enterobacterales; Enterobacteriaceae; Escherichia
MTGDNTLIHSHGINRRDFMKLCAALAATMGLSSKAAAEMAESVTNPQRPPVIWIGAQECT WP_000145410.1
MULTISPECIES: hydrogenase-2 small chain [Proteobacteria] Length: 372\nNumber of predicted TMHs: 1\nExp number of AAs in TMHs: 23.72226\nExp number, first 60 AAs: 0.22038\nTotal prob of N-in: 0.12994\noutside 1 331\nTMhelix 332 354\ninside 355 372

8051 GCF_001420055.1_ASM142005v1 Escherichia coli Proteobacteria; Gammaproteobacteria; Enterobacterales; Enterobacteriaceae; Escherichia
MNNEETFYQAMRRQGVTRRSFLKYCSLAATSLGLGAGMAPKIAWALENKPRIPVVWIHGL WP_001058323.1
MULTISPECIES: hydrogenase-1 small chain [Proteobacteria] Length: 372\nNumber of predicted TMHs: 2\nExp number of AAs in TMHs: 37.2608499999999\nExp number, first 60 AAs: 19.52923\nTotal prob of N-in: 0.94351\nPOSSIBLE N-term signal sequence\ninside 1 19\nTMhelix 20 39\noutside 40 325\nTMhelix 326 348\ninside 349 372

8052 GCF_001440565.1_ASM144056v1 Escherichia coli Proteobacteria; Gammaproteobacteria; Enterobacterales; Enterobacteriaceae; Escherichia
MTGDNTLIHSHGINRRDFMKLCAALAATMGLSSKAAAEMAESVTNPQRPPVIWIGAQECT WP_000145410.1
MULTISPECIES: hydrogenase-2 small chain [Proteobacteria] Length: 372\nNumber of predicted TMHs: 1\nExp number of AAs in TMHs: 23.72226\nExp number, first 60 AAs: 0.22038\nTotal prob of N-in: 0.12994\noutside 1 331\nTMhelix 332 354\ninside 355 372

8053 GCF_001440595.1_ASM144059v1 Escherichia coli Proteobacteria; Gammaproteobacteria; Enterobacterales; Enterobacteriaceae; Escherichia
MTGDNTLIHSHGINRRDFMKLCAALAATMGLSSKAAAEMAESVTNPQRPPVIWIGAQECT WP_000145410.1
MULTISPECIES: hydrogenase-2 small chain [Proteobacteria] Length: 372\nNumber of predicted TMHs: 1\nExp number of AAs in TMHs: 23.72226\nExp number, first 60 AAs: 0.22038\nTotal prob of N-in: 0.12994\noutside 1 331\nTMhelix 332 354\ninside 355 372

8054 GCF_001440655.1_ASM144065v1 Escherichia coli Proteobacteria; Gammaproteobacteria; Enterobacterales; Enterobacteriaceae; Escherichia
MTGDNTLIHSHGINRRDFMKLCAALAATMGLSSKAAAEMAESVTNPQRPPVIWIGAQECT WP_000145410.1
MULTISPECIES: hydrogenase-2 small chain [Proteobacteria] Length: 372\nNumber of predicted TMHs: 1\nExp number of AAs in TMHs: 23.72226\nExp number, first 60 AAs: 0.22038\nTotal prob of N-in: 0.12994\noutside 1 331\nTMhelix 332 354\ninside 355 372

8055 GCF_001440685.1_ASM144068v1 Escherichia coli Proteobacteria; Gammaproteobacteria; Enterobacterales; Enterobacteriaceae; Escherichia
MTGDNTLIHSHGINRRDFMKLCAALAATMGLSSKAAAEMAESVTNPQRPPVIWIGAQECT WP_000145410.1
MULTISPECIES: hydrogenase-2 small chain [Proteobacteria] Length: 372\nNumber of predicted TMHs: 1\nExp number of AAs in TMHs: 23.72226\nExp number, first 60 AAs: 0.22038\nTotal prob of N-in: 0.12994\noutside 1 331\nTMhelix 332 354\ninside 355 372

8056 GCF_001440565.1_ASM144056v1 Escherichia coli Proteobacteria; Gammaproteobacteria; Enterobacterales; Enterobacteriaceae; Escherichia
MNNEETFYQAMRRQGVTRRSFLKSCSLAATSLGLGAGMAPKIAWALENKPRIPVVWIHGL WP_001058318.1
hydrogenase [Escherichia coli] Length: 372\nNumber of predicted TMHs: 1\nExp number of AAs in TMHs: 33.59637\nExp number, first 60 AAs: 15.81976\nTotal prob of N-in: 0.77394\nPOSSIBLE N-term signal sequence\noutside 1 325\nTMhelix 326 348\ninside 349 372

8057 GCF_001440655.1_ASM144065v1 Escherichia coli Proteobacteria; Gammaproteobacteria; Enterobacterales; Enterobacteriaceae; Escherichia
MNNEETFYQAMRRQGVTRRSFLKSCSLAATSLGLGAGMAPKIAWALENKPRIPVVWIHGL WP_001058318.1
hydrogenase [Escherichia coli] Length: 372\nNumber of predicted TMHs: 1\nExp number of AAs in TMHs: 33.59637\nExp number, first 60 AAs: 15.81976\nTotal prob of N-in: 0.77394\nPOSSIBLE N-term signal sequence\noutside 1 325\nTMhelix 326 348\ninside 349 372

8058 GCF_001440595.1_ASM144059v1 Escherichia coli Proteobacteria; Gammaproteobacteria; Enterobacterales; Enterobacteriaceae; Escherichia
MNNEETFYQAMRRQGVTRRSFLKSCSLAATSLGLGAGMAPKIAWALENKPRIPVVWIHGL WP_001058318.1
hydrogenase [Escherichia coli] Length: 372\nNumber of predicted TMHs: 1\nExp number of AAs in TMHs: 33.59637\nExp number, first 60 AAs: 15.81976\nTotal prob of N-in: 0.77394\nPOSSIBLE N-term signal sequence\noutside 1 325\nTMhelix 326 348\ninside 349 372

8059 GCF_001440685.1_ASM144068v1 Escherichia coli Proteobacteria; Gammaproteobacteria; Enterobacterales; Enterobacteriaceae; Escherichia
MNNEETFYQAMRRQGVTRRSFLKSCSLAATSLGLGAGMAPKIAWALENKPRIPVVWIHGL WP_001058318.1
hydrogenase [Escherichia coli] Length: 372\nNumber of predicted TMHs: 1\nExp number of AAs in TMHs: 33.59637\nExp number, first 60 AAs: 15.81976\nTotal prob of N-in: 0.77394\nPOSSIBLE N-term signal sequence\noutside 1 325\nTMhelix 326 348\ninside 349 372

8060 GCF_001420955.1_ASM142095v1 Escherichia coli Proteobacteria; Gammaproteobacteria; Enterobacterales; Enterobacteriaceae; Escherichia
MNNEETFYQAMRRQGVTRRSFLKYCSLAATSLGLGAGMAPKIAWALENKPRIPVVWIHGL WP_001058323.1
MULTISPECIES: hydrogenase-1 small chain [Proteobacteria] Length: 372\nNumber of predicted TMHs: 2\nExp number of AAs in TMHs: 37.2608499999999\nExp number, first 60 AAs: 19.52923\nTotal prob of N-in: 0.94351\nPOSSIBLE N-term signal sequence\ninside 1 19\nTMhelix 20 39\noutside 40 325\nTMhelix 326 348\ninside 349 372

8061 GCF_001420125.1_ASM142012v1 Escherichia coli Proteobacteria; Gammaproteobacteria; Enterobacterales; Enterobacteriaceae; Escherichia
MNNEETFYQAMRRQGVTRRSFLKYCSLAATSLGLGAGMAPKIAWALENKPRIPVVWIHGL WP_001295940.1
MULTISPECIES: hydrogenase [Enterobacteriaceae] Length: 372\nNumber of predicted TMHs: 2\nExp number of AAs in TMHs: 37.2608899999999\nExp number, first 60 AAs: 19.52923\nTotal prob of N-in: 0.94351\nPOSSIBLE N-term signal sequence\ninside 1 19\nTMhelix 20 39\noutside 40 325\nTMhelix 326 348\ninside 349 372

8062 GCF_001420055.1_ASM142005v1 Escherichia coli Proteobacteria; Gammaproteobacteria; Enterobacterales; Enterobacteriaceae; Escherichia
MTGDNTLIHSHGINRRDFMKLCAALATTMGLSSKAAAEMAESVTNPQRPPVIWIGAQECT WP_023149629.1

MULTISPECIES: hydrogenase-2 small chain [Enterobacteriaceae] Length: 372\nNumber of predicted TMHs: 1\nExp number of AAs in TMHs: 23.63603\nExp number, first 60 AAs: 0.12971\nTotal prob of N-in: 0.12608\noutside 1 331\nTMhelix 332 354\ninside 355 372

8063 GCF_001440355.1_ECO5_020 Escherichia coli Proteobacteria; Gammaproteobacteria; Enterobacterales; Enterobacteriaceae; Escherichia MNNEETFYQAMRRQGVTRRSFLKSCSLAATSLGLGAGMAPKIAWALENKPRIPVVWIHGL WP_001058318.1 hydrogenase [Escherichia coli] Length: 372\nNumber of predicted TMHs: 1\nExp number of AAs in TMHs: 33.59637\nExp number, first 60 AAs: 15.81976\nTotal prob of N-in: 0.77394\nPOSSIBLE N-term signal sequence\noutside 1 325\nTMhelix 326 348\ninside 349 372

8064 GCF_001420165.1_ASM142016v1 Escherichia coli Proteobacteria; Gammaproteobacteria; Enterobacterales; Enterobacteriaceae; Escherichia MNNEETFYQAMRRQGVTRRSFLKYCSLAATSLGLGAGMAPKIAWALENKPRIPVVWIHGL WP_001295940.1 MULTISPECIES: hydrogenase [Enterobacteriaceae] Length: 372\nNumber of predicted TMHs: 2\nExp number of AAs in TMHs: 37.2608899999999\nExp number, first 60 AAs: 19.52923\nTotal prob of N-in: 0.94351\nPOSSIBLE N-term signal sequence\ninside 1 19\nTMhelix 20 39\noutside 40 325\nTMhelix 326 348\ninside 349 372

8065 GCF_001440755.1_ASM144075v1 Escherichia coli Proteobacteria; Gammaproteobacteria; Enterobacterales; Enterobacteriaceae; Escherichia MTGDNTLIHSHGINRRDFMKLCAALAATMGLSSKAAAEMAESVTNPQRPPVIWIGAQECT WP_000145410.1 MULTISPECIES: hydrogenase-2 small chain [Proteobacteria] Length: 372\nNumber of predicted TMHs: 1\nExp number of AAs in TMHs: 23.72226\nExp number, first 60 AAs: 0.22038\nTotal prob of N-in: 0.12994\noutside 1 331\nTMhelix 332 354\ninside 355 372

8066 GCF_001440805.1_ASM144080v1 Escherichia coli Proteobacteria; Gammaproteobacteria; Enterobacterales; Enterobacteriaceae; Escherichia MTGDNTLIHSHGINRRDFMKLCAALAATMGLSSKAAAEMAESVTNPQRPPVIWIGAQECT WP_000145410.1 MULTISPECIES: hydrogenase-2 small chain [Proteobacteria] Length: 372\nNumber of predicted TMHs: 1\nExp number of AAs in TMHs: 23.72226\nExp number, first 60 AAs: 0.22038\nTotal prob of N-in: 0.12994\noutside 1 331\nTMhelix 332 354\ninside 355 372

8067 GCF_001440755.1_ASM144075v1 Escherichia coli Proteobacteria; Gammaproteobacteria; Enterobacterales; Enterobacteriaceae; Escherichia MNNEETFYQAMRRQGVTRRSFLKSCSLAATSLGLGAGMAPKIAWALENKPRIPVVWIHGL WP_001058318.1 hydrogenase [Escherichia coli] Length: 372\nNumber of predicted TMHs: 1\nExp number of AAs in TMHs: 33.59637\nExp number, first 60 AAs: 15.81976\nTotal prob of N-in: 0.77394\nPOSSIBLE N-term signal sequence\noutside 1 325\nTMhelix 326 348\ninside 349 372

8068 GCF_001440805.1_ASM144080v1 Escherichia coli Proteobacteria; Gammaproteobacteria; Enterobacterales; Enterobacteriaceae; Escherichia MNNEETFYQAMRRQGVTRRSFLKSCSLAATSLGLGAGMAPKIAWALENKPRIPVVWIHGL WP_001058318.1 hydrogenase [Escherichia coli] Length: 372\nNumber of predicted TMHs: 1\nExp number of AAs in TMHs: 33.59637\nExp number, first 60 AAs: 15.81976\nTotal prob of N-in: 0.77394\nPOSSIBLE N-term signal sequence\noutside 1 325\nTMhelix 326 348\ninside 349 372

8069 GCF_001442595.1_ASM144259v1 Escherichia coli Proteobacteria; Gammaproteobacteria; Enterobacterales; Enterobacteriaceae; Escherichia MTGDNTLIHSHGINRRDFMKLCAALAATMGLSSKAAAEMVESVTNPQRPPVIWIGAQECT WP_000145423.1 hydrogenase 2 small subunit [Escherichia coli] Length: 372\nNumber of predicted TMHs: 1\nExp number of AAs in TMHs: 23.78201\nExp number, first 60 AAs: 0.28355\nTotal prob of N-in: 0.13257\noutside 1 331\nTMhelix 332 354\ninside 355 372

8070 GCF_001443175.1_ASM144317v1 Escherichia coli Proteobacteria; Gammaproteobacteria; Enterobacterales; Enterobacteriaceae; Escherichia MTGDNTLIHSHGINRRDFMKLCAALAATMGLSSKAAAEMAESVTNPQRPPVIWIGAQECT WP_000145410.1 MULTISPECIES: hydrogenase-2 small chain [Proteobacteria] Length: 372\nNumber of predicted TMHs: 1\nExp number of AAs in TMHs: 23.72226\nExp number, first 60 AAs: 0.22038\nTotal prob of N-in: 0.12994\noutside 1 331\nTMhelix 332 354\ninside 355 372

8071 GCF_001446595.1_ASM144659v1 Escherichia coli Proteobacteria; Gammaproteobacteria; Enterobacterales; Enterobacteriaceae; Escherichia
MTGDNTLIHSHGINRRDFMKLCAALAATMGLSSKAAAEMAESVTNPQRPPVIWIGAQECT WP_000145410.1
MULTISPECIES: hydrogenase-2 small chain [Proteobacteria] Length: 372\nNumber of predicted TMHs: 1\nExp number of AAs in TMHs: 23.72226\nExp number, first 60 AAs: 0.22038\nTotal prob of N-in: 0.12994\noutside 1 331\nTMhelix 332 354\ninside 355 372

8072 GCF_001446615.1_ASM144661v1 Escherichia coli Proteobacteria; Gammaproteobacteria; Enterobacterales; Enterobacteriaceae; Escherichia
MTGDNTLIHSHGINRRDFMKLCAALAATMGLSSKAAAEMAESVTNPQRPPVIWIGAQECT WP_000145410.1
MULTISPECIES: hydrogenase-2 small chain [Proteobacteria] Length: 372\nNumber of predicted TMHs: 1\nExp number of AAs in TMHs: 23.72226\nExp number, first 60 AAs: 0.22038\nTotal prob of N-in: 0.12994\noutside 1 331\nTMhelix 332 354\ninside 355 372

8073 GCF_001519145.1_ASM151914v1 Escherichia coli Proteobacteria; Gammaproteobacteria; Enterobacterales; Enterobacteriaceae; Escherichia
MTGDNTLIHSHGINRRDFMKLCAALAATMGLSSKAAAEMAESVTNPQRPPVIWIGAQECT WP_000145410.1
MULTISPECIES: hydrogenase-2 small chain [Proteobacteria] Length: 372\nNumber of predicted TMHs: 1\nExp number of AAs in TMHs: 23.72226\nExp number, first 60 AAs: 0.22038\nTotal prob of N-in: 0.12994\noutside 1 331\nTMhelix 332 354\ninside 355 372

8074 GCF_001443175.1_ASM144317v1 Escherichia coli Proteobacteria; Gammaproteobacteria; Enterobacterales; Enterobacteriaceae; Escherichia
MNNEETFYQAMRRQGVTRRSFLKYCSLAATSLGLGAGMAPKIAWALENKPRIPVVWIHGL WP_001058323.1
MULTISPECIES: hydrogenase-1 small chain [Proteobacteria] Length: 372\nNumber of predicted TMHs: 2\nExp number of AAs in TMHs: 37.2608499999999\nExp number, first 60 AAs: 19.52923\nTotal prob of N-in: 0.94351\nPOSSIBLE N-term signal sequence\ninside 1 19\nTMhelix 20 39\noutside 40 325\nTMhelix 326 348\ninside 349 372

8075 GCF_001455405.1_ASM145540v1 Escherichia coli Proteobacteria; Gammaproteobacteria; Enterobacterales; Enterobacteriaceae; Escherichia
MTGDNTLIHSHGINRRDFMKLCAALAATMGLSSKAAAEMAESVTNPQRPPVIWIGAQECT WP_000145410.1
MULTISPECIES: hydrogenase-2 small chain [Proteobacteria] Length: 372\nNumber of predicted TMHs: 1\nExp number of AAs in TMHs: 23.72226\nExp number, first 60 AAs: 0.22038\nTotal prob of N-in: 0.12994\noutside 1 331\nTMhelix 332 354\ninside 355 372

8076 GCF_001462925.1_ASM146292v1 Escherichia coli Proteobacteria; Gammaproteobacteria; Enterobacterales; Enterobacteriaceae; Escherichia
MTGDNTLIHSHGINRRDFMKLCAALAATMGLSSKAAAEMAESVTNPQRPPVIWIGAQECT WP_000145410.1
MULTISPECIES: hydrogenase-2 small chain [Proteobacteria] Length: 372\nNumber of predicted TMHs: 1\nExp number of AAs in TMHs: 23.72226\nExp number, first 60 AAs: 0.22038\nTotal prob of N-in: 0.12994\noutside 1 331\nTMhelix 332 354\ninside 355 372

8077 GCF_001455025.1_EC222071v1.0 Escherichia coli Proteobacteria; Gammaproteobacteria; Enterobacterales; Enterobacteriaceae; Escherichia
MTGDNTLIHSHGINRRDFMKLCAALAATMGLSSKAAAEMAESVTNPQRPPVIWIGAQECT WP_000145410.1
MULTISPECIES: hydrogenase-2 small chain [Proteobacteria] Length: 372\nNumber of predicted TMHs: 1\nExp number of AAs in TMHs: 23.72226\nExp number, first 60 AAs: 0.22038\nTotal prob of N-in: 0.12994\noutside 1 331\nTMhelix 332 354\ninside 355 372

8078 GCF_001463195.1_ASM146319v1 Escherichia coli Proteobacteria; Gammaproteobacteria; Enterobacterales; Enterobacteriaceae; Escherichia
MTGDNTLIHSHGINRRDFMKLCAALAATMGLSSKAAAEMVESVTNPQRPPVIWIGAQECT WP_000145423.1
hydrogenase 2 small subunit [Escherichia coli] Length: 372\nNumber of predicted TMHs: 1\nExp number of AAs in TMHs: 23.78201\nExp number, first 60 AAs: 0.28355\nTotal prob of N-in: 0.13257\noutside 1 331\nTMhelix 332 354\ninside 355 372

8079 GCF_001462925.1_ASM146292v1 Escherichia coli Proteobacteria; Gammaproteobacteria; Enterobacterales; Enterobacteriaceae; Escherichia
MNNEETFYQAMRRQGVTRRSFLKYCSLAATSLGLGAGMAPKIAWALENKPRIPVVWIHGL WP_001058323.1
MULTISPECIES: hydrogenase-1 small chain [Proteobacteria] Length: 372\nNumber of predicted TMHs:

2\nExp number of AAs in TMHs: 37.2608499999999\nExp number, first 60 AAs:  19.52923\nTotal prob of N-in: 0.94351\nPOSSIBLE N-term signal sequence\ninside   1   19\nTMhelix   20   39\noutside   40   325\nTMhelix   326   348\ninside   349   372

8080 GCF_001455405.1_ASM145540v1 Escherichia coli Proteobacteria; Gammaproteobacteria; Enterobacterales; Enterobacteriaceae; Escherichia
MNNEETFYQAMRRQGVTRRSFLKYCSLAATSLGLGAGMAPKIAWALENKPRIPVVWIHGL WP_001058323.1
MULTISPECIES: hydrogenase-1 small chain [Proteobacteria] Length: 372\nNumber of predicted TMHs: 2\nExp number of AAs in TMHs: 37.2608499999999\nExp number, first 60 AAs:  19.52923\nTotal prob of N-in: 0.94351\nPOSSIBLE N-term signal sequence\ninside   1   19\nTMhelix   20   39\noutside   40   325\nTMhelix   326   348\ninside   349   372

8081 GCF_001442595.1_ASM144259v1 Escherichia coli Proteobacteria; Gammaproteobacteria; Enterobacterales; Enterobacteriaceae; Escherichia
MNNEETFYQAMRRQGVTRRSFLKYCSLAATSLGLGAGMAPKIAWALENKPRIPVVWIHGL WP_001295940.1
MULTISPECIES: hydrogenase [Enterobacteriaceae] Length: 372\nNumber of predicted TMHs:  2\nExp number of AAs in TMHs: 37.2608899999999\nExp number, first 60 AAs:  19.52923\nTotal prob of N-in: 0.94351\nPOSSIBLE N-term signal sequence\ninside   1   19\nTMhelix   20   39\noutside   40   325\nTMhelix   326   348\ninside   349   372

8082 GCF_001455025.1_EC222071v1.0 Escherichia coli Proteobacteria; Gammaproteobacteria; Enterobacterales; Enterobacteriaceae; Escherichia
MNNEETFYQAMRRQGVTRRSFLKYCSLAATSLGLGAGMAPKIAWALENKPRIPVVWIHGL WP_001058323.1
MULTISPECIES: hydrogenase-1 small chain [Proteobacteria] Length: 372\nNumber of predicted TMHs: 2\nExp number of AAs in TMHs: 37.2608499999999\nExp number, first 60 AAs:  19.52923\nTotal prob of N-in: 0.94351\nPOSSIBLE N-term signal sequence\ninside   1   19\nTMhelix   20   39\noutside   40   325\nTMhelix   326   348\ninside   349   372

8083 GCF_001446595.1_ASM144659v1 Escherichia coli Proteobacteria; Gammaproteobacteria; Enterobacterales; Enterobacteriaceae; Escherichia
MNNEETFYQAMRRQGVTRRSFLKYCSLAATSLGLGAGMAPKIAWALENKPRIPVVWIHGL WP_058055433.1
hydrogenase [Escherichia coli] Length: 372\nNumber of predicted TMHs:  2\nExp number of AAs in TMHs: 36.72011\nExp number, first 60 AAs:  19.53447\nTotal prob of N-in:   0.94308\nPOSSIBLE N-term signal sequence\ninside   1   19\nTMhelix   20   39\noutside   40   325\nTMhelix   326   348\ninside   349   372

8084 GCF_001463405.1_ASM146340v1 Escherichia coli Proteobacteria; Gammaproteobacteria; Enterobacterales; Enterobacteriaceae; Escherichia
MTGDNTLIHSHGINRRDFMKLCAALAATMGLSSKAAAEMAESVTNPQRPPVIWIGAQECT WP_000145410.1
MULTISPECIES: hydrogenase-2 small chain [Proteobacteria] Length: 372\nNumber of predicted TMHs: 1\nExp number of AAs in TMHs: 23.72226\nExp number, first 60 AAs:  0.22038\nTotal prob of N-in:   0.12994\noutside   1   331\nTMhelix   332   354\ninside   355   372

8085 GCF_001446615.1_ASM144661v1 Escherichia coli Proteobacteria; Gammaproteobacteria; Enterobacterales; Enterobacteriaceae; Escherichia
MNNEETFYQAMRRQGVTRRSFLKYCSLAATSLGLGAGMAPKIAWALENKPRIPVVWIHGL WP_058055433.1
hydrogenase [Escherichia coli] Length: 372\nNumber of predicted TMHs:  2\nExp number of AAs in TMHs: 36.72011\nExp number, first 60 AAs:  19.53447\nTotal prob of N-in:   0.94308\nPOSSIBLE N-term signal sequence\ninside   1   19\nTMhelix   20   39\noutside   40   325\nTMhelix   326   348\ninside   349   372

8086 GCF_001463535.1_ASM146353v1 Escherichia coli Proteobacteria; Gammaproteobacteria; Enterobacterales; Enterobacteriaceae; Escherichia
MTGDNTLIHSHGINRRDFMKLCAALAATMGLSSKAAAEMAESVTNPQRPPVIWIGAQECT WP_000145410.1
MULTISPECIES: hydrogenase-2 small chain [Proteobacteria] Length: 372\nNumber of predicted TMHs: 1\nExp number of AAs in TMHs: 23.72226\nExp number, first 60 AAs:  0.22038\nTotal prob of N-in:   0.12994\noutside   1   331\nTMhelix   332   354\ninside   355   372

8087 GCF_001484235.1_ASM148423v1 Escherichia coli Proteobacteria; Gammaproteobacteria; Enterobacterales; Enterobacteriaceae; Escherichia
MTGDNTLIHSHGINRRDFMKLCAALAATMGLSSKAAAEMAESVTNPQRPPVIWIGAQECT WP_000145410.1
MULTISPECIES: hydrogenase-2 small chain [Proteobacteria] Length: 372\nNumber of predicted TMHs:

1\nExp number of AAs in TMHs: 23.72226\nExp number, first 60 AAs: 0.22038\nTotal prob of N-in: 0.12994\noutside 1 331\nTMhelix 332 354\ninside 355 372

8088 GCF_001463325.1_ASM146332v1 Escherichia coli Proteobacteria; Gammaproteobacteria; Enterobacterales; Enterobacteriaceae; Escherichia
MNNEETFYQAMRRQGVTRRSFLKYCSLAATSLGLGAGMAPKIAWALENKPRIPVVWIHGL WP_001058323.1
MULTISPECIES: hydrogenase-1 small chain [Proteobacteria] Length: 372\nNumber of predicted TMHs: 2\nExp number of AAs in TMHs: 37.2608499999999\nExp number, first 60 AAs: 19.52923\nTotal prob of N-in: 0.94351\nPOSSIBLE N-term signal sequence\ninside 1 19\nTMhelix 20 39\noutside 40 325\nTMhelix 326 348\ninside 349 372

8089 GCF_001463325.1_ASM146332v1 Escherichia coli Proteobacteria; Gammaproteobacteria; Enterobacterales; Enterobacteriaceae; Escherichia
MTGDNTLIHSHGINRRDFMKLCAALATTMGLSSKAAAEMAESVTNPQRPPVIWIGAQECT WP_023149629.1
MULTISPECIES: hydrogenase-2 small chain [Enterobacteriaceae] Length: 372\nNumber of predicted TMHs: 1\nExp number of AAs in TMHs: 23.63603\nExp number, first 60 AAs: 0.12971\nTotal prob of N-in: 0.12608\noutside 1 331\nTMhelix 332 354\ninside 355 372

8090 GCF_001463195.1_ASM146319v1 Escherichia coli Proteobacteria; Gammaproteobacteria; Enterobacterales; Enterobacteriaceae; Escherichia
MNNEETFYQAMRRQGVTRRSFLKYCSLAATSLGLGAGMAPKIAWALENKPRIPVVWIHGL WP_001295940.1
MULTISPECIES: hydrogenase [Enterobacteriaceae] Length: 372\nNumber of predicted TMHs: 2\nExp number of AAs in TMHs: 37.2608899999999\nExp number, first 60 AAs: 19.52923\nTotal prob of N-in: 0.94351\nPOSSIBLE N-term signal sequence\ninside 1 19\nTMhelix 20 39\noutside 40 325\nTMhelix 326 348\ninside 349 372

8091 GCF_001484265.1_ASM148426v1 Escherichia coli Proteobacteria; Gammaproteobacteria; Enterobacterales; Enterobacteriaceae; Escherichia
MTGDNTLIHSHGINRRDFMKLCAALAATMGLSSKAAAEMAESVTNPQRPPVIWIGAQECT WP_000145410.1
MULTISPECIES: hydrogenase-2 small chain [Proteobacteria] Length: 372\nNumber of predicted TMHs: 1\nExp number of AAs in TMHs: 23.72226\nExp number, first 60 AAs: 0.22038\nTotal prob of N-in: 0.12994\noutside 1 331\nTMhelix 332 354\ninside 355 372

8092 GCF_001484335.1_ASM148433v1 Escherichia coli Proteobacteria; Gammaproteobacteria; Enterobacterales; Enterobacteriaceae; Escherichia
MTGDNTLIHSHGINRRDFMKLCAALAATMGLSSKAAAEMAESVTNPQRPPVIWIGAQECT WP_000145410.1
MULTISPECIES: hydrogenase-2 small chain [Proteobacteria] Length: 372\nNumber of predicted TMHs: 1\nExp number of AAs in TMHs: 23.72226\nExp number, first 60 AAs: 0.22038\nTotal prob of N-in: 0.12994\noutside 1 331\nTMhelix 332 354\ninside 355 372

8093 GCF_001484375.1_ASM148437v1 Escherichia coli Proteobacteria; Gammaproteobacteria; Enterobacterales; Enterobacteriaceae; Escherichia
MTGDNTLIHSHGINRRDFMKLCAALAATMGLSSKAAAEMAESVTNPQRPPVIWIGAQECT WP_000145410.1
MULTISPECIES: hydrogenase-2 small chain [Proteobacteria] Length: 372\nNumber of predicted TMHs: 1\nExp number of AAs in TMHs: 23.72226\nExp number, first 60 AAs: 0.22038\nTotal prob of N-in: 0.12994\noutside 1 331\nTMhelix 332 354\ninside 355 372

8094 GCF_001484415.1_ASM148441v1 Escherichia coli Proteobacteria; Gammaproteobacteria; Enterobacterales; Enterobacteriaceae; Escherichia
MTGDNTLIHSHGINRRDFMKLCAALAATMGLSSKAAAEMAESVTNPQRPPVIWIGAQECT WP_000145410.1
MULTISPECIES: hydrogenase-2 small chain [Proteobacteria] Length: 372\nNumber of predicted TMHs: 1\nExp number of AAs in TMHs: 23.72226\nExp number, first 60 AAs: 0.22038\nTotal prob of N-in: 0.12994\noutside 1 331\nTMhelix 332 354\ninside 355 372

8095 GCF_001485455.1_ASM148545v1 Escherichia coli Proteobacteria; Gammaproteobacteria; Enterobacterales; Enterobacteriaceae; Escherichia
MTGDNTLIHSHGINRRDFMKLCAALAATMGLSSKAAAEMAESVTNPQRPPVIWIGAQECT WP_000145410.1
MULTISPECIES: hydrogenase-2 small chain [Proteobacteria] Length: 372\nNumber of predicted TMHs: 1\nExp number of AAs in TMHs: 23.72226\nExp number, first 60 AAs: 0.22038\nTotal prob of N-in: 0.12994\noutside 1 331\nTMhelix 332 354\ninside 355 372

8096 GCF_001484475.1_ASM148447v1 Escherichia coli Proteobacteria; Gammaproteobacteria; Enterobacterales; Enterobacteriaceae; Escherichia
MTGDNTLIHSHGINRRDFMKLCAALAATMGLSSKAAAEMAESVTNPQRPPVIWIGAQECT WP_000145410.1
MULTISPECIES: hydrogenase-2 small chain [Proteobacteria] Length: 372\nNumber of predicted TMHs: 1\nExp number of AAs in TMHs: 23.72226\nExp number, first 60 AAs: 0.22038\nTotal prob of N-in: 0.12994\noutside 1 331\nTMhelix 332 354\ninside 355 372

8097 GCF_001484335.1_ASM148433v1 Escherichia coli Proteobacteria; Gammaproteobacteria; Enterobacterales; Enterobacteriaceae; Escherichia
MNNEETFYQAMRRQGVTRRSFLKYCSLAATSLGLGAGMAPKIAWALENKPRIPVVWIHGL WP_001058323.1
MULTISPECIES: hydrogenase-1 small chain [Proteobacteria] Length: 372\nNumber of predicted TMHs: 2\nExp number of AAs in TMHs: 37.2608499999999\nExp number, first 60 AAs: 19.52923\nTotal prob of N-in: 0.94351\nPOSSIBLE N-term signal sequence\ninside 1 19\nTMhelix 20 39\noutside 40 325\nTMhelix 326 348\ninside 349 372

8098 GCF_001484265.1_ASM148426v1 Escherichia coli Proteobacteria; Gammaproteobacteria; Enterobacterales; Enterobacteriaceae; Escherichia
MNNEETFYQAMRRQGVTRRSFLKYCSLAATSLGLGAGMAPKIAWALENKPRIPVVWIHGL WP_001058323.1
MULTISPECIES: hydrogenase-1 small chain [Proteobacteria] Length: 372\nNumber of predicted TMHs: 2\nExp number of AAs in TMHs: 37.2608499999999\nExp number, first 60 AAs: 19.52923\nTotal prob of N-in: 0.94351\nPOSSIBLE N-term signal sequence\ninside 1 19\nTMhelix 20 39\noutside 40 325\nTMhelix 326 348\ninside 349 372

8099 GCF_001484235.1_ASM148423v1 Escherichia coli Proteobacteria; Gammaproteobacteria; Enterobacterales; Enterobacteriaceae; Escherichia
MNNEETFYQAMRRQGVTRRSFLKYCSLAATSLGLGAGMAPKIAWALENKPRIPVVWIHGL WP_001717971.1
hydrogenase-1 small chain [Escherichia coli] Length: 372\nNumber of predicted TMHs: 2\nExp number of AAs in TMHs: 38.00708\nExp number, first 60 AAs: 19.5286\nTotal prob of N-in: 0.94354\nPOSSIBLE N-term signal sequence\ninside 1 19\nTMhelix 20 39\noutside 40 325\nTMhelix 326 348\ninside 349 372

8100 GCF_001463535.1_ASM146353v1 Escherichia coli Proteobacteria; Gammaproteobacteria; Enterobacterales; Enterobacteriaceae; Escherichia
MNNEETFYQAMRRQGVTRRSFLKYCSLAATSLGLGAGMAPKIAWALENKPRIPVVWIHGL WP_001295940.1
MULTISPECIES: hydrogenase [Enterobacteriaceae] Length: 372\nNumber of predicted TMHs: 2\nExp number of AAs in TMHs: 37.2608899999999\nExp number, first 60 AAs: 19.52923\nTotal prob of N-in: 0.94351\nPOSSIBLE N-term signal sequence\ninside 1 19\nTMhelix 20 39\noutside 40 325\nTMhelix 326 348\ninside 349 372

8101 GCF_001485455.1_ASM148545v1 Escherichia coli Proteobacteria; Gammaproteobacteria; Enterobacterales; Enterobacteriaceae; Escherichia
MNNEETFYQAMRRQGVTRRSFLKYCSLAATSLGLGAGMAPKIAWALENKPRIPVVWIHGL WP_001058323.1
MULTISPECIES: hydrogenase-1 small chain [Proteobacteria] Length: 372\nNumber of predicted TMHs: 2\nExp number of AAs in TMHs: 37.2608499999999\nExp number, first 60 AAs: 19.52923\nTotal prob of N-in: 0.94351\nPOSSIBLE N-term signal sequence\ninside 1 19\nTMhelix 20 39\noutside 40 325\nTMhelix 326 348\ninside 349 372

8102 GCF_001484375.1_ASM148437v1 Escherichia coli Proteobacteria; Gammaproteobacteria; Enterobacterales; Enterobacteriaceae; Escherichia
MNNEETFYQAMRRQGVTRRSFLKYCSLAATSLGLGAGMAPKIAWALENKPRIPVVWIHGL WP_001058323.1
MULTISPECIES: hydrogenase-1 small chain [Proteobacteria] Length: 372\nNumber of predicted TMHs: 2\nExp number of AAs in TMHs: 37.2608499999999\nExp number, first 60 AAs: 19.52923\nTotal prob of N-in: 0.94351\nPOSSIBLE N-term signal sequence\ninside 1 19\nTMhelix 20 39\noutside 40 325\nTMhelix 326 348\ninside 349 372

8103 GCF_001484415.1_ASM148441v1 Escherichia coli Proteobacteria; Gammaproteobacteria; Enterobacterales; Enterobacteriaceae; Escherichia
MNNEETFYQAMRRQGVTRRSFLKYCSLAATSLGLGAGMAPKIAWALENKPRIPVVWIHGL WP_001058323.1
MULTISPECIES: hydrogenase-1 small chain [Proteobacteria] Length: 372\nNumber of predicted TMHs: 2\nExp number of AAs in TMHs: 37.2608499999999\nExp number, first 60 AAs: 19.52923\nTotal prob of N-in: 0.94351\nPOSSIBLE N-term signal sequence\ninside 1 19\nTMhelix 20 39\noutside 40 325\nTMhelix 326 348\ninside 349 372

8104 GCF_001499595.1_AL505-MS Escherichia coli Proteobacteria; Gammaproteobacteria; Enterobacterales; Enterobacteriaceae; Escherichia MTGDNTLIHSHGINRRDFMKLCAALAATMGLSSKAAAEMAESVTNPQRPPVIWIGAQECT WP_000145410.1 MULTISPECIES: hydrogenase-2 small chain [Proteobacteria] Length: 372\nNumber of predicted TMHs: 1\nExp number of AAs in TMHs: 23.72226\nExp number, first 60 AAs: 0.22038\nTotal prob of N-in: 0.12994\noutside 1 331\nTMhelix 332 354\ninside 355 372

8105 GCF_001484475.1_ASM148447v1 Escherichia coli Proteobacteria; Gammaproteobacteria; Enterobacterales; Enterobacteriaceae; Escherichia MNNEETFYQAMRRQGVTRRSFLKYCSLAATSLGLGAGMAPKIAWALENKPRIPVVWIHGL WP_001717971.1 hydrogenase-1 small chain [Escherichia coli] Length: 372\nNumber of predicted TMHs: 2\nExp number of AAs in TMHs: 38.00708\nExp number, first 60 AAs: 19.5286\nTotal prob of N-in: 0.94354\nPOSSIBLE N-term signal sequence\ninside 1 19\nTMhelix 20 39\noutside 40 325\nTMhelix 326 348\ninside 349 372

8106 GCF_001506145.1_703450 Escherichia coli Proteobacteria; Gammaproteobacteria; Enterobacterales; Enterobacteriaceae; Escherichia MTGDNTLIHSHGINRRDFMKLCAALAATMGLSSKAAAEMAESVTNPQHPPVIWIGAQECT WP_000145403.1 hydrogenase 2 small subunit [Escherichia coli] Length: 372\nNumber of predicted TMHs: 1\nExp number of AAs in TMHs: 23.66365\nExp number, first 60 AAs: 0.1537\nTotal prob of N-in: 0.12709\noutside 1 331\nTMhelix 332 354\ninside 355 372

8107 GCF_001509665.1_ASM150966v1 Escherichia coli Proteobacteria; Gammaproteobacteria; Enterobacterales; Enterobacteriaceae; Escherichia MTGDNTLIHSHGINRRDFMKLCAALAATMGLSSKAAAEMAESVTNPQRPPVIWIGAQECT WP_000145410.1 MULTISPECIES: hydrogenase-2 small chain [Proteobacteria] Length: 372\nNumber of predicted TMHs: 1\nExp number of AAs in TMHs: 23.72226\nExp number, first 60 AAs: 0.22038\nTotal prob of N-in: 0.12994\noutside 1 331\nTMhelix 332 354\ninside 355 372

8108 GCF_001509735.1_ASM150973v1 Escherichia coli Proteobacteria; Gammaproteobacteria; Enterobacterales; Enterobacteriaceae; Escherichia MTGDNTLIHSHGINRRDFMKLCAALAATMGLSSKAAAEMAESVTNPQRPPVIWIGAQECT WP_000145410.1 MULTISPECIES: hydrogenase-2 small chain [Proteobacteria] Length: 372\nNumber of predicted TMHs: 1\nExp number of AAs in TMHs: 23.72226\nExp number, first 60 AAs: 0.22038\nTotal prob of N-in: 0.12994\noutside 1 331\nTMhelix 332 354\ninside 355 372

8109 GCF_001513615.1_ASM151361v1 Escherichia coli Proteobacteria; Gammaproteobacteria; Enterobacterales; Enterobacteriaceae; Escherichia MTGDNTLIHSHGINRRDFMKLCAALAATMGLSSKAAAEMVESVTNPQRPPVIWIGAQECT WP_000145423.1 hydrogenase 2 small subunit [Escherichia coli] Length: 372\nNumber of predicted TMHs: 1\nExp number of AAs in TMHs: 23.78201\nExp number, first 60 AAs: 0.28355\nTotal prob of N-in: 0.13257\noutside 1 331\nTMhelix 332 354\ninside 355 372

8110 GCF_001518355.1_ASM151835v1 Escherichia coli Proteobacteria; Gammaproteobacteria; Enterobacterales; Enterobacteriaceae; Escherichia MTGDNTLIHSHGINRRDFMKLCAALAATMGLSSKAAAEMAESVTNPQRPPVIWIGAQECT WP_000145410.1 MULTISPECIES: hydrogenase-2 small chain [Proteobacteria] Length: 372\nNumber of predicted TMHs: 1\nExp number of AAs in TMHs: 23.72226\nExp number, first 60 AAs: 0.22038\nTotal prob of N-in: 0.12994\noutside 1 331\nTMhelix 332 354\ninside 355 372

8111 GCF_001519125.1_ASM151912v1 Escherichia coli Proteobacteria; Gammaproteobacteria; Enterobacterales; Enterobacteriaceae; Escherichia MTGDNTLIHSHGINRRDFMKLCAALAATMGLSSKAAAEMAESVTNPQRPPVIWIGAQECT WP_000145410.1 MULTISPECIES: hydrogenase-2 small chain [Proteobacteria] Length: 372\nNumber of predicted TMHs: 1\nExp number of AAs in TMHs: 23.72226\nExp number, first 60 AAs: 0.22038\nTotal prob of N-in: 0.12994\noutside 1 331\nTMhelix 332 354\ninside 355 372

8112 GCF_001509665.1_ASM150966v1 Escherichia coli Proteobacteria; Gammaproteobacteria; Enterobacterales; Enterobacteriaceae; Escherichia MNNEETFYQAMRRQGVTRRSFLKYCSLAATSLGLGAGMAPKIAWALENKPRIPVVWIHGL WP_001058323.1 MULTISPECIES: hydrogenase-1 small chain [Proteobacteria] Length: 372\nNumber of predicted TMHs: 2\nExp number of AAs in TMHs: 37.2608499999999\nExp number, first 60 AAs: 19.52923\nTotal prob of N-in:

0.94351\nPOSSIBLE N-term signal sequence\ninside 1 19\nTMhelix 20 39\noutside 40 325\nTMhelix 326 348\ninside 349 372

8113 GCF_001519235.1_ASM151923v1 Escherichia coli Proteobacteria; Gammaproteobacteria; Enterobacterales; Enterobacteriaceae; Escherichia MTGDNTLIHSHGINRRDFMKLCAALAATMGLSSKAAAEMAESVTNPQRPPVIWIGAQECT WP_000145410.1 MULTISPECIES: hydrogenase-2 small chain [Proteobacteria] Length: 372\nNumber of predicted TMHs: 1\nExp number of AAs in TMHs: 23.72226\nExp number, first 60 AAs: 0.22038\nTotal prob of N-in: 0.12994\noutside 1 331\nTMhelix 332 354\ninside 355 372

8114 GCF_001506145.1_703450 Escherichia coli Proteobacteria; Gammaproteobacteria; Enterobacterales; Enterobacteriaceae; Escherichia MNNEETFYQAMRRQGVTRRSFLKYCSLAATSLGLGAGMAPKIAWALENKPRIPVVWIHGL WP_001295940.1 MULTISPECIES: hydrogenase [Enterobacteriaceae] Length: 372\nNumber of predicted TMHs: 2\nExp number of AAs in TMHs: 37.2608899999999\nExp number, first 60 AAs: 19.52923\nTotal prob of N-in: 0.94351\nPOSSIBLE N-term signal sequence\ninside 1 19\nTMhelix 20 39\noutside 40 325\nTMhelix 326 348\ninside 349 372

8115 GCF_001519275.1_ASM151927v1 Escherichia coli Proteobacteria; Gammaproteobacteria; Enterobacterales; Enterobacteriaceae; Escherichia MTGDNTLIHSHGINRRDFMKLCAALAATMGLSSKAAAEMAESVTNPQRPPVIWIGAQECT WP_000145410.1 MULTISPECIES: hydrogenase-2 small chain [Proteobacteria] Length: 372\nNumber of predicted TMHs: 1\nExp number of AAs in TMHs: 23.72226\nExp number, first 60 AAs: 0.22038\nTotal prob of N-in: 0.12994\noutside 1 331\nTMhelix 332 354\ninside 355 372

8116 GCF_001519275.1_ASM151927v1 Escherichia coli Proteobacteria; Gammaproteobacteria; Enterobacterales; Enterobacteriaceae; Escherichia MNNEETFYQAMRRQGVTRRSFLKYCSLAATSLGLGAGMAPKIAWALENKPRIPVVWIHGL WP_001058323.1 MULTISPECIES: hydrogenase-1 small chain [Proteobacteria] Length: 372\nNumber of predicted TMHs: 2\nExp number of AAs in TMHs: 37.2608499999999\nExp number, first 60 AAs: 19.52923\nTotal prob of N-in: 0.94351\nPOSSIBLE N-term signal sequence\ninside 1 19\nTMhelix 20 39\noutside 40 325\nTMhelix 326 348\ninside 349 372

8117 GCF_001509735.1_ASM150973v1 Escherichia coli Proteobacteria; Gammaproteobacteria; Enterobacterales; Enterobacteriaceae; Escherichia MNNEETFYQAMRRQGVTRRSFLKYCSLAATSLGLGAGMAPKIAWALENKPRIPVVWIHGL WP_001058323.1 MULTISPECIES: hydrogenase-1 small chain [Proteobacteria] Length: 372\nNumber of predicted TMHs: 2\nExp number of AAs in TMHs: 37.2608499999999\nExp number, first 60 AAs: 19.52923\nTotal prob of N-in: 0.94351\nPOSSIBLE N-term signal sequence\ninside 1 19\nTMhelix 20 39\noutside 40 325\nTMhelix 326 348\ninside 349 372

8118 GCF_001518355.1_ASM151835v1 Escherichia coli Proteobacteria; Gammaproteobacteria; Enterobacterales; Enterobacteriaceae; Escherichia MNNEETFYQAMRRQGVTRRSFLKYCSLAATSLGLGAGMAPKIAWALENKPRIPVVWIHGL WP_001058323.1 MULTISPECIES: hydrogenase-1 small chain [Proteobacteria] Length: 372\nNumber of predicted TMHs: 2\nExp number of AAs in TMHs: 37.2608499999999\nExp number, first 60 AAs: 19.52923\nTotal prob of N-in: 0.94351\nPOSSIBLE N-term signal sequence\ninside 1 19\nTMhelix 20 39\noutside 40 325\nTMhelix 326 348\ninside 349 372

8119 GCF_001519145.1_ASM151914v1 Escherichia coli Proteobacteria; Gammaproteobacteria; Enterobacterales; Enterobacteriaceae; Escherichia MNNEETFYQAMRRQGVTRRSFLKYCSLAATSLGLGAGMAPKIAWALENKPRIPVVWIHGL WP_001058323.1 MULTISPECIES: hydrogenase-1 small chain [Proteobacteria] Length: 372\nNumber of predicted TMHs: 2\nExp number of AAs in TMHs: 37.2608499999999\nExp number, first 60 AAs: 19.52923\nTotal prob of N-in: 0.94351\nPOSSIBLE N-term signal sequence\ninside 1 19\nTMhelix 20 39\noutside 40 325\nTMhelix 326 348\ninside 349 372

8120 GCF_001519125.1_ASM151912v1 Escherichia coli Proteobacteria; Gammaproteobacteria; Enterobacterales; Enterobacteriaceae; Escherichia MNNEETFYQAMRRQGVTRRSFLKYCSLAATSLGLGAGMAPKIAWALENKPRIPVVWIHGL WP_001058323.1 MULTISPECIES: hydrogenase-1 small chain [Proteobacteria] Length: 372\nNumber of predicted TMHs: 2\nExp number of AAs in TMHs: 37.2608499999999\nExp number, first 60 AAs: 19.52923\nTotal prob of N-in:

0.94351\nPOSSIBLE N-term signal sequence\ninside 1 19\nTMhelix 20 39\noutside 40 325\nTMhelix 326 348\ninside 349 372

8121 GCF_001519335.1_ASM151933v1 Escherichia coli Proteobacteria; Gammaproteobacteria; Enterobacterales; Enterobacteriaceae; Escherichia MTGDNTLIHSHGINRRDFMKLCAALAATMGLSSKAAAEMAESVTNPQRPPVIWIGAQECT WP_000145410.1 MULTISPECIES: hydrogenase-2 small chain [Proteobacteria] Length: 372\nNumber of predicted TMHs: 1\nExp number of AAs in TMHs: 23.72226\nExp number, first 60 AAs: 0.22038\nTotal prob of N-in: 0.12994\noutside 1 331\nTMhelix 332 354\ninside 355 372

8122 GCF_001519365.1_ASM151936v1 Escherichia coli Proteobacteria; Gammaproteobacteria; Enterobacterales; Enterobacteriaceae; Escherichia MTGDNTLIHSHGINRRDFMKLCAALAATMGLSSKAAAEMAESVTNPQRPPVIWIGAQECT WP_000145418.1 hydrogenase 2 small subunit [Escherichia coli] Length: 372\nNumber of predicted TMHs: 1\nExp number of AAs in TMHs: 23.72313\nExp number, first 60 AAs: 0.22034\nTotal prob of N-in: 0.12997\noutside 1 331\nTMhelix 332 354\ninside 355 372

8123 GCF_001519365.1_ASM151936v1 Escherichia coli Proteobacteria; Gammaproteobacteria; Enterobacterales; Enterobacteriaceae; Escherichia MNNEETFYQAMRRQGVTRRSFLKYCSLAATSLGLGAGMAPKIAWALENKPRIPVVWIHGL WP_001058323.1 MULTISPECIES: hydrogenase-1 small chain [Proteobacteria] Length: 372\nNumber of predicted TMHs: 2\nExp number of AAs in TMHs: 37.2608499999999\nExp number, first 60 AAs: 19.52923\nTotal prob of N-in: 0.94351\nPOSSIBLE N-term signal sequence\ninside 1 19\nTMhelix 20 39\noutside 40 325\nTMhelix 326 348\ninside 349 372

8124 GCF_001519335.1_ASM151933v1 Escherichia coli Proteobacteria; Gammaproteobacteria; Enterobacterales; Enterobacteriaceae; Escherichia MNNEETFYQAMRRQGVTRRSFLKYCSLAATSLGLGAGMAPKIAWALENKPRIPVVWIHGL WP_001295940.1 MULTISPECIES: hydrogenase [Enterobacteriaceae] Length: 372\nNumber of predicted TMHs: 2\nExp number of AAs in TMHs: 37.2608899999999\nExp number, first 60 AAs: 19.52923\nTotal prob of N-in: 0.94351\nPOSSIBLE N-term signal sequence\ninside 1 19\nTMhelix 20 39\noutside 40 325\nTMhelix 326 348\ninside 349 372

8125 GCF_001513615.1_ASM151361v1 Escherichia coli Proteobacteria; Gammaproteobacteria; Enterobacterales; Enterobacteriaceae; Escherichia MNNEETFYQAMRRQGVTRRSFLKYCSLAATSLGLGAGMAPKIAWALENKPRIPVVWIHGL WP_001295940.1 MULTISPECIES: hydrogenase [Enterobacteriaceae] Length: 372\nNumber of predicted TMHs: 2\nExp number of AAs in TMHs: 37.2608899999999\nExp number, first 60 AAs: 19.52923\nTotal prob of N-in: 0.94351\nPOSSIBLE N-term signal sequence\ninside 1 19\nTMhelix 20 39\noutside 40 325\nTMhelix 326 348\ninside 349 372

8126 GCF_001519435.1_ASM151943v1 Escherichia coli Proteobacteria; Gammaproteobacteria; Enterobacterales; Enterobacteriaceae; Escherichia MTGDNTLIHSHGINRRDFMKLCAALAATMGLSSKAAAEMAESVTNPQRPPVIWIGAQECT WP_000145418.1 hydrogenase 2 small subunit [Escherichia coli] Length: 372\nNumber of predicted TMHs: 1\nExp number of AAs in TMHs: 23.72313\nExp number, first 60 AAs: 0.22034\nTotal prob of N-in: 0.12997\noutside 1 331\nTMhelix 332 354\ninside 355 372

8127 GCF_001519475.1_ASM151947v1 Escherichia coli Proteobacteria; Gammaproteobacteria; Enterobacterales; Enterobacteriaceae; Escherichia MTGDNTLIHSHGINRRDFMKLCAALAATMGLSSKAAAEMAESVTNPQRPPVIWIGAQECT WP_000145410.1 MULTISPECIES: hydrogenase-2 small chain [Proteobacteria] Length: 372\nNumber of predicted TMHs: 1\nExp number of AAs in TMHs: 23.72226\nExp number, first 60 AAs: 0.22038\nTotal prob of N-in: 0.12994\noutside 1 331\nTMhelix 332 354\ninside 355 372

8128 GCF_001519525.1_ASM151952v1 Escherichia coli Proteobacteria; Gammaproteobacteria; Enterobacterales; Enterobacteriaceae; Escherichia MTGDNTLIHSHGINRRDFMKLCAALAATMGLSSKAAAEMAESVTNPQRPPVIWIGAQECT WP_000145410.1 MULTISPECIES: hydrogenase-2 small chain [Proteobacteria] Length: 372\nNumber of predicted TMHs: 1\nExp number of AAs in TMHs: 23.72226\nExp number, first 60 AAs: 0.22038\nTotal prob of N-in: 0.12994\noutside 1 331\nTMhelix 332 354\ninside 355 372

8129 GCF_001519435.1_ASM151943v1 Escherichia coli Proteobacteria; Gammaproteobacteria; Enterobacterales; Enterobacteriaceae; Escherichia
MNNEETFYQAMRRQGVTRRSFLKYCSLAATSLGLGAGMAPKIAWALENKPRIPVVWIHGL WP_001058323.1
MULTISPECIES: hydrogenase-1 small chain [Proteobacteria] Length: 372\nNumber of predicted TMHs: 2\nExp number of AAs in TMHs: 37.2608499999999\nExp number, first 60 AAs: 19.52923\nTotal prob of N-in: 0.94351\nPOSSIBLE N-term signal sequence\ninside 1 19\nTMhelix 20 39\noutside 40 325\nTMhelix 326 348\ninside 349 372

8130 GCF_001519675.1_ASM151967v1 Escherichia coli Proteobacteria; Gammaproteobacteria; Enterobacterales; Enterobacteriaceae; Escherichia
MTGDNTLIHSHGINRRDFMKLCAALAATMGLSSKAAAEMAESVTNPQRPPVIWIGAQECT WP_000145410.1
MULTISPECIES: hydrogenase-2 small chain [Proteobacteria] Length: 372\nNumber of predicted TMHs: 1\nExp number of AAs in TMHs: 23.72226\nExp number, first 60 AAs: 0.22038\nTotal prob of N-in: 0.12994\noutside 1 331\nTMhelix 332 354\ninside 355 372

8131 GCF_001519575.1_ASM151957v1 Escherichia coli Proteobacteria; Gammaproteobacteria; Enterobacterales; Enterobacteriaceae; Escherichia
MTGDNTLIHSHGINRRDFMKLCAALAATMGLSSKAAAEMAESVTNPQRPPVIWIGAQECT WP_000145410.1
MULTISPECIES: hydrogenase-2 small chain [Proteobacteria] Length: 372\nNumber of predicted TMHs: 1\nExp number of AAs in TMHs: 23.72226\nExp number, first 60 AAs: 0.22038\nTotal prob of N-in: 0.12994\noutside 1 331\nTMhelix 332 354\ninside 355 372

8132 GCF_001519635.1_ASM151963v1 Escherichia coli Proteobacteria; Gammaproteobacteria; Enterobacterales; Enterobacteriaceae; Escherichia
MTGDNTLIHSHGINRRDFMKLCAALAATMGLSSKAAAEMAESVTNPQRPPVIWIGAQECT WP_000145410.1
MULTISPECIES: hydrogenase-2 small chain [Proteobacteria] Length: 372\nNumber of predicted TMHs: 1\nExp number of AAs in TMHs: 23.72226\nExp number, first 60 AAs: 0.22038\nTotal prob of N-in: 0.12994\noutside 1 331\nTMhelix 332 354\ninside 355 372

8133 GCF_001519715.1_ASM151971v1 Escherichia coli Proteobacteria; Gammaproteobacteria; Enterobacterales; Enterobacteriaceae; Escherichia
MTGDNTLIHSHGINRRDFMKLCAALAATMGLSSKAAAEMAESVTNPQRPPVIWIGAQECT WP_000145410.1
MULTISPECIES: hydrogenase-2 small chain [Proteobacteria] Length: 372\nNumber of predicted TMHs: 1\nExp number of AAs in TMHs: 23.72226\nExp number, first 60 AAs: 0.22038\nTotal prob of N-in: 0.12994\noutside 1 331\nTMhelix 332 354\ninside 355 372

8134 GCF_001519775.1_ASM151977v1 Escherichia coli Proteobacteria; Gammaproteobacteria; Enterobacterales; Enterobacteriaceae; Escherichia
MTGDNTLIHSHGINRRDFMKLCAALAATMGLSSKAAAEMAESVTNPQRPPVIWIGAQECT WP_000145410.1
MULTISPECIES: hydrogenase-2 small chain [Proteobacteria] Length: 372\nNumber of predicted TMHs: 1\nExp number of AAs in TMHs: 23.72226\nExp number, first 60 AAs: 0.22038\nTotal prob of N-in: 0.12994\noutside 1 331\nTMhelix 332 354\ninside 355 372

8135 GCF_001519635.1_ASM151963v1 Escherichia coli Proteobacteria; Gammaproteobacteria; Enterobacterales; Enterobacteriaceae; Escherichia
MNNEETFYQAMRRQGVTRRSFLKYCSLAATSLGLGAGMAPKIAWALENKPRIPVVWIHGL WP_001058323.1
MULTISPECIES: hydrogenase-1 small chain [Proteobacteria] Length: 372\nNumber of predicted TMHs: 2\nExp number of AAs in TMHs: 37.2608499999999\nExp number, first 60 AAs: 19.52923\nTotal prob of N-in: 0.94351\nPOSSIBLE N-term signal sequence\ninside 1 19\nTMhelix 20 39\noutside 40 325\nTMhelix 326 348\ninside 349 372

8136 GCF_001519775.1_ASM151977v1 Escherichia coli Proteobacteria; Gammaproteobacteria; Enterobacterales; Enterobacteriaceae; Escherichia
MNNEETFYQAMRRQGVTRRSFLKYCSLAATSLGLGAGMAPKIAWALENKPRIPVVWIHGL WP_001058323.1
MULTISPECIES: hydrogenase-1 small chain [Proteobacteria] Length: 372\nNumber of predicted TMHs: 2\nExp number of AAs in TMHs: 37.2608499999999\nExp number, first 60 AAs: 19.52923\nTotal prob of N-in: 0.94351\nPOSSIBLE N-term signal sequence\ninside 1 19\nTMhelix 20 39\noutside 40 325\nTMhelix 326 348\ninside 349 372

8137 GCF_001519475.1_ASM151947v1 Escherichia coli Proteobacteria; Gammaproteobacteria; Enterobacterales; Enterobacteriaceae; Escherichia

MNNEETFYQAMRRQGVTRRSFLKYCSLAATSLGLGAGMAPKIAWALENKPRIPVVWIHGL WP_001058323.1 MULTISPECIES: hydrogenase-1 small chain [Proteobacteria] Length: 372\nNumber of predicted TMHs: 2\nExp number of AAs in TMHs: 37.2608499999999\nExp number, first 60 AAs: 19.52923\nTotal prob of N-in: 0.94351\nPOSSIBLE N-term signal sequence\ninside 1 19\nTMhelix 20 39\noutside 40 325\nTMhelix 326 348\ninside 349 372

8138 GCF_001519815.1_ASM151981v1 Escherichia coli Proteobacteria; Gammaproteobacteria; Enterobacterales; Enterobacteriaceae; Escherichia MTGDNTLIHSHGINRRDFMKLCAALAATMGLSSKAAAEMAESVTNPQRPPVIWIGAQECT WP_000145410.1 MULTISPECIES: hydrogenase-2 small chain [Proteobacteria] Length: 372\nNumber of predicted TMHs: 1\nExp number of AAs in TMHs: 23.72226\nExp number, first 60 AAs: 0.22038\nTotal prob of N-in: 0.12994\noutside 1 331\nTMhelix 332 354\ninside 355 372

8139 GCF_001519875.1_ASM151987v1 Escherichia coli Proteobacteria; Gammaproteobacteria; Enterobacterales; Enterobacteriaceae; Escherichia MTGDNTLIHSHGINRRDFMKLCAALAATMGLSSKAAAEMAESVTNPQRPPVIWIGAQECT WP_000145410.1 MULTISPECIES: hydrogenase-2 small chain [Proteobacteria] Length: 372\nNumber of predicted TMHs: 1\nExp number of AAs in TMHs: 23.72226\nExp number, first 60 AAs: 0.22038\nTotal prob of N-in: 0.12994\noutside 1 331\nTMhelix 332 354\ninside 355 372

8140 GCF_001519875.1_ASM151987v1 Escherichia coli Proteobacteria; Gammaproteobacteria; Enterobacterales; Enterobacteriaceae; Escherichia MNNEETFYQAMRRQGVTRRSFLKYCSLAATSLGLGAGMAPKIAWALENKPRIPVVWIHGL WP_001058323.1 MULTISPECIES: hydrogenase-1 small chain [Proteobacteria] Length: 372\nNumber of predicted TMHs: 2\nExp number of AAs in TMHs: 37.2608499999999\nExp number, first 60 AAs: 19.52923\nTotal prob of N-in: 0.94351\nPOSSIBLE N-term signal sequence\ninside 1 19\nTMhelix 20 39\noutside 40 325\nTMhelix 326 348\ninside 349 372

8141 GCF_001519815.1_ASM151981v1 Escherichia coli Proteobacteria; Gammaproteobacteria; Enterobacterales; Enterobacteriaceae; Escherichia MNNEETFYQAMRRQGVTRRSFLKYCSLAATSLGLGAGMAPKIAWALENKPRIPVVWIHGL WP_001058323.1 MULTISPECIES: hydrogenase-1 small chain [Proteobacteria] Length: 372\nNumber of predicted TMHs: 2\nExp number of AAs in TMHs: 37.2608499999999\nExp number, first 60 AAs: 19.52923\nTotal prob of N-in: 0.94351\nPOSSIBLE N-term signal sequence\ninside 1 19\nTMhelix 20 39\noutside 40 325\nTMhelix 326 348\ninside 349 372

8142 GCF_001519575.1_ASM151957v1 Escherichia coli Proteobacteria; Gammaproteobacteria; Enterobacterales; Enterobacteriaceae; Escherichia MNNEETFYQAMRRQGVTRRSFLKYCSLAATSLGLGAGMAPKIAWALENKPRIPVVWIHGL WP_001058320.1 hydrogenase [Escherichia coli] Length: 372\nNumber of predicted TMHs: 2\nExp number of AAs in TMHs: 37.26082\nExp number, first 60 AAs: 19.52923\nTotal prob of N-in: 0.94351\nPOSSIBLE N-term signal sequence\ninside 1 19\nTMhelix 20 39\noutside 40 325\nTMhelix 326 348\ninside 349 372

8143 GCF_001519675.1_ASM151967v1 Escherichia coli Proteobacteria; Gammaproteobacteria; Enterobacterales; Enterobacteriaceae; Escherichia MNNEETFYQAMRRQGVTRRSFLKYCSLAATSLGLGAGMAPKIAWALENKPRIPVVWIHGL WP_001058323.1 MULTISPECIES: hydrogenase-1 small chain [Proteobacteria] Length: 372\nNumber of predicted TMHs: 2\nExp number of AAs in TMHs: 37.2608499999999\nExp number, first 60 AAs: 19.52923\nTotal prob of N-in: 0.94351\nPOSSIBLE N-term signal sequence\ninside 1 19\nTMhelix 20 39\noutside 40 325\nTMhelix 326 348\ninside 349 372

8144 GCF_001519915.1_ASM151991v1 Escherichia coli Proteobacteria; Gammaproteobacteria; Enterobacterales; Enterobacteriaceae; Escherichia MTGDNTLIHSHGINRRDFMKLCAALAATMGLSSKAAAEMVESVTNPQRPPVIWIGAQECT WP_000145423.1 hydrogenase 2 small subunit [Escherichia coli] Length: 372\nNumber of predicted TMHs: 1\nExp number of AAs in TMHs: 23.78201\nExp number, first 60 AAs: 0.28355\nTotal prob of N-in: 0.13257\noutside 1 331\nTMhelix 332 354\ninside 355 372

8145 GCF_001519975.1_ASM151997v1 Escherichia coli Proteobacteria; Gammaproteobacteria; Enterobacterales; Enterobacteriaceae; Escherichia MTGDNTLIHSHGINRRDFMKLCAALAATMGLSSKAAAEMAESVTNPQRPPVIWIGAQECT WP_000145410.1

MULTISPECIES: hydrogenase-2 small chain [Proteobacteria]	Length: 372\nNumber of predicted TMHs: 1\nExp number of AAs in TMHs: 23.72226\nExp number, first 60 AAs: 0.22038\nTotal prob of N-in: 0.12994\noutside 1 331\nTMhelix 332 354\ninside 355 372

8146 GCF_001519915.1_ASM151991v1 Escherichia coli Proteobacteria; Gammaproteobacteria; Enterobacterales; Enterobacteriaceae; Escherichia
MNNEETFYQAMRRQGVTRRSFLKYCSLAATSLGLGAGMAPKIAWALENKPRIPVVWIHGL WP_001295940.1
MULTISPECIES: hydrogenase [Enterobacteriaceae] Length: 372\nNumber of predicted TMHs: 2\nExp number of AAs in TMHs: 37.2608899999999\nExp number, first 60 AAs: 19.52923\nTotal prob of N-in: 0.94351\nPOSSIBLE N-term signal sequence\ninside 1 19\nTMhelix 20 39\noutside 40 325\nTMhelix 326 348\ninside 349 372

8147 GCF_001519975.1_ASM151997v1 Escherichia coli Proteobacteria; Gammaproteobacteria; Enterobacterales; Enterobacteriaceae; Escherichia
MNNEETFYQAMRRQGVTRRSFLKYCSLAATSLGLGAGMAPKIAWALENKPRIPVVWIHGL WP_001058323.1
MULTISPECIES: hydrogenase-1 small chain [Proteobacteria] Length: 372\nNumber of predicted TMHs: 2\nExp number of AAs in TMHs: 37.2608499999999\nExp number, first 60 AAs: 19.52923\nTotal prob of N-in: 0.94351\nPOSSIBLE N-term signal sequence\ninside 1 19\nTMhelix 20 39\noutside 40 325\nTMhelix 326 348\ninside 349 372

8148 GCF_001519525.1_ASM151952v1 Escherichia coli Proteobacteria; Gammaproteobacteria; Enterobacterales; Enterobacteriaceae; Escherichia
MNNEETFYQAMRRQGVTRRSFLKYCSLAATSLGLGAGMAPKIAWALENKPRIPVVWIHGL WP_059321607.1
hydrogenase [Escherichia coli] Length: 372\nNumber of predicted TMHs: 2\nExp number of AAs in TMHs: 37.2608299999999\nExp number, first 60 AAs: 19.52923\nTotal prob of N-in: 0.94351\nPOSSIBLE N-term signal sequence\ninside 1 19\nTMhelix 20 39\noutside 40 325\nTMhelix 326 348\ninside 349 372

8149 GCF_001520015.1_ASM152001v1 Escherichia coli Proteobacteria; Gammaproteobacteria; Enterobacterales; Enterobacteriaceae; Escherichia
MTGDNTLIHSHGINRRDFMKLCAALAATMGLSSKAAAEMAESVTNPQRPPVIWIGAQECT WP_000145410.1
MULTISPECIES: hydrogenase-2 small chain [Proteobacteria] Length: 372\nNumber of predicted TMHs: 1\nExp number of AAs in TMHs: 23.72226\nExp number, first 60 AAs: 0.22038\nTotal prob of N-in: 0.12994\noutside 1 331\nTMhelix 332 354\ninside 355 372

8150 GCF_001520075.1_ASM152007v1 Escherichia coli Proteobacteria; Gammaproteobacteria; Enterobacterales; Enterobacteriaceae; Escherichia
MTGDNTLIHSHGINRRDFMKLCAALAATMGLSSKAAAEMVESVTNPQRPPVIWIGAQECT WP_000145423.1
hydrogenase 2 small subunit [Escherichia coli] Length: 372\nNumber of predicted TMHs: 1\nExp number of AAs in TMHs: 23.78201\nExp number, first 60 AAs: 0.28355\nTotal prob of N-in: 0.13257\noutside 1 331\nTMhelix 332 354\ninside 355 372

8151 GCF_001520115.1_ASM152011v1 Escherichia coli Proteobacteria; Gammaproteobacteria; Enterobacterales; Enterobacteriaceae; Escherichia
MTGDNTLIHSHGINRRDFMKLCAALAATMGLSSKAAAEMAESVTNPQRPPVIWIGAQECT WP_000145410.1
MULTISPECIES: hydrogenase-2 small chain [Proteobacteria] Length: 372\nNumber of predicted TMHs: 1\nExp number of AAs in TMHs: 23.72226\nExp number, first 60 AAs: 0.22038\nTotal prob of N-in: 0.12994\noutside 1 331\nTMhelix 332 354\ninside 355 372

8152 GCF_001520175.1_ASM152017v1 Escherichia coli Proteobacteria; Gammaproteobacteria; Enterobacterales; Enterobacteriaceae; Escherichia
MTGDNTLIHSHGINRRDFMKLCAALAATMGLSSKAAAEMAESVTNPQRPPVIWIGAQECT WP_000145410.1
MULTISPECIES: hydrogenase-2 small chain [Proteobacteria] Length: 372\nNumber of predicted TMHs: 1\nExp number of AAs in TMHs: 23.72226\nExp number, first 60 AAs: 0.22038\nTotal prob of N-in: 0.12994\noutside 1 331\nTMhelix 332 354\ninside 355 372

8153 GCF_001520215.1_ASM152021v1 Escherichia coli Proteobacteria; Gammaproteobacteria; Enterobacterales; Enterobacteriaceae; Escherichia
MTGDNTLIHSHGINRRDFMKLCAALAATMGLSSKAAAEMAESVTNPQRPPVIWIGAQECT WP_000145410.1
MULTISPECIES: hydrogenase-2 small chain [Proteobacteria] Length: 372\nNumber of predicted TMHs: 1\nExp number of AAs in TMHs: 23.72226\nExp number, first 60 AAs: 0.22038\nTotal prob of N-in: 0.12994\noutside 1 331\nTMhelix 332 354\ninside 355 372

8154 GCF_001520275.1_ASM152027v1 Escherichia coli Proteobacteria; Gammaproteobacteria; Enterobacterales; Enterobacteriaceae; Escherichia
MTGDNTLIHSHGINRRDFMKLCAALAATMGLSSKAAAEMAESVTNPQRPPVIWIGAQECT WP_000145410.1
MULTISPECIES: hydrogenase-2 small chain [Proteobacteria] Length: 372\nNumber of predicted TMHs: 1\nExp number of AAs in TMHs: 23.72226\nExp number, first 60 AAs: 0.22038\nTotal prob of N-in: 0.12994\noutside 1 331\nTMhelix 332 354\ninside 355 372

8155 GCF_001520315.1_ASM152031v1 Escherichia coli Proteobacteria; Gammaproteobacteria; Enterobacterales; Enterobacteriaceae; Escherichia
MTGDNTLIHSHGINRRDFMKLCAALAATMGLSSKAAAEMAESVTNPQRPPVIWIGAQECT WP_000145410.1
MULTISPECIES: hydrogenase-2 small chain [Proteobacteria] Length: 372\nNumber of predicted TMHs: 1\nExp number of AAs in TMHs: 23.72226\nExp number, first 60 AAs: 0.22038\nTotal prob of N-in: 0.12994\noutside 1 331\nTMhelix 332 354\ninside 355 372

8156 GCF_001520015.1_ASM152001v1 Escherichia coli Proteobacteria; Gammaproteobacteria; Enterobacterales; Enterobacteriaceae; Escherichia
MNNEETFYQAMRRQGVTRRSFLKYCSLAATSLGLGAGMAPKIAWALENKPRIPVVWIHGL WP_001295940.1
MULTISPECIES: hydrogenase [Enterobacteriaceae] Length: 372\nNumber of predicted TMHs: 2\nExp number of AAs in TMHs: 37.2608899999999\nExp number, first 60 AAs: 19.52923\nTotal prob of N-in: 0.94351\nPOSSIBLE N-term signal sequence\ninside 1 19\nTMhelix 20 39\noutside 40 325\nTMhelix 326 348\ninside 349 372

8157 GCF_001520375.1_ASM152037v1 Escherichia coli Proteobacteria; Gammaproteobacteria; Enterobacterales; Enterobacteriaceae; Escherichia
MTGDNTLIHSHGINRRDFMKLCAALAATMGLSSKAAAEMAESVTNPQRPPVIWIGAQECT WP_000145410.1
MULTISPECIES: hydrogenase-2 small chain [Proteobacteria] Length: 372\nNumber of predicted TMHs: 1\nExp number of AAs in TMHs: 23.72226\nExp number, first 60 AAs: 0.22038\nTotal prob of N-in: 0.12994\noutside 1 331\nTMhelix 332 354\ninside 355 372

8158 GCF_001520375.1_ASM152037v1 Escherichia coli Proteobacteria; Gammaproteobacteria; Enterobacterales; Enterobacteriaceae; Escherichia
MNNEETFYQAMRRQGVTRRSFLKYCSLAATSLGLGAGMAPKIAWALENKPRIPVVWIHGL WP_001058323.1
MULTISPECIES: hydrogenase-1 small chain [Proteobacteria] Length: 372\nNumber of predicted TMHs: 2\nExp number of AAs in TMHs: 37.2608499999999\nExp number, first 60 AAs: 19.52923\nTotal prob of N-in: 0.94351\nPOSSIBLE N-term signal sequence\ninside 1 19\nTMhelix 20 39\noutside 40 325\nTMhelix 326 348\ninside 349 372

8159 GCF_001520175.1_ASM152017v1 Escherichia coli Proteobacteria; Gammaproteobacteria; Enterobacterales; Enterobacteriaceae; Escherichia
MNNEETFYQAMRRQGVTRRSFLKYCSLAATSLGLGAGMAPKIAWALENKPRIPVVWIHGL WP_001058323.1
MULTISPECIES: hydrogenase-1 small chain [Proteobacteria] Length: 372\nNumber of predicted TMHs: 2\nExp number of AAs in TMHs: 37.2608499999999\nExp number, first 60 AAs: 19.52923\nTotal prob of N-in: 0.94351\nPOSSIBLE N-term signal sequence\ninside 1 19\nTMhelix 20 39\noutside 40 325\nTMhelix 326 348\ninside 349 372

8160 GCF_001520435.1_ASM152043v1 Escherichia coli Proteobacteria; Gammaproteobacteria; Enterobacterales; Enterobacteriaceae; Escherichia
MTGDNTLIHSHGINRRDFMKLCAALAATMGLSSKAAAEMAESVTNPQRPPVIWIGAQECT WP_000145410.1
MULTISPECIES: hydrogenase-2 small chain [Proteobacteria] Length: 372\nNumber of predicted TMHs: 1\nExp number of AAs in TMHs: 23.72226\nExp number, first 60 AAs: 0.22038\nTotal prob of N-in: 0.12994\noutside 1 331\nTMhelix 332 354\ninside 355 372

8161 GCF_001520435.1_ASM152043v1 Escherichia coli Proteobacteria; Gammaproteobacteria; Enterobacterales; Enterobacteriaceae; Escherichia
MNNEETFYQAMRRQGVTRRSFLKYCSLAATSLGLGAGMAPKIAWALENKPRIPVVWIHGL WP_001058323.1
MULTISPECIES: hydrogenase-1 small chain [Proteobacteria] Length: 372\nNumber of predicted TMHs: 2\nExp number of AAs in TMHs: 37.2608499999999\nExp number, first 60 AAs: 19.52923\nTotal prob of N-in: 0.94351\nPOSSIBLE N-term signal sequence\ninside 1 19\nTMhelix 20 39\noutside 40 325\nTMhelix 326 348\ninside 349 372

8162 GCF_001520315.1_ASM152031v1 Escherichia coli Proteobacteria; Gammaproteobacteria; Enterobacterales; Enterobacteriaceae; Escherichia
MNNEETFYQAMRRQGVTRRSFLKYCSLAATSLGLGAGMAPKIAWALENKPRIPVVWIHGL WP_001058323.1
MULTISPECIES: hydrogenase-1 small chain [Proteobacteria] Length: 372\nNumber of predicted TMHs: 2\nExp number of AAs in TMHs: 37.2608499999999\nExp number, first 60 AAs: 19.52923\nTotal prob of N-in: 0.94351\nPOSSIBLE N-term signal sequence\ninside 1 19\nTMhelix 20 39\noutside 40 325\nTMhelix 326 348\ninside 349 372

8163 GCF_001520405.1_ASM152040v1 Escherichia coli Proteobacteria; Gammaproteobacteria; Enterobacterales; Enterobacteriaceae; Escherichia
MTGDNTLIHSHGINRRDFMKLCAALAATMGLSSKAAAEMVESVTNPQRPPVIWIGAQECT WP_000145423.1
hydrogenase 2 small subunit [Escherichia coli] Length: 372\nNumber of predicted TMHs: 1\nExp number of AAs in TMHs: 23.78201\nExp number, first 60 AAs: 0.28355\nTotal prob of N-in: 0.13257\noutside 1 331\nTMhelix 332 354\ninside 355 372

8164 GCF_001520405.1_ASM152040v1 Escherichia coli Proteobacteria; Gammaproteobacteria; Enterobacterales; Enterobacteriaceae; Escherichia
MNNEETFYQAMRRQGVTRRSFLKYCSLAATSLGLGAGMAPKIAWALENKPRIPVVWIHGL WP_001295940.1
MULTISPECIES: hydrogenase [Enterobacteriaceae] Length: 372\nNumber of predicted TMHs: 2\nExp number of AAs in TMHs: 37.2608899999999\nExp number, first 60 AAs: 19.52923\nTotal prob of N-in: 0.94351\nPOSSIBLE N-term signal sequence\ninside 1 19\nTMhelix 20 39\noutside 40 325\nTMhelix 326 348\ninside 349 372

8165 GCF_001520115.1_ASM152011v1 Escherichia coli Proteobacteria; Gammaproteobacteria; Enterobacterales; Enterobacteriaceae; Escherichia
MNNEETFYQAMRRQGVTRRSFLKYCSLAATSLGLGAGMAPKIAWALENKPRIPVVWIHGL WP_001058323.1
MULTISPECIES: hydrogenase-1 small chain [Proteobacteria] Length: 372\nNumber of predicted TMHs: 2\nExp number of AAs in TMHs: 37.2608499999999\nExp number, first 60 AAs: 19.52923\nTotal prob of N-in: 0.94351\nPOSSIBLE N-term signal sequence\ninside 1 19\nTMhelix 20 39\noutside 40 325\nTMhelix 326 348\ninside 349 372

8166 GCF_001520215.1_ASM152021v1 Escherichia coli Proteobacteria; Gammaproteobacteria; Enterobacterales; Enterobacteriaceae; Escherichia
MNNEETFYQAMRRQGVTRRSFLKYCSLAATSLGLGAGMAPKIAWALENKPRIPVVWIHGL WP_001058323.1
MULTISPECIES: hydrogenase-1 small chain [Proteobacteria] Length: 372\nNumber of predicted TMHs: 2\nExp number of AAs in TMHs: 37.2608499999999\nExp number, first 60 AAs: 19.52923\nTotal prob of N-in: 0.94351\nPOSSIBLE N-term signal sequence\ninside 1 19\nTMhelix 20 39\noutside 40 325\nTMhelix 326 348\ninside 349 372

8167 GCF_001520275.1_ASM152027v1 Escherichia coli Proteobacteria; Gammaproteobacteria; Enterobacterales; Enterobacteriaceae; Escherichia
MNNEETFYQAMRRQGVTRRSFLKYCSLAATSLGLGAGMAPKIAWALENKPRIPVVWIHGL WP_001058323.1
MULTISPECIES: hydrogenase-1 small chain [Proteobacteria] Length: 372\nNumber of predicted TMHs: 2\nExp number of AAs in TMHs: 37.2608499999999\nExp number, first 60 AAs: 19.52923\nTotal prob of N-in: 0.94351\nPOSSIBLE N-term signal sequence\ninside 1 19\nTMhelix 20 39\noutside 40 325\nTMhelix 326 348\ninside 349 372

8168 GCF_001520475.1_ASM152047v1 Escherichia coli Proteobacteria; Gammaproteobacteria; Enterobacterales; Enterobacteriaceae; Escherichia
MTGDNTLIHSHGINRRDFMKLCAALAATMGLSSKAAAEMVESVTNPQRPPVIWIGAQECT WP_000145423.1
hydrogenase 2 small subunit [Escherichia coli] Length: 372\nNumber of predicted TMHs: 1\nExp number of AAs in TMHs: 23.78201\nExp number, first 60 AAs: 0.28355\nTotal prob of N-in: 0.13257\noutside 1 331\nTMhelix 332 354\ninside 355 372

8169 GCF_001520475.1_ASM152047v1 Escherichia coli Proteobacteria; Gammaproteobacteria; Enterobacterales; Enterobacteriaceae; Escherichia
MNNEETFYQAMRRQGVTRRSFLKYCSLAATSLGLGAGMAPKIAWALENKPRIPVVWIHGL WP_001295940.1
MULTISPECIES: hydrogenase [Enterobacteriaceae] Length: 372\nNumber of predicted TMHs: 2\nExp number of AAs in TMHs: 37.2608899999999\nExp number, first 60 AAs: 19.52923\nTotal prob of N-in: 0.94351\nPOSSIBLE N-term signal sequence\ninside 1 19\nTMhelix 20 39\noutside 40 325\nTMhelix 326 348\ninside 349 372

8170 GCF_001520075.1_ASM152007v1 Escherichia coli Proteobacteria; Gammaproteobacteria; Enterobacterales; Enterobacteriaceae; Escherichia
MNNEETFYQAMRRQGVTRRSFLKYCSLAATSLGLGAGMAPKIAWALENKPRIPVVWIHGL WP_001295940.1
MULTISPECIES: hydrogenase [Enterobacteriaceae] Length: 372\nNumber of predicted TMHs: 2\nExp number of AAs in TMHs: 37.2608899999999\nExp number, first 60 AAs: 19.52923\nTotal prob of N-in: 0.94351\nPOSSIBLE N-term signal sequence\ninside 1 19\nTMhelix 20 39\noutside 40 325\nTMhelix 326 348\ninside 349 372

8171 GCF_001520515.1_ASM152051v1 Escherichia coli Proteobacteria; Gammaproteobacteria; Enterobacterales; Enterobacteriaceae; Escherichia
MTGDNTLIHSHGINRRDFMKLCAALAATMGLSSKAAAEMVESVTNPQRPPVIWIGAQECT WP_000145423.1
hydrogenase 2 small subunit [Escherichia coli] Length: 372\nNumber of predicted TMHs: 1\nExp number of AAs in TMHs: 23.78201\nExp number, first 60 AAs: 0.28355\nTotal prob of N-in: 0.13257\noutside 1 331\nTMhelix 332 354\ninside 355 372

8172 GCF_001520575.1_ASM152057v1 Escherichia coli Proteobacteria; Gammaproteobacteria; Enterobacterales; Enterobacteriaceae; Escherichia
MTGDNTLIHSHGINRRDFMKLCAALAATMGLSSKAAAEMAESVTNPQRPPVIWIGAQECT WP_000145410.1
MULTISPECIES: hydrogenase-2 small chain [Proteobacteria] Length: 372\nNumber of predicted TMHs: 1\nExp number of AAs in TMHs: 23.72226\nExp number, first 60 AAs: 0.22038\nTotal prob of N-in: 0.12994\noutside 1 331\nTMhelix 332 354\ninside 355 372

8173 GCF_001520675.1_ASM152067v1 Escherichia coli Proteobacteria; Gammaproteobacteria; Enterobacterales; Enterobacteriaceae; Escherichia
MTGDNTLIHSHGINRRDFMKLCAALAATMGLSSKAAAEMVESVTNPQRPPVIWIGAQECT WP_000145423.1
hydrogenase 2 small subunit [Escherichia coli] Length: 372\nNumber of predicted TMHs: 1\nExp number of AAs in TMHs: 23.78201\nExp number, first 60 AAs: 0.28355\nTotal prob of N-in: 0.13257\noutside 1 331\nTMhelix 332 354\ninside 355 372

8174 GCF_001520615.1_ASM152061v1 Escherichia coli Proteobacteria; Gammaproteobacteria; Enterobacterales; Enterobacteriaceae; Escherichia
MTGDNTLIHSHGINRRDFMKLCAALAATMGLSSKAAAEMVESVTNPQRPPVIWIGAQECT WP_000145423.1
hydrogenase 2 small subunit [Escherichia coli] Length: 372\nNumber of predicted TMHs: 1\nExp number of AAs in TMHs: 23.78201\nExp number, first 60 AAs: 0.28355\nTotal prob of N-in: 0.13257\noutside 1 331\nTMhelix 332 354\ninside 355 372

8175 GCF_001520715.1_ASM152071v1 Escherichia coli Proteobacteria; Gammaproteobacteria; Enterobacterales; Enterobacteriaceae; Escherichia
MTGDNTLIHSHGINRRDFMKLCAALAATMGLSSKAAAEMAESVTNPQRPPVIWIGAQECT WP_000145410.1
MULTISPECIES: hydrogenase-2 small chain [Proteobacteria] Length: 372\nNumber of predicted TMHs: 1\nExp number of AAs in TMHs: 23.72226\nExp number, first 60 AAs: 0.22038\nTotal prob of N-in: 0.12994\noutside 1 331\nTMhelix 332 354\ninside 355 372

8176 GCF_001520575.1_ASM152057v1 Escherichia coli Proteobacteria; Gammaproteobacteria; Enterobacterales; Enterobacteriaceae; Escherichia
MNNEETFYQAMRRQGVTRRSFLKYCSLAATSLGLGAGMAPKIAWALENKPRIPVVWIHGL WP_001058323.1
MULTISPECIES: hydrogenase-1 small chain [Proteobacteria] Length: 372\nNumber of predicted TMHs: 2\nExp number of AAs in TMHs: 37.2608499999999\nExp number, first 60 AAs: 19.52923\nTotal prob of N-in: 0.94351\nPOSSIBLE N-term signal sequence\ninside 1 19\nTMhelix 20 39\noutside 40 325\nTMhelix 326 348\ninside 349 372

8177 GCF_001520775.1_ASM152077v1 Escherichia coli Proteobacteria; Gammaproteobacteria; Enterobacterales; Enterobacteriaceae; Escherichia
MTGDNTLIHSHGINRRDFMKLCAALAATMGLSSKAAAEMAESVTNPQRPPVIWIGAQECT WP_000145410.1
MULTISPECIES: hydrogenase-2 small chain [Proteobacteria] Length: 372\nNumber of predicted TMHs: 1\nExp number of AAs in TMHs: 23.72226\nExp number, first 60 AAs: 0.22038\nTotal prob of N-in: 0.12994\noutside 1 331\nTMhelix 332 354\ninside 355 372

8178 GCF_001520815.1_ASM152081v1 Escherichia coli Proteobacteria; Gammaproteobacteria; Enterobacterales; Enterobacteriaceae; Escherichia
MTGDNTLIHSHGINRRDFMKLCAALAATMGLSSKAAAEMAESVTNPQRPPVIWIGAQECT WP_000145410.1

MULTISPECIES: hydrogenase-2 small chain [Proteobacteria] Length: 372\nNumber of predicted TMHs: 1\nExp number of AAs in TMHs: 23.72226\nExp number, first 60 AAs: 0.22038\nTotal prob of N-in: 0.12994\noutside 1 331\nTMhelix 332 354\ninside 355 372

8179 GCF_001520875.1_ASM152087v1 Escherichia coli Proteobacteria; Gammaproteobacteria; Enterobacterales; Enterobacteriaceae; Escherichia
MTGDNTLIHSHGINRRDFMKLCAALAATMGLSSKAAAEMAESVTNPQRPPVIWIGAQECT WP_000145410.1
MULTISPECIES: hydrogenase-2 small chain [Proteobacteria] Length: 372\nNumber of predicted TMHs: 1\nExp number of AAs in TMHs: 23.72226\nExp number, first 60 AAs: 0.22038\nTotal prob of N-in: 0.12994\noutside 1 331\nTMhelix 332 354\ninside 355 372

8180 GCF_001520915.1_ASM152091v1 Escherichia coli Proteobacteria; Gammaproteobacteria; Enterobacterales; Enterobacteriaceae; Escherichia
MTGDNTLIHSHGINRRDFMKLCAALAATMGLSSKAAAEMAESVTNPQRPPVIWIGAQECT WP_000145410.1
MULTISPECIES: hydrogenase-2 small chain [Proteobacteria] Length: 372\nNumber of predicted TMHs: 1\nExp number of AAs in TMHs: 23.72226\nExp number, first 60 AAs: 0.22038\nTotal prob of N-in: 0.12994\noutside 1 331\nTMhelix 332 354\ninside 355 372

8181 GCF_001520975.1_ASM152097v1 Escherichia coli Proteobacteria; Gammaproteobacteria; Enterobacterales; Enterobacteriaceae; Escherichia
MTGDNTLIHSHGINRRDFMKLCAALAATMGLSSKAAAEMVESVTNPQRPPVIWIGAQECT WP_000145423.1
hydrogenase 2 small subunit [Escherichia coli] Length: 372\nNumber of predicted TMHs: 1\nExp number of AAs in TMHs: 23.78201\nExp number, first 60 AAs: 0.28355\nTotal prob of N-in: 0.13257\noutside 1 331\nTMhelix 332 354\ninside 355 372

8182 GCF_001521015.1_ASM152101v1 Escherichia coli Proteobacteria; Gammaproteobacteria; Enterobacterales; Enterobacteriaceae; Escherichia
MTGDNTLIHSHGINRRDFMKLCAALAATMGLSSKAAAEMAESVTNPQRPPVIWIGAQECT WP_000145410.1
MULTISPECIES: hydrogenase-2 small chain [Proteobacteria] Length: 372\nNumber of predicted TMHs: 1\nExp number of AAs in TMHs: 23.72226\nExp number, first 60 AAs: 0.22038\nTotal prob of N-in: 0.12994\noutside 1 331\nTMhelix 332 354\ninside 355 372

8183 GCF_001520815.1_ASM152081v1 Escherichia coli Proteobacteria; Gammaproteobacteria; Enterobacterales; Enterobacteriaceae; Escherichia
MNNEETFYQAMRRQGVTRRSFLKYCSLAATSLGLGAGMAPKIAWALENKPRIPVVWIHGL WP_001058323.1
MULTISPECIES: hydrogenase-1 small chain [Proteobacteria] Length: 372\nNumber of predicted TMHs: 2\nExp number of AAs in TMHs: 37.2608499999999\nExp number, first 60 AAs: 19.52923\nTotal prob of N-in: 0.94351\nPOSSIBLE N-term signal sequence\ninside 1 19\nTMhelix 20 39\noutside 40 325\nTMhelix 326 348\ninside 349 372

8184 GCF_001521515.1_ASM152151v1 Escherichia coli Proteobacteria; Gammaproteobacteria; Enterobacterales; Enterobacteriaceae; Escherichia
MTGDNTLIHSHGINRRDFMKLCAALAATMGLSSKAAAEMVESVTNPQRPPVIWIGAQECT WP_000145423.1
hydrogenase 2 small subunit [Escherichia coli] Length: 372\nNumber of predicted TMHs: 1\nExp number of AAs in TMHs: 23.78201\nExp number, first 60 AAs: 0.28355\nTotal prob of N-in: 0.13257\noutside 1 331\nTMhelix 332 354\ninside 355 372

8185 GCF_001520715.1_ASM152071v1 Escherichia coli Proteobacteria; Gammaproteobacteria; Enterobacterales; Enterobacteriaceae; Escherichia
MNNEETFYQAMRRQGVTRRSFLKYCSLAATSLGLGAGMAPKIAWALENKPRIPVVWIHGL WP_001058323.1
MULTISPECIES: hydrogenase-1 small chain [Proteobacteria] Length: 372\nNumber of predicted TMHs: 2\nExp number of AAs in TMHs: 37.2608499999999\nExp number, first 60 AAs: 19.52923\nTotal prob of N-in: 0.94351\nPOSSIBLE N-term signal sequence\ninside 1 19\nTMhelix 20 39\noutside 40 325\nTMhelix 326 348\ninside 349 372

8186 GCF_001521075.1_ASM152107v1 Escherichia coli Proteobacteria; Gammaproteobacteria; Enterobacterales; Enterobacteriaceae; Escherichia
MTGDNTLIHSHGINRRDFMKLCAALAATMGLSSKAAAEMAESVTNPQRPPVIWIGAQECT WP_000145410.1
MULTISPECIES: hydrogenase-2 small chain [Proteobacteria] Length: 372\nNumber of predicted TMHs: 1\nExp number of AAs in TMHs: 23.72226\nExp number, first 60 AAs: 0.22038\nTotal prob of N-in: 0.12994\noutside 1 331\nTMhelix 332 354\ninside 355 372

8187 GCF_001521075.1_ASM152107v1 Escherichia coli Proteobacteria; Gammaproteobacteria; Enterobacterales; Enterobacteriaceae; Escherichia
MNNEETFYQAMRRQGVTRRSFLKYCSLAATSLGLGAGMAPKIAWALENKPRIPVVWIHGL WP_059327874.1
hydrogenase [Escherichia coli] Length: 372\nNumber of predicted TMHs: 2\nExp number of AAs in TMHs: 37.666\nExp number, first 60 AAs: 19.52872\nTotal prob of N-in: 0.94374\nPOSSIBLE N-term signal sequence\ninside 1 19\nTMhelix 20 39\noutside 40 325\nTMhelix 326 348\ninside 349 372

8188 GCF_001520515.1_ASM152051v1 Escherichia coli Proteobacteria; Gammaproteobacteria; Enterobacterales; Enterobacteriaceae; Escherichia
MNNEETFYQAMRRQGVTRRSFLKYCSLAATSLGLGAGMAPKIAWALENKPRIPVVWIHGL WP_001295940.1
MULTISPECIES: hydrogenase [Enterobacteriaceae] Length: 372\nNumber of predicted TMHs: 2\nExp number of AAs in TMHs: 37.2608899999999\nExp number, first 60 AAs: 19.52923\nTotal prob of N-in: 0.94351\nPOSSIBLE N-term signal sequence\ninside 1 19\nTMhelix 20 39\noutside 40 325\nTMhelix 326 348\ninside 349 372

8189 GCF_001520675.1_ASM152067v1 Escherichia coli Proteobacteria; Gammaproteobacteria; Enterobacterales; Enterobacteriaceae; Escherichia
MNNEETFYQAMRRQGVTRRSFLKYCSLAATSLGLGAGMAPKIAWALENKPRIPVVWIHGL WP_001295940.1
MULTISPECIES: hydrogenase [Enterobacteriaceae] Length: 372\nNumber of predicted TMHs: 2\nExp number of AAs in TMHs: 37.2608899999999\nExp number, first 60 AAs: 19.52923\nTotal prob of N-in: 0.94351\nPOSSIBLE N-term signal sequence\ninside 1 19\nTMhelix 20 39\noutside 40 325\nTMhelix 326 348\ninside 349 372

8190 GCF_001520915.1_ASM152091v1 Escherichia coli Proteobacteria; Gammaproteobacteria; Enterobacterales; Enterobacteriaceae; Escherichia
MNNEETFYQAMRRQGVTRRSFLKYCSLAATSLGLGAGMAPKIAWALENKPRIPVVWIHGL WP_001058323.1
MULTISPECIES: hydrogenase-1 small chain [Proteobacteria] Length: 372\nNumber of predicted TMHs: 2\nExp number of AAs in TMHs: 37.2608499999999\nExp number, first 60 AAs: 19.52923\nTotal prob of N-in: 0.94351\nPOSSIBLE N-term signal sequence\ninside 1 19\nTMhelix 20 39\noutside 40 325\nTMhelix 326 348\ninside 349 372

8191 GCF_001520615.1_ASM152061v1 Escherichia coli Proteobacteria; Gammaproteobacteria; Enterobacterales; Enterobacteriaceae; Escherichia
MNNEETFYQAMRRQGVTRRSFLKYCSLAATSLGLGAGMAPKIAWALENKPRIPVVWIHGL WP_001295940.1
MULTISPECIES: hydrogenase [Enterobacteriaceae] Length: 372\nNumber of predicted TMHs: 2\nExp number of AAs in TMHs: 37.2608899999999\nExp number, first 60 AAs: 19.52923\nTotal prob of N-in: 0.94351\nPOSSIBLE N-term signal sequence\ninside 1 19\nTMhelix 20 39\noutside 40 325\nTMhelix 326 348\ninside 349 372

8192 GCF_001521015.1_ASM152101v1 Escherichia coli Proteobacteria; Gammaproteobacteria; Enterobacterales; Enterobacteriaceae; Escherichia
MNNEETFYQAMRRQGVTRRSFLKYCSLAATSLGLGAGMAPKIAWALENKPRIPVVWIHGL WP_001058323.1
MULTISPECIES: hydrogenase-1 small chain [Proteobacteria] Length: 372\nNumber of predicted TMHs: 2\nExp number of AAs in TMHs: 37.2608499999999\nExp number, first 60 AAs: 19.52923\nTotal prob of N-in: 0.94351\nPOSSIBLE N-term signal sequence\ninside 1 19\nTMhelix 20 39\noutside 40 325\nTMhelix 326 348\ninside 349 372

8193 GCF_001521115.1_ASM152111v1 Escherichia coli Proteobacteria; Gammaproteobacteria; Enterobacterales; Enterobacteriaceae; Escherichia
MTGDNTLIHSHGINRRDFMKLCAALAATMGLSSKAAAEMAESVTNPQRPPVIWIGAQECT WP_000145418.1
hydrogenase 2 small subunit [Escherichia coli] Length: 372\nNumber of predicted TMHs: 1\nExp number of AAs in TMHs: 23.72313\nExp number, first 60 AAs: 0.22034\nTotal prob of N-in: 0.12997\noutside 1 331\nTMhelix 332 354\ninside 355 372

8194 GCF_001521175.1_ASM152117v1 Escherichia coli Proteobacteria; Gammaproteobacteria; Enterobacterales; Enterobacteriaceae; Escherichia
MTGDNTLIHSHGINRRDFMKLCAALAATMGLSSKAAAEMVESVTNPQRPPVIWIGAQECT WP_000145423.1
hydrogenase 2 small subunit [Escherichia coli] Length: 372\nNumber of predicted TMHs: 1\nExp number of AAs in TMHs: 23.78201\nExp number, first 60 AAs: 0.28355\nTotal prob of N-in: 0.13257\noutside 1 331\nTMhelix 332 354\ninside 355 372

8195 GCF_001521115.1_ASM152111v1 Escherichia coli Proteobacteria; Gammaproteobacteria; Enterobacterales; Enterobacteriaceae; Escherichia
MNNEETFYQAMRRQGVTRRSFLKYCSLAATSLGLGAGMAPKIAWALENKPRIPVVWIHGL WP_021520630.1
hydrogenase-1 small chain [Escherichia coli] Length: 372\nNumber of predicted TMHs: 2\nExp number of AAs in TMHs: 37.8197599999999\nExp number, first 60 AAs: 19.53295\nTotal prob of N-in: 0.94396\nPOSSIBLE N-term signal sequence\ninside 1 19\nTMhelix 20 39\noutside 40 325\nTMhelix 326 348\ninside 349 372

8196 GCF_001521175.1_ASM152117v1 Escherichia coli Proteobacteria; Gammaproteobacteria; Enterobacterales; Enterobacteriaceae; Escherichia
MNNEETFYQAMRRQGVTRRSFLKYCSLAATSLGLGAGMAPKIAWALENKPRIPVVWIHGL WP_001295940.1
MULTISPECIES: hydrogenase [Enterobacteriaceae] Length: 372\nNumber of predicted TMHs: 2\nExp number of AAs in TMHs: 37.2608899999999\nExp number, first 60 AAs: 19.52923\nTotal prob of N-in: 0.94351\nPOSSIBLE N-term signal sequence\ninside 1 19\nTMhelix 20 39\noutside 40 325\nTMhelix 326 348\ninside 349 372

8197 GCF_001520975.1_ASM152097v1 Escherichia coli Proteobacteria; Gammaproteobacteria; Enterobacterales; Enterobacteriaceae; Escherichia
MNNEETFYQAMRRQGVTRRSFLKYCSLAATSLGLGAGMAPKIAWALENKPRIPVVWIHGL WP_001295940.1
MULTISPECIES: hydrogenase [Enterobacteriaceae] Length: 372\nNumber of predicted TMHs: 2\nExp number of AAs in TMHs: 37.2608899999999\nExp number, first 60 AAs: 19.52923\nTotal prob of N-in: 0.94351\nPOSSIBLE N-term signal sequence\ninside 1 19\nTMhelix 20 39\noutside 40 325\nTMhelix 326 348\ninside 349 372

8198 GCF_001521205.1_ASM152120v1 Escherichia coli Proteobacteria; Gammaproteobacteria; Enterobacterales; Enterobacteriaceae; Escherichia
MTGDNTLIHSHGINRRDFMKLCAALAATMGLSSKAAAEMVESVTNPQRPPVIWIGAQECT WP_000145423.1
hydrogenase 2 small subunit [Escherichia coli] Length: 372\nNumber of predicted TMHs: 1\nExp number of AAs in TMHs: 23.78201\nExp number, first 60 AAs: 0.28355\nTotal prob of N-in: 0.13257\noutside 1 331\nTMhelix 332 354\ninside 355 372

8199 GCF_001521215.1_ASM152121v1 Escherichia coli Proteobacteria; Gammaproteobacteria; Enterobacterales; Enterobacteriaceae; Escherichia
MTGDNTLIHSHGINRRDFMKLCAALAATMGLSSKAAAEMAESVTNPQRPPVIWIGAQECT WP_000145418.1
hydrogenase 2 small subunit [Escherichia coli] Length: 372\nNumber of predicted TMHs: 1\nExp number of AAs in TMHs: 23.72313\nExp number, first 60 AAs: 0.22034\nTotal prob of N-in: 0.12997\noutside 1 331\nTMhelix 332 354\ninside 355 372

8200 GCF_001521225.1_ASM152122v1 Escherichia coli Proteobacteria; Gammaproteobacteria; Enterobacterales; Enterobacteriaceae; Escherichia
MTGDNTLIHSHGINRRDFMKLCAALAATMGLSSKAAAEMAESVTNPQRPPVIWIGAQECT WP_000145410.1
MULTISPECIES: hydrogenase-2 small chain [Proteobacteria] Length: 372\nNumber of predicted TMHs: 1\nExp number of AAs in TMHs: 23.72226\nExp number, first 60 AAs: 0.22038\nTotal prob of N-in: 0.12994\noutside 1 331\nTMhelix 332 354\ninside 355 372

8201 GCF_001521415.1_ASM152141v1 Escherichia coli Proteobacteria; Gammaproteobacteria; Enterobacterales; Enterobacteriaceae; Escherichia
MTGDNTLIHSHGINRRDFMKLCAALAATMGLSSKAAAEMAESVTNPQRPPVIWIGAQECT WP_000145410.1
MULTISPECIES: hydrogenase-2 small chain [Proteobacteria] Length: 372\nNumber of predicted TMHs: 1\nExp number of AAs in TMHs: 23.72226\nExp number, first 60 AAs: 0.22038\nTotal prob of N-in: 0.12994\noutside 1 331\nTMhelix 332 354\ninside 355 372

8202 GCF_001521355.1_ASM152135v1 Escherichia coli Proteobacteria; Gammaproteobacteria; Enterobacterales; Enterobacteriaceae; Escherichia
MTGDNTLIHSHGINRRDFMKLCAALAATMGLSSKAAAEMAESVTNPQRPPVIWIGAQECT WP_000145410.1
MULTISPECIES: hydrogenase-2 small chain [Proteobacteria] Length: 372\nNumber of predicted TMHs: 1\nExp number of AAs in TMHs: 23.72226\nExp number, first 60 AAs: 0.22038\nTotal prob of N-in: 0.12994\noutside 1 331\nTMhelix 332 354\ninside 355 372

8203 GCF_001521315.1_ASM152131v1 Escherichia coli Proteobacteria; Gammaproteobacteria; Enterobacterales; Enterobacteriaceae; Escherichia
MTGDNTLIHSHGINRRDFMKLCAALAATMGLSSKAAAEMAESVTNPQRPPVIWIGAQECT WP_000145410.1

MULTISPECIES: hydrogenase-2 small chain [Proteobacteria] Length: 372\nNumber of predicted TMHs: 1\nExp number of AAs in TMHs: 23.72226\nExp number, first 60 AAs: 0.22038\nTotal prob of N-in: 0.12994\noutside 1 331\nTMhelix 332 354\ninside 355 372

8204 GCF_001521455.1_ASM152145v1 Escherichia coli Proteobacteria; Gammaproteobacteria; Enterobacterales; Enterobacteriaceae; Escherichia
MTGDNTLIHSHGINRRDFMKLCAALAATMGLSSKAAAEMAESVTNPQRPPVIWIGAQECT WP_000145410.1
MULTISPECIES: hydrogenase-2 small chain [Proteobacteria] Length: 372\nNumber of predicted TMHs: 1\nExp number of AAs in TMHs: 23.72226\nExp number, first 60 AAs: 0.22038\nTotal prob of N-in: 0.12994\noutside 1 331\nTMhelix 332 354\ninside 355 372

8205 GCF_001521415.1_ASM152141v1 Escherichia coli Proteobacteria; Gammaproteobacteria; Enterobacterales; Enterobacteriaceae; Escherichia
MNNEETFYQAMRRQGVTRRSFLKYCSLAATSLGLGAGMAPKIAWALENKPRIPVVWIHGL WP_001058323.1
MULTISPECIES: hydrogenase-1 small chain [Proteobacteria] Length: 372\nNumber of predicted TMHs: 2\nExp number of AAs in TMHs: 37.2608499999999\nExp number, first 60 AAs: 19.52923\nTotal prob of N-in: 0.94351\nPOSSIBLE N-term signal sequence\ninside 1 19\nTMhelix 20 39\noutside 40 325\nTMhelix 326 348\ninside 349 372

8206 GCF_001521225.1_ASM152122v1 Escherichia coli Proteobacteria; Gammaproteobacteria; Enterobacterales; Enterobacteriaceae; Escherichia
MNNEETFYQAMRRQGVTRRSFLKYCSLAATSLGLGAGMAPKIAWALENKPRIPVVWIHGL WP_001058323.1
MULTISPECIES: hydrogenase-1 small chain [Proteobacteria] Length: 372\nNumber of predicted TMHs: 2\nExp number of AAs in TMHs: 37.2608499999999\nExp number, first 60 AAs: 19.52923\nTotal prob of N-in: 0.94351\nPOSSIBLE N-term signal sequence\ninside 1 19\nTMhelix 20 39\noutside 40 325\nTMhelix 326 348\ninside 349 372

8207 GCF_001521205.1_ASM152120v1 Escherichia coli Proteobacteria; Gammaproteobacteria; Enterobacterales; Enterobacteriaceae; Escherichia
MNNEETFYQAMRRQGVTRRSFLKYCSLAATSLGLGAGMAPKIAWALENKPRIPVVWIHGL WP_001295940.1
MULTISPECIES: hydrogenase [Enterobacteriaceae] Length: 372\nNumber of predicted TMHs: 2\nExp number of AAs in TMHs: 37.2608899999999\nExp number, first 60 AAs: 19.52923\nTotal prob of N-in: 0.94351\nPOSSIBLE N-term signal sequence\ninside 1 19\nTMhelix 20 39\noutside 40 325\nTMhelix 326 348\ninside 349 372

8208 GCF_001521555.1_ASM152155v1 Escherichia coli Proteobacteria; Gammaproteobacteria; Enterobacterales; Enterobacteriaceae; Escherichia
MTGDNTLIHSHGINRRDFMKLCAALAATMGLSSKAAAEMAESVTNPQRPPVIWIGAQECT WP_000145410.1
MULTISPECIES: hydrogenase-2 small chain [Proteobacteria] Length: 372\nNumber of predicted TMHs: 1\nExp number of AAs in TMHs: 23.72226\nExp number, first 60 AAs: 0.22038\nTotal prob of N-in: 0.12994\noutside 1 331\nTMhelix 332 354\ninside 355 372

8209 GCF_001521555.1_ASM152155v1 Escherichia coli Proteobacteria; Gammaproteobacteria; Enterobacterales; Enterobacteriaceae; Escherichia
MNNEETFYQAMRRQGVTRRSFLKYCSLAATSLGLGAGMAPKIAWALENKPRIPVVWIHGL WP_001058323.1
MULTISPECIES: hydrogenase-1 small chain [Proteobacteria] Length: 372\nNumber of predicted TMHs: 2\nExp number of AAs in TMHs: 37.2608499999999\nExp number, first 60 AAs: 19.52923\nTotal prob of N-in: 0.94351\nPOSSIBLE N-term signal sequence\ninside 1 19\nTMhelix 20 39\noutside 40 325\nTMhelix 326 348\ninside 349 372

8210 GCF_001521315.1_ASM152131v1 Escherichia coli Proteobacteria; Gammaproteobacteria; Enterobacterales; Enterobacteriaceae; Escherichia
MNNEETFYQAMRRQGVTRRSFLKYCSLAATSLGLGAGMAPKIAWALENKPRIPVVWIHGL WP_001058323.1
MULTISPECIES: hydrogenase-1 small chain [Proteobacteria] Length: 372\nNumber of predicted TMHs: 2\nExp number of AAs in TMHs: 37.2608499999999\nExp number, first 60 AAs: 19.52923\nTotal prob of N-in: 0.94351\nPOSSIBLE N-term signal sequence\ninside 1 19\nTMhelix 20 39\noutside 40 325\nTMhelix 326 348\ninside 349 372

8211 GCF_001521355.1_ASM152135v1 Escherichia coli Proteobacteria; Gammaproteobacteria; Enterobacterales; Enterobacteriaceae; Escherichia
MNNEETFYQAMRRQGVTRRSFLKYCSLAATSLGLGAGMAPKIAWALENKPRIPVVWIHGL WP_001058323.1

MULTISPECIES: hydrogenase-1 small chain [Proteobacteria] Length: 372\nNumber of predicted TMHs: 2\nExp number of AAs in TMHs: 37.2608499999999\nExp number, first 60 AAs: 19.52923\nTotal prob of N-in: 0.94351\nPOSSIBLE N-term signal sequence\ninside 1 19\nTMhelix 20 39\noutside 40 325\nTMhelix 326 348\ninside 349 372

8212 GCF_001521215.1_ASM152121v1 Escherichia coli Proteobacteria; Gammaproteobacteria; Enterobacterales; Enterobacteriaceae; Escherichia
MNNEETFYQAMRRQGVTRRSFLKYCSLAATSLGLGAGMAPKIAWALENKPRIPVVWIHGL WP_021520630.1
hydrogenase-1 small chain [Escherichia coli] Length: 372\nNumber of predicted TMHs: 2\nExp number of AAs in TMHs: 37.8197599999999\nExp number, first 60 AAs: 19.53295\nTotal prob of N-in: 0.94396\nPOSSIBLE N-term signal sequence\ninside 1 19\nTMhelix 20 39\noutside 40 325\nTMhelix 326 348\ninside 349 372

8213 GCF_001521615.1_ASM152161v1 Escherichia coli Proteobacteria; Gammaproteobacteria; Enterobacterales; Enterobacteriaceae; Escherichia
MTGDNTLIHSHGINRRDFMKLCAALAATMGLSSKAAAEMVESVTNPQRPPVIWIGAQECT WP_000145423.1
hydrogenase 2 small subunit [Escherichia coli] Length: 372\nNumber of predicted TMHs: 1\nExp number of AAs in TMHs: 23.78201\nExp number, first 60 AAs: 0.28355\nTotal prob of N-in: 0.13257\noutside 1 331\nTMhelix 332 354\ninside 355 372

8214 GCF_001521655.1_ASM152165v1 Escherichia coli Proteobacteria; Gammaproteobacteria; Enterobacterales; Enterobacteriaceae; Escherichia
MTGDNTLIHSHGINRRDFMKLCAALAATMGLSSKAAAEMVESVTNPQRPPVIWIGAQECT WP_000145423.1
hydrogenase 2 small subunit [Escherichia coli] Length: 372\nNumber of predicted TMHs: 1\nExp number of AAs in TMHs: 23.78201\nExp number, first 60 AAs: 0.28355\nTotal prob of N-in: 0.13257\noutside 1 331\nTMhelix 332 354\ninside 355 372

8215 GCF_001524885.1_ASM152488v1 Escherichia coli Proteobacteria; Gammaproteobacteria; Enterobacterales; Enterobacteriaceae; Escherichia
MTGDNTLIHSHGINRRDFMKLCAALAATMGLSSKAAAEMAESVTNPQRPPVIWIGAQECT WP_000145410.1
MULTISPECIES: hydrogenase-2 small chain [Proteobacteria] Length: 372\nNumber of predicted TMHs: 1\nExp number of AAs in TMHs: 23.72226\nExp number, first 60 AAs: 0.22038\nTotal prob of N-in: 0.12994\noutside 1 331\nTMhelix 332 354\ninside 355 372

8216 GCF_001521615.1_ASM152161v1 Escherichia coli Proteobacteria; Gammaproteobacteria; Enterobacterales; Enterobacteriaceae; Escherichia
MNNEETFYQAMRRQGVTRRSFLKYCSLAATSLGLGAGMAPKIAWALENKPRIPVVWIHGL WP_001295940.1
MULTISPECIES: hydrogenase [Enterobacteriaceae] Length: 372\nNumber of predicted TMHs: 2\nExp number of AAs in TMHs: 37.2608899999999\nExp number, first 60 AAs: 19.52923\nTotal prob of N-in: 0.94351\nPOSSIBLE N-term signal sequence\ninside 1 19\nTMhelix 20 39\noutside 40 325\nTMhelix 326 348\ninside 349 372

8217 GCF_001521655.1_ASM152165v1 Escherichia coli Proteobacteria; Gammaproteobacteria; Enterobacterales; Enterobacteriaceae; Escherichia
MNNEETFYQAMRRQGVTRRSFLKYCSLAATSLGLGAGMAPKIAWALENKPRIPVVWIHGL WP_001295940.1
MULTISPECIES: hydrogenase [Enterobacteriaceae] Length: 372\nNumber of predicted TMHs: 2\nExp number of AAs in TMHs: 37.2608899999999\nExp number, first 60 AAs: 19.52923\nTotal prob of N-in: 0.94351\nPOSSIBLE N-term signal sequence\ninside 1 19\nTMhelix 20 39\noutside 40 325\nTMhelix 326 348\ninside 349 372

8218 GCF_001521515.1_ASM152151v1 Escherichia coli Proteobacteria; Gammaproteobacteria; Enterobacterales; Enterobacteriaceae; Escherichia
MNNEETFYQAMRRQGVTRRSFLKYCSLAATSLGLGAGMAPKIAWALENKPRIPVVWIHGL WP_001295940.1
MULTISPECIES: hydrogenase [Enterobacteriaceae] Length: 372\nNumber of predicted TMHs: 2\nExp number of AAs in TMHs: 37.2608899999999\nExp number, first 60 AAs: 19.52923\nTotal prob of N-in: 0.94351\nPOSSIBLE N-term signal sequence\ninside 1 19\nTMhelix 20 39\noutside 40 325\nTMhelix 326 348\ninside 349 372

8219 GCF_001524905.1_ASM152490v1 Escherichia coli Proteobacteria; Gammaproteobacteria; Enterobacterales; Enterobacteriaceae; Escherichia
MTGDNTLIHSHGINRRDFMKLCAALAATMGLSSKAAAEMAESVTNPQRPPVIWIGAQECT WP_000145410.1
MULTISPECIES: hydrogenase-2 small chain [Proteobacteria] Length: 372\nNumber of predicted TMHs:

1\nExp number of AAs in TMHs: 23.72226\nExp number, first 60 AAs: 0.22038\nTotal prob of N-in: 0.12994\noutside 1 331\nTMhelix 332 354\ninside 355 372

8220 GCF_001555635.1_ASM155563v1 Escherichia coli Proteobacteria; Gammaproteobacteria; Enterobacterales; Enterobacteriaceae; Escherichia MTGDNTLIHSHGINRRDFMKLCAALAATMGLSSKAAAEMVESVTNPQRPPVIWIGAQECT WP_000145423.1 hydrogenase 2 small subunit [Escherichia coli] Length: 372\nNumber of predicted TMHs: 1\nExp number of AAs in TMHs: 23.78201\nExp number, first 60 AAs: 0.28355\nTotal prob of N-in: 0.13257\noutside 1 331\nTMhelix 332 354\ninside 355 372

8221 GCF_001561025.1_ASM156102v1 Escherichia coli Proteobacteria; Gammaproteobacteria; Enterobacterales; Enterobacteriaceae; Escherichia MTGDNTLIHSHGINRRDFMKLCAALAATMGLSSKAAAEMVESVTNPQRPPVIWIGAQECT WP_000145423.1 hydrogenase 2 small subunit [Escherichia coli] Length: 372\nNumber of predicted TMHs: 1\nExp number of AAs in TMHs: 23.78201\nExp number, first 60 AAs: 0.28355\nTotal prob of N-in: 0.13257\noutside 1 331\nTMhelix 332 354\ninside 355 372

8222 GCF_001561035.1_ASM156103v1 Escherichia coli Proteobacteria; Gammaproteobacteria; Enterobacterales; Enterobacteriaceae; Escherichia MTGDNTLIHSHGINRRDFMKLCAALAATMGLSSKAAAEMVESVTNPQRPPVIWIGAQECT WP_000145423.1 hydrogenase 2 small subunit [Escherichia coli] Length: 372\nNumber of predicted TMHs: 1\nExp number of AAs in TMHs: 23.78201\nExp number, first 60 AAs: 0.28355\nTotal prob of N-in: 0.13257\noutside 1 331\nTMhelix 332 354\ninside 355 372

8223 GCF_001561095.1_ASM156109v1 Escherichia coli Proteobacteria; Gammaproteobacteria; Enterobacterales; Enterobacteriaceae; Escherichia MTGDNTLIHSHGINRRDFMKLCAALAATMGLSSKAAAEMVESVTNPQRPPVIWIGAQECT WP_000145423.1 hydrogenase 2 small subunit [Escherichia coli] Length: 372\nNumber of predicted TMHs: 1\nExp number of AAs in TMHs: 23.78201\nExp number, first 60 AAs: 0.28355\nTotal prob of N-in: 0.13257\noutside 1 331\nTMhelix 332 354\ninside 355 372

8224 GCF_001561035.1_ASM156103v1 Escherichia coli Proteobacteria; Gammaproteobacteria; Enterobacterales; Enterobacteriaceae; Escherichia MNNEETFYQAMRRQGVTRRSFLKYCSLAATSLGLGAGMAPKIAWALENKPRIPVVWIHGL WP_001295940.1 MULTISPECIES: hydrogenase [Enterobacteriaceae] Length: 372\nNumber of predicted TMHs: 2\nExp number of AAs in TMHs: 37.2608899999999\nExp number, first 60 AAs: 19.52923\nTotal prob of N-in: 0.94351\nPOSSIBLE N-term signal sequence\ninside 1 19\nTMhelix 20 39\noutside 40 325\nTMhelix 326 348\ninside 349 372

8225 GCF_001561095.1_ASM156109v1 Escherichia coli Proteobacteria; Gammaproteobacteria; Enterobacterales; Enterobacteriaceae; Escherichia MNNEETFYQAMRRQGVTRRSFLKYCSLAATSLGLGAGMAPKIAWALENKPRIPVVWIHGL WP_001295940.1 MULTISPECIES: hydrogenase [Enterobacteriaceae] Length: 372\nNumber of predicted TMHs: 2\nExp number of AAs in TMHs: 37.2608899999999\nExp number, first 60 AAs: 19.52923\nTotal prob of N-in: 0.94351\nPOSSIBLE N-term signal sequence\ninside 1 19\nTMhelix 20 39\noutside 40 325\nTMhelix 326 348\ninside 349 372

8226 GCF_001561115.1_ASM156111v1 Escherichia coli Proteobacteria; Gammaproteobacteria; Enterobacterales; Enterobacteriaceae; Escherichia MTGDNTLIHSHGINRRDFMKLCAALAATMGLSSKAAAEMVESVTNPQRPPVIWIGAQECT WP_000145423.1 hydrogenase 2 small subunit [Escherichia coli] Length: 372\nNumber of predicted TMHs: 1\nExp number of AAs in TMHs: 23.78201\nExp number, first 60 AAs: 0.28355\nTotal prob of N-in: 0.13257\noutside 1 331\nTMhelix 332 354\ninside 355 372

8227 GCF_001561175.1_ASM156117v1 Escherichia coli Proteobacteria; Gammaproteobacteria; Enterobacterales; Enterobacteriaceae; Escherichia MTGDNTLIHSHGINRRDFMKLCAALAATMGLSSKAAAEMAESVTNPQRPPVIWIGAQECT WP_000145410.1 MULTISPECIES: hydrogenase-2 small chain [Proteobacteria] Length: 372\nNumber of predicted TMHs: 1\nExp number of AAs in TMHs: 23.72226\nExp number, first 60 AAs: 0.22038\nTotal prob of N-in: 0.12994\noutside 1 331\nTMhelix 332 354\ninside 355 372

8228	GCF_001561115.1_ASM156111v1	Escherichia coli	Proteobacteria; Gammaproteobacteria; Enterobacterales; Enterobacteriaceae; Escherichia
MNNEETFYQAMRRQGVTRRSFLKYCSLAATSLGLGAGMAPKIAWALENKPRIPVVWIHGL	WP_001295940.1
MULTISPECIES: hydrogenase [Enterobacteriaceae]	Length: 372\nNumber of predicted TMHs:  2\nExp number of AAs in TMHs: 37.2608899999999\nExp number, first 60 AAs:  19.52923\nTotal prob of N-in: 0.94351\nPOSSIBLE N-term signal sequence\ninside   1   19\nTMhelix   20   39\noutside   40   325\nTMhelix   326 348\ninside   349   372

8229	GCF_001561175.1_ASM156117v1	Escherichia coli	Proteobacteria; Gammaproteobacteria; Enterobacterales; Enterobacteriaceae; Escherichia
MNNEETFYQAMRRQGVTRRSFLKYCSLAATSLGLGAGMAPKIAWALENKPRIPVVWIHGL	WP_001295940.1
MULTISPECIES: hydrogenase [Enterobacteriaceae]	Length: 372\nNumber of predicted TMHs:  2\nExp number of AAs in TMHs: 37.2608899999999\nExp number, first 60 AAs:  19.52923\nTotal prob of N-in: 0.94351\nPOSSIBLE N-term signal sequence\ninside   1   19\nTMhelix   20   39\noutside   40   325\nTMhelix   326 348\ninside   349   372

8230	GCF_001561215.1_ASM156121v1	Escherichia coli	Proteobacteria; Gammaproteobacteria; Enterobacterales; Enterobacteriaceae; Escherichia
MTGDNTLIHSHGINRRDFMKLCAALAATMGLSSKAAAEMVESVTNPQRPPVIWIGAQECT	WP_000145423.1
hydrogenase 2 small subunit [Escherichia coli]	Length: 372\nNumber of predicted TMHs:  1\nExp number of AAs in TMHs: 23.78201\nExp number, first 60 AAs:  0.28355\nTotal prob of N-in:   0.13257\noutside   1   331\nTMhelix 332   354\ninside   355   372

8231	GCF_001561215.1_ASM156121v1	Escherichia coli	Proteobacteria; Gammaproteobacteria; Enterobacterales; Enterobacteriaceae; Escherichia
MNNEETFYQAMRRQGVTRRSFLKYCSLAATSLGLGAGMAPKIAWALENKPRIPVVWIHGL	WP_001295940.1
MULTISPECIES: hydrogenase [Enterobacteriaceae]	Length: 372\nNumber of predicted TMHs:  2\nExp number of AAs in TMHs: 37.2608899999999\nExp number, first 60 AAs:  19.52923\nTotal prob of N-in: 0.94351\nPOSSIBLE N-term signal sequence\ninside   1   19\nTMhelix   20   39\noutside   40   325\nTMhelix   326 348\ninside   349   372

8232	GCF_001555635.1_ASM155563v1	Escherichia coli	Proteobacteria; Gammaproteobacteria; Enterobacterales; Enterobacteriaceae; Escherichia
MNNEETFYQAMRRQGVTRRSFLKYCSLAATSLGLGAGMAPKIAWALENKPRIPVVWIHGL	WP_001295940.1
MULTISPECIES: hydrogenase [Enterobacteriaceae]	Length: 372\nNumber of predicted TMHs:  2\nExp number of AAs in TMHs: 37.2608899999999\nExp number, first 60 AAs:  19.52923\nTotal prob of N-in: 0.94351\nPOSSIBLE N-term signal sequence\ninside   1   19\nTMhelix   20   39\noutside   40   325\nTMhelix   326 348\ninside   349   372

8233	GCF_001561025.1_ASM156102v1	Escherichia coli	Proteobacteria; Gammaproteobacteria; Enterobacterales; Enterobacteriaceae; Escherichia
MNNEETFYQAMRRQGVTRRSFLKYCSLAATSLGLGAGMAPKIAWALENKPRIPVVWIHGL	WP_001295940.1
MULTISPECIES: hydrogenase [Enterobacteriaceae]	Length: 372\nNumber of predicted TMHs:  2\nExp number of AAs in TMHs: 37.2608899999999\nExp number, first 60 AAs:  19.52923\nTotal prob of N-in: 0.94351\nPOSSIBLE N-term signal sequence\ninside   1   19\nTMhelix   20   39\noutside   40   325\nTMhelix   326 348\ninside   349   372

8234	GCF_001561335.1_ASM156133v1	Escherichia coli	Proteobacteria; Gammaproteobacteria; Enterobacterales; Enterobacteriaceae; Escherichia
MTGDNTLIHSHGINRRDFMKLCAALAATMGLSSKAAAEMAESVTNPQRPPVIWIGAQECT	WP_000145410.1
MULTISPECIES: hydrogenase-2 small chain [Proteobacteria]	Length: 372\nNumber of predicted TMHs: 1\nExp number of AAs in TMHs: 23.72226\nExp number, first 60 AAs:  0.22038\nTotal prob of N-in:   0.12994\noutside   1 331\nTMhelix   332   354\ninside   355   372

8235	GCF_001561365.1_ASM156136v1	Escherichia coli	Proteobacteria; Gammaproteobacteria; Enterobacterales; Enterobacteriaceae; Escherichia
MTGDNTLIHSHGINRRDFMKLCAALAATMGLSSKAAAEMVESVTNPQRPPVIWIGAQECT	WP_000145423.1
hydrogenase 2 small subunit [Escherichia coli]	Length: 372\nNumber of predicted TMHs:  1\nExp number of AAs in TMHs: 23.78201\nExp number, first 60 AAs:  0.28355\nTotal prob of N-in:   0.13257\noutside   1   331\nTMhelix 332   354\ninside   355   372

8236 GCF_001561435.1_ASM156143v1 Escherichia coli Proteobacteria; Gammaproteobacteria; Enterobacterales; Enterobacteriaceae; Escherichia
MTGDNTLIHSHGINRRDFMKLCAALAATMGLSSKAAAEMAESVTNPQRPPVIWIGAQECT WP_000145410.1
MULTISPECIES: hydrogenase-2 small chain [Proteobacteria] Length: 372\nNumber of predicted TMHs: 1\nExp number of AAs in TMHs: 23.72226\nExp number, first 60 AAs: 0.22038\nTotal prob of N-in: 0.12994\noutside 1 331\nTMhelix 332 354\ninside 355 372

8237 GCF_001561675.1_ASM156167v1 Escherichia coli Proteobacteria; Gammaproteobacteria; Enterobacterales; Enterobacteriaceae; Escherichia
MTGDNTLIHSHGINRRDFMKLCAALAATMGLSSKAAAEMAESVTNPQRPPVIWIGAQECT WP_000145417.1
MULTISPECIES: hydrogenase-2 small chain [Escherichia] Length: 372\nNumber of predicted TMHs: 1\nExp number of AAs in TMHs: 23.72512\nExp number, first 60 AAs: 0.22034\nTotal prob of N-in: 0.13000\noutside 1 331\nTMhelix 332 354\ninside 355 372

8238 GCF_001561455.1_ASM156145v1 Escherichia coli Proteobacteria; Gammaproteobacteria; Enterobacterales; Enterobacteriaceae; Escherichia
MTGDNTLIHSHGINRRDFMKLCAALAATMGLSSKAAAEMAESVTNPQRPPVIWIGAQECT WP_000145410.1
MULTISPECIES: hydrogenase-2 small chain [Proteobacteria] Length: 372\nNumber of predicted TMHs: 1\nExp number of AAs in TMHs: 23.72226\nExp number, first 60 AAs: 0.22038\nTotal prob of N-in: 0.12994\noutside 1 331\nTMhelix 332 354\ninside 355 372

8239 GCF_001561505.1_ASM156150v1 Escherichia coli Proteobacteria; Gammaproteobacteria; Enterobacterales; Enterobacteriaceae; Escherichia
MTGDNTLIHSHGINRRDFMKLCAALAATMGLSSKAAAEMVESVTNPQRPPVIWIGAQECT WP_000145423.1
hydrogenase 2 small subunit [Escherichia coli] Length: 372\nNumber of predicted TMHs: 1\nExp number of AAs in TMHs: 23.78201\nExp number, first 60 AAs: 0.28355\nTotal prob of N-in: 0.13257\noutside 1 331\nTMhelix 332 354\ninside 355 372

8240 GCF_001561605.1_ASM156160v1 Escherichia coli Proteobacteria; Gammaproteobacteria; Enterobacterales; Enterobacteriaceae; Escherichia
MTGDNTLIHSHGINRRDFMKLCAALAATMGLSSKAAAEMVESVTNPQRPPVIWIGAQECT WP_000145423.1
hydrogenase 2 small subunit [Escherichia coli] Length: 372\nNumber of predicted TMHs: 1\nExp number of AAs in TMHs: 23.78201\nExp number, first 60 AAs: 0.28355\nTotal prob of N-in: 0.13257\noutside 1 331\nTMhelix 332 354\ninside 355 372

8241 GCF_001561575.1_ASM156157v1 Escherichia coli Proteobacteria; Gammaproteobacteria; Enterobacterales; Enterobacteriaceae; Escherichia
MTGDNTLIHSHGINRRDFMKLCAALAATMGLSSKAAAEMAESVTNPQRPPVIWIGAQECT WP_000145410.1
MULTISPECIES: hydrogenase-2 small chain [Proteobacteria] Length: 372\nNumber of predicted TMHs: 1\nExp number of AAs in TMHs: 23.72226\nExp number, first 60 AAs: 0.22038\nTotal prob of N-in: 0.12994\noutside 1 331\nTMhelix 332 354\ninside 355 372

8242 GCF_001561705.1_ASM156170v1 Escherichia coli Proteobacteria; Gammaproteobacteria; Enterobacterales; Enterobacteriaceae; Escherichia
MTGDNTLIHSHGINRRDFMKLCAALAATMGLSSKAAAEMAESVTNPQRPPVIWIGAQECT WP_000145410.1
MULTISPECIES: hydrogenase-2 small chain [Proteobacteria] Length: 372\nNumber of predicted TMHs: 1\nExp number of AAs in TMHs: 23.72226\nExp number, first 60 AAs: 0.22038\nTotal prob of N-in: 0.12994\noutside 1 331\nTMhelix 332 354\ninside 355 372

8243 GCF_001561275.1_ASM156127v1 Escherichia coli Proteobacteria; Gammaproteobacteria; Enterobacterales; Enterobacteriaceae; Escherichia
MNNEETFYQAMRRQGVTRRSFLKYCSLAATSLGLGAGMAPKIAWALENKPRIPVVWIHGL WP_001295940.1
MULTISPECIES: hydrogenase [Enterobacteriaceae] Length: 372\nNumber of predicted TMHs: 2\nExp number of AAs in TMHs: 37.2608899999999\nExp number, first 60 AAs: 19.52923\nTotal prob of N-in: 0.94351\nPOSSIBLE N-term signal sequence\ninside 1 19\nTMhelix 20 39\noutside 40 325\nTMhelix 326 348\ninside 349 372

8244 GCF_001561575.1_ASM156157v1 Escherichia coli Proteobacteria; Gammaproteobacteria; Enterobacterales; Enterobacteriaceae; Escherichia
MNNEETFYQAMRRQGVTRRSFLKYCSLAATSLGLGAGMAPKIAWALENKPRIPVVWIHGL WP_001058323.1
MULTISPECIES: hydrogenase-1 small chain [Proteobacteria] Length: 372\nNumber of predicted TMHs:

2\nExp number of AAs in TMHs: 37.2608499999999\nExp number, first 60 AAs: 19.52923\nTotal prob of N-in: 0.94351\nPOSSIBLE N-term signal sequence\ninside 1 19\nTMhelix 20 39\noutside 40 325\nTMhelix 326 348\ninside 349 372

8245 GCF_001561705.1_ASM156170v1 Escherichia coli Proteobacteria; Gammaproteobacteria; Enterobacterales; Enterobacteriaceae; Escherichia MNNEETFYQAMRRQGVTRRSFLKYCSLAATSLGLGAGMAPKIAWALENKPRIPVVWIHGL WP_001295940.1 MULTISPECIES: hydrogenase [Enterobacteriaceae] Length: 372\nNumber of predicted TMHs: 2\nExp number of AAs in TMHs: 37.2608899999999\nExp number, first 60 AAs: 19.52923\nTotal prob of N-in: 0.94351\nPOSSIBLE N-term signal sequence\ninside 1 19\nTMhelix 20 39\noutside 40 325\nTMhelix 326 348\ninside 349 372

8246 GCF_001561675.1_ASM156167v1 Escherichia coli Proteobacteria; Gammaproteobacteria; Enterobacterales; Enterobacteriaceae; Escherichia MNNEETFYQAMRRQGVTRRSFLKYCSLAATSLGLGAGMAPKIAWALENKPRIPVVWIHGL WP_001516139.1 MULTISPECIES: hydrogenase [Escherichia] Length: 372\nNumber of predicted TMHs: 2\nExp number of AAs in TMHs: 37.98718\nExp number, first 60 AAs: 19.52876\nTotal prob of N-in: 0.94354\nPOSSIBLE N-term signal sequence\ninside 1 19\nTMhelix 20 39\noutside 40 325\nTMhelix 326 348\ninside 349 372

8247 GCF_001561335.1_ASM156133v1 Escherichia coli Proteobacteria; Gammaproteobacteria; Enterobacterales; Enterobacteriaceae; Escherichia MNNEETFYQAMRRQGVTRRSFLKYCSLAATSLGLGAGMAPKIAWALENKPRIPVVWIHGL WP_001295940.1 MULTISPECIES: hydrogenase [Enterobacteriaceae] Length: 372\nNumber of predicted TMHs: 2\nExp number of AAs in TMHs: 37.2608899999999\nExp number, first 60 AAs: 19.52923\nTotal prob of N-in: 0.94351\nPOSSIBLE N-term signal sequence\ninside 1 19\nTMhelix 20 39\noutside 40 325\nTMhelix 326 348\ninside 349 372

8248 GCF_001561765.1_ASM156176v1 Escherichia coli Proteobacteria; Gammaproteobacteria; Enterobacterales; Enterobacteriaceae; Escherichia MTGDNTLIHSHGINRRDFMKLCAALAATMGLSSKAAAEMVESVTNPQRPPVIWIGAQECT WP_000145423.1 hydrogenase 2 small subunit [Escherichia coli] Length: 372\nNumber of predicted TMHs: 1\nExp number of AAs in TMHs: 23.78201\nExp number, first 60 AAs: 0.28355\nTotal prob of N-in: 0.13257\noutside 1 331\nTMhelix 332 354\ninside 355 372

8249 GCF_001561765.1_ASM156176v1 Escherichia coli Proteobacteria; Gammaproteobacteria; Enterobacterales; Enterobacteriaceae; Escherichia MNNEETFYQAMRRQGVTRRSFLKYCSLAATSLGLGAGMAPKIAWALENKPRIPVVWIHGL WP_001295940.1 MULTISPECIES: hydrogenase [Enterobacteriaceae] Length: 372\nNumber of predicted TMHs: 2\nExp number of AAs in TMHs: 37.2608899999999\nExp number, first 60 AAs: 19.52923\nTotal prob of N-in: 0.94351\nPOSSIBLE N-term signal sequence\ninside 1 19\nTMhelix 20 39\noutside 40 325\nTMhelix 326 348\ninside 349 372

8250 GCF_001561365.1_ASM156136v1 Escherichia coli Proteobacteria; Gammaproteobacteria; Enterobacterales; Enterobacteriaceae; Escherichia MNNEETFYQAMRRQGVTRRSFLKYCSLAATSLGLGAGMAPKIAWALENKPRIPVVWIHGL WP_001295940.1 MULTISPECIES: hydrogenase [Enterobacteriaceae] Length: 372\nNumber of predicted TMHs: 2\nExp number of AAs in TMHs: 37.2608899999999\nExp number, first 60 AAs: 19.52923\nTotal prob of N-in: 0.94351\nPOSSIBLE N-term signal sequence\ninside 1 19\nTMhelix 20 39\noutside 40 325\nTMhelix 326 348\ninside 349 372

8251 GCF_001561815.1_ASM156181v1 Escherichia coli Proteobacteria; Gammaproteobacteria; Enterobacterales; Enterobacteriaceae; Escherichia MTGDNTLIHSHGINRRDFMKLCAALAATMGLSSKAAAEMVESVTNPQRPPVIWIGAQECT WP_000145423.1 hydrogenase 2 small subunit [Escherichia coli] Length: 372\nNumber of predicted TMHs: 1\nExp number of AAs in TMHs: 23.78201\nExp number, first 60 AAs: 0.28355\nTotal prob of N-in: 0.13257\noutside 1 331\nTMhelix 332 354\ninside 355 372

8252 GCF_001561815.1_ASM156181v1 Escherichia coli Proteobacteria; Gammaproteobacteria; Enterobacterales; Enterobacteriaceae; Escherichia MNNEETFYQAMRRQGVTRRSFLKYCSLAATSLGLGAGMAPKIAWALENKPRIPVVWIHGL WP_001295940.1 MULTISPECIES: hydrogenase [Enterobacteriaceae] Length: 372\nNumber of predicted TMHs: 2\nExp

number of AAs in TMHs: 37.2608899999999\nExp number, first 60 AAs: 19.52923\nTotal prob of N-in: 0.94351\nPOSSIBLE N-term signal sequence\ninside 1 19\nTMhelix 20 39\noutside 40 325\nTMhelix 326 348\ninside 349 372

8253 GCF_001561435.1_ASM156143v1 Escherichia coli Proteobacteria; Gammaproteobacteria; Enterobacterales; Enterobacteriaceae; Escherichia
MNNEETFYQAMRRQGVTRRSFLKYCSLAATSLGLGAGMAPKIAWALENKPRIPVVWIHGL WP_001295940.1
MULTISPECIES: hydrogenase [Enterobacteriaceae] Length: 372\nNumber of predicted TMHs: 2\nExp number of AAs in TMHs: 37.2608899999999\nExp number, first 60 AAs: 19.52923\nTotal prob of N-in: 0.94351\nPOSSIBLE N-term signal sequence\ninside 1 19\nTMhelix 20 39\noutside 40 325\nTMhelix 326 348\ninside 349 372

8254 GCF_001561865.1_ASM156186v1 Escherichia coli Proteobacteria; Gammaproteobacteria; Enterobacterales; Enterobacteriaceae; Escherichia
MTGDNTLIHSHGINRRDFMKLCAALAATMGLSSKAAAEMVESVTNPQRPPVIWIGAQECT WP_000145423.1
hydrogenase 2 small subunit [Escherichia coli] Length: 372\nNumber of predicted TMHs: 1\nExp number of AAs in TMHs: 23.78201\nExp number, first 60 AAs: 0.28355\nTotal prob of N-in: 0.13257\noutside 1 331\nTMhelix 332 354\ninside 355 372

8255 GCF_001561865.1_ASM156186v1 Escherichia coli Proteobacteria; Gammaproteobacteria; Enterobacterales; Enterobacteriaceae; Escherichia
MNNEETFYQAMRRQGVTRRSFLKYCSLAATSLGLGAGMAPKIAWALENKPRIPVVWIHGL WP_001295940.1
MULTISPECIES: hydrogenase [Enterobacteriaceae] Length: 372\nNumber of predicted TMHs: 2\nExp number of AAs in TMHs: 37.2608899999999\nExp number, first 60 AAs: 19.52923\nTotal prob of N-in: 0.94351\nPOSSIBLE N-term signal sequence\ninside 1 19\nTMhelix 20 39\noutside 40 325\nTMhelix 326 348\ninside 349 372

8256 GCF_001561455.1_ASM156145v1 Escherichia coli Proteobacteria; Gammaproteobacteria; Enterobacterales; Enterobacteriaceae; Escherichia
MNNEETFYQAMRRQGVTRRSFLKYCSLAATSLGLGAGMAPKIAWALENKPRIPVVWIHGL WP_001295940.1
MULTISPECIES: hydrogenase [Enterobacteriaceae] Length: 372\nNumber of predicted TMHs: 2\nExp number of AAs in TMHs: 37.2608899999999\nExp number, first 60 AAs: 19.52923\nTotal prob of N-in: 0.94351\nPOSSIBLE N-term signal sequence\ninside 1 19\nTMhelix 20 39\noutside 40 325\nTMhelix 326 348\ninside 349 372

8257 GCF_001561605.1_ASM156160v1 Escherichia coli Proteobacteria; Gammaproteobacteria; Enterobacterales; Enterobacteriaceae; Escherichia
MNNEETFYQAMRRQGVTRRSFLKYCSLAATSLGLGAGMAPKIAWALENKPRIPVVWIHGL WP_001295940.1
MULTISPECIES: hydrogenase [Enterobacteriaceae] Length: 372\nNumber of predicted TMHs: 2\nExp number of AAs in TMHs: 37.2608899999999\nExp number, first 60 AAs: 19.52923\nTotal prob of N-in: 0.94351\nPOSSIBLE N-term signal sequence\ninside 1 19\nTMhelix 20 39\noutside 40 325\nTMhelix 326 348\ninside 349 372

8258 GCF_001561505.1_ASM156150v1 Escherichia coli Proteobacteria; Gammaproteobacteria; Enterobacterales; Enterobacteriaceae; Escherichia
MNNEETFYQAMRRQGVTRRSFLKYCSLAATSLGLGAGMAPKIAWALENKPRIPVVWIHGL WP_001295940.1
MULTISPECIES: hydrogenase [Enterobacteriaceae] Length: 372\nNumber of predicted TMHs: 2\nExp number of AAs in TMHs: 37.2608899999999\nExp number, first 60 AAs: 19.52923\nTotal prob of N-in: 0.94351\nPOSSIBLE N-term signal sequence\ninside 1 19\nTMhelix 20 39\noutside 40 325\nTMhelix 326 348\ninside 349 372

8259 GCF_001561895.1_ASM156189v1 Escherichia coli Proteobacteria; Gammaproteobacteria; Enterobacterales; Enterobacteriaceae; Escherichia
MTGDNTLIHSHGINRRDFMKLCAALAATMGLSSKAAAEMVESVTNPQRPPVIWIGAQECT WP_000145423.1
hydrogenase 2 small subunit [Escherichia coli] Length: 372\nNumber of predicted TMHs: 1\nExp number of AAs in TMHs: 23.78201\nExp number, first 60 AAs: 0.28355\nTotal prob of N-in: 0.13257\noutside 1 331\nTMhelix 332 354\ninside 355 372

8260 GCF_001562345.1_ASM156234v1 Escherichia coli Proteobacteria; Gammaproteobacteria; Enterobacterales; Enterobacteriaceae; Escherichia
MTGDNTLIHSHGINRRDFMKLCAALAATMGLSSKAAAEMVESVTNPQRPPVIWIGAQECT WP_000145423.1

hydrogenase 2 small subunit [Escherichia coli] Length: 372\nNumber of predicted TMHs: 1\nExp number of AAs in TMHs: 23.78201\nExp number, first 60 AAs: 0.28355\nTotal prob of N-in: 0.13257\noutside 1 331\nTMhelix 332 354\ninside 355 372

8261 GCF_001562385.1_ASM156238v1 Escherichia coli Proteobacteria; Gammaproteobacteria; Enterobacterales; Enterobacteriaceae; Escherichia
MTGDNTLIHSHGINRRDFMKLCAALAATMGLSSKAAAEMVESVTNPQRPPVIWIGAQECT WP_000145423.1
hydrogenase 2 small subunit [Escherichia coli] Length: 372\nNumber of predicted TMHs: 1\nExp number of AAs in TMHs: 23.78201\nExp number, first 60 AAs: 0.28355\nTotal prob of N-in: 0.13257\noutside 1 331\nTMhelix 332 354\ninside 355 372

8262 GCF_001562705.1_ASM156270v1 Escherichia coli Proteobacteria; Gammaproteobacteria; Enterobacterales; Enterobacteriaceae; Escherichia
MTGDNTLIHSHGINRRDFMKLCAALAATMGLSSKAAAEMAESVTNPQRPPVIWIGAQECT WP_000145410.1
MULTISPECIES: hydrogenase-2 small chain [Proteobacteria] Length: 372\nNumber of predicted TMHs: 1\nExp number of AAs in TMHs: 23.72226\nExp number, first 60 AAs: 0.22038\nTotal prob of N-in: 0.12994\noutside 1 331\nTMhelix 332 354\ninside 355 372

8263 GCF_001562655.1_ASM156265v1 Escherichia coli Proteobacteria; Gammaproteobacteria; Enterobacterales; Enterobacteriaceae; Escherichia
MTGDNTLIHSHGINRRDFMKLCAALAATMGLSSKAAAEMAESVTNPQRPPVIWIGAQECT WP_000145410.1
MULTISPECIES: hydrogenase-2 small chain [Proteobacteria] Length: 372\nNumber of predicted TMHs: 1\nExp number of AAs in TMHs: 23.72226\nExp number, first 60 AAs: 0.22038\nTotal prob of N-in: 0.12994\noutside 1 331\nTMhelix 332 354\ninside 355 372

8264 GCF_001562835.1_ASM156283v1 Escherichia coli Proteobacteria; Gammaproteobacteria; Enterobacterales; Enterobacteriaceae; Escherichia
MTGDNTLIHSHGINRRDFMKLCAALAATMGLSSKAAAEMAESVTNPQRPPVIWIGAQECT WP_000145410.1
MULTISPECIES: hydrogenase-2 small chain [Proteobacteria] Length: 372\nNumber of predicted TMHs: 1\nExp number of AAs in TMHs: 23.72226\nExp number, first 60 AAs: 0.22038\nTotal prob of N-in: 0.12994\noutside 1 331\nTMhelix 332 354\ninside 355 372

8265 GCF_001563395.1_ASM156339v1 Escherichia coli Proteobacteria; Gammaproteobacteria; Enterobacterales; Enterobacteriaceae; Escherichia
MTGDNTLIHSHGINRRDFMKLCAALAATMGLSSKAAAEMVESVTNPQRPPVIWIGAQECT WP_000145423.1
hydrogenase 2 small subunit [Escherichia coli] Length: 372\nNumber of predicted TMHs: 1\nExp number of AAs in TMHs: 23.78201\nExp number, first 60 AAs: 0.28355\nTotal prob of N-in: 0.13257\noutside 1 331\nTMhelix 332 354\ninside 355 372

8266 GCF_001562765.1_ASM156276v1 Escherichia coli Proteobacteria; Gammaproteobacteria; Enterobacterales; Enterobacteriaceae; Escherichia
MTGDNTLIHSHGINRRDFMKLCAALAATMGLSSKAAAEMAESVTNPQRPPVIWIGAQECT WP_000145410.1
MULTISPECIES: hydrogenase-2 small chain [Proteobacteria] Length: 372\nNumber of predicted TMHs: 1\nExp number of AAs in TMHs: 23.72226\nExp number, first 60 AAs: 0.22038\nTotal prob of N-in: 0.12994\noutside 1 331\nTMhelix 332 354\ninside 355 372

8267 GCF_001563415.1_ASM156341v1 Escherichia coli Proteobacteria; Gammaproteobacteria; Enterobacterales; Enterobacteriaceae; Escherichia
MTGDNTLIHSHGINRRDFMKLCAALAATMGLSSKAAAEMVESVTNPQRPPVIWIGAQECT WP_000145423.1
hydrogenase 2 small subunit [Escherichia coli] Length: 372\nNumber of predicted TMHs: 1\nExp number of AAs in TMHs: 23.78201\nExp number, first 60 AAs: 0.28355\nTotal prob of N-in: 0.13257\noutside 1 331\nTMhelix 332 354\ninside 355 372

8268 GCF_001562765.1_ASM156276v1 Escherichia coli Proteobacteria; Gammaproteobacteria; Enterobacterales; Enterobacteriaceae; Escherichia
MNNEETFYQAMRRQGVTRRSFLKYCSLAATSLGLGAGMAPKIAWALENKPRIPVVWIHGL WP_001058323.1
MULTISPECIES: hydrogenase-1 small chain [Proteobacteria] Length: 372\nNumber of predicted TMHs: 2\nExp number of AAs in TMHs: 37.2608499999999\nExp number, first 60 AAs: 19.52923\nTotal prob of N-in: 0.94351\nPOSSIBLE N-term signal sequence\ninside 1 19\nTMhelix 20 39\noutside 40 325\nTMhelix 326 348\ninside 349 372

8269 GCF_001563415.1_ASM156341v1 Escherichia coli Proteobacteria; Gammaproteobacteria; Enterobacterales; Enterobacteriaceae; Escherichia
MNNEETFYQAMRRQGVTRRSFLKYCSLAATSLGLGAGMAPKIAWALENKPRIPVVWIHGL WP_001295940.1
MULTISPECIES: hydrogenase [Enterobacteriaceae] Length: 372\nNumber of predicted TMHs: 2\nExp number of AAs in TMHs: 37.2608899999999\nExp number, first 60 AAs: 19.52923\nTotal prob of N-in: 0.94351\nPOSSIBLE N-term signal sequence\ninside 1 19\nTMhelix 20 39\noutside 40 325\nTMhelix 326 348\ninside 349 372

8270 GCF_001562655.1_ASM156265v1 Escherichia coli Proteobacteria; Gammaproteobacteria; Enterobacterales; Enterobacteriaceae; Escherichia
MNNEETFYQAMRRQGVTRRSFLKYCSLAATSLGLGAGMAPKIAWALENKPRIPVVWIHGL WP_001058323.1
MULTISPECIES: hydrogenase-1 small chain [Proteobacteria] Length: 372\nNumber of predicted TMHs: 2\nExp number of AAs in TMHs: 37.2608499999999\nExp number, first 60 AAs: 19.52923\nTotal prob of N-in: 0.94351\nPOSSIBLE N-term signal sequence\ninside 1 19\nTMhelix 20 39\noutside 40 325\nTMhelix 326 348\ninside 349 372

8271 GCF_001562705.1_ASM156270v1 Escherichia coli Proteobacteria; Gammaproteobacteria; Enterobacterales; Enterobacteriaceae; Escherichia
MNNEETFYQAMRRQGVTRRSFLKYCSLAATSLGLGAGMAPKIAWALENKPRIPVVWIHGL WP_001058323.1
MULTISPECIES: hydrogenase-1 small chain [Proteobacteria] Length: 372\nNumber of predicted TMHs: 2\nExp number of AAs in TMHs: 37.2608499999999\nExp number, first 60 AAs: 19.52923\nTotal prob of N-in: 0.94351\nPOSSIBLE N-term signal sequence\ninside 1 19\nTMhelix 20 39\noutside 40 325\nTMhelix 326 348\ninside 349 372

8272 GCF_001562835.1_ASM156283v1 Escherichia coli Proteobacteria; Gammaproteobacteria; Enterobacterales; Enterobacteriaceae; Escherichia
MNNEETFYQAMRRQGVTRRSFLKYCSLAATSLGLGAGMAPKIAWALENKPRIPVVWIHGL WP_001058323.1
MULTISPECIES: hydrogenase-1 small chain [Proteobacteria] Length: 372\nNumber of predicted TMHs: 2\nExp number of AAs in TMHs: 37.2608499999999\nExp number, first 60 AAs: 19.52923\nTotal prob of N-in: 0.94351\nPOSSIBLE N-term signal sequence\ninside 1 19\nTMhelix 20 39\noutside 40 325\nTMhelix 326 348\ninside 349 372

8273 GCF_001566675.1_ASM156667v1 Escherichia coli Proteobacteria; Gammaproteobacteria; Enterobacterales; Enterobacteriaceae; Escherichia
MTGDNTLIHSHGINRRDFMKLCAALAATMGLSSKAAAEMVESVTNPQRPPVIWIGAQECT WP_000145423.1
hydrogenase 2 small subunit [Escherichia coli] Length: 372\nNumber of predicted TMHs: 1\nExp number of AAs in TMHs: 23.78201\nExp number, first 60 AAs: 0.28355\nTotal prob of N-in: 0.13257\noutside 1 331\nTMhelix 332 354\ninside 355 372

8274 GCF_001571505.1_ASM157150v1 Escherichia coli Proteobacteria; Gammaproteobacteria; Enterobacterales; Enterobacteriaceae; Escherichia
MTGDNTLIHSHGINRRDFMKLCAALAATMGLSSKAAAEMAESVTNPQRPPVIWIGAQECT WP_000145410.1
MULTISPECIES: hydrogenase-2 small chain [Proteobacteria] Length: 372\nNumber of predicted TMHs: 1\nExp number of AAs in TMHs: 23.72226\nExp number, first 60 AAs: 0.22038\nTotal prob of N-in: 0.12994\noutside 1 331\nTMhelix 332 354\ninside 355 372

8275 GCF_001571505.1_ASM157150v1 Escherichia coli Proteobacteria; Gammaproteobacteria; Enterobacterales; Enterobacteriaceae; Escherichia
MNNEETFYQAMRRQGVTRRSFLKYCSLAATSLGLGAGMAPKIAWALENKPRIPVVWIHGL WP_001058323.1
MULTISPECIES: hydrogenase-1 small chain [Proteobacteria] Length: 372\nNumber of predicted TMHs: 2\nExp number of AAs in TMHs: 37.2608499999999\nExp number, first 60 AAs: 19.52923\nTotal prob of N-in: 0.94351\nPOSSIBLE N-term signal sequence\ninside 1 19\nTMhelix 20 39\noutside 40 325\nTMhelix 326 348\ninside 349 372

8276 GCF_001566675.1_ASM156667v1 Escherichia coli Proteobacteria; Gammaproteobacteria; Enterobacterales; Enterobacteriaceae; Escherichia
MNNEETFYQAMRRQGVTRRSFLKYCSLAATSLGLGAGMAPKIAWALENKPRIPVVWIHGL WP_001295940.1
MULTISPECIES: hydrogenase [Enterobacteriaceae] Length: 372\nNumber of predicted TMHs: 2\nExp number of AAs in TMHs: 37.2608899999999\nExp number, first 60 AAs: 19.52923\nTotal prob of N-in: 0.94351\nPOSSIBLE N-term signal sequence\ninside 1 19\nTMhelix 20 39\noutside 40 325\nTMhelix 326 348\ninside 349 372

8277 GCF_001561895.1_ASM156189v1 Escherichia coli Proteobacteria; Gammaproteobacteria; Enterobacterales; Enterobacteriaceae; Escherichia
MNNEETFYQAMRRQGVTRRSFLKYCSLAATSLGLGAGMAPKIAWALENKPRIPVVWIHGL WP_001295940.1
MULTISPECIES: hydrogenase [Enterobacteriaceae] Length: 372\nNumber of predicted TMHs: 2\nExp number of AAs in TMHs: 37.2608899999999\nExp number, first 60 AAs: 19.52923\nTotal prob of N-in: 0.94351\nPOSSIBLE N-term signal sequence\ninside 1 19\nTMhelix 20 39\noutside 40 325\nTMhelix 326 348\ninside 349 372

8278 GCF_001562345.1_ASM156234v1 Escherichia coli Proteobacteria; Gammaproteobacteria; Enterobacterales; Enterobacteriaceae; Escherichia
MNNEETFYQAMRRQGVTRRSFLKYCSLAATSLGLGAGMAPKIAWALENKPRIPVVWIHGL WP_001295940.1
MULTISPECIES: hydrogenase [Enterobacteriaceae] Length: 372\nNumber of predicted TMHs: 2\nExp number of AAs in TMHs: 37.2608899999999\nExp number, first 60 AAs: 19.52923\nTotal prob of N-in: 0.94351\nPOSSIBLE N-term signal sequence\ninside 1 19\nTMhelix 20 39\noutside 40 325\nTMhelix 326 348\ninside 349 372

8279 GCF_001562385.1_ASM156238v1 Escherichia coli Proteobacteria; Gammaproteobacteria; Enterobacterales; Enterobacteriaceae; Escherichia
MNNEETFYQAMRRQGVTRRSFLKYCSLAATSLGLGAGMAPKIAWALENKPRIPVVWIHGL WP_001295940.1
MULTISPECIES: hydrogenase [Enterobacteriaceae] Length: 372\nNumber of predicted TMHs: 2\nExp number of AAs in TMHs: 37.2608899999999\nExp number, first 60 AAs: 19.52923\nTotal prob of N-in: 0.94351\nPOSSIBLE N-term signal sequence\ninside 1 19\nTMhelix 20 39\noutside 40 325\nTMhelix 326 348\ninside 349 372

8280 GCF_001563395.1_ASM156339v1 Escherichia coli Proteobacteria; Gammaproteobacteria; Enterobacterales; Enterobacteriaceae; Escherichia
MNNEETFYQAMRRQGVTRRSFLKYCSLAATSLGLGAGMAPKIAWALENKPRIPVVWIHGL WP_001295940.1
MULTISPECIES: hydrogenase [Enterobacteriaceae] Length: 372\nNumber of predicted TMHs: 2\nExp number of AAs in TMHs: 37.2608899999999\nExp number, first 60 AAs: 19.52923\nTotal prob of N-in: 0.94351\nPOSSIBLE N-term signal sequence\ninside 1 19\nTMhelix 20 39\noutside 40 325\nTMhelix 326 348\ninside 349 372

8281 GCF_001571685.1_ASM157168v1 Escherichia coli Proteobacteria; Gammaproteobacteria; Enterobacterales; Enterobacteriaceae; Escherichia
MTGDNTLIHSHGINRRDFMKLCAALAATMGLSSKAAAEMAESVTNPQRPPVIWIGAQECT WP_000145410.1
MULTISPECIES: hydrogenase-2 small chain [Proteobacteria] Length: 372\nNumber of predicted TMHs: 1\nExp number of AAs in TMHs: 23.72226\nExp number, first 60 AAs: 0.22038\nTotal prob of N-in: 0.12994\noutside 1 331\nTMhelix 332 354\ninside 355 372

8282 GCF_001571575.1_ASM157157v1 Escherichia coli Proteobacteria; Gammaproteobacteria; Enterobacterales; Enterobacteriaceae; Escherichia
MTGDNTLIHSHGINRRDFMKLCAALAATMGLSSKAAAEMAESVTNPQRPPVIWIGAQECT WP_000145410.1
MULTISPECIES: hydrogenase-2 small chain [Proteobacteria] Length: 372\nNumber of predicted TMHs: 1\nExp number of AAs in TMHs: 23.72226\nExp number, first 60 AAs: 0.22038\nTotal prob of N-in: 0.12994\noutside 1 331\nTMhelix 332 354\ninside 355 372

8283 GCF_001571665.1_ASM157166v1 Escherichia coli Proteobacteria; Gammaproteobacteria; Enterobacterales; Enterobacteriaceae; Escherichia
MTGDNTLIHSHGINRRDFMKLCAALAATMGLSSKAAAEMAESVTNPQRPPVIWIGAQECT WP_000145410.1
MULTISPECIES: hydrogenase-2 small chain [Proteobacteria] Length: 372\nNumber of predicted TMHs: 1\nExp number of AAs in TMHs: 23.72226\nExp number, first 60 AAs: 0.22038\nTotal prob of N-in: 0.12994\noutside 1 331\nTMhelix 332 354\ninside 355 372

8284 GCF_001571755.1_ASM157175v1 Escherichia coli Proteobacteria; Gammaproteobacteria; Enterobacterales; Enterobacteriaceae; Escherichia
MTGDNTLIHSHGINRRDFMKLCAALAATMGLSSKAAAEMAESVTNPQRPPVIWIGAQECT WP_000145410.1
MULTISPECIES: hydrogenase-2 small chain [Proteobacteria] Length: 372\nNumber of predicted TMHs: 1\nExp number of AAs in TMHs: 23.72226\nExp number, first 60 AAs: 0.22038\nTotal prob of N-in: 0.12994\noutside 1 331\nTMhelix 332 354\ninside 355 372

8285 GCF_001571885.1_ASM157188v1 Escherichia coli Proteobacteria; Gammaproteobacteria; Enterobacterales; Enterobacteriaceae; Escherichia
MTGDNTLIHSHGINRRDFMKLCAALAATMGLSSKAAAEMAESVTNPQRPPVIWIGAQECT WP_000145410.1
MULTISPECIES: hydrogenase-2 small chain [Proteobacteria] Length: 372\nNumber of predicted TMHs: 1\nExp number of AAs in TMHs: 23.72226\nExp number, first 60 AAs: 0.22038\nTotal prob of N-in: 0.12994\noutside 1 331\nTMhelix 332 354\ninside 355 372

8286 GCF_001571835.1_ASM157183v1 Escherichia coli Proteobacteria; Gammaproteobacteria; Enterobacterales; Enterobacteriaceae; Escherichia
MTGDNTLIHSHGINRRDFMKLCAALAATMGLSSKAAAEMAESVTNPQRPPVIWIGAQECT WP_000145410.1
MULTISPECIES: hydrogenase-2 small chain [Proteobacteria] Length: 372\nNumber of predicted TMHs: 1\nExp number of AAs in TMHs: 23.72226\nExp number, first 60 AAs: 0.22038\nTotal prob of N-in: 0.12994\noutside 1 331\nTMhelix 332 354\ninside 355 372

8287 GCF_001571825.1_ASM157182v1 Escherichia coli Proteobacteria; Gammaproteobacteria; Enterobacterales; Enterobacteriaceae; Escherichia
MTGDNTLIHSHGINRRDFMKLCAALAATMGLSSKAAAEMAESVTNPQRPPVIWIGAQECT WP_000145410.1
MULTISPECIES: hydrogenase-2 small chain [Proteobacteria] Length: 372\nNumber of predicted TMHs: 1\nExp number of AAs in TMHs: 23.72226\nExp number, first 60 AAs: 0.22038\nTotal prob of N-in: 0.12994\noutside 1 331\nTMhelix 332 354\ninside 355 372

8288 GCF_001571685.1_ASM157168v1 Escherichia coli Proteobacteria; Gammaproteobacteria; Enterobacterales; Enterobacteriaceae; Escherichia
MNNEETFYQAMRRQGVTRRSFLKYCSLAATSLGLGAGMAPKIAWALENKPRIPVVWIHGL WP_001058323.1
MULTISPECIES: hydrogenase-1 small chain [Proteobacteria] Length: 372\nNumber of predicted TMHs: 2\nExp number of AAs in TMHs: 37.2608499999999\nExp number, first 60 AAs: 19.52923\nTotal prob of N-in: 0.94351\nPOSSIBLE N-term signal sequence\ninside 1 19\nTMhelix 20 39\noutside 40 325\nTMhelix 326 348\ninside 349 372

8289 GCF_001571915.1_ASM157191v1 Escherichia coli Proteobacteria; Gammaproteobacteria; Enterobacterales; Enterobacteriaceae; Escherichia
MTGDNTLIHSHGINRRDFMKLCAALAATMGLSSKAAAEMAESVTNPQRPPVIWIGAQECT WP_000145410.1
MULTISPECIES: hydrogenase-2 small chain [Proteobacteria] Length: 372\nNumber of predicted TMHs: 1\nExp number of AAs in TMHs: 23.72226\nExp number, first 60 AAs: 0.22038\nTotal prob of N-in: 0.12994\noutside 1 331\nTMhelix 332 354\ninside 355 372

8290 GCF_001572605.1_ASM157260v1 Escherichia coli Proteobacteria; Gammaproteobacteria; Enterobacterales; Enterobacteriaceae; Escherichia
MTGDNTLIHSHGINRRDFMKLCAALAATMGLSSKAAAEMAESVTNPQRPPVIWIGAQECT WP_000145410.1
MULTISPECIES: hydrogenase-2 small chain [Proteobacteria] Length: 372\nNumber of predicted TMHs: 1\nExp number of AAs in TMHs: 23.72226\nExp number, first 60 AAs: 0.22038\nTotal prob of N-in: 0.12994\noutside 1 331\nTMhelix 332 354\ninside 355 372

8291 GCF_001571915.1_ASM157191v1 Escherichia coli Proteobacteria; Gammaproteobacteria; Enterobacterales; Enterobacteriaceae; Escherichia
MNNEETFYQAMRRQGVTRRSFLKYCSLAATSLGLGAGMAPKIAWALENKPRIPVVWIHGL WP_001058323.1
MULTISPECIES: hydrogenase-1 small chain [Proteobacteria] Length: 372\nNumber of predicted TMHs: 2\nExp number of AAs in TMHs: 37.2608499999999\nExp number, first 60 AAs: 19.52923\nTotal prob of N-in: 0.94351\nPOSSIBLE N-term signal sequence\ninside 1 19\nTMhelix 20 39\noutside 40 325\nTMhelix 326 348\ninside 349 372

8292 GCF_001571575.1_ASM157157v1 Escherichia coli Proteobacteria; Gammaproteobacteria; Enterobacterales; Enterobacteriaceae; Escherichia
MNNEETFYQAMRRQGVTRRSFLKYCSLAATSLGLGAGMAPKIAWALENKPRIPVVWIHGL WP_001058323.1
MULTISPECIES: hydrogenase-1 small chain [Proteobacteria] Length: 372\nNumber of predicted TMHs: 2\nExp number of AAs in TMHs: 37.2608499999999\nExp number, first 60 AAs: 19.52923\nTotal prob of N-in: 0.94351\nPOSSIBLE N-term signal sequence\ninside 1 19\nTMhelix 20 39\noutside 40 325\nTMhelix 326 348\ninside 349 372

8293 GCF_001571985.1_ASM157198v1 Escherichia coli Proteobacteria; Gammaproteobacteria; Enterobacterales; Enterobacteriaceae; Escherichia

MTGDNTLIHSHGINRRDFMKLCAALAATMGLSSKAAAEMAESVTNPQRPPVIWIGAQECT WP_000145410.1 MULTISPECIES: hydrogenase-2 small chain [Proteobacteria] Length: 372\nNumber of predicted TMHs: 1\nExp number of AAs in TMHs: 23.72226\nExp number, first 60 AAs: 0.22038\nTotal prob of N-in: 0.12994\noutside 1 331\nTMhelix 332 354\ninside 355 372

8294 GCF_001571985.1_ASM157198v1 Escherichia coli Proteobacteria; Gammaproteobacteria; Enterobacterales; Enterobacteriaceae; Escherichia MNNEETFYQAMRRQGVTRRSFLKYCSLAATSLGLGAGMAPKIAWALENKPRIPVVWIHGL WP_001058323.1 MULTISPECIES: hydrogenase-1 small chain [Proteobacteria] Length: 372\nNumber of predicted TMHs: 2\nExp number of AAs in TMHs: 37.2608499999999\nExp number, first 60 AAs: 19.52923\nTotal prob of N-in: 0.94351\nPOSSIBLE N-term signal sequence\ninside 1 19\nTMhelix 20 39\noutside 40 325\nTMhelix 326 348\ninside 349 372

8295 GCF_001571665.1_ASM157166v1 Escherichia coli Proteobacteria; Gammaproteobacteria; Enterobacterales; Enterobacteriaceae; Escherichia MNNEETFYQAMRRQGVTRRSFLKYCSLAATSLGLGAGMAPKIAWALENKPRIPVVWIHGL WP_001058323.1 MULTISPECIES: hydrogenase-1 small chain [Proteobacteria] Length: 372\nNumber of predicted TMHs: 2\nExp number of AAs in TMHs: 37.2608499999999\nExp number, first 60 AAs: 19.52923\nTotal prob of N-in: 0.94351\nPOSSIBLE N-term signal sequence\ninside 1 19\nTMhelix 20 39\noutside 40 325\nTMhelix 326 348\ninside 349 372

8296 GCF_001572005.1_ASM157200v1 Escherichia coli Proteobacteria; Gammaproteobacteria; Enterobacterales; Enterobacteriaceae; Escherichia MTGDNTLIHSHGINRRDFMKLCAALAATMGLSSKAAAEMAESVTNPQRPPVIWIGAQECT WP_000145410.1 MULTISPECIES: hydrogenase-2 small chain [Proteobacteria] Length: 372\nNumber of predicted TMHs: 1\nExp number of AAs in TMHs: 23.72226\nExp number, first 60 AAs: 0.22038\nTotal prob of N-in: 0.12994\noutside 1 331\nTMhelix 332 354\ninside 355 372

8297 GCF_001572145.1_ASM157214v1 Escherichia coli Proteobacteria; Gammaproteobacteria; Enterobacterales; Enterobacteriaceae; Escherichia MTGDNTLIHSHGINRRDFMKLCAALAATMGLSSKAAAEMAESVTNPQRPPVIWIGAQECT WP_000145410.1 MULTISPECIES: hydrogenase-2 small chain [Proteobacteria] Length: 372\nNumber of predicted TMHs: 1\nExp number of AAs in TMHs: 23.72226\nExp number, first 60 AAs: 0.22038\nTotal prob of N-in: 0.12994\noutside 1 331\nTMhelix 332 354\ninside 355 372

8298 GCF_001572005.1_ASM157200v1 Escherichia coli Proteobacteria; Gammaproteobacteria; Enterobacterales; Enterobacteriaceae; Escherichia MNNEETFYQAMRRQGVTRRSFLKYCSLAATSLGLGAGMAPKIAWALENKPRIPVVWIHGL WP_001058323.1 MULTISPECIES: hydrogenase-1 small chain [Proteobacteria] Length: 372\nNumber of predicted TMHs: 2\nExp number of AAs in TMHs: 37.2608499999999\nExp number, first 60 AAs: 19.52923\nTotal prob of N-in: 0.94351\nPOSSIBLE N-term signal sequence\ninside 1 19\nTMhelix 20 39\noutside 40 325\nTMhelix 326 348\ninside 349 372

8299 GCF_001572145.1_ASM157214v1 Escherichia coli Proteobacteria; Gammaproteobacteria; Enterobacterales; Enterobacteriaceae; Escherichia MNNEETFYQAMRRQGVTRRSFLKYCSLAATSLGLGAGMAPKIAWALENKPRIPVVWIHGL WP_001058323.1 MULTISPECIES: hydrogenase-1 small chain [Proteobacteria] Length: 372\nNumber of predicted TMHs: 2\nExp number of AAs in TMHs: 37.2608499999999\nExp number, first 60 AAs: 19.52923\nTotal prob of N-in: 0.94351\nPOSSIBLE N-term signal sequence\ninside 1 19\nTMhelix 20 39\noutside 40 325\nTMhelix 326 348\ninside 349 372

8300 GCF_001571825.1_ASM157182v1 Escherichia coli Proteobacteria; Gammaproteobacteria; Enterobacterales; Enterobacteriaceae; Escherichia MNNEETFYQAMRRQGVTRRSFLKYCSLAATSLGLGAGMAPKIAWALENKPRIPVVWIHGL WP_001058323.1 MULTISPECIES: hydrogenase-1 small chain [Proteobacteria] Length: 372\nNumber of predicted TMHs: 2\nExp number of AAs in TMHs: 37.2608499999999\nExp number, first 60 AAs: 19.52923\nTotal prob of N-in: 0.94351\nPOSSIBLE N-term signal sequence\ninside 1 19\nTMhelix 20 39\noutside 40 325\nTMhelix 326 348\ninside 349 372

8301 GCF_001571835.1_ASM157183v1 Escherichia coli Proteobacteria; Gammaproteobacteria; Enterobacterales; Enterobacteriaceae; Escherichia

MNNEETFYQAMRRQGVTRRSFLKYCSLAATSLGLGAGMAPKIAWALENKPRIPVVWIHGL WP_001058323.1 MULTISPECIES: hydrogenase-1 small chain [Proteobacteria] Length: 372\nNumber of predicted TMHs: 2\nExp number of AAs in TMHs: 37.2608499999999\nExp number, first 60 AAs: 19.52923\nTotal prob of N-in: 0.94351\nPOSSIBLE N-term signal sequence\ninside 1 19\nTMhelix 20 39\noutside 40 325\nTMhelix 326 348\ninside 349 372

8302 GCF_001571885.1_ASM157188v1 Escherichia coli Proteobacteria; Gammaproteobacteria; Enterobacterales; Enterobacteriaceae; Escherichia MNNEETFYQAMRRQGVTRRSFLKYCSLAATSLGLGAGMAPKIAWALENKPRIPVVWIHGL WP_001058323.1 MULTISPECIES: hydrogenase-1 small chain [Proteobacteria] Length: 372\nNumber of predicted TMHs: 2\nExp number of AAs in TMHs: 37.2608499999999\nExp number, first 60 AAs: 19.52923\nTotal prob of N-in: 0.94351\nPOSSIBLE N-term signal sequence\ninside 1 19\nTMhelix 20 39\noutside 40 325\nTMhelix 326 348\ninside 349 372

8303 GCF_001572205.1_ASM157220v1 Escherichia coli Proteobacteria; Gammaproteobacteria; Enterobacterales; Enterobacteriaceae; Escherichia MTGDNTLIHSHGINRRDFMKLCAALAATMGLSSKAAAEMAESVTNPQRPPVIWIGAQECT WP_000145410.1 MULTISPECIES: hydrogenase-2 small chain [Proteobacteria] Length: 372\nNumber of predicted TMHs: 1\nExp number of AAs in TMHs: 23.72226\nExp number, first 60 AAs: 0.22038\nTotal prob of N-in: 0.12994\noutside 1 331\nTMhelix 332 354\ninside 355 372

8304 GCF_001572265.1_ASM157226v1 Escherichia coli Proteobacteria; Gammaproteobacteria; Enterobacterales; Enterobacteriaceae; Escherichia MTGDNTLIHSHGINRRDFMKLCAALAATMGLSSKAAAEMAESVTNPQRPPVIWIGAQECT WP_000145410.1 MULTISPECIES: hydrogenase-2 small chain [Proteobacteria] Length: 372\nNumber of predicted TMHs: 1\nExp number of AAs in TMHs: 23.72226\nExp number, first 60 AAs: 0.22038\nTotal prob of N-in: 0.12994\noutside 1 331\nTMhelix 332 354\ninside 355 372

8305 GCF_001572205.1_ASM157220v1 Escherichia coli Proteobacteria; Gammaproteobacteria; Enterobacterales; Enterobacteriaceae; Escherichia MNNEETFYQAMRRQGVTRRSFLKYCSLAATSLGLGAGMAPKIAWALENKPRIPVVWIHGL WP_001058323.1 MULTISPECIES: hydrogenase-1 small chain [Proteobacteria] Length: 372\nNumber of predicted TMHs: 2\nExp number of AAs in TMHs: 37.2608499999999\nExp number, first 60 AAs: 19.52923\nTotal prob of N-in: 0.94351\nPOSSIBLE N-term signal sequence\ninside 1 19\nTMhelix 20 39\noutside 40 325\nTMhelix 326 348\ninside 349 372

8306 GCF_001572295.1_ASM157229v1 Escherichia coli Proteobacteria; Gammaproteobacteria; Enterobacterales; Enterobacteriaceae; Escherichia MTGDNTLIHSHGINRRDFMKLCAALAATMGLSSKAAAEMAESVTNPQRPPVIWIGAQECT WP_000145410.1 MULTISPECIES: hydrogenase-2 small chain [Proteobacteria] Length: 372\nNumber of predicted TMHs: 1\nExp number of AAs in TMHs: 23.72226\nExp number, first 60 AAs: 0.22038\nTotal prob of N-in: 0.12994\noutside 1 331\nTMhelix 332 354\ninside 355 372

8307 GCF_001572365.1_ASM157236v1 Escherichia coli Proteobacteria; Gammaproteobacteria; Enterobacterales; Enterobacteriaceae; Escherichia MTGDNTLIHSHGINRRDFMKLCAALAATMGLSSKAAAEMAESVTNPQRPPVIWIGAQECT WP_000145410.1 MULTISPECIES: hydrogenase-2 small chain [Proteobacteria] Length: 372\nNumber of predicted TMHs: 1\nExp number of AAs in TMHs: 23.72226\nExp number, first 60 AAs: 0.22038\nTotal prob of N-in: 0.12994\noutside 1 331\nTMhelix 332 354\ninside 355 372

8308 GCF_001572405.1_ASM157240v1 Escherichia coli Proteobacteria; Gammaproteobacteria; Enterobacterales; Enterobacteriaceae; Escherichia MTGDNTLIHSHGINRRDFMKLCAALAATMGLSSKAAAEMAESVTNPQRPPVIWIGAQECT WP_000145410.1 MULTISPECIES: hydrogenase-2 small chain [Proteobacteria] Length: 372\nNumber of predicted TMHs: 1\nExp number of AAs in TMHs: 23.72226\nExp number, first 60 AAs: 0.22038\nTotal prob of N-in: 0.12994\noutside 1 331\nTMhelix 332 354\ninside 355 372

8309 GCF_001572455.1_ASM157245v1 Escherichia coli Proteobacteria; Gammaproteobacteria; Enterobacterales; Enterobacteriaceae; Escherichia MTGDNTLIHSHGINRRDFMKLCAALAATMGLSSKAAAEMAESVTNPQRPPVIWIGAQECT WP_000145410.1 MULTISPECIES: hydrogenase-2 small chain [Proteobacteria] Length: 372\nNumber of predicted TMHs:

1\nExp number of AAs in TMHs: 23.72226\nExp number, first 60 AAs: 0.22038\nTotal prob of N-in: 0.12994\noutside 1 331\nTMhelix 332 354\ninside 355 372

8310 GCF_001572505.1_ASM157250v1 Escherichia coli Proteobacteria; Gammaproteobacteria; Enterobacterales; Enterobacteriaceae; Escherichia
MTGDNTLIHSHGINRRDFMKLCAALAATMGLSSKAAAEMAESVTNPQRPPVIWIGAQECT WP_000145410.1
MULTISPECIES: hydrogenase-2 small chain [Proteobacteria] Length: 372\nNumber of predicted TMHs: 1\nExp number of AAs in TMHs: 23.72226\nExp number, first 60 AAs: 0.22038\nTotal prob of N-in: 0.12994\noutside 1 331\nTMhelix 332 354\ninside 355 372

8311 GCF_001572565.1_ASM157256v1 Escherichia coli Proteobacteria; Gammaproteobacteria; Enterobacterales; Enterobacteriaceae; Escherichia
MTGDNTLIHSHGINRRDFMKLCAALAATMGLSSKAAAEMAESVTNPQRPPVIWIGAQECT WP_000145410.1
MULTISPECIES: hydrogenase-2 small chain [Proteobacteria] Length: 372\nNumber of predicted TMHs: 1\nExp number of AAs in TMHs: 23.72226\nExp number, first 60 AAs: 0.22038\nTotal prob of N-in: 0.12994\noutside 1 331\nTMhelix 332 354\ninside 355 372

8312 GCF_001572455.1_ASM157245v1 Escherichia coli Proteobacteria; Gammaproteobacteria; Enterobacterales; Enterobacteriaceae; Escherichia
MNNEETFYQAMRRQGVTRRSFLKYCSLAATSLGLGAGMAPKIAWALENKPRIPVVWIHGL WP_001058323.1
MULTISPECIES: hydrogenase-1 small chain [Proteobacteria] Length: 372\nNumber of predicted TMHs: 2\nExp number of AAs in TMHs: 37.2608499999999\nExp number, first 60 AAs: 19.52923\nTotal prob of N-in: 0.94351\nPOSSIBLE N-term signal sequence\ninside 1 19\nTMhelix 20 39\noutside 40 325\nTMhelix 326 348\ninside 349 372

8313 GCF_001572565.1_ASM157256v1 Escherichia coli Proteobacteria; Gammaproteobacteria; Enterobacterales; Enterobacteriaceae; Escherichia
MNNEETFYQAMRRQGVTRRSFLKYCSLAATSLGLGAGMAPKIAWALENKPRIPVVWIHGL WP_001058323.1
MULTISPECIES: hydrogenase-1 small chain [Proteobacteria] Length: 372\nNumber of predicted TMHs: 2\nExp number of AAs in TMHs: 37.2608499999999\nExp number, first 60 AAs: 19.52923\nTotal prob of N-in: 0.94351\nPOSSIBLE N-term signal sequence\ninside 1 19\nTMhelix 20 39\noutside 40 325\nTMhelix 326 348\ninside 349 372

8314 GCF_001572505.1_ASM157250v1 Escherichia coli Proteobacteria; Gammaproteobacteria; Enterobacterales; Enterobacteriaceae; Escherichia
MNNEETFYQAMRRQGVTRRSFLKYCSLAATSLGLGAGMAPKIAWALENKPRIPVVWIHGL WP_001058323.1
MULTISPECIES: hydrogenase-1 small chain [Proteobacteria] Length: 372\nNumber of predicted TMHs: 2\nExp number of AAs in TMHs: 37.2608499999999\nExp number, first 60 AAs: 19.52923\nTotal prob of N-in: 0.94351\nPOSSIBLE N-term signal sequence\ninside 1 19\nTMhelix 20 39\noutside 40 325\nTMhelix 326 348\ninside 349 372

8315 GCF_001572265.1_ASM157226v1 Escherichia coli Proteobacteria; Gammaproteobacteria; Enterobacterales; Enterobacteriaceae; Escherichia
MNNEETFYQAMRRQGVTRRSFLKYCSLAATSLGLGAGMAPKIAWALENKPRIPVVWIHGL WP_001058323.1
MULTISPECIES: hydrogenase-1 small chain [Proteobacteria] Length: 372\nNumber of predicted TMHs: 2\nExp number of AAs in TMHs: 37.2608499999999\nExp number, first 60 AAs: 19.52923\nTotal prob of N-in: 0.94351\nPOSSIBLE N-term signal sequence\ninside 1 19\nTMhelix 20 39\noutside 40 325\nTMhelix 326 348\ninside 349 372

8316 GCF_001572295.1_ASM157229v1 Escherichia coli Proteobacteria; Gammaproteobacteria; Enterobacterales; Enterobacteriaceae; Escherichia
MNNEETFYQAMRRQGVTRRSFLKYCSLAATSLGLGAGMAPKIAWALENKPRIPVVWIHGL WP_001058323.1
MULTISPECIES: hydrogenase-1 small chain [Proteobacteria] Length: 372\nNumber of predicted TMHs: 2\nExp number of AAs in TMHs: 37.2608499999999\nExp number, first 60 AAs: 19.52923\nTotal prob of N-in: 0.94351\nPOSSIBLE N-term signal sequence\ninside 1 19\nTMhelix 20 39\noutside 40 325\nTMhelix 326 348\ninside 349 372

8317 GCF_001575275.1_ASM157527v1 Escherichia coli Proteobacteria; Gammaproteobacteria; Enterobacterales; Enterobacteriaceae; Escherichia
MTGDNTLIHSHGINRRDFMKLCAALAATMGLSSKAAAEMAESVTNPQRPPVIWIGAQECT WP_000145410.1
MULTISPECIES: hydrogenase-2 small chain [Proteobacteria] Length: 372\nNumber of predicted TMHs:

1\nExp number of AAs in TMHs: 23.72226\nExp number, first 60 AAs: 0.22038\nTotal prob of N-in: 0.12994\noutside 1 331\nTMhelix 332 354\ninside 355 372

8318 GCF_001575275.1_ASM157527v1 Escherichia coli Proteobacteria; Gammaproteobacteria; Enterobacterales; Enterobacteriaceae; Escherichia MNNEETFYQAMRRQGVTRRSFLKYCSLAATSLGLGAGMAPKIAWALENKPRIPVVWIHGL WP_001058323.1 MULTISPECIES: hydrogenase-1 small chain [Proteobacteria] Length: 372\nNumber of predicted TMHs: 2\nExp number of AAs in TMHs: 37.2608499999999\nExp number, first 60 AAs: 19.52923\nTotal prob of N-in: 0.94351\nPOSSIBLE N-term signal sequence\ninside 1 19\nTMhelix 20 39\noutside 40 325\nTMhelix 326 348\ninside 349 372

8319 GCF_001572365.1_ASM157236v1 Escherichia coli Proteobacteria; Gammaproteobacteria; Enterobacterales; Enterobacteriaceae; Escherichia MNNEETFYQAMRRQGVTRRSFLKYCSLAATSLGLGAGMAPKIAWALENKPRIPVVWIHGL WP_001058323.1 MULTISPECIES: hydrogenase-1 small chain [Proteobacteria] Length: 372\nNumber of predicted TMHs: 2\nExp number of AAs in TMHs: 37.2608499999999\nExp number, first 60 AAs: 19.52923\nTotal prob of N-in: 0.94351\nPOSSIBLE N-term signal sequence\ninside 1 19\nTMhelix 20 39\noutside 40 325\nTMhelix 326 348\ninside 349 372

8320 GCF_001575295.1_ASM157529v1 Escherichia coli Proteobacteria; Gammaproteobacteria; Enterobacterales; Enterobacteriaceae; Escherichia MTGDNTLIHSHGINRRDFMKLCAALAATMGLSSKAAAEMAESVTNPQRPPVIWIGAQECT WP_000145410.1 MULTISPECIES: hydrogenase-2 small chain [Proteobacteria] Length: 372\nNumber of predicted TMHs: 1\nExp number of AAs in TMHs: 23.72226\nExp number, first 60 AAs: 0.22038\nTotal prob of N-in: 0.12994\noutside 1 331\nTMhelix 332 354\ninside 355 372

8321 GCF_001575305.1_ASM157530v1 Escherichia coli Proteobacteria; Gammaproteobacteria; Enterobacterales; Enterobacteriaceae; Escherichia MTGDNTLIHSHGINRRDFMKLCAALAATMGLSSKAAAEMAESVTNPQRPPVIWIGAQECT WP_000145410.1 MULTISPECIES: hydrogenase-2 small chain [Proteobacteria] Length: 372\nNumber of predicted TMHs: 1\nExp number of AAs in TMHs: 23.72226\nExp number, first 60 AAs: 0.22038\nTotal prob of N-in: 0.12994\noutside 1 331\nTMhelix 332 354\ninside 355 372

8322 GCF_001575305.1_ASM157530v1 Escherichia coli Proteobacteria; Gammaproteobacteria; Enterobacterales; Enterobacteriaceae; Escherichia MNNEETFYQAMRRQGVTRRSFLKYCSLAATSLGLGAGMAPKIAWALENKPRIPVVWIHGL WP_001058323.1 MULTISPECIES: hydrogenase-1 small chain [Proteobacteria] Length: 372\nNumber of predicted TMHs: 2\nExp number of AAs in TMHs: 37.2608499999999\nExp number, first 60 AAs: 19.52923\nTotal prob of N-in: 0.94351\nPOSSIBLE N-term signal sequence\ninside 1 19\nTMhelix 20 39\noutside 40 325\nTMhelix 326 348\ninside 349 372

8323 GCF_001575295.1_ASM157529v1 Escherichia coli Proteobacteria; Gammaproteobacteria; Enterobacterales; Enterobacteriaceae; Escherichia MNNEETFYQAMRRQGVTRRSFLKYCSLAATSLGLGAGMAPKIAWALENKPRIPVVWIHGL WP_001058323.1 MULTISPECIES: hydrogenase-1 small chain [Proteobacteria] Length: 372\nNumber of predicted TMHs: 2\nExp number of AAs in TMHs: 37.2608499999999\nExp number, first 60 AAs: 19.52923\nTotal prob of N-in: 0.94351\nPOSSIBLE N-term signal sequence\ninside 1 19\nTMhelix 20 39\noutside 40 325\nTMhelix 326 348\ninside 349 372

8324 GCF_001572405.1_ASM157240v1 Escherichia coli Proteobacteria; Gammaproteobacteria; Enterobacterales; Enterobacteriaceae; Escherichia MNNEETFYQAMRRQGVTRRSFLKYCSLAATSLGLGAGMAPKIAWALENKPRIPVVWIHGL WP_001058323.1 MULTISPECIES: hydrogenase-1 small chain [Proteobacteria] Length: 372\nNumber of predicted TMHs: 2\nExp number of AAs in TMHs: 37.2608499999999\nExp number, first 60 AAs: 19.52923\nTotal prob of N-in: 0.94351\nPOSSIBLE N-term signal sequence\ninside 1 19\nTMhelix 20 39\noutside 40 325\nTMhelix 326 348\ninside 349 372

8325 GCF_001575355.1_ASM157535v1 Escherichia coli Proteobacteria; Gammaproteobacteria; Enterobacterales; Enterobacteriaceae; Escherichia MTGDNTLIHSHGINRRDFMKLCAALAATMGLSSKAAAEMAESVTNPQRPPVIWIGAQECT WP_000145410.1 MULTISPECIES: hydrogenase-2 small chain [Proteobacteria] Length: 372\nNumber of predicted TMHs:

1\nExp number of AAs in TMHs: 23.72226\nExp number, first 60 AAs: 0.22038\nTotal prob of N-in: 0.12994\noutside 1 331\nTMhelix 332 354\ninside 355 372

8326 GCF_001575355.1_ASM157535v1 Escherichia coli Proteobacteria; Gammaproteobacteria; Enterobacterales; Enterobacteriaceae; Escherichia MNNEETFYQAMRRQGVTRRSFLKYCSLAATSLGLGAGMAPKIAWALENKPRIPVVWIHGL WP_001058323.1 MULTISPECIES: hydrogenase-1 small chain [Proteobacteria] Length: 372\nNumber of predicted TMHs: 2\nExp number of AAs in TMHs: 37.2608499999999\nExp number, first 60 AAs: 19.52923\nTotal prob of N-in: 0.94351\nPOSSIBLE N-term signal sequence\ninside 1 19\nTMhelix 20 39\noutside 40 325\nTMhelix 326 348\ninside 349 372

8327 GCF_001572605.1_ASM157260v1 Escherichia coli Proteobacteria; Gammaproteobacteria; Enterobacterales; Enterobacteriaceae; Escherichia MNNEETFYQAMRRQGVTRRSFLKYCSLAATSLGLGAGMAPKIAWALENKPRIPVVWIHGL WP_001058323.1 MULTISPECIES: hydrogenase-1 small chain [Proteobacteria] Length: 372\nNumber of predicted TMHs: 2\nExp number of AAs in TMHs: 37.2608499999999\nExp number, first 60 AAs: 19.52923\nTotal prob of N-in: 0.94351\nPOSSIBLE N-term signal sequence\ninside 1 19\nTMhelix 20 39\noutside 40 325\nTMhelix 326 348\ninside 349 372

8328 GCF_001575385.1_ASM157538v1 Escherichia coli Proteobacteria; Gammaproteobacteria; Enterobacterales; Enterobacteriaceae; Escherichia MTGDNTLIHSHGINRRDFMKLCAALAATMGLSSKAAAEMAESVTNPQRPPVIWIGAQECT WP_000145410.1 MULTISPECIES: hydrogenase-2 small chain [Proteobacteria] Length: 372\nNumber of predicted TMHs: 1\nExp number of AAs in TMHs: 23.72226\nExp number, first 60 AAs: 0.22038\nTotal prob of N-in: 0.12994\noutside 1 331\nTMhelix 332 354\ninside 355 372

8329 GCF_001575435.1_ASM157543v1 Escherichia coli Proteobacteria; Gammaproteobacteria; Enterobacterales; Enterobacteriaceae; Escherichia MTGDNTLIHSHGINRRDFMKLCAALAATMGLSSKAAAEMAESVTNPQRPPVIWIGAQECT WP_000145410.1 MULTISPECIES: hydrogenase-2 small chain [Proteobacteria] Length: 372\nNumber of predicted TMHs: 1\nExp number of AAs in TMHs: 23.72226\nExp number, first 60 AAs: 0.22038\nTotal prob of N-in: 0.12994\noutside 1 331\nTMhelix 332 354\ninside 355 372

8330 GCF_001575535.1_ASM157553v1 Escherichia coli Proteobacteria; Gammaproteobacteria; Enterobacterales; Enterobacteriaceae; Escherichia MTGDNTLIHSHGINRRDFMKLCAALAATMGLSSKAAAEMAESVTNPQRPPVIWIGAQECT WP_000145410.1 MULTISPECIES: hydrogenase-2 small chain [Proteobacteria] Length: 372\nNumber of predicted TMHs: 1\nExp number of AAs in TMHs: 23.72226\nExp number, first 60 AAs: 0.22038\nTotal prob of N-in: 0.12994\noutside 1 331\nTMhelix 332 354\ninside 355 372

8331 GCF_001575625.1_ASM157562v1 Escherichia coli Proteobacteria; Gammaproteobacteria; Enterobacterales; Enterobacteriaceae; Escherichia MTGDNTLIHSHGINRRDFMKLCAALAATMGLSSKAAAEMAESVTNPQRPPVIWIGAQECT WP_000145410.1 MULTISPECIES: hydrogenase-2 small chain [Proteobacteria] Length: 372\nNumber of predicted TMHs: 1\nExp number of AAs in TMHs: 23.72226\nExp number, first 60 AAs: 0.22038\nTotal prob of N-in: 0.12994\noutside 1 331\nTMhelix 332 354\ninside 355 372

8332 GCF_001575475.1_ASM157547v1 Escherichia coli Proteobacteria; Gammaproteobacteria; Enterobacterales; Enterobacteriaceae; Escherichia MTGDNTLIHSHGINRRDFMKLCAALAATMGLSSKAAAEMAESVTNPQRPPVIWIGAQECT WP_000145410.1 MULTISPECIES: hydrogenase-2 small chain [Proteobacteria] Length: 372\nNumber of predicted TMHs: 1\nExp number of AAs in TMHs: 23.72226\nExp number, first 60 AAs: 0.22038\nTotal prob of N-in: 0.12994\noutside 1 331\nTMhelix 332 354\ninside 355 372

8333 GCF_001575595.1_ASM157559v1 Escherichia coli Proteobacteria; Gammaproteobacteria; Enterobacterales; Enterobacteriaceae; Escherichia MTGDNTLIHSHGINRRDFMKLCAALAATMGLSSKAAAEMAESVTNPQRPPVIWIGAQECT WP_000145410.1 MULTISPECIES: hydrogenase-2 small chain [Proteobacteria] Length: 372\nNumber of predicted TMHs: 1\nExp number of AAs in TMHs: 23.72226\nExp number, first 60 AAs: 0.22038\nTotal prob of N-in: 0.12994\noutside 1 331\nTMhelix 332 354\ninside 355 372

8334 GCF_001575715.1_ASM157571v1 Escherichia coli Proteobacteria; Gammaproteobacteria; Enterobacterales; Enterobacteriaceae; Escherichia
MTGDNTLIHSHGINRRDFMKLCAALAATMGLSSKAAAEMAESVTNPQRPPVIWIGAQECT WP_000145410.1
MULTISPECIES: hydrogenase-2 small chain [Proteobacteria] Length: 372\nNumber of predicted TMHs: 1\nExp number of AAs in TMHs: 23.72226\nExp number, first 60 AAs: 0.22038\nTotal prob of N-in: 0.12994\noutside 1 331\nTMhelix 332 354\ninside 355 372

8335 GCF_001575695.1_ASM157569v1 Escherichia coli Proteobacteria; Gammaproteobacteria; Enterobacterales; Enterobacteriaceae; Escherichia
MTGDNTLIHSHGINRRDFMKLCAALAATMGLSSKAAAEMAESVTNPQRPPVIWIGAQECT WP_000145410.1
MULTISPECIES: hydrogenase-2 small chain [Proteobacteria] Length: 372\nNumber of predicted TMHs: 1\nExp number of AAs in TMHs: 23.72226\nExp number, first 60 AAs: 0.22038\nTotal prob of N-in: 0.12994\noutside 1 331\nTMhelix 332 354\ninside 355 372

8336 GCF_001575695.1_ASM157569v1 Escherichia coli Proteobacteria; Gammaproteobacteria; Enterobacterales; Enterobacteriaceae; Escherichia
MNNEETFYQAMRRQGVTRRSFLKYCSLAATSLGLGAGMAPKIAWALENKPRIPVVWIHGL WP_001058323.1
MULTISPECIES: hydrogenase-1 small chain [Proteobacteria] Length: 372\nNumber of predicted TMHs: 2\nExp number of AAs in TMHs: 37.2608499999999\nExp number, first 60 AAs: 19.52923\nTotal prob of N-in: 0.94351\nPOSSIBLE N-term signal sequence\ninside 1 19\nTMhelix 20 39\noutside 40 325\nTMhelix 326 348\ninside 349 372

8337 GCF_001575385.1_ASM157538v1 Escherichia coli Proteobacteria; Gammaproteobacteria; Enterobacterales; Enterobacteriaceae; Escherichia
MNNEETFYQAMRRQGVTRRSFLKYCSLAATSLGLGAGMAPKIAWALENKPRIPVVWIHGL WP_001058323.1
MULTISPECIES: hydrogenase-1 small chain [Proteobacteria] Length: 372\nNumber of predicted TMHs: 2\nExp number of AAs in TMHs: 37.2608499999999\nExp number, first 60 AAs: 19.52923\nTotal prob of N-in: 0.94351\nPOSSIBLE N-term signal sequence\ninside 1 19\nTMhelix 20 39\noutside 40 325\nTMhelix 326 348\ninside 349 372

8338 GCF_001575785.1_ASM157578v1 Escherichia coli Proteobacteria; Gammaproteobacteria; Enterobacterales; Enterobacteriaceae; Escherichia
MTGDNTLIHSHGINRRDFMKLCAALAATMGLSSKAAAEMAESVTNPQRPPVIWIGAQECT WP_000145410.1
MULTISPECIES: hydrogenase-2 small chain [Proteobacteria] Length: 372\nNumber of predicted TMHs: 1\nExp number of AAs in TMHs: 23.72226\nExp number, first 60 AAs: 0.22038\nTotal prob of N-in: 0.12994\noutside 1 331\nTMhelix 332 354\ninside 355 372

8339 GCF_001575785.1_ASM157578v1 Escherichia coli Proteobacteria; Gammaproteobacteria; Enterobacterales; Enterobacteriaceae; Escherichia
MNNEETFYQAMRRQGVTRRSFLKYCSLAATSLGLGAGMAPKIAWALENKPRIPVVWIHGL WP_001058323.1
MULTISPECIES: hydrogenase-1 small chain [Proteobacteria] Length: 372\nNumber of predicted TMHs: 2\nExp number of AAs in TMHs: 37.2608499999999\nExp number, first 60 AAs: 19.52923\nTotal prob of N-in: 0.94351\nPOSSIBLE N-term signal sequence\ninside 1 19\nTMhelix 20 39\noutside 40 325\nTMhelix 326 348\ninside 349 372

8340 GCF_001575435.1_ASM157543v1 Escherichia coli Proteobacteria; Gammaproteobacteria; Enterobacterales; Enterobacteriaceae; Escherichia
MNNEETFYQAMRRQGVTRRSFLKYCSLAATSLGLGAGMAPKIAWALENKPRIPVVWIHGL WP_001058323.1
MULTISPECIES: hydrogenase-1 small chain [Proteobacteria] Length: 372\nNumber of predicted TMHs: 2\nExp number of AAs in TMHs: 37.2608499999999\nExp number, first 60 AAs: 19.52923\nTotal prob of N-in: 0.94351\nPOSSIBLE N-term signal sequence\ninside 1 19\nTMhelix 20 39\noutside 40 325\nTMhelix 326 348\ninside 349 372

8341 GCF_001575535.1_ASM157553v1 Escherichia coli Proteobacteria; Gammaproteobacteria; Enterobacterales; Enterobacteriaceae; Escherichia
MNNEETFYQAMRRQGVTRRSFLKYCSLAATSLGLGAGMAPKIAWALENKPRIPVVWIHGL WP_001058323.1
MULTISPECIES: hydrogenase-1 small chain [Proteobacteria] Length: 372\nNumber of predicted TMHs: 2\nExp number of AAs in TMHs: 37.2608499999999\nExp number, first 60 AAs: 19.52923\nTotal prob of N-in: 0.94351\nPOSSIBLE N-term signal sequence\ninside 1 19\nTMhelix 20 39\noutside 40 325\nTMhelix 326 348\ninside 349 372

8342	GCF_001575885.1_ASM157588v1	Escherichia coli	Proteobacteria; Gammaproteobacteria; Enterobacterales; Enterobacteriaceae; Escherichia
MTGDNTLIHSHGINRRDFMKLCAALAATMGLSSKAAAEMAESVTNPQRPPVIWIGAQECT	WP_000145410.1
MULTISPECIES: hydrogenase-2 small chain [Proteobacteria]	Length: 372\nNumber of predicted TMHs: 1\nExp number of AAs in TMHs: 23.72226\nExp number, first 60 AAs: 0.22038\nTotal prob of N-in: 0.12994\noutside 1 331\nTMhelix 332 354\ninside 355 372

8343	GCF_001575835.1_ASM157583v1	Escherichia coli	Proteobacteria; Gammaproteobacteria; Enterobacterales; Enterobacteriaceae; Escherichia
MTGDNTLIHSHGINRRDFMKLCAALAATMGLSSKAAAEMAESVTNPQRPPVIWIGAQECT	WP_000145410.1
MULTISPECIES: hydrogenase-2 small chain [Proteobacteria]	Length: 372\nNumber of predicted TMHs: 1\nExp number of AAs in TMHs: 23.72226\nExp number, first 60 AAs: 0.22038\nTotal prob of N-in: 0.12994\noutside 1 331\nTMhelix 332 354\ninside 355 372

8344	GCF_001575885.1_ASM157588v1	Escherichia coli	Proteobacteria; Gammaproteobacteria; Enterobacterales; Enterobacteriaceae; Escherichia
MNNEETFYQAMRRQGVTRRSFLKYCSLAATSLGLGAGMAPKIAWALENKPRIPVVWIHGL	WP_001058323.1
MULTISPECIES: hydrogenase-1 small chain [Proteobacteria]	Length: 372\nNumber of predicted TMHs: 2\nExp number of AAs in TMHs: 37.2608499999999\nExp number, first 60 AAs: 19.52923\nTotal prob of N-in: 0.94351\nPOSSIBLE N-term signal sequence\ninside 1 19\nTMhelix 20 39\noutside 40 325\nTMhelix 326 348\ninside 349 372

8345	GCF_001575835.1_ASM157583v1	Escherichia coli	Proteobacteria; Gammaproteobacteria; Enterobacterales; Enterobacteriaceae; Escherichia
MNNEETFYQAMRRQGVTRRSFLKYCSLAATSLGLGAGMAPKIAWALENKPRIPVVWIHGL	WP_001058323.1
MULTISPECIES: hydrogenase-1 small chain [Proteobacteria]	Length: 372\nNumber of predicted TMHs: 2\nExp number of AAs in TMHs: 37.2608499999999\nExp number, first 60 AAs: 19.52923\nTotal prob of N-in: 0.94351\nPOSSIBLE N-term signal sequence\ninside 1 19\nTMhelix 20 39\noutside 40 325\nTMhelix 326 348\ninside 349 372

8346	GCF_001575715.1_ASM157571v1	Escherichia coli	Proteobacteria; Gammaproteobacteria; Enterobacterales; Enterobacteriaceae; Escherichia
MNNEETFYQAMRRQGVTRRSFLKYCSLAATSLGLGAGMAPKIAWALENKPRIPVVWIHGL	WP_001058323.1
MULTISPECIES: hydrogenase-1 small chain [Proteobacteria]	Length: 372\nNumber of predicted TMHs: 2\nExp number of AAs in TMHs: 37.2608499999999\nExp number, first 60 AAs: 19.52923\nTotal prob of N-in: 0.94351\nPOSSIBLE N-term signal sequence\ninside 1 19\nTMhelix 20 39\noutside 40 325\nTMhelix 326 348\ninside 349 372

8347	GCF_001575625.1_ASM157562v1	Escherichia coli	Proteobacteria; Gammaproteobacteria; Enterobacterales; Enterobacteriaceae; Escherichia
MNNEETFYQAMRRQGVTRRSFLKYCSLAATSLGLGAGMAPKIAWALENKPRIPVVWIHGL	WP_001058323.1
MULTISPECIES: hydrogenase-1 small chain [Proteobacteria]	Length: 372\nNumber of predicted TMHs: 2\nExp number of AAs in TMHs: 37.2608499999999\nExp number, first 60 AAs: 19.52923\nTotal prob of N-in: 0.94351\nPOSSIBLE N-term signal sequence\ninside 1 19\nTMhelix 20 39\noutside 40 325\nTMhelix 326 348\ninside 349 372

8348	GCF_001575935.1_ASM157593v1	Escherichia coli	Proteobacteria; Gammaproteobacteria; Enterobacterales; Enterobacteriaceae; Escherichia
MTGDNTLIHSHGINRRDFMKLCAALAATMGLSSKAAAEMAESVTNPQRPPVIWIGAQECT	WP_000145410.1
MULTISPECIES: hydrogenase-2 small chain [Proteobacteria]	Length: 372\nNumber of predicted TMHs: 1\nExp number of AAs in TMHs: 23.72226\nExp number, first 60 AAs: 0.22038\nTotal prob of N-in: 0.12994\noutside 1 331\nTMhelix 332 354\ninside 355 372

8349	GCF_001575935.1_ASM157593v1	Escherichia coli	Proteobacteria; Gammaproteobacteria; Enterobacterales; Enterobacteriaceae; Escherichia
MNNEETFYQAMRRQGVTRRSFLKYCSLAATSLGLGAGMAPKIAWALENKPRIPVVWIHGL	WP_001058323.1
MULTISPECIES: hydrogenase-1 small chain [Proteobacteria]	Length: 372\nNumber of predicted TMHs: 2\nExp number of AAs in TMHs: 37.2608499999999\nExp number, first 60 AAs: 19.52923\nTotal prob of N-in: 0.94351\nPOSSIBLE N-term signal sequence\ninside 1 19\nTMhelix 20 39\noutside 40 325\nTMhelix 326 348\ninside 349 372

8350	GCF_001575595.1_ASM157559v1	Escherichia coli	Proteobacteria; Gammaproteobacteria; Enterobacterales; Enterobacteriaceae; Escherichia
	MNNEETFYQAMRRQGVTRRSFLKYCSLAATSLGLGAGMAPKIAWALENKPRIPVVWIHGL	WP_001058323.1
	MULTISPECIES: hydrogenase-1 small chain [Proteobacteria]	Length: 372\nNumber of predicted TMHs: 2\nExp number of AAs in TMHs: 37.2608499999999\nExp number, first 60 AAs: 19.52923\nTotal prob of N-in: 0.94351\nPOSSIBLE N-term signal sequence\ninside 1 19\nTMhelix 20 39\noutside 40 325\nTMhelix 326 348\ninside 349 372

8351	GCF_001575475.1_ASM157547v1	Escherichia coli	Proteobacteria; Gammaproteobacteria; Enterobacterales; Enterobacteriaceae; Escherichia
	MNNEETFYQAMRRQGVTRRSFLKYCSLAATSLGLGAGMAPKIAWALENKPRIPVVWIHGL	WP_001058323.1
	MULTISPECIES: hydrogenase-1 small chain [Proteobacteria]	Length: 372\nNumber of predicted TMHs: 2\nExp number of AAs in TMHs: 37.2608499999999\nExp number, first 60 AAs: 19.52923\nTotal prob of N-in: 0.94351\nPOSSIBLE N-term signal sequence\ninside 1 19\nTMhelix 20 39\noutside 40 325\nTMhelix 326 348\ninside 349 372

8352	GCF_001575995.1_ASM157599v1	Escherichia coli	Proteobacteria; Gammaproteobacteria; Enterobacterales; Enterobacteriaceae; Escherichia
	MTGDNTLIHSHGINRRDFMKLCAALAATMGLSSKAAAEMAESVTNPQRPPVIWIGAQECT	WP_000145410.1
	MULTISPECIES: hydrogenase-2 small chain [Proteobacteria]	Length: 372\nNumber of predicted TMHs: 1\nExp number of AAs in TMHs: 23.72226\nExp number, first 60 AAs: 0.22038\nTotal prob of N-in: 0.12994\noutside 1 331\nTMhelix 332 354\ninside 355 372

8353	GCF_001576025.1_ASM157602v1	Escherichia coli	Proteobacteria; Gammaproteobacteria; Enterobacterales; Enterobacteriaceae; Escherichia
	MTGDNTLIHSHGINRRDFMKLCAALAATMGLSSKAAAEMAESVTNPQRPPVIWIGAQECT	WP_000145410.1
	MULTISPECIES: hydrogenase-2 small chain [Proteobacteria]	Length: 372\nNumber of predicted TMHs: 1\nExp number of AAs in TMHs: 23.72226\nExp number, first 60 AAs: 0.22038\nTotal prob of N-in: 0.12994\noutside 1 331\nTMhelix 332 354\ninside 355 372

8354	GCF_001576135.1_ASM157613v1	Escherichia coli	Proteobacteria; Gammaproteobacteria; Enterobacterales; Enterobacteriaceae; Escherichia
	MTGDNTLIHSHGINRRDFMKLCAALAATMGLSSKAAAEMAESVTNPQRPPVIWIGAQECT	WP_000145410.1
	MULTISPECIES: hydrogenase-2 small chain [Proteobacteria]	Length: 372\nNumber of predicted TMHs: 1\nExp number of AAs in TMHs: 23.72226\nExp number, first 60 AAs: 0.22038\nTotal prob of N-in: 0.12994\noutside 1 331\nTMhelix 332 354\ninside 355 372

8355	GCF_001576155.1_ASM157615v1	Escherichia coli	Proteobacteria; Gammaproteobacteria; Enterobacterales; Enterobacteriaceae; Escherichia
	MTGDNTLIHSHGINRRDFMKLCAALAATMGLSSKAAAEMAESVTNPQRPPVIWIGAQECT	WP_000145410.1
	MULTISPECIES: hydrogenase-2 small chain [Proteobacteria]	Length: 372\nNumber of predicted TMHs: 1\nExp number of AAs in TMHs: 23.72226\nExp number, first 60 AAs: 0.22038\nTotal prob of N-in: 0.12994\noutside 1 331\nTMhelix 332 354\ninside 355 372

8356	GCF_001576095.1_ASM157609v1	Escherichia coli	Proteobacteria; Gammaproteobacteria; Enterobacterales; Enterobacteriaceae; Escherichia
	MTGDNTLIHSHGINRRDFMKLCAALAATMGLSSKAAAEMAESVTNPQRPPVIWIGAQECT	WP_000145410.1
	MULTISPECIES: hydrogenase-2 small chain [Proteobacteria]	Length: 372\nNumber of predicted TMHs: 1\nExp number of AAs in TMHs: 23.72226\nExp number, first 60 AAs: 0.22038\nTotal prob of N-in: 0.12994\noutside 1 331\nTMhelix 332 354\ninside 355 372

8357	GCF_001576175.1_ASM157617v1	Escherichia coli	Proteobacteria; Gammaproteobacteria; Enterobacterales; Enterobacteriaceae; Escherichia
	MTGDNTLIHSHGINRRDFMKLCAALAATMGLSSKAAAEMAESVTNPQRPPVIWIGAQECT	WP_000145410.1
	MULTISPECIES: hydrogenase-2 small chain [Proteobacteria]	Length: 372\nNumber of predicted TMHs: 1\nExp number of AAs in TMHs: 23.72226\nExp number, first 60 AAs: 0.22038\nTotal prob of N-in: 0.12994\noutside 1 331\nTMhelix 332 354\ninside 355 372

8358	GCF_001576205.1_ASM157620v1	Escherichia coli	Proteobacteria; Gammaproteobacteria; Enterobacterales; Enterobacteriaceae; Escherichia
	MTGDNTLIHSHGINRRDFMKLCAALAATMGLSSKAAAEMAESVTNPQRPPVIWIGAQECT	WP_000145410.1

MULTISPECIES: hydrogenase-2 small chain [Proteobacteria] Length: 372\nNumber of predicted TMHs: 1\nExp number of AAs in TMHs: 23.72226\nExp number, first 60 AAs: 0.22038\nTotal prob of N-in: 0.12994\noutside 1 331\nTMhelix 332 354\ninside 355 372

8359 GCF_001576305.1_ASM157630v1 Escherichia coli Proteobacteria; Gammaproteobacteria; Enterobacterales; Enterobacteriaceae; Escherichia MTGDNTLIHSHGINRRDFMKLCAALAATMGLSSKAAAEMAESVTNPQRPPVIWIGAQECT WP_000145410.1 MULTISPECIES: hydrogenase-2 small chain [Proteobacteria] Length: 372\nNumber of predicted TMHs: 1\nExp number of AAs in TMHs: 23.72226\nExp number, first 60 AAs: 0.22038\nTotal prob of N-in: 0.12994\noutside 1 331\nTMhelix 332 354\ninside 355 372

8360 GCF_001576395.1_ASM157639v1 Escherichia coli Proteobacteria; Gammaproteobacteria; Enterobacterales; Enterobacteriaceae; Escherichia MTGDNTLIHSHGINRRDFMKLCAALAATMGLSSKAAAEMAESVTNPQRPPVIWIGAQECT WP_000145410.1 MULTISPECIES: hydrogenase-2 small chain [Proteobacteria] Length: 372\nNumber of predicted TMHs: 1\nExp number of AAs in TMHs: 23.72226\nExp number, first 60 AAs: 0.22038\nTotal prob of N-in: 0.12994\noutside 1 331\nTMhelix 332 354\ninside 355 372

8361 GCF_001576305.1_ASM157630v1 Escherichia coli Proteobacteria; Gammaproteobacteria; Enterobacterales; Enterobacteriaceae; Escherichia MNNEETFYQAMRRQGVTRRSFLKYCSLAATSLGLGAGMAPKIAWALENKPRIPVVWIHGL WP_001058323.1 MULTISPECIES: hydrogenase-1 small chain [Proteobacteria] Length: 372\nNumber of predicted TMHs: 2\nExp number of AAs in TMHs: 37.2608499999999\nExp number, first 60 AAs: 19.52923\nTotal prob of N-in: 0.94351\nPOSSIBLE N-term signal sequence\ninside 1 19\nTMhelix 20 39\noutside 40 325\nTMhelix 326 348\ninside 349 372

8362 GCF_001576395.1_ASM157639v1 Escherichia coli Proteobacteria; Gammaproteobacteria; Enterobacterales; Enterobacteriaceae; Escherichia MNNEETFYQAMRRQGVTRRSFLKYCSLAATSLGLGAGMAPKIAWALENKPRIPVVWIHGL WP_001058323.1 MULTISPECIES: hydrogenase-1 small chain [Proteobacteria] Length: 372\nNumber of predicted TMHs: 2\nExp number of AAs in TMHs: 37.2608499999999\nExp number, first 60 AAs: 19.52923\nTotal prob of N-in: 0.94351\nPOSSIBLE N-term signal sequence\ninside 1 19\nTMhelix 20 39\noutside 40 325\nTMhelix 326 348\ninside 349 372

8363 GCF_001575995.1_ASM157599v1 Escherichia coli Proteobacteria; Gammaproteobacteria; Enterobacterales; Enterobacteriaceae; Escherichia MNNEETFYQAMRRQGVTRRSFLKYCSLAATSLGLGAGMAPKIAWALENKPRIPVVWIHGL WP_001058323.1 MULTISPECIES: hydrogenase-1 small chain [Proteobacteria] Length: 372\nNumber of predicted TMHs: 2\nExp number of AAs in TMHs: 37.2608499999999\nExp number, first 60 AAs: 19.52923\nTotal prob of N-in: 0.94351\nPOSSIBLE N-term signal sequence\ninside 1 19\nTMhelix 20 39\noutside 40 325\nTMhelix 326 348\ninside 349 372

8364 GCF_001576025.1_ASM157602v1 Escherichia coli Proteobacteria; Gammaproteobacteria; Enterobacterales; Enterobacteriaceae; Escherichia MNNEETFYQAMRRQGVTRRSFLKYCSLAATSLGLGAGMAPKIAWALENKPRIPVVWIHGL WP_001058323.1 MULTISPECIES: hydrogenase-1 small chain [Proteobacteria] Length: 372\nNumber of predicted TMHs: 2\nExp number of AAs in TMHs: 37.2608499999999\nExp number, first 60 AAs: 19.52923\nTotal prob of N-in: 0.94351\nPOSSIBLE N-term signal sequence\ninside 1 19\nTMhelix 20 39\noutside 40 325\nTMhelix 326 348\ninside 349 372

8365 GCF_001576135.1_ASM157613v1 Escherichia coli Proteobacteria; Gammaproteobacteria; Enterobacterales; Enterobacteriaceae; Escherichia MNNEETFYQAMRRQGVTRRSFLKYCSLAATSLGLGAGMAPKIAWALENKPRIPVVWIHGL WP_001058323.1 MULTISPECIES: hydrogenase-1 small chain [Proteobacteria] Length: 372\nNumber of predicted TMHs: 2\nExp number of AAs in TMHs: 37.2608499999999\nExp number, first 60 AAs: 19.52923\nTotal prob of N-in: 0.94351\nPOSSIBLE N-term signal sequence\ninside 1 19\nTMhelix 20 39\noutside 40 325\nTMhelix 326 348\ninside 349 372

8366 GCF_001576325.1_ASM157632v1 Escherichia coli Proteobacteria; Gammaproteobacteria; Enterobacterales; Enterobacteriaceae; Escherichia MTGDNTLIHSHGINRRDFMKLCAALAATMGLSSKAAAEMAESVTNPQRPPVIWIGAQECT WP_000145410.1

MULTISPECIES: hydrogenase-2 small chain [Proteobacteria] Length: 372\nNumber of predicted TMHs: 1\nExp number of AAs in TMHs: 23.72226\nExp number, first 60 AAs: 0.22038\nTotal prob of N-in: 0.12994\noutside 1 331\nTMhelix 332 354\ninside 355 372

8367 GCF_001576325.1_ASM157632v1 Escherichia coli Proteobacteria; Gammaproteobacteria; Enterobacterales; Enterobacteriaceae; Escherichia MNNEETFYQAMRRQGVTRRSFLKYCSLAATSLGLGAGMAPKIAWALENKPRIPVVWIHGL WP_001058323.1 MULTISPECIES: hydrogenase-1 small chain [Proteobacteria] Length: 372\nNumber of predicted TMHs: 2\nExp number of AAs in TMHs: 37.2608499999999\nExp number, first 60 AAs: 19.52923\nTotal prob of N-in: 0.94351\nPOSSIBLE N-term signal sequence\ninside 1 19\nTMhelix 20 39\noutside 40 325\nTMhelix 326 348\ninside 349 372

8368 GCF_001576175.1_ASM157617v1 Escherichia coli Proteobacteria; Gammaproteobacteria; Enterobacterales; Enterobacteriaceae; Escherichia MNNEETFYQAMRRQGVTRRSFLKYCSLAATSLGLGAGMAPKIAWALENKPRIPVVWIHGL WP_001058323.1 MULTISPECIES: hydrogenase-1 small chain [Proteobacteria] Length: 372\nNumber of predicted TMHs: 2\nExp number of AAs in TMHs: 37.2608499999999\nExp number, first 60 AAs: 19.52923\nTotal prob of N-in: 0.94351\nPOSSIBLE N-term signal sequence\ninside 1 19\nTMhelix 20 39\noutside 40 325\nTMhelix 326 348\ninside 349 372

8369 GCF_001576095.1_ASM157609v1 Escherichia coli Proteobacteria; Gammaproteobacteria; Enterobacterales; Enterobacteriaceae; Escherichia MNNEETFYQAMRRQGVTRRSFLKYCSLAATSLGLGAGMAPKIAWALENKPRIPVVWIHGL WP_001058323.1 MULTISPECIES: hydrogenase-1 small chain [Proteobacteria] Length: 372\nNumber of predicted TMHs: 2\nExp number of AAs in TMHs: 37.2608499999999\nExp number, first 60 AAs: 19.52923\nTotal prob of N-in: 0.94351\nPOSSIBLE N-term signal sequence\ninside 1 19\nTMhelix 20 39\noutside 40 325\nTMhelix 326 348\ninside 349 372

8370 GCF_001576155.1_ASM157615v1 Escherichia coli Proteobacteria; Gammaproteobacteria; Enterobacterales; Enterobacteriaceae; Escherichia MNNEETFYQAMRRQGVTRRSFLKYCSLAATSLGLGAGMAPKIAWALENKPRIPVVWIHGL WP_001058323.1 MULTISPECIES: hydrogenase-1 small chain [Proteobacteria] Length: 372\nNumber of predicted TMHs: 2\nExp number of AAs in TMHs: 37.2608499999999\nExp number, first 60 AAs: 19.52923\nTotal prob of N-in: 0.94351\nPOSSIBLE N-term signal sequence\ninside 1 19\nTMhelix 20 39\noutside 40 325\nTMhelix 326 348\ninside 349 372

8371 GCF_001576455.1_ASM157645v1 Escherichia coli Proteobacteria; Gammaproteobacteria; Enterobacterales; Enterobacteriaceae; Escherichia MTGDNTLIHSHGINRRDFMKLCAALAATMGLSSKAAAEMAESVTNPQRPPVIWIGAQECT WP_000145410.1 MULTISPECIES: hydrogenase-2 small chain [Proteobacteria] Length: 372\nNumber of predicted TMHs: 1\nExp number of AAs in TMHs: 23.72226\nExp number, first 60 AAs: 0.22038\nTotal prob of N-in: 0.12994\noutside 1 331\nTMhelix 332 354\ninside 355 372

8372 GCF_001576505.1_ASM157650v1 Escherichia coli Proteobacteria; Gammaproteobacteria; Enterobacterales; Enterobacteriaceae; Escherichia MTGDNTLIHSHGINRRDFMKLCAALAATMGLSSKAAAEMAESVTNPQRPPVIWIGAQECT WP_000145410.1 MULTISPECIES: hydrogenase-2 small chain [Proteobacteria] Length: 372\nNumber of predicted TMHs: 1\nExp number of AAs in TMHs: 23.72226\nExp number, first 60 AAs: 0.22038\nTotal prob of N-in: 0.12994\noutside 1 331\nTMhelix 332 354\ninside 355 372

8373 GCF_001576505.1_ASM157650v1 Escherichia coli Proteobacteria; Gammaproteobacteria; Enterobacterales; Enterobacteriaceae; Escherichia MNNEETFYQAMRRQGVTRRSFLKYCSLAATSLGLGAGMAPKIAWALENKPRIPVVWIHGL WP_001058323.1 MULTISPECIES: hydrogenase-1 small chain [Proteobacteria] Length: 372\nNumber of predicted TMHs: 2\nExp number of AAs in TMHs: 37.2608499999999\nExp number, first 60 AAs: 19.52923\nTotal prob of N-in: 0.94351\nPOSSIBLE N-term signal sequence\ninside 1 19\nTMhelix 20 39\noutside 40 325\nTMhelix 326 348\ninside 349 372

8374 GCF_001576455.1_ASM157645v1 Escherichia coli Proteobacteria; Gammaproteobacteria; Enterobacterales; Enterobacteriaceae; Escherichia MNNEETFYQAMRRQGVTRRSFLKYCSLAATSLGLGAGMAPKIAWALENKPRIPVVWIHGL WP_001058323.1

MULTISPECIES: hydrogenase-1 small chain [Proteobacteria] Length: 372\nNumber of predicted TMHs: 2\nExp number of AAs in TMHs: 37.2608499999999\nExp number, first 60 AAs: 19.52923\nTotal prob of N-in: 0.94351\nPOSSIBLE N-term signal sequence\ninside 1 19\nTMhelix 20 39\noutside 40 325\nTMhelix 326 348\ninside 349 372

8375 GCF_001576205.1_ASM157620v1 Escherichia coli Proteobacteria; Gammaproteobacteria; Enterobacterales; Enterobacteriaceae; Escherichia
MNNEETFYQAMRRQGVTRRSFLKYCSLAATSLGLGAGMAPKIAWALENKPRIPVVWIHGL WP_001058323.1
MULTISPECIES: hydrogenase-1 small chain [Proteobacteria] Length: 372\nNumber of predicted TMHs: 2\nExp number of AAs in TMHs: 37.2608499999999\nExp number, first 60 AAs: 19.52923\nTotal prob of N-in: 0.94351\nPOSSIBLE N-term signal sequence\ninside 1 19\nTMhelix 20 39\noutside 40 325\nTMhelix 326 348\ninside 349 372

8376 GCF_001577325.1_ASM157732v1 Escherichia coli Proteobacteria; Gammaproteobacteria; Enterobacterales; Enterobacteriaceae; Escherichia
MTGDNTLIHSHGINRRDFMKLCAALAATMGLSSKAAAEMAESVTNPQRPPVIWIGAQECT WP_000145410.1
MULTISPECIES: hydrogenase-2 small chain [Proteobacteria] Length: 372\nNumber of predicted TMHs: 1\nExp number of AAs in TMHs: 23.72226\nExp number, first 60 AAs: 0.22038\nTotal prob of N-in: 0.12994\noutside 1 331\nTMhelix 332 354\ninside 355 372

8377 GCF_001596855.1_ASM159685v1 Escherichia coli Proteobacteria; Gammaproteobacteria; Enterobacterales; Enterobacteriaceae; Escherichia
MTGDNTLIHSHGINRRDFMKLCAALAATMGLSSKAAAEMAESVTNPQRPPVIWIGAQECT WP_000145410.1
MULTISPECIES: hydrogenase-2 small chain [Proteobacteria] Length: 372\nNumber of predicted TMHs: 1\nExp number of AAs in TMHs: 23.72226\nExp number, first 60 AAs: 0.22038\nTotal prob of N-in: 0.12994\noutside 1 331\nTMhelix 332 354\ninside 355 372

8378 GCF_001601655.1_ASM160165v1 Escherichia coli Proteobacteria; Gammaproteobacteria; Enterobacterales; Enterobacteriaceae; Escherichia
MTGDNTLIHSHGINRRDFMKLCAALAATMGLSSKAAAEMVESVTNPQRPPVIWIGAQECT WP_000145423.1
hydrogenase 2 small subunit [Escherichia coli] Length: 372\nNumber of predicted TMHs: 1\nExp number of AAs in TMHs: 23.78201\nExp number, first 60 AAs: 0.28355\nTotal prob of N-in: 0.13257\noutside 1 331\nTMhelix 332 354\ninside 355 372

8379 GCF_001606355.1_ASM160635v1 Escherichia coli Proteobacteria; Gammaproteobacteria; Enterobacterales; Enterobacteriaceae; Escherichia
MTGDNTLIHSHGINRRDFMKLCAALAATMGLSSKAAAEMAESVTNPQRPPVIWIGAQECT WP_000145410.1
MULTISPECIES: hydrogenase-2 small chain [Proteobacteria] Length: 372\nNumber of predicted TMHs: 1\nExp number of AAs in TMHs: 23.72226\nExp number, first 60 AAs: 0.22038\nTotal prob of N-in: 0.12994\noutside 1 331\nTMhelix 332 354\ninside 355 372

8380 GCF_001606385.1_ASM160638v1 Escherichia coli Proteobacteria; Gammaproteobacteria; Enterobacterales; Enterobacteriaceae; Escherichia
MTGDNTLIHSHGINRRDFMKLCAALAATMGLSSKAAAEMAESVTNPQRPPVIWIGAQECT WP_000145410.1
MULTISPECIES: hydrogenase-2 small chain [Proteobacteria] Length: 372\nNumber of predicted TMHs: 1\nExp number of AAs in TMHs: 23.72226\nExp number, first 60 AAs: 0.22038\nTotal prob of N-in: 0.12994\noutside 1 331\nTMhelix 332 354\ninside 355 372

8381 GCF_001606445.1_ASM160644v1 Escherichia coli Proteobacteria; Gammaproteobacteria; Enterobacterales; Enterobacteriaceae; Escherichia
MTGDNTLIHSHGINRRDFMKLCAALAATMGLSSKAAAEMAESVTNPQRPPVIWIGAQECT WP_000145410.1
MULTISPECIES: hydrogenase-2 small chain [Proteobacteria] Length: 372\nNumber of predicted TMHs: 1\nExp number of AAs in TMHs: 23.72226\nExp number, first 60 AAs: 0.22038\nTotal prob of N-in: 0.12994\noutside 1 331\nTMhelix 332 354\ninside 355 372

8382 GCF_001606355.1_ASM160635v1 Escherichia coli Proteobacteria; Gammaproteobacteria; Enterobacterales; Enterobacteriaceae; Escherichia
MNNEETFYQAMRRQGVTRRSFLKYCSLAATSLGLGAGMAPKIAWALENKPRIPVVWIHGL WP_001058323.1
MULTISPECIES: hydrogenase-1 small chain [Proteobacteria] Length: 372\nNumber of predicted TMHs: 2\nExp number of AAs in TMHs: 37.2608499999999\nExp number, first 60 AAs: 19.52923\nTotal prob of N-in:

0.94351\nPOSSIBLE N-term signal sequence\ninside 1 19\nTMhelix 20 39\noutside 40 325\nTMhelix 326 348\ninside 349 372

8383 GCF_001606445.1_ASM160644v1 Escherichia coli Proteobacteria; Gammaproteobacteria; Enterobacterales; Enterobacteriaceae; Escherichia
MNNEETFYQAMRRQGVTRRSFLKYCSLAATSLGLGAGMAPKIAWALENKPRIPVVWIHGL WP_001058323.1
MULTISPECIES: hydrogenase-1 small chain [Proteobacteria] Length: 372\nNumber of predicted TMHs: 2\nExp number of AAs in TMHs: 37.2608499999999\nExp number, first 60 AAs: 19.52923\nTotal prob of N-in: 0.94351\nPOSSIBLE N-term signal sequence\ninside 1 19\nTMhelix 20 39\noutside 40 325\nTMhelix 326 348\ninside 349 372

8384 GCF_001601655.1_ASM160165v1 Escherichia coli Proteobacteria; Gammaproteobacteria; Enterobacterales; Enterobacteriaceae; Escherichia
MNNEETFYQAMRRQGVTRRSFLKYCSLAATSLGLGAGMAPKIAWALENKPRIPVVWIHGL WP_001295940.1
MULTISPECIES: hydrogenase [Enterobacteriaceae] Length: 372\nNumber of predicted TMHs: 2\nExp number of AAs in TMHs: 37.2608899999999\nExp number, first 60 AAs: 19.52923\nTotal prob of N-in: 0.94351\nPOSSIBLE N-term signal sequence\ninside 1 19\nTMhelix 20 39\noutside 40 325\nTMhelix 326 348\ninside 349 372

8385 GCF_001596855.1_ASM159685v1 Escherichia coli Proteobacteria; Gammaproteobacteria; Enterobacterales; Enterobacteriaceae; Escherichia
MNNEETFYQAMRRQGVTRRSFLKYCSLAATSLGLGAGMAPKIAWALENKPRIPVVWIHGL WP_001058323.1
MULTISPECIES: hydrogenase-1 small chain [Proteobacteria] Length: 372\nNumber of predicted TMHs: 2\nExp number of AAs in TMHs: 37.2608499999999\nExp number, first 60 AAs: 19.52923\nTotal prob of N-in: 0.94351\nPOSSIBLE N-term signal sequence\ninside 1 19\nTMhelix 20 39\noutside 40 325\nTMhelix 326 348\ninside 349 372

8386 GCF_001606515.1_ASM160651v1 Escherichia coli Proteobacteria; Gammaproteobacteria; Enterobacterales; Enterobacteriaceae; Escherichia
MTGDNTLIHSHGINRRDFMKLCAALAATMGLSSKAAAEMAESVTNPQRPPVIWIGAQECT WP_000145410.1
MULTISPECIES: hydrogenase-2 small chain [Proteobacteria] Length: 372\nNumber of predicted TMHs: 1\nExp number of AAs in TMHs: 23.72226\nExp number, first 60 AAs: 0.22038\nTotal prob of N-in: 0.12994\noutside 1 331\nTMhelix 332 354\ninside 355 372

8387 GCF_001606545.1_ASM160654v1 Escherichia coli Proteobacteria; Gammaproteobacteria; Enterobacterales; Enterobacteriaceae; Escherichia
MTGDNTLIHSHGINRRDFMKLCAALAATMGLSSKAAAEMAESVTNPQRPPVIWIGAQECT WP_000145410.1
MULTISPECIES: hydrogenase-2 small chain [Proteobacteria] Length: 372\nNumber of predicted TMHs: 1\nExp number of AAs in TMHs: 23.72226\nExp number, first 60 AAs: 0.22038\nTotal prob of N-in: 0.12994\noutside 1 331\nTMhelix 332 354\ninside 355 372

8388 GCF_001606515.1_ASM160651v1 Escherichia coli Proteobacteria; Gammaproteobacteria; Enterobacterales; Enterobacteriaceae; Escherichia
MNNEETFYQAMRRQGVTRRSFLKYCSLAATSLGLGAGMAPKIAWALENKPRIPVVWIHGL WP_001058323.1
MULTISPECIES: hydrogenase-1 small chain [Proteobacteria] Length: 372\nNumber of predicted TMHs: 2\nExp number of AAs in TMHs: 37.2608499999999\nExp number, first 60 AAs: 19.52923\nTotal prob of N-in: 0.94351\nPOSSIBLE N-term signal sequence\ninside 1 19\nTMhelix 20 39\noutside 40 325\nTMhelix 326 348\ninside 349 372

8389 GCF_001606545.1_ASM160654v1 Escherichia coli Proteobacteria; Gammaproteobacteria; Enterobacterales; Enterobacteriaceae; Escherichia
MNNEETFYQAMRRQGVTRRSFLKYCSLAATSLGLGAGMAPKIAWALENKPRIPVVWIHGL WP_001295940.1
MULTISPECIES: hydrogenase [Enterobacteriaceae] Length: 372\nNumber of predicted TMHs: 2\nExp number of AAs in TMHs: 37.2608899999999\nExp number, first 60 AAs: 19.52923\nTotal prob of N-in: 0.94351\nPOSSIBLE N-term signal sequence\ninside 1 19\nTMhelix 20 39\noutside 40 325\nTMhelix 326 348\ninside 349 372

8390 GCF_001577325.1_ASM157732v1 Escherichia coli Proteobacteria; Gammaproteobacteria; Enterobacterales; Enterobacteriaceae; Escherichia
MNNEETFYQAMRRQGVTRRSFLKYCSLAATSLGLGAGMAPKIAWALENKPRIPVVWIHGL WP_001295940.1
MULTISPECIES: hydrogenase [Enterobacteriaceae] Length: 372\nNumber of predicted TMHs: 2\nExp

number of AAs in TMHs: 37.2608899999999\nExp number, first 60 AAs: 19.52923\nTotal prob of N-in: 0.94351\nPOSSIBLE N-term signal sequence\ninside 1 19\nTMhelix 20 39\noutside 40 325\nTMhelix 326 348\ninside 349 372

8391 GCF_001606555.1_ASM160655v1 Escherichia coli Proteobacteria; Gammaproteobacteria; Enterobacterales; Enterobacteriaceae; Escherichia MNNEETFYQAMRRQGVTRRSFLKYCSLAATSLGLGAGMAPKIAWALENKPRIPVVWIHGL WP_001058323.1 MULTISPECIES: hydrogenase-1 small chain [Proteobacteria] Length: 372\nNumber of predicted TMHs: 2\nExp number of AAs in TMHs: 37.2608499999999\nExp number, first 60 AAs: 19.52923\nTotal prob of N-in: 0.94351\nPOSSIBLE N-term signal sequence\ninside 1 19\nTMhelix 20 39\noutside 40 325\nTMhelix 326 348\ninside 349 372

8392 GCF_001606555.1_ASM160655v1 Escherichia coli Proteobacteria; Gammaproteobacteria; Enterobacterales; Enterobacteriaceae; Escherichia MTGDNTLIHSHGINRRDFMKLCAALAATMGLSSKAAAEMAESVTNPQRPPVIWIGAQECT WP_024245965.1 hydrogenase 2 small subunit [Escherichia coli] Length: 372\nNumber of predicted TMHs: 1\nExp number of AAs in TMHs: 24.15412\nExp number, first 60 AAs: 0.21967\nTotal prob of N-in: 0.14289\noutside 1 331\nTMhelix 332 354\ninside 355 372

8393 GCF_001606385.1_ASM160638v1 Escherichia coli Proteobacteria; Gammaproteobacteria; Enterobacterales; Enterobacteriaceae; Escherichia MNNEETFYQAMRRQGVTRRSFLKYCSLAATSLGLGAGMAPKIAWALENKPRIPVVWIHGL WP_001295940.1 MULTISPECIES: hydrogenase [Enterobacteriaceae] Length: 372\nNumber of predicted TMHs: 2\nExp number of AAs in TMHs: 37.2608899999999\nExp number, first 60 AAs: 19.52923\nTotal prob of N-in: 0.94351\nPOSSIBLE N-term signal sequence\ninside 1 19\nTMhelix 20 39\noutside 40 325\nTMhelix 326 348\ninside 349 372

8394 GCF_001606705.1_ASM160670v1 Escherichia coli Proteobacteria; Gammaproteobacteria; Enterobacterales; Enterobacteriaceae; Escherichia MTGDNTLIHSHGINRRDFMKLCAALAATMGLSSKAAAEMVESVTNPQRPPVIWIGAQECT WP_000145423.1 hydrogenase 2 small subunit [Escherichia coli] Length: 372\nNumber of predicted TMHs: 1\nExp number of AAs in TMHs: 23.78201\nExp number, first 60 AAs: 0.28355\nTotal prob of N-in: 0.13257\noutside 1 331\nTMhelix 332 354\ninside 355 372

8395 GCF_001607045.1_ASM160704v1 Escherichia coli Proteobacteria; Gammaproteobacteria; Enterobacterales; Enterobacteriaceae; Escherichia MTGDNTLIHSHGINRRDFMKLCAALAATMGLSSKAAAEMVESVTNPQRPPVIWIGAQECT WP_000145423.1 hydrogenase 2 small subunit [Escherichia coli] Length: 372\nNumber of predicted TMHs: 1\nExp number of AAs in TMHs: 23.78201\nExp number, first 60 AAs: 0.28355\nTotal prob of N-in: 0.13257\noutside 1 331\nTMhelix 332 354\ninside 355 372

8396 GCF_001606815.1_ASM160681v1 Escherichia coli Proteobacteria; Gammaproteobacteria; Enterobacterales; Enterobacteriaceae; Escherichia MTGDNTLIHSHGINRRDFMKLCAALAATMGLSSKAAAEMAESVTNPQRPPVIWIGAQECT WP_000145410.1 MULTISPECIES: hydrogenase-2 small chain [Proteobacteria] Length: 372\nNumber of predicted TMHs: 1\nExp number of AAs in TMHs: 23.72226\nExp number, first 60 AAs: 0.22038\nTotal prob of N-in: 0.12994\noutside 1 331\nTMhelix 332 354\ninside 355 372

8397 GCF_001607105.1_ASM160710v1 Escherichia coli Proteobacteria; Gammaproteobacteria; Enterobacterales; Enterobacteriaceae; Escherichia MTGDNTLIHSHGINRRDFMKLCAALAATMGLSSKAAAEMAESVTNPQRPPVIWIGAQECT WP_000145410.1 MULTISPECIES: hydrogenase-2 small chain [Proteobacteria] Length: 372\nNumber of predicted TMHs: 1\nExp number of AAs in TMHs: 23.72226\nExp number, first 60 AAs: 0.22038\nTotal prob of N-in: 0.12994\noutside 1 331\nTMhelix 332 354\ninside 355 372

8398 GCF_001607155.1_ASM160715v1 Escherichia coli Proteobacteria; Gammaproteobacteria; Enterobacterales; Enterobacteriaceae; Escherichia MTGDNTLIHSHGINRRDFMKLCAALAATMGLSSKAAAEMAESVTNPQRPPVIWIGAQECT WP_000145410.1 MULTISPECIES: hydrogenase-2 small chain [Proteobacteria] Length: 372\nNumber of predicted TMHs: 1\nExp number of AAs in TMHs: 23.72226\nExp number, first 60 AAs: 0.22038\nTotal prob of N-in: 0.12994\noutside 1 331\nTMhelix 332 354\ninside 355 372

8399 GCF_001607015.1_ASM160701v1 Escherichia coli Proteobacteria; Gammaproteobacteria; Enterobacterales; Enterobacteriaceae; Escherichia
MTGDNTLIHSHGINRRDFMKLCAALAATMGLSSKAAAEMAESVTNPQRPPVIWIGAQECT WP_000145410.1
MULTISPECIES: hydrogenase-2 small chain [Proteobacteria] Length: 372\nNumber of predicted TMHs: 1\nExp number of AAs in TMHs: 23.72226\nExp number, first 60 AAs: 0.22038\nTotal prob of N-in: 0.12994\noutside 1 331\nTMhelix 332 354\ninside 355 372

8400 GCF_001606755.1_ASM160675v1 Escherichia coli Proteobacteria; Gammaproteobacteria; Enterobacterales; Enterobacteriaceae; Escherichia
MTGDNTLIHSHGINRRDFMKLCAALAATMGLSSKAAAEMAESVTNPQRPPVIWIGAQECT WP_000145410.1
MULTISPECIES: hydrogenase-2 small chain [Proteobacteria] Length: 372\nNumber of predicted TMHs: 1\nExp number of AAs in TMHs: 23.72226\nExp number, first 60 AAs: 0.22038\nTotal prob of N-in: 0.12994\noutside 1 331\nTMhelix 332 354\ninside 355 372

8401 GCF_001606915.1_ASM160691v1 Escherichia coli Proteobacteria; Gammaproteobacteria; Enterobacterales; Enterobacteriaceae; Escherichia
MTGDNTLIHSHGINRRDFMKLCAALAATMGLSSKAAAEMAESVTNPQRPPVIWIGAQECT WP_000145410.1
MULTISPECIES: hydrogenase-2 small chain [Proteobacteria] Length: 372\nNumber of predicted TMHs: 1\nExp number of AAs in TMHs: 23.72226\nExp number, first 60 AAs: 0.22038\nTotal prob of N-in: 0.12994\noutside 1 331\nTMhelix 332 354\ninside 355 372

8402 GCF_001606855.1_ASM160685v1 Escherichia coli Proteobacteria; Gammaproteobacteria; Enterobacterales; Enterobacteriaceae; Escherichia
MNNEETFYQAMRRQGVTRRSFLKYCSLAATSLGLGAGMAPKIAWALENKPRIPVVWIHGL WP_001058323.1
MULTISPECIES: hydrogenase-1 small chain [Proteobacteria] Length: 372\nNumber of predicted TMHs: 2\nExp number of AAs in TMHs: 37.2608499999999\nExp number, first 60 AAs: 19.52923\nTotal prob of N-in: 0.94351\nPOSSIBLE N-term signal sequence\ninside 1 19\nTMhelix 20 39\noutside 40 325\nTMhelix 326 348\ninside 349 372

8403 GCF_001606615.1_ASM160661v1 Escherichia coli Proteobacteria; Gammaproteobacteria; Enterobacterales; Enterobacteriaceae; Escherichia
MTGDNTLIHSHGINRRDFMKLCAALAATMGLSSKAAAEMAESVTNPQRPPVIWIGAQECT WP_000145410.1
MULTISPECIES: hydrogenase-2 small chain [Proteobacteria] Length: 372\nNumber of predicted TMHs: 1\nExp number of AAs in TMHs: 23.72226\nExp number, first 60 AAs: 0.22038\nTotal prob of N-in: 0.12994\noutside 1 331\nTMhelix 332 354\ninside 355 372

8404 GCF_001606955.1_ASM160695v1 Escherichia coli Proteobacteria; Gammaproteobacteria; Enterobacterales; Enterobacteriaceae; Escherichia
MTGDNTLIHSHGINRRDFMKLCAALAATMGLSSKAAAEMAESVTNPQRPPVIWIGAQECT WP_000145410.1
MULTISPECIES: hydrogenase-2 small chain [Proteobacteria] Length: 372\nNumber of predicted TMHs: 1\nExp number of AAs in TMHs: 23.72226\nExp number, first 60 AAs: 0.22038\nTotal prob of N-in: 0.12994\noutside 1 331\nTMhelix 332 354\ninside 355 372

8405 GCF_001606635.1_ASM160663v1 Escherichia coli Proteobacteria; Gammaproteobacteria; Enterobacterales; Enterobacteriaceae; Escherichia
MTGDNTLIHSHGINRRDFMKLCAALAATMGLSSKAAAEMAESVTNPQRPPVIWIGAQECT WP_000145410.1
MULTISPECIES: hydrogenase-2 small chain [Proteobacteria] Length: 372\nNumber of predicted TMHs: 1\nExp number of AAs in TMHs: 23.72226\nExp number, first 60 AAs: 0.22038\nTotal prob of N-in: 0.12994\noutside 1 331\nTMhelix 332 354\ninside 355 372

8406 GCF_001606615.1_ASM160661v1 Escherichia coli Proteobacteria; Gammaproteobacteria; Enterobacterales; Enterobacteriaceae; Escherichia
MNNEETFYQAMRRQGVTRRSFLKYCSLAATSLGLGAGMAPKIAWALENKPRIPVVWIHGL WP_001058323.1
MULTISPECIES: hydrogenase-1 small chain [Proteobacteria] Length: 372\nNumber of predicted TMHs: 2\nExp number of AAs in TMHs: 37.2608499999999\nExp number, first 60 AAs: 19.52923\nTotal prob of N-in: 0.94351\nPOSSIBLE N-term signal sequence\ninside 1 19\nTMhelix 20 39\noutside 40 325\nTMhelix 326 348\ninside 349 372

8407 GCF_001606915.1_ASM160691v1 Escherichia coli Proteobacteria; Gammaproteobacteria; Enterobacterales; Enterobacteriaceae; Escherichia
MNNEETFYQAMRRQGVTRRSFLKYCSLAATSLGLGAGMAPKIAWALENKPRIPVVWIHGL WP_001295940.1

MULTISPECIES: hydrogenase [Enterobacteriaceae] Length: 372\nNumber of predicted TMHs: 2\nExp number of AAs in TMHs: 37.2608899999999\nExp number, first 60 AAs: 19.52923\nTotal prob of N-in: 0.94351\nPOSSIBLE N-term signal sequence\ninside 1 19\nTMhelix 20 39\noutside 40 325\nTMhelix 326 348\ninside 349 372

8408 GCF_001606705.1_ASM160670v1 Escherichia coli Proteobacteria; Gammaproteobacteria; Enterobacterales; Enterobacteriaceae; Escherichia
MNNEETFYQAMRRQGVTRRSFLKYCSLAATSLGLGAGMAPKIAWALENKPRIPVVWIHGL WP_001295940.1
MULTISPECIES: hydrogenase [Enterobacteriaceae] Length: 372\nNumber of predicted TMHs: 2\nExp number of AAs in TMHs: 37.2608899999999\nExp number, first 60 AAs: 19.52923\nTotal prob of N-in: 0.94351\nPOSSIBLE N-term signal sequence\ninside 1 19\nTMhelix 20 39\noutside 40 325\nTMhelix 326 348\ninside 349 372

8409 GCF_001607045.1_ASM160704v1 Escherichia coli Proteobacteria; Gammaproteobacteria; Enterobacterales; Enterobacteriaceae; Escherichia
MNNEETFYQAMRRQGVTRRSFLKYCSLAATSLGLGAGMAPKIAWALENKPRIPVVWIHGL WP_001295940.1
MULTISPECIES: hydrogenase [Enterobacteriaceae] Length: 372\nNumber of predicted TMHs: 2\nExp number of AAs in TMHs: 37.2608899999999\nExp number, first 60 AAs: 19.52923\nTotal prob of N-in: 0.94351\nPOSSIBLE N-term signal sequence\ninside 1 19\nTMhelix 20 39\noutside 40 325\nTMhelix 326 348\ninside 349 372

8410 GCF_001606815.1_ASM160681v1 Escherichia coli Proteobacteria; Gammaproteobacteria; Enterobacterales; Enterobacteriaceae; Escherichia
MNNEETFYQAMRRQGVTRRSFLKYCSLAATSLGLGAGMAPKIAWALENKPRIPVVWIHGL WP_001058323.1
MULTISPECIES: hydrogenase-1 small chain [Proteobacteria] Length: 372\nNumber of predicted TMHs: 2\nExp number of AAs in TMHs: 37.2608499999999\nExp number, first 60 AAs: 19.52923\nTotal prob of N-in: 0.94351\nPOSSIBLE N-term signal sequence\ninside 1 19\nTMhelix 20 39\noutside 40 325\nTMhelix 326 348\ninside 349 372

8411 GCF_001607155.1_ASM160715v1 Escherichia coli Proteobacteria; Gammaproteobacteria; Enterobacterales; Enterobacteriaceae; Escherichia
MNNEETFYQAMRRQGVTRRSFLKYCSLAATSLGLGAGMAPKIAWALENKPRIPVVWIHGL WP_001058323.1
MULTISPECIES: hydrogenase-1 small chain [Proteobacteria] Length: 372\nNumber of predicted TMHs: 2\nExp number of AAs in TMHs: 37.2608499999999\nExp number, first 60 AAs: 19.52923\nTotal prob of N-in: 0.94351\nPOSSIBLE N-term signal sequence\ninside 1 19\nTMhelix 20 39\noutside 40 325\nTMhelix 326 348\ninside 349 372

8412 GCF_001607105.1_ASM160710v1 Escherichia coli Proteobacteria; Gammaproteobacteria; Enterobacterales; Enterobacteriaceae; Escherichia
MNNEETFYQAMRRQGVTRRSFLKYCSLAATSLGLGAGMAPKIAWALENKPRIPVVWIHGL WP_001058323.1
MULTISPECIES: hydrogenase-1 small chain [Proteobacteria] Length: 372\nNumber of predicted TMHs: 2\nExp number of AAs in TMHs: 37.2608499999999\nExp number, first 60 AAs: 19.52923\nTotal prob of N-in: 0.94351\nPOSSIBLE N-term signal sequence\ninside 1 19\nTMhelix 20 39\noutside 40 325\nTMhelix 326 348\ninside 349 372

8413 GCF_001607015.1_ASM160701v1 Escherichia coli Proteobacteria; Gammaproteobacteria; Enterobacterales; Enterobacteriaceae; Escherichia
MNNEETFYQAMRRQGVTRRSFLKYCSLAATSLGLGAGMAPKIAWALENKPRIPVVWIHGL WP_001058323.1
MULTISPECIES: hydrogenase-1 small chain [Proteobacteria] Length: 372\nNumber of predicted TMHs: 2\nExp number of AAs in TMHs: 37.2608499999999\nExp number, first 60 AAs: 19.52923\nTotal prob of N-in: 0.94351\nPOSSIBLE N-term signal sequence\ninside 1 19\nTMhelix 20 39\noutside 40 325\nTMhelix 326 348\ninside 349 372

8414 GCF_001606855.1_ASM160685v1 Escherichia coli Proteobacteria; Gammaproteobacteria; Enterobacterales; Enterobacteriaceae; Escherichia
MTGDNTLIHSHGINRRDFMKLCAALAATMGLSSKAAAEMAESVTNPQRPPVIWIGAQECT WP_024245965.1
hydrogenase 2 small subunit [Escherichia coli] Length: 372\nNumber of predicted TMHs: 1\nExp number of AAs in TMHs: 24.15412\nExp number, first 60 AAs: 0.21967\nTotal prob of N-in: 0.14289\noutside 1 331\nTMhelix 332 354\ninside 355 372

8415 GCF_001606635.1_ASM160663v1 Escherichia coli Proteobacteria; Gammaproteobacteria; Enterobacterales; Enterobacteriaceae; Escherichia
MNNEETFYQAMRRQGVTRRSFLKYCSLAATSLGLGAGMAPKIAWALENKPRIPVVWIHGL WP_001295940.1
MULTISPECIES: hydrogenase [Enterobacteriaceae] Length: 372\nNumber of predicted TMHs: 2\nExp number of AAs in TMHs: 37.2608899999999\nExp number, first 60 AAs: 19.52923\nTotal prob of N-in: 0.94351\nPOSSIBLE N-term signal sequence\ninside 1 19\nTMhelix 20 39\noutside 40 325\nTMhelix 326 348\ninside 349 372

8416 GCF_001606955.1_ASM160695v1 Escherichia coli Proteobacteria; Gammaproteobacteria; Enterobacterales; Enterobacteriaceae; Escherichia
MNNEETFYQAMRRQGVTRRSFLKYCSLAATSLGLGAGMAPKIAWALENKPRIPVVWIHGL WP_001295940.1
MULTISPECIES: hydrogenase [Enterobacteriaceae] Length: 372\nNumber of predicted TMHs: 2\nExp number of AAs in TMHs: 37.2608899999999\nExp number, first 60 AAs: 19.52923\nTotal prob of N-in: 0.94351\nPOSSIBLE N-term signal sequence\ninside 1 19\nTMhelix 20 39\noutside 40 325\nTMhelix 326 348\ninside 349 372

8417 GCF_001606755.1_ASM160675v1 Escherichia coli Proteobacteria; Gammaproteobacteria; Enterobacterales; Enterobacteriaceae; Escherichia
MNNEETFYQAMRRQGVTRRSFLKYCSLAATSLGLGAGMAPKIAWALENKPRIPVVWIHGL WP_062898093.1
hydrogenase [Escherichia coli] Length: 372\nNumber of predicted TMHs: 2\nExp number of AAs in TMHs: 37.5090799999999\nExp number, first 60 AAs: 19.52755\nTotal prob of N-in: 0.94365\nPOSSIBLE N-term signal sequence\ninside 1 19\nTMhelix 20 39\noutside 40 325\nTMhelix 326 348\ninside 349 372

8418 GCF_001607205.1_ASM160720v1 Escherichia coli Proteobacteria; Gammaproteobacteria; Enterobacterales; Enterobacteriaceae; Escherichia
MTGDNTLIHSHGINRRDFMKLCAALAATMGLSSKAAAEMAESVTNPQRPPVIWIGAQECT WP_000145410.1
MULTISPECIES: hydrogenase-2 small chain [Proteobacteria] Length: 372\nNumber of predicted TMHs: 1\nExp number of AAs in TMHs: 23.72226\nExp number, first 60 AAs: 0.22038\nTotal prob of N-in: 0.12994\noutside 1 331\nTMhelix 332 354\ninside 355 372

8419 GCF_001607255.1_ASM160725v1 Escherichia coli Proteobacteria; Gammaproteobacteria; Enterobacterales; Enterobacteriaceae; Escherichia
MTGDNTLIHSHGINRRDFMKLCAALAATMGLSSKAAAEMAESVTNPQRPPVIWIGAQECT WP_000145410.1
MULTISPECIES: hydrogenase-2 small chain [Proteobacteria] Length: 372\nNumber of predicted TMHs: 1\nExp number of AAs in TMHs: 23.72226\nExp number, first 60 AAs: 0.22038\nTotal prob of N-in: 0.12994\noutside 1 331\nTMhelix 332 354\ninside 355 372

8420 GCF_001607515.1_ASM160751v1 Escherichia coli Proteobacteria; Gammaproteobacteria; Enterobacterales; Enterobacteriaceae; Escherichia
MTGDNTLIHSHGINRRDFMKLCAALAATMGLSSKAAAEMVESVTNPQRPPVIWIGAQECT WP_000145423.1
hydrogenase 2 small subunit [Escherichia coli] Length: 372\nNumber of predicted TMHs: 1\nExp number of AAs in TMHs: 23.78201\nExp number, first 60 AAs: 0.28355\nTotal prob of N-in: 0.13257\noutside 1 331\nTMhelix 332 354\ninside 355 372

8421 GCF_001607355.1_ASM160735v1 Escherichia coli Proteobacteria; Gammaproteobacteria; Enterobacterales; Enterobacteriaceae; Escherichia
MTGDNTLIHSHGINRRDFMKLCAALAATMGLSSKAAAEMAESVTNPQRPPVIWIGAQECT WP_000145410.1
MULTISPECIES: hydrogenase-2 small chain [Proteobacteria] Length: 372\nNumber of predicted TMHs: 1\nExp number of AAs in TMHs: 23.72226\nExp number, first 60 AAs: 0.22038\nTotal prob of N-in: 0.12994\noutside 1 331\nTMhelix 332 354\ninside 355 372

8422 GCF_001607295.1_ASM160729v1 Escherichia coli Proteobacteria; Gammaproteobacteria; Enterobacterales; Enterobacteriaceae; Escherichia
MTGDNTLIHSHGINRRDFMKLCAALAATMGLSSKAAAEMAESVTNPQRPPVIWIGAQECT WP_000145410.1
MULTISPECIES: hydrogenase-2 small chain [Proteobacteria] Length: 372\nNumber of predicted TMHs: 1\nExp number of AAs in TMHs: 23.72226\nExp number, first 60 AAs: 0.22038\nTotal prob of N-in: 0.12994\noutside 1 331\nTMhelix 332 354\ninside 355 372

8423 GCF_001607415.1_ASM160741v1 Escherichia coli Proteobacteria; Gammaproteobacteria; Enterobacterales; Enterobacteriaceae; Escherichia
MTGDNTLIHSHGINRRDFMKLCAALAATMGLSSKAAAEMAESVTNPQRPPVIWIGAQECT WP_000145410.1

MULTISPECIES: hydrogenase-2 small chain [Proteobacteria] Length: 372\nNumber of predicted TMHs: 1\nExp number of AAs in TMHs: 23.72226\nExp number, first 60 AAs: 0.22038\nTotal prob of N-in: 0.12994\noutside 1 331\nTMhelix 332 354\ninside 355 372

8424 GCF_001607605.1_ASM160760v1 Escherichia coli Proteobacteria; Gammaproteobacteria; Enterobacterales; Enterobacteriaceae; Escherichia
MTGDNTLIHSHGINRRDFMKLCAALAATMGLSSKAAAEMAESVTNPQRPPVIWIGAQECT WP_000145410.1
MULTISPECIES: hydrogenase-2 small chain [Proteobacteria] Length: 372\nNumber of predicted TMHs: 1\nExp number of AAs in TMHs: 23.72226\nExp number, first 60 AAs: 0.22038\nTotal prob of N-in: 0.12994\noutside 1 331\nTMhelix 332 354\ninside 355 372

8425 GCF_001607365.1_ASM160736v1 Escherichia coli Proteobacteria; Gammaproteobacteria; Enterobacterales; Enterobacteriaceae; Escherichia
MTGDNTLIHSHGINRRDFMKLCAALAATMGLSSKAAAEMAESVTNPQRPPVIWIGAQECT WP_000145410.1
MULTISPECIES: hydrogenase-2 small chain [Proteobacteria] Length: 372\nNumber of predicted TMHs: 1\nExp number of AAs in TMHs: 23.72226\nExp number, first 60 AAs: 0.22038\nTotal prob of N-in: 0.12994\noutside 1 331\nTMhelix 332 354\ninside 355 372

8426 GCF_001607555.1_ASM160755v1 Escherichia coli Proteobacteria; Gammaproteobacteria; Enterobacterales; Enterobacteriaceae; Escherichia
MTGDNTLIHSHGINRRDFMKLCAALAATMGLSSKAAAEMAESVTNPQRPPVIWIGAQECT WP_000145410.1
MULTISPECIES: hydrogenase-2 small chain [Proteobacteria] Length: 372\nNumber of predicted TMHs: 1\nExp number of AAs in TMHs: 23.72226\nExp number, first 60 AAs: 0.22038\nTotal prob of N-in: 0.12994\noutside 1 331\nTMhelix 332 354\ninside 355 372

8427 GCF_001607445.1_ASM160744v1 Escherichia coli Proteobacteria; Gammaproteobacteria; Enterobacterales; Enterobacteriaceae; Escherichia
MTGDNTLIHSHGINRRDFMKLCAALAATMGLSSKAAAEMAESVTNPQRPPVIWIGAQECT WP_000145410.1
MULTISPECIES: hydrogenase-2 small chain [Proteobacteria] Length: 372\nNumber of predicted TMHs: 1\nExp number of AAs in TMHs: 23.72226\nExp number, first 60 AAs: 0.22038\nTotal prob of N-in: 0.12994\noutside 1 331\nTMhelix 332 354\ninside 355 372

8428 GCF_001607715.1_ASM160771v1 Escherichia coli Proteobacteria; Gammaproteobacteria; Enterobacterales; Enterobacteriaceae; Escherichia
MTGDNTLIHSHGINRRDFMKLCAALAATMGLSSKAAAEMAESVTNPQRPPVIWIGAQECT WP_000145410.1
MULTISPECIES: hydrogenase-2 small chain [Proteobacteria] Length: 372\nNumber of predicted TMHs: 1\nExp number of AAs in TMHs: 23.72226\nExp number, first 60 AAs: 0.22038\nTotal prob of N-in: 0.12994\noutside 1 331\nTMhelix 332 354\ninside 355 372

8429 GCF_001607655.1_ASM160765v1 Escherichia coli Proteobacteria; Gammaproteobacteria; Enterobacterales; Enterobacteriaceae; Escherichia
MTGDNTLIHSHGINRRDFMKLCAALAATMGLSSKAAAEMAESVTNPQRPPVIWIGAQECT WP_000145410.1
MULTISPECIES: hydrogenase-2 small chain [Proteobacteria] Length: 372\nNumber of predicted TMHs: 1\nExp number of AAs in TMHs: 23.72226\nExp number, first 60 AAs: 0.22038\nTotal prob of N-in: 0.12994\noutside 1 331\nTMhelix 332 354\ninside 355 372

8430 GCF_001607655.1_ASM160765v1 Escherichia coli Proteobacteria; Gammaproteobacteria; Enterobacterales; Enterobacteriaceae; Escherichia
MNNEETFYQAMRRQGVTRRSFLKYCSLAATSLGLGAGMAPKIAWALENKPRIPVVWIHGL WP_001058323.1
MULTISPECIES: hydrogenase-1 small chain [Proteobacteria] Length: 372\nNumber of predicted TMHs: 2\nExp number of AAs in TMHs: 37.2608499999999\nExp number, first 60 AAs: 19.52923\nTotal prob of N-in: 0.94351\nPOSSIBLE N-term signal sequence\ninside 1 19\nTMhelix 20 39\noutside 40 325\nTMhelix 326 348\ninside 349 372

8431 GCF_001607715.1_ASM160771v1 Escherichia coli Proteobacteria; Gammaproteobacteria; Enterobacterales; Enterobacteriaceae; Escherichia
MNNEETFYQAMRRQGVTRRSFLKYCSLAATSLGLGAGMAPKIAWALENKPRIPVVWIHGL WP_001058323.1
MULTISPECIES: hydrogenase-1 small chain [Proteobacteria] Length: 372\nNumber of predicted TMHs: 2\nExp number of AAs in TMHs: 37.2608499999999\nExp number, first 60 AAs: 19.52923\nTotal prob of N-in: 0.94351\nPOSSIBLE N-term signal sequence\ninside 1 19\nTMhelix 20 39\noutside 40 325\nTMhelix 326 348\ninside 349 372

8432	GCF_001607515.1_ASM160751v1	Escherichia coli	Proteobacteria; Gammaproteobacteria; Enterobacterales; Enterobacteriaceae; Escherichia
MNNEETFYQAMRRQGVTRRSFLKYCSLAATSLGLGAGMAPKIAWALENKPRIPVVWIHGL	WP_001058323.1
MULTISPECIES: hydrogenase-1 small chain [Proteobacteria]	Length: 372\nNumber of predicted TMHs: 2\nExp number of AAs in TMHs: 37.2608499999999\nExp number, first 60 AAs: 19.52923\nTotal prob of N-in: 0.94351\nPOSSIBLE N-term signal sequence\ninside 1 19\nTMhelix 20 39\noutside 40 325\nTMhelix 326 348\ninside 349 372

8433	GCF_001607355.1_ASM160735v1	Escherichia coli	Proteobacteria; Gammaproteobacteria; Enterobacterales; Enterobacteriaceae; Escherichia
MNNEETFYQAMRRQGVTRRSFLKYCSLAATSLGLGAGMAPKIAWALENKPRIPVVWIHGL	WP_001058323.1
MULTISPECIES: hydrogenase-1 small chain [Proteobacteria]	Length: 372\nNumber of predicted TMHs: 2\nExp number of AAs in TMHs: 37.2608499999999\nExp number, first 60 AAs: 19.52923\nTotal prob of N-in: 0.94351\nPOSSIBLE N-term signal sequence\ninside 1 19\nTMhelix 20 39\noutside 40 325\nTMhelix 326 348\ninside 349 372

8434	GCF_001607415.1_ASM160741v1	Escherichia coli	Proteobacteria; Gammaproteobacteria; Enterobacterales; Enterobacteriaceae; Escherichia
MNNEETFYQAMRRQGVTRRSFLKYCSLAATSLGLGAGMAPKIAWALENKPRIPVVWIHGL	WP_001058323.1
MULTISPECIES: hydrogenase-1 small chain [Proteobacteria]	Length: 372\nNumber of predicted TMHs: 2\nExp number of AAs in TMHs: 37.2608499999999\nExp number, first 60 AAs: 19.52923\nTotal prob of N-in: 0.94351\nPOSSIBLE N-term signal sequence\ninside 1 19\nTMhelix 20 39\noutside 40 325\nTMhelix 326 348\ninside 349 372

8435	GCF_001607205.1_ASM160720v1	Escherichia coli	Proteobacteria; Gammaproteobacteria; Enterobacterales; Enterobacteriaceae; Escherichia
MNNEETFYQAMRRQGVTRRSFLKYCSLAATSLGLGAGMAPKIAWALENKPRIPVVWIHGL	WP_001058323.1
MULTISPECIES: hydrogenase-1 small chain [Proteobacteria]	Length: 372\nNumber of predicted TMHs: 2\nExp number of AAs in TMHs: 37.2608499999999\nExp number, first 60 AAs: 19.52923\nTotal prob of N-in: 0.94351\nPOSSIBLE N-term signal sequence\ninside 1 19\nTMhelix 20 39\noutside 40 325\nTMhelix 326 348\ninside 349 372

8436	GCF_001607605.1_ASM160760v1	Escherichia coli	Proteobacteria; Gammaproteobacteria; Enterobacterales; Enterobacteriaceae; Escherichia
MNNEETFYQAMRRQGVTRRSFLKYCSLAATSLGLGAGMAPKIAWALENKPRIPVVWIHGL	WP_001058323.1
MULTISPECIES: hydrogenase-1 small chain [Proteobacteria]	Length: 372\nNumber of predicted TMHs: 2\nExp number of AAs in TMHs: 37.2608499999999\nExp number, first 60 AAs: 19.52923\nTotal prob of N-in: 0.94351\nPOSSIBLE N-term signal sequence\ninside 1 19\nTMhelix 20 39\noutside 40 325\nTMhelix 326 348\ninside 349 372

8437	GCF_001607255.1_ASM160725v1	Escherichia coli	Proteobacteria; Gammaproteobacteria; Enterobacterales; Enterobacteriaceae; Escherichia
MNNEETFYQAMRRQGVTRRSFLKYCSLAATSLGLGAGMAPKIAWALENKPRIPVVWIHGL	WP_001058323.1
MULTISPECIES: hydrogenase-1 small chain [Proteobacteria]	Length: 372\nNumber of predicted TMHs: 2\nExp number of AAs in TMHs: 37.2608499999999\nExp number, first 60 AAs: 19.52923\nTotal prob of N-in: 0.94351\nPOSSIBLE N-term signal sequence\ninside 1 19\nTMhelix 20 39\noutside 40 325\nTMhelix 326 348\ninside 349 372

8438	GCF_001607295.1_ASM160729v1	Escherichia coli	Proteobacteria; Gammaproteobacteria; Enterobacterales; Enterobacteriaceae; Escherichia
MNNEETFYQAMRRQGVTRRSFLKYCSLAATSLGLGAGMAPKIAWALENKPRIPVVWIHGL	WP_001295940.1
MULTISPECIES: hydrogenase [Enterobacteriaceae]	Length: 372\nNumber of predicted TMHs: 2\nExp number of AAs in TMHs: 37.2608899999999\nExp number, first 60 AAs: 19.52923\nTotal prob of N-in: 0.94351\nPOSSIBLE N-term signal sequence\ninside 1 19\nTMhelix 20 39\noutside 40 325\nTMhelix 326 348\ninside 349 372

8439	GCF_001607555.1_ASM160755v1	Escherichia coli	Proteobacteria; Gammaproteobacteria; Enterobacterales; Enterobacteriaceae; Escherichia
MNNEETFYQAMRRQGVTRRSFLKYCSLAATSLGLGAGMAPKIAWALENKPRIPVVWIHGL	WP_001295940.1
MULTISPECIES: hydrogenase [Enterobacteriaceae]	Length: 372\nNumber of predicted TMHs: 2\nExp number of AAs in TMHs: 37.2608899999999\nExp number, first 60 AAs: 19.52923\nTotal prob of N-in:

0.94351\nPOSSIBLE N-term signal sequence\ninside 1 19\nTMhelix 20 39\noutside 40 325\nTMhelix 326 348\ninside 349 372

8440 GCF_001607445.1_ASM160744v1 Escherichia coli Proteobacteria; Gammaproteobacteria; Enterobacterales; Enterobacteriaceae; Escherichia
MNNEETFYQAMRRQGVTRRSFLKYCSLAATSLGLGAGMAPKIAWALENKPRIPVVWIHGL WP_001295940.1
MULTISPECIES: hydrogenase [Enterobacteriaceae] Length: 372\nNumber of predicted TMHs: 2\nExp number of AAs in TMHs: 37.2608899999999\nExp number, first 60 AAs: 19.52923\nTotal prob of N-in: 0.94351\nPOSSIBLE N-term signal sequence\ninside 1 19\nTMhelix 20 39\noutside 40 325\nTMhelix 326 348\ninside 349 372

8441 GCF_001607365.1_ASM160736v1 Escherichia coli Proteobacteria; Gammaproteobacteria; Enterobacterales; Enterobacteriaceae; Escherichia
MNNEETFYQAMRRQGVTRRSFLKYCSLAATSLGLGAGMAPKIAWALENKPRIPVVWIHGL WP_062870334.1
hydrogenase [Escherichia coli] Length: 372\nNumber of predicted TMHs: 2\nExp number of AAs in TMHs: 37.0125199999999\nExp number, first 60 AAs: 19.52914\nTotal prob of N-in: 0.94337\nPOSSIBLE N-term signal sequence\ninside 1 19\nTMhelix 20 39\noutside 40 325\nTMhelix 326 348\ninside 349 372

8442 GCF_001607815.1_ASM160781v1 Escherichia coli Proteobacteria; Gammaproteobacteria; Enterobacterales; Enterobacteriaceae; Escherichia
MTGDNTLIHSHGINRRDFMKLCAALAATMGLSSKAAAEMAESVTNPQRPPVIWIGAQECT WP_000145410.1
MULTISPECIES: hydrogenase-2 small chain [Proteobacteria] Length: 372\nNumber of predicted TMHs: 1\nExp number of AAs in TMHs: 23.72226\nExp number, first 60 AAs: 0.22038\nTotal prob of N-in: 0.12994\noutside 1 331\nTMhelix 332 354\ninside 355 372

8443 GCF_001607755.1_ASM160775v1 Escherichia coli Proteobacteria; Gammaproteobacteria; Enterobacterales; Enterobacteriaceae; Escherichia
MTGDNTLIHSHGINRRDFMKLCAALAATMGLSSKAAAEMAESVTNPQRPPVIWIGAQECT WP_000145410.1
MULTISPECIES: hydrogenase-2 small chain [Proteobacteria] Length: 372\nNumber of predicted TMHs: 1\nExp number of AAs in TMHs: 23.72226\nExp number, first 60 AAs: 0.22038\nTotal prob of N-in: 0.12994\noutside 1 331\nTMhelix 332 354\ninside 355 372

8444 GCF_001608005.1_ASM160800v1 Escherichia coli Proteobacteria; Gammaproteobacteria; Enterobacterales; Enterobacteriaceae; Escherichia
MTGDNTLIHSHGINRRDFMKLCAALAATMGLSSKAAAEMAESVTNPQRPPVIWIGAQECT WP_000145410.1
MULTISPECIES: hydrogenase-2 small chain [Proteobacteria] Length: 372\nNumber of predicted TMHs: 1\nExp number of AAs in TMHs: 23.72226\nExp number, first 60 AAs: 0.22038\nTotal prob of N-in: 0.12994\noutside 1 331\nTMhelix 332 354\ninside 355 372

8445 GCF_001607915.1_ASM160791v1 Escherichia coli Proteobacteria; Gammaproteobacteria; Enterobacterales; Enterobacteriaceae; Escherichia
MTGDNTLIHSHGINRRDFMKLCAALAATMGLSSKAAAEMAESVTNPQRPPVIWIGAQECT WP_000145410.1
MULTISPECIES: hydrogenase-2 small chain [Proteobacteria] Length: 372\nNumber of predicted TMHs: 1\nExp number of AAs in TMHs: 23.72226\nExp number, first 60 AAs: 0.22038\nTotal prob of N-in: 0.12994\noutside 1 331\nTMhelix 332 354\ninside 355 372

8446 GCF_001608055.1_ASM160805v1 Escherichia coli Proteobacteria; Gammaproteobacteria; Enterobacterales; Enterobacteriaceae; Escherichia
MTGDNTLIHSHGINRRDFMKLCAALAATMGLSSKAAAEMAESVTNPQRPPVIWIGAQECT WP_000145410.1
MULTISPECIES: hydrogenase-2 small chain [Proteobacteria] Length: 372\nNumber of predicted TMHs: 1\nExp number of AAs in TMHs: 23.72226\nExp number, first 60 AAs: 0.22038\nTotal prob of N-in: 0.12994\noutside 1 331\nTMhelix 332 354\ninside 355 372

8447 GCF_001607955.1_ASM160795v1 Escherichia coli Proteobacteria; Gammaproteobacteria; Enterobacterales; Enterobacteriaceae; Escherichia
MTGDNTLIHSHGINRRDFMKLCAALAATMGLSSKAAAEMAESVTNPQRPPVIWIGAQECT WP_000145410.1
MULTISPECIES: hydrogenase-2 small chain [Proteobacteria] Length: 372\nNumber of predicted TMHs: 1\nExp number of AAs in TMHs: 23.72226\nExp number, first 60 AAs: 0.22038\nTotal prob of N-in: 0.12994\noutside 1 331\nTMhelix 332 354\ninside 355 372

8448 GCF_001608165.1_ASM160816v1 Escherichia coli Proteobacteria; Gammaproteobacteria; Enterobacterales; Enterobacteriaceae; Escherichia
MTGDNTLIHSHGINRRDFMKLCAALAATMGLSSKAAAEMAESVTNPQRPPVIWIGAQECT WP_000145410.1
MULTISPECIES: hydrogenase-2 small chain [Proteobacteria] Length: 372\nNumber of predicted TMHs: 1\nExp number of AAs in TMHs: 23.72226\nExp number, first 60 AAs: 0.22038\nTotal prob of N-in: 0.12994\noutside 1 331\nTMhelix 332 354\ninside 355 372

8449 GCF_001608155.1_ASM160815v1 Escherichia coli Proteobacteria; Gammaproteobacteria; Enterobacterales; Enterobacteriaceae; Escherichia
MTGDNTLIHSHGINRRDFMKLCAALAATMGLSSKAAAEMAESVTNPQRPPVIWIGAQECT WP_000145410.1
MULTISPECIES: hydrogenase-2 small chain [Proteobacteria] Length: 372\nNumber of predicted TMHs: 1\nExp number of AAs in TMHs: 23.72226\nExp number, first 60 AAs: 0.22038\nTotal prob of N-in: 0.12994\noutside 1 331\nTMhelix 332 354\ninside 355 372

8450 GCF_001608105.1_ASM160810v1 Escherichia coli Proteobacteria; Gammaproteobacteria; Enterobacterales; Enterobacteriaceae; Escherichia
MTGDNTLIHSHGINRRDFMKLCAALAATMGLSSKAAAEMAESVTNPQRPPVIWIGAQECT WP_000145410.1
MULTISPECIES: hydrogenase-2 small chain [Proteobacteria] Length: 372\nNumber of predicted TMHs: 1\nExp number of AAs in TMHs: 23.72226\nExp number, first 60 AAs: 0.22038\nTotal prob of N-in: 0.12994\noutside 1 331\nTMhelix 332 354\ninside 355 372

8451 GCF_001608205.1_ASM160820v1 Escherichia coli Proteobacteria; Gammaproteobacteria; Enterobacterales; Enterobacteriaceae; Escherichia
MTGDNTLIHSHGINRRDFMKLCAALAATMGLSSKAAAEMAESVTNPQRPPVIWIGAQECT WP_000145410.1
MULTISPECIES: hydrogenase-2 small chain [Proteobacteria] Length: 372\nNumber of predicted TMHs: 1\nExp number of AAs in TMHs: 23.72226\nExp number, first 60 AAs: 0.22038\nTotal prob of N-in: 0.12994\noutside 1 331\nTMhelix 332 354\ninside 355 372

8452 GCF_001608245.1_ASM160824v1 Escherichia coli Proteobacteria; Gammaproteobacteria; Enterobacterales; Enterobacteriaceae; Escherichia
MTGDNTLIHSHGINRRDFMKLCAALAATMGLSSKAAAEMAESVTNPQRPPVIWIGAQECT WP_000145410.1
MULTISPECIES: hydrogenase-2 small chain [Proteobacteria] Length: 372\nNumber of predicted TMHs: 1\nExp number of AAs in TMHs: 23.72226\nExp number, first 60 AAs: 0.22038\nTotal prob of N-in: 0.12994\noutside 1 331\nTMhelix 332 354\ninside 355 372

8453 GCF_001607845.1_ASM160784v1 Escherichia coli Proteobacteria; Gammaproteobacteria; Enterobacterales; Enterobacteriaceae; Escherichia
MNNEETFYQAMRRQGVTRRSFLKYCSLAATSLGLGAGMAPKIAWALENKPRIPVVWIHGL WP_001058323.1
MULTISPECIES: hydrogenase-1 small chain [Proteobacteria] Length: 372\nNumber of predicted TMHs: 2\nExp number of AAs in TMHs: 37.2608499999999\nExp number, first 60 AAs: 19.52923\nTotal prob of N-in: 0.94351\nPOSSIBLE N-term signal sequence\ninside 1 19\nTMhelix 20 39\noutside 40 325\nTMhelix 326 348\ninside 349 372

8454 GCF_001608155.1_ASM160815v1 Escherichia coli Proteobacteria; Gammaproteobacteria; Enterobacterales; Enterobacteriaceae; Escherichia
MNNEETFYQAMRRQGVTRRSFLKYCSLAATSLGLGAGMAPKIAWALENKPRIPVVWIHGL WP_001058323.1
MULTISPECIES: hydrogenase-1 small chain [Proteobacteria] Length: 372\nNumber of predicted TMHs: 2\nExp number of AAs in TMHs: 37.2608499999999\nExp number, first 60 AAs: 19.52923\nTotal prob of N-in: 0.94351\nPOSSIBLE N-term signal sequence\ninside 1 19\nTMhelix 20 39\noutside 40 325\nTMhelix 326 348\ninside 349 372

8455 GCF_001608105.1_ASM160810v1 Escherichia coli Proteobacteria; Gammaproteobacteria; Enterobacterales; Enterobacteriaceae; Escherichia
MNNEETFYQAMRRQGVTRRSFLKYCSLAATSLGLGAGMAPKIAWALENKPRIPVVWIHGL WP_001295940.1
MULTISPECIES: hydrogenase [Enterobacteriaceae] Length: 372\nNumber of predicted TMHs: 2\nExp number of AAs in TMHs: 37.2608899999999\nExp number, first 60 AAs: 19.52923\nTotal prob of N-in: 0.94351\nPOSSIBLE N-term signal sequence\ninside 1 19\nTMhelix 20 39\noutside 40 325\nTMhelix 326 348\ninside 349 372

8456 GCF_001607955.1_ASM160795v1 Escherichia coli Proteobacteria; Gammaproteobacteria; Enterobacterales; Enterobacteriaceae; Escherichia

MNNEETFYQAMRRQGVTRRSFLKYCSLAATSLGLGAGMAPKIAWALENKPRIPVVWIHGL WP_001058323.1 MULTISPECIES: hydrogenase-1 small chain [Proteobacteria] Length: 372\nNumber of predicted TMHs: 2\nExp number of AAs in TMHs: 37.2608499999999\nExp number, first 60 AAs: 19.52923\nTotal prob of N-in: 0.94351\nPOSSIBLE N-term signal sequence\ninside 1 19\nTMhelix 20 39\noutside 40 325\nTMhelix 326 348\ninside 349 372

8457 GCF_001608005.1_ASM160800v1 Escherichia coli Proteobacteria; Gammaproteobacteria; Enterobacterales; Enterobacteriaceae; Escherichia MNNEETFYQAMRRQGVTRRSFLKYCSLAATSLGLGAGMAPKIAWALENKPRIPVVWIHGL WP_001058323.1 MULTISPECIES: hydrogenase-1 small chain [Proteobacteria] Length: 372\nNumber of predicted TMHs: 2\nExp number of AAs in TMHs: 37.2608499999999\nExp number, first 60 AAs: 19.52923\nTotal prob of N-in: 0.94351\nPOSSIBLE N-term signal sequence\ninside 1 19\nTMhelix 20 39\noutside 40 325\nTMhelix 326 348\ninside 349 372

8458 GCF_001608165.1_ASM160816v1 Escherichia coli Proteobacteria; Gammaproteobacteria; Enterobacterales; Enterobacteriaceae; Escherichia MNNEETFYQAMRRQGVTRRSFLKYCSLAATSLGLGAGMAPKIAWALENKPRIPVVWIHGL WP_001058323.1 MULTISPECIES: hydrogenase-1 small chain [Proteobacteria] Length: 372\nNumber of predicted TMHs: 2\nExp number of AAs in TMHs: 37.2608499999999\nExp number, first 60 AAs: 19.52923\nTotal prob of N-in: 0.94351\nPOSSIBLE N-term signal sequence\ninside 1 19\nTMhelix 20 39\noutside 40 325\nTMhelix 326 348\ninside 349 372

8459 GCF_001607915.1_ASM160791v1 Escherichia coli Proteobacteria; Gammaproteobacteria; Enterobacterales; Enterobacteriaceae; Escherichia MNNEETFYQAMRRQGVTRRSFLKYCSLAATSLGLGAGMAPKIAWALENKPRIPVVWIHGL WP_001058323.1 MULTISPECIES: hydrogenase-1 small chain [Proteobacteria] Length: 372\nNumber of predicted TMHs: 2\nExp number of AAs in TMHs: 37.2608499999999\nExp number, first 60 AAs: 19.52923\nTotal prob of N-in: 0.94351\nPOSSIBLE N-term signal sequence\ninside 1 19\nTMhelix 20 39\noutside 40 325\nTMhelix 326 348\ninside 349 372

8460 GCF_001607815.1_ASM160781v1 Escherichia coli Proteobacteria; Gammaproteobacteria; Enterobacterales; Enterobacteriaceae; Escherichia MNNEETFYQAMRRQGVTRRSFLKYCSLAATSLGLGAGMAPKIAWALENKPRIPVVWIHGL WP_062876341.1 hydrogenase [Escherichia coli] Length: 372\nNumber of predicted TMHs: 2\nExp number of AAs in TMHs: 37.43626\nExp number, first 60 AAs: 19.52875\nTotal prob of N-in: 0.94361\nPOSSIBLE N-term signal sequence\ninside 1 19\nTMhelix 20 39\noutside 40 325\nTMhelix 326 348\ninside 349 372

8461 GCF_001608055.1_ASM160805v1 Escherichia coli Proteobacteria; Gammaproteobacteria; Enterobacterales; Enterobacteriaceae; Escherichia MNNEETFYQAMRRQGVTRRSFLKYCSLAATSLGLGAGMAPKIAWALENKPRIPVVWIHGL WP_001058323.1 MULTISPECIES: hydrogenase-1 small chain [Proteobacteria] Length: 372\nNumber of predicted TMHs: 2\nExp number of AAs in TMHs: 37.2608499999999\nExp number, first 60 AAs: 19.52923\nTotal prob of N-in: 0.94351\nPOSSIBLE N-term signal sequence\ninside 1 19\nTMhelix 20 39\noutside 40 325\nTMhelix 326 348\ninside 349 372

8462 GCF_001607755.1_ASM160775v1 Escherichia coli Proteobacteria; Gammaproteobacteria; Enterobacterales; Enterobacteriaceae; Escherichia MNNEETFYQAMRRQGVTRRSFLKYCSLAATSLGLGAGMAPKIAWALENKPRIPVVWIHGL WP_001295940.1 MULTISPECIES: hydrogenase [Enterobacteriaceae] Length: 372\nNumber of predicted TMHs: 2\nExp number of AAs in TMHs: 37.2608899999999\nExp number, first 60 AAs: 19.52923\nTotal prob of N-in: 0.94351\nPOSSIBLE N-term signal sequence\ninside 1 19\nTMhelix 20 39\noutside 40 325\nTMhelix 326 348\ninside 349 372

8463 GCF_001608245.1_ASM160824v1 Escherichia coli Proteobacteria; Gammaproteobacteria; Enterobacterales; Enterobacteriaceae; Escherichia MNNEETFYQAMRRQGVTRRSFLKYCSLAATSLGLGAGMAPKIAWALENKPRIPVVWIHGL WP_001058323.1 MULTISPECIES: hydrogenase-1 small chain [Proteobacteria] Length: 372\nNumber of predicted TMHs: 2\nExp number of AAs in TMHs: 37.2608499999999\nExp number, first 60 AAs: 19.52923\nTotal prob of N-in: 0.94351\nPOSSIBLE N-term signal sequence\ninside 1 19\nTMhelix 20 39\noutside 40 325\nTMhelix 326 348\ninside 349 372

8464 GCF_001607845.1_ASM160784v1 Escherichia coli Proteobacteria; Gammaproteobacteria; Enterobacterales; Enterobacteriaceae; Escherichia
MTGDNTLIHSHGINRRDFMKLCAALAATMGLSSKAAAEMAESVTNPQRPPVIWIGAQECT WP_062874794.1
hydrogenase 2 small subunit [Escherichia coli] Length: 372\nNumber of predicted TMHs: 1\nExp number of AAs in TMHs: 24.6459\nExp number, first 60 AAs: 0.21946\nTotal prob of N-in: 0.16283\noutside 1 331\nTMhelix 332 354\ninside 355 372

8465 GCF_001608205.1_ASM160820v1 Escherichia coli Proteobacteria; Gammaproteobacteria; Enterobacterales; Enterobacteriaceae; Escherichia
MNNEETFYQAMRRQGVTRRSFLKYCSLAATSLGLGAGMAPKIAWALENKPRIPVVWIHGL WP_001295940.1
MULTISPECIES: hydrogenase [Enterobacteriaceae] Length: 372\nNumber of predicted TMHs: 2\nExp number of AAs in TMHs: 37.2608899999999\nExp number, first 60 AAs: 19.52923\nTotal prob of N-in: 0.94351\nPOSSIBLE N-term signal sequence\ninside 1 19\nTMhelix 20 39\noutside 40 325\nTMhelix 326 348\ninside 349 372

8466 GCF_001608315.1_ASM160831v1 Escherichia coli Proteobacteria; Gammaproteobacteria; Enterobacterales; Enterobacteriaceae; Escherichia
MTGDNTLIHSHGINRRDFMKLCAALAATMGLSSKAAAEMAESVTNPQRPPVIWIGAQECT WP_000145410.1
MULTISPECIES: hydrogenase-2 small chain [Proteobacteria] Length: 372\nNumber of predicted TMHs: 1\nExp number of AAs in TMHs: 23.72226\nExp number, first 60 AAs: 0.22038\nTotal prob of N-in: 0.12994\noutside 1 331\nTMhelix 332 354\ninside 355 372

8467 GCF_001609765.1_ASM160976v1 Escherichia coli Proteobacteria; Gammaproteobacteria; Enterobacterales; Enterobacteriaceae; Escherichia
MTGDNTLIHSHGINRRDFMKLCAALAATMGLSSKAAAEMAESVTNPQRPPVIWIGAQECT WP_000145410.1
MULTISPECIES: hydrogenase-2 small chain [Proteobacteria] Length: 372\nNumber of predicted TMHs: 1\nExp number of AAs in TMHs: 23.72226\nExp number, first 60 AAs: 0.22038\nTotal prob of N-in: 0.12994\noutside 1 331\nTMhelix 332 354\ninside 355 372

8468 GCF_001609235.1_ASM160923v1 Escherichia coli Proteobacteria; Gammaproteobacteria; Enterobacterales; Enterobacteriaceae; Escherichia
MTGDNTLIHSHGINRRDFMKLCAALAATMGLSSKAAAEMAESVTNPQRPPVIWIGAQECT WP_000145410.1
MULTISPECIES: hydrogenase-2 small chain [Proteobacteria] Length: 372\nNumber of predicted TMHs: 1\nExp number of AAs in TMHs: 23.72226\nExp number, first 60 AAs: 0.22038\nTotal prob of N-in: 0.12994\noutside 1 331\nTMhelix 332 354\ninside 355 372

8469 GCF_001609295.1_ASM160929v1 Escherichia coli Proteobacteria; Gammaproteobacteria; Enterobacterales; Enterobacteriaceae; Escherichia
MTGDNTLIHSHGINRRDFMKLCAALAATMGLSSKAAAEMAESVTNPQRPPVIWIGAQECT WP_000145410.1
MULTISPECIES: hydrogenase-2 small chain [Proteobacteria] Length: 372\nNumber of predicted TMHs: 1\nExp number of AAs in TMHs: 23.72226\nExp number, first 60 AAs: 0.22038\nTotal prob of N-in: 0.12994\noutside 1 331\nTMhelix 332 354\ninside 355 372

8470 GCF_001609335.1_ASM160933v1 Escherichia coli Proteobacteria; Gammaproteobacteria; Enterobacterales; Enterobacteriaceae; Escherichia
MTGDNTLIHSHGINRRDFMKLCAALAATMGLSSKAAAEMAESVTNPQRPPVIWIGAQECT WP_000145410.1
MULTISPECIES: hydrogenase-2 small chain [Proteobacteria] Length: 372\nNumber of predicted TMHs: 1\nExp number of AAs in TMHs: 23.72226\nExp number, first 60 AAs: 0.22038\nTotal prob of N-in: 0.12994\noutside 1 331\nTMhelix 332 354\ninside 355 372

8471 GCF_001609815.1_ASM160981v1 Escherichia coli Proteobacteria; Gammaproteobacteria; Enterobacterales; Enterobacteriaceae; Escherichia
MTGDNTLIHSHGINRRDFMKLCAALAATMGLSSKAAAEMAESVTNPQRPPVIWIGAQECT WP_000145410.1
MULTISPECIES: hydrogenase-2 small chain [Proteobacteria] Length: 372\nNumber of predicted TMHs: 1\nExp number of AAs in TMHs: 23.72226\nExp number, first 60 AAs: 0.22038\nTotal prob of N-in: 0.12994\noutside 1 331\nTMhelix 332 354\ninside 355 372

8472 GCF_001614385.1_ASM161438v1 Escherichia coli Proteobacteria; Gammaproteobacteria; Enterobacterales; Enterobacteriaceae; Escherichia
MTGDNTLIHSHGINRRDFMKLCAALAATMGLSSKAAAEMAESVTNPQRPPVIWIGAQECT WP_000145410.1
MULTISPECIES: hydrogenase-2 small chain [Proteobacteria] Length: 372\nNumber of predicted TMHs:

1\nExp number of AAs in TMHs: 23.72226\nExp number, first 60 AAs: 0.22038\nTotal prob of N-in: 0.12994\noutside 1 331\nTMhelix 332 354\ninside 355 372

8473 GCF_001614485.1_ASM161448v1 Escherichia coli Proteobacteria; Gammaproteobacteria; Enterobacterales; Enterobacteriaceae; Escherichia
MTGDNTLIHSHGINRRDFMKLCAALAATMGLSSKAAAEMAESVTNPQRPPVIWIGAQECT WP_000145410.1
MULTISPECIES: hydrogenase-2 small chain [Proteobacteria] Length: 372\nNumber of predicted TMHs: 1\nExp number of AAs in TMHs: 23.72226\nExp number, first 60 AAs: 0.22038\nTotal prob of N-in: 0.12994\noutside 1 331\nTMhelix 332 354\ninside 355 372

8474 GCF_001612495.1_ASM161249v1 Escherichia coli Proteobacteria; Gammaproteobacteria; Enterobacterales; Enterobacteriaceae; Escherichia
MTGDNTLIHSHGINRRDFMKLCAALAATMGLSSKAAAEMAESVTNPQRPPVIWIGAQECT WP_000145410.1
MULTISPECIES: hydrogenase-2 small chain [Proteobacteria] Length: 372\nNumber of predicted TMHs: 1\nExp number of AAs in TMHs: 23.72226\nExp number, first 60 AAs: 0.22038\nTotal prob of N-in: 0.12994\noutside 1 331\nTMhelix 332 354\ninside 355 372

8475 GCF_001614485.1_ASM161448v1 Escherichia coli Proteobacteria; Gammaproteobacteria; Enterobacterales; Enterobacteriaceae; Escherichia
MNNEETFYQAMRRQGVTRRSFLKYCSLAATSLGLGAGMAPKIAWALENKPRIPVVWIHGL WP_001058323.1
MULTISPECIES: hydrogenase-1 small chain [Proteobacteria] Length: 372\nNumber of predicted TMHs: 2\nExp number of AAs in TMHs: 37.2608499999999\nExp number, first 60 AAs: 19.52923\nTotal prob of N-in: 0.94351\nPOSSIBLE N-term signal sequence\ninside 1 19\nTMhelix 20 39\noutside 40 325\nTMhelix 326 348\ninside 349 372

8476 GCF_001612495.1_ASM161249v1 Escherichia coli Proteobacteria; Gammaproteobacteria; Enterobacterales; Enterobacteriaceae; Escherichia
MNNEETFYQAMRRQGVTRRSFLKYCSLAATSLGLGAGMAPKIAWALENKPRIPVVWIHGL WP_001301613.1
hydrogenase [Escherichia coli] Length: 372\nNumber of predicted TMHs: 2\nExp number of AAs in TMHs: 37.46598\nExp number, first 60 AAs: 19.53139\nTotal prob of N-in: 0.94364\nPOSSIBLE N-term signal sequence\ninside 1 19\nTMhelix 20 39\noutside 40 325\nTMhelix 326 348\ninside 349 372

8477 GCF_001608315.1_ASM160831v1 Escherichia coli Proteobacteria; Gammaproteobacteria; Enterobacterales; Enterobacteriaceae; Escherichia
MNNEETFYQAMRRQGVTRRSFLKYCSLAATSLGLGAGMAPKIAWALENKPRIPVVWIHGL WP_001058323.1
MULTISPECIES: hydrogenase-1 small chain [Proteobacteria] Length: 372\nNumber of predicted TMHs: 2\nExp number of AAs in TMHs: 37.2608499999999\nExp number, first 60 AAs: 19.52923\nTotal prob of N-in: 0.94351\nPOSSIBLE N-term signal sequence\ninside 1 19\nTMhelix 20 39\noutside 40 325\nTMhelix 326 348\ninside 349 372

8478 GCF_001614405.1_ASM161440v1 Escherichia coli Proteobacteria; Gammaproteobacteria; Enterobacterales; Enterobacteriaceae; Escherichia
MTGDNTLIHSHGINRRDFMKLCAALAATMGLSSKAAAEMAESVTNPQRPPVIWIGAQECT WP_000145410.1
MULTISPECIES: hydrogenase-2 small chain [Proteobacteria] Length: 372\nNumber of predicted TMHs: 1\nExp number of AAs in TMHs: 23.72226\nExp number, first 60 AAs: 0.22038\nTotal prob of N-in: 0.12994\noutside 1 331\nTMhelix 332 354\ninside 355 372

8479 GCF_001614405.1_ASM161440v1 Escherichia coli Proteobacteria; Gammaproteobacteria; Enterobacterales; Enterobacteriaceae; Escherichia
MNNEETFYQAMRRQGVTRRSFLKYCSLAATSLGLGAGMAPKIAWALENKPRIPVVWIHGL WP_001058323.1
MULTISPECIES: hydrogenase-1 small chain [Proteobacteria] Length: 372\nNumber of predicted TMHs: 2\nExp number of AAs in TMHs: 37.2608499999999\nExp number, first 60 AAs: 19.52923\nTotal prob of N-in: 0.94351\nPOSSIBLE N-term signal sequence\ninside 1 19\nTMhelix 20 39\noutside 40 325\nTMhelix 326 348\ninside 349 372

8480 GCF_001609765.1_ASM160976v1 Escherichia coli Proteobacteria; Gammaproteobacteria; Enterobacterales; Enterobacteriaceae; Escherichia
MNNEETFYQAMRRQGVTRRSFLKYCSLAATSLGLGAGMAPKIAWALENKPRIPVVWIHGL WP_001058323.1
MULTISPECIES: hydrogenase-1 small chain [Proteobacteria] Length: 372\nNumber of predicted TMHs: 2\nExp number of AAs in TMHs: 37.2608499999999\nExp number, first 60 AAs: 19.52923\nTotal prob of N-in:

0.94351\nPOSSIBLE N-term signal sequence\ninside 1 19\nTMhelix 20 39\noutside 40 325\nTMhelix 326 348\ninside 349 372

8481 GCF_001614535.1_ASM161453v1 Escherichia coli Proteobacteria; Gammaproteobacteria; Enterobacterales; Enterobacteriaceae; Escherichia
MTGDNTLIHSHGINRRDFMKLCAALAATMGLSSKAAAEMAESVTNPQRPPVIWIGAQECT WP_000145410.1
MULTISPECIES: hydrogenase-2 small chain [Proteobacteria] Length: 372\nNumber of predicted TMHs: 1\nExp number of AAs in TMHs: 23.72226\nExp number, first 60 AAs: 0.22038\nTotal prob of N-in: 0.12994\noutside 1 331\nTMhelix 332 354\ninside 355 372

8482 GCF_001614535.1_ASM161453v1 Escherichia coli Proteobacteria; Gammaproteobacteria; Enterobacterales; Enterobacteriaceae; Escherichia
MNNEETFYQAMRRQGVTRRSFLKYCSLAATSLGLGAGMAPKIAWALENKPRIPVVWIHGL WP_001058323.1
MULTISPECIES: hydrogenase-1 small chain [Proteobacteria] Length: 372\nNumber of predicted TMHs: 2\nExp number of AAs in TMHs: 37.2608499999999\nExp number, first 60 AAs: 19.52923\nTotal prob of N-in: 0.94351\nPOSSIBLE N-term signal sequence\ninside 1 19\nTMhelix 20 39\noutside 40 325\nTMhelix 326 348\ninside 349 372

8483 GCF_001609295.1_ASM160929v1 Escherichia coli Proteobacteria; Gammaproteobacteria; Enterobacterales; Enterobacteriaceae; Escherichia
MNNEETFYQAMRRQGVTRRSFLKYCSLAATSLGLGAGMAPKIAWALENKPRIPVVWIHGL WP_001058323.1
MULTISPECIES: hydrogenase-1 small chain [Proteobacteria] Length: 372\nNumber of predicted TMHs: 2\nExp number of AAs in TMHs: 37.2608499999999\nExp number, first 60 AAs: 19.52923\nTotal prob of N-in: 0.94351\nPOSSIBLE N-term signal sequence\ninside 1 19\nTMhelix 20 39\noutside 40 325\nTMhelix 326 348\ninside 349 372

8484 GCF_001609815.1_ASM160981v1 Escherichia coli Proteobacteria; Gammaproteobacteria; Enterobacterales; Enterobacteriaceae; Escherichia
MNNEETFYQAMRRQGVTRRSFLKSCSLAATSLGLGAGMAPKIAWALENKPRIPVVWIHGL WP_001058318.1
hydrogenase [Escherichia coli] Length: 372\nNumber of predicted TMHs: 1\nExp number of AAs in TMHs: 33.59637\nExp number, first 60 AAs: 15.81976\nTotal prob of N-in: 0.77394\nPOSSIBLE N-term signal sequence\noutside 1 325\nTMhelix 326 348\ninside 349 372

8485 GCF_001614385.1_ASM161438v1 Escherichia coli Proteobacteria; Gammaproteobacteria; Enterobacterales; Enterobacteriaceae; Escherichia
MNNEETFYQAMRRQGVTRRSFLKYCSLAATSLGLGAGMAPKIAWALENKPRIPVVWIHGL WP_001058323.1
MULTISPECIES: hydrogenase-1 small chain [Proteobacteria] Length: 372\nNumber of predicted TMHs: 2\nExp number of AAs in TMHs: 37.2608499999999\nExp number, first 60 AAs: 19.52923\nTotal prob of N-in: 0.94351\nPOSSIBLE N-term signal sequence\ninside 1 19\nTMhelix 20 39\noutside 40 325\nTMhelix 326 348\ninside 349 372

8486 GCF_001609235.1_ASM160923v1 Escherichia coli Proteobacteria; Gammaproteobacteria; Enterobacterales; Enterobacteriaceae; Escherichia
MNNEETFYQAMRRQGVTRRSFLKYCSLAATSLGLGAGMAPKIAWALENKPRIPVVWIHGL WP_001295940.1
MULTISPECIES: hydrogenase [Enterobacteriaceae] Length: 372\nNumber of predicted TMHs: 2\nExp number of AAs in TMHs: 37.2608899999999\nExp number, first 60 AAs: 19.52923\nTotal prob of N-in: 0.94351\nPOSSIBLE N-term signal sequence\ninside 1 19\nTMhelix 20 39\noutside 40 325\nTMhelix 326 348\ninside 349 372

8487 GCF_001609335.1_ASM160933v1 Escherichia coli Proteobacteria; Gammaproteobacteria; Enterobacterales; Enterobacteriaceae; Escherichia
MNNEETFYQAMRRQGVTRRSFLKYCSLAATSLGLGAGMAPKIAWALENKPRIPVVWIHGL WP_001295940.1
MULTISPECIES: hydrogenase [Enterobacteriaceae] Length: 372\nNumber of predicted TMHs: 2\nExp number of AAs in TMHs: 37.2608899999999\nExp number, first 60 AAs: 19.52923\nTotal prob of N-in: 0.94351\nPOSSIBLE N-term signal sequence\ninside 1 19\nTMhelix 20 39\noutside 40 325\nTMhelix 326 348\ninside 349 372

8488 GCF_001614575.1_ASM161457v1 Escherichia coli Proteobacteria; Gammaproteobacteria; Enterobacterales; Enterobacteriaceae; Escherichia
MTGDNTLIHSHGINRRDFMKLCAALAATMGLSSKAAAEMAESVTNPQRPPVIWIGAQECT WP_000145410.1
MULTISPECIES: hydrogenase-2 small chain [Proteobacteria] Length: 372\nNumber of predicted TMHs:

1\nExp number of AAs in TMHs: 23.72226\nExp number, first 60 AAs: 0.22038\nTotal prob of N-in: 0.12994\noutside 1 331\nTMhelix 332 354\ninside 355 372

8489 GCF_001614645.1_ASM161464v1 Escherichia coli Proteobacteria; Gammaproteobacteria; Enterobacterales; Enterobacteriaceae; Escherichia
MTGDNTLIHSHGINRRDFMKLCAALAATMGLSSKAAAEMAESVTNPQRPPVIWIGAQECT WP_000145410.1
MULTISPECIES: hydrogenase-2 small chain [Proteobacteria] Length: 372\nNumber of predicted TMHs: 1\nExp number of AAs in TMHs: 23.72226\nExp number, first 60 AAs: 0.22038\nTotal prob of N-in: 0.12994\noutside 1 331\nTMhelix 332 354\ninside 355 372

8490 GCF_001614635.1_ASM161463v1 Escherichia coli Proteobacteria; Gammaproteobacteria; Enterobacterales; Enterobacteriaceae; Escherichia
MTGDNTLIHSHGINRRDFMKLCAALAATMGLSSKAAAEMAESVTNPQRPPVIWIGAQECT WP_000145410.1
MULTISPECIES: hydrogenase-2 small chain [Proteobacteria] Length: 372\nNumber of predicted TMHs: 1\nExp number of AAs in TMHs: 23.72226\nExp number, first 60 AAs: 0.22038\nTotal prob of N-in: 0.12994\noutside 1 331\nTMhelix 332 354\ninside 355 372

8491 GCF_001614695.1_ASM161469v1 Escherichia coli Proteobacteria; Gammaproteobacteria; Enterobacterales; Enterobacteriaceae; Escherichia
MTGDNTLIHSHGINRRDFMKLCAALAATMGLSSKAAAEMAESVTNPQRPPVIWIGAQECT WP_000145410.1
MULTISPECIES: hydrogenase-2 small chain [Proteobacteria] Length: 372\nNumber of predicted TMHs: 1\nExp number of AAs in TMHs: 23.72226\nExp number, first 60 AAs: 0.22038\nTotal prob of N-in: 0.12994\noutside 1 331\nTMhelix 332 354\ninside 355 372

8492 GCF_001614725.1_ASM161472v1 Escherichia coli Proteobacteria; Gammaproteobacteria; Enterobacterales; Enterobacteriaceae; Escherichia
MTGDNTLIHSHGINRRDFMKLCAALAATMGLSSKAAAEMAESVTNPQRPPVIWIGAQECT WP_000145410.1
MULTISPECIES: hydrogenase-2 small chain [Proteobacteria] Length: 372\nNumber of predicted TMHs: 1\nExp number of AAs in TMHs: 23.72226\nExp number, first 60 AAs: 0.22038\nTotal prob of N-in: 0.12994\noutside 1 331\nTMhelix 332 354\ninside 355 372

8493 GCF_001614795.1_ASM161479v1 Escherichia coli Proteobacteria; Gammaproteobacteria; Enterobacterales; Enterobacteriaceae; Escherichia
MTGDNTLIHSHGINRRDFMKLCAALAATMGLSSKAAAEMAESVTNPQRPPVIWIGAQECT WP_000145410.1
MULTISPECIES: hydrogenase-2 small chain [Proteobacteria] Length: 372\nNumber of predicted TMHs: 1\nExp number of AAs in TMHs: 23.72226\nExp number, first 60 AAs: 0.22038\nTotal prob of N-in: 0.12994\noutside 1 331\nTMhelix 332 354\ninside 355 372

8494 GCF_001614825.1_ASM161482v1 Escherichia coli Proteobacteria; Gammaproteobacteria; Enterobacterales; Enterobacteriaceae; Escherichia
MTGDNTLIHSHGINRRDFMKLCAALAATMGLSSKAAAEMAESVTNPQRPPVIWIGAQECT WP_000145410.1
MULTISPECIES: hydrogenase-2 small chain [Proteobacteria] Length: 372\nNumber of predicted TMHs: 1\nExp number of AAs in TMHs: 23.72226\nExp number, first 60 AAs: 0.22038\nTotal prob of N-in: 0.12994\noutside 1 331\nTMhelix 332 354\ninside 355 372

8495 GCF_001614955.1_ASM161495v1 Escherichia coli Proteobacteria; Gammaproteobacteria; Enterobacterales; Enterobacteriaceae; Escherichia
MTGDNTLIHSHGINRRDFMKLCAALAATMGLSSKAAAEMAESVTNPQRPPVIWIGAQECT WP_000145410.1
MULTISPECIES: hydrogenase-2 small chain [Proteobacteria] Length: 372\nNumber of predicted TMHs: 1\nExp number of AAs in TMHs: 23.72226\nExp number, first 60 AAs: 0.22038\nTotal prob of N-in: 0.12994\noutside 1 331\nTMhelix 332 354\ninside 355 372

8496 GCF_001615015.1_ASM161501v1 Escherichia coli Proteobacteria; Gammaproteobacteria; Enterobacterales; Enterobacteriaceae; Escherichia
MTGDNTLIHSHGINRRDFMKLCAALAATMGLSSKAAAEMAESVTNPQRPPVIWIGAQECT WP_000145410.1
MULTISPECIES: hydrogenase-2 small chain [Proteobacteria] Length: 372\nNumber of predicted TMHs: 1\nExp number of AAs in TMHs: 23.72226\nExp number, first 60 AAs: 0.22038\nTotal prob of N-in: 0.12994\noutside 1 331\nTMhelix 332 354\ninside 355 372

8497 GCF_001614885.1_ASM161488v1 Escherichia coli Proteobacteria; Gammaproteobacteria; Enterobacterales; Enterobacteriaceae; Escherichia

MTGDNTLIHSHGINRRDFMKLCAALAATMGLSSKAAAEMAESVTNPQRPPVIWIGAQECT WP_000145410.1 MULTISPECIES: hydrogenase-2 small chain [Proteobacteria] Length: 372\nNumber of predicted TMHs: 1\nExp number of AAs in TMHs: 23.72226\nExp number, first 60 AAs: 0.22038\nTotal prob of N-in: 0.12994\noutside 1 331\nTMhelix 332 354\ninside 355 372

8498 GCF_001614955.1_ASM161495v1 Escherichia coli Proteobacteria; Gammaproteobacteria; Enterobacterales; Enterobacteriaceae; Escherichia MNNEETFYQAMRRQGVTRRSFLKYCSLAATSLGLGAGMAPKIAWALENKPRIPVVWIHGL WP_001058323.1 MULTISPECIES: hydrogenase-1 small chain [Proteobacteria] Length: 372\nNumber of predicted TMHs: 2\nExp number of AAs in TMHs: 37.2608499999999\nExp number, first 60 AAs: 19.52923\nTotal prob of N-in: 0.94351\nPOSSIBLE N-term signal sequence\ninside 1 19\nTMhelix 20 39\noutside 40 325\nTMhelix 326 348\ninside 349 372

8499 GCF_001615045.1_ASM161504v1 Escherichia coli Proteobacteria; Gammaproteobacteria; Enterobacterales; Enterobacteriaceae; Escherichia MTGDNTLIHSHGINRRDFMKLCAALAATMGLSSKAAAEMAESVTNPQRPPVIWIGAQECT WP_000145410.1 MULTISPECIES: hydrogenase-2 small chain [Proteobacteria] Length: 372\nNumber of predicted TMHs: 1\nExp number of AAs in TMHs: 23.72226\nExp number, first 60 AAs: 0.22038\nTotal prob of N-in: 0.12994\noutside 1 331\nTMhelix 332 354\ninside 355 372

8500 GCF_001615115.1_ASM161511v1 Escherichia coli Proteobacteria; Gammaproteobacteria; Enterobacterales; Enterobacteriaceae; Escherichia MTGDNTLIHSHGINRRDFMKLCAALAATMGLSSKAAAEMAESVTNPQRPPVIWIGAQECT WP_000145410.1 MULTISPECIES: hydrogenase-2 small chain [Proteobacteria] Length: 372\nNumber of predicted TMHs: 1\nExp number of AAs in TMHs: 23.72226\nExp number, first 60 AAs: 0.22038\nTotal prob of N-in: 0.12994\noutside 1 331\nTMhelix 332 354\ninside 355 372

8501 GCF_001614885.1_ASM161488v1 Escherichia coli Proteobacteria; Gammaproteobacteria; Enterobacterales; Enterobacteriaceae; Escherichia MNNEETFYQAMRRQGVTRRSFLKYCSLAATSLGLGAGMAPKIAWALENKPRIPVVWIHGL WP_001058320.1 hydrogenase [Escherichia coli] Length: 372\nNumber of predicted TMHs: 2\nExp number of AAs in TMHs: 37.26082\nExp number, first 60 AAs: 19.52923\nTotal prob of N-in: 0.94351\nPOSSIBLE N-term signal sequence\ninside 1 19\nTMhelix 20 39\noutside 40 325\nTMhelix 326 348\ninside 349 372

8502 GCF_001615015.1_ASM161501v1 Escherichia coli Proteobacteria; Gammaproteobacteria; Enterobacterales; Enterobacteriaceae; Escherichia MNNEETFYQAMRRQGVTRRSFLKYCSLAATSLGLGAGMAPKIAWALENKPRIPVVWIHGL WP_001058323.1 MULTISPECIES: hydrogenase-1 small chain [Proteobacteria] Length: 372\nNumber of predicted TMHs: 2\nExp number of AAs in TMHs: 37.2608499999999\nExp number, first 60 AAs: 19.52923\nTotal prob of N-in: 0.94351\nPOSSIBLE N-term signal sequence\ninside 1 19\nTMhelix 20 39\noutside 40 325\nTMhelix 326 348\ninside 349 372

8503 GCF_001614575.1_ASM161457v1 Escherichia coli Proteobacteria; Gammaproteobacteria; Enterobacterales; Enterobacteriaceae; Escherichia MNNEETFYQAMRRQGVTRRSFLKYCSLAATSLGLGAGMAPKIAWALENKPRIPVVWIHGL WP_001058323.1 MULTISPECIES: hydrogenase-1 small chain [Proteobacteria] Length: 372\nNumber of predicted TMHs: 2\nExp number of AAs in TMHs: 37.2608499999999\nExp number, first 60 AAs: 19.52923\nTotal prob of N-in: 0.94351\nPOSSIBLE N-term signal sequence\ninside 1 19\nTMhelix 20 39\noutside 40 325\nTMhelix 326 348\ninside 349 372

8504 GCF_001614725.1_ASM161472v1 Escherichia coli Proteobacteria; Gammaproteobacteria; Enterobacterales; Enterobacteriaceae; Escherichia MNNEETFYQAMRRQGVTRRSFLKYCSLAATSLGLGAGMAPKIAWALENKPRIPVVWIHGL WP_001058323.1 MULTISPECIES: hydrogenase-1 small chain [Proteobacteria] Length: 372\nNumber of predicted TMHs: 2\nExp number of AAs in TMHs: 37.2608499999999\nExp number, first 60 AAs: 19.52923\nTotal prob of N-in: 0.94351\nPOSSIBLE N-term signal sequence\ninside 1 19\nTMhelix 20 39\noutside 40 325\nTMhelix 326 348\ninside 349 372

8505 GCF_001614635.1_ASM161463v1 Escherichia coli Proteobacteria; Gammaproteobacteria; Enterobacterales; Enterobacteriaceae; Escherichia MNNEETFYQAMRRQGVTRRSFLKYCSLAATSLGLGAGMAPKIAWALENKPRIPVVWIHGL WP_001058323.1

MULTISPECIES: hydrogenase-1 small chain [Proteobacteria] Length: 372\nNumber of predicted TMHs: 2\nExp number of AAs in TMHs: 37.2608499999999\nExp number, first 60 AAs: 19.52923\nTotal prob of N-in: 0.94351\nPOSSIBLE N-term signal sequence\ninside 1 19\nTMhelix 20 39\noutside 40 325\nTMhelix 326 348\ninside 349 372

8506 GCF_001614825.1_ASM161482v1 Escherichia coli Proteobacteria; Gammaproteobacteria; Enterobacterales; Enterobacteriaceae; Escherichia
MNNEETFYQAMRRQGVTRRSFLKYCSLAATSLGLGAGMAPKIAWALENKPRIPVVWIHGL WP_001058320.1
hydrogenase [Escherichia coli] Length: 372\nNumber of predicted TMHs: 2\nExp number of AAs in TMHs: 37.26082\nExp number, first 60 AAs: 19.52923\nTotal prob of N-in: 0.94351\nPOSSIBLE N-term signal sequence\ninside 1 19\nTMhelix 20 39\noutside 40 325\nTMhelix 326 348\ninside 349 372

8507 GCF_001614795.1_ASM161479v1 Escherichia coli Proteobacteria; Gammaproteobacteria; Enterobacterales; Enterobacteriaceae; Escherichia
MNNEETFYQAMRRQGVTRRSFLKYCSLAATSLGLGAGMAPKIAWALENKPRIPVVWIHGL WP_001058323.1
MULTISPECIES: hydrogenase-1 small chain [Proteobacteria] Length: 372\nNumber of predicted TMHs: 2\nExp number of AAs in TMHs: 37.2608499999999\nExp number, first 60 AAs: 19.52923\nTotal prob of N-in: 0.94351\nPOSSIBLE N-term signal sequence\ninside 1 19\nTMhelix 20 39\noutside 40 325\nTMhelix 326 348\ninside 349 372

8508 GCF_001615115.1_ASM161511v1 Escherichia coli Proteobacteria; Gammaproteobacteria; Enterobacterales; Enterobacteriaceae; Escherichia
MNNEETFYQAMRRQGVTRRSFLKYCSLAATSLGLGAGMAPKIAWALENKPRIPVVWIHGL WP_001058323.1
MULTISPECIES: hydrogenase-1 small chain [Proteobacteria] Length: 372\nNumber of predicted TMHs: 2\nExp number of AAs in TMHs: 37.2608499999999\nExp number, first 60 AAs: 19.52923\nTotal prob of N-in: 0.94351\nPOSSIBLE N-term signal sequence\ninside 1 19\nTMhelix 20 39\noutside 40 325\nTMhelix 326 348\ninside 349 372

8509 GCF_001614645.1_ASM161464v1 Escherichia coli Proteobacteria; Gammaproteobacteria; Enterobacterales; Enterobacteriaceae; Escherichia
MNNEETFYQAMRRQGVTRRSFLKYCSLAATSLGLGAGMAPKIAWALENKPRIPVVWIHGL WP_001295940.1
MULTISPECIES: hydrogenase [Enterobacteriaceae] Length: 372\nNumber of predicted TMHs: 2\nExp number of AAs in TMHs: 37.2608899999999\nExp number, first 60 AAs: 19.52923\nTotal prob of N-in: 0.94351\nPOSSIBLE N-term signal sequence\ninside 1 19\nTMhelix 20 39\noutside 40 325\nTMhelix 326 348\ninside 349 372

8510 GCF_001615045.1_ASM161504v1 Escherichia coli Proteobacteria; Gammaproteobacteria; Enterobacterales; Enterobacteriaceae; Escherichia
MNNEETFYQAMRRQGVTRRSFLKYCSLAATSLGLGAGMAPKIAWALENKPRIPVVWIHGL WP_001058320.1
hydrogenase [Escherichia coli] Length: 372\nNumber of predicted TMHs: 2\nExp number of AAs in TMHs: 37.26082\nExp number, first 60 AAs: 19.52923\nTotal prob of N-in: 0.94351\nPOSSIBLE N-term signal sequence\ninside 1 19\nTMhelix 20 39\noutside 40 325\nTMhelix 326 348\ninside 349 372

8511 GCF_001614695.1_ASM161469v1 Escherichia coli Proteobacteria; Gammaproteobacteria; Enterobacterales; Enterobacteriaceae; Escherichia
MNNEETFYQAMRRQGVTRRSFLKYCSLAATSLGLGAGMAPKIAWALENKPRIPVVWIHGL WP_001295940.1
MULTISPECIES: hydrogenase [Enterobacteriaceae] Length: 372\nNumber of predicted TMHs: 2\nExp number of AAs in TMHs: 37.2608899999999\nExp number, first 60 AAs: 19.52923\nTotal prob of N-in: 0.94351\nPOSSIBLE N-term signal sequence\ninside 1 19\nTMhelix 20 39\noutside 40 325\nTMhelix 326 348\ninside 349 372

8512 GCF_001615145.1_ASM161514v1 Escherichia coli Proteobacteria; Gammaproteobacteria; Enterobacterales; Enterobacteriaceae; Escherichia
MTGDNTLIHSHGINRRDFMKLCAALAATMGLSSKAAAEMAESVTNPQRPPVIWIGAQECT WP_000145410.1
MULTISPECIES: hydrogenase-2 small chain [Proteobacteria] Length: 372\nNumber of predicted TMHs: 1\nExp number of AAs in TMHs: 23.72226\nExp number, first 60 AAs: 0.22038\nTotal prob of N-in: 0.12994\noutside 1 331\nTMhelix 332 354\ninside 355 372

8513 GCF_001615355.1_ASM161535v1 Escherichia coli Proteobacteria; Gammaproteobacteria; Enterobacterales; Enterobacteriaceae; Escherichia
MTGDNTLIHSHGINRRDFMKLCAALAATMGLSSKAAAEMAESVTNPQRPPVIWIGAQECT WP_000145410.1

MULTISPECIES: hydrogenase-2 small chain [Proteobacteria] Length: 372\nNumber of predicted TMHs: 1\nExp number of AAs in TMHs: 23.72226\nExp number, first 60 AAs: 0.22038\nTotal prob of N-in: 0.12994\noutside 1 331\nTMhelix 332 354\ninside 355 372

8514 GCF_001615305.1_ASM161530v1 Escherichia coli Proteobacteria; Gammaproteobacteria; Enterobacterales; Enterobacteriaceae; Escherichia MTGDNTLIHSHGINRRDFMKLCAALAATMGLSSKAAAEMAESVTNPQRPPVIWIGAQECT WP_000145410.1 MULTISPECIES: hydrogenase-2 small chain [Proteobacteria] Length: 372\nNumber of predicted TMHs: 1\nExp number of AAs in TMHs: 23.72226\nExp number, first 60 AAs: 0.22038\nTotal prob of N-in: 0.12994\noutside 1 331\nTMhelix 332 354\ninside 355 372

8515 GCF_001615235.1_ASM161523v1 Escherichia coli Proteobacteria; Gammaproteobacteria; Enterobacterales; Enterobacteriaceae; Escherichia MTGDNTLIHSHGINRRDFMKLCAALAATMGLSSKAAAEMAESVTNPQRPPVIWIGAQECT WP_000145410.1 MULTISPECIES: hydrogenase-2 small chain [Proteobacteria] Length: 372\nNumber of predicted TMHs: 1\nExp number of AAs in TMHs: 23.72226\nExp number, first 60 AAs: 0.22038\nTotal prob of N-in: 0.12994\noutside 1 331\nTMhelix 332 354\ninside 355 372

8516 GCF_001615475.1_ASM161547v1 Escherichia coli Proteobacteria; Gammaproteobacteria; Enterobacterales; Enterobacteriaceae; Escherichia MTGDNTLIHSHGINRRDFMKLCAALAATMGLSSKAAAEMAESVTNPQRPPVIWIGAQECT WP_000145410.1 MULTISPECIES: hydrogenase-2 small chain [Proteobacteria] Length: 372\nNumber of predicted TMHs: 1\nExp number of AAs in TMHs: 23.72226\nExp number, first 60 AAs: 0.22038\nTotal prob of N-in: 0.12994\noutside 1 331\nTMhelix 332 354\ninside 355 372

8517 GCF_001615205.1_ASM161520v1 Escherichia coli Proteobacteria; Gammaproteobacteria; Enterobacterales; Enterobacteriaceae; Escherichia MTGDNTLIHSHGINRRDFMKLCAALAATMGLSSKAAAEMAESVTNPQRPPVIWIGAQECT WP_000145410.1 MULTISPECIES: hydrogenase-2 small chain [Proteobacteria] Length: 372\nNumber of predicted TMHs: 1\nExp number of AAs in TMHs: 23.72226\nExp number, first 60 AAs: 0.22038\nTotal prob of N-in: 0.12994\noutside 1 331\nTMhelix 332 354\ninside 355 372

8518 GCF_001615455.1_ASM161545v1 Escherichia coli Proteobacteria; Gammaproteobacteria; Enterobacterales; Enterobacteriaceae; Escherichia MTGDNTLIHSHGINRRDFMKLCAALAATMGLSSKAAAEMAESVTNPQRPPVIWIGAQECT WP_000145410.1 MULTISPECIES: hydrogenase-2 small chain [Proteobacteria] Length: 372\nNumber of predicted TMHs: 1\nExp number of AAs in TMHs: 23.72226\nExp number, first 60 AAs: 0.22038\nTotal prob of N-in: 0.12994\noutside 1 331\nTMhelix 332 354\ninside 355 372

8519 GCF_001615495.1_ASM161549v1 Escherichia coli Proteobacteria; Gammaproteobacteria; Enterobacterales; Enterobacteriaceae; Escherichia MTGDNTLIHSHGINRRDFMKLCAALAATMGLSSKAAAEMAESVTNPQRPPVIWIGAQECT WP_000145410.1 MULTISPECIES: hydrogenase-2 small chain [Proteobacteria] Length: 372\nNumber of predicted TMHs: 1\nExp number of AAs in TMHs: 23.72226\nExp number, first 60 AAs: 0.22038\nTotal prob of N-in: 0.12994\noutside 1 331\nTMhelix 332 354\ninside 355 372

8520 GCF_001615615.1_ASM161561v1 Escherichia coli Proteobacteria; Gammaproteobacteria; Enterobacterales; Enterobacteriaceae; Escherichia MTGDNTLIHSHGINRRDFMKLCAALAATMGLSSKAAAEMAESVTNPQRPPVIWIGAQECT WP_000145410.1 MULTISPECIES: hydrogenase-2 small chain [Proteobacteria] Length: 372\nNumber of predicted TMHs: 1\nExp number of AAs in TMHs: 23.72226\nExp number, first 60 AAs: 0.22038\nTotal prob of N-in: 0.12994\noutside 1 331\nTMhelix 332 354\ninside 355 372

8521 GCF_001615545.1_ASM161554v1 Escherichia coli Proteobacteria; Gammaproteobacteria; Enterobacterales; Enterobacteriaceae; Escherichia MTGDNTLIHSHGINRRDFMKLCAALAATMGLSSKAAAEMAESVTNPQRPPVIWIGAQECT WP_000145410.1 MULTISPECIES: hydrogenase-2 small chain [Proteobacteria] Length: 372\nNumber of predicted TMHs: 1\nExp number of AAs in TMHs: 23.72226\nExp number, first 60 AAs: 0.22038\nTotal prob of N-in: 0.12994\noutside 1 331\nTMhelix 332 354\ninside 355 372

8522 GCF_001615145.1_ASM161514v1 Escherichia coli Proteobacteria; Gammaproteobacteria; Enterobacterales; Enterobacteriaceae; Escherichia
MNNEETFYQAMRRQGVTRRSFLKYCSLAATSLGLGAGMAPKIAWALENKPRIPVVWIHGL WP_001058323.1
MULTISPECIES: hydrogenase-1 small chain [Proteobacteria] Length: 372\nNumber of predicted TMHs: 2\nExp number of AAs in TMHs: 37.2608499999999\nExp number, first 60 AAs: 19.52923\nTotal prob of N-in: 0.94351\nPOSSIBLE N-term signal sequence\ninside 1 19\nTMhelix 20 39\noutside 40 325\nTMhelix 326 348\ninside 349 372

8523 GCF_001615305.1_ASM161530v1 Escherichia coli Proteobacteria; Gammaproteobacteria; Enterobacterales; Enterobacteriaceae; Escherichia
MNNEETFYQAMRRQGVTRRSFLKYCSLAATSLGLGAGMAPKIAWALENKPRIPVVWIHGL WP_001058323.1
MULTISPECIES: hydrogenase-1 small chain [Proteobacteria] Length: 372\nNumber of predicted TMHs: 2\nExp number of AAs in TMHs: 37.2608499999999\nExp number, first 60 AAs: 19.52923\nTotal prob of N-in: 0.94351\nPOSSIBLE N-term signal sequence\ninside 1 19\nTMhelix 20 39\noutside 40 325\nTMhelix 326 348\ninside 349 372

8524 GCF_001615405.1_ASM161540v1 Escherichia coli Proteobacteria; Gammaproteobacteria; Enterobacterales; Enterobacteriaceae; Escherichia
MNNEETFYQAMRRQGVTRRSFLKYCSLAATSLGLGAGMAPKIAWALENKPRIPVVWIHGL WP_001058323.1
MULTISPECIES: hydrogenase-1 small chain [Proteobacteria] Length: 372\nNumber of predicted TMHs: 2\nExp number of AAs in TMHs: 37.2608499999999\nExp number, first 60 AAs: 19.52923\nTotal prob of N-in: 0.94351\nPOSSIBLE N-term signal sequence\ninside 1 19\nTMhelix 20 39\noutside 40 325\nTMhelix 326 348\ninside 349 372

8525 GCF_001615615.1_ASM161561v1 Escherichia coli Proteobacteria; Gammaproteobacteria; Enterobacterales; Enterobacteriaceae; Escherichia
MNNEETFYQAMRRQGVTRRSFLKYCSLAATSLGLGAGMAPKIAWALENKPRIPVVWIHGL WP_001058323.1
MULTISPECIES: hydrogenase-1 small chain [Proteobacteria] Length: 372\nNumber of predicted TMHs: 2\nExp number of AAs in TMHs: 37.2608499999999\nExp number, first 60 AAs: 19.52923\nTotal prob of N-in: 0.94351\nPOSSIBLE N-term signal sequence\ninside 1 19\nTMhelix 20 39\noutside 40 325\nTMhelix 326 348\ninside 349 372

8526 GCF_001615455.1_ASM161545v1 Escherichia coli Proteobacteria; Gammaproteobacteria; Enterobacterales; Enterobacteriaceae; Escherichia
MNNEETFYQAMRRQGVTRRSFLKYCSLAATSLGLGAGMAPKIAWALENKPRIPVVWIHGL WP_001058320.1
hydrogenase [Escherichia coli] Length: 372\nNumber of predicted TMHs: 2\nExp number of AAs in TMHs: 37.26082\nExp number, first 60 AAs: 19.52923\nTotal prob of N-in: 0.94351\nPOSSIBLE N-term signal sequence\ninside 1 19\nTMhelix 20 39\noutside 40 325\nTMhelix 326 348\ninside 349 372

8527 GCF_001615645.1_ASM161564v1 Escherichia coli Proteobacteria; Gammaproteobacteria; Enterobacterales; Enterobacteriaceae; Escherichia
MTGDNTLIHSHGINRRDFMKLCAALAATMGLSSKAAAEMAESVTNPQRPPVIWIGAQECT WP_000145410.1
MULTISPECIES: hydrogenase-2 small chain [Proteobacteria] Length: 372\nNumber of predicted TMHs: 1\nExp number of AAs in TMHs: 23.72226\nExp number, first 60 AAs: 0.22038\nTotal prob of N-in: 0.12994\noutside 1 331\nTMhelix 332 354\ninside 355 372

8528 GCF_001615645.1_ASM161564v1 Escherichia coli Proteobacteria; Gammaproteobacteria; Enterobacterales; Enterobacteriaceae; Escherichia
MNNEETFYQAMRRQGVTRRSFLKYCSLAATSLGLGAGMAPKIAWALENKPRIPVVWIHGL WP_001058323.1
MULTISPECIES: hydrogenase-1 small chain [Proteobacteria] Length: 372\nNumber of predicted TMHs: 2\nExp number of AAs in TMHs: 37.2608499999999\nExp number, first 60 AAs: 19.52923\nTotal prob of N-in: 0.94351\nPOSSIBLE N-term signal sequence\ninside 1 19\nTMhelix 20 39\noutside 40 325\nTMhelix 326 348\ninside 349 372

8529 GCF_001615545.1_ASM161554v1 Escherichia coli Proteobacteria; Gammaproteobacteria; Enterobacterales; Enterobacteriaceae; Escherichia
MNNEETFYQAMRRQGVTRRSFLKYCSLAATSLGLGAGMAPKIAWALENKPRIPVVWIHGL WP_001058323.1
MULTISPECIES: hydrogenase-1 small chain [Proteobacteria] Length: 372\nNumber of predicted TMHs: 2\nExp number of AAs in TMHs: 37.2608499999999\nExp number, first 60 AAs: 19.52923\nTotal prob of N-in: 0.94351\nPOSSIBLE N-term signal sequence\ninside 1 19\nTMhelix 20 39\noutside 40 325\nTMhelix 326 348\ninside 349 372

8530 GCF_001615355.1_ASM161535v1 Escherichia coli Proteobacteria; Gammaproteobacteria; Enterobacterales; Enterobacteriaceae; Escherichia
MNNEETFYQAMRRQGVTRRSFLKYCSLAATSLGLGAGMAPKIAWALENKPRIPVVWIHGL WP_001058323.1
MULTISPECIES: hydrogenase-1 small chain [Proteobacteria] Length: 372\nNumber of predicted TMHs: 2\nExp number of AAs in TMHs: 37.2608499999999\nExp number, first 60 AAs: 19.52923\nTotal prob of N-in: 0.94351\nPOSSIBLE N-term signal sequence\ninside 1 19\nTMhelix 20 39\noutside 40 325\nTMhelix 326 348\ninside 349 372

8531 GCF_001615235.1_ASM161523v1 Escherichia coli Proteobacteria; Gammaproteobacteria; Enterobacterales; Enterobacteriaceae; Escherichia
MNNEETFYQAMRRQGVTRRSFLKYCSLAATSLGLGAGMAPKIAWALENKPRIPVVWIHGL WP_001058323.1
MULTISPECIES: hydrogenase-1 small chain [Proteobacteria] Length: 372\nNumber of predicted TMHs: 2\nExp number of AAs in TMHs: 37.2608499999999\nExp number, first 60 AAs: 19.52923\nTotal prob of N-in: 0.94351\nPOSSIBLE N-term signal sequence\ninside 1 19\nTMhelix 20 39\noutside 40 325\nTMhelix 326 348\ninside 349 372

8532 GCF_001615205.1_ASM161520v1 Escherichia coli Proteobacteria; Gammaproteobacteria; Enterobacterales; Enterobacteriaceae; Escherichia
MNNEETFYQAMRRQGVTRRSFLKYCSLAATSLGLGAGMAPKIAWALENKPRIPVVWIHGL WP_001058323.1
MULTISPECIES: hydrogenase-1 small chain [Proteobacteria] Length: 372\nNumber of predicted TMHs: 2\nExp number of AAs in TMHs: 37.2608499999999\nExp number, first 60 AAs: 19.52923\nTotal prob of N-in: 0.94351\nPOSSIBLE N-term signal sequence\ninside 1 19\nTMhelix 20 39\noutside 40 325\nTMhelix 326 348\ninside 349 372

8533 GCF_001615495.1_ASM161549v1 Escherichia coli Proteobacteria; Gammaproteobacteria; Enterobacterales; Enterobacteriaceae; Escherichia
MNNEETFYQAMRRQGVTRRSFLKYCSLAATSLGLGAGMAPKIAWALENKPRIPVVWIHGL WP_001058323.1
MULTISPECIES: hydrogenase-1 small chain [Proteobacteria] Length: 372\nNumber of predicted TMHs: 2\nExp number of AAs in TMHs: 37.2608499999999\nExp number, first 60 AAs: 19.52923\nTotal prob of N-in: 0.94351\nPOSSIBLE N-term signal sequence\ninside 1 19\nTMhelix 20 39\noutside 40 325\nTMhelix 326 348\ninside 349 372

8534 GCF_001615405.1_ASM161540v1 Escherichia coli Proteobacteria; Gammaproteobacteria; Enterobacterales; Enterobacteriaceae; Escherichia
MCXGDNTLIHSHGINRRDFMKLCAALAATMGLSSKAAAEMAESVTNPQRPPVIWIGAQEC WP_074150867.1
hydrogenase 2 small subunit [Escherichia coli] Length: 373\nNumber of predicted TMHs: 1\nExp number of AAs in TMHs: 23.6495\nExp number, first 60 AAs: 0.06148\nTotal prob of N-in: 0.12814\noutside 1 332\nTMhelix 333 355\ninside 356 373

8535 GCF_001615745.1_ASM161574v1 Escherichia coli Proteobacteria; Gammaproteobacteria; Enterobacterales; Enterobacteriaceae; Escherichia
MTGDNTLIHSHGINRRDFMKLCAALAATMGLSSKAAAEMAESVTNPQRPPVIWIGAQECT WP_000145410.1
MULTISPECIES: hydrogenase-2 small chain [Proteobacteria] Length: 372\nNumber of predicted TMHs: 1\nExp number of AAs in TMHs: 23.72226\nExp number, first 60 AAs: 0.22038\nTotal prob of N-in: 0.12994\noutside 1 331\nTMhelix 332 354\ninside 355 372

8536 GCF_001615705.1_ASM161570v1 Escherichia coli Proteobacteria; Gammaproteobacteria; Enterobacterales; Enterobacteriaceae; Escherichia
MTGDNTLIHSHGINRRDFMKLCAALAATMGLSSKAAAEMAESVTNPQRPPVIWIGAQECT WP_000145410.1
MULTISPECIES: hydrogenase-2 small chain [Proteobacteria] Length: 372\nNumber of predicted TMHs: 1\nExp number of AAs in TMHs: 23.72226\nExp number, first 60 AAs: 0.22038\nTotal prob of N-in: 0.12994\noutside 1 331\nTMhelix 332 354\ninside 355 372

8537 GCF_001615805.1_ASM161580v1 Escherichia coli Proteobacteria; Gammaproteobacteria; Enterobacterales; Enterobacteriaceae; Escherichia
MTGDNTLIHSHGINRRDFMKLCAALAATMGLSSKAAAEMAESVTNPQRPPVIWIGAQECT WP_000145410.1
MULTISPECIES: hydrogenase-2 small chain [Proteobacteria] Length: 372\nNumber of predicted TMHs: 1\nExp number of AAs in TMHs: 23.72226\nExp number, first 60 AAs: 0.22038\nTotal prob of N-in: 0.12994\noutside 1 331\nTMhelix 332 354\ninside 355 372

8538 GCF_001615855.1_ASM161585v1 Escherichia coli Proteobacteria; Gammaproteobacteria; Enterobacterales; Enterobacteriaceae; Escherichia
MTGDNTLIHSHGINRRDFMKLCAALAATMGLSSKAAAEMAESVTNPQRPPVIWIGAQECT WP_000145410.1
MULTISPECIES: hydrogenase-2 small chain [Proteobacteria] Length: 372\nNumber of predicted TMHs: 1\nExp number of AAs in TMHs: 23.72226\nExp number, first 60 AAs: 0.22038\nTotal prob of N-in: 0.12994\noutside 1 331\nTMhelix 332 354\ninside 355 372

8539 GCF_001615955.1_ASM161595v1 Escherichia coli Proteobacteria; Gammaproteobacteria; Enterobacterales; Enterobacteriaceae; Escherichia
MTGDNTLIHSHGINRRDFMKLCAALAATMGLSSKAAAEMAESVTNPQRPPVIWIGAQECT WP_000145410.1
MULTISPECIES: hydrogenase-2 small chain [Proteobacteria] Length: 372\nNumber of predicted TMHs: 1\nExp number of AAs in TMHs: 23.72226\nExp number, first 60 AAs: 0.22038\nTotal prob of N-in: 0.12994\noutside 1 331\nTMhelix 332 354\ninside 355 372

8540 GCF_001616045.1_ASM161604v1 Escherichia coli Proteobacteria; Gammaproteobacteria; Enterobacterales; Enterobacteriaceae; Escherichia
MTGDNTLIHSHGINRRDFMKLCAALAATMGLSSKAAAEMAESVTNPQRPPVIWIGAQECT WP_000145410.1
MULTISPECIES: hydrogenase-2 small chain [Proteobacteria] Length: 372\nNumber of predicted TMHs: 1\nExp number of AAs in TMHs: 23.72226\nExp number, first 60 AAs: 0.22038\nTotal prob of N-in: 0.12994\noutside 1 331\nTMhelix 332 354\ninside 355 372

8541 GCF_001616015.1_ASM161601v1 Escherichia coli Proteobacteria; Gammaproteobacteria; Enterobacterales; Enterobacteriaceae; Escherichia
MTGDNTLIHSHGINRRDFMKLCAALAATMGLSSKAAAEMAESVTNPQRPPVIWIGAQECT WP_000145410.1
MULTISPECIES: hydrogenase-2 small chain [Proteobacteria] Length: 372\nNumber of predicted TMHs: 1\nExp number of AAs in TMHs: 23.72226\nExp number, first 60 AAs: 0.22038\nTotal prob of N-in: 0.12994\noutside 1 331\nTMhelix 332 354\ninside 355 372

8542 GCF_001616105.1_ASM161610v1 Escherichia coli Proteobacteria; Gammaproteobacteria; Enterobacterales; Enterobacteriaceae; Escherichia
MTGDNTLIHSHGINRRDFMKLCAALAATMGLSSKAAAEMAESVTNPQRPPVIWIGAQECT WP_000145410.1
MULTISPECIES: hydrogenase-2 small chain [Proteobacteria] Length: 372\nNumber of predicted TMHs: 1\nExp number of AAs in TMHs: 23.72226\nExp number, first 60 AAs: 0.22038\nTotal prob of N-in: 0.12994\noutside 1 331\nTMhelix 332 354\ninside 355 372

8543 GCF_001616105.1_ASM161610v1 Escherichia coli Proteobacteria; Gammaproteobacteria; Enterobacterales; Enterobacteriaceae; Escherichia
MNNEETFYQAMRRQGVTRRSFLKYCSLAATSLGLGAGMAPKIAWALENKPRIPVVWIHGL WP_063078438.1
hydrogenase [Escherichia coli] Length: 372\nNumber of predicted TMHs: 2\nExp number of AAs in TMHs: 37.06316\nExp number, first 60 AAs: 19.52931\nTotal prob of N-in: 0.94340\nPOSSIBLE N-term signal sequence\ninside 1 19\nTMhelix 20 39\noutside 40 325\nTMhelix 326 348\ninside 349 372

8544 GCF_001616655.1_ASM161665v1 Escherichia coli Proteobacteria; Gammaproteobacteria; Enterobacterales; Enterobacteriaceae; Escherichia
MTGDNTLIHSHGINRRDFMKLCAALAATMGLSSKAAAEMAESVTNPQRPPVIWIGAQECT WP_000145410.1
MULTISPECIES: hydrogenase-2 small chain [Proteobacteria] Length: 372\nNumber of predicted TMHs: 1\nExp number of AAs in TMHs: 23.72226\nExp number, first 60 AAs: 0.22038\nTotal prob of N-in: 0.12994\noutside 1 331\nTMhelix 332 354\ninside 355 372

8545 GCF_001615805.1_ASM161580v1 Escherichia coli Proteobacteria; Gammaproteobacteria; Enterobacterales; Enterobacteriaceae; Escherichia
MNNEETFYQAMRRQGVTRRSFLKYCSLAATSLGLGAGMAPKIAWALENKPRIPVVWIHGL WP_001058323.1
MULTISPECIES: hydrogenase-1 small chain [Proteobacteria] Length: 372\nNumber of predicted TMHs: 2\nExp number of AAs in TMHs: 37.2608499999999\nExp number, first 60 AAs: 19.52923\nTotal prob of N-in: 0.94351\nPOSSIBLE N-term signal sequence\ninside 1 19\nTMhelix 20 39\noutside 40 325\nTMhelix 326 348\ninside 349 372

8546 GCF_001616145.1_ASM161614v1 Escherichia coli Proteobacteria; Gammaproteobacteria; Enterobacterales; Enterobacteriaceae; Escherichia
MTGDNTLIHSHGINRRDFMKLCAALAATMGLSSKAAAEMAESVTNPQRPPVIWIGAQECT WP_000145410.1
MULTISPECIES: hydrogenase-2 small chain [Proteobacteria] Length: 372\nNumber of predicted TMHs:

1\nExp number of AAs in TMHs: 23.72226\nExp number, first 60 AAs: 0.22038\nTotal prob of N-in: 0.12994\noutside 1 331\nTMhelix 332 354\ninside 355 372

8547 GCF_001616205.1_ASM161620v1 Escherichia coli Proteobacteria; Gammaproteobacteria; Enterobacterales; Enterobacteriaceae; Escherichia
MTGDNTLIHSHGINRRDFMKLCAALAATMGLSSKAAAEMAESVTNPQRPPVIWIGAQECT WP_000145410.1
MULTISPECIES: hydrogenase-2 small chain [Proteobacteria] Length: 372\nNumber of predicted TMHs: 1\nExp number of AAs in TMHs: 23.72226\nExp number, first 60 AAs: 0.22038\nTotal prob of N-in: 0.12994\noutside 1 331\nTMhelix 332 354\ninside 355 372

8548 GCF_001616145.1_ASM161614v1 Escherichia coli Proteobacteria; Gammaproteobacteria; Enterobacterales; Enterobacteriaceae; Escherichia
MNNEETFYQAMRRQGVTRRSFLKYCSLAATSLGLGAGMAPKIAWALENKPRIPVVWIHGL WP_001058323.1
MULTISPECIES: hydrogenase-1 small chain [Proteobacteria] Length: 372\nNumber of predicted TMHs: 2\nExp number of AAs in TMHs: 37.2608499999999\nExp number, first 60 AAs: 19.52923\nTotal prob of N-in: 0.94351\nPOSSIBLE N-term signal sequence\ninside 1 19\nTMhelix 20 39\noutside 40 325\nTMhelix 326 348\ninside 349 372

8549 GCF_001616205.1_ASM161620v1 Escherichia coli Proteobacteria; Gammaproteobacteria; Enterobacterales; Enterobacteriaceae; Escherichia
MNNEETFYQAMRRQGVTRRSFLKYCSLAATSLGLGAGMAPKIAWALENKPRIPVVWIHGL WP_001058323.1
MULTISPECIES: hydrogenase-1 small chain [Proteobacteria] Length: 372\nNumber of predicted TMHs: 2\nExp number of AAs in TMHs: 37.2608499999999\nExp number, first 60 AAs: 19.52923\nTotal prob of N-in: 0.94351\nPOSSIBLE N-term signal sequence\ninside 1 19\nTMhelix 20 39\noutside 40 325\nTMhelix 326 348\ninside 349 372

8550 GCF_001615745.1_ASM161574v1 Escherichia coli Proteobacteria; Gammaproteobacteria; Enterobacterales; Enterobacteriaceae; Escherichia
MNNEETFYQAMRRQGVTRRSFLKYCSLAATSLGLGAGMAPKIAWALENKPRIPVVWIHGL WP_001058323.1
MULTISPECIES: hydrogenase-1 small chain [Proteobacteria] Length: 372\nNumber of predicted TMHs: 2\nExp number of AAs in TMHs: 37.2608499999999\nExp number, first 60 AAs: 19.52923\nTotal prob of N-in: 0.94351\nPOSSIBLE N-term signal sequence\ninside 1 19\nTMhelix 20 39\noutside 40 325\nTMhelix 326 348\ninside 349 372

8551 GCF_001615705.1_ASM161570v1 Escherichia coli Proteobacteria; Gammaproteobacteria; Enterobacterales; Enterobacteriaceae; Escherichia
MNNEETFYQAMRRQGVTRRSFLKYCSLAATSLGLGAGMAPKIAWALENKPRIPVVWIHGL WP_001058323.1
MULTISPECIES: hydrogenase-1 small chain [Proteobacteria] Length: 372\nNumber of predicted TMHs: 2\nExp number of AAs in TMHs: 37.2608499999999\nExp number, first 60 AAs: 19.52923\nTotal prob of N-in: 0.94351\nPOSSIBLE N-term signal sequence\ninside 1 19\nTMhelix 20 39\noutside 40 325\nTMhelix 326 348\ninside 349 372

8552 GCF_001616045.1_ASM161604v1 Escherichia coli Proteobacteria; Gammaproteobacteria; Enterobacterales; Enterobacteriaceae; Escherichia
MNNEETFYQAMRRQGVTRRSFLKYCSLAATSLGLGAGMAPKIAWALENKPRIPVVWIHGL WP_001058323.1
MULTISPECIES: hydrogenase-1 small chain [Proteobacteria] Length: 372\nNumber of predicted TMHs: 2\nExp number of AAs in TMHs: 37.2608499999999\nExp number, first 60 AAs: 19.52923\nTotal prob of N-in: 0.94351\nPOSSIBLE N-term signal sequence\ninside 1 19\nTMhelix 20 39\noutside 40 325\nTMhelix 326 348\ninside 349 372

8553 GCF_001615855.1_ASM161585v1 Escherichia coli Proteobacteria; Gammaproteobacteria; Enterobacterales; Enterobacteriaceae; Escherichia
MNNEETFYQAMRRQGVTRRSFLKYCSLAATSLGLGAGMAPKIAWALENKPRIPVVWIHGL WP_001058323.1
MULTISPECIES: hydrogenase-1 small chain [Proteobacteria] Length: 372\nNumber of predicted TMHs: 2\nExp number of AAs in TMHs: 37.2608499999999\nExp number, first 60 AAs: 19.52923\nTotal prob of N-in: 0.94351\nPOSSIBLE N-term signal sequence\ninside 1 19\nTMhelix 20 39\noutside 40 325\nTMhelix 326 348\ninside 349 372

8554 GCF_001615915.1_ASM161591v1 Escherichia coli Proteobacteria; Gammaproteobacteria; Enterobacterales; Enterobacteriaceae; Escherichia
MNNEETFYQAMRRQGVTRRSFLKYCSLAATSLGLGAGMAPKIAWALENKPRIPVVWIHGL WP_001058323.1

MULTISPECIES: hydrogenase-1 small chain [Proteobacteria] Length: 372\nNumber of predicted TMHs: 2\nExp number of AAs in TMHs: 37.2608499999999\nExp number, first 60 AAs: 19.52923\nTotal prob of N-in: 0.94351\nPOSSIBLE N-term signal sequence\ninside 1 19\nTMhelix 20 39\noutside 40 325\nTMhelix 326 348\ninside 349 372

8555 GCF_001615915.1_ASM161591v1 Escherichia coli Proteobacteria; Gammaproteobacteria; Enterobacterales; Enterobacteriaceae; Escherichia
MTGDNTLIHSHGINRRDFMKLCAALAATMGLSSKAAAEMAESVTNPQRPPVIWIGAQECT WP_032298447.1
hydrogenase 2 small subunit [Escherichia coli] Length: 372\nNumber of predicted TMHs: 1\nExp number of AAs in TMHs: 23.69807\nExp number, first 60 AAs: 0.221\nTotal prob of N-in: 0.12853\noutside 1 331\nTMhelix 332 354\ninside 355 372

8556 GCF_001615955.1_ASM161595v1 Escherichia coli Proteobacteria; Gammaproteobacteria; Enterobacterales; Enterobacteriaceae; Escherichia
MNNEETFYQAMRRQGVTRRSFLKYCSLAATSLGLGAGMAPKIAWALENKPRIPVVWIHGL WP_001058323.1
MULTISPECIES: hydrogenase-1 small chain [Proteobacteria] Length: 372\nNumber of predicted TMHs: 2\nExp number of AAs in TMHs: 37.2608499999999\nExp number, first 60 AAs: 19.52923\nTotal prob of N-in: 0.94351\nPOSSIBLE N-term signal sequence\ninside 1 19\nTMhelix 20 39\noutside 40 325\nTMhelix 326 348\ninside 349 372

8557 GCF_001616255.1_ASM161625v1 Escherichia coli Proteobacteria; Gammaproteobacteria; Enterobacterales; Enterobacteriaceae; Escherichia
MTGDNTLIHSHGINRRDFMKLCAALAATMGLSSKAAAEMAESVTNPQRPPVIWIGAQECT WP_000145410.1
MULTISPECIES: hydrogenase-2 small chain [Proteobacteria] Length: 372\nNumber of predicted TMHs: 1\nExp number of AAs in TMHs: 23.72226\nExp number, first 60 AAs: 0.22038\nTotal prob of N-in: 0.12994\noutside 1 331\nTMhelix 332 354\ninside 355 372

8558 GCF_001616255.1_ASM161625v1 Escherichia coli Proteobacteria; Gammaproteobacteria; Enterobacterales; Enterobacteriaceae; Escherichia
MNNEETFYQAMRRQGVTRRSFLKYCSLAATSLGLGAGMAPKIAWALENKPRIPVVWIHGL WP_001295940.1
MULTISPECIES: hydrogenase [Enterobacteriaceae] Length: 372\nNumber of predicted TMHs: 2\nExp number of AAs in TMHs: 37.2608899999999\nExp number, first 60 AAs: 19.52923\nTotal prob of N-in: 0.94351\nPOSSIBLE N-term signal sequence\ninside 1 19\nTMhelix 20 39\noutside 40 325\nTMhelix 326 348\ninside 349 372

8559 GCF_001616015.1_ASM161601v1 Escherichia coli Proteobacteria; Gammaproteobacteria; Enterobacterales; Enterobacteriaceae; Escherichia
MNNEETFYQAMRRQGVTRRSFLKYCSLAATSLGLGAGMAPKIAWALENKPRIPVVWIHGL WP_001295940.1
MULTISPECIES: hydrogenase [Enterobacteriaceae] Length: 372\nNumber of predicted TMHs: 2\nExp number of AAs in TMHs: 37.2608899999999\nExp number, first 60 AAs: 19.52923\nTotal prob of N-in: 0.94351\nPOSSIBLE N-term signal sequence\ninside 1 19\nTMhelix 20 39\noutside 40 325\nTMhelix 326 348\ninside 349 372

8560 GCF_001616275.1_ASM161627v1 Escherichia coli Proteobacteria; Gammaproteobacteria; Enterobacterales; Enterobacteriaceae; Escherichia
MTGDNTLIHSHGINRRDFMKLCAALAATMGLSSKAAAEMAESVTNPQRPPVIWIGAQECT WP_000145410.1
MULTISPECIES: hydrogenase-2 small chain [Proteobacteria] Length: 372\nNumber of predicted TMHs: 1\nExp number of AAs in TMHs: 23.72226\nExp number, first 60 AAs: 0.22038\nTotal prob of N-in: 0.12994\noutside 1 331\nTMhelix 332 354\ninside 355 372

8561 GCF_001616305.1_ASM161630v1 Escherichia coli Proteobacteria; Gammaproteobacteria; Enterobacterales; Enterobacteriaceae; Escherichia
MTGDNTLIHSHGINRRDFMKLCAALAATMGLSSKAAAEMAESVTNPQRPPVIWIGAQECT WP_000145410.1
MULTISPECIES: hydrogenase-2 small chain [Proteobacteria] Length: 372\nNumber of predicted TMHs: 1\nExp number of AAs in TMHs: 23.72226\nExp number, first 60 AAs: 0.22038\nTotal prob of N-in: 0.12994\noutside 1 331\nTMhelix 332 354\ninside 355 372

8562 GCF_001616345.1_ASM161634v1 Escherichia coli Proteobacteria; Gammaproteobacteria; Enterobacterales; Enterobacteriaceae; Escherichia
MTGDNTLIHSHGINRRDFMKLCAALAATMGLSSKAAAEMAESVTNPQRPPVIWIGAQECT WP_000145410.1
MULTISPECIES: hydrogenase-2 small chain [Proteobacteria] Length: 372\nNumber of predicted TMHs:

1\nExp number of AAs in TMHs: 23.72226\nExp number, first 60 AAs: 0.22038\nTotal prob of N-in: 0.12994\noutside 1 331\nTMhelix 332 354\ninside 355 372

8563 GCF_001616455.1_ASM161645v1 Escherichia coli Proteobacteria; Gammaproteobacteria; Enterobacterales; Enterobacteriaceae; Escherichia
MTGDNTLIHSHGINRRDFMKLCAALAATMGLSSKAAAEMAESVTNPQRPPVIWIGAQECT WP_000145410.1
MULTISPECIES: hydrogenase-2 small chain [Proteobacteria] Length: 372\nNumber of predicted TMHs: 1\nExp number of AAs in TMHs: 23.72226\nExp number, first 60 AAs: 0.22038\nTotal prob of N-in: 0.12994\noutside 1 331\nTMhelix 332 354\ninside 355 372

8564 GCF_001616515.1_ASM161651v1 Escherichia coli Proteobacteria; Gammaproteobacteria; Enterobacterales; Enterobacteriaceae; Escherichia
MTGDNTLIHSHGINRRDFMKLCAALAATMGLSSKAAAEMAESVTNPQRPPVIWIGAQECT WP_000145410.1
MULTISPECIES: hydrogenase-2 small chain [Proteobacteria] Length: 372\nNumber of predicted TMHs: 1\nExp number of AAs in TMHs: 23.72226\nExp number, first 60 AAs: 0.22038\nTotal prob of N-in: 0.12994\noutside 1 331\nTMhelix 332 354\ninside 355 372

8565 GCF_001616615.1_ASM161661v1 Escherichia coli Proteobacteria; Gammaproteobacteria; Enterobacterales; Enterobacteriaceae; Escherichia
MTGDNTLIHSHGINRRDFMKLCAALAATMGLSSKAAAEMAESVTNPQRPPVIWIGAQECT WP_000145410.1
MULTISPECIES: hydrogenase-2 small chain [Proteobacteria] Length: 372\nNumber of predicted TMHs: 1\nExp number of AAs in TMHs: 23.72226\nExp number, first 60 AAs: 0.22038\nTotal prob of N-in: 0.12994\noutside 1 331\nTMhelix 332 354\ninside 355 372

8566 GCF_001616555.1_ASM161655v1 Escherichia coli Proteobacteria; Gammaproteobacteria; Enterobacterales; Enterobacteriaceae; Escherichia
MTGDNTLIHSHGINRRDFMKLCAALAATMGLSSKAAAEMAESVTNPQRPPVIWIGAQECT WP_000145410.1
MULTISPECIES: hydrogenase-2 small chain [Proteobacteria] Length: 372\nNumber of predicted TMHs: 1\nExp number of AAs in TMHs: 23.72226\nExp number, first 60 AAs: 0.22038\nTotal prob of N-in: 0.12994\noutside 1 331\nTMhelix 332 354\ninside 355 372

8567 GCF_001616715.1_ASM161671v1 Escherichia coli Proteobacteria; Gammaproteobacteria; Enterobacterales; Enterobacteriaceae; Escherichia
MTGDNTLIHSHGINRRDFMKLCAALAATMGLSSKAAAEMAESVTNPQRPPVIWIGAQECT WP_000145410.1
MULTISPECIES: hydrogenase-2 small chain [Proteobacteria] Length: 372\nNumber of predicted TMHs: 1\nExp number of AAs in TMHs: 23.72226\nExp number, first 60 AAs: 0.22038\nTotal prob of N-in: 0.12994\noutside 1 331\nTMhelix 332 354\ninside 355 372

8568 GCF_001616755.1_ASM161675v1 Escherichia coli Proteobacteria; Gammaproteobacteria; Enterobacterales; Enterobacteriaceae; Escherichia
MTGDNTLIHSHGINRRDFMKLCAALAATMGLSSKAAAEMAESVTNPQRPPVIWIGAQECT WP_000145410.1
MULTISPECIES: hydrogenase-2 small chain [Proteobacteria] Length: 372\nNumber of predicted TMHs: 1\nExp number of AAs in TMHs: 23.72226\nExp number, first 60 AAs: 0.22038\nTotal prob of N-in: 0.12994\noutside 1 331\nTMhelix 332 354\ninside 355 372

8569 GCF_001616655.1_ASM161665v1 Escherichia coli Proteobacteria; Gammaproteobacteria; Enterobacterales; Enterobacteriaceae; Escherichia
MNNEETFYQAMRRQGVTRRSFLKYCSLAATSLGLGAGMAPKIAWALENKPRIPVVWIHGL WP_001058323.1
MULTISPECIES: hydrogenase-1 small chain [Proteobacteria] Length: 372\nNumber of predicted TMHs: 2\nExp number of AAs in TMHs: 37.2608499999999\nExp number, first 60 AAs: 19.52923\nTotal prob of N-in: 0.94351\nPOSSIBLE N-term signal sequence\ninside 1 19\nTMhelix 20 39\noutside 40 325\nTMhelix 326 348\ninside 349 372

8570 GCF_001616715.1_ASM161671v1 Escherichia coli Proteobacteria; Gammaproteobacteria; Enterobacterales; Enterobacteriaceae; Escherichia
MNNEETFYQAMRRQGVTRRSFLKYCSLAATSLGLGAGMAPKIAWALENKPRIPVVWIHGL WP_001058323.1
MULTISPECIES: hydrogenase-1 small chain [Proteobacteria] Length: 372\nNumber of predicted TMHs: 2\nExp number of AAs in TMHs: 37.2608499999999\nExp number, first 60 AAs: 19.52923\nTotal prob of N-in: 0.94351\nPOSSIBLE N-term signal sequence\ninside 1 19\nTMhelix 20 39\noutside 40 325\nTMhelix 326 348\ninside 349 372

8571	GCF_001616755.1_ASM161675v1	Escherichia coli	Proteobacteria; Gammaproteobacteria; Enterobacterales; Enterobacteriaceae; Escherichia
MNNEETFYQAMRRQGVTRRSFLKYCSLAATSLGLGAGMAPKIAWALENKPRIPVVWIHGL	WP_001058323.1	MULTISPECIES: hydrogenase-1 small chain [Proteobacteria]	Length: 372\nNumber of predicted TMHs: 2\nExp number of AAs in TMHs: 37.2608499999999\nExp number, first 60 AAs: 19.52923\nTotal prob of N-in: 0.94351\nPOSSIBLE N-term signal sequence\ninside 1 19\nTMhelix 20 39\noutside 40 325\nTMhelix 326 348\ninside 349 372

8572	GCF_001616305.1_ASM161630v1	Escherichia coli	Proteobacteria; Gammaproteobacteria; Enterobacterales; Enterobacteriaceae; Escherichia
MNNEETFYQAMRRQGVTRRSFLKYCSLAATSLGLGAGMAPKIAWALENKPRIPVVWIHGL	WP_001058323.1	MULTISPECIES: hydrogenase-1 small chain [Proteobacteria]	Length: 372\nNumber of predicted TMHs: 2\nExp number of AAs in TMHs: 37.2608499999999\nExp number, first 60 AAs: 19.52923\nTotal prob of N-in: 0.94351\nPOSSIBLE N-term signal sequence\ninside 1 19\nTMhelix 20 39\noutside 40 325\nTMhelix 326 348\ninside 349 372

8573	GCF_001616275.1_ASM161627v1	Escherichia coli	Proteobacteria; Gammaproteobacteria; Enterobacterales; Enterobacteriaceae; Escherichia
MNNEETFYQAMRRQGVTRRSFLKYCSLAATSLGLGAGMAPKIAWALENKPRIPVVWIHGL	WP_001058323.1	MULTISPECIES: hydrogenase-1 small chain [Proteobacteria]	Length: 372\nNumber of predicted TMHs: 2\nExp number of AAs in TMHs: 37.2608499999999\nExp number, first 60 AAs: 19.52923\nTotal prob of N-in: 0.94351\nPOSSIBLE N-term signal sequence\ninside 1 19\nTMhelix 20 39\noutside 40 325\nTMhelix 326 348\ninside 349 372

8574	GCF_001616345.1_ASM161634v1	Escherichia coli	Proteobacteria; Gammaproteobacteria; Enterobacterales; Enterobacteriaceae; Escherichia
MNNEETFYQAMRRQGVTRRSFLKYCSLAATSLGLGAGMAPKIAWALENKPRIPVVWIHGL	WP_001058323.1	MULTISPECIES: hydrogenase-1 small chain [Proteobacteria]	Length: 372\nNumber of predicted TMHs: 2\nExp number of AAs in TMHs: 37.2608499999999\nExp number, first 60 AAs: 19.52923\nTotal prob of N-in: 0.94351\nPOSSIBLE N-term signal sequence\ninside 1 19\nTMhelix 20 39\noutside 40 325\nTMhelix 326 348\ninside 349 372

8575	GCF_001616515.1_ASM161651v1	Escherichia coli	Proteobacteria; Gammaproteobacteria; Enterobacterales; Enterobacteriaceae; Escherichia
MNNEETFYQAMRRQGVTRRSFLKYCSLAATSLGLGAGMAPKIAWALENKPRIPVVWIHGL	WP_001058323.1	MULTISPECIES: hydrogenase-1 small chain [Proteobacteria]	Length: 372\nNumber of predicted TMHs: 2\nExp number of AAs in TMHs: 37.2608499999999\nExp number, first 60 AAs: 19.52923\nTotal prob of N-in: 0.94351\nPOSSIBLE N-term signal sequence\ninside 1 19\nTMhelix 20 39\noutside 40 325\nTMhelix 326 348\ninside 349 372

8576	GCF_001616455.1_ASM161645v1	Escherichia coli	Proteobacteria; Gammaproteobacteria; Enterobacterales; Enterobacteriaceae; Escherichia
MNNEETFYQAMRRQGVTRRSFLKYCSLAATSLGLGAGMAPKIAWALENKPRIPVVWIHGL	WP_001058323.1	MULTISPECIES: hydrogenase-1 small chain [Proteobacteria]	Length: 372\nNumber of predicted TMHs: 2\nExp number of AAs in TMHs: 37.2608499999999\nExp number, first 60 AAs: 19.52923\nTotal prob of N-in: 0.94351\nPOSSIBLE N-term signal sequence\ninside 1 19\nTMhelix 20 39\noutside 40 325\nTMhelix 326 348\ninside 349 372

8577	GCF_001616555.1_ASM161655v1	Escherichia coli	Proteobacteria; Gammaproteobacteria; Enterobacterales; Enterobacteriaceae; Escherichia
MNNEETFYQAMRRQGVTRRSFLKYCSLAATSLGLGAGMAPKIAWALENKPRIPVVWIHGL	WP_001058323.1	MULTISPECIES: hydrogenase-1 small chain [Proteobacteria]	Length: 372\nNumber of predicted TMHs: 2\nExp number of AAs in TMHs: 37.2608499999999\nExp number, first 60 AAs: 19.52923\nTotal prob of N-in: 0.94351\nPOSSIBLE N-term signal sequence\ninside 1 19\nTMhelix 20 39\noutside 40 325\nTMhelix 326 348\ninside 349 372

8578	GCF_001616415.1_ASM161641v1	Escherichia coli	Proteobacteria; Gammaproteobacteria; Enterobacterales; Enterobacteriaceae; Escherichia
MNNEETFYQAMRRQGVTRRSFLKYCSLAATSLGLGAGMAPKIAWALENKPRIPVVWIHGL	WP_001058323.1	MULTISPECIES: hydrogenase-1 small chain [Proteobacteria]	Length: 372\nNumber of predicted TMHs: 2\nExp number of AAs in TMHs: 37.2608499999999\nExp number, first 60 AAs: 19.52923\nTotal prob of N-in:

0.94351\nPOSSIBLE N-term signal sequence\ninside 1 19\nTMhelix 20 39\noutside 40 325\nTMhelix 326 348\ninside 349 372

8579 GCF_001616815.1_ASM161681v1 Escherichia coli Proteobacteria; Gammaproteobacteria; Enterobacterales; Enterobacteriaceae; Escherichia
MTGDNTLIHSHGINRRDFMKLCAALAATMGLSSKAAAEMAESVTNPQRPPVIWIGAQECT WP_000145410.1
MULTISPECIES: hydrogenase-2 small chain [Proteobacteria] Length: 372\nNumber of predicted TMHs: 1\nExp number of AAs in TMHs: 23.72226\nExp number, first 60 AAs: 0.22038\nTotal prob of N-in: 0.12994\noutside 1 331\nTMhelix 332 354\ninside 355 372

8580 GCF_001616415.1_ASM161641v1 Escherichia coli Proteobacteria; Gammaproteobacteria; Enterobacterales; Enterobacteriaceae; Escherichia
MTGDNTLIHSHGINRRDFMKLCAALAATMGLSSKAAAELAESVTNPQRPPVIWIGAQECT WP_063080715.1
hydrogenase 2 small subunit [Escherichia coli] Length: 372\nNumber of predicted TMHs: 1\nExp number of AAs in TMHs: 23.69631\nExp number, first 60 AAs: 0.19388\nTotal prob of N-in: 0.12881\noutside 1 331\nTMhelix 332 354\ninside 355 372

8581 GCF_001616815.1_ASM161681v1 Escherichia coli Proteobacteria; Gammaproteobacteria; Enterobacterales; Enterobacteriaceae; Escherichia
MNNEETFYQAMRRQGVTRRSFLKYCSLAATSLGLGAGMAPKIAWALENKPRIPVVWIHGL WP_001295940.1
MULTISPECIES: hydrogenase [Enterobacteriaceae] Length: 372\nNumber of predicted TMHs: 2\nExp number of AAs in TMHs: 37.2608899999999\nExp number, first 60 AAs: 19.52923\nTotal prob of N-in: 0.94351\nPOSSIBLE N-term signal sequence\ninside 1 19\nTMhelix 20 39\noutside 40 325\nTMhelix 326 348\ninside 349 372

8582 GCF_001616615.1_ASM161661v1 Escherichia coli Proteobacteria; Gammaproteobacteria; Enterobacterales; Enterobacteriaceae; Escherichia
MNNEETFYQAMRRQGVTRRSFLKYCSLAATSLGLGAGMAPKIAWALENKPRIPVVWIHGL WP_001058323.1
MULTISPECIES: hydrogenase-1 small chain [Proteobacteria] Length: 372\nNumber of predicted TMHs: 2\nExp number of AAs in TMHs: 37.2608499999999\nExp number, first 60 AAs: 19.52923\nTotal prob of N-in: 0.94351\nPOSSIBLE N-term signal sequence\ninside 1 19\nTMhelix 20 39\noutside 40 325\nTMhelix 326 348\ninside 349 372

8583 GCF_001616905.1_ASM161690v1 Escherichia coli Proteobacteria; Gammaproteobacteria; Enterobacterales; Enterobacteriaceae; Escherichia
MTGDNTLIHSHGINRRDFMKLCAALAATMGLSSKAAAEMAESVTNPQRPPVIWIGAQECT WP_000145410.1
MULTISPECIES: hydrogenase-2 small chain [Proteobacteria] Length: 372\nNumber of predicted TMHs: 1\nExp number of AAs in TMHs: 23.72226\nExp number, first 60 AAs: 0.22038\nTotal prob of N-in: 0.12994\noutside 1 331\nTMhelix 332 354\ninside 355 372

8584 GCF_001616855.1_ASM161685v1 Escherichia coli Proteobacteria; Gammaproteobacteria; Enterobacterales; Enterobacteriaceae; Escherichia
MTGDNTLIHSHGINRRDFMKLCAALAATMGLSSKAAAEMAESVTNPQRPPVIWIGAQECT WP_000145410.1
MULTISPECIES: hydrogenase-2 small chain [Proteobacteria] Length: 372\nNumber of predicted TMHs: 1\nExp number of AAs in TMHs: 23.72226\nExp number, first 60 AAs: 0.22038\nTotal prob of N-in: 0.12994\noutside 1 331\nTMhelix 332 354\ninside 355 372

8585 GCF_001616955.1_ASM161695v1 Escherichia coli Proteobacteria; Gammaproteobacteria; Enterobacterales; Enterobacteriaceae; Escherichia
MTGDNTLIHSHGINRRDFMKLCAALAATMGLSSKAAAEMAESVTNPQRPPVIWIGAQECT WP_000145410.1
MULTISPECIES: hydrogenase-2 small chain [Proteobacteria] Length: 372\nNumber of predicted TMHs: 1\nExp number of AAs in TMHs: 23.72226\nExp number, first 60 AAs: 0.22038\nTotal prob of N-in: 0.12994\noutside 1 331\nTMhelix 332 354\ninside 355 372

8586 GCF_001617055.1_ASM161705v1 Escherichia coli Proteobacteria; Gammaproteobacteria; Enterobacterales; Enterobacteriaceae; Escherichia
MTGDNTLIHSHGINRRDFMKLCAALAATMGLSSKAAAEMAESVTNPQRPPVIWIGAQECT WP_000145410.1
MULTISPECIES: hydrogenase-2 small chain [Proteobacteria] Length: 372\nNumber of predicted TMHs: 1\nExp number of AAs in TMHs: 23.72226\nExp number, first 60 AAs: 0.22038\nTotal prob of N-in: 0.12994\noutside 1 331\nTMhelix 332 354\ninside 355 372

8587 GCF_001617015.1_ASM161701v1 Escherichia coli Proteobacteria; Gammaproteobacteria; Enterobacterales; Enterobacteriaceae; Escherichia
MTGDNTLIHSHGINRRDFMKLCAALAATMGLSSKAAAEMAESVTNPQRPPVIWIGAQECT WP_000145410.1
MULTISPECIES: hydrogenase-2 small chain [Proteobacteria] Length: 372\nNumber of predicted TMHs: 1\nExp number of AAs in TMHs: 23.72226\nExp number, first 60 AAs: 0.22038\nTotal prob of N-in: 0.12994\noutside 1 331\nTMhelix 332 354\ninside 355 372

8588 GCF_001617095.1_ASM161709v1 Escherichia coli Proteobacteria; Gammaproteobacteria; Enterobacterales; Enterobacteriaceae; Escherichia
MTGDNTLIHSHGINRRDFMKLCAALAATMGLSSKAAAEMAESVTNPQRPPVIWIGAQECT WP_000145410.1
MULTISPECIES: hydrogenase-2 small chain [Proteobacteria] Length: 372\nNumber of predicted TMHs: 1\nExp number of AAs in TMHs: 23.72226\nExp number, first 60 AAs: 0.22038\nTotal prob of N-in: 0.12994\noutside 1 331\nTMhelix 332 354\ninside 355 372

8589 GCF_001617145.1_ASM161714v1 Escherichia coli Proteobacteria; Gammaproteobacteria; Enterobacterales; Enterobacteriaceae; Escherichia
MTGDNTLIHSHGINRRDFMKLCAALAATMGLSSKAAAEMAESVTNPQRPPVIWIGAQECT WP_000145410.1
MULTISPECIES: hydrogenase-2 small chain [Proteobacteria] Length: 372\nNumber of predicted TMHs: 1\nExp number of AAs in TMHs: 23.72226\nExp number, first 60 AAs: 0.22038\nTotal prob of N-in: 0.12994\noutside 1 331\nTMhelix 332 354\ninside 355 372

8590 GCF_001617075.1_ASM161707v1 Escherichia coli Proteobacteria; Gammaproteobacteria; Enterobacterales; Enterobacteriaceae; Escherichia
MTGDNTLIHSHGINRRDFMKLCAALAATMGLSSKAAAEMAESVTNPQRPPVIWIGAQECT WP_000145410.1
MULTISPECIES: hydrogenase-2 small chain [Proteobacteria] Length: 372\nNumber of predicted TMHs: 1\nExp number of AAs in TMHs: 23.72226\nExp number, first 60 AAs: 0.22038\nTotal prob of N-in: 0.12994\noutside 1 331\nTMhelix 332 354\ninside 355 372

8591 GCF_001617195.1_ASM161719v1 Escherichia coli Proteobacteria; Gammaproteobacteria; Enterobacterales; Enterobacteriaceae; Escherichia
MTGDNTLIHSHGINRRDFMKLCAALAATMGLSSKAAAEMAESVTNPQRPPVIWIGAQECT WP_000145410.1
MULTISPECIES: hydrogenase-2 small chain [Proteobacteria] Length: 372\nNumber of predicted TMHs: 1\nExp number of AAs in TMHs: 23.72226\nExp number, first 60 AAs: 0.22038\nTotal prob of N-in: 0.12994\noutside 1 331\nTMhelix 332 354\ninside 355 372

8592 GCF_001617075.1_ASM161707v1 Escherichia coli Proteobacteria; Gammaproteobacteria; Enterobacterales; Enterobacteriaceae; Escherichia
MNNEETFYQAMRRQGVTRRSFLKYCSLAATSLGLGAGMAPKIAWALENKPRIPVVWIHGL WP_001058323.1
MULTISPECIES: hydrogenase-1 small chain [Proteobacteria] Length: 372\nNumber of predicted TMHs: 2\nExp number of AAs in TMHs: 37.2608499999999\nExp number, first 60 AAs: 19.52923\nTotal prob of N-in: 0.94351\nPOSSIBLE N-term signal sequence\ninside 1 19\nTMhelix 20 39\noutside 40 325\nTMhelix 326 348\ninside 349 372

8593 GCF_001617195.1_ASM161719v1 Escherichia coli Proteobacteria; Gammaproteobacteria; Enterobacterales; Enterobacteriaceae; Escherichia
MNNEETFYQAMRRQGVTRRSFLKYCSLAATSLGLGAGMAPKIAWALENKPRIPVVWIHGL WP_001295940.1
MULTISPECIES: hydrogenase [Enterobacteriaceae] Length: 372\nNumber of predicted TMHs: 2\nExp number of AAs in TMHs: 37.2608899999999\nExp number, first 60 AAs: 19.52923\nTotal prob of N-in: 0.94351\nPOSSIBLE N-term signal sequence\ninside 1 19\nTMhelix 20 39\noutside 40 325\nTMhelix 326 348\ninside 349 372

8594 GCF_001616905.1_ASM161690v1 Escherichia coli Proteobacteria; Gammaproteobacteria; Enterobacterales; Enterobacteriaceae; Escherichia
MNNEETFYQAMRRQGVTRRSFLKYCSLAATSLGLGAGMAPKIAWALENKPRIPVVWIHGL WP_001058323.1
MULTISPECIES: hydrogenase-1 small chain [Proteobacteria] Length: 372\nNumber of predicted TMHs: 2\nExp number of AAs in TMHs: 37.2608499999999\nExp number, first 60 AAs: 19.52923\nTotal prob of N-in: 0.94351\nPOSSIBLE N-term signal sequence\ninside 1 19\nTMhelix 20 39\noutside 40 325\nTMhelix 326 348\ninside 349 372

8595 GCF_001617255.1_ASM161725v1 Escherichia coli Proteobacteria; Gammaproteobacteria; Enterobacterales; Enterobacteriaceae; Escherichia

MTGDNTLIHSHGINRRDFMKLCAALAATMGLSSKAAAEMAESVTNPQRPPVIWIGAQECT WP_000145410.1 MULTISPECIES: hydrogenase-2 small chain [Proteobacteria] Length: 372\nNumber of predicted TMHs: 1\nExp number of AAs in TMHs: 23.72226\nExp number, first 60 AAs: 0.22038\nTotal prob of N-in: 0.12994\noutside 1 331\nTMhelix 332 354\ninside 355 372

8596 GCF_001617565.1_ASM161756v1 Escherichia coli Proteobacteria; Gammaproteobacteria; Enterobacterales; Enterobacteriaceae; Escherichia MTGDNTLIHSHGINRRDFMKLCAALAATMGLSSKAAAEMVESVTNPQRPPVIWIGAQECT WP_000145423.1 hydrogenase 2 small subunit [Escherichia coli] Length: 372\nNumber of predicted TMHs: 1\nExp number of AAs in TMHs: 23.78201\nExp number, first 60 AAs: 0.28355\nTotal prob of N-in: 0.13257\noutside 1 331\nTMhelix 332 354\ninside 355 372

8597 GCF_001617255.1_ASM161725v1 Escherichia coli Proteobacteria; Gammaproteobacteria; Enterobacterales; Enterobacteriaceae; Escherichia MNNEETFYQAMRRQGVTRRSFLKYCSLAATSLGLGAGMAPKIAWALENKPRIPVVWIHGL WP_001058323.1 MULTISPECIES: hydrogenase-1 small chain [Proteobacteria] Length: 372\nNumber of predicted TMHs: 2\nExp number of AAs in TMHs: 37.2608499999999\nExp number, first 60 AAs: 19.52923\nTotal prob of N-in: 0.94351\nPOSSIBLE N-term signal sequence\ninside 1 19\nTMhelix 20 39\noutside 40 325\nTMhelix 326 348\ninside 349 372

8598 GCF_001617565.1_ASM161756v1 Escherichia coli Proteobacteria; Gammaproteobacteria; Enterobacterales; Enterobacteriaceae; Escherichia MNNEETFYQAMRRQGVTRRSFLKYCSLAATSLGLGAGMAPKIAWALENKPRIPVVWIHGL WP_001295940.1 MULTISPECIES: hydrogenase [Enterobacteriaceae] Length: 372\nNumber of predicted TMHs: 2\nExp number of AAs in TMHs: 37.2608899999999\nExp number, first 60 AAs: 19.52923\nTotal prob of N-in: 0.94351\nPOSSIBLE N-term signal sequence\ninside 1 19\nTMhelix 20 39\noutside 40 325\nTMhelix 326 348\ninside 349 372

8599 GCF_001616855.1_ASM161685v1 Escherichia coli Proteobacteria; Gammaproteobacteria; Enterobacterales; Enterobacteriaceae; Escherichia MNNEETFYQAMRRQGVTRRSFLKYCSLAATSLGLGAGMAPKIAWALENKPRIPVVWIHGL WP_001058323.1 MULTISPECIES: hydrogenase-1 small chain [Proteobacteria] Length: 372\nNumber of predicted TMHs: 2\nExp number of AAs in TMHs: 37.2608499999999\nExp number, first 60 AAs: 19.52923\nTotal prob of N-in: 0.94351\nPOSSIBLE N-term signal sequence\ninside 1 19\nTMhelix 20 39\noutside 40 325\nTMhelix 326 348\ninside 349 372

8600 GCF_001618365.1_ASM161836v1 Escherichia coli Proteobacteria; Gammaproteobacteria; Enterobacterales; Enterobacteriaceae; Escherichia MTGDNTLIHSHGINRRDFMKLCAALAATMGLSSKAAAEMAESVTNPQRPPVIWIGAQECT WP_000145410.1 MULTISPECIES: hydrogenase-2 small chain [Proteobacteria] Length: 372\nNumber of predicted TMHs: 1\nExp number of AAs in TMHs: 23.72226\nExp number, first 60 AAs: 0.22038\nTotal prob of N-in: 0.12994\noutside 1 331\nTMhelix 332 354\ninside 355 372

8601 GCF_001618365.1_ASM161836v1 Escherichia coli Proteobacteria; Gammaproteobacteria; Enterobacterales; Enterobacteriaceae; Escherichia MNNEETFYQAMRRQGVTRRSFLKYCSLAATSLGLGAGMAPKIAWALENKPRIPVVWIHGL WP_001058323.1 MULTISPECIES: hydrogenase-1 small chain [Proteobacteria] Length: 372\nNumber of predicted TMHs: 2\nExp number of AAs in TMHs: 37.2608499999999\nExp number, first 60 AAs: 19.52923\nTotal prob of N-in: 0.94351\nPOSSIBLE N-term signal sequence\ninside 1 19\nTMhelix 20 39\noutside 40 325\nTMhelix 326 348\ninside 349 372

8602 GCF_001616955.1_ASM161695v1 Escherichia coli Proteobacteria; Gammaproteobacteria; Enterobacterales; Enterobacteriaceae; Escherichia MNNEETFYQAMRRQGVTRRSFLKYCSLAATSLGLGAGMAPKIAWALENKPRIPVVWIHGL WP_001295940.1 MULTISPECIES: hydrogenase [Enterobacteriaceae] Length: 372\nNumber of predicted TMHs: 2\nExp number of AAs in TMHs: 37.2608899999999\nExp number, first 60 AAs: 19.52923\nTotal prob of N-in: 0.94351\nPOSSIBLE N-term signal sequence\ninside 1 19\nTMhelix 20 39\noutside 40 325\nTMhelix 326 348\ninside 349 372

8603 GCF_001617015.1_ASM161701v1 Escherichia coli Proteobacteria; Gammaproteobacteria; Enterobacterales; Enterobacteriaceae; Escherichia

MNNEETFYQAMRRQGVTRRSFLKYCSLAATSLGLGAGMAPKIAWALENKPRIPVVWIHGL WP_001058323.1 MULTISPECIES: hydrogenase-1 small chain [Proteobacteria] Length: 372\nNumber of predicted TMHs: 2\nExp number of AAs in TMHs: 37.2608499999999\nExp number, first 60 AAs: 19.52923\nTotal prob of N-in: 0.94351\nPOSSIBLE N-term signal sequence\ninside 1 19\nTMhelix 20 39\noutside 40 325\nTMhelix 326 348\ninside 349 372

8604 GCF_001617055.1_ASM161705v1 Escherichia coli Proteobacteria; Gammaproteobacteria; Enterobacterales; Enterobacteriaceae; Escherichia MNNEETFYQAMRRQGVTRRSFLKYCSLAATSLGLGAGMAPKIAWALENKPRIPVVWIHGL WP_001295940.1 MULTISPECIES: hydrogenase [Enterobacteriaceae] Length: 372\nNumber of predicted TMHs: 2\nExp number of AAs in TMHs: 37.2608899999999\nExp number, first 60 AAs: 19.52923\nTotal prob of N-in: 0.94351\nPOSSIBLE N-term signal sequence\ninside 1 19\nTMhelix 20 39\noutside 40 325\nTMhelix 326 348\ninside 349 372

8605 GCF_001617145.1_ASM161714v1 Escherichia coli Proteobacteria; Gammaproteobacteria; Enterobacterales; Enterobacteriaceae; Escherichia MNNEETFYQAMRRQGVTRRSFLKYCSLAATSLGLGAGMAPKIAWALENKPRIPVVWIHGL WP_001295940.1 MULTISPECIES: hydrogenase [Enterobacteriaceae] Length: 372\nNumber of predicted TMHs: 2\nExp number of AAs in TMHs: 37.2608899999999\nExp number, first 60 AAs: 19.52923\nTotal prob of N-in: 0.94351\nPOSSIBLE N-term signal sequence\ninside 1 19\nTMhelix 20 39\noutside 40 325\nTMhelix 326 348\ninside 349 372

8606 GCF_001617095.1_ASM161709v1 Escherichia coli Proteobacteria; Gammaproteobacteria; Enterobacterales; Enterobacteriaceae; Escherichia MNNEETFYQAMRRQGVTRRSFLKYCSLAATSLGLGAGMAPKIAWALENKPRIPVVWIHGL WP_001295940.1 MULTISPECIES: hydrogenase [Enterobacteriaceae] Length: 372\nNumber of predicted TMHs: 2\nExp number of AAs in TMHs: 37.2608899999999\nExp number, first 60 AAs: 19.52923\nTotal prob of N-in: 0.94351\nPOSSIBLE N-term signal sequence\ninside 1 19\nTMhelix 20 39\noutside 40 325\nTMhelix 326 348\ninside 349 372

8607 GCF_001619075.1_ASM161907v1 Escherichia coli Proteobacteria; Gammaproteobacteria; Enterobacterales; Enterobacteriaceae; Escherichia MTGDNTLIHSHGINRRDFMKLCAALAATMGLSSKAAAEMAESVTNPQRPPVIWIGAQECT WP_000145410.1 MULTISPECIES: hydrogenase-2 small chain [Proteobacteria] Length: 372\nNumber of predicted TMHs: 1\nExp number of AAs in TMHs: 23.72226\nExp number, first 60 AAs: 0.22038\nTotal prob of N-in: 0.12994\noutside 1 331\nTMhelix 332 354\ninside 355 372

8608 GCF_001619135.1_ASM161913v1 Escherichia coli Proteobacteria; Gammaproteobacteria; Enterobacterales; Enterobacteriaceae; Escherichia MTGDNTLIHSHGINRRDFMKLCAALAATMGLSSKAAAEMAESVTNPQRPPVIWIGAQECT WP_000145410.1 MULTISPECIES: hydrogenase-2 small chain [Proteobacteria] Length: 372\nNumber of predicted TMHs: 1\nExp number of AAs in TMHs: 23.72226\nExp number, first 60 AAs: 0.22038\nTotal prob of N-in: 0.12994\noutside 1 331\nTMhelix 332 354\ninside 355 372

8609 GCF_001619155.1_ASM161915v1 Escherichia coli Proteobacteria; Gammaproteobacteria; Enterobacterales; Enterobacteriaceae; Escherichia MTGDNTLIHSHGINRRDFMKLCAALAATMGLSSKAAAEMAESVTNPQRPPVIWIGAQECT WP_000145410.1 MULTISPECIES: hydrogenase-2 small chain [Proteobacteria] Length: 372\nNumber of predicted TMHs: 1\nExp number of AAs in TMHs: 23.72226\nExp number, first 60 AAs: 0.22038\nTotal prob of N-in: 0.12994\noutside 1 331\nTMhelix 332 354\ninside 355 372

8610 GCF_001620905.1_ASM162090v1 Escherichia coli Proteobacteria; Gammaproteobacteria; Enterobacterales; Enterobacteriaceae; Escherichia MTGDNTLIHSHGINRRDFMKLCAALAATMGLSSKAAAEMVESVTNPQRPPVIWIGAQECT WP_000145423.1 hydrogenase 2 small subunit [Escherichia coli] Length: 372\nNumber of predicted TMHs: 1\nExp number of AAs in TMHs: 23.78201\nExp number, first 60 AAs: 0.28355\nTotal prob of N-in: 0.13257\noutside 1 331\nTMhelix 332 354\ninside 355 372

8611 GCF_001619225.1_ASM161922v1 Escherichia coli Proteobacteria; Gammaproteobacteria; Enterobacterales; Enterobacteriaceae; Escherichia MTGDNTLIHSHGINRRDFMKLCAALAATMGLSSKAAAEMAESVTNPQRPPVIWIGAQECT WP_000145410.1

MULTISPECIES: hydrogenase-2 small chain [Proteobacteria] Length: 372\nNumber of predicted TMHs: 1\nExp number of AAs in TMHs: 23.72226\nExp number, first 60 AAs: 0.22038\nTotal prob of N-in: 0.12994\noutside 1 331\nTMhelix 332 354\ninside 355 372

8612 GCF_001620935.1_ASM162093v1 Escherichia coli Proteobacteria; Gammaproteobacteria; Enterobacterales; Enterobacteriaceae; Escherichia
MTGDNTLIHSHGINRRDFMKLCAALAATMGLSSKAAAEMVESVTNPQRPPVIWIGAQECT WP_000145423.1
hydrogenase 2 small subunit [Escherichia coli] Length: 372\nNumber of predicted TMHs: 1\nExp number of AAs in TMHs: 23.78201\nExp number, first 60 AAs: 0.28355\nTotal prob of N-in: 0.13257\noutside 1 331\nTMhelix 332 354\ninside 355 372

8613 GCF_001621065.1_ASM162106v1 Escherichia coli Proteobacteria; Gammaproteobacteria; Enterobacterales; Enterobacteriaceae; Escherichia
MTGDNTLIHSHGINRRDFMKLCAALAATMGLSSKAAAEMVESVTNPQRPPVIWIGAQECT WP_000145423.1
hydrogenase 2 small subunit [Escherichia coli] Length: 372\nNumber of predicted TMHs: 1\nExp number of AAs in TMHs: 23.78201\nExp number, first 60 AAs: 0.28355\nTotal prob of N-in: 0.13257\noutside 1 331\nTMhelix 332 354\ninside 355 372

8614 GCF_001620995.1_ASM162099v1 Escherichia coli Proteobacteria; Gammaproteobacteria; Enterobacterales; Enterobacteriaceae; Escherichia
MTGDNTLIHSHGINRRDFMKLCAALAATMGLSSKAAAEMAESVTNPQRPPVIWIGAQECT WP_000145410.1
MULTISPECIES: hydrogenase-2 small chain [Proteobacteria] Length: 372\nNumber of predicted TMHs: 1\nExp number of AAs in TMHs: 23.72226\nExp number, first 60 AAs: 0.22038\nTotal prob of N-in: 0.12994\noutside 1 331\nTMhelix 332 354\ninside 355 372

8615 GCF_001621085.1_ASM162108v1 Escherichia coli Proteobacteria; Gammaproteobacteria; Enterobacterales; Enterobacteriaceae; Escherichia
MTGDNTLIHSHGINRRDFMKLCAALAATMGLSSKAAAEMAESVTNPQRPPVIWIGAQECT WP_000145410.1
MULTISPECIES: hydrogenase-2 small chain [Proteobacteria] Length: 372\nNumber of predicted TMHs: 1\nExp number of AAs in TMHs: 23.72226\nExp number, first 60 AAs: 0.22038\nTotal prob of N-in: 0.12994\noutside 1 331\nTMhelix 332 354\ninside 355 372

8616 GCF_001621175.1_ASM162117v1 Escherichia coli Proteobacteria; Gammaproteobacteria; Enterobacterales; Enterobacteriaceae; Escherichia
MTGDNTLIHSHGINRRDFMKLCAALAATMGLSSKAAAEMAESVTNPQRPPVIWIGAQECT WP_000145410.1
MULTISPECIES: hydrogenase-2 small chain [Proteobacteria] Length: 372\nNumber of predicted TMHs: 1\nExp number of AAs in TMHs: 23.72226\nExp number, first 60 AAs: 0.22038\nTotal prob of N-in: 0.12994\noutside 1 331\nTMhelix 332 354\ninside 355 372

8617 GCF_001621165.1_ASM162116v1 Escherichia coli Proteobacteria; Gammaproteobacteria; Enterobacterales; Enterobacteriaceae; Escherichia
MTGDNTLIHSHGINRRDFMKLCAALAATMGLSSKAAAEMAESVTNPQRPPVIWIGAQECT WP_000145410.1
MULTISPECIES: hydrogenase-2 small chain [Proteobacteria] Length: 372\nNumber of predicted TMHs: 1\nExp number of AAs in TMHs: 23.72226\nExp number, first 60 AAs: 0.22038\nTotal prob of N-in: 0.12994\noutside 1 331\nTMhelix 332 354\ninside 355 372

8618 GCF_001620995.1_ASM162099v1 Escherichia coli Proteobacteria; Gammaproteobacteria; Enterobacterales; Enterobacteriaceae; Escherichia
MNNEETFYQAMRRQGVTRRSFLKYCSLAATSLGLGAGMAPKIAWALENKPRIPVVWIHGL WP_001058323.1
MULTISPECIES: hydrogenase-1 small chain [Proteobacteria] Length: 372\nNumber of predicted TMHs: 2\nExp number of AAs in TMHs: 37.2608499999999\nExp number, first 60 AAs: 19.52923\nTotal prob of N-in: 0.94351\nPOSSIBLE N-term signal sequence\ninside 1 19\nTMhelix 20 39\noutside 40 325\nTMhelix 326 348\ninside 349 372

8619 GCF_001621225.1_ASM162122v1 Escherichia coli Proteobacteria; Gammaproteobacteria; Enterobacterales; Enterobacteriaceae; Escherichia
MTGDNTLIHSHGINRRDFMKLCAALAATMGLSSKAAAEMAESVTNPQRPPVIWIGAQECT WP_000145410.1
MULTISPECIES: hydrogenase-2 small chain [Proteobacteria] Length: 372\nNumber of predicted TMHs: 1\nExp number of AAs in TMHs: 23.72226\nExp number, first 60 AAs: 0.22038\nTotal prob of N-in: 0.12994\noutside 1 331\nTMhelix 332 354\ninside 355 372

8620 GCF_001621175.1_ASM162117v1 Escherichia coli Proteobacteria; Gammaproteobacteria; Enterobacterales; Enterobacteriaceae; Escherichia MNNEETFYQAMRRQGVTRRSFLKYCSLAATSLGLGAGMAPKIAWALENKPRIPVVWIHGL WP_001058323.1 MULTISPECIES: hydrogenase-1 small chain [Proteobacteria] Length: 372\nNumber of predicted TMHs: 2\nExp number of AAs in TMHs: 37.2608499999999\nExp number, first 60 AAs: 19.52923\nTotal prob of N-in: 0.94351\nPOSSIBLE N-term signal sequence\ninside 1 19\nTMhelix 20 39\noutside 40 325\nTMhelix 326 348\ninside 349 372

8621 GCF_001621165.1_ASM162116v1 Escherichia coli Proteobacteria; Gammaproteobacteria; Enterobacterales; Enterobacteriaceae; Escherichia MNNEETFYQAMRRQGVTRRSFLKYCSLAATSLGLGAGMAPKIAWALENKPRIPVVWIHGL WP_001058323.1 MULTISPECIES: hydrogenase-1 small chain [Proteobacteria] Length: 372\nNumber of predicted TMHs: 2\nExp number of AAs in TMHs: 37.2608499999999\nExp number, first 60 AAs: 19.52923\nTotal prob of N-in: 0.94351\nPOSSIBLE N-term signal sequence\ninside 1 19\nTMhelix 20 39\noutside 40 325\nTMhelix 326 348\ninside 349 372

8622 GCF_001619075.1_ASM161907v1 Escherichia coli Proteobacteria; Gammaproteobacteria; Enterobacterales; Enterobacteriaceae; Escherichia MNNEETFYQAMRRQGVTRRSFLKYCSLAATSLGLGAGMAPKIAWALENKPRIPVVWIHGL WP_001301613.1 hydrogenase [Escherichia coli] Length: 372\nNumber of predicted TMHs: 2\nExp number of AAs in TMHs: 37.46598\nExp number, first 60 AAs: 19.53139\nTotal prob of N-in: 0.94364\nPOSSIBLE N-term signal sequence\ninside 1 19\nTMhelix 20 39\noutside 40 325\nTMhelix 326 348\ninside 349 372

8623 GCF_001619135.1_ASM161913v1 Escherichia coli Proteobacteria; Gammaproteobacteria; Enterobacterales; Enterobacteriaceae; Escherichia MNNEETFYQAMRRQGVTRRSFLKYCSLAATSLGLGAGMAPKIAWALENKPRIPVVWIHGL WP_001301613.1 hydrogenase [Escherichia coli] Length: 372\nNumber of predicted TMHs: 2\nExp number of AAs in TMHs: 37.46598\nExp number, first 60 AAs: 19.53139\nTotal prob of N-in: 0.94364\nPOSSIBLE N-term signal sequence\ninside 1 19\nTMhelix 20 39\noutside 40 325\nTMhelix 326 348\ninside 349 372

8624 GCF_001620935.1_ASM162093v1 Escherichia coli Proteobacteria; Gammaproteobacteria; Enterobacterales; Enterobacteriaceae; Escherichia MNNEETFYQAMRRQGVTRRSFLKYCSLAATSLGLGAGMAPKIAWALENKPRIPVVWIHGL WP_001295940.1 MULTISPECIES: hydrogenase [Enterobacteriaceae] Length: 372\nNumber of predicted TMHs: 2\nExp number of AAs in TMHs: 37.2608899999999\nExp number, first 60 AAs: 19.52923\nTotal prob of N-in: 0.94351\nPOSSIBLE N-term signal sequence\ninside 1 19\nTMhelix 20 39\noutside 40 325\nTMhelix 326 348\ninside 349 372

8625 GCF_001621065.1_ASM162106v1 Escherichia coli Proteobacteria; Gammaproteobacteria; Enterobacterales; Enterobacteriaceae; Escherichia MNNEETFYQAMRRQGVTRRSFLKYCSLAATSLGLGAGMAPKIAWALENKPRIPVVWIHGL WP_001295940.1 MULTISPECIES: hydrogenase [Enterobacteriaceae] Length: 372\nNumber of predicted TMHs: 2\nExp number of AAs in TMHs: 37.2608899999999\nExp number, first 60 AAs: 19.52923\nTotal prob of N-in: 0.94351\nPOSSIBLE N-term signal sequence\ninside 1 19\nTMhelix 20 39\noutside 40 325\nTMhelix 326 348\ninside 349 372

8626 GCF_001619155.1_ASM161915v1 Escherichia coli Proteobacteria; Gammaproteobacteria; Enterobacterales; Enterobacteriaceae; Escherichia MNNEETFYQAMRRQGVTRRSFLKYCSLAATSLGLGAGMAPKIAWALENKPRIPVVWIHGL WP_001301613.1 hydrogenase [Escherichia coli] Length: 372\nNumber of predicted TMHs: 2\nExp number of AAs in TMHs: 37.46598\nExp number, first 60 AAs: 19.53139\nTotal prob of N-in: 0.94364\nPOSSIBLE N-term signal sequence\ninside 1 19\nTMhelix 20 39\noutside 40 325\nTMhelix 326 348\ninside 349 372

8627 GCF_001620905.1_ASM162090v1 Escherichia coli Proteobacteria; Gammaproteobacteria; Enterobacterales; Enterobacteriaceae; Escherichia MNNEETFYQAMRRQGVTRRSFLKYCSLAATSLGLGAGMAPKIAWALENKPRIPVVWIHGL WP_001295940.1 MULTISPECIES: hydrogenase [Enterobacteriaceae] Length: 372\nNumber of predicted TMHs: 2\nExp number of AAs in TMHs: 37.2608899999999\nExp number, first 60 AAs: 19.52923\nTotal prob of N-in: 0.94351\nPOSSIBLE N-term signal sequence\ninside 1 19\nTMhelix 20 39\noutside 40 325\nTMhelix 326 348\ninside 349 372

8628 GCF_001619225.1_ASM161922v1 Escherichia coli Proteobacteria; Gammaproteobacteria; Enterobacterales; Enterobacteriaceae; Escherichia
MNNEETFYQAMRRQGVTRRSFLKYCSLAATSLGLGAGMAPKIAWALENKPRIPVVWIHGL WP_001301613.1
hydrogenase [Escherichia coli] Length: 372\nNumber of predicted TMHs: 2\nExp number of AAs in TMHs: 37.46598\nExp number, first 60 AAs: 19.53139\nTotal prob of N-in: 0.94364\nPOSSIBLE N-term signal sequence\ninside 1 19\nTMhelix 20 39\noutside 40 325\nTMhelix 326 348\ninside 349 372

8629 GCF_001621225.1_ASM162122v1 Escherichia coli Proteobacteria; Gammaproteobacteria; Enterobacterales; Enterobacteriaceae; Escherichia
MNNEETFYQAMRRQGVTRRSFLKYCSLAATSLGLGAGMAPKIAWALENKPRIPVVWIHGL WP_001058323.1
MULTISPECIES: hydrogenase-1 small chain [Proteobacteria] Length: 372\nNumber of predicted TMHs: 2\nExp number of AAs in TMHs: 37.2608499999999\nExp number, first 60 AAs: 19.52923\nTotal prob of N-in: 0.94351\nPOSSIBLE N-term signal sequence\ninside 1 19\nTMhelix 20 39\noutside 40 325\nTMhelix 326 348\ninside 349 372

8630 GCF_001621325.1_ASM162132v1 Escherichia coli Proteobacteria; Gammaproteobacteria; Enterobacterales; Enterobacteriaceae; Escherichia
MTGDNTLIHSHGINRRDFMKLCAALAATMGLSSKAAAEMVESVTNPQRPPVIWIGAQECT WP_000145423.1
hydrogenase 2 small subunit [Escherichia coli] Length: 372\nNumber of predicted TMHs: 1\nExp number of AAs in TMHs: 23.78201\nExp number, first 60 AAs: 0.28355\nTotal prob of N-in: 0.13257\noutside 1 331\nTMhelix 332 354\ninside 355 372

8631 GCF_001621265.1_ASM162126v1 Escherichia coli Proteobacteria; Gammaproteobacteria; Enterobacterales; Enterobacteriaceae; Escherichia
MTGDNTLIHSHGINRRDFMKLCAALAATMGLSSKAAAEMVESVTNPQRPPVIWIGAQECT WP_000145423.1
hydrogenase 2 small subunit [Escherichia coli] Length: 372\nNumber of predicted TMHs: 1\nExp number of AAs in TMHs: 23.78201\nExp number, first 60 AAs: 0.28355\nTotal prob of N-in: 0.13257\noutside 1 331\nTMhelix 332 354\ninside 355 372

8632 GCF_001621355.1_ASM162135v1 Escherichia coli Proteobacteria; Gammaproteobacteria; Enterobacterales; Enterobacteriaceae; Escherichia
MTGDNTLIHSHGINRRDFMKLCAALAATMGLSSKAAAEMAESVTNPQRPPVIWIGAQECT WP_000145410.1
MULTISPECIES: hydrogenase-2 small chain [Proteobacteria] Length: 372\nNumber of predicted TMHs: 1\nExp number of AAs in TMHs: 23.72226\nExp number, first 60 AAs: 0.22038\nTotal prob of N-in: 0.12994\noutside 1 331\nTMhelix 332 354\ninside 355 372

8633 GCF_001621415.1_ASM162141v1 Escherichia coli Proteobacteria; Gammaproteobacteria; Enterobacterales; Enterobacteriaceae; Escherichia
MTGDNTLIHSHGINRRDFMKLCAALAATMGLSSKAAAEMVESVTNPQRPPVIWIGAQECT WP_000145423.1
hydrogenase 2 small subunit [Escherichia coli] Length: 372\nNumber of predicted TMHs: 1\nExp number of AAs in TMHs: 23.78201\nExp number, first 60 AAs: 0.28355\nTotal prob of N-in: 0.13257\noutside 1 331\nTMhelix 332 354\ninside 355 372

8634 GCF_001621515.1_ASM162151v1 Escherichia coli Proteobacteria; Gammaproteobacteria; Enterobacterales; Enterobacteriaceae; Escherichia
MTGDNTLIHSHGINRRDFMKLCAALAATMGLSSKAAAEMVESVTNPQRPPVIWIGAQECT WP_000145423.1
hydrogenase 2 small subunit [Escherichia coli] Length: 372\nNumber of predicted TMHs: 1\nExp number of AAs in TMHs: 23.78201\nExp number, first 60 AAs: 0.28355\nTotal prob of N-in: 0.13257\noutside 1 331\nTMhelix 332 354\ninside 355 372

8635 GCF_001621465.1_ASM162146v1 Escherichia coli Proteobacteria; Gammaproteobacteria; Enterobacterales; Enterobacteriaceae; Escherichia
MTGDNTLIHSHGINRRDFMKLCAALAATMGLSSKAAAEMVESVTNPQRPPVIWIGAQECT WP_000145423.1
hydrogenase 2 small subunit [Escherichia coli] Length: 372\nNumber of predicted TMHs: 1\nExp number of AAs in TMHs: 23.78201\nExp number, first 60 AAs: 0.28355\nTotal prob of N-in: 0.13257\noutside 1 331\nTMhelix 332 354\ninside 355 372

8636 GCF_001621565.1_ASM162156v1 Escherichia coli Proteobacteria; Gammaproteobacteria; Enterobacterales; Enterobacteriaceae; Escherichia
MTGDNTLIHSHGINRRDFMKLCAALAATMGLSSKAAAEMVESVTNPQRPPVIWIGAQECT WP_000145423.1
hydrogenase 2 small subunit [Escherichia coli] Length: 372\nNumber of predicted TMHs: 1\nExp number of

AAs in TMHs: 23.78201\nExp number, first 60 AAs: 0.28355\nTotal prob of N-in: 0.13257\noutside 1 331\nTMhelix 332 354\ninside 355 372

8637 GCF_001621625.1_ASM162162v1 Escherichia coli Proteobacteria; Gammaproteobacteria; Enterobacterales; Enterobacteriaceae; Escherichia MTGDNTLIHSHGINRRDFMKLCAALAATMGLSSKAAAEMAESVTNPQRPPVIWIGAQECT WP_000145410.1 MULTISPECIES: hydrogenase-2 small chain [Proteobacteria] Length: 372\nNumber of predicted TMHs: 1\nExp number of AAs in TMHs: 23.72226\nExp number, first 60 AAs: 0.22038\nTotal prob of N-in: 0.12994\noutside 1 331\nTMhelix 332 354\ninside 355 372

8638 GCF_001621665.1_ASM162166v1 Escherichia coli Proteobacteria; Gammaproteobacteria; Enterobacterales; Enterobacteriaceae; Escherichia MTGDNTLIHSHGINRRDFMKLCAALAATMGLSSKAAAEMAESVTNPQRPPVIWIGAQECT WP_000145410.1 MULTISPECIES: hydrogenase-2 small chain [Proteobacteria] Length: 372\nNumber of predicted TMHs: 1\nExp number of AAs in TMHs: 23.72226\nExp number, first 60 AAs: 0.22038\nTotal prob of N-in: 0.12994\noutside 1 331\nTMhelix 332 354\ninside 355 372

8639 GCF_001621725.1_ASM162172v1 Escherichia coli Proteobacteria; Gammaproteobacteria; Enterobacterales; Enterobacteriaceae; Escherichia MTGDNTLIHSHGINRRDFMKLCAALAATMGLSSKAAAEMVESVTNPQRPPVIWIGAQECT WP_000145423.1 hydrogenase 2 small subunit [Escherichia coli] Length: 372\nNumber of predicted TMHs: 1\nExp number of AAs in TMHs: 23.78201\nExp number, first 60 AAs: 0.28355\nTotal prob of N-in: 0.13257\noutside 1 331\nTMhelix 332 354\ninside 355 372

8640 GCF_001621755.1_ASM162175v1 Escherichia coli Proteobacteria; Gammaproteobacteria; Enterobacterales; Enterobacteriaceae; Escherichia MTGDNTLIHSHGINRRDFMKLCAALAATMGLSSKAAAEMVESVTNPQRPPVIWIGAQECT WP_000145423.1 hydrogenase 2 small subunit [Escherichia coli] Length: 372\nNumber of predicted TMHs: 1\nExp number of AAs in TMHs: 23.78201\nExp number, first 60 AAs: 0.28355\nTotal prob of N-in: 0.13257\noutside 1 331\nTMhelix 332 354\ninside 355 372

8641 GCF_001621825.1_ASM162182v1 Escherichia coli Proteobacteria; Gammaproteobacteria; Enterobacterales; Enterobacteriaceae; Escherichia MTGDNTLIHSHGINRRDFMKLCAALAATMGLSSKAAAEMVESVTNPQRPPVIWIGAQECT WP_000145423.1 hydrogenase 2 small subunit [Escherichia coli] Length: 372\nNumber of predicted TMHs: 1\nExp number of AAs in TMHs: 23.78201\nExp number, first 60 AAs: 0.28355\nTotal prob of N-in: 0.13257\noutside 1 331\nTMhelix 332 354\ninside 355 372

8642 GCF_001621625.1_ASM162162v1 Escherichia coli Proteobacteria; Gammaproteobacteria; Enterobacterales; Enterobacteriaceae; Escherichia MNNEETFYQAMRRQGVTRRSFLKYCSLAATSLGLGAGMAPKIAWALENKPRIPVVWIHGL WP_001058323.1 MULTISPECIES: hydrogenase-1 small chain [Proteobacteria] Length: 372\nNumber of predicted TMHs: 2\nExp number of AAs in TMHs: 37.2608499999999\nExp number, first 60 AAs: 19.52923\nTotal prob of N-in: 0.94351\nPOSSIBLE N-term signal sequence\ninside 1 19\nTMhelix 20 39\noutside 40 325\nTMhelix 326 348\ninside 349 372

8643 GCF_001621725.1_ASM162172v1 Escherichia coli Proteobacteria; Gammaproteobacteria; Enterobacterales; Enterobacteriaceae; Escherichia MNNEETFYQAMRRQGVTRRSFLKYCSLAATSLGLGAGMAPKIAWALENKPRIPVVWIHGL WP_001295940.1 MULTISPECIES: hydrogenase [Enterobacteriaceae] Length: 372\nNumber of predicted TMHs: 2\nExp number of AAs in TMHs: 37.2608899999999\nExp number, first 60 AAs: 19.52923\nTotal prob of N-in: 0.94351\nPOSSIBLE N-term signal sequence\ninside 1 19\nTMhelix 20 39\noutside 40 325\nTMhelix 326 348\ninside 349 372

8644 GCF_001621355.1_ASM162135v1 Escherichia coli Proteobacteria; Gammaproteobacteria; Enterobacterales; Enterobacteriaceae; Escherichia MNNEETFYQAMRRQGVTRRSFLKYCSLAATSLGLGAGMAPKIAWALENKPRIPVVWIHGL WP_001058323.1 MULTISPECIES: hydrogenase-1 small chain [Proteobacteria] Length: 372\nNumber of predicted TMHs: 2\nExp number of AAs in TMHs: 37.2608499999999\nExp number, first 60 AAs: 19.52923\nTotal prob of N-in: 0.94351\nPOSSIBLE N-term signal sequence\ninside 1 19\nTMhelix 20 39\noutside 40 325\nTMhelix 326 348\ninside 349 372

8645 GCF_001621325.1_ASM162132v1 Escherichia coli Proteobacteria; Gammaproteobacteria; Enterobacterales; Enterobacteriaceae; Escherichia
MNNEETFYQAMRRQGVTRRSFLKYCSLAATSLGLGAGMAPKIAWALENKPRIPVVWIHGL WP_001295940.1
MULTISPECIES: hydrogenase [Enterobacteriaceae] Length: 372\nNumber of predicted TMHs: 2\nExp number of AAs in TMHs: 37.2608899999999\nExp number, first 60 AAs: 19.52923\nTotal prob of N-in: 0.94351\nPOSSIBLE N-term signal sequence\ninside 1 19\nTMhelix 20 39\noutside 40 325\nTMhelix 326 348\ninside 349 372

8646 GCF_001621565.1_ASM162156v1 Escherichia coli Proteobacteria; Gammaproteobacteria; Enterobacterales; Enterobacteriaceae; Escherichia
MNNEETFYQAMRRQGVTRRSFLKYCSLAATSLGLGAGMAPKIAWALENKPRIPVVWIHGL WP_001295940.1
MULTISPECIES: hydrogenase [Enterobacteriaceae] Length: 372\nNumber of predicted TMHs: 2\nExp number of AAs in TMHs: 37.2608899999999\nExp number, first 60 AAs: 19.52923\nTotal prob of N-in: 0.94351\nPOSSIBLE N-term signal sequence\ninside 1 19\nTMhelix 20 39\noutside 40 325\nTMhelix 326 348\ninside 349 372

8647 GCF_001621265.1_ASM162126v1 Escherichia coli Proteobacteria; Gammaproteobacteria; Enterobacterales; Enterobacteriaceae; Escherichia
MNNEETFYQAMRRQGVTRRSFLKYCSLAATSLGLGAGMAPKIAWALENKPRIPVVWIHGL WP_001295940.1
MULTISPECIES: hydrogenase [Enterobacteriaceae] Length: 372\nNumber of predicted TMHs: 2\nExp number of AAs in TMHs: 37.2608899999999\nExp number, first 60 AAs: 19.52923\nTotal prob of N-in: 0.94351\nPOSSIBLE N-term signal sequence\ninside 1 19\nTMhelix 20 39\noutside 40 325\nTMhelix 326 348\ninside 349 372

8648 GCF_001621465.1_ASM162146v1 Escherichia coli Proteobacteria; Gammaproteobacteria; Enterobacterales; Enterobacteriaceae; Escherichia
MNNEETFYQAMRRQGVTRRSFLKYCSLAATSLGLGAGMAPKIAWALENKPRIPVVWIHGL WP_001295940.1
MULTISPECIES: hydrogenase [Enterobacteriaceae] Length: 372\nNumber of predicted TMHs: 2\nExp number of AAs in TMHs: 37.2608899999999\nExp number, first 60 AAs: 19.52923\nTotal prob of N-in: 0.94351\nPOSSIBLE N-term signal sequence\ninside 1 19\nTMhelix 20 39\noutside 40 325\nTMhelix 326 348\ninside 349 372

8649 GCF_001621515.1_ASM162151v1 Escherichia coli Proteobacteria; Gammaproteobacteria; Enterobacterales; Enterobacteriaceae; Escherichia
MNNEETFYQAMRRQGVTRRSFLKYCSLAATSLGLGAGMAPKIAWALENKPRIPVVWIHGL WP_001295940.1
MULTISPECIES: hydrogenase [Enterobacteriaceae] Length: 372\nNumber of predicted TMHs: 2\nExp number of AAs in TMHs: 37.2608899999999\nExp number, first 60 AAs: 19.52923\nTotal prob of N-in: 0.94351\nPOSSIBLE N-term signal sequence\ninside 1 19\nTMhelix 20 39\noutside 40 325\nTMhelix 326 348\ninside 349 372

8650 GCF_001621415.1_ASM162141v1 Escherichia coli Proteobacteria; Gammaproteobacteria; Enterobacterales; Enterobacteriaceae; Escherichia
MNNEETFYQAMRRQGVTRRSFLKYCSLAATSLGLGAGMAPKIAWALENKPRIPVVWIHGL WP_001295940.1
MULTISPECIES: hydrogenase [Enterobacteriaceae] Length: 372\nNumber of predicted TMHs: 2\nExp number of AAs in TMHs: 37.2608899999999\nExp number, first 60 AAs: 19.52923\nTotal prob of N-in: 0.94351\nPOSSIBLE N-term signal sequence\ninside 1 19\nTMhelix 20 39\noutside 40 325\nTMhelix 326 348\ninside 349 372

8651 GCF_001621755.1_ASM162175v1 Escherichia coli Proteobacteria; Gammaproteobacteria; Enterobacterales; Enterobacteriaceae; Escherichia
MNNEETFYQAMRRQGVTRRSFLKYCSLAATSLGLGAGMAPKIAWALENKPRIPVVWIHGL WP_001295940.1
MULTISPECIES: hydrogenase [Enterobacteriaceae] Length: 372\nNumber of predicted TMHs: 2\nExp number of AAs in TMHs: 37.2608899999999\nExp number, first 60 AAs: 19.52923\nTotal prob of N-in: 0.94351\nPOSSIBLE N-term signal sequence\ninside 1 19\nTMhelix 20 39\noutside 40 325\nTMhelix 326 348\ninside 349 372

8652 GCF_001621825.1_ASM162182v1 Escherichia coli Proteobacteria; Gammaproteobacteria; Enterobacterales; Enterobacteriaceae; Escherichia
MNNEETFYQAMRRQGVTRRSFLKYCSLAATSLGLGAGMAPKIAWALENKPRIPVVWIHGL WP_001295940.1
MULTISPECIES: hydrogenase [Enterobacteriaceae] Length: 372\nNumber of predicted TMHs: 2\nExp number of AAs in TMHs: 37.2608899999999\nExp number, first 60 AAs: 19.52923\nTotal prob of N-in:

0.94351\nPOSSIBLE N-term signal sequence\ninside 1 19\nTMhelix 20 39\noutside 40 325\nTMhelix 326 348\ninside 349 372

8653 GCF_001621855.1_ASM162185v1 Escherichia coli Proteobacteria; Gammaproteobacteria; Enterobacterales; Enterobacteriaceae; Escherichia MTGDNTLIHSHGINRRDFMKLCAALAATMGLSSKAAAEMVESVTNPQRPPVIWIGAQECT WP_000145423.1 hydrogenase 2 small subunit [Escherichia coli] Length: 372\nNumber of predicted TMHs: 1\nExp number of AAs in TMHs: 23.78201\nExp number, first 60 AAs: 0.28355\nTotal prob of N-in: 0.13257\noutside 1 331\nTMhelix 332 354\ninside 355 372

8654 GCF_001621995.1_ASM162199v1 Escherichia coli Proteobacteria; Gammaproteobacteria; Enterobacterales; Enterobacteriaceae; Escherichia MTGDNTLIHSHGINRRDFMKLCAALAATMGLSSKAAAEMAESVTNPQRPPVIWIGAQECT WP_000145410.1 MULTISPECIES: hydrogenase-2 small chain [Proteobacteria] Length: 372\nNumber of predicted TMHs: 1\nExp number of AAs in TMHs: 23.72226\nExp number, first 60 AAs: 0.22038\nTotal prob of N-in: 0.12994\noutside 1 331\nTMhelix 332 354\ninside 355 372

8655 GCF_001621975.1_ASM162197v1 Escherichia coli Proteobacteria; Gammaproteobacteria; Enterobacterales; Enterobacteriaceae; Escherichia MTGDNTLIHSHGINRRDFMKLCAALAATMGLSSKAAAEMVESVTNPQRPPVIWIGAQECT WP_000145423.1 hydrogenase 2 small subunit [Escherichia coli] Length: 372\nNumber of predicted TMHs: 1\nExp number of AAs in TMHs: 23.78201\nExp number, first 60 AAs: 0.28355\nTotal prob of N-in: 0.13257\noutside 1 331\nTMhelix 332 354\ninside 355 372

8656 GCF_001621915.1_ASM162191v1 Escherichia coli Proteobacteria; Gammaproteobacteria; Enterobacterales; Enterobacteriaceae; Escherichia MTGDNTLIHSHGINRRDFMKLCAALAATMGLSSKAAAEMAESVTNPQRPPVIWIGAQECT WP_000145410.1 MULTISPECIES: hydrogenase-2 small chain [Proteobacteria] Length: 372\nNumber of predicted TMHs: 1\nExp number of AAs in TMHs: 23.72226\nExp number, first 60 AAs: 0.22038\nTotal prob of N-in: 0.12994\noutside 1 331\nTMhelix 332 354\ninside 355 372

8657 GCF_001622105.1_ASM162210v1 Escherichia coli Proteobacteria; Gammaproteobacteria; Enterobacterales; Enterobacteriaceae; Escherichia MTGDNTLIHSHGINRRDFMKLCAALAATMGLSSKAAAEMAESVTNPQRPPVIWIGAQECT WP_000145410.1 MULTISPECIES: hydrogenase-2 small chain [Proteobacteria] Length: 372\nNumber of predicted TMHs: 1\nExp number of AAs in TMHs: 23.72226\nExp number, first 60 AAs: 0.22038\nTotal prob of N-in: 0.12994\noutside 1 331\nTMhelix 332 354\ninside 355 372

8658 GCF_001621965.1_ASM162196v1 Escherichia coli Proteobacteria; Gammaproteobacteria; Enterobacterales; Enterobacteriaceae; Escherichia MTGDNTLIHSHGINRRDFMKLCAALAATMGLSSKAAAEMVESVTNPQRPPVIWIGAQECT WP_000145423.1 hydrogenase 2 small subunit [Escherichia coli] Length: 372\nNumber of predicted TMHs: 1\nExp number of AAs in TMHs: 23.78201\nExp number, first 60 AAs: 0.28355\nTotal prob of N-in: 0.13257\noutside 1 331\nTMhelix 332 354\ninside 355 372

8659 GCF_001622045.1_ASM162204v1 Escherichia coli Proteobacteria; Gammaproteobacteria; Enterobacterales; Enterobacteriaceae; Escherichia MTGDNTLIHSHGINRRDFMKLCAALAATMGLSSKAAAEMVESVTNPQRPPVIWIGAQECT WP_000145423.1 hydrogenase 2 small subunit [Escherichia coli] Length: 372\nNumber of predicted TMHs: 1\nExp number of AAs in TMHs: 23.78201\nExp number, first 60 AAs: 0.28355\nTotal prob of N-in: 0.13257\noutside 1 331\nTMhelix 332 354\ninside 355 372

8660 GCF_001622135.1_ASM162213v1 Escherichia coli Proteobacteria; Gammaproteobacteria; Enterobacterales; Enterobacteriaceae; Escherichia MTGDNTLIHSHGINRRDFMKLCAALAATMGLSSKAAAEMAESVTNPQRPPVIWIGAQECT WP_000145410.1 MULTISPECIES: hydrogenase-2 small chain [Proteobacteria] Length: 372\nNumber of predicted TMHs: 1\nExp number of AAs in TMHs: 23.72226\nExp number, first 60 AAs: 0.22038\nTotal prob of N-in: 0.12994\noutside 1 331\nTMhelix 332 354\ninside 355 372

8661 GCF_001637645.1_ASM163764v1 Escherichia coli Proteobacteria; Gammaproteobacteria; Enterobacterales; Enterobacteriaceae; Escherichia

MTGDNTLIHSHGINRRDFMKLCAALAATMGLSSKAAAEMAESVTNPQRPPVIWIGAQECT WP_000145410.1 MULTISPECIES: hydrogenase-2 small chain [Proteobacteria] Length: 372\nNumber of predicted TMHs: 1\nExp number of AAs in TMHs: 23.72226\nExp number, first 60 AAs: 0.22038\nTotal prob of N-in: 0.12994\noutside 1 331\nTMhelix 332 354\ninside 355 372

8662 GCF_001637745.1_ASM163774v1 Escherichia coli Proteobacteria; Gammaproteobacteria; Enterobacterales; Enterobacteriaceae; Escherichia MTGDNTLIHSHGINRRDFMKLCAALAATMGLSSKAAAEMAESVTNPQRPPVIWIGAQECT WP_000145410.1 MULTISPECIES: hydrogenase-2 small chain [Proteobacteria] Length: 372\nNumber of predicted TMHs: 1\nExp number of AAs in TMHs: 23.72226\nExp number, first 60 AAs: 0.22038\nTotal prob of N-in: 0.12994\noutside 1 331\nTMhelix 332 354\ninside 355 372

8663 GCF_001637705.1_ASM163770v1 Escherichia coli Proteobacteria; Gammaproteobacteria; Enterobacterales; Enterobacteriaceae; Escherichia MTGDNTLIHSHGINRRDFMKLCAALAATMGLSSKAAAEMAESVTNPQRPPVIWIGAQECT WP_000145410.1 MULTISPECIES: hydrogenase-2 small chain [Proteobacteria] Length: 372\nNumber of predicted TMHs: 1\nExp number of AAs in TMHs: 23.72226\nExp number, first 60 AAs: 0.22038\nTotal prob of N-in: 0.12994\noutside 1 331\nTMhelix 332 354\ninside 355 372

8664 GCF_001637795.1_ASM163779v1 Escherichia coli Proteobacteria; Gammaproteobacteria; Enterobacterales; Enterobacteriaceae; Escherichia MTGDNTLIHSHGINRRDFMKLCAALAATMGLSSKAAAEMAESVTNPQRPPVIWIGAQECT WP_000145410.1 MULTISPECIES: hydrogenase-2 small chain [Proteobacteria] Length: 372\nNumber of predicted TMHs: 1\nExp number of AAs in TMHs: 23.72226\nExp number, first 60 AAs: 0.22038\nTotal prob of N-in: 0.12994\noutside 1 331\nTMhelix 332 354\ninside 355 372

8665 GCF_001637745.1_ASM163774v1 Escherichia coli Proteobacteria; Gammaproteobacteria; Enterobacterales; Enterobacteriaceae; Escherichia MNNEETFYQAMRRQGVTRRSFLKYCSLAATSLGLGAGMAPKIAWALENKPRIPVVWIHGL WP_001058323.1 MULTISPECIES: hydrogenase-1 small chain [Proteobacteria] Length: 372\nNumber of predicted TMHs: 2\nExp number of AAs in TMHs: 37.2608499999999\nExp number, first 60 AAs: 19.52923\nTotal prob of N-in: 0.94351\nPOSSIBLE N-term signal sequence\ninside 1 19\nTMhelix 20 39\noutside 40 325\nTMhelix 326 348\ninside 349 372

8666 GCF_001637645.1_ASM163764v1 Escherichia coli Proteobacteria; Gammaproteobacteria; Enterobacterales; Enterobacteriaceae; Escherichia MNNEETFYQAMRRQGVTRRSFLKYCSLAATSLGLGAGMAPKIAWALENKPRIPVVWIHGL WP_001058323.1 MULTISPECIES: hydrogenase-1 small chain [Proteobacteria] Length: 372\nNumber of predicted TMHs: 2\nExp number of AAs in TMHs: 37.2608499999999\nExp number, first 60 AAs: 19.52923\nTotal prob of N-in: 0.94351\nPOSSIBLE N-term signal sequence\ninside 1 19\nTMhelix 20 39\noutside 40 325\nTMhelix 326 348\ninside 349 372

8667 GCF_001621915.1_ASM162191v1 Escherichia coli Proteobacteria; Gammaproteobacteria; Enterobacterales; Enterobacteriaceae; Escherichia MNNEETFYQAMRRQGVTRRSFLKYCSLAATSLGLGAGMAPKIAWALENKPRIPVVWIHGL WP_001058323.1 MULTISPECIES: hydrogenase-1 small chain [Proteobacteria] Length: 372\nNumber of predicted TMHs: 2\nExp number of AAs in TMHs: 37.2608499999999\nExp number, first 60 AAs: 19.52923\nTotal prob of N-in: 0.94351\nPOSSIBLE N-term signal sequence\ninside 1 19\nTMhelix 20 39\noutside 40 325\nTMhelix 326 348\ninside 349 372

8668 GCF_001621995.1_ASM162199v1 Escherichia coli Proteobacteria; Gammaproteobacteria; Enterobacterales; Enterobacteriaceae; Escherichia MNNEETFYQAMRRQGVTRRSFLKYCSLAATSLGLGAGMAPKIAWALENKPRIPVVWIHGL WP_001058323.1 MULTISPECIES: hydrogenase-1 small chain [Proteobacteria] Length: 372\nNumber of predicted TMHs: 2\nExp number of AAs in TMHs: 37.2608499999999\nExp number, first 60 AAs: 19.52923\nTotal prob of N-in: 0.94351\nPOSSIBLE N-term signal sequence\ninside 1 19\nTMhelix 20 39\noutside 40 325\nTMhelix 326 348\ninside 349 372

8669 GCF_001621855.1_ASM162185v1 Escherichia coli Proteobacteria; Gammaproteobacteria; Enterobacterales; Enterobacteriaceae; Escherichia MNNEETFYQAMRRQGVTRRSFLKYCSLAATSLGLGAGMAPKIAWALENKPRIPVVWIHGL WP_001295940.1

MULTISPECIES: hydrogenase [Enterobacteriaceae] Length: 372\nNumber of predicted TMHs: 2\nExp number of AAs in TMHs: 37.2608899999999\nExp number, first 60 AAs: 19.52923\nTotal prob of N-in: 0.94351\nPOSSIBLE N-term signal sequence\ninside 1 19\nTMhelix 20 39\noutside 40 325\nTMhelix 326 348\ninside 349 372

8670 GCF_001637705.1_ASM163770v1 Escherichia coli Proteobacteria; Gammaproteobacteria; Enterobacterales; Enterobacteriaceae; Escherichia
MNNEETFYQAMRRQGVTRRSFLKYCSLAATSLGLGAGMAPKIAWALENKPRIPVVWIHGL WP_001058323.1
MULTISPECIES: hydrogenase-1 small chain [Proteobacteria] Length: 372\nNumber of predicted TMHs: 2\nExp number of AAs in TMHs: 37.2608499999999\nExp number, first 60 AAs: 19.52923\nTotal prob of N-in: 0.94351\nPOSSIBLE N-term signal sequence\ninside 1 19\nTMhelix 20 39\noutside 40 325\nTMhelix 326 348\ninside 349 372

8671 GCF_001622135.1_ASM162213v1 Escherichia coli Proteobacteria; Gammaproteobacteria; Enterobacterales; Enterobacteriaceae; Escherichia
MNNEETFYQAMRRQGVTRRSFLKYCSLAATSLGLGAGMAPKIAWALENKPRIPVVWIHGL WP_001295940.1
MULTISPECIES: hydrogenase [Enterobacteriaceae] Length: 372\nNumber of predicted TMHs: 2\nExp number of AAs in TMHs: 37.2608899999999\nExp number, first 60 AAs: 19.52923\nTotal prob of N-in: 0.94351\nPOSSIBLE N-term signal sequence\ninside 1 19\nTMhelix 20 39\noutside 40 325\nTMhelix 326 348\ninside 349 372

8672 GCF_001622045.1_ASM162204v1 Escherichia coli Proteobacteria; Gammaproteobacteria; Enterobacterales; Enterobacteriaceae; Escherichia
MNNEETFYQAMRRQGVTRRSFLKYCSLAATSLGLGAGMAPKIAWALENKPRIPVVWIHGL WP_001295940.1
MULTISPECIES: hydrogenase [Enterobacteriaceae] Length: 372\nNumber of predicted TMHs: 2\nExp number of AAs in TMHs: 37.2608899999999\nExp number, first 60 AAs: 19.52923\nTotal prob of N-in: 0.94351\nPOSSIBLE N-term signal sequence\ninside 1 19\nTMhelix 20 39\noutside 40 325\nTMhelix 326 348\ninside 349 372

8673 GCF_001621975.1_ASM162197v1 Escherichia coli Proteobacteria; Gammaproteobacteria; Enterobacterales; Enterobacteriaceae; Escherichia
MNNEETFYQAMRRQGVTRRSFLKYCSLAATSLGLGAGMAPKIAWALENKPRIPVVWIHGL WP_001295940.1
MULTISPECIES: hydrogenase [Enterobacteriaceae] Length: 372\nNumber of predicted TMHs: 2\nExp number of AAs in TMHs: 37.2608899999999\nExp number, first 60 AAs: 19.52923\nTotal prob of N-in: 0.94351\nPOSSIBLE N-term signal sequence\ninside 1 19\nTMhelix 20 39\noutside 40 325\nTMhelix 326 348\ninside 349 372

8674 GCF_001621965.1_ASM162196v1 Escherichia coli Proteobacteria; Gammaproteobacteria; Enterobacterales; Enterobacteriaceae; Escherichia
MNNEETFYQAMRRQGVTRRSFLKYCSLAATSLGLGAGMAPKIAWALENKPRIPVVWIHGL WP_001295940.1
MULTISPECIES: hydrogenase [Enterobacteriaceae] Length: 372\nNumber of predicted TMHs: 2\nExp number of AAs in TMHs: 37.2608899999999\nExp number, first 60 AAs: 19.52923\nTotal prob of N-in: 0.94351\nPOSSIBLE N-term signal sequence\ninside 1 19\nTMhelix 20 39\noutside 40 325\nTMhelix 326 348\ninside 349 372

8675 GCF_001637795.1_ASM163779v1 Escherichia coli Proteobacteria; Gammaproteobacteria; Enterobacterales; Enterobacteriaceae; Escherichia
MNNEETFYQAMRRQGVTRRSFLKYCSLAATSLGLGAGMAPKIAWALENKPRIPVVWIHGL WP_001058323.1
MULTISPECIES: hydrogenase-1 small chain [Proteobacteria] Length: 372\nNumber of predicted TMHs: 2\nExp number of AAs in TMHs: 37.2608499999999\nExp number, first 60 AAs: 19.52923\nTotal prob of N-in: 0.94351\nPOSSIBLE N-term signal sequence\ninside 1 19\nTMhelix 20 39\noutside 40 325\nTMhelix 326 348\ninside 349 372

8676 GCF_001637865.1_ASM163786v1 Escherichia coli Proteobacteria; Gammaproteobacteria; Enterobacterales; Enterobacteriaceae; Escherichia
MTGDNTLIHSHGINRRDFMKLCAALAATMGLSSKAAAEMVESVTNPQRPPVIWIGAQECT WP_000145423.1
hydrogenase 2 small subunit [Escherichia coli] Length: 372\nNumber of predicted TMHs: 1\nExp number of AAs in TMHs: 23.78201\nExp number, first 60 AAs: 0.28355\nTotal prob of N-in: 0.13257\noutside 1 331\nTMhelix 332 354\ninside 355 372

8677 GCF_001637895.1_ASM163789v1 Escherichia coli Proteobacteria; Gammaproteobacteria; Enterobacterales; Enterobacteriaceae; Escherichia
MTGDNTLIHSHGINRRDFMKLCAALAATMGLSSKAAAEMAESVTNPQRPPVIWIGAQECT WP_000145410.1
MULTISPECIES: hydrogenase-2 small chain [Proteobacteria] Length: 372\nNumber of predicted TMHs: 1\nExp number of AAs in TMHs: 23.72226\nExp number, first 60 AAs: 0.22038\nTotal prob of N-in: 0.12994\noutside 1 331\nTMhelix 332 354\ninside 355 372

8678 GCF_001638005.1_ASM163800v1 Escherichia coli Proteobacteria; Gammaproteobacteria; Enterobacterales; Enterobacteriaceae; Escherichia
MTGDNTLIHSHGINRRDFMKLCAALAATMGLSSKAAAEMVESVTNPQRPPVIWIGAQECT WP_000145423.1
hydrogenase 2 small subunit [Escherichia coli] Length: 372\nNumber of predicted TMHs: 1\nExp number of AAs in TMHs: 23.78201\nExp number, first 60 AAs: 0.28355\nTotal prob of N-in: 0.13257\noutside 1 331\nTMhelix 332 354\ninside 355 372

8679 GCF_001637965.1_ASM163796v1 Escherichia coli Proteobacteria; Gammaproteobacteria; Enterobacterales; Enterobacteriaceae; Escherichia
MTGDNTLIHSHGINRRDFMKLCAALAATMGLSSKAAAEMAESVTNPQRPPVIWIGAQECT WP_000145410.1
MULTISPECIES: hydrogenase-2 small chain [Proteobacteria] Length: 372\nNumber of predicted TMHs: 1\nExp number of AAs in TMHs: 23.72226\nExp number, first 60 AAs: 0.22038\nTotal prob of N-in: 0.12994\noutside 1 331\nTMhelix 332 354\ninside 355 372

8680 GCF_001638055.1_ASM163805v1 Escherichia coli Proteobacteria; Gammaproteobacteria; Enterobacterales; Enterobacteriaceae; Escherichia
MTGDNTLIHSHGINRRDFMKLCAALAATMGLSSKAAAEMAESVTNPQRPPVIWIGAQECT WP_000145410.1
MULTISPECIES: hydrogenase-2 small chain [Proteobacteria] Length: 372\nNumber of predicted TMHs: 1\nExp number of AAs in TMHs: 23.72226\nExp number, first 60 AAs: 0.22038\nTotal prob of N-in: 0.12994\noutside 1 331\nTMhelix 332 354\ninside 355 372

8681 GCF_001638075.1_ASM163807v1 Escherichia coli Proteobacteria; Gammaproteobacteria; Enterobacterales; Enterobacteriaceae; Escherichia
MTGDNTLIHSHGINRRDFMKLCAALAATMGLSSKAAAEMAESVTNPQRPPVIWIGAQECT WP_000145410.1
MULTISPECIES: hydrogenase-2 small chain [Proteobacteria] Length: 372\nNumber of predicted TMHs: 1\nExp number of AAs in TMHs: 23.72226\nExp number, first 60 AAs: 0.22038\nTotal prob of N-in: 0.12994\noutside 1 331\nTMhelix 332 354\ninside 355 372

8682 GCF_001638155.1_ASM163815v1 Escherichia coli Proteobacteria; Gammaproteobacteria; Enterobacterales; Enterobacteriaceae; Escherichia
MTGDNTLIHSHGINRRDFMKLCAALAATMGLSSKAAAEMVESVTNPQRPPVIWIGAQECT WP_000145423.1
hydrogenase 2 small subunit [Escherichia coli] Length: 372\nNumber of predicted TMHs: 1\nExp number of AAs in TMHs: 23.78201\nExp number, first 60 AAs: 0.28355\nTotal prob of N-in: 0.13257\noutside 1 331\nTMhelix 332 354\ninside 355 372

8683 GCF_001638205.1_ASM163820v1 Escherichia coli Proteobacteria; Gammaproteobacteria; Enterobacterales; Enterobacteriaceae; Escherichia
MTGDNTLIHSHGINRRDFMKLCAALAATMGLSSKAAAEMVESVTNPQRPPVIWIGAQECT WP_000145423.1
hydrogenase 2 small subunit [Escherichia coli] Length: 372\nNumber of predicted TMHs: 1\nExp number of AAs in TMHs: 23.78201\nExp number, first 60 AAs: 0.28355\nTotal prob of N-in: 0.13257\noutside 1 331\nTMhelix 332 354\ninside 355 372

8684 GCF_001640205.1_ASM164020v1 Escherichia coli Proteobacteria; Gammaproteobacteria; Enterobacterales; Enterobacteriaceae; Escherichia
MTGDNTLIHSHGINRRDFMKLCAALAATMGLSSKAAAEMVESVTNPQRPPVIWIGAQECT WP_000145423.1
hydrogenase 2 small subunit [Escherichia coli] Length: 372\nNumber of predicted TMHs: 1\nExp number of AAs in TMHs: 23.78201\nExp number, first 60 AAs: 0.28355\nTotal prob of N-in: 0.13257\noutside 1 331\nTMhelix 332 354\ninside 355 372

8685 GCF_001640245.1_ASM164024v1 Escherichia coli Proteobacteria; Gammaproteobacteria; Enterobacterales; Enterobacteriaceae; Escherichia
MTGDNTLIHSHGINRRDFMKLCAALAATMGLSSKAAAEMVESVTNPQRPPVIWIGAQECT WP_000145423.1
hydrogenase 2 small subunit [Escherichia coli] Length: 372\nNumber of predicted TMHs: 1\nExp number of

AAs in TMHs: 23.78201\nExp number, first 60 AAs: 0.28355\nTotal prob of N-in: 0.13257\noutside 1 331\nTMhelix 332 354\ninside 355 372

8686 GCF_001641555.1_ASM164155v1 Escherichia coli Proteobacteria; Gammaproteobacteria; Enterobacterales; Enterobacteriaceae; Escherichia
MTGDNTLIHSHGINRRDFMKLCAALAATMGLSSKAAAEMAESVTNPQRPPVIWIGAQECT WP_000145410.1
MULTISPECIES: hydrogenase-2 small chain [Proteobacteria] Length: 372\nNumber of predicted TMHs: 1\nExp number of AAs in TMHs: 23.72226\nExp number, first 60 AAs: 0.22038\nTotal prob of N-in: 0.12994\noutside 1 331\nTMhelix 332 354\ninside 355 372

8687 GCF_001641875.1_ASM164187v1 Escherichia coli Proteobacteria; Gammaproteobacteria; Enterobacterales; Enterobacteriaceae; Escherichia
MTGDNTLIHSHGINRRDFMKLCAALAATMGLSSKAAAEMAESVTNPQRPPVIWIGAQECT WP_000145410.1
MULTISPECIES: hydrogenase-2 small chain [Proteobacteria] Length: 372\nNumber of predicted TMHs: 1\nExp number of AAs in TMHs: 23.72226\nExp number, first 60 AAs: 0.22038\nTotal prob of N-in: 0.12994\noutside 1 331\nTMhelix 332 354\ninside 355 372

8688 GCF_001641555.1_ASM164155v1 Escherichia coli Proteobacteria; Gammaproteobacteria; Enterobacterales; Enterobacteriaceae; Escherichia
MNNEETFYQAMRRQGVTRRSFLKYCSLAATSLGLGAGMAPKIAWALENKPRIPVVWIHGL WP_001058323.1
MULTISPECIES: hydrogenase-1 small chain [Proteobacteria] Length: 372\nNumber of predicted TMHs: 2\nExp number of AAs in TMHs: 37.2608499999999\nExp number, first 60 AAs: 19.52923\nTotal prob of N-in: 0.94351\nPOSSIBLE N-term signal sequence\ninside 1 19\nTMhelix 20 39\noutside 40 325\nTMhelix 326 348\ninside 349 372

8689 GCF_001641875.1_ASM164187v1 Escherichia coli Proteobacteria; Gammaproteobacteria; Enterobacterales; Enterobacteriaceae; Escherichia
MNNEETFYQAMRRQGVTRRSFLKYCSLAATSLGLGAGMAPKIAWALENKPRIPVVWIHGL WP_001058323.1
MULTISPECIES: hydrogenase-1 small chain [Proteobacteria] Length: 372\nNumber of predicted TMHs: 2\nExp number of AAs in TMHs: 37.2608499999999\nExp number, first 60 AAs: 19.52923\nTotal prob of N-in: 0.94351\nPOSSIBLE N-term signal sequence\ninside 1 19\nTMhelix 20 39\noutside 40 325\nTMhelix 326 348\ninside 349 372

8690 GCF_001637895.1_ASM163789v1 Escherichia coli Proteobacteria; Gammaproteobacteria; Enterobacterales; Enterobacteriaceae; Escherichia
MNNEETFYQAMRRQGVTRRSFLKYCSLAATSLGLGAGMAPKIAWALENKPRIPVVWIHGL WP_001058323.1
MULTISPECIES: hydrogenase-1 small chain [Proteobacteria] Length: 372\nNumber of predicted TMHs: 2\nExp number of AAs in TMHs: 37.2608499999999\nExp number, first 60 AAs: 19.52923\nTotal prob of N-in: 0.94351\nPOSSIBLE N-term signal sequence\ninside 1 19\nTMhelix 20 39\noutside 40 325\nTMhelix 326 348\ninside 349 372

8691 GCF_001638075.1_ASM163807v1 Escherichia coli Proteobacteria; Gammaproteobacteria; Enterobacterales; Enterobacteriaceae; Escherichia
MNNEETFYQAMRRQGVTRRSFLKYCSLAATSLGLGAGMAPKIAWALENKPRIPVVWIHGL WP_001058323.1
MULTISPECIES: hydrogenase-1 small chain [Proteobacteria] Length: 372\nNumber of predicted TMHs: 2\nExp number of AAs in TMHs: 37.2608499999999\nExp number, first 60 AAs: 19.52923\nTotal prob of N-in: 0.94351\nPOSSIBLE N-term signal sequence\ninside 1 19\nTMhelix 20 39\noutside 40 325\nTMhelix 326 348\ninside 349 372

8692 GCF_001637865.1_ASM163786v1 Escherichia coli Proteobacteria; Gammaproteobacteria; Enterobacterales; Enterobacteriaceae; Escherichia
MNNEETFYQAMRRQGVTRRSFLKYCSLAATSLGLGAGMAPKIAWALENKPRIPVVWIHGL WP_001295940.1
MULTISPECIES: hydrogenase [Enterobacteriaceae] Length: 372\nNumber of predicted TMHs: 2\nExp number of AAs in TMHs: 37.2608899999999\nExp number, first 60 AAs: 19.52923\nTotal prob of N-in: 0.94351\nPOSSIBLE N-term signal sequence\ninside 1 19\nTMhelix 20 39\noutside 40 325\nTMhelix 326 348\ninside 349 372

8693 GCF_001638055.1_ASM163805v1 Escherichia coli Proteobacteria; Gammaproteobacteria; Enterobacterales; Enterobacteriaceae; Escherichia
MNNEETFYQAMRRQGVTRRSFLKYCSLAATSLGLGAGMAPKIAWALENKPRIPVVWIHGL WP_001295940.1
MULTISPECIES: hydrogenase [Enterobacteriaceae] Length: 372\nNumber of predicted TMHs: 2\nExp

number of AAs in TMHs: 37.2608899999999\nExp number, first 60 AAs: 19.52923\nTotal prob of N-in: 0.94351\nPOSSIBLE N-term signal sequence\ninside 1 19\nTMhelix 20 39\noutside 40 325\nTMhelix 326 348\ninside 349 372

8694 GCF_001637965.1_ASM163796v1 Escherichia coli Proteobacteria; Gammaproteobacteria; Enterobacterales; Enterobacteriaceae; Escherichia MNNEETFYQAMRRQGVTRRSFLKYCSLAATSLGLGAGMAPKIAWALENKPRIPVVWIHGL WP_001295940.1 MULTISPECIES: hydrogenase [Enterobacteriaceae] Length: 372\nNumber of predicted TMHs: 2\nExp number of AAs in TMHs: 37.2608899999999\nExp number, first 60 AAs: 19.52923\nTotal prob of N-in: 0.94351\nPOSSIBLE N-term signal sequence\ninside 1 19\nTMhelix 20 39\noutside 40 325\nTMhelix 326 348\ninside 349 372

8695 GCF_001638005.1_ASM163800v1 Escherichia coli Proteobacteria; Gammaproteobacteria; Enterobacterales; Enterobacteriaceae; Escherichia MNNEETFYQAMRRQGVTRRSFLKYCSLAATSLGLGAGMAPKIAWALENKPRIPVVWIHGL WP_001295940.1 MULTISPECIES: hydrogenase [Enterobacteriaceae] Length: 372\nNumber of predicted TMHs: 2\nExp number of AAs in TMHs: 37.2608899999999\nExp number, first 60 AAs: 19.52923\nTotal prob of N-in: 0.94351\nPOSSIBLE N-term signal sequence\ninside 1 19\nTMhelix 20 39\noutside 40 325\nTMhelix 326 348\ninside 349 372

8696 GCF_001638155.1_ASM163815v1 Escherichia coli Proteobacteria; Gammaproteobacteria; Enterobacterales; Enterobacteriaceae; Escherichia MNNEETFYQAMRRQGVTRRSFLKYCSLAATSLGLGAGMAPKIAWALENKPRIPVVWIHGL WP_001295940.1 MULTISPECIES: hydrogenase [Enterobacteriaceae] Length: 372\nNumber of predicted TMHs: 2\nExp number of AAs in TMHs: 37.2608899999999\nExp number, first 60 AAs: 19.52923\nTotal prob of N-in: 0.94351\nPOSSIBLE N-term signal sequence\ninside 1 19\nTMhelix 20 39\noutside 40 325\nTMhelix 326 348\ninside 349 372

8697 GCF_001638205.1_ASM163820v1 Escherichia coli Proteobacteria; Gammaproteobacteria; Enterobacterales; Enterobacteriaceae; Escherichia MNNEETFYQAMRRQGVTRRSFLKYCSLAATSLGLGAGMAPKIAWALENKPRIPVVWIHGL WP_001295940.1 MULTISPECIES: hydrogenase [Enterobacteriaceae] Length: 372\nNumber of predicted TMHs: 2\nExp number of AAs in TMHs: 37.2608899999999\nExp number, first 60 AAs: 19.52923\nTotal prob of N-in: 0.94351\nPOSSIBLE N-term signal sequence\ninside 1 19\nTMhelix 20 39\noutside 40 325\nTMhelix 326 348\ninside 349 372

8698 GCF_001641935.1_ASM164193v1 Escherichia coli Proteobacteria; Gammaproteobacteria; Enterobacterales; Enterobacteriaceae; Escherichia MTGDNTLIHSHGINRRDFMKLCAALAATMGLSSKAAAEMAESVTNPQRPPVIWIGAQECT WP_000145410.1 MULTISPECIES: hydrogenase-2 small chain [Proteobacteria] Length: 372\nNumber of predicted TMHs: 1\nExp number of AAs in TMHs: 23.72226\nExp number, first 60 AAs: 0.22038\nTotal prob of N-in: 0.12994\noutside 1 331\nTMhelix 332 354\ninside 355 372

8699 GCF_001641935.1_ASM164193v1 Escherichia coli Proteobacteria; Gammaproteobacteria; Enterobacterales; Enterobacteriaceae; Escherichia MNNEETFYQAMRRQGVTRRSFLKYCSLAATSLGLGAGMAPKIAWALENKPRIPVVWIHGL WP_001058323.1 MULTISPECIES: hydrogenase-1 small chain [Proteobacteria] Length: 372\nNumber of predicted TMHs: 2\nExp number of AAs in TMHs: 37.2608499999999\nExp number, first 60 AAs: 19.52923\nTotal prob of N-in: 0.94351\nPOSSIBLE N-term signal sequence\ninside 1 19\nTMhelix 20 39\noutside 40 325\nTMhelix 326 348\ninside 349 372

8700 GCF_001640205.1_ASM164020v1 Escherichia coli Proteobacteria; Gammaproteobacteria; Enterobacterales; Enterobacteriaceae; Escherichia MNNEETFYQAMRRQGVTRRSFLKYCSLAATSLGLGAGMAPKIAWALENKPRIPVVWIHGL WP_001295940.1 MULTISPECIES: hydrogenase [Enterobacteriaceae] Length: 372\nNumber of predicted TMHs: 2\nExp number of AAs in TMHs: 37.2608899999999\nExp number, first 60 AAs: 19.52923\nTotal prob of N-in: 0.94351\nPOSSIBLE N-term signal sequence\ninside 1 19\nTMhelix 20 39\noutside 40 325\nTMhelix 326 348\ninside 349 372

8701 GCF_001640245.1_ASM164024v1 Escherichia coli Proteobacteria; Gammaproteobacteria; Enterobacterales; Enterobacteriaceae; Escherichia

MNNEETFYQAMRRQGVTRRSFLKYCSLAATSLGLGAGMAPKIAWALENKPRIPVVWIHGL WP_001295940.1 MULTISPECIES: hydrogenase [Enterobacteriaceae] Length: 372\nNumber of predicted TMHs: 2\nExp number of AAs in TMHs: 37.2608899999999\nExp number, first 60 AAs: 19.52923\nTotal prob of N-in: 0.94351\nPOSSIBLE N-term signal sequence\ninside 1 19\nTMhelix 20 39\noutside 40 325\nTMhelix 326 348\ninside 349 372

8702 GCF_001644725.1_ASM164472v1 Escherichia coli Proteobacteria; Gammaproteobacteria; Enterobacterales; Enterobacteriaceae; Escherichia MTGDNTLIHSHGINRRDFMKLCAALAATMGLSSKAAAEMAESVTNPQRPPVIWIGAQECT WP_000145410.1 MULTISPECIES: hydrogenase-2 small chain [Proteobacteria] Length: 372\nNumber of predicted TMHs: 1\nExp number of AAs in TMHs: 23.72226\nExp number, first 60 AAs: 0.22038\nTotal prob of N-in: 0.12994\noutside 1 331\nTMhelix 332 354\ninside 355 372

8703 GCF_001645225.1_ASM164522v1 Escherichia coli Proteobacteria; Gammaproteobacteria; Enterobacterales; Enterobacteriaceae; Escherichia MTGDNTLIHSHGINRRDFMKLCAALAATMGLSSKAAAEMAESVTNPQRPPVIWIGAQECT WP_000145410.1 MULTISPECIES: hydrogenase-2 small chain [Proteobacteria] Length: 372\nNumber of predicted TMHs: 1\nExp number of AAs in TMHs: 23.72226\nExp number, first 60 AAs: 0.22038\nTotal prob of N-in: 0.12994\noutside 1 331\nTMhelix 332 354\ninside 355 372

8704 GCF_001647315.1_ASM164731v1 Escherichia coli Proteobacteria; Gammaproteobacteria; Enterobacterales; Enterobacteriaceae; Escherichia MTGDNTLIHSHGINRRDFMKLCAALAATMGLSSKAAAEMVESVTNPQRPPVIWIGAQECT WP_000145423.1 hydrogenase 2 small subunit [Escherichia coli] Length: 372\nNumber of predicted TMHs: 1\nExp number of AAs in TMHs: 23.78201\nExp number, first 60 AAs: 0.28355\nTotal prob of N-in: 0.13257\noutside 1 331\nTMhelix 332 354\ninside 355 372

8705 GCF_001647335.1_ASM164733v1 Escherichia coli Proteobacteria; Gammaproteobacteria; Enterobacterales; Enterobacteriaceae; Escherichia MTGDNTLIHSHGINRRDFMKLCAALAATMGLSSKAAAEMAESVTNPQRPPVIWIGAQECT WP_000145410.1 MULTISPECIES: hydrogenase-2 small chain [Proteobacteria] Length: 372\nNumber of predicted TMHs: 1\nExp number of AAs in TMHs: 23.72226\nExp number, first 60 AAs: 0.22038\nTotal prob of N-in: 0.12994\noutside 1 331\nTMhelix 332 354\ninside 355 372

8706 GCF_001647405.1_ASM164740v1 Escherichia coli Proteobacteria; Gammaproteobacteria; Enterobacterales; Enterobacteriaceae; Escherichia MTGDNTLIHSHGINRRDFMKLCAALAATMGLSSKAAAEMAESVTNPQRPPVIWIGAQECT WP_000145410.1 MULTISPECIES: hydrogenase-2 small chain [Proteobacteria] Length: 372\nNumber of predicted TMHs: 1\nExp number of AAs in TMHs: 23.72226\nExp number, first 60 AAs: 0.22038\nTotal prob of N-in: 0.12994\noutside 1 331\nTMhelix 332 354\ninside 355 372

8707 GCF_001648915.1_ASM164891v1 Escherichia coli Proteobacteria; Gammaproteobacteria; Enterobacterales; Enterobacteriaceae; Escherichia MTGDNTLIHSHGINRRDFMKLCAALAATMGLSSKAAAEMAESVTNPQRPPVIWIGAQECT WP_000145410.1 MULTISPECIES: hydrogenase-2 small chain [Proteobacteria] Length: 372\nNumber of predicted TMHs: 1\nExp number of AAs in TMHs: 23.72226\nExp number, first 60 AAs: 0.22038\nTotal prob of N-in: 0.12994\noutside 1 331\nTMhelix 332 354\ninside 355 372

8708 GCF_001660095.1_ASM166009v1 Escherichia coli Proteobacteria; Gammaproteobacteria; Enterobacterales; Enterobacteriaceae; Escherichia MTGDNTLIHSHGINRRDFMKLCAALAATMGLSSKAAAEMAESVTNPQRPPVIWIGAQECT WP_000145410.1 MULTISPECIES: hydrogenase-2 small chain [Proteobacteria] Length: 372\nNumber of predicted TMHs: 1\nExp number of AAs in TMHs: 23.72226\nExp number, first 60 AAs: 0.22038\nTotal prob of N-in: 0.12994\noutside 1 331\nTMhelix 332 354\ninside 355 372

8709 GCF_001650605.1_ASM165060v1 Escherichia coli Proteobacteria; Gammaproteobacteria; Enterobacterales; Enterobacteriaceae; Escherichia MTGDNTLIHSHGINRRDFMKLCAALAATMGLSSKAAAEMAESVTNPQRPPVIWIGAQECT WP_000145410.1 MULTISPECIES: hydrogenase-2 small chain [Proteobacteria] Length: 372\nNumber of predicted TMHs: 1\nExp number of AAs in TMHs: 23.72226\nExp number, first 60 AAs: 0.22038\nTotal prob of N-in: 0.12994\noutside 1 331\nTMhelix 332 354\ninside 355 372

8710 GCF_001650555.1_ASM165055v1 Escherichia coli Proteobacteria; Gammaproteobacteria; Enterobacterales; Enterobacteriaceae; Escherichia
MTGDNTLIHSHGINRRDFMKLCAALAATMGLSSKAAAEMAESVTNPQRPPVIWIGAQECT WP_000145410.1
MULTISPECIES: hydrogenase-2 small chain [Proteobacteria] Length: 372\nNumber of predicted TMHs: 1\nExp number of AAs in TMHs: 23.72226\nExp number, first 60 AAs: 0.22038\nTotal prob of N-in: 0.12994\noutside 1 331\nTMhelix 332 354\ninside 355 372

8711 GCF_001651615.1_ASM165161v1 Escherichia coli Proteobacteria; Gammaproteobacteria; Enterobacterales; Enterobacteriaceae; Escherichia
MTGDNTLIHSHGINRRDFMKLCAALAATMGLSSKAAAEMAESVTNPQRPPVIWIGAQECT WP_000145410.1
MULTISPECIES: hydrogenase-2 small chain [Proteobacteria] Length: 372\nNumber of predicted TMHs: 1\nExp number of AAs in TMHs: 23.72226\nExp number, first 60 AAs: 0.22038\nTotal prob of N-in: 0.12994\noutside 1 331\nTMhelix 332 354\ninside 355 372

8712 GCF_001651615.1_ASM165161v1 Escherichia coli Proteobacteria; Gammaproteobacteria; Enterobacterales; Enterobacteriaceae; Escherichia
MNNEETFYQAMRRQGVTRRSFLKYCSLAATSLGLGAGMAPKIAWALENKPRIPVVWIHGL WP_001058323.1
MULTISPECIES: hydrogenase-1 small chain [Proteobacteria] Length: 372\nNumber of predicted TMHs: 2\nExp number of AAs in TMHs: 37.2608499999999\nExp number, first 60 AAs: 19.52923\nTotal prob of N-in: 0.94351\nPOSSIBLE N-term signal sequence\ninside 1 19\nTMhelix 20 39\noutside 40 325\nTMhelix 326 348\ninside 349 372

8713 GCF_001644725.1_ASM164472v1 Escherichia coli Proteobacteria; Gammaproteobacteria; Enterobacterales; Enterobacteriaceae; Escherichia
MNNEETFYQAMRRQGVTRRSFLKYCSLAATSLGLGAGMAPKIAWALENKPRIPVVWIHGL WP_001058323.1
MULTISPECIES: hydrogenase-1 small chain [Proteobacteria] Length: 372\nNumber of predicted TMHs: 2\nExp number of AAs in TMHs: 37.2608499999999\nExp number, first 60 AAs: 19.52923\nTotal prob of N-in: 0.94351\nPOSSIBLE N-term signal sequence\ninside 1 19\nTMhelix 20 39\noutside 40 325\nTMhelix 326 348\ninside 349 372

8714 GCF_001645225.1_ASM164522v1 Escherichia coli Proteobacteria; Gammaproteobacteria; Enterobacterales; Enterobacteriaceae; Escherichia
MNNEETFYQAMRRQGVTRRSFLKYCSLAATSLGLGAGMAPKIAWALENKPRIPVVWIHGL WP_001058323.1
MULTISPECIES: hydrogenase-1 small chain [Proteobacteria] Length: 372\nNumber of predicted TMHs: 2\nExp number of AAs in TMHs: 37.2608499999999\nExp number, first 60 AAs: 19.52923\nTotal prob of N-in: 0.94351\nPOSSIBLE N-term signal sequence\ninside 1 19\nTMhelix 20 39\noutside 40 325\nTMhelix 326 348\ninside 349 372

8715 GCF_001648915.1_ASM164891v1 Escherichia coli Proteobacteria; Gammaproteobacteria; Enterobacterales; Enterobacteriaceae; Escherichia
MNNEETFYQAMRRQGVTRRSFLKYCSLAATSLGLGAGMAPKIAWALENKPRIPVVWIHGL WP_001058323.1
MULTISPECIES: hydrogenase-1 small chain [Proteobacteria] Length: 372\nNumber of predicted TMHs: 2\nExp number of AAs in TMHs: 37.2608499999999\nExp number, first 60 AAs: 19.52923\nTotal prob of N-in: 0.94351\nPOSSIBLE N-term signal sequence\ninside 1 19\nTMhelix 20 39\noutside 40 325\nTMhelix 326 348\ninside 349 372

8716 GCF_001650605.1_ASM165060v1 Escherichia coli Proteobacteria; Gammaproteobacteria; Enterobacterales; Enterobacteriaceae; Escherichia
MNNEETFYQAMRRQGVTRRSFLKYCSLAATSLGLGAGMAPKIAWALENKPRIPVVWIHGL WP_001058323.1
MULTISPECIES: hydrogenase-1 small chain [Proteobacteria] Length: 372\nNumber of predicted TMHs: 2\nExp number of AAs in TMHs: 37.2608499999999\nExp number, first 60 AAs: 19.52923\nTotal prob of N-in: 0.94351\nPOSSIBLE N-term signal sequence\ninside 1 19\nTMhelix 20 39\noutside 40 325\nTMhelix 326 348\ninside 349 372

8717 GCF_001647405.1_ASM164740v1 Escherichia coli Proteobacteria; Gammaproteobacteria; Enterobacterales; Enterobacteriaceae; Escherichia
MNNEETFYQAMRRQGVTRRSFLKYCSLAATSLGLGAGMAPKIAWALENKPRIPVVWIHGL WP_001295940.1
MULTISPECIES: hydrogenase [Enterobacteriaceae] Length: 372\nNumber of predicted TMHs: 2\nExp number of AAs in TMHs: 37.2608899999999\nExp number, first 60 AAs: 19.52923\nTotal prob of N-in: 0.94351\nPOSSIBLE N-term signal sequence\ninside 1 19\nTMhelix 20 39\noutside 40 325\nTMhelix 326 348\ninside 349 372

8718 GCF_001647335.1_ASM164733v1 Escherichia coli Proteobacteria; Gammaproteobacteria; Enterobacterales; Enterobacteriaceae; Escherichia
MNNEETFYQAMRRQGVTRRSFLKYCSLAATSLGLGAGMAPKIAWALENKPRIPVVWIHGL WP_001295940.1
MULTISPECIES: hydrogenase [Enterobacteriaceae] Length: 372\nNumber of predicted TMHs: 2\nExp number of AAs in TMHs: 37.2608899999999\nExp number, first 60 AAs: 19.52923\nTotal prob of N-in: 0.94351\nPOSSIBLE N-term signal sequence\ninside 1 19\nTMhelix 20 39\noutside 40 325\nTMhelix 326 348\ninside 349 372

8719 GCF_001647315.1_ASM164731v1 Escherichia coli Proteobacteria; Gammaproteobacteria; Enterobacterales; Enterobacteriaceae; Escherichia
MNNEETFYQAMRRQGVTRRSFLKYCSLAATSLGLGAGMAPKIAWALENKPRIPVVWIHGL WP_001295940.1
MULTISPECIES: hydrogenase [Enterobacteriaceae] Length: 372\nNumber of predicted TMHs: 2\nExp number of AAs in TMHs: 37.2608899999999\nExp number, first 60 AAs: 19.52923\nTotal prob of N-in: 0.94351\nPOSSIBLE N-term signal sequence\ninside 1 19\nTMhelix 20 39\noutside 40 325\nTMhelix 326 348\ninside 349 372

8720 GCF_001651715.1_ASM165171v1 Escherichia coli Proteobacteria; Gammaproteobacteria; Enterobacterales; Enterobacteriaceae; Escherichia
MTGDNTLIHSHGINRRDFMKLCAALAATMGLSSKAAAEMAESVTNPQRPPVIWIGAQECT WP_000145410.1
MULTISPECIES: hydrogenase-2 small chain [Proteobacteria] Length: 372\nNumber of predicted TMHs: 1\nExp number of AAs in TMHs: 23.72226\nExp number, first 60 AAs: 0.22038\nTotal prob of N-in: 0.12994\noutside 1 331\nTMhelix 332 354\ninside 355 372

8721 GCF_001651725.1_ASM165172v1 Escherichia coli Proteobacteria; Gammaproteobacteria; Enterobacterales; Enterobacteriaceae; Escherichia
MTGDNTLIHSHGINRRDFMKLCAALAATMGLSSKAAAEMVESVTNPQRPPVIWIGAQECT WP_000145423.1
hydrogenase 2 small subunit [Escherichia coli] Length: 372\nNumber of predicted TMHs: 1\nExp number of AAs in TMHs: 23.78201\nExp number, first 60 AAs: 0.28355\nTotal prob of N-in: 0.13257\noutside 1 331\nTMhelix 332 354\ninside 355 372

8722 GCF_001651715.1_ASM165171v1 Escherichia coli Proteobacteria; Gammaproteobacteria; Enterobacterales; Enterobacteriaceae; Escherichia
MNNEETFYQAMRRQGVTRRSFLKYCSLAATSLGLGAGMAPKIAWALENKPRIPVVWIHGL WP_001058323.1
MULTISPECIES: hydrogenase-1 small chain [Proteobacteria] Length: 372\nNumber of predicted TMHs: 2\nExp number of AAs in TMHs: 37.2608499999999\nExp number, first 60 AAs: 19.52923\nTotal prob of N-in: 0.94351\nPOSSIBLE N-term signal sequence\ninside 1 19\nTMhelix 20 39\noutside 40 325\nTMhelix 326 348\ninside 349 372

8723 GCF_001651725.1_ASM165172v1 Escherichia coli Proteobacteria; Gammaproteobacteria; Enterobacterales; Enterobacteriaceae; Escherichia
MNNEETFYQAMRRQGVTRRSFLKYCSLAATSLGLGAGMAPKIAWALENKPRIPVVWIHGL WP_001538532.1
hydrogenase-1 small chain [Escherichia coli] Length: 372\nNumber of predicted TMHs: 2\nExp number of AAs in TMHs: 37.27041\nExp number, first 60 AAs: 19.52927\nTotal prob of N-in: 0.94352\nPOSSIBLE N-term signal sequence\ninside 1 19\nTMhelix 20 39\noutside 40 325\nTMhelix 326 348\ninside 349 372

8724 GCF_001652345.1_ASM165234v1 Escherichia coli Proteobacteria; Gammaproteobacteria; Enterobacterales; Enterobacteriaceae; Escherichia
MTGDNTLIHSHGINRRDFMKLCAALAATMGLSSKAAAEMAESVTNPQRPPVIWIGAQECT WP_000145410.1
MULTISPECIES: hydrogenase-2 small chain [Proteobacteria] Length: 372\nNumber of predicted TMHs: 1\nExp number of AAs in TMHs: 23.72226\nExp number, first 60 AAs: 0.22038\nTotal prob of N-in: 0.12994\noutside 1 331\nTMhelix 332 354\ninside 355 372

8725 GCF_001652345.1_ASM165234v1 Escherichia coli Proteobacteria; Gammaproteobacteria; Enterobacterales; Enterobacteriaceae; Escherichia
MNNEETFYQAMRRQGVTRRSFLKYCSLAATSLGLGAGMAPKIAWALENKPRIPVVWIHGL WP_001058323.1
MULTISPECIES: hydrogenase-1 small chain [Proteobacteria] Length: 372\nNumber of predicted TMHs: 2\nExp number of AAs in TMHs: 37.2608499999999\nExp number, first 60 AAs: 19.52923\nTotal prob of N-in: 0.94351\nPOSSIBLE N-term signal sequence\ninside 1 19\nTMhelix 20 39\noutside 40 325\nTMhelix 326 348\ninside 349 372

8726 GCF_001650555.1_ASM165055v1 Escherichia coli Proteobacteria; Gammaproteobacteria; Enterobacterales; Enterobacteriaceae; Escherichia
MNNEETFYQAMRRQGVTRRSFLKYCSLAATSLGLGAGMAPKIAWALENKPRIPVVWIHGL WP_001353311.1
MULTISPECIES: hydrogenase [Escherichia] Length: 372\nNumber of predicted TMHs: 2\nExp number of AAs in TMHs: 37.2638699999999\nExp number, first 60 AAs: 19.52924\nTotal prob of N-in: 0.94351\nPOSSIBLE N-term signal sequence\ninside 1 19\nTMhelix 20 39\noutside 40 325\nTMhelix 326 348\ninside 349 372

8727 GCF_001653535.1_ASM165353v1 Escherichia coli Proteobacteria; Gammaproteobacteria; Enterobacterales; Enterobacteriaceae; Escherichia
MTGDNTLIHSHGINRRDFMKLCAALAATMGLSSKAAAEMAESVTNPQRPPVIWIGAQECT WP_000145410.1
MULTISPECIES: hydrogenase-2 small chain [Proteobacteria] Length: 372\nNumber of predicted TMHs: 1\nExp number of AAs in TMHs: 23.72226\nExp number, first 60 AAs: 0.22038\nTotal prob of N-in: 0.12994\noutside 1 331\nTMhelix 332 354\ninside 355 372

8728 GCF_001654735.1_ASM165473v1 Escherichia coli Proteobacteria; Gammaproteobacteria; Enterobacterales; Enterobacteriaceae; Escherichia
MTGDNTLIHSHGINRRDFMKLCAALAATMGLSSKAAAEMAESVTNPQRPPVIWIGAQECT WP_000145410.1
MULTISPECIES: hydrogenase-2 small chain [Proteobacteria] Length: 372\nNumber of predicted TMHs: 1\nExp number of AAs in TMHs: 23.72226\nExp number, first 60 AAs: 0.22038\nTotal prob of N-in: 0.12994\noutside 1 331\nTMhelix 332 354\ninside 355 372

8729 GCF_001657505.1_ASM165750v1 Escherichia coli Proteobacteria; Gammaproteobacteria; Enterobacterales; Enterobacteriaceae; Escherichia
MTGDNTLIHSHGINRRDFMKLCAALAATMGLSSKAAAEMAESVTNPQRPPVIWIGAQECT WP_000145410.1
MULTISPECIES: hydrogenase-2 small chain [Proteobacteria] Length: 372\nNumber of predicted TMHs: 1\nExp number of AAs in TMHs: 23.72226\nExp number, first 60 AAs: 0.22038\nTotal prob of N-in: 0.12994\noutside 1 331\nTMhelix 332 354\ninside 355 372

8730 GCF_001659815.1_ASM165981v1 Escherichia coli Proteobacteria; Gammaproteobacteria; Enterobacterales; Enterobacteriaceae; Escherichia
MTGDNTLIHSHGINRRDFMKLCAALAATMGLSSKAAAEMAESVTNPQRPPVIWIGAQECT WP_000145410.1
MULTISPECIES: hydrogenase-2 small chain [Proteobacteria] Length: 372\nNumber of predicted TMHs: 1\nExp number of AAs in TMHs: 23.72226\nExp number, first 60 AAs: 0.22038\nTotal prob of N-in: 0.12994\noutside 1 331\nTMhelix 332 354\ninside 355 372

8731 GCF_001653535.1_ASM165353v1 Escherichia coli Proteobacteria; Gammaproteobacteria; Enterobacterales; Enterobacteriaceae; Escherichia
MNNEETFYQAMRRQGVTRRSFLKYCSLAATSLGLGAGMAPKIAWALENKPRIPVVWIHGL WP_001058323.1
MULTISPECIES: hydrogenase-1 small chain [Proteobacteria] Length: 372\nNumber of predicted TMHs: 2\nExp number of AAs in TMHs: 37.2608499999999\nExp number, first 60 AAs: 19.52923\nTotal prob of N-in: 0.94351\nPOSSIBLE N-term signal sequence\ninside 1 19\nTMhelix 20 39\noutside 40 325\nTMhelix 326 348\ninside 349 372

8732 GCF_001660185.1_ASM166018v1 Escherichia coli Proteobacteria; Gammaproteobacteria; Enterobacterales; Enterobacteriaceae; Escherichia
MTGDNTLIHSHGINRRDFMKLCAALAATMGLSSKAAAEMAESVTNPQRPPVIWIGAQECT WP_000145410.1
MULTISPECIES: hydrogenase-2 small chain [Proteobacteria] Length: 372\nNumber of predicted TMHs: 1\nExp number of AAs in TMHs: 23.72226\nExp number, first 60 AAs: 0.22038\nTotal prob of N-in: 0.12994\noutside 1 331\nTMhelix 332 354\ninside 355 372

8733 GCF_001660185.1_ASM166018v1 Escherichia coli Proteobacteria; Gammaproteobacteria; Enterobacterales; Enterobacteriaceae; Escherichia
MNNEETFYQAMRRQGVTRRSFLKYCSLAATSLGLGAGMAPKIAWALENKPRIPVVWIHGL WP_001301613.1
hydrogenase [Escherichia coli] Length: 372\nNumber of predicted TMHs: 2\nExp number of AAs in TMHs: 37.46598\nExp number, first 60 AAs: 19.53139\nTotal prob of N-in: 0.94364\nPOSSIBLE N-term signal sequence\ninside 1 19\nTMhelix 20 39\noutside 40 325\nTMhelix 326 348\ninside 349 372

8734 GCF_001654735.1_ASM165473v1 Escherichia coli Proteobacteria; Gammaproteobacteria; Enterobacterales; Enterobacteriaceae; Escherichia
MNNEETFYQAMRRQGVTRRSFLKYCSLAATSLGLGAGMAPKIAWALENKPRIPVVWIHGL WP_001058323.1
MULTISPECIES: hydrogenase-1 small chain [Proteobacteria] Length: 372\nNumber of predicted TMHs:

2\nExp number of AAs in TMHs: 37.2608499999999\nExp number, first 60 AAs: 19.52923\nTotal prob of N-in: 0.94351\nPOSSIBLE N-term signal sequence\ninside 1 19\nTMhelix 20 39\noutside 40 325\nTMhelix 326 348\ninside 349 372

8735 GCF_001660305.1_ASM166030v1 Escherichia coli Proteobacteria; Gammaproteobacteria; Enterobacterales; Enterobacteriaceae; Escherichia
MTGDNTLIHSHGINRRDFMKLCAALAATMGLSSKAAAEMAESVTNPQRPPVIWIGAQECT WP_000145410.1
MULTISPECIES: hydrogenase-2 small chain [Proteobacteria] Length: 372\nNumber of predicted TMHs: 1\nExp number of AAs in TMHs: 23.72226\nExp number, first 60 AAs: 0.22038\nTotal prob of N-in: 0.12994\noutside 1 331\nTMhelix 332 354\ninside 355 372

8736 GCF_001660245.1_ASM166024v1 Escherichia coli Proteobacteria; Gammaproteobacteria; Enterobacterales; Enterobacteriaceae; Escherichia
MTGDNTLIHSHGINRRDFMKLCAALAATMGLSSKAAAEMAESVTNPQRPPVIWIGAQECT WP_000145410.1
MULTISPECIES: hydrogenase-2 small chain [Proteobacteria] Length: 372\nNumber of predicted TMHs: 1\nExp number of AAs in TMHs: 23.72226\nExp number, first 60 AAs: 0.22038\nTotal prob of N-in: 0.12994\noutside 1 331\nTMhelix 332 354\ninside 355 372

8737 GCF_001660245.1_ASM166024v1 Escherichia coli Proteobacteria; Gammaproteobacteria; Enterobacterales; Enterobacteriaceae; Escherichia
MNNEETFYQAMRRQGVTRRSFLKYCSLAATSLGLGAGMAPKIAWALENKPRIPVVWIHGL WP_001058323.1
MULTISPECIES: hydrogenase-1 small chain [Proteobacteria] Length: 372\nNumber of predicted TMHs: 2\nExp number of AAs in TMHs: 37.2608499999999\nExp number, first 60 AAs: 19.52923\nTotal prob of N-in: 0.94351\nPOSSIBLE N-term signal sequence\ninside 1 19\nTMhelix 20 39\noutside 40 325\nTMhelix 326 348\ninside 349 372

8738 GCF_001660305.1_ASM166030v1 Escherichia coli Proteobacteria; Gammaproteobacteria; Enterobacterales; Enterobacteriaceae; Escherichia
MNNEETFYQAMRRQGVTRRSFLKYCSLAATSLGLGAGMAPKIAWALENKPRIPVVWIHGL WP_001301613.1
hydrogenase [Escherichia coli] Length: 372\nNumber of predicted TMHs: 2\nExp number of AAs in TMHs: 37.46598\nExp number, first 60 AAs: 19.53139\nTotal prob of N-in: 0.94364\nPOSSIBLE N-term signal sequence\ninside 1 19\nTMhelix 20 39\noutside 40 325\nTMhelix 326 348\ninside 349 372

8739 GCF_001657505.1_ASM165750v1 Escherichia coli Proteobacteria; Gammaproteobacteria; Enterobacterales; Enterobacteriaceae; Escherichia
MNNEETFYQAMRRQGVTRRSFLKYCSLAATSLGLGAGMAPKIAWALENKPRIPVVWIHGL WP_001058320.1
hydrogenase [Escherichia coli] Length: 372\nNumber of predicted TMHs: 2\nExp number of AAs in TMHs: 37.26082\nExp number, first 60 AAs: 19.52923\nTotal prob of N-in: 0.94351\nPOSSIBLE N-term signal sequence\ninside 1 19\nTMhelix 20 39\noutside 40 325\nTMhelix 326 348\ninside 349 372

8740 GCF_001659815.1_ASM165981v1 Escherichia coli Proteobacteria; Gammaproteobacteria; Enterobacterales; Enterobacteriaceae; Escherichia
MNNEETFYQAMRRQGVTRRSFLKYCSLAATSLGLGAGMAPKIAWALENKPRIPVVWIHGL WP_001058320.1
hydrogenase [Escherichia coli] Length: 372\nNumber of predicted TMHs: 2\nExp number of AAs in TMHs: 37.26082\nExp number, first 60 AAs: 19.52923\nTotal prob of N-in: 0.94351\nPOSSIBLE N-term signal sequence\ninside 1 19\nTMhelix 20 39\noutside 40 325\nTMhelix 326 348\ninside 349 372

8741 GCF_001660095.1_ASM166009v1 Escherichia coli Proteobacteria; Gammaproteobacteria; Enterobacterales; Enterobacteriaceae; Escherichia
MNNEETFYQAMRRQGVTRRSFLKYCSLAATSLGLGAGMAPKIAWALENKPRIPVVWIHGL WP_001058323.1
MULTISPECIES: hydrogenase-1 small chain [Proteobacteria] Length: 372\nNumber of predicted TMHs: 2\nExp number of AAs in TMHs: 37.2608499999999\nExp number, first 60 AAs: 19.52923\nTotal prob of N-in: 0.94351\nPOSSIBLE N-term signal sequence\ninside 1 19\nTMhelix 20 39\noutside 40 325\nTMhelix 326 348\ninside 349 372

8742 GCF_001660335.1_ASM166033v1 Escherichia coli Proteobacteria; Gammaproteobacteria; Enterobacterales; Enterobacteriaceae; Escherichia
MTGDNTLIHSHGINRRDFMKLCAALAATMGLSSKAAAEMAESVTNPQRPPVIWIGAQECT WP_000145410.1
MULTISPECIES: hydrogenase-2 small chain [Proteobacteria] Length: 372\nNumber of predicted TMHs: 1\nExp number of AAs in TMHs: 23.72226\nExp number, first 60 AAs: 0.22038\nTotal prob of N-in: 0.12994\noutside 1 331\nTMhelix 332 354\ninside 355 372

8743 GCF_001661055.2_ASM166105v2 Escherichia coli Proteobacteria; Gammaproteobacteria; Enterobacterales; Enterobacteriaceae; Escherichia
MTGDNTLIHSHGINRRDFMKLCAALAATMGLSSKAAAEMVESVTNPQRPPVIWIGAQECT WP_000145423.1
hydrogenase 2 small subunit [Escherichia coli] Length: 372\nNumber of predicted TMHs: 1\nExp number of AAs in TMHs: 23.78201\nExp number, first 60 AAs: 0.28355\nTotal prob of N-in: 0.13257\noutside 1 331\nTMhelix 332 354\ninside 355 372

8744 GCF_001660565.1_ASM166056v1 Escherichia coli Proteobacteria; Gammaproteobacteria; Enterobacterales; Enterobacteriaceae; Escherichia
MTGDNTLIHSHGINRRDFMKLCAALAATMGLSSKAAAEMAESVTNPQRPPVIWIGAQECT WP_000145410.1
MULTISPECIES: hydrogenase-2 small chain [Proteobacteria] Length: 372\nNumber of predicted TMHs: 1\nExp number of AAs in TMHs: 23.72226\nExp number, first 60 AAs: 0.22038\nTotal prob of N-in: 0.12994\noutside 1 331\nTMhelix 332 354\ninside 355 372

8745 GCF_001675705.1_ASM167570v1 Escherichia coli Proteobacteria; Gammaproteobacteria; Enterobacterales; Enterobacteriaceae; Escherichia
MTGDNTLIHSHGINRRDFMKLCAALAATMGLSSKAAAEMAESVTNPQRPPVIWIGAQECT WP_000145410.1
MULTISPECIES: hydrogenase-2 small chain [Proteobacteria] Length: 372\nNumber of predicted TMHs: 1\nExp number of AAs in TMHs: 23.72226\nExp number, first 60 AAs: 0.22038\nTotal prob of N-in: 0.12994\noutside 1 331\nTMhelix 332 354\ninside 355 372

8746 GCF_001677475.1_ASM167747v1 Escherichia coli Proteobacteria; Gammaproteobacteria; Enterobacterales; Enterobacteriaceae; Escherichia
MTGDNTLIHSHGINRRDFMKLCAALAATMGLSSKAAAEMAESVTNPQRPPVIWIGAQECT WP_000145410.1
MULTISPECIES: hydrogenase-2 small chain [Proteobacteria] Length: 372\nNumber of predicted TMHs: 1\nExp number of AAs in TMHs: 23.72226\nExp number, first 60 AAs: 0.22038\nTotal prob of N-in: 0.12994\noutside 1 331\nTMhelix 332 354\ninside 355 372

8747 GCF_001672015.1_ASM167201v1 Escherichia coli Proteobacteria; Gammaproteobacteria; Enterobacterales; Enterobacteriaceae; Escherichia
MTGDNTLIHSHGINRRDFMKLCAALAATMGLSSKAAAEMAESVTNPQRPPVIWIGAQECT WP_000145410.1
MULTISPECIES: hydrogenase-2 small chain [Proteobacteria] Length: 372\nNumber of predicted TMHs: 1\nExp number of AAs in TMHs: 23.72226\nExp number, first 60 AAs: 0.22038\nTotal prob of N-in: 0.12994\noutside 1 331\nTMhelix 332 354\ninside 355 372

8748 GCF_001676985.1_ASM167698v1 Escherichia coli Proteobacteria; Gammaproteobacteria; Enterobacterales; Enterobacteriaceae; Escherichia
MTGDNTLIHSHGINRRDFMKLCAALAATMGLSSKAAAEMAESVTNPQRPPVIWIGAQECT WP_000145410.1
MULTISPECIES: hydrogenase-2 small chain [Proteobacteria] Length: 372\nNumber of predicted TMHs: 1\nExp number of AAs in TMHs: 23.72226\nExp number, first 60 AAs: 0.22038\nTotal prob of N-in: 0.12994\noutside 1 331\nTMhelix 332 354\ninside 355 372

8749 GCF_001660335.1_ASM166033v1 Escherichia coli Proteobacteria; Gammaproteobacteria; Enterobacterales; Enterobacteriaceae; Escherichia
MNNEETFYQAMRRQGVTRRSFLKYCSLAATSLGLGAGMAPKIAWALENKPRIPVVWIHGL WP_001058323.1
MULTISPECIES: hydrogenase-1 small chain [Proteobacteria] Length: 372\nNumber of predicted TMHs: 2\nExp number of AAs in TMHs: 37.2608499999999\nExp number, first 60 AAs: 19.52923\nTotal prob of N-in: 0.94351\nPOSSIBLE N-term signal sequence\ninside 1 19\nTMhelix 20 39\noutside 40 325\nTMhelix 326 348\ninside 349 372

8750 GCF_001677495.1_ASM167749v1 Escherichia coli Proteobacteria; Gammaproteobacteria; Enterobacterales; Enterobacteriaceae; Escherichia
MTGDNTLIHSHGINRRDFMKLCAALAATMGLSSKAAAEMAESVTNPQRPPVIWIGAQECT WP_000145410.1
MULTISPECIES: hydrogenase-2 small chain [Proteobacteria] Length: 372\nNumber of predicted TMHs: 1\nExp number of AAs in TMHs: 23.72226\nExp number, first 60 AAs: 0.22038\nTotal prob of N-in: 0.12994\noutside 1 331\nTMhelix 332 354\ninside 355 372

8751 GCF_001677495.1_ASM167749v1 Escherichia coli Proteobacteria; Gammaproteobacteria; Enterobacterales; Enterobacteriaceae; Escherichia
MNNEETFYQAMRRQGVTRRSFLKYCSLAATSLGLGAGMAPKIAWALENKPRIPVVWIHGL WP_001295940.1
MULTISPECIES: hydrogenase [Enterobacteriaceae] Length: 372\nNumber of predicted TMHs: 2\nExp

number of AAs in TMHs: 37.2608899999999\nExp number, first 60 AAs: 19.52923\nTotal prob of N-in: 0.94351\nPOSSIBLE N-term signal sequence\ninside 1 19\nTMhelix 20 39\noutside 40 325\nTMhelix 326 348\ninside 349 372

8752 GCF_001677475.1_ASM167747v1 Escherichia coli Proteobacteria; Gammaproteobacteria; Enterobacterales; Enterobacteriaceae; Escherichia MNNEETFYQAMRRQGVTRRSFLKYCSLAATSLGLGAGMAPKIAWALENKPRIPVVWIHGL WP_001295940.1 MULTISPECIES: hydrogenase [Enterobacteriaceae] Length: 372\nNumber of predicted TMHs: 2\nExp number of AAs in TMHs: 37.2608899999999\nExp number, first 60 AAs: 19.52923\nTotal prob of N-in: 0.94351\nPOSSIBLE N-term signal sequence\ninside 1 19\nTMhelix 20 39\noutside 40 325\nTMhelix 326 348\ninside 349 372

8753 GCF_001660565.1_ASM166056v1 Escherichia coli Proteobacteria; Gammaproteobacteria; Enterobacterales; Enterobacteriaceae; Escherichia MNNEETFYQAMRRQGVTRRSFLKYCSLAATSLGLGAGMAPKIAWALENKPRIPVVWIHGL WP_001058323.1 MULTISPECIES: hydrogenase-1 small chain [Proteobacteria] Length: 372\nNumber of predicted TMHs: 2\nExp number of AAs in TMHs: 37.2608499999999\nExp number, first 60 AAs: 19.52923\nTotal prob of N-in: 0.94351\nPOSSIBLE N-term signal sequence\ninside 1 19\nTMhelix 20 39\noutside 40 325\nTMhelix 326 348\ninside 349 372

8754 GCF_001677545.2_ASM167754v2 Escherichia coli Proteobacteria; Gammaproteobacteria; Enterobacterales; Enterobacteriaceae; Escherichia MTGDNTLIHSHGINRRDFMKLCAALAATMGLSSKAAAEMAESVTNPQRPPVIWIGAQECT WP_000145410.1 MULTISPECIES: hydrogenase-2 small chain [Proteobacteria] Length: 372\nNumber of predicted TMHs: 1\nExp number of AAs in TMHs: 23.72226\nExp number, first 60 AAs: 0.22038\nTotal prob of N-in: 0.12994\noutside 1 331\nTMhelix 332 354\ninside 355 372

8755 GCF_001677565.2_ASM167756v2 Escherichia coli Proteobacteria; Gammaproteobacteria; Enterobacterales; Enterobacteriaceae; Escherichia MTGDNTLIHSHGINRRDFMKLCAALAATMGLSSKAAAEMAESVTNPQRPPVIWIGAQECT WP_000145410.1 MULTISPECIES: hydrogenase-2 small chain [Proteobacteria] Length: 372\nNumber of predicted TMHs: 1\nExp number of AAs in TMHs: 23.72226\nExp number, first 60 AAs: 0.22038\nTotal prob of N-in: 0.12994\noutside 1 331\nTMhelix 332 354\ninside 355 372

8756 GCF_001677635.2_ASM167763v2 Escherichia coli Proteobacteria; Gammaproteobacteria; Enterobacterales; Enterobacteriaceae; Escherichia MTGDNTLIHSHGINRRDFMKLCAALAATMGLSSKAAAEMAESVTNPQRPPVIWIGAQECT WP_000145410.1 MULTISPECIES: hydrogenase-2 small chain [Proteobacteria] Length: 372\nNumber of predicted TMHs: 1\nExp number of AAs in TMHs: 23.72226\nExp number, first 60 AAs: 0.22038\nTotal prob of N-in: 0.12994\noutside 1 331\nTMhelix 332 354\ninside 355 372

8757 GCF_001677635.2_ASM167763v2 Escherichia coli Proteobacteria; Gammaproteobacteria; Enterobacterales; Enterobacteriaceae; Escherichia MNNEETFYQAMRRQGVTRRSFLKYCSLAATSLGLGAGMAPKIAWALENKPRIPVVWIHGL WP_001301613.1 hydrogenase [Escherichia coli] Length: 372\nNumber of predicted TMHs: 2\nExp number of AAs in TMHs: 37.46598\nExp number, first 60 AAs: 19.53139\nTotal prob of N-in: 0.94364\nPOSSIBLE N-term signal sequence\ninside 1 19\nTMhelix 20 39\noutside 40 325\nTMhelix 326 348\ninside 349 372

8758 GCF_001677545.2_ASM167754v2 Escherichia coli Proteobacteria; Gammaproteobacteria; Enterobacterales; Enterobacteriaceae; Escherichia MNNEETFYQAMRRQGVTRRSFLKYCSLAATSLGLGAGMAPKIAWALENKPRIPVVWIHGL WP_001301613.1 hydrogenase [Escherichia coli] Length: 372\nNumber of predicted TMHs: 2\nExp number of AAs in TMHs: 37.46598\nExp number, first 60 AAs: 19.53139\nTotal prob of N-in: 0.94364\nPOSSIBLE N-term signal sequence\ninside 1 19\nTMhelix 20 39\noutside 40 325\nTMhelix 326 348\ninside 349 372

8759 GCF_001677565.2_ASM167756v2 Escherichia coli Proteobacteria; Gammaproteobacteria; Enterobacterales; Enterobacteriaceae; Escherichia MNNEETFYQAMRRQGVTRRSFLKYCSLAATSLGLGAGMAPKIAWALENKPRIPVVWIHGL WP_001301613.1 hydrogenase [Escherichia coli] Length: 372\nNumber of predicted TMHs: 2\nExp number of AAs in TMHs: 37.46598\nExp number, first 60 AAs: 19.53139\nTotal prob of N-in: 0.94364\nPOSSIBLE N-term signal sequence\ninside 1 19\nTMhelix 20 39\noutside 40 325\nTMhelix 326 348\ninside 349 372

8760 GCF_001672015.1_ASM167201v1 Escherichia coli Proteobacteria; Gammaproteobacteria; Enterobacterales; Enterobacteriaceae; Escherichia
MNNEETFYQAMRRQGVTRRSFLKYCSLAATSLGLGAGMAPKIAWALENKPRIPVVWIHGL WP_001058323.1
MULTISPECIES: hydrogenase-1 small chain [Proteobacteria] Length: 372\nNumber of predicted TMHs: 2\nExp number of AAs in TMHs: 37.2608499999999\nExp number, first 60 AAs: 19.52923\nTotal prob of N-in: 0.94351\nPOSSIBLE N-term signal sequence\ninside 1 19\nTMhelix 20 39\noutside 40 325\nTMhelix 326 348\ninside 349 372

8761 GCF_001677695.2_ASM167769v2 Escherichia coli Proteobacteria; Gammaproteobacteria; Enterobacterales; Enterobacteriaceae; Escherichia
MTGDNTLIHSHGINRRDFMKLCAALAATMGLSSKAAAEMAESVTNPQRPPVIWIGAQECT WP_000145410.1
MULTISPECIES: hydrogenase-2 small chain [Proteobacteria] Length: 372\nNumber of predicted TMHs: 1\nExp number of AAs in TMHs: 23.72226\nExp number, first 60 AAs: 0.22038\nTotal prob of N-in: 0.12994\noutside 1 331\nTMhelix 332 354\ninside 355 372

8762 GCF_001677695.2_ASM167769v2 Escherichia coli Proteobacteria; Gammaproteobacteria; Enterobacterales; Enterobacteriaceae; Escherichia
MNNEETFYQAMRRQGVTRRSFLKYCSLAATSLGLGAGMAPKIAWALENKPRIPVVWIHGL WP_001301613.1
hydrogenase [Escherichia coli] Length: 372\nNumber of predicted TMHs: 2\nExp number of AAs in TMHs: 37.46598\nExp number, first 60 AAs: 19.53139\nTotal prob of N-in: 0.94364\nPOSSIBLE N-term signal sequence\ninside 1 19\nTMhelix 20 39\noutside 40 325\nTMhelix 326 348\ninside 349 372

8763 GCF_001661055.2_ASM166105v2 Escherichia coli Proteobacteria; Gammaproteobacteria; Enterobacterales; Enterobacteriaceae; Escherichia
MNNEETFYQAMRRQGVTRRSFLKYCSLAATSLGLGAGMAPKIAWALENKPRIPVVWIHGL WP_001295940.1
MULTISPECIES: hydrogenase [Enterobacteriaceae] Length: 372\nNumber of predicted TMHs: 2\nExp number of AAs in TMHs: 37.2608899999999\nExp number, first 60 AAs: 19.52923\nTotal prob of N-in: 0.94351\nPOSSIBLE N-term signal sequence\ninside 1 19\nTMhelix 20 39\noutside 40 325\nTMhelix 326 348\ninside 349 372

8764 GCF_001676985.1_ASM167698v1 Escherichia coli Proteobacteria; Gammaproteobacteria; Enterobacterales; Enterobacteriaceae; Escherichia
MNNEETFYQAMRRQGVTRRSFLKYCSLAATSLGLGAGMAPKIAWALENKPRIPVVWIHGL WP_001058323.1
MULTISPECIES: hydrogenase-1 small chain [Proteobacteria] Length: 372\nNumber of predicted TMHs: 2\nExp number of AAs in TMHs: 37.2608499999999\nExp number, first 60 AAs: 19.52923\nTotal prob of N-in: 0.94351\nPOSSIBLE N-term signal sequence\ninside 1 19\nTMhelix 20 39\noutside 40 325\nTMhelix 326 348\ninside 349 372

8765 GCF_001675705.1_ASM167570v1 Escherichia coli Proteobacteria; Gammaproteobacteria; Enterobacterales; Enterobacteriaceae; Escherichia
MNNEETFYQAMRRQGVTRRSFLKYCSLAATSLGLGAGMAPKIAWALENKPRIPVVWIHGL WP_001295940.1
MULTISPECIES: hydrogenase [Enterobacteriaceae] Length: 372\nNumber of predicted TMHs: 2\nExp number of AAs in TMHs: 37.2608899999999\nExp number, first 60 AAs: 19.52923\nTotal prob of N-in: 0.94351\nPOSSIBLE N-term signal sequence\ninside 1 19\nTMhelix 20 39\noutside 40 325\nTMhelix 326 348\ninside 349 372

8766 GCF_001677725.2_ASM167772v2 Escherichia coli Proteobacteria; Gammaproteobacteria; Enterobacterales; Enterobacteriaceae; Escherichia
MTGDNTLIHSHGINRRDFMKLCAALAATMGLSSKAAAEMAESVTNPQRPPVIWIGAQECT WP_000145410.1
MULTISPECIES: hydrogenase-2 small chain [Proteobacteria] Length: 372\nNumber of predicted TMHs: 1\nExp number of AAs in TMHs: 23.72226\nExp number, first 60 AAs: 0.22038\nTotal prob of N-in: 0.12994\noutside 1 331\nTMhelix 332 354\ninside 355 372

8767 GCF_001677785.2_ASM167778v2 Escherichia coli Proteobacteria; Gammaproteobacteria; Enterobacterales; Enterobacteriaceae; Escherichia
MTGDNTLIHSHGINRRDFMKLCAALAATMGLSSKAAAEMAESVTNPQRPPVIWIGAQECT WP_000145410.1
MULTISPECIES: hydrogenase-2 small chain [Proteobacteria] Length: 372\nNumber of predicted TMHs: 1\nExp number of AAs in TMHs: 23.72226\nExp number, first 60 AAs: 0.22038\nTotal prob of N-in: 0.12994\noutside 1 331\nTMhelix 332 354\ninside 355 372

8768 GCF_001677855.2_ASM167785v2 Escherichia coli Proteobacteria; Gammaproteobacteria; Enterobacterales; Enterobacteriaceae; Escherichia
MTGDNTLIHSHGINRRDFMKLCAALAATMGLSSKAAAEMAESVTNPQRPPVIWIGAQECT WP_000145410.1
MULTISPECIES: hydrogenase-2 small chain [Proteobacteria] Length: 372\nNumber of predicted TMHs: 1\nExp number of AAs in TMHs: 23.72226\nExp number, first 60 AAs: 0.22038\nTotal prob of N-in: 0.12994\noutside 1 331\nTMhelix 332 354\ninside 355 372

8769 GCF_001677875.2_ASM167787v2 Escherichia coli Proteobacteria; Gammaproteobacteria; Enterobacterales; Enterobacteriaceae; Escherichia
MTGDNTLIHSHGINRRDFMKLCAALAATMGLSSKAAAEMAESVTNPQRPPVIWIGAQECT WP_000145410.1
MULTISPECIES: hydrogenase-2 small chain [Proteobacteria] Length: 372\nNumber of predicted TMHs: 1\nExp number of AAs in TMHs: 23.72226\nExp number, first 60 AAs: 0.22038\nTotal prob of N-in: 0.12994\noutside 1 331\nTMhelix 332 354\ninside 355 372

8770 GCF_001679985.1_ASM167998v1 Escherichia coli Proteobacteria; Gammaproteobacteria; Enterobacterales; Enterobacteriaceae; Escherichia
MTGDNTLIHSHGINRRDFMKLCAALAATMGLSSKAAAEMAESVTNPQRPPVIWIGAQECT WP_000145410.1
MULTISPECIES: hydrogenase-2 small chain [Proteobacteria] Length: 372\nNumber of predicted TMHs: 1\nExp number of AAs in TMHs: 23.72226\nExp number, first 60 AAs: 0.22038\nTotal prob of N-in: 0.12994\noutside 1 331\nTMhelix 332 354\ninside 355 372

8771 GCF_001683435.1_ASM168343v1 Escherichia coli Proteobacteria; Gammaproteobacteria; Enterobacterales; Enterobacteriaceae; Escherichia
MTGDNTLIHSHGINRRDFMKLCAALAATMGLSSKAAAEMAESVTNPQRPPVIWIGAQECT WP_000145410.1
MULTISPECIES: hydrogenase-2 small chain [Proteobacteria] Length: 372\nNumber of predicted TMHs: 1\nExp number of AAs in TMHs: 23.72226\nExp number, first 60 AAs: 0.22038\nTotal prob of N-in: 0.12994\noutside 1 331\nTMhelix 332 354\ninside 355 372

8772 GCF_001683595.1_NGF2 Escherichia coli Proteobacteria; Gammaproteobacteria; Enterobacterales; Enterobacteriaceae; Escherichia MTGDNTLIHSHGINRRDFMKLCAALAATMGLSSKAAAEMVESVTNPQRPPVIWIGAQECT
WP_000145423.1 hydrogenase 2 small subunit [Escherichia coli] Length: 372\nNumber of predicted TMHs: 1\nExp number of AAs in TMHs: 23.78201\nExp number, first 60 AAs: 0.28355\nTotal prob of N-in: 0.13257\noutside 1 331\nTMhelix 332 354\ninside 355 372

8773 GCF_001692785.1_ASM169278v1 Escherichia coli Proteobacteria; Gammaproteobacteria; Enterobacterales; Enterobacteriaceae; Escherichia
MTGDNTLIHSHGINRRDFMKLCAALAATMGLSSKAAAEMVESVTNPQRPPVIWIGAQECT WP_000145423.1
hydrogenase 2 small subunit [Escherichia coli] Length: 372\nNumber of predicted TMHs: 1\nExp number of AAs in TMHs: 23.78201\nExp number, first 60 AAs: 0.28355\nTotal prob of N-in: 0.13257\noutside 1 331\nTMhelix 332 354\ninside 355 372

8774 GCF_001692865.1_ASM169286v1 Escherichia coli Proteobacteria; Gammaproteobacteria; Enterobacterales; Enterobacteriaceae; Escherichia
MTGDNTLIHSHGINRRDFMKLCAALAATMGLSSKAAAEMVESVTNPQRPPVIWIGAQECT WP_000145423.1
hydrogenase 2 small subunit [Escherichia coli] Length: 372\nNumber of predicted TMHs: 1\nExp number of AAs in TMHs: 23.78201\nExp number, first 60 AAs: 0.28355\nTotal prob of N-in: 0.13257\noutside 1 331\nTMhelix 332 354\ninside 355 372

8775 GCF_001692785.1_ASM169278v1 Escherichia coli Proteobacteria; Gammaproteobacteria; Enterobacterales; Enterobacteriaceae; Escherichia
MNNEETFYQAMRRQGVTRRSFLKYCSLAATSLGLGAGMAPKIAWALENKPRIPVVWIHGL WP_001058323.1
MULTISPECIES: hydrogenase-1 small chain [Proteobacteria] Length: 372\nNumber of predicted TMHs: 2\nExp number of AAs in TMHs: 37.2608499999999\nExp number, first 60 AAs: 19.52923\nTotal prob of N-in: 0.94351\nPOSSIBLE N-term signal sequence\ninside 1 19\nTMhelix 20 39\noutside 40 325\nTMhelix 326 348\ninside 349 372

8776 GCF_001692865.1_ASM169286v1 Escherichia coli Proteobacteria; Gammaproteobacteria; Enterobacterales; Enterobacteriaceae; Escherichia
MNNEETFYQAMRRQGVTRRSFLKYCSLAATSLGLGAGMAPKIAWALENKPRIPVVWIHGL WP_001058323.1
MULTISPECIES: hydrogenase-1 small chain [Proteobacteria] Length: 372\nNumber of predicted TMHs: 2\nExp number of AAs in TMHs: 37.2608499999999\nExp number, first 60 AAs: 19.52923\nTotal prob of N-in:

0.94351\nPOSSIBLE N-term signal sequence\ninside 1 19\nTMhelix 20 39\noutside 40 325\nTMhelix 326 348\ninside 349 372

8777 GCF_001683435.1_ASM168343v1 Escherichia coli Proteobacteria; Gammaproteobacteria; Enterobacterales; Enterobacteriaceae; Escherichia MNNEETFYQAMRRQGVTRRSFLKYCSLAATSLGLGAGMAPKIAWALENKPRIPVVWIHGL WP_001058323.1 MULTISPECIES: hydrogenase-1 small chain [Proteobacteria] Length: 372\nNumber of predicted TMHs: 2\nExp number of AAs in TMHs: 37.2608499999999\nExp number, first 60 AAs: 19.52923\nTotal prob of N-in: 0.94351\nPOSSIBLE N-term signal sequence\ninside 1 19\nTMhelix 20 39\noutside 40 325\nTMhelix 326 348\ninside 349 372

8778 GCF_001683595.1_NGF2 Escherichia coli Proteobacteria; Gammaproteobacteria; Enterobacterales; Enterobacteriaceae; Escherichia MNNEETFYQAMRRQGVTRRSFLKYCSLAATSLGLGAGMAPKIAWALENKPRIPVVWIHGL WP_001058323.1 MULTISPECIES: hydrogenase-1 small chain [Proteobacteria] Length: 372\nNumber of predicted TMHs: 2\nExp number of AAs in TMHs: 37.2608499999999\nExp number, first 60 AAs: 19.52923\nTotal prob of N-in: 0.94351\nPOSSIBLE N-term signal sequence\ninside 1 19\nTMhelix 20 39\noutside 40 325\nTMhelix 326 348\ninside 349 372

8779 GCF_001696245.1_ASM169624v1 Escherichia coli Proteobacteria; Gammaproteobacteria; Enterobacterales; Enterobacteriaceae; Escherichia MTGDNTLIHSHGINRRDFMKLCAALAATMGLSSKAAAEMAESVTNPQRPPVIWIGAQECT WP_000145410.1 MULTISPECIES: hydrogenase-2 small chain [Proteobacteria] Length: 372\nNumber of predicted TMHs: 1\nExp number of AAs in TMHs: 23.72226\nExp number, first 60 AAs: 0.22038\nTotal prob of N-in: 0.12994\noutside 1 331\nTMhelix 332 354\ninside 355 372

8780 GCF_001696245.1_ASM169624v1 Escherichia coli Proteobacteria; Gammaproteobacteria; Enterobacterales; Enterobacteriaceae; Escherichia MNNEETFYQAMRRQGVTRRSFLKYCSLAATSLGLGAGMAPKIAWALENKPRIPVVWIHGL WP_001058323.1 MULTISPECIES: hydrogenase-1 small chain [Proteobacteria] Length: 372\nNumber of predicted TMHs: 2\nExp number of AAs in TMHs: 37.2608499999999\nExp number, first 60 AAs: 19.52923\nTotal prob of N-in: 0.94351\nPOSSIBLE N-term signal sequence\ninside 1 19\nTMhelix 20 39\noutside 40 325\nTMhelix 326 348\ninside 349 372

8781 GCF_001677855.2_ASM167785v2 Escherichia coli Proteobacteria; Gammaproteobacteria; Enterobacterales; Enterobacteriaceae; Escherichia MNNEETFYQAMRRQGVTRRSFLKYCSLAATSLGLGAGMAPKIAWALENKPRIPVVWIHGL WP_001301613.1 hydrogenase [Escherichia coli] Length: 372\nNumber of predicted TMHs: 2\nExp number of AAs in TMHs: 37.46598\nExp number, first 60 AAs: 19.53139\nTotal prob of N-in: 0.94364\nPOSSIBLE N-term signal sequence\ninside 1 19\nTMhelix 20 39\noutside 40 325\nTMhelix 326 348\ninside 349 372

8782 GCF_001679985.1_ASM167998v1 Escherichia coli Proteobacteria; Gammaproteobacteria; Enterobacterales; Enterobacteriaceae; Escherichia MNNEETFYQAMRRQGVTRRSFLKYCSLAATSLGLGAGMAPKIAWALENKPRIPVVWIHGL WP_001295940.1 MULTISPECIES: hydrogenase [Enterobacteriaceae] Length: 372\nNumber of predicted TMHs: 2\nExp number of AAs in TMHs: 37.2608899999999\nExp number, first 60 AAs: 19.52923\nTotal prob of N-in: 0.94351\nPOSSIBLE N-term signal sequence\ninside 1 19\nTMhelix 20 39\noutside 40 325\nTMhelix 326 348\ninside 349 372

8783 GCF_001696335.1_ASM169633v1 Escherichia coli Proteobacteria; Gammaproteobacteria; Enterobacterales; Enterobacteriaceae; Escherichia MTGDNTLIHSHGINRRDFMKLCAALAATMGLSSKAAAEMAESVTNPQRPPVIWIGAQECT WP_000145410.1 MULTISPECIES: hydrogenase-2 small chain [Proteobacteria] Length: 372\nNumber of predicted TMHs: 1\nExp number of AAs in TMHs: 23.72226\nExp number, first 60 AAs: 0.22038\nTotal prob of N-in: 0.12994\noutside 1 331\nTMhelix 332 354\ninside 355 372

8784 GCF_001696335.1_ASM169633v1 Escherichia coli Proteobacteria; Gammaproteobacteria; Enterobacterales; Enterobacteriaceae; Escherichia MNNEETFYQAMRRQGVTRRSFLKYCSLAATSLGLGAGMAPKIAWALENKPRIPVVWIHGL WP_001058323.1 MULTISPECIES: hydrogenase-1 small chain [Proteobacteria] Length: 372\nNumber of predicted TMHs: 2\nExp number of AAs in TMHs: 37.2608499999999\nExp number, first 60 AAs: 19.52923\nTotal prob of N-in:

0.94351\nPOSSIBLE N-term signal sequence\ninside 1 19\nTMhelix 20 39\noutside 40 325\nTMhelix 326 348\ninside 349 372

8785 GCF_001677725.2_ASM167772v2 Escherichia coli Proteobacteria; Gammaproteobacteria; Enterobacterales; Enterobacteriaceae; Escherichia MNNEETFYQAMRRQGVTRRSFLKYCSLAATSLGLGAGMAPKIAWALENKPRIPVVWIHGL WP_001301613.1 hydrogenase [Escherichia coli] Length: 372\nNumber of predicted TMHs: 2\nExp number of AAs in TMHs: 37.46598\nExp number, first 60 AAs: 19.53139\nTotal prob of N-in: 0.94364\nPOSSIBLE N-term signal sequence\ninside 1 19\nTMhelix 20 39\noutside 40 325\nTMhelix 326 348\ninside 349 372

8786 GCF_001677785.2_ASM167778v2 Escherichia coli Proteobacteria; Gammaproteobacteria; Enterobacterales; Enterobacteriaceae; Escherichia MNNEETFYQAMRRQGVTRRSFLKYCSLAATSLGLGAGMAPKIAWALENKPRIPVVWIHGL WP_001301613.1 hydrogenase [Escherichia coli] Length: 372\nNumber of predicted TMHs: 2\nExp number of AAs in TMHs: 37.46598\nExp number, first 60 AAs: 19.53139\nTotal prob of N-in: 0.94364\nPOSSIBLE N-term signal sequence\ninside 1 19\nTMhelix 20 39\noutside 40 325\nTMhelix 326 348\ninside 349 372

8787 GCF_001677875.2_ASM167787v2 Escherichia coli Proteobacteria; Gammaproteobacteria; Enterobacterales; Enterobacteriaceae; Escherichia MNNEETFYQAMRRQGVTRRSFLKYCSLAATSLGLGAGMAPKIAWALENKPRIPVVWIHGL WP_001301613.1 hydrogenase [Escherichia coli] Length: 372\nNumber of predicted TMHs: 2\nExp number of AAs in TMHs: 37.46598\nExp number, first 60 AAs: 19.53139\nTotal prob of N-in: 0.94364\nPOSSIBLE N-term signal sequence\ninside 1 19\nTMhelix 20 39\noutside 40 325\nTMhelix 326 348\ninside 349 372

8788 GCF_001696775.1_ASM169677v1 Escherichia coli Proteobacteria; Gammaproteobacteria; Enterobacterales; Enterobacteriaceae; Escherichia MTGDNTLIHSHGINRRDFMKLCAALAATMGLSSKAAAEMAESVTNPQRPPVIWIGAQECT WP_000145410.1 MULTISPECIES: hydrogenase-2 small chain [Proteobacteria] Length: 372\nNumber of predicted TMHs: 1\nExp number of AAs in TMHs: 23.72226\nExp number, first 60 AAs: 0.22038\nTotal prob of N-in: 0.12994\noutside 1 331\nTMhelix 332 354\ninside 355 372

8789 GCF_001700025.1_ASM170002v1 Escherichia coli Proteobacteria; Gammaproteobacteria; Enterobacterales; Enterobacteriaceae; Escherichia MTGDNTLIHSHGINRRDFMKLCAALAATMGLSSKAAAEMAESVTNPQRPPVIWIGAQECT WP_000145410.1 MULTISPECIES: hydrogenase-2 small chain [Proteobacteria] Length: 372\nNumber of predicted TMHs: 1\nExp number of AAs in TMHs: 23.72226\nExp number, first 60 AAs: 0.22038\nTotal prob of N-in: 0.12994\noutside 1 331\nTMhelix 332 354\ninside 355 372

8790 GCF_001693635.1_ASM169363v1 Escherichia coli Proteobacteria; Gammaproteobacteria; Enterobacterales; Enterobacteriaceae; Escherichia MTGDNTLIHSHGINRRDFMKLCAALAATMGLSSKAAAEMAESVTNPQRPPVIWIGAQECT WP_029402609.1 hydrogenase 2 small subunit [Escherichia coli] Length: 372\nNumber of predicted TMHs: 1\nExp number of AAs in TMHs: 23.72225\nExp number, first 60 AAs: 0.22038\nTotal prob of N-in: 0.12994\noutside 1 331\nTMhelix 332 354\ninside 355 372

8791 GCF_001700025.1_ASM170002v1 Escherichia coli Proteobacteria; Gammaproteobacteria; Enterobacterales; Enterobacteriaceae; Escherichia MNNEETFYQAMRRQGVTRRSFLKYCSLAATSLGLGAGMAPKIAWALENKPRIPVVWIHGL WP_001295940.1 MULTISPECIES: hydrogenase [Enterobacteriaceae] Length: 372\nNumber of predicted TMHs: 2\nExp number of AAs in TMHs: 37.2608899999999\nExp number, first 60 AAs: 19.52923\nTotal prob of N-in: 0.94351\nPOSSIBLE N-term signal sequence\ninside 1 19\nTMhelix 20 39\noutside 40 325\nTMhelix 326 348\ninside 349 372

8792 GCF_001704555.1_NCTC86 Escherichia coli Proteobacteria; Gammaproteobacteria; Enterobacterales; Enterobacteriaceae; Escherichia MNNEETFYQAMRRQGVTRRSFLKYCSLAATSLGLGAGMAPKIAWALENKPRIPVVWIHGL WP_001058323.1 MULTISPECIES: hydrogenase-1 small chain [Proteobacteria] Length: 372\nNumber of predicted TMHs: 2\nExp number of AAs in TMHs: 37.2608499999999\nExp number, first 60 AAs: 19.52923\nTotal prob of N-in: 0.94351\nPOSSIBLE N-term signal sequence\ninside 1 19\nTMhelix 20 39\noutside 40 325\nTMhelix 326 348\ninside 349 372

8793 GCF_001713625.1_ASM171362v1 Escherichia coli Proteobacteria; Gammaproteobacteria; Enterobacterales; Enterobacteriaceae; Escherichia MTGDNTLIHSHGINRRDFMKLCAALAATMGLSSKAAAEMAESVTNPQRPPVIWIGAQECT WP_000145410.1 MULTISPECIES: hydrogenase-2 small chain [Proteobacteria] Length: 372\nNumber of predicted TMHs: 1\nExp number of AAs in TMHs: 23.72226\nExp number, first 60 AAs: 0.22038\nTotal prob of N-in: 0.12994\noutside 1 331\nTMhelix 332 354\ninside 355 372

8794 GCF_001713555.1_ASM171355v1 Escherichia coli Proteobacteria; Gammaproteobacteria; Enterobacterales; Enterobacteriaceae; Escherichia MTGDNTLIHSHGINRRDFMKLCAALAATMGLSSKAAAEMAESVTNPQRPPVIWIGAQECT WP_000145410.1 MULTISPECIES: hydrogenase-2 small chain [Proteobacteria] Length: 372\nNumber of predicted TMHs: 1\nExp number of AAs in TMHs: 23.72226\nExp number, first 60 AAs: 0.22038\nTotal prob of N-in: 0.12994\noutside 1 331\nTMhelix 332 354\ninside 355 372

8795 GCF_001713625.1_ASM171362v1 Escherichia coli Proteobacteria; Gammaproteobacteria; Enterobacterales; Enterobacteriaceae; Escherichia MNNEETFYQAMRRQGVTRRSFLKYCSLAATSLGLGAGMAPKIAWALENKPRIPVVWIHGL WP_001058323.1 MULTISPECIES: hydrogenase-1 small chain [Proteobacteria] Length: 372\nNumber of predicted TMHs: 2\nExp number of AAs in TMHs: 37.2608499999999\nExp number, first 60 AAs: 19.52923\nTotal prob of N-in: 0.94351\nPOSSIBLE N-term signal sequence\ninside 1 19\nTMhelix 20 39\noutside 40 325\nTMhelix 326 348\ninside 349 372

8796 GCF_001704555.1_NCTC86 Escherichia coli Proteobacteria; Gammaproteobacteria; Enterobacterales; Enterobacteriaceae; Escherichia MTGDNTLIHSHGINRRDFMKLCAALAATMGLSSKAAAEMAESVTNPQRPPVIWIGAQECT WP_066009678.1 hydrogenase 2 small subunit [Escherichia coli] Length: 372\nNumber of predicted TMHs: 1\nExp number of AAs in TMHs: 23.72258\nExp number, first 60 AAs: 0.22053\nTotal prob of N-in: 0.12995\noutside 1 331\nTMhelix 332 354\ninside 355 372

8797 GCF_001713555.1_ASM171355v1 Escherichia coli Proteobacteria; Gammaproteobacteria; Enterobacterales; Enterobacteriaceae; Escherichia MNNEETFYQAMRRQGVTRRSFLKYCSLAATSLGLGAGMAPKIAWALENKPRIPVVWIHGL WP_001295940.1 MULTISPECIES: hydrogenase [Enterobacteriaceae] Length: 372\nNumber of predicted TMHs: 2\nExp number of AAs in TMHs: 37.2608899999999\nExp number, first 60 AAs: 19.52923\nTotal prob of N-in: 0.94351\nPOSSIBLE N-term signal sequence\ninside 1 19\nTMhelix 20 39\noutside 40 325\nTMhelix 326 348\ninside 349 372

8798 GCF_001700405.1_ASM170040v1 Escherichia coli Proteobacteria; Gammaproteobacteria; Enterobacterales; Enterobacteriaceae; Escherichia MNNEETFYQAMRRQGVTRRSFLKYCSLAATSLGLGAGMAPKIAWALENKPRIPVVWIHGL WP_001058323.1 MULTISPECIES: hydrogenase-1 small chain [Proteobacteria] Length: 372\nNumber of predicted TMHs: 2\nExp number of AAs in TMHs: 37.2608499999999\nExp number, first 60 AAs: 19.52923\nTotal prob of N-in: 0.94351\nPOSSIBLE N-term signal sequence\ninside 1 19\nTMhelix 20 39\noutside 40 325\nTMhelix 326 348\ninside 349 372

8799 GCF_001723505.1_ASM172350v1 Escherichia coli Proteobacteria; Gammaproteobacteria; Enterobacterales; Enterobacteriaceae; Escherichia MTGDNTLIHSHGINRRDFMKLCAALAATMGLSSKAAAEMAESVTNPQRPPVIWIGAQECT WP_000145410.1 MULTISPECIES: hydrogenase-2 small chain [Proteobacteria] Length: 372\nNumber of predicted TMHs: 1\nExp number of AAs in TMHs: 23.72226\nExp number, first 60 AAs: 0.22038\nTotal prob of N-in: 0.12994\noutside 1 331\nTMhelix 332 354\ninside 355 372

8800 GCF_001735705.1_ASM173570v1 Escherichia coli Proteobacteria; Gammaproteobacteria; Enterobacterales; Enterobacteriaceae; Escherichia MTGDNTLIHSHGINRRDFMKLCAALAATMGLSSKAAAEMAESVTNPQRPPVIWIGAQECT WP_000145410.1 MULTISPECIES: hydrogenase-2 small chain [Proteobacteria] Length: 372\nNumber of predicted TMHs: 1\nExp number of AAs in TMHs: 23.72226\nExp number, first 60 AAs: 0.22038\nTotal prob of N-in: 0.12994\noutside 1 331\nTMhelix 332 354\ninside 355 372

8801 GCF_001721225.1_ASM172122v1 Escherichia coli Proteobacteria; Gammaproteobacteria; Enterobacterales; Enterobacteriaceae; Escherichia MNNEETFYQAMRRQGVTRRSFLKYCSLAATSLGLGAGMAPKIAWALENKPRIPVVWIHGL WP_001058323.1

MULTISPECIES: hydrogenase-1 small chain [Proteobacteria] Length: 372\nNumber of predicted TMHs: 2\nExp number of AAs in TMHs: 37.2608499999999\nExp number, first 60 AAs: 19.52923\nTotal prob of N-in: 0.94351\nPOSSIBLE N-term signal sequence\ninside 1 19\nTMhelix 20 39\noutside 40 325\nTMhelix 326 348\ninside 349 372

8802 GCF_001723505.1_ASM172350v1 Escherichia coli Proteobacteria; Gammaproteobacteria; Enterobacterales; Enterobacteriaceae; Escherichia
MNNEETFYQAMRRQGVTRRSFLKYCSLAATSLGLGAGMAPKIAWALENKPRIPVVWIHGL WP_001058323.1
MULTISPECIES: hydrogenase-1 small chain [Proteobacteria] Length: 372\nNumber of predicted TMHs: 2\nExp number of AAs in TMHs: 37.2608499999999\nExp number, first 60 AAs: 19.52923\nTotal prob of N-in: 0.94351\nPOSSIBLE N-term signal sequence\ninside 1 19\nTMhelix 20 39\noutside 40 325\nTMhelix 326 348\ninside 349 372

8803 GCF_001735705.1_ASM173570v1 Escherichia coli Proteobacteria; Gammaproteobacteria; Enterobacterales; Enterobacteriaceae; Escherichia
MNNEETFYQAMRRQGVTRRSFLKYCSLAATSLGLGAGMAPKIAWALENKPRIPVVWIHGL WP_001058323.1
MULTISPECIES: hydrogenase-1 small chain [Proteobacteria] Length: 372\nNumber of predicted TMHs: 2\nExp number of AAs in TMHs: 37.2608499999999\nExp number, first 60 AAs: 19.52923\nTotal prob of N-in: 0.94351\nPOSSIBLE N-term signal sequence\ninside 1 19\nTMhelix 20 39\noutside 40 325\nTMhelix 326 348\ninside 349 372

8804 GCF_001742645.1_ASM174264v1 Escherichia coli Proteobacteria; Gammaproteobacteria; Enterobacterales; Enterobacteriaceae; Escherichia
MNNEETFYQAMRRQGVTRRSFLKYCSLAATSLGLGAGMAPKIAWALENKPRIPVVWIHGL WP_001058323.1
MULTISPECIES: hydrogenase-1 small chain [Proteobacteria] Length: 372\nNumber of predicted TMHs: 2\nExp number of AAs in TMHs: 37.2608499999999\nExp number, first 60 AAs: 19.52923\nTotal prob of N-in: 0.94351\nPOSSIBLE N-term signal sequence\ninside 1 19\nTMhelix 20 39\noutside 40 325\nTMhelix 326 348\ninside 349 372

8805 GCF_001721225.1_ASM172122v1 Escherichia coli Proteobacteria; Gammaproteobacteria; Enterobacterales; Enterobacteriaceae; Escherichia
MTGDNTLIHSHGINRRDFMKLCAALAATMGLSSKAAAEMAESVTNPQRPPVIWIGAQECT WP_069357453.1
hydrogenase 2 small subunit [Escherichia coli] Length: 372\nNumber of predicted TMHs: 1\nExp number of AAs in TMHs: 23.72156\nExp number, first 60 AAs: 0.22038\nTotal prob of N-in: 0.12992\noutside 1 331\nTMhelix 332 354\ninside 355 372

8806 GCF_001742665.1_ASM174266v1 Escherichia coli Proteobacteria; Gammaproteobacteria; Enterobacterales; Enterobacteriaceae; Escherichia
MTGDNTLIHSHGINRRDFMKLCAALAATMGLSSKAAAEMAESVTNPQRPPVIWIGAQECT WP_000145406.1
hydrogenase-2 small chain [Escherichia coli] Length: 372\nNumber of predicted TMHs: 1\nExp number of AAs in TMHs: 23.72251\nExp number, first 60 AAs: 0.22027\nTotal prob of N-in: 0.13006\noutside 1 331\nTMhelix 332 354\ninside 355 372

8807 GCF_001747365.1_ASM174736v2 Escherichia coli Proteobacteria; Gammaproteobacteria; Enterobacterales; Enterobacteriaceae; Escherichia
MTGDNTLIHSHGINRRDFMKLCAALAATMGLSSKAAAEMAESVTNPQRPPVIWIGAQECT WP_000145410.1
MULTISPECIES: hydrogenase-2 small chain [Proteobacteria] Length: 372\nNumber of predicted TMHs: 1\nExp number of AAs in TMHs: 23.72226\nExp number, first 60 AAs: 0.22038\nTotal prob of N-in: 0.12994\noutside 1 331\nTMhelix 332 354\ninside 355 372

8808 GCF_001748605.1_ASM174860v1 Escherichia coli Proteobacteria; Gammaproteobacteria; Enterobacterales; Enterobacteriaceae; Escherichia
MTGDNTLIHSHGINRRDFMKLCAALAATMGLSSKAAAEMAESVTNPQRPPVIWIGAQECT WP_000145410.1
MULTISPECIES: hydrogenase-2 small chain [Proteobacteria] Length: 372\nNumber of predicted TMHs: 1\nExp number of AAs in TMHs: 23.72226\nExp number, first 60 AAs: 0.22038\nTotal prob of N-in: 0.12994\noutside 1 331\nTMhelix 332 354\ninside 355 372

8809 GCF_001747365.1_ASM174736v2 Escherichia coli Proteobacteria; Gammaproteobacteria; Enterobacterales; Enterobacteriaceae; Escherichia
MNNEETFYQAMRRQGVTRRSFLKYCSLAATSLGLGAGMAPKIAWALENKPRIPVVWIHGL WP_001058323.1
MULTISPECIES: hydrogenase-1 small chain [Proteobacteria] Length: 372\nNumber of predicted TMHs:

2\nExp number of AAs in TMHs: 37.2608499999999\nExp number, first 60 AAs: 19.52923\nTotal prob of N-in: 0.94351\nPOSSIBLE N-term signal sequence\ninside 1 19\nTMhelix 20 39\noutside 40 325\nTMhelix 326 348\ninside 349 372

8810 GCF_001742665.1_ASM174266v1 Escherichia coli Proteobacteria; Gammaproteobacteria; Enterobacterales; Enterobacteriaceae; Escherichia
MNNEETFYQAMRRQGVTRRSFLKYCSLAATSLGLGAGMAPKIAWALENKPRIPVVWIHGL WP_001058323.1
MULTISPECIES: hydrogenase-1 small chain [Proteobacteria] Length: 372\nNumber of predicted TMHs: 2\nExp number of AAs in TMHs: 37.2608499999999\nExp number, first 60 AAs: 19.52923\nTotal prob of N-in: 0.94351\nPOSSIBLE N-term signal sequence\ninside 1 19\nTMhelix 20 39\noutside 40 325\nTMhelix 326 348\ninside 349 372

8811 GCF_001748695.1_ASM174869v1 Escherichia coli Proteobacteria; Gammaproteobacteria; Enterobacterales; Enterobacteriaceae; Escherichia
MTGDNTLIHSHGINRRDFMKLCAALAATMGLSSKAAAEMAESVTNPQRPPVIWIGAQECT WP_000145410.1
MULTISPECIES: hydrogenase-2 small chain [Proteobacteria] Length: 372\nNumber of predicted TMHs: 1\nExp number of AAs in TMHs: 23.72226\nExp number, first 60 AAs: 0.22038\nTotal prob of N-in: 0.12994\noutside 1 331\nTMhelix 332 354\ninside 355 372

8812 GCF_001748735.1_ASM174873v1 Escherichia coli Proteobacteria; Gammaproteobacteria; Enterobacterales; Enterobacteriaceae; Escherichia
MNNEETFYQAMRRQGVTRRSFLKYCSLAATSLGLGAGMAPKIAWALENKPRIPVVWIHGL WP_001058323.1
MULTISPECIES: hydrogenase-1 small chain [Proteobacteria] Length: 372\nNumber of predicted TMHs: 2\nExp number of AAs in TMHs: 37.2608499999999\nExp number, first 60 AAs: 19.52923\nTotal prob of N-in: 0.94351\nPOSSIBLE N-term signal sequence\ninside 1 19\nTMhelix 20 39\noutside 40 325\nTMhelix 326 348\ninside 349 372

8813 GCF_001748625.1_ASM174862v1 Escherichia coli Proteobacteria; Gammaproteobacteria; Enterobacterales; Enterobacteriaceae; Escherichia
MNNEETFYQAMRRQGVTRRSFLKYCSLAATSLGLGAGMAPKIAWALENKPRIPVVWIHGL WP_001058323.1
MULTISPECIES: hydrogenase-1 small chain [Proteobacteria] Length: 372\nNumber of predicted TMHs: 2\nExp number of AAs in TMHs: 37.2608499999999\nExp number, first 60 AAs: 19.52923\nTotal prob of N-in: 0.94351\nPOSSIBLE N-term signal sequence\ninside 1 19\nTMhelix 20 39\noutside 40 325\nTMhelix 326 348\ninside 349 372

8814 GCF_001748815.1_ASM174881v1 Escherichia coli Proteobacteria; Gammaproteobacteria; Enterobacterales; Enterobacteriaceae; Escherichia
MTGDNTLIHSHGINRRDFMKLCAALAATMGLSSKAAAEMAESVTNPQRPPVIWIGAQECT WP_000145410.1
MULTISPECIES: hydrogenase-2 small chain [Proteobacteria] Length: 372\nNumber of predicted TMHs: 1\nExp number of AAs in TMHs: 23.72226\nExp number, first 60 AAs: 0.22038\nTotal prob of N-in: 0.12994\noutside 1 331\nTMhelix 332 354\ninside 355 372

8815 GCF_001748695.1_ASM174869v1 Escherichia coli Proteobacteria; Gammaproteobacteria; Enterobacterales; Enterobacteriaceae; Escherichia
MNNEETFYQAMRRQGVTRRSFLKYCSLAATSLGLGAGMAPKIAWALENKPRIPVVWIHGL WP_001295940.1
MULTISPECIES: hydrogenase [Enterobacteriaceae] Length: 372\nNumber of predicted TMHs: 2\nExp number of AAs in TMHs: 37.2608899999999\nExp number, first 60 AAs: 19.52923\nTotal prob of N-in: 0.94351\nPOSSIBLE N-term signal sequence\ninside 1 19\nTMhelix 20 39\noutside 40 325\nTMhelix 326 348\ninside 349 372

8816 GCF_001748965.1_ASM174896v1 Escherichia coli Proteobacteria; Gammaproteobacteria; Enterobacterales; Enterobacteriaceae; Escherichia
MTGDNTLIHSHGINRRDFMKLCAALAATMGLSSKAAAEMAESVTNPQRPPVIWIGAQECT WP_000145410.1
MULTISPECIES: hydrogenase-2 small chain [Proteobacteria] Length: 372\nNumber of predicted TMHs: 1\nExp number of AAs in TMHs: 23.72226\nExp number, first 60 AAs: 0.22038\nTotal prob of N-in: 0.12994\noutside 1 331\nTMhelix 332 354\ninside 355 372

8817 GCF_001748995.1_ASM174899v1 Escherichia coli Proteobacteria; Gammaproteobacteria; Enterobacterales; Enterobacteriaceae; Escherichia
MTGDNTLIHSHGINRRDFMKLCAALAATMGLSSKAAAEMAESVTNPQRPPVIWIGAQECT WP_000145410.1
MULTISPECIES: hydrogenase-2 small chain [Proteobacteria] Length: 372\nNumber of predicted TMHs:

1\nExp number of AAs in TMHs: 23.72226\nExp number, first 60 AAs: 0.22038\nTotal prob of N-in: 0.12994\noutside 1 331\nTMhelix 332 354\ninside 355 372

8818 GCF_001748835.1_ASM174883v1 Escherichia coli Proteobacteria; Gammaproteobacteria; Enterobacterales; Enterobacteriaceae; Escherichia
MTGDNTLIHSHGINRRDFMKLCAALAATMGLSSKAAAEMAESVTNPQRPPVIWIGAQECT WP_000145410.1
MULTISPECIES: hydrogenase-2 small chain [Proteobacteria] Length: 372\nNumber of predicted TMHs: 1\nExp number of AAs in TMHs: 23.72226\nExp number, first 60 AAs: 0.22038\nTotal prob of N-in: 0.12994\noutside 1 331\nTMhelix 332 354\ninside 355 372

8819 GCF_001748905.1_ASM174890v1 Escherichia coli Proteobacteria; Gammaproteobacteria; Enterobacterales; Enterobacteriaceae; Escherichia
MTGDNTLIHSHGINRRDFMKLCAALAATMGLSSKAAAEMAESVTNPQRPPVIWIGAQECT WP_000145410.1
MULTISPECIES: hydrogenase-2 small chain [Proteobacteria] Length: 372\nNumber of predicted TMHs: 1\nExp number of AAs in TMHs: 23.72226\nExp number, first 60 AAs: 0.22038\nTotal prob of N-in: 0.12994\noutside 1 331\nTMhelix 332 354\ninside 355 372

8820 GCF_001749055.1_ASM174905v1 Escherichia coli Proteobacteria; Gammaproteobacteria; Enterobacterales; Enterobacteriaceae; Escherichia
MTGDNTLIHSHGINRRDFMKLCAALAATMGLSSKAAAEMAESVTNPQRPPVIWIGAQECT WP_000145410.1
MULTISPECIES: hydrogenase-2 small chain [Proteobacteria] Length: 372\nNumber of predicted TMHs: 1\nExp number of AAs in TMHs: 23.72226\nExp number, first 60 AAs: 0.22038\nTotal prob of N-in: 0.12994\noutside 1 331\nTMhelix 332 354\ninside 355 372

8821 GCF_001749075.1_ASM174907v1 Escherichia coli Proteobacteria; Gammaproteobacteria; Enterobacterales; Enterobacteriaceae; Escherichia
MTGDNTLIHSHGINRRDFMKLCAALAATMGLSSKAAAEMAESVTNPQRPPVIWIGAQECT WP_000145410.1
MULTISPECIES: hydrogenase-2 small chain [Proteobacteria] Length: 372\nNumber of predicted TMHs: 1\nExp number of AAs in TMHs: 23.72226\nExp number, first 60 AAs: 0.22038\nTotal prob of N-in: 0.12994\noutside 1 331\nTMhelix 332 354\ninside 355 372

8822 GCF_001865905.1_ASM186590v1 Escherichia coli Proteobacteria; Gammaproteobacteria; Enterobacterales; Enterobacteriaceae; Escherichia
MTGDNTLIHSHGINRRDFMKLCAALAATMGLSSKAAAEMAESVTNPQRPPVIWIGAQECT WP_000145410.1
MULTISPECIES: hydrogenase-2 small chain [Proteobacteria] Length: 372\nNumber of predicted TMHs: 1\nExp number of AAs in TMHs: 23.72226\nExp number, first 60 AAs: 0.22038\nTotal prob of N-in: 0.12994\noutside 1 331\nTMhelix 332 354\ninside 355 372

8823 GCF_001748965.1_ASM174896v1 Escherichia coli Proteobacteria; Gammaproteobacteria; Enterobacterales; Enterobacteriaceae; Escherichia
MNNEETFYQAMRRQGVTRRSFLKYCSLAATSLGLGAGMAPKIAWALENKPRIPVVWIHGL WP_001058323.1
MULTISPECIES: hydrogenase-1 small chain [Proteobacteria] Length: 372\nNumber of predicted TMHs: 2\nExp number of AAs in TMHs: 37.2608499999999\nExp number, first 60 AAs: 19.52923\nTotal prob of N-in: 0.94351\nPOSSIBLE N-term signal sequence\ninside 1 19\nTMhelix 20 39\noutside 40 325\nTMhelix 326 348\ninside 349 372

8824 GCF_001748995.1_ASM174899v1 Escherichia coli Proteobacteria; Gammaproteobacteria; Enterobacterales; Enterobacteriaceae; Escherichia
MNNEETFYQAMRRQGVTRRSFLKYCSLAATSLGLGAGMAPKIAWALENKPRIPVVWIHGL WP_001058323.1
MULTISPECIES: hydrogenase-1 small chain [Proteobacteria] Length: 372\nNumber of predicted TMHs: 2\nExp number of AAs in TMHs: 37.2608499999999\nExp number, first 60 AAs: 19.52923\nTotal prob of N-in: 0.94351\nPOSSIBLE N-term signal sequence\ninside 1 19\nTMhelix 20 39\noutside 40 325\nTMhelix 326 348\ninside 349 372

8825 GCF_001749295.1_ASM174929v1 Escherichia coli Proteobacteria; Gammaproteobacteria; Enterobacterales; Enterobacteriaceae; Escherichia
MTGDNTLIHSHGINRRDFMKLCAALAATMGLSSKAAAEMAESVTNPQRPPVIWIGAQECT WP_000145410.1
MULTISPECIES: hydrogenase-2 small chain [Proteobacteria] Length: 372\nNumber of predicted TMHs: 1\nExp number of AAs in TMHs: 23.72226\nExp number, first 60 AAs: 0.22038\nTotal prob of N-in: 0.12994\noutside 1 331\nTMhelix 332 354\ninside 355 372

8826 GCF_001749235.1_ASM174923v1 Escherichia coli Proteobacteria; Gammaproteobacteria; Enterobacterales; Enterobacteriaceae; Escherichia
MTGDNTLIHSHGINRRDFMKLCAALAATMGLSSKAAAEMAESVTNPQRPPVIWIGAQECT WP_000145410.1
MULTISPECIES: hydrogenase-2 small chain [Proteobacteria] Length: 372\nNumber of predicted TMHs: 1\nExp number of AAs in TMHs: 23.72226\nExp number, first 60 AAs: 0.22038\nTotal prob of N-in: 0.12994\noutside 1 331\nTMhelix 332 354\ninside 355 372

8827 GCF_001749325.1_ASM174932v1 Escherichia coli Proteobacteria; Gammaproteobacteria; Enterobacterales; Enterobacteriaceae; Escherichia
MTGDNTLIHSHGINRRDFMKLCAALAATMGLSSKAAAEMAESVTNPQRPPVIWIGAQECT WP_000145410.1
MULTISPECIES: hydrogenase-2 small chain [Proteobacteria] Length: 372\nNumber of predicted TMHs: 1\nExp number of AAs in TMHs: 23.72226\nExp number, first 60 AAs: 0.22038\nTotal prob of N-in: 0.12994\noutside 1 331\nTMhelix 332 354\ninside 355 372

8828 GCF_001749295.1_ASM174929v1 Escherichia coli Proteobacteria; Gammaproteobacteria; Enterobacterales; Enterobacteriaceae; Escherichia
MNNEETFYQAMRRQGVTRRSFLKYCSLAATSLGLGAGMAPKIAWALENKPRIPVVWIHGL WP_001058323.1
MULTISPECIES: hydrogenase-1 small chain [Proteobacteria] Length: 372\nNumber of predicted TMHs: 2\nExp number of AAs in TMHs: 37.2608499999999\nExp number, first 60 AAs: 19.52923\nTotal prob of N-in: 0.94351\nPOSSIBLE N-term signal sequence\ninside 1 19\nTMhelix 20 39\noutside 40 325\nTMhelix 326 348\ninside 349 372

8829 GCF_001749235.1_ASM174923v1 Escherichia coli Proteobacteria; Gammaproteobacteria; Enterobacterales; Enterobacteriaceae; Escherichia
MNNEETFYQAMRRQGVTRRSFLKYCSLAATSLGLGAGMAPKIAWALENKPRIPVVWIHGL WP_001058323.1
MULTISPECIES: hydrogenase-1 small chain [Proteobacteria] Length: 372\nNumber of predicted TMHs: 2\nExp number of AAs in TMHs: 37.2608499999999\nExp number, first 60 AAs: 19.52923\nTotal prob of N-in: 0.94351\nPOSSIBLE N-term signal sequence\ninside 1 19\nTMhelix 20 39\noutside 40 325\nTMhelix 326 348\ninside 349 372

8830 GCF_001749325.1_ASM174932v1 Escherichia coli Proteobacteria; Gammaproteobacteria; Enterobacterales; Enterobacteriaceae; Escherichia
MNNEETFYQAMRRQGVTRRSFLKYCSLAATSLGLGAGMAPKIAWALENKPRIPVVWIHGL WP_001058323.1
MULTISPECIES: hydrogenase-1 small chain [Proteobacteria] Length: 372\nNumber of predicted TMHs: 2\nExp number of AAs in TMHs: 37.2608499999999\nExp number, first 60 AAs: 19.52923\nTotal prob of N-in: 0.94351\nPOSSIBLE N-term signal sequence\ninside 1 19\nTMhelix 20 39\noutside 40 325\nTMhelix 326 348\ninside 349 372

8831 GCF_001748815.1_ASM174881v1 Escherichia coli Proteobacteria; Gammaproteobacteria; Enterobacterales; Enterobacteriaceae; Escherichia
MNNEETFYQAMRRQGVTRRSFLKYCSLAATSLGLGAGMAPKIAWALENKPRIPVVWIHGL WP_069892209.1
hydrogenase [Escherichia coli] Length: 372\nNumber of predicted TMHs: 2\nExp number of AAs in TMHs: 37.2609499999999\nExp number, first 60 AAs: 19.52923\nTotal prob of N-in: 0.94351\nPOSSIBLE N-term signal sequence\ninside 1 19\nTMhelix 20 39\noutside 40 325\nTMhelix 326 348\ninside 349 372

8832 GCF_001749385.1_ASM174938v1 Escherichia coli Proteobacteria; Gammaproteobacteria; Enterobacterales; Enterobacteriaceae; Escherichia
MNNEETFYQAMRRQGVTRRSFLKYCSLAATSLGLGAGMAPKIAWALENKPRIPVVWIHGL WP_001058323.1
MULTISPECIES: hydrogenase-1 small chain [Proteobacteria] Length: 372\nNumber of predicted TMHs: 2\nExp number of AAs in TMHs: 37.2608499999999\nExp number, first 60 AAs: 19.52923\nTotal prob of N-in: 0.94351\nPOSSIBLE N-term signal sequence\ninside 1 19\nTMhelix 20 39\noutside 40 325\nTMhelix 326 348\ninside 349 372

8833 GCF_001749595.1_ASM174959v1 Escherichia coli Proteobacteria; Gammaproteobacteria; Enterobacterales; Enterobacteriaceae; Escherichia
MNNEETFYQAMRRQGVTRRSFLKYCSLAATSLGLGAGMAPKIAWALENKPRIPVVWIHGL WP_001058323.1
MULTISPECIES: hydrogenase-1 small chain [Proteobacteria] Length: 372\nNumber of predicted TMHs: 2\nExp number of AAs in TMHs: 37.2608499999999\nExp number, first 60 AAs: 19.52923\nTotal prob of N-in: 0.94351\nPOSSIBLE N-term signal sequence\ninside 1 19\nTMhelix 20 39\noutside 40 325\nTMhelix 326 348\ninside 349 372

8834 GCF_001749535.1_ASM174953v1 Escherichia coli Proteobacteria; Gammaproteobacteria; Enterobacterales; Enterobacteriaceae; Escherichia
MNNEETFYQAMRRQGVTRRSFLKYCSLAATSLGLGAGMAPKIAWALENKPRIPVVWIHGL WP_001058323.1
MULTISPECIES: hydrogenase-1 small chain [Proteobacteria] Length: 372\nNumber of predicted TMHs: 2\nExp number of AAs in TMHs: 37.2608499999999\nExp number, first 60 AAs: 19.52923\nTotal prob of N-in: 0.94351\nPOSSIBLE N-term signal sequence\ninside 1 19\nTMhelix 20 39\noutside 40 325\nTMhelix 326 348\ninside 349 372

8835 GCF_001749505.1_ASM174950v1 Escherichia coli Proteobacteria; Gammaproteobacteria; Enterobacterales; Enterobacteriaceae; Escherichia
MNNEETFYQAMRRQGVTRRSFLKYCSLAATSLGLGAGMAPKIAWALENKPRIPVVWIHGL WP_001058323.1
MULTISPECIES: hydrogenase-1 small chain [Proteobacteria] Length: 372\nNumber of predicted TMHs: 2\nExp number of AAs in TMHs: 37.2608499999999\nExp number, first 60 AAs: 19.52923\nTotal prob of N-in: 0.94351\nPOSSIBLE N-term signal sequence\ninside 1 19\nTMhelix 20 39\noutside 40 325\nTMhelix 326 348\ninside 349 372

8836 GCF_001749635.1_ASM174963v1 Escherichia coli Proteobacteria; Gammaproteobacteria; Enterobacterales; Enterobacteriaceae; Escherichia
MNNEETFYQAMRRQGVTRRSFLKYCSLAATSLGLGAGMAPKIAWALENKPRIPVVWIHGL WP_001058323.1
MULTISPECIES: hydrogenase-1 small chain [Proteobacteria] Length: 372\nNumber of predicted TMHs: 2\nExp number of AAs in TMHs: 37.2608499999999\nExp number, first 60 AAs: 19.52923\nTotal prob of N-in: 0.94351\nPOSSIBLE N-term signal sequence\ninside 1 19\nTMhelix 20 39\noutside 40 325\nTMhelix 326 348\ninside 349 372

8837 GCF_001816315.1_ASM181631v1 Escherichia coli Proteobacteria; Gammaproteobacteria; Enterobacterales; Enterobacteriaceae; Escherichia
MTGDNTLIHSHGINRRDFMKLCAALAATMGLSSKAAAEMAESVTNPQRPPVIWIGAQECT WP_000145410.1
MULTISPECIES: hydrogenase-2 small chain [Proteobacteria] Length: 372\nNumber of predicted TMHs: 1\nExp number of AAs in TMHs: 23.72226\nExp number, first 60 AAs: 0.22038\nTotal prob of N-in: 0.12994\noutside 1 331\nTMhelix 332 354\ninside 355 372

8838 GCF_001865185.1_ASM186518v1 Escherichia coli Proteobacteria; Gammaproteobacteria; Enterobacterales; Enterobacteriaceae; Escherichia
MTGDNTLIHSHGINRRDFMKLCAALAATMGLSSKAAAEMVESVTNPQRPPVIWIGAQECT WP_000145423.1
hydrogenase 2 small subunit [Escherichia coli] Length: 372\nNumber of predicted TMHs: 1\nExp number of AAs in TMHs: 23.78201\nExp number, first 60 AAs: 0.28355\nTotal prob of N-in: 0.13257\noutside 1 331\nTMhelix 332 354\ninside 355 372

8839 GCF_001758245.1_ASM175824v1 Escherichia coli Proteobacteria; Gammaproteobacteria; Enterobacterales; Enterobacteriaceae; Escherichia
MNNEETFYQAMRRQGVTRRSFLKYCSLAATSLGLGAGMAPKIAWALENKPRIPVVWIHGL WP_001058320.1
hydrogenase [Escherichia coli] Length: 372\nNumber of predicted TMHs: 2\nExp number of AAs in TMHs: 37.26082\nExp number, first 60 AAs: 19.52923\nTotal prob of N-in: 0.94351\nPOSSIBLE N-term signal sequence\ninside 1 19\nTMhelix 20 39\noutside 40 325\nTMhelix 326 348\ninside 349 372

8840 GCF_001865195.1_ASM186519v1 Escherichia coli Proteobacteria; Gammaproteobacteria; Enterobacterales; Enterobacteriaceae; Escherichia
MTGDNTLIHSHGINRRDFMKLCAALAATMGLSSKAAAEMAESVTNPQRPPVIWIGAQECT WP_000145410.1
MULTISPECIES: hydrogenase-2 small chain [Proteobacteria] Length: 372\nNumber of predicted TMHs: 1\nExp number of AAs in TMHs: 23.72226\nExp number, first 60 AAs: 0.22038\nTotal prob of N-in: 0.12994\noutside 1 331\nTMhelix 332 354\ninside 355 372

8841 GCF_001865725.1_ASM186572v1 Escherichia coli Proteobacteria; Gammaproteobacteria; Enterobacterales; Enterobacteriaceae; Escherichia
MTGDNTLIHSHGINRRDFMKLCAALAATMGLSSKAAAEMAESVTNPQRPPVIWIGAQECT WP_000145410.1
MULTISPECIES: hydrogenase-2 small chain [Proteobacteria] Length: 372\nNumber of predicted TMHs: 1\nExp number of AAs in TMHs: 23.72226\nExp number, first 60 AAs: 0.22038\nTotal prob of N-in: 0.12994\noutside 1 331\nTMhelix 332 354\ninside 355 372

8842 GCF_001865975.1_ASM186597v1 Escherichia coli Proteobacteria; Gammaproteobacteria; Enterobacterales; Enterobacteriaceae; Escherichia

MTGDNTLIHSHGINRRDFMKLCAALAATMGLSSKAAAEMAESVTNPQRPPVIWIGAQECT WP_000145410.1 MULTISPECIES: hydrogenase-2 small chain [Proteobacteria] Length: 372\nNumber of predicted TMHs: 1\nExp number of AAs in TMHs: 23.72226\nExp number, first 60 AAs: 0.22038\nTotal prob of N-in: 0.12994\noutside 1 331\nTMhelix 332 354\ninside 355 372

8843 GCF_001867205.1_ASM186720v1 Escherichia coli Proteobacteria; Gammaproteobacteria; Enterobacterales; Enterobacteriaceae; Escherichia MTGDNTLIHSHGINRRDFMKLCAALAATMGLSSKAAAEMAESVTNPQRPPVIWIGAQECT WP_000145410.1 MULTISPECIES: hydrogenase-2 small chain [Proteobacteria] Length: 372\nNumber of predicted TMHs: 1\nExp number of AAs in TMHs: 23.72226\nExp number, first 60 AAs: 0.22038\nTotal prob of N-in: 0.12994\noutside 1 331\nTMhelix 332 354\ninside 355 372

8844 GCF_001881225.1_ASM188122v1 Escherichia coli Proteobacteria; Gammaproteobacteria; Enterobacterales; Enterobacteriaceae; Escherichia MTGDNTLIHSHGINRRDFMKLCAALAATMGLSSKAAAEMAESVTNPQRPPVIWIGAQECT WP_000145410.1 MULTISPECIES: hydrogenase-2 small chain [Proteobacteria] Length: 372\nNumber of predicted TMHs: 1\nExp number of AAs in TMHs: 23.72226\nExp number, first 60 AAs: 0.22038\nTotal prob of N-in: 0.12994\noutside 1 331\nTMhelix 332 354\ninside 355 372

8845 GCF_001865905.1_ASM186590v1 Escherichia coli Proteobacteria; Gammaproteobacteria; Enterobacterales; Enterobacteriaceae; Escherichia MNNEETFYQAMRRQGVTRRSFLKYCSLAATSLGLGAGMAPKIAWALENKPRIPVVWIHGL WP_001058323.1 MULTISPECIES: hydrogenase-1 small chain [Proteobacteria] Length: 372\nNumber of predicted TMHs: 2\nExp number of AAs in TMHs: 37.2608499999999\nExp number, first 60 AAs: 19.52923\nTotal prob of N-in: 0.94351\nPOSSIBLE N-term signal sequence\ninside 1 19\nTMhelix 20 39\noutside 40 325\nTMhelix 326 348\ninside 349 372

8846 GCF_001877805.1_ASM187780v1 Escherichia coli Proteobacteria; Gammaproteobacteria; Enterobacterales; Enterobacteriaceae; Escherichia MTGDNTLIHSHGINRRDFMKLCAALAATMGLSSKAAAEMAESVTNPQRPPVIWIGAQECT WP_000145410.1 MULTISPECIES: hydrogenase-2 small chain [Proteobacteria] Length: 372\nNumber of predicted TMHs: 1\nExp number of AAs in TMHs: 23.72226\nExp number, first 60 AAs: 0.22038\nTotal prob of N-in: 0.12994\noutside 1 331\nTMhelix 332 354\ninside 355 372

8847 GCF_001881075.1_ASM188107v1 Escherichia coli Proteobacteria; Gammaproteobacteria; Enterobacterales; Enterobacteriaceae; Escherichia MTGDNTLIHSHGINRRDFMKLCAALAATMGLSSKAAAEMAESVTNPQRPPVIWIGAQECT WP_000145410.1 MULTISPECIES: hydrogenase-2 small chain [Proteobacteria] Length: 372\nNumber of predicted TMHs: 1\nExp number of AAs in TMHs: 23.72226\nExp number, first 60 AAs: 0.22038\nTotal prob of N-in: 0.12994\noutside 1 331\nTMhelix 332 354\ninside 355 372

8848 GCF_001881125.1_ASM188112v1 Escherichia coli Proteobacteria; Gammaproteobacteria; Enterobacterales; Enterobacteriaceae; Escherichia MTGDNTLIHSHGINRRDFMKLCAALAATMGLSSKAAAEMAESVTNPQRPPVIWIGAQECT WP_000145410.1 MULTISPECIES: hydrogenase-2 small chain [Proteobacteria] Length: 372\nNumber of predicted TMHs: 1\nExp number of AAs in TMHs: 23.72226\nExp number, first 60 AAs: 0.22038\nTotal prob of N-in: 0.12994\noutside 1 331\nTMhelix 332 354\ninside 355 372

8849 GCF_001881165.1_ASM188116v1 Escherichia coli Proteobacteria; Gammaproteobacteria; Enterobacterales; Enterobacteriaceae; Escherichia MTGDNTLIHSHGINRRDFMKLCAALAATMGLSSKAAAEMAESVTNPQRPPVIWIGAQECT WP_000145410.1 MULTISPECIES: hydrogenase-2 small chain [Proteobacteria] Length: 372\nNumber of predicted TMHs: 1\nExp number of AAs in TMHs: 23.72226\nExp number, first 60 AAs: 0.22038\nTotal prob of N-in: 0.12994\noutside 1 331\nTMhelix 332 354\ninside 355 372

8850 GCF_001867205.1_ASM186720v1 Escherichia coli Proteobacteria; Gammaproteobacteria; Enterobacterales; Enterobacteriaceae; Escherichia MNNEETFYQAMRRQGVTRRSFLKYCSLAATSLGLGAGMAPKIAWALENKPRIPVVWIHGL WP_063501981.1 hydrogenase [Escherichia coli] Length: 372\nNumber of predicted TMHs: 2\nExp number of AAs in TMHs: 37.06322\nExp number, first 60 AAs: 19.52932\nTotal prob of N-in: 0.94340\nPOSSIBLE N-term signal sequence\ninside 1 19\nTMhelix 20 39\noutside 40 325\nTMhelix 326 348\ninside 349 372

8851 GCF_001881595.3_ASM188159v3 Escherichia coli Proteobacteria; Gammaproteobacteria; Enterobacterales; Enterobacteriaceae; Escherichia
MNNEETFYQAMRRQGVTRRSFLKYCSLAATSLGLGAGMAPKIAWALENKPRIPVVWIHGL WP_001058323.1
MULTISPECIES: hydrogenase-1 small chain [Proteobacteria] Length: 372\nNumber of predicted TMHs: 2\nExp number of AAs in TMHs: 37.2608499999999\nExp number, first 60 AAs: 19.52923\nTotal prob of N-in: 0.94351\nPOSSIBLE N-term signal sequence\ninside 1 19\nTMhelix 20 39\noutside 40 325\nTMhelix 326 348\ninside 349 372

8852 GCF_001884785.1_ASM188478v1 Escherichia coli Proteobacteria; Gammaproteobacteria; Enterobacterales; Enterobacteriaceae; Escherichia
MTGDNTLIHSHGINRRDFMKLCAALAATMGLSSKAAAEMAESVTNPQRPPVIWIGAQECT WP_000145410.1
MULTISPECIES: hydrogenase-2 small chain [Proteobacteria] Length: 372\nNumber of predicted TMHs: 1\nExp number of AAs in TMHs: 23.72226\nExp number, first 60 AAs: 0.22038\nTotal prob of N-in: 0.12994\noutside 1 331\nTMhelix 332 354\ninside 355 372

8853 GCF_001881505.1_ASM188150v1 Escherichia coli Proteobacteria; Gammaproteobacteria; Enterobacterales; Enterobacteriaceae; Escherichia
MNNEETFYQAMRRQGVTRRSFLKYCSLAATSLGLGAGMAPKIAWALENKPRIPVVWIHGL WP_001295940.1
MULTISPECIES: hydrogenase [Enterobacteriaceae] Length: 372\nNumber of predicted TMHs: 2\nExp number of AAs in TMHs: 37.2608899999999\nExp number, first 60 AAs: 19.52923\nTotal prob of N-in: 0.94351\nPOSSIBLE N-term signal sequence\ninside 1 19\nTMhelix 20 39\noutside 40 325\nTMhelix 326 348\ninside 349 372

8854 GCF_001881645.1_ASM188164v1 Escherichia coli Proteobacteria; Gammaproteobacteria; Enterobacterales; Enterobacteriaceae; Escherichia
MNNEETFYQAMRRQGVTRRSFLKYCSLAATSLGLGAGMAPKIAWALENKPRIPVVWIHGL WP_001295940.1
MULTISPECIES: hydrogenase [Enterobacteriaceae] Length: 372\nNumber of predicted TMHs: 2\nExp number of AAs in TMHs: 37.2608899999999\nExp number, first 60 AAs: 19.52923\nTotal prob of N-in: 0.94351\nPOSSIBLE N-term signal sequence\ninside 1 19\nTMhelix 20 39\noutside 40 325\nTMhelix 326 348\ninside 349 372

8855 GCF_001885555.1_ASM188555v1 Escherichia coli Proteobacteria; Gammaproteobacteria; Enterobacterales; Enterobacteriaceae; Escherichia
MTGDNTLIHSHGINRRDFMKLCAALAATMGLSSKAAAEMAESVTNPQRPPVIWIGAQECT WP_000145410.1
MULTISPECIES: hydrogenase-2 small chain [Proteobacteria] Length: 372\nNumber of predicted TMHs: 1\nExp number of AAs in TMHs: 23.72226\nExp number, first 60 AAs: 0.22038\nTotal prob of N-in: 0.12994\noutside 1 331\nTMhelix 332 354\ninside 355 372

8856 GCF_001885555.1_ASM188555v1 Escherichia coli Proteobacteria; Gammaproteobacteria; Enterobacterales; Enterobacteriaceae; Escherichia
MNNEETFYQAMRRQGVTRRSFLKYCSLAATSLGLGAGMAPKIAWALENKPRIPVVWIHGL WP_001058320.1
hydrogenase [Escherichia coli] Length: 372\nNumber of predicted TMHs: 2\nExp number of AAs in TMHs: 37.26082\nExp number, first 60 AAs: 19.52923\nTotal prob of N-in: 0.94351\nPOSSIBLE N-term signal sequence\ninside 1 19\nTMhelix 20 39\noutside 40 325\nTMhelix 326 348\ninside 349 372

8857 GCF_001881605.1_ASM188160v1 Escherichia coli Proteobacteria; Gammaproteobacteria; Enterobacterales; Enterobacteriaceae; Escherichia
MNNEETFYQAMRRQGVTRRSFLKYCSLAATSLGLGAGMAPKIAWALENKPRIPVVWIHGL WP_001295940.1
MULTISPECIES: hydrogenase [Enterobacteriaceae] Length: 372\nNumber of predicted TMHs: 2\nExp number of AAs in TMHs: 37.2608899999999\nExp number, first 60 AAs: 19.52923\nTotal prob of N-in: 0.94351\nPOSSIBLE N-term signal sequence\ninside 1 19\nTMhelix 20 39\noutside 40 325\nTMhelix 326 348\ninside 349 372

8858 GCF_001884785.1_ASM188478v1 Escherichia coli Proteobacteria; Gammaproteobacteria; Enterobacterales; Enterobacteriaceae; Escherichia
MNNEETFYQAMRRQGVTRRSFLKYCSLAATSLGLGAGMAPKIAWALENKPRIPVVWIHGL WP_001359719.1
hydrogenase [Escherichia coli] Length: 372\nNumber of predicted TMHs: 2\nExp number of AAs in TMHs: 37.2176\nExp number, first 60 AAs: 19.53448\nTotal prob of N-in: 0.94360\nPOSSIBLE N-term signal sequence\ninside 1 19\nTMhelix 20 39\noutside 40 325\nTMhelix 326 348\ninside 349 372

8859 GCF_001886575.1_ASM188657v1 Escherichia coli Proteobacteria; Gammaproteobacteria; Enterobacterales; Enterobacteriaceae; Escherichia
MNNEETFYQAMRRQGVTRRSFLKYCSLAATSLGLGAGMAPKIAWALENKPRIPVVWIHGL WP_001058323.1
MULTISPECIES: hydrogenase-1 small chain [Proteobacteria] Length: 372\nNumber of predicted TMHs: 2\nExp number of AAs in TMHs: 37.2608499999999\nExp number, first 60 AAs: 19.52923\nTotal prob of N-in: 0.94351\nPOSSIBLE N-term signal sequence\ninside 1 19\nTMhelix 20 39\noutside 40 325\nTMhelix 326 348\ninside 349 372

8860 GCF_001886575.1_ASM188657v1 Escherichia coli Proteobacteria; Gammaproteobacteria; Enterobacterales; Enterobacteriaceae; Escherichia
MTGDNTLIHSHGINRRDFMKLCAALATTMGLSSKAAAEMAESVTNPQRPPVIWIGAQECT WP_023149629.1
MULTISPECIES: hydrogenase-2 small chain [Enterobacteriaceae] Length: 372\nNumber of predicted TMHs: 1\nExp number of AAs in TMHs: 23.63603\nExp number, first 60 AAs: 0.12971\nTotal prob of N-in: 0.12608\noutside 1 331\nTMhelix 332 354\ninside 355 372

8861 GCF_001884945.1_ASM188494v1 Escherichia coli Proteobacteria; Gammaproteobacteria; Enterobacterales; Enterobacteriaceae; Escherichia
MTGDNTLIHSHGINRRDFMKLCAALAATMGLSSKAAAEMAESVTNPQRPPVIWIGAQECS WP_061069243.1
hydrogenase 2 small subunit [Escherichia coli] Length: 372\nNumber of predicted TMHs: 1\nExp number of AAs in TMHs: 23.72219\nExp number, first 60 AAs: 0.22034\nTotal prob of N-in: 0.12994\noutside 1 331\nTMhelix 332 354\ninside 355 372

8862 GCF_001890365.1_ASM189036v1 Escherichia coli Proteobacteria; Gammaproteobacteria; Enterobacterales; Enterobacteriaceae; Escherichia
MTGDNTLIHSHGINRRDFMKLCAALAATMGLSSKAAAEMAESVTNPQRPPVIWIGAQECT WP_000145410.1
MULTISPECIES: hydrogenase-2 small chain [Proteobacteria] Length: 372\nNumber of predicted TMHs: 1\nExp number of AAs in TMHs: 23.72226\nExp number, first 60 AAs: 0.22038\nTotal prob of N-in: 0.12994\noutside 1 331\nTMhelix 332 354\ninside 355 372

8863 GCF_001884975.1_ASM188497v1 Escherichia coli Proteobacteria; Gammaproteobacteria; Enterobacterales; Enterobacteriaceae; Escherichia
MNNEETFYQAMRRQGVTRRSFLKYCSLAATSLGLGAGMAPKIAWALENKPRIPVVWIHGL WP_001301613.1
hydrogenase [Escherichia coli] Length: 372\nNumber of predicted TMHs: 2\nExp number of AAs in TMHs: 37.46598\nExp number, first 60 AAs: 19.53139\nTotal prob of N-in: 0.94364\nPOSSIBLE N-term signal sequence\ninside 1 19\nTMhelix 20 39\noutside 40 325\nTMhelix 326 348\ninside 349 372

8864 GCF_001884975.1_ASM188497v1 Escherichia coli Proteobacteria; Gammaproteobacteria; Enterobacterales; Enterobacteriaceae; Escherichia
MTGDNTLIHSHGINRRDFMKLCAALAATMGLSSKAAAEMAESVTNPQRPPVIWIGAQECS WP_061069243.1
hydrogenase 2 small subunit [Escherichia coli] Length: 372\nNumber of predicted TMHs: 1\nExp number of AAs in TMHs: 23.72219\nExp number, first 60 AAs: 0.22034\nTotal prob of N-in: 0.12994\noutside 1 331\nTMhelix 332 354\ninside 355 372

8865 GCF_001891385.1_ASM189138v1 Escherichia coli Proteobacteria; Gammaproteobacteria; Enterobacterales; Enterobacteriaceae; Escherichia
MNNEETFYQAMRRQGVTRRSFLKYCSLAATSLGLGAGMAPKIAWALENKPRIPVVWIHGL WP_001058323.1
MULTISPECIES: hydrogenase-1 small chain [Proteobacteria] Length: 372\nNumber of predicted TMHs: 2\nExp number of AAs in TMHs: 37.2608499999999\nExp number, first 60 AAs: 19.52923\nTotal prob of N-in: 0.94351\nPOSSIBLE N-term signal sequence\ninside 1 19\nTMhelix 20 39\noutside 40 325\nTMhelix 326 348\ninside 349 372

8866 GCF_001890225.1_ASM189022v1 Escherichia coli Proteobacteria; Gammaproteobacteria; Enterobacterales; Enterobacteriaceae; Escherichia
MNNEETFYQAMRRQGVTRRSFLKYCSLAATSLGLGAGMAPKIAWALENKPRIPVVWIHGL WP_001301613.1
hydrogenase [Escherichia coli] Length: 372\nNumber of predicted TMHs: 2\nExp number of AAs in TMHs: 37.46598\nExp number, first 60 AAs: 19.53139\nTotal prob of N-in: 0.94364\nPOSSIBLE N-term signal sequence\ninside 1 19\nTMhelix 20 39\noutside 40 325\nTMhelix 326 348\ninside 349 372

8867 GCF_001890325.1_ASM189032v1 Escherichia coli Proteobacteria; Gammaproteobacteria; Enterobacterales; Enterobacteriaceae; Escherichia
MNNEETFYQAMRRQGVTRRSFLKYCSLAATSLGLGAGMAPKIAWALENKPRIPVVWIHGL WP_001301613.1

hydrogenase [Escherichia coli] Length: 372\nNumber of predicted TMHs: 2\nExp number of AAs in TMHs: 37.46598\nExp number, first 60 AAs: 19.53139\nTotal prob of N-in: 0.94364\nPOSSIBLE N-term signal sequence\ninside 1 19\nTMhelix 20 39\noutside 40 325\nTMhelix 326 348\ninside 349 372

8868 GCF_001890265.1_ASM189026v1 Escherichia coli Proteobacteria; Gammaproteobacteria; Enterobacterales; Enterobacteriaceae; Escherichia MNNEETFYQAMRRQGVTRRSFLKYCSLAATSLGLGAGMAPKIAWALENKPRIPVVWIHGL WP_001301613.1 hydrogenase [Escherichia coli] Length: 372\nNumber of predicted TMHs: 2\nExp number of AAs in TMHs: 37.46598\nExp number, first 60 AAs: 19.53139\nTotal prob of N-in: 0.94364\nPOSSIBLE N-term signal sequence\ninside 1 19\nTMhelix 20 39\noutside 40 325\nTMhelix 326 348\ninside 349 372

8869 GCF_001891455.1_ASM189145v1 Escherichia coli Proteobacteria; Gammaproteobacteria; Enterobacterales; Enterobacteriaceae; Escherichia MTGDNTLIHSHGINRRDFMKLCAALAATMGLSSKAAAEMAESVTNPQRPPVIWIGAQECT WP_000145410.1 MULTISPECIES: hydrogenase-2 small chain [Proteobacteria] Length: 372\nNumber of predicted TMHs: 1\nExp number of AAs in TMHs: 23.72226\nExp number, first 60 AAs: 0.22038\nTotal prob of N-in: 0.12994\noutside 1 331\nTMhelix 332 354\ninside 355 372

8870 GCF_001891465.1_ASM189146v1 Escherichia coli Proteobacteria; Gammaproteobacteria; Enterobacterales; Enterobacteriaceae; Escherichia MTGDNTLIHSHGINRRDFMKLCAALAATMGLSSKAAAEMAESVTNPQRPPVIWIGAQECT WP_000145410.1 MULTISPECIES: hydrogenase-2 small chain [Proteobacteria] Length: 372\nNumber of predicted TMHs: 1\nExp number of AAs in TMHs: 23.72226\nExp number, first 60 AAs: 0.22038\nTotal prob of N-in: 0.12994\noutside 1 331\nTMhelix 332 354\ninside 355 372

8871 GCF_001891535.1_ASM189153v1 Escherichia coli Proteobacteria; Gammaproteobacteria; Enterobacterales; Enterobacteriaceae; Escherichia MTGDNTLIHSHGINRRDFMKLCAALAATMGLSSKAAAEMAESVTNPQRPPVIWIGAQECT WP_000145410.1 MULTISPECIES: hydrogenase-2 small chain [Proteobacteria] Length: 372\nNumber of predicted TMHs: 1\nExp number of AAs in TMHs: 23.72226\nExp number, first 60 AAs: 0.22038\nTotal prob of N-in: 0.12994\noutside 1 331\nTMhelix 332 354\ninside 355 372

8872 GCF_001891705.1_ASM189170v1 Escherichia coli Proteobacteria; Gammaproteobacteria; Enterobacterales; Enterobacteriaceae; Escherichia MTGDNTLIHSHGINRRDFMKLCAALAATMGLSSKAAAEMAESVTNPQRPPVIWIGAQECT WP_000145410.1 MULTISPECIES: hydrogenase-2 small chain [Proteobacteria] Length: 372\nNumber of predicted TMHs: 1\nExp number of AAs in TMHs: 23.72226\nExp number, first 60 AAs: 0.22038\nTotal prob of N-in: 0.12994\noutside 1 331\nTMhelix 332 354\ninside 355 372

8873 GCF_001891785.1_ASM189178v1 Escherichia coli Proteobacteria; Gammaproteobacteria; Enterobacterales; Enterobacteriaceae; Escherichia MTGDNTLIHSHGINRRDFMKLCAALAATMGLSSKAAAEMAESVTNPQRPPVIWIGAQECT WP_000145410.1 MULTISPECIES: hydrogenase-2 small chain [Proteobacteria] Length: 372\nNumber of predicted TMHs: 1\nExp number of AAs in TMHs: 23.72226\nExp number, first 60 AAs: 0.22038\nTotal prob of N-in: 0.12994\noutside 1 331\nTMhelix 332 354\ninside 355 372

8874 GCF_001891615.1_ASM189161v1 Escherichia coli Proteobacteria; Gammaproteobacteria; Enterobacterales; Enterobacteriaceae; Escherichia MNNEETFYQAMRRQGVTRRSFLKYCSLAATSLGLGAGMAPKIAWALENKPRIPVVWIHGL WP_001058323.1 MULTISPECIES: hydrogenase-1 small chain [Proteobacteria] Length: 372\nNumber of predicted TMHs: 2\nExp number of AAs in TMHs: 37.2608499999999\nExp number, first 60 AAs: 19.52923\nTotal prob of N-in: 0.94351\nPOSSIBLE N-term signal sequence\ninside 1 19\nTMhelix 20 39\noutside 40 325\nTMhelix 326 348\ninside 349 372

8875 GCF_001891705.1_ASM189170v1 Escherichia coli Proteobacteria; Gammaproteobacteria; Enterobacterales; Enterobacteriaceae; Escherichia MNNEETFYQAMRRQGVTRRSFLKYCSLAATSLGLGAGMAPKIAWALENKPRIPVVWIHGL WP_001058323.1 MULTISPECIES: hydrogenase-1 small chain [Proteobacteria] Length: 372\nNumber of predicted TMHs: 2\nExp number of AAs in TMHs: 37.2608499999999\nExp number, first 60 AAs: 19.52923\nTotal prob of N-in: 0.94351\nPOSSIBLE N-term signal sequence\ninside 1 19\nTMhelix 20 39\noutside 40 325\nTMhelix 326 348\ninside 349 372

8876	GCF_001891455.1_ASM189145v1	Escherichia coli	Proteobacteria; Gammaproteobacteria; Enterobacterales; Enterobacteriaceae; Escherichia
MNNEETFYQAMRRQGVTRRSFLKYCSLAATSLGLGAGMAPKIAWALENKPRIPVVWIHGL	WP_001295940.1
MULTISPECIES: hydrogenase [Enterobacteriaceae]	Length: 372\nNumber of predicted TMHs: 2\nExp number of AAs in TMHs: 37.2608899999999\nExp number, first 60 AAs: 19.52923\nTotal prob of N-in: 0.94351\nPOSSIBLE N-term signal sequence\ninside 1 19\nTMhelix 20 39\noutside 40 325\nTMhelix 326 348\ninside 349 372

8877	GCF_001891805.1_ASM189180v1	Escherichia coli	Proteobacteria; Gammaproteobacteria; Enterobacterales; Enterobacteriaceae; Escherichia
MTGDNTLIHSHGINRRDFMKLCAALAATMGLSSKAAAEMAESVTNPQRPPVIWIGAQECT	WP_000145410.1
MULTISPECIES: hydrogenase-2 small chain [Proteobacteria]	Length: 372\nNumber of predicted TMHs: 1\nExp number of AAs in TMHs: 23.72226\nExp number, first 60 AAs: 0.22038\nTotal prob of N-in: 0.12994\noutside 1 331\nTMhelix 332 354\ninside 355 372

8878	GCF_001891805.1_ASM189180v1	Escherichia coli	Proteobacteria; Gammaproteobacteria; Enterobacterales; Enterobacteriaceae; Escherichia
MNNEETFYQAMRRQGVTRRSFLKYCSLAATSLGLGAGMAPKIAWALENKPRIPVVWIHGL	WP_001058323.1
MULTISPECIES: hydrogenase-1 small chain [Proteobacteria]	Length: 372\nNumber of predicted TMHs: 2\nExp number of AAs in TMHs: 37.2608499999999\nExp number, first 60 AAs: 19.52923\nTotal prob of N-in: 0.94351\nPOSSIBLE N-term signal sequence\ninside 1 19\nTMhelix 20 39\noutside 40 325\nTMhelix 326 348\ninside 349 372

8879	GCF_001891975.1_ASM189197v1	Escherichia coli	Proteobacteria; Gammaproteobacteria; Enterobacterales; Enterobacteriaceae; Escherichia
MNNEETFYQAMRRQGVTRRSFLKYCSLAATSLGLGAGMAPKIAWALENKPRIPVVWIHGL	WP_001295940.1
MULTISPECIES: hydrogenase [Enterobacteriaceae]	Length: 372\nNumber of predicted TMHs: 2\nExp number of AAs in TMHs: 37.2608899999999\nExp number, first 60 AAs: 19.52923\nTotal prob of N-in: 0.94351\nPOSSIBLE N-term signal sequence\ninside 1 19\nTMhelix 20 39\noutside 40 325\nTMhelix 326 348\ninside 349 372

8880	GCF_001891685.1_ASM189168v1	Escherichia coli	Proteobacteria; Gammaproteobacteria; Enterobacterales; Enterobacteriaceae; Escherichia
MNNEETFYQAMRRQGVTRRSFLKYCSLAATSLGLGAGMAPKIAWALENKPRIPVVWIHGL	WP_001295940.1
MULTISPECIES: hydrogenase [Enterobacteriaceae]	Length: 372\nNumber of predicted TMHs: 2\nExp number of AAs in TMHs: 37.2608899999999\nExp number, first 60 AAs: 19.52923\nTotal prob of N-in: 0.94351\nPOSSIBLE N-term signal sequence\ninside 1 19\nTMhelix 20 39\noutside 40 325\nTMhelix 326 348\ninside 349 372

8881	GCF_001892025.1_ASM189202v1	Escherichia coli	Proteobacteria; Gammaproteobacteria; Enterobacterales; Enterobacteriaceae; Escherichia
MTGDNTLIHSHGINRRDFMKLCAALAATMGLSSKAAAEMAESVTNPQRPPVIWIGAQECT	WP_000145410.1
MULTISPECIES: hydrogenase-2 small chain [Proteobacteria]	Length: 372\nNumber of predicted TMHs: 1\nExp number of AAs in TMHs: 23.72226\nExp number, first 60 AAs: 0.22038\nTotal prob of N-in: 0.12994\noutside 1 331\nTMhelix 332 354\ninside 355 372

8882	GCF_001892055.1_ASM189205v1	Escherichia coli	Proteobacteria; Gammaproteobacteria; Enterobacterales; Enterobacteriaceae; Escherichia
MTGDNTLIHSHGINRRDFMKLCAALAATMGLSSKAAAEMAESVTNPQRPPVIWIGAQECT	WP_000145410.1
MULTISPECIES: hydrogenase-2 small chain [Proteobacteria]	Length: 372\nNumber of predicted TMHs: 1\nExp number of AAs in TMHs: 23.72226\nExp number, first 60 AAs: 0.22038\nTotal prob of N-in: 0.12994\noutside 1 331\nTMhelix 332 354\ninside 355 372

8883	GCF_001892125.1_ASM189212v1	Escherichia coli	Proteobacteria; Gammaproteobacteria; Enterobacterales; Enterobacteriaceae; Escherichia
MTGDNTLIHSHGINRRDFMKLCAALAATMGLSSKAAAEMAESVTNPQRPPVIWIGAQECT	WP_000145410.1
MULTISPECIES: hydrogenase-2 small chain [Proteobacteria]	Length: 372\nNumber of predicted TMHs: 1\nExp number of AAs in TMHs: 23.72226\nExp number, first 60 AAs: 0.22038\nTotal prob of N-in: 0.12994\noutside 1 331\nTMhelix 332 354\ninside 355 372

8884 GCF_001892145.1_ASM189214v1 Escherichia coli Proteobacteria; Gammaproteobacteria; Enterobacterales; Enterobacteriaceae; Escherichia
MTGDNTLIHSHGINRRDFMKLCAALAATMGLSSKAAAEMAESVTNPQRPPVIWIGAQECT WP_000145410.1
MULTISPECIES: hydrogenase-2 small chain [Proteobacteria] Length: 372\nNumber of predicted TMHs: 1\nExp number of AAs in TMHs: 23.72226\nExp number, first 60 AAs: 0.22038\nTotal prob of N-in: 0.12994\noutside 1 331\nTMhelix 332 354\ninside 355 372

8885 GCF_001892215.1_ASM189221v1 Escherichia coli Proteobacteria; Gammaproteobacteria; Enterobacterales; Enterobacteriaceae; Escherichia
MTGDNTLIHSHGINRRDFMKLCAALAATMGLSSKAAAEMAESVTNPQRPPVIWIGAQECT WP_000145410.1
MULTISPECIES: hydrogenase-2 small chain [Proteobacteria] Length: 372\nNumber of predicted TMHs: 1\nExp number of AAs in TMHs: 23.72226\nExp number, first 60 AAs: 0.22038\nTotal prob of N-in: 0.12994\noutside 1 331\nTMhelix 332 354\ninside 355 372

8886 GCF_001892365.1_ASM189236v1 Escherichia coli Proteobacteria; Gammaproteobacteria; Enterobacterales; Enterobacteriaceae; Escherichia
MTGDNTLIHSHGINRRDFMKLCAALAATMGLSSKAAAEMAESVTNPQRPPVIWIGAQECT WP_000145410.1
MULTISPECIES: hydrogenase-2 small chain [Proteobacteria] Length: 372\nNumber of predicted TMHs: 1\nExp number of AAs in TMHs: 23.72226\nExp number, first 60 AAs: 0.22038\nTotal prob of N-in: 0.12994\noutside 1 331\nTMhelix 332 354\ninside 355 372

8887 GCF_001892365.1_ASM189236v1 Escherichia coli Proteobacteria; Gammaproteobacteria; Enterobacterales; Enterobacteriaceae; Escherichia
MNNEETFYQAMRRQGVTRRSFLKYCSLAATSLGLGAGMAPKIAWALENKPRIPVVWIHGL WP_001058323.1
MULTISPECIES: hydrogenase-1 small chain [Proteobacteria] Length: 372\nNumber of predicted TMHs: 2\nExp number of AAs in TMHs: 37.2608499999999\nExp number, first 60 AAs: 19.52923\nTotal prob of N-in: 0.94351\nPOSSIBLE N-term signal sequence\ninside 1 19\nTMhelix 20 39\noutside 40 325\nTMhelix 326 348\ninside 349 372

8888 GCF_001892145.1_ASM189214v1 Escherichia coli Proteobacteria; Gammaproteobacteria; Enterobacterales; Enterobacteriaceae; Escherichia
MNNEETFYQAMRRQGVTRRSFLKYCSLAATSLGLGAGMAPKIAWALENKPRIPVVWIHGL WP_001295940.1
MULTISPECIES: hydrogenase [Enterobacteriaceae] Length: 372\nNumber of predicted TMHs: 2\nExp number of AAs in TMHs: 37.2608899999999\nExp number, first 60 AAs: 19.52923\nTotal prob of N-in: 0.94351\nPOSSIBLE N-term signal sequence\ninside 1 19\nTMhelix 20 39\noutside 40 325\nTMhelix 326 348\ninside 349 372

8889 GCF_001892215.1_ASM189221v1 Escherichia coli Proteobacteria; Gammaproteobacteria; Enterobacterales; Enterobacteriaceae; Escherichia
MNNEETFYQAMRRQGVTRRSFLKYCSLAATSLGLGAGMAPKIAWALENKPRIPVVWIHGL WP_001295940.1
MULTISPECIES: hydrogenase [Enterobacteriaceae] Length: 372\nNumber of predicted TMHs: 2\nExp number of AAs in TMHs: 37.2608899999999\nExp number, first 60 AAs: 19.52923\nTotal prob of N-in: 0.94351\nPOSSIBLE N-term signal sequence\ninside 1 19\nTMhelix 20 39\noutside 40 325\nTMhelix 326 348\ninside 349 372

8890 GCF_001892325.1_ASM189232v1 Escherichia coli Proteobacteria; Gammaproteobacteria; Enterobacterales; Enterobacteriaceae; Escherichia
MNNEETFYQAMRRQGVTRRSFLKYCSLAATSLGLGAGMAPKIAWALENKPRIPVVWIHGL WP_001058323.1
MULTISPECIES: hydrogenase-1 small chain [Proteobacteria] Length: 372\nNumber of predicted TMHs: 2\nExp number of AAs in TMHs: 37.2608499999999\nExp number, first 60 AAs: 19.52923\nTotal prob of N-in: 0.94351\nPOSSIBLE N-term signal sequence\ninside 1 19\nTMhelix 20 39\noutside 40 325\nTMhelix 326 348\ninside 349 372

8891 GCF_001892425.1_ASM189242v1 Escherichia coli Proteobacteria; Gammaproteobacteria; Enterobacterales; Enterobacteriaceae; Escherichia
MTGDNTLIHSHGINRRDFMKLCAALAATMGLSSKAAAEMAESVTNPQRPPVIWIGAQECT WP_000145410.1
MULTISPECIES: hydrogenase-2 small chain [Proteobacteria] Length: 372\nNumber of predicted TMHs: 1\nExp number of AAs in TMHs: 23.72226\nExp number, first 60 AAs: 0.22038\nTotal prob of N-in: 0.12994\noutside 1 331\nTMhelix 332 354\ninside 355 372

8892	GCF_001892515.1_ASM189251v1	Escherichia coli	Proteobacteria; Gammaproteobacteria; Enterobacterales; Enterobacteriaceae; Escherichia
	MTGDNTLIHSHGINRRDFMKLCAALAATMGLSSKAAAEMAESVTNPQRPPVIWIGAQECT	WP_000145410.1
	MULTISPECIES: hydrogenase-2 small chain [Proteobacteria]	Length: 372\nNumber of predicted TMHs: 1\nExp number of AAs in TMHs: 23.72226\nExp number, first 60 AAs: 0.22038\nTotal prob of N-in: 0.12994\noutside 1 331\nTMhelix 332 354\ninside 355 372

8893	GCF_001892755.1_ASM189275v1	Escherichia coli	Proteobacteria; Gammaproteobacteria; Enterobacterales; Enterobacteriaceae; Escherichia
	MNNEETFYQAMRRQGVTRRSFLKYCSLAATSLGLGAGMAPKIAWALENKPRIPVVWIHGL	WP_001295940.1
	MULTISPECIES: hydrogenase [Enterobacteriaceae]	Length: 372\nNumber of predicted TMHs: 2\nExp number of AAs in TMHs: 37.2608899999999\nExp number, first 60 AAs: 19.52923\nTotal prob of N-in: 0.94351\nPOSSIBLE N-term signal sequence\ninside 1 19\nTMhelix 20 39\noutside 40 325\nTMhelix 326 348\ninside 349 372

8894	GCF_001892725.1_ASM189272v1	Escherichia coli	Proteobacteria; Gammaproteobacteria; Enterobacterales; Enterobacteriaceae; Escherichia
	MNNEETFYQAMRRQGVTRRSFLKYCSLAATSLGLGAGMAPKIAWALENKPRIPVVWIHGL	WP_001295940.1
	MULTISPECIES: hydrogenase [Enterobacteriaceae]	Length: 372\nNumber of predicted TMHs: 2\nExp number of AAs in TMHs: 37.2608899999999\nExp number, first 60 AAs: 19.52923\nTotal prob of N-in: 0.94351\nPOSSIBLE N-term signal sequence\ninside 1 19\nTMhelix 20 39\noutside 40 325\nTMhelix 326 348\ninside 349 372

8895	GCF_001892565.1_ASM189256v1	Escherichia coli	Proteobacteria; Gammaproteobacteria; Enterobacterales; Enterobacteriaceae; Escherichia
	MNNEETFYQAMRRQGVTRRSFLKYCSLAATSLGLGAGMAPKIAWALENKPRIPVVWIHGL	WP_001058323.1
	MULTISPECIES: hydrogenase-1 small chain [Proteobacteria]	Length: 372\nNumber of predicted TMHs: 2\nExp number of AAs in TMHs: 37.2608499999999\nExp number, first 60 AAs: 19.52923\nTotal prob of N-in: 0.94351\nPOSSIBLE N-term signal sequence\ninside 1 19\nTMhelix 20 39\noutside 40 325\nTMhelix 326 348\ninside 349 372

8896	GCF_001892515.1_ASM189251v1	Escherichia coli	Proteobacteria; Gammaproteobacteria; Enterobacterales; Enterobacteriaceae; Escherichia
	MNNEETFYQAMRRQGVTRRSFLKYCSLAATSLGLGAGMAPKIAWALENKPRIPVVWIHGL	WP_001058323.1
	MULTISPECIES: hydrogenase-1 small chain [Proteobacteria]	Length: 372\nNumber of predicted TMHs: 2\nExp number of AAs in TMHs: 37.2608499999999\nExp number, first 60 AAs: 19.52923\nTotal prob of N-in: 0.94351\nPOSSIBLE N-term signal sequence\ninside 1 19\nTMhelix 20 39\noutside 40 325\nTMhelix 326 348\ninside 349 372

8897	GCF_001892425.1_ASM189242v1	Escherichia coli	Proteobacteria; Gammaproteobacteria; Enterobacterales; Enterobacteriaceae; Escherichia
	MNNEETFYQAMRRQGVTRRSFLKYCSLAATSLGLGAGMAPKIAWALENKPRIPVVWIHGL	WP_001295940.1
	MULTISPECIES: hydrogenase [Enterobacteriaceae]	Length: 372\nNumber of predicted TMHs: 2\nExp number of AAs in TMHs: 37.2608899999999\nExp number, first 60 AAs: 19.52923\nTotal prob of N-in: 0.94351\nPOSSIBLE N-term signal sequence\ninside 1 19\nTMhelix 20 39\noutside 40 325\nTMhelix 326 348\ninside 349 372

8898	GCF_001892445.1_ASM189244v1	Escherichia coli	Proteobacteria; Gammaproteobacteria; Enterobacterales; Enterobacteriaceae; Escherichia
	MNNEETFYQAMRRQGVTRRSFLKYCSLAATSLGLGAGMAPKIAWALENKPRIPVVWIHGL	WP_001295940.1
	MULTISPECIES: hydrogenase [Enterobacteriaceae]	Length: 372\nNumber of predicted TMHs: 2\nExp number of AAs in TMHs: 37.2608899999999\nExp number, first 60 AAs: 19.52923\nTotal prob of N-in: 0.94351\nPOSSIBLE N-term signal sequence\ninside 1 19\nTMhelix 20 39\noutside 40 325\nTMhelix 326 348\ninside 349 372

8899	GCF_001892795.1_ASM189279v1	Escherichia coli	Proteobacteria; Gammaproteobacteria; Enterobacterales; Enterobacteriaceae; Escherichia
	MTGDNTLIHSHGINRRDFMKLCAALAATMGLSSKAAAEMAESVTNPQRPPVIWIGAQECT	WP_000145410.1
	MULTISPECIES: hydrogenase-2 small chain [Proteobacteria]	Length: 372\nNumber of predicted TMHs: 1\nExp number of AAs in TMHs: 23.72226\nExp number, first 60 AAs: 0.22038\nTotal prob of N-in: 0.12994\noutside 1 331\nTMhelix 332 354\ninside 355 372

8900 GCF_001893105.1_ASM189310v1 Escherichia coli Proteobacteria; Gammaproteobacteria; Enterobacterales; Enterobacteriaceae; Escherichia
MTGDNTLIHSHGINRRDFMKLCAALAATMGLSSKAAAEMAESVTNPQRPPVIWIGAQECT WP_000145410.1
MULTISPECIES: hydrogenase-2 small chain [Proteobacteria] Length: 372\nNumber of predicted TMHs: 1\nExp number of AAs in TMHs: 23.72226\nExp number, first 60 AAs: 0.22038\nTotal prob of N-in: 0.12994\noutside 1 331\nTMhelix 332 354\ninside 355 372

8901 GCF_001893025.1_ASM189302v1 Escherichia coli Proteobacteria; Gammaproteobacteria; Enterobacterales; Enterobacteriaceae; Escherichia
MTGDNTLIHSHGINRRDFMKLCAALAATMGLSSKAAAEMAESVTNPQRPPVIWIGAQECT WP_000145410.1
MULTISPECIES: hydrogenase-2 small chain [Proteobacteria] Length: 372\nNumber of predicted TMHs: 1\nExp number of AAs in TMHs: 23.72226\nExp number, first 60 AAs: 0.22038\nTotal prob of N-in: 0.12994\noutside 1 331\nTMhelix 332 354\ninside 355 372

8902 GCF_001893135.1_ASM189313v1 Escherichia coli Proteobacteria; Gammaproteobacteria; Enterobacterales; Enterobacteriaceae; Escherichia
MTGDNTLIHSHGINRRDFMKLCAALAATMGLSSKAAAEMAESVTNPQRPPVIWIGAQECT WP_000145410.1
MULTISPECIES: hydrogenase-2 small chain [Proteobacteria] Length: 372\nNumber of predicted TMHs: 1\nExp number of AAs in TMHs: 23.72226\nExp number, first 60 AAs: 0.22038\nTotal prob of N-in: 0.12994\noutside 1 331\nTMhelix 332 354\ninside 355 372

8903 GCF_001893135.1_ASM189313v1 Escherichia coli Proteobacteria; Gammaproteobacteria; Enterobacterales; Enterobacteriaceae; Escherichia
MNNEETFYQAMRRQGVTRRSFLKYCSLAATSLGLGAGMAPKIAWALENKPRIPVVWIHGL WP_001058323.1
MULTISPECIES: hydrogenase-1 small chain [Proteobacteria] Length: 372\nNumber of predicted TMHs: 2\nExp number of AAs in TMHs: 37.2608499999999\nExp number, first 60 AAs: 19.52923\nTotal prob of N-in: 0.94351\nPOSSIBLE N-term signal sequence\ninside 1 19\nTMhelix 20 39\noutside 40 325\nTMhelix 326 348\ninside 349 372

8904 GCF_001893165.1_ASM189316v1 Escherichia coli Proteobacteria; Gammaproteobacteria; Enterobacterales; Enterobacteriaceae; Escherichia
MTGDNTLIHSHGINRRDFMKLCAALAATMGLSSKAAAEMAESVTNPQRPPVIWIGAQECT WP_000145410.1
MULTISPECIES: hydrogenase-2 small chain [Proteobacteria] Length: 372\nNumber of predicted TMHs: 1\nExp number of AAs in TMHs: 23.72226\nExp number, first 60 AAs: 0.22038\nTotal prob of N-in: 0.12994\noutside 1 331\nTMhelix 332 354\ninside 355 372

8905 GCF_001893205.1_ASM189320v1 Escherichia coli Proteobacteria; Gammaproteobacteria; Enterobacterales; Enterobacteriaceae; Escherichia
MTGDNTLIHSHGINRRDFMKLCAALAATMGLSSKAAAEMAESVTNPQRPPVIWIGAQECT WP_000145410.1
MULTISPECIES: hydrogenase-2 small chain [Proteobacteria] Length: 372\nNumber of predicted TMHs: 1\nExp number of AAs in TMHs: 23.72226\nExp number, first 60 AAs: 0.22038\nTotal prob of N-in: 0.12994\noutside 1 331\nTMhelix 332 354\ninside 355 372

8906 GCF_001893245.1_ASM189324v1 Escherichia coli Proteobacteria; Gammaproteobacteria; Enterobacterales; Enterobacteriaceae; Escherichia
MTGDNTLIHSHGINRRDFMKLCAALAATMGLSSKAAAEMAESVTNPQRPPVIWIGAQECT WP_000145410.1
MULTISPECIES: hydrogenase-2 small chain [Proteobacteria] Length: 372\nNumber of predicted TMHs: 1\nExp number of AAs in TMHs: 23.72226\nExp number, first 60 AAs: 0.22038\nTotal prob of N-in: 0.12994\noutside 1 331\nTMhelix 332 354\ninside 355 372

8907 GCF_001893775.1_ASM189377v1 Escherichia coli Proteobacteria; Gammaproteobacteria; Enterobacterales; Enterobacteriaceae; Escherichia
MTGDNTLIHSHGINRRDFMKLCAALAATMGLSSKAAAEMAESVTNPQRPPVIWIGAQECT WP_000145410.1
MULTISPECIES: hydrogenase-2 small chain [Proteobacteria] Length: 372\nNumber of predicted TMHs: 1\nExp number of AAs in TMHs: 23.72226\nExp number, first 60 AAs: 0.22038\nTotal prob of N-in: 0.12994\noutside 1 331\nTMhelix 332 354\ninside 355 372

8908 GCF_001893345.1_ASM189334v1 Escherichia coli Proteobacteria; Gammaproteobacteria; Enterobacterales; Enterobacteriaceae; Escherichia
MNNEETFYQAMRRQGVTRRSFLKYCSLAATSLGLGAGMAPKIAWALENKPRIPVVWIHGL WP_001058323.1
MULTISPECIES: hydrogenase-1 small chain [Proteobacteria] Length: 372\nNumber of predicted TMHs:

2\nExp number of AAs in TMHs: 37.2608499999999\nExp number, first 60 AAs: 19.52923\nTotal prob of N-in: 0.94351\nPOSSIBLE N-term signal sequence\ninside 1 19\nTMhelix 20 39\noutside 40 325\nTMhelix 326 348\ninside 349 372

8909 GCF_001893105.1_ASM189310v1 Escherichia coli Proteobacteria; Gammaproteobacteria; Enterobacterales; Enterobacteriaceae; Escherichia MNNEETFYQAMRRQGVTRRSFLKYCSLAATSLGLGAGMAPKIAWALENKSRIPVVWIHGL WP_049595080.1 hydrogenase [Escherichia coli] Length: 372\nNumber of predicted TMHs: 2\nExp number of AAs in TMHs: 37.1898399999999\nExp number, first 60 AAs: 19.44607\nTotal prob of N-in: 0.94159\nPOSSIBLE N-term signal sequence\ninside 1 19\nTMhelix 20 39\noutside 40 325\nTMhelix 326 348\ninside 349 372

8910 GCF_001893385.1_ASM189338v1 Escherichia coli Proteobacteria; Gammaproteobacteria; Enterobacterales; Enterobacteriaceae; Escherichia MTGDNTLIHSHGINRRDFMKLCAALAATMGLSSKAAAEMAESVTNPQRPPVIWIGAQECT WP_000145410.1 MULTISPECIES: hydrogenase-2 small chain [Proteobacteria] Length: 372\nNumber of predicted TMHs: 1\nExp number of AAs in TMHs: 23.72226\nExp number, first 60 AAs: 0.22038\nTotal prob of N-in: 0.12994\noutside 1 331\nTMhelix 332 354\ninside 355 372

8911 GCF_001893445.1_ASM189344v1 Escherichia coli Proteobacteria; Gammaproteobacteria; Enterobacterales; Enterobacteriaceae; Escherichia MTGDNTLIHSHGINRRDFMKLCAALAATMGLSSKAAAEMAESVTNPQRPPVIWIGAQECT WP_000145410.1 MULTISPECIES: hydrogenase-2 small chain [Proteobacteria] Length: 372\nNumber of predicted TMHs: 1\nExp number of AAs in TMHs: 23.72226\nExp number, first 60 AAs: 0.22038\nTotal prob of N-in: 0.12994\noutside 1 331\nTMhelix 332 354\ninside 355 372

8912 GCF_001893475.1_ASM189347v1 Escherichia coli Proteobacteria; Gammaproteobacteria; Enterobacterales; Enterobacteriaceae; Escherichia MTGDNTLIHSHGINRRDFMKLCAALAATMGLSSKAAAEMAESVTNPQRPPVIWIGAQECT WP_000145410.1 MULTISPECIES: hydrogenase-2 small chain [Proteobacteria] Length: 372\nNumber of predicted TMHs: 1\nExp number of AAs in TMHs: 23.72226\nExp number, first 60 AAs: 0.22038\nTotal prob of N-in: 0.12994\noutside 1 331\nTMhelix 332 354\ninside 355 372

8913 GCF_001893535.1_ASM189353v1 Escherichia coli Proteobacteria; Gammaproteobacteria; Enterobacterales; Enterobacteriaceae; Escherichia MTGDNTLIHSHGINRRDFMKLCAALAATMGLSSKAAAEMAESVTNPQRPPVIWIGAQECT WP_000145410.1 MULTISPECIES: hydrogenase-2 small chain [Proteobacteria] Length: 372\nNumber of predicted TMHs: 1\nExp number of AAs in TMHs: 23.72226\nExp number, first 60 AAs: 0.22038\nTotal prob of N-in: 0.12994\noutside 1 331\nTMhelix 332 354\ninside 355 372

8914 GCF_001893385.1_ASM189338v1 Escherichia coli Proteobacteria; Gammaproteobacteria; Enterobacterales; Enterobacteriaceae; Escherichia MNNEETFYQAMRRQGVTRRSFLKYCSLAATSLGLGAGMAPKIAWALENKPRIPVVWIHGL WP_001058323.1 MULTISPECIES: hydrogenase-1 small chain [Proteobacteria] Length: 372\nNumber of predicted TMHs: 2\nExp number of AAs in TMHs: 37.2608499999999\nExp number, first 60 AAs: 19.52923\nTotal prob of N-in: 0.94351\nPOSSIBLE N-term signal sequence\ninside 1 19\nTMhelix 20 39\noutside 40 325\nTMhelix 326 348\ninside 349 372

8915 GCF_001893905.1_ASM189390v1 Escherichia coli Proteobacteria; Gammaproteobacteria; Enterobacterales; Enterobacteriaceae; Escherichia MTGDNTLIHSHGINRRDFMKLCAALAATMGLSSKAAAEMAESVTNPQRPPVIWIGAQECT WP_000145410.1 MULTISPECIES: hydrogenase-2 small chain [Proteobacteria] Length: 372\nNumber of predicted TMHs: 1\nExp number of AAs in TMHs: 23.72226\nExp number, first 60 AAs: 0.22038\nTotal prob of N-in: 0.12994\noutside 1 331\nTMhelix 332 354\ninside 355 372

8916 GCF_001893865.1_ASM189386v1 Escherichia coli Proteobacteria; Gammaproteobacteria; Enterobacterales; Enterobacteriaceae; Escherichia MTGDNTLIHSHGINRRDFMKLCAALAATMGLSSKAAAEMAESVTNPQRPPVIWIGAQECT WP_000145410.1 MULTISPECIES: hydrogenase-2 small chain [Proteobacteria] Length: 372\nNumber of predicted TMHs: 1\nExp number of AAs in TMHs: 23.72226\nExp number, first 60 AAs: 0.22038\nTotal prob of N-in: 0.12994\noutside 1 331\nTMhelix 332 354\ninside 355 372

8917	GCF_001893905.1_ASM189390v1	Escherichia coli	Proteobacteria; Gammaproteobacteria; Enterobacterales; Enterobacteriaceae; Escherichia
MNNEETFYQAMRRQGVTRRSFLKYCSLAATSLGLGAGMAPKIAWALENKPRIPVVWIHGL	WP_001058323.1
MULTISPECIES: hydrogenase-1 small chain [Proteobacteria]	Length: 372\nNumber of predicted TMHs: 2\nExp number of AAs in TMHs: 37.2608499999999\nExp number, first 60 AAs: 19.52923\nTotal prob of N-in: 0.94351\nPOSSIBLE N-term signal sequence\ninside 1 19\nTMhelix 20 39\noutside 40 325\nTMhelix 326 348\ninside 349 372

8918	GCF_001893605.1_ASM189360v1	Escherichia coli	Proteobacteria; Gammaproteobacteria; Enterobacterales; Enterobacteriaceae; Escherichia
MNNEETFYQAMRRQGVTRRSFLKYCSLAATSLGLGAGMAPKIAWALENKPRIPVVWIHGL	WP_001058323.1
MULTISPECIES: hydrogenase-1 small chain [Proteobacteria]	Length: 372\nNumber of predicted TMHs: 2\nExp number of AAs in TMHs: 37.2608499999999\nExp number, first 60 AAs: 19.52923\nTotal prob of N-in: 0.94351\nPOSSIBLE N-term signal sequence\ninside 1 19\nTMhelix 20 39\noutside 40 325\nTMhelix 326 348\ninside 349 372

8919	GCF_001893535.1_ASM189353v1	Escherichia coli	Proteobacteria; Gammaproteobacteria; Enterobacterales; Enterobacteriaceae; Escherichia
MNNEETFYQAMRRQGVTRRSFLKYCSLAATSLGLGAGMAPKIAWALENKPRIPVVWIHGL	WP_001295940.1
MULTISPECIES: hydrogenase [Enterobacteriaceae]	Length: 372\nNumber of predicted TMHs: 2\nExp number of AAs in TMHs: 37.2608899999999\nExp number, first 60 AAs: 19.52923\nTotal prob of N-in: 0.94351\nPOSSIBLE N-term signal sequence\ninside 1 19\nTMhelix 20 39\noutside 40 325\nTMhelix 326 348\ninside 349 372

8920	GCF_001893475.1_ASM189347v1	Escherichia coli	Proteobacteria; Gammaproteobacteria; Enterobacterales; Enterobacteriaceae; Escherichia
MNNEETFYQAMRRQGVTRRSFLKYCSLAATSLGLGAGMAPKIAWALENKPRIPVVWIHGL	WP_063501981.1
hydrogenase [Escherichia coli]	Length: 372\nNumber of predicted TMHs: 2\nExp number of AAs in TMHs: 37.06322\nExp number, first 60 AAs: 19.52932\nTotal prob of N-in: 0.94340\nPOSSIBLE N-term signal sequence\ninside 1 19\nTMhelix 20 39\noutside 40 325\nTMhelix 326 348\ninside 349 372

8921	GCF_001893955.1_ASM189395v1	Escherichia coli	Proteobacteria; Gammaproteobacteria; Enterobacterales; Enterobacteriaceae; Escherichia
MTGDNTLIHSHGINRRDFMKLCAALAATMGLSSKAAAEMAESVTNPQRPPVIWIGAQECT	WP_000145410.1
MULTISPECIES: hydrogenase-2 small chain [Proteobacteria]	Length: 372\nNumber of predicted TMHs: 1\nExp number of AAs in TMHs: 23.72226\nExp number, first 60 AAs: 0.22038\nTotal prob of N-in: 0.12994\noutside 1 331\nTMhelix 332 354\ninside 355 372

8922	GCF_001894065.1_ASM189406v1	Escherichia coli	Proteobacteria; Gammaproteobacteria; Enterobacterales; Enterobacteriaceae; Escherichia
MTGDNTLIHSHGINRRDFMKLCAALAATMGLSSKAAAEMAESVTNPQRPPVIWIGAQECT	WP_000145410.1
MULTISPECIES: hydrogenase-2 small chain [Proteobacteria]	Length: 372\nNumber of predicted TMHs: 1\nExp number of AAs in TMHs: 23.72226\nExp number, first 60 AAs: 0.22038\nTotal prob of N-in: 0.12994\noutside 1 331\nTMhelix 332 354\ninside 355 372

8923	GCF_001894155.1_ASM189415v1	Escherichia coli	Proteobacteria; Gammaproteobacteria; Enterobacterales; Enterobacteriaceae; Escherichia
MTGDNTLIHSHGINRRDFMKLCAALAATMGLSSKAAAEMAESVTNPQRPPVIWIGAQECT	WP_000145410.1
MULTISPECIES: hydrogenase-2 small chain [Proteobacteria]	Length: 372\nNumber of predicted TMHs: 1\nExp number of AAs in TMHs: 23.72226\nExp number, first 60 AAs: 0.22038\nTotal prob of N-in: 0.12994\noutside 1 331\nTMhelix 332 354\ninside 355 372

8924	GCF_001894195.1_ASM189419v1	Escherichia coli	Proteobacteria; Gammaproteobacteria; Enterobacterales; Enterobacteriaceae; Escherichia
MTGDNTLIHSHGINRRDFMKLCAALAATMGLSSKAAAEMAESVTNPQRPPVIWIGAQECT	WP_000145410.1
MULTISPECIES: hydrogenase-2 small chain [Proteobacteria]	Length: 372\nNumber of predicted TMHs: 1\nExp number of AAs in TMHs: 23.72226\nExp number, first 60 AAs: 0.22038\nTotal prob of N-in: 0.12994\noutside 1 331\nTMhelix 332 354\ninside 355 372

8925	GCF_001900315.1_ASM190031v1	Escherichia coli	Proteobacteria; Gammaproteobacteria; Enterobacterales; Enterobacteriaceae; Escherichia

MTGDNTLIHSHGINRRDFMKLCAALAATMGLSSKAAAEMAESVTNPQRPPVIWIGAQECT WP_000145410.1 MULTISPECIES: hydrogenase-2 small chain [Proteobacteria] Length: 372\nNumber of predicted TMHs: 1\nExp number of AAs in TMHs: 23.72226\nExp number, first 60 AAs: 0.22038\nTotal prob of N-in: 0.12994\noutside 1 331\nTMhelix 332 354\ninside 355 372

8926 GCF_001893995.1_ASM189399v1 Escherichia coli Proteobacteria; Gammaproteobacteria; Enterobacterales; Enterobacteriaceae; Escherichia MNNEETFYQAMRRQGVTRRSFLKYCSLAATSLGLGAGMAPKIAWALENKPRIPVVWIHGL WP_001058323.1 MULTISPECIES: hydrogenase-1 small chain [Proteobacteria] Length: 372\nNumber of predicted TMHs: 2\nExp number of AAs in TMHs: 37.2608499999999\nExp number, first 60 AAs: 19.52923\nTotal prob of N-in: 0.94351\nPOSSIBLE N-term signal sequence\ninside 1 19\nTMhelix 20 39\noutside 40 325\nTMhelix 326 348\ninside 349 372

8927 GCF_001894245.1_ASM189424v1 Escherichia coli Proteobacteria; Gammaproteobacteria; Enterobacterales; Enterobacteriaceae; Escherichia MTGDNTLIHSHGINRRDFMKLCAALAATMGLSSKAAAEMAESVTNPQRPPVIWIGAQECT WP_000145410.1 MULTISPECIES: hydrogenase-2 small chain [Proteobacteria] Length: 372\nNumber of predicted TMHs: 1\nExp number of AAs in TMHs: 23.72226\nExp number, first 60 AAs: 0.22038\nTotal prob of N-in: 0.12994\noutside 1 331\nTMhelix 332 354\ninside 355 372

8928 GCF_001894245.1_ASM189424v1 Escherichia coli Proteobacteria; Gammaproteobacteria; Enterobacterales; Enterobacteriaceae; Escherichia MNNEETFYQAMRRQGVTRRSFLKYCSLAATSLGLGAGMAPKIAWALENKPRIPVVWIHGL WP_001058323.1 MULTISPECIES: hydrogenase-1 small chain [Proteobacteria] Length: 372\nNumber of predicted TMHs: 2\nExp number of AAs in TMHs: 37.2608499999999\nExp number, first 60 AAs: 19.52923\nTotal prob of N-in: 0.94351\nPOSSIBLE N-term signal sequence\ninside 1 19\nTMhelix 20 39\noutside 40 325\nTMhelix 326 348\ninside 349 372

8929 GCF_001894305.1_ASM189430v1 Escherichia coli Proteobacteria; Gammaproteobacteria; Enterobacterales; Enterobacteriaceae; Escherichia MNNEETFYQAMRRQGVTRRSFLKYCSLAATSLGLGAGMAPKIAWALENKPRIPVVWIHGL WP_001058323.1 MULTISPECIES: hydrogenase-1 small chain [Proteobacteria] Length: 372\nNumber of predicted TMHs: 2\nExp number of AAs in TMHs: 37.2608499999999\nExp number, first 60 AAs: 19.52923\nTotal prob of N-in: 0.94351\nPOSSIBLE N-term signal sequence\ninside 1 19\nTMhelix 20 39\noutside 40 325\nTMhelix 326 348\ninside 349 372

8930 GCF_001894335.1_ASM189433v1 Escherichia coli Proteobacteria; Gammaproteobacteria; Enterobacterales; Enterobacteriaceae; Escherichia MNNEETFYQAMRRQGVTRRSFLKYCSLAATSLGLGAGMAPKIAWALENKPRIPVVWIHGL WP_001058323.1 MULTISPECIES: hydrogenase-1 small chain [Proteobacteria] Length: 372\nNumber of predicted TMHs: 2\nExp number of AAs in TMHs: 37.2608499999999\nExp number, first 60 AAs: 19.52923\nTotal prob of N-in: 0.94351\nPOSSIBLE N-term signal sequence\ninside 1 19\nTMhelix 20 39\noutside 40 325\nTMhelix 326 348\ninside 349 372

8931 GCF_001894195.1_ASM189419v1 Escherichia coli Proteobacteria; Gammaproteobacteria; Enterobacterales; Enterobacteriaceae; Escherichia MNNEETFYQAMRRQGVTRRSFLKYCSLAATSLGLGAGMAPKIAWALENKPRIPVVWIHGL WP_001058323.1 MULTISPECIES: hydrogenase-1 small chain [Proteobacteria] Length: 372\nNumber of predicted TMHs: 2\nExp number of AAs in TMHs: 37.2608499999999\nExp number, first 60 AAs: 19.52923\nTotal prob of N-in: 0.94351\nPOSSIBLE N-term signal sequence\ninside 1 19\nTMhelix 20 39\noutside 40 325\nTMhelix 326 348\ninside 349 372

8932 GCF_001894495.1_ASM189449v1 Escherichia coli Proteobacteria; Gammaproteobacteria; Enterobacterales; Enterobacteriaceae; Escherichia MTGDNTLIHSHGINRRDFMKLCAALAATMGLSSKAAAEMAESVTNPQRPPVIWIGAQECT WP_000145410.1 MULTISPECIES: hydrogenase-2 small chain [Proteobacteria] Length: 372\nNumber of predicted TMHs: 1\nExp number of AAs in TMHs: 23.72226\nExp number, first 60 AAs: 0.22038\nTotal prob of N-in: 0.12994\noutside 1 331\nTMhelix 332 354\ninside 355 372

8933 GCF_001894495.1_ASM189449v1 Escherichia coli Proteobacteria; Gammaproteobacteria; Enterobacterales; Enterobacteriaceae; Escherichia

MNNEETFYQAMRRQGVTRRSFLKYCSLAATSLGLGAGMAPKIAWALENKPRIPVVWIHGL WP_001295940.1 MULTISPECIES: hydrogenase [Enterobacteriaceae] Length: 372\nNumber of predicted TMHs: 2\nExp number of AAs in TMHs: 37.2608899999999\nExp number, first 60 AAs: 19.52923\nTotal prob of N-in: 0.94351\nPOSSIBLE N-term signal sequence\ninside 1 19\nTMhelix 20 39\noutside 40 325\nTMhelix 326 348\ninside 349 372

8934 GCF_001894155.1_ASM189415v1 Escherichia coli Proteobacteria; Gammaproteobacteria; Enterobacterales; Enterobacteriaceae; Escherichia MNNEETFYQAMRRQGVTRRSFLKYCSLAATSLGLGAGMAPKIAWALENKPRIPVVWIHGL WP_001295940.1 MULTISPECIES: hydrogenase [Enterobacteriaceae] Length: 372\nNumber of predicted TMHs: 2\nExp number of AAs in TMHs: 37.2608899999999\nExp number, first 60 AAs: 19.52923\nTotal prob of N-in: 0.94351\nPOSSIBLE N-term signal sequence\ninside 1 19\nTMhelix 20 39\noutside 40 325\nTMhelix 326 348\ninside 349 372

8935 GCF_001894395.1_ASM189439v1 Escherichia coli Proteobacteria; Gammaproteobacteria; Enterobacterales; Enterobacteriaceae; Escherichia MNNEETFYQAMRRQGVTRRSFLKYCSLAATSLGLGAGMAPKIAWALENKPRIPVVWIHGL WP_001058323.1 MULTISPECIES: hydrogenase-1 small chain [Proteobacteria] Length: 372\nNumber of predicted TMHs: 2\nExp number of AAs in TMHs: 37.2608499999999\nExp number, first 60 AAs: 19.52923\nTotal prob of N-in: 0.94351\nPOSSIBLE N-term signal sequence\ninside 1 19\nTMhelix 20 39\noutside 40 325\nTMhelix 326 348\ninside 349 372

8936 GCF_001894635.1_ASM189463v1 Escherichia coli Proteobacteria; Gammaproteobacteria; Enterobacterales; Enterobacteriaceae; Escherichia MNNEETFYQAMRRQGVTRRSFLKYCSLAATSLGLGAGMAPKIAWALENKPRIPVVWIHGL WP_001058320.1 hydrogenase [Escherichia coli] Length: 372\nNumber of predicted TMHs: 2\nExp number of AAs in TMHs: 37.26082\nExp number, first 60 AAs: 19.52923\nTotal prob of N-in: 0.94351\nPOSSIBLE N-term signal sequence\ninside 1 19\nTMhelix 20 39\noutside 40 325\nTMhelix 326 348\ninside 349 372

8937 GCF_001900415.1_ASM190041v1 Escherichia coli Proteobacteria; Gammaproteobacteria; Enterobacterales; Enterobacteriaceae; Escherichia MTGDNTLIHSHGINRRDFMKLCAALAATMGLSSKAAAEMAESVTNPQRPPVIWIGAQECT WP_000145410.1 MULTISPECIES: hydrogenase-2 small chain [Proteobacteria] Length: 372\nNumber of predicted TMHs: 1\nExp number of AAs in TMHs: 23.72226\nExp number, first 60 AAs: 0.22038\nTotal prob of N-in: 0.12994\noutside 1 331\nTMhelix 332 354\ninside 355 372

8938 GCF_001900455.1_ASM190045v1 Escherichia coli Proteobacteria; Gammaproteobacteria; Enterobacterales; Enterobacteriaceae; Escherichia MTGDNTLIHSHGINRRDFMKLCAALAATMGLSSKAAAEMAESVTNPQRPPVIWIGAQECT WP_000145410.1 MULTISPECIES: hydrogenase-2 small chain [Proteobacteria] Length: 372\nNumber of predicted TMHs: 1\nExp number of AAs in TMHs: 23.72226\nExp number, first 60 AAs: 0.22038\nTotal prob of N-in: 0.12994\noutside 1 331\nTMhelix 332 354\ninside 355 372

8939 GCF_001900375.1_ASM190037v1 Escherichia coli Proteobacteria; Gammaproteobacteria; Enterobacterales; Enterobacteriaceae; Escherichia MTGDNTLIHSHGINRRDFMKLCAALAATMGLSSKAAAEMAESVTNPQRPPVIWIGAQECT WP_000145410.1 MULTISPECIES: hydrogenase-2 small chain [Proteobacteria] Length: 372\nNumber of predicted TMHs: 1\nExp number of AAs in TMHs: 23.72226\nExp number, first 60 AAs: 0.22038\nTotal prob of N-in: 0.12994\noutside 1 331\nTMhelix 332 354\ninside 355 372

8940 GCF_001900415.1_ASM190041v1 Escherichia coli Proteobacteria; Gammaproteobacteria; Enterobacterales; Enterobacteriaceae; Escherichia MNNEETFYQAMRRQGVTRRSFLKYCSLAATSLGLGAGMAPKIAWALENKPRIPVVWIHGL WP_001058323.1 MULTISPECIES: hydrogenase-1 small chain [Proteobacteria] Length: 372\nNumber of predicted TMHs: 2\nExp number of AAs in TMHs: 37.2608499999999\nExp number, first 60 AAs: 19.52923\nTotal prob of N-in: 0.94351\nPOSSIBLE N-term signal sequence\ninside 1 19\nTMhelix 20 39\noutside 40 325\nTMhelix 326 348\ninside 349 372

8941 GCF_001900375.1_ASM190037v1 Escherichia coli Proteobacteria; Gammaproteobacteria; Enterobacterales; Enterobacteriaceae; Escherichia MNNEETFYQAMRRQGVTRRSFLKYCSLAATSLGLGAGMAPKIAWALENKPRIPVVWIHGL WP_001058323.1

MULTISPECIES: hydrogenase-1 small chain [Proteobacteria] Length: 372\nNumber of predicted TMHs: 2\nExp number of AAs in TMHs: 37.2608499999999\nExp number, first 60 AAs: 19.52923\nTotal prob of N-in: 0.94351\nPOSSIBLE N-term signal sequence\ninside 1 19\nTMhelix 20 39\noutside 40 325\nTMhelix 326 348\ninside 349 372

8942 GCF_001900455.1_ASM190045v1 Escherichia coli Proteobacteria; Gammaproteobacteria; Enterobacterales; Enterobacteriaceae; Escherichia
MNNEETFYQAMRRQGVTRRSFLKYCSLAATSLGLGAGMAPKIAWALENKPRIPVVWIHGL WP_001058323.1
MULTISPECIES: hydrogenase-1 small chain [Proteobacteria] Length: 372\nNumber of predicted TMHs: 2\nExp number of AAs in TMHs: 37.2608499999999\nExp number, first 60 AAs: 19.52923\nTotal prob of N-in: 0.94351\nPOSSIBLE N-term signal sequence\ninside 1 19\nTMhelix 20 39\noutside 40 325\nTMhelix 326 348\ninside 349 372

8943 GCF_001899625.1_ASM189962v1 Escherichia coli Proteobacteria; Gammaproteobacteria; Enterobacterales; Enterobacteriaceae; Escherichia
MNNEETFYQAMRRQGVTRRSFLKYCSLAATSLGLGAGMAPKIAWALENKPRIPVVWIHGL WP_001058320.1
hydrogenase [Escherichia coli] Length: 372\nNumber of predicted TMHs: 2\nExp number of AAs in TMHs: 37.26082\nExp number, first 60 AAs: 19.52923\nTotal prob of N-in: 0.94351\nPOSSIBLE N-term signal sequence\ninside 1 19\nTMhelix 20 39\noutside 40 325\nTMhelix 326 348\ninside 349 372

8944 GCF_001900595.1_ASM190059v1 Escherichia coli Proteobacteria; Gammaproteobacteria; Enterobacterales; Enterobacteriaceae; Escherichia
MTGDNTLIHSHGINRRDFMKLCAALAATMGLSSKAAAEMAESVTNPQRPPVIWIGAQECT WP_000145410.1
MULTISPECIES: hydrogenase-2 small chain [Proteobacteria] Length: 372\nNumber of predicted TMHs: 1\nExp number of AAs in TMHs: 23.72226\nExp number, first 60 AAs: 0.22038\nTotal prob of N-in: 0.12994\noutside 1 331\nTMhelix 332 354\ninside 355 372

8945 GCF_001900655.1_ASM190065v1 Escherichia coli Proteobacteria; Gammaproteobacteria; Enterobacterales; Enterobacteriaceae; Escherichia
MTGDNTLIHSHGINRRDFMKLCAALAATMGLSSKAAAEMAESVTNPQRPPVIWIGAQECT WP_000145410.1
MULTISPECIES: hydrogenase-2 small chain [Proteobacteria] Length: 372\nNumber of predicted TMHs: 1\nExp number of AAs in TMHs: 23.72226\nExp number, first 60 AAs: 0.22038\nTotal prob of N-in: 0.12994\noutside 1 331\nTMhelix 332 354\ninside 355 372

8946 GCF_001900775.1_ASM190077v1 Escherichia coli Proteobacteria; Gammaproteobacteria; Enterobacterales; Enterobacteriaceae; Escherichia
MTGDNTLIHSHGINRRDFMKLCAALAATMGLSSKAAAEMAESVTNPQRPPVIWIGAQECT WP_000145410.1
MULTISPECIES: hydrogenase-2 small chain [Proteobacteria] Length: 372\nNumber of predicted TMHs: 1\nExp number of AAs in TMHs: 23.72226\nExp number, first 60 AAs: 0.22038\nTotal prob of N-in: 0.12994\noutside 1 331\nTMhelix 332 354\ninside 355 372

8947 GCF_001900695.1_ASM190069v1 Escherichia coli Proteobacteria; Gammaproteobacteria; Enterobacterales; Enterobacteriaceae; Escherichia
MTGDNTLIHSHGINRRDFMKLCAALAATMGLSSKAAAEMAESVTNPQRPPVIWIGAQECT WP_000145410.1
MULTISPECIES: hydrogenase-2 small chain [Proteobacteria] Length: 372\nNumber of predicted TMHs: 1\nExp number of AAs in TMHs: 23.72226\nExp number, first 60 AAs: 0.22038\nTotal prob of N-in: 0.12994\noutside 1 331\nTMhelix 332 354\ninside 355 372

8948 GCF_001900595.1_ASM190059v1 Escherichia coli Proteobacteria; Gammaproteobacteria; Enterobacterales; Enterobacteriaceae; Escherichia
MNNEETFYQAMRRQGVTRRSFLKYCSLAATSLGLGAGMAPKIAWALENKPRIPVVWIHGL WP_001058323.1
MULTISPECIES: hydrogenase-1 small chain [Proteobacteria] Length: 372\nNumber of predicted TMHs: 2\nExp number of AAs in TMHs: 37.2608499999999\nExp number, first 60 AAs: 19.52923\nTotal prob of N-in: 0.94351\nPOSSIBLE N-term signal sequence\ninside 1 19\nTMhelix 20 39\noutside 40 325\nTMhelix 326 348\ninside 349 372

8949 GCF_001900815.1_ASM190081v1 Escherichia coli Proteobacteria; Gammaproteobacteria; Enterobacterales; Enterobacteriaceae; Escherichia
MTGDNTLIHSHGINRRDFMKLCAALAATMGLSSKAAAEMAESVTNPQRPPVIWIGAQECT WP_000145410.1
MULTISPECIES: hydrogenase-2 small chain [Proteobacteria] Length: 372\nNumber of predicted TMHs:

1\nExp number of AAs in TMHs: 23.72226\nExp number, first 60 AAs: 0.22038\nTotal prob of N-in: 0.12994\noutside 1 331\nTMhelix 332 354\ninside 355 372

8950 GCF_001901105.1_ASM190110v1 Escherichia coli Proteobacteria; Gammaproteobacteria; Enterobacterales; Enterobacteriaceae; Escherichia
MTGDNTLIHSHGINRRDFMKLCAALAATMGLSSKAAAEMAESVTNPQRPPVIWIGAQECT WP_000145410.1
MULTISPECIES: hydrogenase-2 small chain [Proteobacteria] Length: 372\nNumber of predicted TMHs: 1\nExp number of AAs in TMHs: 23.72226\nExp number, first 60 AAs: 0.22038\nTotal prob of N-in: 0.12994\noutside 1 331\nTMhelix 332 354\ninside 355 372

8951 GCF_001901045.1_ASM190104v1 Escherichia coli Proteobacteria; Gammaproteobacteria; Enterobacterales; Enterobacteriaceae; Escherichia
MTGDNTLIHSHGINRRDFMKLCAALAATMGLSSKAAAEMAESVTNPQRPPVIWIGAQECT WP_000145410.1
MULTISPECIES: hydrogenase-2 small chain [Proteobacteria] Length: 372\nNumber of predicted TMHs: 1\nExp number of AAs in TMHs: 23.72226\nExp number, first 60 AAs: 0.22038\nTotal prob of N-in: 0.12994\noutside 1 331\nTMhelix 332 354\ninside 355 372

8952 GCF_001901145.1_ASM190114v1 Escherichia coli Proteobacteria; Gammaproteobacteria; Enterobacterales; Enterobacteriaceae; Escherichia
MTGDNTLIHSHGINRRDFMKLCAALAATMGLSSKAAAEMAESVTNPQRPPVIWIGAQECT WP_000145410.1
MULTISPECIES: hydrogenase-2 small chain [Proteobacteria] Length: 372\nNumber of predicted TMHs: 1\nExp number of AAs in TMHs: 23.72226\nExp number, first 60 AAs: 0.22038\nTotal prob of N-in: 0.12994\noutside 1 331\nTMhelix 332 354\ninside 355 372

8953 GCF_001901005.1_ASM190100v1 Escherichia coli Proteobacteria; Gammaproteobacteria; Enterobacterales; Enterobacteriaceae; Escherichia
MTGDNTLIHSHGINRRDFMKLCAALAATMGLSSKAAAEMAESVTNPQRPPVIWIGAQECT WP_000145410.1
MULTISPECIES: hydrogenase-2 small chain [Proteobacteria] Length: 372\nNumber of predicted TMHs: 1\nExp number of AAs in TMHs: 23.72226\nExp number, first 60 AAs: 0.22038\nTotal prob of N-in: 0.12994\noutside 1 331\nTMhelix 332 354\ninside 355 372

8954 GCF_001901215.1_ASM190121v1 Escherichia coli Proteobacteria; Gammaproteobacteria; Enterobacterales; Enterobacteriaceae; Escherichia
MTGDNTLIHSHGINRRDFMKLCAALAATMGLSSKAAAEMAESVTNPQRPPVIWIGAQECT WP_000145410.1
MULTISPECIES: hydrogenase-2 small chain [Proteobacteria] Length: 372\nNumber of predicted TMHs: 1\nExp number of AAs in TMHs: 23.72226\nExp number, first 60 AAs: 0.22038\nTotal prob of N-in: 0.12994\noutside 1 331\nTMhelix 332 354\ninside 355 372

8955 GCF_001901145.1_ASM190114v1 Escherichia coli Proteobacteria; Gammaproteobacteria; Enterobacterales; Enterobacteriaceae; Escherichia
MNNEETFYQAMRRQGVTRRSFLKYCSLAATSLGLGAGMAPKIAWALENKPRIPVVWIHGL WP_001058323.1
MULTISPECIES: hydrogenase-1 small chain [Proteobacteria] Length: 372\nNumber of predicted TMHs: 2\nExp number of AAs in TMHs: 37.2608499999999\nExp number, first 60 AAs: 19.52923\nTotal prob of N-in: 0.94351\nPOSSIBLE N-term signal sequence\ninside 1 19\nTMhelix 20 39\noutside 40 325\nTMhelix 326 348\ninside 349 372

8956 GCF_001900815.1_ASM190081v1 Escherichia coli Proteobacteria; Gammaproteobacteria; Enterobacterales; Enterobacteriaceae; Escherichia
MNNEETFYQAMRRQGVTRRSFLKYCSLAATSLGLGAGMAPKIAWALENKPRIPVVWIHGL WP_001295940.1
MULTISPECIES: hydrogenase [Enterobacteriaceae] Length: 372\nNumber of predicted TMHs: 2\nExp number of AAs in TMHs: 37.2608899999999\nExp number, first 60 AAs: 19.52923\nTotal prob of N-in: 0.94351\nPOSSIBLE N-term signal sequence\ninside 1 19\nTMhelix 20 39\noutside 40 325\nTMhelix 326 348\ninside 349 372

8957 GCF_001901445.1_ASM190144v1 Escherichia coli Proteobacteria; Gammaproteobacteria; Enterobacterales; Enterobacteriaceae; Escherichia
MNNEETFYQAMRRQGVTRRSFLKYCSLAATSLGLGAGMAPKIAWALENKPRIPVVWIHGL WP_001058323.1
MULTISPECIES: hydrogenase-1 small chain [Proteobacteria] Length: 372\nNumber of predicted TMHs: 2\nExp number of AAs in TMHs: 37.2608499999999\nExp number, first 60 AAs: 19.52923\nTotal prob of N-in: 0.94351\nPOSSIBLE N-term signal sequence\ninside 1 19\nTMhelix 20 39\noutside 40 325\nTMhelix 326 348\ninside 349 372

8958 GCF_001901005.1_ASM190100v1 Escherichia coli Proteobacteria; Gammaproteobacteria; Enterobacterales; Enterobacteriaceae; Escherichia
MNNEETFYQAMRRQGVTRRSFLKYCSLAATSLGLGAGMAPKIAWALENKPRIPVVWIHGL WP_001295940.1
MULTISPECIES: hydrogenase [Enterobacteriaceae] Length: 372\nNumber of predicted TMHs: 2\nExp number of AAs in TMHs: 37.2608899999999\nExp number, first 60 AAs: 19.52923\nTotal prob of N-in: 0.94351\nPOSSIBLE N-term signal sequence\ninside 1 19\nTMhelix 20 39\noutside 40 325\nTMhelix 326 348\ninside 349 372

8959 GCF_001901045.1_ASM190104v1 Escherichia coli Proteobacteria; Gammaproteobacteria; Enterobacterales; Enterobacteriaceae; Escherichia
MNNEETFYQAMRRQGVTRRSFLKYCSLAATSLGLGAGMAPKIAWALENKPRIPVVWIHGL WP_053264900.1
hydrogenase [Escherichia coli] Length: 372\nNumber of predicted TMHs: 2\nExp number of AAs in TMHs: 37.24714\nExp number, first 60 AAs: 19.52921\nTotal prob of N-in: 0.94351\nPOSSIBLE N-term signal sequence\ninside 1 19\nTMhelix 20 39\noutside 40 325\nTMhelix 326 348\ninside 349 372

8960 GCF_001902685.1_ASM190268v1 Escherichia coli Proteobacteria; Gammaproteobacteria; Enterobacterales; Enterobacteriaceae; Escherichia
MTGDNTLIHSHGINRRDFMKLCAALAATMGLSSKAAAEMAESVTNPQRPPVIWIGAQECT WP_000145410.1
MULTISPECIES: hydrogenase-2 small chain [Proteobacteria] Length: 372\nNumber of predicted TMHs: 1\nExp number of AAs in TMHs: 23.72226\nExp number, first 60 AAs: 0.22038\nTotal prob of N-in: 0.12994\noutside 1 331\nTMhelix 332 354\ninside 355 372

8961 GCF_001907315.1_ASM190731v1 Escherichia coli Proteobacteria; Gammaproteobacteria; Enterobacterales; Enterobacteriaceae; Escherichia
MTGDNTLIHSHGINRRDFMKLCAALAATMGLSSKAAAEMAESVTNPQRPPVIWIGAQECT WP_000145410.1
MULTISPECIES: hydrogenase-2 small chain [Proteobacteria] Length: 372\nNumber of predicted TMHs: 1\nExp number of AAs in TMHs: 23.72226\nExp number, first 60 AAs: 0.22038\nTotal prob of N-in: 0.12994\noutside 1 331\nTMhelix 332 354\ninside 355 372

8962 GCF_001907375.1_ASM190737v1 Escherichia coli Proteobacteria; Gammaproteobacteria; Enterobacterales; Enterobacteriaceae; Escherichia
MTGDNTLIHSHGINRRDFMKLCAALAATMGLSSKAAAEMAESVTNPQRPPVIWIGAQECT WP_000145410.1
MULTISPECIES: hydrogenase-2 small chain [Proteobacteria] Length: 372\nNumber of predicted TMHs: 1\nExp number of AAs in TMHs: 23.72226\nExp number, first 60 AAs: 0.22038\nTotal prob of N-in: 0.12994\noutside 1 331\nTMhelix 332 354\ninside 355 372

8963 GCF_001907465.1_ASM190746v1 Escherichia coli Proteobacteria; Gammaproteobacteria; Enterobacterales; Enterobacteriaceae; Escherichia
MTGDNTLIHSHGINRRDFMKLCAALAATMGLSSKAAAEMAESVTNPQRPPVIWIGAQECT WP_000145410.1
MULTISPECIES: hydrogenase-2 small chain [Proteobacteria] Length: 372\nNumber of predicted TMHs: 1\nExp number of AAs in TMHs: 23.72226\nExp number, first 60 AAs: 0.22038\nTotal prob of N-in: 0.12994\noutside 1 331\nTMhelix 332 354\ninside 355 372

8964 GCF_001907565.1_ASM190756v1 Escherichia coli Proteobacteria; Gammaproteobacteria; Enterobacterales; Enterobacteriaceae; Escherichia
MTGDNTLIHSHGINRRDFMKLCAALAATMGLSSKAAAEMAESVTNPQRPPVIWIGAQECT WP_038339805.1
hydrogenase 2 small subunit [Escherichia coli] Length: 372\nNumber of predicted TMHs: 1\nExp number of AAs in TMHs: 23.86829\nExp number, first 60 AAs: 0.22037\nTotal prob of N-in: 0.12535\noutside 1 332\nTMhelix 333 355\ninside 356 372

8965 GCF_001907505.1_ASM190750v1 Escherichia coli Proteobacteria; Gammaproteobacteria; Enterobacterales; Enterobacteriaceae; Escherichia
MNNEETFYQAMRRQGVTRRSFLKYCSLAATSLGLGAGMAPKIAWALENKPRIPVVWIHGL WP_001058323.1
MULTISPECIES: hydrogenase-1 small chain [Proteobacteria] Length: 372\nNumber of predicted TMHs: 2\nExp number of AAs in TMHs: 37.2608499999999\nExp number, first 60 AAs: 19.52923\nTotal prob of N-in: 0.94351\nPOSSIBLE N-term signal sequence\ninside 1 19\nTMhelix 20 39\noutside 40 325\nTMhelix 326 348\ninside 349 372

8966 GCF_001909105.1_ASM190910v1 Escherichia coli Proteobacteria; Gammaproteobacteria; Enterobacterales; Enterobacteriaceae; Escherichia
MTGDNTLIHSHGINRRDFMKLCAALAATMGLSSKAAAEMAESVTNPQRPPVIWIGAQECT WP_000145410.1

MULTISPECIES: hydrogenase-2 small chain [Proteobacteria] Length: 372\nNumber of predicted TMHs: 1\nExp number of AAs in TMHs: 23.72226\nExp number, first 60 AAs: 0.22038\nTotal prob of N-in: 0.12994\noutside 1 331\nTMhelix 332 354\ninside 355 372

8967 GCF_001909105.1_ASM190910v1 Escherichia coli Proteobacteria; Gammaproteobacteria; Enterobacterales; Enterobacteriaceae; Escherichia
MNNEETFYQAMRRQGVTRRSFLKYCSLAATSLGLGAGMAPKIAWALENKPRIPVVWIHGL WP_001058323.1
MULTISPECIES: hydrogenase-1 small chain [Proteobacteria] Length: 372\nNumber of predicted TMHs: 2\nExp number of AAs in TMHs: 37.2608499999999\nExp number, first 60 AAs: 19.52923\nTotal prob of N-in: 0.94351\nPOSSIBLE N-term signal sequence\ninside 1 19\nTMhelix 20 39\noutside 40 325\nTMhelix 326 348\ninside 349 372

8968 GCF_001907395.1_ASM190739v1 Escherichia coli Proteobacteria; Gammaproteobacteria; Enterobacterales; Enterobacteriaceae; Escherichia
MTGDNTLIHSHGINRRDFMKLCAALAATMGLSSKAAAEMAESVTNPQRPPVIWIGAQECT WP_038339805.1
hydrogenase 2 small subunit [Escherichia coli] Length: 372\nNumber of predicted TMHs: 1\nExp number of AAs in TMHs: 23.86829\nExp number, first 60 AAs: 0.22037\nTotal prob of N-in: 0.12535\noutside 1 332\nTMhelix 333 355\ninside 356 372

8969 GCF_001910835.1_ASM191083v1 Escherichia coli Proteobacteria; Gammaproteobacteria; Enterobacterales; Enterobacteriaceae; Escherichia
MTGDNTLIHSHGINRRDFMKLCAALAATMGLSSKAAAEMAESVTNPQRPPVIWIGAQECT WP_000145410.1
MULTISPECIES: hydrogenase-2 small chain [Proteobacteria] Length: 372\nNumber of predicted TMHs: 1\nExp number of AAs in TMHs: 23.72226\nExp number, first 60 AAs: 0.22038\nTotal prob of N-in: 0.12994\noutside 1 331\nTMhelix 332 354\ninside 355 372

8970 GCF_001910855.1_ASM191085v1 Escherichia coli Proteobacteria; Gammaproteobacteria; Enterobacterales; Enterobacteriaceae; Escherichia
MTGDNTLIHSHGINRRDFMKLCAALAATMGLSSKAAAEMAESVTNPQRPPVIWIGAQECT WP_000145410.1
MULTISPECIES: hydrogenase-2 small chain [Proteobacteria] Length: 372\nNumber of predicted TMHs: 1\nExp number of AAs in TMHs: 23.72226\nExp number, first 60 AAs: 0.22038\nTotal prob of N-in: 0.12994\noutside 1 331\nTMhelix 332 354\ninside 355 372

8971 GCF_001911255.1_ASM191125v1 Escherichia coli Proteobacteria; Gammaproteobacteria; Enterobacterales; Enterobacteriaceae; Escherichia
MTGDNTLIHSHGINRRDFMKLCAALAATMGLSSKAAAEMAESVTNPQRPPVIWIGAQECT WP_000145410.1
MULTISPECIES: hydrogenase-2 small chain [Proteobacteria] Length: 372\nNumber of predicted TMHs: 1\nExp number of AAs in TMHs: 23.72226\nExp number, first 60 AAs: 0.22038\nTotal prob of N-in: 0.12994\noutside 1 331\nTMhelix 332 354\ninside 355 372

8972 GCF_001911385.1_ASM191138v1 Escherichia coli Proteobacteria; Gammaproteobacteria; Enterobacterales; Enterobacteriaceae; Escherichia
MTGDNTLIHSHGINRRDFMKLCAALAATMGLSSKAAAEMAESVTNPQRPPVIWIGAQECT WP_000145410.1
MULTISPECIES: hydrogenase-2 small chain [Proteobacteria] Length: 372\nNumber of predicted TMHs: 1\nExp number of AAs in TMHs: 23.72226\nExp number, first 60 AAs: 0.22038\nTotal prob of N-in: 0.12994\noutside 1 331\nTMhelix 332 354\ninside 355 372

8973 GCF_001911085.1_ASM191108v1 Escherichia coli Proteobacteria; Gammaproteobacteria; Enterobacterales; Enterobacteriaceae; Escherichia
MTGDNTLIHSHGINRRDFMKLCAALAATMGLSSKAAAEMAESVTNPQRPPVIWIGAQECT WP_001685818.1
MULTISPECIES: hydrogenase-2 small chain [Enterobacteriaceae] Length: 372\nNumber of predicted TMHs: 1\nExp number of AAs in TMHs: 23.82505\nExp number, first 60 AAs: 0.21974\nTotal prob of N-in: 0.13379\noutside 1 331\nTMhelix 332 354\ninside 355 372

8974 GCF_001911165.1_ASM191116v1 Escherichia coli Proteobacteria; Gammaproteobacteria; Enterobacterales; Enterobacteriaceae; Escherichia
MNNEETFYQAMRRQGVTRRSFLKYCSLAATSLGLGAGMAPKIAWALENKPRIPVVWIHGL WP_001058323.1
MULTISPECIES: hydrogenase-1 small chain [Proteobacteria] Length: 372\nNumber of predicted TMHs: 2\nExp number of AAs in TMHs: 37.2608499999999\nExp number, first 60 AAs: 19.52923\nTotal prob of N-in: 0.94351\nPOSSIBLE N-term signal sequence\ninside 1 19\nTMhelix 20 39\noutside 40 325\nTMhelix 326 348\ninside 349 372

8975 GCF_001911325.1_ASM191132v1 Escherichia coli Proteobacteria; Gammaproteobacteria; Enterobacterales; Enterobacteriaceae; Escherichia
MTGDNTLIHSHGINRRDFMKLCAALAATMGLSSKAAAEMAESVTNPQRPPVIWIGAQECT WP_000145410.1
MULTISPECIES: hydrogenase-2 small chain [Proteobacteria] Length: 372\nNumber of predicted TMHs: 1\nExp number of AAs in TMHs: 23.72226\nExp number, first 60 AAs: 0.22038\nTotal prob of N-in: 0.12994\noutside 1 331\nTMhelix 332 354\ninside 355 372

8976 GCF_001911325.1_ASM191132v1 Escherichia coli Proteobacteria; Gammaproteobacteria; Enterobacterales; Enterobacteriaceae; Escherichia
MNNEETFYQAMRRQGVTRRSFLKYCSLAATSLGLGAGMAPKIAWALENKPRIPVVWIHGL WP_001058323.1
MULTISPECIES: hydrogenase-1 small chain [Proteobacteria] Length: 372\nNumber of predicted TMHs: 2\nExp number of AAs in TMHs: 37.2608499999999\nExp number, first 60 AAs: 19.52923\nTotal prob of N-in: 0.94351\nPOSSIBLE N-term signal sequence\ninside 1 19\nTMhelix 20 39\noutside 40 325\nTMhelix 326 348\ninside 349 372

8977 GCF_001911235.1_ASM191123v1 Escherichia coli Proteobacteria; Gammaproteobacteria; Enterobacterales; Enterobacteriaceae; Escherichia
MNNEETFYQAMRRQGVTRRSFLKYCSLAATSLGLGAGMAPKIAWALENKPRIPVVWIHGL WP_001058323.1
MULTISPECIES: hydrogenase-1 small chain [Proteobacteria] Length: 372\nNumber of predicted TMHs: 2\nExp number of AAs in TMHs: 37.2608499999999\nExp number, first 60 AAs: 19.52923\nTotal prob of N-in: 0.94351\nPOSSIBLE N-term signal sequence\ninside 1 19\nTMhelix 20 39\noutside 40 325\nTMhelix 326 348\ninside 349 372

8978 GCF_001911485.1_ASM191148v1 Escherichia coli Proteobacteria; Gammaproteobacteria; Enterobacterales; Enterobacteriaceae; Escherichia
MTGDNTLIHSHGINRRDFMKLCAALAATMGLSSKAAAEMAESVTNPQRPPVIWIGAQECT WP_000145410.1
MULTISPECIES: hydrogenase-2 small chain [Proteobacteria] Length: 372\nNumber of predicted TMHs: 1\nExp number of AAs in TMHs: 23.72226\nExp number, first 60 AAs: 0.22038\nTotal prob of N-in: 0.12994\noutside 1 331\nTMhelix 332 354\ninside 355 372

8979 GCF_001911485.1_ASM191148v1 Escherichia coli Proteobacteria; Gammaproteobacteria; Enterobacterales; Enterobacteriaceae; Escherichia
MNNEETFYQAMRRQGVTRRSFLKYCSLAATSLGLGAGMAPKIAWALENKPRIPVVWIHGL WP_001058323.1
MULTISPECIES: hydrogenase-1 small chain [Proteobacteria] Length: 372\nNumber of predicted TMHs: 2\nExp number of AAs in TMHs: 37.2608499999999\nExp number, first 60 AAs: 19.52923\nTotal prob of N-in: 0.94351\nPOSSIBLE N-term signal sequence\ninside 1 19\nTMhelix 20 39\noutside 40 325\nTMhelix 326 348\ninside 349 372

8980 GCF_001911255.1_ASM191125v1 Escherichia coli Proteobacteria; Gammaproteobacteria; Enterobacterales; Enterobacteriaceae; Escherichia
MNNEETFYQAMRRQGVTRRSFLKYCSLAATSLGLGAGMAPKIAWALENKPRIPVVWIHGL WP_001058320.1
hydrogenase [Escherichia coli] Length: 372\nNumber of predicted TMHs: 2\nExp number of AAs in TMHs: 37.26082\nExp number, first 60 AAs: 19.52923\nTotal prob of N-in: 0.94351\nPOSSIBLE N-term signal sequence\ninside 1 19\nTMhelix 20 39\noutside 40 325\nTMhelix 326 348\ninside 349 372

8981 GCF_001911385.1_ASM191138v1 Escherichia coli Proteobacteria; Gammaproteobacteria; Enterobacterales; Enterobacteriaceae; Escherichia
MNNEETFYQAMRRQGVTRRSFLKYCSLAATSLGLGAGMAPKIAWALENKPRIPVVWIHGL WP_001058323.1
MULTISPECIES: hydrogenase-1 small chain [Proteobacteria] Length: 372\nNumber of predicted TMHs: 2\nExp number of AAs in TMHs: 37.2608499999999\nExp number, first 60 AAs: 19.52923\nTotal prob of N-in: 0.94351\nPOSSIBLE N-term signal sequence\ninside 1 19\nTMhelix 20 39\noutside 40 325\nTMhelix 326 348\ninside 349 372

8982 GCF_001911505.1_ASM191150v1 Escherichia coli Proteobacteria; Gammaproteobacteria; Enterobacterales; Enterobacteriaceae; Escherichia
MTGDNTLIHSHGINRRDFMKLCAALAATMGLSSKAAAEMAESVTNPQRPPVIWIGAQECT WP_000145410.1
MULTISPECIES: hydrogenase-2 small chain [Proteobacteria] Length: 372\nNumber of predicted TMHs: 1\nExp number of AAs in TMHs: 23.72226\nExp number, first 60 AAs: 0.22038\nTotal prob of N-in: 0.12994\noutside 1 331\nTMhelix 332 354\ninside 355 372

8983 GCF_001911585.1_ASM191158v1 Escherichia coli Proteobacteria; Gammaproteobacteria; Enterobacterales; Enterobacteriaceae; Escherichia
MTGDNTLIHSHGINRRDFMKLCAALAATMGLSSKAAAEMAESVTNPQRPPVIWIGAQECT WP_000145410.1
MULTISPECIES: hydrogenase-2 small chain [Proteobacteria] Length: 372\nNumber of predicted TMHs: 1\nExp number of AAs in TMHs: 23.72226\nExp number, first 60 AAs: 0.22038\nTotal prob of N-in: 0.12994\noutside 1 331\nTMhelix 332 354\ninside 355 372

8984 GCF_001911585.1_ASM191158v1 Escherichia coli Proteobacteria; Gammaproteobacteria; Enterobacterales; Enterobacteriaceae; Escherichia
MNNEETFYQAMRRQGVTRRSFLKYCSLAATSLGLGAGMAPKIAWALENKPRIPVVWIHGL WP_001058323.1
MULTISPECIES: hydrogenase-1 small chain [Proteobacteria] Length: 372\nNumber of predicted TMHs: 2\nExp number of AAs in TMHs: 37.2608499999999\nExp number, first 60 AAs: 19.52923\nTotal prob of N-in: 0.94351\nPOSSIBLE N-term signal sequence\ninside 1 19\nTMhelix 20 39\noutside 40 325\nTMhelix 326 348\ninside 349 372

8985 GCF_001911845.1_ASM191184v1 Escherichia coli Proteobacteria; Gammaproteobacteria; Enterobacterales; Enterobacteriaceae; Escherichia
MTGDNTLIHSHGINRRDFMKLCAALAATMGLSSKAAAEMAESVTNPQRPPVIWIGAQECT WP_000145410.1
MULTISPECIES: hydrogenase-2 small chain [Proteobacteria] Length: 372\nNumber of predicted TMHs: 1\nExp number of AAs in TMHs: 23.72226\nExp number, first 60 AAs: 0.22038\nTotal prob of N-in: 0.12994\noutside 1 331\nTMhelix 332 354\ninside 355 372

8986 GCF_001911725.1_ASM191172v1 Escherichia coli Proteobacteria; Gammaproteobacteria; Enterobacterales; Enterobacteriaceae; Escherichia
MNNEETFYQAMRRQGVTRRSFLKYCSLAATSLGLGAGMAPKIAWALENKPRIPVVWIHGL WP_001058323.1
MULTISPECIES: hydrogenase-1 small chain [Proteobacteria] Length: 372\nNumber of predicted TMHs: 2\nExp number of AAs in TMHs: 37.2608499999999\nExp number, first 60 AAs: 19.52923\nTotal prob of N-in: 0.94351\nPOSSIBLE N-term signal sequence\ninside 1 19\nTMhelix 20 39\noutside 40 325\nTMhelix 326 348\ninside 349 372

8987 GCF_001911855.1_ASM191185v1 Escherichia coli Proteobacteria; Gammaproteobacteria; Enterobacterales; Enterobacteriaceae; Escherichia
MTGDNTLIHSHGINRRDFMKLCAALAATMGLSSKAAAEMAESVTNPQRPPVIWIGAQECT WP_000145410.1
MULTISPECIES: hydrogenase-2 small chain [Proteobacteria] Length: 372\nNumber of predicted TMHs: 1\nExp number of AAs in TMHs: 23.72226\nExp number, first 60 AAs: 0.22038\nTotal prob of N-in: 0.12994\noutside 1 331\nTMhelix 332 354\ninside 355 372

8988 GCF_001911855.1_ASM191185v1 Escherichia coli Proteobacteria; Gammaproteobacteria; Enterobacterales; Enterobacteriaceae; Escherichia
MNNEETFYQAMRRQGVTRRSFLKYCSLAATSLGLGAGMAPKIAWALENKPRIPVVWIHGL WP_001058323.1
MULTISPECIES: hydrogenase-1 small chain [Proteobacteria] Length: 372\nNumber of predicted TMHs: 2\nExp number of AAs in TMHs: 37.2608499999999\nExp number, first 60 AAs: 19.52923\nTotal prob of N-in: 0.94351\nPOSSIBLE N-term signal sequence\ninside 1 19\nTMhelix 20 39\noutside 40 325\nTMhelix 326 348\ninside 349 372

8989 GCF_001911775.1_ASM191177v1 Escherichia coli Proteobacteria; Gammaproteobacteria; Enterobacterales; Enterobacteriaceae; Escherichia
MNNEETFYQAMRRQGVTRRSFLKYCSLAATSLGLGAGMAPKIAWALENKPRIPVVWIHGL WP_001058323.1
MULTISPECIES: hydrogenase-1 small chain [Proteobacteria] Length: 372\nNumber of predicted TMHs: 2\nExp number of AAs in TMHs: 37.2608499999999\nExp number, first 60 AAs: 19.52923\nTotal prob of N-in: 0.94351\nPOSSIBLE N-term signal sequence\ninside 1 19\nTMhelix 20 39\noutside 40 325\nTMhelix 326 348\ninside 349 372

8990 GCF_001911675.1_ASM191167v1 Escherichia coli Proteobacteria; Gammaproteobacteria; Enterobacterales; Enterobacteriaceae; Escherichia
MNNEETFYQAMRRQGVTRRSFLKYCSLAATSLGLGAGMAPKIAWALENKPRIPVVWIHGL WP_001058323.1
MULTISPECIES: hydrogenase-1 small chain [Proteobacteria] Length: 372\nNumber of predicted TMHs: 2\nExp number of AAs in TMHs: 37.2608499999999\nExp number, first 60 AAs: 19.52923\nTotal prob of N-in: 0.94351\nPOSSIBLE N-term signal sequence\ninside 1 19\nTMhelix 20 39\noutside 40 325\nTMhelix 326 348\ninside 349 372

8991 GCF_001911905.1_ASM191190v1 Escherichia coli Proteobacteria; Gammaproteobacteria; Enterobacterales; Enterobacteriaceae; Escherichia
MTGDNTLIHSHGINRRDFMKLCAALAATMGLSSKAAAEMAESVTNPQRPPVIWIGAQECT WP_000145410.1
MULTISPECIES: hydrogenase-2 small chain [Proteobacteria] Length: 372\nNumber of predicted TMHs: 1\nExp number of AAs in TMHs: 23.72226\nExp number, first 60 AAs: 0.22038\nTotal prob of N-in: 0.12994\noutside 1 331\nTMhelix 332 354\ninside 355 372

8992 GCF_001911955.1_ASM191195v1 Escherichia coli Proteobacteria; Gammaproteobacteria; Enterobacterales; Enterobacteriaceae; Escherichia
MTGDNTLIHSHGINRRDFMKLCAALAATMGLSSKAAAEMAESVTNPQRPPVIWIGAQECT WP_000145410.1
MULTISPECIES: hydrogenase-2 small chain [Proteobacteria] Length: 372\nNumber of predicted TMHs: 1\nExp number of AAs in TMHs: 23.72226\nExp number, first 60 AAs: 0.22038\nTotal prob of N-in: 0.12994\noutside 1 331\nTMhelix 332 354\ninside 355 372

8993 GCF_001911985.1_ASM191198v1 Escherichia coli Proteobacteria; Gammaproteobacteria; Enterobacterales; Enterobacteriaceae; Escherichia
MTGDNTLIHSHGINRRDFMKLCAALAATMGLSSKAAAEMAESVTNPQRPPVIWIGAQECT WP_000145410.1
MULTISPECIES: hydrogenase-2 small chain [Proteobacteria] Length: 372\nNumber of predicted TMHs: 1\nExp number of AAs in TMHs: 23.72226\nExp number, first 60 AAs: 0.22038\nTotal prob of N-in: 0.12994\noutside 1 331\nTMhelix 332 354\ninside 355 372

8994 GCF_001911955.1_ASM191195v1 Escherichia coli Proteobacteria; Gammaproteobacteria; Enterobacterales; Enterobacteriaceae; Escherichia
MNNEETFYQAMRRQGVTRRSFLKYCSLAATSLGLGAGMAPKIAWALENKPRIPVVWIHGL WP_001058323.1
MULTISPECIES: hydrogenase-1 small chain [Proteobacteria] Length: 372\nNumber of predicted TMHs: 2\nExp number of AAs in TMHs: 37.2608499999999\nExp number, first 60 AAs: 19.52923\nTotal prob of N-in: 0.94351\nPOSSIBLE N-term signal sequence\ninside 1 19\nTMhelix 20 39\noutside 40 325\nTMhelix 326 348\ninside 349 372

8995 GCF_001911985.1_ASM191198v1 Escherichia coli Proteobacteria; Gammaproteobacteria; Enterobacterales; Enterobacteriaceae; Escherichia
MNNEETFYQAMRRQGVTRRSFLKYCSLAATSLGLGAGMAPKIAWALENKPRIPVVWIHGL WP_001058323.1
MULTISPECIES: hydrogenase-1 small chain [Proteobacteria] Length: 372\nNumber of predicted TMHs: 2\nExp number of AAs in TMHs: 37.2608499999999\nExp number, first 60 AAs: 19.52923\nTotal prob of N-in: 0.94351\nPOSSIBLE N-term signal sequence\ninside 1 19\nTMhelix 20 39\noutside 40 325\nTMhelix 326 348\ninside 349 372

8996 GCF_001911825.1_ASM191182v1 Escherichia coli Proteobacteria; Gammaproteobacteria; Enterobacterales; Enterobacteriaceae; Escherichia
MTGDNTLIHSHGINRRDFMKLCAALAATMGLSSKAAAEMAESVTNPQRPPVIWIGAQECT WP_053918372.1
hydrogenase 2 small subunit [Escherichia coli] Length: 372\nNumber of predicted TMHs: 1\nExp number of AAs in TMHs: 23.73548\nExp number, first 60 AAs: 0.22027\nTotal prob of N-in: 0.13021\noutside 1 331\nTMhelix 332 354\ninside 355 372

8997 GCF_001911925.1_ASM191192v1 Escherichia coli Proteobacteria; Gammaproteobacteria; Enterobacterales; Enterobacteriaceae; Escherichia
MNNEETFYQAMRRQGVTRRSFLKYCSLAATSLGLGAGMAPKIAWALENKPRIPVVWIHGL WP_001058323.1
MULTISPECIES: hydrogenase-1 small chain [Proteobacteria] Length: 372\nNumber of predicted TMHs: 2\nExp number of AAs in TMHs: 37.2608499999999\nExp number, first 60 AAs: 19.52923\nTotal prob of N-in: 0.94351\nPOSSIBLE N-term signal sequence\ninside 1 19\nTMhelix 20 39\noutside 40 325\nTMhelix 326 348\ninside 349 372

8998 GCF_001911935.1_ASM191193v1 Escherichia coli Proteobacteria; Gammaproteobacteria; Enterobacterales; Enterobacteriaceae; Escherichia
MNNEETFYQAMRRQGVTRRSFLKYCSLAATSLGLGAGMAPKIAWALENKPRIPVVWIHGL WP_001058323.1
MULTISPECIES: hydrogenase-1 small chain [Proteobacteria] Length: 372\nNumber of predicted TMHs: 2\nExp number of AAs in TMHs: 37.2608499999999\nExp number, first 60 AAs: 19.52923\nTotal prob of N-in: 0.94351\nPOSSIBLE N-term signal sequence\ninside 1 19\nTMhelix 20 39\noutside 40 325\nTMhelix 326 348\ninside 349 372

8999 GCF_001912005.1_ASM191200v1 Escherichia coli Proteobacteria; Gammaproteobacteria; Enterobacterales; Enterobacteriaceae; Escherichia
MNNEETFYQAMRRQGVTRRSFLKYCSLAATSLGLGAGMAPKIAWALENKPRIPVVWIHGL WP_001058323.1
MULTISPECIES: hydrogenase-1 small chain [Proteobacteria] Length: 372\nNumber of predicted TMHs: 2\nExp number of AAs in TMHs: 37.2608499999999\nExp number, first 60 AAs: 19.52923\nTotal prob of N-in: 0.94351\nPOSSIBLE N-term signal sequence\ninside 1 19\nTMhelix 20 39\noutside 40 325\nTMhelix 326 348\ninside 349 372

9000 GCF_001912145.1_ASM191214v1 Escherichia coli Proteobacteria; Gammaproteobacteria; Enterobacterales; Enterobacteriaceae; Escherichia
MTGDNTLIHSHGINRRDFMKLCAALAATMGLSSKAAAEMAESVTNPQRPPVIWIGAQECT WP_000145410.1
MULTISPECIES: hydrogenase-2 small chain [Proteobacteria] Length: 372\nNumber of predicted TMHs: 1\nExp number of AAs in TMHs: 23.72226\nExp number, first 60 AAs: 0.22038\nTotal prob of N-in: 0.12994\noutside 1 331\nTMhelix 332 354\ninside 355 372

9001 GCF_001912015.1_ASM191201v1 Escherichia coli Proteobacteria; Gammaproteobacteria; Enterobacterales; Enterobacteriaceae; Escherichia
MNNEETFYQAMRRQGVTRRSFLKYCSLAATSLGLGAGMAPKIAWALENKPRIPVVWIHGL WP_001488656.1
hydrogenase-1 small chain [Escherichia coli] Length: 372\nNumber of predicted TMHs: 2\nExp number of AAs in TMHs: 37.0420999999999\nExp number, first 60 AAs: 19.52951\nTotal prob of N-in: 0.94339\nPOSSIBLE N-term signal sequence\ninside 1 19\nTMhelix 20 39\noutside 40 325\nTMhelix 326 348\ninside 349 372

9002 GCF_001912185.1_ASM191218v1 Escherichia coli Proteobacteria; Gammaproteobacteria; Enterobacterales; Enterobacteriaceae; Escherichia
MTGDNTLIHSHGINRRDFMKLCAALAATMGLSSKAAAEMAESVTNPQRPPVIWIGAQECT WP_000145410.1
MULTISPECIES: hydrogenase-2 small chain [Proteobacteria] Length: 372\nNumber of predicted TMHs: 1\nExp number of AAs in TMHs: 23.72226\nExp number, first 60 AAs: 0.22038\nTotal prob of N-in: 0.12994\noutside 1 331\nTMhelix 332 354\ninside 355 372

9003 GCF_001912195.1_ASM191219v1 Escherichia coli Proteobacteria; Gammaproteobacteria; Enterobacterales; Enterobacteriaceae; Escherichia
MTGDNTLIHSHGINRRDFMKLCAALAATMGLSSKAAAEMAESVTNPQRPPVIWIGAQECT WP_000145410.1
MULTISPECIES: hydrogenase-2 small chain [Proteobacteria] Length: 372\nNumber of predicted TMHs: 1\nExp number of AAs in TMHs: 23.72226\nExp number, first 60 AAs: 0.22038\nTotal prob of N-in: 0.12994\noutside 1 331\nTMhelix 332 354\ninside 355 372

9004 GCF_001912015.1_ASM191201v1 Escherichia coli Proteobacteria; Gammaproteobacteria; Enterobacterales; Enterobacteriaceae; Escherichia
MTGDNTLIHSHGINRRDFMKLCAALAATMGLSSKAAAEMAESVTNPQRPPVIWIGAQECT WP_074534817.1
hydrogenase 2 small subunit [Escherichia coli] Length: 372\nNumber of predicted TMHs: 1\nExp number of AAs in TMHs: 23.7182\nExp number, first 60 AAs: 0.22037\nTotal prob of N-in: 0.12983\noutside 1 331\nTMhelix 332 354\ninside 355 372

9005 GCF_001912085.1_ASM191208v1 Escherichia coli Proteobacteria; Gammaproteobacteria; Enterobacterales; Enterobacteriaceae; Escherichia
MNNEETFYQAMRRQGVTRRSFLKYCSLAATSLGLGAGMAPKIAWALENKPRIPVVWIHGL WP_001058320.1
hydrogenase [Escherichia coli] Length: 372\nNumber of predicted TMHs: 2\nExp number of AAs in TMHs: 37.26082\nExp number, first 60 AAs: 19.52923\nTotal prob of N-in: 0.94351\nPOSSIBLE N-term signal sequence\ninside 1 19\nTMhelix 20 39\noutside 40 325\nTMhelix 326 348\ninside 349 372

9006 GCF_001912145.1_ASM191214v1 Escherichia coli Proteobacteria; Gammaproteobacteria; Enterobacterales; Enterobacteriaceae; Escherichia
MNNEETFYQAMRRQGVTRRSFLKYCSLAATSLGLGAGMAPKIAWALENKPRIPVVWIHGL WP_001058323.1
MULTISPECIES: hydrogenase-1 small chain [Proteobacteria] Length: 372\nNumber of predicted TMHs: 2\nExp number of AAs in TMHs: 37.2608499999999\nExp number, first 60 AAs: 19.52923\nTotal prob of N-in: 0.94351\nPOSSIBLE N-term signal sequence\ninside 1 19\nTMhelix 20 39\noutside 40 325\nTMhelix 326 348\ninside 349 372

9007 GCF_001912185.1_ASM191218v1 Escherichia coli Proteobacteria; Gammaproteobacteria; Enterobacterales; Enterobacteriaceae; Escherichia
MNNEETFYQAMRRQGVTRRSFLKYCSLAATSLGLGAGMAPKIAWALENKPRIPVVWIHGL WP_001058320.1

hydrogenase [Escherichia coli]	Length: 372\nNumber of predicted TMHs: 2\nExp number of AAs in TMHs: 37.26082\nExp number, first 60 AAs: 19.52923\nTotal prob of N-in: 0.94351\nPOSSIBLE N-term signal sequence\ninside 1 19\nTMhelix 20 39\noutside 40 325\nTMhelix 326 348\ninside 349 372

9008	GCF_001912195.1_ASM191219v1	Escherichia coli	Proteobacteria; Gammaproteobacteria; Enterobacterales; Enterobacteriaceae; Escherichia	MNNEETFYQAMRRQGVTRRSFLKYCSLAATSLGLGAGMAPKIAWALENKPRIPVVWIHGL	WP_001058323.1	MULTISPECIES: hydrogenase-1 small chain [Proteobacteria]	Length: 372\nNumber of predicted TMHs: 2\nExp number of AAs in TMHs: 37.2608499999999\nExp number, first 60 AAs: 19.52923\nTotal prob of N-in: 0.94351\nPOSSIBLE N-term signal sequence\ninside 1 19\nTMhelix 20 39\noutside 40 325\nTMhelix 326 348\ninside 349 372

9009	GCF_001912265.1_ASM191226v1	Escherichia coli	Proteobacteria; Gammaproteobacteria; Enterobacterales; Enterobacteriaceae; Escherichia	MTGDNTLIHSHGINRRDFMKLCAALAATMGLSSKAAAEMAESVTNPQRPPVIWIGAQECT	WP_000145410.1	MULTISPECIES: hydrogenase-2 small chain [Proteobacteria]	Length: 372\nNumber of predicted TMHs: 1\nExp number of AAs in TMHs: 23.72226\nExp number, first 60 AAs: 0.22038\nTotal prob of N-in: 0.12994\noutside 1 331\nTMhelix 332 354\ninside 355 372

9010	GCF_001912275.1_ASM191227v1	Escherichia coli	Proteobacteria; Gammaproteobacteria; Enterobacterales; Enterobacteriaceae; Escherichia	MTGDNTLIHSHGINRRDFMKLCAALAATMGLSSKAAAEMAESVTNPQRPPVIWIGAQECT	WP_000145410.1	MULTISPECIES: hydrogenase-2 small chain [Proteobacteria]	Length: 372\nNumber of predicted TMHs: 1\nExp number of AAs in TMHs: 23.72226\nExp number, first 60 AAs: 0.22038\nTotal prob of N-in: 0.12994\noutside 1 331\nTMhelix 332 354\ninside 355 372

9011	GCF_001912305.1_ASM191230v1	Escherichia coli	Proteobacteria; Gammaproteobacteria; Enterobacterales; Enterobacteriaceae; Escherichia	MTGDNTLIHSHGINRRDFMKLCAALAATMGLSSKAAAEMAESVTNPQRPPVIWIGAQECT	WP_000145410.1	MULTISPECIES: hydrogenase-2 small chain [Proteobacteria]	Length: 372\nNumber of predicted TMHs: 1\nExp number of AAs in TMHs: 23.72226\nExp number, first 60 AAs: 0.22038\nTotal prob of N-in: 0.12994\noutside 1 331\nTMhelix 332 354\ninside 355 372

9012	GCF_001912315.1_ASM191231v1	Escherichia coli	Proteobacteria; Gammaproteobacteria; Enterobacterales; Enterobacteriaceae; Escherichia	MTGDNTLIHSHGINRRDFMKLCAALAATMGLSSKAAAEMAESVTNPQRPPVIWIGAQECT	WP_000145410.1	MULTISPECIES: hydrogenase-2 small chain [Proteobacteria]	Length: 372\nNumber of predicted TMHs: 1\nExp number of AAs in TMHs: 23.72226\nExp number, first 60 AAs: 0.22038\nTotal prob of N-in: 0.12994\noutside 1 331\nTMhelix 332 354\ninside 355 372

9013	GCF_001912265.1_ASM191226v1	Escherichia coli	Proteobacteria; Gammaproteobacteria; Enterobacterales; Enterobacteriaceae; Escherichia	MNNEETFYQAMRRQGVTRRSFLKYCSLAATSLGLGAGMAPKIAWALENKPRIPVVWIHGL	WP_001295940.1	MULTISPECIES: hydrogenase [Enterobacteriaceae]	Length: 372\nNumber of predicted TMHs: 2\nExp number of AAs in TMHs: 37.2608899999999\nExp number, first 60 AAs: 19.52923\nTotal prob of N-in: 0.94351\nPOSSIBLE N-term signal sequence\ninside 1 19\nTMhelix 20 39\noutside 40 325\nTMhelix 326 348\ninside 349 372

9014	GCF_001912385.1_ASM191238v1	Escherichia coli	Proteobacteria; Gammaproteobacteria; Enterobacterales; Enterobacteriaceae; Escherichia	MTGDNTLIHSHGINRRDFMKLCAALAATMGLSSKAAAEMAESVTNPQRPPVIWIGAQECT	WP_000145410.1	MULTISPECIES: hydrogenase-2 small chain [Proteobacteria]	Length: 372\nNumber of predicted TMHs: 1\nExp number of AAs in TMHs: 23.72226\nExp number, first 60 AAs: 0.22038\nTotal prob of N-in: 0.12994\noutside 1 331\nTMhelix 332 354\ninside 355 372

9015	GCF_001912385.1_ASM191238v1	Escherichia coli	Proteobacteria; Gammaproteobacteria; Enterobacterales; Enterobacteriaceae; Escherichia	MNNEETFYQAMRRQGVTRRSFLKYCSLAATSLGLGAGMAPKIAWALENKPRIPVVWIHGL	WP_001353311.1	MULTISPECIES: hydrogenase [Escherichia]	Length: 372\nNumber of predicted TMHs: 2\nExp number of AAs in TMHs: 37.2638699999999\nExp number, first 60 AAs: 19.52924\nTotal prob of N-in: 0.94351\nPOSSIBLE N-term signal sequence\ninside 1 19\nTMhelix 20 39\noutside 40 325\nTMhelix 326 348\ninside 349 372

9016 GCF_001912315.1_ASM191231v1 Escherichia coli Proteobacteria; Gammaproteobacteria; Enterobacterales; Enterobacteriaceae; Escherichia
MNNEETFYQAMRRQGVTRRSFLKYCSLAATSLGLGAGMAPKIAWALENKPRIPVVWIHGL WP_001058323.1
MULTISPECIES: hydrogenase-1 small chain [Proteobacteria] Length: 372\nNumber of predicted TMHs: 2\nExp number of AAs in TMHs: 37.2608499999999\nExp number, first 60 AAs: 19.52923\nTotal prob of N-in: 0.94351\nPOSSIBLE N-term signal sequence\ninside 1 19\nTMhelix 20 39\noutside 40 325\nTMhelix 326 348\ninside 349 372

9017 GCF_001912425.1_ASM191242v1 Escherichia coli Proteobacteria; Gammaproteobacteria; Enterobacterales; Enterobacteriaceae; Escherichia
MTGDNTLIHSHGINRRDFMKLCAALAATMGLSSKAAAEMAESVTNPQRPPVIWIGAQECT WP_000145410.1
MULTISPECIES: hydrogenase-2 small chain [Proteobacteria] Length: 372\nNumber of predicted TMHs: 1\nExp number of AAs in TMHs: 23.72226\nExp number, first 60 AAs: 0.22038\nTotal prob of N-in: 0.12994\noutside 1 331\nTMhelix 332 354\ninside 355 372

9018 GCF_001912445.1_ASM191244v1 Escherichia coli Proteobacteria; Gammaproteobacteria; Enterobacterales; Enterobacteriaceae; Escherichia
MTGDNTLIHSHGINRRDFMKLCAALAATMGLSSKAAAEMAESVTNPQRPPVIWIGAQECT WP_000145410.1
MULTISPECIES: hydrogenase-2 small chain [Proteobacteria] Length: 372\nNumber of predicted TMHs: 1\nExp number of AAs in TMHs: 23.72226\nExp number, first 60 AAs: 0.22038\nTotal prob of N-in: 0.12994\noutside 1 331\nTMhelix 332 354\ninside 355 372

9019 GCF_001912425.1_ASM191242v1 Escherichia coli Proteobacteria; Gammaproteobacteria; Enterobacterales; Enterobacteriaceae; Escherichia
MNNEETFYQAMRRQGVTRRSFLKYCSLAATSLGLGAGMAPKIAWALENKPRIPVVWIHGL WP_001058323.1
MULTISPECIES: hydrogenase-1 small chain [Proteobacteria] Length: 372\nNumber of predicted TMHs: 2\nExp number of AAs in TMHs: 37.2608499999999\nExp number, first 60 AAs: 19.52923\nTotal prob of N-in: 0.94351\nPOSSIBLE N-term signal sequence\ninside 1 19\nTMhelix 20 39\noutside 40 325\nTMhelix 326 348\ninside 349 372

9020 GCF_001912555.1_ASM191255v1 Escherichia coli Proteobacteria; Gammaproteobacteria; Enterobacterales; Enterobacteriaceae; Escherichia
MTGDNTLIHSHGINRRDFMKLCAALAATMGLSSKAAAEMAESVTNPQRPPVIWIGAQECT WP_000145410.1
MULTISPECIES: hydrogenase-2 small chain [Proteobacteria] Length: 372\nNumber of predicted TMHs: 1\nExp number of AAs in TMHs: 23.72226\nExp number, first 60 AAs: 0.22038\nTotal prob of N-in: 0.12994\noutside 1 331\nTMhelix 332 354\ninside 355 372

9021 GCF_001912475.1_ASM191247v1 Escherichia coli Proteobacteria; Gammaproteobacteria; Enterobacterales; Enterobacteriaceae; Escherichia
MNNEETFYQAMRRQGVTRRSFLKYCSLAATSLGLGAGMAPKIAWALENKPRIPVVWIHGL WP_001058323.1
MULTISPECIES: hydrogenase-1 small chain [Proteobacteria] Length: 372\nNumber of predicted TMHs: 2\nExp number of AAs in TMHs: 37.2608499999999\nExp number, first 60 AAs: 19.52923\nTotal prob of N-in: 0.94351\nPOSSIBLE N-term signal sequence\ninside 1 19\nTMhelix 20 39\noutside 40 325\nTMhelix 326 348\ninside 349 372

9022 GCF_001912605.1_ASM191260v1 Escherichia coli Proteobacteria; Gammaproteobacteria; Enterobacterales; Enterobacteriaceae; Escherichia
MTGDNTLIHSHGINRRDFMKLCAALAATMGLSSKAAAEMAESVTNPQRPPVIWIGAQECT WP_000145410.1
MULTISPECIES: hydrogenase-2 small chain [Proteobacteria] Length: 372\nNumber of predicted TMHs: 1\nExp number of AAs in TMHs: 23.72226\nExp number, first 60 AAs: 0.22038\nTotal prob of N-in: 0.12994\noutside 1 331\nTMhelix 332 354\ninside 355 372

9023 GCF_001912565.1_ASM191256v1 Escherichia coli Proteobacteria; Gammaproteobacteria; Enterobacterales; Enterobacteriaceae; Escherichia
MTGDNTLIHSHGINRRDFMKLCAALAATMGLSSKAAAEMAESVTNPQRPPVIWIGAQECT WP_000145410.1
MULTISPECIES: hydrogenase-2 small chain [Proteobacteria] Length: 372\nNumber of predicted TMHs: 1\nExp number of AAs in TMHs: 23.72226\nExp number, first 60 AAs: 0.22038\nTotal prob of N-in: 0.12994\noutside 1 331\nTMhelix 332 354\ninside 355 372

9024 GCF_001912615.1_ASM191261v1 Escherichia coli Proteobacteria; Gammaproteobacteria; Enterobacterales; Enterobacteriaceae; Escherichia

MTGDNTLIHSHGINRRDFMKLCAALAATMGLSSKAAAEMAESVTNPQRPPVIWIGAQECT WP_000145410.1 MULTISPECIES: hydrogenase-2 small chain [Proteobacteria] Length: 372\nNumber of predicted TMHs: 1\nExp number of AAs in TMHs: 23.72226\nExp number, first 60 AAs: 0.22038\nTotal prob of N-in: 0.12994\noutside 1 331\nTMhelix 332 354\ninside 355 372

9025 GCF_001912605.1_ASM191260v1 Escherichia coli Proteobacteria; Gammaproteobacteria; Enterobacterales; Enterobacteriaceae; Escherichia MNNEETFYQAMRRQGVTRRSFLKYCSLAATSLGLGAGMAPKIAWALENKPRIPVVWIHGL WP_001058323.1 MULTISPECIES: hydrogenase-1 small chain [Proteobacteria] Length: 372\nNumber of predicted TMHs: 2\nExp number of AAs in TMHs: 37.2608499999999\nExp number, first 60 AAs: 19.52923\nTotal prob of N-in: 0.94351\nPOSSIBLE N-term signal sequence\ninside 1 19\nTMhelix 20 39\noutside 40 325\nTMhelix 326 348\ninside 349 372

9026 GCF_001912615.1_ASM191261v1 Escherichia coli Proteobacteria; Gammaproteobacteria; Enterobacterales; Enterobacteriaceae; Escherichia MNNEETFYQAMRRQGVTRRSFLKYCSLAATSLGLGAGMAPKIAWALENKPRIPVVWIHGL WP_001058323.1 MULTISPECIES: hydrogenase-1 small chain [Proteobacteria] Length: 372\nNumber of predicted TMHs: 2\nExp number of AAs in TMHs: 37.2608499999999\nExp number, first 60 AAs: 19.52923\nTotal prob of N-in: 0.94351\nPOSSIBLE N-term signal sequence\ninside 1 19\nTMhelix 20 39\noutside 40 325\nTMhelix 326 348\ninside 349 372

9027 GCF_001912705.1_ASM191270v1 Escherichia coli Proteobacteria; Gammaproteobacteria; Enterobacterales; Enterobacteriaceae; Escherichia MTGDNTLIHSHGINRRDFMKLCAALAATMGLSSKAAAEMAESVTNPQRPPVIWIGAQECT WP_000145410.1 MULTISPECIES: hydrogenase-2 small chain [Proteobacteria] Length: 372\nNumber of predicted TMHs: 1\nExp number of AAs in TMHs: 23.72226\nExp number, first 60 AAs: 0.22038\nTotal prob of N-in: 0.12994\noutside 1 331\nTMhelix 332 354\ninside 355 372

9028 GCF_001912545.1_ASM191254v1 Escherichia coli Proteobacteria; Gammaproteobacteria; Enterobacterales; Enterobacteriaceae; Escherichia MNNEETFYQAMRRQGVTRRSFLKYCSLAATSLGLGAGMAPNIAWALENKPRIPVVWIHGL WP_074523463.1 hydrogenase [Escherichia coli] Length: 372\nNumber of predicted TMHs: 2\nExp number of AAs in TMHs: 39.56696\nExp number, first 60 AAs: 21.8441\nTotal prob of N-in: 0.97537\nPOSSIBLE N-term signal sequence\ninside 1 20\nTMhelix 21 43\noutside 44 325\nTMhelix 326 348\ninside 349 372

9029 GCF_001912545.1_ASM191254v1 Escherichia coli Proteobacteria; Gammaproteobacteria; Enterobacterales; Enterobacteriaceae; Escherichia MTGDNTLIHSHGINRRDFMKLCAALAATMGLSSKAAAEMAESVTNPQRPPVIWIGAQECT WP_074523761.1 hydrogenase 2 small subunit [Escherichia coli] Length: 372\nNumber of predicted TMHs: 1\nExp number of AAs in TMHs: 23.72224\nExp number, first 60 AAs: 0.22037\nTotal prob of N-in: 0.12994\noutside 1 331\nTMhelix 332 354\ninside 355 372

9030 GCF_001912675.1_ASM191267v1 Escherichia coli Proteobacteria; Gammaproteobacteria; Enterobacterales; Enterobacteriaceae; Escherichia MNNEETFYQAMRRQGVTRRSFLKYCSLAATSLGLGAGMAPKIAWALENKPRIPVVWIHGL WP_001058323.1 MULTISPECIES: hydrogenase-1 small chain [Proteobacteria] Length: 372\nNumber of predicted TMHs: 2\nExp number of AAs in TMHs: 37.2608499999999\nExp number, first 60 AAs: 19.52923\nTotal prob of N-in: 0.94351\nPOSSIBLE N-term signal sequence\ninside 1 19\nTMhelix 20 39\noutside 40 325\nTMhelix 326 348\ninside 349 372

9031 GCF_001931785.1_ASM193178v1 Escherichia coli Proteobacteria; Gammaproteobacteria; Enterobacterales; Enterobacteriaceae; Escherichia MTGDNTLIHSHGINRRDFMKLCAALAATMGLSSKAAAEMAESVTNPQRPPVIWIGAQECT WP_000145410.1 MULTISPECIES: hydrogenase-2 small chain [Proteobacteria] Length: 372\nNumber of predicted TMHs: 1\nExp number of AAs in TMHs: 23.72226\nExp number, first 60 AAs: 0.22038\nTotal prob of N-in: 0.12994\noutside 1 331\nTMhelix 332 354\ninside 355 372

9032 GCF_001931785.1_ASM193178v1 Escherichia coli Proteobacteria; Gammaproteobacteria; Enterobacterales; Enterobacteriaceae; Escherichia MNNEETFYQAMRRQGVTRRSFLKYCSLAATSLGLGAGMAPKIAWALENKPRIPVVWIHGL WP_001058323.1 MULTISPECIES: hydrogenase-1 small chain [Proteobacteria] Length: 372\nNumber of predicted TMHs:

2\nExp number of AAs in TMHs: 37.2608499999999\nExp number, first 60 AAs: 19.52923\nTotal prob of N-in: 0.94351\nPOSSIBLE N-term signal sequence\ninside 1 19\nTMhelix 20 39\noutside 40 325\nTMhelix 326 348\ninside 349 372

9033 GCF_001912665.1_ASM191266v1 Escherichia coli Proteobacteria; Gammaproteobacteria; Enterobacterales; Enterobacteriaceae; Escherichia MNNEETFYQAMRRQGVTRRSFLKYCSLAATSLGLGAGMAPKIAWALENKPRIPVVWIHGL WP_001058323.1 MULTISPECIES: hydrogenase-1 small chain [Proteobacteria] Length: 372\nNumber of predicted TMHs: 2\nExp number of AAs in TMHs: 37.2608499999999\nExp number, first 60 AAs: 19.52923\nTotal prob of N-in: 0.94351\nPOSSIBLE N-term signal sequence\ninside 1 19\nTMhelix 20 39\noutside 40 325\nTMhelix 326 348\ninside 349 372

9034 GCF_001936315.1_ASM193631v1 Escherichia coli Proteobacteria; Gammaproteobacteria; Enterobacterales; Enterobacteriaceae; Escherichia MNNEETFYQAMRRQGVTRRSFLKYCSLAATSLGLGAGMAPKIAWALENKPRIPVVWIHGL WP_001058323.1 MULTISPECIES: hydrogenase-1 small chain [Proteobacteria] Length: 372\nNumber of predicted TMHs: 2\nExp number of AAs in TMHs: 37.2608499999999\nExp number, first 60 AAs: 19.52923\nTotal prob of N-in: 0.94351\nPOSSIBLE N-term signal sequence\ninside 1 19\nTMhelix 20 39\noutside 40 325\nTMhelix 326 348\ninside 349 372

9035 GCF_001932515.1_ASM193251v1 Escherichia coli Proteobacteria; Gammaproteobacteria; Enterobacterales; Enterobacteriaceae; Escherichia MNNEETFYQAMRRQGVTRRSFLKYCSLAATSLGLGAGMAPKIAWALENKPRIPVVWIHGL WP_001058323.1 MULTISPECIES: hydrogenase-1 small chain [Proteobacteria] Length: 372\nNumber of predicted TMHs: 2\nExp number of AAs in TMHs: 37.2608499999999\nExp number, first 60 AAs: 19.52923\nTotal prob of N-in: 0.94351\nPOSSIBLE N-term signal sequence\ninside 1 19\nTMhelix 20 39\noutside 40 325\nTMhelix 326 348\ninside 349 372

9036 GCF_001938625.2_ASM193862v2 Escherichia coli Proteobacteria; Gammaproteobacteria; Enterobacterales; Enterobacteriaceae; Escherichia MNNEETFYQAMRRQGVTRRSFLKYCSLAATSLGLGAGMAPKIAWALENKPRIPVVWIHGL WP_001058323.1 MULTISPECIES: hydrogenase-1 small chain [Proteobacteria] Length: 372\nNumber of predicted TMHs: 2\nExp number of AAs in TMHs: 37.2608499999999\nExp number, first 60 AAs: 19.52923\nTotal prob of N-in: 0.94351\nPOSSIBLE N-term signal sequence\ninside 1 19\nTMhelix 20 39\noutside 40 325\nTMhelix 326 348\ninside 349 372

9037 GCF_001937985.1_ASM193798v1 Escherichia coli Proteobacteria; Gammaproteobacteria; Enterobacterales; Enterobacteriaceae; Escherichia MNNEETFYQAMRRQGVTRRSFLKYCSLAATSLGLGAGMAPKIAWALENKPRIPVVWIHGL WP_001058323.1 MULTISPECIES: hydrogenase-1 small chain [Proteobacteria] Length: 372\nNumber of predicted TMHs: 2\nExp number of AAs in TMHs: 37.2608499999999\nExp number, first 60 AAs: 19.52923\nTotal prob of N-in: 0.94351\nPOSSIBLE N-term signal sequence\ninside 1 19\nTMhelix 20 39\noutside 40 325\nTMhelix 326 348\ninside 349 372

9038 GCF_001937255.1_ASM193725v1 Escherichia coli Proteobacteria; Gammaproteobacteria; Enterobacterales; Enterobacteriaceae; Escherichia MNNEETFYQAMRRQGVTRRSFLKYCSLAATSLGLGAGMAPKIAWALENKPRIPVVWIHGL WP_001058323.1 MULTISPECIES: hydrogenase-1 small chain [Proteobacteria] Length: 372\nNumber of predicted TMHs: 2\nExp number of AAs in TMHs: 37.2608499999999\nExp number, first 60 AAs: 19.52923\nTotal prob of N-in: 0.94351\nPOSSIBLE N-term signal sequence\ninside 1 19\nTMhelix 20 39\noutside 40 325\nTMhelix 326 348\ninside 349 372

9039 GCF_001942065.1_ASM194206v1 Escherichia coli Proteobacteria; Gammaproteobacteria; Enterobacterales; Enterobacteriaceae; Escherichia MNNEETFYQAMRRQGVTRRSFLKYCSLAATSLGLGAGMAPKIAWALENKPRIPVVWIHGL WP_001058323.1 MULTISPECIES: hydrogenase-1 small chain [Proteobacteria] Length: 372\nNumber of predicted TMHs: 2\nExp number of AAs in TMHs: 37.2608499999999\nExp number, first 60 AAs: 19.52923\nTotal prob of N-in: 0.94351\nPOSSIBLE N-term signal sequence\ninside 1 19\nTMhelix 20 39\noutside 40 325\nTMhelix 326 348\ninside 349 372

9040 GCF_001940485.1_ASM194048v1 Escherichia coli Proteobacteria; Gammaproteobacteria; Enterobacterales; Enterobacteriaceae; Escherichia
MNNEETFYQAMRRQGVTRRSFLKYCSLAATSLGLGAGMAPKIAWALENKPRIPVVWIHGL WP_001058323.1
MULTISPECIES: hydrogenase-1 small chain [Proteobacteria] Length: 372\nNumber of predicted TMHs: 2\nExp number of AAs in TMHs: 37.2608499999999\nExp number, first 60 AAs: 19.52923\nTotal prob of N-in: 0.94351\nPOSSIBLE N-term signal sequence\ninside 1 19\nTMhelix 20 39\noutside 40 325\nTMhelix 326 348\ninside 349 372

9041 GCF_001942145.1_ASM194214v1 Escherichia coli Proteobacteria; Gammaproteobacteria; Enterobacterales; Enterobacteriaceae; Escherichia
MTGDNTLIHSHGINRRDFMKLCAALAATMGLSSKAAAEMAESVTNPQRPPVIWIGAQECT WP_000145410.1
MULTISPECIES: hydrogenase-2 small chain [Proteobacteria] Length: 372\nNumber of predicted TMHs: 1\nExp number of AAs in TMHs: 23.72226\nExp number, first 60 AAs: 0.22038\nTotal prob of N-in: 0.12994\noutside 1 331\nTMhelix 332 354\ninside 355 372

9042 GCF_001942165.1_ASM194216v1 Escherichia coli Proteobacteria; Gammaproteobacteria; Enterobacterales; Enterobacteriaceae; Escherichia
MTGDNTLIHSHGINRRDFMKLCAALAATMGLSSKAAAEMAESVTNPQRPPVIWIGAQECT WP_000145410.1
MULTISPECIES: hydrogenase-2 small chain [Proteobacteria] Length: 372\nNumber of predicted TMHs: 1\nExp number of AAs in TMHs: 23.72226\nExp number, first 60 AAs: 0.22038\nTotal prob of N-in: 0.12994\noutside 1 331\nTMhelix 332 354\ninside 355 372

9043 GCF_001942145.1_ASM194214v1 Escherichia coli Proteobacteria; Gammaproteobacteria; Enterobacterales; Enterobacteriaceae; Escherichia
MNNEETFYQAMRRQGVTRRSFLKYCSLAATSLGLGAGMAPKIAWALENKPRIPVVWIHGL WP_001058323.1
MULTISPECIES: hydrogenase-1 small chain [Proteobacteria] Length: 372\nNumber of predicted TMHs: 2\nExp number of AAs in TMHs: 37.2608499999999\nExp number, first 60 AAs: 19.52923\nTotal prob of N-in: 0.94351\nPOSSIBLE N-term signal sequence\ninside 1 19\nTMhelix 20 39\noutside 40 325\nTMhelix 326 348\ninside 349 372

9044 GCF_001942165.1_ASM194216v1 Escherichia coli Proteobacteria; Gammaproteobacteria; Enterobacterales; Enterobacteriaceae; Escherichia
MNNEETFYQAMRRQGVTRRSFLKYCSLAATSLGLGAGMAPKIAWALENKPRIPVVWIHGL WP_001058323.1
MULTISPECIES: hydrogenase-1 small chain [Proteobacteria] Length: 372\nNumber of predicted TMHs: 2\nExp number of AAs in TMHs: 37.2608499999999\nExp number, first 60 AAs: 19.52923\nTotal prob of N-in: 0.94351\nPOSSIBLE N-term signal sequence\ninside 1 19\nTMhelix 20 39\noutside 40 325\nTMhelix 326 348\ninside 349 372

9045 GCF_001950735.1_ASM195073v1 Escherichia coli Proteobacteria; Gammaproteobacteria; Enterobacterales; Enterobacteriaceae; Escherichia
MTGDNTLIHSHGINRRDFMKLCAALAATMGLSSKAAAEMAESVTNPQRPPVIWIGAQECT WP_000145410.1
MULTISPECIES: hydrogenase-2 small chain [Proteobacteria] Length: 372\nNumber of predicted TMHs: 1\nExp number of AAs in TMHs: 23.72226\nExp number, first 60 AAs: 0.22038\nTotal prob of N-in: 0.12994\noutside 1 331\nTMhelix 332 354\ninside 355 372

9046 GCF_001950755.1_ASM195075v1 Escherichia coli Proteobacteria; Gammaproteobacteria; Enterobacterales; Enterobacteriaceae; Escherichia
MTGDNTLIHSHGINRRDFMKLCAALAATMGLSSKAAAEMAESVTNPQRPPVIWIGAQECT WP_000145410.1
MULTISPECIES: hydrogenase-2 small chain [Proteobacteria] Length: 372\nNumber of predicted TMHs: 1\nExp number of AAs in TMHs: 23.72226\nExp number, first 60 AAs: 0.22038\nTotal prob of N-in: 0.12994\noutside 1 331\nTMhelix 332 354\ninside 355 372

9047 GCF_001950775.1_ASM195077v1 Escherichia coli Proteobacteria; Gammaproteobacteria; Enterobacterales; Enterobacteriaceae; Escherichia
MTGDNTLIHSHGINRRDFMKLCAALAATMGLSSKAAAEMAESVTNPQRPPVIWIGAQECT WP_000145410.1
MULTISPECIES: hydrogenase-2 small chain [Proteobacteria] Length: 372\nNumber of predicted TMHs: 1\nExp number of AAs in TMHs: 23.72226\nExp number, first 60 AAs: 0.22038\nTotal prob of N-in: 0.12994\noutside 1 331\nTMhelix 332 354\ninside 355 372

9048 GCF_001950715.1_ASM195071v1 Escherichia coli Proteobacteria; Gammaproteobacteria; Enterobacterales; Enterobacteriaceae; Escherichia

MNNEETFYQAMRRQGVTRRSFLKYCSLAATSLGLGAGMAPKIAWALENKPRIPVVWIHGL WP_001058323.1 MULTISPECIES: hydrogenase-1 small chain [Proteobacteria] Length: 372\nNumber of predicted TMHs: 2\nExp number of AAs in TMHs: 37.2608499999999\nExp number, first 60 AAs: 19.52923\nTotal prob of N-in: 0.94351\nPOSSIBLE N-term signal sequence\ninside 1 19\nTMhelix 20 39\noutside 40 325\nTMhelix 326 348\ninside 349 372

9049 GCF_001950755.1_ASM195075v1 Escherichia coli Proteobacteria; Gammaproteobacteria; Enterobacterales; Enterobacteriaceae; Escherichia MNNEETFYQAMRRQGVTRRSFLKYCSLAATSLGLGAGMAPKIAWALENKPRIPVVWIHGL WP_001058323.1 MULTISPECIES: hydrogenase-1 small chain [Proteobacteria] Length: 372\nNumber of predicted TMHs: 2\nExp number of AAs in TMHs: 37.2608499999999\nExp number, first 60 AAs: 19.52923\nTotal prob of N-in: 0.94351\nPOSSIBLE N-term signal sequence\ninside 1 19\nTMhelix 20 39\noutside 40 325\nTMhelix 326 348\ninside 349 372

9050 GCF_001957215.1_ASM195721v1 Escherichia coli Proteobacteria; Gammaproteobacteria; Enterobacterales; Enterobacteriaceae; Escherichia MTGDNTLIHSHGINRRDFMKLCAALAATMGLSSKAAAEMAESVTNPQRPPVIWIGAQECT WP_000145410.1 MULTISPECIES: hydrogenase-2 small chain [Proteobacteria] Length: 372\nNumber of predicted TMHs: 1\nExp number of AAs in TMHs: 23.72226\nExp number, first 60 AAs: 0.22038\nTotal prob of N-in: 0.12994\noutside 1 331\nTMhelix 332 354\ninside 355 372

9051 GCF_001957205.1_ASM195720v1 Escherichia coli Proteobacteria; Gammaproteobacteria; Enterobacterales; Enterobacteriaceae; Escherichia MTGDNTLIHSHGINRRDFMKLCAALAATMGLSSKAAAEMAESVTNPQRPPVIWIGAQECT WP_032298447.1 hydrogenase 2 small subunit [Escherichia coli] Length: 372\nNumber of predicted TMHs: 1\nExp number of AAs in TMHs: 23.69807\nExp number, first 60 AAs: 0.221\nTotal prob of N-in: 0.12853\noutside 1 331\nTMhelix 332 354\ninside 355 372

9052 GCF_001969285.1_ASM196928v1 Escherichia coli Proteobacteria; Gammaproteobacteria; Enterobacterales; Enterobacteriaceae; Escherichia MNNEETFYQAMRRQGVTRRSFLKYCSLAATSLGLGAGMAPKIAWALENKPRIPVVWIHGL WP_063501981.1 hydrogenase [Escherichia coli] Length: 372\nNumber of predicted TMHs: 2\nExp number of AAs in TMHs: 37.06322\nExp number, first 60 AAs: 19.52932\nTotal prob of N-in: 0.94340\nPOSSIBLE N-term signal sequence\ninside 1 19\nTMhelix 20 39\noutside 40 325\nTMhelix 326 348\ninside 349 372

9053 GCF_001982505.1_ASM198250v1 Escherichia coli Proteobacteria; Gammaproteobacteria; Enterobacterales; Enterobacteriaceae; Escherichia MTGDNTLIHSHGINRRDFMKLCAALAATMGLSSKAAAEMAESVTNPQRPPVIWIGAQECT WP_000145410.1 MULTISPECIES: hydrogenase-2 small chain [Proteobacteria] Length: 372\nNumber of predicted TMHs: 1\nExp number of AAs in TMHs: 23.72226\nExp number, first 60 AAs: 0.22038\nTotal prob of N-in: 0.12994\noutside 1 331\nTMhelix 332 354\ninside 355 372

9054 GCF_001982555.1_ASM198255v1 Escherichia coli Proteobacteria; Gammaproteobacteria; Enterobacterales; Enterobacteriaceae; Escherichia MTGDNTLIHSHGINRRDFMKLCAALAATMGLSSKAAAEMAESVTNPQRPPVIWIGAQECT WP_000145410.1 MULTISPECIES: hydrogenase-2 small chain [Proteobacteria] Length: 372\nNumber of predicted TMHs: 1\nExp number of AAs in TMHs: 23.72226\nExp number, first 60 AAs: 0.22038\nTotal prob of N-in: 0.12994\noutside 1 331\nTMhelix 332 354\ninside 355 372

9055 GCF_001982555.1_ASM198255v1 Escherichia coli Proteobacteria; Gammaproteobacteria; Enterobacterales; Enterobacteriaceae; Escherichia MNNEETFYQAMRRQGVTRRSFLKYCSLAATSLGLGAGMAPKIAWALENKPRIPVVWIHGL WP_001058323.1 MULTISPECIES: hydrogenase-1 small chain [Proteobacteria] Length: 372\nNumber of predicted TMHs: 2\nExp number of AAs in TMHs: 37.2608499999999\nExp number, first 60 AAs: 19.52923\nTotal prob of N-in: 0.94351\nPOSSIBLE N-term signal sequence\ninside 1 19\nTMhelix 20 39\noutside 40 325\nTMhelix 326 348\ninside 349 372

9056 GCF_001982505.1_ASM198250v1 Escherichia coli Proteobacteria; Gammaproteobacteria; Enterobacterales; Enterobacteriaceae; Escherichia MNNEETFYQAMRRQGVTRRSFLKYCSLAATSLGLGAGMAPKIAWALENKPRIPVVWIHGL WP_001058323.1 MULTISPECIES: hydrogenase-1 small chain [Proteobacteria] Length: 372\nNumber of predicted TMHs:

2\nExp number of AAs in TMHs: 37.2608499999999\nExp number, first 60 AAs: 19.52923\nTotal prob of N-in: 0.94351\nPOSSIBLE N-term signal sequence\ninside 1 19\nTMhelix 20 39\noutside 40 325\nTMhelix 326 348\ninside 349 372

9057 GCF_001982585.1_ASM198258v1 Escherichia coli Proteobacteria; Gammaproteobacteria; Enterobacterales; Enterobacteriaceae; Escherichia MTGDNTLIHSHGINRRDFMKLCAALAATMGLSSKAAAEMAESVTNPQRPPVIWIGAQECT WP_000145410.1 MULTISPECIES: hydrogenase-2 small chain [Proteobacteria] Length: 372\nNumber of predicted TMHs: 1\nExp number of AAs in TMHs: 23.72226\nExp number, first 60 AAs: 0.22038\nTotal prob of N-in: 0.12994\noutside 1 331\nTMhelix 332 354\ninside 355 372

9058 GCF_001984845.1_ASM198484v1 Escherichia coli Proteobacteria; Gammaproteobacteria; Enterobacterales; Enterobacteriaceae; Escherichia MTGDNTLIHSHGINRRDFMKLCAALAATMGLSSKAAAEMAESVTNPQRPPVIWIGAQECT WP_000145410.1 MULTISPECIES: hydrogenase-2 small chain [Proteobacteria] Length: 372\nNumber of predicted TMHs: 1\nExp number of AAs in TMHs: 23.72226\nExp number, first 60 AAs: 0.22038\nTotal prob of N-in: 0.12994\noutside 1 331\nTMhelix 332 354\ninside 355 372

9059 GCF_001990925.1_ASM199092v1 Escherichia coli Proteobacteria; Gammaproteobacteria; Enterobacterales; Enterobacteriaceae; Escherichia MTGDNTLIHSHGINRRDFMKLCAALAATMGLSSKAAAEMAESVTNPQRPPVIWIGAQECT WP_000145410.1 MULTISPECIES: hydrogenase-2 small chain [Proteobacteria] Length: 372\nNumber of predicted TMHs: 1\nExp number of AAs in TMHs: 23.72226\nExp number, first 60 AAs: 0.22038\nTotal prob of N-in: 0.12994\noutside 1 331\nTMhelix 332 354\ninside 355 372

9060 GCF_001990875.1_ASM199087v1 Escherichia coli Proteobacteria; Gammaproteobacteria; Enterobacterales; Enterobacteriaceae; Escherichia MTGDNTLIHSHGINRRDFMKLCAALAATMGLSSKAAAEMAESVTNPQRPPVIWIGAQECT WP_000145410.1 MULTISPECIES: hydrogenase-2 small chain [Proteobacteria] Length: 372\nNumber of predicted TMHs: 1\nExp number of AAs in TMHs: 23.72226\nExp number, first 60 AAs: 0.22038\nTotal prob of N-in: 0.12994\noutside 1 331\nTMhelix 332 354\ninside 355 372

9061 GCF_001990855.1_ASM199085v1 Escherichia coli Proteobacteria; Gammaproteobacteria; Enterobacterales; Enterobacteriaceae; Escherichia MNNEETFYQAMRRQGVTRRSFLKYCSLAATSLGLGAGMAPKIAWALENKPRIPVVWIHGL WP_001058323.1 MULTISPECIES: hydrogenase-1 small chain [Proteobacteria] Length: 372\nNumber of predicted TMHs: 2\nExp number of AAs in TMHs: 37.2608499999999\nExp number, first 60 AAs: 19.52923\nTotal prob of N-in: 0.94351\nPOSSIBLE N-term signal sequence\ninside 1 19\nTMhelix 20 39\noutside 40 325\nTMhelix 326 348\ninside 349 372

9062 GCF_001990925.1_ASM199092v1 Escherichia coli Proteobacteria; Gammaproteobacteria; Enterobacterales; Enterobacteriaceae; Escherichia MNNEETFYQAMRRQGVTRRSFLKYCSLAATSLGLGAGMAPKIAWALENKPRIPVVWIHGL WP_001058323.1 MULTISPECIES: hydrogenase-1 small chain [Proteobacteria] Length: 372\nNumber of predicted TMHs: 2\nExp number of AAs in TMHs: 37.2608499999999\nExp number, first 60 AAs: 19.52923\nTotal prob of N-in: 0.94351\nPOSSIBLE N-term signal sequence\ninside 1 19\nTMhelix 20 39\noutside 40 325\nTMhelix 326 348\ninside 349 372

9063 GCF_001990545.1_ASM199054v1 Escherichia coli Proteobacteria; Gammaproteobacteria; Enterobacterales; Enterobacteriaceae; Escherichia MNNEETFYQAMRRQGVTRRSFLKYCSLAATSLGLGAGMAPKIAWALENKPRIPVVWIHGL WP_001058323.1 MULTISPECIES: hydrogenase-1 small chain [Proteobacteria] Length: 372\nNumber of predicted TMHs: 2\nExp number of AAs in TMHs: 37.2608499999999\nExp number, first 60 AAs: 19.52923\nTotal prob of N-in: 0.94351\nPOSSIBLE N-term signal sequence\ninside 1 19\nTMhelix 20 39\noutside 40 325\nTMhelix 326 348\ninside 349 372

9064 GCF_001990935.1_ASM199093v1 Escherichia coli Proteobacteria; Gammaproteobacteria; Enterobacterales; Enterobacteriaceae; Escherichia MTGDNTLIHSHGINRRDFMKLCAALAATMGLSSKAAAEMAESVTNPQRPPVIWIGAQECT WP_000145410.1 MULTISPECIES: hydrogenase-2 small chain [Proteobacteria] Length: 372\nNumber of predicted TMHs:

1\nExp number of AAs in TMHs: 23.72226\nExp number, first 60 AAs: 0.22038\nTotal prob of N-in: 0.12994\noutside 1 331\nTMhelix 332 354\ninside 355 372

9065 GCF_001990965.1_ASM199096v1 Escherichia coli Proteobacteria; Gammaproteobacteria; Enterobacterales; Enterobacteriaceae; Escherichia
MTGDNTLIHSHGINRRDFMKLCAALAATMGLSSKAAAEMAESVTNPQRPPVIWIGAQECT WP_000145410.1
MULTISPECIES: hydrogenase-2 small chain [Proteobacteria] Length: 372\nNumber of predicted TMHs: 1\nExp number of AAs in TMHs: 23.72226\nExp number, first 60 AAs: 0.22038\nTotal prob of N-in: 0.12994\noutside 1 331\nTMhelix 332 354\ninside 355 372

9066 GCF_001990845.1_ASM199084v1 Escherichia coli Proteobacteria; Gammaproteobacteria; Enterobacterales; Enterobacteriaceae; Escherichia
MNNEETFYQAMRRQGVTRRSFLKYCSLAATSLGLGAGMAPKIAWALENKPRIPVVWIHGL WP_001295940.1
MULTISPECIES: hydrogenase [Enterobacteriaceae] Length: 372\nNumber of predicted TMHs: 2\nExp number of AAs in TMHs: 37.2608899999999\nExp number, first 60 AAs: 19.52923\nTotal prob of N-in: 0.94351\nPOSSIBLE N-term signal sequence\ninside 1 19\nTMhelix 20 39\noutside 40 325\nTMhelix 326 348\ninside 349 372

9067 GCF_001990865.1_ASM199086v1 Escherichia coli Proteobacteria; Gammaproteobacteria; Enterobacterales; Enterobacteriaceae; Escherichia
MNNEETFYQAMRRQGVTRRSFLKYCSLAATSLGLGAGMAPKIAWALENKPRIPVVWIHGL WP_001295940.1
MULTISPECIES: hydrogenase [Enterobacteriaceae] Length: 372\nNumber of predicted TMHs: 2\nExp number of AAs in TMHs: 37.2608899999999\nExp number, first 60 AAs: 19.52923\nTotal prob of N-in: 0.94351\nPOSSIBLE N-term signal sequence\ninside 1 19\nTMhelix 20 39\noutside 40 325\nTMhelix 326 348\ninside 349 372

9068 GCF_001997045.1_ASM199704v1 Escherichia coli Proteobacteria; Gammaproteobacteria; Enterobacterales; Enterobacteriaceae; Escherichia
MTGDNTLIHSHGINRRDFMKLCAALAATMGLSSKAAAEMAESVTNPQRPPVIWIGAQECT WP_000145410.1
MULTISPECIES: hydrogenase-2 small chain [Proteobacteria] Length: 372\nNumber of predicted TMHs: 1\nExp number of AAs in TMHs: 23.72226\nExp number, first 60 AAs: 0.22038\nTotal prob of N-in: 0.12994\noutside 1 331\nTMhelix 332 354\ninside 355 372

9069 GCF_001997065.1_ASM199706v1 Escherichia coli Proteobacteria; Gammaproteobacteria; Enterobacterales; Enterobacteriaceae; Escherichia
MTGDNTLIHSHGINRRDFMKLCAALAATMGLSSKAAAEMAESVTNPQRPPVIWIGAQECT WP_000145410.1
MULTISPECIES: hydrogenase-2 small chain [Proteobacteria] Length: 372\nNumber of predicted TMHs: 1\nExp number of AAs in TMHs: 23.72226\nExp number, first 60 AAs: 0.22038\nTotal prob of N-in: 0.12994\noutside 1 331\nTMhelix 332 354\ninside 355 372

9070 GCF_001997075.1_ASM199707v1 Escherichia coli Proteobacteria; Gammaproteobacteria; Enterobacterales; Enterobacteriaceae; Escherichia
MTGDNTLIHSHGINRRDFMKLCAALAATMGLSSKAAAEMAESVTNPQRPPVIWIGAQECT WP_000145410.1
MULTISPECIES: hydrogenase-2 small chain [Proteobacteria] Length: 372\nNumber of predicted TMHs: 1\nExp number of AAs in TMHs: 23.72226\nExp number, first 60 AAs: 0.22038\nTotal prob of N-in: 0.12994\noutside 1 331\nTMhelix 332 354\ninside 355 372

9071 GCF_001997405.1_ASM199740v1 Escherichia coli Proteobacteria; Gammaproteobacteria; Enterobacterales; Enterobacteriaceae; Escherichia
MTGDNTLIHSHGINRRDFMKLCAALAATMGLSSKAAAEMAESVTNPQRPPVIWIGAQECT WP_000145410.1
MULTISPECIES: hydrogenase-2 small chain [Proteobacteria] Length: 372\nNumber of predicted TMHs: 1\nExp number of AAs in TMHs: 23.72226\nExp number, first 60 AAs: 0.22038\nTotal prob of N-in: 0.12994\noutside 1 331\nTMhelix 332 354\ninside 355 372

9072 GCF_002001545.1_ASM200154v1 Escherichia coli Proteobacteria; Gammaproteobacteria; Enterobacterales; Enterobacteriaceae; Escherichia
MTGDNTLIHSHGINRRDFMKLCAALAATMGLSSKAAAEMVESVTNPQRPPVIWIGAQECT WP_000145423.1
hydrogenase 2 small subunit [Escherichia coli] Length: 372\nNumber of predicted TMHs: 1\nExp number of AAs in TMHs: 23.78201\nExp number, first 60 AAs: 0.28355\nTotal prob of N-in: 0.13257\noutside 1 331\nTMhelix 332 354\ninside 355 372

9073 GCF_002001595.1_ASM200159v1 Escherichia coli Proteobacteria; Gammaproteobacteria; Enterobacterales; Enterobacteriaceae; Escherichia
MTGDNTLIHSHGINRRDFMKLCAALAATMGLSSKAAAEMAESVTNPQRPPVIWIGAQECT WP_000145410.1
MULTISPECIES: hydrogenase-2 small chain [Proteobacteria] Length: 372\nNumber of predicted TMHs: 1\nExp number of AAs in TMHs: 23.72226\nExp number, first 60 AAs: 0.22038\nTotal prob of N-in: 0.12994\noutside 1 331\nTMhelix 332 354\ninside 355 372

9074 GCF_002001625.1_ASM200162v1 Escherichia coli Proteobacteria; Gammaproteobacteria; Enterobacterales; Enterobacteriaceae; Escherichia
MTGDNTLIHSHGINRRDFMKLCAALAATMGLSSKAAAEMAESVTNPQRPPVIWIGAQECT WP_000145410.1
MULTISPECIES: hydrogenase-2 small chain [Proteobacteria] Length: 372\nNumber of predicted TMHs: 1\nExp number of AAs in TMHs: 23.72226\nExp number, first 60 AAs: 0.22038\nTotal prob of N-in: 0.12994\noutside 1 331\nTMhelix 332 354\ninside 355 372

9075 GCF_002001595.1_ASM200159v1 Escherichia coli Proteobacteria; Gammaproteobacteria; Enterobacterales; Enterobacteriaceae; Escherichia
MNNEETFYQAMRRQGVTRRSFLKYCSLAATSLGLGAGMAPKIAWALENKPRIPVVWIHGL WP_001295940.1
MULTISPECIES: hydrogenase [Enterobacteriaceae] Length: 372\nNumber of predicted TMHs: 2\nExp number of AAs in TMHs: 37.2608899999999\nExp number, first 60 AAs: 19.52923\nTotal prob of N-in: 0.94351\nPOSSIBLE N-term signal sequence\ninside 1 19\nTMhelix 20 39\noutside 40 325\nTMhelix 326 348\ninside 349 372

9076 GCF_002001625.1_ASM200162v1 Escherichia coli Proteobacteria; Gammaproteobacteria; Enterobacterales; Enterobacteriaceae; Escherichia
MNNEETFYQAMRRQGVTRRSFLKYCSLAATSLGLGAGMAPKIAWALENKPRIPVVWIHGL WP_001295940.1
MULTISPECIES: hydrogenase [Enterobacteriaceae] Length: 372\nNumber of predicted TMHs: 2\nExp number of AAs in TMHs: 37.2608899999999\nExp number, first 60 AAs: 19.52923\nTotal prob of N-in: 0.94351\nPOSSIBLE N-term signal sequence\ninside 1 19\nTMhelix 20 39\noutside 40 325\nTMhelix 326 348\ninside 349 372

9077 GCF_002001535.1_ASM200153v1 Escherichia coli Proteobacteria; Gammaproteobacteria; Enterobacterales; Enterobacteriaceae; Escherichia
MNNEETFYQAMRRQGVTRRSFLKYCSLAATSLGLGAGMAPKIAWALENKPRIPVVWIHGL WP_001058323.1
MULTISPECIES: hydrogenase-1 small chain [Proteobacteria] Length: 372\nNumber of predicted TMHs: 2\nExp number of AAs in TMHs: 37.2608499999999\nExp number, first 60 AAs: 19.52923\nTotal prob of N-in: 0.94351\nPOSSIBLE N-term signal sequence\ninside 1 19\nTMhelix 20 39\noutside 40 325\nTMhelix 326 348\ninside 349 372

9078 GCF_001999185.1_ASM199918v1 Escherichia coli Proteobacteria; Gammaproteobacteria; Enterobacterales; Enterobacteriaceae; Escherichia
MNNEETFYQAMRRQGVTRRSFLKYCSLAATSLGLGAGMAPKIAWALENKPRIPVVWIHGL WP_001058323.1
MULTISPECIES: hydrogenase-1 small chain [Proteobacteria] Length: 372\nNumber of predicted TMHs: 2\nExp number of AAs in TMHs: 37.2608499999999\nExp number, first 60 AAs: 19.52923\nTotal prob of N-in: 0.94351\nPOSSIBLE N-term signal sequence\ninside 1 19\nTMhelix 20 39\noutside 40 325\nTMhelix 326 348\ninside 349 372

9079 GCF_002001545.1_ASM200154v1 Escherichia coli Proteobacteria; Gammaproteobacteria; Enterobacterales; Enterobacteriaceae; Escherichia
MNNEETFYQAMRRQGVTRRSFLKYCSLAATSLGLGAGMAPKIAWALENKPRIPVVWIHGL WP_001538532.1
hydrogenase-1 small chain [Escherichia coli] Length: 372\nNumber of predicted TMHs: 2\nExp number of AAs in TMHs: 37.27041\nExp number, first 60 AAs: 19.52927\nTotal prob of N-in: 0.94352\nPOSSIBLE N-term signal sequence\ninside 1 19\nTMhelix 20 39\noutside 40 325\nTMhelix 326 348\ninside 349 372

9080 GCF_002001715.1_ASM200171v1 Escherichia coli Proteobacteria; Gammaproteobacteria; Enterobacterales; Enterobacteriaceae; Escherichia
MTGDNTLIHSHGINRRDFMKLCAALAATMGLSSKAAAEMAESVTNPQRPPVIWIGAQECT WP_000145410.1
MULTISPECIES: hydrogenase-2 small chain [Proteobacteria] Length: 372\nNumber of predicted TMHs: 1\nExp number of AAs in TMHs: 23.72226\nExp number, first 60 AAs: 0.22038\nTotal prob of N-in: 0.12994\noutside 1 331\nTMhelix 332 354\ninside 355 372

9081	GCF_002001745.1_ASM200174v1	Escherichia coli	Proteobacteria; Gammaproteobacteria; Enterobacterales; Enterobacteriaceae; Escherichia
MTGDNTLIHSHGINRRDFMKLCAALAATMGLSSKAAAEMAESVTNPQRPPVIWIGAQECT	WP_000145410.1
MULTISPECIES: hydrogenase-2 small chain [Proteobacteria]	Length: 372\nNumber of predicted TMHs: 1\nExp number of AAs in TMHs: 23.72226\nExp number, first 60 AAs: 0.22038\nTotal prob of N-in: 0.12994\noutside 1 331\nTMhelix 332 354\ninside 355 372

9082	GCF_002001705.1_ASM200170v1	Escherichia coli	Proteobacteria; Gammaproteobacteria; Enterobacterales; Enterobacteriaceae; Escherichia
MNNEETFYQAMRRQGVTRRSFLKYCSLAATSLGLGAGMAPKIAWALENKPRIPVVWIHGL	WP_001058323.1
MULTISPECIES: hydrogenase-1 small chain [Proteobacteria]	Length: 372\nNumber of predicted TMHs: 2\nExp number of AAs in TMHs: 37.2608499999999\nExp number, first 60 AAs: 19.52923\nTotal prob of N-in: 0.94351\nPOSSIBLE N-term signal sequence\ninside 1 19\nTMhelix 20 39\noutside 40 325\nTMhelix 326 348\ninside 349 372

9083	GCF_002001805.1_ASM200180v1	Escherichia coli	Proteobacteria; Gammaproteobacteria; Enterobacterales; Enterobacteriaceae; Escherichia
MTGDNTLIHSHGINRRDFMKLCAALAATMGLSSKAAAEMAESVTNPQRPPVIWIGAQECT	WP_000145410.1
MULTISPECIES: hydrogenase-2 small chain [Proteobacteria]	Length: 372\nNumber of predicted TMHs: 1\nExp number of AAs in TMHs: 23.72226\nExp number, first 60 AAs: 0.22038\nTotal prob of N-in: 0.12994\noutside 1 331\nTMhelix 332 354\ninside 355 372

9084	GCF_002001825.1_ASM200182v1	Escherichia coli	Proteobacteria; Gammaproteobacteria; Enterobacterales; Enterobacteriaceae; Escherichia
MTGDNTLIHSHGINRRDFMKLCAALAATMGLSSKAAAEMAESVTNPQRPPVIWIGAQECT	WP_000145410.1
MULTISPECIES: hydrogenase-2 small chain [Proteobacteria]	Length: 372\nNumber of predicted TMHs: 1\nExp number of AAs in TMHs: 23.72226\nExp number, first 60 AAs: 0.22038\nTotal prob of N-in: 0.12994\noutside 1 331\nTMhelix 332 354\ninside 355 372

9085	GCF_002001765.1_ASM200176v1	Escherichia coli	Proteobacteria; Gammaproteobacteria; Enterobacterales; Enterobacteriaceae; Escherichia
MTGDNTLIHSHGINRRDFMKLCAALAATMGLSSKAAAEMAESVTNPQRPPVIWIGAQECT	WP_000145410.1
MULTISPECIES: hydrogenase-2 small chain [Proteobacteria]	Length: 372\nNumber of predicted TMHs: 1\nExp number of AAs in TMHs: 23.72226\nExp number, first 60 AAs: 0.22038\nTotal prob of N-in: 0.12994\noutside 1 331\nTMhelix 332 354\ninside 355 372

9086	GCF_002001785.1_ASM200178v1	Escherichia coli	Proteobacteria; Gammaproteobacteria; Enterobacterales; Enterobacteriaceae; Escherichia
MTGDNTLIHSHGINRRDFMKLCAALAATMGLSSKAAAEMAESVTNPQRPPVIWIGAQECT	WP_000145410.1
MULTISPECIES: hydrogenase-2 small chain [Proteobacteria]	Length: 372\nNumber of predicted TMHs: 1\nExp number of AAs in TMHs: 23.72226\nExp number, first 60 AAs: 0.22038\nTotal prob of N-in: 0.12994\noutside 1 331\nTMhelix 332 354\ninside 355 372

9087	GCF_002001845.1_ASM200184v1	Escherichia coli	Proteobacteria; Gammaproteobacteria; Enterobacterales; Enterobacteriaceae; Escherichia
MTGDNTLIHSHGINRRDFMKLCAALAATMGLSSKAAAEMAESVTNPQRPPVIWIGAQECT	WP_000145410.1
MULTISPECIES: hydrogenase-2 small chain [Proteobacteria]	Length: 372\nNumber of predicted TMHs: 1\nExp number of AAs in TMHs: 23.72226\nExp number, first 60 AAs: 0.22038\nTotal prob of N-in: 0.12994\noutside 1 331\nTMhelix 332 354\ninside 355 372

9088	GCF_002001965.1_ASM200196v1	Escherichia coli	Proteobacteria; Gammaproteobacteria; Enterobacterales; Enterobacteriaceae; Escherichia
MTGDNTLIHSHGINRRDFMKLCAALAATMGLSSKAAAEMAESVTNPQRPPVIWIGAQECT	WP_000145410.1
MULTISPECIES: hydrogenase-2 small chain [Proteobacteria]	Length: 372\nNumber of predicted TMHs: 1\nExp number of AAs in TMHs: 23.72226\nExp number, first 60 AAs: 0.22038\nTotal prob of N-in: 0.12994\noutside 1 331\nTMhelix 332 354\ninside 355 372

9089	GCF_002001945.1_ASM200194v1	Escherichia coli	Proteobacteria; Gammaproteobacteria; Enterobacterales; Enterobacteriaceae; Escherichia
MTGDNTLIHSHGINRRDFMKLCAALAATMGLSSKAAAEMAESVTNPQRPPVIWIGAQECT	WP_000145410.1
MULTISPECIES: hydrogenase-2 small chain [Proteobacteria]	Length: 372\nNumber of predicted TMHs:

1\nExp number of AAs in TMHs: 23.72226\nExp number, first 60 AAs: 0.22038\nTotal prob of N-in: 0.12994\noutside 1 331\nTMhelix 332 354\ninside 355 372

9090 GCF_002001975.1_ASM200197v1 Escherichia coli Proteobacteria; Gammaproteobacteria; Enterobacterales; Enterobacteriaceae; Escherichia MTGDNTLIHSHGINRRDFMKLCAALAATMGLSSKAAAEMAESVTNPQRPPVIWIGAQECT WP_000145410.1 MULTISPECIES: hydrogenase-2 small chain [Proteobacteria] Length: 372\nNumber of predicted TMHs: 1\nExp number of AAs in TMHs: 23.72226\nExp number, first 60 AAs: 0.22038\nTotal prob of N-in: 0.12994\noutside 1 331\nTMhelix 332 354\ninside 355 372

9091 GCF_002001995.1_ASM200199v1 Escherichia coli Proteobacteria; Gammaproteobacteria; Enterobacterales; Enterobacteriaceae; Escherichia MTGDNTLIHSHGINRRDFMKLCAALAATMGLSSKAAAEMAESVTNPQRPPVIWIGAQECT WP_000145410.1 MULTISPECIES: hydrogenase-2 small chain [Proteobacteria] Length: 372\nNumber of predicted TMHs: 1\nExp number of AAs in TMHs: 23.72226\nExp number, first 60 AAs: 0.22038\nTotal prob of N-in: 0.12994\noutside 1 331\nTMhelix 332 354\ninside 355 372

9092 GCF_002002045.1_ASM200204v1 Escherichia coli Proteobacteria; Gammaproteobacteria; Enterobacterales; Enterobacteriaceae; Escherichia MTGDNTLIHSHGINRRDFMKLCAALAATMGLSSKAAAEMAESVTNPQRPPVIWIGAQECT WP_000145410.1 MULTISPECIES: hydrogenase-2 small chain [Proteobacteria] Length: 372\nNumber of predicted TMHs: 1\nExp number of AAs in TMHs: 23.72226\nExp number, first 60 AAs: 0.22038\nTotal prob of N-in: 0.12994\noutside 1 331\nTMhelix 332 354\ninside 355 372

9093 GCF_002002055.1_ASM200205v1 Escherichia coli Proteobacteria; Gammaproteobacteria; Enterobacterales; Enterobacteriaceae; Escherichia MTGDNTLIHSHGINRRDFMKLCAALAATMGLSSKAAAEMAESVTNPQRPPVIWIGAQECT WP_000145410.1 MULTISPECIES: hydrogenase-2 small chain [Proteobacteria] Length: 372\nNumber of predicted TMHs: 1\nExp number of AAs in TMHs: 23.72226\nExp number, first 60 AAs: 0.22038\nTotal prob of N-in: 0.12994\noutside 1 331\nTMhelix 332 354\ninside 355 372

9094 GCF_002002005.1_ASM200200v1 Escherichia coli Proteobacteria; Gammaproteobacteria; Enterobacterales; Enterobacteriaceae; Escherichia MTGDNTLIHSHGINRRDFMKLCAALAATMGLSSKAAAEMAESVTNPQRPPVIWIGAQECT WP_000145410.1 MULTISPECIES: hydrogenase-2 small chain [Proteobacteria] Length: 372\nNumber of predicted TMHs: 1\nExp number of AAs in TMHs: 23.72226\nExp number, first 60 AAs: 0.22038\nTotal prob of N-in: 0.12994\noutside 1 331\nTMhelix 332 354\ninside 355 372

9095 GCF_002001975.1_ASM200197v1 Escherichia coli Proteobacteria; Gammaproteobacteria; Enterobacterales; Enterobacteriaceae; Escherichia MNNEETFYQAMRRQGVTRRSFLKYCSLAATSLGLGAGMAPKIAWALENKPRIPVVWIHGL WP_001058323.1 MULTISPECIES: hydrogenase-1 small chain [Proteobacteria] Length: 372\nNumber of predicted TMHs: 2\nExp number of AAs in TMHs: 37.2608499999999\nExp number, first 60 AAs: 19.52923\nTotal prob of N-in: 0.94351\nPOSSIBLE N-term signal sequence\ninside 1 19\nTMhelix 20 39\noutside 40 325\nTMhelix 326 348\ninside 349 372

9096 GCF_002002105.1_ASM200210v1 Escherichia coli Proteobacteria; Gammaproteobacteria; Enterobacterales; Enterobacteriaceae; Escherichia MTGDNTLIHSHGINRRDFMKLCAALAATMGLSSKAAAEMAESVTNPQRPPVIWIGAQECT WP_000145410.1 MULTISPECIES: hydrogenase-2 small chain [Proteobacteria] Length: 372\nNumber of predicted TMHs: 1\nExp number of AAs in TMHs: 23.72226\nExp number, first 60 AAs: 0.22038\nTotal prob of N-in: 0.12994\noutside 1 331\nTMhelix 332 354\ninside 355 372

9097 GCF_002002105.1_ASM200210v1 Escherichia coli Proteobacteria; Gammaproteobacteria; Enterobacterales; Enterobacteriaceae; Escherichia MNNEETFYQAMRRQGVTRRSFLKYCSLAATSLGLGAGMAPKIAWALENKPRIPVVWIHGL WP_001058323.1 MULTISPECIES: hydrogenase-1 small chain [Proteobacteria] Length: 372\nNumber of predicted TMHs: 2\nExp number of AAs in TMHs: 37.2608499999999\nExp number, first 60 AAs: 19.52923\nTotal prob of N-in: 0.94351\nPOSSIBLE N-term signal sequence\ninside 1 19\nTMhelix 20 39\noutside 40 325\nTMhelix 326 348\ninside 349 372

9098 GCF_002001945.1_ASM200194v1 Escherichia coli Proteobacteria; Gammaproteobacteria; Enterobacterales; Enterobacteriaceae; Escherichia
MNNEETFYQAMRRQGVTRRSFLKYCSLAATSLGLGAGMAPKIAWALENKPRIPVVWIHGL WP_001295940.1
MULTISPECIES: hydrogenase [Enterobacteriaceae] Length: 372\nNumber of predicted TMHs: 2\nExp number of AAs in TMHs: 37.2608899999999\nExp number, first 60 AAs: 19.52923\nTotal prob of N-in: 0.94351\nPOSSIBLE N-term signal sequence\ninside 1 19\nTMhelix 20 39\noutside 40 325\nTMhelix 326 348\ninside 349 372

9099 GCF_002002145.1_ASM200214v1 Escherichia coli Proteobacteria; Gammaproteobacteria; Enterobacterales; Enterobacteriaceae; Escherichia
MTGDNTLIHSHGINRRDFMKLCAALAATMGLSSKAAAEMAESVTNPQRPPVIWIGAQECT WP_000145410.1
MULTISPECIES: hydrogenase-2 small chain [Proteobacteria] Length: 372\nNumber of predicted TMHs: 1\nExp number of AAs in TMHs: 23.72226\nExp number, first 60 AAs: 0.22038\nTotal prob of N-in: 0.12994\noutside 1 331\nTMhelix 332 354\ninside 355 372

9100 GCF_002002125.1_ASM200212v1 Escherichia coli Proteobacteria; Gammaproteobacteria; Enterobacterales; Enterobacteriaceae; Escherichia
MTGDNTLIHSHGINRRDFMKLCAALAATMGLSSKAAAEMAESVTNPQRPPVIWIGAQECT WP_000145410.1
MULTISPECIES: hydrogenase-2 small chain [Proteobacteria] Length: 372\nNumber of predicted TMHs: 1\nExp number of AAs in TMHs: 23.72226\nExp number, first 60 AAs: 0.22038\nTotal prob of N-in: 0.12994\noutside 1 331\nTMhelix 332 354\ninside 355 372

9101 GCF_002002165.1_ASM200216v1 Escherichia coli Proteobacteria; Gammaproteobacteria; Enterobacterales; Enterobacteriaceae; Escherichia
MTGDNTLIHSHGINRRDFMKLCAALAATMGLSSKAAAEMAESVTNPQRPPVIWIGAQECT WP_000145410.1
MULTISPECIES: hydrogenase-2 small chain [Proteobacteria] Length: 372\nNumber of predicted TMHs: 1\nExp number of AAs in TMHs: 23.72226\nExp number, first 60 AAs: 0.22038\nTotal prob of N-in: 0.12994\noutside 1 331\nTMhelix 332 354\ninside 355 372

9102 GCF_002002145.1_ASM200214v1 Escherichia coli Proteobacteria; Gammaproteobacteria; Enterobacterales; Enterobacteriaceae; Escherichia
MNNEETFYQAMRRQGVTRRSFLKYCSLAATSLGLGAGMAPKIAWALENKPRIPVVWIHGL WP_001058323.1
MULTISPECIES: hydrogenase-1 small chain [Proteobacteria] Length: 372\nNumber of predicted TMHs: 2\nExp number of AAs in TMHs: 37.2608499999999\nExp number, first 60 AAs: 19.52923\nTotal prob of N-in: 0.94351\nPOSSIBLE N-term signal sequence\ninside 1 19\nTMhelix 20 39\noutside 40 325\nTMhelix 326 348\ninside 349 372

9103 GCF_002002235.1_ASM200223v1 Escherichia coli Proteobacteria; Gammaproteobacteria; Enterobacterales; Enterobacteriaceae; Escherichia
MNNEETFYQAMRRQGVTRRSFLKYCSLAATSLGLGAGMAPKIAWALENKPRIPVVWIHGL WP_001058323.1
MULTISPECIES: hydrogenase-1 small chain [Proteobacteria] Length: 372\nNumber of predicted TMHs: 2\nExp number of AAs in TMHs: 37.2608499999999\nExp number, first 60 AAs: 19.52923\nTotal prob of N-in: 0.94351\nPOSSIBLE N-term signal sequence\ninside 1 19\nTMhelix 20 39\noutside 40 325\nTMhelix 326 348\ninside 349 372

9104 GCF_002002165.1_ASM200216v1 Escherichia coli Proteobacteria; Gammaproteobacteria; Enterobacterales; Enterobacteriaceae; Escherichia
MNNEETFYQAMRRQGVTRRSFLKYCSLAATSLGLGAGMAPKIAWALENKPRIPVVWIHGL WP_001058323.1
MULTISPECIES: hydrogenase-1 small chain [Proteobacteria] Length: 372\nNumber of predicted TMHs: 2\nExp number of AAs in TMHs: 37.2608499999999\nExp number, first 60 AAs: 19.52923\nTotal prob of N-in: 0.94351\nPOSSIBLE N-term signal sequence\ninside 1 19\nTMhelix 20 39\noutside 40 325\nTMhelix 326 348\ninside 349 372

9105 GCF_002002175.1_ASM200217v1 Escherichia coli Proteobacteria; Gammaproteobacteria; Enterobacterales; Enterobacteriaceae; Escherichia
MNNEETFYQAMRRQGVTRRSFLKYCSLAATSLGLGAGMAPKIAWALENKPRIPVVWIHGL WP_001058323.1
MULTISPECIES: hydrogenase-1 small chain [Proteobacteria] Length: 372\nNumber of predicted TMHs: 2\nExp number of AAs in TMHs: 37.2608499999999\nExp number, first 60 AAs: 19.52923\nTotal prob of N-in: 0.94351\nPOSSIBLE N-term signal sequence\ninside 1 19\nTMhelix 20 39\noutside 40 325\nTMhelix 326 348\ninside 349 372

9106 GCF_002002315.1_ASM200231v1 Escherichia coli Proteobacteria; Gammaproteobacteria; Enterobacterales; Enterobacteriaceae; Escherichia
MTGDNTLIHSHGINRRDFMKLCAALAATMGLSSKAAAEMAESVTNPQRPPVIWIGAQECT WP_000145410.1
MULTISPECIES: hydrogenase-2 small chain [Proteobacteria] Length: 372\nNumber of predicted TMHs: 1\nExp number of AAs in TMHs: 23.72226\nExp number, first 60 AAs: 0.22038\nTotal prob of N-in: 0.12994\noutside 1 331\nTMhelix 332 354\ninside 355 372

9107 GCF_002002225.1_ASM200222v1 Escherichia coli Proteobacteria; Gammaproteobacteria; Enterobacterales; Enterobacteriaceae; Escherichia
MNNEETFYQAMRRQGVTRRSFLKYCSLAATSLGLGAGMAPKIAWALENKPRIPVVWIHGL WP_001058323.1
MULTISPECIES: hydrogenase-1 small chain [Proteobacteria] Length: 372\nNumber of predicted TMHs: 2\nExp number of AAs in TMHs: 37.2608499999999\nExp number, first 60 AAs: 19.52923\nTotal prob of N-in: 0.94351\nPOSSIBLE N-term signal sequence\ninside 1 19\nTMhelix 20 39\noutside 40 325\nTMhelix 326 348\ninside 349 372

9108 GCF_002002195.1_ASM200219v1 Escherichia coli Proteobacteria; Gammaproteobacteria; Enterobacterales; Enterobacteriaceae; Escherichia
MNNEETFYQAMRRQGVTRRSFLKYCSLAATSLGLGAGMAPKIAWALENKPRIPVVWIHGL WP_001058323.1
MULTISPECIES: hydrogenase-1 small chain [Proteobacteria] Length: 372\nNumber of predicted TMHs: 2\nExp number of AAs in TMHs: 37.2608499999999\nExp number, first 60 AAs: 19.52923\nTotal prob of N-in: 0.94351\nPOSSIBLE N-term signal sequence\ninside 1 19\nTMhelix 20 39\noutside 40 325\nTMhelix 326 348\ninside 349 372

9109 GCF_002002325.1_ASM200232v1 Escherichia coli Proteobacteria; Gammaproteobacteria; Enterobacterales; Enterobacteriaceae; Escherichia
MTGDNTLIHSHGINRRDFMKLCAALAATMGLSSKAAAEMAESVTNPQRPPVIWIGAQECT WP_000145410.1
MULTISPECIES: hydrogenase-2 small chain [Proteobacteria] Length: 372\nNumber of predicted TMHs: 1\nExp number of AAs in TMHs: 23.72226\nExp number, first 60 AAs: 0.22038\nTotal prob of N-in: 0.12994\noutside 1 331\nTMhelix 332 354\ninside 355 372

9110 GCF_002002595.1_ASM200259v1 Escherichia coli Proteobacteria; Gammaproteobacteria; Enterobacterales; Enterobacteriaceae; Escherichia
MTGDNTLIHSHGINRRDFMKLCAALAATMGLSSKAAAEMAESVTNPQRPPVIWIGAQECT WP_000145410.1
MULTISPECIES: hydrogenase-2 small chain [Proteobacteria] Length: 372\nNumber of predicted TMHs: 1\nExp number of AAs in TMHs: 23.72226\nExp number, first 60 AAs: 0.22038\nTotal prob of N-in: 0.12994\noutside 1 331\nTMhelix 332 354\ninside 355 372

9111 GCF_002002685.1_ASM200268v1 Escherichia coli Proteobacteria; Gammaproteobacteria; Enterobacterales; Enterobacteriaceae; Escherichia
MTGDNTLIHSHGINRRDFMKLCAALAATMGLSSKAAAEMAESVTNPQRPPVIWIGAQECT WP_000145410.1
MULTISPECIES: hydrogenase-2 small chain [Proteobacteria] Length: 372\nNumber of predicted TMHs: 1\nExp number of AAs in TMHs: 23.72226\nExp number, first 60 AAs: 0.22038\nTotal prob of N-in: 0.12994\noutside 1 331\nTMhelix 332 354\ninside 355 372

9112 GCF_002002465.1_ASM200246v1 Escherichia coli Proteobacteria; Gammaproteobacteria; Enterobacterales; Enterobacteriaceae; Escherichia
MNNEETFYQAMRRQGVTRRSFLKYCSLAATSLGLGAGMAPKIAWALENKPRIPVVWIHGL WP_001058323.1
MULTISPECIES: hydrogenase-1 small chain [Proteobacteria] Length: 372\nNumber of predicted TMHs: 2\nExp number of AAs in TMHs: 37.2608499999999\nExp number, first 60 AAs: 19.52923\nTotal prob of N-in: 0.94351\nPOSSIBLE N-term signal sequence\ninside 1 19\nTMhelix 20 39\noutside 40 325\nTMhelix 326 348\ninside 349 372

9113 GCF_002002805.1_ASM200280v1 Escherichia coli Proteobacteria; Gammaproteobacteria; Enterobacterales; Enterobacteriaceae; Escherichia
MTGDNTLIHSHGINRRDFMKLCAALAATMGLSSKAAAEMVESVTNPQRPPVIWIGAQECT WP_000145423.1
hydrogenase 2 small subunit [Escherichia coli] Length: 372\nNumber of predicted TMHs: 1\nExp number of AAs in TMHs: 23.78201\nExp number, first 60 AAs: 0.28355\nTotal prob of N-in: 0.13257\noutside 1 331\nTMhelix 332 354\ninside 355 372

9114 GCF_002002815.1_ASM200281v1 Escherichia coli Proteobacteria; Gammaproteobacteria; Enterobacterales; Enterobacteriaceae; Escherichia

MTGDNTLIHSHGINRRDFMKLCAALAATMGLSSKAAAEMAESVTNPQRPPVIWIGAQECT WP_000145410.1 MULTISPECIES: hydrogenase-2 small chain [Proteobacteria] Length: 372\nNumber of predicted TMHs: 1\nExp number of AAs in TMHs: 23.72226\nExp number, first 60 AAs: 0.22038\nTotal prob of N-in: 0.12994\noutside 1 331\nTMhelix 332 354\ninside 355 372

9115 GCF_002002805.1_ASM200280v1 Escherichia coli Proteobacteria; Gammaproteobacteria; Enterobacterales; Enterobacteriaceae; Escherichia MNNEETFYQAMRRQGVTRRSFLKYCSLAATSLGLGAGMAPKIAWALENKPRIPVVWIHGL WP_001538532.1 hydrogenase-1 small chain [Escherichia coli] Length: 372\nNumber of predicted TMHs: 2\nExp number of AAs in TMHs: 37.27041\nExp number, first 60 AAs: 19.52927\nTotal prob of N-in: 0.94352\nPOSSIBLE N-term signal sequence\ninside 1 19\nTMhelix 20 39\noutside 40 325\nTMhelix 326 348\ninside 349 372

9116 GCF_002002815.1_ASM200281v1 Escherichia coli Proteobacteria; Gammaproteobacteria; Enterobacterales; Enterobacteriaceae; Escherichia MNNEETFYQAMRRQGVTRRSFLKYCSLAATSLGLGAGMAPKIAWALENKPRIPVVWIHGL WP_001058323.1 MULTISPECIES: hydrogenase-1 small chain [Proteobacteria] Length: 372\nNumber of predicted TMHs: 2\nExp number of AAs in TMHs: 37.2608499999999\nExp number, first 60 AAs: 19.52923\nTotal prob of N-in: 0.94351\nPOSSIBLE N-term signal sequence\ninside 1 19\nTMhelix 20 39\noutside 40 325\nTMhelix 326 348\ninside 349 372

9117 GCF_002002545.1_ASM200254v1 Escherichia coli Proteobacteria; Gammaproteobacteria; Enterobacterales; Enterobacteriaceae; Escherichia MNNEETFYQAMRRQGVTRRSFLKYCSLAATSLGLGAGMAPKIAWALENKPRIPVVWIHGL WP_001058323.1 MULTISPECIES: hydrogenase-1 small chain [Proteobacteria] Length: 372\nNumber of predicted TMHs: 2\nExp number of AAs in TMHs: 37.2608499999999\nExp number, first 60 AAs: 19.52923\nTotal prob of N-in: 0.94351\nPOSSIBLE N-term signal sequence\ninside 1 19\nTMhelix 20 39\noutside 40 325\nTMhelix 326 348\ninside 349 372

9118 GCF_002006525.1_ASM200652v1 Escherichia coli Proteobacteria; Gammaproteobacteria; Enterobacterales; Enterobacteriaceae; Escherichia MTGDNTLIHSHGINRRDFMKLCAALAATMGLSSKAAAEMAESVTNPQRPPVIWIGAQECT WP_000145410.1 MULTISPECIES: hydrogenase-2 small chain [Proteobacteria] Length: 372\nNumber of predicted TMHs: 1\nExp number of AAs in TMHs: 23.72226\nExp number, first 60 AAs: 0.22038\nTotal prob of N-in: 0.12994\noutside 1 331\nTMhelix 332 354\ninside 355 372

9119 GCF_002002685.1_ASM200268v1 Escherichia coli Proteobacteria; Gammaproteobacteria; Enterobacterales; Enterobacteriaceae; Escherichia MNNEETFYQAMRRQGVTRRSFLKYCSLAATSLGLGAGMAPKIAWALENKPRIPVVWIHGL WP_001058323.1 MULTISPECIES: hydrogenase-1 small chain [Proteobacteria] Length: 372\nNumber of predicted TMHs: 2\nExp number of AAs in TMHs: 37.2608499999999\nExp number, first 60 AAs: 19.52923\nTotal prob of N-in: 0.94351\nPOSSIBLE N-term signal sequence\ninside 1 19\nTMhelix 20 39\noutside 40 325\nTMhelix 326 348\ninside 349 372

9120 GCF_002006865.1_ASM200686v1 Escherichia coli Proteobacteria; Gammaproteobacteria; Enterobacterales; Enterobacteriaceae; Escherichia MTGDNTLIHSHGINRRDFMKLCAALAATMGLSSKAAAEMAESVTNPQRPPVIWIGAQECT WP_000145410.1 MULTISPECIES: hydrogenase-2 small chain [Proteobacteria] Length: 372\nNumber of predicted TMHs: 1\nExp number of AAs in TMHs: 23.72226\nExp number, first 60 AAs: 0.22038\nTotal prob of N-in: 0.12994\noutside 1 331\nTMhelix 332 354\ninside 355 372

9121 GCF_002006965.1_ASM200696v1 Escherichia coli Proteobacteria; Gammaproteobacteria; Enterobacterales; Enterobacteriaceae; Escherichia MTGDNTLIHSHGINRRDFMKLCAALAATMGLSSKAAAEMAESVTNPQRPPVIWIGAQECT WP_000145410.1 MULTISPECIES: hydrogenase-2 small chain [Proteobacteria] Length: 372\nNumber of predicted TMHs: 1\nExp number of AAs in TMHs: 23.72226\nExp number, first 60 AAs: 0.22038\nTotal prob of N-in: 0.12994\noutside 1 331\nTMhelix 332 354\ninside 355 372

9122 GCF_002007165.1_ASM200716v1 Escherichia coli Proteobacteria; Gammaproteobacteria; Enterobacterales; Enterobacteriaceae; Escherichia MTGDNTLIHSHGINRRDFMKLCAALAATMGLSSKAAAEMAESVTNPQRPPVIWIGAQECT WP_000145410.1 MULTISPECIES: hydrogenase-2 small chain [Proteobacteria] Length: 372\nNumber of predicted TMHs:

1\nExp number of AAs in TMHs: 23.72226\nExp number, first 60 AAs: 0.22038\nTotal prob of N-in: 0.12994\noutside 1 331\nTMhelix 332 354\ninside 355 372

9123 GCF_002006965.1_ASM200696v1 Escherichia coli Proteobacteria; Gammaproteobacteria; Enterobacterales; Enterobacteriaceae; Escherichia
MNNEETFYQAMRRQGVTRRSFLKYCSLAATSLGLGAGMAPKIAWALENKPRIPVVWIHGL WP_059321607.1
hydrogenase [Escherichia coli] Length: 372\nNumber of predicted TMHs: 2\nExp number of AAs in TMHs: 37.2608299999999\nExp number, first 60 AAs: 19.52923\nTotal prob of N-in: 0.94351\nPOSSIBLE N-term signal sequence\ninside 1 19\nTMhelix 20 39\noutside 40 325\nTMhelix 326 348\ninside 349 372

9124 GCF_002011945.1_ASM201194v1 Escherichia coli Proteobacteria; Gammaproteobacteria; Enterobacterales; Enterobacteriaceae; Escherichia
MNNEETFYQAMRRQGVTRRSFLKYCSLAATSLGLGAGMAPKIAWALENKPRIPVVWIHGL WP_001295940.1
MULTISPECIES: hydrogenase [Enterobacteriaceae] Length: 372\nNumber of predicted TMHs: 2\nExp number of AAs in TMHs: 37.2608899999999\nExp number, first 60 AAs: 19.52923\nTotal prob of N-in: 0.94351\nPOSSIBLE N-term signal sequence\ninside 1 19\nTMhelix 20 39\noutside 40 325\nTMhelix 326 348\ninside 349 372

9125 GCF_002007165.1_ASM200716v1 Escherichia coli Proteobacteria; Gammaproteobacteria; Enterobacterales; Enterobacteriaceae; Escherichia
MNNEETFYQAMRRQGVTRRSFLKYCSLAATSLGLGAGMAPKIAWALENKPRIPVVWIHGL WP_001058323.1
MULTISPECIES: hydrogenase-1 small chain [Proteobacteria] Length: 372\nNumber of predicted TMHs: 2\nExp number of AAs in TMHs: 37.2608499999999\nExp number, first 60 AAs: 19.52923\nTotal prob of N-in: 0.94351\nPOSSIBLE N-term signal sequence\ninside 1 19\nTMhelix 20 39\noutside 40 325\nTMhelix 326 348\ninside 349 372

9126 GCF_002011965.1_ASM201196v1 Escherichia coli Proteobacteria; Gammaproteobacteria; Enterobacterales; Enterobacteriaceae; Escherichia
MTGDNTLIHSHGINRRDFMKLCAALAATMGLSSKAAAEMVESVTNPQRPPVIWIGAQECT WP_000145423.1
hydrogenase 2 small subunit [Escherichia coli] Length: 372\nNumber of predicted TMHs: 1\nExp number of AAs in TMHs: 23.78201\nExp number, first 60 AAs: 0.28355\nTotal prob of N-in: 0.13257\noutside 1 331\nTMhelix 332 354\ninside 355 372

9127 GCF_002011965.1_ASM201196v1 Escherichia coli Proteobacteria; Gammaproteobacteria; Enterobacterales; Enterobacteriaceae; Escherichia
MNNEETFYQAMRRQGVTRRSFLKYCSLAATSLGLGAGMAPKIAWALENKPRIPVVWIHGL WP_001295940.1
MULTISPECIES: hydrogenase [Enterobacteriaceae] Length: 372\nNumber of predicted TMHs: 2\nExp number of AAs in TMHs: 37.2608899999999\nExp number, first 60 AAs: 19.52923\nTotal prob of N-in: 0.94351\nPOSSIBLE N-term signal sequence\ninside 1 19\nTMhelix 20 39\noutside 40 325\nTMhelix 326 348\ninside 349 372

9128 GCF_002007705.1_ASM200770v1 Escherichia coli Proteobacteria; Gammaproteobacteria; Enterobacterales; Enterobacteriaceae; Escherichia
MNNEETFYQAMRRQGVTRRSFLKYCSLAATSLGLGAGMAPKIAWALENKPRIPVVWIHGL WP_001058323.1
MULTISPECIES: hydrogenase-1 small chain [Proteobacteria] Length: 372\nNumber of predicted TMHs: 2\nExp number of AAs in TMHs: 37.2608499999999\nExp number, first 60 AAs: 19.52923\nTotal prob of N-in: 0.94351\nPOSSIBLE N-term signal sequence\ninside 1 19\nTMhelix 20 39\noutside 40 325\nTMhelix 326 348\ninside 349 372

9129 GCF_002012005.1_ASM201200v1 Escherichia coli Proteobacteria; Gammaproteobacteria; Enterobacterales; Enterobacteriaceae; Escherichia
MTGDNTLIHSHGINRRDFMKLCAALAATMGLSSKAAAEMAESVTNPQRPPVIWIGAQECT WP_000145410.1
MULTISPECIES: hydrogenase-2 small chain [Proteobacteria] Length: 372\nNumber of predicted TMHs: 1\nExp number of AAs in TMHs: 23.72226\nExp number, first 60 AAs: 0.22038\nTotal prob of N-in: 0.12994\noutside 1 331\nTMhelix 332 354\ninside 355 372

9130 GCF_002012025.1_ASM201202v1 Escherichia coli Proteobacteria; Gammaproteobacteria; Enterobacterales; Enterobacteriaceae; Escherichia
MTGDNTLIHSHGINRRDFMKLCAALAATMGLSSKAAAEMVESVTNPQRPPVIWIGAQECT WP_000145423.1
hydrogenase 2 small subunit [Escherichia coli] Length: 372\nNumber of predicted TMHs: 1\nExp number of

AAs in TMHs: 23.78201\nExp number, first 60 AAs:  0.28355\nTotal prob of N-in:   0.13257\noutside   1   331\nTMhelix   332   354\ninside   355   372

9131 GCF_002012105.1_ASM201210v1 Escherichia coli Proteobacteria; Gammaproteobacteria; Enterobacterales; Enterobacteriaceae; Escherichia MTGDNTLIHSHGINRRDFMKLCAALAATMGLSSKAAAEMAESVTNPQRPPVIWIGAQECT WP_000145410.1 MULTISPECIES: hydrogenase-2 small chain [Proteobacteria] Length: 372\nNumber of predicted TMHs: 1\nExp number of AAs in TMHs: 23.72226\nExp number, first 60 AAs:  0.22038\nTotal prob of N-in:   0.12994\noutside   1   331\nTMhelix   332   354\ninside   355   372

9132 GCF_002012105.1_ASM201210v1 Escherichia coli Proteobacteria; Gammaproteobacteria; Enterobacterales; Enterobacteriaceae; Escherichia MNNEETFYQAMRRQGVTRRSFLKYCSLAATSLGLGAGMAPKIAWALENKPRIPVVWIHGL WP_001058323.1 MULTISPECIES: hydrogenase-1 small chain [Proteobacteria] Length: 372\nNumber of predicted TMHs: 2\nExp number of AAs in TMHs: 37.2608499999999\nExp number, first 60 AAs:  19.52923\nTotal prob of N-in: 0.94351\nPOSSIBLE N-term signal sequence\ninside   1   19\nTMhelix   20   39\noutside   40   325\nTMhelix   326   348\ninside   349   372

9133 GCF_002012025.1_ASM201202v1 Escherichia coli Proteobacteria; Gammaproteobacteria; Enterobacterales; Enterobacteriaceae; Escherichia MNNEETFYQAMRRQGVTRRSFLKYCSLAATSLGLGAGMAPKIAWALENKPRIPVVWIHGL WP_001295940.1 MULTISPECIES: hydrogenase [Enterobacteriaceae] Length: 372\nNumber of predicted TMHs:  2\nExp number of AAs in TMHs: 37.2608899999999\nExp number, first 60 AAs:  19.52923\nTotal prob of N-in: 0.94351\nPOSSIBLE N-term signal sequence\ninside   1   19\nTMhelix   20   39\noutside   40   325\nTMhelix   326   348\ninside   349   372

9134 GCF_002012125.1_ASM201212v1 Escherichia coli Proteobacteria; Gammaproteobacteria; Enterobacterales; Enterobacteriaceae; Escherichia MTGDNTLIHSHGINRRDFMKLCAALAATMGLSSKAAAEMAESVTNPQRPPVIWIGAQECT WP_000145410.1 MULTISPECIES: hydrogenase-2 small chain [Proteobacteria] Length: 372\nNumber of predicted TMHs: 1\nExp number of AAs in TMHs: 23.72226\nExp number, first 60 AAs:  0.22038\nTotal prob of N-in:   0.12994\noutside   1   331\nTMhelix   332   354\ninside   355   372

9135 GCF_002012145.1_ASM201214v1 Escherichia coli Proteobacteria; Gammaproteobacteria; Enterobacterales; Enterobacteriaceae; Escherichia MTGDNTLIHSHGINRRDFMKLCAALAATMGLSSKAAAEMAESVTNPQRPPVIWIGAQECT WP_000145410.1 MULTISPECIES: hydrogenase-2 small chain [Proteobacteria] Length: 372\nNumber of predicted TMHs: 1\nExp number of AAs in TMHs: 23.72226\nExp number, first 60 AAs:  0.22038\nTotal prob of N-in:   0.12994\noutside   1   331\nTMhelix   332   354\ninside   355   372

9136 GCF_002012165.1_ASM201216v1 Escherichia coli Proteobacteria; Gammaproteobacteria; Enterobacterales; Enterobacteriaceae; Escherichia MTGDNTLIHSHGINRRDFMKLCAALAATMGLSSKAAAEMVESVTNPQRPPVIWIGAQECT WP_000145423.1 hydrogenase 2 small subunit [Escherichia coli] Length: 372\nNumber of predicted TMHs:  1\nExp number of AAs in TMHs: 23.78201\nExp number, first 60 AAs:  0.28355\nTotal prob of N-in:   0.13257\noutside   1   331\nTMhelix   332   354\ninside   355   372

9137 GCF_002012125.1_ASM201212v1 Escherichia coli Proteobacteria; Gammaproteobacteria; Enterobacterales; Enterobacteriaceae; Escherichia MNNEETFYQAMRRQGVTRRSFLKYCSLAATSLGLGAGMAPKIAWALENKPRIPVVWIHGL WP_001058323.1 MULTISPECIES: hydrogenase-1 small chain [Proteobacteria] Length: 372\nNumber of predicted TMHs: 2\nExp number of AAs in TMHs: 37.2608499999999\nExp number, first 60 AAs:  19.52923\nTotal prob of N-in: 0.94351\nPOSSIBLE N-term signal sequence\ninside   1   19\nTMhelix   20   39\noutside   40   325\nTMhelix   326   348\ninside   349   372

9138 GCF_002012225.1_ASM201222v1 Escherichia coli Proteobacteria; Gammaproteobacteria; Enterobacterales; Enterobacteriaceae; Escherichia MNNEETFYQAMRRQGVTRRSFLKYCSLAATSLGLGAGMAPKIAWALENKPRIPVVWIHGL WP_001058323.1 MULTISPECIES: hydrogenase-1 small chain [Proteobacteria] Length: 372\nNumber of predicted TMHs: 2\nExp number of AAs in TMHs: 37.2608499999999\nExp number, first 60 AAs:  19.52923\nTotal prob of N-in:

0.94351\nPOSSIBLE N-term signal sequence\ninside 1 19\nTMhelix 20 39\noutside 40 325\nTMhelix 326 348\ninside 349 372

9139 GCF_002012245.1_ASM201224v1 Escherichia coli Proteobacteria; Gammaproteobacteria; Enterobacterales; Enterobacteriaceae; Escherichia MTGDNTLIHSHGINRRDFMKLCAALAATMGLSSKAAAEMVESVTNPQRPPVIWIGAQECT WP_000145423.1 hydrogenase 2 small subunit [Escherichia coli] Length: 372\nNumber of predicted TMHs: 1\nExp number of AAs in TMHs: 23.78201\nExp number, first 60 AAs: 0.28355\nTotal prob of N-in: 0.13257\noutside 1 331\nTMhelix 332 354\ninside 355 372

9140 GCF_002012265.1_ASM201226v1 Escherichia coli Proteobacteria; Gammaproteobacteria; Enterobacterales; Enterobacteriaceae; Escherichia MTGDNTLIHSHGINRRDFMKLCAALAATMGLSSKAAAEMVESVTNPQRPPVIWIGAQECT WP_000145423.1 hydrogenase 2 small subunit [Escherichia coli] Length: 372\nNumber of predicted TMHs: 1\nExp number of AAs in TMHs: 23.78201\nExp number, first 60 AAs: 0.28355\nTotal prob of N-in: 0.13257\noutside 1 331\nTMhelix 332 354\ninside 355 372

9141 GCF_002012305.1_ASM201230v1 Escherichia coli Proteobacteria; Gammaproteobacteria; Enterobacterales; Enterobacteriaceae; Escherichia MTGDNTLIHSHGINRRDFMKLCAALAATMGLSSKAAAEMAESVTNPQRPPVIWIGAQECT WP_000145410.1 MULTISPECIES: hydrogenase-2 small chain [Proteobacteria] Length: 372\nNumber of predicted TMHs: 1\nExp number of AAs in TMHs: 23.72226\nExp number, first 60 AAs: 0.22038\nTotal prob of N-in: 0.12994\noutside 1 331\nTMhelix 332 354\ninside 355 372

9142 GCF_002012305.1_ASM201230v1 Escherichia coli Proteobacteria; Gammaproteobacteria; Enterobacterales; Enterobacteriaceae; Escherichia MNNEETFYQAMRRQGVTRRSFLKYCSLAATSLGLGAGMAPKIAWALENKPRIPVVWIHGL WP_001295940.1 MULTISPECIES: hydrogenase [Enterobacteriaceae] Length: 372\nNumber of predicted TMHs: 2\nExp number of AAs in TMHs: 37.2608899999999\nExp number, first 60 AAs: 19.52923\nTotal prob of N-in: 0.94351\nPOSSIBLE N-term signal sequence\ninside 1 19\nTMhelix 20 39\noutside 40 325\nTMhelix 326 348\ninside 349 372

9143 GCF_002012185.1_ASM201218v1 Escherichia coli Proteobacteria; Gammaproteobacteria; Enterobacterales; Enterobacteriaceae; Escherichia MNNEETFYQAMRRQGVTRRSFLKYCSLAATSLGLGAGMAPKIAWALENKPRIPVVWIHGL WP_001058323.1 MULTISPECIES: hydrogenase-1 small chain [Proteobacteria] Length: 372\nNumber of predicted TMHs: 2\nExp number of AAs in TMHs: 37.2608499999999\nExp number, first 60 AAs: 19.52923\nTotal prob of N-in: 0.94351\nPOSSIBLE N-term signal sequence\ninside 1 19\nTMhelix 20 39\noutside 40 325\nTMhelix 326 348\ninside 349 372

9144 GCF_002015205.1_ASM201520v1 Escherichia coli Proteobacteria; Gammaproteobacteria; Enterobacterales; Enterobacteriaceae; Escherichia MTGDNTLIHSHGINRRDFMKLCAALAATMGLSSKAAAEMAESVTNPQRPPVIWIGAQECT WP_000145410.1 MULTISPECIES: hydrogenase-2 small chain [Proteobacteria] Length: 372\nNumber of predicted TMHs: 1\nExp number of AAs in TMHs: 23.72226\nExp number, first 60 AAs: 0.22038\nTotal prob of N-in: 0.12994\noutside 1 331\nTMhelix 332 354\ninside 355 372

9145 GCF_002015265.1_ASM201526v1 Escherichia coli Proteobacteria; Gammaproteobacteria; Enterobacterales; Enterobacteriaceae; Escherichia MNNEETFYQAMRRQGVTRRSFLKYCSLAATSLGLGAGMAPKIAWALENKPRIPVVWIHGL WP_024246676.1 hydrogenase-1 small chain [Escherichia coli] Length: 372\nNumber of predicted TMHs: 2\nExp number of AAs in TMHs: 37.8214899999999\nExp number, first 60 AAs: 19.52326\nTotal prob of N-in: 0.94395\nPOSSIBLE N-term signal sequence\ninside 1 19\nTMhelix 20 39\noutside 40 325\nTMhelix 326 348\ninside 349 372

9146 GCF_002015255.1_ASM201525v1 Escherichia coli Proteobacteria; Gammaproteobacteria; Enterobacterales; Enterobacteriaceae; Escherichia MNNEETFYQAMRRQGVTRRSFLKYCSLAATSLGLGAGMAPKIAWALENKPRIPVVWIHGL WP_024246676.1 hydrogenase-1 small chain [Escherichia coli] Length: 372\nNumber of predicted TMHs: 2\nExp number of AAs in TMHs: 37.8214899999999\nExp number, first 60 AAs: 19.52326\nTotal prob of N-in: 0.94395\nPOSSIBLE N-term signal sequence\ninside 1 19\nTMhelix 20 39\noutside 40 325\nTMhelix 326 348\ninside 349 372

9147 GCF_002012265.1_ASM201226v1 Escherichia coli Proteobacteria; Gammaproteobacteria; Enterobacterales; Enterobacteriaceae; Escherichia
MNNEETFYQAMRRQGVTRRSFLKYCSLAATSLGLGAGMAPKIAWALENKPRIPVVWIHGL WP_001295940.1
MULTISPECIES: hydrogenase [Enterobacteriaceae] Length: 372\nNumber of predicted TMHs: 2\nExp number of AAs in TMHs: 37.2608899999999\nExp number, first 60 AAs: 19.52923\nTotal prob of N-in: 0.94351\nPOSSIBLE N-term signal sequence\ninside 1 19\nTMhelix 20 39\noutside 40 325\nTMhelix 326 348\ninside 349 372

9148 GCF_002015275.1_ASM201527v1 Escherichia coli Proteobacteria; Gammaproteobacteria; Enterobacterales; Enterobacteriaceae; Escherichia
MTGDNTLIHSHGINRRDFMKLCAALAATMGLSSKAAAEMAESVTNPQRPPVIWIGAQECT WP_000145410.1
MULTISPECIES: hydrogenase-2 small chain [Proteobacteria] Length: 372\nNumber of predicted TMHs: 1\nExp number of AAs in TMHs: 23.72226\nExp number, first 60 AAs: 0.22038\nTotal prob of N-in: 0.12994\noutside 1 331\nTMhelix 332 354\ninside 355 372

9149 GCF_002015275.1_ASM201527v1 Escherichia coli Proteobacteria; Gammaproteobacteria; Enterobacterales; Enterobacteriaceae; Escherichia
MNNEETFYQAMRRQGVTRRSFLKYCSLAATSLGLGAGMAPKIAWALENKPRIPVVWIHGL WP_024246676.1
hydrogenase-1 small chain [Escherichia coli] Length: 372\nNumber of predicted TMHs: 2\nExp number of AAs in TMHs: 37.8214899999999\nExp number, first 60 AAs: 19.52326\nTotal prob of N-in: 0.94395\nPOSSIBLE N-term signal sequence\ninside 1 19\nTMhelix 20 39\noutside 40 325\nTMhelix 326 348\ninside 349 372

9150 GCF_002015195.1_ASM201519v1 Escherichia coli Proteobacteria; Gammaproteobacteria; Enterobacterales; Enterobacteriaceae; Escherichia
MNNEETFYQAMRRQGVTRRSFLKYCSLAATSLGLGAGMAPKIAWALENKPRIPVVWIHGL WP_024246676.1
hydrogenase-1 small chain [Escherichia coli] Length: 372\nNumber of predicted TMHs: 2\nExp number of AAs in TMHs: 37.8214899999999\nExp number, first 60 AAs: 19.52326\nTotal prob of N-in: 0.94395\nPOSSIBLE N-term signal sequence\ninside 1 19\nTMhelix 20 39\noutside 40 325\nTMhelix 326 348\ninside 349 372

9151 GCF_002015315.1_ASM201531v1 Escherichia coli Proteobacteria; Gammaproteobacteria; Enterobacterales; Enterobacteriaceae; Escherichia
MTGDNTLIHSHGINRRDFMKLCAALAATMGLSSKAAAEMAESVTNPQRPPVIWIGAQECT WP_000145410.1
MULTISPECIES: hydrogenase-2 small chain [Proteobacteria] Length: 372\nNumber of predicted TMHs: 1\nExp number of AAs in TMHs: 23.72226\nExp number, first 60 AAs: 0.22038\nTotal prob of N-in: 0.12994\noutside 1 331\nTMhelix 332 354\ninside 355 372

9152 GCF_002015315.1_ASM201531v1 Escherichia coli Proteobacteria; Gammaproteobacteria; Enterobacterales; Enterobacteriaceae; Escherichia
MNNEETFYQAMRRQGVTRRSFLKYCSLAATSLGLGAGMAPKIAWALENKPRIPVVWIHGL WP_001058323.1
MULTISPECIES: hydrogenase-1 small chain [Proteobacteria] Length: 372\nNumber of predicted TMHs: 2\nExp number of AAs in TMHs: 37.2608499999999\nExp number, first 60 AAs: 19.52923\nTotal prob of N-in: 0.94351\nPOSSIBLE N-term signal sequence\ninside 1 19\nTMhelix 20 39\noutside 40 325\nTMhelix 326 348\ninside 349 372

9153 GCF_002015465.1_ASM201546v1 Escherichia coli Proteobacteria; Gammaproteobacteria; Enterobacterales; Enterobacteriaceae; Escherichia
MTGDNTLIHSHGINRRDFMKLCAALAATMGLSSKAAAEMAESVTNPQRPPVIWIGAQECT WP_000145410.1
MULTISPECIES: hydrogenase-2 small chain [Proteobacteria] Length: 372\nNumber of predicted TMHs: 1\nExp number of AAs in TMHs: 23.72226\nExp number, first 60 AAs: 0.22038\nTotal prob of N-in: 0.12994\noutside 1 331\nTMhelix 332 354\ninside 355 372

9154 GCF_002015465.1_ASM201546v1 Escherichia coli Proteobacteria; Gammaproteobacteria; Enterobacterales; Enterobacteriaceae; Escherichia
MNNEETFYQAMRRQGVTRRSFLKYCSLAATSLGLGAGMAPKIAWALENKPRIPVVWIHGL WP_001295940.1
MULTISPECIES: hydrogenase [Enterobacteriaceae] Length: 372\nNumber of predicted TMHs: 2\nExp number of AAs in TMHs: 37.2608899999999\nExp number, first 60 AAs: 19.52923\nTotal prob of N-in: 0.94351\nPOSSIBLE N-term signal sequence\ninside 1 19\nTMhelix 20 39\noutside 40 325\nTMhelix 326 348\ninside 349 372

9155 GCF_002015325.1_ASM201532v1 Escherichia coli Proteobacteria; Gammaproteobacteria; Enterobacterales; Enterobacteriaceae; Escherichia

MNNEETFYQAMRRQGVTRRSFLKYCSLAATSLGLGAGMAPKIAWALENKPRIPVVWIHGL WP_024246676.1 hydrogenase-1 small chain [Escherichia coli] Length: 372\nNumber of predicted TMHs: 2\nExp number of AAs in TMHs: 37.8214899999999\nExp number, first 60 AAs: 19.52326\nTotal prob of N-in: 0.94395\nPOSSIBLE N-term signal sequence\ninside 1 19\nTMhelix 20 39\noutside 40 325\nTMhelix 326 348\ninside 349 372

9156 GCF_002015355.1_ASM201535v1 Escherichia coli Proteobacteria; Gammaproteobacteria; Enterobacterales; Enterobacteriaceae; Escherichia MNNEETFYQAMRRQGVTRRSFLKYCSLAATSLGLGAGMAPKIAWALENKPRIPVVWIHGL WP_024246676.1 hydrogenase-1 small chain [Escherichia coli] Length: 372\nNumber of predicted TMHs: 2\nExp number of AAs in TMHs: 37.8214899999999\nExp number, first 60 AAs: 19.52326\nTotal prob of N-in: 0.94395\nPOSSIBLE N-term signal sequence\ninside 1 19\nTMhelix 20 39\noutside 40 325\nTMhelix 326 348\ninside 349 372

9157 GCF_002015365.1_ASM201536v1 Escherichia coli Proteobacteria; Gammaproteobacteria; Enterobacterales; Enterobacteriaceae; Escherichia MNNEETFYQAMRRQGVTRRSFLKYCSLAATSLGLGAGMAPKIAWALENKPRIPVVWIHGL WP_024246676.1 hydrogenase-1 small chain [Escherichia coli] Length: 372\nNumber of predicted TMHs: 2\nExp number of AAs in TMHs: 37.8214899999999\nExp number, first 60 AAs: 19.52326\nTotal prob of N-in: 0.94395\nPOSSIBLE N-term signal sequence\ninside 1 19\nTMhelix 20 39\noutside 40 325\nTMhelix 326 348\ninside 349 372

9158 GCF_002015375.1_ASM201537v1 Escherichia coli Proteobacteria; Gammaproteobacteria; Enterobacterales; Enterobacteriaceae; Escherichia MNNEETFYQAMRRQGVTRRSFLKYCSLAATSLGLGAGMAPKIAWALENKPRIPVVWIHGL WP_024246676.1 hydrogenase-1 small chain [Escherichia coli] Length: 372\nNumber of predicted TMHs: 2\nExp number of AAs in TMHs: 37.8214899999999\nExp number, first 60 AAs: 19.52326\nTotal prob of N-in: 0.94395\nPOSSIBLE N-term signal sequence\ninside 1 19\nTMhelix 20 39\noutside 40 325\nTMhelix 326 348\ninside 349 372

9159 GCF_002015435.1_ASM201543v1 Escherichia coli Proteobacteria; Gammaproteobacteria; Enterobacterales; Enterobacteriaceae; Escherichia MNNEETFYQAMRRQGVTRRSFLKYCSLAATSLGLGAGMAPKIAWALENKPRIPVVWIHGL WP_001295940.1 MULTISPECIES: hydrogenase [Enterobacteriaceae] Length: 372\nNumber of predicted TMHs: 2\nExp number of AAs in TMHs: 37.2608899999999\nExp number, first 60 AAs: 19.52923\nTotal prob of N-in: 0.94351\nPOSSIBLE N-term signal sequence\ninside 1 19\nTMhelix 20 39\noutside 40 325\nTMhelix 326 348\ninside 349 372

9160 GCF_002015475.1_ASM201547v1 Escherichia coli Proteobacteria; Gammaproteobacteria; Enterobacterales; Enterobacteriaceae; Escherichia MTGDNTLIHSHGINRRDFMKLCAALAATMGLSSKAAAEMAESVTNPQRPPVIWIGAQECT WP_000145410.1 MULTISPECIES: hydrogenase-2 small chain [Proteobacteria] Length: 372\nNumber of predicted TMHs: 1\nExp number of AAs in TMHs: 23.72226\nExp number, first 60 AAs: 0.22038\nTotal prob of N-in: 0.12994\noutside 1 331\nTMhelix 332 354\ninside 355 372

9161 GCF_002015515.1_ASM201551v1 Escherichia coli Proteobacteria; Gammaproteobacteria; Enterobacterales; Enterobacteriaceae; Escherichia MTGDNTLIHSHGINRRDFMKLCAALAATMGLSSKAAAEMAESVTNPQRPPVIWIGAQECT WP_000145410.1 MULTISPECIES: hydrogenase-2 small chain [Proteobacteria] Length: 372\nNumber of predicted TMHs: 1\nExp number of AAs in TMHs: 23.72226\nExp number, first 60 AAs: 0.22038\nTotal prob of N-in: 0.12994\noutside 1 331\nTMhelix 332 354\ninside 355 372

9162 GCF_002015475.1_ASM201547v1 Escherichia coli Proteobacteria; Gammaproteobacteria; Enterobacterales; Enterobacteriaceae; Escherichia MNNEETFYQAMRRQGVTRRSFLKYCSLAATSLGLGAGMAPKIAWALENKPRIPVVWIHGL WP_001295940.1 MULTISPECIES: hydrogenase [Enterobacteriaceae] Length: 372\nNumber of predicted TMHs: 2\nExp number of AAs in TMHs: 37.2608899999999\nExp number, first 60 AAs: 19.52923\nTotal prob of N-in: 0.94351\nPOSSIBLE N-term signal sequence\ninside 1 19\nTMhelix 20 39\noutside 40 325\nTMhelix 326 348\ninside 349 372

9163 GCF_002015515.1_ASM201551v1 Escherichia coli Proteobacteria; Gammaproteobacteria; Enterobacterales; Enterobacteriaceae; Escherichia MNNEETFYQAMRRQGVTRRSFLKYCSLAATSLGLGAGMAPKIAWALENKPRIPVVWIHGL WP_001295940.1 MULTISPECIES: hydrogenase [Enterobacteriaceae] Length: 372\nNumber of predicted TMHs: 2\nExp number of AAs in TMHs: 37.2608899999999\nExp number, first 60 AAs: 19.52923\nTotal prob of N-in:

0.94351\nPOSSIBLE N-term signal sequence\ninside 1 19\nTMhelix 20 39\noutside 40 325\nTMhelix 326 348\ninside 349 372

9164 GCF_002015385.1_ASM201538v1 Escherichia coli Proteobacteria; Gammaproteobacteria; Enterobacterales; Enterobacteriaceae; Escherichia
MNNEETFYQAMRRQGVTRRSFLKYCSLAATSLGLGAGMAPKIAWALENKPRIPVVWIHGL WP_024246676.1
hydrogenase-1 small chain [Escherichia coli] Length: 372\nNumber of predicted TMHs: 2\nExp number of AAs in TMHs: 37.8214899999999\nExp number, first 60 AAs: 19.52326\nTotal prob of N-in: 0.94395\nPOSSIBLE N-term signal sequence\ninside 1 19\nTMhelix 20 39\noutside 40 325\nTMhelix 326 348\ninside 349 372

9165 GCF_002015535.1_ASM201553v1 Escherichia coli Proteobacteria; Gammaproteobacteria; Enterobacterales; Enterobacteriaceae; Escherichia
MTGDNTLIHSHGINRRDFMKLCAALAATMGLSSKAAAEMAESVTNPQRPPVIWIGAQECT WP_000145410.1
MULTISPECIES: hydrogenase-2 small chain [Proteobacteria] Length: 372\nNumber of predicted TMHs: 1\nExp number of AAs in TMHs: 23.72226\nExp number, first 60 AAs: 0.22038\nTotal prob of N-in: 0.12994\noutside 1 331\nTMhelix 332 354\ninside 355 372

9166 GCF_002015675.1_ASM201567v1 Escherichia coli Proteobacteria; Gammaproteobacteria; Enterobacterales; Enterobacteriaceae; Escherichia
MNNEETFYQAMRRQGVTRRSFLKYCSLAATSLGLGAGMAPKIAWALENKPRIPVVWIHGL WP_001295940.1
MULTISPECIES: hydrogenase [Enterobacteriaceae] Length: 372\nNumber of predicted TMHs: 2\nExp number of AAs in TMHs: 37.2608899999999\nExp number, first 60 AAs: 19.52923\nTotal prob of N-in: 0.94351\nPOSSIBLE N-term signal sequence\ninside 1 19\nTMhelix 20 39\noutside 40 325\nTMhelix 326 348\ninside 349 372

9167 GCF_002015695.1_ASM201569v1 Escherichia coli Proteobacteria; Gammaproteobacteria; Enterobacterales; Enterobacteriaceae; Escherichia
MNNEETFYQAMRRQGVTRRSFLKYCSLAATSLGLGAGMAPKIAWALENKPRIPVVWIHGL WP_001295940.1
MULTISPECIES: hydrogenase [Enterobacteriaceae] Length: 372\nNumber of predicted TMHs: 2\nExp number of AAs in TMHs: 37.2608899999999\nExp number, first 60 AAs: 19.52923\nTotal prob of N-in: 0.94351\nPOSSIBLE N-term signal sequence\ninside 1 19\nTMhelix 20 39\noutside 40 325\nTMhelix 326 348\ninside 349 372

9168 GCF_002015535.1_ASM201553v1 Escherichia coli Proteobacteria; Gammaproteobacteria; Enterobacterales; Enterobacteriaceae; Escherichia
MNNEETFYQAMRRQGVTRRSFLKYCSLAATSLGLGAGMAPKIAWALENKPRIPVVWIHGL WP_001295940.1
MULTISPECIES: hydrogenase [Enterobacteriaceae] Length: 372\nNumber of predicted TMHs: 2\nExp number of AAs in TMHs: 37.2608899999999\nExp number, first 60 AAs: 19.52923\nTotal prob of N-in: 0.94351\nPOSSIBLE N-term signal sequence\ninside 1 19\nTMhelix 20 39\noutside 40 325\nTMhelix 326 348\ninside 349 372

9169 GCF_002015545.1_ASM201554v1 Escherichia coli Proteobacteria; Gammaproteobacteria; Enterobacterales; Enterobacteriaceae; Escherichia
MNNEETFYQAMRRQGVTRRSFLKYCSLAATSLGLGAGMAPKIAWALENKPRIPVVWIHGL WP_001295940.1
MULTISPECIES: hydrogenase [Enterobacteriaceae] Length: 372\nNumber of predicted TMHs: 2\nExp number of AAs in TMHs: 37.2608899999999\nExp number, first 60 AAs: 19.52923\nTotal prob of N-in: 0.94351\nPOSSIBLE N-term signal sequence\ninside 1 19\nTMhelix 20 39\noutside 40 325\nTMhelix 326 348\ninside 349 372

9170 GCF_002015625.1_ASM201562v1 Escherichia coli Proteobacteria; Gammaproteobacteria; Enterobacterales; Enterobacteriaceae; Escherichia
MNNEETFYQAMRRQGVTRRSFLKYCSLAATSLGLGAGMAPKIAWALENKPRIPVVWIHGL WP_001295940.1
MULTISPECIES: hydrogenase [Enterobacteriaceae] Length: 372\nNumber of predicted TMHs: 2\nExp number of AAs in TMHs: 37.2608899999999\nExp number, first 60 AAs: 19.52923\nTotal prob of N-in: 0.94351\nPOSSIBLE N-term signal sequence\ninside 1 19\nTMhelix 20 39\noutside 40 325\nTMhelix 326 348\ninside 349 372

9171 GCF_002015555.1_ASM201555v1 Escherichia coli Proteobacteria; Gammaproteobacteria; Enterobacterales; Enterobacteriaceae; Escherichia
MNNEETFYQAMRRQGVTRRSFLKYCSLAATSLGLGAGMAPKIAWALENKPRIPVVWIHGL WP_001295940.1
MULTISPECIES: hydrogenase [Enterobacteriaceae] Length: 372\nNumber of predicted TMHs: 2\nExp

number of AAs in TMHs: 37.2608899999999\nExp number, first 60 AAs:  19.52923\nTotal prob of N-in:   0.94351\nPOSSIBLE N-term signal sequence\ninside   1   19\nTMhelix   20   39\noutside   40   325\nTMhelix   326   348\ninside   349   372

9172 GCF_002015705.1_ASM201570v1 Escherichia coli Proteobacteria; Gammaproteobacteria; Enterobacterales; Enterobacteriaceae; Escherichia
MTGDNTLIHSHGINRRDFMKLCAALAATMGLSSKAAAEMAESVTNPQRPPVIWIGAQECT WP_000145410.1
MULTISPECIES: hydrogenase-2 small chain [Proteobacteria] Length: 372\nNumber of predicted TMHs:  1\nExp number of AAs in TMHs: 23.72226\nExp number, first 60 AAs:  0.22038\nTotal prob of N-in:   0.12994\noutside   1   331\nTMhelix   332   354\ninside   355   372

9173 GCF_002015825.1_ASM201582v1 Escherichia coli Proteobacteria; Gammaproteobacteria; Enterobacterales; Enterobacteriaceae; Escherichia
MTGDNTLIHSHGINRRDFMKLCAALAATMGLSSKAAAEMAESVTNPQRPPVIWIGAQECT WP_000145410.1
MULTISPECIES: hydrogenase-2 small chain [Proteobacteria] Length: 372\nNumber of predicted TMHs:  1\nExp number of AAs in TMHs: 23.72226\nExp number, first 60 AAs:  0.22038\nTotal prob of N-in:   0.12994\noutside   1   331\nTMhelix   332   354\ninside   355   372

9174 GCF_002015875.1_ASM201587v1 Escherichia coli Proteobacteria; Gammaproteobacteria; Enterobacterales; Enterobacteriaceae; Escherichia
MTGDNTLIHSHGINRRDFMKLCAALAATMGLSSKAAAEMAESVTNPQRPPVIWIGAQECT WP_000145410.1
MULTISPECIES: hydrogenase-2 small chain [Proteobacteria] Length: 372\nNumber of predicted TMHs:  1\nExp number of AAs in TMHs: 23.72226\nExp number, first 60 AAs:  0.22038\nTotal prob of N-in:   0.12994\noutside   1   331\nTMhelix   332   354\ninside   355   372

9175 GCF_002015855.1_ASM201585v1 Escherichia coli Proteobacteria; Gammaproteobacteria; Enterobacterales; Enterobacteriaceae; Escherichia
MTGDNTLIHSHGINRRDFMKLCAALAATMGLSSKAAAEMAESVTNPQRPPVIWIGAQECT WP_000145410.1
MULTISPECIES: hydrogenase-2 small chain [Proteobacteria] Length: 372\nNumber of predicted TMHs:  1\nExp number of AAs in TMHs: 23.72226\nExp number, first 60 AAs:  0.22038\nTotal prob of N-in:   0.12994\noutside   1   331\nTMhelix   332   354\ninside   355   372

9176 GCF_002016155.1_ASM201615v1 Escherichia coli Proteobacteria; Gammaproteobacteria; Enterobacterales; Enterobacteriaceae; Escherichia
MTGDNTLIHSHGINRRDFMKLCAALAATMGLSSKAAAEMAESVTNPQRPPVIWIGAQECT WP_000145410.1
MULTISPECIES: hydrogenase-2 small chain [Proteobacteria] Length: 372\nNumber of predicted TMHs:  1\nExp number of AAs in TMHs: 23.72226\nExp number, first 60 AAs:  0.22038\nTotal prob of N-in:   0.12994\noutside   1   331\nTMhelix   332   354\ninside   355   372

9177 GCF_002015735.1_ASM201573v1 Escherichia coli Proteobacteria; Gammaproteobacteria; Enterobacterales; Enterobacteriaceae; Escherichia
MNNEETFYQAMRRQGVTRRSFLKYCSLAATSLGLGAGMAPKIAWALENKPRIPVVWIHGL WP_001295940.1
MULTISPECIES: hydrogenase [Enterobacteriaceae] Length: 372\nNumber of predicted TMHs:  2\nExp number of AAs in TMHs: 37.2608899999999\nExp number, first 60 AAs:  19.52923\nTotal prob of N-in:   0.94351\nPOSSIBLE N-term signal sequence\ninside   1   19\nTMhelix   20   39\noutside   40   325\nTMhelix   326   348\ninside   349   372

9178 GCF_002015885.1_ASM201588v1 Escherichia coli Proteobacteria; Gammaproteobacteria; Enterobacterales; Enterobacteriaceae; Escherichia
MTGDNTLIHSHGINRRDFMKLCAALAATMGLSSKAAAEMAESVTNPQRPPVIWIGAQECT WP_000145410.1
MULTISPECIES: hydrogenase-2 small chain [Proteobacteria] Length: 372\nNumber of predicted TMHs:  1\nExp number of AAs in TMHs: 23.72226\nExp number, first 60 AAs:  0.22038\nTotal prob of N-in:   0.12994\noutside   1   331\nTMhelix   332   354\ninside   355   372

9179 GCF_002015755.1_ASM201575v1 Escherichia coli Proteobacteria; Gammaproteobacteria; Enterobacterales; Enterobacteriaceae; Escherichia
MNNEETFYQAMRRQGVTRRSFLKYCSLAATSLGLGAGMAPKIAWALENKPRIPVVWIHGL WP_001295940.1
MULTISPECIES: hydrogenase [Enterobacteriaceae] Length: 372\nNumber of predicted TMHs:  2\nExp number of AAs in TMHs: 37.2608899999999\nExp number, first 60 AAs:  19.52923\nTotal prob of N-in:   0.94351\nPOSSIBLE N-term signal sequence\ninside   1   19\nTMhelix   20   39\noutside   40   325\nTMhelix   326   348\ninside   349   372

9180 GCF_002015945.1_ASM201594v1 Escherichia coli Proteobacteria; Gammaproteobacteria; Enterobacterales; Enterobacteriaceae; Escherichia
MTGDNTLIHSHGINRRDFMKLCAALAATMGLSSKAAAEMAESVTNPQRPPVIWIGAQECT WP_000145410.1
MULTISPECIES: hydrogenase-2 small chain [Proteobacteria] Length: 372\nNumber of predicted TMHs: 1\nExp number of AAs in TMHs: 23.72226\nExp number, first 60 AAs: 0.22038\nTotal prob of N-in: 0.12994\noutside 1 331\nTMhelix 332 354\ninside 355 372

9181 GCF_002015945.1_ASM201594v1 Escherichia coli Proteobacteria; Gammaproteobacteria; Enterobacterales; Enterobacteriaceae; Escherichia
MNNEETFYQAMRRQGVTRRSFLKYCSLAATSLGLGAGMAPKIAWALENKPRIPVVWIHGL WP_001295940.1
MULTISPECIES: hydrogenase [Enterobacteriaceae] Length: 372\nNumber of predicted TMHs: 2\nExp number of AAs in TMHs: 37.2608899999999\nExp number, first 60 AAs: 19.52923\nTotal prob of N-in: 0.94351\nPOSSIBLE N-term signal sequence\ninside 1 19\nTMhelix 20 39\noutside 40 325\nTMhelix 326 348\ninside 349 372

9182 GCF_002015825.1_ASM201582v1 Escherichia coli Proteobacteria; Gammaproteobacteria; Enterobacterales; Enterobacteriaceae; Escherichia
MNNEETFYQAMRRQGVTRRSFLKYCSLAATSLGLGAGMAPKIAWALENKPRIPVVWIHGL WP_001295940.1
MULTISPECIES: hydrogenase [Enterobacteriaceae] Length: 372\nNumber of predicted TMHs: 2\nExp number of AAs in TMHs: 37.2608899999999\nExp number, first 60 AAs: 19.52923\nTotal prob of N-in: 0.94351\nPOSSIBLE N-term signal sequence\ninside 1 19\nTMhelix 20 39\noutside 40 325\nTMhelix 326 348\ninside 349 372

9183 GCF_002015875.1_ASM201587v1 Escherichia coli Proteobacteria; Gammaproteobacteria; Enterobacterales; Enterobacteriaceae; Escherichia
MNNEETFYQAMRRQGVTRRSFLKYCSLAATSLGLGAGMAPKIAWALENKPRIPVVWIHGL WP_001295940.1
MULTISPECIES: hydrogenase [Enterobacteriaceae] Length: 372\nNumber of predicted TMHs: 2\nExp number of AAs in TMHs: 37.2608899999999\nExp number, first 60 AAs: 19.52923\nTotal prob of N-in: 0.94351\nPOSSIBLE N-term signal sequence\ninside 1 19\nTMhelix 20 39\noutside 40 325\nTMhelix 326 348\ninside 349 372

9184 GCF_002015855.1_ASM201585v1 Escherichia coli Proteobacteria; Gammaproteobacteria; Enterobacterales; Enterobacteriaceae; Escherichia
MNNEETFYQAMRRQGVTRRSFLKYCSLAATSLGLGAGMAPKIAWALENKPRIPVVWIHGL WP_001295940.1
MULTISPECIES: hydrogenase [Enterobacteriaceae] Length: 372\nNumber of predicted TMHs: 2\nExp number of AAs in TMHs: 37.2608899999999\nExp number, first 60 AAs: 19.52923\nTotal prob of N-in: 0.94351\nPOSSIBLE N-term signal sequence\ninside 1 19\nTMhelix 20 39\noutside 40 325\nTMhelix 326 348\ninside 349 372

9185 GCF_002015965.1_ASM201596v1 Escherichia coli Proteobacteria; Gammaproteobacteria; Enterobacterales; Enterobacteriaceae; Escherichia
MTGDNTLIHSHGINRRDFMKLCAALAATMGLSSKAAAEMAESVTNPQRPPVIWIGAQECT WP_000145410.1
MULTISPECIES: hydrogenase-2 small chain [Proteobacteria] Length: 372\nNumber of predicted TMHs: 1\nExp number of AAs in TMHs: 23.72226\nExp number, first 60 AAs: 0.22038\nTotal prob of N-in: 0.12994\noutside 1 331\nTMhelix 332 354\ninside 355 372

9186 GCF_002015965.1_ASM201596v1 Escherichia coli Proteobacteria; Gammaproteobacteria; Enterobacterales; Enterobacteriaceae; Escherichia
MNNEETFYQAMRRQGVTRRSFLKYCSLAATSLGLGAGMAPKIAWALENKPRIPVVWIHGL WP_001295940.1
MULTISPECIES: hydrogenase [Enterobacteriaceae] Length: 372\nNumber of predicted TMHs: 2\nExp number of AAs in TMHs: 37.2608899999999\nExp number, first 60 AAs: 19.52923\nTotal prob of N-in: 0.94351\nPOSSIBLE N-term signal sequence\ninside 1 19\nTMhelix 20 39\noutside 40 325\nTMhelix 326 348\ninside 349 372

9187 GCF_002016025.1_ASM201602v1 Escherichia coli Proteobacteria; Gammaproteobacteria; Enterobacterales; Enterobacteriaceae; Escherichia
MTGDNTLIHSHGINRRDFMKLCAALAATMGLSSKAAAEMAESVTNPQRPPVIWIGAQECT WP_000145410.1
MULTISPECIES: hydrogenase-2 small chain [Proteobacteria] Length: 372\nNumber of predicted TMHs: 1\nExp number of AAs in TMHs: 23.72226\nExp number, first 60 AAs: 0.22038\nTotal prob of N-in: 0.12994\noutside 1 331\nTMhelix 332 354\ninside 355 372

9188 GCF_002016065.1_ASM201606v1 Escherichia coli Proteobacteria; Gammaproteobacteria; Enterobacterales; Enterobacteriaceae; Escherichia
MTGDNTLIHSHGINRRDFMKLCAALAATMGLSSKAAAEMAESVTNPQRPPVIWIGAQECT WP_000145410.1
MULTISPECIES: hydrogenase-2 small chain [Proteobacteria] Length: 372\nNumber of predicted TMHs: 1\nExp number of AAs in TMHs: 23.72226\nExp number, first 60 AAs: 0.22038\nTotal prob of N-in: 0.12994\noutside 1 331\nTMhelix 332 354\ninside 355 372

9189 GCF_002016075.1_ASM201607v1 Escherichia coli Proteobacteria; Gammaproteobacteria; Enterobacterales; Enterobacteriaceae; Escherichia
MTGDNTLIHSHGINRRDFMKLCAALAATMGLSSKAAAEMAESVTNPQRPPVIWIGAQECT WP_000145410.1
MULTISPECIES: hydrogenase-2 small chain [Proteobacteria] Length: 372\nNumber of predicted TMHs: 1\nExp number of AAs in TMHs: 23.72226\nExp number, first 60 AAs: 0.22038\nTotal prob of N-in: 0.12994\noutside 1 331\nTMhelix 332 354\ninside 355 372

9190 GCF_002016085.1_ASM201608v1 Escherichia coli Proteobacteria; Gammaproteobacteria; Enterobacterales; Enterobacteriaceae; Escherichia
MTGDNTLIHSHGINRRDFMKLCAALAATMGLSSKAAAEMAESVTNPQRPPVIWIGAQECT WP_000145410.1
MULTISPECIES: hydrogenase-2 small chain [Proteobacteria] Length: 372\nNumber of predicted TMHs: 1\nExp number of AAs in TMHs: 23.72226\nExp number, first 60 AAs: 0.22038\nTotal prob of N-in: 0.12994\noutside 1 331\nTMhelix 332 354\ninside 355 372

9191 GCF_002016135.1_ASM201613v1 Escherichia coli Proteobacteria; Gammaproteobacteria; Enterobacterales; Enterobacteriaceae; Escherichia
MTGDNTLIHSHGINRRDFMKLCAALAATMGLSSKAAAEMAESVTNPQRPPVIWIGAQECT WP_000145410.1
MULTISPECIES: hydrogenase-2 small chain [Proteobacteria] Length: 372\nNumber of predicted TMHs: 1\nExp number of AAs in TMHs: 23.72226\nExp number, first 60 AAs: 0.22038\nTotal prob of N-in: 0.12994\noutside 1 331\nTMhelix 332 354\ninside 355 372

9192 GCF_002016165.1_ASM201616v1 Escherichia coli Proteobacteria; Gammaproteobacteria; Enterobacterales; Enterobacteriaceae; Escherichia
MTGDNTLIHSHGINRRDFMKLCAALAATMGLSSKAAAEMAESVTNPQRPPVIWIGAQECT WP_000145410.1
MULTISPECIES: hydrogenase-2 small chain [Proteobacteria] Length: 372\nNumber of predicted TMHs: 1\nExp number of AAs in TMHs: 23.72226\nExp number, first 60 AAs: 0.22038\nTotal prob of N-in: 0.12994\noutside 1 331\nTMhelix 332 354\ninside 355 372

9193 GCF_002016165.1_ASM201616v1 Escherichia coli Proteobacteria; Gammaproteobacteria; Enterobacterales; Enterobacteriaceae; Escherichia
MNNEETFYQAMRRQGVTRRSFLKYCSLAATSLGLGAGMAPKIAWALENKPRIPVVWIHGL WP_001295940.1
MULTISPECIES: hydrogenase [Enterobacteriaceae] Length: 372\nNumber of predicted TMHs: 2\nExp number of AAs in TMHs: 37.2608899999999\nExp number, first 60 AAs: 19.52923\nTotal prob of N-in: 0.94351\nPOSSIBLE N-term signal sequence\ninside 1 19\nTMhelix 20 39\noutside 40 325\nTMhelix 326 348\ninside 349 372

9194 GCF_002016285.1_ASM201628v1 Escherichia coli Proteobacteria; Gammaproteobacteria; Enterobacterales; Enterobacteriaceae; Escherichia
MTGDNTLIHSHGINRRDFMKLCAALAATMGLSSKAAAEMAESVTNPQRPPVIWIGAQECT WP_000145410.1
MULTISPECIES: hydrogenase-2 small chain [Proteobacteria] Length: 372\nNumber of predicted TMHs: 1\nExp number of AAs in TMHs: 23.72226\nExp number, first 60 AAs: 0.22038\nTotal prob of N-in: 0.12994\noutside 1 331\nTMhelix 332 354\ninside 355 372

9195 GCF_002016135.1_ASM201613v1 Escherichia coli Proteobacteria; Gammaproteobacteria; Enterobacterales; Enterobacteriaceae; Escherichia
MNNEETFYQAMRRQGVTRRSFLKYCSLAATSLGLGAGMAPKIAWALENKPRIPVVWIHGL WP_001295940.1
MULTISPECIES: hydrogenase [Enterobacteriaceae] Length: 372\nNumber of predicted TMHs: 2\nExp number of AAs in TMHs: 37.2608899999999\nExp number, first 60 AAs: 19.52923\nTotal prob of N-in: 0.94351\nPOSSIBLE N-term signal sequence\ninside 1 19\nTMhelix 20 39\noutside 40 325\nTMhelix 326 348\ninside 349 372

9196 GCF_002016085.1_ASM201608v1 Escherichia coli Proteobacteria; Gammaproteobacteria; Enterobacterales; Enterobacteriaceae; Escherichia
MNNEETFYQAMRRQGVTRRSFLKYCSLAATSLGLGAGMAPKIAWALENKPRIPVVWIHGL WP_001295940.1

MULTISPECIES: hydrogenase [Enterobacteriaceae] Length: 372\nNumber of predicted TMHs: 2\nExp number of AAs in TMHs: 37.2608899999999\nExp number, first 60 AAs: 19.52923\nTotal prob of N-in: 0.94351\nPOSSIBLE N-term signal sequence\ninside 1 19\nTMhelix 20 39\noutside 40 325\nTMhelix 326 348\ninside 349 372

9197 GCF_002016155.1_ASM201615v1 Escherichia coli Proteobacteria; Gammaproteobacteria; Enterobacterales; Enterobacteriaceae; Escherichia
MNNEETFYQAMRRQGVTRRSFLKYCSLAATSLGLGAGMAPKIAWALENKPRIPVVWIHGL WP_001295940.1
MULTISPECIES: hydrogenase [Enterobacteriaceae] Length: 372\nNumber of predicted TMHs: 2\nExp number of AAs in TMHs: 37.2608899999999\nExp number, first 60 AAs: 19.52923\nTotal prob of N-in: 0.94351\nPOSSIBLE N-term signal sequence\ninside 1 19\nTMhelix 20 39\noutside 40 325\nTMhelix 326 348\ninside 349 372

9198 GCF_002016205.1_ASM201620v1 Escherichia coli Proteobacteria; Gammaproteobacteria; Enterobacterales; Enterobacteriaceae; Escherichia
MTGDNTLIHSHGINRRDFMKLCAALAATMGLSSKAAAEMAESVTNPQRPPVIWIGAQECT WP_000145410.1
MULTISPECIES: hydrogenase-2 small chain [Proteobacteria] Length: 372\nNumber of predicted TMHs: 1\nExp number of AAs in TMHs: 23.72226\nExp number, first 60 AAs: 0.22038\nTotal prob of N-in: 0.12994\noutside 1 331\nTMhelix 332 354\ninside 355 372

9199 GCF_002016215.1_ASM201621v1 Escherichia coli Proteobacteria; Gammaproteobacteria; Enterobacterales; Enterobacteriaceae; Escherichia
MTGDNTLIHSHGINRRDFMKLCAALAATMGLSSKAAAEMAESVTNPQRPPVIWIGAQECT WP_000145410.1
MULTISPECIES: hydrogenase-2 small chain [Proteobacteria] Length: 372\nNumber of predicted TMHs: 1\nExp number of AAs in TMHs: 23.72226\nExp number, first 60 AAs: 0.22038\nTotal prob of N-in: 0.12994\noutside 1 331\nTMhelix 332 354\ninside 355 372

9200 GCF_002016275.1_ASM201627v1 Escherichia coli Proteobacteria; Gammaproteobacteria; Enterobacterales; Enterobacteriaceae; Escherichia
MTGDNTLIHSHGINRRDFMKLCAALAATMGLSSKAAAEMAESVTNPQRPPVIWIGAQECT WP_000145410.1
MULTISPECIES: hydrogenase-2 small chain [Proteobacteria] Length: 372\nNumber of predicted TMHs: 1\nExp number of AAs in TMHs: 23.72226\nExp number, first 60 AAs: 0.22038\nTotal prob of N-in: 0.12994\noutside 1 331\nTMhelix 332 354\ninside 355 372

9201 GCF_002016225.1_ASM201622v1 Escherichia coli Proteobacteria; Gammaproteobacteria; Enterobacterales; Enterobacteriaceae; Escherichia
MTGDNTLIHSHGINRRDFMKLCAALAATMGLSSKAAAEMAESVTNPQRPPVIWIGAQECT WP_000145410.1
MULTISPECIES: hydrogenase-2 small chain [Proteobacteria] Length: 372\nNumber of predicted TMHs: 1\nExp number of AAs in TMHs: 23.72226\nExp number, first 60 AAs: 0.22038\nTotal prob of N-in: 0.12994\noutside 1 331\nTMhelix 332 354\ninside 355 372

9202 GCF_002016375.1_ASM201637v1 Escherichia coli Proteobacteria; Gammaproteobacteria; Enterobacterales; Enterobacteriaceae; Escherichia
MNNEETFYQAMRRQGVTRRSFLKYCSLAATSLGLGAGMAPKIAWALENKPRIPVVWIHGL WP_001058323.1
MULTISPECIES: hydrogenase-1 small chain [Proteobacteria] Length: 372\nNumber of predicted TMHs: 2\nExp number of AAs in TMHs: 37.2608499999999\nExp number, first 60 AAs: 19.52923\nTotal prob of N-in: 0.94351\nPOSSIBLE N-term signal sequence\ninside 1 19\nTMhelix 20 39\noutside 40 325\nTMhelix 326 348\ninside 349 372

9203 GCF_002016405.1_ASM201640v1 Escherichia coli Proteobacteria; Gammaproteobacteria; Enterobacterales; Enterobacteriaceae; Escherichia
MNNEETFYQAMRRQGVTRRSFLKYCSLAATSLGLGAGMAPKIAWALENKPRIPVVWIHGL WP_001058323.1
MULTISPECIES: hydrogenase-1 small chain [Proteobacteria] Length: 372\nNumber of predicted TMHs: 2\nExp number of AAs in TMHs: 37.2608499999999\nExp number, first 60 AAs: 19.52923\nTotal prob of N-in: 0.94351\nPOSSIBLE N-term signal sequence\ninside 1 19\nTMhelix 20 39\noutside 40 325\nTMhelix 326 348\ninside 349 372

9204 GCF_002016205.1_ASM201620v1 Escherichia coli Proteobacteria; Gammaproteobacteria; Enterobacterales; Enterobacteriaceae; Escherichia
MNNEETFYQAMRRQGVTRRSFLKYCSLAATSLGLGAGMAPKIAWALENKPRIPVVWIHGL WP_001295940.1
MULTISPECIES: hydrogenase [Enterobacteriaceae] Length: 372\nNumber of predicted TMHs: 2\nExp

number of AAs in TMHs: 37.2608899999999\nExp number, first 60 AAs: 19.52923\nTotal prob of N-in: 0.94351\nPOSSIBLE N-term signal sequence\ninside 1 19\nTMhelix 20 39\noutside 40 325\nTMhelix 326 348\ninside 349 372

9205 GCF_002016215.1_ASM201621v1 Escherichia coli Proteobacteria; Gammaproteobacteria; Enterobacterales; Enterobacteriaceae; Escherichia
MNNEETFYQAMRRQGVTRRSFLKYCSLAATSLGLGAGMAPKIAWALENKPRIPVVWIHGL WP_001295940.1
MULTISPECIES: hydrogenase [Enterobacteriaceae] Length: 372\nNumber of predicted TMHs: 2\nExp number of AAs in TMHs: 37.2608899999999\nExp number, first 60 AAs: 19.52923\nTotal prob of N-in: 0.94351\nPOSSIBLE N-term signal sequence\ninside 1 19\nTMhelix 20 39\noutside 40 325\nTMhelix 326 348\ninside 349 372

9206 GCF_002016275.1_ASM201627v1 Escherichia coli Proteobacteria; Gammaproteobacteria; Enterobacterales; Enterobacteriaceae; Escherichia
MNNEETFYQAMRRQGVTRRSFLKYCSLAATSLGLGAGMAPKIAWALENKPRIPVVWIHGL WP_001295940.1
MULTISPECIES: hydrogenase [Enterobacteriaceae] Length: 372\nNumber of predicted TMHs: 2\nExp number of AAs in TMHs: 37.2608899999999\nExp number, first 60 AAs: 19.52923\nTotal prob of N-in: 0.94351\nPOSSIBLE N-term signal sequence\ninside 1 19\nTMhelix 20 39\noutside 40 325\nTMhelix 326 348\ninside 349 372

9207 GCF_002016305.1_ASM201630v1 Escherichia coli Proteobacteria; Gammaproteobacteria; Enterobacterales; Enterobacteriaceae; Escherichia
MNNEETFYQAMRRQGVTRRSFLKYCSLAATSLGLGAGMAPKIAWALENKPRIPVVWIHGL WP_001058323.1
MULTISPECIES: hydrogenase-1 small chain [Proteobacteria] Length: 372\nNumber of predicted TMHs: 2\nExp number of AAs in TMHs: 37.2608499999999\nExp number, first 60 AAs: 19.52923\nTotal prob of N-in: 0.94351\nPOSSIBLE N-term signal sequence\ninside 1 19\nTMhelix 20 39\noutside 40 325\nTMhelix 326 348\ninside 349 372

9208 GCF_002016285.1_ASM201628v1 Escherichia coli Proteobacteria; Gammaproteobacteria; Enterobacterales; Enterobacteriaceae; Escherichia
MNNEETFYQAMRRQGVTRRSFLKYCSLAATSLGLGAGMAPKIAWALENKPRIPVVWIHGL WP_001295940.1
MULTISPECIES: hydrogenase [Enterobacteriaceae] Length: 372\nNumber of predicted TMHs: 2\nExp number of AAs in TMHs: 37.2608899999999\nExp number, first 60 AAs: 19.52923\nTotal prob of N-in: 0.94351\nPOSSIBLE N-term signal sequence\ninside 1 19\nTMhelix 20 39\noutside 40 325\nTMhelix 326 348\ninside 349 372

9209 GCF_002016435.1_ASM201643v1 Escherichia coli Proteobacteria; Gammaproteobacteria; Enterobacterales; Enterobacteriaceae; Escherichia
MNNEETFYQAMRRQGVTRRSFLKYCSLAATSLGLGAGMAPKIAWALENKPRIPVVWIHGL WP_001058323.1
MULTISPECIES: hydrogenase-1 small chain [Proteobacteria] Length: 372\nNumber of predicted TMHs: 2\nExp number of AAs in TMHs: 37.2608499999999\nExp number, first 60 AAs: 19.52923\nTotal prob of N-in: 0.94351\nPOSSIBLE N-term signal sequence\ninside 1 19\nTMhelix 20 39\noutside 40 325\nTMhelix 326 348\ninside 349 372

9210 GCF_002018425.1_ASM201842v1 Escherichia coli Proteobacteria; Gammaproteobacteria; Enterobacterales; Enterobacteriaceae; Escherichia
MTGDNTLIHSHGINRRDFMKLCAALAATMGLSSKAAAEMAESVTNPQRPPVIWIGAQECT WP_000145410.1
MULTISPECIES: hydrogenase-2 small chain [Proteobacteria] Length: 372\nNumber of predicted TMHs: 1\nExp number of AAs in TMHs: 23.72226\nExp number, first 60 AAs: 0.22038\nTotal prob of N-in: 0.12994\noutside 1 331\nTMhelix 332 354\ninside 355 372

9211 GCF_002016545.1_ASM201654v1 Escherichia coli Proteobacteria; Gammaproteobacteria; Enterobacterales; Enterobacteriaceae; Escherichia
MTGDNTLIHSHGINRRDFMKLCAALAATMGLSSKAAAEMAESVTNPQRPPVIWIGAQECT WP_000145410.1
MULTISPECIES: hydrogenase-2 small chain [Proteobacteria] Length: 372\nNumber of predicted TMHs: 1\nExp number of AAs in TMHs: 23.72226\nExp number, first 60 AAs: 0.22038\nTotal prob of N-in: 0.12994\noutside 1 331\nTMhelix 332 354\ninside 355 372

9212 GCF_002016535.1_ASM201653v1 Escherichia coli Proteobacteria; Gammaproteobacteria; Enterobacterales; Enterobacteriaceae; Escherichia
MTGDNTLIHSHGINRRDFMKLCAALAATMGLSSKAAAEMAESVTNPQRPPVIWIGAQECT WP_000145410.1

MULTISPECIES: hydrogenase-2 small chain [Proteobacteria] Length: 372\nNumber of predicted TMHs: 1\nExp number of AAs in TMHs: 23.72226\nExp number, first 60 AAs: 0.22038\nTotal prob of N-in: 0.12994\noutside 1 331\nTMhelix 332 354\ninside 355 372

9213 GCF_002018795.1_ASM201879v1 Escherichia coli Proteobacteria; Gammaproteobacteria; Enterobacterales; Enterobacteriaceae; Escherichia
MTGDNTLIHSHGINRRDFMKLCAALAATMGLSSKAAAEMAESVTNPQRPPVIWIGAQECT WP_000145410.1
MULTISPECIES: hydrogenase-2 small chain [Proteobacteria] Length: 372\nNumber of predicted TMHs: 1\nExp number of AAs in TMHs: 23.72226\nExp number, first 60 AAs: 0.22038\nTotal prob of N-in: 0.12994\noutside 1 331\nTMhelix 332 354\ninside 355 372

9214 GCF_002018815.1_ASM201881v1 Escherichia coli Proteobacteria; Gammaproteobacteria; Enterobacterales; Enterobacteriaceae; Escherichia
MTGDNTLIHSHGINRRDFMKLCAALAATMGLSSKAAAEMAESVTNPQRPPVIWIGAQECT WP_000145410.1
MULTISPECIES: hydrogenase-2 small chain [Proteobacteria] Length: 372\nNumber of predicted TMHs: 1\nExp number of AAs in TMHs: 23.72226\nExp number, first 60 AAs: 0.22038\nTotal prob of N-in: 0.12994\noutside 1 331\nTMhelix 332 354\ninside 355 372

9215 GCF_002018425.1_ASM201842v1 Escherichia coli Proteobacteria; Gammaproteobacteria; Enterobacterales; Enterobacteriaceae; Escherichia
MNNEETFYQAMRRQGVTRRSFLKYCSLAATSLGLGAGMAPKIAWALENKPRIPVVWIHGL WP_001058323.1
MULTISPECIES: hydrogenase-1 small chain [Proteobacteria] Length: 372\nNumber of predicted TMHs: 2\nExp number of AAs in TMHs: 37.2608499999999\nExp number, first 60 AAs: 19.52923\nTotal prob of N-in: 0.94351\nPOSSIBLE N-term signal sequence\ninside 1 19\nTMhelix 20 39\noutside 40 325\nTMhelix 326 348\ninside 349 372

9216 GCF_002016545.1_ASM201654v1 Escherichia coli Proteobacteria; Gammaproteobacteria; Enterobacterales; Enterobacteriaceae; Escherichia
MNNEETFYQAMRRQGVTRRSFLKYCSLAATSLGLGAGMAPKIAWALENKPRIPVVWIHGL WP_001058323.1
MULTISPECIES: hydrogenase-1 small chain [Proteobacteria] Length: 372\nNumber of predicted TMHs: 2\nExp number of AAs in TMHs: 37.2608499999999\nExp number, first 60 AAs: 19.52923\nTotal prob of N-in: 0.94351\nPOSSIBLE N-term signal sequence\ninside 1 19\nTMhelix 20 39\noutside 40 325\nTMhelix 326 348\ninside 349 372

9217 GCF_002016535.1_ASM201653v1 Escherichia coli Proteobacteria; Gammaproteobacteria; Enterobacterales; Enterobacteriaceae; Escherichia
MNNEETFYQAMRRQGVTRRSFLKYCSLAATSLGLGAGMAPKIAWALENKPRIPVVWIHGL WP_001058323.1
MULTISPECIES: hydrogenase-1 small chain [Proteobacteria] Length: 372\nNumber of predicted TMHs: 2\nExp number of AAs in TMHs: 37.2608499999999\nExp number, first 60 AAs: 19.52923\nTotal prob of N-in: 0.94351\nPOSSIBLE N-term signal sequence\ninside 1 19\nTMhelix 20 39\noutside 40 325\nTMhelix 326 348\ninside 349 372

9218 GCF_002018795.1_ASM201879v1 Escherichia coli Proteobacteria; Gammaproteobacteria; Enterobacterales; Enterobacteriaceae; Escherichia
MNNEETFYQAMRRQGVTRRSFLKYCSLAATSLGLGAGMAPKIAWALENKPRIPVVWIHGL WP_001058323.1
MULTISPECIES: hydrogenase-1 small chain [Proteobacteria] Length: 372\nNumber of predicted TMHs: 2\nExp number of AAs in TMHs: 37.2608499999999\nExp number, first 60 AAs: 19.52923\nTotal prob of N-in: 0.94351\nPOSSIBLE N-term signal sequence\ninside 1 19\nTMhelix 20 39\noutside 40 325\nTMhelix 326 348\ninside 349 372

9219 GCF_002024905.1_ASM202490v1 Escherichia coli Proteobacteria; Gammaproteobacteria; Enterobacterales; Enterobacteriaceae; Escherichia
MTGDNTLIHSHGINRRDFMKLCAALAATMGLSSKAAAEMAESVTNPQRPPVIWIGAQECT WP_000145410.1
MULTISPECIES: hydrogenase-2 small chain [Proteobacteria] Length: 372\nNumber of predicted TMHs: 1\nExp number of AAs in TMHs: 23.72226\nExp number, first 60 AAs: 0.22038\nTotal prob of N-in: 0.12994\noutside 1 331\nTMhelix 332 354\ninside 355 372

9220 GCF_000782255.1_ASM78225v1 Escherichia coli Proteobacteria; Gammaproteobacteria; Enterobacterales; Enterobacteriaceae; Escherichia
MTGDNTLIHSHGINRRDFMKLCAALAATMGLSSKAAAEMVESVTNPQRPPVIWIGAQECT WP_000145423.1
hydrogenase 2 small subunit [Escherichia coli] Length: 372\nNumber of predicted TMHs: 1\nExp number of

AAs in TMHs: 23.78201\nExp number, first 60 AAs: 0.28355\nTotal prob of N-in: 0.13257\noutside 1 331\nTMhelix 332 354\ninside 355 372

9221 GCF_000781415.1_ASM78141v1 Escherichia coli Proteobacteria; Gammaproteobacteria; Enterobacterales; Enterobacteriaceae; Escherichia
MTGDNTLIHSHGINRRDFMKLCAALAATMGLSSKAAAEMAESVTNPQRPPVIWIGAQECT WP_000145410.1
MULTISPECIES: hydrogenase-2 small chain [Proteobacteria] Length: 372\nNumber of predicted TMHs: 1\nExp number of AAs in TMHs: 23.72226\nExp number, first 60 AAs: 0.22038\nTotal prob of N-in: 0.12994\noutside 1 331\nTMhelix 332 354\ninside 355 372

9222 GCF_000781415.1_ASM78141v1 Escherichia coli Proteobacteria; Gammaproteobacteria; Enterobacterales; Enterobacteriaceae; Escherichia
MNNEETFYQAMRRQGVTRRSFLKYCSLAATSLGLGAGMAPKIAWALENKPRIPVVWIHGL WP_001058323.1
MULTISPECIES: hydrogenase-1 small chain [Proteobacteria] Length: 372\nNumber of predicted TMHs: 2\nExp number of AAs in TMHs: 37.2608499999999\nExp number, first 60 AAs: 19.52923\nTotal prob of N-in: 0.94351\nPOSSIBLE N-term signal sequence\ninside 1 19\nTMhelix 20 39\noutside 40 325\nTMhelix 326 348\ninside 349 372

9223 GCF_000781515.1_ASM78151v1 Escherichia coli Proteobacteria; Gammaproteobacteria; Enterobacterales; Enterobacteriaceae; Escherichia
MTGDNTLIHSHGINRRDFMKLCAALAATMGLSSKAAAEMVESVTNPQRPPVIWIGAQECT WP_000145423.1
hydrogenase 2 small subunit [Escherichia coli] Length: 372\nNumber of predicted TMHs: 1\nExp number of AAs in TMHs: 23.78201\nExp number, first 60 AAs: 0.28355\nTotal prob of N-in: 0.13257\noutside 1 331\nTMhelix 332 354\ninside 355 372

9224 GCF_000781515.1_ASM78151v1 Escherichia coli Proteobacteria; Gammaproteobacteria; Enterobacterales; Enterobacteriaceae; Escherichia
MNNEETFYQAMRRQGVTRRSFLKYCSLAATSLGLGAGMAPKIAWALENKPRIPVVWIHGL WP_001295940.1
MULTISPECIES: hydrogenase [Enterobacteriaceae] Length: 372\nNumber of predicted TMHs: 2\nExp number of AAs in TMHs: 37.2608899999999\nExp number, first 60 AAs: 19.52923\nTotal prob of N-in: 0.94351\nPOSSIBLE N-term signal sequence\ninside 1 19\nTMhelix 20 39\noutside 40 325\nTMhelix 326 348\ninside 349 372

9225 GCF_000781595.1_ASM78159v1 Escherichia coli Proteobacteria; Gammaproteobacteria; Enterobacterales; Enterobacteriaceae; Escherichia
MTGDNTLIHSHGINRRDFMKLCAALAATMGLSSKAAAEMAESVTNPQRPPVIWIGAQECT WP_000145410.1
MULTISPECIES: hydrogenase-2 small chain [Proteobacteria] Length: 372\nNumber of predicted TMHs: 1\nExp number of AAs in TMHs: 23.72226\nExp number, first 60 AAs: 0.22038\nTotal prob of N-in: 0.12994\noutside 1 331\nTMhelix 332 354\ninside 355 372

9226 GCF_000781595.1_ASM78159v1 Escherichia coli Proteobacteria; Gammaproteobacteria; Enterobacterales; Enterobacteriaceae; Escherichia
MNNEETFYQAMRRQGVTRRSFLKYCSLAATSLGLGAGMAPKIAWALENKPRIPVVWIHGL WP_001058323.1
MULTISPECIES: hydrogenase-1 small chain [Proteobacteria] Length: 372\nNumber of predicted TMHs: 2\nExp number of AAs in TMHs: 37.2608499999999\nExp number, first 60 AAs: 19.52923\nTotal prob of N-in: 0.94351\nPOSSIBLE N-term signal sequence\ninside 1 19\nTMhelix 20 39\noutside 40 325\nTMhelix 326 348\ninside 349 372

9227 GCF_000781635.1_ASM78163v1 Escherichia coli Proteobacteria; Gammaproteobacteria; Enterobacterales; Enterobacteriaceae; Escherichia
MTGDNTLIHSHGINRRDFMKLCAALAATMGLSSKAAAEMAESVTNPQRPPVIWIGAQECT WP_000145410.1
MULTISPECIES: hydrogenase-2 small chain [Proteobacteria] Length: 372\nNumber of predicted TMHs: 1\nExp number of AAs in TMHs: 23.72226\nExp number, first 60 AAs: 0.22038\nTotal prob of N-in: 0.12994\noutside 1 331\nTMhelix 332 354\ninside 355 372

9228 GCF_000781635.1_ASM78163v1 Escherichia coli Proteobacteria; Gammaproteobacteria; Enterobacterales; Enterobacteriaceae; Escherichia
MNNEETFYQAMRRQGVTRRSFLKYCSLAATSLGLGAGMAPKIAWALENKPRIPVVWIHGL WP_001058323.1
MULTISPECIES: hydrogenase-1 small chain [Proteobacteria] Length: 372\nNumber of predicted TMHs: 2\nExp number of AAs in TMHs: 37.2608499999999\nExp number, first 60 AAs: 19.52923\nTotal prob of N-in:

0.94351\nPOSSIBLE N-term signal sequence\ninside 1 19\nTMhelix 20 39\noutside 40 325\nTMhelix 326 348\ninside 349 372

9229 GCF_000781695.1_ASM78169v1 Escherichia coli Proteobacteria; Gammaproteobacteria; Enterobacterales; Enterobacteriaceae; Escherichia
MTGDNTLIHSHGINRRDFMKLCAALAATMGLSSKAAAEMVESVTNPQRPPVIWIGAQECT WP_000145423.1
hydrogenase 2 small subunit [Escherichia coli] Length: 372\nNumber of predicted TMHs: 1\nExp number of AAs in TMHs: 23.78201\nExp number, first 60 AAs: 0.28355\nTotal prob of N-in: 0.13257\noutside 1 331\nTMhelix 332 354\ninside 355 372

9230 GCF_000781695.1_ASM78169v1 Escherichia coli Proteobacteria; Gammaproteobacteria; Enterobacterales; Enterobacteriaceae; Escherichia
MNNEETFYQAMRRQGVTRRSFLKYCSLAATSLGLGAGMAPKIAWALENKPRIPVVWIHGL WP_001295940.1
MULTISPECIES: hydrogenase [Enterobacteriaceae] Length: 372\nNumber of predicted TMHs: 2\nExp number of AAs in TMHs: 37.2608899999999\nExp number, first 60 AAs: 19.52923\nTotal prob of N-in: 0.94351\nPOSSIBLE N-term signal sequence\ninside 1 19\nTMhelix 20 39\noutside 40 325\nTMhelix 326 348\ninside 349 372

9231 GCF_000781775.1_ASM78177v1 Escherichia coli Proteobacteria; Gammaproteobacteria; Enterobacterales; Enterobacteriaceae; Escherichia
MTGDNTLIHSHGINRRDFMKLCAALAATMGLSSKAAAEMAESVTNPQRPPVIWIGAQECT WP_000145410.1
MULTISPECIES: hydrogenase-2 small chain [Proteobacteria] Length: 372\nNumber of predicted TMHs: 1\nExp number of AAs in TMHs: 23.72226\nExp number, first 60 AAs: 0.22038\nTotal prob of N-in: 0.12994\noutside 1 331\nTMhelix 332 354\ninside 355 372

9232 GCF_000781775.1_ASM78177v1 Escherichia coli Proteobacteria; Gammaproteobacteria; Enterobacterales; Enterobacteriaceae; Escherichia
MNNEETFYQAMRRQGVTRRSFLKYCSLAATSLGLGAGMAPKIAWALENKPRIPVVWIHGL WP_001295940.1
MULTISPECIES: hydrogenase [Enterobacteriaceae] Length: 372\nNumber of predicted TMHs: 2\nExp number of AAs in TMHs: 37.2608899999999\nExp number, first 60 AAs: 19.52923\nTotal prob of N-in: 0.94351\nPOSSIBLE N-term signal sequence\ninside 1 19\nTMhelix 20 39\noutside 40 325\nTMhelix 326 348\ninside 349 372

9233 GCF_000781855.1_ASM78185v1 Escherichia coli Proteobacteria; Gammaproteobacteria; Enterobacterales; Enterobacteriaceae; Escherichia
MNNEETFYQAMRRQGVTRRSFLKYCSLAATSLGLGAGMAPKIAWALENKPRIPVVWIHGL WP_001058323.1
MULTISPECIES: hydrogenase-1 small chain [Proteobacteria] Length: 372\nNumber of predicted TMHs: 2\nExp number of AAs in TMHs: 37.2608499999999\nExp number, first 60 AAs: 19.52923\nTotal prob of N-in: 0.94351\nPOSSIBLE N-term signal sequence\ninside 1 19\nTMhelix 20 39\noutside 40 325\nTMhelix 326 348\ninside 349 372

9234 GCF_000781915.1_ASM78191v1 Escherichia coli Proteobacteria; Gammaproteobacteria; Enterobacterales; Enterobacteriaceae; Escherichia
MTGDNTLIHSHGINRRDFMKLCAALAATMGLSSKAAAEMAESVTNPQRPPVIWIGAQECT WP_000145410.1
MULTISPECIES: hydrogenase-2 small chain [Proteobacteria] Length: 372\nNumber of predicted TMHs: 1\nExp number of AAs in TMHs: 23.72226\nExp number, first 60 AAs: 0.22038\nTotal prob of N-in: 0.12994\noutside 1 331\nTMhelix 332 354\ninside 355 372

9235 GCF_000781915.1_ASM78191v1 Escherichia coli Proteobacteria; Gammaproteobacteria; Enterobacterales; Enterobacteriaceae; Escherichia
MNNEETFYQAMRRQGVTRRSFLKYCSLAATSLGLGAGMAPKIAWALENKPRIPVVWIHGL WP_001058323.1
MULTISPECIES: hydrogenase-1 small chain [Proteobacteria] Length: 372\nNumber of predicted TMHs: 2\nExp number of AAs in TMHs: 37.2608499999999\nExp number, first 60 AAs: 19.52923\nTotal prob of N-in: 0.94351\nPOSSIBLE N-term signal sequence\ninside 1 19\nTMhelix 20 39\noutside 40 325\nTMhelix 326 348\ninside 349 372

9236 GCF_000781955.1_ASM78195v1 Escherichia coli Proteobacteria; Gammaproteobacteria; Enterobacterales; Enterobacteriaceae; Escherichia
MTGDNTLIHSHGINRRDFMKLCAALAATMGLSSKAAAEMAESVTNPQRPPVIWIGAQECT WP_000145410.1
MULTISPECIES: hydrogenase-2 small chain [Proteobacteria] Length: 372\nNumber of predicted TMHs:

1\nExp number of AAs in TMHs: 23.72226\nExp number, first 60 AAs: 0.22038\nTotal prob of N-in: 0.12994\noutside 1 331\nTMhelix 332 354\ninside 355 372

9237 GCF_000781955.1_ASM78195v1 Escherichia coli Proteobacteria; Gammaproteobacteria; Enterobacterales; Enterobacteriaceae; Escherichia
MNNEETFYQAMRRQGVTRRSFLKYCSLAATSLGLGAGMAPKIAWALENKPRIPVVWIHGL WP_001058323.1
MULTISPECIES: hydrogenase-1 small chain [Proteobacteria] Length: 372\nNumber of predicted TMHs: 2\nExp number of AAs in TMHs: 37.2608499999999\nExp number, first 60 AAs: 19.52923\nTotal prob of N-in: 0.94351\nPOSSIBLE N-term signal sequence\ninside 1 19\nTMhelix 20 39\noutside 40 325\nTMhelix 326 348\ninside 349 372

9238 GCF_000782155.1_ASM78215v1 Escherichia coli Proteobacteria; Gammaproteobacteria; Enterobacterales; Enterobacteriaceae; Escherichia
MTGDNTLIHSHGINRRDFMKLCAALAATMGLSSKAAAEMAESVTNPQRPPVIWIGAQECT WP_000145410.1
MULTISPECIES: hydrogenase-2 small chain [Proteobacteria] Length: 372\nNumber of predicted TMHs: 1\nExp number of AAs in TMHs: 23.72226\nExp number, first 60 AAs: 0.22038\nTotal prob of N-in: 0.12994\noutside 1 331\nTMhelix 332 354\ninside 355 372

9239 GCF_000782155.1_ASM78215v1 Escherichia coli Proteobacteria; Gammaproteobacteria; Enterobacterales; Enterobacteriaceae; Escherichia
MNNEETFYQAMRRQGVTRRSFLKYCSLAATSLGLGAGMAPKIAWALENKPRIPVVWIHGL WP_001058323.1
MULTISPECIES: hydrogenase-1 small chain [Proteobacteria] Length: 372\nNumber of predicted TMHs: 2\nExp number of AAs in TMHs: 37.2608499999999\nExp number, first 60 AAs: 19.52923\nTotal prob of N-in: 0.94351\nPOSSIBLE N-term signal sequence\ninside 1 19\nTMhelix 20 39\noutside 40 325\nTMhelix 326 348\ninside 349 372

9240 GCF_000782215.1_ASM78221v1 Escherichia coli Proteobacteria; Gammaproteobacteria; Enterobacterales; Enterobacteriaceae; Escherichia
MTGDNTLIHSHGINRRDFMKLCAALAATMGLSSKAAAEMAESVTNPQRPPVIWIGAQECT WP_000145410.1
MULTISPECIES: hydrogenase-2 small chain [Proteobacteria] Length: 372\nNumber of predicted TMHs: 1\nExp number of AAs in TMHs: 23.72226\nExp number, first 60 AAs: 0.22038\nTotal prob of N-in: 0.12994\noutside 1 331\nTMhelix 332 354\ninside 355 372

9241 GCF_001058555.1_ASM105855v1 Escherichia coli Proteobacteria; Gammaproteobacteria; Enterobacterales; Enterobacteriaceae; Escherichia
MTGDNTLIHSHGINRRDFMKLCAALAATMGLSSKAAAEMAESVTNPQRPPVIWIGAQECT WP_000145410.1
MULTISPECIES: hydrogenase-2 small chain [Proteobacteria] Length: 372\nNumber of predicted TMHs: 1\nExp number of AAs in TMHs: 23.72226\nExp number, first 60 AAs: 0.22038\nTotal prob of N-in: 0.12994\noutside 1 331\nTMhelix 332 354\ninside 355 372

9242 GCF_001058555.1_ASM105855v1 Escherichia coli Proteobacteria; Gammaproteobacteria; Enterobacterales; Enterobacteriaceae; Escherichia
MNNEETFYQAMRRQGVTRRSFLKYCSLAATSLGLGAGMAPKIAWALENKPRIPVVWIHGL WP_001058323.1
MULTISPECIES: hydrogenase-1 small chain [Proteobacteria] Length: 372\nNumber of predicted TMHs: 2\nExp number of AAs in TMHs: 37.2608499999999\nExp number, first 60 AAs: 19.52923\nTotal prob of N-in: 0.94351\nPOSSIBLE N-term signal sequence\ninside 1 19\nTMhelix 20 39\noutside 40 325\nTMhelix 326 348\ninside 349 372

9243 GCF_000782375.1_ASM78237v1 Escherichia coli Proteobacteria; Gammaproteobacteria; Enterobacterales; Enterobacteriaceae; Escherichia
MTGDNTLIHSHGINRRDFMKLCAALAATMGLSSKAAAEMAESVTNPQRPPVIWIGAQECT WP_000145410.1
MULTISPECIES: hydrogenase-2 small chain [Proteobacteria] Length: 372\nNumber of predicted TMHs: 1\nExp number of AAs in TMHs: 23.72226\nExp number, first 60 AAs: 0.22038\nTotal prob of N-in: 0.12994\noutside 1 331\nTMhelix 332 354\ninside 355 372

9244 GCF_000782375.1_ASM78237v1 Escherichia coli Proteobacteria; Gammaproteobacteria; Enterobacterales; Enterobacteriaceae; Escherichia
MNNEETFYQAMRRQGVTRRSFLKYCSLAATSLGLGAGMAPKIAWALENKPRIPVVWIHGL WP_001058323.1
MULTISPECIES: hydrogenase-1 small chain [Proteobacteria] Length: 372\nNumber of predicted TMHs: 2\nExp number of AAs in TMHs: 37.2608499999999\nExp number, first 60 AAs: 19.52923\nTotal prob of N-in:

0.94351\nPOSSIBLE N-term signal sequence\ninside 1 19\nTMhelix 20 39\noutside 40 325\nTMhelix 326 348\ninside 349 372

9245 GCF_000968905.2_ASM96890v2 Escherichia fergusonii Proteobacteria; Gammaproteobacteria; Enterobacterales; Enterobacteriaceae; Escherichia MNNEETFYQAMRRQGVTRRSFLKYCSLAATSLGLGAGMAPKIAWALENKPRIPVVWIHGL WP_046077437.1 hydrogenase [Escherichia fergusonii] Length: 372\nNumber of predicted TMHs: 2\nExp number of AAs in TMHs: 37.08355\nExp number, first 60 AAs: 19.52938\nTotal prob of N-in: 0.94341\nPOSSIBLE N-term signal sequence\ninside 1 19\nTMhelix 20 39\noutside 40 325\nTMhelix 326 348\ninside 349 372

9246 GCF_001471755.1_ASM147175v1 Escherichia fergusonii Proteobacteria; Gammaproteobacteria; Enterobacterales; Enterobacteriaceae; Escherichia MTGDNTLIHSHGINRRDFMKLCAALAATMGLSSKAAAEMAESVTNPQRPPVIWIGAQECT WP_000145410.1 MULTISPECIES: hydrogenase-2 small chain [Proteobacteria] Length: 372\nNumber of predicted TMHs: 1\nExp number of AAs in TMHs: 23.72226\nExp number, first 60 AAs: 0.22038\nTotal prob of N-in: 0.12994\noutside 1 331\nTMhelix 332 354\ninside 355 372

9247 GCF_000968895.2_ASM96889v2 Escherichia fergusonii Proteobacteria; Gammaproteobacteria; Enterobacterales; Enterobacteriaceae; Escherichia MNNEETFYQAMRRQGVTRRSFLKYCSLAATSLGLGAGMAPKIAWALENKPRIPVVWIHGL WP_001295940.1 MULTISPECIES: hydrogenase [Enterobacteriaceae] Length: 372\nNumber of predicted TMHs: 2\nExp number of AAs in TMHs: 37.2608899999999\nExp number, first 60 AAs: 19.52923\nTotal prob of N-in: 0.94351\nPOSSIBLE N-term signal sequence\ninside 1 19\nTMhelix 20 39\noutside 40 325\nTMhelix 326 348\ninside 349 372

9248 GCF_000968885.2_ASM96888v2 Escherichia fergusonii Proteobacteria; Gammaproteobacteria; Enterobacterales; Enterobacteriaceae; Escherichia MNNEETFYQAMRRQGVTRRSFLKYCSLAATSLGLGAGMAPKIAWALENKPRIPVVWIHGL WP_002431613.1 hydrogenase-1 small chain [Escherichia fergusonii] Length: 372\nNumber of predicted TMHs: 2\nExp number of AAs in TMHs: 37.25693\nExp number, first 60 AAs: 19.52926\nTotal prob of N-in: 0.94351\nPOSSIBLE N-term signal sequence\ninside 1 19\nTMhelix 20 39\noutside 40 325\nTMhelix 326 348\ninside 349 372

9249 GCF_001471755.1_ASM147175v1 Escherichia fergusonii Proteobacteria; Gammaproteobacteria; Enterobacterales; Enterobacteriaceae; Escherichia MNNEETFYQAMRRQGVTRRSFLKYCSLAATSLGLGAGMAPKIAWALENKPRIPVVWIHGL WP_001058323.1 MULTISPECIES: hydrogenase-1 small chain [Proteobacteria] Length: 372\nNumber of predicted TMHs: 2\nExp number of AAs in TMHs: 37.2608499999999\nExp number, first 60 AAs: 19.52923\nTotal prob of N-in: 0.94351\nPOSSIBLE N-term signal sequence\ninside 1 19\nTMhelix 20 39\noutside 40 325\nTMhelix 326 348\ninside 349 372

9250 GCF_000968885.2_ASM96888v2 Escherichia fergusonii Proteobacteria; Gammaproteobacteria; Enterobacterales; Enterobacteriaceae; Escherichia MTGDNTLDNSHGINRRDFMKLCAALAATMGLSSKAAAEMAESVANPQRPPVIWIGAQECT WP_000145402.1 hydrogenase 2 small subunit [Escherichia fergusonii] Length: 372\nNumber of predicted TMHs: 1\nExp number of AAs in TMHs: 24.63122\nExp number, first 60 AAs: 0.32267\nTotal prob of N-in: 0.18195\noutside 1 331\nTMhelix 332 354\ninside 355 372

9251 GCF_000968895.2_ASM96889v2 Escherichia fergusonii Proteobacteria; Gammaproteobacteria; Enterobacterales; Enterobacteriaceae; Escherichia MTGDNTLDNSHGINRRDFMKLCAALAATMGLSSKAAAEMAESVANPQRPPVIWIGAQECT WP_000145402.1 hydrogenase 2 small subunit [Escherichia fergusonii] Length: 372\nNumber of predicted TMHs: 1\nExp number of AAs in TMHs: 24.63122\nExp number, first 60 AAs: 0.32267\nTotal prob of N-in: 0.18195\noutside 1 331\nTMhelix 332 354\ninside 355 372

9252 GCF_000968905.2_ASM96890v2 Escherichia fergusonii Proteobacteria; Gammaproteobacteria; Enterobacterales; Enterobacteriaceae; Escherichia MTGDNTLDNSHGINRRDFMKLCAALAATMGLSSKAAAEMAESVANPQRPPVIWIGAQECT WP_000145402.1 hydrogenase 2 small subunit [Escherichia fergusonii] Length: 372\nNumber of predicted TMHs: 1\nExp number of AAs in TMHs: 24.63122\nExp number, first 60 AAs: 0.32267\nTotal prob of N-in: 0.18195\noutside 1 331\nTMhelix 332 354\ninside 355 372

9253 GCF_000807695.1_ASM80769v3 Escherichia marmotae Proteobacteria; Gammaproteobacteria; Enterobacterales; Enterobacteriaceae; Escherichia
MNNEETFYQAMRRQGVTRRSFLKYCSLAATSLGLGAGMAPKIAWALENKPRIPVVWIHGL WP_038355281.1
hydrogenase [Escherichia marmotae] Length: 372\nNumber of predicted TMHs: 2\nExp number of AAs in TMHs: 38.3313\nExp number, first 60 AAs: 19.52893\nTotal prob of N-in: 0.94355\nPOSSIBLE N-term signal sequence\ninside 1 19\nTMhelix 20 39\noutside 40 325\nTMhelix 326 348\ninside 349 372

9254 GCF_000807695.1_ASM80769v3 Escherichia marmotae Proteobacteria; Gammaproteobacteria; Enterobacterales; Enterobacteriaceae; Escherichia
MTGDNTLIHSHGINRRDFMKLCAALAATMGLSSKAAAEMAESVTNPQRPPVIWIGAQECT WP_000145417.1
MULTISPECIES: hydrogenase-2 small chain [Escherichia] Length: 372\nNumber of predicted TMHs: 1\nExp number of AAs in TMHs: 23.72512\nExp number, first 60 AAs: 0.22034\nTotal prob of N-in: 0.13000\noutside 1 331\nTMhelix 332 354\ninside 355 372

9255 GCF_000159895.2_Escherichia_sp_1_1_43_V2 Escherichia sp. 1_1_43 Proteobacteria; Gammaproteobacteria; Enterobacterales; Enterobacteriaceae; Escherichia
MNNEETFYQAMRRQGVTRRSFLKYCSLAATSLGLGAGMAPKIAWALENKPRIPVVWIHGL WP_001058323.1
MULTISPECIES: hydrogenase-1 small chain [Proteobacteria] Length: 372\nNumber of predicted TMHs: 2\nExp number of AAs in TMHs: 37.2608499999999\nExp number, first 60 AAs: 19.52923\nTotal prob of N-in: 0.94351\nPOSSIBLE N-term signal sequence\ninside 1 19\nTMhelix 20 39\noutside 40 325\nTMhelix 326 348\ninside 349 372

9256 GCF_000159895.2_Escherichia_sp_1_1_43_V2 Escherichia sp. 1_1_43 Proteobacteria; Gammaproteobacteria; Enterobacterales; Enterobacteriaceae; Escherichia
MTGDNTLIHSHGINRRDFMKLCAALAATMGLSSKAAAEMAESVTNPQRPPVIWIGAQECT WP_000145410.1
MULTISPECIES: hydrogenase-2 small chain [Proteobacteria] Length: 372\nNumber of predicted TMHs: 1\nExp number of AAs in TMHs: 23.72226\nExp number, first 60 AAs: 0.22038\nTotal prob of N-in: 0.12994\noutside 1 331\nTMhelix 332 354\ninside 355 372

9257 GCF_000157115.2_Escherichia_sp_3_2_53FAA_V2 Escherichia sp. 3_2_53FAA Proteobacteria; Gammaproteobacteria; Enterobacterales; Enterobacteriaceae; Escherichia
MTGDNTLIHSHGINRRDFMKLCAALAATMGLSSKAAAEMAESVTNPQRPPVIWIGAQECT WP_000145410.1
MULTISPECIES: hydrogenase-2 small chain [Proteobacteria] Length: 372\nNumber of predicted TMHs: 1\nExp number of AAs in TMHs: 23.72226\nExp number, first 60 AAs: 0.22038\nTotal prob of N-in: 0.12994\noutside 1 331\nTMhelix 332 354\ninside 355 372

9258 GCF_000158415.2_Escherichia_sp_4_1_40B_V2 Escherichia sp. 4_1_40B Proteobacteria; Gammaproteobacteria; Enterobacterales; Enterobacteriaceae; Escherichia
MNNEETFYQAMRRQGVTRRSFLKYCSLAATSLGLGAGMAPKIAWALENKPRIPVVWIHGL WP_001058323.1
MULTISPECIES: hydrogenase-1 small chain [Proteobacteria] Length: 372\nNumber of predicted TMHs: 2\nExp number of AAs in TMHs: 37.2608499999999\nExp number, first 60 AAs: 19.52923\nTotal prob of N-in: 0.94351\nPOSSIBLE N-term signal sequence\ninside 1 19\nTMhelix 20 39\noutside 40 325\nTMhelix 326 348\ninside 349 372

9259 GCF_000158415.2_Escherichia_sp_4_1_40B_V2 Escherichia sp. 4_1_40B Proteobacteria; Gammaproteobacteria; Enterobacterales; Enterobacteriaceae; Escherichia
MTGDNTLIHSHGINRRDFMKLCAALAATMGLSSKAAAEMAESVTNPQRPPVIWIGAQECT WP_000145410.1
MULTISPECIES: hydrogenase-2 small chain [Proteobacteria] Length: 372\nNumber of predicted TMHs: 1\nExp number of AAs in TMHs: 23.72226\nExp number, first 60 AAs: 0.22038\nTotal prob of N-in: 0.12994\noutside 1 331\nTMhelix 332 354\ninside 355 372

9260 GCF_001660175.1_ASM166017v1 Escherichia sp. B1147 Proteobacteria; Gammaproteobacteria; Enterobacterales; Enterobacteriaceae; Escherichia
MTGDNTLIHSHGINRRDFMKLCAALAATMGLSSKAAAEMAESVTNPQRPPVIWIGAQECT WP_000145410.1
MULTISPECIES: hydrogenase-2 small chain [Proteobacteria] Length: 372\nNumber of predicted TMHs: 1\nExp number of AAs in TMHs: 23.72226\nExp number, first 60 AAs: 0.22038\nTotal prob of N-in: 0.12994\noutside 1 331\nTMhelix 332 354\ninside 355 372

9261 GCF_001660175.1_ASM166017v1 Escherichia sp. B1147 Proteobacteria; Gammaproteobacteria; Enterobacterales; Enterobacteriaceae; Escherichia
MNNEETFYQAMRRQGVTRRSFLKYCSLAATSLGLGAGMAPKIAWALENKPRIPVVWIHGL WP_064528419.1

hydrogenase [Escherichia sp. B1147] Length: 372\nNumber of predicted TMHs: 2\nExp number of AAs in TMHs: 38.04888\nExp number, first 60 AAs: 19.5298\nTotal prob of N-in: 0.94352\nPOSSIBLE N-term signal sequence\ninside 1 19\nTMhelix 20 39\noutside 40 325\nTMhelix 326 348\ninside 349 372

9262 GCF_000350705.1_Esch_coli_KTE11_V1 Escherichia sp. KTE11 Proteobacteria; Gammaproteobacteria; Enterobacterales; Enterobacteriaceae; Escherichia MNNEETFYQAMRRQGVTRRSFLKYCSLAATSLGLGAGMAPKIAWALENKPRIPVVWIHGL WP_001516139.1 MULTISPECIES: hydrogenase [Escherichia] Length: 372\nNumber of predicted TMHs: 2\nExp number of AAs in TMHs: 37.98718\nExp number, first 60 AAs: 19.52876\nTotal prob of N-in: 0.94354\nPOSSIBLE N-term signal sequence\ninside 1 19\nTMhelix 20 39\noutside 40 325\nTMhelix 326 348\ninside 349 372

9263 GCF_000350705.1_Esch_coli_KTE11_V1 Escherichia sp. KTE11 Proteobacteria; Gammaproteobacteria; Enterobacterales; Enterobacteriaceae; Escherichia MTGDNTLIHSHGINRRDFMKLCAALAATMGLSSKAAAEMAESVTNPQRPPVIWIGAQECT WP_000145417.1 MULTISPECIES: hydrogenase-2 small chain [Escherichia] Length: 372\nNumber of predicted TMHs: 1\nExp number of AAs in TMHs: 23.72512\nExp number, first 60 AAs: 0.22034\nTotal prob of N-in: 0.13000\noutside 1 331\nTMhelix 332 354\ninside 355 372

9264 GCF_000407765.1_Esch_coli_KTE114_V1 Escherichia sp. KTE114 Proteobacteria; Gammaproteobacteria; Enterobacterales; Enterobacteriaceae; Escherichia MTGDNTLIHSHGINRRDFMKLCAALAATMGLSSKAAAEMAESVTNPQRPPVIWIGAQECT WP_000145412.1 MULTISPECIES: hydrogenase-2 small chain [Escherichia] Length: 372\nNumber of predicted TMHs: 1\nExp number of AAs in TMHs: 23.72209\nExp number, first 60 AAs: 0.22038\nTotal prob of N-in: 0.12994\noutside 1 331\nTMhelix 332 354\ninside 355 372

9265 GCF_000407765.1_Esch_coli_KTE114_V1 Escherichia sp. KTE114 Proteobacteria; Gammaproteobacteria; Enterobacterales; Enterobacteriaceae; Escherichia MNNEETFYQAMRRQGVTRRSFLKYCSLAATSLGLGAGMAPKIAWALENKPRIPVVWIHGL WP_010344442.1 MULTISPECIES: hydrogenase [Escherichia] Length: 372\nNumber of predicted TMHs: 2\nExp number of AAs in TMHs: 38.04874\nExp number, first 60 AAs: 19.5298\nTotal prob of N-in: 0.94352\nPOSSIBLE N-term signal sequence\ninside 1 19\nTMhelix 20 39\noutside 40 325\nTMhelix 326 348\ninside 349 372

9266 GCF_000408805.1_Esch_coli_KTE159_V1 Escherichia sp. KTE159 Proteobacteria; Gammaproteobacteria; Enterobacterales; Enterobacteriaceae; Escherichia MNNEETFYQAMRRQGVTRRSFLKYCSLAATSLGLGAGMAPKIAWALENKPRIPVVWIHGL WP_001516139.1 MULTISPECIES: hydrogenase [Escherichia] Length: 372\nNumber of predicted TMHs: 2\nExp number of AAs in TMHs: 37.98718\nExp number, first 60 AAs: 19.52876\nTotal prob of N-in: 0.94354\nPOSSIBLE N-term signal sequence\ninside 1 19\nTMhelix 20 39\noutside 40 325\nTMhelix 326 348\ninside 349 372

9267 GCF_000408805.1_Esch_coli_KTE159_V1 Escherichia sp. KTE159 Proteobacteria; Gammaproteobacteria; Enterobacterales; Enterobacteriaceae; Escherichia MTGDNTLIHSHGINRRDFMKLCAALAATMGLSSKAAAEMAESVTNPQRPPVIWIGAQECT WP_000145417.1 MULTISPECIES: hydrogenase-2 small chain [Escherichia] Length: 372\nNumber of predicted TMHs: 1\nExp number of AAs in TMHs: 23.72512\nExp number, first 60 AAs: 0.22034\nTotal prob of N-in: 0.13000\noutside 1 331\nTMhelix 332 354\ninside 355 372

9268 GCF_000408845.1_Esch_coli_KTE172_V1 Escherichia sp. KTE172 Proteobacteria; Gammaproteobacteria; Enterobacterales; Enterobacteriaceae; Escherichia MTGDNTLIHSHGINRRDFMKLCAALAATMGLSSKAAAEMAESVTNPQRPPVIWIGAQECT WP_000145410.1 MULTISPECIES: hydrogenase-2 small chain [Proteobacteria] Length: 372\nNumber of predicted TMHs: 1\nExp number of AAs in TMHs: 23.72226\nExp number, first 60 AAs: 0.22038\nTotal prob of N-in: 0.12994\noutside 1 331\nTMhelix 332 354\ninside 355 372

9269 GCF_000408845.1_Esch_coli_KTE172_V1 Escherichia sp. KTE172 Proteobacteria; Gammaproteobacteria; Enterobacterales; Enterobacteriaceae; Escherichia MNNEETFYQAMRRQGVTRRSFLKYCSLAATSLGLGAGMAPKIAWALENKPRIPVVWIHGL WP_001058323.1 MULTISPECIES: hydrogenase-1 small chain [Proteobacteria] Length: 372\nNumber of predicted TMHs: 2\nExp number of AAs in TMHs: 37.2608499999999\nExp number, first 60 AAs: 19.52923\nTotal prob of N-in: 0.94351\nPOSSIBLE N-term signal sequence\ninside 1 19\nTMhelix 20 39\noutside 40 325\nTMhelix 326 348\ninside 349 372

9270	GCF_000407925.1_Esch_coli_KTE31_V1	Escherichia sp. KTE31	Proteobacteria; Gammaproteobacteria; Enterobacterales; Enterobacteriaceae; Escherichia
MNNEETFYQAMRRQGVTRRSFLKYCSLAATSLGLGAGMAPKIAWALENKPRIPVVWIHGL	WP_010344442.1
MULTISPECIES: hydrogenase [Escherichia]	Length: 372\nNumber of predicted TMHs: 2\nExp number of AAs in TMHs: 38.04874\nExp number, first 60 AAs: 19.5298\nTotal prob of N-in: 0.94352\nPOSSIBLE N-term signal sequence\ninside 1 19\nTMhelix 20 39\noutside 40 325\nTMhelix 326 348\ninside 349 372

9271	GCF_000407925.1_Esch_coli_KTE31_V1	Escherichia sp. KTE31	Proteobacteria; Gammaproteobacteria; Enterobacterales; Enterobacteriaceae; Escherichia
MTGDNTLIHSHGINRRDFMKLCAALAATMGLSSKAAAEMAESVTNPQRPPVIWIGAQECT	WP_000145410.1
MULTISPECIES: hydrogenase-2 small chain [Proteobacteria]	Length: 372\nNumber of predicted TMHs: 1\nExp number of AAs in TMHs: 23.72226\nExp number, first 60 AAs: 0.22038\nTotal prob of N-in: 0.12994\noutside 1 331\nTMhelix 332 354\ninside 355 372

9272	GCF_000408325.1_Esch_coli_KTE52_V1	Escherichia sp. KTE52	Proteobacteria; Gammaproteobacteria; Enterobacterales; Enterobacteriaceae; Escherichia
MNNEETFYQAMRRQGVTRRSFLKYCSLAATSLGLGAGMAPKIAWALENKPRIPVVWIHGL	WP_001516139.1
MULTISPECIES: hydrogenase [Escherichia]	Length: 372\nNumber of predicted TMHs: 2\nExp number of AAs in TMHs: 37.98718\nExp number, first 60 AAs: 19.52876\nTotal prob of N-in: 0.94354\nPOSSIBLE N-term signal sequence\ninside 1 19\nTMhelix 20 39\noutside 40 325\nTMhelix 326 348\ninside 349 372

9273	GCF_000408325.1_Esch_coli_KTE52_V1	Escherichia sp. KTE52	Proteobacteria; Gammaproteobacteria; Enterobacterales; Enterobacteriaceae; Escherichia
MTGDNTLIHSHGINRRDFMKLCAALAATMGLSSKAAAEMAESVTNPQRPPVIWIGAQECT	WP_000145417.1
MULTISPECIES: hydrogenase-2 small chain [Escherichia]	Length: 372\nNumber of predicted TMHs: 1\nExp number of AAs in TMHs: 23.72512\nExp number, first 60 AAs: 0.22034\nTotal prob of N-in: 0.13000\noutside 1 331\nTMhelix 332 354\ninside 355 372

9274	GCF_000408525.1_Esch_coli_KTE96_V1	Escherichia sp. KTE96	Proteobacteria; Gammaproteobacteria; Enterobacterales; Enterobacteriaceae; Escherichia
MNNEETFYQAMRRQGVTRRSFLKYCSLAATSLGLGAGMAPKIAWALENKPRIPVVWIHGL	WP_001516139.1
MULTISPECIES: hydrogenase [Escherichia]	Length: 372\nNumber of predicted TMHs: 2\nExp number of AAs in TMHs: 37.98718\nExp number, first 60 AAs: 19.52876\nTotal prob of N-in: 0.94354\nPOSSIBLE N-term signal sequence\ninside 1 19\nTMhelix 20 39\noutside 40 325\nTMhelix 326 348\ninside 349 372

9275	GCF_000408525.1_Esch_coli_KTE96_V1	Escherichia sp. KTE96	Proteobacteria; Gammaproteobacteria; Enterobacterales; Enterobacteriaceae; Escherichia
MTGDNTLIHSHGINRRDFMKLCAALAATMGLSSKAAAEMAESVTNPQRPPVIWIGAQECT	WP_000145417.1
MULTISPECIES: hydrogenase-2 small chain [Escherichia]	Length: 372\nNumber of predicted TMHs: 1\nExp number of AAs in TMHs: 23.72512\nExp number, first 60 AAs: 0.22034\nTotal prob of N-in: 0.13000\noutside 1 331\nTMhelix 332 354\ninside 355 372

9276	GCF_000208465.1_ASM20846v2	Escherichia sp. TW09231	Proteobacteria; Gammaproteobacteria; Enterobacterales; Enterobacteriaceae; Escherichia
MTGDNTLIHSHGINRRDFMKLCAALAATMGLSSKAAAEMAESVTNPQRPPVIWIGAQECT	WP_000145410.1
MULTISPECIES: hydrogenase-2 small chain [Proteobacteria]	Length: 372\nNumber of predicted TMHs: 1\nExp number of AAs in TMHs: 23.72226\nExp number, first 60 AAs: 0.22038\nTotal prob of N-in: 0.12994\noutside 1 331\nTMhelix 332 354\ninside 355 372

9277	GCF_000208445.1_ASM20844v2	Escherichia sp. TW09276	Proteobacteria; Gammaproteobacteria; Enterobacterales; Enterobacteriaceae; Escherichia
MTGDNTLIHSHGINRRDFMKLCAALAATMGLSSKAAAEMAESVTNPQRPPVIWIGAQECT	WP_000145412.1
MULTISPECIES: hydrogenase-2 small chain [Escherichia]	Length: 372\nNumber of predicted TMHs: 1\nExp number of AAs in TMHs: 23.72209\nExp number, first 60 AAs: 0.22038\nTotal prob of N-in: 0.12994\noutside 1 331\nTMhelix 332 354\ninside 355 372

9278	GCF_000208445.1_ASM20844v2	Escherichia sp. TW09276	Proteobacteria; Gammaproteobacteria; Enterobacterales; Enterobacteriaceae; Escherichia
MNNEETFYQAMRRQGVTRRSFLKYCSLAATSLGLGAGMAPKIAWALENKPRIPVVWIHGL	WP_010344442.1
MULTISPECIES: hydrogenase [Escherichia]	Length: 372\nNumber of predicted TMHs: 2\nExp number of

AAs in TMHs: 38.04874\nExp number, first 60 AAs: 19.5298\nTotal prob of N-in: 0.94352\nPOSSIBLE N-term signal sequence\ninside 1 19\nTMhelix 20 39\noutside 40 325\nTMhelix 326 348\ninside 349 372

9279 GCF_000208565.1_ASM20856v2 Escherichia sp. TW09308 Proteobacteria; Gammaproteobacteria; Enterobacterales; Enterobacteriaceae; Escherichia MTGDNTLIHSHGINRRDFMKLCAALAATMGLSSKAAAEMAESVTNPQRPPVIWIGAQECT WP_000145417.1 MULTISPECIES: hydrogenase-2 small chain [Escherichia] Length: 372\nNumber of predicted TMHs: 1\nExp number of AAs in TMHs: 23.72512\nExp number, first 60 AAs: 0.22034\nTotal prob of N-in: 0.13000\noutside 1 331\nTMhelix 332 354\ninside 355 372

9280 GCF_000208565.1_ASM20856v2 Escherichia sp. TW09308 Proteobacteria; Gammaproteobacteria; Enterobacterales; Enterobacteriaceae; Escherichia MNNEETFYQAMRRQGVTRRSFLKYCSLAATSLGLGAGMAPKIAWALENKPRIPVVWIHGL WP_001516139.1 MULTISPECIES: hydrogenase [Escherichia] Length: 372\nNumber of predicted TMHs: 2\nExp number of AAs in TMHs: 37.98718\nExp number, first 60 AAs: 19.52876\nTotal prob of N-in: 0.94354\nPOSSIBLE N-term signal sequence\ninside 1 19\nTMhelix 20 39\noutside 40 325\nTMhelix 326 348\ninside 349 372

9281 GCF_000208545.1_ASM20854v2 Escherichia sp. TW10509 Proteobacteria; Gammaproteobacteria; Enterobacterales; Enterobacteriaceae; Escherichia MNNEETFYQAMRRQGVTRRSFLKYCSLAATSLGLGAGMAPKIAWALENKPRIPVVWIHGL WP_001353311.1 MULTISPECIES: hydrogenase [Escherichia] Length: 372\nNumber of predicted TMHs: 2\nExp number of AAs in TMHs: 37.2638699999999\nExp number, first 60 AAs: 19.52924\nTotal prob of N-in: 0.94351\nPOSSIBLE N-term signal sequence\ninside 1 19\nTMhelix 20 39\noutside 40 325\nTMhelix 326 348\ninside 349 372

9282 GCF_000208545.1_ASM20854v2 Escherichia sp. TW10509 Proteobacteria; Gammaproteobacteria; Enterobacterales; Enterobacteriaceae; Escherichia MTGDNTLIHSHGINRRDFMKLCAALAATMGLSSKAAAEMAESVTNPQRPPVIWIGAQECT WP_000145410.1 MULTISPECIES: hydrogenase-2 small chain [Proteobacteria] Length: 372\nNumber of predicted TMHs: 1\nExp number of AAs in TMHs: 23.72226\nExp number, first 60 AAs: 0.22038\nTotal prob of N-in: 0.12994\noutside 1 331\nTMhelix 332 354\ninside 355 372

9283 GCF_000208525.1_ASM20852v2 Escherichia sp. TW14182 Proteobacteria; Gammaproteobacteria; Enterobacterales; Enterobacteriaceae; Escherichia MNNEETFYQAMRRQGVTRRSFLKYCSLAATSLGLGAGMAPKIAWALENKPRIPVVWIHGL WP_010344442.1 MULTISPECIES: hydrogenase [Escherichia] Length: 372\nNumber of predicted TMHs: 2\nExp number of AAs in TMHs: 38.04874\nExp number, first 60 AAs: 19.5298\nTotal prob of N-in: 0.94352\nPOSSIBLE N-term signal sequence\ninside 1 19\nTMhelix 20 39\noutside 40 325\nTMhelix 326 348\ninside 349 372

9284 GCF_000208525.1_ASM20852v2 Escherichia sp. TW14182 Proteobacteria; Gammaproteobacteria; Enterobacterales; Enterobacteriaceae; Escherichia MTGDNTLIHSHGINRRDFMKLCAALAATMGLSSKAAAEMAESVTNPQRPPVIWIGAQECT WP_000145416.1 hydrogenase 2 small subunit [Escherichia sp. TW14182] Length: 372\nNumber of predicted TMHs: 1\nExp number of AAs in TMHs: 23.72226\nExp number, first 60 AAs: 0.22038\nTotal prob of N-in: 0.12994\noutside 1 331\nTMhelix 332 354\ninside 355 372

9285 GCF_000208485.1_ASM20848v2 Escherichia sp. TW15838 Proteobacteria; Gammaproteobacteria; Enterobacterales; Enterobacteriaceae; Escherichia MTGDNTLIHSHGINRRDFMKLCAALAATMGLSSKAAAEMAESVTNPQRPPVIWIGAQECT WP_000145410.1 MULTISPECIES: hydrogenase-2 small chain [Proteobacteria] Length: 372\nNumber of predicted TMHs: 1\nExp number of AAs in TMHs: 23.72226\nExp number, first 60 AAs: 0.22038\nTotal prob of N-in: 0.12994\noutside 1 331\nTMhelix 332 354\ninside 355 372

9286 GCF_000208485.1_ASM20848v2 Escherichia sp. TW15838 Proteobacteria; Gammaproteobacteria; Enterobacterales; Enterobacteriaceae; Escherichia MNNEETFYQAMRRQGVTRRSFLKYCSLAATSLGLGAGMAPKIAWALENKPRIPVVWIHGL WP_001353311.1 MULTISPECIES: hydrogenase [Escherichia] Length: 372\nNumber of predicted TMHs: 2\nExp number of AAs in TMHs: 37.2638699999999\nExp number, first 60 AAs: 19.52924\nTotal prob of N-in: 0.94351\nPOSSIBLE N-term signal sequence\ninside 1 19\nTMhelix 20 39\noutside 40 325\nTMhelix 326 348\ninside 349 372

9287 GCF_002109845.1_ASM210984v1 Escherichia albertii B156 Proteobacteria; Gammaproteobacteria; Enterobacterales; Enterobacteriaceae; Escherichia; Escherichia albertii

MNNEETFYQSMRRQGVTRRSFLKYCSLAATSLGLGAGMAPKIAWALENKPRIPVVWIHGL WP_002461900.1 MULTISPECIES: hydrogenase-1 small chain [Escherichia] Length: 372\nNumber of predicted TMHs: 2\nExp number of AAs in TMHs: 37.1074899999999\nExp number, first 60 AAs: 19.37835\nTotal prob of N-in: 0.93614\nPOSSIBLE N-term signal sequence\ninside 1 19\nTMhelix 20 39\noutside 40 325\nTMhelix 326 348\ninside 349 372

9288 GCF_002109845.1_ASM210984v1 Escherichia albertii B156 Proteobacteria; Gammaproteobacteria; Enterobacterales; Enterobacteriaceae; Escherichia; Escherichia albertii MTGDNTLIHSHGINRRDFMKLCAALAATMGLSSKAAAEMAESVTNPQRPPVIWIGAQECT WP_000145410.1 MULTISPECIES: hydrogenase-2 small chain [Proteobacteria] Length: 372\nNumber of predicted TMHs: 1\nExp number of AAs in TMHs: 23.72226\nExp number, first 60 AAs: 0.22038\nTotal prob of N-in: 0.12994\noutside 1 331\nTMhelix 332 354\ninside 355 372

9289 GCF_000512125.1_ASM51212v1 Escherichia albertii KF1 Proteobacteria; Gammaproteobacteria; Enterobacterales; Enterobacteriaceae; Escherichia; Escherichia albertii MTGDNTLIHSHGINRRDFMKLCAALAATMGLSSKAAAEMAESVTNPQRPPVIWIGAQECT WP_025238349.1 hydrogenase 2 small subunit [Escherichia albertii] Length: 372\nNumber of predicted TMHs: 1\nExp number of AAs in TMHs: 23.72042\nExp number, first 60 AAs: 0.22025\nTotal prob of N-in: 0.12992\noutside 1 331\nTMhelix 332 354\ninside 355 372

9290 GCF_000512125.1_ASM51212v1 Escherichia albertii KF1 Proteobacteria; Gammaproteobacteria; Enterobacterales; Enterobacteriaceae; Escherichia; Escherichia albertii MNNEETFYQSMRRQGVTRRSFLKYCSLAATSLGLGAGMAPKIAWALENKPRIPVVWIHGL WP_025237223.1 hydrogenase [Escherichia albertii] Length: 372\nNumber of predicted TMHs: 2\nExp number of AAs in TMHs: 36.9974299999999\nExp number, first 60 AAs: 19.37688\nTotal prob of N-in: 0.93606\nPOSSIBLE N-term signal sequence\ninside 1 19\nTMhelix 20 39\noutside 40 325\nTMhelix 326 348\ninside 349 372

9291 GCF_000759775.1_ASM75977v1 Escherichia albertii NBRC 107761 Proteobacteria; Gammaproteobacteria; Enterobacterales; Enterobacteriaceae; Escherichia; Escherichia albertii MNNEETFYQSMRRQGVTRRSFLKYCSLAATSLGLGAGMAPKIAWALENKPRIPVVWIHGL WP_002461900.1 MULTISPECIES: hydrogenase-1 small chain [Escherichia] Length: 372\nNumber of predicted TMHs: 2\nExp number of AAs in TMHs: 37.1074899999999\nExp number, first 60 AAs: 19.37835\nTotal prob of N-in: 0.93614\nPOSSIBLE N-term signal sequence\ninside 1 19\nTMhelix 20 39\noutside 40 325\nTMhelix 326 348\ninside 349 372

9292 GCF_000759775.1_ASM75977v1 Escherichia albertii NBRC 107761 Proteobacteria; Gammaproteobacteria; Enterobacterales; Enterobacteriaceae; Escherichia; Escherichia albertii MTGDNTLIHSHGINRRDFMKLCAALAATMGLSSKAAAEMAESVTNPQRPPVIWIGAQECT WP_000145410.1 MULTISPECIES: hydrogenase-2 small chain [Proteobacteria] Length: 372\nNumber of predicted TMHs: 1\nExp number of AAs in TMHs: 23.72226\nExp number, first 60 AAs: 0.22038\nTotal prob of N-in: 0.12994\noutside 1 331\nTMhelix 332 354\ninside 355 372

9293 GCF_000155105.1_ASM15510v1 Escherichia albertii TW07627 Proteobacteria; Gammaproteobacteria; Enterobacterales; Enterobacteriaceae; Escherichia; Escherichia albertii MNNEETFYQSMRRQGVTRRSFLKYCSLAATSLGLGAGMAPKIAWALENKPRIPVVWIHGL WP_002461900.1 MULTISPECIES: hydrogenase-1 small chain [Escherichia] Length: 372\nNumber of predicted TMHs: 2\nExp number of AAs in TMHs: 37.1074899999999\nExp number, first 60 AAs: 19.37835\nTotal prob of N-in: 0.93614\nPOSSIBLE N-term signal sequence\ninside 1 19\nTMhelix 20 39\noutside 40 325\nTMhelix 326 348\ninside 349 372

9294 GCF_000155105.1_ASM15510v1 Escherichia albertii TW07627 Proteobacteria; Gammaproteobacteria; Enterobacterales; Enterobacteriaceae; Escherichia; Escherichia albertii MTGDNTLIHSHGINRRDFMKLCAALAATMGLSSKAAAEMAESVTNPQRPPVIWIGAQECT WP_000145410.1 MULTISPECIES: hydrogenase-2 small chain [Proteobacteria] Length: 372\nNumber of predicted TMHs: 1\nExp number of AAs in TMHs: 23.72226\nExp number, first 60 AAs: 0.22038\nTotal prob of N-in: 0.12994\noutside 1 331\nTMhelix 332 354\ninside 355 372

9295 GCF_000208425.1_ASM20842v2 Escherichia albertii TW08933 Proteobacteria; Gammaproteobacteria; Enterobacterales; Enterobacteriaceae; Escherichia; Escherichia albertii MNNEETFYQSMRRQGVTRRSFLKYCSLAATSLGLGAGMAPKIAWALENKPRIPVVWIHGL WP_010318334.1 hydrogenase-1 small chain [Escherichia albertii] Length: 372\nNumber of predicted TMHs: 2\nExp number of

AAs in TMHs: 36.9963899999999\nExp number, first 60 AAs:  19.37682\nTotal prob of N-in:   0.93606\nPOSSIBLE N-term signal sequence\ninside   1   19\nTMhelix   20   39\noutside   40   325\nTMhelix   326   348\ninside   349   372

9296	GCF_000208425.1_ASM20842v2	Escherichia albertii TW08933 Proteobacteria; Gammaproteobacteria; Enterobacterales; Enterobacteriaceae; Escherichia; Escherichia albertii
	MTGDNTLIHSHGINRRDFMKLCAALAATMGLSSKAAAEMAESVTNPQRPPVIWIGAQECT	WP_000145422.1
	hydrogenase 2 small subunit [Escherichia albertii]	Length: 372\nNumber of predicted TMHs:  1\nExp number of AAs in TMHs: 23.69552\nExp number, first 60 AAs:  0.2204\nTotal prob of N-in:   0.13100\noutside   1   331\nTMhelix   332   354\ninside   355   372

9297	GCF_000208505.1_ASM20850v2	Escherichia albertii TW15818 Proteobacteria; Gammaproteobacteria; Enterobacterales; Enterobacteriaceae; Escherichia; Escherichia albertii
	MTGDNTLIHSHGINRRDFMKLCAALAATMGLSSKAAAEMAESVTNPQRPPVIWIGAQECT	WP_000145410.1
	MULTISPECIES: hydrogenase-2 small chain [Proteobacteria]	Length: 372\nNumber of predicted TMHs:  1\nExp number of AAs in TMHs: 23.72226\nExp number, first 60 AAs:  0.22038\nTotal prob of N-in:   0.12994\noutside   1   331\nTMhelix   332   354\ninside   355   372

9298	GCF_000208505.1_ASM20850v2	Escherichia albertii TW15818 Proteobacteria; Gammaproteobacteria; Enterobacterales; Enterobacteriaceae; Escherichia; Escherichia albertii
	MNNEETFYQSMRRQGVTRRSFLKYCSLAATSLGLGAGMAPKIAWALENKPRIPVVWIHGL	WP_002461900.1
	MULTISPECIES: hydrogenase-1 small chain [Escherichia]	Length: 372\nNumber of predicted TMHs:  2\nExp number of AAs in TMHs: 37.1074899999999\nExp number, first 60 AAs:  19.37835\nTotal prob of N-in:   0.93614\nPOSSIBLE N-term signal sequence\ninside   1   19\nTMhelix   20   39\noutside   40   325\nTMhelix   326   348\ninside   349   372

9299	GCF_000303255.2_ASM30325v2	Escherichia coli 0.1288	Proteobacteria; Gammaproteobacteria; Enterobacterales; Enterobacteriaceae; Escherichia; Escherichia coli
	MTGDNTLIHSHGINRRDFMKLCAALAATMGLSSKAAAEMAESVTNPQRPPVIWIGAQECT	WP_000145410.1
	MULTISPECIES: hydrogenase-2 small chain [Proteobacteria]	Length: 372\nNumber of predicted TMHs:  1\nExp number of AAs in TMHs: 23.72226\nExp number, first 60 AAs:  0.22038\nTotal prob of N-in:   0.12994\noutside   1   331\nTMhelix   332   354\ninside   355   372

9300	GCF_000303255.2_ASM30325v2	Escherichia coli 0.1288	Proteobacteria; Gammaproteobacteria; Enterobacterales; Enterobacteriaceae; Escherichia; Escherichia coli
	MNNEETFYQAMRRQGVTRRSFLKYCSLAATSLGLGAGMAPKIAWALENKPRIPVVWIHGL	WP_001498899.1
	hydrogenase-1 small chain [Escherichia coli]	Length: 372\nNumber of predicted TMHs:  2\nExp number of AAs in TMHs: 39.34459\nExp number, first 60 AAs:  19.53328\nTotal prob of N-in:   0.94350\nPOSSIBLE N-term signal sequence\ninside   1   19\nTMhelix   20   39\noutside   40   325\nTMhelix   326   348\ninside   349   372

9301	GCF_000303235.2_ASM30323v2	Escherichia coli 0.1304	Proteobacteria; Gammaproteobacteria; Enterobacterales; Enterobacteriaceae; Escherichia; Escherichia coli
	MTGDNTLIHSHGINRRDFMKLCAALAATMGLSSKAAAEMAESVTNPQRPPVIWIGAQECT	WP_000145410.1
	MULTISPECIES: hydrogenase-2 small chain [Proteobacteria]	Length: 372\nNumber of predicted TMHs:  1\nExp number of AAs in TMHs: 23.72226\nExp number, first 60 AAs:  0.22038\nTotal prob of N-in:   0.12994\noutside   1   331\nTMhelix   332   354\ninside   355   372

9302	GCF_000303235.2_ASM30323v2	Escherichia coli 0.1304	Proteobacteria; Gammaproteobacteria; Enterobacterales; Enterobacteriaceae; Escherichia; Escherichia coli
	MNNEETFYQAMRRQGVTRRSFLKYCSLAATSLGLGAGMAPKIAWALENKPRIPVVWIHGL	WP_001301613.1
	hydrogenase [Escherichia coli]	Length: 372\nNumber of predicted TMHs:  2\nExp number of AAs in TMHs: 37.46598\nExp number, first 60 AAs:  19.53139\nTotal prob of N-in:   0.94364\nPOSSIBLE N-term signal sequence\ninside   1   19\nTMhelix   20   39\noutside   40   325\nTMhelix   326   348\ninside   349   372

9303	GCF_000303655.2_ASM30365v2	Escherichia coli 07798	Proteobacteria; Gammaproteobacteria; Enterobacterales; Enterobacteriaceae; Escherichia; Escherichia coli
	MNNEETFYQAMRRQGVTRRSFLKYCSLAATSLGLGAGMAPKIAWALENKPRIPVVWIHGL	WP_001058323.1
	MULTISPECIES: hydrogenase-1 small chain [Proteobacteria]	Length: 372\nNumber of predicted TMHs:  2\nExp number of AAs in TMHs: 37.2608499999999\nExp number, first 60 AAs:  19.52923\nTotal prob of N-in:   0.94351\nPOSSIBLE N-term signal sequence\ninside   1   19\nTMhelix   20   39\noutside   40   325\nTMhelix   326   348\ninside   349   372

9304 GCF_000303655.2_ASM30365v2 Escherichia coli 07798 Proteobacteria; Gammaproteobacteria; Enterobacterales; Enterobacteriaceae; Escherichia; Escherichia coli
MTGDNTLIHSHGINRRDFMKLCAALAATMGLSSKAAAEMAESVTNPQRPPVIWIGAQECT WP_000145406.1
hydrogenase-2 small chain [Escherichia coli] Length: 372\nNumber of predicted TMHs: 1\nExp number of AAs in TMHs: 23.72251\nExp number, first 60 AAs: 0.22027\nTotal prob of N-in: 0.13006\noutside 1 331\nTMhelix 332 354\ninside 355 372

9305 GCF_000462305.2_ASM46230v2 Escherichia coli 08BKT055439 Proteobacteria; Gammaproteobacteria; Enterobacterales; Enterobacteriaceae; Escherichia; Escherichia coli
MNNEETFYQAMRRQGVTRRSFLKYCSLAATSLGLGAGMAPKIAWALENKPRIPVVWIHGL WP_001301613.1
hydrogenase [Escherichia coli] Length: 372\nNumber of predicted TMHs: 2\nExp number of AAs in TMHs: 37.46598\nExp number, first 60 AAs: 19.53139\nTotal prob of N-in: 0.94364\nPOSSIBLE N-term signal sequence\ninside 1 19\nTMhelix 20 39\noutside 40 325\nTMhelix 326 348\ninside 349 372

9306 GCF_000462305.2_ASM46230v2 Escherichia coli 08BKT055439 Proteobacteria; Gammaproteobacteria; Enterobacterales; Enterobacteriaceae; Escherichia; Escherichia coli
MTGDNTLIHSHGINRRDFMKLCAALAATMGLSSKAAAEMAESVTNPQRPPVIWIGAQECT WP_000145410.1
MULTISPECIES: hydrogenase-2 small chain [Proteobacteria] Length: 372\nNumber of predicted TMHs: 1\nExp number of AAs in TMHs: 23.72226\nExp number, first 60 AAs: 0.22038\nTotal prob of N-in: 0.12994\noutside 1 331\nTMhelix 332 354\ninside 355 372

9307 GCF_000462325.2_ASM46232v2 Escherichia coli 08BKT77219 Proteobacteria; Gammaproteobacteria; Enterobacterales; Enterobacteriaceae; Escherichia; Escherichia coli
MNNEETFYQAMRRQGVTRRSFLKYCSLAATSLGLGAGMAPKIAWALENKPRIPVVWIHGL WP_001301613.1
hydrogenase [Escherichia coli] Length: 372\nNumber of predicted TMHs: 2\nExp number of AAs in TMHs: 37.46598\nExp number, first 60 AAs: 19.53139\nTotal prob of N-in: 0.94364\nPOSSIBLE N-term signal sequence\ninside 1 19\nTMhelix 20 39\noutside 40 325\nTMhelix 326 348\ninside 349 372

9308 GCF_000462325.2_ASM46232v2 Escherichia coli 08BKT77219 Proteobacteria; Gammaproteobacteria; Enterobacterales; Enterobacteriaceae; Escherichia; Escherichia coli
MTGDNTLIHSHGINRRDFMKLCAALAATMGLSSKAAAEMAESVTNPQRPPVIWIGAQECT WP_000145410.1
MULTISPECIES: hydrogenase-2 small chain [Proteobacteria] Length: 372\nNumber of predicted TMHs: 1\nExp number of AAs in TMHs: 23.72226\nExp number, first 60 AAs: 0.22038\nTotal prob of N-in: 0.12994\noutside 1 331\nTMhelix 332 354\ninside 355 372

9309 GCF_000462345.2_ASM46234v2 Escherichia coli 09BKT024447 Proteobacteria; Gammaproteobacteria; Enterobacterales; Enterobacteriaceae; Escherichia; Escherichia coli
MTGDNTLIHSHGINRRDFMKLCAALAATMGLSSKAAAEMAESVTNPQRPPVIWIGAQECT WP_000145410.1
MULTISPECIES: hydrogenase-2 small chain [Proteobacteria] Length: 372\nNumber of predicted TMHs: 1\nExp number of AAs in TMHs: 23.72226\nExp number, first 60 AAs: 0.22038\nTotal prob of N-in: 0.12994\noutside 1 331\nTMhelix 332 354\ninside 355 372

9310 GCF_000462345.2_ASM46234v2 Escherichia coli 09BKT024447 Proteobacteria; Gammaproteobacteria; Enterobacterales; Enterobacteriaceae; Escherichia; Escherichia coli
MNNEETFYQAMRRQGVTRRSFLKYCSLAATSLGLGAGMAPKIAWALENKPRIPVVWIHGL WP_001301613.1
hydrogenase [Escherichia coli] Length: 372\nNumber of predicted TMHs: 2\nExp number of AAs in TMHs: 37.46598\nExp number, first 60 AAs: 19.53139\nTotal prob of N-in: 0.94364\nPOSSIBLE N-term signal sequence\ninside 1 19\nTMhelix 20 39\noutside 40 325\nTMhelix 326 348\ninside 349 372

9311 GCF_000461935.2_ASM46193v2 Escherichia coli 09BKT076207 Proteobacteria; Gammaproteobacteria; Enterobacterales; Enterobacteriaceae; Escherichia; Escherichia coli
MNNEETFYQAMRRQGVTRRSFLKYCSLAATSLGLGAGMAPKIAWALENKPRIPVVWIHGL WP_001301613.1
hydrogenase [Escherichia coli] Length: 372\nNumber of predicted TMHs: 2\nExp number of AAs in TMHs: 37.46598\nExp number, first 60 AAs: 19.53139\nTotal prob of N-in: 0.94364\nPOSSIBLE N-term signal sequence\ninside 1 19\nTMhelix 20 39\noutside 40 325\nTMhelix 326 348\ninside 349 372

9312 GCF_000461935.2_ASM46193v2 Escherichia coli 09BKT076207 Proteobacteria; Gammaproteobacteria; Enterobacterales; Enterobacteriaceae; Escherichia; Escherichia coli
MTGDNTLIHSHGINRRDFMKLCAALAATMGLSSKAAAEMAESVTNPQRPPVIWIGAQECT WP_000145410.1
MULTISPECIES: hydrogenase-2 small chain [Proteobacteria] Length: 372\nNumber of predicted TMHs:

1\nExp number of AAs in TMHs: 23.72226\nExp number, first 60 AAs: 0.22038\nTotal prob of N-in: 0.12994\noutside 1 331\nTMhelix 332 354\ninside 355 372

9313 GCF_000340455.2_ASM34045v2 Escherichia coli 09BKT078844 Proteobacteria; Gammaproteobacteria; Enterobacterales; Enterobacteriaceae; Escherichia; Escherichia coli
MNNEETFYQAMRRQGVTRRSFLKYCSLAATSLGLGAGMAPKIAWALENKPRIPVVWIHGL WP_001301613.1
hydrogenase [Escherichia coli] Length: 372\nNumber of predicted TMHs: 2\nExp number of AAs in TMHs: 37.46598\nExp number, first 60 AAs: 19.53139\nTotal prob of N-in: 0.94364\nPOSSIBLE N-term signal sequence\ninside 1 19\nTMhelix 20 39\noutside 40 325\nTMhelix 326 348\ninside 349 372

9314 GCF_000340455.2_ASM34045v2 Escherichia coli 09BKT078844 Proteobacteria; Gammaproteobacteria; Enterobacterales; Enterobacteriaceae; Escherichia; Escherichia coli
MTGDNTLIHSHGINRRDFMKLCAALAATMGLSSKAAAEMAESVTNPQRPPVIWIGAQECT WP_000145410.1
MULTISPECIES: hydrogenase-2 small chain [Proteobacteria] Length: 372\nNumber of predicted TMHs: 1\nExp number of AAs in TMHs: 23.72226\nExp number, first 60 AAs: 0.22038\nTotal prob of N-in: 0.12994\noutside 1 331\nTMhelix 332 354\ninside 355 372

9315 GCF_001265615.1_100329 Escherichia coli 100329 Proteobacteria; Gammaproteobacteria; Enterobacterales; Enterobacteriaceae; Escherichia; Escherichia coli
MTGDNTLIHSHGINRRDFMKLCAALAATMGLSSKAAAEMAESVTNPQRPPVIWIGAQECT WP_000145410.1
MULTISPECIES: hydrogenase-2 small chain [Proteobacteria] Length: 372\nNumber of predicted TMHs: 1\nExp number of AAs in TMHs: 23.72226\nExp number, first 60 AAs: 0.22038\nTotal prob of N-in: 0.12994\noutside 1 331\nTMhelix 332 354\ninside 355 372

9316 GCF_001265615.1_100329 Escherichia coli 100329 Proteobacteria; Gammaproteobacteria; Enterobacterales; Enterobacteriaceae; Escherichia; Escherichia coli
MNNEETFYQAMRRQGVTRRSFLKYCSLAATSLGLGAGMAPKIAWALENKPRIPVVWIHGL WP_052935581.1
hydrogenase [Escherichia coli] Length: 372\nNumber of predicted TMHs: 2\nExp number of AAs in TMHs: 37.2610899999999\nExp number, first 60 AAs: 19.52923\nTotal prob of N-in: 0.94351\nPOSSIBLE N-term signal sequence\ninside 1 19\nTMhelix 20 39\noutside 40 325\nTMhelix 326 348\ninside 349 372

9317 GCF_000304855.2_ASM30485v2 Escherichia coli 10.0821 Proteobacteria; Gammaproteobacteria; Enterobacterales; Enterobacteriaceae; Escherichia; Escherichia coli
MNNEETFYQAMRRQGVTRRSFLKYCSLAATSLGLGAGMAPKIAWALENKPRIPVVWIHGL WP_001301613.1
hydrogenase [Escherichia coli] Length: 372\nNumber of predicted TMHs: 2\nExp number of AAs in TMHs: 37.46598\nExp number, first 60 AAs: 19.53139\nTotal prob of N-in: 0.94364\nPOSSIBLE N-term signal sequence\ninside 1 19\nTMhelix 20 39\noutside 40 325\nTMhelix 326 348\ninside 349 372

9318 GCF_000304855.2_ASM30485v2 Escherichia coli 10.0821 Proteobacteria; Gammaproteobacteria; Enterobacterales; Enterobacteriaceae; Escherichia; Escherichia coli
MTGDNTLIHSHGINRRDFMKLCAALAATMGLSSKAAAEMAESVTNPQRPPVIWIGAQECT WP_000145410.1
MULTISPECIES: hydrogenase-2 small chain [Proteobacteria] Length: 372\nNumber of predicted TMHs: 1\nExp number of AAs in TMHs: 23.72226\nExp number, first 60 AAs: 0.22038\nTotal prob of N-in: 0.12994\noutside 1 331\nTMhelix 332 354\ninside 355 372

9319 GCF_000305415.2_ASM30541v2 Escherichia coli 10.0833 Proteobacteria; Gammaproteobacteria; Enterobacterales; Enterobacteriaceae; Escherichia; Escherichia coli
MNNEETFYQAMRRQGVTRRSFLKYCSLAATSLGLGAGMAPKIAWALENKPRIPVVWIHGL WP_001301613.1
hydrogenase [Escherichia coli] Length: 372\nNumber of predicted TMHs: 2\nExp number of AAs in TMHs: 37.46598\nExp number, first 60 AAs: 19.53139\nTotal prob of N-in: 0.94364\nPOSSIBLE N-term signal sequence\ninside 1 19\nTMhelix 20 39\noutside 40 325\nTMhelix 326 348\ninside 349 372

9320 GCF_000305415.2_ASM30541v2 Escherichia coli 10.0833 Proteobacteria; Gammaproteobacteria; Enterobacterales; Enterobacteriaceae; Escherichia; Escherichia coli
MTGDNTLIHSHGINRRDFMKLCAALAATMGLSSKAAAEMAESVTNPQRPPVIWIGAQECT WP_000145410.1
MULTISPECIES: hydrogenase-2 small chain [Proteobacteria] Length: 372\nNumber of predicted TMHs: 1\nExp number of AAs in TMHs: 23.72226\nExp number, first 60 AAs: 0.22038\nTotal prob of N-in: 0.12994\noutside 1 331\nTMhelix 332 354\ninside 355 372

9321 GCF_000305435.2_ASM30543v2 Escherichia coli 10.0869 Proteobacteria; Gammaproteobacteria; Enterobacterales; Enterobacteriaceae; Escherichia; Escherichia coli

MIGDNTLIHSHGINRRDFMKLCAALAATMGLSSKAAAEMAESVTNPQRPPVIWIGAQECT WP_000578724.1 hydrogenase 2 small subunit [Escherichia coli] Length: 372\nNumber of predicted TMHs: 1\nExp number of AAs in TMHs: 23.05601\nExp number, first 60 AAs: 0.15431\nTotal prob of N-in: 0.09116\noutside 1 331\nTMhelix 332 354\ninside 355 372

9322 GCF_000305435.2_ASM30543v2 Escherichia coli 10.0869 Proteobacteria; Gammaproteobacteria; Enterobacterales; Enterobacteriaceae; Escherichia; Escherichia coli MNNEETFYQAMRRQGVTRRSFLKYCSLAATSLGLGAGMAPKIAWALENKPRIPVVWIHGL WP_001301613.1 hydrogenase [Escherichia coli] Length: 372\nNumber of predicted TMHs: 2\nExp number of AAs in TMHs: 37.46598\nExp number, first 60 AAs: 19.53139\nTotal prob of N-in: 0.94364\nPOSSIBLE N-term signal sequence\ninside 1 19\nTMhelix 20 39\noutside 40 325\nTMhelix 326 348\ninside 349 372

9323 GCF_000168095.1_ASM16809v1 Escherichia coli 101-1 Proteobacteria; Gammaproteobacteria; Enterobacterales; Enterobacteriaceae; Escherichia; Escherichia coli MNNEETFYQAMRRQGVTRRSFLKYCSLAATSLGLGAGMAPKIAWALENKPRIPVVWIHGL WP_001058323.1 MULTISPECIES: hydrogenase-1 small chain [Proteobacteria] Length: 372\nNumber of predicted TMHs: 2\nExp number of AAs in TMHs: 37.2608499999999\nExp number, first 60 AAs: 19.52923\nTotal prob of N-in: 0.94351\nPOSSIBLE N-term signal sequence\ninside 1 19\nTMhelix 20 39\noutside 40 325\nTMhelix 326 348\ninside 349 372

9324 GCF_000168095.1_ASM16809v1 Escherichia coli 101-1 Proteobacteria; Gammaproteobacteria; Enterobacterales; Enterobacteriaceae; Escherichia; Escherichia coli MTGDNTLIHSHGINRRDFMKLCAALAATMGLSSKAAAEMAESVTNPQRPPVIWIGAQECT WP_000145410.1 MULTISPECIES: hydrogenase-2 small chain [Proteobacteria] Length: 372\nNumber of predicted TMHs: 1\nExp number of AAs in TMHs: 23.72226\nExp number, first 60 AAs: 0.22038\nTotal prob of N-in: 0.12994\noutside 1 331\nTMhelix 332 354\ninside 355 372

9325 GCF_001265235.1_103385 Escherichia coli 103385 Proteobacteria; Gammaproteobacteria; Enterobacterales; Enterobacteriaceae; Escherichia; Escherichia coli MNNEETFYQAMRRQGVTRRSFLKYCSLAATSLGLGAGMAPKIAWALENKPRIPVVWIHGL WP_001058323.1 MULTISPECIES: hydrogenase-1 small chain [Proteobacteria] Length: 372\nNumber of predicted TMHs: 2\nExp number of AAs in TMHs: 37.2608499999999\nExp number, first 60 AAs: 19.52923\nTotal prob of N-in: 0.94351\nPOSSIBLE N-term signal sequence\ninside 1 19\nTMhelix 20 39\noutside 40 325\nTMhelix 326 348\ninside 349 372

9326 GCF_001265235.1_103385 Escherichia coli 103385 Proteobacteria; Gammaproteobacteria; Enterobacterales; Enterobacteriaceae; Escherichia; Escherichia coli MTGDNTLIHSHGINRRDFMKLCAALAATMGLSSKAAAEMAESVTNPQRPPVIWIGAQECT WP_000145410.1 MULTISPECIES: hydrogenase-2 small chain [Proteobacteria] Length: 372\nNumber of predicted TMHs: 1\nExp number of AAs in TMHs: 23.72226\nExp number, first 60 AAs: 0.22038\nTotal prob of N-in: 0.12994\noutside 1 331\nTMhelix 332 354\ninside 355 372

9327 GCF_001265255.1_103573 Escherichia coli 103573 Proteobacteria; Gammaproteobacteria; Enterobacterales; Enterobacteriaceae; Escherichia; Escherichia coli MNNEETFYQAMRRQGVTRRSFLKYCSLAATSLGLGAGMAPKIAWALENKPRIPVVWIHGL WP_001058323.1 MULTISPECIES: hydrogenase-1 small chain [Proteobacteria] Length: 372\nNumber of predicted TMHs: 2\nExp number of AAs in TMHs: 37.2608499999999\nExp number, first 60 AAs: 19.52923\nTotal prob of N-in: 0.94351\nPOSSIBLE N-term signal sequence\ninside 1 19\nTMhelix 20 39\noutside 40 325\nTMhelix 326 348\ninside 349 372

9328 GCF_001265255.1_103573 Escherichia coli 103573 Proteobacteria; Gammaproteobacteria; Enterobacterales; Enterobacteriaceae; Escherichia; Escherichia coli MTGDNTLIHSHGINRRDFMKLCAALAATMGLSSKAAAEMAESVTNPQRPPVIWIGAQECT WP_000145410.1 MULTISPECIES: hydrogenase-2 small chain [Proteobacteria] Length: 372\nNumber of predicted TMHs: 1\nExp number of AAs in TMHs: 23.72226\nExp number, first 60 AAs: 0.22038\nTotal prob of N-in: 0.12994\noutside 1 331\nTMhelix 332 354\ninside 355 372

9329 GCF_000503655.1_Ecoli1047 Escherichia coli 1047 Proteobacteria; Gammaproteobacteria; Enterobacterales; Enterobacteriaceae; Escherichia; Escherichia coli MTGDNTLIHSHGINRRDFMKLCAALAATMGLSSKAAAEMAESVTNPQRPPVIWIGAQECT WP_000145410.1 MULTISPECIES: hydrogenase-2 small chain [Proteobacteria] Length: 372\nNumber of predicted TMHs:

1\nExp number of AAs in TMHs: 23.72226\nExp number, first 60 AAs: 0.22038\nTotal prob of N-in: 0.12994\noutside 1 331\nTMhelix 332 354\ninside 355 372

9330 GCF_000503655.1_Ecoli1047 Escherichia coli 1047 Proteobacteria; Gammaproteobacteria; Enterobacterales; Enterobacteriaceae; Escherichia; Escherichia coli MNNEETFYQAMRRQGVTRRSFLKYCSLAATSLGLGAGMAPKIAWALENKPRIPVVWIHGL WP_001058323.1 MULTISPECIES: hydrogenase-1 small chain [Proteobacteria] Length: 372\nNumber of predicted TMHs: 2\nExp number of AAs in TMHs: 37.2608499999999\nExp number, first 60 AAs: 19.52923\nTotal prob of N-in: 0.94351\nPOSSIBLE N-term signal sequence\ninside 1 19\nTMhelix 20 39\noutside 40 325\nTMhelix 326 348\ninside 349 372

9331 GCF_000699305.1_10810 Escherichia coli 10810 Proteobacteria; Gammaproteobacteria; Enterobacterales; Enterobacteriaceae; Escherichia; Escherichia coli MTGDNTLIHSHGINRRDFMKLCAALAATMGLSSKAAAEMVESVTNPQRPPVIWIGAQECT WP_000145423.1 hydrogenase 2 small subunit [Escherichia coli] Length: 372\nNumber of predicted TMHs: 1\nExp number of AAs in TMHs: 23.78201\nExp number, first 60 AAs: 0.28355\nTotal prob of N-in: 0.13257\noutside 1 331\nTMhelix 332 354\ninside 355 372

9332 GCF_000699305.1_10810 Escherichia coli 10810 Proteobacteria; Gammaproteobacteria; Enterobacterales; Enterobacteriaceae; Escherichia; Escherichia coli MNNEETFYQAMRRQGVTRRSFLKYCSLAATSLGLGAGMAPKIAWALENKPRIPVVWIHGL WP_001295940.1 MULTISPECIES: hydrogenase [Enterobacteriaceae] Length: 372\nNumber of predicted TMHs: 2\nExp number of AAs in TMHs: 37.2608899999999\nExp number, first 60 AAs: 19.52923\nTotal prob of N-in: 0.94351\nPOSSIBLE N-term signal sequence\ninside 1 19\nTMhelix 20 39\noutside 40 325\nTMhelix 326 348\ninside 349 372

9333 GCF_000485615.1_ASM48561v1 Escherichia coli 110957 Proteobacteria; Gammaproteobacteria; Enterobacterales; Enterobacteriaceae; Escherichia; Escherichia coli MNNEETFYQAMRRQGVTRRSFLKYCSLAATSLGLGAGMAPKIAWALENKPRIPVVWIHGL WP_001058323.1 MULTISPECIES: hydrogenase-1 small chain [Proteobacteria] Length: 372\nNumber of predicted TMHs: 2\nExp number of AAs in TMHs: 37.2608499999999\nExp number, first 60 AAs: 19.52923\nTotal prob of N-in: 0.94351\nPOSSIBLE N-term signal sequence\ninside 1 19\nTMhelix 20 39\noutside 40 325\nTMhelix 326 348\ninside 349 372

9334 GCF_000485615.1_ASM48561v1 Escherichia coli 110957 Proteobacteria; Gammaproteobacteria; Enterobacterales; Enterobacteriaceae; Escherichia; Escherichia coli MTGDNTLIHSHGINRRDFMKLCAALAATMGLSSKAAAEMAESVTNPQRPPVIWIGAQECT WP_000145410.1 MULTISPECIES: hydrogenase-2 small chain [Proteobacteria] Length: 372\nNumber of predicted TMHs: 1\nExp number of AAs in TMHs: 23.72226\nExp number, first 60 AAs: 0.22038\nTotal prob of N-in: 0.12994\noutside 1 331\nTMhelix 332 354\ninside 355 372

9335 GCF_000601315.1_ASM60131v1 Escherichia coli 1-110-08_S1_C1 Proteobacteria; Gammaproteobacteria; Enterobacterales; Enterobacteriaceae; Escherichia; Escherichia coli MNNEETFYQAMRRQGVTRRSFLKYCSLAATSLGLGAGMAPKIAWALENKPRIPVVWIHGL WP_001058323.1 MULTISPECIES: hydrogenase-1 small chain [Proteobacteria] Length: 372\nNumber of predicted TMHs: 2\nExp number of AAs in TMHs: 37.2608499999999\nExp number, first 60 AAs: 19.52923\nTotal prob of N-in: 0.94351\nPOSSIBLE N-term signal sequence\ninside 1 19\nTMhelix 20 39\noutside 40 325\nTMhelix 326 348\ninside 349 372

9336 GCF_000601315.1_ASM60131v1 Escherichia coli 1-110-08_S1_C1 Proteobacteria; Gammaproteobacteria; Enterobacterales; Enterobacteriaceae; Escherichia; Escherichia coli MTGDNTLIHSHGINRRDFMKLCAALAATMGLSSKAAAEMAESVTNPQRPPVIWIGAQECT WP_000145410.1 MULTISPECIES: hydrogenase-2 small chain [Proteobacteria] Length: 372\nNumber of predicted TMHs: 1\nExp number of AAs in TMHs: 23.72226\nExp number, first 60 AAs: 0.22038\nTotal prob of N-in: 0.12994\noutside 1 331\nTMhelix 332 354\ninside 355 372

9337 GCF_000601275.1_ASM60127v1 Escherichia coli 1-110-08_S1_C2 Proteobacteria; Gammaproteobacteria; Enterobacterales; Enterobacteriaceae; Escherichia; Escherichia coli MTGDNTLIHSHGINRRDFMKLCAALAATMGLSSKAAAEMAESVTNPQRPPVIWIGAQECT WP_000145410.1 MULTISPECIES: hydrogenase-2 small chain [Proteobacteria] Length: 372\nNumber of predicted TMHs:

1\nExp number of AAs in TMHs: 23.72226\nExp number, first 60 AAs: 0.22038\nTotal prob of N-in: 0.12994\noutside 1 331\nTMhelix 332 354\ninside 355 372

9338 GCF_000601275.1_ASM60127v1 Escherichia coli 1-110-08_S1_C2 Proteobacteria; Gammaproteobacteria; Enterobacterales; Enterobacteriaceae; Escherichia; Escherichia coli MNNEETFYQAMRRQGVTRRSFLKYCSLAATSLGLGAGMAPKIAWALENKPRIPVVWIHGL WP_001058323.1 MULTISPECIES: hydrogenase-1 small chain [Proteobacteria] Length: 372\nNumber of predicted TMHs: 2\nExp number of AAs in TMHs: 37.2608499999999\nExp number, first 60 AAs: 19.52923\nTotal prob of N-in: 0.94351\nPOSSIBLE N-term signal sequence\ninside 1 19\nTMhelix 20 39\noutside 40 325\nTMhelix 326 348\ninside 349 372

9339 GCF_000601295.1_ASM60129v1 Escherichia coli 1-110-08_S1_C3 Proteobacteria; Gammaproteobacteria; Enterobacterales; Enterobacteriaceae; Escherichia; Escherichia coli MNNEETFYQAMRRQGVTRRSFLKYCSLAATSLGLGAGMAPKIAWALENKPRIPVVWIHGL WP_001058323.1 MULTISPECIES: hydrogenase-1 small chain [Proteobacteria] Length: 372\nNumber of predicted TMHs: 2\nExp number of AAs in TMHs: 37.2608499999999\nExp number, first 60 AAs: 19.52923\nTotal prob of N-in: 0.94351\nPOSSIBLE N-term signal sequence\ninside 1 19\nTMhelix 20 39\noutside 40 325\nTMhelix 326 348\ninside 349 372

9340 GCF_000601295.1_ASM60129v1 Escherichia coli 1-110-08_S1_C3 Proteobacteria; Gammaproteobacteria; Enterobacterales; Enterobacteriaceae; Escherichia; Escherichia coli MTGDNTLIHSHGINRRDFMKLCAALAATMGLSSKAAAEMAESVTNPQRPPVIWIGAQECT WP_000145410.1 MULTISPECIES: hydrogenase-2 small chain [Proteobacteria] Length: 372\nNumber of predicted TMHs: 1\nExp number of AAs in TMHs: 23.72226\nExp number, first 60 AAs: 0.22038\nTotal prob of N-in: 0.12994\noutside 1 331\nTMhelix 332 354\ninside 355 372

9341 GCF_000601155.1_ASM60115v1 Escherichia coli 1-110-08_S3_C1 Proteobacteria; Gammaproteobacteria; Enterobacterales; Enterobacteriaceae; Escherichia; Escherichia coli MTGDNTLIHSHGINRRDFMKLCAALAATMGLSSKAAAEMAESVTNPQRPPVIWIGAQECT WP_000145410.1 MULTISPECIES: hydrogenase-2 small chain [Proteobacteria] Length: 372\nNumber of predicted TMHs: 1\nExp number of AAs in TMHs: 23.72226\nExp number, first 60 AAs: 0.22038\nTotal prob of N-in: 0.12994\noutside 1 331\nTMhelix 332 354\ninside 355 372

9342 GCF_000601155.1_ASM60115v1 Escherichia coli 1-110-08_S3_C1 Proteobacteria; Gammaproteobacteria; Enterobacterales; Enterobacteriaceae; Escherichia; Escherichia coli MNNEETFYQAMRRQGVTRRSFLKYCSLAATSLGLGAGMAPKIAWALENKPRIPVVWIHGL WP_001058323.1 MULTISPECIES: hydrogenase-1 small chain [Proteobacteria] Length: 372\nNumber of predicted TMHs: 2\nExp number of AAs in TMHs: 37.2608499999999\nExp number, first 60 AAs: 19.52923\nTotal prob of N-in: 0.94351\nPOSSIBLE N-term signal sequence\ninside 1 19\nTMhelix 20 39\noutside 40 325\nTMhelix 326 348\ninside 349 372

9343 GCF_000601175.1_ASM60117v1 Escherichia coli 1-110-08_S3_C2 Proteobacteria; Gammaproteobacteria; Enterobacterales; Enterobacteriaceae; Escherichia; Escherichia coli MTGDNTLIHSHGINRRDFMKLCAALAATMGLSSKAAAEMAESVTNPQRPPVIWIGAQECT WP_000145410.1 MULTISPECIES: hydrogenase-2 small chain [Proteobacteria] Length: 372\nNumber of predicted TMHs: 1\nExp number of AAs in TMHs: 23.72226\nExp number, first 60 AAs: 0.22038\nTotal prob of N-in: 0.12994\noutside 1 331\nTMhelix 332 354\ninside 355 372

9344 GCF_000601175.1_ASM60117v1 Escherichia coli 1-110-08_S3_C2 Proteobacteria; Gammaproteobacteria; Enterobacterales; Enterobacteriaceae; Escherichia; Escherichia coli MNNEETFYQAMRRQGVTRRSFLKYCSLAATSLGLGAGMAPKIAWALENKPRIPVVWIHGL WP_001058323.1 MULTISPECIES: hydrogenase-1 small chain [Proteobacteria] Length: 372\nNumber of predicted TMHs: 2\nExp number of AAs in TMHs: 37.2608499999999\nExp number, first 60 AAs: 19.52923\nTotal prob of N-in: 0.94351\nPOSSIBLE N-term signal sequence\ninside 1 19\nTMhelix 20 39\noutside 40 325\nTMhelix 326 348\ninside 349 372

9345 GCF_000601335.1_ASM60133v1 Escherichia coli 1-110-08_S3_C3 Proteobacteria; Gammaproteobacteria; Enterobacterales; Enterobacteriaceae; Escherichia; Escherichia coli MTGDNTLIHSHGINRRDFMKLCAALAATMGLSSKAAAEMAESVTNPQRPPVIWIGAQECT WP_000145410.1 MULTISPECIES: hydrogenase-2 small chain [Proteobacteria] Length: 372\nNumber of predicted TMHs:

1\nExp number of AAs in TMHs: 23.72226\nExp number, first 60 AAs: 0.22038\nTotal prob of N-in: 0.12994\noutside 1 331\nTMhelix 332 354\ninside 355 372

9346 GCF_000601335.1_ASM60133v1 Escherichia coli 1-110-08_S3_C3 Proteobacteria; Gammaproteobacteria; Enterobacterales; Enterobacteriaceae; Escherichia; Escherichia coli
MNNEETFYQAMRRQGVTRRSFLKYCSLAATSLGLGAGMAPKIAWALENKPRIPVVWIHGL WP_001058323.1
MULTISPECIES: hydrogenase-1 small chain [Proteobacteria] Length: 372\nNumber of predicted TMHs: 2\nExp number of AAs in TMHs: 37.2608499999999\nExp number, first 60 AAs: 19.52923\nTotal prob of N-in: 0.94351\nPOSSIBLE N-term signal sequence\ninside 1 19\nTMhelix 20 39\noutside 40 325\nTMhelix 326 348\ninside 349 372

9347 GCF_000601355.1_ASM60135v1 Escherichia coli 1-110-08_S4_C1 Proteobacteria; Gammaproteobacteria; Enterobacterales; Enterobacteriaceae; Escherichia; Escherichia coli
MTGDNTLIHSHGINRRDFMKLCAALAATMGLSSKAAAEMAESVTNPQRPPVIWIGAQECT WP_000145410.1
MULTISPECIES: hydrogenase-2 small chain [Proteobacteria] Length: 372\nNumber of predicted TMHs: 1\nExp number of AAs in TMHs: 23.72226\nExp number, first 60 AAs: 0.22038\nTotal prob of N-in: 0.12994\noutside 1 331\nTMhelix 332 354\ninside 355 372

9348 GCF_000601355.1_ASM60135v1 Escherichia coli 1-110-08_S4_C1 Proteobacteria; Gammaproteobacteria; Enterobacterales; Enterobacteriaceae; Escherichia; Escherichia coli
MNNEETFYQAMRRQGVTRRSFLKYCSLAATSLGLGAGMAPKIAWALENKPRIPVVWIHGL WP_001058323.1
MULTISPECIES: hydrogenase-1 small chain [Proteobacteria] Length: 372\nNumber of predicted TMHs: 2\nExp number of AAs in TMHs: 37.2608499999999\nExp number, first 60 AAs: 19.52923\nTotal prob of N-in: 0.94351\nPOSSIBLE N-term signal sequence\ninside 1 19\nTMhelix 20 39\noutside 40 325\nTMhelix 326 348\ninside 349 372

9349 GCF_000601215.1_ASM60121v1 Escherichia coli 1-110-08_S4_C2 Proteobacteria; Gammaproteobacteria; Enterobacterales; Enterobacteriaceae; Escherichia; Escherichia coli
MNNEETFYQAMRRQGVTRRSFLKYCSLAATSLGLGAGMAPKIAWALENKPRIPVVWIHGL WP_001058323.1
MULTISPECIES: hydrogenase-1 small chain [Proteobacteria] Length: 372\nNumber of predicted TMHs: 2\nExp number of AAs in TMHs: 37.2608499999999\nExp number, first 60 AAs: 19.52923\nTotal prob of N-in: 0.94351\nPOSSIBLE N-term signal sequence\ninside 1 19\nTMhelix 20 39\noutside 40 325\nTMhelix 326 348\ninside 349 372

9350 GCF_000601215.1_ASM60121v1 Escherichia coli 1-110-08_S4_C2 Proteobacteria; Gammaproteobacteria; Enterobacterales; Enterobacteriaceae; Escherichia; Escherichia coli
MTGDNTLIHSHGINRRDFMKLCAALAATMGLSSKAAAEMAESVTNPQRPPVIWIGAQECT WP_000145410.1
MULTISPECIES: hydrogenase-2 small chain [Proteobacteria] Length: 372\nNumber of predicted TMHs: 1\nExp number of AAs in TMHs: 23.72226\nExp number, first 60 AAs: 0.22038\nTotal prob of N-in: 0.12994\noutside 1 331\nTMhelix 332 354\ninside 355 372

9351 GCF_000601235.1_ASM60123v1 Escherichia coli 1-110-08_S4_C3 Proteobacteria; Gammaproteobacteria; Enterobacterales; Enterobacteriaceae; Escherichia; Escherichia coli
MTGDNTLIHSHGINRRDFMKLCAALAATMGLSSKAAAEMAESVTNPQRPPVIWIGAQECT WP_000145410.1
MULTISPECIES: hydrogenase-2 small chain [Proteobacteria] Length: 372\nNumber of predicted TMHs: 1\nExp number of AAs in TMHs: 23.72226\nExp number, first 60 AAs: 0.22038\nTotal prob of N-in: 0.12994\noutside 1 331\nTMhelix 332 354\ninside 355 372

9352 GCF_000601235.1_ASM60123v1 Escherichia coli 1-110-08_S4_C3 Proteobacteria; Gammaproteobacteria; Enterobacterales; Enterobacteriaceae; Escherichia; Escherichia coli
MNNEETFYQAMRRQGVTRRSFLKYCSLAATSLGLGAGMAPKIAWALENKPRIPVVWIHGL WP_001058323.1
MULTISPECIES: hydrogenase-1 small chain [Proteobacteria] Length: 372\nNumber of predicted TMHs: 2\nExp number of AAs in TMHs: 37.2608499999999\nExp number, first 60 AAs: 19.52923\nTotal prob of N-in: 0.94351\nPOSSIBLE N-term signal sequence\ninside 1 19\nTMhelix 20 39\noutside 40 325\nTMhelix 326 348\ninside 349 372

9353 GCF_000699385.1_11117 Escherichia coli 11117 Proteobacteria; Gammaproteobacteria; Enterobacterales; Enterobacteriaceae; Escherichia; Escherichia coli
MTGDNTLIHSHGINRRDFMKLCAALAATMGLSSKAAAEMVESVTNPQRPPVIWIGAQECT WP_000145423.1
hydrogenase 2 small subunit [Escherichia coli] Length: 372\nNumber of predicted TMHs: 1\nExp number of

AAs in TMHs: 23.78201\nExp number, first 60 AAs: 0.28355\nTotal prob of N-in: 0.13257\noutside 1 331\nTMhelix 332 354\ninside 355 372

9354 GCF_000699385.1_11117 Escherichia coli 11117 Proteobacteria; Gammaproteobacteria; Enterobacterales; Enterobacteriaceae; Escherichia; Escherichia coli MNNEETFYQAMRRQGVTRRSFLKYCSLAATSLGLGAGMAPKIAWALENKPRIPVVWIHGL WP_001295940.1 MULTISPECIES: hydrogenase [Enterobacteriaceae] Length: 372\nNumber of predicted TMHs: 2\nExp number of AAs in TMHs: 37.2608899999999\nExp number, first 60 AAs: 19.52923\nTotal prob of N-in: 0.94351\nPOSSIBLE N-term signal sequence\ninside 1 19\nTMhelix 20 39\noutside 40 325\nTMhelix 326 348\ninside 349 372

9355 GCF_000418795.1_E_coli112469215-isolate1-1.0Escherichia coli 112469215-isolate1 Proteobacteria; Gammaproteobacteria; Enterobacterales; Enterobacteriaceae; Escherichia; Escherichia coli MNNEETFYQAMRRQGVTRRSFLKYCSLAATSLGLGAGMAPKIAWALENKPRIPVVWIHGL WP_001058323.1 MULTISPECIES: hydrogenase-1 small chain [Proteobacteria] Length: 372\nNumber of predicted TMHs: 2\nExp number of AAs in TMHs: 37.2608499999999\nExp number, first 60 AAs: 19.52923\nTotal prob of N-in: 0.94351\nPOSSIBLE N-term signal sequence\ninside 1 19\nTMhelix 20 39\noutside 40 325\nTMhelix 326 348\ninside 349 372

9356 GCF_000418795.1_E_coli112469215-isolate1-1.0Escherichia coli 112469215-isolate1 Proteobacteria; Gammaproteobacteria; Enterobacterales; Enterobacteriaceae; Escherichia; Escherichia coli MTGDNTLIHSHGINRRDFMKLCAALAATMGLSSKAAAEMAESVTNPQRPPVIWIGAQECT WP_000145410.1 MULTISPECIES: hydrogenase-2 small chain [Proteobacteria] Length: 372\nNumber of predicted TMHs: 1\nExp number of AAs in TMHs: 23.72226\nExp number, first 60 AAs: 0.22038\nTotal prob of N-in: 0.12994\noutside 1 331\nTMhelix 332 354\ninside 355 372

9357 GCF_000418775.1_E_coli112469218-isolate1-1.0Escherichia coli 112469218-isolate1 Proteobacteria; Gammaproteobacteria; Enterobacterales; Enterobacteriaceae; Escherichia; Escherichia coli MNNEETFYQAMRRQGVTRRSFLKYCSLAATSLGLGAGMAPKIAWALENKPRIPVVWIHGL WP_001058323.1 MULTISPECIES: hydrogenase-1 small chain [Proteobacteria] Length: 372\nNumber of predicted TMHs: 2\nExp number of AAs in TMHs: 37.2608499999999\nExp number, first 60 AAs: 19.52923\nTotal prob of N-in: 0.94351\nPOSSIBLE N-term signal sequence\ninside 1 19\nTMhelix 20 39\noutside 40 325\nTMhelix 326 348\ninside 349 372

9358 GCF_000418775.1_E_coli112469218-isolate1-1.0Escherichia coli 112469218-isolate1 Proteobacteria; Gammaproteobacteria; Enterobacterales; Enterobacteriaceae; Escherichia; Escherichia coli MTGDNTLIHSHGINRRDFMKLCAALAATMGLSSKAAAEMAESVTNPQRPPVIWIGAQECT WP_000145410.1 MULTISPECIES: hydrogenase-2 small chain [Proteobacteria] Length: 372\nNumber of predicted TMHs: 1\nExp number of AAs in TMHs: 23.72226\nExp number, first 60 AAs: 0.22038\nTotal prob of N-in: 0.12994\noutside 1 331\nTMhelix 332 354\ninside 355 372

9359 GCF_000485635.1_ASM48563v1 Escherichia coli 113290 Proteobacteria; Gammaproteobacteria; Enterobacterales; Enterobacteriaceae; Escherichia; Escherichia coli MTGDNTLIHSHGINRRDFMKLCAALAATMGLSSKAAAEMAESVTNPQRPPVIWIGAQECT WP_000145410.1 MULTISPECIES: hydrogenase-2 small chain [Proteobacteria] Length: 372\nNumber of predicted TMHs: 1\nExp number of AAs in TMHs: 23.72226\nExp number, first 60 AAs: 0.22038\nTotal prob of N-in: 0.12994\noutside 1 331\nTMhelix 332 354\ninside 355 372

9360 GCF_000485635.1_ASM48563v1 Escherichia coli 113290 Proteobacteria; Gammaproteobacteria; Enterobacterales; Enterobacteriaceae; Escherichia; Escherichia coli MNNEETFYQAMRRQGVTRRSFLKYCSLAATSLGLGAGMAPKIAWALENKPRIPVVWIHGL WP_001058323.1 MULTISPECIES: hydrogenase-1 small chain [Proteobacteria] Length: 372\nNumber of predicted TMHs: 2\nExp number of AAs in TMHs: 37.2608499999999\nExp number, first 60 AAs: 19.52923\nTotal prob of N-in: 0.94351\nPOSSIBLE N-term signal sequence\ninside 1 19\nTMhelix 20 39\noutside 40 325\nTMhelix 326 348\ninside 349 372

9361 GCF_000488295.1_ASM48829v1 Escherichia coli 113302 Proteobacteria; Gammaproteobacteria; Enterobacterales; Enterobacteriaceae; Escherichia; Escherichia coli MTGDNTLIHSHGINRRDFMKLCAALAATMGLSSKAAAEMAESVTNPQRPPVIWIGAQECT WP_000145410.1 MULTISPECIES: hydrogenase-2 small chain [Proteobacteria] Length: 372\nNumber of predicted TMHs:

1\nExp number of AAs in TMHs: 23.72226\nExp number, first 60 AAs: 0.22038\nTotal prob of N-in: 0.12994\noutside 1 331\nTMhelix 332 354\ninside 355 372

9362 GCF_000488295.1_ASM48829v1 Escherichia coli 113302 Proteobacteria; Gammaproteobacteria; Enterobacterales; Enterobacteriaceae; Escherichia; Escherichia coli
MNNEETFYQAMRRQGVTRRSFLKYCSLAATSLGLGAGMAPKIAWALENKPRIPVVWIHGL WP_001295940.1
MULTISPECIES: hydrogenase [Enterobacteriaceae] Length: 372\nNumber of predicted TMHs: 2\nExp number of AAs in TMHs: 37.2608899999999\nExp number, first 60 AAs: 19.52923\nTotal prob of N-in: 0.94351\nPOSSIBLE N-term signal sequence\ninside 1 19\nTMhelix 20 39\noutside 40 325\nTMhelix 326 348\ninside 349 372

9363 GCF_000485655.1_ASM48565v1 Escherichia coli 113303 Proteobacteria; Gammaproteobacteria; Enterobacterales; Enterobacteriaceae; Escherichia; Escherichia coli
MNNEETFYQAMRRQGVTRRSFLKYCSLAATSLGLGAGMAPKIAWALENKPRIPVVWIHGL WP_001058323.1
MULTISPECIES: hydrogenase-1 small chain [Proteobacteria] Length: 372\nNumber of predicted TMHs: 2\nExp number of AAs in TMHs: 37.2608499999999\nExp number, first 60 AAs: 19.52923\nTotal prob of N-in: 0.94351\nPOSSIBLE N-term signal sequence\ninside 1 19\nTMhelix 20 39\noutside 40 325\nTMhelix 326 348\ninside 349 372

9364 GCF_000485655.1_ASM48565v1 Escherichia coli 113303 Proteobacteria; Gammaproteobacteria; Enterobacterales; Enterobacteriaceae; Escherichia; Escherichia coli
MTGDNTLIHSHGINRRDFMKLCAALAATMGLSSKAAAEMAESVTNPQRPPVIWIGAQECT WP_000145410.1
MULTISPECIES: hydrogenase-2 small chain [Proteobacteria] Length: 372\nNumber of predicted TMHs: 1\nExp number of AAs in TMHs: 23.72226\nExp number, first 60 AAs: 0.22038\nTotal prob of N-in: 0.12994\noutside 1 331\nTMhelix 332 354\ninside 355 372

9365 GCF_000601255.1_ASM60125v1 Escherichia coli 1-176-05_S1_C1 Proteobacteria; Gammaproteobacteria; Enterobacterales; Enterobacteriaceae; Escherichia; Escherichia coli
MNNEETFYQAMRRQGVTRRSFLKYCSLAATSLGLGAGMAPKIAWALENKPRIPVVWIHGL WP_001058323.1
MULTISPECIES: hydrogenase-1 small chain [Proteobacteria] Length: 372\nNumber of predicted TMHs: 2\nExp number of AAs in TMHs: 37.2608499999999\nExp number, first 60 AAs: 19.52923\nTotal prob of N-in: 0.94351\nPOSSIBLE N-term signal sequence\ninside 1 19\nTMhelix 20 39\noutside 40 325\nTMhelix 326 348\ninside 349 372

9366 GCF_000601255.1_ASM60125v1 Escherichia coli 1-176-05_S1_C1 Proteobacteria; Gammaproteobacteria; Enterobacterales; Enterobacteriaceae; Escherichia; Escherichia coli
MTGDNTLIHSHGINRRDFMKLCAALAATMGLSSKAAAEMAESVTNPQRPPVIWIGAQECT WP_000145410.1
MULTISPECIES: hydrogenase-2 small chain [Proteobacteria] Length: 372\nNumber of predicted TMHs: 1\nExp number of AAs in TMHs: 23.72226\nExp number, first 60 AAs: 0.22038\nTotal prob of N-in: 0.12994\noutside 1 331\nTMhelix 332 354\ninside 355 372

9367 GCF_000627905.1_ASM62790v1 Escherichia coli 1-176-05_S1_C2 Proteobacteria; Gammaproteobacteria; Enterobacterales; Enterobacteriaceae; Escherichia; Escherichia coli
MNNEETFYQAMRRQGVTRRSFLKYCSLAATSLGLGAGMAPKIAWALENKPRIPVVWIHGL WP_001058323.1
MULTISPECIES: hydrogenase-1 small chain [Proteobacteria] Length: 372\nNumber of predicted TMHs: 2\nExp number of AAs in TMHs: 37.2608499999999\nExp number, first 60 AAs: 19.52923\nTotal prob of N-in: 0.94351\nPOSSIBLE N-term signal sequence\ninside 1 19\nTMhelix 20 39\noutside 40 325\nTMhelix 326 348\ninside 349 372

9368 GCF_000627905.1_ASM62790v1 Escherichia coli 1-176-05_S1_C2 Proteobacteria; Gammaproteobacteria; Enterobacterales; Enterobacteriaceae; Escherichia; Escherichia coli
MTGDNTLIHSHGINRRDFMKLCAALAATMGLSSKAAAEMAESVTNPQRPPVIWIGAQECT WP_000145410.1
MULTISPECIES: hydrogenase-2 small chain [Proteobacteria] Length: 372\nNumber of predicted TMHs: 1\nExp number of AAs in TMHs: 23.72226\nExp number, first 60 AAs: 0.22038\nTotal prob of N-in: 0.12994\noutside 1 331\nTMhelix 332 354\ninside 355 372

9369 GCF_000627945.1_ASM62794v1 Escherichia coli 1-176-05_S1_C3 Proteobacteria; Gammaproteobacteria; Enterobacterales; Enterobacteriaceae; Escherichia; Escherichia coli
MTGDNTLIHSHGINRRDFMKLCAALAATMGLSSKAAAEMAESVTNPQRPPVIWIGAQECT WP_000145410.1
MULTISPECIES: hydrogenase-2 small chain [Proteobacteria] Length: 372\nNumber of predicted TMHs:

1\nExp number of AAs in TMHs: 23.72226\nExp number, first 60 AAs: 0.22038\nTotal prob of N-in: 0.12994\noutside 1 331\nTMhelix 332 354\ninside 355 372

9370 GCF_000627945.1_ASM62794v1 Escherichia coli 1-176-05_S1_C3 Proteobacteria; Gammaproteobacteria; Enterobacterales; Enterobacteriaceae; Escherichia; Escherichia coli MNNEETFYQAMRRQGVTRRSFLKYCSLAATSLGLGAGMAPKIAWALENKPRIPVVWIHGL WP_001058323.1 MULTISPECIES: hydrogenase-1 small chain [Proteobacteria] Length: 372\nNumber of predicted TMHs: 2\nExp number of AAs in TMHs: 37.2608499999999\nExp number, first 60 AAs: 19.52923\nTotal prob of N-in: 0.94351\nPOSSIBLE N-term signal sequence\ninside 1 19\nTMhelix 20 39\noutside 40 325\nTMhelix 326 348\ninside 349 372

9371 GCF_000601135.1_ASM60113v1 Escherichia coli 1-176-05_S3_C1 Proteobacteria; Gammaproteobacteria; Enterobacterales; Enterobacteriaceae; Escherichia; Escherichia coli MTGDNTLIHSHGINRRDFMKLCAALAATMGLSSKAAAEMAESVTNPQRPPVIWIGAQECT WP_000145410.1 MULTISPECIES: hydrogenase-2 small chain [Proteobacteria] Length: 372\nNumber of predicted TMHs: 1\nExp number of AAs in TMHs: 23.72226\nExp number, first 60 AAs: 0.22038\nTotal prob of N-in: 0.12994\noutside 1 331\nTMhelix 332 354\ninside 355 372

9372 GCF_000601135.1_ASM60113v1 Escherichia coli 1-176-05_S3_C1 Proteobacteria; Gammaproteobacteria; Enterobacterales; Enterobacteriaceae; Escherichia; Escherichia coli MNNEETFYQAMRRQGVTRRSFLKYCSLAATSLGLGAGMAPKIAWALENKPRIPVVWIHGL WP_001058323.1 MULTISPECIES: hydrogenase-1 small chain [Proteobacteria] Length: 372\nNumber of predicted TMHs: 2\nExp number of AAs in TMHs: 37.2608499999999\nExp number, first 60 AAs: 19.52923\nTotal prob of N-in: 0.94351\nPOSSIBLE N-term signal sequence\ninside 1 19\nTMhelix 20 39\noutside 40 325\nTMhelix 326 348\ninside 349 372

9373 GCF_000601195.1_ASM60119v1 Escherichia coli 1-176-05_S3_C2 Proteobacteria; Gammaproteobacteria; Enterobacterales; Enterobacteriaceae; Escherichia; Escherichia coli MNNEETFYQAMRRQGVTRRSFLKYCSLAATSLGLGAGMAPKIAWALENKPRIPVVWIHGL WP_010344442.1 MULTISPECIES: hydrogenase [Escherichia] Length: 372\nNumber of predicted TMHs: 2\nExp number of AAs in TMHs: 38.04874\nExp number, first 60 AAs: 19.5298\nTotal prob of N-in: 0.94352\nPOSSIBLE N-term signal sequence\ninside 1 19\nTMhelix 20 39\noutside 40 325\nTMhelix 326 348\ninside 349 372

9374 GCF_000601195.1_ASM60119v1 Escherichia coli 1-176-05_S3_C2 Proteobacteria; Gammaproteobacteria; Enterobacterales; Enterobacteriaceae; Escherichia; Escherichia coli MTGDNTLIHSHGINRRDFMKLCAALAATMGLSSKAAAEMAESVTNPQRPPVIWIGAQECT WP_032226272.1 hydrogenase 2 small subunit [Escherichia coli] Length: 372\nNumber of predicted TMHs: 1\nExp number of AAs in TMHs: 23.45172\nExp number, first 60 AAs: 0.22095\nTotal prob of N-in: 0.12250\noutside 1 332\nTMhelix 333 355\ninside 356 372

9375 GCF_000627315.1_ASM62731v1 Escherichia coli 1-176-05_S4_C1 Proteobacteria; Gammaproteobacteria; Enterobacterales; Enterobacteriaceae; Escherichia; Escherichia coli MNNEETFYQAMRRQGVTRRSFLKYCSLAATSLGLGAGMAPKIAWALENKPRIPVVWIHGL WP_001058323.1 MULTISPECIES: hydrogenase-1 small chain [Proteobacteria] Length: 372\nNumber of predicted TMHs: 2\nExp number of AAs in TMHs: 37.2608499999999\nExp number, first 60 AAs: 19.52923\nTotal prob of N-in: 0.94351\nPOSSIBLE N-term signal sequence\ninside 1 19\nTMhelix 20 39\noutside 40 325\nTMhelix 326 348\ninside 349 372

9376 GCF_000627315.1_ASM62731v1 Escherichia coli 1-176-05_S4_C1 Proteobacteria; Gammaproteobacteria; Enterobacterales; Enterobacteriaceae; Escherichia; Escherichia coli MTGDNTLIHSHGINRRDFMKLCAALAATMGLSSKAAAEMAESVTNPQRPPVIWIGAQECT WP_000145410.1 MULTISPECIES: hydrogenase-2 small chain [Proteobacteria] Length: 372\nNumber of predicted TMHs: 1\nExp number of AAs in TMHs: 23.72226\nExp number, first 60 AAs: 0.22038\nTotal prob of N-in: 0.12994\noutside 1 331\nTMhelix 332 354\ninside 355 372

9377 GCF_000687005.1_ASM68700v1 Escherichia coli 1-176-05_S4_C2 Proteobacteria; Gammaproteobacteria; Enterobacterales; Enterobacteriaceae; Escherichia; Escherichia coli MNNEETFYQAMRRQGVTRRSFLKYCSLAATSLGLGAGMAPKIAWALENKPRIPVVWIHGL WP_001058323.1 MULTISPECIES: hydrogenase-1 small chain [Proteobacteria] Length: 372\nNumber of predicted TMHs: 2\nExp number of AAs in TMHs: 37.2608499999999\nExp number, first 60 AAs: 19.52923\nTotal prob of N-in:

0.94351\nPOSSIBLE N-term signal sequence\ninside 1 19\nTMhelix 20 39\noutside 40 325\nTMhelix 326 348\ninside 349 372

9378 GCF_000687005.1_ASM68700v1 Escherichia coli 1-176-05_S4_C2 Proteobacteria; Gammaproteobacteria; Enterobacterales; Enterobacteriaceae; Escherichia; Escherichia coli MTGDNTLIHSHGINRRDFMKLCAALAATMGLSSKAAAEMAESVTNPQRPPVIWIGAQECT WP_000145410.1 MULTISPECIES: hydrogenase-2 small chain [Proteobacteria] Length: 372\nNumber of predicted TMHs: 1\nExp number of AAs in TMHs: 23.72226\nExp number, first 60 AAs: 0.22038\nTotal prob of N-in: 0.12994\noutside 1 331\nTMhelix 332 354\ninside 355 372

9379 GCF_000627135.1_ASM62713v1 Escherichia coli 1-176-05_S4_C3 Proteobacteria; Gammaproteobacteria; Enterobacterales; Enterobacteriaceae; Escherichia; Escherichia coli MNNEETFYQAMRRQGVTRRSFLKYCSLAATSLGLGAGMAPKIAWALENKPRIPVVWIHGL WP_001058323.1 MULTISPECIES: hydrogenase-1 small chain [Proteobacteria] Length: 372\nNumber of predicted TMHs: 2\nExp number of AAs in TMHs: 37.2608499999999\nExp number, first 60 AAs: 19.52923\nTotal prob of N-in: 0.94351\nPOSSIBLE N-term signal sequence\ninside 1 19\nTMhelix 20 39\noutside 40 325\nTMhelix 326 348\ninside 349 372

9380 GCF_000627135.1_ASM62713v1 Escherichia coli 1-176-05_S4_C3 Proteobacteria; Gammaproteobacteria; Enterobacterales; Enterobacteriaceae; Escherichia; Escherichia coli MTGDNTLIHSHGINRRDFMKLCAALAATMGLSSKAAAEMAESVTNPQRPPVIWIGAQECT WP_000145410.1 MULTISPECIES: hydrogenase-2 small chain [Proteobacteria] Length: 372\nNumber of predicted TMHs: 1\nExp number of AAs in TMHs: 23.72226\nExp number, first 60 AAs: 0.22038\nTotal prob of N-in: 0.12994\noutside 1 331\nTMhelix 332 354\ninside 355 372

9381 GCF_000627855.1_ASM62785v1 Escherichia coli 1-182-04_S1_C1 Proteobacteria; Gammaproteobacteria; Enterobacterales; Enterobacteriaceae; Escherichia; Escherichia coli MNNEETFYQAMRRQGVTRRSFLKYCSLAATSLGLGAGMAPKIAWALENKPRIPVVWIHGL WP_001058320.1 hydrogenase [Escherichia coli] Length: 372\nNumber of predicted TMHs: 2\nExp number of AAs in TMHs: 37.26082\nExp number, first 60 AAs: 19.52923\nTotal prob of N-in: 0.94351\nPOSSIBLE N-term signal sequence\ninside 1 19\nTMhelix 20 39\noutside 40 325\nTMhelix 326 348\ninside 349 372

9382 GCF_000627855.1_ASM62785v1 Escherichia coli 1-182-04_S1_C1 Proteobacteria; Gammaproteobacteria; Enterobacterales; Enterobacteriaceae; Escherichia; Escherichia coli MTGDNTLIHSHGINRRDFMKLCAALAATMGLSSKAAAEMAESVTNPQRPPVIWIGAQECT WP_000145410.1 MULTISPECIES: hydrogenase-2 small chain [Proteobacteria] Length: 372\nNumber of predicted TMHs: 1\nExp number of AAs in TMHs: 23.72226\nExp number, first 60 AAs: 0.22038\nTotal prob of N-in: 0.12994\noutside 1 331\nTMhelix 332 354\ninside 355 372

9383 GCF_000687045.1_ASM68704v1 Escherichia coli 1-182-04_S1_C2 Proteobacteria; Gammaproteobacteria; Enterobacterales; Enterobacteriaceae; Escherichia; Escherichia coli MNNEETFYQAMRRQGVTRRSFLKYCSLAATSLGLGAGMAPKIAWALENKPRIPVVWIHGL WP_001058320.1 hydrogenase [Escherichia coli] Length: 372\nNumber of predicted TMHs: 2\nExp number of AAs in TMHs: 37.26082\nExp number, first 60 AAs: 19.52923\nTotal prob of N-in: 0.94351\nPOSSIBLE N-term signal sequence\ninside 1 19\nTMhelix 20 39\noutside 40 325\nTMhelix 326 348\ninside 349 372

9384 GCF_000687045.1_ASM68704v1 Escherichia coli 1-182-04_S1_C2 Proteobacteria; Gammaproteobacteria; Enterobacterales; Enterobacteriaceae; Escherichia; Escherichia coli MTGDNTLIHSHGINRRDFMKLCAALAATMGLSSKAAAEMAESVTNPQRPPVIWIGAQECT WP_000145410.1 MULTISPECIES: hydrogenase-2 small chain [Proteobacteria] Length: 372\nNumber of predicted TMHs: 1\nExp number of AAs in TMHs: 23.72226\nExp number, first 60 AAs: 0.22038\nTotal prob of N-in: 0.12994\noutside 1 331\nTMhelix 332 354\ninside 355 372

9385 GCF_000627795.1_ASM62779v1 Escherichia coli 1-182-04_S1_C3 Proteobacteria; Gammaproteobacteria; Enterobacterales; Enterobacteriaceae; Escherichia; Escherichia coli MTGDNTLIHSHGINRRDFMKLCAALAATMGLSSKAAAEMAESVTNPQRPPVIWIGAQECT WP_000145410.1 MULTISPECIES: hydrogenase-2 small chain [Proteobacteria] Length: 372\nNumber of predicted TMHs: 1\nExp number of AAs in TMHs: 23.72226\nExp number, first 60 AAs: 0.22038\nTotal prob of N-in: 0.12994\noutside 1 331\nTMhelix 332 354\ninside 355 372

9386 GCF_000627795.1_ASM62779v1 Escherichia coli 1-182-04_S1_C3 Proteobacteria; Gammaproteobacteria; Enterobacterales; Enterobacteriaceae; Escherichia; Escherichia coli MNNEETFYQAMRRQGVTRRSFLKYCSLAATSLGLGAGMAPKIAWALENKPRIPVVWIHGL WP_001058320.1 hydrogenase [Escherichia coli] Length: 372\nNumber of predicted TMHs: 2\nExp number of AAs in TMHs: 37.26082\nExp number, first 60 AAs: 19.52923\nTotal prob of N-in: 0.94351\nPOSSIBLE N-term signal sequence\ninside 1 19\nTMhelix 20 39\noutside 40 325\nTMhelix 326 348\ninside 349 372

9387 GCF_000627965.1_ASM62796v1 Escherichia coli 1-182-04_S3_C1 Proteobacteria; Gammaproteobacteria; Enterobacterales; Enterobacteriaceae; Escherichia; Escherichia coli MNNEETFYQAMRRQGVTRRSFLKYCSLAATSLGLGAGMAPKIAWALENKPRIPVVWIHGL WP_001058323.1 MULTISPECIES: hydrogenase-1 small chain [Proteobacteria] Length: 372\nNumber of predicted TMHs: 2\nExp number of AAs in TMHs: 37.2608499999999\nExp number, first 60 AAs: 19.52923\nTotal prob of N-in: 0.94351\nPOSSIBLE N-term signal sequence\ninside 1 19\nTMhelix 20 39\noutside 40 325\nTMhelix 326 348\ninside 349 372

9388 GCF_000627965.1_ASM62796v1 Escherichia coli 1-182-04_S3_C1 Proteobacteria; Gammaproteobacteria; Enterobacterales; Enterobacteriaceae; Escherichia; Escherichia coli MTGDNTLIHSHGINRRDFMKLCAALAATMGLSSKAAAEMAESVTNPQRPPVIWIGAQECT WP_000145410.1 MULTISPECIES: hydrogenase-2 small chain [Proteobacteria] Length: 372\nNumber of predicted TMHs: 1\nExp number of AAs in TMHs: 23.72226\nExp number, first 60 AAs: 0.22038\nTotal prob of N-in: 0.12994\noutside 1 331\nTMhelix 332 354\ninside 355 372

9389 GCF_000627455.1_ASM62745v1 Escherichia coli 1-182-04_S3_C2 Proteobacteria; Gammaproteobacteria; Enterobacterales; Enterobacteriaceae; Escherichia; Escherichia coli MTGDNTLIHSHGINRRDFMKLCAALAATMGLSSKAAAEMAESVTNPQRPPVIWIGAQECT WP_000145410.1 MULTISPECIES: hydrogenase-2 small chain [Proteobacteria] Length: 372\nNumber of predicted TMHs: 1\nExp number of AAs in TMHs: 23.72226\nExp number, first 60 AAs: 0.22038\nTotal prob of N-in: 0.12994\noutside 1 331\nTMhelix 332 354\ninside 355 372

9390 GCF_000627455.1_ASM62745v1 Escherichia coli 1-182-04_S3_C2 Proteobacteria; Gammaproteobacteria; Enterobacterales; Enterobacteriaceae; Escherichia; Escherichia coli MNNEETFYQAMRRQGVTRRSFLKYCSLAATSLGLGAGMAPKIAWALENKPRIPVVWIHGL WP_001058323.1 MULTISPECIES: hydrogenase-1 small chain [Proteobacteria] Length: 372\nNumber of predicted TMHs: 2\nExp number of AAs in TMHs: 37.2608499999999\nExp number, first 60 AAs: 19.52923\nTotal prob of N-in: 0.94351\nPOSSIBLE N-term signal sequence\ninside 1 19\nTMhelix 20 39\noutside 40 325\nTMhelix 326 348\ninside 349 372

9391 GCF_000627475.1_ASM62747v1 Escherichia coli 1-182-04_S3_C3 Proteobacteria; Gammaproteobacteria; Enterobacterales; Enterobacteriaceae; Escherichia; Escherichia coli MNNEETFYQAMRRQGVTRRSFLKYCSLAATSLGLGAGMAPKIAWALENKPRIPVVWIHGL WP_001058323.1 MULTISPECIES: hydrogenase-1 small chain [Proteobacteria] Length: 372\nNumber of predicted TMHs: 2\nExp number of AAs in TMHs: 37.2608499999999\nExp number, first 60 AAs: 19.52923\nTotal prob of N-in: 0.94351\nPOSSIBLE N-term signal sequence\ninside 1 19\nTMhelix 20 39\noutside 40 325\nTMhelix 326 348\ninside 349 372

9392 GCF_000627475.1_ASM62747v1 Escherichia coli 1-182-04_S3_C3 Proteobacteria; Gammaproteobacteria; Enterobacterales; Enterobacteriaceae; Escherichia; Escherichia coli MTGDNTLIHSHGINRRDFMKLCAALAATMGLSSKAAAEMAESVTNPQRPPVIWIGAQECT WP_000145410.1 MULTISPECIES: hydrogenase-2 small chain [Proteobacteria] Length: 372\nNumber of predicted TMHs: 1\nExp number of AAs in TMHs: 23.72226\nExp number, first 60 AAs: 0.22038\nTotal prob of N-in: 0.12994\noutside 1 331\nTMhelix 332 354\ninside 355 372

9393 GCF_000627175.1_ASM62717v1 Escherichia coli 1-182-04_S4_C1 Proteobacteria; Gammaproteobacteria; Enterobacterales; Enterobacteriaceae; Escherichia; Escherichia coli MNNEETFYQAMRRQGVTRRSFLKYCSLAATSLGLGAGMAPKIAWALENKPRIPVVWIHGL WP_001058323.1 MULTISPECIES: hydrogenase-1 small chain [Proteobacteria] Length: 372\nNumber of predicted TMHs: 2\nExp number of AAs in TMHs: 37.2608499999999\nExp number, first 60 AAs: 19.52923\nTotal prob of N-in: 0.94351\nPOSSIBLE N-term signal sequence\ninside 1 19\nTMhelix 20 39\noutside 40 325\nTMhelix 326 348\ninside 349 372

9394 GCF_000627175.1_ASM62717v1 Escherichia coli 1-182-04_S4_C1 Proteobacteria; Gammaproteobacteria; Enterobacterales; Enterobacteriaceae; Escherichia; Escherichia coli
MTGDNTLIHSHGINRRDFMKLCAALAATMGLSSKAAAEMAESVTNPQRPPVIWIGAQECT WP_000145410.1
MULTISPECIES: hydrogenase-2 small chain [Proteobacteria] Length: 372\nNumber of predicted TMHs: 1\nExp number of AAs in TMHs: 23.72226\nExp number, first 60 AAs: 0.22038\nTotal prob of N-in: 0.12994\noutside 1 331\nTMhelix 332 354\ninside 355 372

9395 GCF_000627215.1_ASM62721v1 Escherichia coli 1-182-04_S4_C2 Proteobacteria; Gammaproteobacteria; Enterobacterales; Enterobacteriaceae; Escherichia; Escherichia coli
MNNEETFYQAMRRQGVTRRSFLKYCSLAATSLGLGAGMAPKIAWALENKPRIPVVWIHGL WP_001058323.1
MULTISPECIES: hydrogenase-1 small chain [Proteobacteria] Length: 372\nNumber of predicted TMHs: 2\nExp number of AAs in TMHs: 37.2608499999999\nExp number, first 60 AAs: 19.52923\nTotal prob of N-in: 0.94351\nPOSSIBLE N-term signal sequence\ninside 1 19\nTMhelix 20 39\noutside 40 325\nTMhelix 326 348\ninside 349 372

9396 GCF_000627215.1_ASM62721v1 Escherichia coli 1-182-04_S4_C2 Proteobacteria; Gammaproteobacteria; Enterobacterales; Enterobacteriaceae; Escherichia; Escherichia coli
MTGDNTLIHSHGINRRDFMKLCAALAATMGLSSKAAAEMAESVTNPQRPPVIWIGAQECT WP_000145410.1
MULTISPECIES: hydrogenase-2 small chain [Proteobacteria] Length: 372\nNumber of predicted TMHs: 1\nExp number of AAs in TMHs: 23.72226\nExp number, first 60 AAs: 0.22038\nTotal prob of N-in: 0.12994\noutside 1 331\nTMhelix 332 354\ninside 355 372

9397 GCF_000627095.1_ASM62709v1 Escherichia coli 1-182-04_S4_C3 Proteobacteria; Gammaproteobacteria; Enterobacterales; Enterobacteriaceae; Escherichia; Escherichia coli
MNNEETFYQAMRRQGVTRRSFLKYCSLAATSLGLGAGMAPKIAWALENKPRIPVVWIHGL WP_001058323.1
MULTISPECIES: hydrogenase-1 small chain [Proteobacteria] Length: 372\nNumber of predicted TMHs: 2\nExp number of AAs in TMHs: 37.2608499999999\nExp number, first 60 AAs: 19.52923\nTotal prob of N-in: 0.94351\nPOSSIBLE N-term signal sequence\ninside 1 19\nTMhelix 20 39\noutside 40 325\nTMhelix 326 348\ninside 349 372

9398 GCF_000627095.1_ASM62709v1 Escherichia coli 1-182-04_S4_C3 Proteobacteria; Gammaproteobacteria; Enterobacterales; Enterobacteriaceae; Escherichia; Escherichia coli
MTGDNTLIHSHGINRRDFMKLCAALAATMGLSSKAAAEMAESVTNPQRPPVIWIGAQECT WP_000145410.1
MULTISPECIES: hydrogenase-2 small chain [Proteobacteria] Length: 372\nNumber of predicted TMHs: 1\nExp number of AAs in TMHs: 23.72226\nExp number, first 60 AAs: 0.22038\nTotal prob of N-in: 0.12994\noutside 1 331\nTMhelix 332 354\ninside 355 372

9399 GCF_000194415.1_ASM19441v2 Escherichia coli 1.2264 Proteobacteria; Gammaproteobacteria; Enterobacterales; Enterobacteriaceae; Escherichia; Escherichia coli
MNNEETFYQAMRRQGVTRRSFLKYCSLAATSLGLGAGMAPKIAWALENKPRIPVVWIHGL WP_001058323.1
MULTISPECIES: hydrogenase-1 small chain [Proteobacteria] Length: 372\nNumber of predicted TMHs: 2\nExp number of AAs in TMHs: 37.2608499999999\nExp number, first 60 AAs: 19.52923\nTotal prob of N-in: 0.94351\nPOSSIBLE N-term signal sequence\ninside 1 19\nTMhelix 20 39\noutside 40 325\nTMhelix 326 348\ninside 349 372

9400 GCF_000194415.1_ASM19441v2 Escherichia coli 1.2264 Proteobacteria; Gammaproteobacteria; Enterobacterales; Enterobacteriaceae; Escherichia; Escherichia coli
MTGDNTLIHSHGINRRDFMKLCAALAATMGLSSKAAAEMAESVTNPQRPPVIWIGAQECT WP_000145410.1
MULTISPECIES: hydrogenase-2 small chain [Proteobacteria] Length: 372\nNumber of predicted TMHs: 1\nExp number of AAs in TMHs: 23.72226\nExp number, first 60 AAs: 0.22038\nTotal prob of N-in: 0.12994\noutside 1 331\nTMhelix 332 354\ninside 355 372

9401 GCF_000503695.1_Ecoli1240 Escherichia coli 1240 Proteobacteria; Gammaproteobacteria; Enterobacterales; Enterobacteriaceae; Escherichia; Escherichia coli
MNNEETFYQAMRRQGVTRRSFLKYCSLAATSLGLGAGMAPKIAWALENKPRIPVVWIHGL WP_001058323.1
MULTISPECIES: hydrogenase-1 small chain [Proteobacteria] Length: 372\nNumber of predicted TMHs: 2\nExp number of AAs in TMHs: 37.2608499999999\nExp number, first 60 AAs: 19.52923\nTotal prob of N-in: 0.94351\nPOSSIBLE N-term signal sequence\ninside 1 19\nTMhelix 20 39\noutside 40 325\nTMhelix 326 348\ninside 349 372

9402 GCF_000503695.1_Ecoli1240 Escherichia coli 1240 Proteobacteria; Gammaproteobacteria; Enterobacterales; Enterobacteriaceae; Escherichia; Escherichia coli
MTGDNTLIHSHGINRRDFMKLCAALAATMGLSSKAAAEMAESVTNPQRPPVIWIGAQECT WP_000145410.1
MULTISPECIES: hydrogenase-2 small chain [Proteobacteria] Length: 372\nNumber of predicted TMHs: 1\nExp number of AAs in TMHs: 23.72226\nExp number, first 60 AAs: 0.22038\nTotal prob of N-in: 0.12994\noutside 1 331\nTMhelix 332 354\ninside 355 372

9403 GCF_000704245.1_ASM70424v1 Escherichia coli 1-250-04_S1_C1 Proteobacteria; Gammaproteobacteria; Enterobacterales; Enterobacteriaceae; Escherichia; Escherichia coli
MNNEETFYQAMRRQGVTRRSFLKYCSLAATSLGLGAGMAPKIAWALENKPRIPVVWIHGL WP_001058323.1
MULTISPECIES: hydrogenase-1 small chain [Proteobacteria] Length: 372\nNumber of predicted TMHs: 2\nExp number of AAs in TMHs: 37.2608499999999\nExp number, first 60 AAs: 19.52923\nTotal prob of N-in: 0.94351\nPOSSIBLE N-term signal sequence\ninside 1 19\nTMhelix 20 39\noutside 40 325\nTMhelix 326 348\ninside 349 372

9404 GCF_000704245.1_ASM70424v1 Escherichia coli 1-250-04_S1_C1 Proteobacteria; Gammaproteobacteria; Enterobacterales; Enterobacteriaceae; Escherichia; Escherichia coli
MTGDNTLIHSHGINRRDFMKLCAALAATMGLSSKAAAEMAESVTNPQRPPVIWIGAQECT WP_000145410.1
MULTISPECIES: hydrogenase-2 small chain [Proteobacteria] Length: 372\nNumber of predicted TMHs: 1\nExp number of AAs in TMHs: 23.72226\nExp number, first 60 AAs: 0.22038\nTotal prob of N-in: 0.12994\noutside 1 331\nTMhelix 332 354\ninside 355 372

9405 GCF_000704265.1_ASM70426v1 Escherichia coli 1-250-04_S1_C2 Proteobacteria; Gammaproteobacteria; Enterobacterales; Enterobacteriaceae; Escherichia; Escherichia coli
MNNEETFYQAMRRQGVTRRSFLKYCSLAATSLGLGAGMAPKIAWALENKPRIPVVWIHGL WP_001058323.1
MULTISPECIES: hydrogenase-1 small chain [Proteobacteria] Length: 372\nNumber of predicted TMHs: 2\nExp number of AAs in TMHs: 37.2608499999999\nExp number, first 60 AAs: 19.52923\nTotal prob of N-in: 0.94351\nPOSSIBLE N-term signal sequence\ninside 1 19\nTMhelix 20 39\noutside 40 325\nTMhelix 326 348\ninside 349 372

9406 GCF_000704265.1_ASM70426v1 Escherichia coli 1-250-04_S1_C2 Proteobacteria; Gammaproteobacteria; Enterobacterales; Enterobacteriaceae; Escherichia; Escherichia coli
MTGDNTLIHSHGINRRDFMKLCAALAATMGLSSKAAAEMAESVTNPQRPPVIWIGAQECT WP_000145410.1
MULTISPECIES: hydrogenase-2 small chain [Proteobacteria] Length: 372\nNumber of predicted TMHs: 1\nExp number of AAs in TMHs: 23.72226\nExp number, first 60 AAs: 0.22038\nTotal prob of N-in: 0.12994\noutside 1 331\nTMhelix 332 354\ninside 355 372

9407 GCF_000627815.1_ASM62781v1 Escherichia coli 1-250-04_S1_C3 Proteobacteria; Gammaproteobacteria; Enterobacterales; Enterobacteriaceae; Escherichia; Escherichia coli
MTGDNTLIHSHGINRRDFMKLCAALAATMGLSSKAAAEMAESVTNPQRPPVIWIGAQECT WP_000145410.1
MULTISPECIES: hydrogenase-2 small chain [Proteobacteria] Length: 372\nNumber of predicted TMHs: 1\nExp number of AAs in TMHs: 23.72226\nExp number, first 60 AAs: 0.22038\nTotal prob of N-in: 0.12994\noutside 1 331\nTMhelix 332 354\ninside 355 372

9408 GCF_000627815.1_ASM62781v1 Escherichia coli 1-250-04_S1_C3 Proteobacteria; Gammaproteobacteria; Enterobacterales; Enterobacteriaceae; Escherichia; Escherichia coli
MNNEETFYQAMRRQGVTRRSFLKYCSLAATSLGLGAGMAPKIAWALENKPRIPVVWIHGL WP_001058323.1
MULTISPECIES: hydrogenase-1 small chain [Proteobacteria] Length: 372\nNumber of predicted TMHs: 2\nExp number of AAs in TMHs: 37.2608499999999\nExp number, first 60 AAs: 19.52923\nTotal prob of N-in: 0.94351\nPOSSIBLE N-term signal sequence\ninside 1 19\nTMhelix 20 39\noutside 40 325\nTMhelix 326 348\ninside 349 372

9409 GCF_000627435.1_ASM62743v1 Escherichia coli 1-250-04_S3_C1 Proteobacteria; Gammaproteobacteria; Enterobacterales; Enterobacteriaceae; Escherichia; Escherichia coli
MNNEETFYQAMRRQGVTRRSFLKYCSLAATSLGLGAGMAPKIAWALENKPRIPVVWIHGL WP_032280669.1
hydrogenase [Escherichia coli] Length: 372\nNumber of predicted TMHs: 2\nExp number of AAs in TMHs: 37.2609299999999\nExp number, first 60 AAs: 19.52923\nTotal prob of N-in: 0.94351\nPOSSIBLE N-term signal sequence\ninside 1 19\nTMhelix 20 39\noutside 40 325\nTMhelix 326 348\ninside 349 372

9410 GCF_000627435.1_ASM62743v1 Escherichia coli 1-250-04_S3_C1 Proteobacteria; Gammaproteobacteria; Enterobacterales; Enterobacteriaceae; Escherichia; Escherichia coli

MTGDNTLIHSHGINRRDFMKLCAALAATMGLSSKAAAEMAESVTNPQRPPVIWIGAQECT WP_000145410.1 MULTISPECIES: hydrogenase-2 small chain [Proteobacteria] Length: 372\nNumber of predicted TMHs: 1\nExp number of AAs in TMHs: 23.72226\nExp number, first 60 AAs: 0.22038\nTotal prob of N-in: 0.12994\noutside 1 331\nTMhelix 332 354\ninside 355 372

9411 GCF_000713815.1_ASM71381v1 Escherichia coli 1-250-04_S3_C2 Proteobacteria; Gammaproteobacteria; Enterobacterales; Enterobacteriaceae; Escherichia; Escherichia coli MTGDNTLIHSHGINRRDFMKLCAALAATMGLSSKAAAEMAESVTNPQRPPVIWIGAQECT WP_000145410.1 MULTISPECIES: hydrogenase-2 small chain [Proteobacteria] Length: 372\nNumber of predicted TMHs: 1\nExp number of AAs in TMHs: 23.72226\nExp number, first 60 AAs: 0.22038\nTotal prob of N-in: 0.12994\noutside 1 331\nTMhelix 332 354\ninside 355 372

9412 GCF_000713815.1_ASM71381v1 Escherichia coli 1-250-04_S3_C2 Proteobacteria; Gammaproteobacteria; Enterobacterales; Enterobacteriaceae; Escherichia; Escherichia coli MNNEETFYQAMRRQGVTRRSFLKYCSLAATSLGLGAGMAPKIAWALENKPRIPVVWIHGL WP_032280669.1 hydrogenase [Escherichia coli] Length: 372\nNumber of predicted TMHs: 2\nExp number of AAs in TMHs: 37.2609299999999\nExp number, first 60 AAs: 19.52923\nTotal prob of N-in: 0.94351\nPOSSIBLE N-term signal sequence\ninside 1 19\nTMhelix 20 39\noutside 40 325\nTMhelix 326 348\ninside 349 372

9413 GCF_000627155.1_ASM62715v1 Escherichia coli 1-250-04_S4_C1 Proteobacteria; Gammaproteobacteria; Enterobacterales; Enterobacteriaceae; Escherichia; Escherichia coli MTGDNTLIHSHGINRRDFMKLCAALAATMGLSSKAAAEMAESVTNPQRPPVIWIGAQECT WP_000145410.1 MULTISPECIES: hydrogenase-2 small chain [Proteobacteria] Length: 372\nNumber of predicted TMHs: 1\nExp number of AAs in TMHs: 23.72226\nExp number, first 60 AAs: 0.22038\nTotal prob of N-in: 0.12994\noutside 1 331\nTMhelix 332 354\ninside 355 372

9414 GCF_000627155.1_ASM62715v1 Escherichia coli 1-250-04_S4_C1 Proteobacteria; Gammaproteobacteria; Enterobacterales; Enterobacteriaceae; Escherichia; Escherichia coli MNNEETFYQAMRRQGVTRRSFLKYCSLAATSLGLGAGMAPKIAWALENKPRIPVVWIHGL WP_001058323.1 MULTISPECIES: hydrogenase-1 small chain [Proteobacteria] Length: 372\nNumber of predicted TMHs: 2\nExp number of AAs in TMHs: 37.2608499999999\nExp number, first 60 AAs: 19.52923\nTotal prob of N-in: 0.94351\nPOSSIBLE N-term signal sequence\ninside 1 19\nTMhelix 20 39\noutside 40 325\nTMhelix 326 348\ninside 349 372

9415 GCF_000627235.1_ASM62723v1 Escherichia coli 1-250-04_S4_C2 Proteobacteria; Gammaproteobacteria; Enterobacterales; Enterobacteriaceae; Escherichia; Escherichia coli MTGDNTLIHSHGINRRDFMKLCAALAATMGLSSKAAAEMAESVTNPQRPPVIWIGAQECT WP_000145410.1 MULTISPECIES: hydrogenase-2 small chain [Proteobacteria] Length: 372\nNumber of predicted TMHs: 1\nExp number of AAs in TMHs: 23.72226\nExp number, first 60 AAs: 0.22038\nTotal prob of N-in: 0.12994\noutside 1 331\nTMhelix 332 354\ninside 355 372

9416 GCF_000627235.1_ASM62723v1 Escherichia coli 1-250-04_S4_C2 Proteobacteria; Gammaproteobacteria; Enterobacterales; Enterobacteriaceae; Escherichia; Escherichia coli MNNEETFYQAMRRQGVTRRSFLKYCSLAATSLGLGAGMAPKIAWALENKPRIPVVWIHGL WP_001058323.1 MULTISPECIES: hydrogenase-1 small chain [Proteobacteria] Length: 372\nNumber of predicted TMHs: 2\nExp number of AAs in TMHs: 37.2608499999999\nExp number, first 60 AAs: 19.52923\nTotal prob of N-in: 0.94351\nPOSSIBLE N-term signal sequence\ninside 1 19\nTMhelix 20 39\noutside 40 325\nTMhelix 326 348\ninside 349 372

9417 GCF_000194175.1_ASM19417v2 Escherichia coli 1.2741 Proteobacteria; Gammaproteobacteria; Enterobacterales; Enterobacteriaceae; Escherichia; Escherichia coli MNNEETFYQAMRRQGVTRRSFLKYCSLAATSLGLGAGMAPKIAWALENKPRIPVVWIHGL WP_001353311.1 MULTISPECIES: hydrogenase [Escherichia] Length: 372\nNumber of predicted TMHs: 2\nExp number of AAs in TMHs: 37.2638699999999\nExp number, first 60 AAs: 19.52924\nTotal prob of N-in: 0.94351\nPOSSIBLE N-term signal sequence\ninside 1 19\nTMhelix 20 39\noutside 40 325\nTMhelix 326 348\ninside 349 372

9418 GCF_000194175.1_ASM19417v2 Escherichia coli 1.2741 Proteobacteria; Gammaproteobacteria; Enterobacterales; Enterobacteriaceae; Escherichia; Escherichia coli MTGDNTLIHSHGINRRDFMKLCAALAATMGLSSKAAAEMAESVTNPQRPPVIWIGAQECT WP_000145410.1 MULTISPECIES: hydrogenase-2 small chain [Proteobacteria] Length: 372\nNumber of predicted TMHs:

1\nExp number of AAs in TMHs: 23.72226\nExp number, first 60 AAs: 0.22038\nTotal prob of N-in: 0.12994\noutside 1 331\nTMhelix 332 354\ninside 355 372

9419 GCF_000829985.1_ASM82998v1 Escherichia coli 1303 Proteobacteria; Gammaproteobacteria; Enterobacterales; Enterobacteriaceae; Escherichia; Escherichia coli
MTGDNTLIHSHGINRRDFMKLCAALAATMGLSSKAAAEMAESVTNPQRPPVIWIGAQECT WP_000145410.1
MULTISPECIES: hydrogenase-2 small chain [Proteobacteria] Length: 372\nNumber of predicted TMHs: 1\nExp number of AAs in TMHs: 23.72226\nExp number, first 60 AAs: 0.22038\nTotal prob of N-in: 0.12994\noutside 1 331\nTMhelix 332 354\ninside 355 372

9420 GCF_000829985.1_ASM82998v1 Escherichia coli 1303 Proteobacteria; Gammaproteobacteria; Enterobacterales; Enterobacteriaceae; Escherichia; Escherichia coli
MNNEETFYQAMRRQGVTRRSFLKYCSLAATSLGLGAGMAPKIAWALENKPRIPVVWIHGL WP_001058323.1
MULTISPECIES: hydrogenase-1 small chain [Proteobacteria] Length: 372\nNumber of predicted TMHs: 2\nExp number of AAs in TMHs: 37.2608499999999\nExp number, first 60 AAs: 19.52923\nTotal prob of N-in: 0.94351\nPOSSIBLE N-term signal sequence\ninside 1 19\nTMhelix 20 39\noutside 40 325\nTMhelix 326 348\ninside 349 372

9421 GCF_000503715.1_Ecoli1350 Escherichia coli 1350 Proteobacteria; Gammaproteobacteria; Enterobacterales; Enterobacteriaceae; Escherichia; Escherichia coli
MNNEETFYQAMRRQGVTRRSFLKYCSLAATSLGLGAGMAPKIAWALENKPRIPVVWIHGL WP_001058323.1
MULTISPECIES: hydrogenase-1 small chain [Proteobacteria] Length: 372\nNumber of predicted TMHs: 2\nExp number of AAs in TMHs: 37.2608499999999\nExp number, first 60 AAs: 19.52923\nTotal prob of N-in: 0.94351\nPOSSIBLE N-term signal sequence\ninside 1 19\nTMhelix 20 39\noutside 40 325\nTMhelix 326 348\ninside 349 372

9422 GCF_000503715.1_Ecoli1350 Escherichia coli 1350 Proteobacteria; Gammaproteobacteria; Enterobacterales; Enterobacteriaceae; Escherichia; Escherichia coli
MTGDNTLIHSHGINRRDFMKLCAALAATMGLSSKAAAEMAESVTNPQRPPVIWIGAQECT WP_000145410.1
MULTISPECIES: hydrogenase-2 small chain [Proteobacteria] Length: 372\nNumber of predicted TMHs: 1\nExp number of AAs in TMHs: 23.72226\nExp number, first 60 AAs: 0.22038\nTotal prob of N-in: 0.12994\noutside 1 331\nTMhelix 332 354\ninside 355 372

9423 GCF_000503275.1_Ecoli1365 Escherichia coli 1365 Proteobacteria; Gammaproteobacteria; Enterobacterales; Enterobacteriaceae; Escherichia; Escherichia coli
MTGDNTLIHSHGINRRDFMKLCAALAATMGLSSKAAAEMAESVTNPQRPPVIWIGAQECT WP_000145410.1
MULTISPECIES: hydrogenase-2 small chain [Proteobacteria] Length: 372\nNumber of predicted TMHs: 1\nExp number of AAs in TMHs: 23.72226\nExp number, first 60 AAs: 0.22038\nTotal prob of N-in: 0.12994\noutside 1 331\nTMhelix 332 354\ninside 355 372

9424 GCF_000503275.1_Ecoli1365 Escherichia coli 1365 Proteobacteria; Gammaproteobacteria; Enterobacterales; Enterobacteriaceae; Escherichia; Escherichia coli
MNNEETFYQAMRRQGVTRRSFLKYCSLAATSLGLGAGMAPKIAWALENKPRIPVVWIHGL WP_001058323.1
MULTISPECIES: hydrogenase-1 small chain [Proteobacteria] Length: 372\nNumber of predicted TMHs: 2\nExp number of AAs in TMHs: 37.2608499999999\nExp number, first 60 AAs: 19.52923\nTotal prob of N-in: 0.94351\nPOSSIBLE N-term signal sequence\ninside 1 19\nTMhelix 20 39\noutside 40 325\nTMhelix 326 348\ninside 349 372

9425 GCF_000703865.1_ASM70386v1 Escherichia coli 1-392-07_S1_C1 Proteobacteria; Gammaproteobacteria; Enterobacterales; Enterobacteriaceae; Escherichia; Escherichia coli
MNNEETFYQAMRRQGVTRRSFLKYCSLAATSLGLGAGMAPKIAWALENKPRIPVVWIHGL WP_001058323.1
MULTISPECIES: hydrogenase-1 small chain [Proteobacteria] Length: 372\nNumber of predicted TMHs: 2\nExp number of AAs in TMHs: 37.2608499999999\nExp number, first 60 AAs: 19.52923\nTotal prob of N-in: 0.94351\nPOSSIBLE N-term signal sequence\ninside 1 19\nTMhelix 20 39\noutside 40 325\nTMhelix 326 348\ninside 349 372

9426 GCF_000703865.1_ASM70386v1 Escherichia coli 1-392-07_S1_C1 Proteobacteria; Gammaproteobacteria; Enterobacterales; Enterobacteriaceae; Escherichia; Escherichia coli
MTGDNTLIHSHGINRRDFMKLCAALAATMGLSSKAAAEMAESVTNPQRPPVIWIGAQECT WP_000145410.1
MULTISPECIES: hydrogenase-2 small chain [Proteobacteria] Length: 372\nNumber of predicted TMHs:

1\nExp number of AAs in TMHs: 23.72226\nExp number, first 60 AAs: 0.22038\nTotal prob of N-in: 0.12994\noutside 1 331\nTMhelix 332 354\ninside 355 372

9427 GCF_000703885.1_ASM70388v1 Escherichia coli 1-392-07_S1_C2 Proteobacteria; Gammaproteobacteria; Enterobacterales; Enterobacteriaceae; Escherichia; Escherichia coli MNNEETFYQAMRRQGVTRRSFLKYCSLAATSLGLGAGMAPKIAWALENKPRIPVVWIHGL WP_001058323.1 MULTISPECIES: hydrogenase-1 small chain [Proteobacteria] Length: 372\nNumber of predicted TMHs: 2\nExp number of AAs in TMHs: 37.2608499999999\nExp number, first 60 AAs: 19.52923\nTotal prob of N-in: 0.94351\nPOSSIBLE N-term signal sequence\ninside 1 19\nTMhelix 20 39\noutside 40 325\nTMhelix 326 348\ninside 349 372

9428 GCF_000703885.1_ASM70388v1 Escherichia coli 1-392-07_S1_C2 Proteobacteria; Gammaproteobacteria; Enterobacterales; Enterobacteriaceae; Escherichia; Escherichia coli MTGDNTLIHSHGINRRDFMKLCAALAATMGLSSKAAAEMAESVTNPQRPPVIWIGAQECT WP_000145410.1 MULTISPECIES: hydrogenase-2 small chain [Proteobacteria] Length: 372\nNumber of predicted TMHs: 1\nExp number of AAs in TMHs: 23.72226\nExp number, first 60 AAs: 0.22038\nTotal prob of N-in: 0.12994\noutside 1 331\nTMhelix 332 354\ninside 355 372

9429 GCF_000703545.1_ASM70354v1 Escherichia coli 1-392-07_S3_C1 Proteobacteria; Gammaproteobacteria; Enterobacterales; Enterobacteriaceae; Escherichia; Escherichia coli MNNEETFYQAMRRQGVTRRSFLKYCSLAATSLGLGAGMAPKIAWALENKPRIPVVWIHGL WP_001058323.1 MULTISPECIES: hydrogenase-1 small chain [Proteobacteria] Length: 372\nNumber of predicted TMHs: 2\nExp number of AAs in TMHs: 37.2608499999999\nExp number, first 60 AAs: 19.52923\nTotal prob of N-in: 0.94351\nPOSSIBLE N-term signal sequence\ninside 1 19\nTMhelix 20 39\noutside 40 325\nTMhelix 326 348\ninside 349 372

9430 GCF_000703545.1_ASM70354v1 Escherichia coli 1-392-07_S3_C1 Proteobacteria; Gammaproteobacteria; Enterobacterales; Enterobacteriaceae; Escherichia; Escherichia coli MTGDNTLIHSHGINRRDFMKLCAALAATMGLSSKAAAEMAESVTNPQRPPVIWIGAQECT WP_000145410.1 MULTISPECIES: hydrogenase-2 small chain [Proteobacteria] Length: 372\nNumber of predicted TMHs: 1\nExp number of AAs in TMHs: 23.72226\nExp number, first 60 AAs: 0.22038\nTotal prob of N-in: 0.12994\noutside 1 331\nTMhelix 332 354\ninside 355 372

9431 GCF_000703785.1_ASM70378v1 Escherichia coli 1-392-07_S3_C2 Proteobacteria; Gammaproteobacteria; Enterobacterales; Enterobacteriaceae; Escherichia; Escherichia coli MTGDNTLIHSHGINRRDFMKLCAALAATMGLSSKAAAEMAESVTNPQRPPVIWIGAQECT WP_000145410.1 MULTISPECIES: hydrogenase-2 small chain [Proteobacteria] Length: 372\nNumber of predicted TMHs: 1\nExp number of AAs in TMHs: 23.72226\nExp number, first 60 AAs: 0.22038\nTotal prob of N-in: 0.12994\noutside 1 331\nTMhelix 332 354\ninside 355 372

9432 GCF_000703785.1_ASM70378v1 Escherichia coli 1-392-07_S3_C2 Proteobacteria; Gammaproteobacteria; Enterobacterales; Enterobacteriaceae; Escherichia; Escherichia coli MNNEETFYQAMRRQGVTRRSFLKYCSLAATSLGLGAGMAPKIAWALENKPRIPVVWIHGL WP_001058323.1 MULTISPECIES: hydrogenase-1 small chain [Proteobacteria] Length: 372\nNumber of predicted TMHs: 2\nExp number of AAs in TMHs: 37.2608499999999\nExp number, first 60 AAs: 19.52923\nTotal prob of N-in: 0.94351\nPOSSIBLE N-term signal sequence\ninside 1 19\nTMhelix 20 39\noutside 40 325\nTMhelix 326 348\ninside 349 372

9433 GCF_000627295.1_ASM62729v1 Escherichia coli 1-392-07_S3_C3 Proteobacteria; Gammaproteobacteria; Enterobacterales; Enterobacteriaceae; Escherichia; Escherichia coli MNNEETFYQAMRRQGVTRRSFLKYCSLAATSLGLGAGMAPKIAWALENKPRIPVVWIHGL WP_001058323.1 MULTISPECIES: hydrogenase-1 small chain [Proteobacteria] Length: 372\nNumber of predicted TMHs: 2\nExp number of AAs in TMHs: 37.2608499999999\nExp number, first 60 AAs: 19.52923\nTotal prob of N-in: 0.94351\nPOSSIBLE N-term signal sequence\ninside 1 19\nTMhelix 20 39\noutside 40 325\nTMhelix 326 348\ninside 349 372

9434 GCF_000627295.1_ASM62729v1 Escherichia coli 1-392-07_S3_C3 Proteobacteria; Gammaproteobacteria; Enterobacterales; Enterobacteriaceae; Escherichia; Escherichia coli MTGDNTLIHSHGINRRDFMKLCAALAATMGLSSKAAAEMAESVTNPQRPPVIWIGAQECT WP_000145410.1 MULTISPECIES: hydrogenase-2 small chain [Proteobacteria] Length: 372\nNumber of predicted TMHs:

1\nExp number of AAs in TMHs: 23.72226\nExp number, first 60 AAs: 0.22038\nTotal prob of N-in: 0.12994\noutside 1 331\nTMhelix 332 354\ninside 355 372

9435 GCF_000713675.1_ASM71367v1 Escherichia coli 1-392-07_S4_C1 Proteobacteria; Gammaproteobacteria; Enterobacterales; Enterobacteriaceae; Escherichia; Escherichia coli
MTGDNTLIHSHGINRRDFMKLCAALAATMGLSSKAAAEMAESVTNPQRPPVIWIGAQECT WP_000145410.1
MULTISPECIES: hydrogenase-2 small chain [Proteobacteria] Length: 372\nNumber of predicted TMHs: 1\nExp number of AAs in TMHs: 23.72226\nExp number, first 60 AAs: 0.22038\nTotal prob of N-in: 0.12994\noutside 1 331\nTMhelix 332 354\ninside 355 372

9436 GCF_000713675.1_ASM71367v1 Escherichia coli 1-392-07_S4_C1 Proteobacteria; Gammaproteobacteria; Enterobacterales; Enterobacteriaceae; Escherichia; Escherichia coli
MNNEETFYQAMRRQGVTRRSFLKYCSLAATSLGLGAGMAPKIAWALENKPRIPVVWIHGL WP_001058323.1
MULTISPECIES: hydrogenase-1 small chain [Proteobacteria] Length: 372\nNumber of predicted TMHs: 2\nExp number of AAs in TMHs: 37.2608499999999\nExp number, first 60 AAs: 19.52923\nTotal prob of N-in: 0.94351\nPOSSIBLE N-term signal sequence\ninside 1 19\nTMhelix 20 39\noutside 40 325\nTMhelix 326 348\ninside 349 372

9437 GCF_000627255.1_ASM62725v1 Escherichia coli 1-392-07_S4_C2 Proteobacteria; Gammaproteobacteria; Enterobacterales; Enterobacteriaceae; Escherichia; Escherichia coli
MTGDNTLIHSHGINRRDFMKLCAALAATMGLSSKAAAEMAESVTNPQRPPVIWIGAQECT WP_000145410.1
MULTISPECIES: hydrogenase-2 small chain [Proteobacteria] Length: 372\nNumber of predicted TMHs: 1\nExp number of AAs in TMHs: 23.72226\nExp number, first 60 AAs: 0.22038\nTotal prob of N-in: 0.12994\noutside 1 331\nTMhelix 332 354\ninside 355 372

9438 GCF_000627255.1_ASM62725v1 Escherichia coli 1-392-07_S4_C2 Proteobacteria; Gammaproteobacteria; Enterobacterales; Enterobacteriaceae; Escherichia; Escherichia coli
MNNEETFYQAMRRQGVTRRSFLKYCSLAATSLGLGAGMAPKIAWALENKPRIPVVWIHGL WP_001058320.1
hydrogenase [Escherichia coli] Length: 372\nNumber of predicted TMHs: 2\nExp number of AAs in TMHs: 37.26082\nExp number, first 60 AAs: 19.52923\nTotal prob of N-in: 0.94351\nPOSSIBLE N-term signal sequence\ninside 1 19\nTMhelix 20 39\noutside 40 325\nTMhelix 326 348\ninside 349 372

9439 GCF_000713615.1_ASM71361v1 Escherichia coli 1-392-07_S4_C3 Proteobacteria; Gammaproteobacteria; Enterobacterales; Enterobacteriaceae; Escherichia; Escherichia coli
MNNEETFYQAMRRQGVTRRSFLKYCSLAATSLGLGAGMAPKIAWALENKPRIPVVWIHGL WP_001058323.1
MULTISPECIES: hydrogenase-1 small chain [Proteobacteria] Length: 372\nNumber of predicted TMHs: 2\nExp number of AAs in TMHs: 37.2608499999999\nExp number, first 60 AAs: 19.52923\nTotal prob of N-in: 0.94351\nPOSSIBLE N-term signal sequence\ninside 1 19\nTMhelix 20 39\noutside 40 325\nTMhelix 326 348\ninside 349 372

9440 GCF_000713615.1_ASM71361v1 Escherichia coli 1-392-07_S4_C3 Proteobacteria; Gammaproteobacteria; Enterobacterales; Enterobacteriaceae; Escherichia; Escherichia coli
MTGDNTLIHSHGINRRDFMKLCAALAATMGLSSKAAAEMAESVTNPQRPPVIWIGAQECT WP_000145410.1
MULTISPECIES: hydrogenase-2 small chain [Proteobacteria] Length: 372\nNumber of predicted TMHs: 1\nExp number of AAs in TMHs: 23.72226\nExp number, first 60 AAs: 0.22038\nTotal prob of N-in: 0.12994\noutside 1 331\nTMhelix 332 354\ninside 355 372

9441 GCF_000511425.1_Ecoli148 Escherichia coli 148 Proteobacteria; Gammaproteobacteria; Enterobacterales; Enterobacteriaceae; Escherichia; Escherichia coli
MTGDNTLIHSHGINRRDFMKLCAALAATMGLSSKAAAEMAESVTNPQRPPVIWIGAQECT WP_000145410.1
MULTISPECIES: hydrogenase-2 small chain [Proteobacteria] Length: 372\nNumber of predicted TMHs: 1\nExp number of AAs in TMHs: 23.72226\nExp number, first 60 AAs: 0.22038\nTotal prob of N-in: 0.12994\noutside 1 331\nTMhelix 332 354\ninside 355 372

9442 GCF_000511425.1_Ecoli148 Escherichia coli 148 Proteobacteria; Gammaproteobacteria; Enterobacterales; Enterobacteriaceae; Escherichia; Escherichia coli
MNNEETFYQAMRRQGVTRRSFLKYCSLAATSLGLGAGMAPKIAWALENKPRIPVVWIHGL WP_001058323.1
MULTISPECIES: hydrogenase-1 small chain [Proteobacteria] Length: 372\nNumber of predicted TMHs: 2\nExp number of AAs in TMHs: 37.2608499999999\nExp number, first 60 AAs: 19.52923\nTotal prob of N-in: 0.94351\nPOSSIBLE N-term signal sequence\ninside 1 19\nTMhelix 20 39\noutside 40 325\nTMhelix 326 348\ninside 349 372

9443 GCF_000462225.2_ASM46222v2 Escherichia coli 14A Proteobacteria; Gammaproteobacteria; Enterobacterales; Enterobacteriaceae; Escherichia; Escherichia coli
MNNEETFYQAMRRQGVTRRSFLKYCSLAATSLGLGAGMAPKIAWALENKPRIPVVWIHGL WP_001301613.1
hydrogenase [Escherichia coli] Length: 372\nNumber of predicted TMHs: 2\nExp number of AAs in TMHs: 37.46598\nExp number, first 60 AAs: 19.53139\nTotal prob of N-in: 0.94364\nPOSSIBLE N-term signal sequence\ninside 1 19\nTMhelix 20 39\noutside 40 325\nTMhelix 326 348\ninside 349 372

9444 GCF_000462225.2_ASM46222v2 Escherichia coli 14A Proteobacteria; Gammaproteobacteria; Enterobacterales; Enterobacteriaceae; Escherichia; Escherichia coli
MTGDNTLIHSHGINRRDFMKLCAALAATMGLSSKAAAEMAESVTNPQRPPVIWIGAQECT WP_000145410.1
MULTISPECIES: hydrogenase-2 small chain [Proteobacteria] Length: 372\nNumber of predicted TMHs: 1\nExp number of AAs in TMHs: 23.72226\nExp number, first 60 AAs: 0.22038\nTotal prob of N-in: 0.12994\noutside 1 331\nTMhelix 332 354\ninside 355 372

9445 GCF_000354915.1_ASM35491v1 Escherichia coli 174750 Proteobacteria; Gammaproteobacteria; Enterobacterales; Enterobacteriaceae; Escherichia; Escherichia coli
MNNEETFYQAMRRQGVTRRSFLKYCSLAATSLGLGAGMAPKIAWALENKPRIPVVWIHGL WP_001058323.1
MULTISPECIES: hydrogenase-1 small chain [Proteobacteria] Length: 372\nNumber of predicted TMHs: 2\nExp number of AAs in TMHs: 37.2608499999999\nExp number, first 60 AAs: 19.52923\nTotal prob of N-in: 0.94351\nPOSSIBLE N-term signal sequence\ninside 1 19\nTMhelix 20 39\noutside 40 325\nTMhelix 326 348\ninside 349 372

9446 GCF_000354915.1_ASM35491v1 Escherichia coli 174750 Proteobacteria; Gammaproteobacteria; Enterobacterales; Enterobacteriaceae; Escherichia; Escherichia coli
MTGDNTLIHSHGINRRDFMKLCAALAATMGLSSKAAAEMAESVTNPQRPPVIWIGAQECT WP_000145410.1
MULTISPECIES: hydrogenase-2 small chain [Proteobacteria] Length: 372\nNumber of predicted TMHs: 1\nExp number of AAs in TMHs: 23.72226\nExp number, first 60 AAs: 0.22038\nTotal prob of N-in: 0.12994\noutside 1 331\nTMhelix 332 354\ninside 355 372

9447 GCF_000356105.2_ASM35610v2 Escherichia coli 174900 Proteobacteria; Gammaproteobacteria; Enterobacterales; Enterobacteriaceae; Escherichia; Escherichia coli
MTGDNTLIHSHGINRRDFMKLCAALAATMGLSSKAAAEMAESVTNPQRPPVIWIGAQECT WP_000145410.1
MULTISPECIES: hydrogenase-2 small chain [Proteobacteria] Length: 372\nNumber of predicted TMHs: 1\nExp number of AAs in TMHs: 23.72226\nExp number, first 60 AAs: 0.22038\nTotal prob of N-in: 0.12994\noutside 1 331\nTMhelix 332 354\ninside 355 372

9448 GCF_000356105.2_ASM35610v2 Escherichia coli 174900 Proteobacteria; Gammaproteobacteria; Enterobacterales; Enterobacteriaceae; Escherichia; Escherichia coli
MNNEETFYQAMRRQGVTRRSFLKYCSLAATSLGLGAGMAPKIAWALENKPRIPVVWIHGL WP_001295940.1
MULTISPECIES: hydrogenase [Enterobacteriaceae] Length: 372\nNumber of predicted TMHs: 2\nExp number of AAs in TMHs: 37.2608899999999\nExp number, first 60 AAs: 19.52923\nTotal prob of N-in: 0.94351\nPOSSIBLE N-term signal sequence\ninside 1 19\nTMhelix 20 39\noutside 40 325\nTMhelix 326 348\ninside 349 372

9449 GCF_000358895.1_ASM35889v1 Escherichia coli 178200 Proteobacteria; Gammaproteobacteria; Enterobacterales; Enterobacteriaceae; Escherichia; Escherichia coli
MNNEETFYQAMRRQGVTRRSFLKYCSLAATSLGLGAGMAPKIAWALENKPRIPVVWIHGL WP_001058320.1
hydrogenase [Escherichia coli] Length: 372\nNumber of predicted TMHs: 2\nExp number of AAs in TMHs: 37.26082\nExp number, first 60 AAs: 19.52923\nTotal prob of N-in: 0.94351\nPOSSIBLE N-term signal sequence\ninside 1 19\nTMhelix 20 39\noutside 40 325\nTMhelix 326 348\ninside 349 372

9450 GCF_000358895.1_ASM35889v1 Escherichia coli 178200 Proteobacteria; Gammaproteobacteria; Enterobacterales; Enterobacteriaceae; Escherichia; Escherichia coli
MTGDNTLIHSHGINRRDFMKLCAALAATMGLSSKAAAEMAESVTNPQRPPVIWIGAQECT WP_000145410.1
MULTISPECIES: hydrogenase-2 small chain [Proteobacteria] Length: 372\nNumber of predicted TMHs: 1\nExp number of AAs in TMHs: 23.72226\nExp number, first 60 AAs: 0.22038\nTotal prob of N-in: 0.12994\noutside 1 331\nTMhelix 332 354\ninside 355 372

9451 GCF_000358755.2_ASM35875v2 Escherichia coli 178850 Proteobacteria; Gammaproteobacteria; Enterobacterales; Enterobacteriaceae; Escherichia; Escherichia coli
MNNEETFYQAMRRQGVTRRSFLKYCSLAATSLGLGAGMAPKIAWALENKPRIPVVWIHGL WP_001058320.1

hydrogenase [Escherichia coli] Length: 372\nNumber of predicted TMHs: 2\nExp number of AAs in TMHs: 37.26082\nExp number, first 60 AAs: 19.52923\nTotal prob of N-in: 0.94351\nPOSSIBLE N-term signal sequence\ninside 1 19\nTMhelix 20 39\noutside 40 325\nTMhelix 326 348\ninside 349 372

9452 GCF_000358755.2_ASM35875v2 Escherichia coli 178850 Proteobacteria; Gammaproteobacteria; Enterobacterales; Enterobacteriaceae; Escherichia; Escherichia coli MTGDNTLIHSHGINRRDFMKLCAALAATMGLSSKAAAEMAESVTNPQRPPVIWIGAQECT WP_000145410.1 MULTISPECIES: hydrogenase-2 small chain [Proteobacteria] Length: 372\nNumber of predicted TMHs: 1\nExp number of AAs in TMHs: 23.72226\nExp number, first 60 AAs: 0.22038\nTotal prob of N-in: 0.12994\noutside 1 331\nTMhelix 332 354\ninside 355 372

9453 GCF_000356345.2_ASM35634v2 Escherichia coli 178900 Proteobacteria; Gammaproteobacteria; Enterobacterales; Enterobacteriaceae; Escherichia; Escherichia coli MTGDNTLIHSHGINRRDFMKLCAALAAPMGLSSKAAAEMAESVTNPQRPPVIWIGAQECT WP_001384145.1 hydrogenase-2 small chain [Escherichia coli] Length: 372\nNumber of predicted TMHs: 1\nExp number of AAs in TMHs: 23.58872\nExp number, first 60 AAs: 0.07992\nTotal prob of N-in: 0.12396\noutside 1 331\nTMhelix 332 354\ninside 355 372

9454 GCF_000356345.2_ASM35634v2 Escherichia coli 178900 Proteobacteria; Gammaproteobacteria; Enterobacterales; Enterobacteriaceae; Escherichia; Escherichia coli MNNEETFYQAMRRQGVTRRSFLKYCSLAATSLGLGAGMAPKIAWALENKPRIPVVWIHGL WP_001058323.1 MULTISPECIES: hydrogenase-1 small chain [Proteobacteria] Length: 372\nNumber of predicted TMHs: 2\nExp number of AAs in TMHs: 37.2608499999999\nExp number, first 60 AAs: 19.52923\nTotal prob of N-in: 0.94351\nPOSSIBLE N-term signal sequence\ninside 1 19\nTMhelix 20 39\noutside 40 325\nTMhelix 326 348\ninside 349 372

9455 GCF_000356905.1_ASM35690v1 Escherichia coli 179100 Proteobacteria; Gammaproteobacteria; Enterobacterales; Enterobacteriaceae; Escherichia; Escherichia coli MNNEETFYQAMRRQGVTRRSFLKYCSLAATSLGLGAGMAPKIAWALENKPRIPVVWIHGL WP_001058323.1 MULTISPECIES: hydrogenase-1 small chain [Proteobacteria] Length: 372\nNumber of predicted TMHs: 2\nExp number of AAs in TMHs: 37.2608499999999\nExp number, first 60 AAs: 19.52923\nTotal prob of N-in: 0.94351\nPOSSIBLE N-term signal sequence\ninside 1 19\nTMhelix 20 39\noutside 40 325\nTMhelix 326 348\ninside 349 372

9456 GCF_000356905.1_ASM35690v1 Escherichia coli 179100 Proteobacteria; Gammaproteobacteria; Enterobacterales; Enterobacteriaceae; Escherichia; Escherichia coli MTGDNTLIHSHGINRRDFMKLCAALAATMGLSSKAAAEMAESVTNPQRPPVIWIGAQECT WP_000145410.1 MULTISPECIES: hydrogenase-2 small chain [Proteobacteria] Length: 372\nNumber of predicted TMHs: 1\nExp number of AAs in TMHs: 23.72226\nExp number, first 60 AAs: 0.22038\nTotal prob of N-in: 0.12994\noutside 1 331\nTMhelix 332 354\ninside 355 372

9457 GCF_000356365.2_ASM35636v2 Escherichia coli 179550 Proteobacteria; Gammaproteobacteria; Enterobacterales; Enterobacteriaceae; Escherichia; Escherichia coli MTGDNTLIHSHGINRRDFMKLCVALAATMGLSSKAAAEMAESVTNPQRPPVIWIGAQECT WP_000145425.1 hydrogenase 2 small subunit [Escherichia coli] Length: 372\nNumber of predicted TMHs: 1\nExp number of AAs in TMHs: 23.94354\nExp number, first 60 AAs: 0.45405\nTotal prob of N-in: 0.13997\noutside 1 331\nTMhelix 332 354\ninside 355 372

9458 GCF_000356365.2_ASM35636v2 Escherichia coli 179550 Proteobacteria; Gammaproteobacteria; Enterobacterales; Enterobacteriaceae; Escherichia; Escherichia coli MNNEETFYQAMRRQGVTRRSFLKYCSLAATSLGLGAGMAPKIAWALENKPRIPVVWIHGL WP_001295940.1 MULTISPECIES: hydrogenase [Enterobacteriaceae] Length: 372\nNumber of predicted TMHs: 2\nExp number of AAs in TMHs: 37.2608899999999\nExp number, first 60 AAs: 19.52923\nTotal prob of N-in: 0.94351\nPOSSIBLE N-term signal sequence\ninside 1 19\nTMhelix 20 39\noutside 40 325\nTMhelix 326 348\ninside 349 372

9459 GCF_000356045.2_ASM35604v2 Escherichia coli 180050 Proteobacteria; Gammaproteobacteria; Enterobacterales; Enterobacteriaceae; Escherichia; Escherichia coli MNNEETFYQAMRRQGVTRRSFLKYCSLAATSLGLGAGMAPKIAWALENKPRIPVVWIHGL WP_001058323.1 MULTISPECIES: hydrogenase-1 small chain [Proteobacteria] Length: 372\nNumber of predicted TMHs: 2\nExp number of AAs in TMHs: 37.2608499999999\nExp number, first 60 AAs: 19.52923\nTotal prob of N-in:

0.94351\nPOSSIBLE N-term signal sequence\ninside 1 19\nTMhelix 20 39\noutside 40 325\nTMhelix 326 348\ninside 349 372

9460 GCF_000356045.2_ASM35604v2 Escherichia coli 180050 Proteobacteria; Gammaproteobacteria; Enterobacterales; Enterobacteriaceae; Escherichia; Escherichia coli
MTGDNPLIHSHGINRRDFMKLCAALAAPMGLSSKAAAEMAESVTNPQRPPVIWIGAQECT WP_024168075.1
hydrogenase 2 small subunit [Escherichia coli] Length: 372\nNumber of predicted TMHs: 1\nExp number of AAs in TMHs: 22.53411\nExp number, first 60 AAs: 0.04002\nTotal prob of N-in: 0.06079\noutside 1 331\nTMhelix 332 354\ninside 355 372

9461 GCF_000356305.2_ASM35630v2 Escherichia coli 180200 Proteobacteria; Gammaproteobacteria; Enterobacterales; Enterobacteriaceae; Escherichia; Escherichia coli
MTGDNTLIHSHGINRRDFMKLCVALAATMGLSSKAAAEMAESVTNPQRPPVIWIGAQECT WP_000145425.1
hydrogenase 2 small subunit [Escherichia coli] Length: 372\nNumber of predicted TMHs: 1\nExp number of AAs in TMHs: 23.94354\nExp number, first 60 AAs: 0.45405\nTotal prob of N-in: 0.13997\noutside 1 331\nTMhelix 332 354\ninside 355 372

9462 GCF_000356305.2_ASM35630v2 Escherichia coli 180200 Proteobacteria; Gammaproteobacteria; Enterobacterales; Enterobacteriaceae; Escherichia; Escherichia coli
MNNEETFYQAMRRQGVTRRSFLKYCSLAATSLGLGAGMAPKIAWALENKPRIPVVWIHGL WP_001295940.1
MULTISPECIES: hydrogenase [Enterobacteriaceae] Length: 372\nNumber of predicted TMHs: 2\nExp number of AAs in TMHs: 37.2608899999999\nExp number, first 60 AAs: 19.52923\nTotal prob of N-in: 0.94351\nPOSSIBLE N-term signal sequence\ninside 1 19\nTMhelix 20 39\noutside 40 325\nTMhelix 326 348\ninside 349 372

9463 GCF_000354835.2_ASM35483v2 Escherichia coli 180600 Proteobacteria; Gammaproteobacteria; Enterobacterales; Enterobacteriaceae; Escherichia; Escherichia coli
MTGDNTLIHSHGINRRDFMKLCAALAATMGLSSKAAAEMAESVTNPQRPPVIWIGAQECT WP_000145410.1
MULTISPECIES: hydrogenase-2 small chain [Proteobacteria] Length: 372\nNumber of predicted TMHs: 1\nExp number of AAs in TMHs: 23.72226\nExp number, first 60 AAs: 0.22038\nTotal prob of N-in: 0.12994\noutside 1 331\nTMhelix 332 354\ninside 355 372

9464 GCF_000354835.2_ASM35483v2 Escherichia coli 180600 Proteobacteria; Gammaproteobacteria; Enterobacterales; Enterobacteriaceae; Escherichia; Escherichia coli
MNNEETFYQAMRRQGVTRRSFLKYCSLAATSLGLGAGMAPKIAWALENKPRIPVVWIHGL WP_001058320.1
hydrogenase [Escherichia coli] Length: 372\nNumber of predicted TMHs: 2\nExp number of AAs in TMHs: 37.26082\nExp number, first 60 AAs: 19.52923\nTotal prob of N-in: 0.94351\nPOSSIBLE N-term signal sequence\ninside 1 19\nTMhelix 20 39\noutside 40 325\nTMhelix 326 348\ninside 349 372

9465 GCF_000356125.1_ASM35612v1 Escherichia coli 199900.1 Proteobacteria; Gammaproteobacteria; Enterobacterales; Enterobacteriaceae; Escherichia; Escherichia coli
MTGDNTLIHSHGINRRDFMKLCAALAATMGLSSKAAAEMAESVTNPQRPPVIWIGAQECT WP_000145410.1
MULTISPECIES: hydrogenase-2 small chain [Proteobacteria] Length: 372\nNumber of predicted TMHs: 1\nExp number of AAs in TMHs: 23.72226\nExp number, first 60 AAs: 0.22038\nTotal prob of N-in: 0.12994\noutside 1 331\nTMhelix 332 354\ninside 355 372

9466 GCF_000503535.1_Ecoli1A Escherichia coli 1A Proteobacteria; Gammaproteobacteria; Enterobacterales; Enterobacteriaceae; Escherichia; Escherichia coli
MNNEETFYQAMRRQGVTRRSFLKYCSLAATSLGLGAGMAPKIAWALENKPRIPVVWIHGL WP_001058320.1
hydrogenase [Escherichia coli] Length: 372\nNumber of predicted TMHs: 2\nExp number of AAs in TMHs: 37.26082\nExp number, first 60 AAs: 19.52923\nTotal prob of N-in: 0.94351\nPOSSIBLE N-term signal sequence\ninside 1 19\nTMhelix 20 39\noutside 40 325\nTMhelix 326 348\ninside 349 372

9467 GCF_000503535.1_Ecoli1A Escherichia coli 1A Proteobacteria; Gammaproteobacteria; Enterobacterales; Enterobacteriaceae; Escherichia; Escherichia coli
MTGDNTLIHSHGINRRDFMKLCAALAATMGLSSKAAAEMAESVTNPQRPPVIWIGAQECT WP_000145410.1
MULTISPECIES: hydrogenase-2 small chain [Proteobacteria] Length: 372\nNumber of predicted TMHs: 1\nExp number of AAs in TMHs: 23.72226\nExp number, first 60 AAs: 0.22038\nTotal prob of N-in: 0.12994\noutside 1 331\nTMhelix 332 354\ninside 355 372

9468 GCF_000627835.1_ASM62783v1 Escherichia coli 2-005-03_S1_C1 Proteobacteria; Gammaproteobacteria; Enterobacterales; Enterobacteriaceae; Escherichia; Escherichia coli
MNNEETFYQAMRRQGVTRRSFLKYCSLAATSLGLGAGMAPKIAWALENKPRIPVVWIHGL WP_001058320.1
hydrogenase [Escherichia coli] Length: 372\nNumber of predicted TMHs: 2\nExp number of AAs in TMHs: 37.26082\nExp number, first 60 AAs: 19.52923\nTotal prob of N-in: 0.94351\nPOSSIBLE N-term signal sequence\ninside 1 19\nTMhelix 20 39\noutside 40 325\nTMhelix 326 348\ninside 349 372

9469 GCF_000627835.1_ASM62783v1 Escherichia coli 2-005-03_S1_C1 Proteobacteria; Gammaproteobacteria; Enterobacterales; Enterobacteriaceae; Escherichia; Escherichia coli
MTGDNTLIHSHGINRRDFMKLCAALAATMGLSSKAAAEMAESVTNPQRPPVIWIGAQECT WP_000145410.1
MULTISPECIES: hydrogenase-2 small chain [Proteobacteria] Length: 372\nNumber of predicted TMHs: 1\nExp number of AAs in TMHs: 23.72226\nExp number, first 60 AAs: 0.22038\nTotal prob of N-in: 0.12994\noutside 1 331\nTMhelix 332 354\ninside 355 372

9470 GCF_000627985.1_ASM62798v1 Escherichia coli 2-005-03_S1_C2 Proteobacteria; Gammaproteobacteria; Enterobacterales; Enterobacteriaceae; Escherichia; Escherichia coli
MNNEETFYQAMRRQGVTRRSFLKYCSLAATSLGLGAGMAPKIAWALENKPRIPVVWIHGL WP_001058320.1
hydrogenase [Escherichia coli] Length: 372\nNumber of predicted TMHs: 2\nExp number of AAs in TMHs: 37.26082\nExp number, first 60 AAs: 19.52923\nTotal prob of N-in: 0.94351\nPOSSIBLE N-term signal sequence\ninside 1 19\nTMhelix 20 39\noutside 40 325\nTMhelix 326 348\ninside 349 372

9471 GCF_000627985.1_ASM62798v1 Escherichia coli 2-005-03_S1_C2 Proteobacteria; Gammaproteobacteria; Enterobacterales; Enterobacteriaceae; Escherichia; Escherichia coli
MTGDNTLIHSHGINRRDFMKLCAALAATMGLSSKAAAEMAESVTNPQRPPVIWIGAQECT WP_000145410.1
MULTISPECIES: hydrogenase-2 small chain [Proteobacteria] Length: 372\nNumber of predicted TMHs: 1\nExp number of AAs in TMHs: 23.72226\nExp number, first 60 AAs: 0.22038\nTotal prob of N-in: 0.12994\noutside 1 331\nTMhelix 332 354\ninside 355 372

9472 GCF_000627925.1_ASM62792v1 Escherichia coli 2-005-03_S1_C3 Proteobacteria; Gammaproteobacteria; Enterobacterales; Enterobacteriaceae; Escherichia; Escherichia coli
MNNEETFYQAMRRQGVTRRSFLKYCSLAATSLGLGAGMAPKIAWALENKPRIPVVWIHGL WP_001058320.1
hydrogenase [Escherichia coli] Length: 372\nNumber of predicted TMHs: 2\nExp number of AAs in TMHs: 37.26082\nExp number, first 60 AAs: 19.52923\nTotal prob of N-in: 0.94351\nPOSSIBLE N-term signal sequence\ninside 1 19\nTMhelix 20 39\noutside 40 325\nTMhelix 326 348\ninside 349 372

9473 GCF_000627925.1_ASM62792v1 Escherichia coli 2-005-03_S1_C3 Proteobacteria; Gammaproteobacteria; Enterobacterales; Enterobacteriaceae; Escherichia; Escherichia coli
MTGDNTLIHSHGINRRDFMKLCAALAATMGLSSKAAAEMAESVTNPQRPPVIWIGAQECT WP_000145410.1
MULTISPECIES: hydrogenase-2 small chain [Proteobacteria] Length: 372\nNumber of predicted TMHs: 1\nExp number of AAs in TMHs: 23.72226\nExp number, first 60 AAs: 0.22038\nTotal prob of N-in: 0.12994\noutside 1 331\nTMhelix 332 354\ninside 355 372

9474 GCF_000703805.1_ASM70380v1 Escherichia coli 2-005-03_S3_C1 Proteobacteria; Gammaproteobacteria; Enterobacterales; Enterobacteriaceae; Escherichia; Escherichia coli
MTGDNTLIHSHGINRRDFMKLCAALAATMGLSSKAAAEMAESVTNPQRPPVIWIGAQECT WP_000145410.1
MULTISPECIES: hydrogenase-2 small chain [Proteobacteria] Length: 372\nNumber of predicted TMHs: 1\nExp number of AAs in TMHs: 23.72226\nExp number, first 60 AAs: 0.22038\nTotal prob of N-in: 0.12994\noutside 1 331\nTMhelix 332 354\ninside 355 372

9475 GCF_000703805.1_ASM70380v1 Escherichia coli 2-005-03_S3_C1 Proteobacteria; Gammaproteobacteria; Enterobacterales; Enterobacteriaceae; Escherichia; Escherichia coli
MNNEETFYQAMRRQGVTRRSFLKYCSLAATSLGLGAGMAPKIAWALENKPRIPVVWIHGL WP_001058323.1
MULTISPECIES: hydrogenase-1 small chain [Proteobacteria] Length: 372\nNumber of predicted TMHs: 2\nExp number of AAs in TMHs: 37.2608499999999\nExp number, first 60 AAs: 19.52923\nTotal prob of N-in: 0.94351\nPOSSIBLE N-term signal sequence\ninside 1 19\nTMhelix 20 39\noutside 40 325\nTMhelix 326 348\ninside 349 372

9476 GCF_000687105.1_ASM68710v1 Escherichia coli 2-005-03_S3_C2 Proteobacteria; Gammaproteobacteria; Enterobacterales; Enterobacteriaceae; Escherichia; Escherichia coli
MNNEETFYQAMRRQGVTRRSFLKYCSLAATSLGLGAGMAPKIAWALENKPRIPVVWIHGL WP_001058323.1
MULTISPECIES: hydrogenase-1 small chain [Proteobacteria] Length: 372\nNumber of predicted TMHs:

2\nExp number of AAs in TMHs: 37.2608499999999\nExp number, first 60 AAs: 19.52923\nTotal prob of N-in: 0.94351\nPOSSIBLE N-term signal sequence\ninside 1 19\nTMhelix 20 39\noutside 40 325\nTMhelix 326 348\ninside 349 372

9477 GCF_000687105.1_ASM68710v1 Escherichia coli 2-005-03_S3_C2 Proteobacteria; Gammaproteobacteria; Enterobacterales; Enterobacteriaceae; Escherichia; Escherichia coli MTGDNTLIHSHGINRRDFMKLCAALAATMGLSSKAAAEMAESVTNPQRPPVIWIGAQECT WP_000145410.1 MULTISPECIES: hydrogenase-2 small chain [Proteobacteria] Length: 372\nNumber of predicted TMHs: 1\nExp number of AAs in TMHs: 23.72226\nExp number, first 60 AAs: 0.22038\nTotal prob of N-in: 0.12994\noutside 1 331\nTMhelix 332 354\ninside 355 372

9478 GCF_000703825.1_ASM70382v1 Escherichia coli 2-005-03_S3_C3 Proteobacteria; Gammaproteobacteria; Enterobacterales; Enterobacteriaceae; Escherichia; Escherichia coli MNNEETFYQAMRRQGVTRRSFLKYCSLAATSLGLGAGMAPKIAWALENKPRIPVVWIHGL WP_001058323.1 MULTISPECIES: hydrogenase-1 small chain [Proteobacteria] Length: 372\nNumber of predicted TMHs: 2\nExp number of AAs in TMHs: 37.2608499999999\nExp number, first 60 AAs: 19.52923\nTotal prob of N-in: 0.94351\nPOSSIBLE N-term signal sequence\ninside 1 19\nTMhelix 20 39\noutside 40 325\nTMhelix 326 348\ninside 349 372

9479 GCF_000703825.1_ASM70382v1 Escherichia coli 2-005-03_S3_C3 Proteobacteria; Gammaproteobacteria; Enterobacterales; Enterobacteriaceae; Escherichia; Escherichia coli MTGDNTLIHSHGINRRDFMKLCAALAATMGLSSKAAAEMAESVTNPQRPPVIWIGAQECT WP_000145410.1 MULTISPECIES: hydrogenase-2 small chain [Proteobacteria] Length: 372\nNumber of predicted TMHs: 1\nExp number of AAs in TMHs: 23.72226\nExp number, first 60 AAs: 0.22038\nTotal prob of N-in: 0.12994\noutside 1 331\nTMhelix 332 354\ninside 355 372

9480 GCF_000703845.1_ASM70384v1 Escherichia coli 2-005-03_S4_C1 Proteobacteria; Gammaproteobacteria; Enterobacterales; Enterobacteriaceae; Escherichia; Escherichia coli MNNEETFYQAMRRQGVTRRSFLKYCSLAATSLGLGAGMAPKIAWALENKPRIPVVWIHGL WP_001058323.1 MULTISPECIES: hydrogenase-1 small chain [Proteobacteria] Length: 372\nNumber of predicted TMHs: 2\nExp number of AAs in TMHs: 37.2608499999999\nExp number, first 60 AAs: 19.52923\nTotal prob of N-in: 0.94351\nPOSSIBLE N-term signal sequence\ninside 1 19\nTMhelix 20 39\noutside 40 325\nTMhelix 326 348\ninside 349 372

9481 GCF_000703845.1_ASM70384v1 Escherichia coli 2-005-03_S4_C1 Proteobacteria; Gammaproteobacteria; Enterobacterales; Enterobacteriaceae; Escherichia; Escherichia coli MTGDNTLIHSHGINRRDFMKLCAALAATMGLSSKAAAEMAESVTNPQRPPVIWIGAQECT WP_000145410.1 MULTISPECIES: hydrogenase-2 small chain [Proteobacteria] Length: 372\nNumber of predicted TMHs: 1\nExp number of AAs in TMHs: 23.72226\nExp number, first 60 AAs: 0.22038\nTotal prob of N-in: 0.12994\noutside 1 331\nTMhelix 332 354\ninside 355 372

9482 GCF_000627195.1_ASM62719v1 Escherichia coli 2-005-03_S4_C2 Proteobacteria; Gammaproteobacteria; Enterobacterales; Enterobacteriaceae; Escherichia; Escherichia coli MTGDNTLIHSHGINRRDFMKLCAALAATMGLSSKAAAEMAESVTNPQRPPVIWIGAQECT WP_000145418.1 hydrogenase 2 small subunit [Escherichia coli] Length: 372\nNumber of predicted TMHs: 1\nExp number of AAs in TMHs: 23.72313\nExp number, first 60 AAs: 0.22034\nTotal prob of N-in: 0.12997\noutside 1 331\nTMhelix 332 354\ninside 355 372

9483 GCF_000627195.1_ASM62719v1 Escherichia coli 2-005-03_S4_C2 Proteobacteria; Gammaproteobacteria; Enterobacterales; Enterobacteriaceae; Escherichia; Escherichia coli MNNEETFYQAMRRQGVTRRSFLKYCSLAATSLGLGAGMAPKIAWALENKPRIPVVWIHGL WP_001295940.1 MULTISPECIES: hydrogenase [Enterobacteriaceae] Length: 372\nNumber of predicted TMHs: 2\nExp number of AAs in TMHs: 37.2608899999999\nExp number, first 60 AAs: 19.52923\nTotal prob of N-in: 0.94351\nPOSSIBLE N-term signal sequence\ninside 1 19\nTMhelix 20 39\noutside 40 325\nTMhelix 326 348\ninside 349 372

9484 GCF_000627075.1_ASM62707v1 Escherichia coli 2-005-03_S4_C3 Proteobacteria; Gammaproteobacteria; Enterobacterales; Enterobacteriaceae; Escherichia; Escherichia coli MNNEETFYQAMRRQGVTRRSFLKYCSLAATSLGLGAGMAPKIAWALENKPRIPVVWIHGL WP_001295940.1 MULTISPECIES: hydrogenase [Enterobacteriaceae] Length: 372\nNumber of predicted TMHs: 2\nExp number of AAs in TMHs: 37.2608899999999\nExp number, first 60 AAs: 19.52923\nTotal prob of N-in:

0.94351\nPOSSIBLE N-term signal sequence\ninside 1 19\nTMhelix 20 39\noutside 40 325\nTMhelix 326 348\ninside 349 372

9485 GCF_000627075.1_ASM62707v1 Escherichia coli 2-005-03_S4_C3 Proteobacteria; Gammaproteobacteria; Enterobacterales; Enterobacteriaceae; Escherichia; Escherichia coli MTGDNTLIHSHGINRRDFMKLCAALAATMGLSSKAAAEMAESVTNPQRPPVIWIGAQECT WP_000145418.1 hydrogenase 2 small subunit [Escherichia coli] Length: 372\nNumber of predicted TMHs: 1\nExp number of AAs in TMHs: 23.72313\nExp number, first 60 AAs: 0.22034\nTotal prob of N-in: 0.12997\noutside 1 331\nTMhelix 332 354\ninside 355 372

9486 GCF_000687125.1_ASM68712v1 Escherichia coli 2-011-08_S1_C1 Proteobacteria; Gammaproteobacteria; Enterobacterales; Enterobacteriaceae; Escherichia; Escherichia coli MNNEETFYQAMRRQGVTRRSFLKYCSLAATSLGLGAGMAPKIAWALENKPRIPVVWIHGL WP_001353311.1 MULTISPECIES: hydrogenase [Escherichia] Length: 372\nNumber of predicted TMHs: 2\nExp number of AAs in TMHs: 37.2638699999999\nExp number, first 60 AAs: 19.52924\nTotal prob of N-in: 0.94351\nPOSSIBLE N-term signal sequence\ninside 1 19\nTMhelix 20 39\noutside 40 325\nTMhelix 326 348\ninside 349 372

9487 GCF_000687125.1_ASM68712v1 Escherichia coli 2-011-08_S1_C1 Proteobacteria; Gammaproteobacteria; Enterobacterales; Enterobacteriaceae; Escherichia; Escherichia coli MTGDNTLIHSHGINRRDFMKLCAALAATMGLSSKAAAEMAESVTNPQRPPVIWIGAQECT WP_000145404.1 hydrogenase-2 small chain [Escherichia coli] Length: 372\nNumber of predicted TMHs: 1\nExp number of AAs in TMHs: 23.72251\nExp number, first 60 AAs: 0.2205\nTotal prob of N-in: 0.12994\noutside 1 331\nTMhelix 332 354\ninside 355 372

9488 GCF_000627885.1_ASM62788v1 Escherichia coli 2-011-08_S1_C2 Proteobacteria; Gammaproteobacteria; Enterobacterales; Enterobacteriaceae; Escherichia; Escherichia coli MTGDNTLIHSHGINRRDFMKLCAALAATMGLSSKAAAEMAESVTNPQRPPVIWIGAQECT WP_000145410.1 MULTISPECIES: hydrogenase-2 small chain [Proteobacteria] Length: 372\nNumber of predicted TMHs: 1\nExp number of AAs in TMHs: 23.72226\nExp number, first 60 AAs: 0.22038\nTotal prob of N-in: 0.12994\noutside 1 331\nTMhelix 332 354\ninside 355 372

9489 GCF_000627885.1_ASM62788v1 Escherichia coli 2-011-08_S1_C2 Proteobacteria; Gammaproteobacteria; Enterobacterales; Enterobacteriaceae; Escherichia; Escherichia coli MNNEETFYQAMRRQGVTRRSFLKYCSLAATSLGLGAGMAPKIAWALENKPRIPVVWIHGL WP_001058323.1 MULTISPECIES: hydrogenase-1 small chain [Proteobacteria] Length: 372\nNumber of predicted TMHs: 2\nExp number of AAs in TMHs: 37.2608499999999\nExp number, first 60 AAs: 19.52923\nTotal prob of N-in: 0.94351\nPOSSIBLE N-term signal sequence\ninside 1 19\nTMhelix 20 39\noutside 40 325\nTMhelix 326 348\ninside 349 372

9490 GCF_000700165.1_ASM70016v1 Escherichia coli 2-011-08_S1_C3 Proteobacteria; Gammaproteobacteria; Enterobacterales; Enterobacteriaceae; Escherichia; Escherichia coli MNNEETFYQAMRRQGVTRRSFLKYCSLAATSLGLGAGMAPKIAWALENKPRIPVVWIHGL WP_001058323.1 MULTISPECIES: hydrogenase-1 small chain [Proteobacteria] Length: 372\nNumber of predicted TMHs: 2\nExp number of AAs in TMHs: 37.2608499999999\nExp number, first 60 AAs: 19.52923\nTotal prob of N-in: 0.94351\nPOSSIBLE N-term signal sequence\ninside 1 19\nTMhelix 20 39\noutside 40 325\nTMhelix 326 348\ninside 349 372

9491 GCF_000700165.1_ASM70016v1 Escherichia coli 2-011-08_S1_C3 Proteobacteria; Gammaproteobacteria; Enterobacterales; Enterobacteriaceae; Escherichia; Escherichia coli MTGDNTLIHSHGINRRDFMKLCAALAATMGLSSKAAAEMAESVTNPQRPPVIWIGAQECT WP_000145410.1 MULTISPECIES: hydrogenase-2 small chain [Proteobacteria] Length: 372\nNumber of predicted TMHs: 1\nExp number of AAs in TMHs: 23.72226\nExp number, first 60 AAs: 0.22038\nTotal prob of N-in: 0.12994\noutside 1 331\nTMhelix 332 354\ninside 355 372

9492 GCF_000700505.1_ASM70050v1 Escherichia coli 2-011-08_S3_C1 Proteobacteria; Gammaproteobacteria; Enterobacterales; Enterobacteriaceae; Escherichia; Escherichia coli MNNEETFYQAMRRQGVTRRSFLKYCSLAATSLGLGAGMAPKIAWALENKPRIPVVWIHGL WP_001058323.1 MULTISPECIES: hydrogenase-1 small chain [Proteobacteria] Length: 372\nNumber of predicted TMHs: 2\nExp number of AAs in TMHs: 37.2608499999999\nExp number, first 60 AAs: 19.52923\nTotal prob of N-in: 0.94351\nPOSSIBLE N-term signal sequence\ninside 1 19\nTMhelix 20 39\noutside 40 325\nTMhelix 326 348\ninside 349 372

9493 GCF_000700505.1_ASM70050v1 Escherichia coli 2-011-08_S3_C1 Proteobacteria; Gammaproteobacteria; Enterobacterales; Enterobacteriaceae; Escherichia; Escherichia coli MTGDNTLIHSHGINRRDFMKLCAALAATMGLSSKAAAEMAESVTNPQRPPVIWIGAQECT WP_000145410.1 MULTISPECIES: hydrogenase-2 small chain [Proteobacteria] Length: 372\nNumber of predicted TMHs: 1\nExp number of AAs in TMHs: 23.72226\nExp number, first 60 AAs: 0.22038\nTotal prob of N-in: 0.12994\noutside 1 331\nTMhelix 332 354\ninside 355 372

9494 GCF_000687025.1_ASM68702v1 Escherichia coli 2-011-08_S3_C2 Proteobacteria; Gammaproteobacteria; Enterobacterales; Enterobacteriaceae; Escherichia; Escherichia coli MNNEETFYQAMRRQGVTRRSFLKYCSLAATSLGLGAGMAPKIAWALENKPRIPVVWIHGL WP_001058323.1 MULTISPECIES: hydrogenase-1 small chain [Proteobacteria] Length: 372\nNumber of predicted TMHs: 2\nExp number of AAs in TMHs: 37.2608499999999\nExp number, first 60 AAs: 19.52923\nTotal prob of N-in: 0.94351\nPOSSIBLE N-term signal sequence\ninside 1 19\nTMhelix 20 39\noutside 40 325\nTMhelix 326 348\ninside 349 372

9495 GCF_000687025.1_ASM68702v1 Escherichia coli 2-011-08_S3_C2 Proteobacteria; Gammaproteobacteria; Enterobacterales; Enterobacteriaceae; Escherichia; Escherichia coli MTGDNTLIHSHGINRRDFMKLCAALAATMGLSSKAAAEMAESVTNPQRPPVIWIGAQECT WP_000145410.1 MULTISPECIES: hydrogenase-2 small chain [Proteobacteria] Length: 372\nNumber of predicted TMHs: 1\nExp number of AAs in TMHs: 23.72226\nExp number, first 60 AAs: 0.22038\nTotal prob of N-in: 0.12994\noutside 1 331\nTMhelix 332 354\ninside 355 372

9496 GCF_000687065.1_ASM68706v1 Escherichia coli 2-011-08_S3_C3 Proteobacteria; Gammaproteobacteria; Enterobacterales; Enterobacteriaceae; Escherichia; Escherichia coli MTGDNTLIHSHGINRRDFMKLCAALAATMGLSSKAAAEMAESVTNPQRPPVIWIGAQECT WP_000145410.1 MULTISPECIES: hydrogenase-2 small chain [Proteobacteria] Length: 372\nNumber of predicted TMHs: 1\nExp number of AAs in TMHs: 23.72226\nExp number, first 60 AAs: 0.22038\nTotal prob of N-in: 0.12994\noutside 1 331\nTMhelix 332 354\ninside 355 372

9497 GCF_000687065.1_ASM68706v1 Escherichia coli 2-011-08_S3_C3 Proteobacteria; Gammaproteobacteria; Enterobacterales; Enterobacteriaceae; Escherichia; Escherichia coli MNNEETFYQAMRRQGVTRRSFLKYCSLAATSLGLGAGMAPKIAWALENKPRIPVVWIHGL WP_001058323.1 MULTISPECIES: hydrogenase-1 small chain [Proteobacteria] Length: 372\nNumber of predicted TMHs: 2\nExp number of AAs in TMHs: 37.2608499999999\nExp number, first 60 AAs: 19.52923\nTotal prob of N-in: 0.94351\nPOSSIBLE N-term signal sequence\ninside 1 19\nTMhelix 20 39\noutside 40 325\nTMhelix 326 348\ninside 349 372

9498 GCF_000700585.1_ASM70058v1 Escherichia coli 2-011-08_S4_C1 Proteobacteria; Gammaproteobacteria; Enterobacterales; Enterobacteriaceae; Escherichia; Escherichia coli MTGDNTLIHSHGINRRDFMKLCAALAATMGLSSKAAAEMAESVTNPQRPPVIWIGAQECT WP_000145410.1 MULTISPECIES: hydrogenase-2 small chain [Proteobacteria] Length: 372\nNumber of predicted TMHs: 1\nExp number of AAs in TMHs: 23.72226\nExp number, first 60 AAs: 0.22038\nTotal prob of N-in: 0.12994\noutside 1 331\nTMhelix 332 354\ninside 355 372

9499 GCF_000700585.1_ASM70058v1 Escherichia coli 2-011-08_S4_C1 Proteobacteria; Gammaproteobacteria; Enterobacterales; Enterobacteriaceae; Escherichia; Escherichia coli MNNEETFYQAMRRQGVTRRSFLKYCSLAATSLGLGAGMAPKIAWALENKPRIPVVWIHGL WP_001058323.1 MULTISPECIES: hydrogenase-1 small chain [Proteobacteria] Length: 372\nNumber of predicted TMHs: 2\nExp number of AAs in TMHs: 37.2608499999999\nExp number, first 60 AAs: 19.52923\nTotal prob of N-in: 0.94351\nPOSSIBLE N-term signal sequence\ninside 1 19\nTMhelix 20 39\noutside 40 325\nTMhelix 326 348\ninside 349 372

9500 GCF_000700525.1_ASM70052v1 Escherichia coli 2-011-08_S4_C3 Proteobacteria; Gammaproteobacteria; Enterobacterales; Enterobacteriaceae; Escherichia; Escherichia coli MTGDNTLIHSHGINRRDFMKLCAALAATMGLSSKAAAEMAESVTNPQRPPVIWIGAQECT WP_000145410.1 MULTISPECIES: hydrogenase-2 small chain [Proteobacteria] Length: 372\nNumber of predicted TMHs: 1\nExp number of AAs in TMHs: 23.72226\nExp number, first 60 AAs: 0.22038\nTotal prob of N-in: 0.12994\noutside 1 331\nTMhelix 332 354\ninside 355 372

9501 GCF_000700525.1_ASM70052v1 Escherichia coli 2-011-08_S4_C3 Proteobacteria; Gammaproteobacteria; Enterobacterales; Enterobacteriaceae; Escherichia; Escherichia coli

MNNEETFYQAMRRQGVTRRSFLKYCSLAATSLGLGAGMAPKIAWALENKPRIPVVWIHGL WP_001058323.1 MULTISPECIES: hydrogenase-1 small chain [Proteobacteria] Length: 372\nNumber of predicted TMHs: 2\nExp number of AAs in TMHs: 37.2608499999999\nExp number, first 60 AAs: 19.52923\nTotal prob of N-in: 0.94351\nPOSSIBLE N-term signal sequence\ninside 1 19\nTMhelix 20 39\noutside 40 325\nTMhelix 326 348\ninside 349 372

9502 GCF_000356745.2_ASM35674v2 Escherichia coli 201600.1 Proteobacteria; Gammaproteobacteria; Enterobacterales; Enterobacteriaceae; Escherichia; Escherichia coli MTGDNTLIHSHGINRRDFMKLCAALAATMGLSSKAAAEMAESVTNPQRPPVIWIGAQECT WP_000145410.1 MULTISPECIES: hydrogenase-2 small chain [Proteobacteria] Length: 372\nNumber of predicted TMHs: 1\nExp number of AAs in TMHs: 23.72226\nExp number, first 60 AAs: 0.22038\nTotal prob of N-in: 0.12994\noutside 1 331\nTMhelix 332 354\ninside 355 372

9503 GCF_000356745.2_ASM35674v2 Escherichia coli 201600.1 Proteobacteria; Gammaproteobacteria; Enterobacterales; Enterobacteriaceae; Escherichia; Escherichia coli MNNEETFYQAMRRQGVTRRSFLKYCSLAATSLGLGAGMAPKIAWALENKPRIPVVWIHGL WP_001058323.1 MULTISPECIES: hydrogenase-1 small chain [Proteobacteria] Length: 372\nNumber of predicted TMHs: 2\nExp number of AAs in TMHs: 37.2608499999999\nExp number, first 60 AAs: 19.52923\nTotal prob of N-in: 0.94351\nPOSSIBLE N-term signal sequence\ninside 1 19\nTMhelix 20 39\noutside 40 325\nTMhelix 326 348\ninside 349 372

9504 GCF_000687085.1_ASM68708v1 Escherichia coli 2-052-05_S1_C1 Proteobacteria; Gammaproteobacteria; Enterobacterales; Enterobacteriaceae; Escherichia; Escherichia coli MTGDNTLIHSHGINRRDFMKLCAALAATMGLSSKAAAEMAESVTNPQRPPVIWIGAQECT WP_000145410.1 MULTISPECIES: hydrogenase-2 small chain [Proteobacteria] Length: 372\nNumber of predicted TMHs: 1\nExp number of AAs in TMHs: 23.72226\nExp number, first 60 AAs: 0.22038\nTotal prob of N-in: 0.12994\noutside 1 331\nTMhelix 332 354\ninside 355 372

9505 GCF_000687085.1_ASM68708v1 Escherichia coli 2-052-05_S1_C1 Proteobacteria; Gammaproteobacteria; Enterobacterales; Enterobacteriaceae; Escherichia; Escherichia coli MNNEETFYQAMRRQGVTRRSFLKYCSLAATSLGLGAGMAPKIAWALENKPRIPVVWIHGL WP_001058320.1 hydrogenase [Escherichia coli] Length: 372\nNumber of predicted TMHs: 2\nExp number of AAs in TMHs: 37.26082\nExp number, first 60 AAs: 19.52923\nTotal prob of N-in: 0.94351\nPOSSIBLE N-term signal sequence\ninside 1 19\nTMhelix 20 39\noutside 40 325\nTMhelix 326 348\ninside 349 372

9506 GCF_000700565.1_ASM70056v1 Escherichia coli 2-052-05_S1_C3 Proteobacteria; Gammaproteobacteria; Enterobacterales; Enterobacteriaceae; Escherichia; Escherichia coli MTGDNTLIHSHGINRRDFMKLCAALAATMGLSSKAAAEMAESVTNPQRPPVIWIGAQECT WP_000145410.1 MULTISPECIES: hydrogenase-2 small chain [Proteobacteria] Length: 372\nNumber of predicted TMHs: 1\nExp number of AAs in TMHs: 23.72226\nExp number, first 60 AAs: 0.22038\nTotal prob of N-in: 0.12994\noutside 1 331\nTMhelix 332 354\ninside 355 372

9507 GCF_000700565.1_ASM70056v1 Escherichia coli 2-052-05_S1_C3 Proteobacteria; Gammaproteobacteria; Enterobacterales; Enterobacteriaceae; Escherichia; Escherichia coli MNNEETFYQAMRRQGVTRRSFLKYCSLAATSLGLGAGMAPKIAWALENKPRIPVVWIHGL WP_001058320.1 hydrogenase [Escherichia coli] Length: 372\nNumber of predicted TMHs: 2\nExp number of AAs in TMHs: 37.26082\nExp number, first 60 AAs: 19.52923\nTotal prob of N-in: 0.94351\nPOSSIBLE N-term signal sequence\ninside 1 19\nTMhelix 20 39\noutside 40 325\nTMhelix 326 348\ninside 349 372

9508 GCF_000700605.1_ASM70060v1 Escherichia coli 2-052-05_S3_C1 Proteobacteria; Gammaproteobacteria; Enterobacterales; Enterobacteriaceae; Escherichia; Escherichia coli MNNEETFYQAMRRQGVTRRSFLKYCSLAATSLGLGAGMAPKIAWALENKPRIPVVWIHGL WP_001058323.1 MULTISPECIES: hydrogenase-1 small chain [Proteobacteria] Length: 372\nNumber of predicted TMHs: 2\nExp number of AAs in TMHs: 37.2608499999999\nExp number, first 60 AAs: 19.52923\nTotal prob of N-in: 0.94351\nPOSSIBLE N-term signal sequence\ninside 1 19\nTMhelix 20 39\noutside 40 325\nTMhelix 326 348\ninside 349 372

9509 GCF_000700605.1_ASM70060v1 Escherichia coli 2-052-05_S3_C1 Proteobacteria; Gammaproteobacteria; Enterobacterales; Enterobacteriaceae; Escherichia; Escherichia coli MTGDNTLIHSHGINRRDFMKLCAALAATMGLSSKAAAEMAESVTNPQRPPVIWIGAQECT WP_000145410.1 MULTISPECIES: hydrogenase-2 small chain [Proteobacteria] Length: 372\nNumber of predicted TMHs:

1\nExp number of AAs in TMHs: 23.72226\nExp number, first 60 AAs: 0.22038\nTotal prob of N-in: 0.12994\noutside 1 331\nTMhelix 332 354\ninside 355 372

9510 GCF_000713655.1_ASM71365v1 Escherichia coli 2-052-05_S3_C2 Proteobacteria; Gammaproteobacteria; Enterobacterales; Enterobacteriaceae; Escherichia; Escherichia coli MTGDNTLIHSHGINRRDFMKLCAALAATMGLSSKAAAEMAESVTNPQRPPVIWIGAQECT WP_000145410.1 MULTISPECIES: hydrogenase-2 small chain [Proteobacteria] Length: 372\nNumber of predicted TMHs: 1\nExp number of AAs in TMHs: 23.72226\nExp number, first 60 AAs: 0.22038\nTotal prob of N-in: 0.12994\noutside 1 331\nTMhelix 332 354\ninside 355 372

9511 GCF_000713655.1_ASM71365v1 Escherichia coli 2-052-05_S3_C2 Proteobacteria; Gammaproteobacteria; Enterobacterales; Enterobacteriaceae; Escherichia; Escherichia coli MNNEETFYQAMRRQGVTRRSFLKYCSLAATSLGLGAGMAPKIAWALENKPRIPVVWIHGL WP_001058323.1 MULTISPECIES: hydrogenase-1 small chain [Proteobacteria] Length: 372\nNumber of predicted TMHs: 2\nExp number of AAs in TMHs: 37.2608499999999\nExp number, first 60 AAs: 19.52923\nTotal prob of N-in: 0.94351\nPOSSIBLE N-term signal sequence\ninside 1 19\nTMhelix 20 39\noutside 40 325\nTMhelix 326 348\ninside 349 372

9512 GCF_000703605.1_ASM70360v1 Escherichia coli 2-052-05_S3_C3 Proteobacteria; Gammaproteobacteria; Enterobacterales; Enterobacteriaceae; Escherichia; Escherichia coli MNNEETFYQAMRRQGVTRRSFLKYCSLAATSLGLGAGMAPKIAWALENKPRIPVVWIHGL WP_001058323.1 MULTISPECIES: hydrogenase-1 small chain [Proteobacteria] Length: 372\nNumber of predicted TMHs: 2\nExp number of AAs in TMHs: 37.2608499999999\nExp number, first 60 AAs: 19.52923\nTotal prob of N-in: 0.94351\nPOSSIBLE N-term signal sequence\ninside 1 19\nTMhelix 20 39\noutside 40 325\nTMhelix 326 348\ninside 349 372

9513 GCF_000703605.1_ASM70360v1 Escherichia coli 2-052-05_S3_C3 Proteobacteria; Gammaproteobacteria; Enterobacterales; Enterobacteriaceae; Escherichia; Escherichia coli MTGDNTLIHSHGINRRDFMKLCAALAATMGLSSKAAAEMAESVTNPQRPPVIWIGAQECT WP_000145410.1 MULTISPECIES: hydrogenase-2 small chain [Proteobacteria] Length: 372\nNumber of predicted TMHs: 1\nExp number of AAs in TMHs: 23.72226\nExp number, first 60 AAs: 0.22038\nTotal prob of N-in: 0.12994\noutside 1 331\nTMhelix 332 354\ninside 355 372

9514 GCF_000700545.1_ASM70054v1 Escherichia coli 2-052-05_S4_C1 Proteobacteria; Gammaproteobacteria; Enterobacterales; Enterobacteriaceae; Escherichia; Escherichia coli MTGDNTLIHSHGINRRDFMKLCAALAATMGLSSKAAAEMAESVTNPQRPPVIWIGAQECT WP_000145418.1 hydrogenase 2 small subunit [Escherichia coli] Length: 372\nNumber of predicted TMHs: 1\nExp number of AAs in TMHs: 23.72313\nExp number, first 60 AAs: 0.22034\nTotal prob of N-in: 0.12997\noutside 1 331\nTMhelix 332 354\ninside 355 372

9515 GCF_000700545.1_ASM70054v1 Escherichia coli 2-052-05_S4_C1 Proteobacteria; Gammaproteobacteria; Enterobacterales; Enterobacteriaceae; Escherichia; Escherichia coli MNNEETFYQAMRRQGVTRRSFLKYCSLAATSLGLGAGMAPKIAWALENKPRIPVVWIHGL WP_001295940.1 MULTISPECIES: hydrogenase [Enterobacteriaceae] Length: 372\nNumber of predicted TMHs: 2\nExp number of AAs in TMHs: 37.2608899999999\nExp number, first 60 AAs: 19.52923\nTotal prob of N-in: 0.94351\nPOSSIBLE N-term signal sequence\ninside 1 19\nTMhelix 20 39\noutside 40 325\nTMhelix 326 348\ninside 349 372

9516 GCF_000703385.1_ASM70338v1 Escherichia coli 2-052-05_S4_C2 Proteobacteria; Gammaproteobacteria; Enterobacterales; Enterobacteriaceae; Escherichia; Escherichia coli MNNEETFYQAMRRQGVTRRSFLKYCSLAATSLGLGAGMAPKIAWALENKPRIPVVWIHGL WP_001058323.1 MULTISPECIES: hydrogenase-1 small chain [Proteobacteria] Length: 372\nNumber of predicted TMHs: 2\nExp number of AAs in TMHs: 37.2608499999999\nExp number, first 60 AAs: 19.52923\nTotal prob of N-in: 0.94351\nPOSSIBLE N-term signal sequence\ninside 1 19\nTMhelix 20 39\noutside 40 325\nTMhelix 326 348\ninside 349 372

9517 GCF_000703385.1_ASM70338v1 Escherichia coli 2-052-05_S4_C2 Proteobacteria; Gammaproteobacteria; Enterobacterales; Enterobacteriaceae; Escherichia; Escherichia coli MTGDNTLIHSHGINRRDFMKLCAALAATMGLSSKAAAEMAESVTNPQRPPVIWIGAQECT WP_000145410.1 MULTISPECIES: hydrogenase-2 small chain [Proteobacteria] Length: 372\nNumber of predicted TMHs:

1\nExp number of AAs in TMHs: 23.72226\nExp number, first 60 AAs: 0.22038\nTotal prob of N-in: 0.12994\noutside 1 331\nTMhelix 332 354\ninside 355 372

9518 GCF_000703445.1_ASM70344v1 Escherichia coli 2-052-05_S4_C3 Proteobacteria; Gammaproteobacteria; Enterobacterales; Enterobacteriaceae; Escherichia; Escherichia coli MTGDNTLIHSHGINRRDFMKLCAALAATMGLSSKAAAEMAESVTNPQRPPVIWIGAQECT WP_000145410.1 MULTISPECIES: hydrogenase-2 small chain [Proteobacteria] Length: 372\nNumber of predicted TMHs: 1\nExp number of AAs in TMHs: 23.72226\nExp number, first 60 AAs: 0.22038\nTotal prob of N-in: 0.12994\noutside 1 331\nTMhelix 332 354\ninside 355 372

9519 GCF_000703445.1_ASM70344v1 Escherichia coli 2-052-05_S4_C3 Proteobacteria; Gammaproteobacteria; Enterobacterales; Enterobacteriaceae; Escherichia; Escherichia coli MNNEETFYQAMRRQGVTRRSFLKYCSLAATSLGLGAGMAPKIAWALENKPRIPVVWIHGL WP_001058323.1 MULTISPECIES: hydrogenase-1 small chain [Proteobacteria] Length: 372\nNumber of predicted TMHs: 2\nExp number of AAs in TMHs: 37.2608499999999\nExp number, first 60 AAs: 19.52923\nTotal prob of N-in: 0.94351\nPOSSIBLE N-term signal sequence\ninside 1 19\nTMhelix 20 39\noutside 40 325\nTMhelix 326 348\ninside 349 372

9520 GCF_000703505.1_ASM70350v1 Escherichia coli 2-156-04_S1_C3 Proteobacteria; Gammaproteobacteria; Enterobacterales; Enterobacteriaceae; Escherichia; Escherichia coli MTGDNTLIHSHGINRRDFMKLCAALAATMGLSSKAAAEMAESVTNPQRPPVIWIGAQECT WP_000145410.1 MULTISPECIES: hydrogenase-2 small chain [Proteobacteria] Length: 372\nNumber of predicted TMHs: 1\nExp number of AAs in TMHs: 23.72226\nExp number, first 60 AAs: 0.22038\nTotal prob of N-in: 0.12994\noutside 1 331\nTMhelix 332 354\ninside 355 372

9521 GCF_000703505.1_ASM70350v1 Escherichia coli 2-156-04_S1_C3 Proteobacteria; Gammaproteobacteria; Enterobacterales; Enterobacteriaceae; Escherichia; Escherichia coli MNNEETFYQAMRRQGVTRRSFLKYCSLAATSLGLGAGMAPKIAWALENKPRIPVVWIHGL WP_001058323.1 MULTISPECIES: hydrogenase-1 small chain [Proteobacteria] Length: 372\nNumber of predicted TMHs: 2\nExp number of AAs in TMHs: 37.2608499999999\nExp number, first 60 AAs: 19.52923\nTotal prob of N-in: 0.94351\nPOSSIBLE N-term signal sequence\ninside 1 19\nTMhelix 20 39\noutside 40 325\nTMhelix 326 348\ninside 349 372

9522 GCF_000703525.1_ASM70352v1 Escherichia coli 2-156-04_S3_C1 Proteobacteria; Gammaproteobacteria; Enterobacterales; Enterobacteriaceae; Escherichia; Escherichia coli MTGDNTLIHSHGINRRDFMKLCAALAATMGLSSKAAAEMAESVTNPQRPPVIWIGAQECT WP_000145410.1 MULTISPECIES: hydrogenase-2 small chain [Proteobacteria] Length: 372\nNumber of predicted TMHs: 1\nExp number of AAs in TMHs: 23.72226\nExp number, first 60 AAs: 0.22038\nTotal prob of N-in: 0.12994\noutside 1 331\nTMhelix 332 354\ninside 355 372

9523 GCF_000703525.1_ASM70352v1 Escherichia coli 2-156-04_S3_C1 Proteobacteria; Gammaproteobacteria; Enterobacterales; Enterobacteriaceae; Escherichia; Escherichia coli MNNEETFYQAMRRQGVTRRSFLKYCSLAATSLGLGAGMAPKIAWALENKPRIPVVWIHGL WP_001295940.1 MULTISPECIES: hydrogenase [Enterobacteriaceae] Length: 372\nNumber of predicted TMHs: 2\nExp number of AAs in TMHs: 37.2608899999999\nExp number, first 60 AAs: 19.52923\nTotal prob of N-in: 0.94351\nPOSSIBLE N-term signal sequence\ninside 1 19\nTMhelix 20 39\noutside 40 325\nTMhelix 326 348\ninside 349 372

9524 GCF_000703725.1_ASM70372v1 Escherichia coli 2-156-04_S3_C2 Proteobacteria; Gammaproteobacteria; Enterobacterales; Enterobacteriaceae; Escherichia; Escherichia coli MNNEETFYQAMRRQGVTRRSFLKYCSLAATSLGLGAGMAPKIAWALENKPRIPVVWIHGL WP_001353311.1 MULTISPECIES: hydrogenase [Escherichia] Length: 372\nNumber of predicted TMHs: 2\nExp number of AAs in TMHs: 37.2638699999999\nExp number, first 60 AAs: 19.52924\nTotal prob of N-in: 0.94351\nPOSSIBLE N-term signal sequence\ninside 1 19\nTMhelix 20 39\noutside 40 325\nTMhelix 326 348\ninside 349 372

9525 GCF_000703725.1_ASM70372v1 Escherichia coli 2-156-04_S3_C2 Proteobacteria; Gammaproteobacteria; Enterobacterales; Enterobacteriaceae; Escherichia; Escherichia coli MTGDNTLIHSHGINRRDFMKLCAALAATMGLSSKAAAEMAESVTNPQRPPVIWIGAQECT WP_000145410.1 MULTISPECIES: hydrogenase-2 small chain [Proteobacteria] Length: 372\nNumber of predicted TMHs: 1\nExp number of AAs in TMHs: 23.72226\nExp number, first 60 AAs: 0.22038\nTotal prob of N-in: 0.12994\noutside 1 331\nTMhelix 332 354\ninside 355 372

9526 GCF_000703745.1_ASM70374v1 Escherichia coli 2-156-04_S3_C3 Proteobacteria; Gammaproteobacteria; Enterobacterales; Enterobacteriaceae; Escherichia; Escherichia coli MTGDNTLIHSHGINRRDFMKLCAALAATMGLSSKAAAEMAESVTNPQRPPVIWIGAQECT WP_000145410.1 MULTISPECIES: hydrogenase-2 small chain [Proteobacteria] Length: 372\nNumber of predicted TMHs: 1\nExp number of AAs in TMHs: 23.72226\nExp number, first 60 AAs: 0.22038\nTotal prob of N-in: 0.12994\noutside 1 331\nTMhelix 332 354\ninside 355 372

9527 GCF_000703745.1_ASM70374v1 Escherichia coli 2-156-04_S3_C3 Proteobacteria; Gammaproteobacteria; Enterobacterales; Enterobacteriaceae; Escherichia; Escherichia coli MNNEETFYQAMRRQGVTRRSFLKYCSLAATSLGLGAGMAPKIAWALENKPRIPVVWIHGL WP_001058323.1 MULTISPECIES: hydrogenase-1 small chain [Proteobacteria] Length: 372\nNumber of predicted TMHs: 2\nExp number of AAs in TMHs: 37.2608499999999\nExp number, first 60 AAs: 19.52923\nTotal prob of N-in: 0.94351\nPOSSIBLE N-term signal sequence\ninside 1 19\nTMhelix 20 39\noutside 40 325\nTMhelix 326 348\ninside 349 372

9528 GCF_000703665.1_ASM70366v1 Escherichia coli 2-156-04_S4_C1 Proteobacteria; Gammaproteobacteria; Enterobacterales; Enterobacteriaceae; Escherichia; Escherichia coli MTGDNTLIHSHGINRRDFMKLCAALAATMGLSSKAAAEMAESVTNPQRPPVIWIGAQECT WP_032181345.1 hydrogenase 2 small subunit [Escherichia coli] Length: 372\nNumber of predicted TMHs: 1\nExp number of AAs in TMHs: 23.73477\nExp number, first 60 AAs: 0.22039\nTotal prob of N-in: 0.13690\noutside 1 332\nTMhelix 333 355\ninside 356 372

9529 GCF_000703665.1_ASM70366v1 Escherichia coli 2-156-04_S4_C1 Proteobacteria; Gammaproteobacteria; Enterobacterales; Enterobacteriaceae; Escherichia; Escherichia coli MNNEETFYQAMRRQGVTRRSFLKYCSLAATSLGLGAGMAPKIAWALENKPRIPVVWIHGL WP_004999783.1 MULTISPECIES: hydrogenase [Enterobacteriaceae] Length: 372\nNumber of predicted TMHs: 2\nExp number of AAs in TMHs: 37.2630799999999\nExp number, first 60 AAs: 19.52922\nTotal prob of N-in: 0.94351\nPOSSIBLE N-term signal sequence\ninside 1 19\nTMhelix 20 39\noutside 40 325\nTMhelix 326 348\ninside 349 372

9530 GCF_000704025.1_ASM70402v1 Escherichia coli 2-156-04_S4_C2 Proteobacteria; Gammaproteobacteria; Enterobacterales; Enterobacteriaceae; Escherichia; Escherichia coli MTGDNTLIHSHGINRRDFMKLCAALAATMGLSSKAAAEMAESVTNPQRPPVIWIGAQECT WP_000145410.1 MULTISPECIES: hydrogenase-2 small chain [Proteobacteria] Length: 372\nNumber of predicted TMHs: 1\nExp number of AAs in TMHs: 23.72226\nExp number, first 60 AAs: 0.22038\nTotal prob of N-in: 0.12994\noutside 1 331\nTMhelix 332 354\ninside 355 372

9531 GCF_000704025.1_ASM70402v1 Escherichia coli 2-156-04_S4_C2 Proteobacteria; Gammaproteobacteria; Enterobacterales; Enterobacteriaceae; Escherichia; Escherichia coli MNNEETFYQAMRRQGVTRRSFLKYCSLAATSLGLGAGMAPKIAWALENKPRIPVVWIHGL WP_001058323.1 MULTISPECIES: hydrogenase-1 small chain [Proteobacteria] Length: 372\nNumber of predicted TMHs: 2\nExp number of AAs in TMHs: 37.2608499999999\nExp number, first 60 AAs: 19.52923\nTotal prob of N-in: 0.94351\nPOSSIBLE N-term signal sequence\ninside 1 19\nTMhelix 20 39\noutside 40 325\nTMhelix 326 348\ninside 349 372

9532 GCF_000704105.1_ASM70410v1 Escherichia coli 2-156-04_S4_C3 Proteobacteria; Gammaproteobacteria; Enterobacterales; Enterobacteriaceae; Escherichia; Escherichia coli MNNEETFYQAMRRQGVTRRSFLKYCSLAATSLGLGAGMAPKIAWALENKPRIPVVWIHGL WP_001058323.1 MULTISPECIES: hydrogenase-1 small chain [Proteobacteria] Length: 372\nNumber of predicted TMHs: 2\nExp number of AAs in TMHs: 37.2608499999999\nExp number, first 60 AAs: 19.52923\nTotal prob of N-in: 0.94351\nPOSSIBLE N-term signal sequence\ninside 1 19\nTMhelix 20 39\noutside 40 325\nTMhelix 326 348\ninside 349 372

9533 GCF_000704105.1_ASM70410v1 Escherichia coli 2-156-04_S4_C3 Proteobacteria; Gammaproteobacteria; Enterobacterales; Enterobacteriaceae; Escherichia; Escherichia coli MTGDNTLIHSHGINRRDFMKLCAALAATMGLSSKAAAEMAESVTNPQRPPVIWIGAQECT WP_032170173.1 hydrogenase 2 small subunit [Escherichia coli] Length: 372\nNumber of predicted TMHs: 1\nExp number of AAs in TMHs: 23.72138\nExp number, first 60 AAs: 0.22038\nTotal prob of N-in: 0.12992\noutside 1 331\nTMhelix 332 354\ninside 355 372

9534	GCF_000703425.1_ASM70342v1	Escherichia coli 2-177-06_S1_C1	Proteobacteria; Gammaproteobacteria; Enterobacterales; Enterobacteriaceae; Escherichia; Escherichia coli
	MTGDNTLIHSHGINRRDFMKLCAALAATMGLSSKAAAEMAESVTNPQRPPVIWIGAQECT	WP_000145410.1
	MULTISPECIES: hydrogenase-2 small chain [Proteobacteria]	Length: 372\nNumber of predicted TMHs: 1\nExp number of AAs in TMHs: 23.72226\nExp number, first 60 AAs: 0.22038\nTotal prob of N-in: 0.12994\noutside 1 331\nTMhelix 332 354\ninside 355 372

9535	GCF_000703425.1_ASM70342v1	Escherichia coli 2-177-06_S1_C1	Proteobacteria; Gammaproteobacteria; Enterobacterales; Enterobacteriaceae; Escherichia; Escherichia coli
	MNNEETFYQAMRRQGVTRRSFLKYCSLAATSLGLGAGMAPKIAWALENKPRIPVVWIHGL	WP_001058320.1
	hydrogenase [Escherichia coli]	Length: 372\nNumber of predicted TMHs: 2\nExp number of AAs in TMHs: 37.26082\nExp number, first 60 AAs: 19.52923\nTotal prob of N-in: 0.94351\nPOSSIBLE N-term signal sequence\ninside 1 19\nTMhelix 20 39\noutside 40 325\nTMhelix 326 348\ninside 349 372

9536	GCF_000703705.1_ASM70370v1	Escherichia coli 2-177-06_S1_C2	Proteobacteria; Gammaproteobacteria; Enterobacterales; Enterobacteriaceae; Escherichia; Escherichia coli
	MNNEETFYQAMRRQGVTRRSFLKYCSLAATSLGLGAGMAPKIAWALENKPRIPVVWIHGL	WP_001058320.1
	hydrogenase [Escherichia coli]	Length: 372\nNumber of predicted TMHs: 2\nExp number of AAs in TMHs: 37.26082\nExp number, first 60 AAs: 19.52923\nTotal prob of N-in: 0.94351\nPOSSIBLE N-term signal sequence\ninside 1 19\nTMhelix 20 39\noutside 40 325\nTMhelix 326 348\ninside 349 372

9537	GCF_000703705.1_ASM70370v1	Escherichia coli 2-177-06_S1_C2	Proteobacteria; Gammaproteobacteria; Enterobacterales; Enterobacteriaceae; Escherichia; Escherichia coli
	MTGDNTLIHSHGINRRDFMKLCAALAATMGLSSKAAAEMAESVTNPQRPPVIWIGAQECT	WP_000145410.1
	MULTISPECIES: hydrogenase-2 small chain [Proteobacteria]	Length: 372\nNumber of predicted TMHs: 1\nExp number of AAs in TMHs: 23.72226\nExp number, first 60 AAs: 0.22038\nTotal prob of N-in: 0.12994\noutside 1 331\nTMhelix 332 354\ninside 355 372

9538	GCF_000703685.1_ASM70368v1	Escherichia coli 2-177-06_S1_C3	Proteobacteria; Gammaproteobacteria; Enterobacterales; Enterobacteriaceae; Escherichia; Escherichia coli
	MNNEETFYQAMRRQGVTRRSFLKYCSLAATSLGLGAGMAPKIAWALENKPRIPVVWIHGL	WP_001058320.1
	hydrogenase [Escherichia coli]	Length: 372\nNumber of predicted TMHs: 2\nExp number of AAs in TMHs: 37.26082\nExp number, first 60 AAs: 19.52923\nTotal prob of N-in: 0.94351\nPOSSIBLE N-term signal sequence\ninside 1 19\nTMhelix 20 39\noutside 40 325\nTMhelix 326 348\ninside 349 372

9539	GCF_000703685.1_ASM70368v1	Escherichia coli 2-177-06_S1_C3	Proteobacteria; Gammaproteobacteria; Enterobacterales; Enterobacteriaceae; Escherichia; Escherichia coli
	MTGDNTLIHSHGINRRDFMKLCAALAATMGLSSKAAAEMAESVTNPQRPPVIWIGAQECT	WP_000145410.1
	MULTISPECIES: hydrogenase-2 small chain [Proteobacteria]	Length: 372\nNumber of predicted TMHs: 1\nExp number of AAs in TMHs: 23.72226\nExp number, first 60 AAs: 0.22038\nTotal prob of N-in: 0.12994\noutside 1 331\nTMhelix 332 354\ninside 355 372

9540	GCF_000703645.1_ASM70364v1	Escherichia coli 2-177-06_S3_C1	Proteobacteria; Gammaproteobacteria; Enterobacterales; Enterobacteriaceae; Escherichia; Escherichia coli
	MTGDNTLIHSHGINRRDFMKLCAALAATMGLSSKAAAEMAESVTNPQRPPVIWIGAQECT	WP_000145410.1
	MULTISPECIES: hydrogenase-2 small chain [Proteobacteria]	Length: 372\nNumber of predicted TMHs: 1\nExp number of AAs in TMHs: 23.72226\nExp number, first 60 AAs: 0.22038\nTotal prob of N-in: 0.12994\noutside 1 331\nTMhelix 332 354\ninside 355 372

9541	GCF_000703645.1_ASM70364v1	Escherichia coli 2-177-06_S3_C1	Proteobacteria; Gammaproteobacteria; Enterobacterales; Enterobacteriaceae; Escherichia; Escherichia coli
	MNNEETFYQAMRRQGVTRRSFLKYCSLAATSLGLGAGMAPKIAWALENKPRIPVVWIHGL	WP_001058323.1
	MULTISPECIES: hydrogenase-1 small chain [Proteobacteria]	Length: 372\nNumber of predicted TMHs: 2\nExp number of AAs in TMHs: 37.2608499999999\nExp number, first 60 AAs: 19.52923\nTotal prob of N-in: 0.94351\nPOSSIBLE N-term signal sequence\ninside 1 19\nTMhelix 20 39\noutside 40 325\nTMhelix 326 348\ninside 349 372

9542	GCF_000704005.1_ASM70400v1	Escherichia coli 2-177-06_S3_C2	Proteobacteria; Gammaproteobacteria; Enterobacterales; Enterobacteriaceae; Escherichia; Escherichia coli
	MNNEETFYQAMRRQGVTRRSFLKYCSLAATSLGLGAGMAPKIAWALENKPRIPVVWIHGL	WP_001058323.1
	MULTISPECIES: hydrogenase-1 small chain [Proteobacteria]	Length: 372\nNumber of predicted TMHs:

2\nExp number of AAs in TMHs: 37.2608499999999\nExp number, first 60 AAs: 19.52923\nTotal prob of N-in: 0.94351\nPOSSIBLE N-term signal sequence\ninside 1 19\nTMhelix 20 39\noutside 40 325\nTMhelix 326 348\ninside 349 372

9543 GCF_000704005.1_ASM70400v1 Escherichia coli 2-177-06_S3_C2 Proteobacteria; Gammaproteobacteria; Enterobacterales; Enterobacteriaceae; Escherichia; Escherichia coli MTGDNTLIHSHGINRRDFMKLCAALAATMGLSSKAAAEMAESVTNPQRPPVIWIGAQECT WP_000145410.1 MULTISPECIES: hydrogenase-2 small chain [Proteobacteria] Length: 372\nNumber of predicted TMHs: 1\nExp number of AAs in TMHs: 23.72226\nExp number, first 60 AAs: 0.22038\nTotal prob of N-in: 0.12994\noutside 1 331\nTMhelix 332 354\ninside 355 372

9544 GCF_000713745.1_ASM71374v1 Escherichia coli 2-177-06_S3_C3 Proteobacteria; Gammaproteobacteria; Enterobacterales; Enterobacteriaceae; Escherichia; Escherichia coli MNNEETFYQAMRRQGVTRRSFLKYCSLAATSLGLGAGMAPKIAWALENKPRIPVVWIHGL WP_001058323.1 MULTISPECIES: hydrogenase-1 small chain [Proteobacteria] Length: 372\nNumber of predicted TMHs: 2\nExp number of AAs in TMHs: 37.2608499999999\nExp number, first 60 AAs: 19.52923\nTotal prob of N-in: 0.94351\nPOSSIBLE N-term signal sequence\ninside 1 19\nTMhelix 20 39\noutside 40 325\nTMhelix 326 348\ninside 349 372

9545 GCF_000713745.1_ASM71374v1 Escherichia coli 2-177-06_S3_C3 Proteobacteria; Gammaproteobacteria; Enterobacterales; Enterobacteriaceae; Escherichia; Escherichia coli MTGDNTLIHSHGINRRDFMKLCAALAATMGLSSKAAAEMAESVTNPQRPPVIWIGAQECT WP_000145410.1 MULTISPECIES: hydrogenase-2 small chain [Proteobacteria] Length: 372\nNumber of predicted TMHs: 1\nExp number of AAs in TMHs: 23.72226\nExp number, first 60 AAs: 0.22038\nTotal prob of N-in: 0.12994\noutside 1 331\nTMhelix 332 354\ninside 355 372

9546 GCF_000704225.1_ASM70422v1 Escherichia coli 2-177-06_S4_C1 Proteobacteria; Gammaproteobacteria; Enterobacterales; Enterobacteriaceae; Escherichia; Escherichia coli MTGDNTLIHSHGINRRDFMKLCAALAATMGLSSKAAAEMAESVTNPQRPPVIWIGAQECT WP_000145410.1 MULTISPECIES: hydrogenase-2 small chain [Proteobacteria] Length: 372\nNumber of predicted TMHs: 1\nExp number of AAs in TMHs: 23.72226\nExp number, first 60 AAs: 0.22038\nTotal prob of N-in: 0.12994\noutside 1 331\nTMhelix 332 354\ninside 355 372

9547 GCF_000704225.1_ASM70422v1 Escherichia coli 2-177-06_S4_C1 Proteobacteria; Gammaproteobacteria; Enterobacterales; Enterobacteriaceae; Escherichia; Escherichia coli MNNEETFYQAMRRQGVTRRSFLKYCSLAATSLGLGAGMAPKIAWALENKPRIPVVWIHGL WP_001058323.1 MULTISPECIES: hydrogenase-1 small chain [Proteobacteria] Length: 372\nNumber of predicted TMHs: 2\nExp number of AAs in TMHs: 37.2608499999999\nExp number, first 60 AAs: 19.52923\nTotal prob of N-in: 0.94351\nPOSSIBLE N-term signal sequence\ninside 1 19\nTMhelix 20 39\noutside 40 325\nTMhelix 326 348\ninside 349 372

9548 GCF_000703625.1_ASM70362v1 Escherichia coli 2-177-06_S4_C2 Proteobacteria; Gammaproteobacteria; Enterobacterales; Enterobacteriaceae; Escherichia; Escherichia coli MTGDNTLIHSHGINRRDFMKLCAALAATMGLSSKAAAEMAESVTNPQRPPVIWIGAQECT WP_000145410.1 MULTISPECIES: hydrogenase-2 small chain [Proteobacteria] Length: 372\nNumber of predicted TMHs: 1\nExp number of AAs in TMHs: 23.72226\nExp number, first 60 AAs: 0.22038\nTotal prob of N-in: 0.12994\noutside 1 331\nTMhelix 332 354\ninside 355 372

9549 GCF_000703625.1_ASM70362v1 Escherichia coli 2-177-06_S4_C2 Proteobacteria; Gammaproteobacteria; Enterobacterales; Enterobacteriaceae; Escherichia; Escherichia coli MNNEETFYQAMRRQGVTRRSFLKYCSLAATSLGLGAGMAPKIAWALENKPRIPVVWIHGL WP_001058323.1 MULTISPECIES: hydrogenase-1 small chain [Proteobacteria] Length: 372\nNumber of predicted TMHs: 2\nExp number of AAs in TMHs: 37.2608499999999\nExp number, first 60 AAs: 19.52923\nTotal prob of N-in: 0.94351\nPOSSIBLE N-term signal sequence\ninside 1 19\nTMhelix 20 39\noutside 40 325\nTMhelix 326 348\ninside 349 372

9550 GCF_000703565.1_ASM70356v1 Escherichia coli 2-177-06_S4_C3 Proteobacteria; Gammaproteobacteria; Enterobacterales; Enterobacteriaceae; Escherichia; Escherichia coli MTGDNTLIHSHGINRRDFMKLCAALAATMGLSSKAAAEMAESVTNPQRPPVIWIGAQECT WP_000145410.1 MULTISPECIES: hydrogenase-2 small chain [Proteobacteria] Length: 372\nNumber of predicted TMHs:

1\nExp number of AAs in TMHs: 23.72226\nExp number, first 60 AAs: 0.22038\nTotal prob of N-in: 0.12994\noutside 1 331\nTMhelix 332 354\ninside 355 372

9551 GCF_000703565.1_ASM70356v1 Escherichia coli 2-177-06_S4_C3 Proteobacteria; Gammaproteobacteria; Enterobacterales; Enterobacteriaceae; Escherichia; Escherichia coli MNNEETFYQAMRRQGVTRRSFLKYCSLAATSLGLGAGMAPKIAWALENKPRIPVVWIHGL WP_001058323.1 MULTISPECIES: hydrogenase-1 small chain [Proteobacteria] Length: 372\nNumber of predicted TMHs: 2\nExp number of AAs in TMHs: 37.2608499999999\nExp number, first 60 AAs: 19.52923\nTotal prob of N-in: 0.94351\nPOSSIBLE N-term signal sequence\ninside 1 19\nTMhelix 20 39\noutside 40 325\nTMhelix 326 348\ninside 349 372

9552 GCF_000703905.1_ASM70390v1 Escherichia coli 2-210-07_S1_C2 Proteobacteria; Gammaproteobacteria; Enterobacterales; Enterobacteriaceae; Escherichia; Escherichia coli MTGDNTLIHSHGINRRDFMKLCAALAATMGLSSKAAAEMAESVTNPQRPPVIWIGAQECT WP_000145410.1 MULTISPECIES: hydrogenase-2 small chain [Proteobacteria] Length: 372\nNumber of predicted TMHs: 1\nExp number of AAs in TMHs: 23.72226\nExp number, first 60 AAs: 0.22038\nTotal prob of N-in: 0.12994\noutside 1 331\nTMhelix 332 354\ninside 355 372

9553 GCF_000703905.1_ASM70390v1 Escherichia coli 2-210-07_S1_C2 Proteobacteria; Gammaproteobacteria; Enterobacterales; Enterobacteriaceae; Escherichia; Escherichia coli MNNEETFYQAMRRQGVTRRSFLKYCSLAATSLGLGAGMAPKIAWALENKPRIPVVWIHGL WP_001058323.1 MULTISPECIES: hydrogenase-1 small chain [Proteobacteria] Length: 372\nNumber of predicted TMHs: 2\nExp number of AAs in TMHs: 37.2608499999999\nExp number, first 60 AAs: 19.52923\nTotal prob of N-in: 0.94351\nPOSSIBLE N-term signal sequence\ninside 1 19\nTMhelix 20 39\noutside 40 325\nTMhelix 326 348\ninside 349 372

9554 GCF_000703765.1_ASM70376v1 Escherichia coli 2-210-07_S1_C3 Proteobacteria; Gammaproteobacteria; Enterobacterales; Enterobacteriaceae; Escherichia; Escherichia coli MTGDNTLIHSHGINRRDFMKLCAALAATMGLSSKAAAEMAESVTNPQRPPVIWIGAQECT WP_000145410.1 MULTISPECIES: hydrogenase-2 small chain [Proteobacteria] Length: 372\nNumber of predicted TMHs: 1\nExp number of AAs in TMHs: 23.72226\nExp number, first 60 AAs: 0.22038\nTotal prob of N-in: 0.12994\noutside 1 331\nTMhelix 332 354\ninside 355 372

9555 GCF_000703765.1_ASM70376v1 Escherichia coli 2-210-07_S1_C3 Proteobacteria; Gammaproteobacteria; Enterobacterales; Enterobacteriaceae; Escherichia; Escherichia coli MNNEETFYQAMRRQGVTRRSFLKYCSLAATSLGLGAGMAPKIAWALENKPRIPVVWIHGL WP_001058323.1 MULTISPECIES: hydrogenase-1 small chain [Proteobacteria] Length: 372\nNumber of predicted TMHs: 2\nExp number of AAs in TMHs: 37.2608499999999\nExp number, first 60 AAs: 19.52923\nTotal prob of N-in: 0.94351\nPOSSIBLE N-term signal sequence\ninside 1 19\nTMhelix 20 39\noutside 40 325\nTMhelix 326 348\ninside 349 372

9556 GCF_000704205.1_ASM70420v1 Escherichia coli 2-210-07_S3_C1 Proteobacteria; Gammaproteobacteria; Enterobacterales; Enterobacteriaceae; Escherichia; Escherichia coli MTGDNTLIHSHGINRRDFMKLCAALAATMGLSSKAAAEMAESVTNPQRPPVIWIGAQECT WP_000145410.1 MULTISPECIES: hydrogenase-2 small chain [Proteobacteria] Length: 372\nNumber of predicted TMHs: 1\nExp number of AAs in TMHs: 23.72226\nExp number, first 60 AAs: 0.22038\nTotal prob of N-in: 0.12994\noutside 1 331\nTMhelix 332 354\ninside 355 372

9557 GCF_000703945.1_ASM70394v1 Escherichia coli 2-210-07_S3_C2 Proteobacteria; Gammaproteobacteria; Enterobacterales; Enterobacteriaceae; Escherichia; Escherichia coli MTGDNTLIHSHGINRRDFMKLCAALAATMGLSSKAAAEMAESVTNPQRPPVIWIGAQECT WP_001685818.1 MULTISPECIES: hydrogenase-2 small chain [Enterobacteriaceae] Length: 372\nNumber of predicted TMHs: 1\nExp number of AAs in TMHs: 23.82505\nExp number, first 60 AAs: 0.21974\nTotal prob of N-in: 0.13379\noutside 1 331\nTMhelix 332 354\ninside 355 372

9558 GCF_000703945.1_ASM70394v1 Escherichia coli 2-210-07_S3_C2 Proteobacteria; Gammaproteobacteria; Enterobacterales; Enterobacteriaceae; Escherichia; Escherichia coli MNNEETFYQAMRRQGVTRRSFLKYCSLAATSLGLGAGMAPKIAWALENKPRIPVVWIHGL WP_001058320.1 hydrogenase [Escherichia coli] Length: 372\nNumber of predicted TMHs: 2\nExp number of AAs in TMHs: 37.26082\nExp number, first 60 AAs: 19.52923\nTotal prob of N-in: 0.94351\nPOSSIBLE N-term signal sequence\ninside 1 19\nTMhelix 20 39\noutside 40 325\nTMhelix 326 348\ninside 349 372

9559 GCF_000703585.1_ASM70358v1 Escherichia coli 2-210-07_S3_C3 Proteobacteria; Gammaproteobacteria; Enterobacterales; Enterobacteriaceae; Escherichia; Escherichia coli
MTGDNTLIHSHGINRRDFMKLCAALAATMGLSSKAAAEMAESVTNPQRPPVIWIGAQECT WP_000145410.1
MULTISPECIES: hydrogenase-2 small chain [Proteobacteria] Length: 372\nNumber of predicted TMHs: 1\nExp number of AAs in TMHs: 23.72226\nExp number, first 60 AAs: 0.22038\nTotal prob of N-in: 0.12994\noutside 1 331\nTMhelix 332 354\ninside 355 372

9560 GCF_000703585.1_ASM70358v1 Escherichia coli 2-210-07_S3_C3 Proteobacteria; Gammaproteobacteria; Enterobacterales; Enterobacteriaceae; Escherichia; Escherichia coli
MNNEETFYQAMRRQGVTRRSFLKYCSLAATSLGLGAGMAPKIAWALENKPRIPVVWIHGL WP_001058323.1
MULTISPECIES: hydrogenase-1 small chain [Proteobacteria] Length: 372\nNumber of predicted TMHs: 2\nExp number of AAs in TMHs: 37.2608499999999\nExp number, first 60 AAs: 19.52923\nTotal prob of N-in: 0.94351\nPOSSIBLE N-term signal sequence\ninside 1 19\nTMhelix 20 39\noutside 40 325\nTMhelix 326 348\ninside 349 372

9561 GCF_000703925.1_ASM70392v1 Escherichia coli 2-210-07_S4_C1 Proteobacteria; Gammaproteobacteria; Enterobacterales; Enterobacteriaceae; Escherichia; Escherichia coli
MTGDNTLIHSHGINRRDFMKLCAALAATMGLSSKAAAEMAESVTNPQRPPVIWIGAQECT WP_000145410.1
MULTISPECIES: hydrogenase-2 small chain [Proteobacteria] Length: 372\nNumber of predicted TMHs: 1\nExp number of AAs in TMHs: 23.72226\nExp number, first 60 AAs: 0.22038\nTotal prob of N-in: 0.12994\noutside 1 331\nTMhelix 332 354\ninside 355 372

9562 GCF_000703925.1_ASM70392v1 Escherichia coli 2-210-07_S4_C1 Proteobacteria; Gammaproteobacteria; Enterobacterales; Enterobacteriaceae; Escherichia; Escherichia coli
MNNEETFYQAMRRQGVTRRSFLKYCSLAATSLGLGAGMAPKIAWALENKPRIPVVWIHGL WP_001058320.1
hydrogenase [Escherichia coli] Length: 372\nNumber of predicted TMHs: 2\nExp number of AAs in TMHs: 37.26082\nExp number, first 60 AAs: 19.52923\nTotal prob of N-in: 0.94351\nPOSSIBLE N-term signal sequence\ninside 1 19\nTMhelix 20 39\noutside 40 325\nTMhelix 326 348\ninside 349 372

9563 GCF_000704185.1_ASM70418v1 Escherichia coli 2-210-07_S4_C2 Proteobacteria; Gammaproteobacteria; Enterobacterales; Enterobacteriaceae; Escherichia; Escherichia coli
MNNEETFYQAMRRQGVTRRSFLKYCSLAATSLGLGAGMAPKIAWALENKPRIPVVWIHGL WP_001058323.1
MULTISPECIES: hydrogenase-1 small chain [Proteobacteria] Length: 372\nNumber of predicted TMHs: 2\nExp number of AAs in TMHs: 37.2608499999999\nExp number, first 60 AAs: 19.52923\nTotal prob of N-in: 0.94351\nPOSSIBLE N-term signal sequence\ninside 1 19\nTMhelix 20 39\noutside 40 325\nTMhelix 326 348\ninside 349 372

9564 GCF_000704185.1_ASM70418v1 Escherichia coli 2-210-07_S4_C2 Proteobacteria; Gammaproteobacteria; Enterobacterales; Enterobacteriaceae; Escherichia; Escherichia coli
MTGDNTLIHSHGINRRDFMKLCAALAATMGLSSKAAAEMAESVTNPQRPPVIWIGAQECT WP_032170173.1
hydrogenase 2 small subunit [Escherichia coli] Length: 372\nNumber of predicted TMHs: 1\nExp number of AAs in TMHs: 23.72138\nExp number, first 60 AAs: 0.22038\nTotal prob of N-in: 0.12992\noutside 1 331\nTMhelix 332 354\ninside 355 372

9565 GCF_000703985.1_ASM70398v1 Escherichia coli 2-210-07_S4_C3 Proteobacteria; Gammaproteobacteria; Enterobacterales; Enterobacteriaceae; Escherichia; Escherichia coli
MNNEETFYQAMRRQGVTRRSFLKYCSLAATSLGLGAGMAPKIAWALENKPRIPVVWIHGL WP_001058323.1
MULTISPECIES: hydrogenase-1 small chain [Proteobacteria] Length: 372\nNumber of predicted TMHs: 2\nExp number of AAs in TMHs: 37.2608499999999\nExp number, first 60 AAs: 19.52923\nTotal prob of N-in: 0.94351\nPOSSIBLE N-term signal sequence\ninside 1 19\nTMhelix 20 39\noutside 40 325\nTMhelix 326 348\ninside 349 372

9566 GCF_000703985.1_ASM70398v1 Escherichia coli 2-210-07_S4_C3 Proteobacteria; Gammaproteobacteria; Enterobacterales; Enterobacteriaceae; Escherichia; Escherichia coli
MTGDNTLIHSHGINRRDFMKLCAALAATMGLSSKAAAEMAESVTNPQRPPVIWIGAQECT WP_032170173.1
hydrogenase 2 small subunit [Escherichia coli] Length: 372\nNumber of predicted TMHs: 1\nExp number of AAs in TMHs: 23.72138\nExp number, first 60 AAs: 0.22038\nTotal prob of N-in: 0.12992\noutside 1 331\nTMhelix 332 354\ninside 355 372

9567 GCF_000703965.1_ASM70396v1 Escherichia coli 2-222-05_S1_C1 Proteobacteria; Gammaproteobacteria; Enterobacterales; Enterobacteriaceae; Escherichia; Escherichia coli

MNNEETFYQAMRRQGVTRRSFLKYCSLAATSLGLGAGMAPKIAWALENKPRIPVVWIHGL WP_001295940.1 MULTISPECIES: hydrogenase [Enterobacteriaceae] Length: 372\nNumber of predicted TMHs: 2\nExp number of AAs in TMHs: 37.2608899999999\nExp number, first 60 AAs: 19.52923\nTotal prob of N-in: 0.94351\nPOSSIBLE N-term signal sequence\ninside 1 19\nTMhelix 20 39\noutside 40 325\nTMhelix 326 348\ninside 349 372

9568 GCF_000703965.1_ASM70396v1 Escherichia coli 2-222-05_S1_C1 Proteobacteria; Gammaproteobacteria; Enterobacterales; Enterobacteriaceae; Escherichia; Escherichia coli MTGDNTLIHSHGINRRDFMKLCAALAATMGLSSKAAAEMAESVTNPQRPPVIWIGAQECT WP_000145410.1 MULTISPECIES: hydrogenase-2 small chain [Proteobacteria] Length: 372\nNumber of predicted TMHs: 1\nExp number of AAs in TMHs: 23.72226\nExp number, first 60 AAs: 0.22038\nTotal prob of N-in: 0.12994\noutside 1 331\nTMhelix 332 354\ninside 355 372

9569 GCF_000704125.1_ASM70412v1 Escherichia coli 2-222-05_S1_C2 Proteobacteria; Gammaproteobacteria; Enterobacterales; Enterobacteriaceae; Escherichia; Escherichia coli MTGDNTLIHSHGINRRDFMKLCAALAATMGLSSKAAAEMAESVTNPQRPPVIWIGAQECT WP_000145410.1 MULTISPECIES: hydrogenase-2 small chain [Proteobacteria] Length: 372\nNumber of predicted TMHs: 1\nExp number of AAs in TMHs: 23.72226\nExp number, first 60 AAs: 0.22038\nTotal prob of N-in: 0.12994\noutside 1 331\nTMhelix 332 354\ninside 355 372

9570 GCF_000704125.1_ASM70412v1 Escherichia coli 2-222-05_S1_C2 Proteobacteria; Gammaproteobacteria; Enterobacterales; Enterobacteriaceae; Escherichia; Escherichia coli MNNEETFYQAMRRQGVTRRSFLKYCSLAATSLGLGAGMAPKIAWALENKPRIPVVWIHGL WP_001295940.1 MULTISPECIES: hydrogenase [Enterobacteriaceae] Length: 372\nNumber of predicted TMHs: 2\nExp number of AAs in TMHs: 37.2608899999999\nExp number, first 60 AAs: 19.52923\nTotal prob of N-in: 0.94351\nPOSSIBLE N-term signal sequence\ninside 1 19\nTMhelix 20 39\noutside 40 325\nTMhelix 326 348\ninside 349 372

9571 GCF_000704285.1_ASM70428v1 Escherichia coli 2-222-05_S1_C3 Proteobacteria; Gammaproteobacteria; Enterobacterales; Enterobacteriaceae; Escherichia; Escherichia coli MNNEETFYQAMRRQGVTRRSFLKYCSLAATSLGLGAGMAPKIAWALENKPRIPVVWIHGL WP_001295940.1 MULTISPECIES: hydrogenase [Enterobacteriaceae] Length: 372\nNumber of predicted TMHs: 2\nExp number of AAs in TMHs: 37.2608899999999\nExp number, first 60 AAs: 19.52923\nTotal prob of N-in: 0.94351\nPOSSIBLE N-term signal sequence\ninside 1 19\nTMhelix 20 39\noutside 40 325\nTMhelix 326 348\ninside 349 372

9572 GCF_000704285.1_ASM70428v1 Escherichia coli 2-222-05_S1_C3 Proteobacteria; Gammaproteobacteria; Enterobacterales; Enterobacteriaceae; Escherichia; Escherichia coli MTGDNTLIHSHGINRRDFMKLCAALAATMGLSSKAAAEMAESVTNPQRPPVIWIGAQECT WP_000145410.1 MULTISPECIES: hydrogenase-2 small chain [Proteobacteria] Length: 372\nNumber of predicted TMHs: 1\nExp number of AAs in TMHs: 23.72226\nExp number, first 60 AAs: 0.22038\nTotal prob of N-in: 0.12994\noutside 1 331\nTMhelix 332 354\ninside 355 372

9573 GCF_000713695.1_ASM71369v1 Escherichia coli 2-222-05_S3_C1 Proteobacteria; Gammaproteobacteria; Enterobacterales; Enterobacteriaceae; Escherichia; Escherichia coli MTGDNTLIHSHGINRRDFMKLCAALAATMGLSSKAAAEMAESVTNPQRPPVIWIGAQECT WP_000145410.1 MULTISPECIES: hydrogenase-2 small chain [Proteobacteria] Length: 372\nNumber of predicted TMHs: 1\nExp number of AAs in TMHs: 23.72226\nExp number, first 60 AAs: 0.22038\nTotal prob of N-in: 0.12994\noutside 1 331\nTMhelix 332 354\ninside 355 372

9574 GCF_000713695.1_ASM71369v1 Escherichia coli 2-222-05_S3_C1 Proteobacteria; Gammaproteobacteria; Enterobacterales; Enterobacteriaceae; Escherichia; Escherichia coli MNNEETFYQAMRRQGVTRRSFLKYCSLAATSLGLGAGMAPKIAWALENKPRIPVVWIHGL WP_001058323.1 MULTISPECIES: hydrogenase-1 small chain [Proteobacteria] Length: 372\nNumber of predicted TMHs: 2\nExp number of AAs in TMHs: 37.2608499999999\nExp number, first 60 AAs: 19.52923\nTotal prob of N-in: 0.94351\nPOSSIBLE N-term signal sequence\ninside 1 19\nTMhelix 20 39\noutside 40 325\nTMhelix 326 348\ninside 349 372

9575 GCF_000713735.1_ASM71373v1 Escherichia coli 2-222-05_S3_C2 Proteobacteria; Gammaproteobacteria; Enterobacterales; Enterobacteriaceae; Escherichia; Escherichia coli MNNEETFYQAMRRQGVTRRSFLKYCSLAATSLGLGAGMAPKIAWALENKPRIPVVWIHGL WP_001058323.1

MULTISPECIES: hydrogenase-1 small chain [Proteobacteria] Length: 372\nNumber of predicted TMHs: 2\nExp number of AAs in TMHs: 37.2608499999999\nExp number, first 60 AAs: 19.52923\nTotal prob of N-in: 0.94351\nPOSSIBLE N-term signal sequence\ninside 1 19\nTMhelix 20 39\noutside 40 325\nTMhelix 326 348\ninside 349 372

9576 GCF_000713735.1_ASM71373v1 Escherichia coli 2-222-05_S3_C2 Proteobacteria; Gammaproteobacteria; Enterobacterales; Enterobacteriaceae; Escherichia; Escherichia coli MTGDNTLIHSHGINRRDFMKLCAALAATMGLSSKAAAEMAESVTNPQRPPVIWIGAQECT WP_000145410.1 MULTISPECIES: hydrogenase-2 small chain [Proteobacteria] Length: 372\nNumber of predicted TMHs: 1\nExp number of AAs in TMHs: 23.72226\nExp number, first 60 AAs: 0.22038\nTotal prob of N-in: 0.12994\noutside 1 331\nTMhelix 332 354\ninside 355 372

9577 GCF_000704145.1_ASM70414v1 Escherichia coli 2-222-05_S3_C3 Proteobacteria; Gammaproteobacteria; Enterobacterales; Enterobacteriaceae; Escherichia; Escherichia coli MNNEETFYQAMRRQGVTRRSFLKYCSLAATSLGLGAGMAPKIAWALENKPRIPVVWIHGL WP_001058323.1 MULTISPECIES: hydrogenase-1 small chain [Proteobacteria] Length: 372\nNumber of predicted TMHs: 2\nExp number of AAs in TMHs: 37.2608499999999\nExp number, first 60 AAs: 19.52923\nTotal prob of N-in: 0.94351\nPOSSIBLE N-term signal sequence\ninside 1 19\nTMhelix 20 39\noutside 40 325\nTMhelix 326 348\ninside 349 372

9578 GCF_000704145.1_ASM70414v1 Escherichia coli 2-222-05_S3_C3 Proteobacteria; Gammaproteobacteria; Enterobacterales; Enterobacteriaceae; Escherichia; Escherichia coli MTGDNTLIHSHGINRRDFMKLCAALAATMGLSSKAAAEMAESVTNPQRPPVIWIGAQECT WP_000145410.1 MULTISPECIES: hydrogenase-2 small chain [Proteobacteria] Length: 372\nNumber of predicted TMHs: 1\nExp number of AAs in TMHs: 23.72226\nExp number, first 60 AAs: 0.22038\nTotal prob of N-in: 0.12994\noutside 1 331\nTMhelix 332 354\ninside 355 372

9579 GCF_000713275.1_ASM71327v1 Escherichia coli 2-222-05_S4_C1 Proteobacteria; Gammaproteobacteria; Enterobacterales; Enterobacteriaceae; Escherichia; Escherichia coli MNNEETFYQAMRRQGVTRRSFLKYCSLAATSLGLGAGMAPKIAWALENKPRIPVVWIHGL WP_001058323.1 MULTISPECIES: hydrogenase-1 small chain [Proteobacteria] Length: 372\nNumber of predicted TMHs: 2\nExp number of AAs in TMHs: 37.2608499999999\nExp number, first 60 AAs: 19.52923\nTotal prob of N-in: 0.94351\nPOSSIBLE N-term signal sequence\ninside 1 19\nTMhelix 20 39\noutside 40 325\nTMhelix 326 348\ninside 349 372

9580 GCF_000713275.1_ASM71327v1 Escherichia coli 2-222-05_S4_C1 Proteobacteria; Gammaproteobacteria; Enterobacterales; Enterobacteriaceae; Escherichia; Escherichia coli MTGDNTLIHSHGINRRDFMKLCAALAATMGLSSKAAAEMAESVTNPQRPPVIWIGAQECT WP_032170173.1 hydrogenase 2 small subunit [Escherichia coli] Length: 372\nNumber of predicted TMHs: 1\nExp number of AAs in TMHs: 23.72138\nExp number, first 60 AAs: 0.22038\nTotal prob of N-in: 0.12992\noutside 1 331\nTMhelix 332 354\ninside 355 372

9581 GCF_000704165.1_ASM70416v1 Escherichia coli 2-222-05_S4_C2 Proteobacteria; Gammaproteobacteria; Enterobacterales; Enterobacteriaceae; Escherichia; Escherichia coli MTGDNTLIHSHGINRRDFMKLCAALAATMGLSSKAAAEMAESVTNPQRPPVIWIGAQECT WP_000145410.1 MULTISPECIES: hydrogenase-2 small chain [Proteobacteria] Length: 372\nNumber of predicted TMHs: 1\nExp number of AAs in TMHs: 23.72226\nExp number, first 60 AAs: 0.22038\nTotal prob of N-in: 0.12994\noutside 1 331\nTMhelix 332 354\ninside 355 372

9582 GCF_000704165.1_ASM70416v1 Escherichia coli 2-222-05_S4_C2 Proteobacteria; Gammaproteobacteria; Enterobacterales; Enterobacteriaceae; Escherichia; Escherichia coli MNNEETFYQAMRRQGVTRRSFLKYCSLAATSLGLGAGMAPKIAWALENKPRIPVVWIHGL WP_001058323.1 MULTISPECIES: hydrogenase-1 small chain [Proteobacteria] Length: 372\nNumber of predicted TMHs: 2\nExp number of AAs in TMHs: 37.2608499999999\nExp number, first 60 AAs: 19.52923\nTotal prob of N-in: 0.94351\nPOSSIBLE N-term signal sequence\ninside 1 19\nTMhelix 20 39\noutside 40 325\nTMhelix 326 348\ninside 349 372

9583 GCF_000713785.1_ASM71378v1 Escherichia coli 2-222-05_S4_C3 Proteobacteria; Gammaproteobacteria; Enterobacterales; Enterobacteriaceae; Escherichia; Escherichia coli MTGDNTLIHSHGINRRDFMKLCAALAATMGLSSKAAAEMAESVTNPQRPPVIWIGAQECT WP_000145410.1 MULTISPECIES: hydrogenase-2 small chain [Proteobacteria] Length: 372\nNumber of predicted TMHs:

1\nExp number of AAs in TMHs: 23.72226\nExp number, first 60 AAs: 0.22038\nTotal prob of N-in: 0.12994\noutside 1 331\nTMhelix 332 354\ninside 355 372

9584 GCF_000713785.1_ASM71378v1 Escherichia coli 2-222-05_S4_C3 Proteobacteria; Gammaproteobacteria; Enterobacterales; Enterobacteriaceae; Escherichia; Escherichia coli MNNEETFYQAMRRQGVTRRSFLKYCSLAATSLGLGAGMAPKIAWALENKPRIPVVWIHGL WP_001058323.1 MULTISPECIES: hydrogenase-1 small chain [Proteobacteria] Length: 372\nNumber of predicted TMHs: 2\nExp number of AAs in TMHs: 37.2608499999999\nExp number, first 60 AAs: 19.52923\nTotal prob of N-in: 0.94351\nPOSSIBLE N-term signal sequence\ninside 1 19\nTMhelix 20 39\noutside 40 325\nTMhelix 326 348\ninside 349 372

9585 GCF_000711475.1_ASM71147v1 Escherichia coli 2-316-03_S1_C1 Proteobacteria; Gammaproteobacteria; Enterobacterales; Enterobacteriaceae; Escherichia; Escherichia coli MNNEETFYQAMRRQGVTRRSFLKYCSLAATSLGLGAGMAPKIAWALENKPRIPVVWIHGL WP_001058323.1 MULTISPECIES: hydrogenase-1 small chain [Proteobacteria] Length: 372\nNumber of predicted TMHs: 2\nExp number of AAs in TMHs: 37.2608499999999\nExp number, first 60 AAs: 19.52923\nTotal prob of N-in: 0.94351\nPOSSIBLE N-term signal sequence\ninside 1 19\nTMhelix 20 39\noutside 40 325\nTMhelix 326 348\ninside 349 372

9586 GCF_000711475.1_ASM71147v1 Escherichia coli 2-316-03_S1_C1 Proteobacteria; Gammaproteobacteria; Enterobacterales; Enterobacteriaceae; Escherichia; Escherichia coli MTGDNTLIHSHGINRRDFMKLCAALAATMGLSSKAAAEMAESVTNPQRPPVIWIGAQECT WP_000145410.1 MULTISPECIES: hydrogenase-2 small chain [Proteobacteria] Length: 372\nNumber of predicted TMHs: 1\nExp number of AAs in TMHs: 23.72226\nExp number, first 60 AAs: 0.22038\nTotal prob of N-in: 0.12994\noutside 1 331\nTMhelix 332 354\ninside 355 372

9587 GCF_000711415.1_ASM71141v1 Escherichia coli 2-316-03_S1_C2 Proteobacteria; Gammaproteobacteria; Enterobacterales; Enterobacteriaceae; Escherichia; Escherichia coli MNNEETFYQAMRRQGVTRRSFLKYCSLAATSLGLGAGMAPKIAWALENKPRIPVVWIHGL WP_001058323.1 MULTISPECIES: hydrogenase-1 small chain [Proteobacteria] Length: 372\nNumber of predicted TMHs: 2\nExp number of AAs in TMHs: 37.2608499999999\nExp number, first 60 AAs: 19.52923\nTotal prob of N-in: 0.94351\nPOSSIBLE N-term signal sequence\ninside 1 19\nTMhelix 20 39\noutside 40 325\nTMhelix 326 348\ninside 349 372

9588 GCF_000711415.1_ASM71141v1 Escherichia coli 2-316-03_S1_C2 Proteobacteria; Gammaproteobacteria; Enterobacterales; Enterobacteriaceae; Escherichia; Escherichia coli MTGDNTLIHSHGINRRDFMKLCAALAATMGLSSKAAAEMAESVTNPQRPPVIWIGAQECT WP_000145410.1 MULTISPECIES: hydrogenase-2 small chain [Proteobacteria] Length: 372\nNumber of predicted TMHs: 1\nExp number of AAs in TMHs: 23.72226\nExp number, first 60 AAs: 0.22038\nTotal prob of N-in: 0.12994\noutside 1 331\nTMhelix 332 354\ninside 355 372

9589 GCF_000704045.1_ASM70404v1 Escherichia coli 2-316-03_S3_C1 Proteobacteria; Gammaproteobacteria; Enterobacterales; Enterobacteriaceae; Escherichia; Escherichia coli MTGDNTLIHSHGINRRDFMKLCAALAATMGLSSKAAAEMAESVTNPQRPPVIWIGAQECT WP_000145410.1 MULTISPECIES: hydrogenase-2 small chain [Proteobacteria] Length: 372\nNumber of predicted TMHs: 1\nExp number of AAs in TMHs: 23.72226\nExp number, first 60 AAs: 0.22038\nTotal prob of N-in: 0.12994\noutside 1 331\nTMhelix 332 354\ninside 355 372

9590 GCF_000704045.1_ASM70404v1 Escherichia coli 2-316-03_S3_C1 Proteobacteria; Gammaproteobacteria; Enterobacterales; Enterobacteriaceae; Escherichia; Escherichia coli MNNEETFYQAMRRQGVTRRSFLKYCSLAATSLGLGAGMAPKIAWALENKPRIPVVWIHGL WP_001058323.1 MULTISPECIES: hydrogenase-1 small chain [Proteobacteria] Length: 372\nNumber of predicted TMHs: 2\nExp number of AAs in TMHs: 37.2608499999999\nExp number, first 60 AAs: 19.52923\nTotal prob of N-in: 0.94351\nPOSSIBLE N-term signal sequence\ninside 1 19\nTMhelix 20 39\noutside 40 325\nTMhelix 326 348\ninside 349 372

9591 GCF_000704065.1_ASM70406v1 Escherichia coli 2-316-03_S3_C2 Proteobacteria; Gammaproteobacteria; Enterobacterales; Enterobacteriaceae; Escherichia; Escherichia coli MNNEETFYQAMRRQGVTRRSFLKYCSLAATSLGLGAGMAPKIAWALENKPRIPVVWIHGL WP_001058323.1 MULTISPECIES: hydrogenase-1 small chain [Proteobacteria] Length: 372\nNumber of predicted TMHs: 2\nExp number of AAs in TMHs: 37.2608499999999\nExp number, first 60 AAs: 19.52923\nTotal prob of N-in:

0.94351\nPOSSIBLE N-term signal sequence\ninside 1 19\nTMhelix 20 39\noutside 40 325\nTMhelix 326 348\ninside 349 372

9592 GCF_000704065.1_ASM70406v1 Escherichia coli 2-316-03_S3_C2 Proteobacteria; Gammaproteobacteria; Enterobacterales; Enterobacteriaceae; Escherichia; Escherichia coli MTGDNTLIHSHGINRRDFMKLCAALAATMGLSSKAAAEMAESVTNPQRPPVIWIGAQECT WP_000145410.1 MULTISPECIES: hydrogenase-2 small chain [Proteobacteria] Length: 372\nNumber of predicted TMHs: 1\nExp number of AAs in TMHs: 23.72226\nExp number, first 60 AAs: 0.22038\nTotal prob of N-in: 0.12994\noutside 1 331\nTMhelix 332 354\ninside 355 372

9593 GCF_000704085.1_ASM70408v1 Escherichia coli 2-316-03_S3_C3 Proteobacteria; Gammaproteobacteria; Enterobacterales; Enterobacteriaceae; Escherichia; Escherichia coli MTGDNTLIHSHGINRRDFMKLCAALAATMGLSSKAAAEMAESVTNPQRPPVIWIGAQECT WP_000145410.1 MULTISPECIES: hydrogenase-2 small chain [Proteobacteria] Length: 372\nNumber of predicted TMHs: 1\nExp number of AAs in TMHs: 23.72226\nExp number, first 60 AAs: 0.22038\nTotal prob of N-in: 0.12994\noutside 1 331\nTMhelix 332 354\ninside 355 372

9594 GCF_000704085.1_ASM70408v1 Escherichia coli 2-316-03_S3_C3 Proteobacteria; Gammaproteobacteria; Enterobacterales; Enterobacteriaceae; Escherichia; Escherichia coli MNNEETFYQAMRRQGVTRRSFLKYCSLAATSLGLGAGMAPKIAWALENKPRIPVVWIHGL WP_001058323.1 MULTISPECIES: hydrogenase-1 small chain [Proteobacteria] Length: 372\nNumber of predicted TMHs: 2\nExp number of AAs in TMHs: 37.2608499999999\nExp number, first 60 AAs: 19.52923\nTotal prob of N-in: 0.94351\nPOSSIBLE N-term signal sequence\ninside 1 19\nTMhelix 20 39\noutside 40 325\nTMhelix 326 348\ninside 349 372

9595 GCF_000704305.1_ASM70430v1 Escherichia coli 2-316-03_S4_C1 Proteobacteria; Gammaproteobacteria; Enterobacterales; Enterobacteriaceae; Escherichia; Escherichia coli MTGDNTLIHSHGINRRDFMKLCAALAATMGLSSKAAAEMAESVTNPQRPPVIWIGAQECT WP_000145410.1 MULTISPECIES: hydrogenase-2 small chain [Proteobacteria] Length: 372\nNumber of predicted TMHs: 1\nExp number of AAs in TMHs: 23.72226\nExp number, first 60 AAs: 0.22038\nTotal prob of N-in: 0.12994\noutside 1 331\nTMhelix 332 354\ninside 355 372

9596 GCF_000704305.1_ASM70430v1 Escherichia coli 2-316-03_S4_C1 Proteobacteria; Gammaproteobacteria; Enterobacterales; Enterobacteriaceae; Escherichia; Escherichia coli MNNEETFYQAMRRQGVTRRSFLKYCSLAATSLGLGAGMAPKIAWALENKPRIPVVWIHGL WP_001058323.1 MULTISPECIES: hydrogenase-1 small chain [Proteobacteria] Length: 372\nNumber of predicted TMHs: 2\nExp number of AAs in TMHs: 37.2608499999999\nExp number, first 60 AAs: 19.52923\nTotal prob of N-in: 0.94351\nPOSSIBLE N-term signal sequence\ninside 1 19\nTMhelix 20 39\noutside 40 325\nTMhelix 326 348\ninside 349 372

9597 GCF_000704325.1_ASM70432v1 Escherichia coli 2-316-03_S4_C2 Proteobacteria; Gammaproteobacteria; Enterobacterales; Enterobacteriaceae; Escherichia; Escherichia coli MNNEETFYQAMRRQGVTRRSFLKYCSLAATSLGLGAGMAPKIAWALENKPRIPVVWIHGL WP_001058323.1 MULTISPECIES: hydrogenase-1 small chain [Proteobacteria] Length: 372\nNumber of predicted TMHs: 2\nExp number of AAs in TMHs: 37.2608499999999\nExp number, first 60 AAs: 19.52923\nTotal prob of N-in: 0.94351\nPOSSIBLE N-term signal sequence\ninside 1 19\nTMhelix 20 39\noutside 40 325\nTMhelix 326 348\ninside 349 372

9598 GCF_000704325.1_ASM70432v1 Escherichia coli 2-316-03_S4_C2 Proteobacteria; Gammaproteobacteria; Enterobacterales; Enterobacteriaceae; Escherichia; Escherichia coli MTGDNTLIHSHGINRRDFMKLCAALAATMGLSSKAAAEMAESVTNPQRPPVIWIGAQECT WP_000145410.1 MULTISPECIES: hydrogenase-2 small chain [Proteobacteria] Length: 372\nNumber of predicted TMHs: 1\nExp number of AAs in TMHs: 23.72226\nExp number, first 60 AAs: 0.22038\nTotal prob of N-in: 0.12994\noutside 1 331\nTMhelix 332 354\ninside 355 372

9599 GCF_000704345.1_ASM70434v1 Escherichia coli 2-316-03_S4_C3 Proteobacteria; Gammaproteobacteria; Enterobacterales; Enterobacteriaceae; Escherichia; Escherichia coli MNNEETFYQAMRRQGVTRRSFLKYCSLAATSLGLGAGMAPKIAWALENKPRIPVVWIHGL WP_001295940.1 MULTISPECIES: hydrogenase [Enterobacteriaceae] Length: 372\nNumber of predicted TMHs: 2\nExp number of AAs in TMHs: 37.2608899999999\nExp number, first 60 AAs: 19.52923\nTotal prob of N-in:

0.94351\nPOSSIBLE N-term signal sequence\ninside 1 19\nTMhelix 20 39\noutside 40 325\nTMhelix 326 348\ninside 349 372

9600 GCF_000704345.1_ASM70434v1 Escherichia coli 2-316-03_S4_C3 Proteobacteria; Gammaproteobacteria; Enterobacterales; Enterobacteriaceae; Escherichia; Escherichia coli MTGDNTLIHSHGINRRDFMKLCAALAATMGLSSKAAAEMAESVTNPQRPPVIWIGAQECT WP_000145410.1 MULTISPECIES: hydrogenase-2 small chain [Proteobacteria] Length: 372\nNumber of predicted TMHs: 1\nExp number of AAs in TMHs: 23.72226\nExp number, first 60 AAs: 0.22038\nTotal prob of N-in: 0.12994\noutside 1 331\nTMhelix 332 354\ninside 355 372

9601 GCF_000183005.1_ASM18300v1 Escherichia coli 2362-75 Proteobacteria; Gammaproteobacteria; Enterobacterales; Enterobacteriaceae; Escherichia; Escherichia coli MTGDNTLIHSHGINRRDFMKLCAALAATMGLSSKAAAEMAESVTNPQRPPVIWIGAQECT WP_000145410.1 MULTISPECIES: hydrogenase-2 small chain [Proteobacteria] Length: 372\nNumber of predicted TMHs: 1\nExp number of AAs in TMHs: 23.72226\nExp number, first 60 AAs: 0.22038\nTotal prob of N-in: 0.12994\noutside 1 331\nTMhelix 332 354\ninside 355 372

9602 GCF_000183005.1_ASM18300v1 Escherichia coli 2362-75 Proteobacteria; Gammaproteobacteria; Enterobacterales; Enterobacteriaceae; Escherichia; Escherichia coli MNNEETFYQAMRRQGVTRRSFLKYCSLAATSLGLGAGMAPKIAWALENKPRIPVVWIHGL WP_001058323.1 MULTISPECIES: hydrogenase-1 small chain [Proteobacteria] Length: 372\nNumber of predicted TMHs: 2\nExp number of AAs in TMHs: 37.2608499999999\nExp number, first 60 AAs: 19.52923\nTotal prob of N-in: 0.94351\nPOSSIBLE N-term signal sequence\ninside 1 19\nTMhelix 20 39\noutside 40 325\nTMhelix 326 348\ninside 349 372

9603 GCF_000194535.1_ASM19453v2 Escherichia coli 2.3916 Proteobacteria; Gammaproteobacteria; Enterobacterales; Enterobacteriaceae; Escherichia; Escherichia coli MTGDNTLIHSHGINRRDFMKLCAALAATMGLSSKAAAEMAESVTNPQRPPVIWIGAQECT WP_000145410.1 MULTISPECIES: hydrogenase-2 small chain [Proteobacteria] Length: 372\nNumber of predicted TMHs: 1\nExp number of AAs in TMHs: 23.72226\nExp number, first 60 AAs: 0.22038\nTotal prob of N-in: 0.12994\noutside 1 331\nTMhelix 332 354\ninside 355 372

9604 GCF_000194535.1_ASM19453v2 Escherichia coli 2.3916 Proteobacteria; Gammaproteobacteria; Enterobacterales; Enterobacteriaceae; Escherichia; Escherichia coli MNNEETFYQAMRRQGVTRRSFLKYCSLAATSLGLGAGMAPKIAWALENKPRIPVVWIHGL WP_001058323.1 MULTISPECIES: hydrogenase-1 small chain [Proteobacteria] Length: 372\nNumber of predicted TMHs: 2\nExp number of AAs in TMHs: 37.2608499999999\nExp number, first 60 AAs: 19.52923\nTotal prob of N-in: 0.94351\nPOSSIBLE N-term signal sequence\ninside 1 19\nTMhelix 20 39\noutside 40 325\nTMhelix 326 348\ninside 349 372

9605 GCF_000194555.1_ASM19455v2 Escherichia coli 2.4168 Proteobacteria; Gammaproteobacteria; Enterobacterales; Enterobacteriaceae; Escherichia; Escherichia coli MNNEETFYQAMRRQGVTRRSFLKYCSLAATSLGLGAGMAPKIAWALENKPRIPVVWIHGL WP_001058323.1 MULTISPECIES: hydrogenase-1 small chain [Proteobacteria] Length: 372\nNumber of predicted TMHs: 2\nExp number of AAs in TMHs: 37.2608499999999\nExp number, first 60 AAs: 19.52923\nTotal prob of N-in: 0.94351\nPOSSIBLE N-term signal sequence\ninside 1 19\nTMhelix 20 39\noutside 40 325\nTMhelix 326 348\ninside 349 372

9606 GCF_000194555.1_ASM19455v2 Escherichia coli 2.4168 Proteobacteria; Gammaproteobacteria; Enterobacterales; Enterobacteriaceae; Escherichia; Escherichia coli MTGDNTLIHSHGINRRDFMKLCAALAATMGLSSKAAAEMAESVTNPQRPPVIWIGAQECT WP_000145410.1 MULTISPECIES: hydrogenase-2 small chain [Proteobacteria] Length: 372\nNumber of predicted TMHs: 1\nExp number of AAs in TMHs: 23.72226\nExp number, first 60 AAs: 0.22038\nTotal prob of N-in: 0.12994\noutside 1 331\nTMhelix 332 354\ninside 355 372

9607 GCF_000704925.1_ASM70492v1 Escherichia coli 2-427-07_S1_C1 Proteobacteria; Gammaproteobacteria; Enterobacterales; Enterobacteriaceae; Escherichia; Escherichia coli MNNEETFYQAMRRQGVTRRSFLKYCSLAATSLGLGAGMAPKIAWALENKPRIPVVWIHGL WP_001058323.1 MULTISPECIES: hydrogenase-1 small chain [Proteobacteria] Length: 372\nNumber of predicted TMHs: 2\nExp number of AAs in TMHs: 37.2608499999999\nExp number, first 60 AAs: 19.52923\nTotal prob of N-in:

0.94351\nPOSSIBLE N-term signal sequence\ninside 1 19\nTMhelix 20 39\noutside 40 325\nTMhelix 326 348\ninside 349 372

9608 GCF_000704925.1_ASM70492v1 Escherichia coli 2-427-07_S1_C1 Proteobacteria; Gammaproteobacteria; Enterobacterales; Enterobacteriaceae; Escherichia; Escherichia coli MTGDNTLIHSHGINRRDFMKLCAALAATMGLSSKAAAEMAESVTNPQRPPVIWIGAQECT WP_000145410.1 MULTISPECIES: hydrogenase-2 small chain [Proteobacteria] Length: 372\nNumber of predicted TMHs: 1\nExp number of AAs in TMHs: 23.72226\nExp number, first 60 AAs: 0.22038\nTotal prob of N-in: 0.12994\noutside 1 331\nTMhelix 332 354\ninside 355 372

9609 GCF_000704365.1_ASM70436v1 Escherichia coli 2-427-07_S1_C2 Proteobacteria; Gammaproteobacteria; Enterobacterales; Enterobacteriaceae; Escherichia; Escherichia coli MTGDNTLIHSHGINRRDFMKLCAALAATMGLSSKAAAEMAESVTNPQRPPVIWIGAQECT WP_000145410.1 MULTISPECIES: hydrogenase-2 small chain [Proteobacteria] Length: 372\nNumber of predicted TMHs: 1\nExp number of AAs in TMHs: 23.72226\nExp number, first 60 AAs: 0.22038\nTotal prob of N-in: 0.12994\noutside 1 331\nTMhelix 332 354\ninside 355 372

9610 GCF_000704365.1_ASM70436v1 Escherichia coli 2-427-07_S1_C2 Proteobacteria; Gammaproteobacteria; Enterobacterales; Enterobacteriaceae; Escherichia; Escherichia coli MNNEETFYQAMRRQGVTRRSFLKYCSLAATSLGLGAGMAPKIAWALENKPRIPVVWIHGL WP_001058323.1 MULTISPECIES: hydrogenase-1 small chain [Proteobacteria] Length: 372\nNumber of predicted TMHs: 2\nExp number of AAs in TMHs: 37.2608499999999\nExp number, first 60 AAs: 19.52923\nTotal prob of N-in: 0.94351\nPOSSIBLE N-term signal sequence\ninside 1 19\nTMhelix 20 39\noutside 40 325\nTMhelix 326 348\ninside 349 372

9611 GCF_000704625.1_ASM70462v1 Escherichia coli 2-427-07_S1_C3 Proteobacteria; Gammaproteobacteria; Enterobacterales; Enterobacteriaceae; Escherichia; Escherichia coli MNNEETFYQAMRRQGVTRRSFLKYCSLAATSLGLGAGMAPKIAWALENKPRIPVVWIHGL WP_001058323.1 MULTISPECIES: hydrogenase-1 small chain [Proteobacteria] Length: 372\nNumber of predicted TMHs: 2\nExp number of AAs in TMHs: 37.2608499999999\nExp number, first 60 AAs: 19.52923\nTotal prob of N-in: 0.94351\nPOSSIBLE N-term signal sequence\ninside 1 19\nTMhelix 20 39\noutside 40 325\nTMhelix 326 348\ninside 349 372

9612 GCF_000704625.1_ASM70462v1 Escherichia coli 2-427-07_S1_C3 Proteobacteria; Gammaproteobacteria; Enterobacterales; Enterobacteriaceae; Escherichia; Escherichia coli MTGDNTLIHSHGINRRDFMKLCAALAATMGLSSKAAAEMAESVTNPQRPPVIWIGAQECT WP_000145410.1 MULTISPECIES: hydrogenase-2 small chain [Proteobacteria] Length: 372\nNumber of predicted TMHs: 1\nExp number of AAs in TMHs: 23.72226\nExp number, first 60 AAs: 0.22038\nTotal prob of N-in: 0.12994\noutside 1 331\nTMhelix 332 354\ninside 355 372

9613 GCF_000704385.1_ASM70438v1 Escherichia coli 2-427-07_S3_C1 Proteobacteria; Gammaproteobacteria; Enterobacterales; Enterobacteriaceae; Escherichia; Escherichia coli MNNEETFYQAMRRQGVTRRSFLKYCSLAATSLGLGAGMAPKIAWALENKPRIPVVWIHGL WP_001295940.1 MULTISPECIES: hydrogenase [Enterobacteriaceae] Length: 372\nNumber of predicted TMHs: 2\nExp number of AAs in TMHs: 37.2608899999999\nExp number, first 60 AAs: 19.52923\nTotal prob of N-in: 0.94351\nPOSSIBLE N-term signal sequence\ninside 1 19\nTMhelix 20 39\noutside 40 325\nTMhelix 326 348\ninside 349 372

9614 GCF_000704385.1_ASM70438v1 Escherichia coli 2-427-07_S3_C1 Proteobacteria; Gammaproteobacteria; Enterobacterales; Enterobacteriaceae; Escherichia; Escherichia coli MTGDNTLIHSHGINRRDFMKLCAALAATMGLSSKAAAEMAESVTNPQRPPVIWIGAQECT WP_000145410.1 MULTISPECIES: hydrogenase-2 small chain [Proteobacteria] Length: 372\nNumber of predicted TMHs: 1\nExp number of AAs in TMHs: 23.72226\nExp number, first 60 AAs: 0.22038\nTotal prob of N-in: 0.12994\noutside 1 331\nTMhelix 332 354\ninside 355 372

9615 GCF_000704405.1_ASM70440v1 Escherichia coli 2-427-07_S3_C3 Proteobacteria; Gammaproteobacteria; Enterobacterales; Enterobacteriaceae; Escherichia; Escherichia coli MNNEETFYQAMRRQGVTRRSFLKYCSLAATSLGLGAGMAPKIAWALENKPRIPVVWIHGL WP_001295940.1 MULTISPECIES: hydrogenase [Enterobacteriaceae] Length: 372\nNumber of predicted TMHs: 2\nExp number of AAs in TMHs: 37.2608899999999\nExp number, first 60 AAs: 19.52923\nTotal prob of N-in:

0.94351\nPOSSIBLE N-term signal sequence\ninside 1 19\nTMhelix 20 39\noutside 40 325\nTMhelix 326 348\ninside 349 372

9616 GCF_000704405.1_ASM70440v1 Escherichia coli 2-427-07_S3_C3 Proteobacteria; Gammaproteobacteria; Enterobacterales; Enterobacteriaceae; Escherichia; Escherichia coli MTGDNTLIHSHGINRRDFMKLCAALAATMGLSSKAAAEMAESVTNPQRPPVIWIGAQECT WP_000145410.1 MULTISPECIES: hydrogenase-2 small chain [Proteobacteria] Length: 372\nNumber of predicted TMHs: 1\nExp number of AAs in TMHs: 23.72226\nExp number, first 60 AAs: 0.22038\nTotal prob of N-in: 0.12994\noutside 1 331\nTMhelix 332 354\ninside 355 372

9617 GCF_000704425.1_ASM70442v1 Escherichia coli 2-427-07_S4_C1 Proteobacteria; Gammaproteobacteria; Enterobacterales; Enterobacteriaceae; Escherichia; Escherichia coli MTGDNTLIHSHGINRRDFMKLCAALAATMGLSSKAAAEMAESVTNPQRPPVIWIGAQECT WP_000145410.1 MULTISPECIES: hydrogenase-2 small chain [Proteobacteria] Length: 372\nNumber of predicted TMHs: 1\nExp number of AAs in TMHs: 23.72226\nExp number, first 60 AAs: 0.22038\nTotal prob of N-in: 0.12994\noutside 1 331\nTMhelix 332 354\ninside 355 372

9618 GCF_000704425.1_ASM70442v1 Escherichia coli 2-427-07_S4_C1 Proteobacteria; Gammaproteobacteria; Enterobacterales; Enterobacteriaceae; Escherichia; Escherichia coli MNNEETFYQAMRRQGVTRRSFLKYCSLAATSLGLGAGMAPKIAWALENKPRIPVVWIHGL WP_001058323.1 MULTISPECIES: hydrogenase-1 small chain [Proteobacteria] Length: 372\nNumber of predicted TMHs: 2\nExp number of AAs in TMHs: 37.2608499999999\nExp number, first 60 AAs: 19.52923\nTotal prob of N-in: 0.94351\nPOSSIBLE N-term signal sequence\ninside 1 19\nTMhelix 20 39\noutside 40 325\nTMhelix 326 348\ninside 349 372

9619 GCF_000704445.1_ASM70444v1 Escherichia coli 2-427-07_S4_C2 Proteobacteria; Gammaproteobacteria; Enterobacterales; Enterobacteriaceae; Escherichia; Escherichia coli MNNEETFYQAMRRQGVTRRSFLKYCSLAATSLGLGAGMAPKIAWALENKPRIPVVWIHGL WP_001295940.1 MULTISPECIES: hydrogenase [Enterobacteriaceae] Length: 372\nNumber of predicted TMHs: 2\nExp number of AAs in TMHs: 37.2608899999999\nExp number, first 60 AAs: 19.52923\nTotal prob of N-in: 0.94351\nPOSSIBLE N-term signal sequence\ninside 1 19\nTMhelix 20 39\noutside 40 325\nTMhelix 326 348\ninside 349 372

9620 GCF_000704445.1_ASM70444v1 Escherichia coli 2-427-07_S4_C2 Proteobacteria; Gammaproteobacteria; Enterobacterales; Enterobacteriaceae; Escherichia; Escherichia coli MTGDNTLIHSHGINRRDFMKLCAALAATMGLSSKAAAEMVESVTNPQRPPVIWIGAQECT WP_000145423.1 hydrogenase 2 small subunit [Escherichia coli] Length: 372\nNumber of predicted TMHs: 1\nExp number of AAs in TMHs: 23.78201\nExp number, first 60 AAs: 0.28355\nTotal prob of N-in: 0.13257\noutside 1 331\nTMhelix 332 354\ninside 355 372

9621 GCF_000711355.1_ASM71135v1 Escherichia coli 2-427-07_S4_C3 Proteobacteria; Gammaproteobacteria; Enterobacterales; Enterobacteriaceae; Escherichia; Escherichia coli MTGDNTLIHSHGINRRDFMKLCAALAATMGLSSKAAAEMAESVTNPQRPPVIWIGAQECT WP_000145410.1 MULTISPECIES: hydrogenase-2 small chain [Proteobacteria] Length: 372\nNumber of predicted TMHs: 1\nExp number of AAs in TMHs: 23.72226\nExp number, first 60 AAs: 0.22038\nTotal prob of N-in: 0.12994\noutside 1 331\nTMhelix 332 354\ninside 355 372

9622 GCF_000711355.1_ASM71135v1 Escherichia coli 2-427-07_S4_C3 Proteobacteria; Gammaproteobacteria; Enterobacterales; Enterobacteriaceae; Escherichia; Escherichia coli MNNEETFYQAMRRQGVTRRSFLKYCSLAATSLGLGAGMAPKIAWALENKPRIPVVWIHGL WP_001058323.1 MULTISPECIES: hydrogenase-1 small chain [Proteobacteria] Length: 372\nNumber of predicted TMHs: 2\nExp number of AAs in TMHs: 37.2608499999999\nExp number, first 60 AAs: 19.52923\nTotal prob of N-in: 0.94351\nPOSSIBLE N-term signal sequence\ninside 1 19\nTMhelix 20 39\noutside 40 325\nTMhelix 326 348\ninside 349 372

9623 GCF_000713825.1_ASM71382v1 Escherichia coli 2-460-02_S1_C1 Proteobacteria; Gammaproteobacteria; Enterobacterales; Enterobacteriaceae; Escherichia; Escherichia coli MTGDNTLIHSHGINRRDFMKLCAALAATMGLSSKAAAEMAESVTNPQRPPVIWIGAQECT WP_000145410.1 MULTISPECIES: hydrogenase-2 small chain [Proteobacteria] Length: 372\nNumber of predicted TMHs: 1\nExp number of AAs in TMHs: 23.72226\nExp number, first 60 AAs: 0.22038\nTotal prob of N-in: 0.12994\noutside 1 331\nTMhelix 332 354\ninside 355 372

9624	GCF_000713825.1_ASM71382v1	Escherichia coli 2-460-02_S1_C1	Proteobacteria; Gammaproteobacteria; Enterobacterales; Enterobacteriaceae; Escherichia; Escherichia coli
MNNEETFYQAMRRQGVTRRSFLKYCSLAATSLGLGAGMAPKIAWALENKPRIPVVWIHGL	WP_001058323.1
MULTISPECIES: hydrogenase-1 small chain [Proteobacteria]	Length: 372\nNumber of predicted TMHs: 2\nExp number of AAs in TMHs: 37.2608499999999\nExp number, first 60 AAs: 19.52923\nTotal prob of N-in: 0.94351\nPOSSIBLE N-term signal sequence\ninside 1 19\nTMhelix 20 39\noutside 40 325\nTMhelix 326 348\ninside 349 372

9625	GCF_000713865.1_ASM71386v1	Escherichia coli 2-460-02_S1_C2	Proteobacteria; Gammaproteobacteria; Enterobacterales; Enterobacteriaceae; Escherichia; Escherichia coli
MTGDNTLIHSHGINRRDFMKLCAALAATMGLSSKAAAEMVESVTNPQRPPVIWIGAQECT	WP_000145423.1
hydrogenase 2 small subunit [Escherichia coli]	Length: 372\nNumber of predicted TMHs: 1\nExp number of AAs in TMHs: 23.78201\nExp number, first 60 AAs: 0.28355\nTotal prob of N-in: 0.13257\noutside 1 331\nTMhelix 332 354\ninside 355 372

9626	GCF_000713865.1_ASM71386v1	Escherichia coli 2-460-02_S1_C2	Proteobacteria; Gammaproteobacteria; Enterobacterales; Enterobacteriaceae; Escherichia; Escherichia coli
MNNEETFYQAMRRQGVTRRSFLKYCSLAATSLGLGAGMAPKIAWALENKPRIPVVWIHGL	WP_001295940.1
MULTISPECIES: hydrogenase [Enterobacteriaceae]	Length: 372\nNumber of predicted TMHs: 2\nExp number of AAs in TMHs: 37.2608899999999\nExp number, first 60 AAs: 19.52923\nTotal prob of N-in: 0.94351\nPOSSIBLE N-term signal sequence\ninside 1 19\nTMhelix 20 39\noutside 40 325\nTMhelix 326 348\ninside 349 372

9627	GCF_000711485.1_ASM71148v1	Escherichia coli 2-460-02_S1_C3	Proteobacteria; Gammaproteobacteria; Enterobacterales; Enterobacteriaceae; Escherichia; Escherichia coli
MNNEETFYQAMRRQGVTRRSFLKYCSLAATSLGLGAGMAPKIAWALENKPRIPVVWIHGL	WP_001295940.1
MULTISPECIES: hydrogenase [Enterobacteriaceae]	Length: 372\nNumber of predicted TMHs: 2\nExp number of AAs in TMHs: 37.2608899999999\nExp number, first 60 AAs: 19.52923\nTotal prob of N-in: 0.94351\nPOSSIBLE N-term signal sequence\ninside 1 19\nTMhelix 20 39\noutside 40 325\nTMhelix 326 348\ninside 349 372

9628	GCF_000711485.1_ASM71148v1	Escherichia coli 2-460-02_S1_C3	Proteobacteria; Gammaproteobacteria; Enterobacterales; Enterobacteriaceae; Escherichia; Escherichia coli
MTGDNTLIHSHGINRRDFMKLCAALAATMGLSSKAAAEMVESVTNPQRPPVIWIGAQECT	WP_000145423.1
hydrogenase 2 small subunit [Escherichia coli]	Length: 372\nNumber of predicted TMHs: 1\nExp number of AAs in TMHs: 23.78201\nExp number, first 60 AAs: 0.28355\nTotal prob of N-in: 0.13257\noutside 1 331\nTMhelix 332 354\ninside 355 372

9629	GCF_000704465.1_ASM70446v1	Escherichia coli 2-460-02_S3_C1	Proteobacteria; Gammaproteobacteria; Enterobacterales; Enterobacteriaceae; Escherichia; Escherichia coli
MNNEETFYQAMRRQGVTRRSFLKYCSLAATSLGLGAGMAPKIAWALENKPRIPVVWIHGL	WP_001058323.1
MULTISPECIES: hydrogenase-1 small chain [Proteobacteria]	Length: 372\nNumber of predicted TMHs: 2\nExp number of AAs in TMHs: 37.2608499999999\nExp number, first 60 AAs: 19.52923\nTotal prob of N-in: 0.94351\nPOSSIBLE N-term signal sequence\ninside 1 19\nTMhelix 20 39\noutside 40 325\nTMhelix 326 348\ninside 349 372

9630	GCF_000704465.1_ASM70446v1	Escherichia coli 2-460-02_S3_C1	Proteobacteria; Gammaproteobacteria; Enterobacterales; Enterobacteriaceae; Escherichia; Escherichia coli
MTGDNTLIHSHGINRRDFMKLCAALAATMGLSSKAAAEMAESVTNPQRPPVIWIGAQECT	WP_000145410.1
MULTISPECIES: hydrogenase-2 small chain [Proteobacteria]	Length: 372\nNumber of predicted TMHs: 1\nExp number of AAs in TMHs: 23.72226\nExp number, first 60 AAs: 0.22038\nTotal prob of N-in: 0.12994\noutside 1 331\nTMhelix 332 354\ninside 355 372

9631	GCF_000704485.1_ASM70448v1	Escherichia coli 2-460-02_S3_C2	Proteobacteria; Gammaproteobacteria; Enterobacterales; Enterobacteriaceae; Escherichia; Escherichia coli
MTGDNTLIHSHGINRRDFMKLCAALAATMGLSSKAAAEMAESVTNPQRPPVIWIGAQECT	WP_000145410.1
MULTISPECIES: hydrogenase-2 small chain [Proteobacteria]	Length: 372\nNumber of predicted TMHs: 1\nExp number of AAs in TMHs: 23.72226\nExp number, first 60 AAs: 0.22038\nTotal prob of N-in: 0.12994\noutside 1 331\nTMhelix 332 354\ninside 355 372

9632 GCF_000704485.1_ASM70448v1 Escherichia coli 2-460-02_S3_C2 Proteobacteria; Gammaproteobacteria; Enterobacterales; Enterobacteriaceae; Escherichia; Escherichia coli MNNEETFYQAMRRQGVTRRSFLKYCSLAATSLGLGAGMAPKIAWALENKPRIPVVWIHGL WP_001058323.1 MULTISPECIES: hydrogenase-1 small chain [Proteobacteria] Length: 372\nNumber of predicted TMHs: 2\nExp number of AAs in TMHs: 37.2608499999999\nExp number, first 60 AAs: 19.52923\nTotal prob of N-in: 0.94351\nPOSSIBLE N-term signal sequence\ninside 1 19\nTMhelix 20 39\noutside 40 325\nTMhelix 326 348\ninside 349 372

9633 GCF_000704505.1_ASM70450v1 Escherichia coli 2-460-02_S3_C3 Proteobacteria; Gammaproteobacteria; Enterobacterales; Enterobacteriaceae; Escherichia; Escherichia coli MTGDNTLIHSHGINRRDFMKLCAALAATMGLSSKAAAEMAESVTNPQRPPVIWIGAQECT WP_000145410.1 MULTISPECIES: hydrogenase-2 small chain [Proteobacteria] Length: 372\nNumber of predicted TMHs: 1\nExp number of AAs in TMHs: 23.72226\nExp number, first 60 AAs: 0.22038\nTotal prob of N-in: 0.12994\noutside 1 331\nTMhelix 332 354\ninside 355 372

9634 GCF_000704505.1_ASM70450v1 Escherichia coli 2-460-02_S3_C3 Proteobacteria; Gammaproteobacteria; Enterobacterales; Enterobacteriaceae; Escherichia; Escherichia coli MNNEETFYQAMRRQGVTRRSFLKYCSLAATSLGLGAGMAPKIAWALENKPRIPVVWIHGL WP_001058323.1 MULTISPECIES: hydrogenase-1 small chain [Proteobacteria] Length: 372\nNumber of predicted TMHs: 2\nExp number of AAs in TMHs: 37.2608499999999\nExp number, first 60 AAs: 19.52923\nTotal prob of N-in: 0.94351\nPOSSIBLE N-term signal sequence\ninside 1 19\nTMhelix 20 39\noutside 40 325\nTMhelix 326 348\ninside 349 372

9635 GCF_000711565.1_ASM71156v1 Escherichia coli 2-460-02_S4_C1 Proteobacteria; Gammaproteobacteria; Enterobacterales; Enterobacteriaceae; Escherichia; Escherichia coli MTGDNTLIHSHGINRRDFMKLCAALAATMGLSSKAAAEMAESVTNPQRPPVIWIGAQECT WP_000145410.1 MULTISPECIES: hydrogenase-2 small chain [Proteobacteria] Length: 372\nNumber of predicted TMHs: 1\nExp number of AAs in TMHs: 23.72226\nExp number, first 60 AAs: 0.22038\nTotal prob of N-in: 0.12994\noutside 1 331\nTMhelix 332 354\ninside 355 372

9636 GCF_000711565.1_ASM71156v1 Escherichia coli 2-460-02_S4_C1 Proteobacteria; Gammaproteobacteria; Enterobacterales; Enterobacteriaceae; Escherichia; Escherichia coli MNNEETFYQAMRRQGVTRRSFLKYCSLAATSLGLGAGMAPKIAWALENKPRIPVVWIHGL WP_001058323.1 MULTISPECIES: hydrogenase-1 small chain [Proteobacteria] Length: 372\nNumber of predicted TMHs: 2\nExp number of AAs in TMHs: 37.2608499999999\nExp number, first 60 AAs: 19.52923\nTotal prob of N-in: 0.94351\nPOSSIBLE N-term signal sequence\ninside 1 19\nTMhelix 20 39\noutside 40 325\nTMhelix 326 348\ninside 349 372

9637 GCF_000704525.1_ASM70452v1 Escherichia coli 2-460-02_S4_C2 Proteobacteria; Gammaproteobacteria; Enterobacterales; Enterobacteriaceae; Escherichia; Escherichia coli MNNEETFYQAMRRQGVTRRSFLKYCSLAATSLGLGAGMAPKIAWALENKPRIPVVWIHGL WP_001058323.1 MULTISPECIES: hydrogenase-1 small chain [Proteobacteria] Length: 372\nNumber of predicted TMHs: 2\nExp number of AAs in TMHs: 37.2608499999999\nExp number, first 60 AAs: 19.52923\nTotal prob of N-in: 0.94351\nPOSSIBLE N-term signal sequence\ninside 1 19\nTMhelix 20 39\noutside 40 325\nTMhelix 326 348\ninside 349 372

9638 GCF_000704525.1_ASM70452v1 Escherichia coli 2-460-02_S4_C2 Proteobacteria; Gammaproteobacteria; Enterobacterales; Enterobacteriaceae; Escherichia; Escherichia coli MTGDNTLIHSHGINRRDFMKLCAALAATMGLSSKAAAEMAESVTNPQRPPVIWIGAQECT WP_000145410.1 MULTISPECIES: hydrogenase-2 small chain [Proteobacteria] Length: 372\nNumber of predicted TMHs: 1\nExp number of AAs in TMHs: 23.72226\nExp number, first 60 AAs: 0.22038\nTotal prob of N-in: 0.12994\noutside 1 331\nTMhelix 332 354\ninside 355 372

9639 GCF_000704545.1_ASM70454v1 Escherichia coli 2-460-02_S4_C3 Proteobacteria; Gammaproteobacteria; Enterobacterales; Enterobacteriaceae; Escherichia; Escherichia coli MTGDNTLIHSHGINRRDFMKLCAALAATMGLSSKAAAEMAESVTNPQRPPVIWIGAQECT WP_000145410.1 MULTISPECIES: hydrogenase-2 small chain [Proteobacteria] Length: 372\nNumber of predicted TMHs: 1\nExp number of AAs in TMHs: 23.72226\nExp number, first 60 AAs: 0.22038\nTotal prob of N-in: 0.12994\noutside 1 331\nTMhelix 332 354\ninside 355 372

9640 GCF_000704545.1_ASM70454v1 Escherichia coli 2-460-02_S4_C3 Proteobacteria; Gammaproteobacteria; Enterobacterales; Enterobacteriaceae; Escherichia; Escherichia coli MNNEETFYQAMRRQGVTRRSFLKYCSLAATSLGLGAGMAPKIAWALENKPRIPVVWIHGL WP_001058323.1 MULTISPECIES: hydrogenase-1 small chain [Proteobacteria] Length: 372\nNumber of predicted TMHs: 2\nExp number of AAs in TMHs: 37.2608499999999\nExp number, first 60 AAs: 19.52923\nTotal prob of N-in: 0.94351\nPOSSIBLE N-term signal sequence\ninside 1 19\nTMhelix 20 39\noutside 40 325\nTMhelix 326 348\ninside 349 372

9641 GCF_000704565.1_ASM70456v1 Escherichia coli 2-474-04_S1_C1 Proteobacteria; Gammaproteobacteria; Enterobacterales; Enterobacteriaceae; Escherichia; Escherichia coli MNNEETFYQAMRRQGVTRRSFLKYCSLAATSLGLGAGMAPKIAWALENKPRIPVVWIHGL WP_001058323.1 MULTISPECIES: hydrogenase-1 small chain [Proteobacteria] Length: 372\nNumber of predicted TMHs: 2\nExp number of AAs in TMHs: 37.2608499999999\nExp number, first 60 AAs: 19.52923\nTotal prob of N-in: 0.94351\nPOSSIBLE N-term signal sequence\ninside 1 19\nTMhelix 20 39\noutside 40 325\nTMhelix 326 348\ninside 349 372

9642 GCF_000704565.1_ASM70456v1 Escherichia coli 2-474-04_S1_C1 Proteobacteria; Gammaproteobacteria; Enterobacterales; Enterobacteriaceae; Escherichia; Escherichia coli MTGDNTLIHSHGINRRDFMKLCAALAATMGLSSKAAAEMAESVTNPQRPPVIWIGAQECT WP_000145410.1 MULTISPECIES: hydrogenase-2 small chain [Proteobacteria] Length: 372\nNumber of predicted TMHs: 1\nExp number of AAs in TMHs: 23.72226\nExp number, first 60 AAs: 0.22038\nTotal prob of N-in: 0.12994\noutside 1 331\nTMhelix 332 354\ninside 355 372

9643 GCF_000704645.1_ASM70464v1 Escherichia coli 2-474-04_S1_C2 Proteobacteria; Gammaproteobacteria; Enterobacterales; Enterobacteriaceae; Escherichia; Escherichia coli MNNEETFYQAMRRQGVTRRSFLKYCSLAATSLGLGAGMAPKIAWALENKPRIPVVWIHGL WP_001058323.1 MULTISPECIES: hydrogenase-1 small chain [Proteobacteria] Length: 372\nNumber of predicted TMHs: 2\nExp number of AAs in TMHs: 37.2608499999999\nExp number, first 60 AAs: 19.52923\nTotal prob of N-in: 0.94351\nPOSSIBLE N-term signal sequence\ninside 1 19\nTMhelix 20 39\noutside 40 325\nTMhelix 326 348\ninside 349 372

9644 GCF_000704645.1_ASM70464v1 Escherichia coli 2-474-04_S1_C2 Proteobacteria; Gammaproteobacteria; Enterobacterales; Enterobacteriaceae; Escherichia; Escherichia coli MTGDNTLIHSHGINRRDFMKLCAALAATMGLSSKAAAEMAESVTNPQRPPVIWIGAQECT WP_000145410.1 MULTISPECIES: hydrogenase-2 small chain [Proteobacteria] Length: 372\nNumber of predicted TMHs: 1\nExp number of AAs in TMHs: 23.72226\nExp number, first 60 AAs: 0.22038\nTotal prob of N-in: 0.12994\noutside 1 331\nTMhelix 332 354\ninside 355 372

9645 GCF_000704585.1_ASM70458v1 Escherichia coli 2-474-04_S3_C1 Proteobacteria; Gammaproteobacteria; Enterobacterales; Enterobacteriaceae; Escherichia; Escherichia coli MNNEETFYQAMRRQGVTRRSFLKYCSLAATSLGLGAGMAPKIAWALENKPRIPVVWIHGL WP_001058323.1 MULTISPECIES: hydrogenase-1 small chain [Proteobacteria] Length: 372\nNumber of predicted TMHs: 2\nExp number of AAs in TMHs: 37.2608499999999\nExp number, first 60 AAs: 19.52923\nTotal prob of N-in: 0.94351\nPOSSIBLE N-term signal sequence\ninside 1 19\nTMhelix 20 39\noutside 40 325\nTMhelix 326 348\ninside 349 372

9646 GCF_000704585.1_ASM70458v1 Escherichia coli 2-474-04_S3_C1 Proteobacteria; Gammaproteobacteria; Enterobacterales; Enterobacteriaceae; Escherichia; Escherichia coli MTGDNTLIHSHGINRRDFMKLCAALAATMGLSSKAAAEMAESVTNPQRPPVIWIGAQECT WP_000145410.1 MULTISPECIES: hydrogenase-2 small chain [Proteobacteria] Length: 372\nNumber of predicted TMHs: 1\nExp number of AAs in TMHs: 23.72226\nExp number, first 60 AAs: 0.22038\nTotal prob of N-in: 0.12994\noutside 1 331\nTMhelix 332 354\ninside 355 372

9647 GCF_000704605.1_ASM70460v1 Escherichia coli 2-474-04_S3_C2 Proteobacteria; Gammaproteobacteria; Enterobacterales; Enterobacteriaceae; Escherichia; Escherichia coli MTGDNTLIHSHGINRRDFMKLCAALAATMGLSSKAAAEMAESVTNPQRPPVIWIGAQECT WP_000145410.1 MULTISPECIES: hydrogenase-2 small chain [Proteobacteria] Length: 372\nNumber of predicted TMHs: 1\nExp number of AAs in TMHs: 23.72226\nExp number, first 60 AAs: 0.22038\nTotal prob of N-in: 0.12994\noutside 1 331\nTMhelix 332 354\ninside 355 372

9648 GCF_000704605.1_ASM70460v1 Escherichia coli 2-474-04_S3_C2 Proteobacteria; Gammaproteobacteria; Enterobacterales; Enterobacteriaceae; Escherichia; Escherichia coli MNNEETFYQAMRRQGVTRRSFLKYCSLAATSLGLGAGMAPKIAWALENKPRIPVVWIHGL WP_001058323.1 MULTISPECIES: hydrogenase-1 small chain [Proteobacteria] Length: 372\nNumber of predicted TMHs: 2\nExp number of AAs in TMHs: 37.2608499999999\nExp number, first 60 AAs: 19.52923\nTotal prob of N-in: 0.94351\nPOSSIBLE N-term signal sequence\ninside 1 19\nTMhelix 20 39\noutside 40 325\nTMhelix 326 348\ninside 349 372

9649 GCF_000704665.1_ASM70466v1 Escherichia coli 2-474-04_S3_C3 Proteobacteria; Gammaproteobacteria; Enterobacterales; Enterobacteriaceae; Escherichia; Escherichia coli MNNEETFYQAMRRQGVTRRSFLKYCSLAATSLGLGAGMAPKIAWALENKPRIPVVWIHGL WP_001058323.1 MULTISPECIES: hydrogenase-1 small chain [Proteobacteria] Length: 372\nNumber of predicted TMHs: 2\nExp number of AAs in TMHs: 37.2608499999999\nExp number, first 60 AAs: 19.52923\nTotal prob of N-in: 0.94351\nPOSSIBLE N-term signal sequence\ninside 1 19\nTMhelix 20 39\noutside 40 325\nTMhelix 326 348\ninside 349 372

9650 GCF_000704665.1_ASM70466v1 Escherichia coli 2-474-04_S3_C3 Proteobacteria; Gammaproteobacteria; Enterobacterales; Enterobacteriaceae; Escherichia; Escherichia coli MTGDNTLIHSHGINRRDFMKLCAALAATMGLSSKAAAEMAESVTNPQRPPVIWIGAQECT WP_000145410.1 MULTISPECIES: hydrogenase-2 small chain [Proteobacteria] Length: 372\nNumber of predicted TMHs: 1\nExp number of AAs in TMHs: 23.72226\nExp number, first 60 AAs: 0.22038\nTotal prob of N-in: 0.12994\noutside 1 331\nTMhelix 332 354\ninside 355 372

9651 GCF_000713685.1_ASM71368v1 Escherichia coli 2-474-04_S4_C1 Proteobacteria; Gammaproteobacteria; Enterobacterales; Enterobacteriaceae; Escherichia; Escherichia coli MTGDNTLIHSHGINRRDFMKLCAALAATMGLSSKAAAEMAESVTNPQRPPVIWIGAQECT WP_000145410.1 MULTISPECIES: hydrogenase-2 small chain [Proteobacteria] Length: 372\nNumber of predicted TMHs: 1\nExp number of AAs in TMHs: 23.72226\nExp number, first 60 AAs: 0.22038\nTotal prob of N-in: 0.12994\noutside 1 331\nTMhelix 332 354\ninside 355 372

9652 GCF_000713685.1_ASM71368v1 Escherichia coli 2-474-04_S4_C1 Proteobacteria; Gammaproteobacteria; Enterobacterales; Enterobacteriaceae; Escherichia; Escherichia coli MNNEETFYQAMRRQGVTRRSFLKYCSLAATSLGLGAGMAPKIAWALENKPRIPVVWIHGL WP_001295940.1 MULTISPECIES: hydrogenase [Enterobacteriaceae] Length: 372\nNumber of predicted TMHs: 2\nExp number of AAs in TMHs: 37.2608899999999\nExp number, first 60 AAs: 19.52923\nTotal prob of N-in: 0.94351\nPOSSIBLE N-term signal sequence\ninside 1 19\nTMhelix 20 39\noutside 40 325\nTMhelix 326 348\ninside 349 372

9653 GCF_000704705.1_ASM70470v1 Escherichia coli 2-474-04_S4_C2 Proteobacteria; Gammaproteobacteria; Enterobacterales; Enterobacteriaceae; Escherichia; Escherichia coli MTGDNTLIHSHGINRRDFMKLCAALAATMGLSSKAAAEMAESVTNPQRPPVIWIGAQECT WP_000145410.1 MULTISPECIES: hydrogenase-2 small chain [Proteobacteria] Length: 372\nNumber of predicted TMHs: 1\nExp number of AAs in TMHs: 23.72226\nExp number, first 60 AAs: 0.22038\nTotal prob of N-in: 0.12994\noutside 1 331\nTMhelix 332 354\ninside 355 372

9654 GCF_000704705.1_ASM70470v1 Escherichia coli 2-474-04_S4_C2 Proteobacteria; Gammaproteobacteria; Enterobacterales; Enterobacteriaceae; Escherichia; Escherichia coli MNNEETFYQAMRRQGVTRRSFLKYCSLAATSLGLGAGMAPKIAWALENKPRIPVVWIHGL WP_001295940.1 MULTISPECIES: hydrogenase [Enterobacteriaceae] Length: 372\nNumber of predicted TMHs: 2\nExp number of AAs in TMHs: 37.2608899999999\nExp number, first 60 AAs: 19.52923\nTotal prob of N-in: 0.94351\nPOSSIBLE N-term signal sequence\ninside 1 19\nTMhelix 20 39\noutside 40 325\nTMhelix 326 348\ninside 349 372

9655 GCF_000704725.1_ASM70472v1 Escherichia coli 2-474-04_S4_C3 Proteobacteria; Gammaproteobacteria; Enterobacterales; Enterobacteriaceae; Escherichia; Escherichia coli MTGDNTLIHSHGINRRDFMKLCAALAATMGLSSKAAAEMAESVTNPQRPPVIWIGAQECT WP_000145410.1 MULTISPECIES: hydrogenase-2 small chain [Proteobacteria] Length: 372\nNumber of predicted TMHs: 1\nExp number of AAs in TMHs: 23.72226\nExp number, first 60 AAs: 0.22038\nTotal prob of N-in: 0.12994\noutside 1 331\nTMhelix 332 354\ninside 355 372

9656	GCF_000704725.1_ASM70472v1	Escherichia coli 2-474-04_S4_C3	Proteobacteria; Gammaproteobacteria; Enterobacterales; Enterobacteriaceae; Escherichia; Escherichia coli
MNNEETFYQAMRRQGVTRRSFLKYCSLAATSLGLGAGMAPKIAWALENKPRIPVVWIHGL	WP_001295940.1
MULTISPECIES: hydrogenase [Enterobacteriaceae]	Length: 372\nNumber of predicted TMHs: 2\nExp number of AAs in TMHs: 37.2608899999999\nExp number, first 60 AAs: 19.52923\nTotal prob of N-in: 0.94351\nPOSSIBLE N-term signal sequence\ninside 1 19\nTMhelix 20 39\noutside 40 325\nTMhelix 326 348\ninside 349 372

9657	GCF_000225065.1_ASM22506v2	Escherichia coli 2534-86	Proteobacteria; Gammaproteobacteria; Enterobacterales; Enterobacteriaceae; Escherichia; Escherichia coli
MTGDNTLIHSHGINRRDFMKLCAALAATMGLSSKAAAEMAESVTNPQRPPVIWIGAQECT	WP_000145410.1
MULTISPECIES: hydrogenase-2 small chain [Proteobacteria]	Length: 372\nNumber of predicted TMHs: 1\nExp number of AAs in TMHs: 23.72226\nExp number, first 60 AAs: 0.22038\nTotal prob of N-in: 0.12994\noutside 1 331\nTMhelix 332 354\ninside 355 372

9658	GCF_000225065.1_ASM22506v2	Escherichia coli 2534-86	Proteobacteria; Gammaproteobacteria; Enterobacterales; Enterobacteriaceae; Escherichia; Escherichia coli
MNNEETFYQAMRRQGVTRRSFLKYCSLAATSLGLGAGMAPKIAWALENKPRIPVVWIHGL	WP_001058323.1
MULTISPECIES: hydrogenase-1 small chain [Proteobacteria]	Length: 372\nNumber of predicted TMHs: 2\nExp number of AAs in TMHs: 37.2608499999999\nExp number, first 60 AAs: 19.52923\nTotal prob of N-in: 0.94351\nPOSSIBLE N-term signal sequence\ninside 1 19\nTMhelix 20 39\noutside 40 325\nTMhelix 326 348\ninside 349 372

9659	GCF_000355155.2_ASM35515v2	Escherichia coli 2719100	Proteobacteria; Gammaproteobacteria; Enterobacterales; Enterobacteriaceae; Escherichia; Escherichia coli
MTGDNTLIHSHGINRRDFMKLCAALAATMGLSSKAAAEMAESVTNPQRPPVIWIGAQECT	WP_000145410.1
MULTISPECIES: hydrogenase-2 small chain [Proteobacteria]	Length: 372\nNumber of predicted TMHs: 1\nExp number of AAs in TMHs: 23.72226\nExp number, first 60 AAs: 0.22038\nTotal prob of N-in: 0.12994\noutside 1 331\nTMhelix 332 354\ninside 355 372

9660	GCF_000355155.2_ASM35515v2	Escherichia coli 2719100	Proteobacteria; Gammaproteobacteria; Enterobacterales; Enterobacteriaceae; Escherichia; Escherichia coli
MNNEETFYQAMRRQGVTRRSFLKYCSLAATSLGLGAGMAPKIAWALENKPRIPVVWIHGL	WP_001058323.1
MULTISPECIES: hydrogenase-1 small chain [Proteobacteria]	Length: 372\nNumber of predicted TMHs: 2\nExp number of AAs in TMHs: 37.2608499999999\nExp number, first 60 AAs: 19.52923\nTotal prob of N-in: 0.94351\nPOSSIBLE N-term signal sequence\ninside 1 19\nTMhelix 20 39\noutside 40 325\nTMhelix 326 348\ninside 349 372

9661	GCF_000355175.1_ASM35517v1	Escherichia coli 2720900	Proteobacteria; Gammaproteobacteria; Enterobacterales; Enterobacteriaceae; Escherichia; Escherichia coli
MNNEETFYQAMRRQGVTRRSFLKYCSLAATSLGLGAGMAPKIAWALENKPRIPVVWIHGL	WP_001295940.1
MULTISPECIES: hydrogenase [Enterobacteriaceae]	Length: 372\nNumber of predicted TMHs: 2\nExp number of AAs in TMHs: 37.2608899999999\nExp number, first 60 AAs: 19.52923\nTotal prob of N-in: 0.94351\nPOSSIBLE N-term signal sequence\ninside 1 19\nTMhelix 20 39\noutside 40 325\nTMhelix 326 348\ninside 349 372

9662	GCF_000355175.1_ASM35517v1	Escherichia coli 2720900	Proteobacteria; Gammaproteobacteria; Enterobacterales; Enterobacteriaceae; Escherichia; Escherichia coli
MTGDNTLIHSHGINRRDFMKLCAALAATMGLSSKAAAEMAESVTNPQRPPVIWIGAQECT	WP_000145410.1
MULTISPECIES: hydrogenase-2 small chain [Proteobacteria]	Length: 372\nNumber of predicted TMHs: 1\nExp number of AAs in TMHs: 23.72226\nExp number, first 60 AAs: 0.22038\nTotal prob of N-in: 0.12994\noutside 1 331\nTMhelix 332 354\ninside 355 372

9663	GCF_000356185.2_ASM35618v2	Escherichia coli 2722950	Proteobacteria; Gammaproteobacteria; Enterobacterales; Enterobacteriaceae; Escherichia; Escherichia coli
MNNEETFYQAMRRQGVTRRSFLKYCSLAATSLGLGAGMAPKIAWALENKPRIPVVWIHGL	WP_001058323.1
MULTISPECIES: hydrogenase-1 small chain [Proteobacteria]	Length: 372\nNumber of predicted TMHs: 2\nExp number of AAs in TMHs: 37.2608499999999\nExp number, first 60 AAs: 19.52923\nTotal prob of N-in: 0.94351\nPOSSIBLE N-term signal sequence\ninside 1 19\nTMhelix 20 39\noutside 40 325\nTMhelix 326 348\ninside 349 372

9664	GCF_000356185.2_ASM35618v2	Escherichia coli 2722950	Proteobacteria; Gammaproteobacteria; Enterobacterales; Enterobacteriaceae; Escherichia; Escherichia coli
MTGDNTLIHSHGINRRDFMKLCAALAATMGLSSKAAAEMAESVTNPQRPPVIWIGAQECT	WP_000145410.1
MULTISPECIES: hydrogenase-2 small chain [Proteobacteria]	Length: 372\nNumber of predicted TMHs: 1\nExp number of AAs in TMHs: 23.72226\nExp number, first 60 AAs: 0.22038\nTotal prob of N-in: 0.12994\noutside 1 331\nTMhelix 332 354\ninside 355 372

9665	GCF_000356625.2_ASM35662v2	Escherichia coli 2726800	Proteobacteria; Gammaproteobacteria; Enterobacterales; Enterobacteriaceae; Escherichia; Escherichia coli
MNNEETFYQAMRRQGVTRRSFLKYCSLAATSLGLGAGMAPKIAWALENKPRIPVVWIHGL	WP_001058323.1
MULTISPECIES: hydrogenase-1 small chain [Proteobacteria]	Length: 372\nNumber of predicted TMHs: 2\nExp number of AAs in TMHs: 37.2608499999999\nExp number, first 60 AAs: 19.52923\nTotal prob of N-in: 0.94351\nPOSSIBLE N-term signal sequence\ninside 1 19\nTMhelix 20 39\noutside 40 325\nTMhelix 326 348\ninside 349 372

9666	GCF_000356625.2_ASM35662v2	Escherichia coli 2726800	Proteobacteria; Gammaproteobacteria; Enterobacterales; Enterobacteriaceae; Escherichia; Escherichia coli
MTGDNTLIHSHGINRRDFMKLCAALAATMGLSTKAAAEMAESVTNPQRPPVIWIGAQECT	WP_001507912.1
hydrogenase-2 small chain [Escherichia coli]	Length: 372\nNumber of predicted TMHs: 1\nExp number of AAs in TMHs: 23.86013\nExp number, first 60 AAs: 0.3669\nTotal prob of N-in: 0.13628\noutside 1 331\nTMhelix 332 354\ninside 355 372

9667	GCF_000356145.2_ASM35614v2	Escherichia coli 2726950	Proteobacteria; Gammaproteobacteria; Enterobacterales; Enterobacteriaceae; Escherichia; Escherichia coli
MNNEETFYQAMRRQGVTRRSFLKYCSLAATSLGLGAGMAPKIAWALENKPRIPVVWIHGL	WP_001058323.1
MULTISPECIES: hydrogenase-1 small chain [Proteobacteria]	Length: 372\nNumber of predicted TMHs: 2\nExp number of AAs in TMHs: 37.2608499999999\nExp number, first 60 AAs: 19.52923\nTotal prob of N-in: 0.94351\nPOSSIBLE N-term signal sequence\ninside 1 19\nTMhelix 20 39\noutside 40 325\nTMhelix 326 348\ninside 349 372

9668	GCF_000356145.2_ASM35614v2	Escherichia coli 2726950	Proteobacteria; Gammaproteobacteria; Enterobacterales; Enterobacteriaceae; Escherichia; Escherichia coli
MTGDNTLIHSHGINRRDFMKLCAALAATMGLSSKAAAEMAESVTNPQRPPVIWIGAQECT	WP_001655824.1
hydrogenase-2 small chain [Escherichia coli]	Length: 372\nNumber of predicted TMHs: 1\nExp number of AAs in TMHs: 23.72049\nExp number, first 60 AAs: 0.22038\nTotal prob of N-in: 0.12989\noutside 1 331\nTMhelix 332 354\ninside 355 372

9669	GCF_000356065.2_ASM35606v2	Escherichia coli 2729250	Proteobacteria; Gammaproteobacteria; Enterobacterales; Enterobacteriaceae; Escherichia; Escherichia coli
MTGDNTLIHSHGINRRDFMKLCAALAATMGLSSKAAAEMAESVTNPQRPPVIWIGAQECT	WP_000145410.1
MULTISPECIES: hydrogenase-2 small chain [Proteobacteria]	Length: 372\nNumber of predicted TMHs: 1\nExp number of AAs in TMHs: 23.72226\nExp number, first 60 AAs: 0.22038\nTotal prob of N-in: 0.12994\noutside 1 331\nTMhelix 332 354\ninside 355 372

9670	GCF_000356065.2_ASM35606v2	Escherichia coli 2729250	Proteobacteria; Gammaproteobacteria; Enterobacterales; Enterobacteriaceae; Escherichia; Escherichia coli
MNNEETFYKAMRRQGVTRRSFLKYCSLAATSLGLGAGMAPKIVWALENKPRIPVVWIHGL	WP_001736230.1
hydrogenase-1 small chain [Escherichia coli]	Length: 372\nNumber of predicted TMHs: 2\nExp number of AAs in TMHs: 37.81688\nExp number, first 60 AAs: 20.05665\nTotal prob of N-in: 0.95630\nPOSSIBLE N-term signal sequence\ninside 1 20\nTMhelix 21 43\noutside 44 325\nTMhelix 326 348\ninside 349 372

9671	GCF_000356325.2_ASM35632v2	Escherichia coli 2730350	Proteobacteria; Gammaproteobacteria; Enterobacterales; Enterobacteriaceae; Escherichia; Escherichia coli
MNNEETFYQAMRRQGVTRRSFLKYCSLAATSLGLGAGMAPKIAWALENKPRIPVVWIHGL	WP_001058320.1
hydrogenase [Escherichia coli]	Length: 372\nNumber of predicted TMHs: 2\nExp number of AAs in TMHs: 37.26082\nExp number, first 60 AAs: 19.52923\nTotal prob of N-in: 0.94351\nPOSSIBLE N-term signal sequence\ninside 1 19\nTMhelix 20 39\noutside 40 325\nTMhelix 326 348\ninside 349 372

9672	GCF_000356325.2_ASM35632v2	Escherichia coli 2730350	Proteobacteria; Gammaproteobacteria; Enterobacterales; Enterobacteriaceae; Escherichia; Escherichia coli
MTGDNTLIHSHGINRRDFMKLCAALAATMGLSSKAAAEMAESVTNPQRPPVIWIGAQECT	WP_000145410.1

MULTISPECIES: hydrogenase-2 small chain [Proteobacteria] Length: 372\nNumber of predicted TMHs: 1\nExp number of AAs in TMHs: 23.72226\nExp number, first 60 AAs: 0.22038\nTotal prob of N-in: 0.12994\noutside 1 331\nTMhelix 332 354\ninside 355 372

9673 GCF_000356205.2_ASM35620v2 Escherichia coli 2730450 Proteobacteria; Gammaproteobacteria; Enterobacterales; Enterobacteriaceae; Escherichia; Escherichia coli MNNEETFYQAMRRQGVTRRSFLKYCSLAATSLGLGAGMAPKIAWALENKPRIPVVWIHGL WP_001058323.1 MULTISPECIES: hydrogenase-1 small chain [Proteobacteria] Length: 372\nNumber of predicted TMHs: 2\nExp number of AAs in TMHs: 37.2608499999999\nExp number, first 60 AAs: 19.52923\nTotal prob of N-in: 0.94351\nPOSSIBLE N-term signal sequence\ninside 1 19\nTMhelix 20 39\noutside 40 325\nTMhelix 326 348\ninside 349 372

9674 GCF_000356205.2_ASM35620v2 Escherichia coli 2730450 Proteobacteria; Gammaproteobacteria; Enterobacterales; Enterobacteriaceae; Escherichia; Escherichia coli MTGDNTLIHSHGINRRDFMKLCAALAATMGLSSKAAAEMAESVTNPQRPPVIWIGAQECT WP_000145410.1 MULTISPECIES: hydrogenase-2 small chain [Proteobacteria] Length: 372\nNumber of predicted TMHs: 1\nExp number of AAs in TMHs: 23.72226\nExp number, first 60 AAs: 0.22038\nTotal prob of N-in: 0.12994\noutside 1 331\nTMhelix 332 354\ninside 355 372

9675 GCF_000354895.2_ASM35489v2 Escherichia coli 2731150 Proteobacteria; Gammaproteobacteria; Enterobacterales; Enterobacteriaceae; Escherichia; Escherichia coli MNNEETFYQAMRRQGVTRRSFLKYCSLAATSLGLGAGMAPKIAWALENKPRIPVVWIHGL WP_001058323.1 MULTISPECIES: hydrogenase-1 small chain [Proteobacteria] Length: 372\nNumber of predicted TMHs: 2\nExp number of AAs in TMHs: 37.2608499999999\nExp number, first 60 AAs: 19.52923\nTotal prob of N-in: 0.94351\nPOSSIBLE N-term signal sequence\ninside 1 19\nTMhelix 20 39\noutside 40 325\nTMhelix 326 348\ninside 349 372

9676 GCF_000354895.2_ASM35489v2 Escherichia coli 2731150 Proteobacteria; Gammaproteobacteria; Enterobacterales; Enterobacteriaceae; Escherichia; Escherichia coli MTGDNTLIHSHGINRRDFMKLCAALAATMGLSSKAAAEMAESVTNPQRPPVIWIGAQECT WP_000145410.1 MULTISPECIES: hydrogenase-2 small chain [Proteobacteria] Length: 372\nNumber of predicted TMHs: 1\nExp number of AAs in TMHs: 23.72226\nExp number, first 60 AAs: 0.22038\nTotal prob of N-in: 0.12994\noutside 1 331\nTMhelix 332 354\ninside 355 372

9677 GCF_000356925.2_ASM35692v2 Escherichia coli 2733950 Proteobacteria; Gammaproteobacteria; Enterobacterales; Enterobacteriaceae; Escherichia; Escherichia coli MNNEETFYQAMRRQGVTRRSFLKYCSLAATSLGLGAGMAPKIAWALENKPRIPVVWIHGL WP_001058323.1 MULTISPECIES: hydrogenase-1 small chain [Proteobacteria] Length: 372\nNumber of predicted TMHs: 2\nExp number of AAs in TMHs: 37.2608499999999\nExp number, first 60 AAs: 19.52923\nTotal prob of N-in: 0.94351\nPOSSIBLE N-term signal sequence\ninside 1 19\nTMhelix 20 39\noutside 40 325\nTMhelix 326 348\ninside 349 372

9678 GCF_000356925.2_ASM35692v2 Escherichia coli 2733950 Proteobacteria; Gammaproteobacteria; Enterobacterales; Enterobacteriaceae; Escherichia; Escherichia coli MTGDNTLIHSHGINRRDFMKLCAALAATMGLSSKAAAEMAESVTNPQRPPVIWIGAQECT WP_001506281.1 hydrogenase-2 small chain [Escherichia coli] Length: 372\nNumber of predicted TMHs: 1\nExp number of AAs in TMHs: 23.72225\nExp number, first 60 AAs: 0.22038\nTotal prob of N-in: 0.12994\noutside 1 331\nTMhelix 332 354\ninside 355 372

9679 GCF_000356085.2_ASM35608v2 Escherichia coli 2735000 Proteobacteria; Gammaproteobacteria; Enterobacterales; Enterobacteriaceae; Escherichia; Escherichia coli MTGDNPLIHSHGINRRDFMKLCVALAATMGLSSKAAAEMAESVTNPQRPPVIWIGAQECT WP_001766310.1 hydrogenase-2 small chain [Escherichia coli] Length: 372\nNumber of predicted TMHs: 1\nExp number of AAs in TMHs: 22.72313\nExp number, first 60 AAs: 0.22827\nTotal prob of N-in: 0.06927\noutside 1 331\nTMhelix 332 354\ninside 355 372

9680 GCF_000356085.2_ASM35608v2 Escherichia coli 2735000 Proteobacteria; Gammaproteobacteria; Enterobacterales; Enterobacteriaceae; Escherichia; Escherichia coli MNNEETFYQAMRRQGVTRRSFLKYCSLAATSLGLGAGMAPKIAWALENKPRIPVVWIHGL WP_001295940.1 MULTISPECIES: hydrogenase [Enterobacteriaceae] Length: 372\nNumber of predicted TMHs: 2\nExp number of AAs in TMHs: 37.2608899999999\nExp number, first 60 AAs: 19.52923\nTotal prob of N-in:

0.94351\nPOSSIBLE N-term signal sequence\ninside 1 19\nTMhelix 20 39\noutside 40 325\nTMhelix 326 348\ninside 349 372

9681 GCF_000356225.2_ASM35622v2 Escherichia coli 2741950 Proteobacteria; Gammaproteobacteria; Enterobacterales; Enterobacteriaceae; Escherichia; Escherichia coli
MNNEETFYQAMRRQGVTRRSFLKYCSLAATSLGLGAGMAPKIAWALENKPRIPVVWIHGL WP_001058323.1
MULTISPECIES: hydrogenase-1 small chain [Proteobacteria] Length: 372\nNumber of predicted TMHs: 2\nExp number of AAs in TMHs: 37.2608499999999\nExp number, first 60 AAs: 19.52923\nTotal prob of N-in: 0.94351\nPOSSIBLE N-term signal sequence\ninside 1 19\nTMhelix 20 39\noutside 40 325\nTMhelix 326 348\ninside 349 372

9682 GCF_000356225.2_ASM35622v2 Escherichia coli 2741950 Proteobacteria; Gammaproteobacteria; Enterobacterales; Enterobacteriaceae; Escherichia; Escherichia coli
MTGDNTLIHSHGINRRDFMKLCAALAATMGLSSKAAAEMAESVTNPQRPPVIWIGAQECT WP_000145410.1
MULTISPECIES: hydrogenase-2 small chain [Proteobacteria] Length: 372\nNumber of predicted TMHs: 1\nExp number of AAs in TMHs: 23.72226\nExp number, first 60 AAs: 0.22038\nTotal prob of N-in: 0.12994\noutside 1 331\nTMhelix 332 354\ninside 355 372

9683 GCF_000354775.2_ASM35477v2 Escherichia coli 2747800 Proteobacteria; Gammaproteobacteria; Enterobacterales; Enterobacteriaceae; Escherichia; Escherichia coli
MNNEETFYQAMRRQGVTRRSFLKYCSLAATSLGLGAGMAPKIVWALENKPRIPVVWIHGL WP_001398876.1
hydrogenase-1 small chain [Escherichia coli] Length: 372\nNumber of predicted TMHs: 2\nExp number of AAs in TMHs: 37.60268\nExp number, first 60 AAs: 19.84677\nTotal prob of N-in: 0.94621\nPOSSIBLE N-term signal sequence\ninside 1 20\nTMhelix 21 43\noutside 44 325\nTMhelix 326 348\ninside 349 372

9684 GCF_000354775.2_ASM35477v2 Escherichia coli 2747800 Proteobacteria; Gammaproteobacteria; Enterobacterales; Enterobacteriaceae; Escherichia; Escherichia coli
MTGDNTLIHSHGINRRDFMKLCAALAATMGLSSKAAAEMAESVTNPQRPPVIWIGAQECT WP_000145410.1
MULTISPECIES: hydrogenase-2 small chain [Proteobacteria] Length: 372\nNumber of predicted TMHs: 1\nExp number of AAs in TMHs: 23.72226\nExp number, first 60 AAs: 0.22038\nTotal prob of N-in: 0.12994\noutside 1 331\nTMhelix 332 354\ninside 355 372

9685 GCF_000354875.1_ASM35487v1 Escherichia coli 2749250 Proteobacteria; Gammaproteobacteria; Enterobacterales; Enterobacteriaceae; Escherichia; Escherichia coli
MTGDNTLIHSHGINRRDFMKLCAALAATMGLSSKAAAEMAESVTNPQRPPVIWIGAQECT WP_000145410.1
MULTISPECIES: hydrogenase-2 small chain [Proteobacteria] Length: 372\nNumber of predicted TMHs: 1\nExp number of AAs in TMHs: 23.72226\nExp number, first 60 AAs: 0.22038\nTotal prob of N-in: 0.12994\noutside 1 331\nTMhelix 332 354\ninside 355 372

9686 GCF_000354875.1_ASM35487v1 Escherichia coli 2749250 Proteobacteria; Gammaproteobacteria; Enterobacterales; Enterobacteriaceae; Escherichia; Escherichia coli
MNNEETFYQAMRRQGVTRRSFLKYCSLAATSLGLGAGMAPKIVWALENKPRIPVVWIHGL WP_001398876.1
hydrogenase-1 small chain [Escherichia coli] Length: 372\nNumber of predicted TMHs: 2\nExp number of AAs in TMHs: 37.60268\nExp number, first 60 AAs: 19.84677\nTotal prob of N-in: 0.94621\nPOSSIBLE N-term signal sequence\ninside 1 20\nTMhelix 21 43\noutside 44 325\nTMhelix 326 348\ninside 349 372

9687 GCF_000355395.2_ASM35539v2 Escherichia coli 2756500 Proteobacteria; Gammaproteobacteria; Enterobacterales; Enterobacteriaceae; Escherichia; Escherichia coli
MTGDNTLIHSHGINRRDFMKLCVALAATMGLSSKAAAEMAESVTNPQRPPVIWIGAQECT WP_000145425.1
hydrogenase 2 small subunit [Escherichia coli] Length: 372\nNumber of predicted TMHs: 1\nExp number of AAs in TMHs: 23.94354\nExp number, first 60 AAs: 0.45405\nTotal prob of N-in: 0.13997\noutside 1 331\nTMhelix 332 354\ninside 355 372

9688 GCF_000355395.2_ASM35539v2 Escherichia coli 2756500 Proteobacteria; Gammaproteobacteria; Enterobacterales; Enterobacteriaceae; Escherichia; Escherichia coli
MNNEETFYQAMRRQGVTRRSFLKYCSLAATSLGLGAGMAPKIAWALENKPRIPVVWIHGL WP_001295940.1
MULTISPECIES: hydrogenase [Enterobacteriaceae] Length: 372\nNumber of predicted TMHs: 2\nExp number of AAs in TMHs: 37.2608899999999\nExp number, first 60 AAs: 19.52923\nTotal prob of N-in: 0.94351\nPOSSIBLE N-term signal sequence\ninside 1 19\nTMhelix 20 39\noutside 40 325\nTMhelix 326 348\ninside 349 372

9689 GCF_000355535.1_ASM35553v1 Escherichia coli 2762100 Proteobacteria; Gammaproteobacteria; Enterobacterales; Enterobacteriaceae; Escherichia; Escherichia coli
MTGDNPLIHSHGINRRDFMKLCAALAATMGLSSKAAAEMAESVTNPQRPPVIWIGAQECT WP_001461571.1
hydrogenase-2 small chain [Escherichia coli] Length: 372\nNumber of predicted TMHs: 1\nExp number of AAs in TMHs: 22.60469\nExp number, first 60 AAs: 0.1106\nTotal prob of N-in: 0.06393\noutside 1 331\nTMhelix 332 354\ninside 355 372

9690 GCF_000355535.1_ASM35553v1 Escherichia coli 2762100 Proteobacteria; Gammaproteobacteria; Enterobacterales; Enterobacteriaceae; Escherichia; Escherichia coli
MNNEETFYQAMRRQGVTRRSFLKYCSLAATSLGLGAGMAPKIAWALENKPRIPVVWIHGL WP_001425448.1
hydrogenase-1 small chain [Escherichia coli] Length: 372\nNumber of predicted TMHs: 2\nExp number of AAs in TMHs: 37.14002\nExp number, first 60 AAs: 19.53586\nTotal prob of N-in: 0.94339\nPOSSIBLE N-term signal sequence\ninside 1 19\nTMhelix 20 39\noutside 40 325\nTMhelix 326 348\ninside 349 372

9691 GCF_000355615.2_ASM35561v2 Escherichia coli 2770900 Proteobacteria; Gammaproteobacteria; Enterobacterales; Enterobacteriaceae; Escherichia; Escherichia coli
MTGDNTLIHSHGINRRDFMKLCAALAATMGLSSKAAAEMAESVTNPQRPPVIWIGAQECT WP_000145410.1
MULTISPECIES: hydrogenase-2 small chain [Proteobacteria] Length: 372\nNumber of predicted TMHs: 1\nExp number of AAs in TMHs: 23.72226\nExp number, first 60 AAs: 0.22038\nTotal prob of N-in: 0.12994\noutside 1 331\nTMhelix 332 354\ninside 355 372

9692 GCF_000355615.2_ASM35561v2 Escherichia coli 2770900 Proteobacteria; Gammaproteobacteria; Enterobacterales; Enterobacteriaceae; Escherichia; Escherichia coli
MNNEETFYQAMRRQGVTRRSFLKYCSLAATSLGLGAGMAPKIVWALENKPRIPVVWIHGL WP_001398876.1
hydrogenase-1 small chain [Escherichia coli] Length: 372\nNumber of predicted TMHs: 2\nExp number of AAs in TMHs: 37.60268\nExp number, first 60 AAs: 19.84677\nTotal prob of N-in: 0.94621\nPOSSIBLE N-term signal sequence\ninside 1 20\nTMhelix 21 43\noutside 44 325\nTMhelix 326 348\ninside 349 372

9693 GCF_000355555.2_ASM35555v2 Escherichia coli 2780750 Proteobacteria; Gammaproteobacteria; Enterobacterales; Enterobacteriaceae; Escherichia; Escherichia coli
MNNEETFYQAMRRQGVTRRSFLKYCSLAATSLGLGAGMAPKIAWALENKPRIPVVWIHGL WP_001058323.1
MULTISPECIES: hydrogenase-1 small chain [Proteobacteria] Length: 372\nNumber of predicted TMHs: 2\nExp number of AAs in TMHs: 37.2608499999999\nExp number, first 60 AAs: 19.52923\nTotal prob of N-in: 0.94351\nPOSSIBLE N-term signal sequence\ninside 1 19\nTMhelix 20 39\noutside 40 325\nTMhelix 326 348\ninside 349 372

9694 GCF_000355555.2_ASM35555v2 Escherichia coli 2780750 Proteobacteria; Gammaproteobacteria; Enterobacterales; Enterobacteriaceae; Escherichia; Escherichia coli
MTGDNTLIHSHGINRRDFMKLCAALAATMGLSSKAAAEMAESVTNPQRPPVIWIGAQECT WP_000145410.1
MULTISPECIES: hydrogenase-2 small chain [Proteobacteria] Length: 372\nNumber of predicted TMHs: 1\nExp number of AAs in TMHs: 23.72226\nExp number, first 60 AAs: 0.22038\nTotal prob of N-in: 0.12994\noutside 1 331\nTMhelix 332 354\ninside 355 372

9695 GCF_000355575.2_ASM35557v2 Escherichia coli 2785200 Proteobacteria; Gammaproteobacteria; Enterobacterales; Enterobacteriaceae; Escherichia; Escherichia coli
MNNEETFYQAMRRQGVTRRSFLKYCSLAATSLGLGAGMAPKIAWALENKPRIPVVWIHGL WP_001058323.1
MULTISPECIES: hydrogenase-1 small chain [Proteobacteria] Length: 372\nNumber of predicted TMHs: 2\nExp number of AAs in TMHs: 37.2608499999999\nExp number, first 60 AAs: 19.52923\nTotal prob of N-in: 0.94351\nPOSSIBLE N-term signal sequence\ninside 1 19\nTMhelix 20 39\noutside 40 325\nTMhelix 326 348\ninside 349 372

9696 GCF_000355575.2_ASM35557v2 Escherichia coli 2785200 Proteobacteria; Gammaproteobacteria; Enterobacterales; Enterobacteriaceae; Escherichia; Escherichia coli
MTGDNTLIHSHGINRRDFMKLCAALAATMGLSSKAAAEMAESVTNPQRPPVIWIGAQECT WP_000145410.1
MULTISPECIES: hydrogenase-2 small chain [Proteobacteria] Length: 372\nNumber of predicted TMHs: 1\nExp number of AAs in TMHs: 23.72226\nExp number, first 60 AAs: 0.22038\nTotal prob of N-in: 0.12994\noutside 1 331\nTMhelix 332 354\ninside 355 372

9697 GCF_000355595.1_ASM35559v1 Escherichia coli 2788150 Proteobacteria; Gammaproteobacteria; Enterobacterales; Enterobacteriaceae; Escherichia; Escherichia coli
MTGDNPLIHSHGINRRDFMKLCAALAATMGLSSKAAAEMAESVTNPQRPPVIWIGAQECT WP_001461571.1

hydrogenase-2 small chain [Escherichia coli]	Length: 372\nNumber of predicted TMHs:  1\nExp number of AAs in TMHs: 22.60469\nExp number, first 60 AAs:  0.1106\nTotal prob of N-in:   0.06393\noutside   1   331\nTMhelix   332   354\ninside   355   372

9698	GCF_000355595.1_ASM35559v1	Escherichia coli 2788150	Proteobacteria; Gammaproteobacteria; Enterobacterales; Enterobacteriaceae; Escherichia; Escherichia coli
MNNEETFYQAMRRQGVTRRSFLKYCSLAATSLGLGAGMAPKIAWALENKPRIPVVWIHGL	WP_001058323.1
MULTISPECIES: hydrogenase-1 small chain [Proteobacteria]	Length: 372\nNumber of predicted TMHs: 2\nExp number of AAs in TMHs: 37.2608499999999\nExp number, first 60 AAs:  19.52923\nTotal prob of N-in: 0.94351\nPOSSIBLE N-term signal sequence\ninside   1   19\nTMhelix   20   39\noutside   40   325\nTMhelix   326   348\ninside   349   372

9699	GCF_000503575.1_Ecoli27A	Escherichia coli 27A	Proteobacteria; Gammaproteobacteria; Enterobacterales; Enterobacteriaceae; Escherichia; Escherichia coli
MTGDNTLIHSHGINRRDFMKLCAALAATMGLSSKAAAEMAESVTNPQRPPVIWIGAQECT	WP_000145410.1
MULTISPECIES: hydrogenase-2 small chain [Proteobacteria]	Length: 372\nNumber of predicted TMHs: 1\nExp number of AAs in TMHs: 23.72226\nExp number, first 60 AAs:  0.22038\nTotal prob of N-in:   0.12994\noutside   1   331\nTMhelix   332   354\ninside   355   372

9700	GCF_000503575.1_Ecoli27A	Escherichia coli 27A	Proteobacteria; Gammaproteobacteria; Enterobacterales; Enterobacteriaceae; Escherichia; Escherichia coli
MNNEETFYQAMRRQGVTRRSFLKYCSLAATSLGLGAGMAPKIAWALENKPRIPVVWIHGL	WP_001058320.1
hydrogenase [Escherichia coli]	Length: 372\nNumber of predicted TMHs:  2\nExp number of AAs in TMHs: 37.26082\nExp number, first 60 AAs:  19.52923\nTotal prob of N-in:   0.94351\nPOSSIBLE N-term signal sequence\ninside   1   19\nTMhelix   20   39\noutside   40   325\nTMhelix   326   348\ninside   349   372

9701	GCF_000355415.2_ASM35541v2	Escherichia coli 2845350	Proteobacteria; Gammaproteobacteria; Enterobacterales; Enterobacteriaceae; Escherichia; Escherichia coli
MTGDNPLIHSHGINRRDFMKLCAALAATMGLSSKAAAEMAESVTNPQRPPVIWIGAQECT	WP_001461571.1
hydrogenase-2 small chain [Escherichia coli]	Length: 372\nNumber of predicted TMHs:  1\nExp number of AAs in TMHs: 22.60469\nExp number, first 60 AAs:  0.1106\nTotal prob of N-in:   0.06393\noutside   1   331\nTMhelix   332   354\ninside   355   372

9702	GCF_000355415.2_ASM35541v2	Escherichia coli 2845350	Proteobacteria; Gammaproteobacteria; Enterobacterales; Enterobacteriaceae; Escherichia; Escherichia coli
MNNEETFYQAMRRQGVTRRSFLKYCSLAATSLGLGAGMAPKIAWALENKPRIPVVWIHGL	WP_001058323.1
MULTISPECIES: hydrogenase-1 small chain [Proteobacteria]	Length: 372\nNumber of predicted TMHs: 2\nExp number of AAs in TMHs: 37.2608499999999\nExp number, first 60 AAs:  19.52923\nTotal prob of N-in: 0.94351\nPOSSIBLE N-term signal sequence\ninside   1   19\nTMhelix   20   39\noutside   40   325\nTMhelix   326   348\ninside   349   372

9703	GCF_000355435.2_ASM35543v2	Escherichia coli 2845650	Proteobacteria; Gammaproteobacteria; Enterobacterales; Enterobacteriaceae; Escherichia; Escherichia coli
MTGDNPLIHSHGINRRDFMKLCAALAATMGLSSKAAAEMAESVTNPQRPPVIWIGAQECT	WP_001461571.1
hydrogenase-2 small chain [Escherichia coli]	Length: 372\nNumber of predicted TMHs:  1\nExp number of AAs in TMHs: 22.60469\nExp number, first 60 AAs:  0.1106\nTotal prob of N-in:   0.06393\noutside   1   331\nTMhelix   332   354\ninside   355   372

9704	GCF_000355435.2_ASM35543v2	Escherichia coli 2845650	Proteobacteria; Gammaproteobacteria; Enterobacterales; Enterobacteriaceae; Escherichia; Escherichia coli
MNNEETFYQAMRRQGVTRRSFLKYCSLAATSLGLGAGMAPKIAWALENKPRIPVVWIHGL	WP_001717971.1
hydrogenase-1 small chain [Escherichia coli]	Length: 372\nNumber of predicted TMHs:  2\nExp number of AAs in TMHs: 38.00708\nExp number, first 60 AAs:  19.5286\nTotal prob of N-in:   0.94354\nPOSSIBLE N-term signal sequence\ninside   1   19\nTMhelix   20   39\noutside   40   325\nTMhelix   326   348\ninside   349   372

9705	GCF_000356685.2_ASM35668v2	Escherichia coli 2846750	Proteobacteria; Gammaproteobacteria; Enterobacterales; Enterobacteriaceae; Escherichia; Escherichia coli
MNNEETFYQAMRRQGVTRRSFLKYCSLAATSLGLGAGMAPKIAWALENKPRIPVVWIHGL	WP_001058323.1
MULTISPECIES: hydrogenase-1 small chain [Proteobacteria]	Length: 372\nNumber of predicted TMHs: 2\nExp number of AAs in TMHs: 37.2608499999999\nExp number, first 60 AAs:  19.52923\nTotal prob of N-in:

0.94351\nPOSSIBLE N-term signal sequence\ninside 1 19\nTMhelix 20 39\noutside 40 325\nTMhelix 326 348\ninside 349 372

9706 GCF_000356685.2_ASM35668v2 Escherichia coli 2846750 Proteobacteria; Gammaproteobacteria; Enterobacterales; Enterobacteriaceae; Escherichia; Escherichia coli MTGDNPLIHSHGINRRDFMKLCAALAATMGLSSKAAAEMAESVTNPQRPPVIWIGAQECT WP_001461571.1 hydrogenase-2 small chain [Escherichia coli] Length: 372\nNumber of predicted TMHs: 1\nExp number of AAs in TMHs: 22.60469\nExp number, first 60 AAs: 0.1106\nTotal prob of N-in: 0.06393\noutside 1 331\nTMhelix 332 354\ninside 355 372

9707 GCF_000355375.2_ASM35537v2 Escherichia coli 2848050 Proteobacteria; Gammaproteobacteria; Enterobacterales; Enterobacteriaceae; Escherichia; Escherichia coli MTGDNTLIHSHGINRRDFMKLCAALAATMGLSSKAAAEMAESVTNPQRPPVIWIGAQECT WP_000145410.1 MULTISPECIES: hydrogenase-2 small chain [Proteobacteria] Length: 372\nNumber of predicted TMHs: 1\nExp number of AAs in TMHs: 23.72226\nExp number, first 60 AAs: 0.22038\nTotal prob of N-in: 0.12994\noutside 1 331\nTMhelix 332 354\ninside 355 372

9708 GCF_000355375.2_ASM35537v2 Escherichia coli 2848050 Proteobacteria; Gammaproteobacteria; Enterobacterales; Enterobacteriaceae; Escherichia; Escherichia coli MNNEETFYQAMRRQGVTRRSFLKYCSLAATSLGLGAGMAPKIAWALENKPRIPVVWIHGL WP_001058323.1 MULTISPECIES: hydrogenase-1 small chain [Proteobacteria] Length: 372\nNumber of predicted TMHs: 2\nExp number of AAs in TMHs: 37.2608499999999\nExp number, first 60 AAs: 19.52923\nTotal prob of N-in: 0.94351\nPOSSIBLE N-term signal sequence\ninside 1 19\nTMhelix 20 39\noutside 40 325\nTMhelix 326 348\ninside 349 372

9709 GCF_000355055.1_ASM35505v1 Escherichia coli 2850400 Proteobacteria; Gammaproteobacteria; Enterobacterales; Enterobacteriaceae; Escherichia; Escherichia coli MTGDNTLIHSHGINRRDFMKLCVALAATMGLSSKAAAEMAESVTNPQRPPVIWIGAQECT WP_000145425.1 hydrogenase 2 small subunit [Escherichia coli] Length: 372\nNumber of predicted TMHs: 1\nExp number of AAs in TMHs: 23.94354\nExp number, first 60 AAs: 0.45405\nTotal prob of N-in: 0.13997\noutside 1 331\nTMhelix 332 354\ninside 355 372

9710 GCF_000355055.1_ASM35505v1 Escherichia coli 2850400 Proteobacteria; Gammaproteobacteria; Enterobacterales; Enterobacteriaceae; Escherichia; Escherichia coli MNNEETFYQAMRRQGVTRRSFLKYCSLAATSLGLGAGMAPKIAWALENKPRIPVVWIHGL WP_001295940.1 MULTISPECIES: hydrogenase [Enterobacteriaceae] Length: 372\nNumber of predicted TMHs: 2\nExp number of AAs in TMHs: 37.2608899999999\nExp number, first 60 AAs: 19.52923\nTotal prob of N-in: 0.94351\nPOSSIBLE N-term signal sequence\ninside 1 19\nTMhelix 20 39\noutside 40 325\nTMhelix 326 348\ninside 349 372

9711 GCF_000355335.2_ASM35533v2 Escherichia coli 2850750 Proteobacteria; Gammaproteobacteria; Enterobacterales; Enterobacteriaceae; Escherichia; Escherichia coli MNNEETFYQAMRRQGVTRRSFLKYCSLAATSLGLGAGMAPKIAWALENKPRIPVVWIHGL WP_001295940.1 MULTISPECIES: hydrogenase [Enterobacteriaceae] Length: 372\nNumber of predicted TMHs: 2\nExp number of AAs in TMHs: 37.2608899999999\nExp number, first 60 AAs: 19.52923\nTotal prob of N-in: 0.94351\nPOSSIBLE N-term signal sequence\ninside 1 19\nTMhelix 20 39\noutside 40 325\nTMhelix 326 348\ninside 349 372

9712 GCF_000355335.2_ASM35533v2 Escherichia coli 2850750 Proteobacteria; Gammaproteobacteria; Enterobacterales; Enterobacteriaceae; Escherichia; Escherichia coli MTGDNTLIHSHGINRRDFMKLCVALAATMGLSSKAAAEMAESVTNPQRPPVIWIGAQECT WP_000145425.1 hydrogenase 2 small subunit [Escherichia coli] Length: 372\nNumber of predicted TMHs: 1\nExp number of AAs in TMHs: 23.94354\nExp number, first 60 AAs: 0.45405\nTotal prob of N-in: 0.13997\noutside 1 331\nTMhelix 332 354\ninside 355 372

9713 GCF_000355015.2_ASM35501v2 Escherichia coli 2851500 Proteobacteria; Gammaproteobacteria; Enterobacterales; Enterobacteriaceae; Escherichia; Escherichia coli MNNEETFYQAMRRQGVTRRSFLKYCSLAATSLGLGAGMAPKIAWALENKPRIPVVWIHGL WP_001295940.1 MULTISPECIES: hydrogenase [Enterobacteriaceae] Length: 372\nNumber of predicted TMHs: 2\nExp number of AAs in TMHs: 37.2608899999999\nExp number, first 60 AAs: 19.52923\nTotal prob of N-in:

0.94351\nPOSSIBLE N-term signal sequence\ninside 1 19\nTMhelix 20 39\noutside 40 325\nTMhelix 326 348\ninside 349 372

9714 GCF_000355015.2_ASM35501v2 Escherichia coli 2851500 Proteobacteria; Gammaproteobacteria; Enterobacterales; Enterobacteriaceae; Escherichia; Escherichia coli
MTGDNPLIHSHGINRRDFMKLCVALAATMGLSSKAAAEMAESVTNPQRPPVIWIGAQECT WP_001713585.1
hydrogenase-2 small chain [Escherichia coli] Length: 372\nNumber of predicted TMHs: 1\nExp number of AAs in TMHs: 22.72435\nExp number, first 60 AAs: 0.22827\nTotal prob of N-in: 0.06929\noutside 1 331\nTMhelix 332 354\ninside 355 372

9715 GCF_000355355.1_ASM35535v1 Escherichia coli 2853500 Proteobacteria; Gammaproteobacteria; Enterobacterales; Enterobacteriaceae; Escherichia; Escherichia coli
MNNEETFYQAMRRQGVTRRSFLKYCSLAATSLGLGAGMAPKIAWALENKPRIPVVWIHGL WP_001295940.1
MULTISPECIES: hydrogenase [Enterobacteriaceae] Length: 372\nNumber of predicted TMHs: 2\nExp number of AAs in TMHs: 37.2608899999999\nExp number, first 60 AAs: 19.52923\nTotal prob of N-in: 0.94351\nPOSSIBLE N-term signal sequence\ninside 1 19\nTMhelix 20 39\noutside 40 325\nTMhelix 326 348\ninside 349 372

9716 GCF_000355355.1_ASM35535v1 Escherichia coli 2853500 Proteobacteria; Gammaproteobacteria; Enterobacterales; Enterobacteriaceae; Escherichia; Escherichia coli
MTGDNTLIHSHGINRRDFMKLCVALAATMGLSSKAAAEMAESVTNPQRPPVIWIGAQECT WP_000145425.1
hydrogenase 2 small subunit [Escherichia coli] Length: 372\nNumber of predicted TMHs: 1\nExp number of AAs in TMHs: 23.94354\nExp number, first 60 AAs: 0.45405\nTotal prob of N-in: 0.13997\noutside 1 331\nTMhelix 332 354\ninside 355 372

9717 GCF_000356845.2_ASM35684v2 Escherichia coli 2854350 Proteobacteria; Gammaproteobacteria; Enterobacterales; Enterobacteriaceae; Escherichia; Escherichia coli
MTGDNTLIHSHGINRRDFMKLCAALAATMGLSTKAAAEMAESVTNPQRPPVIWIGAQECT WP_001507912.1
hydrogenase-2 small chain [Escherichia coli] Length: 372\nNumber of predicted TMHs: 1\nExp number of AAs in TMHs: 23.86013\nExp number, first 60 AAs: 0.3669\nTotal prob of N-in: 0.13628\noutside 1 331\nTMhelix 332 354\ninside 355 372

9718 GCF_000356845.2_ASM35684v2 Escherichia coli 2854350 Proteobacteria; Gammaproteobacteria; Enterobacterales; Enterobacteriaceae; Escherichia; Escherichia coli
MNNEETFYQAMRRQGVTRRSFLKYCSLAATSLGLGAGMAPKIAWALENKPRIPVVWIHGL WP_001058323.1
MULTISPECIES: hydrogenase-1 small chain [Proteobacteria] Length: 372\nNumber of predicted TMHs: 2\nExp number of AAs in TMHs: 37.2608499999999\nExp number, first 60 AAs: 19.52923\nTotal prob of N-in: 0.94351\nPOSSIBLE N-term signal sequence\ninside 1 19\nTMhelix 20 39\noutside 40 325\nTMhelix 326 348\ninside 349 372

9719 GCF_000354995.2_ASM35499v2 Escherichia coli 2860050 Proteobacteria; Gammaproteobacteria; Enterobacterales; Enterobacteriaceae; Escherichia; Escherichia coli
MTGDNTLIHSHGINRRDFMKLCAALAATMGLSSKAAAEMAESVTNPQRPPVIWIGAQECT WP_000145410.1
MULTISPECIES: hydrogenase-2 small chain [Proteobacteria] Length: 372\nNumber of predicted TMHs: 1\nExp number of AAs in TMHs: 23.72226\nExp number, first 60 AAs: 0.22038\nTotal prob of N-in: 0.12994\noutside 1 331\nTMhelix 332 354\ninside 355 372

9720 GCF_000356445.2_ASM35644v2 Escherichia coli 2860650 Proteobacteria; Gammaproteobacteria; Enterobacterales; Enterobacteriaceae; Escherichia; Escherichia coli
MNNEETFYQAMRRQGVTRRSFLKYCSLAATSLGLGAGMAPKIVWALENKPRIPVVWIHGL WP_001398876.1
hydrogenase-1 small chain [Escherichia coli] Length: 372\nNumber of predicted TMHs: 2\nExp number of AAs in TMHs: 37.60268\nExp number, first 60 AAs: 19.84677\nTotal prob of N-in: 0.94621\nPOSSIBLE N-term signal sequence\ninside 1 20\nTMhelix 21 43\noutside 44 325\nTMhelix 326 348\ninside 349 372

9721 GCF_000356445.2_ASM35644v2 Escherichia coli 2860650 Proteobacteria; Gammaproteobacteria; Enterobacterales; Enterobacteriaceae; Escherichia; Escherichia coli
MTGDNTLIHSHGINRRDFMKLCAALAATMGLSSKAAAEMAESVTNPQRPPVIWIGAQECT WP_000145410.1
MULTISPECIES: hydrogenase-2 small chain [Proteobacteria] Length: 372\nNumber of predicted TMHs: 1\nExp number of AAs in TMHs: 23.72226\nExp number, first 60 AAs: 0.22038\nTotal prob of N-in: 0.12994\noutside 1 331\nTMhelix 332 354\ninside 355 372

9722 GCF_000355035.2_ASM35503v2 Escherichia coli 2861200 Proteobacteria; Gammaproteobacteria; Enterobacterales; Enterobacteriaceae; Escherichia; Escherichia coli
MTGDNPLIHSHGINRRDFMKLCAALAATMGLSSKAAAEMAESVTNPQRPPVIWIGAQECT WP_001647491.1
hydrogenase-2 small chain [Escherichia coli] Length: 372\nNumber of predicted TMHs: 1\nExp number of AAs in TMHs: 22.60351\nExp number, first 60 AAs: 0.1106\nTotal prob of N-in: 0.06390\noutside 1 331\nTMhelix 332 354\ninside 355 372

9723 GCF_000355035.2_ASM35503v2 Escherichia coli 2861200 Proteobacteria; Gammaproteobacteria; Enterobacterales; Enterobacteriaceae; Escherichia; Escherichia coli
MNNEETFYQAMRRQGVTRRSFLKYCSLAATSLGLGAGMAPKIAWALENKPRIPVVWIHGL WP_001425448.1
hydrogenase-1 small chain [Escherichia coli] Length: 372\nNumber of predicted TMHs: 2\nExp number of AAs in TMHs: 37.14002\nExp number, first 60 AAs: 19.53586\nTotal prob of N-in: 0.94339\nPOSSIBLE N-term signal sequence\ninside 1 19\nTMhelix 20 39\noutside 40 325\nTMhelix 326 348\ninside 349 372

9724 GCF_000356465.2_ASM35646v2 Escherichia coli 2862600 Proteobacteria; Gammaproteobacteria; Enterobacterales; Enterobacteriaceae; Escherichia; Escherichia coli
MTGDNTLIHSHGINRRDFMKLCAALAATMGLSSKAAAEMAESVTNPQRPPVIWIGAQECT WP_000145410.1
MULTISPECIES: hydrogenase-2 small chain [Proteobacteria] Length: 372\nNumber of predicted TMHs: 1\nExp number of AAs in TMHs: 23.72226\nExp number, first 60 AAs: 0.22038\nTotal prob of N-in: 0.12994\noutside 1 331\nTMhelix 332 354\ninside 355 372

9725 GCF_000356465.2_ASM35646v2 Escherichia coli 2862600 Proteobacteria; Gammaproteobacteria; Enterobacterales; Enterobacteriaceae; Escherichia; Escherichia coli
MNNEETFYQAMRRQGVTRRSFLKYCSLAATSLGLGAGMAPKIAWALENKPRIPVVWIHGL WP_001058323.1
MULTISPECIES: hydrogenase-1 small chain [Proteobacteria] Length: 372\nNumber of predicted TMHs: 2\nExp number of AAs in TMHs: 37.2608499999999\nExp number, first 60 AAs: 19.52923\nTotal prob of N-in: 0.94351\nPOSSIBLE N-term signal sequence\ninside 1 19\nTMhelix 20 39\noutside 40 325\nTMhelix 326 348\ninside 349 372

9726 GCF_000356485.2_ASM35648v2 Escherichia coli 2864350 Proteobacteria; Gammaproteobacteria; Enterobacterales; Enterobacteriaceae; Escherichia; Escherichia coli
MNNEETFYQAMRRQGVTRRSFLKYCSLAATSLGLGAGMAPKIAWALENKPRIPVVWIHGL WP_001058323.1
MULTISPECIES: hydrogenase-1 small chain [Proteobacteria] Length: 372\nNumber of predicted TMHs: 2\nExp number of AAs in TMHs: 37.2608499999999\nExp number, first 60 AAs: 19.52923\nTotal prob of N-in: 0.94351\nPOSSIBLE N-term signal sequence\ninside 1 19\nTMhelix 20 39\noutside 40 325\nTMhelix 326 348\ninside 349 372

9727 GCF_000356485.2_ASM35648v2 Escherichia coli 2864350 Proteobacteria; Gammaproteobacteria; Enterobacterales; Enterobacteriaceae; Escherichia; Escherichia coli
MTGDNTLIHSHGINRRDFMKLCAALAATMGLSSKAAAEMAESVTNPQRPPVIWIGAQECT WP_000145410.1
MULTISPECIES: hydrogenase-2 small chain [Proteobacteria] Length: 372\nNumber of predicted TMHs: 1\nExp number of AAs in TMHs: 23.72226\nExp number, first 60 AAs: 0.22038\nTotal prob of N-in: 0.12994\noutside 1 331\nTMhelix 332 354\ninside 355 372

9728 GCF_000354975.1_ASM35497v1 Escherichia coli 2865200 Proteobacteria; Gammaproteobacteria; Enterobacterales; Enterobacteriaceae; Escherichia; Escherichia coli
MNNEETFYQAMRRQGVTRRSFLKYCSLAATSLGLGAGMAPKIAWALENKPRIPVVWIHGL WP_001058323.1
MULTISPECIES: hydrogenase-1 small chain [Proteobacteria] Length: 372\nNumber of predicted TMHs: 2\nExp number of AAs in TMHs: 37.2608499999999\nExp number, first 60 AAs: 19.52923\nTotal prob of N-in: 0.94351\nPOSSIBLE N-term signal sequence\ninside 1 19\nTMhelix 20 39\noutside 40 325\nTMhelix 326 348\ninside 349 372

9729 GCF_000354975.1_ASM35497v1 Escherichia coli 2865200 Proteobacteria; Gammaproteobacteria; Enterobacterales; Enterobacteriaceae; Escherichia; Escherichia coli
MTGDNTLIHSHGINRRDFMKLCAALAATMGLSSKAAAEMAESVTNPQRPPVIWIGAQECT WP_000145410.1
MULTISPECIES: hydrogenase-2 small chain [Proteobacteria] Length: 372\nNumber of predicted TMHs: 1\nExp number of AAs in TMHs: 23.72226\nExp number, first 60 AAs: 0.22038\nTotal prob of N-in: 0.12994\noutside 1 331\nTMhelix 332 354\ninside 355 372

9730 GCF_000356505.2_ASM35650v2 Escherichia coli 2866350 Proteobacteria; Gammaproteobacteria; Enterobacterales; Enterobacteriaceae; Escherichia; Escherichia coli

MNNEETFYQAMRRQGVTRRSFLKYCSLAATSLGLGAGMAPKIVWALENKPRIPVVWIHGL WP_001398876.1 hydrogenase-1 small chain [Escherichia coli] Length: 372\nNumber of predicted TMHs: 2\nExp number of AAs in TMHs: 37.60268\nExp number, first 60 AAs: 19.84677\nTotal prob of N-in: 0.94621\nPOSSIBLE N-term signal sequence\ninside 1 20\nTMhelix 21 43\noutside 44 325\nTMhelix 326 348\ninside 349 372

9731 GCF_000356505.2_ASM35650v2 Escherichia coli 2866350 Proteobacteria; Gammaproteobacteria; Enterobacterales; Enterobacteriaceae; Escherichia; Escherichia coli MTGDNTLIHSHGINRRDFMKLCAALAATMGLSSKAAAEMAESVTNPQRPPVIWIGAQECT WP_000145410.1 MULTISPECIES: hydrogenase-2 small chain [Proteobacteria] Length: 372\nNumber of predicted TMHs: 1\nExp number of AAs in TMHs: 23.72226\nExp number, first 60 AAs: 0.22038\nTotal prob of N-in: 0.12994\noutside 1 331\nTMhelix 332 354\ninside 355 372

9732 GCF_000354935.2_ASM35493v2 Escherichia coli 2866450 Proteobacteria; Gammaproteobacteria; Enterobacterales; Enterobacteriaceae; Escherichia; Escherichia coli MTGDNTLIHSHGINRRDFMKLCVALAATMGLSSKAAAEMAESVTNPQRPPVIWIGAQECT WP_000145425.1 hydrogenase 2 small subunit [Escherichia coli] Length: 372\nNumber of predicted TMHs: 1\nExp number of AAs in TMHs: 23.94354\nExp number, first 60 AAs: 0.45405\nTotal prob of N-in: 0.13997\noutside 1 331\nTMhelix 332 354\ninside 355 372

9733 GCF_000354935.2_ASM35493v2 Escherichia coli 2866450 Proteobacteria; Gammaproteobacteria; Enterobacterales; Enterobacteriaceae; Escherichia; Escherichia coli MNNEETFYQAMRRQGVTRRSFLKYCSLAATSLGLGAGMAPKIAWALENKPRIPVVWIHGL WP_001295940.1 MULTISPECIES: hydrogenase [Enterobacteriaceae] Length: 372\nNumber of predicted TMHs: 2\nExp number of AAs in TMHs: 37.2608899999999\nExp number, first 60 AAs: 19.52923\nTotal prob of N-in: 0.94351\nPOSSIBLE N-term signal sequence\ninside 1 19\nTMhelix 20 39\noutside 40 325\nTMhelix 326 348\ninside 349 372

9734 GCF_000354955.1_ASM35495v1 Escherichia coli 2866550 Proteobacteria; Gammaproteobacteria; Enterobacterales; Enterobacteriaceae; Escherichia; Escherichia coli MTGDNTLIHSHGINRRDFMKLCVALAATMGLSSKAAAEMAESVTNPQRPPVIWIGAQECT WP_000145425.1 hydrogenase 2 small subunit [Escherichia coli] Length: 372\nNumber of predicted TMHs: 1\nExp number of AAs in TMHs: 23.94354\nExp number, first 60 AAs: 0.45405\nTotal prob of N-in: 0.13997\noutside 1 331\nTMhelix 332 354\ninside 355 372

9735 GCF_000354955.1_ASM35495v1 Escherichia coli 2866550 Proteobacteria; Gammaproteobacteria; Enterobacterales; Enterobacteriaceae; Escherichia; Escherichia coli MNNEETFYQAMRRQGVTRRSFLKYCSLAATSLGLGAGMAPKIAWALENKPRIPVVWIHGL WP_001295940.1 MULTISPECIES: hydrogenase [Enterobacteriaceae] Length: 372\nNumber of predicted TMHs: 2\nExp number of AAs in TMHs: 37.2608899999999\nExp number, first 60 AAs: 19.52923\nTotal prob of N-in: 0.94351\nPOSSIBLE N-term signal sequence\ninside 1 19\nTMhelix 20 39\noutside 40 325\nTMhelix 326 348\ninside 349 372

9736 GCF_000354755.2_ASM35475v2 Escherichia coli 2866750 Proteobacteria; Gammaproteobacteria; Enterobacterales; Enterobacteriaceae; Escherichia; Escherichia coli MTGDNTLIHSHGINRRDFMKLCVALAATMGLSSKAAAEMAESVTNPQRPPVIWIGAQECT WP_000145425.1 hydrogenase 2 small subunit [Escherichia coli] Length: 372\nNumber of predicted TMHs: 1\nExp number of AAs in TMHs: 23.94354\nExp number, first 60 AAs: 0.45405\nTotal prob of N-in: 0.13997\noutside 1 331\nTMhelix 332 354\ninside 355 372

9737 GCF_000354755.2_ASM35475v2 Escherichia coli 2866750 Proteobacteria; Gammaproteobacteria; Enterobacterales; Enterobacteriaceae; Escherichia; Escherichia coli MNNEETFYQAMRRQGVTRRSFLKYCSLAATSLGLGAGMAPKIAWALENKPRIPVVWIHGL WP_001295940.1 MULTISPECIES: hydrogenase [Enterobacteriaceae] Length: 372\nNumber of predicted TMHs: 2\nExp number of AAs in TMHs: 37.2608899999999\nExp number, first 60 AAs: 19.52923\nTotal prob of N-in: 0.94351\nPOSSIBLE N-term signal sequence\ninside 1 19\nTMhelix 20 39\noutside 40 325\nTMhelix 326 348\ninside 349 372

9738 GCF_000354815.2_ASM35481v2 Escherichia coli 2867750 Proteobacteria; Gammaproteobacteria; Enterobacterales; Enterobacteriaceae; Escherichia; Escherichia coli MTGDNTLIHSHGINRRDFMKLCVALAATMGLSSKAAAEMAESVTNPQRPPVIWIGAQECT WP_000145425.1 hydrogenase 2 small subunit [Escherichia coli] Length: 372\nNumber of predicted TMHs: 1\nExp number of

AAs in TMHs: 23.94354\nExp number, first 60 AAs: 0.45405\nTotal prob of N-in: 0.13997\noutside 1 331\nTMhelix 332 354\ninside 355 372

9739 GCF_000354815.2_ASM35481v2 Escherichia coli 2867750 Proteobacteria; Gammaproteobacteria; Enterobacterales; Enterobacteriaceae; Escherichia; Escherichia coli MNNEETFYQAMRRQGVTRRSFLKYCSLAATSLGLGAGMAPKIAWALENKPRIPVVWIHGL WP_001295940.1 MULTISPECIES: hydrogenase [Enterobacteriaceae] Length: 372\nNumber of predicted TMHs: 2\nExp number of AAs in TMHs: 37.2608899999999\nExp number, first 60 AAs: 19.52923\nTotal prob of N-in: 0.94351\nPOSSIBLE N-term signal sequence\ninside 1 19\nTMhelix 20 39\noutside 40 325\nTMhelix 326 348\ninside 349 372

9740 GCF_000355455.2_ASM35545v2 Escherichia coli 2871950 Proteobacteria; Gammaproteobacteria; Enterobacterales; Enterobacteriaceae; Escherichia; Escherichia coli MNNEETFYQAMRRQGVTRRSFLKYCSLAATSLGLGAGMAPKIAWALENKPRIPVVWIHGL WP_001058323.1 MULTISPECIES: hydrogenase-1 small chain [Proteobacteria] Length: 372\nNumber of predicted TMHs: 2\nExp number of AAs in TMHs: 37.2608499999999\nExp number, first 60 AAs: 19.52923\nTotal prob of N-in: 0.94351\nPOSSIBLE N-term signal sequence\ninside 1 19\nTMhelix 20 39\noutside 40 325\nTMhelix 326 348\ninside 349 372

9741 GCF_000355455.2_ASM35545v2 Escherichia coli 2871950 Proteobacteria; Gammaproteobacteria; Enterobacterales; Enterobacteriaceae; Escherichia; Escherichia coli MTGDNPLIHSHGINRRDFMKLCAALAATMGLSSKAAAEMAESVTNPQRPPVIWIGAQECT WP_001461571.1 hydrogenase-2 small chain [Escherichia coli] Length: 372\nNumber of predicted TMHs: 1\nExp number of AAs in TMHs: 22.60469\nExp number, first 60 AAs: 0.1106\nTotal prob of N-in: 0.06393\noutside 1 331\nTMhelix 332 354\ninside 355 372

9742 GCF_000355475.2_ASM35547v2 Escherichia coli 2872000 Proteobacteria; Gammaproteobacteria; Enterobacterales; Enterobacteriaceae; Escherichia; Escherichia coli MNNEETFYQAMRRQGVTRRSFLKYCSLAATSLGLGAGMAPKIAWALENKPRIPVVWIHGL WP_001058323.1 MULTISPECIES: hydrogenase-1 small chain [Proteobacteria] Length: 372\nNumber of predicted TMHs: 2\nExp number of AAs in TMHs: 37.2608499999999\nExp number, first 60 AAs: 19.52923\nTotal prob of N-in: 0.94351\nPOSSIBLE N-term signal sequence\ninside 1 19\nTMhelix 20 39\noutside 40 325\nTMhelix 326 348\ninside 349 372

9743 GCF_000355475.2_ASM35547v2 Escherichia coli 2872000 Proteobacteria; Gammaproteobacteria; Enterobacterales; Enterobacteriaceae; Escherichia; Escherichia coli MTGDNTLIHSHGINRRDFMKLCAALAATMGLSSKAAAEMAESVTNPQRPPVIWIGAQECT WP_000145410.1 MULTISPECIES: hydrogenase-2 small chain [Proteobacteria] Length: 372\nNumber of predicted TMHs: 1\nExp number of AAs in TMHs: 23.72226\nExp number, first 60 AAs: 0.22038\nTotal prob of N-in: 0.12994\noutside 1 331\nTMhelix 332 354\ninside 355 372

9744 GCF_000354455.1_ASM35445v1 Escherichia coli 2872800 Proteobacteria; Gammaproteobacteria; Enterobacterales; Enterobacteriaceae; Escherichia; Escherichia coli MTGDNTLIHSHGINRRDFMKLCVALAATMGLSSKAAAEMAESVTNPQRPPVIWIGAQECT WP_000145425.1 hydrogenase 2 small subunit [Escherichia coli] Length: 372\nNumber of predicted TMHs: 1\nExp number of AAs in TMHs: 23.94354\nExp number, first 60 AAs: 0.45405\nTotal prob of N-in: 0.13997\noutside 1 331\nTMhelix 332 354\ninside 355 372

9745 GCF_000354455.1_ASM35445v1 Escherichia coli 2872800 Proteobacteria; Gammaproteobacteria; Enterobacterales; Enterobacteriaceae; Escherichia; Escherichia coli MNNEETFYQAMRRQGVTRRSFLKYCSLAATSLGLGAGMAPKIAWALENKPRIPVVWIHGL WP_001295940.1 MULTISPECIES: hydrogenase [Enterobacteriaceae] Length: 372\nNumber of predicted TMHs: 2\nExp number of AAs in TMHs: 37.2608899999999\nExp number, first 60 AAs: 19.52923\nTotal prob of N-in: 0.94351\nPOSSIBLE N-term signal sequence\ninside 1 19\nTMhelix 20 39\noutside 40 325\nTMhelix 326 348\ninside 349 372

9746 GCF_000354275.2_ASM35427v2 Escherichia coli 2875000 Proteobacteria; Gammaproteobacteria; Enterobacterales; Enterobacteriaceae; Escherichia; Escherichia coli MTGDNTLIHSHGINRRDFMKLCVALAATMGLSSKAAAEMAESVTNPQRPPVIWIGAQECT WP_000145425.1 hydrogenase 2 small subunit [Escherichia coli] Length: 372\nNumber of predicted TMHs: 1\nExp number of

AAs in TMHs: 23.94354\nExp number, first 60 AAs:  0.45405\nTotal prob of N-in:   0.13997\noutside   1   331\nTMhelix 332   354\ninside   355   372

9747 GCF_000354275.2_ASM35427v2 Escherichia coli 2875000 Proteobacteria; Gammaproteobacteria; Enterobacterales; Enterobacteriaceae; Escherichia; Escherichia coli
MNNEETFYQAMRRQGVTRRSFLKYCSLAATSLGLGAGMAPKIAWALENKPRIPVVWIHGL WP_001295940.1
MULTISPECIES: hydrogenase [Enterobacteriaceae] Length: 372\nNumber of predicted TMHs:  2\nExp number of AAs in TMHs: 37.2608899999999\nExp number, first 60 AAs:  19.52923\nTotal prob of N-in: 0.94351\nPOSSIBLE N-term signal sequence\ninside   1   19\nTMhelix   20   39\noutside   40   325\nTMhelix   326 348\ninside   349   372

9748 GCF_000356245.1_ASM35624v1 Escherichia coli 2875150 Proteobacteria; Gammaproteobacteria; Enterobacterales; Enterobacteriaceae; Escherichia; Escherichia coli
MNNEETFYQAMRRQGVTRRSFLKYCSLAATSLGLGAGMAPKIAWALENKPRIPVVWIHGL WP_001404960.1
hydrogenase-1 small chain [Escherichia coli] Length: 372\nNumber of predicted TMHs:  2\nExp number of AAs in TMHs: 37.25632\nExp number, first 60 AAs:  19.52904\nTotal prob of N-in:   0.94351\nPOSSIBLE N-term signal sequence\ninside   1   19\nTMhelix   20   39\noutside   40   325\nTMhelix   326   348\ninside   349   372

9749 GCF_000356245.1_ASM35624v1 Escherichia coli 2875150 Proteobacteria; Gammaproteobacteria; Enterobacterales; Enterobacteriaceae; Escherichia; Escherichia coli
MTGDNTLIHSHGINRRDFMKLCAALAATMGLSSKAAAEMAESVTNPQRPPVIWIGAQECT WP_000145410.1
MULTISPECIES: hydrogenase-2 small chain [Proteobacteria] Length: 372\nNumber of predicted TMHs: 1\nExp number of AAs in TMHs: 23.72226\nExp number, first 60 AAs:  0.22038\nTotal prob of N-in:   0.12994\noutside   1 331\nTMhelix   332   354\ninside   355   372

9750 GCF_000462245.2_ASM46224v2 Escherichia coli 2886-75 Proteobacteria; Gammaproteobacteria; Enterobacterales; Enterobacteriaceae; Escherichia; Escherichia coli
MTGDNTLIHSHGINRRDFMKLCAALAATMGLSSKAAAEMAESVTNPQRPPVIWIGAQECT WP_000145410.1
MULTISPECIES: hydrogenase-2 small chain [Proteobacteria] Length: 372\nNumber of predicted TMHs: 1\nExp number of AAs in TMHs: 23.72226\nExp number, first 60 AAs:  0.22038\nTotal prob of N-in:   0.12994\noutside   1 331\nTMhelix   332   354\ninside   355   372

9751 GCF_000462245.2_ASM46224v2 Escherichia coli 2886-75 Proteobacteria; Gammaproteobacteria; Enterobacterales; Enterobacteriaceae; Escherichia; Escherichia coli
MNNEETFYQAMRRQGVTRRSFLKYCSLAATSLGLGAGMAPKIAWALENKPRIPVVWIHGL WP_001301613.1
hydrogenase [Escherichia coli] Length: 372\nNumber of predicted TMHs:  2\nExp number of AAs in TMHs: 37.46598\nExp number, first 60 AAs:  19.53139\nTotal prob of N-in:   0.94364\nPOSSIBLE N-term signal sequence\ninside   1   19\nTMhelix   20   39\noutside   40   325\nTMhelix   326   348\ninside   349   372

9752 GCF_001265295.1_300059 Escherichia coli 300059 Proteobacteria; Gammaproteobacteria; Enterobacterales; Enterobacteriaceae; Escherichia; Escherichia coli
MTGDNTLIHSHGINRRDFMKLCAALAATMGLSSKAAAEMAESVTNPQRPPVIWIGAQECT WP_000145410.1
MULTISPECIES: hydrogenase-2 small chain [Proteobacteria] Length: 372\nNumber of predicted TMHs: 1\nExp number of AAs in TMHs: 23.72226\nExp number, first 60 AAs:  0.22038\nTotal prob of N-in:   0.12994\noutside   1 331\nTMhelix   332   354\ninside   355   372

9753 GCF_001265295.1_300059 Escherichia coli 300059 Proteobacteria; Gammaproteobacteria; Enterobacterales; Enterobacteriaceae; Escherichia; Escherichia coli
MNNEETFYQAMRRQGVTRRSFLKYCSLAATSLGLGAGMAPKIAWALENKPRIPVVWIHGL WP_001058323.1
MULTISPECIES: hydrogenase-1 small chain [Proteobacteria] Length: 372\nNumber of predicted TMHs: 2\nExp number of AAs in TMHs: 37.2608499999999\nExp number, first 60 AAs:  19.52923\nTotal prob of N-in: 0.94351\nPOSSIBLE N-term signal sequence\ninside   1   19\nTMhelix   20   39\noutside   40   325\nTMhelix   326 348\ninside   349   372

9754 GCF_000194665.1_ASM19466v2 Escherichia coli 3003 Proteobacteria; Gammaproteobacteria; Enterobacterales; Enterobacteriaceae; Escherichia; Escherichia coli
MTGDNTLIHSHGINRRDFMKLCAALAATMGLSSKAAAEMAESVTNPQRPPVIWIGAQECT WP_000145410.1
MULTISPECIES: hydrogenase-2 small chain [Proteobacteria] Length: 372\nNumber of predicted TMHs: 1\nExp number of AAs in TMHs: 23.72226\nExp number, first 60 AAs:  0.22038\nTotal prob of N-in:   0.12994\noutside   1 331\nTMhelix   332   354\ninside   355   372

9755 GCF_000194665.1_ASM19466v2 Escherichia coli 3003 Proteobacteria; Gammaproteobacteria; Enterobacterales; Enterobacteriaceae; Escherichia; Escherichia coli
MNNEETFYQAMRRQGVTRRSFLKYCSLAATSLGLGAGMAPKIAWALENKPRIPVVWIHGL WP_001058323.1
MULTISPECIES: hydrogenase-1 small chain [Proteobacteria] Length: 372\nNumber of predicted TMHs: 2\nExp number of AAs in TMHs: 37.2608499999999\nExp number, first 60 AAs: 19.52923\nTotal prob of N-in: 0.94351\nPOSSIBLE N-term signal sequence\ninside 1 19\nTMhelix 20 39\noutside 40 325\nTMhelix 326 348\ninside 349 372

9756 GCF_000304135.2_ASM30413v2 Escherichia coli 3006 Proteobacteria; Gammaproteobacteria; Enterobacterales; Enterobacteriaceae; Escherichia; Escherichia coli
MTGDNTLIHSHGINRRDFMKLCAALAATMGLSSKAAAEMAESVTNPQRPPVIWIGAQECT WP_000145410.1
MULTISPECIES: hydrogenase-2 small chain [Proteobacteria] Length: 372\nNumber of predicted TMHs: 1\nExp number of AAs in TMHs: 23.72226\nExp number, first 60 AAs: 0.22038\nTotal prob of N-in: 0.12994\noutside 1 331\nTMhelix 332 354\ninside 355 372

9757 GCF_000304135.2_ASM30413v2 Escherichia coli 3006 Proteobacteria; Gammaproteobacteria; Enterobacterales; Enterobacteriaceae; Escherichia; Escherichia coli
MNNEETFYQAMRRQGVTRRSFLKYCSLAATSLGLGAGMAPKIAWALENKPRIPVVWIHGL WP_001058323.1
MULTISPECIES: hydrogenase-1 small chain [Proteobacteria] Length: 372\nNumber of predicted TMHs: 2\nExp number of AAs in TMHs: 37.2608499999999\nExp number, first 60 AAs: 19.52923\nTotal prob of N-in: 0.94351\nPOSSIBLE N-term signal sequence\ninside 1 19\nTMhelix 20 39\noutside 40 325\nTMhelix 326 348\ninside 349 372

9758 GCF_000704685.1_ASM70468v1 Escherichia coli 3-020-07_S1_C1 Proteobacteria; Gammaproteobacteria; Enterobacterales; Enterobacteriaceae; Escherichia; Escherichia coli
MTGDNTLIHSHGINRRDFMKLCAALAATMGLSSKAAAEMAESVTNPQRPPVIWIGAQECT WP_000145410.1
MULTISPECIES: hydrogenase-2 small chain [Proteobacteria] Length: 372\nNumber of predicted TMHs: 1\nExp number of AAs in TMHs: 23.72226\nExp number, first 60 AAs: 0.22038\nTotal prob of N-in: 0.12994\noutside 1 331\nTMhelix 332 354\ninside 355 372

9759 GCF_000704685.1_ASM70468v1 Escherichia coli 3-020-07_S1_C1 Proteobacteria; Gammaproteobacteria; Enterobacterales; Enterobacteriaceae; Escherichia; Escherichia coli
MNNEETFYQAMRRQGVTRRSFLKYCSLAATSLGLGAGMAPKIAWALENKPRIPVVWIHGL WP_001058323.1
MULTISPECIES: hydrogenase-1 small chain [Proteobacteria] Length: 372\nNumber of predicted TMHs: 2\nExp number of AAs in TMHs: 37.2608499999999\nExp number, first 60 AAs: 19.52923\nTotal prob of N-in: 0.94351\nPOSSIBLE N-term signal sequence\ninside 1 19\nTMhelix 20 39\noutside 40 325\nTMhelix 326 348\ninside 349 372

9760 GCF_000704745.1_ASM70474v1 Escherichia coli 3-020-07_S1_C2 Proteobacteria; Gammaproteobacteria; Enterobacterales; Enterobacteriaceae; Escherichia; Escherichia coli
MNNEETFYQAMRRQGVTRRSFLKYCSLAATSLGLGAGMAPKIAWALENKPRIPVVWIHGL WP_001058323.1
MULTISPECIES: hydrogenase-1 small chain [Proteobacteria] Length: 372\nNumber of predicted TMHs: 2\nExp number of AAs in TMHs: 37.2608499999999\nExp number, first 60 AAs: 19.52923\nTotal prob of N-in: 0.94351\nPOSSIBLE N-term signal sequence\ninside 1 19\nTMhelix 20 39\noutside 40 325\nTMhelix 326 348\ninside 349 372

9761 GCF_000704745.1_ASM70474v1 Escherichia coli 3-020-07_S1_C2 Proteobacteria; Gammaproteobacteria; Enterobacterales; Enterobacteriaceae; Escherichia; Escherichia coli
MTGDNTLIHSHGINRRDFMKLCAALAATMGLSSKAAAEMAESVTNPQRPPVIWIGAQECT WP_000145410.1
MULTISPECIES: hydrogenase-2 small chain [Proteobacteria] Length: 372\nNumber of predicted TMHs: 1\nExp number of AAs in TMHs: 23.72226\nExp number, first 60 AAs: 0.22038\nTotal prob of N-in: 0.12994\noutside 1 331\nTMhelix 332 354\ninside 355 372

9762 GCF_000704765.1_ASM70476v1 Escherichia coli 3-020-07_S1_C3 Proteobacteria; Gammaproteobacteria; Enterobacterales; Enterobacteriaceae; Escherichia; Escherichia coli
MNNEETFYQAMRRQGVTRRSFLKYCSLAATSLGLGAGMAPKIAWALENKPRIPVVWIHGL WP_001058323.1
MULTISPECIES: hydrogenase-1 small chain [Proteobacteria] Length: 372\nNumber of predicted TMHs: 2\nExp number of AAs in TMHs: 37.2608499999999\nExp number, first 60 AAs: 19.52923\nTotal prob of N-in: 0.94351\nPOSSIBLE N-term signal sequence\ninside 1 19\nTMhelix 20 39\noutside 40 325\nTMhelix 326 348\ninside 349 372

9763 GCF_000704765.1_ASM70476v1 Escherichia coli 3-020-07_S1_C3 Proteobacteria; Gammaproteobacteria; Enterobacterales; Enterobacteriaceae; Escherichia; Escherichia coli MTGDNTLIHSHGINRRDFMKLCAALAATMGLSSKAAAEMAESVTNPQRPPVIWIGAQECT WP_000145410.1 MULTISPECIES: hydrogenase-2 small chain [Proteobacteria] Length: 372\nNumber of predicted TMHs: 1\nExp number of AAs in TMHs: 23.72226\nExp number, first 60 AAs: 0.22038\nTotal prob of N-in: 0.12994\noutside 1 331\nTMhelix 332 354\ninside 355 372

9764 GCF_000704785.1_ASM70478v1 Escherichia coli 3-020-07_S3_C1 Proteobacteria; Gammaproteobacteria; Enterobacterales; Enterobacteriaceae; Escherichia; Escherichia coli MTGDNTLIHSHGINRRDFMKLCAALAATMGLSSKAAAEMAESVTNPQRPPVIWIGAQECT WP_000145410.1 MULTISPECIES: hydrogenase-2 small chain [Proteobacteria] Length: 372\nNumber of predicted TMHs: 1\nExp number of AAs in TMHs: 23.72226\nExp number, first 60 AAs: 0.22038\nTotal prob of N-in: 0.12994\noutside 1 331\nTMhelix 332 354\ninside 355 372

9765 GCF_000704785.1_ASM70478v1 Escherichia coli 3-020-07_S3_C1 Proteobacteria; Gammaproteobacteria; Enterobacterales; Enterobacteriaceae; Escherichia; Escherichia coli MNNEETFYQAMRRQGVTRRSFLKYCSLAATSLGLGAGMAPKIAWALENKPRIPVVWIHGL WP_001058323.1 MULTISPECIES: hydrogenase-1 small chain [Proteobacteria] Length: 372\nNumber of predicted TMHs: 2\nExp number of AAs in TMHs: 37.2608499999999\nExp number, first 60 AAs: 19.52923\nTotal prob of N-in: 0.94351\nPOSSIBLE N-term signal sequence\ninside 1 19\nTMhelix 20 39\noutside 40 325\nTMhelix 326 348\ninside 349 372

9766 GCF_000711515.1_ASM71151v1 Escherichia coli 3-020-07_S3_C2 Proteobacteria; Gammaproteobacteria; Enterobacterales; Enterobacteriaceae; Escherichia; Escherichia coli MTGDNTLIHSHGINRRDFMKLCAALAATMGLSSKAAAEMAESVTNPQRPPVIWIGAQECT WP_000145410.1 MULTISPECIES: hydrogenase-2 small chain [Proteobacteria] Length: 372\nNumber of predicted TMHs: 1\nExp number of AAs in TMHs: 23.72226\nExp number, first 60 AAs: 0.22038\nTotal prob of N-in: 0.12994\noutside 1 331\nTMhelix 332 354\ninside 355 372

9767 GCF_000711515.1_ASM71151v1 Escherichia coli 3-020-07_S3_C2 Proteobacteria; Gammaproteobacteria; Enterobacterales; Enterobacteriaceae; Escherichia; Escherichia coli MNNEETFYQAMRRQGVTRRSFLKYCSLAATSLGLGAGMAPKIAWALENKPRIPVVWIHGL WP_001058323.1 MULTISPECIES: hydrogenase-1 small chain [Proteobacteria] Length: 372\nNumber of predicted TMHs: 2\nExp number of AAs in TMHs: 37.2608499999999\nExp number, first 60 AAs: 19.52923\nTotal prob of N-in: 0.94351\nPOSSIBLE N-term signal sequence\ninside 1 19\nTMhelix 20 39\noutside 40 325\nTMhelix 326 348\ninside 349 372

9768 GCF_000711555.1_ASM71155v1 Escherichia coli 3-020-07_S4_C1 Proteobacteria; Gammaproteobacteria; Enterobacterales; Enterobacteriaceae; Escherichia; Escherichia coli MTGDNTLIHSHGINRRDFMKLCAALAATMGLSSKAAAEMAESVTNPQRPPVIWIGAQECT WP_000145410.1 MULTISPECIES: hydrogenase-2 small chain [Proteobacteria] Length: 372\nNumber of predicted TMHs: 1\nExp number of AAs in TMHs: 23.72226\nExp number, first 60 AAs: 0.22038\nTotal prob of N-in: 0.12994\noutside 1 331\nTMhelix 332 354\ninside 355 372

9769 GCF_000711555.1_ASM71155v1 Escherichia coli 3-020-07_S4_C1 Proteobacteria; Gammaproteobacteria; Enterobacterales; Enterobacteriaceae; Escherichia; Escherichia coli MNNEETFYQAMRRQGVTRRSFLKYCSLAATSLGLGAGMAPKIAWALENKPRIPVVWIHGL WP_001295940.1 MULTISPECIES: hydrogenase [Enterobacteriaceae] Length: 372\nNumber of predicted TMHs: 2\nExp number of AAs in TMHs: 37.2608899999999\nExp number, first 60 AAs: 19.52923\nTotal prob of N-in: 0.94351\nPOSSIBLE N-term signal sequence\ninside 1 19\nTMhelix 20 39\noutside 40 325\nTMhelix 326 348\ninside 349 372

9770 GCF_000704805.1_ASM70480v1 Escherichia coli 3-020-07_S4_C2 Proteobacteria; Gammaproteobacteria; Enterobacterales; Enterobacteriaceae; Escherichia; Escherichia coli MNNEETFYQAMRRQGVTRRSFLKYCSLAATSLGLGAGMAPKIAWALENKPRIPVVWIHGL WP_001058323.1 MULTISPECIES: hydrogenase-1 small chain [Proteobacteria] Length: 372\nNumber of predicted TMHs: 2\nExp number of AAs in TMHs: 37.2608499999999\nExp number, first 60 AAs: 19.52923\nTotal prob of N-in: 0.94351\nPOSSIBLE N-term signal sequence\ninside 1 19\nTMhelix 20 39\noutside 40 325\nTMhelix 326 348\ninside 349 372

9771 GCF_000704805.1_ASM70480v1 Escherichia coli 3-020-07_S4_C2 Proteobacteria; Gammaproteobacteria; Enterobacterales; Enterobacteriaceae; Escherichia; Escherichia coli
MTGDNTLIHSHGINRRDFMKLCAALAATMGLSSKAAAEMAESVTNPQRPPVIWIGAQECT WP_000145410.1
MULTISPECIES: hydrogenase-2 small chain [Proteobacteria] Length: 372\nNumber of predicted TMHs: 1\nExp number of AAs in TMHs: 23.72226\nExp number, first 60 AAs: 0.22038\nTotal prob of N-in: 0.12994\noutside 1 331\nTMhelix 332 354\ninside 355 372

9772 GCF_000704825.1_ASM70482v1 Escherichia coli 3-020-07_S4_C3 Proteobacteria; Gammaproteobacteria; Enterobacterales; Enterobacteriaceae; Escherichia; Escherichia coli
MTGDNTLIHSHGINRRDFMKLCAALAATMGLSSKAAAEMAESVTNPQRPPVIWIGAQECT WP_000145410.1
MULTISPECIES: hydrogenase-2 small chain [Proteobacteria] Length: 372\nNumber of predicted TMHs: 1\nExp number of AAs in TMHs: 23.72226\nExp number, first 60 AAs: 0.22038\nTotal prob of N-in: 0.12994\noutside 1 331\nTMhelix 332 354\ninside 355 372

9773 GCF_000704825.1_ASM70482v1 Escherichia coli 3-020-07_S4_C3 Proteobacteria; Gammaproteobacteria; Enterobacterales; Enterobacteriaceae; Escherichia; Escherichia coli
MNNEETFYQAMRRQGVTRRSFLKYCSLAATSLGLGAGMAPKIAWALENKPRIPVVWIHGL WP_001058323.1
MULTISPECIES: hydrogenase-1 small chain [Proteobacteria] Length: 372\nNumber of predicted TMHs: 2\nExp number of AAs in TMHs: 37.2608499999999\nExp number, first 60 AAs: 19.52923\nTotal prob of N-in: 0.94351\nPOSSIBLE N-term signal sequence\ninside 1 19\nTMhelix 20 39\noutside 40 325\nTMhelix 326 348\ninside 349 372

9774 GCF_001265345.1_302053 Escherichia coli 302053 Proteobacteria; Gammaproteobacteria; Enterobacterales; Enterobacteriaceae; Escherichia; Escherichia coli
MNNEETFYQAMRRQGVTRRSFLKYCSLAATSLGLGAGMAPKIAWALENKPRIPVVWIHGL WP_001631259.1
hydrogenase-1 small chain [Escherichia coli] Length: 372\nNumber of predicted TMHs: 2\nExp number of AAs in TMHs: 37.14002\nExp number, first 60 AAs: 19.53586\nTotal prob of N-in: 0.94339\nPOSSIBLE N-term signal sequence\ninside 1 19\nTMhelix 20 39\noutside 40 325\nTMhelix 326 348\ninside 349 372

9775 GCF_000704845.1_ASM70484v1 Escherichia coli 3-073-06_S1_C1 Proteobacteria; Gammaproteobacteria; Enterobacterales; Enterobacteriaceae; Escherichia; Escherichia coli
MNNEETFYQAMRRQGVTRRSFLKYCSLAATSLGLGAGMAPKIAWALENKPRIPVVWIHGL WP_001058323.1
MULTISPECIES: hydrogenase-1 small chain [Proteobacteria] Length: 372\nNumber of predicted TMHs: 2\nExp number of AAs in TMHs: 37.2608499999999\nExp number, first 60 AAs: 19.52923\nTotal prob of N-in: 0.94351\nPOSSIBLE N-term signal sequence\ninside 1 19\nTMhelix 20 39\noutside 40 325\nTMhelix 326 348\ninside 349 372

9776 GCF_000704845.1_ASM70484v1 Escherichia coli 3-073-06_S1_C1 Proteobacteria; Gammaproteobacteria; Enterobacterales; Enterobacteriaceae; Escherichia; Escherichia coli
MTGDNTLIHSHGINRRDFMKLCAALAATMGLSSKAAAEMAESVTNPQRPPVIWIGAQECT WP_000145406.1
hydrogenase-2 small chain [Escherichia coli] Length: 372\nNumber of predicted TMHs: 1\nExp number of AAs in TMHs: 23.72251\nExp number, first 60 AAs: 0.22027\nTotal prob of N-in: 0.13006\noutside 1 331\nTMhelix 332 354\ninside 355 372

9777 GCF_000704865.1_ASM70486v1 Escherichia coli 3-073-06_S1_C2 Proteobacteria; Gammaproteobacteria; Enterobacterales; Enterobacteriaceae; Escherichia; Escherichia coli
MNNEETFYQAMRRQGVTRRSFLKYCSLAATSLGLGAGMAPKIAWALENKPRIPVVWIHGL WP_001058323.1
MULTISPECIES: hydrogenase-1 small chain [Proteobacteria] Length: 372\nNumber of predicted TMHs: 2\nExp number of AAs in TMHs: 37.2608499999999\nExp number, first 60 AAs: 19.52923\nTotal prob of N-in: 0.94351\nPOSSIBLE N-term signal sequence\ninside 1 19\nTMhelix 20 39\noutside 40 325\nTMhelix 326 348\ninside 349 372

9778 GCF_000704865.1_ASM70486v1 Escherichia coli 3-073-06_S1_C2 Proteobacteria; Gammaproteobacteria; Enterobacterales; Enterobacteriaceae; Escherichia; Escherichia coli
MTGDNTLIHSHGINRRDFMKLCAALAATMGLSSKAAAEMAESVTNPQRPPVIWIGAQECT WP_000145410.1
MULTISPECIES: hydrogenase-2 small chain [Proteobacteria] Length: 372\nNumber of predicted TMHs: 1\nExp number of AAs in TMHs: 23.72226\nExp number, first 60 AAs: 0.22038\nTotal prob of N-in: 0.12994\noutside 1 331\nTMhelix 332 354\ninside 355 372

9779 GCF_000704885.1_ASM70488v1 Escherichia coli 3-073-06_S3_C1 Proteobacteria; Gammaproteobacteria; Enterobacterales; Enterobacteriaceae; Escherichia; Escherichia coli

MNNEETFYQAMRRQGVTRRSFLKYCSLAATSLGLGAGMAPKIAWALENKPRIPVVWIHGL WP_001058323.1 MULTISPECIES: hydrogenase-1 small chain [Proteobacteria] Length: 372\nNumber of predicted TMHs: 2\nExp number of AAs in TMHs: 37.2608499999999\nExp number, first 60 AAs: 19.52923\nTotal prob of N-in: 0.94351\nPOSSIBLE N-term signal sequence\ninside 1 19\nTMhelix 20 39\noutside 40 325\nTMhelix 326 348\ninside 349 372

9780 GCF_000704885.1_ASM70488v1 Escherichia coli 3-073-06_S3_C1 Proteobacteria; Gammaproteobacteria; Enterobacterales; Enterobacteriaceae; Escherichia; Escherichia coli MTGDNTLIHSHGINRRDFMKLCAALAATMGLSSKAAAEMAESVTNPQRPPVIWIGAQECT WP_000145410.1 MULTISPECIES: hydrogenase-2 small chain [Proteobacteria] Length: 372\nNumber of predicted TMHs: 1\nExp number of AAs in TMHs: 23.72226\nExp number, first 60 AAs: 0.22038\nTotal prob of N-in: 0.12994\noutside 1 331\nTMhelix 332 354\ninside 355 372

9781 GCF_000704905.1_ASM70490v1 Escherichia coli 3-073-06_S3_C2 Proteobacteria; Gammaproteobacteria; Enterobacterales; Enterobacteriaceae; Escherichia; Escherichia coli MNNEETFYQAMRRQGVTRRSFLKYCSLAATSLGLGAGMAPKIAWALENKPRIPVVWIHGL WP_001058323.1 MULTISPECIES: hydrogenase-1 small chain [Proteobacteria] Length: 372\nNumber of predicted TMHs: 2\nExp number of AAs in TMHs: 37.2608499999999\nExp number, first 60 AAs: 19.52923\nTotal prob of N-in: 0.94351\nPOSSIBLE N-term signal sequence\ninside 1 19\nTMhelix 20 39\noutside 40 325\nTMhelix 326 348\ninside 349 372

9782 GCF_000704905.1_ASM70490v1 Escherichia coli 3-073-06_S3_C2 Proteobacteria; Gammaproteobacteria; Enterobacterales; Enterobacteriaceae; Escherichia; Escherichia coli MTGDNTLIHSHGINRRDFMKLCAALAATMGLSSKAAAEMAESVTNPQRPPVIWIGAQECT WP_000145410.1 MULTISPECIES: hydrogenase-2 small chain [Proteobacteria] Length: 372\nNumber of predicted TMHs: 1\nExp number of AAs in TMHs: 23.72226\nExp number, first 60 AAs: 0.22038\nTotal prob of N-in: 0.12994\noutside 1 331\nTMhelix 332 354\ninside 355 372

9783 GCF_000700485.1_ASM70048v1 Escherichia coli 3-073-06_S4_C1 Proteobacteria; Gammaproteobacteria; Enterobacterales; Enterobacteriaceae; Escherichia; Escherichia coli MTGDNTLIHSHGINRRDFMKLCAALAATMGLSSKAAAEMAESVTNPQRPPVIWIGAQECT WP_000145410.1 MULTISPECIES: hydrogenase-2 small chain [Proteobacteria] Length: 372\nNumber of predicted TMHs: 1\nExp number of AAs in TMHs: 23.72226\nExp number, first 60 AAs: 0.22038\nTotal prob of N-in: 0.12994\noutside 1 331\nTMhelix 332 354\ninside 355 372

9784 GCF_000700485.1_ASM70048v1 Escherichia coli 3-073-06_S4_C1 Proteobacteria; Gammaproteobacteria; Enterobacterales; Enterobacteriaceae; Escherichia; Escherichia coli MNNEETFYQAMRRQGVTRRSFLKYCSLAATSLGLGAGMAPKIAWALENKPRIPVVWIHGL WP_001058323.1 MULTISPECIES: hydrogenase-1 small chain [Proteobacteria] Length: 372\nNumber of predicted TMHs: 2\nExp number of AAs in TMHs: 37.2608499999999\nExp number, first 60 AAs: 19.52923\nTotal prob of N-in: 0.94351\nPOSSIBLE N-term signal sequence\ninside 1 19\nTMhelix 20 39\noutside 40 325\nTMhelix 326 348\ninside 349 372

9785 GCF_000704945.1_ASM70494v1 Escherichia coli 3-073-06_S4_C2 Proteobacteria; Gammaproteobacteria; Enterobacterales; Enterobacteriaceae; Escherichia; Escherichia coli MNNEETFYQAMRRQGVTRRSFLKYCSLAATSLGLGAGMAPKIAWALENKPRIPVVWIHGL WP_032267472.1 hydrogenase-1 small chain [Escherichia coli] Length: 372\nNumber of predicted TMHs: 2\nExp number of AAs in TMHs: 36.9217\nExp number, first 60 AAs: 19.53335\nTotal prob of N-in: 0.94324\nPOSSIBLE N-term signal sequence\ninside 1 19\nTMhelix 20 39\noutside 40 325\nTMhelix 326 348\ninside 349 372

9786 GCF_000704945.1_ASM70494v1 Escherichia coli 3-073-06_S4_C2 Proteobacteria; Gammaproteobacteria; Enterobacterales; Enterobacteriaceae; Escherichia; Escherichia coli MTGDNTLIHSHGINRRDFMKLCAALAATMGLSSKAAAEMAESVTNPQRPPVIWIGAQECT WP_000145410.1 MULTISPECIES: hydrogenase-2 small chain [Proteobacteria] Length: 372\nNumber of predicted TMHs: 1\nExp number of AAs in TMHs: 23.72226\nExp number, first 60 AAs: 0.22038\nTotal prob of N-in: 0.12994\noutside 1 331\nTMhelix 332 354\ninside 355 372

9787 GCF_000704965.1_ASM70496v1 Escherichia coli 3-073-06_S4_C3 Proteobacteria; Gammaproteobacteria; Enterobacterales; Enterobacteriaceae; Escherichia; Escherichia coli MNNEETFYQAMRRQGVTRRSFLKYCSLAATSLGLGAGMAPKIAWALENKPRIPVVWIHGL WP_001058323.1 MULTISPECIES: hydrogenase-1 small chain [Proteobacteria] Length: 372\nNumber of predicted TMHs:

2\nExp number of AAs in TMHs: 37.2608499999999\nExp number, first 60 AAs:  19.52923\nTotal prob of N-in:  0.94351\nPOSSIBLE N-term signal sequence\ninside   1   19\nTMhelix   20   39\noutside   40   325\nTMhelix   326   348\ninside   349   372

9788 GCF_000704965.1_ASM70496v1 Escherichia coli 3-073-06_S4_C3 Proteobacteria; Gammaproteobacteria; Enterobacterales; Enterobacteriaceae; Escherichia; Escherichia coli MTGDNTLIHSHGINRRDFMKLCAALAATMGLSSKAAAEMAESVTNPQRPPVIWIGAQECT WP_000145410.1 MULTISPECIES: hydrogenase-2 small chain [Proteobacteria] Length: 372\nNumber of predicted TMHs: 1\nExp number of AAs in TMHs: 23.72226\nExp number, first 60 AAs:  0.22038\nTotal prob of N-in:   0.12994\noutside   1   331\nTMhelix   332   354\ninside   355   372

9789 GCF_000700125.1_ASM70012v1 Escherichia coli 3-105-05_S1_C1 Proteobacteria; Gammaproteobacteria; Enterobacterales; Enterobacteriaceae; Escherichia; Escherichia coli MTGDNTLIHSHGINRRDFMKLCAALAATMGLSSKAAAEMAESVTNPQRPPVIWIGAQECT WP_000145410.1 MULTISPECIES: hydrogenase-2 small chain [Proteobacteria] Length: 372\nNumber of predicted TMHs: 1\nExp number of AAs in TMHs: 23.72226\nExp number, first 60 AAs:  0.22038\nTotal prob of N-in:   0.12994\noutside   1   331\nTMhelix   332   354\ninside   355   372

9790 GCF_000700125.1_ASM70012v1 Escherichia coli 3-105-05_S1_C1 Proteobacteria; Gammaproteobacteria; Enterobacterales; Enterobacteriaceae; Escherichia; Escherichia coli MNNEETFYQAMRRQGVTRRSFLKYCSLAATSLGLGAGMAPKIAWALENKPRIPVVWIHGL WP_001058323.1 MULTISPECIES: hydrogenase-1 small chain [Proteobacteria] Length: 372\nNumber of predicted TMHs: 2\nExp number of AAs in TMHs: 37.2608499999999\nExp number, first 60 AAs:  19.52923\nTotal prob of N-in:  0.94351\nPOSSIBLE N-term signal sequence\ninside   1   19\nTMhelix   20   39\noutside   40   325\nTMhelix   326   348\ninside   349   372

9791 GCF_000704985.1_ASM70498v1 Escherichia coli 3-105-05_S1_C2 Proteobacteria; Gammaproteobacteria; Enterobacterales; Enterobacteriaceae; Escherichia; Escherichia coli MTGDNTLIHSHGINRRDFMKLCAALAATMGLSSKAAAEMAESVTNPQRPPVIWIGAQECT WP_000145410.1 MULTISPECIES: hydrogenase-2 small chain [Proteobacteria] Length: 372\nNumber of predicted TMHs: 1\nExp number of AAs in TMHs: 23.72226\nExp number, first 60 AAs:  0.22038\nTotal prob of N-in:   0.12994\noutside   1   331\nTMhelix   332   354\ninside   355   372

9792 GCF_000704985.1_ASM70498v1 Escherichia coli 3-105-05_S1_C2 Proteobacteria; Gammaproteobacteria; Enterobacterales; Enterobacteriaceae; Escherichia; Escherichia coli MNNEETFYQAMRRQGVTRRSFLKYCSLAATSLGLGAGMAPKIAWALENKPRIPVVWIHGL WP_001058323.1 MULTISPECIES: hydrogenase-1 small chain [Proteobacteria] Length: 372\nNumber of predicted TMHs: 2\nExp number of AAs in TMHs: 37.2608499999999\nExp number, first 60 AAs:  19.52923\nTotal prob of N-in:  0.94351\nPOSSIBLE N-term signal sequence\ninside   1   19\nTMhelix   20   39\noutside   40   325\nTMhelix   326   348\ninside   349   372

9793 GCF_000705005.1_ASM70500v1 Escherichia coli 3-105-05_S1_C3 Proteobacteria; Gammaproteobacteria; Enterobacterales; Enterobacteriaceae; Escherichia; Escherichia coli MNNEETFYQAMRRQGVTRRSFLKYCSLAATSLGLGAGMAPKIAWALENKPRIPVVWIHGL WP_001058323.1 MULTISPECIES: hydrogenase-1 small chain [Proteobacteria] Length: 372\nNumber of predicted TMHs: 2\nExp number of AAs in TMHs: 37.2608499999999\nExp number, first 60 AAs:  19.52923\nTotal prob of N-in:  0.94351\nPOSSIBLE N-term signal sequence\ninside   1   19\nTMhelix   20   39\noutside   40   325\nTMhelix   326   348\ninside   349   372

9794 GCF_000705005.1_ASM70500v1 Escherichia coli 3-105-05_S1_C3 Proteobacteria; Gammaproteobacteria; Enterobacterales; Enterobacteriaceae; Escherichia; Escherichia coli MTGDNTLIHSHGINRRDFMKLCAALAATMGLSSKAAAEMAESVTNPQRPPVIWIGAQECT WP_000145410.1 MULTISPECIES: hydrogenase-2 small chain [Proteobacteria] Length: 372\nNumber of predicted TMHs: 1\nExp number of AAs in TMHs: 23.72226\nExp number, first 60 AAs:  0.22038\nTotal prob of N-in:   0.12994\noutside   1   331\nTMhelix   332   354\ninside   355   372

9795 GCF_000700185.1_ASM70018v1 Escherichia coli 3-105-05_S3_C1 Proteobacteria; Gammaproteobacteria; Enterobacterales; Enterobacteriaceae; Escherichia; Escherichia coli MTGDNTLIHSHGINRRDFMKLCAALAATMGLSSKAAAEMAESVTNPQRPPVIWIGAQECT WP_000145410.1 MULTISPECIES: hydrogenase-2 small chain [Proteobacteria] Length: 372\nNumber of predicted TMHs:

1\nExp number of AAs in TMHs: 23.72226\nExp number, first 60 AAs: 0.22038\nTotal prob of N-in: 0.12994\noutside 1 331\nTMhelix 332 354\ninside 355 372

9796 GCF_000700185.1_ASM70018v1 Escherichia coli 3-105-05_S3_C1 Proteobacteria; Gammaproteobacteria; Enterobacterales; Enterobacteriaceae; Escherichia; Escherichia coli
MNNEETFYQAMRRQGVTRRSFLKYCSLAATSLGLGAGMAPKIAWALENKPRIPVVWIHGL WP_001058323.1
MULTISPECIES: hydrogenase-1 small chain [Proteobacteria] Length: 372\nNumber of predicted TMHs: 2\nExp number of AAs in TMHs: 37.2608499999999\nExp number, first 60 AAs: 19.52923\nTotal prob of N-in: 0.94351\nPOSSIBLE N-term signal sequence\ninside 1 19\nTMhelix 20 39\noutside 40 325\nTMhelix 326 348\ninside 349 372

9797 GCF_000700205.1_ASM70020v1 Escherichia coli 3-105-05_S3_C2 Proteobacteria; Gammaproteobacteria; Enterobacterales; Enterobacteriaceae; Escherichia; Escherichia coli
MNNEETFYQAMRRQGVTRRSFLKYCSLAATSLGLGAGMAPKIAWALENKPRIPVVWIHGL WP_001058323.1
MULTISPECIES: hydrogenase-1 small chain [Proteobacteria] Length: 372\nNumber of predicted TMHs: 2\nExp number of AAs in TMHs: 37.2608499999999\nExp number, first 60 AAs: 19.52923\nTotal prob of N-in: 0.94351\nPOSSIBLE N-term signal sequence\ninside 1 19\nTMhelix 20 39\noutside 40 325\nTMhelix 326 348\ninside 349 372

9798 GCF_000700205.1_ASM70020v1 Escherichia coli 3-105-05_S3_C2 Proteobacteria; Gammaproteobacteria; Enterobacterales; Enterobacteriaceae; Escherichia; Escherichia coli
MTGDNTLIHSHGINRRDFMKLCAALAATMGLSSKAAAEMAESVTNPQRPPVIWIGAQECT WP_000145410.1
MULTISPECIES: hydrogenase-2 small chain [Proteobacteria] Length: 372\nNumber of predicted TMHs: 1\nExp number of AAs in TMHs: 23.72226\nExp number, first 60 AAs: 0.22038\nTotal prob of N-in: 0.12994\noutside 1 331\nTMhelix 332 354\ninside 355 372

9799 GCF_000700385.1_ASM70038v1 Escherichia coli 3-105-05_S3_C3 Proteobacteria; Gammaproteobacteria; Enterobacterales; Enterobacteriaceae; Escherichia; Escherichia coli
MTGDNTLIHSHGINRRDFMKLCAALAATMGLSSKAAAEMAESVTNPQRPPVIWIGAQECT WP_000145410.1
MULTISPECIES: hydrogenase-2 small chain [Proteobacteria] Length: 372\nNumber of predicted TMHs: 1\nExp number of AAs in TMHs: 23.72226\nExp number, first 60 AAs: 0.22038\nTotal prob of N-in: 0.12994\noutside 1 331\nTMhelix 332 354\ninside 355 372

9800 GCF_000700385.1_ASM70038v1 Escherichia coli 3-105-05_S3_C3 Proteobacteria; Gammaproteobacteria; Enterobacterales; Enterobacteriaceae; Escherichia; Escherichia coli
MNNEETFYQAMRRQGVTRRSFLKYCSLAATSLGLGAGMAPKIAWALENKPRIPVVWIHGL WP_001058323.1
MULTISPECIES: hydrogenase-1 small chain [Proteobacteria] Length: 372\nNumber of predicted TMHs: 2\nExp number of AAs in TMHs: 37.2608499999999\nExp number, first 60 AAs: 19.52923\nTotal prob of N-in: 0.94351\nPOSSIBLE N-term signal sequence\ninside 1 19\nTMhelix 20 39\noutside 40 325\nTMhelix 326 348\ninside 349 372

9801 GCF_000700325.1_ASM70032v1 Escherichia coli 3-105-05_S4_C1 Proteobacteria; Gammaproteobacteria; Enterobacterales; Enterobacteriaceae; Escherichia; Escherichia coli
MNNEETFYQAMRRQGVTRRSFLKYCSLAATSLGLGAGMAPKIAWALENKPRIPVVWIHGL WP_001058323.1
MULTISPECIES: hydrogenase-1 small chain [Proteobacteria] Length: 372\nNumber of predicted TMHs: 2\nExp number of AAs in TMHs: 37.2608499999999\nExp number, first 60 AAs: 19.52923\nTotal prob of N-in: 0.94351\nPOSSIBLE N-term signal sequence\ninside 1 19\nTMhelix 20 39\noutside 40 325\nTMhelix 326 348\ninside 349 372

9802 GCF_000700325.1_ASM70032v1 Escherichia coli 3-105-05_S4_C1 Proteobacteria; Gammaproteobacteria; Enterobacterales; Enterobacteriaceae; Escherichia; Escherichia coli
MTGDNTLIHSHGINRRDFMKLCAALAATMGLSSKAAAEMAESVTNPQRPPVIWIGAQECT WP_000145410.1
MULTISPECIES: hydrogenase-2 small chain [Proteobacteria] Length: 372\nNumber of predicted TMHs: 1\nExp number of AAs in TMHs: 23.72226\nExp number, first 60 AAs: 0.22038\nTotal prob of N-in: 0.12994\noutside 1 331\nTMhelix 332 354\ninside 355 372

9803 GCF_000700145.1_ASM70014v1 Escherichia coli 3-105-05_S4_C2 Proteobacteria; Gammaproteobacteria; Enterobacterales; Enterobacteriaceae; Escherichia; Escherichia coli
MNNEETFYQAMRRQGVTRRSFLKYCSLAATSLGLGAGMAPKIAWALENKPRIPVVWIHGL WP_001058323.1
MULTISPECIES: hydrogenase-1 small chain [Proteobacteria] Length: 372\nNumber of predicted TMHs: 2\nExp number of AAs in TMHs: 37.2608499999999\nExp number, first 60 AAs: 19.52923\nTotal prob of N-in:

0.94351\nPOSSIBLE N-term signal sequence\ninside 1 19\nTMhelix 20 39\noutside 40 325\nTMhelix 326 348\ninside 349 372

9804 GCF_000700145.1_ASM70014v1 Escherichia coli 3-105-05_S4_C2 Proteobacteria; Gammaproteobacteria; Enterobacterales; Enterobacteriaceae; Escherichia; Escherichia coli
MTGDNTLIHSHGINRRDFMKLCAALAATMGLSSKAAAEMAESVTNPQRPPVIWIGAQECT WP_000145410.1
MULTISPECIES: hydrogenase-2 small chain [Proteobacteria] Length: 372\nNumber of predicted TMHs: 1\nExp number of AAs in TMHs: 23.72226\nExp number, first 60 AAs: 0.22038\nTotal prob of N-in: 0.12994\noutside 1 331\nTMhelix 332 354\ninside 355 372

9805 GCF_000700065.1_ASM70006v1 Escherichia coli 3-105-05_S4_C3 Proteobacteria; Gammaproteobacteria; Enterobacterales; Enterobacteriaceae; Escherichia; Escherichia coli
MTGDNTLIHSHGINRRDFMKLCAALAATMGLSSKAAAEMAESVTNPQRPPVIWIGAQECT WP_000145410.1
MULTISPECIES: hydrogenase-2 small chain [Proteobacteria] Length: 372\nNumber of predicted TMHs: 1\nExp number of AAs in TMHs: 23.72226\nExp number, first 60 AAs: 0.22038\nTotal prob of N-in: 0.12994\noutside 1 331\nTMhelix 332 354\ninside 355 372

9806 GCF_000700065.1_ASM70006v1 Escherichia coli 3-105-05_S4_C3 Proteobacteria; Gammaproteobacteria; Enterobacterales; Enterobacteriaceae; Escherichia; Escherichia coli
MNNEETFYQAMRRQGVTRRSFLKYCSLAATSLGLGAGMAPKIAWALENKPRIPVVWIHGL WP_001058323.1
MULTISPECIES: hydrogenase-1 small chain [Proteobacteria] Length: 372\nNumber of predicted TMHs: 2\nExp number of AAs in TMHs: 37.2608499999999\nExp number, first 60 AAs: 19.52923\nTotal prob of N-in: 0.94351\nPOSSIBLE N-term signal sequence\ninside 1 19\nTMhelix 20 39\noutside 40 325\nTMhelix 326 348\ninside 349 372

9807 GCF_000503315.1_Ecoli320 Escherichia coli 320Proteobacteria; Gammaproteobacteria; Enterobacterales; Enterobacteriaceae; Escherichia; Escherichia coli
MTGDNTLIHSHGINRRDFMKLCAALAATMGLSSKAAAEMAESVTNPQRPPVIWIGAQECT WP_000145410.1
MULTISPECIES: hydrogenase-2 small chain [Proteobacteria] Length: 372\nNumber of predicted TMHs: 1\nExp number of AAs in TMHs: 23.72226\nExp number, first 60 AAs: 0.22038\nTotal prob of N-in: 0.12994\noutside 1 331\nTMhelix 332 354\ninside 355 372

9808 GCF_000503315.1_Ecoli320 Escherichia coli 320Proteobacteria; Gammaproteobacteria; Enterobacterales; Enterobacteriaceae; Escherichia; Escherichia coli
MNNEETFYQAMRRQGVTRRSFLKYCSLAATSLGLGAGMAPKIAWALENKPRIPVVWIHGL WP_001058323.1
MULTISPECIES: hydrogenase-1 small chain [Proteobacteria] Length: 372\nNumber of predicted TMHs: 2\nExp number of AAs in TMHs: 37.2608499999999\nExp number, first 60 AAs: 19.52923\nTotal prob of N-in: 0.94351\nPOSSIBLE N-term signal sequence\ninside 1 19\nTMhelix 20 39\noutside 40 325\nTMhelix 326 348\ninside 349 372

9809 GCF_000194645.1_ASM19464v2 Escherichia coli 3.2303 Proteobacteria; Gammaproteobacteria; Enterobacterales; Enterobacteriaceae; Escherichia; Escherichia coli
MTGDNTLIHSHGINRRDFMKLCAALAATMGLSSKAAAEMAESVTNPQRPPVIWIGAQECT WP_000145410.1
MULTISPECIES: hydrogenase-2 small chain [Proteobacteria] Length: 372\nNumber of predicted TMHs: 1\nExp number of AAs in TMHs: 23.72226\nExp number, first 60 AAs: 0.22038\nTotal prob of N-in: 0.12994\noutside 1 331\nTMhelix 332 354\ninside 355 372

9810 GCF_000194645.1_ASM19464v2 Escherichia coli 3.2303 Proteobacteria; Gammaproteobacteria; Enterobacterales; Enterobacteriaceae; Escherichia; Escherichia coli
MNNEETFYQAMRRQGVTRRSFLKYCSLAATSLGLGAGMAPKIAWALENKPRIPVVWIHGL WP_001058323.1
MULTISPECIES: hydrogenase-1 small chain [Proteobacteria] Length: 372\nNumber of predicted TMHs: 2\nExp number of AAs in TMHs: 37.2608499999999\nExp number, first 60 AAs: 19.52923\nTotal prob of N-in: 0.94351\nPOSSIBLE N-term signal sequence\ninside 1 19\nTMhelix 20 39\noutside 40 325\nTMhelix 326 348\ninside 349 372

9811 GCF_000215205.1_ASM21520v2 Escherichia coli 3.2608 Proteobacteria; Gammaproteobacteria; Enterobacterales; Enterobacteriaceae; Escherichia; Escherichia coli
MNNEETFYQAMRRQGVTRRSFLKYCSLAATSLGLGAGMAPKIAWALENKPRIPVVWIHGL WP_001295940.1
MULTISPECIES: hydrogenase [Enterobacteriaceae] Length: 372\nNumber of predicted TMHs: 2\nExp number of AAs in TMHs: 37.2608899999999\nExp number, first 60 AAs: 19.52923\nTotal prob of N-in:

0.94351\nPOSSIBLE N-term signal sequence\ninside 1 19\nTMhelix 20 39\noutside 40 325\nTMhelix 326 348\ninside 349 372

9812 GCF_000215205.1_ASM21520v2 Escherichia coli 3.2608 Proteobacteria; Gammaproteobacteria; Enterobacterales; Enterobacteriaceae; Escherichia; Escherichia coli MTGDNTLIHSHGINRRDFMKLCAALAATMGLSSKAAAEMAESVTNPQRPPVIWIGAQECT WP_000145410.1 MULTISPECIES: hydrogenase-2 small chain [Proteobacteria] Length: 372\nNumber of predicted TMHs: 1\nExp number of AAs in TMHs: 23.72226\nExp number, first 60 AAs: 0.22038\nTotal prob of N-in: 0.12994\noutside 1 331\nTMhelix 332 354\ninside 355 372

9813 GCF_000700405.1_ASM70040v1 Escherichia coli 3-267-03_S1_C1 Proteobacteria; Gammaproteobacteria; Enterobacterales; Enterobacteriaceae; Escherichia; Escherichia coli MNNEETFYQAMRRQGVTRRSFLKYCSLAATSLGLGAGMAPKIAWALENKPRIPVVWIHGL WP_001058323.1 MULTISPECIES: hydrogenase-1 small chain [Proteobacteria] Length: 372\nNumber of predicted TMHs: 2\nExp number of AAs in TMHs: 37.2608499999999\nExp number, first 60 AAs: 19.52923\nTotal prob of N-in: 0.94351\nPOSSIBLE N-term signal sequence\ninside 1 19\nTMhelix 20 39\noutside 40 325\nTMhelix 326 348\ninside 349 372

9814 GCF_000700405.1_ASM70040v1 Escherichia coli 3-267-03_S1_C1 Proteobacteria; Gammaproteobacteria; Enterobacterales; Enterobacteriaceae; Escherichia; Escherichia coli MTGDNTLIHSHGINRRDFMKLCAALAATMGLSSKAAAEMAESVTNPQRPPVIWIGAQECT WP_000145410.1 MULTISPECIES: hydrogenase-2 small chain [Proteobacteria] Length: 372\nNumber of predicted TMHs: 1\nExp number of AAs in TMHs: 23.72226\nExp number, first 60 AAs: 0.22038\nTotal prob of N-in: 0.12994\noutside 1 331\nTMhelix 332 354\ninside 355 372

9815 GCF_000700365.1_ASM70036v1 Escherichia coli 3-267-03_S1_C2 Proteobacteria; Gammaproteobacteria; Enterobacterales; Enterobacteriaceae; Escherichia; Escherichia coli MNNEETFYQAMRRQGVTRRSFLKYCSLAATSLGLGAGMAPKIAWALENKPRIPVVWIHGL WP_001058323.1 MULTISPECIES: hydrogenase-1 small chain [Proteobacteria] Length: 372\nNumber of predicted TMHs: 2\nExp number of AAs in TMHs: 37.2608499999999\nExp number, first 60 AAs: 19.52923\nTotal prob of N-in: 0.94351\nPOSSIBLE N-term signal sequence\ninside 1 19\nTMhelix 20 39\noutside 40 325\nTMhelix 326 348\ninside 349 372

9816 GCF_000700365.1_ASM70036v1 Escherichia coli 3-267-03_S1_C2 Proteobacteria; Gammaproteobacteria; Enterobacterales; Enterobacteriaceae; Escherichia; Escherichia coli MTGDNTLIHSHGINRRDFMKLCAALAATMGLSSKAAAEMAESVTNPQRPPVIWIGAQECT WP_000145410.1 MULTISPECIES: hydrogenase-2 small chain [Proteobacteria] Length: 372\nNumber of predicted TMHs: 1\nExp number of AAs in TMHs: 23.72226\nExp number, first 60 AAs: 0.22038\nTotal prob of N-in: 0.12994\noutside 1 331\nTMhelix 332 354\ninside 355 372

9817 GCF_000700085.1_ASM70008v1 Escherichia coli 3-267-03_S1_C3 Proteobacteria; Gammaproteobacteria; Enterobacterales; Enterobacteriaceae; Escherichia; Escherichia coli MTGDNTLIHSHGINRRDFMKLCAALAATMGLSSKAAAEMAESVTNPQRPPVIWIGAQECT WP_000145410.1 MULTISPECIES: hydrogenase-2 small chain [Proteobacteria] Length: 372\nNumber of predicted TMHs: 1\nExp number of AAs in TMHs: 23.72226\nExp number, first 60 AAs: 0.22038\nTotal prob of N-in: 0.12994\noutside 1 331\nTMhelix 332 354\ninside 355 372

9818 GCF_000700085.1_ASM70008v1 Escherichia coli 3-267-03_S1_C3 Proteobacteria; Gammaproteobacteria; Enterobacterales; Enterobacteriaceae; Escherichia; Escherichia coli MNNEETFYQAMRRQGVTRRSFLKYCSLAATSLGLGAGMAPKIAWALENKPRIPVVWIHGL WP_001058323.1 MULTISPECIES: hydrogenase-1 small chain [Proteobacteria] Length: 372\nNumber of predicted TMHs: 2\nExp number of AAs in TMHs: 37.2608499999999\nExp number, first 60 AAs: 19.52923\nTotal prob of N-in: 0.94351\nPOSSIBLE N-term signal sequence\ninside 1 19\nTMhelix 20 39\noutside 40 325\nTMhelix 326 348\ninside 349 372

9819 GCF_000700105.1_ASM70010v1 Escherichia coli 3-267-03_S3_C1 Proteobacteria; Gammaproteobacteria; Enterobacterales; Enterobacteriaceae; Escherichia; Escherichia coli MNNEETFYQAMRRQGVTRRSFLKYCSLAATSLGLGAGMAPKIAWALENKPRIPVVWIHGL WP_001058323.1 MULTISPECIES: hydrogenase-1 small chain [Proteobacteria] Length: 372\nNumber of predicted TMHs: 2\nExp number of AAs in TMHs: 37.2608499999999\nExp number, first 60 AAs: 19.52923\nTotal prob of N-in:

0.94351\nPOSSIBLE N-term signal sequence\ninside 1 19\nTMhelix 20 39\noutside 40 325\nTMhelix 326 348\ninside 349 372

9820 GCF_000700105.1_ASM70010v1 Escherichia coli 3-267-03_S3_C1 Proteobacteria; Gammaproteobacteria; Enterobacterales; Enterobacteriaceae; Escherichia; Escherichia coli MTGDNTLIHSHGINRRDFMKLCAALAATMGLSSKAAAEMAESVTNPQRPPVIWIGAQECT WP_000145410.1 MULTISPECIES: hydrogenase-2 small chain [Proteobacteria] Length: 372\nNumber of predicted TMHs: 1\nExp number of AAs in TMHs: 23.72226\nExp number, first 60 AAs: 0.22038\nTotal prob of N-in: 0.12994\noutside 1 331\nTMhelix 332 354\ninside 355 372

9821 GCF_000700345.1_ASM70034v1 Escherichia coli 3-267-03_S3_C2 Proteobacteria; Gammaproteobacteria; Enterobacterales; Enterobacteriaceae; Escherichia; Escherichia coli MTGDNTLIHSHGINRRDFMKLCAALAATMGLSSKAAAEMAESVTNPQRPPVIWIGAQECT WP_000145410.1 MULTISPECIES: hydrogenase-2 small chain [Proteobacteria] Length: 372\nNumber of predicted TMHs: 1\nExp number of AAs in TMHs: 23.72226\nExp number, first 60 AAs: 0.22038\nTotal prob of N-in: 0.12994\noutside 1 331\nTMhelix 332 354\ninside 355 372

9822 GCF_000700345.1_ASM70034v1 Escherichia coli 3-267-03_S3_C2 Proteobacteria; Gammaproteobacteria; Enterobacterales; Enterobacteriaceae; Escherichia; Escherichia coli MNNEETFYQAMRRQGVTRRSFLKYCSLAATSLGLGAGMAPKIAWALENKPRIPVVWIHGL WP_001295940.1 MULTISPECIES: hydrogenase [Enterobacteriaceae] Length: 372\nNumber of predicted TMHs: 2\nExp number of AAs in TMHs: 37.2608899999999\nExp number, first 60 AAs: 19.52923\nTotal prob of N-in: 0.94351\nPOSSIBLE N-term signal sequence\ninside 1 19\nTMhelix 20 39\noutside 40 325\nTMhelix 326 348\ninside 349 372

9823 GCF_000711595.1_ASM71159v1 Escherichia coli 3-267-03_S4_C1 Proteobacteria; Gammaproteobacteria; Enterobacterales; Enterobacteriaceae; Escherichia; Escherichia coli MTGDNTLIHSHGINRRDFMKLCAALAATMGLSSKAAAEMAESVTNPQRPPVIWIGAQECT WP_000145410.1 MULTISPECIES: hydrogenase-2 small chain [Proteobacteria] Length: 372\nNumber of predicted TMHs: 1\nExp number of AAs in TMHs: 23.72226\nExp number, first 60 AAs: 0.22038\nTotal prob of N-in: 0.12994\noutside 1 331\nTMhelix 332 354\ninside 355 372

9824 GCF_000711595.1_ASM71159v1 Escherichia coli 3-267-03_S4_C1 Proteobacteria; Gammaproteobacteria; Enterobacterales; Enterobacteriaceae; Escherichia; Escherichia coli MNNEETFYQAMRRQGVTRRSFLKYCSLAATSLGLGAGMAPKIAWALENKPRIPVVWIHGL WP_001058323.1 MULTISPECIES: hydrogenase-1 small chain [Proteobacteria] Length: 372\nNumber of predicted TMHs: 2\nExp number of AAs in TMHs: 37.2608499999999\nExp number, first 60 AAs: 19.52923\nTotal prob of N-in: 0.94351\nPOSSIBLE N-term signal sequence\ninside 1 19\nTMhelix 20 39\noutside 40 325\nTMhelix 326 348\ninside 349 372

9825 GCF_000700445.1_ASM70044v1 Escherichia coli 3-267-03_S4_C2 Proteobacteria; Gammaproteobacteria; Enterobacterales; Enterobacteriaceae; Escherichia; Escherichia coli MNNEETFYQAMRRQGVTRRSFLKYCSLAATSLGLGAGMAPKIAWALENKPRIPVVWIHGL WP_001295940.1 MULTISPECIES: hydrogenase [Enterobacteriaceae] Length: 372\nNumber of predicted TMHs: 2\nExp number of AAs in TMHs: 37.2608899999999\nExp number, first 60 AAs: 19.52923\nTotal prob of N-in: 0.94351\nPOSSIBLE N-term signal sequence\ninside 1 19\nTMhelix 20 39\noutside 40 325\nTMhelix 326 348\ninside 349 372

9826 GCF_000700445.1_ASM70044v1 Escherichia coli 3-267-03_S4_C2 Proteobacteria; Gammaproteobacteria; Enterobacterales; Enterobacteriaceae; Escherichia; Escherichia coli MTGDNTLIHSHGINRRDFMKLCAALAATMGLSSKAAAEMAESVTNPQRPPVIWIGAQECT WP_000145410.1 MULTISPECIES: hydrogenase-2 small chain [Proteobacteria] Length: 372\nNumber of predicted TMHs: 1\nExp number of AAs in TMHs: 23.72226\nExp number, first 60 AAs: 0.22038\nTotal prob of N-in: 0.12994\noutside 1 331\nTMhelix 332 354\ninside 355 372

9827 GCF_000503435.1_Ecoli328 Escherichia coli 328Proteobacteria; Gammaproteobacteria; Enterobacterales; Enterobacteriaceae; Escherichia; Escherichia coli MTGDNTLIHSHGINRRDFMKLCAALAATMGLSSKAAAEMAESVTNPQRPPVIWIGAQECT WP_000145410.1 MULTISPECIES: hydrogenase-2 small chain [Proteobacteria] Length: 372\nNumber of predicted TMHs: 1\nExp number of AAs in TMHs: 23.72226\nExp number, first 60 AAs: 0.22038\nTotal prob of N-in: 0.12994\noutside 1 331\nTMhelix 332 354\ninside 355 372

9828 GCF_000503435.1_Ecoli328 Escherichia coli 328 Proteobacteria; Gammaproteobacteria; Enterobacterales; Enterobacteriaceae; Escherichia; Escherichia coli
MNNEETFYQAMRRQGVTRRSFLKYCSLAATSLGLGAGMAPKIAWALENKPRIPVVWIHGL WP_001058323.1
MULTISPECIES: hydrogenase-1 small chain [Proteobacteria] Length: 372\nNumber of predicted TMHs: 2\nExp number of AAs in TMHs: 37.2608499999999\nExp number, first 60 AAs: 19.52923\nTotal prob of N-in: 0.94351\nPOSSIBLE N-term signal sequence\ninside 1 19\nTMhelix 20 39\noutside 40 325\nTMhelix 326 348\ninside 349 372

9829 GCF_000700645.1_ASM70064v1 Escherichia coli 3-373-03_S1_C1 Proteobacteria; Gammaproteobacteria; Enterobacterales; Enterobacteriaceae; Escherichia; Escherichia coli
MNNEETFYQAMRRQGVTRRSFLKYCSLAATSLGLGAGMAPKIAWALENKPRIPVVWIHGL WP_001058323.1
MULTISPECIES: hydrogenase-1 small chain [Proteobacteria] Length: 372\nNumber of predicted TMHs: 2\nExp number of AAs in TMHs: 37.2608499999999\nExp number, first 60 AAs: 19.52923\nTotal prob of N-in: 0.94351\nPOSSIBLE N-term signal sequence\ninside 1 19\nTMhelix 20 39\noutside 40 325\nTMhelix 326 348\ninside 349 372

9830 GCF_000700645.1_ASM70064v1 Escherichia coli 3-373-03_S1_C1 Proteobacteria; Gammaproteobacteria; Enterobacterales; Enterobacteriaceae; Escherichia; Escherichia coli
MTGDNTLIHSHGINRRDFMKLCAALAATMGLSSKAAAEMAESVTNPQRPPVIWIGAQECT WP_000145410.1
MULTISPECIES: hydrogenase-2 small chain [Proteobacteria] Length: 372\nNumber of predicted TMHs: 1\nExp number of AAs in TMHs: 23.72226\nExp number, first 60 AAs: 0.22038\nTotal prob of N-in: 0.12994\noutside 1 331\nTMhelix 332 354\ninside 355 372

9831 GCF_000700265.1_ASM70026v1 Escherichia coli 3-373-03_S1_C2 Proteobacteria; Gammaproteobacteria; Enterobacterales; Enterobacteriaceae; Escherichia; Escherichia coli
MNNEETFYQAMRRQGVTRRSFLKYCSLAATSLGLGAGMAPKIAWALENKPRIPVVWIHGL WP_001058323.1
MULTISPECIES: hydrogenase-1 small chain [Proteobacteria] Length: 372\nNumber of predicted TMHs: 2\nExp number of AAs in TMHs: 37.2608499999999\nExp number, first 60 AAs: 19.52923\nTotal prob of N-in: 0.94351\nPOSSIBLE N-term signal sequence\ninside 1 19\nTMhelix 20 39\noutside 40 325\nTMhelix 326 348\ninside 349 372

9832 GCF_000700265.1_ASM70026v1 Escherichia coli 3-373-03_S1_C2 Proteobacteria; Gammaproteobacteria; Enterobacterales; Enterobacteriaceae; Escherichia; Escherichia coli
MTGDNTLIHSHGINRRDFMKLCAALAATMGLSSKAAAEMAESVTNPQRPPVIWIGAQECT WP_000145410.1
MULTISPECIES: hydrogenase-2 small chain [Proteobacteria] Length: 372\nNumber of predicted TMHs: 1\nExp number of AAs in TMHs: 23.72226\nExp number, first 60 AAs: 0.22038\nTotal prob of N-in: 0.12994\noutside 1 331\nTMhelix 332 354\ninside 355 372

9833 GCF_000700625.1_ASM70062v1 Escherichia coli 3-373-03_S1_C3 Proteobacteria; Gammaproteobacteria; Enterobacterales; Enterobacteriaceae; Escherichia; Escherichia coli
MNNEETFYQAMRRQGVTRRSFLKYCSLAATSLGLGAGMAPKIAWALENKPRIPVVWIHGL WP_001058323.1
MULTISPECIES: hydrogenase-1 small chain [Proteobacteria] Length: 372\nNumber of predicted TMHs: 2\nExp number of AAs in TMHs: 37.2608499999999\nExp number, first 60 AAs: 19.52923\nTotal prob of N-in: 0.94351\nPOSSIBLE N-term signal sequence\ninside 1 19\nTMhelix 20 39\noutside 40 325\nTMhelix 326 348\ninside 349 372

9834 GCF_000700625.1_ASM70062v1 Escherichia coli 3-373-03_S1_C3 Proteobacteria; Gammaproteobacteria; Enterobacterales; Enterobacteriaceae; Escherichia; Escherichia coli
MTGDNTLIHSHGINRRDFMKLCAALAATMGLSSKAAAEMAESVTNPQRPPVIWIGAQECT WP_000145410.1
MULTISPECIES: hydrogenase-2 small chain [Proteobacteria] Length: 372\nNumber of predicted TMHs: 1\nExp number of AAs in TMHs: 23.72226\nExp number, first 60 AAs: 0.22038\nTotal prob of N-in: 0.12994\noutside 1 331\nTMhelix 332 354\ninside 355 372

9835 GCF_000700225.1_ASM70022v1 Escherichia coli 3-373-03_S3_C1 Proteobacteria; Gammaproteobacteria; Enterobacterales; Enterobacteriaceae; Escherichia; Escherichia coli
MNNEETFYQAMRRQGVTRRSFLKYCSLAATSLGLGAGMAPKIAWALENKPRIPVVWIHGL WP_001058323.1
MULTISPECIES: hydrogenase-1 small chain [Proteobacteria] Length: 372\nNumber of predicted TMHs: 2\nExp number of AAs in TMHs: 37.2608499999999\nExp number, first 60 AAs: 19.52923\nTotal prob of N-in: 0.94351\nPOSSIBLE N-term signal sequence\ninside 1 19\nTMhelix 20 39\noutside 40 325\nTMhelix 326 348\ninside 349 372

9836 GCF_000700225.1_ASM70022v1 Escherichia coli 3-373-03_S3_C1 Proteobacteria; Gammaproteobacteria; Enterobacterales; Enterobacteriaceae; Escherichia; Escherichia coli MTGDNTLIHSHGINRRDFMKLCAALAATMGLSSKAAAEMAESVTNPQRPPVIWIGAQECT WP_000145410.1 MULTISPECIES: hydrogenase-2 small chain [Proteobacteria] Length: 372\nNumber of predicted TMHs: 1\nExp number of AAs in TMHs: 23.72226\nExp number, first 60 AAs: 0.22038\nTotal prob of N-in: 0.12994\noutside 1 331\nTMhelix 332 354\ninside 355 372

9837 GCF_000700425.1_ASM70042v1 Escherichia coli 3-373-03_S3_C2 Proteobacteria; Gammaproteobacteria; Enterobacterales; Enterobacteriaceae; Escherichia; Escherichia coli MTGDNTLIHSHGINRRDFMKLCAALAATMGLSSKAAAEMAESVTNPQRPPVIWIGAQECT WP_000145410.1 MULTISPECIES: hydrogenase-2 small chain [Proteobacteria] Length: 372\nNumber of predicted TMHs: 1\nExp number of AAs in TMHs: 23.72226\nExp number, first 60 AAs: 0.22038\nTotal prob of N-in: 0.12994\noutside 1 331\nTMhelix 332 354\ninside 355 372

9838 GCF_000700425.1_ASM70042v1 Escherichia coli 3-373-03_S3_C2 Proteobacteria; Gammaproteobacteria; Enterobacterales; Enterobacteriaceae; Escherichia; Escherichia coli MNNEETFYQAMRRQGVTRRSFLKYCSLAATSLGLGAGMAPKIAWALENKPRIPVVWIHGL WP_001058323.1 MULTISPECIES: hydrogenase-1 small chain [Proteobacteria] Length: 372\nNumber of predicted TMHs: 2\nExp number of AAs in TMHs: 37.2608499999999\nExp number, first 60 AAs: 19.52923\nTotal prob of N-in: 0.94351\nPOSSIBLE N-term signal sequence\ninside 1 19\nTMhelix 20 39\noutside 40 325\nTMhelix 326 348\ninside 349 372

9839 GCF_000700245.1_ASM70024v1 Escherichia coli 3-373-03_S3_C3 Proteobacteria; Gammaproteobacteria; Enterobacterales; Enterobacteriaceae; Escherichia; Escherichia coli MNNEETFYQAMRRQGVTRRSFLKYCSLAATSLGLGAGMAPKIAWALENKPRIPVVWIHGL WP_001058323.1 MULTISPECIES: hydrogenase-1 small chain [Proteobacteria] Length: 372\nNumber of predicted TMHs: 2\nExp number of AAs in TMHs: 37.2608499999999\nExp number, first 60 AAs: 19.52923\nTotal prob of N-in: 0.94351\nPOSSIBLE N-term signal sequence\ninside 1 19\nTMhelix 20 39\noutside 40 325\nTMhelix 326 348\ninside 349 372

9840 GCF_000700245.1_ASM70024v1 Escherichia coli 3-373-03_S3_C3 Proteobacteria; Gammaproteobacteria; Enterobacterales; Enterobacteriaceae; Escherichia; Escherichia coli MTGDNTLIHSHGINRRDFMKLCAALAATMGLSSKAAAEMAESVTNPQRPPVIWIGAQECT WP_000145410.1 MULTISPECIES: hydrogenase-2 small chain [Proteobacteria] Length: 372\nNumber of predicted TMHs: 1\nExp number of AAs in TMHs: 23.72226\nExp number, first 60 AAs: 0.22038\nTotal prob of N-in: 0.12994\noutside 1 331\nTMhelix 332 354\ninside 355 372

9841 GCF_000700665.1_ASM70066v1 Escherichia coli 3-373-03_S4_C1 Proteobacteria; Gammaproteobacteria; Enterobacterales; Enterobacteriaceae; Escherichia; Escherichia coli MTGDNTLIHSHGINRRDFMKLCAALAATMGLSSKAAAEMAESVTNPQRPPVIWIGAQECT WP_000145410.1 MULTISPECIES: hydrogenase-2 small chain [Proteobacteria] Length: 372\nNumber of predicted TMHs: 1\nExp number of AAs in TMHs: 23.72226\nExp number, first 60 AAs: 0.22038\nTotal prob of N-in: 0.12994\noutside 1 331\nTMhelix 332 354\ninside 355 372

9842 GCF_000700665.1_ASM70066v1 Escherichia coli 3-373-03_S4_C1 Proteobacteria; Gammaproteobacteria; Enterobacterales; Enterobacteriaceae; Escherichia; Escherichia coli MNNEETFYQAMRRQGVTRRSFLKYCSLAATSLGLGAGMAPKIAWALENKPRIPVVWIHGL WP_001058323.1 MULTISPECIES: hydrogenase-1 small chain [Proteobacteria] Length: 372\nNumber of predicted TMHs: 2\nExp number of AAs in TMHs: 37.2608499999999\nExp number, first 60 AAs: 19.52923\nTotal prob of N-in: 0.94351\nPOSSIBLE N-term signal sequence\ninside 1 19\nTMhelix 20 39\noutside 40 325\nTMhelix 326 348\ninside 349 372

9843 GCF_000700465.1_ASM70046v1 Escherichia coli 3-373-03_S4_C2 Proteobacteria; Gammaproteobacteria; Enterobacterales; Enterobacteriaceae; Escherichia; Escherichia coli MTGDNTLIHSHGINRRDFMKLCAALAATMGLSSKAAAEMAESVTNPQRPPVIWIGAQECT WP_000145410.1 MULTISPECIES: hydrogenase-2 small chain [Proteobacteria] Length: 372\nNumber of predicted TMHs: 1\nExp number of AAs in TMHs: 23.72226\nExp number, first 60 AAs: 0.22038\nTotal prob of N-in: 0.12994\noutside 1 331\nTMhelix 332 354\ninside 355 372

9844 GCF_000700465.1_ASM70046v1 Escherichia coli 3-373-03_S4_C2 Proteobacteria; Gammaproteobacteria; Enterobacterales; Enterobacteriaceae; Escherichia; Escherichia coli

MNNEETFYQAMRRQGVTRRSFLKYCSLAATSLGLGAGMAPKIAWALENKPRIPVVWIHGL WP_001058323.1 MULTISPECIES: hydrogenase-1 small chain [Proteobacteria] Length: 372\nNumber of predicted TMHs: 2\nExp number of AAs in TMHs: 37.2608499999999\nExp number, first 60 AAs: 19.52923\nTotal prob of N-in: 0.94351\nPOSSIBLE N-term signal sequence\ninside 1 19\nTMhelix 20 39\noutside 40 325\nTMhelix 326 348\ninside 349 372

9845 GCF_000714065.1_ASM71406v1 Escherichia coli 3-373-03_S4_C3 Proteobacteria; Gammaproteobacteria; Enterobacterales; Enterobacteriaceae; Escherichia; Escherichia coli MTGDNTLIHSHGINRRDFMKLCAALAATMGLSSKAAAEMAESVTNPQRPPVIWIGAQECT WP_000145410.1 MULTISPECIES: hydrogenase-2 small chain [Proteobacteria] Length: 372\nNumber of predicted TMHs: 1\nExp number of AAs in TMHs: 23.72226\nExp number, first 60 AAs: 0.22038\nTotal prob of N-in: 0.12994\noutside 1 331\nTMhelix 332 354\ninside 355 372

9846 GCF_000714065.1_ASM71406v1 Escherichia coli 3-373-03_S4_C3 Proteobacteria; Gammaproteobacteria; Enterobacterales; Enterobacteriaceae; Escherichia; Escherichia coli MNNEETFYQAMRRQGVTRRSFLKYCSLAATSLGLGAGMAPKIAWALENKPRIPVVWIHGL WP_001058323.1 MULTISPECIES: hydrogenase-1 small chain [Proteobacteria] Length: 372\nNumber of predicted TMHs: 2\nExp number of AAs in TMHs: 37.2608499999999\nExp number, first 60 AAs: 19.52923\nTotal prob of N-in: 0.94351\nPOSSIBLE N-term signal sequence\ninside 1 19\nTMhelix 20 39\noutside 40 325\nTMhelix 326 348\ninside 349 372

9847 GCF_000215285.1_ASM21528v2 Escherichia coli 3.3884 Proteobacteria; Gammaproteobacteria; Enterobacterales; Enterobacteriaceae; Escherichia; Escherichia coli MTGDNTLIHSHGINRRDFMKLCAALAATMGLSSKAAAEMAESVTNPQRPPVIWIGAQECT WP_000145410.1 MULTISPECIES: hydrogenase-2 small chain [Proteobacteria] Length: 372\nNumber of predicted TMHs: 1\nExp number of AAs in TMHs: 23.72226\nExp number, first 60 AAs: 0.22038\nTotal prob of N-in: 0.12994\noutside 1 331\nTMhelix 332 354\ninside 355 372

9848 GCF_000215285.1_ASM21528v2 Escherichia coli 3.3884 Proteobacteria; Gammaproteobacteria; Enterobacterales; Enterobacteriaceae; Escherichia; Escherichia coli MNNEETFYQAMRRQGVTRRSFLKYCSLAATSLGLGAGMAPKIAWALENKPRIPVVWIHGL WP_001058323.1 MULTISPECIES: hydrogenase-1 small chain [Proteobacteria] Length: 372\nNumber of predicted TMHs: 2\nExp number of AAs in TMHs: 37.2608499999999\nExp number, first 60 AAs: 19.52923\nTotal prob of N-in: 0.94351\nPOSSIBLE N-term signal sequence\ninside 1 19\nTMhelix 20 39\noutside 40 325\nTMhelix 326 348\ninside 349 372

9849 GCF_000184765.1_ASM18476v2 Escherichia coli 3431 Proteobacteria; Gammaproteobacteria; Enterobacterales; Enterobacteriaceae; Escherichia; Escherichia coli MNNEETFYQAMRRQGVTRRSFLKYCSLAATSLGLGAGMAPKIAWALENKPRIPVVWIHGL WP_001058323.1 MULTISPECIES: hydrogenase-1 small chain [Proteobacteria] Length: 372\nNumber of predicted TMHs: 2\nExp number of AAs in TMHs: 37.2608499999999\nExp number, first 60 AAs: 19.52923\nTotal prob of N-in: 0.94351\nPOSSIBLE N-term signal sequence\ninside 1 19\nTMhelix 20 39\noutside 40 325\nTMhelix 326 348\ninside 349 372

9850 GCF_000184765.1_ASM18476v2 Escherichia coli 3431 Proteobacteria; Gammaproteobacteria; Enterobacterales; Enterobacteriaceae; Escherichia; Escherichia coli MTGDNTLIHSHGINRRDFMKLCAALAATMGLSSKAAAEMAESVTNPQRPPVIWIGAQECT WP_000145410.1 MULTISPECIES: hydrogenase-2 small chain [Proteobacteria] Length: 372\nNumber of predicted TMHs: 1\nExp number of AAs in TMHs: 23.72226\nExp number, first 60 AAs: 0.22038\nTotal prob of N-in: 0.12994\noutside 1 331\nTMhelix 332 354\ninside 355 372

9851 GCF_000700285.1_ASM70028v1 Escherichia coli 3-475-03_S1_C1 Proteobacteria; Gammaproteobacteria; Enterobacterales; Enterobacteriaceae; Escherichia; Escherichia coli MTGDNTLIHSHGINRRDFMKLCAALAATMGLSSKAAAEMAESVTNPQRPPVIWIGAQECT WP_000145410.1 MULTISPECIES: hydrogenase-2 small chain [Proteobacteria] Length: 372\nNumber of predicted TMHs: 1\nExp number of AAs in TMHs: 23.72226\nExp number, first 60 AAs: 0.22038\nTotal prob of N-in: 0.12994\noutside 1 331\nTMhelix 332 354\ninside 355 372

9852 GCF_000700285.1_ASM70028v1 Escherichia coli 3-475-03_S1_C1 Proteobacteria; Gammaproteobacteria; Enterobacterales; Enterobacteriaceae; Escherichia; Escherichia coli MNNEETFYQAMRRQGVTRRSFLKYCSLAATSLGLGAGMAPKIAWALENKPRIPVVWIHGL WP_001058323.1

MULTISPECIES: hydrogenase-1 small chain [Proteobacteria] Length: 372\nNumber of predicted TMHs: 2\nExp number of AAs in TMHs: 37.2608499999999\nExp number, first 60 AAs: 19.52923\nTotal prob of N-in: 0.94351\nPOSSIBLE N-term signal sequence\ninside 1 19\nTMhelix 20 39\noutside 40 325\nTMhelix 326 348\ninside 349 372

9853 GCF_000713905.1_ASM71390v1 Escherichia coli 3-475-03_S1_C2 Proteobacteria; Gammaproteobacteria; Enterobacterales; Enterobacteriaceae; Escherichia; Escherichia coli MTGDNTLIHSHGINRRDFMKLCAALAATMGLSSKAAAEMAESVTNPQRPPVIWIGAQECT WP_000145410.1 MULTISPECIES: hydrogenase-2 small chain [Proteobacteria] Length: 372\nNumber of predicted TMHs: 1\nExp number of AAs in TMHs: 23.72226\nExp number, first 60 AAs: 0.22038\nTotal prob of N-in: 0.12994\noutside 1 331\nTMhelix 332 354\ninside 355 372

9854 GCF_000713905.1_ASM71390v1 Escherichia coli 3-475-03_S1_C2 Proteobacteria; Gammaproteobacteria; Enterobacterales; Enterobacteriaceae; Escherichia; Escherichia coli MNNEETFYQAMRRQGVTRRSFLKYCSLAATSLGLGAGMAPKIAWALENKPRIPVVWIHGL WP_001058320.1 hydrogenase [Escherichia coli] Length: 372\nNumber of predicted TMHs: 2\nExp number of AAs in TMHs: 37.26082\nExp number, first 60 AAs: 19.52923\nTotal prob of N-in: 0.94351\nPOSSIBLE N-term signal sequence\ninside 1 19\nTMhelix 20 39\noutside 40 325\nTMhelix 326 348\ninside 349 372

9855 GCF_000713895.1_ASM71389v1 Escherichia coli 3-475-03_S3_C1 Proteobacteria; Gammaproteobacteria; Enterobacterales; Enterobacteriaceae; Escherichia; Escherichia coli MTGDNTLIHSHGINRRDFMKLCAALAATMGLSSKAAAEMAESVTNPQRPPVIWIGAQECT WP_000145410.1 MULTISPECIES: hydrogenase-2 small chain [Proteobacteria] Length: 372\nNumber of predicted TMHs: 1\nExp number of AAs in TMHs: 23.72226\nExp number, first 60 AAs: 0.22038\nTotal prob of N-in: 0.12994\noutside 1 331\nTMhelix 332 354\ninside 355 372

9856 GCF_000713935.1_ASM71393v1 Escherichia coli 3-475-03_S3_C2 Proteobacteria; Gammaproteobacteria; Enterobacterales; Enterobacteriaceae; Escherichia; Escherichia coli MTGDNTLIHSHGINRRDFMKLCAALAATMGLSSKAAAEMAESVTNPQRPPVIWIGAQECT WP_000145410.1 MULTISPECIES: hydrogenase-2 small chain [Proteobacteria] Length: 372\nNumber of predicted TMHs: 1\nExp number of AAs in TMHs: 23.72226\nExp number, first 60 AAs: 0.22038\nTotal prob of N-in: 0.12994\noutside 1 331\nTMhelix 332 354\ninside 355 372

9857 GCF_000713935.1_ASM71393v1 Escherichia coli 3-475-03_S3_C2 Proteobacteria; Gammaproteobacteria; Enterobacterales; Enterobacteriaceae; Escherichia; Escherichia coli MNNEETFYQAMRRQGVTRRSFLKYCSLAATSLGLGAGMAPKIAWALENKPRIPVVWIHGL WP_001058323.1 MULTISPECIES: hydrogenase-1 small chain [Proteobacteria] Length: 372\nNumber of predicted TMHs: 2\nExp number of AAs in TMHs: 37.2608499999999\nExp number, first 60 AAs: 19.52923\nTotal prob of N-in: 0.94351\nPOSSIBLE N-term signal sequence\ninside 1 19\nTMhelix 20 39\noutside 40 325\nTMhelix 326 348\ninside 349 372

9858 GCF_000700305.1_ASM70030v1 Escherichia coli 3-475-03_S4_C1 Proteobacteria; Gammaproteobacteria; Enterobacterales; Enterobacteriaceae; Escherichia; Escherichia coli MNNEETFYQAMRRQGVTRRSFLKYCSLAATSLGLGAGMAPKIAWALENKPRIPVVWIHGL WP_001058323.1 MULTISPECIES: hydrogenase-1 small chain [Proteobacteria] Length: 372\nNumber of predicted TMHs: 2\nExp number of AAs in TMHs: 37.2608499999999\nExp number, first 60 AAs: 19.52923\nTotal prob of N-in: 0.94351\nPOSSIBLE N-term signal sequence\ninside 1 19\nTMhelix 20 39\noutside 40 325\nTMhelix 326 348\ninside 349 372

9859 GCF_000700305.1_ASM70030v1 Escherichia coli 3-475-03_S4_C1 Proteobacteria; Gammaproteobacteria; Enterobacterales; Enterobacteriaceae; Escherichia; Escherichia coli MTGDNTLIHSHGINRRDFMKLCAALAATMGLSSKAAAEMAESVTNPQRPPVIWIGAQECT WP_000145410.1 MULTISPECIES: hydrogenase-2 small chain [Proteobacteria] Length: 372\nNumber of predicted TMHs: 1\nExp number of AAs in TMHs: 23.72226\nExp number, first 60 AAs: 0.22038\nTotal prob of N-in: 0.12994\noutside 1 331\nTMhelix 332 354\ninside 355 372

9860 GCF_000700685.1_ASM70068v1 Escherichia coli 3-475-03_S4_C2 Proteobacteria; Gammaproteobacteria; Enterobacterales; Enterobacteriaceae; Escherichia; Escherichia coli MNNEETFYQAMRRQGVTRRSFLKYCSLAATSLGLGAGMAPKIAWALENKPRIPVVWIHGL WP_001058323.1 MULTISPECIES: hydrogenase-1 small chain [Proteobacteria] Length: 372\nNumber of predicted TMHs: 2\nExp number of AAs in TMHs: 37.2608499999999\nExp number, first 60 AAs: 19.52923\nTotal prob of N-in:

0.94351\nPOSSIBLE N-term signal sequence\ninside 1 19\nTMhelix 20 39\noutside 40 325\nTMhelix 326 348\ninside 349 372

9861 GCF_000700685.1_ASM70068v1 Escherichia coli 3-475-03_S4_C2 Proteobacteria; Gammaproteobacteria; Enterobacterales; Enterobacteriaceae; Escherichia; Escherichia coli MTGDNTLIHSHGINRRDFMKLCAALAATMGLSSKAAAEMAESVTNPQRPPVIWIGAQECT WP_000145410.1 MULTISPECIES: hydrogenase-2 small chain [Proteobacteria] Length: 372\nNumber of predicted TMHs: 1\nExp number of AAs in TMHs: 23.72226\nExp number, first 60 AAs: 0.22038\nTotal prob of N-in: 0.12994\noutside 1 331\nTMhelix 332 354\ninside 355 372

9862 GCF_000304815.2_ASM30481v2 Escherichia coli 3.4870 Proteobacteria; Gammaproteobacteria; Enterobacterales; Enterobacteriaceae; Escherichia; Escherichia coli MTGDNTLIHSHGINRRDFMKLCAALAATMGLSSKAAAEMAESVTNPQRPPVIWIGAQECT WP_000145410.1 MULTISPECIES: hydrogenase-2 small chain [Proteobacteria] Length: 372\nNumber of predicted TMHs: 1\nExp number of AAs in TMHs: 23.72226\nExp number, first 60 AAs: 0.22038\nTotal prob of N-in: 0.12994\noutside 1 331\nTMhelix 332 354\ninside 355 372

9863 GCF_000304815.2_ASM30481v2 Escherichia coli 3.4870 Proteobacteria; Gammaproteobacteria; Enterobacterales; Enterobacteriaceae; Escherichia; Escherichia coli MNNEETFYQAMRRQGVTRRSFLKYCSLAATSLGLGAGMAPKIAWALENKPRIPVVWIHGL WP_001301613.1 hydrogenase [Escherichia coli] Length: 372\nNumber of predicted TMHs: 2\nExp number of AAs in TMHs: 37.46598\nExp number, first 60 AAs: 19.53139\nTotal prob of N-in: 0.94364\nPOSSIBLE N-term signal sequence\ninside 1 19\nTMhelix 20 39\noutside 40 325\nTMhelix 326 348\ninside 349 372

9864 GCF_000503295.1_Ecoli38.16 Escherichia coli 38.16 Proteobacteria; Gammaproteobacteria; Enterobacterales; Enterobacteriaceae; Escherichia; Escherichia coli MTGDNTLIHSHGINRRDFMKLCAALAATMGLSSKAAAEMAESVTNPQRPPVIWIGAQECT WP_000145410.1 MULTISPECIES: hydrogenase-2 small chain [Proteobacteria] Length: 372\nNumber of predicted TMHs: 1\nExp number of AAs in TMHs: 23.72226\nExp number, first 60 AAs: 0.22038\nTotal prob of N-in: 0.12994\noutside 1 331\nTMhelix 332 354\ninside 355 372

9865 GCF_000503295.1_Ecoli38.16 Escherichia coli 38.16 Proteobacteria; Gammaproteobacteria; Enterobacterales; Enterobacteriaceae; Escherichia; Escherichia coli MNNEETFYQAMRRQGVTRRSFLKYCSLAATSLGLGAGMAPKIAWALENKPRIPVVWIHGL WP_001058323.1 MULTISPECIES: hydrogenase-1 small chain [Proteobacteria] Length: 372\nNumber of predicted TMHs: 2\nExp number of AAs in TMHs: 37.2608499999999\nExp number, first 60 AAs: 19.52923\nTotal prob of N-in: 0.94351\nPOSSIBLE N-term signal sequence\ninside 1 19\nTMhelix 20 39\noutside 40 325\nTMhelix 326 348\ninside 349 372

9866 GCF_000503355.1_Ecoli38.27 Escherichia coli 38.27 Proteobacteria; Gammaproteobacteria; Enterobacterales; Enterobacteriaceae; Escherichia; Escherichia coli MTGDNTLIHSHGINRRDFMKLCAALAATMGLSSKAAAEMAESVTNPQRPPVIWIGAQECT WP_000145410.1 MULTISPECIES: hydrogenase-2 small chain [Proteobacteria] Length: 372\nNumber of predicted TMHs: 1\nExp number of AAs in TMHs: 23.72226\nExp number, first 60 AAs: 0.22038\nTotal prob of N-in: 0.12994\noutside 1 331\nTMhelix 332 354\ninside 355 372

9867 GCF_000503355.1_Ecoli38.27 Escherichia coli 38.27 Proteobacteria; Gammaproteobacteria; Enterobacterales; Enterobacteriaceae; Escherichia; Escherichia coli MNNEETFYQAMRRQGVTRRSFLKYCSLAATSLGLGAGMAPKIAWALENKPRIPVVWIHGL WP_001058323.1 MULTISPECIES: hydrogenase-1 small chain [Proteobacteria] Length: 372\nNumber of predicted TMHs: 2\nExp number of AAs in TMHs: 37.2608499999999\nExp number, first 60 AAs: 19.52923\nTotal prob of N-in: 0.94351\nPOSSIBLE N-term signal sequence\ninside 1 19\nTMhelix 20 39\noutside 40 325\nTMhelix 326 348\ninside 349 372

9868 GCF_000503375.1_Ecoli38.34 Escherichia coli 38.34 Proteobacteria; Gammaproteobacteria; Enterobacterales; Enterobacteriaceae; Escherichia; Escherichia coli MTGDNTLIHSHGINRRDFMKLCAALAATMGLSSKAAAEMAESVTNPQRPPVIWIGAQECT WP_000145410.1 MULTISPECIES: hydrogenase-2 small chain [Proteobacteria] Length: 372\nNumber of predicted TMHs: 1\nExp number of AAs in TMHs: 23.72226\nExp number, first 60 AAs: 0.22038\nTotal prob of N-in: 0.12994\noutside 1 331\nTMhelix 332 354\ninside 355 372

9869 GCF_000503375.1_Ecoli38.34 Escherichia coli 38.34 Proteobacteria; Gammaproteobacteria; Enterobacterales; Enterobacteriaceae; Escherichia; Escherichia coli
MNNEETFYQAMRRQGVTRRSFLKYCSLAATSLGLGAGMAPKIAWALENKPRIPVVWIHGL WP_001058323.1
MULTISPECIES: hydrogenase-1 small chain [Proteobacteria] Length: 372\nNumber of predicted TMHs: 2\nExp number of AAs in TMHs: 37.2608499999999\nExp number, first 60 AAs: 19.52923\nTotal prob of N-in: 0.94351\nPOSSIBLE N-term signal sequence\ninside 1 19\nTMhelix 20 39\noutside 40 325\nTMhelix 326 348\ninside 349 372

9870 GCF_000503675.1_Ecoli38.52 Escherichia coli 38.52 Proteobacteria; Gammaproteobacteria; Enterobacterales; Enterobacteriaceae; Escherichia; Escherichia coli
MTGDNTLIHSHGINRRDFMKLCAALAATMGLSSKAAAEMAESVTNPQRPPVIWIGAQECT WP_000145410.1
MULTISPECIES: hydrogenase-2 small chain [Proteobacteria] Length: 372\nNumber of predicted TMHs: 1\nExp number of AAs in TMHs: 23.72226\nExp number, first 60 AAs: 0.22038\nTotal prob of N-in: 0.12994\noutside 1 331\nTMhelix 332 354\ninside 355 372

9871 GCF_000503675.1_Ecoli38.52 Escherichia coli 38.52 Proteobacteria; Gammaproteobacteria; Enterobacterales; Enterobacteriaceae; Escherichia; Escherichia coli
MNNEETFYQAMRRQGVTRRSFLKYCSLAATSLGLGAGMAPKIAWALENKPRIPVVWIHGL WP_001058323.1
MULTISPECIES: hydrogenase-1 small chain [Proteobacteria] Length: 372\nNumber of predicted TMHs: 2\nExp number of AAs in TMHs: 37.2608499999999\nExp number, first 60 AAs: 19.52923\nTotal prob of N-in: 0.94351\nPOSSIBLE N-term signal sequence\ninside 1 19\nTMhelix 20 39\noutside 40 325\nTMhelix 326 348\ninside 349 372

9872 GCF_001265465.1_401140 Escherichia coli 401140 Proteobacteria; Gammaproteobacteria; Enterobacterales; Enterobacteriaceae; Escherichia; Escherichia coli
MTGDNTLIHSHGINRRDFMKLCAALAATMGLSSKAAAEMAESVTNPQRPPVIWIGAQECT WP_000145410.1
MULTISPECIES: hydrogenase-2 small chain [Proteobacteria] Length: 372\nNumber of predicted TMHs: 1\nExp number of AAs in TMHs: 23.72226\nExp number, first 60 AAs: 0.22038\nTotal prob of N-in: 0.12994\noutside 1 331\nTMhelix 332 354\ninside 355 372

9873 GCF_001265465.1_401140 Escherichia coli 401140 Proteobacteria; Gammaproteobacteria; Enterobacterales; Enterobacteriaceae; Escherichia; Escherichia coli
MNNEETFYQAMRRQGVTRRSFLKYCSLAATSLGLGAGMAPKIAWALENKPRIPVVWIHGL WP_001058323.1
MULTISPECIES: hydrogenase-1 small chain [Proteobacteria] Length: 372\nNumber of predicted TMHs: 2\nExp number of AAs in TMHs: 37.2608499999999\nExp number, first 60 AAs: 19.52923\nTotal prob of N-in: 0.94351\nPOSSIBLE N-term signal sequence\ninside 1 19\nTMhelix 20 39\noutside 40 325\nTMhelix 326 348\ninside 349 372

9874 GCF_001265545.1_402290 Escherichia coli 402290 Proteobacteria; Gammaproteobacteria; Enterobacterales; Enterobacteriaceae; Escherichia; Escherichia coli
MNNEETFYQAMRRQGVTRRSFLKYCSLAATSLGLGAGMAPKIAWALENKPRIPVVWIHGL WP_001058323.1
MULTISPECIES: hydrogenase-1 small chain [Proteobacteria] Length: 372\nNumber of predicted TMHs: 2\nExp number of AAs in TMHs: 37.2608499999999\nExp number, first 60 AAs: 19.52923\nTotal prob of N-in: 0.94351\nPOSSIBLE N-term signal sequence\ninside 1 19\nTMhelix 20 39\noutside 40 325\nTMhelix 326 348\ninside 349 372

9875 GCF_001265545.1_402290 Escherichia coli 402290 Proteobacteria; Gammaproteobacteria; Enterobacterales; Enterobacteriaceae; Escherichia; Escherichia coli
MTGDNTLIHSHGINRRDFMKLCAALAATMGLSSKAAAEMAESVTNPQRPPVIWIGAQECT WP_000145410.1
MULTISPECIES: hydrogenase-2 small chain [Proteobacteria] Length: 372\nNumber of predicted TMHs: 1\nExp number of AAs in TMHs: 23.72226\nExp number, first 60 AAs: 0.22038\nTotal prob of N-in: 0.12994\noutside 1 331\nTMhelix 332 354\ninside 355 372

9876 GCF_000194335.1_ASM19433v2 Escherichia coli 4.0522 Proteobacteria; Gammaproteobacteria; Enterobacterales; Enterobacteriaceae; Escherichia; Escherichia coli
MNNEETFYQAMRRQGVTRRSFLKYCSLAATSLGLGAGMAPKIAWALENKPRIPVVWIHGL WP_001058323.1
MULTISPECIES: hydrogenase-1 small chain [Proteobacteria] Length: 372\nNumber of predicted TMHs: 2\nExp number of AAs in TMHs: 37.2608499999999\nExp number, first 60 AAs: 19.52923\nTotal prob of N-in: 0.94351\nPOSSIBLE N-term signal sequence\ninside 1 19\nTMhelix 20 39\noutside 40 325\nTMhelix 326 348\ninside 349 372

9877 GCF_000194335.1_ASM19433v2 Escherichia coli 4.0522 Proteobacteria; Gammaproteobacteria; Enterobacterales; Enterobacteriaceae; Escherichia; Escherichia coli
MTGDNTLIHSHGINRRDFMKLCAALAATMGLSSKAAAEMAESVTNPQRPPVIWIGAQECT WP_000145410.1
MULTISPECIES: hydrogenase-2 small chain [Proteobacteria] Length: 372\nNumber of predicted TMHs: 1\nExp number of AAs in TMHs: 23.72226\nExp number, first 60 AAs: 0.22038\nTotal prob of N-in: 0.12994\noutside 1 331\nTMhelix 332 354\ninside 355 372

9878 GCF_000194495.1_ASM19449v2 Escherichia coli 4.0967 Proteobacteria; Gammaproteobacteria; Enterobacterales; Enterobacteriaceae; Escherichia; Escherichia coli
MTGDNTLIHSHGINRRDFMKLCAALAATMGLSSKAAAEMAESVTNPQRPPVIWIGAQECT WP_000145410.1
MULTISPECIES: hydrogenase-2 small chain [Proteobacteria] Length: 372\nNumber of predicted TMHs: 1\nExp number of AAs in TMHs: 23.72226\nExp number, first 60 AAs: 0.22038\nTotal prob of N-in: 0.12994\noutside 1 331\nTMhelix 332 354\ninside 355 372

9879 GCF_000194495.1_ASM19449v2 Escherichia coli 4.0967 Proteobacteria; Gammaproteobacteria; Enterobacterales; Enterobacteriaceae; Escherichia; Escherichia coli
MNNEETFYQAMRRQGVTRRSFLKYCSLAATSLGLGAGMAPKIAWALENKPRIPVVWIHGL WP_001295940.1
MULTISPECIES: hydrogenase [Enterobacteriaceae] Length: 372\nNumber of predicted TMHs: 2\nExp number of AAs in TMHs: 37.2608899999999\nExp number, first 60 AAs: 19.52923\nTotal prob of N-in: 0.94351\nPOSSIBLE N-term signal sequence\ninside 1 19\nTMhelix 20 39\noutside 40 325\nTMhelix 326 348\ninside 349 372

9880 GCF_000700705.1_ASM70070v1 Escherichia coli 4-203-08_S1_C1 Proteobacteria; Gammaproteobacteria; Enterobacterales; Enterobacteriaceae; Escherichia; Escherichia coli
MTGDNTLIHSHGINRRDFMKLCAALAATMGLSSKAAAEMAESVTNPQRPPVIWIGAQECT WP_000145410.1
MULTISPECIES: hydrogenase-2 small chain [Proteobacteria] Length: 372\nNumber of predicted TMHs: 1\nExp number of AAs in TMHs: 23.72226\nExp number, first 60 AAs: 0.22038\nTotal prob of N-in: 0.12994\noutside 1 331\nTMhelix 332 354\ninside 355 372

9881 GCF_000700705.1_ASM70070v1 Escherichia coli 4-203-08_S1_C1 Proteobacteria; Gammaproteobacteria; Enterobacterales; Enterobacteriaceae; Escherichia; Escherichia coli
MNNEETFYQAMRRQGVTRRSFLKYCSLAATSLGLGAGMAPKIAWALENKPRIPVVWIHGL WP_001058323.1
MULTISPECIES: hydrogenase-1 small chain [Proteobacteria] Length: 372\nNumber of predicted TMHs: 2\nExp number of AAs in TMHs: 37.2608499999999\nExp number, first 60 AAs: 19.52923\nTotal prob of N-in: 0.94351\nPOSSIBLE N-term signal sequence\ninside 1 19\nTMhelix 20 39\noutside 40 325\nTMhelix 326 348\ninside 349 372

9882 GCF_000713945.1_ASM71394v1 Escherichia coli 4-203-08_S1_C2 Proteobacteria; Gammaproteobacteria; Enterobacterales; Enterobacteriaceae; Escherichia; Escherichia coli
MNNEETFYQAMRRQGVTRRSFLKYCSLAATSLGLGAGMAPKIAWALENKPRIPVVWIHGL WP_001058323.1
MULTISPECIES: hydrogenase-1 small chain [Proteobacteria] Length: 372\nNumber of predicted TMHs: 2\nExp number of AAs in TMHs: 37.2608499999999\nExp number, first 60 AAs: 19.52923\nTotal prob of N-in: 0.94351\nPOSSIBLE N-term signal sequence\ninside 1 19\nTMhelix 20 39\noutside 40 325\nTMhelix 326 348\ninside 349 372

9883 GCF_000713945.1_ASM71394v1 Escherichia coli 4-203-08_S1_C2 Proteobacteria; Gammaproteobacteria; Enterobacterales; Enterobacteriaceae; Escherichia; Escherichia coli
MTGDNTLIHSHGINRRDFMKLCAALAATMGLSSKAAAEMAESVTNPQRPPVIWIGAQECT WP_000145410.1
MULTISPECIES: hydrogenase-2 small chain [Proteobacteria] Length: 372\nNumber of predicted TMHs: 1\nExp number of AAs in TMHs: 23.72226\nExp number, first 60 AAs: 0.22038\nTotal prob of N-in: 0.12994\noutside 1 331\nTMhelix 332 354\ninside 355 372

9884 GCF_000713975.1_ASM71397v1 Escherichia coli 4-203-08_S1_C3 Proteobacteria; Gammaproteobacteria; Enterobacterales; Enterobacteriaceae; Escherichia; Escherichia coli
MTGDNTLIHSHGINRRDFMKLCAALAATMGLSSKAAAEMAESVTNPQRPPVIWIGAQECT WP_000145410.1
MULTISPECIES: hydrogenase-2 small chain [Proteobacteria] Length: 372\nNumber of predicted TMHs: 1\nExp number of AAs in TMHs: 23.72226\nExp number, first 60 AAs: 0.22038\nTotal prob of N-in: 0.12994\noutside 1 331\nTMhelix 332 354\ninside 355 372

9885 GCF_000713975.1_ASM71397v1 Escherichia coli 4-203-08_S1_C3 Proteobacteria; Gammaproteobacteria; Enterobacterales; Enterobacteriaceae; Escherichia; Escherichia coli

MNNEETFYQAMRRQGVTRRSFLKYCSLAATSLGLGAGMAPKIAWALENKPRIPVVWIHGL WP_001058323.1 MULTISPECIES: hydrogenase-1 small chain [Proteobacteria] Length: 372\nNumber of predicted TMHs: 2\nExp number of AAs in TMHs: 37.2608499999999\nExp number, first 60 AAs: 19.52923\nTotal prob of N-in: 0.94351\nPOSSIBLE N-term signal sequence\ninside 1 19\nTMhelix 20 39\noutside 40 325\nTMhelix 326 348\ninside 349 372

9886 GCF_000714025.1_ASM71402v1 Escherichia coli 4-203-08_S3_C1 Proteobacteria; Gammaproteobacteria; Enterobacterales; Enterobacteriaceae; Escherichia; Escherichia coli MTGDNTLIHSHGINRRDFMKLCAALAATMGLSSKAAAEMAESVTNPQRPPVIWIGAQECT WP_000145410.1 MULTISPECIES: hydrogenase-2 small chain [Proteobacteria] Length: 372\nNumber of predicted TMHs: 1\nExp number of AAs in TMHs: 23.72226\nExp number, first 60 AAs: 0.22038\nTotal prob of N-in: 0.12994\noutside 1 331\nTMhelix 332 354\ninside 355 372

9887 GCF_000714025.1_ASM71402v1 Escherichia coli 4-203-08_S3_C1 Proteobacteria; Gammaproteobacteria; Enterobacterales; Enterobacteriaceae; Escherichia; Escherichia coli MNNEETFYQAMRRQGVTRRSFLKYCSLAATSLGLGAGMAPKIAWALENKPRIPVVWIHGL WP_001058323.1 MULTISPECIES: hydrogenase-1 small chain [Proteobacteria] Length: 372\nNumber of predicted TMHs: 2\nExp number of AAs in TMHs: 37.2608499999999\nExp number, first 60 AAs: 19.52923\nTotal prob of N-in: 0.94351\nPOSSIBLE N-term signal sequence\ninside 1 19\nTMhelix 20 39\noutside 40 325\nTMhelix 326 348\ninside 349 372

9888 GCF_000714055.1_ASM71405v1 Escherichia coli 4-203-08_S3_C2 Proteobacteria; Gammaproteobacteria; Enterobacterales; Enterobacteriaceae; Escherichia; Escherichia coli MNNEETFYQAMRRQGVTRRSFLKYCSLAATSLGLGAGMAPKIAWALENKPRIPVVWIHGL WP_001058323.1 MULTISPECIES: hydrogenase-1 small chain [Proteobacteria] Length: 372\nNumber of predicted TMHs: 2\nExp number of AAs in TMHs: 37.2608499999999\nExp number, first 60 AAs: 19.52923\nTotal prob of N-in: 0.94351\nPOSSIBLE N-term signal sequence\ninside 1 19\nTMhelix 20 39\noutside 40 325\nTMhelix 326 348\ninside 349 372

9889 GCF_000714055.1_ASM71405v1 Escherichia coli 4-203-08_S3_C2 Proteobacteria; Gammaproteobacteria; Enterobacterales; Enterobacteriaceae; Escherichia; Escherichia coli MTGDNTLIHSHGINRRDFMKLCAALAATMGLSSKAAAEMAESVTNPQRPPVIWIGAQECT WP_000145410.1 MULTISPECIES: hydrogenase-2 small chain [Proteobacteria] Length: 372\nNumber of predicted TMHs: 1\nExp number of AAs in TMHs: 23.72226\nExp number, first 60 AAs: 0.22038\nTotal prob of N-in: 0.12994\noutside 1 331\nTMhelix 332 354\ninside 355 372

9890 GCF_000713985.1_ASM71398v1 Escherichia coli 4-203-08_S3_C3 Proteobacteria; Gammaproteobacteria; Enterobacterales; Enterobacteriaceae; Escherichia; Escherichia coli MTGDNTLIHSHGINRRDFMKLCAALAATMGLSSKAAAEMAESVTNPQRPPVIWIGAQECT WP_000145410.1 MULTISPECIES: hydrogenase-2 small chain [Proteobacteria] Length: 372\nNumber of predicted TMHs: 1\nExp number of AAs in TMHs: 23.72226\nExp number, first 60 AAs: 0.22038\nTotal prob of N-in: 0.12994\noutside 1 331\nTMhelix 332 354\ninside 355 372

9891 GCF_000713985.1_ASM71398v1 Escherichia coli 4-203-08_S3_C3 Proteobacteria; Gammaproteobacteria; Enterobacterales; Enterobacteriaceae; Escherichia; Escherichia coli MNNEETFYQAMRRQGVTRRSFLKYCSLAATSLGLGAGMAPKIAWALENKPRIPVVWIHGL WP_001058323.1 MULTISPECIES: hydrogenase-1 small chain [Proteobacteria] Length: 372\nNumber of predicted TMHs: 2\nExp number of AAs in TMHs: 37.2608499999999\nExp number, first 60 AAs: 19.52923\nTotal prob of N-in: 0.94351\nPOSSIBLE N-term signal sequence\ninside 1 19\nTMhelix 20 39\noutside 40 325\nTMhelix 326 348\ninside 349 372

9892 GCF_000714015.1_ASM71401v1 Escherichia coli 4-203-08_S4_C2 Proteobacteria; Gammaproteobacteria; Enterobacterales; Enterobacteriaceae; Escherichia; Escherichia coli MNNEETFYQAMRRQGVTRRSFLKYCSLAATSLGLGAGMAPKIAWALENKPRIPVVWIHGL WP_001058323.1 MULTISPECIES: hydrogenase-1 small chain [Proteobacteria] Length: 372\nNumber of predicted TMHs: 2\nExp number of AAs in TMHs: 37.2608499999999\nExp number, first 60 AAs: 19.52923\nTotal prob of N-in: 0.94351\nPOSSIBLE N-term signal sequence\ninside 1 19\nTMhelix 20 39\noutside 40 325\nTMhelix 326 348\ninside 349 372

9893 GCF_000714015.1_ASM71401v1 Escherichia coli 4-203-08_S4_C2 Proteobacteria; Gammaproteobacteria; Enterobacterales; Enterobacteriaceae; Escherichia; Escherichia coli

MTGDNTLIHSHGINRRDFMKLCAALAATMGLSSKAAAEMAESVTNPQRPPVIWIGAQECT WP_000145410.1 MULTISPECIES: hydrogenase-2 small chain [Proteobacteria] Length: 372\nNumber of predicted TMHs: 1\nExp number of AAs in TMHs: 23.72226\nExp number, first 60 AAs: 0.22038\nTotal prob of N-in: 0.12994\noutside 1 331\nTMhelix 332 354\ninside 355 372

9894 GCF_000700725.1_ASM70072v1 Escherichia coli 4-203-08_S4_C3 Proteobacteria; Gammaproteobacteria; Enterobacterales; Enterobacteriaceae; Escherichia; Escherichia coli MTGDNTLIHSHGINRRDFMKLCAALAATMGLSSKAAAEMAESVTNPQRPPVIWIGAQECT WP_000145410.1 MULTISPECIES: hydrogenase-2 small chain [Proteobacteria] Length: 372\nNumber of predicted TMHs: 1\nExp number of AAs in TMHs: 23.72226\nExp number, first 60 AAs: 0.22038\nTotal prob of N-in: 0.12994\noutside 1 331\nTMhelix 332 354\ninside 355 372

9895 GCF_000700725.1_ASM70072v1 Escherichia coli 4-203-08_S4_C3 Proteobacteria; Gammaproteobacteria; Enterobacterales; Enterobacteriaceae; Escherichia; Escherichia coli MNNEETFYQAMRRQGVTRRSFLKYCSLAATSLGLGAGMAPKIAWALENKPRIPVVWIHGL WP_001058323.1 MULTISPECIES: hydrogenase-1 small chain [Proteobacteria] Length: 372\nNumber of predicted TMHs: 2\nExp number of AAs in TMHs: 37.2608499999999\nExp number, first 60 AAs: 19.52923\nTotal prob of N-in: 0.94351\nPOSSIBLE N-term signal sequence\ninside 1 19\nTMhelix 20 39\noutside 40 325\nTMhelix 326 348\ninside 349 372

9896 GCF_000503455.1_Ecoli435 Escherichia coli 435 Proteobacteria; Gammaproteobacteria; Enterobacterales; Enterobacteriaceae; Escherichia; Escherichia coli MTGDNTLIHSHGINRRDFMKLCAALAATMGLSSKAAAEMAESVTNPQRPPVIWIGAQECT WP_000145410.1 MULTISPECIES: hydrogenase-2 small chain [Proteobacteria] Length: 372\nNumber of predicted TMHs: 1\nExp number of AAs in TMHs: 23.72226\nExp number, first 60 AAs: 0.22038\nTotal prob of N-in: 0.12994\noutside 1 331\nTMhelix 332 354\ninside 355 372

9897 GCF_000503455.1_Ecoli435 Escherichia coli 435 Proteobacteria; Gammaproteobacteria; Enterobacterales; Enterobacteriaceae; Escherichia; Escherichia coli MNNEETFYQAMRRQGVTRRSFLKYCSLAATSLGLGAGMAPKIAWALENKPRIPVVWIHGL WP_001058323.1 MULTISPECIES: hydrogenase-1 small chain [Proteobacteria] Length: 372\nNumber of predicted TMHs: 2\nExp number of AAs in TMHs: 37.2608499999999\nExp number, first 60 AAs: 19.52923\nTotal prob of N-in: 0.94351\nPOSSIBLE N-term signal sequence\ninside 1 19\nTMhelix 20 39\noutside 40 325\nTMhelix 326 348\ninside 349 372

9898 GCF_000699265.1_4541-1 Escherichia coli 4541-1 Proteobacteria; Gammaproteobacteria; Enterobacterales; Enterobacteriaceae; Escherichia; Escherichia coli MTGDNTLIHSHGINRRDFMKLCAALAATMGLSSKAAAEMAESVTNPQRPPVIWIGAQECT WP_000145410.1 MULTISPECIES: hydrogenase-2 small chain [Proteobacteria] Length: 372\nNumber of predicted TMHs: 1\nExp number of AAs in TMHs: 23.72226\nExp number, first 60 AAs: 0.22038\nTotal prob of N-in: 0.12994\noutside 1 331\nTMhelix 332 354\ninside 355 372

9899 GCF_000699265.1_4541-1 Escherichia coli 4541-1 Proteobacteria; Gammaproteobacteria; Enterobacterales; Enterobacteriaceae; Escherichia; Escherichia coli MNNEETFYQAMRRQGVTRRSFLKYCSLAATSLGLGAGMAPKIAWALENKPRIPVVWIHGL WP_001058323.1 MULTISPECIES: hydrogenase-1 small chain [Proteobacteria] Length: 372\nNumber of predicted TMHs: 2\nExp number of AAs in TMHs: 37.2608499999999\nExp number, first 60 AAs: 19.52923\nTotal prob of N-in: 0.94351\nPOSSIBLE N-term signal sequence\ninside 1 19\nTMhelix 20 39\noutside 40 325\nTMhelix 326 348\ninside 349 372

9900 GCF_000699285.1_4552-1 Escherichia coli 4552-1 Proteobacteria; Gammaproteobacteria; Enterobacterales; Enterobacteriaceae; Escherichia; Escherichia coli MTGDNTLIHSHGINRRDFMKLCAALAATMGLSSKAAAEMAESVTNPQRPPVIWIGAQECT WP_000145410.1 MULTISPECIES: hydrogenase-2 small chain [Proteobacteria] Length: 372\nNumber of predicted TMHs: 1\nExp number of AAs in TMHs: 23.72226\nExp number, first 60 AAs: 0.22038\nTotal prob of N-in: 0.12994\noutside 1 331\nTMhelix 332 354\ninside 355 372

9901 GCF_000699285.1_4552-1 Escherichia coli 4552-1 Proteobacteria; Gammaproteobacteria; Enterobacterales; Enterobacteriaceae; Escherichia; Escherichia coli MNNEETFYQAMRRQGVTRRSFLKYCSLAATSLGLGAGMAPKIAWALENKPRIPVVWIHGL WP_001058323.1 MULTISPECIES: hydrogenase-1 small chain [Proteobacteria] Length: 372\nNumber of predicted TMHs:

2\nExp number of AAs in TMHs: 37.2608499999999\nExp number, first 60 AAs: 19.52923\nTotal prob of N-in: 0.94351\nPOSSIBLE N-term signal sequence\ninside 1 19\nTMhelix 20 39\noutside 40 325\nTMhelix 326 348\ninside 349 372

9902 GCF_000215145.1_ASM21514v2 Escherichia coli 5.0588 Proteobacteria; Gammaproteobacteria; Enterobacterales; Enterobacteriaceae; Escherichia; Escherichia coli MNNEETFYQAMRRQGVTRRSFLKYCSLAATSLGLGAGMAPKIAWALENKPRIPVVWIHGL WP_001295940.1 MULTISPECIES: hydrogenase [Enterobacteriaceae] Length: 372\nNumber of predicted TMHs: 2\nExp number of AAs in TMHs: 37.2608899999999\nExp number, first 60 AAs: 19.52923\nTotal prob of N-in: 0.94351\nPOSSIBLE N-term signal sequence\ninside 1 19\nTMhelix 20 39\noutside 40 325\nTMhelix 326 348\ninside 349 372

9903 GCF_000215145.1_ASM21514v2 Escherichia coli 5.0588 Proteobacteria; Gammaproteobacteria; Enterobacterales; Enterobacteriaceae; Escherichia; Escherichia coli MTGDNTLIHSHGINRRDFMKLCAALAATMGLSSKAAAEMAESVTNPQRPPVIWIGAQECT WP_000145410.1 MULTISPECIES: hydrogenase-2 small chain [Proteobacteria] Length: 372\nNumber of predicted TMHs: 1\nExp number of AAs in TMHs: 23.72226\nExp number, first 60 AAs: 0.22038\nTotal prob of N-in: 0.12994\noutside 1 331\nTMhelix 332 354\ninside 355 372

9904 GCF_000194395.1_ASM19439v2 Escherichia coli 5.0959 Proteobacteria; Gammaproteobacteria; Enterobacterales; Enterobacteriaceae; Escherichia; Escherichia coli MTGDNTLIHSHGINRRDFMKLCAALAATMGLSSKAAAEMAESVTNPQRPPVIWIGAQECT WP_000145410.1 MULTISPECIES: hydrogenase-2 small chain [Proteobacteria] Length: 372\nNumber of predicted TMHs: 1\nExp number of AAs in TMHs: 23.72226\nExp number, first 60 AAs: 0.22038\nTotal prob of N-in: 0.12994\noutside 1 331\nTMhelix 332 354\ninside 355 372

9905 GCF_000714215.1_ASM71421v1 Escherichia coli 5-172-05_S1_C3 Proteobacteria; Gammaproteobacteria; Enterobacterales; Enterobacteriaceae; Escherichia; Escherichia coli MTGDNTLIHSHGINRRDFMKLCAALAATMGLSSKAAAEMAESVTNPQRPPVIWIGAQECT WP_032298447.1 hydrogenase 2 small subunit [Escherichia coli] Length: 372\nNumber of predicted TMHs: 1\nExp number of AAs in TMHs: 23.69807\nExp number, first 60 AAs: 0.221\nTotal prob of N-in: 0.12853\noutside 1 331\nTMhelix 332 354\ninside 355 372

9906 GCF_000714175.1_ASM71417v1 Escherichia coli 5-172-05_S3_C1 Proteobacteria; Gammaproteobacteria; Enterobacterales; Enterobacteriaceae; Escherichia; Escherichia coli MTGDNTLIHSHGINRRDFMKLCAALAATMGLSSKAAAEMAESVTNPQRPPVIWIGAQECT WP_000145410.1 MULTISPECIES: hydrogenase-2 small chain [Proteobacteria] Length: 372\nNumber of predicted TMHs: 1\nExp number of AAs in TMHs: 23.72226\nExp number, first 60 AAs: 0.22038\nTotal prob of N-in: 0.12994\noutside 1 331\nTMhelix 332 354\ninside 355 372

9907 GCF_000714175.1_ASM71417v1 Escherichia coli 5-172-05_S3_C1 Proteobacteria; Gammaproteobacteria; Enterobacterales; Enterobacteriaceae; Escherichia; Escherichia coli MNNEETFYQAMRRQGVTRRSFLKYCSLAATSLGLGAGMAPKIAWALENKPRIPVVWIHGL WP_001058323.1 MULTISPECIES: hydrogenase-1 small chain [Proteobacteria] Length: 372\nNumber of predicted TMHs: 2\nExp number of AAs in TMHs: 37.2608499999999\nExp number, first 60 AAs: 19.52923\nTotal prob of N-in: 0.94351\nPOSSIBLE N-term signal sequence\ninside 1 19\nTMhelix 20 39\noutside 40 325\nTMhelix 326 348\ninside 349 372

9908 GCF_000714135.1_ASM71413v1 Escherichia coli 5-172-05_S3_C3 Proteobacteria; Gammaproteobacteria; Enterobacterales; Enterobacteriaceae; Escherichia; Escherichia coli MNNEETFYQAMRRQGVTRRSFLKYCSLAATSLGLGAGMAPKIAWALENKPRIPVVWIHGL WP_001058323.1 MULTISPECIES: hydrogenase-1 small chain [Proteobacteria] Length: 372\nNumber of predicted TMHs: 2\nExp number of AAs in TMHs: 37.2608499999999\nExp number, first 60 AAs: 19.52923\nTotal prob of N-in: 0.94351\nPOSSIBLE N-term signal sequence\ninside 1 19\nTMhelix 20 39\noutside 40 325\nTMhelix 326 348\ninside 349 372

9909 GCF_000714135.1_ASM71413v1 Escherichia coli 5-172-05_S3_C3 Proteobacteria; Gammaproteobacteria; Enterobacterales; Enterobacteriaceae; Escherichia; Escherichia coli MTGDNTLIHSHGINRRDFMKLCAALAATMGLSSKAAAEMAESVTNPQRPPVIWIGAQECT WP_000145410.1 MULTISPECIES: hydrogenase-2 small chain [Proteobacteria] Length: 372\nNumber of predicted TMHs:

1\nExp number of AAs in TMHs: 23.72226\nExp number, first 60 AAs: 0.22038\nTotal prob of N-in: 0.12994\noutside 1 331\nTMhelix 332 354\ninside 355 372

9910 GCF_000714145.1_ASM71414v1 Escherichia coli 5-172-05_S4_C1 Proteobacteria; Gammaproteobacteria; Enterobacterales; Enterobacteriaceae; Escherichia; Escherichia coli MTGDNTLIHSHGINRRDFMKLCAALAATMGLSSKAAAEMAESVTNPQRPPVIWIGAQECT WP_000145410.1 MULTISPECIES: hydrogenase-2 small chain [Proteobacteria] Length: 372\nNumber of predicted TMHs: 1\nExp number of AAs in TMHs: 23.72226\nExp number, first 60 AAs: 0.22038\nTotal prob of N-in: 0.12994\noutside 1 331\nTMhelix 332 354\ninside 355 372

9911 GCF_000714145.1_ASM71414v1 Escherichia coli 5-172-05_S4_C1 Proteobacteria; Gammaproteobacteria; Enterobacterales; Enterobacteriaceae; Escherichia; Escherichia coli MNNEETFYQAMRRQGVTRRSFLKYCSLAATSLGLGAGMAPKIAWALENKPRIPVVWIHGL WP_001058323.1 MULTISPECIES: hydrogenase-1 small chain [Proteobacteria] Length: 372\nNumber of predicted TMHs: 2\nExp number of AAs in TMHs: 37.2608499999999\nExp number, first 60 AAs: 19.52923\nTotal prob of N-in: 0.94351\nPOSSIBLE N-term signal sequence\ninside 1 19\nTMhelix 20 39\noutside 40 325\nTMhelix 326 348\ninside 349 372

9912 GCF_000714105.1_ASM71410v1 Escherichia coli 5-172-05_S4_C2 Proteobacteria; Gammaproteobacteria; Enterobacterales; Enterobacteriaceae; Escherichia; Escherichia coli MNNEETFYQAMRRQGVTRRSFLKYCSLAATSLGLGAGMAPKIAWALENKPRIPVVWIHGL WP_001058323.1 MULTISPECIES: hydrogenase-1 small chain [Proteobacteria] Length: 372\nNumber of predicted TMHs: 2\nExp number of AAs in TMHs: 37.2608499999999\nExp number, first 60 AAs: 19.52923\nTotal prob of N-in: 0.94351\nPOSSIBLE N-term signal sequence\ninside 1 19\nTMhelix 20 39\noutside 40 325\nTMhelix 326 348\ninside 349 372

9913 GCF_000714105.1_ASM71410v1 Escherichia coli 5-172-05_S4_C2 Proteobacteria; Gammaproteobacteria; Enterobacterales; Enterobacteriaceae; Escherichia; Escherichia coli MTGDNTLIHSHGINRRDFMKLCAALAATMGLSSKAAAEMAESVTNPQRPPVIWIGAQECT WP_000145410.1 MULTISPECIES: hydrogenase-2 small chain [Proteobacteria] Length: 372\nNumber of predicted TMHs: 1\nExp number of AAs in TMHs: 23.72226\nExp number, first 60 AAs: 0.22038\nTotal prob of N-in: 0.12994\noutside 1 331\nTMhelix 332 354\ninside 355 372

9914 GCF_000714225.1_ASM71422v1 Escherichia coli 5-172-05_S4_C3 Proteobacteria; Gammaproteobacteria; Enterobacterales; Enterobacteriaceae; Escherichia; Escherichia coli MNNEETFYQAMRRQGVTRRSFLKYCSLAATSLGLGAGMAPKIAWALENKPRIPVVWIHGL WP_001058323.1 MULTISPECIES: hydrogenase-1 small chain [Proteobacteria] Length: 372\nNumber of predicted TMHs: 2\nExp number of AAs in TMHs: 37.2608499999999\nExp number, first 60 AAs: 19.52923\nTotal prob of N-in: 0.94351\nPOSSIBLE N-term signal sequence\ninside 1 19\nTMhelix 20 39\noutside 40 325\nTMhelix 326 348\ninside 349 372

9915 GCF_000714225.1_ASM71422v1 Escherichia coli 5-172-05_S4_C3 Proteobacteria; Gammaproteobacteria; Enterobacterales; Enterobacteriaceae; Escherichia; Escherichia coli MTGDNTLIHSHGINRRDFMKLCAALAATMGLSSKAAAEMAESVTNPQRPPVIWIGAQECT WP_000145410.1 MULTISPECIES: hydrogenase-2 small chain [Proteobacteria] Length: 372\nNumber of predicted TMHs: 1\nExp number of AAs in TMHs: 23.72226\nExp number, first 60 AAs: 0.22038\nTotal prob of N-in: 0.12994\noutside 1 331\nTMhelix 332 354\ninside 355 372

9916 GCF_000304875.2_ASM30487v2 Escherichia coli 5.2239 Proteobacteria; Gammaproteobacteria; Enterobacterales; Enterobacteriaceae; Escherichia; Escherichia coli MTGDNTLIHSHGINRRDFMKLCAALAATMGLSSKAAAEMAESVTNPQRPPVIWIGAQECT WP_000145410.1 MULTISPECIES: hydrogenase-2 small chain [Proteobacteria] Length: 372\nNumber of predicted TMHs: 1\nExp number of AAs in TMHs: 23.72226\nExp number, first 60 AAs: 0.22038\nTotal prob of N-in: 0.12994\noutside 1 331\nTMhelix 332 354\ninside 355 372

9917 GCF_000304875.2_ASM30487v2 Escherichia coli 5.2239 Proteobacteria; Gammaproteobacteria; Enterobacterales; Enterobacteriaceae; Escherichia; Escherichia coli MNNEETFYQAMRRQGVTRRSFLKYCSLAATSLGLGAGMAPKIAWALENKPRIPVVWIHGL WP_001301613.1 hydrogenase [Escherichia coli] Length: 372\nNumber of predicted TMHs: 2\nExp number of AAs in TMHs: 37.46598\nExp number, first 60 AAs: 19.53139\nTotal prob of N-in: 0.94364\nPOSSIBLE N-term signal sequence\ninside 1 19\nTMhelix 20 39\noutside 40 325\nTMhelix 326 348\ninside 349 372

9918 GCF_000013305.1_ASM1330v1 Escherichia coli 536 Proteobacteria; Gammaproteobacteria; Enterobacterales; Enterobacteriaceae; Escherichia; Escherichia coli
MTGDNTLIHSHGINRRDFMKLCAALAATMGLSSKAAAEMAESVTNPQRPPVIWIGAQECT WP_000145405.1
hydrogenase 2 small subunit [Escherichia coli] Length: 372\nNumber of predicted TMHs: 1\nExp number of AAs in TMHs: 24.53687\nExp number, first 60 AAs: 0.21619\nTotal prob of N-in: 0.16244\noutside 1 331\nTMhelix 332 354\ninside 355 372

9919 GCF_000013305.1_ASM1330v1 Escherichia coli 536 Proteobacteria; Gammaproteobacteria; Enterobacterales; Enterobacteriaceae; Escherichia; Escherichia coli
MNNEETFYQAMRRQGVTRRSFLKYCSLAATSLGLGAGMAPKIAWALENKPRIPVVWIHGL WP_001058323.1
MULTISPECIES: hydrogenase-1 small chain [Proteobacteria] Length: 372\nNumber of predicted TMHs: 2\nExp number of AAs in TMHs: 37.2608499999999\nExp number, first 60 AAs: 19.52923\nTotal prob of N-in: 0.94351\nPOSSIBLE N-term signal sequence\ninside 1 19\nTMhelix 20 39\noutside 40 325\nTMhelix 326 348\ninside 349 372

9920 GCF_000167915.2_ASM16791v2 Escherichia coli 53638 Proteobacteria; Gammaproteobacteria; Enterobacterales; Enterobacteriaceae; Escherichia; Escherichia coli
MTGDNTLIHSHGINRRDFMKLCAALAATMGLSSKAAAEMAESVTNPQRPPVIWIGAQECT WP_000145410.1
MULTISPECIES: hydrogenase-2 small chain [Proteobacteria] Length: 372\nNumber of predicted TMHs: 1\nExp number of AAs in TMHs: 23.72226\nExp number, first 60 AAs: 0.22038\nTotal prob of N-in: 0.12994\noutside 1 331\nTMhelix 332 354\ninside 355 372

9921 GCF_000167915.2_ASM16791v2 Escherichia coli 53638 Proteobacteria; Gammaproteobacteria; Enterobacterales; Enterobacteriaceae; Escherichia; Escherichia coli
MNNEETFYQAMRRQGVTRRSFLKYCSLAATSLGLGAGMAPKIAWALENKPRIPVVWIHGL WP_001058323.1
MULTISPECIES: hydrogenase-1 small chain [Proteobacteria] Length: 372\nNumber of predicted TMHs: 2\nExp number of AAs in TMHs: 37.2608499999999\nExp number, first 60 AAs: 19.52923\nTotal prob of N-in: 0.94351\nPOSSIBLE N-term signal sequence\ninside 1 19\nTMhelix 20 39\noutside 40 325\nTMhelix 326 348\ninside 349 372

9922 GCF_000714255.1_ASM71425v1 Escherichia coli 5-366-08_S1_C1 Proteobacteria; Gammaproteobacteria; Enterobacterales; Enterobacteriaceae; Escherichia; Escherichia coli
MTGDNTLIHSHGINRRDFMKLCAALAATMGLSSKAAAEMAESVTNPQRPPVIWIGAQECT WP_000145410.1
MULTISPECIES: hydrogenase-2 small chain [Proteobacteria] Length: 372\nNumber of predicted TMHs: 1\nExp number of AAs in TMHs: 23.72226\nExp number, first 60 AAs: 0.22038\nTotal prob of N-in: 0.12994\noutside 1 331\nTMhelix 332 354\ninside 355 372

9923 GCF_000714255.1_ASM71425v1 Escherichia coli 5-366-08_S1_C1 Proteobacteria; Gammaproteobacteria; Enterobacterales; Enterobacteriaceae; Escherichia; Escherichia coli
MNNEETFYQAMRRQGVTRRSFLKYCSLAATSLGLGAGMAPKIAWALENKPRIPVVWIHGL WP_001058323.1
MULTISPECIES: hydrogenase-1 small chain [Proteobacteria] Length: 372\nNumber of predicted TMHs: 2\nExp number of AAs in TMHs: 37.2608499999999\nExp number, first 60 AAs: 19.52923\nTotal prob of N-in: 0.94351\nPOSSIBLE N-term signal sequence\ninside 1 19\nTMhelix 20 39\noutside 40 325\nTMhelix 326 348\ninside 349 372

9924 GCF_000711605.1_ASM71160v1 Escherichia coli 5-366-08_S1_C3 Proteobacteria; Gammaproteobacteria; Enterobacterales; Enterobacteriaceae; Escherichia; Escherichia coli
MNNEETFYQAMRRQGVTRRSFLKYCSLAATSLGLGAGMAPKIAWALENKPRIPVVWIHGL WP_001058323.1
MULTISPECIES: hydrogenase-1 small chain [Proteobacteria] Length: 372\nNumber of predicted TMHs: 2\nExp number of AAs in TMHs: 37.2608499999999\nExp number, first 60 AAs: 19.52923\nTotal prob of N-in: 0.94351\nPOSSIBLE N-term signal sequence\ninside 1 19\nTMhelix 20 39\noutside 40 325\nTMhelix 326 348\ninside 349 372

9925 GCF_000711605.1_ASM71160v1 Escherichia coli 5-366-08_S1_C3 Proteobacteria; Gammaproteobacteria; Enterobacterales; Enterobacteriaceae; Escherichia; Escherichia coli
MTGDNTLIHSHGINRRDFMKLCAALAATMGLSSKAAAEMAESVTNPQRPPVIWIGAQECT WP_000145410.1
MULTISPECIES: hydrogenase-2 small chain [Proteobacteria] Length: 372\nNumber of predicted TMHs: 1\nExp number of AAs in TMHs: 23.72226\nExp number, first 60 AAs: 0.22038\nTotal prob of N-in: 0.12994\noutside 1 331\nTMhelix 332 354\ninside 355 372

9926 GCF_000714335.1_ASM71433v1 Escherichia coli 5-366-08_S3_C1 Proteobacteria; Gammaproteobacteria; Enterobacterales; Enterobacteriaceae; Escherichia; Escherichia coli MTGDNTLIHSHGINRRDFMKLCAALAATMGLSSKAAAEMAESVTNPQRPPVIWIGAQECT WP_000145410.1 MULTISPECIES: hydrogenase-2 small chain [Proteobacteria] Length: 372\nNumber of predicted TMHs: 1\nExp number of AAs in TMHs: 23.72226\nExp number, first 60 AAs: 0.22038\nTotal prob of N-in: 0.12994\noutside 1 331\nTMhelix 332 354\ninside 355 372

9927 GCF_000714335.1_ASM71433v1 Escherichia coli 5-366-08_S3_C1 Proteobacteria; Gammaproteobacteria; Enterobacterales; Enterobacteriaceae; Escherichia; Escherichia coli MNNEETFYQAMRRQGVTRRSFLKYCSLAATSLGLGAGMAPKIAWALENKPRIPVVWIHGL WP_001058323.1 MULTISPECIES: hydrogenase-1 small chain [Proteobacteria] Length: 372\nNumber of predicted TMHs: 2\nExp number of AAs in TMHs: 37.2608499999999\nExp number, first 60 AAs: 19.52923\nTotal prob of N-in: 0.94351\nPOSSIBLE N-term signal sequence\ninside 1 19\nTMhelix 20 39\noutside 40 325\nTMhelix 326 348\ninside 349 372

9928 GCF_000714295.1_ASM71429v1 Escherichia coli 5-366-08_S3_C2 Proteobacteria; Gammaproteobacteria; Enterobacterales; Enterobacteriaceae; Escherichia; Escherichia coli MNNEETFYQAMRRQGVTRRSFLKYCSLAATSLGLGAGMAPKIAWALENKPRIPVVWIHGL WP_001295940.1 MULTISPECIES: hydrogenase [Enterobacteriaceae] Length: 372\nNumber of predicted TMHs: 2\nExp number of AAs in TMHs: 37.2608899999999\nExp number, first 60 AAs: 19.52923\nTotal prob of N-in: 0.94351\nPOSSIBLE N-term signal sequence\ninside 1 19\nTMhelix 20 39\noutside 40 325\nTMhelix 326 348\ninside 349 372

9929 GCF_000714295.1_ASM71429v1 Escherichia coli 5-366-08_S3_C2 Proteobacteria; Gammaproteobacteria; Enterobacterales; Enterobacteriaceae; Escherichia; Escherichia coli MTGDNTLIHSHGINRRDFMKLCAALAATMGLSSKAAAEMAESVTNPQRPPVIWIGAQECT WP_000145410.1 MULTISPECIES: hydrogenase-2 small chain [Proteobacteria] Length: 372\nNumber of predicted TMHs: 1\nExp number of AAs in TMHs: 23.72226\nExp number, first 60 AAs: 0.22038\nTotal prob of N-in: 0.12994\noutside 1 331\nTMhelix 332 354\ninside 355 372

9930 GCF_000714265.1_ASM71426v1 Escherichia coli 5-366-08_S3_C3 Proteobacteria; Gammaproteobacteria; Enterobacterales; Enterobacteriaceae; Escherichia; Escherichia coli MTGDNTLIHSHGINRRDFMKLCAALAATMGLSSKAAAEMAESVTNPQRPPVIWIGAQECT WP_000145410.1 MULTISPECIES: hydrogenase-2 small chain [Proteobacteria] Length: 372\nNumber of predicted TMHs: 1\nExp number of AAs in TMHs: 23.72226\nExp number, first 60 AAs: 0.22038\nTotal prob of N-in: 0.12994\noutside 1 331\nTMhelix 332 354\ninside 355 372

9931 GCF_000714265.1_ASM71426v1 Escherichia coli 5-366-08_S3_C3 Proteobacteria; Gammaproteobacteria; Enterobacterales; Enterobacteriaceae; Escherichia; Escherichia coli MNNEETFYQAMRRQGVTRRSFLKYCSLAATSLGLGAGMAPKIAWALENKPRIPVVWIHGL WP_001058323.1 MULTISPECIES: hydrogenase-1 small chain [Proteobacteria] Length: 372\nNumber of predicted TMHs: 2\nExp number of AAs in TMHs: 37.2608499999999\nExp number, first 60 AAs: 19.52923\nTotal prob of N-in: 0.94351\nPOSSIBLE N-term signal sequence\ninside 1 19\nTMhelix 20 39\noutside 40 325\nTMhelix 326 348\ninside 349 372

9932 GCF_000713855.1_ASM71385v1 Escherichia coli 5-366-08_S4_C1 Proteobacteria; Gammaproteobacteria; Enterobacterales; Enterobacteriaceae; Escherichia; Escherichia coli MTGDNTLIHSHGINRRDFMKLCAALAATMGLSSKAAAEMAESVTNPQRPPVIWIGAQECT WP_000145410.1 MULTISPECIES: hydrogenase-2 small chain [Proteobacteria] Length: 372\nNumber of predicted TMHs: 1\nExp number of AAs in TMHs: 23.72226\nExp number, first 60 AAs: 0.22038\nTotal prob of N-in: 0.12994\noutside 1 331\nTMhelix 332 354\ninside 355 372

9933 GCF_000713855.1_ASM71385v1 Escherichia coli 5-366-08_S4_C1 Proteobacteria; Gammaproteobacteria; Enterobacterales; Enterobacteriaceae; Escherichia; Escherichia coli MNNEETFYQAMRRQGVTRRSFLKYCSLAATSLGLGAGMAPKIAWALENKPRIPVVWIHGL WP_001058323.1 MULTISPECIES: hydrogenase-1 small chain [Proteobacteria] Length: 372\nNumber of predicted TMHs: 2\nExp number of AAs in TMHs: 37.2608499999999\nExp number, first 60 AAs: 19.52923\nTotal prob of N-in: 0.94351\nPOSSIBLE N-term signal sequence\ninside 1 19\nTMhelix 20 39\noutside 40 325\nTMhelix 326 348\ninside 349 372

9934 GCF_000714095.1_ASM71409v1 Escherichia coli 5-366-08_S4_C2 Proteobacteria; Gammaproteobacteria; Enterobacterales; Enterobacteriaceae; Escherichia; Escherichia coli MTGDNTLIHSHGINRRDFMKLCAALAATMGLSSKAAAEMAESVTNPQRPPVIWIGAQECT WP_000145410.1 MULTISPECIES: hydrogenase-2 small chain [Proteobacteria] Length: 372\nNumber of predicted TMHs: 1\nExp number of AAs in TMHs: 23.72226\nExp number, first 60 AAs: 0.22038\nTotal prob of N-in: 0.12994\noutside 1 331\nTMhelix 332 354\ninside 355 372

9935 GCF_000714095.1_ASM71409v1 Escherichia coli 5-366-08_S4_C2 Proteobacteria; Gammaproteobacteria; Enterobacterales; Enterobacteriaceae; Escherichia; Escherichia coli MNNEETFYQAMRRQGVTRRSFLKYCSLAATSLGLGAGMAPKIAWALENKPRIPVVWIHGL WP_001058323.1 MULTISPECIES: hydrogenase-1 small chain [Proteobacteria] Length: 372\nNumber of predicted TMHs: 2\nExp number of AAs in TMHs: 37.2608499999999\nExp number, first 60 AAs: 19.52923\nTotal prob of N-in: 0.94351\nPOSSIBLE N-term signal sequence\ninside 1 19\nTMhelix 20 39\noutside 40 325\nTMhelix 326 348\ninside 349 372

9936 GCF_000503395.1_Ecoli53A Escherichia coli 53A Proteobacteria; Gammaproteobacteria; Enterobacterales; Enterobacteriaceae; Escherichia; Escherichia coli MNNEETFYQAMRRQGVTRRSFLKYCSLAATSLGLGAGMAPKIAWALENKPRIPVVWIHGL WP_001058323.1 MULTISPECIES: hydrogenase-1 small chain [Proteobacteria] Length: 372\nNumber of predicted TMHs: 2\nExp number of AAs in TMHs: 37.2608499999999\nExp number, first 60 AAs: 19.52923\nTotal prob of N-in: 0.94351\nPOSSIBLE N-term signal sequence\ninside 1 19\nTMhelix 20 39\noutside 40 325\nTMhelix 326 348\ninside 349 372

9937 GCF_000503395.1_Ecoli53A Escherichia coli 53A Proteobacteria; Gammaproteobacteria; Enterobacterales; Enterobacteriaceae; Escherichia; Escherichia coli MTGDNTLIHSHGINRRDFMKLCAALAATMGLSSKAAAEMAESVTNPQRPPVIWIGAQECT WP_000145410.1 MULTISPECIES: hydrogenase-2 small chain [Proteobacteria] Length: 372\nNumber of predicted TMHs: 1\nExp number of AAs in TMHs: 23.72226\nExp number, first 60 AAs: 0.22038\nTotal prob of N-in: 0.12994\noutside 1 331\nTMhelix 332 354\ninside 355 372

9938 GCF_000503595.1_Ecoli53C Escherichia coli 53CProteobacteria; Gammaproteobacteria; Enterobacterales; Enterobacteriaceae; Escherichia; Escherichia coli MTGDNTLIHSHGINRRDFMKLCAALAATMGLSSKAAAEMAESVTNPQRPPVIWIGAQECT WP_000145410.1 MULTISPECIES: hydrogenase-2 small chain [Proteobacteria] Length: 372\nNumber of predicted TMHs: 1\nExp number of AAs in TMHs: 23.72226\nExp number, first 60 AAs: 0.22038\nTotal prob of N-in: 0.12994\noutside 1 331\nTMhelix 332 354\ninside 355 372

9939 GCF_000831715.1_ASM83171v1 Escherichia coli 53CProteobacteria; Gammaproteobacteria; Enterobacterales; Enterobacteriaceae; Escherichia; Escherichia coli MTGDNTLIHSHGINRRDFMKLCAALAATMGLSSKAAAEMAESVTNPQRPPVIWIGAQECT WP_000145410.1 MULTISPECIES: hydrogenase-2 small chain [Proteobacteria] Length: 372\nNumber of predicted TMHs: 1\nExp number of AAs in TMHs: 23.72226\nExp number, first 60 AAs: 0.22038\nTotal prob of N-in: 0.12994\noutside 1 331\nTMhelix 332 354\ninside 355 372

9940 GCF_000831715.1_ASM83171v1 Escherichia coli 53CProteobacteria; Gammaproteobacteria; Enterobacterales; Enterobacteriaceae; Escherichia; Escherichia coli MNNEETFYQAMRRQGVTRRSFLKYCSLAATSLGLGAGMAPKIAWALENKPRIPVVWIHGL WP_001058323.1 MULTISPECIES: hydrogenase-1 small chain [Proteobacteria] Length: 372\nNumber of predicted TMHs: 2\nExp number of AAs in TMHs: 37.2608499999999\nExp number, first 60 AAs: 19.52923\nTotal prob of N-in: 0.94351\nPOSSIBLE N-term signal sequence\ninside 1 19\nTMhelix 20 39\noutside 40 325\nTMhelix 326 348\ninside 349 372

9941 GCF_000503595.1_Ecoli53C Escherichia coli 53CProteobacteria; Gammaproteobacteria; Enterobacterales; Enterobacteriaceae; Escherichia; Escherichia coli MNNEETFYQAMRRQGVTRRSFLKYCSLAATSLGLGAGMAPKIAWALENKPRIPVVWIHGL WP_001058323.1 MULTISPECIES: hydrogenase-1 small chain [Proteobacteria] Length: 372\nNumber of predicted TMHs: 2\nExp number of AAs in TMHs: 37.2608499999999\nExp number, first 60 AAs: 19.52923\nTotal prob of N-in: 0.94351\nPOSSIBLE N-term signal sequence\ninside 1 19\nTMhelix 20 39\noutside 40 325\nTMhelix 326 348\ninside 349 372

9942 GCF_000264215.1_ASM26421v1 Escherichia coli 541-1 Proteobacteria; Gammaproteobacteria; Enterobacterales; Enterobacteriaceae; Escherichia; Escherichia coli
MNNEETFYQAMRRQGVTRRSFLKYCSLAATSLGLGAGMAPKIAWALENKPRIPVVWIHGL WP_001058323.1
MULTISPECIES: hydrogenase-1 small chain [Proteobacteria] Length: 372\nNumber of predicted TMHs: 2\nExp number of AAs in TMHs: 37.2608499999999\nExp number, first 60 AAs: 19.52923\nTotal prob of N-in: 0.94351\nPOSSIBLE N-term signal sequence\ninside 1 19\nTMhelix 20 39\noutside 40 325\nTMhelix 326 348\ninside 349 372

9943 GCF_000264215.1_ASM26421v1 Escherichia coli 541-1 Proteobacteria; Gammaproteobacteria; Enterobacterales; Enterobacteriaceae; Escherichia; Escherichia coli
MTGDNTLIHSHGINRRDFMKLCAALAATMGLSSKAAAEMAESVTNPQRPPVIWIGAQECT WP_000145410.1
MULTISPECIES: hydrogenase-2 small chain [Proteobacteria] Length: 372\nNumber of predicted TMHs: 1\nExp number of AAs in TMHs: 23.72226\nExp number, first 60 AAs: 0.22038\nTotal prob of N-in: 0.12994\noutside 1 331\nTMhelix 332 354\ninside 355 372

9944 GCF_000264115.1_ASM26411v1 Escherichia coli 541-15 Proteobacteria; Gammaproteobacteria; Enterobacterales; Enterobacteriaceae; Escherichia; Escherichia coli
MTGDNTLIHSHGINRRDFMKLCAALAATMGLSSKAAAEMAESVTNPQRPPVIWIGAQECT WP_000145410.1
MULTISPECIES: hydrogenase-2 small chain [Proteobacteria] Length: 372\nNumber of predicted TMHs: 1\nExp number of AAs in TMHs: 23.72226\nExp number, first 60 AAs: 0.22038\nTotal prob of N-in: 0.12994\noutside 1 331\nTMhelix 332 354\ninside 355 372

9945 GCF_000264115.1_ASM26411v1 Escherichia coli 541-15 Proteobacteria; Gammaproteobacteria; Enterobacterales; Enterobacteriaceae; Escherichia; Escherichia coli
MNNEETFYQAMRRQGVTRRSFLKYCSLAATSLGLGAGMAPKIAWALENKPRIPVVWIHGL WP_001058323.1
MULTISPECIES: hydrogenase-1 small chain [Proteobacteria] Length: 372\nNumber of predicted TMHs: 2\nExp number of AAs in TMHs: 37.2608499999999\nExp number, first 60 AAs: 19.52923\nTotal prob of N-in: 0.94351\nPOSSIBLE N-term signal sequence\ninside 1 19\nTMhelix 20 39\noutside 40 325\nTMhelix 326 348\ninside 349 372

9946 GCF_000304055.2_ASM30405v2 Escherichia coli 5412 Proteobacteria; Gammaproteobacteria; Enterobacterales; Enterobacteriaceae; Escherichia; Escherichia coli
MTGDNTLIHSHGINRRDFMKLCAALAATMGLSSKAAAEMAESVTNPQRPPVIWIGAQECT WP_000145410.1
MULTISPECIES: hydrogenase-2 small chain [Proteobacteria] Length: 372\nNumber of predicted TMHs: 1\nExp number of AAs in TMHs: 23.72226\nExp number, first 60 AAs: 0.22038\nTotal prob of N-in: 0.12994\noutside 1 331\nTMhelix 332 354\ninside 355 372

9947 GCF_000304055.2_ASM30405v2 Escherichia coli 5412 Proteobacteria; Gammaproteobacteria; Enterobacterales; Enterobacteriaceae; Escherichia; Escherichia coli
MNNEETFYQAMRRQGVTRRSFLKYCSLAATSLGLGAGMAPKIAWALENKPRIPVVWIHGL WP_001301613.1
hydrogenase [Escherichia coli] Length: 372\nNumber of predicted TMHs: 2\nExp number of AAs in TMHs: 37.46598\nExp number, first 60 AAs: 19.53139\nTotal prob of N-in: 0.94364\nPOSSIBLE N-term signal sequence\ninside 1 19\nTMhelix 20 39\noutside 40 325\nTMhelix 326 348\ninside 349 372

9948 GCF_000026245.1_ASM2624v1 Escherichia coli 55989 Proteobacteria; Gammaproteobacteria; Enterobacterales; Enterobacteriaceae; Escherichia; Escherichia coli
MTGDNTLIHSHGINRRDFMKLCAALAATMGLSSKAAAEMAESVTNPQRPPVIWIGAQECT WP_000145410.1
MULTISPECIES: hydrogenase-2 small chain [Proteobacteria] Length: 372\nNumber of predicted TMHs: 1\nExp number of AAs in TMHs: 23.72226\nExp number, first 60 AAs: 0.22038\nTotal prob of N-in: 0.12994\noutside 1 331\nTMhelix 332 354\ninside 355 372

9949 GCF_000026245.1_ASM2624v1 Escherichia coli 55989 Proteobacteria; Gammaproteobacteria; Enterobacterales; Enterobacteriaceae; Escherichia; Escherichia coli
MNNEETFYQAMRRQGVTRRSFLKYCSLAATSLGLGAGMAPKIAWALENKPRIPVVWIHGL WP_001058323.1
MULTISPECIES: hydrogenase-1 small chain [Proteobacteria] Length: 372\nNumber of predicted TMHs: 2\nExp number of AAs in TMHs: 37.2608499999999\nExp number, first 60 AAs: 19.52923\nTotal prob of N-in: 0.94351\nPOSSIBLE N-term signal sequence\ninside 1 19\nTMhelix 20 39\noutside 40 325\nTMhelix 326 348\ninside 349 372

9950 GCF_000264135.1_ASM26413v1 Escherichia coli 576-1 Proteobacteria; Gammaproteobacteria; Enterobacterales; Enterobacteriaceae; Escherichia; Escherichia coli

MTGDNTLIHSHGINRRDFMKLCAALAATMGLSSKAAAEMAESVTNPQRPPVIWIGAQECT WP_000145410.1 MULTISPECIES: hydrogenase-2 small chain [Proteobacteria] Length: 372\nNumber of predicted TMHs: 1\nExp number of AAs in TMHs: 23.72226\nExp number, first 60 AAs: 0.22038\nTotal prob of N-in: 0.12994\noutside 1 331\nTMhelix 332 354\ninside 355 372

9951 GCF_000264135.1_ASM26413v1 Escherichia coli 576-1 Proteobacteria; Gammaproteobacteria; Enterobacterales; Enterobacteriaceae; Escherichia; Escherichia coli MNNEETFYQAMRRQGVTRRSFLKYCSLAATSLGLGAGMAPKIAWALENKPRIPVVWIHGL WP_001058323.1 MULTISPECIES: hydrogenase-1 small chain [Proteobacteria] Length: 372\nNumber of predicted TMHs: 2\nExp number of AAs in TMHs: 37.2608499999999\nExp number, first 60 AAs: 19.52923\nTotal prob of N-in: 0.94351\nPOSSIBLE N-term signal sequence\ninside 1 19\nTMhelix 20 39\noutside 40 325\nTMhelix 326 348\ninside 349 372

9952 GCF_000303995.2_ASM30399v2 Escherichia coli 5905 Proteobacteria; Gammaproteobacteria; Enterobacterales; Enterobacteriaceae; Escherichia; Escherichia coli MNNEETFYQAMRRQGVTRRSFLKYCSLAATSLGLGAGMAPKIAWALENKPRIPVVWIHGL WP_001359719.1 hydrogenase [Escherichia coli] Length: 372\nNumber of predicted TMHs: 2\nExp number of AAs in TMHs: 37.2176\nExp number, first 60 AAs: 19.53448\nTotal prob of N-in: 0.94360\nPOSSIBLE N-term signal sequence\ninside 1 19\nTMhelix 20 39\noutside 40 325\nTMhelix 326 348\ninside 349 372

9953 GCF_000303995.2_ASM30399v2 Escherichia coli 5905 Proteobacteria; Gammaproteobacteria; Enterobacterales; Enterobacteriaceae; Escherichia; Escherichia coli MTGDNTLIHSHGINRRDFMKLCAALAATMGLSSKAAAEMAESVTNPQRPPVIWIGAQECT WP_000145410.1 MULTISPECIES: hydrogenase-2 small chain [Proteobacteria] Length: 372\nNumber of predicted TMHs: 1\nExp number of AAs in TMHs: 23.72226\nExp number, first 60 AAs: 0.22038\nTotal prob of N-in: 0.12994\noutside 1 331\nTMhelix 332 354\ninside 355 372

9954 GCF_000503475.1_Ecoli597 Escherichia coli 597 Proteobacteria; Gammaproteobacteria; Enterobacterales; Enterobacteriaceae; Escherichia; Escherichia coli MTGDNTLIHSHGINRRDFMKLCAALAATMGLSSKAAAEMAESVTNPQRPPVIWIGAQECT WP_000145410.1 MULTISPECIES: hydrogenase-2 small chain [Proteobacteria] Length: 372\nNumber of predicted TMHs: 1\nExp number of AAs in TMHs: 23.72226\nExp number, first 60 AAs: 0.22038\nTotal prob of N-in: 0.12994\noutside 1 331\nTMhelix 332 354\ninside 355 372

9955 GCF_000305155.2_ASM30515v2 Escherichia coli 6.0172 Proteobacteria; Gammaproteobacteria; Enterobacterales; Enterobacteriaceae; Escherichia; Escherichia coli MTGDNTLIHSHGINRRDFMKLCAALAATMGLSSKAAAEMAESVTNPQRPPVIWIGAQECT WP_000145410.1 MULTISPECIES: hydrogenase-2 small chain [Proteobacteria] Length: 372\nNumber of predicted TMHs: 1\nExp number of AAs in TMHs: 23.72226\nExp number, first 60 AAs: 0.22038\nTotal prob of N-in: 0.12994\noutside 1 331\nTMhelix 332 354\ninside 355 372

9956 GCF_000305155.2_ASM30515v2 Escherichia coli 6.0172 Proteobacteria; Gammaproteobacteria; Enterobacterales; Enterobacteriaceae; Escherichia; Escherichia coli MNNEETFYQAMRRQGVTRRSFLKYCSLAATSLGLGAGMAPKIAWALENKPRIPVVWIHGL WP_001301613.1 hydrogenase [Escherichia coli] Length: 372\nNumber of predicted TMHs: 2\nExp number of AAs in TMHs: 37.46598\nExp number, first 60 AAs: 19.53139\nTotal prob of N-in: 0.94364\nPOSSIBLE N-term signal sequence\ninside 1 19\nTMhelix 20 39\noutside 40 325\nTMhelix 326 348\ninside 349 372

9957 GCF_000503495.1_Ecoli606 Escherichia coli 606 Proteobacteria; Gammaproteobacteria; Enterobacterales; Enterobacteriaceae; Escherichia; Escherichia coli MTGDNTLIHSHGINRRDFMKLCAALAATMGLSSKAAAEMAESVTNPQRPPVIWIGAQECT WP_029402609.1 hydrogenase 2 small subunit [Escherichia coli] Length: 372\nNumber of predicted TMHs: 1\nExp number of AAs in TMHs: 23.72225\nExp number, first 60 AAs: 0.22038\nTotal prob of N-in: 0.12994\noutside 1 331\nTMhelix 332 354\ninside 355 372

9958 GCF_000503495.1_Ecoli606 Escherichia coli 606 Proteobacteria; Gammaproteobacteria; Enterobacterales; Enterobacteriaceae; Escherichia; Escherichia coli MNNEETFYQAMRRQGVTRRSFLKYCSLAATSLGLGAGMAPKIAWALENKPRIPVVWIHGL WP_001295940.1 MULTISPECIES: hydrogenase [Enterobacteriaceae] Length: 372\nNumber of predicted TMHs: 2\nExp number of AAs in TMHs: 37.2608899999999\nExp number, first 60 AAs: 19.52923\nTotal prob of N-in:

0.94351\nPOSSIBLE N-term signal sequence\ninside 1 19\nTMhelix 20 39\noutside 40 325\nTMhelix 326 348\ninside 349 372

9959 GCF_000714185.1_ASM71418v1 Escherichia coli 6-175-07_S1_C1 Proteobacteria; Gammaproteobacteria; Enterobacterales; Enterobacteriaceae; Escherichia; Escherichia coli MNNEETFYQAMRRQGVTRRSFLKYCSLAATSLGLGAGMAPKIAWALENKPRIPVVWIHGL WP_001295940.1 MULTISPECIES: hydrogenase [Enterobacteriaceae] Length: 372\nNumber of predicted TMHs: 2\nExp number of AAs in TMHs: 37.2608899999999\nExp number, first 60 AAs: 19.52923\nTotal prob of N-in: 0.94351\nPOSSIBLE N-term signal sequence\ninside 1 19\nTMhelix 20 39\noutside 40 325\nTMhelix 326 348\ninside 349 372

9960 GCF_000714185.1_ASM71418v1 Escherichia coli 6-175-07_S1_C1 Proteobacteria; Gammaproteobacteria; Enterobacterales; Enterobacteriaceae; Escherichia; Escherichia coli MTGDNTLIHSHGINRRDFMKLCAALAATMGLSSKAAAEMAESVTNPQRPPVIWIGAQECT WP_000145410.1 MULTISPECIES: hydrogenase-2 small chain [Proteobacteria] Length: 372\nNumber of predicted TMHs: 1\nExp number of AAs in TMHs: 23.72226\nExp number, first 60 AAs: 0.22038\nTotal prob of N-in: 0.12994\noutside 1 331\nTMhelix 332 354\ninside 355 372

9961 GCF_000711375.1_ASM71137v1 Escherichia coli 6-175-07_S1_C2 Proteobacteria; Gammaproteobacteria; Enterobacterales; Enterobacteriaceae; Escherichia; Escherichia coli MNNEETFYQAMRRQGVTRRSFLKYCSLAATSLGLGAGMAPKIAWALENKPRIPVVWIHGL WP_001058323.1 MULTISPECIES: hydrogenase-1 small chain [Proteobacteria] Length: 372\nNumber of predicted TMHs: 2\nExp number of AAs in TMHs: 37.2608499999999\nExp number, first 60 AAs: 19.52923\nTotal prob of N-in: 0.94351\nPOSSIBLE N-term signal sequence\ninside 1 19\nTMhelix 20 39\noutside 40 325\nTMhelix 326 348\ninside 349 372

9962 GCF_000711375.1_ASM71137v1 Escherichia coli 6-175-07_S1_C2 Proteobacteria; Gammaproteobacteria; Enterobacterales; Enterobacteriaceae; Escherichia; Escherichia coli MTGDNTLIHSHGINRRDFMKLCAALAATMGLSSKAAAEMAESVTNPQRPPVIWIGAQECT WP_000145410.1 MULTISPECIES: hydrogenase-2 small chain [Proteobacteria] Length: 372\nNumber of predicted TMHs: 1\nExp number of AAs in TMHs: 23.72226\nExp number, first 60 AAs: 0.22038\nTotal prob of N-in: 0.12994\noutside 1 331\nTMhelix 332 354\ninside 355 372

9963 GCF_000713135.1_ASM71313v1 Escherichia coli 6-175-07_S1_C3 Proteobacteria; Gammaproteobacteria; Enterobacterales; Enterobacteriaceae; Escherichia; Escherichia coli MTGDNTLIHSHGINRRDFMKLCAALAATMGLSSKAAAEMAESVTNPQRPPVIWIGAQECT WP_000145410.1 MULTISPECIES: hydrogenase-2 small chain [Proteobacteria] Length: 372\nNumber of predicted TMHs: 1\nExp number of AAs in TMHs: 23.72226\nExp number, first 60 AAs: 0.22038\nTotal prob of N-in: 0.12994\noutside 1 331\nTMhelix 332 354\ninside 355 372

9964 GCF_000713135.1_ASM71313v1 Escherichia coli 6-175-07_S1_C3 Proteobacteria; Gammaproteobacteria; Enterobacterales; Enterobacteriaceae; Escherichia; Escherichia coli MNNEETFYQAMRRQGVTRRSFLKYCSLAATSLGLGAGMAPKIAWALENKPRIPVVWIHGL WP_001058323.1 MULTISPECIES: hydrogenase-1 small chain [Proteobacteria] Length: 372\nNumber of predicted TMHs: 2\nExp number of AAs in TMHs: 37.2608499999999\nExp number, first 60 AAs: 19.52923\nTotal prob of N-in: 0.94351\nPOSSIBLE N-term signal sequence\ninside 1 19\nTMhelix 20 39\noutside 40 325\nTMhelix 326 348\ninside 349 372

9965 GCF_000714345.1_ASM71434v1 Escherichia coli 6-175-07_S3_C1 Proteobacteria; Gammaproteobacteria; Enterobacterales; Enterobacteriaceae; Escherichia; Escherichia coli MNNEETFYQAMRRQGVTRRSFLKYCSLAATSLGLGAGMAPKIAWALENKPRIPVVWIHGL WP_001058323.1 MULTISPECIES: hydrogenase-1 small chain [Proteobacteria] Length: 372\nNumber of predicted TMHs: 2\nExp number of AAs in TMHs: 37.2608499999999\nExp number, first 60 AAs: 19.52923\nTotal prob of N-in: 0.94351\nPOSSIBLE N-term signal sequence\ninside 1 19\nTMhelix 20 39\noutside 40 325\nTMhelix 326 348\ninside 349 372

9966 GCF_000714345.1_ASM71434v1 Escherichia coli 6-175-07_S3_C1 Proteobacteria; Gammaproteobacteria; Enterobacterales; Enterobacteriaceae; Escherichia; Escherichia coli MTGDNTLIHSHGINRRDFMKLCAALAATMGLSSKAAAEMAESVTNPQRPPVIWIGAQECT WP_000145410.1 MULTISPECIES: hydrogenase-2 small chain [Proteobacteria] Length: 372\nNumber of predicted TMHs:

1\nExp number of AAs in TMHs: 23.72226\nExp number, first 60 AAs: 0.22038\nTotal prob of N-in: 0.12994\noutside 1 331\nTMhelix 332 354\ninside 355 372

9967 GCF_000711525.1_ASM71152v1 Escherichia coli 6-175-07_S3_C2 Proteobacteria; Gammaproteobacteria; Enterobacterales; Enterobacteriaceae; Escherichia; Escherichia coli MNNEETFYQAMRRQGVTRRSFLKYCSLAATSLGLGAGMAPKIAWALENKPRIPVVWIHGL WP_001058323.1 MULTISPECIES: hydrogenase-1 small chain [Proteobacteria] Length: 372\nNumber of predicted TMHs: 2\nExp number of AAs in TMHs: 37.2608499999999\nExp number, first 60 AAs: 19.52923\nTotal prob of N-in: 0.94351\nPOSSIBLE N-term signal sequence\ninside 1 19\nTMhelix 20 39\noutside 40 325\nTMhelix 326 348\ninside 349 372

9968 GCF_000711525.1_ASM71152v1 Escherichia coli 6-175-07_S3_C2 Proteobacteria; Gammaproteobacteria; Enterobacterales; Enterobacteriaceae; Escherichia; Escherichia coli MTGDNTLIHSHGINRRDFMKLCAALAATMGLSSKAAAEMAESVTNPQRPPVIWIGAQECT WP_000145410.1 MULTISPECIES: hydrogenase-2 small chain [Proteobacteria] Length: 372\nNumber of predicted TMHs: 1\nExp number of AAs in TMHs: 23.72226\nExp number, first 60 AAs: 0.22038\nTotal prob of N-in: 0.12994\noutside 1 331\nTMhelix 332 354\ninside 355 372

9969 GCF_000714375.1_ASM71437v1 Escherichia coli 6-175-07_S3_C3 Proteobacteria; Gammaproteobacteria; Enterobacterales; Enterobacteriaceae; Escherichia; Escherichia coli MNNEETFYQAMRRQGVTRRSFLKYCSLAATSLGLGAGMAPKIAWALENKPRIPVVWIHGL WP_001058323.1 MULTISPECIES: hydrogenase-1 small chain [Proteobacteria] Length: 372\nNumber of predicted TMHs: 2\nExp number of AAs in TMHs: 37.2608499999999\nExp number, first 60 AAs: 19.52923\nTotal prob of N-in: 0.94351\nPOSSIBLE N-term signal sequence\ninside 1 19\nTMhelix 20 39\noutside 40 325\nTMhelix 326 348\ninside 349 372

9970 GCF_000714375.1_ASM71437v1 Escherichia coli 6-175-07_S3_C3 Proteobacteria; Gammaproteobacteria; Enterobacterales; Enterobacteriaceae; Escherichia; Escherichia coli MTGDNTLIHSHGINRRDFMKLCAALAATMGLSSKAAAEMAESVTNPQRPPVIWIGAQECT WP_000145410.1 MULTISPECIES: hydrogenase-2 small chain [Proteobacteria] Length: 372\nNumber of predicted TMHs: 1\nExp number of AAs in TMHs: 23.72226\nExp number, first 60 AAs: 0.22038\nTotal prob of N-in: 0.12994\noutside 1 331\nTMhelix 332 354\ninside 355 372

9971 GCF_000714385.1_ASM71438v1 Escherichia coli 6-175-07_S4_C1 Proteobacteria; Gammaproteobacteria; Enterobacterales; Enterobacteriaceae; Escherichia; Escherichia coli MTGDNTLIHSHGINRRDFMKLCAALAATMGLSSKAAAEMAESVTNPQRPPVIWIGAQECT WP_000145410.1 MULTISPECIES: hydrogenase-2 small chain [Proteobacteria] Length: 372\nNumber of predicted TMHs: 1\nExp number of AAs in TMHs: 23.72226\nExp number, first 60 AAs: 0.22038\nTotal prob of N-in: 0.12994\noutside 1 331\nTMhelix 332 354\ninside 355 372

9972 GCF_000714385.1_ASM71438v1 Escherichia coli 6-175-07_S4_C1 Proteobacteria; Gammaproteobacteria; Enterobacterales; Enterobacteriaceae; Escherichia; Escherichia coli MNNEETFYQAMRRQGVTRRSFLKYCSLAATSLGLGAGMAPKIAWALENKPRIPVVWIHGL WP_001058323.1 MULTISPECIES: hydrogenase-1 small chain [Proteobacteria] Length: 372\nNumber of predicted TMHs: 2\nExp number of AAs in TMHs: 37.2608499999999\nExp number, first 60 AAs: 19.52923\nTotal prob of N-in: 0.94351\nPOSSIBLE N-term signal sequence\ninside 1 19\nTMhelix 20 39\noutside 40 325\nTMhelix 326 348\ninside 349 372

9973 GCF_000714415.1_ASM71441v1 Escherichia coli 6-175-07_S4_C2 Proteobacteria; Gammaproteobacteria; Enterobacterales; Enterobacteriaceae; Escherichia; Escherichia coli MNNEETFYQAMRRQGVTRRSFLKYCSLAATSLGLGAGMAPKIAWALENKPRIPVVWIHGL WP_001058323.1 MULTISPECIES: hydrogenase-1 small chain [Proteobacteria] Length: 372\nNumber of predicted TMHs: 2\nExp number of AAs in TMHs: 37.2608499999999\nExp number, first 60 AAs: 19.52923\nTotal prob of N-in: 0.94351\nPOSSIBLE N-term signal sequence\ninside 1 19\nTMhelix 20 39\noutside 40 325\nTMhelix 326 348\ninside 349 372

9974 GCF_000714415.1_ASM71441v1 Escherichia coli 6-175-07_S4_C2 Proteobacteria; Gammaproteobacteria; Enterobacterales; Enterobacteriaceae; Escherichia; Escherichia coli MTGDNTLIHSHGINRRDFMKLCAALAATMGLSSKAAAEMAESVTNPQRPPVIWIGAQECT WP_000145410.1 MULTISPECIES: hydrogenase-2 small chain [Proteobacteria] Length: 372\nNumber of predicted TMHs:

1\nExp number of AAs in TMHs: 23.72226\nExp number, first 60 AAs: 0.22038\nTotal prob of N-in: 0.12994\noutside 1 331\nTMhelix 332 354\ninside 355 372

9975 GCF_000713175.1_ASM71317v1 Escherichia coli 6-175-07_S4_C3 Proteobacteria; Gammaproteobacteria; Enterobacterales; Enterobacteriaceae; Escherichia; Escherichia coli MNNEETFYQAMRRQGVTRRSFLKYCSLAATSLGLGAGMAPKIAWALENKPRIPVVWIHGL WP_001058323.1 MULTISPECIES: hydrogenase-1 small chain [Proteobacteria] Length: 372\nNumber of predicted TMHs: 2\nExp number of AAs in TMHs: 37.2608499999999\nExp number, first 60 AAs: 19.52923\nTotal prob of N-in: 0.94351\nPOSSIBLE N-term signal sequence\ninside 1 19\nTMhelix 20 39\noutside 40 325\nTMhelix 326 348\ninside 349 372

9976 GCF_000713175.1_ASM71317v1 Escherichia coli 6-175-07_S4_C3 Proteobacteria; Gammaproteobacteria; Enterobacterales; Enterobacteriaceae; Escherichia; Escherichia coli MTGDNTLIHSHGINRRDFMKLCAALAATMGLSSKAAAEMAESVTNPQRPPVIWIGAQECT WP_000145410.1 MULTISPECIES: hydrogenase-2 small chain [Proteobacteria] Length: 372\nNumber of predicted TMHs: 1\nExp number of AAs in TMHs: 23.72226\nExp number, first 60 AAs: 0.22038\nTotal prob of N-in: 0.12994\noutside 1 331\nTMhelix 332 354\ninside 355 372

9977 GCF_000713335.1_ASM71333v1 Escherichia coli 6-319-05_S1_C1 Proteobacteria; Gammaproteobacteria; Enterobacterales; Enterobacteriaceae; Escherichia; Escherichia coli MNNEETFYQAMRRQGVTRRSFLKYCSLAATSLGLGAGMAPKIAWALENKPRIPVVWIHGL WP_001058323.1 MULTISPECIES: hydrogenase-1 small chain [Proteobacteria] Length: 372\nNumber of predicted TMHs: 2\nExp number of AAs in TMHs: 37.2608499999999\nExp number, first 60 AAs: 19.52923\nTotal prob of N-in: 0.94351\nPOSSIBLE N-term signal sequence\ninside 1 19\nTMhelix 20 39\noutside 40 325\nTMhelix 326 348\ninside 349 372

9978 GCF_000713335.1_ASM71333v1 Escherichia coli 6-319-05_S1_C1 Proteobacteria; Gammaproteobacteria; Enterobacterales; Enterobacteriaceae; Escherichia; Escherichia coli MTGDNTLIHSHGINRRDFMKLCAALAATMGLSSKAAAEMAESVTNPQRPPVIWIGAQECT WP_000145410.1 MULTISPECIES: hydrogenase-2 small chain [Proteobacteria] Length: 372\nNumber of predicted TMHs: 1\nExp number of AAs in TMHs: 23.72226\nExp number, first 60 AAs: 0.22038\nTotal prob of N-in: 0.12994\noutside 1 331\nTMhelix 332 354\ninside 355 372

9979 GCF_000713025.1_ASM71302v1 Escherichia coli 6-319-05_S1_C2 Proteobacteria; Gammaproteobacteria; Enterobacterales; Enterobacteriaceae; Escherichia; Escherichia coli MNNEETFYQAMRRQGVTRRSFLKYCSLAATSLGLGAGMAPKIAWALENKPRIPVVWIHGL WP_001058323.1 MULTISPECIES: hydrogenase-1 small chain [Proteobacteria] Length: 372\nNumber of predicted TMHs: 2\nExp number of AAs in TMHs: 37.2608499999999\nExp number, first 60 AAs: 19.52923\nTotal prob of N-in: 0.94351\nPOSSIBLE N-term signal sequence\ninside 1 19\nTMhelix 20 39\noutside 40 325\nTMhelix 326 348\ninside 349 372

9980 GCF_000713025.1_ASM71302v1 Escherichia coli 6-319-05_S1_C2 Proteobacteria; Gammaproteobacteria; Enterobacterales; Enterobacteriaceae; Escherichia; Escherichia coli MTGDNTLIHSHGINRRDFMKLCAALAATMGLSSKAAAEMAESVTNPQRPPVIWIGAQECT WP_000145410.1 MULTISPECIES: hydrogenase-2 small chain [Proteobacteria] Length: 372\nNumber of predicted TMHs: 1\nExp number of AAs in TMHs: 23.72226\nExp number, first 60 AAs: 0.22038\nTotal prob of N-in: 0.12994\noutside 1 331\nTMhelix 332 354\ninside 355 372

9981 GCF_000713015.1_ASM71301v1 Escherichia coli 6-319-05_S1_C3 Proteobacteria; Gammaproteobacteria; Enterobacterales; Enterobacteriaceae; Escherichia; Escherichia coli MNNEETFYQAMRRQGVTRRSFLKYCSLAATSLGLGAGMAPKIAWALENKPRIPVVWIHGL WP_001058323.1 MULTISPECIES: hydrogenase-1 small chain [Proteobacteria] Length: 372\nNumber of predicted TMHs: 2\nExp number of AAs in TMHs: 37.2608499999999\nExp number, first 60 AAs: 19.52923\nTotal prob of N-in: 0.94351\nPOSSIBLE N-term signal sequence\ninside 1 19\nTMhelix 20 39\noutside 40 325\nTMhelix 326 348\ninside 349 372

9982 GCF_000713015.1_ASM71301v1 Escherichia coli 6-319-05_S1_C3 Proteobacteria; Gammaproteobacteria; Enterobacterales; Enterobacteriaceae; Escherichia; Escherichia coli MTGDNTLIHSHGINRRDFMKLCAALAATMGLSSKAAAEMAESVTNPQRPPVIWIGAQECT WP_000145410.1 MULTISPECIES: hydrogenase-2 small chain [Proteobacteria] Length: 372\nNumber of predicted TMHs:

1\nExp number of AAs in TMHs: 23.72226\nExp number, first 60 AAs: 0.22038\nTotal prob of N-in: 0.12994\noutside 1 331\nTMhelix 332 354\ninside 355 372

9983 GCF_000713045.1_ASM71304v1 Escherichia coli 6-319-05_S3_C1 Proteobacteria; Gammaproteobacteria; Enterobacterales; Enterobacteriaceae; Escherichia; Escherichia coli
MNNEETFYQAMRRQGVTRRSFLKYCSLAATSLGLGAGMAPKIAWALENKPRIPVVWIHGL WP_001058323.1
MULTISPECIES: hydrogenase-1 small chain [Proteobacteria] Length: 372\nNumber of predicted TMHs: 2\nExp number of AAs in TMHs: 37.2608499999999\nExp number, first 60 AAs: 19.52923\nTotal prob of N-in: 0.94351\nPOSSIBLE N-term signal sequence\ninside 1 19\nTMhelix 20 39\noutside 40 325\nTMhelix 326 348\ninside 349 372

9984 GCF_000713045.1_ASM71304v1 Escherichia coli 6-319-05_S3_C1 Proteobacteria; Gammaproteobacteria; Enterobacterales; Enterobacteriaceae; Escherichia; Escherichia coli
MTGDNTLIHSHGINRRDFMKLCAALAATMGLSSKAAAEMAESVTNPQRPPVIWIGAQECT WP_000145410.1
MULTISPECIES: hydrogenase-2 small chain [Proteobacteria] Length: 372\nNumber of predicted TMHs: 1\nExp number of AAs in TMHs: 23.72226\nExp number, first 60 AAs: 0.22038\nTotal prob of N-in: 0.12994\noutside 1 331\nTMhelix 332 354\ninside 355 372

9985 GCF_000713035.1_ASM71303v1 Escherichia coli 6-319-05_S3_C2 Proteobacteria; Gammaproteobacteria; Enterobacterales; Enterobacteriaceae; Escherichia; Escherichia coli
MTGDNTLIHSHGINRRDFMKLCAALAATMGLSSKAAAEMAESVTNPQRPPVIWIGAQECT WP_000145410.1
MULTISPECIES: hydrogenase-2 small chain [Proteobacteria] Length: 372\nNumber of predicted TMHs: 1\nExp number of AAs in TMHs: 23.72226\nExp number, first 60 AAs: 0.22038\nTotal prob of N-in: 0.12994\noutside 1 331\nTMhelix 332 354\ninside 355 372

9986 GCF_000713035.1_ASM71303v1 Escherichia coli 6-319-05_S3_C2 Proteobacteria; Gammaproteobacteria; Enterobacterales; Enterobacteriaceae; Escherichia; Escherichia coli
MNNEETFYQAMRRQGVTRRSFLKYCSLAATSLGLGAGMAPKIAWALENKPRIPVVWIHGL WP_001058323.1
MULTISPECIES: hydrogenase-1 small chain [Proteobacteria] Length: 372\nNumber of predicted TMHs: 2\nExp number of AAs in TMHs: 37.2608499999999\nExp number, first 60 AAs: 19.52923\nTotal prob of N-in: 0.94351\nPOSSIBLE N-term signal sequence\ninside 1 19\nTMhelix 20 39\noutside 40 325\nTMhelix 326 348\ninside 349 372

9987 GCF_000713185.1_ASM71318v1 Escherichia coli 6-319-05_S3_C3 Proteobacteria; Gammaproteobacteria; Enterobacterales; Enterobacteriaceae; Escherichia; Escherichia coli
MTGDNTLIHSHGINRRDFMKLCAALAATMGLSSKAAAEMAESVTNPQRPPVIWIGAQECT WP_000145410.1
MULTISPECIES: hydrogenase-2 small chain [Proteobacteria] Length: 372\nNumber of predicted TMHs: 1\nExp number of AAs in TMHs: 23.72226\nExp number, first 60 AAs: 0.22038\nTotal prob of N-in: 0.12994\noutside 1 331\nTMhelix 332 354\ninside 355 372

9988 GCF_000713185.1_ASM71318v1 Escherichia coli 6-319-05_S3_C3 Proteobacteria; Gammaproteobacteria; Enterobacterales; Enterobacteriaceae; Escherichia; Escherichia coli
MNNEETFYQAMRRQGVTRRSFLKYCSLAATSLGLGAGMAPKIAWALENKPRIPVVWIHGL WP_001058323.1
MULTISPECIES: hydrogenase-1 small chain [Proteobacteria] Length: 372\nNumber of predicted TMHs: 2\nExp number of AAs in TMHs: 37.2608499999999\nExp number, first 60 AAs: 19.52923\nTotal prob of N-in: 0.94351\nPOSSIBLE N-term signal sequence\ninside 1 19\nTMhelix 20 39\noutside 40 325\nTMhelix 326 348\ninside 349 372

9989 GCF_000713095.1_ASM71309v1 Escherichia coli 6-319-05_S4_C2 Proteobacteria; Gammaproteobacteria; Enterobacterales; Enterobacteriaceae; Escherichia; Escherichia coli
MTGDNTLIHSHGINRRDFMKLCAALAATMGLSSKAAAEMAESVTNPQRPPVIWIGAQECT WP_000145410.1
MULTISPECIES: hydrogenase-2 small chain [Proteobacteria] Length: 372\nNumber of predicted TMHs: 1\nExp number of AAs in TMHs: 23.72226\nExp number, first 60 AAs: 0.22038\nTotal prob of N-in: 0.12994\noutside 1 331\nTMhelix 332 354\ninside 355 372

9990 GCF_000713095.1_ASM71309v1 Escherichia coli 6-319-05_S4_C2 Proteobacteria; Gammaproteobacteria; Enterobacterales; Enterobacteriaceae; Escherichia; Escherichia coli
MNNEETFYQAMRRQGVTRRSFLKYCSLAATSLGLGAGMAPKIAWALENKPRIPVVWIHGL WP_001058323.1
MULTISPECIES: hydrogenase-1 small chain [Proteobacteria] Length: 372\nNumber of predicted TMHs: 2\nExp number of AAs in TMHs: 37.2608499999999\nExp number, first 60 AAs: 19.52923\nTotal prob of N-in:

0.94351\nPOSSIBLE N-term signal sequence\ninside 1 19\nTMhelix 20 39\noutside 40 325\nTMhelix 326 348\ninside 349 372

9991 GCF_000713115.1_ASM71311v1 Escherichia coli 6-319-05_S4_C3 Proteobacteria; Gammaproteobacteria; Enterobacterales; Enterobacteriaceae; Escherichia; Escherichia coli MTGDNTLIHSHGINRRDFMKLCAALAATMGLSSKAAAEMAESVTNPQRPPVIWIGAQECT WP_000145410.1 MULTISPECIES: hydrogenase-2 small chain [Proteobacteria] Length: 372\nNumber of predicted TMHs: 1\nExp number of AAs in TMHs: 23.72226\nExp number, first 60 AAs: 0.22038\nTotal prob of N-in: 0.12994\noutside 1 331\nTMhelix 332 354\ninside 355 372

9992 GCF_000713115.1_ASM71311v1 Escherichia coli 6-319-05_S4_C3 Proteobacteria; Gammaproteobacteria; Enterobacterales; Enterobacteriaceae; Escherichia; Escherichia coli MNNEETFYQAMRRQGVTRRSFLKYCSLAATSLGLGAGMAPKIAWALENKPRIPVVWIHGL WP_001058323.1 MULTISPECIES: hydrogenase-1 small chain [Proteobacteria] Length: 372\nNumber of predicted TMHs: 2\nExp number of AAs in TMHs: 37.2608499999999\nExp number, first 60 AAs: 19.52923\nTotal prob of N-in: 0.94351\nPOSSIBLE N-term signal sequence\ninside 1 19\nTMhelix 20 39\noutside 40 325\nTMhelix 326 348\ninside 349 372

9993 GCF_000713105.1_ASM71310v1 Escherichia coli 6-537-08_S1_C1 Proteobacteria; Gammaproteobacteria; Enterobacterales; Enterobacteriaceae; Escherichia; Escherichia coli MTGDNTLIHSHGINRRDFMKLCAALAATMGLSSKAAAEMAESVTNPQRPPVIWIGAQECT WP_000145410.1 MULTISPECIES: hydrogenase-2 small chain [Proteobacteria] Length: 372\nNumber of predicted TMHs: 1\nExp number of AAs in TMHs: 23.72226\nExp number, first 60 AAs: 0.22038\nTotal prob of N-in: 0.12994\noutside 1 331\nTMhelix 332 354\ninside 355 372

9994 GCF_000713105.1_ASM71310v1 Escherichia coli 6-537-08_S1_C1 Proteobacteria; Gammaproteobacteria; Enterobacterales; Enterobacteriaceae; Escherichia; Escherichia coli MNNEETFYQAMRRQGVTRRSFLKYCSLAATSLGLGAGMAPKIAWALENKPRIPVVWIHGL WP_001058323.1 MULTISPECIES: hydrogenase-1 small chain [Proteobacteria] Length: 372\nNumber of predicted TMHs: 2\nExp number of AAs in TMHs: 37.2608499999999\nExp number, first 60 AAs: 19.52923\nTotal prob of N-in: 0.94351\nPOSSIBLE N-term signal sequence\ninside 1 19\nTMhelix 20 39\noutside 40 325\nTMhelix 326 348\ninside 349 372

9995 GCF_000713505.1_ASM71350v1 Escherichia coli 6-537-08_S1_C2 Proteobacteria; Gammaproteobacteria; Enterobacterales; Enterobacteriaceae; Escherichia; Escherichia coli MNNEETFYQAMRRQGVTRRSFLKYCSLAATSLGLGAGMAPKIAWALENKPRIPVVWIHGL WP_001058323.1 MULTISPECIES: hydrogenase-1 small chain [Proteobacteria] Length: 372\nNumber of predicted TMHs: 2\nExp number of AAs in TMHs: 37.2608499999999\nExp number, first 60 AAs: 19.52923\nTotal prob of N-in: 0.94351\nPOSSIBLE N-term signal sequence\ninside 1 19\nTMhelix 20 39\noutside 40 325\nTMhelix 326 348\ninside 349 372

9996 GCF_000713505.1_ASM71350v1 Escherichia coli 6-537-08_S1_C2 Proteobacteria; Gammaproteobacteria; Enterobacterales; Enterobacteriaceae; Escherichia; Escherichia coli MTGDNTLIHSHGINRRDFMKLCAALAATMGLSSKAAAEMAESVTNPQRPPVIWIGAQECT WP_000145410.1 MULTISPECIES: hydrogenase-2 small chain [Proteobacteria] Length: 372\nNumber of predicted TMHs: 1\nExp number of AAs in TMHs: 23.72226\nExp number, first 60 AAs: 0.22038\nTotal prob of N-in: 0.12994\noutside 1 331\nTMhelix 332 354\ninside 355 372

9997 GCF_000713575.1_ASM71357v1 Escherichia coli 6-537-08_S1_C3 Proteobacteria; Gammaproteobacteria; Enterobacterales; Enterobacteriaceae; Escherichia; Escherichia coli MNNEETFYQAMRRQGVTRRSFLKYCSLAATSLGLGAGMAPKIAWALENKPRIPVVWIHGL WP_001058323.1 MULTISPECIES: hydrogenase-1 small chain [Proteobacteria] Length: 372\nNumber of predicted TMHs: 2\nExp number of AAs in TMHs: 37.2608499999999\nExp number, first 60 AAs: 19.52923\nTotal prob of N-in: 0.94351\nPOSSIBLE N-term signal sequence\ninside 1 19\nTMhelix 20 39\noutside 40 325\nTMhelix 326 348\ninside 349 372

9998 GCF_000713575.1_ASM71357v1 Escherichia coli 6-537-08_S1_C3 Proteobacteria; Gammaproteobacteria; Enterobacterales; Enterobacteriaceae; Escherichia; Escherichia coli MTGDNTLIHSHGINRRDFMKLCAALAATMGLSSKAAAEMAESVTNPQRPPVIWIGAQECT WP_000145410.1 MULTISPECIES: hydrogenase-2 small chain [Proteobacteria] Length: 372\nNumber of predicted TMHs:

1\nExp number of AAs in TMHs: 23.72226\nExp number, first 60 AAs:  0.22038\nTotal prob of N-in:   0.12994\noutside   1   331\nTMhelix   332   354\ninside   355   372

9999 GCF_000713265.1_ASM71326v1 Escherichia coli 6-537-08_S3_C1 Proteobacteria; Gammaproteobacteria; Enterobacterales; Enterobacteriaceae; Escherichia; Escherichia coli MNNEETFYQAMRRQGVTRRSFLKYCSLAATSLGLGAGMAPKIAWALENKPRIPVVWIHGL WP_001295940.1 MULTISPECIES: hydrogenase [Enterobacteriaceae] Length: 372\nNumber of predicted TMHs:  2\nExp number of AAs in TMHs: 37.2608899999999\nExp number, first 60 AAs:  19.52923\nTotal prob of N-in:   0.94351\nPOSSIBLE N-term signal sequence\ninside   1   19\nTMhelix   20   39\noutside   40   325\nTMhelix   326   348\ninside   349   372

10000 GCF_000713265.1_ASM71326v1 Escherichia coli 6-537-08_S3_C1 Proteobacteria; Gammaproteobacteria; Enterobacterales; Enterobacteriaceae; Escherichia; Escherichia coli MTGDNTLIHSHGINRRDFMKLCAALAATMGLSSKAAAEMAESVTNPQRPPVIWIGAQECT WP_000145410.1 MULTISPECIES: hydrogenase-2 small chain [Proteobacteria] Length: 372\nNumber of predicted TMHs:  1\nExp number of AAs in TMHs: 23.72226\nExp number, first 60 AAs:  0.22038\nTotal prob of N-in:   0.12994\noutside   1   331\nTMhelix   332   354\ninside   355   372

10001 GCF_000713415.1_ASM71341v1 Escherichia coli 6-537-08_S3_C2 Proteobacteria; Gammaproteobacteria; Enterobacterales; Enterobacteriaceae; Escherichia; Escherichia coli MNNEETFYQAMRRQGVTRRSFLKYCSLAATSLGLGAGMAPKIAWALENKPRIPVVWIHGL WP_001295940.1 MULTISPECIES: hydrogenase [Enterobacteriaceae] Length: 372\nNumber of predicted TMHs:  2\nExp number of AAs in TMHs: 37.2608899999999\nExp number, first 60 AAs:  19.52923\nTotal prob of N-in:   0.94351\nPOSSIBLE N-term signal sequence\ninside   1   19\nTMhelix   20   39\noutside   40   325\nTMhelix   326   348\ninside   349   372

10002 GCF_000713415.1_ASM71341v1 Escherichia coli 6-537-08_S3_C2 Proteobacteria; Gammaproteobacteria; Enterobacterales; Enterobacteriaceae; Escherichia; Escherichia coli MTGDNTLIHSHGINRRDFMKLCAALAATMGLSSKAAAEMAESVTNPQRPPVIWIGAQECT WP_000145410.1 MULTISPECIES: hydrogenase-2 small chain [Proteobacteria] Length: 372\nNumber of predicted TMHs:  1\nExp number of AAs in TMHs: 23.72226\nExp number, first 60 AAs:  0.22038\nTotal prob of N-in:   0.12994\noutside   1   331\nTMhelix   332   354\ninside   355   372

10003 GCF_000713315.1_ASM71331v1 Escherichia coli 6-537-08_S3_C3 Proteobacteria; Gammaproteobacteria; Enterobacterales; Enterobacteriaceae; Escherichia; Escherichia coli MTGDNTLIHSHGINRRDFMKLCAALAATMGLSSKAAAEMAESVTNPQRPPVIWIGAQECT WP_000145410.1 MULTISPECIES: hydrogenase-2 small chain [Proteobacteria] Length: 372\nNumber of predicted TMHs:  1\nExp number of AAs in TMHs: 23.72226\nExp number, first 60 AAs:  0.22038\nTotal prob of N-in:   0.12994\noutside   1   331\nTMhelix   332   354\ninside   355   372

10004 GCF_000713315.1_ASM71331v1 Escherichia coli 6-537-08_S3_C3 Proteobacteria; Gammaproteobacteria; Enterobacterales; Enterobacteriaceae; Escherichia; Escherichia coli MNNEETFYQAMRRQGVTRRSFLKYCSLAATSLGLGAGMAPKIAWALENKPRIPVVWIHGL WP_001295940.1 MULTISPECIES: hydrogenase [Enterobacteriaceae] Length: 372\nNumber of predicted TMHs:  2\nExp number of AAs in TMHs: 37.2608899999999\nExp number, first 60 AAs:  19.52923\nTotal prob of N-in:   0.94351\nPOSSIBLE N-term signal sequence\ninside   1   19\nTMhelix   20   39\noutside   40   325\nTMhelix   326   348\ninside   349   372

10005 GCF_000713255.1_ASM71325v1 Escherichia coli 6-537-08_S4_C1 Proteobacteria; Gammaproteobacteria; Enterobacterales; Enterobacteriaceae; Escherichia; Escherichia coli MTGDNTLIHSHGINRRDFMKLCAALAATMGLSSKAAAEMAESVTNPQRPPVIWIGAQECT WP_000145410.1 MULTISPECIES: hydrogenase-2 small chain [Proteobacteria] Length: 372\nNumber of predicted TMHs:  1\nExp number of AAs in TMHs: 23.72226\nExp number, first 60 AAs:  0.22038\nTotal prob of N-in:   0.12994\noutside   1   331\nTMhelix   332   354\ninside   355   372

10006 GCF_000713255.1_ASM71325v1 Escherichia coli 6-537-08_S4_C1 Proteobacteria; Gammaproteobacteria; Enterobacterales; Enterobacteriaceae; Escherichia; Escherichia coli MNNEETFYQAMRRQGVTRRSFLKYCSLAATSLGLGAGMAPKIAWALENKPRIPVVWIHGL WP_001058323.1 MULTISPECIES: hydrogenase-1 small chain [Proteobacteria] Length: 372\nNumber of predicted TMHs:  2\nExp number of AAs in TMHs: 37.2608499999999\nExp number, first 60 AAs:  19.52923\nTotal prob of N-in:

0.94351\nPOSSIBLE N-term signal sequence\ninside 1 19\nTMhelix 20 39\noutside 40 325\nTMhelix 326 348\ninside 349 372

10007 GCF_000713605.1_ASM71360v1 Escherichia coli 6-537-08_S4_C2 Proteobacteria; Gammaproteobacteria; Enterobacterales; Enterobacteriaceae; Escherichia; Escherichia coli MTGDNTLIHSHGINRRDFMKLCAALAATMGLSSKAAAEMAESVTNPQRPPVIWIGAQECT WP_000145410.1 MULTISPECIES: hydrogenase-2 small chain [Proteobacteria] Length: 372\nNumber of predicted TMHs: 1\nExp number of AAs in TMHs: 23.72226\nExp number, first 60 AAs: 0.22038\nTotal prob of N-in: 0.12994\noutside 1 331\nTMhelix 332 354\ninside 355 372

10008 GCF_000713605.1_ASM71360v1 Escherichia coli 6-537-08_S4_C2 Proteobacteria; Gammaproteobacteria; Enterobacterales; Enterobacteriaceae; Escherichia; Escherichia coli MNNEETFYQAMRRQGVTRRSFLKYCSLAATSLGLGAGMAPKIAWALENKPRIPVVWIHGL WP_001058323.1 MULTISPECIES: hydrogenase-1 small chain [Proteobacteria] Length: 372\nNumber of predicted TMHs: 2\nExp number of AAs in TMHs: 37.2608499999999\nExp number, first 60 AAs: 19.52923\nTotal prob of N-in: 0.94351\nPOSSIBLE N-term signal sequence\ninside 1 19\nTMhelix 20 39\noutside 40 325\nTMhelix 326 348\ninside 349 372

10009 GCF_000503515.1_Ecoli668 Escherichia coli 668 Proteobacteria; Gammaproteobacteria; Enterobacterales; Enterobacteriaceae; Escherichia; Escherichia coli MNNEETFYQAMRRQGVTRRSFLKYCSLAATSLGLGAGMAPKIAWALENKPRIPVVWIHGL WP_001058323.1 MULTISPECIES: hydrogenase-1 small chain [Proteobacteria] Length: 372\nNumber of predicted TMHs: 2\nExp number of AAs in TMHs: 37.2608499999999\nExp number, first 60 AAs: 19.52923\nTotal prob of N-in: 0.94351\nPOSSIBLE N-term signal sequence\ninside 1 19\nTMhelix 20 39\noutside 40 325\nTMhelix 326 348\ninside 349 372

10010 GCF_000503515.1_Ecoli668 Escherichia coli 668 Proteobacteria; Gammaproteobacteria; Enterobacterales; Enterobacteriaceae; Escherichia; Escherichia coli MTGDNTLIHSHGINRRDFMKLCAALAATMGLSSKAAAEMAESVTNPQRPPVIWIGAQECT WP_000145410.1 MULTISPECIES: hydrogenase-2 small chain [Proteobacteria] Length: 372\nNumber of predicted TMHs: 1\nExp number of AAs in TMHs: 23.72226\nExp number, first 60 AAs: 0.22038\nTotal prob of N-in: 0.12994\noutside 1 331\nTMhelix 332 354\ninside 355 372

10011 GCF_000503335.1_Ecoli681 Escherichia coli 681 Proteobacteria; Gammaproteobacteria; Enterobacterales; Enterobacteriaceae; Escherichia; Escherichia coli MNNEETFYQAMRRQGVTRRSFLKYCSLAATSLGLGAGMAPKIAWALENKPRIPVVWIHGL WP_001058323.1 MULTISPECIES: hydrogenase-1 small chain [Proteobacteria] Length: 372\nNumber of predicted TMHs: 2\nExp number of AAs in TMHs: 37.2608499999999\nExp number, first 60 AAs: 19.52923\nTotal prob of N-in: 0.94351\nPOSSIBLE N-term signal sequence\ninside 1 19\nTMhelix 20 39\noutside 40 325\nTMhelix 326 348\ninside 349 372

10012 GCF_000503335.1_Ecoli681 Escherichia coli 681 Proteobacteria; Gammaproteobacteria; Enterobacterales; Enterobacteriaceae; Escherichia; Escherichia coli MTGDNTLIHSHGINRRDFMKLCAALAATMGLSSKAAAEMAESVTNPQRPPVIWIGAQECT WP_000145410.1 MULTISPECIES: hydrogenase-2 small chain [Proteobacteria] Length: 372\nNumber of predicted TMHs: 1\nExp number of AAs in TMHs: 23.72226\nExp number, first 60 AAs: 0.22038\nTotal prob of N-in: 0.12994\noutside 1 331\nTMhelix 332 354\ninside 355 372

10013 GCF_000713195.1_ASM71319v1 Escherichia coli 7-233-03_S1_C2 Proteobacteria; Gammaproteobacteria; Enterobacterales; Enterobacteriaceae; Escherichia; Escherichia coli MTGDNTLIHSHGINRRDFMKLCAALAATMGLSSKAAAEMAESVTNPQRPPVIWIGAQECT WP_000145410.1 MULTISPECIES: hydrogenase-2 small chain [Proteobacteria] Length: 372\nNumber of predicted TMHs: 1\nExp number of AAs in TMHs: 23.72226\nExp number, first 60 AAs: 0.22038\nTotal prob of N-in: 0.12994\noutside 1 331\nTMhelix 332 354\ninside 355 372

10014 GCF_000713195.1_ASM71319v1 Escherichia coli 7-233-03_S1_C2 Proteobacteria; Gammaproteobacteria; Enterobacterales; Enterobacteriaceae; Escherichia; Escherichia coli MNNEETFYQAMRRQGVTRRSFLKYCSLAATSLGLGAGMAPKIAWALENKPRIPVVWIHGL WP_001058323.1 MULTISPECIES: hydrogenase-1 small chain [Proteobacteria] Length: 372\nNumber of predicted TMHs: 2\nExp number of AAs in TMHs: 37.2608499999999\nExp number, first 60 AAs: 19.52923\nTotal prob of N-in:

0.94351\nPOSSIBLE N-term signal sequence\ninside 1 19\nTMhelix 20 39\noutside 40 325\nTMhelix 326 348\ninside 349 372

10015 GCF_000713345.1_ASM71334v1 Escherichia coli 7-233-03_S1_C3 Proteobacteria; Gammaproteobacteria; Enterobacterales; Enterobacteriaceae; Escherichia; Escherichia coli MTGDNTLIHSHGINRRDFMKLCAALAATMGLSSKAAAEMAESVTNPQRPPVIWIGAQECT WP_000145410.1 MULTISPECIES: hydrogenase-2 small chain [Proteobacteria] Length: 372\nNumber of predicted TMHs: 1\nExp number of AAs in TMHs: 23.72226\nExp number, first 60 AAs: 0.22038\nTotal prob of N-in: 0.12994\noutside 1 331\nTMhelix 332 354\ninside 355 372

10016 GCF_000713345.1_ASM71334v1 Escherichia coli 7-233-03_S1_C3 Proteobacteria; Gammaproteobacteria; Enterobacterales; Enterobacteriaceae; Escherichia; Escherichia coli MNNEETFYQAMRRQGVTRRSFLKYCSLAATSLGLGAGMAPKIAWALENKPRIPVVWIHGL WP_001058323.1 MULTISPECIES: hydrogenase-1 small chain [Proteobacteria] Length: 372\nNumber of predicted TMHs: 2\nExp number of AAs in TMHs: 37.2608499999999\nExp number, first 60 AAs: 19.52923\nTotal prob of N-in: 0.94351\nPOSSIBLE N-term signal sequence\ninside 1 19\nTMhelix 20 39\noutside 40 325\nTMhelix 326 348\ninside 349 372

10017 GCF_000713235.1_ASM71323v1 Escherichia coli 7-233-03_S3_C1 Proteobacteria; Gammaproteobacteria; Enterobacterales; Enterobacteriaceae; Escherichia; Escherichia coli MTGDNTLIHSHGINRRDFMKLCAALAATMGLSSKAAAEMAESVTNPQRPPVIWIGAQECT WP_000145410.1 MULTISPECIES: hydrogenase-2 small chain [Proteobacteria] Length: 372\nNumber of predicted TMHs: 1\nExp number of AAs in TMHs: 23.72226\nExp number, first 60 AAs: 0.22038\nTotal prob of N-in: 0.12994\noutside 1 331\nTMhelix 332 354\ninside 355 372

10018 GCF_000713235.1_ASM71323v1 Escherichia coli 7-233-03_S3_C1 Proteobacteria; Gammaproteobacteria; Enterobacterales; Enterobacteriaceae; Escherichia; Escherichia coli MNNEETFYQAMRRQGVTRRSFLKYCSLAATSLGLGAGMAPKIAWALENKPRIPVVWIHGL WP_001058323.1 MULTISPECIES: hydrogenase-1 small chain [Proteobacteria] Length: 372\nNumber of predicted TMHs: 2\nExp number of AAs in TMHs: 37.2608499999999\nExp number, first 60 AAs: 19.52923\nTotal prob of N-in: 0.94351\nPOSSIBLE N-term signal sequence\ninside 1 19\nTMhelix 20 39\noutside 40 325\nTMhelix 326 348\ninside 349 372

10019 GCF_000713435.1_ASM71343v1 Escherichia coli 7-233-03_S3_C2 Proteobacteria; Gammaproteobacteria; Enterobacterales; Enterobacteriaceae; Escherichia; Escherichia coli MTGDNTLIHSHGINRRDFMKLCAALAATMGLSSKAAAEMAESVTNPQRPPVIWIGAQECT WP_000145410.1 MULTISPECIES: hydrogenase-2 small chain [Proteobacteria] Length: 372\nNumber of predicted TMHs: 1\nExp number of AAs in TMHs: 23.72226\nExp number, first 60 AAs: 0.22038\nTotal prob of N-in: 0.12994\noutside 1 331\nTMhelix 332 354\ninside 355 372

10020 GCF_000713435.1_ASM71343v1 Escherichia coli 7-233-03_S3_C2 Proteobacteria; Gammaproteobacteria; Enterobacterales; Enterobacteriaceae; Escherichia; Escherichia coli MNNEETFYQAMRRQGVTRRSFLKYCSLAATSLGLGAGMAPKIAWALENKPRIPVVWIHGL WP_001058323.1 MULTISPECIES: hydrogenase-1 small chain [Proteobacteria] Length: 372\nNumber of predicted TMHs: 2\nExp number of AAs in TMHs: 37.2608499999999\nExp number, first 60 AAs: 19.52923\nTotal prob of N-in: 0.94351\nPOSSIBLE N-term signal sequence\ninside 1 19\nTMhelix 20 39\noutside 40 325\nTMhelix 326 348\ninside 349 372

10021 GCF_000713355.1_ASM71335v1 Escherichia coli 7-233-03_S3_C3 Proteobacteria; Gammaproteobacteria; Enterobacterales; Enterobacteriaceae; Escherichia; Escherichia coli MNNEETFYQAMRRQGVTRRSFLKYCSLAATSLGLGAGMAPKIAWALENKPRIPVVWIHGL WP_001058323.1 MULTISPECIES: hydrogenase-1 small chain [Proteobacteria] Length: 372\nNumber of predicted TMHs: 2\nExp number of AAs in TMHs: 37.2608499999999\nExp number, first 60 AAs: 19.52923\nTotal prob of N-in: 0.94351\nPOSSIBLE N-term signal sequence\ninside 1 19\nTMhelix 20 39\noutside 40 325\nTMhelix 326 348\ninside 349 372

10022 GCF_000713355.1_ASM71335v1 Escherichia coli 7-233-03_S3_C3 Proteobacteria; Gammaproteobacteria; Enterobacterales; Enterobacteriaceae; Escherichia; Escherichia coli MTGDNTLIHSHGINRRDFMKLCAALAATMGLSSKAAAEMAESVTNPQRPPVIWIGAQECT WP_000145410.1 MULTISPECIES: hydrogenase-2 small chain [Proteobacteria] Length: 372\nNumber of predicted TMHs:

1\nExp number of AAs in TMHs: 23.72226\nExp number, first 60 AAs: 0.22038\nTotal prob of N-in: 0.12994\noutside 1 331\nTMhelix 332 354\ninside 355 372

10023 GCF_000713525.1_ASM71352v1 Escherichia coli 7-233-03_S4_C1 Proteobacteria; Gammaproteobacteria; Enterobacterales; Enterobacteriaceae; Escherichia; Escherichia coli MTGDNTLIHSHGINRRDFMKLCAALAATMGLSSKAAAEMAESVTNPQRPPVIWIGAQECT WP_000145410.1 MULTISPECIES: hydrogenase-2 small chain [Proteobacteria] Length: 372\nNumber of predicted TMHs: 1\nExp number of AAs in TMHs: 23.72226\nExp number, first 60 AAs: 0.22038\nTotal prob of N-in: 0.12994\noutside 1 331\nTMhelix 332 354\ninside 355 372

10024 GCF_000713525.1_ASM71352v1 Escherichia coli 7-233-03_S4_C1 Proteobacteria; Gammaproteobacteria; Enterobacterales; Enterobacteriaceae; Escherichia; Escherichia coli MNNEETFYQAMRRQGVTRRSFLKYCSLAATSLGLGAGMAPKIAWALENKPRIPVVWIHGL WP_032292164.1 hydrogenase-1 small chain [Escherichia coli] Length: 372\nNumber of predicted TMHs: 2\nExp number of AAs in TMHs: 37.50401\nExp number, first 60 AAs: 19.52955\nTotal prob of N-in: 0.94349\nPOSSIBLE N-term signal sequence\ninside 1 19\nTMhelix 20 39\noutside 40 325\nTMhelix 326 348\ninside 349 372

10025 GCF_000713395.1_ASM71339v1 Escherichia coli 7-233-03_S4_C2 Proteobacteria; Gammaproteobacteria; Enterobacterales; Enterobacteriaceae; Escherichia; Escherichia coli MTGDNTLIHSHGINRRDFMKLCAALAATMGLSSKAAAEMAESVTNPQRPPVIWIGAQECT WP_000145410.1 MULTISPECIES: hydrogenase-2 small chain [Proteobacteria] Length: 372\nNumber of predicted TMHs: 1\nExp number of AAs in TMHs: 23.72226\nExp number, first 60 AAs: 0.22038\nTotal prob of N-in: 0.12994\noutside 1 331\nTMhelix 332 354\ninside 355 372

10026 GCF_000713395.1_ASM71339v1 Escherichia coli 7-233-03_S4_C2 Proteobacteria; Gammaproteobacteria; Enterobacterales; Enterobacteriaceae; Escherichia; Escherichia coli MNNEETFYQAMRRQGVTRRSFLKYCSLAATSLGLGAGMAPKIAWALENKPRIPVVWIHGL WP_001058323.1 MULTISPECIES: hydrogenase-1 small chain [Proteobacteria] Length: 372\nNumber of predicted TMHs: 2\nExp number of AAs in TMHs: 37.2608499999999\nExp number, first 60 AAs: 19.52923\nTotal prob of N-in: 0.94351\nPOSSIBLE N-term signal sequence\ninside 1 19\nTMhelix 20 39\noutside 40 325\nTMhelix 326 348\ninside 349 372

10027 GCF_000713535.1_ASM71353v1 Escherichia coli 7-233-03_S4_C3 Proteobacteria; Gammaproteobacteria; Enterobacterales; Enterobacteriaceae; Escherichia; Escherichia coli MTGDNTLIHSHGINRRDFMKLCAALAATMGLSSKAAAEMAESVTNPQRPPVIWIGAQECT WP_000145410.1 MULTISPECIES: hydrogenase-2 small chain [Proteobacteria] Length: 372\nNumber of predicted TMHs: 1\nExp number of AAs in TMHs: 23.72226\nExp number, first 60 AAs: 0.22038\nTotal prob of N-in: 0.12994\noutside 1 331\nTMhelix 332 354\ninside 355 372

10028 GCF_000713535.1_ASM71353v1 Escherichia coli 7-233-03_S4_C3 Proteobacteria; Gammaproteobacteria; Enterobacterales; Enterobacteriaceae; Escherichia; Escherichia coli MNNEETFYQAMRRQGVTRRSFLKYCSLAATSLGLGAGMAPKIAWALENKPRIPVVWIHGL WP_032292164.1 hydrogenase-1 small chain [Escherichia coli] Length: 372\nNumber of predicted TMHs: 2\nExp number of AAs in TMHs: 37.50401\nExp number, first 60 AAs: 19.52955\nTotal prob of N-in: 0.94349\nPOSSIBLE N-term signal sequence\ninside 1 19\nTMhelix 20 39\noutside 40 325\nTMhelix 326 348\ninside 349 372

10029 GCF_000264155.1_ASM26415v1 Escherichia coli 75 Proteobacteria; Gammaproteobacteria; Enterobacterales; Enterobacteriaceae; Escherichia; Escherichia coli MTGDNTLIHSHGINRRDFMKLCAALAATMGLSSKAAAEMAESVTNPQRPPVIWIGAQECT WP_000145410.1 MULTISPECIES: hydrogenase-2 small chain [Proteobacteria] Length: 372\nNumber of predicted TMHs: 1\nExp number of AAs in TMHs: 23.72226\nExp number, first 60 AAs: 0.22038\nTotal prob of N-in: 0.12994\noutside 1 331\nTMhelix 332 354\ninside 355 372

10030 GCF_000264155.1_ASM26415v1 Escherichia coli 75 Proteobacteria; Gammaproteobacteria; Enterobacterales; Enterobacteriaceae; Escherichia; Escherichia coli MNNEETFYQAMRRQGVTRRSFLKYCSLAATSLGLGAGMAPKIAWALENKPRIPVVWIHGL WP_001058323.1 MULTISPECIES: hydrogenase-1 small chain [Proteobacteria] Length: 372\nNumber of predicted TMHs: 2\nExp number of AAs in TMHs: 37.2608499999999\nExp number, first 60 AAs: 19.52923\nTotal prob of N-in: 0.94351\nPOSSIBLE N-term signal sequence\ninside 1 19\nTMhelix 20 39\noutside 40 325\nTMhelix 326 348\ninside 349 372

10031 GCF_000699365.1_7996-1 Escherichia coli 7996-1 Proteobacteria; Gammaproteobacteria; Enterobacterales; Enterobacteriaceae; Escherichia; Escherichia coli
MTGDNTLIHSHGINRRDFMKLCAALAATMGLSSKAAAEMAESVTNPQRPPVIWIGAQECT WP_000145410.1
MULTISPECIES: hydrogenase-2 small chain [Proteobacteria] Length: 372\nNumber of predicted TMHs: 1\nExp number of AAs in TMHs: 23.72226\nExp number, first 60 AAs: 0.22038\nTotal prob of N-in: 0.12994\noutside 1 331\nTMhelix 332 354\ninside 355 372

10032 GCF_000699365.1_7996-1 Escherichia coli 7996-1 Proteobacteria; Gammaproteobacteria; Enterobacterales; Enterobacteriaceae; Escherichia; Escherichia coli
MNNEETFYQAMRRQGVTRRSFLKYCSLAATSLGLGAGMAPKIAWALENKPRIPVVWIHGL WP_001058323.1
MULTISPECIES: hydrogenase-1 small chain [Proteobacteria] Length: 372\nNumber of predicted TMHs: 2\nExp number of AAs in TMHs: 37.2608499999999\nExp number, first 60 AAs: 19.52923\nTotal prob of N-in: 0.94351\nPOSSIBLE N-term signal sequence\ninside 1 19\nTMhelix 20 39\noutside 40 325\nTMhelix 326 348\ninside 349 372

10033 GCF_000304835.2_ASM30483v2 Escherichia coli 8.0416 Proteobacteria; Gammaproteobacteria; Enterobacterales; Enterobacteriaceae; Escherichia; Escherichia coli
MTGDNTLIHSHGINRRDFMKLCAALAATMGLSSKAAAEMAESVTNPQRPPVIWIGAQECT WP_000145410.1
MULTISPECIES: hydrogenase-2 small chain [Proteobacteria] Length: 372\nNumber of predicted TMHs: 1\nExp number of AAs in TMHs: 23.72226\nExp number, first 60 AAs: 0.22038\nTotal prob of N-in: 0.12994\noutside 1 331\nTMhelix 332 354\ninside 355 372

10034 GCF_000304835.2_ASM30483v2 Escherichia coli 8.0416 Proteobacteria; Gammaproteobacteria; Enterobacterales; Enterobacteriaceae; Escherichia; Escherichia coli
MNNEETFYQAMRRQGVTRRSFLKYCSLAATSLGLGAGMAPKIAWALENKPRIPVVWIHGL WP_001301613.1
hydrogenase [Escherichia coli] Length: 372\nNumber of predicted TMHs: 2\nExp number of AAs in TMHs: 37.46598\nExp number, first 60 AAs: 19.53139\nTotal prob of N-in: 0.94364\nPOSSIBLE N-term signal sequence\ninside 1 19\nTMhelix 20 39\noutside 40 325\nTMhelix 326 348\ninside 349 372

10035 GCF_000305175.1_ASM30517v1 Escherichia coli 8.0566 Proteobacteria; Gammaproteobacteria; Enterobacterales; Enterobacteriaceae; Escherichia; Escherichia coli
MTGDNTLIHSHGINRRDFMKLCAALAATMGLSSKAAAEMAESVTNPQRPPVIWIGAQECT WP_000145410.1
MULTISPECIES: hydrogenase-2 small chain [Proteobacteria] Length: 372\nNumber of predicted TMHs: 1\nExp number of AAs in TMHs: 23.72226\nExp number, first 60 AAs: 0.22038\nTotal prob of N-in: 0.12994\noutside 1 331\nTMhelix 332 354\ninside 355 372

10036 GCF_000305355.1_ASM30535v1 Escherichia coli 8.0569 Proteobacteria; Gammaproteobacteria; Enterobacterales; Enterobacteriaceae; Escherichia; Escherichia coli
MTGDNTLIHSHGINRRDFMKLCAALAATMGLSSKAAAEMAESVTNPQRPPVIWIGAQECT WP_000145410.1
MULTISPECIES: hydrogenase-2 small chain [Proteobacteria] Length: 372\nNumber of predicted TMHs: 1\nExp number of AAs in TMHs: 23.72226\nExp number, first 60 AAs: 0.22038\nTotal prob of N-in: 0.12994\noutside 1 331\nTMhelix 332 354\ninside 355 372

10037 GCF_000305375.2_ASM30537v2 Escherichia coli 8.0586 Proteobacteria; Gammaproteobacteria; Enterobacterales; Enterobacteriaceae; Escherichia; Escherichia coli
MTGDNTLIHSHGINRRDFMKLCAALAATMGLSSKAAAEMAESVTNPQRPPVIWIGAQECT WP_000145410.1
MULTISPECIES: hydrogenase-2 small chain [Proteobacteria] Length: 372\nNumber of predicted TMHs: 1\nExp number of AAs in TMHs: 23.72226\nExp number, first 60 AAs: 0.22038\nTotal prob of N-in: 0.12994\noutside 1 331\nTMhelix 332 354\ninside 355 372

10038 GCF_000305375.2_ASM30537v2 Escherichia coli 8.0586 Proteobacteria; Gammaproteobacteria; Enterobacterales; Enterobacteriaceae; Escherichia; Escherichia coli
MNNEETFYQAMRRQGVTRRSFLKYCSLAATSLGLGAGMAPKIAWALENKPRIPVVWIHGL WP_001301613.1
hydrogenase [Escherichia coli] Length: 372\nNumber of predicted TMHs: 2\nExp number of AAs in TMHs: 37.46598\nExp number, first 60 AAs: 19.53139\nTotal prob of N-in: 0.94364\nPOSSIBLE N-term signal sequence\ninside 1 19\nTMhelix 20 39\noutside 40 325\nTMhelix 326 348\ninside 349 372

10039 GCF_000305395.2_ASM30539v2 Escherichia coli 8.2524 Proteobacteria; Gammaproteobacteria; Enterobacterales; Enterobacteriaceae; Escherichia; Escherichia coli
MTGDNTLIHSHGINRRDFMKLCAALAATMGLSSKAAAEMAESVTNPQRPPVIWIGAQECT WP_000145410.1
MULTISPECIES: hydrogenase-2 small chain [Proteobacteria] Length: 372\nNumber of predicted TMHs:

1\nExp number of AAs in TMHs: 23.72226\nExp number, first 60 AAs: 0.22038\nTotal prob of N-in: 0.12994\noutside 1 331\nTMhelix 332 354\ninside 355 372

10040 GCF_000305395.2_ASM30539v2 Escherichia coli 8.2524 Proteobacteria; Gammaproteobacteria; Enterobacterales; Enterobacteriaceae; Escherichia; Escherichia coli MNNEETFYQAMRRQGVTRRSFLKYCSLAATSLGLGAGMAPKIAWALENKPRIPVVWIHGL WP_001301613.1 hydrogenase [Escherichia coli] Length: 372\nNumber of predicted TMHs: 2\nExp number of AAs in TMHs: 37.46598\nExp number, first 60 AAs: 19.53139\nTotal prob of N-in: 0.94364\nPOSSIBLE N-term signal sequence\ninside 1 19\nTMhelix 20 39\noutside 40 325\nTMhelix 326 348\ninside 349 372

10041 GCF_000159295.1_ASM15929v1 Escherichia coli 83972 Proteobacteria; Gammaproteobacteria; Enterobacterales; Enterobacteriaceae; Escherichia; Escherichia coli MNNEETFYQAMRRQGVTRRSFLKYCSLAATSLGLGAGMAPKIAWALENKPRIPVVWIHGL WP_001058323.1 MULTISPECIES: hydrogenase-1 small chain [Proteobacteria] Length: 372\nNumber of predicted TMHs: 2\nExp number of AAs in TMHs: 37.2608499999999\nExp number, first 60 AAs: 19.52923\nTotal prob of N-in: 0.94351\nPOSSIBLE N-term signal sequence\ninside 1 19\nTMhelix 20 39\noutside 40 325\nTMhelix 326 348\ninside 349 372

10042 GCF_000159295.1_ASM15929v1 Escherichia coli 83972 Proteobacteria; Gammaproteobacteria; Enterobacterales; Enterobacteriaceae; Escherichia; Escherichia coli MTGDNTLIHSHGINRRDFMKLCAALAATMGLSSKAAAEMAESVTNPQRPPVIWIGAQECT WP_000145410.1 MULTISPECIES: hydrogenase-2 small chain [Proteobacteria] Length: 372\nNumber of predicted TMHs: 1\nExp number of AAs in TMHs: 23.72226\nExp number, first 60 AAs: 0.22038\nTotal prob of N-in: 0.12994\noutside 1 331\nTMhelix 332 354\ninside 355 372

10043 GCF_000713425.1_ASM71342v1 Escherichia coli 8-415-05_S1_C1 Proteobacteria; Gammaproteobacteria; Enterobacterales; Enterobacteriaceae; Escherichia; Escherichia coli MNNEETFYQAMRRQGVTRRSFLKYCSLAATSLGLGAGMAPKIAWALENKPRIPVVWIHGL WP_001058323.1 MULTISPECIES: hydrogenase-1 small chain [Proteobacteria] Length: 372\nNumber of predicted TMHs: 2\nExp number of AAs in TMHs: 37.2608499999999\nExp number, first 60 AAs: 19.52923\nTotal prob of N-in: 0.94351\nPOSSIBLE N-term signal sequence\ninside 1 19\nTMhelix 20 39\noutside 40 325\nTMhelix 326 348\ninside 349 372

10044 GCF_000713425.1_ASM71342v1 Escherichia coli 8-415-05_S1_C1 Proteobacteria; Gammaproteobacteria; Enterobacterales; Enterobacteriaceae; Escherichia; Escherichia coli MTGDNTLIHSHGINRRDFMKLCAALAATMGLSSKAAAEMAESVTNPQRPPVIWIGAQECT WP_000145410.1 MULTISPECIES: hydrogenase-2 small chain [Proteobacteria] Length: 372\nNumber of predicted TMHs: 1\nExp number of AAs in TMHs: 23.72226\nExp number, first 60 AAs: 0.22038\nTotal prob of N-in: 0.12994\noutside 1 331\nTMhelix 332 354\ninside 355 372

10045 GCF_000713775.1_ASM71377v1 Escherichia coli 8-415-05_S1_C2 Proteobacteria; Gammaproteobacteria; Enterobacterales; Enterobacteriaceae; Escherichia; Escherichia coli MTGDNTLIHSHGINRRDFMKLCAALAATMGLSSKAAAEMAESVTNPQRPPVIWIGAQECT WP_000145410.1 MULTISPECIES: hydrogenase-2 small chain [Proteobacteria] Length: 372\nNumber of predicted TMHs: 1\nExp number of AAs in TMHs: 23.72226\nExp number, first 60 AAs: 0.22038\nTotal prob of N-in: 0.12994\noutside 1 331\nTMhelix 332 354\ninside 355 372

10046 GCF_000713775.1_ASM71377v1 Escherichia coli 8-415-05_S1_C2 Proteobacteria; Gammaproteobacteria; Enterobacterales; Enterobacteriaceae; Escherichia; Escherichia coli MNNEETFYQAMRRQGVTRRSFLKYCSLAATSLGLGAGMAPKIAWALENKPRIPVVWIHGL WP_001058323.1 MULTISPECIES: hydrogenase-1 small chain [Proteobacteria] Length: 372\nNumber of predicted TMHs: 2\nExp number of AAs in TMHs: 37.2608499999999\nExp number, first 60 AAs: 19.52923\nTotal prob of N-in: 0.94351\nPOSSIBLE N-term signal sequence\ninside 1 19\nTMhelix 20 39\noutside 40 325\nTMhelix 326 348\ninside 349 372

10047 GCF_000713495.1_ASM71349v1 Escherichia coli 8-415-05_S3_C1 Proteobacteria; Gammaproteobacteria; Enterobacterales; Enterobacteriaceae; Escherichia; Escherichia coli MTGDNTLIHSHGINRRDFMKLCAALAATMGLSSKAAAEMAESVTNPQRPPVIWIGAQECT WP_000145410.1 MULTISPECIES: hydrogenase-2 small chain [Proteobacteria] Length: 372\nNumber of predicted TMHs: 1\nExp number of AAs in TMHs: 23.72226\nExp number, first 60 AAs: 0.22038\nTotal prob of N-in: 0.12994\noutside 1 331\nTMhelix 332 354\ninside 355 372

10048 GCF_000713495.1_ASM71349v1 Escherichia coli 8-415-05_S3_C1 Proteobacteria; Gammaproteobacteria; Enterobacterales; Enterobacteriaceae; Escherichia; Escherichia coli
MNNEETFYQAMRRQGVTRRSFLKYCSLAATSLGLGAGMAPKIAWALENKPRIPVVWIHGL WP_001058323.1
MULTISPECIES: hydrogenase-1 small chain [Proteobacteria] Length: 372\nNumber of predicted TMHs: 2\nExp number of AAs in TMHs: 37.2608499999999\nExp number, first 60 AAs: 19.52923\nTotal prob of N-in: 0.94351\nPOSSIBLE N-term signal sequence\ninside 1 19\nTMhelix 20 39\noutside 40 325\nTMhelix 326 348\ninside 349 372

10049 GCF_000713585.1_ASM71358v1 Escherichia coli 8-415-05_S3_C2 Proteobacteria; Gammaproteobacteria; Enterobacterales; Enterobacteriaceae; Escherichia; Escherichia coli
MNNEETFYQAMRRQGVTRRSFLKYCSLAATSLGLGAGMAPKIAWALENKPRIPVVWIHGL WP_001058323.1
MULTISPECIES: hydrogenase-1 small chain [Proteobacteria] Length: 372\nNumber of predicted TMHs: 2\nExp number of AAs in TMHs: 37.2608499999999\nExp number, first 60 AAs: 19.52923\nTotal prob of N-in: 0.94351\nPOSSIBLE N-term signal sequence\ninside 1 19\nTMhelix 20 39\noutside 40 325\nTMhelix 326 348\ninside 349 372

10050 GCF_000713585.1_ASM71358v1 Escherichia coli 8-415-05_S3_C2 Proteobacteria; Gammaproteobacteria; Enterobacterales; Enterobacteriaceae; Escherichia; Escherichia coli
MTGDNTLIHSHGINRRDFMKLCAALAATMGLSSKAAAEMAESVTNPQRPPVIWIGAQECT WP_000145410.1
MULTISPECIES: hydrogenase-2 small chain [Proteobacteria] Length: 372\nNumber of predicted TMHs: 1\nExp number of AAs in TMHs: 23.72226\nExp number, first 60 AAs: 0.22038\nTotal prob of N-in: 0.12994\noutside 1 331\nTMhelix 332 354\ninside 355 372

10051 GCF_000713455.1_ASM71345v1 Escherichia coli 8-415-05_S3_C3 Proteobacteria; Gammaproteobacteria; Enterobacterales; Enterobacteriaceae; Escherichia; Escherichia coli
MNNEETFYQAMRRQGVTRRSFLKYCSLAATSLGLGAGMAPKIAWALENKPRIPVVWIHGL WP_001058323.1
MULTISPECIES: hydrogenase-1 small chain [Proteobacteria] Length: 372\nNumber of predicted TMHs: 2\nExp number of AAs in TMHs: 37.2608499999999\nExp number, first 60 AAs: 19.52923\nTotal prob of N-in: 0.94351\nPOSSIBLE N-term signal sequence\ninside 1 19\nTMhelix 20 39\noutside 40 325\nTMhelix 326 348\ninside 349 372

10052 GCF_000713455.1_ASM71345v1 Escherichia coli 8-415-05_S3_C3 Proteobacteria; Gammaproteobacteria; Enterobacterales; Enterobacteriaceae; Escherichia; Escherichia coli
MTGDNTLIHSHGINRRDFMKLCAALAATMGLSSKAAAEMAESVTNPQRPPVIWIGAQECT WP_000145410.1
MULTISPECIES: hydrogenase-2 small chain [Proteobacteria] Length: 372\nNumber of predicted TMHs: 1\nExp number of AAs in TMHs: 23.72226\nExp number, first 60 AAs: 0.22038\nTotal prob of N-in: 0.12994\noutside 1 331\nTMhelix 332 354\ninside 355 372

10053 GCF_000711455.1_ASM71145v1 Escherichia coli 8-415-05_S4_C1 Proteobacteria; Gammaproteobacteria; Enterobacterales; Enterobacteriaceae; Escherichia; Escherichia coli
MTGDNTLIHSHGINRRDFMKLCAALAATMGLSSKAAAEMAESVTNPQRPPVIWIGAQECT WP_000145410.1
MULTISPECIES: hydrogenase-2 small chain [Proteobacteria] Length: 372\nNumber of predicted TMHs: 1\nExp number of AAs in TMHs: 23.72226\nExp number, first 60 AAs: 0.22038\nTotal prob of N-in: 0.12994\noutside 1 331\nTMhelix 332 354\ninside 355 372

10054 GCF_000711455.1_ASM71145v1 Escherichia coli 8-415-05_S4_C1 Proteobacteria; Gammaproteobacteria; Enterobacterales; Enterobacteriaceae; Escherichia; Escherichia coli
MNNEETFYQAMRRQGVTRRSFLKYCSLAATSLGLGAGMAPKIAWALENKPRIPVVWIHGL WP_001058323.1
MULTISPECIES: hydrogenase-1 small chain [Proteobacteria] Length: 372\nNumber of predicted TMHs: 2\nExp number of AAs in TMHs: 37.2608499999999\nExp number, first 60 AAs: 19.52923\nTotal prob of N-in: 0.94351\nPOSSIBLE N-term signal sequence\ninside 1 19\nTMhelix 20 39\noutside 40 325\nTMhelix 326 348\ninside 349 372

10055 GCF_000711365.1_ASM71136v1 Escherichia coli 8-415-05_S4_C2 Proteobacteria; Gammaproteobacteria; Enterobacterales; Enterobacteriaceae; Escherichia; Escherichia coli
MTGDNTLIHSHGINRRDFMKLCAALAATMGLSSKAAAEMAESVTNPQRPPVIWIGAQECT WP_000145410.1
MULTISPECIES: hydrogenase-2 small chain [Proteobacteria] Length: 372\nNumber of predicted TMHs: 1\nExp number of AAs in TMHs: 23.72226\nExp number, first 60 AAs: 0.22038\nTotal prob of N-in: 0.12994\noutside 1 331\nTMhelix 332 354\ninside 355 372

10056 GCF_000711365.1_ASM71136v1 Escherichia coli 8-415-05_S4_C2 Proteobacteria; Gammaproteobacteria; Enterobacterales; Enterobacteriaceae; Escherichia; Escherichia coli
MNNEETFYQAMRRQGVTRRSFLKYCSLAATSLGLGAGMAPKIAWALENKPRIPVVWIHGL WP_001058323.1
MULTISPECIES: hydrogenase-1 small chain [Proteobacteria] Length: 372\nNumber of predicted TMHs: 2\nExp number of AAs in TMHs: 37.2608499999999\nExp number, first 60 AAs: 19.52923\nTotal prob of N-in: 0.94351\nPOSSIBLE N-term signal sequence\ninside 1 19\nTMhelix 20 39\noutside 40 325\nTMhelix 326 348\ninside 349 372

10057 GCF_000711435.1_ASM71143v1 Escherichia coli 8-415-05_S4_C3 Proteobacteria; Gammaproteobacteria; Enterobacterales; Enterobacteriaceae; Escherichia; Escherichia coli
MNNEETFYQAMRRQGVTRRSFLKYCSLAATSLGLGAGMAPKIAWALENKPRIPVVWIHGL WP_001058323.1
MULTISPECIES: hydrogenase-1 small chain [Proteobacteria] Length: 372\nNumber of predicted TMHs: 2\nExp number of AAs in TMHs: 37.2608499999999\nExp number, first 60 AAs: 19.52923\nTotal prob of N-in: 0.94351\nPOSSIBLE N-term signal sequence\ninside 1 19\nTMhelix 20 39\noutside 40 325\nTMhelix 326 348\ninside 349 372

10058 GCF_000711435.1_ASM71143v1 Escherichia coli 8-415-05_S4_C3 Proteobacteria; Gammaproteobacteria; Enterobacterales; Enterobacteriaceae; Escherichia; Escherichia coli
MTGDNTLIHSHGINRRDFMKLCAALAATMGLSSKAAAEMAESVTNPQRPPVIWIGAQECT WP_000145410.1
MULTISPECIES: hydrogenase-2 small chain [Proteobacteria] Length: 372\nNumber of predicted TMHs: 1\nExp number of AAs in TMHs: 23.72226\nExp number, first 60 AAs: 0.22038\nTotal prob of N-in: 0.12994\noutside 1 331\nTMhelix 332 354\ninside 355 372

10059 GCF_000503415.1_Ecoli85B Escherichia coli 85BProteobacteria; Gammaproteobacteria; Enterobacterales; Enterobacteriaceae; Escherichia; Escherichia coli
MNNEETFYQAMRRQGVTRRSFLKYCSLAATSLGLGAGMAPKIAWALENKPRIPVVWIHGL WP_001058323.1
MULTISPECIES: hydrogenase-1 small chain [Proteobacteria] Length: 372\nNumber of predicted TMHs: 2\nExp number of AAs in TMHs: 37.2608499999999\nExp number, first 60 AAs: 19.52923\nTotal prob of N-in: 0.94351\nPOSSIBLE N-term signal sequence\ninside 1 19\nTMhelix 20 39\noutside 40 325\nTMhelix 326 348\ninside 349 372

10060 GCF_000503415.1_Ecoli85B Escherichia coli 85BProteobacteria; Gammaproteobacteria; Enterobacterales; Enterobacteriaceae; Escherichia; Escherichia coli
MTGDNTLIHSHGINRRDFMKLCAALAATMGLSSKAAAEMAESVTNPQRPPVIWIGAQECT WP_000145410.1
MULTISPECIES: hydrogenase-2 small chain [Proteobacteria] Length: 372\nNumber of predicted TMHs: 1\nExp number of AAs in TMHs: 23.72226\nExp number, first 60 AAs: 0.22038\nTotal prob of N-in: 0.12994\noutside 1 331\nTMhelix 332 354\ninside 355 372

10061 GCF_000478215.1_ASM47821v1 Escherichia coli 8624 Proteobacteria; Gammaproteobacteria; Enterobacterales; Enterobacteriaceae; Escherichia; Escherichia coli
MNNEETFYQAMRRQGVTRRSFLKYCSLAATSLGLGAGMAPKIAWALENKPRIPVVWIHGL WP_001301613.1
hydrogenase [Escherichia coli] Length: 372\nNumber of predicted TMHs: 2\nExp number of AAs in TMHs: 37.46598\nExp number, first 60 AAs: 19.53139\nTotal prob of N-in: 0.94364\nPOSSIBLE N-term signal sequence\ninside 1 19\nTMhelix 20 39\noutside 40 325\nTMhelix 326 348\ninside 349 372

10062 GCF_000478215.1_ASM47821v1 Escherichia coli 8624 Proteobacteria; Gammaproteobacteria; Enterobacterales; Enterobacteriaceae; Escherichia; Escherichia coli
MTGDNTLIHSHGINRRDFMKLCAALAATMGLSSKAAAEMAESVTNPQRPPVIWIGAQECT WP_000145410.1
MULTISPECIES: hydrogenase-2 small chain [Proteobacteria] Length: 372\nNumber of predicted TMHs: 1\nExp number of AAs in TMHs: 23.72226\nExp number, first 60 AAs: 0.22038\nTotal prob of N-in: 0.12994\noutside 1 331\nTMhelix 332 354\ninside 355 372

10063 GCF_000503615.1_Ecoli87A Escherichia coli 87A Proteobacteria; Gammaproteobacteria; Enterobacterales; Enterobacteriaceae; Escherichia; Escherichia coli
MNNEETFYQAMRRQGVTRRSFLKYCSLAATSLGLGAGMAPKIAWALENKPRIPVVWIHGL WP_001058323.1
MULTISPECIES: hydrogenase-1 small chain [Proteobacteria] Length: 372\nNumber of predicted TMHs: 2\nExp number of AAs in TMHs: 37.2608499999999\nExp number, first 60 AAs: 19.52923\nTotal prob of N-in: 0.94351\nPOSSIBLE N-term signal sequence\ninside 1 19\nTMhelix 20 39\noutside 40 325\nTMhelix 326 348\ninside 349 372

10064 GCF_000503615.1_Ecoli87A Escherichia coli 87A Proteobacteria; Gammaproteobacteria; Enterobacterales; Enterobacteriaceae; Escherichia; Escherichia coli
MTGDNTLIHSHGINRRDFMKLCAALAATMGLSSKAAAEMAESVTNPQRPPVIWIGAQECT WP_000145410.1
MULTISPECIES: hydrogenase-2 small chain [Proteobacteria] Length: 372\nNumber of predicted TMHs: 1\nExp number of AAs in TMHs: 23.72226\nExp number, first 60 AAs: 0.22038\nTotal prob of N-in: 0.12994\noutside 1 331\nTMhelix 332 354\ninside 355 372

10065 GCF_000305455.2_ASM30545v2 Escherichia coli 88.0221 Proteobacteria; Gammaproteobacteria; Enterobacterales; Enterobacteriaceae; Escherichia; Escherichia coli
MNNEETFYQAMRRQGVTRRSFLKYCSLAATSLGLGAGMAPKIAWALENKPRIPVVWIHGL WP_001301613.1
hydrogenase [Escherichia coli] Length: 372\nNumber of predicted TMHs: 2\nExp number of AAs in TMHs: 37.46598\nExp number, first 60 AAs: 19.53139\nTotal prob of N-in: 0.94364\nPOSSIBLE N-term signal sequence\ninside 1 19\nTMhelix 20 39\noutside 40 325\nTMhelix 326 348\ninside 349 372

10066 GCF_000305455.2_ASM30545v2 Escherichia coli 88.0221 Proteobacteria; Gammaproteobacteria; Enterobacterales; Enterobacteriaceae; Escherichia; Escherichia coli
MTGDNTLIHSHGINRRDFMKLCAALAATMGLSSKAAAEMAESVTNPQRPPVIWIGAQECT WP_000145410.1
MULTISPECIES: hydrogenase-2 small chain [Proteobacteria] Length: 372\nNumber of predicted TMHs: 1\nExp number of AAs in TMHs: 23.72226\nExp number, first 60 AAs: 0.22038\nTotal prob of N-in: 0.12994\noutside 1 331\nTMhelix 332 354\ninside 355 372

10067 GCF_000316345.2_ASM31634v2 Escherichia coli 88.1042 Proteobacteria; Gammaproteobacteria; Enterobacterales; Enterobacteriaceae; Escherichia; Escherichia coli
MNNEETFYQAMRRQGVTRRSFLKYCSLAATSLGLGAGMAPKIAWALENKPRIPVVWIHGL WP_001301613.1
hydrogenase [Escherichia coli] Length: 372\nNumber of predicted TMHs: 2\nExp number of AAs in TMHs: 37.46598\nExp number, first 60 AAs: 19.53139\nTotal prob of N-in: 0.94364\nPOSSIBLE N-term signal sequence\ninside 1 19\nTMhelix 20 39\noutside 40 325\nTMhelix 326 348\ninside 349 372

10068 GCF_000316345.2_ASM31634v2 Escherichia coli 88.1042 Proteobacteria; Gammaproteobacteria; Enterobacterales; Enterobacteriaceae; Escherichia; Escherichia coli
MTGDNTLIHSHGINRRDFMKLCAALAATMGLSSKAAAEMAESVTNPQRPPVIWIGAQECT WP_000145410.1
MULTISPECIES: hydrogenase-2 small chain [Proteobacteria] Length: 372\nNumber of predicted TMHs: 1\nExp number of AAs in TMHs: 23.72226\nExp number, first 60 AAs: 0.22038\nTotal prob of N-in: 0.12994\noutside 1 331\nTMhelix 332 354\ninside 355 372

10069 GCF_000316325.2_ASM31632v2 Escherichia coli 88.1467 Proteobacteria; Gammaproteobacteria; Enterobacterales; Enterobacteriaceae; Escherichia; Escherichia coli
MTGDNTLIHSHGINRRDFMKLCAALAATMGLSSKAAAEMAESVTNPQRPPVIWIGAQECT WP_000145410.1
MULTISPECIES: hydrogenase-2 small chain [Proteobacteria] Length: 372\nNumber of predicted TMHs: 1\nExp number of AAs in TMHs: 23.72226\nExp number, first 60 AAs: 0.22038\nTotal prob of N-in: 0.12994\noutside 1 331\nTMhelix 332 354\ninside 355 372

10070 GCF_000316325.2_ASM31632v2 Escherichia coli 88.1467 Proteobacteria; Gammaproteobacteria; Enterobacterales; Enterobacteriaceae; Escherichia; Escherichia coli
MNNEETFYQAMRRQGVTRRSFLKYCSLAATSLGLGAGMAPKIAWALENKPRIPVVWIHGL WP_001301613.1
hydrogenase [Escherichia coli] Length: 372\nNumber of predicted TMHs: 2\nExp number of AAs in TMHs: 37.46598\nExp number, first 60 AAs: 19.53139\nTotal prob of N-in: 0.94364\nPOSSIBLE N-term signal sequence\ninside 1 19\nTMhelix 20 39\noutside 40 325\nTMhelix 326 348\ninside 349 372

10071 GCF_000316365.2_ASM31636v2 Escherichia coli 89.0511 Proteobacteria; Gammaproteobacteria; Enterobacterales; Enterobacteriaceae; Escherichia; Escherichia coli
MTGDNTLIHSHGINRRDFMKLCAALAATMGLSSKAAAEMAESVTNPQRPPVIWIGAQECT WP_000145410.1
MULTISPECIES: hydrogenase-2 small chain [Proteobacteria] Length: 372\nNumber of predicted TMHs: 1\nExp number of AAs in TMHs: 23.72226\nExp number, first 60 AAs: 0.22038\nTotal prob of N-in: 0.12994\noutside 1 331\nTMhelix 332 354\ninside 355 372

10072 GCF_000316365.2_ASM31636v2 Escherichia coli 89.0511 Proteobacteria; Gammaproteobacteria; Enterobacterales; Enterobacteriaceae; Escherichia; Escherichia coli
MNNEETFYQAMRRQGVTRRSFLKYCSLAATSLGLGAGMAPKIAWALENKPRIPVVWIHGL WP_001301613.1
hydrogenase [Escherichia coli] Length: 372\nNumber of predicted TMHs: 2\nExp number of AAs in

TMHs: 37.46598\nExp number, first 60 AAs: 19.53139\nTotal prob of N-in: 0.94364\nPOSSIBLE N-term signal sequence\ninside 1 19\nTMhelix 20 39\noutside 40 325\nTMhelix 326 348\ninside 349 372

10073 GCF_000503255.1_Ecoli897 Escherichia coli 897 Proteobacteria; Gammaproteobacteria; Enterobacterales; Enterobacteriaceae; Escherichia; Escherichia coli
MNNEETFYQAMRRQGVTRRSFLKYCSLAATSLGLGAGMAPKIAWALENKPRIPVVWIHGL WP_001058323.1
MULTISPECIES: hydrogenase-1 small chain [Proteobacteria] Length: 372\nNumber of predicted TMHs: 2\nExp number of AAs in TMHs: 37.2608499999999\nExp number, first 60 AAs: 19.52923\nTotal prob of N-in: 0.94351\nPOSSIBLE N-term signal sequence\ninside 1 19\nTMhelix 20 39\noutside 40 325\nTMhelix 326 348\ninside 349 372

10074 GCF_000503255.1_Ecoli897 Escherichia coli 897 Proteobacteria; Gammaproteobacteria; Enterobacterales; Enterobacteriaceae; Escherichia; Escherichia coli
MTGDNTLIHSHGINRRDFMKLCAALAATMGLSSKAAAEMAESVTNPQRPPVIWIGAQECT WP_000145410.1
MULTISPECIES: hydrogenase-2 small chain [Proteobacteria] Length: 372\nNumber of predicted TMHs: 1\nExp number of AAs in TMHs: 23.72226\nExp number, first 60 AAs: 0.22038\nTotal prob of N-in: 0.12994\noutside 1 331\nTMhelix 332 354\ninside 355 372

10075 GCF_000316405.2_ASM31640v2 Escherichia coli 90.0039 Proteobacteria; Gammaproteobacteria; Enterobacterales; Enterobacteriaceae; Escherichia; Escherichia coli
MTGDNTLIHSHGINRRDFMKLCAALAATMGLSSKAAAEMAESVTNPQRPPVIWIGAQECT WP_000145410.1
MULTISPECIES: hydrogenase-2 small chain [Proteobacteria] Length: 372\nNumber of predicted TMHs: 1\nExp number of AAs in TMHs: 23.72226\nExp number, first 60 AAs: 0.22038\nTotal prob of N-in: 0.12994\noutside 1 331\nTMhelix 332 354\ninside 355 372

10076 GCF_000316405.2_ASM31640v2 Escherichia coli 90.0039 Proteobacteria; Gammaproteobacteria; Enterobacterales; Enterobacteriaceae; Escherichia; Escherichia coli
MNNEETFYQAMRRQGVTRRSFLKYCSLAATSLGLGAGMAPKIAWALENKPRIPVVWIHGL WP_001301613.1
hydrogenase [Escherichia coli] Length: 372\nNumber of predicted TMHs: 2\nExp number of AAs in TMHs: 37.46598\nExp number, first 60 AAs: 19.53139\nTotal prob of N-in: 0.94364\nPOSSIBLE N-term signal sequence\ninside 1 19\nTMhelix 20 39\noutside 40 325\nTMhelix 326 348\ninside 349 372

10077 GCF_000316385.2_ASM31638v2 Escherichia coli 90.0091 Proteobacteria; Gammaproteobacteria; Enterobacterales; Enterobacteriaceae; Escherichia; Escherichia coli
MTGDNTLIHSHGINRRDFMKLCAALAATMGLSSKAAAEMAESVTNPQRPPVIWIGAQECT WP_000145410.1
MULTISPECIES: hydrogenase-2 small chain [Proteobacteria] Length: 372\nNumber of predicted TMHs: 1\nExp number of AAs in TMHs: 23.72226\nExp number, first 60 AAs: 0.22038\nTotal prob of N-in: 0.12994\noutside 1 331\nTMhelix 332 354\ninside 355 372

10078 GCF_000316385.2_ASM31638v2 Escherichia coli 90.0091 Proteobacteria; Gammaproteobacteria; Enterobacterales; Enterobacteriaceae; Escherichia; Escherichia coli
MNNEETFYQAMRRQGVTRRSFLKYCSLAATSLGLGAGMAPKIAWALENKPRIPVVWIHGL WP_001301613.1
hydrogenase [Escherichia coli] Length: 372\nNumber of predicted TMHs: 2\nExp number of AAs in TMHs: 37.46598\nExp number, first 60 AAs: 19.53139\nTotal prob of N-in: 0.94364\nPOSSIBLE N-term signal sequence\ninside 1 19\nTMhelix 20 39\noutside 40 325\nTMhelix 326 348\ninside 349 372

10079 GCF_000194725.1_ASM19472v2 Escherichia coli 900105 (10e) Proteobacteria; Gammaproteobacteria; Enterobacterales; Enterobacteriaceae; Escherichia; Escherichia coli
MTGDNTLIHSHGINRRDFMKLCAALAATMGLSSKAAAEMAESVTNPQRPPVIWIGAQECT WP_000145410.1
MULTISPECIES: hydrogenase-2 small chain [Proteobacteria] Length: 372\nNumber of predicted TMHs: 1\nExp number of AAs in TMHs: 23.72226\nExp number, first 60 AAs: 0.22038\nTotal prob of N-in: 0.12994\noutside 1 331\nTMhelix 332 354\ninside 355 372

10080 GCF_000194725.1_ASM19472v2 Escherichia coli 900105 (10e) Proteobacteria; Gammaproteobacteria; Enterobacterales; Enterobacteriaceae; Escherichia; Escherichia coli
MNNEETFYQAMRRQGVTRRSFLKYCSLAATSLGLGAGMAPKIAWALENKPRIPVVWIHGL WP_001058323.1
MULTISPECIES: hydrogenase-1 small chain [Proteobacteria] Length: 372\nNumber of predicted TMHs: 2\nExp number of AAs in TMHs: 37.2608499999999\nExp number, first 60 AAs: 19.52923\nTotal prob of N-in: 0.94351\nPOSSIBLE N-term signal sequence\ninside 1 19\nTMhelix 20 39\noutside 40 325\nTMhelix 326 348\ninside 349 372

10081 GCF_000316425.2_ASM31642v2 Escherichia coli 90.2281 Proteobacteria; Gammaproteobacteria; Enterobacterales; Enterobacteriaceae; Escherichia; Escherichia coli
MNNEETFYQAMRRQGVTRRSFLKYCSLAATSLGLGAGMAPKIAWALENKPRIPVVWIHGL WP_001301613.1
hydrogenase [Escherichia coli] Length: 372\nNumber of predicted TMHs: 2\nExp number of AAs in TMHs: 37.46598\nExp number, first 60 AAs: 19.53139\nTotal prob of N-in: 0.94364\nPOSSIBLE N-term signal sequence\ninside 1 19\nTMhelix 20 39\noutside 40 325\nTMhelix 326 348\ninside 349 372

10082 GCF_000316425.2_ASM31642v2 Escherichia coli 90.2281 Proteobacteria; Gammaproteobacteria; Enterobacterales; Enterobacteriaceae; Escherichia; Escherichia coli
MTGDNTLIHSHGINRRDFMKLCAALAATMGLSSKAAAEMAESVTNPQRPPVIWIGAQECT WP_000145410.1
MULTISPECIES: hydrogenase-2 small chain [Proteobacteria] Length: 372\nNumber of predicted TMHs: 1\nExp number of AAs in TMHs: 23.72226\nExp number, first 60 AAs: 0.22038\nTotal prob of N-in: 0.12994\noutside 1 331\nTMhelix 332 354\ninside 355 372

10083 GCF_000485675.1_ASM48567v1 Escherichia coli 907357 Proteobacteria; Gammaproteobacteria; Enterobacterales; Enterobacteriaceae; Escherichia; Escherichia coli
MNNEETFYQAMRRQGVTRRSFLKYCSLAATSLGLGAGMAPKIAWALENKPRIPVVWIHGL WP_001295940.1
MULTISPECIES: hydrogenase [Enterobacteriaceae] Length: 372\nNumber of predicted TMHs: 2\nExp number of AAs in TMHs: 37.2608899999999\nExp number, first 60 AAs: 19.52923\nTotal prob of N-in: 0.94351\nPOSSIBLE N-term signal sequence\ninside 1 19\nTMhelix 20 39\noutside 40 325\nTMhelix 326 348\ninside 349 372

10084 GCF_000485675.1_ASM48567v1 Escherichia coli 907357 Proteobacteria; Gammaproteobacteria; Enterobacterales; Enterobacteriaceae; Escherichia; Escherichia coli
MTGDNTLIHSHGINRRDFMKLCAALAATMGLSSKAAAEMAESVTNPQRPPVIWIGAQECT WP_000145410.1
MULTISPECIES: hydrogenase-2 small chain [Proteobacteria] Length: 372\nNumber of predicted TMHs: 1\nExp number of AAs in TMHs: 23.72226\nExp number, first 60 AAs: 0.22038\nTotal prob of N-in: 0.12994\noutside 1 331\nTMhelix 332 354\ninside 355 372

10085 GCF_000488315.1_ASM48831v1 Escherichia coli 907391 Proteobacteria; Gammaproteobacteria; Enterobacterales; Enterobacteriaceae; Escherichia; Escherichia coli
MNNEETFYQAMRRQGVTRRSFLKYCSLAATSLGLGAGMAPKIAWALENKPRIPVVWIHGL WP_001058323.1
MULTISPECIES: hydrogenase-1 small chain [Proteobacteria] Length: 372\nNumber of predicted TMHs: 2\nExp number of AAs in TMHs: 37.2608499999999\nExp number, first 60 AAs: 19.52923\nTotal prob of N-in: 0.94351\nPOSSIBLE N-term signal sequence\ninside 1 19\nTMhelix 20 39\noutside 40 325\nTMhelix 326 348\ninside 349 372

10086 GCF_000488315.1_ASM48831v1 Escherichia coli 907391 Proteobacteria; Gammaproteobacteria; Enterobacterales; Enterobacteriaceae; Escherichia; Escherichia coli
MTGDNTLIHSHGINRRDFMKLCAALAATMGLSSKAAAEMAESVTNPQRPPVIWIGAQECT WP_000145410.1
MULTISPECIES: hydrogenase-2 small chain [Proteobacteria] Length: 372\nNumber of predicted TMHs: 1\nExp number of AAs in TMHs: 23.72226\nExp number, first 60 AAs: 0.22038\nTotal prob of N-in: 0.12994\noutside 1 331\nTMhelix 332 354\ninside 355 372

10087 GCF_000488335.1_ASM48833v1 Escherichia coli 907446 Proteobacteria; Gammaproteobacteria; Enterobacterales; Enterobacteriaceae; Escherichia; Escherichia coli
MNNEETFYQAMRRQGVTRRSFLKYCSLAATSLGLGAGMAPKIAWALENKPRIPVVWIHGL WP_001058323.1
MULTISPECIES: hydrogenase-1 small chain [Proteobacteria] Length: 372\nNumber of predicted TMHs: 2\nExp number of AAs in TMHs: 37.2608499999999\nExp number, first 60 AAs: 19.52923\nTotal prob of N-in: 0.94351\nPOSSIBLE N-term signal sequence\ninside 1 19\nTMhelix 20 39\noutside 40 325\nTMhelix 326 348\ninside 349 372

10088 GCF_000488335.1_ASM48833v1 Escherichia coli 907446 Proteobacteria; Gammaproteobacteria; Enterobacterales; Enterobacteriaceae; Escherichia; Escherichia coli
MTGDNTLIHSHGINRRDFMKLCAALAATMGLSSKAAAEMAESVTNPQRPPVIWIGAQECT WP_000145410.1
MULTISPECIES: hydrogenase-2 small chain [Proteobacteria] Length: 372\nNumber of predicted TMHs: 1\nExp number of AAs in TMHs: 23.72226\nExp number, first 60 AAs: 0.22038\nTotal prob of N-in: 0.12994\noutside 1 331\nTMhelix 332 354\ninside 355 372

10089 GCF_000488355.1_ASM48835v1 Escherichia coli 907672 Proteobacteria; Gammaproteobacteria; Enterobacterales; Enterobacteriaceae; Escherichia; Escherichia coli

MNNEETFYQAMRRQGVTRRSFLKYCSLAATSLGLGAGMAPKIAWALENKPRIPVVWIHGL WP_001058323.1 MULTISPECIES: hydrogenase-1 small chain [Proteobacteria] Length: 372\nNumber of predicted TMHs: 2\nExp number of AAs in TMHs: 37.2608499999999\nExp number, first 60 AAs: 19.52923\nTotal prob of N-in: 0.94351\nPOSSIBLE N-term signal sequence\ninside 1 19\nTMhelix 20 39\noutside 40 325\nTMhelix 326 348\ninside 349 372

10090 GCF_000488355.1_ASM48835v1 Escherichia coli 907672 Proteobacteria; Gammaproteobacteria; Enterobacterales; Enterobacteriaceae; Escherichia; Escherichia coli MTGDNTLIHSHGINRRDFMKLCAALAATMGLSSKAAAEMAESVTNPQRPPVIWIGAQECT WP_000145410.1 MULTISPECIES: hydrogenase-2 small chain [Proteobacteria] Length: 372\nNumber of predicted TMHs: 1\nExp number of AAs in TMHs: 23.72226\nExp number, first 60 AAs: 0.22038\nTotal prob of N-in: 0.12994\noutside 1 331\nTMhelix 332 354\ninside 355 372

10091 GCF_000488375.1_ASM48837v1 Escherichia coli 907700 Proteobacteria; Gammaproteobacteria; Enterobacterales; Enterobacteriaceae; Escherichia; Escherichia coli MNNEETFYQAMRRQGVTRRSFLKYCSLAATSLGLGAGMAPKIAWALENKPRIPVVWIHGL WP_001295940.1 MULTISPECIES: hydrogenase [Enterobacteriaceae] Length: 372\nNumber of predicted TMHs: 2\nExp number of AAs in TMHs: 37.2608899999999\nExp number, first 60 AAs: 19.52923\nTotal prob of N-in: 0.94351\nPOSSIBLE N-term signal sequence\ninside 1 19\nTMhelix 20 39\noutside 40 325\nTMhelix 326 348\ninside 349 372

10092 GCF_000488375.1_ASM48837v1 Escherichia coli 907700 Proteobacteria; Gammaproteobacteria; Enterobacterales; Enterobacteriaceae; Escherichia; Escherichia coli MTGDNTLIHSHGINRRDFMKLCAALAATMGLSSKAAAEMVESVTNPQRPPVIWIGAQECT WP_000145423.1 hydrogenase 2 small subunit [Escherichia coli] Length: 372\nNumber of predicted TMHs: 1\nExp number of AAs in TMHs: 23.78201\nExp number, first 60 AAs: 0.28355\nTotal prob of N-in: 0.13257\noutside 1 331\nTMhelix 332 354\ninside 355 372

10093 GCF_000488395.1_ASM48839v1 Escherichia coli 907701 Proteobacteria; Gammaproteobacteria; Enterobacterales; Enterobacteriaceae; Escherichia; Escherichia coli MNNEETFYQAMRRQGVTRRSFLKYCSLAATSLGLGAGMAPKIAWALENKPRIPVVWIHGL WP_001295940.1 MULTISPECIES: hydrogenase [Enterobacteriaceae] Length: 372\nNumber of predicted TMHs: 2\nExp number of AAs in TMHs: 37.2608899999999\nExp number, first 60 AAs: 19.52923\nTotal prob of N-in: 0.94351\nPOSSIBLE N-term signal sequence\ninside 1 19\nTMhelix 20 39\noutside 40 325\nTMhelix 326 348\ninside 349 372

10094 GCF_000488395.1_ASM48839v1 Escherichia coli 907701 Proteobacteria; Gammaproteobacteria; Enterobacterales; Enterobacteriaceae; Escherichia; Escherichia coli MTGDNTLIHSHGINRRDFMKLCAALAATMGLSSKAAAEMVESVTNPQRPPVIWIGAQECT WP_000145423.1 hydrogenase 2 small subunit [Escherichia coli] Length: 372\nNumber of predicted TMHs: 1\nExp number of AAs in TMHs: 23.78201\nExp number, first 60 AAs: 0.28355\nTotal prob of N-in: 0.13257\noutside 1 331\nTMhelix 332 354\ninside 355 372

10095 GCF_000488415.1_ASM48841v1 Escherichia coli 907710 Proteobacteria; Gammaproteobacteria; Enterobacterales; Enterobacteriaceae; Escherichia; Escherichia coli MTGDNTLIHSHGINRRDFMKLCAALAATMGLSSKAAAEMAESVTNPQRPPVIWIGAQECT WP_000145410.1 MULTISPECIES: hydrogenase-2 small chain [Proteobacteria] Length: 372\nNumber of predicted TMHs: 1\nExp number of AAs in TMHs: 23.72226\nExp number, first 60 AAs: 0.22038\nTotal prob of N-in: 0.12994\noutside 1 331\nTMhelix 332 354\ninside 355 372

10096 GCF_000488415.1_ASM48841v1 Escherichia coli 907710 Proteobacteria; Gammaproteobacteria; Enterobacterales; Enterobacteriaceae; Escherichia; Escherichia coli MNNEETFYQAMRRQGVTRRSFLKYCSLAATSLGLGAGMAPKIAWALENKPRIPVVWIHGL WP_001295940.1 MULTISPECIES: hydrogenase [Enterobacteriaceae] Length: 372\nNumber of predicted TMHs: 2\nExp number of AAs in TMHs: 37.2608899999999\nExp number, first 60 AAs: 19.52923\nTotal prob of N-in: 0.94351\nPOSSIBLE N-term signal sequence\ninside 1 19\nTMhelix 20 39\noutside 40 325\nTMhelix 326 348\ninside 349 372

10097 GCF_000485695.1_ASM48569v1 Escherichia coli 907713 Proteobacteria; Gammaproteobacteria; Enterobacterales; Enterobacteriaceae; Escherichia; Escherichia coli MTGDNTLIHSHGINRRDFMKLCAALAATMGLSSKAAAEMAESVTNPQRPPVIWIGAQECT WP_000145410.1

MULTISPECIES: hydrogenase-2 small chain [Proteobacteria] Length: 372\nNumber of predicted TMHs: 1\nExp number of AAs in TMHs: 23.72226\nExp number, first 60 AAs: 0.22038\nTotal prob of N-in: 0.12994\noutside 1 331\nTMhelix 332 354\ninside 355 372

10098 GCF_000485695.1_ASM48569v1 Escherichia coli 907713 Proteobacteria; Gammaproteobacteria; Enterobacterales; Enterobacteriaceae; Escherichia; Escherichia coli
MNNEETFYQAMRRQGVTRRSFLKYCSLAATSLGLGAGMAPKIAWALENKPRIPVVWIHGL WP_001058323.1
MULTISPECIES: hydrogenase-1 small chain [Proteobacteria] Length: 372\nNumber of predicted TMHs: 2\nExp number of AAs in TMHs: 37.2608499999999\nExp number, first 60 AAs: 19.52923\nTotal prob of N-in: 0.94351\nPOSSIBLE N-term signal sequence\ninside 1 19\nTMhelix 20 39\noutside 40 325\nTMhelix 326 348\ninside 349 372

10099 GCF_000488435.1_ASM48843v1 Escherichia coli 907715 Proteobacteria; Gammaproteobacteria; Enterobacterales; Enterobacteriaceae; Escherichia; Escherichia coli
MNNEETFYQAMRRQGVTRRSFLKYCSLAATSLGLGAGMAPKIAWALENKPRIPVVWIHGL WP_001058323.1
MULTISPECIES: hydrogenase-1 small chain [Proteobacteria] Length: 372\nNumber of predicted TMHs: 2\nExp number of AAs in TMHs: 37.2608499999999\nExp number, first 60 AAs: 19.52923\nTotal prob of N-in: 0.94351\nPOSSIBLE N-term signal sequence\ninside 1 19\nTMhelix 20 39\noutside 40 325\nTMhelix 326 348\ninside 349 372

10100 GCF_000488435.1_ASM48843v1 Escherichia coli 907715 Proteobacteria; Gammaproteobacteria; Enterobacterales; Enterobacteriaceae; Escherichia; Escherichia coli
MTGDNTLIHSHGINRRDFMKLCAALAATMGLSSKAAAEMAESVTNPQRPPVIWIGAQECT WP_000145410.1
MULTISPECIES: hydrogenase-2 small chain [Proteobacteria] Length: 372\nNumber of predicted TMHs: 1\nExp number of AAs in TMHs: 23.72226\nExp number, first 60 AAs: 0.22038\nTotal prob of N-in: 0.12994\noutside 1 331\nTMhelix 332 354\ninside 355 372

10101 GCF_000485715.1_ASM48571v1 Escherichia coli 907779 Proteobacteria; Gammaproteobacteria; Enterobacterales; Enterobacteriaceae; Escherichia; Escherichia coli
MNNEETFYQAMRRQGVTRRSFLKYCSLAATSLGLGAGMAPKIAWALENKPRIPVVWIHGL WP_001295940.1
MULTISPECIES: hydrogenase [Enterobacteriaceae] Length: 372\nNumber of predicted TMHs: 2\nExp number of AAs in TMHs: 37.2608899999999\nExp number, first 60 AAs: 19.52923\nTotal prob of N-in: 0.94351\nPOSSIBLE N-term signal sequence\ninside 1 19\nTMhelix 20 39\noutside 40 325\nTMhelix 326 348\ninside 349 372

10102 GCF_000485715.1_ASM48571v1 Escherichia coli 907779 Proteobacteria; Gammaproteobacteria; Enterobacterales; Enterobacteriaceae; Escherichia; Escherichia coli
MTGDNTLIHSHGINRRDFMKLCAALAATMGLSSKAAAEMVESVTNPQRPPVIWIGAQECT WP_000145423.1
hydrogenase 2 small subunit [Escherichia coli] Length: 372\nNumber of predicted TMHs: 1\nExp number of AAs in TMHs: 23.78201\nExp number, first 60 AAs: 0.28355\nTotal prob of N-in: 0.13257\noutside 1 331\nTMhelix 332 354\ninside 355 372

10103 GCF_000488455.1_ASM48845v1 Escherichia coli 907889 Proteobacteria; Gammaproteobacteria; Enterobacterales; Enterobacteriaceae; Escherichia; Escherichia coli
MNNEETFYQAMRRQGVTRRSFLKYCSLAATSLGLGAGMAPKIAWALENKPRIPVVWIHGL WP_001058323.1
MULTISPECIES: hydrogenase-1 small chain [Proteobacteria] Length: 372\nNumber of predicted TMHs: 2\nExp number of AAs in TMHs: 37.2608499999999\nExp number, first 60 AAs: 19.52923\nTotal prob of N-in: 0.94351\nPOSSIBLE N-term signal sequence\ninside 1 19\nTMhelix 20 39\noutside 40 325\nTMhelix 326 348\ninside 349 372

10104 GCF_000488455.1_ASM48845v1 Escherichia coli 907889 Proteobacteria; Gammaproteobacteria; Enterobacterales; Enterobacteriaceae; Escherichia; Escherichia coli
MTGDNTLIHSHGINRRDFMKLCAALAATMGLSSKAAAEMAESVTNPQRPPVIWIGAQECT WP_000145410.1
MULTISPECIES: hydrogenase-2 small chain [Proteobacteria] Length: 372\nNumber of predicted TMHs: 1\nExp number of AAs in TMHs: 23.72226\nExp number, first 60 AAs: 0.22038\nTotal prob of N-in: 0.12994\noutside 1 331\nTMhelix 332 354\ninside 355 372

10105 GCF_000488475.1_ASM48847v1 Escherichia coli 907892 Proteobacteria; Gammaproteobacteria; Enterobacterales; Enterobacteriaceae; Escherichia; Escherichia coli
MTGDNTLIHSHGINRRDFMKLCAALAATMGLSSKAAAEMAESVTNPQRPPVIWIGAQECT WP_000145410.1
MULTISPECIES: hydrogenase-2 small chain [Proteobacteria] Length: 372\nNumber of predicted TMHs:

1\nExp number of AAs in TMHs: 23.72226\nExp number, first 60 AAs: 0.22038\nTotal prob of N-in: 0.12994\noutside 1 331\nTMhelix 332 354\ninside 355 372

10106 GCF_000488475.1_ASM48847v1 Escherichia coli 907892 Proteobacteria; Gammaproteobacteria; Enterobacterales; Enterobacteriaceae; Escherichia; Escherichia coli MNNEETFYQAMRRQGVTRRSFLKYCSLAATSLGLGAGMAPKIAWALENKPRIPVVWIHGL WP_001058323.1 MULTISPECIES: hydrogenase-1 small chain [Proteobacteria] Length: 372\nNumber of predicted TMHs: 2\nExp number of AAs in TMHs: 37.2608499999999\nExp number, first 60 AAs: 19.52923\nTotal prob of N-in: 0.94351\nPOSSIBLE N-term signal sequence\ninside 1 19\nTMhelix 20 39\noutside 40 325\nTMhelix 326 348\ninside 349 372

10107 GCF_000488495.1_ASM48849v1 Escherichia coli 908519 Proteobacteria; Gammaproteobacteria; Enterobacterales; Enterobacteriaceae; Escherichia; Escherichia coli MTGDNTLIHSHGINRRDFMKLCAALAATMGLSSKAAAEMAESVTNPQRPPVIWIGAQECT WP_000145410.1 MULTISPECIES: hydrogenase-2 small chain [Proteobacteria] Length: 372\nNumber of predicted TMHs: 1\nExp number of AAs in TMHs: 23.72226\nExp number, first 60 AAs: 0.22038\nTotal prob of N-in: 0.12994\noutside 1 331\nTMhelix 332 354\ninside 355 372

10108 GCF_000488495.1_ASM48849v1 Escherichia coli 908519 Proteobacteria; Gammaproteobacteria; Enterobacterales; Enterobacteriaceae; Escherichia; Escherichia coli MNNEETFYQAMRRQGVTRRSFLKYCSLAATSLGLGAGMAPKIAWALENKPRIPVVWIHGL WP_001058323.1 MULTISPECIES: hydrogenase-1 small chain [Proteobacteria] Length: 372\nNumber of predicted TMHs: 2\nExp number of AAs in TMHs: 37.2608499999999\nExp number, first 60 AAs: 19.52923\nTotal prob of N-in: 0.94351\nPOSSIBLE N-term signal sequence\ninside 1 19\nTMhelix 20 39\noutside 40 325\nTMhelix 326 348\ninside 349 372

10109 GCF_000488515.1_ASM48851v1 Escherichia coli 908521 Proteobacteria; Gammaproteobacteria; Enterobacterales; Enterobacteriaceae; Escherichia; Escherichia coli MNNEETFYQAMRRQGVTRRSFLKYCSLAATSLGLGAGMAPKIAWALENKPRIPVVWIHGL WP_001058323.1 MULTISPECIES: hydrogenase-1 small chain [Proteobacteria] Length: 372\nNumber of predicted TMHs: 2\nExp number of AAs in TMHs: 37.2608499999999\nExp number, first 60 AAs: 19.52923\nTotal prob of N-in: 0.94351\nPOSSIBLE N-term signal sequence\ninside 1 19\nTMhelix 20 39\noutside 40 325\nTMhelix 326 348\ninside 349 372

10110 GCF_000488515.1_ASM48851v1 Escherichia coli 908521 Proteobacteria; Gammaproteobacteria; Enterobacterales; Enterobacteriaceae; Escherichia; Escherichia coli MTGDNTLIHSHGINRRDFMKLCAALAATMGLSSKAAAEMAESVTNPQRPPVIWIGAQECT WP_000145410.1 MULTISPECIES: hydrogenase-2 small chain [Proteobacteria] Length: 372\nNumber of predicted TMHs: 1\nExp number of AAs in TMHs: 23.72226\nExp number, first 60 AAs: 0.22038\nTotal prob of N-in: 0.12994\noutside 1 331\nTMhelix 332 354\ninside 355 372

10111 GCF_000488535.1_ASM48853v1 Escherichia coli 908522 Proteobacteria; Gammaproteobacteria; Enterobacterales; Enterobacteriaceae; Escherichia; Escherichia coli MNNEETFYQAMRRQGVTRRSFLKYCSLAATSLGLGAGMAPKIAWALENKPRIPVVWIHGL WP_001058323.1 MULTISPECIES: hydrogenase-1 small chain [Proteobacteria] Length: 372\nNumber of predicted TMHs: 2\nExp number of AAs in TMHs: 37.2608499999999\nExp number, first 60 AAs: 19.52923\nTotal prob of N-in: 0.94351\nPOSSIBLE N-term signal sequence\ninside 1 19\nTMhelix 20 39\noutside 40 325\nTMhelix 326 348\ninside 349 372

10112 GCF_000488535.1_ASM48853v1 Escherichia coli 908522 Proteobacteria; Gammaproteobacteria; Enterobacterales; Enterobacteriaceae; Escherichia; Escherichia coli MTGDNTLIHSHGINRRDFMKLCAALAATMGLSSKAAAEMAESVTNPQRPPVIWIGAQECT WP_000145410.1 MULTISPECIES: hydrogenase-2 small chain [Proteobacteria] Length: 372\nNumber of predicted TMHs: 1\nExp number of AAs in TMHs: 23.72226\nExp number, first 60 AAs: 0.22038\nTotal prob of N-in: 0.12994\noutside 1 331\nTMhelix 332 354\ninside 355 372

10113 GCF_000488555.1_ASM48855v1 Escherichia coli 908524 Proteobacteria; Gammaproteobacteria; Enterobacterales; Enterobacteriaceae; Escherichia; Escherichia coli MTGDNTLIHSHGINRRDFMKLCAALAATMGLSSKAAAEMVESVTNPQRPPVIWIGAQECT WP_000145423.1 hydrogenase 2 small subunit [Escherichia coli] Length: 372\nNumber of predicted TMHs: 1\nExp number of

AAs in TMHs: 23.78201\nExp number, first 60 AAs:  0.28355\nTotal prob of N-in:   0.13257\noutside   1   331\nTMhelix   332   354\ninside   355   372

10114	GCF_000488555.1_ASM48855v1	Escherichia coli 908524	Proteobacteria; Gammaproteobacteria; Enterobacterales; Enterobacteriaceae; Escherichia; Escherichia coli
MNNEETFYQAMRRQGVTRRSFLKYCSLAATSLGLGAGMAPKIAWALENKPRIPVVWIHGL	WP_001295940.1
MULTISPECIES: hydrogenase [Enterobacteriaceae]	Length: 372\nNumber of predicted TMHs:  2\nExp number of AAs in TMHs: 37.2608899999999\nExp number, first 60 AAs:  19.52923\nTotal prob of N-in:   0.94351\nPOSSIBLE N-term signal sequence\ninside   1   19\nTMhelix   20   39\noutside   40   325\nTMhelix   326   348\ninside   349   372

10115	GCF_000488575.1_ASM48857v1	Escherichia coli 908525	Proteobacteria; Gammaproteobacteria; Enterobacterales; Enterobacteriaceae; Escherichia; Escherichia coli
MNNEETFYQAMRRQGVTRRSFLKYCSLAATSLGLGAGMAPKIAWALENKPRIPVVWIHGL	WP_001058323.1
MULTISPECIES: hydrogenase-1 small chain [Proteobacteria]	Length: 372\nNumber of predicted TMHs: 2\nExp number of AAs in TMHs: 37.2608499999999\nExp number, first 60 AAs:  19.52923\nTotal prob of N-in:   0.94351\nPOSSIBLE N-term signal sequence\ninside   1   19\nTMhelix   20   39\noutside   40   325\nTMhelix   326   348\ninside   349   372

10116	GCF_000488575.1_ASM48857v1	Escherichia coli 908525	Proteobacteria; Gammaproteobacteria; Enterobacterales; Enterobacteriaceae; Escherichia; Escherichia coli
MTGDNTLIHSHGINRRDFMKLCAALAATMGLSSKAAAEMAESVTNPQRPPVIWIGAQECT	WP_000145410.1
MULTISPECIES: hydrogenase-2 small chain [Proteobacteria]	Length: 372\nNumber of predicted TMHs: 1\nExp number of AAs in TMHs: 23.72226\nExp number, first 60 AAs:  0.22038\nTotal prob of N-in:   0.12994\noutside   1   331\nTMhelix   332   354\ninside   355   372

10117	GCF_000488595.1_ASM48859v1	Escherichia coli 908541	Proteobacteria; Gammaproteobacteria; Enterobacterales; Enterobacteriaceae; Escherichia; Escherichia coli
MTGDNTLIHSHGINRRDFMKLCAALAATMGLSSKAAAEMAESVTNPQRPPVIWIGAQECT	WP_000145410.1
MULTISPECIES: hydrogenase-2 small chain [Proteobacteria]	Length: 372\nNumber of predicted TMHs: 1\nExp number of AAs in TMHs: 23.72226\nExp number, first 60 AAs:  0.22038\nTotal prob of N-in:   0.12994\noutside   1   331\nTMhelix   332   354\ninside   355   372

10118	GCF_000488595.1_ASM48859v1	Escherichia coli 908541	Proteobacteria; Gammaproteobacteria; Enterobacterales; Enterobacteriaceae; Escherichia; Escherichia coli
MNNEETFYQAMRRQGVTRRSFLKYCSLAATSLGLGAGMAPKIAWALENKPRIPVVWIHGL	WP_001058323.1
MULTISPECIES: hydrogenase-1 small chain [Proteobacteria]	Length: 372\nNumber of predicted TMHs: 2\nExp number of AAs in TMHs: 37.2608499999999\nExp number, first 60 AAs:  19.52923\nTotal prob of N-in:   0.94351\nPOSSIBLE N-term signal sequence\ninside   1   19\nTMhelix   20   39\noutside   40   325\nTMhelix   326   348\ninside   349   372

10119	GCF_000488615.1_ASM48861v1	Escherichia coli 908555	Proteobacteria; Gammaproteobacteria; Enterobacterales; Enterobacteriaceae; Escherichia; Escherichia coli
MTGDNTLIHSHGINRRDFMKLCAALAATMGLSSKAAAEMAESVTNPQRPPVIWIGAQECT	WP_000145410.1
MULTISPECIES: hydrogenase-2 small chain [Proteobacteria]	Length: 372\nNumber of predicted TMHs: 1\nExp number of AAs in TMHs: 23.72226\nExp number, first 60 AAs:  0.22038\nTotal prob of N-in:   0.12994\noutside   1   331\nTMhelix   332   354\ninside   355   372

10120	GCF_000488615.1_ASM48861v1	Escherichia coli 908555	Proteobacteria; Gammaproteobacteria; Enterobacterales; Enterobacteriaceae; Escherichia; Escherichia coli
MNNEETFYQAMRRQGVTRRSFLKYCSLAATSLGLGAGMAPKIAWALENKPRIPVVWIHGL	WP_001058320.1
hydrogenase [Escherichia coli]	Length: 372\nNumber of predicted TMHs:  2\nExp number of AAs in TMHs: 37.26082\nExp number, first 60 AAs:  19.52923\nTotal prob of N-in:   0.94351\nPOSSIBLE N-term signal sequence\ninside   1   19\nTMhelix   20   39\noutside   40   325\nTMhelix   326   348\ninside   349   372

10121	GCF_000488635.1_ASM48863v1	Escherichia coli 908573	Proteobacteria; Gammaproteobacteria; Enterobacterales; Enterobacteriaceae; Escherichia; Escherichia coli
MNNEETFYQAMRRQGVTRRSFLKYCSLAATSLGLGAGMAPKIAWALENKPRIPVVWIHGL	WP_001058323.1
MULTISPECIES: hydrogenase-1 small chain [Proteobacteria]	Length: 372\nNumber of predicted TMHs: 2\nExp number of AAs in TMHs: 37.2608499999999\nExp number, first 60 AAs:  19.52923\nTotal prob of N-in:

0.94351\nPOSSIBLE N-term signal sequence\ninside 1 19\nTMhelix 20 39\noutside 40 325\nTMhelix 326 348\ninside 349 372

10122 GCF_000488655.1_ASM48865v1 Escherichia coli 908585 Proteobacteria; Gammaproteobacteria; Enterobacterales; Enterobacteriaceae; Escherichia; Escherichia coli
MTGDNTLIHSHGINRRDFMKLCAALAATMGLSSKAAAEMAESVTNPQRPPVIWIGAQECT WP_000145410.1
MULTISPECIES: hydrogenase-2 small chain [Proteobacteria] Length: 372\nNumber of predicted TMHs: 1\nExp number of AAs in TMHs: 23.72226\nExp number, first 60 AAs: 0.22038\nTotal prob of N-in: 0.12994\noutside 1 331\nTMhelix 332 354\ninside 355 372

10123 GCF_000488655.1_ASM48865v1 Escherichia coli 908585 Proteobacteria; Gammaproteobacteria; Enterobacterales; Enterobacteriaceae; Escherichia; Escherichia coli
MNNEETFYQAMRRQGVTRRSFLKYCSLAATSLGLGAGMAPKIAWALENKPRIPVVWIHGL WP_001058323.1
MULTISPECIES: hydrogenase-1 small chain [Proteobacteria] Length: 372\nNumber of predicted TMHs: 2\nExp number of AAs in TMHs: 37.2608499999999\nExp number, first 60 AAs: 19.52923\nTotal prob of N-in: 0.94351\nPOSSIBLE N-term signal sequence\ninside 1 19\nTMhelix 20 39\noutside 40 325\nTMhelix 326 348\ninside 349 372

10124 GCF_000488675.1_ASM48867v1 Escherichia coli 908616 Proteobacteria; Gammaproteobacteria; Enterobacterales; Enterobacteriaceae; Escherichia; Escherichia coli
MTGDNTLIHSHGINRRDFMKLCAALAATMGLSSKAAAEMAESVTNPQRPPVIWIGAQECT WP_000145410.1
MULTISPECIES: hydrogenase-2 small chain [Proteobacteria] Length: 372\nNumber of predicted TMHs: 1\nExp number of AAs in TMHs: 23.72226\nExp number, first 60 AAs: 0.22038\nTotal prob of N-in: 0.12994\noutside 1 331\nTMhelix 332 354\ninside 355 372

10125 GCF_000488675.1_ASM48867v1 Escherichia coli 908616 Proteobacteria; Gammaproteobacteria; Enterobacterales; Enterobacteriaceae; Escherichia; Escherichia coli
MNNEETFYQAMRRQGVTRRSFLKYCSLAATSLGLGAGMAPKIAWALENKPRIPVVWIHGL WP_001058323.1
MULTISPECIES: hydrogenase-1 small chain [Proteobacteria] Length: 372\nNumber of predicted TMHs: 2\nExp number of AAs in TMHs: 37.2608499999999\nExp number, first 60 AAs: 19.52923\nTotal prob of N-in: 0.94351\nPOSSIBLE N-term signal sequence\ninside 1 19\nTMhelix 20 39\noutside 40 325\nTMhelix 326 348\ninside 349 372

10126 GCF_000488695.1_ASM48869v1 Escherichia coli 908624 Proteobacteria; Gammaproteobacteria; Enterobacterales; Enterobacteriaceae; Escherichia; Escherichia coli
MNNEETFYQAMRRQGVTRRSFLKYCSLAATSLGLGAGMAPKIAWALENKPRIPVVWIHGL WP_001295940.1
MULTISPECIES: hydrogenase [Enterobacteriaceae] Length: 372\nNumber of predicted TMHs: 2\nExp number of AAs in TMHs: 37.2608899999999\nExp number, first 60 AAs: 19.52923\nTotal prob of N-in: 0.94351\nPOSSIBLE N-term signal sequence\ninside 1 19\nTMhelix 20 39\noutside 40 325\nTMhelix 326 348\ninside 349 372

10127 GCF_000488695.1_ASM48869v1 Escherichia coli 908624 Proteobacteria; Gammaproteobacteria; Enterobacterales; Enterobacteriaceae; Escherichia; Escherichia coli
MTGDNTLIHSHGINRRDFMKLCAALAATMGLSSKAAAEMVESVTNPQRPPVIWIGAQECT WP_000145423.1
hydrogenase 2 small subunit [Escherichia coli] Length: 372\nNumber of predicted TMHs: 1\nExp number of AAs in TMHs: 23.78201\nExp number, first 60 AAs: 0.28355\nTotal prob of N-in: 0.13257\noutside 1 331\nTMhelix 332 354\ninside 355 372

10128 GCF_000488715.1_ASM48871v1 Escherichia coli 908632 Proteobacteria; Gammaproteobacteria; Enterobacterales; Enterobacteriaceae; Escherichia; Escherichia coli
MNNEETFYQAMRRQGVTRRSFLKYCSLAATSLGLGAGMAPKIAWALENKPRIPVVWIHGL WP_001295940.1
MULTISPECIES: hydrogenase [Enterobacteriaceae] Length: 372\nNumber of predicted TMHs: 2\nExp number of AAs in TMHs: 37.2608899999999\nExp number, first 60 AAs: 19.52923\nTotal prob of N-in: 0.94351\nPOSSIBLE N-term signal sequence\ninside 1 19\nTMhelix 20 39\noutside 40 325\nTMhelix 326 348\ninside 349 372

10129 GCF_000488715.1_ASM48871v1 Escherichia coli 908632 Proteobacteria; Gammaproteobacteria; Enterobacterales; Enterobacteriaceae; Escherichia; Escherichia coli
MTGDNTLIHSHGINRRDFMKLCAALAATMGLSSKAAAEMVESVTNPQRPPVIWIGAQECT WP_000145423.1
hydrogenase 2 small subunit [Escherichia coli] Length: 372\nNumber of predicted TMHs: 1\nExp number of

AAs in TMHs: 23.78201\nExp number, first 60 AAs: 0.28355\nTotal prob of N-in: 0.13257\noutside 1 331\nTMhelix 332 354\ninside 355 372

10130 GCF_000488735.1_ASM48873v1 Escherichia coli 908658 Proteobacteria; Gammaproteobacteria; Enterobacterales; Enterobacteriaceae; Escherichia; Escherichia coli
MNNEETFYQAMRRQGVTRRSFLKYCSLAATSLGLGAGMAPKIAWALENKPRIPVVWIHGL WP_001058323.1
MULTISPECIES: hydrogenase-1 small chain [Proteobacteria] Length: 372\nNumber of predicted TMHs: 2\nExp number of AAs in TMHs: 37.2608499999999\nExp number, first 60 AAs: 19.52923\nTotal prob of N-in: 0.94351\nPOSSIBLE N-term signal sequence\ninside 1 19\nTMhelix 20 39\noutside 40 325\nTMhelix 326 348\ninside 349 372

10131 GCF_000488735.1_ASM48873v1 Escherichia coli 908658 Proteobacteria; Gammaproteobacteria; Enterobacterales; Enterobacteriaceae; Escherichia; Escherichia coli
MTGDNTLIHSHGINRRDFMKLCAALATTMGLSSKAAAEMAESVTNPQRPPVIWIGAQECT WP_023149629.1
MULTISPECIES: hydrogenase-2 small chain [Enterobacteriaceae] Length: 372\nNumber of predicted TMHs: 1\nExp number of AAs in TMHs: 23.63603\nExp number, first 60 AAs: 0.12971\nTotal prob of N-in: 0.12608\noutside 1 331\nTMhelix 332 354\ninside 355 372

10132 GCF_000488755.1_ASM48875v1 Escherichia coli 908675 Proteobacteria; Gammaproteobacteria; Enterobacterales; Enterobacteriaceae; Escherichia; Escherichia coli
MTGDNTLIHSHGINRRDFMKLCAALAATMGLSSKAAAEMAESVTNPQRPPVIWIGAQECT WP_000145410.1
MULTISPECIES: hydrogenase-2 small chain [Proteobacteria] Length: 372\nNumber of predicted TMHs: 1\nExp number of AAs in TMHs: 23.72226\nExp number, first 60 AAs: 0.22038\nTotal prob of N-in: 0.12994\noutside 1 331\nTMhelix 332 354\ninside 355 372

10133 GCF_000488775.1_ASM48877v1 Escherichia coli 908691 Proteobacteria; Gammaproteobacteria; Enterobacterales; Enterobacteriaceae; Escherichia; Escherichia coli
MNNEETFYQAMRRQGVTRRSFLKYCSLAATSLGLGAGMAPKIAWALENKPRIPVVWIHGL WP_001295940.1
MULTISPECIES: hydrogenase [Enterobacteriaceae] Length: 372\nNumber of predicted TMHs: 2\nExp number of AAs in TMHs: 37.2608899999999\nExp number, first 60 AAs: 19.52923\nTotal prob of N-in: 0.94351\nPOSSIBLE N-term signal sequence\ninside 1 19\nTMhelix 20 39\noutside 40 325\nTMhelix 326 348\ninside 349 372

10134 GCF_000488775.1_ASM48877v1 Escherichia coli 908691 Proteobacteria; Gammaproteobacteria; Enterobacterales; Enterobacteriaceae; Escherichia; Escherichia coli
MTGDNTLIHSHGINRRDFMKLCAALAATMGLSSKAAAEMVESVTNPQRPPVIWIGAQECT WP_000145423.1
hydrogenase 2 small subunit [Escherichia coli] Length: 372\nNumber of predicted TMHs: 1\nExp number of AAs in TMHs: 23.78201\nExp number, first 60 AAs: 0.28355\nTotal prob of N-in: 0.13257\noutside 1 331\nTMhelix 332 354\ninside 355 372

10135 GCF_000485735.1_ASM48573v1 Escherichia coli 909945-2 Proteobacteria; Gammaproteobacteria; Enterobacterales; Enterobacteriaceae; Escherichia; Escherichia coli
MTGDNTLIHSHGINRRDFMKLCAALAATMGLSSKAAAEMAESVTNPQRPPVIWIGAQECT WP_000145410.1
MULTISPECIES: hydrogenase-2 small chain [Proteobacteria] Length: 372\nNumber of predicted TMHs: 1\nExp number of AAs in TMHs: 23.72226\nExp number, first 60 AAs: 0.22038\nTotal prob of N-in: 0.12994\noutside 1 331\nTMhelix 332 354\ninside 355 372

10136 GCF_000485735.1_ASM48573v1 Escherichia coli 909945-2 Proteobacteria; Gammaproteobacteria; Enterobacterales; Enterobacteriaceae; Escherichia; Escherichia coli
MNNEETFYQAMRRQGVTRRSFLKYCSLAATSLGLGAGMAPKIAWALENKPRIPVVWIHGL WP_001058323.1
MULTISPECIES: hydrogenase-1 small chain [Proteobacteria] Length: 372\nNumber of predicted TMHs: 2\nExp number of AAs in TMHs: 37.2608499999999\nExp number, first 60 AAs: 19.52923\nTotal prob of N-in: 0.94351\nPOSSIBLE N-term signal sequence\ninside 1 19\nTMhelix 20 39\noutside 40 325\nTMhelix 326 348\ninside 349 372

10137 GCF_000488795.1_ASM48879v1 Escherichia coli 910096-2 Proteobacteria; Gammaproteobacteria; Enterobacterales; Enterobacteriaceae; Escherichia; Escherichia coli
MNNEETFYQAMRRQGVTRRSFLKYCSLAATSLGLGAGMAPKIAWALENKPRIPVVWIHGL WP_001538532.1
hydrogenase-1 small chain [Escherichia coli] Length: 372\nNumber of predicted TMHs: 2\nExp number of AAs in TMHs: 37.27041\nExp number, first 60 AAs: 19.52927\nTotal prob of N-in: 0.94352\nPOSSIBLE N-term signal sequence\ninside 1 19\nTMhelix 20 39\noutside 40 325\nTMhelix 326 348\ninside 349 372

10138 GCF_000488795.1_ASM48879v1 Escherichia coli 910096-2 Proteobacteria; Gammaproteobacteria; Enterobacterales; Enterobacteriaceae; Escherichia; Escherichia coli
MTGDNTLIHSHGINRRDFMKLCAALAATMGLSSKAAAEMVESVTNPQRPPVIWIGAQECT WP_000145423.1
hydrogenase 2 small subunit [Escherichia coli] Length: 372\nNumber of predicted TMHs: 1\nExp number of AAs in TMHs: 23.78201\nExp number, first 60 AAs: 0.28355\nTotal prob of N-in: 0.13257\noutside 1 331\nTMhelix 332 354\ninside 355 372

10139 GCF_000194475.2_ASM19447v2 Escherichia coli 9.1649 Proteobacteria; Gammaproteobacteria; Enterobacterales; Enterobacteriaceae; Escherichia; Escherichia coli
MTGDNTLIHSHGINRRDFMKLCAALAATMGLSSKAAAEMVESVTNPQRPPVIWIGAQECT WP_000145423.1
hydrogenase 2 small subunit [Escherichia coli] Length: 372\nNumber of predicted TMHs: 1\nExp number of AAs in TMHs: 23.78201\nExp number, first 60 AAs: 0.28355\nTotal prob of N-in: 0.13257\noutside 1 331\nTMhelix 332 354\ninside 355 372

10140 GCF_000194475.2_ASM19447v2 Escherichia coli 9.1649 Proteobacteria; Gammaproteobacteria; Enterobacterales; Enterobacteriaceae; Escherichia; Escherichia coli
MNNEETFYQAMRRQGVTRRSFLKYCSLAATSLGLGAGMAPKIAWALENKPRIPVVWIHGL WP_001058323.1
MULTISPECIES: hydrogenase-1 small chain [Proteobacteria] Length: 372\nNumber of predicted TMHs: 2\nExp number of AAs in TMHs: 37.2608499999999\nExp number, first 60 AAs: 19.52923\nTotal prob of N-in: 0.94351\nPOSSIBLE N-term signal sequence\ninside 1 19\nTMhelix 20 39\noutside 40 325\nTMhelix 326 348\ninside 349 372

10141 GCF_000267945.2_ASM26794v2 Escherichia coli 93-001 Proteobacteria; Gammaproteobacteria; Enterobacterales; Enterobacteriaceae; Escherichia; Escherichia coli
MNNEETFYQAMRRQGVTRRSFLKYCSLAATSLGLGAGMAPKIAWALENKPRIPVVWIHGL WP_001301613.1
hydrogenase [Escherichia coli] Length: 372\nNumber of predicted TMHs: 2\nExp number of AAs in TMHs: 37.46598\nExp number, first 60 AAs: 19.53139\nTotal prob of N-in: 0.94364\nPOSSIBLE N-term signal sequence\ninside 1 19\nTMhelix 20 39\noutside 40 325\nTMhelix 326 348\ninside 349 372

10142 GCF_000267945.2_ASM26794v2 Escherichia coli 93-001 Proteobacteria; Gammaproteobacteria; Enterobacterales; Enterobacteriaceae; Escherichia; Escherichia coli
MTGDNTLIHSHGINRRDFMKLCAALAATMGLSSKAAAEMAESVTNPQRPPVIWIGAQECT WP_000145410.1
MULTISPECIES: hydrogenase-2 small chain [Proteobacteria] Length: 372\nNumber of predicted TMHs: 1\nExp number of AAs in TMHs: 23.72226\nExp number, first 60 AAs: 0.22038\nTotal prob of N-in: 0.12994\noutside 1 331\nTMhelix 332 354\ninside 355 372

10143 GCF_000316445.2_ASM31644v2 Escherichia coli 93.0055 Proteobacteria; Gammaproteobacteria; Enterobacterales; Enterobacteriaceae; Escherichia; Escherichia coli
MNNEETFYQAMRRQGVTRRSFLKYCSLAATSLGLGAGMAPKIAWALENKPRIPVVWIHGL WP_001301613.1
hydrogenase [Escherichia coli] Length: 372\nNumber of predicted TMHs: 2\nExp number of AAs in TMHs: 37.46598\nExp number, first 60 AAs: 19.53139\nTotal prob of N-in: 0.94364\nPOSSIBLE N-term signal sequence\ninside 1 19\nTMhelix 20 39\noutside 40 325\nTMhelix 326 348\ninside 349 372

10144 GCF_000316445.2_ASM31644v2 Escherichia coli 93.0055 Proteobacteria; Gammaproteobacteria; Enterobacterales; Enterobacteriaceae; Escherichia; Escherichia coli
MTGDNTLIHSHGINRRDFMKLCAALAATMGLSSKAAAEMAESVTNPQRPPVIWIGAQECT WP_000145410.1
MULTISPECIES: hydrogenase-2 small chain [Proteobacteria] Length: 372\nNumber of predicted TMHs: 1\nExp number of AAs in TMHs: 23.72226\nExp number, first 60 AAs: 0.22038\nTotal prob of N-in: 0.12994\noutside 1 331\nTMhelix 332 354\ninside 355 372

10145 GCF_000316465.1_ASM31646v1 Escherichia coli 93.0056 Proteobacteria; Gammaproteobacteria; Enterobacterales; Enterobacteriaceae; Escherichia; Escherichia coli
MTGDNTLIHSHGINRRDFMKLCAALAATMGLSSKAAAEMAESVTNPQRPPVIWIGAQECT WP_000145410.1
MULTISPECIES: hydrogenase-2 small chain [Proteobacteria] Length: 372\nNumber of predicted TMHs: 1\nExp number of AAs in TMHs: 23.72226\nExp number, first 60 AAs: 0.22038\nTotal prob of N-in: 0.12994\noutside 1 331\nTMhelix 332 354\ninside 355 372

10146 GCF_000316465.1_ASM31646v1 Escherichia coli 93.0056 Proteobacteria; Gammaproteobacteria; Enterobacterales; Enterobacteriaceae; Escherichia; Escherichia coli
MNNEETFYQAMRRQGVTRRSFLKYCSLAATSLGLGAGMAPKIAWALENKPRIPVVWIHGL WP_001301613.1
hydrogenase [Escherichia coli] Length: 372\nNumber of predicted TMHs: 2\nExp number of AAs in

TMHs: 37.46598\nExp number, first 60 AAs: 19.53139\nTotal prob of N-in: 0.94364\nPOSSIBLE N-term signal sequence\ninside 1 19\nTMhelix 20 39\noutside 40 325\nTMhelix 326 348\ninside 349 372

10147 GCF_000215225.1_ASM21522v2 Escherichia coli 93.0624 Proteobacteria; Gammaproteobacteria; Enterobacterales; Enterobacteriaceae; Escherichia; Escherichia coli MNNEETFYQAMRRQGVTRRSFLKYCSLAATSLGLGAGMAPKIAWALENKPRIPVVWIHGL WP_001295940.1 MULTISPECIES: hydrogenase [Enterobacteriaceae] Length: 372\nNumber of predicted TMHs: 2\nExp number of AAs in TMHs: 37.2608899999999\nExp number, first 60 AAs: 19.52923\nTotal prob of N-in: 0.94351\nPOSSIBLE N-term signal sequence\ninside 1 19\nTMhelix 20 39\noutside 40 325\nTMhelix 326 348\ninside 349 372

10148 GCF_000215225.1_ASM21522v2 Escherichia coli 93.0624 Proteobacteria; Gammaproteobacteria; Enterobacterales; Enterobacteriaceae; Escherichia; Escherichia coli MTGDNTLIHSHGINRRDFMKLCAALAATMGLSSKAAAEMAESVTNPQRPPVIWIGAQECT WP_000145410.1 MULTISPECIES: hydrogenase-2 small chain [Proteobacteria] Length: 372\nNumber of predicted TMHs: 1\nExp number of AAs in TMHs: 23.72226\nExp number, first 60 AAs: 0.22038\nTotal prob of N-in: 0.12994\noutside 1 331\nTMhelix 332 354\ninside 355 372

10149 GCF_000316485.1_ASM31648v1 Escherichia coli 94.0618 Proteobacteria; Gammaproteobacteria; Enterobacterales; Enterobacteriaceae; Escherichia; Escherichia coli MNNEETFYQAMRRQGVTRRSFLKYCSLAATSLGLGAGMAPKIAWALENKPRIPVVWIHGL WP_001301613.1 hydrogenase [Escherichia coli] Length: 372\nNumber of predicted TMHs: 2\nExp number of AAs in TMHs: 37.46598\nExp number, first 60 AAs: 19.53139\nTotal prob of N-in: 0.94364\nPOSSIBLE N-term signal sequence\ninside 1 19\nTMhelix 20 39\noutside 40 325\nTMhelix 326 348\ninside 349 372

10150 GCF_000316485.1_ASM31648v1 Escherichia coli 94.0618 Proteobacteria; Gammaproteobacteria; Enterobacterales; Enterobacteriaceae; Escherichia; Escherichia coli MTGDNTLIHSHGINRRDFMKLCAALAATMGLSSKAAAEMAESVTNPQRPPVIWIGAQECT WP_000145410.1 MULTISPECIES: hydrogenase-2 small chain [Proteobacteria] Length: 372\nNumber of predicted TMHs: 1\nExp number of AAs in TMHs: 23.72226\nExp number, first 60 AAs: 0.22038\nTotal prob of N-in: 0.12994\noutside 1 331\nTMhelix 332 354\ninside 355 372

10151 GCF_000316845.2_ASM31684v2 Escherichia coli 95.0183 Proteobacteria; Gammaproteobacteria; Enterobacterales; Enterobacteriaceae; Escherichia; Escherichia coli MNNEETFYQAMRRQGVTRRSFLKYCSLAATSLGLGAGMAPKIAWALENKPRIPVVWIHGL WP_001301613.1 hydrogenase [Escherichia coli] Length: 372\nNumber of predicted TMHs: 2\nExp number of AAs in TMHs: 37.46598\nExp number, first 60 AAs: 19.53139\nTotal prob of N-in: 0.94364\nPOSSIBLE N-term signal sequence\ninside 1 19\nTMhelix 20 39\noutside 40 325\nTMhelix 326 348\ninside 349 372

10152 GCF_000316845.2_ASM31684v2 Escherichia coli 95.0183 Proteobacteria; Gammaproteobacteria; Enterobacterales; Enterobacteriaceae; Escherichia; Escherichia coli MTGDNTLIHSHGINRRDFMKLCAALAATMGLSSKAAAEMAESVTNPQRPPVIWIGAQECT WP_000145410.1 MULTISPECIES: hydrogenase-2 small chain [Proteobacteria] Length: 372\nNumber of predicted TMHs: 1\nExp number of AAs in TMHs: 23.72226\nExp number, first 60 AAs: 0.22038\nTotal prob of N-in: 0.12994\noutside 1 331\nTMhelix 332 354\ninside 355 372

10153 GCF_000194255.1_ASM19425v1 Escherichia coli 95.0941 Proteobacteria; Gammaproteobacteria; Enterobacterales; Enterobacteriaceae; Escherichia; Escherichia coli MTGDNTLIHSHGINRRDFMKLCAALAATMGLSSKAAAEMAESVTNPQRPPVIWIGAQECT WP_000145410.1 MULTISPECIES: hydrogenase-2 small chain [Proteobacteria] Length: 372\nNumber of predicted TMHs: 1\nExp number of AAs in TMHs: 23.72226\nExp number, first 60 AAs: 0.22038\nTotal prob of N-in: 0.12994\noutside 1 331\nTMhelix 332 354\ninside 355 372

10154 GCF_000194255.1_ASM19425v1 Escherichia coli 95.0941 Proteobacteria; Gammaproteobacteria; Enterobacterales; Enterobacteriaceae; Escherichia; Escherichia coli MNNEETFYQAMRRQGVTRRSFLKYCSLAATSLGLGAGMAPKIAWALENKPRIPVVWIHGL WP_001058323.1 MULTISPECIES: hydrogenase-1 small chain [Proteobacteria] Length: 372\nNumber of predicted TMHs: 2\nExp number of AAs in TMHs: 37.2608499999999\nExp number, first 60 AAs: 19.52923\nTotal prob of N-in: 0.94351\nPOSSIBLE N-term signal sequence\ninside 1 19\nTMhelix 20 39\noutside 40 325\nTMhelix 326 348\ninside 349 372

10155 GCF_000316885.2_ASM31688v2 Escherichia coli 95.0943 Proteobacteria; Gammaproteobacteria; Enterobacterales; Enterobacteriaceae; Escherichia; Escherichia coli
MTGDNTLIHSHGINRRDFMKLCAALAATMGLSSKAAAEMAESVTNPQRPPVIWIGAQECT WP_000145410.1
MULTISPECIES: hydrogenase-2 small chain [Proteobacteria] Length: 372\nNumber of predicted TMHs: 1\nExp number of AAs in TMHs: 23.72226\nExp number, first 60 AAs: 0.22038\nTotal prob of N-in: 0.12994\noutside 1 331\nTMhelix 332 354\ninside 355 372

10156 GCF_000316885.2_ASM31688v2 Escherichia coli 95.0943 Proteobacteria; Gammaproteobacteria; Enterobacterales; Enterobacteriaceae; Escherichia; Escherichia coli
MNNEETFYQAMRRQGVTRRSFLKYCSLAATSLGLGAGMAPKIAWALENKPRIPVVWIHGL WP_001301613.1
hydrogenase [Escherichia coli] Length: 372\nNumber of predicted TMHs: 2\nExp number of AAs in TMHs: 37.46598\nExp number, first 60 AAs: 19.53139\nTotal prob of N-in: 0.94364\nPOSSIBLE N-term signal sequence\ninside 1 19\nTMhelix 20 39\noutside 40 325\nTMhelix 326 348\ninside 349 372

10157 GCF_000316865.2_ASM31686v2 Escherichia coli 95.1288 Proteobacteria; Gammaproteobacteria; Enterobacterales; Enterobacteriaceae; Escherichia; Escherichia coli
MTGDNTLIHSHGINRRDFMKLCAALAATMGLSSKAAAEMAESVTNPQRPPVIWIGAQECT WP_000145410.1
MULTISPECIES: hydrogenase-2 small chain [Proteobacteria] Length: 372\nNumber of predicted TMHs: 1\nExp number of AAs in TMHs: 23.72226\nExp number, first 60 AAs: 0.22038\nTotal prob of N-in: 0.12994\noutside 1 331\nTMhelix 332 354\ninside 355 372

10158 GCF_000478705.1_95JB1 Escherichia coli 95JB1 Proteobacteria; Gammaproteobacteria; Enterobacterales; Enterobacteriaceae; Escherichia; Escherichia coli
MNNEETFYQAMRRQGVTRRSFLKYCSLAATSLGLGAGMAPKIAWALENKPRIPVVWIHGL WP_001058323.1
MULTISPECIES: hydrogenase-1 small chain [Proteobacteria] Length: 372\nNumber of predicted TMHs: 2\nExp number of AAs in TMHs: 37.2608499999999\nExp number, first 60 AAs: 19.52923\nTotal prob of N-in: 0.94351\nPOSSIBLE N-term signal sequence\ninside 1 19\nTMhelix 20 39\noutside 40 325\nTMhelix 326 348\ninside 349 372

10159 GCF_000478705.1_95JB1 Escherichia coli 95JB1 Proteobacteria; Gammaproteobacteria; Enterobacterales; Enterobacteriaceae; Escherichia; Escherichia coli
MTGDNTLIHSHGINRRDFMKLCAALAATMGLSSKAAAEMAESVTNPQRPPVIWIGAQECT WP_000145410.1
MULTISPECIES: hydrogenase-2 small chain [Proteobacteria] Length: 372\nNumber of predicted TMHs: 1\nExp number of AAs in TMHs: 23.72226\nExp number, first 60 AAs: 0.22038\nTotal prob of N-in: 0.12994\noutside 1 331\nTMhelix 332 354\ninside 355 372

10160 GCF_000467695.1_95NR1 Escherichia coli 95NR1 Proteobacteria; Gammaproteobacteria; Enterobacterales; Enterobacteriaceae; Escherichia; Escherichia coli
MNNEETFYQAMRRQGVTRRSFLKYCSLAATSLGLGAGMAPKIAWALENKPRIPVVWIHGL WP_001058323.1
MULTISPECIES: hydrogenase-1 small chain [Proteobacteria] Length: 372\nNumber of predicted TMHs: 2\nExp number of AAs in TMHs: 37.2608499999999\nExp number, first 60 AAs: 19.52923\nTotal prob of N-in: 0.94351\nPOSSIBLE N-term signal sequence\ninside 1 19\nTMhelix 20 39\noutside 40 325\nTMhelix 326 348\ninside 349 372

10161 GCF_000467695.1_95NR1 Escherichia coli 95NR1 Proteobacteria; Gammaproteobacteria; Enterobacterales; Enterobacteriaceae; Escherichia; Escherichia coli
MTGDNTLIHSHGINRRDFMKLCAALAATMGLSSKAAAEMAESVTNPQRPPVIWIGAQECT WP_000145410.1
MULTISPECIES: hydrogenase-2 small chain [Proteobacteria] Length: 372\nNumber of predicted TMHs: 1\nExp number of AAs in TMHs: 23.72226\nExp number, first 60 AAs: 0.22038\nTotal prob of N-in: 0.12994\noutside 1 331\nTMhelix 332 354\ninside 355 372

10162 GCF_000316765.2_ASM31676v2 Escherichia coli 96.0107 Proteobacteria; Gammaproteobacteria; Enterobacterales; Enterobacteriaceae; Escherichia; Escherichia coli
MTGDNTLIHSHGINRRDFMKLCAALAATMGLSSKAAAEMAESVTNPQRPPVIWIGAQECT WP_000145410.1
MULTISPECIES: hydrogenase-2 small chain [Proteobacteria] Length: 372\nNumber of predicted TMHs: 1\nExp number of AAs in TMHs: 23.72226\nExp number, first 60 AAs: 0.22038\nTotal prob of N-in: 0.12994\noutside 1 331\nTMhelix 332 354\ninside 355 372

10163 GCF_000316765.2_ASM31676v2 Escherichia coli 96.0107 Proteobacteria; Gammaproteobacteria; Enterobacterales; Enterobacteriaceae; Escherichia; Escherichia coli
MNNEETFYQAMRRQGVTRRSFLKYCSLAATSLGLGAGMAPKIAWALENKPRIPVVWIHGL WP_001301613.1

hydrogenase [Escherichia coli] Length: 372\nNumber of predicted TMHs: 2\nExp number of AAs in TMHs: 37.46598\nExp number, first 60 AAs: 19.53139\nTotal prob of N-in: 0.94364\nPOSSIBLE N-term signal sequence\ninside 1 19\nTMhelix 20 39\noutside 40 325\nTMhelix 326 348\ninside 349 372

10164 GCF_000318425.2_ASM31842v2 Escherichia coli 96.0109 Proteobacteria; Gammaproteobacteria; Enterobacterales; Enterobacteriaceae; Escherichia; Escherichia coli MNNEETFYQAMRRQGVTRRSFLKYCSLAATSLGLGAGMAPKIAWALENKPRIPVVWIHGL WP_001301613.1 hydrogenase [Escherichia coli] Length: 372\nNumber of predicted TMHs: 2\nExp number of AAs in TMHs: 37.46598\nExp number, first 60 AAs: 19.53139\nTotal prob of N-in: 0.94364\nPOSSIBLE N-term signal sequence\ninside 1 19\nTMhelix 20 39\noutside 40 325\nTMhelix 326 348\ninside 349 372

10165 GCF_000318425.2_ASM31842v2 Escherichia coli 96.0109 Proteobacteria; Gammaproteobacteria; Enterobacterales; Enterobacteriaceae; Escherichia; Escherichia coli MTGDNTLIHSHGINRRDFMKLCAALAATMGLSSKAAAEMAESVTNPQRPPVIWIGAQECT WP_000145410.1 MULTISPECIES: hydrogenase-2 small chain [Proteobacteria] Length: 372\nNumber of predicted TMHs: 1\nExp number of AAs in TMHs: 23.72226\nExp number, first 60 AAs: 0.22038\nTotal prob of N-in: 0.12994\noutside 1 331\nTMhelix 332 354\ninside 355 372

10166 GCF_000316825.2_ASM31682v2 Escherichia coli 96.0427 Proteobacteria; Gammaproteobacteria; Enterobacterales; Enterobacteriaceae; Escherichia; Escherichia coli MNNEETFYQAMRRQGVTRRSFLKYCSLAATSLGLGAGMAPKIAWALENKPRIPVVWIHGL WP_001301613.1 hydrogenase [Escherichia coli] Length: 372\nNumber of predicted TMHs: 2\nExp number of AAs in TMHs: 37.46598\nExp number, first 60 AAs: 19.53139\nTotal prob of N-in: 0.94364\nPOSSIBLE N-term signal sequence\ninside 1 19\nTMhelix 20 39\noutside 40 325\nTMhelix 326 348\ninside 349 372

10167 GCF_000316825.2_ASM31682v2 Escherichia coli 96.0427 Proteobacteria; Gammaproteobacteria; Enterobacterales; Enterobacteriaceae; Escherichia; Escherichia coli MTGDNTLIHSHGINRRDFMKLCAALAATMGLSSKAAAEMAESVTNPQRPPVIWIGAQECT WP_000145410.1 MULTISPECIES: hydrogenase-2 small chain [Proteobacteria] Length: 372\nNumber of predicted TMHs: 1\nExp number of AAs in TMHs: 23.72226\nExp number, first 60 AAs: 0.22038\nTotal prob of N-in: 0.12994\noutside 1 331\nTMhelix 332 354\ninside 355 372

10168 GCF_000316905.2_ASM31690v2 Escherichia coli 96.0428 Proteobacteria; Gammaproteobacteria; Enterobacterales; Enterobacteriaceae; Escherichia; Escherichia coli MNNEETFYQAMRRQGVTRRSFLKYCSLAATSLGLGAGMAPKIAWALENKPRIPVVWIHGL WP_001301613.1 hydrogenase [Escherichia coli] Length: 372\nNumber of predicted TMHs: 2\nExp number of AAs in TMHs: 37.46598\nExp number, first 60 AAs: 19.53139\nTotal prob of N-in: 0.94364\nPOSSIBLE N-term signal sequence\ninside 1 19\nTMhelix 20 39\noutside 40 325\nTMhelix 326 348\ninside 349 372

10169 GCF_000316905.2_ASM31690v2 Escherichia coli 96.0428 Proteobacteria; Gammaproteobacteria; Enterobacterales; Enterobacteriaceae; Escherichia; Escherichia coli MTGDNTLIHSHGINRRDFMKLCAALAATMGLSSKAAAEMAESVTNPQRPPVIWIGAQECT WP_000145410.1 MULTISPECIES: hydrogenase-2 small chain [Proteobacteria] Length: 372\nNumber of predicted TMHs: 1\nExp number of AAs in TMHs: 23.72226\nExp number, first 60 AAs: 0.22038\nTotal prob of N-in: 0.12994\noutside 1 331\nTMhelix 332 354\ninside 355 372

10170 GCF_000215185.1_ASM21518v2 Escherichia coli 96.0497 Proteobacteria; Gammaproteobacteria; Enterobacterales; Enterobacteriaceae; Escherichia; Escherichia coli MTGDNTLIHSHGINRRDFMKLCAALAATMGLSSKAAAEMAESVTNPQRPPVIWIGAQECT WP_000145410.1 MULTISPECIES: hydrogenase-2 small chain [Proteobacteria] Length: 372\nNumber of predicted TMHs: 1\nExp number of AAs in TMHs: 23.72226\nExp number, first 60 AAs: 0.22038\nTotal prob of N-in: 0.12994\noutside 1 331\nTMhelix 332 354\ninside 355 372

10171 GCF_000215185.1_ASM21518v2 Escherichia coli 96.0497 Proteobacteria; Gammaproteobacteria; Enterobacterales; Enterobacteriaceae; Escherichia; Escherichia coli MNNEETFYQAMRRQGVTRRSFLKYCSLAATSLGLGAGMAPKIAWALENKPRIPVVWIHGL WP_001295940.1 MULTISPECIES: hydrogenase [Enterobacteriaceae] Length: 372\nNumber of predicted TMHs: 2\nExp number of AAs in TMHs: 37.2608899999999\nExp number, first 60 AAs: 19.52923\nTotal prob of N-in: 0.94351\nPOSSIBLE N-term signal sequence\ninside 1 19\nTMhelix 20 39\noutside 40 325\nTMhelix 326 348\ninside 349 372

10172 GCF_000316805.2_ASM31680v2 Escherichia coli 96.0932 Proteobacteria; Gammaproteobacteria; Enterobacterales; Enterobacteriaceae; Escherichia; Escherichia coli
MNNEETFYQAMRRQGVTRRSFLKYCSLAATSLGLGAGMAPKIAWALENKPRIPVVWIHGL WP_001301613.1
hydrogenase [Escherichia coli] Length: 372\nNumber of predicted TMHs: 2\nExp number of AAs in TMHs: 37.46598\nExp number, first 60 AAs: 19.53139\nTotal prob of N-in: 0.94364\nPOSSIBLE N-term signal sequence\ninside 1 19\nTMhelix 20 39\noutside 40 325\nTMhelix 326 348\ninside 349 372

10173 GCF_000316805.2_ASM31680v2 Escherichia coli 96.0932 Proteobacteria; Gammaproteobacteria; Enterobacterales; Enterobacteriaceae; Escherichia; Escherichia coli
MTGDNTLIHSHGINRRDFMKLCAALAATMGLSSKAAAEMAESVTNPQRPPVIWIGAQECT WP_000145410.1
MULTISPECIES: hydrogenase-2 small chain [Proteobacteria] Length: 372\nNumber of predicted TMHs: 1\nExp number of AAs in TMHs: 23.72226\nExp number, first 60 AAs: 0.22038\nTotal prob of N-in: 0.12994\noutside 1 331\nTMhelix 332 354\ninside 355 372

10174 GCF_000316785.2_ASM31678v2 Escherichia coli 96.0939 Proteobacteria; Gammaproteobacteria; Enterobacterales; Enterobacteriaceae; Escherichia; Escherichia coli
MTGDNTLIHSHGINRRDFMKLCAALAATMGLSSKAAAEMAESVTNPQRPPVIWIGAQECT WP_000145410.1
MULTISPECIES: hydrogenase-2 small chain [Proteobacteria] Length: 372\nNumber of predicted TMHs: 1\nExp number of AAs in TMHs: 23.72226\nExp number, first 60 AAs: 0.22038\nTotal prob of N-in: 0.12994\noutside 1 331\nTMhelix 332 354\ninside 355 372

10175 GCF_000316785.2_ASM31678v2 Escherichia coli 96.0939 Proteobacteria; Gammaproteobacteria; Enterobacterales; Enterobacteriaceae; Escherichia; Escherichia coli
MNNEETFYQAMRRQGVTRRSFLKYCSLAATSLGLGAGMAPKIAWALENKPRIPVVWIHGL WP_001301613.1
hydrogenase [Escherichia coli] Length: 372\nNumber of predicted TMHs: 2\nExp number of AAs in TMHs: 37.46598\nExp number, first 60 AAs: 19.53139\nTotal prob of N-in: 0.94364\nPOSSIBLE N-term signal sequence\ninside 1 19\nTMhelix 20 39\noutside 40 325\nTMhelix 326 348\ninside 349 372

10176 GCF_000215245.1_ASM21524v2 Escherichia coli 96.154 Proteobacteria; Gammaproteobacteria; Enterobacterales; Enterobacteriaceae; Escherichia; Escherichia coli
MTGDNTLIHSHGINRRDFMKLCAALAATMGLSSKAAAEMAESVTNPQRPPVIWIGAQECT WP_000145410.1
MULTISPECIES: hydrogenase-2 small chain [Proteobacteria] Length: 372\nNumber of predicted TMHs: 1\nExp number of AAs in TMHs: 23.72226\nExp number, first 60 AAs: 0.22038\nTotal prob of N-in: 0.12994\noutside 1 331\nTMhelix 332 354\ninside 355 372

10177 GCF_000215245.1_ASM21524v2 Escherichia coli 96.154 Proteobacteria; Gammaproteobacteria; Enterobacterales; Enterobacteriaceae; Escherichia; Escherichia coli
MNNEETFYQAMRRQGVTRRSFLKYCSLAATSLGLGAGMAPKIAWALENKPRIPVVWIHGL WP_001058323.1
MULTISPECIES: hydrogenase-1 small chain [Proteobacteria] Length: 372\nNumber of predicted TMHs: 2\nExp number of AAs in TMHs: 37.2608499999999\nExp number, first 60 AAs: 19.52923\nTotal prob of N-in: 0.94351\nPOSSIBLE N-term signal sequence\ninside 1 19\nTMhelix 20 39\noutside 40 325\nTMhelix 326 348\ninside 349 372

10178 GCF_000316745.2_ASM31674v2 Escherichia coli 97.0003 Proteobacteria; Gammaproteobacteria; Enterobacterales; Enterobacteriaceae; Escherichia; Escherichia coli
MTGDNTLIHSHGINRRDFMKLCAALAATMGLSSKAAAEMAESVTNPQRPPVIWIGAQECT WP_000145410.1
MULTISPECIES: hydrogenase-2 small chain [Proteobacteria] Length: 372\nNumber of predicted TMHs: 1\nExp number of AAs in TMHs: 23.72226\nExp number, first 60 AAs: 0.22038\nTotal prob of N-in: 0.12994\noutside 1 331\nTMhelix 332 354\ninside 355 372

10179 GCF_000316745.2_ASM31674v2 Escherichia coli 97.0003 Proteobacteria; Gammaproteobacteria; Enterobacterales; Enterobacteriaceae; Escherichia; Escherichia coli
MNNEETFYQAMRRQGVTRRSFLKYCSLAATSLGLGAGMAPKIAWALENKPRIPVVWIHGL WP_001301613.1
hydrogenase [Escherichia coli] Length: 372\nNumber of predicted TMHs: 2\nExp number of AAs in TMHs: 37.46598\nExp number, first 60 AAs: 19.53139\nTotal prob of N-in: 0.94364\nPOSSIBLE N-term signal sequence\ninside 1 19\nTMhelix 20 39\noutside 40 325\nTMhelix 326 348\ninside 349 372

10180 GCF_000316505.2_ASM31650v2 Escherichia coli 97.0007 Proteobacteria; Gammaproteobacteria; Enterobacterales; Enterobacteriaceae; Escherichia; Escherichia coli
MNNEETFYQAMRRQGVTRRSFLKYCSLAATSLGLGAGMAPKIAWALENKPRIPVVWIHGL WP_001301613.1
hydrogenase [Escherichia coli] Length: 372\nNumber of predicted TMHs: 2\nExp number of AAs in

TMHs: 37.46598\nExp number, first 60 AAs:  19.53139\nTotal prob of N-in:   0.94364\nPOSSIBLE N-term signal sequence\ninside   1   19\nTMhelix   20   39\noutside   40   325\nTMhelix   326   348\ninside   349   372

10181 GCF_000316505.2_ASM31650v2 Escherichia coli 97.0007 Proteobacteria; Gammaproteobacteria; Enterobacterales; Enterobacteriaceae; Escherichia; Escherichia coli
MTGDNTLIHSHGINRRDFMKLCAALAATMGLSSKAAAEMAESVTNPQRPPVIWIGAQECT WP_000145410.1
MULTISPECIES: hydrogenase-2 small chain [Proteobacteria] Length: 372\nNumber of predicted TMHs: 1\nExp number of AAs in TMHs: 23.72226\nExp number, first 60 AAs:  0.22038\nTotal prob of N-in:   0.12994\noutside   1   331\nTMhelix   332   354\ninside   355   372

10182 GCF_000318445.2_ASM31844v2 Escherichia coli 97.0010 Proteobacteria; Gammaproteobacteria; Enterobacterales; Enterobacteriaceae; Escherichia; Escherichia coli
MNNEETFYQAMRRQGVTRRSFLKYCSLAATSLGLGAGMAPKIAWALENKPRIPVVWIHGL WP_001301613.1
hydrogenase [Escherichia coli] Length: 372\nNumber of predicted TMHs:  2\nExp number of AAs in TMHs: 37.46598\nExp number, first 60 AAs:  19.53139\nTotal prob of N-in:   0.94364\nPOSSIBLE N-term signal sequence\ninside   1   19\nTMhelix   20   39\noutside   40   325\nTMhelix   326   348\ninside   349   372

10183 GCF_000318445.2_ASM31844v2 Escherichia coli 97.0010 Proteobacteria; Gammaproteobacteria; Enterobacterales; Enterobacteriaceae; Escherichia; Escherichia coli
MTGDNTLIHSHGINRRDFMKLCAALAATMGLSSKAAAEMAESVTNPQRPPVIWIGAQECT WP_000145410.1
MULTISPECIES: hydrogenase-2 small chain [Proteobacteria] Length: 372\nNumber of predicted TMHs: 1\nExp number of AAs in TMHs: 23.72226\nExp number, first 60 AAs:  0.22038\nTotal prob of N-in:   0.12994\noutside   1   331\nTMhelix   332   354\ninside   355   372

10184 GCF_000194215.1_ASM19421v2 Escherichia coli 97.0246 Proteobacteria; Gammaproteobacteria; Enterobacterales; Enterobacteriaceae; Escherichia; Escherichia coli
MTGDNTLIHSHGINRRDFMKLCAALAATMGLSSKAAAEMAESVTNPQRPPVIWIGAQECT WP_000145410.1
MULTISPECIES: hydrogenase-2 small chain [Proteobacteria] Length: 372\nNumber of predicted TMHs: 1\nExp number of AAs in TMHs: 23.72226\nExp number, first 60 AAs:  0.22038\nTotal prob of N-in:   0.12994\noutside   1   331\nTMhelix   332   354\ninside   355   372

10185 GCF_000194215.1_ASM19421v2 Escherichia coli 97.0246 Proteobacteria; Gammaproteobacteria; Enterobacterales; Enterobacteriaceae; Escherichia; Escherichia coli
MNNEETFYQAMRRQGVTRRSFLKSCSLAATSLGLGAGMAPKIAWALENKPRIPVVWIHGL WP_001058318.1
hydrogenase [Escherichia coli] Length: 372\nNumber of predicted TMHs:  1\nExp number of AAs in TMHs: 33.59637\nExp number, first 60 AAs:  15.81976\nTotal prob of N-in:   0.77394\nPOSSIBLE N-term signal sequence\noutside   1   325\nTMhelix   326   348\ninside   349   372

10186 GCF_000194235.1_ASM19423v2 Escherichia coli 97.0259 Proteobacteria; Gammaproteobacteria; Enterobacterales; Enterobacteriaceae; Escherichia; Escherichia coli
MTGDNTLIHSHGINRRDFMKLCAALAATMGLSSKAAAEMAESVTNPQRPPVIWIGAQECT WP_000145410.1
MULTISPECIES: hydrogenase-2 small chain [Proteobacteria] Length: 372\nNumber of predicted TMHs: 1\nExp number of AAs in TMHs: 23.72226\nExp number, first 60 AAs:  0.22038\nTotal prob of N-in:   0.12994\noutside   1   331\nTMhelix   332   354\ninside   355   372

10187 GCF_000194235.1_ASM19423v2 Escherichia coli 97.0259 Proteobacteria; Gammaproteobacteria; Enterobacterales; Enterobacteriaceae; Escherichia; Escherichia coli
MNNEETFYQAMRRQGVTRRSFLKYCSLAATSLGLGAGMAPKIAWALENKPRIPVVWIHGL WP_001058323.1
MULTISPECIES: hydrogenase-1 small chain [Proteobacteria] Length: 372\nNumber of predicted TMHs: 2\nExp number of AAs in TMHs: 37.2608499999999\nExp number, first 60 AAs:  19.52923\nTotal prob of N-in:   0.94351\nPOSSIBLE N-term signal sequence\ninside   1   19\nTMhelix   20   39\noutside   40   325\nTMhelix   326   348\ninside   349   372

10188 GCF_000194295.2_ASM19429v2 Escherichia coli 97.0264 Proteobacteria; Gammaproteobacteria; Enterobacterales; Enterobacteriaceae; Escherichia; Escherichia coli
MNNEETFYQAMRRQGVTRRSFLKYCSLAATSLGLGAGMAPKIAWALENKPRIPVVWIHGL WP_001058323.1
MULTISPECIES: hydrogenase-1 small chain [Proteobacteria] Length: 372\nNumber of predicted TMHs: 2\nExp number of AAs in TMHs: 37.2608499999999\nExp number, first 60 AAs:  19.52923\nTotal prob of N-in:   0.94351\nPOSSIBLE N-term signal sequence\ninside   1   19\nTMhelix   20   39\noutside   40   325\nTMhelix   326   348\ninside   349   372

10189	GCF_000194295.2_ASM19429v2	Escherichia coli 97.0264	Proteobacteria; Gammaproteobacteria; Enterobacterales; Enterobacteriaceae; Escherichia; Escherichia coli
MTGDNTLIHSHGINRRDFMKLCAALAATMGLSSKAAAEMAESVTNPQRPPVIWIGAQECT	WP_000145410.1
MULTISPECIES: hydrogenase-2 small chain [Proteobacteria]	Length: 372\nNumber of predicted TMHs: 1\nExp number of AAs in TMHs: 23.72226\nExp number, first 60 AAs: 0.22038\nTotal prob of N-in: 0.12994\noutside 1 331\nTMhelix 332 354\ninside 355 372

10190	GCF_000316725.2_ASM31672v2	Escherichia coli 97.1742	Proteobacteria; Gammaproteobacteria; Enterobacterales; Enterobacteriaceae; Escherichia; Escherichia coli
MTGDNTLIHSHGINRRDFMKLCAALAATMGLSSKAAAEMAESVTNPQRPPVIWIGAQECT	WP_000145410.1
MULTISPECIES: hydrogenase-2 small chain [Proteobacteria]	Length: 372\nNumber of predicted TMHs: 1\nExp number of AAs in TMHs: 23.72226\nExp number, first 60 AAs: 0.22038\nTotal prob of N-in: 0.12994\noutside 1 331\nTMhelix 332 354\ninside 355 372

10191	GCF_000316725.2_ASM31672v2	Escherichia coli 97.1742	Proteobacteria; Gammaproteobacteria; Enterobacterales; Enterobacteriaceae; Escherichia; Escherichia coli
MNNEETFYQAMRRQGVTRRSFLKYCSLAATSLGLGAGMAPKIAWALENKPRIPVVWIHGL	WP_001301613.1
hydrogenase [Escherichia coli]	Length: 372\nNumber of predicted TMHs: 2\nExp number of AAs in TMHs: 37.46598\nExp number, first 60 AAs: 19.53139\nTotal prob of N-in: 0.94364\nPOSSIBLE N-term signal sequence\ninside 1 19\nTMhelix 20 39\noutside 40 325\nTMhelix 326 348\ninside 349 372

10192	GCF_000335215.2_ASM33521v2	Escherichia coli 99.0670	Proteobacteria; Gammaproteobacteria; Enterobacterales; Enterobacteriaceae; Escherichia; Escherichia coli
MNNEETFYQAMRRQGVTRRSFLKYCSLAATSLGLGAGMAPKIAWALENKPRIPVVWIHGL	WP_001301613.1
hydrogenase [Escherichia coli]	Length: 372\nNumber of predicted TMHs: 2\nExp number of AAs in TMHs: 37.46598\nExp number, first 60 AAs: 19.53139\nTotal prob of N-in: 0.94364\nPOSSIBLE N-term signal sequence\ninside 1 19\nTMhelix 20 39\noutside 40 325\nTMhelix 326 348\ninside 349 372

10193	GCF_000335215.2_ASM33521v2	Escherichia coli 99.0670	Proteobacteria; Gammaproteobacteria; Enterobacterales; Enterobacteriaceae; Escherichia; Escherichia coli
MTGDNTLIHSHGINRRDFMKLCAALAATMGLSSKAAAEMAESVTNPQRPPVIWIGAQECT	WP_000145410.1
MULTISPECIES: hydrogenase-2 small chain [Proteobacteria]	Length: 372\nNumber of predicted TMHs: 1\nExp number of AAs in TMHs: 23.72226\nExp number, first 60 AAs: 0.22038\nTotal prob of N-in: 0.12994\noutside 1 331\nTMhelix 332 354\ninside 355 372

10194	GCF_000316705.2_ASM31670v2	Escherichia coli 99.0672	Proteobacteria; Gammaproteobacteria; Enterobacterales; Enterobacteriaceae; Escherichia; Escherichia coli
MNNEETFYQAMRRQGVTRRSFLKYCSLAATSLGLGAGMAPKIAWALENKPRIPVVWIHGL	WP_001301613.1
hydrogenase [Escherichia coli]	Length: 372\nNumber of predicted TMHs: 2\nExp number of AAs in TMHs: 37.46598\nExp number, first 60 AAs: 19.53139\nTotal prob of N-in: 0.94364\nPOSSIBLE N-term signal sequence\ninside 1 19\nTMhelix 20 39\noutside 40 325\nTMhelix 326 348\ninside 349 372

10195	GCF_000316705.2_ASM31670v2	Escherichia coli 99.0672	Proteobacteria; Gammaproteobacteria; Enterobacterales; Enterobacteriaceae; Escherichia; Escherichia coli
MTGDNTLIHSHGINRRDFMKLCAALAATMGLSSKAAAEMAESVTNPQRPPVIWIGAQECT	WP_000145410.1
MULTISPECIES: hydrogenase-2 small chain [Proteobacteria]	Length: 372\nNumber of predicted TMHs: 1\nExp number of AAs in TMHs: 23.72226\nExp number, first 60 AAs: 0.22038\nTotal prob of N-in: 0.12994\noutside 1 331\nTMhelix 332 354\ninside 355 372

10196	GCF_000316545.2_ASM31654v2	Escherichia coli 99.0678	Proteobacteria; Gammaproteobacteria; Enterobacterales; Enterobacteriaceae; Escherichia; Escherichia coli
MNNEETFYQAMRRQGVTRRSFLKYCSLAATSLGLGAGMAPKIAWALENKPRIPVVWIHGL	WP_001301613.1
hydrogenase [Escherichia coli]	Length: 372\nNumber of predicted TMHs: 2\nExp number of AAs in TMHs: 37.46598\nExp number, first 60 AAs: 19.53139\nTotal prob of N-in: 0.94364\nPOSSIBLE N-term signal sequence\ninside 1 19\nTMhelix 20 39\noutside 40 325\nTMhelix 326 348\ninside 349 372

10197	GCF_000316545.2_ASM31654v2	Escherichia coli 99.0678	Proteobacteria; Gammaproteobacteria; Enterobacterales; Enterobacteriaceae; Escherichia; Escherichia coli
MTGDNTLIHSHGINRRDFMKLCAALAATMGLSSKAAAEMAESVTNPQRPPVIWIGAQECT	WP_000145410.1
MULTISPECIES: hydrogenase-2 small chain [Proteobacteria]	Length: 372\nNumber of predicted TMHs:

1\nExp number of AAs in TMHs: 23.72226\nExp number, first 60 AAs: 0.22038\nTotal prob of N-in: 0.12994\noutside 1 331\nTMhelix 332 354\ninside 355 372

10198 GCF_000316565.2_ASM31656v2 Escherichia coli 99.0713 Proteobacteria; Gammaproteobacteria; Enterobacterales; Enterobacteriaceae; Escherichia; Escherichia coli MTGDNTLIHSHGINRRDFMKLCAALAATMGLSSKAAAEMAESVTNPQRPPVIWIGAQECT WP_000145410.1 MULTISPECIES: hydrogenase-2 small chain [Proteobacteria] Length: 372\nNumber of predicted TMHs: 1\nExp number of AAs in TMHs: 23.72226\nExp number, first 60 AAs: 0.22038\nTotal prob of N-in: 0.12994\noutside 1 331\nTMhelix 332 354\ninside 355 372

10199 GCF_000316565.2_ASM31656v2 Escherichia coli 99.0713 Proteobacteria; Gammaproteobacteria; Enterobacterales; Enterobacteriaceae; Escherichia; Escherichia coli MNNEETFYQAMRRQGVTRRSFLKYCSLAATSLGLGAGMAPKIAWALENKPRIPVVWIHGL WP_001301613.1 hydrogenase [Escherichia coli] Length: 372\nNumber of predicted TMHs: 2\nExp number of AAs in TMHs: 37.46598\nExp number, first 60 AAs: 19.53139\nTotal prob of N-in: 0.94364\nPOSSIBLE N-term signal sequence\ninside 1 19\nTMhelix 20 39\noutside 40 325\nTMhelix 326 348\ninside 349 372

10200 GCF_000194435.1_ASM19443v2 Escherichia coli 99.0741 Proteobacteria; Gammaproteobacteria; Enterobacterales; Enterobacteriaceae; Escherichia; Escherichia coli MTGDNTLIHSHGINRRDFMKLCAALAATMGLSSKAAAEMAESVTNPQRPPVIWIGAQECT WP_000145410.1 MULTISPECIES: hydrogenase-2 small chain [Proteobacteria] Length: 372\nNumber of predicted TMHs: 1\nExp number of AAs in TMHs: 23.72226\nExp number, first 60 AAs: 0.22038\nTotal prob of N-in: 0.12994\noutside 1 331\nTMhelix 332 354\ninside 355 372

10201 GCF_000194435.1_ASM19443v2 Escherichia coli 99.0741 Proteobacteria; Gammaproteobacteria; Enterobacterales; Enterobacteriaceae; Escherichia; Escherichia coli MNNEETFYQAMRRQGVTRRSFLKYCSLAATSLGLGAGMAPKIAWALENKPRIPVVWIHGL WP_001058323.1 MULTISPECIES: hydrogenase-1 small chain [Proteobacteria] Length: 372\nNumber of predicted TMHs: 2\nExp number of AAs in TMHs: 37.2608499999999\nExp number, first 60 AAs: 19.52923\nTotal prob of N-in: 0.94351\nPOSSIBLE N-term signal sequence\ninside 1 19\nTMhelix 20 39\noutside 40 325\nTMhelix 326 348\ninside 349 372

10202 GCF_000335235.2_ASM33523v2 Escherichia coli 99.0814 Proteobacteria; Gammaproteobacteria; Enterobacterales; Enterobacteriaceae; Escherichia; Escherichia coli MTGDNTLIHSHGINRRDFMKLCAALAATMGLSSKAAAEMAESVTNPQRPPVIWIGAQECT WP_001685818.1 MULTISPECIES: hydrogenase-2 small chain [Enterobacteriaceae] Length: 372\nNumber of predicted TMHs: 1\nExp number of AAs in TMHs: 23.82505\nExp number, first 60 AAs: 0.21974\nTotal prob of N-in: 0.13379\noutside 1 331\nTMhelix 332 354\ninside 355 372

10203 GCF_000335235.2_ASM33523v2 Escherichia coli 99.0814 Proteobacteria; Gammaproteobacteria; Enterobacterales; Enterobacteriaceae; Escherichia; Escherichia coli MNNEETFYQAMRRQGVTRRSFLKYCSLAATSLGLGAGMAPKIAWALENKPRIPVVWIHGL WP_001301613.1 hydrogenase [Escherichia coli] Length: 372\nNumber of predicted TMHs: 2\nExp number of AAs in TMHs: 37.46598\nExp number, first 60 AAs: 19.53139\nTotal prob of N-in: 0.94364\nPOSSIBLE N-term signal sequence\ninside 1 19\nTMhelix 20 39\noutside 40 325\nTMhelix 326 348\ninside 349 372

10204 GCF_000334975.2_ASM33497v2 Escherichia coli 99.0815 Proteobacteria; Gammaproteobacteria; Enterobacterales; Enterobacteriaceae; Escherichia; Escherichia coli MTGDNTLIHSHGINRRDFMKLCAALAATMGLSSKAAAEMAESVTNPQRPPVIWIGAQECT WP_001685818.1 MULTISPECIES: hydrogenase-2 small chain [Enterobacteriaceae] Length: 372\nNumber of predicted TMHs: 1\nExp number of AAs in TMHs: 23.82505\nExp number, first 60 AAs: 0.21974\nTotal prob of N-in: 0.13379\noutside 1 331\nTMhelix 332 354\ninside 355 372

10205 GCF_000334975.2_ASM33497v2 Escherichia coli 99.0815 Proteobacteria; Gammaproteobacteria; Enterobacterales; Enterobacteriaceae; Escherichia; Escherichia coli MNNEETFYQAMRRQGVTRRSFLKYCSLAATSLGLGAGMAPKIAWALENKPRIPVVWIHGL WP_001301613.1 hydrogenase [Escherichia coli] Length: 372\nNumber of predicted TMHs: 2\nExp number of AAs in TMHs: 37.46598\nExp number, first 60 AAs: 19.53139\nTotal prob of N-in: 0.94364\nPOSSIBLE N-term signal sequence\ninside 1 19\nTMhelix 20 39\noutside 40 325\nTMhelix 326 348\ninside 349 372

10206 GCF_000334995.2_ASM33499v2 Escherichia coli 99.0816 Proteobacteria; Gammaproteobacteria; Enterobacterales; Enterobacteriaceae; Escherichia; Escherichia coli MNNEETFYQAMRRQGVTRRSFLKYCSLAATSLGLGAGMAPKIAWALENKPRIPVVWIHGL WP_001301613.1 hydrogenase [Escherichia coli] Length: 372\nNumber of predicted TMHs: 2\nExp number of AAs in TMHs: 37.46598\nExp number, first 60 AAs: 19.53139\nTotal prob of N-in: 0.94364\nPOSSIBLE N-term signal sequence\ninside 1 19\nTMhelix 20 39\noutside 40 325\nTMhelix 326 348\ninside 349 372

10207 GCF_000334995.2_ASM33499v2 Escherichia coli 99.0816 Proteobacteria; Gammaproteobacteria; Enterobacterales; Enterobacteriaceae; Escherichia; Escherichia coli MTGDNTLIHSHGINRRDFMKLCAALAATMGLSSKAAAEMAESVTNPQRPPVIWIGAQECT WP_001685818.1 MULTISPECIES: hydrogenase-2 small chain [Enterobacteriaceae] Length: 372\nNumber of predicted TMHs: 1\nExp number of AAs in TMHs: 23.82505\nExp number, first 60 AAs: 0.21974\nTotal prob of N-in: 0.13379\noutside 1 331\nTMhelix 332 354\ninside 355 372

10208 GCF_000335255.2_ASM33525v2 Escherichia coli 99.0839 Proteobacteria; Gammaproteobacteria; Enterobacterales; Enterobacteriaceae; Escherichia; Escherichia coli MTGDNTLIHSHGINRRDFMKLCAALAATMGLSSKAAAEMAESVTNPQRPPVIWIGAQECT WP_000145410.1 MULTISPECIES: hydrogenase-2 small chain [Proteobacteria] Length: 372\nNumber of predicted TMHs: 1\nExp number of AAs in TMHs: 23.72226\nExp number, first 60 AAs: 0.22038\nTotal prob of N-in: 0.12994\noutside 1 331\nTMhelix 332 354\ninside 355 372

10209 GCF_000335255.2_ASM33525v2 Escherichia coli 99.0839 Proteobacteria; Gammaproteobacteria; Enterobacterales; Enterobacteriaceae; Escherichia; Escherichia coli MNNEETFYQAMRRQGVTRRSFLKYCSLAATSLGLGAGMAPKIAWALENKPRIPVVWIHGL WP_001301613.1 hydrogenase [Escherichia coli] Length: 372\nNumber of predicted TMHs: 2\nExp number of AAs in TMHs: 37.46598\nExp number, first 60 AAs: 19.53139\nTotal prob of N-in: 0.94364\nPOSSIBLE N-term signal sequence\ninside 1 19\nTMhelix 20 39\noutside 40 325\nTMhelix 326 348\ninside 349 372

10210 GCF_000335275.2_ASM33527v2 Escherichia coli 99.0848 Proteobacteria; Gammaproteobacteria; Enterobacterales; Enterobacteriaceae; Escherichia; Escherichia coli MTGDNTLIHSHGINRRDFMKLCAALAATMGLSSKAAAEMAESVTNPQRPPVIWIGAQECT WP_000145410.1 MULTISPECIES: hydrogenase-2 small chain [Proteobacteria] Length: 372\nNumber of predicted TMHs: 1\nExp number of AAs in TMHs: 23.72226\nExp number, first 60 AAs: 0.22038\nTotal prob of N-in: 0.12994\noutside 1 331\nTMhelix 332 354\ninside 355 372

10211 GCF_000335275.2_ASM33527v2 Escherichia coli 99.0848 Proteobacteria; Gammaproteobacteria; Enterobacterales; Enterobacteriaceae; Escherichia; Escherichia coli MNNEETFYQAMRRQGVTRRSFLKYCSLAATSLGLGAGMAPKIAWALENKPRIPVVWIHGL WP_001301613.1 hydrogenase [Escherichia coli] Length: 372\nNumber of predicted TMHs: 2\nExp number of AAs in TMHs: 37.46598\nExp number, first 60 AAs: 19.53139\nTotal prob of N-in: 0.94364\nPOSSIBLE N-term signal sequence\ninside 1 19\nTMhelix 20 39\noutside 40 325\nTMhelix 326 348\ninside 349 372

10212 GCF_000335295.2_ASM33529v2 Escherichia coli 99.1753 Proteobacteria; Gammaproteobacteria; Enterobacterales; Enterobacteriaceae; Escherichia; Escherichia coli MNNEETFYQAMRRQGVTRRSFLKYCSLAATSLGLGAGMAPKIAWALENKPRIPVVWIHGL WP_001301613.1 hydrogenase [Escherichia coli] Length: 372\nNumber of predicted TMHs: 2\nExp number of AAs in TMHs: 37.46598\nExp number, first 60 AAs: 19.53139\nTotal prob of N-in: 0.94364\nPOSSIBLE N-term signal sequence\ninside 1 19\nTMhelix 20 39\noutside 40 325\nTMhelix 326 348\ninside 349 372

10213 GCF_000335295.2_ASM33529v2 Escherichia coli 99.1753 Proteobacteria; Gammaproteobacteria; Enterobacterales; Enterobacteriaceae; Escherichia; Escherichia coli MTGDNTLIHSHGINRRDFMKLCAALAATMGLSSKAAAEMAESVTNPQRPPVIWIGAQECT WP_000145410.1 MULTISPECIES: hydrogenase-2 small chain [Proteobacteria] Length: 372\nNumber of predicted TMHs: 1\nExp number of AAs in TMHs: 23.72226\nExp number, first 60 AAs: 0.22038\nTotal prob of N-in: 0.12994\noutside 1 331\nTMhelix 332 354\ninside 355 372

10214 GCF_000335155.2_ASM33515v2 Escherichia coli 99.1762 Proteobacteria; Gammaproteobacteria; Enterobacterales; Enterobacteriaceae; Escherichia; Escherichia coli MTGDNTLIHSHGINRRDFMKLCAALAATMGLSSKAAAEMAESVTNPQRPPVIWIGAQECT WP_000145410.1 MULTISPECIES: hydrogenase-2 small chain [Proteobacteria] Length: 372\nNumber of predicted TMHs:

1\nExp number of AAs in TMHs: 23.72226\nExp number, first 60 AAs: 0.22038\nTotal prob of N-in: 0.12994\noutside 1 331\nTMhelix 332 354\ninside 355 372

10215 GCF_000335155.2_ASM33515v2 Escherichia coli 99.1762 Proteobacteria; Gammaproteobacteria; Enterobacterales; Enterobacteriaceae; Escherichia; Escherichia coli MNNEETFYQAMRRQGVTRRSFLKYCSLAATSLGLGAGMAPKIAWALENKPRIPVVWIHGL WP_001301613.1 hydrogenase [Escherichia coli] Length: 372\nNumber of predicted TMHs: 2\nExp number of AAs in TMHs: 37.46598\nExp number, first 60 AAs: 19.53139\nTotal prob of N-in: 0.94364\nPOSSIBLE N-term signal sequence\ninside 1 19\nTMhelix 20 39\noutside 40 325\nTMhelix 326 348\ninside 349 372

10216 GCF_000335315.2_ASM33531v2 Escherichia coli 99.1775 Proteobacteria; Gammaproteobacteria; Enterobacterales; Enterobacteriaceae; Escherichia; Escherichia coli MNNEETFYQAMRRQGVTRRSFLKYCSLAATSLGLGAGMAPKIAWALENKPRIPVVWIHGL WP_001301613.1 hydrogenase [Escherichia coli] Length: 372\nNumber of predicted TMHs: 2\nExp number of AAs in TMHs: 37.46598\nExp number, first 60 AAs: 19.53139\nTotal prob of N-in: 0.94364\nPOSSIBLE N-term signal sequence\ninside 1 19\nTMhelix 20 39\noutside 40 325\nTMhelix 326 348\ninside 349 372

10217 GCF_000335315.2_ASM33531v2 Escherichia coli 99.1775 Proteobacteria; Gammaproteobacteria; Enterobacterales; Enterobacteriaceae; Escherichia; Escherichia coli MTGDNTLIHSHGINRRDFMKLCAALAATMGLSSKAAAEMAESVTNPQRPPVIWIGAQECT WP_000145410.1 MULTISPECIES: hydrogenase-2 small chain [Proteobacteria] Length: 372\nNumber of predicted TMHs: 1\nExp number of AAs in TMHs: 23.72226\nExp number, first 60 AAs: 0.22038\nTotal prob of N-in: 0.12994\noutside 1 331\nTMhelix 332 354\ninside 355 372

10218 GCF_000335135.1_ASM33513v1 Escherichia coli 99.1781 Proteobacteria; Gammaproteobacteria; Enterobacterales; Enterobacteriaceae; Escherichia; Escherichia coli MNNEETFYQAMRRQGVTRRSFLKYCSLAATSLGLGAGMAPKIAWALENKPRIPVVWIHGL WP_001301613.1 hydrogenase [Escherichia coli] Length: 372\nNumber of predicted TMHs: 2\nExp number of AAs in TMHs: 37.46598\nExp number, first 60 AAs: 19.53139\nTotal prob of N-in: 0.94364\nPOSSIBLE N-term signal sequence\ninside 1 19\nTMhelix 20 39\noutside 40 325\nTMhelix 326 348\ninside 349 372

10219 GCF_000335135.1_ASM33513v1 Escherichia coli 99.1781 Proteobacteria; Gammaproteobacteria; Enterobacterales; Enterobacteriaceae; Escherichia; Escherichia coli MTGDNTLIHSHGINRRDFMKLCAALAATMGLSSKAAAEMAESVTNPQRPPVIWIGAQECT WP_000145410.1 MULTISPECIES: hydrogenase-2 small chain [Proteobacteria] Length: 372\nNumber of predicted TMHs: 1\nExp number of AAs in TMHs: 23.72226\nExp number, first 60 AAs: 0.22038\nTotal prob of N-in: 0.12994\noutside 1 331\nTMhelix 332 354\ninside 355 372

10220 GCF_000335015.2_ASM33501v2 Escherichia coli 99.1793 Proteobacteria; Gammaproteobacteria; Enterobacterales; Enterobacteriaceae; Escherichia; Escherichia coli MNNEETFYQAMRRQGVTRRSFLKYCSLAATSLGLGAGMAPKIAWALENKPRIPVVWIHGL WP_001301613.1 hydrogenase [Escherichia coli] Length: 372\nNumber of predicted TMHs: 2\nExp number of AAs in TMHs: 37.46598\nExp number, first 60 AAs: 19.53139\nTotal prob of N-in: 0.94364\nPOSSIBLE N-term signal sequence\ninside 1 19\nTMhelix 20 39\noutside 40 325\nTMhelix 326 348\ninside 349 372

10221 GCF_000335015.2_ASM33501v2 Escherichia coli 99.1793 Proteobacteria; Gammaproteobacteria; Enterobacterales; Enterobacteriaceae; Escherichia; Escherichia coli MTGDNTLIHSHGINRRDFMKLCAALAATMGLSSKAAAEMAESVTNPQRPPVIWIGAQECT WP_000145410.1 MULTISPECIES: hydrogenase-2 small chain [Proteobacteria] Length: 372\nNumber of predicted TMHs: 1\nExp number of AAs in TMHs: 23.72226\nExp number, first 60 AAs: 0.22038\nTotal prob of N-in: 0.12994\noutside 1 331\nTMhelix 332 354\ninside 355 372

10222 GCF_000335035.2_ASM33503v2 Escherichia coli 99.1805 Proteobacteria; Gammaproteobacteria; Enterobacterales; Enterobacteriaceae; Escherichia; Escherichia coli MNNEETFYQAMRRQGVTRRSFLKYCSLAATSLGLGAGMAPKIAWALENKPRIPVVWIHGL WP_001301613.1 hydrogenase [Escherichia coli] Length: 372\nNumber of predicted TMHs: 2\nExp number of AAs in TMHs: 37.46598\nExp number, first 60 AAs: 19.53139\nTotal prob of N-in: 0.94364\nPOSSIBLE N-term signal sequence\ninside 1 19\nTMhelix 20 39\noutside 40 325\nTMhelix 326 348\ninside 349 372

10223 GCF_000335035.2_ASM33503v2 Escherichia coli 99.1805 Proteobacteria; Gammaproteobacteria; Enterobacterales; Enterobacteriaceae; Escherichia; Escherichia coli

MTGDNTLIHSHGINRRDFMKLCAALAATMGLSSKAAAEMAESVTNPQRPPVIWIGAQECT WP_000145410.1 MULTISPECIES: hydrogenase-2 small chain [Proteobacteria] Length: 372\nNumber of predicted TMHs: 1\nExp number of AAs in TMHs: 23.72226\nExp number, first 60 AAs: 0.22038\nTotal prob of N-in: 0.12994\noutside 1 331\nTMhelix 332 354\ninside 355 372

10224 GCF_000503555.1_Ecoli9B Escherichia coli 9B Proteobacteria; Gammaproteobacteria; Enterobacterales; Enterobacteriaceae; Escherichia; Escherichia coli MTGDNTLIHSHGINRRDFMKLCAALAATMGLSSKAAAEMAESVTNPQRPPVIWIGAQECT WP_000145410.1 MULTISPECIES: hydrogenase-2 small chain [Proteobacteria] Length: 372\nNumber of predicted TMHs: 1\nExp number of AAs in TMHs: 23.72226\nExp number, first 60 AAs: 0.22038\nTotal prob of N-in: 0.12994\noutside 1 331\nTMhelix 332 354\ninside 355 372

10225 GCF_000503555.1_Ecoli9B Escherichia coli 9B Proteobacteria; Gammaproteobacteria; Enterobacterales; Enterobacteriaceae; Escherichia; Escherichia coli MNNEETFYQAMRRQGVTRRSFLKYCSLAATSLGLGAGMAPKIAWALENKPRIPVVWIHGL WP_001058323.1 MULTISPECIES: hydrogenase-1 small chain [Proteobacteria] Length: 372\nNumber of predicted TMHs: 2\nExp number of AAs in TMHs: 37.2608499999999\nExp number, first 60 AAs: 19.52923\nTotal prob of N-in: 0.94351\nPOSSIBLE N-term signal sequence\ninside 1 19\nTMhelix 20 39\noutside 40 325\nTMhelix 326 348\ninside 349 372

10226 GCF_000488815.1_ASM48881v1 Escherichia coli A25922R Proteobacteria; Gammaproteobacteria; Enterobacterales; Enterobacteriaceae; Escherichia; Escherichia coli MTGDNTLIHSHGINRRDFMKLCAALAATMGLSSKAAAEMAESVTNPQRPPVIWIGAQECT WP_000145410.1 MULTISPECIES: hydrogenase-2 small chain [Proteobacteria] Length: 372\nNumber of predicted TMHs: 1\nExp number of AAs in TMHs: 23.72226\nExp number, first 60 AAs: 0.22038\nTotal prob of N-in: 0.12994\noutside 1 331\nTMhelix 332 354\ninside 355 372

10227 GCF_000488815.1_ASM48881v1 Escherichia coli A25922R Proteobacteria; Gammaproteobacteria; Enterobacterales; Enterobacteriaceae; Escherichia; Escherichia coli MNNEETFYQAMRRQGVTRRSFLKYCSLAATSLGLGAGMAPKIAWALENKPRIPVVWIHGL WP_001058323.1 MULTISPECIES: hydrogenase-1 small chain [Proteobacteria] Length: 372\nNumber of predicted TMHs: 2\nExp number of AAs in TMHs: 37.2608499999999\nExp number, first 60 AAs: 19.52923\nTotal prob of N-in: 0.94351\nPOSSIBLE N-term signal sequence\ninside 1 19\nTMhelix 20 39\noutside 40 325\nTMhelix 326 348\ninside 349 372

10228 GCF_000488835.1_ASM48883v1 Escherichia coli A35218R Proteobacteria; Gammaproteobacteria; Enterobacterales; Enterobacteriaceae; Escherichia; Escherichia coli MNNEETFYQAMRRQGVTRRSFLKYCSLAATSLGLGAGMAPKIAWALENKPRIPVVWIHGL WP_001058323.1 MULTISPECIES: hydrogenase-1 small chain [Proteobacteria] Length: 372\nNumber of predicted TMHs: 2\nExp number of AAs in TMHs: 37.2608499999999\nExp number, first 60 AAs: 19.52923\nTotal prob of N-in: 0.94351\nPOSSIBLE N-term signal sequence\ninside 1 19\nTMhelix 20 39\noutside 40 325\nTMhelix 326 348\ninside 349 372

10229 GCF_000488835.1_ASM48883v1 Escherichia coli A35218R Proteobacteria; Gammaproteobacteria; Enterobacterales; Enterobacteriaceae; Escherichia; Escherichia coli MTGDNTLIHSHGINRRDFMKLCAALAATMGLSSKAAAEMAESVTNPQRPPVIWIGAQECT WP_000145410.1 MULTISPECIES: hydrogenase-2 small chain [Proteobacteria] Length: 372\nNumber of predicted TMHs: 1\nExp number of AAs in TMHs: 23.72226\nExp number, first 60 AAs: 0.22038\nTotal prob of N-in: 0.12994\noutside 1 331\nTMhelix 332 354\ninside 355 372

10230 GCF_000211395.1_ASM21139v1 Escherichia coli AA86 Proteobacteria; Gammaproteobacteria; Enterobacterales; Enterobacteriaceae; Escherichia; Escherichia coli MTGDNTLIHSHGINRRDFMKLCAALAATMGLSSKAAAEMAESVTNPQRPPVIWIGAQECT WP_000145410.1 MULTISPECIES: hydrogenase-2 small chain [Proteobacteria] Length: 372\nNumber of predicted TMHs: 1\nExp number of AAs in TMHs: 23.72226\nExp number, first 60 AAs: 0.22038\nTotal prob of N-in: 0.12994\noutside 1 331\nTMhelix 332 354\ninside 355 372

10231 GCF_000211395.1_ASM21139v1 Escherichia coli AA86 Proteobacteria; Gammaproteobacteria; Enterobacterales; Enterobacteriaceae; Escherichia; Escherichia coli MNNEETFYQAMRRQGVTRRSFLKYCSLAATSLGLGAGMAPKIAWALENKPRIPVVWIHGL WP_001058323.1 MULTISPECIES: hydrogenase-1 small chain [Proteobacteria] Length: 372\nNumber of predicted TMHs:

2\nExp number of AAs in TMHs: 37.2608499999999\nExp number, first 60 AAs: 19.52923\nTotal prob of N-in: 0.94351\nPOSSIBLE N-term signal sequence\ninside 1 19\nTMhelix 20 39\noutside 40 325\nTMhelix 326 348\ninside 349 372

10232 GCF_000418755.2_E_coliAB42410445-isolate1-1.0 Escherichia coli AB42410445-isolate1 Proteobacteria; Gammaproteobacteria; Enterobacterales; Enterobacteriaceae; Escherichia; Escherichia coli MNNEETFYQAMRRQGVTRRSFLKYCSLAATSLGLGAGMAPKIAWALENKPRIPVVWIHGL WP_001058323.1 MULTISPECIES: hydrogenase-1 small chain [Proteobacteria] Length: 372\nNumber of predicted TMHs: 2\nExp number of AAs in TMHs: 37.2608499999999\nExp number, first 60 AAs: 19.52923\nTotal prob of N-in: 0.94351\nPOSSIBLE N-term signal sequence\ninside 1 19\nTMhelix 20 39\noutside 40 325\nTMhelix 326 348\ninside 349 372

10233 GCF_000418755.2_E_coliAB42410445-isolate1-1.0 Escherichia coli AB42410445-isolate1 Proteobacteria; Gammaproteobacteria; Enterobacterales; Enterobacteriaceae; Escherichia; Escherichia coli MTGDNTLIHSHGINRRDFMKLCAALAATMGLSSKAAAEMAESVTNPQRPPVIWIGAQECT WP_000145410.1 MULTISPECIES: hydrogenase-2 small chain [Proteobacteria] Length: 372\nNumber of predicted TMHs: 1\nExp number of AAs in TMHs: 23.72226\nExp number, first 60 AAs: 0.22038\nTotal prob of N-in: 0.12994\noutside 1 331\nTMhelix 332 354\ninside 355 372

10234 GCF_000418735.2_E_coliAB42554418-isolate1-1.0 Escherichia coli AB42554418-isolate1 Proteobacteria; Gammaproteobacteria; Enterobacterales; Enterobacteriaceae; Escherichia; Escherichia coli MTGDNTLIHSHGINRRDFMKLCAALAATMGLSSKAAAEMAESVTNPQRPPVIWIGAQECT WP_000145410.1 MULTISPECIES: hydrogenase-2 small chain [Proteobacteria] Length: 372\nNumber of predicted TMHs: 1\nExp number of AAs in TMHs: 23.72226\nExp number, first 60 AAs: 0.22038\nTotal prob of N-in: 0.12994\noutside 1 331\nTMhelix 332 354\ninside 355 372

10235 GCF_000418735.2_E_coliAB42554418-isolate1-1.0 Escherichia coli AB42554418-isolate1 Proteobacteria; Gammaproteobacteria; Enterobacterales; Enterobacteriaceae; Escherichia; Escherichia coli MNNEETFYQAMRRQGVTRRSFLKYCSLAATSLGLGAGMAPKIAWALENKPRIPVVWIHGL WP_001058323.1 MULTISPECIES: hydrogenase-1 small chain [Proteobacteria] Length: 372\nNumber of predicted TMHs: 2\nExp number of AAs in TMHs: 37.2608499999999\nExp number, first 60 AAs: 19.52923\nTotal prob of N-in: 0.94351\nPOSSIBLE N-term signal sequence\ninside 1 19\nTMhelix 20 39\noutside 40 325\nTMhelix 326 348\ninside 349 372

10236 GCF_000418715.1_E_coliAB42602061-isolate1-1.0 Escherichia coli AB42602061-isolate1 Proteobacteria; Gammaproteobacteria; Enterobacterales; Enterobacteriaceae; Escherichia; Escherichia coli MNNEETFYQAMRRQGVTRRSFLKYCSLAATSLGLGAGMAPKIAWALENKPRIPVVWIHGL WP_001058323.1 MULTISPECIES: hydrogenase-1 small chain [Proteobacteria] Length: 372\nNumber of predicted TMHs: 2\nExp number of AAs in TMHs: 37.2608499999999\nExp number, first 60 AAs: 19.52923\nTotal prob of N-in: 0.94351\nPOSSIBLE N-term signal sequence\ninside 1 19\nTMhelix 20 39\noutside 40 325\nTMhelix 326 348\ninside 349 372

10237 GCF_000418715.1_E_coliAB42602061-isolate1-1.0 Escherichia coli AB42602061-isolate1 Proteobacteria; Gammaproteobacteria; Enterobacterales; Enterobacteriaceae; Escherichia; Escherichia coli MTGDNTLIHSHGINRRDFMKLCAALAATMGLSSKAAAEMAESVTNPQRPPVIWIGAQECT WP_000145410.1 MULTISPECIES: hydrogenase-2 small chain [Proteobacteria] Length: 372\nNumber of predicted TMHs: 1\nExp number of AAs in TMHs: 23.72226\nExp number, first 60 AAs: 0.22038\nTotal prob of N-in: 0.12994\noutside 1 331\nTMhelix 332 354\ninside 355 372

10238 GCF_000418695.2_E_coliAB43739056-isolate1-1.0 Escherichia coli AB43739056-isolate1 Proteobacteria; Gammaproteobacteria; Enterobacterales; Enterobacteriaceae; Escherichia; Escherichia coli MNNEETFYQAMRRQGVTRRSFLKYCSLAATSLGLGAGMAPKIAWALENKPRIPVVWIHGL WP_001058323.1 MULTISPECIES: hydrogenase-1 small chain [Proteobacteria] Length: 372\nNumber of predicted TMHs: 2\nExp number of AAs in TMHs: 37.2608499999999\nExp number, first 60 AAs: 19.52923\nTotal prob of N-in: 0.94351\nPOSSIBLE N-term signal sequence\ninside 1 19\nTMhelix 20 39\noutside 40 325\nTMhelix 326 348\ninside 349 372

10239 GCF_000418695.2_E_coliAB43739056-isolate1-1.0 Escherichia coli AB43739056-isolate1 Proteobacteria; Gammaproteobacteria; Enterobacterales; Enterobacteriaceae; Escherichia; Escherichia coli MTGDNTLIHSHGINRRDFMKLCAALAATMGLSSKAAAEMAESVTNPQRPPVIWIGAQECT WP_000145410.1 MULTISPECIES: hydrogenase-2 small chain [Proteobacteria] Length: 372\nNumber of predicted TMHs:

1\nExp number of AAs in TMHs: 23.72226\nExp number, first 60 AAs: 0.22038\nTotal prob of N-in: 0.12994\noutside 1 331\nTMhelix 332 354\ninside 355 372

10240 GCF_000148365.1_ASM14836v1 Escherichia coli ABU 83972 Proteobacteria; Gammaproteobacteria; Enterobacterales; Enterobacteriaceae; Escherichia; Escherichia coli
MNNEETFYQAMRRQGVTRRSFLKYCSLAATSLGLGAGMAPKIAWALENKPRIPVVWIHGL WP_001058323.1
MULTISPECIES: hydrogenase-1 small chain [Proteobacteria] Length: 372\nNumber of predicted TMHs: 2\nExp number of AAs in TMHs: 37.2608499999999\nExp number, first 60 AAs: 19.52923\nTotal prob of N-in: 0.94351\nPOSSIBLE N-term signal sequence\ninside 1 19\nTMhelix 20 39\noutside 40 325\nTMhelix 326 348\ninside 349 372

10241 GCF_000148365.1_ASM14836v1 Escherichia coli ABU 83972 Proteobacteria; Gammaproteobacteria; Enterobacterales; Enterobacteriaceae; Escherichia; Escherichia coli
MTGDNTLIHSHGINRRDFMKLCAALAATMGLSSKAAAEMAESVTNPQRPPVIWIGAQECT WP_000145410.1
MULTISPECIES: hydrogenase-2 small chain [Proteobacteria] Length: 372\nNumber of predicted TMHs: 1\nExp number of AAs in TMHs: 23.72226\nExp number, first 60 AAs: 0.22038\nTotal prob of N-in: 0.12994\noutside 1 331\nTMhelix 332 354\ninside 355 372

10242 GCF_001051135.1_ASM105113v1 Escherichia coli ACN001 Proteobacteria; Gammaproteobacteria; Enterobacterales; Enterobacteriaceae; Escherichia; Escherichia coli
MNNEETFYQAMRRQGVTRRSFLKYCSLAATSLGLGAGMAPKIAWALENKPRIPVVWIHGL WP_001058320.1
hydrogenase [Escherichia coli] Length: 372\nNumber of predicted TMHs: 2\nExp number of AAs in TMHs: 37.26082\nExp number, first 60 AAs: 19.52923\nTotal prob of N-in: 0.94351\nPOSSIBLE N-term signal sequence\ninside 1 19\nTMhelix 20 39\noutside 40 325\nTMhelix 326 348\ninside 349 372

10243 GCF_001051135.1_ASM105113v1 Escherichia coli ACN001 Proteobacteria; Gammaproteobacteria; Enterobacterales; Enterobacteriaceae; Escherichia; Escherichia coli
MTGDNTLIHSHGINRRDFMKLCAALAATMGLSSKAAAEMAESVTNPQRPPVIWIGAQECT WP_000145410.1
MULTISPECIES: hydrogenase-2 small chain [Proteobacteria] Length: 372\nNumber of predicted TMHs: 1\nExp number of AAs in TMHs: 23.72226\nExp number, first 60 AAs: 0.22038\nTotal prob of N-in: 0.12994\noutside 1 331\nTMhelix 332 354\ninside 355 372

10244 GCF_000304255.1_ECAD30.1Escherichia coli AD30 Proteobacteria; Gammaproteobacteria; Enterobacterales; Enterobacteriaceae; Escherichia; Escherichia coli
MNNEETFYQAMRRQGVTRRSFLKYCSLAATSLGLGAGMAPKIAWALENKPRIPVVWIHGL WP_001058323.1
MULTISPECIES: hydrogenase-1 small chain [Proteobacteria] Length: 372\nNumber of predicted TMHs: 2\nExp number of AAs in TMHs: 37.2608499999999\nExp number, first 60 AAs: 19.52923\nTotal prob of N-in: 0.94351\nPOSSIBLE N-term signal sequence\ninside 1 19\nTMhelix 20 39\noutside 40 325\nTMhelix 326 348\ninside 349 372

10245 GCF_000304255.1_ECAD30.1Escherichia coli AD30 Proteobacteria; Gammaproteobacteria; Enterobacterales; Enterobacteriaceae; Escherichia; Escherichia coli
MTGDNTLIHSHGINRRDFMKLCAALAATMGLSSKAAAEMAESVTNPQRPPVIWIGAQECT WP_000145410.1
MULTISPECIES: hydrogenase-2 small chain [Proteobacteria] Length: 372\nNumber of predicted TMHs: 1\nExp number of AAs in TMHs: 23.72226\nExp number, first 60 AAs: 0.22038\nTotal prob of N-in: 0.12994\noutside 1 331\nTMhelix 332 354\ninside 355 372

10246 GCF_001244915.1_1.ECAD30.1 Escherichia coli AD30 Proteobacteria; Gammaproteobacteria; Enterobacterales; Enterobacteriaceae; Escherichia; Escherichia coli
MTGDNTLIHSHGINRRDFMKLCAALAATMGLSSKAAAEMAESVTNPQRPPVIWIGAQECT WP_000145410.1
MULTISPECIES: hydrogenase-2 small chain [Proteobacteria] Length: 372\nNumber of predicted TMHs: 1\nExp number of AAs in TMHs: 23.72226\nExp number, first 60 AAs: 0.22038\nTotal prob of N-in: 0.12994\noutside 1 331\nTMhelix 332 354\ninside 355 372

10247 GCF_001244915.1_1.ECAD30.1 Escherichia coli AD30 Proteobacteria; Gammaproteobacteria; Enterobacterales; Enterobacteriaceae; Escherichia; Escherichia coli
MNNEETFYQAMRRQGVTRRSFLKYCSLAATSLGLGAGMAPKIAWALENKPRIPVVWIHGL WP_001058323.1
MULTISPECIES: hydrogenase-1 small chain [Proteobacteria] Length: 372\nNumber of predicted TMHs: 2\nExp number of AAs in TMHs: 37.2608499999999\nExp number, first 60 AAs: 19.52923\nTotal prob of N-in: 0.94351\nPOSSIBLE N-term signal sequence\ninside 1 19\nTMhelix 20 39\noutside 40 325\nTMhelix 326 348\ninside 349 372

10248 GCF_000259135.1_ECAI27.hybrid.1 Escherichia coli AI27 Proteobacteria; Gammaproteobacteria; Enterobacterales; Enterobacteriaceae; Escherichia; Escherichia coli
MNNEETFYQAMRRQGVTRRSFLKYCSLAATSLGLGAGMAPKIAWALENKPRIPVVWIHGL WP_001058323.1
MULTISPECIES: hydrogenase-1 small chain [Proteobacteria] Length: 372\nNumber of predicted TMHs: 2\nExp number of AAs in TMHs: 37.2608499999999\nExp number, first 60 AAs: 19.52923\nTotal prob of N-in: 0.94351\nPOSSIBLE N-term signal sequence\ninside 1 19\nTMhelix 20 39\noutside 40 325\nTMhelix 326 348\ninside 349 372

10249 GCF_000259135.1_ECAI27.hybrid.1 Escherichia coli AI27 Proteobacteria; Gammaproteobacteria; Enterobacterales; Enterobacteriaceae; Escherichia; Escherichia coli
MTGDNTLIHSHGINRRDFMKLCAALAATMGLSSKAAAEMAESVTNPQRPPVIWIGAQECT WP_000145410.1
MULTISPECIES: hydrogenase-2 small chain [Proteobacteria] Length: 372\nNumber of predicted TMHs: 1\nExp number of AAs in TMHs: 23.72226\nExp number, first 60 AAs: 0.22038\nTotal prob of N-in: 0.12994\noutside 1 331\nTMhelix 332 354\ninside 355 372

10250 GCF_000813165.1_ASM81316v1 Escherichia coli APEC IMT5155 Proteobacteria; Gammaproteobacteria; Enterobacterales; Enterobacteriaceae; Escherichia; Escherichia coli
MTGDNTLIHSHGINRRDFMKLCAALAATMGLSSKAAAEMAESVTNPQRPPVIWIGAQECT WP_000145410.1
MULTISPECIES: hydrogenase-2 small chain [Proteobacteria] Length: 372\nNumber of predicted TMHs: 1\nExp number of AAs in TMHs: 23.72226\nExp number, first 60 AAs: 0.22038\nTotal prob of N-in: 0.12994\noutside 1 331\nTMhelix 332 354\ninside 355 372

10251 GCF_000014845.1_ASM1484v1 Escherichia coli APEC O1 Proteobacteria; Gammaproteobacteria; Enterobacterales; Enterobacteriaceae; Escherichia; Escherichia coli
MTGDNTLIHSHGINRRDFMKLCAALAATMGLSSKAAAEMAESVTNPQRPPVIWIGAQECT WP_000145410.1
MULTISPECIES: hydrogenase-2 small chain [Proteobacteria] Length: 372\nNumber of predicted TMHs: 1\nExp number of AAs in TMHs: 23.72226\nExp number, first 60 AAs: 0.22038\nTotal prob of N-in: 0.12994\noutside 1 331\nTMhelix 332 354\ninside 355 372

10252 GCF_001021615.1_APECO18 Escherichia coli APEC O18 Proteobacteria; Gammaproteobacteria; Enterobacterales; Enterobacteriaceae; Escherichia; Escherichia coli
MTGDNTLIHSHGINRRDFMKLCAALAATMGLSSKAAAEMAESVTNPQRPPVIWIGAQECT WP_000145410.1
MULTISPECIES: hydrogenase-2 small chain [Proteobacteria] Length: 372\nNumber of predicted TMHs: 1\nExp number of AAs in TMHs: 23.72226\nExp number, first 60 AAs: 0.22038\nTotal prob of N-in: 0.12994\noutside 1 331\nTMhelix 332 354\ninside 355 372

10253 GCF_001620375.1_ASM162037v1 Escherichia coli APEC O2 Proteobacteria; Gammaproteobacteria; Enterobacterales; Enterobacteriaceae; Escherichia; Escherichia coli
MTGDNTLIHSHGINRRDFMKLCAALAATMGLSSKAAAEMVESVTNPQRPPVIWIGAQECT WP_000145423.1
hydrogenase 2 small subunit [Escherichia coli] Length: 372\nNumber of predicted TMHs: 1\nExp number of AAs in TMHs: 23.78201\nExp number, first 60 AAs: 0.28355\nTotal prob of N-in: 0.13257\noutside 1 331\nTMhelix 332 354\ninside 355 372

10254 GCF_001620375.1_ASM162037v1 Escherichia coli APEC O2 Proteobacteria; Gammaproteobacteria; Enterobacterales; Enterobacteriaceae; Escherichia; Escherichia coli
MNNEETFYQAMRRQGVTRRSFLKYCSLAATSLGLGAGMAPKIAWALENKPRIPVVWIHGL WP_001058323.1
MULTISPECIES: hydrogenase-1 small chain [Proteobacteria] Length: 372\nNumber of predicted TMHs: 2\nExp number of AAs in TMHs: 37.2608499999999\nExp number, first 60 AAs: 19.52923\nTotal prob of N-in: 0.94351\nPOSSIBLE N-term signal sequence\ninside 1 19\nTMhelix 20 39\noutside 40 325\nTMhelix 326 348\ninside 349 372

10255 GCF_001021635.1_ASM102163v1 Escherichia coli APEC O2-211 Proteobacteria; Gammaproteobacteria; Enterobacterales; Enterobacteriaceae; Escherichia; Escherichia coli
MNNEETFYQAMRRQGVTRRSFLKYCSLAATSLGLGAGMAPKIAWALENKPRIPVVWIHGL WP_001058323.1
MULTISPECIES: hydrogenase-1 small chain [Proteobacteria] Length: 372\nNumber of predicted TMHs: 2\nExp number of AAs in TMHs: 37.2608499999999\nExp number, first 60 AAs: 19.52923\nTotal prob of N-in: 0.94351\nPOSSIBLE N-term signal sequence\ninside 1 19\nTMhelix 20 39\noutside 40 325\nTMhelix 326 348\ninside 349 372

10256 GCF_001021635.1_ASM102163v1 Escherichia coli APEC O2-211 Proteobacteria; Gammaproteobacteria; Enterobacterales; Enterobacteriaceae; Escherichia; Escherichia coli

MTGDNTLIHSHGINRRDFMKLCAALAATMGLSSKAAAEMAESVTNPQRPPVIWIGAQECT WP_000145410.1 MULTISPECIES: hydrogenase-2 small chain [Proteobacteria] Length: 372\nNumber of predicted TMHs: 1\nExp number of AAs in TMHs: 23.72226\nExp number, first 60 AAs: 0.22038\nTotal prob of N-in: 0.12994\noutside 1 331\nTMhelix 332 354\ninside 355 372

10257 GCF_000332755.1_ASM33275v1 Escherichia coli APEC O78 Proteobacteria; Gammaproteobacteria; Enterobacterales; Enterobacteriaceae; Escherichia; Escherichia coli MTGDNTLIHSHGINRRDFMKLCAALAATMGLSSKAAAEMAESVTNPQRPPVIWIGAQECT WP_000145410.1 MULTISPECIES: hydrogenase-2 small chain [Proteobacteria] Length: 372\nNumber of predicted TMHs: 1\nExp number of AAs in TMHs: 23.72226\nExp number, first 60 AAs: 0.22038\nTotal prob of N-in: 0.12994\noutside 1 331\nTMhelix 332 354\ninside 355 372

10258 GCF_000332755.1_ASM33275v1 Escherichia coli APEC O78 Proteobacteria; Gammaproteobacteria; Enterobacterales; Enterobacteriaceae; Escherichia; Escherichia coli MNNEETFYQAMRRQGVTRRSFLKYCSLAATSLGLGAGMAPKIAWALENKPRIPVVWIHGL WP_001058320.1 hydrogenase [Escherichia coli] Length: 372\nNumber of predicted TMHs: 2\nExp number of AAs in TMHs: 37.26082\nExp number, first 60 AAs: 19.52923\nTotal prob of N-in: 0.94351\nPOSSIBLE N-term signal sequence\ninside 1 19\nTMhelix 20 39\noutside 40 325\nTMhelix 326 348\ninside 349 372

10259 GCF_000304095.2_ASM30409v2 Escherichia coli ARS4.2123 Proteobacteria; Gammaproteobacteria; Enterobacterales; Enterobacteriaceae; Escherichia; Escherichia coli MNNEETFYQAMRRQGVTRRSFLKYCSLAATSLGLGAGMAPKIAWALENKPRIPVVWIHGL WP_001058323.1 MULTISPECIES: hydrogenase-1 small chain [Proteobacteria] Length: 372\nNumber of predicted TMHs: 2\nExp number of AAs in TMHs: 37.2608499999999\nExp number, first 60 AAs: 19.52923\nTotal prob of N-in: 0.94351\nPOSSIBLE N-term signal sequence\ninside 1 19\nTMhelix 20 39\noutside 40 325\nTMhelix 326 348\ninside 349 372

10260 GCF_000304095.2_ASM30409v2 Escherichia coli ARS4.2123 Proteobacteria; Gammaproteobacteria; Enterobacterales; Enterobacteriaceae; Escherichia; Escherichia coli MTGDNTLIHSHGINRRDFMKLCAALAATMGLSSKAAAEMAESVTNPQRPPVIWIGAQECT WP_000145410.1 MULTISPECIES: hydrogenase-2 small chain [Proteobacteria] Length: 372\nNumber of predicted TMHs: 1\nExp number of AAs in TMHs: 23.72226\nExp number, first 60 AAs: 0.22038\nTotal prob of N-in: 0.12994\noutside 1 331\nTMhelix 332 354\ninside 355 372

10261 GCF_000401755.1_Escherichia_coli_ATCC_25922 Escherichia coli ATCC 25922 Proteobacteria; Gammaproteobacteria; Enterobacterales; Enterobacteriaceae; Escherichia; Escherichia coli MNNEETFYQAMRRQGVTRRSFLKYCSLAATSLGLGAGMAPKIAWALENKPRIPVVWIHGL WP_001058323.1 MULTISPECIES: hydrogenase-1 small chain [Proteobacteria] Length: 372\nNumber of predicted TMHs: 2\nExp number of AAs in TMHs: 37.2608499999999\nExp number, first 60 AAs: 19.52923\nTotal prob of N-in: 0.94351\nPOSSIBLE N-term signal sequence\ninside 1 19\nTMhelix 20 39\noutside 40 325\nTMhelix 326 348\ninside 349 372

10262 GCF_000743255.1_ASM74325v1 Escherichia coli ATCC 25922 Proteobacteria; Gammaproteobacteria; Enterobacterales; Enterobacteriaceae; Escherichia; Escherichia coli MNNEETFYQAMRRQGVTRRSFLKYCSLAATSLGLGAGMAPKIAWALENKPRIPVVWIHGL WP_001058323.1 MULTISPECIES: hydrogenase-1 small chain [Proteobacteria] Length: 372\nNumber of predicted TMHs: 2\nExp number of AAs in TMHs: 37.2608499999999\nExp number, first 60 AAs: 19.52923\nTotal prob of N-in: 0.94351\nPOSSIBLE N-term signal sequence\ninside 1 19\nTMhelix 20 39\noutside 40 325\nTMhelix 326 348\ninside 349 372

10263 GCF_000743255.1_ASM74325v1 Escherichia coli ATCC 25922 Proteobacteria; Gammaproteobacteria; Enterobacterales; Enterobacteriaceae; Escherichia; Escherichia coli MTGDNTLIHSHGINRRDFMKLCAALAATMGLSSKAAAEMAESVTNPQRPPVIWIGAQECT WP_000145410.1 MULTISPECIES: hydrogenase-2 small chain [Proteobacteria] Length: 372\nNumber of predicted TMHs: 1\nExp number of AAs in TMHs: 23.72226\nExp number, first 60 AAs: 0.22038\nTotal prob of N-in: 0.12994\noutside 1 331\nTMhelix 332 354\ninside 355 372

10264 GCF_000401755.1_Escherichia_coli_ATCC_25922 Escherichia coli ATCC 25922 Proteobacteria; Gammaproteobacteria; Enterobacterales; Enterobacteriaceae; Escherichia; Escherichia coli MTGDNTLIHSHGINRRDFMKLCAALAATMGLSSKAAAEMAESVTNPQRPPVIWIGAQECT WP_000145410.1 MULTISPECIES: hydrogenase-2 small chain [Proteobacteria] Length: 372\nNumber of predicted TMHs:

1\nExp number of AAs in TMHs: 23.72226\nExp number, first 60 AAs: 0.22038\nTotal prob of N-in: 0.12994\noutside 1 331\nTMhelix 332 354\ninside 355 372

10265 GCF_000473725.2_O199v4 Escherichia coli ATCC 35150 Proteobacteria; Gammaproteobacteria; Enterobacterales; Enterobacteriaceae; Escherichia; Escherichia coli
MTGDNTLIHSHGINRRDFMKLCAALAATMGLSSKAAAEMAESVTNPQRPPVIWIGAQECT WP_000145410.1
MULTISPECIES: hydrogenase-2 small chain [Proteobacteria] Length: 372\nNumber of predicted TMHs: 1\nExp number of AAs in TMHs: 23.72226\nExp number, first 60 AAs: 0.22038\nTotal prob of N-in: 0.12994\noutside 1 331\nTMhelix 332 354\ninside 355 372

10266 GCF_000473725.2_O199v4 Escherichia coli ATCC 35150 Proteobacteria; Gammaproteobacteria; Enterobacterales; Enterobacteriaceae; Escherichia; Escherichia coli
MNNEETFYQAMRRQGVTRRSFLKYCSLAATSLGLGAGMAPKIAWALENKPRIPVVWIHGL WP_001058323.1
MULTISPECIES: hydrogenase-1 small chain [Proteobacteria] Length: 372\nNumber of predicted TMHs: 2\nExp number of AAs in TMHs: 37.2608499999999\nExp number, first 60 AAs: 19.52923\nTotal prob of N-in: 0.94351\nPOSSIBLE N-term signal sequence\ninside 1 19\nTMhelix 20 39\noutside 40 325\nTMhelix 326 348\ninside 349 372

10267 GCF_000496345.2_O455v2 Escherichia coli ATCC BAA-2192 Proteobacteria; Gammaproteobacteria; Enterobacterales; Enterobacteriaceae; Escherichia; Escherichia coli
MNNEETFYQAMRRQGVTRRSFLKYCSLAATSLGLGAGMAPKIAWALENKPRIPVVWIHGL WP_001301613.1
hydrogenase [Escherichia coli] Length: 372\nNumber of predicted TMHs: 2\nExp number of AAs in TMHs: 37.46598\nExp number, first 60 AAs: 19.53139\nTotal prob of N-in: 0.94364\nPOSSIBLE N-term signal sequence\ninside 1 19\nTMhelix 20 39\noutside 40 325\nTMhelix 326 348\ninside 349 372

10268 GCF_000496345.2_O455v2 Escherichia coli ATCC BAA-2192 Proteobacteria; Gammaproteobacteria; Enterobacterales; Enterobacteriaceae; Escherichia; Escherichia coli
MTGDNTLIHSHGINRRDFMKLCAALAATMGLSSKAAAEMAESVTNPQRPPVIWIGAQECT WP_000145410.1
MULTISPECIES: hydrogenase-2 small chain [Proteobacteria] Length: 372\nNumber of predicted TMHs: 1\nExp number of AAs in TMHs: 23.72226\nExp number, first 60 AAs: 0.22038\nTotal prob of N-in: 0.12994\noutside 1 331\nTMhelix 332 354\ninside 355 372

10269 GCF_000496365.2_Q456v4 Escherichia coli ATCC BAA-2193 Proteobacteria; Gammaproteobacteria; Enterobacterales; Enterobacteriaceae; Escherichia; Escherichia coli
MTGDNTLIHSHGINRRDFMKLCAALAATMGLSSKAAAEMAESVTNPQRPPVIWIGAQECT WP_000145410.1
MULTISPECIES: hydrogenase-2 small chain [Proteobacteria] Length: 372\nNumber of predicted TMHs: 1\nExp number of AAs in TMHs: 23.72226\nExp number, first 60 AAs: 0.22038\nTotal prob of N-in: 0.12994\noutside 1 331\nTMhelix 332 354\ninside 355 372

10270 GCF_000496365.2_Q456v4 Escherichia coli ATCC BAA-2193 Proteobacteria; Gammaproteobacteria; Enterobacterales; Enterobacteriaceae; Escherichia; Escherichia coli
MNNEETFYQAMRRQGVTRRSFLKYCSLAATSLGLGAGMAPKIAWALENKPRIPVVWIHGL WP_001058323.1
MULTISPECIES: hydrogenase-1 small chain [Proteobacteria] Length: 372\nNumber of predicted TMHs: 2\nExp number of AAs in TMHs: 37.2608499999999\nExp number, first 60 AAs: 19.52923\nTotal prob of N-in: 0.94351\nPOSSIBLE N-term signal sequence\ninside 1 19\nTMhelix 20 39\noutside 40 325\nTMhelix 326 348\ninside 349 372

10271 GCF_000508365.1_Q457 Escherichia coli ATCC BAA-2196 Proteobacteria; Gammaproteobacteria; Enterobacterales; Enterobacteriaceae; Escherichia; Escherichia coli
MTGDNTLIHSHGINRRDFMKLCAALAATMGLSSKAAAEMAESVTNPQRPPVIWIGAQECT WP_000145410.1
MULTISPECIES: hydrogenase-2 small chain [Proteobacteria] Length: 372\nNumber of predicted TMHs: 1\nExp number of AAs in TMHs: 23.72226\nExp number, first 60 AAs: 0.22038\nTotal prob of N-in: 0.12994\noutside 1 331\nTMhelix 332 354\ninside 355 372

10272 GCF_000508365.1_Q457 Escherichia coli ATCC BAA-2196 Proteobacteria; Gammaproteobacteria; Enterobacterales; Enterobacteriaceae; Escherichia; Escherichia coli
MNNEETFYQAMRRQGVTRRSFLKYCSLAATSLGLGAGMAPKIAWALENKPRIPVVWIHGL WP_001058323.1
MULTISPECIES: hydrogenase-1 small chain [Proteobacteria] Length: 372\nNumber of predicted TMHs: 2\nExp number of AAs in TMHs: 37.2608499999999\nExp number, first 60 AAs: 19.52923\nTotal prob of N-in: 0.94351\nPOSSIBLE N-term signal sequence\ninside 1 19\nTMhelix 20 39\noutside 40 325\nTMhelix 326 348\ninside 349 372

10273 GCF_000506825.1_Q458v4 Escherichia coli ATCC BAA-2209 Proteobacteria; Gammaproteobacteria; Enterobacterales; Enterobacteriaceae; Escherichia; Escherichia coli
MNNEETFYQAMRRQGVTRRSFLKYCSLAATSLGLGAGMAPKIAWALENKPRIPVVWIHGL WP_001058323.1
MULTISPECIES: hydrogenase-1 small chain [Proteobacteria] Length: 372\nNumber of predicted TMHs: 2\nExp number of AAs in TMHs: 37.2608499999999\nExp number, first 60 AAs: 19.52923\nTotal prob of N-in: 0.94351\nPOSSIBLE N-term signal sequence\ninside 1 19\nTMhelix 20 39\noutside 40 325\nTMhelix 326 348\ninside 349 372

10274 GCF_000506825.1_Q458v4 Escherichia coli ATCC BAA-2209 Proteobacteria; Gammaproteobacteria; Enterobacterales; Enterobacteriaceae; Escherichia; Escherichia coli
MTGDNTLIHSHGINRRDFMKLCAALAATMGLSSKAAAEMAESVTNPQRPPVIWIGAQECT WP_000145410.1
MULTISPECIES: hydrogenase-2 small chain [Proteobacteria] Length: 372\nNumber of predicted TMHs: 1\nExp number of AAs in TMHs: 23.72226\nExp number, first 60 AAs: 0.22038\nTotal prob of N-in: 0.12994\noutside 1 331\nTMhelix 332 354\ninside 355 372

10275 GCF_000506845.1_Q459v3 Escherichia coli ATCC BAA-2215 Proteobacteria; Gammaproteobacteria; Enterobacterales; Enterobacteriaceae; Escherichia; Escherichia coli
MTGDNTLIHSHGINRRDFMKLCAALAATMGLSSKAAAEMAESVTNPQRPPVIWIGAQECT WP_000145410.1
MULTISPECIES: hydrogenase-2 small chain [Proteobacteria] Length: 372\nNumber of predicted TMHs: 1\nExp number of AAs in TMHs: 23.72226\nExp number, first 60 AAs: 0.22038\nTotal prob of N-in: 0.12994\noutside 1 331\nTMhelix 332 354\ninside 355 372

10276 GCF_000506845.1_Q459v3 Escherichia coli ATCC BAA-2215 Proteobacteria; Gammaproteobacteria; Enterobacterales; Enterobacteriaceae; Escherichia; Escherichia coli
MNNEETFYQAMRRQGVTRRSFLKYCSLAATSLGLGAGIAPKIAWALENKPRIPVVWIHGL WP_023982790.1
hydrogenase [Escherichia coli] Length: 372\nNumber of predicted TMHs: 2\nExp number of AAs in TMHs: 37.8310099999999\nExp number, first 60 AAs: 20.15722\nTotal prob of N-in: 0.95896\nPOSSIBLE N-term signal sequence\ninside 1 20\nTMhelix 21 43\noutside 44 325\nTMhelix 326 348\ninside 349 372

10277 GCF_000508385.1_Q460 Escherichia coli ATCC BAA-2219 Proteobacteria; Gammaproteobacteria; Enterobacterales; Enterobacteriaceae; Escherichia; Escherichia coli
MTGDNTLIHSHGINRRDFMKLCAALAATMGLSSKAAAEMAESVTNPQRPPVIWIGAQECT WP_000145410.1
MULTISPECIES: hydrogenase-2 small chain [Proteobacteria] Length: 372\nNumber of predicted TMHs: 1\nExp number of AAs in TMHs: 23.72226\nExp number, first 60 AAs: 0.22038\nTotal prob of N-in: 0.12994\noutside 1 331\nTMhelix 332 354\ninside 355 372

10278 GCF_001559635.1_ASM155963v1 Escherichia coli B Proteobacteria; Gammaproteobacteria; Enterobacterales; Enterobacteriaceae; Escherichia; Escherichia coli
MNNEETFYQAMRRQGVTRRSFLKYCSLAATSLGLGAGMAPKIAWALENKPRIPVVWIHGL WP_001058323.1
MULTISPECIES: hydrogenase-1 small chain [Proteobacteria] Length: 372\nNumber of predicted TMHs: 2\nExp number of AAs in TMHs: 37.2608499999999\nExp number, first 60 AAs: 19.52923\nTotal prob of N-in: 0.94351\nPOSSIBLE N-term signal sequence\ninside 1 19\nTMhelix 20 39\noutside 40 325\nTMhelix 326 348\ninside 349 372

10279 GCF_001559635.1_ASM155963v1 Escherichia coli B Proteobacteria; Gammaproteobacteria; Enterobacterales; Enterobacteriaceae; Escherichia; Escherichia coli
MTGDNTLIHSHGINRRDFMKLCAALAATMGLSSKAAAEMAESVTNPQRPPVIWIGAQECT WP_000145410.1
MULTISPECIES: hydrogenase-2 small chain [Proteobacteria] Length: 372\nNumber of predicted TMHs: 1\nExp number of AAs in TMHs: 23.72226\nExp number, first 60 AAs: 0.22038\nTotal prob of N-in: 0.12994\noutside 1 331\nTMhelix 332 354\ninside 355 372

10280 GCF_001559615.2_ASM155961v2 Escherichia coli B Proteobacteria; Gammaproteobacteria; Enterobacterales; Enterobacteriaceae; Escherichia; Escherichia coli
MNNEETFYQAMRRQGVTRRSFLKYCSLAATSLGLGAGMAPKIAWALENKPRIPVVWIHGL WP_001058323.1
MULTISPECIES: hydrogenase-1 small chain [Proteobacteria] Length: 372\nNumber of predicted TMHs: 2\nExp number of AAs in TMHs: 37.2608499999999\nExp number, first 60 AAs: 19.52923\nTotal prob of N-in: 0.94351\nPOSSIBLE N-term signal sequence\ninside 1 19\nTMhelix 20 39\noutside 40 325\nTMhelix 326 348\ninside 349 372

10281 GCF_001559615.2_ASM155961v2 Escherichia coli B Proteobacteria; Gammaproteobacteria; Enterobacterales; Enterobacteriaceae; Escherichia; Escherichia coli

MTGDNTLIHSHGINRRDFMKLCAALAATMGLSSKAAAEMAESVTNPQRPPVIWIGAQECT WP_000145410.1 MULTISPECIES: hydrogenase-2 small chain [Proteobacteria] Length: 372\nNumber of predicted TMHs: 1\nExp number of AAs in TMHs: 23.72226\nExp number, first 60 AAs: 0.22038\nTotal prob of N-in: 0.12994\noutside 1 331\nTMhelix 332 354\ninside 355 372

10282 GCF_000163155.1_ASM16315v1 Escherichia coli B088 Proteobacteria; Gammaproteobacteria; Enterobacterales; Enterobacteriaceae; Escherichia; Escherichia coli MNNEETFYQAMRRQGVTRRSFLKYCSLAATSLGLGAGMAPKIAWALENKPRIPVVWIHGL WP_001295940.1 MULTISPECIES: hydrogenase [Enterobacteriaceae] Length: 372\nNumber of predicted TMHs: 2\nExp number of AAs in TMHs: 37.2608899999999\nExp number, first 60 AAs: 19.52923\nTotal prob of N-in: 0.94351\nPOSSIBLE N-term signal sequence\ninside 1 19\nTMhelix 20 39\noutside 40 325\nTMhelix 326 348\ninside 349 372

10283 GCF_000163155.1_ASM16315v1 Escherichia coli B088 Proteobacteria; Gammaproteobacteria; Enterobacterales; Enterobacteriaceae; Escherichia; Escherichia coli MTGDNTLIHSHGINRRDFMKLCAALAATMGLSSKAAAEMAESVTNPQRPPVIWIGAQECT WP_000145410.1 MULTISPECIES: hydrogenase-2 small chain [Proteobacteria] Length: 372\nNumber of predicted TMHs: 1\nExp number of AAs in TMHs: 23.72226\nExp number, first 60 AAs: 0.22038\nTotal prob of N-in: 0.12994\noutside 1 331\nTMhelix 332 354\ninside 355 372

10284 GCF_000242015.1_Esch_coli_B093_V1 Escherichia coli B093 Proteobacteria; Gammaproteobacteria; Enterobacterales; Enterobacteriaceae; Escherichia; Escherichia coli MNNEETFYQAMRRQGVTRRSFLKYCSLAATSLGLGAGMAPKIAWALENKPRIPVVWIHGL WP_001058323.1 MULTISPECIES: hydrogenase-1 small chain [Proteobacteria] Length: 372\nNumber of predicted TMHs: 2\nExp number of AAs in TMHs: 37.2608499999999\nExp number, first 60 AAs: 19.52923\nTotal prob of N-in: 0.94351\nPOSSIBLE N-term signal sequence\ninside 1 19\nTMhelix 20 39\noutside 40 325\nTMhelix 326 348\ninside 349 372

10285 GCF_000242015.1_Esch_coli_B093_V1 Escherichia coli B093 Proteobacteria; Gammaproteobacteria; Enterobacterales; Enterobacteriaceae; Escherichia; Escherichia coli MTGDNTLIHSHGINRRDFMKLCAALAATMGLSSKAAAEMAESVTNPQRPPVIWIGAQECT WP_000145410.1 MULTISPECIES: hydrogenase-2 small chain [Proteobacteria] Length: 372\nNumber of predicted TMHs: 1\nExp number of AAs in TMHs: 23.72226\nExp number, first 60 AAs: 0.22038\nTotal prob of N-in: 0.12994\noutside 1 331\nTMhelix 332 354\ninside 355 372

10286 GCF_000461955.2_ASM46195v2 Escherichia coli B102 Proteobacteria; Gammaproteobacteria; Enterobacterales; Enterobacteriaceae; Escherichia; Escherichia coli MNNEETFYQAMRRQGVTRRSFLKYCSLAATSLGLGAGMAPKIAWALENKPRIPVVWIHGL WP_001301613.1 hydrogenase [Escherichia coli] Length: 372\nNumber of predicted TMHs: 2\nExp number of AAs in TMHs: 37.46598\nExp number, first 60 AAs: 19.53139\nTotal prob of N-in: 0.94364\nPOSSIBLE N-term signal sequence\ninside 1 19\nTMhelix 20 39\noutside 40 325\nTMhelix 326 348\ninside 349 372

10287 GCF_000461955.2_ASM46195v2 Escherichia coli B102 Proteobacteria; Gammaproteobacteria; Enterobacterales; Enterobacteriaceae; Escherichia; Escherichia coli MTGDNTLIHSHGINRRDFMKLCAALAATMGLSSKAAAEMAESVTNPQRPPVIWIGAQECT WP_000145410.1 MULTISPECIES: hydrogenase-2 small chain [Proteobacteria] Length: 372\nNumber of predicted TMHs: 1\nExp number of AAs in TMHs: 23.72226\nExp number, first 60 AAs: 0.22038\nTotal prob of N-in: 0.12994\noutside 1 331\nTMhelix 332 354\ninside 355 372

10288 GCF_000462485.2_ASM46248v2 Escherichia coli B103 Proteobacteria; Gammaproteobacteria; Enterobacterales; Enterobacteriaceae; Escherichia; Escherichia coli MTGDNTLIHSHGINRRDFMKLCAALAATMGLSSKAAAEMAESVTNPQRPPVIWIGAQECT WP_000145410.1 MULTISPECIES: hydrogenase-2 small chain [Proteobacteria] Length: 372\nNumber of predicted TMHs: 1\nExp number of AAs in TMHs: 23.72226\nExp number, first 60 AAs: 0.22038\nTotal prob of N-in: 0.12994\noutside 1 331\nTMhelix 332 354\ninside 355 372

10289 GCF_000462485.2_ASM46248v2 Escherichia coli B103 Proteobacteria; Gammaproteobacteria; Enterobacterales; Enterobacteriaceae; Escherichia; Escherichia coli MNNEETFYQAMRRQGVTRRSFLKYCSLAATSLGLGAGMAPKIAWALENKPRIPVVWIHGL WP_001301613.1 hydrogenase [Escherichia coli] Length: 372\nNumber of predicted TMHs: 2\nExp number of AAs in

TMHs: 37.46598\nExp number, first 60 AAs: 19.53139\nTotal prob of N-in: 0.94364\nPOSSIBLE N-term signal sequence\ninside 1 19\nTMhelix 20 39\noutside 40 325\nTMhelix 326 348\ninside 349 372

10290 GCF_000462505.2_ASM46250v2 Escherichia coli B104 Proteobacteria; Gammaproteobacteria; Enterobacterales; Enterobacteriaceae; Escherichia; Escherichia coli MTGDNTLIHSHGINRRDFMKLCAALAATMGLSSKAAAEMAESVTNPQRPPVIWIGAQECT WP_000145410.1 MULTISPECIES: hydrogenase-2 small chain [Proteobacteria] Length: 372\nNumber of predicted TMHs: 1\nExp number of AAs in TMHs: 23.72226\nExp number, first 60 AAs: 0.22038\nTotal prob of N-in: 0.12994\noutside 1 331\nTMhelix 332 354\ninside 355 372

10291 GCF_000462505.2_ASM46250v2 Escherichia coli B104 Proteobacteria; Gammaproteobacteria; Enterobacterales; Enterobacteriaceae; Escherichia; Escherichia coli MNNEETFYQAMRRQGVTRRSFLKYCSLAATSLGLGAGMAPKIAWALENKPRIPVVWIHGL WP_001301613.1 hydrogenase [Escherichia coli] Length: 372\nNumber of predicted TMHs: 2\nExp number of AAs in TMHs: 37.46598\nExp number, first 60 AAs: 19.53139\nTotal prob of N-in: 0.94364\nPOSSIBLE N-term signal sequence\ninside 1 19\nTMhelix 20 39\noutside 40 325\nTMhelix 326 348\ninside 349 372

10292 GCF_000462525.2_ASM46252v2 Escherichia coli B105 Proteobacteria; Gammaproteobacteria; Enterobacterales; Enterobacteriaceae; Escherichia; Escherichia coli MTGDNTLIHSHGINRRDFMKLCAALAATMGLSSKAAAEMAESVTNPQRPPVIWIGAQECT WP_000145410.1 MULTISPECIES: hydrogenase-2 small chain [Proteobacteria] Length: 372\nNumber of predicted TMHs: 1\nExp number of AAs in TMHs: 23.72226\nExp number, first 60 AAs: 0.22038\nTotal prob of N-in: 0.12994\noutside 1 331\nTMhelix 332 354\ninside 355 372

10293 GCF_000462525.2_ASM46252v2 Escherichia coli B105 Proteobacteria; Gammaproteobacteria; Enterobacterales; Enterobacteriaceae; Escherichia; Escherichia coli MNNEETFYQAMRRQGVTRRSFLKYCSLAATSLGLGAGMAPKIAWALENKPRIPVVWIHGL WP_001301613.1 hydrogenase [Escherichia coli] Length: 372\nNumber of predicted TMHs: 2\nExp number of AAs in TMHs: 37.46598\nExp number, first 60 AAs: 19.53139\nTotal prob of N-in: 0.94364\nPOSSIBLE N-term signal sequence\ninside 1 19\nTMhelix 20 39\noutside 40 325\nTMhelix 326 348\ninside 349 372

10294 GCF_000462545.2_ASM46254v2 Escherichia coli B106 Proteobacteria; Gammaproteobacteria; Enterobacterales; Enterobacteriaceae; Escherichia; Escherichia coli MTGDNTLIHSHGINRRDFMKLCAALAATMGLSSKAAAEMAESVTNPQRPPVIWIGAQECT WP_000145410.1 MULTISPECIES: hydrogenase-2 small chain [Proteobacteria] Length: 372\nNumber of predicted TMHs: 1\nExp number of AAs in TMHs: 23.72226\nExp number, first 60 AAs: 0.22038\nTotal prob of N-in: 0.12994\noutside 1 331\nTMhelix 332 354\ninside 355 372

10295 GCF_000462545.2_ASM46254v2 Escherichia coli B106 Proteobacteria; Gammaproteobacteria; Enterobacterales; Enterobacteriaceae; Escherichia; Escherichia coli MNNEETFYQAMRRQGVTRRSFLKYCSLAATSLGLGAGMAPKIAWALENKPRIPVVWIHGL WP_001301613.1 hydrogenase [Escherichia coli] Length: 372\nNumber of predicted TMHs: 2\nExp number of AAs in TMHs: 37.46598\nExp number, first 60 AAs: 19.53139\nTotal prob of N-in: 0.94364\nPOSSIBLE N-term signal sequence\ninside 1 19\nTMhelix 20 39\noutside 40 325\nTMhelix 326 348\ninside 349 372

10296 GCF_000461995.2_ASM46199v2 Escherichia coli B107 Proteobacteria; Gammaproteobacteria; Enterobacterales; Enterobacteriaceae; Escherichia; Escherichia coli MTGDNTLIHSHGINRRDFMKLCAALAATMGLSSKAAAEMAESVTNPQRPPVIWIGAQECT WP_000145410.1 MULTISPECIES: hydrogenase-2 small chain [Proteobacteria] Length: 372\nNumber of predicted TMHs: 1\nExp number of AAs in TMHs: 23.72226\nExp number, first 60 AAs: 0.22038\nTotal prob of N-in: 0.12994\noutside 1 331\nTMhelix 332 354\ninside 355 372

10297 GCF_000461995.2_ASM46199v2 Escherichia coli B107 Proteobacteria; Gammaproteobacteria; Enterobacterales; Enterobacteriaceae; Escherichia; Escherichia coli MNNEETFYQAMRRQGVTRRSFLKYCSLAATSLGLGAGMAPKIAWALENKPRIPVVWIHGL WP_001301613.1 hydrogenase [Escherichia coli] Length: 372\nNumber of predicted TMHs: 2\nExp number of AAs in TMHs: 37.46598\nExp number, first 60 AAs: 19.53139\nTotal prob of N-in: 0.94364\nPOSSIBLE N-term signal sequence\ninside 1 19\nTMhelix 20 39\noutside 40 325\nTMhelix 326 348\ninside 349 372

10298 GCF_002109615.1_ASM210961v1 Escherichia coli B108 Proteobacteria; Gammaproteobacteria; Enterobacterales; Enterobacteriaceae; Escherichia; Escherichia coli

MTGDNTLIHSHGINRRDFMKLCAALAATMGLSSKAAAEMAESVTNPQRPPVIWIGAQECT WP_000145410.1 MULTISPECIES: hydrogenase-2 small chain [Proteobacteria] Length: 372\nNumber of predicted TMHs: 1\nExp number of AAs in TMHs: 23.72226\nExp number, first 60 AAs: 0.22038\nTotal prob of N-in: 0.12994\noutside 1 331\nTMhelix 332 354\ninside 355 372

10299 GCF_002109615.1_ASM210961v1 Escherichia coli B108 Proteobacteria; Gammaproteobacteria; Enterobacterales; Enterobacteriaceae; Escherichia; Escherichia coli MNNEETFYQAMRRQGVTRRSFLKYCSLAATSLGLGAGMAPKIAWALENKPRIPVVWIHGL WP_001058323.1 MULTISPECIES: hydrogenase-1 small chain [Proteobacteria] Length: 372\nNumber of predicted TMHs: 2\nExp number of AAs in TMHs: 37.2608499999999\nExp number, first 60 AAs: 19.52923\nTotal prob of N-in: 0.94351\nPOSSIBLE N-term signal sequence\ninside 1 19\nTMhelix 20 39\noutside 40 325\nTMhelix 326 348\ninside 349 372

10300 GCF_000462565.2_ASM46256v2 Escherichia coli B108 Proteobacteria; Gammaproteobacteria; Enterobacterales; Enterobacteriaceae; Escherichia; Escherichia coli MTGDNTLIHSHGINRRDFMKLCAALAATMGLSSKAAAEMAESVTNPQRPPVIWIGAQECT WP_000145410.1 MULTISPECIES: hydrogenase-2 small chain [Proteobacteria] Length: 372\nNumber of predicted TMHs: 1\nExp number of AAs in TMHs: 23.72226\nExp number, first 60 AAs: 0.22038\nTotal prob of N-in: 0.12994\noutside 1 331\nTMhelix 332 354\ninside 355 372

10301 GCF_000462565.2_ASM46256v2 Escherichia coli B108 Proteobacteria; Gammaproteobacteria; Enterobacterales; Enterobacteriaceae; Escherichia; Escherichia coli MNNEETFYQAMRRQGVTRRSFLKYCSLAATSLGLGAGMAPKIAWALENKPRIPVVWIHGL WP_001301613.1 hydrogenase [Escherichia coli] Length: 372\nNumber of predicted TMHs: 2\nExp number of AAs in TMHs: 37.46598\nExp number, first 60 AAs: 19.53139\nTotal prob of N-in: 0.94364\nPOSSIBLE N-term signal sequence\ninside 1 19\nTMhelix 20 39\noutside 40 325\nTMhelix 326 348\ninside 349 372

10302 GCF_000462585.2_ASM46258v2 Escherichia coli B109 Proteobacteria; Gammaproteobacteria; Enterobacterales; Enterobacteriaceae; Escherichia; Escherichia coli MNNEETFYQAMRRQGVTRRSFLKYCSLAATSLGLGAGMAPKIAWALENKPRIPVVWIHGL WP_001301613.1 hydrogenase [Escherichia coli] Length: 372\nNumber of predicted TMHs: 2\nExp number of AAs in TMHs: 37.46598\nExp number, first 60 AAs: 19.53139\nTotal prob of N-in: 0.94364\nPOSSIBLE N-term signal sequence\ninside 1 19\nTMhelix 20 39\noutside 40 325\nTMhelix 326 348\ninside 349 372

10303 GCF_000462585.2_ASM46258v2 Escherichia coli B109 Proteobacteria; Gammaproteobacteria; Enterobacterales; Enterobacteriaceae; Escherichia; Escherichia coli MTGDNTLIHSHGINRRDFMKLCAALAATMGLSSKAAAEMAESVTNPQRPPVIWIGAQECT WP_000145410.1 MULTISPECIES: hydrogenase-2 small chain [Proteobacteria] Length: 372\nNumber of predicted TMHs: 1\nExp number of AAs in TMHs: 23.72226\nExp number, first 60 AAs: 0.22038\nTotal prob of N-in: 0.12994\noutside 1 331\nTMhelix 332 354\ninside 355 372

10304 GCF_000462605.2_ASM46260v2 Escherichia coli B112 Proteobacteria; Gammaproteobacteria; Enterobacterales; Enterobacteriaceae; Escherichia; Escherichia coli MTGDNTLIHSHGINRRDFMKLCAALAATMGLSSKAAAEMAESVTNPQRPPVIWIGAQECT WP_000145410.1 MULTISPECIES: hydrogenase-2 small chain [Proteobacteria] Length: 372\nNumber of predicted TMHs: 1\nExp number of AAs in TMHs: 23.72226\nExp number, first 60 AAs: 0.22038\nTotal prob of N-in: 0.12994\noutside 1 331\nTMhelix 332 354\ninside 355 372

10305 GCF_000462605.2_ASM46260v2 Escherichia coli B112 Proteobacteria; Gammaproteobacteria; Enterobacterales; Enterobacteriaceae; Escherichia; Escherichia coli MNNEETFYQAMRRQGVTRRSFLKYCSLAATSLGLGAGMAPKIAWALENKPRIPVVWIHGL WP_001301613.1 hydrogenase [Escherichia coli] Length: 372\nNumber of predicted TMHs: 2\nExp number of AAs in TMHs: 37.46598\nExp number, first 60 AAs: 19.53139\nTotal prob of N-in: 0.94364\nPOSSIBLE N-term signal sequence\ninside 1 19\nTMhelix 20 39\noutside 40 325\nTMhelix 326 348\ninside 349 372

10306 GCF_000462625.2_ASM46262v2 Escherichia coli B113 Proteobacteria; Gammaproteobacteria; Enterobacterales; Enterobacteriaceae; Escherichia; Escherichia coli MNNEETFYQAMRRQGVTRRSFLKYCSLAATSLGLGAGMAPKIAWALENKPRIPVVWIHGL WP_001301613.1 hydrogenase [Escherichia coli] Length: 372\nNumber of predicted TMHs: 2\nExp number of AAs in TMHs: 37.46598\nExp number, first 60 AAs: 19.53139\nTotal prob of N-in: 0.94364\nPOSSIBLE N-term signal sequence\ninside 1 19\nTMhelix 20 39\noutside 40 325\nTMhelix 326 348\ninside 349 372

10307 GCF_000462625.2_ASM46262v2 Escherichia coli B113 Proteobacteria; Gammaproteobacteria; Enterobacterales; Enterobacteriaceae; Escherichia; Escherichia coli
MTGDNTLIHSHGINRRDFMKLCAALAATMGLSSKAAAEMAESVTNPQRPPVIWIGAQECT WP_000145410.1
MULTISPECIES: hydrogenase-2 small chain [Proteobacteria] Length: 372\nNumber of predicted TMHs: 1\nExp number of AAs in TMHs: 23.72226\nExp number, first 60 AAs: 0.22038\nTotal prob of N-in: 0.12994\noutside 1 331\nTMhelix 332 354\ninside 355 372

10308 GCF_000462645.2_ASM46264v2 Escherichia coli B114 Proteobacteria; Gammaproteobacteria; Enterobacterales; Enterobacteriaceae; Escherichia; Escherichia coli
MTGDNTLIHSHGINRRDFMKLCAALAATMGLSSKAAAEMAESVTNPQRPPVIWIGAQECT WP_000145410.1
MULTISPECIES: hydrogenase-2 small chain [Proteobacteria] Length: 372\nNumber of predicted TMHs: 1\nExp number of AAs in TMHs: 23.72226\nExp number, first 60 AAs: 0.22038\nTotal prob of N-in: 0.12994\noutside 1 331\nTMhelix 332 354\ninside 355 372

10309 GCF_000462645.2_ASM46264v2 Escherichia coli B114 Proteobacteria; Gammaproteobacteria; Enterobacterales; Enterobacteriaceae; Escherichia; Escherichia coli
MNNEETFYQAMRRQGVTRRSFLKYCSLAATSLGLGAGMAPKIAWALENKPRIPVVWIHGL WP_001301613.1
hydrogenase [Escherichia coli] Length: 372\nNumber of predicted TMHs: 2\nExp number of AAs in TMHs: 37.46598\nExp number, first 60 AAs: 19.53139\nTotal prob of N-in: 0.94364\nPOSSIBLE N-term signal sequence\ninside 1 19\nTMhelix 20 39\noutside 40 325\nTMhelix 326 348\ninside 349 372

10310 GCF_000462185.2_ASM46218v2 Escherichia coli B15Proteobacteria; Gammaproteobacteria; Enterobacterales; Enterobacteriaceae; Escherichia; Escherichia coli
MTGDNTLIHSHGINRRDFMKLCAALAATMGLSSKAAAEMAESVTNPQRPPVIWIGAQECT WP_000145410.1
MULTISPECIES: hydrogenase-2 small chain [Proteobacteria] Length: 372\nNumber of predicted TMHs: 1\nExp number of AAs in TMHs: 23.72226\nExp number, first 60 AAs: 0.22038\nTotal prob of N-in: 0.12994\noutside 1 331\nTMhelix 332 354\ninside 355 372

10311 GCF_000462185.2_ASM46218v2 Escherichia coli B15Proteobacteria; Gammaproteobacteria; Enterobacterales; Enterobacteriaceae; Escherichia; Escherichia coli
MNNEETFYQAMRRQGVTRRSFLKYCSLAATSLGLGAGMAPKIAWALENKPRIPVVWIHGL WP_001301613.1
hydrogenase [Escherichia coli] Length: 372\nNumber of predicted TMHs: 2\nExp number of AAs in TMHs: 37.46598\nExp number, first 60 AAs: 19.53139\nTotal prob of N-in: 0.94364\nPOSSIBLE N-term signal sequence\ninside 1 19\nTMhelix 20 39\noutside 40 325\nTMhelix 326 348\ninside 349 372

10312 GCF_000462665.2_ASM46266v2 Escherichia coli B17Proteobacteria; Gammaproteobacteria; Enterobacterales; Enterobacteriaceae; Escherichia; Escherichia coli
MTGDNTLIHSHGINRRDFMKLCAALAATMGLSSKAAAEMAESVTNPQRPPVIWIGAQECT WP_000145410.1
MULTISPECIES: hydrogenase-2 small chain [Proteobacteria] Length: 372\nNumber of predicted TMHs: 1\nExp number of AAs in TMHs: 23.72226\nExp number, first 60 AAs: 0.22038\nTotal prob of N-in: 0.12994\noutside 1 331\nTMhelix 332 354\ninside 355 372

10313 GCF_000462665.2_ASM46266v2 Escherichia coli B17Proteobacteria; Gammaproteobacteria; Enterobacterales; Enterobacteriaceae; Escherichia; Escherichia coli
MNNEETFYQAMRRQGVTRRSFLKYCSLAATSLGLGAGMAPKIAWALENKPRIPVVWIHGL WP_001301613.1
hydrogenase [Escherichia coli] Length: 372\nNumber of predicted TMHs: 2\nExp number of AAs in TMHs: 37.46598\nExp number, first 60 AAs: 19.53139\nTotal prob of N-in: 0.94364\nPOSSIBLE N-term signal sequence\ninside 1 19\nTMhelix 20 39\noutside 40 325\nTMhelix 326 348\ninside 349 372

10314 GCF_000167895.2_ASM16789v2 Escherichia coli B171 Proteobacteria; Gammaproteobacteria; Enterobacterales; Enterobacteriaceae; Escherichia; Escherichia coli
MNNEETFYQAMRRQGVTRRSFLKYCSLAATSLGLGAGMAPKIAWALENKPRIPVVWIHGL WP_001295940.1
MULTISPECIES: hydrogenase [Enterobacteriaceae] Length: 372\nNumber of predicted TMHs: 2\nExp number of AAs in TMHs: 37.2608899999999\nExp number, first 60 AAs: 19.52923\nTotal prob of N-in: 0.94351\nPOSSIBLE N-term signal sequence\ninside 1 19\nTMhelix 20 39\noutside 40 325\nTMhelix 326 348\ninside 349 372

10315 GCF_000167895.2_ASM16789v2 Escherichia coli B171 Proteobacteria; Gammaproteobacteria; Enterobacterales; Enterobacteriaceae; Escherichia; Escherichia coli
MTGDNTLIHSHGINRRDFMKLCAALAATMGLSSKAAAEMAESVTNPQRPPVIWIGAQECT WP_000145410.1
MULTISPECIES: hydrogenase-2 small chain [Proteobacteria] Length: 372\nNumber of predicted TMHs:

1\nExp number of AAs in TMHs: 23.72226\nExp number, first 60 AAs: 0.22038\nTotal prob of N-in: 0.12994\noutside 1 331\nTMhelix 332 354\ninside 355 372

10316 GCF_002109815.1_ASM210981v1 Escherichia coli B175 Proteobacteria; Gammaproteobacteria; Enterobacterales; Enterobacteriaceae; Escherichia; Escherichia coli MTGDNTLIHSHGINRRDFMKLCAALAATMGLSSKAAAEMAESVTNPQRPPVIWIGAQECT WP_000145410.1 MULTISPECIES: hydrogenase-2 small chain [Proteobacteria] Length: 372\nNumber of predicted TMHs: 1\nExp number of AAs in TMHs: 23.72226\nExp number, first 60 AAs: 0.22038\nTotal prob of N-in: 0.12994\noutside 1 331\nTMhelix 332 354\ninside 355 372

10317 GCF_002109815.1_ASM210981v1 Escherichia coli B175 Proteobacteria; Gammaproteobacteria; Enterobacterales; Enterobacteriaceae; Escherichia; Escherichia coli MNNEETFYQAMRRQGVTRRSFLKYCSLAATSLGLGAGMAPKIAWALENKPRIPVVWIHGL WP_001058323.1 MULTISPECIES: hydrogenase-1 small chain [Proteobacteria] Length: 372\nNumber of predicted TMHs: 2\nExp number of AAs in TMHs: 37.2608499999999\nExp number, first 60 AAs: 19.52923\nTotal prob of N-in: 0.94351\nPOSSIBLE N-term signal sequence\ninside 1 19\nTMhelix 20 39\noutside 40 325\nTMhelix 326 348\ninside 349 372

10318 GCF_000163175.1_ASM16317v1 Escherichia coli B185 Proteobacteria; Gammaproteobacteria; Enterobacterales; Enterobacteriaceae; Escherichia; Escherichia coli MNNEETFYQAMRRQGVTRRSFLKYCSLAATSLGLGAGMAPKIAWALENKPRIPVVWIHGL WP_001058323.1 MULTISPECIES: hydrogenase-1 small chain [Proteobacteria] Length: 372\nNumber of predicted TMHs: 2\nExp number of AAs in TMHs: 37.2608499999999\nExp number, first 60 AAs: 19.52923\nTotal prob of N-in: 0.94351\nPOSSIBLE N-term signal sequence\ninside 1 19\nTMhelix 20 39\noutside 40 325\nTMhelix 326 348\ninside 349 372

10319 GCF_000163175.1_ASM16317v1 Escherichia coli B185 Proteobacteria; Gammaproteobacteria; Enterobacterales; Enterobacteriaceae; Escherichia; Escherichia coli MTGDNTLIHSHGINRRDFMKLCAALAATMGLSSKAAAEMAESVTNPQRPPVIWIGAQECT WP_000145410.1 MULTISPECIES: hydrogenase-2 small chain [Proteobacteria] Length: 372\nNumber of predicted TMHs: 1\nExp number of AAs in TMHs: 23.72226\nExp number, first 60 AAs: 0.22038\nTotal prob of N-in: 0.12994\noutside 1 331\nTMhelix 332 354\ninside 355 372

10320 GCF_000462025.2_ASM46202v2 Escherichia coli B26-1 Proteobacteria; Gammaproteobacteria; Enterobacterales; Enterobacteriaceae; Escherichia; Escherichia coli MNNEETFYQAMRRQGVTRRSFLKYCSLAATSLGLGAGMAPKIAWALENKPRIPVVWIHGL WP_001301613.1 hydrogenase [Escherichia coli] Length: 372\nNumber of predicted TMHs: 2\nExp number of AAs in TMHs: 37.46598\nExp number, first 60 AAs: 19.53139\nTotal prob of N-in: 0.94364\nPOSSIBLE N-term signal sequence\ninside 1 19\nTMhelix 20 39\noutside 40 325\nTMhelix 326 348\ninside 349 372

10321 GCF_000462025.2_ASM46202v2 Escherichia coli B26-1 Proteobacteria; Gammaproteobacteria; Enterobacterales; Enterobacteriaceae; Escherichia; Escherichia coli MTGDNTLIHSHGINRRDFMKLCAALAATMGLSSKAAAEMAESVTNPQRPPVIWIGAQECT WP_000145410.1 MULTISPECIES: hydrogenase-2 small chain [Proteobacteria] Length: 372\nNumber of predicted TMHs: 1\nExp number of AAs in TMHs: 23.72226\nExp number, first 60 AAs: 0.22038\nTotal prob of N-in: 0.12994\noutside 1 331\nTMhelix 332 354\ninside 355 372

10322 GCF_000461915.2_ASM46191v2 Escherichia coli B26-2 Proteobacteria; Gammaproteobacteria; Enterobacterales; Enterobacteriaceae; Escherichia; Escherichia coli MTGDNTLIHSHGINRRDFMKLCAALAATMGLSSKAAAEMAESVTNPQRPPVIWIGAQECT WP_000145410.1 MULTISPECIES: hydrogenase-2 small chain [Proteobacteria] Length: 372\nNumber of predicted TMHs: 1\nExp number of AAs in TMHs: 23.72226\nExp number, first 60 AAs: 0.22038\nTotal prob of N-in: 0.12994\noutside 1 331\nTMhelix 332 354\ninside 355 372

10323 GCF_000461915.2_ASM46191v2 Escherichia coli B26-2 Proteobacteria; Gammaproteobacteria; Enterobacterales; Enterobacteriaceae; Escherichia; Escherichia coli MNNEETFYQAMRRQGVTRRSFLKYCSLAATSLGLGAGMAPKIAWALENKPRIPVVWIHGL WP_001301613.1 hydrogenase [Escherichia coli] Length: 372\nNumber of predicted TMHs: 2\nExp number of AAs in TMHs: 37.46598\nExp number, first 60 AAs: 19.53139\nTotal prob of N-in: 0.94364\nPOSSIBLE N-term signal sequence\ninside 1 19\nTMhelix 20 39\noutside 40 325\nTMhelix 326 348\ninside 349 372

10324	GCF_000462045.2_ASM46204v2	Escherichia coli B28-1	Proteobacteria; Gammaproteobacteria; Enterobacterales; Enterobacteriaceae; Escherichia; Escherichia coli
MTGDNTLIHSHGINRRDFMKLCAALAATMGLSSKAAAEMAESVTNPQRPPVIWIGAQECT	WP_000145410.1
MULTISPECIES: hydrogenase-2 small chain [Proteobacteria]	Length: 372\nNumber of predicted TMHs: 1\nExp number of AAs in TMHs: 23.72226\nExp number, first 60 AAs: 0.22038\nTotal prob of N-in: 0.12994\noutside 1 331\nTMhelix 332 354\ninside 355 372

10325	GCF_000462045.2_ASM46204v2	Escherichia coli B28-1	Proteobacteria; Gammaproteobacteria; Enterobacterales; Enterobacteriaceae; Escherichia; Escherichia coli
MNNEETFYQAMRRQGVTRRSFLKYCSLAATSLGLGAGMAPKIAWALENKPRIPVVWIHGL	WP_001301613.1
hydrogenase [Escherichia coli]	Length: 372\nNumber of predicted TMHs: 2\nExp number of AAs in TMHs: 37.46598\nExp number, first 60 AAs: 19.53139\nTotal prob of N-in: 0.94364\nPOSSIBLE N-term signal sequence\ninside 1 19\nTMhelix 20 39\noutside 40 325\nTMhelix 326 348\ninside 349 372

10326	GCF_000477495.2_ASM47749v2	Escherichia coli B28-2	Proteobacteria; Gammaproteobacteria; Enterobacterales; Enterobacteriaceae; Escherichia; Escherichia coli
MTGDNTLIHSHGINRRDFMKLCAALAATMGLSSKAAAEMAESVTNPQRPPVIWIGAQECT	WP_000145410.1
MULTISPECIES: hydrogenase-2 small chain [Proteobacteria]	Length: 372\nNumber of predicted TMHs: 1\nExp number of AAs in TMHs: 23.72226\nExp number, first 60 AAs: 0.22038\nTotal prob of N-in: 0.12994\noutside 1 331\nTMhelix 332 354\ninside 355 372

10327	GCF_000477495.2_ASM47749v2	Escherichia coli B28-2	Proteobacteria; Gammaproteobacteria; Enterobacterales; Enterobacteriaceae; Escherichia; Escherichia coli
MNNEETFYQAMRRQGVTRRSFLKYCSLAATSLGLGAGMAPKIAWALENKPRIPVVWIHGL	WP_001301613.1
hydrogenase [Escherichia coli]	Length: 372\nNumber of predicted TMHs: 2\nExp number of AAs in TMHs: 37.46598\nExp number, first 60 AAs: 19.53139\nTotal prob of N-in: 0.94364\nPOSSIBLE N-term signal sequence\ninside 1 19\nTMhelix 20 39\noutside 40 325\nTMhelix 326 348\ninside 349 372

10328	GCF_000461975.2_ASM46197v2	Escherichia coli B29-1	Proteobacteria; Gammaproteobacteria; Enterobacterales; Enterobacteriaceae; Escherichia; Escherichia coli
MNNEETFYQAMRRQGVTRRSFLKYCSLAATSLGLGAGMAPKIAWALENKPRIPVVWIHGL	WP_001301613.1
hydrogenase [Escherichia coli]	Length: 372\nNumber of predicted TMHs: 2\nExp number of AAs in TMHs: 37.46598\nExp number, first 60 AAs: 19.53139\nTotal prob of N-in: 0.94364\nPOSSIBLE N-term signal sequence\ninside 1 19\nTMhelix 20 39\noutside 40 325\nTMhelix 326 348\ninside 349 372

10329	GCF_000461975.2_ASM46197v2	Escherichia coli B29-1	Proteobacteria; Gammaproteobacteria; Enterobacterales; Enterobacteriaceae; Escherichia; Escherichia coli
MTGDNTLIHSHGINRRDFMKLCAALAATMGLSSKAAAEMAESVTNPQRPPVIWIGAQECT	WP_000145410.1
MULTISPECIES: hydrogenase-2 small chain [Proteobacteria]	Length: 372\nNumber of predicted TMHs: 1\nExp number of AAs in TMHs: 23.72226\nExp number, first 60 AAs: 0.22038\nTotal prob of N-in: 0.12994\noutside 1 331\nTMhelix 332 354\ninside 355 372

10330	GCF_000462105.2_ASM46210v2	Escherichia coli B29-2	Proteobacteria; Gammaproteobacteria; Enterobacterales; Enterobacteriaceae; Escherichia; Escherichia coli
MTGDNTLIHSHGINRRDFMKLCAALAATMGLSSKAAAEMAESVTNPQRPPVIWIGAQECT	WP_000145410.1
MULTISPECIES: hydrogenase-2 small chain [Proteobacteria]	Length: 372\nNumber of predicted TMHs: 1\nExp number of AAs in TMHs: 23.72226\nExp number, first 60 AAs: 0.22038\nTotal prob of N-in: 0.12994\noutside 1 331\nTMhelix 332 354\ninside 355 372

10331	GCF_000462105.2_ASM46210v2	Escherichia coli B29-2	Proteobacteria; Gammaproteobacteria; Enterobacterales; Enterobacteriaceae; Escherichia; Escherichia coli
MNNEETFYQAMRRQGVTRRSFLKYCSLAATSLGLGAGMAPKIAWALENKPRIPVVWIHGL	WP_001301613.1
hydrogenase [Escherichia coli]	Length: 372\nNumber of predicted TMHs: 2\nExp number of AAs in TMHs: 37.46598\nExp number, first 60 AAs: 19.53139\nTotal prob of N-in: 0.94364\nPOSSIBLE N-term signal sequence\ninside 1 19\nTMhelix 20 39\noutside 40 325\nTMhelix 326 348\ninside 349 372

10332	GCF_000163195.1_ASM16319v1	Escherichia coli B354	Proteobacteria; Gammaproteobacteria; Enterobacterales; Enterobacteriaceae; Escherichia; Escherichia coli
MTGDNTLIHSHGINRRDFMKLCAALAATMGLSSKAAAEMAESVTNPQRPPVIWIGAQECT	WP_000145410.1
MULTISPECIES: hydrogenase-2 small chain [Proteobacteria]	Length: 372\nNumber of predicted TMHs:

1\nExp number of AAs in TMHs: 23.72226\nExp number, first 60 AAs: 0.22038\nTotal prob of N-in: 0.12994\noutside 1 331\nTMhelix 332 354\ninside 355 372

10333 GCF_000163195.1_ASM16319v1 Escherichia coli B354 Proteobacteria; Gammaproteobacteria; Enterobacterales; Enterobacteriaceae; Escherichia; Escherichia coli
MNNEETFYQAMRRQGVTRRSFLKYCSLAATSLGLGAGMAPKIAWALENKPRIPVVWIHGL WP_001058323.1
MULTISPECIES: hydrogenase-1 small chain [Proteobacteria] Length: 372\nNumber of predicted TMHs: 2\nExp number of AAs in TMHs: 37.2608499999999\nExp number, first 60 AAs: 19.52923\nTotal prob of N-in: 0.94351\nPOSSIBLE N-term signal sequence\ninside 1 19\nTMhelix 20 39\noutside 40 325\nTMhelix 326 348\ninside 349 372

10334 GCF_000462125.2_ASM46212v2 Escherichia coli B36-1 Proteobacteria; Gammaproteobacteria; Enterobacterales; Enterobacteriaceae; Escherichia; Escherichia coli
MNNEETFYQAMRRQGVTRRSFLKYCSLAATSLGLGAGMAPKIAWALENKPRIPVVWIHGL WP_001301613.1
hydrogenase [Escherichia coli] Length: 372\nNumber of predicted TMHs: 2\nExp number of AAs in TMHs: 37.46598\nExp number, first 60 AAs: 19.53139\nTotal prob of N-in: 0.94364\nPOSSIBLE N-term signal sequence\ninside 1 19\nTMhelix 20 39\noutside 40 325\nTMhelix 326 348\ninside 349 372

10335 GCF_000462125.2_ASM46212v2 Escherichia coli B36-1 Proteobacteria; Gammaproteobacteria; Enterobacterales; Enterobacteriaceae; Escherichia; Escherichia coli
MTGDNTLIHSHGINRRDFMKLCAALAATMGLSSKAAAEMAESVTNPQRPPVIWIGAQECT WP_000145410.1
MULTISPECIES: hydrogenase-2 small chain [Proteobacteria] Length: 372\nNumber of predicted TMHs: 1\nExp number of AAs in TMHs: 23.72226\nExp number, first 60 AAs: 0.22038\nTotal prob of N-in: 0.12994\noutside 1 331\nTMhelix 332 354\ninside 355 372

10336 GCF_000462065.2_ASM46206v2 Escherichia coli B36-2 Proteobacteria; Gammaproteobacteria; Enterobacterales; Enterobacteriaceae; Escherichia; Escherichia coli
MNNEETFYQAMRRQGVTRRSFLKYCSLAATSLGLGAGMAPKIAWALENKPRIPVVWIHGL WP_001301613.1
hydrogenase [Escherichia coli] Length: 372\nNumber of predicted TMHs: 2\nExp number of AAs in TMHs: 37.46598\nExp number, first 60 AAs: 19.53139\nTotal prob of N-in: 0.94364\nPOSSIBLE N-term signal sequence\ninside 1 19\nTMhelix 20 39\noutside 40 325\nTMhelix 326 348\ninside 349 372

10337 GCF_000462065.2_ASM46206v2 Escherichia coli B36-2 Proteobacteria; Gammaproteobacteria; Enterobacterales; Enterobacteriaceae; Escherichia; Escherichia coli
MTGDNTLIHSHGINRRDFMKLCAALAATMGLSSKAAAEMAESVTNPQRPPVIWIGAQECT WP_000145410.1
MULTISPECIES: hydrogenase-2 small chain [Proteobacteria] Length: 372\nNumber of predicted TMHs: 1\nExp number of AAs in TMHs: 23.72226\nExp number, first 60 AAs: 0.22038\nTotal prob of N-in: 0.12994\noutside 1 331\nTMhelix 332 354\ninside 355 372

10338 GCF_002110145.1_ASM211014v1 Escherichia coli B367 Proteobacteria; Gammaproteobacteria; Enterobacterales; Enterobacteriaceae; Escherichia; Escherichia coli
MTGDNTLIHSHGINRRDFMKLCAALAATMGLSSKAAAEMAESVTNPQRPPVIWIGAQECT WP_000145410.1
MULTISPECIES: hydrogenase-2 small chain [Proteobacteria] Length: 372\nNumber of predicted TMHs: 1\nExp number of AAs in TMHs: 23.72226\nExp number, first 60 AAs: 0.22038\nTotal prob of N-in: 0.12994\noutside 1 331\nTMhelix 332 354\ninside 355 372

10339 GCF_002110145.1_ASM211014v1 Escherichia coli B367 Proteobacteria; Gammaproteobacteria; Enterobacterales; Enterobacteriaceae; Escherichia; Escherichia coli
MNNEETFYQAMRRQGVTRRSFLKYCSLAATSLGLGAGMAPKIAWALENKPRIPVVWIHGL WP_001058323.1
MULTISPECIES: hydrogenase-1 small chain [Proteobacteria] Length: 372\nNumber of predicted TMHs: 2\nExp number of AAs in TMHs: 37.2608499999999\nExp number, first 60 AAs: 19.52923\nTotal prob of N-in: 0.94351\nPOSSIBLE N-term signal sequence\ninside 1 19\nTMhelix 20 39\noutside 40 325\nTMhelix 326 348\ninside 349 372

10340 GCF_000462685.2_ASM46268v2 Escherichia coli B40-1 Proteobacteria; Gammaproteobacteria; Enterobacterales; Enterobacteriaceae; Escherichia; Escherichia coli
MNNEETFYQAMRRQGVTRRSFLKYCSLAATSLGLGAGMAPKIAWALENKPRIPVVWIHGL WP_001301613.1
hydrogenase [Escherichia coli] Length: 372\nNumber of predicted TMHs: 2\nExp number of AAs in TMHs: 37.46598\nExp number, first 60 AAs: 19.53139\nTotal prob of N-in: 0.94364\nPOSSIBLE N-term signal sequence\ninside 1 19\nTMhelix 20 39\noutside 40 325\nTMhelix 326 348\ninside 349 372

10341	GCF_000462685.2_ASM46268v2	Escherichia coli B40-1	Proteobacteria; Gammaproteobacteria; Enterobacterales; Enterobacteriaceae; Escherichia; Escherichia coli
	MTGDNTLIHSHGINRRDFMKLCAALAATMGLSSKAAAEMAESVTNPQRPPVIWIGAQECT	WP_000145410.1
	MULTISPECIES: hydrogenase-2 small chain [Proteobacteria]	Length: 372\nNumber of predicted TMHs: 1\nExp number of AAs in TMHs: 23.72226\nExp number, first 60 AAs: 0.22038\nTotal prob of N-in: 0.12994\noutside 1 331\nTMhelix 332 354\ninside 355 372

10342	GCF_000462705.2_ASM46270v2	Escherichia coli B40-2	Proteobacteria; Gammaproteobacteria; Enterobacterales; Enterobacteriaceae; Escherichia; Escherichia coli
	MTGDNTLIHSHGINRRDFMKLCAALAATMGLSSKAAAEMAESVTNPQRPPVIWIGAQECT	WP_000145410.1
	MULTISPECIES: hydrogenase-2 small chain [Proteobacteria]	Length: 372\nNumber of predicted TMHs: 1\nExp number of AAs in TMHs: 23.72226\nExp number, first 60 AAs: 0.22038\nTotal prob of N-in: 0.12994\noutside 1 331\nTMhelix 332 354\ninside 355 372

10343	GCF_000462705.2_ASM46270v2	Escherichia coli B40-2	Proteobacteria; Gammaproteobacteria; Enterobacterales; Enterobacteriaceae; Escherichia; Escherichia coli
	MNNEETFYQAMRRQGVTRRSFLKYCSLAATSLGLGAGMAPKIAWALENKPRIPVVWIHGL	WP_001301613.1
	hydrogenase [Escherichia coli]	Length: 372\nNumber of predicted TMHs: 2\nExp number of AAs in TMHs: 37.46598\nExp number, first 60 AAs: 19.53139\nTotal prob of N-in: 0.94364\nPOSSIBLE N-term signal sequence\ninside 1 19\nTMhelix 20 39\noutside 40 325\nTMhelix 326 348\ninside 349 372

10344	GCF_000194705.1_ASM19470v2	Escherichia coli B41Proteobacteria; Gammaproteobacteria; Enterobacterales; Enterobacteriaceae; Escherichia; Escherichia coli
	MTGDNTLIHSHGINRRDFMKLCAALAATMGLSSKAAAEMAESVTNPQRPPVIWIGAQECT	WP_000145410.1
	MULTISPECIES: hydrogenase-2 small chain [Proteobacteria]	Length: 372\nNumber of predicted TMHs: 1\nExp number of AAs in TMHs: 23.72226\nExp number, first 60 AAs: 0.22038\nTotal prob of N-in: 0.12994\noutside 1 331\nTMhelix 332 354\ninside 355 372

10345	GCF_000194705.1_ASM19470v2	Escherichia coli B41Proteobacteria; Gammaproteobacteria; Enterobacterales; Enterobacteriaceae; Escherichia; Escherichia coli
	MNNEETFYQAMRRQGVTRRSFLKYCSLAATSLGLGAGMAPKIAWALENKPRIPVVWIHGL	WP_001058323.1
	MULTISPECIES: hydrogenase-1 small chain [Proteobacteria]	Length: 372\nNumber of predicted TMHs: 2\nExp number of AAs in TMHs: 37.2608499999999\nExp number, first 60 AAs: 19.52923\nTotal prob of N-in: 0.94351\nPOSSIBLE N-term signal sequence\ninside 1 19\nTMhelix 20 39\noutside 40 325\nTMhelix 326 348\ninside 349 372

10346	GCF_000462725.2_ASM46272v2	Escherichia coli B49-2	Proteobacteria; Gammaproteobacteria; Enterobacterales; Enterobacteriaceae; Escherichia; Escherichia coli
	MTGDNTLIHSHGINRRDFMKLCAALAATMGLSSKAAAEMAESVTNPQRPPVIWIGAQECT	WP_000145410.1
	MULTISPECIES: hydrogenase-2 small chain [Proteobacteria]	Length: 372\nNumber of predicted TMHs: 1\nExp number of AAs in TMHs: 23.72226\nExp number, first 60 AAs: 0.22038\nTotal prob of N-in: 0.12994\noutside 1 331\nTMhelix 332 354\ninside 355 372

10347	GCF_000462725.2_ASM46272v2	Escherichia coli B49-2	Proteobacteria; Gammaproteobacteria; Enterobacterales; Enterobacteriaceae; Escherichia; Escherichia coli
	MNNEETFYQAMRRQGVTRRSFLKYCSLAATSLGLGAGMAPKIAWALENKPRIPVVWIHGL	WP_001301613.1
	hydrogenase [Escherichia coli]	Length: 372\nNumber of predicted TMHs: 2\nExp number of AAs in TMHs: 37.46598\nExp number, first 60 AAs: 19.53139\nTotal prob of N-in: 0.94364\nPOSSIBLE N-term signal sequence\ninside 1 19\nTMhelix 20 39\noutside 40 325\nTMhelix 326 348\ninside 349 372

10348	GCF_000462745.2_ASM46274v2	Escherichia coli B5-2	Proteobacteria; Gammaproteobacteria; Enterobacterales; Enterobacteriaceae; Escherichia; Escherichia coli
	MTGDNTLIHSHGINRRDFMKLCAALAATMGLSSKAAAEMAESVTNPQRPPVIWIGAQECT	WP_000145410.1
	MULTISPECIES: hydrogenase-2 small chain [Proteobacteria]	Length: 372\nNumber of predicted TMHs: 1\nExp number of AAs in TMHs: 23.72226\nExp number, first 60 AAs: 0.22038\nTotal prob of N-in: 0.12994\noutside 1 331\nTMhelix 332 354\ninside 355 372

10349	GCF_000462745.2_ASM46274v2	Escherichia coli B5-2	Proteobacteria; Gammaproteobacteria; Enterobacterales; Enterobacteriaceae; Escherichia; Escherichia coli
	MNNEETFYQAMRRQGVTRRSFLKYCSLAATSLGLGAGMAPKIAWALENKPRIPVVWIHGL	WP_001301613.1
	hydrogenase [Escherichia coli]	Length: 372\nNumber of predicted TMHs: 2\nExp number of AAs in

TMHs: 37.46598\nExp number, first 60 AAs: 19.53139\nTotal prob of N-in: 0.94364\nPOSSIBLE N-term signal sequence\ninside 1 19\nTMhelix 20 39\noutside 40 325\nTMhelix 326 348\ninside 349 372

10350 GCF_002110045.1_ASM211004v1 Escherichia coli B574 Proteobacteria; Gammaproteobacteria; Enterobacterales; Enterobacteriaceae; Escherichia; Escherichia coli MNNEETFYQAMRRQGVTRRSFLKYCSLAATSLGLGAGMAPKIAWALENKPRIPVVWIHGL WP_001058323.1 MULTISPECIES: hydrogenase-1 small chain [Proteobacteria] Length: 372\nNumber of predicted TMHs: 2\nExp number of AAs in TMHs: 37.2608499999999\nExp number, first 60 AAs: 19.52923\nTotal prob of N-in: 0.94351\nPOSSIBLE N-term signal sequence\ninside 1 19\nTMhelix 20 39\noutside 40 325\nTMhelix 326 348\ninside 349 372

10351 GCF_002110045.1_ASM211004v1 Escherichia coli B574 Proteobacteria; Gammaproteobacteria; Enterobacterales; Enterobacteriaceae; Escherichia; Escherichia coli MTGDNTLIHSHGINRRDFMKLCAALAATMGLSSKAAAEMAESVTNPQRPPVIWIGAQECT WP_000145410.1 MULTISPECIES: hydrogenase-2 small chain [Proteobacteria] Length: 372\nNumber of predicted TMHs: 1\nExp number of AAs in TMHs: 23.72226\nExp number, first 60 AAs: 0.22038\nTotal prob of N-in: 0.12994\noutside 1 331\nTMhelix 332 354\ninside 355 372

10352 GCF_002109655.1_ASM210965v1 Escherichia coli B671 Proteobacteria; Gammaproteobacteria; Enterobacterales; Enterobacteriaceae; Escherichia; Escherichia coli MTGDNTLIHSHGINRRDFMKLCAALAATMGLSSKAAAEMAESVTNPQRPPVIWIGAQECT WP_000145410.1 MULTISPECIES: hydrogenase-2 small chain [Proteobacteria] Length: 372\nNumber of predicted TMHs: 1\nExp number of AAs in TMHs: 23.72226\nExp number, first 60 AAs: 0.22038\nTotal prob of N-in: 0.12994\noutside 1 331\nTMhelix 332 354\ninside 355 372

10353 GCF_002109655.1_ASM210965v1 Escherichia coli B671 Proteobacteria; Gammaproteobacteria; Enterobacterales; Enterobacteriaceae; Escherichia; Escherichia coli MNNEETFYQAMRRQGVTRRSFLKYCSLAATSLGLGAGMAPKIAWALENKPRIPVVWIHGL WP_001058323.1 MULTISPECIES: hydrogenase-1 small chain [Proteobacteria] Length: 372\nNumber of predicted TMHs: 2\nExp number of AAs in TMHs: 37.2608499999999\nExp number, first 60 AAs: 19.52923\nTotal prob of N-in: 0.94351\nPOSSIBLE N-term signal sequence\ninside 1 19\nTMhelix 20 39\noutside 40 325\nTMhelix 326 348\ninside 349 372

10354 GCF_000462085.2_ASM46208v2 Escherichia coli B7-1 Proteobacteria; Gammaproteobacteria; Enterobacterales; Enterobacteriaceae; Escherichia; Escherichia coli MNNEETFYQAMRRQGVTRRSFLKYCSLAATSLGLGAGMAPKIAWALENKPRIPVVWIHGL WP_001301613.1 hydrogenase [Escherichia coli] Length: 372\nNumber of predicted TMHs: 2\nExp number of AAs in TMHs: 37.46598\nExp number, first 60 AAs: 19.53139\nTotal prob of N-in: 0.94364\nPOSSIBLE N-term signal sequence\ninside 1 19\nTMhelix 20 39\noutside 40 325\nTMhelix 326 348\ninside 349 372

10355 GCF_000462085.2_ASM46208v2 Escherichia coli B7-1 Proteobacteria; Gammaproteobacteria; Enterobacterales; Enterobacteriaceae; Escherichia; Escherichia coli MTGDNTLIHSHGINRRDFMKLCAALAATMGLSSKAAAEMAESVTNPQRPPVIWIGAQECT WP_000145410.1 MULTISPECIES: hydrogenase-2 small chain [Proteobacteria] Length: 372\nNumber of predicted TMHs: 1\nExp number of AAs in TMHs: 23.72226\nExp number, first 60 AAs: 0.22038\nTotal prob of N-in: 0.12994\noutside 1 331\nTMhelix 332 354\ninside 355 372

10356 GCF_000462405.2_ASM46240v2 Escherichia coli B7-2 Proteobacteria; Gammaproteobacteria; Enterobacterales; Enterobacteriaceae; Escherichia; Escherichia coli MTGDNTLIHSHGINRRDFMKLCAALAATMGLSSKAAAEMAESVTNPQRPPVIWIGAQECT WP_000145410.1 MULTISPECIES: hydrogenase-2 small chain [Proteobacteria] Length: 372\nNumber of predicted TMHs: 1\nExp number of AAs in TMHs: 23.72226\nExp number, first 60 AAs: 0.22038\nTotal prob of N-in: 0.12994\noutside 1 331\nTMhelix 332 354\ninside 355 372

10357 GCF_000462405.2_ASM46240v2 Escherichia coli B7-2 Proteobacteria; Gammaproteobacteria; Enterobacterales; Enterobacteriaceae; Escherichia; Escherichia coli MNNEETFYQAMRRQGVTRRSFLKYCSLAATSLGLGAGMAPKIAWALENKPRIPVVWIHGL WP_001301613.1 hydrogenase [Escherichia coli] Length: 372\nNumber of predicted TMHs: 2\nExp number of AAs in TMHs: 37.46598\nExp number, first 60 AAs: 19.53139\nTotal prob of N-in: 0.94364\nPOSSIBLE N-term signal sequence\ninside 1 19\nTMhelix 20 39\noutside 40 325\nTMhelix 326 348\ninside 349 372

10358 GCF_000261385.1_Esch_coli_B799_V1 Escherichia coli B799 Proteobacteria; Gammaproteobacteria; Enterobacterales; Enterobacteriaceae; Escherichia; Escherichia coli
MNNEETFYQAMRRQGVTRRSFLKYCSLAATSLGLGAGMAPKIAWALENKPRIPVVWIHGL WP_001058323.1
MULTISPECIES: hydrogenase-1 small chain [Proteobacteria] Length: 372\nNumber of predicted TMHs: 2\nExp number of AAs in TMHs: 37.2608499999999\nExp number, first 60 AAs: 19.52923\nTotal prob of N-in: 0.94351\nPOSSIBLE N-term signal sequence\ninside 1 19\nTMhelix 20 39\noutside 40 325\nTMhelix 326 348\ninside 349 372

10359 GCF_000261385.1_Esch_coli_B799_V1 Escherichia coli B799 Proteobacteria; Gammaproteobacteria; Enterobacterales; Enterobacteriaceae; Escherichia; Escherichia coli
MTGDNTLIHSHGINRRDFMKLCAALAATMGLSSKAAAEMAESVTNPQRPPVIWIGAQECT WP_000145410.1
MULTISPECIES: hydrogenase-2 small chain [Proteobacteria] Length: 372\nNumber of predicted TMHs: 1\nExp number of AAs in TMHs: 23.72226\nExp number, first 60 AAs: 0.22038\nTotal prob of N-in: 0.12994\noutside 1 331\nTMhelix 332 354\ninside 355 372

10360 GCF_000167815.1_ASM16781v1 Escherichia coli B7A Proteobacteria; Gammaproteobacteria; Enterobacterales; Enterobacteriaceae; Escherichia; Escherichia coli
MTGDNTLIHSHGINRRDFMKLCAALAATMGLSSKAAAEMAESVTNPQRPPVIWIGAQECT WP_000145410.1
MULTISPECIES: hydrogenase-2 small chain [Proteobacteria] Length: 372\nNumber of predicted TMHs: 1\nExp number of AAs in TMHs: 23.72226\nExp number, first 60 AAs: 0.22038\nTotal prob of N-in: 0.12994\noutside 1 331\nTMhelix 332 354\ninside 355 372

10361 GCF_000167815.1_ASM16781v1 Escherichia coli B7A Proteobacteria; Gammaproteobacteria; Enterobacterales; Enterobacteriaceae; Escherichia; Escherichia coli
MNNEETFYQAMRRQGVTRRSFLKYCSLAATSLGLGAGMAPKIAWALENKPRIPVVWIHGL WP_001058323.1
MULTISPECIES: hydrogenase-1 small chain [Proteobacteria] Length: 372\nNumber of predicted TMHs: 2\nExp number of AAs in TMHs: 37.2608499999999\nExp number, first 60 AAs: 19.52923\nTotal prob of N-in: 0.94351\nPOSSIBLE N-term signal sequence\ninside 1 19\nTMhelix 20 39\noutside 40 325\nTMhelix 326 348\ninside 349 372

10362 GCF_000462765.2_ASM46276v2 Escherichia coli B83Proteobacteria; Gammaproteobacteria; Enterobacterales; Enterobacteriaceae; Escherichia; Escherichia coli
MTGDNTLIHSHGINRRDFMKLCAALAATMGLSSKAAAEMAESVTNPQRPPVIWIGAQECT WP_000145410.1
MULTISPECIES: hydrogenase-2 small chain [Proteobacteria] Length: 372\nNumber of predicted TMHs: 1\nExp number of AAs in TMHs: 23.72226\nExp number, first 60 AAs: 0.22038\nTotal prob of N-in: 0.12994\noutside 1 331\nTMhelix 332 354\ninside 355 372

10363 GCF_000462765.2_ASM46276v2 Escherichia coli B83Proteobacteria; Gammaproteobacteria; Enterobacterales; Enterobacteriaceae; Escherichia; Escherichia coli
MNNEETFYQAMRRQGVTRRSFLKYCSLAATSLGLGAGMAPKIAWALENKPRIPVVWIHGL WP_001301613.1
hydrogenase [Escherichia coli] Length: 372\nNumber of predicted TMHs: 2\nExp number of AAs in TMHs: 37.46598\nExp number, first 60 AAs: 19.53139\nTotal prob of N-in: 0.94364\nPOSSIBLE N-term signal sequence\ninside 1 19\nTMhelix 20 39\noutside 40 325\nTMhelix 326 348\ninside 349 372

10364 GCF_000462785.2_ASM46278v2 Escherichia coli B84Proteobacteria; Gammaproteobacteria; Enterobacterales; Enterobacteriaceae; Escherichia; Escherichia coli
MTGDNTLIHSHGINRRDFMKLCAALAATMGLSSKAAAEMAESVTNPQRPPVIWIGAQECT WP_000145410.1
MULTISPECIES: hydrogenase-2 small chain [Proteobacteria] Length: 372\nNumber of predicted TMHs: 1\nExp number of AAs in TMHs: 23.72226\nExp number, first 60 AAs: 0.22038\nTotal prob of N-in: 0.12994\noutside 1 331\nTMhelix 332 354\ninside 355 372

10365 GCF_000462785.2_ASM46278v2 Escherichia coli B84Proteobacteria; Gammaproteobacteria; Enterobacterales; Enterobacteriaceae; Escherichia; Escherichia coli
MNNEETFYQAMRRQGVTRRSFLKYCSLAATSLGLGAGMAPKIAWALENKPRIPVVWIHGL WP_001301613.1
hydrogenase [Escherichia coli] Length: 372\nNumber of predicted TMHs: 2\nExp number of AAs in TMHs: 37.46598\nExp number, first 60 AAs: 19.53139\nTotal prob of N-in: 0.94364\nPOSSIBLE N-term signal sequence\ninside 1 19\nTMhelix 20 39\noutside 40 325\nTMhelix 326 348\ninside 349 372

10366 GCF_000462805.2_ASM46280v2 Escherichia coli B85Proteobacteria; Gammaproteobacteria; Enterobacterales; Enterobacteriaceae; Escherichia; Escherichia coli
MNNEETFYQAMRRQGVTRRSFLKYCSLAATSLGLGAGMAPKIAWALENKPRIPVVWIHGL WP_001301613.1

hydrogenase [Escherichia coli] Length: 372\nNumber of predicted TMHs: 2\nExp number of AAs in TMHs: 37.46598\nExp number, first 60 AAs: 19.53139\nTotal prob of N-in: 0.94364\nPOSSIBLE N-term signal sequence\ninside 1 19\nTMhelix 20 39\noutside 40 325\nTMhelix 326 348\ninside 349 372

10367 GCF_000462805.2_ASM46280v2 Escherichia coli B85Proteobacteria; Gammaproteobacteria; Enterobacterales; Enterobacteriaceae; Escherichia; Escherichia coli MTGDNTLIHSHGINRRDFMKLCAALAATMGLSSKAAAEMAESVTNPQRPPVIWIGAQECT WP_000145410.1 MULTISPECIES: hydrogenase-2 small chain [Proteobacteria] Length: 372\nNumber of predicted TMHs: 1\nExp number of AAs in TMHs: 23.72226\nExp number, first 60 AAs: 0.22038\nTotal prob of N-in: 0.12994\noutside 1 331\nTMhelix 332 354\ninside 355 372

10368 GCF_000462825.2_ASM46282v2 Escherichia coli B86Proteobacteria; Gammaproteobacteria; Enterobacterales; Enterobacteriaceae; Escherichia; Escherichia coli MTGDNTLIHSHGINRRDFMKLCAALAATMGLSSKAAAEMAESVTNPQRPPVIWIGAQECT WP_000145410.1 MULTISPECIES: hydrogenase-2 small chain [Proteobacteria] Length: 372\nNumber of predicted TMHs: 1\nExp number of AAs in TMHs: 23.72226\nExp number, first 60 AAs: 0.22038\nTotal prob of N-in: 0.12994\noutside 1 331\nTMhelix 332 354\ninside 355 372

10369 GCF_000462825.2_ASM46282v2 Escherichia coli B86Proteobacteria; Gammaproteobacteria; Enterobacterales; Enterobacteriaceae; Escherichia; Escherichia coli MNNEETFYQAMRRQGVTRRSFLKYCSLAATSLGLGAGMAPKIAWALENKPRIPVVWIHGL WP_001301613.1 hydrogenase [Escherichia coli] Length: 372\nNumber of predicted TMHs: 2\nExp number of AAs in TMHs: 37.46598\nExp number, first 60 AAs: 19.53139\nTotal prob of N-in: 0.94364\nPOSSIBLE N-term signal sequence\ninside 1 19\nTMhelix 20 39\noutside 40 325\nTMhelix 326 348\ninside 349 372

10370 GCF_000462845.2_ASM46284v2 Escherichia coli B89Proteobacteria; Gammaproteobacteria; Enterobacterales; Enterobacteriaceae; Escherichia; Escherichia coli MNNEETFYQAMRRQGVTRRSFLKYCSLAATSLGLGAGMAPKIAWALENKPRIPVVWIHGL WP_001301613.1 hydrogenase [Escherichia coli] Length: 372\nNumber of predicted TMHs: 2\nExp number of AAs in TMHs: 37.46598\nExp number, first 60 AAs: 19.53139\nTotal prob of N-in: 0.94364\nPOSSIBLE N-term signal sequence\ninside 1 19\nTMhelix 20 39\noutside 40 325\nTMhelix 326 348\ninside 349 372

10371 GCF_000462845.2_ASM46284v2 Escherichia coli B89Proteobacteria; Gammaproteobacteria; Enterobacterales; Enterobacteriaceae; Escherichia; Escherichia coli MTGDNTLIHSHGINRRDFMKLCAALAATMGLSSKAAAEMAESVTNPQRPPVIWIGAQECT WP_000145410.1 MULTISPECIES: hydrogenase-2 small chain [Proteobacteria] Length: 372\nNumber of predicted TMHs: 1\nExp number of AAs in TMHs: 23.72226\nExp number, first 60 AAs: 0.22038\nTotal prob of N-in: 0.12994\noutside 1 331\nTMhelix 332 354\ninside 355 372

10372 GCF_000462865.2_ASM46286v2 Escherichia coli B90Proteobacteria; Gammaproteobacteria; Enterobacterales; Enterobacteriaceae; Escherichia; Escherichia coli MTGDNTLIHSHGINRRDFMKLCAALAATMGLSSKAAAEMAESVTNPQRPPVIWIGAQECT WP_000145410.1 MULTISPECIES: hydrogenase-2 small chain [Proteobacteria] Length: 372\nNumber of predicted TMHs: 1\nExp number of AAs in TMHs: 23.72226\nExp number, first 60 AAs: 0.22038\nTotal prob of N-in: 0.12994\noutside 1 331\nTMhelix 332 354\ninside 355 372

10373 GCF_000462865.2_ASM46286v2 Escherichia coli B90Proteobacteria; Gammaproteobacteria; Enterobacterales; Enterobacteriaceae; Escherichia; Escherichia coli MNNEETFYQAMRRQGVTRRSFLKYCSLAATSLGLGAGMAPKIAWALENKPRIPVVWIHGL WP_001301613.1 hydrogenase [Escherichia coli] Length: 372\nNumber of predicted TMHs: 2\nExp number of AAs in TMHs: 37.46598\nExp number, first 60 AAs: 19.53139\nTotal prob of N-in: 0.94364\nPOSSIBLE N-term signal sequence\ninside 1 19\nTMhelix 20 39\noutside 40 325\nTMhelix 326 348\ninside 349 372

10374 GCF_002110105.1_ASM211010v1 Escherichia coli B921 Proteobacteria; Gammaproteobacteria; Enterobacterales; Enterobacteriaceae; Escherichia; Escherichia coli MTGDNTLIHSHGINRRDFMKLCAALAATMGLSSKAAAEMAESVTNPQRPPVIWIGAQECT WP_000145410.1 MULTISPECIES: hydrogenase-2 small chain [Proteobacteria] Length: 372\nNumber of predicted TMHs: 1\nExp number of AAs in TMHs: 23.72226\nExp number, first 60 AAs: 0.22038\nTotal prob of N-in: 0.12994\noutside 1 331\nTMhelix 332 354\ninside 355 372

10375 GCF_002110105.1_ASM211010v1 Escherichia coli B921 Proteobacteria; Gammaproteobacteria; Enterobacterales; Enterobacteriaceae; Escherichia; Escherichia coli
MNNEETFYQAMRRQGVTRRSFLKYCSLAATSLGLGAGMAPKIAWALENKPRIPVVWIHGL WP_001058323.1
MULTISPECIES: hydrogenase-1 small chain [Proteobacteria] Length: 372\nNumber of predicted TMHs: 2\nExp number of AAs in TMHs: 37.2608499999999\nExp number, first 60 AAs: 19.52923\nTotal prob of N-in: 0.94351\nPOSSIBLE N-term signal sequence\ninside 1 19\nTMhelix 20 39\noutside 40 325\nTMhelix 326 348\ninside 349 372

10376 GCF_000462425.2_ASM46242v2 Escherichia coli B93Proteobacteria; Gammaproteobacteria; Enterobacterales; Enterobacteriaceae; Escherichia; Escherichia coli
MTGDNTLIHSHGINRRDFMKLCAALAATMGLSSKAAAEMAESVTNPQRPPVIWIGAQECT WP_000145410.1
MULTISPECIES: hydrogenase-2 small chain [Proteobacteria] Length: 372\nNumber of predicted TMHs: 1\nExp number of AAs in TMHs: 23.72226\nExp number, first 60 AAs: 0.22038\nTotal prob of N-in: 0.12994\noutside 1 331\nTMhelix 332 354\ninside 355 372

10377 GCF_000462425.2_ASM46242v2 Escherichia coli B93Proteobacteria; Gammaproteobacteria; Enterobacterales; Enterobacteriaceae; Escherichia; Escherichia coli
MNNEETFYQAMRRQGVTRRSFLKYCSLAATSLGLGAGMAPKIAWALENKPRIPVVWIHGL WP_001301613.1
hydrogenase [Escherichia coli] Length: 372\nNumber of predicted TMHs: 2\nExp number of AAs in TMHs: 37.46598\nExp number, first 60 AAs: 19.53139\nTotal prob of N-in: 0.94364\nPOSSIBLE N-term signal sequence\ninside 1 19\nTMhelix 20 39\noutside 40 325\nTMhelix 326 348\ninside 349 372

10378 GCF_000462445.2_ASM46244v2 Escherichia coli B94Proteobacteria; Gammaproteobacteria; Enterobacterales; Enterobacteriaceae; Escherichia; Escherichia coli
MNNEETFYQAMRRQGVTRRSFLKYCSLAATSLGLGAGMAPKIAWALENKPRIPVVWIHGL WP_001301613.1
hydrogenase [Escherichia coli] Length: 372\nNumber of predicted TMHs: 2\nExp number of AAs in TMHs: 37.46598\nExp number, first 60 AAs: 19.53139\nTotal prob of N-in: 0.94364\nPOSSIBLE N-term signal sequence\ninside 1 19\nTMhelix 20 39\noutside 40 325\nTMhelix 326 348\ninside 349 372

10379 GCF_000462445.2_ASM46244v2 Escherichia coli B94Proteobacteria; Gammaproteobacteria; Enterobacterales; Enterobacteriaceae; Escherichia; Escherichia coli
MTGDNTLIHSHGINRRDFMKLCAALAATMGLSSKAAAEMAESVTNPQRPPVIWIGAQECT WP_000145410.1
MULTISPECIES: hydrogenase-2 small chain [Proteobacteria] Length: 372\nNumber of predicted TMHs: 1\nExp number of AAs in TMHs: 23.72226\nExp number, first 60 AAs: 0.22038\nTotal prob of N-in: 0.12994\noutside 1 331\nTMhelix 332 354\ninside 355 372

10380 GCF_000462465.2_ASM46246v2 Escherichia coli B95Proteobacteria; Gammaproteobacteria; Enterobacterales; Enterobacteriaceae; Escherichia; Escherichia coli
MTGDNTLIHSHGINRRDFMKLCAALAATMGLSSKAAAEMAESVTNPQRPPVIWIGAQECT WP_000145410.1
MULTISPECIES: hydrogenase-2 small chain [Proteobacteria] Length: 372\nNumber of predicted TMHs: 1\nExp number of AAs in TMHs: 23.72226\nExp number, first 60 AAs: 0.22038\nTotal prob of N-in: 0.12994\noutside 1 331\nTMhelix 332 354\ninside 355 372

10381 GCF_000462465.2_ASM46246v2 Escherichia coli B95Proteobacteria; Gammaproteobacteria; Enterobacterales; Enterobacteriaceae; Escherichia; Escherichia coli
MNNEETFYQAMRRQGVTRRSFLKYCSLAATSLGLGAGMAPKIAWALENKPRIPVVWIHGL WP_001301613.1
hydrogenase [Escherichia coli] Length: 372\nNumber of predicted TMHs: 2\nExp number of AAs in TMHs: 37.46598\nExp number, first 60 AAs: 19.53139\nTotal prob of N-in: 0.94364\nPOSSIBLE N-term signal sequence\ninside 1 19\nTMhelix 20 39\noutside 40 325\nTMhelix 326 348\ninside 349 372

10382 GCF_000355495.2_ASM35549v2 Escherichia coli BCE001_MS16 Proteobacteria; Gammaproteobacteria; Enterobacterales; Enterobacteriaceae; Escherichia; Escherichia coli
MTGDNTLIHSHGINRRDFMKLCAALAATMGLSSKAAAEMAESVTNPQRPPVIWIGAQECT WP_000145410.1
MULTISPECIES: hydrogenase-2 small chain [Proteobacteria] Length: 372\nNumber of predicted TMHs: 1\nExp number of AAs in TMHs: 23.72226\nExp number, first 60 AAs: 0.22038\nTotal prob of N-in: 0.12994\noutside 1 331\nTMhelix 332 354\ninside 355 372

10383 GCF_000355495.2_ASM35549v2 Escherichia coli BCE001_MS16 Proteobacteria; Gammaproteobacteria; Enterobacterales; Enterobacteriaceae; Escherichia; Escherichia coli
MNNEETFYQAMRRQGVTRRSFLKYCSLAATSLGLGAGMAPKIAWALENKPRIPVVWIHGL WP_001058323.1
MULTISPECIES: hydrogenase-1 small chain [Proteobacteria] Length: 372\nNumber of predicted TMHs:

2\nExp number of AAs in TMHs: 37.2608499999999\nExp number, first 60 AAs: 19.52923\nTotal prob of N-in: 0.94351\nPOSSIBLE N-term signal sequence\ninside 1 19\nTMhelix 20 39\noutside 40 325\nTMhelix 326 348\ninside 349 372

10384 GCF_000354295.1_ASM35429v1 Escherichia coli BCE002_MS12 Proteobacteria; Gammaproteobacteria; Enterobacterales; Enterobacteriaceae; Escherichia; Escherichia coli MTGDNTLIHSHGINRRDFMKLCAALAATMGLSSKAAAEMAESVTNPQRPPVIWIGAQECT WP_000145410.1 MULTISPECIES: hydrogenase-2 small chain [Proteobacteria] Length: 372\nNumber of predicted TMHs: 1\nExp number of AAs in TMHs: 23.72226\nExp number, first 60 AAs: 0.22038\nTotal prob of N-in: 0.12994\noutside 1 331\nTMhelix 332 354\ninside 355 372

10385 GCF_000354295.1_ASM35429v1 Escherichia coli BCE002_MS12 Proteobacteria; Gammaproteobacteria; Enterobacterales; Enterobacteriaceae; Escherichia; Escherichia coli MNNEETFYQAMRRQGVTRRSFLKYCSLAATSLGLGAGMAPKIAWALENKPRIPVVWIHGL WP_001058323.1 MULTISPECIES: hydrogenase-1 small chain [Proteobacteria] Length: 372\nNumber of predicted TMHs: 2\nExp number of AAs in TMHs: 37.2608499999999\nExp number, first 60 AAs: 19.52923\nTotal prob of N-in: 0.94351\nPOSSIBLE N-term signal sequence\ninside 1 19\nTMhelix 20 39\noutside 40 325\nTMhelix 326 348\ninside 349 372

10386 GCF_000356885.2_ASM35688v2 Escherichia coli BCE006_MS-23 Proteobacteria; Gammaproteobacteria; Enterobacterales; Enterobacteriaceae; Escherichia; Escherichia coli MTGDNTLIHSHGINRRDFMKLCAALAATMGLSSKAAAEMAESVTNPQRPPVIWIGAQECT WP_001509018.1 hydrogenase-2 small chain [Escherichia coli] Length: 372\nNumber of predicted TMHs: 1\nExp number of AAs in TMHs: 23.72887\nExp number, first 60 AAs: 0.22038\nTotal prob of N-in: 0.13011\noutside 1 331\nTMhelix 332 354\ninside 355 372

10387 GCF_000356885.2_ASM35688v2 Escherichia coli BCE006_MS-23 Proteobacteria; Gammaproteobacteria; Enterobacterales; Enterobacteriaceae; Escherichia; Escherichia coli MNNEETFYQAMRRQGVTRRSFLKYCSLAATSLGLGAGMAPKIAWALENKPRIPVVWIHGL WP_001058323.1 MULTISPECIES: hydrogenase-1 small chain [Proteobacteria] Length: 372\nNumber of predicted TMHs: 2\nExp number of AAs in TMHs: 37.2608499999999\nExp number, first 60 AAs: 19.52923\nTotal prob of N-in: 0.94351\nPOSSIBLE N-term signal sequence\ninside 1 19\nTMhelix 20 39\noutside 40 325\nTMhelix 326 348\ninside 349 372

10388 GCF_000356705.1_ASM35670v1 Escherichia coli BCE007_MS-11 Proteobacteria; Gammaproteobacteria; Enterobacterales; Enterobacteriaceae; Escherichia; Escherichia coli MNNEETFYQAMRRQGVTRRSFLKYCSLAATSLGLGAGMAPKIAWALENKPRIPVVWIHGL WP_001058323.1 MULTISPECIES: hydrogenase-1 small chain [Proteobacteria] Length: 372\nNumber of predicted TMHs: 2\nExp number of AAs in TMHs: 37.2608499999999\nExp number, first 60 AAs: 19.52923\nTotal prob of N-in: 0.94351\nPOSSIBLE N-term signal sequence\ninside 1 19\nTMhelix 20 39\noutside 40 325\nTMhelix 326 348\ninside 349 372

10389 GCF_000356705.1_ASM35670v1 Escherichia coli BCE007_MS-11 Proteobacteria; Gammaproteobacteria; Enterobacterales; Enterobacteriaceae; Escherichia; Escherichia coli MTGDNTLIHSHGINRRDFMKLCAALAATMGLSSKAAAEMAESVTNPQRPPVIWIGAQECT WP_000145410.1 MULTISPECIES: hydrogenase-2 small chain [Proteobacteria] Length: 372\nNumber of predicted TMHs: 1\nExp number of AAs in TMHs: 23.72226\nExp number, first 60 AAs: 0.22038\nTotal prob of N-in: 0.12994\noutside 1 331\nTMhelix 332 354\ninside 355 372

10390 GCF_000356265.2_ASM35626v2 Escherichia coli BCE008_MS-01 Proteobacteria; Gammaproteobacteria; Enterobacterales; Enterobacteriaceae; Escherichia; Escherichia coli MNNEETFYQAMRRQGVTRRSFLKYCSLAATSLGLGAGMAPKIAWALENKPRIPVVWIHGL WP_001058323.1 MULTISPECIES: hydrogenase-1 small chain [Proteobacteria] Length: 372\nNumber of predicted TMHs: 2\nExp number of AAs in TMHs: 37.2608499999999\nExp number, first 60 AAs: 19.52923\nTotal prob of N-in: 0.94351\nPOSSIBLE N-term signal sequence\ninside 1 19\nTMhelix 20 39\noutside 40 325\nTMhelix 326 348\ninside 349 372

10391 GCF_000356265.2_ASM35626v2 Escherichia coli BCE008_MS-01 Proteobacteria; Gammaproteobacteria; Enterobacterales; Enterobacteriaceae; Escherichia; Escherichia coli MTGDNTLIHSHGINRRDFMKLCAALAATMGLSSKAAAEMAESVTNPQRPPVIWIGAQECT WP_000145410.1 MULTISPECIES: hydrogenase-2 small chain [Proteobacteria] Length: 372\nNumber of predicted TMHs:

1\nExp number of AAs in TMHs: 23.72226\nExp number, first 60 AAs: 0.22038\nTotal prob of N-in: 0.12994\noutside 1 331\nTMhelix 332 354\ninside 355 372

10392 GCF_000356725.1_ASM35672v1 Escherichia coli BCE008_MS-13 Proteobacteria; Gammaproteobacteria; Enterobacterales; Enterobacteriaceae; Escherichia; Escherichia coli MNNEETFYQAMRRQGVTRRSFLKYCSLAATSLGLGAGMAPKIAWALENKPRIPVVWIHGL WP_001058323.1 MULTISPECIES: hydrogenase-1 small chain [Proteobacteria] Length: 372\nNumber of predicted TMHs: 2\nExp number of AAs in TMHs: 37.2608499999999\nExp number, first 60 AAs: 19.52923\nTotal prob of N-in: 0.94351\nPOSSIBLE N-term signal sequence\ninside 1 19\nTMhelix 20 39\noutside 40 325\nTMhelix 326 348\ninside 349 372

10393 GCF_000356725.1_ASM35672v1 Escherichia coli BCE008_MS-13 Proteobacteria; Gammaproteobacteria; Enterobacterales; Enterobacteriaceae; Escherichia; Escherichia coli MTGDNTLIHSHGINRRDFMKLCAALAATMGLSSKAAAEMAESVTNPQRPPVIWIGAQECT WP_000145410.1 MULTISPECIES: hydrogenase-2 small chain [Proteobacteria] Length: 372\nNumber of predicted TMHs: 1\nExp number of AAs in TMHs: 23.72226\nExp number, first 60 AAs: 0.22038\nTotal prob of N-in: 0.12994\noutside 1 331\nTMhelix 332 354\ninside 355 372

10394 GCF_000356285.2_ASM35628v2 Escherichia coli BCE011_MS-01 Proteobacteria; Gammaproteobacteria; Enterobacterales; Enterobacteriaceae; Escherichia; Escherichia coli MTGDNTLIHSHGINRRDFMKLCAALAATMGLSSKAAAEMAESVTNPQRPPVIWIGAQECT WP_000145410.1 MULTISPECIES: hydrogenase-2 small chain [Proteobacteria] Length: 372\nNumber of predicted TMHs: 1\nExp number of AAs in TMHs: 23.72226\nExp number, first 60 AAs: 0.22038\nTotal prob of N-in: 0.12994\noutside 1 331\nTMhelix 332 354\ninside 355 372

10395 GCF_000356285.2_ASM35628v2 Escherichia coli BCE011_MS-01 Proteobacteria; Gammaproteobacteria; Enterobacterales; Enterobacteriaceae; Escherichia; Escherichia coli MNNEETFYQAMRRQGVTRRSFLKYCSLAATSLGLGAGMAPKIAWALENKPRIPVVWIHGL WP_001058323.1 MULTISPECIES: hydrogenase-1 small chain [Proteobacteria] Length: 372\nNumber of predicted TMHs: 2\nExp number of AAs in TMHs: 37.2608499999999\nExp number, first 60 AAs: 19.52923\nTotal prob of N-in: 0.94351\nPOSSIBLE N-term signal sequence\ninside 1 19\nTMhelix 20 39\noutside 40 325\nTMhelix 326 348\ninside 349 372

10396 GCF_000354315.2_ASM35431v2 Escherichia coli BCE019_MS-13 Proteobacteria; Gammaproteobacteria; Enterobacterales; Enterobacteriaceae; Escherichia; Escherichia coli MTGDNTLIHSHGINRRDFMKLCAALAATMGLSSKAAAEMAESVTNPQRPPVIWIGAQECT WP_000145410.1 MULTISPECIES: hydrogenase-2 small chain [Proteobacteria] Length: 372\nNumber of predicted TMHs: 1\nExp number of AAs in TMHs: 23.72226\nExp number, first 60 AAs: 0.22038\nTotal prob of N-in: 0.12994\noutside 1 331\nTMhelix 332 354\ninside 355 372

10397 GCF_000354315.2_ASM35431v2 Escherichia coli BCE019_MS-13 Proteobacteria; Gammaproteobacteria; Enterobacterales; Enterobacteriaceae; Escherichia; Escherichia coli MNNEETFYQAMRRQGVTRRSFLKYCSLAATSLGLGAGMAPKIAWALENKPRIPVVWIHGL WP_001058323.1 MULTISPECIES: hydrogenase-1 small chain [Proteobacteria] Length: 372\nNumber of predicted TMHs: 2\nExp number of AAs in TMHs: 37.2608499999999\nExp number, first 60 AAs: 19.52923\nTotal prob of N-in: 0.94351\nPOSSIBLE N-term signal sequence\ninside 1 19\nTMhelix 20 39\noutside 40 325\nTMhelix 326 348\ninside 349 372

10398 GCF_000356385.1_ASM35638v1 Escherichia coli BCE030_MS-09 Proteobacteria; Gammaproteobacteria; Enterobacterales; Enterobacteriaceae; Escherichia; Escherichia coli MNNEETFYQAMRRQGVTRRSFLKYCSLAATSLGLGAGMAPKIAWALENKPRIPVVWIHGL WP_001058323.1 MULTISPECIES: hydrogenase-1 small chain [Proteobacteria] Length: 372\nNumber of predicted TMHs: 2\nExp number of AAs in TMHs: 37.2608499999999\nExp number, first 60 AAs: 19.52923\nTotal prob of N-in: 0.94351\nPOSSIBLE N-term signal sequence\ninside 1 19\nTMhelix 20 39\noutside 40 325\nTMhelix 326 348\ninside 349 372

10399 GCF_000356385.1_ASM35638v1 Escherichia coli BCE030_MS-09 Proteobacteria; Gammaproteobacteria; Enterobacterales; Enterobacteriaceae; Escherichia; Escherichia coli MTGDNTLIHSHGINRRDFMKLCAALAATMGLSSKAAAEMAESVTNPQRPPVIWIGAQECT WP_000145410.1 MULTISPECIES: hydrogenase-2 small chain [Proteobacteria] Length: 372\nNumber of predicted TMHs:

1\nExp number of AAs in TMHs: 23.72226\nExp number, first 60 AAs:  0.22038\nTotal prob of N-in:   0.12994\noutside   1   331\nTMhelix   332   354\ninside   355   372

10400 GCF_000356405.2_ASM35640v2 Escherichia coli BCE032_MS-12 Proteobacteria; Gammaproteobacteria; Enterobacterales; Enterobacteriaceae; Escherichia; Escherichia coli
MTGDNTLIHSHGINRRDFMKLCAALAATMGLSSKAAAEMAESVTNPQRPPVIWIGAQECT WP_000145410.1
MULTISPECIES: hydrogenase-2 small chain [Proteobacteria] Length: 372\nNumber of predicted TMHs: 1\nExp number of AAs in TMHs: 23.72226\nExp number, first 60 AAs:  0.22038\nTotal prob of N-in:   0.12994\noutside   1   331\nTMhelix   332   354\ninside   355   372

10401 GCF_000356405.2_ASM35640v2 Escherichia coli BCE032_MS-12 Proteobacteria; Gammaproteobacteria; Enterobacterales; Enterobacteriaceae; Escherichia; Escherichia coli
MNNEETFYQAMRRQGVTRRSFLKYCSLAATSLGLGAGMAPKIAWALENKPRIPVVWIHGL WP_001058323.1
MULTISPECIES: hydrogenase-1 small chain [Proteobacteria] Length: 372\nNumber of predicted TMHs: 2\nExp number of AAs in TMHs: 37.2608499999999\nExp number, first 60 AAs:  19.52923\nTotal prob of N-in: 0.94351\nPOSSIBLE N-term signal sequence\ninside   1   19\nTMhelix   20   39\noutside   40   325\nTMhelix   326   348\ninside   349   372

10402 GCF_000354335.2_ASM35433v2 Escherichia coli BCE034_MS-14 Proteobacteria; Gammaproteobacteria; Enterobacterales; Enterobacteriaceae; Escherichia; Escherichia coli
MNNEETFYQAMRRQGVTRRSFLKYCSLAATSLGLGAGMAPKIAWALENKPRIPVVWIHGL WP_001295940.1
MULTISPECIES: hydrogenase [Enterobacteriaceae] Length: 372\nNumber of predicted TMHs: 2\nExp number of AAs in TMHs: 37.2608899999999\nExp number, first 60 AAs:  19.52923\nTotal prob of N-in: 0.94351\nPOSSIBLE N-term signal sequence\ninside   1   19\nTMhelix   20   39\noutside   40   325\nTMhelix   326   348\ninside   349   372

10403 GCF_000354335.2_ASM35433v2 Escherichia coli BCE034_MS-14 Proteobacteria; Gammaproteobacteria; Enterobacterales; Enterobacteriaceae; Escherichia; Escherichia coli
MTGDNTLIHSHGINRRDFMKLCAALAATMGLSSKAAAEMAESVTNPQRPPVIWIGAQECT WP_000145410.1
MULTISPECIES: hydrogenase-2 small chain [Proteobacteria] Length: 372\nNumber of predicted TMHs: 1\nExp number of AAs in TMHs: 23.72226\nExp number, first 60 AAs:  0.22038\nTotal prob of N-in:   0.12994\noutside   1   331\nTMhelix   332   354\ninside   355   372

10404 GCF_000462365.2_ASM46236v2 Escherichia coli Bd5610_99 Proteobacteria; Gammaproteobacteria; Enterobacterales; Enterobacteriaceae; Escherichia; Escherichia coli
MNNEETFYQAMRRQGVTRRSFLKYCSLAATSLGLGAGMAPKIAWALENKPRIPVVWIHGL WP_001301613.1
hydrogenase [Escherichia coli] Length: 372\nNumber of predicted TMHs: 2\nExp number of AAs in TMHs: 37.46598\nExp number, first 60 AAs:  19.53139\nTotal prob of N-in:   0.94364\nPOSSIBLE N-term signal sequence\ninside   1   19\nTMhelix   20   39\noutside   40   325\nTMhelix   326   348\ninside   349   372

10405 GCF_000462365.2_ASM46236v2 Escherichia coli Bd5610_99 Proteobacteria; Gammaproteobacteria; Enterobacterales; Enterobacteriaceae; Escherichia; Escherichia coli
MTGDNTLIHSHGINRRDFMKLCAALAATMGLSSKAAAEMAESVTNPQRPPVIWIGAQECT WP_000145410.1
MULTISPECIES: hydrogenase-2 small chain [Proteobacteria] Length: 372\nNumber of predicted TMHs: 1\nExp number of AAs in TMHs: 23.72226\nExp number, first 60 AAs:  0.22038\nTotal prob of N-in:   0.12994\noutside   1   331\nTMhelix   332   354\ninside   355   372

10406 GCF_000522245.1_Esch_coli_BIDMC_15_V2 Escherichia coli BIDMC 15 Proteobacteria; Gammaproteobacteria; Enterobacterales; Enterobacteriaceae; Escherichia; Escherichia coli
MTGDNTLIHSHGINRRDFMKLCAALAATMGLSSKAAAEMAESVTNPQRPPVIWIGAQECT WP_000145410.1
MULTISPECIES: hydrogenase-2 small chain [Proteobacteria] Length: 372\nNumber of predicted TMHs: 1\nExp number of AAs in TMHs: 23.72226\nExp number, first 60 AAs:  0.22038\nTotal prob of N-in:   0.12994\noutside   1   331\nTMhelix   332   354\ninside   355   372

10407 GCF_000522245.1_Esch_coli_BIDMC_15_V2 Escherichia coli BIDMC 15 Proteobacteria; Gammaproteobacteria; Enterobacterales; Enterobacteriaceae; Escherichia; Escherichia coli
MNNEETFYQAMRRQGVTRRSFLKYCSLAATSLGLGAGMAPKIAWALENKPRIPVVWIHGL WP_001058323.1
MULTISPECIES: hydrogenase-1 small chain [Proteobacteria] Length: 372\nNumber of predicted TMHs: 2\nExp number of AAs in TMHs: 37.2608499999999\nExp number, first 60 AAs:  19.52923\nTotal prob of N-in: 0.94351\nPOSSIBLE N-term signal sequence\ninside   1   19\nTMhelix   20   39\noutside   40   325\nTMhelix   326   348\ninside   349   372

10408 GCF_000522225.1_Esch_coli_BIDMC_17A_V2 Escherichia coli BIDMC 17A Proteobacteria; Gammaproteobacteria; Enterobacterales; Enterobacteriaceae; Escherichia; Escherichia coli MNNEETFYQAMRRQGVTRRSFLKYCSLAATSLGLGAGMAPKIAWALENKPRIPVVWIHGL WP_001058323.1 MULTISPECIES: hydrogenase-1 small chain [Proteobacteria] Length: 372\nNumber of predicted TMHs: 2\nExp number of AAs in TMHs: 37.2608499999999\nExp number, first 60 AAs: 19.52923\nTotal prob of N-in: 0.94351\nPOSSIBLE N-term signal sequence\ninside 1 19\nTMhelix 20 39\noutside 40 325\nTMhelix 326 348\ninside 349 372

10409 GCF_000522225.1_Esch_coli_BIDMC_17A_V2 Escherichia coli BIDMC 17A Proteobacteria; Gammaproteobacteria; Enterobacterales; Enterobacteriaceae; Escherichia; Escherichia coli MTGDNTLIHSHGINRRDFMKLCAALAATMGLSSKAAAEMAESVTNPQRPPVIWIGAQECT WP_000145410.1 MULTISPECIES: hydrogenase-2 small chain [Proteobacteria] Length: 372\nNumber of predicted TMHs: 1\nExp number of AAs in TMHs: 23.72226\nExp number, first 60 AAs: 0.22038\nTotal prob of N-in: 0.12994\noutside 1 331\nTMhelix 332 354\ninside 355 372

10410 GCF_000522205.1_Esch_coli_BIDMC_17B_V2 Escherichia coli BIDMC 17B Proteobacteria; Gammaproteobacteria; Enterobacterales; Enterobacteriaceae; Escherichia; Escherichia coli MNNEETFYQAMRRQGVTRRSFLKYCSLAATSLGLGAGMAPKIAWALENKPRIPVVWIHGL WP_001058323.1 MULTISPECIES: hydrogenase-1 small chain [Proteobacteria] Length: 372\nNumber of predicted TMHs: 2\nExp number of AAs in TMHs: 37.2608499999999\nExp number, first 60 AAs: 19.52923\nTotal prob of N-in: 0.94351\nPOSSIBLE N-term signal sequence\ninside 1 19\nTMhelix 20 39\noutside 40 325\nTMhelix 326 348\ninside 349 372

10411 GCF_000522205.1_Esch_coli_BIDMC_17B_V2 Escherichia coli BIDMC 17B Proteobacteria; Gammaproteobacteria; Enterobacterales; Enterobacteriaceae; Escherichia; Escherichia coli MTGDNTLIHSHGINRRDFMKLCAALAATMGLSSKAAAEMAESVTNPQRPPVIWIGAQECT WP_000145410.1 MULTISPECIES: hydrogenase-2 small chain [Proteobacteria] Length: 372\nNumber of predicted TMHs: 1\nExp number of AAs in TMHs: 23.72226\nExp number, first 60 AAs: 0.22038\nTotal prob of N-in: 0.12994\noutside 1 331\nTMhelix 332 354\ninside 355 372

10412 GCF_000522185.1_Esch_coli_BIDMC_19A_V2 Escherichia coli BIDMC 19A Proteobacteria; Gammaproteobacteria; Enterobacterales; Enterobacteriaceae; Escherichia; Escherichia coli MNNEETFYQAMRRQGVTRRSFLKYCSLAATSLGLGAGMAPKIAWALENKPRIPVVWIHGL WP_001058323.1 MULTISPECIES: hydrogenase-1 small chain [Proteobacteria] Length: 372\nNumber of predicted TMHs: 2\nExp number of AAs in TMHs: 37.2608499999999\nExp number, first 60 AAs: 19.52923\nTotal prob of N-in: 0.94351\nPOSSIBLE N-term signal sequence\ninside 1 19\nTMhelix 20 39\noutside 40 325\nTMhelix 326 348\ninside 349 372

10413 GCF_000522185.1_Esch_coli_BIDMC_19A_V2 Escherichia coli BIDMC 19A Proteobacteria; Gammaproteobacteria; Enterobacterales; Enterobacteriaceae; Escherichia; Escherichia coli MTGDNTLIHSHGINRRDFMKLCAALAATMGLSSKAAAEMAESVTNPQRPPVIWIGAQECT WP_000145410.1 MULTISPECIES: hydrogenase-2 small chain [Proteobacteria] Length: 372\nNumber of predicted TMHs: 1\nExp number of AAs in TMHs: 23.72226\nExp number, first 60 AAs: 0.22038\nTotal prob of N-in: 0.12994\noutside 1 331\nTMhelix 332 354\ninside 355 372

10414 GCF_000522165.1_Esch_coli_BIDMC_19B_V2 Escherichia coli BIDMC 19B Proteobacteria; Gammaproteobacteria; Enterobacterales; Enterobacteriaceae; Escherichia; Escherichia coli MNNEETFYQAMRRQGVTRRSFLKYCSLAATSLGLGAGMAPKIAWALENKPRIPVVWIHGL WP_001058323.1 MULTISPECIES: hydrogenase-1 small chain [Proteobacteria] Length: 372\nNumber of predicted TMHs: 2\nExp number of AAs in TMHs: 37.2608499999999\nExp number, first 60 AAs: 19.52923\nTotal prob of N-in: 0.94351\nPOSSIBLE N-term signal sequence\ninside 1 19\nTMhelix 20 39\noutside 40 325\nTMhelix 326 348\ninside 349 372

10415 GCF_000522165.1_Esch_coli_BIDMC_19B_V2 Escherichia coli BIDMC 19B Proteobacteria; Gammaproteobacteria; Enterobacterales; Enterobacteriaceae; Escherichia; Escherichia coli MTGDNTLIHSHGINRRDFMKLCAALAATMGLSSKAAAEMAESVTNPQRPPVIWIGAQECT WP_000145410.1 MULTISPECIES: hydrogenase-2 small chain [Proteobacteria] Length: 372\nNumber of predicted TMHs: 1\nExp number of AAs in TMHs: 23.72226\nExp number, first 60 AAs: 0.22038\nTotal prob of N-in: 0.12994\noutside 1 331\nTMhelix 332 354\ninside 355 372

10416 GCF_000474825.1_Esch_coli_BIDMC_19C_V1_PacBio Escherichia coli BIDMC 19C Proteobacteria; Gammaproteobacteria; Enterobacterales; Enterobacteriaceae; Escherichia; Escherichia coli MNNEETFYQAMRRQGVTRRSFLKYCSLAATSLGLGAGMAPKIAWALENKPRIPVVWIHGL WP_001058323.1 MULTISPECIES: hydrogenase-1 small chain [Proteobacteria] Length: 372\nNumber of predicted TMHs: 2\nExp number of AAs in TMHs: 37.2608499999999\nExp number, first 60 AAs: 19.52923\nTotal prob of N-in: 0.94351\nPOSSIBLE N-term signal sequence\ninside 1 19\nTMhelix 20 39\noutside 40 325\nTMhelix 326 348\ninside 349 372

10417 GCF_000474825.1_Esch_coli_BIDMC_19C_V1_PacBio Escherichia coli BIDMC 19C Proteobacteria; Gammaproteobacteria; Enterobacterales; Enterobacteriaceae; Escherichia; Escherichia coli MTGDNTLIHSHGINRRDFMKLCAALAATMGLSSKAAAEMAESVTNPQRPPVIWIGAQECT WP_000145410.1 MULTISPECIES: hydrogenase-2 small chain [Proteobacteria] Length: 372\nNumber of predicted TMHs: 1\nExp number of AAs in TMHs: 23.72226\nExp number, first 60 AAs: 0.22038\nTotal prob of N-in: 0.12994\noutside 1 331\nTMhelix 332 354\ninside 355 372

10418 GCF_000522145.1_Esch_coli_BIDMC_20A_V2 Escherichia coli BIDMC 20A Proteobacteria; Gammaproteobacteria; Enterobacterales; Enterobacteriaceae; Escherichia; Escherichia coli MTGDNTLIHSHGINRRDFMKLCAALAATMGLSSKAAAEMVESVTNPQRPPVIWIGAQECT WP_000145423.1 hydrogenase 2 small subunit [Escherichia coli] Length: 372\nNumber of predicted TMHs: 1\nExp number of AAs in TMHs: 23.78201\nExp number, first 60 AAs: 0.28355\nTotal prob of N-in: 0.13257\noutside 1 331\nTMhelix 332 354\ninside 355 372

10419 GCF_000522145.1_Esch_coli_BIDMC_20A_V2 Escherichia coli BIDMC 20A Proteobacteria; Gammaproteobacteria; Enterobacterales; Enterobacteriaceae; Escherichia; Escherichia coli MNNEETFYQAMRRQGVTRRSFLKYCSLAATSLGLGAGMAPKIAWALENKPRIPVVWIHGL WP_001295940.1 MULTISPECIES: hydrogenase [Enterobacteriaceae] Length: 372\nNumber of predicted TMHs: 2\nExp number of AAs in TMHs: 37.2608899999999\nExp number, first 60 AAs: 19.52923\nTotal prob of N-in: 0.94351\nPOSSIBLE N-term signal sequence\ninside 1 19\nTMhelix 20 39\noutside 40 325\nTMhelix 326 348\ninside 349 372

10420 GCF_000522125.1_Esch_coli_BIDMC_20B_V1 Escherichia coli BIDMC 20B Proteobacteria; Gammaproteobacteria; Enterobacterales; Enterobacteriaceae; Escherichia; Escherichia coli MNNEETFYQAMRRQGVTRRSFLKYCSLAATSLGLGAGMAPKIAWALENKPRIPVVWIHGL WP_001295940.1 MULTISPECIES: hydrogenase [Enterobacteriaceae] Length: 372\nNumber of predicted TMHs: 2\nExp number of AAs in TMHs: 37.2608899999999\nExp number, first 60 AAs: 19.52923\nTotal prob of N-in: 0.94351\nPOSSIBLE N-term signal sequence\ninside 1 19\nTMhelix 20 39\noutside 40 325\nTMhelix 326 348\ninside 349 372

10421 GCF_000522125.1_Esch_coli_BIDMC_20B_V1 Escherichia coli BIDMC 20B Proteobacteria; Gammaproteobacteria; Enterobacterales; Enterobacteriaceae; Escherichia; Escherichia coli MTGDNTLIHSHGINRRDFMKLCAALAATMGLSSKAAAEMVESVTNPQRPPVIWIGAQECT WP_000145423.1 hydrogenase 2 small subunit [Escherichia coli] Length: 372\nNumber of predicted TMHs: 1\nExp number of AAs in TMHs: 23.78201\nExp number, first 60 AAs: 0.28355\nTotal prob of N-in: 0.13257\noutside 1 331\nTMhelix 332 354\ninside 355 372

10422 GCF_000522305.1_Esch_coli_BIDMC_2B_V2 Escherichia coli BIDMC 2B Proteobacteria; Gammaproteobacteria; Enterobacterales; Enterobacteriaceae; Escherichia; Escherichia coli MTGDNTLIHSHGINRRDFMKLCAALAATMGLSSKAAAEMAESVTNPQRPPVIWIGAQECT WP_000145410.1 MULTISPECIES: hydrogenase-2 small chain [Proteobacteria] Length: 372\nNumber of predicted TMHs: 1\nExp number of AAs in TMHs: 23.72226\nExp number, first 60 AAs: 0.22038\nTotal prob of N-in: 0.12994\noutside 1 331\nTMhelix 332 354\ninside 355 372

10423 GCF_000522305.1_Esch_coli_BIDMC_2B_V2 Escherichia coli BIDMC 2B Proteobacteria; Gammaproteobacteria; Enterobacterales; Enterobacteriaceae; Escherichia; Escherichia coli MNNEETFYQAMRRQGVTRRSFLKYCSLAATSLGLGAGMAPKIAWALENKPRIPVVWIHGL WP_001058323.1 MULTISPECIES: hydrogenase-1 small chain [Proteobacteria] Length: 372\nNumber of predicted TMHs: 2\nExp number of AAs in TMHs: 37.2608499999999\nExp number, first 60 AAs: 19.52923\nTotal prob of N-in: 0.94351\nPOSSIBLE N-term signal sequence\ninside 1 19\nTMhelix 20 39\noutside 40 325\nTMhelix 326 348\ninside 349 372

10424 GCF_000522285.1_Esch_coli_BIDMC_3_V2 Escherichia coli BIDMC 3 Proteobacteria; Gammaproteobacteria; Enterobacterales; Enterobacteriaceae; Escherichia; Escherichia coli
MTGDNTLIHSHGINRRDFMKLCAALAATMGLSSKAAAEMAESVTNPQRPPVIWIGAQECT WP_000145410.1
MULTISPECIES: hydrogenase-2 small chain [Proteobacteria] Length: 372\nNumber of predicted TMHs: 1\nExp number of AAs in TMHs: 23.72226\nExp number, first 60 AAs: 0.22038\nTotal prob of N-in: 0.12994\noutside 1 331\nTMhelix 332 354\ninside 355 372

10425 GCF_000522285.1_Esch_coli_BIDMC_3_V2 Escherichia coli BIDMC 3 Proteobacteria; Gammaproteobacteria; Enterobacterales; Enterobacteriaceae; Escherichia; Escherichia coli
MNNEETFYQAMRRQGVTRRSFLKYCSLAATSLGLGAGMAPKIAWALENKPRIPVVWIHGL WP_001058323.1
MULTISPECIES: hydrogenase-1 small chain [Proteobacteria] Length: 372\nNumber of predicted TMHs: 2\nExp number of AAs in TMHs: 37.2608499999999\nExp number, first 60 AAs: 19.52923\nTotal prob of N-in: 0.94351\nPOSSIBLE N-term signal sequence\ninside 1 19\nTMhelix 20 39\noutside 40 325\nTMhelix 326 348\ninside 349 372

10426 GCF_000492275.1_Esch_coli_BIDMC_37_V1 Escherichia coli BIDMC 37 Proteobacteria; Gammaproteobacteria; Enterobacterales; Enterobacteriaceae; Escherichia; Escherichia coli
MNNEETFYQAMRRQGVTRRSFLKYCSLAATSLGLGAGMAPKIAWALENKPRIPVVWIHGL WP_001058323.1
MULTISPECIES: hydrogenase-1 small chain [Proteobacteria] Length: 372\nNumber of predicted TMHs: 2\nExp number of AAs in TMHs: 37.2608499999999\nExp number, first 60 AAs: 19.52923\nTotal prob of N-in: 0.94351\nPOSSIBLE N-term signal sequence\ninside 1 19\nTMhelix 20 39\noutside 40 325\nTMhelix 326 348\ninside 349 372

10427 GCF_000492275.1_Esch_coli_BIDMC_37_V1 Escherichia coli BIDMC 37 Proteobacteria; Gammaproteobacteria; Enterobacterales; Enterobacteriaceae; Escherichia; Escherichia coli
MTGDNTLIHSHGINRRDFMKLCAALAATMGLSSKAAAEMAESVTNPQRPPVIWIGAQECT WP_000145410.1
MULTISPECIES: hydrogenase-2 small chain [Proteobacteria] Length: 372\nNumber of predicted TMHs: 1\nExp number of AAs in TMHs: 23.72226\nExp number, first 60 AAs: 0.22038\nTotal prob of N-in: 0.12994\noutside 1 331\nTMhelix 332 354\ninside 355 372

10428 GCF_000492255.1_Esch_coli_BIDMC_38_V1 Escherichia coli BIDMC 38 Proteobacteria; Gammaproteobacteria; Enterobacterales; Enterobacteriaceae; Escherichia; Escherichia coli
MTGDNTLIHSHGINRRDFMKLCAALAATMGLSSKAAAEMVESVTNPQRPPVIWIGAQECT WP_000145423.1
hydrogenase 2 small subunit [Escherichia coli] Length: 372\nNumber of predicted TMHs: 1\nExp number of AAs in TMHs: 23.78201\nExp number, first 60 AAs: 0.28355\nTotal prob of N-in: 0.13257\noutside 1 331\nTMhelix 332 354\ninside 355 372

10429 GCF_000492255.1_Esch_coli_BIDMC_38_V1 Escherichia coli BIDMC 38 Proteobacteria; Gammaproteobacteria; Enterobacterales; Enterobacteriaceae; Escherichia; Escherichia coli
MNNEETFYQAMRRQGVTRRSFLKYCSLAATSLGLGAGMAPKIAWALENKPRIPVVWIHGL WP_001295940.1
MULTISPECIES: hydrogenase [Enterobacteriaceae] Length: 372\nNumber of predicted TMHs: 2\nExp number of AAs in TMHs: 37.2608899999999\nExp number, first 60 AAs: 19.52923\nTotal prob of N-in: 0.94351\nPOSSIBLE N-term signal sequence\ninside 1 19\nTMhelix 20 39\noutside 40 325\nTMhelix 326 348\ninside 349 372

10430 GCF_000492235.1_Esch_coli_BIDMC_39_V1 Escherichia coli BIDMC 39 Proteobacteria; Gammaproteobacteria; Enterobacterales; Enterobacteriaceae; Escherichia; Escherichia coli
MTGDNTLIHSHGINRRDFMKLCAALAATMGLSSKAAAEMAESVTNPQRPPVIWIGAQECT WP_000145410.1
MULTISPECIES: hydrogenase-2 small chain [Proteobacteria] Length: 372\nNumber of predicted TMHs: 1\nExp number of AAs in TMHs: 23.72226\nExp number, first 60 AAs: 0.22038\nTotal prob of N-in: 0.12994\noutside 1 331\nTMhelix 332 354\ninside 355 372

10431 GCF_000492235.1_Esch_coli_BIDMC_39_V1 Escherichia coli BIDMC 39 Proteobacteria; Gammaproteobacteria; Enterobacterales; Enterobacteriaceae; Escherichia; Escherichia coli
MNNEETFYQAMRRQGVTRRSFLKYCSLAATSLGLGAGMAPKIAWAQENKPRIPVVWIHGL WP_023281173.1
hydrogenase-1 small chain [Escherichia coli] Length: 372\nNumber of predicted TMHs: 2\nExp number of AAs in TMHs: 36.61267\nExp number, first 60 AAs: 18.81809\nTotal prob of N-in: 0.91316\nPOSSIBLE N-term signal sequence\ninside 1 19\nTMhelix 20 39\noutside 40 325\nTMhelix 326 348\ninside 349 372

10432 GCF_000522105.1_Esch_coli_BIDMC_43a_V1 Escherichia coli BIDMC 43a Proteobacteria; Gammaproteobacteria; Enterobacterales; Enterobacteriaceae; Escherichia; Escherichia coli

MNNEETFYQAMRRQGVTRRSFLKYCSLAATSLGLGAGMAPKIAWALENKPRIPVVWIHGL WP_001058323.1 MULTISPECIES: hydrogenase-1 small chain [Proteobacteria] Length: 372\nNumber of predicted TMHs: 2\nExp number of AAs in TMHs: 37.2608499999999\nExp number, first 60 AAs: 19.52923\nTotal prob of N-in: 0.94351\nPOSSIBLE N-term signal sequence\ninside 1 19\nTMhelix 20 39\noutside 40 325\nTMhelix 326 348\ninside 349 372

10433 GCF_000522105.1_Esch_coli_BIDMC_43a_V1 Escherichia coli BIDMC 43a Proteobacteria; Gammaproteobacteria; Enterobacterales; Enterobacteriaceae; Escherichia; Escherichia coli MTGDNTLIHSHGINRRDFMKLCAALAATMGLSSKAAAEMAESVTNPQRPPVIWIGAQECT WP_000145410.1 MULTISPECIES: hydrogenase-2 small chain [Proteobacteria] Length: 372\nNumber of predicted TMHs: 1\nExp number of AAs in TMHs: 23.72226\nExp number, first 60 AAs: 0.22038\nTotal prob of N-in: 0.12994\noutside 1 331\nTMhelix 332 354\ninside 355 372

10434 GCF_000522085.1_Esch_coli_BIDMC_43b_V1 Escherichia coli BIDMC 43b Proteobacteria; Gammaproteobacteria; Enterobacterales; Enterobacteriaceae; Escherichia; Escherichia coli MNNEETFYQAMRRQGVTRRSFLKYCSLAATSLGLGAGMAPKIAWALENKPRIPVVWIHGL WP_001058323.1 MULTISPECIES: hydrogenase-1 small chain [Proteobacteria] Length: 372\nNumber of predicted TMHs: 2\nExp number of AAs in TMHs: 37.2608499999999\nExp number, first 60 AAs: 19.52923\nTotal prob of N-in: 0.94351\nPOSSIBLE N-term signal sequence\ninside 1 19\nTMhelix 20 39\noutside 40 325\nTMhelix 326 348\ninside 349 372

10435 GCF_000522085.1_Esch_coli_BIDMC_43b_V1 Escherichia coli BIDMC 43b Proteobacteria; Gammaproteobacteria; Enterobacterales; Enterobacteriaceae; Escherichia; Escherichia coli MTGDNTLIHSHGINRRDFMKLCAALAATMGLSSKAAAEMAESVTNPQRPPVIWIGAQECT WP_000145410.1 MULTISPECIES: hydrogenase-2 small chain [Proteobacteria] Length: 372\nNumber of predicted TMHs: 1\nExp number of AAs in TMHs: 23.72226\nExp number, first 60 AAs: 0.22038\nTotal prob of N-in: 0.12994\noutside 1 331\nTMhelix 332 354\ninside 355 372

10436 GCF_000522385.1_Esch_coli_BIDMC_49a_V1 Escherichia coli BIDMC 49a Proteobacteria; Gammaproteobacteria; Enterobacterales; Enterobacteriaceae; Escherichia; Escherichia coli MTGDNTLIHSHGINRRDFMKLCAALAATMGLSSKAAAEMAESVTNPQRPPVIWIGAQECT WP_000145410.1 MULTISPECIES: hydrogenase-2 small chain [Proteobacteria] Length: 372\nNumber of predicted TMHs: 1\nExp number of AAs in TMHs: 23.72226\nExp number, first 60 AAs: 0.22038\nTotal prob of N-in: 0.12994\noutside 1 331\nTMhelix 332 354\ninside 355 372

10437 GCF_000522365.1_Esch_coli_BIDMC_49b_V1 Escherichia coli BIDMC 49b Proteobacteria; Gammaproteobacteria; Enterobacterales; Enterobacteriaceae; Escherichia; Escherichia coli MTGDNTLIHSHGINRRDFMKLCAALAATMGLSSKAAAEMAESVTNPQRPPVIWIGAQECT WP_000145410.1 MULTISPECIES: hydrogenase-2 small chain [Proteobacteria] Length: 372\nNumber of predicted TMHs: 1\nExp number of AAs in TMHs: 23.72226\nExp number, first 60 AAs: 0.22038\nTotal prob of N-in: 0.12994\noutside 1 331\nTMhelix 332 354\ninside 355 372

10438 GCF_000692375.1_Esch_coli_BIDMC_58_V1 Escherichia coli BIDMC 58 Proteobacteria; Gammaproteobacteria; Enterobacterales; Enterobacteriaceae; Escherichia; Escherichia coli MTGDNTLIHSHGINRRDFMKLCAALAATMGLSSKAAAEMVESVTNPQRPPVIWIGAQECT WP_000145423.1 hydrogenase 2 small subunit [Escherichia coli] Length: 372\nNumber of predicted TMHs: 1\nExp number of AAs in TMHs: 23.78201\nExp number, first 60 AAs: 0.28355\nTotal prob of N-in: 0.13257\noutside 1 331\nTMhelix 332 354\ninside 355 372

10439 GCF_000692375.1_Esch_coli_BIDMC_58_V1 Escherichia coli BIDMC 58 Proteobacteria; Gammaproteobacteria; Enterobacterales; Enterobacteriaceae; Escherichia; Escherichia coli MNNEETFYQAMRRQGVTRRSFLKYCSLAATSLGLGAGMAPKIAWALENKPRIPVVWIHGL WP_001295940.1 MULTISPECIES: hydrogenase [Enterobacteriaceae] Length: 372\nNumber of predicted TMHs: 2\nExp number of AAs in TMHs: 37.2608899999999\nExp number, first 60 AAs: 19.52923\nTotal prob of N-in: 0.94351\nPOSSIBLE N-term signal sequence\ninside 1 19\nTMhelix 20 39\noutside 40 325\nTMhelix 326 348\ninside 349 372

10440 GCF_000692395.1_Esch_coli_BIDMC_59_V1 Escherichia coli BIDMC 59 Proteobacteria; Gammaproteobacteria; Enterobacterales; Enterobacteriaceae; Escherichia; Escherichia coli MNNEETFYQAMRRQGVTRRSFLKYCSLAATSLGLGAGMAPKIAWALENKPRIPVVWIHGL WP_001058323.1 MULTISPECIES: hydrogenase-1 small chain [Proteobacteria] Length: 372\nNumber of predicted TMHs:

2\nExp number of AAs in TMHs: 37.2608499999999\nExp number, first 60 AAs: 19.52923\nTotal prob of N-in: 0.94351\nPOSSIBLE N-term signal sequence\ninside 1 19\nTMhelix 20 39\noutside 40 325\nTMhelix 326 348\ninside 349 372

10441 GCF_000692395.1_Esch_coli_BIDMC_59_V1 Escherichia coli BIDMC 59 Proteobacteria; Gammaproteobacteria; Enterobacterales; Enterobacteriaceae; Escherichia; Escherichia coli MTGDNTLIHSHGINRRDFMKLCAALAATMGLSSKAAAEMAESVTNPQRPPVIWIGAQECT WP_000145410.1 MULTISPECIES: hydrogenase-2 small chain [Proteobacteria] Length: 372\nNumber of predicted TMHs: 1\nExp number of AAs in TMHs: 23.72226\nExp number, first 60 AAs: 0.22038\nTotal prob of N-in: 0.12994\noutside 1 331\nTMhelix 332 354\ninside 355 372

10442 GCF_000522405.1_Esch_coli_BIDMC_6_V2 Escherichia coli BIDMC 6 Proteobacteria; Gammaproteobacteria; Enterobacterales; Enterobacteriaceae; Escherichia; Escherichia coli MNNEETFYQAMRRQGVTRRSFLKYCSLAATSLGLGAGMAPKIAWALENKPRIPVVWIHGL WP_001058323.1 MULTISPECIES: hydrogenase-1 small chain [Proteobacteria] Length: 372\nNumber of predicted TMHs: 2\nExp number of AAs in TMHs: 37.2608499999999\nExp number, first 60 AAs: 19.52923\nTotal prob of N-in: 0.94351\nPOSSIBLE N-term signal sequence\ninside 1 19\nTMhelix 20 39\noutside 40 325\nTMhelix 326 348\ninside 349 372

10443 GCF_000522405.1_Esch_coli_BIDMC_6_V2 Escherichia coli BIDMC 6 Proteobacteria; Gammaproteobacteria; Enterobacterales; Enterobacteriaceae; Escherichia; Escherichia coli MTGDNTLIHSHGINRRDFMKLCAALAATMGLSSKAAAEMAESVTNPQRPPVIWIGAQECT WP_000145410.1 MULTISPECIES: hydrogenase-2 small chain [Proteobacteria] Length: 372\nNumber of predicted TMHs: 1\nExp number of AAs in TMHs: 23.72226\nExp number, first 60 AAs: 0.22038\nTotal prob of N-in: 0.12994\noutside 1 331\nTMhelix 332 354\ninside 355 372

10444 GCF_000692415.1_Esch_coli_BIDMC_62_V1 Escherichia coli BIDMC 62 Proteobacteria; Gammaproteobacteria; Enterobacterales; Enterobacteriaceae; Escherichia; Escherichia coli MTGDNTLIHSHGINRRDFMKLCAALAATMGLSSKAAAEMVESVTNPQRPPVIWIGAQECT WP_000145423.1 hydrogenase 2 small subunit [Escherichia coli] Length: 372\nNumber of predicted TMHs: 1\nExp number of AAs in TMHs: 23.78201\nExp number, first 60 AAs: 0.28355\nTotal prob of N-in: 0.13257\noutside 1 331\nTMhelix 332 354\ninside 355 372

10445 GCF_000692415.1_Esch_coli_BIDMC_62_V1 Escherichia coli BIDMC 62 Proteobacteria; Gammaproteobacteria; Enterobacterales; Enterobacteriaceae; Escherichia; Escherichia coli MNNEETFYQAMRRQGVTRRSFLKYCSLAATSLGLGAGMAPKIAWALENKPRIPVVWIHGL WP_001295940.1 MULTISPECIES: hydrogenase [Enterobacteriaceae] Length: 372\nNumber of predicted TMHs: 2\nExp number of AAs in TMHs: 37.2608899999999\nExp number, first 60 AAs: 19.52923\nTotal prob of N-in: 0.94351\nPOSSIBLE N-term signal sequence\ninside 1 19\nTMhelix 20 39\noutside 40 325\nTMhelix 326 348\ninside 349 372

10446 GCF_000692435.1_Esch_coli_BIDMC_63_V1 Escherichia coli BIDMC 63 Proteobacteria; Gammaproteobacteria; Enterobacterales; Enterobacteriaceae; Escherichia; Escherichia coli MNNEETFYQAMRRQGVTRRSFLKYCSLAATSLGLGAGMAPKIAWALENKPRIPVVWIHGL WP_001295940.1 MULTISPECIES: hydrogenase [Enterobacteriaceae] Length: 372\nNumber of predicted TMHs: 2\nExp number of AAs in TMHs: 37.2608899999999\nExp number, first 60 AAs: 19.52923\nTotal prob of N-in: 0.94351\nPOSSIBLE N-term signal sequence\ninside 1 19\nTMhelix 20 39\noutside 40 325\nTMhelix 326 348\ninside 349 372

10447 GCF_000692435.1_Esch_coli_BIDMC_63_V1 Escherichia coli BIDMC 63 Proteobacteria; Gammaproteobacteria; Enterobacterales; Enterobacteriaceae; Escherichia; Escherichia coli MTGDNTLIHSHGINRRDFMKLCAALAATMGLSSKAAAEMVESVTNPQRPPVIWIGAQECT WP_000145423.1 hydrogenase 2 small subunit [Escherichia coli] Length: 372\nNumber of predicted TMHs: 1\nExp number of AAs in TMHs: 23.78201\nExp number, first 60 AAs: 0.28355\nTotal prob of N-in: 0.13257\noutside 1 331\nTMhelix 332 354\ninside 355 372

10448 GCF_000692455.1_Esch_coli_BIDMC_64_V1 Escherichia coli BIDMC 64 Proteobacteria; Gammaproteobacteria; Enterobacterales; Enterobacteriaceae; Escherichia; Escherichia coli MNNEETFYQAMRRQGVTRRSFLKYCSLAATSLGLGAGMAPKIAWALENKPRIPVVWIHGL WP_001295940.1 MULTISPECIES: hydrogenase [Enterobacteriaceae] Length: 372\nNumber of predicted TMHs: 2\nExp number of AAs in TMHs: 37.2608899999999\nExp number, first 60 AAs: 19.52923\nTotal prob of N-in:

0.94351\nPOSSIBLE N-term signal sequence\ninside 1 19\nTMhelix 20 39\noutside 40 325\nTMhelix 326 348\ninside 349 372

10449 GCF_000692455.1_Esch_coli_BIDMC_64_V1 Escherichia coli BIDMC 64 Proteobacteria; Gammaproteobacteria; Enterobacterales; Enterobacteriaceae; Escherichia; Escherichia coli MTGDNTLIHSHGINRRDFMKLCAALAATMGLSSKAAAEMVESVTNPQRPPVIWIGAQECT WP_000145423.1 hydrogenase 2 small subunit [Escherichia coli] Length: 372\nNumber of predicted TMHs: 1\nExp number of AAs in TMHs: 23.78201\nExp number, first 60 AAs: 0.28355\nTotal prob of N-in: 0.13257\noutside 1 331\nTMhelix 332 354\ninside 355 372

10450 GCF_000692475.1_Esch_coli_BIDMC_65_V1 Escherichia coli BIDMC 65 Proteobacteria; Gammaproteobacteria; Enterobacterales; Enterobacteriaceae; Escherichia; Escherichia coli MTGDNTLIHSHGINRRDFMKLCAALAATMGLSSKAAAEMAESVTNPQRPPVIWIGAQECT WP_000145410.1 MULTISPECIES: hydrogenase-2 small chain [Proteobacteria] Length: 372\nNumber of predicted TMHs: 1\nExp number of AAs in TMHs: 23.72226\nExp number, first 60 AAs: 0.22038\nTotal prob of N-in: 0.12994\noutside 1 331\nTMhelix 332 354\ninside 355 372

10451 GCF_000692495.1_Esch_coli_BIDMC_70_V1 Escherichia coli BIDMC 70 Proteobacteria; Gammaproteobacteria; Enterobacterales; Enterobacteriaceae; Escherichia; Escherichia coli MNNEETFYQAMRRQGVTRRSFLKYCSLAATSLGLGAGMAPKIAWALENKPRIPVVWIHGL WP_001295940.1 MULTISPECIES: hydrogenase [Enterobacteriaceae] Length: 372\nNumber of predicted TMHs: 2\nExp number of AAs in TMHs: 37.2608899999999\nExp number, first 60 AAs: 19.52923\nTotal prob of N-in: 0.94351\nPOSSIBLE N-term signal sequence\ninside 1 19\nTMhelix 20 39\noutside 40 325\nTMhelix 326 348\ninside 349 372

10452 GCF_000692495.1_Esch_coli_BIDMC_70_V1 Escherichia coli BIDMC 70 Proteobacteria; Gammaproteobacteria; Enterobacterales; Enterobacteriaceae; Escherichia; Escherichia coli MTGDNTLIHSHGINRRDFMKLCAALAATMGLSSKAAAEMVESVTNPQRPPVIWIGAQECT WP_000145423.1 hydrogenase 2 small subunit [Escherichia coli] Length: 372\nNumber of predicted TMHs: 1\nExp number of AAs in TMHs: 23.78201\nExp number, first 60 AAs: 0.28355\nTotal prob of N-in: 0.13257\noutside 1 331\nTMhelix 332 354\ninside 355 372

10453 GCF_000692515.1_Esch_coli_BIDMC_71_V1 Escherichia coli BIDMC 71 Proteobacteria; Gammaproteobacteria; Enterobacterales; Enterobacteriaceae; Escherichia; Escherichia coli MTGDNTLIHSHGINRRDFMKLCAALAATMGLSSKAAAEMAESVTNPQRPPVIWIGAQECT WP_000145410.1 MULTISPECIES: hydrogenase-2 small chain [Proteobacteria] Length: 372\nNumber of predicted TMHs: 1\nExp number of AAs in TMHs: 23.72226\nExp number, first 60 AAs: 0.22038\nTotal prob of N-in: 0.12994\noutside 1 331\nTMhelix 332 354\ninside 355 372

10454 GCF_000692515.1_Esch_coli_BIDMC_71_V1 Escherichia coli BIDMC 71 Proteobacteria; Gammaproteobacteria; Enterobacterales; Enterobacteriaceae; Escherichia; Escherichia coli MNNEETFYQAMRRQGVTRRSFLKYCSLAATSLGLGAGMAPKIAWALENKPRIPVVWIHGL WP_001058323.1 MULTISPECIES: hydrogenase-1 small chain [Proteobacteria] Length: 372\nNumber of predicted TMHs: 2\nExp number of AAs in TMHs: 37.2608499999999\nExp number, first 60 AAs: 19.52923\nTotal prob of N-in: 0.94351\nPOSSIBLE N-term signal sequence\ninside 1 19\nTMhelix 20 39\noutside 40 325\nTMhelix 326 348\ninside 349 372

10455 GCF_000692535.1_Esch_coli_BIDMC_72_V1 Escherichia coli BIDMC 72 Proteobacteria; Gammaproteobacteria; Enterobacterales; Enterobacteriaceae; Escherichia; Escherichia coli MTGDNTLIHSHGINRRDFMKLCAALAATMGLSSKAAAEMVESVTNPQRPPVIWIGAQECT WP_000145423.1 hydrogenase 2 small subunit [Escherichia coli] Length: 372\nNumber of predicted TMHs: 1\nExp number of AAs in TMHs: 23.78201\nExp number, first 60 AAs: 0.28355\nTotal prob of N-in: 0.13257\noutside 1 331\nTMhelix 332 354\ninside 355 372

10456 GCF_000692535.1_Esch_coli_BIDMC_72_V1 Escherichia coli BIDMC 72 Proteobacteria; Gammaproteobacteria; Enterobacterales; Enterobacteriaceae; Escherichia; Escherichia coli MNNEETFYQAMRRQGVTRRSFLKYCSLAATSLGLGAGMAPKIAWALENKPRIPVVWIHGL WP_001538532.1 hydrogenase-1 small chain [Escherichia coli] Length: 372\nNumber of predicted TMHs: 2\nExp number of AAs in TMHs: 37.27041\nExp number, first 60 AAs: 19.52927\nTotal prob of N-in: 0.94352\nPOSSIBLE N-term signal sequence\ninside 1 19\nTMhelix 20 39\noutside 40 325\nTMhelix 326 348\ninside 349 372

10457 GCF_000692555.1_Esch_coli_BIDMC_73_V1 Escherichia coli BIDMC 73 Proteobacteria; Gammaproteobacteria; Enterobacterales; Enterobacteriaceae; Escherichia; Escherichia coli
MNNEETFYQAMRRQGVTRRSFLKYCSLAATSLGLGAGMAPKIAWALENKPRIPVVWIHGL WP_001538532.1
hydrogenase-1 small chain [Escherichia coli] Length: 372\nNumber of predicted TMHs: 2\nExp number of AAs in TMHs: 37.27041\nExp number, first 60 AAs: 19.52927\nTotal prob of N-in: 0.94352\nPOSSIBLE N-term signal sequence\ninside 1 19\nTMhelix 20 39\noutside 40 325\nTMhelix 326 348\ninside 349 372

10458 GCF_000692555.1_Esch_coli_BIDMC_73_V1 Escherichia coli BIDMC 73 Proteobacteria; Gammaproteobacteria; Enterobacterales; Enterobacteriaceae; Escherichia; Escherichia coli
MTGDNTLIHSHGINRRDFMKLCAALAATMGLSSKAAAEMVESVTNPQRPPVIWIGAQECT WP_000145423.1
hydrogenase 2 small subunit [Escherichia coli] Length: 372\nNumber of predicted TMHs: 1\nExp number of AAs in TMHs: 23.78201\nExp number, first 60 AAs: 0.28355\nTotal prob of N-in: 0.13257\noutside 1 331\nTMhelix 332 354\ninside 355 372

10459 GCF_000692575.1_Esch_coli_BIDMC_74_V1 Escherichia coli BIDMC 74 Proteobacteria; Gammaproteobacteria; Enterobacterales; Enterobacteriaceae; Escherichia; Escherichia coli
MTGDNTLIHSHGINRRDFMKLCAALAATMGLSSKAAAEMAESVTNPQRPPVIWIGAQECT WP_000145410.1
MULTISPECIES: hydrogenase-2 small chain [Proteobacteria] Length: 372\nNumber of predicted TMHs: 1\nExp number of AAs in TMHs: 23.72226\nExp number, first 60 AAs: 0.22038\nTotal prob of N-in: 0.12994\noutside 1 331\nTMhelix 332 354\ninside 355 372

10460 GCF_000692575.1_Esch_coli_BIDMC_74_V1 Escherichia coli BIDMC 74 Proteobacteria; Gammaproteobacteria; Enterobacterales; Enterobacteriaceae; Escherichia; Escherichia coli
MNNEETFYQAMRRQGVTRRSFLKYCSLAATSLGLGAGMAPKIAWALENKPRIPVVWIHGL WP_001058323.1
MULTISPECIES: hydrogenase-1 small chain [Proteobacteria] Length: 372\nNumber of predicted TMHs: 2\nExp number of AAs in TMHs: 37.2608499999999\nExp number, first 60 AAs: 19.52923\nTotal prob of N-in: 0.94351\nPOSSIBLE N-term signal sequence\ninside 1 19\nTMhelix 20 39\noutside 40 325\nTMhelix 326 348\ninside 349 372

10461 GCF_000692595.1_Esch_coli_BIDMC_75_V1 Escherichia coli BIDMC 75 Proteobacteria; Gammaproteobacteria; Enterobacterales; Enterobacteriaceae; Escherichia; Escherichia coli
MTGDNTLIHSHGINRRDFMKLCAALAATMGLSSKAAAEMVESVTNPQRPPVIWIGAQECT WP_000145423.1
hydrogenase 2 small subunit [Escherichia coli] Length: 372\nNumber of predicted TMHs: 1\nExp number of AAs in TMHs: 23.78201\nExp number, first 60 AAs: 0.28355\nTotal prob of N-in: 0.13257\noutside 1 331\nTMhelix 332 354\ninside 355 372

10462 GCF_000692595.1_Esch_coli_BIDMC_75_V1 Escherichia coli BIDMC 75 Proteobacteria; Gammaproteobacteria; Enterobacterales; Enterobacteriaceae; Escherichia; Escherichia coli
MNNEETFYQAMRRQGVTRRSFLKYCSLAATSLGLGAGMAPKIAWALENKPRIPVVWIHGL WP_001295940.1
MULTISPECIES: hydrogenase [Enterobacteriaceae] Length: 372\nNumber of predicted TMHs: 2\nExp number of AAs in TMHs: 37.2608899999999\nExp number, first 60 AAs: 19.52923\nTotal prob of N-in: 0.94351\nPOSSIBLE N-term signal sequence\ninside 1 19\nTMhelix 20 39\noutside 40 325\nTMhelix 326 348\ninside 349 372

10463 GCF_000692615.1_Esch_coli_BIDMC_76_V1 Escherichia coli BIDMC 76 Proteobacteria; Gammaproteobacteria; Enterobacterales; Enterobacteriaceae; Escherichia; Escherichia coli
MTGDNTLIHSHGINRRDFMKLCAALAATMGLSSKAAAEMVESVTNPQRPPVIWIGAQECT WP_000145423.1
hydrogenase 2 small subunit [Escherichia coli] Length: 372\nNumber of predicted TMHs: 1\nExp number of AAs in TMHs: 23.78201\nExp number, first 60 AAs: 0.28355\nTotal prob of N-in: 0.13257\noutside 1 331\nTMhelix 332 354\ninside 355 372

10464 GCF_000692615.1_Esch_coli_BIDMC_76_V1 Escherichia coli BIDMC 76 Proteobacteria; Gammaproteobacteria; Enterobacterales; Enterobacteriaceae; Escherichia; Escherichia coli
MNNEETFYQAMRRQGVTRRSFLKYCSLAATSLGLGAGMAPKIAWALENKPRIPVVWIHGL WP_001295940.1
MULTISPECIES: hydrogenase [Enterobacteriaceae] Length: 372\nNumber of predicted TMHs: 2\nExp number of AAs in TMHs: 37.2608899999999\nExp number, first 60 AAs: 19.52923\nTotal prob of N-in: 0.94351\nPOSSIBLE N-term signal sequence\ninside 1 19\nTMhelix 20 39\noutside 40 325\nTMhelix 326 348\ninside 349 372

10465 GCF_000692635.1_Esch_coli_BIDMC_77_V1 Escherichia coli BIDMC 77 Proteobacteria; Gammaproteobacteria; Enterobacterales; Enterobacteriaceae; Escherichia; Escherichia coli

MNNEETFYQAMRRQGVTRRSFLKYCSLAATSLGLGAGMAPKIAWALENKPRIPVVWIHGL WP_001058323.1 MULTISPECIES: hydrogenase-1 small chain [Proteobacteria] Length: 372\nNumber of predicted TMHs: 2\nExp number of AAs in TMHs: 37.2608499999999\nExp number, first 60 AAs: 19.52923\nTotal prob of N-in: 0.94351\nPOSSIBLE N-term signal sequence\ninside 1 19\nTMhelix 20 39\noutside 40 325\nTMhelix 326 348\ninside 349 372

10466 GCF_000692635.1_Esch_coli_BIDMC_77_V1 Escherichia coli BIDMC 77 Proteobacteria; Gammaproteobacteria; Enterobacterales; Enterobacteriaceae; Escherichia; Escherichia coli MTGDNTLIHSHGINRRDFMKLCAALAATMGLSSKAAAEMAESVTNPQRPPVIWIGAQECT WP_000145410.1 MULTISPECIES: hydrogenase-2 small chain [Proteobacteria] Length: 372\nNumber of predicted TMHs: 1\nExp number of AAs in TMHs: 23.72226\nExp number, first 60 AAs: 0.22038\nTotal prob of N-in: 0.12994\noutside 1 331\nTMhelix 332 354\ninside 355 372

10467 GCF_000692655.1_Esch_coli_BIDMC_78_V1 Escherichia coli BIDMC 78 Proteobacteria; Gammaproteobacteria; Enterobacterales; Enterobacteriaceae; Escherichia; Escherichia coli MTGDNTLIHSHGINRRDFMKLCAALAATMGLSSKAAAEMVESVTNPQRPPVIWIGAQECT WP_000145423.1 hydrogenase 2 small subunit [Escherichia coli] Length: 372\nNumber of predicted TMHs: 1\nExp number of AAs in TMHs: 23.78201\nExp number, first 60 AAs: 0.28355\nTotal prob of N-in: 0.13257\noutside 1 331\nTMhelix 332 354\ninside 355 372

10468 GCF_000692655.1_Esch_coli_BIDMC_78_V1 Escherichia coli BIDMC 78 Proteobacteria; Gammaproteobacteria; Enterobacterales; Enterobacteriaceae; Escherichia; Escherichia coli MNNEETFYQAMRRQGVTRRSFLKYCSLAATSLGLGAGMAPKIAWALENKPRIPVVWIHGL WP_001295940.1 MULTISPECIES: hydrogenase [Enterobacteriaceae] Length: 372\nNumber of predicted TMHs: 2\nExp number of AAs in TMHs: 37.2608899999999\nExp number, first 60 AAs: 19.52923\nTotal prob of N-in: 0.94351\nPOSSIBLE N-term signal sequence\ninside 1 19\nTMhelix 20 39\noutside 40 325\nTMhelix 326 348\ninside 349 372

10469 GCF_000692675.1_Esch_coli_BIDMC_79_V1 Escherichia coli BIDMC 79 Proteobacteria; Gammaproteobacteria; Enterobacterales; Enterobacteriaceae; Escherichia; Escherichia coli MTGDNTLIHSHGINRRDFMKLCAALAATMGLSSKAAAEMVESVTNPQRPPVIWIGAQECT WP_000145423.1 hydrogenase 2 small subunit [Escherichia coli] Length: 372\nNumber of predicted TMHs: 1\nExp number of AAs in TMHs: 23.78201\nExp number, first 60 AAs: 0.28355\nTotal prob of N-in: 0.13257\noutside 1 331\nTMhelix 332 354\ninside 355 372

10470 GCF_000692675.1_Esch_coli_BIDMC_79_V1 Escherichia coli BIDMC 79 Proteobacteria; Gammaproteobacteria; Enterobacterales; Enterobacteriaceae; Escherichia; Escherichia coli MNNEETFYQAMRRQGVTRRSFLKYCSLAATSLGLGAGMAPKIAWALENKPRIPVVWIHGL WP_001295940.1 MULTISPECIES: hydrogenase [Enterobacteriaceae] Length: 372\nNumber of predicted TMHs: 2\nExp number of AAs in TMHs: 37.2608899999999\nExp number, first 60 AAs: 19.52923\nTotal prob of N-in: 0.94351\nPOSSIBLE N-term signal sequence\ninside 1 19\nTMhelix 20 39\noutside 40 325\nTMhelix 326 348\ninside 349 372

10471 GCF_000633675.1_Esch_coli_BIDMC_82_V1 Escherichia coli BIDMC 82 Proteobacteria; Gammaproteobacteria; Enterobacterales; Enterobacteriaceae; Escherichia; Escherichia coli MNNEETFYQAMRRQGVTRRSFLKYCSLAATSLGLGAGMAPKIAWALENKPRIPVVWIHGL WP_001058323.1 MULTISPECIES: hydrogenase-1 small chain [Proteobacteria] Length: 372\nNumber of predicted TMHs: 2\nExp number of AAs in TMHs: 37.2608499999999\nExp number, first 60 AAs: 19.52923\nTotal prob of N-in: 0.94351\nPOSSIBLE N-term signal sequence\ninside 1 19\nTMhelix 20 39\noutside 40 325\nTMhelix 326 348\ninside 349 372

10472 GCF_000633675.1_Esch_coli_BIDMC_82_V1 Escherichia coli BIDMC 82 Proteobacteria; Gammaproteobacteria; Enterobacterales; Enterobacteriaceae; Escherichia; Escherichia coli MTGDNTLIHSHGINRRDFMKLCAALAATMGLSSKAAAEMAESVTNPQRPPVIWIGAQECT WP_000145410.1 MULTISPECIES: hydrogenase-2 small chain [Proteobacteria] Length: 372\nNumber of predicted TMHs: 1\nExp number of AAs in TMHs: 23.72226\nExp number, first 60 AAs: 0.22038\nTotal prob of N-in: 0.12994\noutside 1 331\nTMhelix 332 354\ninside 355 372

10473 GCF_000633655.1_Esch_coli_BIDMC_83_V1 Escherichia coli BIDMC 83 Proteobacteria; Gammaproteobacteria; Enterobacterales; Enterobacteriaceae; Escherichia; Escherichia coli MTGDNTLIHSHGINRRDFMKLCAALAATMGLSSKAAAEMAESVTNPQRPPVIWIGAQECT WP_000145410.1

MULTISPECIES: hydrogenase-2 small chain [Proteobacteria] Length: 372\nNumber of predicted TMHs: 1\nExp number of AAs in TMHs: 23.72226\nExp number, first 60 AAs: 0.22038\nTotal prob of N-in: 0.12994\noutside 1 331\nTMhelix 332 354\ninside 355 372

10474 GCF_000633655.1_Esch_coli_BIDMC_83_V1 Escherichia coli BIDMC 83 Proteobacteria; Gammaproteobacteria; Enterobacterales; Enterobacteriaceae; Escherichia; Escherichia coli MNNEETFYQAMRRQGVTRRSFLKYCSLAATSLGLGAGMAPKIAWALENKPRIPVVWIHGL WP_001058323.1 MULTISPECIES: hydrogenase-1 small chain [Proteobacteria] Length: 372\nNumber of predicted TMHs: 2\nExp number of AAs in TMHs: 37.2608499999999\nExp number, first 60 AAs: 19.52923\nTotal prob of N-in: 0.94351\nPOSSIBLE N-term signal sequence\ninside 1 19\nTMhelix 20 39\noutside 40 325\nTMhelix 326 348\ninside 349 372

10475 GCF_000522265.1_Esch_coli_BIDMC_9_V2 Escherichia coli BIDMC 9 Proteobacteria; Gammaproteobacteria; Enterobacterales; Enterobacteriaceae; Escherichia; Escherichia coli MTGDNTLIHSHGINRRDFMKLCAALAATMGLSSKAAAEMAESVTNPQRPPVIWIGAQECT WP_000145410.1 MULTISPECIES: hydrogenase-2 small chain [Proteobacteria] Length: 372\nNumber of predicted TMHs: 1\nExp number of AAs in TMHs: 23.72226\nExp number, first 60 AAs: 0.22038\nTotal prob of N-in: 0.12994\noutside 1 331\nTMhelix 332 354\ninside 355 372

10476 GCF_000522265.1_Esch_coli_BIDMC_9_V2 Escherichia coli BIDMC 9 Proteobacteria; Gammaproteobacteria; Enterobacterales; Enterobacteriaceae; Escherichia; Escherichia coli MNNEETFYQAMRRQGVTRRSFLKYCSLAATSLGLGAGMAPKIAWALENKPRIPVVWIHGL WP_001058323.1 MULTISPECIES: hydrogenase-1 small chain [Proteobacteria] Length: 372\nNumber of predicted TMHs: 2\nExp number of AAs in TMHs: 37.2608499999999\nExp number, first 60 AAs: 19.52923\nTotal prob of N-in: 0.94351\nPOSSIBLE N-term signal sequence\ninside 1 19\nTMhelix 20 39\noutside 40 325\nTMhelix 326 348\ninside 349 372

10477 GCF_000022665.1_ASM2266v1 Escherichia coli BL21(DE3) Proteobacteria; Gammaproteobacteria; Enterobacterales; Enterobacteriaceae; Escherichia; Escherichia coli MTGDNTLIHSHGINRRDFMKLCAALAATMGLSSKAAAEMAESVTNPQRPPVIWIGAQECT WP_000145410.1 MULTISPECIES: hydrogenase-2 small chain [Proteobacteria] Length: 372\nNumber of predicted TMHs: 1\nExp number of AAs in TMHs: 23.72226\nExp number, first 60 AAs: 0.22038\nTotal prob of N-in: 0.12994\noutside 1 331\nTMhelix 332 354\ninside 355 372

10478 GCF_000022665.1_ASM2266v1 Escherichia coli BL21(DE3) Proteobacteria; Gammaproteobacteria; Enterobacterales; Enterobacteriaceae; Escherichia; Escherichia coli MNNEETFYQAMRRQGVTRRSFLKYCSLAATSLGLGAGMAPKIAWALENKPRIPVVWIHGL WP_001058323.1 MULTISPECIES: hydrogenase-1 small chain [Proteobacteria] Length: 372\nNumber of predicted TMHs: 2\nExp number of AAs in TMHs: 37.2608499999999\nExp number, first 60 AAs: 19.52923\nTotal prob of N-in: 0.94351\nPOSSIBLE N-term signal sequence\ninside 1 19\nTMhelix 20 39\noutside 40 325\nTMhelix 326 348\ninside 349 372

10479 GCF_000009565.1_ASM956v1 Escherichia coli BL21(DE3) Proteobacteria; Gammaproteobacteria; Enterobacterales; Enterobacteriaceae; Escherichia; Escherichia coli MTGDNTLIHSHGINRRDFMKLCAALAATMGLSSKAAAEMAESVTNPQRPPVIWIGAQECT WP_000145410.1 MULTISPECIES: hydrogenase-2 small chain [Proteobacteria] Length: 372\nNumber of predicted TMHs: 1\nExp number of AAs in TMHs: 23.72226\nExp number, first 60 AAs: 0.22038\nTotal prob of N-in: 0.12994\noutside 1 331\nTMhelix 332 354\ninside 355 372

10480 GCF_000009565.1_ASM956v1 Escherichia coli BL21(DE3) Proteobacteria; Gammaproteobacteria; Enterobacterales; Enterobacteriaceae; Escherichia; Escherichia coli MNNEETFYQAMRRQGVTRRSFLKYCSLAATSLGLGAGMAPKIAWALENKPRIPVVWIHGL WP_001058323.1 MULTISPECIES: hydrogenase-1 small chain [Proteobacteria] Length: 372\nNumber of predicted TMHs: 2\nExp number of AAs in TMHs: 37.2608499999999\nExp number, first 60 AAs: 19.52923\nTotal prob of N-in: 0.94351\nPOSSIBLE N-term signal sequence\ninside 1 19\nTMhelix 20 39\noutside 40 325\nTMhelix 326 348\ninside 349 372

10481 GCF_000474845.1_Esch_coli_BWH_24_V1_PacBio Escherichia coli BWH 24 Proteobacteria; Gammaproteobacteria; Enterobacterales; Enterobacteriaceae; Escherichia; Escherichia coli MNNEETFYQAMRRQGVTRRSFLKYCSLAATSLGLGAGMAPKIAWALENKPRIPVVWIHGL WP_001295940.1 MULTISPECIES: hydrogenase [Enterobacteriaceae] Length: 372\nNumber of predicted TMHs: 2\nExp

number of AAs in TMHs: 37.2608899999999\nExp number, first 60 AAs: 19.52923\nTotal prob of N-in: 0.94351\nPOSSIBLE N-term signal sequence\ninside 1 19\nTMhelix 20 39\noutside 40 325\nTMhelix 326 348\ninside 349 372

10482 GCF_000474845.1_Esch_coli_BWH_24_V1_PacBio Escherichia coli BWH 24 Proteobacteria; Gammaproteobacteria; Enterobacterales; Enterobacteriaceae; Escherichia; Escherichia coli MTGDNTLIHSHGINRRDFMKLCAALAATMGLSSKAAAEMVESVTNPQRPPVIWIGAQECT WP_000145423.1 hydrogenase 2 small subunit [Escherichia coli] Length: 372\nNumber of predicted TMHs: 1\nExp number of AAs in TMHs: 23.78201\nExp number, first 60 AAs: 0.28355\nTotal prob of N-in: 0.13257\noutside 1 331\nTMhelix 332 354\ninside 355 372

10483 GCF_000492655.1_Esch_coli_BWH_32_V1 Escherichia coli BWH 32 Proteobacteria; Gammaproteobacteria; Enterobacterales; Enterobacteriaceae; Escherichia; Escherichia coli MNNEETFYQAMRRQGVTRRSFLKYCSLAATSLGLGAGMAPKIAWALENKPRIPVVWIHGL WP_001058323.1 MULTISPECIES: hydrogenase-1 small chain [Proteobacteria] Length: 372\nNumber of predicted TMHs: 2\nExp number of AAs in TMHs: 37.2608499999999\nExp number, first 60 AAs: 19.52923\nTotal prob of N-in: 0.94351\nPOSSIBLE N-term signal sequence\ninside 1 19\nTMhelix 20 39\noutside 40 325\nTMhelix 326 348\ninside 349 372

10484 GCF_000492655.1_Esch_coli_BWH_32_V1 Escherichia coli BWH 32 Proteobacteria; Gammaproteobacteria; Enterobacterales; Enterobacteriaceae; Escherichia; Escherichia coli MTGDNTLIHSHGINRRDFMKLCAALAATMGLSSKAAAEMAESVTNPQRPPVIWIGAQECT WP_000145410.1 MULTISPECIES: hydrogenase-2 small chain [Proteobacteria] Length: 372\nNumber of predicted TMHs: 1\nExp number of AAs in TMHs: 23.72226\nExp number, first 60 AAs: 0.22038\nTotal prob of N-in: 0.12994\noutside 1 331\nTMhelix 332 354\ninside 355 372

10485 GCF_000522345.1_Esch_coli_BWH_34_V1 Escherichia coli BWH 34 Proteobacteria; Gammaproteobacteria; Enterobacterales; Enterobacteriaceae; Escherichia; Escherichia coli MNNEETFYQAMRRQGVTRRSFLKYCSLAATSLGLGAGMAPKIAWALENKPRIPVVWIHGL WP_001295940.1 MULTISPECIES: hydrogenase [Enterobacteriaceae] Length: 372\nNumber of predicted TMHs: 2\nExp number of AAs in TMHs: 37.2608899999999\nExp number, first 60 AAs: 19.52923\nTotal prob of N-in: 0.94351\nPOSSIBLE N-term signal sequence\ninside 1 19\nTMhelix 20 39\noutside 40 325\nTMhelix 326 348\ninside 349 372

10486 GCF_000522345.1_Esch_coli_BWH_34_V1 Escherichia coli BWH 34 Proteobacteria; Gammaproteobacteria; Enterobacterales; Enterobacteriaceae; Escherichia; Escherichia coli MTGDNTLIHSHGINRRDFMKLCAALAATMGLSSKAAAEMVESVTNPQRPPVIWIGAQECT WP_000145423.1 hydrogenase 2 small subunit [Escherichia coli] Length: 372\nNumber of predicted TMHs: 1\nExp number of AAs in TMHs: 23.78201\nExp number, first 60 AAs: 0.28355\nTotal prob of N-in: 0.13257\noutside 1 331\nTMhelix 332 354\ninside 355 372

10487 GCF_000522325.1_Esch_coli_BWH_40_V1 Escherichia coli BWH 40 Proteobacteria; Gammaproteobacteria; Enterobacterales; Enterobacteriaceae; Escherichia; Escherichia coli MNNEETFYQAMRRQGVTRRSFLKYCSLAATSLGLGAGMAPKIAWALENKPRIPVVWIHGL WP_001058323.1 MULTISPECIES: hydrogenase-1 small chain [Proteobacteria] Length: 372\nNumber of predicted TMHs: 2\nExp number of AAs in TMHs: 37.2608499999999\nExp number, first 60 AAs: 19.52923\nTotal prob of N-in: 0.94351\nPOSSIBLE N-term signal sequence\ninside 1 19\nTMhelix 20 39\noutside 40 325\nTMhelix 326 348\ninside 349 372

10488 GCF_000522325.1_Esch_coli_BWH_40_V1 Escherichia coli BWH 40 Proteobacteria; Gammaproteobacteria; Enterobacterales; Enterobacteriaceae; Escherichia; Escherichia coli MTGDNTLIHSHGINRRDFMKLCAALAATMGLSSKAAAEMAESVTNPQRPPVIWIGAQECT WP_000145410.1 MULTISPECIES: hydrogenase-2 small chain [Proteobacteria] Length: 372\nNumber of predicted TMHs: 1\nExp number of AAs in TMHs: 23.72226\nExp number, first 60 AAs: 0.22038\nTotal prob of N-in: 0.12994\noutside 1 331\nTMhelix 332 354\ninside 355 372

10489 GCF_002092895.1_ASM209289v1 Escherichia coli C Proteobacteria; Gammaproteobacteria; Enterobacterales; Enterobacteriaceae; Escherichia; Escherichia coli MTGDNTLIHSHGINRRDFMKLCAALAATMGLSSKAAAEMAESVTNPQRPPVIWIGAQECT WP_000145410.1 MULTISPECIES: hydrogenase-2 small chain [Proteobacteria] Length: 372\nNumber of predicted TMHs:

1\nExp number of AAs in TMHs: 23.72226\nExp number, first 60 AAs: 0.22038\nTotal prob of N-in: 0.12994\noutside 1 331\nTMhelix 332 354\ninside 355 372

10490 GCF_002079225.1_ASM207922v1 Escherichia coli C Proteobacteria; Gammaproteobacteria; Enterobacterales; Enterobacteriaceae; Escherichia; Escherichia coli MNNEETFYQAMRRQGVTRRSFLKYCSLAATSLGLGAGMAPKIAWALENKPRIPVVWIHGL WP_001058323.1 MULTISPECIES: hydrogenase-1 small chain [Proteobacteria] Length: 372\nNumber of predicted TMHs: 2\nExp number of AAs in TMHs: 37.2608499999999\nExp number, first 60 AAs: 19.52923\nTotal prob of N-in: 0.94351\nPOSSIBLE N-term signal sequence\ninside 1 19\nTMhelix 20 39\noutside 40 325\nTMhelix 326 348\ninside 349 372

10491 GCF_002092895.1_ASM209289v1 Escherichia coli C Proteobacteria; Gammaproteobacteria; Enterobacterales; Enterobacteriaceae; Escherichia; Escherichia coli MNNEETFYQAMRRQGVTRRSFLKYCSLAATSLGLGAGMAPKIAWALENKPRIPVVWIHGL WP_001058323.1 MULTISPECIES: hydrogenase-1 small chain [Proteobacteria] Length: 372\nNumber of predicted TMHs: 2\nExp number of AAs in TMHs: 37.2608499999999\nExp number, first 60 AAs: 19.52923\nTotal prob of N-in: 0.94351\nPOSSIBLE N-term signal sequence\ninside 1 19\nTMhelix 20 39\noutside 40 325\nTMhelix 326 348\ninside 349 372

10492 GCF_002079225.1_ASM207922v1 Escherichia coli C Proteobacteria; Gammaproteobacteria; Enterobacterales; Enterobacteriaceae; Escherichia; Escherichia coli MTGDNTLIHSHGINRRDFMKLCAALAATMGLSSKAAAEMAESVTNPQRPPVIWIGAQECT WP_000145410.1 MULTISPECIES: hydrogenase-2 small chain [Proteobacteria] Length: 372\nNumber of predicted TMHs: 1\nExp number of AAs in TMHs: 23.72226\nExp number, first 60 AAs: 0.22038\nTotal prob of N-in: 0.12994\noutside 1 331\nTMhelix 332 354\ninside 355 372

10493 GCF_000446115.2_ASM44611v2 Escherichia coli C1214_90 Proteobacteria; Gammaproteobacteria; Enterobacterales; Enterobacteriaceae; Escherichia; Escherichia coli MNNEETFYQAMRRQGVTRRSFLKYCSLAATSLGLGAGMAPKIAWALENKPRIPVVWIHGL WP_001058323.1 MULTISPECIES: hydrogenase-1 small chain [Proteobacteria] Length: 372\nNumber of predicted TMHs: 2\nExp number of AAs in TMHs: 37.2608499999999\nExp number, first 60 AAs: 19.52923\nTotal prob of N-in: 0.94351\nPOSSIBLE N-term signal sequence\ninside 1 19\nTMhelix 20 39\noutside 40 325\nTMhelix 326 348\ninside 349 372

10494 GCF_000446115.2_ASM44611v2 Escherichia coli C1214_90 Proteobacteria; Gammaproteobacteria; Enterobacterales; Enterobacteriaceae; Escherichia; Escherichia coli MTGDNTLIHSHGINRRDFMKLCAALAATMGLSSKAAAEMAESVTNPQRPPVIWIGAQECT WP_000145410.1 MULTISPECIES: hydrogenase-2 small chain [Proteobacteria] Length: 372\nNumber of predicted TMHs: 1\nExp number of AAs in TMHs: 23.72226\nExp number, first 60 AAs: 0.22038\nTotal prob of N-in: 0.12994\noutside 1 331\nTMhelix 332 354\ninside 355 372

10495 GCF_000446095.2_ASM44609v2 Escherichia coli C1244_91 Proteobacteria; Gammaproteobacteria; Enterobacterales; Enterobacteriaceae; Escherichia; Escherichia coli MNNEETFYQAMRRQGVTRRSFLKYCSLAATSLGLGAGMAPKIAWALENKPRIPVVWIHGL WP_001359719.1 hydrogenase [Escherichia coli] Length: 372\nNumber of predicted TMHs: 2\nExp number of AAs in TMHs: 37.2176\nExp number, first 60 AAs: 19.53448\nTotal prob of N-in: 0.94360\nPOSSIBLE N-term signal sequence\ninside 1 19\nTMhelix 20 39\noutside 40 325\nTMhelix 326 348\ninside 349 372

10496 GCF_000446095.2_ASM44609v2 Escherichia coli C1244_91 Proteobacteria; Gammaproteobacteria; Enterobacterales; Enterobacteriaceae; Escherichia; Escherichia coli MTGDNTLIHSHGINRRDFMKLCAALAATMGLSSKAAAEMAESVTNPQRPPVIWIGAQECT WP_000145410.1 MULTISPECIES: hydrogenase-2 small chain [Proteobacteria] Length: 372\nNumber of predicted TMHs: 1\nExp number of AAs in TMHs: 23.72226\nExp number, first 60 AAs: 0.22038\nTotal prob of N-in: 0.12994\noutside 1 331\nTMhelix 332 354\ninside 355 372

10497 GCF_000446075.2_ASM44607v2 Escherichia coli C12_92 Proteobacteria; Gammaproteobacteria; Enterobacterales; Enterobacteriaceae; Escherichia; Escherichia coli MTGDNTLIHSHGINRRDFMKLCAALAATMGLSSKAAAEMAESVTNPQRPPVIWIGAQECT WP_000145410.1 MULTISPECIES: hydrogenase-2 small chain [Proteobacteria] Length: 372\nNumber of predicted TMHs: 1\nExp number of AAs in TMHs: 23.72226\nExp number, first 60 AAs: 0.22038\nTotal prob of N-in: 0.12994\noutside 1 331\nTMhelix 332 354\ninside 355 372

10498 GCF_000446075.2_ASM44607v2 Escherichia coli C12_92 Proteobacteria; Gammaproteobacteria; Enterobacterales; Enterobacteriaceae; Escherichia; Escherichia coli
MNNEETFYQAMRRQGVTRRSFLKYCSLAATSLGLGAGMAPKIAWALENKPRIPVVWIHGL WP_001359719.1
hydrogenase [Escherichia coli] Length: 372\nNumber of predicted TMHs: 2\nExp number of AAs in TMHs: 37.2176\nExp number, first 60 AAs: 19.53448\nTotal prob of N-in: 0.94360\nPOSSIBLE N-term signal sequence\ninside 1 19\nTMhelix 20 39\noutside 40 325\nTMhelix 326 348\ninside 349 372

10499 GCF_000446135.2_ASM44613v2 Escherichia coli C154_11 Proteobacteria; Gammaproteobacteria; Enterobacterales; Enterobacteriaceae; Escherichia; Escherichia coli
MNNEETFYQAMRRQGVTRRSFLKYCSLAATSLGLGAGMAPKIAWALENKPRIPVVWIHGL WP_001359719.1
hydrogenase [Escherichia coli] Length: 372\nNumber of predicted TMHs: 2\nExp number of AAs in TMHs: 37.2176\nExp number, first 60 AAs: 19.53448\nTotal prob of N-in: 0.94360\nPOSSIBLE N-term signal sequence\ninside 1 19\nTMhelix 20 39\noutside 40 325\nTMhelix 326 348\ninside 349 372

10500 GCF_000446155.2_ASM44615v2 Escherichia coli C155_11 Proteobacteria; Gammaproteobacteria; Enterobacterales; Enterobacteriaceae; Escherichia; Escherichia coli
MTGDNTLIHSHGINRRDFMKLCAALAATMGLSSKAAAEMAESVTNPQRPPVIWIGAQECT WP_000145410.1
MULTISPECIES: hydrogenase-2 small chain [Proteobacteria] Length: 372\nNumber of predicted TMHs: 1\nExp number of AAs in TMHs: 23.72226\nExp number, first 60 AAs: 0.22038\nTotal prob of N-in: 0.12994\noutside 1 331\nTMhelix 332 354\ninside 355 372

10501 GCF_000446155.2_ASM44615v2 Escherichia coli C155_11 Proteobacteria; Gammaproteobacteria; Enterobacterales; Enterobacteriaceae; Escherichia; Escherichia coli
MNNEETFYQAMRRQGVTRRSFLKYCSLAATSLGLGAGMAPKIAWALENKPRIPVVWIHGL WP_001058323.1
MULTISPECIES: hydrogenase-1 small chain [Proteobacteria] Length: 372\nNumber of predicted TMHs: 2\nExp number of AAs in TMHs: 37.2608499999999\nExp number, first 60 AAs: 19.52923\nTotal prob of N-in: 0.94351\nPOSSIBLE N-term signal sequence\ninside 1 19\nTMhelix 20 39\noutside 40 325\nTMhelix 326 348\ninside 349 372

10502 GCF_000446175.2_ASM44617v2 Escherichia coli C157_11 Proteobacteria; Gammaproteobacteria; Enterobacterales; Enterobacteriaceae; Escherichia; Escherichia coli
MNNEETFYQAMRRQGVTRRSFLKYCSLAATSLGLGAGMAPKIAWALENKPRIPVVWIHGL WP_001359719.1
hydrogenase [Escherichia coli] Length: 372\nNumber of predicted TMHs: 2\nExp number of AAs in TMHs: 37.2176\nExp number, first 60 AAs: 19.53448\nTotal prob of N-in: 0.94360\nPOSSIBLE N-term signal sequence\ninside 1 19\nTMhelix 20 39\noutside 40 325\nTMhelix 326 348\ninside 349 372

10503 GCF_000446175.2_ASM44617v2 Escherichia coli C157_11 Proteobacteria; Gammaproteobacteria; Enterobacterales; Enterobacteriaceae; Escherichia; Escherichia coli
MTGDNTLIHSHGINRRDFMKLCAALAATMGLSSKAAAEMAESVTNPQRPPVIWIGAQECT WP_000145410.1
MULTISPECIES: hydrogenase-2 small chain [Proteobacteria] Length: 372\nNumber of predicted TMHs: 1\nExp number of AAs in TMHs: 23.72226\nExp number, first 60 AAs: 0.22038\nTotal prob of N-in: 0.12994\noutside 1 331\nTMhelix 332 354\ninside 355 372

10504 GCF_000446195.2_ASM44619v2 Escherichia coli C161_11 Proteobacteria; Gammaproteobacteria; Enterobacterales; Enterobacteriaceae; Escherichia; Escherichia coli
MNNEETFYQAMRRQGVTRRSFLKYCSLAATSLGLGAGMAPKIAWALENKPRIPVVWIHGL WP_001058323.1
MULTISPECIES: hydrogenase-1 small chain [Proteobacteria] Length: 372\nNumber of predicted TMHs: 2\nExp number of AAs in TMHs: 37.2608499999999\nExp number, first 60 AAs: 19.52923\nTotal prob of N-in: 0.94351\nPOSSIBLE N-term signal sequence\ninside 1 19\nTMhelix 20 39\noutside 40 325\nTMhelix 326 348\ninside 349 372

10505 GCF_000446195.2_ASM44619v2 Escherichia coli C161_11 Proteobacteria; Gammaproteobacteria; Enterobacterales; Enterobacteriaceae; Escherichia; Escherichia coli
MTGDNTLIHSHGINRRDFMKLCAALAATMGLSSKAAAEMAESVTNPQRPPVIWIGAQECT WP_000145410.1
MULTISPECIES: hydrogenase-2 small chain [Proteobacteria] Length: 372\nNumber of predicted TMHs: 1\nExp number of AAs in TMHs: 23.72226\nExp number, first 60 AAs: 0.22038\nTotal prob of N-in: 0.12994\noutside 1 331\nTMhelix 332 354\ninside 355 372

10506 GCF_000446265.2_ASM44626v2 Escherichia coli C166_11 Proteobacteria; Gammaproteobacteria; Enterobacterales; Enterobacteriaceae; Escherichia; Escherichia coli
MNNEETFYQAMRRQGVTRRSFLKYCSLAATSLGLGAGMAPKIAWALENKPRIPVVWIHGL WP_001058323.1

MULTISPECIES: hydrogenase-1 small chain [Proteobacteria] Length: 372\nNumber of predicted TMHs: 2\nExp number of AAs in TMHs: 37.2608499999999\nExp number, first 60 AAs: 19.52923\nTotal prob of N-in: 0.94351\nPOSSIBLE N-term signal sequence\ninside 1 19\nTMhelix 20 39\noutside 40 325\nTMhelix 326 348\ninside 349 372

10507 GCF_000446265.2_ASM44626v2 Escherichia coli C166_11 Proteobacteria; Gammaproteobacteria; Enterobacterales; Enterobacteriaceae; Escherichia; Escherichia coli
MTGDNTLIHSHGINRRDFMKLCAALAATMGLSSKAAAEMAESVTNPQRPPVIWIGAQECT WP_000145410.1
MULTISPECIES: hydrogenase-2 small chain [Proteobacteria] Length: 372\nNumber of predicted TMHs: 1\nExp number of AAs in TMHs: 23.72226\nExp number, first 60 AAs: 0.22038\nTotal prob of N-in: 0.12994\noutside 1 331\nTMhelix 332 354\ninside 355 372

10508 GCF_000447145.2_ASM44714v2 Escherichia coli C170_11 Proteobacteria; Gammaproteobacteria; Enterobacterales; Enterobacteriaceae; Escherichia; Escherichia coli
MTGDNTLIHSHGINRRDFMKLCAALAATMGLSSKAAAEMAESVTNPQRPPVIWIGAQECT WP_000145410.1
MULTISPECIES: hydrogenase-2 small chain [Proteobacteria] Length: 372\nNumber of predicted TMHs: 1\nExp number of AAs in TMHs: 23.72226\nExp number, first 60 AAs: 0.22038\nTotal prob of N-in: 0.12994\noutside 1 331\nTMhelix 332 354\ninside 355 372

10509 GCF_000447145.2_ASM44714v2 Escherichia coli C170_11 Proteobacteria; Gammaproteobacteria; Enterobacterales; Enterobacteriaceae; Escherichia; Escherichia coli
MNNEETFYQAMRRQGVTRRSFLKYCSLAATSLGLGAGMAPKIAWALENKPRIPVVWIHGL WP_001295940.1
MULTISPECIES: hydrogenase [Enterobacteriaceae] Length: 372\nNumber of predicted TMHs: 2\nExp number of AAs in TMHs: 37.2608899999999\nExp number, first 60 AAs: 19.52923\nTotal prob of N-in: 0.94351\nPOSSIBLE N-term signal sequence\ninside 1 19\nTMhelix 20 39\noutside 40 325\nTMhelix 326 348\ninside 349 372

10510 GCF_000446225.2_ASM44622v2 Escherichia coli C213_10 Proteobacteria; Gammaproteobacteria; Enterobacterales; Enterobacteriaceae; Escherichia; Escherichia coli
MNNEETFYQAMRRQGVTRRSFLKYCSLAATSLGLGAGMAPKIAWALENKPRIPVVWIHGL WP_001058323.1
MULTISPECIES: hydrogenase-1 small chain [Proteobacteria] Length: 372\nNumber of predicted TMHs: 2\nExp number of AAs in TMHs: 37.2608499999999\nExp number, first 60 AAs: 19.52923\nTotal prob of N-in: 0.94351\nPOSSIBLE N-term signal sequence\ninside 1 19\nTMhelix 20 39\noutside 40 325\nTMhelix 326 348\ninside 349 372

10511 GCF_000446225.2_ASM44622v2 Escherichia coli C213_10 Proteobacteria; Gammaproteobacteria; Enterobacterales; Enterobacteriaceae; Escherichia; Escherichia coli
MTGDNTLIHSHGINRRDFMKLCAALAATMGLSSKAAAEMAESVTNPQRPPVIWIGAQECT WP_000145410.1
MULTISPECIES: hydrogenase-2 small chain [Proteobacteria] Length: 372\nNumber of predicted TMHs: 1\nExp number of AAs in TMHs: 23.72226\nExp number, first 60 AAs: 0.22038\nTotal prob of N-in: 0.12994\noutside 1 331\nTMhelix 332 354\ninside 355 372

10512 GCF_000446285.2_ASM44628v2 Escherichia coli C2139_99 Proteobacteria; Gammaproteobacteria; Enterobacterales; Enterobacteriaceae; Escherichia; Escherichia coli
MTGDNTLIHSHGINRRDFMKLCAALAATMGLSSKAAAEMAESVTNPQRPPVIWIGAQECT WP_000145410.1
MULTISPECIES: hydrogenase-2 small chain [Proteobacteria] Length: 372\nNumber of predicted TMHs: 1\nExp number of AAs in TMHs: 23.72226\nExp number, first 60 AAs: 0.22038\nTotal prob of N-in: 0.12994\noutside 1 331\nTMhelix 332 354\ninside 355 372

10513 GCF_000446285.2_ASM44628v2 Escherichia coli C2139_99 Proteobacteria; Gammaproteobacteria; Enterobacterales; Enterobacteriaceae; Escherichia; Escherichia coli
MNNEETFYQAMRRQGVTRRSFLKYCSLAATSLGLGAGMAPKIAWALENKPRIPVVWIHGL WP_001058323.1
MULTISPECIES: hydrogenase-1 small chain [Proteobacteria] Length: 372\nNumber of predicted TMHs: 2\nExp number of AAs in TMHs: 37.2608499999999\nExp number, first 60 AAs: 19.52923\nTotal prob of N-in: 0.94351\nPOSSIBLE N-term signal sequence\ninside 1 19\nTMhelix 20 39\noutside 40 325\nTMhelix 326 348\ninside 349 372

10514 GCF_000446305.2_ASM44630v2 Escherichia coli C238_91 Proteobacteria; Gammaproteobacteria; Enterobacterales; Enterobacteriaceae; Escherichia; Escherichia coli
MTGDNTLIHSHGINRRDFMKLCAALAATMGLSSKAAAEMAESVTNPQRPPVIWIGAQECT WP_000145410.1
MULTISPECIES: hydrogenase-2 small chain [Proteobacteria] Length: 372\nNumber of predicted TMHs:

1\nExp number of AAs in TMHs: 23.72226\nExp number, first 60 AAs: 0.22038\nTotal prob of N-in: 0.12994\noutside 1 331\nTMhelix 332 354\ninside 355 372

10515 GCF_000446305.2_ASM44630v2 Escherichia coli C238_91 Proteobacteria; Gammaproteobacteria; Enterobacterales; Enterobacteriaceae; Escherichia; Escherichia coli MNNEETFYQAMRRQGVTRRSFLKYCSLAATSLGLGAGMAPKIAWALENKPRIPVVWIHGL WP_001058323.1 MULTISPECIES: hydrogenase-1 small chain [Proteobacteria] Length: 372\nNumber of predicted TMHs: 2\nExp number of AAs in TMHs: 37.2608499999999\nExp number, first 60 AAs: 19.52923\nTotal prob of N-in: 0.94351\nPOSSIBLE N-term signal sequence\ninside 1 19\nTMhelix 20 39\noutside 40 325\nTMhelix 326 348\ninside 349 372

10516 GCF_000446325.2_ASM44632v2 Escherichia coli C260_92 Proteobacteria; Gammaproteobacteria; Enterobacterales; Enterobacteriaceae; Escherichia; Escherichia coli MNNEETFYQAMRRQGVTRRSFLKYCSLAATSLGLGAGMAPKIAWALENKPRIPVVWIHGL WP_001058323.1 MULTISPECIES: hydrogenase-1 small chain [Proteobacteria] Length: 372\nNumber of predicted TMHs: 2\nExp number of AAs in TMHs: 37.2608499999999\nExp number, first 60 AAs: 19.52923\nTotal prob of N-in: 0.94351\nPOSSIBLE N-term signal sequence\ninside 1 19\nTMhelix 20 39\noutside 40 325\nTMhelix 326 348\ninside 349 372

10517 GCF_000446325.2_ASM44632v2 Escherichia coli C260_92 Proteobacteria; Gammaproteobacteria; Enterobacterales; Enterobacteriaceae; Escherichia; Escherichia coli MTGDNTLIHSHGINRRDFMKLCAALAATMGLSSKAAAEMAESVTNPQRPPVIWIGAQECT WP_000145410.1 MULTISPECIES: hydrogenase-2 small chain [Proteobacteria] Length: 372\nNumber of predicted TMHs: 1\nExp number of AAs in TMHs: 23.72226\nExp number, first 60 AAs: 0.22038\nTotal prob of N-in: 0.12994\noutside 1 331\nTMhelix 332 354\ninside 355 372

10518 GCF_000446345.2_ASM44634v2 Escherichia coli C262_10 Proteobacteria; Gammaproteobacteria; Enterobacterales; Enterobacteriaceae; Escherichia; Escherichia coli MTGDNTLIHSHGINRRDFMKLCAALAATMGLSSKAAAEMAESVTNPQRPPVIWIGAQECT WP_000145410.1 MULTISPECIES: hydrogenase-2 small chain [Proteobacteria] Length: 372\nNumber of predicted TMHs: 1\nExp number of AAs in TMHs: 23.72226\nExp number, first 60 AAs: 0.22038\nTotal prob of N-in: 0.12994\noutside 1 331\nTMhelix 332 354\ninside 355 372

10519 GCF_000446345.2_ASM44634v2 Escherichia coli C262_10 Proteobacteria; Gammaproteobacteria; Enterobacterales; Enterobacteriaceae; Escherichia; Escherichia coli MNNEETFYQAMRRQGVTRRSFLKYCSLAATSLGLGAGMAPKIAWALENKPRIPVVWIHGL WP_024225128.1 hydrogenase-1 small chain [Escherichia coli] Length: 372\nNumber of predicted TMHs: 2\nExp number of AAs in TMHs: 38.67234\nExp number, first 60 AAs: 19.53155\nTotal prob of N-in: 0.94349\nPOSSIBLE N-term signal sequence\ninside 1 19\nTMhelix 20 39\noutside 40 325\nTMhelix 326 348\ninside 349 372

10520 GCF_000446365.2_ASM44636v2 Escherichia coli C283_09 Proteobacteria; Gammaproteobacteria; Enterobacterales; Enterobacteriaceae; Escherichia; Escherichia coli MNNEETFYQAMRRQGVTRRSFLKYCSLAATSLGLGAGMAPKIAWALENKPRIPVVWIHGL WP_001058323.1 MULTISPECIES: hydrogenase-1 small chain [Proteobacteria] Length: 372\nNumber of predicted TMHs: 2\nExp number of AAs in TMHs: 37.2608499999999\nExp number, first 60 AAs: 19.52923\nTotal prob of N-in: 0.94351\nPOSSIBLE N-term signal sequence\ninside 1 19\nTMhelix 20 39\noutside 40 325\nTMhelix 326 348\ninside 349 372

10521 GCF_000446365.2_ASM44636v2 Escherichia coli C283_09 Proteobacteria; Gammaproteobacteria; Enterobacterales; Enterobacteriaceae; Escherichia; Escherichia coli MTGDNTLIHSHGINRRDFMKLCAALAATMGLSSKAAAEMAESVTNPQRPPVIWIGAQECT WP_000145410.1 MULTISPECIES: hydrogenase-2 small chain [Proteobacteria] Length: 372\nNumber of predicted TMHs: 1\nExp number of AAs in TMHs: 23.72226\nExp number, first 60 AAs: 0.22038\nTotal prob of N-in: 0.12994\noutside 1 331\nTMhelix 332 354\ninside 355 372

10522 GCF_000446385.2_ASM44638v2 Escherichia coli C295_10 Proteobacteria; Gammaproteobacteria; Enterobacterales; Enterobacteriaceae; Escherichia; Escherichia coli MTGDNTLIHSHGINRRDFMKLCAALAATMGLSSKAAAEMAESVTNPQRPPVIWIGAQECT WP_000145410.1 MULTISPECIES: hydrogenase-2 small chain [Proteobacteria] Length: 372\nNumber of predicted TMHs: 1\nExp number of AAs in TMHs: 23.72226\nExp number, first 60 AAs: 0.22038\nTotal prob of N-in: 0.12994\noutside 1 331\nTMhelix 332 354\ninside 355 372

10523 GCF_000446385.2_ASM44638v2 Escherichia coli C295_10 Proteobacteria; Gammaproteobacteria; Enterobacterales; Enterobacteriaceae; Escherichia; Escherichia coli
MNNEETFYQAMRRQGVTRRSFLKYCSLAATSLGLGAGMAPKIAWALENKPRIPVVWIHGL WP_001591878.1
hydrogenase-1 small chain [Escherichia coli] Length: 372\nNumber of predicted TMHs: 2\nExp number of AAs in TMHs: 37.25867\nExp number, first 60 AAs: 19.52849\nTotal prob of N-in: 0.94350\nPOSSIBLE N-term signal sequence\ninside 1 19\nTMhelix 20 39\noutside 40 325\nTMhelix 326 348\ninside 349 372

10524 GCF_000446405.2_ASM44640v2 Escherichia coli C341_10 Proteobacteria; Gammaproteobacteria; Enterobacterales; Enterobacteriaceae; Escherichia; Escherichia coli
MNNEETFYQAMRRQGVTRRSFLKYCSLAATSLGLGAGMAPKIAWALENKPRIPVVWIHGL WP_001295940.1
MULTISPECIES: hydrogenase [Enterobacteriaceae] Length: 372\nNumber of predicted TMHs: 2\nExp number of AAs in TMHs: 37.2608899999999\nExp number, first 60 AAs: 19.52923\nTotal prob of N-in: 0.94351\nPOSSIBLE N-term signal sequence\ninside 1 19\nTMhelix 20 39\noutside 40 325\nTMhelix 326 348\ninside 349 372

10525 GCF_000446405.2_ASM44640v2 Escherichia coli C341_10 Proteobacteria; Gammaproteobacteria; Enterobacterales; Enterobacteriaceae; Escherichia; Escherichia coli
MTGDNTLIHSHGINRRDFMKLCAALAATMGLSSKAAAEMAESVTNPQRPPVIWIGAQECT WP_000145410.1
MULTISPECIES: hydrogenase-2 small chain [Proteobacteria] Length: 372\nNumber of predicted TMHs: 1\nExp number of AAs in TMHs: 23.72226\nExp number, first 60 AAs: 0.22038\nTotal prob of N-in: 0.12994\noutside 1 331\nTMhelix 332 354\ninside 355 372

10526 GCF_000446425.2_ASM44642v2 Escherichia coli C343_08 Proteobacteria; Gammaproteobacteria; Enterobacterales; Enterobacteriaceae; Escherichia; Escherichia coli
MNNEETFYQAMRRQGVTRRSFLKYCSLAATSLGLGAGMAPKIAWALENKPRIPVVWIHGL WP_001058323.1
MULTISPECIES: hydrogenase-1 small chain [Proteobacteria] Length: 372\nNumber of predicted TMHs: 2\nExp number of AAs in TMHs: 37.2608499999999\nExp number, first 60 AAs: 19.52923\nTotal prob of N-in: 0.94351\nPOSSIBLE N-term signal sequence\ninside 1 19\nTMhelix 20 39\noutside 40 325\nTMhelix 326 348\ninside 349 372

10527 GCF_000446425.2_ASM44642v2 Escherichia coli C343_08 Proteobacteria; Gammaproteobacteria; Enterobacterales; Enterobacteriaceae; Escherichia; Escherichia coli
MTGDNTLIHSHGINRRDFMKLCAALAATMGLSSKAAAEMAESVTNPQRPPVIWIGAQECT WP_000145410.1
MULTISPECIES: hydrogenase-2 small chain [Proteobacteria] Length: 372\nNumber of predicted TMHs: 1\nExp number of AAs in TMHs: 23.72226\nExp number, first 60 AAs: 0.22038\nTotal prob of N-in: 0.12994\noutside 1 331\nTMhelix 332 354\ninside 355 372

10528 GCF_000354355.2_ASM35435v2 Escherichia coli C-34666 Proteobacteria; Gammaproteobacteria; Enterobacterales; Enterobacteriaceae; Escherichia; Escherichia coli
MNNEETFYQAMRRQGVTRRSFLKYCSLAATSLGLGAGMAPKIAWALENKPRIPVVWIHGL WP_001058320.1
hydrogenase [Escherichia coli] Length: 372\nNumber of predicted TMHs: 2\nExp number of AAs in TMHs: 37.26082\nExp number, first 60 AAs: 19.52923\nTotal prob of N-in: 0.94351\nPOSSIBLE N-term signal sequence\ninside 1 19\nTMhelix 20 39\noutside 40 325\nTMhelix 326 348\ninside 349 372

10529 GCF_000354355.2_ASM35435v2 Escherichia coli C-34666 Proteobacteria; Gammaproteobacteria; Enterobacterales; Enterobacteriaceae; Escherichia; Escherichia coli
MTGDNTLIHSHGINRRDFMKLCAALAATMGLSSKAAAEMAESVTNPQRPPVIWIGAQECT WP_000145410.1
MULTISPECIES: hydrogenase-2 small chain [Proteobacteria] Length: 372\nNumber of predicted TMHs: 1\nExp number of AAs in TMHs: 23.72226\nExp number, first 60 AAs: 0.22038\nTotal prob of N-in: 0.12994\noutside 1 331\nTMhelix 332 354\ninside 355 372

10530 GCF_000446445.2_ASM44644v2 Escherichia coli C347_93 Proteobacteria; Gammaproteobacteria; Enterobacterales; Enterobacteriaceae; Escherichia; Escherichia coli
MNNEETFYQAMRRQGVTRRSFLKYCSLAATSLGLGAGMAPKIAWALENKPRIPVVWIHGL WP_024228890.1
hydrogenase [Escherichia coli] Length: 372\nNumber of predicted TMHs: 2\nExp number of AAs in TMHs: 38.3063\nExp number, first 60 AAs: 19.52635\nTotal prob of N-in: 0.94360\nPOSSIBLE N-term signal sequence\ninside 1 19\nTMhelix 20 39\noutside 40 325\nTMhelix 326 348\ninside 349 372

10531 GCF_000446445.2_ASM44644v2 Escherichia coli C347_93 Proteobacteria; Gammaproteobacteria; Enterobacterales; Enterobacteriaceae; Escherichia; Escherichia coli
MTGDNTLIHSHGINRRDFMKLCAALAATMGLSSKAAAEMAESVTNPQRPPVIWIGAQECT WP_024228528.1

hydrogenase 2 small subunit [Escherichia coli] Length: 372\nNumber of predicted TMHs: 1\nExp number of AAs in TMHs: 23.61909\nExp number, first 60 AAs: 0.22137\nTotal prob of N-in: 0.12572\noutside 1 331\nTMhelix 332 354\ninside 355 372

10532 GCF_000446465.2_ASM44646v2 Escherichia coli C353_09 Proteobacteria; Gammaproteobacteria; Enterobacterales; Enterobacteriaceae; Escherichia; Escherichia coli
MTGDNTLIHSHGINRRDFMKLCAALAATMGLSSKAAAEMAESVTNPQRPPVIWIGAQECT WP_000145410.1
MULTISPECIES: hydrogenase-2 small chain [Proteobacteria] Length: 372\nNumber of predicted TMHs: 1\nExp number of AAs in TMHs: 23.72226\nExp number, first 60 AAs: 0.22038\nTotal prob of N-in: 0.12994\noutside 1 331\nTMhelix 332 354\ninside 355 372

10533 GCF_000446465.2_ASM44646v2 Escherichia coli C353_09 Proteobacteria; Gammaproteobacteria; Enterobacterales; Enterobacteriaceae; Escherichia; Escherichia coli
MNNEETFYQAMRRQGVTRRSFLKYCSLAATSLGLGAGMAPKIAWALENKPRIPVVWIHGL WP_001058323.1
MULTISPECIES: hydrogenase-1 small chain [Proteobacteria] Length: 372\nNumber of predicted TMHs: 2\nExp number of AAs in TMHs: 37.2608499999999\nExp number, first 60 AAs: 19.52923\nTotal prob of N-in: 0.94351\nPOSSIBLE N-term signal sequence\ninside 1 19\nTMhelix 20 39\noutside 40 325\nTMhelix 326 348\ninside 349 372

10534 GCF_000446485.2_ASM44648v2 Escherichia coli C354_03B Proteobacteria; Gammaproteobacteria; Enterobacterales; Enterobacteriaceae; Escherichia; Escherichia coli
MNNEETFYQAMRRQGVTRRSFLKYCSLAATSLGLGAGMAPKIAWALENKPRIPVVWIHGL WP_001301613.1
hydrogenase [Escherichia coli] Length: 372\nNumber of predicted TMHs: 2\nExp number of AAs in TMHs: 37.46598\nExp number, first 60 AAs: 19.53139\nTotal prob of N-in: 0.94364\nPOSSIBLE N-term signal sequence\ninside 1 19\nTMhelix 20 39\noutside 40 325\nTMhelix 326 348\ninside 349 372

10535 GCF_000446485.2_ASM44648v2 Escherichia coli C354_03B Proteobacteria; Gammaproteobacteria; Enterobacterales; Enterobacteriaceae; Escherichia; Escherichia coli
MTGDNTLIHSHGINRRDFMKLCAALAATMGLSSKAAAEMAESVTNPQRPPVIWIGAQECT WP_000145410.1
MULTISPECIES: hydrogenase-2 small chain [Proteobacteria] Length: 372\nNumber of predicted TMHs: 1\nExp number of AAs in TMHs: 23.72226\nExp number, first 60 AAs: 0.22038\nTotal prob of N-in: 0.12994\noutside 1 331\nTMhelix 332 354\ninside 355 372

10536 GCF_000446505.2_ASM44650v2 Escherichia coli C40_11 Proteobacteria; Gammaproteobacteria; Enterobacterales; Enterobacteriaceae; Escherichia; Escherichia coli
MNNEETFYQAMRRQGVTRRSFLKYCSLAATSLGLGAGMAPKIAWALENKPRIPVVWIHGL WP_001058323.1
MULTISPECIES: hydrogenase-1 small chain [Proteobacteria] Length: 372\nNumber of predicted TMHs: 2\nExp number of AAs in TMHs: 37.2608499999999\nExp number, first 60 AAs: 19.52923\nTotal prob of N-in: 0.94351\nPOSSIBLE N-term signal sequence\ninside 1 19\nTMhelix 20 39\noutside 40 325\nTMhelix 326 348\ninside 349 372

10537 GCF_000446505.2_ASM44650v2 Escherichia coli C40_11 Proteobacteria; Gammaproteobacteria; Enterobacterales; Enterobacteriaceae; Escherichia; Escherichia coli
MTGDNTLIHSHGINRRDFMKLCAALAATMGLSSKAAAEMAESVTNPQRPPVIWIGAQECT WP_000145410.1
MULTISPECIES: hydrogenase-2 small chain [Proteobacteria] Length: 372\nNumber of predicted TMHs: 1\nExp number of AAs in TMHs: 23.72226\nExp number, first 60 AAs: 0.22038\nTotal prob of N-in: 0.12994\noutside 1 331\nTMhelix 332 354\ninside 355 372

10538 GCF_000446525.2_ASM44652v2 Escherichia coli C418_89 Proteobacteria; Gammaproteobacteria; Enterobacterales; Enterobacteriaceae; Escherichia; Escherichia coli
MTGDNTLIHSHGINRRDFMKLCAALAATMGLSSKAAAEMVESVTNPQRPPVIWIGAQECT WP_000145423.1
hydrogenase 2 small subunit [Escherichia coli] Length: 372\nNumber of predicted TMHs: 1\nExp number of AAs in TMHs: 23.78201\nExp number, first 60 AAs: 0.28355\nTotal prob of N-in: 0.13257\noutside 1 331\nTMhelix 332 354\ninside 355 372

10539 GCF_000446525.2_ASM44652v2 Escherichia coli C418_89 Proteobacteria; Gammaproteobacteria; Enterobacterales; Enterobacteriaceae; Escherichia; Escherichia coli
MNNEETFYQAMRRQGVTRRSFLKYCSLAATSLGLGAGMAPKIAWALENKPRIPVVWIHGL WP_001295940.1
MULTISPECIES: hydrogenase [Enterobacteriaceae] Length: 372\nNumber of predicted TMHs: 2\nExp number of AAs in TMHs: 37.2608899999999\nExp number, first 60 AAs: 19.52923\nTotal prob of N-in:

0.94351\nPOSSIBLE N-term signal sequence\ninside 1 19\nTMhelix 20 39\noutside 40 325\nTMhelix 326 348\ninside 349 372

10540 GCF_000446545.2_ASM44654v2 Escherichia coli C458_10 Proteobacteria; Gammaproteobacteria; Enterobacterales; Enterobacteriaceae; Escherichia; Escherichia coli MNNEETFYQAMRRQGVTRRSFLKYCSLAATSLGLGAGMAPKIAWALENKPRIPVVWIHGL WP_001058320.1 hydrogenase [Escherichia coli] Length: 372\nNumber of predicted TMHs: 2\nExp number of AAs in TMHs: 37.26082\nExp number, first 60 AAs: 19.52923\nTotal prob of N-in: 0.94351\nPOSSIBLE N-term signal sequence\ninside 1 19\nTMhelix 20 39\noutside 40 325\nTMhelix 326 348\ninside 349 372

10541 GCF_000446545.2_ASM44654v2 Escherichia coli C458_10 Proteobacteria; Gammaproteobacteria; Enterobacterales; Enterobacteriaceae; Escherichia; Escherichia coli MTGDNTLIHSHGINRRDFMKLCAALAATMGLSSKAAAEMAESVTNPQRPPVIWIGAQECT WP_000145410.1 MULTISPECIES: hydrogenase-2 small chain [Proteobacteria] Length: 372\nNumber of predicted TMHs: 1\nExp number of AAs in TMHs: 23.72226\nExp number, first 60 AAs: 0.22038\nTotal prob of N-in: 0.12994\noutside 1 331\nTMhelix 332 354\ninside 355 372

10542 GCF_000446565.2_ASM44656v2 Escherichia coli C488_07 Proteobacteria; Gammaproteobacteria; Enterobacterales; Enterobacteriaceae; Escherichia; Escherichia coli MNNEETFYQAMRRQGVTRRSFLKYCSLAATSLGLGAGMAPKIAWALENKPRIPVVWIHGL WP_001295940.1 MULTISPECIES: hydrogenase [Enterobacteriaceae] Length: 372\nNumber of predicted TMHs: 2\nExp number of AAs in TMHs: 37.2608899999999\nExp number, first 60 AAs: 19.52923\nTotal prob of N-in: 0.94351\nPOSSIBLE N-term signal sequence\ninside 1 19\nTMhelix 20 39\noutside 40 325\nTMhelix 326 348\ninside 349 372

10543 GCF_000446565.2_ASM44656v2 Escherichia coli C488_07 Proteobacteria; Gammaproteobacteria; Enterobacterales; Enterobacteriaceae; Escherichia; Escherichia coli MTGDNTLIHSHGINRRDFMKLCAALAATMGLSSKAAAEMAESVTNPQRPPVIWIGAQECT WP_000145410.1 MULTISPECIES: hydrogenase-2 small chain [Proteobacteria] Length: 372\nNumber of predicted TMHs: 1\nExp number of AAs in TMHs: 23.72226\nExp number, first 60 AAs: 0.22038\nTotal prob of N-in: 0.12994\noutside 1 331\nTMhelix 332 354\ninside 355 372

10544 GCF_000446585.2_ASM44658v2 Escherichia coli C496_10 Proteobacteria; Gammaproteobacteria; Enterobacterales; Enterobacteriaceae; Escherichia; Escherichia coli MTGDNTLIHSHGINRRDFMKLCAALAATMGLSSKAAAEMAESVTNPQRPPVIWIGAQECT WP_000145410.1 MULTISPECIES: hydrogenase-2 small chain [Proteobacteria] Length: 372\nNumber of predicted TMHs: 1\nExp number of AAs in TMHs: 23.72226\nExp number, first 60 AAs: 0.22038\nTotal prob of N-in: 0.12994\noutside 1 331\nTMhelix 332 354\ninside 355 372

10545 GCF_000446585.2_ASM44658v2 Escherichia coli C496_10 Proteobacteria; Gammaproteobacteria; Enterobacterales; Enterobacteriaceae; Escherichia; Escherichia coli MNNEETFYQAMRRQGVTRRSFLKYCSLAATSLGLGAGMAPKIAWALENKPRIPVVWIHGL WP_001058323.1 MULTISPECIES: hydrogenase-1 small chain [Proteobacteria] Length: 372\nNumber of predicted TMHs: 2\nExp number of AAs in TMHs: 37.2608499999999\nExp number, first 60 AAs: 19.52923\nTotal prob of N-in: 0.94351\nPOSSIBLE N-term signal sequence\ninside 1 19\nTMhelix 20 39\noutside 40 325\nTMhelix 326 348\ninside 349 372

10546 GCF_000446605.2_ASM44660v2 Escherichia coli C497_10 Proteobacteria; Gammaproteobacteria; Enterobacterales; Enterobacteriaceae; Escherichia; Escherichia coli MTGDNTLIHSHGINRRDFMKLCAALAATMGLSSKAAAEMAESVTNPQRPPVIWIGAQECT WP_000145410.1 MULTISPECIES: hydrogenase-2 small chain [Proteobacteria] Length: 372\nNumber of predicted TMHs: 1\nExp number of AAs in TMHs: 23.72226\nExp number, first 60 AAs: 0.22038\nTotal prob of N-in: 0.12994\noutside 1 331\nTMhelix 332 354\ninside 355 372

10547 GCF_000446605.2_ASM44660v2 Escherichia coli C497_10 Proteobacteria; Gammaproteobacteria; Enterobacterales; Enterobacteriaceae; Escherichia; Escherichia coli MNNEETFYQAMRRQGVTRRSFLKYCSLAATSLGLGAGMAPKIAWALENKPRIPVVWIHGL WP_001295940.1 MULTISPECIES: hydrogenase [Enterobacteriaceae] Length: 372\nNumber of predicted TMHs: 2\nExp number of AAs in TMHs: 37.2608899999999\nExp number, first 60 AAs: 19.52923\nTotal prob of N-in: 0.94351\nPOSSIBLE N-term signal sequence\ninside 1 19\nTMhelix 20 39\noutside 40 325\nTMhelix 326 348\ninside 349 372

10548 GCF_000446625.2_ASM44662v2 Escherichia coli C527_94 Proteobacteria; Gammaproteobacteria; Enterobacterales; Enterobacteriaceae; Escherichia; Escherichia coli
MNNEETFYQAMRRQGVTRRSFLKYCSLAATSLGLGAGMAPKIAWALENKPRIPVVWIHGL WP_001058323.1
MULTISPECIES: hydrogenase-1 small chain [Proteobacteria] Length: 372\nNumber of predicted TMHs: 2\nExp number of AAs in TMHs: 37.2608499999999\nExp number, first 60 AAs: 19.52923\nTotal prob of N-in: 0.94351\nPOSSIBLE N-term signal sequence\ninside 1 19\nTMhelix 20 39\noutside 40 325\nTMhelix 326 348\ninside 349 372

10549 GCF_000446625.2_ASM44662v2 Escherichia coli C527_94 Proteobacteria; Gammaproteobacteria; Enterobacterales; Enterobacteriaceae; Escherichia; Escherichia coli
MTGDNTLIHSHGINRRDFMKLCAALAATMGLSSKAAAEMAESVTNPQRPPVIWIGAQECT WP_000145410.1
MULTISPECIES: hydrogenase-2 small chain [Proteobacteria] Length: 372\nNumber of predicted TMHs: 1\nExp number of AAs in TMHs: 23.72226\nExp number, first 60 AAs: 0.22038\nTotal prob of N-in: 0.12994\noutside 1 331\nTMhelix 332 354\ninside 355 372

10550 GCF_000446645.2_ASM44664v2 Escherichia coli C581_05 Proteobacteria; Gammaproteobacteria; Enterobacterales; Enterobacteriaceae; Escherichia; Escherichia coli
MTGDNTLIHSHGINRRDFMKLCAALAATMGLSSKAAAEMAESVTNPQRPPVIWIGAQECT WP_000145410.1
MULTISPECIES: hydrogenase-2 small chain [Proteobacteria] Length: 372\nNumber of predicted TMHs: 1\nExp number of AAs in TMHs: 23.72226\nExp number, first 60 AAs: 0.22038\nTotal prob of N-in: 0.12994\noutside 1 331\nTMhelix 332 354\ninside 355 372

10551 GCF_000446645.2_ASM44664v2 Escherichia coli C581_05 Proteobacteria; Gammaproteobacteria; Enterobacterales; Enterobacteriaceae; Escherichia; Escherichia coli
MNNEETFYQAMRRQGVTRRSFLKYCSLAATSLGLGAGMAPKIAWALENKPRIPVVWIHGL WP_001058323.1
MULTISPECIES: hydrogenase-1 small chain [Proteobacteria] Length: 372\nNumber of predicted TMHs: 2\nExp number of AAs in TMHs: 37.2608499999999\nExp number, first 60 AAs: 19.52923\nTotal prob of N-in: 0.94351\nPOSSIBLE N-term signal sequence\ninside 1 19\nTMhelix 20 39\noutside 40 325\nTMhelix 326 348\ninside 349 372

10552 GCF_000446665.2_ASM44666v2 Escherichia coli C58_11 Proteobacteria; Gammaproteobacteria; Enterobacterales; Enterobacteriaceae; Escherichia; Escherichia coli
MNNEETFYQAMRRQGVTRRSFLKYCSLAATSLGLGAGMAPKIAWALENKPRIPVVWIHGL WP_001058323.1
MULTISPECIES: hydrogenase-1 small chain [Proteobacteria] Length: 372\nNumber of predicted TMHs: 2\nExp number of AAs in TMHs: 37.2608499999999\nExp number, first 60 AAs: 19.52923\nTotal prob of N-in: 0.94351\nPOSSIBLE N-term signal sequence\ninside 1 19\nTMhelix 20 39\noutside 40 325\nTMhelix 326 348\ninside 349 372

10553 GCF_000446665.2_ASM44666v2 Escherichia coli C58_11 Proteobacteria; Gammaproteobacteria; Enterobacterales; Enterobacteriaceae; Escherichia; Escherichia coli
MTGDNTLIHSHGINRRDFMKLCAALAATMGLSSKAAAEMAESVTNPQRPPVIWIGAQECT WP_000145410.1
MULTISPECIES: hydrogenase-2 small chain [Proteobacteria] Length: 372\nNumber of predicted TMHs: 1\nExp number of AAs in TMHs: 23.72226\nExp number, first 60 AAs: 0.22038\nTotal prob of N-in: 0.12994\noutside 1 331\nTMhelix 332 354\ninside 355 372

10554 GCF_000446685.2_ASM44668v2 Escherichia coli C586_05 Proteobacteria; Gammaproteobacteria; Enterobacterales; Enterobacteriaceae; Escherichia; Escherichia coli
MNNEETFYQAMRRQGVTRRSFLKYCSLAATSLGLGAGMAPKIAWALENKPRIPVVWIHGL WP_001058323.1
MULTISPECIES: hydrogenase-1 small chain [Proteobacteria] Length: 372\nNumber of predicted TMHs: 2\nExp number of AAs in TMHs: 37.2608499999999\nExp number, first 60 AAs: 19.52923\nTotal prob of N-in: 0.94351\nPOSSIBLE N-term signal sequence\ninside 1 19\nTMhelix 20 39\noutside 40 325\nTMhelix 326 348\ninside 349 372

10555 GCF_000446685.2_ASM44668v2 Escherichia coli C586_05 Proteobacteria; Gammaproteobacteria; Enterobacterales; Enterobacteriaceae; Escherichia; Escherichia coli
MTGDNTLIHSHGINRRDFMKLCAALAATMGLSSKAAAEMAESVTNPQRPPVIWIGAQECT WP_000145410.1
MULTISPECIES: hydrogenase-2 small chain [Proteobacteria] Length: 372\nNumber of predicted TMHs: 1\nExp number of AAs in TMHs: 23.72226\nExp number, first 60 AAs: 0.22038\nTotal prob of N-in: 0.12994\noutside 1 331\nTMhelix 332 354\ninside 355 372

10556 GCF_000410655.2_ASM41065v2 Escherichia coli C639_08 Proteobacteria; Gammaproteobacteria; Enterobacterales; Enterobacteriaceae; Escherichia; Escherichia coli
MTGDNTLIHSHGINRRDFMKLCAALAATMGLSSKAAAEMAESVTNPQRPPVIWIGAQECT WP_000145410.1
MULTISPECIES: hydrogenase-2 small chain [Proteobacteria] Length: 372\nNumber of predicted TMHs: 1\nExp number of AAs in TMHs: 23.72226\nExp number, first 60 AAs: 0.22038\nTotal prob of N-in: 0.12994\noutside 1 331\nTMhelix 332 354\ninside 355 372

10557 GCF_000410655.2_ASM41065v2 Escherichia coli C639_08 Proteobacteria; Gammaproteobacteria; Enterobacterales; Enterobacteriaceae; Escherichia; Escherichia coli
MNNEETFYQAMRRQGVTRRSFLKYCSLAATSLGLGAGMAPKIAWALENKPRIPVVWIHGL WP_001058323.1
MULTISPECIES: hydrogenase-1 small chain [Proteobacteria] Length: 372\nNumber of predicted TMHs: 2\nExp number of AAs in TMHs: 37.2608499999999\nExp number, first 60 AAs: 19.52923\nTotal prob of N-in: 0.94351\nPOSSIBLE N-term signal sequence\ninside 1 19\nTMhelix 20 39\noutside 40 325\nTMhelix 326 348\ninside 349 372

10558 GCF_000446705.2_ASM44670v2 Escherichia coli C652_10 Proteobacteria; Gammaproteobacteria; Enterobacterales; Enterobacteriaceae; Escherichia; Escherichia coli
MNNEETFYQAMRRQGVTRRSFLKYCSLAATSLGLGAGMAPKIAWALENKPRIPVVWIHGL WP_001295940.1
MULTISPECIES: hydrogenase [Enterobacteriaceae] Length: 372\nNumber of predicted TMHs: 2\nExp number of AAs in TMHs: 37.2608899999999\nExp number, first 60 AAs: 19.52923\nTotal prob of N-in: 0.94351\nPOSSIBLE N-term signal sequence\ninside 1 19\nTMhelix 20 39\noutside 40 325\nTMhelix 326 348\ninside 349 372

10559 GCF_000446705.2_ASM44670v2 Escherichia coli C652_10 Proteobacteria; Gammaproteobacteria; Enterobacterales; Enterobacteriaceae; Escherichia; Escherichia coli
MTGDNTLIHSHGINRRDFMKLCAALAATMGLSSKAAAEMAESVTNPQRPPVIWIGAQECT WP_000145410.1
MULTISPECIES: hydrogenase-2 small chain [Proteobacteria] Length: 372\nNumber of predicted TMHs: 1\nExp number of AAs in TMHs: 23.72226\nExp number, first 60 AAs: 0.22038\nTotal prob of N-in: 0.12994\noutside 1 331\nTMhelix 332 354\ninside 355 372

10560 GCF_000446725.2_ASM44672v2 Escherichia coli C654_09 Proteobacteria; Gammaproteobacteria; Enterobacterales; Enterobacteriaceae; Escherichia; Escherichia coli
MNNEETFYQAMRRQGVTRRSFLKYCSLAATSLGLGAGMAPKIAWALENKPRIPVVWIHGL WP_001058323.1
MULTISPECIES: hydrogenase-1 small chain [Proteobacteria] Length: 372\nNumber of predicted TMHs: 2\nExp number of AAs in TMHs: 37.2608499999999\nExp number, first 60 AAs: 19.52923\nTotal prob of N-in: 0.94351\nPOSSIBLE N-term signal sequence\ninside 1 19\nTMhelix 20 39\noutside 40 325\nTMhelix 326 348\ninside 349 372

10561 GCF_000446725.2_ASM44672v2 Escherichia coli C654_09 Proteobacteria; Gammaproteobacteria; Enterobacterales; Enterobacteriaceae; Escherichia; Escherichia coli
MTGDNTLIHSHGINRRDFMKLCAALAATMGLSSKAAAEMAESVTNPQRPPVIWIGAQECT WP_000145410.1
MULTISPECIES: hydrogenase-2 small chain [Proteobacteria] Length: 372\nNumber of predicted TMHs: 1\nExp number of AAs in TMHs: 23.72226\nExp number, first 60 AAs: 0.22038\nTotal prob of N-in: 0.12994\noutside 1 331\nTMhelix 332 354\ninside 355 372

10562 GCF_000818965.1_CLC_assembly_of_Escherichia_coli_C691-71 Escherichia coli C691-71 (14b)
Proteobacteria; Gammaproteobacteria; Enterobacterales; Enterobacteriaceae; Escherichia; Escherichia coli
MTGDNTLIHSHGINRRDFMKLCAALAATMGLSSKAAAEMAESVTNPQRPPVIWIGAQECT WP_000145410.1
MULTISPECIES: hydrogenase-2 small chain [Proteobacteria] Length: 372\nNumber of predicted TMHs: 1\nExp number of AAs in TMHs: 23.72226\nExp number, first 60 AAs: 0.22038\nTotal prob of N-in: 0.12994\noutside 1 331\nTMhelix 332 354\ninside 355 372

10563 GCF_000818965.1_CLC_assembly_of_Escherichia_coli_C691-71 Escherichia coli C691-71 (14b)
Proteobacteria; Gammaproteobacteria; Enterobacterales; Enterobacteriaceae; Escherichia; Escherichia coli
MNNEETFYQAMRRQGVTRRSFLKYCSLAATSLGLGAGMAPKIAWALENKPRIPVVWIHGL WP_001058323.1
MULTISPECIES: hydrogenase-1 small chain [Proteobacteria] Length: 372\nNumber of predicted TMHs: 2\nExp number of AAs in TMHs: 37.2608499999999\nExp number, first 60 AAs: 19.52923\nTotal prob of N-in: 0.94351\nPOSSIBLE N-term signal sequence\ninside 1 19\nTMhelix 20 39\noutside 40 325\nTMhelix 326 348\ninside 349 372

10564 GCF_000446745.2_ASM44674v2 Escherichia coli C717_10 Proteobacteria; Gammaproteobacteria; Enterobacterales; Enterobacteriaceae; Escherichia; Escherichia coli
MTGDNTLIHSHGINRRDFMKLCAALAATMGLSSKAAAEMAESVTNPQRPPVIWIGAQECT WP_000145410.1
MULTISPECIES: hydrogenase-2 small chain [Proteobacteria] Length: 372\nNumber of predicted TMHs: 1\nExp number of AAs in TMHs: 23.72226\nExp number, first 60 AAs: 0.22038\nTotal prob of N-in: 0.12994\noutside 1 331\nTMhelix 332 354\ninside 355 372

10565 GCF_000446745.2_ASM44674v2 Escherichia coli C717_10 Proteobacteria; Gammaproteobacteria; Enterobacterales; Enterobacteriaceae; Escherichia; Escherichia coli
MNNEETFYQAMRRQGVTRRSFLKYCSLAATSLGLGAGMAPKIAWALENKPRIPVVWIHGL WP_024237484.1
hydrogenase-1 small chain [Escherichia coli] Length: 372\nNumber of predicted TMHs: 2\nExp number of AAs in TMHs: 37.2691999999999\nExp number, first 60 AAs: 19.53117\nTotal prob of N-in: 0.94352\nPOSSIBLE N-term signal sequence\ninside 1 19\nTMhelix 20 39\noutside 40 325\nTMhelix 326 348\ninside 349 372

10566 GCF_000446765.2_ASM44676v2 Escherichia coli C725_88 Proteobacteria; Gammaproteobacteria; Enterobacterales; Enterobacteriaceae; Escherichia; Escherichia coli
MNNEETFYQAMRRQGVTRRSFLKYCSLAATSLGLGAGMAPKIAWALENKPRIPVVWIHGL WP_001058323.1
MULTISPECIES: hydrogenase-1 small chain [Proteobacteria] Length: 372\nNumber of predicted TMHs: 2\nExp number of AAs in TMHs: 37.2608499999999\nExp number, first 60 AAs: 19.52923\nTotal prob of N-in: 0.94351\nPOSSIBLE N-term signal sequence\ninside 1 19\nTMhelix 20 39\noutside 40 325\nTMhelix 326 348\ninside 349 372

10567 GCF_000446765.2_ASM44676v2 Escherichia coli C725_88 Proteobacteria; Gammaproteobacteria; Enterobacterales; Enterobacteriaceae; Escherichia; Escherichia coli
MTGDNTLIHSHGINRRDFMKLCAALAATMGLSSKAAAEMAESVTNPQRPPVIWIGAQECT WP_000145410.1
MULTISPECIES: hydrogenase-2 small chain [Proteobacteria] Length: 372\nNumber of predicted TMHs: 1\nExp number of AAs in TMHs: 23.72226\nExp number, first 60 AAs: 0.22038\nTotal prob of N-in: 0.12994\noutside 1 331\nTMhelix 332 354\ninside 355 372

10568 GCF_000446785.2_ASM44678v2 Escherichia coli C732_98 Proteobacteria; Gammaproteobacteria; Enterobacterales; Enterobacteriaceae; Escherichia; Escherichia coli
MNNEETFYQAMRRQGVTRRSFLKYCSLAATSLGLGAGMAPKIAWALENKPRIPVVWIHGL WP_001058323.1
MULTISPECIES: hydrogenase-1 small chain [Proteobacteria] Length: 372\nNumber of predicted TMHs: 2\nExp number of AAs in TMHs: 37.2608499999999\nExp number, first 60 AAs: 19.52923\nTotal prob of N-in: 0.94351\nPOSSIBLE N-term signal sequence\ninside 1 19\nTMhelix 20 39\noutside 40 325\nTMhelix 326 348\ninside 349 372

10569 GCF_000446785.2_ASM44678v2 Escherichia coli C732_98 Proteobacteria; Gammaproteobacteria; Enterobacterales; Enterobacteriaceae; Escherichia; Escherichia coli
MTGDNTLIHSHGINRRDFMKLCAALAATMGLSSKAAAEMAESVTNPQRPPVIWIGAQECT WP_000145410.1
MULTISPECIES: hydrogenase-2 small chain [Proteobacteria] Length: 372\nNumber of predicted TMHs: 1\nExp number of AAs in TMHs: 23.72226\nExp number, first 60 AAs: 0.22038\nTotal prob of N-in: 0.12994\noutside 1 331\nTMhelix 332 354\ninside 355 372

10570 GCF_000446805.2_ASM44680v2 Escherichia coli C743_03 Proteobacteria; Gammaproteobacteria; Enterobacterales; Enterobacteriaceae; Escherichia; Escherichia coli
MNNEETFYQAMRRQGVTRRSFLKYCSLAATSLGLGAGMAPKIAWALENKPRIPVVWIHGL WP_001295940.1
MULTISPECIES: hydrogenase [Enterobacteriaceae] Length: 372\nNumber of predicted TMHs: 2\nExp number of AAs in TMHs: 37.2608899999999\nExp number, first 60 AAs: 19.52923\nTotal prob of N-in: 0.94351\nPOSSIBLE N-term signal sequence\ninside 1 19\nTMhelix 20 39\noutside 40 325\nTMhelix 326 348\ninside 349 372

10571 GCF_000446805.2_ASM44680v2 Escherichia coli C743_03 Proteobacteria; Gammaproteobacteria; Enterobacterales; Enterobacteriaceae; Escherichia; Escherichia coli
MTGDNTLIHSHGINRRDFMKLCAALAATMGLSSKAAAEMAESVTNPQRPPVIWIGAQECT WP_000145410.1
MULTISPECIES: hydrogenase-2 small chain [Proteobacteria] Length: 372\nNumber of predicted TMHs: 1\nExp number of AAs in TMHs: 23.72226\nExp number, first 60 AAs: 0.22038\nTotal prob of N-in: 0.12994\noutside 1 331\nTMhelix 332 354\ninside 355 372

10572 GCF_000446825.2_ASM44682v2 Escherichia coli C751_03 Proteobacteria; Gammaproteobacteria; Enterobacterales; Enterobacteriaceae; Escherichia; Escherichia coli

MTGDNTLIHSHGINRRDFMKLCAALAATMGLSSKAAAEMAESVTNPQRPPVIWIGAQECT WP_000145410.1 MULTISPECIES: hydrogenase-2 small chain [Proteobacteria] Length: 372\nNumber of predicted TMHs: 1\nExp number of AAs in TMHs: 23.72226\nExp number, first 60 AAs: 0.22038\nTotal prob of N-in: 0.12994\noutside 1 331\nTMhelix 332 354\ninside 355 372

10573 GCF_000446825.2_ASM44682v2 Escherichia coli C751_03 Proteobacteria; Gammaproteobacteria; Enterobacterales; Enterobacteriaceae; Escherichia; Escherichia coli MNNEETFYQAMRRQGVTRRSFLKYCSLAATSLGLGAGMAPKIAWALENKPRIPVVWIHGL WP_001058323.1 MULTISPECIES: hydrogenase-1 small chain [Proteobacteria] Length: 372\nNumber of predicted TMHs: 2\nExp number of AAs in TMHs: 37.2608499999999\nExp number, first 60 AAs: 19.52923\nTotal prob of N-in: 0.94351\nPOSSIBLE N-term signal sequence\ninside 1 19\nTMhelix 20 39\noutside 40 325\nTMhelix 326 348\ninside 349 372

10574 GCF_000446845.2_ASM44684v2 Escherichia coli C78_09C Proteobacteria; Gammaproteobacteria; Enterobacterales; Enterobacteriaceae; Escherichia; Escherichia coli MTGDNTLIHSHGINRRDFMKLCAALAATMGLSSKAAAEMAESVTNPQRPPVIWIGAQECT WP_000145410.1 MULTISPECIES: hydrogenase-2 small chain [Proteobacteria] Length: 372\nNumber of predicted TMHs: 1\nExp number of AAs in TMHs: 23.72226\nExp number, first 60 AAs: 0.22038\nTotal prob of N-in: 0.12994\noutside 1 331\nTMhelix 332 354\ninside 355 372

10575 GCF_000446845.2_ASM44684v2 Escherichia coli C78_09C Proteobacteria; Gammaproteobacteria; Enterobacterales; Enterobacteriaceae; Escherichia; Escherichia coli MNNEETFYQAMRRQGVTRRSFLKYCSLAATSLGLGAGMAPKIAWALENKPRIPVVWIHGL WP_001058323.1 MULTISPECIES: hydrogenase-1 small chain [Proteobacteria] Length: 372\nNumber of predicted TMHs: 2\nExp number of AAs in TMHs: 37.2608499999999\nExp number, first 60 AAs: 19.52923\nTotal prob of N-in: 0.94351\nPOSSIBLE N-term signal sequence\ninside 1 19\nTMhelix 20 39\noutside 40 325\nTMhelix 326 348\ninside 349 372

10576 GCF_000446865.2_ASM44686v2 Escherichia coli C79_08 Proteobacteria; Gammaproteobacteria; Enterobacterales; Enterobacteriaceae; Escherichia; Escherichia coli MNNEETFYQAMRRQGVTRRSFLKYCSLAATSLGLGAGMAPKIAWALENKPRIPVVWIHGL WP_001058323.1 MULTISPECIES: hydrogenase-1 small chain [Proteobacteria] Length: 372\nNumber of predicted TMHs: 2\nExp number of AAs in TMHs: 37.2608499999999\nExp number, first 60 AAs: 19.52923\nTotal prob of N-in: 0.94351\nPOSSIBLE N-term signal sequence\ninside 1 19\nTMhelix 20 39\noutside 40 325\nTMhelix 326 348\ninside 349 372

10577 GCF_000446865.2_ASM44686v2 Escherichia coli C79_08 Proteobacteria; Gammaproteobacteria; Enterobacterales; Enterobacteriaceae; Escherichia; Escherichia coli MTGDNTLIHSHGINRRDFMKLCAALAATMGLSSKAAAEMAESVTNPQRPPVIWIGAQECT WP_000145410.1 MULTISPECIES: hydrogenase-2 small chain [Proteobacteria] Length: 372\nNumber of predicted TMHs: 1\nExp number of AAs in TMHs: 23.72226\nExp number, first 60 AAs: 0.22038\nTotal prob of N-in: 0.12994\noutside 1 331\nTMhelix 332 354\ninside 355 372

10578 GCF_000446885.2_ASM44688v2 Escherichia coli C792_92 Proteobacteria; Gammaproteobacteria; Enterobacterales; Enterobacteriaceae; Escherichia; Escherichia coli MTGDNTLIHSHGINRRDFMKLCAALAATMGLSSKAAAEMAESVTNPQRPPVIWIGAQECT WP_000145410.1 MULTISPECIES: hydrogenase-2 small chain [Proteobacteria] Length: 372\nNumber of predicted TMHs: 1\nExp number of AAs in TMHs: 23.72226\nExp number, first 60 AAs: 0.22038\nTotal prob of N-in: 0.12994\noutside 1 331\nTMhelix 332 354\ninside 355 372

10579 GCF_000446885.2_ASM44688v2 Escherichia coli C792_92 Proteobacteria; Gammaproteobacteria; Enterobacterales; Enterobacteriaceae; Escherichia; Escherichia coli MNNEETFYQAMRRQGVTRRSFLKYCSLAATSLGLGAGMAPKIAWALENKPRIPVVWIHGL WP_001058323.1 MULTISPECIES: hydrogenase-1 small chain [Proteobacteria] Length: 372\nNumber of predicted TMHs: 2\nExp number of AAs in TMHs: 37.2608499999999\nExp number, first 60 AAs: 19.52923\nTotal prob of N-in: 0.94351\nPOSSIBLE N-term signal sequence\ninside 1 19\nTMhelix 20 39\noutside 40 325\nTMhelix 326 348\ninside 349 372

10580 GCF_000446905.2_ASM44690v2 Escherichia coli C796_10 Proteobacteria; Gammaproteobacteria; Enterobacterales; Enterobacteriaceae; Escherichia; Escherichia coli MTGDNTLIHSHGINRRDFMKLCAALAATMGLSSKAAAEMAESVTNPQRPPVIWIGAQECT WP_000145410.1

MULTISPECIES: hydrogenase-2 small chain [Proteobacteria] Length: 372\nNumber of predicted TMHs: 1\nExp number of AAs in TMHs: 23.72226\nExp number, first 60 AAs: 0.22038\nTotal prob of N-in: 0.12994\noutside 1 331\nTMhelix 332 354\ninside 355 372

10581 GCF_000446905.2_ASM44690v2 Escherichia coli C796_10 Proteobacteria; Gammaproteobacteria; Enterobacterales; Enterobacteriaceae; Escherichia; Escherichia coli
MNNEETFYQAMRRQGVTRRSFLKYCSLAATSLGLGAGMAPKIAWALENKPRIPVVWIHGL WP_001295940.1
MULTISPECIES: hydrogenase [Enterobacteriaceae] Length: 372\nNumber of predicted TMHs: 2\nExp number of AAs in TMHs: 37.2608899999999\nExp number, first 60 AAs: 19.52923\nTotal prob of N-in: 0.94351\nPOSSIBLE N-term signal sequence\ninside 1 19\nTMhelix 20 39\noutside 40 325\nTMhelix 326 348\ninside 349 372

10582 GCF_000446925.2_ASM44692v2 Escherichia coli C799_92 Proteobacteria; Gammaproteobacteria; Enterobacterales; Enterobacteriaceae; Escherichia; Escherichia coli
MTGDNTLIHSHGINRRDFMKLCAALAATMGLSSKAAAEMAESVTNPQRPPVIWIGAQECT WP_000145410.1
MULTISPECIES: hydrogenase-2 small chain [Proteobacteria] Length: 372\nNumber of predicted TMHs: 1\nExp number of AAs in TMHs: 23.72226\nExp number, first 60 AAs: 0.22038\nTotal prob of N-in: 0.12994\noutside 1 331\nTMhelix 332 354\ninside 355 372

10583 GCF_000446925.2_ASM44692v2 Escherichia coli C799_92 Proteobacteria; Gammaproteobacteria; Enterobacterales; Enterobacteriaceae; Escherichia; Escherichia coli
MNNEETFYQAMRRQGVTRRSFLKYCSLAATSLGLGAGMAPKIAWALENKPRIPVVWIHGL WP_001058323.1
MULTISPECIES: hydrogenase-1 small chain [Proteobacteria] Length: 372\nNumber of predicted TMHs: 2\nExp number of AAs in TMHs: 37.2608499999999\nExp number, first 60 AAs: 19.52923\nTotal prob of N-in: 0.94351\nPOSSIBLE N-term signal sequence\ninside 1 19\nTMhelix 20 39\noutside 40 325\nTMhelix 326 348\ninside 349 372

10584 GCF_000446945.2_ASM44694v2 Escherichia coli C80_08 Proteobacteria; Gammaproteobacteria; Enterobacterales; Enterobacteriaceae; Escherichia; Escherichia coli
MTGDNTLIHSHGINRRDFMKLCAALAATMGLSSKAAAEMAESVTNPQRPPVIWIGAQECT WP_000145410.1
MULTISPECIES: hydrogenase-2 small chain [Proteobacteria] Length: 372\nNumber of predicted TMHs: 1\nExp number of AAs in TMHs: 23.72226\nExp number, first 60 AAs: 0.22038\nTotal prob of N-in: 0.12994\noutside 1 331\nTMhelix 332 354\ninside 355 372

10585 GCF_000446945.2_ASM44694v2 Escherichia coli C80_08 Proteobacteria; Gammaproteobacteria; Enterobacterales; Enterobacteriaceae; Escherichia; Escherichia coli
MNNEETFYQAMRRQGVTRRSFLKYCSLAATSLGLGAGMAPKIAWALENKPRIPVVWIHGL WP_001058323.1
MULTISPECIES: hydrogenase-1 small chain [Proteobacteria] Length: 372\nNumber of predicted TMHs: 2\nExp number of AAs in TMHs: 37.2608499999999\nExp number, first 60 AAs: 19.52923\nTotal prob of N-in: 0.94351\nPOSSIBLE N-term signal sequence\ninside 1 19\nTMhelix 20 39\noutside 40 325\nTMhelix 326 348\ninside 349 372

10586 GCF_000446965.2_ASM44696v2 Escherichia coli C807_09 Proteobacteria; Gammaproteobacteria; Enterobacterales; Enterobacteriaceae; Escherichia; Escherichia coli
MNNEETFYQAMRRQGVTRRSFLKYCSLAATSLGLGAGMAPKIAWALENKPRIPVVWIHGL WP_001058323.1
MULTISPECIES: hydrogenase-1 small chain [Proteobacteria] Length: 372\nNumber of predicted TMHs: 2\nExp number of AAs in TMHs: 37.2608499999999\nExp number, first 60 AAs: 19.52923\nTotal prob of N-in: 0.94351\nPOSSIBLE N-term signal sequence\ninside 1 19\nTMhelix 20 39\noutside 40 325\nTMhelix 326 348\ninside 349 372

10587 GCF_000446965.2_ASM44696v2 Escherichia coli C807_09 Proteobacteria; Gammaproteobacteria; Enterobacterales; Enterobacteriaceae; Escherichia; Escherichia coli
MTGDNTLIHSHGINRRDFMKLCAALAATMGLSSKAAAEMAESVTNPQRPPVIWIGAQECT WP_000145410.1
MULTISPECIES: hydrogenase-2 small chain [Proteobacteria] Length: 372\nNumber of predicted TMHs: 1\nExp number of AAs in TMHs: 23.72226\nExp number, first 60 AAs: 0.22038\nTotal prob of N-in: 0.12994\noutside 1 331\nTMhelix 332 354\ninside 355 372

10588 GCF_000446985.2_ASM44698v2 Escherichia coli C82_11 Proteobacteria; Gammaproteobacteria; Enterobacterales; Enterobacteriaceae; Escherichia; Escherichia coli
MNNEETFYQAMRRQGVTRRSFLKYCSLAATSLGLGAGMAPKIAWALENKPRIPVVWIHGL WP_001295940.1
MULTISPECIES: hydrogenase [Enterobacteriaceae] Length: 372\nNumber of predicted TMHs: 2\nExp

number of AAs in TMHs: 37.2608899999999\nExp number, first 60 AAs: 19.52923\nTotal prob of N-in: 0.94351\nPOSSIBLE N-term signal sequence\ninside 1 19\nTMhelix 20 39\noutside 40 325\nTMhelix 326 348\ninside 349 372

10589 GCF_000446985.2_ASM44698v2 Escherichia coli C82_11 Proteobacteria; Gammaproteobacteria; Enterobacterales; Enterobacteriaceae; Escherichia; Escherichia coli MTGDNTLIHSHGINRRDFMKLCAALAATMGLSSKAAAEMAESVTNPQRPPVIWIGAQECT WP_000145410.1 MULTISPECIES: hydrogenase-2 small chain [Proteobacteria] Length: 372\nNumber of predicted TMHs: 1\nExp number of AAs in TMHs: 23.72226\nExp number, first 60 AAs: 0.22038\nTotal prob of N-in: 0.12994\noutside 1 331\nTMhelix 332 354\ninside 355 372

10590 GCF_000447005.2_ASM44700v2 Escherichia coli C824_10 Proteobacteria; Gammaproteobacteria; Enterobacterales; Enterobacteriaceae; Escherichia; Escherichia coli MTGDNTLIHSHGINRRDFMKLCAALAATMGLSSKAAAEMAESVTNPQRPPVIWIGAQECT WP_000145410.1 MULTISPECIES: hydrogenase-2 small chain [Proteobacteria] Length: 372\nNumber of predicted TMHs: 1\nExp number of AAs in TMHs: 23.72226\nExp number, first 60 AAs: 0.22038\nTotal prob of N-in: 0.12994\noutside 1 331\nTMhelix 332 354\ninside 355 372

10591 GCF_000447005.2_ASM44700v2 Escherichia coli C824_10 Proteobacteria; Gammaproteobacteria; Enterobacterales; Enterobacteriaceae; Escherichia; Escherichia coli MNNEETFYQAMRRQGVTRRSFLKYCSLAATSLGLGAGMAPKIAWALENKPRIPVVWIHGL WP_001058323.1 MULTISPECIES: hydrogenase-1 small chain [Proteobacteria] Length: 372\nNumber of predicted TMHs: 2\nExp number of AAs in TMHs: 37.2608499999999\nExp number, first 60 AAs: 19.52923\nTotal prob of N-in: 0.94351\nPOSSIBLE N-term signal sequence\ninside 1 19\nTMhelix 20 39\noutside 40 325\nTMhelix 326 348\ninside 349 372

10592 GCF_000447025.2_ASM44702v2 Escherichia coli C842_97 Proteobacteria; Gammaproteobacteria; Enterobacterales; Enterobacteriaceae; Escherichia; Escherichia coli MTGDNTLIHSHGINRRDFMKLCAALAATMGLSSKAAAEMAESVTNPQRPPVIWIGAQECT WP_000145410.1 MULTISPECIES: hydrogenase-2 small chain [Proteobacteria] Length: 372\nNumber of predicted TMHs: 1\nExp number of AAs in TMHs: 23.72226\nExp number, first 60 AAs: 0.22038\nTotal prob of N-in: 0.12994\noutside 1 331\nTMhelix 332 354\ninside 355 372

10593 GCF_000447025.2_ASM44702v2 Escherichia coli C842_97 Proteobacteria; Gammaproteobacteria; Enterobacterales; Enterobacteriaceae; Escherichia; Escherichia coli MNNEETFYQAMRRQGVTRRSFLKYCSLAATSLGLGAGMAPKIAWALENKPRIPVVWIHGL WP_001301613.1 hydrogenase [Escherichia coli] Length: 372\nNumber of predicted TMHs: 2\nExp number of AAs in TMHs: 37.46598\nExp number, first 60 AAs: 19.53139\nTotal prob of N-in: 0.94364\nPOSSIBLE N-term signal sequence\ninside 1 19\nTMhelix 20 39\noutside 40 325\nTMhelix 326 348\ninside 349 372

10594 GCF_000410675.2_ASM41067v2 Escherichia coli C844_97 Proteobacteria; Gammaproteobacteria; Enterobacterales; Enterobacteriaceae; Escherichia; Escherichia coli MNNEETFYQAMRRQGVTRRSFLKYCSLAATSLGLGAGMAPKIAWALENKPRIPVVWIHGL WP_001058323.1 MULTISPECIES: hydrogenase-1 small chain [Proteobacteria] Length: 372\nNumber of predicted TMHs: 2\nExp number of AAs in TMHs: 37.2608499999999\nExp number, first 60 AAs: 19.52923\nTotal prob of N-in: 0.94351\nPOSSIBLE N-term signal sequence\ninside 1 19\nTMhelix 20 39\noutside 40 325\nTMhelix 326 348\ninside 349 372

10595 GCF_000410675.2_ASM41067v2 Escherichia coli C844_97 Proteobacteria; Gammaproteobacteria; Enterobacterales; Enterobacteriaceae; Escherichia; Escherichia coli MTGDNTLIHSHGINRRDFMKLCAALAATMGLSSKAAAEMAESVTNPQRPPVIWIGAQECT WP_000145410.1 MULTISPECIES: hydrogenase-2 small chain [Proteobacteria] Length: 372\nNumber of predicted TMHs: 1\nExp number of AAs in TMHs: 23.72226\nExp number, first 60 AAs: 0.22038\nTotal prob of N-in: 0.12994\noutside 1 331\nTMhelix 332 354\ninside 355 372

10596 GCF_000447045.2_ASM44704v2 Escherichia coli C87_11 Proteobacteria; Gammaproteobacteria; Enterobacterales; Enterobacteriaceae; Escherichia; Escherichia coli MTGDNTLIHSHGINRRDFMKLCAALAATMGLSSKAAAEMAESVTNPQRPPVIWIGAQECT WP_000145410.1 MULTISPECIES: hydrogenase-2 small chain [Proteobacteria] Length: 372\nNumber of predicted TMHs: 1\nExp number of AAs in TMHs: 23.72226\nExp number, first 60 AAs: 0.22038\nTotal prob of N-in: 0.12994\noutside 1 331\nTMhelix 332 354\ninside 355 372

10597 GCF_000447045.2_ASM44704v2 Escherichia coli C87_11 Proteobacteria; Gammaproteobacteria; Enterobacterales; Enterobacteriaceae; Escherichia; Escherichia coli
MNNEETFYQAMRRQGVTRRSFLKYCSLAATSLGLGAGMAPKIAWALENKPRIPVVWIHGL WP_001058323.1
MULTISPECIES: hydrogenase-1 small chain [Proteobacteria] Length: 372\nNumber of predicted TMHs: 2\nExp number of AAs in TMHs: 37.2608499999999\nExp number, first 60 AAs: 19.52923\nTotal prob of N-in: 0.94351\nPOSSIBLE N-term signal sequence\ninside 1 19\nTMhelix 20 39\noutside 40 325\nTMhelix 326 348\ninside 349 372

10598 GCF_000447065.2_ASM44706v2 Escherichia coli C887_10 Proteobacteria; Gammaproteobacteria; Enterobacterales; Enterobacteriaceae; Escherichia; Escherichia coli
MNNEETFYQAMRRQGVTRRSFLKYCSLAATSLGLGAGMAPKIAWALENKPRIPVVWIHGL WP_001058323.1
MULTISPECIES: hydrogenase-1 small chain [Proteobacteria] Length: 372\nNumber of predicted TMHs: 2\nExp number of AAs in TMHs: 37.2608499999999\nExp number, first 60 AAs: 19.52923\nTotal prob of N-in: 0.94351\nPOSSIBLE N-term signal sequence\ninside 1 19\nTMhelix 20 39\noutside 40 325\nTMhelix 326 348\ninside 349 372

10599 GCF_000447065.2_ASM44706v2 Escherichia coli C887_10 Proteobacteria; Gammaproteobacteria; Enterobacterales; Enterobacteriaceae; Escherichia; Escherichia coli
MTGDNTLIHSHGINRRDFMKLCAALAATMGLSSKAAAEMAESVTNPQRPPVIWIGAQECT WP_024245965.1
hydrogenase 2 small subunit [Escherichia coli] Length: 372\nNumber of predicted TMHs: 1\nExp number of AAs in TMHs: 24.15412\nExp number, first 60 AAs: 0.21967\nTotal prob of N-in: 0.14289\noutside 1 331\nTMhelix 332 354\ninside 355 372

10600 GCF_000447085.2_ASM44708v2 Escherichia coli C900_01 Proteobacteria; Gammaproteobacteria; Enterobacterales; Enterobacteriaceae; Escherichia; Escherichia coli
MTGDNTLIHSHGINRRDFMKLCAALAATMGLSSKAAAEMAESVTNPQRPPVIWIGAQECT WP_000145410.1
MULTISPECIES: hydrogenase-2 small chain [Proteobacteria] Length: 372\nNumber of predicted TMHs: 1\nExp number of AAs in TMHs: 23.72226\nExp number, first 60 AAs: 0.22038\nTotal prob of N-in: 0.12994\noutside 1 331\nTMhelix 332 354\ninside 355 372

10601 GCF_000447085.2_ASM44708v2 Escherichia coli C900_01 Proteobacteria; Gammaproteobacteria; Enterobacterales; Enterobacteriaceae; Escherichia; Escherichia coli
MNNEETFYQAMRRQGVTRRSFLKYCSLAATSLGLGAGMAPKIAWALENKPRIPVVWIHGL WP_024246676.1
hydrogenase-1 small chain [Escherichia coli] Length: 372\nNumber of predicted TMHs: 2\nExp number of AAs in TMHs: 37.8214899999999\nExp number, first 60 AAs: 19.52326\nTotal prob of N-in: 0.94395\nPOSSIBLE N-term signal sequence\ninside 1 19\nTMhelix 20 39\noutside 40 325\nTMhelix 326 348\ninside 349 372

10602 GCF_000447105.2_ASM44710v2 Escherichia coli C93_11 Proteobacteria; Gammaproteobacteria; Enterobacterales; Enterobacteriaceae; Escherichia; Escherichia coli
MTGDNTLIHSHGINRRDFMKLCAALAATMGLSSKAAAEMAESVTNPQRPPVIWIGAQECT WP_000145406.1
hydrogenase-2 small chain [Escherichia coli] Length: 372\nNumber of predicted TMHs: 1\nExp number of AAs in TMHs: 23.72251\nExp number, first 60 AAs: 0.22027\nTotal prob of N-in: 0.13006\noutside 1 331\nTMhelix 332 354\ninside 355 372

10603 GCF_000447105.2_ASM44710v2 Escherichia coli C93_11 Proteobacteria; Gammaproteobacteria; Enterobacterales; Enterobacteriaceae; Escherichia; Escherichia coli
MNNEETFYQAMRRQGVTRRSFLKYCSLAATSLGLGAGMAPKIAWALENKPRIPVVWIHGL WP_001058323.1
MULTISPECIES: hydrogenase-1 small chain [Proteobacteria] Length: 372\nNumber of predicted TMHs: 2\nExp number of AAs in TMHs: 37.2608499999999\nExp number, first 60 AAs: 19.52923\nTotal prob of N-in: 0.94351\nPOSSIBLE N-term signal sequence\ninside 1 19\nTMhelix 20 39\noutside 40 325\nTMhelix 326 348\ninside 349 372

10604 GCF_000447125.2_ASM44712v2 Escherichia coli C9_92 Proteobacteria; Gammaproteobacteria; Enterobacterales; Enterobacteriaceae; Escherichia; Escherichia coli
MNNEETFYQAMRRQGVTRRSFLKYCSLAATSLGLGAGMAPKIAWALENKPRIPVVWIHGL WP_001058323.1
MULTISPECIES: hydrogenase-1 small chain [Proteobacteria] Length: 372\nNumber of predicted TMHs: 2\nExp number of AAs in TMHs: 37.2608499999999\nExp number, first 60 AAs: 19.52923\nTotal prob of N-in: 0.94351\nPOSSIBLE N-term signal sequence\ninside 1 19\nTMhelix 20 39\noutside 40 325\nTMhelix 326 348\ninside 349 372

10605 GCF_000447125.2_ASM44712v2 Escherichia coli C9_92 Proteobacteria; Gammaproteobacteria; Enterobacterales; Enterobacteriaceae; Escherichia; Escherichia coli
MTGDNTLIHSHGINRRDFMKLCAALAATMGLSSKAAAEMAESVTNPQRPPVIWIGAQECT WP_000145410.1
MULTISPECIES: hydrogenase-2 small chain [Proteobacteria] Length: 372\nNumber of predicted TMHs: 1\nExp number of AAs in TMHs: 23.72226\nExp number, first 60 AAs: 0.22038\nTotal prob of N-in: 0.12994\noutside 1 331\nTMhelix 332 354\ninside 355 372

10606 GCF_000304015.2_ASM30401v2 Escherichia coli CB7326 Proteobacteria; Gammaproteobacteria; Enterobacterales; Enterobacteriaceae; Escherichia; Escherichia coli
MTGDNTLIHSHGINRRDFMKLCAALAATMGLSSKAAAEMAESVTNPQRPPVIWIGAQECT WP_000145410.1
MULTISPECIES: hydrogenase-2 small chain [Proteobacteria] Length: 372\nNumber of predicted TMHs: 1\nExp number of AAs in TMHs: 23.72226\nExp number, first 60 AAs: 0.22038\nTotal prob of N-in: 0.12994\noutside 1 331\nTMhelix 332 354\ninside 355 372

10607 GCF_000304015.2_ASM30401v2 Escherichia coli CB7326 Proteobacteria; Gammaproteobacteria; Enterobacterales; Enterobacteriaceae; Escherichia; Escherichia coli
MNNEETFYQAMRRQGVTRRSFLKYCSLAATSLGLGAGMAPKIAWALENKPRIPVVWIHGL WP_001301613.1
hydrogenase [Escherichia coli] Length: 372\nNumber of predicted TMHs: 2\nExp number of AAs in TMHs: 37.46598\nExp number, first 60 AAs: 19.53139\nTotal prob of N-in: 0.94364\nPOSSIBLE N-term signal sequence\ninside 1 19\nTMhelix 20 39\noutside 40 325\nTMhelix 326 348\ninside 349 372

10608 GCF_000496975.1_CE418_1 Escherichia coli CE418 Proteobacteria; Gammaproteobacteria; Enterobacterales; Enterobacteriaceae; Escherichia; Escherichia coli
MTGDNTLIHSHGINRRDFMKLCAALAATMGLSSKAAAEMAESVTNPQRPPVIWIGAQECT WP_000145410.1
MULTISPECIES: hydrogenase-2 small chain [Proteobacteria] Length: 372\nNumber of predicted TMHs: 1\nExp number of AAs in TMHs: 23.72226\nExp number, first 60 AAs: 0.22038\nTotal prob of N-in: 0.12994\noutside 1 331\nTMhelix 332 354\ninside 355 372

10609 GCF_000496995.1_CE516_1 Escherichia coli CE516 Proteobacteria; Gammaproteobacteria; Enterobacterales; Enterobacteriaceae; Escherichia; Escherichia coli
MNNEETFYQAMRRQGVTRRSFLKYCSLAATSLGLGAGMAPKIAWALENKPRIPVVWIHGL WP_001058323.1
MULTISPECIES: hydrogenase-1 small chain [Proteobacteria] Length: 372\nNumber of predicted TMHs: 2\nExp number of AAs in TMHs: 37.2608499999999\nExp number, first 60 AAs: 19.52923\nTotal prob of N-in: 0.94351\nPOSSIBLE N-term signal sequence\ninside 1 19\nTMhelix 20 39\noutside 40 325\nTMhelix 326 348\ninside 349 372

10610 GCF_000496995.1_CE516_1 Escherichia coli CE516 Proteobacteria; Gammaproteobacteria; Enterobacterales; Enterobacteriaceae; Escherichia; Escherichia coli
MTGDNTLIHSHGINRRDFMKLCAALAATMGLSSKAAAEMAESVTNPQRPPVIWIGAQECT WP_000145410.1
MULTISPECIES: hydrogenase-2 small chain [Proteobacteria] Length: 372\nNumber of predicted TMHs: 1\nExp number of AAs in TMHs: 23.72226\nExp number, first 60 AAs: 0.22038\nTotal prob of N-in: 0.12994\noutside 1 331\nTMhelix 332 354\ninside 355 372

10611 GCF_000497015.1_CE549_1 Escherichia coli CE549 Proteobacteria; Gammaproteobacteria; Enterobacterales; Enterobacteriaceae; Escherichia; Escherichia coli
MNNEETFYQAMRRQGVTRRSFLKYCSLAATSLGLGAGMAPKIAWALENKPRIPVVWIHGL WP_001058323.1
MULTISPECIES: hydrogenase-1 small chain [Proteobacteria] Length: 372\nNumber of predicted TMHs: 2\nExp number of AAs in TMHs: 37.2608499999999\nExp number, first 60 AAs: 19.52923\nTotal prob of N-in: 0.94351\nPOSSIBLE N-term signal sequence\ninside 1 19\nTMhelix 20 39\noutside 40 325\nTMhelix 326 348\ninside 349 372

10612 GCF_000497015.1_CE549_1 Escherichia coli CE549 Proteobacteria; Gammaproteobacteria; Enterobacterales; Enterobacteriaceae; Escherichia; Escherichia coli
MTGDNTLIHSHGINRRDFMKLCAALAATMGLSSKAAAEMAESVTNPQRPPVIWIGAQECT WP_000145410.1
MULTISPECIES: hydrogenase-2 small chain [Proteobacteria] Length: 372\nNumber of predicted TMHs: 1\nExp number of AAs in TMHs: 23.72226\nExp number, first 60 AAs: 0.22038\nTotal prob of N-in: 0.12994\noutside 1 331\nTMhelix 332 354\ninside 355 372

10613 GCF_000007445.1_ASM744v1 Escherichia coli CFT073 Proteobacteria; Gammaproteobacteria; Enterobacterales; Enterobacteriaceae; Escherichia; Escherichia coli
MTGDNTLIHSHGINRRDFMKLCAALAATMGLSSKAAAEMAESVTNPQRPPVIWIGAQECT WP_000145410.1

MULTISPECIES: hydrogenase-2 small chain [Proteobacteria] Length: 372\nNumber of predicted TMHs: 1\nExp number of AAs in TMHs: 23.72226\nExp number, first 60 AAs: 0.22038\nTotal prob of N-in: 0.12994\noutside 1 331\nTMhelix 332 354\ninside 355 372

10614 GCF_000007445.1_ASM744v1 Escherichia coli CFT073 Proteobacteria; Gammaproteobacteria; Enterobacterales; Enterobacteriaceae; Escherichia; Escherichia coli MNNEETFYQAMRRQGVTRRSFLKYCSLAATSLGLGAGMAPKIAWALENKPRIPVVWIHGL WP_001058323.1 MULTISPECIES: hydrogenase-1 small chain [Proteobacteria] Length: 372\nNumber of predicted TMHs: 2\nExp number of AAs in TMHs: 37.2608499999999\nExp number, first 60 AAs: 19.52923\nTotal prob of N-in: 0.94351\nPOSSIBLE N-term signal sequence\ninside 1 19\nTMhelix 20 39\noutside 40 325\nTMhelix 326 348\ninside 349 372

10615 GCF_000307205.1_ASM30720v1 Escherichia coli chi7122 Proteobacteria; Gammaproteobacteria; Enterobacterales; Enterobacteriaceae; Escherichia; Escherichia coli MTGDNTLIHSHGINRRDFMKLCAALAATMGLSSKAAAEMAESVTNPQRPPVIWIGAQECT WP_000145410.1 MULTISPECIES: hydrogenase-2 small chain [Proteobacteria] Length: 372\nNumber of predicted TMHs: 1\nExp number of AAs in TMHs: 23.72226\nExp number, first 60 AAs: 0.22038\nTotal prob of N-in: 0.12994\noutside 1 331\nTMhelix 332 354\ninside 355 372

10616 GCF_000307205.1_ASM30720v1 Escherichia coli chi7122 Proteobacteria; Gammaproteobacteria; Enterobacterales; Enterobacteriaceae; Escherichia; Escherichia coli MNNEETFYQAMRRQGVTRRSFLKYCSLAATSLGLGAGMAPKIAWALENKPRIPVVWIHGL WP_001058320.1 hydrogenase [Escherichia coli] Length: 372\nNumber of predicted TMHs: 2\nExp number of AAs in TMHs: 37.26082\nExp number, first 60 AAs: 19.52923\nTotal prob of N-in: 0.94351\nPOSSIBLE N-term signal sequence\ninside 1 19\nTMhelix 20 39\noutside 40 325\nTMhelix 326 348\ninside 349 372

10617 GCF_000692695.1_Esch_coli_CHS_68_V1 Escherichia coli CHS 68 Proteobacteria; Gammaproteobacteria; Enterobacterales; Enterobacteriaceae; Escherichia; Escherichia coli MTGDNTLIHSHGINRRDFMKLCAALAATMGLSSKAAAEMVESVTNPQRPPVIWIGAQECT WP_000145423.1 hydrogenase 2 small subunit [Escherichia coli] Length: 372\nNumber of predicted TMHs: 1\nExp number of AAs in TMHs: 23.78201\nExp number, first 60 AAs: 0.28355\nTotal prob of N-in: 0.13257\noutside 1 331\nTMhelix 332 354\ninside 355 372

10618 GCF_000692695.1_Esch_coli_CHS_68_V1 Escherichia coli CHS 68 Proteobacteria; Gammaproteobacteria; Enterobacterales; Enterobacteriaceae; Escherichia; Escherichia coli MNNEETFYQAMRRQGVTRRSFLKYCSLAATSLGLGAGMAPKIAWALENKPRIPVVWIHGL WP_001295940.1 MULTISPECIES: hydrogenase [Enterobacteriaceae] Length: 372\nNumber of predicted TMHs: 2\nExp number of AAs in TMHs: 37.2608899999999\nExp number, first 60 AAs: 19.52923\nTotal prob of N-in: 0.94351\nPOSSIBLE N-term signal sequence\ninside 1 19\nTMhelix 20 39\noutside 40 325\nTMhelix 326 348\ninside 349 372

10619 GCF_000692715.1_Esch_coli_CHS_69_V1 Escherichia coli CHS 69 Proteobacteria; Gammaproteobacteria; Enterobacterales; Enterobacteriaceae; Escherichia; Escherichia coli MTGDNTLIHSHGINRRDFMKLCAALAATMGLSSKAAAEMVESVTNPQRPPVIWIGAQECT WP_000145423.1 hydrogenase 2 small subunit [Escherichia coli] Length: 372\nNumber of predicted TMHs: 1\nExp number of AAs in TMHs: 23.78201\nExp number, first 60 AAs: 0.28355\nTotal prob of N-in: 0.13257\noutside 1 331\nTMhelix 332 354\ninside 355 372

10620 GCF_000692715.1_Esch_coli_CHS_69_V1 Escherichia coli CHS 69 Proteobacteria; Gammaproteobacteria; Enterobacterales; Enterobacteriaceae; Escherichia; Escherichia coli MNNEETFYQAMRRQGVTRRSFLKYCSLAATSLGLGAGMAPKIAWALENKPRIPVVWIHGL WP_001295940.1 MULTISPECIES: hydrogenase [Enterobacteriaceae] Length: 372\nNumber of predicted TMHs: 2\nExp number of AAs in TMHs: 37.2608899999999\nExp number, first 60 AAs: 19.52923\nTotal prob of N-in: 0.94351\nPOSSIBLE N-term signal sequence\ninside 1 19\nTMhelix 20 39\noutside 40 325\nTMhelix 326 348\ninside 349 372

10621 GCF_000692735.1_Esch_coli_CHS_77_V1 Escherichia coli CHS 77 Proteobacteria; Gammaproteobacteria; Enterobacterales; Enterobacteriaceae; Escherichia; Escherichia coli MTGDNTLIHSHGINRRDFMKLCAALAATMGLSSKAAAEMAESVTNPQRPPVIWIGAQECT WP_000145410.1 MULTISPECIES: hydrogenase-2 small chain [Proteobacteria] Length: 372\nNumber of predicted TMHs:

1\nExp number of AAs in TMHs: 23.72226\nExp number, first 60 AAs: 0.22038\nTotal prob of N-in: 0.12994\noutside 1 331\nTMhelix 332 354\ninside 355 372

10622 GCF_000692735.1_Esch_coli_CHS_77_V1 Escherichia coli CHS 77 Proteobacteria; Gammaproteobacteria; Enterobacterales; Enterobacteriaceae; Escherichia; Escherichia coli MNNEETFYQAMRRQGVTRRSFLKYCSLAATSLGLGAGMAPKIAWALENKPRIPVVWIHGL WP_001058323.1 MULTISPECIES: hydrogenase-1 small chain [Proteobacteria] Length: 372\nNumber of predicted TMHs: 2\nExp number of AAs in TMHs: 37.2608499999999\nExp number, first 60 AAs: 19.52923\nTotal prob of N-in: 0.94351\nPOSSIBLE N-term signal sequence\ninside 1 19\nTMhelix 20 39\noutside 40 325\nTMhelix 326 348\ninside 349 372

10623 GCF_000233675.1_ASM23367v2 Escherichia coli cloneA_i1 Proteobacteria; Gammaproteobacteria; Enterobacterales; Enterobacteriaceae; Escherichia; Escherichia coli MTGDNTLIHSHGINRRDFMKLCAALAATMGLSSKAAAEMAESVTNPQRPPVIWIGAQECT WP_000145415.1 hydrogenase 2 small subunit [Escherichia coli] Length: 372\nNumber of predicted TMHs: 1\nExp number of AAs in TMHs: 23.72233\nExp number, first 60 AAs: 0.22038\nTotal prob of N-in: 0.12994\noutside 1 331\nTMhelix 332 354\ninside 355 372

10624 GCF_000264235.1_ASM26423v1 Escherichia coli CUMT8 Proteobacteria; Gammaproteobacteria; Enterobacterales; Enterobacteriaceae; Escherichia; Escherichia coli MNNEETFYQAMRRQGVTRRSFLKYCSLAATSLGLGAGMAPKIAWALENKPRIPVVWIHGL WP_001295940.1 MULTISPECIES: hydrogenase [Enterobacteriaceae] Length: 372\nNumber of predicted TMHs: 2\nExp number of AAs in TMHs: 37.2608899999999\nExp number, first 60 AAs: 19.52923\nTotal prob of N-in: 0.94351\nPOSSIBLE N-term signal sequence\ninside 1 19\nTMhelix 20 39\noutside 40 325\nTMhelix 326 348\ninside 349 372

10625 GCF_000264235.1_ASM26423v1 Escherichia coli CUMT8 Proteobacteria; Gammaproteobacteria; Enterobacterales; Enterobacteriaceae; Escherichia; Escherichia coli MTGDNTLIHSHGINRRDFMKLCAALAATMGLSSKAAAEMAESVTNPQRPPVIWIGAQECT WP_000145410.1 MULTISPECIES: hydrogenase-2 small chain [Proteobacteria] Length: 372\nNumber of predicted TMHs: 1\nExp number of AAs in TMHs: 23.72226\nExp number, first 60 AAs: 0.22038\nTotal prob of N-in: 0.12994\noutside 1 331\nTMhelix 332 354\ninside 355 372

10626 GCF_000731355.1_V1 Escherichia coli D6-113.11 Proteobacteria; Gammaproteobacteria; Enterobacterales; Enterobacteriaceae; Escherichia; Escherichia coli MNNEETFYQAMRRQGVTRRSFLKYCSLAATSLGLGAGMAPKIAWALENKPRIPVVWIHGL WP_001058323.1 MULTISPECIES: hydrogenase-1 small chain [Proteobacteria] Length: 372\nNumber of predicted TMHs: 2\nExp number of AAs in TMHs: 37.2608499999999\nExp number, first 60 AAs: 19.52923\nTotal prob of N-in: 0.94351\nPOSSIBLE N-term signal sequence\ninside 1 19\nTMhelix 20 39\noutside 40 325\nTMhelix 326 348\ninside 349 372

10627 GCF_000731355.1_V1 Escherichia coli D6-113.11 Proteobacteria; Gammaproteobacteria; Enterobacterales; Enterobacteriaceae; Escherichia; Escherichia coli MTGDNTLIHSHGINRRDFMKLCAALAATMGLSSKAAAEMAESVTNPQRPPVIWIGAQECT WP_000145410.1 MULTISPECIES: hydrogenase-2 small chain [Proteobacteria] Length: 372\nNumber of predicted TMHs: 1\nExp number of AAs in TMHs: 23.72226\nExp number, first 60 AAs: 0.22038\nTotal prob of N-in: 0.12994\noutside 1 331\nTMhelix 332 354\ninside 355 372

10628 GCF_000723325.1_V1 Escherichia coli D6-117.29 Proteobacteria; Gammaproteobacteria; Enterobacterales; Enterobacteriaceae; Escherichia; Escherichia coli MNNEETFYQAMRRQGVTRRSFLKYCSLAATSLGLGAGMAPKIAWALENKPRIPVVWIHGL WP_001058323.1 MULTISPECIES: hydrogenase-1 small chain [Proteobacteria] Length: 372\nNumber of predicted TMHs: 2\nExp number of AAs in TMHs: 37.2608499999999\nExp number, first 60 AAs: 19.52923\nTotal prob of N-in: 0.94351\nPOSSIBLE N-term signal sequence\ninside 1 19\nTMhelix 20 39\noutside 40 325\nTMhelix 326 348\ninside 349 372

10629 GCF_000723325.1_V1 Escherichia coli D6-117.29 Proteobacteria; Gammaproteobacteria; Enterobacterales; Enterobacteriaceae; Escherichia; Escherichia coli MTGDNTLIHSHGINRRDFMKLCAALAATMGLSSKAAAEMAESVTNPQRPPVIWIGAQECT WP_000145410.1 MULTISPECIES: hydrogenase-2 small chain [Proteobacteria] Length: 372\nNumber of predicted TMHs:

1\nExp number of AAs in TMHs: 23.72226\nExp number, first 60 AAs: 0.22038\nTotal prob of N-in: 0.12994\noutside 1 331\nTMhelix 332 354\ninside 355 372

10630 GCF_000158395.1_ASM15839v1 Escherichia coli D9 Proteobacteria; Gammaproteobacteria; Enterobacterales; Enterobacteriaceae; Escherichia; Escherichia coli
MNNEETFYQAMRRQGVTRRSFLKYCSLAATSLGLGAGMAPKIAWALENKPRIPVVWIHGL WP_001058323.1
MULTISPECIES: hydrogenase-1 small chain [Proteobacteria] Length: 372\nNumber of predicted TMHs: 2\nExp number of AAs in TMHs: 37.2608499999999\nExp number, first 60 AAs: 19.52923\nTotal prob of N-in: 0.94351\nPOSSIBLE N-term signal sequence\ninside 1 19\nTMhelix 20 39\noutside 40 325\nTMhelix 326 348\ninside 349 372

10631 GCF_000158395.1_ASM15839v1 Escherichia coli D9 Proteobacteria; Gammaproteobacteria; Enterobacterales; Enterobacteriaceae; Escherichia; Escherichia coli
MTGDNTLIHSHGINRRDFMKLCAALAATMGLSSKAAAEMAESVTNPQRPPVIWIGAQECT WP_000145410.1
MULTISPECIES: hydrogenase-2 small chain [Proteobacteria] Length: 372\nNumber of predicted TMHs: 1\nExp number of AAs in TMHs: 23.72226\nExp number, first 60 AAs: 0.22038\nTotal prob of N-in: 0.12994\noutside 1 331\nTMhelix 332 354\ninside 355 372

10632 GCF_000250075.1_ASM25007v2 Escherichia coli DEC10E Proteobacteria; Gammaproteobacteria; Enterobacterales; Enterobacteriaceae; Escherichia; Escherichia coli
MNNEETFYQAMRRQGVTRRSFLKYCSLAATSLGLGAGMAPKIAWALENKPRIPVVWIHGL WP_001058323.1
MULTISPECIES: hydrogenase-1 small chain [Proteobacteria] Length: 372\nNumber of predicted TMHs: 2\nExp number of AAs in TMHs: 37.2608499999999\nExp number, first 60 AAs: 19.52923\nTotal prob of N-in: 0.94351\nPOSSIBLE N-term signal sequence\ninside 1 19\nTMhelix 20 39\noutside 40 325\nTMhelix 326 348\ninside 349 372

10633 GCF_000250075.1_ASM25007v2 Escherichia coli DEC10E Proteobacteria; Gammaproteobacteria; Enterobacterales; Enterobacteriaceae; Escherichia; Escherichia coli
MTGDNTLIHSHGINRRDFMKLCAALAATMGLSSKAAAEMAESVTNPQRPPVIWIGAQECT WP_000145410.1
MULTISPECIES: hydrogenase-2 small chain [Proteobacteria] Length: 372\nNumber of predicted TMHs: 1\nExp number of AAs in TMHs: 23.72226\nExp number, first 60 AAs: 0.22038\nTotal prob of N-in: 0.12994\noutside 1 331\nTMhelix 332 354\ninside 355 372

10634 GCF_000250095.1_ASM25009v2 Escherichia coli DEC10F Proteobacteria; Gammaproteobacteria; Enterobacterales; Enterobacteriaceae; Escherichia; Escherichia coli
MTGDNTLIHSHGINRRDFMKLCAALAATMGLSSKAAAEMAESVTNPQRPPVIWIGAQECT WP_000145410.1
MULTISPECIES: hydrogenase-2 small chain [Proteobacteria] Length: 372\nNumber of predicted TMHs: 1\nExp number of AAs in TMHs: 23.72226\nExp number, first 60 AAs: 0.22038\nTotal prob of N-in: 0.12994\noutside 1 331\nTMhelix 332 354\ninside 355 372

10635 GCF_000250095.1_ASM25009v2 Escherichia coli DEC10F Proteobacteria; Gammaproteobacteria; Enterobacterales; Enterobacteriaceae; Escherichia; Escherichia coli
MNNEETFYQAMRRQGVTRRSFLKYCSLAATSLGLGAGMAPKIAWALENKPRIPVVWIHGL WP_001058323.1
MULTISPECIES: hydrogenase-1 small chain [Proteobacteria] Length: 372\nNumber of predicted TMHs: 2\nExp number of AAs in TMHs: 37.2608499999999\nExp number, first 60 AAs: 19.52923\nTotal prob of N-in: 0.94351\nPOSSIBLE N-term signal sequence\ninside 1 19\nTMhelix 20 39\noutside 40 325\nTMhelix 326 348\ninside 349 372

10636 GCF_000250115.1_ASM25011v2 Escherichia coli DEC11A Proteobacteria; Gammaproteobacteria; Enterobacterales; Enterobacteriaceae; Escherichia; Escherichia coli
MNNEETFYQAMRRQGVTRRSFLKYCSLAATSLGLGAGMAPKIAWALENKPRIPVVWIHGL WP_001295940.1
MULTISPECIES: hydrogenase [Enterobacteriaceae] Length: 372\nNumber of predicted TMHs: 2\nExp number of AAs in TMHs: 37.2608899999999\nExp number, first 60 AAs: 19.52923\nTotal prob of N-in: 0.94351\nPOSSIBLE N-term signal sequence\ninside 1 19\nTMhelix 20 39\noutside 40 325\nTMhelix 326 348\ninside 349 372

10637 GCF_000250115.1_ASM25011v2 Escherichia coli DEC11A Proteobacteria; Gammaproteobacteria; Enterobacterales; Enterobacteriaceae; Escherichia; Escherichia coli
MTGDNTLIHSHGINRRDFMKLCAALAATMGLSSKAAAEMAESVTNPQRPPVIWIGAQECT WP_000145410.1
MULTISPECIES: hydrogenase-2 small chain [Proteobacteria] Length: 372\nNumber of predicted TMHs:

1\nExp number of AAs in TMHs: 23.72226\nExp number, first 60 AAs: 0.22038\nTotal prob of N-in: 0.12994\noutside 1 331\nTMhelix 332 354\ninside 355 372

10638 GCF_000250135.1_ASM25013v2 Escherichia coli DEC11B Proteobacteria; Gammaproteobacteria; Enterobacterales; Enterobacteriaceae; Escherichia; Escherichia coli
MTGDNTLIHSHGINRRDFMKLCAALAATMGLSSKAAAEMAESVTNPQRPPVIWIGAQECT WP_000145410.1
MULTISPECIES: hydrogenase-2 small chain [Proteobacteria] Length: 372\nNumber of predicted TMHs: 1\nExp number of AAs in TMHs: 23.72226\nExp number, first 60 AAs: 0.22038\nTotal prob of N-in: 0.12994\noutside 1 331\nTMhelix 332 354\ninside 355 372

10639 GCF_000250135.1_ASM25013v2 Escherichia coli DEC11B Proteobacteria; Gammaproteobacteria; Enterobacterales; Enterobacteriaceae; Escherichia; Escherichia coli
MNNEETFYQAMRRQGVTRRSFLKYCSLAATSLGLGAGMAPKIAWALENKPRIPVVWIHGL WP_001295940.1
MULTISPECIES: hydrogenase [Enterobacteriaceae] Length: 372\nNumber of predicted TMHs: 2\nExp number of AAs in TMHs: 37.2608899999999\nExp number, first 60 AAs: 19.52923\nTotal prob of N-in: 0.94351\nPOSSIBLE N-term signal sequence\ninside 1 19\nTMhelix 20 39\noutside 40 325\nTMhelix 326 348\ninside 349 372

10640 GCF_000250175.1_ASM25017v2 Escherichia coli DEC11D Proteobacteria; Gammaproteobacteria; Enterobacterales; Enterobacteriaceae; Escherichia; Escherichia coli
MNNEETFYQAMRRQGVTRRSFLKYCSLAATSLGLGAGMAPKIAWALENKPRIPVVWIHGL WP_001295940.1
MULTISPECIES: hydrogenase [Enterobacteriaceae] Length: 372\nNumber of predicted TMHs: 2\nExp number of AAs in TMHs: 37.2608899999999\nExp number, first 60 AAs: 19.52923\nTotal prob of N-in: 0.94351\nPOSSIBLE N-term signal sequence\ninside 1 19\nTMhelix 20 39\noutside 40 325\nTMhelix 326 348\ninside 349 372

10641 GCF_000250175.1_ASM25017v2 Escherichia coli DEC11D Proteobacteria; Gammaproteobacteria; Enterobacterales; Enterobacteriaceae; Escherichia; Escherichia coli
MTGDNTLIHSHGINRRDFMKLCAALAATMGLSSKAAAEMAESVTNPQRPPVIWIGAQECT WP_000145410.1
MULTISPECIES: hydrogenase-2 small chain [Proteobacteria] Length: 372\nNumber of predicted TMHs: 1\nExp number of AAs in TMHs: 23.72226\nExp number, first 60 AAs: 0.22038\nTotal prob of N-in: 0.12994\noutside 1 331\nTMhelix 332 354\ninside 355 372

10642 GCF_000250195.1_ASM25019v2 Escherichia coli DEC11E Proteobacteria; Gammaproteobacteria; Enterobacterales; Enterobacteriaceae; Escherichia; Escherichia coli
MNNEETFYQAMRRQGVTRRSFLKYCSLAATSLGLGAGMAPKIAWALENKPRIPVVWIHGL WP_001295940.1
MULTISPECIES: hydrogenase [Enterobacteriaceae] Length: 372\nNumber of predicted TMHs: 2\nExp number of AAs in TMHs: 37.2608899999999\nExp number, first 60 AAs: 19.52923\nTotal prob of N-in: 0.94351\nPOSSIBLE N-term signal sequence\ninside 1 19\nTMhelix 20 39\noutside 40 325\nTMhelix 326 348\ninside 349 372

10643 GCF_000250195.1_ASM25019v2 Escherichia coli DEC11E Proteobacteria; Gammaproteobacteria; Enterobacterales; Enterobacteriaceae; Escherichia; Escherichia coli
MTGDNTLIHSHGINRRDFMKLCAALAATMGLSSKAAAEMAESVTNPQRPPVIWIGAQECT WP_000145410.1
MULTISPECIES: hydrogenase-2 small chain [Proteobacteria] Length: 372\nNumber of predicted TMHs: 1\nExp number of AAs in TMHs: 23.72226\nExp number, first 60 AAs: 0.22038\nTotal prob of N-in: 0.12994\noutside 1 331\nTMhelix 332 354\ninside 355 372

10644 GCF_000250215.1_ASM25021v2 Escherichia coli DEC12A Proteobacteria; Gammaproteobacteria; Enterobacterales; Enterobacteriaceae; Escherichia; Escherichia coli
MTGDNTLIHSHGINRRDFMKLCAALAATMGLSSKAAAEMAESVTNPQRPPVIWIGAQECT WP_000145419.1
hydrogenase 2 small subunit [Escherichia coli] Length: 372\nNumber of predicted TMHs: 1\nExp number of AAs in TMHs: 25.59455\nExp number, first 60 AAs: 0.21473\nTotal prob of N-in: 0.19917\noutside 1 331\nTMhelix 332 354\ninside 355 372

10645 GCF_000250215.1_ASM25021v2 Escherichia coli DEC12A Proteobacteria; Gammaproteobacteria; Enterobacterales; Enterobacteriaceae; Escherichia; Escherichia coli
MNNEETFYQAMRRQGVTRRSFLKYCSLAATSLGLGAGMAPKIAWALENKPRIPVVWIHGL WP_001295940.1
MULTISPECIES: hydrogenase [Enterobacteriaceae] Length: 372\nNumber of predicted TMHs: 2\nExp number of AAs in TMHs: 37.2608899999999\nExp number, first 60 AAs: 19.52923\nTotal prob of N-in:

0.94351\nPOSSIBLE N-term signal sequence\ninside 1 19\nTMhelix 20 39\noutside 40 325\nTMhelix 326 348\ninside 349 372

10646 GCF_000250235.1_ASM25023v2 Escherichia coli DEC12B Proteobacteria; Gammaproteobacteria; Enterobacterales; Enterobacteriaceae; Escherichia; Escherichia coli
MNNEETFYQAMRRQGVTRRSFLKYCSLAATSLGLGAGMAPKIAWALENKPRIPVVWIHGL WP_001295940.1
MULTISPECIES: hydrogenase [Enterobacteriaceae] Length: 372\nNumber of predicted TMHs: 2\nExp number of AAs in TMHs: 37.2608899999999\nExp number, first 60 AAs: 19.52923\nTotal prob of N-in: 0.94351\nPOSSIBLE N-term signal sequence\ninside 1 19\nTMhelix 20 39\noutside 40 325\nTMhelix 326 348\ninside 349 372

10647 GCF_000250235.1_ASM25023v2 Escherichia coli DEC12B Proteobacteria; Gammaproteobacteria; Enterobacterales; Enterobacteriaceae; Escherichia; Escherichia coli
MTGDNTLIHSHGINRRDFMKLCAALAATMGLSSKAAAEMAESVTNPQRPPVIWIGAQECT WP_000145410.1
MULTISPECIES: hydrogenase-2 small chain [Proteobacteria] Length: 372\nNumber of predicted TMHs: 1\nExp number of AAs in TMHs: 23.72226\nExp number, first 60 AAs: 0.22038\nTotal prob of N-in: 0.12994\noutside 1 331\nTMhelix 332 354\ninside 355 372

10648 GCF_000250275.1_ASM25027v2 Escherichia coli DEC12D Proteobacteria; Gammaproteobacteria; Enterobacterales; Enterobacteriaceae; Escherichia; Escherichia coli
MTGDNTLIHSHGINRRDFMKLCAALAATMGLSSKAAAEMAESVTNPQRPPVIWIGAQECT WP_000145410.1
MULTISPECIES: hydrogenase-2 small chain [Proteobacteria] Length: 372\nNumber of predicted TMHs: 1\nExp number of AAs in TMHs: 23.72226\nExp number, first 60 AAs: 0.22038\nTotal prob of N-in: 0.12994\noutside 1 331\nTMhelix 332 354\ninside 355 372

10649 GCF_000250275.1_ASM25027v2 Escherichia coli DEC12D Proteobacteria; Gammaproteobacteria; Enterobacterales; Enterobacteriaceae; Escherichia; Escherichia coli
MNNEETFYQAMRRQGVTRRSFLKYCSLAATSLGLGAGMAPKIAWALENKPRIPVVWIHGL WP_001295940.1
MULTISPECIES: hydrogenase [Enterobacteriaceae] Length: 372\nNumber of predicted TMHs: 2\nExp number of AAs in TMHs: 37.2608899999999\nExp number, first 60 AAs: 19.52923\nTotal prob of N-in: 0.94351\nPOSSIBLE N-term signal sequence\ninside 1 19\nTMhelix 20 39\noutside 40 325\nTMhelix 326 348\ninside 349 372

10650 GCF_000250295.1_ASM25029v2 Escherichia coli DEC12E Proteobacteria; Gammaproteobacteria; Enterobacterales; Enterobacteriaceae; Escherichia; Escherichia coli
MNNEETFYQAMRRQGVTRRSFLKYCSLAATSLGLGAGMAPKIAWALENKPRIPVVWIHGL WP_001437382.1
hydrogenase [Escherichia coli] Length: 372\nNumber of predicted TMHs: 2\nExp number of AAs in TMHs: 37.27324\nExp number, first 60 AAs: 19.52926\nTotal prob of N-in: 0.94351\nPOSSIBLE N-term signal sequence\ninside 1 19\nTMhelix 20 39\noutside 40 325\nTMhelix 326 348\ninside 349 372

10651 GCF_000250295.1_ASM25029v2 Escherichia coli DEC12E Proteobacteria; Gammaproteobacteria; Enterobacterales; Enterobacteriaceae; Escherichia; Escherichia coli
MTGDNTLIHSHGINRRDFMKLCAALAATMGLSSKAAAEMAESVTNPQRPPVIWIGAQECT WP_000145410.1
MULTISPECIES: hydrogenase-2 small chain [Proteobacteria] Length: 372\nNumber of predicted TMHs: 1\nExp number of AAs in TMHs: 23.72226\nExp number, first 60 AAs: 0.22038\nTotal prob of N-in: 0.12994\noutside 1 331\nTMhelix 332 354\ninside 355 372

10652 GCF_000250315.1_ASM25031v2 Escherichia coli DEC13A Proteobacteria; Gammaproteobacteria; Enterobacterales; Enterobacteriaceae; Escherichia; Escherichia coli
MNNEETFYQAMRRQGVTRRSFLKYCSLAATSLGLGAGMAPKIAWALENKPRIPVVWIHGL WP_001058323.1
MULTISPECIES: hydrogenase-1 small chain [Proteobacteria] Length: 372\nNumber of predicted TMHs: 2\nExp number of AAs in TMHs: 37.2608499999999\nExp number, first 60 AAs: 19.52923\nTotal prob of N-in: 0.94351\nPOSSIBLE N-term signal sequence\ninside 1 19\nTMhelix 20 39\noutside 40 325\nTMhelix 326 348\ninside 349 372

10653 GCF_000250315.1_ASM25031v2 Escherichia coli DEC13A Proteobacteria; Gammaproteobacteria; Enterobacterales; Enterobacteriaceae; Escherichia; Escherichia coli
MTGDNTLIHSHGINRRDFMKLCAALAATMGLSSKAAAEMAESVTNPQRPPVIWIGAQECT WP_000145410.1
MULTISPECIES: hydrogenase-2 small chain [Proteobacteria] Length: 372\nNumber of predicted TMHs: 1\nExp number of AAs in TMHs: 23.72226\nExp number, first 60 AAs: 0.22038\nTotal prob of N-in: 0.12994\noutside 1 331\nTMhelix 332 354\ninside 355 372

10654 GCF_000250335.1_ASM25033v2 Escherichia coli DEC13B Proteobacteria; Gammaproteobacteria; Enterobacterales; Enterobacteriaceae; Escherichia; Escherichia coli
MNNEETFYQAMRRQGVTRRSFLKYCSLAATSLGLGAGMAPKIAWALENKPRIPVVWIHGL WP_001058323.1
MULTISPECIES: hydrogenase-1 small chain [Proteobacteria] Length: 372\nNumber of predicted TMHs: 2\nExp number of AAs in TMHs: 37.2608499999999\nExp number, first 60 AAs: 19.52923\nTotal prob of N-in: 0.94351\nPOSSIBLE N-term signal sequence\ninside 1 19\nTMhelix 20 39\noutside 40 325\nTMhelix 326 348\ninside 349 372

10655 GCF_000250335.1_ASM25033v2 Escherichia coli DEC13B Proteobacteria; Gammaproteobacteria; Enterobacterales; Enterobacteriaceae; Escherichia; Escherichia coli
MTGDNTLIHSHGINRRDFMKLCAALAATMGLSSKAAAEMAESVTNPQRPPVIWIGAQECT WP_000145410.1
MULTISPECIES: hydrogenase-2 small chain [Proteobacteria] Length: 372\nNumber of predicted TMHs: 1\nExp number of AAs in TMHs: 23.72226\nExp number, first 60 AAs: 0.22038\nTotal prob of N-in: 0.12994\noutside 1 331\nTMhelix 332 354\ninside 355 372

10656 GCF_000250355.1_ASM25035v2 Escherichia coli DEC13C Proteobacteria; Gammaproteobacteria; Enterobacterales; Enterobacteriaceae; Escherichia; Escherichia coli
MNNEETFYQAMRRQGVTRRSFLKYCSLAATSLGLGAGMAPKIAWALENKPRIPVVWIHGL WP_001058323.1
MULTISPECIES: hydrogenase-1 small chain [Proteobacteria] Length: 372\nNumber of predicted TMHs: 2\nExp number of AAs in TMHs: 37.2608499999999\nExp number, first 60 AAs: 19.52923\nTotal prob of N-in: 0.94351\nPOSSIBLE N-term signal sequence\ninside 1 19\nTMhelix 20 39\noutside 40 325\nTMhelix 326 348\ninside 349 372

10657 GCF_000250355.1_ASM25035v2 Escherichia coli DEC13C Proteobacteria; Gammaproteobacteria; Enterobacterales; Enterobacteriaceae; Escherichia; Escherichia coli
MTGDNTLIHSHGINRRDFMKLCAALAATMGLSSKAAAEMAESVTNPQRPPVIWIGAQECT WP_000145410.1
MULTISPECIES: hydrogenase-2 small chain [Proteobacteria] Length: 372\nNumber of predicted TMHs: 1\nExp number of AAs in TMHs: 23.72226\nExp number, first 60 AAs: 0.22038\nTotal prob of N-in: 0.12994\noutside 1 331\nTMhelix 332 354\ninside 355 372

10658 GCF_000250375.1_ASM25037v2 Escherichia coli DEC13D Proteobacteria; Gammaproteobacteria; Enterobacterales; Enterobacteriaceae; Escherichia; Escherichia coli
MTGDNTLIHSHGINRRDFMKLCAALAATMGLSSKAAAEMAESVTNPQRPPVIWIGAQECT WP_000145410.1
MULTISPECIES: hydrogenase-2 small chain [Proteobacteria] Length: 372\nNumber of predicted TMHs: 1\nExp number of AAs in TMHs: 23.72226\nExp number, first 60 AAs: 0.22038\nTotal prob of N-in: 0.12994\noutside 1 331\nTMhelix 332 354\ninside 355 372

10659 GCF_000250375.1_ASM25037v2 Escherichia coli DEC13D Proteobacteria; Gammaproteobacteria; Enterobacterales; Enterobacteriaceae; Escherichia; Escherichia coli
MNNEETFYQAMRRQGVTRRSFLKYCSLAATSLGLGAGMAPKIAWALENKPRIPVVWIHGL WP_001058323.1
MULTISPECIES: hydrogenase-1 small chain [Proteobacteria] Length: 372\nNumber of predicted TMHs: 2\nExp number of AAs in TMHs: 37.2608499999999\nExp number, first 60 AAs: 19.52923\nTotal prob of N-in: 0.94351\nPOSSIBLE N-term signal sequence\ninside 1 19\nTMhelix 20 39\noutside 40 325\nTMhelix 326 348\ninside 349 372

10660 GCF_000250395.1_ASM25039v2 Escherichia coli DEC13E Proteobacteria; Gammaproteobacteria; Enterobacterales; Enterobacteriaceae; Escherichia; Escherichia coli
MNNEETFYQAMRRQGVTRRSFLKYCSLAATSLGLGAGMAPKIAWALENKPRIPVVWIHGL WP_001058323.1
MULTISPECIES: hydrogenase-1 small chain [Proteobacteria] Length: 372\nNumber of predicted TMHs: 2\nExp number of AAs in TMHs: 37.2608499999999\nExp number, first 60 AAs: 19.52923\nTotal prob of N-in: 0.94351\nPOSSIBLE N-term signal sequence\ninside 1 19\nTMhelix 20 39\noutside 40 325\nTMhelix 326 348\ninside 349 372

10661 GCF_000250395.1_ASM25039v2 Escherichia coli DEC13E Proteobacteria; Gammaproteobacteria; Enterobacterales; Enterobacteriaceae; Escherichia; Escherichia coli
MTGDNTLIHSHGINRRDFMKLCAALAATMGLSSKAAAEMAESVTNPQRPPVIWIGAQECT WP_000145410.1
MULTISPECIES: hydrogenase-2 small chain [Proteobacteria] Length: 372\nNumber of predicted TMHs: 1\nExp number of AAs in TMHs: 23.72226\nExp number, first 60 AAs: 0.22038\nTotal prob of N-in: 0.12994\noutside 1 331\nTMhelix 332 354\ninside 355 372

10662 GCF_000250415.1_ASM25041v2 Escherichia coli DEC14A Proteobacteria; Gammaproteobacteria; Enterobacterales; Enterobacteriaceae; Escherichia; Escherichia coli
MTGDNTLIHSHGINRRDFMKLCAALAATMGLSSKAAAEMAESVTNPQRPPVIWIGAQECT WP_000145410.1
MULTISPECIES: hydrogenase-2 small chain [Proteobacteria] Length: 372\nNumber of predicted TMHs: 1\nExp number of AAs in TMHs: 23.72226\nExp number, first 60 AAs: 0.22038\nTotal prob of N-in: 0.12994\noutside 1 331\nTMhelix 332 354\ninside 355 372

10663 GCF_000250415.1_ASM25041v2 Escherichia coli DEC14A Proteobacteria; Gammaproteobacteria; Enterobacterales; Enterobacteriaceae; Escherichia; Escherichia coli
MNNEETFYQAMRRQGVTRRSFLKYCSLAATSLGLGAGMAPKIAWALENKPRIPVVWIHGL WP_001058323.1
MULTISPECIES: hydrogenase-1 small chain [Proteobacteria] Length: 372\nNumber of predicted TMHs: 2\nExp number of AAs in TMHs: 37.2608499999999\nExp number, first 60 AAs: 19.52923\nTotal prob of N-in: 0.94351\nPOSSIBLE N-term signal sequence\ninside 1 19\nTMhelix 20 39\noutside 40 325\nTMhelix 326 348\ninside 349 372

10664 GCF_000250435.1_ASM25043v2 Escherichia coli DEC14B Proteobacteria; Gammaproteobacteria; Enterobacterales; Enterobacteriaceae; Escherichia; Escherichia coli
MNNEETFYQAMRRQGVTRRSFLKYCSLAATSLGLGAGMAPKIAWALENKPRIPVVWIHGL WP_001058323.1
MULTISPECIES: hydrogenase-1 small chain [Proteobacteria] Length: 372\nNumber of predicted TMHs: 2\nExp number of AAs in TMHs: 37.2608499999999\nExp number, first 60 AAs: 19.52923\nTotal prob of N-in: 0.94351\nPOSSIBLE N-term signal sequence\ninside 1 19\nTMhelix 20 39\noutside 40 325\nTMhelix 326 348\ninside 349 372

10665 GCF_000250435.1_ASM25043v2 Escherichia coli DEC14B Proteobacteria; Gammaproteobacteria; Enterobacterales; Enterobacteriaceae; Escherichia; Escherichia coli
MTGDNTLIHSHGINRRDFMKLCAALAATMGLSSKAAAEMAESVTNPQRPPVIWIGAQECT WP_000145410.1
MULTISPECIES: hydrogenase-2 small chain [Proteobacteria] Length: 372\nNumber of predicted TMHs: 1\nExp number of AAs in TMHs: 23.72226\nExp number, first 60 AAs: 0.22038\nTotal prob of N-in: 0.12994\noutside 1 331\nTMhelix 332 354\ninside 355 372

10666 GCF_000250455.1_ASM25045v2 Escherichia coli DEC14C Proteobacteria; Gammaproteobacteria; Enterobacterales; Enterobacteriaceae; Escherichia; Escherichia coli
MNNEETFYQAMRRQGVTRRSFLKYCSLAAASLGLGAGMAPKIAWALENKPRIPVVWIHGL WP_001439901.1
hydrogenase-1 small chain [Escherichia coli] Length: 372\nNumber of predicted TMHs: 2\nExp number of AAs in TMHs: 37.74488\nExp number, first 60 AAs: 20.0186\nTotal prob of N-in: 0.96589\nPOSSIBLE N-term signal sequence\ninside 1 19\nTMhelix 20 39\noutside 40 325\nTMhelix 326 348\ninside 349 372

10667 GCF_000250455.1_ASM25045v2 Escherichia coli DEC14C Proteobacteria; Gammaproteobacteria; Enterobacterales; Enterobacteriaceae; Escherichia; Escherichia coli
MTGDNTLIHSHGINRRDFMKLCAALAATMGLSSKAAAEMAESVTNPQRPPVIWIGAQECT WP_000145410.1
MULTISPECIES: hydrogenase-2 small chain [Proteobacteria] Length: 372\nNumber of predicted TMHs: 1\nExp number of AAs in TMHs: 23.72226\nExp number, first 60 AAs: 0.22038\nTotal prob of N-in: 0.12994\noutside 1 331\nTMhelix 332 354\ninside 355 372

10668 GCF_000250475.1_ASM25047v2 Escherichia coli DEC14D Proteobacteria; Gammaproteobacteria; Enterobacterales; Enterobacteriaceae; Escherichia; Escherichia coli
MTGDNTLIHSHGINRRDFMKLCAALAATMGLSSKAAAEMAESVTNPQRPPVIWIGAQECT WP_000145410.1
MULTISPECIES: hydrogenase-2 small chain [Proteobacteria] Length: 372\nNumber of predicted TMHs: 1\nExp number of AAs in TMHs: 23.72226\nExp number, first 60 AAs: 0.22038\nTotal prob of N-in: 0.12994\noutside 1 331\nTMhelix 332 354\ninside 355 372

10669 GCF_000250475.1_ASM25047v2 Escherichia coli DEC14D Proteobacteria; Gammaproteobacteria; Enterobacterales; Enterobacteriaceae; Escherichia; Escherichia coli
MNNEETFYQAMRRQGVTRRSFLKYCSLAATSLGLGAGMAPKIAWALENKPRIPVVWIHGL WP_001058323.1
MULTISPECIES: hydrogenase-1 small chain [Proteobacteria] Length: 372\nNumber of predicted TMHs: 2\nExp number of AAs in TMHs: 37.2608499999999\nExp number, first 60 AAs: 19.52923\nTotal prob of N-in: 0.94351\nPOSSIBLE N-term signal sequence\ninside 1 19\nTMhelix 20 39\noutside 40 325\nTMhelix 326 348\ninside 349 372

10670 GCF_000250495.1_ASM25049v2 Escherichia coli DEC15A Proteobacteria; Gammaproteobacteria; Enterobacterales; Enterobacteriaceae; Escherichia; Escherichia coli

MTGDNTLIHSHGINRRDFMKLCAALAATMGLSSKAAAEMAESVTNPQRPPVIWIGAQECT WP_000145410.1 MULTISPECIES: hydrogenase-2 small chain [Proteobacteria] Length: 372\nNumber of predicted TMHs: 1\nExp number of AAs in TMHs: 23.72226\nExp number, first 60 AAs: 0.22038\nTotal prob of N-in: 0.12994\noutside 1 331\nTMhelix 332 354\ninside 355 372

10671 GCF_000250495.1_ASM25049v2 Escherichia coli DEC15A Proteobacteria; Gammaproteobacteria; Enterobacterales; Enterobacteriaceae; Escherichia; Escherichia coli MNNEETFYQAMRRQGVTRRSFLKYCSLAATSLGLGAGMAPKIAWALENKPRIPVVWIHGL WP_001058320.1 hydrogenase [Escherichia coli] Length: 372\nNumber of predicted TMHs: 2\nExp number of AAs in TMHs: 37.26082\nExp number, first 60 AAs: 19.52923\nTotal prob of N-in: 0.94351\nPOSSIBLE N-term signal sequence\ninside 1 19\nTMhelix 20 39\noutside 40 325\nTMhelix 326 348\ninside 349 372

10672 GCF_000250515.1_ASM25051v2 Escherichia coli DEC15B Proteobacteria; Gammaproteobacteria; Enterobacterales; Enterobacteriaceae; Escherichia; Escherichia coli MTGDNTLIHSHGINRRDFMKLCAALAATMGLSSKAAAEMAESVTNPQRPPVIWIGAQECT WP_000145410.1 MULTISPECIES: hydrogenase-2 small chain [Proteobacteria] Length: 372\nNumber of predicted TMHs: 1\nExp number of AAs in TMHs: 23.72226\nExp number, first 60 AAs: 0.22038\nTotal prob of N-in: 0.12994\noutside 1 331\nTMhelix 332 354\ninside 355 372

10673 GCF_000250515.1_ASM25051v2 Escherichia coli DEC15B Proteobacteria; Gammaproteobacteria; Enterobacterales; Enterobacteriaceae; Escherichia; Escherichia coli MNNEETFYQAMRRQGVTRRSFLKYCSLAATSLGLGAGMAPKIAWALENKPRIPVVWIHGL WP_001058320.1 hydrogenase [Escherichia coli] Length: 372\nNumber of predicted TMHs: 2\nExp number of AAs in TMHs: 37.26082\nExp number, first 60 AAs: 19.52923\nTotal prob of N-in: 0.94351\nPOSSIBLE N-term signal sequence\ninside 1 19\nTMhelix 20 39\noutside 40 325\nTMhelix 326 348\ninside 349 372

10674 GCF_000250535.1_ASM25053v2 Escherichia coli DEC15C Proteobacteria; Gammaproteobacteria; Enterobacterales; Enterobacteriaceae; Escherichia; Escherichia coli MTGDNTLIHSHGINRRDFMKLCAALAATMGLSSKAAAEMAESVTNPQRPPVIWIGAQECT WP_000145410.1 MULTISPECIES: hydrogenase-2 small chain [Proteobacteria] Length: 372\nNumber of predicted TMHs: 1\nExp number of AAs in TMHs: 23.72226\nExp number, first 60 AAs: 0.22038\nTotal prob of N-in: 0.12994\noutside 1 331\nTMhelix 332 354\ninside 355 372

10675 GCF_000250535.1_ASM25053v2 Escherichia coli DEC15C Proteobacteria; Gammaproteobacteria; Enterobacterales; Enterobacteriaceae; Escherichia; Escherichia coli MNNEETFYQAMRRQGVTRRSFLKYCSLAATSLGLGAGMAPKIAWALENKPRIPVVWIHGL WP_001058320.1 hydrogenase [Escherichia coli] Length: 372\nNumber of predicted TMHs: 2\nExp number of AAs in TMHs: 37.26082\nExp number, first 60 AAs: 19.52923\nTotal prob of N-in: 0.94351\nPOSSIBLE N-term signal sequence\ninside 1 19\nTMhelix 20 39\noutside 40 325\nTMhelix 326 348\ninside 349 372

10676 GCF_000250555.1_ASM25055v2 Escherichia coli DEC15D Proteobacteria; Gammaproteobacteria; Enterobacterales; Enterobacteriaceae; Escherichia; Escherichia coli MTGDNTLIHSHGINRRDFMKLCAALAATMGLSSKAAAEMAESVTNPQRPPVIWIGAQECT WP_000145410.1 MULTISPECIES: hydrogenase-2 small chain [Proteobacteria] Length: 372\nNumber of predicted TMHs: 1\nExp number of AAs in TMHs: 23.72226\nExp number, first 60 AAs: 0.22038\nTotal prob of N-in: 0.12994\noutside 1 331\nTMhelix 332 354\ninside 355 372

10677 GCF_000250555.1_ASM25055v2 Escherichia coli DEC15D Proteobacteria; Gammaproteobacteria; Enterobacterales; Enterobacteriaceae; Escherichia; Escherichia coli MNNEETFYQAMRRQGVTRRSFLKYCSLAATSLGLGAGMAPKIAWALENKPRIPVVWIHGL WP_001058320.1 hydrogenase [Escherichia coli] Length: 372\nNumber of predicted TMHs: 2\nExp number of AAs in TMHs: 37.26082\nExp number, first 60 AAs: 19.52923\nTotal prob of N-in: 0.94351\nPOSSIBLE N-term signal sequence\ninside 1 19\nTMhelix 20 39\noutside 40 325\nTMhelix 326 348\ninside 349 372

10678 GCF_000250575.1_ASM25057v2 Escherichia coli DEC15E Proteobacteria; Gammaproteobacteria; Enterobacterales; Enterobacteriaceae; Escherichia; Escherichia coli MNNEETFYQAMRRQGVTRRSFLKYCSLAATSLGLGAGMAPKIAWALENKPRIPVVWIHGL WP_001058320.1 hydrogenase [Escherichia coli] Length: 372\nNumber of predicted TMHs: 2\nExp number of AAs in TMHs: 37.26082\nExp number, first 60 AAs: 19.52923\nTotal prob of N-in: 0.94351\nPOSSIBLE N-term signal sequence\ninside 1 19\nTMhelix 20 39\noutside 40 325\nTMhelix 326 348\ninside 349 372

10679 GCF_000250575.1_ASM25057v2 Escherichia coli DEC15E Proteobacteria; Gammaproteobacteria; Enterobacterales; Enterobacteriaceae; Escherichia; Escherichia coli
MTGDNTLIHSHGINRRDFMKLCAALAATMGLSSKAAAEMAESVTNPQRPPVIWIGAQECT WP_000145410.1
MULTISPECIES: hydrogenase-2 small chain [Proteobacteria] Length: 372\nNumber of predicted TMHs: 1\nExp number of AAs in TMHs: 23.72226\nExp number, first 60 AAs: 0.22038\nTotal prob of N-in: 0.12994\noutside 1 331\nTMhelix 332 354\ninside 355 372

10680 GCF_000249055.1_ASM24905v2 Escherichia coli DEC1A Proteobacteria; Gammaproteobacteria; Enterobacterales; Enterobacteriaceae; Escherichia; Escherichia coli
MTGDNTLIHSHGINRRDFMKLCAALAATMGLSSKAAAEMAESVTNPQRPPVIWIGAQECT WP_000145410.1
MULTISPECIES: hydrogenase-2 small chain [Proteobacteria] Length: 372\nNumber of predicted TMHs: 1\nExp number of AAs in TMHs: 23.72226\nExp number, first 60 AAs: 0.22038\nTotal prob of N-in: 0.12994\noutside 1 331\nTMhelix 332 354\ninside 355 372

10681 GCF_000249055.1_ASM24905v2 Escherichia coli DEC1A Proteobacteria; Gammaproteobacteria; Enterobacterales; Enterobacteriaceae; Escherichia; Escherichia coli
MNNEETFYQAMRRQGVTRRSFLKYCSLAATSLGLGAGMAPKIAWALENKPRIPVVWIHGL WP_001058323.1
MULTISPECIES: hydrogenase-1 small chain [Proteobacteria] Length: 372\nNumber of predicted TMHs: 2\nExp number of AAs in TMHs: 37.2608499999999\nExp number, first 60 AAs: 19.52923\nTotal prob of N-in: 0.94351\nPOSSIBLE N-term signal sequence\ninside 1 19\nTMhelix 20 39\noutside 40 325\nTMhelix 326 348\ninside 349 372

10682 GCF_000249075.1_ASM24907v2 Escherichia coli DEC1B Proteobacteria; Gammaproteobacteria; Enterobacterales; Enterobacteriaceae; Escherichia; Escherichia coli
MTGDNTLIHSHGINRRDFMKLCAALAATMGLSSKAAAEMAESVTNPQRPPVIWIGAQECT WP_000145410.1
MULTISPECIES: hydrogenase-2 small chain [Proteobacteria] Length: 372\nNumber of predicted TMHs: 1\nExp number of AAs in TMHs: 23.72226\nExp number, first 60 AAs: 0.22038\nTotal prob of N-in: 0.12994\noutside 1 331\nTMhelix 332 354\ninside 355 372

10683 GCF_000249075.1_ASM24907v2 Escherichia coli DEC1B Proteobacteria; Gammaproteobacteria; Enterobacterales; Enterobacteriaceae; Escherichia; Escherichia coli
MNNEETFYQAMRRQGVTRRSFLKYCSLAATSLGLGAGMAPKIAWALENKPRIPVVWIHGL WP_001058323.1
MULTISPECIES: hydrogenase-1 small chain [Proteobacteria] Length: 372\nNumber of predicted TMHs: 2\nExp number of AAs in TMHs: 37.2608499999999\nExp number, first 60 AAs: 19.52923\nTotal prob of N-in: 0.94351\nPOSSIBLE N-term signal sequence\ninside 1 19\nTMhelix 20 39\noutside 40 325\nTMhelix 326 348\ninside 349 372

10684 GCF_000249095.1_ASM24909v2 Escherichia coli DEC1C Proteobacteria; Gammaproteobacteria; Enterobacterales; Enterobacteriaceae; Escherichia; Escherichia coli
MNNEETFYQAMRRQGVTRRSFLKYCSLAATSLGLGAGMAPKIAWALENKPRIPVVWIHGL WP_001058323.1
MULTISPECIES: hydrogenase-1 small chain [Proteobacteria] Length: 372\nNumber of predicted TMHs: 2\nExp number of AAs in TMHs: 37.2608499999999\nExp number, first 60 AAs: 19.52923\nTotal prob of N-in: 0.94351\nPOSSIBLE N-term signal sequence\ninside 1 19\nTMhelix 20 39\noutside 40 325\nTMhelix 326 348\ninside 349 372

10685 GCF_000249095.1_ASM24909v2 Escherichia coli DEC1C Proteobacteria; Gammaproteobacteria; Enterobacterales; Enterobacteriaceae; Escherichia; Escherichia coli
MTGDNTLIHSHGINRRDFMKLCAALAATMGLSSKAAAEMAESVTNPQRPPVIWIGAQECT WP_000145410.1
MULTISPECIES: hydrogenase-2 small chain [Proteobacteria] Length: 372\nNumber of predicted TMHs: 1\nExp number of AAs in TMHs: 23.72226\nExp number, first 60 AAs: 0.22038\nTotal prob of N-in: 0.12994\noutside 1 331\nTMhelix 332 354\ninside 355 372

10686 GCF_000249115.1_ASM24911v2 Escherichia coli DEC1D Proteobacteria; Gammaproteobacteria; Enterobacterales; Enterobacteriaceae; Escherichia; Escherichia coli
MTGDNTLIHSHGINRRDFMKLCAALAATMGLSSKAAAEMAESVTNPQRPPVIWIGAQECT WP_000145410.1
MULTISPECIES: hydrogenase-2 small chain [Proteobacteria] Length: 372\nNumber of predicted TMHs: 1\nExp number of AAs in TMHs: 23.72226\nExp number, first 60 AAs: 0.22038\nTotal prob of N-in: 0.12994\noutside 1 331\nTMhelix 332 354\ninside 355 372

10687 GCF_000249115.1_ASM24911v2 Escherichia coli DEC1D Proteobacteria; Gammaproteobacteria; Enterobacterales; Enterobacteriaceae; Escherichia; Escherichia coli

MNNEETFYQAMRRQGVTRRSFLKYCSLAATSLGLGAGMAPKIAWALENKPRIPVVWIHGL WP_001058323.1 MULTISPECIES: hydrogenase-1 small chain [Proteobacteria] Length: 372\nNumber of predicted TMHs: 2\nExp number of AAs in TMHs: 37.2608499999999\nExp number, first 60 AAs: 19.52923\nTotal prob of N-in: 0.94351\nPOSSIBLE N-term signal sequence\ninside 1 19\nTMhelix 20 39\noutside 40 325\nTMhelix 326 348\ninside 349 372

10688 GCF_000249135.1_ASM24913v2 Escherichia coli DEC1E Proteobacteria; Gammaproteobacteria; Enterobacterales; Enterobacteriaceae; Escherichia; Escherichia coli MTGDNTLIHSHGINRRDFMKLCAALAATMGLSSKAAAEMAESVTNPQRPPVIWIGAQECT WP_000145410.1 MULTISPECIES: hydrogenase-2 small chain [Proteobacteria] Length: 372\nNumber of predicted TMHs: 1\nExp number of AAs in TMHs: 23.72226\nExp number, first 60 AAs: 0.22038\nTotal prob of N-in: 0.12994\noutside 1 331\nTMhelix 332 354\ninside 355 372

10689 GCF_000249135.1_ASM24913v2 Escherichia coli DEC1E Proteobacteria; Gammaproteobacteria; Enterobacterales; Enterobacteriaceae; Escherichia; Escherichia coli MNNEETFYQAMRRQGVTRRSFLKYCSLAATSLGLGAGMAPKIAWALENKPRIPVVWIHGL WP_001058323.1 MULTISPECIES: hydrogenase-1 small chain [Proteobacteria] Length: 372\nNumber of predicted TMHs: 2\nExp number of AAs in TMHs: 37.2608499999999\nExp number, first 60 AAs: 19.52923\nTotal prob of N-in: 0.94351\nPOSSIBLE N-term signal sequence\ninside 1 19\nTMhelix 20 39\noutside 40 325\nTMhelix 326 348\ninside 349 372

10690 GCF_000249175.1_ASM24917v2 Escherichia coli DEC2B Proteobacteria; Gammaproteobacteria; Enterobacterales; Enterobacteriaceae; Escherichia; Escherichia coli MNNEETFYQAMRRQGVTRRSFLKYCSLAATSLGLGAGMAPKIAWALENKPRIPVVWIHGL WP_001058323.1 MULTISPECIES: hydrogenase-1 small chain [Proteobacteria] Length: 372\nNumber of predicted TMHs: 2\nExp number of AAs in TMHs: 37.2608499999999\nExp number, first 60 AAs: 19.52923\nTotal prob of N-in: 0.94351\nPOSSIBLE N-term signal sequence\ninside 1 19\nTMhelix 20 39\noutside 40 325\nTMhelix 326 348\ninside 349 372

10691 GCF_000249175.1_ASM24917v2 Escherichia coli DEC2B Proteobacteria; Gammaproteobacteria; Enterobacterales; Enterobacteriaceae; Escherichia; Escherichia coli MTGDNTLIHSHGINRRDFMKLCAALAATMGLSSKAAAEMAESVTNPQRPPVIWIGAQECT WP_000145410.1 MULTISPECIES: hydrogenase-2 small chain [Proteobacteria] Length: 372\nNumber of predicted TMHs: 1\nExp number of AAs in TMHs: 23.72226\nExp number, first 60 AAs: 0.22038\nTotal prob of N-in: 0.12994\noutside 1 331\nTMhelix 332 354\ninside 355 372

10692 GCF_000249195.1_ASM24919v2 Escherichia coli DEC2C Proteobacteria; Gammaproteobacteria; Enterobacterales; Enterobacteriaceae; Escherichia; Escherichia coli MNNEETFYQAMRRQGVTRRSFLKYCSLAATSLGLGAGMAPKIAWALENKPRIPVVWIHGL WP_001058323.1 MULTISPECIES: hydrogenase-1 small chain [Proteobacteria] Length: 372\nNumber of predicted TMHs: 2\nExp number of AAs in TMHs: 37.2608499999999\nExp number, first 60 AAs: 19.52923\nTotal prob of N-in: 0.94351\nPOSSIBLE N-term signal sequence\ninside 1 19\nTMhelix 20 39\noutside 40 325\nTMhelix 326 348\ninside 349 372

10693 GCF_000249195.1_ASM24919v2 Escherichia coli DEC2C Proteobacteria; Gammaproteobacteria; Enterobacterales; Enterobacteriaceae; Escherichia; Escherichia coli MTGDNTLIHSHGINRRDFMKLCAALAATMGLSSKAAAEMAESVTNPQRPPVIWIGAQECT WP_000145410.1 MULTISPECIES: hydrogenase-2 small chain [Proteobacteria] Length: 372\nNumber of predicted TMHs: 1\nExp number of AAs in TMHs: 23.72226\nExp number, first 60 AAs: 0.22038\nTotal prob of N-in: 0.12994\noutside 1 331\nTMhelix 332 354\ninside 355 372

10694 GCF_000249215.1_ASM24921v2 Escherichia coli DEC2D Proteobacteria; Gammaproteobacteria; Enterobacterales; Enterobacteriaceae; Escherichia; Escherichia coli MTGDNTLIHSHGINRRDFMKLCAALAATMGLSSKAAAEMAESVTNPQRPPVIWIGAQECT WP_000145410.1 MULTISPECIES: hydrogenase-2 small chain [Proteobacteria] Length: 372\nNumber of predicted TMHs: 1\nExp number of AAs in TMHs: 23.72226\nExp number, first 60 AAs: 0.22038\nTotal prob of N-in: 0.12994\noutside 1 331\nTMhelix 332 354\ninside 355 372

10695 GCF_000249215.1_ASM24921v2 Escherichia coli DEC2D Proteobacteria; Gammaproteobacteria; Enterobacterales; Enterobacteriaceae; Escherichia; Escherichia coli MNNEETFYQAMRRQGVTRRSFLKYCSLAATSLGLGAGMAPKIAWALENKPRIPVVWIHGL WP_001058323.1

MULTISPECIES: hydrogenase-1 small chain [Proteobacteria] Length: 372\nNumber of predicted TMHs: 2\nExp number of AAs in TMHs: 37.2608499999999\nExp number, first 60 AAs: 19.52923\nTotal prob of N-in: 0.94351\nPOSSIBLE N-term signal sequence\ninside 1 19\nTMhelix 20 39\noutside 40 325\nTMhelix 326 348\ninside 349 372

10696 GCF_000249235.1_ASM24923v2 Escherichia coli DEC2E Proteobacteria; Gammaproteobacteria; Enterobacterales; Enterobacteriaceae; Escherichia; Escherichia coli MTGDNTLIHSHGINRRDFMKLCAALAATMGLSSKAAAEMAESVTNPQRPPVIWIGAQECT WP_000145410.1 MULTISPECIES: hydrogenase-2 small chain [Proteobacteria] Length: 372\nNumber of predicted TMHs: 1\nExp number of AAs in TMHs: 23.72226\nExp number, first 60 AAs: 0.22038\nTotal prob of N-in: 0.12994\noutside 1 331\nTMhelix 332 354\ninside 355 372

10697 GCF_000249235.1_ASM24923v2 Escherichia coli DEC2E Proteobacteria; Gammaproteobacteria; Enterobacterales; Enterobacteriaceae; Escherichia; Escherichia coli MNNEETFYQAMRRQGVTRRSFLKYCSLAATSLGLGAGMAPKIAWALENKPRIPVVWIHGL WP_001058323.1 MULTISPECIES: hydrogenase-1 small chain [Proteobacteria] Length: 372\nNumber of predicted TMHs: 2\nExp number of AAs in TMHs: 37.2608499999999\nExp number, first 60 AAs: 19.52923\nTotal prob of N-in: 0.94351\nPOSSIBLE N-term signal sequence\ninside 1 19\nTMhelix 20 39\noutside 40 325\nTMhelix 326 348\ninside 349 372

10698 GCF_000249255.1_ASM24925v2 Escherichia coli DEC3A Proteobacteria; Gammaproteobacteria; Enterobacterales; Enterobacteriaceae; Escherichia; Escherichia coli MTGDNTLIHSHGINRRDFMKLCAALAATMGLSSKAAAEMAESVTNPQRPPVIWIGAQECT WP_000145410.1 MULTISPECIES: hydrogenase-2 small chain [Proteobacteria] Length: 372\nNumber of predicted TMHs: 1\nExp number of AAs in TMHs: 23.72226\nExp number, first 60 AAs: 0.22038\nTotal prob of N-in: 0.12994\noutside 1 331\nTMhelix 332 354\ninside 355 372

10699 GCF_000249255.1_ASM24925v2 Escherichia coli DEC3A Proteobacteria; Gammaproteobacteria; Enterobacterales; Enterobacteriaceae; Escherichia; Escherichia coli MNNEETFYQAMRRQGVTRRSFLKYCSLAATSLGLGAGMAPKIAWALENKPRIPVVWIHGL WP_001301613.1 hydrogenase [Escherichia coli] Length: 372\nNumber of predicted TMHs: 2\nExp number of AAs in TMHs: 37.46598\nExp number, first 60 AAs: 19.53139\nTotal prob of N-in: 0.94364\nPOSSIBLE N-term signal sequence\ninside 1 19\nTMhelix 20 39\noutside 40 325\nTMhelix 326 348\ninside 349 372

10700 GCF_000249275.1_ASM24927v2 Escherichia coli DEC3B Proteobacteria; Gammaproteobacteria; Enterobacterales; Enterobacteriaceae; Escherichia; Escherichia coli MNNEETFYQAMRRQGVTRRSFLKYCSLAATSLGLGAGMAPKIAWALENKPRIPVVWIHGL WP_001301613.1 hydrogenase [Escherichia coli] Length: 372\nNumber of predicted TMHs: 2\nExp number of AAs in TMHs: 37.46598\nExp number, first 60 AAs: 19.53139\nTotal prob of N-in: 0.94364\nPOSSIBLE N-term signal sequence\ninside 1 19\nTMhelix 20 39\noutside 40 325\nTMhelix 326 348\ninside 349 372

10701 GCF_000249275.1_ASM24927v2 Escherichia coli DEC3B Proteobacteria; Gammaproteobacteria; Enterobacterales; Enterobacteriaceae; Escherichia; Escherichia coli MTGDNTLIHSHGINRRDFMKLCAALAATMGLSSKAAAEMAESVTNPQRPPVIWIGAQECT WP_000145410.1 MULTISPECIES: hydrogenase-2 small chain [Proteobacteria] Length: 372\nNumber of predicted TMHs: 1\nExp number of AAs in TMHs: 23.72226\nExp number, first 60 AAs: 0.22038\nTotal prob of N-in: 0.12994\noutside 1 331\nTMhelix 332 354\ninside 355 372

10702 GCF_000249335.1_ASM24933v2 Escherichia coli DEC3E Proteobacteria; Gammaproteobacteria; Enterobacterales; Enterobacteriaceae; Escherichia; Escherichia coli MNNEETFYQAMRRQGVTRRSFLKYCSLAATSLGLGAGMAPKIAWALENKPRIPVVWIHGL WP_001301613.1 hydrogenase [Escherichia coli] Length: 372\nNumber of predicted TMHs: 2\nExp number of AAs in TMHs: 37.46598\nExp number, first 60 AAs: 19.53139\nTotal prob of N-in: 0.94364\nPOSSIBLE N-term signal sequence\ninside 1 19\nTMhelix 20 39\noutside 40 325\nTMhelix 326 348\ninside 349 372

10703 GCF_000249335.1_ASM24933v2 Escherichia coli DEC3E Proteobacteria; Gammaproteobacteria; Enterobacterales; Enterobacteriaceae; Escherichia; Escherichia coli MTGDNTLIHSHGINRRDFMKLCAALAATMGLSSKAAAEMAESVTNPQRPPVIWIGAQECT WP_000145410.1 MULTISPECIES: hydrogenase-2 small chain [Proteobacteria] Length: 372\nNumber of predicted TMHs: 1\nExp number of AAs in TMHs: 23.72226\nExp number, first 60 AAs: 0.22038\nTotal prob of N-in: 0.12994\noutside 1 331\nTMhelix 332 354\ninside 355 372

10704 GCF_000249435.1_ASM24943v2 Escherichia coli DEC4D Proteobacteria; Gammaproteobacteria; Enterobacterales; Enterobacteriaceae; Escherichia; Escherichia coli
MTGDNTLIHSHGINRRDFMKLCAALAATMGLSSKAAAEMAESVTNPQRPPVIWIGAQECT WP_000145410.1
MULTISPECIES: hydrogenase-2 small chain [Proteobacteria] Length: 372\nNumber of predicted TMHs: 1\nExp number of AAs in TMHs: 23.72226\nExp number, first 60 AAs: 0.22038\nTotal prob of N-in: 0.12994\noutside 1 331\nTMhelix 332 354\ninside 355 372

10705 GCF_000249435.1_ASM24943v2 Escherichia coli DEC4D Proteobacteria; Gammaproteobacteria; Enterobacterales; Enterobacteriaceae; Escherichia; Escherichia coli
MNNEETFYQAMRRQGVTRRSFLKYCSLAATSLGLGAGMAPKIAWALENKPRIPVVWIHGL WP_001301613.1
hydrogenase [Escherichia coli] Length: 372\nNumber of predicted TMHs: 2\nExp number of AAs in TMHs: 37.46598\nExp number, first 60 AAs: 19.53139\nTotal prob of N-in: 0.94364\nPOSSIBLE N-term signal sequence\ninside 1 19\nTMhelix 20 39\noutside 40 325\nTMhelix 326 348\ninside 349 372

10706 GCF_000249455.1_ASM24945v2 Escherichia coli DEC4E Proteobacteria; Gammaproteobacteria; Enterobacterales; Enterobacteriaceae; Escherichia; Escherichia coli
MNNEETFYQAMRRQGVTRRSFLKYCSLAATSLGLGAGMAPKIAWALENKPRIPVVWIHGL WP_001301613.1
hydrogenase [Escherichia coli] Length: 372\nNumber of predicted TMHs: 2\nExp number of AAs in TMHs: 37.46598\nExp number, first 60 AAs: 19.53139\nTotal prob of N-in: 0.94364\nPOSSIBLE N-term signal sequence\ninside 1 19\nTMhelix 20 39\noutside 40 325\nTMhelix 326 348\ninside 349 372

10707 GCF_000249455.1_ASM24945v2 Escherichia coli DEC4E Proteobacteria; Gammaproteobacteria; Enterobacterales; Enterobacteriaceae; Escherichia; Escherichia coli
MTGDNTLIHSHGINRRDFMKLCAALAATMGLSSKAAAEMAESVTNPQRPPVIWIGAQECT WP_000145410.1
MULTISPECIES: hydrogenase-2 small chain [Proteobacteria] Length: 372\nNumber of predicted TMHs: 1\nExp number of AAs in TMHs: 23.72226\nExp number, first 60 AAs: 0.22038\nTotal prob of N-in: 0.12994\noutside 1 331\nTMhelix 332 354\ninside 355 372

10708 GCF_000249495.1_ASM24949v2 Escherichia coli DEC5A Proteobacteria; Gammaproteobacteria; Enterobacterales; Enterobacteriaceae; Escherichia; Escherichia coli
MTGDNTLIHSHGINRRDFMKLCAALAATMGLSSKAAAEMAESVTNPQRPPVIWIGAQECT WP_000145410.1
MULTISPECIES: hydrogenase-2 small chain [Proteobacteria] Length: 372\nNumber of predicted TMHs: 1\nExp number of AAs in TMHs: 23.72226\nExp number, first 60 AAs: 0.22038\nTotal prob of N-in: 0.12994\noutside 1 331\nTMhelix 332 354\ninside 355 372

10709 GCF_000249495.1_ASM24949v2 Escherichia coli DEC5A Proteobacteria; Gammaproteobacteria; Enterobacterales; Enterobacteriaceae; Escherichia; Escherichia coli
MNNEETFYQAMRRQGVTRRSFLKYCSLAATSLGLGAGMAPKIAWALENKPRIPVVWIHGL WP_001359719.1
hydrogenase [Escherichia coli] Length: 372\nNumber of predicted TMHs: 2\nExp number of AAs in TMHs: 37.2176\nExp number, first 60 AAs: 19.53448\nTotal prob of N-in: 0.94360\nPOSSIBLE N-term signal sequence\ninside 1 19\nTMhelix 20 39\noutside 40 325\nTMhelix 326 348\ninside 349 372

10710 GCF_000249515.1_ASM24951v2 Escherichia coli DEC5B Proteobacteria; Gammaproteobacteria; Enterobacterales; Enterobacteriaceae; Escherichia; Escherichia coli
MNNEETFYQAMRRQGVTRRSFLKYCSLAATSLGLGAGMAPKIAWALENKPRIPVVWIHGL WP_001301613.1
hydrogenase [Escherichia coli] Length: 372\nNumber of predicted TMHs: 2\nExp number of AAs in TMHs: 37.46598\nExp number, first 60 AAs: 19.53139\nTotal prob of N-in: 0.94364\nPOSSIBLE N-term signal sequence\ninside 1 19\nTMhelix 20 39\noutside 40 325\nTMhelix 326 348\ninside 349 372

10711 GCF_000249515.1_ASM24951v2 Escherichia coli DEC5B Proteobacteria; Gammaproteobacteria; Enterobacterales; Enterobacteriaceae; Escherichia; Escherichia coli
MTGDNTLIHSHGINRRDFMKLCAALAATMGLSSKAAAEMAESVTNPQRPPVIWIGAQECT WP_000145410.1
MULTISPECIES: hydrogenase-2 small chain [Proteobacteria] Length: 372\nNumber of predicted TMHs: 1\nExp number of AAs in TMHs: 23.72226\nExp number, first 60 AAs: 0.22038\nTotal prob of N-in: 0.12994\noutside 1 331\nTMhelix 332 354\ninside 355 372

10712 GCF_000249535.1_ASM24953v2 Escherichia coli DEC5C Proteobacteria; Gammaproteobacteria; Enterobacterales; Enterobacteriaceae; Escherichia; Escherichia coli
MNNEETFYQAMRRQGVTRRSFLKYCSLAATSLGLGAGMAPKIAWALENKPRIPVVWIHGL WP_001359719.1
hydrogenase [Escherichia coli] Length: 372\nNumber of predicted TMHs: 2\nExp number of AAs in

TMHs: 37.2176\nExp number, first 60 AAs:  19.53448\nTotal prob of N-in:   0.94360\nPOSSIBLE N-term signal sequence\ninside   1   19\nTMhelix   20   39\noutside   40   325\nTMhelix   326   348\ninside   349   372

10713 GCF_000249535.1_ASM24953v2 Escherichia coli DEC5C Proteobacteria; Gammaproteobacteria; Enterobacterales; Enterobacteriaceae; Escherichia; Escherichia coli
MTGDNTLIHSHGINRRDFMKLCAALAATMGLSSKAAAEMAESVTNPQRPPVIWIGAQECT WP_000145410.1
MULTISPECIES: hydrogenase-2 small chain [Proteobacteria] Length: 372\nNumber of predicted TMHs: 1\nExp number of AAs in TMHs: 23.72226\nExp number, first 60 AAs:  0.22038\nTotal prob of N-in:   0.12994\noutside   1   331\nTMhelix   332   354\ninside   355   372

10714 GCF_000249555.1_ASM24955v2 Escherichia coli DEC5D Proteobacteria; Gammaproteobacteria; Enterobacterales; Enterobacteriaceae; Escherichia; Escherichia coli
MTGDNTLIHSHGINRRDFMKLCAALAATMGLSSKAAAEMAESVTNPQRPPVIWIGAQECT WP_000145410.1
MULTISPECIES: hydrogenase-2 small chain [Proteobacteria] Length: 372\nNumber of predicted TMHs: 1\nExp number of AAs in TMHs: 23.72226\nExp number, first 60 AAs:  0.22038\nTotal prob of N-in:   0.12994\noutside   1   331\nTMhelix   332   354\ninside   355   372

10715 GCF_000249555.1_ASM24955v2 Escherichia coli DEC5D Proteobacteria; Gammaproteobacteria; Enterobacterales; Enterobacteriaceae; Escherichia; Escherichia coli
MNNEETFYQAMRRQGVTRRSFLKYCSLAATSLGLGAGMAPKIAWALENKPRIPVVWIHGL WP_001359719.1
hydrogenase [Escherichia coli] Length: 372\nNumber of predicted TMHs:  2\nExp number of AAs in TMHs: 37.2176\nExp number, first 60 AAs:  19.53448\nTotal prob of N-in:   0.94360\nPOSSIBLE N-term signal sequence\ninside   1   19\nTMhelix   20   39\noutside   40   325\nTMhelix   326   348\ninside   349   372

10716 GCF_000249575.1_ASM24957v2 Escherichia coli DEC5E Proteobacteria; Gammaproteobacteria; Enterobacterales; Enterobacteriaceae; Escherichia; Escherichia coli
MNNEETFYQAMRRQGVTRRSFLKYCSLAATSLGLGAGMAPKIAWALENKPRIPVVWIHGL WP_001359719.1
hydrogenase [Escherichia coli] Length: 372\nNumber of predicted TMHs:  2\nExp number of AAs in TMHs: 37.2176\nExp number, first 60 AAs:  19.53448\nTotal prob of N-in:   0.94360\nPOSSIBLE N-term signal sequence\ninside   1   19\nTMhelix   20   39\noutside   40   325\nTMhelix   326   348\ninside   349   372

10717 GCF_000249575.1_ASM24957v2 Escherichia coli DEC5E Proteobacteria; Gammaproteobacteria; Enterobacterales; Enterobacteriaceae; Escherichia; Escherichia coli
MTGDNTLIHSHGINRRDFMKLCAALAATMGLSSKAAAEMAESVTNPQRPPVIWIGAQECT WP_000145410.1
MULTISPECIES: hydrogenase-2 small chain [Proteobacteria] Length: 372\nNumber of predicted TMHs: 1\nExp number of AAs in TMHs: 23.72226\nExp number, first 60 AAs:  0.22038\nTotal prob of N-in:   0.12994\noutside   1   331\nTMhelix   332   354\ninside   355   372

10718 GCF_000249595.1_ASM24959v2 Escherichia coli DEC6A Proteobacteria; Gammaproteobacteria; Enterobacterales; Enterobacteriaceae; Escherichia; Escherichia coli
MTGDNTLIHSHGINRRDFMKLCAALAATMGLSSKAAAEMAESVTNPQRPPVIWIGAQECT WP_000145410.1
MULTISPECIES: hydrogenase-2 small chain [Proteobacteria] Length: 372\nNumber of predicted TMHs: 1\nExp number of AAs in TMHs: 23.72226\nExp number, first 60 AAs:  0.22038\nTotal prob of N-in:   0.12994\noutside   1   331\nTMhelix   332   354\ninside   355   372

10719 GCF_000249595.1_ASM24959v2 Escherichia coli DEC6A Proteobacteria; Gammaproteobacteria; Enterobacterales; Enterobacteriaceae; Escherichia; Escherichia coli
MNNEETFYQAMRRQGVTRRSFLKYCSLAATSLGLGAGMAPKIAWALENKPRIPVVWIHGL WP_001058323.1
MULTISPECIES: hydrogenase-1 small chain [Proteobacteria] Length: 372\nNumber of predicted TMHs: 2\nExp number of AAs in TMHs: 37.2608499999999\nExp number, first 60 AAs:  19.52923\nTotal prob of N-in:   0.94351\nPOSSIBLE N-term signal sequence\ninside   1   19\nTMhelix   20   39\noutside   40   325\nTMhelix   326   348\ninside   349   372

10720 GCF_000249675.1_ASM24967v2 Escherichia coli DEC6E Proteobacteria; Gammaproteobacteria; Enterobacterales; Enterobacteriaceae; Escherichia; Escherichia coli
MTGDNTLIHSHGINRRDFMKLCAALAATMGLSSKAAAEMAESVTNPQRPPVIWIGAQECT WP_000145410.1
MULTISPECIES: hydrogenase-2 small chain [Proteobacteria] Length: 372\nNumber of predicted TMHs: 1\nExp number of AAs in TMHs: 23.72226\nExp number, first 60 AAs:  0.22038\nTotal prob of N-in:   0.12994\noutside   1   331\nTMhelix   332   354\ninside   355   372

10721 GCF_000249675.1_ASM24967v2 Escherichia coli DEC6E Proteobacteria; Gammaproteobacteria; Enterobacterales; Enterobacteriaceae; Escherichia; Escherichia coli
MNNEETFYQAMRRQGVTRRSFLKYCSLAATSLGLGAGMAPKIAWALENKPRIPVVWIHGL WP_001058323.1
MULTISPECIES: hydrogenase-1 small chain [Proteobacteria] Length: 372\nNumber of predicted TMHs: 2\nExp number of AAs in TMHs: 37.2608499999999\nExp number, first 60 AAs: 19.52923\nTotal prob of N-in: 0.94351\nPOSSIBLE N-term signal sequence\ninside 1 19\nTMhelix 20 39\noutside 40 325\nTMhelix 326 348\ninside 349 372

10722 GCF_000249695.1_ASM24969v2 Escherichia coli DEC7A Proteobacteria; Gammaproteobacteria; Enterobacterales; Enterobacteriaceae; Escherichia; Escherichia coli
MNNEETFYQAMRRQGVTRRSFLKYCSLAATSLGLGAGMAPKIAWALENKPRIPVVWIHGL WP_001058320.1
hydrogenase [Escherichia coli] Length: 372\nNumber of predicted TMHs: 2\nExp number of AAs in TMHs: 37.26082\nExp number, first 60 AAs: 19.52923\nTotal prob of N-in: 0.94351\nPOSSIBLE N-term signal sequence\ninside 1 19\nTMhelix 20 39\noutside 40 325\nTMhelix 326 348\ninside 349 372

10723 GCF_000249695.1_ASM24969v2 Escherichia coli DEC7A Proteobacteria; Gammaproteobacteria; Enterobacterales; Enterobacteriaceae; Escherichia; Escherichia coli
MTGDNTLIHSHGINRRDFMKLCAALAATMGLSSKAAAEMAESVTNPQRPPVIWIGAQECT WP_000145410.1
MULTISPECIES: hydrogenase-2 small chain [Proteobacteria] Length: 372\nNumber of predicted TMHs: 1\nExp number of AAs in TMHs: 23.72226\nExp number, first 60 AAs: 0.22038\nTotal prob of N-in: 0.12994\noutside 1 331\nTMhelix 332 354\ninside 355 372

10724 GCF_000249715.1_ASM24971v2 Escherichia coli DEC7B Proteobacteria; Gammaproteobacteria; Enterobacterales; Enterobacteriaceae; Escherichia; Escherichia coli
MTGDNTLIHSHGINRRDFMKLCAALAATMGLSSKAAAEMAESVTNPQRPPVIWIGAQECT WP_000145410.1
MULTISPECIES: hydrogenase-2 small chain [Proteobacteria] Length: 372\nNumber of predicted TMHs: 1\nExp number of AAs in TMHs: 23.72226\nExp number, first 60 AAs: 0.22038\nTotal prob of N-in: 0.12994\noutside 1 331\nTMhelix 332 354\ninside 355 372

10725 GCF_000249715.1_ASM24971v2 Escherichia coli DEC7B Proteobacteria; Gammaproteobacteria; Enterobacterales; Enterobacteriaceae; Escherichia; Escherichia coli
MNNEETFYQAMRRQGVTRRSFLKYCSLAATSLGLGAGMAPKIAWALENKPRIPVVWIHGL WP_001058323.1
MULTISPECIES: hydrogenase-1 small chain [Proteobacteria] Length: 372\nNumber of predicted TMHs: 2\nExp number of AAs in TMHs: 37.2608499999999\nExp number, first 60 AAs: 19.52923\nTotal prob of N-in: 0.94351\nPOSSIBLE N-term signal sequence\ninside 1 19\nTMhelix 20 39\noutside 40 325\nTMhelix 326 348\ninside 349 372

10726 GCF_000249735.1_ASM24973v2 Escherichia coli DEC7C Proteobacteria; Gammaproteobacteria; Enterobacterales; Enterobacteriaceae; Escherichia; Escherichia coli
MTGDNTLIHSHGINRRDFMKLCAALAATMGLSSKAAAEMAESVTNPQRPPVIWIGAQECT WP_000145410.1
MULTISPECIES: hydrogenase-2 small chain [Proteobacteria] Length: 372\nNumber of predicted TMHs: 1\nExp number of AAs in TMHs: 23.72226\nExp number, first 60 AAs: 0.22038\nTotal prob of N-in: 0.12994\noutside 1 331\nTMhelix 332 354\ninside 355 372

10727 GCF_000249735.1_ASM24973v2 Escherichia coli DEC7C Proteobacteria; Gammaproteobacteria; Enterobacterales; Enterobacteriaceae; Escherichia; Escherichia coli
MNNEETFYQAMRRQGVTRRSFLKYCSLAATSLGLGAGMAPKIAWALENKPRIPVVWIHGL WP_001058320.1
hydrogenase [Escherichia coli] Length: 372\nNumber of predicted TMHs: 2\nExp number of AAs in TMHs: 37.26082\nExp number, first 60 AAs: 19.52923\nTotal prob of N-in: 0.94351\nPOSSIBLE N-term signal sequence\ninside 1 19\nTMhelix 20 39\noutside 40 325\nTMhelix 326 348\ninside 349 372

10728 GCF_000249755.1_ASM24975v2 Escherichia coli DEC7D Proteobacteria; Gammaproteobacteria; Enterobacterales; Enterobacteriaceae; Escherichia; Escherichia coli
MNNEETFYQAMRRQGVTRRSFLKYCSLAATSLGLGAGMAPKIAWALENKPRIPVVWIHGL WP_001058320.1
hydrogenase [Escherichia coli] Length: 372\nNumber of predicted TMHs: 2\nExp number of AAs in TMHs: 37.26082\nExp number, first 60 AAs: 19.52923\nTotal prob of N-in: 0.94351\nPOSSIBLE N-term signal sequence\ninside 1 19\nTMhelix 20 39\noutside 40 325\nTMhelix 326 348\ninside 349 372

10729 GCF_000249755.1_ASM24975v2 Escherichia coli DEC7D Proteobacteria; Gammaproteobacteria; Enterobacterales; Enterobacteriaceae; Escherichia; Escherichia coli
MTGDNTLIHSHGINRRDFMKLCAALAATMGLSSKAAAEMAESVTNPQRPPVIWIGAQECT WP_000145410.1

MULTISPECIES: hydrogenase-2 small chain [Proteobacteria] Length: 372\nNumber of predicted TMHs: 1\nExp number of AAs in TMHs: 23.72226\nExp number, first 60 AAs: 0.22038\nTotal prob of N-in: 0.12994\noutside 1 331\nTMhelix 332 354\ninside 355 372

10730 GCF_000249775.1_ASM24977v2 Escherichia coli DEC7E Proteobacteria; Gammaproteobacteria; Enterobacterales; Enterobacteriaceae; Escherichia; Escherichia coli
MNNEETFYQAMRRQGVTRRSFLKYCSLAATSLGLGAGMAPKIAWALENKPRIPVVWIHGL WP_001058320.1
hydrogenase [Escherichia coli] Length: 372\nNumber of predicted TMHs: 2\nExp number of AAs in TMHs: 37.26082\nExp number, first 60 AAs: 19.52923\nTotal prob of N-in: 0.94351\nPOSSIBLE N-term signal sequence\ninside 1 19\nTMhelix 20 39\noutside 40 325\nTMhelix 326 348\ninside 349 372

10731 GCF_000249775.1_ASM24977v2 Escherichia coli DEC7E Proteobacteria; Gammaproteobacteria; Enterobacterales; Enterobacteriaceae; Escherichia; Escherichia coli
MTGDNTLIHSHGINRRDFMKLCAALAATMGLSSKAAAEMAESVTNPQRPPVIWIGAQECT WP_000145410.1
MULTISPECIES: hydrogenase-2 small chain [Proteobacteria] Length: 372\nNumber of predicted TMHs: 1\nExp number of AAs in TMHs: 23.72226\nExp number, first 60 AAs: 0.22038\nTotal prob of N-in: 0.12994\noutside 1 331\nTMhelix 332 354\ninside 355 372

10732 GCF_000249855.1_ASM24985v2 Escherichia coli DEC8D Proteobacteria; Gammaproteobacteria; Enterobacterales; Enterobacteriaceae; Escherichia; Escherichia coli
MTGDNTLIHSHGINRRDFMKLCAALAATMGLSSKAAAEMAESVTNPQRPPVIWIGAQECT WP_000145410.1
MULTISPECIES: hydrogenase-2 small chain [Proteobacteria] Length: 372\nNumber of predicted TMHs: 1\nExp number of AAs in TMHs: 23.72226\nExp number, first 60 AAs: 0.22038\nTotal prob of N-in: 0.12994\noutside 1 331\nTMhelix 332 354\ninside 355 372

10733 GCF_000249855.1_ASM24985v2 Escherichia coli DEC8D Proteobacteria; Gammaproteobacteria; Enterobacterales; Enterobacteriaceae; Escherichia; Escherichia coli
MNNEETFYQAMRRQGVTRRSFLKYCSLAATSLGLGAGMAPKIAWALENKPRIPVVWIHGL WP_001058323.1
MULTISPECIES: hydrogenase-1 small chain [Proteobacteria] Length: 372\nNumber of predicted TMHs: 2\nExp number of AAs in TMHs: 37.2608499999999\nExp number, first 60 AAs: 19.52923\nTotal prob of N-in: 0.94351\nPOSSIBLE N-term signal sequence\ninside 1 19\nTMhelix 20 39\noutside 40 325\nTMhelix 326 348\ninside 349 372

10734 GCF_000249875.1_ASM24987v2 Escherichia coli DEC8E Proteobacteria; Gammaproteobacteria; Enterobacterales; Enterobacteriaceae; Escherichia; Escherichia coli
MNNEETFYQAMRRQGVTRRSFLKYCSLAATSLGLGAGMAPKIAWALENKPRIPVVWIHGL WP_001058323.1
MULTISPECIES: hydrogenase-1 small chain [Proteobacteria] Length: 372\nNumber of predicted TMHs: 2\nExp number of AAs in TMHs: 37.2608499999999\nExp number, first 60 AAs: 19.52923\nTotal prob of N-in: 0.94351\nPOSSIBLE N-term signal sequence\ninside 1 19\nTMhelix 20 39\noutside 40 325\nTMhelix 326 348\ninside 349 372

10735 GCF_000249875.1_ASM24987v2 Escherichia coli DEC8E Proteobacteria; Gammaproteobacteria; Enterobacterales; Enterobacteriaceae; Escherichia; Escherichia coli
MTGDNTLIHSHGINRRDFMKLCAALAATMGLSSKAAAEMAESVTNPQRPPVIWIGAQECT WP_000145410.1
MULTISPECIES: hydrogenase-2 small chain [Proteobacteria] Length: 372\nNumber of predicted TMHs: 1\nExp number of AAs in TMHs: 23.72226\nExp number, first 60 AAs: 0.22038\nTotal prob of N-in: 0.12994\noutside 1 331\nTMhelix 332 354\ninside 355 372

10736 GCF_000249895.1_ASM24989v2 Escherichia coli DEC9A Proteobacteria; Gammaproteobacteria; Enterobacterales; Enterobacteriaceae; Escherichia; Escherichia coli
MNNEETFYQAMRRQGVTRRSFLKYCSLAATSLGLGAGMAPKIAWALENKPRIPVVWIHGL WP_001058323.1
MULTISPECIES: hydrogenase-1 small chain [Proteobacteria] Length: 372\nNumber of predicted TMHs: 2\nExp number of AAs in TMHs: 37.2608499999999\nExp number, first 60 AAs: 19.52923\nTotal prob of N-in: 0.94351\nPOSSIBLE N-term signal sequence\ninside 1 19\nTMhelix 20 39\noutside 40 325\nTMhelix 326 348\ninside 349 372

10737 GCF_000249895.1_ASM24989v2 Escherichia coli DEC9A Proteobacteria; Gammaproteobacteria; Enterobacterales; Enterobacteriaceae; Escherichia; Escherichia coli
MTGDNTLIHSHGINRRDFMKLCAALAATMGLSSKAAAEMAESVTNPQRPPVIWIGAQECT WP_000145410.1
MULTISPECIES: hydrogenase-2 small chain [Proteobacteria] Length: 372\nNumber of predicted TMHs:

1\nExp number of AAs in TMHs: 23.72226\nExp number, first 60 AAs: 0.22038\nTotal prob of N-in: 0.12994\noutside 1 331\nTMhelix 332 354\ninside 355 372

10738 GCF_000249915.1_ASM24991v2 Escherichia coli DEC9B Proteobacteria; Gammaproteobacteria; Enterobacterales; Enterobacteriaceae; Escherichia; Escherichia coli
MNNEETFYQAMRRQGVTRRSFLKYCSLAATSLGLGAGMAPKIAWALENKPRIPVVWIHGL WP_001058323.1
MULTISPECIES: hydrogenase-1 small chain [Proteobacteria] Length: 372\nNumber of predicted TMHs: 2\nExp number of AAs in TMHs: 37.2608499999999\nExp number, first 60 AAs: 19.52923\nTotal prob of N-in: 0.94351\nPOSSIBLE N-term signal sequence\ninside 1 19\nTMhelix 20 39\noutside 40 325\nTMhelix 326 348\ninside 349 372

10739 GCF_000249915.1_ASM24991v2 Escherichia coli DEC9B Proteobacteria; Gammaproteobacteria; Enterobacterales; Enterobacteriaceae; Escherichia; Escherichia coli
MTGDNTLIHSHGINRRDFMKLCAALAATMGLSSKAAAEMAESVTNPQRPPVIWIGAQECT WP_000145410.1
MULTISPECIES: hydrogenase-2 small chain [Proteobacteria] Length: 372\nNumber of predicted TMHs: 1\nExp number of AAs in TMHs: 23.72226\nExp number, first 60 AAs: 0.22038\nTotal prob of N-in: 0.12994\noutside 1 331\nTMhelix 332 354\ninside 355 372

10740 GCF_000249935.1_ASM24993v2 Escherichia coli DEC9C Proteobacteria; Gammaproteobacteria; Enterobacterales; Enterobacteriaceae; Escherichia; Escherichia coli
MTGDNTLIHSHGINRRDFMKLCAALAATMGLSSKAAAEMAESVTNPQRPPVIWIGAQECT WP_000145410.1
MULTISPECIES: hydrogenase-2 small chain [Proteobacteria] Length: 372\nNumber of predicted TMHs: 1\nExp number of AAs in TMHs: 23.72226\nExp number, first 60 AAs: 0.22038\nTotal prob of N-in: 0.12994\noutside 1 331\nTMhelix 332 354\ninside 355 372

10741 GCF_000249935.1_ASM24993v2 Escherichia coli DEC9C Proteobacteria; Gammaproteobacteria; Enterobacterales; Enterobacteriaceae; Escherichia; Escherichia coli
MNNEETFYQAMRRQGVTRRSFLKYCSLAATSLGLGAGMAPKIAWALENKPRIPVVWIHGL WP_001058323.1
MULTISPECIES: hydrogenase-1 small chain [Proteobacteria] Length: 372\nNumber of predicted TMHs: 2\nExp number of AAs in TMHs: 37.2608499999999\nExp number, first 60 AAs: 19.52923\nTotal prob of N-in: 0.94351\nPOSSIBLE N-term signal sequence\ninside 1 19\nTMhelix 20 39\noutside 40 325\nTMhelix 326 348\ninside 349 372

10742 GCF_000249955.1_ASM24995v2 Escherichia coli DEC9D Proteobacteria; Gammaproteobacteria; Enterobacterales; Enterobacteriaceae; Escherichia; Escherichia coli
MTGDNTLIHSHGINRRDFMKLCAALAATMGLSSKAAAEMAESVTNPQRPPVIWIGAQECT WP_000145410.1
MULTISPECIES: hydrogenase-2 small chain [Proteobacteria] Length: 372\nNumber of predicted TMHs: 1\nExp number of AAs in TMHs: 23.72226\nExp number, first 60 AAs: 0.22038\nTotal prob of N-in: 0.12994\noutside 1 331\nTMhelix 332 354\ninside 355 372

10743 GCF_000249955.1_ASM24995v2 Escherichia coli DEC9D Proteobacteria; Gammaproteobacteria; Enterobacterales; Enterobacteriaceae; Escherichia; Escherichia coli
MNNEETFYQAMRRQGVTRRSFLKYCSLAATSLGLGAGMAPKIAWALENKPRIPVVWIHGL WP_001058323.1
MULTISPECIES: hydrogenase-1 small chain [Proteobacteria] Length: 372\nNumber of predicted TMHs: 2\nExp number of AAs in TMHs: 37.2608499999999\nExp number, first 60 AAs: 19.52923\nTotal prob of N-in: 0.94351\nPOSSIBLE N-term signal sequence\ninside 1 19\nTMhelix 20 39\noutside 40 325\nTMhelix 326 348\ninside 349 372

10744 GCF_000249975.1_ASM24997v2 Escherichia coli DEC9E Proteobacteria; Gammaproteobacteria; Enterobacterales; Enterobacteriaceae; Escherichia; Escherichia coli
MTGDNTLIHSHGINRRDFMKLCAALAATMGLSSKAAAEMAESVTNPQRPPVIWIGAQECT WP_000145410.1
MULTISPECIES: hydrogenase-2 small chain [Proteobacteria] Length: 372\nNumber of predicted TMHs: 1\nExp number of AAs in TMHs: 23.72226\nExp number, first 60 AAs: 0.22038\nTotal prob of N-in: 0.12994\noutside 1 331\nTMhelix 332 354\ninside 355 372

10745 GCF_000249975.1_ASM24997v2 Escherichia coli DEC9E Proteobacteria; Gammaproteobacteria; Enterobacterales; Enterobacteriaceae; Escherichia; Escherichia coli
MNNEETFYQAMRRQGVTRRSFLKYCSLAATSLGLGAGMAPKIAWALENKPRIPVVWIHGL WP_001058323.1
MULTISPECIES: hydrogenase-1 small chain [Proteobacteria] Length: 372\nNumber of predicted TMHs: 2\nExp number of AAs in TMHs: 37.2608499999999\nExp number, first 60 AAs: 19.52923\nTotal prob of N-in:

0.94351\nPOSSIBLE N-term signal sequence\ninside 1 19\nTMhelix 20 39\noutside 40 325\nTMhelix 326 348\ninside 349 372

10746 GCF_000270105.1_ASM27010v1 Escherichia coli DH1 Proteobacteria; Gammaproteobacteria; Enterobacterales; Enterobacteriaceae; Escherichia; Escherichia coli MNNEETFYQAMRRQGVTRRSFLKYCSLAATSLGLGAGMAPKIAWALENKPRIPVVWIHGL WP_001058323.1 MULTISPECIES: hydrogenase-1 small chain [Proteobacteria] Length: 372\nNumber of predicted TMHs: 2\nExp number of AAs in TMHs: 37.2608499999999\nExp number, first 60 AAs: 19.52923\nTotal prob of N-in: 0.94351\nPOSSIBLE N-term signal sequence\ninside 1 19\nTMhelix 20 39\noutside 40 325\nTMhelix 326 348\ninside 349 372

10747 GCF_000023365.1_ASM2336v1 Escherichia coli DH1 Proteobacteria; Gammaproteobacteria; Enterobacterales; Enterobacteriaceae; Escherichia; Escherichia coli MNNEETFYQAMRRQGVTRRSFLKYCSLAATSLGLGAGMAPKIAWALENKPRIPVVWIHGL WP_001058323.1 MULTISPECIES: hydrogenase-1 small chain [Proteobacteria] Length: 372\nNumber of predicted TMHs: 2\nExp number of AAs in TMHs: 37.2608499999999\nExp number, first 60 AAs: 19.52923\nTotal prob of N-in: 0.94351\nPOSSIBLE N-term signal sequence\ninside 1 19\nTMhelix 20 39\noutside 40 325\nTMhelix 326 348\ninside 349 372

10748 GCF_000270105.1_ASM27010v1 Escherichia coli DH1 Proteobacteria; Gammaproteobacteria; Enterobacterales; Enterobacteriaceae; Escherichia; Escherichia coli MTGDNTLIHSHGINRRDFMKLCAALAATMGLSSKAAAEMAESVTNPQRPPVIWIGAQECT WP_000145410.1 MULTISPECIES: hydrogenase-2 small chain [Proteobacteria] Length: 372\nNumber of predicted TMHs: 1\nExp number of AAs in TMHs: 23.72226\nExp number, first 60 AAs: 0.22038\nTotal prob of N-in: 0.12994\noutside 1 331\nTMhelix 332 354\ninside 355 372

10749 GCF_000023365.1_ASM2336v1 Escherichia coli DH1 Proteobacteria; Gammaproteobacteria; Enterobacterales; Enterobacteriaceae; Escherichia; Escherichia coli MTGDNTLIHSHGINRRDFMKLCAALAATMGLSSKAAAEMAESVTNPQRPPVIWIGAQECT WP_000145410.1 MULTISPECIES: hydrogenase-2 small chain [Proteobacteria] Length: 372\nNumber of predicted TMHs: 1\nExp number of AAs in TMHs: 23.72226\nExp number, first 60 AAs: 0.22038\nTotal prob of N-in: 0.12994\noutside 1 331\nTMhelix 332 354\ninside 355 372

10750 GCF_000982435.1_ASM98243v1 Escherichia coli DH5[alpha] Proteobacteria; Gammaproteobacteria; Enterobacterales; Enterobacteriaceae; Escherichia; Escherichia coli MNNEETFYQAMRRQGVTRRSFLKYCSLAATSLGLGAGMAPKIAWALENKPRIPVVWIHGL WP_001058323.1 MULTISPECIES: hydrogenase-1 small chain [Proteobacteria] Length: 372\nNumber of predicted TMHs: 2\nExp number of AAs in TMHs: 37.2608499999999\nExp number, first 60 AAs: 19.52923\nTotal prob of N-in: 0.94351\nPOSSIBLE N-term signal sequence\ninside 1 19\nTMhelix 20 39\noutside 40 325\nTMhelix 326 348\ninside 349 372

10751 GCF_000982435.1_ASM98243v1 Escherichia coli DH5[alpha] Proteobacteria; Gammaproteobacteria; Enterobacterales; Enterobacteriaceae; Escherichia; Escherichia coli MTGDNTLIHSHGINRRDFMKLCAALAATMGLSSKAAAEMAESVTNPQRPPVIWIGAQECT WP_000145410.1 MULTISPECIES: hydrogenase-2 small chain [Proteobacteria] Length: 372\nNumber of predicted TMHs: 1\nExp number of AAs in TMHs: 23.72226\nExp number, first 60 AAs: 0.22038\nTotal prob of N-in: 0.12994\noutside 1 331\nTMhelix 332 354\ninside 355 372

10752 GCF_000734955.1_ASM73495v1 Escherichia coli DSM 30083 = JCM 1649 = ATCC 11775 Proteobacteria; Gammaproteobacteria; Enterobacterales; Enterobacteriaceae; Escherichia; Escherichia coli MTGDNTLIHSHGINRRDFMKLCAALAATMGLSSKAAAEMAESVTNPQRPPVIWIGAQECT WP_000145410.1 MULTISPECIES: hydrogenase-2 small chain [Proteobacteria] Length: 372\nNumber of predicted TMHs: 1\nExp number of AAs in TMHs: 23.72226\nExp number, first 60 AAs: 0.22038\nTotal prob of N-in: 0.12994\noutside 1 331\nTMhelix 332 354\ninside 355 372

10753 GCF_000690815.1_ASM69081v1 Escherichia coli DSM 30083 = JCM 1649 = ATCC 11775 Proteobacteria; Gammaproteobacteria; Enterobacterales; Enterobacteriaceae; Escherichia; Escherichia coli MTGDNTLIHSHGINRRDFMKLCAALAATMGLSSKAAAEMAESVTNPQRPPVIWIGAQECT WP_000145410.1 MULTISPECIES: hydrogenase-2 small chain [Proteobacteria] Length: 372\nNumber of predicted TMHs: 1\nExp number of AAs in TMHs: 23.72226\nExp number, first 60 AAs: 0.22038\nTotal prob of N-in: 0.12994\noutside 1 331\nTMhelix 332 354\ninside 355 372

10754 GCF_002109745.1_ASM210974v1 Escherichia coli E1002 Proteobacteria; Gammaproteobacteria; Enterobacterales; Enterobacteriaceae; Escherichia; Escherichia coli
MTGDNTLIHSHGINRRDFMKLCAALAATMGLSSKAAAEMAESVTNPQRPPVIWIGAQECT WP_000145410.1
MULTISPECIES: hydrogenase-2 small chain [Proteobacteria] Length: 372\nNumber of predicted TMHs: 1\nExp number of AAs in TMHs: 23.72226\nExp number, first 60 AAs: 0.22038\nTotal prob of N-in: 0.12994\noutside 1 331\nTMhelix 332 354\ninside 355 372

10755 GCF_002109745.1_ASM210974v1 Escherichia coli E1002 Proteobacteria; Gammaproteobacteria; Enterobacterales; Enterobacteriaceae; Escherichia; Escherichia coli
MNNEETFYQAMRRQGVTRRSFLKYCSLAATSLGLGAGMAPKIAWALENKPRIPVVWIHGL WP_001058323.1
MULTISPECIES: hydrogenase-1 small chain [Proteobacteria] Length: 372\nNumber of predicted TMHs: 2\nExp number of AAs in TMHs: 37.2608499999999\nExp number, first 60 AAs: 19.52923\nTotal prob of N-in: 0.94351\nPOSSIBLE N-term signal sequence\ninside 1 19\nTMhelix 20 39\noutside 40 325\nTMhelix 326 348\ninside 349 372

10756 GCF_000241995.1_Esch_coli_E101_V1 Escherichia coli E101 Proteobacteria; Gammaproteobacteria; Enterobacterales; Enterobacteriaceae; Escherichia; Escherichia coli
MTGDNTLIHSHGINRRDFMKLCAALAATMGLSSKAAAEMAESVTNPQRPPVIWIGAQECT WP_000145410.1
MULTISPECIES: hydrogenase-2 small chain [Proteobacteria] Length: 372\nNumber of predicted TMHs: 1\nExp number of AAs in TMHs: 23.72226\nExp number, first 60 AAs: 0.22038\nTotal prob of N-in: 0.12994\noutside 1 331\nTMhelix 332 354\ninside 355 372

10757 GCF_000241995.1_Esch_coli_E101_V1 Escherichia coli E101 Proteobacteria; Gammaproteobacteria; Enterobacterales; Enterobacteriaceae; Escherichia; Escherichia coli
MNNEETFYQAMRRQGVTRRSFLKYCSLAATSLGLGAGMAPKIAWALENKPRIPVVWIHGL WP_001058323.1
MULTISPECIES: hydrogenase-1 small chain [Proteobacteria] Length: 372\nNumber of predicted TMHs: 2\nExp number of AAs in TMHs: 37.2608499999999\nExp number, first 60 AAs: 19.52923\nTotal prob of N-in: 0.94351\nPOSSIBLE N-term signal sequence\ninside 1 19\nTMhelix 20 39\noutside 40 325\nTMhelix 326 348\ninside 349 372

10758 GCF_000167875.1_ASM16787v1 Escherichia coli E110019 Proteobacteria; Gammaproteobacteria; Enterobacterales; Enterobacteriaceae; Escherichia; Escherichia coli
MTGDNTLIHSHGINRRDFMKLCAALAATMGLSSKAAAEMAESVTNPQRPPVIWIGAQECT WP_000145410.1
MULTISPECIES: hydrogenase-2 small chain [Proteobacteria] Length: 372\nNumber of predicted TMHs: 1\nExp number of AAs in TMHs: 23.72226\nExp number, first 60 AAs: 0.22038\nTotal prob of N-in: 0.12994\noutside 1 331\nTMhelix 332 354\ninside 355 372

10759 GCF_000167875.1_ASM16787v1 Escherichia coli E110019 Proteobacteria; Gammaproteobacteria; Enterobacterales; Enterobacteriaceae; Escherichia; Escherichia coli
MNNEETFYQAMRRQGVTRRSFLKYCSLAATSLGLGAGMAPKIAWALENKPRIPVVWIHGL WP_001058323.1
MULTISPECIES: hydrogenase-1 small chain [Proteobacteria] Length: 372\nNumber of predicted TMHs: 2\nExp number of AAs in TMHs: 37.2608499999999\nExp number, first 60 AAs: 19.52923\nTotal prob of N-in: 0.94351\nPOSSIBLE N-term signal sequence\ninside 1 19\nTMhelix 20 39\noutside 40 325\nTMhelix 326 348\ninside 349 372

10760 GCF_002109705.1_ASM210970v1 Escherichia coli E1114 Proteobacteria; Gammaproteobacteria; Enterobacterales; Enterobacteriaceae; Escherichia; Escherichia coli
MTGDNTLIHSHGINRRDFMKLCAALAATMGLSSKAAAEMAESVTNPQRPPVIWIGAQECT WP_000145410.1
MULTISPECIES: hydrogenase-2 small chain [Proteobacteria] Length: 372\nNumber of predicted TMHs: 1\nExp number of AAs in TMHs: 23.72226\nExp number, first 60 AAs: 0.22038\nTotal prob of N-in: 0.12994\noutside 1 331\nTMhelix 332 354\ninside 355 372

10761 GCF_002109705.1_ASM210970v1 Escherichia coli E1114 Proteobacteria; Gammaproteobacteria; Enterobacterales; Enterobacteriaceae; Escherichia; Escherichia coli
MNNEETFYQAMRRQGVTRRSFLKYCSLAATSLGLGAGMAPKIAWALENKPRIPVVWIHGL WP_001058323.1
MULTISPECIES: hydrogenase-1 small chain [Proteobacteria] Length: 372\nNumber of predicted TMHs: 2\nExp number of AAs in TMHs: 37.2608499999999\nExp number, first 60 AAs: 19.52923\nTotal prob of N-in: 0.94351\nPOSSIBLE N-term signal sequence\ninside 1 19\nTMhelix 20 39\noutside 40 325\nTMhelix 326 348\ninside 349 372

10762 GCF_002109985.1_ASM210998v1 Escherichia coli E1118 Proteobacteria; Gammaproteobacteria; Enterobacterales; Enterobacteriaceae; Escherichia; Escherichia coli
MNNEETFYQAMRRQGVTRRSFLKYCSLAATSLGLGAGMAPKIAWALENKPRIPVVWIHGL WP_001516139.1
MULTISPECIES: hydrogenase [Escherichia] Length: 372\nNumber of predicted TMHs: 2\nExp number of AAs in TMHs: 37.98718\nExp number, first 60 AAs: 19.52876\nTotal prob of N-in: 0.94354\nPOSSIBLE N-term signal sequence\ninside 1 19\nTMhelix 20 39\noutside 40 325\nTMhelix 326 348\ninside 349 372

10763 GCF_002109985.1_ASM210998v1 Escherichia coli E1118 Proteobacteria; Gammaproteobacteria; Enterobacterales; Enterobacteriaceae; Escherichia; Escherichia coli
MTGDNTLIHSHGINRRDFMKLCAALAATMGLSSKAAAEMAESVTNPQRPPVIWIGAQECT WP_000145417.1
MULTISPECIES: hydrogenase-2 small chain [Escherichia] Length: 372\nNumber of predicted TMHs: 1\nExp number of AAs in TMHs: 23.72512\nExp number, first 60 AAs: 0.22034\nTotal prob of N-in: 0.13000\noutside 1 331\nTMhelix 332 354\ninside 355 372

10764 GCF_000622285.1_ASM62228v2 Escherichia coli E1140 Proteobacteria; Gammaproteobacteria; Enterobacterales; Enterobacteriaceae; Escherichia; Escherichia coli
MTGDNTLIHSHGINRRDFMKLCAALAATMGLSSKAAAEMAESVTNPQRPPVIWIGAQECT WP_000145410.1
MULTISPECIES: hydrogenase-2 small chain [Proteobacteria] Length: 372\nNumber of predicted TMHs: 1\nExp number of AAs in TMHs: 23.72226\nExp number, first 60 AAs: 0.22038\nTotal prob of N-in: 0.12994\noutside 1 331\nTMhelix 332 354\ninside 355 372

10765 GCF_000622285.1_ASM62228v2 Escherichia coli E1140 Proteobacteria; Gammaproteobacteria; Enterobacterales; Enterobacteriaceae; Escherichia; Escherichia coli
MNNEETFYQAMRRQGVTRRSFLKYCSLAATSLGLGAGMAPKIAWALENKPRIPVVWIHGL WP_001058323.1
MULTISPECIES: hydrogenase-1 small chain [Proteobacteria] Length: 372\nNumber of predicted TMHs: 2\nExp number of AAs in TMHs: 37.2608499999999\nExp number, first 60 AAs: 19.52923\nTotal prob of N-in: 0.94351\nPOSSIBLE N-term signal sequence\ninside 1 19\nTMhelix 20 39\noutside 40 325\nTMhelix 326 348\ninside 349 372

10766 GCF_000190795.1_ASM19079v1 Escherichia coli E1167 Proteobacteria; Gammaproteobacteria; Enterobacterales; Enterobacteriaceae; Escherichia; Escherichia coli
MNNEETFYQAMRRQGVTRRSFLKYCSLAATSLGLGAGMAPKIAWALENKPRIPVVWIHGL WP_001058323.1
MULTISPECIES: hydrogenase-1 small chain [Proteobacteria] Length: 372\nNumber of predicted TMHs: 2\nExp number of AAs in TMHs: 37.2608499999999\nExp number, first 60 AAs: 19.52923\nTotal prob of N-in: 0.94351\nPOSSIBLE N-term signal sequence\ninside 1 19\nTMhelix 20 39\noutside 40 325\nTMhelix 326 348\ninside 349 372

10767 GCF_000190795.1_ASM19079v1 Escherichia coli E1167 Proteobacteria; Gammaproteobacteria; Enterobacterales; Enterobacteriaceae; Escherichia; Escherichia coli
MTGDNTLIHSHGINRRDFMKLCAALAATMGLSSKAAAEMAESVTNPQRPPVIWIGAQECT WP_000145410.1
MULTISPECIES: hydrogenase-2 small chain [Proteobacteria] Length: 372\nNumber of predicted TMHs: 1\nExp number of AAs in TMHs: 23.72226\nExp number, first 60 AAs: 0.22038\nTotal prob of N-in: 0.12994\noutside 1 331\nTMhelix 332 354\ninside 355 372

10768 GCF_000188775.1_ASM18877v2 Escherichia coli E128010 Proteobacteria; Gammaproteobacteria; Enterobacterales; Enterobacteriaceae; Escherichia; Escherichia coli
MNNEETFYQAMRRQGVTRRSFLKYCSLAATSLGLGAGMAPKIAWALENKPRIPVVWIHGL WP_001295940.1
MULTISPECIES: hydrogenase [Enterobacteriaceae] Length: 372\nNumber of predicted TMHs: 2\nExp number of AAs in TMHs: 37.2608899999999\nExp number, first 60 AAs: 19.52923\nTotal prob of N-in: 0.94351\nPOSSIBLE N-term signal sequence\ninside 1 19\nTMhelix 20 39\noutside 40 325\nTMhelix 326 348\ninside 349 372

10769 GCF_000188775.1_ASM18877v2 Escherichia coli E128010 Proteobacteria; Gammaproteobacteria; Enterobacterales; Enterobacteriaceae; Escherichia; Escherichia coli
MTGDNTLIHSHGINRRDFMKLCAALAATMGLSSKAAAEMAESVTNPQRPPVIWIGAQECT WP_000145410.1
MULTISPECIES: hydrogenase-2 small chain [Proteobacteria] Length: 372\nNumber of predicted TMHs: 1\nExp number of AAs in TMHs: 23.72226\nExp number, first 60 AAs: 0.22038\nTotal prob of N-in: 0.12994\noutside 1 331\nTMhelix 332 354\ninside 355 372

10770 GCF_000190815.1_ASM19081v1 Escherichia coli E1520 Proteobacteria; Gammaproteobacteria; Enterobacterales; Enterobacteriaceae; Escherichia; Escherichia coli

MTGDNTLIHSHGINRRDFMKLCAALAATMGLSSKAAAEMAESVTNPQRPPVIWIGAQECT WP_000145410.1 MULTISPECIES: hydrogenase-2 small chain [Proteobacteria] Length: 372\nNumber of predicted TMHs: 1\nExp number of AAs in TMHs: 23.72226\nExp number, first 60 AAs: 0.22038\nTotal prob of N-in: 0.12994\noutside 1 331\nTMhelix 332 354\ninside 355 372

10771 GCF_000190815.1_ASM19081v1 Escherichia coli E1520 Proteobacteria; Gammaproteobacteria; Enterobacterales; Enterobacteriaceae; Escherichia; Escherichia coli MNNEETFYQAMRRQGVTRRSFLKYCSLAATSLGLGAGMAPKIAWALENKPRIPVVWIHGL WP_001058323.1 MULTISPECIES: hydrogenase-1 small chain [Proteobacteria] Length: 372\nNumber of predicted TMHs: 2\nExp number of AAs in TMHs: 37.2608499999999\nExp number, first 60 AAs: 19.52923\nTotal prob of N-in: 0.94351\nPOSSIBLE N-term signal sequence\ninside 1 19\nTMhelix 20 39\noutside 40 325\nTMhelix 326 348\ninside 349 372

10772 GCF_000622265.1_CLC_6.5.1_of_E._coli_genome Escherichia coli E1728 Proteobacteria; Gammaproteobacteria; Enterobacterales; Enterobacteriaceae; Escherichia; Escherichia coli MTGDNTLIHSHGINRRDFMKLCAALAATMGLSSKAAAEMAESVTNPQRPPVIWIGAQECT WP_000145410.1 MULTISPECIES: hydrogenase-2 small chain [Proteobacteria] Length: 372\nNumber of predicted TMHs: 1\nExp number of AAs in TMHs: 23.72226\nExp number, first 60 AAs: 0.22038\nTotal prob of N-in: 0.12994\noutside 1 331\nTMhelix 332 354\ninside 355 372

10773 GCF_000622265.1_CLC_6.5.1_of_E._coli_genome Escherichia coli E1728 Proteobacteria; Gammaproteobacteria; Enterobacterales; Enterobacteriaceae; Escherichia; Escherichia coli MNNEETFYQAMRRQGVTRRSFLKYCSLAATSLGLGAGMAPKIAWALENKPRIPVVWIHGL WP_001058323.1 MULTISPECIES: hydrogenase-1 small chain [Proteobacteria] Length: 372\nNumber of predicted TMHs: 2\nExp number of AAs in TMHs: 37.2608499999999\nExp number, first 60 AAs: 19.52923\nTotal prob of N-in: 0.94351\nPOSSIBLE N-term signal sequence\ninside 1 19\nTMhelix 20 39\noutside 40 325\nTMhelix 326 348\ninside 349 372

10774 GCF_000512625.1_E1777_2 Escherichia coli E1777 Proteobacteria; Gammaproteobacteria; Enterobacterales; Enterobacteriaceae; Escherichia; Escherichia coli MTGDNTLIHSHGINRRDFMKLCAALAATMGLSSKAAAEMAESVTNPQRPPVIWIGAQECT WP_000145410.1 MULTISPECIES: hydrogenase-2 small chain [Proteobacteria] Length: 372\nNumber of predicted TMHs: 1\nExp number of AAs in TMHs: 23.72226\nExp number, first 60 AAs: 0.22038\nTotal prob of N-in: 0.12994\noutside 1 331\nTMhelix 332 354\ninside 355 372

10775 GCF_000512625.1_E1777_2 Escherichia coli E1777 Proteobacteria; Gammaproteobacteria; Enterobacterales; Enterobacteriaceae; Escherichia; Escherichia coli MNNEETFYQAMRRQGVTRRSFLKYCSLAATSLGLGAGMAPKIVWALENKPRIPVVWIHGL WP_001398876.1 hydrogenase-1 small chain [Escherichia coli] Length: 372\nNumber of predicted TMHs: 2\nExp number of AAs in TMHs: 37.60268\nExp number, first 60 AAs: 19.84677\nTotal prob of N-in: 0.94621\nPOSSIBLE N-term signal sequence\ninside 1 20\nTMhelix 21 43\noutside 44 325\nTMhelix 326 348\ninside 349 372

10776 GCF_000415485.1_E2265_1 Escherichia coli E2265 Proteobacteria; Gammaproteobacteria; Enterobacterales; Enterobacteriaceae; Escherichia; Escherichia coli MNNEETFYQAMRRQGVTRRSFLKYCSLAATSLGLGAGMAPKIVWALENKPRIPVVWIHGL WP_001398876.1 hydrogenase-1 small chain [Escherichia coli] Length: 372\nNumber of predicted TMHs: 2\nExp number of AAs in TMHs: 37.60268\nExp number, first 60 AAs: 19.84677\nTotal prob of N-in: 0.94621\nPOSSIBLE N-term signal sequence\ninside 1 20\nTMhelix 21 43\noutside 44 325\nTMhelix 326 348\ninside 349 372

10777 GCF_000415485.1_E2265_1 Escherichia coli E2265 Proteobacteria; Gammaproteobacteria; Enterobacterales; Enterobacteriaceae; Escherichia; Escherichia coli MTGDNTLIHSHGINRRDFMKLCAALAATMGLSSKAAAEMAESVTNPQRPPVIWIGAQECT WP_032190500.1 hydrogenase 2 small subunit [Escherichia coli] Length: 405\nNumber of predicted TMHs: 1\nExp number of AAs in TMHs: 34.53127\nExp number, first 60 AAs: 0.3257\nTotal prob of N-in: 0.21863\noutside 1 364\nTMhelix 365 387\ninside 388 405

10778 GCF_002110065.1_ASM211006v1 Escherichia coli E267 Proteobacteria; Gammaproteobacteria; Enterobacterales; Enterobacteriaceae; Escherichia; Escherichia coli MTGDNTLIHSHGINRRDFMKLCAALAATMGLSSKAAAEMAESVTNPQRPPVIWIGAQECT WP_000145410.1 MULTISPECIES: hydrogenase-2 small chain [Proteobacteria] Length: 372\nNumber of predicted TMHs:

1\nExp number of AAs in TMHs: 23.72226\nExp number, first 60 AAs: 0.22038\nTotal prob of N-in: 0.12994\noutside 1 331\nTMhelix 332 354\ninside 355 372

10779 GCF_002110065.1_ASM211006v1 Escherichia coli E267 Proteobacteria; Gammaproteobacteria; Enterobacterales; Enterobacteriaceae; Escherichia; Escherichia coli
MNNEETFYQAMRRQGVTRRSFLKYCSLAATSLGLGAGMAPKIAWALENKPRIPVVWIHGL WP_001058323.1
MULTISPECIES: hydrogenase-1 small chain [Proteobacteria] Length: 372\nNumber of predicted TMHs: 2\nExp number of AAs in TMHs: 37.2608499999999\nExp number, first 60 AAs: 19.52923\nTotal prob of N-in: 0.94351\nPOSSIBLE N-term signal sequence\ninside 1 19\nTMhelix 20 39\noutside 40 325\nTMhelix 326 348\ninside 349 372

10780 GCF_000190835.1_ASM19083v1 Escherichia coli E482 Proteobacteria; Gammaproteobacteria; Enterobacterales; Enterobacteriaceae; Escherichia; Escherichia coli
MTGDNTLIHSHGINRRDFMKLCAALAATMGLSSKAAAEMAESVTNPQRPPVIWIGAQECT WP_000145410.1
MULTISPECIES: hydrogenase-2 small chain [Proteobacteria] Length: 372\nNumber of predicted TMHs: 1\nExp number of AAs in TMHs: 23.72226\nExp number, first 60 AAs: 0.22038\nTotal prob of N-in: 0.12994\noutside 1 331\nTMhelix 332 354\ninside 355 372

10781 GCF_000190835.1_ASM19083v1 Escherichia coli E482 Proteobacteria; Gammaproteobacteria; Enterobacterales; Enterobacteriaceae; Escherichia; Escherichia coli
MNNEETFYQAMRRQGVTRRSFLKYCSLAATSLGLGAGMAPKIAWALENKPRIPVVWIHGL WP_001058323.1
MULTISPECIES: hydrogenase-1 small chain [Proteobacteria] Length: 372\nNumber of predicted TMHs: 2\nExp number of AAs in TMHs: 37.2608499999999\nExp number, first 60 AAs: 19.52923\nTotal prob of N-in: 0.94351\nPOSSIBLE N-term signal sequence\ninside 1 19\nTMhelix 20 39\noutside 40 325\nTMhelix 326 348\ninside 349 372

10782 GCF_002109725.1_ASM210972v1 Escherichia coli E560 Proteobacteria; Gammaproteobacteria; Enterobacterales; Enterobacteriaceae; Escherichia; Escherichia coli
MNNEETFYQAMRRQGVTRRSFLKYCSLAATSLGLGAGMAPKIAWALENKPRIPVVWIHGL WP_001058323.1
MULTISPECIES: hydrogenase-1 small chain [Proteobacteria] Length: 372\nNumber of predicted TMHs: 2\nExp number of AAs in TMHs: 37.2608499999999\nExp number, first 60 AAs: 19.52923\nTotal prob of N-in: 0.94351\nPOSSIBLE N-term signal sequence\ninside 1 19\nTMhelix 20 39\noutside 40 325\nTMhelix 326 348\ninside 349 372

10783 GCF_002109725.1_ASM210972v1 Escherichia coli E560 Proteobacteria; Gammaproteobacteria; Enterobacterales; Enterobacteriaceae; Escherichia; Escherichia coli
MTGDNTLIHSHGINRRDFMKLCAALAATMGLSSKAAAEMAESVTNPQRPPVIWIGAQECT WP_000145410.1
MULTISPECIES: hydrogenase-2 small chain [Proteobacteria] Length: 372\nNumber of predicted TMHs: 1\nExp number of AAs in TMHs: 23.72226\nExp number, first 60 AAs: 0.22038\nTotal prob of N-in: 0.12994\noutside 1 331\nTMhelix 332 354\ninside 355 372

10784 GCF_002109965.1_ASM210996v1 Escherichia coli E704 Proteobacteria; Gammaproteobacteria; Enterobacterales; Enterobacteriaceae; Escherichia; Escherichia coli
MTGDNTLIHSHGINRRDFMKLCAALAATMGLSSKAAAEMAESVTNPQRPPVIWIGAQECT WP_000145410.1
MULTISPECIES: hydrogenase-2 small chain [Proteobacteria] Length: 372\nNumber of predicted TMHs: 1\nExp number of AAs in TMHs: 23.72226\nExp number, first 60 AAs: 0.22038\nTotal prob of N-in: 0.12994\noutside 1 331\nTMhelix 332 354\ninside 355 372

10785 GCF_002109965.1_ASM210996v1 Escherichia coli E704 Proteobacteria; Gammaproteobacteria; Enterobacterales; Enterobacteriaceae; Escherichia; Escherichia coli
MNNEETFYQAMRRQGVTRRSFLKYCSLAATSLGLGAGMAPKIAWALENKPRIPVVWIHGL WP_001058323.1
MULTISPECIES: hydrogenase-1 small chain [Proteobacteria] Length: 372\nNumber of predicted TMHs: 2\nExp number of AAs in TMHs: 37.2608499999999\nExp number, first 60 AAs: 19.52923\nTotal prob of N-in: 0.94351\nPOSSIBLE N-term signal sequence\ninside 1 19\nTMhelix 20 39\noutside 40 325\nTMhelix 326 348\ninside 349 372

10786 GCF_000564835.2_EC096_10.1 Escherichia coli EC096/10 Proteobacteria; Gammaproteobacteria; Enterobacterales; Enterobacteriaceae; Escherichia; Escherichia coli
MTGDNTLIHSHGINRRDFMKLCAALAATMGLSSKAAAEMAESVTNPQRPPVIWIGAQECT WP_000145410.1
MULTISPECIES: hydrogenase-2 small chain [Proteobacteria] Length: 372\nNumber of predicted TMHs:

1\nExp number of AAs in TMHs: 23.72226\nExp number, first 60 AAs: 0.22038\nTotal prob of N-in: 0.12994\noutside 1 331\nTMhelix 332 354\ninside 355 372

10787 GCF_000564835.2_EC096_10.1 Escherichia coli EC096/10 Proteobacteria; Gammaproteobacteria; Enterobacterales; Enterobacteriaceae; Escherichia; Escherichia coli MNNEETFYQAMRRQGVTRRSFLKYCSLAATSLGLGAGMAPKIAWALENKPRIPVVWIHGL WP_001058323.1 MULTISPECIES: hydrogenase-1 small chain [Proteobacteria] Length: 372\nNumber of predicted TMHs: 2\nExp number of AAs in TMHs: 37.2608499999999\nExp number, first 60 AAs: 19.52923\nTotal prob of N-in: 0.94351\nPOSSIBLE N-term signal sequence\ninside 1 19\nTMhelix 20 39\noutside 40 325\nTMhelix 326 348\ninside 349 372

10788 GCF_000267045.1_ASM26704v1 Escherichia coli EC1734 Proteobacteria; Gammaproteobacteria; Enterobacterales; Enterobacteriaceae; Escherichia; Escherichia coli MNNEETFYQAMRRQGVTRRSFLKYCSLAATSLGLGAGMAPKIAWALENKPRIPVVWIHGL WP_001301613.1 hydrogenase [Escherichia coli] Length: 372\nNumber of predicted TMHs: 2\nExp number of AAs in TMHs: 37.46598\nExp number, first 60 AAs: 19.53139\nTotal prob of N-in: 0.94364\nPOSSIBLE N-term signal sequence\ninside 1 19\nTMhelix 20 39\noutside 40 325\nTMhelix 326 348\ninside 349 372

10789 GCF_000267045.1_ASM26704v1 Escherichia coli EC1734 Proteobacteria; Gammaproteobacteria; Enterobacterales; Enterobacteriaceae; Escherichia; Escherichia coli MTGDNTLIHSHGINRRDFMKLCAALAATMGLSSKAAAEMAESVTNPQRPPVIWIGAQECT WP_000145410.1 MULTISPECIES: hydrogenase-2 small chain [Proteobacteria] Length: 372\nNumber of predicted TMHs: 1\nExp number of AAs in TMHs: 23.72226\nExp number, first 60 AAs: 0.22038\nTotal prob of N-in: 0.12994\noutside 1 331\nTMhelix 332 354\ninside 355 372

10790 GCF_000303615.2_ASM30361v2 Escherichia coli EC1735 Proteobacteria; Gammaproteobacteria; Enterobacterales; Enterobacteriaceae; Escherichia; Escherichia coli MTGDNTLIHSHGINRRDFMKLCAALAATMGLSSKAAAEMAESVTNPQRPPVIWIGAQECT WP_000145410.1 MULTISPECIES: hydrogenase-2 small chain [Proteobacteria] Length: 372\nNumber of predicted TMHs: 1\nExp number of AAs in TMHs: 23.72226\nExp number, first 60 AAs: 0.22038\nTotal prob of N-in: 0.12994\noutside 1 331\nTMhelix 332 354\ninside 355 372

10791 GCF_000303615.2_ASM30361v2 Escherichia coli EC1735 Proteobacteria; Gammaproteobacteria; Enterobacterales; Enterobacteriaceae; Escherichia; Escherichia coli MNNEETFYQAMRRQGVTRRSFLKYCSLAATSLGLGAGMAPKIAWALENKPRIPVVWIHGL WP_001301613.1 hydrogenase [Escherichia coli] Length: 372\nNumber of predicted TMHs: 2\nExp number of AAs in TMHs: 37.46598\nExp number, first 60 AAs: 19.53139\nTotal prob of N-in: 0.94364\nPOSSIBLE N-term signal sequence\ninside 1 19\nTMhelix 20 39\noutside 40 325\nTMhelix 326 348\ninside 349 372

10792 GCF_000303595.2_ASM30359v2 Escherichia coli EC1736 Proteobacteria; Gammaproteobacteria; Enterobacterales; Enterobacteriaceae; Escherichia; Escherichia coli MNNEETFYQAMRRQGVTRRSFLKYCSLAATSLGLGAGMAPKIAWALENKPRIPVVWIHGL WP_001301613.1 hydrogenase [Escherichia coli] Length: 372\nNumber of predicted TMHs: 2\nExp number of AAs in TMHs: 37.46598\nExp number, first 60 AAs: 19.53139\nTotal prob of N-in: 0.94364\nPOSSIBLE N-term signal sequence\ninside 1 19\nTMhelix 20 39\noutside 40 325\nTMhelix 326 348\ninside 349 372

10793 GCF_000303595.2_ASM30359v2 Escherichia coli EC1736 Proteobacteria; Gammaproteobacteria; Enterobacterales; Enterobacteriaceae; Escherichia; Escherichia coli MTGDNTLIHSHGINRRDFMKLCAALAATMGLSSKAAAEMAESVTNPQRPPVIWIGAQECT WP_000145410.1 MULTISPECIES: hydrogenase-2 small chain [Proteobacteria] Length: 372\nNumber of predicted TMHs: 1\nExp number of AAs in TMHs: 23.72226\nExp number, first 60 AAs: 0.22038\nTotal prob of N-in: 0.12994\noutside 1 331\nTMhelix 332 354\ninside 355 372

10794 GCF_000303575.2_ASM30357v2 Escherichia coli EC1737 Proteobacteria; Gammaproteobacteria; Enterobacterales; Enterobacteriaceae; Escherichia; Escherichia coli MNNEETFYQAMRRQGVTRRSFLKYCSLAATSLGLGAGMAPKIAWALENKPRIPVVWIHGL WP_001301613.1 hydrogenase [Escherichia coli] Length: 372\nNumber of predicted TMHs: 2\nExp number of AAs in TMHs: 37.46598\nExp number, first 60 AAs: 19.53139\nTotal prob of N-in: 0.94364\nPOSSIBLE N-term signal sequence\ninside 1 19\nTMhelix 20 39\noutside 40 325\nTMhelix 326 348\ninside 349 372

10795 GCF_000303575.2_ASM30357v2 Escherichia coli EC1737 Proteobacteria; Gammaproteobacteria; Enterobacterales; Enterobacteriaceae; Escherichia; Escherichia coli
MTGDNTLIHSHGINRRDFMKLCAALAATMGLSSKAAAEMAESVTNPQRPPVIWIGAQECT WP_000145410.1
MULTISPECIES: hydrogenase-2 small chain [Proteobacteria] Length: 372\nNumber of predicted TMHs: 1\nExp number of AAs in TMHs: 23.72226\nExp number, first 60 AAs: 0.22038\nTotal prob of N-in: 0.12994\noutside 1 331\nTMhelix 332 354\ninside 355 372

10796 GCF_000267025.1_ASM26702v1 Escherichia coli EC1738 Proteobacteria; Gammaproteobacteria; Enterobacterales; Enterobacteriaceae; Escherichia; Escherichia coli
MTGDNTLIHSHGINRRDFMKLCAALAATMGLSSKAAAEMAESVTNPQRPPVIWIGAQECT WP_000145410.1
MULTISPECIES: hydrogenase-2 small chain [Proteobacteria] Length: 372\nNumber of predicted TMHs: 1\nExp number of AAs in TMHs: 23.72226\nExp number, first 60 AAs: 0.22038\nTotal prob of N-in: 0.12994\noutside 1 331\nTMhelix 332 354\ninside 355 372

10797 GCF_000267025.1_ASM26702v1 Escherichia coli EC1738 Proteobacteria; Gammaproteobacteria; Enterobacterales; Enterobacteriaceae; Escherichia; Escherichia coli
MNNEETFYQAMRRQGVTRRSFLKYCSLAATSLGLGAGMAPKIAWALENKPRIPVVWIHGL WP_001301613.1
hydrogenase [Escherichia coli] Length: 372\nNumber of predicted TMHs: 2\nExp number of AAs in TMHs: 37.46598\nExp number, first 60 AAs: 19.53139\nTotal prob of N-in: 0.94364\nPOSSIBLE N-term signal sequence\ninside 1 19\nTMhelix 20 39\noutside 40 325\nTMhelix 326 348\ninside 349 372

10798 GCF_000267225.2_ASM26722v2 Escherichia coli EC1845 Proteobacteria; Gammaproteobacteria; Enterobacterales; Enterobacteriaceae; Escherichia; Escherichia coli
MTGDNTLIHSHGINRRDFMKLCAALAATMGLSSKAAAEMAESVTNPQRPPVIWIGAQECT WP_000145410.1
MULTISPECIES: hydrogenase-2 small chain [Proteobacteria] Length: 372\nNumber of predicted TMHs: 1\nExp number of AAs in TMHs: 23.72226\nExp number, first 60 AAs: 0.22038\nTotal prob of N-in: 0.12994\noutside 1 331\nTMhelix 332 354\ninside 355 372

10799 GCF_000267225.2_ASM26722v2 Escherichia coli EC1845 Proteobacteria; Gammaproteobacteria; Enterobacterales; Enterobacteriaceae; Escherichia; Escherichia coli
MNNEETFYQAMRRQGVTRRSFLKYCSLAATSLGLGAGMAPKIAWALENKPRIPVVWIHGL WP_001301613.1
hydrogenase [Escherichia coli] Length: 372\nNumber of predicted TMHs: 2\nExp number of AAs in TMHs: 37.46598\nExp number, first 60 AAs: 19.53139\nTotal prob of N-in: 0.94364\nPOSSIBLE N-term signal sequence\ninside 1 19\nTMhelix 20 39\noutside 40 325\nTMhelix 326 348\ninside 349 372

10800 GCF_000303555.2_ASM30355v2 Escherichia coli EC1846 Proteobacteria; Gammaproteobacteria; Enterobacterales; Enterobacteriaceae; Escherichia; Escherichia coli
MNNEETFYQAMRRQGVTRRSFLKYCSLAATSLGLGAGMAPKIAWALENKPRIPVVWIHGL WP_001301613.1
hydrogenase [Escherichia coli] Length: 372\nNumber of predicted TMHs: 2\nExp number of AAs in TMHs: 37.46598\nExp number, first 60 AAs: 19.53139\nTotal prob of N-in: 0.94364\nPOSSIBLE N-term signal sequence\ninside 1 19\nTMhelix 20 39\noutside 40 325\nTMhelix 326 348\ninside 349 372

10801 GCF_000303555.2_ASM30355v2 Escherichia coli EC1846 Proteobacteria; Gammaproteobacteria; Enterobacterales; Enterobacteriaceae; Escherichia; Escherichia coli
MTGDNTLIHSHGINRRDFMKLCAALAATMGLSSKAAAEMAESVTNPQRPPVIWIGAQECT WP_000145410.1
MULTISPECIES: hydrogenase-2 small chain [Proteobacteria] Length: 372\nNumber of predicted TMHs: 1\nExp number of AAs in TMHs: 23.72226\nExp number, first 60 AAs: 0.22038\nTotal prob of N-in: 0.12994\noutside 1 331\nTMhelix 332 354\ninside 355 372

10802 GCF_000303535.2_ASM30353v2 Escherichia coli EC1847 Proteobacteria; Gammaproteobacteria; Enterobacterales; Enterobacteriaceae; Escherichia; Escherichia coli
MNNEETFYQAMRRQGVTRRSFLKYCSLAATSLGLGAGMAPKIAWALENKPRIPVVWIHGL WP_001301613.1
hydrogenase [Escherichia coli] Length: 372\nNumber of predicted TMHs: 2\nExp number of AAs in TMHs: 37.46598\nExp number, first 60 AAs: 19.53139\nTotal prob of N-in: 0.94364\nPOSSIBLE N-term signal sequence\ninside 1 19\nTMhelix 20 39\noutside 40 325\nTMhelix 326 348\ninside 349 372

10803 GCF_000303535.2_ASM30353v2 Escherichia coli EC1847 Proteobacteria; Gammaproteobacteria; Enterobacterales; Enterobacteriaceae; Escherichia; Escherichia coli
MTGDNTLIHSHGINRRDFMKLCAALAATMGLSSKAAAEMAESVTNPQRPPVIWIGAQECT WP_000145410.1
MULTISPECIES: hydrogenase-2 small chain [Proteobacteria] Length: 372\nNumber of predicted TMHs:

1\nExp number of AAs in TMHs: 23.72226\nExp number, first 60 AAs: 0.22038\nTotal prob of N-in: 0.12994\noutside 1 331\nTMhelix 332 354\ninside 355 372

10804 GCF_000303515.2_ASM30351v2 Escherichia coli EC1848 Proteobacteria; Gammaproteobacteria; Enterobacterales; Enterobacteriaceae; Escherichia; Escherichia coli
MNNEETFYQAMRRQGVTRRSFLKYCSLAATSLGLGAGMAPKIAWALENKPRIPVVWIHGL WP_001301613.1
hydrogenase [Escherichia coli] Length: 372\nNumber of predicted TMHs: 2\nExp number of AAs in TMHs: 37.46598\nExp number, first 60 AAs: 19.53139\nTotal prob of N-in: 0.94364\nPOSSIBLE N-term signal sequence\ninside 1 19\nTMhelix 20 39\noutside 40 325\nTMhelix 326 348\ninside 349 372

10805 GCF_000303515.2_ASM30351v2 Escherichia coli EC1848 Proteobacteria; Gammaproteobacteria; Enterobacterales; Enterobacteriaceae; Escherichia; Escherichia coli
MTGDNTLIHSHGINRRDFMKLCAALAATMGLSSKAAAEMAESVTNPQRPPVIWIGAQECT WP_000145410.1
MULTISPECIES: hydrogenase-2 small chain [Proteobacteria] Length: 372\nNumber of predicted TMHs: 1\nExp number of AAs in TMHs: 23.72226\nExp number, first 60 AAs: 0.22038\nTotal prob of N-in: 0.12994\noutside 1 331\nTMhelix 332 354\ninside 355 372

10806 GCF_000303495.2_ASM30349v2 Escherichia coli EC1849 Proteobacteria; Gammaproteobacteria; Enterobacterales; Enterobacteriaceae; Escherichia; Escherichia coli
MNNEETFYQAMRRQGVTRRSFLKYCSLAATSLGLGAGMAPKIAWALENKPRIPVVWIHGL WP_001301613.1
hydrogenase [Escherichia coli] Length: 372\nNumber of predicted TMHs: 2\nExp number of AAs in TMHs: 37.46598\nExp number, first 60 AAs: 19.53139\nTotal prob of N-in: 0.94364\nPOSSIBLE N-term signal sequence\ninside 1 19\nTMhelix 20 39\noutside 40 325\nTMhelix 326 348\ninside 349 372

10807 GCF_000303495.2_ASM30349v2 Escherichia coli EC1849 Proteobacteria; Gammaproteobacteria; Enterobacterales; Enterobacteriaceae; Escherichia; Escherichia coli
MTGDNTLIHSHGINRRDFMKLCAALAATMGLSSKAAAEMAESVTNPQRPPVIWIGAQECT WP_000145410.1
MULTISPECIES: hydrogenase-2 small chain [Proteobacteria] Length: 372\nNumber of predicted TMHs: 1\nExp number of AAs in TMHs: 23.72226\nExp number, first 60 AAs: 0.22038\nTotal prob of N-in: 0.12994\noutside 1 331\nTMhelix 332 354\ninside 355 372

10808 GCF_000303475.2_ASM30347v2 Escherichia coli EC1850 Proteobacteria; Gammaproteobacteria; Enterobacterales; Enterobacteriaceae; Escherichia; Escherichia coli
MTGDNTLIHSHGINRRDFMKLCAALAATMGLSSKAAAEMAESVTNPQRPPVIWIGAQECT WP_000145410.1
MULTISPECIES: hydrogenase-2 small chain [Proteobacteria] Length: 372\nNumber of predicted TMHs: 1\nExp number of AAs in TMHs: 23.72226\nExp number, first 60 AAs: 0.22038\nTotal prob of N-in: 0.12994\noutside 1 331\nTMhelix 332 354\ninside 355 372

10809 GCF_000303475.2_ASM30347v2 Escherichia coli EC1850 Proteobacteria; Gammaproteobacteria; Enterobacterales; Enterobacteriaceae; Escherichia; Escherichia coli
MNNEETFYQAMRRQGVTRRSFLKYCSLAATSLGLGAGMAPKIAWALENKPRIPVVWIHGL WP_001301613.1
hydrogenase [Escherichia coli] Length: 372\nNumber of predicted TMHs: 2\nExp number of AAs in TMHs: 37.46598\nExp number, first 60 AAs: 19.53139\nTotal prob of N-in: 0.94364\nPOSSIBLE N-term signal sequence\ninside 1 19\nTMhelix 20 39\noutside 40 325\nTMhelix 326 348\ninside 349 372

10810 GCF_000303455.2_ASM30345v2 Escherichia coli EC1856 Proteobacteria; Gammaproteobacteria; Enterobacterales; Enterobacteriaceae; Escherichia; Escherichia coli
MTGDNTLIHSHGINRRDFMKLCAALAATMGLSSKAAAEMAESVTNPQRPPVIWIGAQECT WP_000145410.1
MULTISPECIES: hydrogenase-2 small chain [Proteobacteria] Length: 372\nNumber of predicted TMHs: 1\nExp number of AAs in TMHs: 23.72226\nExp number, first 60 AAs: 0.22038\nTotal prob of N-in: 0.12994\noutside 1 331\nTMhelix 332 354\ninside 355 372

10811 GCF_000303455.2_ASM30345v2 Escherichia coli EC1856 Proteobacteria; Gammaproteobacteria; Enterobacterales; Enterobacteriaceae; Escherichia; Escherichia coli
MNNEETFYQAMRRQGVTRRSFLKYCSLAATSLGLGAGMAPKIAWALENKPRIPVVWIHGL WP_001301613.1
hydrogenase [Escherichia coli] Length: 372\nNumber of predicted TMHs: 2\nExp number of AAs in TMHs: 37.46598\nExp number, first 60 AAs: 19.53139\nTotal prob of N-in: 0.94364\nPOSSIBLE N-term signal sequence\ninside 1 19\nTMhelix 20 39\noutside 40 325\nTMhelix 326 348\ninside 349 372

10812 GCF_000303435.2_ASM30343v2 Escherichia coli EC1862 Proteobacteria; Gammaproteobacteria; Enterobacterales; Enterobacteriaceae; Escherichia; Escherichia coli

MTGDNTLIHSHGINRRDFMKLCAALAATMGLSSKAAAEMAESVTNPQRPPVIWIGAQECT WP_000145410.1 MULTISPECIES: hydrogenase-2 small chain [Proteobacteria] Length: 372\nNumber of predicted TMHs: 1\nExp number of AAs in TMHs: 23.72226\nExp number, first 60 AAs: 0.22038\nTotal prob of N-in: 0.12994\noutside 1 331\nTMhelix 332 354\ninside 355 372

10813 GCF_000303435.2_ASM30343v2 Escherichia coli EC1862 Proteobacteria; Gammaproteobacteria; Enterobacterales; Enterobacteriaceae; Escherichia; Escherichia coli MNNEETFYQAMRRQGVTRRSFLKYCSLAATSLGLGAGMAPKIAWALENKPRIPVVWIHGL WP_001301613.1 hydrogenase [Escherichia coli] Length: 372\nNumber of predicted TMHs: 2\nExp number of AAs in TMHs: 37.46598\nExp number, first 60 AAs: 19.53139\nTotal prob of N-in: 0.94364\nPOSSIBLE N-term signal sequence\ninside 1 19\nTMhelix 20 39\noutside 40 325\nTMhelix 326 348\ninside 349 372

10814 GCF_000267205.2_ASM26720v2 Escherichia coli EC1863 Proteobacteria; Gammaproteobacteria; Enterobacterales; Enterobacteriaceae; Escherichia; Escherichia coli MNNEETFYQAMRRQGVTRRSFLKYCSLAATSLGLGAGMAPKIAWALENKPRIPVVWIHGL WP_001301613.1 hydrogenase [Escherichia coli] Length: 372\nNumber of predicted TMHs: 2\nExp number of AAs in TMHs: 37.46598\nExp number, first 60 AAs: 19.53139\nTotal prob of N-in: 0.94364\nPOSSIBLE N-term signal sequence\ninside 1 19\nTMhelix 20 39\noutside 40 325\nTMhelix 326 348\ninside 349 372

10815 GCF_000267205.2_ASM26720v2 Escherichia coli EC1863 Proteobacteria; Gammaproteobacteria; Enterobacterales; Enterobacteriaceae; Escherichia; Escherichia coli MTGDNTLIHSHGINRRDFMKLCAALAATMGLSSKAAAEMAESVTNPQRPPVIWIGAQECT WP_000145410.1 MULTISPECIES: hydrogenase-2 small chain [Proteobacteria] Length: 372\nNumber of predicted TMHs: 1\nExp number of AAs in TMHs: 23.72226\nExp number, first 60 AAs: 0.22038\nTotal prob of N-in: 0.12994\noutside 1 331\nTMhelix 332 354\ninside 355 372

10816 GCF_000303415.2_ASM30341v2 Escherichia coli EC1864 Proteobacteria; Gammaproteobacteria; Enterobacterales; Enterobacteriaceae; Escherichia; Escherichia coli MNNEETFYQAMRRQGVTRRSFLKYCSLAATSLGLGAGMAPKIAWALENKPRIPVVWIHGL WP_001301613.1 hydrogenase [Escherichia coli] Length: 372\nNumber of predicted TMHs: 2\nExp number of AAs in TMHs: 37.46598\nExp number, first 60 AAs: 19.53139\nTotal prob of N-in: 0.94364\nPOSSIBLE N-term signal sequence\ninside 1 19\nTMhelix 20 39\noutside 40 325\nTMhelix 326 348\ninside 349 372

10817 GCF_000303415.2_ASM30341v2 Escherichia coli EC1864 Proteobacteria; Gammaproteobacteria; Enterobacterales; Enterobacteriaceae; Escherichia; Escherichia coli MTGDNTLIHSHGINRRDFMKLCAALAATMGLSSKAAAEMAESVTNPQRPPVIWIGAQECT WP_000145410.1 MULTISPECIES: hydrogenase-2 small chain [Proteobacteria] Length: 372\nNumber of predicted TMHs: 1\nExp number of AAs in TMHs: 23.72226\nExp number, first 60 AAs: 0.22038\nTotal prob of N-in: 0.12994\noutside 1 331\nTMhelix 332 354\ninside 355 372

10818 GCF_000303395.2_ASM30339v2 Escherichia coli EC1865 Proteobacteria; Gammaproteobacteria; Enterobacterales; Enterobacteriaceae; Escherichia; Escherichia coli MNNEETFYQAMRRQGVTRRSFLKYCSLAATSLGLGAGMAPKIAWALENKPRIPVVWIHGL WP_001058323.1 MULTISPECIES: hydrogenase-1 small chain [Proteobacteria] Length: 372\nNumber of predicted TMHs: 2\nExp number of AAs in TMHs: 37.2608499999999\nExp number, first 60 AAs: 19.52923\nTotal prob of N-in: 0.94351\nPOSSIBLE N-term signal sequence\ninside 1 19\nTMhelix 20 39\noutside 40 325\nTMhelix 326 348\ninside 349 372

10819 GCF_000303395.2_ASM30339v2 Escherichia coli EC1865 Proteobacteria; Gammaproteobacteria; Enterobacterales; Enterobacteriaceae; Escherichia; Escherichia coli MTGDNTLIHSHGINRRDFMKLCAALAATMGLSSKAAAEMAESVTNPQRPPVIWIGAQECT WP_000145410.1 MULTISPECIES: hydrogenase-2 small chain [Proteobacteria] Length: 372\nNumber of predicted TMHs: 1\nExp number of AAs in TMHs: 23.72226\nExp number, first 60 AAs: 0.22038\nTotal prob of N-in: 0.12994\noutside 1 331\nTMhelix 332 354\ninside 355 372

10820 GCF_000303375.2_ASM30337v2 Escherichia coli EC1866 Proteobacteria; Gammaproteobacteria; Enterobacterales; Enterobacteriaceae; Escherichia; Escherichia coli MTGDNTLIHSHGINRRDFMKLCAALAATMGLSSKAAAEMAESVTNPQRPPVIWIGAQECT WP_000145410.1 MULTISPECIES: hydrogenase-2 small chain [Proteobacteria] Length: 372\nNumber of predicted TMHs: 1\nExp number of AAs in TMHs: 23.72226\nExp number, first 60 AAs: 0.22038\nTotal prob of N-in: 0.12994\noutside 1 331\nTMhelix 332 354\ninside 355 372

10821 GCF_000303375.2_ASM30337v2 Escherichia coli EC1866 Proteobacteria; Gammaproteobacteria; Enterobacterales; Enterobacteriaceae; Escherichia; Escherichia coli
MNNEETFYQAMRRQGVTRRSFLKYCSLAATSLGLGAGMAPKIAWALENKPRIPVVWIHGL WP_001301613.1
hydrogenase [Escherichia coli] Length: 372\nNumber of predicted TMHs: 2\nExp number of AAs in TMHs: 37.46598\nExp number, first 60 AAs: 19.53139\nTotal prob of N-in: 0.94364\nPOSSIBLE N-term signal sequence\ninside 1 19\nTMhelix 20 39\noutside 40 325\nTMhelix 326 348\ninside 349 372

10822 GCF_000303355.2_ASM30335v2 Escherichia coli EC1868 Proteobacteria; Gammaproteobacteria; Enterobacterales; Enterobacteriaceae; Escherichia; Escherichia coli
MNNEETFYQAMRRQGVTRRSFLKYCSLAATSLGLGAGMAPKIAWALENKPRIPVVWIHGL WP_001301613.1
hydrogenase [Escherichia coli] Length: 372\nNumber of predicted TMHs: 2\nExp number of AAs in TMHs: 37.46598\nExp number, first 60 AAs: 19.53139\nTotal prob of N-in: 0.94364\nPOSSIBLE N-term signal sequence\ninside 1 19\nTMhelix 20 39\noutside 40 325\nTMhelix 326 348\ninside 349 372

10823 GCF_000303355.2_ASM30335v2 Escherichia coli EC1868 Proteobacteria; Gammaproteobacteria; Enterobacterales; Enterobacteriaceae; Escherichia; Escherichia coli
MTGDNTLIHSHGINRRDFMKLCAALAATMGLSSKAAAEMAESVTNPQRPPVIWIGAQECT WP_000145410.1
MULTISPECIES: hydrogenase-2 small chain [Proteobacteria] Length: 372\nNumber of predicted TMHs: 1\nExp number of AAs in TMHs: 23.72226\nExp number, first 60 AAs: 0.22038\nTotal prob of N-in: 0.12994\noutside 1 331\nTMhelix 332 354\ninside 355 372

10824 GCF_000303335.2_ASM30333v2 Escherichia coli EC1869 Proteobacteria; Gammaproteobacteria; Enterobacterales; Enterobacteriaceae; Escherichia; Escherichia coli
MTGDNTLIHSHGINRRDFMKLCAALAATMGLSSKAAAEMAESVTNPQRPPVIWIGAQECT WP_000145410.1
MULTISPECIES: hydrogenase-2 small chain [Proteobacteria] Length: 372\nNumber of predicted TMHs: 1\nExp number of AAs in TMHs: 23.72226\nExp number, first 60 AAs: 0.22038\nTotal prob of N-in: 0.12994\noutside 1 331\nTMhelix 332 354\ninside 355 372

10825 GCF_000303335.2_ASM30333v2 Escherichia coli EC1869 Proteobacteria; Gammaproteobacteria; Enterobacterales; Enterobacteriaceae; Escherichia; Escherichia coli
MNNEETFYQAMRRQGVTRRSFLKYCSLAATSLGLGAGMAPKIAWALENKPRIPVVWIHGL WP_001301613.1
hydrogenase [Escherichia coli] Length: 372\nNumber of predicted TMHs: 2\nExp number of AAs in TMHs: 37.46598\nExp number, first 60 AAs: 19.53139\nTotal prob of N-in: 0.94364\nPOSSIBLE N-term signal sequence\ninside 1 19\nTMhelix 20 39\noutside 40 325\nTMhelix 326 348\ninside 349 372

10826 GCF_000303315.2_ASM30331v2 Escherichia coli EC1870 Proteobacteria; Gammaproteobacteria; Enterobacterales; Enterobacteriaceae; Escherichia; Escherichia coli
MNNEETFYQAMRRQGVTRRSFLKYCSLAATSLGLGAGMAPKIAWALENKPRIPVVWIHGL WP_001301613.1
hydrogenase [Escherichia coli] Length: 372\nNumber of predicted TMHs: 2\nExp number of AAs in TMHs: 37.46598\nExp number, first 60 AAs: 19.53139\nTotal prob of N-in: 0.94364\nPOSSIBLE N-term signal sequence\ninside 1 19\nTMhelix 20 39\noutside 40 325\nTMhelix 326 348\ninside 349 372

10827 GCF_000303315.2_ASM30331v2 Escherichia coli EC1870 Proteobacteria; Gammaproteobacteria; Enterobacterales; Enterobacteriaceae; Escherichia; Escherichia coli
MTGDNTLIHSHGINRRDFMKLCAALAATMGLSSKAAAEMAESVTNPQRPPVIWIGAQECT WP_000145410.1
MULTISPECIES: hydrogenase-2 small chain [Proteobacteria] Length: 372\nNumber of predicted TMHs: 1\nExp number of AAs in TMHs: 23.72226\nExp number, first 60 AAs: 0.22038\nTotal prob of N-in: 0.12994\noutside 1 331\nTMhelix 332 354\ninside 355 372

10828 GCF_000297235.3_EC302_04.1 Escherichia coli EC302/04 Proteobacteria; Gammaproteobacteria; Enterobacterales; Enterobacteriaceae; Escherichia; Escherichia coli
MTGDNTLIHSHGINRRDFMKLCAALAATMGLSSKAAAEMAESVTNPQRPPVIWIGAQECT WP_000145410.1
MULTISPECIES: hydrogenase-2 small chain [Proteobacteria] Length: 372\nNumber of predicted TMHs: 1\nExp number of AAs in TMHs: 23.72226\nExp number, first 60 AAs: 0.22038\nTotal prob of N-in: 0.12994\noutside 1 331\nTMhelix 332 354\ninside 355 372

10829 GCF_000297235.3_EC302_04.1 Escherichia coli EC302/04 Proteobacteria; Gammaproteobacteria; Enterobacterales; Enterobacteriaceae; Escherichia; Escherichia coli
MNNEETFYQAMRRQGVTRRSFLKYCSLAATSLGLGAGMAPKIAWALENKPRIPVVWIHGL WP_001058323.1
MULTISPECIES: hydrogenase-1 small chain [Proteobacteria] Length: 372\nNumber of predicted TMHs: 2\nExp number of AAs in TMHs: 37.2608499999999\nExp number, first 60 AAs: 19.52923\nTotal prob of N-in:

0.94351\nPOSSIBLE N-term signal sequence\ninside 1 19\nTMhelix 20 39\noutside 40 325\nTMhelix 326 348\ninside 349 372

10830 GCF_000267345.2_ASM26734v2 Escherichia coli EC4013 Proteobacteria; Gammaproteobacteria; Enterobacterales; Enterobacteriaceae; Escherichia; Escherichia coli
MTGDNTLIHSHGINRRDFMKLCAALAATMGLSSKAAAEMAESVTNPQRPPVIWIGAQECT WP_000145410.1
MULTISPECIES: hydrogenase-2 small chain [Proteobacteria] Length: 372\nNumber of predicted TMHs: 1\nExp number of AAs in TMHs: 23.72226\nExp number, first 60 AAs: 0.22038\nTotal prob of N-in: 0.12994\noutside 1 331\nTMhelix 332 354\ninside 355 372

10831 GCF_000267345.2_ASM26734v2 Escherichia coli EC4013 Proteobacteria; Gammaproteobacteria; Enterobacterales; Enterobacteriaceae; Escherichia; Escherichia coli
MNNEETFYQAMRRQGVTRRSFLKYCSLAATSLGLGAGMAPKIAWALENKPRIPVVWIHGL WP_001301613.1
hydrogenase [Escherichia coli] Length: 372\nNumber of predicted TMHs: 2\nExp number of AAs in TMHs: 37.46598\nExp number, first 60 AAs: 19.53139\nTotal prob of N-in: 0.94364\nPOSSIBLE N-term signal sequence\ninside 1 19\nTMhelix 20 39\noutside 40 325\nTMhelix 326 348\ninside 349 372

10832 GCF_000193995.1_ASM19399v2 Escherichia coli EC4100B Proteobacteria; Gammaproteobacteria; Enterobacterales; Enterobacteriaceae; Escherichia; Escherichia coli
MNNEETFYQAMRRQGVTRRSFLKYCSLAATSLGLGAGMAPKIAWALENKPRIPVVWIHGL WP_001058323.1
MULTISPECIES: hydrogenase-1 small chain [Proteobacteria] Length: 372\nNumber of predicted TMHs: 2\nExp number of AAs in TMHs: 37.2608499999999\nExp number, first 60 AAs: 19.52923\nTotal prob of N-in: 0.94351\nPOSSIBLE N-term signal sequence\ninside 1 19\nTMhelix 20 39\noutside 40 325\nTMhelix 326 348\ninside 349 372

10833 GCF_000267445.2_ASM26744v2 Escherichia coli EC4196 Proteobacteria; Gammaproteobacteria; Enterobacterales; Enterobacteriaceae; Escherichia; Escherichia coli
MTGDNTLIHSHGINRRDFMKLCAALAATMGLSSKAAAEMAESVTNPQRPPVIWIGAQECT WP_000145410.1
MULTISPECIES: hydrogenase-2 small chain [Proteobacteria] Length: 372\nNumber of predicted TMHs: 1\nExp number of AAs in TMHs: 23.72226\nExp number, first 60 AAs: 0.22038\nTotal prob of N-in: 0.12994\noutside 1 331\nTMhelix 332 354\ninside 355 372

10834 GCF_000267445.2_ASM26744v2 Escherichia coli EC4196 Proteobacteria; Gammaproteobacteria; Enterobacterales; Enterobacteriaceae; Escherichia; Escherichia coli
MNNEETFYQAMRRQGVTRRSFLKYCSLAATSLGLGAGMAPKIAWALENKPRIPVVWIHGL WP_001301613.1
hydrogenase [Escherichia coli] Length: 372\nNumber of predicted TMHs: 2\nExp number of AAs in TMHs: 37.46598\nExp number, first 60 AAs: 19.53139\nTotal prob of N-in: 0.94364\nPOSSIBLE N-term signal sequence\ninside 1 19\nTMhelix 20 39\noutside 40 325\nTMhelix 326 348\ninside 349 372

10835 GCF_000267465.2_ASM26746v2 Escherichia coli EC4203 Proteobacteria; Gammaproteobacteria; Enterobacterales; Enterobacteriaceae; Escherichia; Escherichia coli
MNNEETFYQAMRRQGVTRRSFLKYCSLAATSLGLGAGMAPKIAWALENKPRIPVVWIHGL WP_001301613.1
hydrogenase [Escherichia coli] Length: 372\nNumber of predicted TMHs: 2\nExp number of AAs in TMHs: 37.46598\nExp number, first 60 AAs: 19.53139\nTotal prob of N-in: 0.94364\nPOSSIBLE N-term signal sequence\ninside 1 19\nTMhelix 20 39\noutside 40 325\nTMhelix 326 348\ninside 349 372

10836 GCF_000267465.2_ASM26746v2 Escherichia coli EC4203 Proteobacteria; Gammaproteobacteria; Enterobacterales; Enterobacteriaceae; Escherichia; Escherichia coli
MTGDNTLIHSHGINRRDFMKLCAALAATMGLSSKAAAEMAESVTNPQRPPVIWIGAQECT WP_000145410.1
MULTISPECIES: hydrogenase-2 small chain [Proteobacteria] Length: 372\nNumber of predicted TMHs: 1\nExp number of AAs in TMHs: 23.72226\nExp number, first 60 AAs: 0.22038\nTotal prob of N-in: 0.12994\noutside 1 331\nTMhelix 332 354\ninside 355 372

10837 GCF_000267325.1_ASM26732v1 Escherichia coli EC4402 Proteobacteria; Gammaproteobacteria; Enterobacterales; Enterobacteriaceae; Escherichia; Escherichia coli
MTGDNTLIHSHGINRRDFMKLCAALAATMGLSSKAAAEMAESVTNPQRPPVIWIGAQECT WP_000145410.1
MULTISPECIES: hydrogenase-2 small chain [Proteobacteria] Length: 372\nNumber of predicted TMHs: 1\nExp number of AAs in TMHs: 23.72226\nExp number, first 60 AAs: 0.22038\nTotal prob of N-in: 0.12994\noutside 1 331\nTMhelix 332 354\ninside 355 372

10838 GCF_000267325.1_ASM26732v1 Escherichia coli EC4402 Proteobacteria; Gammaproteobacteria; Enterobacterales; Enterobacteriaceae; Escherichia; Escherichia coli
MNNEETFYQAMRRQGVTRRSFLKYCSLAATSLGLGAGMAPKIAWALENKPRIPVVWIHGL WP_001301613.1
hydrogenase [Escherichia coli] Length: 372\nNumber of predicted TMHs: 2\nExp number of AAs in TMHs: 37.46598\nExp number, first 60 AAs: 19.53139\nTotal prob of N-in: 0.94364\nPOSSIBLE N-term signal sequence\ninside 1 19\nTMhelix 20 39\noutside 40 325\nTMhelix 326 348\ninside 349 372

10839 GCF_000267385.2_ASM26738v2 Escherichia coli EC4421 Proteobacteria; Gammaproteobacteria; Enterobacterales; Enterobacteriaceae; Escherichia; Escherichia coli
MNNEETFYQAMRRQGVTRRSFLKYCSLAATSLGLGAGMAPKIAWALENKPRIPVVWIHGL WP_001301613.1
hydrogenase [Escherichia coli] Length: 372\nNumber of predicted TMHs: 2\nExp number of AAs in TMHs: 37.46598\nExp number, first 60 AAs: 19.53139\nTotal prob of N-in: 0.94364\nPOSSIBLE N-term signal sequence\ninside 1 19\nTMhelix 20 39\noutside 40 325\nTMhelix 326 348\ninside 349 372

10840 GCF_000267385.2_ASM26738v2 Escherichia coli EC4421 Proteobacteria; Gammaproteobacteria; Enterobacterales; Enterobacteriaceae; Escherichia; Escherichia coli
MTGDNTLIHSHGINRRDFMKLCAALAATMGLSSKAAAEMAESVTNPQRPPVIWIGAQECT WP_000145410.1
MULTISPECIES: hydrogenase-2 small chain [Proteobacteria] Length: 372\nNumber of predicted TMHs: 1\nExp number of AAs in TMHs: 23.72226\nExp number, first 60 AAs: 0.22038\nTotal prob of N-in: 0.12994\noutside 1 331\nTMhelix 332 354\ninside 355 372

10841 GCF_000267365.2_ASM26736v2 Escherichia coli EC4422 Proteobacteria; Gammaproteobacteria; Enterobacterales; Enterobacteriaceae; Escherichia; Escherichia coli
MTGDNTLIHSHGINRRDFMKLCAALAATMGLSSKAAAEMAESVTNPQRPPVIWIGAQECT WP_000145410.1
MULTISPECIES: hydrogenase-2 small chain [Proteobacteria] Length: 372\nNumber of predicted TMHs: 1\nExp number of AAs in TMHs: 23.72226\nExp number, first 60 AAs: 0.22038\nTotal prob of N-in: 0.12994\noutside 1 331\nTMhelix 332 354\ninside 355 372

10842 GCF_000267365.2_ASM26736v2 Escherichia coli EC4422 Proteobacteria; Gammaproteobacteria; Enterobacterales; Enterobacteriaceae; Escherichia; Escherichia coli
MNNEETFYQAMRRQGVTRRSFLKYCSLAATSLGLGAGMAPKIAWALENKPRIPVVWIHGL WP_001301613.1
hydrogenase [Escherichia coli] Length: 372\nNumber of predicted TMHs: 2\nExp number of AAs in TMHs: 37.46598\nExp number, first 60 AAs: 19.53139\nTotal prob of N-in: 0.94364\nPOSSIBLE N-term signal sequence\ninside 1 19\nTMhelix 20 39\noutside 40 325\nTMhelix 326 348\ninside 349 372

10843 GCF_000267285.2_ASM26728v2 Escherichia coli EC4436 Proteobacteria; Gammaproteobacteria; Enterobacterales; Enterobacteriaceae; Escherichia; Escherichia coli
MNNEETFYQAMRRQGVTRRSFLKYCSLAATSLGLGAGMAPKIAWALENKPRIPVVWIHGL WP_001301613.1
hydrogenase [Escherichia coli] Length: 372\nNumber of predicted TMHs: 2\nExp number of AAs in TMHs: 37.46598\nExp number, first 60 AAs: 19.53139\nTotal prob of N-in: 0.94364\nPOSSIBLE N-term signal sequence\ninside 1 19\nTMhelix 20 39\noutside 40 325\nTMhelix 326 348\ninside 349 372

10844 GCF_000267285.2_ASM26728v2 Escherichia coli EC4436 Proteobacteria; Gammaproteobacteria; Enterobacterales; Enterobacteriaceae; Escherichia; Escherichia coli
MTGDNTLIHSHGINRRDFMKLCAALAATMGLSSKAAAEMAESVTNPQRPPVIWIGAQECT WP_000145410.1
MULTISPECIES: hydrogenase-2 small chain [Proteobacteria] Length: 372\nNumber of predicted TMHs: 1\nExp number of AAs in TMHs: 23.72226\nExp number, first 60 AAs: 0.22038\nTotal prob of N-in: 0.12994\noutside 1 331\nTMhelix 332 354\ninside 355 372

10845 GCF_000267265.2_ASM26726v2 Escherichia coli EC4437 Proteobacteria; Gammaproteobacteria; Enterobacterales; Enterobacteriaceae; Escherichia; Escherichia coli
MNNEETFYQAMRRQGVTRRSFLKYCSLAATSLGLGAGMAPKIAWALENKPRIPVVWIHGL WP_001301613.1
hydrogenase [Escherichia coli] Length: 372\nNumber of predicted TMHs: 2\nExp number of AAs in TMHs: 37.46598\nExp number, first 60 AAs: 19.53139\nTotal prob of N-in: 0.94364\nPOSSIBLE N-term signal sequence\ninside 1 19\nTMhelix 20 39\noutside 40 325\nTMhelix 326 348\ninside 349 372

10846 GCF_000267265.2_ASM26726v2 Escherichia coli EC4437 Proteobacteria; Gammaproteobacteria; Enterobacterales; Enterobacteriaceae; Escherichia; Escherichia coli
MTGDNTLIHSHGINRRDFMKLCAALAATMGLSSKAAAEMAESVTNPQRPPVIWIGAQECT WP_000145410.1
MULTISPECIES: hydrogenase-2 small chain [Proteobacteria] Length: 372\nNumber of predicted TMHs:

1\nExp number of AAs in TMHs: 23.72226\nExp number, first 60 AAs: 0.22038\nTotal prob of N-in: 0.12994\noutside 1 331\nTMhelix 332 354\ninside 355 372

10847 GCF_000267305.2_ASM26730v2 Escherichia coli EC4439 Proteobacteria; Gammaproteobacteria; Enterobacterales; Enterobacteriaceae; Escherichia; Escherichia coli
MTGDNTLIHSHGINRRDFMKLCAALAATMGLSSKAAAEMAESVTNPQRPPVIWIGAQECT WP_000145410.1
MULTISPECIES: hydrogenase-2 small chain [Proteobacteria] Length: 372\nNumber of predicted TMHs: 1\nExp number of AAs in TMHs: 23.72226\nExp number, first 60 AAs: 0.22038\nTotal prob of N-in: 0.12994\noutside 1 331\nTMhelix 332 354\ninside 355 372

10848 GCF_000267305.2_ASM26730v2 Escherichia coli EC4439 Proteobacteria; Gammaproteobacteria; Enterobacterales; Enterobacteriaceae; Escherichia; Escherichia coli
MNNEETFYQAMRRQGVTRRSFLKYCSLAATSLGLGAGMAPKIAWALENKPRIPVVWIHGL WP_001301613.1
hydrogenase [Escherichia coli] Length: 372\nNumber of predicted TMHs: 2\nExp number of AAs in TMHs: 37.46598\nExp number, first 60 AAs: 19.53139\nTotal prob of N-in: 0.94364\nPOSSIBLE N-term signal sequence\ninside 1 19\nTMhelix 20 39\noutside 40 325\nTMhelix 326 348\ninside 349 372

10849 GCF_000267245.2_ASM26724v2 Escherichia coli EC4448 Proteobacteria; Gammaproteobacteria; Enterobacterales; Enterobacteriaceae; Escherichia; Escherichia coli
MNNEETFYQAMRRQGVTRRSFLKYCSLAATSLGLGAGMAPKIAWALENKPRIPVVWIHGL WP_001301613.1
hydrogenase [Escherichia coli] Length: 372\nNumber of predicted TMHs: 2\nExp number of AAs in TMHs: 37.46598\nExp number, first 60 AAs: 19.53139\nTotal prob of N-in: 0.94364\nPOSSIBLE N-term signal sequence\ninside 1 19\nTMhelix 20 39\noutside 40 325\nTMhelix 326 348\ninside 349 372

10850 GCF_000267245.2_ASM26724v2 Escherichia coli EC4448 Proteobacteria; Gammaproteobacteria; Enterobacterales; Enterobacteriaceae; Escherichia; Escherichia coli
MTGDNTLIHSHGINRRDFMKLCAALAATMGLSSKAAAEMAESVTNPQRPPVIWIGAQECT WP_000145410.1
MULTISPECIES: hydrogenase-2 small chain [Proteobacteria] Length: 372\nNumber of predicted TMHs: 1\nExp number of AAs in TMHs: 23.72226\nExp number, first 60 AAs: 0.22038\nTotal prob of N-in: 0.12994\noutside 1 331\nTMhelix 332 354\ninside 355 372

10851 GCF_000304035.2_ASM30403v2 Escherichia coli EC96038 Proteobacteria; Gammaproteobacteria; Enterobacterales; Enterobacteriaceae; Escherichia; Escherichia coli
MNNEETFYQAMRRQGVTRRSFLKYCSLAATSLGLGAGMAPKIAWALENKPRIPVVWIHGL WP_001301613.1
hydrogenase [Escherichia coli] Length: 372\nNumber of predicted TMHs: 2\nExp number of AAs in TMHs: 37.46598\nExp number, first 60 AAs: 19.53139\nTotal prob of N-in: 0.94364\nPOSSIBLE N-term signal sequence\ninside 1 19\nTMhelix 20 39\noutside 40 325\nTMhelix 326 348\ninside 349 372

10852 GCF_000304035.2_ASM30403v2 Escherichia coli EC96038 Proteobacteria; Gammaproteobacteria; Enterobacterales; Enterobacteriaceae; Escherichia; Escherichia coli
MIGDNTLIHSHGINRRDFMKLCAALAATMGLSSKAAAEMAESVTNPQRPPVIWIGAQECT WP_000578723.1
hydrogenase 2 small subunit [Escherichia coli] Length: 372\nNumber of predicted TMHs: 1\nExp number of AAs in TMHs: 23.05603\nExp number, first 60 AAs: 0.15431\nTotal prob of N-in: 0.09116\noutside 1 331\nTMhelix 332 354\ninside 355 372

10853 GCF_000831565.1_ASM83156v1 Escherichia coli ECC-1470 Proteobacteria; Gammaproteobacteria; Enterobacterales; Enterobacteriaceae; Escherichia; Escherichia coli
MNNEETFYQAMRRQGVTRRSFLKYCSLAATSLGLGAGMAPKIAWALENKPRIPVVWIHGL WP_001295940.1
MULTISPECIES: hydrogenase [Enterobacteriaceae] Length: 372\nNumber of predicted TMHs: 2\nExp number of AAs in TMHs: 37.2608899999999\nExp number, first 60 AAs: 19.52923\nTotal prob of N-in: 0.94351\nPOSSIBLE N-term signal sequence\ninside 1 19\nTMhelix 20 39\noutside 40 325\nTMhelix 326 348\ninside 349 372

10854 GCF_000831565.1_ASM83156v1 Escherichia coli ECC-1470 Proteobacteria; Gammaproteobacteria; Enterobacterales; Enterobacteriaceae; Escherichia; Escherichia coli
MTGDNTLIHSHGINRRDFMKLCAALAATMGLSSKAAAEMAESVTNPQRPPVIWIGAQECT WP_000145410.1
MULTISPECIES: hydrogenase-2 small chain [Proteobacteria] Length: 372\nNumber of predicted TMHs: 1\nExp number of AAs in TMHs: 23.72226\nExp number, first 60 AAs: 0.22038\nTotal prob of N-in: 0.12994\noutside 1 331\nTMhelix 332 354\ninside 355 372

10855 GCF_000498215.1_ASM49821v1 Escherichia coli ECC-1470 Proteobacteria; Gammaproteobacteria; Enterobacterales; Enterobacteriaceae; Escherichia; Escherichia coli
MTGDNTLIHSHGINRRDFMKLCAALAATMGLSSKAAAEMAESVTNPQRPPVIWIGAQECT WP_000145410.1
MULTISPECIES: hydrogenase-2 small chain [Proteobacteria] Length: 372\nNumber of predicted TMHs: 1\nExp number of AAs in TMHs: 23.72226\nExp number, first 60 AAs: 0.22038\nTotal prob of N-in: 0.12994\noutside 1 331\nTMhelix 332 354\ninside 355 372

10856 GCF_000498215.1_ASM49821v1 Escherichia coli ECC-1470 Proteobacteria; Gammaproteobacteria; Enterobacterales; Enterobacteriaceae; Escherichia; Escherichia coli
MNNEETFYQAMRRQGVTRRSFLKYCSLAATSLGLGAGMAPKIAWALENKPRIPVVWIHGL WP_001295940.1
MULTISPECIES: hydrogenase [Enterobacteriaceae] Length: 372\nNumber of predicted TMHs: 2\nExp number of AAs in TMHs: 37.2608899999999\nExp number, first 60 AAs: 19.52923\nTotal prob of N-in: 0.94351\nPOSSIBLE N-term signal sequence\ninside 1 19\nTMhelix 20 39\noutside 40 325\nTMhelix 326 348\ninside 349 372

10857 GCF_000498235.1_ASM49823v1 Escherichia coli ECC-Z Proteobacteria; Gammaproteobacteria; Enterobacterales; Enterobacteriaceae; Escherichia; Escherichia coli
MNNEETFYQAMRRQGVTRRSFLKYCSLAATSLGLGAGMAPKIAWALENKPRIPVVWIHGL WP_001058323.1
MULTISPECIES: hydrogenase-1 small chain [Proteobacteria] Length: 372\nNumber of predicted TMHs: 2\nExp number of AAs in TMHs: 37.2608499999999\nExp number, first 60 AAs: 19.52923\nTotal prob of N-in: 0.94351\nPOSSIBLE N-term signal sequence\ninside 1 19\nTMhelix 20 39\noutside 40 325\nTMhelix 326 348\ninside 349 372

10858 GCF_000498235.1_ASM49823v1 Escherichia coli ECC-Z Proteobacteria; Gammaproteobacteria; Enterobacterales; Enterobacteriaceae; Escherichia; Escherichia coli
MTGDNTLIHSHGINRRDFMKLCAALAATMGLSSKAAAEMAESVTNPQRPPVIWIGAQECT WP_000145410.1
MULTISPECIES: hydrogenase-2 small chain [Proteobacteria] Length: 372\nNumber of predicted TMHs: 1\nExp number of AAs in TMHs: 23.72226\nExp number, first 60 AAs: 0.22038\nTotal prob of N-in: 0.12994\noutside 1 331\nTMhelix 332 354\ninside 355 372

10859 GCF_000026305.1_ASM2630v1 Escherichia coli ED1a Proteobacteria; Gammaproteobacteria; Enterobacterales; Enterobacteriaceae; Escherichia; Escherichia coli
MTGDNTLIHSHGINRRDFMKLCAALAATMGLSSKAAAEMAESVTNPQRPPVIWIGAQECT WP_000145418.1
hydrogenase 2 small subunit [Escherichia coli] Length: 372\nNumber of predicted TMHs: 1\nExp number of AAs in TMHs: 23.72313\nExp number, first 60 AAs: 0.22034\nTotal prob of N-in: 0.12997\noutside 1 331\nTMhelix 332 354\ninside 355 372

10860 GCF_000026305.1_ASM2630v1 Escherichia coli ED1a Proteobacteria; Gammaproteobacteria; Enterobacterales; Enterobacteriaceae; Escherichia; Escherichia coli
MNNEETFYQAMRRQGVTRRSFLKYCSLAATSLGLGAGMAPKIAWALENKPRIPVVWIHGL WP_001295940.1
MULTISPECIES: hydrogenase [Enterobacteriaceae] Length: 372\nNumber of predicted TMHs: 2\nExp number of AAs in TMHs: 37.2608899999999\nExp number, first 60 AAs: 19.52923\nTotal prob of N-in: 0.94351\nPOSSIBLE N-term signal sequence\ninside 1 19\nTMhelix 20 39\noutside 40 325\nTMhelix 326 348\ninside 349 372

10861 GCF_000355195.2_ASM35519v2 Escherichia coli Envira 10/1 Proteobacteria; Gammaproteobacteria; Enterobacterales; Enterobacteriaceae; Escherichia; Escherichia coli
MNNEETFYQAMRRQGVTRRSFLKYCSLAATSLGLGAGMAPKIAWALENKPRIPVVWIHGL WP_001058323.1
MULTISPECIES: hydrogenase-1 small chain [Proteobacteria] Length: 372\nNumber of predicted TMHs: 2\nExp number of AAs in TMHs: 37.2608499999999\nExp number, first 60 AAs: 19.52923\nTotal prob of N-in: 0.94351\nPOSSIBLE N-term signal sequence\ninside 1 19\nTMhelix 20 39\noutside 40 325\nTMhelix 326 348\ninside 349 372

10862 GCF_000355195.2_ASM35519v2 Escherichia coli Envira 10/1 Proteobacteria; Gammaproteobacteria; Enterobacterales; Enterobacteriaceae; Escherichia; Escherichia coli
MTGDNTLIHSHGINRRDFMKLCAALAATMGLSSKAAAEMAESVTNPQRPPVIWIGAQECT WP_000145410.1
MULTISPECIES: hydrogenase-2 small chain [Proteobacteria] Length: 372\nNumber of predicted TMHs: 1\nExp number of AAs in TMHs: 23.72226\nExp number, first 60 AAs: 0.22038\nTotal prob of N-in: 0.12994\noutside 1 331\nTMhelix 332 354\ninside 355 372

10863 GCF_000355215.2_ASM35521v2 Escherichia coli Envira 8/11 Proteobacteria; Gammaproteobacteria; Enterobacterales; Enterobacteriaceae; Escherichia; Escherichia coli
MNNEETFYQAMRRQGVTRRSFLKYCSLAATSLGLGAGMAPKIAWALENKPRIPVVWIHGL WP_001058323.1
MULTISPECIES: hydrogenase-1 small chain [Proteobacteria] Length: 372\nNumber of predicted TMHs: 2\nExp number of AAs in TMHs: 37.2608499999999\nExp number, first 60 AAs: 19.52923\nTotal prob of N-in: 0.94351\nPOSSIBLE N-term signal sequence\ninside 1 19\nTMhelix 20 39\noutside 40 325\nTMhelix 326 348\ninside 349 372

10864 GCF_000355215.2_ASM35521v2 Escherichia coli Envira 8/11 Proteobacteria; Gammaproteobacteria; Enterobacterales; Enterobacteriaceae; Escherichia; Escherichia coli
MTGDNTLIHSHGINRRDFMKLCAALAATMGLSSKAAAEMAESVTNPQRPPVIWIGAQECT WP_000145410.1
MULTISPECIES: hydrogenase-2 small chain [Proteobacteria] Length: 372\nNumber of predicted TMHs: 1\nExp number of AAs in TMHs: 23.72226\nExp number, first 60 AAs: 0.22038\nTotal prob of N-in: 0.12994\noutside 1 331\nTMhelix 332 354\ninside 355 372

10865 GCF_002024105.1_ASM202410v1 Escherichia coli EPEC 32/73 Proteobacteria; Gammaproteobacteria; Enterobacterales; Enterobacteriaceae; Escherichia; Escherichia coli
MTGDNTLIHSHGINRRDFMKLCAALAATMGLSSKAAAEMAESVTNPQHPPVIWIGAQECT WP_000145403.1
hydrogenase 2 small subunit [Escherichia coli] Length: 372\nNumber of predicted TMHs: 1\nExp number of AAs in TMHs: 23.66365\nExp number, first 60 AAs: 0.1537\nTotal prob of N-in: 0.12709\noutside 1 331\nTMhelix 332 354\ninside 355 372

10866 GCF_002024105.1_ASM202410v1 Escherichia coli EPEC 32/73 Proteobacteria; Gammaproteobacteria; Enterobacterales; Enterobacteriaceae; Escherichia; Escherichia coli
MNNEETFYQAMRRQGVTRRSFLKYCSLAATSLGLGAGMAPKIAWALENKPRIPVVWIHGL WP_001295940.1
MULTISPECIES: hydrogenase [Enterobacteriaceae] Length: 372\nNumber of predicted TMHs: 2\nExp number of AAs in TMHs: 37.2608899999999\nExp number, first 60 AAs: 19.52923\nTotal prob of N-in: 0.94351\nPOSSIBLE N-term signal sequence\ninside 1 19\nTMhelix 20 39\noutside 40 325\nTMhelix 326 348\ninside 349 372

10867 GCF_000268205.1_ASM26820v1 Escherichia coli EPEC C342-62 Proteobacteria; Gammaproteobacteria; Enterobacterales; Enterobacteriaceae; Escherichia; Escherichia coli
MNNEETFYQAMRRQGVTRRSFLKYCSLAATSLGLGAGMAPKIAWALENKPRIPVVWIHGL WP_001295940.1
MULTISPECIES: hydrogenase [Enterobacteriaceae] Length: 372\nNumber of predicted TMHs: 2\nExp number of AAs in TMHs: 37.2608899999999\nExp number, first 60 AAs: 19.52923\nTotal prob of N-in: 0.94351\nPOSSIBLE N-term signal sequence\ninside 1 19\nTMhelix 20 39\noutside 40 325\nTMhelix 326 348\ninside 349 372

10868 GCF_000268205.1_ASM26820v1 Escherichia coli EPEC C342-62 Proteobacteria; Gammaproteobacteria; Enterobacterales; Enterobacteriaceae; Escherichia; Escherichia coli
MTGDNTLIHSHGINRRDFMKLCAALAATMGLSSKAAAEMAESVTNPQHPPVIWIGAQECT WP_000145403.1
hydrogenase 2 small subunit [Escherichia coli] Length: 372\nNumber of predicted TMHs: 1\nExp number of AAs in TMHs: 23.66365\nExp number, first 60 AAs: 0.1537\nTotal prob of N-in: 0.12709\noutside 1 331\nTMhelix 332 354\ninside 355 372

10869 GCF_000210475.1_ASM21047v1 Escherichia coli ETEC H10407 Proteobacteria; Gammaproteobacteria; Enterobacterales; Enterobacteriaceae; Escherichia; Escherichia coli
MTGDNTLIHSHGINRRDFMKLCAALAATMGLSSKAAAEMAESVTNPQRPPVIWIGAQECT WP_000145410.1
MULTISPECIES: hydrogenase-2 small chain [Proteobacteria] Length: 372\nNumber of predicted TMHs: 1\nExp number of AAs in TMHs: 23.72226\nExp number, first 60 AAs: 0.22038\nTotal prob of N-in: 0.12994\noutside 1 331\nTMhelix 332 354\ninside 355 372

10870 GCF_000210475.1_ASM21047v1 Escherichia coli ETEC H10407 Proteobacteria; Gammaproteobacteria; Enterobacterales; Enterobacteriaceae; Escherichia; Escherichia coli
MNNEETFYQAMRRQGVTRRSFLKYCSLAATSLGLGAGMAPKIAWALENKPRIPVVWIHGL WP_014639883.1
hydrogenase [Escherichia coli] Length: 372\nNumber of predicted TMHs: 2\nExp number of AAs in TMHs: 37.26086\nExp number, first 60 AAs: 19.52923\nTotal prob of N-in: 0.94351\nPOSSIBLE N-term signal sequence\ninside 1 19\nTMhelix 20 39\noutside 40 325\nTMhelix 326 348\ninside 349 372

10871 GCF_000167835.1_ASM16783v1 Escherichia coli F11 Proteobacteria; Gammaproteobacteria; Enterobacterales; Enterobacteriaceae; Escherichia; Escherichia coli

MTGDNTLIHSHGINRRDFMKLCAALAATMGLSSKAAAEMAESVTNPQRPPVIWIGAQECT WP_000145410.1 MULTISPECIES: hydrogenase-2 small chain [Proteobacteria] Length: 372\nNumber of predicted TMHs: 1\nExp number of AAs in TMHs: 23.72226\nExp number, first 60 AAs: 0.22038\nTotal prob of N-in: 0.12994\noutside 1 331\nTMhelix 332 354\ninside 355 372

10872 GCF_000167835.1_ASM16783v1 Escherichia coli F11 Proteobacteria; Gammaproteobacteria; Enterobacterales; Enterobacteriaceae; Escherichia; Escherichia coli MNNEETFYQAMRRQGVTRRSFLKYCSLAATSLGLGAGMAPKIAWALENKPRIPVVWIHGL WP_001058323.1 MULTISPECIES: hydrogenase-1 small chain [Proteobacteria] Length: 372\nNumber of predicted TMHs: 2\nExp number of AAs in TMHs: 37.2608499999999\nExp number, first 60 AAs: 19.52923\nTotal prob of N-in: 0.94351\nPOSSIBLE N-term signal sequence\ninside 1 19\nTMhelix 20 39\noutside 40 325\nTMhelix 326 348\ninside 349 372

10873 GCF_000511465.1_EcoliFAH1 Escherichia coli FAH1 Proteobacteria; Gammaproteobacteria; Enterobacterales; Enterobacteriaceae; Escherichia; Escherichia coli MTGDNTLIHSHGINRRDFMKLCAALAATMGLSSKAAAEMAESVTNPQRPPVIWIGAQECT WP_000145410.1 MULTISPECIES: hydrogenase-2 small chain [Proteobacteria] Length: 372\nNumber of predicted TMHs: 1\nExp number of AAs in TMHs: 23.72226\nExp number, first 60 AAs: 0.22038\nTotal prob of N-in: 0.12994\noutside 1 331\nTMhelix 332 354\ninside 355 372

10874 GCF_000511465.1_EcoliFAH1 Escherichia coli FAH1 Proteobacteria; Gammaproteobacteria; Enterobacterales; Enterobacteriaceae; Escherichia; Escherichia coli MNNEETFYQAMRRQGVTRRSFLKYCSLAATSLGLGAGMAPKIAWALENKPRIPVVWIHGL WP_001058323.1 MULTISPECIES: hydrogenase-1 small chain [Proteobacteria] Length: 372\nNumber of predicted TMHs: 2\nExp number of AAs in TMHs: 37.2608499999999\nExp number, first 60 AAs: 19.52923\nTotal prob of N-in: 0.94351\nPOSSIBLE N-term signal sequence\ninside 1 19\nTMhelix 20 39\noutside 40 325\nTMhelix 326 348\ninside 349 372

10875 GCF_000511445.1_EcoliFAH2 Escherichia coli FAH2 Proteobacteria; Gammaproteobacteria; Enterobacterales; Enterobacteriaceae; Escherichia; Escherichia coli MTGDNTLIHSHGINRRDFMKLCAALAATMGLSSKAAAEMAESVTNPQRPPVIWIGAQECT WP_000145410.1 MULTISPECIES: hydrogenase-2 small chain [Proteobacteria] Length: 372\nNumber of predicted TMHs: 1\nExp number of AAs in TMHs: 23.72226\nExp number, first 60 AAs: 0.22038\nTotal prob of N-in: 0.12994\noutside 1 331\nTMhelix 332 354\ninside 355 372

10876 GCF_000511445.1_EcoliFAH2 Escherichia coli FAH2 Proteobacteria; Gammaproteobacteria; Enterobacterales; Enterobacteriaceae; Escherichia; Escherichia coli MNNEETFYQAMRRQGVTRRSFLKYCSLAATSLGLGAGMAPKIAWALENKPRIPVVWIHGL WP_001058323.1 MULTISPECIES: hydrogenase-1 small chain [Proteobacteria] Length: 372\nNumber of predicted TMHs: 2\nExp number of AAs in TMHs: 37.2608499999999\nExp number, first 60 AAs: 19.52923\nTotal prob of N-in: 0.94351\nPOSSIBLE N-term signal sequence\ninside 1 19\nTMhelix 20 39\noutside 40 325\nTMhelix 326 348\ninside 349 372

10877 GCF_000765435.1_ASM76543v1 Escherichia coli FAP1 Proteobacteria; Gammaproteobacteria; Enterobacterales; Enterobacteriaceae; Escherichia; Escherichia coli MTGDNTLIHSHGINRRDFMKLCAALAATMGLSSKAAAEMAESVTNPQRPPVIWIGAQECT WP_000145410.1 MULTISPECIES: hydrogenase-2 small chain [Proteobacteria] Length: 372\nNumber of predicted TMHs: 1\nExp number of AAs in TMHs: 23.72226\nExp number, first 60 AAs: 0.22038\nTotal prob of N-in: 0.12994\noutside 1 331\nTMhelix 332 354\ninside 355 372

10878 GCF_000511485.1_EcoliFAP1 Escherichia coli FAP1 Proteobacteria; Gammaproteobacteria; Enterobacterales; Enterobacteriaceae; Escherichia; Escherichia coli MTGDNTLIHSHGINRRDFMKLCAALAATMGLSSKAAAEMAESVTNPQRPPVIWIGAQECT WP_000145410.1 MULTISPECIES: hydrogenase-2 small chain [Proteobacteria] Length: 372\nNumber of predicted TMHs: 1\nExp number of AAs in TMHs: 23.72226\nExp number, first 60 AAs: 0.22038\nTotal prob of N-in: 0.12994\noutside 1 331\nTMhelix 332 354\ninside 355 372

10879 GCF_000765435.1_ASM76543v1 Escherichia coli FAP1 Proteobacteria; Gammaproteobacteria; Enterobacterales; Enterobacteriaceae; Escherichia; Escherichia coli MNNEETFYQAMRRQGVTRRSFLKYCSLAATSLGLGAGMAPKIAWALENKPRIPVVWIHGL WP_001058323.1 MULTISPECIES: hydrogenase-1 small chain [Proteobacteria] Length: 372\nNumber of predicted TMHs:

2\nExp number of AAs in TMHs: 37.2608499999999\nExp number, first 60 AAs: 19.52923\nTotal prob of N-in: 0.94351\nPOSSIBLE N-term signal sequence\ninside 1 19\nTMhelix 20 39\noutside 40 325\nTMhelix 326 348\ninside 349 372

10880 GCF_000511485.1_EcoliFAP1 Escherichia coli FAP1 Proteobacteria; Gammaproteobacteria; Enterobacterales; Enterobacteriaceae; Escherichia; Escherichia coli MNNEETFYQAMRRQGVTRRSFLKYCSLAATSLGLGAGMAPKIAWALENKPRIPVVWIHGL WP_001058323.1 MULTISPECIES: hydrogenase-1 small chain [Proteobacteria] Length: 372\nNumber of predicted TMHs: 2\nExp number of AAs in TMHs: 37.2608499999999\nExp number, first 60 AAs: 19.52923\nTotal prob of N-in: 0.94351\nPOSSIBLE N-term signal sequence\ninside 1 19\nTMhelix 20 39\noutside 40 325\nTMhelix 326 348\ninside 349 372

10881 GCF_000511505.1_EcoliFAP2 Escherichia coli FAP2 Proteobacteria; Gammaproteobacteria; Enterobacterales; Enterobacteriaceae; Escherichia; Escherichia coli MTGDNTLIHSHGINRRDFMKLCAALAATMGLSSKAAAEMAESVTNPQRPPVIWIGAQECT WP_000145410.1 MULTISPECIES: hydrogenase-2 small chain [Proteobacteria] Length: 372\nNumber of predicted TMHs: 1\nExp number of AAs in TMHs: 23.72226\nExp number, first 60 AAs: 0.22038\nTotal prob of N-in: 0.12994\noutside 1 331\nTMhelix 332 354\ninside 355 372

10882 GCF_000511505.1_EcoliFAP2 Escherichia coli FAP2 Proteobacteria; Gammaproteobacteria; Enterobacterales; Enterobacteriaceae; Escherichia; Escherichia coli MNNEETFYQAMRRQGVTRRSFLKYCSLAATSLGLGAGMAPKIAWALENKPRIPVVWIHGL WP_001058323.1 MULTISPECIES: hydrogenase-1 small chain [Proteobacteria] Length: 372\nNumber of predicted TMHs: 2\nExp number of AAs in TMHs: 37.2608499999999\nExp number, first 60 AAs: 19.52923\nTotal prob of N-in: 0.94351\nPOSSIBLE N-term signal sequence\ninside 1 19\nTMhelix 20 39\noutside 40 325\nTMhelix 326 348\ninside 349 372

10883 GCF_000503635.1_EcoliFBH1 Escherichia coli FBH1 Proteobacteria; Gammaproteobacteria; Enterobacterales; Enterobacteriaceae; Escherichia; Escherichia coli MTGDNTLIHSHGINRRDFMKLCAALAATMGLSSKAAAEMAESVTNPQRPPVIWIGAQECT WP_029487472.1 hydrogenase 2 small subunit [Escherichia coli] Length: 372\nNumber of predicted TMHs: 1\nExp number of AAs in TMHs: 22.56987\nExp number, first 60 AAs: 0.23233\nTotal prob of N-in: 0.08221\noutside 1 331\nTMhelix 332 354\ninside 355 372

10884 GCF_000503635.1_EcoliFBH1 Escherichia coli FBH1 Proteobacteria; Gammaproteobacteria; Enterobacterales; Enterobacteriaceae; Escherichia; Escherichia coli MNNEETFYQAMRRQGVTRRSFLKYCSLAATSLGLGAGMAPKIAWALENKPRIPVVWIHGL WP_001058323.1 MULTISPECIES: hydrogenase-1 small chain [Proteobacteria] Length: 372\nNumber of predicted TMHs: 2\nExp number of AAs in TMHs: 37.2608499999999\nExp number, first 60 AAs: 19.52923\nTotal prob of N-in: 0.94351\nPOSSIBLE N-term signal sequence\ninside 1 19\nTMhelix 20 39\noutside 40 325\nTMhelix 326 348\ninside 349 372

10885 GCF_000511525.1_EcoliFBP1 Escherichia coli FBP1 Proteobacteria; Gammaproteobacteria; Enterobacterales; Enterobacteriaceae; Escherichia; Escherichia coli MNNEETFYQAMRRQGVTRRSFLKYCSLAATSLGLGAGMAPKIAWALENKPRIPVVWIHGL WP_001058323.1 MULTISPECIES: hydrogenase-1 small chain [Proteobacteria] Length: 372\nNumber of predicted TMHs: 2\nExp number of AAs in TMHs: 37.2608499999999\nExp number, first 60 AAs: 19.52923\nTotal prob of N-in: 0.94351\nPOSSIBLE N-term signal sequence\ninside 1 19\nTMhelix 20 39\noutside 40 325\nTMhelix 326 348\ninside 349 372

10886 GCF_000511525.1_EcoliFBP1 Escherichia coli FBP1 Proteobacteria; Gammaproteobacteria; Enterobacterales; Enterobacteriaceae; Escherichia; Escherichia coli MTGDNTLIHSHGINRRDFMKLCAALAATMGLSSKAAAEMAESVTNPQRPPVIWIGAQECT WP_000145410.1 MULTISPECIES: hydrogenase-2 small chain [Proteobacteria] Length: 372\nNumber of predicted TMHs: 1\nExp number of AAs in TMHs: 23.72226\nExp number, first 60 AAs: 0.22038\nTotal prob of N-in: 0.12994\noutside 1 331\nTMhelix 332 354\ninside 355 372

10887 GCF_000511545.1_EcoliFCH1 Escherichia coli FCH1 Proteobacteria; Gammaproteobacteria; Enterobacterales; Enterobacteriaceae; Escherichia; Escherichia coli MNNEETFYQAMRRQGVTRRSFLKYCSLAATSLGLGAGMAPKIAWALENKPRIPVVWIHGL WP_001058323.1 MULTISPECIES: hydrogenase-1 small chain [Proteobacteria] Length: 372\nNumber of predicted TMHs:

2\nExp number of AAs in TMHs: 37.2608499999999\nExp number, first 60 AAs: 19.52923\nTotal prob of N-in: 0.94351\nPOSSIBLE N-term signal sequence\ninside 1 19\nTMhelix 20 39\noutside 40 325\nTMhelix 326 348\ninside 349 372

10888 GCF_000511545.1_EcoliFCH1 Escherichia coli FCH1 Proteobacteria; Gammaproteobacteria; Enterobacterales; Enterobacteriaceae; Escherichia; Escherichia coli MTGDNTLIHSHGINRRDFMKLCAALAATMGLSSKAAAEMAESVTNPQRPPVIWIGAQECT WP_000145410.1 MULTISPECIES: hydrogenase-2 small chain [Proteobacteria] Length: 372\nNumber of predicted TMHs: 1\nExp number of AAs in TMHs: 23.72226\nExp number, first 60 AAs: 0.22038\nTotal prob of N-in: 0.12994\noutside 1 331\nTMhelix 332 354\ninside 355 372

10889 GCF_000511565.1_EcoliFCP1 Escherichia coli FCP1 Proteobacteria; Gammaproteobacteria; Enterobacterales; Enterobacteriaceae; Escherichia; Escherichia coli MTGDNTLIHSHGINRRDFMKLCAALAATMGLSSKAAAEMAESVTNPQRPPVIWIGAQECT WP_000145410.1 MULTISPECIES: hydrogenase-2 small chain [Proteobacteria] Length: 372\nNumber of predicted TMHs: 1\nExp number of AAs in TMHs: 23.72226\nExp number, first 60 AAs: 0.22038\nTotal prob of N-in: 0.12994\noutside 1 331\nTMhelix 332 354\ninside 355 372

10890 GCF_000511565.1_EcoliFCP1 Escherichia coli FCP1 Proteobacteria; Gammaproteobacteria; Enterobacterales; Enterobacteriaceae; Escherichia; Escherichia coli MNNEETFYQAMRRQGVTRRSFLKYCSLAATSLGLGAGMAPKIAWALENKPRIPVVWIHGL WP_001058323.1 MULTISPECIES: hydrogenase-1 small chain [Proteobacteria] Length: 372\nNumber of predicted TMHs: 2\nExp number of AAs in TMHs: 37.2608499999999\nExp number, first 60 AAs: 19.52923\nTotal prob of N-in: 0.94351\nPOSSIBLE N-term signal sequence\ninside 1 19\nTMhelix 20 39\noutside 40 325\nTMhelix 326 348\ninside 349 372

10891 GCF_000303755.2_ASM30375v2 Escherichia coli FDA504 Proteobacteria; Gammaproteobacteria; Enterobacterales; Enterobacteriaceae; Escherichia; Escherichia coli MNNEETFYQAMRRQGVTRRSFLKYCSLAATSLGLGAGMAPKIAWALENKPRIPVVWIHGL WP_001301613.1 hydrogenase [Escherichia coli] Length: 372\nNumber of predicted TMHs: 2\nExp number of AAs in TMHs: 37.46598\nExp number, first 60 AAs: 19.53139\nTotal prob of N-in: 0.94364\nPOSSIBLE N-term signal sequence\ninside 1 19\nTMhelix 20 39\noutside 40 325\nTMhelix 326 348\ninside 349 372

10892 GCF_000303755.2_ASM30375v2 Escherichia coli FDA504 Proteobacteria; Gammaproteobacteria; Enterobacterales; Enterobacteriaceae; Escherichia; Escherichia coli MTGDNTLIHSHGINRRDFMKLCAALAATMGLSSKAAAEMAESVTNPQRPPVIWIGAQECT WP_000145410.1 MULTISPECIES: hydrogenase-2 small chain [Proteobacteria] Length: 372\nNumber of predicted TMHs: 1\nExp number of AAs in TMHs: 23.72226\nExp number, first 60 AAs: 0.22038\nTotal prob of N-in: 0.12994\noutside 1 331\nTMhelix 332 354\ninside 355 372

10893 GCF_000267005.2_ASM26700v2 Escherichia coli FDA505 Proteobacteria; Gammaproteobacteria; Enterobacterales; Enterobacteriaceae; Escherichia; Escherichia coli MNNEETFYQAMRRQGVTRRSFLKYCSLAATSLGLGAGMAPKIAWALENKPRIPVVWIHGL WP_001301613.1 hydrogenase [Escherichia coli] Length: 372\nNumber of predicted TMHs: 2\nExp number of AAs in TMHs: 37.46598\nExp number, first 60 AAs: 19.53139\nTotal prob of N-in: 0.94364\nPOSSIBLE N-term signal sequence\ninside 1 19\nTMhelix 20 39\noutside 40 325\nTMhelix 326 348\ninside 349 372

10894 GCF_000267005.2_ASM26700v2 Escherichia coli FDA505 Proteobacteria; Gammaproteobacteria; Enterobacterales; Enterobacteriaceae; Escherichia; Escherichia coli MTGDNTLIHSHGINRRDFMKLCAALAATMGLSSKAAAEMAESVTNPQRPPVIWIGAQECT WP_000145410.1 MULTISPECIES: hydrogenase-2 small chain [Proteobacteria] Length: 372\nNumber of predicted TMHs: 1\nExp number of AAs in TMHs: 23.72226\nExp number, first 60 AAs: 0.22038\nTotal prob of N-in: 0.12994\noutside 1 331\nTMhelix 332 354\ninside 355 372

10895 GCF_000303715.2_ASM30371v2 Escherichia coli FDA506 Proteobacteria; Gammaproteobacteria; Enterobacterales; Enterobacteriaceae; Escherichia; Escherichia coli MTGDNTLIHSHGINRRDFMKLCAALAATMGLSSKAAAEMAESVTNPQRPPVIWIGAQECT WP_000145410.1 MULTISPECIES: hydrogenase-2 small chain [Proteobacteria] Length: 372\nNumber of predicted TMHs: 1\nExp number of AAs in TMHs: 23.72226\nExp number, first 60 AAs: 0.22038\nTotal prob of N-in: 0.12994\noutside 1 331\nTMhelix 332 354\ninside 355 372

10896 GCF_000303715.2_ASM30371v2 Escherichia coli FDA506 Proteobacteria; Gammaproteobacteria; Enterobacterales; Enterobacteriaceae; Escherichia; Escherichia coli
MNNEETFYQAMRRQGVTRRSFLKYCSLAATSLGLGAGMAPKIAWALENKPRIPVVWIHGL WP_001301613.1
hydrogenase [Escherichia coli] Length: 372\nNumber of predicted TMHs: 2\nExp number of AAs in TMHs: 37.46598\nExp number, first 60 AAs: 19.53139\nTotal prob of N-in: 0.94364\nPOSSIBLE N-term signal sequence\ninside 1 19\nTMhelix 20 39\noutside 40 325\nTMhelix 326 348\ninside 349 372

10897 GCF_000303735.2_ASM30373v2 Escherichia coli FDA507 Proteobacteria; Gammaproteobacteria; Enterobacterales; Enterobacteriaceae; Escherichia; Escherichia coli
MTGDNTLIHSHGINRRDFMKLCAALAATMGLSSKAAAEMAESVTNPQRPPVIWIGAQECT WP_000145410.1
MULTISPECIES: hydrogenase-2 small chain [Proteobacteria] Length: 372\nNumber of predicted TMHs: 1\nExp number of AAs in TMHs: 23.72226\nExp number, first 60 AAs: 0.22038\nTotal prob of N-in: 0.12994\noutside 1 331\nTMhelix 332 354\ninside 355 372

10898 GCF_000303735.2_ASM30373v2 Escherichia coli FDA507 Proteobacteria; Gammaproteobacteria; Enterobacterales; Enterobacteriaceae; Escherichia; Escherichia coli
MNNEETFYQAMRRQGVTRRSFLKYCSLAATSLGLGAGMAPKIAWALENKPRIPVVWIHGL WP_001301613.1
hydrogenase [Escherichia coli] Length: 372\nNumber of predicted TMHs: 2\nExp number of AAs in TMHs: 37.46598\nExp number, first 60 AAs: 19.53139\nTotal prob of N-in: 0.94364\nPOSSIBLE N-term signal sequence\ninside 1 19\nTMhelix 20 39\noutside 40 325\nTMhelix 326 348\ninside 349 372

10899 GCF_000267485.2_ASM26748v2 Escherichia coli FDA517 Proteobacteria; Gammaproteobacteria; Enterobacterales; Enterobacteriaceae; Escherichia; Escherichia coli
MNNEETFYQAMRRQGVTRRSFLKYCSLAATSLGLGAGMAPKIAWALENKPRIPVVWIHGL WP_001301613.1
hydrogenase [Escherichia coli] Length: 372\nNumber of predicted TMHs: 2\nExp number of AAs in TMHs: 37.46598\nExp number, first 60 AAs: 19.53139\nTotal prob of N-in: 0.94364\nPOSSIBLE N-term signal sequence\ninside 1 19\nTMhelix 20 39\noutside 40 325\nTMhelix 326 348\ninside 349 372

10900 GCF_000267485.2_ASM26748v2 Escherichia coli FDA517 Proteobacteria; Gammaproteobacteria; Enterobacterales; Enterobacteriaceae; Escherichia; Escherichia coli
MTGDNTLIHSHGINRRDFMKLCAALAATMGLSSKAAAEMAESVTNPQRPPVIWIGAQECT WP_000145410.1
MULTISPECIES: hydrogenase-2 small chain [Proteobacteria] Length: 372\nNumber of predicted TMHs: 1\nExp number of AAs in TMHs: 23.72226\nExp number, first 60 AAs: 0.22038\nTotal prob of N-in: 0.12994\noutside 1 331\nTMhelix 332 354\ninside 355 372

10901 GCF_000267525.2_ASM26752v2 Escherichia coli FRIK1985 Proteobacteria; Gammaproteobacteria; Enterobacterales; Enterobacteriaceae; Escherichia; Escherichia coli
MTGDNTLIHSHGINRRDFMKLCAALAATMGLSSKAAAEMAESVTNPQRPPVIWIGAQECT WP_000145410.1
MULTISPECIES: hydrogenase-2 small chain [Proteobacteria] Length: 372\nNumber of predicted TMHs: 1\nExp number of AAs in TMHs: 23.72226\nExp number, first 60 AAs: 0.22038\nTotal prob of N-in: 0.12994\noutside 1 331\nTMhelix 332 354\ninside 355 372

10902 GCF_000267525.2_ASM26752v2 Escherichia coli FRIK1985 Proteobacteria; Gammaproteobacteria; Enterobacterales; Enterobacteriaceae; Escherichia; Escherichia coli
MNNEETFYQAMRRQGVTRRSFLKYCSLAATSLGLGAGMAPKIAWALENKPRIPVVWIHGL WP_001301613.1
hydrogenase [Escherichia coli] Length: 372\nNumber of predicted TMHs: 2\nExp number of AAs in TMHs: 37.46598\nExp number, first 60 AAs: 19.53139\nTotal prob of N-in: 0.94364\nPOSSIBLE N-term signal sequence\ninside 1 19\nTMhelix 20 39\noutside 40 325\nTMhelix 326 348\ninside 349 372

10903 GCF_000267965.2_ASM26796v2 Escherichia coli FRIK1990 Proteobacteria; Gammaproteobacteria; Enterobacterales; Enterobacteriaceae; Escherichia; Escherichia coli
MNNEETFYQAMRRQGVTRRSFLKYCSLAATSLGLGAGMAPKIAWALENKPRIPVVWIHGL WP_001301613.1
hydrogenase [Escherichia coli] Length: 372\nNumber of predicted TMHs: 2\nExp number of AAs in TMHs: 37.46598\nExp number, first 60 AAs: 19.53139\nTotal prob of N-in: 0.94364\nPOSSIBLE N-term signal sequence\ninside 1 19\nTMhelix 20 39\noutside 40 325\nTMhelix 326 348\ninside 349 372

10904 GCF_000267965.2_ASM26796v2 Escherichia coli FRIK1990 Proteobacteria; Gammaproteobacteria; Enterobacterales; Enterobacteriaceae; Escherichia; Escherichia coli
MTGDNTLIHSHGINRRDFMKLCAALAATMGLSSKAAAEMAESVTNPQRPPVIWIGAQECT WP_000145410.1
MULTISPECIES: hydrogenase-2 small chain [Proteobacteria] Length: 372\nNumber of predicted TMHs:

1\nExp number of AAs in TMHs: 23.72226\nExp number, first 60 AAs: 0.22038\nTotal prob of N-in: 0.12994\noutside 1 331\nTMhelix 332 354\ninside 355 372

10905 GCF_000267505.2_ASM26750v2 Escherichia coli FRIK1996 Proteobacteria; Gammaproteobacteria; Enterobacterales; Enterobacteriaceae; Escherichia; Escherichia coli
MTGDNTLIHSHGINRRDFMKLCAALAATMGLSSKAAAEMAESVTNPQRPPVIWIGAQECT WP_000145410.1
MULTISPECIES: hydrogenase-2 small chain [Proteobacteria] Length: 372\nNumber of predicted TMHs: 1\nExp number of AAs in TMHs: 23.72226\nExp number, first 60 AAs: 0.22038\nTotal prob of N-in: 0.12994\noutside 1 331\nTMhelix 332 354\ninside 355 372

10906 GCF_000267505.2_ASM26750v2 Escherichia coli FRIK1996 Proteobacteria; Gammaproteobacteria; Enterobacterales; Enterobacteriaceae; Escherichia; Escherichia coli
MNNEETFYQAMRRQGVTRRSFLKYCSLAATSLGLGAGMAPKIAWALENKPRIPVVWIHGL WP_001301613.1
hydrogenase [Escherichia coli] Length: 372\nNumber of predicted TMHs: 2\nExp number of AAs in TMHs: 37.46598\nExp number, first 60 AAs: 19.53139\nTotal prob of N-in: 0.94364\nPOSSIBLE N-term signal sequence\ninside 1 19\nTMhelix 20 39\noutside 40 325\nTMhelix 326 348\ninside 349 372

10907 GCF_000303795.2_ASM30379v2 Escherichia coli FRIK1997 Proteobacteria; Gammaproteobacteria; Enterobacterales; Enterobacteriaceae; Escherichia; Escherichia coli
MTGDNTLIHSHGINRRDFMKLCAALAATMGLSSKAAAEMAESVTNPQRPPVIWIGAQECT WP_000145410.1
MULTISPECIES: hydrogenase-2 small chain [Proteobacteria] Length: 372\nNumber of predicted TMHs: 1\nExp number of AAs in TMHs: 23.72226\nExp number, first 60 AAs: 0.22038\nTotal prob of N-in: 0.12994\noutside 1 331\nTMhelix 332 354\ninside 355 372

10908 GCF_000303795.2_ASM30379v2 Escherichia coli FRIK1997 Proteobacteria; Gammaproteobacteria; Enterobacterales; Enterobacteriaceae; Escherichia; Escherichia coli
MNNEETFYQAMRRQGVTRRSFLKYCSLAATSLGLGAGMAPKIAWALENKPRIPVVWIHGL WP_001301613.1
hydrogenase [Escherichia coli] Length: 372\nNumber of predicted TMHs: 2\nExp number of AAs in TMHs: 37.46598\nExp number, first 60 AAs: 19.53139\nTotal prob of N-in: 0.94364\nPOSSIBLE N-term signal sequence\ninside 1 19\nTMhelix 20 39\noutside 40 325\nTMhelix 326 348\ninside 349 372

10909 GCF_000303775.2_ASM30377v2 Escherichia coli FRIK1999 Proteobacteria; Gammaproteobacteria; Enterobacterales; Enterobacteriaceae; Escherichia; Escherichia coli
MTGDNTLIHSHGINRRDFMKLCAALAATMGLSSKAAAEMAESVTNPQRPPVIWIGAQECT WP_000145410.1
MULTISPECIES: hydrogenase-2 small chain [Proteobacteria] Length: 372\nNumber of predicted TMHs: 1\nExp number of AAs in TMHs: 23.72226\nExp number, first 60 AAs: 0.22038\nTotal prob of N-in: 0.12994\noutside 1 331\nTMhelix 332 354\ninside 355 372

10910 GCF_000303775.2_ASM30377v2 Escherichia coli FRIK1999 Proteobacteria; Gammaproteobacteria; Enterobacterales; Enterobacteriaceae; Escherichia; Escherichia coli
MNNEETFYQAMRRQGVTRRSFLKYCSLAATSLGLGAGMAPKIAWALENKPRIPVVWIHGL WP_001301613.1
hydrogenase [Escherichia coli] Length: 372\nNumber of predicted TMHs: 2\nExp number of AAs in TMHs: 37.46598\nExp number, first 60 AAs: 19.53139\nTotal prob of N-in: 0.94364\nPOSSIBLE N-term signal sequence\ninside 1 19\nTMhelix 20 39\noutside 40 325\nTMhelix 326 348\ninside 349 372

10911 GCF_000303855.2_ASM30385v2 Escherichia coli FRIK2001 Proteobacteria; Gammaproteobacteria; Enterobacterales; Enterobacteriaceae; Escherichia; Escherichia coli
MNNEETFYQAMRRQGVTRRSFLKYCSLAATSLGLGAGMAPKIAWALENKPRIPVVWIHGL WP_001301613.1
hydrogenase [Escherichia coli] Length: 372\nNumber of predicted TMHs: 2\nExp number of AAs in TMHs: 37.46598\nExp number, first 60 AAs: 19.53139\nTotal prob of N-in: 0.94364\nPOSSIBLE N-term signal sequence\ninside 1 19\nTMhelix 20 39\noutside 40 325\nTMhelix 326 348\ninside 349 372

10912 GCF_000303855.2_ASM30385v2 Escherichia coli FRIK2001 Proteobacteria; Gammaproteobacteria; Enterobacterales; Enterobacteriaceae; Escherichia; Escherichia coli
MTGDNTLIHSHGINRRDFMKLCAALAATMGLSSKAAAEMAESVTNPQRPPVIWIGAQECT WP_000145410.1
MULTISPECIES: hydrogenase-2 small chain [Proteobacteria] Length: 372\nNumber of predicted TMHs: 1\nExp number of AAs in TMHs: 23.72226\nExp number, first 60 AAs: 0.22038\nTotal prob of N-in: 0.12994\noutside 1 331\nTMhelix 332 354\ninside 355 372

10913 GCF_000303275.2_ASM30327v2 Escherichia coli FRIK523 Proteobacteria; Gammaproteobacteria; Enterobacterales; Enterobacteriaceae; Escherichia; Escherichia coli

MTGDNTLIHSHGINRRDFMKLCAALAATMGLSSKAAAEMAESVTNPQRPPVIWIGAQECT WP_000145410.1 MULTISPECIES: hydrogenase-2 small chain [Proteobacteria] Length: 372\nNumber of predicted TMHs: 1\nExp number of AAs in TMHs: 23.72226\nExp number, first 60 AAs: 0.22038\nTotal prob of N-in: 0.12994\noutside 1 331\nTMhelix 332 354\ninside 355 372

10914 GCF_000303275.2_ASM30327v2 Escherichia coli FRIK523 Proteobacteria; Gammaproteobacteria; Enterobacterales; Enterobacteriaceae; Escherichia; Escherichia coli MNNEETFYQAMRRQGVTRRSFLKYCSLAATSLGLGAGMAPKIAWALENKPRIPVVWIHGL WP_001301613.1 hydrogenase [Escherichia coli] Length: 372\nNumber of predicted TMHs: 2\nExp number of AAs in TMHs: 37.46598\nExp number, first 60 AAs: 19.53139\nTotal prob of N-in: 0.94364\nPOSSIBLE N-term signal sequence\ninside 1 19\nTMhelix 20 39\noutside 40 325\nTMhelix 326 348\ninside 349 372

10915 GCF_000302715.1_ASM30271v1 Escherichia coli FRIK920 Proteobacteria; Gammaproteobacteria; Enterobacterales; Enterobacteriaceae; Escherichia; Escherichia coli MTGDNTLIHSHGINRRDFMKLCAALAATMGLSSKAAAEMAESVTNPQRPPVIWIGAQECT WP_000145410.1 MULTISPECIES: hydrogenase-2 small chain [Proteobacteria] Length: 372\nNumber of predicted TMHs: 1\nExp number of AAs in TMHs: 23.72226\nExp number, first 60 AAs: 0.22038\nTotal prob of N-in: 0.12994\noutside 1 331\nTMhelix 332 354\ninside 355 372

10916 GCF_000302715.1_ASM30271v1 Escherichia coli FRIK920 Proteobacteria; Gammaproteobacteria; Enterobacterales; Enterobacteriaceae; Escherichia; Escherichia coli MNNEETFYQAMRRQGVTRRSFLKYCSLAATSLGLGAGMAPKIAWALENKPRIPVVWIHGL WP_001301613.1 hydrogenase [Escherichia coli] Length: 372\nNumber of predicted TMHs: 2\nExp number of AAs in TMHs: 37.46598\nExp number, first 60 AAs: 19.53139\nTotal prob of N-in: 0.94364\nPOSSIBLE N-term signal sequence\ninside 1 19\nTMhelix 20 39\noutside 40 325\nTMhelix 326 348\ninside 349 372

10917 GCF_000163215.1_ASM16321v1 Escherichia coli FVEC1302 Proteobacteria; Gammaproteobacteria; Enterobacterales; Enterobacteriaceae; Escherichia; Escherichia coli MTGDNTLIHSHGINRRDFMKLCAALAATMGLSSKAAAEMAESVTNPQRPPVIWIGAQECT WP_000145410.1 MULTISPECIES: hydrogenase-2 small chain [Proteobacteria] Length: 372\nNumber of predicted TMHs: 1\nExp number of AAs in TMHs: 23.72226\nExp number, first 60 AAs: 0.22038\nTotal prob of N-in: 0.12994\noutside 1 331\nTMhelix 332 354\ninside 355 372

10918 GCF_000163215.1_ASM16321v1 Escherichia coli FVEC1302 Proteobacteria; Gammaproteobacteria; Enterobacterales; Enterobacteriaceae; Escherichia; Escherichia coli MNNEETFYQAMRRQGVTRRSFLKYCSLAATSLGLGAGMAPKIAWALENKPRIPVVWIHGL WP_001058323.1 MULTISPECIES: hydrogenase-1 small chain [Proteobacteria] Length: 372\nNumber of predicted TMHs: 2\nExp number of AAs in TMHs: 37.2608499999999\nExp number, first 60 AAs: 19.52923\nTotal prob of N-in: 0.94351\nPOSSIBLE N-term signal sequence\ninside 1 19\nTMhelix 20 39\noutside 40 325\nTMhelix 326 348\ninside 349 372

10919 GCF_000163235.1_ASM16323v1 Escherichia coli FVEC1412 Proteobacteria; Gammaproteobacteria; Enterobacterales; Enterobacteriaceae; Escherichia; Escherichia coli MNNEETFYQAMRRQGVTRRSFLKYCSLAATSLGLGAGMAPKIAWALENKPRIPVVWIHGL WP_001058323.1 MULTISPECIES: hydrogenase-1 small chain [Proteobacteria] Length: 372\nNumber of predicted TMHs: 2\nExp number of AAs in TMHs: 37.2608499999999\nExp number, first 60 AAs: 19.52923\nTotal prob of N-in: 0.94351\nPOSSIBLE N-term signal sequence\ninside 1 19\nTMhelix 20 39\noutside 40 325\nTMhelix 326 348\ninside 349 372

10920 GCF_000163235.1_ASM16323v1 Escherichia coli FVEC1412 Proteobacteria; Gammaproteobacteria; Enterobacterales; Enterobacteriaceae; Escherichia; Escherichia coli MTGDNTLIHSHGINRRDFMKLCAALAATMGLSSKAAAEMAESVTNPQRPPVIWIGAQECT WP_000145410.1 MULTISPECIES: hydrogenase-2 small chain [Proteobacteria] Length: 372\nNumber of predicted TMHs: 1\nExp number of AAs in TMHs: 23.72226\nExp number, first 60 AAs: 0.22038\nTotal prob of N-in: 0.12994\noutside 1 331\nTMhelix 332 354\ninside 355 372

10921 GCF_002109565.1_ASM210956v1 Escherichia coli FVEC1465 Proteobacteria; Gammaproteobacteria; Enterobacterales; Enterobacteriaceae; Escherichia; Escherichia coli MNNEETFYQAMRRQGVTRRSFLKYCSLAATSLGLGAGMAPKIAWALENKPRIPVVWIHGL WP_001058323.1 MULTISPECIES: hydrogenase-1 small chain [Proteobacteria] Length: 372\nNumber of predicted TMHs: 2\nExp number of AAs in TMHs: 37.2608499999999\nExp number, first 60 AAs: 19.52923\nTotal prob of N-in:

0.94351\nPOSSIBLE N-term signal sequence\ninside 1 19\nTMhelix 20 39\noutside 40 325\nTMhelix 326 348\ninside 349 372

10922 GCF_002109565.1_ASM210956v1 Escherichia coli FVEC1465 Proteobacteria; Gammaproteobacteria; Enterobacterales; Enterobacteriaceae; Escherichia; Escherichia coli MTGDNTLIHSHGINRRDFMKLCAALAATMGLSSKAAAEMAESVTNPQRPPVIWIGAQECT WP_000145410.1 MULTISPECIES: hydrogenase-2 small chain [Proteobacteria] Length: 372\nNumber of predicted TMHs: 1\nExp number of AAs in TMHs: 23.72226\nExp number, first 60 AAs: 0.22038\nTotal prob of N-in: 0.12994\noutside 1 331\nTMhelix 332 354\ninside 355 372

10923 GCF_000768425.1_ASM76842v1 Escherichia coli G3/10 Proteobacteria; Gammaproteobacteria; Enterobacterales; Enterobacteriaceae; Escherichia; Escherichia coli MTGDNTLIHSHGINRRDFMKLCAALAATMGLSSKAAAEMAESVTNPQRPPVIWIGAQECT WP_000145410.1 MULTISPECIES: hydrogenase-2 small chain [Proteobacteria] Length: 372\nNumber of predicted TMHs: 1\nExp number of AAs in TMHs: 23.72226\nExp number, first 60 AAs: 0.22038\nTotal prob of N-in: 0.12994\noutside 1 331\nTMhelix 332 354\ninside 355 372

10924 GCF_000768425.1_ASM76842v1 Escherichia coli G3/10 Proteobacteria; Gammaproteobacteria; Enterobacterales; Enterobacteriaceae; Escherichia; Escherichia coli MNNEETFYQAMRRQGVTRRSFLKYCSLAATSLGLGAGMAPKIAWALENKPRIPVVWIHGL WP_001058323.1 MULTISPECIES: hydrogenase-1 small chain [Proteobacteria] Length: 372\nNumber of predicted TMHs: 2\nExp number of AAs in TMHs: 37.2608499999999\nExp number, first 60 AAs: 19.52923\nTotal prob of N-in: 0.94351\nPOSSIBLE N-term signal sequence\ninside 1 19\nTMhelix 20 39\noutside 40 325\nTMhelix 326 348\ninside 349 372

10925 GCF_000225165.1_ASM22516v2 Escherichia coli G58-1 Proteobacteria; Gammaproteobacteria; Enterobacterales; Enterobacteriaceae; Escherichia; Escherichia coli MTGDNTLIHSHGINRRDFMKLCAALAATMGLSSKAAAEMAESVTNPQRPPVIWIGAQECT WP_000145410.1 MULTISPECIES: hydrogenase-2 small chain [Proteobacteria] Length: 372\nNumber of predicted TMHs: 1\nExp number of AAs in TMHs: 23.72226\nExp number, first 60 AAs: 0.22038\nTotal prob of N-in: 0.12994\noutside 1 331\nTMhelix 332 354\ninside 355 372

10926 GCF_000225165.1_ASM22516v2 Escherichia coli G58-1 Proteobacteria; Gammaproteobacteria; Enterobacterales; Enterobacteriaceae; Escherichia; Escherichia coli MNNEETFYQAMRRQGVTRRSFLKYCSLAATSLGLGAGMAPKIAWALENKPRIPVVWIHGL WP_001058323.1 MULTISPECIES: hydrogenase-1 small chain [Proteobacteria] Length: 372\nNumber of predicted TMHs: 2\nExp number of AAs in TMHs: 37.2608499999999\nExp number, first 60 AAs: 19.52923\nTotal prob of N-in: 0.94351\nPOSSIBLE N-term signal sequence\ninside 1 19\nTMhelix 20 39\noutside 40 325\nTMhelix 326 348\ninside 349 372

10927 GCF_000500835.1_GSK2022 Escherichia coli GSK2022 Proteobacteria; Gammaproteobacteria; Enterobacterales; Enterobacteriaceae; Escherichia; Escherichia coli MTGDNTLIHSHGINRRDFMKLCAALAATMGLSSKAAAEMVESVTNPQRPPVIWIGAQECT WP_000145423.1 hydrogenase 2 small subunit [Escherichia coli] Length: 372\nNumber of predicted TMHs: 1\nExp number of AAs in TMHs: 23.78201\nExp number, first 60 AAs: 0.28355\nTotal prob of N-in: 0.13257\noutside 1 331\nTMhelix 332 354\ninside 355 372

10928 GCF_000500835.1_GSK2022 Escherichia coli GSK2022 Proteobacteria; Gammaproteobacteria; Enterobacterales; Enterobacteriaceae; Escherichia; Escherichia coli MNNEETFYQAMRRQGVTRRSFLKYCSLAATSLGLGAGMAPKIAWALENKPRIPVVWIHGL WP_001058323.1 MULTISPECIES: hydrogenase-1 small chain [Proteobacteria] Length: 372\nNumber of predicted TMHs: 2\nExp number of AAs in TMHs: 37.2608499999999\nExp number, first 60 AAs: 19.52923\nTotal prob of N-in: 0.94351\nPOSSIBLE N-term signal sequence\ninside 1 19\nTMhelix 20 39\noutside 40 325\nTMhelix 326 348\ninside 349 372

10929 GCF_000500855.1_GSK2023 Escherichia coli GSK2023 Proteobacteria; Gammaproteobacteria; Enterobacterales; Enterobacteriaceae; Escherichia; Escherichia coli MNNEETFYQAMRRQGVTRRSFLKYCSLAATSLGLGAGMAPKIAWALENKPRIPVVWIHGL WP_001058323.1 MULTISPECIES: hydrogenase-1 small chain [Proteobacteria] Length: 372\nNumber of predicted TMHs: 2\nExp number of AAs in TMHs: 37.2608499999999\nExp number, first 60 AAs: 19.52923\nTotal prob of N-in:

0.94351\nPOSSIBLE N-term signal sequence\ninside 1 19\nTMhelix 20 39\noutside 40 325\nTMhelix 326 348\ninside 349 372

10930 GCF_000500855.1_GSK2023 Escherichia coli GSK2023 Proteobacteria; Gammaproteobacteria; Enterobacterales; Enterobacteriaceae; Escherichia; Escherichia coli
MTGDNTLIHSHGINRRDFMKLCAALAATMGLSSKAAAEMVESVTNPQRPPVIWIGAQECT WP_000145423.1
hydrogenase 2 small subunit [Escherichia coli] Length: 372\nNumber of predicted TMHs: 1\nExp number of AAs in TMHs: 23.78201\nExp number, first 60 AAs: 0.28355\nTotal prob of N-in: 0.13257\noutside 1 331\nTMhelix 332 354\ninside 355 372

10931 GCF_000500875.1_GSK2024 Escherichia coli GSK2024 Proteobacteria; Gammaproteobacteria; Enterobacterales; Enterobacteriaceae; Escherichia; Escherichia coli
MNNEETFYQAMRRQGVTRRSFLKYCSLAATSLGLGAGMAPKIAWALENKPRIPVVWIHGL WP_001058323.1
MULTISPECIES: hydrogenase-1 small chain [Proteobacteria] Length: 372\nNumber of predicted TMHs: 2\nExp number of AAs in TMHs: 37.2608499999999\nExp number, first 60 AAs: 19.52923\nTotal prob of N-in: 0.94351\nPOSSIBLE N-term signal sequence\ninside 1 19\nTMhelix 20 39\noutside 40 325\nTMhelix 326 348\ninside 349 372

10932 GCF_000500875.1_GSK2024 Escherichia coli GSK2024 Proteobacteria; Gammaproteobacteria; Enterobacterales; Enterobacteriaceae; Escherichia; Escherichia coli
MTGDNTLIHSHGINRRDFMKLCAALAATMGLSSKAAAEMVESVTNPQRPPVIWIGAQECT WP_000145423.1
hydrogenase 2 small subunit [Escherichia coli] Length: 372\nNumber of predicted TMHs: 1\nExp number of AAs in TMHs: 23.78201\nExp number, first 60 AAs: 0.28355\nTotal prob of N-in: 0.13257\noutside 1 331\nTMhelix 332 354\ninside 355 372

10933 GCF_000500895.1_GSK202B Escherichia coli GSK202B Proteobacteria; Gammaproteobacteria; Enterobacterales; Enterobacteriaceae; Escherichia; Escherichia coli
MNNEETFYQAMRRQGVTRRSFLKYCSLAATSLGLGAGMAPKIAWALENKPRIPVVWIHGL WP_001058323.1
MULTISPECIES: hydrogenase-1 small chain [Proteobacteria] Length: 372\nNumber of predicted TMHs: 2\nExp number of AAs in TMHs: 37.2608499999999\nExp number, first 60 AAs: 19.52923\nTotal prob of N-in: 0.94351\nPOSSIBLE N-term signal sequence\ninside 1 19\nTMhelix 20 39\noutside 40 325\nTMhelix 326 348\ninside 349 372

10934 GCF_000500895.1_GSK202B Escherichia coli GSK202B Proteobacteria; Gammaproteobacteria; Enterobacterales; Enterobacteriaceae; Escherichia; Escherichia coli
MTGDNTLIHSHGINRRDFMKLCAALAATMGLSSKAAAEMVESVTNPQRPPVIWIGAQECT WP_000145423.1
hydrogenase 2 small subunit [Escherichia coli] Length: 372\nNumber of predicted TMHs: 1\nExp number of AAs in TMHs: 23.78201\nExp number, first 60 AAs: 0.28355\nTotal prob of N-in: 0.13257\noutside 1 331\nTMhelix 332 354\ninside 355 372

10935 GCF_000500815.1_GSK202U Escherichia coli GSK202U Proteobacteria; Gammaproteobacteria; Enterobacterales; Enterobacteriaceae; Escherichia; Escherichia coli
MTGDNTLIHSHGINRRDFMKLCAALAATMGLSSKAAAEMVESVTNPQRPPVIWIGAQECT WP_000145423.1
hydrogenase 2 small subunit [Escherichia coli] Length: 372\nNumber of predicted TMHs: 1\nExp number of AAs in TMHs: 23.78201\nExp number, first 60 AAs: 0.28355\nTotal prob of N-in: 0.13257\noutside 1 331\nTMhelix 332 354\ninside 355 372

10936 GCF_000500815.1_GSK202U Escherichia coli GSK202U Proteobacteria; Gammaproteobacteria; Enterobacterales; Enterobacteriaceae; Escherichia; Escherichia coli
MNNEETFYQAMRRQGVTRRSFLKYCSLAATSLGLGAGMAPKIAWALENKPRIPVVWIHGL WP_001058323.1
MULTISPECIES: hydrogenase-1 small chain [Proteobacteria] Length: 372\nNumber of predicted TMHs: 2\nExp number of AAs in TMHs: 37.2608499999999\nExp number, first 60 AAs: 19.52923\nTotal prob of N-in: 0.94351\nPOSSIBLE N-term signal sequence\ninside 1 19\nTMhelix 20 39\noutside 40 325\nTMhelix 326 348\ninside 349 372

10937 GCF_002110085.1_ASM211008v1 Escherichia coli H001 Proteobacteria; Gammaproteobacteria; Enterobacterales; Enterobacteriaceae; Escherichia; Escherichia coli
MTGDNTLIHSHGINRRDFMKLCAALAATMGLSSKAAAEMAESVTNPQRPPVIWIGAQECT WP_000145410.1
MULTISPECIES: hydrogenase-2 small chain [Proteobacteria] Length: 372\nNumber of predicted TMHs: 1\nExp number of AAs in TMHs: 23.72226\nExp number, first 60 AAs: 0.22038\nTotal prob of N-in: 0.12994\noutside 1 331\nTMhelix 332 354\ninside 355 372

10938 GCF_002110085.1_ASM211008v1 Escherichia coli H001 Proteobacteria; Gammaproteobacteria; Enterobacterales; Enterobacteriaceae; Escherichia; Escherichia coli
MNNEETFYQAMRRQGVTRRSFLKYCSLAATSLGLGAGMAPKIAWALENKPRIPVVWIHGL WP_001058323.1
MULTISPECIES: hydrogenase-1 small chain [Proteobacteria] Length: 372\nNumber of predicted TMHs: 2\nExp number of AAs in TMHs: 37.2608499999999\nExp number, first 60 AAs: 19.52923\nTotal prob of N-in: 0.94351\nPOSSIBLE N-term signal sequence\ninside 1 19\nTMhelix 20 39\noutside 40 325\nTMhelix 326 348\ninside 349 372

10939 GCF_000190855.1_ASM19085v1 Escherichia coli H120 Proteobacteria; Gammaproteobacteria; Enterobacterales; Enterobacteriaceae; Escherichia; Escherichia coli
MNNEETFYQAMRRQGVTRRSFLKYCSLAATSLGLGAGMAPKIAWALENKPRIPVVWIHGL WP_001295940.1
MULTISPECIES: hydrogenase [Enterobacteriaceae] Length: 372\nNumber of predicted TMHs: 2\nExp number of AAs in TMHs: 37.2608899999999\nExp number, first 60 AAs: 19.52923\nTotal prob of N-in: 0.94351\nPOSSIBLE N-term signal sequence\ninside 1 19\nTMhelix 20 39\noutside 40 325\nTMhelix 326 348\ninside 349 372

10940 GCF_000190855.1_ASM19085v1 Escherichia coli H120 Proteobacteria; Gammaproteobacteria; Enterobacterales; Enterobacteriaceae; Escherichia; Escherichia coli
MTGDNTLIHSHGINRRDFMKLCAALAATMGLSSKAAAEMAESVTNPQRPPVIWIGAQECT WP_000145410.1
MULTISPECIES: hydrogenase-2 small chain [Proteobacteria] Length: 372\nNumber of predicted TMHs: 1\nExp number of AAs in TMHs: 23.72226\nExp number, first 60 AAs: 0.22038\nTotal prob of N-in: 0.12994\noutside 1 331\nTMhelix 332 354\ninside 355 372

10941 GCF_002110555.1_ASM211055v1 Escherichia coli H223 Proteobacteria; Gammaproteobacteria; Enterobacterales; Enterobacteriaceae; Escherichia; Escherichia coli
MNNEETFYQAMRRQGVTRRSFLKYCSLAATSLGLGAGMAPKIAWALENKPRIPVVWIHGL WP_001058323.1
MULTISPECIES: hydrogenase-1 small chain [Proteobacteria] Length: 372\nNumber of predicted TMHs: 2\nExp number of AAs in TMHs: 37.2608499999999\nExp number, first 60 AAs: 19.52923\nTotal prob of N-in: 0.94351\nPOSSIBLE N-term signal sequence\ninside 1 19\nTMhelix 20 39\noutside 40 325\nTMhelix 326 348\ninside 349 372

10942 GCF_002110555.1_ASM211055v1 Escherichia coli H223 Proteobacteria; Gammaproteobacteria; Enterobacterales; Enterobacteriaceae; Escherichia; Escherichia coli
MTGDNTLIHSHGINRRDFMKLCAALAATMGLSSKAAAEMVESVTNPQRPPVIWIGAQECT WP_000145423.1
hydrogenase 2 small subunit [Escherichia coli] Length: 372\nNumber of predicted TMHs: 1\nExp number of AAs in TMHs: 23.78201\nExp number, first 60 AAs: 0.28355\nTotal prob of N-in: 0.13257\noutside 1 331\nTMhelix 332 354\ninside 355 372

10943 GCF_000190895.1_ASM19089v1 Escherichia coli H252 Proteobacteria; Gammaproteobacteria; Enterobacterales; Enterobacteriaceae; Escherichia; Escherichia coli
MTGDNTLIHSHGINRRDFMKLCAALAATMGLSSKAAAEMAESVTNPQRPPVIWIGAQECT WP_000145410.1
MULTISPECIES: hydrogenase-2 small chain [Proteobacteria] Length: 372\nNumber of predicted TMHs: 1\nExp number of AAs in TMHs: 23.72226\nExp number, first 60 AAs: 0.22038\nTotal prob of N-in: 0.12994\noutside 1 331\nTMhelix 332 354\ninside 355 372

10944 GCF_000190915.1_ASM19091v1 Escherichia coli H263 Proteobacteria; Gammaproteobacteria; Enterobacterales; Enterobacteriaceae; Escherichia; Escherichia coli
MTGDNTLIHSHGINRRDFMKLCAALAATMGLSSKAAAEMAESVTNPQRPPVIWIGAQECT WP_000145410.1
MULTISPECIES: hydrogenase-2 small chain [Proteobacteria] Length: 372\nNumber of predicted TMHs: 1\nExp number of AAs in TMHs: 23.72226\nExp number, first 60 AAs: 0.22038\nTotal prob of N-in: 0.12994\noutside 1 331\nTMhelix 332 354\ninside 355 372

10945 GCF_002109775.1_ASM210977v1 Escherichia coli H296 Proteobacteria; Gammaproteobacteria; Enterobacterales; Enterobacteriaceae; Escherichia; Escherichia coli
MTGDNTLIHSHGINRRDFMKLCAALAATMGLSSKAAAEMAESVTNPQRPPVIWIGAQECT WP_000145410.1
MULTISPECIES: hydrogenase-2 small chain [Proteobacteria] Length: 372\nNumber of predicted TMHs: 1\nExp number of AAs in TMHs: 23.72226\nExp number, first 60 AAs: 0.22038\nTotal prob of N-in: 0.12994\noutside 1 331\nTMhelix 332 354\ninside 355 372

10946 GCF_000176695.2_ASM17669v2 Escherichia coli H299 Proteobacteria; Gammaproteobacteria; Enterobacterales; Enterobacteriaceae; Escherichia; Escherichia coli

MNNEETFYQAMRRQGVTRRSFLKYCSLAATSLGLGAGMAPKIAWALENKPRIPVVWIHGL WP_001058323.1 MULTISPECIES: hydrogenase-1 small chain [Proteobacteria] Length: 372\nNumber of predicted TMHs: 2\nExp number of AAs in TMHs: 37.2608499999999\nExp number, first 60 AAs: 19.52923\nTotal prob of N-in: 0.94351\nPOSSIBLE N-term signal sequence\ninside 1 19\nTMhelix 20 39\noutside 40 325\nTMhelix 326 348\ninside 349 372

10947 GCF_000176695.2_ASM17669v2 Escherichia coli H299 Proteobacteria; Gammaproteobacteria; Enterobacterales; Enterobacteriaceae; Escherichia; Escherichia coli MTGDNTLIHSHGINRRDFMKLCAALAATMGLSSKAAAEMAESVTNPQRPPVIWIGAQECT WP_000145410.1 MULTISPECIES: hydrogenase-2 small chain [Proteobacteria] Length: 372\nNumber of predicted TMHs: 1\nExp number of AAs in TMHs: 23.72226\nExp number, first 60 AAs: 0.22038\nTotal prob of N-in: 0.12994\noutside 1 331\nTMhelix 332 354\ninside 355 372

10948 GCF_000215165.1_ASM21516v1 Escherichia coli H30 Proteobacteria; Gammaproteobacteria; Enterobacterales; Enterobacteriaceae; Escherichia; Escherichia coli MNNEETFYQAMRRQGVTRRSFLKYCSLAATSLGLGAGMAPKIAWALENKPRIPVVWIHGL WP_001301613.1 hydrogenase [Escherichia coli] Length: 372\nNumber of predicted TMHs: 2\nExp number of AAs in TMHs: 37.46598\nExp number, first 60 AAs: 19.53139\nTotal prob of N-in: 0.94364\nPOSSIBLE N-term signal sequence\ninside 1 19\nTMhelix 20 39\noutside 40 325\nTMhelix 326 348\ninside 349 372

10949 GCF_000215165.1_ASM21516v1 Escherichia coli H30 Proteobacteria; Gammaproteobacteria; Enterobacterales; Enterobacteriaceae; Escherichia; Escherichia coli MTGDNTLIHSHGINRRDFMKLCAALAATMGLSSKAAAEMAESVTNPQRPPVIWIGAQECT WP_000145410.1 MULTISPECIES: hydrogenase-2 small chain [Proteobacteria] Length: 372\nNumber of predicted TMHs: 1\nExp number of AAs in TMHs: 23.72226\nExp number, first 60 AAs: 0.22038\nTotal prob of N-in: 0.12994\noutside 1 331\nTMhelix 332 354\ninside 355 372

10950 GCF_002109765.1_ASM210976v1 Escherichia coli H305 Proteobacteria; Gammaproteobacteria; Enterobacterales; Enterobacteriaceae; Escherichia; Escherichia coli MTGDNTLIHSHGINRRDFMKLCAALAATMGLSSKAAAEMAESVTNPQRPPVIWIGAQECT WP_000145410.1 MULTISPECIES: hydrogenase-2 small chain [Proteobacteria] Length: 372\nNumber of predicted TMHs: 1\nExp number of AAs in TMHs: 23.72226\nExp number, first 60 AAs: 0.22038\nTotal prob of N-in: 0.12994\noutside 1 331\nTMhelix 332 354\ninside 355 372

10951 GCF_002110115.1_ASM211011v1 Escherichia coli H378 Proteobacteria; Gammaproteobacteria; Enterobacterales; Enterobacteriaceae; Escherichia; Escherichia coli MNNEETFYQAMRRQGVTRRSFLKYCSLAATSLGLGAGMAPKIAWALENKPRIPVVWIHGL WP_001058323.1 MULTISPECIES: hydrogenase-1 small chain [Proteobacteria] Length: 372\nNumber of predicted TMHs: 2\nExp number of AAs in TMHs: 37.2608499999999\nExp number, first 60 AAs: 19.52923\nTotal prob of N-in: 0.94351\nPOSSIBLE N-term signal sequence\ninside 1 19\nTMhelix 20 39\noutside 40 325\nTMhelix 326 348\ninside 349 372

10952 GCF_002110115.1_ASM211011v1 Escherichia coli H378 Proteobacteria; Gammaproteobacteria; Enterobacterales; Enterobacteriaceae; Escherichia; Escherichia coli MTGDNTLIHSHGINRRDFMKLCAALAATMGLSSKAAAEMAESVTNPQRPPVIWIGAQECT WP_000145410.1 MULTISPECIES: hydrogenase-2 small chain [Proteobacteria] Length: 372\nNumber of predicted TMHs: 1\nExp number of AAs in TMHs: 23.72226\nExp number, first 60 AAs: 0.22038\nTotal prob of N-in: 0.12994\noutside 1 331\nTMhelix 332 354\ninside 355 372

10953 GCF_002110225.1_ASM211022v1 Escherichia coli H383 Proteobacteria; Gammaproteobacteria; Enterobacterales; Enterobacteriaceae; Escherichia; Escherichia coli MNNEETFYQAMRRQGVTRRSFLKYCSLAATSLGLGAGMAPKIAWALENKPRIPVVWIHGL WP_001058323.1 MULTISPECIES: hydrogenase-1 small chain [Proteobacteria] Length: 372\nNumber of predicted TMHs: 2\nExp number of AAs in TMHs: 37.2608499999999\nExp number, first 60 AAs: 19.52923\nTotal prob of N-in: 0.94351\nPOSSIBLE N-term signal sequence\ninside 1 19\nTMhelix 20 39\noutside 40 325\nTMhelix 326 348\ninside 349 372

10954 GCF_002110225.1_ASM211022v1 Escherichia coli H383 Proteobacteria; Gammaproteobacteria; Enterobacterales; Enterobacteriaceae; Escherichia; Escherichia coli MTGDNTLIHSHGINRRDFMKLCAALAATMGLSSKAAAEMAESVTNPQRPPVIWIGAQECT WP_000145410.1 MULTISPECIES: hydrogenase-2 small chain [Proteobacteria] Length: 372\nNumber of predicted TMHs:

1\nExp number of AAs in TMHs: 23.72226\nExp number, first 60 AAs: 0.22038\nTotal prob of N-in: 0.12994\noutside 1 331\nTMhelix 332 354\ninside 355 372

10955 GCF_002110545.1_ASM211054v1 Escherichia coli H386 Proteobacteria; Gammaproteobacteria; Enterobacterales; Enterobacteriaceae; Escherichia; Escherichia coli MTGDNTLIHSHGINRRDFMKLCAALAATMGLSSKAAAEMAESVTNPQRPPVIWIGAQECT WP_000145410.1 MULTISPECIES: hydrogenase-2 small chain [Proteobacteria] Length: 372\nNumber of predicted TMHs: 1\nExp number of AAs in TMHs: 23.72226\nExp number, first 60 AAs: 0.22038\nTotal prob of N-in: 0.12994\noutside 1 331\nTMhelix 332 354\ninside 355 372

10956 GCF_002110545.1_ASM211054v1 Escherichia coli H386 Proteobacteria; Gammaproteobacteria; Enterobacterales; Enterobacteriaceae; Escherichia; Escherichia coli MNNEETFYQAMRRQGVTRRSFLKYCSLAATSLGLGAGMAPKIAWALENKPRIPVVWIHGL WP_001058323.1 MULTISPECIES: hydrogenase-1 small chain [Proteobacteria] Length: 372\nNumber of predicted TMHs: 2\nExp number of AAs in TMHs: 37.2608499999999\nExp number, first 60 AAs: 19.52923\nTotal prob of N-in: 0.94351\nPOSSIBLE N-term signal sequence\ninside 1 19\nTMhelix 20 39\noutside 40 325\nTMhelix 326 348\ninside 349 372

10957 GCF_000241975.1_Esch_coli_H397_V1 Escherichia coli H397 Proteobacteria; Gammaproteobacteria; Enterobacterales; Enterobacteriaceae; Escherichia; Escherichia coli MTGDNTLIHSHGINRRDFMKLCAALAATMGLSSKAAAEMAESVTNPQRPPVIWIGAQECT WP_000145410.1 MULTISPECIES: hydrogenase-2 small chain [Proteobacteria] Length: 372\nNumber of predicted TMHs: 1\nExp number of AAs in TMHs: 23.72226\nExp number, first 60 AAs: 0.22038\nTotal prob of N-in: 0.12994\noutside 1 331\nTMhelix 332 354\ninside 355 372

10958 GCF_002109685.1_ASM210968v1 Escherichia coli H413 Proteobacteria; Gammaproteobacteria; Enterobacterales; Enterobacteriaceae; Escherichia; Escherichia coli MTGDNTLIHSHGINRRDFMKLCAALAATMGLSSKAAAEMAESVTNPQRPPVIWIGAQECT WP_000145410.1 MULTISPECIES: hydrogenase-2 small chain [Proteobacteria] Length: 372\nNumber of predicted TMHs: 1\nExp number of AAs in TMHs: 23.72226\nExp number, first 60 AAs: 0.22038\nTotal prob of N-in: 0.12994\noutside 1 331\nTMhelix 332 354\ninside 355 372

10959 GCF_002110265.1_ASM211026v1 Escherichia coli H420 Proteobacteria; Gammaproteobacteria; Enterobacterales; Enterobacteriaceae; Escherichia; Escherichia coli MNNEETFYQAMRRQGVTRRSFLKYCSLAATSLGLGAGMAPKIAWALENKPRIPVVWIHGL WP_001058323.1 MULTISPECIES: hydrogenase-1 small chain [Proteobacteria] Length: 372\nNumber of predicted TMHs: 2\nExp number of AAs in TMHs: 37.2608499999999\nExp number, first 60 AAs: 19.52923\nTotal prob of N-in: 0.94351\nPOSSIBLE N-term signal sequence\ninside 1 19\nTMhelix 20 39\noutside 40 325\nTMhelix 326 348\ninside 349 372

10960 GCF_002110265.1_ASM211026v1 Escherichia coli H420 Proteobacteria; Gammaproteobacteria; Enterobacterales; Enterobacteriaceae; Escherichia; Escherichia coli MTGDNTLIHSHGINRRDFMKLCAALAATMGLSSKAAAEMAESVTNPQRPPVIWIGAQECT WP_000145410.1 MULTISPECIES: hydrogenase-2 small chain [Proteobacteria] Length: 372\nNumber of predicted TMHs: 1\nExp number of AAs in TMHs: 23.72226\nExp number, first 60 AAs: 0.22038\nTotal prob of N-in: 0.12994\noutside 1 331\nTMhelix 332 354\ninside 355 372

10961 GCF_002110275.1_ASM211027v1 Escherichia coli H454 Proteobacteria; Gammaproteobacteria; Enterobacterales; Enterobacteriaceae; Escherichia; Escherichia coli MTGDNTLIHSHGINRRDFMKLCAALAATMGLSSKAAAEMAESVTNPQRPPVIWIGAQECT WP_000145410.1 MULTISPECIES: hydrogenase-2 small chain [Proteobacteria] Length: 372\nNumber of predicted TMHs: 1\nExp number of AAs in TMHs: 23.72226\nExp number, first 60 AAs: 0.22038\nTotal prob of N-in: 0.12994\noutside 1 331\nTMhelix 332 354\ninside 355 372

10962 GCF_002110275.1_ASM211027v1 Escherichia coli H454 Proteobacteria; Gammaproteobacteria; Enterobacterales; Enterobacteriaceae; Escherichia; Escherichia coli MNNEETFYQAMRRQGVTRRSFLKYCSLAATSLGLGAGMAPKIAWALENKPRIPVVWIHGL WP_001058323.1 MULTISPECIES: hydrogenase-1 small chain [Proteobacteria] Length: 372\nNumber of predicted TMHs: 2\nExp number of AAs in TMHs: 37.2608499999999\nExp number, first 60 AAs: 19.52923\nTotal prob of N-in: 0.94351\nPOSSIBLE N-term signal sequence\ninside 1 19\nTMhelix 20 39\noutside 40 325\nTMhelix 326 348\ninside 349 372

10963 GCF_002110165.1_ASM211016v1 Escherichia coli H461 Proteobacteria; Gammaproteobacteria; Enterobacterales; Enterobacteriaceae; Escherichia; Escherichia coli
MTGDNTLIHSHGINRRDFMKLCAALAATMGLSSKAAAEMAESVTNPQRPPVIWIGAQECT WP_000145410.1
MULTISPECIES: hydrogenase-2 small chain [Proteobacteria] Length: 372\nNumber of predicted TMHs: 1\nExp number of AAs in TMHs: 23.72226\nExp number, first 60 AAs: 0.22038\nTotal prob of N-in: 0.12994\noutside 1 331\nTMhelix 332 354\ninside 355 372

10964 GCF_000191015.1_ASM19101v1 Escherichia coli H489 Proteobacteria; Gammaproteobacteria; Enterobacterales; Enterobacteriaceae; Escherichia; Escherichia coli
MNNEETFYQAMRRQGVTRRSFLKYCSLAATSLGLGAGMAPKIAWALENKPRIPVVWIHGL WP_001058323.1
MULTISPECIES: hydrogenase-1 small chain [Proteobacteria] Length: 372\nNumber of predicted TMHs: 2\nExp number of AAs in TMHs: 37.2608499999999\nExp number, first 60 AAs: 19.52923\nTotal prob of N-in: 0.94351\nPOSSIBLE N-term signal sequence\ninside 1 19\nTMhelix 20 39\noutside 40 325\nTMhelix 326 348\ninside 349 372

10965 GCF_000191015.1_ASM19101v1 Escherichia coli H489 Proteobacteria; Gammaproteobacteria; Enterobacterales; Enterobacteriaceae; Escherichia; Escherichia coli
MTGDNTLIHSHGINRRDFMKLCAALAATMGLSSKAAAEMAESVTNPQRPPVIWIGAQECT WP_000145410.1
MULTISPECIES: hydrogenase-2 small chain [Proteobacteria] Length: 372\nNumber of predicted TMHs: 1\nExp number of AAs in TMHs: 23.72226\nExp number, first 60 AAs: 0.22038\nTotal prob of N-in: 0.12994\noutside 1 331\nTMhelix 332 354\ninside 355 372

10966 GCF_000242035.1_Esch_coli_H494_V1 Escherichia coli H494 Proteobacteria; Gammaproteobacteria; Enterobacterales; Enterobacteriaceae; Escherichia; Escherichia coli
MTGDNTLIHSHGINRRDFMKLCAALAATMGLSSKAAAEMAESVTNPQRPPVIWIGAQECT WP_000145410.1
MULTISPECIES: hydrogenase-2 small chain [Proteobacteria] Length: 372\nNumber of predicted TMHs: 1\nExp number of AAs in TMHs: 23.72226\nExp number, first 60 AAs: 0.22038\nTotal prob of N-in: 0.12994\noutside 1 331\nTMhelix 332 354\ninside 355 372

10967 GCF_000242035.1_Esch_coli_H494_V1 Escherichia coli H494 Proteobacteria; Gammaproteobacteria; Enterobacterales; Enterobacteriaceae; Escherichia; Escherichia coli
MNNEETFYQAMRRQGVTRRSFLKYCSLAATSLGLGAGMAPKIAWALENKPRIPVVWIHGL WP_001058323.1
MULTISPECIES: hydrogenase-1 small chain [Proteobacteria] Length: 372\nNumber of predicted TMHs: 2\nExp number of AAs in TMHs: 37.2608499999999\nExp number, first 60 AAs: 19.52923\nTotal prob of N-in: 0.94351\nPOSSIBLE N-term signal sequence\ninside 1 19\nTMhelix 20 39\noutside 40 325\nTMhelix 326 348\ninside 349 372

10968 GCF_002109645.1_ASM210964v1 Escherichia coli H588 Proteobacteria; Gammaproteobacteria; Enterobacterales; Enterobacteriaceae; Escherichia; Escherichia coli
MTGDNTLIHSHGINRRDFMKLCAALAATMGLSSKAAAEMAESVTNPQRPPVIWIGAQECT WP_000145410.1
MULTISPECIES: hydrogenase-2 small chain [Proteobacteria] Length: 372\nNumber of predicted TMHs: 1\nExp number of AAs in TMHs: 23.72226\nExp number, first 60 AAs: 0.22038\nTotal prob of N-in: 0.12994\noutside 1 331\nTMhelix 332 354\ninside 355 372

10969 GCF_000176675.2_ASM17667v2 Escherichia coli H591 Proteobacteria; Gammaproteobacteria; Enterobacterales; Enterobacteriaceae; Escherichia; Escherichia coli
MNNEETFYQAMRRQGVTRRSFLKYCSLAATSLGLGAGMAPKIAWALENKPRIPVVWIHGL WP_001058323.1
MULTISPECIES: hydrogenase-1 small chain [Proteobacteria] Length: 372\nNumber of predicted TMHs: 2\nExp number of AAs in TMHs: 37.2608499999999\nExp number, first 60 AAs: 19.52923\nTotal prob of N-in: 0.94351\nPOSSIBLE N-term signal sequence\ninside 1 19\nTMhelix 20 39\noutside 40 325\nTMhelix 326 348\ninside 349 372

10970 GCF_000176675.2_ASM17667v2 Escherichia coli H591 Proteobacteria; Gammaproteobacteria; Enterobacterales; Enterobacteriaceae; Escherichia; Escherichia coli
MTGDNTLIHSHGINRRDFMKLCAALAATMGLSSKAAAEMAESVTNPQRPPVIWIGAQECT WP_000145410.1
MULTISPECIES: hydrogenase-2 small chain [Proteobacteria] Length: 372\nNumber of predicted TMHs: 1\nExp number of AAs in TMHs: 23.72226\nExp number, first 60 AAs: 0.22038\nTotal prob of N-in: 0.12994\noutside 1 331\nTMhelix 332 354\ninside 355 372

10971 GCF_002110245.1_ASM211024v1 Escherichia coli H605 Proteobacteria; Gammaproteobacteria; Enterobacterales; Enterobacteriaceae; Escherichia; Escherichia coli

MNNEETFYQAMRRQGVTRRSFLKYCSLAATSLGLGAGMAPKIAWALENKPRIPVVWIHGL WP_010344442.1 MULTISPECIES: hydrogenase [Escherichia] Length: 372\nNumber of predicted TMHs: 2\nExp number of AAs in TMHs: 38.04874\nExp number, first 60 AAs: 19.5298\nTotal prob of N-in: 0.94352\nPOSSIBLE N-term signal sequence\ninside 1 19\nTMhelix 20 39\noutside 40 325\nTMhelix 326 348\ninside 349 372

10972 GCF_002110245.1_ASM211024v1 Escherichia coli H605 Proteobacteria; Gammaproteobacteria; Enterobacterales; Enterobacteriaceae; Escherichia; Escherichia coli MTGDNTLIHSHGINRRDFMKLCAALAATMGLSSKAAAEMAESVTNPQRPPVIWIGAQECT WP_000145410.1 MULTISPECIES: hydrogenase-2 small chain [Proteobacteria] Length: 372\nNumber of predicted TMHs: 1\nExp number of AAs in TMHs: 23.72226\nExp number, first 60 AAs: 0.22038\nTotal prob of N-in: 0.12994\noutside 1 331\nTMhelix 332 354\ninside 355 372

10973 GCF_002109805.1_ASM210980v1 Escherichia coli H617 Proteobacteria; Gammaproteobacteria; Enterobacterales; Enterobacteriaceae; Escherichia; Escherichia coli MNNEETFYQAMRRQGVTRRSFLKYCSLAATSLGLGAGMAPKIAWALENKPRIPVVWIHGL WP_001058323.1 MULTISPECIES: hydrogenase-1 small chain [Proteobacteria] Length: 372\nNumber of predicted TMHs: 2\nExp number of AAs in TMHs: 37.2608499999999\nExp number, first 60 AAs: 19.52923\nTotal prob of N-in: 0.94351\nPOSSIBLE N-term signal sequence\ninside 1 19\nTMhelix 20 39\noutside 40 325\nTMhelix 326 348\ninside 349 372

10974 GCF_002109805.1_ASM210980v1 Escherichia coli H617 Proteobacteria; Gammaproteobacteria; Enterobacterales; Enterobacteriaceae; Escherichia; Escherichia coli MTGDNTLIHSHGINRRDFMKLCAALAATMGLSSKAAAEMAESVTNPQRPPVIWIGAQECT WP_000145410.1 MULTISPECIES: hydrogenase-2 small chain [Proteobacteria] Length: 372\nNumber of predicted TMHs: 1\nExp number of AAs in TMHs: 23.72226\nExp number, first 60 AAs: 0.22038\nTotal prob of N-in: 0.12994\noutside 1 331\nTMhelix 332 354\ninside 355 372

10975 GCF_000261405.1_Esch_coli_H730_V1 Escherichia coli H730 Proteobacteria; Gammaproteobacteria; Enterobacterales; Enterobacteriaceae; Escherichia; Escherichia coli MNNEETFYQAMRRQGVTRRSFLKYCSLAATSLGLGAGMAPKIAWALENKPRIPVVWIHGL WP_001058323.1 MULTISPECIES: hydrogenase-1 small chain [Proteobacteria] Length: 372\nNumber of predicted TMHs: 2\nExp number of AAs in TMHs: 37.2608499999999\nExp number, first 60 AAs: 19.52923\nTotal prob of N-in: 0.94351\nPOSSIBLE N-term signal sequence\ninside 1 19\nTMhelix 20 39\noutside 40 325\nTMhelix 326 348\ninside 349 372

10976 GCF_000261405.1_Esch_coli_H730_V1 Escherichia coli H730 Proteobacteria; Gammaproteobacteria; Enterobacterales; Enterobacteriaceae; Escherichia; Escherichia coli MTGDNTLIHSHGINRRDFMKLCAALAATMGLSSKAAAEMAESVTNPQRPPVIWIGAQECT WP_000145410.1 MULTISPECIES: hydrogenase-2 small chain [Proteobacteria] Length: 372\nNumber of predicted TMHs: 1\nExp number of AAs in TMHs: 23.72226\nExp number, first 60 AAs: 0.22038\nTotal prob of N-in: 0.12994\noutside 1 331\nTMhelix 332 354\ninside 355 372

10977 GCF_000176535.2_ASM17653v2 Escherichia coli H736 Proteobacteria; Gammaproteobacteria; Enterobacterales; Enterobacteriaceae; Escherichia; Escherichia coli MTGDNTLIHSHGINRRDFMKLCAALAATMGLSSKAAAEMAESVTNPQRPPVIWIGAQECT WP_000145410.1 MULTISPECIES: hydrogenase-2 small chain [Proteobacteria] Length: 372\nNumber of predicted TMHs: 1\nExp number of AAs in TMHs: 23.72226\nExp number, first 60 AAs: 0.22038\nTotal prob of N-in: 0.12994\noutside 1 331\nTMhelix 332 354\ninside 355 372

10978 GCF_000176535.2_ASM17653v2 Escherichia coli H736 Proteobacteria; Gammaproteobacteria; Enterobacterales; Enterobacteriaceae; Escherichia; Escherichia coli MNNEETFYQAMRRQGVTRRSFLKYCSLAATSLGLGAGMAPKIAWALENKPRIPVVWIHGL WP_001058323.1 MULTISPECIES: hydrogenase-1 small chain [Proteobacteria] Length: 372\nNumber of predicted TMHs: 2\nExp number of AAs in TMHs: 37.2608499999999\nExp number, first 60 AAs: 19.52923\nTotal prob of N-in: 0.94351\nPOSSIBLE N-term signal sequence\ninside 1 19\nTMhelix 20 39\noutside 40 325\nTMhelix 326 348\ninside 349 372

10979 GCF_000387805.2_ASM38780v2 Escherichia coli HM26 Proteobacteria; Gammaproteobacteria; Enterobacterales; Enterobacteriaceae; Escherichia; Escherichia coli MNNEETFYQAMRRQGVTRRSFLKYCSLAATSLGLGAGMAPKIAWALENKPRIPVVWIHGL WP_001058323.1 MULTISPECIES: hydrogenase-1 small chain [Proteobacteria] Length: 372\nNumber of predicted TMHs:

2\nExp number of AAs in TMHs: 37.2608499999999\nExp number, first 60 AAs: 19.52923\nTotal prob of N-in: 0.94351\nPOSSIBLE N-term signal sequence\ninside 1 19\nTMhelix 20 39\noutside 40 325\nTMhelix 326 348\ninside 349 372

10980 GCF_000387805.2_ASM38780v2 Escherichia coli HM26 Proteobacteria; Gammaproteobacteria; Enterobacterales; Enterobacteriaceae; Escherichia; Escherichia coli MTGDNTLIHSHGINRRDFMKLCAALAATMGLSSKAAAEMAESVTNPQRPPVIWIGAQECT WP_000145410.1 MULTISPECIES: hydrogenase-2 small chain [Proteobacteria] Length: 372\nNumber of predicted TMHs: 1\nExp number of AAs in TMHs: 23.72226\nExp number, first 60 AAs: 0.22038\nTotal prob of N-in: 0.12994\noutside 1 331\nTMhelix 332 354\ninside 355 372

10981 GCF_000387825.2_ASM38782v2 Escherichia coli HM27 Proteobacteria; Gammaproteobacteria; Enterobacterales; Enterobacteriaceae; Escherichia; Escherichia coli MTGDNTLIHSHGINRRDFMKLCAALAATMGLSSKAAAEMAESVTNPQRPPVIWIGAQECT WP_000145410.1 MULTISPECIES: hydrogenase-2 small chain [Proteobacteria] Length: 372\nNumber of predicted TMHs: 1\nExp number of AAs in TMHs: 23.72226\nExp number, first 60 AAs: 0.22038\nTotal prob of N-in: 0.12994\noutside 1 331\nTMhelix 332 354\ninside 355 372

10982 GCF_000387825.2_ASM38782v2 Escherichia coli HM27 Proteobacteria; Gammaproteobacteria; Enterobacterales; Enterobacteriaceae; Escherichia; Escherichia coli MNNEETFYQAMRRQGVTRRSFLKYCSLAATSLGLGAGMAPKIAWALENKPRIPVVWIHGL WP_001058323.1 MULTISPECIES: hydrogenase-1 small chain [Proteobacteria] Length: 372\nNumber of predicted TMHs: 2\nExp number of AAs in TMHs: 37.2608499999999\nExp number, first 60 AAs: 19.52923\nTotal prob of N-in: 0.94351\nPOSSIBLE N-term signal sequence\ninside 1 19\nTMhelix 20 39\noutside 40 325\nTMhelix 326 348\ninside 349 372

10983 GCF_000387845.2_ASM38784v2 Escherichia coli HM46 Proteobacteria; Gammaproteobacteria; Enterobacterales; Enterobacteriaceae; Escherichia; Escherichia coli MTGDNTLIHSHGINRRDFMKLCAALAATMGLSSKAAAEMAESVTNPQRPPVIWIGAQECT WP_000145410.1 MULTISPECIES: hydrogenase-2 small chain [Proteobacteria] Length: 372\nNumber of predicted TMHs: 1\nExp number of AAs in TMHs: 23.72226\nExp number, first 60 AAs: 0.22038\nTotal prob of N-in: 0.12994\noutside 1 331\nTMhelix 332 354\ninside 355 372

10984 GCF_000387845.2_ASM38784v2 Escherichia coli HM46 Proteobacteria; Gammaproteobacteria; Enterobacterales; Enterobacteriaceae; Escherichia; Escherichia coli MNNEETFYQAMRRQGVTRRSFLKYCSLAATSLGLGAGMAPKIAWALENKPRIPVVWIHGL WP_001058323.1 MULTISPECIES: hydrogenase-1 small chain [Proteobacteria] Length: 372\nNumber of predicted TMHs: 2\nExp number of AAs in TMHs: 37.2608499999999\nExp number, first 60 AAs: 19.52923\nTotal prob of N-in: 0.94351\nPOSSIBLE N-term signal sequence\ninside 1 19\nTMhelix 20 39\noutside 40 325\nTMhelix 326 348\ninside 349 372

10985 GCF_000285375.1_ASM28537v1 Escherichia coli HM605 Proteobacteria; Gammaproteobacteria; Enterobacterales; Enterobacteriaceae; Escherichia; Escherichia coli MTGDNTLIHSHGINRRDFMKLCAALAATMGLSSKAAAEMAESVTNPQRPPVIWIGAQECT WP_000145410.1 MULTISPECIES: hydrogenase-2 small chain [Proteobacteria] Length: 372\nNumber of predicted TMHs: 1\nExp number of AAs in TMHs: 23.72226\nExp number, first 60 AAs: 0.22038\nTotal prob of N-in: 0.12994\noutside 1 331\nTMhelix 332 354\ninside 355 372

10986 GCF_000264175.1_ASM26417v1 Escherichia coli HM605 Proteobacteria; Gammaproteobacteria; Enterobacterales; Enterobacteriaceae; Escherichia; Escherichia coli MTGDNTLIHSHGINRRDFMKLCAALAATMGLSSKAAAEMAESVTNPQRPPVIWIGAQECT WP_000145410.1 MULTISPECIES: hydrogenase-2 small chain [Proteobacteria] Length: 372\nNumber of predicted TMHs: 1\nExp number of AAs in TMHs: 23.72226\nExp number, first 60 AAs: 0.22038\nTotal prob of N-in: 0.12994\noutside 1 331\nTMhelix 332 354\ninside 355 372

10987 GCF_000387785.2_ASM38778v2 Escherichia coli HM65 Proteobacteria; Gammaproteobacteria; Enterobacterales; Enterobacteriaceae; Escherichia; Escherichia coli MNNEETFYQAMRRQGVTRRSFLKYCSLAATSLGLGAGMAPKIAWALENKPRIPVVWIHGL WP_001058323.1 MULTISPECIES: hydrogenase-1 small chain [Proteobacteria] Length: 372\nNumber of predicted TMHs: 2\nExp number of AAs in TMHs: 37.2608499999999\nExp number, first 60 AAs: 19.52923\nTotal prob of N-in:

0.94351\nPOSSIBLE N-term signal sequence\ninside 1 19\nTMhelix 20 39\noutside 40 325\nTMhelix 326 348\ninside 349 372

10988 GCF_000387785.2_ASM38778v2 Escherichia coli HM65 Proteobacteria; Gammaproteobacteria; Enterobacterales; Enterobacteriaceae; Escherichia; Escherichia coli MTGDNTLIHSHGINRRDFMKLCAALAATMGLSSKAAAEMVESVTNPQRPPVIWIGAQECT WP_000145423.1 hydrogenase 2 small subunit [Escherichia coli] Length: 372\nNumber of predicted TMHs: 1\nExp number of AAs in TMHs: 23.78201\nExp number, first 60 AAs: 0.28355\nTotal prob of N-in: 0.13257\noutside 1 331\nTMhelix 332 354\ninside 355 372

10989 GCF_000387865.2_ASM38786v2 Escherichia coli HM69 Proteobacteria; Gammaproteobacteria; Enterobacterales; Enterobacteriaceae; Escherichia; Escherichia coli MTGDNTLIHSHGINRRDFMKLCAALAATMGLSSKAAAEMAESVTNPQRPPVIWIGAQECT WP_000145410.1 MULTISPECIES: hydrogenase-2 small chain [Proteobacteria] Length: 372\nNumber of predicted TMHs: 1\nExp number of AAs in TMHs: 23.72226\nExp number, first 60 AAs: 0.22038\nTotal prob of N-in: 0.12994\noutside 1 331\nTMhelix 332 354\ninside 355 372

10990 GCF_000387865.2_ASM38786v2 Escherichia coli HM69 Proteobacteria; Gammaproteobacteria; Enterobacterales; Enterobacteriaceae; Escherichia; Escherichia coli MNNEETFYQAMRRQGVTRRSFLKYCSLAATSLGLGAGMAPKIAWALENKPRIPVVWIHGL WP_001058323.1 MULTISPECIES: hydrogenase-1 small chain [Proteobacteria] Length: 372\nNumber of predicted TMHs: 2\nExp number of AAs in TMHs: 37.2608499999999\nExp number, first 60 AAs: 19.52923\nTotal prob of N-in: 0.94351\nPOSSIBLE N-term signal sequence\ninside 1 19\nTMhelix 20 39\noutside 40 325\nTMhelix 326 348\ninside 349 372

10991 GCF_000017765.1_ASM1776v1 Escherichia coli HS Proteobacteria; Gammaproteobacteria; Enterobacterales; Enterobacteriaceae; Escherichia; Escherichia coli MTGDNTLIHSHGINRRDFMKLCAALAATMGLSSKAAAEMAESVTNPQRPPVIWIGAQECT WP_000145410.1 MULTISPECIES: hydrogenase-2 small chain [Proteobacteria] Length: 372\nNumber of predicted TMHs: 1\nExp number of AAs in TMHs: 23.72226\nExp number, first 60 AAs: 0.22038\nTotal prob of N-in: 0.12994\noutside 1 331\nTMhelix 332 354\ninside 355 372

10992 GCF_000017765.1_ASM1776v1 Escherichia coli HS Proteobacteria; Gammaproteobacteria; Enterobacterales; Enterobacteriaceae; Escherichia; Escherichia coli MNNEETFYQAMRRQGVTRRSFLKYCSLAATSLGLGAGMAPKIAWALENKPRIPVVWIHGL WP_001058323.1 MULTISPECIES: hydrogenase-1 small chain [Proteobacteria] Length: 372\nNumber of predicted TMHs: 2\nExp number of AAs in TMHs: 37.2608499999999\nExp number, first 60 AAs: 19.52923\nTotal prob of N-in: 0.94351\nPOSSIBLE N-term signal sequence\ninside 1 19\nTMhelix 20 39\noutside 40 325\nTMhelix 326 348\ninside 349 372

10993 GCF_000457405.1_Esch_coli_HVH_100_4-2850729_V1 Escherichia coli HVH 100 (4-2850729) Proteobacteria; Gammaproteobacteria; Enterobacterales; Enterobacteriaceae; Escherichia; Escherichia coli MNNEETFYQAMRRQGVTRRSFLKYCSLAATSLGLGAGMAPKIAWALENKPRIPVVWIHGL WP_001058323.1 MULTISPECIES: hydrogenase-1 small chain [Proteobacteria] Length: 372\nNumber of predicted TMHs: 2\nExp number of AAs in TMHs: 37.2608499999999\nExp number, first 60 AAs: 19.52923\nTotal prob of N-in: 0.94351\nPOSSIBLE N-term signal sequence\ninside 1 19\nTMhelix 20 39\noutside 40 325\nTMhelix 326 348\ninside 349 372

10994 GCF_000457405.1_Esch_coli_HVH_100_4-2850729_V1 Escherichia coli HVH 100 (4-2850729) Proteobacteria; Gammaproteobacteria; Enterobacterales; Enterobacteriaceae; Escherichia; Escherichia coli MTGDNTLIHSHGINRRDFMKLCAALAATMGLSSKAAAEMAESVTNPQRPPVIWIGAQECT WP_000145410.1 MULTISPECIES: hydrogenase-2 small chain [Proteobacteria] Length: 372\nNumber of predicted TMHs: 1\nExp number of AAs in TMHs: 23.72226\nExp number, first 60 AAs: 0.22038\nTotal prob of N-in: 0.12994\noutside 1 331\nTMhelix 332 354\ninside 355 372

10995 GCF_000465155.1_Esch_coli_HVH_102_4-6906788_V1 Escherichia coli HVH 102 (4-6906788) Proteobacteria; Gammaproteobacteria; Enterobacterales; Enterobacteriaceae; Escherichia; Escherichia coli MTGDNTLIHSHGINRRDFMKLCAALAATMGLSSKAAAEMAESVTNPQRPPVIWIGAQECT WP_000145410.1 MULTISPECIES: hydrogenase-2 small chain [Proteobacteria] Length: 372\nNumber of predicted TMHs: 1\nExp number of AAs in TMHs: 23.72226\nExp number, first 60 AAs: 0.22038\nTotal prob of N-in: 0.12994\noutside 1 331\nTMhelix 332 354\ninside 355 372

10996 GCF_000457435.1_Esch_coli_HVH_103_4-5904188_V1 Escherichia coli HVH 103 (4-5904188) Proteobacteria; Gammaproteobacteria; Enterobacterales; Enterobacteriaceae; Escherichia; Escherichia coli MTGDNTLIHSHGINRRDFMKLCAALAATMGLSSKAAAEMAESVTNPQRPPVIWIGAQECT WP_000145410.1 MULTISPECIES: hydrogenase-2 small chain [Proteobacteria] Length: 372\nNumber of predicted TMHs: 1\nExp number of AAs in TMHs: 23.72226\nExp number, first 60 AAs: 0.22038\nTotal prob of N-in: 0.12994\noutside 1 331\nTMhelix 332 354\ninside 355 372

10997 GCF_000457435.1_Esch_coli_HVH_103_4-5904188_V1 Escherichia coli HVH 103 (4-5904188) Proteobacteria; Gammaproteobacteria; Enterobacterales; Enterobacteriaceae; Escherichia; Escherichia coli MNNEETFYQAMRRQGVTRRSFLKYCSLAATSLGLGAGMAPKIAWALENKPRIPVVWIHGL WP_001058323.1 MULTISPECIES: hydrogenase-1 small chain [Proteobacteria] Length: 372\nNumber of predicted TMHs: 2\nExp number of AAs in TMHs: 37.2608499999999\nExp number, first 60 AAs: 19.52923\nTotal prob of N-in: 0.94351\nPOSSIBLE N-term signal sequence\ninside 1 19\nTMhelix 20 39\noutside 40 325\nTMhelix 326 348\ninside 349 372

10998 GCF_000457455.1_Esch_coli_HVH_104_4-6977960_V1 Escherichia coli HVH 104 (4-6977960) Proteobacteria; Gammaproteobacteria; Enterobacterales; Enterobacteriaceae; Escherichia; Escherichia coli MTGDNTLIHSHGINRRDFMKLCAALAATMGLSSKAAAEMAESVTNPQRPPVIWIGAQECT WP_000145410.1 MULTISPECIES: hydrogenase-2 small chain [Proteobacteria] Length: 372\nNumber of predicted TMHs: 1\nExp number of AAs in TMHs: 23.72226\nExp number, first 60 AAs: 0.22038\nTotal prob of N-in: 0.12994\noutside 1 331\nTMhelix 332 354\ninside 355 372

10999 GCF_000456165.1_Esch_coli_HVH_10_4-6832164_V1 Escherichia coli HVH 10 (4-6832164) Proteobacteria; Gammaproteobacteria; Enterobacterales; Enterobacteriaceae; Escherichia; Escherichia coli MTGDNTLIHSHGINRRDFMKLCAALAATMGLSSKAAAEMAESVTNPQRPPVIWIGAQECT WP_000145410.1 MULTISPECIES: hydrogenase-2 small chain [Proteobacteria] Length: 372\nNumber of predicted TMHs: 1\nExp number of AAs in TMHs: 23.72226\nExp number, first 60 AAs: 0.22038\nTotal prob of N-in: 0.12994\noutside 1 331\nTMhelix 332 354\ninside 355 372

11000 GCF_000456165.1_Esch_coli_HVH_10_4-6832164_V1 Escherichia coli HVH 10 (4-6832164) Proteobacteria; Gammaproteobacteria; Enterobacterales; Enterobacteriaceae; Escherichia; Escherichia coli MNNEETFYQAMRRQGVTRRSFLKYCSLAATSLGLGAGMAPKIAWALENKPRIPVVWIHGL WP_001058323.1 MULTISPECIES: hydrogenase-1 small chain [Proteobacteria] Length: 372\nNumber of predicted TMHs: 2\nExp number of AAs in TMHs: 37.2608499999999\nExp number, first 60 AAs: 19.52923\nTotal prob of N-in: 0.94351\nPOSSIBLE N-term signal sequence\ninside 1 19\nTMhelix 20 39\noutside 40 325\nTMhelix 326 348\ninside 349 372

11001 GCF_000457475.1_Esch_coli_HVH_106_4-6881831_V1 Escherichia coli HVH 106 (4-6881831) Proteobacteria; Gammaproteobacteria; Enterobacterales; Enterobacteriaceae; Escherichia; Escherichia coli MTGDNTLIHSHGINRRDFMKLCAALAATMGLSSKAAAEMAESVTNPQRPPVIWIGAQECT WP_000145410.1 MULTISPECIES: hydrogenase-2 small chain [Proteobacteria] Length: 372\nNumber of predicted TMHs: 1\nExp number of AAs in TMHs: 23.72226\nExp number, first 60 AAs: 0.22038\nTotal prob of N-in: 0.12994\noutside 1 331\nTMhelix 332 354\ninside 355 372

11002 GCF_000457475.1_Esch_coli_HVH_106_4-6881831_V1 Escherichia coli HVH 106 (4-6881831) Proteobacteria; Gammaproteobacteria; Enterobacterales; Enterobacteriaceae; Escherichia; Escherichia coli MNNEETFYQAMRRQGVTRRSFLKYCSLAATSLGLGAGMAPKIAWALENKPRIPVVWIHGL WP_001058323.1 MULTISPECIES: hydrogenase-1 small chain [Proteobacteria] Length: 372\nNumber of predicted TMHs: 2\nExp number of AAs in TMHs: 37.2608499999999\nExp number, first 60 AAs: 19.52923\nTotal prob of N-in: 0.94351\nPOSSIBLE N-term signal sequence\ninside 1 19\nTMhelix 20 39\noutside 40 325\nTMhelix 326 348\ninside 349 372

11003 GCF_000457495.1_Esch_coli_HVH_107_4-5860571_V1 Escherichia coli HVH 107 (4-5860571) Proteobacteria; Gammaproteobacteria; Enterobacterales; Enterobacteriaceae; Escherichia; Escherichia coli MNNEETFYQAMRRQGVTRRSFLKYCSLAATSLGLGAGMAPKIAWALENKPRIPVVWIHGL WP_001538532.1 hydrogenase-1 small chain [Escherichia coli] Length: 372\nNumber of predicted TMHs: 2\nExp number of AAs in TMHs: 37.27041\nExp number, first 60 AAs: 19.52927\nTotal prob of N-in: 0.94352\nPOSSIBLE N-term signal sequence\ninside 1 19\nTMhelix 20 39\noutside 40 325\nTMhelix 326 348\ninside 349 372

11004 GCF_000457495.1_Esch_coli_HVH_107_4-5860571_V1 Escherichia coli HVH 107 (4-5860571) Proteobacteria; Gammaproteobacteria; Enterobacterales; Enterobacteriaceae; Escherichia; Escherichia coli

MTGDNTLIHSHGINRRDFMKLCAALAATMGLSSKAAAEMVESVTNPQRPPVIWIGAQECT WP_000145423.1 hydrogenase 2 small subunit [Escherichia coli] Length: 372\nNumber of predicted TMHs: 1\nExp number of AAs in TMHs: 23.78201\nExp number, first 60 AAs: 0.28355\nTotal prob of N-in: 0.13257\noutside 1 331\nTMhelix 332 354\ninside 355 372

11005 GCF_000495095.1_Esch_coli_HVH_108_4-6924867_V1 Escherichia coli HVH 108 (4-6924867) Proteobacteria; Gammaproteobacteria; Enterobacterales; Enterobacteriaceae; Escherichia; Escherichia coli MNNEETFYQAMRRQGVTRRSFLKYCSLAATSLGLGAGMAPKIAWALENKPRIPVVWIHGL WP_001295940.1 MULTISPECIES: hydrogenase [Enterobacteriaceae] Length: 372\nNumber of predicted TMHs: 2\nExp number of AAs in TMHs: 37.2608899999999\nExp number, first 60 AAs: 19.52923\nTotal prob of N-in: 0.94351\nPOSSIBLE N-term signal sequence\ninside 1 19\nTMhelix 20 39\noutside 40 325\nTMhelix 326 348\ninside 349 372

11006 GCF_000495095.1_Esch_coli_HVH_108_4-6924867_V1 Escherichia coli HVH 108 (4-6924867) Proteobacteria; Gammaproteobacteria; Enterobacterales; Enterobacteriaceae; Escherichia; Escherichia coli MTGDNTLIHSHGINRRDFMKLCAALAATMGLSSKAAAEMAESVTNPQRPPVIWIGAQECT WP_000145410.1 MULTISPECIES: hydrogenase-2 small chain [Proteobacteria] Length: 372\nNumber of predicted TMHs: 1\nExp number of AAs in TMHs: 23.72226\nExp number, first 60 AAs: 0.22038\nTotal prob of N-in: 0.12994\noutside 1 331\nTMhelix 332 354\ninside 355 372

11007 GCF_000457515.1_Esch_coli_HVH_109_4-6977162_V1 Escherichia coli HVH 109 (4-6977162) Proteobacteria; Gammaproteobacteria; Enterobacterales; Enterobacteriaceae; Escherichia; Escherichia coli MTGDNTLIHSHGINRRDFMKLCAALAATMGLSSKAAAEMAESVTNPQRPPVIWIGAQECT WP_000145410.1 MULTISPECIES: hydrogenase-2 small chain [Proteobacteria] Length: 372\nNumber of predicted TMHs: 1\nExp number of AAs in TMHs: 23.72226\nExp number, first 60 AAs: 0.22038\nTotal prob of N-in: 0.12994\noutside 1 331\nTMhelix 332 354\ninside 355 372

11008 GCF_000457515.1_Esch_coli_HVH_109_4-6977162_V1 Escherichia coli HVH 109 (4-6977162) Proteobacteria; Gammaproteobacteria; Enterobacterales; Enterobacteriaceae; Escherichia; Escherichia coli MNNEETFYQAMRRQGVTRRSFLKYCSLAATSLGLGAGMAPKIAWALENKPRIPVVWIHGL WP_001058323.1 MULTISPECIES: hydrogenase-1 small chain [Proteobacteria] Length: 372\nNumber of predicted TMHs: 2\nExp number of AAs in TMHs: 37.2608499999999\nExp number, first 60 AAs: 19.52923\nTotal prob of N-in: 0.94351\nPOSSIBLE N-term signal sequence\ninside 1 19\nTMhelix 20 39\noutside 40 325\nTMhelix 326 348\ninside 349 372

11009 GCF_000457535.1_Esch_coli_HVH_110_4-6978754_V1 Escherichia coli HVH 110 (4-6978754) Proteobacteria; Gammaproteobacteria; Enterobacterales; Enterobacteriaceae; Escherichia; Escherichia coli MNNEETFYQAMRRQGVTRRSFLKYCSLAATSLGLGAGMAPKIAWALENKPRIPVVWIHGL WP_001058323.1 MULTISPECIES: hydrogenase-1 small chain [Proteobacteria] Length: 372\nNumber of predicted TMHs: 2\nExp number of AAs in TMHs: 37.2608499999999\nExp number, first 60 AAs: 19.52923\nTotal prob of N-in: 0.94351\nPOSSIBLE N-term signal sequence\ninside 1 19\nTMhelix 20 39\noutside 40 325\nTMhelix 326 348\ninside 349 372

11010 GCF_000457535.1_Esch_coli_HVH_110_4-6978754_V1 Escherichia coli HVH 110 (4-6978754) Proteobacteria; Gammaproteobacteria; Enterobacterales; Enterobacteriaceae; Escherichia; Escherichia coli MTGDNTLIHSHGINRRDFMKLCAALAATMGLSSKAAAEMAESVTNPQRPPVIWIGAQECT WP_000145410.1 MULTISPECIES: hydrogenase-2 small chain [Proteobacteria] Length: 372\nNumber of predicted TMHs: 1\nExp number of AAs in TMHs: 23.72226\nExp number, first 60 AAs: 0.22038\nTotal prob of N-in: 0.12994\noutside 1 331\nTMhelix 332 354\ninside 355 372

11011 GCF_000457555.1_Esch_coli_HVH_111_4-7039018_V1 Escherichia coli HVH 111 (4-7039018) Proteobacteria; Gammaproteobacteria; Enterobacterales; Enterobacteriaceae; Escherichia; Escherichia coli MNNEETFYQAMRRQGVTRRSFLKYCSLAATSLGLGAGMAPKIAWALENKPRIPVVWIHGL WP_001058323.1 MULTISPECIES: hydrogenase-1 small chain [Proteobacteria] Length: 372\nNumber of predicted TMHs: 2\nExp number of AAs in TMHs: 37.2608499999999\nExp number, first 60 AAs: 19.52923\nTotal prob of N-in: 0.94351\nPOSSIBLE N-term signal sequence\ninside 1 19\nTMhelix 20 39\noutside 40 325\nTMhelix 326 348\ninside 349 372

11012 GCF_000457555.1_Esch_coli_HVH_111_4-7039018_V1 Escherichia coli HVH 111 (4-7039018) Proteobacteria; Gammaproteobacteria; Enterobacterales; Enterobacteriaceae; Escherichia; Escherichia coli MTGDNTLIHSHGINRRDFMKLCAALAATMGLSSKAAAEMAESVTNPQRPPVIWIGAQECT WP_000145410.1

MULTISPECIES: hydrogenase-2 small chain [Proteobacteria] Length: 372\nNumber of predicted TMHs: 1\nExp number of AAs in TMHs: 23.72226\nExp number, first 60 AAs: 0.22038\nTotal prob of N-in: 0.12994\noutside 1 331\nTMhelix 332 354\ninside 355 372

11013 GCF_000457575.1_Esch_coli_HVH_112_4-5987253_V1 Escherichia coli HVH 112 (4-5987253) Proteobacteria; Gammaproteobacteria; Enterobacterales; Enterobacteriaceae; Escherichia; Escherichia coli MNNEETFYQAMRRQGVTRRSFLKYCSLAATSLGLGAGMAPKIAWALENKPRIPVVWIHGL WP_001058323.1 MULTISPECIES: hydrogenase-1 small chain [Proteobacteria] Length: 372\nNumber of predicted TMHs: 2\nExp number of AAs in TMHs: 37.2608499999999\nExp number, first 60 AAs: 19.52923\nTotal prob of N-in: 0.94351\nPOSSIBLE N-term signal sequence\ninside 1 19\nTMhelix 20 39\noutside 40 325\nTMhelix 326 348\ninside 349 372

11014 GCF_000457575.1_Esch_coli_HVH_112_4-5987253_V1 Escherichia coli HVH 112 (4-5987253) Proteobacteria; Gammaproteobacteria; Enterobacterales; Enterobacteriaceae; Escherichia; Escherichia coli MTGDNTLIHSHGINRRDFMKLCAALAATMGLSSKAAAEMAESVTNPQRPPVIWIGAQECT WP_000145410.1 MULTISPECIES: hydrogenase-2 small chain [Proteobacteria] Length: 372\nNumber of predicted TMHs: 1\nExp number of AAs in TMHs: 23.72226\nExp number, first 60 AAs: 0.22038\nTotal prob of N-in: 0.12994\noutside 1 331\nTMhelix 332 354\ninside 355 372

11015 GCF_000457595.1_Esch_coli_HVH_113_4-7535473_V1 Escherichia coli HVH 113 (4-7535473) Proteobacteria; Gammaproteobacteria; Enterobacterales; Enterobacteriaceae; Escherichia; Escherichia coli MNNEETFYQAMRRQGVTRRSFLKYCSLAATSLGLGAGMAPKIAWALENKPRIPVVWIHGL WP_001058323.1 MULTISPECIES: hydrogenase-1 small chain [Proteobacteria] Length: 372\nNumber of predicted TMHs: 2\nExp number of AAs in TMHs: 37.2608499999999\nExp number, first 60 AAs: 19.52923\nTotal prob of N-in: 0.94351\nPOSSIBLE N-term signal sequence\ninside 1 19\nTMhelix 20 39\noutside 40 325\nTMhelix 326 348\ninside 349 372

11016 GCF_000457595.1_Esch_coli_HVH_113_4-7535473_V1 Escherichia coli HVH 113 (4-7535473) Proteobacteria; Gammaproteobacteria; Enterobacterales; Enterobacteriaceae; Escherichia; Escherichia coli MTGDNTLIHSHGINRRDFMKLCAALAATMGLSSKAAAEMAESVTNPQRPPVIWIGAQECT WP_000145410.1 MULTISPECIES: hydrogenase-2 small chain [Proteobacteria] Length: 372\nNumber of predicted TMHs: 1\nExp number of AAs in TMHs: 23.72226\nExp number, first 60 AAs: 0.22038\nTotal prob of N-in: 0.12994\noutside 1 331\nTMhelix 332 354\ninside 355 372

11017 GCF_000457615.1_Esch_coli_HVH_114_4-7037740_V1 Escherichia coli HVH 114 (4-7037740) Proteobacteria; Gammaproteobacteria; Enterobacterales; Enterobacteriaceae; Escherichia; Escherichia coli MNNEETFYQAMRRQGVTRRSFLKYCSLAATSLGLGAGMAPKIAWALENKPRIPVVWIHGL WP_001058323.1 MULTISPECIES: hydrogenase-1 small chain [Proteobacteria] Length: 372\nNumber of predicted TMHs: 2\nExp number of AAs in TMHs: 37.2608499999999\nExp number, first 60 AAs: 19.52923\nTotal prob of N-in: 0.94351\nPOSSIBLE N-term signal sequence\ninside 1 19\nTMhelix 20 39\noutside 40 325\nTMhelix 326 348\ninside 349 372

11018 GCF_000457615.1_Esch_coli_HVH_114_4-7037740_V1 Escherichia coli HVH 114 (4-7037740) Proteobacteria; Gammaproteobacteria; Enterobacterales; Enterobacteriaceae; Escherichia; Escherichia coli MTGDNTLIHSHGINRRDFMKLCAALAATMGLSSKAAAEMAESVTNPQRPPVIWIGAQECT WP_000145410.1 MULTISPECIES: hydrogenase-2 small chain [Proteobacteria] Length: 372\nNumber of predicted TMHs: 1\nExp number of AAs in TMHs: 23.72226\nExp number, first 60 AAs: 0.22038\nTotal prob of N-in: 0.12994\noutside 1 331\nTMhelix 332 354\ninside 355 372

11019 GCF_000457655.1_Esch_coli_HVH_115_4-4465989_V1 Escherichia coli HVH 115 (4-4465989) Proteobacteria; Gammaproteobacteria; Enterobacterales; Enterobacteriaceae; Escherichia; Escherichia coli MNNEETFYQAMRRQGVTRRSFLKYCSLAATSLGLGAGMAPKIAWALENKPRIPVVWIHGL WP_001058323.1 MULTISPECIES: hydrogenase-1 small chain [Proteobacteria] Length: 372\nNumber of predicted TMHs: 2\nExp number of AAs in TMHs: 37.2608499999999\nExp number, first 60 AAs: 19.52923\nTotal prob of N-in: 0.94351\nPOSSIBLE N-term signal sequence\ninside 1 19\nTMhelix 20 39\noutside 40 325\nTMhelix 326 348\ninside 349 372

11020 GCF_000457655.1_Esch_coli_HVH_115_4-4465989_V1 Escherichia coli HVH 115 (4-4465989) Proteobacteria; Gammaproteobacteria; Enterobacterales; Enterobacteriaceae; Escherichia; Escherichia coli MTGDNTLIHSHGINRRDFMKLCAALAATMGLSSKAAAEMAESVTNPQRPPVIWIGAQECT WP_000145410.1 MULTISPECIES: hydrogenase-2 small chain [Proteobacteria] Length: 372\nNumber of predicted TMHs:

1\nExp number of AAs in TMHs: 23.72226\nExp number, first 60 AAs: 0.22038\nTotal prob of N-in: 0.12994\noutside 1 331\nTMhelix 332 354\ninside 355 372

11021 GCF_000457635.1_Esch_coli_HVH_115_4-4465997_V1 Escherichia coli HVH 115 (4-4465997) Proteobacteria; Gammaproteobacteria; Enterobacterales; Enterobacteriaceae; Escherichia; Escherichia coli MTGDNTLIHSHGINRRDFMKLCAALAATMGLSSKAAAEMAESVTNPQRPPVIWIGAQECT WP_000145410.1 MULTISPECIES: hydrogenase-2 small chain [Proteobacteria] Length: 372\nNumber of predicted TMHs: 1\nExp number of AAs in TMHs: 23.72226\nExp number, first 60 AAs: 0.22038\nTotal prob of N-in: 0.12994\noutside 1 331\nTMhelix 332 354\ninside 355 372

11022 GCF_000457635.1_Esch_coli_HVH_115_4-4465997_V1 Escherichia coli HVH 115 (4-4465997) Proteobacteria; Gammaproteobacteria; Enterobacterales; Enterobacteriaceae; Escherichia; Escherichia coli MNNEETFYQAMRRQGVTRRSFLKYCSLAATSLGLGAGMAPKIAWALENKPRIPVVWIHGL WP_001058323.1 MULTISPECIES: hydrogenase-1 small chain [Proteobacteria] Length: 372\nNumber of predicted TMHs: 2\nExp number of AAs in TMHs: 37.2608499999999\nExp number, first 60 AAs: 19.52923\nTotal prob of N-in: 0.94351\nPOSSIBLE N-term signal sequence\ninside 1 19\nTMhelix 20 39\noutside 40 325\nTMhelix 326 348\ninside 349 372

11023 GCF_000457675.1_Esch_coli_HVH_116_4-6879942_V1 Escherichia coli HVH 116 (4-6879942) Proteobacteria; Gammaproteobacteria; Enterobacterales; Enterobacteriaceae; Escherichia; Escherichia coli MTGDNTLIHSHGINRRDFMKLCAALAATMGLSSKAAAEMAESVTNPQRPPVIWIGAQECT WP_000145410.1 MULTISPECIES: hydrogenase-2 small chain [Proteobacteria] Length: 372\nNumber of predicted TMHs: 1\nExp number of AAs in TMHs: 23.72226\nExp number, first 60 AAs: 0.22038\nTotal prob of N-in: 0.12994\noutside 1 331\nTMhelix 332 354\ninside 355 372

11024 GCF_000457675.1_Esch_coli_HVH_116_4-6879942_V1 Escherichia coli HVH 116 (4-6879942) Proteobacteria; Gammaproteobacteria; Enterobacterales; Enterobacteriaceae; Escherichia; Escherichia coli MNNEETFYQAMRRQGVTRRSFLKYCSLAATSLGLGAGMAPKIAWALENKPRIPVVWIHGL WP_001058323.1 MULTISPECIES: hydrogenase-1 small chain [Proteobacteria] Length: 372\nNumber of predicted TMHs: 2\nExp number of AAs in TMHs: 37.2608499999999\nExp number, first 60 AAs: 19.52923\nTotal prob of N-in: 0.94351\nPOSSIBLE N-term signal sequence\ninside 1 19\nTMhelix 20 39\noutside 40 325\nTMhelix 326 348\ninside 349 372

11025 GCF_000457695.1_Esch_coli_HVH_117_4-6857191_V1 Escherichia coli HVH 117 (4-6857191) Proteobacteria; Gammaproteobacteria; Enterobacterales; Enterobacteriaceae; Escherichia; Escherichia coli MTGDNTLIHSHGINRRDFMKLCAALAATMGLSSKAAAEMAESVTNPQRPPVIWIGAQECT WP_000145410.1 MULTISPECIES: hydrogenase-2 small chain [Proteobacteria] Length: 372\nNumber of predicted TMHs: 1\nExp number of AAs in TMHs: 23.72226\nExp number, first 60 AAs: 0.22038\nTotal prob of N-in: 0.12994\noutside 1 331\nTMhelix 332 354\ninside 355 372

11026 GCF_000457695.1_Esch_coli_HVH_117_4-6857191_V1 Escherichia coli HVH 117 (4-6857191) Proteobacteria; Gammaproteobacteria; Enterobacterales; Enterobacteriaceae; Escherichia; Escherichia coli MNNEETFYQAMRRQGVTRRSFLKYCSLAATSLGLGAGMAPKIAWALENKPRIPVVWIHGL WP_001058323.1 MULTISPECIES: hydrogenase-1 small chain [Proteobacteria] Length: 372\nNumber of predicted TMHs: 2\nExp number of AAs in TMHs: 37.2608499999999\nExp number, first 60 AAs: 19.52923\nTotal prob of N-in: 0.94351\nPOSSIBLE N-term signal sequence\ninside 1 19\nTMhelix 20 39\noutside 40 325\nTMhelix 326 348\ninside 349 372

11027 GCF_000457715.1_Esch_coli_HVH_118_4-7345399_V1 Escherichia coli HVH 118 (4-7345399) Proteobacteria; Gammaproteobacteria; Enterobacterales; Enterobacteriaceae; Escherichia; Escherichia coli MTGDNTLIHSHGINRRDFMKLCAALAATMGLSSKAAAEMAESVTNPQRPPVIWIGAQECT WP_000145410.1 MULTISPECIES: hydrogenase-2 small chain [Proteobacteria] Length: 372\nNumber of predicted TMHs: 1\nExp number of AAs in TMHs: 23.72226\nExp number, first 60 AAs: 0.22038\nTotal prob of N-in: 0.12994\noutside 1 331\nTMhelix 332 354\ninside 355 372

11028 GCF_000457735.1_Esch_coli_HVH_119_4-6879578_V1 Escherichia coli HVH 119 (4-6879578) Proteobacteria; Gammaproteobacteria; Enterobacterales; Enterobacteriaceae; Escherichia; Escherichia coli MTGDNTLIHSHGINRRDFMKLCAALAATMGLSSKAAAEMAESVTNPQRPPVIWIGAQECT WP_000145410.1 MULTISPECIES: hydrogenase-2 small chain [Proteobacteria] Length: 372\nNumber of predicted TMHs: 1\nExp number of AAs in TMHs: 23.72226\nExp number, first 60 AAs: 0.22038\nTotal prob of N-in: 0.12994\noutside 1 331\nTMhelix 332 354\ninside 355 372

11029 GCF_000457735.1_Esch_coli_HVH_119_4-6879578_V1 Escherichia coli HVH 119 (4-6879578) Proteobacteria; Gammaproteobacteria; Enterobacterales; Enterobacteriaceae; Escherichia; Escherichia coli MNNEETFYQAMRRQGVTRRSFLKYCSLAATSLGLGAGMAPKIAWALENKPRIPVVWIHGL WP_001058323.1 MULTISPECIES: hydrogenase-1 small chain [Proteobacteria] Length: 372\nNumber of predicted TMHs: 2\nExp number of AAs in TMHs: 37.2608499999999\nExp number, first 60 AAs: 19.52923\nTotal prob of N-in: 0.94351\nPOSSIBLE N-term signal sequence\ninside 1 19\nTMhelix 20 39\noutside 40 325\nTMhelix 326 348\ninside 349 372

11030 GCF_000457755.1_Esch_coli_HVH_120_4-6978681_V1 Escherichia coli HVH 120 (4-6978681) Proteobacteria; Gammaproteobacteria; Enterobacterales; Enterobacteriaceae; Escherichia; Escherichia coli MTGDNTLIHSHGINRRDFMKLCAALAATMGLSSKAAAEMAESVTNPQRPPVIWIGAQECT WP_000145410.1 MULTISPECIES: hydrogenase-2 small chain [Proteobacteria] Length: 372\nNumber of predicted TMHs: 1\nExp number of AAs in TMHs: 23.72226\nExp number, first 60 AAs: 0.22038\nTotal prob of N-in: 0.12994\noutside 1 331\nTMhelix 332 354\ninside 355 372

11031 GCF_000457755.1_Esch_coli_HVH_120_4-6978681_V1 Escherichia coli HVH 120 (4-6978681) Proteobacteria; Gammaproteobacteria; Enterobacterales; Enterobacteriaceae; Escherichia; Escherichia coli MNNEETFYQAMRRQGVTRRSFLKYCSLAATSLGLGAGMAPKIAWALENKPRIPVVWIHGL WP_001058323.1 MULTISPECIES: hydrogenase-1 small chain [Proteobacteria] Length: 372\nNumber of predicted TMHs: 2\nExp number of AAs in TMHs: 37.2608499999999\nExp number, first 60 AAs: 19.52923\nTotal prob of N-in: 0.94351\nPOSSIBLE N-term signal sequence\ninside 1 19\nTMhelix 20 39\noutside 40 325\nTMhelix 326 348\ninside 349 372

11032 GCF_000457775.1_Esch_coli_HVH_121_4-6877826_V1 Escherichia coli HVH 121 (4-6877826) Proteobacteria; Gammaproteobacteria; Enterobacterales; Enterobacteriaceae; Escherichia; Escherichia coli MNNEETFYQAMRRQGVTRRSFLKYCSLAATSLGLGAGMAPKIAWALENKPRIPVVWIHGL WP_001058320.1 hydrogenase [Escherichia coli] Length: 372\nNumber of predicted TMHs: 2\nExp number of AAs in TMHs: 37.26082\nExp number, first 60 AAs: 19.52923\nTotal prob of N-in: 0.94351\nPOSSIBLE N-term signal sequence\ninside 1 19\nTMhelix 20 39\noutside 40 325\nTMhelix 326 348\ninside 349 372

11033 GCF_000457775.1_Esch_coli_HVH_121_4-6877826_V1 Escherichia coli HVH 121 (4-6877826) Proteobacteria; Gammaproteobacteria; Enterobacterales; Enterobacteriaceae; Escherichia; Escherichia coli MTGDNTLIHSHGINRRDFMKLCAALAATMGLSSKAAAEMAESVTNPQRPPVIWIGAQECT WP_000145410.1 MULTISPECIES: hydrogenase-2 small chain [Proteobacteria] Length: 372\nNumber of predicted TMHs: 1\nExp number of AAs in TMHs: 23.72226\nExp number, first 60 AAs: 0.22038\nTotal prob of N-in: 0.12994\noutside 1 331\nTMhelix 332 354\ninside 355 372

11034 GCF_000457795.1_Esch_coli_HVH_122_4-6851606_V1 Escherichia coli HVH 122 (4-6851606) Proteobacteria; Gammaproteobacteria; Enterobacterales; Enterobacteriaceae; Escherichia; Escherichia coli MTGDNTLIHSHGINRRDFMKLCAALAATMGLSSKAAAEMAESVTNPQRPPVIWIGAQECT WP_000145410.1 MULTISPECIES: hydrogenase-2 small chain [Proteobacteria] Length: 372\nNumber of predicted TMHs: 1\nExp number of AAs in TMHs: 23.72226\nExp number, first 60 AAs: 0.22038\nTotal prob of N-in: 0.12994\noutside 1 331\nTMhelix 332 354\ninside 355 372

11035 GCF_000457795.1_Esch_coli_HVH_122_4-6851606_V1 Escherichia coli HVH 122 (4-6851606) Proteobacteria; Gammaproteobacteria; Enterobacterales; Enterobacteriaceae; Escherichia; Escherichia coli MNNEETFYQAMRRQGVTRRSFLKYCSLAATSLGLGAGMAPKIAWALENKPRIPVVWIHGL WP_001058323.1 MULTISPECIES: hydrogenase-1 small chain [Proteobacteria] Length: 372\nNumber of predicted TMHs: 2\nExp number of AAs in TMHs: 37.2608499999999\nExp number, first 60 AAs: 19.52923\nTotal prob of N-in: 0.94351\nPOSSIBLE N-term signal sequence\ninside 1 19\nTMhelix 20 39\noutside 40 325\nTMhelix 326 348\ninside 349 372

11036 GCF_000494955.1_Esch_coli_HVH_12_4-7653042_V1 Escherichia coli HVH 12 (4-7653042) Proteobacteria; Gammaproteobacteria; Enterobacterales; Enterobacteriaceae; Escherichia; Escherichia coli MTGDNTLIHSHGINRRDFMKLCAALAATMGLSSKAAAEMAESVTNPQRPPVIWIGAQECT WP_000145410.1 MULTISPECIES: hydrogenase-2 small chain [Proteobacteria] Length: 372\nNumber of predicted TMHs: 1\nExp number of AAs in TMHs: 23.72226\nExp number, first 60 AAs: 0.22038\nTotal prob of N-in: 0.12994\noutside 1 331\nTMhelix 332 354\ninside 355 372

11037 GCF_000457815.1_Esch_coli_HVH_125_4-2634716_V1 Escherichia coli HVH 125 (4-2634716) Proteobacteria; Gammaproteobacteria; Enterobacterales; Enterobacteriaceae; Escherichia; Escherichia coli

MNNEETFYQAMRRQGVTRRSFLKYCSLAATSLGLGAGMAPKIAWALENKPRIPVVWIHGL WP_001058323.1 MULTISPECIES: hydrogenase-1 small chain [Proteobacteria] Length: 372\nNumber of predicted TMHs: 2\nExp number of AAs in TMHs: 37.2608499999999\nExp number, first 60 AAs: 19.52923\nTotal prob of N-in: 0.94351\nPOSSIBLE N-term signal sequence\ninside 1 19\nTMhelix 20 39\noutside 40 325\nTMhelix 326 348\ninside 349 372

11038 GCF_000457815.1_Esch_coli_HVH_125_4-2634716_V1 Escherichia coli HVH 125 (4-2634716) Proteobacteria; Gammaproteobacteria; Enterobacterales; Enterobacteriaceae; Escherichia; Escherichia coli MTGDNTLIHSHGINRRDFMKLCAALAATMGLSSKAAAEMAESVTNPQRPPVIWIGAQECT WP_000145410.1 MULTISPECIES: hydrogenase-2 small chain [Proteobacteria] Length: 372\nNumber of predicted TMHs: 1\nExp number of AAs in TMHs: 23.72226\nExp number, first 60 AAs: 0.22038\nTotal prob of N-in: 0.12994\noutside 1 331\nTMhelix 332 354\ninside 355 372

11039 GCF_000457835.1_Esch_coli_HVH_126_4-6034225_V1 Escherichia coli HVH 126 (4-6034225) Proteobacteria; Gammaproteobacteria; Enterobacterales; Enterobacteriaceae; Escherichia; Escherichia coli MTGDNTLIHSHGINRRDFMKLCAALAATMGLSSKAAAEMAESVTNPQRPPVIWIGAQECT WP_000145410.1 MULTISPECIES: hydrogenase-2 small chain [Proteobacteria] Length: 372\nNumber of predicted TMHs: 1\nExp number of AAs in TMHs: 23.72226\nExp number, first 60 AAs: 0.22038\nTotal prob of N-in: 0.12994\noutside 1 331\nTMhelix 332 354\ninside 355 372

11040 GCF_000457855.1_Esch_coli_HVH_127_4-7303629_V1 Escherichia coli HVH 127 (4-7303629) Proteobacteria; Gammaproteobacteria; Enterobacterales; Enterobacteriaceae; Escherichia; Escherichia coli MTGDNTLIHSHGINRRDFMKLCAALAATMGLSSKAAAEMAESVTNPQRPPVIWIGAQECT WP_000145410.1 MULTISPECIES: hydrogenase-2 small chain [Proteobacteria] Length: 372\nNumber of predicted TMHs: 1\nExp number of AAs in TMHs: 23.72226\nExp number, first 60 AAs: 0.22038\nTotal prob of N-in: 0.12994\noutside 1 331\nTMhelix 332 354\ninside 355 372

11041 GCF_000457875.1_Esch_coli_HVH_128_4-7030436_V1 Escherichia coli HVH 128 (4-7030436) Proteobacteria; Gammaproteobacteria; Enterobacterales; Enterobacteriaceae; Escherichia; Escherichia coli MNNEETFYQAMRRQGVTRRSFLKYCSLAATSLGLGAGMAPKIAWALENKPRIPVVWIHGL WP_001538532.1 hydrogenase-1 small chain [Escherichia coli] Length: 372\nNumber of predicted TMHs: 2\nExp number of AAs in TMHs: 37.27041\nExp number, first 60 AAs: 19.52927\nTotal prob of N-in: 0.94352\nPOSSIBLE N-term signal sequence\ninside 1 19\nTMhelix 20 39\noutside 40 325\nTMhelix 326 348\ninside 349 372

11042 GCF_000457875.1_Esch_coli_HVH_128_4-7030436_V1 Escherichia coli HVH 128 (4-7030436) Proteobacteria; Gammaproteobacteria; Enterobacterales; Enterobacteriaceae; Escherichia; Escherichia coli MTGDNTLIHSHGINRRDFMKLCAALAATMGLSSKAAAEMVESVTNPQRPPVIWIGAQECT WP_000145423.1 hydrogenase 2 small subunit [Escherichia coli] Length: 372\nNumber of predicted TMHs: 1\nExp number of AAs in TMHs: 23.78201\nExp number, first 60 AAs: 0.28355\nTotal prob of N-in: 0.13257\noutside 1 331\nTMhelix 332 354\ninside 355 372

11043 GCF_000457895.1_Esch_coli_HVH_130_4-7036876_V1 Escherichia coli HVH 130 (4-7036876) Proteobacteria; Gammaproteobacteria; Enterobacterales; Enterobacteriaceae; Escherichia; Escherichia coli MTGDNTLIHSHGINRRDFMKLCAALAATMGLSSKAAAEMAESVTNPQRPPVIWIGAQECT WP_000145410.1 MULTISPECIES: hydrogenase-2 small chain [Proteobacteria] Length: 372\nNumber of predicted TMHs: 1\nExp number of AAs in TMHs: 23.72226\nExp number, first 60 AAs: 0.22038\nTotal prob of N-in: 0.12994\noutside 1 331\nTMhelix 332 354\ninside 355 372

11044 GCF_000457895.1_Esch_coli_HVH_130_4-7036876_V1 Escherichia coli HVH 130 (4-7036876) Proteobacteria; Gammaproteobacteria; Enterobacterales; Enterobacteriaceae; Escherichia; Escherichia coli MNNEETFYQAMRRQGVTRRSFLKYCSLAATSLGLGAGMAPKIAWALENKPRIPVVWIHGL WP_001058323.1 MULTISPECIES: hydrogenase-1 small chain [Proteobacteria] Length: 372\nNumber of predicted TMHs: 2\nExp number of AAs in TMHs: 37.2608499999999\nExp number, first 60 AAs: 19.52923\nTotal prob of N-in: 0.94351\nPOSSIBLE N-term signal sequence\ninside 1 19\nTMhelix 20 39\noutside 40 325\nTMhelix 326 348\ninside 349 372

11045 GCF_000457915.1_Esch_coli_HVH_132_4-6876862_V1 Escherichia coli HVH 132 (4-6876862) Proteobacteria; Gammaproteobacteria; Enterobacterales; Enterobacteriaceae; Escherichia; Escherichia coli MTGDNTLIHSHGINRRDFMKLCAALAATMGLSSKAAAEMAESVTNPQRPPVIWIGAQECT WP_000145410.1 MULTISPECIES: hydrogenase-2 small chain [Proteobacteria] Length: 372\nNumber of predicted TMHs:

1\nExp number of AAs in TMHs: 23.72226\nExp number, first 60 AAs: 0.22038\nTotal prob of N-in: 0.12994\noutside 1 331\nTMhelix 332 354\ninside 355 372

11046 GCF_000457915.1_Esch_coli_HVH_132_4-6876862_V1 Escherichia coli HVH 132 (4-6876862) Proteobacteria; Gammaproteobacteria; Enterobacterales; Enterobacteriaceae; Escherichia; Escherichia coli MNNEETFYQAMRRQGVTRRSFLKYCSLAATSLGLGAGMAPKIAWALENKPRIPVVWIHGL WP_001058323.1 MULTISPECIES: hydrogenase-1 small chain [Proteobacteria] Length: 372\nNumber of predicted TMHs: 2\nExp number of AAs in TMHs: 37.2608499999999\nExp number, first 60 AAs: 19.52923\nTotal prob of N-in: 0.94351\nPOSSIBLE N-term signal sequence\ninside 1 19\nTMhelix 20 39\noutside 40 325\nTMhelix 326 348\ninside 349 372

11047 GCF_000457935.1_Esch_coli_HVH_133_4-4466519_V1 Escherichia coli HVH 133 (4-4466519) Proteobacteria; Gammaproteobacteria; Enterobacterales; Enterobacteriaceae; Escherichia; Escherichia coli MTGDNTLIHSHGINRRDFMKLCAALAATMGLSSKAAAEMAESVTNPQRPPVIWIGAQECT WP_000145410.1 MULTISPECIES: hydrogenase-2 small chain [Proteobacteria] Length: 372\nNumber of predicted TMHs: 1\nExp number of AAs in TMHs: 23.72226\nExp number, first 60 AAs: 0.22038\nTotal prob of N-in: 0.12994\noutside 1 331\nTMhelix 332 354\ninside 355 372

11048 GCF_000457935.1_Esch_coli_HVH_133_4-4466519_V1 Escherichia coli HVH 133 (4-4466519) Proteobacteria; Gammaproteobacteria; Enterobacterales; Enterobacteriaceae; Escherichia; Escherichia coli MNNEETFYQAMRRQGVTRRSFLKYCSLAATSLGLGAGMAPKIAWALENKPRIPVVWIHGL WP_001058323.1 MULTISPECIES: hydrogenase-1 small chain [Proteobacteria] Length: 372\nNumber of predicted TMHs: 2\nExp number of AAs in TMHs: 37.2608499999999\nExp number, first 60 AAs: 19.52923\nTotal prob of N-in: 0.94351\nPOSSIBLE N-term signal sequence\ninside 1 19\nTMhelix 20 39\noutside 40 325\nTMhelix 326 348\ninside 349 372

11049 GCF_000457955.1_Esch_coli_HVH_134_4-6073441_V1 Escherichia coli HVH 134 (4-6073441) Proteobacteria; Gammaproteobacteria; Enterobacterales; Enterobacteriaceae; Escherichia; Escherichia coli MNNEETFYQAMRRQGVTRRSFLKYCSLAATSLGLGAGMAPKIAWALENKPRIPVVWIHGL WP_001058323.1 MULTISPECIES: hydrogenase-1 small chain [Proteobacteria] Length: 372\nNumber of predicted TMHs: 2\nExp number of AAs in TMHs: 37.2608499999999\nExp number, first 60 AAs: 19.52923\nTotal prob of N-in: 0.94351\nPOSSIBLE N-term signal sequence\ninside 1 19\nTMhelix 20 39\noutside 40 325\nTMhelix 326 348\ninside 349 372

11050 GCF_000457955.1_Esch_coli_HVH_134_4-6073441_V1 Escherichia coli HVH 134 (4-6073441) Proteobacteria; Gammaproteobacteria; Enterobacterales; Enterobacteriaceae; Escherichia; Escherichia coli MTGDNTLIHSHGINRRDFMKLCAALAATMGLSSKAAAEMAESVTNPQRPPVIWIGAQECT WP_000145410.1 MULTISPECIES: hydrogenase-2 small chain [Proteobacteria] Length: 372\nNumber of predicted TMHs: 1\nExp number of AAs in TMHs: 23.72226\nExp number, first 60 AAs: 0.22038\nTotal prob of N-in: 0.12994\noutside 1 331\nTMhelix 332 354\ninside 355 372

11051 GCF_000456185.1_Esch_coli_HVH_13_4-7634056_V1 Escherichia coli HVH 13 (4-7634056) Proteobacteria; Gammaproteobacteria; Enterobacterales; Enterobacteriaceae; Escherichia; Escherichia coli MNNEETFYQAMRRQGVTRRSFLKYCSLAATSLGLGAGMAPKIAWALENKPRIPVVWIHGL WP_001058323.1 MULTISPECIES: hydrogenase-1 small chain [Proteobacteria] Length: 372\nNumber of predicted TMHs: 2\nExp number of AAs in TMHs: 37.2608499999999\nExp number, first 60 AAs: 19.52923\nTotal prob of N-in: 0.94351\nPOSSIBLE N-term signal sequence\ninside 1 19\nTMhelix 20 39\noutside 40 325\nTMhelix 326 348\ninside 349 372

11052 GCF_000456185.1_Esch_coli_HVH_13_4-7634056_V1 Escherichia coli HVH 13 (4-7634056) Proteobacteria; Gammaproteobacteria; Enterobacterales; Enterobacteriaceae; Escherichia; Escherichia coli MTGDNTLIHSHGINRRDFMKLCAALAATMGLSSKAAAEMAESVTNPQRPPVIWIGAQECT WP_000145410.1 MULTISPECIES: hydrogenase-2 small chain [Proteobacteria] Length: 372\nNumber of predicted TMHs: 1\nExp number of AAs in TMHs: 23.72226\nExp number, first 60 AAs: 0.22038\nTotal prob of N-in: 0.12994\noutside 1 331\nTMhelix 332 354\ninside 355 372

11053 GCF_000457975.1_Esch_coli_HVH_135_4-4449320_V1 Escherichia coli HVH 135 (4-4449320) Proteobacteria; Gammaproteobacteria; Enterobacterales; Enterobacteriaceae; Escherichia; Escherichia coli MTGDNTLIHSHGINRRDFMKLCAALAATMGLSSKAAAEMAESVTNPQRPPVIWIGAQECT WP_000145410.1 MULTISPECIES: hydrogenase-2 small chain [Proteobacteria] Length: 372\nNumber of predicted TMHs:

1\nExp number of AAs in TMHs: 23.72226\nExp number, first 60 AAs: 0.22038\nTotal prob of N-in: 0.12994\noutside 1 331\nTMhelix 332 354\ninside 355 372

11054 GCF_000457975.1_Esch_coli_HVH_135_4-4449320_V1 Escherichia coli HVH 135 (4-4449320) Proteobacteria; Gammaproteobacteria; Enterobacterales; Enterobacteriaceae; Escherichia; Escherichia coli MNNEETFYQAMRRQGVTRRSFLKYCSLAATSLGLGAGMAPKIAWALENKPRIPVVWIHGL WP_001058323.1 MULTISPECIES: hydrogenase-1 small chain [Proteobacteria] Length: 372\nNumber of predicted TMHs: 2\nExp number of AAs in TMHs: 37.2608499999999\nExp number, first 60 AAs: 19.52923\nTotal prob of N-in: 0.94351\nPOSSIBLE N-term signal sequence\ninside 1 19\nTMhelix 20 39\noutside 40 325\nTMhelix 326 348\ninside 349 372

11055 GCF_000494995.1_Esch_coli_HVH_136_4-5970458_V1 Escherichia coli HVH 136 (4-5970458) Proteobacteria; Gammaproteobacteria; Enterobacterales; Enterobacteriaceae; Escherichia; Escherichia coli MNNEETFYQAMRRQGVTRRSFLKYCSLAATSLGLGAGMAPKIAWALENKPRIPVVWIHGL WP_001058323.1 MULTISPECIES: hydrogenase-1 small chain [Proteobacteria] Length: 372\nNumber of predicted TMHs: 2\nExp number of AAs in TMHs: 37.2608499999999\nExp number, first 60 AAs: 19.52923\nTotal prob of N-in: 0.94351\nPOSSIBLE N-term signal sequence\ninside 1 19\nTMhelix 20 39\noutside 40 325\nTMhelix 326 348\ninside 349 372

11056 GCF_000494995.1_Esch_coli_HVH_136_4-5970458_V1 Escherichia coli HVH 136 (4-5970458) Proteobacteria; Gammaproteobacteria; Enterobacterales; Enterobacteriaceae; Escherichia; Escherichia coli MTGDNTLIHSHGINRRDFMKLCAALAATMGLSSKAAAEMAESVTNPQRPPVIWIGAQECT WP_000145410.1 MULTISPECIES: hydrogenase-2 small chain [Proteobacteria] Length: 372\nNumber of predicted TMHs: 1\nExp number of AAs in TMHs: 23.72226\nExp number, first 60 AAs: 0.22038\nTotal prob of N-in: 0.12994\noutside 1 331\nTMhelix 332 354\ninside 355 372

11057 GCF_000457995.1_Esch_coli_HVH_137_4-2124971_V1 Escherichia coli HVH 137 (4-2124971) Proteobacteria; Gammaproteobacteria; Enterobacterales; Enterobacteriaceae; Escherichia; Escherichia coli MTGDNTLIHSHGINRRDFMKLCAALAATMGLSSKAAAEMAESVTNPQRPPVIWIGAQECT WP_000145410.1 MULTISPECIES: hydrogenase-2 small chain [Proteobacteria] Length: 372\nNumber of predicted TMHs: 1\nExp number of AAs in TMHs: 23.72226\nExp number, first 60 AAs: 0.22038\nTotal prob of N-in: 0.12994\noutside 1 331\nTMhelix 332 354\ninside 355 372

11058 GCF_000458015.1_Esch_coli_HVH_138_4-6066704_V1 Escherichia coli HVH 138 (4-6066704) Proteobacteria; Gammaproteobacteria; Enterobacterales; Enterobacteriaceae; Escherichia; Escherichia coli MTGDNTLIHSHGINRRDFMKLCAALAATMGLSSKAAAEMAESVTNPQRPPVIWIGAQECT WP_000145410.1 MULTISPECIES: hydrogenase-2 small chain [Proteobacteria] Length: 372\nNumber of predicted TMHs: 1\nExp number of AAs in TMHs: 23.72226\nExp number, first 60 AAs: 0.22038\nTotal prob of N-in: 0.12994\noutside 1 331\nTMhelix 332 354\ninside 355 372

11059 GCF_000458015.1_Esch_coli_HVH_138_4-6066704_V1 Escherichia coli HVH 138 (4-6066704) Proteobacteria; Gammaproteobacteria; Enterobacterales; Enterobacteriaceae; Escherichia; Escherichia coli MNNEETFYQAMRRQGVTRRSFLKYCSLAATSLGLGAGMAPKIAWALENKPRIPVVWIHGL WP_001058323.1 MULTISPECIES: hydrogenase-1 small chain [Proteobacteria] Length: 372\nNumber of predicted TMHs: 2\nExp number of AAs in TMHs: 37.2608499999999\nExp number, first 60 AAs: 19.52923\nTotal prob of N-in: 0.94351\nPOSSIBLE N-term signal sequence\ninside 1 19\nTMhelix 20 39\noutside 40 325\nTMhelix 326 348\ninside 349 372

11060 GCF_000458035.1_Esch_coli_HVH_139_4-3192644_V1 Escherichia coli HVH 139 (4-3192644) Proteobacteria; Gammaproteobacteria; Enterobacterales; Enterobacteriaceae; Escherichia; Escherichia coli MNNEETFYQAMRRQGVTRRSFLKYCSLAATSLGLGAGMAPKIAWALENKPRIPVVWIHGL WP_001295940.1 MULTISPECIES: hydrogenase [Enterobacteriaceae] Length: 372\nNumber of predicted TMHs: 2\nExp number of AAs in TMHs: 37.2608899999999\nExp number, first 60 AAs: 19.52923\nTotal prob of N-in: 0.94351\nPOSSIBLE N-term signal sequence\ninside 1 19\nTMhelix 20 39\noutside 40 325\nTMhelix 326 348\ninside 349 372

11061 GCF_000458035.1_Esch_coli_HVH_139_4-3192644_V1 Escherichia coli HVH 139 (4-3192644) Proteobacteria; Gammaproteobacteria; Enterobacterales; Enterobacteriaceae; Escherichia; Escherichia coli MTGDNTLIHSHGINRRDFMKLCAALAATMGLSSKAAAEMAESVTNPQRPPVIWIGAQECT WP_000145410.1 MULTISPECIES: hydrogenase-2 small chain [Proteobacteria] Length: 372\nNumber of predicted TMHs:

1\nExp number of AAs in TMHs: 23.72226\nExp number, first 60 AAs:  0.22038\nTotal prob of N-in:   0.12994\noutside   1   331\nTMhelix   332   354\ninside   355   372

11062 GCF_000458055.1_Esch_coli_HVH_140_4-5894387_V1 Escherichia coli HVH 140 (4-5894387) Proteobacteria; Gammaproteobacteria; Enterobacterales; Enterobacteriaceae; Escherichia; Escherichia coli MNNEETFYQAMRRQGVTRRSFLKYCSLAATSLGLGAGMAPKIAWALENKPRIPVVWIHGL WP_001058323.1 MULTISPECIES: hydrogenase-1 small chain [Proteobacteria] Length: 372\nNumber of predicted TMHs: 2\nExp number of AAs in TMHs: 37.2608499999999\nExp number, first 60 AAs:  19.52923\nTotal prob of N-in: 0.94351\nPOSSIBLE N-term signal sequence\ninside   1   19\nTMhelix   20   39\noutside   40   325\nTMhelix   326   348\ninside   349   372

11063 GCF_000458055.1_Esch_coli_HVH_140_4-5894387_V1 Escherichia coli HVH 140 (4-5894387) Proteobacteria; Gammaproteobacteria; Enterobacterales; Enterobacteriaceae; Escherichia; Escherichia coli MTGDNTLIHSHGINRRDFMKLCAALAATMGLSSKAAAEMAESVTNPQRPPVIWIGAQECT WP_000145410.1 MULTISPECIES: hydrogenase-2 small chain [Proteobacteria] Length: 372\nNumber of predicted TMHs: 1\nExp number of AAs in TMHs: 23.72226\nExp number, first 60 AAs:  0.22038\nTotal prob of N-in:   0.12994\noutside   1   331\nTMhelix   332   354\ninside   355   372

11064 GCF_000458075.1_Esch_coli_HVH_141_4-5995973_V1 Escherichia coli HVH 141 (4-5995973) Proteobacteria; Gammaproteobacteria; Enterobacterales; Enterobacteriaceae; Escherichia; Escherichia coli MTGDNTLIHSHGINRRDFMKLCAALAATMGLSSKAAAEMAESVTNPQRPPVIWIGAQECT WP_000145410.1 MULTISPECIES: hydrogenase-2 small chain [Proteobacteria] Length: 372\nNumber of predicted TMHs: 1\nExp number of AAs in TMHs: 23.72226\nExp number, first 60 AAs:  0.22038\nTotal prob of N-in:   0.12994\noutside   1   331\nTMhelix   332   354\ninside   355   372

11065 GCF_000458075.1_Esch_coli_HVH_141_4-5995973_V1 Escherichia coli HVH 141 (4-5995973) Proteobacteria; Gammaproteobacteria; Enterobacterales; Enterobacteriaceae; Escherichia; Escherichia coli MNNEETFYQAMRRQGVTRRSFLKYCSLAATSLGLGAGMAPKIAWALENKPRIPVVWIHGL WP_001295940.1 MULTISPECIES: hydrogenase [Enterobacteriaceae] Length: 372\nNumber of predicted TMHs:  2\nExp number of AAs in TMHs: 37.2608899999999\nExp number, first 60 AAs:  19.52923\nTotal prob of N-in: 0.94351\nPOSSIBLE N-term signal sequence\ninside   1   19\nTMhelix   20   39\noutside   40   325\nTMhelix   326   348\ninside   349   372

11066 GCF_000458095.1_Esch_coli_HVH_142_4-5627451_V1 Escherichia coli HVH 142 (4-5627451) Proteobacteria; Gammaproteobacteria; Enterobacterales; Enterobacteriaceae; Escherichia; Escherichia coli MNNEETFYQAMRRQGVTRRSFLKYCSLAATSLGLGAGMAPKIAWALENKPRIPVVWIHGL WP_001058323.1 MULTISPECIES: hydrogenase-1 small chain [Proteobacteria] Length: 372\nNumber of predicted TMHs: 2\nExp number of AAs in TMHs: 37.2608499999999\nExp number, first 60 AAs:  19.52923\nTotal prob of N-in: 0.94351\nPOSSIBLE N-term signal sequence\ninside   1   19\nTMhelix   20   39\noutside   40   325\nTMhelix   326   348\ninside   349   372

11067 GCF_000458095.1_Esch_coli_HVH_142_4-5627451_V1 Escherichia coli HVH 142 (4-5627451) Proteobacteria; Gammaproteobacteria; Enterobacterales; Enterobacteriaceae; Escherichia; Escherichia coli MTGDNTLIHSHGINRRDFMKLCAALAATMGLSSKAAAEMAESVTNPQRPPVIWIGAQECT WP_000145410.1 MULTISPECIES: hydrogenase-2 small chain [Proteobacteria] Length: 372\nNumber of predicted TMHs: 1\nExp number of AAs in TMHs: 23.72226\nExp number, first 60 AAs:  0.22038\nTotal prob of N-in:   0.12994\noutside   1   331\nTMhelix   332   354\ninside   355   372

11068 GCF_000458115.1_Esch_coli_HVH_143_4-5674999_V1 Escherichia coli HVH 143 (4-5674999) Proteobacteria; Gammaproteobacteria; Enterobacterales; Enterobacteriaceae; Escherichia; Escherichia coli MTGDNTLIHSHGINRRDFMKLCAALAATMGLSSKAAAEMAESVTNPQRPPVIWIGAQECT WP_000145410.1 MULTISPECIES: hydrogenase-2 small chain [Proteobacteria] Length: 372\nNumber of predicted TMHs: 1\nExp number of AAs in TMHs: 23.72226\nExp number, first 60 AAs:  0.22038\nTotal prob of N-in:   0.12994\noutside   1   331\nTMhelix   332   354\ninside   355   372

11069 GCF_000458115.1_Esch_coli_HVH_143_4-5674999_V1 Escherichia coli HVH 143 (4-5674999) Proteobacteria; Gammaproteobacteria; Enterobacterales; Enterobacteriaceae; Escherichia; Escherichia coli MNNEETFYQAMRRQGVTRRSFLKYCSLAATSLGLGAGMAPKIAWALENKPRIPVVWIHGL WP_001058323.1 MULTISPECIES: hydrogenase-1 small chain [Proteobacteria] Length: 372\nNumber of predicted TMHs: 2\nExp number of AAs in TMHs: 37.2608499999999\nExp number, first 60 AAs:  19.52923\nTotal prob of N-in:

0.94351\nPOSSIBLE N-term signal sequence\ninside 1 19\nTMhelix 20 39\noutside 40 325\nTMhelix 326 348\ninside 349 372

11070 GCF_000458135.1_Esch_coli_HVH_144_4-4451937_V1 Escherichia coli HVH 144 (4-4451937) Proteobacteria; Gammaproteobacteria; Enterobacterales; Enterobacteriaceae; Escherichia; Escherichia coli MTGDNTLIHSHGINRRDFMKLCAALAATMGLSSKAAAEMAESVTNPQRPPVIWIGAQECT WP_000145410.1 MULTISPECIES: hydrogenase-2 small chain [Proteobacteria] Length: 372\nNumber of predicted TMHs: 1\nExp number of AAs in TMHs: 23.72226\nExp number, first 60 AAs: 0.22038\nTotal prob of N-in: 0.12994\noutside 1 331\nTMhelix 332 354\ninside 355 372

11071 GCF_000458135.1_Esch_coli_HVH_144_4-4451937_V1 Escherichia coli HVH 144 (4-4451937) Proteobacteria; Gammaproteobacteria; Enterobacterales; Enterobacteriaceae; Escherichia; Escherichia coli MNNEETFYQAMRRQGVTRRSFLKYCSLAATSLGLGAGMAPKIAWALENKPRIPVVWIHGL WP_001058323.1 MULTISPECIES: hydrogenase-1 small chain [Proteobacteria] Length: 372\nNumber of predicted TMHs: 2\nExp number of AAs in TMHs: 37.2608499999999\nExp number, first 60 AAs: 19.52923\nTotal prob of N-in: 0.94351\nPOSSIBLE N-term signal sequence\ninside 1 19\nTMhelix 20 39\noutside 40 325\nTMhelix 326 348\ninside 349 372

11072 GCF_000458155.1_Esch_coli_HVH_145_4-5672112_V1 Escherichia coli HVH 145 (4-5672112) Proteobacteria; Gammaproteobacteria; Enterobacterales; Enterobacteriaceae; Escherichia; Escherichia coli MTGDNTLIHSHGINRRDFMKLCAALAATMGLSSKAAAEMAESVTNPQRPPVIWIGAQECT WP_000145410.1 MULTISPECIES: hydrogenase-2 small chain [Proteobacteria] Length: 372\nNumber of predicted TMHs: 1\nExp number of AAs in TMHs: 23.72226\nExp number, first 60 AAs: 0.22038\nTotal prob of N-in: 0.12994\noutside 1 331\nTMhelix 332 354\ninside 355 372

11073 GCF_000458155.1_Esch_coli_HVH_145_4-5672112_V1 Escherichia coli HVH 145 (4-5672112) Proteobacteria; Gammaproteobacteria; Enterobacterales; Enterobacteriaceae; Escherichia; Escherichia coli MNNEETFYQAMRRQGVTRRSFLKYCSLAATSLGLGAGMAPKIAWALENKPRIPVVWIHGL WP_001058323.1 MULTISPECIES: hydrogenase-1 small chain [Proteobacteria] Length: 372\nNumber of predicted TMHs: 2\nExp number of AAs in TMHs: 37.2608499999999\nExp number, first 60 AAs: 19.52923\nTotal prob of N-in: 0.94351\nPOSSIBLE N-term signal sequence\ninside 1 19\nTMhelix 20 39\noutside 40 325\nTMhelix 326 348\ninside 349 372

11074 GCF_000458175.1_Esch_coli_HVH_146_4-3189767_V1 Escherichia coli HVH 146 (4-3189767) Proteobacteria; Gammaproteobacteria; Enterobacterales; Enterobacteriaceae; Escherichia; Escherichia coli MTGDNTLIHSHGINRRDFMKLCAALAATMGLSSKAAAEMAESVTNPQRPPVIWIGAQECT WP_000145410.1 MULTISPECIES: hydrogenase-2 small chain [Proteobacteria] Length: 372\nNumber of predicted TMHs: 1\nExp number of AAs in TMHs: 23.72226\nExp number, first 60 AAs: 0.22038\nTotal prob of N-in: 0.12994\noutside 1 331\nTMhelix 332 354\ninside 355 372

11075 GCF_000458175.1_Esch_coli_HVH_146_4-3189767_V1 Escherichia coli HVH 146 (4-3189767) Proteobacteria; Gammaproteobacteria; Enterobacterales; Enterobacteriaceae; Escherichia; Escherichia coli MNNEETFYQAMRRQGVTRRSFLKYCSLAATSLGLGAGMAPKIAWALENKPRIPVVWIHGL WP_001295940.1 MULTISPECIES: hydrogenase [Enterobacteriaceae] Length: 372\nNumber of predicted TMHs: 2\nExp number of AAs in TMHs: 37.2608899999999\nExp number, first 60 AAs: 19.52923\nTotal prob of N-in: 0.94351\nPOSSIBLE N-term signal sequence\ninside 1 19\nTMhelix 20 39\noutside 40 325\nTMhelix 326 348\ninside 349 372

11076 GCF_000456005.1_Esch_coli_HVH_1_4-6876161_V1 Escherichia coli HVH 1 (4-6876161) Proteobacteria; Gammaproteobacteria; Enterobacterales; Enterobacteriaceae; Escherichia; Escherichia coli MTGDNTLIHSHGINRRDFMKLCAALAATMGLSSKAAAEMAESVTNPQRPPVIWIGAQECT WP_000145410.1 MULTISPECIES: hydrogenase-2 small chain [Proteobacteria] Length: 372\nNumber of predicted TMHs: 1\nExp number of AAs in TMHs: 23.72226\nExp number, first 60 AAs: 0.22038\nTotal prob of N-in: 0.12994\noutside 1 331\nTMhelix 332 354\ninside 355 372

11077 GCF_000458195.1_Esch_coli_HVH_147_4-5893887_V1 Escherichia coli HVH 147 (4-5893887) Proteobacteria; Gammaproteobacteria; Enterobacterales; Enterobacteriaceae; Escherichia; Escherichia coli MNNEETFYQAMRRQGVTRRSFLKYCSLAATSLGLGAGMAPKIAWALENKPRIPVVWIHGL WP_001058323.1 MULTISPECIES: hydrogenase-1 small chain [Proteobacteria] Length: 372\nNumber of predicted TMHs: 2\nExp number of AAs in TMHs: 37.2608499999999\nExp number, first 60 AAs: 19.52923\nTotal prob of N-in:

0.94351\nPOSSIBLE N-term signal sequence\ninside 1 19\nTMhelix 20 39\noutside 40 325\nTMhelix 326 348\ninside 349 372

11078 GCF_000458195.1_Esch_coli_HVH_147_4-5893887_V1 Escherichia coli HVH 147 (4-5893887) Proteobacteria; Gammaproteobacteria; Enterobacterales; Enterobacteriaceae; Escherichia; Escherichia coli MTGDNTLIHSHGINRRDFMKLCAALAATMGLSSKAAAEMAESVTNPQRPPVIWIGAQECT WP_000145410.1 MULTISPECIES: hydrogenase-2 small chain [Proteobacteria] Length: 372\nNumber of predicted TMHs: 1\nExp number of AAs in TMHs: 23.72226\nExp number, first 60 AAs: 0.22038\nTotal prob of N-in: 0.12994\noutside 1 331\nTMhelix 332 354\ninside 355 372

11079 GCF_000495015.1_Esch_coli_HVH_148_4-3192490_V1 Escherichia coli HVH 148 (4-3192490) Proteobacteria; Gammaproteobacteria; Enterobacterales; Enterobacteriaceae; Escherichia; Escherichia coli MTGDNTLIHSHGINRRDFMKLCAALAATMGLSSKAAAEMAESVTNPQRPPVIWIGAQECT WP_000145410.1 MULTISPECIES: hydrogenase-2 small chain [Proteobacteria] Length: 372\nNumber of predicted TMHs: 1\nExp number of AAs in TMHs: 23.72226\nExp number, first 60 AAs: 0.22038\nTotal prob of N-in: 0.12994\noutside 1 331\nTMhelix 332 354\ninside 355 372

11080 GCF_000458215.1_Esch_coli_HVH_149_4-4451880_V1 Escherichia coli HVH 149 (4-4451880) Proteobacteria; Gammaproteobacteria; Enterobacterales; Enterobacteriaceae; Escherichia; Escherichia coli MNNEETFYQAMRRQGVTRRSFLKYCSLAATSLGLGAGMAPKIAWALENKPRIPVVWIHGL WP_001058323.1 MULTISPECIES: hydrogenase-1 small chain [Proteobacteria] Length: 372\nNumber of predicted TMHs: 2\nExp number of AAs in TMHs: 37.2608499999999\nExp number, first 60 AAs: 19.52923\nTotal prob of N-in: 0.94351\nPOSSIBLE N-term signal sequence\ninside 1 19\nTMhelix 20 39\noutside 40 325\nTMhelix 326 348\ninside 349 372

11081 GCF_000458215.1_Esch_coli_HVH_149_4-4451880_V1 Escherichia coli HVH 149 (4-4451880) Proteobacteria; Gammaproteobacteria; Enterobacterales; Enterobacteriaceae; Escherichia; Escherichia coli MTGDNTLIHSHGINRRDFMKLCAALAATMGLSSKAAAEMAESVTNPQRPPVIWIGAQECT WP_000145410.1 MULTISPECIES: hydrogenase-2 small chain [Proteobacteria] Length: 372\nNumber of predicted TMHs: 1\nExp number of AAs in TMHs: 23.72226\nExp number, first 60 AAs: 0.22038\nTotal prob of N-in: 0.12994\noutside 1 331\nTMhelix 332 354\ninside 355 372

11082 GCF_000458235.1_Esch_coli_HVH_150_4-3258106_V1 Escherichia coli HVH 150 (4-3258106) Proteobacteria; Gammaproteobacteria; Enterobacterales; Enterobacteriaceae; Escherichia; Escherichia coli MNNEETFYQAMRRQGVTRRSFLKYCSLAATSLGLGAGMAPKIAWALENKPRIPVVWIHGL WP_001058320.1 hydrogenase [Escherichia coli] Length: 372\nNumber of predicted TMHs: 2\nExp number of AAs in TMHs: 37.26082\nExp number, first 60 AAs: 19.52923\nTotal prob of N-in: 0.94351\nPOSSIBLE N-term signal sequence\ninside 1 19\nTMhelix 20 39\noutside 40 325\nTMhelix 326 348\ninside 349 372

11083 GCF_000458235.1_Esch_coli_HVH_150_4-3258106_V1 Escherichia coli HVH 150 (4-3258106) Proteobacteria; Gammaproteobacteria; Enterobacterales; Enterobacteriaceae; Escherichia; Escherichia coli MTGDNTLIHSHGINRRDFMKLCAALAATMGLSSKAAAEMAESVTNPQRPPVIWIGAQECT WP_000145410.1 MULTISPECIES: hydrogenase-2 small chain [Proteobacteria] Length: 372\nNumber of predicted TMHs: 1\nExp number of AAs in TMHs: 23.72226\nExp number, first 60 AAs: 0.22038\nTotal prob of N-in: 0.12994\noutside 1 331\nTMhelix 332 354\ninside 355 372

11084 GCF_000458255.1_Esch_coli_HVH_151_4-5755573_V1 Escherichia coli HVH 151 (4-5755573) Proteobacteria; Gammaproteobacteria; Enterobacterales; Enterobacteriaceae; Escherichia; Escherichia coli MNNEETFYQAMRRQGVTRRSFLKYCSLAATSLGLGAGMAPKIAWALENKPRIPVVWIHGL WP_001058323.1 MULTISPECIES: hydrogenase-1 small chain [Proteobacteria] Length: 372\nNumber of predicted TMHs: 2\nExp number of AAs in TMHs: 37.2608499999999\nExp number, first 60 AAs: 19.52923\nTotal prob of N-in: 0.94351\nPOSSIBLE N-term signal sequence\ninside 1 19\nTMhelix 20 39\noutside 40 325\nTMhelix 326 348\ninside 349 372

11085 GCF_000458255.1_Esch_coli_HVH_151_4-5755573_V1 Escherichia coli HVH 151 (4-5755573) Proteobacteria; Gammaproteobacteria; Enterobacterales; Enterobacteriaceae; Escherichia; Escherichia coli MTGDNTLIHSHGINRRDFMKLCAALAATMGLSSKAAAEMAESVTNPQRPPVIWIGAQECT WP_000145410.1 MULTISPECIES: hydrogenase-2 small chain [Proteobacteria] Length: 372\nNumber of predicted TMHs: 1\nExp number of AAs in TMHs: 23.72226\nExp number, first 60 AAs: 0.22038\nTotal prob of N-in: 0.12994\noutside 1 331\nTMhelix 332 354\ninside 355 372

11086 GCF_000495035.1_Esch_coli_HVH_152_4-3447545_V1 Escherichia coli HVH 152 (4-3447545)
Proteobacteria; Gammaproteobacteria; Enterobacterales; Enterobacteriaceae; Escherichia; Escherichia coli
MTGDNTLIHSHGINRRDFMKLCAALAATMGLSSKAAAEMAESVTNPQRPPVIWIGAQECT WP_000145410.1
MULTISPECIES: hydrogenase-2 small chain [Proteobacteria] Length: 372\nNumber of predicted TMHs: 1\nExp number of AAs in TMHs: 23.72226\nExp number, first 60 AAs: 0.22038\nTotal prob of N-in: 0.12994\noutside 1 331\nTMhelix 332 354\ninside 355 372

11087 GCF_000495035.1_Esch_coli_HVH_152_4-3447545_V1 Escherichia coli HVH 152 (4-3447545)
Proteobacteria; Gammaproteobacteria; Enterobacterales; Enterobacteriaceae; Escherichia; Escherichia coli
MNNEETFYQAMRRQGVTRRSFLKYCSLAATSLGLGAGMAPKIAWALENKPRIPVVWIHGL WP_001058323.1
MULTISPECIES: hydrogenase-1 small chain [Proteobacteria] Length: 372\nNumber of predicted TMHs: 2\nExp number of AAs in TMHs: 37.2608499999999\nExp number, first 60 AAs: 19.52923\nTotal prob of N-in: 0.94351\nPOSSIBLE N-term signal sequence\ninside 1 19\nTMhelix 20 39\noutside 40 325\nTMhelix 326 348\ninside 349 372

11088 GCF_000458275.1_Esch_coli_HVH_153_3-9344314_V1 Escherichia coli HVH 153 (3-9344314)
Proteobacteria; Gammaproteobacteria; Enterobacterales; Enterobacteriaceae; Escherichia; Escherichia coli
MNNEETFYQAMRRQGVTRRSFLKYCSLAATSLGLGAGMAPKIAWALENKPRIPVVWIHGL WP_021520630.1
hydrogenase-1 small chain [Escherichia coli] Length: 372\nNumber of predicted TMHs: 2\nExp number of AAs in TMHs: 37.8197599999999\nExp number, first 60 AAs: 19.53295\nTotal prob of N-in: 0.94396\nPOSSIBLE N-term signal sequence\ninside 1 19\nTMhelix 20 39\noutside 40 325\nTMhelix 326 348\ninside 349 372

11089 GCF_000458275.1_Esch_coli_HVH_153_3-9344314_V1 Escherichia coli HVH 153 (3-9344314)
Proteobacteria; Gammaproteobacteria; Enterobacterales; Enterobacteriaceae; Escherichia; Escherichia coli
MTGDNTLIHSHGINRRDFMKLCAALAATMGLSSKAAAEMAESVTNPQRPPVIWIGAQECT WP_000145410.1
MULTISPECIES: hydrogenase-2 small chain [Proteobacteria] Length: 372\nNumber of predicted TMHs: 1\nExp number of AAs in TMHs: 23.72226\nExp number, first 60 AAs: 0.22038\nTotal prob of N-in: 0.12994\noutside 1 331\nTMhelix 332 354\ninside 355 372

11090 GCF_000458295.1_Esch_coli_HVH_154_4-5636698_V1 Escherichia coli HVH 154 (4-5636698)
Proteobacteria; Gammaproteobacteria; Enterobacterales; Enterobacteriaceae; Escherichia; Escherichia coli
MTGDNTLIHSHGINRRDFMKLCAALAATMGLSSKAAAEMAESVTNPQRPPVIWIGAQECT WP_000145410.1
MULTISPECIES: hydrogenase-2 small chain [Proteobacteria] Length: 372\nNumber of predicted TMHs: 1\nExp number of AAs in TMHs: 23.72226\nExp number, first 60 AAs: 0.22038\nTotal prob of N-in: 0.12994\noutside 1 331\nTMhelix 332 354\ninside 355 372

11091 GCF_000458295.1_Esch_coli_HVH_154_4-5636698_V1 Escherichia coli HVH 154 (4-5636698)
Proteobacteria; Gammaproteobacteria; Enterobacterales; Enterobacteriaceae; Escherichia; Escherichia coli
MNNEETFYQAMRRQGVTRRSFLKYCSLAATSLGLGAGMAPKIAWALENKPRIPVVWIHGL WP_001295940.1
MULTISPECIES: hydrogenase [Enterobacteriaceae] Length: 372\nNumber of predicted TMHs: 2\nExp number of AAs in TMHs: 37.2608899999999\nExp number, first 60 AAs: 19.52923\nTotal prob of N-in: 0.94351\nPOSSIBLE N-term signal sequence\ninside 1 19\nTMhelix 20 39\noutside 40 325\nTMhelix 326 348\ninside 349 372

11092 GCF_000458315.1_Esch_coli_HVH_155_4-4509048_V1 Escherichia coli HVH 155 (4-4509048)
Proteobacteria; Gammaproteobacteria; Enterobacterales; Enterobacteriaceae; Escherichia; Escherichia coli
MNNEETFYQAMRRQGVTRRSFLKYCSLAATSLGLGAGMAPKIAWALENKPRIPVVWIHGL WP_001058323.1
MULTISPECIES: hydrogenase-1 small chain [Proteobacteria] Length: 372\nNumber of predicted TMHs: 2\nExp number of AAs in TMHs: 37.2608499999999\nExp number, first 60 AAs: 19.52923\nTotal prob of N-in: 0.94351\nPOSSIBLE N-term signal sequence\ninside 1 19\nTMhelix 20 39\noutside 40 325\nTMhelix 326 348\ninside 349 372

11093 GCF_000458315.1_Esch_coli_HVH_155_4-4509048_V1 Escherichia coli HVH 155 (4-4509048)
Proteobacteria; Gammaproteobacteria; Enterobacterales; Enterobacteriaceae; Escherichia; Escherichia coli
MTGDNTLIHSHGINRRDFMKLCAALAATMGLSSKAAAEMAESVTNPQRPPVIWIGAQECT WP_000145410.1
MULTISPECIES: hydrogenase-2 small chain [Proteobacteria] Length: 372\nNumber of predicted TMHs: 1\nExp number of AAs in TMHs: 23.72226\nExp number, first 60 AAs: 0.22038\nTotal prob of N-in: 0.12994\noutside 1 331\nTMhelix 332 354\ninside 355 372

11094 GCF_000458335.1_Esch_coli_HVH_156_4-3206505_V1 Escherichia coli HVH 156 (4-3206505)
Proteobacteria; Gammaproteobacteria; Enterobacterales; Enterobacteriaceae; Escherichia; Escherichia coli

MNNEETFYQAMRRQGVTRRSFLKYCSLAATSLGLGAGMAPKIAWALENKPRIPVVWIHGL WP_001058323.1 MULTISPECIES: hydrogenase-1 small chain [Proteobacteria] Length: 372\nNumber of predicted TMHs: 2\nExp number of AAs in TMHs: 37.2608499999999\nExp number, first 60 AAs: 19.52923\nTotal prob of N-in: 0.94351\nPOSSIBLE N-term signal sequence\ninside 1 19\nTMhelix 20 39\noutside 40 325\nTMhelix 326 348\ninside 349 372

11095 GCF_000458335.1_Esch_coli_HVH_156_4-3206505_V1 Escherichia coli HVH 156 (4-3206505) Proteobacteria; Gammaproteobacteria; Enterobacterales; Enterobacteriaceae; Escherichia; Escherichia coli MTGDNTLIHSHGINRRDFMKLCAALAATMGLSSKAAAEMAESVTNPQRPPVIWIGAQECT WP_000145410.1 MULTISPECIES: hydrogenase-2 small chain [Proteobacteria] Length: 372\nNumber of predicted TMHs: 1\nExp number of AAs in TMHs: 23.72226\nExp number, first 60 AAs: 0.22038\nTotal prob of N-in: 0.12994\noutside 1 331\nTMhelix 332 354\ninside 355 372

11096 GCF_000458355.1_Esch_coli_HVH_157_4-3406229_V1 Escherichia coli HVH 157 (4-3406229) Proteobacteria; Gammaproteobacteria; Enterobacterales; Enterobacteriaceae; Escherichia; Escherichia coli MNNEETFYQAMRRQGVTRRSFLKYCSLAATSLGLGAGMAPKIAWALENKPRIPVVWIHGL WP_001058323.1 MULTISPECIES: hydrogenase-1 small chain [Proteobacteria] Length: 372\nNumber of predicted TMHs: 2\nExp number of AAs in TMHs: 37.2608499999999\nExp number, first 60 AAs: 19.52923\nTotal prob of N-in: 0.94351\nPOSSIBLE N-term signal sequence\ninside 1 19\nTMhelix 20 39\noutside 40 325\nTMhelix 326 348\ninside 349 372

11097 GCF_000458355.1_Esch_coli_HVH_157_4-3406229_V1 Escherichia coli HVH 157 (4-3406229) Proteobacteria; Gammaproteobacteria; Enterobacterales; Enterobacteriaceae; Escherichia; Escherichia coli MTGDNTLIHSHGINRRDFMKLCAALAATMGLSSKAAAEMAESVTNPQRPPVIWIGAQECT WP_000145410.1 MULTISPECIES: hydrogenase-2 small chain [Proteobacteria] Length: 372\nNumber of predicted TMHs: 1\nExp number of AAs in TMHs: 23.72226\nExp number, first 60 AAs: 0.22038\nTotal prob of N-in: 0.12994\noutside 1 331\nTMhelix 332 354\ninside 355 372

11098 GCF_000458375.1_Esch_coli_HVH_158_4-3224287_V1 Escherichia coli HVH 158 (4-3224287) Proteobacteria; Gammaproteobacteria; Enterobacterales; Enterobacteriaceae; Escherichia; Escherichia coli MTGDNTLIHSHGINRRDFMKLCAALAATMGLSSKAAAEMAESVTNPQRPPVIWIGAQECT WP_000145418.1 hydrogenase 2 small subunit [Escherichia coli] Length: 372\nNumber of predicted TMHs: 1\nExp number of AAs in TMHs: 23.72313\nExp number, first 60 AAs: 0.22034\nTotal prob of N-in: 0.12997\noutside 1 331\nTMhelix 332 354\ninside 355 372

11099 GCF_000458375.1_Esch_coli_HVH_158_4-3224287_V1 Escherichia coli HVH 158 (4-3224287) Proteobacteria; Gammaproteobacteria; Enterobacterales; Enterobacteriaceae; Escherichia; Escherichia coli MNNEETFYQAMRRQGVTRRSFLKYCSLAATSLGLGAGMAPKIAWALENKPRIPVVWIHGL WP_001058323.1 MULTISPECIES: hydrogenase-1 small chain [Proteobacteria] Length: 372\nNumber of predicted TMHs: 2\nExp number of AAs in TMHs: 37.2608499999999\nExp number, first 60 AAs: 19.52923\nTotal prob of N-in: 0.94351\nPOSSIBLE N-term signal sequence\ninside 1 19\nTMhelix 20 39\noutside 40 325\nTMhelix 326 348\ninside 349 372

11100 GCF_000458395.1_Esch_coli_HVH_159_4-5818141_V1 Escherichia coli HVH 159 (4-5818141) Proteobacteria; Gammaproteobacteria; Enterobacterales; Enterobacteriaceae; Escherichia; Escherichia coli MTGDNTLIHSHGINRRDFMKLCAALAATMGLSSKAAAEMAESVTNPQRPPVIWIGAQECT WP_000145410.1 MULTISPECIES: hydrogenase-2 small chain [Proteobacteria] Length: 372\nNumber of predicted TMHs: 1\nExp number of AAs in TMHs: 23.72226\nExp number, first 60 AAs: 0.22038\nTotal prob of N-in: 0.12994\noutside 1 331\nTMhelix 332 354\ninside 355 372

11101 GCF_000458395.1_Esch_coli_HVH_159_4-5818141_V1 Escherichia coli HVH 159 (4-5818141) Proteobacteria; Gammaproteobacteria; Enterobacterales; Enterobacteriaceae; Escherichia; Escherichia coli MNNEETFYQAMRRQGVTRRSFLKYCSLAATSLGLGAGMAPKIAWALENKPRIPVVWIHGL WP_001058323.1 MULTISPECIES: hydrogenase-1 small chain [Proteobacteria] Length: 372\nNumber of predicted TMHs: 2\nExp number of AAs in TMHs: 37.2608499999999\nExp number, first 60 AAs: 19.52923\nTotal prob of N-in: 0.94351\nPOSSIBLE N-term signal sequence\ninside 1 19\nTMhelix 20 39\noutside 40 325\nTMhelix 326 348\ninside 349 372

11102 GCF_000458415.1_Esch_coli_HVH_160_4-5695937_V1 Escherichia coli HVH 160 (4-5695937) Proteobacteria; Gammaproteobacteria; Enterobacterales; Enterobacteriaceae; Escherichia; Escherichia coli MTGDNTLIHSHGINRRDFMKLCAALAATMGLSSKAAAEMAESVTNPQRPPVIWIGAQECT WP_000145410.1

MULTISPECIES: hydrogenase-2 small chain [Proteobacteria] Length: 372\nNumber of predicted TMHs: 1\nExp number of AAs in TMHs: 23.72226\nExp number, first 60 AAs: 0.22038\nTotal prob of N-in: 0.12994\noutside 1 331\nTMhelix 332 354\ninside 355 372

11103 GCF_000458415.1_Esch_coli_HVH_160_4-5695937_V1 Escherichia coli HVH 160 (4-5695937) Proteobacteria; Gammaproteobacteria; Enterobacterales; Enterobacteriaceae; Escherichia; Escherichia coli MNNEETFYQAMRRQGVTRRSFLKYCSLAATSLGLGAGMAPKIAWALENKPRIPVVWIHGL WP_001058323.1 MULTISPECIES: hydrogenase-1 small chain [Proteobacteria] Length: 372\nNumber of predicted TMHs: 2\nExp number of AAs in TMHs: 37.2608499999999\nExp number, first 60 AAs: 19.52923\nTotal prob of N-in: 0.94351\nPOSSIBLE N-term signal sequence\ninside 1 19\nTMhelix 20 39\noutside 40 325\nTMhelix 326 348\ninside 349 372

11104 GCF_000458435.1_Esch_coli_HVH_161_4-3119890_V1 Escherichia coli HVH 161 (4-3119890) Proteobacteria; Gammaproteobacteria; Enterobacterales; Enterobacteriaceae; Escherichia; Escherichia coli MNNEETFYQAMRRQGVTRRSFLKYCSLAATSLGLGAGMAPKIAWALENKPRIPVVWIHGL WP_001058323.1 MULTISPECIES: hydrogenase-1 small chain [Proteobacteria] Length: 372\nNumber of predicted TMHs: 2\nExp number of AAs in TMHs: 37.2608499999999\nExp number, first 60 AAs: 19.52923\nTotal prob of N-in: 0.94351\nPOSSIBLE N-term signal sequence\ninside 1 19\nTMhelix 20 39\noutside 40 325\nTMhelix 326 348\ninside 349 372

11105 GCF_000458435.1_Esch_coli_HVH_161_4-3119890_V1 Escherichia coli HVH 161 (4-3119890) Proteobacteria; Gammaproteobacteria; Enterobacterales; Enterobacteriaceae; Escherichia; Escherichia coli MTGDNTLIHSHGINRRDFMKLCAALAATMGLSSKAAAEMAESVTNPQRPPVIWIGAQECT WP_000145410.1 MULTISPECIES: hydrogenase-2 small chain [Proteobacteria] Length: 372\nNumber of predicted TMHs: 1\nExp number of AAs in TMHs: 23.72226\nExp number, first 60 AAs: 0.22038\nTotal prob of N-in: 0.12994\noutside 1 331\nTMhelix 332 354\ninside 355 372

11106 GCF_000458455.1_Esch_coli_HVH_162_4-5627982_V1 Escherichia coli HVH 162 (4-5627982) Proteobacteria; Gammaproteobacteria; Enterobacterales; Enterobacteriaceae; Escherichia; Escherichia coli MNNEETFYQAMRRQGVTRRSFLKYCSLAATSLGLGAGMAPKIAWALENKPRIPVVWIHGL WP_001058323.1 MULTISPECIES: hydrogenase-1 small chain [Proteobacteria] Length: 372\nNumber of predicted TMHs: 2\nExp number of AAs in TMHs: 37.2608499999999\nExp number, first 60 AAs: 19.52923\nTotal prob of N-in: 0.94351\nPOSSIBLE N-term signal sequence\ninside 1 19\nTMhelix 20 39\noutside 40 325\nTMhelix 326 348\ninside 349 372

11107 GCF_000458455.1_Esch_coli_HVH_162_4-5627982_V1 Escherichia coli HVH 162 (4-5627982) Proteobacteria; Gammaproteobacteria; Enterobacterales; Enterobacteriaceae; Escherichia; Escherichia coli MTGDNTLIHSHGINRRDFMKLCAALAATMGLSSKAAAEMAESVTNPQRPPVIWIGAQECT WP_000145410.1 MULTISPECIES: hydrogenase-2 small chain [Proteobacteria] Length: 372\nNumber of predicted TMHs: 1\nExp number of AAs in TMHs: 23.72226\nExp number, first 60 AAs: 0.22038\nTotal prob of N-in: 0.12994\noutside 1 331\nTMhelix 332 354\ninside 355 372

11108 GCF_000458475.1_Esch_coli_HVH_163_4-4697553_V1 Escherichia coli HVH 163 (4-4697553) Proteobacteria; Gammaproteobacteria; Enterobacterales; Enterobacteriaceae; Escherichia; Escherichia coli MNNEETFYQAMRRQGVTRRSFLKYCSLAATSLGLGAGMAPKIAWALENKPRIPVVWIHGL WP_001058323.1 MULTISPECIES: hydrogenase-1 small chain [Proteobacteria] Length: 372\nNumber of predicted TMHs: 2\nExp number of AAs in TMHs: 37.2608499999999\nExp number, first 60 AAs: 19.52923\nTotal prob of N-in: 0.94351\nPOSSIBLE N-term signal sequence\ninside 1 19\nTMhelix 20 39\noutside 40 325\nTMhelix 326 348\ninside 349 372

11109 GCF_000458475.1_Esch_coli_HVH_163_4-4697553_V1 Escherichia coli HVH 163 (4-4697553) Proteobacteria; Gammaproteobacteria; Enterobacterales; Enterobacteriaceae; Escherichia; Escherichia coli MTGDNTLIHSHGINRRDFMKLCAALAATMGLSSKAAAEMAESVTNPQRPPVIWIGAQECT WP_000145410.1 MULTISPECIES: hydrogenase-2 small chain [Proteobacteria] Length: 372\nNumber of predicted TMHs: 1\nExp number of AAs in TMHs: 23.72226\nExp number, first 60 AAs: 0.22038\nTotal prob of N-in: 0.12994\noutside 1 331\nTMhelix 332 354\ninside 355 372

11110 GCF_000458495.1_Esch_coli_HVH_164_4-5953081_V1 Escherichia coli HVH 164 (4-5953081) Proteobacteria; Gammaproteobacteria; Enterobacterales; Enterobacteriaceae; Escherichia; Escherichia coli MNNEETFYQAMRRQGVTRRSFLKYCSLAATSLGLGAGMAPKIAWALENKPRIPVVWIHGL WP_001058323.1 MULTISPECIES: hydrogenase-1 small chain [Proteobacteria] Length: 372\nNumber of predicted TMHs:

2\nExp number of AAs in TMHs: 37.2608499999999\nExp number, first 60 AAs: 19.52923\nTotal prob of N-in: 0.94351\nPOSSIBLE N-term signal sequence\ninside 1 19\nTMhelix 20 39\noutside 40 325\nTMhelix 326 348\ninside 349 372

11111 GCF_000458495.1_Esch_coli_HVH_164_4-5953081_V1 Escherichia coli HVH 164 (4-5953081) Proteobacteria; Gammaproteobacteria; Enterobacterales; Enterobacteriaceae; Escherichia; Escherichia coli MTGDNTLIHSHGINRRDFMKLCAALAATMGLSSKAAAEMAESVTNPQRPPVIWIGAQECT WP_000145410.1 MULTISPECIES: hydrogenase-2 small chain [Proteobacteria] Length: 372\nNumber of predicted TMHs: 1\nExp number of AAs in TMHs: 23.72226\nExp number, first 60 AAs: 0.22038\nTotal prob of N-in: 0.12994\noutside 1 331\nTMhelix 332 354\ninside 355 372

11112 GCF_000456205.1_Esch_coli_HVH_16_4-7649002_V1 Escherichia coli HVH 16 (4-7649002) Proteobacteria; Gammaproteobacteria; Enterobacterales; Enterobacteriaceae; Escherichia; Escherichia coli MTGDNTLIHSHGINRRDFMKLCAALAATMGLSSKAAAEMAESVTNPQRPPVIWIGAQECT WP_000145410.1 MULTISPECIES: hydrogenase-2 small chain [Proteobacteria] Length: 372\nNumber of predicted TMHs: 1\nExp number of AAs in TMHs: 23.72226\nExp number, first 60 AAs: 0.22038\nTotal prob of N-in: 0.12994\noutside 1 331\nTMhelix 332 354\ninside 355 372

11113 GCF_000456205.1_Esch_coli_HVH_16_4-7649002_V1 Escherichia coli HVH 16 (4-7649002) Proteobacteria; Gammaproteobacteria; Enterobacterales; Enterobacteriaceae; Escherichia; Escherichia coli MNNEETFYQAMRRQGVTRRSFLKYCSLAATSLGLGAGMAPKIAWALENKPRIPVVWIHGL WP_001058323.1 MULTISPECIES: hydrogenase-1 small chain [Proteobacteria] Length: 372\nNumber of predicted TMHs: 2\nExp number of AAs in TMHs: 37.2608499999999\nExp number, first 60 AAs: 19.52923\nTotal prob of N-in: 0.94351\nPOSSIBLE N-term signal sequence\ninside 1 19\nTMhelix 20 39\noutside 40 325\nTMhelix 326 348\ninside 349 372

11114 GCF_000458515.1_Esch_coli_HVH_167_4-6073565_V1 Escherichia coli HVH 167 (4-6073565) Proteobacteria; Gammaproteobacteria; Enterobacterales; Enterobacteriaceae; Escherichia; Escherichia coli MNNEETFYQAMRRQGVTRRSFLKYCSLAATSLGLGAGMAPKIAWALENKPRIPVVWIHGL WP_001058323.1 MULTISPECIES: hydrogenase-1 small chain [Proteobacteria] Length: 372\nNumber of predicted TMHs: 2\nExp number of AAs in TMHs: 37.2608499999999\nExp number, first 60 AAs: 19.52923\nTotal prob of N-in: 0.94351\nPOSSIBLE N-term signal sequence\ninside 1 19\nTMhelix 20 39\noutside 40 325\nTMhelix 326 348\ninside 349 372

11115 GCF_000458515.1_Esch_coli_HVH_167_4-6073565_V1 Escherichia coli HVH 167 (4-6073565) Proteobacteria; Gammaproteobacteria; Enterobacterales; Enterobacteriaceae; Escherichia; Escherichia coli MTGDNTLIHSHGINRRDFMKLCAALAATMGLSSKAAAEMAESVTNPQRPPVIWIGAQECT WP_000145410.1 MULTISPECIES: hydrogenase-2 small chain [Proteobacteria] Length: 372\nNumber of predicted TMHs: 1\nExp number of AAs in TMHs: 23.72226\nExp number, first 60 AAs: 0.22038\nTotal prob of N-in: 0.12994\noutside 1 331\nTMhelix 332 354\ninside 355 372

11116 GCF_000458535.1_Esch_coli_HVH_169_4-1075578_V1 Escherichia coli HVH 169 (4-1075578) Proteobacteria; Gammaproteobacteria; Enterobacterales; Enterobacteriaceae; Escherichia; Escherichia coli MNNEETFYQAMRRQGVTRRSFLKYCSLAATSLGLGAGMAPKIAWALENKPRIPVVWIHGL WP_001058323.1 MULTISPECIES: hydrogenase-1 small chain [Proteobacteria] Length: 372\nNumber of predicted TMHs: 2\nExp number of AAs in TMHs: 37.2608499999999\nExp number, first 60 AAs: 19.52923\nTotal prob of N-in: 0.94351\nPOSSIBLE N-term signal sequence\ninside 1 19\nTMhelix 20 39\noutside 40 325\nTMhelix 326 348\ninside 349 372

11117 GCF_000458535.1_Esch_coli_HVH_169_4-1075578_V1 Escherichia coli HVH 169 (4-1075578) Proteobacteria; Gammaproteobacteria; Enterobacterales; Enterobacteriaceae; Escherichia; Escherichia coli MTGDNTLIHSHGINRRDFMKLCAALAATMGLSSKAAAEMAESVTNPQRPPVIWIGAQECT WP_000145410.1 MULTISPECIES: hydrogenase-2 small chain [Proteobacteria] Length: 372\nNumber of predicted TMHs: 1\nExp number of AAs in TMHs: 23.72226\nExp number, first 60 AAs: 0.22038\nTotal prob of N-in: 0.12994\noutside 1 331\nTMhelix 332 354\ninside 355 372

11118 GCF_000458555.1_Esch_coli_HVH_170_4-3026949_V1 Escherichia coli HVH 170 (4-3026949) Proteobacteria; Gammaproteobacteria; Enterobacterales; Enterobacteriaceae; Escherichia; Escherichia coli MTGDNTLIHSHGINRRDFMKLCAALAATMGLSSKAAAEMAESVTNPQRPPVIWIGAQECT WP_000145410.1 MULTISPECIES: hydrogenase-2 small chain [Proteobacteria] Length: 372\nNumber of predicted TMHs:

1\nExp number of AAs in TMHs: 23.72226\nExp number, first 60 AAs:  0.22038\nTotal prob of N-in:   0.12994\noutside   1   331\nTMhelix   332   354\ninside   355   372

11119 GCF_000458575.1_Esch_coli_HVH_171_4-3191958_V1 Escherichia coli HVH 171 (4-3191958) Proteobacteria; Gammaproteobacteria; Enterobacterales; Enterobacteriaceae; Escherichia; Escherichia coli MTGDNTLIHSHGINRRDFMKLCAALAATMGLSSKAAAEMAESVTNPQRPPVIWIGAQECT WP_000145410.1 MULTISPECIES: hydrogenase-2 small chain [Proteobacteria] Length: 372\nNumber of predicted TMHs: 1\nExp number of AAs in TMHs: 23.72226\nExp number, first 60 AAs:  0.22038\nTotal prob of N-in:   0.12994\noutside   1   331\nTMhelix   332   354\ninside   355   372

11120 GCF_000458575.1_Esch_coli_HVH_171_4-3191958_V1 Escherichia coli HVH 171 (4-3191958) Proteobacteria; Gammaproteobacteria; Enterobacterales; Enterobacteriaceae; Escherichia; Escherichia coli MNNEETFYQAMRRQGVTRRSFLKYCSLAATSLGLGAGMAPKIAWALENKPRIPVVWIHGL WP_001058323.1 MULTISPECIES: hydrogenase-1 small chain [Proteobacteria] Length: 372\nNumber of predicted TMHs: 2\nExp number of AAs in TMHs: 37.2608499999999\nExp number, first 60 AAs:  19.52923\nTotal prob of N-in: 0.94351\nPOSSIBLE N-term signal sequence\ninside   1   19\nTMhelix   20   39\noutside   40   325\nTMhelix   326   348\ninside   349   372

11121 GCF_000458605.1_Esch_coli_HVH_172_4-3248542_V1 Escherichia coli HVH 172 (4-3248542) Proteobacteria; Gammaproteobacteria; Enterobacterales; Enterobacteriaceae; Escherichia; Escherichia coli MTGDNTLIHSHGINRRDFMKLCAALAATMGLSSKAAAEMAESVTNPQRPPVIWIGAQECT WP_000145410.1 MULTISPECIES: hydrogenase-2 small chain [Proteobacteria] Length: 372\nNumber of predicted TMHs: 1\nExp number of AAs in TMHs: 23.72226\nExp number, first 60 AAs:  0.22038\nTotal prob of N-in:   0.12994\noutside   1   331\nTMhelix   332   354\ninside   355   372

11122 GCF_000458605.1_Esch_coli_HVH_172_4-3248542_V1 Escherichia coli HVH 172 (4-3248542) Proteobacteria; Gammaproteobacteria; Enterobacterales; Enterobacteriaceae; Escherichia; Escherichia coli MNNEETFYQAMRRQGVTRRSFLKYCSLAATSLGLGAGMAPKIAWALENKPRIPVVWIHGL WP_001058323.1 MULTISPECIES: hydrogenase-1 small chain [Proteobacteria] Length: 372\nNumber of predicted TMHs: 2\nExp number of AAs in TMHs: 37.2608499999999\nExp number, first 60 AAs:  19.52923\nTotal prob of N-in: 0.94351\nPOSSIBLE N-term signal sequence\ninside   1   19\nTMhelix   20   39\noutside   40   325\nTMhelix   326   348\ninside   349   372

11123 GCF_000458625.1_Esch_coli_HVH_173_3-9175482_V1 Escherichia coli HVH 173 (3-9175482) Proteobacteria; Gammaproteobacteria; Enterobacterales; Enterobacteriaceae; Escherichia; Escherichia coli MTGDNTLIHSHGINRRDFMKLCAALAATMGLSSKAAAEMAESVTNPQRPPVIWIGAQECT WP_000145410.1 MULTISPECIES: hydrogenase-2 small chain [Proteobacteria] Length: 372\nNumber of predicted TMHs: 1\nExp number of AAs in TMHs: 23.72226\nExp number, first 60 AAs:  0.22038\nTotal prob of N-in:   0.12994\noutside   1   331\nTMhelix   332   354\ninside   355   372

11124 GCF_000458625.1_Esch_coli_HVH_173_3-9175482_V1 Escherichia coli HVH 173 (3-9175482) Proteobacteria; Gammaproteobacteria; Enterobacterales; Enterobacteriaceae; Escherichia; Escherichia coli MNNEETFYQAMRRQGVTRRSFLKYCSLAATSLGLGAGMAPKIAWALENKPRIPVVWIHGL WP_001058323.1 MULTISPECIES: hydrogenase-1 small chain [Proteobacteria] Length: 372\nNumber of predicted TMHs: 2\nExp number of AAs in TMHs: 37.2608499999999\nExp number, first 60 AAs:  19.52923\nTotal prob of N-in: 0.94351\nPOSSIBLE N-term signal sequence\ninside   1   19\nTMhelix   20   39\noutside   40   325\nTMhelix   326   348\ninside   349   372

11125 GCF_000456225.1_Esch_coli_HVH_17_4-7473087_V1 Escherichia coli HVH 17 (4-7473087) Proteobacteria; Gammaproteobacteria; Enterobacterales; Enterobacteriaceae; Escherichia; Escherichia coli MNNEETFYQAMRRQGVTRRSFLKYCSLAATSLGLGAGMAPKIAWALENKPRIPVVWIHGL WP_001295940.1 MULTISPECIES: hydrogenase [Enterobacteriaceae] Length: 372\nNumber of predicted TMHs:  2\nExp number of AAs in TMHs: 37.2608899999999\nExp number, first 60 AAs:  19.52923\nTotal prob of N-in: 0.94351\nPOSSIBLE N-term signal sequence\ninside   1   19\nTMhelix   20   39\noutside   40   325\nTMhelix   326   348\ninside   349   372

11126 GCF_000456225.1_Esch_coli_HVH_17_4-7473087_V1 Escherichia coli HVH 17 (4-7473087) Proteobacteria; Gammaproteobacteria; Enterobacterales; Enterobacteriaceae; Escherichia; Escherichia coli MTGDNTLIHSHGINRRDFMKLCAALAATMGLSSKAAAEMAESVTNPQRPPVIWIGAQECT WP_000145410.1 MULTISPECIES: hydrogenase-2 small chain [Proteobacteria] Length: 372\nNumber of predicted TMHs:

1\nExp number of AAs in TMHs: 23.72226\nExp number, first 60 AAs: 0.22038\nTotal prob of N-in: 0.12994\noutside 1 331\nTMhelix 332 354\ninside 355 372

11127 GCF_000458645.1_Esch_coli_HVH_175_4-3405184_V1 Escherichia coli HVH 175 (4-3405184) Proteobacteria; Gammaproteobacteria; Enterobacterales; Enterobacteriaceae; Escherichia; Escherichia coli MTGDNTLIHSHGINRRDFMKLCAALAATMGLSSKAAAEMAESVTNPQRPPVIWIGAQECT WP_000145418.1 hydrogenase 2 small subunit [Escherichia coli] Length: 372\nNumber of predicted TMHs: 1\nExp number of AAs in TMHs: 23.72313\nExp number, first 60 AAs: 0.22034\nTotal prob of N-in: 0.12997\noutside 1 331\nTMhelix 332 354\ninside 355 372

11128 GCF_000458645.1_Esch_coli_HVH_175_4-3405184_V1 Escherichia coli HVH 175 (4-3405184) Proteobacteria; Gammaproteobacteria; Enterobacterales; Enterobacteriaceae; Escherichia; Escherichia coli MNNEETFYQAMRRQGVTRRSFLKYCSLAATSLGLGAGMAPKIAWALENKPRIPVVWIHGL WP_001058323.1 MULTISPECIES: hydrogenase-1 small chain [Proteobacteria] Length: 372\nNumber of predicted TMHs: 2\nExp number of AAs in TMHs: 37.2608499999999\nExp number, first 60 AAs: 19.52923\nTotal prob of N-in: 0.94351\nPOSSIBLE N-term signal sequence\ninside 1 19\nTMhelix 20 39\noutside 40 325\nTMhelix 326 348\ninside 349 372

11129 GCF_000458665.1_Esch_coli_HVH_176_4-3428664_V1 Escherichia coli HVH 176 (4-3428664) Proteobacteria; Gammaproteobacteria; Enterobacterales; Enterobacteriaceae; Escherichia; Escherichia coli MTGDNTLIHSHGINRRDFMKLCAALAATMGLSSKAAAEMVESVTNPQRPPVIWIGAQECT WP_000145423.1 hydrogenase 2 small subunit [Escherichia coli] Length: 372\nNumber of predicted TMHs: 1\nExp number of AAs in TMHs: 23.78201\nExp number, first 60 AAs: 0.28355\nTotal prob of N-in: 0.13257\noutside 1 331\nTMhelix 332 354\ninside 355 372

11130 GCF_000458665.1_Esch_coli_HVH_176_4-3428664_V1 Escherichia coli HVH 176 (4-3428664) Proteobacteria; Gammaproteobacteria; Enterobacterales; Enterobacteriaceae; Escherichia; Escherichia coli MNNEETFYQAMRRQGVTRRSFLKYCSLAATSLGLGAGMAPKIAWALENKPRIPVVWIHGL WP_001295940.1 MULTISPECIES: hydrogenase [Enterobacteriaceae] Length: 372\nNumber of predicted TMHs: 2\nExp number of AAs in TMHs: 37.2608899999999\nExp number, first 60 AAs: 19.52923\nTotal prob of N-in: 0.94351\nPOSSIBLE N-term signal sequence\ninside 1 19\nTMhelix 20 39\noutside 40 325\nTMhelix 326 348\ninside 349 372

11131 GCF_000507645.1_Esch_coli_HVH_177_4-2876612_V1 Escherichia coli HVH 177 (4-2876612) Proteobacteria; Gammaproteobacteria; Enterobacterales; Enterobacteriaceae; Escherichia; Escherichia coli MNNEETFYQAMRRQGVTRRSFLKYCSLAATSLGLGAGMAPKIAWALENKPRIPVVWIHGL WP_001295940.1 MULTISPECIES: hydrogenase [Enterobacteriaceae] Length: 372\nNumber of predicted TMHs: 2\nExp number of AAs in TMHs: 37.2608899999999\nExp number, first 60 AAs: 19.52923\nTotal prob of N-in: 0.94351\nPOSSIBLE N-term signal sequence\ninside 1 19\nTMhelix 20 39\noutside 40 325\nTMhelix 326 348\ninside 349 372

11132 GCF_000507645.1_Esch_coli_HVH_177_4-2876612_V1 Escherichia coli HVH 177 (4-2876612) Proteobacteria; Gammaproteobacteria; Enterobacterales; Enterobacteriaceae; Escherichia; Escherichia coli MTGDNTLIHSHGINRRDFMKLCAALAATMGLSSKAAAEMAESVTNPQRPPVIWIGAQECT WP_000145410.1 MULTISPECIES: hydrogenase-2 small chain [Proteobacteria] Length: 372\nNumber of predicted TMHs: 1\nExp number of AAs in TMHs: 23.72226\nExp number, first 60 AAs: 0.22038\nTotal prob of N-in: 0.12994\noutside 1 331\nTMhelix 332 354\ninside 355 372

11133 GCF_000495055.1_Esch_coli_HVH_178_4-3189163_V1 Escherichia coli HVH 178 (4-3189163) Proteobacteria; Gammaproteobacteria; Enterobacterales; Enterobacteriaceae; Escherichia; Escherichia coli MTGDNTLIHSHGINRRDFMKLCAALAATMGLSSKAAAEMAESVTNPQRPPVIWIGAQECT WP_000145410.1 MULTISPECIES: hydrogenase-2 small chain [Proteobacteria] Length: 372\nNumber of predicted TMHs: 1\nExp number of AAs in TMHs: 23.72226\nExp number, first 60 AAs: 0.22038\nTotal prob of N-in: 0.12994\noutside 1 331\nTMhelix 332 354\ninside 355 372

11134 GCF_000458685.1_Esch_coli_HVH_180_4-3051617_V1 Escherichia coli HVH 180 (4-3051617) Proteobacteria; Gammaproteobacteria; Enterobacterales; Enterobacteriaceae; Escherichia; Escherichia coli MTGDNTLIHSHGINRRDFMKLCAALAATMGLSSKAAAEMAESVTNPQRPPVIWIGAQECT WP_000145410.1 MULTISPECIES: hydrogenase-2 small chain [Proteobacteria] Length: 372\nNumber of predicted TMHs: 1\nExp number of AAs in TMHs: 23.72226\nExp number, first 60 AAs: 0.22038\nTotal prob of N-in: 0.12994\noutside 1 331\nTMhelix 332 354\ninside 355 372

11135 GCF_000458705.1_Esch_coli_HVH_182_4-0985554_V1 Escherichia coli HVH 182 (4-0985554) Proteobacteria; Gammaproteobacteria; Enterobacterales; Enterobacteriaceae; Escherichia; Escherichia coli MTGDNTLIHSHGINRRDFMKLCAALAATMGLSSKAAAEMAESVTNPQRPPVIWIGAQECT WP_000145410.1 MULTISPECIES: hydrogenase-2 small chain [Proteobacteria] Length: 372\nNumber of predicted TMHs: 1\nExp number of AAs in TMHs: 23.72226\nExp number, first 60 AAs: 0.22038\nTotal prob of N-in: 0.12994\noutside 1 331\nTMhelix 332 354\ninside 355 372

11136 GCF_000458705.1_Esch_coli_HVH_182_4-0985554_V1 Escherichia coli HVH 182 (4-0985554) Proteobacteria; Gammaproteobacteria; Enterobacterales; Enterobacteriaceae; Escherichia; Escherichia coli MNNEETFYQAMRRQGVTRRSFLKYCSLAATSLGLGAGMAPKIAWALENKPRIPVVWIHGL WP_001058323.1 MULTISPECIES: hydrogenase-1 small chain [Proteobacteria] Length: 372\nNumber of predicted TMHs: 2\nExp number of AAs in TMHs: 37.2608499999999\nExp number, first 60 AAs: 19.52923\nTotal prob of N-in: 0.94351\nPOSSIBLE N-term signal sequence\ninside 1 19\nTMhelix 20 39\noutside 40 325\nTMhelix 326 348\ninside 349 372

11137 GCF_000458725.1_Esch_coli_HVH_183_4-3205932_V1 Escherichia coli HVH 183 (4-3205932) Proteobacteria; Gammaproteobacteria; Enterobacterales; Enterobacteriaceae; Escherichia; Escherichia coli MNNEETFYQAMRRQGVTRRSFLKYCSLAATSLGLGAGMAPKIAWALENKPRIPVVWIHGL WP_001058323.1 MULTISPECIES: hydrogenase-1 small chain [Proteobacteria] Length: 372\nNumber of predicted TMHs: 2\nExp number of AAs in TMHs: 37.2608499999999\nExp number, first 60 AAs: 19.52923\nTotal prob of N-in: 0.94351\nPOSSIBLE N-term signal sequence\ninside 1 19\nTMhelix 20 39\noutside 40 325\nTMhelix 326 348\ninside 349 372

11138 GCF_000458725.1_Esch_coli_HVH_183_4-3205932_V1 Escherichia coli HVH 183 (4-3205932) Proteobacteria; Gammaproteobacteria; Enterobacterales; Enterobacteriaceae; Escherichia; Escherichia coli MTGDNTLIHSHGINRRDFMKLCAALAATMGLSSKAAAEMAESVTNPQRPPVIWIGAQECT WP_000145410.1 MULTISPECIES: hydrogenase-2 small chain [Proteobacteria] Length: 372\nNumber of predicted TMHs: 1\nExp number of AAs in TMHs: 23.72226\nExp number, first 60 AAs: 0.22038\nTotal prob of N-in: 0.12994\noutside 1 331\nTMhelix 332 354\ninside 355 372

11139 GCF_000458745.1_Esch_coli_HVH_184_4-3343286_V1 Escherichia coli HVH 184 (4-3343286) Proteobacteria; Gammaproteobacteria; Enterobacterales; Enterobacteriaceae; Escherichia; Escherichia coli MNNEETFYQAMRRQGVTRRSFLKYCSLAATSLGLGAGMAPKIAWALENKPRIPVVWIHGL WP_001058323.1 MULTISPECIES: hydrogenase-1 small chain [Proteobacteria] Length: 372\nNumber of predicted TMHs: 2\nExp number of AAs in TMHs: 37.2608499999999\nExp number, first 60 AAs: 19.52923\nTotal prob of N-in: 0.94351\nPOSSIBLE N-term signal sequence\ninside 1 19\nTMhelix 20 39\noutside 40 325\nTMhelix 326 348\ninside 349 372

11140 GCF_000458745.1_Esch_coli_HVH_184_4-3343286_V1 Escherichia coli HVH 184 (4-3343286) Proteobacteria; Gammaproteobacteria; Enterobacterales; Enterobacteriaceae; Escherichia; Escherichia coli MTGDNTLIHSHGINRRDFMKLCAALAATMGLSSKAAAEMAESVTNPQRPPVIWIGAQECT WP_000145418.1 hydrogenase 2 small subunit [Escherichia coli] Length: 372\nNumber of predicted TMHs: 1\nExp number of AAs in TMHs: 23.72313\nExp number, first 60 AAs: 0.22034\nTotal prob of N-in: 0.12997\noutside 1 331\nTMhelix 332 354\ninside 355 372

11141 GCF_000456245.1_Esch_coli_HVH_18_4-8589585_V1 Escherichia coli HVH 18 (4-8589585) Proteobacteria; Gammaproteobacteria; Enterobacterales; Enterobacteriaceae; Escherichia; Escherichia coli MTGDNTLIHSHGINRRDFMKLCAALAATMGLSSKAAAEMAESVTNPQRPPVIWIGAQECT WP_000145410.1 MULTISPECIES: hydrogenase-2 small chain [Proteobacteria] Length: 372\nNumber of predicted TMHs: 1\nExp number of AAs in TMHs: 23.72226\nExp number, first 60 AAs: 0.22038\nTotal prob of N-in: 0.12994\noutside 1 331\nTMhelix 332 354\ninside 355 372

11142 GCF_000456245.1_Esch_coli_HVH_18_4-8589585_V1 Escherichia coli HVH 18 (4-8589585) Proteobacteria; Gammaproteobacteria; Enterobacterales; Enterobacteriaceae; Escherichia; Escherichia coli MNNEETFYQAMRRQGVTRRSFLKYCSLAATSLGLGAGMAPKIAWALENKPRIPVVWIHGL WP_001058320.1 hydrogenase [Escherichia coli] Length: 372\nNumber of predicted TMHs: 2\nExp number of AAs in TMHs: 37.26082\nExp number, first 60 AAs: 19.52923\nTotal prob of N-in: 0.94351\nPOSSIBLE N-term signal sequence\ninside 1 19\nTMhelix 20 39\noutside 40 325\nTMhelix 326 348\ninside 349 372

11143 GCF_000458765.1_Esch_coli_HVH_185_4-2876639_V1 Escherichia coli HVH 185 (4-2876639) Proteobacteria; Gammaproteobacteria; Enterobacterales; Enterobacteriaceae; Escherichia; Escherichia coli

MNNEETFYQAMRRQGVTRRSFLKYCSLAATSLGLGAGMAPKIAWALENKPRIPVVWIHGL WP_001058323.1 MULTISPECIES: hydrogenase-1 small chain [Proteobacteria] Length: 372\nNumber of predicted TMHs: 2\nExp number of AAs in TMHs: 37.2608499999999\nExp number, first 60 AAs: 19.52923\nTotal prob of N-in: 0.94351\nPOSSIBLE N-term signal sequence\ninside 1 19\nTMhelix 20 39\noutside 40 325\nTMhelix 326 348\ninside 349 372

11144 GCF_000458765.1_Esch_coli_HVH_185_4-2876639_V1 Escherichia coli HVH 185 (4-2876639) Proteobacteria; Gammaproteobacteria; Enterobacterales; Enterobacteriaceae; Escherichia; Escherichia coli MTGDNTLIHSHGINRRDFMKLCAALAATMGLSSKAAAEMAESVTNPQRPPVIWIGAQECT WP_000145410.1 MULTISPECIES: hydrogenase-2 small chain [Proteobacteria] Length: 372\nNumber of predicted TMHs: 1\nExp number of AAs in TMHs: 23.72226\nExp number, first 60 AAs: 0.22038\nTotal prob of N-in: 0.12994\noutside 1 331\nTMhelix 332 354\ninside 355 372

11145 GCF_000458785.1_Esch_coli_HVH_186_4-3405044_V1 Escherichia coli HVH 186 (4-3405044) Proteobacteria; Gammaproteobacteria; Enterobacterales; Enterobacteriaceae; Escherichia; Escherichia coli MNNEETFYQAMRRQGVTRRSFLKYCSLAATSLGLGAGMAPKIAWALENKPRIPVVWIHGL WP_001295940.1 MULTISPECIES: hydrogenase [Enterobacteriaceae] Length: 372\nNumber of predicted TMHs: 2\nExp number of AAs in TMHs: 37.2608899999999\nExp number, first 60 AAs: 19.52923\nTotal prob of N-in: 0.94351\nPOSSIBLE N-term signal sequence\ninside 1 19\nTMhelix 20 39\noutside 40 325\nTMhelix 326 348\ninside 349 372

11146 GCF_000458785.1_Esch_coli_HVH_186_4-3405044_V1 Escherichia coli HVH 186 (4-3405044) Proteobacteria; Gammaproteobacteria; Enterobacterales; Enterobacteriaceae; Escherichia; Escherichia coli MTGDNTLIHSHGINRRDFMKLCAALAATMGLSSKAAAEMAESVTNPQRPPVIWIGAQECT WP_000145410.1 MULTISPECIES: hydrogenase-2 small chain [Proteobacteria] Length: 372\nNumber of predicted TMHs: 1\nExp number of AAs in TMHs: 23.72226\nExp number, first 60 AAs: 0.22038\nTotal prob of N-in: 0.12994\noutside 1 331\nTMhelix 332 354\ninside 355 372

11147 GCF_000458805.1_Esch_coli_HVH_187_4-4471660_V1 Escherichia coli HVH 187 (4-4471660) Proteobacteria; Gammaproteobacteria; Enterobacterales; Enterobacteriaceae; Escherichia; Escherichia coli MTGDNTLIHSHGINRRDFMKLCAALAATMGLSSKAAAEMAESVTNPQRPPVIWIGAQECT WP_000145410.1 MULTISPECIES: hydrogenase-2 small chain [Proteobacteria] Length: 372\nNumber of predicted TMHs: 1\nExp number of AAs in TMHs: 23.72226\nExp number, first 60 AAs: 0.22038\nTotal prob of N-in: 0.12994\noutside 1 331\nTMhelix 332 354\ninside 355 372

11148 GCF_000458805.1_Esch_coli_HVH_187_4-4471660_V1 Escherichia coli HVH 187 (4-4471660) Proteobacteria; Gammaproteobacteria; Enterobacterales; Enterobacteriaceae; Escherichia; Escherichia coli MNNEETFYQAMRRQGVTRRSFLKYCSLAATSLGLGAGMAPKIAWALENKPRIPVVWIHGL WP_001295940.1 MULTISPECIES: hydrogenase [Enterobacteriaceae] Length: 372\nNumber of predicted TMHs: 2\nExp number of AAs in TMHs: 37.2608899999999\nExp number, first 60 AAs: 19.52923\nTotal prob of N-in: 0.94351\nPOSSIBLE N-term signal sequence\ninside 1 19\nTMhelix 20 39\noutside 40 325\nTMhelix 326 348\ninside 349 372

11149 GCF_000458825.1_Esch_coli_HVH_188_4-2356988_V1 Escherichia coli HVH 188 (4-2356988) Proteobacteria; Gammaproteobacteria; Enterobacterales; Enterobacteriaceae; Escherichia; Escherichia coli MNNEETFYQAMRRQGVTRRSFLKYCSLAATSLGLGAGMAPKIAWALENKPRIPVVWIHGL WP_001058323.1 MULTISPECIES: hydrogenase-1 small chain [Proteobacteria] Length: 372\nNumber of predicted TMHs: 2\nExp number of AAs in TMHs: 37.2608499999999\nExp number, first 60 AAs: 19.52923\nTotal prob of N-in: 0.94351\nPOSSIBLE N-term signal sequence\ninside 1 19\nTMhelix 20 39\noutside 40 325\nTMhelix 326 348\ninside 349 372

11150 GCF_000458825.1_Esch_coli_HVH_188_4-2356988_V1 Escherichia coli HVH 188 (4-2356988) Proteobacteria; Gammaproteobacteria; Enterobacterales; Enterobacteriaceae; Escherichia; Escherichia coli MTGDNTLIHSHGINRRDFMKLCAALAATMGLSSKAAAEMAESVTNPQRPPVIWIGAQECT WP_000145410.1 MULTISPECIES: hydrogenase-2 small chain [Proteobacteria] Length: 372\nNumber of predicted TMHs: 1\nExp number of AAs in TMHs: 23.72226\nExp number, first 60 AAs: 0.22038\nTotal prob of N-in: 0.12994\noutside 1 331\nTMhelix 332 354\ninside 355 372

11151 GCF_000458845.1_Esch_coli_HVH_189_4-3220125_V1 Escherichia coli HVH 189 (4-3220125) Proteobacteria; Gammaproteobacteria; Enterobacterales; Enterobacteriaceae; Escherichia; Escherichia coli MTGDNTLIHSHGINRRDFMKLCAALAATMGLSSKAAAEMAESVTNPQRPPVIWIGAQECT WP_000145410.1

MULTISPECIES: hydrogenase-2 small chain [Proteobacteria] Length: 372\nNumber of predicted TMHs: 1\nExp number of AAs in TMHs: 23.72226\nExp number, first 60 AAs: 0.22038\nTotal prob of N-in: 0.12994\noutside 1 331\nTMhelix 332 354\ninside 355 372

11152 GCF_000458845.1_Esch_coli_HVH_189_4-3220125_V1 Escherichia coli HVH 189 (4-3220125) Proteobacteria; Gammaproteobacteria; Enterobacterales; Enterobacteriaceae; Escherichia; Escherichia coli MNNEETFYQAMRRQGVTRRSFLKYCSLAATSLGLGAGMAPKIAWALENKPRIPVVWIHGL WP_001058323.1 MULTISPECIES: hydrogenase-1 small chain [Proteobacteria] Length: 372\nNumber of predicted TMHs: 2\nExp number of AAs in TMHs: 37.2608499999999\nExp number, first 60 AAs: 19.52923\nTotal prob of N-in: 0.94351\nPOSSIBLE N-term signal sequence\ninside 1 19\nTMhelix 20 39\noutside 40 325\nTMhelix 326 348\ninside 349 372

11153 GCF_000465135.1_Esch_coli_HVH_190_4-3255514_V1 Escherichia coli HVH 190 (4-3255514) Proteobacteria; Gammaproteobacteria; Enterobacterales; Enterobacteriaceae; Escherichia; Escherichia coli MTGDNTLIHSHGINRRDFMKLCAALAATMGLSSKAAAEMAESVTNPQRPPVIWIGAQECT WP_000145410.1 MULTISPECIES: hydrogenase-2 small chain [Proteobacteria] Length: 372\nNumber of predicted TMHs: 1\nExp number of AAs in TMHs: 23.72226\nExp number, first 60 AAs: 0.22038\nTotal prob of N-in: 0.12994\noutside 1 331\nTMhelix 332 354\ninside 355 372

11154 GCF_000465135.1_Esch_coli_HVH_190_4-3255514_V1 Escherichia coli HVH 190 (4-3255514) Proteobacteria; Gammaproteobacteria; Enterobacterales; Enterobacteriaceae; Escherichia; Escherichia coli MNNEETFYQAMRRQGVTRRSFLKYCSLAATSLGLGAGMAPKIAWALENKPRIPVVWIHGL WP_001295940.1 MULTISPECIES: hydrogenase [Enterobacteriaceae] Length: 372\nNumber of predicted TMHs: 2\nExp number of AAs in TMHs: 37.2608899999999\nExp number, first 60 AAs: 19.52923\nTotal prob of N-in: 0.94351\nPOSSIBLE N-term signal sequence\ninside 1 19\nTMhelix 20 39\noutside 40 325\nTMhelix 326 348\ninside 349 372

11155 GCF_000458875.1_Esch_coli_HVH_191_3-9341900_V1 Escherichia coli HVH 191 (3-9341900) Proteobacteria; Gammaproteobacteria; Enterobacterales; Enterobacteriaceae; Escherichia; Escherichia coli MTGDNTLIHSHGINRRDFMKLCAALAATMGLSSKAAAEMAESVTNPQRPPVIWIGAQECT WP_000145410.1 MULTISPECIES: hydrogenase-2 small chain [Proteobacteria] Length: 372\nNumber of predicted TMHs: 1\nExp number of AAs in TMHs: 23.72226\nExp number, first 60 AAs: 0.22038\nTotal prob of N-in: 0.12994\noutside 1 331\nTMhelix 332 354\ninside 355 372

11156 GCF_000458895.1_Esch_coli_HVH_192_4-3054470_V1 Escherichia coli HVH 192 (4-3054470) Proteobacteria; Gammaproteobacteria; Enterobacterales; Enterobacteriaceae; Escherichia; Escherichia coli MTGDNTLIHSHGINRRDFMKLCAALAATMGLSSKAAAEMAESVTNPQRPPVIWIGAQECT WP_000145410.1 MULTISPECIES: hydrogenase-2 small chain [Proteobacteria] Length: 372\nNumber of predicted TMHs: 1\nExp number of AAs in TMHs: 23.72226\nExp number, first 60 AAs: 0.22038\nTotal prob of N-in: 0.12994\noutside 1 331\nTMhelix 332 354\ninside 355 372

11157 GCF_000458915.1_Esch_coli_HVH_193_4-3331423_V1 Escherichia coli HVH 193 (4-3331423) Proteobacteria; Gammaproteobacteria; Enterobacterales; Enterobacteriaceae; Escherichia; Escherichia coli MTGDNTLIHSHGINRRDFMKLCAALAATMGLSSKAAAEMAESVTNPQRPPVIWIGAQECT WP_000145410.1 MULTISPECIES: hydrogenase-2 small chain [Proteobacteria] Length: 372\nNumber of predicted TMHs: 1\nExp number of AAs in TMHs: 23.72226\nExp number, first 60 AAs: 0.22038\nTotal prob of N-in: 0.12994\noutside 1 331\nTMhelix 332 354\ninside 355 372

11158 GCF_000458915.1_Esch_coli_HVH_193_4-3331423_V1 Escherichia coli HVH 193 (4-3331423) Proteobacteria; Gammaproteobacteria; Enterobacterales; Enterobacteriaceae; Escherichia; Escherichia coli MNNEETFYQAMRRQGVTRRSFLKYCSLAATSLGLGAGMAPKIAWALENKPRIPVVWIHGL WP_001058323.1 MULTISPECIES: hydrogenase-1 small chain [Proteobacteria] Length: 372\nNumber of predicted TMHs: 2\nExp number of AAs in TMHs: 37.2608499999999\nExp number, first 60 AAs: 19.52923\nTotal prob of N-in: 0.94351\nPOSSIBLE N-term signal sequence\ninside 1 19\nTMhelix 20 39\noutside 40 325\nTMhelix 326 348\ninside 349 372

11159 GCF_000458935.1_Esch_coli_HVH_194_4-2356805_V1 Escherichia coli HVH 194 (4-2356805) Proteobacteria; Gammaproteobacteria; Enterobacterales; Enterobacteriaceae; Escherichia; Escherichia coli MTGDNTLIHSHGINRRDFMKLCAALAATMGLSSKAAAEMAESVTNPQRPPVIWIGAQECT WP_000145418.1 hydrogenase 2 small subunit [Escherichia coli] Length: 372\nNumber of predicted TMHs: 1\nExp number of

AAs in TMHs: 23.72313\nExp number, first 60 AAs:  0.22034\nTotal prob of N-in:   0.12997\noutside   1   331\nTMhelix   332   354\ninside   355   372

11160 GCF_000458935.1_Esch_coli_HVH_194_4-2356805_V1 Escherichia coli HVH 194 (4-2356805) Proteobacteria; Gammaproteobacteria; Enterobacterales; Enterobacteriaceae; Escherichia; Escherichia coli MNNEETFYQAMRRQGVTRRSFLKYCSLAATSLGLGAGMAPKIAWALENKPRIPVVWIHGL WP_001058323.1 MULTISPECIES: hydrogenase-1 small chain [Proteobacteria] Length: 372\nNumber of predicted TMHs: 2\nExp number of AAs in TMHs: 37.2608499999999\nExp number, first 60 AAs:  19.52923\nTotal prob of N-in: 0.94351\nPOSSIBLE N-term signal sequence\ninside   1   19\nTMhelix   20   39\noutside   40   325\nTMhelix   326 348\ninside   349   372

11161 GCF_000456265.1_Esch_coli_HVH_19_4-7154984_V1 Escherichia coli HVH 19 (4-7154984) Proteobacteria; Gammaproteobacteria; Enterobacterales; Enterobacteriaceae; Escherichia; Escherichia coli MTGDNTLIHSHGINRRDFMKLCAALAATMGLSSKAAAEMAESVTNPQRPPVIWIGAQECT WP_000145410.1 MULTISPECIES: hydrogenase-2 small chain [Proteobacteria] Length: 372\nNumber of predicted TMHs: 1\nExp number of AAs in TMHs: 23.72226\nExp number, first 60 AAs:  0.22038\nTotal prob of N-in:   0.12994\noutside   1 331\nTMhelix   332   354\ninside   355   372

11162 GCF_000458955.1_Esch_coli_HVH_195_3-7155360_V1 Escherichia coli HVH 195 (3-7155360) Proteobacteria; Gammaproteobacteria; Enterobacterales; Enterobacteriaceae; Escherichia; Escherichia coli MNNEETFYQAMRRQGVTRRSFLKYCSLAATSLGLGAGMAPKIAWALENKPRIPVVWIHGL WP_001295940.1 MULTISPECIES: hydrogenase [Enterobacteriaceae] Length: 372\nNumber of predicted TMHs:  2\nExp number of AAs in TMHs: 37.2608899999999\nExp number, first 60 AAs:  19.52923\nTotal prob of N-in: 0.94351\nPOSSIBLE N-term signal sequence\ninside   1   19\nTMhelix   20   39\noutside   40   325\nTMhelix   326 348\ninside   349   372

11163 GCF_000458955.1_Esch_coli_HVH_195_3-7155360_V1 Escherichia coli HVH 195 (3-7155360) Proteobacteria; Gammaproteobacteria; Enterobacterales; Enterobacteriaceae; Escherichia; Escherichia coli MTGDNTLIHSHGINRRDFMKLCAALAATMGLSSKAAAEMAESVTNPQRPPVIWIGAQECT WP_000145410.1 MULTISPECIES: hydrogenase-2 small chain [Proteobacteria] Length: 372\nNumber of predicted TMHs: 1\nExp number of AAs in TMHs: 23.72226\nExp number, first 60 AAs:  0.22038\nTotal prob of N-in:   0.12994\noutside   1 331\nTMhelix   332   354\ninside   355   372

11164 GCF_000458975.1_Esch_coli_HVH_196_4-4530470_V1 Escherichia coli HVH 196 (4-4530470) Proteobacteria; Gammaproteobacteria; Enterobacterales; Enterobacteriaceae; Escherichia; Escherichia coli MTGDNTLIHSHGINRRDFMKLCAALAATMGLSSKAAAEMAESVTNPQRPPVIWIGAQECT WP_000145410.1 MULTISPECIES: hydrogenase-2 small chain [Proteobacteria] Length: 372\nNumber of predicted TMHs: 1\nExp number of AAs in TMHs: 23.72226\nExp number, first 60 AAs:  0.22038\nTotal prob of N-in:   0.12994\noutside   1 331\nTMhelix   332   354\ninside   355   372

11165 GCF_000458995.1_Esch_coli_HVH_197_4-4466217_V1 Escherichia coli HVH 197 (4-4466217) Proteobacteria; Gammaproteobacteria; Enterobacterales; Enterobacteriaceae; Escherichia; Escherichia coli MTGDNTLIHSHGINRRDFMKLCAALAATMGLSSKAAAEMAESVTNPQRPPVIWIGAQECT WP_000145410.1 MULTISPECIES: hydrogenase-2 small chain [Proteobacteria] Length: 372\nNumber of predicted TMHs: 1\nExp number of AAs in TMHs: 23.72226\nExp number, first 60 AAs:  0.22038\nTotal prob of N-in:   0.12994\noutside   1 331\nTMhelix   332   354\ninside   355   372

11166 GCF_000458995.1_Esch_coli_HVH_197_4-4466217_V1 Escherichia coli HVH 197 (4-4466217) Proteobacteria; Gammaproteobacteria; Enterobacterales; Enterobacteriaceae; Escherichia; Escherichia coli MNNEETFYQAMRRQGVTRRSFLKYCSLAATSLGLGAGMAPKIAWALENKPRIPVVWIHGL WP_001058323.1 MULTISPECIES: hydrogenase-1 small chain [Proteobacteria] Length: 372\nNumber of predicted TMHs: 2\nExp number of AAs in TMHs: 37.2608499999999\nExp number, first 60 AAs:  19.52923\nTotal prob of N-in: 0.94351\nPOSSIBLE N-term signal sequence\ninside   1   19\nTMhelix   20   39\noutside   40   325\nTMhelix   326 348\ninside   349   372

11167 GCF_000459015.1_Esch_coli_HVH_198_4-3206106_V1 Escherichia coli HVH 198 (4-3206106) Proteobacteria; Gammaproteobacteria; Enterobacterales; Enterobacteriaceae; Escherichia; Escherichia coli MTGDNTLIHSHGINRRDFMKLCAALAATMGLSSKAAAEMAESVTNPQRPPVIWIGAQECT WP_000145410.1 MULTISPECIES: hydrogenase-2 small chain [Proteobacteria] Length: 372\nNumber of predicted TMHs: 1\nExp number of AAs in TMHs: 23.72226\nExp number, first 60 AAs:  0.22038\nTotal prob of N-in:   0.12994\noutside   1 331\nTMhelix   332   354\ninside   355   372

11168 GCF_000459015.1_Esch_coli_HVH_198_4-3206106_V1 Escherichia coli HVH 198 (4-3206106) Proteobacteria; Gammaproteobacteria; Enterobacterales; Enterobacteriaceae; Escherichia; Escherichia coli MNNEETFYQAMRRQGVTRRSFLKYCSLAATSLGLGAGMAPKIAWALENKPRIPVVWIHGL WP_001058323.1 MULTISPECIES: hydrogenase-1 small chain [Proteobacteria] Length: 372\nNumber of predicted TMHs: 2\nExp number of AAs in TMHs: 37.2608499999999\nExp number, first 60 AAs: 19.52923\nTotal prob of N-in: 0.94351\nPOSSIBLE N-term signal sequence\ninside 1 19\nTMhelix 20 39\noutside 40 325\nTMhelix 326 348\ninside 349 372

11169 GCF_000459035.1_Esch_coli_HVH_199_4-5670322_V1 Escherichia coli HVH 199 (4-5670322) Proteobacteria; Gammaproteobacteria; Enterobacterales; Enterobacteriaceae; Escherichia; Escherichia coli MTGDNTLIHSHGINRRDFMKLCAALAATMGLSSKAAAEMAESVTNPQRPPVIWIGAQECT WP_000145410.1 MULTISPECIES: hydrogenase-2 small chain [Proteobacteria] Length: 372\nNumber of predicted TMHs: 1\nExp number of AAs in TMHs: 23.72226\nExp number, first 60 AAs: 0.22038\nTotal prob of N-in: 0.12994\noutside 1 331\nTMhelix 332 354\ninside 355 372

11170 GCF_000459055.1_Esch_coli_HVH_200_4-4449924_V1 Escherichia coli HVH 200 (4-4449924) Proteobacteria; Gammaproteobacteria; Enterobacterales; Enterobacteriaceae; Escherichia; Escherichia coli MTGDNTLIHSHGINRRDFMKLCAALAATMGLSSKAAAEMAESVTNPQRPPVIWIGAQECT WP_000145410.1 MULTISPECIES: hydrogenase-2 small chain [Proteobacteria] Length: 372\nNumber of predicted TMHs: 1\nExp number of AAs in TMHs: 23.72226\nExp number, first 60 AAs: 0.22038\nTotal prob of N-in: 0.12994\noutside 1 331\nTMhelix 332 354\ninside 355 372

11171 GCF_000459055.1_Esch_coli_HVH_200_4-4449924_V1 Escherichia coli HVH 200 (4-4449924) Proteobacteria; Gammaproteobacteria; Enterobacterales; Enterobacteriaceae; Escherichia; Escherichia coli MNNEETFYQAMRRQGVTRRSFLKYCSLAATSLGLGAGMAPKIAWALENKPRIPVVWIHGL WP_001058320.1 hydrogenase [Escherichia coli] Length: 372\nNumber of predicted TMHs: 2\nExp number of AAs in TMHs: 37.26082\nExp number, first 60 AAs: 19.52923\nTotal prob of N-in: 0.94351\nPOSSIBLE N-term signal sequence\ninside 1 19\nTMhelix 20 39\noutside 40 325\nTMhelix 326 348\ninside 349 372

11172 GCF_000459075.1_Esch_coli_HVH_201_4-4459431_V1 Escherichia coli HVH 201 (4-4459431) Proteobacteria; Gammaproteobacteria; Enterobacterales; Enterobacteriaceae; Escherichia; Escherichia coli MTGDNTLIHSHGINRRDFMKLCAALAATMGLSSKAAAEMAESVTNPQRPPVIWIGAQECT WP_000145410.1 MULTISPECIES: hydrogenase-2 small chain [Proteobacteria] Length: 372\nNumber of predicted TMHs: 1\nExp number of AAs in TMHs: 23.72226\nExp number, first 60 AAs: 0.22038\nTotal prob of N-in: 0.12994\noutside 1 331\nTMhelix 332 354\ninside 355 372

11173 GCF_000459095.1_Esch_coli_HVH_202_4-3163997_V1 Escherichia coli HVH 202 (4-3163997) Proteobacteria; Gammaproteobacteria; Enterobacterales; Enterobacteriaceae; Escherichia; Escherichia coli MNNEETFYQAMRRQGVTRRSFLKYCSLAATSLGLGAGMAPKIAWALENKPRIPVVWIHGL WP_001295940.1 MULTISPECIES: hydrogenase [Enterobacteriaceae] Length: 372\nNumber of predicted TMHs: 2\nExp number of AAs in TMHs: 37.2608899999999\nExp number, first 60 AAs: 19.52923\nTotal prob of N-in: 0.94351\nPOSSIBLE N-term signal sequence\ninside 1 19\nTMhelix 20 39\noutside 40 325\nTMhelix 326 348\ninside 349 372

11174 GCF_000459095.1_Esch_coli_HVH_202_4-3163997_V1 Escherichia coli HVH 202 (4-3163997) Proteobacteria; Gammaproteobacteria; Enterobacterales; Enterobacteriaceae; Escherichia; Escherichia coli MTGDNTLIHSHGINRRDFMKLCAALAATMGLSSKAAAEMAESVTNPQRPPVIWIGAQECT WP_000145410.1 MULTISPECIES: hydrogenase-2 small chain [Proteobacteria] Length: 372\nNumber of predicted TMHs: 1\nExp number of AAs in TMHs: 23.72226\nExp number, first 60 AAs: 0.22038\nTotal prob of N-in: 0.12994\noutside 1 331\nTMhelix 332 354\ninside 355 372

11175 GCF_000459115.1_Esch_coli_HVH_203_4-3126218_V1 Escherichia coli HVH 203 (4-3126218) Proteobacteria; Gammaproteobacteria; Enterobacterales; Enterobacteriaceae; Escherichia; Escherichia coli MTGDNTLIHSHGINRRDFMKLCAALAATMGLSSKAAAEMAESVTNPQRPPVIWIGAQECT WP_000145410.1 MULTISPECIES: hydrogenase-2 small chain [Proteobacteria] Length: 372\nNumber of predicted TMHs: 1\nExp number of AAs in TMHs: 23.72226\nExp number, first 60 AAs: 0.22038\nTotal prob of N-in: 0.12994\noutside 1 331\nTMhelix 332 354\ninside 355 372

11176 GCF_000459135.1_Esch_coli_HVH_204_4-3112802_V1 Escherichia coli HVH 204 (4-3112802) Proteobacteria; Gammaproteobacteria; Enterobacterales; Enterobacteriaceae; Escherichia; Escherichia coli MNNEETFYQAMRRQGVTRRSFLKYCSLAATSLGLGAGMAPKIAWALENKPRIPVVWIHGL WP_001058323.1

MULTISPECIES: hydrogenase-1 small chain [Proteobacteria] Length: 372\nNumber of predicted TMHs: 2\nExp number of AAs in TMHs: 37.2608499999999\nExp number, first 60 AAs: 19.52923\nTotal prob of N-in: 0.94351\nPOSSIBLE N-term signal sequence\ninside 1 19\nTMhelix 20 39\noutside 40 325\nTMhelix 326 348\ninside 349 372

11177 GCF_000459135.1_Esch_coli_HVH_204_4-3112802_V1 Escherichia coli HVH 204 (4-3112802) Proteobacteria; Gammaproteobacteria; Enterobacterales; Enterobacteriaceae; Escherichia; Escherichia coli MTGDNTLIHSHGINRRDFMKLCAALAATMGLSSKAAAEMAESVTNPQRPPVIWIGAQECT WP_000145410.1 MULTISPECIES: hydrogenase-2 small chain [Proteobacteria] Length: 372\nNumber of predicted TMHs: 1\nExp number of AAs in TMHs: 23.72226\nExp number, first 60 AAs: 0.22038\nTotal prob of N-in: 0.12994\noutside 1 331\nTMhelix 332 354\ninside 355 372

11178 GCF_000456285.1_Esch_coli_HVH_20_4-5865042_V1 Escherichia coli HVH 20 (4-5865042) Proteobacteria; Gammaproteobacteria; Enterobacterales; Enterobacteriaceae; Escherichia; Escherichia coli MTGDNTLIHSHGINRRDFMKLCAALAATMGLSSKAAAEMAESVTNPQRPPVIWIGAQECT WP_000145410.1 MULTISPECIES: hydrogenase-2 small chain [Proteobacteria] Length: 372\nNumber of predicted TMHs: 1\nExp number of AAs in TMHs: 23.72226\nExp number, first 60 AAs: 0.22038\nTotal prob of N-in: 0.12994\noutside 1 331\nTMhelix 332 354\ninside 355 372

11179 GCF_000456285.1_Esch_coli_HVH_20_4-5865042_V1 Escherichia coli HVH 20 (4-5865042) Proteobacteria; Gammaproteobacteria; Enterobacterales; Enterobacteriaceae; Escherichia; Escherichia coli MNNEETFYQAMRRQGVTRRSFLKYCSLAATSLGLGAGMAPKIAWALENKPRIPVVWIHGL WP_001058323.1 MULTISPECIES: hydrogenase-1 small chain [Proteobacteria] Length: 372\nNumber of predicted TMHs: 2\nExp number of AAs in TMHs: 37.2608499999999\nExp number, first 60 AAs: 19.52923\nTotal prob of N-in: 0.94351\nPOSSIBLE N-term signal sequence\ninside 1 19\nTMhelix 20 39\noutside 40 325\nTMhelix 326 348\ninside 349 372

11180 GCF_000459155.1_Esch_coli_HVH_205_4-3094677_V1 Escherichia coli HVH 205 (4-3094677) Proteobacteria; Gammaproteobacteria; Enterobacterales; Enterobacteriaceae; Escherichia; Escherichia coli MNNEETFYQAMRRQGVTRRSFLKYCSLAATSLGLGAGMAPKIAWALENKPRIPVVWIHGL WP_001295940.1 MULTISPECIES: hydrogenase [Enterobacteriaceae] Length: 372\nNumber of predicted TMHs: 2\nExp number of AAs in TMHs: 37.2608899999999\nExp number, first 60 AAs: 19.52923\nTotal prob of N-in: 0.94351\nPOSSIBLE N-term signal sequence\ninside 1 19\nTMhelix 20 39\noutside 40 325\nTMhelix 326 348\ninside 349 372

11181 GCF_000459155.1_Esch_coli_HVH_205_4-3094677_V1 Escherichia coli HVH 205 (4-3094677) Proteobacteria; Gammaproteobacteria; Enterobacterales; Enterobacteriaceae; Escherichia; Escherichia coli MTGDNTLIHSHGINRRDFMKLCAALAATMGLSSKAAAEMVESVTNPQRPPVIWIGAQECT WP_000145423.1 hydrogenase 2 small subunit [Escherichia coli] Length: 372\nNumber of predicted TMHs: 1\nExp number of AAs in TMHs: 23.78201\nExp number, first 60 AAs: 0.28355\nTotal prob of N-in: 0.13257\noutside 1 331\nTMhelix 332 354\ninside 355 372

11182 GCF_000459175.1_Esch_coli_HVH_206_4-3128229_V1 Escherichia coli HVH 206 (4-3128229) Proteobacteria; Gammaproteobacteria; Enterobacterales; Enterobacteriaceae; Escherichia; Escherichia coli MTGDNTLIHSHGINRRDFMKLCAALAATMGLSSKAAAEMAESVTNPQRPPVIWIGAQECT WP_000145410.1 MULTISPECIES: hydrogenase-2 small chain [Proteobacteria] Length: 372\nNumber of predicted TMHs: 1\nExp number of AAs in TMHs: 23.72226\nExp number, first 60 AAs: 0.22038\nTotal prob of N-in: 0.12994\noutside 1 331\nTMhelix 332 354\ninside 355 372

11183 GCF_000459175.1_Esch_coli_HVH_206_4-3128229_V1 Escherichia coli HVH 206 (4-3128229) Proteobacteria; Gammaproteobacteria; Enterobacterales; Enterobacteriaceae; Escherichia; Escherichia coli MNNEETFYQAMRRQGVTRRSFLKYCSLAATSLGLGAGMAPKIAWALENKPRIPVVWIHGL WP_001058323.1 MULTISPECIES: hydrogenase-1 small chain [Proteobacteria] Length: 372\nNumber of predicted TMHs: 2\nExp number of AAs in TMHs: 37.2608499999999\nExp number, first 60 AAs: 19.52923\nTotal prob of N-in: 0.94351\nPOSSIBLE N-term signal sequence\ninside 1 19\nTMhelix 20 39\noutside 40 325\nTMhelix 326 348\ninside 349 372

11184 GCF_000459195.1_Esch_coli_HVH_207_4-3113221_V1 Escherichia coli HVH 207 (4-3113221) Proteobacteria; Gammaproteobacteria; Enterobacterales; Enterobacteriaceae; Escherichia; Escherichia coli MTGDNTLIHSHGINRRDFMKLCAALAATMGLSSKAAAEMAESVTNPQRPPVIWIGAQECT WP_000145410.1 MULTISPECIES: hydrogenase-2 small chain [Proteobacteria] Length: 372\nNumber of predicted TMHs:

1\nExp number of AAs in TMHs: 23.72226\nExp number, first 60 AAs: 0.22038\nTotal prob of N-in: 0.12994\noutside 1 331\nTMhelix 332 354\ninside 355 372

11185 GCF_000459195.1_Esch_coli_HVH_207_4-3113221_V1 Escherichia coli HVH 207 (4-3113221) Proteobacteria; Gammaproteobacteria; Enterobacterales; Enterobacteriaceae; Escherichia; Escherichia coli MNNEETFYQAMRRQGVTRRSFLKYCSLAATSLGLGAGMAPKIAWALENKPRIPVVWIHGL WP_001058323.1 MULTISPECIES: hydrogenase-1 small chain [Proteobacteria] Length: 372\nNumber of predicted TMHs: 2\nExp number of AAs in TMHs: 37.2608499999999\nExp number, first 60 AAs: 19.52923\nTotal prob of N-in: 0.94351\nPOSSIBLE N-term signal sequence\ninside 1 19\nTMhelix 20 39\noutside 40 325\nTMhelix 326 348\ninside 349 372

11186 GCF_000459215.1_Esch_coli_HVH_208_4-3112292_V1 Escherichia coli HVH 208 (4-3112292) Proteobacteria; Gammaproteobacteria; Enterobacterales; Enterobacteriaceae; Escherichia; Escherichia coli MTGDNTLIHSHGINRRDFMKLCAALAATMGLSSKAAAEMAESVTNPQRPPVIWIGAQECT WP_000145410.1 MULTISPECIES: hydrogenase-2 small chain [Proteobacteria] Length: 372\nNumber of predicted TMHs: 1\nExp number of AAs in TMHs: 23.72226\nExp number, first 60 AAs: 0.22038\nTotal prob of N-in: 0.12994\noutside 1 331\nTMhelix 332 354\ninside 355 372

11187 GCF_000459215.1_Esch_coli_HVH_208_4-3112292_V1 Escherichia coli HVH 208 (4-3112292) Proteobacteria; Gammaproteobacteria; Enterobacterales; Enterobacteriaceae; Escherichia; Escherichia coli MNNEETFYQAMRRQGVTRRSFLKYCSLAATSLGLGAGMAPKIAWALENKPRIPVVWIHGL WP_001058323.1 MULTISPECIES: hydrogenase-1 small chain [Proteobacteria] Length: 372\nNumber of predicted TMHs: 2\nExp number of AAs in TMHs: 37.2608499999999\nExp number, first 60 AAs: 19.52923\nTotal prob of N-in: 0.94351\nPOSSIBLE N-term signal sequence\ninside 1 19\nTMhelix 20 39\noutside 40 325\nTMhelix 326 348\ninside 349 372

11188 GCF_000459235.1_Esch_coli_HVH_209_4-3062651_V1 Escherichia coli HVH 209 (4-3062651) Proteobacteria; Gammaproteobacteria; Enterobacterales; Enterobacteriaceae; Escherichia; Escherichia coli MTGDNTLIHSHGINRRDFMKLCAALAATMGLSSKAAAEMAESVTNPQRPPVIWIGAQECT WP_000145410.1 MULTISPECIES: hydrogenase-2 small chain [Proteobacteria] Length: 372\nNumber of predicted TMHs: 1\nExp number of AAs in TMHs: 23.72226\nExp number, first 60 AAs: 0.22038\nTotal prob of N-in: 0.12994\noutside 1 331\nTMhelix 332 354\ninside 355 372

11189 GCF_000459235.1_Esch_coli_HVH_209_4-3062651_V1 Escherichia coli HVH 209 (4-3062651) Proteobacteria; Gammaproteobacteria; Enterobacterales; Enterobacteriaceae; Escherichia; Escherichia coli MNNEETFYQAMRRQGVTRRSFLKYCSLAATSLGLGAGMAPKIAWALENKPRIPVVWIHGL WP_001058323.1 MULTISPECIES: hydrogenase-1 small chain [Proteobacteria] Length: 372\nNumber of predicted TMHs: 2\nExp number of AAs in TMHs: 37.2608499999999\nExp number, first 60 AAs: 19.52923\nTotal prob of N-in: 0.94351\nPOSSIBLE N-term signal sequence\ninside 1 19\nTMhelix 20 39\noutside 40 325\nTMhelix 326 348\ninside 349 372

11190 GCF_000459255.1_Esch_coli_HVH_210_4-3042480_V1 Escherichia coli HVH 210 (4-3042480) Proteobacteria; Gammaproteobacteria; Enterobacterales; Enterobacteriaceae; Escherichia; Escherichia coli MTGDNTLIHSHGINRRDFMKLCAALAATMGLSSKAAAEMAESVTNPQRPPVIWIGAQECT WP_000145410.1 MULTISPECIES: hydrogenase-2 small chain [Proteobacteria] Length: 372\nNumber of predicted TMHs: 1\nExp number of AAs in TMHs: 23.72226\nExp number, first 60 AAs: 0.22038\nTotal prob of N-in: 0.12994\noutside 1 331\nTMhelix 332 354\ninside 355 372

11191 GCF_000459275.1_Esch_coli_HVH_211_4-3041891_V1 Escherichia coli HVH 211 (4-3041891) Proteobacteria; Gammaproteobacteria; Enterobacterales; Enterobacteriaceae; Escherichia; Escherichia coli MTGDNTLIHSHGINRRDFMKLCAALAATMGLSSKAAAEMAESVTNPQRPPVIWIGAQECT WP_000145410.1 MULTISPECIES: hydrogenase-2 small chain [Proteobacteria] Length: 372\nNumber of predicted TMHs: 1\nExp number of AAs in TMHs: 23.72226\nExp number, first 60 AAs: 0.22038\nTotal prob of N-in: 0.12994\noutside 1 331\nTMhelix 332 354\ninside 355 372

11192 GCF_000459295.1_Esch_coli_HVH_212_3-9305343_V1 Escherichia coli HVH 212 (3-9305343) Proteobacteria; Gammaproteobacteria; Enterobacterales; Enterobacteriaceae; Escherichia; Escherichia coli MTGDNTLIHSHGINRRDFMKLCAALAATMGLSSKAAAEMAESVTNPQRPPVIWIGAQECT WP_000145410.1 MULTISPECIES: hydrogenase-2 small chain [Proteobacteria] Length: 372\nNumber of predicted TMHs: 1\nExp number of AAs in TMHs: 23.72226\nExp number, first 60 AAs: 0.22038\nTotal prob of N-in: 0.12994\noutside 1 331\nTMhelix 332 354\ninside 355 372

11193 GCF_000459295.1_Esch_coli_HVH_212_3-9305343_V1 Escherichia coli HVH 212 (3-9305343) Proteobacteria; Gammaproteobacteria; Enterobacterales; Enterobacteriaceae; Escherichia; Escherichia coli MNNEETFYQAMRRQGVTRRSFLKYCSLAATSLGLGAGMAPKIAWALENKPRIPVVWIHGL WP_001058323.1 MULTISPECIES: hydrogenase-1 small chain [Proteobacteria] Length: 372\nNumber of predicted TMHs: 2\nExp number of AAs in TMHs: 37.2608499999999\nExp number, first 60 AAs: 19.52923\nTotal prob of N-in: 0.94351\nPOSSIBLE N-term signal sequence\ninside 1 19\nTMhelix 20 39\noutside 40 325\nTMhelix 326 348\ninside 349 372

11194 GCF_000459315.1_Esch_coli_HVH_213_4-3042928_V1 Escherichia coli HVH 213 (4-3042928) Proteobacteria; Gammaproteobacteria; Enterobacterales; Enterobacteriaceae; Escherichia; Escherichia coli MNNEETFYQAMRRQGVTRRSFLKYCSLAATSLGLGAGMAPKIAWALENKPRIPVVWIHGL WP_001538532.1 hydrogenase-1 small chain [Escherichia coli] Length: 372\nNumber of predicted TMHs: 2\nExp number of AAs in TMHs: 37.27041\nExp number, first 60 AAs: 19.52927\nTotal prob of N-in: 0.94352\nPOSSIBLE N-term signal sequence\ninside 1 19\nTMhelix 20 39\noutside 40 325\nTMhelix 326 348\ninside 349 372

11195 GCF_000459315.1_Esch_coli_HVH_213_4-3042928_V1 Escherichia coli HVH 213 (4-3042928) Proteobacteria; Gammaproteobacteria; Enterobacterales; Enterobacteriaceae; Escherichia; Escherichia coli MTGDNTLIHSHGINRRDFMKLCAALAATMGLSSKAAAEMVESVTNPQRPPVIWIGAQECT WP_000145423.1 hydrogenase 2 small subunit [Escherichia coli] Length: 372\nNumber of predicted TMHs: 1\nExp number of AAs in TMHs: 23.78201\nExp number, first 60 AAs: 0.28355\nTotal prob of N-in: 0.13257\noutside 1 331\nTMhelix 332 354\ninside 355 372

11196 GCF_000507665.1_Esch_coli_HVH_214_4-3062198_V1 Escherichia coli HVH 214 (4-3062198) Proteobacteria; Gammaproteobacteria; Enterobacterales; Enterobacteriaceae; Escherichia; Escherichia coli MTGDNTLIHSHGINRRDFMKLCAALAATMGLSSKAAAEMAESVTNPQRPPVIWIGAQECT WP_000145410.1 MULTISPECIES: hydrogenase-2 small chain [Proteobacteria] Length: 372\nNumber of predicted TMHs: 1\nExp number of AAs in TMHs: 23.72226\nExp number, first 60 AAs: 0.22038\nTotal prob of N-in: 0.12994\noutside 1 331\nTMhelix 332 354\ninside 355 372

11197 GCF_000456305.1_Esch_coli_HVH_21_4-4517873_V1 Escherichia coli HVH 21 (4-4517873) Proteobacteria; Gammaproteobacteria; Enterobacterales; Enterobacteriaceae; Escherichia; Escherichia coli MNNEETFYQAMRRQGVTRRSFLKYCSLAATSLGLGAGMAPKIAWALENKPRIPVVWIHGL WP_001058323.1 MULTISPECIES: hydrogenase-1 small chain [Proteobacteria] Length: 372\nNumber of predicted TMHs: 2\nExp number of AAs in TMHs: 37.2608499999999\nExp number, first 60 AAs: 19.52923\nTotal prob of N-in: 0.94351\nPOSSIBLE N-term signal sequence\ninside 1 19\nTMhelix 20 39\noutside 40 325\nTMhelix 326 348\ninside 349 372

11198 GCF_000456305.1_Esch_coli_HVH_21_4-4517873_V1 Escherichia coli HVH 21 (4-4517873) Proteobacteria; Gammaproteobacteria; Enterobacterales; Enterobacteriaceae; Escherichia; Escherichia coli MTGDNTLIHSHGINRRDFMKLCAALAATMGLSSKAAAEMAESVTNPQRPPVIWIGAQECT WP_000145410.1 MULTISPECIES: hydrogenase-2 small chain [Proteobacteria] Length: 372\nNumber of predicted TMHs: 1\nExp number of AAs in TMHs: 23.72226\nExp number, first 60 AAs: 0.22038\nTotal prob of N-in: 0.12994\noutside 1 331\nTMhelix 332 354\ninside 355 372

11199 GCF_000459335.1_Esch_coli_HVH_215_4-3008371_V1 Escherichia coli HVH 215 (4-3008371) Proteobacteria; Gammaproteobacteria; Enterobacterales; Enterobacteriaceae; Escherichia; Escherichia coli MNNEETFYQAMRRQGVTRRSFLKYCSLAATSLGLGAGMAPKIAWALENKPRIPVVWIHGL WP_001058323.1 MULTISPECIES: hydrogenase-1 small chain [Proteobacteria] Length: 372\nNumber of predicted TMHs: 2\nExp number of AAs in TMHs: 37.2608499999999\nExp number, first 60 AAs: 19.52923\nTotal prob of N-in: 0.94351\nPOSSIBLE N-term signal sequence\ninside 1 19\nTMhelix 20 39\noutside 40 325\nTMhelix 326 348\ninside 349 372

11200 GCF_000459335.1_Esch_coli_HVH_215_4-3008371_V1 Escherichia coli HVH 215 (4-3008371) Proteobacteria; Gammaproteobacteria; Enterobacterales; Enterobacteriaceae; Escherichia; Escherichia coli MTGDNTLIHSHGINRRDFMKLCAALAATMGLSSKAAAEMAESVTNPQRPPVIWIGAQECT WP_000145410.1 MULTISPECIES: hydrogenase-2 small chain [Proteobacteria] Length: 372\nNumber of predicted TMHs: 1\nExp number of AAs in TMHs: 23.72226\nExp number, first 60 AAs: 0.22038\nTotal prob of N-in: 0.12994\noutside 1 331\nTMhelix 332 354\ninside 355 372

11201 GCF_000459355.1_Esch_coli_HVH_216_4-3042952_V1 Escherichia coli HVH 216 (4-3042952) Proteobacteria; Gammaproteobacteria; Enterobacterales; Enterobacteriaceae; Escherichia; Escherichia coli

MTGDNTLIHSHGINRRDFMKLCAALAATMGLSSKAAAEMAESVTNPQRPPVIWIGAQECT WP_000145410.1 MULTISPECIES: hydrogenase-2 small chain [Proteobacteria] Length: 372\nNumber of predicted TMHs: 1\nExp number of AAs in TMHs: 23.72226\nExp number, first 60 AAs: 0.22038\nTotal prob of N-in: 0.12994\noutside 1 331\nTMhelix 332 354\ninside 355 372

11202 GCF_000459355.1_Esch_coli_HVH_216_4-3042952_V1 Escherichia coli HVH 216 (4-3042952) Proteobacteria; Gammaproteobacteria; Enterobacterales; Enterobacteriaceae; Escherichia; Escherichia coli MNNEETFYQAMRRQGVTRRSFLKYCSLAATSLGLGAGMAPKIAWALENKPRIPVVWIHGL WP_021545218.1 hydrogenase-1 small chain [Escherichia coli] Length: 372\nNumber of predicted TMHs: 2\nExp number of AAs in TMHs: 37.2587199999999\nExp number, first 60 AAs: 19.52855\nTotal prob of N-in: 0.94351\nPOSSIBLE N-term signal sequence\ninside 1 19\nTMhelix 20 39\noutside 40 325\nTMhelix 326 348\ninside 349 372

11203 GCF_000459375.1_Esch_coli_HVH_217_4-1022806_V1 Escherichia coli HVH 217 (4-1022806) Proteobacteria; Gammaproteobacteria; Enterobacterales; Enterobacteriaceae; Escherichia; Escherichia coli MTGDNTLIHSHGINRRDFMKLCAALAATMGLSSKAAAEMAESVTNPQRPPVIWIGAQECT WP_000145410.1 MULTISPECIES: hydrogenase-2 small chain [Proteobacteria] Length: 372\nNumber of predicted TMHs: 1\nExp number of AAs in TMHs: 23.72226\nExp number, first 60 AAs: 0.22038\nTotal prob of N-in: 0.12994\noutside 1 331\nTMhelix 332 354\ninside 355 372

11204 GCF_000459395.1_Esch_coli_HVH_218_4-4500903_V1 Escherichia coli HVH 218 (4-4500903) Proteobacteria; Gammaproteobacteria; Enterobacterales; Enterobacteriaceae; Escherichia; Escherichia coli MTGDNTLIHSHGINRRDFMKLCAALAATMGLSSKAAAEMAESVTNPQRPPVIWIGAQECT WP_000145410.1 MULTISPECIES: hydrogenase-2 small chain [Proteobacteria] Length: 372\nNumber of predicted TMHs: 1\nExp number of AAs in TMHs: 23.72226\nExp number, first 60 AAs: 0.22038\nTotal prob of N-in: 0.12994\noutside 1 331\nTMhelix 332 354\ninside 355 372

11205 GCF_000459395.1_Esch_coli_HVH_218_4-4500903_V1 Escherichia coli HVH 218 (4-4500903) Proteobacteria; Gammaproteobacteria; Enterobacterales; Enterobacteriaceae; Escherichia; Escherichia coli MNNEETFYQAMRRQGVTRRSFLKYCSLAATSLGLGAGMAPKIAWALENKPRIPVVWIHGL WP_001058323.1 MULTISPECIES: hydrogenase-1 small chain [Proteobacteria] Length: 372\nNumber of predicted TMHs: 2\nExp number of AAs in TMHs: 37.2608499999999\nExp number, first 60 AAs: 19.52923\nTotal prob of N-in: 0.94351\nPOSSIBLE N-term signal sequence\ninside 1 19\nTMhelix 20 39\noutside 40 325\nTMhelix 326 348\ninside 349 372

11206 GCF_000459415.1_Esch_coli_HVH_220_4-5876842_V1 Escherichia coli HVH 220 (4-5876842) Proteobacteria; Gammaproteobacteria; Enterobacterales; Enterobacteriaceae; Escherichia; Escherichia coli MTGDNTLIHSHGINRRDFMKLCAALAATMGLSSKAAAEMAESVTNPQRPPVIWIGAQECT WP_000145410.1 MULTISPECIES: hydrogenase-2 small chain [Proteobacteria] Length: 372\nNumber of predicted TMHs: 1\nExp number of AAs in TMHs: 23.72226\nExp number, first 60 AAs: 0.22038\nTotal prob of N-in: 0.12994\noutside 1 331\nTMhelix 332 354\ninside 355 372

11207 GCF_000459415.1_Esch_coli_HVH_220_4-5876842_V1 Escherichia coli HVH 220 (4-5876842) Proteobacteria; Gammaproteobacteria; Enterobacterales; Enterobacteriaceae; Escherichia; Escherichia coli MNNEETFYQAMRRQGVTRRSFLKYCSLAATSLGLGAGMAPKIAWALENKPRIPVVWIHGL WP_001058323.1 MULTISPECIES: hydrogenase-1 small chain [Proteobacteria] Length: 372\nNumber of predicted TMHs: 2\nExp number of AAs in TMHs: 37.2608499999999\nExp number, first 60 AAs: 19.52923\nTotal prob of N-in: 0.94351\nPOSSIBLE N-term signal sequence\ninside 1 19\nTMhelix 20 39\noutside 40 325\nTMhelix 326 348\ninside 349 372

11208 GCF_000459435.1_Esch_coli_HVH_221_4-3136817_V1 Escherichia coli HVH 221 (4-3136817) Proteobacteria; Gammaproteobacteria; Enterobacterales; Enterobacteriaceae; Escherichia; Escherichia coli MTGDNTLIHSHGINRRDFMKLCAALAATMGLSSKAAAEMAESVTNPQRPPVIWIGAQECT WP_000145410.1 MULTISPECIES: hydrogenase-2 small chain [Proteobacteria] Length: 372\nNumber of predicted TMHs: 1\nExp number of AAs in TMHs: 23.72226\nExp number, first 60 AAs: 0.22038\nTotal prob of N-in: 0.12994\noutside 1 331\nTMhelix 332 354\ninside 355 372

11209 GCF_000459435.1_Esch_coli_HVH_221_4-3136817_V1 Escherichia coli HVH 221 (4-3136817) Proteobacteria; Gammaproteobacteria; Enterobacterales; Enterobacteriaceae; Escherichia; Escherichia coli MNNEETFYQAMRRQGVTRRSFLKYCSLAATSLGLGAGMAPKIAWALENKPRIPVVWIHGL WP_001058323.1 MULTISPECIES: hydrogenase-1 small chain [Proteobacteria] Length: 372\nNumber of predicted TMHs: 2\nExp number of AAs in TMHs: 37.2608499999999\nExp number, first 60 AAs: 19.52923\nTotal prob of N-in:

0.94351\nPOSSIBLE N-term signal sequence\ninside 1 19\nTMhelix 20 39\noutside 40 325\nTMhelix 326 348\ninside 349 372

11210 GCF_000459455.1_Esch_coli_HVH_222_4-2977443_V1 Escherichia coli HVH 222 (4-2977443) Proteobacteria; Gammaproteobacteria; Enterobacterales; Enterobacteriaceae; Escherichia; Escherichia coli MTGDNTLIHSHGINRRDFMKLCAALAATMGLSSKAAAEMAESVTNPQRPPVIWIGAQECT WP_000145410.1 MULTISPECIES: hydrogenase-2 small chain [Proteobacteria] Length: 372\nNumber of predicted TMHs: 1\nExp number of AAs in TMHs: 23.72226\nExp number, first 60 AAs: 0.22038\nTotal prob of N-in: 0.12994\noutside 1 331\nTMhelix 332 354\ninside 355 372

11211 GCF_000459475.1_Esch_coli_HVH_223_4-2976528_V1 Escherichia coli HVH 223 (4-2976528) Proteobacteria; Gammaproteobacteria; Enterobacterales; Enterobacteriaceae; Escherichia; Escherichia coli MNNEETFYQAMRRQGVTRRSFLKYCSLAATSLGLGAGMAPKIAWALENKPRIPVVWIHGL WP_001058323.1 MULTISPECIES: hydrogenase-1 small chain [Proteobacteria] Length: 372\nNumber of predicted TMHs: 2\nExp number of AAs in TMHs: 37.2608499999999\nExp number, first 60 AAs: 19.52923\nTotal prob of N-in: 0.94351\nPOSSIBLE N-term signal sequence\ninside 1 19\nTMhelix 20 39\noutside 40 325\nTMhelix 326 348\ninside 349 372

11212 GCF_000459475.1_Esch_coli_HVH_223_4-2976528_V1 Escherichia coli HVH 223 (4-2976528) Proteobacteria; Gammaproteobacteria; Enterobacterales; Enterobacteriaceae; Escherichia; Escherichia coli MTGDNTLIHSHGINRRDFMKLCAALAATMGLSSKAAAEMAESVTNPQRPPVIWIGAQECT WP_000145410.1 MULTISPECIES: hydrogenase-2 small chain [Proteobacteria] Length: 372\nNumber of predicted TMHs: 1\nExp number of AAs in TMHs: 23.72226\nExp number, first 60 AAs: 0.22038\nTotal prob of N-in: 0.12994\noutside 1 331\nTMhelix 332 354\ninside 355 372

11213 GCF_000456325.1_Esch_coli_HVH_22_4-2258986_V1 Escherichia coli HVH 22 (4-2258986) Proteobacteria; Gammaproteobacteria; Enterobacterales; Enterobacteriaceae; Escherichia; Escherichia coli MTGDNTLIHSHGINRRDFMKLCAALAATMGLSSKAAAEMAESVTNPQRPPVIWIGAQECT WP_000145410.1 MULTISPECIES: hydrogenase-2 small chain [Proteobacteria] Length: 372\nNumber of predicted TMHs: 1\nExp number of AAs in TMHs: 23.72226\nExp number, first 60 AAs: 0.22038\nTotal prob of N-in: 0.12994\noutside 1 331\nTMhelix 332 354\ninside 355 372

11214 GCF_000456325.1_Esch_coli_HVH_22_4-2258986_V1 Escherichia coli HVH 22 (4-2258986) Proteobacteria; Gammaproteobacteria; Enterobacterales; Enterobacteriaceae; Escherichia; Escherichia coli MNNEETFYQAMRRQGVTRRSFLKYCSLAATSLGLGAGMAPKIAWALENKPRIPVVWIHGL WP_001058323.1 MULTISPECIES: hydrogenase-1 small chain [Proteobacteria] Length: 372\nNumber of predicted TMHs: 2\nExp number of AAs in TMHs: 37.2608499999999\nExp number, first 60 AAs: 19.52923\nTotal prob of N-in: 0.94351\nPOSSIBLE N-term signal sequence\ninside 1 19\nTMhelix 20 39\noutside 40 325\nTMhelix 326 348\ninside 349 372

11215 GCF_000459495.1_Esch_coli_HVH_225_4-1273116_V1 Escherichia coli HVH 225 (4-1273116) Proteobacteria; Gammaproteobacteria; Enterobacterales; Enterobacteriaceae; Escherichia; Escherichia coli MTGDNTLIHSHGINRRDFMKLCAALAATMGLSSKAAAEMVESVTNPQRPPVIWIGAQECT WP_000145423.1 hydrogenase 2 small subunit [Escherichia coli] Length: 372\nNumber of predicted TMHs: 1\nExp number of AAs in TMHs: 23.78201\nExp number, first 60 AAs: 0.28355\nTotal prob of N-in: 0.13257\noutside 1 331\nTMhelix 332 354\ninside 355 372

11216 GCF_000459495.1_Esch_coli_HVH_225_4-1273116_V1 Escherichia coli HVH 225 (4-1273116) Proteobacteria; Gammaproteobacteria; Enterobacterales; Enterobacteriaceae; Escherichia; Escherichia coli MNNEETFYQAMRRQGVTRRSFLKYCSLAATSLGLGAGMAPKIAWALENKPRIPVVWIHGL WP_001538532.1 hydrogenase-1 small chain [Escherichia coli] Length: 372\nNumber of predicted TMHs: 2\nExp number of AAs in TMHs: 37.27041\nExp number, first 60 AAs: 19.52927\nTotal prob of N-in: 0.94352\nPOSSIBLE N-term signal sequence\ninside 1 19\nTMhelix 20 39\noutside 40 325\nTMhelix 326 348\ninside 349 372

11217 GCF_000459515.1_Esch_coli_HVH_227_4-2277670_V1 Escherichia coli HVH 227 (4-2277670) Proteobacteria; Gammaproteobacteria; Enterobacterales; Enterobacteriaceae; Escherichia; Escherichia coli MNNEETFYQAMRRQGVTRRSFLKYCSLAATSLGLGAGMAPKIAWALENKPRIPVVWIHGL WP_001058323.1 MULTISPECIES: hydrogenase-1 small chain [Proteobacteria] Length: 372\nNumber of predicted TMHs: 2\nExp number of AAs in TMHs: 37.2608499999999\nExp number, first 60 AAs: 19.52923\nTotal prob of N-in: 0.94351\nPOSSIBLE N-term signal sequence\ninside 1 19\nTMhelix 20 39\noutside 40 325\nTMhelix 326 348\ninside 349 372

11218 GCF_000459515.1_Esch_coli_HVH_227_4-2277670_V1 Escherichia coli HVH 227 (4-2277670) Proteobacteria; Gammaproteobacteria; Enterobacterales; Enterobacteriaceae; Escherichia; Escherichia coli MTGDNTLIHSHGINRRDFMKLCAALAATMGLSSKAAAEMAESVTNPQRPPVIWIGAQECT WP_000145410.1 MULTISPECIES: hydrogenase-2 small chain [Proteobacteria] Length: 372\nNumber of predicted TMHs: 1\nExp number of AAs in TMHs: 23.72226\nExp number, first 60 AAs: 0.22038\nTotal prob of N-in: 0.12994\noutside 1 331\nTMhelix 332 354\ninside 355 372

11219 GCF_000459535.1_Esch_coli_HVH_228_4-7787030_V1 Escherichia coli HVH 228 (4-7787030) Proteobacteria; Gammaproteobacteria; Enterobacterales; Enterobacteriaceae; Escherichia; Escherichia coli MTGDNTLIHSHGINRRDFMKLCAALAATMGLSSKAAAEMAESVTNPQRPPVIWIGAQECT WP_000145410.1 MULTISPECIES: hydrogenase-2 small chain [Proteobacteria] Length: 372\nNumber of predicted TMHs: 1\nExp number of AAs in TMHs: 23.72226\nExp number, first 60 AAs: 0.22038\nTotal prob of N-in: 0.12994\noutside 1 331\nTMhelix 332 354\ninside 355 372

11220 GCF_000459535.1_Esch_coli_HVH_228_4-7787030_V1 Escherichia coli HVH 228 (4-7787030) Proteobacteria; Gammaproteobacteria; Enterobacterales; Enterobacteriaceae; Escherichia; Escherichia coli MNNEETFYQAMRRQGVTRRSFLKYCSLAATSLGLGAGMAPKIAWALENKPRIPVVWIHGL WP_001058323.1 MULTISPECIES: hydrogenase-1 small chain [Proteobacteria] Length: 372\nNumber of predicted TMHs: 2\nExp number of AAs in TMHs: 37.2608499999999\nExp number, first 60 AAs: 19.52923\nTotal prob of N-in: 0.94351\nPOSSIBLE N-term signal sequence\ninside 1 19\nTMhelix 20 39\noutside 40 325\nTMhelix 326 348\ninside 349 372

11221 GCF_000507605.1_Esch_coli_HVH_23_4-6066488_V1 Escherichia coli HVH 23 (4-6066488) Proteobacteria; Gammaproteobacteria; Enterobacterales; Enterobacteriaceae; Escherichia; Escherichia coli MNNEETFYQAMRRQGVTRRSFLKYCSLAATSLGLGAGMAPKIAWALENKPRIPVVWIHGL WP_001058323.1 MULTISPECIES: hydrogenase-1 small chain [Proteobacteria] Length: 372\nNumber of predicted TMHs: 2\nExp number of AAs in TMHs: 37.2608499999999\nExp number, first 60 AAs: 19.52923\nTotal prob of N-in: 0.94351\nPOSSIBLE N-term signal sequence\ninside 1 19\nTMhelix 20 39\noutside 40 325\nTMhelix 326 348\ninside 349 372

11222 GCF_000507605.1_Esch_coli_HVH_23_4-6066488_V1 Escherichia coli HVH 23 (4-6066488) Proteobacteria; Gammaproteobacteria; Enterobacterales; Enterobacteriaceae; Escherichia; Escherichia coli MTGDNTLIHSHGINRRDFMKLCAALAATMGLSSKAAAEMAESVTNPQRPPVIWIGAQECT WP_000145410.1 MULTISPECIES: hydrogenase-2 small chain [Proteobacteria] Length: 372\nNumber of predicted TMHs: 1\nExp number of AAs in TMHs: 23.72226\nExp number, first 60 AAs: 0.22038\nTotal prob of N-in: 0.12994\noutside 1 331\nTMhelix 332 354\ninside 355 372

11223 GCF_000456345.1_Esch_coli_HVH_24_4-5985145_V1 Escherichia coli HVH 24 (4-5985145) Proteobacteria; Gammaproteobacteria; Enterobacterales; Enterobacteriaceae; Escherichia; Escherichia coli MNNEETFYQAMRRQGVTRRSFLKYCSLAATSLGLGAGMAPKIAWALENKPRIPVVWIHGL WP_001058323.1 MULTISPECIES: hydrogenase-1 small chain [Proteobacteria] Length: 372\nNumber of predicted TMHs: 2\nExp number of AAs in TMHs: 37.2608499999999\nExp number, first 60 AAs: 19.52923\nTotal prob of N-in: 0.94351\nPOSSIBLE N-term signal sequence\ninside 1 19\nTMhelix 20 39\noutside 40 325\nTMhelix 326 348\ninside 349 372

11224 GCF_000456345.1_Esch_coli_HVH_24_4-5985145_V1 Escherichia coli HVH 24 (4-5985145) Proteobacteria; Gammaproteobacteria; Enterobacterales; Enterobacteriaceae; Escherichia; Escherichia coli MTGDNTLIHSHGINRRDFMKLCAALAATMGLSSKAAAEMVESVTNPRRPPVIWIGAQECT WP_021516469.1 hydrogenase-2 small chain [Escherichia coli] Length: 372\nNumber of predicted TMHs: 1\nExp number of AAs in TMHs: 23.91885\nExp number, first 60 AAs: 0.43255\nTotal prob of N-in: 0.13883\noutside 1 331\nTMhelix 332 354\ninside 355 372

11225 GCF_000456025.1_Esch_coli_HVH_2_4-6943160_V1 Escherichia coli HVH 2 (4-6943160) Proteobacteria; Gammaproteobacteria; Enterobacterales; Enterobacteriaceae; Escherichia; Escherichia coli MNNEETFYQAMRRQGVTRRSFLKYCSLAATSLGLGAGMAPKIAWALENKPRIPVVWIHGL WP_001058323.1 MULTISPECIES: hydrogenase-1 small chain [Proteobacteria] Length: 372\nNumber of predicted TMHs: 2\nExp number of AAs in TMHs: 37.2608499999999\nExp number, first 60 AAs: 19.52923\nTotal prob of N-in: 0.94351\nPOSSIBLE N-term signal sequence\ninside 1 19\nTMhelix 20 39\noutside 40 325\nTMhelix 326 348\ninside 349 372

11226 GCF_000456025.1_Esch_coli_HVH_2_4-6943160_V1 Escherichia coli HVH 2 (4-6943160) Proteobacteria; Gammaproteobacteria; Enterobacterales; Enterobacteriaceae; Escherichia; Escherichia coli MTGDNTLIHSHGINRRDFMKLCAALAATMGLSSKAAAEMAESVTNPQRPPVIWIGAQECT WP_000145410.1 MULTISPECIES: hydrogenase-2 small chain [Proteobacteria] Length: 372\nNumber of predicted TMHs: 1\nExp number of AAs in TMHs: 23.72226\nExp number, first 60 AAs: 0.22038\nTotal prob of N-in: 0.12994\noutside 1 331\nTMhelix 332 354\ninside 355 372

11227 GCF_000456365.1_Esch_coli_HVH_25_4-5851939_V1 Escherichia coli HVH 25 (4-5851939) Proteobacteria; Gammaproteobacteria; Enterobacterales; Enterobacteriaceae; Escherichia; Escherichia coli MNNEETFYQAMRRQGVTRRSFLKYCSLAATSLGLGAGMAPKIAWALENKPRIPVVWIHGL WP_001058323.1 MULTISPECIES: hydrogenase-1 small chain [Proteobacteria] Length: 372\nNumber of predicted TMHs: 2\nExp number of AAs in TMHs: 37.2608499999999\nExp number, first 60 AAs: 19.52923\nTotal prob of N-in: 0.94351\nPOSSIBLE N-term signal sequence\ninside 1 19\nTMhelix 20 39\noutside 40 325\nTMhelix 326 348\ninside 349 372

11228 GCF_000456365.1_Esch_coli_HVH_25_4-5851939_V1 Escherichia coli HVH 25 (4-5851939) Proteobacteria; Gammaproteobacteria; Enterobacterales; Enterobacteriaceae; Escherichia; Escherichia coli MTGDNTLIHSHGINRRDFMKLCAALAATMGLSSKAAAEMAESVTNPQRPPVIWIGAQECT WP_000145410.1 MULTISPECIES: hydrogenase-2 small chain [Proteobacteria] Length: 372\nNumber of predicted TMHs: 1\nExp number of AAs in TMHs: 23.72226\nExp number, first 60 AAs: 0.22038\nTotal prob of N-in: 0.12994\noutside 1 331\nTMhelix 332 354\ninside 355 372

11229 GCF_000456385.1_Esch_coli_HVH_26_4-5703913_V1 Escherichia coli HVH 26 (4-5703913) Proteobacteria; Gammaproteobacteria; Enterobacterales; Enterobacteriaceae; Escherichia; Escherichia coli MTGDNTLIHSHGINRRDFMKLCAALAATMGLSSKAAAEMVESVTNPQRPPVIWIGAQECT WP_000145423.1 hydrogenase 2 small subunit [Escherichia coli] Length: 372\nNumber of predicted TMHs: 1\nExp number of AAs in TMHs: 23.78201\nExp number, first 60 AAs: 0.28355\nTotal prob of N-in: 0.13257\noutside 1 331\nTMhelix 332 354\ninside 355 372

11230 GCF_000456385.1_Esch_coli_HVH_26_4-5703913_V1 Escherichia coli HVH 26 (4-5703913) Proteobacteria; Gammaproteobacteria; Enterobacterales; Enterobacteriaceae; Escherichia; Escherichia coli MNNEETFYQAMRRQGVTRRSFLKYCSLAATSLGLGAGMAPKIAWALENKPRIPVVWIHGL WP_001538532.1 hydrogenase-1 small chain [Escherichia coli] Length: 372\nNumber of predicted TMHs: 2\nExp number of AAs in TMHs: 37.27041\nExp number, first 60 AAs: 19.52927\nTotal prob of N-in: 0.94352\nPOSSIBLE N-term signal sequence\ninside 1 19\nTMhelix 20 39\noutside 40 325\nTMhelix 326 348\ninside 349 372

11231 GCF_000456405.1_Esch_coli_HVH_27_4-7449267_V1 Escherichia coli HVH 27 (4-7449267) Proteobacteria; Gammaproteobacteria; Enterobacterales; Enterobacteriaceae; Escherichia; Escherichia coli MNNEETFYQAMRRQGVTRRSFLKYCSLAATSLGLGAGMAPKIAWALENKPRIPVVWIHGL WP_001058323.1 MULTISPECIES: hydrogenase-1 small chain [Proteobacteria] Length: 372\nNumber of predicted TMHs: 2\nExp number of AAs in TMHs: 37.2608499999999\nExp number, first 60 AAs: 19.52923\nTotal prob of N-in: 0.94351\nPOSSIBLE N-term signal sequence\ninside 1 19\nTMhelix 20 39\noutside 40 325\nTMhelix 326 348\ninside 349 372

11232 GCF_000456405.1_Esch_coli_HVH_27_4-7449267_V1 Escherichia coli HVH 27 (4-7449267) Proteobacteria; Gammaproteobacteria; Enterobacterales; Enterobacteriaceae; Escherichia; Escherichia coli MTGDNTLIHSHGINRRDFMKLCAALAATMGLSSKAAAEMAESVTNPQRPPVIWIGAQECT WP_000145410.1 MULTISPECIES: hydrogenase-2 small chain [Proteobacteria] Length: 372\nNumber of predicted TMHs: 1\nExp number of AAs in TMHs: 23.72226\nExp number, first 60 AAs: 0.22038\nTotal prob of N-in: 0.12994\noutside 1 331\nTMhelix 332 354\ninside 355 372

11233 GCF_000456425.1_Esch_coli_HVH_28_4-0907367_V1 Escherichia coli HVH 28 (4-0907367) Proteobacteria; Gammaproteobacteria; Enterobacterales; Enterobacteriaceae; Escherichia; Escherichia coli MNNEETFYQAMRRQGVTRRSFLKYCSLAATSLGLGAGMAPKIAWALENKPRIPVVWIHGL WP_001058323.1 MULTISPECIES: hydrogenase-1 small chain [Proteobacteria] Length: 372\nNumber of predicted TMHs: 2\nExp number of AAs in TMHs: 37.2608499999999\nExp number, first 60 AAs: 19.52923\nTotal prob of N-in: 0.94351\nPOSSIBLE N-term signal sequence\ninside 1 19\nTMhelix 20 39\noutside 40 325\nTMhelix 326 348\ninside 349 372

11234 GCF_000456425.1_Esch_coli_HVH_28_4-0907367_V1 Escherichia coli HVH 28 (4-0907367) Proteobacteria; Gammaproteobacteria; Enterobacterales; Enterobacteriaceae; Escherichia; Escherichia coli

MTGDNTLIHSHGINRRDFMKLCAALAATMGLSSKAAAEMAESVTNPQRPPVIWIGAQECT WP_000145410.1 MULTISPECIES: hydrogenase-2 small chain [Proteobacteria] Length: 372\nNumber of predicted TMHs: 1\nExp number of AAs in TMHs: 23.72226\nExp number, first 60 AAs: 0.22038\nTotal prob of N-in: 0.12994\noutside 1 331\nTMhelix 332 354\ninside 355 372

11235 GCF_000456445.1_Esch_coli_HVH_29_4-3418073_V1 Escherichia coli HVH 29 (4-3418073) Proteobacteria; Gammaproteobacteria; Enterobacterales; Enterobacteriaceae; Escherichia; Escherichia coli MNNEETFYQAMRRQGVTRRSFLKYCSLAATSLGLGAGMAPKIAWALENKPRIPVVWIHGL WP_001058323.1 MULTISPECIES: hydrogenase-1 small chain [Proteobacteria] Length: 372\nNumber of predicted TMHs: 2\nExp number of AAs in TMHs: 37.2608499999999\nExp number, first 60 AAs: 19.52923\nTotal prob of N-in: 0.94351\nPOSSIBLE N-term signal sequence\ninside 1 19\nTMhelix 20 39\noutside 40 325\nTMhelix 326 348\ninside 349 372

11236 GCF_000456445.1_Esch_coli_HVH_29_4-3418073_V1 Escherichia coli HVH 29 (4-3418073) Proteobacteria; Gammaproteobacteria; Enterobacterales; Enterobacteriaceae; Escherichia; Escherichia coli MTGDNTLIHSHGINRRDFMKLCAALAATMGLSSKAAAEMAESVTNPQRPPVIWIGAQECT WP_000145410.1 MULTISPECIES: hydrogenase-2 small chain [Proteobacteria] Length: 372\nNumber of predicted TMHs: 1\nExp number of AAs in TMHs: 23.72226\nExp number, first 60 AAs: 0.22038\nTotal prob of N-in: 0.12994\noutside 1 331\nTMhelix 332 354\ninside 355 372

11237 GCF_000456465.1_Esch_coli_HVH_30_4-2661829_V1 Escherichia coli HVH 30 (4-2661829) Proteobacteria; Gammaproteobacteria; Enterobacterales; Enterobacteriaceae; Escherichia; Escherichia coli MTGDNTLIHSHGINRRDFMKLCAALAATMGLSSKAAAEMAESVTNPQRPPVIWIGAQECT WP_000145410.1 MULTISPECIES: hydrogenase-2 small chain [Proteobacteria] Length: 372\nNumber of predicted TMHs: 1\nExp number of AAs in TMHs: 23.72226\nExp number, first 60 AAs: 0.22038\nTotal prob of N-in: 0.12994\noutside 1 331\nTMhelix 332 354\ninside 355 372

11238 GCF_000456485.1_Esch_coli_HVH_31_4-2602156_V1 Escherichia coli HVH 31 (4-2602156) Proteobacteria; Gammaproteobacteria; Enterobacterales; Enterobacteriaceae; Escherichia; Escherichia coli MNNEETFYQAMRRQGVTRRSFLKYCSLAATSLGLGAGMAPKIAWALENKPRIPVVWIHGL WP_001058323.1 MULTISPECIES: hydrogenase-1 small chain [Proteobacteria] Length: 372\nNumber of predicted TMHs: 2\nExp number of AAs in TMHs: 37.2608499999999\nExp number, first 60 AAs: 19.52923\nTotal prob of N-in: 0.94351\nPOSSIBLE N-term signal sequence\ninside 1 19\nTMhelix 20 39\noutside 40 325\nTMhelix 326 348\ninside 349 372

11239 GCF_000456485.1_Esch_coli_HVH_31_4-2602156_V1 Escherichia coli HVH 31 (4-2602156) Proteobacteria; Gammaproteobacteria; Enterobacterales; Enterobacteriaceae; Escherichia; Escherichia coli MTGDNTLIHSHGINRRDFMKLCAALAATMGLSSKAAAEMAESVTNPQRPPVIWIGAQECT WP_000145410.1 MULTISPECIES: hydrogenase-2 small chain [Proteobacteria] Length: 372\nNumber of predicted TMHs: 1\nExp number of AAs in TMHs: 23.72226\nExp number, first 60 AAs: 0.22038\nTotal prob of N-in: 0.12994\noutside 1 331\nTMhelix 332 354\ninside 355 372

11240 GCF_000456505.1_Esch_coli_HVH_32_4-3773988_V1 Escherichia coli HVH 32 (4-3773988) Proteobacteria; Gammaproteobacteria; Enterobacterales; Enterobacteriaceae; Escherichia; Escherichia coli MTGDNTLIHSHGINRRDFMKLCAALAATMGLSSKAAAEMAESVTNPQRPPVIWIGAQECT WP_000145410.1 MULTISPECIES: hydrogenase-2 small chain [Proteobacteria] Length: 372\nNumber of predicted TMHs: 1\nExp number of AAs in TMHs: 23.72226\nExp number, first 60 AAs: 0.22038\nTotal prob of N-in: 0.12994\noutside 1 331\nTMhelix 332 354\ninside 355 372

11241 GCF_000456525.1_Esch_coli_HVH_33_4-2174936_V1 Escherichia coli HVH 33 (4-2174936) Proteobacteria; Gammaproteobacteria; Enterobacterales; Enterobacteriaceae; Escherichia; Escherichia coli MNNEETFYQAMRRQGVTRRSFLKYCSLAATSLGLGAGMAPKIAWALENKPRIPVVWIHGL WP_001058320.1 hydrogenase [Escherichia coli] Length: 372\nNumber of predicted TMHs: 2\nExp number of AAs in TMHs: 37.26082\nExp number, first 60 AAs: 19.52923\nTotal prob of N-in: 0.94351\nPOSSIBLE N-term signal sequence\ninside 1 19\nTMhelix 20 39\noutside 40 325\nTMhelix 326 348\ninside 349 372

11242 GCF_000456525.1_Esch_coli_HVH_33_4-2174936_V1 Escherichia coli HVH 33 (4-2174936) Proteobacteria; Gammaproteobacteria; Enterobacterales; Enterobacteriaceae; Escherichia; Escherichia coli MTGDNTLIHSHGINRRDFMKLCAALAATMGLSSKAAAEMAESVTNPQRPPVIWIGAQECT WP_000145410.1 MULTISPECIES: hydrogenase-2 small chain [Proteobacteria] Length: 372\nNumber of predicted TMHs:

1\nExp number of AAs in TMHs: 23.72226\nExp number, first 60 AAs: 0.22038\nTotal prob of N-in: 0.12994\noutside 1 331\nTMhelix 332 354\ninside 355 372

11243 GCF_000456045.1_Esch_coli_HVH_3_4-7276001_V1 Escherichia coli HVH 3 (4-7276001) Proteobacteria; Gammaproteobacteria; Enterobacterales; Enterobacteriaceae; Escherichia; Escherichia coli MTGDNTLIHSHGINRRDFMKLCAALAATMGLSSKAAAEMAESVTNPQRPPVIWIGAQECT WP_000145410.1 MULTISPECIES: hydrogenase-2 small chain [Proteobacteria] Length: 372\nNumber of predicted TMHs: 1\nExp number of AAs in TMHs: 23.72226\nExp number, first 60 AAs: 0.22038\nTotal prob of N-in: 0.12994\noutside 1 331\nTMhelix 332 354\ninside 355 372

11244 GCF_000456545.1_Esch_coli_HVH_35_4-2962667_V1 Escherichia coli HVH 35 (4-2962667) Proteobacteria; Gammaproteobacteria; Enterobacterales; Enterobacteriaceae; Escherichia; Escherichia coli MTGDNTLIHSHGINRRDFMKLCAALAATMGLSSKAAAEMAESVTNPQRPPVIWIGAQECT WP_000145410.1 MULTISPECIES: hydrogenase-2 small chain [Proteobacteria] Length: 372\nNumber of predicted TMHs: 1\nExp number of AAs in TMHs: 23.72226\nExp number, first 60 AAs: 0.22038\nTotal prob of N-in: 0.12994\noutside 1 331\nTMhelix 332 354\ninside 355 372

11245 GCF_000494935.1_Esch_coli_HVH_36_4-5675286_V1 Escherichia coli HVH 36 (4-5675286) Proteobacteria; Gammaproteobacteria; Enterobacterales; Enterobacteriaceae; Escherichia; Escherichia coli MNNEETFYQAMRRQGVTRRSFLKYCSLAATSLGLGAGMAPKIAWALENKPRIPVVWIHGL WP_001058323.1 MULTISPECIES: hydrogenase-1 small chain [Proteobacteria] Length: 372\nNumber of predicted TMHs: 2\nExp number of AAs in TMHs: 37.2608499999999\nExp number, first 60 AAs: 19.52923\nTotal prob of N-in: 0.94351\nPOSSIBLE N-term signal sequence\ninside 1 19\nTMhelix 20 39\noutside 40 325\nTMhelix 326 348\ninside 349 372

11246 GCF_000494935.1_Esch_coli_HVH_36_4-5675286_V1 Escherichia coli HVH 36 (4-5675286) Proteobacteria; Gammaproteobacteria; Enterobacterales; Enterobacteriaceae; Escherichia; Escherichia coli MTGDNTLIHSHGINRRDFMKLCAALAATMGLSSKAAAEMAESVTNPQRPPVIWIGAQECT WP_000145410.1 MULTISPECIES: hydrogenase-2 small chain [Proteobacteria] Length: 372\nNumber of predicted TMHs: 1\nExp number of AAs in TMHs: 23.72226\nExp number, first 60 AAs: 0.22038\nTotal prob of N-in: 0.12994\noutside 1 331\nTMhelix 332 354\ninside 355 372

11247 GCF_000456565.1_Esch_coli_HVH_37_4-2773848_V1 Escherichia coli HVH 37 (4-2773848) Proteobacteria; Gammaproteobacteria; Enterobacterales; Enterobacteriaceae; Escherichia; Escherichia coli MNNEETFYQAMRRQGVTRRSFLKYCSLAATSLGLGAGMAPKIAWALENKPRIPVVWIHGL WP_001058323.1 MULTISPECIES: hydrogenase-1 small chain [Proteobacteria] Length: 372\nNumber of predicted TMHs: 2\nExp number of AAs in TMHs: 37.2608499999999\nExp number, first 60 AAs: 19.52923\nTotal prob of N-in: 0.94351\nPOSSIBLE N-term signal sequence\ninside 1 19\nTMhelix 20 39\noutside 40 325\nTMhelix 326 348\ninside 349 372

11248 GCF_000456565.1_Esch_coli_HVH_37_4-2773848_V1 Escherichia coli HVH 37 (4-2773848) Proteobacteria; Gammaproteobacteria; Enterobacterales; Enterobacteriaceae; Escherichia; Escherichia coli MTGDNTLIHSHGINRRDFMKLCAALAATMGLSSKAAAEMAESVTNPQRPPVIWIGAQECT WP_000145410.1 MULTISPECIES: hydrogenase-2 small chain [Proteobacteria] Length: 372\nNumber of predicted TMHs: 1\nExp number of AAs in TMHs: 23.72226\nExp number, first 60 AAs: 0.22038\nTotal prob of N-in: 0.12994\noutside 1 331\nTMhelix 332 354\ninside 355 372

11249 GCF_000456585.1_Esch_coli_HVH_38_4-2774682_V1 Escherichia coli HVH 38 (4-2774682) Proteobacteria; Gammaproteobacteria; Enterobacterales; Enterobacteriaceae; Escherichia; Escherichia coli MTGDNTLIHSHGINRRDFMKLCAALAATMGLSSKAAAEMVESVTNPQRPPVIWIGAQECT WP_000145423.1 hydrogenase 2 small subunit [Escherichia coli] Length: 372\nNumber of predicted TMHs: 1\nExp number of AAs in TMHs: 23.78201\nExp number, first 60 AAs: 0.28355\nTotal prob of N-in: 0.13257\noutside 1 331\nTMhelix 332 354\ninside 355 372

11250 GCF_000456585.1_Esch_coli_HVH_38_4-2774682_V1 Escherichia coli HVH 38 (4-2774682) Proteobacteria; Gammaproteobacteria; Enterobacterales; Enterobacteriaceae; Escherichia; Escherichia coli MNNEETFYQAMRRQGVTRRSFLKYCSLAATSLGLGAGMAPKIAWALENKPRIPVVWIHGL WP_001058323.1 MULTISPECIES: hydrogenase-1 small chain [Proteobacteria] Length: 372\nNumber of predicted TMHs: 2\nExp number of AAs in TMHs: 37.2608499999999\nExp number, first 60 AAs: 19.52923\nTotal prob of N-in: 0.94351\nPOSSIBLE N-term signal sequence\ninside 1 19\nTMhelix 20 39\noutside 40 325\nTMhelix 326 348\ninside 349 372

11251 GCF_000456605.1_Esch_coli_HVH_39_4-2679949_V1 Escherichia coli HVH 39 (4-2679949) Proteobacteria; Gammaproteobacteria; Enterobacterales; Enterobacteriaceae; Escherichia; Escherichia coli MTGDNTLIHSHGINRRDFMKLCAALAATMGLSSKAAAEMAESVTNPQRPPVIWIGAQECT WP_021519494.1 hydrogenase-2 small chain [Escherichia coli] Length: 372\nNumber of predicted TMHs: 1\nExp number of AAs in TMHs: 23.72331\nExp number, first 60 AAs: 0.22039\nTotal prob of N-in: 0.12995\noutside 1 331\nTMhelix 332 354\ninside 355 372

11252 GCF_000456605.1_Esch_coli_HVH_39_4-2679949_V1 Escherichia coli HVH 39 (4-2679949) Proteobacteria; Gammaproteobacteria; Enterobacterales; Enterobacteriaceae; Escherichia; Escherichia coli MNNEETFYQAMRRQGVTRRSFLKYCSLAATSLGLGAGMAPKIAWALENKPRIPVVWIHGL WP_001058323.1 MULTISPECIES: hydrogenase-1 small chain [Proteobacteria] Length: 372\nNumber of predicted TMHs: 2\nExp number of AAs in TMHs: 37.2608499999999\nExp number, first 60 AAs: 19.52923\nTotal prob of N-in: 0.94351\nPOSSIBLE N-term signal sequence\ninside 1 19\nTMhelix 20 39\noutside 40 325\nTMhelix 326 348\ninside 349 372

11253 GCF_000456625.1_Esch_coli_HVH_40_4-1219782_V1 Escherichia coli HVH 40 (4-1219782) Proteobacteria; Gammaproteobacteria; Enterobacterales; Enterobacteriaceae; Escherichia; Escherichia coli MNNEETFYQAMRRQGVTRRSFLKYCSLAATSLGLGAGMAPKIAWALENKPRIPVVWIHGL WP_001058323.1 MULTISPECIES: hydrogenase-1 small chain [Proteobacteria] Length: 372\nNumber of predicted TMHs: 2\nExp number of AAs in TMHs: 37.2608499999999\nExp number, first 60 AAs: 19.52923\nTotal prob of N-in: 0.94351\nPOSSIBLE N-term signal sequence\ninside 1 19\nTMhelix 20 39\noutside 40 325\nTMhelix 326 348\ninside 349 372

11254 GCF_000456625.1_Esch_coli_HVH_40_4-1219782_V1 Escherichia coli HVH 40 (4-1219782) Proteobacteria; Gammaproteobacteria; Enterobacterales; Enterobacteriaceae; Escherichia; Escherichia coli MTGDNTLIHSHGINRRDFMKLCAALAATMGLSSKAAAEMAESVTNPQRPPVIWIGAQECT WP_000145410.1 MULTISPECIES: hydrogenase-2 small chain [Proteobacteria] Length: 372\nNumber of predicted TMHs: 1\nExp number of AAs in TMHs: 23.72226\nExp number, first 60 AAs: 0.22038\nTotal prob of N-in: 0.12994\noutside 1 331\nTMhelix 332 354\ninside 355 372

11255 GCF_000456645.1_Esch_coli_HVH_41_4-2677849_V1 Escherichia coli HVH 41 (4-2677849) Proteobacteria; Gammaproteobacteria; Enterobacterales; Enterobacteriaceae; Escherichia; Escherichia coli MTGDNTLIHSHGINRRDFMKLCAALAATMGLSSKAAAEMAESVTNPQRPPVIWIGAQECT WP_000145410.1 MULTISPECIES: hydrogenase-2 small chain [Proteobacteria] Length: 372\nNumber of predicted TMHs: 1\nExp number of AAs in TMHs: 23.72226\nExp number, first 60 AAs: 0.22038\nTotal prob of N-in: 0.12994\noutside 1 331\nTMhelix 332 354\ninside 355 372

11256 GCF_000456645.1_Esch_coli_HVH_41_4-2677849_V1 Escherichia coli HVH 41 (4-2677849) Proteobacteria; Gammaproteobacteria; Enterobacterales; Enterobacteriaceae; Escherichia; Escherichia coli MNNEETFYQAMRRQGVTRRSFLKYCSLAATSLGLGAGMAPKIAWALENKPRIPVVWIHGL WP_001058323.1 MULTISPECIES: hydrogenase-1 small chain [Proteobacteria] Length: 372\nNumber of predicted TMHs: 2\nExp number of AAs in TMHs: 37.2608499999999\nExp number, first 60 AAs: 19.52923\nTotal prob of N-in: 0.94351\nPOSSIBLE N-term signal sequence\ninside 1 19\nTMhelix 20 39\noutside 40 325\nTMhelix 326 348\ninside 349 372

11257 GCF_000456665.1_Esch_coli_HVH_42_4-2100061_V1 Escherichia coli HVH 42 (4-2100061) Proteobacteria; Gammaproteobacteria; Enterobacterales; Enterobacteriaceae; Escherichia; Escherichia coli MTGDNTLIHSHGINRRDFMKLCAALAATMGLSSKAAAEMAESVTNPQRPPVIWIGAQECT WP_000145410.1 MULTISPECIES: hydrogenase-2 small chain [Proteobacteria] Length: 372\nNumber of predicted TMHs: 1\nExp number of AAs in TMHs: 23.72226\nExp number, first 60 AAs: 0.22038\nTotal prob of N-in: 0.12994\noutside 1 331\nTMhelix 332 354\ninside 355 372

11258 GCF_000456685.1_Esch_coli_HVH_43_4-2173468_V1 Escherichia coli HVH 43 (4-2173468) Proteobacteria; Gammaproteobacteria; Enterobacterales; Enterobacteriaceae; Escherichia; Escherichia coli MNNEETFYQAMRRQGVTRRSFLKYCSLAATSLGLGAGMAPKIAWALENKPRIPVVWIHGL WP_001058323.1 MULTISPECIES: hydrogenase-1 small chain [Proteobacteria] Length: 372\nNumber of predicted TMHs: 2\nExp number of AAs in TMHs: 37.2608499999999\nExp number, first 60 AAs: 19.52923\nTotal prob of N-in: 0.94351\nPOSSIBLE N-term signal sequence\ninside 1 19\nTMhelix 20 39\noutside 40 325\nTMhelix 326 348\ninside 349 372

11259 GCF_000456685.1_Esch_coli_HVH_43_4-2173468_V1 Escherichia coli HVH 43 (4-2173468) Proteobacteria; Gammaproteobacteria; Enterobacterales; Enterobacteriaceae; Escherichia; Escherichia coli MTGDNTLIHSHGINRRDFMKLCAALAATMGLSSKAAAEMAESVTNPQRPPVIWIGAQECT WP_000145410.1 MULTISPECIES: hydrogenase-2 small chain [Proteobacteria] Length: 372\nNumber of predicted TMHs: 1\nExp number of AAs in TMHs: 23.72226\nExp number, first 60 AAs: 0.22038\nTotal prob of N-in: 0.12994\noutside 1 331\nTMhelix 332 354\ninside 355 372

11260 GCF_000456705.1_Esch_coli_HVH_44_4-2298570_V1 Escherichia coli HVH 44 (4-2298570) Proteobacteria; Gammaproteobacteria; Enterobacterales; Enterobacteriaceae; Escherichia; Escherichia coli MTGDNTLIHSHGINRRDFMKLCAALAATMGLSSKAAAEMAESVTNPQRPPVIWIGAQECT WP_000145410.1 MULTISPECIES: hydrogenase-2 small chain [Proteobacteria] Length: 372\nNumber of predicted TMHs: 1\nExp number of AAs in TMHs: 23.72226\nExp number, first 60 AAs: 0.22038\nTotal prob of N-in: 0.12994\noutside 1 331\nTMhelix 332 354\ninside 355 372

11261 GCF_000456705.1_Esch_coli_HVH_44_4-2298570_V1 Escherichia coli HVH 44 (4-2298570) Proteobacteria; Gammaproteobacteria; Enterobacterales; Enterobacteriaceae; Escherichia; Escherichia coli MNNEETFYQAMRRQGVTRRSFLKYCSLAATSLGLGAGMAPKIAWALENKPRIPVVWIHGL WP_001058323.1 MULTISPECIES: hydrogenase-1 small chain [Proteobacteria] Length: 372\nNumber of predicted TMHs: 2\nExp number of AAs in TMHs: 37.2608499999999\nExp number, first 60 AAs: 19.52923\nTotal prob of N-in: 0.94351\nPOSSIBLE N-term signal sequence\ninside 1 19\nTMhelix 20 39\noutside 40 325\nTMhelix 326 348\ninside 349 372

11262 GCF_000456065.1_Esch_coli_HVH_4_4-7276109_V1 Escherichia coli HVH 4 (4-7276109) Proteobacteria; Gammaproteobacteria; Enterobacterales; Enterobacteriaceae; Escherichia; Escherichia coli MTGDNTLIHSHGINRRDFMKLCAALAATMGLSSKAAAEMAESVTNPQRPPVIWIGAQECT WP_000145410.1 MULTISPECIES: hydrogenase-2 small chain [Proteobacteria] Length: 372\nNumber of predicted TMHs: 1\nExp number of AAs in TMHs: 23.72226\nExp number, first 60 AAs: 0.22038\nTotal prob of N-in: 0.12994\noutside 1 331\nTMhelix 332 354\ninside 355 372

11263 GCF_000456065.1_Esch_coli_HVH_4_4-7276109_V1 Escherichia coli HVH 4 (4-7276109) Proteobacteria; Gammaproteobacteria; Enterobacterales; Enterobacteriaceae; Escherichia; Escherichia coli MNNEETFYQAMRRQGVTRRSFLKYCSLAATSLGLGAGMAPKIAWALENKPRIPVVWIHGL WP_001058323.1 MULTISPECIES: hydrogenase-1 small chain [Proteobacteria] Length: 372\nNumber of predicted TMHs: 2\nExp number of AAs in TMHs: 37.2608499999999\nExp number, first 60 AAs: 19.52923\nTotal prob of N-in: 0.94351\nPOSSIBLE N-term signal sequence\ninside 1 19\nTMhelix 20 39\noutside 40 325\nTMhelix 326 348\ninside 349 372

11264 GCF_000456725.1_Esch_coli_HVH_45_4-3129918_V1 Escherichia coli HVH 45 (4-3129918) Proteobacteria; Gammaproteobacteria; Enterobacterales; Enterobacteriaceae; Escherichia; Escherichia coli MNNEETFYQAMRRQGVTRRSFLKYCSLAATSLGLGAGMAPKIAWALENKPRIPVVWIHGL WP_001058323.1 MULTISPECIES: hydrogenase-1 small chain [Proteobacteria] Length: 372\nNumber of predicted TMHs: 2\nExp number of AAs in TMHs: 37.2608499999999\nExp number, first 60 AAs: 19.52923\nTotal prob of N-in: 0.94351\nPOSSIBLE N-term signal sequence\ninside 1 19\nTMhelix 20 39\noutside 40 325\nTMhelix 326 348\ninside 349 372

11265 GCF_000456725.1_Esch_coli_HVH_45_4-3129918_V1 Escherichia coli HVH 45 (4-3129918) Proteobacteria; Gammaproteobacteria; Enterobacterales; Enterobacteriaceae; Escherichia; Escherichia coli MTGDNTLIHSHGINRRDFMKLCAALAATMGLSSKAAAEMAESVTNPQRPPVIWIGAQECT WP_000145410.1 MULTISPECIES: hydrogenase-2 small chain [Proteobacteria] Length: 372\nNumber of predicted TMHs: 1\nExp number of AAs in TMHs: 23.72226\nExp number, first 60 AAs: 0.22038\nTotal prob of N-in: 0.12994\noutside 1 331\nTMhelix 332 354\ninside 355 372

11266 GCF_000456745.1_Esch_coli_HVH_46_4-2758776_V1 Escherichia coli HVH 46 (4-2758776) Proteobacteria; Gammaproteobacteria; Enterobacterales; Enterobacteriaceae; Escherichia; Escherichia coli MNNEETFYQAMRRQGVTRRSFLKYCSLAATSLGLGAGMAPKIAWALENKPRIPVVWIHGL WP_021520630.1 hydrogenase-1 small chain [Escherichia coli] Length: 372\nNumber of predicted TMHs: 2\nExp number of AAs in TMHs: 37.8197599999999\nExp number, first 60 AAs: 19.53295\nTotal prob of N-in: 0.94396\nPOSSIBLE N-term signal sequence\ninside 1 19\nTMhelix 20 39\noutside 40 325\nTMhelix 326 348\ninside 349 372

11267 GCF_000456745.1_Esch_coli_HVH_46_4-2758776_V1 Escherichia coli HVH 46 (4-2758776) Proteobacteria; Gammaproteobacteria; Enterobacterales; Enterobacteriaceae; Escherichia; Escherichia coli

MTGDNTLIHSHGINRRDFMKLCAALAATMGLSSKAAAEMAESVTNPQRPPVIWIGAQECT WP_000145410.1 MULTISPECIES: hydrogenase-2 small chain [Proteobacteria] Length: 372\nNumber of predicted TMHs: 1\nExp number of AAs in TMHs: 23.72226\nExp number, first 60 AAs: 0.22038\nTotal prob of N-in: 0.12994\noutside 1 331\nTMhelix 332 354\ninside 355 372

11268 GCF_000456765.1_Esch_coli_HVH_48_4-2658593_V1 Escherichia coli HVH 48 (4-2658593) Proteobacteria; Gammaproteobacteria; Enterobacterales; Enterobacteriaceae; Escherichia; Escherichia coli MTGDNTLIHSHGINRRDFMKLCAALAATMGLSSKAAAEMAESVTNPQRPPVIWIGAQECT WP_000145410.1 MULTISPECIES: hydrogenase-2 small chain [Proteobacteria] Length: 372\nNumber of predicted TMHs: 1\nExp number of AAs in TMHs: 23.72226\nExp number, first 60 AAs: 0.22038\nTotal prob of N-in: 0.12994\noutside 1 331\nTMhelix 332 354\ninside 355 372

11269 GCF_000488015.1_Esch_coli_HVH_50_4-2593475_V1 Escherichia coli HVH 50 (4-2593475) Proteobacteria; Gammaproteobacteria; Enterobacterales; Enterobacteriaceae; Escherichia; Escherichia coli MNNEETFYQAMRRQGVTRRSFLKYCSLAATSLGLGAGMAPKIAWALENKPRIPVVWIHGL WP_001058323.1 MULTISPECIES: hydrogenase-1 small chain [Proteobacteria] Length: 372\nNumber of predicted TMHs: 2\nExp number of AAs in TMHs: 37.2608499999999\nExp number, first 60 AAs: 19.52923\nTotal prob of N-in: 0.94351\nPOSSIBLE N-term signal sequence\ninside 1 19\nTMhelix 20 39\noutside 40 325\nTMhelix 326 348\ninside 349 372

11270 GCF_000488015.1_Esch_coli_HVH_50_4-2593475_V1 Escherichia coli HVH 50 (4-2593475) Proteobacteria; Gammaproteobacteria; Enterobacterales; Enterobacteriaceae; Escherichia; Escherichia coli MTGDNTLIHSHGINRRDFMKLCAALAATMGLSSKAAAEMAESVTNPQRPPVIWIGAQECT WP_000145410.1 MULTISPECIES: hydrogenase-2 small chain [Proteobacteria] Length: 372\nNumber of predicted TMHs: 1\nExp number of AAs in TMHs: 23.72226\nExp number, first 60 AAs: 0.22038\nTotal prob of N-in: 0.12994\noutside 1 331\nTMhelix 332 354\ninside 355 372

11271 GCF_000488015.1_Esch_coli_HVH_50_4-2593475_V1 Escherichia coli HVH 50 (4-2593475) Proteobacteria; Gammaproteobacteria; Enterobacterales; Enterobacteriaceae; Escherichia; Escherichia coli MKGVLEKCTVEDFNYISATLDSYVSFTNDKRRKTLLSAYQNNPALHAELISLIDTQIKYF WP_001515206.1 MULTISPECIES: hypothetical protein [Enterobacteriaceae] Length: 249\nNumber of predicted TMHs: 2\nExp number of AAs in TMHs: 39.68696\nExp number, first 60 AAs: 0.01961\nTotal prob of N-in: 0.09053\noutside 1 157\nTMhelix 158 180\ninside 181 200\nTMhelix 201 220\noutside 221 249

11272 GCF_000456785.1_Esch_coli_HVH_51_4-2172526_V1 Escherichia coli HVH 51 (4-2172526) Proteobacteria; Gammaproteobacteria; Enterobacterales; Enterobacteriaceae; Escherichia; Escherichia coli MTGDNTLIHSHGINRRDFMKLCAALAATMGLSSKAAAEMAESVTNPQRPPVIWIGAQECT WP_000145410.1 MULTISPECIES: hydrogenase-2 small chain [Proteobacteria] Length: 372\nNumber of predicted TMHs: 1\nExp number of AAs in TMHs: 23.72226\nExp number, first 60 AAs: 0.22038\nTotal prob of N-in: 0.12994\noutside 1 331\nTMhelix 332 354\ninside 355 372

11273 GCF_000456785.1_Esch_coli_HVH_51_4-2172526_V1 Escherichia coli HVH 51 (4-2172526) Proteobacteria; Gammaproteobacteria; Enterobacterales; Enterobacteriaceae; Escherichia; Escherichia coli MNNEETFYQAMRRQGVTRRSFLKYCSLAATSLGLGAGMAPKIAWALENKPRIPVVWIHGL WP_001058323.1 MULTISPECIES: hydrogenase-1 small chain [Proteobacteria] Length: 372\nNumber of predicted TMHs: 2\nExp number of AAs in TMHs: 37.2608499999999\nExp number, first 60 AAs: 19.52923\nTotal prob of N-in: 0.94351\nPOSSIBLE N-term signal sequence\ninside 1 19\nTMhelix 20 39\noutside 40 325\nTMhelix 326 348\ninside 349 372

11274 GCF_000456805.1_Esch_coli_HVH_53_4-0631051_V1 Escherichia coli HVH 53 (4-0631051) Proteobacteria; Gammaproteobacteria; Enterobacterales; Enterobacteriaceae; Escherichia; Escherichia coli MTGDNTLIHSHGINRRDFMKLCAALAATMGLSSKAAAEMAESVTNPQRPPVIWIGAQECT WP_000145410.1 MULTISPECIES: hydrogenase-2 small chain [Proteobacteria] Length: 372\nNumber of predicted TMHs: 1\nExp number of AAs in TMHs: 23.72226\nExp number, first 60 AAs: 0.22038\nTotal prob of N-in: 0.12994\noutside 1 331\nTMhelix 332 354\ninside 355 372

11275 GCF_000456805.1_Esch_coli_HVH_53_4-0631051_V1 Escherichia coli HVH 53 (4-0631051) Proteobacteria; Gammaproteobacteria; Enterobacterales; Enterobacteriaceae; Escherichia; Escherichia coli MNNEETFYQAMRRQGVTRRSFLKYCSLAATSLGLGAGMAPKIAWALENKPRIPVVWIHGL WP_001295940.1 MULTISPECIES: hydrogenase [Enterobacteriaceae] Length: 372\nNumber of predicted TMHs: 2\nExp number of AAs in TMHs: 37.2608899999999\nExp number, first 60 AAs: 19.52923\nTotal prob of N-in:

0.94351\nPOSSIBLE N-term signal sequence\ninside 1 19\nTMhelix 20 39\noutside 40 325\nTMhelix 326 348\ninside 349 372

11276 GCF_000456085.1_Esch_coli_HVH_5_4-7148410_V1 Escherichia coli HVH 5 (4-7148410) Proteobacteria; Gammaproteobacteria; Enterobacterales; Enterobacteriaceae; Escherichia; Escherichia coli MTGDNTLIHSHGINRRDFMKLCAALAATMGLSSKAAAEMAESVTNPQRPPVIWIGAQECT WP_000145410.1 MULTISPECIES: hydrogenase-2 small chain [Proteobacteria] Length: 372\nNumber of predicted TMHs: 1\nExp number of AAs in TMHs: 23.72226\nExp number, first 60 AAs: 0.22038\nTotal prob of N-in: 0.12994\noutside 1 331\nTMhelix 332 354\ninside 355 372

11277 GCF_000456825.1_Esch_coli_HVH_55_4-2646161_V1 Escherichia coli HVH 55 (4-2646161) Proteobacteria; Gammaproteobacteria; Enterobacterales; Enterobacteriaceae; Escherichia; Escherichia coli MNNEETFYQAMRRQGVTRRSFLKYCSLAATSLGLGAGMAPKIAWALENKPRIPVVWIHGL WP_001538532.1 hydrogenase-1 small chain [Escherichia coli] Length: 372\nNumber of predicted TMHs: 2\nExp number of AAs in TMHs: 37.27041\nExp number, first 60 AAs: 19.52927\nTotal prob of N-in: 0.94352\nPOSSIBLE N-term signal sequence\ninside 1 19\nTMhelix 20 39\noutside 40 325\nTMhelix 326 348\ninside 349 372

11278 GCF_000456825.1_Esch_coli_HVH_55_4-2646161_V1 Escherichia coli HVH 55 (4-2646161) Proteobacteria; Gammaproteobacteria; Enterobacterales; Enterobacteriaceae; Escherichia; Escherichia coli MTGDNTLIHSHGINRRDFMKLCAALAATMGLSSKAAAEMVESVTNPQRPPVIWIGAQECT WP_000145423.1 hydrogenase 2 small subunit [Escherichia coli] Length: 372\nNumber of predicted TMHs: 1\nExp number of AAs in TMHs: 23.78201\nExp number, first 60 AAs: 0.28355\nTotal prob of N-in: 0.13257\noutside 1 331\nTMhelix 332 354\ninside 355 372

11279 GCF_000456845.1_Esch_coli_HVH_56_4-2153033_V1 Escherichia coli HVH 56 (4-2153033) Proteobacteria; Gammaproteobacteria; Enterobacterales; Enterobacteriaceae; Escherichia; Escherichia coli MNNEETFYQAMRRQGVTRRSFLKYCSLAATSLGLGAGMAPKIAWALENKPRIPVVWIHGL WP_001058323.1 MULTISPECIES: hydrogenase-1 small chain [Proteobacteria] Length: 372\nNumber of predicted TMHs: 2\nExp number of AAs in TMHs: 37.2608499999999\nExp number, first 60 AAs: 19.52923\nTotal prob of N-in: 0.94351\nPOSSIBLE N-term signal sequence\ninside 1 19\nTMhelix 20 39\noutside 40 325\nTMhelix 326 348\ninside 349 372

11280 GCF_000456845.1_Esch_coli_HVH_56_4-2153033_V1 Escherichia coli HVH 56 (4-2153033) Proteobacteria; Gammaproteobacteria; Enterobacterales; Enterobacteriaceae; Escherichia; Escherichia coli MTGDNTLIHSHGINRRDFMKLCAALAATMGLSSKAAAEMAESVTNPQRPPVIWIGAQECT WP_000145410.1 MULTISPECIES: hydrogenase-2 small chain [Proteobacteria] Length: 372\nNumber of predicted TMHs: 1\nExp number of AAs in TMHs: 23.72226\nExp number, first 60 AAs: 0.22038\nTotal prob of N-in: 0.12994\noutside 1 331\nTMhelix 332 354\ninside 355 372

11281 GCF_000456865.1_Esch_coli_HVH_58_4-2839709_V1 Escherichia coli HVH 58 (4-2839709) Proteobacteria; Gammaproteobacteria; Enterobacterales; Enterobacteriaceae; Escherichia; Escherichia coli MTGDNTLIHSHGINRRDFMKLCAALAATMGLSSKAAAEMAESVTNPQRPPVIWIGAQECT WP_000145410.1 MULTISPECIES: hydrogenase-2 small chain [Proteobacteria] Length: 372\nNumber of predicted TMHs: 1\nExp number of AAs in TMHs: 23.72226\nExp number, first 60 AAs: 0.22038\nTotal prob of N-in: 0.12994\noutside 1 331\nTMhelix 332 354\ninside 355 372

11282 GCF_000456865.1_Esch_coli_HVH_58_4-2839709_V1 Escherichia coli HVH 58 (4-2839709) Proteobacteria; Gammaproteobacteria; Enterobacterales; Enterobacteriaceae; Escherichia; Escherichia coli MNNEETFYQAMRRQGVTRRSFLKYCSLAATSLGLGAGMAPKIAWALENKPRIPVVWIHGL WP_001058323.1 MULTISPECIES: hydrogenase-1 small chain [Proteobacteria] Length: 372\nNumber of predicted TMHs: 2\nExp number of AAs in TMHs: 37.2608499999999\nExp number, first 60 AAs: 19.52923\nTotal prob of N-in: 0.94351\nPOSSIBLE N-term signal sequence\ninside 1 19\nTMhelix 20 39\noutside 40 325\nTMhelix 326 348\ninside 349 372

11283 GCF_000456885.1_Esch_coli_HVH_59_4-1119338_V1 Escherichia coli HVH 59 (4-1119338) Proteobacteria; Gammaproteobacteria; Enterobacterales; Enterobacteriaceae; Escherichia; Escherichia coli MTGDNTLIHSHGINRRDFMKLCAALAATMGLSSKAAAEMAESVTNPQRPPVIWIGAQECT WP_000145410.1 MULTISPECIES: hydrogenase-2 small chain [Proteobacteria] Length: 372\nNumber of predicted TMHs: 1\nExp number of AAs in TMHs: 23.72226\nExp number, first 60 AAs: 0.22038\nTotal prob of N-in: 0.12994\noutside 1 331\nTMhelix 332 354\ninside 355 372

11284 GCF_000456905.1_Esch_coli_HVH_61_4-2736020_V1 Escherichia coli HVH 61 (4-2736020) Proteobacteria; Gammaproteobacteria; Enterobacterales; Enterobacteriaceae; Escherichia; Escherichia coli MNNEETFYQAMRRQGVTRRSFLKYCSLAATSLGLGAGMAPKIAWALENKPRIPVVWIHGL WP_001058323.1 MULTISPECIES: hydrogenase-1 small chain [Proteobacteria] Length: 372\nNumber of predicted TMHs: 2\nExp number of AAs in TMHs: 37.2608499999999\nExp number, first 60 AAs: 19.52923\nTotal prob of N-in: 0.94351\nPOSSIBLE N-term signal sequence\ninside 1 19\nTMhelix 20 39\noutside 40 325\nTMhelix 326 348\ninside 349 372

11285 GCF_000456905.1_Esch_coli_HVH_61_4-2736020_V1 Escherichia coli HVH 61 (4-2736020) Proteobacteria; Gammaproteobacteria; Enterobacterales; Enterobacteriaceae; Escherichia; Escherichia coli MTGDNTLIHSHGINRRDFMKLCAALAATMGLSSKAAAEMAESVTNPQRPPVIWIGAQECT WP_000145410.1 MULTISPECIES: hydrogenase-2 small chain [Proteobacteria] Length: 372\nNumber of predicted TMHs: 1\nExp number of AAs in TMHs: 23.72226\nExp number, first 60 AAs: 0.22038\nTotal prob of N-in: 0.12994\noutside 1 331\nTMhelix 332 354\ninside 355 372

11286 GCF_000456925.1_Esch_coli_HVH_63_4-2542528_V1 Escherichia coli HVH 63 (4-2542528) Proteobacteria; Gammaproteobacteria; Enterobacterales; Enterobacteriaceae; Escherichia; Escherichia coli MNNEETFYQAMRRQGVTRRSFLKYCSLAATSLGLGAGMAPKIAWALENKPRIPVVWIHGL WP_001058323.1 MULTISPECIES: hydrogenase-1 small chain [Proteobacteria] Length: 372\nNumber of predicted TMHs: 2\nExp number of AAs in TMHs: 37.2608499999999\nExp number, first 60 AAs: 19.52923\nTotal prob of N-in: 0.94351\nPOSSIBLE N-term signal sequence\ninside 1 19\nTMhelix 20 39\noutside 40 325\nTMhelix 326 348\ninside 349 372

11287 GCF_000456925.1_Esch_coli_HVH_63_4-2542528_V1 Escherichia coli HVH 63 (4-2542528) Proteobacteria; Gammaproteobacteria; Enterobacterales; Enterobacteriaceae; Escherichia; Escherichia coli MTGDNTLIHSHGINRRDFMKLCAALAATMGLSSKAAAEMAESVTNPQRPPVIWIGAQECT WP_000145410.1 MULTISPECIES: hydrogenase-2 small chain [Proteobacteria] Length: 372\nNumber of predicted TMHs: 1\nExp number of AAs in TMHs: 23.72226\nExp number, first 60 AAs: 0.22038\nTotal prob of N-in: 0.12994\noutside 1 331\nTMhelix 332 354\ninside 355 372

11288 GCF_000456105.1_Esch_coli_HVH_6_3-8296502_V1 Escherichia coli HVH 6 (3-8296502) Proteobacteria; Gammaproteobacteria; Enterobacterales; Enterobacteriaceae; Escherichia; Escherichia coli MTGDNTLIHSHGINRRDFMKLCAALAATMGLSSKAAAEMAESVTNPQRPPVIWIGAQECT WP_000145410.1 MULTISPECIES: hydrogenase-2 small chain [Proteobacteria] Length: 372\nNumber of predicted TMHs: 1\nExp number of AAs in TMHs: 23.72226\nExp number, first 60 AAs: 0.22038\nTotal prob of N-in: 0.12994\noutside 1 331\nTMhelix 332 354\ninside 355 372

11289 GCF_000456105.1_Esch_coli_HVH_6_3-8296502_V1 Escherichia coli HVH 6 (3-8296502) Proteobacteria; Gammaproteobacteria; Enterobacterales; Enterobacteriaceae; Escherichia; Escherichia coli MNNEETFYQAMRRQGVTRRSFLKYCSLAATSLGLGAGMAPKIAWALENKPRIPVVWIHGL WP_001058323.1 MULTISPECIES: hydrogenase-1 small chain [Proteobacteria] Length: 372\nNumber of predicted TMHs: 2\nExp number of AAs in TMHs: 37.2608499999999\nExp number, first 60 AAs: 19.52923\nTotal prob of N-in: 0.94351\nPOSSIBLE N-term signal sequence\ninside 1 19\nTMhelix 20 39\noutside 40 325\nTMhelix 326 348\ninside 349 372

11290 GCF_000456945.1_Esch_coli_HVH_65_4-2262045_V1 Escherichia coli HVH 65 (4-2262045) Proteobacteria; Gammaproteobacteria; Enterobacterales; Enterobacteriaceae; Escherichia; Escherichia coli MTGDNTLIHSHGINRRDFMKLCAALAATMGLSSKAAAEMAESVTNPQRPPVIWIGAQECT WP_000145410.1 MULTISPECIES: hydrogenase-2 small chain [Proteobacteria] Length: 372\nNumber of predicted TMHs: 1\nExp number of AAs in TMHs: 23.72226\nExp number, first 60 AAs: 0.22038\nTotal prob of N-in: 0.12994\noutside 1 331\nTMhelix 332 354\ninside 355 372

11291 GCF_000456945.1_Esch_coli_HVH_65_4-2262045_V1 Escherichia coli HVH 65 (4-2262045) Proteobacteria; Gammaproteobacteria; Enterobacterales; Enterobacteriaceae; Escherichia; Escherichia coli MNNEETFYQAMRRQGVTRRSFLKYCSLAATSLGLGAGMAPKIAWALENKPRIPVVWIHGL WP_001058323.1 MULTISPECIES: hydrogenase-1 small chain [Proteobacteria] Length: 372\nNumber of predicted TMHs: 2\nExp number of AAs in TMHs: 37.2608499999999\nExp number, first 60 AAs: 19.52923\nTotal prob of N-in: 0.94351\nPOSSIBLE N-term signal sequence\ninside 1 19\nTMhelix 20 39\noutside 40 325\nTMhelix 326 348\ninside 349 372

11292 GCF_000456965.1_Esch_coli_HVH_68_4-0888028_V1 Escherichia coli HVH 68 (4-0888028) Proteobacteria; Gammaproteobacteria; Enterobacterales; Enterobacteriaceae; Escherichia; Escherichia coli MTGDNTLIHSHGINRRDFMKLCAALAATMGLSSKAAAEMAESVTNPQRPPVIWIGAQECT WP_000145410.1 MULTISPECIES: hydrogenase-2 small chain [Proteobacteria] Length: 372\nNumber of predicted TMHs: 1\nExp number of AAs in TMHs: 23.72226\nExp number, first 60 AAs: 0.22038\nTotal prob of N-in: 0.12994\noutside 1 331\nTMhelix 332 354\ninside 355 372

11293 GCF_000456965.1_Esch_coli_HVH_68_4-0888028_V1 Escherichia coli HVH 68 (4-0888028) Proteobacteria; Gammaproteobacteria; Enterobacterales; Enterobacteriaceae; Escherichia; Escherichia coli MNNEETFYQAMRRQGVTRRSFLKYCSLAATSLGLGAGMAPKIAWALENKPRIPVVWIHGL WP_001058323.1 MULTISPECIES: hydrogenase-1 small chain [Proteobacteria] Length: 372\nNumber of predicted TMHs: 2\nExp number of AAs in TMHs: 37.2608499999999\nExp number, first 60 AAs: 19.52923\nTotal prob of N-in: 0.94351\nPOSSIBLE N-term signal sequence\ninside 1 19\nTMhelix 20 39\noutside 40 325\nTMhelix 326 348\ninside 349 372

11294 GCF_000456985.1_Esch_coli_HVH_69_4-2837072_V1 Escherichia coli HVH 69 (4-2837072) Proteobacteria; Gammaproteobacteria; Enterobacterales; Enterobacteriaceae; Escherichia; Escherichia coli MNNEETFYQAMRRQGVTRRSFLKYCSLAATSLGLGAGMAPKIAWALENKPRIPVVWIHGL WP_001058323.1 MULTISPECIES: hydrogenase-1 small chain [Proteobacteria] Length: 372\nNumber of predicted TMHs: 2\nExp number of AAs in TMHs: 37.2608499999999\nExp number, first 60 AAs: 19.52923\nTotal prob of N-in: 0.94351\nPOSSIBLE N-term signal sequence\ninside 1 19\nTMhelix 20 39\noutside 40 325\nTMhelix 326 348\ninside 349 372

11295 GCF_000456985.1_Esch_coli_HVH_69_4-2837072_V1 Escherichia coli HVH 69 (4-2837072) Proteobacteria; Gammaproteobacteria; Enterobacterales; Enterobacteriaceae; Escherichia; Escherichia coli MTGDNTLIHSHGINRRDFMKLCAALAATMGLSSKAAAEMAESVTNPQRPPVIWIGAQECT WP_000145410.1 MULTISPECIES: hydrogenase-2 small chain [Proteobacteria] Length: 372\nNumber of predicted TMHs: 1\nExp number of AAs in TMHs: 23.72226\nExp number, first 60 AAs: 0.22038\nTotal prob of N-in: 0.12994\noutside 1 331\nTMhelix 332 354\ninside 355 372

11296 GCF_000457005.1_Esch_coli_HVH_70_4-2963531_V1 Escherichia coli HVH 70 (4-2963531) Proteobacteria; Gammaproteobacteria; Enterobacterales; Enterobacteriaceae; Escherichia; Escherichia coli MNNEETFYQAMRRQGVTRRSFLKYCSLAATSLGLGAGMAPKIAWALENKPRIPVVWIHGL WP_001058323.1 MULTISPECIES: hydrogenase-1 small chain [Proteobacteria] Length: 372\nNumber of predicted TMHs: 2\nExp number of AAs in TMHs: 37.2608499999999\nExp number, first 60 AAs: 19.52923\nTotal prob of N-in: 0.94351\nPOSSIBLE N-term signal sequence\ninside 1 19\nTMhelix 20 39\noutside 40 325\nTMhelix 326 348\ninside 349 372

11297 GCF_000457005.1_Esch_coli_HVH_70_4-2963531_V1 Escherichia coli HVH 70 (4-2963531) Proteobacteria; Gammaproteobacteria; Enterobacterales; Enterobacteriaceae; Escherichia; Escherichia coli MTGDNTLIHSHGINRRDFMKLCAALAATMGLSSKAAAEMAESVTNPQRPPVIWIGAQECT WP_000145418.1 hydrogenase 2 small subunit [Escherichia coli] Length: 372\nNumber of predicted TMHs: 1\nExp number of AAs in TMHs: 23.72313\nExp number, first 60 AAs: 0.22034\nTotal prob of N-in: 0.12997\noutside 1 331\nTMhelix 332 354\ninside 355 372

11298 GCF_000457025.1_Esch_coli_HVH_73_4-2393174_V1 Escherichia coli HVH 73 (4-2393174) Proteobacteria; Gammaproteobacteria; Enterobacterales; Enterobacteriaceae; Escherichia; Escherichia coli MTGDNTLIHSHGINRRDFMKLCAALAATMGLSSKAAAEMAESVTNPQRPPVIWIGAQECT WP_000145410.1 MULTISPECIES: hydrogenase-2 small chain [Proteobacteria] Length: 372\nNumber of predicted TMHs: 1\nExp number of AAs in TMHs: 23.72226\nExp number, first 60 AAs: 0.22038\nTotal prob of N-in: 0.12994\noutside 1 331\nTMhelix 332 354\ninside 355 372

11299 GCF_000457045.1_Esch_coli_HVH_74_4-1034782_V1 Escherichia coli HVH 74 (4-1034782) Proteobacteria; Gammaproteobacteria; Enterobacterales; Enterobacteriaceae; Escherichia; Escherichia coli MNNEETFYQAMRRQGVTRRSFLKYCSLAATSLGLGAGMAPKIAWALENKPRIPVVWIHGL WP_001058323.1 MULTISPECIES: hydrogenase-1 small chain [Proteobacteria] Length: 372\nNumber of predicted TMHs: 2\nExp number of AAs in TMHs: 37.2608499999999\nExp number, first 60 AAs: 19.52923\nTotal prob of N-in: 0.94351\nPOSSIBLE N-term signal sequence\ninside 1 19\nTMhelix 20 39\noutside 40 325\nTMhelix 326 348\ninside 349 372

11300 GCF_000457045.1_Esch_coli_HVH_74_4-1034782_V1 Escherichia coli HVH 74 (4-1034782) Proteobacteria; Gammaproteobacteria; Enterobacterales; Enterobacteriaceae; Escherichia; Escherichia coli MTGDNTLIHSHGINRRDFMKLCAALAATMGLSSKAAAEMAESVTNPQRPPVIWIGAQECT WP_000145410.1 MULTISPECIES: hydrogenase-2 small chain [Proteobacteria] Length: 372\nNumber of predicted TMHs: 1\nExp number of AAs in TMHs: 23.72226\nExp number, first 60 AAs: 0.22038\nTotal prob of N-in: 0.12994\noutside 1 331\nTMhelix 332 354\ninside 355 372

11301 GCF_000456125.1_Esch_coli_HVH_7_4-7315031_V1 Escherichia coli HVH 7 (4-7315031) Proteobacteria; Gammaproteobacteria; Enterobacterales; Enterobacteriaceae; Escherichia; Escherichia coli MNNEETFYQAMRRQGVTRRSFLKYCSLAATSLGLGAGMAPKIAWALENKPRIPVVWIHGL WP_001058323.1 MULTISPECIES: hydrogenase-1 small chain [Proteobacteria] Length: 372\nNumber of predicted TMHs: 2\nExp number of AAs in TMHs: 37.2608499999999\nExp number, first 60 AAs: 19.52923\nTotal prob of N-in: 0.94351\nPOSSIBLE N-term signal sequence\ninside 1 19\nTMhelix 20 39\noutside 40 325\nTMhelix 326 348\ninside 349 372

11302 GCF_000456125.1_Esch_coli_HVH_7_4-7315031_V1 Escherichia coli HVH 7 (4-7315031) Proteobacteria; Gammaproteobacteria; Enterobacterales; Enterobacteriaceae; Escherichia; Escherichia coli MTGDNTLIHSHGINRRDFMKLCAALAATMGLSSKAAAEMAESVTNPQRPPVIWIGAQECT WP_000145410.1 MULTISPECIES: hydrogenase-2 small chain [Proteobacteria] Length: 372\nNumber of predicted TMHs: 1\nExp number of AAs in TMHs: 23.72226\nExp number, first 60 AAs: 0.22038\nTotal prob of N-in: 0.12994\noutside 1 331\nTMhelix 332 354\ninside 355 372

11303 GCF_000457065.1_Esch_coli_HVH_76_4-2538717_V1 Escherichia coli HVH 76 (4-2538717) Proteobacteria; Gammaproteobacteria; Enterobacterales; Enterobacteriaceae; Escherichia; Escherichia coli MTGDNTLIHSHGINRRDFMKLCAALAATMGLSSKAAAEMAESVTNPQRPPVIWIGAQECT WP_000145410.1 MULTISPECIES: hydrogenase-2 small chain [Proteobacteria] Length: 372\nNumber of predicted TMHs: 1\nExp number of AAs in TMHs: 23.72226\nExp number, first 60 AAs: 0.22038\nTotal prob of N-in: 0.12994\noutside 1 331\nTMhelix 332 354\ninside 355 372

11304 GCF_000457085.1_Esch_coli_HVH_77_4-2605759_V1 Escherichia coli HVH 77 (4-2605759) Proteobacteria; Gammaproteobacteria; Enterobacterales; Enterobacteriaceae; Escherichia; Escherichia coli MTGDNTLIHSHGINRRDFMKLCAALAATMGLSSKAAAEMAESVTNPQRPPVIWIGAQECT WP_000145410.1 MULTISPECIES: hydrogenase-2 small chain [Proteobacteria] Length: 372\nNumber of predicted TMHs: 1\nExp number of AAs in TMHs: 23.72226\nExp number, first 60 AAs: 0.22038\nTotal prob of N-in: 0.12994\noutside 1 331\nTMhelix 332 354\ninside 355 372

11305 GCF_000457085.1_Esch_coli_HVH_77_4-2605759_V1 Escherichia coli HVH 77 (4-2605759) Proteobacteria; Gammaproteobacteria; Enterobacterales; Enterobacteriaceae; Escherichia; Escherichia coli MNNEETFYQAMRRQGVTRRSFLKYCSLAATSLGLGAGMAPKIAWALENKPRIPVVWIHGL WP_001058323.1 MULTISPECIES: hydrogenase-1 small chain [Proteobacteria] Length: 372\nNumber of predicted TMHs: 2\nExp number of AAs in TMHs: 37.2608499999999\nExp number, first 60 AAs: 19.52923\nTotal prob of N-in: 0.94351\nPOSSIBLE N-term signal sequence\ninside 1 19\nTMhelix 20 39\noutside 40 325\nTMhelix 326 348\ninside 349 372

11306 GCF_000457105.1_Esch_coli_HVH_78_4-2735946_V1 Escherichia coli HVH 78 (4-2735946) Proteobacteria; Gammaproteobacteria; Enterobacterales; Enterobacteriaceae; Escherichia; Escherichia coli MNNEETFYQAMRRQGVTRRSFLKYCSLAATSLGLGAGMAPKIAWALENKPRIPVVWIHGL WP_001058323.1 MULTISPECIES: hydrogenase-1 small chain [Proteobacteria] Length: 372\nNumber of predicted TMHs: 2\nExp number of AAs in TMHs: 37.2608499999999\nExp number, first 60 AAs: 19.52923\nTotal prob of N-in: 0.94351\nPOSSIBLE N-term signal sequence\ninside 1 19\nTMhelix 20 39\noutside 40 325\nTMhelix 326 348\ninside 349 372

11307 GCF_000457105.1_Esch_coli_HVH_78_4-2735946_V1 Escherichia coli HVH 78 (4-2735946) Proteobacteria; Gammaproteobacteria; Enterobacterales; Enterobacteriaceae; Escherichia; Escherichia coli MTGDNTLIHSHGINRRDFMKLCAALAATMGLSSKAAAEMAESVTNPQRPPVIWIGAQECT WP_000145410.1 MULTISPECIES: hydrogenase-2 small chain [Proteobacteria] Length: 372\nNumber of predicted TMHs: 1\nExp number of AAs in TMHs: 23.72226\nExp number, first 60 AAs: 0.22038\nTotal prob of N-in: 0.12994\noutside 1 331\nTMhelix 332 354\ninside 355 372

11308 GCF_000457125.1_Esch_coli_HVH_79_4-2512823_V1 Escherichia coli HVH 79 (4-2512823) Proteobacteria; Gammaproteobacteria; Enterobacterales; Enterobacteriaceae; Escherichia; Escherichia coli

MTGDNTLIHSHGINRRDFMKLCAALAATMGLSSKAAAEMAESVTNPQRPPVIWIGAQECT WP_000145410.1 MULTISPECIES: hydrogenase-2 small chain [Proteobacteria] Length: 372\nNumber of predicted TMHs: 1\nExp number of AAs in TMHs: 23.72226\nExp number, first 60 AAs: 0.22038\nTotal prob of N-in: 0.12994\noutside 1 331\nTMhelix 332 354\ninside 355 372

11309 GCF_000457125.1_Esch_coli_HVH_79_4-2512823_V1 Escherichia coli HVH 79 (4-2512823) Proteobacteria; Gammaproteobacteria; Enterobacterales; Enterobacteriaceae; Escherichia; Escherichia coli MNNEETFYQAMRRQGVTRRSFLKYCSLAATSLGLGAGMAPKIAWALENKPRIPVVWIHGL WP_001058323.1 MULTISPECIES: hydrogenase-1 small chain [Proteobacteria] Length: 372\nNumber of predicted TMHs: 2\nExp number of AAs in TMHs: 37.2608499999999\nExp number, first 60 AAs: 19.52923\nTotal prob of N-in: 0.94351\nPOSSIBLE N-term signal sequence\ninside 1 19\nTMhelix 20 39\noutside 40 325\nTMhelix 326 348\ninside 349 372

11310 GCF_000457145.1_Esch_coli_HVH_80_4-2428830_V1 Escherichia coli HVH 80 (4-2428830) Proteobacteria; Gammaproteobacteria; Enterobacterales; Enterobacteriaceae; Escherichia; Escherichia coli MTGDNTLIHSHGINRRDFMKLCAALAATMGLSSKAAAEMAESVTNPQRPPVIWIGAQECT WP_000145410.1 MULTISPECIES: hydrogenase-2 small chain [Proteobacteria] Length: 372\nNumber of predicted TMHs: 1\nExp number of AAs in TMHs: 23.72226\nExp number, first 60 AAs: 0.22038\nTotal prob of N-in: 0.12994\noutside 1 331\nTMhelix 332 354\ninside 355 372

11311 GCF_000457165.1_Esch_coli_HVH_82_4-2209276_V1 Escherichia coli HVH 82 (4-2209276) Proteobacteria; Gammaproteobacteria; Enterobacterales; Enterobacteriaceae; Escherichia; Escherichia coli MTGDNTLIHSHGINRRGFMKLCAALAATMGLSSKAAAEMAESVTNPQRPPVIWIGAQECT WP_021525990.1 hydrogenase-2 small chain [Escherichia coli] Length: 372\nNumber of predicted TMHs: 1\nExp number of AAs in TMHs: 23.90966\nExp number, first 60 AAs: 0.40466\nTotal prob of N-in: 0.13702\noutside 1 331\nTMhelix 332 354\ninside 355 372

11312 GCF_000457165.1_Esch_coli_HVH_82_4-2209276_V1 Escherichia coli HVH 82 (4-2209276) Proteobacteria; Gammaproteobacteria; Enterobacterales; Enterobacteriaceae; Escherichia; Escherichia coli MNNEETFYQAMRRQGVTRRSFLKYCSLAATSLGLGAGMAPKIAWALENKPRIPVVWIHGL WP_001058320.1 hydrogenase [Escherichia coli] Length: 372\nNumber of predicted TMHs: 2\nExp number of AAs in TMHs: 37.26082\nExp number, first 60 AAs: 19.52923\nTotal prob of N-in: 0.94351\nPOSSIBLE N-term signal sequence\ninside 1 19\nTMhelix 20 39\noutside 40 325\nTMhelix 326 348\ninside 349 372

11313 GCF_000507625.1_Esch_coli_HVH_83_4-2051087_V1 Escherichia coli HVH 83 (4-2051087) Proteobacteria; Gammaproteobacteria; Enterobacterales; Enterobacteriaceae; Escherichia; Escherichia coli MNNEETFYQAMRRQGVTRRSFLKYCSLAATSLGLGAGMAPKIAWALENKPRIPVVWIHGL WP_001058323.1 MULTISPECIES: hydrogenase-1 small chain [Proteobacteria] Length: 372\nNumber of predicted TMHs: 2\nExp number of AAs in TMHs: 37.2608499999999\nExp number, first 60 AAs: 19.52923\nTotal prob of N-in: 0.94351\nPOSSIBLE N-term signal sequence\ninside 1 19\nTMhelix 20 39\noutside 40 325\nTMhelix 326 348\ninside 349 372

11314 GCF_000507625.1_Esch_coli_HVH_83_4-2051087_V1 Escherichia coli HVH 83 (4-2051087) Proteobacteria; Gammaproteobacteria; Enterobacterales; Enterobacteriaceae; Escherichia; Escherichia coli MTGDNTLIHSHGINRRDFMKLCAALAATMGLSSKAAAEMAESVTNPQRPPVIWIGAQECT WP_000145410.1 MULTISPECIES: hydrogenase-2 small chain [Proteobacteria] Length: 372\nNumber of predicted TMHs: 1\nExp number of AAs in TMHs: 23.72226\nExp number, first 60 AAs: 0.22038\nTotal prob of N-in: 0.12994\noutside 1 331\nTMhelix 332 354\ninside 355 372

11315 GCF_000457185.1_Esch_coli_HVH_84_4-1021478_V1 Escherichia coli HVH 84 (4-1021478) Proteobacteria; Gammaproteobacteria; Enterobacterales; Enterobacteriaceae; Escherichia; Escherichia coli MTGDNTLIHSHGINRRDFMKLCAALAATMGLSSKAAAEMVESVTNPQRPPVIWIGAQECT WP_000145423.1 hydrogenase 2 small subunit [Escherichia coli] Length: 372\nNumber of predicted TMHs: 1\nExp number of AAs in TMHs: 23.78201\nExp number, first 60 AAs: 0.28355\nTotal prob of N-in: 0.13257\noutside 1 331\nTMhelix 332 354\ninside 355 372

11316 GCF_000457185.1_Esch_coli_HVH_84_4-1021478_V1 Escherichia coli HVH 84 (4-1021478) Proteobacteria; Gammaproteobacteria; Enterobacterales; Enterobacteriaceae; Escherichia; Escherichia coli MNNEETFYQAMRRQGVTRRSFLKYCSLAATSLGLGAGMAPKIAWALENKPRIPVVWIHGL WP_001295940.1 MULTISPECIES: hydrogenase [Enterobacteriaceae] Length: 372\nNumber of predicted TMHs: 2\nExp number of AAs in TMHs: 37.2608899999999\nExp number, first 60 AAs: 19.52923\nTotal prob of N-in:

0.94351\nPOSSIBLE N-term signal sequence\ninside 1 19\nTMhelix 20 39\noutside 40 325\nTMhelix 326 348\ninside 349 372

11317 GCF_000457205.1_Esch_coli_HVH_85_4-0792144_V1 Escherichia coli HVH 85 (4-0792144) Proteobacteria; Gammaproteobacteria; Enterobacterales; Enterobacteriaceae; Escherichia; Escherichia coli MNNEETFYQAMRRQGVTRRSFLKYCSLAATSLGLGAGMAPKIAWALENKPRIPVVWIHGL WP_001058323.1 MULTISPECIES: hydrogenase-1 small chain [Proteobacteria] Length: 372\nNumber of predicted TMHs: 2\nExp number of AAs in TMHs: 37.2608499999999\nExp number, first 60 AAs: 19.52923\nTotal prob of N-in: 0.94351\nPOSSIBLE N-term signal sequence\ninside 1 19\nTMhelix 20 39\noutside 40 325\nTMhelix 326 348\ninside 349 372

11318 GCF_000457205.1_Esch_coli_HVH_85_4-0792144_V1 Escherichia coli HVH 85 (4-0792144) Proteobacteria; Gammaproteobacteria; Enterobacterales; Enterobacteriaceae; Escherichia; Escherichia coli MTGDNTLIHSHGINRRDFMKLCAALAATMGLSSKAAAEMAESVTNPQRPPVIWIGAQECT WP_000145410.1 MULTISPECIES: hydrogenase-2 small chain [Proteobacteria] Length: 372\nNumber of predicted TMHs: 1\nExp number of AAs in TMHs: 23.72226\nExp number, first 60 AAs: 0.22038\nTotal prob of N-in: 0.12994\noutside 1 331\nTMhelix 332 354\ninside 355 372

11319 GCF_000494975.1_Esch_coli_HVH_86_4-7026218_V1 Escherichia coli HVH 86 (4-7026218) Proteobacteria; Gammaproteobacteria; Enterobacterales; Enterobacteriaceae; Escherichia; Escherichia coli MTGDNTLIHSHGINRRDFMKLCAALAATMGLSSKAAAEMAESVTNPQRPPVIWIGAQECT WP_000145410.1 MULTISPECIES: hydrogenase-2 small chain [Proteobacteria] Length: 372\nNumber of predicted TMHs: 1\nExp number of AAs in TMHs: 23.72226\nExp number, first 60 AAs: 0.22038\nTotal prob of N-in: 0.12994\noutside 1 331\nTMhelix 332 354\ninside 355 372

11320 GCF_000494975.1_Esch_coli_HVH_86_4-7026218_V1 Escherichia coli HVH 86 (4-7026218) Proteobacteria; Gammaproteobacteria; Enterobacterales; Enterobacteriaceae; Escherichia; Escherichia coli MNNEETFYQAMRRQGVTRRSFLKYCSLAATSLGLGAGMAPKIAWALENKPRIPVVWIHGL WP_001058323.1 MULTISPECIES: hydrogenase-1 small chain [Proteobacteria] Length: 372\nNumber of predicted TMHs: 2\nExp number of AAs in TMHs: 37.2608499999999\nExp number, first 60 AAs: 19.52923\nTotal prob of N-in: 0.94351\nPOSSIBLE N-term signal sequence\ninside 1 19\nTMhelix 20 39\noutside 40 325\nTMhelix 326 348\ninside 349 372

11321 GCF_000457225.1_Esch_coli_HVH_87_4-5977630_V1 Escherichia coli HVH 87 (4-5977630) Proteobacteria; Gammaproteobacteria; Enterobacterales; Enterobacteriaceae; Escherichia; Escherichia coli MNNEETFYQAMRRQGVTRRSFLKYCSLAATSLGLGAGMAPKIAWALENKPRIPVVWIHGL WP_001058323.1 MULTISPECIES: hydrogenase-1 small chain [Proteobacteria] Length: 372\nNumber of predicted TMHs: 2\nExp number of AAs in TMHs: 37.2608499999999\nExp number, first 60 AAs: 19.52923\nTotal prob of N-in: 0.94351\nPOSSIBLE N-term signal sequence\ninside 1 19\nTMhelix 20 39\noutside 40 325\nTMhelix 326 348\ninside 349 372

11322 GCF_000457225.1_Esch_coli_HVH_87_4-5977630_V1 Escherichia coli HVH 87 (4-5977630) Proteobacteria; Gammaproteobacteria; Enterobacterales; Enterobacteriaceae; Escherichia; Escherichia coli MTGDNTLIHSHGINRRDFMKLCAALAATMGLSSKAAAEMAESVTNPQRPPVIWIGAQECT WP_000145410.1 MULTISPECIES: hydrogenase-2 small chain [Proteobacteria] Length: 372\nNumber of predicted TMHs: 1\nExp number of AAs in TMHs: 23.72226\nExp number, first 60 AAs: 0.22038\nTotal prob of N-in: 0.12994\noutside 1 331\nTMhelix 332 354\ninside 355 372

11323 GCF_000457245.1_Esch_coli_HVH_88_4-5854636_V1 Escherichia coli HVH 88 (4-5854636) Proteobacteria; Gammaproteobacteria; Enterobacterales; Enterobacteriaceae; Escherichia; Escherichia coli MTGDNTLIHSHGINRRDFMKLCAALAATMGLSSKAAAEMAESVTNPQRPPVIWIGAQECT WP_000145410.1 MULTISPECIES: hydrogenase-2 small chain [Proteobacteria] Length: 372\nNumber of predicted TMHs: 1\nExp number of AAs in TMHs: 23.72226\nExp number, first 60 AAs: 0.22038\nTotal prob of N-in: 0.12994\noutside 1 331\nTMhelix 332 354\ninside 355 372

11324 GCF_000457245.1_Esch_coli_HVH_88_4-5854636_V1 Escherichia coli HVH 88 (4-5854636) Proteobacteria; Gammaproteobacteria; Enterobacterales; Enterobacteriaceae; Escherichia; Escherichia coli MNNEETFYQAMRRQGVTRRSFLKYCSLAATSLGLGAGMAPKIAWALENKPRIPVVWIHGL WP_001058323.1 MULTISPECIES: hydrogenase-1 small chain [Proteobacteria] Length: 372\nNumber of predicted TMHs: 2\nExp number of AAs in TMHs: 37.2608499999999\nExp number, first 60 AAs: 19.52923\nTotal prob of N-in:

0.94351\nPOSSIBLE N-term signal sequence\ninside 1 19\nTMhelix 20 39\noutside 40 325\nTMhelix 326 348\ninside 349 372

11325 GCF_000457265.1_Esch_coli_HVH_89_4-5885604_V1 Escherichia coli HVH 89 (4-5885604) Proteobacteria; Gammaproteobacteria; Enterobacterales; Enterobacteriaceae; Escherichia; Escherichia coli MNNEETFYQAMRRQGVTRRSFLKYCSLAATSLGLGTGMAPKIAWALENKPRIPVVWIHGL WP_021527074.1 hydrogenase-1 small chain [Escherichia coli] Length: 372\nNumber of predicted TMHs: 2\nExp number of AAs in TMHs: 36.47672\nExp number, first 60 AAs: 18.73629\nTotal prob of N-in: 0.90684\nPOSSIBLE N-term signal sequence\ninside 1 19\nTMhelix 20 39\noutside 40 325\nTMhelix 326 348\ninside 349 372

11326 GCF_000457265.1_Esch_coli_HVH_89_4-5885604_V1 Escherichia coli HVH 89 (4-5885604) Proteobacteria; Gammaproteobacteria; Enterobacterales; Enterobacteriaceae; Escherichia; Escherichia coli MTGDNTLIHSHGINRRDFMKLCAALAATMGLSSKAAAEMAESVTNPQRPPVIWIGAQECT WP_000145410.1 MULTISPECIES: hydrogenase-2 small chain [Proteobacteria] Length: 372\nNumber of predicted TMHs: 1\nExp number of AAs in TMHs: 23.72226\nExp number, first 60 AAs: 0.22038\nTotal prob of N-in: 0.12994\noutside 1 331\nTMhelix 332 354\ninside 355 372

11327 GCF_000457285.1_Esch_coli_HVH_90_4-3191362_V1 Escherichia coli HVH 90 (4-3191362) Proteobacteria; Gammaproteobacteria; Enterobacterales; Enterobacteriaceae; Escherichia; Escherichia coli MNNEETFYQAMRRQGVTRRSFLKYCSLAATSLGLGAGMAPKIAWALENKPRIPVVWIHGL WP_001295940.1 MULTISPECIES: hydrogenase [Enterobacteriaceae] Length: 372\nNumber of predicted TMHs: 2\nExp number of AAs in TMHs: 37.2608899999999\nExp number, first 60 AAs: 19.52923\nTotal prob of N-in: 0.94351\nPOSSIBLE N-term signal sequence\ninside 1 19\nTMhelix 20 39\noutside 40 325\nTMhelix 326 348\ninside 349 372

11328 GCF_000457285.1_Esch_coli_HVH_90_4-3191362_V1 Escherichia coli HVH 90 (4-3191362) Proteobacteria; Gammaproteobacteria; Enterobacterales; Enterobacteriaceae; Escherichia; Escherichia coli MTGDNTLIHSHGINRRDFMKLCAALAATMGLSSKAAAEMAESVTNPQRPPVIWIGAQECT WP_000145410.1 MULTISPECIES: hydrogenase-2 small chain [Proteobacteria] Length: 372\nNumber of predicted TMHs: 1\nExp number of AAs in TMHs: 23.72226\nExp number, first 60 AAs: 0.22038\nTotal prob of N-in: 0.12994\noutside 1 331\nTMhelix 332 354\ninside 355 372

11329 GCF_000457305.1_Esch_coli_HVH_91_4-4638751_V1 Escherichia coli HVH 91 (4-4638751) Proteobacteria; Gammaproteobacteria; Enterobacterales; Enterobacteriaceae; Escherichia; Escherichia coli MTGDNTLIHSHGINRRDFMKLCAALAATMGLSSKAAAEMAESVTNPQRPPVIWIGAQECT WP_000145410.1 MULTISPECIES: hydrogenase-2 small chain [Proteobacteria] Length: 372\nNumber of predicted TMHs: 1\nExp number of AAs in TMHs: 23.72226\nExp number, first 60 AAs: 0.22038\nTotal prob of N-in: 0.12994\noutside 1 331\nTMhelix 332 354\ninside 355 372

11330 GCF_000457305.1_Esch_coli_HVH_91_4-4638751_V1 Escherichia coli HVH 91 (4-4638751) Proteobacteria; Gammaproteobacteria; Enterobacterales; Enterobacteriaceae; Escherichia; Escherichia coli MNNEETFYQAMRRQGVTRRSFLKYCSLAATSLGLGAGMAPKIAWALENKPRIPVVWIHGL WP_001058323.1 MULTISPECIES: hydrogenase-1 small chain [Proteobacteria] Length: 372\nNumber of predicted TMHs: 2\nExp number of AAs in TMHs: 37.2608499999999\nExp number, first 60 AAs: 19.52923\nTotal prob of N-in: 0.94351\nPOSSIBLE N-term signal sequence\ninside 1 19\nTMhelix 20 39\noutside 40 325\nTMhelix 326 348\ninside 349 372

11331 GCF_000457325.1_Esch_coli_HVH_92_4-5930790_V1 Escherichia coli HVH 92 (4-5930790) Proteobacteria; Gammaproteobacteria; Enterobacterales; Enterobacteriaceae; Escherichia; Escherichia coli MNNEETFYQAMRRQGVTRRSFLKYCSLAATSLGLGAGMAPKIAWALENKPRIPVVWIHGL WP_001058323.1 MULTISPECIES: hydrogenase-1 small chain [Proteobacteria] Length: 372\nNumber of predicted TMHs: 2\nExp number of AAs in TMHs: 37.2608499999999\nExp number, first 60 AAs: 19.52923\nTotal prob of N-in: 0.94351\nPOSSIBLE N-term signal sequence\ninside 1 19\nTMhelix 20 39\noutside 40 325\nTMhelix 326 348\ninside 349 372

11332 GCF_000457325.1_Esch_coli_HVH_92_4-5930790_V1 Escherichia coli HVH 92 (4-5930790) Proteobacteria; Gammaproteobacteria; Enterobacterales; Enterobacteriaceae; Escherichia; Escherichia coli MTGDNTLIHSHGINRRDFMKLCAALAATMGLSSKAAAEMAESVTNPQRPPVIWIGAQECT WP_000145410.1 MULTISPECIES: hydrogenase-2 small chain [Proteobacteria] Length: 372\nNumber of predicted TMHs: 1\nExp number of AAs in TMHs: 23.72226\nExp number, first 60 AAs: 0.22038\nTotal prob of N-in: 0.12994\noutside 1 331\nTMhelix 332 354\ninside 355 372

11333	GCF_000456145.1_Esch_coli_HVH_9_4-6942539_V1	Escherichia coli HVH 9 (4-6942539)
Proteobacteria; Gammaproteobacteria; Enterobacterales; Enterobacteriaceae; Escherichia; Escherichia coli
MTGDNTLIHSHGINRRDFMKLCAALAATMGLSSKAAAEMVESVTNPQRPPVIWIGAQECT	WP_000145423.1
hydrogenase 2 small subunit [Escherichia coli]	Length: 372\nNumber of predicted TMHs: 1\nExp number of AAs in TMHs: 23.78201\nExp number, first 60 AAs: 0.28355\nTotal prob of N-in: 0.13257\noutside 1 331\nTMhelix 332 354\ninside 355 372

11334	GCF_000456145.1_Esch_coli_HVH_9_4-6942539_V1	Escherichia coli HVH 9 (4-6942539)
Proteobacteria; Gammaproteobacteria; Enterobacterales; Enterobacteriaceae; Escherichia; Escherichia coli
MNNEETFYQAMRRQGVTRRSFLKYCSLAATSLGLGAGMAPKIAWALENKPRIPVVWIHGL	WP_001058323.1
MULTISPECIES: hydrogenase-1 small chain [Proteobacteria]	Length: 372\nNumber of predicted TMHs: 2\nExp number of AAs in TMHs: 37.2608499999999\nExp number, first 60 AAs: 19.52923\nTotal prob of N-in: 0.94351\nPOSSIBLE N-term signal sequence\ninside 1 19\nTMhelix 20 39\noutside 40 325\nTMhelix 326 348\ninside 349 372

11335	GCF_000457345.1_Esch_coli_HVH_95_4-6074464_V1	Escherichia coli HVH 95 (4-6074464)
Proteobacteria; Gammaproteobacteria; Enterobacterales; Enterobacteriaceae; Escherichia; Escherichia coli
MNNEETFYQAMRRQGVTRRSFLKYCSLAATSLGLGAGMAPKIAWALENKPRIPVVWIHGL	WP_001058323.1
MULTISPECIES: hydrogenase-1 small chain [Proteobacteria]	Length: 372\nNumber of predicted TMHs: 2\nExp number of AAs in TMHs: 37.2608499999999\nExp number, first 60 AAs: 19.52923\nTotal prob of N-in: 0.94351\nPOSSIBLE N-term signal sequence\ninside 1 19\nTMhelix 20 39\noutside 40 325\nTMhelix 326 348\ninside 349 372

11336	GCF_000457345.1_Esch_coli_HVH_95_4-6074464_V1	Escherichia coli HVH 95 (4-6074464)
Proteobacteria; Gammaproteobacteria; Enterobacterales; Enterobacteriaceae; Escherichia; Escherichia coli
MTGDNTLIHSHGINRRDFMKLCAALAATMGLSSKAAAEMAESVTNPQRPPVIWIGAQECT	WP_000145410.1
MULTISPECIES: hydrogenase-2 small chain [Proteobacteria]	Length: 372\nNumber of predicted TMHs: 1\nExp number of AAs in TMHs: 23.72226\nExp number, first 60 AAs: 0.22038\nTotal prob of N-in: 0.12994\noutside 1 331\nTMhelix 332 354\ninside 355 372

11337	GCF_000457385.1_Esch_coli_HVH_96_4-5934869_V1	Escherichia coli HVH 96 (4-5934869)
Proteobacteria; Gammaproteobacteria; Enterobacterales; Enterobacteriaceae; Escherichia; Escherichia coli
MNNEETFYQAMRRQGVTRRSFLKYCSLAATSLGLGAGMAPKIAWALENKPRIPVVWIHGL	WP_001058323.1
MULTISPECIES: hydrogenase-1 small chain [Proteobacteria]	Length: 372\nNumber of predicted TMHs: 2\nExp number of AAs in TMHs: 37.2608499999999\nExp number, first 60 AAs: 19.52923\nTotal prob of N-in: 0.94351\nPOSSIBLE N-term signal sequence\ninside 1 19\nTMhelix 20 39\noutside 40 325\nTMhelix 326 348\ninside 349 372

11338	GCF_000457385.1_Esch_coli_HVH_96_4-5934869_V1	Escherichia coli HVH 96 (4-5934869)
Proteobacteria; Gammaproteobacteria; Enterobacterales; Enterobacteriaceae; Escherichia; Escherichia coli
MTGDNTLIHSHGINRRDFMKLCAALAATMGLSSKAAAEMAESVTNPQRPPVIWIGAQECT	WP_000145410.1
MULTISPECIES: hydrogenase-2 small chain [Proteobacteria]	Length: 372\nNumber of predicted TMHs: 1\nExp number of AAs in TMHs: 23.72226\nExp number, first 60 AAs: 0.22038\nTotal prob of N-in: 0.12994\noutside 1 331\nTMhelix 332 354\ninside 355 372

11339	GCF_000488035.1_Esch_coli_HVH_98_4-5799287_V1	Escherichia coli HVH 98 (4-5799287)
Proteobacteria; Gammaproteobacteria; Enterobacterales; Enterobacteriaceae; Escherichia; Escherichia coli
MTGDNTLIHSHGINRRDFMKLCAALAATMGLSSKAAAEMAESVTNPQRPPVIWIGAQECT	WP_000145410.1
MULTISPECIES: hydrogenase-2 small chain [Proteobacteria]	Length: 372\nNumber of predicted TMHs: 1\nExp number of AAs in TMHs: 23.72226\nExp number, first 60 AAs: 0.22038\nTotal prob of N-in: 0.12994\noutside 1 331\nTMhelix 332 354\ninside 355 372

11340	GCF_000488035.1_Esch_coli_HVH_98_4-5799287_V1	Escherichia coli HVH 98 (4-5799287)
Proteobacteria; Gammaproteobacteria; Enterobacterales; Enterobacteriaceae; Escherichia; Escherichia coli
MNNEETFYQAMRRQGVTRRSFLKYCSLAATSLGLGAGMAPKIAWALENKPRIPVVWIHGL	WP_001295940.1
MULTISPECIES: hydrogenase [Enterobacteriaceae]	Length: 372\nNumber of predicted TMHs: 2\nExp number of AAs in TMHs: 37.2608899999999\nExp number, first 60 AAs: 19.52923\nTotal prob of N-in: 0.94351\nPOSSIBLE N-term signal sequence\ninside 1 19\nTMhelix 20 39\noutside 40 325\nTMhelix 326 348\ninside 349 372

11341	GCF_000026265.1_ASM2626v1	Escherichia coli IAI1	Proteobacteria; Gammaproteobacteria; Enterobacterales; Enterobacteriaceae; Escherichia; Escherichia coli
MTGDNTLIHSHGINRRDFMKLCAALAATMGLSSKAAAEMAESVTNPQRPPVIWIGAQECT	WP_000145410.1
MULTISPECIES: hydrogenase-2 small chain [Proteobacteria]	Length: 372\nNumber of predicted TMHs: 1\nExp number of AAs in TMHs: 23.72226\nExp number, first 60 AAs: 0.22038\nTotal prob of N-in: 0.12994\noutside 1 331\nTMhelix 332 354\ninside 355 372

11342	GCF_000026265.1_ASM2626v1	Escherichia coli IAI1	Proteobacteria; Gammaproteobacteria; Enterobacterales; Enterobacteriaceae; Escherichia; Escherichia coli
MNNEETFYQAMRRQGVTRRSFLKYCSLAATSLGLGAGMAPKIAWALENKPRIPVVWIHGL	WP_001295940.1
MULTISPECIES: hydrogenase [Enterobacteriaceae]	Length: 372\nNumber of predicted TMHs: 2\nExp number of AAs in TMHs: 37.2608899999999\nExp number, first 60 AAs: 19.52923\nTotal prob of N-in: 0.94351\nPOSSIBLE N-term signal sequence\ninside 1 19\nTMhelix 20 39\noutside 40 325\nTMhelix 326 348\ninside 349 372

11343	GCF_000026345.1_ASM2634v1	Escherichia coli IAI39	Proteobacteria; Gammaproteobacteria; Enterobacterales; Enterobacteriaceae; Escherichia; Escherichia coli
MTGDNTLIHSHGINRRDFMKLCAALAATMGLSSKAAAEMAESVTNPQRPPVIWIGAQECT	YP_002409400.1
hydrogenase 2 small subunit [Escherichia coli IAI39]	Length: 372\nNumber of predicted TMHs: 1\nExp number of AAs in TMHs: 23.72226\nExp number, first 60 AAs: 0.22038\nTotal prob of N-in: 0.12994\noutside 1 331\nTMhelix 332 354\ninside 355 372

11344	GCF_000026345.1_ASM2634v1	Escherichia coli IAI39	Proteobacteria; Gammaproteobacteria; Enterobacterales; Enterobacteriaceae; Escherichia; Escherichia coli
MNNEETFYQAMRRQGVTRRSFLKYCSLAATSLGLGAGMAPKIAWALENKPRIPVVWIHGL	YP_002408137.1
hydrogenase 1, small subunit [Escherichia coli IAI39]	Length: 372\nNumber of predicted TMHs: 2\nExp number of AAs in TMHs: 37.2608899999999\nExp number, first 60 AAs: 19.52923\nTotal prob of N-in: 0.94351\nPOSSIBLE N-term signal sequence\ninside 1 19\nTMhelix 20 39\noutside 40 325\nTMhelix 326 348\ninside 349 372

11345	GCF_000025745.1_ASM2574v1	Escherichia coli IHE3034	Proteobacteria; Gammaproteobacteria; Enterobacterales; Enterobacteriaceae; Escherichia; Escherichia coli
MTGDNTLIHSHGINRRDFMKLCAALAATMGLSSKAAAEMAESVTNPQRPPVIWIGAQECT	WP_000145410.1
MULTISPECIES: hydrogenase-2 small chain [Proteobacteria]	Length: 372\nNumber of predicted TMHs: 1\nExp number of AAs in TMHs: 23.72226\nExp number, first 60 AAs: 0.22038\nTotal prob of N-in: 0.12994\noutside 1 331\nTMhelix 332 354\ninside 355 372

11346	GCF_000414155.2_IMT8073_mix_1	Escherichia coli IMT8073	Proteobacteria; Gammaproteobacteria; Enterobacterales; Enterobacteriaceae; Escherichia; Escherichia coli
MTGDNTLIHSHGINRRDFMKLCAALAATMGLSSKAAAEMAESVTNPQRPPVIWIGAQECT	WP_000145410.1
MULTISPECIES: hydrogenase-2 small chain [Proteobacteria]	Length: 372\nNumber of predicted TMHs: 1\nExp number of AAs in TMHs: 23.72226\nExp number, first 60 AAs: 0.22038\nTotal prob of N-in: 0.12994\noutside 1 331\nTMhelix 332 354\ninside 355 372

11347	GCF_000414155.2_IMT8073_mix_1	Escherichia coli IMT8073	Proteobacteria; Gammaproteobacteria; Enterobacterales; Enterobacteriaceae; Escherichia; Escherichia coli
MNNEETFYQAMRRQGVTRRSFLKYCSLAATSLGLGAGMAPKIAWALENKPRIPVVWIHGL	WP_001058323.1
MULTISPECIES: hydrogenase-1 small chain [Proteobacteria]	Length: 372\nNumber of predicted TMHs: 2\nExp number of AAs in TMHs: 37.2608499999999\nExp number, first 60 AAs: 19.52923\nTotal prob of N-in: 0.94351\nPOSSIBLE N-term signal sequence\ninside 1 19\nTMhelix 20 39\noutside 40 325\nTMhelix 326 348\ninside 349 372

11348	GCF_000513455.1_IS1v1	Escherichia coli IS1	Proteobacteria; Gammaproteobacteria; Enterobacterales; Enterobacteriaceae; Escherichia; Escherichia coli
MNNEETFYQAMRRQGVTRRSFLKYCSLAATSLGLGAGMAPKIAWALENKPRIPVVWIHGL	WP_001058323.1
MULTISPECIES: hydrogenase-1 small chain [Proteobacteria]	Length: 372\nNumber of predicted TMHs: 2\nExp number of AAs in TMHs: 37.2608499999999\nExp number, first 60 AAs: 19.52923\nTotal prob of N-in: 0.94351\nPOSSIBLE N-term signal sequence\ninside 1 19\nTMhelix 20 39\noutside 40 325\nTMhelix 326 348\ninside 349 372

11349 GCF_000529355.1_IS25v1 Escherichia coli IS25 Proteobacteria; Gammaproteobacteria; Enterobacterales; Enterobacteriaceae; Escherichia; Escherichia coli
MTGDNTLIHSHGINRRDFMKLCAALAATMGLSSKAAAEMAESVTNPQRPPVIWIGAQECT WP_000145410.1
MULTISPECIES: hydrogenase-2 small chain [Proteobacteria] Length: 372\nNumber of predicted TMHs: 1\nExp number of AAs in TMHs: 23.72226\nExp number, first 60 AAs: 0.22038\nTotal prob of N-in: 0.12994\noutside 1 331\nTMhelix 332 354\ninside 355 372

11350 GCF_000529355.1_IS25v1 Escherichia coli IS25 Proteobacteria; Gammaproteobacteria; Enterobacterales; Enterobacteriaceae; Escherichia; Escherichia coli
MNNEETFYQAMRRQGVTRRSFLKYCSLAATSLGLGAGMAPKIAWALENKPRIPVVWIHGL WP_001058323.1
MULTISPECIES: hydrogenase-1 small chain [Proteobacteria] Length: 372\nNumber of predicted TMHs: 2\nExp number of AAs in TMHs: 37.2608499999999\nExp number, first 60 AAs: 19.52923\nTotal prob of N-in: 0.94351\nPOSSIBLE N-term signal sequence\ninside 1 19\nTMhelix 20 39\noutside 40 325\nTMhelix 326 348\ninside 349 372

11351 GCF_000529975.1_IS29v1 Escherichia coli IS29 Proteobacteria; Gammaproteobacteria; Enterobacterales; Enterobacteriaceae; Escherichia; Escherichia coli
MTGDNTLIHSHGINRRDFMKLCAALAATMGLSSKAAAEMVESVTNPQRPPVIWIGAQECT WP_000145423.1
hydrogenase 2 small subunit [Escherichia coli] Length: 372\nNumber of predicted TMHs: 1\nExp number of AAs in TMHs: 23.78201\nExp number, first 60 AAs: 0.28355\nTotal prob of N-in: 0.13257\noutside 1 331\nTMhelix 332 354\ninside 355 372

11352 GCF_000529975.1_IS29v1 Escherichia coli IS29 Proteobacteria; Gammaproteobacteria; Enterobacterales; Enterobacteriaceae; Escherichia; Escherichia coli
MNNEETFYQAMRRQGVTRRSFLKYCSLAATSLGLGAGMAPKIAWALENKPRIPVVWIHGL WP_001295940.1
MULTISPECIES: hydrogenase [Enterobacteriaceae] Length: 372\nNumber of predicted TMHs: 2\nExp number of AAs in TMHs: 37.2608899999999\nExp number, first 60 AAs: 19.52923\nTotal prob of N-in: 0.94351\nPOSSIBLE N-term signal sequence\ninside 1 19\nTMhelix 20 39\noutside 40 325\nTMhelix 326 348\ninside 349 372

11353 GCF_000513375.1_IS5 Escherichia coli IS5 Proteobacteria; Gammaproteobacteria; Enterobacterales; Enterobacteriaceae; Escherichia; Escherichia coli
MNNEETFYQAMRRQGVTRRSFLKYCSLAATSLGLGAGMAPKIAWALENKPRIPVVWIHGL WP_001058323.1
MULTISPECIES: hydrogenase-1 small chain [Proteobacteria] Length: 372\nNumber of predicted TMHs: 2\nExp number of AAs in TMHs: 37.2608499999999\nExp number, first 60 AAs: 19.52923\nTotal prob of N-in: 0.94351\nPOSSIBLE N-term signal sequence\ninside 1 19\nTMhelix 20 39\noutside 40 325\nTMhelix 326 348\ninside 349 372

11354 GCF_000513375.1_IS5 Escherichia coli IS5 Proteobacteria; Gammaproteobacteria; Enterobacterales; Enterobacteriaceae; Escherichia; Escherichia coli
MTGDNTLIHSHGINRRDFMKLCAALAATMGLSSKAAAEMAESVTNPQRPPVIWIGAQECT WP_000145410.1
MULTISPECIES: hydrogenase-2 small chain [Proteobacteria] Length: 372\nNumber of predicted TMHs: 1\nExp number of AAs in TMHs: 23.72226\nExp number, first 60 AAs: 0.22038\nTotal prob of N-in: 0.12994\noutside 1 331\nTMhelix 332 354\ninside 355 372

11355 GCF_000689315.1_IS9 Escherichia coli IS9 Proteobacteria; Gammaproteobacteria; Enterobacterales; Enterobacteriaceae; Escherichia; Escherichia coli
MNNEETFYQAMRRQGVTRRSFLKYCSLAATSLGLGAGMAPKIAWALENKPRIPVVWIHGL WP_001295940.1
MULTISPECIES: hydrogenase [Enterobacteriaceae] Length: 372\nNumber of predicted TMHs: 2\nExp number of AAs in TMHs: 37.2608899999999\nExp number, first 60 AAs: 19.52923\nTotal prob of N-in: 0.94351\nPOSSIBLE N-term signal sequence\ninside 1 19\nTMhelix 20 39\noutside 40 325\nTMhelix 326 348\ninside 349 372

11356 GCF_000689315.1_IS9 Escherichia coli IS9 Proteobacteria; Gammaproteobacteria; Enterobacterales; Enterobacteriaceae; Escherichia; Escherichia coli
MTGDNTLIHSHGINRRDFMKLCAALAATMGLSSKAAAEMAESVTNPQRPPVIWIGAQECT WP_000145410.1
MULTISPECIES: hydrogenase-2 small chain [Proteobacteria] Length: 372\nNumber of predicted TMHs: 1\nExp number of AAs in TMHs: 23.72226\nExp number, first 60 AAs: 0.22038\nTotal prob of N-in: 0.12994\noutside 1 331\nTMhelix 332 354\ninside 355 372

11357 GCF_000258865.1_ASM25886v1 Escherichia coli J53 Proteobacteria; Gammaproteobacteria; Enterobacterales; Enterobacteriaceae; Escherichia; Escherichia coli
MTGDNTLIHSHGINRRDFMKLCAALAATMGLSSKAAAEMAESVTNPQRPPVIWIGAQECT WP_000145410.1
MULTISPECIES: hydrogenase-2 small chain [Proteobacteria] Length: 372\nNumber of predicted TMHs: 1\nExp number of AAs in TMHs: 23.72226\nExp number, first 60 AAs: 0.22038\nTotal prob of N-in: 0.12994\noutside 1 331\nTMhelix 332 354\ninside 355 372

11358 GCF_000258865.1_ASM25886v1 Escherichia coli J53 Proteobacteria; Gammaproteobacteria; Enterobacterales; Enterobacteriaceae; Escherichia; Escherichia coli
MNNEETFYQAMRRQGVTRRSFLKYCSLAATSLGLGAGMAPKIAWALENKPRIPVVWIHGL WP_001058323.1
MULTISPECIES: hydrogenase-1 small chain [Proteobacteria] Length: 372\nNumber of predicted TMHs: 2\nExp number of AAs in TMHs: 37.2608499999999\nExp number, first 60 AAs: 19.52923\nTotal prob of N-in: 0.94351\nPOSSIBLE N-term signal sequence\ninside 1 19\nTMhelix 20 39\noutside 40 325\nTMhelix 326 348\ninside 349 372

11359 GCF_000295775.2_ASM29577v2 Escherichia coli J96 Proteobacteria; Gammaproteobacteria; Enterobacterales; Enterobacteriaceae; Escherichia; Escherichia coli
MTGDNTLIHSHGINRRDFMKLCAALAATMGLSSKAAAEMAESVTNPQRPPVIWIGAQECT WP_000145410.1
MULTISPECIES: hydrogenase-2 small chain [Proteobacteria] Length: 372\nNumber of predicted TMHs: 1\nExp number of AAs in TMHs: 23.72226\nExp number, first 60 AAs: 0.22038\nTotal prob of N-in: 0.12994\noutside 1 331\nTMhelix 332 354\ninside 355 372

11360 GCF_000295775.2_ASM29577v2 Escherichia coli J96 Proteobacteria; Gammaproteobacteria; Enterobacterales; Enterobacteriaceae; Escherichia; Escherichia coli
MNNEETFYQAMRRQGVTRRSFLKYCSLAATSLGLGAGMAPKIAWALENKPRIPVVWIHGL WP_001058323.1
MULTISPECIES: hydrogenase-1 small chain [Proteobacteria] Length: 372\nNumber of predicted TMHs: 2\nExp number of AAs in TMHs: 37.2608499999999\nExp number, first 60 AAs: 19.52923\nTotal prob of N-in: 0.94351\nPOSSIBLE N-term signal sequence\ninside 1 19\nTMhelix 20 39\noutside 40 325\nTMhelix 326 348\ninside 349 372

11361 GCF_000194355.1_ASM19435v2 Escherichia coli JB1-95 Proteobacteria; Gammaproteobacteria; Enterobacterales; Enterobacteriaceae; Escherichia; Escherichia coli
MNNEETFYQAMRRQGVTRRSFLKYCSLAATSLGLGAGMAPKIAWALENKPRIPVVWIHGL WP_001058323.1
MULTISPECIES: hydrogenase-1 small chain [Proteobacteria] Length: 372\nNumber of predicted TMHs: 2\nExp number of AAs in TMHs: 37.2608499999999\nExp number, first 60 AAs: 19.52923\nTotal prob of N-in: 0.94351\nPOSSIBLE N-term signal sequence\ninside 1 19\nTMhelix 20 39\noutside 40 325\nTMhelix 326 348\ninside 349 372

11362 GCF_000194355.1_ASM19435v2 Escherichia coli JB1-95 Proteobacteria; Gammaproteobacteria; Enterobacterales; Enterobacteriaceae; Escherichia; Escherichia coli
MTGDNTLIHSHGINRRDFMKLCAALAATMGLSSKAAAEMAESVTNPQRPPVIWIGAQECT WP_000145410.1
MULTISPECIES: hydrogenase-2 small chain [Proteobacteria] Length: 372\nNumber of predicted TMHs: 1\nExp number of AAs in TMHs: 23.72226\nExp number, first 60 AAs: 0.22038\nTotal prob of N-in: 0.12994\noutside 1 331\nTMhelix 332 354\ninside 355 372

11363 GCF_000493755.1_ASM49375v1 Escherichia coli JJ1886 Proteobacteria; Gammaproteobacteria; Enterobacterales; Enterobacteriaceae; Escherichia; Escherichia coli
MNNEETFYQAMRRQGVTRRSFLKYCSLAATSLGLGAGMAPKIAWALENKPRIPVVWIHGL WP_001295940.1
MULTISPECIES: hydrogenase [Enterobacteriaceae] Length: 372\nNumber of predicted TMHs: 2\nExp number of AAs in TMHs: 37.2608899999999\nExp number, first 60 AAs: 19.52923\nTotal prob of N-in: 0.94351\nPOSSIBLE N-term signal sequence\ninside 1 19\nTMhelix 20 39\noutside 40 325\nTMhelix 326 348\ninside 349 372

11364 GCF_000493755.1_ASM49375v1 Escherichia coli JJ1886 Proteobacteria; Gammaproteobacteria; Enterobacterales; Enterobacteriaceae; Escherichia; Escherichia coli
MTGDNTLIHSHGINRRDFMKLCAALAATMGLSSKAAAEMVESVTNPQRPPVIWIGAQECT WP_000145423.1
hydrogenase 2 small subunit [Escherichia coli] Length: 372\nNumber of predicted TMHs: 1\nExp number of AAs in TMHs: 23.78201\nExp number, first 60 AAs: 0.28355\nTotal prob of N-in: 0.13257\noutside 1 331\nTMhelix 332 354\ninside 355 372

11365 GCF_001593565.1_ASM159356v1 Escherichia coli JJ1887 Proteobacteria; Gammaproteobacteria; Enterobacterales; Enterobacteriaceae; Escherichia; Escherichia coli
MNNEETFYQAMRRQGVTRRSFLKYCSLAATSLGLGAGMAPKIAWALENKPRIPVVWIHGL WP_001295940.1
MULTISPECIES: hydrogenase [Enterobacteriaceae] Length: 372\nNumber of predicted TMHs: 2\nExp number of AAs in TMHs: 37.2608899999999\nExp number, first 60 AAs: 19.52923\nTotal prob of N-in: 0.94351\nPOSSIBLE N-term signal sequence\ninside 1 19\nTMhelix 20 39\noutside 40 325\nTMhelix 326 348\ninside 349 372

11366 GCF_001593565.1_ASM159356v1 Escherichia coli JJ1887 Proteobacteria; Gammaproteobacteria; Enterobacterales; Enterobacteriaceae; Escherichia; Escherichia coli
MTGDNTLIHSHGINRRDFMKLCAALAATMGLSSKAAAEMVESVTNPQRPPVIWIGAQECT WP_000145423.1
hydrogenase 2 small subunit [Escherichia coli] Length: 372\nNumber of predicted TMHs: 1\nExp number of AAs in TMHs: 23.78201\nExp number, first 60 AAs: 0.28355\nTotal prob of N-in: 0.13257\noutside 1 331\nTMhelix 332 354\ninside 355 372

11367 GCF_000809145.1_ASM80914v1 Escherichia coli JT5 Proteobacteria; Gammaproteobacteria; Enterobacterales; Enterobacteriaceae; Escherichia; Escherichia coli
MNNEETFYQAMRRQGVTRRSFLKYCSLAATSLGLGAGMAPKIAWALENKPRIPVVWIHGL WP_001295940.1
MULTISPECIES: hydrogenase [Enterobacteriaceae] Length: 372\nNumber of predicted TMHs: 2\nExp number of AAs in TMHs: 37.2608899999999\nExp number, first 60 AAs: 19.52923\nTotal prob of N-in: 0.94351\nPOSSIBLE N-term signal sequence\ninside 1 19\nTMhelix 20 39\noutside 40 325\nTMhelix 326 348\ninside 349 372

11368 GCF_000809145.1_ASM80914v1 Escherichia coli JT5 Proteobacteria; Gammaproteobacteria; Enterobacterales; Enterobacteriaceae; Escherichia; Escherichia coli
MTGDNTLIHSHGINRRDFMKLCAALAATMGLSSKAAAEMAESVTNPQRPPVIWIGAQECT WP_000145410.1
MULTISPECIES: hydrogenase-2 small chain [Proteobacteria] Length: 372\nNumber of predicted TMHs: 1\nExp number of AAs in TMHs: 23.72226\nExp number, first 60 AAs: 0.22038\nTotal prob of N-in: 0.12994\noutside 1 331\nTMhelix 332 354\ninside 355 372

11369 GCF_000355235.2_ASM35523v2 Escherichia coli Jurua 18/11 Proteobacteria; Gammaproteobacteria; Enterobacterales; Enterobacteriaceae; Escherichia; Escherichia coli
MNNEETFYQAMRRQGVTRRSFLKYCSLAATSLGLGAGMAPKIAWALENKPRIPVVWIHGL WP_001058320.1
hydrogenase [Escherichia coli] Length: 372\nNumber of predicted TMHs: 2\nExp number of AAs in TMHs: 37.26082\nExp number, first 60 AAs: 19.52923\nTotal prob of N-in: 0.94351\nPOSSIBLE N-term signal sequence\ninside 1 19\nTMhelix 20 39\noutside 40 325\nTMhelix 326 348\ninside 349 372

11370 GCF_000355235.2_ASM35523v2 Escherichia coli Jurua 18/11 Proteobacteria; Gammaproteobacteria; Enterobacterales; Enterobacteriaceae; Escherichia; Escherichia coli
MTGDNTLIHSHGINRRDFMKLCAALAATMGLSSKAAAEMAESVTNPQRPPVIWIGAQECT WP_000145410.1
MULTISPECIES: hydrogenase-2 small chain [Proteobacteria] Length: 372\nNumber of predicted TMHs: 1\nExp number of AAs in TMHs: 23.72226\nExp number, first 60 AAs: 0.22038\nTotal prob of N-in: 0.12994\noutside 1 331\nTMhelix 332 354\ninside 355 372

11371 GCF_000355255.2_ASM35525v2 Escherichia coli Jurua 20/10 Proteobacteria; Gammaproteobacteria; Enterobacterales; Enterobacteriaceae; Escherichia; Escherichia coli
MTGDNTLIHSHGINRRDFMKLCVALAATMGLSSKAAAEMAESVTNPQRPPVIWIGAQECT WP_000145425.1
hydrogenase 2 small subunit [Escherichia coli] Length: 372\nNumber of predicted TMHs: 1\nExp number of AAs in TMHs: 23.94354\nExp number, first 60 AAs: 0.45405\nTotal prob of N-in: 0.13997\noutside 1 331\nTMhelix 332 354\ninside 355 372

11372 GCF_000355255.2_ASM35525v2 Escherichia coli Jurua 20/10 Proteobacteria; Gammaproteobacteria; Enterobacterales; Enterobacteriaceae; Escherichia; Escherichia coli
MNNEETFYQAMRRQGVTRRSFLKYCSLAATSLGLGAGMAPKIAWALENKPRIPVVWIHGL WP_001295940.1
MULTISPECIES: hydrogenase [Enterobacteriaceae] Length: 372\nNumber of predicted TMHs: 2\nExp number of AAs in TMHs: 37.2608899999999\nExp number, first 60 AAs: 19.52923\nTotal prob of N-in: 0.94351\nPOSSIBLE N-term signal sequence\ninside 1 19\nTMhelix 20 39\noutside 40 325\nTMhelix 326 348\ninside 349 372

11373 GCF_000607285.1_Esch_coli_K02_V1 Escherichia coli K02Proteobacteria; Gammaproteobacteria; Enterobacterales; Enterobacteriaceae; Escherichia; Escherichia coli

MNNEETFYQAMRRQGVTRRSFLKYCSLAATSLGLGAGMAPKIAWALENKPRIPVVWIHGL WP_001058323.1 MULTISPECIES: hydrogenase-1 small chain [Proteobacteria] Length: 372\nNumber of predicted TMHs: 2\nExp number of AAs in TMHs: 37.2608499999999\nExp number, first 60 AAs: 19.52923\nTotal prob of N-in: 0.94351\nPOSSIBLE N-term signal sequence\ninside 1 19\nTMhelix 20 39\noutside 40 325\nTMhelix 326 348\ninside 349 372

11374 GCF_000607285.1_Esch_coli_K02_V1 Escherichia coli K02 Proteobacteria; Gammaproteobacteria; Enterobacterales; Enterobacteriaceae; Escherichia; Escherichia coli MTGDNTLIHSHGINRRDFMKLCAALAATMGLSSKAAAEMVESVTNPQRPPVIWIGAQECT WP_000145423.1 hydrogenase 2 small subunit [Escherichia coli] Length: 372\nNumber of predicted TMHs: 1\nExp number of AAs in TMHs: 23.78201\nExp number, first 60 AAs: 0.28355\nTotal prob of N-in: 0.13257\noutside 1 331\nTMhelix 332 354\ninside 355 372

11375 GCF_000471505.1_E_coliK1-1.0 Escherichia coli K1 Proteobacteria; Gammaproteobacteria; Enterobacterales; Enterobacteriaceae; Escherichia; Escherichia coli MTGDNTLIHSHGINRRDFMKLCAALAATMGLSSKAAAEMAESVTNPQRPPVIWIGAQECT WP_000145410.1 MULTISPECIES: hydrogenase-2 small chain [Proteobacteria] Length: 372\nNumber of predicted TMHs: 1\nExp number of AAs in TMHs: 23.72226\nExp number, first 60 AAs: 0.22038\nTotal prob of N-in: 0.12994\noutside 1 331\nTMhelix 332 354\ninside 355 372

11376 GCF_000471505.1_E_coliK1-1.0 Escherichia coli K1 Proteobacteria; Gammaproteobacteria; Enterobacterales; Enterobacteriaceae; Escherichia; Escherichia coli MNNEETFYQAMRRQGVTRRSFLKYCSLAATSLGLGAGMAPKIAWALENKPRIPVVWIHGL WP_001301613.1 hydrogenase [Escherichia coli] Length: 372\nNumber of predicted TMHs: 2\nExp number of AAs in TMHs: 37.46598\nExp number, first 60 AAs: 19.53139\nTotal prob of N-in: 0.94364\nPOSSIBLE N-term signal sequence\ninside 1 19\nTMhelix 20 39\noutside 40 325\nTMhelix 326 348\ninside 349 372

11377 GCF_000742555.1_E_coli_k1_929 Escherichia coli K1 Proteobacteria; Gammaproteobacteria; Enterobacterales; Enterobacteriaceae; Escherichia; Escherichia coli MTGDNTLIHSHGINRRDFMKLCAALAATMGLSSKAAAEMAESVTNPQRPPVIWIGAQECT WP_000145410.1 MULTISPECIES: hydrogenase-2 small chain [Proteobacteria] Length: 372\nNumber of predicted TMHs: 1\nExp number of AAs in TMHs: 23.72226\nExp number, first 60 AAs: 0.22038\nTotal prob of N-in: 0.12994\noutside 1 331\nTMhelix 332 354\ninside 355 372

11378 GCF_000742575.1_E_coli_k1_943 Escherichia coli K1 Proteobacteria; Gammaproteobacteria; Enterobacterales; Enterobacteriaceae; Escherichia; Escherichia coli MTGDNTLIHSHGINRRDFMKLCAALAATMGLSSKAAAEMAESVTNPQRPPVIWIGAQECT WP_000145410.1 MULTISPECIES: hydrogenase-2 small chain [Proteobacteria] Length: 372\nNumber of predicted TMHs: 1\nExp number of AAs in TMHs: 23.72226\nExp number, first 60 AAs: 0.22038\nTotal prob of N-in: 0.12994\noutside 1 331\nTMhelix 332 354\ninside 355 372

11379 GCF_000742505.1_E_coli_k1_910 Escherichia coli K1 Proteobacteria; Gammaproteobacteria; Enterobacterales; Enterobacteriaceae; Escherichia; Escherichia coli MTGDNTLIHSHGINRRDFMKLCAALAATMGLSSKAAAEMAESVTNPQRPPVIWIGAQECT WP_000145410.1 MULTISPECIES: hydrogenase-2 small chain [Proteobacteria] Length: 372\nNumber of predicted TMHs: 1\nExp number of AAs in TMHs: 23.72226\nExp number, first 60 AAs: 0.22038\nTotal prob of N-in: 0.12994\noutside 1 331\nTMhelix 332 354\ninside 355 372

11380 GCF_000974505.1_ASM97450v1 Escherichia coli K-12 Proteobacteria; Gammaproteobacteria; Enterobacterales; Enterobacteriaceae; Escherichia; Escherichia coli MTGDNTLIHSHGINRRDFMKLCAALAATMGLSSKAAAEMAESVTNPQRPPVIWIGAQECT WP_000145410.1 MULTISPECIES: hydrogenase-2 small chain [Proteobacteria] Length: 372\nNumber of predicted TMHs: 1\nExp number of AAs in TMHs: 23.72226\nExp number, first 60 AAs: 0.22038\nTotal prob of N-in: 0.12994\noutside 1 331\nTMhelix 332 354\ninside 355 372

11381 GCF_001021005.2_ASM102100v2 Escherichia coli K-12 Proteobacteria; Gammaproteobacteria; Enterobacterales; Enterobacteriaceae; Escherichia; Escherichia coli MTGDNTLIHSHGINRRDFMKLCAALAATMGLSSKAAAEMAESVTNPQRPPVIWIGAQECT WP_000145410.1 MULTISPECIES: hydrogenase-2 small chain [Proteobacteria] Length: 372\nNumber of predicted TMHs: 1\nExp number of AAs in TMHs: 23.72226\nExp number, first 60 AAs: 0.22038\nTotal prob of N-in: 0.12994\noutside 1 331\nTMhelix 332 354\ninside 355 372

11382 GCF_001021005.2_ASM102100v2 Escherichia coli K-12 Proteobacteria; Gammaproteobacteria; Enterobacterales; Enterobacteriaceae; Escherichia; Escherichia coli
MNNEETFYQAMRRQGVTRRSFLKYCSLAATSLGLGAGMAPKIAWALENKPRIPVVWIHGL WP_001058323.1
MULTISPECIES: hydrogenase-1 small chain [Proteobacteria] Length: 372\nNumber of predicted TMHs: 2\nExp number of AAs in TMHs: 37.2608499999999\nExp number, first 60 AAs: 19.52923\nTotal prob of N-in: 0.94351\nPOSSIBLE N-term signal sequence\ninside 1 19\nTMhelix 20 39\noutside 40 325\nTMhelix 326 348\ninside 349 372

11383 GCF_001559675.1_ASM155967v1 Escherichia coli K-12 Proteobacteria; Gammaproteobacteria; Enterobacterales; Enterobacteriaceae; Escherichia; Escherichia coli
MTGDNTLIHSHGINRRDFMKLCAALAATMGLSSKAAAEMAESVTNPQRPPVIWIGAQECT WP_000145410.1
MULTISPECIES: hydrogenase-2 small chain [Proteobacteria] Length: 372\nNumber of predicted TMHs: 1\nExp number of AAs in TMHs: 23.72226\nExp number, first 60 AAs: 0.22038\nTotal prob of N-in: 0.12994\noutside 1 331\nTMhelix 332 354\ninside 355 372

11384 GCF_000974465.1_ASM97446v1 Escherichia coli K-12 Proteobacteria; Gammaproteobacteria; Enterobacterales; Enterobacteriaceae; Escherichia; Escherichia coli
MTGDNTLIHSHGINRRDFMKLCAALAATMGLSSKAAAEMAESVTNPQRPPVIWIGAQECT WP_000145410.1
MULTISPECIES: hydrogenase-2 small chain [Proteobacteria] Length: 372\nNumber of predicted TMHs: 1\nExp number of AAs in TMHs: 23.72226\nExp number, first 60 AAs: 0.22038\nTotal prob of N-in: 0.12994\noutside 1 331\nTMhelix 332 354\ninside 355 372

11385 GCF_000974465.1_ASM97446v1 Escherichia coli K-12 Proteobacteria; Gammaproteobacteria; Enterobacterales; Enterobacteriaceae; Escherichia; Escherichia coli
MNNEETFYQAMRRQGVTRRSFLKYCSLAATSLGLGAGMAPKIAWALENKPRIPVVWIHGL WP_001058323.1
MULTISPECIES: hydrogenase-1 small chain [Proteobacteria] Length: 372\nNumber of predicted TMHs: 2\nExp number of AAs in TMHs: 37.2608499999999\nExp number, first 60 AAs: 19.52923\nTotal prob of N-in: 0.94351\nPOSSIBLE N-term signal sequence\ninside 1 19\nTMhelix 20 39\noutside 40 325\nTMhelix 326 348\ninside 349 372

11386 GCF_000974535.1_ASM97453v1 Escherichia coli K-12 Proteobacteria; Gammaproteobacteria; Enterobacterales; Enterobacteriaceae; Escherichia; Escherichia coli
MTGDNTLIHSHGINRRDFMKLCAALAATMGLSSKAAAEMAESVTNPQRPPVIWIGAQECT WP_000145410.1
MULTISPECIES: hydrogenase-2 small chain [Proteobacteria] Length: 372\nNumber of predicted TMHs: 1\nExp number of AAs in TMHs: 23.72226\nExp number, first 60 AAs: 0.22038\nTotal prob of N-in: 0.12994\noutside 1 331\nTMhelix 332 354\ninside 355 372

11387 GCF_000974535.1_ASM97453v1 Escherichia coli K-12 Proteobacteria; Gammaproteobacteria; Enterobacterales; Enterobacteriaceae; Escherichia; Escherichia coli
MNNEETFYQAMRRQGVTRRSFLKYCSLAATSLGLGAGMAPKIAWALENKPRIPVVWIHGL WP_001058323.1
MULTISPECIES: hydrogenase-1 small chain [Proteobacteria] Length: 372\nNumber of predicted TMHs: 2\nExp number of AAs in TMHs: 37.2608499999999\nExp number, first 60 AAs: 19.52923\nTotal prob of N-in: 0.94351\nPOSSIBLE N-term signal sequence\ninside 1 19\nTMhelix 20 39\noutside 40 325\nTMhelix 326 348\ninside 349 372

11388 GCF_000974825.1_ASM97482v1 Escherichia coli K-12 Proteobacteria; Gammaproteobacteria; Enterobacterales; Enterobacteriaceae; Escherichia; Escherichia coli
MTGDNTLIHSHGINRRDFMKLCAALAATMGLSSKAAAEMAESVTNPQRPPVIWIGAQECT WP_000145410.1
MULTISPECIES: hydrogenase-2 small chain [Proteobacteria] Length: 372\nNumber of predicted TMHs: 1\nExp number of AAs in TMHs: 23.72226\nExp number, first 60 AAs: 0.22038\nTotal prob of N-in: 0.12994\noutside 1 331\nTMhelix 332 354\ninside 355 372

11389 GCF_000974825.1_ASM97482v1 Escherichia coli K-12 Proteobacteria; Gammaproteobacteria; Enterobacterales; Enterobacteriaceae; Escherichia; Escherichia coli
MNNEETFYQAMRRQGVTRRSFLKYCSLAATSLGLGAGMAPKIAWALENKPRIPVVWIHGL WP_001058323.1
MULTISPECIES: hydrogenase-1 small chain [Proteobacteria] Length: 372\nNumber of predicted TMHs: 2\nExp number of AAs in TMHs: 37.2608499999999\nExp number, first 60 AAs: 19.52923\nTotal prob of N-in: 0.94351\nPOSSIBLE N-term signal sequence\ninside 1 19\nTMhelix 20 39\noutside 40 325\nTMhelix 326 348\ninside 349 372

11390 GCF_000974865.1_ASM97486v1 Escherichia coli K-12 Proteobacteria; Gammaproteobacteria; Enterobacterales; Enterobacteriaceae; Escherichia; Escherichia coli MNNEETFYQAMRRQGVTRRSFLKYCSLAATSLGLGAGMAPKIAWALENKPRIPVVWIHGL WP_001058323.1 MULTISPECIES: hydrogenase-1 small chain [Proteobacteria] Length: 372\nNumber of predicted TMHs: 2\nExp number of AAs in TMHs: 37.2608499999999\nExp number, first 60 AAs: 19.52923\nTotal prob of N-in: 0.94351\nPOSSIBLE N-term signal sequence\ninside 1 19\nTMhelix 20 39\noutside 40 325\nTMhelix 326 348\ninside 349 372

11391 GCF_001559675.1_ASM155967v1 Escherichia coli K-12 Proteobacteria; Gammaproteobacteria; Enterobacterales; Enterobacteriaceae; Escherichia; Escherichia coli MNNEETFYQAMRRQGVTRRSFLKYCSLAATSLGLGAGMAPKIAWALENKPRIPVVWIHGL WP_001058323.1 MULTISPECIES: hydrogenase-1 small chain [Proteobacteria] Length: 372\nNumber of predicted TMHs: 2\nExp number of AAs in TMHs: 37.2608499999999\nExp number, first 60 AAs: 19.52923\nTotal prob of N-in: 0.94351\nPOSSIBLE N-term signal sequence\ninside 1 19\nTMhelix 20 39\noutside 40 325\nTMhelix 326 348\ninside 349 372

11392 GCF_000974865.1_ASM97486v1 Escherichia coli K-12 Proteobacteria; Gammaproteobacteria; Enterobacterales; Enterobacteriaceae; Escherichia; Escherichia coli MTGDNTLIHSHGINRRDFMKLCAALAATMGLSSKAAAEMAESVTNPQRPPVIWIGAQECT WP_000145410.1 MULTISPECIES: hydrogenase-2 small chain [Proteobacteria] Length: 372\nNumber of predicted TMHs: 1\nExp number of AAs in TMHs: 23.72226\nExp number, first 60 AAs: 0.22038\nTotal prob of N-in: 0.12994\noutside 1 331\nTMhelix 332 354\ninside 355 372

11393 GCF_000800765.1_ASM80076v1 Escherichia coli K-12 Proteobacteria; Gammaproteobacteria; Enterobacterales; Enterobacteriaceae; Escherichia; Escherichia coli MTGDNTLIHSHGINRRDFMKLCAALAATMGLSSKAAAEMAESVTNPQRPPVIWIGAQECT WP_000145410.1 MULTISPECIES: hydrogenase-2 small chain [Proteobacteria] Length: 372\nNumber of predicted TMHs: 1\nExp number of AAs in TMHs: 23.72226\nExp number, first 60 AAs: 0.22038\nTotal prob of N-in: 0.12994\noutside 1 331\nTMhelix 332 354\ninside 355 372

11394 GCF_000800765.1_ASM80076v1 Escherichia coli K-12 Proteobacteria; Gammaproteobacteria; Enterobacterales; Enterobacteriaceae; Escherichia; Escherichia coli MNNEETFYQAMRRQGVTRRSFLKYCSLAATSLGLGAGMAPKIAWALENKPRIPVVWIHGL WP_001058323.1 MULTISPECIES: hydrogenase-1 small chain [Proteobacteria] Length: 372\nNumber of predicted TMHs: 2\nExp number of AAs in TMHs: 37.2608499999999\nExp number, first 60 AAs: 19.52923\nTotal prob of N-in: 0.94351\nPOSSIBLE N-term signal sequence\ninside 1 19\nTMhelix 20 39\noutside 40 325\nTMhelix 326 348\ninside 349 372

11395 GCF_000981485.1_EcoliK12AG100 Escherichia coli K-12 Proteobacteria; Gammaproteobacteria; Enterobacterales; Enterobacteriaceae; Escherichia; Escherichia coli MNNEETFYQAMRRQGVTRRSFLKYCSLAATSLGLGAGMAPKIAWALENKPRIPVVWIHGL WP_001058323.1 MULTISPECIES: hydrogenase-1 small chain [Proteobacteria] Length: 372\nNumber of predicted TMHs: 2\nExp number of AAs in TMHs: 37.2608499999999\nExp number, first 60 AAs: 19.52923\nTotal prob of N-in: 0.94351\nPOSSIBLE N-term signal sequence\ninside 1 19\nTMhelix 20 39\noutside 40 325\nTMhelix 326 348\ninside 349 372

11396 GCF_001020945.2_ASM102094v2 Escherichia coli K-12 Proteobacteria; Gammaproteobacteria; Enterobacterales; Enterobacteriaceae; Escherichia; Escherichia coli MNNEETFYQAMRRQGVTRRSFLKYCSLAATSLGLGAGMAPKIAWALENKPRIPVVWIHGL WP_001058323.1 MULTISPECIES: hydrogenase-1 small chain [Proteobacteria] Length: 372\nNumber of predicted TMHs: 2\nExp number of AAs in TMHs: 37.2608499999999\nExp number, first 60 AAs: 19.52923\nTotal prob of N-in: 0.94351\nPOSSIBLE N-term signal sequence\ninside 1 19\nTMhelix 20 39\noutside 40 325\nTMhelix 326 348\ninside 349 372

11397 GCF_000981485.1_EcoliK12AG100 Escherichia coli K-12 Proteobacteria; Gammaproteobacteria; Enterobacterales; Enterobacteriaceae; Escherichia; Escherichia coli MTGDNTLIHSHGINRRDFMKLCAALAATMGLSSKAAAEMAESVTNPQRPPVIWIGAQECT WP_000145410.1 MULTISPECIES: hydrogenase-2 small chain [Proteobacteria] Length: 372\nNumber of predicted TMHs: 1\nExp number of AAs in TMHs: 23.72226\nExp number, first 60 AAs: 0.22038\nTotal prob of N-in: 0.12994\noutside 1 331\nTMhelix 332 354\ninside 355 372

11398 GCF_001020945.2_ASM102094v2 Escherichia coli K-12 Proteobacteria; Gammaproteobacteria; Enterobacterales; Enterobacteriaceae; Escherichia; Escherichia coli
MTGDNTLIHSHGINRRDFMKLCAALAATMGLSSKAAAEMAESVTNPQRPPVIWIGAQECT WP_000145410.1
MULTISPECIES: hydrogenase-2 small chain [Proteobacteria] Length: 372\nNumber of predicted TMHs: 1\nExp number of AAs in TMHs: 23.72226\nExp number, first 60 AAs: 0.22038\nTotal prob of N-in: 0.12994\noutside 1 331\nTMhelix 332 354\ninside 355 372

11399 GCF_001559655.1_ASM155965v1 Escherichia coli K-12 Proteobacteria; Gammaproteobacteria; Enterobacterales; Enterobacteriaceae; Escherichia; Escherichia coli
MTGDNTLIHSHGINRRDFMKLCAALAATMGLSSKAAAEMAESVTNPQRPPVIWIGAQECT WP_000145410.1
MULTISPECIES: hydrogenase-2 small chain [Proteobacteria] Length: 372\nNumber of predicted TMHs: 1\nExp number of AAs in TMHs: 23.72226\nExp number, first 60 AAs: 0.22038\nTotal prob of N-in: 0.12994\noutside 1 331\nTMhelix 332 354\ninside 355 372

11400 GCF_001559655.1_ASM155965v1 Escherichia coli K-12 Proteobacteria; Gammaproteobacteria; Enterobacterales; Enterobacteriaceae; Escherichia; Escherichia coli
MNNEETFYQAMRRQGVTRRSFLKYCSLAATSLGLGAGMAPKIAWALENKPRIPVVWIHGL WP_001058323.1
MULTISPECIES: hydrogenase-1 small chain [Proteobacteria] Length: 372\nNumber of predicted TMHs: 2\nExp number of AAs in TMHs: 37.2608499999999\nExp number, first 60 AAs: 19.52923\nTotal prob of N-in: 0.94351\nPOSSIBLE N-term signal sequence\ninside 1 19\nTMhelix 20 39\noutside 40 325\nTMhelix 326 348\ninside 349 372

11401 GCF_000974885.1_ASM97488v1 Escherichia coli K-12 Proteobacteria; Gammaproteobacteria; Enterobacterales; Enterobacteriaceae; Escherichia; Escherichia coli
MTGDNTLIHSHGINRRDFMKLCAALAATMGLSSKAAAEMAESVTNPQRPPVIWIGAQECT WP_000145410.1
MULTISPECIES: hydrogenase-2 small chain [Proteobacteria] Length: 372\nNumber of predicted TMHs: 1\nExp number of AAs in TMHs: 23.72226\nExp number, first 60 AAs: 0.22038\nTotal prob of N-in: 0.12994\noutside 1 331\nTMhelix 332 354\ninside 355 372

11402 GCF_000974505.1_ASM97450v1 Escherichia coli K-12 Proteobacteria; Gammaproteobacteria; Enterobacterales; Enterobacteriaceae; Escherichia; Escherichia coli
MNNEETFYQAMRRQGVTRRSFLKYCSLAATSLGLGAGMAPKIAWALENKPRIPVVWIHGL WP_001058323.1
MULTISPECIES: hydrogenase-1 small chain [Proteobacteria] Length: 372\nNumber of predicted TMHs: 2\nExp number of AAs in TMHs: 37.2608499999999\nExp number, first 60 AAs: 19.52923\nTotal prob of N-in: 0.94351\nPOSSIBLE N-term signal sequence\ninside 1 19\nTMhelix 20 39\noutside 40 325\nTMhelix 326 348\ninside 349 372

11403 GCF_000974885.1_ASM97488v1 Escherichia coli K-12 Proteobacteria; Gammaproteobacteria; Enterobacterales; Enterobacteriaceae; Escherichia; Escherichia coli
MNNEETFYQAMRRQGVTRRSFLKYCSLAATSLGLGAGMAPKIAWALENKPRIPVVWIHGL WP_001058323.1
MULTISPECIES: hydrogenase-1 small chain [Proteobacteria] Length: 372\nNumber of predicted TMHs: 2\nExp number of AAs in TMHs: 37.2608499999999\nExp number, first 60 AAs: 19.52923\nTotal prob of N-in: 0.94351\nPOSSIBLE N-term signal sequence\ninside 1 19\nTMhelix 20 39\noutside 40 325\nTMhelix 326 348\ninside 349 372

11404 GCF_000974405.1_ASM97440v1 Escherichia coli K-12 Proteobacteria; Gammaproteobacteria; Enterobacterales; Enterobacteriaceae; Escherichia; Escherichia coli
MTGDNTLIHSHGINRRDFMKLCAALAATMGLSSKAAAEMAESVTNPQRPPVIWIGAQECT WP_000145410.1
MULTISPECIES: hydrogenase-2 small chain [Proteobacteria] Length: 372\nNumber of predicted TMHs: 1\nExp number of AAs in TMHs: 23.72226\nExp number, first 60 AAs: 0.22038\nTotal prob of N-in: 0.12994\noutside 1 331\nTMhelix 332 354\ninside 355 372

11405 GCF_000974405.1_ASM97440v1 Escherichia coli K-12 Proteobacteria; Gammaproteobacteria; Enterobacterales; Enterobacteriaceae; Escherichia; Escherichia coli
MNNEETFYQAMRRQGVTRRSFLKYCSLAATSLGLGAGMAPKIAWALENKPRIPVVWIHGL WP_001058323.1
MULTISPECIES: hydrogenase-1 small chain [Proteobacteria] Length: 372\nNumber of predicted TMHs: 2\nExp number of AAs in TMHs: 37.2608499999999\nExp number, first 60 AAs: 19.52923\nTotal prob of N-in: 0.94351\nPOSSIBLE N-term signal sequence\ninside 1 19\nTMhelix 20 39\noutside 40 325\nTMhelix 326 348\ninside 349 372

11406 GCF_000974575.1_ASM97457v1 Escherichia coli K-12 Proteobacteria; Gammaproteobacteria; Enterobacterales; Enterobacteriaceae; Escherichia; Escherichia coli
MTGDNTLIHSHGINRRDFMKLCAALAATMGLSSKAAAEMAESVTNPQRPPVIWIGAQECT WP_000145410.1
MULTISPECIES: hydrogenase-2 small chain [Proteobacteria] Length: 372\nNumber of predicted TMHs: 1\nExp number of AAs in TMHs: 23.72226\nExp number, first 60 AAs: 0.22038\nTotal prob of N-in: 0.12994\noutside 1 331\nTMhelix 332 354\ninside 355 372

11407 GCF_000974575.1_ASM97457v1 Escherichia coli K-12 Proteobacteria; Gammaproteobacteria; Enterobacterales; Enterobacteriaceae; Escherichia; Escherichia coli
MNNEETFYQAMRRQGVTRRSFLKYCSLAATSLGLGAGMAPKIAWALENKPRIPVVWIHGL WP_001058323.1
MULTISPECIES: hydrogenase-1 small chain [Proteobacteria] Length: 372\nNumber of predicted TMHs: 2\nExp number of AAs in TMHs: 37.2608499999999\nExp number, first 60 AAs: 19.52923\nTotal prob of N-in: 0.94351\nPOSSIBLE N-term signal sequence\ninside 1 19\nTMhelix 20 39\noutside 40 325\nTMhelix 326 348\ninside 349 372

11408 GCF_000471525.1_E_coliK2-1.0 Escherichia coli K2 Proteobacteria; Gammaproteobacteria; Enterobacterales; Enterobacteriaceae; Escherichia; Escherichia coli
MNNEETFYQAMRRQGVTRRSFLKYCSLAATSLGLGAGMAPKIAWALENKPRIPVVWIHGL WP_001301613.1
hydrogenase [Escherichia coli] Length: 372\nNumber of predicted TMHs: 2\nExp number of AAs in TMHs: 37.46598\nExp number, first 60 AAs: 19.53139\nTotal prob of N-in: 0.94364\nPOSSIBLE N-term signal sequence\ninside 1 19\nTMhelix 20 39\noutside 40 325\nTMhelix 326 348\ninside 349 372

11409 GCF_000471525.1_E_coliK2-1.0 Escherichia coli K2 Proteobacteria; Gammaproteobacteria; Enterobacterales; Enterobacteriaceae; Escherichia; Escherichia coli
MTGDNTLIHSHGINRRDFMKLCAALAATMGLSSKAAAEMAESVTNPQRPPVIWIGAQECT WP_000145410.1
MULTISPECIES: hydrogenase-2 small chain [Proteobacteria] Length: 372\nNumber of predicted TMHs: 1\nExp number of AAs in TMHs: 23.72226\nExp number, first 60 AAs: 0.22038\nTotal prob of N-in: 0.12994\noutside 1 331\nTMhelix 332 354\ninside 355 372

11410 GCF_001441345.1_ASM144134v1 Escherichia coli K71Proteobacteria; Gammaproteobacteria; Enterobacterales; Enterobacteriaceae; Escherichia; Escherichia coli
MTGDNTLIHSHGINRRDFMKLCAALAATMGLSSKAAAEMAESVTNPQRPPVIWIGAQECT WP_000145410.1
MULTISPECIES: hydrogenase-2 small chain [Proteobacteria] Length: 372\nNumber of predicted TMHs: 1\nExp number of AAs in TMHs: 23.72226\nExp number, first 60 AAs: 0.22038\nTotal prob of N-in: 0.12994\noutside 1 331\nTMhelix 332 354\ninside 355 372

11411 GCF_001441345.1_ASM144134v1 Escherichia coli K71Proteobacteria; Gammaproteobacteria; Enterobacterales; Enterobacteriaceae; Escherichia; Escherichia coli
MNNEETFYQAMRRQGVTRRSFLKYCSLAATSLGLGAGMAPKIAWALENKPRIPVVWIHGL WP_001058323.1
MULTISPECIES: hydrogenase-1 small chain [Proteobacteria] Length: 372\nNumber of predicted TMHs: 2\nExp number of AAs in TMHs: 37.2608499999999\nExp number, first 60 AAs: 19.52923\nTotal prob of N-in: 0.94351\nPOSSIBLE N-term signal sequence\ninside 1 19\nTMhelix 20 39\noutside 40 325\nTMhelix 326 348\ninside 349 372

11412 GCF_000264095.1_ASM26409v1 Escherichia coli KD1 Proteobacteria; Gammaproteobacteria; Enterobacterales; Enterobacteriaceae; Escherichia; Escherichia coli
MTGDNTLIHSHGINRRDFMKLCAALAATMGLSSKAAAEMAESVTNPQRPPVIWIGAQECT WP_000145410.1
MULTISPECIES: hydrogenase-2 small chain [Proteobacteria] Length: 372\nNumber of predicted TMHs: 1\nExp number of AAs in TMHs: 23.72226\nExp number, first 60 AAs: 0.22038\nTotal prob of N-in: 0.12994\noutside 1 331\nTMhelix 332 354\ninside 355 372

11413 GCF_000264095.1_ASM26409v1 Escherichia coli KD1 Proteobacteria; Gammaproteobacteria; Enterobacterales; Enterobacteriaceae; Escherichia; Escherichia coli
MNNEETFYQAMRRQGVTRRSFLKYCSLAATSLGLGAGMAPKIAWALENKPRIPVVWIHGL WP_001058323.1
MULTISPECIES: hydrogenase-1 small chain [Proteobacteria] Length: 372\nNumber of predicted TMHs: 2\nExp number of AAs in TMHs: 37.2608499999999\nExp number, first 60 AAs: 19.52923\nTotal prob of N-in: 0.94351\nPOSSIBLE N-term signal sequence\ninside 1 19\nTMhelix 20 39\noutside 40 325\nTMhelix 326 348\ninside 349 372

11414 GCF_000264195.1_ASM26419v1 Escherichia coli KD2 Proteobacteria; Gammaproteobacteria; Enterobacterales; Enterobacteriaceae; Escherichia; Escherichia coli

MNNEETFYQAMRRQGVTRRSFLKYCSLAATSLGLGAGMAPKIAWALENKPRIPVVWIHGL WP_001058323.1 MULTISPECIES: hydrogenase-1 small chain [Proteobacteria] Length: 372\nNumber of predicted TMHs: 2\nExp number of AAs in TMHs: 37.2608499999999\nExp number, first 60 AAs: 19.52923\nTotal prob of N-in: 0.94351\nPOSSIBLE N-term signal sequence\ninside 1 19\nTMhelix 20 39\noutside 40 325\nTMhelix 326 348\ninside 349 372

11415 GCF_000264195.1_ASM26419v1 Escherichia coli KD2 Proteobacteria; Gammaproteobacteria; Enterobacterales; Enterobacteriaceae; Escherichia; Escherichia coli MTGDNTLIHSHGINRRDFMKLCAALAATMGLSSKAAAEMAESVTNPQRPPVIWIGAQECT WP_000145410.1 MULTISPECIES: hydrogenase-2 small chain [Proteobacteria] Length: 372\nNumber of predicted TMHs: 1\nExp number of AAs in TMHs: 23.72226\nExp number, first 60 AAs: 0.22038\nTotal prob of N-in: 0.12994\noutside 1 331\nTMhelix 332 354\ninside 355 372

11416 GCF_000725305.1_ASM72530v1 Escherichia coli KLY Proteobacteria; Gammaproteobacteria; Enterobacterales; Enterobacteriaceae; Escherichia; Escherichia coli MNNEETFYQAMRRQGVTRRSFLKYCSLAATSLGLGAGMAPKIAWALENKPRIPVVWIHGL WP_001058323.1 MULTISPECIES: hydrogenase-1 small chain [Proteobacteria] Length: 372\nNumber of predicted TMHs: 2\nExp number of AAs in TMHs: 37.2608499999999\nExp number, first 60 AAs: 19.52923\nTotal prob of N-in: 0.94351\nPOSSIBLE N-term signal sequence\ninside 1 19\nTMhelix 20 39\noutside 40 325\nTMhelix 326 348\ninside 349 372

11417 GCF_000725305.1_ASM72530v1 Escherichia coli KLY Proteobacteria; Gammaproteobacteria; Enterobacterales; Enterobacteriaceae; Escherichia; Escherichia coli MTGDNTLIHSHGINRRDFMKLCAALAATMGLSSKAAAEMAESVTNPQRPPVIWIGAQECT WP_000145410.1 MULTISPECIES: hydrogenase-2 small chain [Proteobacteria] Length: 372\nNumber of predicted TMHs: 1\nExp number of AAs in TMHs: 23.72226\nExp number, first 60 AAs: 0.22038\nTotal prob of N-in: 0.12994\noutside 1 331\nTMhelix 332 354\ninside 355 372

11418 GCF_000258025.1_ASM25802v1 Escherichia coli KO11FL Proteobacteria; Gammaproteobacteria; Enterobacterales; Enterobacteriaceae; Escherichia; Escherichia coli MTGDNTLIHSHGINRRDFMKLCAALAATMGLSSKAAAEMAESVTNPQRPPVIWIGAQECT WP_000145410.1 MULTISPECIES: hydrogenase-2 small chain [Proteobacteria] Length: 372\nNumber of predicted TMHs: 1\nExp number of AAs in TMHs: 23.72226\nExp number, first 60 AAs: 0.22038\nTotal prob of N-in: 0.12994\noutside 1 331\nTMhelix 332 354\ninside 355 372

11419 GCF_000147855.2_ASM14785v3 Escherichia coli KO11FL Proteobacteria; Gammaproteobacteria; Enterobacterales; Enterobacteriaceae; Escherichia; Escherichia coli MTGDNTLIHSHGINRRDFMKLCAALAATMGLSSKAAAEMAESVTNPQRPPVIWIGAQECT WP_000145410.1 MULTISPECIES: hydrogenase-2 small chain [Proteobacteria] Length: 372\nNumber of predicted TMHs: 1\nExp number of AAs in TMHs: 23.72226\nExp number, first 60 AAs: 0.22038\nTotal prob of N-in: 0.12994\noutside 1 331\nTMhelix 332 354\ninside 355 372

11420 GCF_000147855.2_ASM14785v3 Escherichia coli KO11FL Proteobacteria; Gammaproteobacteria; Enterobacterales; Enterobacteriaceae; Escherichia; Escherichia coli MNNEETFYQAMRRQGVTRRSFLKYCSLAATSLGLGAGMAPKIAWALENKPRIPVVWIHGL WP_001349947.1 hydrogenase [Escherichia coli] Length: 372\nNumber of predicted TMHs: 2\nExp number of AAs in TMHs: 37.2615299999999\nExp number, first 60 AAs: 19.52923\nTotal prob of N-in: 0.94351\nPOSSIBLE N-term signal sequence\ninside 1 19\nTMhelix 20 39\noutside 40 325\nTMhelix 326 348\ninside 349 372

11421 GCF_000258025.1_ASM25802v1 Escherichia coli KO11FL Proteobacteria; Gammaproteobacteria; Enterobacterales; Enterobacteriaceae; Escherichia; Escherichia coli MNNEETFYQAMRRQGVTRRSFLKYCSLAATSLGLGAGMAPKIAWALENKPRIPVVWIHGL WP_001349947.1 hydrogenase [Escherichia coli] Length: 372\nNumber of predicted TMHs: 2\nExp number of AAs in TMHs: 37.2615299999999\nExp number, first 60 AAs: 19.52923\nTotal prob of N-in: 0.94351\nPOSSIBLE N-term signal sequence\ninside 1 19\nTMhelix 20 39\noutside 40 325\nTMhelix 326 348\ninside 349 372

11422 GCF_000459595.1_Esch_coli_KOEGE_10_25a_V1 Escherichia coli KOEGE 10 (25a) Proteobacteria; Gammaproteobacteria; Enterobacterales; Enterobacteriaceae; Escherichia; Escherichia coli MTGDNTLIHSHGINRRDFMKLCAALAATMGLSSKAAAEMAESVTNPQRPPVIWIGAQECT WP_000145418.1 hydrogenase 2 small subunit [Escherichia coli] Length: 372\nNumber of predicted TMHs: 1\nExp number of

AAs in TMHs: 23.72313\nExp number, first 60 AAs:  0.22034\nTotal prob of N-in:   0.12997\noutside   1   331\nTMhelix   332   354\ninside   355   372

11423 GCF_000459595.1_Esch_coli_KOEGE_10_25a_V1 Escherichia coli KOEGE 10 (25a) Proteobacteria; Gammaproteobacteria; Enterobacterales; Enterobacteriaceae; Escherichia; Escherichia coli MNNEETFYQAMRRQGVTRRSFLKYCSLAATSLGLGAGMAPKIAWALENKPRIPVVWIHGL WP_001058323.1 MULTISPECIES: hydrogenase-1 small chain [Proteobacteria] Length: 372\nNumber of predicted TMHs: 2\nExp number of AAs in TMHs: 37.2608499999999\nExp number, first 60 AAs:  19.52923\nTotal prob of N-in: 0.94351\nPOSSIBLE N-term signal sequence\ninside   1   19\nTMhelix   20   39\noutside   40   325\nTMhelix   326   348\ninside   349   372

11424 GCF_000459915.1_Esch_coli_KOEGE_118_317a_V1 Escherichia coli KOEGE 118 (317a) Proteobacteria; Gammaproteobacteria; Enterobacterales; Enterobacteriaceae; Escherichia; Escherichia coli MNNEETFYQAMRRQGVTRRSFLKYCSLAATSLGLGAGMAPKIAWALENKPRIPVVWIHGL WP_001058323.1 MULTISPECIES: hydrogenase-1 small chain [Proteobacteria] Length: 372\nNumber of predicted TMHs: 2\nExp number of AAs in TMHs: 37.2608499999999\nExp number, first 60 AAs:  19.52923\nTotal prob of N-in: 0.94351\nPOSSIBLE N-term signal sequence\ninside   1   19\nTMhelix   20   39\noutside   40   325\nTMhelix   326   348\ninside   349   372

11425 GCF_000459915.1_Esch_coli_KOEGE_118_317a_V1 Escherichia coli KOEGE 118 (317a) Proteobacteria; Gammaproteobacteria; Enterobacterales; Enterobacteriaceae; Escherichia; Escherichia coli MTGDNTLIHSHGINRRDFMKLCAALAATMGLSSKAAAEMAESVTNPQRPPVIWIGAQECT WP_000145410.1 MULTISPECIES: hydrogenase-2 small chain [Proteobacteria] Length: 372\nNumber of predicted TMHs: 1\nExp number of AAs in TMHs: 23.72226\nExp number, first 60 AAs:  0.22038\nTotal prob of N-in:   0.12994\noutside   1   331\nTMhelix   332   354\ninside   355   372

11426 GCF_000459935.1_Esch_coli_KOEGE_131_358a_V1 Escherichia coli KOEGE 131 (358a) Proteobacteria; Gammaproteobacteria; Enterobacterales; Enterobacteriaceae; Escherichia; Escherichia coli MTGDNTLIHSHGINRRDFMKLCAALAATMGLSSKAAAEMAESVTNPQRPPVIWIGAQECT WP_000145410.1 MULTISPECIES: hydrogenase-2 small chain [Proteobacteria] Length: 372\nNumber of predicted TMHs: 1\nExp number of AAs in TMHs: 23.72226\nExp number, first 60 AAs:  0.22038\nTotal prob of N-in:   0.12994\noutside   1   331\nTMhelix   332   354\ninside   355   372

11427 GCF_000459935.1_Esch_coli_KOEGE_131_358a_V1 Escherichia coli KOEGE 131 (358a) Proteobacteria; Gammaproteobacteria; Enterobacterales; Enterobacteriaceae; Escherichia; Escherichia coli MNNEETFYQAMRRQGVTRRSFLKYCSLAATSLGLGAGMAPKIAWALENKPRIPVVWIHGL WP_001058320.1 hydrogenase [Escherichia coli] Length: 372\nNumber of predicted TMHs:  2\nExp number of AAs in TMHs: 37.26082\nExp number, first 60 AAs:  19.52923\nTotal prob of N-in:   0.94351\nPOSSIBLE N-term signal sequence\ninside   1   19\nTMhelix   20   39\noutside   40   325\nTMhelix   326   348\ninside   349   372

11428 GCF_000459615.1_Esch_coli_KOEGE_30_63a_V1 Escherichia coli KOEGE 30 (63a) Proteobacteria; Gammaproteobacteria; Enterobacterales; Enterobacteriaceae; Escherichia; Escherichia coli MNNEETFYQAMRRQGVTRRSFLKYCSLAATSLGLGAGMAPKIAWALENKPRIPVVWIHGL WP_001058323.1 MULTISPECIES: hydrogenase-1 small chain [Proteobacteria] Length: 372\nNumber of predicted TMHs: 2\nExp number of AAs in TMHs: 37.2608499999999\nExp number, first 60 AAs:  19.52923\nTotal prob of N-in: 0.94351\nPOSSIBLE N-term signal sequence\ninside   1   19\nTMhelix   20   39\noutside   40   325\nTMhelix   326   348\ninside   349   372

11429 GCF_000459615.1_Esch_coli_KOEGE_30_63a_V1 Escherichia coli KOEGE 30 (63a) Proteobacteria; Gammaproteobacteria; Enterobacterales; Enterobacteriaceae; Escherichia; Escherichia coli MTGDNTLIHSHGINRRDFMKLCAALAATMGLSSKAAAEMAESVTNPQRPPVIWIGAQECT WP_000145410.1 MULTISPECIES: hydrogenase-2 small chain [Proteobacteria] Length: 372\nNumber of predicted TMHs: 1\nExp number of AAs in TMHs: 23.72226\nExp number, first 60 AAs:  0.22038\nTotal prob of N-in:   0.12994\noutside   1   331\nTMhelix   332   354\ninside   355   372

11430 GCF_000459635.1_Esch_coli_KOEGE_32_66a_V1 Escherichia coli KOEGE 32 (66a) Proteobacteria; Gammaproteobacteria; Enterobacterales; Enterobacteriaceae; Escherichia; Escherichia coli MTGDNTLIHSHGINRRDFMKLCAALAATMGLSSKAAAEMAESVTNPQRPPVIWIGAQECT WP_000145410.1 MULTISPECIES: hydrogenase-2 small chain [Proteobacteria] Length: 372\nNumber of predicted TMHs: 1\nExp number of AAs in TMHs: 23.72226\nExp number, first 60 AAs:  0.22038\nTotal prob of N-in:   0.12994\noutside   1   331\nTMhelix   332   354\ninside   355   372

11431 GCF_000459655.1_Esch_coli_KOEGE_33_68a_V1 Escherichia coli KOEGE 33 (68a) Proteobacteria; Gammaproteobacteria; Enterobacterales; Enterobacteriaceae; Escherichia; Escherichia coli MTGDNTLIHSHGINRRDFMKLCAALAATMGLSSKAAAEMAESVTNPQRPPVIWIGAQECT WP_000145410.1 MULTISPECIES: hydrogenase-2 small chain [Proteobacteria] Length: 372\nNumber of predicted TMHs: 1\nExp number of AAs in TMHs: 23.72226\nExp number, first 60 AAs: 0.22038\nTotal prob of N-in: 0.12994\noutside 1 331\nTMhelix 332 354\ninside 355 372

11432 GCF_000459655.1_Esch_coli_KOEGE_33_68a_V1 Escherichia coli KOEGE 33 (68a) Proteobacteria; Gammaproteobacteria; Enterobacterales; Enterobacteriaceae; Escherichia; Escherichia coli MNNEETFYQAMRRQGVTRRSFLKYCSLAATSLGLGAGMAPKIAWALENKPRIPVVWIHGL WP_001058323.1 MULTISPECIES: hydrogenase-1 small chain [Proteobacteria] Length: 372\nNumber of predicted TMHs: 2\nExp number of AAs in TMHs: 37.2608499999999\nExp number, first 60 AAs: 19.52923\nTotal prob of N-in: 0.94351\nPOSSIBLE N-term signal sequence\ninside 1 19\nTMhelix 20 39\noutside 40 325\nTMhelix 326 348\ninside 349 372

11433 GCF_000459555.1_Esch_coli_KOEGE_3_4a_V1 Escherichia coli KOEGE 3 (4a) Proteobacteria; Gammaproteobacteria; Enterobacterales; Enterobacteriaceae; Escherichia; Escherichia coli MTGDNTLIHSHGINRRDFMKLCAALAATMGLSSKAAAEMAESVTNPQRPPVIWIGAQECT WP_000145410.1 MULTISPECIES: hydrogenase-2 small chain [Proteobacteria] Length: 372\nNumber of predicted TMHs: 1\nExp number of AAs in TMHs: 23.72226\nExp number, first 60 AAs: 0.22038\nTotal prob of N-in: 0.12994\noutside 1 331\nTMhelix 332 354\ninside 355 372

11434 GCF_000459555.1_Esch_coli_KOEGE_3_4a_V1 Escherichia coli KOEGE 3 (4a) Proteobacteria; Gammaproteobacteria; Enterobacterales; Enterobacteriaceae; Escherichia; Escherichia coli MNNEETFYQAMRRQGVTRRSFLKYCSLAATSLGLGAGMAPKIAWALENKPRIPVVWIHGL WP_001058323.1 MULTISPECIES: hydrogenase-1 small chain [Proteobacteria] Length: 372\nNumber of predicted TMHs: 2\nExp number of AAs in TMHs: 37.2608499999999\nExp number, first 60 AAs: 19.52923\nTotal prob of N-in: 0.94351\nPOSSIBLE N-term signal sequence\ninside 1 19\nTMhelix 20 39\noutside 40 325\nTMhelix 326 348\ninside 349 372

11435 GCF_000459675.1_Esch_coli_KOEGE_40_102a_V1 Escherichia coli KOEGE 40 (102a) Proteobacteria; Gammaproteobacteria; Enterobacterales; Enterobacteriaceae; Escherichia; Escherichia coli MNNEETFYQAMRRQGVTRRSFLKYCSLAATSLGLGAGMAPKIAWALENKPRIPVVWIHGL WP_001058323.1 MULTISPECIES: hydrogenase-1 small chain [Proteobacteria] Length: 372\nNumber of predicted TMHs: 2\nExp number of AAs in TMHs: 37.2608499999999\nExp number, first 60 AAs: 19.52923\nTotal prob of N-in: 0.94351\nPOSSIBLE N-term signal sequence\ninside 1 19\nTMhelix 20 39\noutside 40 325\nTMhelix 326 348\ninside 349 372

11436 GCF_000459675.1_Esch_coli_KOEGE_40_102a_V1 Escherichia coli KOEGE 40 (102a) Proteobacteria; Gammaproteobacteria; Enterobacterales; Enterobacteriaceae; Escherichia; Escherichia coli MTGDNTLIHSHGINRRDFMKLCAALAATMGLSSKAAAEMAESVTNPQRPPVIWIGAQECT WP_000145410.1 MULTISPECIES: hydrogenase-2 small chain [Proteobacteria] Length: 372\nNumber of predicted TMHs: 1\nExp number of AAs in TMHs: 23.72226\nExp number, first 60 AAs: 0.22038\nTotal prob of N-in: 0.12994\noutside 1 331\nTMhelix 332 354\ninside 355 372

11437 GCF_000459695.1_Esch_coli_KOEGE_43_105a_V1 Escherichia coli KOEGE 43 (105a) Proteobacteria; Gammaproteobacteria; Enterobacterales; Enterobacteriaceae; Escherichia; Escherichia coli MNNEETFYQAMRRQGVTRRSFLKYCSLAATSLGLGAGMAPKIAWALENKPRIPVVWIHGL WP_001058323.1 MULTISPECIES: hydrogenase-1 small chain [Proteobacteria] Length: 372\nNumber of predicted TMHs: 2\nExp number of AAs in TMHs: 37.2608499999999\nExp number, first 60 AAs: 19.52923\nTotal prob of N-in: 0.94351\nPOSSIBLE N-term signal sequence\ninside 1 19\nTMhelix 20 39\noutside 40 325\nTMhelix 326 348\ninside 349 372

11438 GCF_000459695.1_Esch_coli_KOEGE_43_105a_V1 Escherichia coli KOEGE 43 (105a) Proteobacteria; Gammaproteobacteria; Enterobacterales; Enterobacteriaceae; Escherichia; Escherichia coli MTGDNTLIHSHGINRRDFMKLCAALAATMGLSSKAAAEMAESVTNPQRPPVIWIGAQECT WP_000145410.1 MULTISPECIES: hydrogenase-2 small chain [Proteobacteria] Length: 372\nNumber of predicted TMHs: 1\nExp number of AAs in TMHs: 23.72226\nExp number, first 60 AAs: 0.22038\nTotal prob of N-in: 0.12994\noutside 1 331\nTMhelix 332 354\ninside 355 372

11439 GCF_000459715.1_Esch_coli_KOEGE_44_106a_V1 Escherichia coli KOEGE 44 (106a) Proteobacteria; Gammaproteobacteria; Enterobacterales; Enterobacteriaceae; Escherichia; Escherichia coli MTGDNTLIHSHGINRRDFMKLCAALAATMGLSSKAAAEMAESVTNPQRPPVIWIGAQECT WP_000145410.1 MULTISPECIES: hydrogenase-2 small chain [Proteobacteria] Length: 372\nNumber of predicted TMHs: 1\nExp number of AAs in TMHs: 23.72226\nExp number, first 60 AAs: 0.22038\nTotal prob of N-in: 0.12994\noutside 1 331\nTMhelix 332 354\ninside 355 372

11440 GCF_000459715.1_Esch_coli_KOEGE_44_106a_V1 Escherichia coli KOEGE 44 (106a) Proteobacteria; Gammaproteobacteria; Enterobacterales; Enterobacteriaceae; Escherichia; Escherichia coli MNNEETFYQAMRRQGVTRRSFLKYCSLAATSLGLGAGMAPKIAWALENKPRIPVVWIHGL WP_001058323.1 MULTISPECIES: hydrogenase-1 small chain [Proteobacteria] Length: 372\nNumber of predicted TMHs: 2\nExp number of AAs in TMHs: 37.2608499999999\nExp number, first 60 AAs: 19.52923\nTotal prob of N-in: 0.94351\nPOSSIBLE N-term signal sequence\ninside 1 19\nTMhelix 20 39\noutside 40 325\nTMhelix 326 348\ninside 349 372

11441 GCF_000459735.1_Esch_coli_KOEGE_56_169a_V1 Escherichia coli KOEGE 56 (169a) Proteobacteria; Gammaproteobacteria; Enterobacterales; Enterobacteriaceae; Escherichia; Escherichia coli MNNEETFYQAMRRQGVTRRSFLKYCSLAATSLGLGAGMAPKIAWALENKPRIPVVWIHGL WP_001058323.1 MULTISPECIES: hydrogenase-1 small chain [Proteobacteria] Length: 372\nNumber of predicted TMHs: 2\nExp number of AAs in TMHs: 37.2608499999999\nExp number, first 60 AAs: 19.52923\nTotal prob of N-in: 0.94351\nPOSSIBLE N-term signal sequence\ninside 1 19\nTMhelix 20 39\noutside 40 325\nTMhelix 326 348\ninside 349 372

11442 GCF_000459735.1_Esch_coli_KOEGE_56_169a_V1 Escherichia coli KOEGE 56 (169a) Proteobacteria; Gammaproteobacteria; Enterobacterales; Enterobacteriaceae; Escherichia; Escherichia coli MTGDNTLIHSHGINRRDFMKLCAALAATMGLSSKAAAEMAESVTNPQRPPVIWIGAQECT WP_000145410.1 MULTISPECIES: hydrogenase-2 small chain [Proteobacteria] Length: 372\nNumber of predicted TMHs: 1\nExp number of AAs in TMHs: 23.72226\nExp number, first 60 AAs: 0.22038\nTotal prob of N-in: 0.12994\noutside 1 331\nTMhelix 332 354\ninside 355 372

11443 GCF_000459755.1_Esch_coli_KOEGE_58_171a_V1 Escherichia coli KOEGE 58 (171a) Proteobacteria; Gammaproteobacteria; Enterobacterales; Enterobacteriaceae; Escherichia; Escherichia coli MTGDNTLIHSHGINRRDFMKLCAALAATMGLSSKAAAEMAESVTNPQRPPVIWIGAQECT WP_000145410.1 MULTISPECIES: hydrogenase-2 small chain [Proteobacteria] Length: 372\nNumber of predicted TMHs: 1\nExp number of AAs in TMHs: 23.72226\nExp number, first 60 AAs: 0.22038\nTotal prob of N-in: 0.12994\noutside 1 331\nTMhelix 332 354\ninside 355 372

11444 GCF_000459755.1_Esch_coli_KOEGE_58_171a_V1 Escherichia coli KOEGE 58 (171a) Proteobacteria; Gammaproteobacteria; Enterobacterales; Enterobacteriaceae; Escherichia; Escherichia coli MNNEETFYQAMRRQGVTRRSFLKYCSLAATSLGLGAGMAPKIAWALENKPRIPVVWIHGL WP_001058323.1 MULTISPECIES: hydrogenase-1 small chain [Proteobacteria] Length: 372\nNumber of predicted TMHs: 2\nExp number of AAs in TMHs: 37.2608499999999\nExp number, first 60 AAs: 19.52923\nTotal prob of N-in: 0.94351\nPOSSIBLE N-term signal sequence\ninside 1 19\nTMhelix 20 39\noutside 40 325\nTMhelix 326 348\ninside 349 372

11445 GCF_000459775.1_Esch_coli_KOEGE_61_174a_V1 Escherichia coli KOEGE 61 (174a) Proteobacteria; Gammaproteobacteria; Enterobacterales; Enterobacteriaceae; Escherichia; Escherichia coli MTGDNTLIHSHGINRRDFMKLCAALAATMGLSSKAAAEMAESVTNPQRPPVIWIGAQECT WP_000145410.1 MULTISPECIES: hydrogenase-2 small chain [Proteobacteria] Length: 372\nNumber of predicted TMHs: 1\nExp number of AAs in TMHs: 23.72226\nExp number, first 60 AAs: 0.22038\nTotal prob of N-in: 0.12994\noutside 1 331\nTMhelix 332 354\ninside 355 372

11446 GCF_000459775.1_Esch_coli_KOEGE_61_174a_V1 Escherichia coli KOEGE 61 (174a) Proteobacteria; Gammaproteobacteria; Enterobacterales; Enterobacteriaceae; Escherichia; Escherichia coli MNNEETFYQAMRRQGVTRRSFLKYCSLAATSLGLGAGMAPKIAWALENKPRIPVVWIHGL WP_001058323.1 MULTISPECIES: hydrogenase-1 small chain [Proteobacteria] Length: 372\nNumber of predicted TMHs: 2\nExp number of AAs in TMHs: 37.2608499999999\nExp number, first 60 AAs: 19.52923\nTotal prob of N-in: 0.94351\nPOSSIBLE N-term signal sequence\ninside 1 19\nTMhelix 20 39\noutside 40 325\nTMhelix 326 348\ninside 349 372

11447 GCF_000459795.1_Esch_coli_KOEGE_62_175a_V1 Escherichia coli KOEGE 62 (175a) Proteobacteria; Gammaproteobacteria; Enterobacterales; Enterobacteriaceae; Escherichia; Escherichia coli MNNEETFYQAMRRQGVTRRSFLKYCSLAATSLGLGAGMAPKIAWALENKPRIPVVWIHGL WP_001058323.1 MULTISPECIES: hydrogenase-1 small chain [Proteobacteria] Length: 372\nNumber of predicted TMHs: 2\nExp number of AAs in TMHs: 37.2608499999999\nExp number, first 60 AAs: 19.52923\nTotal prob of N-in: 0.94351\nPOSSIBLE N-term signal sequence\ninside 1 19\nTMhelix 20 39\noutside 40 325\nTMhelix 326 348\ninside 349 372

11448 GCF_000459795.1_Esch_coli_KOEGE_62_175a_V1 Escherichia coli KOEGE 62 (175a) Proteobacteria; Gammaproteobacteria; Enterobacterales; Enterobacteriaceae; Escherichia; Escherichia coli MTGDNTLIHSHGINRRDFMKLCAALAATMGLSSKAAAEMAESVTNPQRPPVIWIGAQECT WP_000145410.1 MULTISPECIES: hydrogenase-2 small chain [Proteobacteria] Length: 372\nNumber of predicted TMHs: 1\nExp number of AAs in TMHs: 23.72226\nExp number, first 60 AAs: 0.22038\nTotal prob of N-in: 0.12994\noutside 1 331\nTMhelix 332 354\ninside 355 372

11449 GCF_000459815.1_Esch_coli_KOEGE_68_182a_V1 Escherichia coli KOEGE 68 (182a) Proteobacteria; Gammaproteobacteria; Enterobacterales; Enterobacteriaceae; Escherichia; Escherichia coli MNNEETFYQAMRRQGVTRRSFLKYCSLAATSLGLGAGMAPKIAWALENKPRIPVVWIHGL WP_001058323.1 MULTISPECIES: hydrogenase-1 small chain [Proteobacteria] Length: 372\nNumber of predicted TMHs: 2\nExp number of AAs in TMHs: 37.2608499999999\nExp number, first 60 AAs: 19.52923\nTotal prob of N-in: 0.94351\nPOSSIBLE N-term signal sequence\ninside 1 19\nTMhelix 20 39\noutside 40 325\nTMhelix 326 348\ninside 349 372

11450 GCF_000459815.1_Esch_coli_KOEGE_68_182a_V1 Escherichia coli KOEGE 68 (182a) Proteobacteria; Gammaproteobacteria; Enterobacterales; Enterobacteriaceae; Escherichia; Escherichia coli MTGDNTLIHSHGINRRDFMKLCAALAATMGLSSKAAAEMAESVTNPQRPPVIWIGAQECT WP_000145410.1 MULTISPECIES: hydrogenase-2 small chain [Proteobacteria] Length: 372\nNumber of predicted TMHs: 1\nExp number of AAs in TMHs: 23.72226\nExp number, first 60 AAs: 0.22038\nTotal prob of N-in: 0.12994\noutside 1 331\nTMhelix 332 354\ninside 355 372

11451 GCF_000459835.1_Esch_coli_KOEGE_70_185a_V1 Escherichia coli KOEGE 70 (185a) Proteobacteria; Gammaproteobacteria; Enterobacterales; Enterobacteriaceae; Escherichia; Escherichia coli MTGDNTLIHSHGINRRDFMKLCAALAATMGLSSKAAAEMAESVTNPQRPPVIWIGAQECT WP_000145410.1 MULTISPECIES: hydrogenase-2 small chain [Proteobacteria] Length: 372\nNumber of predicted TMHs: 1\nExp number of AAs in TMHs: 23.72226\nExp number, first 60 AAs: 0.22038\nTotal prob of N-in: 0.12994\noutside 1 331\nTMhelix 332 354\ninside 355 372

11452 GCF_000459835.1_Esch_coli_KOEGE_70_185a_V1 Escherichia coli KOEGE 70 (185a) Proteobacteria; Gammaproteobacteria; Enterobacterales; Enterobacteriaceae; Escherichia; Escherichia coli MNNEETFYQAMRRQGVTRRSFLKYCSLAATSLGLGAGMAPKIAWALENKPRIPVVWIHGL WP_001058323.1 MULTISPECIES: hydrogenase-1 small chain [Proteobacteria] Length: 372\nNumber of predicted TMHs: 2\nExp number of AAs in TMHs: 37.2608499999999\nExp number, first 60 AAs: 19.52923\nTotal prob of N-in: 0.94351\nPOSSIBLE N-term signal sequence\ninside 1 19\nTMhelix 20 39\noutside 40 325\nTMhelix 326 348\ninside 349 372

11453 GCF_000459855.1_Esch_coli_KOEGE_71_186a_V1 Escherichia coli KOEGE 71 (186a) Proteobacteria; Gammaproteobacteria; Enterobacterales; Enterobacteriaceae; Escherichia; Escherichia coli MNNEETFYQAMRRQGVTRRSFLKYCSLAATSLGLGAGMAPKIAWALENKPRIPVVWIHGL WP_010344442.1 MULTISPECIES: hydrogenase [Escherichia] Length: 372\nNumber of predicted TMHs: 2\nExp number of AAs in TMHs: 38.04874\nExp number, first 60 AAs: 19.5298\nTotal prob of N-in: 0.94352\nPOSSIBLE N-term signal sequence\ninside 1 19\nTMhelix 20 39\noutside 40 325\nTMhelix 326 348\ninside 349 372

11454 GCF_000459855.1_Esch_coli_KOEGE_71_186a_V1 Escherichia coli KOEGE 71 (186a) Proteobacteria; Gammaproteobacteria; Enterobacterales; Enterobacteriaceae; Escherichia; Escherichia coli MTGDNTLIHSHGINRRDFMKLCAALAATMGLSSKAAAEMAESVTNPQRPPVIWIGAQECT WP_021551423.1 hydrogenase-2 small chain [Escherichia coli] Length: 372\nNumber of predicted TMHs: 1\nExp number of AAs in TMHs: 23.72222\nExp number, first 60 AAs: 0.22038\nTotal prob of N-in: 0.12994\noutside 1 331\nTMhelix 332 354\ninside 355 372

11455 GCF_000459575.1_Esch_coli_KOEGE_7_16a_V1 Escherichia coli KOEGE 7 (16a) Proteobacteria; Gammaproteobacteria; Enterobacterales; Enterobacteriaceae; Escherichia; Escherichia coli

MNNEETFYQAMRRQGVTRRSFLKYCSLAATSLGLGAGMAPKIAWALENKPRIPVVWIHGL WP_001058323.1 MULTISPECIES: hydrogenase-1 small chain [Proteobacteria] Length: 372\nNumber of predicted TMHs: 2\nExp number of AAs in TMHs: 37.2608499999999\nExp number, first 60 AAs: 19.52923\nTotal prob of N-in: 0.94351\nPOSSIBLE N-term signal sequence\ninside 1 19\nTMhelix 20 39\noutside 40 325\nTMhelix 326 348\ninside 349 372

11456 GCF_000459575.1_Esch_coli_KOEGE_7_16a_V1 Escherichia coli KOEGE 7 (16a) Proteobacteria; Gammaproteobacteria; Enterobacterales; Enterobacteriaceae; Escherichia; Escherichia coli MTGDNTLIHSHGINRRDFMKLCAALAATMGLSSKAAAEMAESVTNPQRPPVIWIGAQECT WP_000145410.1 MULTISPECIES: hydrogenase-2 small chain [Proteobacteria] Length: 372\nNumber of predicted TMHs: 1\nExp number of AAs in TMHs: 23.72226\nExp number, first 60 AAs: 0.22038\nTotal prob of N-in: 0.12994\noutside 1 331\nTMhelix 332 354\ninside 355 372

11457 GCF_000459875.1_Esch_coli_KOEGE_73_195a_V1 Escherichia coli KOEGE 73 (195a) Proteobacteria; Gammaproteobacteria; Enterobacterales; Enterobacteriaceae; Escherichia; Escherichia coli MNNEETFYQAMRRQGVTRRSFLKYCSLAATSLGLGAGMAPKIAWALENKPRIPVVWIHGL WP_001058323.1 MULTISPECIES: hydrogenase-1 small chain [Proteobacteria] Length: 372\nNumber of predicted TMHs: 2\nExp number of AAs in TMHs: 37.2608499999999\nExp number, first 60 AAs: 19.52923\nTotal prob of N-in: 0.94351\nPOSSIBLE N-term signal sequence\ninside 1 19\nTMhelix 20 39\noutside 40 325\nTMhelix 326 348\ninside 349 372

11458 GCF_000459875.1_Esch_coli_KOEGE_73_195a_V1 Escherichia coli KOEGE 73 (195a) Proteobacteria; Gammaproteobacteria; Enterobacterales; Enterobacteriaceae; Escherichia; Escherichia coli MTGDNTLIHSHGINRRDFMKLCAALAATMGLSSKAAAEMAESVTNPQRPPVIWIGAQECT WP_000145410.1 MULTISPECIES: hydrogenase-2 small chain [Proteobacteria] Length: 372\nNumber of predicted TMHs: 1\nExp number of AAs in TMHs: 23.72226\nExp number, first 60 AAs: 0.22038\nTotal prob of N-in: 0.12994\noutside 1 331\nTMhelix 332 354\ninside 355 372

11459 GCF_000459895.1_Esch_coli_KOEGE_77_202a_V1 Escherichia coli KOEGE 77 (202a) Proteobacteria; Gammaproteobacteria; Enterobacterales; Enterobacteriaceae; Escherichia; Escherichia coli MNNEETFYQAMRRQGVTRRSFLKYCSLAATSLGLGAGMAPKIAWALENKPRIPVVWIHGL WP_001058323.1 MULTISPECIES: hydrogenase-1 small chain [Proteobacteria] Length: 372\nNumber of predicted TMHs: 2\nExp number of AAs in TMHs: 37.2608499999999\nExp number, first 60 AAs: 19.52923\nTotal prob of N-in: 0.94351\nPOSSIBLE N-term signal sequence\ninside 1 19\nTMhelix 20 39\noutside 40 325\nTMhelix 326 348\ninside 349 372

11460 GCF_000459895.1_Esch_coli_KOEGE_77_202a_V1 Escherichia coli KOEGE 77 (202a) Proteobacteria; Gammaproteobacteria; Enterobacterales; Enterobacteriaceae; Escherichia; Escherichia coli MTGDNTLIHSHGINRRDFMKLCAALAATMGLSSKAAAEMAESVTNPQRPPVIWIGAQECT WP_000145410.1 MULTISPECIES: hydrogenase-2 small chain [Proteobacteria] Length: 372\nNumber of predicted TMHs: 1\nExp number of AAs in TMHs: 23.72226\nExp number, first 60 AAs: 0.22038\nTotal prob of N-in: 0.12994\noutside 1 331\nTMhelix 332 354\ninside 355 372

11461 GCF_001291365.1_KRX Escherichia coli KRX Proteobacteria; Gammaproteobacteria; Enterobacterales; Enterobacteriaceae; Escherichia; Escherichia coli MNNEETFYQAMRRQGVTRRSFLKYCSLAATSLGLGAGMAPKIAWALENKPRIPVVWIHGL WP_001058323.1 MULTISPECIES: hydrogenase-1 small chain [Proteobacteria] Length: 372\nNumber of predicted TMHs: 2\nExp number of AAs in TMHs: 37.2608499999999\nExp number, first 60 AAs: 19.52923\nTotal prob of N-in: 0.94351\nPOSSIBLE N-term signal sequence\ninside 1 19\nTMhelix 20 39\noutside 40 325\nTMhelix 326 348\ninside 349 372

11462 GCF_001291365.1_KRX Escherichia coli KRX Proteobacteria; Gammaproteobacteria; Enterobacterales; Enterobacteriaceae; Escherichia; Escherichia coli MTGDNTLIHSHGINRRDFMKLCAALAATMGLSSKAAAEMAESVTNPQRPPVIWIGAQECT WP_000145410.1 MULTISPECIES: hydrogenase-2 small chain [Proteobacteria] Length: 372\nNumber of predicted TMHs: 1\nExp number of AAs in TMHs: 23.72226\nExp number, first 60 AAs: 0.22038\nTotal prob of N-in: 0.12994\noutside 1 331\nTMhelix 332 354\ninside 355 372

11463 GCF_000407665.1_Esch_coli_KTE1_V1 Escherichia coli KTE1 Proteobacteria; Gammaproteobacteria; Enterobacterales; Enterobacteriaceae; Escherichia; Escherichia coli MTGDNTLIHSHGINRRDFMKLCAALAATMGLSSKAAAEMAESVTNPQRPPVIWIGAQECT WP_000145410.1

MULTISPECIES: hydrogenase-2 small chain [Proteobacteria] Length: 372\nNumber of predicted TMHs: 1\nExp number of AAs in TMHs: 23.72226\nExp number, first 60 AAs: 0.22038\nTotal prob of N-in: 0.12994\noutside 1 331\nTMhelix 332 354\ninside 355 372

11464 GCF_000407665.1_Esch_coli_KTE1_V1 Escherichia coli KTE1 Proteobacteria; Gammaproteobacteria; Enterobacterales; Enterobacteriaceae; Escherichia; Escherichia coli MNNEETFYQAMRRQGVTRRSFLKYCSLAATSLGLGAGMAPKIAWALENKPRIPVVWIHGL WP_001058323.1 MULTISPECIES: hydrogenase-1 small chain [Proteobacteria] Length: 372\nNumber of predicted TMHs: 2\nExp number of AAs in TMHs: 37.2608499999999\nExp number, first 60 AAs: 19.52923\nTotal prob of N-in: 0.94351\nPOSSIBLE N-term signal sequence\ninside 1 19\nTMhelix 20 39\noutside 40 325\nTMhelix 326 348\ninside 349 372

11465 GCF_000350685.1_Esch_coli_KTE10_V1 Escherichia coli KTE10 Proteobacteria; Gammaproteobacteria; Enterobacterales; Enterobacteriaceae; Escherichia; Escherichia coli MKGVLEKCTVEDFNYISATLDSYVSFTNDKRRKTLLSAYQNNPALHAELISLIDTQIKYF WP_001515206.1 MULTISPECIES: hypothetical protein [Enterobacteriaceae] Length: 249\nNumber of predicted TMHs: 2\nExp number of AAs in TMHs: 39.68696\nExp number, first 60 AAs: 0.01961\nTotal prob of N-in: 0.09053\noutside 1 157\nTMhelix 158 180\ninside 181 200\nTMhelix 201 220\noutside 221 249

11466 GCF_000350685.1_Esch_coli_KTE10_V1 Escherichia coli KTE10 Proteobacteria; Gammaproteobacteria; Enterobacterales; Enterobacteriaceae; Escherichia; Escherichia coli MTGDNTLIHSHGINRRDFMKLCAALAATMGLSSKAAAEMAESVTNPQRPPVIWIGAQECT WP_000145410.1 MULTISPECIES: hydrogenase-2 small chain [Proteobacteria] Length: 372\nNumber of predicted TMHs: 1\nExp number of AAs in TMHs: 23.72226\nExp number, first 60 AAs: 0.22038\nTotal prob of N-in: 0.12994\noutside 1 331\nTMhelix 332 354\ninside 355 372

11467 GCF_000350685.1_Esch_coli_KTE10_V1 Escherichia coli KTE10 Proteobacteria; Gammaproteobacteria; Enterobacterales; Enterobacteriaceae; Escherichia; Escherichia coli MNNEETFYQAMRRQGVTRRSFLKYCSLAATSLGLGAGMAPKIAWALENKPRIPVVWIHGL WP_001058323.1 MULTISPECIES: hydrogenase-1 small chain [Proteobacteria] Length: 372\nNumber of predicted TMHs: 2\nExp number of AAs in TMHs: 37.2608499999999\nExp number, first 60 AAs: 19.52923\nTotal prob of N-in: 0.94351\nPOSSIBLE N-term signal sequence\ninside 1 19\nTMhelix 20 39\noutside 40 325\nTMhelix 326 348\ninside 349 372

11468 GCF_000408565.1_Esch_coli_KTE100_V1 Escherichia coli KTE100 Proteobacteria; Gammaproteobacteria; Enterobacterales; Enterobacteriaceae; Escherichia; Escherichia coli MNNEETFYQAMRRQGVTRRSFLKYCSLAATSLGLGAGMAPKIAWALENKPRIPVVWIHGL WP_001058323.1 MULTISPECIES: hydrogenase-1 small chain [Proteobacteria] Length: 372\nNumber of predicted TMHs: 2\nExp number of AAs in TMHs: 37.2608499999999\nExp number, first 60 AAs: 19.52923\nTotal prob of N-in: 0.94351\nPOSSIBLE N-term signal sequence\ninside 1 19\nTMhelix 20 39\noutside 40 325\nTMhelix 326 348\ninside 349 372

11469 GCF_000408565.1_Esch_coli_KTE100_V1 Escherichia coli KTE100 Proteobacteria; Gammaproteobacteria; Enterobacterales; Enterobacteriaceae; Escherichia; Escherichia coli MTGDNTLIHSHGINRRDFMKLCAALAATMGLSSKAAAEMAESVTNPQRPPVIWIGAQECT WP_000145410.1 MULTISPECIES: hydrogenase-2 small chain [Proteobacteria] Length: 372\nNumber of predicted TMHs: 1\nExp number of AAs in TMHs: 23.72226\nExp number, first 60 AAs: 0.22038\nTotal prob of N-in: 0.12994\noutside 1 331\nTMhelix 332 354\ninside 355 372

11470 GCF_000352505.1_Esch_coli_KTE101_V1 Escherichia coli KTE101 Proteobacteria; Gammaproteobacteria; Enterobacterales; Enterobacteriaceae; Escherichia; Escherichia coli MTGDNTLIHSHGINRRDFMKLCAALAATMGLSSKAAAEMAESVTNPQRPPVIWIGAQECT WP_000145410.1 MULTISPECIES: hydrogenase-2 small chain [Proteobacteria] Length: 372\nNumber of predicted TMHs: 1\nExp number of AAs in TMHs: 23.72226\nExp number, first 60 AAs: 0.22038\nTotal prob of N-in: 0.12994\noutside 1 331\nTMhelix 332 354\ninside 355 372

11471 GCF_000352505.1_Esch_coli_KTE101_V1 Escherichia coli KTE101 Proteobacteria; Gammaproteobacteria; Enterobacterales; Enterobacteriaceae; Escherichia; Escherichia coli MNNEETFYQAMRRQGVTRRSFLKYCSLAATSLGLGAGMAPKIAWALENKPRIPVVWIHGL WP_001058323.1 MULTISPECIES: hydrogenase-1 small chain [Proteobacteria] Length: 372\nNumber of predicted TMHs: 2\nExp number of AAs in TMHs: 37.2608499999999\nExp number, first 60 AAs: 19.52923\nTotal prob of N-in:

0.94351\nPOSSIBLE N-term signal sequence\ninside 1 19\nTMhelix 20 39\noutside 40 325\nTMhelix 326 348\ninside 349 372

11472 GCF_000408585.1_Esch_coli_KTE102_V1 Escherichia coli KTE102 Proteobacteria; Gammaproteobacteria; Enterobacterales; Enterobacteriaceae; Escherichia; Escherichia coli MNNEETFYQAMRRQGVTRRSFLKYCSLAATSLGLGAGMAPKIAWALENKPRIPVVWIHGL WP_001058323.1 MULTISPECIES: hydrogenase-1 small chain [Proteobacteria] Length: 372\nNumber of predicted TMHs: 2\nExp number of AAs in TMHs: 37.2608499999999\nExp number, first 60 AAs: 19.52923\nTotal prob of N-in: 0.94351\nPOSSIBLE N-term signal sequence\ninside 1 19\nTMhelix 20 39\noutside 40 325\nTMhelix 326 348\ninside 349 372

11473 GCF_000408585.1_Esch_coli_KTE102_V1 Escherichia coli KTE102 Proteobacteria; Gammaproteobacteria; Enterobacterales; Enterobacteriaceae; Escherichia; Escherichia coli MTGDNTLIHSHGINRRDFMKLCAALAATMGLSSKAAAEMAESVTNPQRPPVIWIGAQECT WP_000145410.1 MULTISPECIES: hydrogenase-2 small chain [Proteobacteria] Length: 372\nNumber of predicted TMHs: 1\nExp number of AAs in TMHs: 23.72226\nExp number, first 60 AAs: 0.22038\nTotal prob of N-in: 0.12994\noutside 1 331\nTMhelix 332 354\ninside 355 372

11474 GCF_000408605.1_Esch_coli_KTE103_V1 Escherichia coli KTE103 Proteobacteria; Gammaproteobacteria; Enterobacterales; Enterobacteriaceae; Escherichia; Escherichia coli MTGDNTLIHSHGINRRDFMKLCAALAATMGLSSKAAAEMAESVTNPQRPPVIWIGAQECT WP_016245522.1 hydrogenase-2 small chain [Escherichia coli] Length: 372\nNumber of predicted TMHs: 1\nExp number of AAs in TMHs: 22.46122\nExp number, first 60 AAs: 0.22133\nTotal prob of N-in: 0.14101\noutside 1 331\nTMhelix 332 354\ninside 355 372

11475 GCF_000408605.1_Esch_coli_KTE103_V1 Escherichia coli KTE103 Proteobacteria; Gammaproteobacteria; Enterobacterales; Enterobacteriaceae; Escherichia; Escherichia coli MNNEETFYQAMRRQGVTRRSFLKYCSLAATSLGLGAGMAPKIAWALENKPRIPVVWIHGL WP_001058323.1 MULTISPECIES: hydrogenase-1 small chain [Proteobacteria] Length: 372\nNumber of predicted TMHs: 2\nExp number of AAs in TMHs: 37.2608499999999\nExp number, first 60 AAs: 19.52923\nTotal prob of N-in: 0.94351\nPOSSIBLE N-term signal sequence\ninside 1 19\nTMhelix 20 39\noutside 40 325\nTMhelix 326 348\ninside 349 372

11476 GCF_000353185.1_Esch_coli_KTE104_V1 Escherichia coli KTE104 Proteobacteria; Gammaproteobacteria; Enterobacterales; Enterobacteriaceae; Escherichia; Escherichia coli MNNEETFYQAMRRQGVTRRSFLKYCSLAATSLGLGAGMAPKIAWALENKPRIPVVWIHGL WP_001058323.1 MULTISPECIES: hydrogenase-1 small chain [Proteobacteria] Length: 372\nNumber of predicted TMHs: 2\nExp number of AAs in TMHs: 37.2608499999999\nExp number, first 60 AAs: 19.52923\nTotal prob of N-in: 0.94351\nPOSSIBLE N-term signal sequence\ninside 1 19\nTMhelix 20 39\noutside 40 325\nTMhelix 326 348\ninside 349 372

11477 GCF_000353185.1_Esch_coli_KTE104_V1 Escherichia coli KTE104 Proteobacteria; Gammaproteobacteria; Enterobacterales; Enterobacteriaceae; Escherichia; Escherichia coli MTGDNTLIHSHGINRRDFMKLCAALAATMGLSSKAAAEMAESVTNPQRPPVIWIGAQECT WP_000145410.1 MULTISPECIES: hydrogenase-2 small chain [Proteobacteria] Length: 372\nNumber of predicted TMHs: 1\nExp number of AAs in TMHs: 23.72226\nExp number, first 60 AAs: 0.22038\nTotal prob of N-in: 0.12994\noutside 1 331\nTMhelix 332 354\ninside 355 372

11478 GCF_000326145.1_Esch_coli_KTE105_V1 Escherichia coli KTE105 Proteobacteria; Gammaproteobacteria; Enterobacterales; Enterobacteriaceae; Escherichia; Escherichia coli MNNEETFYQAMRRQGVTRRSFLKYCSLAATSLGLGAGMAPKIAWALENKPRIPVVWIHGL WP_001058323.1 MULTISPECIES: hydrogenase-1 small chain [Proteobacteria] Length: 372\nNumber of predicted TMHs: 2\nExp number of AAs in TMHs: 37.2608499999999\nExp number, first 60 AAs: 19.52923\nTotal prob of N-in: 0.94351\nPOSSIBLE N-term signal sequence\ninside 1 19\nTMhelix 20 39\noutside 40 325\nTMhelix 326 348\ninside 349 372

11479 GCF_000326145.1_Esch_coli_KTE105_V1 Escherichia coli KTE105 Proteobacteria; Gammaproteobacteria; Enterobacterales; Enterobacteriaceae; Escherichia; Escherichia coli MTGDNTLIHSHGINRRDFMKLCAALAATMGLSSKAAAEMAESVTNPQRPPVIWIGAQECT WP_000145410.1 MULTISPECIES: hydrogenase-2 small chain [Proteobacteria] Length: 372\nNumber of predicted TMHs:

1\nExp number of AAs in TMHs: 23.72226\nExp number, first 60 AAs: 0.22038\nTotal prob of N-in: 0.12994\noutside 1 331\nTMhelix 332 354\ninside 355 372

11480 GCF_000326165.1_Esch_coli_KTE106_V1 Escherichia coli KTE106 Proteobacteria; Gammaproteobacteria; Enterobacterales; Enterobacteriaceae; Escherichia; Escherichia coli MNNEETFYQAMRRQGVTRRSFLKYCSLAATSLGLGAGMAPKIAWALENKPRIPVVWIHGL WP_001058323.1 MULTISPECIES: hydrogenase-1 small chain [Proteobacteria] Length: 372\nNumber of predicted TMHs: 2\nExp number of AAs in TMHs: 37.2608499999999\nExp number, first 60 AAs: 19.52923\nTotal prob of N-in: 0.94351\nPOSSIBLE N-term signal sequence\ninside 1 19\nTMhelix 20 39\noutside 40 325\nTMhelix 326 348\ninside 349 372

11481 GCF_000326165.1_Esch_coli_KTE106_V1 Escherichia coli KTE106 Proteobacteria; Gammaproteobacteria; Enterobacterales; Enterobacteriaceae; Escherichia; Escherichia coli MTGDNTLIHSHGINRRDFMKLCAALAATMGLSSKAAAEMAESVTNPQRPPVIWIGAQECT WP_000145410.1 MULTISPECIES: hydrogenase-2 small chain [Proteobacteria] Length: 372\nNumber of predicted TMHs: 1\nExp number of AAs in TMHs: 23.72226\nExp number, first 60 AAs: 0.22038\nTotal prob of N-in: 0.12994\noutside 1 331\nTMhelix 332 354\ninside 355 372

11482 GCF_000408625.1_Esch_coli_KTE107_V1 Escherichia coli KTE107 Proteobacteria; Gammaproteobacteria; Enterobacterales; Enterobacteriaceae; Escherichia; Escherichia coli MNNEETFYQAMRRQGVTRRSFLKYCSLAATSLGLGAGMAPKIAWALENKPRIPVVWIHGL WP_001058323.1 MULTISPECIES: hydrogenase-1 small chain [Proteobacteria] Length: 372\nNumber of predicted TMHs: 2\nExp number of AAs in TMHs: 37.2608499999999\nExp number, first 60 AAs: 19.52923\nTotal prob of N-in: 0.94351\nPOSSIBLE N-term signal sequence\ninside 1 19\nTMhelix 20 39\noutside 40 325\nTMhelix 326 348\ninside 349 372

11483 GCF_000408625.1_Esch_coli_KTE107_V1 Escherichia coli KTE107 Proteobacteria; Gammaproteobacteria; Enterobacterales; Enterobacteriaceae; Escherichia; Escherichia coli MTGDNTLIHSHGINRRDFMKLCAALAATMGLSSKAAAEMAESVTNPQRPPVIWIGAQECT WP_000145410.1 MULTISPECIES: hydrogenase-2 small chain [Proteobacteria] Length: 372\nNumber of predicted TMHs: 1\nExp number of AAs in TMHs: 23.72226\nExp number, first 60 AAs: 0.22038\nTotal prob of N-in: 0.12994\noutside 1 331\nTMhelix 332 354\ninside 355 372

11484 GCF_000408645.1_Esch_coli_KTE108_V1 Escherichia coli KTE108 Proteobacteria; Gammaproteobacteria; Enterobacterales; Enterobacteriaceae; Escherichia; Escherichia coli MNNEETFYQAMRRQGVTRRSFLKYCSLAATSLGLGAGMAPKIAWALENKPRIPVVWIHGL WP_001058323.1 MULTISPECIES: hydrogenase-1 small chain [Proteobacteria] Length: 372\nNumber of predicted TMHs: 2\nExp number of AAs in TMHs: 37.2608499999999\nExp number, first 60 AAs: 19.52923\nTotal prob of N-in: 0.94351\nPOSSIBLE N-term signal sequence\ninside 1 19\nTMhelix 20 39\noutside 40 325\nTMhelix 326 348\ninside 349 372

11485 GCF_000408645.1_Esch_coli_KTE108_V1 Escherichia coli KTE108 Proteobacteria; Gammaproteobacteria; Enterobacterales; Enterobacteriaceae; Escherichia; Escherichia coli MTGDNTLIHSHGINRRDFMKLCAALAATMGLSSKAAAEMAESVTNPQRPPVIWIGAQECT WP_000145410.1 MULTISPECIES: hydrogenase-2 small chain [Proteobacteria] Length: 372\nNumber of predicted TMHs: 1\nExp number of AAs in TMHs: 23.72226\nExp number, first 60 AAs: 0.22038\nTotal prob of N-in: 0.12994\noutside 1 331\nTMhelix 332 354\ninside 355 372

11486 GCF_000326185.1_Esch_coli_KTE109_V1 Escherichia coli KTE109 Proteobacteria; Gammaproteobacteria; Enterobacterales; Enterobacteriaceae; Escherichia; Escherichia coli MNNEETFYQAMRRQGVTRRSFLKYCSLAATSLGLGAGMAPKIAWALENKPRIPVVWIHGL WP_001058323.1 MULTISPECIES: hydrogenase-1 small chain [Proteobacteria] Length: 372\nNumber of predicted TMHs: 2\nExp number of AAs in TMHs: 37.2608499999999\nExp number, first 60 AAs: 19.52923\nTotal prob of N-in: 0.94351\nPOSSIBLE N-term signal sequence\ninside 1 19\nTMhelix 20 39\noutside 40 325\nTMhelix 326 348\ninside 349 372

11487 GCF_000326185.1_Esch_coli_KTE109_V1 Escherichia coli KTE109 Proteobacteria; Gammaproteobacteria; Enterobacterales; Enterobacteriaceae; Escherichia; Escherichia coli MTGDNTLIHSHGINRRDFMKLCAALAATMGLSSKAAAEMVESVTNPQRPPVIWIGAQECT WP_000145423.1 hydrogenase 2 small subunit [Escherichia coli] Length: 372\nNumber of predicted TMHs: 1\nExp number of

AAs in TMHs: 23.78201\nExp number, first 60 AAs: 0.28355\nTotal prob of N-in: 0.13257\noutside 1 331\nTMhelix 332 354\ninside 355 372

11488 GCF_000351865.1_Esch_coli_KTE111_V1 Escherichia coli KTE111 Proteobacteria; Gammaproteobacteria; Enterobacterales; Enterobacteriaceae; Escherichia; Escherichia coli MNNEETFYQAMRRQGVTRRSFLKYCSLAATSLGLGAGMAPKIAWALENKPRIPVVWIHGL WP_001058323.1 MULTISPECIES: hydrogenase-1 small chain [Proteobacteria] Length: 372\nNumber of predicted TMHs: 2\nExp number of AAs in TMHs: 37.2608499999999\nExp number, first 60 AAs: 19.52923\nTotal prob of N-in: 0.94351\nPOSSIBLE N-term signal sequence\ninside 1 19\nTMhelix 20 39\noutside 40 325\nTMhelix 326 348\ninside 349 372

11489 GCF_000351865.1_Esch_coli_KTE111_V1 Escherichia coli KTE111 Proteobacteria; Gammaproteobacteria; Enterobacterales; Enterobacteriaceae; Escherichia; Escherichia coli MTGDNTLIHSHGINRRDFMKLCAALAATMGLSSKAAAEMAESVTNPQRPPVIWIGAQECT WP_000145410.1 MULTISPECIES: hydrogenase-2 small chain [Proteobacteria] Length: 372\nNumber of predicted TMHs: 1\nExp number of AAs in TMHs: 23.72226\nExp number, first 60 AAs: 0.22038\nTotal prob of N-in: 0.12994\noutside 1 331\nTMhelix 332 354\ninside 355 372

11490 GCF_000326565.1_Esch_coli_KTE112_V1 Escherichia coli KTE112 Proteobacteria; Gammaproteobacteria; Enterobacterales; Enterobacteriaceae; Escherichia; Escherichia coli MTGDNTLIHSHGINRRDFMKLCAALAATMGLSSKAAAEMAESVTNPQRPPVIWIGAQECT WP_000145410.1 MULTISPECIES: hydrogenase-2 small chain [Proteobacteria] Length: 372\nNumber of predicted TMHs: 1\nExp number of AAs in TMHs: 23.72226\nExp number, first 60 AAs: 0.22038\nTotal prob of N-in: 0.12994\noutside 1 331\nTMhelix 332 354\ninside 355 372

11491 GCF_000326565.1_Esch_coli_KTE112_V1 Escherichia coli KTE112 Proteobacteria; Gammaproteobacteria; Enterobacterales; Enterobacteriaceae; Escherichia; Escherichia coli MNNEETFYQAMRRQGVTRRSFLKYCSLAATSLGLGAGMAPKIAWALENKPRIPVVWIHGL WP_001058323.1 MULTISPECIES: hydrogenase-1 small chain [Proteobacteria] Length: 372\nNumber of predicted TMHs: 2\nExp number of AAs in TMHs: 37.2608499999999\nExp number, first 60 AAs: 19.52923\nTotal prob of N-in: 0.94351\nPOSSIBLE N-term signal sequence\ninside 1 19\nTMhelix 20 39\noutside 40 325\nTMhelix 326 348\ninside 349 372

11492 GCF_000326585.1_Esch_coli_KTE113_V1 Escherichia coli KTE113 Proteobacteria; Gammaproteobacteria; Enterobacterales; Enterobacteriaceae; Escherichia; Escherichia coli MTGDNTLIHSHGINRRDFMKLCAALAATMGLSSKAAAEMVESVTNPQRPPVIWIGAQECT WP_000145423.1 hydrogenase 2 small subunit [Escherichia coli] Length: 372\nNumber of predicted TMHs: 1\nExp number of AAs in TMHs: 23.78201\nExp number, first 60 AAs: 0.28355\nTotal prob of N-in: 0.13257\noutside 1 331\nTMhelix 332 354\ninside 355 372

11493 GCF_000326585.1_Esch_coli_KTE113_V1 Escherichia coli KTE113 Proteobacteria; Gammaproteobacteria; Enterobacterales; Enterobacteriaceae; Escherichia; Escherichia coli MNNEETFYQAMRRQGVTRRSFLKYCSLAATSLGLGAGMAPKIAWALENKPRIPVVWIHGL WP_001058323.1 MULTISPECIES: hydrogenase-1 small chain [Proteobacteria] Length: 372\nNumber of predicted TMHs: 2\nExp number of AAs in TMHs: 37.2608499999999\nExp number, first 60 AAs: 19.52923\nTotal prob of N-in: 0.94351\nPOSSIBLE N-term signal sequence\ninside 1 19\nTMhelix 20 39\noutside 40 325\nTMhelix 326 348\ninside 349 372

11494 GCF_000352525.1_Esch_coli_KTE115_V1 Escherichia coli KTE115 Proteobacteria; Gammaproteobacteria; Enterobacterales; Enterobacteriaceae; Escherichia; Escherichia coli MNNEETFYQAMRRQGVTRRSFLKYCSLAATSLGLGAGMAPKIAWALENKPRIPVVWIHGL WP_001058323.1 MULTISPECIES: hydrogenase-1 small chain [Proteobacteria] Length: 372\nNumber of predicted TMHs: 2\nExp number of AAs in TMHs: 37.2608499999999\nExp number, first 60 AAs: 19.52923\nTotal prob of N-in: 0.94351\nPOSSIBLE N-term signal sequence\ninside 1 19\nTMhelix 20 39\noutside 40 325\nTMhelix 326 348\ninside 349 372

11495 GCF_000352525.1_Esch_coli_KTE115_V1 Escherichia coli KTE115 Proteobacteria; Gammaproteobacteria; Enterobacterales; Enterobacteriaceae; Escherichia; Escherichia coli MTGDNTLIHSHGINRRDFMKLCAALAATMGLSSKAAAEMAESVTNPQRPPVIWIGAQECT WP_000145410.1 MULTISPECIES: hydrogenase-2 small chain [Proteobacteria] Length: 372\nNumber of predicted TMHs:

1\nExp number of AAs in TMHs: 23.72226\nExp number, first 60 AAs: 0.22038\nTotal prob of N-in: 0.12994\noutside 1 331\nTMhelix 332 354\ninside 355 372

11496 GCF_000351885.1_Esch_coli_KTE116_V1 Escherichia coli KTE116 Proteobacteria; Gammaproteobacteria; Enterobacterales; Enterobacteriaceae; Escherichia; Escherichia coli MNNEETFYQAMRRQGVTRRSFLKYCSLAATSLGLGAGMAPKIAWALENKPRIPVVWIHGL WP_001058323.1 MULTISPECIES: hydrogenase-1 small chain [Proteobacteria] Length: 372\nNumber of predicted TMHs: 2\nExp number of AAs in TMHs: 37.2608499999999\nExp number, first 60 AAs: 19.52923\nTotal prob of N-in: 0.94351\nPOSSIBLE N-term signal sequence\ninside 1 19\nTMhelix 20 39\noutside 40 325\nTMhelix 326 348\ninside 349 372

11497 GCF_000351885.1_Esch_coli_KTE116_V1 Escherichia coli KTE116 Proteobacteria; Gammaproteobacteria; Enterobacterales; Enterobacteriaceae; Escherichia; Escherichia coli MTGDNTLIHSHGINRRDFMKLCAALAATMGLSSKAAAEMAESVTNPQRPPVIWIGAQECT WP_000145410.1 MULTISPECIES: hydrogenase-2 small chain [Proteobacteria] Length: 372\nNumber of predicted TMHs: 1\nExp number of AAs in TMHs: 23.72226\nExp number, first 60 AAs: 0.22038\nTotal prob of N-in: 0.12994\noutside 1 331\nTMhelix 332 354\ninside 355 372

11498 GCF_000326205.1_Esch_coli_KTE117_V1 Escherichia coli KTE117 Proteobacteria; Gammaproteobacteria; Enterobacterales; Enterobacteriaceae; Escherichia; Escherichia coli MNNEETFYQAMRRQGVTRRSFLKYCSLAATSLGLGAGMAPKIAWALENKPRIPVVWIHGL WP_001058323.1 MULTISPECIES: hydrogenase-1 small chain [Proteobacteria] Length: 372\nNumber of predicted TMHs: 2\nExp number of AAs in TMHs: 37.2608499999999\nExp number, first 60 AAs: 19.52923\nTotal prob of N-in: 0.94351\nPOSSIBLE N-term signal sequence\ninside 1 19\nTMhelix 20 39\noutside 40 325\nTMhelix 326 348\ninside 349 372

11499 GCF_000326205.1_Esch_coli_KTE117_V1 Escherichia coli KTE117 Proteobacteria; Gammaproteobacteria; Enterobacterales; Enterobacteriaceae; Escherichia; Escherichia coli MTGDNTLIHSHGINRRDFMKLCAALAATMGLSSKAAAEMAESVTNPQRPPVIWIGAQECT WP_000145410.1 MULTISPECIES: hydrogenase-2 small chain [Proteobacteria] Length: 372\nNumber of predicted TMHs: 1\nExp number of AAs in TMHs: 23.72226\nExp number, first 60 AAs: 0.22038\nTotal prob of N-in: 0.12994\noutside 1 331\nTMhelix 332 354\ninside 355 372

11500 GCF_000352545.1_Esch_coli_KTE118_V1 Escherichia coli KTE118 Proteobacteria; Gammaproteobacteria; Enterobacterales; Enterobacteriaceae; Escherichia; Escherichia coli MTGDNTLIHSHGINRRDFMKLCAALAATMGLSSKAAAEMAESVTNPQRPPVIWIGAQECT WP_000145410.1 MULTISPECIES: hydrogenase-2 small chain [Proteobacteria] Length: 372\nNumber of predicted TMHs: 1\nExp number of AAs in TMHs: 23.72226\nExp number, first 60 AAs: 0.22038\nTotal prob of N-in: 0.12994\noutside 1 331\nTMhelix 332 354\ninside 355 372

11501 GCF_000351905.1_Esch_coli_KTE119_V1 Escherichia coli KTE119 Proteobacteria; Gammaproteobacteria; Enterobacterales; Enterobacteriaceae; Escherichia; Escherichia coli MTGDNTLIHSHGINRRDFMKLCAALAATMGLSSKAAAEMAESVTNPQRPPVIWIGAQECT WP_000145410.1 MULTISPECIES: hydrogenase-2 small chain [Proteobacteria] Length: 372\nNumber of predicted TMHs: 1\nExp number of AAs in TMHs: 23.72226\nExp number, first 60 AAs: 0.22038\nTotal prob of N-in: 0.12994\noutside 1 331\nTMhelix 332 354\ninside 355 372

11502 GCF_000351905.1_Esch_coli_KTE119_V1 Escherichia coli KTE119 Proteobacteria; Gammaproteobacteria; Enterobacterales; Enterobacteriaceae; Escherichia; Escherichia coli MNNEETFYQAMRRQGVTRRSFLKYCSLAATSLGLGAGMAPKIAWALENKPRIPVVWIHGL WP_001058323.1 MULTISPECIES: hydrogenase-1 small chain [Proteobacteria] Length: 372\nNumber of predicted TMHs: 2\nExp number of AAs in TMHs: 37.2608499999999\nExp number, first 60 AAs: 19.52923\nTotal prob of N-in: 0.94351\nPOSSIBLE N-term signal sequence\ninside 1 19\nTMhelix 20 39\noutside 40 325\nTMhelix 326 348\ninside 349 372

11503 GCF_000350725.1_Esch_coli_KTE12_V1 Escherichia coli KTE12 Proteobacteria; Gammaproteobacteria; Enterobacterales; Enterobacteriaceae; Escherichia; Escherichia coli MNNEETFYQAMRRQGVTRRSFLKYCSLAATSLGLGAGMAPKIAWALENKPRIPVVWIHGL WP_001058323.1 MULTISPECIES: hydrogenase-1 small chain [Proteobacteria] Length: 372\nNumber of predicted TMHs: 2\nExp number of AAs in TMHs: 37.2608499999999\nExp number, first 60 AAs: 19.52923\nTotal prob of N-in:

0.94351\nPOSSIBLE N-term signal sequence\ninside 1 19\nTMhelix 20 39\noutside 40 325\nTMhelix 326 348\ninside 349 372

11504 GCF_000350725.1_Esch_coli_KTE12_V1 Escherichia coli KTE12 Proteobacteria; Gammaproteobacteria; Enterobacterales; Enterobacteriaceae; Escherichia; Escherichia coli MTGDNTLIHSHGINRRDFMKLCAALAATMGLSSKAAAEMAESVTNPQRPPVIWIGAQECT WP_000145410.1 MULTISPECIES: hydrogenase-2 small chain [Proteobacteria] Length: 372\nNumber of predicted TMHs: 1\nExp number of AAs in TMHs: 23.72226\nExp number, first 60 AAs: 0.22038\nTotal prob of N-in: 0.12994\noutside 1 331\nTMhelix 332 354\ninside 355 372

11505 GCF_000326925.1_Esch_coli_KTE120_V1 Escherichia coli KTE120 Proteobacteria; Gammaproteobacteria; Enterobacterales; Enterobacteriaceae; Escherichia; Escherichia coli MNNEETFYQAMRRQGVTRRSFLKYCSLAATSLGLGAGMAPKIAWALENKPRIPVVWIHGL WP_001058323.1 MULTISPECIES: hydrogenase-1 small chain [Proteobacteria] Length: 372\nNumber of predicted TMHs: 2\nExp number of AAs in TMHs: 37.2608499999999\nExp number, first 60 AAs: 19.52923\nTotal prob of N-in: 0.94351\nPOSSIBLE N-term signal sequence\ninside 1 19\nTMhelix 20 39\noutside 40 325\nTMhelix 326 348\ninside 349 372

11506 GCF_000326925.1_Esch_coli_KTE120_V1 Escherichia coli KTE120 Proteobacteria; Gammaproteobacteria; Enterobacterales; Enterobacteriaceae; Escherichia; Escherichia coli MTGDNTLIHSHGINRRDFMKLCAALAATMGLSSKAAAEMAESVTNPQRPPVIWIGAQECT WP_000145410.1 MULTISPECIES: hydrogenase-2 small chain [Proteobacteria] Length: 372\nNumber of predicted TMHs: 1\nExp number of AAs in TMHs: 23.72226\nExp number, first 60 AAs: 0.22038\nTotal prob of N-in: 0.12994\noutside 1 331\nTMhelix 332 354\ninside 355 372

11507 GCF_000408665.1_Esch_coli_KTE121_V1 Escherichia coli KTE121 Proteobacteria; Gammaproteobacteria; Enterobacterales; Enterobacteriaceae; Escherichia; Escherichia coli MNNEETFYQAMRRQGVTRRSFLKYCSLAATSLGLGAGMAPKIAWALENKPRIPVVWIHGL WP_001058323.1 MULTISPECIES: hydrogenase-1 small chain [Proteobacteria] Length: 372\nNumber of predicted TMHs: 2\nExp number of AAs in TMHs: 37.2608499999999\nExp number, first 60 AAs: 19.52923\nTotal prob of N-in: 0.94351\nPOSSIBLE N-term signal sequence\ninside 1 19\nTMhelix 20 39\noutside 40 325\nTMhelix 326 348\ninside 349 372

11508 GCF_000408665.1_Esch_coli_KTE121_V1 Escherichia coli KTE121 Proteobacteria; Gammaproteobacteria; Enterobacterales; Enterobacteriaceae; Escherichia; Escherichia coli MTGDNTLIHSHGINRRDFMKLCAALAATMGLSSKAAAEMAESVTNPQRPPVIWIGAQECT WP_000145410.1 MULTISPECIES: hydrogenase-2 small chain [Proteobacteria] Length: 372\nNumber of predicted TMHs: 1\nExp number of AAs in TMHs: 23.72226\nExp number, first 60 AAs: 0.22038\nTotal prob of N-in: 0.12994\noutside 1 331\nTMhelix 332 354\ninside 355 372

11509 GCF_000326945.1_Esch_coli_KTE122_V1 Escherichia coli KTE122 Proteobacteria; Gammaproteobacteria; Enterobacterales; Enterobacteriaceae; Escherichia; Escherichia coli MNNEETFYQAMRRQGVTRRSFLKYCSLAATSLGLGAGMAPKIAWALENKPRIPVVWIHGL WP_001058323.1 MULTISPECIES: hydrogenase-1 small chain [Proteobacteria] Length: 372\nNumber of predicted TMHs: 2\nExp number of AAs in TMHs: 37.2608499999999\nExp number, first 60 AAs: 19.52923\nTotal prob of N-in: 0.94351\nPOSSIBLE N-term signal sequence\ninside 1 19\nTMhelix 20 39\noutside 40 325\nTMhelix 326 348\ninside 349 372

11510 GCF_000326945.1_Esch_coli_KTE122_V1 Escherichia coli KTE122 Proteobacteria; Gammaproteobacteria; Enterobacterales; Enterobacteriaceae; Escherichia; Escherichia coli MTGDNTLIHSHGINRRDFMKLCAALAATMGLSSKAAAEMAESVTNPQRPPVIWIGAQECT WP_000145410.1 MULTISPECIES: hydrogenase-2 small chain [Proteobacteria] Length: 372\nNumber of predicted TMHs: 1\nExp number of AAs in TMHs: 23.72226\nExp number, first 60 AAs: 0.22038\nTotal prob of N-in: 0.12994\noutside 1 331\nTMhelix 332 354\ninside 355 372

11511 GCF_000352565.1_Esch_coli_KTE123_V1 Escherichia coli KTE123 Proteobacteria; Gammaproteobacteria; Enterobacterales; Enterobacteriaceae; Escherichia; Escherichia coli MTGDNTLIHSHGINRRDFMKLCAALAATMGLSSKAAAEMAESVTNPQRPPVIWIGAQECT WP_000145410.1 MULTISPECIES: hydrogenase-2 small chain [Proteobacteria] Length: 372\nNumber of predicted TMHs: 1\nExp number of AAs in TMHs: 23.72226\nExp number, first 60 AAs: 0.22038\nTotal prob of N-in: 0.12994\noutside 1 331\nTMhelix 332 354\ninside 355 372

11512 GCF_000326225.1_Esch_coli_KTE124_V1 Escherichia coli KTE124 Proteobacteria; Gammaproteobacteria; Enterobacterales; Enterobacteriaceae; Escherichia; Escherichia coli MTGDNTLIHSHGINRRDFMKLCAALAATMGLSSKAAAEMAESVTNPQRPPVIWIGAQECT WP_000145410.1 MULTISPECIES: hydrogenase-2 small chain [Proteobacteria] Length: 372\nNumber of predicted TMHs: 1\nExp number of AAs in TMHs: 23.72226\nExp number, first 60 AAs: 0.22038\nTotal prob of N-in: 0.12994\noutside 1 331\nTMhelix 332 354\ninside 355 372

11513 GCF_000326225.1_Esch_coli_KTE124_V1 Escherichia coli KTE124 Proteobacteria; Gammaproteobacteria; Enterobacterales; Enterobacteriaceae; Escherichia; Escherichia coli MNNEETFYQAMRRQGVTRRSFLKYCSLAATSLGLGAGMAPKIAWALENKPRIPVVWIHGL WP_001058323.1 MULTISPECIES: hydrogenase-1 small chain [Proteobacteria] Length: 372\nNumber of predicted TMHs: 2\nExp number of AAs in TMHs: 37.2608499999999\nExp number, first 60 AAs: 19.52923\nTotal prob of N-in: 0.94351\nPOSSIBLE N-term signal sequence\ninside 1 19\nTMhelix 20 39\noutside 40 325\nTMhelix 326 348\ninside 349 372

11514 GCF_000326965.1_Esch_coli_KTE125_V1 Escherichia coli KTE125 Proteobacteria; Gammaproteobacteria; Enterobacterales; Enterobacteriaceae; Escherichia; Escherichia coli MTGDNTLIHSHGINRRDFMKLCAALAATMGLSSKAAAEMAESVTNPQRPPVIWIGAQECT WP_000145410.1 MULTISPECIES: hydrogenase-2 small chain [Proteobacteria] Length: 372\nNumber of predicted TMHs: 1\nExp number of AAs in TMHs: 23.72226\nExp number, first 60 AAs: 0.22038\nTotal prob of N-in: 0.12994\noutside 1 331\nTMhelix 332 354\ninside 355 372

11515 GCF_000326965.1_Esch_coli_KTE125_V1 Escherichia coli KTE125 Proteobacteria; Gammaproteobacteria; Enterobacterales; Enterobacteriaceae; Escherichia; Escherichia coli MNNEETFYQAMRRQGVTRRSFLKYCSLAATSLGLGAGMAPKIAWALENKPRIPVVWIHGL WP_001058323.1 MULTISPECIES: hydrogenase-1 small chain [Proteobacteria] Length: 372\nNumber of predicted TMHs: 2\nExp number of AAs in TMHs: 37.2608499999999\nExp number, first 60 AAs: 19.52923\nTotal prob of N-in: 0.94351\nPOSSIBLE N-term signal sequence\ninside 1 19\nTMhelix 20 39\noutside 40 325\nTMhelix 326 348\ninside 349 372

11516 GCF_000408685.1_Esch_coli_KTE126_V1 Escherichia coli KTE126 Proteobacteria; Gammaproteobacteria; Enterobacterales; Enterobacteriaceae; Escherichia; Escherichia coli MNNEETFYQAMRRQGVTRRSFLKYCSLAATSLGLGAGMAPKIAWALENKPRIPVVWIHGL WP_001058323.1 MULTISPECIES: hydrogenase-1 small chain [Proteobacteria] Length: 372\nNumber of predicted TMHs: 2\nExp number of AAs in TMHs: 37.2608499999999\nExp number, first 60 AAs: 19.52923\nTotal prob of N-in: 0.94351\nPOSSIBLE N-term signal sequence\ninside 1 19\nTMhelix 20 39\noutside 40 325\nTMhelix 326 348\ninside 349 372

11517 GCF_000408685.1_Esch_coli_KTE126_V1 Escherichia coli KTE126 Proteobacteria; Gammaproteobacteria; Enterobacterales; Enterobacteriaceae; Escherichia; Escherichia coli MTGDNTLIHSHGINRRDFMKLCAALAATMGLSSKAAAEMAESVTNPQRPPVIWIGAQECT WP_000145410.1 MULTISPECIES: hydrogenase-2 small chain [Proteobacteria] Length: 372\nNumber of predicted TMHs: 1\nExp number of AAs in TMHs: 23.72226\nExp number, first 60 AAs: 0.22038\nTotal prob of N-in: 0.12994\noutside 1 331\nTMhelix 332 354\ninside 355 372

11518 GCF_000408705.1_Esch_coli_KTE127_V1 Escherichia coli KTE127 Proteobacteria; Gammaproteobacteria; Enterobacterales; Enterobacteriaceae; Escherichia; Escherichia coli MTGDNTLIHSHGINRRDFMKLCAALAATMGLSSKAAAEMAESVTNPQRPPVIWIGAQECT WP_000145410.1 MULTISPECIES: hydrogenase-2 small chain [Proteobacteria] Length: 372\nNumber of predicted TMHs: 1\nExp number of AAs in TMHs: 23.72226\nExp number, first 60 AAs: 0.22038\nTotal prob of N-in: 0.12994\noutside 1 331\nTMhelix 332 354\ninside 355 372

11519 GCF_000408705.1_Esch_coli_KTE127_V1 Escherichia coli KTE127 Proteobacteria; Gammaproteobacteria; Enterobacterales; Enterobacteriaceae; Escherichia; Escherichia coli MNNEETFYQAMRRQGVTRRSFLKYCSLAATSLGLGAGMAPKIAWALENKPRIPVVWIHGL WP_001058323.1 MULTISPECIES: hydrogenase-1 small chain [Proteobacteria] Length: 372\nNumber of predicted TMHs: 2\nExp number of AAs in TMHs: 37.2608499999999\nExp number, first 60 AAs: 19.52923\nTotal prob of N-in: 0.94351\nPOSSIBLE N-term signal sequence\ninside 1 19\nTMhelix 20 39\noutside 40 325\nTMhelix 326 348\ninside 349 372

11520 GCF_000326245.1_Esch_coli_KTE128_V1 Escherichia coli KTE128 Proteobacteria; Gammaproteobacteria; Enterobacterales; Enterobacteriaceae; Escherichia; Escherichia coli MTGDNTLIHSHGINRRDFMKLCAALAATMGLSSKAAAEMAESVTNPQRPPVIWIGAQECT WP_000145410.1 MULTISPECIES: hydrogenase-2 small chain [Proteobacteria] Length: 372\nNumber of predicted TMHs: 1\nExp number of AAs in TMHs: 23.72226\nExp number, first 60 AAs: 0.22038\nTotal prob of N-in: 0.12994\noutside 1 331\nTMhelix 332 354\ninside 355 372

11521 GCF_000326245.1_Esch_coli_KTE128_V1 Escherichia coli KTE128 Proteobacteria; Gammaproteobacteria; Enterobacterales; Enterobacteriaceae; Escherichia; Escherichia coli MNNEETFYQAMRRQGVTRRSFLKYCSLAATSLGLGAGMAPKIAWALENKPRIPVVWIHGL WP_001058323.1 MULTISPECIES: hydrogenase-1 small chain [Proteobacteria] Length: 372\nNumber of predicted TMHs: 2\nExp number of AAs in TMHs: 37.2608499999999\nExp number, first 60 AAs: 19.52923\nTotal prob of N-in: 0.94351\nPOSSIBLE N-term signal sequence\ninside 1 19\nTMhelix 20 39\noutside 40 325\nTMhelix 326 348\ninside 349 372

11522 GCF_000326265.1_Esch_coli_KTE129_V1 Escherichia coli KTE129 Proteobacteria; Gammaproteobacteria; Enterobacterales; Enterobacteriaceae; Escherichia; Escherichia coli MTGDNTLIHSHGINRRDFMKLCAALAATMGLSSKAAAEMVESVTNPQRPPVIWIGAQECT WP_000145423.1 hydrogenase 2 small subunit [Escherichia coli] Length: 372\nNumber of predicted TMHs: 1\nExp number of AAs in TMHs: 23.78201\nExp number, first 60 AAs: 0.28355\nTotal prob of N-in: 0.13257\noutside 1 331\nTMhelix 332 354\ninside 355 372

11523 GCF_000326265.1_Esch_coli_KTE129_V1 Escherichia coli KTE129 Proteobacteria; Gammaproteobacteria; Enterobacterales; Enterobacteriaceae; Escherichia; Escherichia coli MNNEETFYQAMRRQGVTRRSFLKYCSLAATSLGLGAGMAPKIAWALENKPRIPVVWIHGL WP_001538532.1 hydrogenase-1 small chain [Escherichia coli] Length: 372\nNumber of predicted TMHs: 2\nExp number of AAs in TMHs: 37.27041\nExp number, first 60 AAs: 19.52927\nTotal prob of N-in: 0.94352\nPOSSIBLE N-term signal sequence\ninside 1 19\nTMhelix 20 39\noutside 40 325\nTMhelix 326 348\ninside 349 372

11524 GCF_000407725.1_Esch_coli_KTE13_V1 Escherichia coli KTE13 Proteobacteria; Gammaproteobacteria; Enterobacterales; Enterobacteriaceae; Escherichia; Escherichia coli MTGDNTLIHSHGINRRDFMKLCAALAATMGLSSKAAAEMAESVTNPQRPPVIWIGAQECT WP_000145410.1 MULTISPECIES: hydrogenase-2 small chain [Proteobacteria] Length: 372\nNumber of predicted TMHs: 1\nExp number of AAs in TMHs: 23.72226\nExp number, first 60 AAs: 0.22038\nTotal prob of N-in: 0.12994\noutside 1 331\nTMhelix 332 354\ninside 355 372

11525 GCF_000407725.1_Esch_coli_KTE13_V1 Escherichia coli KTE13 Proteobacteria; Gammaproteobacteria; Enterobacterales; Enterobacteriaceae; Escherichia; Escherichia coli MNNEETFYQAMRRQGVTRRSFLKYCSLAATSLGLGAGMAPKIAWALENKPRIPVVWIHGL WP_001058323.1 MULTISPECIES: hydrogenase-1 small chain [Proteobacteria] Length: 372\nNumber of predicted TMHs: 2\nExp number of AAs in TMHs: 37.2608499999999\nExp number, first 60 AAs: 19.52923\nTotal prob of N-in: 0.94351\nPOSSIBLE N-term signal sequence\ninside 1 19\nTMhelix 20 39\noutside 40 325\nTMhelix 326 348\ninside 349 372

11526 GCF_000408725.1_Esch_coli_KTE130_V1 Escherichia coli KTE130 Proteobacteria; Gammaproteobacteria; Enterobacterales; Enterobacteriaceae; Escherichia; Escherichia coli MNNEETFYQAMRRQGVTRRSFLKYCSLAATSLGLGAGMAPKIAWALENKPRIPVVWIHGL WP_001058323.1 MULTISPECIES: hydrogenase-1 small chain [Proteobacteria] Length: 372\nNumber of predicted TMHs: 2\nExp number of AAs in TMHs: 37.2608499999999\nExp number, first 60 AAs: 19.52923\nTotal prob of N-in: 0.94351\nPOSSIBLE N-term signal sequence\ninside 1 19\nTMhelix 20 39\noutside 40 325\nTMhelix 326 348\ninside 349 372

11527 GCF_000408725.1_Esch_coli_KTE130_V1 Escherichia coli KTE130 Proteobacteria; Gammaproteobacteria; Enterobacterales; Enterobacteriaceae; Escherichia; Escherichia coli MTGDNTLIHSHGINRRDFMKLCAALAATMGLSSKAAAEMAESVTNPQRPPVIWIGAQECT WP_000145410.1 MULTISPECIES: hydrogenase-2 small chain [Proteobacteria] Length: 372\nNumber of predicted TMHs: 1\nExp number of AAs in TMHs: 23.72226\nExp number, first 60 AAs: 0.22038\nTotal prob of N-in: 0.12994\noutside 1 331\nTMhelix 332 354\ninside 355 372

11528 GCF_000326285.1_Esch_coli_KTE131_V1 Escherichia coli KTE131 Proteobacteria; Gammaproteobacteria; Enterobacterales; Enterobacteriaceae; Escherichia; Escherichia coli

MNNEETFYQAMRRQGVTRRSFLKYCSLAATSLGLGAGMAPKIAWALENKPRIPVVWIHGL WP_001058323.1 MULTISPECIES: hydrogenase-1 small chain [Proteobacteria] Length: 372\nNumber of predicted TMHs: 2\nExp number of AAs in TMHs: 37.2608499999999\nExp number, first 60 AAs: 19.52923\nTotal prob of N-in: 0.94351\nPOSSIBLE N-term signal sequence\ninside 1 19\nTMhelix 20 39\noutside 40 325\nTMhelix 326 348\ninside 349 372

11529 GCF_000326285.1_Esch_coli_KTE131_V1 Escherichia coli KTE131 Proteobacteria; Gammaproteobacteria; Enterobacterales; Enterobacteriaceae; Escherichia; Escherichia coli MTGDNTLIHSHGINRRDFMKLCAALAATMGLSSKAAAEMAESVTNPQRPPVIWIGAQECT WP_000145410.1 MULTISPECIES: hydrogenase-2 small chain [Proteobacteria] Length: 372\nNumber of predicted TMHs: 1\nExp number of AAs in TMHs: 23.72226\nExp number, first 60 AAs: 0.22038\nTotal prob of N-in: 0.12994\noutside 1 331\nTMhelix 332 354\ninside 355 372

11530 GCF_000408745.1_Esch_coli_KTE132_V1 Escherichia coli KTE132 Proteobacteria; Gammaproteobacteria; Enterobacterales; Enterobacteriaceae; Escherichia; Escherichia coli MNNEETFYQAMRRQGVTRRSFLKYCSLAATSLGLGAGMAPKIAWALENKPRIPVVWIHGL WP_001058323.1 MULTISPECIES: hydrogenase-1 small chain [Proteobacteria] Length: 372\nNumber of predicted TMHs: 2\nExp number of AAs in TMHs: 37.2608499999999\nExp number, first 60 AAs: 19.52923\nTotal prob of N-in: 0.94351\nPOSSIBLE N-term signal sequence\ninside 1 19\nTMhelix 20 39\noutside 40 325\nTMhelix 326 348\ninside 349 372

11531 GCF_000408745.1_Esch_coli_KTE132_V1 Escherichia coli KTE132 Proteobacteria; Gammaproteobacteria; Enterobacterales; Enterobacteriaceae; Escherichia; Escherichia coli MTGDNTLIHSHGINRRDFMKLCAALAATMGLSSKAAAEMAESVTNPQRPPVIWIGAQECT WP_000145410.1 MULTISPECIES: hydrogenase-2 small chain [Proteobacteria] Length: 372\nNumber of predicted TMHs: 1\nExp number of AAs in TMHs: 23.72226\nExp number, first 60 AAs: 0.22038\nTotal prob of N-in: 0.12994\noutside 1 331\nTMhelix 332 354\ninside 355 372

11532 GCF_000326305.1_Esch_coli_KTE133_V1 Escherichia coli KTE133 Proteobacteria; Gammaproteobacteria; Enterobacterales; Enterobacteriaceae; Escherichia; Escherichia coli MTGDNTLIHSHGINRRDFMKLCAALAATMGLSSKAAAEMVESVTNPQRPPVIWIGAQECT WP_000145423.1 hydrogenase 2 small subunit [Escherichia coli] Length: 372\nNumber of predicted TMHs: 1\nExp number of AAs in TMHs: 23.78201\nExp number, first 60 AAs: 0.28355\nTotal prob of N-in: 0.13257\noutside 1 331\nTMhelix 332 354\ninside 355 372

11533 GCF_000326305.1_Esch_coli_KTE133_V1 Escherichia coli KTE133 Proteobacteria; Gammaproteobacteria; Enterobacterales; Enterobacteriaceae; Escherichia; Escherichia coli MNNEETFYQAMRRQGVTRRSFLKYCSLAATSLGLGAGMAPKIAWALENKPRIPVVWIHGL WP_001058323.1 MULTISPECIES: hydrogenase-1 small chain [Proteobacteria] Length: 372\nNumber of predicted TMHs: 2\nExp number of AAs in TMHs: 37.2608499999999\nExp number, first 60 AAs: 19.52923\nTotal prob of N-in: 0.94351\nPOSSIBLE N-term signal sequence\ninside 1 19\nTMhelix 20 39\noutside 40 325\nTMhelix 326 348\ninside 349 372

11534 GCF_000408765.1_Esch_coli_KTE134_V1 Escherichia coli KTE134 Proteobacteria; Gammaproteobacteria; Enterobacterales; Enterobacteriaceae; Escherichia; Escherichia coli MNNEETFYQAMRRQGVTRRSFLKYCSLAATSLGLGAGMAPKIAWALENKPRIPVVWIHGL WP_001058323.1 MULTISPECIES: hydrogenase-1 small chain [Proteobacteria] Length: 372\nNumber of predicted TMHs: 2\nExp number of AAs in TMHs: 37.2608499999999\nExp number, first 60 AAs: 19.52923\nTotal prob of N-in: 0.94351\nPOSSIBLE N-term signal sequence\ninside 1 19\nTMhelix 20 39\noutside 40 325\nTMhelix 326 348\ninside 349 372

11535 GCF_000408765.1_Esch_coli_KTE134_V1 Escherichia coli KTE134 Proteobacteria; Gammaproteobacteria; Enterobacterales; Enterobacteriaceae; Escherichia; Escherichia coli MTGDNTLIHSHGINRRDFMKLCAALAATMGLSSKAAAEMAESVTNPQRPPVIWIGAQECT WP_000145410.1 MULTISPECIES: hydrogenase-2 small chain [Proteobacteria] Length: 372\nNumber of predicted TMHs: 1\nExp number of AAs in TMHs: 23.72226\nExp number, first 60 AAs: 0.22038\nTotal prob of N-in: 0.12994\noutside 1 331\nTMhelix 332 354\ninside 355 372

11536 GCF_000352585.1_Esch_coli_KTE135_V1 Escherichia coli KTE135 Proteobacteria; Gammaproteobacteria; Enterobacterales; Enterobacteriaceae; Escherichia; Escherichia coli MNNEETFYQAMRRQGVTRRSFLKYCSLAATSLGLGAGMAPKIAWALENKPRIPVVWIHGL WP_001058323.1

MULTISPECIES: hydrogenase-1 small chain [Proteobacteria] Length: 372\nNumber of predicted TMHs: 2\nExp number of AAs in TMHs: 37.2608499999999\nExp number, first 60 AAs: 19.52923\nTotal prob of N-in: 0.94351\nPOSSIBLE N-term signal sequence\ninside 1 19\nTMhelix 20 39\noutside 40 325\nTMhelix 326 348\ninside 349 372

11537 GCF_000352585.1_Esch_coli_KTE135_V1 Escherichia coli KTE135 Proteobacteria; Gammaproteobacteria; Enterobacterales; Enterobacteriaceae; Escherichia; Escherichia coli MTGDNTLIHSHGINRRDFMKLCAALAATMGLSSKAAAEMAESVTNPQRPPVIWIGAQECT WP_000145410.1 MULTISPECIES: hydrogenase-2 small chain [Proteobacteria] Length: 372\nNumber of predicted TMHs: 1\nExp number of AAs in TMHs: 23.72226\nExp number, first 60 AAs: 0.22038\nTotal prob of N-in: 0.12994\noutside 1 331\nTMhelix 332 354\ninside 355 372

11538 GCF_000352605.1_Esch_coli_KTE136_V1 Escherichia coli KTE136 Proteobacteria; Gammaproteobacteria; Enterobacterales; Enterobacteriaceae; Escherichia; Escherichia coli MNNEETFYQAMRRQGVTRRSFLKYCSLAATSLGLGAGMAPKIAWALENKPRIPVVWIHGL WP_001058323.1 MULTISPECIES: hydrogenase-1 small chain [Proteobacteria] Length: 372\nNumber of predicted TMHs: 2\nExp number of AAs in TMHs: 37.2608499999999\nExp number, first 60 AAs: 19.52923\nTotal prob of N-in: 0.94351\nPOSSIBLE N-term signal sequence\ninside 1 19\nTMhelix 20 39\noutside 40 325\nTMhelix 326 348\ninside 349 372

11539 GCF_000352605.1_Esch_coli_KTE136_V1 Escherichia coli KTE136 Proteobacteria; Gammaproteobacteria; Enterobacterales; Enterobacteriaceae; Escherichia; Escherichia coli MTGDNTLIHSHGINRRDFMKLCAALAATMGLSSKAAAEMAESVTNPQRPPVIWIGAQECT WP_000145410.1 MULTISPECIES: hydrogenase-2 small chain [Proteobacteria] Length: 372\nNumber of predicted TMHs: 1\nExp number of AAs in TMHs: 23.72226\nExp number, first 60 AAs: 0.22038\nTotal prob of N-in: 0.12994\noutside 1 331\nTMhelix 332 354\ninside 355 372

11540 GCF_000326325.1_Esch_coli_KTE137_V1 Escherichia coli KTE137 Proteobacteria; Gammaproteobacteria; Enterobacterales; Enterobacteriaceae; Escherichia; Escherichia coli MTGDNTLIHSHGINRRDFMKLCAALAATMGLSSKAAAEMAESVTNPQRPPVIWIGAQECT WP_000145410.1 MULTISPECIES: hydrogenase-2 small chain [Proteobacteria] Length: 372\nNumber of predicted TMHs: 1\nExp number of AAs in TMHs: 23.72226\nExp number, first 60 AAs: 0.22038\nTotal prob of N-in: 0.12994\noutside 1 331\nTMhelix 332 354\ninside 355 372

11541 GCF_000326325.1_Esch_coli_KTE137_V1 Escherichia coli KTE137 Proteobacteria; Gammaproteobacteria; Enterobacterales; Enterobacteriaceae; Escherichia; Escherichia coli MNNEETFYQAMRRQGVTRRSFLKYCSLAATSLGLGAGMAPKIAWALENKPRIPVVWIHGL WP_001058323.1 MULTISPECIES: hydrogenase-1 small chain [Proteobacteria] Length: 372\nNumber of predicted TMHs: 2\nExp number of AAs in TMHs: 37.2608499999999\nExp number, first 60 AAs: 19.52923\nTotal prob of N-in: 0.94351\nPOSSIBLE N-term signal sequence\ninside 1 19\nTMhelix 20 39\noutside 40 325\nTMhelix 326 348\ninside 349 372

11542 GCF_000326685.1_Esch_coli_KTE138_V1 Escherichia coli KTE138 Proteobacteria; Gammaproteobacteria; Enterobacterales; Enterobacteriaceae; Escherichia; Escherichia coli MTGDNTLIHSHGINRRDFMKLCAALAATMGLSSKAAAEMAESVTNPQRPPVIWIGAQECT WP_000145410.1 MULTISPECIES: hydrogenase-2 small chain [Proteobacteria] Length: 372\nNumber of predicted TMHs: 1\nExp number of AAs in TMHs: 23.72226\nExp number, first 60 AAs: 0.22038\nTotal prob of N-in: 0.12994\noutside 1 331\nTMhelix 332 354\ninside 355 372

11543 GCF_000326685.1_Esch_coli_KTE138_V1 Escherichia coli KTE138 Proteobacteria; Gammaproteobacteria; Enterobacterales; Enterobacteriaceae; Escherichia; Escherichia coli MNNEETFYQAMRRQGVTRRSFLKYCSLAATSLGLGAGMAPKIAWALENKPRIPVVWIHGL WP_001295940.1 MULTISPECIES: hydrogenase [Enterobacteriaceae] Length: 372\nNumber of predicted TMHs: 2\nExp number of AAs in TMHs: 37.2608899999999\nExp number, first 60 AAs: 19.52923\nTotal prob of N-in: 0.94351\nPOSSIBLE N-term signal sequence\ninside 1 19\nTMhelix 20 39\noutside 40 325\nTMhelix 326 348\ninside 349 372

11544 GCF_000326345.1_Esch_coli_KTE139_V1 Escherichia coli KTE139 Proteobacteria; Gammaproteobacteria; Enterobacterales; Enterobacteriaceae; Escherichia; Escherichia coli MTGDNTLIHSHGINRRDFMKLCAALAATMGLSSKAAAEMAESVTNPQRPPVIWIGAQECT WP_000145410.1 MULTISPECIES: hydrogenase-2 small chain [Proteobacteria] Length: 372\nNumber of predicted TMHs:

1\nExp number of AAs in TMHs: 23.72226\nExp number, first 60 AAs: 0.22038\nTotal prob of N-in: 0.12994\noutside 1 331\nTMhelix 332 354\ninside 355 372

11545 GCF_000326345.1_Esch_coli_KTE139_V1 Escherichia coli KTE139 Proteobacteria; Gammaproteobacteria; Enterobacterales; Enterobacteriaceae; Escherichia; Escherichia coli MNNEETFYQAMRRQGVTRRSFLKYCSLAATSLGLGAGMAPKIAWALENKPRIPVVWIHGL WP_001058323.1 MULTISPECIES: hydrogenase-1 small chain [Proteobacteria] Length: 372\nNumber of predicted TMHs: 2\nExp number of AAs in TMHs: 37.2608499999999\nExp number, first 60 AAs: 19.52923\nTotal prob of N-in: 0.94351\nPOSSIBLE N-term signal sequence\ninside 1 19\nTMhelix 20 39\noutside 40 325\nTMhelix 326 348\ninside 349 372

11546 GCF_000407805.1_Esch_coli_KTE14_V1 Escherichia coli KTE14 Proteobacteria; Gammaproteobacteria; Enterobacterales; Enterobacteriaceae; Escherichia; Escherichia coli MNNEETFYQAMRRQGVTRRSFLKYCSLAATSLGLGAGMAPKIAWALENKPRIPVVWIHGL WP_001058323.1 MULTISPECIES: hydrogenase-1 small chain [Proteobacteria] Length: 372\nNumber of predicted TMHs: 2\nExp number of AAs in TMHs: 37.2608499999999\nExp number, first 60 AAs: 19.52923\nTotal prob of N-in: 0.94351\nPOSSIBLE N-term signal sequence\ninside 1 19\nTMhelix 20 39\noutside 40 325\nTMhelix 326 348\ninside 349 372

11547 GCF_000407805.1_Esch_coli_KTE14_V1 Escherichia coli KTE14 Proteobacteria; Gammaproteobacteria; Enterobacterales; Enterobacteriaceae; Escherichia; Escherichia coli MTGDNTLIHSHGINRRDFMKLCAALAATMGLSSKAAAEMAESVTNPQRPPVIWIGAQECT WP_000145410.1 MULTISPECIES: hydrogenase-2 small chain [Proteobacteria] Length: 372\nNumber of predicted TMHs: 1\nExp number of AAs in TMHs: 23.72226\nExp number, first 60 AAs: 0.22038\nTotal prob of N-in: 0.12994\noutside 1 331\nTMhelix 332 354\ninside 355 372

11548 GCF_000352625.1_Esch_coli_KTE140_V1 Escherichia coli KTE140 Proteobacteria; Gammaproteobacteria; Enterobacterales; Enterobacteriaceae; Escherichia; Escherichia coli MTGDNTLIHSHGINRRDFMKLCAALAATMGLSSKAAAEMAESVTNPQRPPVIWIGAQECT WP_000145410.1 MULTISPECIES: hydrogenase-2 small chain [Proteobacteria] Length: 372\nNumber of predicted TMHs: 1\nExp number of AAs in TMHs: 23.72226\nExp number, first 60 AAs: 0.22038\nTotal prob of N-in: 0.12994\noutside 1 331\nTMhelix 332 354\ninside 355 372

11549 GCF_000352625.1_Esch_coli_KTE140_V1 Escherichia coli KTE140 Proteobacteria; Gammaproteobacteria; Enterobacterales; Enterobacteriaceae; Escherichia; Escherichia coli MNNEETFYQAMRRQGVTRRSFLKYCSLAATSLGLGAGMAPKIAWALENKPRIPVVWIHGL WP_001058323.1 MULTISPECIES: hydrogenase-1 small chain [Proteobacteria] Length: 372\nNumber of predicted TMHs: 2\nExp number of AAs in TMHs: 37.2608499999999\nExp number, first 60 AAs: 19.52923\nTotal prob of N-in: 0.94351\nPOSSIBLE N-term signal sequence\ninside 1 19\nTMhelix 20 39\noutside 40 325\nTMhelix 326 348\ninside 349 372

11550 GCF_000352645.1_Esch_coli_KTE141_V1 Escherichia coli KTE141 Proteobacteria; Gammaproteobacteria; Enterobacterales; Enterobacteriaceae; Escherichia; Escherichia coli MTGDNTLIHSHGINRRDFMKLCAALAATMGLSSKAAAEMAESVTNPQRPPVIWIGAQECT WP_000145410.1 MULTISPECIES: hydrogenase-2 small chain [Proteobacteria] Length: 372\nNumber of predicted TMHs: 1\nExp number of AAs in TMHs: 23.72226\nExp number, first 60 AAs: 0.22038\nTotal prob of N-in: 0.12994\noutside 1 331\nTMhelix 332 354\ninside 355 372

11551 GCF_000352645.1_Esch_coli_KTE141_V1 Escherichia coli KTE141 Proteobacteria; Gammaproteobacteria; Enterobacterales; Enterobacteriaceae; Escherichia; Escherichia coli MNNEETFYQAMRRQGVTRRSFLKYCSLAATSLGLGAGMAPKIAWALENKPRIPVVWIHGL WP_001058323.1 MULTISPECIES: hydrogenase-1 small chain [Proteobacteria] Length: 372\nNumber of predicted TMHs: 2\nExp number of AAs in TMHs: 37.2608499999999\nExp number, first 60 AAs: 19.52923\nTotal prob of N-in: 0.94351\nPOSSIBLE N-term signal sequence\ninside 1 19\nTMhelix 20 39\noutside 40 325\nTMhelix 326 348\ninside 349 372

11552 GCF_000351925.1_Esch_coli_KTE142_V1 Escherichia coli KTE142 Proteobacteria; Gammaproteobacteria; Enterobacterales; Enterobacteriaceae; Escherichia; Escherichia coli MTGDNTLIHSHGINRRDFMKLCAALAATMGLSSKAAAEMAESVTNPQRPPVIWIGAQECT WP_000145410.1 MULTISPECIES: hydrogenase-2 small chain [Proteobacteria] Length: 372\nNumber of predicted TMHs:

1\nExp number of AAs in TMHs: 23.72226\nExp number, first 60 AAs: 0.22038\nTotal prob of N-in: 0.12994\noutside 1 331\nTMhelix 332 354\ninside 355 372

11553 GCF_000351925.1_Esch_coli_KTE142_V1 Escherichia coli KTE142 Proteobacteria; Gammaproteobacteria; Enterobacterales; Enterobacteriaceae; Escherichia; Escherichia coli MNNEETFYQAMRRQGVTRRSFLKYCSLAATSLGLGAGMAPKIAWALENKPRIPVVWIHGL WP_001058323.1 MULTISPECIES: hydrogenase-1 small chain [Proteobacteria] Length: 372\nNumber of predicted TMHs: 2\nExp number of AAs in TMHs: 37.2608499999999\nExp number, first 60 AAs: 19.52923\nTotal prob of N-in: 0.94351\nPOSSIBLE N-term signal sequence\ninside 1 19\nTMhelix 20 39\noutside 40 325\nTMhelix 326 348\ninside 349 372

11554 GCF_000351945.1_Esch_coli_KTE143_V1 Escherichia coli KTE143 Proteobacteria; Gammaproteobacteria; Enterobacterales; Enterobacteriaceae; Escherichia; Escherichia coli MTGDNTLIHSHGINRRDFMKLCAALAATMGLSSKAAAEMAESVTNPQRPPVIWIGAQECT WP_000145410.1 MULTISPECIES: hydrogenase-2 small chain [Proteobacteria] Length: 372\nNumber of predicted TMHs: 1\nExp number of AAs in TMHs: 23.72226\nExp number, first 60 AAs: 0.22038\nTotal prob of N-in: 0.12994\noutside 1 331\nTMhelix 332 354\ninside 355 372

11555 GCF_000351945.1_Esch_coli_KTE143_V1 Escherichia coli KTE143 Proteobacteria; Gammaproteobacteria; Enterobacterales; Enterobacteriaceae; Escherichia; Escherichia coli MNNEETFYQAMRRQGVTRRSFLKYCSLAATSLGLGAGMAPKIAWALENKPRIPVVWIHGL WP_001058323.1 MULTISPECIES: hydrogenase-1 small chain [Proteobacteria] Length: 372\nNumber of predicted TMHs: 2\nExp number of AAs in TMHs: 37.2608499999999\nExp number, first 60 AAs: 19.52923\nTotal prob of N-in: 0.94351\nPOSSIBLE N-term signal sequence\ninside 1 19\nTMhelix 20 39\noutside 40 325\nTMhelix 326 348\ninside 349 372

11556 GCF_000352665.1_Esch_coli_KTE144_V1 Escherichia coli KTE144 Proteobacteria; Gammaproteobacteria; Enterobacterales; Enterobacteriaceae; Escherichia; Escherichia coli MTGDNTLIHSHGINRRDFMKLCAALAATMGLSSKAAAEMAESVTNPQRPPVIWIGAQECT WP_000145410.1 MULTISPECIES: hydrogenase-2 small chain [Proteobacteria] Length: 372\nNumber of predicted TMHs: 1\nExp number of AAs in TMHs: 23.72226\nExp number, first 60 AAs: 0.22038\nTotal prob of N-in: 0.12994\noutside 1 331\nTMhelix 332 354\ninside 355 372

11557 GCF_000352665.1_Esch_coli_KTE144_V1 Escherichia coli KTE144 Proteobacteria; Gammaproteobacteria; Enterobacterales; Enterobacteriaceae; Escherichia; Escherichia coli MNNEETFYQAMRRQGVTRRSFLKYCSLAATSLGLGAGMAPKIAWALENKPRIPVVWIHGL WP_001599658.1 hydrogenase-1 small chain [Escherichia coli] Length: 372\nNumber of predicted TMHs: 2\nExp number of AAs in TMHs: 37.2619299999999\nExp number, first 60 AAs: 19.52918\nTotal prob of N-in: 0.94351\nPOSSIBLE N-term signal sequence\ninside 1 19\nTMhelix 20 39\noutside 40 325\nTMhelix 326 348\ninside 349 372

11558 GCF_000326705.1_Esch_coli_KTE145_V1 Escherichia coli KTE145 Proteobacteria; Gammaproteobacteria; Enterobacterales; Enterobacteriaceae; Escherichia; Escherichia coli MNNEETFYQAMRRQGVTRRSFLKYCSLAATSLGLGAGMAPKIAWALENKPRIPVVWIHGL WP_001058323.1 MULTISPECIES: hydrogenase-1 small chain [Proteobacteria] Length: 372\nNumber of predicted TMHs: 2\nExp number of AAs in TMHs: 37.2608499999999\nExp number, first 60 AAs: 19.52923\nTotal prob of N-in: 0.94351\nPOSSIBLE N-term signal sequence\ninside 1 19\nTMhelix 20 39\noutside 40 325\nTMhelix 326 348\ninside 349 372

11559 GCF_000326705.1_Esch_coli_KTE145_V1 Escherichia coli KTE145 Proteobacteria; Gammaproteobacteria; Enterobacterales; Enterobacteriaceae; Escherichia; Escherichia coli MTGDNTLIHSHGINRRDFMKLCAALAATMGLSSKAAAEMVESVTNPQRPPVIWIGAQECT WP_000145423.1 hydrogenase 2 small subunit [Escherichia coli] Length: 372\nNumber of predicted TMHs: 1\nExp number of AAs in TMHs: 23.78201\nExp number, first 60 AAs: 0.28355\nTotal prob of N-in: 0.13257\noutside 1 331\nTMhelix 332 354\ninside 355 372

11560 GCF_000352685.1_Esch_coli_KTE146_V1 Escherichia coli KTE146 Proteobacteria; Gammaproteobacteria; Enterobacterales; Enterobacteriaceae; Escherichia; Escherichia coli MTGDNTLIHSHGINRRDFMKLCAALAATMGLSSKAAAEMAESVTNPQRPPVIWIGAQECT WP_000145410.1 MULTISPECIES: hydrogenase-2 small chain [Proteobacteria] Length: 372\nNumber of predicted TMHs: 1\nExp number of AAs in TMHs: 23.72226\nExp number, first 60 AAs: 0.22038\nTotal prob of N-in: 0.12994\noutside 1 331\nTMhelix 332 354\ninside 355 372

11561 GCF_000352685.1_Esch_coli_KTE146_V1 Escherichia coli KTE146 Proteobacteria; Gammaproteobacteria; Enterobacterales; Enterobacteriaceae; Escherichia; Escherichia coli MNNEETFYQAMRRQGVTRRSFLKYCSLAATSLGLGAGMAPKIAWALENKPRIPVVWIHGL WP_001058323.1 MULTISPECIES: hydrogenase-1 small chain [Proteobacteria] Length: 372\nNumber of predicted TMHs: 2\nExp number of AAs in TMHs: 37.2608499999999\nExp number, first 60 AAs: 19.52923\nTotal prob of N-in: 0.94351\nPOSSIBLE N-term signal sequence\ninside 1 19\nTMhelix 20 39\noutside 40 325\nTMhelix 326 348\ninside 349 372

11562 GCF_000352705.1_Esch_coli_KTE147_V1 Escherichia coli KTE147 Proteobacteria; Gammaproteobacteria; Enterobacterales; Enterobacteriaceae; Escherichia; Escherichia coli MNNEETFYQAMRRQGVTRRSFLKYCSLAATSLGLGAGMAPKIAWALENKPRIPVVWIHGL WP_001058323.1 MULTISPECIES: hydrogenase-1 small chain [Proteobacteria] Length: 372\nNumber of predicted TMHs: 2\nExp number of AAs in TMHs: 37.2608499999999\nExp number, first 60 AAs: 19.52923\nTotal prob of N-in: 0.94351\nPOSSIBLE N-term signal sequence\ninside 1 19\nTMhelix 20 39\noutside 40 325\nTMhelix 326 348\ninside 349 372

11563 GCF_000352705.1_Esch_coli_KTE147_V1 Escherichia coli KTE147 Proteobacteria; Gammaproteobacteria; Enterobacterales; Enterobacteriaceae; Escherichia; Escherichia coli MTGDNTLIHSHGINRRDFMKLCAALAATMGLSSKAAAEMAESVTNPQRPPVIWIGAQECT WP_000145410.1 MULTISPECIES: hydrogenase-2 small chain [Proteobacteria] Length: 372\nNumber of predicted TMHs: 1\nExp number of AAs in TMHs: 23.72226\nExp number, first 60 AAs: 0.22038\nTotal prob of N-in: 0.12994\noutside 1 331\nTMhelix 332 354\ninside 355 372

11564 GCF_000326365.1_Esch_coli_KTE148_V1 Escherichia coli KTE148 Proteobacteria; Gammaproteobacteria; Enterobacterales; Enterobacteriaceae; Escherichia; Escherichia coli MTGDNTLIHSHGINRRDFMKLCAALAATMGLSSKAAAEMAESVTNPQRPPVIWIGAQECT WP_000145410.1 MULTISPECIES: hydrogenase-2 small chain [Proteobacteria] Length: 372\nNumber of predicted TMHs: 1\nExp number of AAs in TMHs: 23.72226\nExp number, first 60 AAs: 0.22038\nTotal prob of N-in: 0.12994\noutside 1 331\nTMhelix 332 354\ninside 355 372

11565 GCF_000326365.1_Esch_coli_KTE148_V1 Escherichia coli KTE148 Proteobacteria; Gammaproteobacteria; Enterobacterales; Enterobacteriaceae; Escherichia; Escherichia coli MNNEETFYQAMRRQGVTRRSFLKYCSLAATSLGLGAGMAPKIAWALENKPRIPVVWIHGL WP_001058323.1 MULTISPECIES: hydrogenase-1 small chain [Proteobacteria] Length: 372\nNumber of predicted TMHs: 2\nExp number of AAs in TMHs: 37.2608499999999\nExp number, first 60 AAs: 19.52923\nTotal prob of N-in: 0.94351\nPOSSIBLE N-term signal sequence\ninside 1 19\nTMhelix 20 39\noutside 40 325\nTMhelix 326 348\ninside 349 372

11566 GCF_000350745.1_Esch_coli_KTE15_V1 Escherichia coli KTE15 Proteobacteria; Gammaproteobacteria; Enterobacterales; Enterobacteriaceae; Escherichia; Escherichia coli MTGDNTLIHSHGINRRDFMKLCAALAATMGLSSKAAAEMVESVTNPQRPPVIWIGAQECT WP_000145423.1 hydrogenase 2 small subunit [Escherichia coli] Length: 372\nNumber of predicted TMHs: 1\nExp number of AAs in TMHs: 23.78201\nExp number, first 60 AAs: 0.28355\nTotal prob of N-in: 0.13257\noutside 1 331\nTMhelix 332 354\ninside 355 372

11567 GCF_000350745.1_Esch_coli_KTE15_V1 Escherichia coli KTE15 Proteobacteria; Gammaproteobacteria; Enterobacterales; Enterobacteriaceae; Escherichia; Escherichia coli MNNEETFYQAMRRQGVTRRSFLKYCSLAATSLGLGAGMAPKIAWALENKPRIPVVWIHGL WP_001058323.1 MULTISPECIES: hydrogenase-1 small chain [Proteobacteria] Length: 372\nNumber of predicted TMHs: 2\nExp number of AAs in TMHs: 37.2608499999999\nExp number, first 60 AAs: 19.52923\nTotal prob of N-in: 0.94351\nPOSSIBLE N-term signal sequence\ninside 1 19\nTMhelix 20 39\noutside 40 325\nTMhelix 326 348\ninside 349 372

11568 GCF_000326725.1_Esch_coli_KTE150_V1 Escherichia coli KTE150 Proteobacteria; Gammaproteobacteria; Enterobacterales; Enterobacteriaceae; Escherichia; Escherichia coli MNNEETFYQAMRRQGVTRRSFLKYCSLAATSLGLGAGMAPKIAWALENKPRIPVVWIHGL WP_001058323.1 MULTISPECIES: hydrogenase-1 small chain [Proteobacteria] Length: 372\nNumber of predicted TMHs: 2\nExp number of AAs in TMHs: 37.2608499999999\nExp number, first 60 AAs: 19.52923\nTotal prob of N-in: 0.94351\nPOSSIBLE N-term signal sequence\ninside 1 19\nTMhelix 20 39\noutside 40 325\nTMhelix 326 348\ninside 349 372

11569 GCF_000326725.1_Esch_coli_KTE150_V1 Escherichia coli KTE150 Proteobacteria; Gammaproteobacteria; Enterobacterales; Enterobacteriaceae; Escherichia; Escherichia coli MTGDNTLIHSHGINRRDFMKLCAALAATMGLSSKAAAEMAESVTNPQRPPVIWIGAQECT WP_000145410.1 MULTISPECIES: hydrogenase-2 small chain [Proteobacteria] Length: 372\nNumber of predicted TMHs: 1\nExp number of AAs in TMHs: 23.72226\nExp number, first 60 AAs: 0.22038\nTotal prob of N-in: 0.12994\noutside 1 331\nTMhelix 332 354\ninside 355 372

11570 GCF_000326385.1_Esch_coli_KTE153_V1 Escherichia coli KTE153 Proteobacteria; Gammaproteobacteria; Enterobacterales; Enterobacteriaceae; Escherichia; Escherichia coli MTGDNTLIHSHGINRRDFMKLCAALAATMGLSSKAAAEMAESVTNPQRPPVIWIGAQECT WP_000145410.1 MULTISPECIES: hydrogenase-2 small chain [Proteobacteria] Length: 372\nNumber of predicted TMHs: 1\nExp number of AAs in TMHs: 23.72226\nExp number, first 60 AAs: 0.22038\nTotal prob of N-in: 0.12994\noutside 1 331\nTMhelix 332 354\ninside 355 372

11571 GCF_000326385.1_Esch_coli_KTE153_V1 Escherichia coli KTE153 Proteobacteria; Gammaproteobacteria; Enterobacterales; Enterobacteriaceae; Escherichia; Escherichia coli MNNEETFYQAMRRQGVTRRSFLKYCSLAATSLGLGAGMAPKIAWALENKPRIPVVWIHGL WP_001058323.1 MULTISPECIES: hydrogenase-1 small chain [Proteobacteria] Length: 372\nNumber of predicted TMHs: 2\nExp number of AAs in TMHs: 37.2608499999999\nExp number, first 60 AAs: 19.52923\nTotal prob of N-in: 0.94351\nPOSSIBLE N-term signal sequence\ninside 1 19\nTMhelix 20 39\noutside 40 325\nTMhelix 326 348\ninside 349 372

11572 GCF_000352725.1_Esch_coli_KTE154_V1 Escherichia coli KTE154 Proteobacteria; Gammaproteobacteria; Enterobacterales; Enterobacteriaceae; Escherichia; Escherichia coli MNNEETFYQAMRRQGVTRRSFLKYCSLAATSLGLGAGMAPKIAWALENKPRIPVVWIHGL WP_001058323.1 MULTISPECIES: hydrogenase-1 small chain [Proteobacteria] Length: 372\nNumber of predicted TMHs: 2\nExp number of AAs in TMHs: 37.2608499999999\nExp number, first 60 AAs: 19.52923\nTotal prob of N-in: 0.94351\nPOSSIBLE N-term signal sequence\ninside 1 19\nTMhelix 20 39\noutside 40 325\nTMhelix 326 348\ninside 349 372

11573 GCF_000352725.1_Esch_coli_KTE154_V1 Escherichia coli KTE154 Proteobacteria; Gammaproteobacteria; Enterobacterales; Enterobacteriaceae; Escherichia; Escherichia coli MTGDNTLIHSHGINRRDFMKLCAALAATMGLSSKAAAEMAESVTNPQRPPVIWIGAQECT WP_000145410.1 MULTISPECIES: hydrogenase-2 small chain [Proteobacteria] Length: 372\nNumber of predicted TMHs: 1\nExp number of AAs in TMHs: 23.72226\nExp number, first 60 AAs: 0.22038\nTotal prob of N-in: 0.12994\noutside 1 331\nTMhelix 332 354\ninside 355 372

11574 GCF_000408785.1_Esch_coli_KTE155_V1 Escherichia coli KTE155 Proteobacteria; Gammaproteobacteria; Enterobacterales; Enterobacteriaceae; Escherichia; Escherichia coli MTGDNTLIHSHGINRRDFMKLCAALAATMGLSSKAAAEMAESVTNPQRPPVIWIGAQECT WP_000145410.1 MULTISPECIES: hydrogenase-2 small chain [Proteobacteria] Length: 372\nNumber of predicted TMHs: 1\nExp number of AAs in TMHs: 23.72226\nExp number, first 60 AAs: 0.22038\nTotal prob of N-in: 0.12994\noutside 1 331\nTMhelix 332 354\ninside 355 372

11575 GCF_000408785.1_Esch_coli_KTE155_V1 Escherichia coli KTE155 Proteobacteria; Gammaproteobacteria; Enterobacterales; Enterobacteriaceae; Escherichia; Escherichia coli MNNEETFYQAMRRQGVTRRSFLKYCSLAATSLGLGAGMAPKIAWALENKPRIPVVWIHGL WP_001058323.1 MULTISPECIES: hydrogenase-1 small chain [Proteobacteria] Length: 372\nNumber of predicted TMHs: 2\nExp number of AAs in TMHs: 37.2608499999999\nExp number, first 60 AAs: 19.52923\nTotal prob of N-in: 0.94351\nPOSSIBLE N-term signal sequence\ninside 1 19\nTMhelix 20 39\noutside 40 325\nTMhelix 326 348\ninside 349 372

11576 GCF_000351965.1_Esch_coli_KTE156_V1 Escherichia coli KTE156 Proteobacteria; Gammaproteobacteria; Enterobacterales; Enterobacteriaceae; Escherichia; Escherichia coli MNNEETFYQAMRRQGVTRRSFLKYCSLAATSLGLGAGMAPKIAWALENKPRIPVVWIHGL WP_001058323.1 MULTISPECIES: hydrogenase-1 small chain [Proteobacteria] Length: 372\nNumber of predicted TMHs: 2\nExp number of AAs in TMHs: 37.2608499999999\nExp number, first 60 AAs: 19.52923\nTotal prob of N-in: 0.94351\nPOSSIBLE N-term signal sequence\ninside 1 19\nTMhelix 20 39\noutside 40 325\nTMhelix 326 348\ninside 349 372

11577 GCF_000351965.1_Esch_coli_KTE156_V1 Escherichia coli KTE156 Proteobacteria; Gammaproteobacteria; Enterobacterales; Enterobacteriaceae; Escherichia; Escherichia coli MTGDNTLIHSHGINRRDFMKLCAALAATMGLSSKAAAEMAESVTNPQRPPVIWIGAQECT WP_000145410.1 MULTISPECIES: hydrogenase-2 small chain [Proteobacteria] Length: 372\nNumber of predicted TMHs: 1\nExp number of AAs in TMHs: 23.72226\nExp number, first 60 AAs: 0.22038\nTotal prob of N-in: 0.12994\noutside 1 331\nTMhelix 332 354\ninside 355 372

11578 GCF_000326745.1_Esch_coli_KTE157_V1 Escherichia coli KTE157 Proteobacteria; Gammaproteobacteria; Enterobacterales; Enterobacteriaceae; Escherichia; Escherichia coli MTGDNTLIHSHGINRRDFMKLCAALAATMGLSSKAAAEMVESVTNPQRPPVIWIGAQECT WP_000145423.1 hydrogenase 2 small subunit [Escherichia coli] Length: 372\nNumber of predicted TMHs: 1\nExp number of AAs in TMHs: 23.78201\nExp number, first 60 AAs: 0.28355\nTotal prob of N-in: 0.13257\noutside 1 331\nTMhelix 332 354\ninside 355 372

11579 GCF_000326745.1_Esch_coli_KTE157_V1 Escherichia coli KTE157 Proteobacteria; Gammaproteobacteria; Enterobacterales; Enterobacteriaceae; Escherichia; Escherichia coli MNNEETFYQAMRRQGVTRRSFLKYCSLAATSLGLGAGMAPKIAWALENKPRIPVVWIHGL WP_001538532.1 hydrogenase-1 small chain [Escherichia coli] Length: 372\nNumber of predicted TMHs: 2\nExp number of AAs in TMHs: 37.27041\nExp number, first 60 AAs: 19.52927\nTotal prob of N-in: 0.94352\nPOSSIBLE N-term signal sequence\ninside 1 19\nTMhelix 20 39\noutside 40 325\nTMhelix 326 348\ninside 349 372

11580 GCF_000352745.1_Esch_coli_KTE158_V1 Escherichia coli KTE158 Proteobacteria; Gammaproteobacteria; Enterobacterales; Enterobacteriaceae; Escherichia; Escherichia coli MNNEETFYQAMRRQGVTRRSFLKYCSLAATSLGLGAGMAPKIAWALENKPRIPVVWIHGL WP_001058323.1 MULTISPECIES: hydrogenase-1 small chain [Proteobacteria] Length: 372\nNumber of predicted TMHs: 2\nExp number of AAs in TMHs: 37.2608499999999\nExp number, first 60 AAs: 19.52923\nTotal prob of N-in: 0.94351\nPOSSIBLE N-term signal sequence\ninside 1 19\nTMhelix 20 39\noutside 40 325\nTMhelix 326 348\ninside 349 372

11581 GCF_000352745.1_Esch_coli_KTE158_V1 Escherichia coli KTE158 Proteobacteria; Gammaproteobacteria; Enterobacterales; Enterobacteriaceae; Escherichia; Escherichia coli MTGDNTLIHSHGINRRDFMKLCAALAATMGLSSKAAAEMAESVTNPQRPPVIWIGAQECT WP_000145410.1 MULTISPECIES: hydrogenase-2 small chain [Proteobacteria] Length: 372\nNumber of predicted TMHs: 1\nExp number of AAs in TMHs: 23.72226\nExp number, first 60 AAs: 0.22038\nTotal prob of N-in: 0.12994\noutside 1 331\nTMhelix 332 354\ninside 355 372

11582 GCF_000350765.1_Esch_coli_KTE16_V1 Escherichia coli KTE16 Proteobacteria; Gammaproteobacteria; Enterobacterales; Enterobacteriaceae; Escherichia; Escherichia coli MTGDNTLIHSHGINRRDFMKLCAALAATMGLSSKAAAEMVESVTNPQRPPVIWIGAQECT WP_000145423.1 hydrogenase 2 small subunit [Escherichia coli] Length: 372\nNumber of predicted TMHs: 1\nExp number of AAs in TMHs: 23.78201\nExp number, first 60 AAs: 0.28355\nTotal prob of N-in: 0.13257\noutside 1 331\nTMhelix 332 354\ninside 355 372

11583 GCF_000350765.1_Esch_coli_KTE16_V1 Escherichia coli KTE16 Proteobacteria; Gammaproteobacteria; Enterobacterales; Enterobacteriaceae; Escherichia; Escherichia coli MNNEETFYQAMRRQGVTRRSFLKYCSLAATSLGLGAGMAPKIAWALENKPRIPVVWIHGL WP_001058323.1 MULTISPECIES: hydrogenase-1 small chain [Proteobacteria] Length: 372\nNumber of predicted TMHs: 2\nExp number of AAs in TMHs: 37.2608499999999\nExp number, first 60 AAs: 19.52923\nTotal prob of N-in: 0.94351\nPOSSIBLE N-term signal sequence\ninside 1 19\nTMhelix 20 39\noutside 40 325\nTMhelix 326 348\ninside 349 372

11584 GCF_000326405.1_Esch_coli_KTE160_V1 Escherichia coli KTE160 Proteobacteria; Gammaproteobacteria; Enterobacterales; Enterobacteriaceae; Escherichia; Escherichia coli MTGDNTLIHSHGINRRDFMKLCAALAATMGLSSKAAAEMAESVTNPQRPPVIWIGAQECT WP_000145410.1 MULTISPECIES: hydrogenase-2 small chain [Proteobacteria] Length: 372\nNumber of predicted TMHs: 1\nExp number of AAs in TMHs: 23.72226\nExp number, first 60 AAs: 0.22038\nTotal prob of N-in: 0.12994\noutside 1 331\nTMhelix 332 354\ninside 355 372

11585 GCF_000326405.1_Esch_coli_KTE160_V1 Escherichia coli KTE160 Proteobacteria; Gammaproteobacteria; Enterobacterales; Enterobacteriaceae; Escherichia; Escherichia coli MNNEETFYQAMRRQGVTRRSFLKYCSLAATSLGLGAGMAPKIAWALENKPRIPVVWIHGL WP_001058323.1

MULTISPECIES: hydrogenase-1 small chain [Proteobacteria] Length: 372\nNumber of predicted TMHs: 2\nExp number of AAs in TMHs: 37.2608499999999\nExp number, first 60 AAs: 19.52923\nTotal prob of N-in: 0.94351\nPOSSIBLE N-term signal sequence\ninside 1 19\nTMhelix 20 39\noutside 40 325\nTMhelix 326 348\ninside 349 372

11586 GCF_000351985.1_Esch_coli_KTE161_V1 Escherichia coli KTE161 Proteobacteria; Gammaproteobacteria; Enterobacterales; Enterobacteriaceae; Escherichia; Escherichia coli MTGDNTLIHSHGINRRDFMKLCAALAATMGLSSKAAAEMAESVTNPQRPPVIWIGAQECT WP_000145410.1 MULTISPECIES: hydrogenase-2 small chain [Proteobacteria] Length: 372\nNumber of predicted TMHs: 1\nExp number of AAs in TMHs: 23.72226\nExp number, first 60 AAs: 0.22038\nTotal prob of N-in: 0.12994\noutside 1 331\nTMhelix 332 354\ninside 355 372

11587 GCF_000351985.1_Esch_coli_KTE161_V1 Escherichia coli KTE161 Proteobacteria; Gammaproteobacteria; Enterobacterales; Enterobacteriaceae; Escherichia; Escherichia coli MNNEETFYQAMRRQGVTRRSFLKYCSLAATSLGLGAGMAPKIAWALENKPRIPVVWIHGL WP_001058323.1 MULTISPECIES: hydrogenase-1 small chain [Proteobacteria] Length: 372\nNumber of predicted TMHs: 2\nExp number of AAs in TMHs: 37.2608499999999\nExp number, first 60 AAs: 19.52923\nTotal prob of N-in: 0.94351\nPOSSIBLE N-term signal sequence\ninside 1 19\nTMhelix 20 39\noutside 40 325\nTMhelix 326 348\ninside 349 372

11588 GCF_000352005.1_Esch_coli_KTE162_V1 Escherichia coli KTE162 Proteobacteria; Gammaproteobacteria; Enterobacterales; Enterobacteriaceae; Escherichia; Escherichia coli MTGDNTLIHSHGINRRDFMKLCAALAATMGLSSKAAAEMAESVTNPQRPPVIWIGAQECT WP_000145410.1 MULTISPECIES: hydrogenase-2 small chain [Proteobacteria] Length: 372\nNumber of predicted TMHs: 1\nExp number of AAs in TMHs: 23.72226\nExp number, first 60 AAs: 0.22038\nTotal prob of N-in: 0.12994\noutside 1 331\nTMhelix 332 354\ninside 355 372

11589 GCF_000352005.1_Esch_coli_KTE162_V1 Escherichia coli KTE162 Proteobacteria; Gammaproteobacteria; Enterobacterales; Enterobacteriaceae; Escherichia; Escherichia coli MNNEETFYQAMRRQGVTRRSFLKYCSLAATSLGLGAGMAPKIAWALENKPRIPVVWIHGL WP_001058323.1 MULTISPECIES: hydrogenase-1 small chain [Proteobacteria] Length: 372\nNumber of predicted TMHs: 2\nExp number of AAs in TMHs: 37.2608499999999\nExp number, first 60 AAs: 19.52923\nTotal prob of N-in: 0.94351\nPOSSIBLE N-term signal sequence\ninside 1 19\nTMhelix 20 39\noutside 40 325\nTMhelix 326 348\ninside 349 372

11590 GCF_000326425.1_Esch_coli_KTE163_V1 Escherichia coli KTE163 Proteobacteria; Gammaproteobacteria; Enterobacterales; Enterobacteriaceae; Escherichia; Escherichia coli MNNEETFYQAMRRQGVTRRSFLKYCSLAATSLGLGAGMAPKIAWALENKPRIPVVWIHGL WP_001058320.1 hydrogenase [Escherichia coli] Length: 372\nNumber of predicted TMHs: 2\nExp number of AAs in TMHs: 37.26082\nExp number, first 60 AAs: 19.52923\nTotal prob of N-in: 0.94351\nPOSSIBLE N-term signal sequence\ninside 1 19\nTMhelix 20 39\noutside 40 325\nTMhelix 326 348\ninside 349 372

11591 GCF_000326425.1_Esch_coli_KTE163_V1 Escherichia coli KTE163 Proteobacteria; Gammaproteobacteria; Enterobacterales; Enterobacteriaceae; Escherichia; Escherichia coli MTGDNTLIHSHGINRRDFMKLCAALAATMGLSSKAAAEMAESVTNPQRPPVIWIGAQECT WP_000145410.1 MULTISPECIES: hydrogenase-2 small chain [Proteobacteria] Length: 372\nNumber of predicted TMHs: 1\nExp number of AAs in TMHs: 23.72226\nExp number, first 60 AAs: 0.22038\nTotal prob of N-in: 0.12994\noutside 1 331\nTMhelix 332 354\ninside 355 372

11592 GCF_000352765.1_Esch_coli_KTE165_V1 Escherichia coli KTE165 Proteobacteria; Gammaproteobacteria; Enterobacterales; Enterobacteriaceae; Escherichia; Escherichia coli MTGDNTLIHSHGINRRDFMKLCAALAATMGLSSKAAAEMAESVTNPQRPPVIWIGAQECT WP_000145410.1 MULTISPECIES: hydrogenase-2 small chain [Proteobacteria] Length: 372\nNumber of predicted TMHs: 1\nExp number of AAs in TMHs: 23.72226\nExp number, first 60 AAs: 0.22038\nTotal prob of N-in: 0.12994\noutside 1 331\nTMhelix 332 354\ninside 355 372

11593 GCF_000352765.1_Esch_coli_KTE165_V1 Escherichia coli KTE165 Proteobacteria; Gammaproteobacteria; Enterobacterales; Enterobacteriaceae; Escherichia; Escherichia coli MNNEETFYQAMRRQGVTRRSFLKYCSLAATSLGLGAGMAPKIAWALENKPRIPVVWIHGL WP_001295940.1 MULTISPECIES: hydrogenase [Enterobacteriaceae] Length: 372\nNumber of predicted TMHs: 2\nExp number of AAs in TMHs: 37.2608899999999\nExp number, first 60 AAs: 19.52923\nTotal prob of N-in:

0.94351\nPOSSIBLE N-term signal sequence\ninside 1 19\nTMhelix 20 39\noutside 40 325\nTMhelix 326 348\ninside 349 372

11594 GCF_000326765.1_Esch_coli_KTE166_V1 Escherichia coli KTE166 Proteobacteria; Gammaproteobacteria; Enterobacterales; Enterobacteriaceae; Escherichia; Escherichia coli MNNEETFYQAMRRQGVTRRSFLKYCSLAATSLGLGAGMAPKIAWALENKPRIPVVWIHGL WP_001058320.1 hydrogenase [Escherichia coli] Length: 372\nNumber of predicted TMHs: 2\nExp number of AAs in TMHs: 37.26082\nExp number, first 60 AAs: 19.52923\nTotal prob of N-in: 0.94351\nPOSSIBLE N-term signal sequence\ninside 1 19\nTMhelix 20 39\noutside 40 325\nTMhelix 326 348\ninside 349 372

11595 GCF_000326765.1_Esch_coli_KTE166_V1 Escherichia coli KTE166 Proteobacteria; Gammaproteobacteria; Enterobacterales; Enterobacteriaceae; Escherichia; Escherichia coli MTGDNTLIHSHGINRRDFMKLCAALAATMGLSSKAAAEMAESVTNPQRPPVIWIGAQECT WP_000145410.1 MULTISPECIES: hydrogenase-2 small chain [Proteobacteria] Length: 372\nNumber of predicted TMHs: 1\nExp number of AAs in TMHs: 23.72226\nExp number, first 60 AAs: 0.22038\nTotal prob of N-in: 0.12994\noutside 1 331\nTMhelix 332 354\ninside 355 372

11596 GCF_000326905.1_Esch_coli_KTE167_V1 Escherichia coli KTE167 Proteobacteria; Gammaproteobacteria; Enterobacterales; Enterobacteriaceae; Escherichia; Escherichia coli MTGDNTLIHSHGINRRDFMKLCAALAATMGLSSKAAAEMAESVTNPQRPPVIWIGAQECT WP_000145410.1 MULTISPECIES: hydrogenase-2 small chain [Proteobacteria] Length: 372\nNumber of predicted TMHs: 1\nExp number of AAs in TMHs: 23.72226\nExp number, first 60 AAs: 0.22038\nTotal prob of N-in: 0.12994\noutside 1 331\nTMhelix 332 354\ninside 355 372

11597 GCF_000326905.1_Esch_coli_KTE167_V1 Escherichia coli KTE167 Proteobacteria; Gammaproteobacteria; Enterobacterales; Enterobacteriaceae; Escherichia; Escherichia coli MNNEETFYQAMRRQGVTRRSFLKYCSLAATSLGLGAGMAPKIAWALENKPRIPVVWIHGL WP_001058323.1 MULTISPECIES: hydrogenase-1 small chain [Proteobacteria] Length: 372\nNumber of predicted TMHs: 2\nExp number of AAs in TMHs: 37.2608499999999\nExp number, first 60 AAs: 19.52923\nTotal prob of N-in: 0.94351\nPOSSIBLE N-term signal sequence\ninside 1 19\nTMhelix 20 39\noutside 40 325\nTMhelix 326 348\ninside 349 372

11598 GCF_000326445.1_Esch_coli_KTE168_V1 Escherichia coli KTE168 Proteobacteria; Gammaproteobacteria; Enterobacterales; Enterobacteriaceae; Escherichia; Escherichia coli MNNEETFYQAMRRQGVTRRSFLKYCSLAATSLGLGAGMAPKIAWALENKPRIPVVWIHGL WP_001058323.1 MULTISPECIES: hydrogenase-1 small chain [Proteobacteria] Length: 372\nNumber of predicted TMHs: 2\nExp number of AAs in TMHs: 37.2608499999999\nExp number, first 60 AAs: 19.52923\nTotal prob of N-in: 0.94351\nPOSSIBLE N-term signal sequence\ninside 1 19\nTMhelix 20 39\noutside 40 325\nTMhelix 326 348\ninside 349 372

11599 GCF_000326445.1_Esch_coli_KTE168_V1 Escherichia coli KTE168 Proteobacteria; Gammaproteobacteria; Enterobacterales; Enterobacteriaceae; Escherichia; Escherichia coli MTGDNTLIHSHGINRRDFMKLCAALAATMGLSSKAAAEMVESVTNPQRPPVIWIGAQECT WP_000145423.1 hydrogenase 2 small subunit [Escherichia coli] Length: 372\nNumber of predicted TMHs: 1\nExp number of AAs in TMHs: 23.78201\nExp number, first 60 AAs: 0.28355\nTotal prob of N-in: 0.13257\noutside 1 331\nTMhelix 332 354\ninside 355 372

11600 GCF_000352025.1_Esch_coli_KTE169_V1 Escherichia coli KTE169 Proteobacteria; Gammaproteobacteria; Enterobacterales; Enterobacteriaceae; Escherichia; Escherichia coli MNNEETFYQAMRRQGVTRRSFLKYCSLAATSLGLGAGMAPKIAWALENKPRIPVVWIHGL WP_001058323.1 MULTISPECIES: hydrogenase-1 small chain [Proteobacteria] Length: 372\nNumber of predicted TMHs: 2\nExp number of AAs in TMHs: 37.2608499999999\nExp number, first 60 AAs: 19.52923\nTotal prob of N-in: 0.94351\nPOSSIBLE N-term signal sequence\ninside 1 19\nTMhelix 20 39\noutside 40 325\nTMhelix 326 348\ninside 349 372

11601 GCF_000352025.1_Esch_coli_KTE169_V1 Escherichia coli KTE169 Proteobacteria; Gammaproteobacteria; Enterobacterales; Enterobacteriaceae; Escherichia; Escherichia coli MTGDNTLIHSHGINRRDFMKLCAALAATMGLSSKAAAEMVESVTNPQRPPVIWIGAQECT WP_000145423.1 hydrogenase 2 small subunit [Escherichia coli] Length: 372\nNumber of predicted TMHs: 1\nExp number of AAs in TMHs: 23.78201\nExp number, first 60 AAs: 0.28355\nTotal prob of N-in: 0.13257\noutside 1 331\nTMhelix 332 354\ninside 355 372

11602 GCF_000352125.1_Esch_coli_KTE17_V1 Escherichia coli KTE17 Proteobacteria; Gammaproteobacteria; Enterobacterales; Enterobacteriaceae; Escherichia; Escherichia coli MNNEETFYQAMRRQGVTRRSFLKYCSLAATSLGLGAGMAPKIAWALENKPRIPVVWIHGL WP_001058323.1 MULTISPECIES: hydrogenase-1 small chain [Proteobacteria] Length: 372\nNumber of predicted TMHs: 2\nExp number of AAs in TMHs: 37.2608499999999\nExp number, first 60 AAs: 19.52923\nTotal prob of N-in: 0.94351\nPOSSIBLE N-term signal sequence\ninside 1 19\nTMhelix 20 39\noutside 40 325\nTMhelix 326 348\ninside 349 372

11603 GCF_000352125.1_Esch_coli_KTE17_V1 Escherichia coli KTE17 Proteobacteria; Gammaproteobacteria; Enterobacterales; Enterobacteriaceae; Escherichia; Escherichia coli MTGDNTLIHSHGINRRDFMKLCAALAATMGLSSKAAAEMAESVTNPQRPPVIWIGAQECT WP_000145410.1 MULTISPECIES: hydrogenase-2 small chain [Proteobacteria] Length: 372\nNumber of predicted TMHs: 1\nExp number of AAs in TMHs: 23.72226\nExp number, first 60 AAs: 0.22038\nTotal prob of N-in: 0.12994\noutside 1 331\nTMhelix 332 354\ninside 355 372

11604 GCF_000408825.1_Esch_coli_KTE170_V1 Escherichia coli KTE170 Proteobacteria; Gammaproteobacteria; Enterobacterales; Enterobacteriaceae; Escherichia; Escherichia coli MTGDNTLIHSHGINRRDFMKLCAALAATMGLSSKAAAEMAESVTNPQRPPVIWIGAQECT WP_000145410.1 MULTISPECIES: hydrogenase-2 small chain [Proteobacteria] Length: 372\nNumber of predicted TMHs: 1\nExp number of AAs in TMHs: 23.72226\nExp number, first 60 AAs: 0.22038\nTotal prob of N-in: 0.12994\noutside 1 331\nTMhelix 332 354\ninside 355 372

11605 GCF_000408825.1_Esch_coli_KTE170_V1 Escherichia coli KTE170 Proteobacteria; Gammaproteobacteria; Enterobacterales; Enterobacteriaceae; Escherichia; Escherichia coli MNNEETFYQAMRRQGVTRRSFLKYCSLAATSLGLGAGMAPKIAWALENKPRIPVVWIHGL WP_001058323.1 MULTISPECIES: hydrogenase-1 small chain [Proteobacteria] Length: 372\nNumber of predicted TMHs: 2\nExp number of AAs in TMHs: 37.2608499999999\nExp number, first 60 AAs: 19.52923\nTotal prob of N-in: 0.94351\nPOSSIBLE N-term signal sequence\ninside 1 19\nTMhelix 20 39\noutside 40 325\nTMhelix 326 348\ninside 349 372

11606 GCF_000352045.1_Esch_coli_KTE171_V1 Escherichia coli KTE171 Proteobacteria; Gammaproteobacteria; Enterobacterales; Enterobacteriaceae; Escherichia; Escherichia coli MTGDNTLIHSHGINRRDFMKLCAALAATMGLSSKAAAEMAESVTNPQRPPVIWIGAQECT WP_000145410.1 MULTISPECIES: hydrogenase-2 small chain [Proteobacteria] Length: 372\nNumber of predicted TMHs: 1\nExp number of AAs in TMHs: 23.72226\nExp number, first 60 AAs: 0.22038\nTotal prob of N-in: 0.12994\noutside 1 331\nTMhelix 332 354\ninside 355 372

11607 GCF_000352045.1_Esch_coli_KTE171_V1 Escherichia coli KTE171 Proteobacteria; Gammaproteobacteria; Enterobacterales; Enterobacteriaceae; Escherichia; Escherichia coli MNNEETFYQAMRRQGVTRRSFLKYCSLAATSLGLGAGMAPKIAWALENKPRIPVVWIHGL WP_001058323.1 MULTISPECIES: hydrogenase-1 small chain [Proteobacteria] Length: 372\nNumber of predicted TMHs: 2\nExp number of AAs in TMHs: 37.2608499999999\nExp number, first 60 AAs: 19.52923\nTotal prob of N-in: 0.94351\nPOSSIBLE N-term signal sequence\ninside 1 19\nTMhelix 20 39\noutside 40 325\nTMhelix 326 348\ninside 349 372

11608 GCF_000352845.1_Esch_coli_KTE173_V1 Escherichia coli KTE173 Proteobacteria; Gammaproteobacteria; Enterobacterales; Enterobacteriaceae; Escherichia; Escherichia coli MNNEETFYQAMRRQGVTRRSFLKYCSLAATSLGLGAGMAPKIAWALENKPRIPVVWIHGL WP_001058323.1 MULTISPECIES: hydrogenase-1 small chain [Proteobacteria] Length: 372\nNumber of predicted TMHs: 2\nExp number of AAs in TMHs: 37.2608499999999\nExp number, first 60 AAs: 19.52923\nTotal prob of N-in: 0.94351\nPOSSIBLE N-term signal sequence\ninside 1 19\nTMhelix 20 39\noutside 40 325\nTMhelix 326 348\ninside 349 372

11609 GCF_000352845.1_Esch_coli_KTE173_V1 Escherichia coli KTE173 Proteobacteria; Gammaproteobacteria; Enterobacterales; Enterobacteriaceae; Escherichia; Escherichia coli MTGDNTLIHSHGINRRDFMKLCAALAATMGLSSKAAAEMAESVTNPQRPPVIWIGAQECT WP_000145410.1 MULTISPECIES: hydrogenase-2 small chain [Proteobacteria] Length: 372\nNumber of predicted TMHs: 1\nExp number of AAs in TMHs: 23.72226\nExp number, first 60 AAs: 0.22038\nTotal prob of N-in: 0.12994\noutside 1 331\nTMhelix 332 354\ninside 355 372

11610 GCF_000326605.1_Esch_coli_KTE174_V1 Escherichia coli KTE174 Proteobacteria; Gammaproteobacteria; Enterobacterales; Enterobacteriaceae; Escherichia; Escherichia coli MNNEETFYQAMRRQGVTRRSFLKYCSLAATSLGLGAGMAPKIAWALENKPRIPVVWIHGL WP_001058323.1 MULTISPECIES: hydrogenase-1 small chain [Proteobacteria] Length: 372\nNumber of predicted TMHs: 2\nExp number of AAs in TMHs: 37.2608499999999\nExp number, first 60 AAs: 19.52923\nTotal prob of N-in: 0.94351\nPOSSIBLE N-term signal sequence\ninside 1 19\nTMhelix 20 39\noutside 40 325\nTMhelix 326 348\ninside 349 372

11611 GCF_000326605.1_Esch_coli_KTE174_V1 Escherichia coli KTE174 Proteobacteria; Gammaproteobacteria; Enterobacterales; Enterobacteriaceae; Escherichia; Escherichia coli MTGDNTLIHSHGINRRDFMKLCAALAATMGLSSKAAAEMAESVTNPQRPPVIWIGAQECT WP_000145410.1 MULTISPECIES: hydrogenase-2 small chain [Proteobacteria] Length: 372\nNumber of predicted TMHs: 1\nExp number of AAs in TMHs: 23.72226\nExp number, first 60 AAs: 0.22038\nTotal prob of N-in: 0.12994\noutside 1 331\nTMhelix 332 354\ninside 355 372

11612 GCF_000352865.1_Esch_coli_KTE175_V1 Escherichia coli KTE175 Proteobacteria; Gammaproteobacteria; Enterobacterales; Enterobacteriaceae; Escherichia; Escherichia coli MNNEETFYQAMRRQGVTRRSFLKYCSLAATSLGLGAGMAPKIAWALENKPRIPVVWIHGL WP_001058323.1 MULTISPECIES: hydrogenase-1 small chain [Proteobacteria] Length: 372\nNumber of predicted TMHs: 2\nExp number of AAs in TMHs: 37.2608499999999\nExp number, first 60 AAs: 19.52923\nTotal prob of N-in: 0.94351\nPOSSIBLE N-term signal sequence\ninside 1 19\nTMhelix 20 39\noutside 40 325\nTMhelix 326 348\ninside 349 372

11613 GCF_000352865.1_Esch_coli_KTE175_V1 Escherichia coli KTE175 Proteobacteria; Gammaproteobacteria; Enterobacterales; Enterobacteriaceae; Escherichia; Escherichia coli MTGDNTLIHSHGINRRDFMKLCAALAATMGLSSKAAAEMAESVTNPQRPPVIWIGAQECT WP_000145410.1 MULTISPECIES: hydrogenase-2 small chain [Proteobacteria] Length: 372\nNumber of predicted TMHs: 1\nExp number of AAs in TMHs: 23.72226\nExp number, first 60 AAs: 0.22038\nTotal prob of N-in: 0.12994\noutside 1 331\nTMhelix 332 354\ninside 355 372

11614 GCF_000326465.1_Esch_coli_KTE176_V1 Escherichia coli KTE176 Proteobacteria; Gammaproteobacteria; Enterobacterales; Enterobacteriaceae; Escherichia; Escherichia coli MTGDNTLIHSHGINRRDFMKLCAALAATMGLSSKAAAEMAESVTNPQRPPVIWIGAQECT WP_000145410.1 MULTISPECIES: hydrogenase-2 small chain [Proteobacteria] Length: 372\nNumber of predicted TMHs: 1\nExp number of AAs in TMHs: 23.72226\nExp number, first 60 AAs: 0.22038\nTotal prob of N-in: 0.12994\noutside 1 331\nTMhelix 332 354\ninside 355 372

11615 GCF_000326785.1_Esch_coli_KTE177_V1 Escherichia coli KTE177 Proteobacteria; Gammaproteobacteria; Enterobacterales; Enterobacteriaceae; Escherichia; Escherichia coli MTGDNTLIHSHGINRRDFMKLCAALAATMGLSSKAAAEMAESVTNPQRPPVIWIGAQECT WP_000145410.1 MULTISPECIES: hydrogenase-2 small chain [Proteobacteria] Length: 372\nNumber of predicted TMHs: 1\nExp number of AAs in TMHs: 23.72226\nExp number, first 60 AAs: 0.22038\nTotal prob of N-in: 0.12994\noutside 1 331\nTMhelix 332 354\ninside 355 372

11616 GCF_000326785.1_Esch_coli_KTE177_V1 Escherichia coli KTE177 Proteobacteria; Gammaproteobacteria; Enterobacterales; Enterobacteriaceae; Escherichia; Escherichia coli MNNEETFYQAMRRQGVTRRSFLKYCSLAATSLGLGAGMAPKIAWALENKPRIPVVWIHGL WP_001058323.1 MULTISPECIES: hydrogenase-1 small chain [Proteobacteria] Length: 372\nNumber of predicted TMHs: 2\nExp number of AAs in TMHs: 37.2608499999999\nExp number, first 60 AAs: 19.52923\nTotal prob of N-in: 0.94351\nPOSSIBLE N-term signal sequence\ninside 1 19\nTMhelix 20 39\noutside 40 325\nTMhelix 326 348\ninside 349 372

11617 GCF_000350905.1_Esch_coli_KTE178_V1 Escherichia coli KTE178 Proteobacteria; Gammaproteobacteria; Enterobacterales; Enterobacteriaceae; Escherichia; Escherichia coli MNNEETFYQAMRRQGVTRRSFLKYCSLAATSLGLGAGMAPKIAWALENKPRIPVVWIHGL WP_001295940.1 MULTISPECIES: hydrogenase [Enterobacteriaceae] Length: 372\nNumber of predicted TMHs: 2\nExp number of AAs in TMHs: 37.2608899999999\nExp number, first 60 AAs: 19.52923\nTotal prob of N-in: 0.94351\nPOSSIBLE N-term signal sequence\ninside 1 19\nTMhelix 20 39\noutside 40 325\nTMhelix 326 348\ninside 349 372

11618 GCF_000350905.1_Esch_coli_KTE178_V1 Escherichia coli KTE178 Proteobacteria; Gammaproteobacteria; Enterobacterales; Enterobacteriaceae; Escherichia; Escherichia coli
MTGDNTLIHSHGINRRDFMKLCAALAATMGLSSKAAAEMAESVTNPQRPPVIWIGAQECT WP_000145410.1
MULTISPECIES: hydrogenase-2 small chain [Proteobacteria] Length: 372\nNumber of predicted TMHs: 1\nExp number of AAs in TMHs: 23.72226\nExp number, first 60 AAs: 0.22038\nTotal prob of N-in: 0.12994\noutside 1 331\nTMhelix 332 354\ninside 355 372

11619 GCF_000326485.1_Esch_coli_KTE179_V1 Escherichia coli KTE179 Proteobacteria; Gammaproteobacteria; Enterobacterales; Enterobacteriaceae; Escherichia; Escherichia coli
MTGDNTLIHSHGINRRDFMKLCAALAATMGLSSKAAAEMAESVTNPQRPPVIWIGAQECT WP_000145410.1
MULTISPECIES: hydrogenase-2 small chain [Proteobacteria] Length: 372\nNumber of predicted TMHs: 1\nExp number of AAs in TMHs: 23.72226\nExp number, first 60 AAs: 0.22038\nTotal prob of N-in: 0.12994\noutside 1 331\nTMhelix 332 354\ninside 355 372

11620 GCF_000352145.1_Esch_coli_KTE18_V1 Escherichia coli KTE18 Proteobacteria; Gammaproteobacteria; Enterobacterales; Enterobacteriaceae; Escherichia; Escherichia coli
MTGDNTLIHSHGINRRDFMKLCAALAATMGLSSKAAAEMAESVTNPQRPPVIWIGAQECT WP_000145410.1
MULTISPECIES: hydrogenase-2 small chain [Proteobacteria] Length: 372\nNumber of predicted TMHs: 1\nExp number of AAs in TMHs: 23.72226\nExp number, first 60 AAs: 0.22038\nTotal prob of N-in: 0.12994\noutside 1 331\nTMhelix 332 354\ninside 355 372

11621 GCF_000352145.1_Esch_coli_KTE18_V1 Escherichia coli KTE18 Proteobacteria; Gammaproteobacteria; Enterobacterales; Enterobacteriaceae; Escherichia; Escherichia coli
MNNEETFYQAMRRQGVTRRSFLKYCSLAATSLGLGAGMAPKIAWALENKPRIPVVWIHGL WP_001058323.1
MULTISPECIES: hydrogenase-1 small chain [Proteobacteria] Length: 372\nNumber of predicted TMHs: 2\nExp number of AAs in TMHs: 37.2608499999999\nExp number, first 60 AAs: 19.52923\nTotal prob of N-in: 0.94351\nPOSSIBLE N-term signal sequence\ninside 1 19\nTMhelix 20 39\noutside 40 325\nTMhelix 326 348\ninside 349 372

11622 GCF_000326805.1_Esch_coli_KTE180_V1 Escherichia coli KTE180 Proteobacteria; Gammaproteobacteria; Enterobacterales; Enterobacteriaceae; Escherichia; Escherichia coli
MTGDNTLIHSHGINRRDFMKLCAALAATMGLSSKAAAEMAESVTNPQRPPVIWIGAQECT WP_000145410.1
MULTISPECIES: hydrogenase-2 small chain [Proteobacteria] Length: 372\nNumber of predicted TMHs: 1\nExp number of AAs in TMHs: 23.72226\nExp number, first 60 AAs: 0.22038\nTotal prob of N-in: 0.12994\noutside 1 331\nTMhelix 332 354\ninside 355 372

11623 GCF_000350925.1_Esch_coli_KTE181_V1 Escherichia coli KTE181 Proteobacteria; Gammaproteobacteria; Enterobacterales; Enterobacteriaceae; Escherichia; Escherichia coli
MNNEETFYQAMRRQGVTRRSFLKYCSLAATSLGLGAGMAPKIAWALENKPRIPVVWIHGL WP_001058323.1
MULTISPECIES: hydrogenase-1 small chain [Proteobacteria] Length: 372\nNumber of predicted TMHs: 2\nExp number of AAs in TMHs: 37.2608499999999\nExp number, first 60 AAs: 19.52923\nTotal prob of N-in: 0.94351\nPOSSIBLE N-term signal sequence\ninside 1 19\nTMhelix 20 39\noutside 40 325\nTMhelix 326 348\ninside 349 372

11624 GCF_000350925.1_Esch_coli_KTE181_V1 Escherichia coli KTE181 Proteobacteria; Gammaproteobacteria; Enterobacterales; Enterobacteriaceae; Escherichia; Escherichia coli
MTGDNTLIHSHGINRRDFMKLCAALAATMGLSSKAAAEMAESVTNPQRPPVIWIGAQECT WP_000145410.1
MULTISPECIES: hydrogenase-2 small chain [Proteobacteria] Length: 372\nNumber of predicted TMHs: 1\nExp number of AAs in TMHs: 23.72226\nExp number, first 60 AAs: 0.22038\nTotal prob of N-in: 0.12994\noutside 1 331\nTMhelix 332 354\ninside 355 372

11625 GCF_000408065.1_Esch_coli_KTE182_V1 Escherichia coli KTE182 Proteobacteria; Gammaproteobacteria; Enterobacterales; Enterobacteriaceae; Escherichia; Escherichia coli
MTGDNTLIHSHGINRRDFMKLCAALAATMGLSSKAAAEMVESVTNPQRPPVIWIGAQECT WP_000145423.1
hydrogenase 2 small subunit [Escherichia coli] Length: 372\nNumber of predicted TMHs: 1\nExp number of AAs in TMHs: 23.78201\nExp number, first 60 AAs: 0.28355\nTotal prob of N-in: 0.13257\noutside 1 331\nTMhelix 332 354\ninside 355 372

11626 GCF_000408065.1_Esch_coli_KTE182_V1 Escherichia coli KTE182 Proteobacteria; Gammaproteobacteria; Enterobacterales; Enterobacteriaceae; Escherichia; Escherichia coli
MNNEETFYQAMRRQGVTRRSFLKYCSLAATSLGLGAGMAPKIAWALENKPRIPVVWIHGL WP_001538532.1

hydrogenase-1 small chain [Escherichia coli]	Length: 372\nNumber of predicted TMHs:  2\nExp number of AAs in TMHs: 37.27041\nExp number, first 60 AAs:  19.52927\nTotal prob of N-in:   0.94352\nPOSSIBLE N-term signal sequence\ninside   1   19\nTMhelix   20   39\noutside   40   325\nTMhelix   326   348\ninside   349   372

11627	GCF_000352905.1_Esch_coli_KTE183_V1	Escherichia coli KTE183	Proteobacteria; Gammaproteobacteria; Enterobacterales; Enterobacteriaceae; Escherichia; Escherichia coli	MNNEETFYQAMRRQGVTRRSFLKYCSLAATSLGLGAGMAPKIAWALENKPRIPVVWIHGL	WP_001058323.1	MULTISPECIES: hydrogenase-1 small chain [Proteobacteria]	Length: 372\nNumber of predicted TMHs: 2\nExp number of AAs in TMHs: 37.2608499999999\nExp number, first 60 AAs:  19.52923\nTotal prob of N-in: 0.94351\nPOSSIBLE N-term signal sequence\ninside   1   19\nTMhelix   20   39\noutside   40   325\nTMhelix   326 348\ninside   349   372

11628	GCF_000352905.1_Esch_coli_KTE183_V1	Escherichia coli KTE183	Proteobacteria; Gammaproteobacteria; Enterobacterales; Enterobacteriaceae; Escherichia; Escherichia coli	MTGDNTLIHSHGINRRDFMKLCAALAATMGLSSKAAAEMVESVTNPQRPPVIWIGAQECT	WP_000145423.1	hydrogenase 2 small subunit [Escherichia coli]	Length: 372\nNumber of predicted TMHs:  1\nExp number of AAs in TMHs: 23.78201\nExp number, first 60 AAs:  0.28355\nTotal prob of N-in:   0.13257\noutside   1   331\nTMhelix 332   354\ninside   355   372

11629	GCF_000352885.1_Esch_coli_KTE184_V1	Escherichia coli KTE184	Proteobacteria; Gammaproteobacteria; Enterobacterales; Enterobacteriaceae; Escherichia; Escherichia coli	MNNEETFYQAMRRQGVTRRSFLKYCSLAATSLGLGAGMAPKIAWALENKPRIPVVWIHGL	WP_001295940.1	MULTISPECIES: hydrogenase [Enterobacteriaceae]	Length: 372\nNumber of predicted TMHs:  2\nExp number of AAs in TMHs: 37.2608899999999\nExp number, first 60 AAs:  19.52923\nTotal prob of N-in: 0.94351\nPOSSIBLE N-term signal sequence\ninside   1   19\nTMhelix   20   39\noutside   40   325\nTMhelix   326 348\ninside   349   372

11630	GCF_000352885.1_Esch_coli_KTE184_V1	Escherichia coli KTE184	Proteobacteria; Gammaproteobacteria; Enterobacterales; Enterobacteriaceae; Escherichia; Escherichia coli	MTGDNTLIHSHGINRRDFMKLCAALAATMGLSSKAAAEMAESVTNPQRPPVIWIGAQECT	WP_000145410.1	MULTISPECIES: hydrogenase-2 small chain [Proteobacteria]	Length: 372\nNumber of predicted TMHs: 1\nExp number of AAs in TMHs: 23.72226\nExp number, first 60 AAs:  0.22038\nTotal prob of N-in:   0.12994\noutside   1 331\nTMhelix   332   354\ninside   355   372

11631	GCF_000408085.1_Esch_coli_KTE185_V1	Escherichia coli KTE185	Proteobacteria; Gammaproteobacteria; Enterobacterales; Enterobacteriaceae; Escherichia; Escherichia coli	MTGDNTLIHSHGINRRDFMKLCAALAATMGLSSKAAAEMAESVTNPQRPPVIWIGAQECT	WP_000145410.1	MULTISPECIES: hydrogenase-2 small chain [Proteobacteria]	Length: 372\nNumber of predicted TMHs: 1\nExp number of AAs in TMHs: 23.72226\nExp number, first 60 AAs:  0.22038\nTotal prob of N-in:   0.12994\noutside   1 331\nTMhelix   332   354\ninside   355   372

11632	GCF_000408085.1_Esch_coli_KTE185_V1	Escherichia coli KTE185	Proteobacteria; Gammaproteobacteria; Enterobacterales; Enterobacteriaceae; Escherichia; Escherichia coli	MNNEETFYQAMRRQGVTRRSFLKYCSLAATSLGLGAGMAPKIAWALENKPRIPVVWIHGL	WP_001058323.1	MULTISPECIES: hydrogenase-1 small chain [Proteobacteria]	Length: 372\nNumber of predicted TMHs: 2\nExp number of AAs in TMHs: 37.2608499999999\nExp number, first 60 AAs:  19.52923\nTotal prob of N-in: 0.94351\nPOSSIBLE N-term signal sequence\ninside   1   19\nTMhelix   20   39\noutside   40   325\nTMhelix   326 348\ninside   349   372

11633	GCF_000408105.1_Esch_coli_KTE186_V1	Escherichia coli KTE186	Proteobacteria; Gammaproteobacteria; Enterobacterales; Enterobacteriaceae; Escherichia; Escherichia coli	MNNEETFYQAMRRQGVTRRSFLKYCSLAATSLGLGAGMAPKIAWALENKPRIPVVWIHGL	WP_001295940.1	MULTISPECIES: hydrogenase [Enterobacteriaceae]	Length: 372\nNumber of predicted TMHs:  2\nExp number of AAs in TMHs: 37.2608899999999\nExp number, first 60 AAs:  19.52923\nTotal prob of N-in: 0.94351\nPOSSIBLE N-term signal sequence\ninside   1   19\nTMhelix   20   39\noutside   40   325\nTMhelix   326 348\ninside   349   372

11634	GCF_000408105.1_Esch_coli_KTE186_V1	Escherichia coli KTE186	Proteobacteria; Gammaproteobacteria; Enterobacterales; Enterobacteriaceae; Escherichia; Escherichia coli	MTGDNTLIHSHGINRRDFMKLCAALAATMGLSSKAAAEMAESVTNPQRPPVIWIGAQECT	WP_000145418.1	hydrogenase 2 small subunit [Escherichia coli]	Length: 372\nNumber of predicted TMHs:  1\nExp number of

AAs in TMHs: 23.72313\nExp number, first 60 AAs: 0.22034\nTotal prob of N-in: 0.12997\noutside 1 331\nTMhelix 332 354\ninside 355 372

11635 GCF_000350945.1_Esch_coli_KTE187_V1 Escherichia coli KTE187 Proteobacteria; Gammaproteobacteria; Enterobacterales; Enterobacteriaceae; Escherichia; Escherichia coli MNNEETFYQAMRRQGVTRRSFLKYCSLAATSLGLGAGMAPKIAWALENKPRIPVVWIHGL WP_001058323.1 MULTISPECIES: hydrogenase-1 small chain [Proteobacteria] Length: 372\nNumber of predicted TMHs: 2\nExp number of AAs in TMHs: 37.2608499999999\nExp number, first 60 AAs: 19.52923\nTotal prob of N-in: 0.94351\nPOSSIBLE N-term signal sequence\ninside 1 19\nTMhelix 20 39\noutside 40 325\nTMhelix 326 348\ninside 349 372

11636 GCF_000350945.1_Esch_coli_KTE187_V1 Escherichia coli KTE187 Proteobacteria; Gammaproteobacteria; Enterobacterales; Enterobacteriaceae; Escherichia; Escherichia coli MTGDNTLIHSHGINRRDFMKLCAALAATMGLSSKAAAEMAESVTNPQRPPVIWIGAQECT WP_000145410.1 MULTISPECIES: hydrogenase-2 small chain [Proteobacteria] Length: 372\nNumber of predicted TMHs: 1\nExp number of AAs in TMHs: 23.72226\nExp number, first 60 AAs: 0.22038\nTotal prob of N-in: 0.12994\noutside 1 331\nTMhelix 332 354\ninside 355 372

11637 GCF_000350965.1_Esch_coli_KTE188_V1 Escherichia coli KTE188 Proteobacteria; Gammaproteobacteria; Enterobacterales; Enterobacteriaceae; Escherichia; Escherichia coli MNNEETFYQAMRRQGVTRRSFLKYCSLAATSLGLGAGMAPKIAWALENKPRIPVVWIHGL WP_001058323.1 MULTISPECIES: hydrogenase-1 small chain [Proteobacteria] Length: 372\nNumber of predicted TMHs: 2\nExp number of AAs in TMHs: 37.2608499999999\nExp number, first 60 AAs: 19.52923\nTotal prob of N-in: 0.94351\nPOSSIBLE N-term signal sequence\ninside 1 19\nTMhelix 20 39\noutside 40 325\nTMhelix 326 348\ninside 349 372

11638 GCF_000350965.1_Esch_coli_KTE188_V1 Escherichia coli KTE188 Proteobacteria; Gammaproteobacteria; Enterobacterales; Enterobacteriaceae; Escherichia; Escherichia coli MTGDNTLIHSHGINRRDFMKLCAALAATMGLSSKAAAEMAESVTNPQRPPVIWIGAQECT WP_000145418.1 hydrogenase 2 small subunit [Escherichia coli] Length: 372\nNumber of predicted TMHs: 1\nExp number of AAs in TMHs: 23.72313\nExp number, first 60 AAs: 0.22034\nTotal prob of N-in: 0.12997\noutside 1 331\nTMhelix 332 354\ninside 355 372

11639 GCF_000350985.1_Esch_coli_KTE189_V1 Escherichia coli KTE189 Proteobacteria; Gammaproteobacteria; Enterobacterales; Enterobacteriaceae; Escherichia; Escherichia coli MTGDNTLIHSHGINRRDFMKLCAALAATMGLSSKAAAEMAESVTNPQRPPVIWIGAQECT WP_000145410.1 MULTISPECIES: hydrogenase-2 small chain [Proteobacteria] Length: 372\nNumber of predicted TMHs: 1\nExp number of AAs in TMHs: 23.72226\nExp number, first 60 AAs: 0.22038\nTotal prob of N-in: 0.12994\noutside 1 331\nTMhelix 332 354\ninside 355 372

11640 GCF_000350985.1_Esch_coli_KTE189_V1 Escherichia coli KTE189 Proteobacteria; Gammaproteobacteria; Enterobacterales; Enterobacteriaceae; Escherichia; Escherichia coli MNNEETFYQAMRRQGVTRRSFLKYCSLAATSLGLGAGMAPKIAWALENKPRIPVVWIHGL WP_001058323.1 MULTISPECIES: hydrogenase-1 small chain [Proteobacteria] Length: 372\nNumber of predicted TMHs: 2\nExp number of AAs in TMHs: 37.2608499999999\nExp number, first 60 AAs: 19.52923\nTotal prob of N-in: 0.94351\nPOSSIBLE N-term signal sequence\ninside 1 19\nTMhelix 20 39\noutside 40 325\nTMhelix 326 348\ninside 349 372

11641 GCF_000407825.1_Esch_coli_KTE19_V1 Escherichia coli KTE19 Proteobacteria; Gammaproteobacteria; Enterobacterales; Enterobacteriaceae; Escherichia; Escherichia coli MNNEETFYQAMRRQGVTRRSFLKYCSLAATSLGLGAGMAPKIAWALENKPRIPVVWIHGL WP_001058323.1 MULTISPECIES: hydrogenase-1 small chain [Proteobacteria] Length: 372\nNumber of predicted TMHs: 2\nExp number of AAs in TMHs: 37.2608499999999\nExp number, first 60 AAs: 19.52923\nTotal prob of N-in: 0.94351\nPOSSIBLE N-term signal sequence\ninside 1 19\nTMhelix 20 39\noutside 40 325\nTMhelix 326 348\ninside 349 372

11642 GCF_000407825.1_Esch_coli_KTE19_V1 Escherichia coli KTE19 Proteobacteria; Gammaproteobacteria; Enterobacterales; Enterobacteriaceae; Escherichia; Escherichia coli MTGDNTLIHSHGINRRDFMKLCAALAATMGLSSKAAAEMVESVTNPQRPPVIWIGAQECT WP_000145423.1 hydrogenase 2 small subunit [Escherichia coli] Length: 372\nNumber of predicted TMHs: 1\nExp number of

AAs in TMHs: 23.78201\nExp number, first 60 AAs:  0.28355\nTotal prob of N-in:   0.13257\noutside   1   331\nTMhelix 332   354\ninside   355   372

11643 GCF_000352825.1_Esch_coli_KTE190_V1 Escherichia coli KTE190 Proteobacteria; Gammaproteobacteria; Enterobacterales; Enterobacteriaceae; Escherichia; Escherichia coli MTGDNTLIHSHGINRRDFMKLCAALAATMGLSSKAAAEMAESVTNPQRPPVIWIGAQECT WP_000145410.1 MULTISPECIES: hydrogenase-2 small chain [Proteobacteria] Length: 372\nNumber of predicted TMHs: 1\nExp number of AAs in TMHs: 23.72226\nExp number, first 60 AAs:  0.22038\nTotal prob of N-in:   0.12994\noutside   1 331\nTMhelix   332   354\ninside   355   372

11644 GCF_000352825.1_Esch_coli_KTE190_V1 Escherichia coli KTE190 Proteobacteria; Gammaproteobacteria; Enterobacterales; Enterobacteriaceae; Escherichia; Escherichia coli MNNEETFYQAMRRQGVTRRSFLKYCSLAATSLGLGAGMAPKIAWALENKPRIPVVWIHGL WP_001058323.1 MULTISPECIES: hydrogenase-1 small chain [Proteobacteria] Length: 372\nNumber of predicted TMHs: 2\nExp number of AAs in TMHs: 37.2608499999999\nExp number, first 60 AAs:  19.52923\nTotal prob of N-in: 0.94351\nPOSSIBLE N-term signal sequence\ninside   1   19\nTMhelix   20   39\noutside   40   325\nTMhelix   326 348\ninside   349   372

11645 GCF_000351005.1_Esch_coli_KTE191_V1 Escherichia coli KTE191 Proteobacteria; Gammaproteobacteria; Enterobacterales; Enterobacteriaceae; Escherichia; Escherichia coli MTGDNTLIHSHGINRRDFMKLCAALAATMGLSSKAAAEMAESVTNPQRPPVIWIGAQECT WP_000145410.1 MULTISPECIES: hydrogenase-2 small chain [Proteobacteria] Length: 372\nNumber of predicted TMHs: 1\nExp number of AAs in TMHs: 23.72226\nExp number, first 60 AAs:  0.22038\nTotal prob of N-in:   0.12994\noutside   1 331\nTMhelix   332   354\ninside   355   372

11646 GCF_000351005.1_Esch_coli_KTE191_V1 Escherichia coli KTE191 Proteobacteria; Gammaproteobacteria; Enterobacterales; Enterobacteriaceae; Escherichia; Escherichia coli MNNEETFYQAMRRQGVTRRSFLKYCSLAATSLGLGAGMAPKIAWALENKPRIPVVWIHGL WP_001058323.1 MULTISPECIES: hydrogenase-1 small chain [Proteobacteria] Length: 372\nNumber of predicted TMHs: 2\nExp number of AAs in TMHs: 37.2608499999999\nExp number, first 60 AAs:  19.52923\nTotal prob of N-in: 0.94351\nPOSSIBLE N-term signal sequence\ninside   1   19\nTMhelix   20   39\noutside   40   325\nTMhelix   326 348\ninside   349   372

11647 GCF_000352785.1_Esch_coli_KTE192_V1 Escherichia coli KTE192 Proteobacteria; Gammaproteobacteria; Enterobacterales; Enterobacteriaceae; Escherichia; Escherichia coli MNNEETFYQAMRRQGVTRRSFLKYCSLAATSLGLGAGMAPKIAWALENKPRIPVVWIHGL WP_001058323.1 MULTISPECIES: hydrogenase-1 small chain [Proteobacteria] Length: 372\nNumber of predicted TMHs: 2\nExp number of AAs in TMHs: 37.2608499999999\nExp number, first 60 AAs:  19.52923\nTotal prob of N-in: 0.94351\nPOSSIBLE N-term signal sequence\ninside   1   19\nTMhelix   20   39\noutside   40   325\nTMhelix   326 348\ninside   349   372

11648 GCF_000352785.1_Esch_coli_KTE192_V1 Escherichia coli KTE192 Proteobacteria; Gammaproteobacteria; Enterobacterales; Enterobacteriaceae; Escherichia; Escherichia coli MTGDNTLIHSHGINRRDFMKLCAALAATMGLSSKAAAEMVESVTNPQRPPVIWIGAQECT WP_000145423.1 hydrogenase 2 small subunit [Escherichia coli] Length: 372\nNumber of predicted TMHs:  1\nExp number of AAs in TMHs: 23.78201\nExp number, first 60 AAs:  0.28355\nTotal prob of N-in:   0.13257\noutside   1   331\nTMhelix 332   354\ninside   355   372

11649 GCF_000351025.1_Esch_coli_KTE193_V1 Escherichia coli KTE193 Proteobacteria; Gammaproteobacteria; Enterobacterales; Enterobacteriaceae; Escherichia; Escherichia coli MTGDNTLIHSHGINRRDFMKLCAALAATMGLSSKAAAEMAESVTNPQRPPVIWIGAQECT WP_000145410.1 MULTISPECIES: hydrogenase-2 small chain [Proteobacteria] Length: 372\nNumber of predicted TMHs: 1\nExp number of AAs in TMHs: 23.72226\nExp number, first 60 AAs:  0.22038\nTotal prob of N-in:   0.12994\noutside   1 331\nTMhelix   332   354\ninside   355   372

11650 GCF_000351025.1_Esch_coli_KTE193_V1 Escherichia coli KTE193 Proteobacteria; Gammaproteobacteria; Enterobacterales; Enterobacteriaceae; Escherichia; Escherichia coli MNNEETFYQAMRRQGVTRRSFLKYCSLAATSLGLGAGMAPKIAWALENKPRIPVVWIHGL WP_001058323.1 MULTISPECIES: hydrogenase-1 small chain [Proteobacteria] Length: 372\nNumber of predicted TMHs: 2\nExp number of AAs in TMHs: 37.2608499999999\nExp number, first 60 AAs:  19.52923\nTotal prob of N-in:

0.94351\nPOSSIBLE N-term signal sequence\ninside 1 19\nTMhelix 20 39\noutside 40 325\nTMhelix 326 348\ninside 349 372

11651 GCF_000352805.1_Esch_coli_KTE194_V1 Escherichia coli KTE194 Proteobacteria; Gammaproteobacteria; Enterobacterales; Enterobacteriaceae; Escherichia; Escherichia coli MTGDNTLIHSHGINRRDFMKLCAALAATMGLSSKAAAEMVESVTNPQRPPVIWIGAQECT WP_000145423.1 hydrogenase 2 small subunit [Escherichia coli] Length: 372\nNumber of predicted TMHs: 1\nExp number of AAs in TMHs: 23.78201\nExp number, first 60 AAs: 0.28355\nTotal prob of N-in: 0.13257\noutside 1 331\nTMhelix 332 354\ninside 355 372

11652 GCF_000352805.1_Esch_coli_KTE194_V1 Escherichia coli KTE194 Proteobacteria; Gammaproteobacteria; Enterobacterales; Enterobacteriaceae; Escherichia; Escherichia coli MNNEETFYQAMRRQGVTRRSFLKYCSLAATSLGLGAGMAPKIAWALENKPRIPVVWIHGL WP_001058323.1 MULTISPECIES: hydrogenase-1 small chain [Proteobacteria] Length: 372\nNumber of predicted TMHs: 2\nExp number of AAs in TMHs: 37.2608499999999\nExp number, first 60 AAs: 19.52923\nTotal prob of N-in: 0.94351\nPOSSIBLE N-term signal sequence\ninside 1 19\nTMhelix 20 39\noutside 40 325\nTMhelix 326 348\ninside 349 372

11653 GCF_000408125.1_Esch_coli_KTE195_V1 Escherichia coli KTE195 Proteobacteria; Gammaproteobacteria; Enterobacterales; Enterobacteriaceae; Escherichia; Escherichia coli MNNEETFYQAMRRQGVTRRSFLKYCSLAATSLGLGAGMAPKIAWALENKPRIPVVWIHGL WP_001058323.1 MULTISPECIES: hydrogenase-1 small chain [Proteobacteria] Length: 372\nNumber of predicted TMHs: 2\nExp number of AAs in TMHs: 37.2608499999999\nExp number, first 60 AAs: 19.52923\nTotal prob of N-in: 0.94351\nPOSSIBLE N-term signal sequence\ninside 1 19\nTMhelix 20 39\noutside 40 325\nTMhelix 326 348\ninside 349 372

11654 GCF_000408125.1_Esch_coli_KTE195_V1 Escherichia coli KTE195 Proteobacteria; Gammaproteobacteria; Enterobacterales; Enterobacteriaceae; Escherichia; Escherichia coli MTGDNTLIHSHGINRRDFMKLCAALAATMGLSSKAAAEMAESVTNPQRPPVIWIGAQECT WP_000145410.1 MULTISPECIES: hydrogenase-2 small chain [Proteobacteria] Length: 372\nNumber of predicted TMHs: 1\nExp number of AAs in TMHs: 23.72226\nExp number, first 60 AAs: 0.22038\nTotal prob of N-in: 0.12994\noutside 1 331\nTMhelix 332 354\ninside 355 372

11655 GCF_000352925.1_Esch_coli_KTE196_V1 Escherichia coli KTE196 Proteobacteria; Gammaproteobacteria; Enterobacterales; Enterobacteriaceae; Escherichia; Escherichia coli MNNEETFYQAMRRQGVTRRSFLKYCSLAATSLGLGAGMAPKIAWALENKPRIPVVWIHGL WP_001058323.1 MULTISPECIES: hydrogenase-1 small chain [Proteobacteria] Length: 372\nNumber of predicted TMHs: 2\nExp number of AAs in TMHs: 37.2608499999999\nExp number, first 60 AAs: 19.52923\nTotal prob of N-in: 0.94351\nPOSSIBLE N-term signal sequence\ninside 1 19\nTMhelix 20 39\noutside 40 325\nTMhelix 326 348\ninside 349 372

11656 GCF_000352925.1_Esch_coli_KTE196_V1 Escherichia coli KTE196 Proteobacteria; Gammaproteobacteria; Enterobacterales; Enterobacteriaceae; Escherichia; Escherichia coli MTGDNTLIHSHGINRRDFMKLCAALAATMGLSSKAAAEMAESVTNPQRPPVIWIGAQECT WP_000145410.1 MULTISPECIES: hydrogenase-2 small chain [Proteobacteria] Length: 372\nNumber of predicted TMHs: 1\nExp number of AAs in TMHs: 23.72226\nExp number, first 60 AAs: 0.22038\nTotal prob of N-in: 0.12994\noutside 1 331\nTMhelix 332 354\ninside 355 372

11657 GCF_000352945.1_Esch_coli_KTE197_V1 Escherichia coli KTE197 Proteobacteria; Gammaproteobacteria; Enterobacterales; Enterobacteriaceae; Escherichia; Escherichia coli MNNEETFYQAMRRQGVTRRSFLKYCSLAATSLGLGAGMAPKIAWALENKPRIPVVWIHGL WP_001058323.1 MULTISPECIES: hydrogenase-1 small chain [Proteobacteria] Length: 372\nNumber of predicted TMHs: 2\nExp number of AAs in TMHs: 37.2608499999999\nExp number, first 60 AAs: 19.52923\nTotal prob of N-in: 0.94351\nPOSSIBLE N-term signal sequence\ninside 1 19\nTMhelix 20 39\noutside 40 325\nTMhelix 326 348\ninside 349 372

11658 GCF_000352945.1_Esch_coli_KTE197_V1 Escherichia coli KTE197 Proteobacteria; Gammaproteobacteria; Enterobacterales; Enterobacteriaceae; Escherichia; Escherichia coli MTGDNTLIHSHGINRRDFMKLCAALAATMGLSSKAAAEMAESVTNPQRPPVIWIGAQECT WP_000145410.1 MULTISPECIES: hydrogenase-2 small chain [Proteobacteria] Length: 372\nNumber of predicted TMHs:

1\nExp number of AAs in TMHs: 23.72226\nExp number, first 60 AAs: 0.22038\nTotal prob of N-in: 0.12994\noutside 1 331\nTMhelix 332 354\ninside 355 372

11659 GCF_000408145.1_Esch_coli_KTE198_V1 Escherichia coli KTE198 Proteobacteria; Gammaproteobacteria; Enterobacterales; Enterobacteriaceae; Escherichia; Escherichia coli MTGDNTLIHSHGINRRDFMKLCAALAATMGLSSKAAAEMAESVTNPQRPPVIWIGAQECT WP_000145410.1 MULTISPECIES: hydrogenase-2 small chain [Proteobacteria] Length: 372\nNumber of predicted TMHs: 1\nExp number of AAs in TMHs: 23.72226\nExp number, first 60 AAs: 0.22038\nTotal prob of N-in: 0.12994\noutside 1 331\nTMhelix 332 354\ninside 355 372

11660 GCF_000408145.1_Esch_coli_KTE198_V1 Escherichia coli KTE198 Proteobacteria; Gammaproteobacteria; Enterobacterales; Enterobacteriaceae; Escherichia; Escherichia coli MNNEETFYQAMRRQGVTRRSFLKYCSLAATSLGLGAGMAPKIAWALENKPRIPVVWIHGL WP_001058323.1 MULTISPECIES: hydrogenase-1 small chain [Proteobacteria] Length: 372\nNumber of predicted TMHs: 2\nExp number of AAs in TMHs: 37.2608499999999\nExp number, first 60 AAs: 19.52923\nTotal prob of N-in: 0.94351\nPOSSIBLE N-term signal sequence\ninside 1 19\nTMhelix 20 39\noutside 40 325\nTMhelix 326 348\ninside 349 372

11661 GCF_000408165.1_Esch_coli_KTE199_V1 Escherichia coli KTE199 Proteobacteria; Gammaproteobacteria; Enterobacterales; Enterobacteriaceae; Escherichia; Escherichia coli MNNEETFYQAMRRQGVTRRSFLKYCSLAATSLGLGAGMAPKIAWALENKPRIPVVWIHGL WP_001058323.1 MULTISPECIES: hydrogenase-1 small chain [Proteobacteria] Length: 372\nNumber of predicted TMHs: 2\nExp number of AAs in TMHs: 37.2608499999999\nExp number, first 60 AAs: 19.52923\nTotal prob of N-in: 0.94351\nPOSSIBLE N-term signal sequence\ninside 1 19\nTMhelix 20 39\noutside 40 325\nTMhelix 326 348\ninside 349 372

11662 GCF_000408165.1_Esch_coli_KTE199_V1 Escherichia coli KTE199 Proteobacteria; Gammaproteobacteria; Enterobacterales; Enterobacteriaceae; Escherichia; Escherichia coli MTGDNTLIHSHGINRRDFMKLCAALAATMGLSSKAAAEMAESVTNPQRPPVIWIGAQECT WP_000145410.1 MULTISPECIES: hydrogenase-2 small chain [Proteobacteria] Length: 372\nNumber of predicted TMHs: 1\nExp number of AAs in TMHs: 23.72226\nExp number, first 60 AAs: 0.22038\nTotal prob of N-in: 0.12994\noutside 1 331\nTMhelix 332 354\ninside 355 372

11663 GCF_000350625.1_Esch_coli_KTE2_V1 Escherichia coli KTE2 Proteobacteria; Gammaproteobacteria; Enterobacterales; Enterobacteriaceae; Escherichia; Escherichia coli MTGDNTLIHSHGINRRDFMKLCAALAATMGLSSKAAAEMAESVTNPQRPPVIWIGAQECT WP_000145410.1 MULTISPECIES: hydrogenase-2 small chain [Proteobacteria] Length: 372\nNumber of predicted TMHs: 1\nExp number of AAs in TMHs: 23.72226\nExp number, first 60 AAs: 0.22038\nTotal prob of N-in: 0.12994\noutside 1 331\nTMhelix 332 354\ninside 355 372

11664 GCF_000350625.1_Esch_coli_KTE2_V1 Escherichia coli KTE2 Proteobacteria; Gammaproteobacteria; Enterobacterales; Enterobacteriaceae; Escherichia; Escherichia coli MNNEETFYQAMRRQGVTRRSFLKYCSLAATSLGLGAGMAPKIAWALENKPRIPVVWIHGL WP_001058323.1 MULTISPECIES: hydrogenase-1 small chain [Proteobacteria] Length: 372\nNumber of predicted TMHs: 2\nExp number of AAs in TMHs: 37.2608499999999\nExp number, first 60 AAs: 19.52923\nTotal prob of N-in: 0.94351\nPOSSIBLE N-term signal sequence\ninside 1 19\nTMhelix 20 39\noutside 40 325\nTMhelix 326 348\ninside 349 372

11665 GCF_000407845.1_Esch_coli_KTE20_V1 Escherichia coli KTE20 Proteobacteria; Gammaproteobacteria; Enterobacterales; Enterobacteriaceae; Escherichia; Escherichia coli MNNEETFYQAMRRQGVTRRSFLKYCSLAATSLGLGAGMAPKIAWALENKPRIPVVWIHGL WP_001058323.1 MULTISPECIES: hydrogenase-1 small chain [Proteobacteria] Length: 372\nNumber of predicted TMHs: 2\nExp number of AAs in TMHs: 37.2608499999999\nExp number, first 60 AAs: 19.52923\nTotal prob of N-in: 0.94351\nPOSSIBLE N-term signal sequence\ninside 1 19\nTMhelix 20 39\noutside 40 325\nTMhelix 326 348\ninside 349 372

11666 GCF_000407845.1_Esch_coli_KTE20_V1 Escherichia coli KTE20 Proteobacteria; Gammaproteobacteria; Enterobacterales; Enterobacteriaceae; Escherichia; Escherichia coli MTGDNTLIHSHGINRRDFMKLCAALAATMGLSSKAAAEMAESVTNPQRPPVIWIGAQECT WP_000145410.1 MULTISPECIES: hydrogenase-2 small chain [Proteobacteria] Length: 372\nNumber of predicted TMHs:

1\nExp number of AAs in TMHs: 23.72226\nExp number, first 60 AAs: 0.22038\nTotal prob of N-in: 0.12994\noutside 1 331\nTMhelix 332 354\ninside 355 372

11667 GCF_000408185.1_Esch_coli_KTE200_V1 Escherichia coli KTE200 Proteobacteria; Gammaproteobacteria; Enterobacterales; Enterobacteriaceae; Escherichia; Escherichia coli MNNEETFYQAMRRQGVTRRSFLKYCSLAATSLGLGAGMAPKIAWALENKPRIPVVWIHGL WP_001058323.1 MULTISPECIES: hydrogenase-1 small chain [Proteobacteria] Length: 372\nNumber of predicted TMHs: 2\nExp number of AAs in TMHs: 37.2608499999999\nExp number, first 60 AAs: 19.52923\nTotal prob of N-in: 0.94351\nPOSSIBLE N-term signal sequence\ninside 1 19\nTMhelix 20 39\noutside 40 325\nTMhelix 326 348\ninside 349 372

11668 GCF_000408185.1_Esch_coli_KTE200_V1 Escherichia coli KTE200 Proteobacteria; Gammaproteobacteria; Enterobacterales; Enterobacteriaceae; Escherichia; Escherichia coli MTGDNTLIHSHGINRRDFMKLCAALAATMGLSSKAAAEMAESVTNPQRPPVIWIGAQECT WP_000145410.1 MULTISPECIES: hydrogenase-2 small chain [Proteobacteria] Length: 372\nNumber of predicted TMHs: 1\nExp number of AAs in TMHs: 23.72226\nExp number, first 60 AAs: 0.22038\nTotal prob of N-in: 0.12994\noutside 1 331\nTMhelix 332 354\ninside 355 372

11669 GCF_000351045.1_Esch_coli_KTE201_V1 Escherichia coli KTE201 Proteobacteria; Gammaproteobacteria; Enterobacterales; Enterobacteriaceae; Escherichia; Escherichia coli MTGDNTLIHSHGINRRDFMKLCAALAATMGLSSKAAAEMAESVTNPQRPPVIWIGAQECT WP_000145410.1 MULTISPECIES: hydrogenase-2 small chain [Proteobacteria] Length: 372\nNumber of predicted TMHs: 1\nExp number of AAs in TMHs: 23.72226\nExp number, first 60 AAs: 0.22038\nTotal prob of N-in: 0.12994\noutside 1 331\nTMhelix 332 354\ninside 355 372

11670 GCF_000351045.1_Esch_coli_KTE201_V1 Escherichia coli KTE201 Proteobacteria; Gammaproteobacteria; Enterobacterales; Enterobacteriaceae; Escherichia; Escherichia coli MNNEETFYQAMRRQGVTRRSFLKYCSLAATSLGLGAGMAPKIAWALENKPRIPVVWIHGL WP_001058323.1 MULTISPECIES: hydrogenase-1 small chain [Proteobacteria] Length: 372\nNumber of predicted TMHs: 2\nExp number of AAs in TMHs: 37.2608499999999\nExp number, first 60 AAs: 19.52923\nTotal prob of N-in: 0.94351\nPOSSIBLE N-term signal sequence\ninside 1 19\nTMhelix 20 39\noutside 40 325\nTMhelix 326 348\ninside 349 372

11671 GCF_000352965.1_Esch_coli_KTE202_V1 Escherichia coli KTE202 Proteobacteria; Gammaproteobacteria; Enterobacterales; Enterobacteriaceae; Escherichia; Escherichia coli MTGDNTLIHSHGINRRDFMKLCAALAATMGLSSKAAAEMAESVTNPQRPPVIWIGAQECT WP_000145410.1 MULTISPECIES: hydrogenase-2 small chain [Proteobacteria] Length: 372\nNumber of predicted TMHs: 1\nExp number of AAs in TMHs: 23.72226\nExp number, first 60 AAs: 0.22038\nTotal prob of N-in: 0.12994\noutside 1 331\nTMhelix 332 354\ninside 355 372

11672 GCF_000352965.1_Esch_coli_KTE202_V1 Escherichia coli KTE202 Proteobacteria; Gammaproteobacteria; Enterobacterales; Enterobacteriaceae; Escherichia; Escherichia coli MNNEETFYQAMRRQGVTRRSFLKYCSLAATSLGLGAGMAPKIAWALENKPRIPVVWIHGL WP_001058323.1 MULTISPECIES: hydrogenase-1 small chain [Proteobacteria] Length: 372\nNumber of predicted TMHs: 2\nExp number of AAs in TMHs: 37.2608499999999\nExp number, first 60 AAs: 19.52923\nTotal prob of N-in: 0.94351\nPOSSIBLE N-term signal sequence\ninside 1 19\nTMhelix 20 39\noutside 40 325\nTMhelix 326 348\ninside 349 372

11673 GCF_000352985.1_Esch_coli_KTE203_V1 Escherichia coli KTE203 Proteobacteria; Gammaproteobacteria; Enterobacterales; Enterobacteriaceae; Escherichia; Escherichia coli MTGDNTLIHSHGINRRDFMKLCAALAATMGLSSKAAAEMAESVTNPQRPPVIWIGAQECT WP_000145410.1 MULTISPECIES: hydrogenase-2 small chain [Proteobacteria] Length: 372\nNumber of predicted TMHs: 1\nExp number of AAs in TMHs: 23.72226\nExp number, first 60 AAs: 0.22038\nTotal prob of N-in: 0.12994\noutside 1 331\nTMhelix 332 354\ninside 355 372

11674 GCF_000352985.1_Esch_coli_KTE203_V1 Escherichia coli KTE203 Proteobacteria; Gammaproteobacteria; Enterobacterales; Enterobacteriaceae; Escherichia; Escherichia coli MNNEETFYQAMRRQGVTRRSFLKYCSLAATSLGLGAGMAPKIAWALENKPRIPVVWIHGL WP_001295940.1 MULTISPECIES: hydrogenase [Enterobacteriaceae] Length: 372\nNumber of predicted TMHs: 2\nExp number of AAs in TMHs: 37.2608899999999\nExp number, first 60 AAs: 19.52923\nTotal prob of N-in:

0.94351\nPOSSIBLE N-term signal sequence\ninside 1 19\nTMhelix 20 39\noutside 40 325\nTMhelix 326 348\ninside 349 372

11675 GCF_000351065.1_Esch_coli_KTE204_V1 Escherichia coli KTE204 Proteobacteria; Gammaproteobacteria; Enterobacterales; Enterobacteriaceae; Escherichia; Escherichia coli MTGDNTLIHSHGINRRDFMKLCAALAATMGLSSKAAAEMAESVTNPQRPPVIWIGAQECT WP_000145410.1 MULTISPECIES: hydrogenase-2 small chain [Proteobacteria] Length: 372\nNumber of predicted TMHs: 1\nExp number of AAs in TMHs: 23.72226\nExp number, first 60 AAs: 0.22038\nTotal prob of N-in: 0.12994\noutside 1 331\nTMhelix 332 354\ninside 355 372

11676 GCF_000351065.1_Esch_coli_KTE204_V1 Escherichia coli KTE204 Proteobacteria; Gammaproteobacteria; Enterobacterales; Enterobacteriaceae; Escherichia; Escherichia coli MNNEETFYQAMRRQGVTRRSFLKYCSLAATSLGLGAGMAPKIAWALENKPRIPVVWIHGL WP_001058323.1 MULTISPECIES: hydrogenase-1 small chain [Proteobacteria] Length: 372\nNumber of predicted TMHs: 2\nExp number of AAs in TMHs: 37.2608499999999\nExp number, first 60 AAs: 19.52923\nTotal prob of N-in: 0.94351\nPOSSIBLE N-term signal sequence\ninside 1 19\nTMhelix 20 39\noutside 40 325\nTMhelix 326 348\ninside 349 372

11677 GCF_000351085.1_Esch_coli_KTE205_V1 Escherichia coli KTE205 Proteobacteria; Gammaproteobacteria; Enterobacterales; Enterobacteriaceae; Escherichia; Escherichia coli MTGDNTLIHSHGINRRDFMKLCAALAATMGLSSKAAAEMVESVTNPQRPPVIWIGAQECT WP_000145423.1 hydrogenase 2 small subunit [Escherichia coli] Length: 372\nNumber of predicted TMHs: 1\nExp number of AAs in TMHs: 23.78201\nExp number, first 60 AAs: 0.28355\nTotal prob of N-in: 0.13257\noutside 1 331\nTMhelix 332 354\ninside 355 372

11678 GCF_000351085.1_Esch_coli_KTE205_V1 Escherichia coli KTE205 Proteobacteria; Gammaproteobacteria; Enterobacterales; Enterobacteriaceae; Escherichia; Escherichia coli MNNEETFYQAMRRQGVTRRSFLKYCSLAATSLGLGAGMAPKIAWALENKPRIPVVWIHGL WP_001538532.1 hydrogenase-1 small chain [Escherichia coli] Length: 372\nNumber of predicted TMHs: 2\nExp number of AAs in TMHs: 37.27041\nExp number, first 60 AAs: 19.52927\nTotal prob of N-in: 0.94352\nPOSSIBLE N-term signal sequence\ninside 1 19\nTMhelix 20 39\noutside 40 325\nTMhelix 326 348\ninside 349 372

11679 GCF_000351105.1_Esch_coli_KTE206_V1 Escherichia coli KTE206 Proteobacteria; Gammaproteobacteria; Enterobacterales; Enterobacteriaceae; Escherichia; Escherichia coli MNNEETFYQAMRRQGVTRRSFLKYCSLAATSLGLGAGMAPKIAWALENKPRIPVVWIHGL WP_001058323.1 MULTISPECIES: hydrogenase-1 small chain [Proteobacteria] Length: 372\nNumber of predicted TMHs: 2\nExp number of AAs in TMHs: 37.2608499999999\nExp number, first 60 AAs: 19.52923\nTotal prob of N-in: 0.94351\nPOSSIBLE N-term signal sequence\ninside 1 19\nTMhelix 20 39\noutside 40 325\nTMhelix 326 348\ninside 349 372

11680 GCF_000351105.1_Esch_coli_KTE206_V1 Escherichia coli KTE206 Proteobacteria; Gammaproteobacteria; Enterobacterales; Enterobacteriaceae; Escherichia; Escherichia coli MTGDNTLIHSHGINRRDFMKLCAALAATMGLSSKAAAEMAESVTNPQRPPVIWIGAQECT WP_000145410.1 MULTISPECIES: hydrogenase-2 small chain [Proteobacteria] Length: 372\nNumber of predicted TMHs: 1\nExp number of AAs in TMHs: 23.72226\nExp number, first 60 AAs: 0.22038\nTotal prob of N-in: 0.12994\noutside 1 331\nTMhelix 332 354\ninside 355 372

11681 GCF_000353005.1_Esch_coli_KTE207_V1 Escherichia coli KTE207 Proteobacteria; Gammaproteobacteria; Enterobacterales; Enterobacteriaceae; Escherichia; Escherichia coli MTGDNTLIHSHGINRRDFMKLCAALAATMGLSSKAAAEMVESVTNPQRPPVIWIGAQECT WP_000145423.1 hydrogenase 2 small subunit [Escherichia coli] Length: 372\nNumber of predicted TMHs: 1\nExp number of AAs in TMHs: 23.78201\nExp number, first 60 AAs: 0.28355\nTotal prob of N-in: 0.13257\noutside 1 331\nTMhelix 332 354\ninside 355 372

11682 GCF_000353005.1_Esch_coli_KTE207_V1 Escherichia coli KTE207 Proteobacteria; Gammaproteobacteria; Enterobacterales; Enterobacteriaceae; Escherichia; Escherichia coli MNNEETFYQAMRRQGVTRRSFLKYCSLAATSLGLGAGMAPKIAWALENKPRIPVVWIHGL WP_001058323.1 MULTISPECIES: hydrogenase-1 small chain [Proteobacteria] Length: 372\nNumber of predicted TMHs: 2\nExp number of AAs in TMHs: 37.2608499999999\nExp number, first 60 AAs: 19.52923\nTotal prob of N-in: 0.94351\nPOSSIBLE N-term signal sequence\ninside 1 19\nTMhelix 20 39\noutside 40 325\nTMhelix 326 348\ninside 349 372

11683 GCF_000351125.1_Esch_coli_KTE208_V1 Escherichia coli KTE208 Proteobacteria; Gammaproteobacteria; Enterobacterales; Enterobacteriaceae; Escherichia; Escherichia coli
MNNEETFYQAMRRQGVTRRSFLKYCSLAATSLGLGAGMAPKIAWALENKPRIPVVWIHGL WP_001058323.1
MULTISPECIES: hydrogenase-1 small chain [Proteobacteria] Length: 372\nNumber of predicted TMHs: 2\nExp number of AAs in TMHs: 37.2608499999999\nExp number, first 60 AAs: 19.52923\nTotal prob of N-in: 0.94351\nPOSSIBLE N-term signal sequence\ninside 1 19\nTMhelix 20 39\noutside 40 325\nTMhelix 326 348\ninside 349 372

11684 GCF_000351125.1_Esch_coli_KTE208_V1 Escherichia coli KTE208 Proteobacteria; Gammaproteobacteria; Enterobacterales; Enterobacteriaceae; Escherichia; Escherichia coli
MTGDNTLIHSHGINRRDFMKLCAALAATMGLSSKAAAEMAESVTNPQRPPVIWIGAQECT WP_000145410.1
MULTISPECIES: hydrogenase-2 small chain [Proteobacteria] Length: 372\nNumber of predicted TMHs: 1\nExp number of AAs in TMHs: 23.72226\nExp number, first 60 AAs: 0.22038\nTotal prob of N-in: 0.12994\noutside 1 331\nTMhelix 332 354\ninside 355 372

11685 GCF_000353025.1_Esch_coli_KTE209_V1 Escherichia coli KTE209 Proteobacteria; Gammaproteobacteria; Enterobacterales; Enterobacteriaceae; Escherichia; Escherichia coli
MTGDNTLIHSHGINRRDFMKLCAALAATMGLSSKAAAEMAESVTNPQRPPVIWIGAQECT WP_000145410.1
MULTISPECIES: hydrogenase-2 small chain [Proteobacteria] Length: 372\nNumber of predicted TMHs: 1\nExp number of AAs in TMHs: 23.72226\nExp number, first 60 AAs: 0.22038\nTotal prob of N-in: 0.12994\noutside 1 331\nTMhelix 332 354\ninside 355 372

11686 GCF_000353025.1_Esch_coli_KTE209_V1 Escherichia coli KTE209 Proteobacteria; Gammaproteobacteria; Enterobacterales; Enterobacteriaceae; Escherichia; Escherichia coli
MNNEETFYQAMRRQGVTRRSFLKYCSLAATSLGLGAGMAPKIAWALENKPRIPVVWIHGL WP_001058323.1
MULTISPECIES: hydrogenase-1 small chain [Proteobacteria] Length: 372\nNumber of predicted TMHs: 2\nExp number of AAs in TMHs: 37.2608499999999\nExp number, first 60 AAs: 19.52923\nTotal prob of N-in: 0.94351\nPOSSIBLE N-term signal sequence\ninside 1 19\nTMhelix 20 39\noutside 40 325\nTMhelix 326 348\ninside 349 372

11687 GCF_000350785.1_Esch_coli_KTE21_V1 Escherichia coli KTE21 Proteobacteria; Gammaproteobacteria; Enterobacterales; Enterobacteriaceae; Escherichia; Escherichia coli
MNNEETFYQAMRRQGVTRRSFLKYCSLAATSLGLGAGMAPKIAWALENKPRIPVVWIHGL WP_001058323.1
MULTISPECIES: hydrogenase-1 small chain [Proteobacteria] Length: 372\nNumber of predicted TMHs: 2\nExp number of AAs in TMHs: 37.2608499999999\nExp number, first 60 AAs: 19.52923\nTotal prob of N-in: 0.94351\nPOSSIBLE N-term signal sequence\ninside 1 19\nTMhelix 20 39\noutside 40 325\nTMhelix 326 348\ninside 349 372

11688 GCF_000350785.1_Esch_coli_KTE21_V1 Escherichia coli KTE21 Proteobacteria; Gammaproteobacteria; Enterobacterales; Enterobacteriaceae; Escherichia; Escherichia coli
MTGDNTLIHSHGINRRDFMKLCAALAATMGLSSKAAAEMAESVTNPQRPPVIWIGAQECT WP_000145410.1
MULTISPECIES: hydrogenase-2 small chain [Proteobacteria] Length: 372\nNumber of predicted TMHs: 1\nExp number of AAs in TMHs: 23.72226\nExp number, first 60 AAs: 0.22038\nTotal prob of N-in: 0.12994\noutside 1 331\nTMhelix 332 354\ninside 355 372

11689 GCF_000351145.1_Esch_coli_KTE210_V1 Escherichia coli KTE210 Proteobacteria; Gammaproteobacteria; Enterobacterales; Enterobacteriaceae; Escherichia; Escherichia coli
MNNEETFYQAMRRQGVTRRSFLKYCSLAATSLGLGAGMAPKIAWALENKPRIPVVWIHGL WP_001058323.1
MULTISPECIES: hydrogenase-1 small chain [Proteobacteria] Length: 372\nNumber of predicted TMHs: 2\nExp number of AAs in TMHs: 37.2608499999999\nExp number, first 60 AAs: 19.52923\nTotal prob of N-in: 0.94351\nPOSSIBLE N-term signal sequence\ninside 1 19\nTMhelix 20 39\noutside 40 325\nTMhelix 326 348\ninside 349 372

11690 GCF_000351145.1_Esch_coli_KTE210_V1 Escherichia coli KTE210 Proteobacteria; Gammaproteobacteria; Enterobacterales; Enterobacteriaceae; Escherichia; Escherichia coli
MTGDNTLIHSHGINRRDFMKLCAALAATMGLSSKAAAEMAESVTNPQRPPVIWIGAQECT WP_000145410.1
MULTISPECIES: hydrogenase-2 small chain [Proteobacteria] Length: 372\nNumber of predicted TMHs: 1\nExp number of AAs in TMHs: 23.72226\nExp number, first 60 AAs: 0.22038\nTotal prob of N-in: 0.12994\noutside 1 331\nTMhelix 332 354\ninside 355 372

11691 GCF_000353045.1_Esch_coli_KTE211_V1 Escherichia coli KTE211 Proteobacteria; Gammaproteobacteria; Enterobacterales; Enterobacteriaceae; Escherichia; Escherichia coli MNNEETFYQAMRRQGVTRRSFLKYCSLAATSLGLGAGMAPKIAWALENKPRIPVVWIHGL WP_001058323.1 MULTISPECIES: hydrogenase-1 small chain [Proteobacteria] Length: 372\nNumber of predicted TMHs: 2\nExp number of AAs in TMHs: 37.2608499999999\nExp number, first 60 AAs: 19.52923\nTotal prob of N-in: 0.94351\nPOSSIBLE N-term signal sequence\ninside 1 19\nTMhelix 20 39\noutside 40 325\nTMhelix 326 348\ninside 349 372

11692 GCF_000353045.1_Esch_coli_KTE211_V1 Escherichia coli KTE211 Proteobacteria; Gammaproteobacteria; Enterobacterales; Enterobacteriaceae; Escherichia; Escherichia coli MTGDNTLIHSHGINRRDFMKLCAALAATMGLSSKAAAEMAESVTNPQRPPVIWIGAQECT WP_000145410.1 MULTISPECIES: hydrogenase-2 small chain [Proteobacteria] Length: 372\nNumber of predicted TMHs: 1\nExp number of AAs in TMHs: 23.72226\nExp number, first 60 AAs: 0.22038\nTotal prob of N-in: 0.12994\noutside 1 331\nTMhelix 332 354\ninside 355 372

11693 GCF_000351165.1_Esch_coli_KTE212_V1 Escherichia coli KTE212 Proteobacteria; Gammaproteobacteria; Enterobacterales; Enterobacteriaceae; Escherichia; Escherichia coli MNNEETFYQAMRRQGVTRRSFLKYCSLAATSLGLGAGMAPKIAWALENKPRIPVVWIHGL WP_001058323.1 MULTISPECIES: hydrogenase-1 small chain [Proteobacteria] Length: 372\nNumber of predicted TMHs: 2\nExp number of AAs in TMHs: 37.2608499999999\nExp number, first 60 AAs: 19.52923\nTotal prob of N-in: 0.94351\nPOSSIBLE N-term signal sequence\ninside 1 19\nTMhelix 20 39\noutside 40 325\nTMhelix 326 348\ninside 349 372

11694 GCF_000351165.1_Esch_coli_KTE212_V1 Escherichia coli KTE212 Proteobacteria; Gammaproteobacteria; Enterobacterales; Enterobacteriaceae; Escherichia; Escherichia coli MTGDNTLIHSHGINRRDFMKLCAALAATMGLSSKAAAEMAESVTNPQRPPVIWIGAQECT WP_000145410.1 MULTISPECIES: hydrogenase-2 small chain [Proteobacteria] Length: 372\nNumber of predicted TMHs: 1\nExp number of AAs in TMHs: 23.72226\nExp number, first 60 AAs: 0.22038\nTotal prob of N-in: 0.12994\noutside 1 331\nTMhelix 332 354\ninside 355 372

11695 GCF_000351185.1_Esch_coli_KTE213_V1 Escherichia coli KTE213 Proteobacteria; Gammaproteobacteria; Enterobacterales; Enterobacteriaceae; Escherichia; Escherichia coli MTGDNTLIHSHGINRRDFMKLCAALAATMGLSSKAAAEMAESVTNPQRPPVIWIGAQECT WP_000145410.1 MULTISPECIES: hydrogenase-2 small chain [Proteobacteria] Length: 372\nNumber of predicted TMHs: 1\nExp number of AAs in TMHs: 23.72226\nExp number, first 60 AAs: 0.22038\nTotal prob of N-in: 0.12994\noutside 1 331\nTMhelix 332 354\ninside 355 372

11696 GCF_000351185.1_Esch_coli_KTE213_V1 Escherichia coli KTE213 Proteobacteria; Gammaproteobacteria; Enterobacterales; Enterobacteriaceae; Escherichia; Escherichia coli MNNEETFYQAMRRQGVTRRSFLKYCSLAATSLGLGAGMAPKIAWALENKPRIPVVWIHGL WP_001058323.1 MULTISPECIES: hydrogenase-1 small chain [Proteobacteria] Length: 372\nNumber of predicted TMHs: 2\nExp number of AAs in TMHs: 37.2608499999999\nExp number, first 60 AAs: 19.52923\nTotal prob of N-in: 0.94351\nPOSSIBLE N-term signal sequence\ninside 1 19\nTMhelix 20 39\noutside 40 325\nTMhelix 326 348\ninside 349 372

11697 GCF_000351205.1_Esch_coli_KTE214_V1 Escherichia coli KTE214 Proteobacteria; Gammaproteobacteria; Enterobacterales; Enterobacteriaceae; Escherichia; Escherichia coli MTGDNTLIHSHGINRRDFMKLCAALAATMGLSSKAAAEMAESVTNPQRPPVIWIGAQECT WP_000145410.1 MULTISPECIES: hydrogenase-2 small chain [Proteobacteria] Length: 372\nNumber of predicted TMHs: 1\nExp number of AAs in TMHs: 23.72226\nExp number, first 60 AAs: 0.22038\nTotal prob of N-in: 0.12994\noutside 1 331\nTMhelix 332 354\ninside 355 372

11698 GCF_000351205.1_Esch_coli_KTE214_V1 Escherichia coli KTE214 Proteobacteria; Gammaproteobacteria; Enterobacterales; Enterobacteriaceae; Escherichia; Escherichia coli MNNEETFYQAMRRQGVTRRSFLKYCSLAATSLGLGAGMAPKIAWALENKPRIPVVWIHGL WP_001058323.1 MULTISPECIES: hydrogenase-1 small chain [Proteobacteria] Length: 372\nNumber of predicted TMHs: 2\nExp number of AAs in TMHs: 37.2608499999999\nExp number, first 60 AAs: 19.52923\nTotal prob of N-in: 0.94351\nPOSSIBLE N-term signal sequence\ninside 1 19\nTMhelix 20 39\noutside 40 325\nTMhelix 326 348\ninside 349 372

11699	GCF_000353065.1_Esch_coli_KTE215_V1	Escherichia coli KTE215	Proteobacteria; Gammaproteobacteria; Enterobacterales; Enterobacteriaceae; Escherichia; Escherichia coli
MNNEETFYQAMRRQGVTRRSFLKYCSLAATSLGLGAGMAPKIAWALENKPRIPVVWIHGL	WP_001058323.1
MULTISPECIES: hydrogenase-1 small chain [Proteobacteria]	Length: 372\nNumber of predicted TMHs: 2\nExp number of AAs in TMHs: 37.2608499999999\nExp number, first 60 AAs: 19.52923\nTotal prob of N-in: 0.94351\nPOSSIBLE N-term signal sequence\ninside 1 19\nTMhelix 20 39\noutside 40 325\nTMhelix 326 348\ninside 349 372

11700	GCF_000353065.1_Esch_coli_KTE215_V1	Escherichia coli KTE215	Proteobacteria; Gammaproteobacteria; Enterobacterales; Enterobacteriaceae; Escherichia; Escherichia coli
MTGDNTLIHSHGINRRDFMKLCAALAATMGLSSKAAAEMVESVTNPQRPPVIWIGAQECT	WP_000145423.1
hydrogenase 2 small subunit [Escherichia coli]	Length: 372\nNumber of predicted TMHs: 1\nExp number of AAs in TMHs: 23.78201\nExp number, first 60 AAs: 0.28355\nTotal prob of N-in: 0.13257\noutside 1 331\nTMhelix 332 354\ninside 355 372

11701	GCF_000351225.1_Esch_coli_KTE216_V1	Escherichia coli KTE216	Proteobacteria; Gammaproteobacteria; Enterobacterales; Enterobacteriaceae; Escherichia; Escherichia coli
MNNEETFYQAMRRQGVTRRSFLKYCSLAATSLGLGAGMAPKIAWALENKPRIPVVWIHGL	WP_001058323.1
MULTISPECIES: hydrogenase-1 small chain [Proteobacteria]	Length: 372\nNumber of predicted TMHs: 2\nExp number of AAs in TMHs: 37.2608499999999\nExp number, first 60 AAs: 19.52923\nTotal prob of N-in: 0.94351\nPOSSIBLE N-term signal sequence\ninside 1 19\nTMhelix 20 39\noutside 40 325\nTMhelix 326 348\ninside 349 372

11702	GCF_000351225.1_Esch_coli_KTE216_V1	Escherichia coli KTE216	Proteobacteria; Gammaproteobacteria; Enterobacterales; Enterobacteriaceae; Escherichia; Escherichia coli
MTGDNTLIHSHGINRRDFMKLCAALAATMGLSSKAAAEMAESVTNPQRPPVIWIGAQECT	WP_000145410.1
MULTISPECIES: hydrogenase-2 small chain [Proteobacteria]	Length: 372\nNumber of predicted TMHs: 1\nExp number of AAs in TMHs: 23.72226\nExp number, first 60 AAs: 0.22038\nTotal prob of N-in: 0.12994\noutside 1 331\nTMhelix 332 354\ninside 355 372

11703	GCF_000353085.1_Esch_coli_KTE217_V1	Escherichia coli KTE217	Proteobacteria; Gammaproteobacteria; Enterobacterales; Enterobacteriaceae; Escherichia; Escherichia coli
MNNEETFYQAMRRQGVTRRSFLKYCSLAATSLGLGAGMAPKIAWALENKPRIPVVWIHGL	WP_001058323.1
MULTISPECIES: hydrogenase-1 small chain [Proteobacteria]	Length: 372\nNumber of predicted TMHs: 2\nExp number of AAs in TMHs: 37.2608499999999\nExp number, first 60 AAs: 19.52923\nTotal prob of N-in: 0.94351\nPOSSIBLE N-term signal sequence\ninside 1 19\nTMhelix 20 39\noutside 40 325\nTMhelix 326 348\ninside 349 372

11704	GCF_000353085.1_Esch_coli_KTE217_V1	Escherichia coli KTE217	Proteobacteria; Gammaproteobacteria; Enterobacterales; Enterobacteriaceae; Escherichia; Escherichia coli
MTGDNTLIHSHGINRRDFMKLCAALAATMGLSSKAAAEMAESVTNPQRPPVIWIGAQECT	WP_000145410.1
MULTISPECIES: hydrogenase-2 small chain [Proteobacteria]	Length: 372\nNumber of predicted TMHs: 1\nExp number of AAs in TMHs: 23.72226\nExp number, first 60 AAs: 0.22038\nTotal prob of N-in: 0.12994\noutside 1 331\nTMhelix 332 354\ninside 355 372

11705	GCF_000353105.1_Esch_coli_KTE218_V1	Escherichia coli KTE218	Proteobacteria; Gammaproteobacteria; Enterobacterales; Enterobacteriaceae; Escherichia; Escherichia coli
MNNEETFYQAMRRQGVTRRSFLKYCSLAATSLGLGAGMAPKIAWALENKPRIPVVWIHGL	WP_001058323.1
MULTISPECIES: hydrogenase-1 small chain [Proteobacteria]	Length: 372\nNumber of predicted TMHs: 2\nExp number of AAs in TMHs: 37.2608499999999\nExp number, first 60 AAs: 19.52923\nTotal prob of N-in: 0.94351\nPOSSIBLE N-term signal sequence\ninside 1 19\nTMhelix 20 39\noutside 40 325\nTMhelix 326 348\ninside 349 372

11706	GCF_000353105.1_Esch_coli_KTE218_V1	Escherichia coli KTE218	Proteobacteria; Gammaproteobacteria; Enterobacterales; Enterobacteriaceae; Escherichia; Escherichia coli
MTGDNTLIHSHGINRRDFMKLCAALAATMGLSSKAAAEMAESVTNPQRPPVIWIGAQECT	WP_000145410.1
MULTISPECIES: hydrogenase-2 small chain [Proteobacteria]	Length: 372\nNumber of predicted TMHs: 1\nExp number of AAs in TMHs: 23.72226\nExp number, first 60 AAs: 0.22038\nTotal prob of N-in: 0.12994\noutside 1 331\nTMhelix 332 354\ninside 355 372

11707 GCF_000408205.1_Esch_coli_KTE219_V1 Escherichia coli KTE219 Proteobacteria; Gammaproteobacteria; Enterobacterales; Enterobacteriaceae; Escherichia; Escherichia coli MNNEETFYQAMRRQGVTRRSFLKYCSLAATSLGLGAGMAPKIAWALENKPRIPVVWIHGL WP_001538532.1 hydrogenase-1 small chain [Escherichia coli] Length: 372\nNumber of predicted TMHs: 2\nExp number of AAs in TMHs: 37.27041\nExp number, first 60 AAs: 19.52927\nTotal prob of N-in: 0.94352\nPOSSIBLE N-term signal sequence\ninside 1 19\nTMhelix 20 39\noutside 40 325\nTMhelix 326 348\ninside 349 372

11708 GCF_000408205.1_Esch_coli_KTE219_V1 Escherichia coli KTE219 Proteobacteria; Gammaproteobacteria; Enterobacterales; Enterobacteriaceae; Escherichia; Escherichia coli MTGDNTLIHSHGINRRDFMKLCAALAATMGLSSKAAAEMVESVTNPQRPPVIWIGAQECT WP_000145423.1 hydrogenase 2 small subunit [Escherichia coli] Length: 372\nNumber of predicted TMHs: 1\nExp number of AAs in TMHs: 23.78201\nExp number, first 60 AAs: 0.28355\nTotal prob of N-in: 0.13257\noutside 1 331\nTMhelix 332 354\ninside 355 372

11709 GCF_000352265.1_Esch_coli_KTE22_V1 Escherichia coli KTE22 Proteobacteria; Gammaproteobacteria; Enterobacterales; Enterobacteriaceae; Escherichia; Escherichia coli MTGDNTLIHSHGINRRDFMKLCAALAATMGLSSKAAAEMAESVTNPQRPPVIWIGAQECT WP_000145410.1 MULTISPECIES: hydrogenase-2 small chain [Proteobacteria] Length: 372\nNumber of predicted TMHs: 1\nExp number of AAs in TMHs: 23.72226\nExp number, first 60 AAs: 0.22038\nTotal prob of N-in: 0.12994\noutside 1 331\nTMhelix 332 354\ninside 355 372

11710 GCF_000351245.1_Esch_coli_KTE220_V1 Escherichia coli KTE220 Proteobacteria; Gammaproteobacteria; Enterobacterales; Enterobacteriaceae; Escherichia; Escherichia coli MNNEETFYQAMRRQGVTRRSFLKYCSLAATSLGLGAGMAPKIAWALENKPRIPVVWIHGL WP_001058323.1 MULTISPECIES: hydrogenase-1 small chain [Proteobacteria] Length: 372\nNumber of predicted TMHs: 2\nExp number of AAs in TMHs: 37.2608499999999\nExp number, first 60 AAs: 19.52923\nTotal prob of N-in: 0.94351\nPOSSIBLE N-term signal sequence\ninside 1 19\nTMhelix 20 39\noutside 40 325\nTMhelix 326 348\ninside 349 372

11711 GCF_000351245.1_Esch_coli_KTE220_V1 Escherichia coli KTE220 Proteobacteria; Gammaproteobacteria; Enterobacterales; Enterobacteriaceae; Escherichia; Escherichia coli MTGDNTLIHSHGINRRDFMKLCAALAATMGLSSKAAAEMAESVTNPQRPPVIWIGAQECT WP_000145410.1 MULTISPECIES: hydrogenase-2 small chain [Proteobacteria] Length: 372\nNumber of predicted TMHs: 1\nExp number of AAs in TMHs: 23.72226\nExp number, first 60 AAs: 0.22038\nTotal prob of N-in: 0.12994\noutside 1 331\nTMhelix 332 354\ninside 355 372

11712 GCF_000408225.1_Esch_coli_KTE221_V1 Escherichia coli KTE221 Proteobacteria; Gammaproteobacteria; Enterobacterales; Enterobacteriaceae; Escherichia; Escherichia coli MTGDNTLIHSHGINRRDFMKLCAALAATMGLSSKAAAEMAESVTNPQRPPVIWIGAQECT WP_000145410.1 MULTISPECIES: hydrogenase-2 small chain [Proteobacteria] Length: 372\nNumber of predicted TMHs: 1\nExp number of AAs in TMHs: 23.72226\nExp number, first 60 AAs: 0.22038\nTotal prob of N-in: 0.12994\noutside 1 331\nTMhelix 332 354\ninside 355 372

11713 GCF_000408225.1_Esch_coli_KTE221_V1 Escherichia coli KTE221 Proteobacteria; Gammaproteobacteria; Enterobacterales; Enterobacteriaceae; Escherichia; Escherichia coli MNNEETFYQAMRRQGVTRRSFLKYCSLAATSLGLGAGMAPKIAWALENKPRIPVVWIHGL WP_001058320.1 hydrogenase [Escherichia coli] Length: 372\nNumber of predicted TMHs: 2\nExp number of AAs in TMHs: 37.26082\nExp number, first 60 AAs: 19.52923\nTotal prob of N-in: 0.94351\nPOSSIBLE N-term signal sequence\ninside 1 19\nTMhelix 20 39\noutside 40 325\nTMhelix 326 348\ninside 349 372

11714 GCF_000408245.1_Esch_coli_KTE222_V1 Escherichia coli KTE222 Proteobacteria; Gammaproteobacteria; Enterobacterales; Enterobacteriaceae; Escherichia; Escherichia coli MNNEETFYQAMRRQGVTRRSFLKYCSLAATSLGLGAGMAPKIAWALENKPRIPVVWIHGL WP_001058320.1 hydrogenase [Escherichia coli] Length: 372\nNumber of predicted TMHs: 2\nExp number of AAs in TMHs: 37.26082\nExp number, first 60 AAs: 19.52923\nTotal prob of N-in: 0.94351\nPOSSIBLE N-term signal sequence\ninside 1 19\nTMhelix 20 39\noutside 40 325\nTMhelix 326 348\ninside 349 372

11715 GCF_000408245.1_Esch_coli_KTE222_V1 Escherichia coli KTE222 Proteobacteria; Gammaproteobacteria; Enterobacterales; Enterobacteriaceae; Escherichia; Escherichia coli MTGDNTLIHSHGINRRDFMKLCAALAATMGLSSKAAAEMAESVTNPQRPPVIWIGAQECT WP_000145410.1 MULTISPECIES: hydrogenase-2 small chain [Proteobacteria] Length: 372\nNumber of predicted TMHs:

1\nExp number of AAs in TMHs: 23.72226\nExp number, first 60 AAs: 0.22038\nTotal prob of N-in: 0.12994\noutside 1 331\nTMhelix 332 354\ninside 355 372

11716 GCF_000353125.1_Esch_coli_KTE223_V1 Escherichia coli KTE223 Proteobacteria; Gammaproteobacteria; Enterobacterales; Enterobacteriaceae; Escherichia; Escherichia coli
MNNEETFYQAMRRQGVTRRSFLKYCSLAATSLGLGAGMAPKIAWALENKPRIPVVWIHGL WP_001058323.1
MULTISPECIES: hydrogenase-1 small chain [Proteobacteria] Length: 372\nNumber of predicted TMHs: 2\nExp number of AAs in TMHs: 37.2608499999999\nExp number, first 60 AAs: 19.52923\nTotal prob of N-in: 0.94351\nPOSSIBLE N-term signal sequence\ninside 1 19\nTMhelix 20 39\noutside 40 325\nTMhelix 326 348\ninside 349 372

11717 GCF_000353125.1_Esch_coli_KTE223_V1 Escherichia coli KTE223 Proteobacteria; Gammaproteobacteria; Enterobacterales; Enterobacteriaceae; Escherichia; Escherichia coli
MTGDNTLIHSHGINRRDFMKLCAALAATMGLSSKAAAEMAESVTNPQRPPVIWIGAQECT WP_000145410.1
MULTISPECIES: hydrogenase-2 small chain [Proteobacteria] Length: 372\nNumber of predicted TMHs: 1\nExp number of AAs in TMHs: 23.72226\nExp number, first 60 AAs: 0.22038\nTotal prob of N-in: 0.12994\noutside 1 331\nTMhelix 332 354\ninside 355 372

11718 GCF_000351265.1_Esch_coli_KTE224_V1 Escherichia coli KTE224 Proteobacteria; Gammaproteobacteria; Enterobacterales; Enterobacteriaceae; Escherichia; Escherichia coli
MNNEETFYQAMRRQGVTRRSFLKYCSLAATSLGLGAGMAPKIAWALENKPRIPVVWIHGL WP_001058323.1
MULTISPECIES: hydrogenase-1 small chain [Proteobacteria] Length: 372\nNumber of predicted TMHs: 2\nExp number of AAs in TMHs: 37.2608499999999\nExp number, first 60 AAs: 19.52923\nTotal prob of N-in: 0.94351\nPOSSIBLE N-term signal sequence\ninside 1 19\nTMhelix 20 39\noutside 40 325\nTMhelix 326 348\ninside 349 372

11719 GCF_000351265.1_Esch_coli_KTE224_V1 Escherichia coli KTE224 Proteobacteria; Gammaproteobacteria; Enterobacterales; Enterobacteriaceae; Escherichia; Escherichia coli
MTGDNTLIHSHGINRRDFMKLCAALAATMGLSSKAAAEMVESVTNPQRPPVIWIGAQECT WP_000145423.1
hydrogenase 2 small subunit [Escherichia coli] Length: 372\nNumber of predicted TMHs: 1\nExp number of AAs in TMHs: 23.78201\nExp number, first 60 AAs: 0.28355\nTotal prob of N-in: 0.13257\noutside 1 331\nTMhelix 332 354\ninside 355 372

11720 GCF_000408265.1_Esch_coli_KTE225_V1 Escherichia coli KTE225 Proteobacteria; Gammaproteobacteria; Enterobacterales; Enterobacteriaceae; Escherichia; Escherichia coli
MNNEETFYQAMRRQGVTRRSFLKYCSLAATSLGLGAGMAPKIAWALENKPRIPVVWIHGL WP_001058323.1
MULTISPECIES: hydrogenase-1 small chain [Proteobacteria] Length: 372\nNumber of predicted TMHs: 2\nExp number of AAs in TMHs: 37.2608499999999\nExp number, first 60 AAs: 19.52923\nTotal prob of N-in: 0.94351\nPOSSIBLE N-term signal sequence\ninside 1 19\nTMhelix 20 39\noutside 40 325\nTMhelix 326 348\ninside 349 372

11721 GCF_000408265.1_Esch_coli_KTE225_V1 Escherichia coli KTE225 Proteobacteria; Gammaproteobacteria; Enterobacterales; Enterobacteriaceae; Escherichia; Escherichia coli
MTGDNTLIHSHGINRRDFMKLCAALAATMGLSSKAAAEMAESVTNPQRPPVIWIGAQECT WP_000145410.1
MULTISPECIES: hydrogenase-2 small chain [Proteobacteria] Length: 372\nNumber of predicted TMHs: 1\nExp number of AAs in TMHs: 23.72226\nExp number, first 60 AAs: 0.22038\nTotal prob of N-in: 0.12994\noutside 1 331\nTMhelix 332 354\ninside 355 372

11722 GCF_000408285.1_Esch_coli_KTE226_V1 Escherichia coli KTE226 Proteobacteria; Gammaproteobacteria; Enterobacterales; Enterobacteriaceae; Escherichia; Escherichia coli
MTGDNTLIHSHGINRRDFMKLCAALAATMGLSSKAAAEMAESVTNPQRPPVIWIGAQECT WP_000145410.1
MULTISPECIES: hydrogenase-2 small chain [Proteobacteria] Length: 372\nNumber of predicted TMHs: 1\nExp number of AAs in TMHs: 23.72226\nExp number, first 60 AAs: 0.22038\nTotal prob of N-in: 0.12994\noutside 1 331\nTMhelix 332 354\ninside 355 372

11723 GCF_000408285.1_Esch_coli_KTE226_V1 Escherichia coli KTE226 Proteobacteria; Gammaproteobacteria; Enterobacterales; Enterobacteriaceae; Escherichia; Escherichia coli
MNNEETFYQAMRRQGVTRRSFLKYCSLAATSLGLGAGMAPKIAWALENKPRIPVVWIHGL WP_001058323.1
MULTISPECIES: hydrogenase-1 small chain [Proteobacteria] Length: 372\nNumber of predicted TMHs: 2\nExp number of AAs in TMHs: 37.2608499999999\nExp number, first 60 AAs: 19.52923\nTotal prob of N-in:

0.94351\nPOSSIBLE N-term signal sequence\ninside 1 19\nTMhelix 20 39\noutside 40 325\nTMhelix 326 348\ninside 349 372

11724 GCF_000353145.1_Esch_coli_KTE227_V1 Escherichia coli KTE227 Proteobacteria; Gammaproteobacteria; Enterobacterales; Enterobacteriaceae; Escherichia; Escherichia coli MTGDNTLIHSHGINRRDFMKLCAALAATMGLSSKAAAEMAESVTNPQRPPVIWIGAQECT WP_000145410.1 MULTISPECIES: hydrogenase-2 small chain [Proteobacteria] Length: 372\nNumber of predicted TMHs: 1\nExp number of AAs in TMHs: 23.72226\nExp number, first 60 AAs: 0.22038\nTotal prob of N-in: 0.12994\noutside 1 331\nTMhelix 332 354\ninside 355 372

11725 GCF_000351285.1_Esch_coli_KTE228_V1 Escherichia coli KTE228 Proteobacteria; Gammaproteobacteria; Enterobacterales; Enterobacteriaceae; Escherichia; Escherichia coli MNNEETFYQAMRRQGVTRRSFLKYCSLAATSLGLGAGMAPKIAWALENKPRIPVVWIHGL WP_001058323.1 MULTISPECIES: hydrogenase-1 small chain [Proteobacteria] Length: 372\nNumber of predicted TMHs: 2\nExp number of AAs in TMHs: 37.2608499999999\nExp number, first 60 AAs: 19.52923\nTotal prob of N-in: 0.94351\nPOSSIBLE N-term signal sequence\ninside 1 19\nTMhelix 20 39\noutside 40 325\nTMhelix 326 348\ninside 349 372

11726 GCF_000351285.1_Esch_coli_KTE228_V1 Escherichia coli KTE228 Proteobacteria; Gammaproteobacteria; Enterobacterales; Enterobacteriaceae; Escherichia; Escherichia coli MTGDNTLIHSHGINRRDFMKLCAALAATMGLSSKAAAEMAESVTNPQRPPVIWIGAQECT WP_000145410.1 MULTISPECIES: hydrogenase-2 small chain [Proteobacteria] Length: 372\nNumber of predicted TMHs: 1\nExp number of AAs in TMHs: 23.72226\nExp number, first 60 AAs: 0.22038\nTotal prob of N-in: 0.12994\noutside 1 331\nTMhelix 332 354\ninside 355 372

11727 GCF_000353165.1_Esch_coli_KTE229_V1 Escherichia coli KTE229 Proteobacteria; Gammaproteobacteria; Enterobacterales; Enterobacteriaceae; Escherichia; Escherichia coli MTGDNTLIHSHGINRRDFMKLCAALAATMGLSSKAAAEMAESVTNPQRPPVIWIGAQECT WP_000145410.1 MULTISPECIES: hydrogenase-2 small chain [Proteobacteria] Length: 372\nNumber of predicted TMHs: 1\nExp number of AAs in TMHs: 23.72226\nExp number, first 60 AAs: 0.22038\nTotal prob of N-in: 0.12994\noutside 1 331\nTMhelix 332 354\ninside 355 372

11728 GCF_000352205.1_Esch_coli_KTE23_V1 Escherichia coli KTE23 Proteobacteria; Gammaproteobacteria; Enterobacterales; Enterobacteriaceae; Escherichia; Escherichia coli MNNEETFYQAMRRQGVTRRSFLKYCSLAATSLGLGAGMAPKIAWALENKPRIPVVWIHGL WP_001058323.1 MULTISPECIES: hydrogenase-1 small chain [Proteobacteria] Length: 372\nNumber of predicted TMHs: 2\nExp number of AAs in TMHs: 37.2608499999999\nExp number, first 60 AAs: 19.52923\nTotal prob of N-in: 0.94351\nPOSSIBLE N-term signal sequence\ninside 1 19\nTMhelix 20 39\noutside 40 325\nTMhelix 326 348\ninside 349 372

11729 GCF_000352205.1_Esch_coli_KTE23_V1 Escherichia coli KTE23 Proteobacteria; Gammaproteobacteria; Enterobacterales; Enterobacteriaceae; Escherichia; Escherichia coli MTGDNTLIHSHGINRRDFMKLCAALAATMGLSSKAAAEMAESVTNPQRPPVIWIGAQECT WP_000145410.1 MULTISPECIES: hydrogenase-2 small chain [Proteobacteria] Length: 372\nNumber of predicted TMHs: 1\nExp number of AAs in TMHs: 23.72226\nExp number, first 60 AAs: 0.22038\nTotal prob of N-in: 0.12994\noutside 1 331\nTMhelix 332 354\ninside 355 372

11730 GCF_000351305.1_Esch_coli_KTE230_V1 Escherichia coli KTE230 Proteobacteria; Gammaproteobacteria; Enterobacterales; Enterobacteriaceae; Escherichia; Escherichia coli MNNEETFYQAMRRQGVTRRSFLKYCSLAATSLGLGAGMAPKIAWALENKPRIPVVWIHGL WP_001058323.1 MULTISPECIES: hydrogenase-1 small chain [Proteobacteria] Length: 372\nNumber of predicted TMHs: 2\nExp number of AAs in TMHs: 37.2608499999999\nExp number, first 60 AAs: 19.52923\nTotal prob of N-in: 0.94351\nPOSSIBLE N-term signal sequence\ninside 1 19\nTMhelix 20 39\noutside 40 325\nTMhelix 326 348\ninside 349 372

11731 GCF_000351305.1_Esch_coli_KTE230_V1 Escherichia coli KTE230 Proteobacteria; Gammaproteobacteria; Enterobacterales; Enterobacteriaceae; Escherichia; Escherichia coli MTGDNTLIHSHGINRRDFMKLCAALAATMGLSSKAAAEMAESVTNPQRPPVIWIGAQECT WP_000145418.1 hydrogenase 2 small subunit [Escherichia coli] Length: 372\nNumber of predicted TMHs: 1\nExp number of AAs in TMHs: 23.72313\nExp number, first 60 AAs: 0.22034\nTotal prob of N-in: 0.12997\noutside 1 331\nTMhelix 332 354\ninside 355 372

11732 GCF_000407785.1_Esch_coli_KTE231_V1 Escherichia coli KTE231 Proteobacteria; Gammaproteobacteria; Enterobacterales; Enterobacteriaceae; Escherichia; Escherichia coli MTGDNTLIHSHGINRRDFMKLCAALAATMGLSSKAAAEMAESVTNPQRPPVIWIGAQECT WP_000145410.1 MULTISPECIES: hydrogenase-2 small chain [Proteobacteria] Length: 372\nNumber of predicted TMHs: 1\nExp number of AAs in TMHs: 23.72226\nExp number, first 60 AAs: 0.22038\nTotal prob of N-in: 0.12994\noutside 1 331\nTMhelix 332 354\ninside 355 372

11733 GCF_000407785.1_Esch_coli_KTE231_V1 Escherichia coli KTE231 Proteobacteria; Gammaproteobacteria; Enterobacterales; Enterobacteriaceae; Escherichia; Escherichia coli MNNEETFYQAMRRQGVTRRSFLKYCSLAATSLGLGAGMAPKIAWALENKPRIPVVWIHGL WP_001058323.1 MULTISPECIES: hydrogenase-1 small chain [Proteobacteria] Length: 372\nNumber of predicted TMHs: 2\nExp number of AAs in TMHs: 37.2608499999999\nExp number, first 60 AAs: 19.52923\nTotal prob of N-in: 0.94351\nPOSSIBLE N-term signal sequence\ninside 1 19\nTMhelix 20 39\noutside 40 325\nTMhelix 326 348\ninside 349 372

11734 GCF_000326825.1_Esch_coli_KTE232_V1 Escherichia coli KTE232 Proteobacteria; Gammaproteobacteria; Enterobacterales; Enterobacteriaceae; Escherichia; Escherichia coli MTGDNTLIHSHGINRRDFMKLCAALAATMGLSSKAAAEMAESVTNPQRPPVIWIGAQECT WP_000145410.1 MULTISPECIES: hydrogenase-2 small chain [Proteobacteria] Length: 372\nNumber of predicted TMHs: 1\nExp number of AAs in TMHs: 23.72226\nExp number, first 60 AAs: 0.22038\nTotal prob of N-in: 0.12994\noutside 1 331\nTMhelix 332 354\ninside 355 372

11735 GCF_000326825.1_Esch_coli_KTE232_V1 Escherichia coli KTE232 Proteobacteria; Gammaproteobacteria; Enterobacterales; Enterobacteriaceae; Escherichia; Escherichia coli MNNEETFYQAMRRQGVTRRSFLKYCSLAATSLGLGAGMAPKIAWALENKPRIPVVWIHGL WP_001058323.1 MULTISPECIES: hydrogenase-1 small chain [Proteobacteria] Length: 372\nNumber of predicted TMHs: 2\nExp number of AAs in TMHs: 37.2608499999999\nExp number, first 60 AAs: 19.52923\nTotal prob of N-in: 0.94351\nPOSSIBLE N-term signal sequence\ninside 1 19\nTMhelix 20 39\noutside 40 325\nTMhelix 326 348\ninside 349 372

11736 GCF_000351325.1_Esch_coli_KTE233_V1 Escherichia coli KTE233 Proteobacteria; Gammaproteobacteria; Enterobacterales; Enterobacteriaceae; Escherichia; Escherichia coli MNNEETFYQAMRRQGVTRRSFLKYCSLAATSLGLGAGMAPKIAWALENKPRIPVVWIHGL WP_001058323.1 MULTISPECIES: hydrogenase-1 small chain [Proteobacteria] Length: 372\nNumber of predicted TMHs: 2\nExp number of AAs in TMHs: 37.2608499999999\nExp number, first 60 AAs: 19.52923\nTotal prob of N-in: 0.94351\nPOSSIBLE N-term signal sequence\ninside 1 19\nTMhelix 20 39\noutside 40 325\nTMhelix 326 348\ninside 349 372

11737 GCF_000351325.1_Esch_coli_KTE233_V1 Escherichia coli KTE233 Proteobacteria; Gammaproteobacteria; Enterobacterales; Enterobacteriaceae; Escherichia; Escherichia coli MTGDNTLIHSHGINRRDFMKLCAALAATMGLSSKAAAEMAESVTNPQRPPVIWIGAQECT WP_000145410.1 MULTISPECIES: hydrogenase-2 small chain [Proteobacteria] Length: 372\nNumber of predicted TMHs: 1\nExp number of AAs in TMHs: 23.72226\nExp number, first 60 AAs: 0.22038\nTotal prob of N-in: 0.12994\noutside 1 331\nTMhelix 332 354\ninside 355 372

11738 GCF_000351345.1_Esch_coli_KTE234_V1 Escherichia coli KTE234 Proteobacteria; Gammaproteobacteria; Enterobacterales; Enterobacteriaceae; Escherichia; Escherichia coli MTGDNTLIHSHGINRRDFMKLCAALAATMGLSSKAAAEMAESVTNPQRPPVIWIGAQECT WP_000145410.1 MULTISPECIES: hydrogenase-2 small chain [Proteobacteria] Length: 372\nNumber of predicted TMHs: 1\nExp number of AAs in TMHs: 23.72226\nExp number, first 60 AAs: 0.22038\nTotal prob of N-in: 0.12994\noutside 1 331\nTMhelix 332 354\ninside 355 372

11739 GCF_000351345.1_Esch_coli_KTE234_V1 Escherichia coli KTE234 Proteobacteria; Gammaproteobacteria; Enterobacterales; Enterobacteriaceae; Escherichia; Escherichia coli MNNEETFYQAMRRQGVTRRSFLKYCSLAATSLGLGAGMAPKIAWALENKPRIPVVWIHGL WP_001058323.1 MULTISPECIES: hydrogenase-1 small chain [Proteobacteria] Length: 372\nNumber of predicted TMHs: 2\nExp number of AAs in TMHs: 37.2608499999999\nExp number, first 60 AAs: 19.52923\nTotal prob of N-in: 0.94351\nPOSSIBLE N-term signal sequence\ninside 1 19\nTMhelix 20 39\noutside 40 325\nTMhelix 326 348\ninside 349 372

11740 GCF_000351365.1_Esch_coli_KTE235_V1 Escherichia coli KTE235 Proteobacteria; Gammaproteobacteria; Enterobacterales; Enterobacteriaceae; Escherichia; Escherichia coli MNNEETFYQAMRRQGVTRRSFLKYCSLAATSLGLGAGMAPKIAWALENKPRIPVVWIHGL WP_001058323.1 MULTISPECIES: hydrogenase-1 small chain [Proteobacteria] Length: 372\nNumber of predicted TMHs: 2\nExp number of AAs in TMHs: 37.2608499999999\nExp number, first 60 AAs: 19.52923\nTotal prob of N-in: 0.94351\nPOSSIBLE N-term signal sequence\ninside 1 19\nTMhelix 20 39\noutside 40 325\nTMhelix 326 348\ninside 349 372

11741 GCF_000351365.1_Esch_coli_KTE235_V1 Escherichia coli KTE235 Proteobacteria; Gammaproteobacteria; Enterobacterales; Enterobacteriaceae; Escherichia; Escherichia coli MTGDNTLIHSHGINRRDFMKLCAALAATMGLSSKAAAEMAESVTNPQRPPVIWIGAQECT WP_000145410.1 MULTISPECIES: hydrogenase-2 small chain [Proteobacteria] Length: 372\nNumber of predicted TMHs: 1\nExp number of AAs in TMHs: 23.72226\nExp number, first 60 AAs: 0.22038\nTotal prob of N-in: 0.12994\noutside 1 331\nTMhelix 332 354\ninside 355 372

11742 GCF_000351385.1_Esch_coli_KTE236_V1 Escherichia coli KTE236 Proteobacteria; Gammaproteobacteria; Enterobacterales; Enterobacteriaceae; Escherichia; Escherichia coli MTGDNTLIHSHGINRRDFMKLCAALAATMGLSSKAAAEMAESVTNPQRPPVIWIGAQECT WP_000145410.1 MULTISPECIES: hydrogenase-2 small chain [Proteobacteria] Length: 372\nNumber of predicted TMHs: 1\nExp number of AAs in TMHs: 23.72226\nExp number, first 60 AAs: 0.22038\nTotal prob of N-in: 0.12994\noutside 1 331\nTMhelix 332 354\ninside 355 372

11743 GCF_000351385.1_Esch_coli_KTE236_V1 Escherichia coli KTE236 Proteobacteria; Gammaproteobacteria; Enterobacterales; Enterobacteriaceae; Escherichia; Escherichia coli MNNEETFYQAMRRQGVTRRSFLKYCSLAATSLGLGAGMAPKIAWALENKPRIPVVWIHGL WP_001058323.1 MULTISPECIES: hydrogenase-1 small chain [Proteobacteria] Length: 372\nNumber of predicted TMHs: 2\nExp number of AAs in TMHs: 37.2608499999999\nExp number, first 60 AAs: 19.52923\nTotal prob of N-in: 0.94351\nPOSSIBLE N-term signal sequence\ninside 1 19\nTMhelix 20 39\noutside 40 325\nTMhelix 326 348\ninside 349 372

11744 GCF_000351405.1_Esch_coli_KTE237_V1 Escherichia coli KTE237 Proteobacteria; Gammaproteobacteria; Enterobacterales; Enterobacteriaceae; Escherichia; Escherichia coli MTGDNTLIHSHGINRRDFMKLCAALAATMGLSSKAAAEMAESVTNPQRPPVIWIGAQECT WP_000145410.1 MULTISPECIES: hydrogenase-2 small chain [Proteobacteria] Length: 372\nNumber of predicted TMHs: 1\nExp number of AAs in TMHs: 23.72226\nExp number, first 60 AAs: 0.22038\nTotal prob of N-in: 0.12994\noutside 1 331\nTMhelix 332 354\ninside 355 372

11745 GCF_000351405.1_Esch_coli_KTE237_V1 Escherichia coli KTE237 Proteobacteria; Gammaproteobacteria; Enterobacterales; Enterobacteriaceae; Escherichia; Escherichia coli MNNEETFYQAMRRQGVTRRSFLKYCSLAATSLGLGAGMAPKIAWALENKPRIPVVWIHGL WP_001058323.1 MULTISPECIES: hydrogenase-1 small chain [Proteobacteria] Length: 372\nNumber of predicted TMHs: 2\nExp number of AAs in TMHs: 37.2608499999999\nExp number, first 60 AAs: 19.52923\nTotal prob of N-in: 0.94351\nPOSSIBLE N-term signal sequence\ninside 1 19\nTMhelix 20 39\noutside 40 325\nTMhelix 326 348\ninside 349 372

11746 GCF_000407865.1_Esch_coli_KTE24_V1 Escherichia coli KTE24 Proteobacteria; Gammaproteobacteria; Enterobacterales; Enterobacteriaceae; Escherichia; Escherichia coli MTGDNTLIHSHGINRRDFMKLCAALAATMGLSSKAAAEMAESVTNPQRPPVIWIGAQECT WP_000145410.1 MULTISPECIES: hydrogenase-2 small chain [Proteobacteria] Length: 372\nNumber of predicted TMHs: 1\nExp number of AAs in TMHs: 23.72226\nExp number, first 60 AAs: 0.22038\nTotal prob of N-in: 0.12994\noutside 1 331\nTMhelix 332 354\ninside 355 372

11747 GCF_000407865.1_Esch_coli_KTE24_V1 Escherichia coli KTE24 Proteobacteria; Gammaproteobacteria; Enterobacterales; Enterobacteriaceae; Escherichia; Escherichia coli MNNEETFYQAMRRQGVTRRSFLKYCSLAATSLGLGAGMAPKIAWALENKPRIPVVWIHGL WP_001058323.1 MULTISPECIES: hydrogenase-1 small chain [Proteobacteria] Length: 372\nNumber of predicted TMHs: 2\nExp number of AAs in TMHs: 37.2608499999999\nExp number, first 60 AAs: 19.52923\nTotal prob of N-in: 0.94351\nPOSSIBLE N-term signal sequence\ninside 1 19\nTMhelix 20 39\noutside 40 325\nTMhelix 326 348\ninside 349 372

11748 GCF_000408305.1_Esch_coli_KTE240_V1 Escherichia coli KTE240 Proteobacteria; Gammaproteobacteria; Enterobacterales; Enterobacteriaceae; Escherichia; Escherichia coli
MTGDNTLIHSHGINRRDFMKLCAALAATMGLSSKAAAEMAESVTNPQRPPVIWIGAQECT WP_000145410.1
MULTISPECIES: hydrogenase-2 small chain [Proteobacteria] Length: 372\nNumber of predicted TMHs: 1\nExp number of AAs in TMHs: 23.72226\nExp number, first 60 AAs: 0.22038\nTotal prob of N-in: 0.12994\noutside 1 331\nTMhelix 332 354\ninside 355 372

11749 GCF_000350805.1_Esch_coli_KTE25_V1 Escherichia coli KTE25 Proteobacteria; Gammaproteobacteria; Enterobacterales; Enterobacteriaceae; Escherichia; Escherichia coli
MTGDNTLIHSHGINRRDFMKLCAALAATMGLSSKAAAEMAESVTNPQRPPVIWIGAQECT WP_000145410.1
MULTISPECIES: hydrogenase-2 small chain [Proteobacteria] Length: 372\nNumber of predicted TMHs: 1\nExp number of AAs in TMHs: 23.72226\nExp number, first 60 AAs: 0.22038\nTotal prob of N-in: 0.12994\noutside 1 331\nTMhelix 332 354\ninside 355 372

11750 GCF_000350805.1_Esch_coli_KTE25_V1 Escherichia coli KTE25 Proteobacteria; Gammaproteobacteria; Enterobacterales; Enterobacteriaceae; Escherichia; Escherichia coli
MNNEETFYQAMRRQGVTRRSFLKYCSLAATSLGLGAGMAPKIAWALENKPRIPVVWIHGL WP_001058323.1
MULTISPECIES: hydrogenase-1 small chain [Proteobacteria] Length: 372\nNumber of predicted TMHs: 2\nExp number of AAs in TMHs: 37.2608499999999\nExp number, first 60 AAs: 19.52923\nTotal prob of N-in: 0.94351\nPOSSIBLE N-term signal sequence\ninside 1 19\nTMhelix 20 39\noutside 40 325\nTMhelix 326 348\ninside 349 372

11751 GCF_000350825.1_Esch_coli_KTE26_V1 Escherichia coli KTE26 Proteobacteria; Gammaproteobacteria; Enterobacterales; Enterobacteriaceae; Escherichia; Escherichia coli
MNNEETFYQAMRRQGVTRRSFLKYCSLAATSLGLGAGMAPKIAWALENKPRIPVVWIHGL WP_001058323.1
MULTISPECIES: hydrogenase-1 small chain [Proteobacteria] Length: 372\nNumber of predicted TMHs: 2\nExp number of AAs in TMHs: 37.2608499999999\nExp number, first 60 AAs: 19.52923\nTotal prob of N-in: 0.94351\nPOSSIBLE N-term signal sequence\ninside 1 19\nTMhelix 20 39\noutside 40 325\nTMhelix 326 348\ninside 349 372

11752 GCF_000350825.1_Esch_coli_KTE26_V1 Escherichia coli KTE26 Proteobacteria; Gammaproteobacteria; Enterobacterales; Enterobacteriaceae; Escherichia; Escherichia coli
MTGDNTLIHSHGINRRDFMKLCAALAATMGLSSKAAAEMAESVTNPQRPPVIWIGAQECT WP_000145410.1
MULTISPECIES: hydrogenase-2 small chain [Proteobacteria] Length: 372\nNumber of predicted TMHs: 1\nExp number of AAs in TMHs: 23.72226\nExp number, first 60 AAs: 0.22038\nTotal prob of N-in: 0.12994\noutside 1 331\nTMhelix 332 354\ninside 355 372

11753 GCF_000407885.1_Esch_coli_KTE27_V1 Escherichia coli KTE27 Proteobacteria; Gammaproteobacteria; Enterobacterales; Enterobacteriaceae; Escherichia; Escherichia coli
MTGDNTLIHSHGINRRDFMKLCAALAATMGLSSKAAAEMAESVTNPQRPPVIWIGAQECT WP_000145410.1
MULTISPECIES: hydrogenase-2 small chain [Proteobacteria] Length: 372\nNumber of predicted TMHs: 1\nExp number of AAs in TMHs: 23.72226\nExp number, first 60 AAs: 0.22038\nTotal prob of N-in: 0.12994\noutside 1 331\nTMhelix 332 354\ninside 355 372

11754 GCF_000350845.1_Esch_coli_KTE28_V1 Escherichia coli KTE28 Proteobacteria; Gammaproteobacteria; Enterobacterales; Enterobacteriaceae; Escherichia; Escherichia coli
MNNEETFYQAMRRQGVTRRSFLKYCSLAATSLGLGAGMAPKIAWALENKPRIPVVWIHGL WP_001295940.1
MULTISPECIES: hydrogenase [Enterobacteriaceae] Length: 372\nNumber of predicted TMHs: 2\nExp number of AAs in TMHs: 37.2608899999999\nExp number, first 60 AAs: 19.52923\nTotal prob of N-in: 0.94351\nPOSSIBLE N-term signal sequence\ninside 1 19\nTMhelix 20 39\noutside 40 325\nTMhelix 326 348\ninside 349 372

11755 GCF_000350845.1_Esch_coli_KTE28_V1 Escherichia coli KTE28 Proteobacteria; Gammaproteobacteria; Enterobacterales; Enterobacteriaceae; Escherichia; Escherichia coli
MTGDNTLIHSHGINRRDFMKLCAALAATMGLSSKAAAEMAESVTNPQRPPVIWIGAQECT WP_000145410.1
MULTISPECIES: hydrogenase-2 small chain [Proteobacteria] Length: 372\nNumber of predicted TMHs: 1\nExp number of AAs in TMHs: 23.72226\nExp number, first 60 AAs: 0.22038\nTotal prob of N-in: 0.12994\noutside 1 331\nTMhelix 332 354\ninside 355 372

11756 GCF_000352245.1_Esch_coli_KTE29_V1 Escherichia coli KTE29 Proteobacteria; Gammaproteobacteria; Enterobacterales; Enterobacteriaceae; Escherichia; Escherichia coli

MNNEETFYQAMRRQGVTRRSFLKYCSLAATSLGLGAGMAPKIAWALENKPRIPVVWIHGL WP_001058323.1 MULTISPECIES: hydrogenase-1 small chain [Proteobacteria] Length: 372\nNumber of predicted TMHs: 2\nExp number of AAs in TMHs: 37.2608499999999\nExp number, first 60 AAs: 19.52923\nTotal prob of N-in: 0.94351\nPOSSIBLE N-term signal sequence\ninside 1 19\nTMhelix 20 39\noutside 40 325\nTMhelix 326 348\ninside 349 372

11757 GCF_000352245.1_Esch_coli_KTE29_V1 Escherichia coli KTE29 Proteobacteria; Gammaproteobacteria; Enterobacterales; Enterobacteriaceae; Escherichia; Escherichia coli MTGDNTLIHSHGINRRDFMKLCAALAATMGLSSKAAAEMAESVTNPQRPPVIWIGAQECT WP_000145410.1 MULTISPECIES: hydrogenase-2 small chain [Proteobacteria] Length: 372\nNumber of predicted TMHs: 1\nExp number of AAs in TMHs: 23.72226\nExp number, first 60 AAs: 0.22038\nTotal prob of N-in: 0.12994\noutside 1 331\nTMhelix 332 354\ninside 355 372

11758 GCF_000407685.1_Esch_coli_KTE3_V1 Escherichia coli KTE3 Proteobacteria; Gammaproteobacteria; Enterobacterales; Enterobacteriaceae; Escherichia; Escherichia coli MTGDNTLIHSHGINRRDFMKLCAALAATMGLSSKAAAEMAESVTNPQRPPVIWIGAQECT WP_000145410.1 MULTISPECIES: hydrogenase-2 small chain [Proteobacteria] Length: 372\nNumber of predicted TMHs: 1\nExp number of AAs in TMHs: 23.72226\nExp number, first 60 AAs: 0.22038\nTotal prob of N-in: 0.12994\noutside 1 331\nTMhelix 332 354\ninside 355 372

11759 GCF_000398885.1_Esch_coli_KTE33_V1 Escherichia coli KTE33 Proteobacteria; Gammaproteobacteria; Enterobacterales; Enterobacteriaceae; Escherichia; Escherichia coli MTGDNTLIHSHGINRRDFMKLCAALAATMGLSSKAAAEMAESVTNPQRPPVIWIGAQECT WP_000145410.1 MULTISPECIES: hydrogenase-2 small chain [Proteobacteria] Length: 372\nNumber of predicted TMHs: 1\nExp number of AAs in TMHs: 23.72226\nExp number, first 60 AAs: 0.22038\nTotal prob of N-in: 0.12994\noutside 1 331\nTMhelix 332 354\ninside 355 372

11760 GCF_000398885.1_Esch_coli_KTE33_V1 Escherichia coli KTE33 Proteobacteria; Gammaproteobacteria; Enterobacterales; Enterobacteriaceae; Escherichia; Escherichia coli MNNEETFYQAMRRQGVTRRSFLKYCSLAATSLGLGAGMAPKIAWALENKPRIPVVWIHGL WP_010344442.1 MULTISPECIES: hydrogenase [Escherichia] Length: 372\nNumber of predicted TMHs: 2\nExp number of AAs in TMHs: 38.04874\nExp number, first 60 AAs: 19.5298\nTotal prob of N-in: 0.94352\nPOSSIBLE N-term signal sequence\ninside 1 19\nTMhelix 20 39\noutside 40 325\nTMhelix 326 348\ninside 349 372

11761 GCF_000407945.1_Esch_coli_KTE34_V1 Escherichia coli KTE34 Proteobacteria; Gammaproteobacteria; Enterobacterales; Enterobacteriaceae; Escherichia; Escherichia coli MTGDNTLIHSHGINRRDFMKLCAALAATMGLSSKAAAEMAESVTNPQRPPVIWIGAQECT WP_000145410.1 MULTISPECIES: hydrogenase-2 small chain [Proteobacteria] Length: 372\nNumber of predicted TMHs: 1\nExp number of AAs in TMHs: 23.72226\nExp number, first 60 AAs: 0.22038\nTotal prob of N-in: 0.12994\noutside 1 331\nTMhelix 332 354\ninside 355 372

11762 GCF_000407945.1_Esch_coli_KTE34_V1 Escherichia coli KTE34 Proteobacteria; Gammaproteobacteria; Enterobacterales; Enterobacteriaceae; Escherichia; Escherichia coli MNNEETFYQAMRRQGVTRRSFLKYCSLAATSLGLGAGMAPKIAWALENKPRIPVVWIHGL WP_001058323.1 MULTISPECIES: hydrogenase-1 small chain [Proteobacteria] Length: 372\nNumber of predicted TMHs: 2\nExp number of AAs in TMHs: 37.2608499999999\nExp number, first 60 AAs: 19.52923\nTotal prob of N-in: 0.94351\nPOSSIBLE N-term signal sequence\ninside 1 19\nTMhelix 20 39\noutside 40 325\nTMhelix 326 348\ninside 349 372

11763 GCF_000407745.1_Esch_coli_KTE35_V1 Escherichia coli KTE35 Proteobacteria; Gammaproteobacteria; Enterobacterales; Enterobacteriaceae; Escherichia; Escherichia coli MNNEETFYQAMRRQGVTRRSFLKYCSLAATSLGLGAGMAPKIAWALENKPRIPVVWIHGL WP_001058323.1 MULTISPECIES: hydrogenase-1 small chain [Proteobacteria] Length: 372\nNumber of predicted TMHs: 2\nExp number of AAs in TMHs: 37.2608499999999\nExp number, first 60 AAs: 19.52923\nTotal prob of N-in: 0.94351\nPOSSIBLE N-term signal sequence\ninside 1 19\nTMhelix 20 39\noutside 40 325\nTMhelix 326 348\ninside 349 372

11764 GCF_000407745.1_Esch_coli_KTE35_V1 Escherichia coli KTE35 Proteobacteria; Gammaproteobacteria; Enterobacterales; Enterobacteriaceae; Escherichia; Escherichia coli MTGDNTLIHSHGINRRDFMKLCAALAATMGLSSKAAAEMAESVTNPQRPPVIWIGAQECT WP_000145410.1 MULTISPECIES: hydrogenase-2 small chain [Proteobacteria] Length: 372\nNumber of predicted TMHs:

1\nExp number of AAs in TMHs: 23.72226\nExp number, first 60 AAs: 0.22038\nTotal prob of N-in: 0.12994\noutside 1 331\nTMhelix 332 354\ninside 355 372

11765 GCF_000407965.1_Esch_coli_KTE36_V1 Escherichia coli KTE36 Proteobacteria; Gammaproteobacteria; Enterobacterales; Enterobacteriaceae; Escherichia; Escherichia coli MNNEETFYQAMRRQGVTRRSFLKYCSLAATSLGLGAGMAPKIAWALENKPRIPVVWIHGL WP_001058323.1 MULTISPECIES: hydrogenase-1 small chain [Proteobacteria] Length: 372\nNumber of predicted TMHs: 2\nExp number of AAs in TMHs: 37.2608499999999\nExp number, first 60 AAs: 19.52923\nTotal prob of N-in: 0.94351\nPOSSIBLE N-term signal sequence\ninside 1 19\nTMhelix 20 39\noutside 40 325\nTMhelix 326 348\ninside 349 372

11766 GCF_000407965.1_Esch_coli_KTE36_V1 Escherichia coli KTE36 Proteobacteria; Gammaproteobacteria; Enterobacterales; Enterobacteriaceae; Escherichia; Escherichia coli MTGDNTLIHSHGINRRDFMKLCAALAATMGLSSKAAAEMAESVTNPQRPPVIWIGAQECT WP_000145410.1 MULTISPECIES: hydrogenase-2 small chain [Proteobacteria] Length: 372\nNumber of predicted TMHs: 1\nExp number of AAs in TMHs: 23.72226\nExp number, first 60 AAs: 0.22038\nTotal prob of N-in: 0.12994\noutside 1 331\nTMhelix 332 354\ninside 355 372

11767 GCF_000407985.1_Esch_coli_KTE37_V1 Escherichia coli KTE37 Proteobacteria; Gammaproteobacteria; Enterobacterales; Enterobacteriaceae; Escherichia; Escherichia coli MTGDNTLIHSHGINRRDFMKLCAALAATMGLSSKAAAEMAESVTNPQRPPVIWIGAQECT WP_000145410.1 MULTISPECIES: hydrogenase-2 small chain [Proteobacteria] Length: 372\nNumber of predicted TMHs: 1\nExp number of AAs in TMHs: 23.72226\nExp number, first 60 AAs: 0.22038\nTotal prob of N-in: 0.12994\noutside 1 331\nTMhelix 332 354\ninside 355 372

11768 GCF_000407985.1_Esch_coli_KTE37_V1 Escherichia coli KTE37 Proteobacteria; Gammaproteobacteria; Enterobacterales; Enterobacteriaceae; Escherichia; Escherichia coli MNNEETFYQAMRRQGVTRRSFLKYCSLAATSLGLGAGMAPKIAWALENKPRIPVVWIHGL WP_001058323.1 MULTISPECIES: hydrogenase-1 small chain [Proteobacteria] Length: 372\nNumber of predicted TMHs: 2\nExp number of AAs in TMHs: 37.2608499999999\nExp number, first 60 AAs: 19.52923\nTotal prob of N-in: 0.94351\nPOSSIBLE N-term signal sequence\ninside 1 19\nTMhelix 20 39\noutside 40 325\nTMhelix 326 348\ninside 349 372

11769 GCF_000408005.1_Esch_coli_KTE38_V1 Escherichia coli KTE38 Proteobacteria; Gammaproteobacteria; Enterobacterales; Enterobacteriaceae; Escherichia; Escherichia coli MNNEETFYQAMRRQGVTRRSFLKYCSLAATSLGLGAGMAPKIAWALENKPRIPVVWIHGL WP_001058323.1 MULTISPECIES: hydrogenase-1 small chain [Proteobacteria] Length: 372\nNumber of predicted TMHs: 2\nExp number of AAs in TMHs: 37.2608499999999\nExp number, first 60 AAs: 19.52923\nTotal prob of N-in: 0.94351\nPOSSIBLE N-term signal sequence\ninside 1 19\nTMhelix 20 39\noutside 40 325\nTMhelix 326 348\ninside 349 372

11770 GCF_000408005.1_Esch_coli_KTE38_V1 Escherichia coli KTE38 Proteobacteria; Gammaproteobacteria; Enterobacterales; Enterobacteriaceae; Escherichia; Escherichia coli MTGDNTLIHSHGINRRDFMKLCAALAATMGLSSKAAAEMAESVTNPQRPPVIWIGAQECT WP_000145410.1 MULTISPECIES: hydrogenase-2 small chain [Proteobacteria] Length: 372\nNumber of predicted TMHs: 1\nExp number of AAs in TMHs: 23.72226\nExp number, first 60 AAs: 0.22038\nTotal prob of N-in: 0.12994\noutside 1 331\nTMhelix 332 354\ninside 355 372

11771 GCF_000350865.1_Esch_coli_KTE39_V1 Escherichia coli KTE39 Proteobacteria; Gammaproteobacteria; Enterobacterales; Enterobacteriaceae; Escherichia; Escherichia coli MNNEETFYQAMRRQGVTRRSFLKYCSLAATSLGLGAGMAPKIAWALENKPRIPVVWIHGL WP_001058323.1 MULTISPECIES: hydrogenase-1 small chain [Proteobacteria] Length: 372\nNumber of predicted TMHs: 2\nExp number of AAs in TMHs: 37.2608499999999\nExp number, first 60 AAs: 19.52923\nTotal prob of N-in: 0.94351\nPOSSIBLE N-term signal sequence\ninside 1 19\nTMhelix 20 39\noutside 40 325\nTMhelix 326 348\ninside 349 372

11772 GCF_000350865.1_Esch_coli_KTE39_V1 Escherichia coli KTE39 Proteobacteria; Gammaproteobacteria; Enterobacterales; Enterobacteriaceae; Escherichia; Escherichia coli MTGDNTLIHSHGINRRDFMKLCAALAATMGLSSKAAAEMAESVTNPQRPPVIWIGAQECT WP_000145410.1 MULTISPECIES: hydrogenase-2 small chain [Proteobacteria] Length: 372\nNumber of predicted TMHs:

1\nExp number of AAs in TMHs: 23.72226\nExp number, first 60 AAs: 0.22038\nTotal prob of N-in: 0.12994\noutside 1 331\nTMhelix 332 354\ninside 355 372

11773 GCF_000350645.1_Esch_coli_KTE4_V1 Escherichia coli KTE4 Proteobacteria; Gammaproteobacteria; Enterobacterales; Enterobacteriaceae; Escherichia; Escherichia coli MTGDNTLIHSHGINRRDFMKLCAALAATMGLSSKAAAEMAESVTNPQRPPVIWIGAQECT WP_000145410.1 MULTISPECIES: hydrogenase-2 small chain [Proteobacteria] Length: 372\nNumber of predicted TMHs: 1\nExp number of AAs in TMHs: 23.72226\nExp number, first 60 AAs: 0.22038\nTotal prob of N-in: 0.12994\noutside 1 331\nTMhelix 332 354\ninside 355 372

11774 GCF_000408025.1_Esch_coli_KTE40_V1 Escherichia coli KTE40 Proteobacteria; Gammaproteobacteria; Enterobacterales; Enterobacteriaceae; Escherichia; Escherichia coli MNNEETFYQAMRRQGVTRRSFLKYCSLAATSLGLGAGMAPKIAWALENKPRIPVVWIHGL WP_001058323.1 MULTISPECIES: hydrogenase-1 small chain [Proteobacteria] Length: 372\nNumber of predicted TMHs: 2\nExp number of AAs in TMHs: 37.2608499999999\nExp number, first 60 AAs: 19.52923\nTotal prob of N-in: 0.94351\nPOSSIBLE N-term signal sequence\ninside 1 19\nTMhelix 20 39\noutside 40 325\nTMhelix 326 348\ninside 349 372

11775 GCF_000408025.1_Esch_coli_KTE40_V1 Escherichia coli KTE40 Proteobacteria; Gammaproteobacteria; Enterobacterales; Enterobacteriaceae; Escherichia; Escherichia coli MTGDNTLIHSHGINRRDFMKLCAALAATMGLSSKAAAEMAESVTNPQRPPVIWIGAQECT WP_000145410.1 MULTISPECIES: hydrogenase-2 small chain [Proteobacteria] Length: 372\nNumber of predicted TMHs: 1\nExp number of AAs in TMHs: 23.72226\nExp number, first 60 AAs: 0.22038\nTotal prob of N-in: 0.12994\noutside 1 331\nTMhelix 332 354\ninside 355 372

11776 GCF_000408045.1_Esch_coli_KTE41_V1 Escherichia coli KTE41 Proteobacteria; Gammaproteobacteria; Enterobacterales; Enterobacteriaceae; Escherichia; Escherichia coli MNNEETFYQAMRRQGVTRRSFLKYCSLAATSLGLGAGMAPKIAWALENKPRIPVVWIHGL WP_001058323.1 MULTISPECIES: hydrogenase-1 small chain [Proteobacteria] Length: 372\nNumber of predicted TMHs: 2\nExp number of AAs in TMHs: 37.2608499999999\nExp number, first 60 AAs: 19.52923\nTotal prob of N-in: 0.94351\nPOSSIBLE N-term signal sequence\ninside 1 19\nTMhelix 20 39\noutside 40 325\nTMhelix 326 348\ninside 349 372

11777 GCF_000408045.1_Esch_coli_KTE41_V1 Escherichia coli KTE41 Proteobacteria; Gammaproteobacteria; Enterobacterales; Enterobacteriaceae; Escherichia; Escherichia coli MTGDNTLIHSHGINRRDFMKLCAALAATMGLSSKAAAEMAESVTNPQRPPVIWIGAQECT WP_000145410.1 MULTISPECIES: hydrogenase-2 small chain [Proteobacteria] Length: 372\nNumber of predicted TMHs: 1\nExp number of AAs in TMHs: 23.72226\nExp number, first 60 AAs: 0.22038\nTotal prob of N-in: 0.12994\noutside 1 331\nTMhelix 332 354\ninside 355 372

11778 GCF_000352185.1_Esch_coli_KTE42_V1 Escherichia coli KTE42 Proteobacteria; Gammaproteobacteria; Enterobacterales; Enterobacteriaceae; Escherichia; Escherichia coli MNNEETFYQAMRRQGVTRRSFLKYCSLAATSLGLGAGMAPKIAWALENKPRIPVVWIHGL WP_001058323.1 MULTISPECIES: hydrogenase-1 small chain [Proteobacteria] Length: 372\nNumber of predicted TMHs: 2\nExp number of AAs in TMHs: 37.2608499999999\nExp number, first 60 AAs: 19.52923\nTotal prob of N-in: 0.94351\nPOSSIBLE N-term signal sequence\ninside 1 19\nTMhelix 20 39\noutside 40 325\nTMhelix 326 348\ninside 349 372

11779 GCF_000352185.1_Esch_coli_KTE42_V1 Escherichia coli KTE42 Proteobacteria; Gammaproteobacteria; Enterobacterales; Enterobacteriaceae; Escherichia; Escherichia coli MTGDNTLIHSHGINRRDFMKLCAALAATMGLSSKAAAEMAESVTNPQRPPVIWIGAQECT WP_000145410.1 MULTISPECIES: hydrogenase-2 small chain [Proteobacteria] Length: 372\nNumber of predicted TMHs: 1\nExp number of AAs in TMHs: 23.72226\nExp number, first 60 AAs: 0.22038\nTotal prob of N-in: 0.12994\noutside 1 331\nTMhelix 332 354\ninside 355 372

11780 GCF_000352225.1_Esch_coli_KTE43_V1 Escherichia coli KTE43 Proteobacteria; Gammaproteobacteria; Enterobacterales; Enterobacteriaceae; Escherichia; Escherichia coli MTGDNTLIHSHGINRRDFMKLCAALAATMGLSSKAAAEMVESVTNPQRPPVIWIGAQECT WP_000145423.1 hydrogenase 2 small subunit [Escherichia coli] Length: 372\nNumber of predicted TMHs: 1\nExp number of AAs in TMHs: 23.78201\nExp number, first 60 AAs: 0.28355\nTotal prob of N-in: 0.13257\noutside 1 331\nTMhelix 332 354\ninside 355 372

11781 GCF_000352225.1_Esch_coli_KTE43_V1 Escherichia coli KTE43 Proteobacteria; Gammaproteobacteria; Enterobacterales; Enterobacteriaceae; Escherichia; Escherichia coli MNNEETFYQAMRRQGVTRRSFLKYCSLAATSLGLGAGMAPKIAWALENKPRIPVVWIHGL WP_001058323.1 MULTISPECIES: hydrogenase-1 small chain [Proteobacteria] Length: 372\nNumber of predicted TMHs: 2\nExp number of AAs in TMHs: 37.2608499999999\nExp number, first 60 AAs: 19.52923\nTotal prob of N-in: 0.94351\nPOSSIBLE N-term signal sequence\ninside 1 19\nTMhelix 20 39\noutside 40 325\nTMhelix 326 348\ninside 349 372

11782 GCF_000350885.1_Esch_coli_KTE44_V1 Escherichia coli KTE44 Proteobacteria; Gammaproteobacteria; Enterobacterales; Enterobacteriaceae; Escherichia; Escherichia coli MNNEETFYQAMRRQGVTRRSFLKYCSLAATSLGLGAGMAPKIAWALENKPRIPVVWIHGL WP_001058323.1 MULTISPECIES: hydrogenase-1 small chain [Proteobacteria] Length: 372\nNumber of predicted TMHs: 2\nExp number of AAs in TMHs: 37.2608499999999\nExp number, first 60 AAs: 19.52923\nTotal prob of N-in: 0.94351\nPOSSIBLE N-term signal sequence\ninside 1 19\nTMhelix 20 39\noutside 40 325\nTMhelix 326 348\ninside 349 372

11783 GCF_000350885.1_Esch_coli_KTE44_V1 Escherichia coli KTE44 Proteobacteria; Gammaproteobacteria; Enterobacterales; Enterobacteriaceae; Escherichia; Escherichia coli MTGDNTLIHSHGINRRDFMKLCAALAATMGLSSKAAAEMAESVTNPQRPPVIWIGAQECT WP_000145410.1 MULTISPECIES: hydrogenase-2 small chain [Proteobacteria] Length: 372\nNumber of predicted TMHs: 1\nExp number of AAs in TMHs: 23.72226\nExp number, first 60 AAs: 0.22038\nTotal prob of N-in: 0.12994\noutside 1 331\nTMhelix 332 354\ninside 355 372

11784 GCF_000352165.1_Esch_coli_KTE45_V1 Escherichia coli KTE45 Proteobacteria; Gammaproteobacteria; Enterobacterales; Enterobacteriaceae; Escherichia; Escherichia coli MNNEETFYQAMRRQGVTRRSFLKYCSLAATSLGLGAGMAPKIAWALENKPRIPVVWIHGL WP_001058323.1 MULTISPECIES: hydrogenase-1 small chain [Proteobacteria] Length: 372\nNumber of predicted TMHs: 2\nExp number of AAs in TMHs: 37.2608499999999\nExp number, first 60 AAs: 19.52923\nTotal prob of N-in: 0.94351\nPOSSIBLE N-term signal sequence\ninside 1 19\nTMhelix 20 39\noutside 40 325\nTMhelix 326 348\ninside 349 372

11785 GCF_000352165.1_Esch_coli_KTE45_V1 Escherichia coli KTE45 Proteobacteria; Gammaproteobacteria; Enterobacterales; Enterobacteriaceae; Escherichia; Escherichia coli MTGDNTLIHSHGINRRDFMKLCAALAATMGLSSKAAAEMAESVTNPQRPPVIWIGAQECT WP_000145410.1 MULTISPECIES: hydrogenase-2 small chain [Proteobacteria] Length: 372\nNumber of predicted TMHs: 1\nExp number of AAs in TMHs: 23.72226\nExp number, first 60 AAs: 0.22038\nTotal prob of N-in: 0.12994\noutside 1 331\nTMhelix 332 354\ninside 355 372

11786 GCF_000352285.1_Esch_coli_KTE46_V1 Escherichia coli KTE46 Proteobacteria; Gammaproteobacteria; Enterobacterales; Enterobacteriaceae; Escherichia; Escherichia coli MTGDNTLIHSHGINRRDFMKLCAALAATMGLSSKAAAEMAESVTNPQRPPVIWIGAQECT WP_000145410.1 MULTISPECIES: hydrogenase-2 small chain [Proteobacteria] Length: 372\nNumber of predicted TMHs: 1\nExp number of AAs in TMHs: 23.72226\nExp number, first 60 AAs: 0.22038\nTotal prob of N-in: 0.12994\noutside 1 331\nTMhelix 332 354\ninside 355 372

11787 GCF_000352285.1_Esch_coli_KTE46_V1 Escherichia coli KTE46 Proteobacteria; Gammaproteobacteria; Enterobacterales; Enterobacteriaceae; Escherichia; Escherichia coli MNNEETFYQAMRRQGVTRRSFLKYCSLAATSLGLGAGMAPKIAWALENKPRIPVVWIHGL WP_001058323.1 MULTISPECIES: hydrogenase-1 small chain [Proteobacteria] Length: 372\nNumber of predicted TMHs: 2\nExp number of AAs in TMHs: 37.2608499999999\nExp number, first 60 AAs: 19.52923\nTotal prob of N-in: 0.94351\nPOSSIBLE N-term signal sequence\ninside 1 19\nTMhelix 20 39\noutside 40 325\nTMhelix 326 348\ninside 349 372

11788 GCF_000351425.1_Esch_coli_KTE47_V1 Escherichia coli KTE47 Proteobacteria; Gammaproteobacteria; Enterobacterales; Enterobacteriaceae; Escherichia; Escherichia coli MTGDNTLIHSHGINRRDFMKLCAALAATMGLSSKAAAEMAESVTNPQRPPVIWIGAQECT WP_000145410.1 MULTISPECIES: hydrogenase-2 small chain [Proteobacteria] Length: 372\nNumber of predicted TMHs: 1\nExp number of AAs in TMHs: 23.72226\nExp number, first 60 AAs: 0.22038\nTotal prob of N-in: 0.12994\noutside 1 331\nTMhelix 332 354\ninside 355 372

11789 GCF_000351425.1_Esch_coli_KTE47_V1 Escherichia coli KTE47 Proteobacteria; Gammaproteobacteria; Enterobacterales; Enterobacteriaceae; Escherichia; Escherichia coli MNNEETFYQAMRRQGVTRRSFLKYCSLAATSLGLGAGMAPKIAWALENKPRIPVVWIHGL WP_001058323.1 MULTISPECIES: hydrogenase-1 small chain [Proteobacteria] Length: 372\nNumber of predicted TMHs: 2\nExp number of AAs in TMHs: 37.2608499999999\nExp number, first 60 AAs: 19.52923\nTotal prob of N-in: 0.94351\nPOSSIBLE N-term signal sequence\ninside 1 19\nTMhelix 20 39\noutside 40 325\nTMhelix 326 348\ninside 349 372

11790 GCF_000352305.1_Esch_coli_KTE48_V1 Escherichia coli KTE48 Proteobacteria; Gammaproteobacteria; Enterobacterales; Enterobacteriaceae; Escherichia; Escherichia coli MNNEETFYQAMRRQGVTRRSFLKYCSLAATSLGLGAGMAPKIAWALENKPRIPVVWIHGL WP_001058323.1 MULTISPECIES: hydrogenase-1 small chain [Proteobacteria] Length: 372\nNumber of predicted TMHs: 2\nExp number of AAs in TMHs: 37.2608499999999\nExp number, first 60 AAs: 19.52923\nTotal prob of N-in: 0.94351\nPOSSIBLE N-term signal sequence\ninside 1 19\nTMhelix 20 39\noutside 40 325\nTMhelix 326 348\ninside 349 372

11791 GCF_000352305.1_Esch_coli_KTE48_V1 Escherichia coli KTE48 Proteobacteria; Gammaproteobacteria; Enterobacterales; Enterobacteriaceae; Escherichia; Escherichia coli MTGDNTLIHSHGINRRDFMKLCAALAATMGLSSKAAAEMAESVTNPQRPPVIWIGAQECT WP_000145410.1 MULTISPECIES: hydrogenase-2 small chain [Proteobacteria] Length: 372\nNumber of predicted TMHs: 1\nExp number of AAs in TMHs: 23.72226\nExp number, first 60 AAs: 0.22038\nTotal prob of N-in: 0.12994\noutside 1 331\nTMhelix 332 354\ninside 355 372

11792 GCF_000351445.1_Esch_coli_KTE49_V1 Escherichia coli KTE49 Proteobacteria; Gammaproteobacteria; Enterobacterales; Enterobacteriaceae; Escherichia; Escherichia coli MNNEETFYQAMRRQGVTRRSFLKYCSLAATSLGLGAGMAPKIAWALENKPRIPVVWIHGL WP_001295940.1 MULTISPECIES: hydrogenase [Enterobacteriaceae] Length: 372\nNumber of predicted TMHs: 2\nExp number of AAs in TMHs: 37.2608899999999\nExp number, first 60 AAs: 19.52923\nTotal prob of N-in: 0.94351\nPOSSIBLE N-term signal sequence\ninside 1 19\nTMhelix 20 39\noutside 40 325\nTMhelix 326 348\ninside 349 372

11793 GCF_000351445.1_Esch_coli_KTE49_V1 Escherichia coli KTE49 Proteobacteria; Gammaproteobacteria; Enterobacterales; Enterobacteriaceae; Escherichia; Escherichia coli MTGDNTLIHSHGINRRDFMKLCAALAATMGLSSKAAAEMVESVTNPQRPPVIWIGAQECT WP_000145423.1 hydrogenase 2 small subunit [Escherichia coli] Length: 372\nNumber of predicted TMHs: 1\nExp number of AAs in TMHs: 23.78201\nExp number, first 60 AAs: 0.28355\nTotal prob of N-in: 0.13257\noutside 1 331\nTMhelix 332 354\ninside 355 372

11794 GCF_000350665.1_Esch_coli_KTE5_V1 Escherichia coli KTE5 Proteobacteria; Gammaproteobacteria; Enterobacterales; Enterobacteriaceae; Escherichia; Escherichia coli MTGDNTLIHSHGINRRDFMKLCAALAATMGLSSKAAAEMAESVTNPQRPPVIWIGAQECT WP_000145410.1 MULTISPECIES: hydrogenase-2 small chain [Proteobacteria] Length: 372\nNumber of predicted TMHs: 1\nExp number of AAs in TMHs: 23.72226\nExp number, first 60 AAs: 0.22038\nTotal prob of N-in: 0.12994\noutside 1 331\nTMhelix 332 354\ninside 355 372

11795 GCF_000352325.1_Esch_coli_KTE50_V1 Escherichia coli KTE50 Proteobacteria; Gammaproteobacteria; Enterobacterales; Enterobacteriaceae; Escherichia; Escherichia coli MTGDNTLIHSHGINRRDFMKLCAALAATMGLSSKAAAEMAESVTNPQRPPVIWIGAQECT WP_000145410.1 MULTISPECIES: hydrogenase-2 small chain [Proteobacteria] Length: 372\nNumber of predicted TMHs: 1\nExp number of AAs in TMHs: 23.72226\nExp number, first 60 AAs: 0.22038\nTotal prob of N-in: 0.12994\noutside 1 331\nTMhelix 332 354\ninside 355 372

11796 GCF_000352325.1_Esch_coli_KTE50_V1 Escherichia coli KTE50 Proteobacteria; Gammaproteobacteria; Enterobacterales; Enterobacteriaceae; Escherichia; Escherichia coli MNNEETFYQAMRRQGVTRRSFLKYCSLAATSLGLGAGMAPKIAWALENKPRIPVVWIHGL WP_001058323.1 MULTISPECIES: hydrogenase-1 small chain [Proteobacteria] Length: 372\nNumber of predicted TMHs: 2\nExp number of AAs in TMHs: 37.2608499999999\nExp number, first 60 AAs: 19.52923\nTotal prob of N-in: 0.94351\nPOSSIBLE N-term signal sequence\ninside 1 19\nTMhelix 20 39\noutside 40 325\nTMhelix 326 348\ninside 349 372

11797 GCF_000351465.1_Esch_coli_KTE51_V1 Escherichia coli KTE51 Proteobacteria; Gammaproteobacteria; Enterobacterales; Enterobacteriaceae; Escherichia; Escherichia coli MTGDNTLIHSHGINRRDFMKLCAALAATMGLSSKAAAEMAESVTNPQRPPVIWIGAQECT WP_000145410.1 MULTISPECIES: hydrogenase-2 small chain [Proteobacteria] Length: 372\nNumber of predicted TMHs: 1\nExp number of AAs in TMHs: 23.72226\nExp number, first 60 AAs: 0.22038\nTotal prob of N-in: 0.12994\noutside 1 331\nTMhelix 332 354\ninside 355 372

11798 GCF_000351465.1_Esch_coli_KTE51_V1 Escherichia coli KTE51 Proteobacteria; Gammaproteobacteria; Enterobacterales; Enterobacteriaceae; Escherichia; Escherichia coli MNNEETFYQAMRRQGVTRRSFLKYCSLAATSLGLGAGMAPKIAWALENKPRIPVVWIHGL WP_001058323.1 MULTISPECIES: hydrogenase-1 small chain [Proteobacteria] Length: 372\nNumber of predicted TMHs: 2\nExp number of AAs in TMHs: 37.2608499999999\nExp number, first 60 AAs: 19.52923\nTotal prob of N-in: 0.94351\nPOSSIBLE N-term signal sequence\ninside 1 19\nTMhelix 20 39\noutside 40 325\nTMhelix 326 348\ninside 349 372

11799 GCF_000351485.1_Esch_coli_KTE53_V1 Escherichia coli KTE53 Proteobacteria; Gammaproteobacteria; Enterobacterales; Enterobacteriaceae; Escherichia; Escherichia coli MNNEETFYQAMRRQGVTRRSFLKYCSLAATSLGLGAGMAPKIAWALENKPRIPVVWIHGL WP_001058323.1 MULTISPECIES: hydrogenase-1 small chain [Proteobacteria] Length: 372\nNumber of predicted TMHs: 2\nExp number of AAs in TMHs: 37.2608499999999\nExp number, first 60 AAs: 19.52923\nTotal prob of N-in: 0.94351\nPOSSIBLE N-term signal sequence\ninside 1 19\nTMhelix 20 39\noutside 40 325\nTMhelix 326 348\ninside 349 372

11800 GCF_000351485.1_Esch_coli_KTE53_V1 Escherichia coli KTE53 Proteobacteria; Gammaproteobacteria; Enterobacterales; Enterobacteriaceae; Escherichia; Escherichia coli MTGDNTLIHSHGINRRDFMKLCAALAATMGLSSKAAAEMAESVTNPQRPPVIWIGAQECT WP_000145410.1 MULTISPECIES: hydrogenase-2 small chain [Proteobacteria] Length: 372\nNumber of predicted TMHs: 1\nExp number of AAs in TMHs: 23.72226\nExp number, first 60 AAs: 0.22038\nTotal prob of N-in: 0.12994\noutside 1 331\nTMhelix 332 354\ninside 355 372

11801 GCF_000352345.1_Esch_coli_KTE54_V1 Escherichia coli KTE54 Proteobacteria; Gammaproteobacteria; Enterobacterales; Enterobacteriaceae; Escherichia; Escherichia coli MTGDNTLIHSHGINRRDFMKLCAALAATMGLSSKAAAEMAESVTNPQRPPVIWIGAQECT WP_000145410.1 MULTISPECIES: hydrogenase-2 small chain [Proteobacteria] Length: 372\nNumber of predicted TMHs: 1\nExp number of AAs in TMHs: 23.72226\nExp number, first 60 AAs: 0.22038\nTotal prob of N-in: 0.12994\noutside 1 331\nTMhelix 332 354\ninside 355 372

11802 GCF_000352345.1_Esch_coli_KTE54_V1 Escherichia coli KTE54 Proteobacteria; Gammaproteobacteria; Enterobacterales; Enterobacteriaceae; Escherichia; Escherichia coli MNNEETFYQAMRRQGVTRRSFLKYCSLAATSLGLGAGMAPKIAWALENKPRIPVVWIHGL WP_001058323.1 MULTISPECIES: hydrogenase-1 small chain [Proteobacteria] Length: 372\nNumber of predicted TMHs: 2\nExp number of AAs in TMHs: 37.2608499999999\nExp number, first 60 AAs: 19.52923\nTotal prob of N-in: 0.94351\nPOSSIBLE N-term signal sequence\ninside 1 19\nTMhelix 20 39\noutside 40 325\nTMhelix 326 348\ninside 349 372

11803 GCF_000351505.1_Esch_coli_KTE55_V1 Escherichia coli KTE55 Proteobacteria; Gammaproteobacteria; Enterobacterales; Enterobacteriaceae; Escherichia; Escherichia coli MTGDNTLIHSHGINRRDFMKLCAALAATMGLSSKAAAEMAESVTNPQRPPVIWIGAQECT WP_000145410.1 MULTISPECIES: hydrogenase-2 small chain [Proteobacteria] Length: 372\nNumber of predicted TMHs: 1\nExp number of AAs in TMHs: 23.72226\nExp number, first 60 AAs: 0.22038\nTotal prob of N-in: 0.12994\noutside 1 331\nTMhelix 332 354\ninside 355 372

11804 GCF_000351525.1_Esch_coli_KTE56_V1 Escherichia coli KTE56 Proteobacteria; Gammaproteobacteria; Enterobacterales; Enterobacteriaceae; Escherichia; Escherichia coli MNNEETFYQAMRRQGVTRRSFLKYCSLAATSLGLGAGMAPKIAWALENKPRIPVVWIHGL WP_001058323.1 MULTISPECIES: hydrogenase-1 small chain [Proteobacteria] Length: 372\nNumber of predicted TMHs: 2\nExp number of AAs in TMHs: 37.2608499999999\nExp number, first 60 AAs: 19.52923\nTotal prob of N-in: 0.94351\nPOSSIBLE N-term signal sequence\ninside 1 19\nTMhelix 20 39\noutside 40 325\nTMhelix 326 348\ninside 349 372

11805 GCF_000351525.1_Esch_coli_KTE56_V1 Escherichia coli KTE56 Proteobacteria; Gammaproteobacteria; Enterobacterales; Enterobacteriaceae; Escherichia; Escherichia coli MTGDNTLIHSHGINRRDFMKLCAALAATMGLSSKAAAEMAESVTNPQRPPVIWIGAQECT WP_000145410.1 MULTISPECIES: hydrogenase-2 small chain [Proteobacteria] Length: 372\nNumber of predicted TMHs: 1\nExp number of AAs in TMHs: 23.72226\nExp number, first 60 AAs: 0.22038\nTotal prob of N-in: 0.12994\noutside 1 331\nTMhelix 332 354\ninside 355 372

11806 GCF_000351545.1_Esch_coli_KTE57_V1 Escherichia coli KTE57 Proteobacteria; Gammaproteobacteria; Enterobacterales; Enterobacteriaceae; Escherichia; Escherichia coli MTGDNTLIHSHGINRRDFMKLCAALAATMGLSSKAAAEMVESVTNPQRPPVIWIGAQECT WP_001556793.1 hydrogenase-2 small chain [Escherichia coli] Length: 372\nNumber of predicted TMHs: 1\nExp number of AAs in TMHs: 23.78201\nExp number, first 60 AAs: 0.28355\nTotal prob of N-in: 0.13257\noutside 1 331\nTMhelix 332 354\ninside 355 372

11807 GCF_000351545.1_Esch_coli_KTE57_V1 Escherichia coli KTE57 Proteobacteria; Gammaproteobacteria; Enterobacterales; Enterobacteriaceae; Escherichia; Escherichia coli MNNEETFYQAMRRQGVTRRSFLKYCSLAATSLGLGAGMAPKIAWALENKPRIPVVWIHGL WP_001538532.1 hydrogenase-1 small chain [Escherichia coli] Length: 372\nNumber of predicted TMHs: 2\nExp number of AAs in TMHs: 37.27041\nExp number, first 60 AAs: 19.52927\nTotal prob of N-in: 0.94352\nPOSSIBLE N-term signal sequence\ninside 1 19\nTMhelix 20 39\noutside 40 325\nTMhelix 326 348\ninside 349 372

11808 GCF_000351565.1_Esch_coli_KTE58_V1 Escherichia coli KTE58 Proteobacteria; Gammaproteobacteria; Enterobacterales; Enterobacteriaceae; Escherichia; Escherichia coli MTGDNTLIHSHGINRRDFMKLCAALAATMGLSSKAAAEMAESVTNPQRPPVIWIGAQECT WP_000145410.1 MULTISPECIES: hydrogenase-2 small chain [Proteobacteria] Length: 372\nNumber of predicted TMHs: 1\nExp number of AAs in TMHs: 23.72226\nExp number, first 60 AAs: 0.22038\nTotal prob of N-in: 0.12994\noutside 1 331\nTMhelix 332 354\ninside 355 372

11809 GCF_000352365.1_Esch_coli_KTE59_V1 Escherichia coli KTE59 Proteobacteria; Gammaproteobacteria; Enterobacterales; Enterobacteriaceae; Escherichia; Escherichia coli MTGDNTLIHSHGINRRDFMKLCAALAATMGLSSKAAAEMAESVTNPQRPPVIWIGAQECT WP_000145410.1 MULTISPECIES: hydrogenase-2 small chain [Proteobacteria] Length: 372\nNumber of predicted TMHs: 1\nExp number of AAs in TMHs: 23.72226\nExp number, first 60 AAs: 0.22038\nTotal prob of N-in: 0.12994\noutside 1 331\nTMhelix 332 354\ninside 355 372

11810 GCF_000352065.1_Esch_coli_KTE6_V1 Escherichia coli KTE6 Proteobacteria; Gammaproteobacteria; Enterobacterales; Enterobacteriaceae; Escherichia; Escherichia coli MTGDNTLIHSHGINRRDFMKLCAALAATMGLSSKAAAEMAESVTNPQRPPVIWIGAQECT WP_000145410.1 MULTISPECIES: hydrogenase-2 small chain [Proteobacteria] Length: 372\nNumber of predicted TMHs: 1\nExp number of AAs in TMHs: 23.72226\nExp number, first 60 AAs: 0.22038\nTotal prob of N-in: 0.12994\noutside 1 331\nTMhelix 332 354\ninside 355 372

11811 GCF_000352065.1_Esch_coli_KTE6_V1 Escherichia coli KTE6 Proteobacteria; Gammaproteobacteria; Enterobacterales; Enterobacteriaceae; Escherichia; Escherichia coli MNNEETFYQAMRRQGVTRRSFLKYCSLAATSLGLGAGMAPKIAWALENKPRIPVVWIHGL WP_001295940.1 MULTISPECIES: hydrogenase [Enterobacteriaceae] Length: 372\nNumber of predicted TMHs: 2\nExp number of AAs in TMHs: 37.2608899999999\nExp number, first 60 AAs: 19.52923\nTotal prob of N-in: 0.94351\nPOSSIBLE N-term signal sequence\ninside 1 19\nTMhelix 20 39\noutside 40 325\nTMhelix 326 348\ninside 349 372

11812 GCF_000351585.1_Esch_coli_KTE60_V1 Escherichia coli KTE60 Proteobacteria; Gammaproteobacteria; Enterobacterales; Enterobacteriaceae; Escherichia; Escherichia coli MNNEETFYQAMRRQGVTRRSFLKYCSLAATSLGLGAGMAPKIAWALENKPRIPVVWIHGL WP_001058323.1 MULTISPECIES: hydrogenase-1 small chain [Proteobacteria] Length: 372\nNumber of predicted TMHs: 2\nExp number of AAs in TMHs: 37.2608499999999\nExp number, first 60 AAs: 19.52923\nTotal prob of N-in: 0.94351\nPOSSIBLE N-term signal sequence\ninside 1 19\nTMhelix 20 39\noutside 40 325\nTMhelix 326 348\ninside 349 372

11813 GCF_000351585.1_Esch_coli_KTE60_V1 Escherichia coli KTE60 Proteobacteria; Gammaproteobacteria; Enterobacterales; Enterobacteriaceae; Escherichia; Escherichia coli MTGDNTLIHSHGINRRDFMKLCAALAATMGLSSKAAAEMAESVTNPQRPPVIWIGAQECT WP_000145410.1

MULTISPECIES: hydrogenase-2 small chain [Proteobacteria] Length: 372\nNumber of predicted TMHs: 1\nExp number of AAs in TMHs: 23.72226\nExp number, first 60 AAs: 0.22038\nTotal prob of N-in: 0.12994\noutside 1 331\nTMhelix 332 354\ninside 355 372

11814 GCF_000408345.1_Esch_coli_KTE61_V1 Escherichia coli KTE61 Proteobacteria; Gammaproteobacteria; Enterobacterales; Enterobacteriaceae; Escherichia; Escherichia coli MTGDNTLIHSHGINRRDFMKLCAALAATMGLSSKAAAEMAESVTNPQRPPVIWIGAQECT WP_000145410.1 MULTISPECIES: hydrogenase-2 small chain [Proteobacteria] Length: 372\nNumber of predicted TMHs: 1\nExp number of AAs in TMHs: 23.72226\nExp number, first 60 AAs: 0.22038\nTotal prob of N-in: 0.12994\noutside 1 331\nTMhelix 332 354\ninside 355 372

11815 GCF_000408345.1_Esch_coli_KTE61_V1 Escherichia coli KTE61 Proteobacteria; Gammaproteobacteria; Enterobacterales; Enterobacteriaceae; Escherichia; Escherichia coli MNNEETFYQAMRRQGVTRRSFLKYCSLAATSLGLGAGMAPKIAWALENKPRIPVVWIHGL WP_001058323.1 MULTISPECIES: hydrogenase-1 small chain [Proteobacteria] Length: 372\nNumber of predicted TMHs: 2\nExp number of AAs in TMHs: 37.2608499999999\nExp number, first 60 AAs: 19.52923\nTotal prob of N-in: 0.94351\nPOSSIBLE N-term signal sequence\ninside 1 19\nTMhelix 20 39\noutside 40 325\nTMhelix 326 348\ninside 349 372

11816 GCF_000351605.1_Esch_coli_KTE62_V1 Escherichia coli KTE62 Proteobacteria; Gammaproteobacteria; Enterobacterales; Enterobacteriaceae; Escherichia; Escherichia coli MTGDNTLIHSHGINRRDFMKLCAALAATMGLSSKAAAEMAESVTNPQRPPVIWIGAQECT WP_000145410.1 MULTISPECIES: hydrogenase-2 small chain [Proteobacteria] Length: 372\nNumber of predicted TMHs: 1\nExp number of AAs in TMHs: 23.72226\nExp number, first 60 AAs: 0.22038\nTotal prob of N-in: 0.12994\noutside 1 331\nTMhelix 332 354\ninside 355 372

11817 GCF_000352385.1_Esch_coli_KTE63_V1 Escherichia coli KTE63 Proteobacteria; Gammaproteobacteria; Enterobacterales; Enterobacteriaceae; Escherichia; Escherichia coli MTGDNTLIHSHGINRRDFMKLCAALAATMGLSSKAAAEMAESVTNPQRPPVIWIGAQECT WP_000145410.1 MULTISPECIES: hydrogenase-2 small chain [Proteobacteria] Length: 372\nNumber of predicted TMHs: 1\nExp number of AAs in TMHs: 23.72226\nExp number, first 60 AAs: 0.22038\nTotal prob of N-in: 0.12994\noutside 1 331\nTMhelix 332 354\ninside 355 372

11818 GCF_000352385.1_Esch_coli_KTE63_V1 Escherichia coli KTE63 Proteobacteria; Gammaproteobacteria; Enterobacterales; Enterobacteriaceae; Escherichia; Escherichia coli MNNEETFYQAMRRQGVTRRSFLKYCSLAATSLGLGAGMAPKIAWALENKPRIPVVWIHGL WP_001058323.1 MULTISPECIES: hydrogenase-1 small chain [Proteobacteria] Length: 372\nNumber of predicted TMHs: 2\nExp number of AAs in TMHs: 37.2608499999999\nExp number, first 60 AAs: 19.52923\nTotal prob of N-in: 0.94351\nPOSSIBLE N-term signal sequence\ninside 1 19\nTMhelix 20 39\noutside 40 325\nTMhelix 326 348\ninside 349 372

11819 GCF_000408365.1_Esch_coli_KTE64_V1 Escherichia coli KTE64 Proteobacteria; Gammaproteobacteria; Enterobacterales; Enterobacteriaceae; Escherichia; Escherichia coli MNNEETFYQAMRRQGVTRRSFLKYCSLAATSLGLGAGMAPKIAWALENKPRIPVVWIHGL WP_001058323.1 MULTISPECIES: hydrogenase-1 small chain [Proteobacteria] Length: 372\nNumber of predicted TMHs: 2\nExp number of AAs in TMHs: 37.2608499999999\nExp number, first 60 AAs: 19.52923\nTotal prob of N-in: 0.94351\nPOSSIBLE N-term signal sequence\ninside 1 19\nTMhelix 20 39\noutside 40 325\nTMhelix 326 348\ninside 349 372

11820 GCF_000408365.1_Esch_coli_KTE64_V1 Escherichia coli KTE64 Proteobacteria; Gammaproteobacteria; Enterobacterales; Enterobacteriaceae; Escherichia; Escherichia coli MTGDNTLIHSHGINRRDFMKLCAALAATMGLSSKAAAEMAESVTNPQRPPVIWIGAQECT WP_000145410.1 MULTISPECIES: hydrogenase-2 small chain [Proteobacteria] Length: 372\nNumber of predicted TMHs: 1\nExp number of AAs in TMHs: 23.72226\nExp number, first 60 AAs: 0.22038\nTotal prob of N-in: 0.12994\noutside 1 331\nTMhelix 332 354\ninside 355 372

11821 GCF_000352405.1_Esch_coli_KTE65_V1 Escherichia coli KTE65 Proteobacteria; Gammaproteobacteria; Enterobacterales; Enterobacteriaceae; Escherichia; Escherichia coli MTGDNTLIHSHGINRRDFMKLCAALAATMGLSSKAAAEMAESVTNPQRPPVIWIGAQECT WP_000145410.1 MULTISPECIES: hydrogenase-2 small chain [Proteobacteria] Length: 372\nNumber of predicted TMHs:

1\nExp number of AAs in TMHs: 23.72226\nExp number, first 60 AAs: 0.22038\nTotal prob of N-in: 0.12994\noutside 1 331\nTMhelix 332 354\ninside 355 372

11822 GCF_000351625.1_Esch_coli_KTE66_V1 Escherichia coli KTE66 Proteobacteria; Gammaproteobacteria; Enterobacterales; Enterobacteriaceae; Escherichia; Escherichia coli
MTGDNTLIHSHGINRRDFMKLCAALAATMGLSSKAAAEMAESVTNPQRPPVIWIGAQECT WP_000145410.1
MULTISPECIES: hydrogenase-2 small chain [Proteobacteria] Length: 372\nNumber of predicted TMHs: 1\nExp number of AAs in TMHs: 23.72226\nExp number, first 60 AAs: 0.22038\nTotal prob of N-in: 0.12994\noutside 1 331\nTMhelix 332 354\ninside 355 372

11823 GCF_000351625.1_Esch_coli_KTE66_V1 Escherichia coli KTE66 Proteobacteria; Gammaproteobacteria; Enterobacterales; Enterobacteriaceae; Escherichia; Escherichia coli
MNNEETFYQAMRRQGVTRRSFLKYCSLAATSLGLGAGMAPKIAWALENKPRIPVVWIHGL WP_001058323.1
MULTISPECIES: hydrogenase-1 small chain [Proteobacteria] Length: 372\nNumber of predicted TMHs: 2\nExp number of AAs in TMHs: 37.2608499999999\nExp number, first 60 AAs: 19.52923\nTotal prob of N-in: 0.94351\nPOSSIBLE N-term signal sequence\ninside 1 19\nTMhelix 20 39\noutside 40 325\nTMhelix 326 348\ninside 349 372

11824 GCF_000351645.1_Esch_coli_KTE67_V1 Escherichia coli KTE67 Proteobacteria; Gammaproteobacteria; Enterobacterales; Enterobacteriaceae; Escherichia; Escherichia coli
MTGDNTLIHSHGINRRDFMKLCAALAATMGLSSKAAAEMAESVTNPQRPPVIWIGAQECT WP_000145410.1
MULTISPECIES: hydrogenase-2 small chain [Proteobacteria] Length: 372\nNumber of predicted TMHs: 1\nExp number of AAs in TMHs: 23.72226\nExp number, first 60 AAs: 0.22038\nTotal prob of N-in: 0.12994\noutside 1 331\nTMhelix 332 354\ninside 355 372

11825 GCF_000351645.1_Esch_coli_KTE67_V1 Escherichia coli KTE67 Proteobacteria; Gammaproteobacteria; Enterobacterales; Enterobacteriaceae; Escherichia; Escherichia coli
MNNEETFYQAMRRQGVTRRSFLKYCSLAATSLGLGAGMAPKIAWALENKPRIPVVWIHGL WP_001058323.1
MULTISPECIES: hydrogenase-1 small chain [Proteobacteria] Length: 372\nNumber of predicted TMHs: 2\nExp number of AAs in TMHs: 37.2608499999999\nExp number, first 60 AAs: 19.52923\nTotal prob of N-in: 0.94351\nPOSSIBLE N-term signal sequence\ninside 1 19\nTMhelix 20 39\noutside 40 325\nTMhelix 326 348\ninside 349 372

11826 GCF_000408385.1_Esch_coli_KTE68_V1 Escherichia coli KTE68 Proteobacteria; Gammaproteobacteria; Enterobacterales; Enterobacteriaceae; Escherichia; Escherichia coli
MTGDNTLIHSHGINRRDFMKLCAALAATMGLSSKAAAEMAESVTNPQRPPVIWIGAQECT WP_000145410.1
MULTISPECIES: hydrogenase-2 small chain [Proteobacteria] Length: 372\nNumber of predicted TMHs: 1\nExp number of AAs in TMHs: 23.72226\nExp number, first 60 AAs: 0.22038\nTotal prob of N-in: 0.12994\noutside 1 331\nTMhelix 332 354\ninside 355 372

11827 GCF_000408385.1_Esch_coli_KTE68_V1 Escherichia coli KTE68 Proteobacteria; Gammaproteobacteria; Enterobacterales; Enterobacteriaceae; Escherichia; Escherichia coli
MNNEETFYQAMRRQGVTRRSFLKYCSLAATSLGLGAGMAPKIAWALENKPRIPVVWIHGL WP_001058323.1
MULTISPECIES: hydrogenase-1 small chain [Proteobacteria] Length: 372\nNumber of predicted TMHs: 2\nExp number of AAs in TMHs: 37.2608499999999\nExp number, first 60 AAs: 19.52923\nTotal prob of N-in: 0.94351\nPOSSIBLE N-term signal sequence\ninside 1 19\nTMhelix 20 39\noutside 40 325\nTMhelix 326 348\ninside 349 372

11828 GCF_000408405.1_Esch_coli_KTE69_V1 Escherichia coli KTE69 Proteobacteria; Gammaproteobacteria; Enterobacterales; Enterobacteriaceae; Escherichia; Escherichia coli
MTGDNTLIHSHGINRRDFMKLCAALAATMGLSSKAAAEMAESVTNPQRPPVIWIGAQECT WP_000145410.1
MULTISPECIES: hydrogenase-2 small chain [Proteobacteria] Length: 372\nNumber of predicted TMHs: 1\nExp number of AAs in TMHs: 23.72226\nExp number, first 60 AAs: 0.22038\nTotal prob of N-in: 0.12994\noutside 1 331\nTMhelix 332 354\ninside 355 372

11829 GCF_000408405.1_Esch_coli_KTE69_V1 Escherichia coli KTE69 Proteobacteria; Gammaproteobacteria; Enterobacterales; Enterobacteriaceae; Escherichia; Escherichia coli
MNNEETFYQAMRRQGVTRRSFLKYCSLAATSLGLGAGMAPKIAWALENKPRIPVVWIHGL WP_001058323.1
MULTISPECIES: hydrogenase-1 small chain [Proteobacteria] Length: 372\nNumber of predicted TMHs: 2\nExp number of AAs in TMHs: 37.2608499999999\nExp number, first 60 AAs: 19.52923\nTotal prob of N-in:

0.94351\nPOSSIBLE N-term signal sequence\ninside 1 19\nTMhelix 20 39\noutside 40 325\nTMhelix 326 348\ninside 349 372

11830 GCF_000407705.1_Esch_coli_KTE7_V1 Escherichia coli KTE7 Proteobacteria; Gammaproteobacteria; Enterobacterales; Enterobacteriaceae; Escherichia; Escherichia coli
MTGDNTLIHSHGINRRDFMKLCAALAATMGLSSKAAAEMAESVTNPQRPPVIWIGAQECT WP_000145410.1
MULTISPECIES: hydrogenase-2 small chain [Proteobacteria] Length: 372\nNumber of predicted TMHs: 1\nExp number of AAs in TMHs: 23.72226\nExp number, first 60 AAs: 0.22038\nTotal prob of N-in: 0.12994\noutside 1 331\nTMhelix 332 354\ninside 355 372

11831 GCF_000408425.1_Esch_coli_KTE70_V1 Escherichia coli KTE70 Proteobacteria; Gammaproteobacteria; Enterobacterales; Enterobacteriaceae; Escherichia; Escherichia coli
MNNEETFYQAMRRQGVTRRSFLKYCSLAATSLGLGAGMAPKIAWALENKPRIPVVWIHGL WP_001058323.1
MULTISPECIES: hydrogenase-1 small chain [Proteobacteria] Length: 372\nNumber of predicted TMHs: 2\nExp number of AAs in TMHs: 37.2608499999999\nExp number, first 60 AAs: 19.52923\nTotal prob of N-in: 0.94351\nPOSSIBLE N-term signal sequence\ninside 1 19\nTMhelix 20 39\noutside 40 325\nTMhelix 326 348\ninside 349 372

11832 GCF_000408425.1_Esch_coli_KTE70_V1 Escherichia coli KTE70 Proteobacteria; Gammaproteobacteria; Enterobacterales; Enterobacteriaceae; Escherichia; Escherichia coli
MTGDNTLIHSHGINRRDFMKLCAALAATMGLSSKAAAEMAESVTNPQRPPVIWIGAQECT WP_000145410.1
MULTISPECIES: hydrogenase-2 small chain [Proteobacteria] Length: 372\nNumber of predicted TMHs: 1\nExp number of AAs in TMHs: 23.72226\nExp number, first 60 AAs: 0.22038\nTotal prob of N-in: 0.12994\noutside 1 331\nTMhelix 332 354\ninside 355 372

11833 GCF_000408445.1_Esch_coli_KTE71_V1 Escherichia coli KTE71 Proteobacteria; Gammaproteobacteria; Enterobacterales; Enterobacteriaceae; Escherichia; Escherichia coli
MTGDNTLIHSHGINRRDFMKLCAALAATMGLSSKAAAEMAESVTNPQRPPVIWIGAQECT WP_000145410.1
MULTISPECIES: hydrogenase-2 small chain [Proteobacteria] Length: 372\nNumber of predicted TMHs: 1\nExp number of AAs in TMHs: 23.72226\nExp number, first 60 AAs: 0.22038\nTotal prob of N-in: 0.12994\noutside 1 331\nTMhelix 332 354\ninside 355 372

11834 GCF_000408445.1_Esch_coli_KTE71_V1 Escherichia coli KTE71 Proteobacteria; Gammaproteobacteria; Enterobacterales; Enterobacteriaceae; Escherichia; Escherichia coli
MNNEETFYQAMRRQGVTRRSFLKYCSLAATSLGLGAGMAPKIAWALENKPRIPVVWIHGL WP_001058323.1
MULTISPECIES: hydrogenase-1 small chain [Proteobacteria] Length: 372\nNumber of predicted TMHs: 2\nExp number of AAs in TMHs: 37.2608499999999\nExp number, first 60 AAs: 19.52923\nTotal prob of N-in: 0.94351\nPOSSIBLE N-term signal sequence\ninside 1 19\nTMhelix 20 39\noutside 40 325\nTMhelix 326 348\ninside 349 372

11835 GCF_000351665.1_Esch_coli_KTE72_V1 Escherichia coli KTE72 Proteobacteria; Gammaproteobacteria; Enterobacterales; Enterobacteriaceae; Escherichia; Escherichia coli
MTGDNTLIHSHGINRRDFMKLCAALAATMGLSSKAAAEMVESVTNPQRPPVIWIGAQECT WP_000145423.1
hydrogenase 2 small subunit [Escherichia coli] Length: 372\nNumber of predicted TMHs: 1\nExp number of AAs in TMHs: 23.78201\nExp number, first 60 AAs: 0.28355\nTotal prob of N-in: 0.13257\noutside 1 331\nTMhelix 332 354\ninside 355 372

11836 GCF_000351665.1_Esch_coli_KTE72_V1 Escherichia coli KTE72 Proteobacteria; Gammaproteobacteria; Enterobacterales; Enterobacteriaceae; Escherichia; Escherichia coli
MNNEETFYQAMRRQGVTRRSFLKYCSLAATSLGLGAGMAPKIAWALENKPRIPVVWIHGL WP_001058323.1
MULTISPECIES: hydrogenase-1 small chain [Proteobacteria] Length: 372\nNumber of predicted TMHs: 2\nExp number of AAs in TMHs: 37.2608499999999\nExp number, first 60 AAs: 19.52923\nTotal prob of N-in: 0.94351\nPOSSIBLE N-term signal sequence\ninside 1 19\nTMhelix 20 39\noutside 40 325\nTMhelix 326 348\ninside 349 372

11837 GCF_000408465.1_Esch_coli_KTE73_V1 Escherichia coli KTE73 Proteobacteria; Gammaproteobacteria; Enterobacterales; Enterobacteriaceae; Escherichia; Escherichia coli
MNNEETFYQAMRRQGVTRRSFLKYCSLAATSLGLGAGMAPKIAWALENKPRIPVVWIHGL WP_001058323.1
MULTISPECIES: hydrogenase-1 small chain [Proteobacteria] Length: 372\nNumber of predicted TMHs: 2\nExp number of AAs in TMHs: 37.2608499999999\nExp number, first 60 AAs: 19.52923\nTotal prob of N-in:

0.94351\nPOSSIBLE N-term signal sequence\ninside 1 19\nTMhelix 20 39\noutside 40 325\nTMhelix 326 348\ninside 349 372

11838 GCF_000408465.1_Esch_coli_KTE73_V1 Escherichia coli KTE73 Proteobacteria; Gammaproteobacteria; Enterobacterales; Enterobacteriaceae; Escherichia; Escherichia coli MTGDNTLIHSHGINRRDFMKLCAALAATMGLSSKAAAEMAESVTNPQRPPVIWIGAQECT WP_000145410.1 MULTISPECIES: hydrogenase-2 small chain [Proteobacteria] Length: 372\nNumber of predicted TMHs: 1\nExp number of AAs in TMHs: 23.72226\nExp number, first 60 AAs: 0.22038\nTotal prob of N-in: 0.12994\noutside 1 331\nTMhelix 332 354\ninside 355 372

11839 GCF_000408485.1_Esch_coli_KTE74_V1 Escherichia coli KTE74 Proteobacteria; Gammaproteobacteria; Enterobacterales; Enterobacteriaceae; Escherichia; Escherichia coli MNNEETFYQAMRRQGVTRRSFLKYCSLAATSLGLGAGMAPKIAWALENKPRIPVVWIHGL WP_001058323.1 MULTISPECIES: hydrogenase-1 small chain [Proteobacteria] Length: 372\nNumber of predicted TMHs: 2\nExp number of AAs in TMHs: 37.2608499999999\nExp number, first 60 AAs: 19.52923\nTotal prob of N-in: 0.94351\nPOSSIBLE N-term signal sequence\ninside 1 19\nTMhelix 20 39\noutside 40 325\nTMhelix 326 348\ninside 349 372

11840 GCF_000408485.1_Esch_coli_KTE74_V1 Escherichia coli KTE74 Proteobacteria; Gammaproteobacteria; Enterobacterales; Enterobacteriaceae; Escherichia; Escherichia coli MTGDNTLIHSHGINRRDFMKLCAALAATMGLSSKAAAEMAESVTNPQRPPVIWIGAQECT WP_000145410.1 MULTISPECIES: hydrogenase-2 small chain [Proteobacteria] Length: 372\nNumber of predicted TMHs: 1\nExp number of AAs in TMHs: 23.72226\nExp number, first 60 AAs: 0.22038\nTotal prob of N-in: 0.12994\noutside 1 331\nTMhelix 332 354\ninside 355 372

11841 GCF_000351685.1_Esch_coli_KTE75_V1 Escherichia coli KTE75 Proteobacteria; Gammaproteobacteria; Enterobacterales; Enterobacteriaceae; Escherichia; Escherichia coli MNNEETFYQAMRRQGVTRRSFLKYCSLAATSLGLGAGMAPKIAWALENKPRIPVVWIHGL WP_001563066.1 hydrogenase-1 small chain [Escherichia coli] Length: 372\nNumber of predicted TMHs: 2\nExp number of AAs in TMHs: 37.39588\nExp number, first 60 AAs: 19.52908\nTotal prob of N-in: 0.94358\nPOSSIBLE N-term signal sequence\ninside 1 19\nTMhelix 20 39\noutside 40 325\nTMhelix 326 348\ninside 349 372

11842 GCF_000351685.1_Esch_coli_KTE75_V1 Escherichia coli KTE75 Proteobacteria; Gammaproteobacteria; Enterobacterales; Enterobacteriaceae; Escherichia; Escherichia coli MTGDNTLIHSHGINRRDFMKLCAALAATMGLSSKAAAEMAESVTNPQRPPVIWIGAQECT WP_000145410.1 MULTISPECIES: hydrogenase-2 small chain [Proteobacteria] Length: 372\nNumber of predicted TMHs: 1\nExp number of AAs in TMHs: 23.72226\nExp number, first 60 AAs: 0.22038\nTotal prob of N-in: 0.12994\noutside 1 331\nTMhelix 332 354\ninside 355 372

11843 GCF_000351705.1_Esch_coli_KTE76_V1 Escherichia coli KTE76 Proteobacteria; Gammaproteobacteria; Enterobacterales; Enterobacteriaceae; Escherichia; Escherichia coli MNNEETFYQAMRRQGVTRRSFLKYCSLAATSLGLGAGMAPKIAWALENKPRIPVVWIHGL WP_001058323.1 MULTISPECIES: hydrogenase-1 small chain [Proteobacteria] Length: 372\nNumber of predicted TMHs: 2\nExp number of AAs in TMHs: 37.2608499999999\nExp number, first 60 AAs: 19.52923\nTotal prob of N-in: 0.94351\nPOSSIBLE N-term signal sequence\ninside 1 19\nTMhelix 20 39\noutside 40 325\nTMhelix 326 348\ninside 349 372

11844 GCF_000351705.1_Esch_coli_KTE76_V1 Escherichia coli KTE76 Proteobacteria; Gammaproteobacteria; Enterobacterales; Enterobacteriaceae; Escherichia; Escherichia coli MTGDNTLIHSHGINRRDFMKLCAALAATMGLSSKAAAEMAESVTNPQRPPVIWIGAQECT WP_000145410.1 MULTISPECIES: hydrogenase-2 small chain [Proteobacteria] Length: 372\nNumber of predicted TMHs: 1\nExp number of AAs in TMHs: 23.72226\nExp number, first 60 AAs: 0.22038\nTotal prob of N-in: 0.12994\noutside 1 331\nTMhelix 332 354\ninside 355 372

11845 GCF_000351725.1_Esch_coli_KTE77_V1 Escherichia coli KTE77 Proteobacteria; Gammaproteobacteria; Enterobacterales; Enterobacteriaceae; Escherichia; Escherichia coli MNNEETFYQAMRRQGVTRRSFLKYCSLAATSLGLGAGMAPKIAWALENKPRIPVVWIHGL WP_001058323.1 MULTISPECIES: hydrogenase-1 small chain [Proteobacteria] Length: 372\nNumber of predicted TMHs: 2\nExp number of AAs in TMHs: 37.2608499999999\nExp number, first 60 AAs: 19.52923\nTotal prob of N-in: 0.94351\nPOSSIBLE N-term signal sequence\ninside 1 19\nTMhelix 20 39\noutside 40 325\nTMhelix 326 348\ninside 349 372

11846 GCF_000351725.1_Esch_coli_KTE77_V1 Escherichia coli KTE77 Proteobacteria; Gammaproteobacteria; Enterobacterales; Enterobacteriaceae; Escherichia; Escherichia coli MTGDNTLIHSHGINRRDFMKLCAALAATMGLSSKAAAEMAESVTNPQRPPVIWIGAQECT WP_000145410.1 MULTISPECIES: hydrogenase-2 small chain [Proteobacteria] Length: 372\nNumber of predicted TMHs: 1\nExp number of AAs in TMHs: 23.72226\nExp number, first 60 AAs: 0.22038\nTotal prob of N-in: 0.12994\noutside 1 331\nTMhelix 332 354\ninside 355 372

11847 GCF_000352425.1_Esch_coli_KTE78_V1 Escherichia coli KTE78 Proteobacteria; Gammaproteobacteria; Enterobacterales; Enterobacteriaceae; Escherichia; Escherichia coli MNNEETFYQAMRRQGVTRRSFLKYCSLAATSLGLGAGMAPKIAWALENKPRIPVVWIHGL WP_001058323.1 MULTISPECIES: hydrogenase-1 small chain [Proteobacteria] Length: 372\nNumber of predicted TMHs: 2\nExp number of AAs in TMHs: 37.2608499999999\nExp number, first 60 AAs: 19.52923\nTotal prob of N-in: 0.94351\nPOSSIBLE N-term signal sequence\ninside 1 19\nTMhelix 20 39\noutside 40 325\nTMhelix 326 348\ninside 349 372

11848 GCF_000352425.1_Esch_coli_KTE78_V1 Escherichia coli KTE78 Proteobacteria; Gammaproteobacteria; Enterobacterales; Enterobacteriaceae; Escherichia; Escherichia coli MTGDNTLIHSHGINRRDFMKLCAALAATMGLSSKAAAEMAESVTNPQRPPVIWIGAQECT WP_000145410.1 MULTISPECIES: hydrogenase-2 small chain [Proteobacteria] Length: 372\nNumber of predicted TMHs: 1\nExp number of AAs in TMHs: 23.72226\nExp number, first 60 AAs: 0.22038\nTotal prob of N-in: 0.12994\noutside 1 331\nTMhelix 332 354\ninside 355 372

11849 GCF_000352445.1_Esch_coli_KTE79_V1 Escherichia coli KTE79 Proteobacteria; Gammaproteobacteria; Enterobacterales; Enterobacteriaceae; Escherichia; Escherichia coli MTGDNTLIHSHGINRRDFMKLCAALAATMGLSSKAAAEMAESVTNPQRPPVIWIGAQECT WP_000145410.1 MULTISPECIES: hydrogenase-2 small chain [Proteobacteria] Length: 372\nNumber of predicted TMHs: 1\nExp number of AAs in TMHs: 23.72226\nExp number, first 60 AAs: 0.22038\nTotal prob of N-in: 0.12994\noutside 1 331\nTMhelix 332 354\ninside 355 372

11850 GCF_000352445.1_Esch_coli_KTE79_V1 Escherichia coli KTE79 Proteobacteria; Gammaproteobacteria; Enterobacterales; Enterobacteriaceae; Escherichia; Escherichia coli MNNEETFYQAMRRQGVTRRSFLKYCSLAATSLGLGAGMAPKIAWALENKPRIPVVWIHGL WP_001058323.1 MULTISPECIES: hydrogenase-1 small chain [Proteobacteria] Length: 372\nNumber of predicted TMHs: 2\nExp number of AAs in TMHs: 37.2608499999999\nExp number, first 60 AAs: 19.52923\nTotal prob of N-in: 0.94351\nPOSSIBLE N-term signal sequence\ninside 1 19\nTMhelix 20 39\noutside 40 325\nTMhelix 326 348\ninside 349 372

11851 GCF_000352085.1_Esch_coli_KTE8_V1 Escherichia coli KTE8 Proteobacteria; Gammaproteobacteria; Enterobacterales; Enterobacteriaceae; Escherichia; Escherichia coli MTGDNTLIHSHGINRRDFMKLCAALAATMGLSSKAAAEMAESVTNPQRPPVIWIGAQECT WP_000145410.1 MULTISPECIES: hydrogenase-2 small chain [Proteobacteria] Length: 372\nNumber of predicted TMHs: 1\nExp number of AAs in TMHs: 23.72226\nExp number, first 60 AAs: 0.22038\nTotal prob of N-in: 0.12994\noutside 1 331\nTMhelix 332 354\ninside 355 372

11852 GCF_000352085.1_Esch_coli_KTE8_V1 Escherichia coli KTE8 Proteobacteria; Gammaproteobacteria; Enterobacterales; Enterobacteriaceae; Escherichia; Escherichia coli MNNEETFYQAMRRQGVTRRSFLKYCSLAATSLGLGAGMAPKIAWALENKPRIPVVWIHGL WP_001058323.1 MULTISPECIES: hydrogenase-1 small chain [Proteobacteria] Length: 372\nNumber of predicted TMHs: 2\nExp number of AAs in TMHs: 37.2608499999999\nExp number, first 60 AAs: 19.52923\nTotal prob of N-in: 0.94351\nPOSSIBLE N-term signal sequence\ninside 1 19\nTMhelix 20 39\noutside 40 325\nTMhelix 326 348\ninside 349 372

11853 GCF_000351745.1_Esch_coli_KTE80_V1 Escherichia coli KTE80 Proteobacteria; Gammaproteobacteria; Enterobacterales; Enterobacteriaceae; Escherichia; Escherichia coli MNNEETFYQAMRRQGVTRRSFLKYCSLAATSLGLGAGMAPKIAWALENKPRIPVVWIHGL WP_001058323.1 MULTISPECIES: hydrogenase-1 small chain [Proteobacteria] Length: 372\nNumber of predicted TMHs: 2\nExp number of AAs in TMHs: 37.2608499999999\nExp number, first 60 AAs: 19.52923\nTotal prob of N-in: 0.94351\nPOSSIBLE N-term signal sequence\ninside 1 19\nTMhelix 20 39\noutside 40 325\nTMhelix 326 348\ninside 349 372

11854 GCF_000351745.1_Esch_coli_KTE80_V1 Escherichia coli KTE80 Proteobacteria; Gammaproteobacteria; Enterobacterales; Enterobacteriaceae; Escherichia; Escherichia coli
MTGDNTLIHSHGINRRDFMKLCAALAATMGLSSKAAAEMAESVTNPQRPPVIWIGAQECT WP_000145410.1
MULTISPECIES: hydrogenase-2 small chain [Proteobacteria] Length: 372\nNumber of predicted TMHs: 1\nExp number of AAs in TMHs: 23.72226\nExp number, first 60 AAs: 0.22038\nTotal prob of N-in: 0.12994\noutside 1 331\nTMhelix 332 354\ninside 355 372

11855 GCF_000351765.1_Esch_coli_KTE81_V1 Escherichia coli KTE81 Proteobacteria; Gammaproteobacteria; Enterobacterales; Enterobacteriaceae; Escherichia; Escherichia coli
MNNEETFYQAMRRQGVTRRSFLKYCSLAATSLGLGAGMAPKIAWALENKPRIPVVWIHGL WP_001058323.1
MULTISPECIES: hydrogenase-1 small chain [Proteobacteria] Length: 372\nNumber of predicted TMHs: 2\nExp number of AAs in TMHs: 37.2608499999999\nExp number, first 60 AAs: 19.52923\nTotal prob of N-in: 0.94351\nPOSSIBLE N-term signal sequence\ninside 1 19\nTMhelix 20 39\noutside 40 325\nTMhelix 326 348\ninside 349 372

11856 GCF_000351765.1_Esch_coli_KTE81_V1 Escherichia coli KTE81 Proteobacteria; Gammaproteobacteria; Enterobacterales; Enterobacteriaceae; Escherichia; Escherichia coli
MTGDNTLIHSHGINRRDFMKLCAALAATMGLSSKAAAEMAESVTNPQRPPVIWIGAQECT WP_000145410.1
MULTISPECIES: hydrogenase-2 small chain [Proteobacteria] Length: 372\nNumber of predicted TMHs: 1\nExp number of AAs in TMHs: 23.72226\nExp number, first 60 AAs: 0.22038\nTotal prob of N-in: 0.12994\noutside 1 331\nTMhelix 332 354\ninside 355 372

11857 GCF_000326505.1_Esch_coli_KTE82_V1 Escherichia coli KTE82 Proteobacteria; Gammaproteobacteria; Enterobacterales; Enterobacteriaceae; Escherichia; Escherichia coli
MTGDNTLIHSHGINRRDFMKLCAALAATMGLSSKAAAEMAESVTNPQRPPVIWIGAQECT WP_000145410.1
MULTISPECIES: hydrogenase-2 small chain [Proteobacteria] Length: 372\nNumber of predicted TMHs: 1\nExp number of AAs in TMHs: 23.72226\nExp number, first 60 AAs: 0.22038\nTotal prob of N-in: 0.12994\noutside 1 331\nTMhelix 332 354\ninside 355 372

11858 GCF_000326505.1_Esch_coli_KTE82_V1 Escherichia coli KTE82 Proteobacteria; Gammaproteobacteria; Enterobacterales; Enterobacteriaceae; Escherichia; Escherichia coli
MNNEETFYQAMRRQGVTRRSFLKYCSLAATSLGLGAGMAPKIAWALENKPRIPVVWIHGL WP_001058323.1
MULTISPECIES: hydrogenase-1 small chain [Proteobacteria] Length: 372\nNumber of predicted TMHs: 2\nExp number of AAs in TMHs: 37.2608499999999\nExp number, first 60 AAs: 19.52923\nTotal prob of N-in: 0.94351\nPOSSIBLE N-term signal sequence\ninside 1 19\nTMhelix 20 39\noutside 40 325\nTMhelix 326 348\ninside 349 372

11859 GCF_000351785.1_Esch_coli_KTE83_V1 Escherichia coli KTE83 Proteobacteria; Gammaproteobacteria; Enterobacterales; Enterobacteriaceae; Escherichia; Escherichia coli
MNNEETFYQAMRRQGVTRRSFLKYCSLAATSLGLGAGMAPKIAWALENKPRIPVVWIHGL WP_001058323.1
MULTISPECIES: hydrogenase-1 small chain [Proteobacteria] Length: 372\nNumber of predicted TMHs: 2\nExp number of AAs in TMHs: 37.2608499999999\nExp number, first 60 AAs: 19.52923\nTotal prob of N-in: 0.94351\nPOSSIBLE N-term signal sequence\ninside 1 19\nTMhelix 20 39\noutside 40 325\nTMhelix 326 348\ninside 349 372

11860 GCF_000351785.1_Esch_coli_KTE83_V1 Escherichia coli KTE83 Proteobacteria; Gammaproteobacteria; Enterobacterales; Enterobacteriaceae; Escherichia; Escherichia coli
MTGDNTLIHSHGINRRDFMKLCAALAATMGLSSKAAAEMAESVTNPQRPPVIWIGAQECT WP_000145410.1
MULTISPECIES: hydrogenase-2 small chain [Proteobacteria] Length: 372\nNumber of predicted TMHs: 1\nExp number of AAs in TMHs: 23.72226\nExp number, first 60 AAs: 0.22038\nTotal prob of N-in: 0.12994\noutside 1 331\nTMhelix 332 354\ninside 355 372

11861 GCF_000352465.1_Esch_coli_KTE84_V1 Escherichia coli KTE84 Proteobacteria; Gammaproteobacteria; Enterobacterales; Enterobacteriaceae; Escherichia; Escherichia coli
MNNEETFYQAMRRQGVTRRSFLKYCSLAATSLGLGAGMAPKIAWALENKPRIPVVWIHGL WP_001591878.1
hydrogenase-1 small chain [Escherichia coli] Length: 372\nNumber of predicted TMHs: 2\nExp number of AAs in TMHs: 37.25867\nExp number, first 60 AAs: 19.52849\nTotal prob of N-in: 0.94350\nPOSSIBLE N-term signal sequence\ninside 1 19\nTMhelix 20 39\noutside 40 325\nTMhelix 326 348\ninside 349 372

11862 GCF_000352465.1_Esch_coli_KTE84_V1 Escherichia coli KTE84 Proteobacteria; Gammaproteobacteria; Enterobacterales; Enterobacteriaceae; Escherichia; Escherichia coli

MTGDNTLIHSHGINRRDFMKLCAALAATMGLSSKAAAEMAESVTNPQRPPVIWIGAQECT WP_000145410.1 MULTISPECIES: hydrogenase-2 small chain [Proteobacteria] Length: 372\nNumber of predicted TMHs: 1\nExp number of AAs in TMHs: 23.72226\nExp number, first 60 AAs: 0.22038\nTotal prob of N-in: 0.12994\noutside 1 331\nTMhelix 332 354\ninside 355 372

11863 GCF_000326885.1_Esch_coli_KTE85_V1 Escherichia coli KTE85 Proteobacteria; Gammaproteobacteria; Enterobacterales; Enterobacteriaceae; Escherichia; Escherichia coli MTGDNTLIHSHGINRRDFMKLCAALAATMGLSSKAAAEMVESVTNPQRPPVIWIGAQECT WP_000145423.1 hydrogenase 2 small subunit [Escherichia coli] Length: 372\nNumber of predicted TMHs: 1\nExp number of AAs in TMHs: 23.78201\nExp number, first 60 AAs: 0.28355\nTotal prob of N-in: 0.13257\noutside 1 331\nTMhelix 332 354\ninside 355 372

11864 GCF_000326885.1_Esch_coli_KTE85_V1 Escherichia coli KTE85 Proteobacteria; Gammaproteobacteria; Enterobacterales; Enterobacteriaceae; Escherichia; Escherichia coli MNNEETFYQAMRRQGVTRRSFLKYCSLAATSLGLGAGMAPKIAWALENKPRIPVVWIHGL WP_001538532.1 hydrogenase-1 small chain [Escherichia coli] Length: 372\nNumber of predicted TMHs: 2\nExp number of AAs in TMHs: 37.27041\nExp number, first 60 AAs: 19.52927\nTotal prob of N-in: 0.94352\nPOSSIBLE N-term signal sequence\ninside 1 19\nTMhelix 20 39\noutside 40 325\nTMhelix 326 348\ninside 349 372

11865 GCF_000351805.1_Esch_coli_KTE86_V1 Escherichia coli KTE86 Proteobacteria; Gammaproteobacteria; Enterobacterales; Enterobacteriaceae; Escherichia; Escherichia coli MNNEETFYQAMRRQGVTRRSFLKYCSLAATSLGLGAGMAPKIAWALENKPRIPVVWIHGL WP_001058323.1 MULTISPECIES: hydrogenase-1 small chain [Proteobacteria] Length: 372\nNumber of predicted TMHs: 2\nExp number of AAs in TMHs: 37.2608499999999\nExp number, first 60 AAs: 19.52923\nTotal prob of N-in: 0.94351\nPOSSIBLE N-term signal sequence\ninside 1 19\nTMhelix 20 39\noutside 40 325\nTMhelix 326 348\ninside 349 372

11866 GCF_000351805.1_Esch_coli_KTE86_V1 Escherichia coli KTE86 Proteobacteria; Gammaproteobacteria; Enterobacterales; Enterobacteriaceae; Escherichia; Escherichia coli MTGDNTLIHSHGINRRDFMKLCAALAATMGLSSKAAAEMVESVTNPQRPPVIWIGAQECT WP_000145423.1 hydrogenase 2 small subunit [Escherichia coli] Length: 372\nNumber of predicted TMHs: 1\nExp number of AAs in TMHs: 23.78201\nExp number, first 60 AAs: 0.28355\nTotal prob of N-in: 0.13257\noutside 1 331\nTMhelix 332 354\ninside 355 372

11867 GCF_000351825.1_Esch_coli_KTE87_V1 Escherichia coli KTE87 Proteobacteria; Gammaproteobacteria; Enterobacterales; Enterobacteriaceae; Escherichia; Escherichia coli MNNEETFYQAMRRQGVTRRSFLKYCSLAATSLGLGAGMAPKIAWALENKPRIPVVWIHGL WP_001058323.1 MULTISPECIES: hydrogenase-1 small chain [Proteobacteria] Length: 372\nNumber of predicted TMHs: 2\nExp number of AAs in TMHs: 37.2608499999999\nExp number, first 60 AAs: 19.52923\nTotal prob of N-in: 0.94351\nPOSSIBLE N-term signal sequence\ninside 1 19\nTMhelix 20 39\noutside 40 325\nTMhelix 326 348\ninside 349 372

11868 GCF_000351825.1_Esch_coli_KTE87_V1 Escherichia coli KTE87 Proteobacteria; Gammaproteobacteria; Enterobacterales; Enterobacteriaceae; Escherichia; Escherichia coli MTGDNTLIHSHGINRRDFMKLCAALAATMGLSSKAAAEMAESVTNPQRPPVIWIGAQECT WP_000145410.1 MULTISPECIES: hydrogenase-2 small chain [Proteobacteria] Length: 372\nNumber of predicted TMHs: 1\nExp number of AAs in TMHs: 23.72226\nExp number, first 60 AAs: 0.22038\nTotal prob of N-in: 0.12994\noutside 1 331\nTMhelix 332 354\ninside 355 372

11869 GCF_000326625.1_Esch_coli_KTE88_V1 Escherichia coli KTE88 Proteobacteria; Gammaproteobacteria; Enterobacterales; Enterobacteriaceae; Escherichia; Escherichia coli MNNEETFYQAMRRQGVTRRSFLKYCSLAATSLGLGAGMAPKIAWALENKPRIPVVWIHGL WP_001058323.1 MULTISPECIES: hydrogenase-1 small chain [Proteobacteria] Length: 372\nNumber of predicted TMHs: 2\nExp number of AAs in TMHs: 37.2608499999999\nExp number, first 60 AAs: 19.52923\nTotal prob of N-in: 0.94351\nPOSSIBLE N-term signal sequence\ninside 1 19\nTMhelix 20 39\noutside 40 325\nTMhelix 326 348\ninside 349 372

11870 GCF_000326625.1_Esch_coli_KTE88_V1 Escherichia coli KTE88 Proteobacteria; Gammaproteobacteria; Enterobacterales; Enterobacteriaceae; Escherichia; Escherichia coli MTGDNTLIHSHGINRRDFMKLCAALAATMGLSSKAAAEMVESVTNPQRPPVIWIGAQECT WP_000145423.1 hydrogenase 2 small subunit [Escherichia coli] Length: 372\nNumber of predicted TMHs: 1\nExp number of

AAs in TMHs: 23.78201\nExp number, first 60 AAs: 0.28355\nTotal prob of N-in: 0.13257\noutside 1 331\nTMhelix 332 354\ninside 355 372

11871 GCF_000408505.1_Esch_coli_KTE89_V1 Escherichia coli KTE89 Proteobacteria; Gammaproteobacteria; Enterobacterales; Enterobacteriaceae; Escherichia; Escherichia coli MNNEETFYQAMRRQGVTRRSFLKYCSLAATSLGLGAGMAPKIAWALENKPRIPVVWIHGL WP_001058323.1 MULTISPECIES: hydrogenase-1 small chain [Proteobacteria] Length: 372\nNumber of predicted TMHs: 2\nExp number of AAs in TMHs: 37.2608499999999\nExp number, first 60 AAs: 19.52923\nTotal prob of N-in: 0.94351\nPOSSIBLE N-term signal sequence\ninside 1 19\nTMhelix 20 39\noutside 40 325\nTMhelix 326 348\ninside 349 372

11872 GCF_000408505.1_Esch_coli_KTE89_V1 Escherichia coli KTE89 Proteobacteria; Gammaproteobacteria; Enterobacterales; Enterobacteriaceae; Escherichia; Escherichia coli MTGDNTLIHSHGINRRDFMKLCAALAATMGLSSKAAAEMVESVTNPQRPPVIWIGAQECT WP_000145423.1 hydrogenase 2 small subunit [Escherichia coli] Length: 372\nNumber of predicted TMHs: 1\nExp number of AAs in TMHs: 23.78201\nExp number, first 60 AAs: 0.28355\nTotal prob of N-in: 0.13257\noutside 1 331\nTMhelix 332 354\ninside 355 372

11873 GCF_000352105.1_Esch_coli_KTE9_V1 Escherichia coli KTE9 Proteobacteria; Gammaproteobacteria; Enterobacterales; Enterobacteriaceae; Escherichia; Escherichia coli MNNEETFYQAMRRQGVTRRSFLKYCSLAATSLGLGAGMAPKIAWALENKPRIPVVWIHGL WP_001058323.1 MULTISPECIES: hydrogenase-1 small chain [Proteobacteria] Length: 372\nNumber of predicted TMHs: 2\nExp number of AAs in TMHs: 37.2608499999999\nExp number, first 60 AAs: 19.52923\nTotal prob of N-in: 0.94351\nPOSSIBLE N-term signal sequence\ninside 1 19\nTMhelix 20 39\noutside 40 325\nTMhelix 326 348\ninside 349 372

11874 GCF_000352105.1_Esch_coli_KTE9_V1 Escherichia coli KTE9 Proteobacteria; Gammaproteobacteria; Enterobacterales; Enterobacteriaceae; Escherichia; Escherichia coli MTGDNTLIHSHGINRRDFMKLCAALAATMGLSSKAAAEMAESVTNPQRPPVIWIGAQECT WP_000145410.1 MULTISPECIES: hydrogenase-2 small chain [Proteobacteria] Length: 372\nNumber of predicted TMHs: 1\nExp number of AAs in TMHs: 23.72226\nExp number, first 60 AAs: 0.22038\nTotal prob of N-in: 0.12994\noutside 1 331\nTMhelix 332 354\ninside 355 372

11875 GCF_000326645.1_Esch_coli_KTE90_V1 Escherichia coli KTE90 Proteobacteria; Gammaproteobacteria; Enterobacterales; Enterobacteriaceae; Escherichia; Escherichia coli MTGDNTLIHSHGINRRDFMKLCAALAATMGLSSKAAAEMAESVTNPQRPPVIWIGAQECT WP_000145410.1 MULTISPECIES: hydrogenase-2 small chain [Proteobacteria] Length: 372\nNumber of predicted TMHs: 1\nExp number of AAs in TMHs: 23.72226\nExp number, first 60 AAs: 0.22038\nTotal prob of N-in: 0.12994\noutside 1 331\nTMhelix 332 354\ninside 355 372

11876 GCF_000326645.1_Esch_coli_KTE90_V1 Escherichia coli KTE90 Proteobacteria; Gammaproteobacteria; Enterobacterales; Enterobacteriaceae; Escherichia; Escherichia coli MNNEETFYQAMRRQGVTRRSFLKYCSLAATSLGLGAGMAPKIAWALENKPRIPVVWIHGL WP_001295940.1 MULTISPECIES: hydrogenase [Enterobacteriaceae] Length: 372\nNumber of predicted TMHs: 2\nExp number of AAs in TMHs: 37.2608899999999\nExp number, first 60 AAs: 19.52923\nTotal prob of N-in: 0.94351\nPOSSIBLE N-term signal sequence\ninside 1 19\nTMhelix 20 39\noutside 40 325\nTMhelix 326 348\ninside 349 372

11877 GCF_000352485.1_Esch_coli_KTE91_V1 Escherichia coli KTE91 Proteobacteria; Gammaproteobacteria; Enterobacterales; Enterobacteriaceae; Escherichia; Escherichia coli MNNEETFYQAMRRQGVTRRSFLKYCSLAATSLGLGAGMAPKIAWALENKPRIPVVWIHGL WP_001295940.1 MULTISPECIES: hydrogenase [Enterobacteriaceae] Length: 372\nNumber of predicted TMHs: 2\nExp number of AAs in TMHs: 37.2608899999999\nExp number, first 60 AAs: 19.52923\nTotal prob of N-in: 0.94351\nPOSSIBLE N-term signal sequence\ninside 1 19\nTMhelix 20 39\noutside 40 325\nTMhelix 326 348\ninside 349 372

11878 GCF_000352485.1_Esch_coli_KTE91_V1 Escherichia coli KTE91 Proteobacteria; Gammaproteobacteria; Enterobacterales; Enterobacteriaceae; Escherichia; Escherichia coli MTGDNTLIHSHGINRRDFMKLCAALAATMGLSSKAAAEMAESVTNPQRPPVIWIGAQECT WP_000145410.1 MULTISPECIES: hydrogenase-2 small chain [Proteobacteria] Length: 372\nNumber of predicted TMHs:

1\nExp number of AAs in TMHs: 23.72226\nExp number, first 60 AAs: 0.22038\nTotal prob of N-in: 0.12994\noutside 1 331\nTMhelix 332 354\ninside 355 372

11879 GCF_000351845.1_Esch_coli_KTE93_V1 Escherichia coli KTE93 Proteobacteria; Gammaproteobacteria; Enterobacterales; Enterobacteriaceae; Escherichia; Escherichia coli MTGDNTLIHSHGINRRDFMKLCAALAATMGLSSKAAAEMVESVTNPQRPPVIWIGAQECT WP_000145423.1 hydrogenase 2 small subunit [Escherichia coli] Length: 372\nNumber of predicted TMHs: 1\nExp number of AAs in TMHs: 23.78201\nExp number, first 60 AAs: 0.28355\nTotal prob of N-in: 0.13257\noutside 1 331\nTMhelix 332 354\ninside 355 372

11880 GCF_000351845.1_Esch_coli_KTE93_V1 Escherichia coli KTE93 Proteobacteria; Gammaproteobacteria; Enterobacterales; Enterobacteriaceae; Escherichia; Escherichia coli MNNEETFYQAMRRQGVTRRSFLKYCSLAATSLGLGAGMAPKIAWALENKPRIPVVWIHGL WP_001058323.1 MULTISPECIES: hydrogenase-1 small chain [Proteobacteria] Length: 372\nNumber of predicted TMHs: 2\nExp number of AAs in TMHs: 37.2608499999999\nExp number, first 60 AAs: 19.52923\nTotal prob of N-in: 0.94351\nPOSSIBLE N-term signal sequence\ninside 1 19\nTMhelix 20 39\noutside 40 325\nTMhelix 326 348\ninside 349 372

11881 GCF_000326525.1_Esch_coli_KTE94_V1 Escherichia coli KTE94 Proteobacteria; Gammaproteobacteria; Enterobacterales; Enterobacteriaceae; Escherichia; Escherichia coli MTGDNTLIHSHGINRRDFMKLCAALAATMGLSSKAAAEMAESVTNPQRPPVIWIGAQECT WP_000145410.1 MULTISPECIES: hydrogenase-2 small chain [Proteobacteria] Length: 372\nNumber of predicted TMHs: 1\nExp number of AAs in TMHs: 23.72226\nExp number, first 60 AAs: 0.22038\nTotal prob of N-in: 0.12994\noutside 1 331\nTMhelix 332 354\ninside 355 372

11882 GCF_000326525.1_Esch_coli_KTE94_V1 Escherichia coli KTE94 Proteobacteria; Gammaproteobacteria; Enterobacterales; Enterobacteriaceae; Escherichia; Escherichia coli MNNEETFYQAMRRQGVTRRSFLKYCSLAATSLGLGAGMAPKIAWALENKPRIPVVWIHGL WP_001058323.1 MULTISPECIES: hydrogenase-1 small chain [Proteobacteria] Length: 372\nNumber of predicted TMHs: 2\nExp number of AAs in TMHs: 37.2608499999999\nExp number, first 60 AAs: 19.52923\nTotal prob of N-in: 0.94351\nPOSSIBLE N-term signal sequence\ninside 1 19\nTMhelix 20 39\noutside 40 325\nTMhelix 326 348\ninside 349 372

11883 GCF_000326845.1_Esch_coli_KTE95_V1 Escherichia coli KTE95 Proteobacteria; Gammaproteobacteria; Enterobacterales; Enterobacteriaceae; Escherichia; Escherichia coli MTGDNTLIHSHGINRRDFMKLCAALAATMGLSSKAAAEMAESVTNPQRPPVIWIGAQECT WP_000145410.1 MULTISPECIES: hydrogenase-2 small chain [Proteobacteria] Length: 372\nNumber of predicted TMHs: 1\nExp number of AAs in TMHs: 23.72226\nExp number, first 60 AAs: 0.22038\nTotal prob of N-in: 0.12994\noutside 1 331\nTMhelix 332 354\ninside 355 372

11884 GCF_000326845.1_Esch_coli_KTE95_V1 Escherichia coli KTE95 Proteobacteria; Gammaproteobacteria; Enterobacterales; Enterobacteriaceae; Escherichia; Escherichia coli MNNEETFYQAMRRQGVTRRSFLKYCSLAATSLGLGAGMAPKIAWALENKPRIPVVWIHGL WP_001058323.1 MULTISPECIES: hydrogenase-1 small chain [Proteobacteria] Length: 372\nNumber of predicted TMHs: 2\nExp number of AAs in TMHs: 37.2608499999999\nExp number, first 60 AAs: 19.52923\nTotal prob of N-in: 0.94351\nPOSSIBLE N-term signal sequence\ninside 1 19\nTMhelix 20 39\noutside 40 325\nTMhelix 326 348\ninside 349 372

11885 GCF_000326545.1_Esch_coli_KTE97_V1 Escherichia coli KTE97 Proteobacteria; Gammaproteobacteria; Enterobacterales; Enterobacteriaceae; Escherichia; Escherichia coli MNNEETFYQAMRRQGVTRRSFLKYCSLAATSLGLGAGMAPKIAWALENKPRIPVVWIHGL WP_001058323.1 MULTISPECIES: hydrogenase-1 small chain [Proteobacteria] Length: 372\nNumber of predicted TMHs: 2\nExp number of AAs in TMHs: 37.2608499999999\nExp number, first 60 AAs: 19.52923\nTotal prob of N-in: 0.94351\nPOSSIBLE N-term signal sequence\ninside 1 19\nTMhelix 20 39\noutside 40 325\nTMhelix 326 348\ninside 349 372

11886 GCF_000326545.1_Esch_coli_KTE97_V1 Escherichia coli KTE97 Proteobacteria; Gammaproteobacteria; Enterobacterales; Enterobacteriaceae; Escherichia; Escherichia coli MTGDNTLIHSHGINRRDFMKLCAALAATMGLSSKAAAEMAESVTNPQRPPVIWIGAQECT WP_000145410.1 MULTISPECIES: hydrogenase-2 small chain [Proteobacteria] Length: 372\nNumber of predicted TMHs:

1\nExp number of AAs in TMHs: 23.72226\nExp number, first 60 AAs: 0.22038\nTotal prob of N-in: 0.12994\noutside 1 331\nTMhelix 332 354\ninside 355 372

11887 GCF_000408545.1_Esch_coli_KTE98_V1 Escherichia coli KTE98 Proteobacteria; Gammaproteobacteria; Enterobacterales; Enterobacteriaceae; Escherichia; Escherichia coli MTGDNTLIHSHGINRRDFMKLCAALAATMGLSSKAAAEMAESVTNPQRPPVIWIGAQECT WP_000145410.1 MULTISPECIES: hydrogenase-2 small chain [Proteobacteria] Length: 372\nNumber of predicted TMHs: 1\nExp number of AAs in TMHs: 23.72226\nExp number, first 60 AAs: 0.22038\nTotal prob of N-in: 0.12994\noutside 1 331\nTMhelix 332 354\ninside 355 372

11888 GCF_000408545.1_Esch_coli_KTE98_V1 Escherichia coli KTE98 Proteobacteria; Gammaproteobacteria; Enterobacterales; Enterobacteriaceae; Escherichia; Escherichia coli MNNEETFYQAMRRQGVTRRSFLKYCSLAATSLGLGAGMAPKIAWALENKPRIPVVWIHGL WP_001058323.1 MULTISPECIES: hydrogenase-1 small chain [Proteobacteria] Length: 372\nNumber of predicted TMHs: 2\nExp number of AAs in TMHs: 37.2608499999999\nExp number, first 60 AAs: 19.52923\nTotal prob of N-in: 0.94351\nPOSSIBLE N-term signal sequence\ninside 1 19\nTMhelix 20 39\noutside 40 325\nTMhelix 326 348\ninside 349 372

11889 GCF_000326665.1_Esch_coli_KTE99_V1 Escherichia coli KTE99 Proteobacteria; Gammaproteobacteria; Enterobacterales; Enterobacteriaceae; Escherichia; Escherichia coli MNNEETFYQAMRRQGVTRRSFLKYCSLAATSLGLGAGMAPKIAWALENKPRIPVVWIHGL WP_001058323.1 MULTISPECIES: hydrogenase-1 small chain [Proteobacteria] Length: 372\nNumber of predicted TMHs: 2\nExp number of AAs in TMHs: 37.2608499999999\nExp number, first 60 AAs: 19.52923\nTotal prob of N-in: 0.94351\nPOSSIBLE N-term signal sequence\ninside 1 19\nTMhelix 20 39\noutside 40 325\nTMhelix 326 348\ninside 349 372

11890 GCF_000326665.1_Esch_coli_KTE99_V1 Escherichia coli KTE99 Proteobacteria; Gammaproteobacteria; Enterobacterales; Enterobacteriaceae; Escherichia; Escherichia coli MTGDNTLIHSHGINRRDFMKLCAALAATMGLSSKAAAEMAESVTNPQRPPVIWIGAQECT WP_000145418.1 hydrogenase 2 small subunit [Escherichia coli] Length: 372\nNumber of predicted TMHs: 1\nExp number of AAs in TMHs: 23.72313\nExp number, first 60 AAs: 0.22034\nTotal prob of N-in: 0.12997\noutside 1 331\nTMhelix 332 354\ninside 355 372

11891 GCF_000506505.1_WGS_Escherichia_coli_EC10 Escherichia coli LAU-EC10 Proteobacteria; Gammaproteobacteria; Enterobacterales; Enterobacteriaceae; Escherichia; Escherichia coli MNNEETFYQAMRRQGVTRRSFLKYCSLAATSLGLGAGMAPKIAWALENKPRIPVVWIHGL WP_001058323.1 MULTISPECIES: hydrogenase-1 small chain [Proteobacteria] Length: 372\nNumber of predicted TMHs: 2\nExp number of AAs in TMHs: 37.2608499999999\nExp number, first 60 AAs: 19.52923\nTotal prob of N-in: 0.94351\nPOSSIBLE N-term signal sequence\ninside 1 19\nTMhelix 20 39\noutside 40 325\nTMhelix 326 348\ninside 349 372

11892 GCF_000506505.1_WGS_Escherichia_coli_EC10 Escherichia coli LAU-EC10 Proteobacteria; Gammaproteobacteria; Enterobacterales; Enterobacteriaceae; Escherichia; Escherichia coli MTGDNTLIHSHGINRRDFMKLCAALAATMGLSSKAAAEMAESVTNPQRPPVIWIGAQECT WP_000145410.1 MULTISPECIES: hydrogenase-2 small chain [Proteobacteria] Length: 372\nNumber of predicted TMHs: 1\nExp number of AAs in TMHs: 23.72226\nExp number, first 60 AAs: 0.22038\nTotal prob of N-in: 0.12994\noutside 1 331\nTMhelix 332 354\ninside 355 372

11893 GCF_000498795.2_LAU-EC2 Escherichia coli LAU-EC2 Proteobacteria; Gammaproteobacteria; Enterobacterales; Enterobacteriaceae; Escherichia; Escherichia coli MNNEETFYQAMRRQGVTRRSFLKYCSLAATSLGLGAGMAPKIAWALENKPRIPVVWIHGL WP_001295940.1 MULTISPECIES: hydrogenase [Enterobacteriaceae] Length: 372\nNumber of predicted TMHs: 2\nExp number of AAs in TMHs: 37.2608899999999\nExp number, first 60 AAs: 19.52923\nTotal prob of N-in: 0.94351\nPOSSIBLE N-term signal sequence\ninside 1 19\nTMhelix 20 39\noutside 40 325\nTMhelix 326 348\ninside 349 372

11894 GCF_000498795.2_LAU-EC2 Escherichia coli LAU-EC2 Proteobacteria; Gammaproteobacteria; Enterobacterales; Enterobacteriaceae; Escherichia; Escherichia coli MTGDNTLIHSHGINRRDFMKLCAALAATMGLSSKAAAEMVESVTNPQRPPVIWIGAQECT WP_000145423.1 hydrogenase 2 small subunit [Escherichia coli] Length: 372\nNumber of predicted TMHs: 1\nExp number of

AAs in TMHs: 23.78201\nExp number, first 60 AAs:  0.28355\nTotal prob of N-in:   0.13257\noutside   1   331\nTMhelix   332   354\ninside   355   372

11895 GCF_000498815.2_EC3 Escherichia coli LAU-EC3 Proteobacteria; Gammaproteobacteria; Enterobacterales; Enterobacteriaceae; Escherichia; Escherichia coli
MNNEETFYQAMRRQGVTRRSFLKYCSLAATSLGLGAGMAPKIAWALENKPRIPVVWIHGL WP_001295940.1
MULTISPECIES: hydrogenase [Enterobacteriaceae] Length: 372\nNumber of predicted TMHs:  2\nExp number of AAs in TMHs: 37.2608899999999\nExp number, first 60 AAs:  19.52923\nTotal prob of N-in:   0.94351\nPOSSIBLE N-term signal sequence\ninside   1   19\nTMhelix   20   39\noutside   40   325\nTMhelix   326   348\ninside   349   372

11896 GCF_000498815.2_EC3 Escherichia coli LAU-EC3 Proteobacteria; Gammaproteobacteria; Enterobacterales; Enterobacteriaceae; Escherichia; Escherichia coli
MTGDNTLIHSHGINRRDFMKLCAALAATMGLSSKAAAEMVESVTNPQRPPVIWIGAQECT WP_000145423.1
hydrogenase 2 small subunit [Escherichia coli] Length: 372\nNumber of predicted TMHs:  1\nExp number of AAs in TMHs: 23.78201\nExp number, first 60 AAs:  0.28355\nTotal prob of N-in:   0.13257\noutside   1   331\nTMhelix   332   354\ninside   355   372

11897 GCF_000498835.2_LAU-EC4 Escherichia coli LAU-EC4 Proteobacteria; Gammaproteobacteria; Enterobacterales; Enterobacteriaceae; Escherichia; Escherichia coli
MNNEETFYQAMRRQGVTRRSFLKYCSLAATSLGLGAGMAPKIAWALENKPRIPVVWIHGL WP_001058323.1
MULTISPECIES: hydrogenase-1 small chain [Proteobacteria] Length: 372\nNumber of predicted TMHs:  2\nExp number of AAs in TMHs: 37.2608499999999\nExp number, first 60 AAs:  19.52923\nTotal prob of N-in:   0.94351\nPOSSIBLE N-term signal sequence\ninside   1   19\nTMhelix   20   39\noutside   40   325\nTMhelix   326   348\ninside   349   372

11898 GCF_000498835.2_LAU-EC4 Escherichia coli LAU-EC4 Proteobacteria; Gammaproteobacteria; Enterobacterales; Enterobacteriaceae; Escherichia; Escherichia coli
MTGDNTLIHSHGINRRDFMKLCAALAATMGLSSKAAAEMAESVTNPQRPPVIWIGAQECT WP_000145410.1
MULTISPECIES: hydrogenase-2 small chain [Proteobacteria] Length: 372\nNumber of predicted TMHs:  1\nExp number of AAs in TMHs: 23.72226\nExp number, first 60 AAs:  0.22038\nTotal prob of N-in:   0.12994\noutside   1   331\nTMhelix   332   354\ninside   355   372

11899 GCF_000498875.2_LAU-EC5 Escherichia coli LAU-EC5 Proteobacteria; Gammaproteobacteria; Enterobacterales; Enterobacteriaceae; Escherichia; Escherichia coli
MTGDNTLIHSHGINRRDFMKLCAALAATMGLSSKAAAEMAESVTNPQRPPVIWIGAQECT WP_000145410.1
MULTISPECIES: hydrogenase-2 small chain [Proteobacteria] Length: 372\nNumber of predicted TMHs:  1\nExp number of AAs in TMHs: 23.72226\nExp number, first 60 AAs:  0.22038\nTotal prob of N-in:   0.12994\noutside   1   331\nTMhelix   332   354\ninside   355   372

11900 GCF_000498875.2_LAU-EC5 Escherichia coli LAU-EC5 Proteobacteria; Gammaproteobacteria; Enterobacterales; Enterobacteriaceae; Escherichia; Escherichia coli
MNNEETFYQAMRRQGVTRRSFLKYCSLAATSLGLGAGMAPKIAWALENKPRIPVVWIHGL WP_001058323.1
MULTISPECIES: hydrogenase-1 small chain [Proteobacteria] Length: 372\nNumber of predicted TMHs:  2\nExp number of AAs in TMHs: 37.2608499999999\nExp number, first 60 AAs:  19.52923\nTotal prob of N-in:   0.94351\nPOSSIBLE N-term signal sequence\ninside   1   19\nTMhelix   20   39\noutside   40   325\nTMhelix   326   348\ninside   349   372

11901 GCF_000506445.2_WGS_Escherichia_coli_EC6 Escherichia coli LAU-EC6 Proteobacteria; Gammaproteobacteria; Enterobacterales; Enterobacteriaceae; Escherichia; Escherichia coli
MNNEETFYQAMRRQGVTRRSFLKYCSLAATSLGLGAGMAPKIAWALENKPRIPVVWIHGL WP_001058323.1
MULTISPECIES: hydrogenase-1 small chain [Proteobacteria] Length: 372\nNumber of predicted TMHs:  2\nExp number of AAs in TMHs: 37.2608499999999\nExp number, first 60 AAs:  19.52923\nTotal prob of N-in:   0.94351\nPOSSIBLE N-term signal sequence\ninside   1   19\nTMhelix   20   39\noutside   40   325\nTMhelix   326   348\ninside   349   372

11902 GCF_000506445.2_WGS_Escherichia_coli_EC6 Escherichia coli LAU-EC6 Proteobacteria; Gammaproteobacteria; Enterobacterales; Enterobacteriaceae; Escherichia; Escherichia coli
MTGDNTLIHSHGINRRDFMKLCAALAATMGLSSKAAAEMAESVTNPQRPPVIWIGAQECT WP_000145410.1
MULTISPECIES: hydrogenase-2 small chain [Proteobacteria] Length: 372\nNumber of predicted TMHs:

1\nExp number of AAs in TMHs: 23.72226\nExp number, first 60 AAs:  0.22038\nTotal prob of N-in:   0.12994\noutside   1   331\nTMhelix   332   354\ninside   355   372

11903 GCF_000506465.2_WGS_Escherichia_coli_EC7 Escherichia coli LAU-EC7 Proteobacteria; Gammaproteobacteria; Enterobacterales; Enterobacteriaceae; Escherichia; Escherichia coli MNNEETFYQAMRRQGVTRRSFLKYCSLAATSLGLGAGMAPKIAWALENKPRIPVVWIHGL WP_001295940.1 MULTISPECIES: hydrogenase [Enterobacteriaceae] Length: 372\nNumber of predicted TMHs:  2\nExp number of AAs in TMHs: 37.2608899999999\nExp number, first 60 AAs:  19.52923\nTotal prob of N-in:   0.94351\nPOSSIBLE N-term signal sequence\ninside   1   19\nTMhelix   20   39\noutside   40   325\nTMhelix   326   348\ninside   349   372

11904 GCF_000506465.2_WGS_Escherichia_coli_EC7 Escherichia coli LAU-EC7 Proteobacteria; Gammaproteobacteria; Enterobacterales; Enterobacteriaceae; Escherichia; Escherichia coli MTGDNTLIHSHGINRRDFMKLCAALAATMGLSSKAAAEMVESVTNPQRPPVIWIGAQECT WP_000145423.1 hydrogenase 2 small subunit [Escherichia coli] Length: 372\nNumber of predicted TMHs:  1\nExp number of AAs in TMHs: 23.78201\nExp number, first 60 AAs:  0.28355\nTotal prob of N-in:   0.13257\noutside   1   331\nTMhelix   332   354\ninside   355   372

11905 GCF_000506585.2_WGS_Escherichia_coli_EC8 Escherichia coli LAU-EC8 Proteobacteria; Gammaproteobacteria; Enterobacterales; Enterobacteriaceae; Escherichia; Escherichia coli MNNEETFYQAMRRQGVTRRSFLKYCSLAATSLGLGAGMAPKIAWALENKPRIPVVWIHGL WP_001058323.1 MULTISPECIES: hydrogenase-1 small chain [Proteobacteria] Length: 372\nNumber of predicted TMHs:  2\nExp number of AAs in TMHs: 37.2608499999999\nExp number, first 60 AAs:  19.52923\nTotal prob of N-in:   0.94351\nPOSSIBLE N-term signal sequence\ninside   1   19\nTMhelix   20   39\noutside   40   325\nTMhelix   326   348\ninside   349   372

11906 GCF_000506585.2_WGS_Escherichia_coli_EC8 Escherichia coli LAU-EC8 Proteobacteria; Gammaproteobacteria; Enterobacterales; Enterobacteriaceae; Escherichia; Escherichia coli MTGDNTLIHSHGINRRDFMKLCAALAATMGLSSKAAAEMAESVTNPQRPPVIWIGAQECT WP_000145410.1 MULTISPECIES: hydrogenase-2 small chain [Proteobacteria] Length: 372\nNumber of predicted TMHs:  1\nExp number of AAs in TMHs: 23.72226\nExp number, first 60 AAs:  0.22038\nTotal prob of N-in:   0.12994\noutside   1   331\nTMhelix   332   354\ninside   355   372

11907 GCF_000506485.2_WGS_Escherichia_coli_EC9 Escherichia coli LAU-EC9 Proteobacteria; Gammaproteobacteria; Enterobacterales; Enterobacteriaceae; Escherichia; Escherichia coli MTGDNTLIHSHGINRRDFMKLCAALAATMGLSSKAAAEMAESVTNPQRPPVIWIGAQECT WP_000145410.1 MULTISPECIES: hydrogenase-2 small chain [Proteobacteria] Length: 372\nNumber of predicted TMHs:  1\nExp number of AAs in TMHs: 23.72226\nExp number, first 60 AAs:  0.22038\nTotal prob of N-in:   0.12994\noutside   1   331\nTMhelix   332   354\ninside   355   372

11908 GCF_000506485.2_WGS_Escherichia_coli_EC9 Escherichia coli LAU-EC9 Proteobacteria; Gammaproteobacteria; Enterobacterales; Enterobacteriaceae; Escherichia; Escherichia coli MNNEETFYQAMRRQGVTRRSFLKYCSLAATSLGLGAGMAPKIAWALENKPRIPVVWIHGL WP_001058323.1 MULTISPECIES: hydrogenase-1 small chain [Proteobacteria] Length: 372\nNumber of predicted TMHs:  2\nExp number of AAs in TMHs: 37.2608499999999\nExp number, first 60 AAs:  19.52923\nTotal prob of N-in:   0.94351\nPOSSIBLE N-term signal sequence\ninside   1   19\nTMhelix   20   39\noutside   40   325\nTMhelix   326   348\ninside   349   372

11909 GCF_000259695.1_ASM25969v1 Escherichia coli LCT-EC106 Proteobacteria; Gammaproteobacteria; Enterobacterales; Enterobacteriaceae; Escherichia; Escherichia coli MTGDNTLIHSHGINRRDFMKLCAALAATMGLSSKAAAEMAESVTNPQRPPVIWIGAQECT WP_000145410.1 MULTISPECIES: hydrogenase-2 small chain [Proteobacteria] Length: 372\nNumber of predicted TMHs:  1\nExp number of AAs in TMHs: 23.72226\nExp number, first 60 AAs:  0.22038\nTotal prob of N-in:   0.12994\noutside   1   331\nTMhelix   332   354\ninside   355   372

11910 GCF_000259695.1_ASM25969v1 Escherichia coli LCT-EC106 Proteobacteria; Gammaproteobacteria; Enterobacterales; Enterobacteriaceae; Escherichia; Escherichia coli MNNEETFYQAMRRQGVTRRSFLKYCSLAATSLGLGAGMAPKIAWALENKPRIPVVWIHGL WP_001058323.1 MULTISPECIES: hydrogenase-1 small chain [Proteobacteria] Length: 372\nNumber of predicted TMHs:  2\nExp number of AAs in TMHs: 37.2608499999999\nExp number, first 60 AAs:  19.52923\nTotal prob of N-in:

0.94351\nPOSSIBLE N-term signal sequence\ninside 1 19\nTMhelix 20 39\noutside 40 325\nTMhelix 326 348\ninside 349 372

11911 GCF_000331615.1_genome_assembly_of_LCT-EC52_strain_of_E.coli Escherichia coli LCT-EC52 Proteobacteria; Gammaproteobacteria; Enterobacterales; Enterobacteriaceae; Escherichia; Escherichia coli MNNEETFYQAMRRQGVTRRSFLKYCSLAATSLGLGAGMAPKIAWALENKPRIPVVWIHGL WP_001058323.1 MULTISPECIES: hydrogenase-1 small chain [Proteobacteria] Length: 372\nNumber of predicted TMHs: 2\nExp number of AAs in TMHs: 37.2608499999999\nExp number, first 60 AAs: 19.52923\nTotal prob of N-in: 0.94351\nPOSSIBLE N-term signal sequence\ninside 1 19\nTMhelix 20 39\noutside 40 325\nTMhelix 326 348\ninside 349 372

11912 GCF_000331615.1_genome_assembly_of_LCT-EC52_strain_of_E.coli Escherichia coli LCT-EC52 Proteobacteria; Gammaproteobacteria; Enterobacterales; Enterobacteriaceae; Escherichia; Escherichia coli MTGDNTLIHSHGINRRDFMKLCAALAATMGLSSKAAAEMAESVTNPQRPPVIWIGAQECT WP_000145410.1 MULTISPECIES: hydrogenase-2 small chain [Proteobacteria] Length: 372\nNumber of predicted TMHs: 1\nExp number of AAs in TMHs: 23.72226\nExp number, first 60 AAs: 0.22038\nTotal prob of N-in: 0.12994\noutside 1 331\nTMhelix 332 354\ninside 355 372

11913 GCF_000317395.1_genome_assembly_of_LCT-EC59_strain_of_E.coli Escherichia coli LCT-EC59 Proteobacteria; Gammaproteobacteria; Enterobacterales; Enterobacteriaceae; Escherichia; Escherichia coli MTGDNTLIHSHGINRRDFMKLCAALAATMGLSSKAAAEMAESVTNPQRPPVIWIGAQECT WP_000145410.1 MULTISPECIES: hydrogenase-2 small chain [Proteobacteria] Length: 372\nNumber of predicted TMHs: 1\nExp number of AAs in TMHs: 23.72226\nExp number, first 60 AAs: 0.22038\nTotal prob of N-in: 0.12994\noutside 1 331\nTMhelix 332 354\ninside 355 372

11914 GCF_000317395.1_genome_assembly_of_LCT-EC59_strain_of_E.coli Escherichia coli LCT-EC59 Proteobacteria; Gammaproteobacteria; Enterobacterales; Enterobacteriaceae; Escherichia; Escherichia coli MNNEETFYQAMRRQGVTRRSFLKYCSLAATSLGLGAGMAPKIAWALENKPRIPVVWIHGL WP_001058323.1 MULTISPECIES: hydrogenase-1 small chain [Proteobacteria] Length: 372\nNumber of predicted TMHs: 2\nExp number of AAs in TMHs: 37.2608499999999\nExp number, first 60 AAs: 19.52923\nTotal prob of N-in: 0.94351\nPOSSIBLE N-term signal sequence\ninside 1 19\nTMhelix 20 39\noutside 40 325\nTMhelix 326 348\ninside 349 372

11915 GCF_000284495.1_ASM28449v1 Escherichia coli LF82 Proteobacteria; Gammaproteobacteria; Enterobacterales; Enterobacteriaceae; Escherichia; Escherichia coli MNNEETFYQAMRRQGVTRRSFLKYCSLAATSLGLGAGMAPKIAWALENKPRIPVVWIHGL WP_001058323.1 MULTISPECIES: hydrogenase-1 small chain [Proteobacteria] Length: 372\nNumber of predicted TMHs: 2\nExp number of AAs in TMHs: 37.2608499999999\nExp number, first 60 AAs: 19.52923\nTotal prob of N-in: 0.94351\nPOSSIBLE N-term signal sequence\ninside 1 19\nTMhelix 20 39\noutside 40 325\nTMhelix 326 348\ninside 349 372

11916 GCF_000284495.1_ASM28449v1 Escherichia coli LF82 Proteobacteria; Gammaproteobacteria; Enterobacterales; Enterobacteriaceae; Escherichia; Escherichia coli MTGDNTLIHSHGINRRDFMKLCAALAATMGLSSKAAAEMAESVTNPQRPPVIWIGAQECT WP_000145418.1 hydrogenase 2 small subunit [Escherichia coli] Length: 372\nNumber of predicted TMHs: 1\nExp number of AAs in TMHs: 23.72313\nExp number, first 60 AAs: 0.22034\nTotal prob of N-in: 0.12997\noutside 1 331\nTMhelix 332 354\ninside 355 372

11917 GCF_000586515.1_E.coliLR09v1 Escherichia coli LR09 Proteobacteria; Gammaproteobacteria; Enterobacterales; Enterobacteriaceae; Escherichia; Escherichia coli MNNEETFYQAMRRQGVTRRSFLKYCSLAATSLGLGAGMAPKIAWALENKPRIPVVWIHGL WP_001058323.1 MULTISPECIES: hydrogenase-1 small chain [Proteobacteria] Length: 372\nNumber of predicted TMHs: 2\nExp number of AAs in TMHs: 37.2608499999999\nExp number, first 60 AAs: 19.52923\nTotal prob of N-in: 0.94351\nPOSSIBLE N-term signal sequence\ninside 1 19\nTMhelix 20 39\noutside 40 325\nTMhelix 326 348\ninside 349 372

11918 GCF_000586515.1_E.coliLR09v1 Escherichia coli LR09 Proteobacteria; Gammaproteobacteria; Enterobacterales; Enterobacteriaceae; Escherichia; Escherichia coli MTGDNTLIHSHGINRRDFMKLCAALAATMGLSSKAAAEMAESVTNPQRPPVIWIGAQECT WP_000145410.1 MULTISPECIES: hydrogenase-2 small chain [Proteobacteria] Length: 372\nNumber of predicted TMHs:

1\nExp number of AAs in TMHs: 23.72226\nExp number, first 60 AAs: 0.22038\nTotal prob of N-in: 0.12994\noutside 1 331\nTMhelix 332 354\ninside 355 372

11919 GCF_000468515.1_ASM46851v1 Escherichia coli LY180 Proteobacteria; Gammaproteobacteria; Enterobacterales; Enterobacteriaceae; Escherichia; Escherichia coli
MTGDNTLIHSHGINRRDFMKLCAALAATMGLSSKAAAEMAESVTNPQRPPVIWIGAQECT WP_000145410.1
MULTISPECIES: hydrogenase-2 small chain [Proteobacteria] Length: 372\nNumber of predicted TMHs: 1\nExp number of AAs in TMHs: 23.72226\nExp number, first 60 AAs: 0.22038\nTotal prob of N-in: 0.12994\noutside 1 331\nTMhelix 332 354\ninside 355 372

11920 GCF_000468515.1_ASM46851v1 Escherichia coli LY180 Proteobacteria; Gammaproteobacteria; Enterobacterales; Enterobacteriaceae; Escherichia; Escherichia coli
MNNEETFYQAMRRQGVTRRSFLKYCSLAATSLGLGAGMAPKIAWALENKPRIPVVWIHGL WP_001349947.1
hydrogenase [Escherichia coli] Length: 372\nNumber of predicted TMHs: 2\nExp number of AAs in TMHs: 37.2615299999999\nExp number, first 60 AAs: 19.52923\nTotal prob of N-in: 0.94351\nPOSSIBLE N-term signal sequence\ninside 1 19\nTMhelix 20 39\noutside 40 325\nTMhelix 326 348\ninside 349 372

11921 GCF_002109605.1_ASM210960v1 Escherichia coli M056 Proteobacteria; Gammaproteobacteria; Enterobacterales; Enterobacteriaceae; Escherichia; Escherichia coli
MNNEETFYQAMRRQGVTRRSFLKYCSLAATSLGLGAGMAPKIAWALENKPRIPVVWIHGL WP_001058323.1
MULTISPECIES: hydrogenase-1 small chain [Proteobacteria] Length: 372\nNumber of predicted TMHs: 2\nExp number of AAs in TMHs: 37.2608499999999\nExp number, first 60 AAs: 19.52923\nTotal prob of N-in: 0.94351\nPOSSIBLE N-term signal sequence\ninside 1 19\nTMhelix 20 39\noutside 40 325\nTMhelix 326 348\ninside 349 372

11922 GCF_002109605.1_ASM210960v1 Escherichia coli M056 Proteobacteria; Gammaproteobacteria; Enterobacterales; Enterobacteriaceae; Escherichia; Escherichia coli
MTGDNTLIHSHGINRRDFMKLCAALAATMGLSSKAAAEMAESVTNPQRPPVIWIGAQECT WP_000145410.1
MULTISPECIES: hydrogenase-2 small chain [Proteobacteria] Length: 372\nNumber of predicted TMHs: 1\nExp number of AAs in TMHs: 23.72226\nExp number, first 60 AAs: 0.22038\nTotal prob of N-in: 0.12994\noutside 1 331\nTMhelix 332 354\ninside 355 372

11923 GCF_000466625.1_E_coliM1-1.0 Escherichia coli M1 Proteobacteria; Gammaproteobacteria; Enterobacterales; Enterobacteriaceae; Escherichia; Escherichia coli
MTGDNTLIHSHGINRRDFMKLCAALAATMGLSSKAAAEMAESVTNPQRPPVIWIGAQECT WP_000145410.1
MULTISPECIES: hydrogenase-2 small chain [Proteobacteria] Length: 372\nNumber of predicted TMHs: 1\nExp number of AAs in TMHs: 23.72226\nExp number, first 60 AAs: 0.22038\nTotal prob of N-in: 0.12994\noutside 1 331\nTMhelix 332 354\ninside 355 372

11924 GCF_000466625.1_E_coliM1-1.0 Escherichia coli M1 Proteobacteria; Gammaproteobacteria; Enterobacterales; Enterobacteriaceae; Escherichia; Escherichia coli
MNNEETFYQAMRRQGVTRRSFLKYCSLAATSLGLGAGMAPKIAWALENKPRIPVVWIHGL WP_001301613.1
hydrogenase [Escherichia coli] Length: 372\nNumber of predicted TMHs: 2\nExp number of AAs in TMHs: 37.46598\nExp number, first 60 AAs: 19.53139\nTotal prob of N-in: 0.94364\nPOSSIBLE N-term signal sequence\ninside 1 19\nTMhelix 20 39\noutside 40 325\nTMhelix 326 348\ninside 349 372

11925 GCF_000471225.1_E_coliM10-1.0 Escherichia coli M10 Proteobacteria; Gammaproteobacteria; Enterobacterales; Enterobacteriaceae; Escherichia; Escherichia coli
MTGDNTLIHSHGINRRDFMKLCAALAATMGLSSKAAAEMAESVTNPQRPPVIWIGAQECT WP_000145410.1
MULTISPECIES: hydrogenase-2 small chain [Proteobacteria] Length: 372\nNumber of predicted TMHs: 1\nExp number of AAs in TMHs: 23.72226\nExp number, first 60 AAs: 0.22038\nTotal prob of N-in: 0.12994\noutside 1 331\nTMhelix 332 354\ninside 355 372

11926 GCF_000471225.1_E_coliM10-1.0 Escherichia coli M10 Proteobacteria; Gammaproteobacteria; Enterobacterales; Enterobacteriaceae; Escherichia; Escherichia coli
MNNEETFYQAMRRQGVTRRSFLKYCSLAATSLGLGAGMAPKIAWALENKPRIPVVWIHGL WP_024253159.1
hydrogenase [Escherichia coli] Length: 372\nNumber of predicted TMHs: 2\nExp number of AAs in TMHs: 37.56907\nExp number, first 60 AAs: 19.53147\nTotal prob of N-in: 0.94370\nPOSSIBLE N-term signal sequence\ninside 1 19\nTMhelix 20 39\noutside 40 325\nTMhelix 326 348\ninside 349 372

11927 GCF_000471245.1_E_coliM11-1.0 Escherichia coli M11 Proteobacteria; Gammaproteobacteria; Enterobacterales; Enterobacteriaceae; Escherichia; Escherichia coli
MTGDNTLIHSHGINRRDFMKLCAALAATMGLSSKAAAEMAESVTNPQRPPVIWIGAQECT WP_000145410.1
MULTISPECIES: hydrogenase-2 small chain [Proteobacteria] Length: 372\nNumber of predicted TMHs: 1\nExp number of AAs in TMHs: 23.72226\nExp number, first 60 AAs: 0.22038\nTotal prob of N-in: 0.12994\noutside 1 331\nTMhelix 332 354\ninside 355 372

11928 GCF_000471245.1_E_coliM11-1.0 Escherichia coli M11 Proteobacteria; Gammaproteobacteria; Enterobacterales; Enterobacteriaceae; Escherichia; Escherichia coli
MNNEETFYQAMRRQGVTRRSFLKYCSLAATSLGLGAGMAPKIAWALENKPRIPVVWIHGL WP_001301613.1
hydrogenase [Escherichia coli] Length: 372\nNumber of predicted TMHs: 2\nExp number of AAs in TMHs: 37.46598\nExp number, first 60 AAs: 19.53139\nTotal prob of N-in: 0.94364\nPOSSIBLE N-term signal sequence\ninside 1 19\nTMhelix 20 39\noutside 40 325\nTMhelix 326 348\ninside 349 372

11929 GCF_001186315.1_Esch_coli_M114_V2 Escherichia coli M114 Proteobacteria; Gammaproteobacteria; Enterobacterales; Enterobacteriaceae; Escherichia; Escherichia coli
MNNEETFYQAMRRQGVTRRSFLKYCSLAATSLGLGAGMAPKIAWALENKPRIPVVWIHGL WP_001058323.1
MULTISPECIES: hydrogenase-1 small chain [Proteobacteria] Length: 372\nNumber of predicted TMHs: 2\nExp number of AAs in TMHs: 37.2608499999999\nExp number, first 60 AAs: 19.52923\nTotal prob of N-in: 0.94351\nPOSSIBLE N-term signal sequence\ninside 1 19\nTMhelix 20 39\noutside 40 325\nTMhelix 326 348\ninside 349 372

11930 GCF_001186315.1_Esch_coli_M114_V2 Escherichia coli M114 Proteobacteria; Gammaproteobacteria; Enterobacterales; Enterobacteriaceae; Escherichia; Escherichia coli
MTGDNTLIHSHGINRRDFMKLCAALAATMGLSSKAAAEMAESVTNPQRPPVIWIGAQECT WP_000145410.1
MULTISPECIES: hydrogenase-2 small chain [Proteobacteria] Length: 372\nNumber of predicted TMHs: 1\nExp number of AAs in TMHs: 23.72226\nExp number, first 60 AAs: 0.22038\nTotal prob of N-in: 0.12994\noutside 1 331\nTMhelix 332 354\ninside 355 372

11931 GCF_000471265.1_E_coliM12-1.0 Escherichia coli M12 Proteobacteria; Gammaproteobacteria; Enterobacterales; Enterobacteriaceae; Escherichia; Escherichia coli
MTGDNTLIHSHGINRRDFMKLCAALAATMGLSSKAAAEMAESVTNPQRPPVIWIGAQECT WP_000145410.1
MULTISPECIES: hydrogenase-2 small chain [Proteobacteria] Length: 372\nNumber of predicted TMHs: 1\nExp number of AAs in TMHs: 23.72226\nExp number, first 60 AAs: 0.22038\nTotal prob of N-in: 0.12994\noutside 1 331\nTMhelix 332 354\ninside 355 372

11932 GCF_000471265.1_E_coliM12-1.0 Escherichia coli M12 Proteobacteria; Gammaproteobacteria; Enterobacterales; Enterobacteriaceae; Escherichia; Escherichia coli
MNNEETFYQAMRRQGVTRRSFLKYCSLAATSLGLGAGMAPKIAWALENKPRIPVVWIHGL WP_001301613.1
hydrogenase [Escherichia coli] Length: 372\nNumber of predicted TMHs: 2\nExp number of AAs in TMHs: 37.46598\nExp number, first 60 AAs: 19.53139\nTotal prob of N-in: 0.94364\nPOSSIBLE N-term signal sequence\ninside 1 19\nTMhelix 20 39\noutside 40 325\nTMhelix 326 348\ninside 349 372

11933 GCF_000471285.1_E_coliM13-1.0 Escherichia coli M13 Proteobacteria; Gammaproteobacteria; Enterobacterales; Enterobacteriaceae; Escherichia; Escherichia coli
MNNEETFYQAMRRQGVTRRSFLKYCSLAATSLGLGAGMAPKIAWALENKPRIPVVWIHGL WP_001301613.1
hydrogenase [Escherichia coli] Length: 372\nNumber of predicted TMHs: 2\nExp number of AAs in TMHs: 37.46598\nExp number, first 60 AAs: 19.53139\nTotal prob of N-in: 0.94364\nPOSSIBLE N-term signal sequence\ninside 1 19\nTMhelix 20 39\noutside 40 325\nTMhelix 326 348\ninside 349 372

11934 GCF_000471285.1_E_coliM13-1.0 Escherichia coli M13 Proteobacteria; Gammaproteobacteria; Enterobacterales; Enterobacteriaceae; Escherichia; Escherichia coli
MTGDNTLIHSHGINRRDFMKLCAALAATMGLSSKAAAEMAESVTNPQRPPVIWIGAQECT WP_000145410.1
MULTISPECIES: hydrogenase-2 small chain [Proteobacteria] Length: 372\nNumber of predicted TMHs: 1\nExp number of AAs in TMHs: 23.72226\nExp number, first 60 AAs: 0.22038\nTotal prob of N-in: 0.12994\noutside 1 331\nTMhelix 332 354\ninside 355 372

11935 GCF_000471305.1_E_coliM14-1.0 Escherichia coli M14 Proteobacteria; Gammaproteobacteria; Enterobacterales; Enterobacteriaceae; Escherichia; Escherichia coli
MNNEETFYQAMRRQGVTRRSFLKYCSLAATSLGLGAGMAPKIAWALENKPRIPVVWIHGL WP_024254239.1
hydrogenase [Escherichia coli] Length: 372\nNumber of predicted TMHs: 2\nExp number of AAs in

TMHs: 37.30566\nExp number, first 60 AAs: 19.53326\nTotal prob of N-in: 0.94355\nPOSSIBLE N-term signal sequence\ninside 1 19\nTMhelix 20 39\noutside 40 325\nTMhelix 326 348\ninside 349 372

11936 GCF_000471305.1_E_coliM14-1.0 Escherichia coli M14 Proteobacteria; Gammaproteobacteria; Enterobacterales; Enterobacteriaceae; Escherichia; Escherichia coli
MTGDNTLIHSHGINRRDFMKLCAALAATMGLSSKAAAEMAESVTNPQRPPVIWIGAQECT WP_000145410.1
MULTISPECIES: hydrogenase-2 small chain [Proteobacteria] Length: 372\nNumber of predicted TMHs: 1\nExp number of AAs in TMHs: 23.72226\nExp number, first 60 AAs: 0.22038\nTotal prob of N-in: 0.12994\noutside 1 331\nTMhelix 332 354\ninside 355 372

11937 GCF_000471325.1_E_coliM15-1.0 Escherichia coli M15 Proteobacteria; Gammaproteobacteria; Enterobacterales; Enterobacteriaceae; Escherichia; Escherichia coli
MTGDNTLIHSHGINRRDFMKLCAALAATMGLSSKAAAEMAESVTNPQRPPVIWIGAQECT WP_000145410.1
MULTISPECIES: hydrogenase-2 small chain [Proteobacteria] Length: 372\nNumber of predicted TMHs: 1\nExp number of AAs in TMHs: 23.72226\nExp number, first 60 AAs: 0.22038\nTotal prob of N-in: 0.12994\noutside 1 331\nTMhelix 332 354\ninside 355 372

11938 GCF_000471325.1_E_coliM15-1.0 Escherichia coli M15 Proteobacteria; Gammaproteobacteria; Enterobacterales; Enterobacteriaceae; Escherichia; Escherichia coli
MNNEETFYQAMRRQGVTRRSFLKYCSLAATSLGLGAGMAPKIAWALENKPRIPVVWIHGL WP_001301613.1
hydrogenase [Escherichia coli] Length: 372\nNumber of predicted TMHs: 2\nExp number of AAs in TMHs: 37.46598\nExp number, first 60 AAs: 19.53139\nTotal prob of N-in: 0.94364\nPOSSIBLE N-term signal sequence\ninside 1 19\nTMhelix 20 39\noutside 40 325\nTMhelix 326 348\ninside 349 372

11939 GCF_000471345.1_E_coliM16-1.0 Escherichia coli M16 Proteobacteria; Gammaproteobacteria; Enterobacterales; Enterobacteriaceae; Escherichia; Escherichia coli
MNNEETFYQAMRRQGVTRRSFLKYCSLAATSLGLGAGMAPKIAWALENKPRIPVVWIHGL WP_001301613.1
hydrogenase [Escherichia coli] Length: 372\nNumber of predicted TMHs: 2\nExp number of AAs in TMHs: 37.46598\nExp number, first 60 AAs: 19.53139\nTotal prob of N-in: 0.94364\nPOSSIBLE N-term signal sequence\ninside 1 19\nTMhelix 20 39\noutside 40 325\nTMhelix 326 348\ninside 349 372

11940 GCF_000471345.1_E_coliM16-1.0 Escherichia coli M16 Proteobacteria; Gammaproteobacteria; Enterobacterales; Enterobacteriaceae; Escherichia; Escherichia coli
MTGDNTLIHSHGINRRDFMKLCAALAATMGLSSKAAAEMAESVTNPQRPPVIWIGAQECT WP_000145410.1
MULTISPECIES: hydrogenase-2 small chain [Proteobacteria] Length: 372\nNumber of predicted TMHs: 1\nExp number of AAs in TMHs: 23.72226\nExp number, first 60 AAs: 0.22038\nTotal prob of N-in: 0.12994\noutside 1 331\nTMhelix 332 354\ninside 355 372

11941 GCF_000471365.1_E_coliM17-1.0 Escherichia coli M17 Proteobacteria; Gammaproteobacteria; Enterobacterales; Enterobacteriaceae; Escherichia; Escherichia coli
MTGDNTLIHSHGINRRDFMKLCAALAATMGLSSKAAAEMAESVTNPQRPPVIWIGAQECT WP_000145410.1
MULTISPECIES: hydrogenase-2 small chain [Proteobacteria] Length: 372\nNumber of predicted TMHs: 1\nExp number of AAs in TMHs: 23.72226\nExp number, first 60 AAs: 0.22038\nTotal prob of N-in: 0.12994\noutside 1 331\nTMhelix 332 354\ninside 355 372

11942 GCF_000471365.1_E_coliM17-1.0 Escherichia coli M17 Proteobacteria; Gammaproteobacteria; Enterobacterales; Enterobacteriaceae; Escherichia; Escherichia coli
MNNEETFYQAMRRQGVTRRSFLKYCSLAATSLGLGAGMAPKIAWALENKPRIPVVWIHGL WP_001301613.1
hydrogenase [Escherichia coli] Length: 372\nNumber of predicted TMHs: 2\nExp number of AAs in TMHs: 37.46598\nExp number, first 60 AAs: 19.53139\nTotal prob of N-in: 0.94364\nPOSSIBLE N-term signal sequence\ninside 1 19\nTMhelix 20 39\noutside 40 325\nTMhelix 326 348\ninside 349 372

11943 GCF_000471385.1_E_coliM18-1.0 Escherichia coli M18 Proteobacteria; Gammaproteobacteria; Enterobacterales; Enterobacteriaceae; Escherichia; Escherichia coli
MTGDNTLIHSHGINRRDFMKLCAALAATMGLSSKAAAEMAESVTNPQRPPVIWIGAQECT WP_000145410.1
MULTISPECIES: hydrogenase-2 small chain [Proteobacteria] Length: 372\nNumber of predicted TMHs: 1\nExp number of AAs in TMHs: 23.72226\nExp number, first 60 AAs: 0.22038\nTotal prob of N-in: 0.12994\noutside 1 331\nTMhelix 332 354\ninside 355 372

11944 GCF_000471385.1_E_coliM18-1.0 Escherichia coli M18 Proteobacteria; Gammaproteobacteria; Enterobacterales; Enterobacteriaceae; Escherichia; Escherichia coli

MNNEETFYQAMRRQGVTRRSFLKYCSLAATSLGLGAGMAPKIAWALENKPRIPVVWIHGL WP_001301613.1 hydrogenase [Escherichia coli] Length: 372\nNumber of predicted TMHs: 2\nExp number of AAs in TMHs: 37.46598\nExp number, first 60 AAs: 19.53139\nTotal prob of N-in: 0.94364\nPOSSIBLE N-term signal sequence\ninside 1 19\nTMhelix 20 39\noutside 40 325\nTMhelix 326 348\ninside 349 372

11945 GCF_000471405.1_E_coliM19-1.0 Escherichia coli M19 Proteobacteria; Gammaproteobacteria; Enterobacterales; Enterobacteriaceae; Escherichia; Escherichia coli MTGDNTLIHSHGINRRDFMKLCAALAATMGLSSKAAAEMAESVTNPQRPPVIWIGAQECT WP_000145410.1 MULTISPECIES: hydrogenase-2 small chain [Proteobacteria] Length: 372\nNumber of predicted TMHs: 1\nExp number of AAs in TMHs: 23.72226\nExp number, first 60 AAs: 0.22038\nTotal prob of N-in: 0.12994\noutside 1 331\nTMhelix 332 354\ninside 355 372

11946 GCF_000471405.1_E_coliM19-1.0 Escherichia coli M19 Proteobacteria; Gammaproteobacteria; Enterobacterales; Enterobacteriaceae; Escherichia; Escherichia coli MNNEETFYQAMRRQGVTRRSFLKYCSLAATSLGLGAGMAPKIAWALENKPRIPVVWIHGL WP_001301613.1 hydrogenase [Escherichia coli] Length: 372\nNumber of predicted TMHs: 2\nExp number of AAs in TMHs: 37.46598\nExp number, first 60 AAs: 19.53139\nTotal prob of N-in: 0.94364\nPOSSIBLE N-term signal sequence\ninside 1 19\nTMhelix 20 39\noutside 40 325\nTMhelix 326 348\ninside 349 372

11947 GCF_000471065.1_E_coliM2-1.0 Escherichia coli M2 Proteobacteria; Gammaproteobacteria; Enterobacterales; Enterobacteriaceae; Escherichia; Escherichia coli MTGDNTLIHSHGINRRDFMKLCAALAATMGLSSKAAAEMAESVTNPQRPPVIWIGAQECT WP_000145410.1 MULTISPECIES: hydrogenase-2 small chain [Proteobacteria] Length: 372\nNumber of predicted TMHs: 1\nExp number of AAs in TMHs: 23.72226\nExp number, first 60 AAs: 0.22038\nTotal prob of N-in: 0.12994\noutside 1 331\nTMhelix 332 354\ninside 355 372

11948 GCF_000471065.1_E_coliM2-1.0 Escherichia coli M2 Proteobacteria; Gammaproteobacteria; Enterobacterales; Enterobacteriaceae; Escherichia; Escherichia coli MNNEETFYQAMRRQGVTRRSFLKYCSLAATSLGLGAGMAPKIAWALENKPRIPVVWIHGL WP_001301613.1 hydrogenase [Escherichia coli] Length: 372\nNumber of predicted TMHs: 2\nExp number of AAs in TMHs: 37.46598\nExp number, first 60 AAs: 19.53139\nTotal prob of N-in: 0.94364\nPOSSIBLE N-term signal sequence\ninside 1 19\nTMhelix 20 39\noutside 40 325\nTMhelix 326 348\ninside 349 372

11949 GCF_000471425.1_E_coliM20-1.0 Escherichia coli M20 Proteobacteria; Gammaproteobacteria; Enterobacterales; Enterobacteriaceae; Escherichia; Escherichia coli MTGDNTLIHSHGINRRDFMKLCAALAATMGLSSKAAAEMAESVTNPQRPPVIWIGAQECT WP_000145410.1 MULTISPECIES: hydrogenase-2 small chain [Proteobacteria] Length: 372\nNumber of predicted TMHs: 1\nExp number of AAs in TMHs: 23.72226\nExp number, first 60 AAs: 0.22038\nTotal prob of N-in: 0.12994\noutside 1 331\nTMhelix 332 354\ninside 355 372

11950 GCF_000471425.1_E_coliM20-1.0 Escherichia coli M20 Proteobacteria; Gammaproteobacteria; Enterobacterales; Enterobacteriaceae; Escherichia; Escherichia coli MNNEETFYQAMRRQGVTRRSFLKYCSLAATSLGLGAGMAPKIAWALENKPRIPVVWIHGL WP_001301613.1 hydrogenase [Escherichia coli] Length: 372\nNumber of predicted TMHs: 2\nExp number of AAs in TMHs: 37.46598\nExp number, first 60 AAs: 19.53139\nTotal prob of N-in: 0.94364\nPOSSIBLE N-term signal sequence\ninside 1 19\nTMhelix 20 39\noutside 40 325\nTMhelix 326 348\ninside 349 372

11951 GCF_000471445.1_E_coliM21-1.0 Escherichia coli M21 Proteobacteria; Gammaproteobacteria; Enterobacterales; Enterobacteriaceae; Escherichia; Escherichia coli MTGDNTLIHSHGINRRDFMKLCAALAATMGLSSKAAAEMAESVTNPQRPPVIWIGAQECT WP_000145410.1 MULTISPECIES: hydrogenase-2 small chain [Proteobacteria] Length: 372\nNumber of predicted TMHs: 1\nExp number of AAs in TMHs: 23.72226\nExp number, first 60 AAs: 0.22038\nTotal prob of N-in: 0.12994\noutside 1 331\nTMhelix 332 354\ninside 355 372

11952 GCF_000471445.1_E_coliM21-1.0 Escherichia coli M21 Proteobacteria; Gammaproteobacteria; Enterobacterales; Enterobacteriaceae; Escherichia; Escherichia coli MNNEETFYQAMRRQGVTRRSFLKYCSLAATSLGLGAGMAPKIAWALENKPRIPVVWIHGL WP_001301613.1 hydrogenase [Escherichia coli] Length: 372\nNumber of predicted TMHs: 2\nExp number of AAs in TMHs: 37.46598\nExp number, first 60 AAs: 19.53139\nTotal prob of N-in: 0.94364\nPOSSIBLE N-term signal sequence\ninside 1 19\nTMhelix 20 39\noutside 40 325\nTMhelix 326 348\ninside 349 372

11953 GCF_000471465.1_E_coliM22-1.0 Escherichia coli M22 Proteobacteria; Gammaproteobacteria; Enterobacterales; Enterobacteriaceae; Escherichia; Escherichia coli
MNNEETFYQAMRRQGVTRRSFLKYCSLAATSLGLGAGMAPKIAWALENKPRIPVVWIHGL WP_001301613.1
hydrogenase [Escherichia coli] Length: 372\nNumber of predicted TMHs: 2\nExp number of AAs in TMHs: 37.46598\nExp number, first 60 AAs: 19.53139\nTotal prob of N-in: 0.94364\nPOSSIBLE N-term signal sequence\ninside 1 19\nTMhelix 20 39\noutside 40 325\nTMhelix 326 348\ninside 349 372

11954 GCF_000471465.1_E_coliM22-1.0 Escherichia coli M22 Proteobacteria; Gammaproteobacteria; Enterobacterales; Enterobacteriaceae; Escherichia; Escherichia coli
MTGDNTLIHSHGINRRDFMKLCAALAATMGLSSKAAAEMAESVTNPQRPPVIWIGAQECT WP_000145410.1
MULTISPECIES: hydrogenase-2 small chain [Proteobacteria] Length: 372\nNumber of predicted TMHs: 1\nExp number of AAs in TMHs: 23.72226\nExp number, first 60 AAs: 0.22038\nTotal prob of N-in: 0.12994\noutside 1 331\nTMhelix 332 354\ninside 355 372

11955 GCF_000471485.1_E_coliM23-1.0 Escherichia coli M23 Proteobacteria; Gammaproteobacteria; Enterobacterales; Enterobacteriaceae; Escherichia; Escherichia coli
MTGDNTLIHSHGINRRDFMKLCAALAATMGLSSKAAAEMAESVTNPQRPPVIWIGAQECT WP_000145410.1
MULTISPECIES: hydrogenase-2 small chain [Proteobacteria] Length: 372\nNumber of predicted TMHs: 1\nExp number of AAs in TMHs: 23.72226\nExp number, first 60 AAs: 0.22038\nTotal prob of N-in: 0.12994\noutside 1 331\nTMhelix 332 354\ninside 355 372

11956 GCF_000471485.1_E_coliM23-1.0 Escherichia coli M23 Proteobacteria; Gammaproteobacteria; Enterobacterales; Enterobacteriaceae; Escherichia; Escherichia coli
MNNEETFYQAMRRQGVTRRSFLKYCSLAATSLGLGAGMAPKIAWALENKPRIPVVWIHGL WP_001301613.1
hydrogenase [Escherichia coli] Length: 372\nNumber of predicted TMHs: 2\nExp number of AAs in TMHs: 37.46598\nExp number, first 60 AAs: 19.53139\nTotal prob of N-in: 0.94364\nPOSSIBLE N-term signal sequence\ninside 1 19\nTMhelix 20 39\noutside 40 325\nTMhelix 326 348\ninside 349 372

11957 GCF_000471085.2_E_coliM3-1.0 Escherichia coli M3 Proteobacteria; Gammaproteobacteria; Enterobacterales; Enterobacteriaceae; Escherichia; Escherichia coli
MTGDNTLIHSHGINRRDFMKLCAALAATMGLSSKAAAEMAESVTNPQRPPVIWIGAQECT WP_000145410.1
MULTISPECIES: hydrogenase-2 small chain [Proteobacteria] Length: 372\nNumber of predicted TMHs: 1\nExp number of AAs in TMHs: 23.72226\nExp number, first 60 AAs: 0.22038\nTotal prob of N-in: 0.12994\noutside 1 331\nTMhelix 332 354\ninside 355 372

11958 GCF_000471085.2_E_coliM3-1.0 Escherichia coli M3 Proteobacteria; Gammaproteobacteria; Enterobacterales; Enterobacteriaceae; Escherichia; Escherichia coli
MNNEETFYQAMRRQGVTRRSFLKYCSLAATSLGLGAGMAPKIAWALENKPRIPVVWIHGL WP_001301613.1
hydrogenase [Escherichia coli] Length: 372\nNumber of predicted TMHs: 2\nExp number of AAs in TMHs: 37.46598\nExp number, first 60 AAs: 19.53139\nTotal prob of N-in: 0.94364\nPOSSIBLE N-term signal sequence\ninside 1 19\nTMhelix 20 39\noutside 40 325\nTMhelix 326 348\ninside 349 372

11959 GCF_002149955.1_ASM214995v1 Escherichia coli M4 Proteobacteria; Gammaproteobacteria; Enterobacterales; Enterobacteriaceae; Escherichia; Escherichia coli
MTGDNTLIHSHGINRRDFMKLCAALAATMGLSSKAAAEMAESVTNPQRPPVIWIGAQECT WP_000145410.1
MULTISPECIES: hydrogenase-2 small chain [Proteobacteria] Length: 372\nNumber of predicted TMHs: 1\nExp number of AAs in TMHs: 23.72226\nExp number, first 60 AAs: 0.22038\nTotal prob of N-in: 0.12994\noutside 1 331\nTMhelix 332 354\ninside 355 372

11960 GCF_002149955.1_ASM214995v1 Escherichia coli M4 Proteobacteria; Gammaproteobacteria; Enterobacterales; Enterobacteriaceae; Escherichia; Escherichia coli
MNNEETFYQAMRRQGVTRRSFLKYCSLAATSLGLGAGMAPKIAWALENKPRIPVVWIHGL WP_001058323.1
MULTISPECIES: hydrogenase-1 small chain [Proteobacteria] Length: 372\nNumber of predicted TMHs: 2\nExp number of AAs in TMHs: 37.2608499999999\nExp number, first 60 AAs: 19.52923\nTotal prob of N-in: 0.94351\nPOSSIBLE N-term signal sequence\ninside 1 19\nTMhelix 20 39\noutside 40 325\nTMhelix 326 348\ninside 349 372

11961 GCF_000471105.1_E_coliM4-1.0 Escherichia coli M4 Proteobacteria; Gammaproteobacteria; Enterobacterales; Enterobacteriaceae; Escherichia; Escherichia coli
MNNEETFYQAMRRQGVTRRSFLKYCSLAATSLGLGAGMAPKIAWALENKPRIPVVWIHGL WP_001301613.1
hydrogenase [Escherichia coli] Length: 372\nNumber of predicted TMHs: 2\nExp number of AAs in

TMHs: 37.46598\nExp number, first 60 AAs:  19.53139\nTotal prob of N-in:   0.94364\nPOSSIBLE N-term signal sequence\ninside   1   19\nTMhelix   20   39\noutside   40   325\nTMhelix   326   348\ninside   349   372

11962 GCF_000471105.1_E_coliM4-1.0 Escherichia coli M4 Proteobacteria; Gammaproteobacteria; Enterobacterales; Enterobacteriaceae; Escherichia; Escherichia coli
MTGDNTLIHSHGINRRDFMKLCAALAATMGLSSKAAAEMAESVTNPQRPPVIWIGAQECT WP_000145410.1
MULTISPECIES: hydrogenase-2 small chain [Proteobacteria] Length: 372\nNumber of predicted TMHs: 1\nExp number of AAs in TMHs: 23.72226\nExp number, first 60 AAs:  0.22038\nTotal prob of N-in:   0.12994\noutside   1   331\nTMhelix   332   354\ninside   355   372

11963 GCF_000471125.1_E_coliM5-1.0 Escherichia coli M5 Proteobacteria; Gammaproteobacteria; Enterobacterales; Enterobacteriaceae; Escherichia; Escherichia coli
MNNEETFYQAMRRQGVTRRSFLKYCSLAATSLGLGAGMAPKIAWALENKPRIPVVWIHGL WP_001301613.1
hydrogenase [Escherichia coli] Length: 372\nNumber of predicted TMHs:  2\nExp number of AAs in TMHs: 37.46598\nExp number, first 60 AAs:  19.53139\nTotal prob of N-in:   0.94364\nPOSSIBLE N-term signal sequence\ninside   1   19\nTMhelix   20   39\noutside   40   325\nTMhelix   326   348\ninside   349   372

11964 GCF_000471125.1_E_coliM5-1.0 Escherichia coli M5 Proteobacteria; Gammaproteobacteria; Enterobacterales; Enterobacteriaceae; Escherichia; Escherichia coli
MTGDNTLIHSHGINRRDFMKLCAALAATMGLSSKAAAEMAESVTNPQRPPVIWIGAQECT WP_000145410.1
MULTISPECIES: hydrogenase-2 small chain [Proteobacteria] Length: 372\nNumber of predicted TMHs: 1\nExp number of AAs in TMHs: 23.72226\nExp number, first 60 AAs:  0.22038\nTotal prob of N-in:   0.12994\noutside   1   331\nTMhelix   332   354\ninside   355   372

11965 GCF_000471145.1_E_coliM6-1.0 Escherichia coli M6 Proteobacteria; Gammaproteobacteria; Enterobacterales; Enterobacteriaceae; Escherichia; Escherichia coli
MTGDNTLIHSHGINRRDFMKLCAALAATMGLSSKAAAEMAESVTNPQRPPVIWIGAQECT WP_000145410.1
MULTISPECIES: hydrogenase-2 small chain [Proteobacteria] Length: 372\nNumber of predicted TMHs: 1\nExp number of AAs in TMHs: 23.72226\nExp number, first 60 AAs:  0.22038\nTotal prob of N-in:   0.12994\noutside   1   331\nTMhelix   332   354\ninside   355   372

11966 GCF_000471145.1_E_coliM6-1.0 Escherichia coli M6 Proteobacteria; Gammaproteobacteria; Enterobacterales; Enterobacteriaceae; Escherichia; Escherichia coli
MNNEETFYQAMRRQGVTRRSFLKYCSLAATSLGLGAGMAPKIAWALENKPRIPVVWIHGL WP_001301613.1
hydrogenase [Escherichia coli] Length: 372\nNumber of predicted TMHs:  2\nExp number of AAs in TMHs: 37.46598\nExp number, first 60 AAs:  19.53139\nTotal prob of N-in:   0.94364\nPOSSIBLE N-term signal sequence\ninside   1   19\nTMhelix   20   39\noutside   40   325\nTMhelix   326   348\ninside   349   372

11967 GCF_000176555.2_ASM17655v2 Escherichia coli M605 Proteobacteria; Gammaproteobacteria; Enterobacterales; Enterobacteriaceae; Escherichia; Escherichia coli
MTGDNTLIHSHGINRRDFMKLCAALAATMGLSSKAAAEMAESVTNPQRPPVIWIGAQECT WP_000145410.1
MULTISPECIES: hydrogenase-2 small chain [Proteobacteria] Length: 372\nNumber of predicted TMHs: 1\nExp number of AAs in TMHs: 23.72226\nExp number, first 60 AAs:  0.22038\nTotal prob of N-in:   0.12994\noutside   1   331\nTMhelix   332   354\ninside   355   372

11968 GCF_000176555.2_ASM17655v2 Escherichia coli M605 Proteobacteria; Gammaproteobacteria; Enterobacterales; Enterobacteriaceae; Escherichia; Escherichia coli
MNNEETFYQAMRRQGVTRRSFLKYCSLAATSLGLGAGMAPKIAWALENKPRIPVVWIHGL WP_001058323.1
MULTISPECIES: hydrogenase-1 small chain [Proteobacteria] Length: 372\nNumber of predicted TMHs: 2\nExp number of AAs in TMHs: 37.2608499999999\nExp number, first 60 AAs:  19.52923\nTotal prob of N-in:   0.94351\nPOSSIBLE N-term signal sequence\ninside   1   19\nTMhelix   20   39\noutside   40   325\nTMhelix   326   348\ninside   349   372

11969 GCF_000471165.1_E_coliM7-1.0 Escherichia coli M7 Proteobacteria; Gammaproteobacteria; Enterobacterales; Enterobacteriaceae; Escherichia; Escherichia coli
MTGDNTLIHSHGINRRDFMKLCAALAATMGLSSKAAAEMAESVTNPQRPPVIWIGAQECT WP_000145410.1
MULTISPECIES: hydrogenase-2 small chain [Proteobacteria] Length: 372\nNumber of predicted TMHs: 1\nExp number of AAs in TMHs: 23.72226\nExp number, first 60 AAs:  0.22038\nTotal prob of N-in:   0.12994\noutside   1   331\nTMhelix   332   354\ninside   355   372

11970 GCF_000471165.1_E_coliM7-1.0 Escherichia coli M7 Proteobacteria; Gammaproteobacteria; Enterobacterales; Enterobacteriaceae; Escherichia; Escherichia coli
MNNEETFYQAMRRQGVTRRSFLKYCSLAATSLGLGAGMAPKIAWALENKPRIPVVWIHGL WP_024251880.1
hydrogenase [Escherichia coli] Length: 372\nNumber of predicted TMHs: 2\nExp number of AAs in TMHs: 37.17638\nExp number, first 60 AAs: 19.53124\nTotal prob of N-in: 0.94348\nPOSSIBLE N-term signal sequence\ninside 1 19\nTMhelix 20 39\noutside 40 325\nTMhelix 326 348\ninside 349 372

11971 GCF_000176575.2_ASM17657v2 Escherichia coli M718 Proteobacteria; Gammaproteobacteria; Enterobacterales; Enterobacteriaceae; Escherichia; Escherichia coli
MNNEETFYQAMRRQGVTRRSFLKYCSLAATSLGLGAGMAPKIAWALENKPRIPVVWIHGL WP_001058323.1
MULTISPECIES: hydrogenase-1 small chain [Proteobacteria] Length: 372\nNumber of predicted TMHs: 2\nExp number of AAs in TMHs: 37.2608499999999\nExp number, first 60 AAs: 19.52923\nTotal prob of N-in: 0.94351\nPOSSIBLE N-term signal sequence\ninside 1 19\nTMhelix 20 39\noutside 40 325\nTMhelix 326 348\ninside 349 372

11972 GCF_000176575.2_ASM17657v2 Escherichia coli M718 Proteobacteria; Gammaproteobacteria; Enterobacterales; Enterobacteriaceae; Escherichia; Escherichia coli
MTGDNTLIHSHGINRRDFMKLCAALAATMGLSSKAAAEMAESVTNPQRPPVIWIGAQECT WP_000145410.1
MULTISPECIES: hydrogenase-2 small chain [Proteobacteria] Length: 372\nNumber of predicted TMHs: 1\nExp number of AAs in TMHs: 23.72226\nExp number, first 60 AAs: 0.22038\nTotal prob of N-in: 0.12994\noutside 1 331\nTMhelix 332 354\ninside 355 372

11973 GCF_000471185.1_E_coliM8-1.0 Escherichia coli M8 Proteobacteria; Gammaproteobacteria; Enterobacterales; Enterobacteriaceae; Escherichia; Escherichia coli
MTGDNTLIHSHGINRRDFMKLCAALAATMGLSSKAAAEMAESVTNPQRPPVIWIGAQECT WP_000145410.1
MULTISPECIES: hydrogenase-2 small chain [Proteobacteria] Length: 372\nNumber of predicted TMHs: 1\nExp number of AAs in TMHs: 23.72226\nExp number, first 60 AAs: 0.22038\nTotal prob of N-in: 0.12994\noutside 1 331\nTMhelix 332 354\ninside 355 372

11974 GCF_000471185.1_E_coliM8-1.0 Escherichia coli M8 Proteobacteria; Gammaproteobacteria; Enterobacterales; Enterobacteriaceae; Escherichia; Escherichia coli
MNNEETFYQAMRRQGVTRRSFLKYCSLAATSLGLGAGMAPKIAWALENKPRIPVVWIHGL WP_001301613.1
hydrogenase [Escherichia coli] Length: 372\nNumber of predicted TMHs: 2\nExp number of AAs in TMHs: 37.46598\nExp number, first 60 AAs: 19.53139\nTotal prob of N-in: 0.94364\nPOSSIBLE N-term signal sequence\ninside 1 19\nTMhelix 20 39\noutside 40 325\nTMhelix 326 348\ninside 349 372

11975 GCF_000190955.1_ASM19095v1 Escherichia coli M863 Proteobacteria; Gammaproteobacteria; Enterobacterales; Enterobacteriaceae; Escherichia; Escherichia coli
MTGDNTLIHSHGINRRDFMKLCAALAATMGLSSKAAAEMAESVTNPQRPPVIWIGAQECT WP_000145410.1
MULTISPECIES: hydrogenase-2 small chain [Proteobacteria] Length: 372\nNumber of predicted TMHs: 1\nExp number of AAs in TMHs: 23.72226\nExp number, first 60 AAs: 0.22038\nTotal prob of N-in: 0.12994\noutside 1 331\nTMhelix 332 354\ninside 355 372

11976 GCF_000190955.1_ASM19095v1 Escherichia coli M863 Proteobacteria; Gammaproteobacteria; Enterobacterales; Enterobacteriaceae; Escherichia; Escherichia coli
MNNEETFYQAMRRQGVTRRSFLKYCSLAATSLGLGAGMAPKIAWALENKPRIPVVWIHGL WP_001353311.1
MULTISPECIES: hydrogenase [Escherichia] Length: 372\nNumber of predicted TMHs: 2\nExp number of AAs in TMHs: 37.2638699999999\nExp number, first 60 AAs: 19.52924\nTotal prob of N-in: 0.94351\nPOSSIBLE N-term signal sequence\ninside 1 19\nTMhelix 20 39\noutside 40 325\nTMhelix 326 348\ninside 349 372

11977 GCF_000471205.1_E_coliM9-1.0 Escherichia coli M9 Proteobacteria; Gammaproteobacteria; Enterobacterales; Enterobacteriaceae; Escherichia; Escherichia coli
MNNEETFYQAMRRQGVTRRSFLKYCSLAATSLGLGAGMAPKIAWALENKPRIPVVWIHGL WP_001301613.1
hydrogenase [Escherichia coli] Length: 372\nNumber of predicted TMHs: 2\nExp number of AAs in TMHs: 37.46598\nExp number, first 60 AAs: 19.53139\nTotal prob of N-in: 0.94364\nPOSSIBLE N-term signal sequence\ninside 1 19\nTMhelix 20 39\noutside 40 325\nTMhelix 326 348\ninside 349 372

11978 GCF_000471205.1_E_coliM9-1.0 Escherichia coli M9 Proteobacteria; Gammaproteobacteria; Enterobacterales; Enterobacteriaceae; Escherichia; Escherichia coli
MTGDNTLIHSHGINRRDFMKLCAALAATMGLSSKAAAEMAESVTNPQRPPVIWIGAQECT WP_000145410.1
MULTISPECIES: hydrogenase-2 small chain [Proteobacteria] Length: 372\nNumber of predicted TMHs:

1\nExp number of AAs in TMHs: 23.72226\nExp number, first 60 AAs: 0.22038\nTotal prob of N-in: 0.12994\noutside 1 331\nTMhelix 332 354\ninside 355 372

11979 GCF_000261145.1_Esch_coli_M919_V2 Escherichia coli M919 Proteobacteria; Gammaproteobacteria; Enterobacterales; Enterobacteriaceae; Escherichia; Escherichia coli MNNEETFYQAMRRQGVTRRSFLKYCSLAATSLGLGAGMAPKIAWALENKPRIPVVWIHGL WP_001058323.1 MULTISPECIES: hydrogenase-1 small chain [Proteobacteria] Length: 372\nNumber of predicted TMHs: 2\nExp number of AAs in TMHs: 37.2608499999999\nExp number, first 60 AAs: 19.52923\nTotal prob of N-in: 0.94351\nPOSSIBLE N-term signal sequence\ninside 1 19\nTMhelix 20 39\noutside 40 325\nTMhelix 326 348\ninside 349 372

11980 GCF_000261145.1_Esch_coli_M919_V2 Escherichia coli M919 Proteobacteria; Gammaproteobacteria; Enterobacterales; Enterobacteriaceae; Escherichia; Escherichia coli MTGDNTLIHSHGINRRDFMKLCAALAATMGLSSKAAAEMAESVTNPQRPPVIWIGAQECT WP_000145420.1 hydrogenase-2 small chain [Escherichia coli] Length: 372\nNumber of predicted TMHs: 1\nExp number of AAs in TMHs: 22.88747\nExp number, first 60 AAs: 0.22401\nTotal prob of N-in: 0.09738\noutside 1 331\nTMhelix 332 354\ninside 355 372

11981 GCF_000303975.2_ASM30397v2 Escherichia coli MA6 Proteobacteria; Gammaproteobacteria; Enterobacterales; Enterobacteriaceae; Escherichia; Escherichia coli MTGDNTLIHSHGINRRDFMKLCAALAATMGLSSKAAAEMAESVTNPQRPPVIWIGAQECT WP_000145410.1 MULTISPECIES: hydrogenase-2 small chain [Proteobacteria] Length: 372\nNumber of predicted TMHs: 1\nExp number of AAs in TMHs: 23.72226\nExp number, first 60 AAs: 0.22038\nTotal prob of N-in: 0.12994\noutside 1 331\nTMhelix 332 354\ninside 355 372

11982 GCF_000303975.2_ASM30397v2 Escherichia coli MA6 Proteobacteria; Gammaproteobacteria; Enterobacterales; Enterobacteriaceae; Escherichia; Escherichia coli MNNEETFYQAMRRQGVTRRSFLKYCSLAATSLGLGAGMAPKIAWALENKPRIPVVWIHGL WP_001301613.1 hydrogenase [Escherichia coli] Length: 372\nNumber of predicted TMHs: 2\nExp number of AAs in TMHs: 37.46598\nExp number, first 60 AAs: 19.53139\nTotal prob of N-in: 0.94364\nPOSSIBLE N-term signal sequence\ninside 1 19\nTMhelix 20 39\noutside 40 325\nTMhelix 326 348\ninside 349 372

11983 GCF_000418675.1_E_coliMC19-1.0 Escherichia coli MC19 Proteobacteria; Gammaproteobacteria; Enterobacterales; Enterobacteriaceae; Escherichia; Escherichia coli MNNEETFYQAMRRQGVTRRSFLKYCSLAATSLGLGAGMAPKIAWALENKPRIPVVWIHGL WP_001058323.1 MULTISPECIES: hydrogenase-1 small chain [Proteobacteria] Length: 372\nNumber of predicted TMHs: 2\nExp number of AAs in TMHs: 37.2608499999999\nExp number, first 60 AAs: 19.52923\nTotal prob of N-in: 0.94351\nPOSSIBLE N-term signal sequence\ninside 1 19\nTMhelix 20 39\noutside 40 325\nTMhelix 326 348\ninside 349 372

11984 GCF_000418675.1_E_coliMC19-1.0 Escherichia coli MC19 Proteobacteria; Gammaproteobacteria; Enterobacterales; Enterobacteriaceae; Escherichia; Escherichia coli MTGDNTLIHSHGINRRDFMKLCAALAATMGLSSKAAAEMAESVTNPQRPPVIWIGAQECT WP_000145410.1 MULTISPECIES: hydrogenase-2 small chain [Proteobacteria] Length: 372\nNumber of predicted TMHs: 1\nExp number of AAs in TMHs: 23.72226\nExp number, first 60 AAs: 0.22038\nTotal prob of N-in: 0.12994\noutside 1 331\nTMhelix 332 354\ninside 355 372

11985 GCF_000418655.1_E_coliMC21-1.0 Escherichia coli MC21 Proteobacteria; Gammaproteobacteria; Enterobacterales; Enterobacteriaceae; Escherichia; Escherichia coli MTGDNTLIHSHGINRRDFMKLCAALAATMGLSSKAAAEMAESVTNPQRPPVIWIGAQECT WP_000145410.1 MULTISPECIES: hydrogenase-2 small chain [Proteobacteria] Length: 372\nNumber of predicted TMHs: 1\nExp number of AAs in TMHs: 23.72226\nExp number, first 60 AAs: 0.22038\nTotal prob of N-in: 0.12994\noutside 1 331\nTMhelix 332 354\ninside 355 372

11986 GCF_000418655.1_E_coliMC21-1.0 Escherichia coli MC21 Proteobacteria; Gammaproteobacteria; Enterobacterales; Enterobacteriaceae; Escherichia; Escherichia coli MNNEETFYQAMRRQGVTRRSFLKYCSLAATSLGLGAGMAPKIAWALENKPRIPVVWIHGL WP_001058323.1 MULTISPECIES: hydrogenase-1 small chain [Proteobacteria] Length: 372\nNumber of predicted TMHs: 2\nExp number of AAs in TMHs: 37.2608499999999\nExp number, first 60 AAs: 19.52923\nTotal prob of N-in: 0.94351\nPOSSIBLE N-term signal sequence\ninside 1 19\nTMhelix 20 39\noutside 40 325\nTMhelix 326 348\ninside 349 372

11987 GCF_000418635.1_E_coliMC23-1.0 Escherichia coli MC23 Proteobacteria; Gammaproteobacteria; Enterobacterales; Enterobacteriaceae; Escherichia; Escherichia coli
MNNEETFYQAMRRQGVTRRSFLKYCSLAATSLGLGAGMAPKIAWALENKPRIPVVWIHGL WP_001058323.1
MULTISPECIES: hydrogenase-1 small chain [Proteobacteria] Length: 372\nNumber of predicted TMHs: 2\nExp number of AAs in TMHs: 37.2608499999999\nExp number, first 60 AAs: 19.52923\nTotal prob of N-in: 0.94351\nPOSSIBLE N-term signal sequence\ninside 1 19\nTMhelix 20 39\noutside 40 325\nTMhelix 326 348\ninside 349 372

11988 GCF_000418635.1_E_coliMC23-1.0 Escherichia coli MC23 Proteobacteria; Gammaproteobacteria; Enterobacterales; Enterobacteriaceae; Escherichia; Escherichia coli
MTGDNTLIHSHGINRRDFMKLCAALAATMGLSSKAAAEMAESVTNPQRPPVIWIGAQECT WP_000145410.1
MULTISPECIES: hydrogenase-2 small chain [Proteobacteria] Length: 372\nNumber of predicted TMHs: 1\nExp number of AAs in TMHs: 23.72226\nExp number, first 60 AAs: 0.22038\nTotal prob of N-in: 0.12994\noutside 1 331\nTMhelix 332 354\ninside 355 372

11989 GCF_000418615.2_E_coliMC6002-1.0 Escherichia coli MC6002 Proteobacteria; Gammaproteobacteria; Enterobacterales; Enterobacteriaceae; Escherichia; Escherichia coli
MTGDNTLIHSHGINRRDFMKLCAALAATMGLSSKAAAEMAESVTNPQRPPVIWIGAQECT WP_000145410.1
MULTISPECIES: hydrogenase-2 small chain [Proteobacteria] Length: 372\nNumber of predicted TMHs: 1\nExp number of AAs in TMHs: 23.72226\nExp number, first 60 AAs: 0.22038\nTotal prob of N-in: 0.12994\noutside 1 331\nTMhelix 332 354\ninside 355 372

11990 GCF_000418615.2_E_coliMC6002-1.0 Escherichia coli MC6002 Proteobacteria; Gammaproteobacteria; Enterobacterales; Enterobacteriaceae; Escherichia; Escherichia coli
MNNEETFYQAMRRQGVTRRSFLKYCSLAATSLGLGAGMAPKIAWALENKPRIPVVWIHGL WP_001058323.1
MULTISPECIES: hydrogenase-1 small chain [Proteobacteria] Length: 372\nNumber of predicted TMHs: 2\nExp number of AAs in TMHs: 37.2608499999999\nExp number, first 60 AAs: 19.52923\nTotal prob of N-in: 0.94351\nPOSSIBLE N-term signal sequence\ninside 1 19\nTMhelix 20 39\noutside 40 325\nTMhelix 326 348\ninside 349 372

11991 GCF_000418595.1_E_coliMC6003-1.0 Escherichia coli MC6003 Proteobacteria; Gammaproteobacteria; Enterobacterales; Enterobacteriaceae; Escherichia; Escherichia coli
MTGDNTLIHSHGINRRDFMKLCAALAATMGLSSKAAAEMAESVTNPQRPPVIWIGAQECT WP_000145410.1
MULTISPECIES: hydrogenase-2 small chain [Proteobacteria] Length: 372\nNumber of predicted TMHs: 1\nExp number of AAs in TMHs: 23.72226\nExp number, first 60 AAs: 0.22038\nTotal prob of N-in: 0.12994\noutside 1 331\nTMhelix 332 354\ninside 355 372

11992 GCF_000418595.1_E_coliMC6003-1.0 Escherichia coli MC6003 Proteobacteria; Gammaproteobacteria; Enterobacterales; Enterobacteriaceae; Escherichia; Escherichia coli
MNNEETFYQAMRRQGVTRRSFLKYCSLAATSLGLGAGMAPKIAWALENKPRIPVVWIHGL WP_001058323.1
MULTISPECIES: hydrogenase-1 small chain [Proteobacteria] Length: 372\nNumber of predicted TMHs: 2\nExp number of AAs in TMHs: 37.2608499999999\nExp number, first 60 AAs: 19.52923\nTotal prob of N-in: 0.94351\nPOSSIBLE N-term signal sequence\ninside 1 19\nTMhelix 20 39\noutside 40 325\nTMhelix 326 348\ninside 349 372

11993 GCF_000692755.1_Esch_coli_MGH_57_V1 Escherichia coli MGH 57 Proteobacteria; Gammaproteobacteria; Enterobacterales; Enterobacteriaceae; Escherichia; Escherichia coli
MNNEETFYQAMRRQGVTRRSFLKYCSLAATSLGLGAGMAPKIAWALENKPRIPVVWIHGL WP_001295940.1
MULTISPECIES: hydrogenase [Enterobacteriaceae] Length: 372\nNumber of predicted TMHs: 2\nExp number of AAs in TMHs: 37.2608899999999\nExp number, first 60 AAs: 19.52923\nTotal prob of N-in: 0.94351\nPOSSIBLE N-term signal sequence\ninside 1 19\nTMhelix 20 39\noutside 40 325\nTMhelix 326 348\ninside 349 372

11994 GCF_000692755.1_Esch_coli_MGH_57_V1 Escherichia coli MGH 57 Proteobacteria; Gammaproteobacteria; Enterobacterales; Enterobacteriaceae; Escherichia; Escherichia coli
MTGDNTLIHSHGINRRDFMKLCAALAATMGLSSKAAAEMAESVTNPQRPPVIWIGAQECT WP_000145410.1
MULTISPECIES: hydrogenase-2 small chain [Proteobacteria] Length: 372\nNumber of predicted TMHs: 1\nExp number of AAs in TMHs: 23.72226\nExp number, first 60 AAs: 0.22038\nTotal prob of N-in: 0.12994\noutside 1 331\nTMhelix 332 354\ninside 355 372

11995 GCF_000692775.1_Esch_coli_MGH_58_V1 Escherichia coli MGH 58 Proteobacteria; Gammaproteobacteria; Enterobacterales; Enterobacteriaceae; Escherichia; Escherichia coli
MNNEETFYQAMRRQGVTRRSFLKYCSLAATSLGLGAGMAPKIAWALENKPRIPVVWIHGL WP_001058323.1
MULTISPECIES: hydrogenase-1 small chain [Proteobacteria] Length: 372\nNumber of predicted TMHs: 2\nExp number of AAs in TMHs: 37.2608499999999\nExp number, first 60 AAs: 19.52923\nTotal prob of N-in: 0.94351\nPOSSIBLE N-term signal sequence\ninside 1 19\nTMhelix 20 39\noutside 40 325\nTMhelix 326 348\ninside 349 372

11996 GCF_000692775.1_Esch_coli_MGH_58_V1 Escherichia coli MGH 58 Proteobacteria; Gammaproteobacteria; Enterobacterales; Enterobacteriaceae; Escherichia; Escherichia coli
MTGDNTLIHSHGINRRDFMKLCAALAATMGLSSKAAAEMAESVTNPQRPPVIWIGAQECT WP_000145410.1
MULTISPECIES: hydrogenase-2 small chain [Proteobacteria] Length: 372\nNumber of predicted TMHs: 1\nExp number of AAs in TMHs: 23.72226\nExp number, first 60 AAs: 0.22038\nTotal prob of N-in: 0.12994\noutside 1 331\nTMhelix 332 354\ninside 355 372

11997 GCF_000355275.2_ASM35527v2 Escherichia coli MP020940.1 Proteobacteria; Gammaproteobacteria; Enterobacterales; Enterobacteriaceae; Escherichia; Escherichia coli
MNNEETFYQAMRRQGVTRRSFLKYCSLAATSLGLGAGMAPKIAWALENKPRIPVVWIHGL WP_001295940.1
MULTISPECIES: hydrogenase [Enterobacteriaceae] Length: 372\nNumber of predicted TMHs: 2\nExp number of AAs in TMHs: 37.2608899999999\nExp number, first 60 AAs: 19.52923\nTotal prob of N-in: 0.94351\nPOSSIBLE N-term signal sequence\ninside 1 19\nTMhelix 20 39\noutside 40 325\nTMhelix 326 348\ninside 349 372

11998 GCF_000355275.2_ASM35527v2 Escherichia coli MP020940.1 Proteobacteria; Gammaproteobacteria; Enterobacterales; Enterobacteriaceae; Escherichia; Escherichia coli
MTGDNTLIHSHGINRRDFMKLCAALAATMGLSSKAAAEMAESVTNPQRPPVIWIGAQECT WP_000145410.1
MULTISPECIES: hydrogenase-2 small chain [Proteobacteria] Length: 372\nNumber of predicted TMHs: 1\nExp number of AAs in TMHs: 23.72226\nExp number, first 60 AAs: 0.22038\nTotal prob of N-in: 0.12994\noutside 1 331\nTMhelix 332 354\ninside 355 372

11999 GCF_000358145.2_ASM35814v2 Escherichia coli MP020980.1 Proteobacteria; Gammaproteobacteria; Enterobacterales; Enterobacteriaceae; Escherichia; Escherichia coli
MTGDNTLIHSHGINRRDFMKLCAALAATMGLSSKAAAEMAESVTNPQRPPVIWIGAQECT WP_000145410.1
MULTISPECIES: hydrogenase-2 small chain [Proteobacteria] Length: 372\nNumber of predicted TMHs: 1\nExp number of AAs in TMHs: 23.72226\nExp number, first 60 AAs: 0.22038\nTotal prob of N-in: 0.12994\noutside 1 331\nTMhelix 332 354\ninside 355 372

12000 GCF_000358145.2_ASM35814v2 Escherichia coli MP020980.1 Proteobacteria; Gammaproteobacteria; Enterobacterales; Enterobacteriaceae; Escherichia; Escherichia coli
MNNEETFYQAMRRQGVTRRSFLKYCSLAATSLGLGAGMAPKIAWALENKPRIPVVWIHGL WP_001058323.1
MULTISPECIES: hydrogenase-1 small chain [Proteobacteria] Length: 372\nNumber of predicted TMHs: 2\nExp number of AAs in TMHs: 37.2608499999999\nExp number, first 60 AAs: 19.52923\nTotal prob of N-in: 0.94351\nPOSSIBLE N-term signal sequence\ninside 1 19\nTMhelix 20 39\noutside 40 325\nTMhelix 326 348\ninside 349 372

12001 GCF_000355295.1_ASM35529v1 Escherichia coli MP020980.2 Proteobacteria; Gammaproteobacteria; Enterobacterales; Enterobacteriaceae; Escherichia; Escherichia coli
MNNEETFYQAMRRQGVTRRSFLKYCSLAATSLGLGAGMAPKIAWALENKPRIPVVWIHGL WP_001058323.1
MULTISPECIES: hydrogenase-1 small chain [Proteobacteria] Length: 372\nNumber of predicted TMHs: 2\nExp number of AAs in TMHs: 37.2608499999999\nExp number, first 60 AAs: 19.52923\nTotal prob of N-in: 0.94351\nPOSSIBLE N-term signal sequence\ninside 1 19\nTMhelix 20 39\noutside 40 325\nTMhelix 326 348\ninside 349 372

12002 GCF_000355295.1_ASM35529v1 Escherichia coli MP020980.2 Proteobacteria; Gammaproteobacteria; Enterobacterales; Enterobacteriaceae; Escherichia; Escherichia coli
MTGDNTLIHSHGINRRDFMKLCAALAATMGLSSKAAAEMAESVTNPQRPPVIWIGAQECT WP_000145410.1
MULTISPECIES: hydrogenase-2 small chain [Proteobacteria] Length: 372\nNumber of predicted TMHs: 1\nExp number of AAs in TMHs: 23.72226\nExp number, first 60 AAs: 0.22038\nTotal prob of N-in: 0.12994\noutside 1 331\nTMhelix 332 354\ninside 355 372

12003 GCF_000355515.2_ASM35551v2 Escherichia coli MP021017.1 Proteobacteria; Gammaproteobacteria; Enterobacterales; Enterobacteriaceae; Escherichia; Escherichia coli
MNNEETFYQAMRRQGVTRRSFLKYCSLAATSLGLGAGMAPKIAWALENKPRIPVVWIHGL WP_001058323.1
MULTISPECIES: hydrogenase-1 small chain [Proteobacteria] Length: 372\nNumber of predicted TMHs: 2\nExp number of AAs in TMHs: 37.2608499999999\nExp number, first 60 AAs: 19.52923\nTotal prob of N-in: 0.94351\nPOSSIBLE N-term signal sequence\ninside 1 19\nTMhelix 20 39\noutside 40 325\nTMhelix 326 348\ninside 349 372

12004 GCF_000355515.2_ASM35551v2 Escherichia coli MP021017.1 Proteobacteria; Gammaproteobacteria; Enterobacterales; Enterobacteriaceae; Escherichia; Escherichia coli
MTGDNTLIHSHGINRRDFMKLCAALAATMGLSSKAAAEMAESVTNPQRPPVIWIGAQECT WP_000145410.1
MULTISPECIES: hydrogenase-2 small chain [Proteobacteria] Length: 372\nNumber of predicted TMHs: 1\nExp number of AAs in TMHs: 23.72226\nExp number, first 60 AAs: 0.22038\nTotal prob of N-in: 0.12994\noutside 1 331\nTMhelix 332 354\ninside 355 372

12005 GCF_000354375.2_ASM35437v2 Escherichia coli MP021017.10 Proteobacteria; Gammaproteobacteria; Enterobacterales; Enterobacteriaceae; Escherichia; Escherichia coli
MNNEETFYQAMRRQGVTRRSFLKYCSLAATSLGLGAGMAPKIAWALENKPRIPVVWIHGL WP_001058323.1
MULTISPECIES: hydrogenase-1 small chain [Proteobacteria] Length: 372\nNumber of predicted TMHs: 2\nExp number of AAs in TMHs: 37.2608499999999\nExp number, first 60 AAs: 19.52923\nTotal prob of N-in: 0.94351\nPOSSIBLE N-term signal sequence\ninside 1 19\nTMhelix 20 39\noutside 40 325\nTMhelix 326 348\ninside 349 372

12006 GCF_000354375.2_ASM35437v2 Escherichia coli MP021017.10 Proteobacteria; Gammaproteobacteria; Enterobacterales; Enterobacteriaceae; Escherichia; Escherichia coli
MTGDNTLIHSHGINRRDFMKLCAALAATMGLSSKAAAEMAESVTNPQRPPVIWIGAQECT WP_000145410.1
MULTISPECIES: hydrogenase-2 small chain [Proteobacteria] Length: 372\nNumber of predicted TMHs: 1\nExp number of AAs in TMHs: 23.72226\nExp number, first 60 AAs: 0.22038\nTotal prob of N-in: 0.12994\noutside 1 331\nTMhelix 332 354\ninside 355 372

12007 GCF_000354415.2_ASM35441v2 Escherichia coli MP021017.11 Proteobacteria; Gammaproteobacteria; Enterobacterales; Enterobacteriaceae; Escherichia; Escherichia coli
MTGDNTLIHSHGINRRDFMKLCAALAATMGLSSKAAAEMAESVTNPQRPPVIWIGAQECT WP_000145410.1
MULTISPECIES: hydrogenase-2 small chain [Proteobacteria] Length: 372\nNumber of predicted TMHs: 1\nExp number of AAs in TMHs: 23.72226\nExp number, first 60 AAs: 0.22038\nTotal prob of N-in: 0.12994\noutside 1 331\nTMhelix 332 354\ninside 355 372

12008 GCF_000354415.2_ASM35441v2 Escherichia coli MP021017.11 Proteobacteria; Gammaproteobacteria; Enterobacterales; Enterobacteriaceae; Escherichia; Escherichia coli
MNNEETFYQAMRRQGVTRRSFLKYCSLAATSLGLGAGMAPKIAWALENKPRIPVVWIHGL WP_001058323.1
MULTISPECIES: hydrogenase-1 small chain [Proteobacteria] Length: 372\nNumber of predicted TMHs: 2\nExp number of AAs in TMHs: 37.2608499999999\nExp number, first 60 AAs: 19.52923\nTotal prob of N-in: 0.94351\nPOSSIBLE N-term signal sequence\ninside 1 19\nTMhelix 20 39\noutside 40 325\nTMhelix 326 348\ninside 349 372

12009 GCF_000354395.1_ASM35439v1 Escherichia coli MP021017.12 Proteobacteria; Gammaproteobacteria; Enterobacterales; Enterobacteriaceae; Escherichia; Escherichia coli
MNNEETFYQAMRRQGVTRRSFLKYCSLAATSLGLGAGMAPKIAWALENKPRIPVVWIHGL WP_001058323.1
MULTISPECIES: hydrogenase-1 small chain [Proteobacteria] Length: 372\nNumber of predicted TMHs: 2\nExp number of AAs in TMHs: 37.2608499999999\nExp number, first 60 AAs: 19.52923\nTotal prob of N-in: 0.94351\nPOSSIBLE N-term signal sequence\ninside 1 19\nTMhelix 20 39\noutside 40 325\nTMhelix 326 348\ninside 349 372

12010 GCF_000354395.1_ASM35439v1 Escherichia coli MP021017.12 Proteobacteria; Gammaproteobacteria; Enterobacterales; Enterobacteriaceae; Escherichia; Escherichia coli
MTGDNTLIHSHGINRRDFMKLCAALAATMGLSSKAAAEMAESVTNPQRPPVIWIGAQECT WP_000145410.1
MULTISPECIES: hydrogenase-2 small chain [Proteobacteria] Length: 372\nNumber of predicted TMHs: 1\nExp number of AAs in TMHs: 23.72226\nExp number, first 60 AAs: 0.22038\nTotal prob of N-in: 0.12994\noutside 1 331\nTMhelix 332 354\ninside 355 372

12011 GCF_000354435.2_ASM35443v2 Escherichia coli MP021017.2 Proteobacteria; Gammaproteobacteria; Enterobacterales; Enterobacteriaceae; Escherichia; Escherichia coli
MNNEETFYQAMRRQGVTRRSFLKYCSLAATSLGLGAGMAPKIAWALENKPRIPVVWIHGL WP_001058323.1
MULTISPECIES: hydrogenase-1 small chain [Proteobacteria] Length: 372\nNumber of predicted TMHs: 2\nExp number of AAs in TMHs: 37.2608499999999\nExp number, first 60 AAs: 19.52923\nTotal prob of N-in: 0.94351\nPOSSIBLE N-term signal sequence\ninside 1 19\nTMhelix 20 39\noutside 40 325\nTMhelix 326 348\ninside 349 372

12012 GCF_000354435.2_ASM35443v2 Escherichia coli MP021017.2 Proteobacteria; Gammaproteobacteria; Enterobacterales; Enterobacteriaceae; Escherichia; Escherichia coli
MTGDNPLIHSHGINRRDFMKLCAALAATMGLSSKAAAEMAESVTNPQRPPVIWIGAQECT WP_001461571.1
hydrogenase-2 small chain [Escherichia coli] Length: 372\nNumber of predicted TMHs: 1\nExp number of AAs in TMHs: 22.60469\nExp number, first 60 AAs: 0.1106\nTotal prob of N-in: 0.06393\noutside 1 331\nTMhelix 332 354\ninside 355 372

12013 GCF_000354475.2_ASM35447v2 Escherichia coli MP021017.3 Proteobacteria; Gammaproteobacteria; Enterobacterales; Enterobacteriaceae; Escherichia; Escherichia coli
MTGDNPLIHSHGINRRDFMKLCAALAATMGLSSKAAAEMAESVTNPQRPPVIWIGAQECT WP_001690065.1
hydrogenase-2 small chain [Escherichia coli] Length: 372\nNumber of predicted TMHs: 1\nExp number of AAs in TMHs: 22.60602\nExp number, first 60 AAs: 0.11062\nTotal prob of N-in: 0.06395\noutside 1 331\nTMhelix 332 354\ninside 355 372

12014 GCF_000354475.2_ASM35447v2 Escherichia coli MP021017.3 Proteobacteria; Gammaproteobacteria; Enterobacterales; Enterobacteriaceae; Escherichia; Escherichia coli
MNNEETFYQAMRRQGVTRRSFLKYCSLAATSLGLGAGMAPKIAWALENKPRIPVVWIHGL WP_001058323.1
MULTISPECIES: hydrogenase-1 small chain [Proteobacteria] Length: 372\nNumber of predicted TMHs: 2\nExp number of AAs in TMHs: 37.2608499999999\nExp number, first 60 AAs: 19.52923\nTotal prob of N-in: 0.94351\nPOSSIBLE N-term signal sequence\ninside 1 19\nTMhelix 20 39\noutside 40 325\nTMhelix 326 348\ninside 349 372

12015 GCF_000354495.2_ASM35449v2 Escherichia coli MP021017.4 Proteobacteria; Gammaproteobacteria; Enterobacterales; Enterobacteriaceae; Escherichia; Escherichia coli
MNNEETFYQAMRRQGVTRRSFLKYCSLAATSLGLGAGMAPKIAWALENKPRIPVVWIHGL WP_001058323.1
MULTISPECIES: hydrogenase-1 small chain [Proteobacteria] Length: 372\nNumber of predicted TMHs: 2\nExp number of AAs in TMHs: 37.2608499999999\nExp number, first 60 AAs: 19.52923\nTotal prob of N-in: 0.94351\nPOSSIBLE N-term signal sequence\ninside 1 19\nTMhelix 20 39\noutside 40 325\nTMhelix 326 348\ninside 349 372

12016 GCF_000354495.2_ASM35449v2 Escherichia coli MP021017.4 Proteobacteria; Gammaproteobacteria; Enterobacterales; Enterobacteriaceae; Escherichia; Escherichia coli
MTGDNTLIHSHGINRRDFMKLCAALAATMGLSSKAAAEMAESVTNPQRPPVIWIGAQECT WP_000145410.1
MULTISPECIES: hydrogenase-2 small chain [Proteobacteria] Length: 372\nNumber of predicted TMHs: 1\nExp number of AAs in TMHs: 23.72226\nExp number, first 60 AAs: 0.22038\nTotal prob of N-in: 0.12994\noutside 1 331\nTMhelix 332 354\ninside 355 372

12017 GCF_000354515.2_ASM35451v2 Escherichia coli MP021017.5 Proteobacteria; Gammaproteobacteria; Enterobacterales; Enterobacteriaceae; Escherichia; Escherichia coli
MNNEETFYQAMRRQGVTRRSFLKYCSLAATSLGLGAGMAPKIAWALENKPRIPVVWIHGL WP_001058323.1
MULTISPECIES: hydrogenase-1 small chain [Proteobacteria] Length: 372\nNumber of predicted TMHs: 2\nExp number of AAs in TMHs: 37.2608499999999\nExp number, first 60 AAs: 19.52923\nTotal prob of N-in: 0.94351\nPOSSIBLE N-term signal sequence\ninside 1 19\nTMhelix 20 39\noutside 40 325\nTMhelix 326 348\ninside 349 372

12018 GCF_000354515.2_ASM35451v2 Escherichia coli MP021017.5 Proteobacteria; Gammaproteobacteria; Enterobacterales; Enterobacteriaceae; Escherichia; Escherichia coli
MTGDNTLIHSHGINRRDFMKLCAALAATMGLSSKAAAEMAESVTNPQRPPVIWIGAQECT WP_000145410.1
MULTISPECIES: hydrogenase-2 small chain [Proteobacteria] Length: 372\nNumber of predicted TMHs: 1\nExp number of AAs in TMHs: 23.72226\nExp number, first 60 AAs: 0.22038\nTotal prob of N-in: 0.12994\noutside 1 331\nTMhelix 332 354\ninside 355 372

12019 GCF_000354535.2_ASM35453v2 Escherichia coli MP021017.6 Proteobacteria; Gammaproteobacteria; Enterobacterales; Enterobacteriaceae; Escherichia; Escherichia coli
MTGDNTLIHSHGINRRDFMKLCAALAATMGLSSKAAAEMAESVTNPQRPPVIWIGAQECT WP_000145410.1
MULTISPECIES: hydrogenase-2 small chain [Proteobacteria] Length: 372\nNumber of predicted TMHs: 1\nExp number of AAs in TMHs: 23.72226\nExp number, first 60 AAs: 0.22038\nTotal prob of N-in: 0.12994\noutside 1 331\nTMhelix 332 354\ninside 355 372

12020 GCF_000354535.2_ASM35453v2 Escherichia coli MP021017.6 Proteobacteria; Gammaproteobacteria; Enterobacterales; Enterobacteriaceae; Escherichia; Escherichia coli
MNNEETFYQAMRRQGVTRRSFLKYCSLAATSLGLGAGMAPKIAWALENKPRIPVVWIHGL WP_001058323.1
MULTISPECIES: hydrogenase-1 small chain [Proteobacteria] Length: 372\nNumber of predicted TMHs: 2\nExp number of AAs in TMHs: 37.2608499999999\nExp number, first 60 AAs: 19.52923\nTotal prob of N-in: 0.94351\nPOSSIBLE N-term signal sequence\ninside 1 19\nTMhelix 20 39\noutside 40 325\nTMhelix 326 348\ninside 349 372

12021 GCF_000354555.2_ASM35455v2 Escherichia coli MP021017.9 Proteobacteria; Gammaproteobacteria; Enterobacterales; Enterobacteriaceae; Escherichia; Escherichia coli
MTGDNTLIHSHGINRRDFMKLCAALAATMGLSSKAAAEMAESVTNPQRPPVIWIGAQECT WP_000145410.1
MULTISPECIES: hydrogenase-2 small chain [Proteobacteria] Length: 372\nNumber of predicted TMHs: 1\nExp number of AAs in TMHs: 23.72226\nExp number, first 60 AAs: 0.22038\nTotal prob of N-in: 0.12994\noutside 1 331\nTMhelix 332 354\ninside 355 372

12022 GCF_000354555.2_ASM35455v2 Escherichia coli MP021017.9 Proteobacteria; Gammaproteobacteria; Enterobacterales; Enterobacteriaceae; Escherichia; Escherichia coli
MNNEETFYQAMRRQGVTRRSFLKYCSLAATSLGLGAGMAPKIAWALENKPRIPVVWIHGL WP_001058323.1
MULTISPECIES: hydrogenase-1 small chain [Proteobacteria] Length: 372\nNumber of predicted TMHs: 2\nExp number of AAs in TMHs: 37.2608499999999\nExp number, first 60 AAs: 19.52923\nTotal prob of N-in: 0.94351\nPOSSIBLE N-term signal sequence\ninside 1 19\nTMhelix 20 39\noutside 40 325\nTMhelix 326 348\ninside 349 372

12023 GCF_000354575.2_ASM35457v2 Escherichia coli MP021552.11 Proteobacteria; Gammaproteobacteria; Enterobacterales; Enterobacteriaceae; Escherichia; Escherichia coli
MNNEETFYQAMRRQGVTRRSFLKYCSLAATSLGLGAGMAPKIAWALENKPRIPVVWIHGL WP_001058320.1
hydrogenase [Escherichia coli] Length: 372\nNumber of predicted TMHs: 2\nExp number of AAs in TMHs: 37.26082\nExp number, first 60 AAs: 19.52923\nTotal prob of N-in: 0.94351\nPOSSIBLE N-term signal sequence\ninside 1 19\nTMhelix 20 39\noutside 40 325\nTMhelix 326 348\ninside 349 372

12024 GCF_000354575.2_ASM35457v2 Escherichia coli MP021552.11 Proteobacteria; Gammaproteobacteria; Enterobacterales; Enterobacteriaceae; Escherichia; Escherichia coli
MTGDNTLIHSHGINRRDFMKLCAALAATMGLSSKAAAEMAESVTNPQRPPVIWIGAQECT WP_000145410.1
MULTISPECIES: hydrogenase-2 small chain [Proteobacteria] Length: 372\nNumber of predicted TMHs: 1\nExp number of AAs in TMHs: 23.72226\nExp number, first 60 AAs: 0.22038\nTotal prob of N-in: 0.12994\noutside 1 331\nTMhelix 332 354\ninside 355 372

12025 GCF_000354595.1_ASM35459v1 Escherichia coli MP021552.12 Proteobacteria; Gammaproteobacteria; Enterobacterales; Enterobacteriaceae; Escherichia; Escherichia coli
MNNEETFYQAMRRQGVTRRSFLKYCSLAATSLGLGAGMAPKIAWALENKPRIPVVWIHGL WP_001058320.1
hydrogenase [Escherichia coli] Length: 372\nNumber of predicted TMHs: 2\nExp number of AAs in TMHs: 37.26082\nExp number, first 60 AAs: 19.52923\nTotal prob of N-in: 0.94351\nPOSSIBLE N-term signal sequence\ninside 1 19\nTMhelix 20 39\noutside 40 325\nTMhelix 326 348\ninside 349 372

12026 GCF_000354595.1_ASM35459v1 Escherichia coli MP021552.12 Proteobacteria; Gammaproteobacteria; Enterobacterales; Enterobacteriaceae; Escherichia; Escherichia coli
MTGDNTLIHSHGINRRDFMKLCAALAATMGLSSKAAAEMAESVTNPQRPPVIWIGAQECT WP_000145410.1
MULTISPECIES: hydrogenase-2 small chain [Proteobacteria] Length: 372\nNumber of predicted TMHs: 1\nExp number of AAs in TMHs: 23.72226\nExp number, first 60 AAs: 0.22038\nTotal prob of N-in: 0.12994\noutside 1 331\nTMhelix 332 354\ninside 355 372

12027 GCF_000354615.2_ASM35461v2 Escherichia coli MP021552.7 Proteobacteria; Gammaproteobacteria; Enterobacterales; Enterobacteriaceae; Escherichia; Escherichia coli
MNNEETFYQAMRRQGVTRRSFLKYCSLAATSLGLGAGMAPKIAWALENKPRIPVVWIHGL WP_001058320.1

hydrogenase [Escherichia coli]	Length: 372\nNumber of predicted TMHs: 2\nExp number of AAs in TMHs: 37.26082\nExp number, first 60 AAs: 19.52923\nTotal prob of N-in: 0.94351\nPOSSIBLE N-term signal sequence\ninside 1 19\nTMhelix 20 39\noutside 40 325\nTMhelix 326 348\ninside 349 372

12028 GCF_000354615.2_ASM35461v2 Escherichia coli MP021552.7 Proteobacteria; Gammaproteobacteria; Enterobacterales; Enterobacteriaceae; Escherichia; Escherichia coli MTGDNTLIHSHGINRRDFMKLCAALAATMGLSSKAAAEMAESVTNPQRPPVIWIGAQECT WP_000145410.1 MULTISPECIES: hydrogenase-2 small chain [Proteobacteria] Length: 372\nNumber of predicted TMHs: 1\nExp number of AAs in TMHs: 23.72226\nExp number, first 60 AAs: 0.22038\nTotal prob of N-in: 0.12994\noutside 1 331\nTMhelix 332 354\ninside 355 372

12029 GCF_000354735.2_ASM35473v2 Escherichia coli MP021552.8 Proteobacteria; Gammaproteobacteria; Enterobacterales; Enterobacteriaceae; Escherichia; Escherichia coli MNNEETFYQAMRRQGVTRRSFLKYCSLAATSLGLGAGMAPKIAWALENKPRIPVVWIHGL WP_001058320.1 hydrogenase [Escherichia coli] Length: 372\nNumber of predicted TMHs: 2\nExp number of AAs in TMHs: 37.26082\nExp number, first 60 AAs: 19.52923\nTotal prob of N-in: 0.94351\nPOSSIBLE N-term signal sequence\ninside 1 19\nTMhelix 20 39\noutside 40 325\nTMhelix 326 348\ninside 349 372

12030 GCF_000354735.2_ASM35473v2 Escherichia coli MP021552.8 Proteobacteria; Gammaproteobacteria; Enterobacterales; Enterobacteriaceae; Escherichia; Escherichia coli MTGDNTLIHSHGINRRDFMKLCAALAATMGLSSKAAAEMAESVTNPQRPPVIWIGAQECT WP_000145410.1 MULTISPECIES: hydrogenase-2 small chain [Proteobacteria] Length: 372\nNumber of predicted TMHs: 1\nExp number of AAs in TMHs: 23.72226\nExp number, first 60 AAs: 0.22038\nTotal prob of N-in: 0.12994\noutside 1 331\nTMhelix 332 354\ninside 355 372

12031 GCF_000354795.2_ASM35479v2 Escherichia coli MP021561.2 Proteobacteria; Gammaproteobacteria; Enterobacterales; Enterobacteriaceae; Escherichia; Escherichia coli MNNEETFYQAMRRQGVTRRSFLKYCSLAATSLGLGAGMAPKIAWALENKPRIPVVWIHGL WP_001058323.1 MULTISPECIES: hydrogenase-1 small chain [Proteobacteria] Length: 372\nNumber of predicted TMHs: 2\nExp number of AAs in TMHs: 37.2608499999999\nExp number, first 60 AAs: 19.52923\nTotal prob of N-in: 0.94351\nPOSSIBLE N-term signal sequence\ninside 1 19\nTMhelix 20 39\noutside 40 325\nTMhelix 326 348\ninside 349 372

12032 GCF_000354795.2_ASM35479v2 Escherichia coli MP021561.2 Proteobacteria; Gammaproteobacteria; Enterobacterales; Enterobacteriaceae; Escherichia; Escherichia coli MTGDNTLIHSHGINRRDFMKLCAALAATMGLSSKAAAEMAESVTNPQRPPVIWIGAQECT WP_000145410.1 MULTISPECIES: hydrogenase-2 small chain [Proteobacteria] Length: 372\nNumber of predicted TMHs: 1\nExp number of AAs in TMHs: 23.72226\nExp number, first 60 AAs: 0.22038\nTotal prob of N-in: 0.12994\noutside 1 331\nTMhelix 332 354\ninside 355 372

12033 GCF_000356425.2_ASM35642v2 Escherichia coli MP021561.3 Proteobacteria; Gammaproteobacteria; Enterobacterales; Enterobacteriaceae; Escherichia; Escherichia coli MTGDNTLIHSHGINRRDFMKLCAALAATMGLSSKAAAEMAESVTNPQRPPVIWIGAQECT WP_000145410.1 MULTISPECIES: hydrogenase-2 small chain [Proteobacteria] Length: 372\nNumber of predicted TMHs: 1\nExp number of AAs in TMHs: 23.72226\nExp number, first 60 AAs: 0.22038\nTotal prob of N-in: 0.12994\noutside 1 331\nTMhelix 332 354\ninside 355 372

12034 GCF_000356425.2_ASM35642v2 Escherichia coli MP021561.3 Proteobacteria; Gammaproteobacteria; Enterobacterales; Enterobacteriaceae; Escherichia; Escherichia coli MNNEETFYQAMRRQGVTRRSFLKYCSLAATSLGLGAGMAPKIAWALENKPRIPVVWIHGL WP_001058323.1 MULTISPECIES: hydrogenase-1 small chain [Proteobacteria] Length: 372\nNumber of predicted TMHs: 2\nExp number of AAs in TMHs: 37.2608499999999\nExp number, first 60 AAs: 19.52923\nTotal prob of N-in: 0.94351\nPOSSIBLE N-term signal sequence\ninside 1 19\nTMhelix 20 39\noutside 40 325\nTMhelix 326 348\ninside 349 372

12035 GCF_000354855.1_ASM35485v1 Escherichia coli MP021566.1 Proteobacteria; Gammaproteobacteria; Enterobacterales; Enterobacteriaceae; Escherichia; Escherichia coli MTGDNTLIHSHGINRRDFMKLCAALAATMGLSSKAAAEMAESVTNPQRPPVIWIGAQECT WP_000145410.1 MULTISPECIES: hydrogenase-2 small chain [Proteobacteria] Length: 372\nNumber of predicted TMHs: 1\nExp number of AAs in TMHs: 23.72226\nExp number, first 60 AAs: 0.22038\nTotal prob of N-in: 0.12994\noutside 1 331\nTMhelix 332 354\ninside 355 372

12036 GCF_000354855.1_ASM35485v1 Escherichia coli MP021566.1 Proteobacteria; Gammaproteobacteria; Enterobacterales; Enterobacteriaceae; Escherichia; Escherichia coli
MNNEETFYQAMRRQGVTRRSFLKYCSLAATSLGLGAGMAPKIAWALENKPRIPVVWIHGL WP_001058323.1
MULTISPECIES: hydrogenase-1 small chain [Proteobacteria] Length: 372\nNumber of predicted TMHs: 2\nExp number of AAs in TMHs: 37.2608499999999\nExp number, first 60 AAs: 19.52923\nTotal prob of N-in: 0.94351\nPOSSIBLE N-term signal sequence\ninside 1 19\nTMhelix 20 39\noutside 40 325\nTMhelix 326 348\ninside 349 372

12037 GCF_000576655.1_MP1_1.0 Escherichia coli MP1 Proteobacteria; Gammaproteobacteria; Enterobacterales; Enterobacteriaceae; Escherichia; Escherichia coli
MNNEETFYQAMRRQGVTRRSFLKYCSLAATSLGLGAGMAPKIAWALENKPRIPVVWIHGL WP_001058323.1
MULTISPECIES: hydrogenase-1 small chain [Proteobacteria] Length: 372\nNumber of predicted TMHs: 2\nExp number of AAs in TMHs: 37.2608499999999\nExp number, first 60 AAs: 19.52923\nTotal prob of N-in: 0.94351\nPOSSIBLE N-term signal sequence\ninside 1 19\nTMhelix 20 39\noutside 40 325\nTMhelix 326 348\ninside 349 372

12038 GCF_000576655.1_MP1_1.0 Escherichia coli MP1 Proteobacteria; Gammaproteobacteria; Enterobacterales; Enterobacteriaceae; Escherichia; Escherichia coli
MTGDNTLIHSHGINRRDFMKLCAALAATMGLSSKAAAEMVESVTNPQRPPVIWIGAQECT WP_000145423.1
hydrogenase 2 small subunit [Escherichia coli] Length: 372\nNumber of predicted TMHs: 1\nExp number of AAs in TMHs: 23.78201\nExp number, first 60 AAs: 0.28355\nTotal prob of N-in: 0.13257\noutside 1 331\nTMhelix 332 354\ninside 355 372

12039 GCF_000982035.1_ASM98203v1 Escherichia coli MRSN 10204 Proteobacteria; Gammaproteobacteria; Enterobacterales; Enterobacteriaceae; Escherichia; Escherichia coli
MTGDNTLIHSHGINRRDFMKLCAALAATMGLSSKAAAEMAESVTNPQRPPVIWIGAQECT WP_000145418.1
hydrogenase 2 small subunit [Escherichia coli] Length: 372\nNumber of predicted TMHs: 1\nExp number of AAs in TMHs: 23.72313\nExp number, first 60 AAs: 0.22034\nTotal prob of N-in: 0.12997\noutside 1 331\nTMhelix 332 354\ninside 355 372

12040 GCF_000982035.1_ASM98203v1 Escherichia coli MRSN 10204 Proteobacteria; Gammaproteobacteria; Enterobacterales; Enterobacteriaceae; Escherichia; Escherichia coli
MNNEETFYQAMRRQGVTRRSFLKYCSLAATSLGLGAGMAPKIAWALENKPRIPVVWIHGL WP_001058323.1
MULTISPECIES: hydrogenase-1 small chain [Proteobacteria] Length: 372\nNumber of predicted TMHs: 2\nExp number of AAs in TMHs: 37.2608499999999\nExp number, first 60 AAs: 19.52923\nTotal prob of N-in: 0.94351\nPOSSIBLE N-term signal sequence\ninside 1 19\nTMhelix 20 39\noutside 40 325\nTMhelix 326 348\ninside 349 372

12041 GCF_000179175.1_ASM17917v1 Escherichia coli MS 107-1 Proteobacteria; Gammaproteobacteria; Enterobacterales; Enterobacteriaceae; Escherichia; Escherichia coli
MTGDNTLIHSHGINRRDFMKLCAALAATMGLSSKAAAEMAESVTNPQRPPVIWIGAQECT WP_000145410.1
MULTISPECIES: hydrogenase-2 small chain [Proteobacteria] Length: 372\nNumber of predicted TMHs: 1\nExp number of AAs in TMHs: 23.72226\nExp number, first 60 AAs: 0.22038\nTotal prob of N-in: 0.12994\noutside 1 331\nTMhelix 332 354\ninside 355 372

12042 GCF_000179175.1_ASM17917v1 Escherichia coli MS 107-1 Proteobacteria; Gammaproteobacteria; Enterobacterales; Enterobacteriaceae; Escherichia; Escherichia coli
MNNEETFYQAMRRQGVTRRSFLKYCSLAATSLGLGAGMAPKIAWALENKPRIPVVWIHGL WP_001295940.1
MULTISPECIES: hydrogenase [Enterobacteriaceae] Length: 372\nNumber of predicted TMHs: 2\nExp number of AAs in TMHs: 37.2608899999999\nExp number, first 60 AAs: 19.52923\nTotal prob of N-in: 0.94351\nPOSSIBLE N-term signal sequence\ninside 1 19\nTMhelix 20 39\noutside 40 325\nTMhelix 326 348\ninside 349 372

12043 GCF_000164415.1_ASM16441v1 Escherichia coli MS 110-3 Proteobacteria; Gammaproteobacteria; Enterobacterales; Enterobacteriaceae; Escherichia; Escherichia coli
MTGDNTLIHSHGINRRDFMKLCAALAATMGLSSKAAAEMAESVTNPQRPPVIWIGAQECT WP_000145410.1
MULTISPECIES: hydrogenase-2 small chain [Proteobacteria] Length: 372\nNumber of predicted TMHs: 1\nExp number of AAs in TMHs: 23.72226\nExp number, first 60 AAs: 0.22038\nTotal prob of N-in: 0.12994\noutside 1 331\nTMhelix 332 354\ninside 355 372

12044 GCF_000164235.1_ASM16423v1 Escherichia coli MS 115-1 Proteobacteria; Gammaproteobacteria; Enterobacterales; Enterobacteriaceae; Escherichia; Escherichia coli
MTGDNTLIHSHGINRRDFMKLCAALAATMGLSSKAAAEMAESVTNPQRPPVIWIGAQECT WP_000145410.1
MULTISPECIES: hydrogenase-2 small chain [Proteobacteria] Length: 372\nNumber of predicted TMHs: 1\nExp number of AAs in TMHs: 23.72226\nExp number, first 60 AAs: 0.22038\nTotal prob of N-in: 0.12994\noutside 1 331\nTMhelix 332 354\ninside 355 372

12045 GCF_000164235.1_ASM16423v1 Escherichia coli MS 115-1 Proteobacteria; Gammaproteobacteria; Enterobacterales; Enterobacteriaceae; Escherichia; Escherichia coli
MNNEETFYQAMRRQGVTRRSFLKYCSLAATSLGLGAGMAPKIAWALENKPRIPVVWIHGL WP_001058323.1
MULTISPECIES: hydrogenase-1 small chain [Proteobacteria] Length: 372\nNumber of predicted TMHs: 2\nExp number of AAs in TMHs: 37.2608499999999\nExp number, first 60 AAs: 19.52923\nTotal prob of N-in: 0.94351\nPOSSIBLE N-term signal sequence\ninside 1 19\nTMhelix 20 39\noutside 40 325\nTMhelix 326 348\ninside 349 372

12046 GCF_000164475.1_ASM16447v1 Escherichia coli MS 116-1 Proteobacteria; Gammaproteobacteria; Enterobacterales; Enterobacteriaceae; Escherichia; Escherichia coli
MTGDNTLIHSHGINRRDFMKLCAALAATMGLSSKAAAEMAESVTNPQRPPVIWIGAQECT WP_000145410.1
MULTISPECIES: hydrogenase-2 small chain [Proteobacteria] Length: 372\nNumber of predicted TMHs: 1\nExp number of AAs in TMHs: 23.72226\nExp number, first 60 AAs: 0.22038\nTotal prob of N-in: 0.12994\noutside 1 331\nTMhelix 332 354\ninside 355 372

12047 GCF_000164475.1_ASM16447v1 Escherichia coli MS 116-1 Proteobacteria; Gammaproteobacteria; Enterobacterales; Enterobacteriaceae; Escherichia; Escherichia coli
MNNEETFYQAMRRQGVTRRSFLKYCSLAATSLGLGAGMAPKIAWALENKPRIPVVWIHGL WP_001058323.1
MULTISPECIES: hydrogenase-1 small chain [Proteobacteria] Length: 372\nNumber of predicted TMHs: 2\nExp number of AAs in TMHs: 37.2608499999999\nExp number, first 60 AAs: 19.52923\nTotal prob of N-in: 0.94351\nPOSSIBLE N-term signal sequence\ninside 1 19\nTMhelix 20 39\noutside 40 325\nTMhelix 326 348\ninside 349 372

12048 GCF_000164375.1_ASM16437v1 Escherichia coli MS 117-3 Proteobacteria; Gammaproteobacteria; Enterobacterales; Enterobacteriaceae; Escherichia; Escherichia coli
MTGDNTLIHSHGINRRDFMKLCAALAATMGLSSKAAAEMAESVTNPQRPPVIWIGAQECT WP_000145410.1
MULTISPECIES: hydrogenase-2 small chain [Proteobacteria] Length: 372\nNumber of predicted TMHs: 1\nExp number of AAs in TMHs: 23.72226\nExp number, first 60 AAs: 0.22038\nTotal prob of N-in: 0.12994\noutside 1 331\nTMhelix 332 354\ninside 355 372

12049 GCF_000164375.1_ASM16437v1 Escherichia coli MS 117-3 Proteobacteria; Gammaproteobacteria; Enterobacterales; Enterobacteriaceae; Escherichia; Escherichia coli
MNNEETFYQAMRRQGVTRRSFLKYCSLAATSLGLGAGMAPKIAWALENKPRIPVVWIHGL WP_001295940.1
MULTISPECIES: hydrogenase [Enterobacteriaceae] Length: 372\nNumber of predicted TMHs: 2\nExp number of AAs in TMHs: 37.2608899999999\nExp number, first 60 AAs: 19.52923\nTotal prob of N-in: 0.94351\nPOSSIBLE N-term signal sequence\ninside 1 19\nTMhelix 20 39\noutside 40 325\nTMhelix 326 348\ninside 349 372

12050 GCF_000179155.1_ASM17915v1 Escherichia coli MS 119-7 Proteobacteria; Gammaproteobacteria; Enterobacterales; Enterobacteriaceae; Escherichia; Escherichia coli
MNNEETFYQAMRRQGVTRRSFLKYCSLAATSLGLGAGMAPKIAWALENKPRIPVVWIHGL WP_001058323.1
MULTISPECIES: hydrogenase-1 small chain [Proteobacteria] Length: 372\nNumber of predicted TMHs: 2\nExp number of AAs in TMHs: 37.2608499999999\nExp number, first 60 AAs: 19.52923\nTotal prob of N-in: 0.94351\nPOSSIBLE N-term signal sequence\ninside 1 19\nTMhelix 20 39\noutside 40 325\nTMhelix 326 348\ninside 349 372

12051 GCF_000179155.1_ASM17915v1 Escherichia coli MS 119-7 Proteobacteria; Gammaproteobacteria; Enterobacterales; Enterobacteriaceae; Escherichia; Escherichia coli
MTGDNTLIHSHGINRRDFMKLCAALAATMGLSSKAAAEMAESVTNPQRPPVIWIGAQECT WP_000145410.1
MULTISPECIES: hydrogenase-2 small chain [Proteobacteria] Length: 372\nNumber of predicted TMHs: 1\nExp number of AAs in TMHs: 23.72226\nExp number, first 60 AAs: 0.22038\nTotal prob of N-in: 0.12994\noutside 1 331\nTMhelix 332 354\ninside 355 372

12052 GCF_000179135.1_ASM17913v1 Escherichia coli MS 124-1 Proteobacteria; Gammaproteobacteria; Enterobacterales; Enterobacteriaceae; Escherichia; Escherichia coli
MTGDNTLIHSHGINRRDFMKLCAALAATMGLSSKAAAEMAESVTNPQRPPVIWIGAQECT WP_000145410.1
MULTISPECIES: hydrogenase-2 small chain [Proteobacteria] Length: 372\nNumber of predicted TMHs: 1\nExp number of AAs in TMHs: 23.72226\nExp number, first 60 AAs: 0.22038\nTotal prob of N-in: 0.12994\noutside 1 331\nTMhelix 332 354\ninside 355 372

12053 GCF_000179135.1_ASM17913v1 Escherichia coli MS 124-1 Proteobacteria; Gammaproteobacteria; Enterobacterales; Enterobacteriaceae; Escherichia; Escherichia coli
MNNEETFYQAMRRQGVTRRSFLKYCSLAATSLGLGAGMAPKIAWALENKPRIPVVWIHGL WP_001058320.1
hydrogenase [Escherichia coli] Length: 372\nNumber of predicted TMHs: 2\nExp number of AAs in TMHs: 37.26082\nExp number, first 60 AAs: 19.52923\nTotal prob of N-in: 0.94351\nPOSSIBLE N-term signal sequence\ninside 1 19\nTMhelix 20 39\noutside 40 325\nTMhelix 326 348\ninside 349 372

12054 GCF_000179115.1_ASM17911v1 Escherichia coli MS 145-7 Proteobacteria; Gammaproteobacteria; Enterobacterales; Enterobacteriaceae; Escherichia; Escherichia coli
MNNEETFYQAMRRQGVTRRSFLKYCSLAATSLGLGAGMAPKIAWALENKPRIPVVWIHGL WP_001058323.1
MULTISPECIES: hydrogenase-1 small chain [Proteobacteria] Length: 372\nNumber of predicted TMHs: 2\nExp number of AAs in TMHs: 37.2608499999999\nExp number, first 60 AAs: 19.52923\nTotal prob of N-in: 0.94351\nPOSSIBLE N-term signal sequence\ninside 1 19\nTMhelix 20 39\noutside 40 325\nTMhelix 326 348\ninside 349 372

12055 GCF_000179115.1_ASM17911v1 Escherichia coli MS 145-7 Proteobacteria; Gammaproteobacteria; Enterobacterales; Enterobacteriaceae; Escherichia; Escherichia coli
MTGDNTLIHSHGINRRDFMKLCAALAATMGLSSKAAAEMAESVTNPQRPPVIWIGAQECT WP_000145410.1
MULTISPECIES: hydrogenase-2 small chain [Proteobacteria] Length: 372\nNumber of predicted TMHs: 1\nExp number of AAs in TMHs: 23.72226\nExp number, first 60 AAs: 0.22038\nTotal prob of N-in: 0.12994\noutside 1 331\nTMhelix 332 354\ninside 355 372

12056 GCF_000164275.1_ASM16427v1 Escherichia coli MS 146-1 Proteobacteria; Gammaproteobacteria; Enterobacterales; Enterobacteriaceae; Escherichia; Escherichia coli
MTGDNTLIHSHGINRRDFMKLCAALAATMGLSSKAAAEMAESVTNPQRPPVIWIGAQECT WP_000145410.1
MULTISPECIES: hydrogenase-2 small chain [Proteobacteria] Length: 372\nNumber of predicted TMHs: 1\nExp number of AAs in TMHs: 23.72226\nExp number, first 60 AAs: 0.22038\nTotal prob of N-in: 0.12994\noutside 1 331\nTMhelix 332 354\ninside 355 372

12057 GCF_000164275.1_ASM16427v1 Escherichia coli MS 146-1 Proteobacteria; Gammaproteobacteria; Enterobacterales; Enterobacteriaceae; Escherichia; Escherichia coli
MNNEETFYQAMRRQGVTRRSFLKYCSLAATSLGLGAGMAPKIAWALENKPRIPVVWIHGL WP_001058323.1
MULTISPECIES: hydrogenase-1 small chain [Proteobacteria] Length: 372\nNumber of predicted TMHs: 2\nExp number of AAs in TMHs: 37.2608499999999\nExp number, first 60 AAs: 19.52923\nTotal prob of N-in: 0.94351\nPOSSIBLE N-term signal sequence\ninside 1 19\nTMhelix 20 39\noutside 40 325\nTMhelix 326 348\ninside 349 372

12058 GCF_000164435.1_ASM16443v1 Escherichia coli MS 153-1 Proteobacteria; Gammaproteobacteria; Enterobacterales; Enterobacteriaceae; Escherichia; Escherichia coli
MNNEETFYQAMRRQGVTRRSFLKYCSLAATSLGLGAGMAPKIAWALENKPRIPVVWIHGL WP_001332755.1
hydrogenase [Escherichia coli] Length: 372\nNumber of predicted TMHs: 2\nExp number of AAs in TMHs: 37.2640999999999\nExp number, first 60 AAs: 19.52809\nTotal prob of N-in: 0.94350\nPOSSIBLE N-term signal sequence\ninside 1 19\nTMhelix 20 39\noutside 40 325\nTMhelix 326 348\ninside 349 372

12059 GCF_000164435.1_ASM16443v1 Escherichia coli MS 153-1 Proteobacteria; Gammaproteobacteria; Enterobacterales; Enterobacteriaceae; Escherichia; Escherichia coli
MTGDNTLIHSHGINRRDFMKLCAALAATMGLSSKAAAEMAESVTNPQRPPVIWIGAQECT WP_000145410.1
MULTISPECIES: hydrogenase-2 small chain [Proteobacteria] Length: 372\nNumber of predicted TMHs: 1\nExp number of AAs in TMHs: 23.72226\nExp number, first 60 AAs: 0.22038\nTotal prob of N-in: 0.12994\noutside 1 331\nTMhelix 332 354\ninside 355 372

12060 GCF_000164495.1_ASM16449v1 Escherichia coli MS 16-3 Proteobacteria; Gammaproteobacteria; Enterobacterales; Enterobacteriaceae; Escherichia; Escherichia coli
MNNEETFYQAMRRQGVTRRSFLKYCSLAATSLGLGAGMAPKIAWALENKPRIPVVWIHGL WP_001058323.1

MULTISPECIES: hydrogenase-1 small chain [Proteobacteria] Length: 372\nNumber of predicted TMHs: 2\nExp number of AAs in TMHs: 37.2608499999999\nExp number, first 60 AAs: 19.52923\nTotal prob of N-in: 0.94351\nPOSSIBLE N-term signal sequence\ninside 1 19\nTMhelix 20 39\noutside 40 325\nTMhelix 326 348\ninside 349 372

12061 GCF_000164495.1_ASM16449v1 Escherichia coli MS 16-3 Proteobacteria; Gammaproteobacteria; Enterobacterales; Enterobacteriaceae; Escherichia; Escherichia coli
MTGDNTLIHSHGINRRDFMKLCAALAATMGLSSKAAAEMVESVTNPQRPPVIWIGAQECT WP_000145423.1
hydrogenase 2 small subunit [Escherichia coli] Length: 372\nNumber of predicted TMHs: 1\nExp number of AAs in TMHs: 23.78201\nExp number, first 60 AAs: 0.28355\nTotal prob of N-in: 0.13257\noutside 1 331\nTMhelix 332 354\ninside 355 372

12062 GCF_000164515.1_ASM16451v1 Escherichia coli MS 175-1 Proteobacteria; Gammaproteobacteria; Enterobacterales; Enterobacteriaceae; Escherichia; Escherichia coli
MTGDNTLIHSHGINRRDFMKLCAALAATMGLSSKAAAEMAESVTNPQRPPVIWIGAQECT WP_000145410.1
MULTISPECIES: hydrogenase-2 small chain [Proteobacteria] Length: 372\nNumber of predicted TMHs: 1\nExp number of AAs in TMHs: 23.72226\nExp number, first 60 AAs: 0.22038\nTotal prob of N-in: 0.12994\noutside 1 331\nTMhelix 332 354\ninside 355 372

12063 GCF_000164515.1_ASM16451v1 Escherichia coli MS 175-1 Proteobacteria; Gammaproteobacteria; Enterobacterales; Enterobacteriaceae; Escherichia; Escherichia coli
MNNEETFYQAMRRQGVTRRSFLKYCSLAATSLGLGAGMAPKIAWALENKPRIPVVWIHGL WP_001058323.1
MULTISPECIES: hydrogenase-1 small chain [Proteobacteria] Length: 372\nNumber of predicted TMHs: 2\nExp number of AAs in TMHs: 37.2608499999999\nExp number, first 60 AAs: 19.52923\nTotal prob of N-in: 0.94351\nPOSSIBLE N-term signal sequence\ninside 1 19\nTMhelix 20 39\noutside 40 325\nTMhelix 326 348\ninside 349 372

12064 GCF_000164255.1_ASM16425v1 Escherichia coli MS 182-1 Proteobacteria; Gammaproteobacteria; Enterobacterales; Enterobacteriaceae; Escherichia; Escherichia coli
MTGDNTLIHSHGINRRDFMKLCAALAATMGLSSKAAAEMAESVTNPQRPPVIWIGAQECT WP_000145411.1
hydrogenase-2 small chain [Escherichia coli] Length: 372\nNumber of predicted TMHs: 1\nExp number of AAs in TMHs: 23.70577\nExp number, first 60 AAs: 0.22039\nTotal prob of N-in: 0.12981\noutside 1 331\nTMhelix 332 354\ninside 355 372

12065 GCF_000164255.1_ASM16425v1 Escherichia coli MS 182-1 Proteobacteria; Gammaproteobacteria; Enterobacterales; Enterobacteriaceae; Escherichia; Escherichia coli
MNNEETFYQAMRRQGVTRRSFLKYCSLAATSLGLGAGMAPKIAWALENKPRIPVVWIHGL WP_001058323.1
MULTISPECIES: hydrogenase-1 small chain [Proteobacteria] Length: 372\nNumber of predicted TMHs: 2\nExp number of AAs in TMHs: 37.2608499999999\nExp number, first 60 AAs: 19.52923\nTotal prob of N-in: 0.94351\nPOSSIBLE N-term signal sequence\ninside 1 19\nTMhelix 20 39\noutside 40 325\nTMhelix 326 348\ninside 349 372

12066 GCF_000164575.1_ASM16457v1 Escherichia coli MS 185-1 Proteobacteria; Gammaproteobacteria; Enterobacterales; Enterobacteriaceae; Escherichia; Escherichia coli
MTGDNTLIHSHGINRRDFMKLCAALAATMGLSSKAAAEMAESVTNPQRPPVIWIGAQECT WP_000145410.1
MULTISPECIES: hydrogenase-2 small chain [Proteobacteria] Length: 372\nNumber of predicted TMHs: 1\nExp number of AAs in TMHs: 23.72226\nExp number, first 60 AAs: 0.22038\nTotal prob of N-in: 0.12994\noutside 1 331\nTMhelix 332 354\ninside 355 372

12067 GCF_000164575.1_ASM16457v1 Escherichia coli MS 185-1 Proteobacteria; Gammaproteobacteria; Enterobacterales; Enterobacteriaceae; Escherichia; Escherichia coli
MNNEETFYQAMRRQGVTRRSFLKYCSLAATSLGLGAGMAPKIAWALENKPRIPVVWIHGL WP_001058323.1
MULTISPECIES: hydrogenase-1 small chain [Proteobacteria] Length: 372\nNumber of predicted TMHs: 2\nExp number of AAs in TMHs: 37.2608499999999\nExp number, first 60 AAs: 19.52923\nTotal prob of N-in: 0.94351\nPOSSIBLE N-term signal sequence\ninside 1 19\nTMhelix 20 39\noutside 40 325\nTMhelix 326 348\ninside 349 372

12068 GCF_000164335.1_ASM16433v1 Escherichia coli MS 187-1 Proteobacteria; Gammaproteobacteria; Enterobacterales; Enterobacteriaceae; Escherichia; Escherichia coli
MTGDNTLIHSHGINRRDFMKLCAALAATMGLSSKAAAEMAESVTNPQRPPVIWIGAQECT WP_000145410.1
MULTISPECIES: hydrogenase-2 small chain [Proteobacteria] Length: 372\nNumber of predicted TMHs:

1\nExp number of AAs in TMHs: 23.72226\nExp number, first 60 AAs: 0.22038\nTotal prob of N-in: 0.12994\noutside 1 331\nTMhelix 332 354\ninside 355 372

12069 GCF_000164335.1_ASM16433v1 Escherichia coli MS 187-1 Proteobacteria; Gammaproteobacteria; Enterobacterales; Enterobacteriaceae; Escherichia; Escherichia coli
MNNEETFYQAMRRQGVTRRSFLKYCSLAATSLGLGAGMAPKIAWALENKPRIPVVWIHGL WP_001058323.1
MULTISPECIES: hydrogenase-1 small chain [Proteobacteria] Length: 372\nNumber of predicted TMHs: 2\nExp number of AAs in TMHs: 37.2608499999999\nExp number, first 60 AAs: 19.52923\nTotal prob of N-in: 0.94351\nPOSSIBLE N-term signal sequence\ninside 1 19\nTMhelix 20 39\noutside 40 325\nTMhelix 326 348\ninside 349 372

12070 GCF_000164555.1_ASM16455v1 Escherichia coli MS 196-1 Proteobacteria; Gammaproteobacteria; Enterobacterales; Enterobacteriaceae; Escherichia; Escherichia coli
MNNEETFYQAMRRQGVTRRSFLKYCSLAATSLGLGAGMAPKIAWALENKPRIPVVWIHGL WP_001058323.1
MULTISPECIES: hydrogenase-1 small chain [Proteobacteria] Length: 372\nNumber of predicted TMHs: 2\nExp number of AAs in TMHs: 37.2608499999999\nExp number, first 60 AAs: 19.52923\nTotal prob of N-in: 0.94351\nPOSSIBLE N-term signal sequence\ninside 1 19\nTMhelix 20 39\noutside 40 325\nTMhelix 326 348\ninside 349 372

12071 GCF_000164555.1_ASM16455v1 Escherichia coli MS 196-1 Proteobacteria; Gammaproteobacteria; Enterobacterales; Enterobacteriaceae; Escherichia; Escherichia coli
MTGDNTLIHSHGINRRDFMKLCAALAATMGLSSKAAAEMAESVTNPQRPPVIWIGAQECT WP_000145410.1
MULTISPECIES: hydrogenase-2 small chain [Proteobacteria] Length: 372\nNumber of predicted TMHs: 1\nExp number of AAs in TMHs: 23.72226\nExp number, first 60 AAs: 0.22038\nTotal prob of N-in: 0.12994\noutside 1 331\nTMhelix 332 354\ninside 355 372

12072 GCF_000164195.1_ASM16419v1 Escherichia coli MS 198-1 Proteobacteria; Gammaproteobacteria; Enterobacterales; Enterobacteriaceae; Escherichia; Escherichia coli
MNNEETFYQAMRRQGVTRRSFLKYCSLAATSLGLGAGMAPKIAWALENKPRIPVVWIHGL WP_001058323.1
MULTISPECIES: hydrogenase-1 small chain [Proteobacteria] Length: 372\nNumber of predicted TMHs: 2\nExp number of AAs in TMHs: 37.2608499999999\nExp number, first 60 AAs: 19.52923\nTotal prob of N-in: 0.94351\nPOSSIBLE N-term signal sequence\ninside 1 19\nTMhelix 20 39\noutside 40 325\nTMhelix 326 348\ninside 349 372

12073 GCF_000164195.1_ASM16419v1 Escherichia coli MS 198-1 Proteobacteria; Gammaproteobacteria; Enterobacterales; Enterobacteriaceae; Escherichia; Escherichia coli
MTGDNTLIHSHGINRRDFMKLCAALAATMGLSSKAAAEMAESVTNPQRPPVIWIGAQECT WP_000145410.1
MULTISPECIES: hydrogenase-2 small chain [Proteobacteria] Length: 372\nNumber of predicted TMHs: 1\nExp number of AAs in TMHs: 23.72226\nExp number, first 60 AAs: 0.22038\nTotal prob of N-in: 0.12994\noutside 1 331\nTMhelix 332 354\ninside 355 372

12074 GCF_000164535.1_ASM16453v1 Escherichia coli MS 200-1 Proteobacteria; Gammaproteobacteria; Enterobacterales; Enterobacteriaceae; Escherichia; Escherichia coli
MTGDNTLIHSHGINRRDFMKLCAALAATMGLSSKAAAEMAESVTNPQRPPVIWIGAQECT WP_000145410.1
MULTISPECIES: hydrogenase-2 small chain [Proteobacteria] Length: 372\nNumber of predicted TMHs: 1\nExp number of AAs in TMHs: 23.72226\nExp number, first 60 AAs: 0.22038\nTotal prob of N-in: 0.12994\noutside 1 331\nTMhelix 332 354\ninside 355 372

12075 GCF_000164535.1_ASM16453v1 Escherichia coli MS 200-1 Proteobacteria; Gammaproteobacteria; Enterobacterales; Enterobacteriaceae; Escherichia; Escherichia coli
MNNEETFYQAMRRQGVTRRSFLKYCSLAATSLGLGAGMAPKIAWALENKPRIPVVWIHGL WP_001058323.1
MULTISPECIES: hydrogenase-1 small chain [Proteobacteria] Length: 372\nNumber of predicted TMHs: 2\nExp number of AAs in TMHs: 37.2608499999999\nExp number, first 60 AAs: 19.52923\nTotal prob of N-in: 0.94351\nPOSSIBLE N-term signal sequence\ninside 1 19\nTMhelix 20 39\noutside 40 325\nTMhelix 326 348\ninside 349 372

12076 GCF_000164355.1_ASM16435v1 Escherichia coli MS 21-1 Proteobacteria; Gammaproteobacteria; Enterobacterales; Enterobacteriaceae; Escherichia; Escherichia coli
MNNEETFYQAMRRQGVTRRSFLKYCSLAATSLGLGAGMAPKIAWALENKPRIPVVWIHGL WP_001058323.1
MULTISPECIES: hydrogenase-1 small chain [Proteobacteria] Length: 372\nNumber of predicted TMHs: 2\nExp number of AAs in TMHs: 37.2608499999999\nExp number, first 60 AAs: 19.52923\nTotal prob of N-in:

0.94351\nPOSSIBLE N-term signal sequence\ninside 1 19\nTMhelix 20 39\noutside 40 325\nTMhelix 326 348\ninside 349 372

12077 GCF_000164355.1_ASM16435v1 Escherichia coli MS 21-1 Proteobacteria; Gammaproteobacteria; Enterobacterales; Enterobacteriaceae; Escherichia; Escherichia coli
MTGDNTLIHSHGINRRDFMKLCAALAATMGLSSKAAAEMAESVTNPQRPPVIWIGAQECT WP_000145410.1
MULTISPECIES: hydrogenase-2 small chain [Proteobacteria] Length: 372\nNumber of predicted TMHs: 1\nExp number of AAs in TMHs: 23.72226\nExp number, first 60 AAs: 0.22038\nTotal prob of N-in: 0.12994\noutside 1 331\nTMhelix 332 354\ninside 355 372

12078 GCF_000164295.1_ASM16429v1 Escherichia coli MS 45-1 Proteobacteria; Gammaproteobacteria; Enterobacterales; Enterobacteriaceae; Escherichia; Escherichia coli
MNNEETFYQAMRRQGVTRRSFLKYCSLAATSLGLGAGMAPKIAWALENKPRIPVVWIHGL WP_001058323.1
MULTISPECIES: hydrogenase-1 small chain [Proteobacteria] Length: 372\nNumber of predicted TMHs: 2\nExp number of AAs in TMHs: 37.2608499999999\nExp number, first 60 AAs: 19.52923\nTotal prob of N-in: 0.94351\nPOSSIBLE N-term signal sequence\ninside 1 19\nTMhelix 20 39\noutside 40 325\nTMhelix 326 348\ninside 349 372

12079 GCF_000164295.1_ASM16429v1 Escherichia coli MS 45-1 Proteobacteria; Gammaproteobacteria; Enterobacterales; Enterobacteriaceae; Escherichia; Escherichia coli
MTGDNTLIHSHGINRRDFMKLCAALAATMGLSSKAAAEMAESVTNPQRPPVIWIGAQECT WP_000145410.1
MULTISPECIES: hydrogenase-2 small chain [Proteobacteria] Length: 372\nNumber of predicted TMHs: 1\nExp number of AAs in TMHs: 23.72226\nExp number, first 60 AAs: 0.22038\nTotal prob of N-in: 0.12994\noutside 1 331\nTMhelix 332 354\ninside 355 372

12080 GCF_000701145.1_EcMS4991.0 Escherichia coli MS499 Proteobacteria; Gammaproteobacteria; Enterobacterales; Enterobacteriaceae; Escherichia; Escherichia coli
MTGDNTLIHSHGINRRDFMKLCAALAATMGLSSKAAAEMAESVTNPQRPPVIWIGAQECT WP_000145410.1
MULTISPECIES: hydrogenase-2 small chain [Proteobacteria] Length: 372\nNumber of predicted TMHs: 1\nExp number of AAs in TMHs: 23.72226\nExp number, first 60 AAs: 0.22038\nTotal prob of N-in: 0.12994\noutside 1 331\nTMhelix 332 354\ninside 355 372

12081 GCF_000164615.1_ASM16461v1 Escherichia coli MS 57-2 Proteobacteria; Gammaproteobacteria; Enterobacterales; Enterobacteriaceae; Escherichia; Escherichia coli
MTGDNTLIHSHGINRRDFMKLCAALAATMGLSSKAAAEMAESVTNPQRPPVIWIGAQECT WP_000145410.1
MULTISPECIES: hydrogenase-2 small chain [Proteobacteria] Length: 372\nNumber of predicted TMHs: 1\nExp number of AAs in TMHs: 23.72226\nExp number, first 60 AAs: 0.22038\nTotal prob of N-in: 0.12994\noutside 1 331\nTMhelix 332 354\ninside 355 372

12082 GCF_000164615.1_ASM16461v1 Escherichia coli MS 57-2 Proteobacteria; Gammaproteobacteria; Enterobacterales; Enterobacteriaceae; Escherichia; Escherichia coli
MNNEETFYQAMRRQGVTRRSFLKYCSLAATSLGLGAGMAPKIAWALENKPRIPVVWIHGL WP_001058323.1
MULTISPECIES: hydrogenase-1 small chain [Proteobacteria] Length: 372\nNumber of predicted TMHs: 2\nExp number of AAs in TMHs: 37.2608499999999\nExp number, first 60 AAs: 19.52923\nTotal prob of N-in: 0.94351\nPOSSIBLE N-term signal sequence\ninside 1 19\nTMhelix 20 39\noutside 40 325\nTMhelix 326 348\ninside 349 372

12083 GCF_000164595.1_ASM16459v1 Escherichia coli MS 60-1 Proteobacteria; Gammaproteobacteria; Enterobacterales; Enterobacteriaceae; Escherichia; Escherichia coli
MNNEETFYQAMRRQGVTRRSFLKYCSLAATSLGLGAGMAPKIAWALENKPRIPVVWIHGL WP_001058323.1
MULTISPECIES: hydrogenase-1 small chain [Proteobacteria] Length: 372\nNumber of predicted TMHs: 2\nExp number of AAs in TMHs: 37.2608499999999\nExp number, first 60 AAs: 19.52923\nTotal prob of N-in: 0.94351\nPOSSIBLE N-term signal sequence\ninside 1 19\nTMhelix 20 39\noutside 40 325\nTMhelix 326 348\ninside 349 372

12084 GCF_000164595.1_ASM16459v1 Escherichia coli MS 60-1 Proteobacteria; Gammaproteobacteria; Enterobacterales; Enterobacteriaceae; Escherichia; Escherichia coli
MTGDNTLIHSHGINRRDFMKLCAALAATMGLSSKAAAEMAESVTNPQRPPVIWIGAQECT WP_000145410.1
MULTISPECIES: hydrogenase-2 small chain [Proteobacteria] Length: 372\nNumber of predicted TMHs: 1\nExp number of AAs in TMHs: 23.72226\nExp number, first 60 AAs: 0.22038\nTotal prob of N-in: 0.12994\noutside 1 331\nTMhelix 332 354\ninside 355 372

12085 GCF_000164315.1_ASM16431v1 Escherichia coli MS 69-1 Proteobacteria; Gammaproteobacteria; Enterobacterales; Enterobacteriaceae; Escherichia; Escherichia coli
MNNEETFYQAMRRQGVTRRSFLKYCSLAATSLGLGAGMAPKIAWALENKPRIPVVWIHGL WP_001058323.1
MULTISPECIES: hydrogenase-1 small chain [Proteobacteria] Length: 372\nNumber of predicted TMHs: 2\nExp number of AAs in TMHs: 37.2608499999999\nExp number, first 60 AAs: 19.52923\nTotal prob of N-in: 0.94351\nPOSSIBLE N-term signal sequence\ninside 1 19\nTMhelix 20 39\noutside 40 325\nTMhelix 326 348\ninside 349 372

12086 GCF_000164315.1_ASM16431v1 Escherichia coli MS 69-1 Proteobacteria; Gammaproteobacteria; Enterobacterales; Enterobacteriaceae; Escherichia; Escherichia coli
MTGDNTLIHSHGINRRDFMKLCAALAATMGLSSKAAAEMAESVTNPQRPPVIWIGAQECT WP_000145410.1
MULTISPECIES: hydrogenase-2 small chain [Proteobacteria] Length: 372\nNumber of predicted TMHs: 1\nExp number of AAs in TMHs: 23.72226\nExp number, first 60 AAs: 0.22038\nTotal prob of N-in: 0.12994\noutside 1 331\nTMhelix 332 354\ninside 355 372

12087 GCF_000164455.1_ASM16445v1 Escherichia coli MS 78-1 Proteobacteria; Gammaproteobacteria; Enterobacterales; Enterobacteriaceae; Escherichia; Escherichia coli
MTGDNTLIHSHGINRRDFMKLCAALAATMGLSSKAAAEMAESVTNPQRPPVIWIGAQECT WP_000145404.1
hydrogenase-2 small chain [Escherichia coli] Length: 372\nNumber of predicted TMHs: 1\nExp number of AAs in TMHs: 23.72251\nExp number, first 60 AAs: 0.2205\nTotal prob of N-in: 0.12994\noutside 1 331\nTMhelix 332 354\ninside 355 372

12088 GCF_000164455.1_ASM16445v1 Escherichia coli MS 78-1 Proteobacteria; Gammaproteobacteria; Enterobacterales; Enterobacteriaceae; Escherichia; Escherichia coli
MNNEETFYQAMRRQGVTRRSFLKYCSLAATSLGLGAGMAPKIAWALENKPRIPVVWIHGL WP_001058323.1
MULTISPECIES: hydrogenase-1 small chain [Proteobacteria] Length: 372\nNumber of predicted TMHs: 2\nExp number of AAs in TMHs: 37.2608499999999\nExp number, first 60 AAs: 19.52923\nTotal prob of N-in: 0.94351\nPOSSIBLE N-term signal sequence\ninside 1 19\nTMhelix 20 39\noutside 40 325\nTMhelix 326 348\ninside 349 372

12089 GCF_000179095.1_ASM17909v1 Escherichia coli MS 79-10 Proteobacteria; Gammaproteobacteria; Enterobacterales; Enterobacteriaceae; Escherichia; Escherichia coli
MTGDNTLIHSHGINRRDFMKLCAALAATMGLSSKAAAEMAESVTNPQRPPVIWIGAQECT WP_000145410.1
MULTISPECIES: hydrogenase-2 small chain [Proteobacteria] Length: 372\nNumber of predicted TMHs: 1\nExp number of AAs in TMHs: 23.72226\nExp number, first 60 AAs: 0.22038\nTotal prob of N-in: 0.12994\noutside 1 331\nTMhelix 332 354\ninside 355 372

12090 GCF_000179095.1_ASM17909v1 Escherichia coli MS 79-10 Proteobacteria; Gammaproteobacteria; Enterobacterales; Enterobacteriaceae; Escherichia; Escherichia coli
MNNEETFYQAMRRQGVTRRSFLKYCSLAATSLGLGAGMAPKIAWALENKPRIPVVWIHGL WP_001295940.1
MULTISPECIES: hydrogenase [Enterobacteriaceae] Length: 372\nNumber of predicted TMHs: 2\nExp number of AAs in TMHs: 37.2608899999999\nExp number, first 60 AAs: 19.52923\nTotal prob of N-in: 0.94351\nPOSSIBLE N-term signal sequence\ninside 1 19\nTMhelix 20 39\noutside 40 325\nTMhelix 326 348\ninside 349 372

12091 GCF_000164215.1_ASM16421v1 Escherichia coli MS 84-1 Proteobacteria; Gammaproteobacteria; Enterobacterales; Enterobacteriaceae; Escherichia; Escherichia coli
MNNEETFYQAMRRQGVTRRSFLKYCSLAATSLGLGAGMAPKIAWALENKPRIPVVWIHGL WP_001058320.1
hydrogenase [Escherichia coli] Length: 372\nNumber of predicted TMHs: 2\nExp number of AAs in TMHs: 37.26082\nExp number, first 60 AAs: 19.52923\nTotal prob of N-in: 0.94351\nPOSSIBLE N-term signal sequence\ninside 1 19\nTMhelix 20 39\noutside 40 325\nTMhelix 326 348\ninside 349 372

12092 GCF_000164215.1_ASM16421v1 Escherichia coli MS 84-1 Proteobacteria; Gammaproteobacteria; Enterobacterales; Enterobacteriaceae; Escherichia; Escherichia coli
MTGDNTLIHSHGINRRDFMKLCAALAATMGLSSKAAAEMAESVTNPQRPPVIWIGAQECT WP_000145410.1
MULTISPECIES: hydrogenase-2 small chain [Proteobacteria] Length: 372\nNumber of predicted TMHs: 1\nExp number of AAs in TMHs: 23.72226\nExp number, first 60 AAs: 0.22038\nTotal prob of N-in: 0.12994\noutside 1 331\nTMhelix 332 354\ninside 355 372

12093 GCF_000179075.1_ASM17907v1 Escherichia coli MS 85-1 Proteobacteria; Gammaproteobacteria; Enterobacterales; Enterobacteriaceae; Escherichia; Escherichia coli

MTGDNTLIHSHGINRRDFMKLCAALAATMGLSSKAAAEMAESVTNPQRPPVIWIGAQECT WP_000145410.1 MULTISPECIES: hydrogenase-2 small chain [Proteobacteria] Length: 372\nNumber of predicted TMHs: 1\nExp number of AAs in TMHs: 23.72226\nExp number, first 60 AAs: 0.22038\nTotal prob of N-in: 0.12994\noutside 1 331\nTMhelix 332 354\ninside 355 372

12094 GCF_000179075.1_ASM17907v1 Escherichia coli MS 85-1 Proteobacteria; Gammaproteobacteria; Enterobacterales; Enterobacteriaceae; Escherichia; Escherichia coli MNNEETFYQAMRRQGVTRRSFLKYCSLAATSLGLGAGMAPKIAWALENKPRIPVVWIHGL WP_001058320.1 hydrogenase [Escherichia coli] Length: 372\nNumber of predicted TMHs: 2\nExp number of AAs in TMHs: 37.26082\nExp number, first 60 AAs: 19.52923\nTotal prob of N-in: 0.94351\nPOSSIBLE N-term signal sequence\ninside 1 19\nTMhelix 20 39\noutside 40 325\nTMhelix 326 348\ninside 349 372

12095 GCF_000303635.2_ASM30363v2 Escherichia coli N1 Proteobacteria; Gammaproteobacteria; Enterobacterales; Enterobacteriaceae; Escherichia; Escherichia coli MNNEETFYQAMRRQGVTRRSFLKYCSLAATSLGLGAGMAPKIAWALENKPRIPVVWIHGL WP_001295940.1 MULTISPECIES: hydrogenase [Enterobacteriaceae] Length: 372\nNumber of predicted TMHs: 2\nExp number of AAs in TMHs: 37.2608899999999\nExp number, first 60 AAs: 19.52923\nTotal prob of N-in: 0.94351\nPOSSIBLE N-term signal sequence\ninside 1 19\nTMhelix 20 39\noutside 40 325\nTMhelix 326 348\ninside 349 372

12096 GCF_000303635.2_ASM30363v2 Escherichia coli N1 Proteobacteria; Gammaproteobacteria; Enterobacterales; Enterobacteriaceae; Escherichia; Escherichia coli MTGDNTLIHSHGINRRDFMKLCAALAATMGLSSKAAAEMAESVTNPQRPPVIWIGAQECT WP_000145410.1 MULTISPECIES: hydrogenase-2 small chain [Proteobacteria] Length: 372\nNumber of predicted TMHs: 1\nExp number of AAs in TMHs: 23.72226\nExp number, first 60 AAs: 0.22038\nTotal prob of N-in: 0.12994\noutside 1 331\nTMhelix 332 354\ninside 355 372

12097 GCF_001970075.1_ASM197007v1 Escherichia coli N36254PS Proteobacteria; Gammaproteobacteria; Enterobacterales; Enterobacteriaceae; Escherichia; Escherichia coli MTGDNTLIHSHGINRRDFMKLCAALAATMGLSSKAAAEMAESVTNPQRPPVIWIGAQECT WP_000145410.1 MULTISPECIES: hydrogenase-2 small chain [Proteobacteria] Length: 372\nNumber of predicted TMHs: 1\nExp number of AAs in TMHs: 23.72226\nExp number, first 60 AAs: 0.22038\nTotal prob of N-in: 0.12994\noutside 1 331\nTMhelix 332 354\ninside 355 372

12098 GCF_001970075.1_ASM197007v1 Escherichia coli N36254PS Proteobacteria; Gammaproteobacteria; Enterobacterales; Enterobacteriaceae; Escherichia; Escherichia coli MNNEETFYQAMRRQGVTRRSFLKYCSLAATSLGLGAGMAPKIAWALENKPRIPVVWIHGL WP_001295940.1 MULTISPECIES: hydrogenase [Enterobacteriaceae] Length: 372\nNumber of predicted TMHs: 2\nExp number of AAs in TMHs: 37.2608899999999\nExp number, first 60 AAs: 19.52923\nTotal prob of N-in: 0.94351\nPOSSIBLE N-term signal sequence\ninside 1 19\nTMhelix 20 39\noutside 40 325\nTMhelix 326 348\ninside 349 372

12099 GCF_001970105.1_ASM197010v1 Escherichia coli N36410PS Proteobacteria; Gammaproteobacteria; Enterobacterales; Enterobacteriaceae; Escherichia; Escherichia coli MTGDNTLIHSHGINRRDFMKLCAALAATMGLSSKAAAEMAESVTNPQRPPVIWIGAQECT WP_000145410.1 MULTISPECIES: hydrogenase-2 small chain [Proteobacteria] Length: 372\nNumber of predicted TMHs: 1\nExp number of AAs in TMHs: 23.72226\nExp number, first 60 AAs: 0.22038\nTotal prob of N-in: 0.12994\noutside 1 331\nTMhelix 332 354\ninside 355 372

12100 GCF_001970105.1_ASM197010v1 Escherichia coli N36410PS Proteobacteria; Gammaproteobacteria; Enterobacterales; Enterobacteriaceae; Escherichia; Escherichia coli MNNEETFYQAMRRQGVTRRSFLKYCSLAATSLGLGAGMAPKIAWALENKPRIPVVWIHGL WP_001058323.1 MULTISPECIES: hydrogenase-1 small chain [Proteobacteria] Length: 372\nNumber of predicted TMHs: 2\nExp number of AAs in TMHs: 37.2608499999999\nExp number, first 60 AAs: 19.52923\nTotal prob of N-in: 0.94351\nPOSSIBLE N-term signal sequence\ninside 1 19\nTMhelix 20 39\noutside 40 325\nTMhelix 326 348\ninside 349 372

12101 GCF_001970035.1_ASM197003v1 Escherichia coli N37058PS Proteobacteria; Gammaproteobacteria; Enterobacterales; Enterobacteriaceae; Escherichia; Escherichia coli MNNEETFYQAMRRQGVTRRSFLKYCSLAATSLGLGAGMAPKIAWALENKPRIPVVWIHGL WP_001058323.1 MULTISPECIES: hydrogenase-1 small chain [Proteobacteria] Length: 372\nNumber of predicted TMHs:

2\nExp number of AAs in TMHs: 37.2608499999999\nExp number, first 60 AAs:  19.52923\nTotal prob of N-in:   0.94351\nPOSSIBLE N-term signal sequence\ninside   1   19\nTMhelix   20   39\noutside   40   325\nTMhelix   326   348\ninside   349   372

12102 GCF_001970035.1_ASM197003v1 Escherichia coli N37058PS Proteobacteria; Gammaproteobacteria; Enterobacterales; Enterobacteriaceae; Escherichia; Escherichia coli
MTGDNTLIHSHGINRRDFMKLCAALAATMGLSSKAAAEMAESVTNPQRPPVIWIGAQECT WP_000145410.1
MULTISPECIES: hydrogenase-2 small chain [Proteobacteria] Length: 372\nNumber of predicted TMHs: 1\nExp number of AAs in TMHs: 23.72226\nExp number, first 60 AAs:  0.22038\nTotal prob of N-in:   0.12994\noutside   1   331\nTMhelix   332   354\ninside   355   372

12103 GCF_001970095.1_ASM197009v1 Escherichia coli N37122PS Proteobacteria; Gammaproteobacteria; Enterobacterales; Enterobacteriaceae; Escherichia; Escherichia coli
MTGDNTLIHSHGINRRDFMKLCAALAATMGLSSKAAAEMAESVTNPQRPPVIWIGAQECT WP_000145410.1
MULTISPECIES: hydrogenase-2 small chain [Proteobacteria] Length: 372\nNumber of predicted TMHs: 1\nExp number of AAs in TMHs: 23.72226\nExp number, first 60 AAs:  0.22038\nTotal prob of N-in:   0.12994\noutside   1   331\nTMhelix   332   354\ninside   355   372

12104 GCF_001970095.1_ASM197009v1 Escherichia coli N37122PS Proteobacteria; Gammaproteobacteria; Enterobacterales; Enterobacteriaceae; Escherichia; Escherichia coli
MNNEETFYQAMRRQGVTRRSFLKYCSLAATSLGLGAGMAPKIAWALENKPRIPVVWIHGL WP_001058323.1
MULTISPECIES: hydrogenase-1 small chain [Proteobacteria] Length: 372\nNumber of predicted TMHs: 2\nExp number of AAs in TMHs: 37.2608499999999\nExp number, first 60 AAs:  19.52923\nTotal prob of N-in:   0.94351\nPOSSIBLE N-term signal sequence\ninside   1   19\nTMhelix   20   39\noutside   40   325\nTMhelix   326   348\ninside   349   372

12105 GCF_001970045.1_ASM197004v1 Escherichia coli N40513 Proteobacteria; Gammaproteobacteria; Enterobacterales; Enterobacteriaceae; Escherichia; Escherichia coli
MNNEETFYQAMRRQGVTRRSFLKYCSLAATSLGLGAGMAPKIAWALENKPRIPVVWIHGL WP_001058323.1
MULTISPECIES: hydrogenase-1 small chain [Proteobacteria] Length: 372\nNumber of predicted TMHs: 2\nExp number of AAs in TMHs: 37.2608499999999\nExp number, first 60 AAs:  19.52923\nTotal prob of N-in:   0.94351\nPOSSIBLE N-term signal sequence\ninside   1   19\nTMhelix   20   39\noutside   40   325\nTMhelix   326   348\ninside   349   372

12106 GCF_001970045.1_ASM197004v1 Escherichia coli N40513 Proteobacteria; Gammaproteobacteria; Enterobacterales; Enterobacteriaceae; Escherichia; Escherichia coli
MTGDNTLIHSHGINRRDFMKLCAALAATMGLSSKAAAEMAESVTNPQRPPVIWIGAQECT WP_000145410.1
MULTISPECIES: hydrogenase-2 small chain [Proteobacteria] Length: 372\nNumber of predicted TMHs: 1\nExp number of AAs in TMHs: 23.72226\nExp number, first 60 AAs:  0.22038\nTotal prob of N-in:   0.12994\noutside   1   331\nTMhelix   332   354\ninside   355   372

12107 GCF_000214765.2_ASM21476v3 Escherichia coli NA114 Proteobacteria; Gammaproteobacteria; Enterobacterales; Enterobacteriaceae; Escherichia; Escherichia coli
MNNEETFYQAMRRQGVTRRSFLKYCSLAATSLGLGAGMAPKIAWALENKPRIPVVWIHGL WP_001295940.1
MULTISPECIES: hydrogenase [Enterobacteriaceae] Length: 372\nNumber of predicted TMHs:  2\nExp number of AAs in TMHs: 37.2608899999999\nExp number, first 60 AAs:  19.52923\nTotal prob of N-in:   0.94351\nPOSSIBLE N-term signal sequence\ninside   1   19\nTMhelix   20   39\noutside   40   325\nTMhelix   326   348\ninside   349   372

12108 GCF_000214765.2_ASM21476v3 Escherichia coli NA114 Proteobacteria; Gammaproteobacteria; Enterobacterales; Enterobacteriaceae; Escherichia; Escherichia coli
MTGDNTLIHSHGINRRDFMKLCAALAATMGLSSKAAAEMVESVTNPQRPPVIWIGAQECT WP_000145423.1
hydrogenase 2 small subunit [Escherichia coli] Length: 372\nNumber of predicted TMHs:  1\nExp number of AAs in TMHs: 23.78201\nExp number, first 60 AAs:  0.28355\nTotal prob of N-in:   0.13257\noutside   1   331\nTMhelix   332   354\ninside   355   372

12109 GCF_000982875.1_ASM98287v1 Escherichia coli NB8 Proteobacteria; Gammaproteobacteria; Enterobacterales; Enterobacteriaceae; Escherichia; Escherichia coli
MTGDNTLIHSHGINRRDFMKLCAALAATMGLSSKAAAEMVESVTNPQRPPVIWIGAQECT WP_000145423.1
hydrogenase 2 small subunit [Escherichia coli] Length: 372\nNumber of predicted TMHs:  1\nExp number of

AAs in TMHs: 23.78201\nExp number, first 60 AAs:  0.28355\nTotal prob of N-in:   0.13257\noutside   1   331\nTMhelix   332   354\ninside   355   372

12110 GCF_000982875.1_ASM98287v1 Escherichia coli NB8 Proteobacteria; Gammaproteobacteria; Enterobacterales; Enterobacteriaceae; Escherichia; Escherichia coli
MNNEETFYQAMRRQGVTRRSFLKYCSLAATSLGLGAGMAPKIAWALENKPRIPVVWIHGL WP_001295940.1
MULTISPECIES: hydrogenase [Enterobacteriaceae] Length: 372\nNumber of predicted TMHs:  2\nExp number of AAs in TMHs: 37.2608899999999\nExp number, first 60 AAs:  19.52923\nTotal prob of N-in:   0.94351\nPOSSIBLE N-term signal sequence\ninside   1   19\nTMhelix   20   39\noutside   40   325\nTMhelix   326   348\ninside   349   372

12111 GCF_000179795.1_ASM17979v1 Escherichia coli NC101 Proteobacteria; Gammaproteobacteria; Enterobacterales; Enterobacteriaceae; Escherichia; Escherichia coli
MTGDNTLIHSHGINRRDFMKLCAALAATMGLSSKAAAEMVESVTNPQRPPVIWIGAQECT WP_000145423.1
hydrogenase 2 small subunit [Escherichia coli] Length: 372\nNumber of predicted TMHs:  1\nExp number of AAs in TMHs: 23.78201\nExp number, first 60 AAs:  0.28355\nTotal prob of N-in:   0.13257\noutside   1   331\nTMhelix   332   354\ninside   355   372

12112 GCF_000179795.1_ASM17979v1 Escherichia coli NC101 Proteobacteria; Gammaproteobacteria; Enterobacterales; Enterobacteriaceae; Escherichia; Escherichia coli
MNNEETFYQAMRRQGVTRRSFLKYCSLAATSLGLGAGMAPKIAWALENKPRIPVVWIHGL WP_001058323.1
MULTISPECIES: hydrogenase-1 small chain [Proteobacteria] Length: 372\nNumber of predicted TMHs:  2\nExp number of AAs in TMHs: 37.2608499999999\nExp number, first 60 AAs:  19.52923\nTotal prob of N-in:   0.94351\nPOSSIBLE N-term signal sequence\ninside   1   19\nTMhelix   20   39\noutside   40   325\nTMhelix   326   348\ninside   349   372

12113 GCF_000259385.1_NCCP15647_1.0 Escherichia coli NCCP15647 Proteobacteria; Gammaproteobacteria; Enterobacterales; Enterobacteriaceae; Escherichia; Escherichia coli
MTGDNTLIHSHGINRRDFMKLCAALAATMGLSSKAAAEMAESVTNPQRPPVIWIGAQECT WP_000145410.1
MULTISPECIES: hydrogenase-2 small chain [Proteobacteria] Length: 372\nNumber of predicted TMHs:  1\nExp number of AAs in TMHs: 23.72226\nExp number, first 60 AAs:  0.22038\nTotal prob of N-in:   0.12994\noutside   1   331\nTMhelix   332   354\ninside   355   372

12114 GCF_000259385.1_NCCP15647_1.0 Escherichia coli NCCP15647 Proteobacteria; Gammaproteobacteria; Enterobacterales; Enterobacteriaceae; Escherichia; Escherichia coli
MNNEETFYQAMRRQGVTRRSFLKYCSLAATSLGLGAGMAPKIAWALENKPRIPVVWIHGL WP_001295940.1
MULTISPECIES: hydrogenase [Enterobacteriaceae] Length: 372\nNumber of predicted TMHs:  2\nExp number of AAs in TMHs: 37.2608899999999\nExp number, first 60 AAs:  19.52923\nTotal prob of N-in:   0.94351\nPOSSIBLE N-term signal sequence\ninside   1   19\nTMhelix   20   39\noutside   40   325\nTMhelix   326   348\ninside   349   372

12115 GCF_001466755.1_ASM146675v1 Escherichia coli NCCP 15653 Proteobacteria; Gammaproteobacteria; Enterobacterales; Enterobacteriaceae; Escherichia; Escherichia coli
MNNEETFYQAMRRQGVTRRSFLKYCSLAATSLGLGAGMAPKIAWALENKPRIPVVWIHGL WP_001058323.1
MULTISPECIES: hydrogenase-1 small chain [Proteobacteria] Length: 372\nNumber of predicted TMHs:  2\nExp number of AAs in TMHs: 37.2608499999999\nExp number, first 60 AAs:  19.52923\nTotal prob of N-in:   0.94351\nPOSSIBLE N-term signal sequence\ninside   1   19\nTMhelix   20   39\noutside   40   325\nTMhelix   326   348\ninside   349   372

12116 GCF_001466755.1_ASM146675v1 Escherichia coli NCCP 15653 Proteobacteria; Gammaproteobacteria; Enterobacterales; Enterobacteriaceae; Escherichia; Escherichia coli
MTGDNTLIHSHGINRRDFMKLCAALAATMGLSSKAAAEMAESVTNPQRPPVIWIGAQECT WP_000145410.1
MULTISPECIES: hydrogenase-2 small chain [Proteobacteria] Length: 372\nNumber of predicted TMHs:  1\nExp number of AAs in TMHs: 23.72226\nExp number, first 60 AAs:  0.22038\nTotal prob of N-in:   0.12994\noutside   1   331\nTMhelix   332   354\ninside   355   372

12117 GCF_001467005.1_ASM146700v1 Escherichia coli NCCP 15655 Proteobacteria; Gammaproteobacteria; Enterobacterales; Enterobacteriaceae; Escherichia; Escherichia coli
MTGDNTLIHSHGINRRDFMKLCAALAATMGLSSKAAAEMAESVTNPQRPPVIWIGAQECT WP_000145410.1
MULTISPECIES: hydrogenase-2 small chain [Proteobacteria] Length: 372\nNumber of predicted TMHs:

1\nExp number of AAs in TMHs: 23.72226\nExp number, first 60 AAs: 0.22038\nTotal prob of N-in: 0.12994\noutside 1 331\nTMhelix 332 354\ninside 355 372

12118 GCF_001467005.1_ASM146700v1 Escherichia coli NCCP 15655 Proteobacteria; Gammaproteobacteria; Enterobacterales; Enterobacteriaceae; Escherichia; Escherichia coli
MNNEETFYQAMRRQGVTRRSFLKYCSLAATSLGLGAGMAPKIAWALENKPRIPVVWIHGL WP_001058323.1
MULTISPECIES: hydrogenase-1 small chain [Proteobacteria] Length: 372\nNumber of predicted TMHs: 2\nExp number of AAs in TMHs: 37.2608499999999\nExp number, first 60 AAs: 19.52923\nTotal prob of N-in: 0.94351\nPOSSIBLE N-term signal sequence\ninside 1 19\nTMhelix 20 39\noutside 40 325\nTMhelix 326 348\ninside 349 372

12119 GCF_001466815.1_ASM146681v1 Escherichia coli NCCP 15656 Proteobacteria; Gammaproteobacteria; Enterobacterales; Enterobacteriaceae; Escherichia; Escherichia coli
MTGDNTLIHSHGINRRDFMKLCAALAATMGLSSKAAAEMAESVTNPQRPPVIWIGAQECT WP_000145410.1
MULTISPECIES: hydrogenase-2 small chain [Proteobacteria] Length: 372\nNumber of predicted TMHs: 1\nExp number of AAs in TMHs: 23.72226\nExp number, first 60 AAs: 0.22038\nTotal prob of N-in: 0.12994\noutside 1 331\nTMhelix 332 354\ninside 355 372

12120 GCF_001466815.1_ASM146681v1 Escherichia coli NCCP 15656 Proteobacteria; Gammaproteobacteria; Enterobacterales; Enterobacteriaceae; Escherichia; Escherichia coli
MNNEETFYQAMRRQGVTRRSFLKYCSLAATSLGLGAGMAPKIAWALENKPRIPVVWIHGL WP_001058323.1
MULTISPECIES: hydrogenase-1 small chain [Proteobacteria] Length: 372\nNumber of predicted TMHs: 2\nExp number of AAs in TMHs: 37.2608499999999\nExp number, first 60 AAs: 19.52923\nTotal prob of N-in: 0.94351\nPOSSIBLE N-term signal sequence\ninside 1 19\nTMhelix 20 39\noutside 40 325\nTMhelix 326 348\ninside 349 372

12121 GCF_000258785.1_EscCol1.0 Escherichia coli NCCP15657 Proteobacteria; Gammaproteobacteria; Enterobacterales; Enterobacteriaceae; Escherichia; Escherichia coli
MNNEETFYQAMRRQGVTRRSFLKYCSLAATSLGLGAGMAPKIAWALENKPRIPVVWIHGL WP_001295940.1
MULTISPECIES: hydrogenase [Enterobacteriaceae] Length: 372\nNumber of predicted TMHs: 2\nExp number of AAs in TMHs: 37.2608899999999\nExp number, first 60 AAs: 19.52923\nTotal prob of N-in: 0.94351\nPOSSIBLE N-term signal sequence\ninside 1 19\nTMhelix 20 39\noutside 40 325\nTMhelix 326 348\ninside 349 372

12122 GCF_000258785.1_EscCol1.0 Escherichia coli NCCP15657 Proteobacteria; Gammaproteobacteria; Enterobacterales; Enterobacteriaceae; Escherichia; Escherichia coli
MTGDNTLIHSHGINRRDFMKLCAALAATMGLSSKAAAEMAESVTNPQRPPVIWIGAQECT WP_000145410.1
MULTISPECIES: hydrogenase-2 small chain [Proteobacteria] Length: 372\nNumber of predicted TMHs: 1\nExp number of AAs in TMHs: 23.72226\nExp number, first 60 AAs: 0.22038\nTotal prob of N-in: 0.12994\noutside 1 331\nTMhelix 332 354\ninside 355 372

12123 GCF_000260475.1_NCCP15658v1.0 Escherichia coli NCCP15658 Proteobacteria; Gammaproteobacteria; Enterobacterales; Enterobacteriaceae; Escherichia; Escherichia coli
MNNEETFYQAMRRQGVTRRSFLKYCSLAATSLGLGAGMAPKIAWALENKPRIPVVWIHGL WP_001301613.1
hydrogenase [Escherichia coli] Length: 372\nNumber of predicted TMHs: 2\nExp number of AAs in TMHs: 37.46598\nExp number, first 60 AAs: 19.53139\nTotal prob of N-in: 0.94364\nPOSSIBLE N-term signal sequence\ninside 1 19\nTMhelix 20 39\noutside 40 325\nTMhelix 326 348\ninside 349 372

12124 GCF_000260475.1_NCCP15658v1.0 Escherichia coli NCCP15658 Proteobacteria; Gammaproteobacteria; Enterobacterales; Enterobacteriaceae; Escherichia; Escherichia coli
MTGDNTLIHSHGINRRDFMKLCAALAATMGLSSKAAAEMAESVTNPQRPPVIWIGAQECT WP_000145410.1
MULTISPECIES: hydrogenase-2 small chain [Proteobacteria] Length: 372\nNumber of predicted TMHs: 1\nExp number of AAs in TMHs: 23.72226\nExp number, first 60 AAs: 0.22038\nTotal prob of N-in: 0.12994\noutside 1 331\nTMhelix 332 354\ninside 355 372

12125 GCF_000766965.1_CREcoli50110 Escherichia coli NCTC 50110 Proteobacteria; Gammaproteobacteria; Enterobacterales; Enterobacteriaceae; Escherichia; Escherichia coli
MTGDNTLIHSHGINRRDFMKLCAALAATMGLSSKAAAEMAESVTNPQRPPVIWIGAQECT WP_000145410.1
MULTISPECIES: hydrogenase-2 small chain [Proteobacteria] Length: 372\nNumber of predicted TMHs: 1\nExp number of AAs in TMHs: 23.72226\nExp number, first 60 AAs: 0.22038\nTotal prob of N-in: 0.12994\noutside 1 331\nTMhelix 332 354\ninside 355 372

12126 GCF_000766965.1_CREcoli50110 Escherichia coli NCTC 50110 Proteobacteria; Gammaproteobacteria; Enterobacterales; Enterobacteriaceae; Escherichia; Escherichia coli
MNNEETFYQAMRRQGVTRRSFLKYCSLAATSLGLGAGMAPKIAWALENKPRIPVVWIHGL WP_001058323.1
MULTISPECIES: hydrogenase-1 small chain [Proteobacteria] Length: 372\nNumber of predicted TMHs: 2\nExp number of AAs in TMHs: 37.2608499999999\nExp number, first 60 AAs: 19.52923\nTotal prob of N-in: 0.94351\nPOSSIBLE N-term signal sequence\ninside 1 19\nTMhelix 20 39\noutside 40 325\nTMhelix 326 348\ninside 349 372

12127 GCF_000303835.2_ASM30383v2 Escherichia coli NE037 Proteobacteria; Gammaproteobacteria; Enterobacterales; Enterobacteriaceae; Escherichia; Escherichia coli
MNNEETFYQAMRRQGVTRRSFLKYCSLAATSLGLGAGMAPKIAWALENKPRIPVVWIHGL WP_001301613.1
hydrogenase [Escherichia coli] Length: 372\nNumber of predicted TMHs: 2\nExp number of AAs in TMHs: 37.46598\nExp number, first 60 AAs: 19.53139\nTotal prob of N-in: 0.94364\nPOSSIBLE N-term signal sequence\ninside 1 19\nTMhelix 20 39\noutside 40 325\nTMhelix 326 348\ninside 349 372

12128 GCF_000303835.2_ASM30383v2 Escherichia coli NE037 Proteobacteria; Gammaproteobacteria; Enterobacterales; Enterobacteriaceae; Escherichia; Escherichia coli
MTGDNTLIHSHGINRRDFMKLCAALAATMGLSSKAAAEMAESVTNPQRPPVIWIGAQECT WP_000145410.1
MULTISPECIES: hydrogenase-2 small chain [Proteobacteria] Length: 372\nNumber of predicted TMHs: 1\nExp number of AAs in TMHs: 23.72226\nExp number, first 60 AAs: 0.22038\nTotal prob of N-in: 0.12994\noutside 1 331\nTMhelix 332 354\ninside 355 372

12129 GCF_000303295.2_ASM30329v2 Escherichia coli NE098 Proteobacteria; Gammaproteobacteria; Enterobacterales; Enterobacteriaceae; Escherichia; Escherichia coli
MNNEETFYQAMRRQGVTRRSFLKYCSLAATSLGLGAGMAPKIAWALENKPRIPVVWIHGL WP_001301613.1
hydrogenase [Escherichia coli] Length: 372\nNumber of predicted TMHs: 2\nExp number of AAs in TMHs: 37.46598\nExp number, first 60 AAs: 19.53139\nTotal prob of N-in: 0.94364\nPOSSIBLE N-term signal sequence\ninside 1 19\nTMhelix 20 39\noutside 40 325\nTMhelix 326 348\ninside 349 372

12130 GCF_000303295.2_ASM30329v2 Escherichia coli NE098 Proteobacteria; Gammaproteobacteria; Enterobacterales; Enterobacteriaceae; Escherichia; Escherichia coli
MTGDNTLIHSHGINRRDFMKLCAALAATMGLSSKAAAEMAESVTNPQRPPVIWIGAQECT WP_000145410.1
MULTISPECIES: hydrogenase-2 small chain [Proteobacteria] Length: 372\nNumber of predicted TMHs: 1\nExp number of AAs in TMHs: 23.72226\nExp number, first 60 AAs: 0.22038\nTotal prob of N-in: 0.12994\noutside 1 331\nTMhelix 332 354\ninside 355 372

12131 GCF_000303815.2_ASM30381v2 Escherichia coli NE1487 Proteobacteria; Gammaproteobacteria; Enterobacterales; Enterobacteriaceae; Escherichia; Escherichia coli
MNNEETFYQAMRRQGVTRRSFLKYCSLAATSLGLGAGMAPKIAWALENKPRIPVVWIHGL WP_001301613.1
hydrogenase [Escherichia coli] Length: 372\nNumber of predicted TMHs: 2\nExp number of AAs in TMHs: 37.46598\nExp number, first 60 AAs: 19.53139\nTotal prob of N-in: 0.94364\nPOSSIBLE N-term signal sequence\ninside 1 19\nTMhelix 20 39\noutside 40 325\nTMhelix 326 348\ninside 349 372

12132 GCF_000303815.2_ASM30381v2 Escherichia coli NE1487 Proteobacteria; Gammaproteobacteria; Enterobacterales; Enterobacteriaceae; Escherichia; Escherichia coli
MTGDNTLIHSHGINRRDFMKLCAALAATMGLSSKAAAEMAESVTNPQRPPVIWIGAQECT WP_000145410.1
MULTISPECIES: hydrogenase-2 small chain [Proteobacteria] Length: 372\nNumber of predicted TMHs: 1\nExp number of AAs in TMHs: 23.72226\nExp number, first 60 AAs: 0.22038\nTotal prob of N-in: 0.12994\noutside 1 331\nTMhelix 332 354\ninside 355 372

12133 GCF_000333215.1_ASM33321v1 Escherichia coli Nissle 1917 Proteobacteria; Gammaproteobacteria; Enterobacterales; Enterobacteriaceae; Escherichia; Escherichia coli
MTGDNTLIHSHGINRRDFMKLCAALAATMGLSSKAAAEMAESVTNPQRPPVIWIGAQECT WP_000145410.1
MULTISPECIES: hydrogenase-2 small chain [Proteobacteria] Length: 372\nNumber of predicted TMHs: 1\nExp number of AAs in TMHs: 23.72226\nExp number, first 60 AAs: 0.22038\nTotal prob of N-in: 0.12994\noutside 1 331\nTMhelix 332 354\ninside 355 372

12134 GCF_000714595.1_ASM71459v1 Escherichia coli Nissle 1917 Proteobacteria; Gammaproteobacteria; Enterobacterales; Enterobacteriaceae; Escherichia; Escherichia coli
MTGDNTLIHSHGINRRDFMKLCAALAATMGLSSKAAAEMAESVTNPQRPPVIWIGAQECT WP_000145410.1
MULTISPECIES: hydrogenase-2 small chain [Proteobacteria] Length: 372\nNumber of predicted TMHs:

1\nExp number of AAs in TMHs: 23.72226\nExp number, first 60 AAs: 0.22038\nTotal prob of N-in: 0.12994\noutside 1 331\nTMhelix 332 354\ninside 355 372

12135 GCF_000333215.1_ASM33321v1 Escherichia coli Nissle 1917 Proteobacteria; Gammaproteobacteria; Enterobacterales; Enterobacteriaceae; Escherichia; Escherichia coli
MNNEETFYQAMRRQGVTRRSFLKYCSLAATSLGLGAGMAPKIAWALENKPRIPVVWIHGL WP_001058323.1
MULTISPECIES: hydrogenase-1 small chain [Proteobacteria] Length: 372\nNumber of predicted TMHs: 2\nExp number of AAs in TMHs: 37.2608499999999\nExp number, first 60 AAs: 19.52923\nTotal prob of N-in: 0.94351\nPOSSIBLE N-term signal sequence\ninside 1 19\nTMhelix 20 39\noutside 40 325\nTMhelix 326 348\ninside 349 372

12136 GCF_000714595.1_ASM71459v1 Escherichia coli Nissle 1917 Proteobacteria; Gammaproteobacteria; Enterobacterales; Enterobacteriaceae; Escherichia; Escherichia coli
MNNEETFYQAMRRQGVTRRSFLKYCSLAATSLGLGAGMAPKIAWALENKPRIPVVWIHGL WP_001058323.1
MULTISPECIES: hydrogenase-1 small chain [Proteobacteria] Length: 372\nNumber of predicted TMHs: 2\nExp number of AAs in TMHs: 37.2608499999999\nExp number, first 60 AAs: 19.52923\nTotal prob of N-in: 0.94351\nPOSSIBLE N-term signal sequence\ninside 1 19\nTMhelix 20 39\noutside 40 325\nTMhelix 326 348\ninside 349 372

12137 GCF_002024865.1_ASM202486v1 Escherichia coli NU14 Proteobacteria; Gammaproteobacteria; Enterobacterales; Enterobacteriaceae; Escherichia; Escherichia coli
MTGDNTLIHSHGINRRDFMKLCAALAATMGLSSKAAAEMAESVTNPQRPPVIWIGAQECT WP_000145410.1
MULTISPECIES: hydrogenase-2 small chain [Proteobacteria] Length: 372\nNumber of predicted TMHs: 1\nExp number of AAs in TMHs: 23.72226\nExp number, first 60 AAs: 0.22038\nTotal prob of N-in: 0.12994\noutside 1 331\nTMhelix 332 354\ninside 355 372

12138 GCF_000340235.1_O8 Escherichia coli O08 Proteobacteria; Gammaproteobacteria; Enterobacterales; Enterobacteriaceae; Escherichia; Escherichia coli
MTGDNTLIHSHGINRRDFMKLCAALAATMGLSSKAAAEMAESVTNPQRPPVIWIGAQECT WP_000145410.1
MULTISPECIES: hydrogenase-2 small chain [Proteobacteria] Length: 372\nNumber of predicted TMHs: 1\nExp number of AAs in TMHs: 23.72226\nExp number, first 60 AAs: 0.22038\nTotal prob of N-in: 0.12994\noutside 1 331\nTMhelix 332 354\ninside 355 372

12139 GCF_000340235.1_O8 Escherichia coli O08 Proteobacteria; Gammaproteobacteria; Enterobacterales; Enterobacteriaceae; Escherichia; Escherichia coli
MNNEETFYQAMRRQGVTRRSFLKYCSLAATSLGLGAGMAPKIAWALENKPRIPVVWIHGL WP_001058323.1
MULTISPECIES: hydrogenase-1 small chain [Proteobacteria] Length: 372\nNumber of predicted TMHs: 2\nExp number of AAs in TMHs: 37.2608499999999\nExp number, first 60 AAs: 19.52923\nTotal prob of N-in: 0.94351\nPOSSIBLE N-term signal sequence\ninside 1 19\nTMhelix 20 39\noutside 40 325\nTMhelix 326 348\ninside 349 372

12140 GCF_002209105.1_ASM220910v1 Escherichia coli O104:H4 Proteobacteria; Gammaproteobacteria; Enterobacterales; Enterobacteriaceae; Escherichia; Escherichia coli
MTGDNTLIHSHGINRRDFMKLCAALAATMGLSSKAAAEMAESVTNPQRPPVIWIGAQECT WP_000145410.1
MULTISPECIES: hydrogenase-2 small chain [Proteobacteria] Length: 372\nNumber of predicted TMHs: 1\nExp number of AAs in TMHs: 23.72226\nExp number, first 60 AAs: 0.22038\nTotal prob of N-in: 0.12994\noutside 1 331\nTMhelix 332 354\ninside 355 372

12141 GCF_002209105.1_ASM220910v1 Escherichia coli O104:H4 Proteobacteria; Gammaproteobacteria; Enterobacterales; Enterobacteriaceae; Escherichia; Escherichia coli
MNNEETFYQAMRRQGVTRRSFLKYCSLAATSLGLGAGMAPKIAWALENKPRIPVVWIHGL WP_001058323.1
MULTISPECIES: hydrogenase-1 small chain [Proteobacteria] Length: 372\nNumber of predicted TMHs: 2\nExp number of AAs in TMHs: 37.2608499999999\nExp number, first 60 AAs: 19.52923\nTotal prob of N-in: 0.94351\nPOSSIBLE N-term signal sequence\ninside 1 19\nTMhelix 20 39\noutside 40 325\nTMhelix 326 348\ninside 349 372

12142 GCF_002156555.1_ASM215655v1 Escherichia coli O112ac:H19 Proteobacteria; Gammaproteobacteria; Enterobacterales; Enterobacteriaceae; Escherichia; Escherichia coli
MTGDNTLIHSHGINRRDFMKLCAALAATMGLSSKAAAEMAESVTNPQRPPVIWIGAQECT WP_000145410.1
MULTISPECIES: hydrogenase-2 small chain [Proteobacteria] Length: 372\nNumber of predicted TMHs:

1\nExp number of AAs in TMHs: 23.72226\nExp number, first 60 AAs: 0.22038\nTotal prob of N-in: 0.12994\noutside 1 331\nTMhelix 332 354\ninside 355 372

12143 GCF_001592525.1_ASM159252v1 Escherichia coli O119:H6 Proteobacteria; Gammaproteobacteria; Enterobacterales; Enterobacteriaceae; Escherichia; Escherichia coli
MTGDNTLIHSHGINRRDFMKLCAALAATMGLSSKAAAEMAESVTNPQRPPVIWIGAQECT WP_000145410.1
MULTISPECIES: hydrogenase-2 small chain [Proteobacteria] Length: 372\nNumber of predicted TMHs: 1\nExp number of AAs in TMHs: 23.72226\nExp number, first 60 AAs: 0.22038\nTotal prob of N-in: 0.12994\noutside 1 331\nTMhelix 332 354\ninside 355 372

12144 GCF_001592525.1_ASM159252v1 Escherichia coli O119:H6 Proteobacteria; Gammaproteobacteria; Enterobacterales; Enterobacteriaceae; Escherichia; Escherichia coli
MNNEETFYQAMRRQGVTRRSFLKYCSLAATSLGLGAGMAPKIAWALENKPRIPVVWIHGL WP_001058323.1
MULTISPECIES: hydrogenase-1 small chain [Proteobacteria] Length: 372\nNumber of predicted TMHs: 2\nExp number of AAs in TMHs: 37.2608499999999\nExp number, first 60 AAs: 19.52923\nTotal prob of N-in: 0.94351\nPOSSIBLE N-term signal sequence\ninside 1 19\nTMhelix 20 39\noutside 40 325\nTMhelix 326 348\ninside 349 372

12145 GCF_001592505.1_ASM159250v1 Escherichia coli O119:H6 Proteobacteria; Gammaproteobacteria; Enterobacterales; Enterobacteriaceae; Escherichia; Escherichia coli
MNNEETFYQAMRRQGVTRRSFLKYCSLAATSLGLGAGMAPKIAWALENKPRIPVVWIHGL WP_001058323.1
MULTISPECIES: hydrogenase-1 small chain [Proteobacteria] Length: 372\nNumber of predicted TMHs: 2\nExp number of AAs in TMHs: 37.2608499999999\nExp number, first 60 AAs: 19.52923\nTotal prob of N-in: 0.94351\nPOSSIBLE N-term signal sequence\ninside 1 19\nTMhelix 20 39\noutside 40 325\nTMhelix 326 348\ninside 349 372

12146 GCF_001592545.1_ASM159254v1 Escherichia coli O119:H6 Proteobacteria; Gammaproteobacteria; Enterobacterales; Enterobacteriaceae; Escherichia; Escherichia coli
MNNEETFYQAMRRQGVTRRSFLKYCSLAATSLGLGAGMAPKIAWALENKPRIPVVWIHGL WP_001058323.1
MULTISPECIES: hydrogenase-1 small chain [Proteobacteria] Length: 372\nNumber of predicted TMHs: 2\nExp number of AAs in TMHs: 37.2608499999999\nExp number, first 60 AAs: 19.52923\nTotal prob of N-in: 0.94351\nPOSSIBLE N-term signal sequence\ninside 1 19\nTMhelix 20 39\noutside 40 325\nTMhelix 326 348\ninside 349 372

12147 GCF_001592505.1_ASM159250v1 Escherichia coli O119:H6 Proteobacteria; Gammaproteobacteria; Enterobacterales; Enterobacteriaceae; Escherichia; Escherichia coli
MTGDNTLIHSHGINRRDFMKLCAALAATMGLSSKAAAEMAESVTNPQRPPVIWIGAQECT WP_000145410.1
MULTISPECIES: hydrogenase-2 small chain [Proteobacteria] Length: 372\nNumber of predicted TMHs: 1\nExp number of AAs in TMHs: 23.72226\nExp number, first 60 AAs: 0.22038\nTotal prob of N-in: 0.12994\noutside 1 331\nTMhelix 332 354\ninside 355 372

12148 GCF_001592545.1_ASM159254v1 Escherichia coli O119:H6 Proteobacteria; Gammaproteobacteria; Enterobacterales; Enterobacteriaceae; Escherichia; Escherichia coli
MTGDNTLIHSHGINRRDFMKLCAALAATMGLSSKAAAEMAESVTNPQRPPVIWIGAQECT WP_000145410.1
MULTISPECIES: hydrogenase-2 small chain [Proteobacteria] Length: 372\nNumber of predicted TMHs: 1\nExp number of AAs in TMHs: 23.72226\nExp number, first 60 AAs: 0.22038\nTotal prob of N-in: 0.12994\noutside 1 331\nTMhelix 332 354\ninside 355 372

12149 GCF_001650295.1_ASM165029v1 Escherichia coli O157 Proteobacteria; Gammaproteobacteria; Enterobacterales; Enterobacteriaceae; Escherichia; Escherichia coli
MTGDNTLIHSHGINRRDFMKLCAALAATMGLSSKAAAEMAESVTNPQRPPVIWIGAQECT WP_000145410.1
MULTISPECIES: hydrogenase-2 small chain [Proteobacteria] Length: 372\nNumber of predicted TMHs: 1\nExp number of AAs in TMHs: 23.72226\nExp number, first 60 AAs: 0.22038\nTotal prob of N-in: 0.12994\noutside 1 331\nTMhelix 332 354\ninside 355 372

12150 GCF_001650295.1_ASM165029v1 Escherichia coli O157 Proteobacteria; Gammaproteobacteria; Enterobacterales; Enterobacteriaceae; Escherichia; Escherichia coli
MNNEETFYQAMRRQGVTRRSFLKYCSLAATSLGLGAGMAPKIAWALENKPRIPVVWIHGL WP_001301613.1
hydrogenase [Escherichia coli] Length: 372\nNumber of predicted TMHs: 2\nExp number of AAs in TMHs: 37.46598\nExp number, first 60 AAs: 19.53139\nTotal prob of N-in: 0.94364\nPOSSIBLE N-term signal sequence\ninside 1 19\nTMhelix 20 39\noutside 40 325\nTMhelix 326 348\ninside 349 372

12151 GCF_002206405.1_ASM220640v1 Escherichia coli O157 Proteobacteria; Gammaproteobacteria; Enterobacterales; Enterobacteriaceae; Escherichia; Escherichia coli
MTGDNTLIHSHGINRRDFMKLCAALAATMGLSSKAAAEMAESVTNPQRPPVIWIGAQECT WP_000145410.1
MULTISPECIES: hydrogenase-2 small chain [Proteobacteria] Length: 372\nNumber of predicted TMHs: 1\nExp number of AAs in TMHs: 23.72226\nExp number, first 60 AAs: 0.22038\nTotal prob of N-in: 0.12994\noutside 1 331\nTMhelix 332 354\ninside 355 372

12152 GCF_002208865.1_ASM220886v1 Escherichia coli O157 Proteobacteria; Gammaproteobacteria; Enterobacterales; Enterobacteriaceae; Escherichia; Escherichia coli
MTGDNTLIHSHGINRRDFMKLCAALAATMGLSSKAAAEMAESVTNPQRPPVIWIGAQECT WP_000145410.1
MULTISPECIES: hydrogenase-2 small chain [Proteobacteria] Length: 372\nNumber of predicted TMHs: 1\nExp number of AAs in TMHs: 23.72226\nExp number, first 60 AAs: 0.22038\nTotal prob of N-in: 0.12994\noutside 1 331\nTMhelix 332 354\ninside 355 372

12153 GCF_001650275.1_ASM165027v1 Escherichia coli O157 Proteobacteria; Gammaproteobacteria; Enterobacterales; Enterobacteriaceae; Escherichia; Escherichia coli
MTGDNTLIHSHGINRRDFMKLCAALAATMGLSSKAAAEMAESVTNPQRPPVIWIGAQECT WP_000145410.1
MULTISPECIES: hydrogenase-2 small chain [Proteobacteria] Length: 372\nNumber of predicted TMHs: 1\nExp number of AAs in TMHs: 23.72226\nExp number, first 60 AAs: 0.22038\nTotal prob of N-in: 0.12994\noutside 1 331\nTMhelix 332 354\ninside 355 372

12154 GCF_001650275.1_ASM165027v1 Escherichia coli O157 Proteobacteria; Gammaproteobacteria; Enterobacterales; Enterobacteriaceae; Escherichia; Escherichia coli
MNNEETFYQAMRRQGVTRRSFLKYCSLAATSLGLGAGMAPKIAWALENKPRIPVVWIHGL WP_001301613.1
hydrogenase [Escherichia coli] Length: 372\nNumber of predicted TMHs: 2\nExp number of AAs in TMHs: 37.46598\nExp number, first 60 AAs: 19.53139\nTotal prob of N-in: 0.94364\nPOSSIBLE N-term signal sequence\ninside 1 19\nTMhelix 20 39\noutside 40 325\nTMhelix 326 348\ninside 349 372

12155 GCF_002206405.1_ASM220640v1 Escherichia coli O157 Proteobacteria; Gammaproteobacteria; Enterobacterales; Enterobacteriaceae; Escherichia; Escherichia coli
MNNEETFYQAMRRQGVTRRSFLKYCSLAATSLGLGAGMAPKIAWALENKPRIPVVWIHGL WP_001301613.1
hydrogenase [Escherichia coli] Length: 372\nNumber of predicted TMHs: 2\nExp number of AAs in TMHs: 37.46598\nExp number, first 60 AAs: 19.53139\nTotal prob of N-in: 0.94364\nPOSSIBLE N-term signal sequence\ninside 1 19\nTMhelix 20 39\noutside 40 325\nTMhelix 326 348\ninside 349 372

12156 GCF_002208865.1_ASM220886v1 Escherichia coli O157 Proteobacteria; Gammaproteobacteria; Enterobacterales; Enterobacteriaceae; Escherichia; Escherichia coli
MNNEETFYQAMRRQGVTRRSFLKYCSLAATSLGLGAGMAPKIAWALENKPRIPVVWIHGL WP_001301613.1
hydrogenase [Escherichia coli] Length: 372\nNumber of predicted TMHs: 2\nExp number of AAs in TMHs: 37.46598\nExp number, first 60 AAs: 19.53139\nTotal prob of N-in: 0.94364\nPOSSIBLE N-term signal sequence\ninside 1 19\nTMhelix 20 39\noutside 40 325\nTMhelix 326 348\ninside 349 372

12157 GCF_000827105.1_ASM82710v1 Escherichia coli O157:H16 Proteobacteria; Gammaproteobacteria; Enterobacterales; Enterobacteriaceae; Escherichia; Escherichia coli
MTGDNTLIHSHGINRRDFMKLCAALAATMGLSSKAAAEMAESVTNPQRPPVIWIGAQECT WP_000145410.1
MULTISPECIES: hydrogenase-2 small chain [Proteobacteria] Length: 372\nNumber of predicted TMHs: 1\nExp number of AAs in TMHs: 23.72226\nExp number, first 60 AAs: 0.22038\nTotal prob of N-in: 0.12994\noutside 1 331\nTMhelix 332 354\ninside 355 372

12158 GCF_000827105.1_ASM82710v1 Escherichia coli O157:H16 Proteobacteria; Gammaproteobacteria; Enterobacterales; Enterobacteriaceae; Escherichia; Escherichia coli
MNNEETFYQAMRRQGVTRRSFLKYCSLAATSLGLGAGMAPKIAWALENKPRIPVVWIHGL WP_001058323.1
MULTISPECIES: hydrogenase-1 small chain [Proteobacteria] Length: 372\nNumber of predicted TMHs: 2\nExp number of AAs in TMHs: 37.2608499999999\nExp number, first 60 AAs: 19.52923\nTotal prob of N-in: 0.94351\nPOSSIBLE N-term signal sequence\ninside 1 19\nTMhelix 20 39\noutside 40 325\nTMhelix 326 348\ninside 349 372

12159 GCF_001953095.1_ASM195309v1 Escherichia coli O157:H43 Proteobacteria; Gammaproteobacteria; Enterobacterales; Enterobacteriaceae; Escherichia; Escherichia coli
MTGDNTLIHSHGINRRDFMKLCAALAATMGLSSKAAAEMAESVTNPQRPPVIWIGAQECT WP_000145410.1
MULTISPECIES: hydrogenase-2 small chain [Proteobacteria] Length: 372\nNumber of predicted TMHs:

1\nExp number of AAs in TMHs: 23.72226\nExp number, first 60 AAs: 0.22038\nTotal prob of N-in: 0.12994\noutside 1 331\nTMhelix 332 354\ninside 355 372

12160 GCF_001953095.1_ASM195309v1 Escherichia coli O157:H43 Proteobacteria; Gammaproteobacteria; Enterobacterales; Enterobacteriaceae; Escherichia; Escherichia coli
MNNEETFYQAMRRQGVTRRSFLKYCSLAATSLGLGAGMAPKIAWALENKPRIPVVWIHGL WP_001058323.1
MULTISPECIES: hydrogenase-1 small chain [Proteobacteria] Length: 372\nNumber of predicted TMHs: 2\nExp number of AAs in TMHs: 37.2608499999999\nExp number, first 60 AAs: 19.52923\nTotal prob of N-in: 0.94351\nPOSSIBLE N-term signal sequence\ninside 1 19\nTMhelix 20 39\noutside 40 325\nTMhelix 326 348\ninside 349 372

12161 GCF_001516935.2_ASM151693v2 Escherichia coli O157:H7 Proteobacteria; Gammaproteobacteria; Enterobacterales; Enterobacteriaceae; Escherichia; Escherichia coli
MTGDNTLIHSHGINRRDFMKLCAALAATMGLSSKAAAEMAESVTNPQRPPVIWIGAQECT WP_000145410.1
MULTISPECIES: hydrogenase-2 small chain [Proteobacteria] Length: 372\nNumber of predicted TMHs: 1\nExp number of AAs in TMHs: 23.72226\nExp number, first 60 AAs: 0.22038\nTotal prob of N-in: 0.12994\noutside 1 331\nTMhelix 332 354\ninside 355 372

12162 GCF_002027605.1_ASM202760v1 Escherichia coli O157:H7 Proteobacteria; Gammaproteobacteria; Enterobacterales; Enterobacteriaceae; Escherichia; Escherichia coli
MTGDNTLIHSHGINRRDFMKLCAALAATMGLSSKAAAEMAESVTNPQRPPVIWIGAQECT WP_000145410.1
MULTISPECIES: hydrogenase-2 small chain [Proteobacteria] Length: 372\nNumber of predicted TMHs: 1\nExp number of AAs in TMHs: 23.72226\nExp number, first 60 AAs: 0.22038\nTotal prob of N-in: 0.12994\noutside 1 331\nTMhelix 332 354\ninside 355 372

12163 GCF_001006425.1_ASM100642v1 Escherichia coli O157:H7 Proteobacteria; Gammaproteobacteria; Enterobacterales; Enterobacteriaceae; Escherichia; Escherichia coli
MNNEETFYQAMRRQGVTRRSFLKYCSLAATSLGLGAGMAPKIAWALENKPRIPVVWIHGL WP_001301613.1
hydrogenase [Escherichia coli] Length: 372\nNumber of predicted TMHs: 2\nExp number of AAs in TMHs: 37.46598\nExp number, first 60 AAs: 19.53139\nTotal prob of N-in: 0.94364\nPOSSIBLE N-term signal sequence\ninside 1 19\nTMhelix 20 39\noutside 40 325\nTMhelix 326 348\ninside 349 372

12164 GCF_001432195.1_WS4678 Escherichia coli O157:H7 Proteobacteria; Gammaproteobacteria; Enterobacterales; Enterobacteriaceae; Escherichia; Escherichia coli
MTGDNTLIHSHGINRRDFMKLCAALAATMGLSSKAAAEMAESVTNPQRPPVIWIGAQECT WP_000145410.1
MULTISPECIES: hydrogenase-2 small chain [Proteobacteria] Length: 372\nNumber of predicted TMHs: 1\nExp number of AAs in TMHs: 23.72226\nExp number, first 60 AAs: 0.22038\nTotal prob of N-in: 0.12994\noutside 1 331\nTMhelix 332 354\ninside 355 372

12165 GCF_001516935.2_ASM151693v2 Escherichia coli O157:H7 Proteobacteria; Gammaproteobacteria; Enterobacterales; Enterobacteriaceae; Escherichia; Escherichia coli
MNNEETFYQAMRRQGVTRRSFLKYCSLAATSLGLGAGMAPKIAWALENKPRIPVVWIHGL WP_001301613.1
hydrogenase [Escherichia coli] Length: 372\nNumber of predicted TMHs: 2\nExp number of AAs in TMHs: 37.46598\nExp number, first 60 AAs: 19.53139\nTotal prob of N-in: 0.94364\nPOSSIBLE N-term signal sequence\ninside 1 19\nTMhelix 20 39\noutside 40 325\nTMhelix 326 348\ninside 349 372

12166 GCF_002027605.1_ASM202760v1 Escherichia coli O157:H7 Proteobacteria; Gammaproteobacteria; Enterobacterales; Enterobacteriaceae; Escherichia; Escherichia coli
MNNEETFYQAMRRQGVTRRSFLKYCSLAATSLGLGAGMAPKIAWALENKPRIPVVWIHGL WP_001301613.1
hydrogenase [Escherichia coli] Length: 372\nNumber of predicted TMHs: 2\nExp number of AAs in TMHs: 37.46598\nExp number, first 60 AAs: 19.53139\nTotal prob of N-in: 0.94364\nPOSSIBLE N-term signal sequence\ninside 1 19\nTMhelix 20 39\noutside 40 325\nTMhelix 326 348\ninside 349 372

12167 GCF_002027645.1_ASM202764v1 Escherichia coli O157:H7 Proteobacteria; Gammaproteobacteria; Enterobacterales; Enterobacteriaceae; Escherichia; Escherichia coli
MNNEETFYQAMRRQGVTRRSFLKYCSLAATSLGLGAGMAPKIAWALENKPRIPVVWIHGL WP_001301613.1
hydrogenase [Escherichia coli] Length: 372\nNumber of predicted TMHs: 2\nExp number of AAs in TMHs: 37.46598\nExp number, first 60 AAs: 19.53139\nTotal prob of N-in: 0.94364\nPOSSIBLE N-term signal sequence\ninside 1 19\nTMhelix 20 39\noutside 40 325\nTMhelix 326 348\ninside 349 372

12168 GCF_002027685.1_ASM202768v1 Escherichia coli O157:H7 Proteobacteria; Gammaproteobacteria; Enterobacterales; Enterobacteriaceae; Escherichia; Escherichia coli
MNNEETFYQAMRRQGVTRRSFLKYCSLAATSLGLGAGMAPKIAWALENKPRIPVVWIHGL WP_001301613.1
hydrogenase [Escherichia coli] Length: 372\nNumber of predicted TMHs: 2\nExp number of AAs in TMHs: 37.46598\nExp number, first 60 AAs: 19.53139\nTotal prob of N-in: 0.94364\nPOSSIBLE N-term signal sequence\ninside 1 19\nTMhelix 20 39\noutside 40 325\nTMhelix 326 348\ninside 349 372

12169 GCF_001939185.1_ASM193918v1 Escherichia coli O157:H7 Proteobacteria; Gammaproteobacteria; Enterobacterales; Enterobacteriaceae; Escherichia; Escherichia coli
MTGDNTLIHSHGINRRDFMKLCAALAATMGLSSKAAAEMAESVTNPQRPPVIWIGAQECT WP_000145410.1
MULTISPECIES: hydrogenase-2 small chain [Proteobacteria] Length: 372\nNumber of predicted TMHs: 1\nExp number of AAs in TMHs: 23.72226\nExp number, first 60 AAs: 0.22038\nTotal prob of N-in: 0.12994\noutside 1 331\nTMhelix 332 354\ninside 355 372

12170 GCF_001651925.1_ASM165192v1 Escherichia coli O157:H7 Proteobacteria; Gammaproteobacteria; Enterobacterales; Enterobacteriaceae; Escherichia; Escherichia coli
MTGDNTLIHSHGINRRDFMKLCAALAATMGLSSKAAAEMAESVTNPQRPPVIWIGAQECT WP_000145410.1
MULTISPECIES: hydrogenase-2 small chain [Proteobacteria] Length: 372\nNumber of predicted TMHs: 1\nExp number of AAs in TMHs: 23.72226\nExp number, first 60 AAs: 0.22038\nTotal prob of N-in: 0.12994\noutside 1 331\nTMhelix 332 354\ninside 355 372

12171 GCF_001432195.1_WS4678 Escherichia coli O157:H7 Proteobacteria; Gammaproteobacteria; Enterobacterales; Enterobacteriaceae; Escherichia; Escherichia coli
MNNEETFYQAMRRQGVTRRSFLKYCSLAATSLGLGAGMAPKIAWALENKPRIPVVWIHGL WP_001301613.1
hydrogenase [Escherichia coli] Length: 372\nNumber of predicted TMHs: 2\nExp number of AAs in TMHs: 37.46598\nExp number, first 60 AAs: 19.53139\nTotal prob of N-in: 0.94364\nPOSSIBLE N-term signal sequence\ninside 1 19\nTMhelix 20 39\noutside 40 325\nTMhelix 326 348\ninside 349 372

12172 GCF_001939185.1_ASM193918v1 Escherichia coli O157:H7 Proteobacteria; Gammaproteobacteria; Enterobacterales; Enterobacteriaceae; Escherichia; Escherichia coli
MNNEETFYQAMRRQGVTRRSFLKYCSLAATSLGLGAGMAPKIAWALENKPRIPVVWIHGL WP_001301613.1
hydrogenase [Escherichia coli] Length: 372\nNumber of predicted TMHs: 2\nExp number of AAs in TMHs: 37.46598\nExp number, first 60 AAs: 19.53139\nTotal prob of N-in: 0.94364\nPOSSIBLE N-term signal sequence\ninside 1 19\nTMhelix 20 39\noutside 40 325\nTMhelix 326 348\ninside 349 372

12173 GCF_001753445.1_ASM175344v1 Escherichia coli O157:H7 Proteobacteria; Gammaproteobacteria; Enterobacterales; Enterobacteriaceae; Escherichia; Escherichia coli
MTGDNTLIHSHGINRRDFMKLCAALAATMGLSSKAAAEMAESVTNPQRPPVIWIGAQECT WP_000145410.1
MULTISPECIES: hydrogenase-2 small chain [Proteobacteria] Length: 372\nNumber of predicted TMHs: 1\nExp number of AAs in TMHs: 23.72226\nExp number, first 60 AAs: 0.22038\nTotal prob of N-in: 0.12994\noutside 1 331\nTMhelix 332 354\ninside 355 372

12174 GCF_001753445.1_ASM175344v1 Escherichia coli O157:H7 Proteobacteria; Gammaproteobacteria; Enterobacterales; Enterobacteriaceae; Escherichia; Escherichia coli
MNNEETFYQAMRRQGVTRRSFLKYCSLAATSLGLGAGMAPKIAWALENKPRIPVVWIHGL WP_001301613.1
hydrogenase [Escherichia coli] Length: 372\nNumber of predicted TMHs: 2\nExp number of AAs in TMHs: 37.46598\nExp number, first 60 AAs: 19.53139\nTotal prob of N-in: 0.94364\nPOSSIBLE N-term signal sequence\ninside 1 19\nTMhelix 20 39\noutside 40 325\nTMhelix 326 348\ninside 349 372

12175 GCF_001651925.1_ASM165192v1 Escherichia coli O157:H7 Proteobacteria; Gammaproteobacteria; Enterobacterales; Enterobacteriaceae; Escherichia; Escherichia coli
MNNEETFYQAMRRQGVTRRSFLKYCSLAATSLGLGAGMAPKIAWALENKPRIPVVWIHGL WP_001301613.1
hydrogenase [Escherichia coli] Length: 372\nNumber of predicted TMHs: 2\nExp number of AAs in TMHs: 37.46598\nExp number, first 60 AAs: 19.53139\nTotal prob of N-in: 0.94364\nPOSSIBLE N-term signal sequence\ninside 1 19\nTMhelix 20 39\noutside 40 325\nTMhelix 326 348\ninside 349 372

12176 GCF_001006425.1_ASM100642v1 Escherichia coli O157:H7 Proteobacteria; Gammaproteobacteria; Enterobacterales; Enterobacteriaceae; Escherichia; Escherichia coli
MTGDNTLIHSHGINRRDFMKLCAALAATMGLSSKAAAEMAESVTNPQRPPVIWIGAQECT WP_000145410.1
MULTISPECIES: hydrogenase-2 small chain [Proteobacteria] Length: 372\nNumber of predicted TMHs:

1\nExp number of AAs in TMHs: 23.72226\nExp number, first 60 AAs: 0.22038\nTotal prob of N-in: 0.12994\noutside 1 331\nTMhelix 332 354\ninside 355 372

12177 GCF_001440045.1_ASM144004v1 Escherichia coli O157:H7 Proteobacteria; Gammaproteobacteria; Enterobacterales; Enterobacteriaceae; Escherichia; Escherichia coli
MTGDNTLIHSHGINRRDFMKLCAALAATMGLSSKAAAEMAESVTNPQRPPVIWIGAQECT WP_000145410.1
MULTISPECIES: hydrogenase-2 small chain [Proteobacteria] Length: 372\nNumber of predicted TMHs: 1\nExp number of AAs in TMHs: 23.72226\nExp number, first 60 AAs: 0.22038\nTotal prob of N-in: 0.12994\noutside 1 331\nTMhelix 332 354\ninside 355 372

12178 GCF_001753505.1_ASM175350v1 Escherichia coli O157:H7 Proteobacteria; Gammaproteobacteria; Enterobacterales; Enterobacteriaceae; Escherichia; Escherichia coli
MNNEETFYQAMRRQGVTRRSFLKYCSLAATSLGLGAGMAPKIAWALENKPRIPVVWIHGL WP_001301613.1
hydrogenase [Escherichia coli] Length: 372\nNumber of predicted TMHs: 2\nExp number of AAs in TMHs: 37.46598\nExp number, first 60 AAs: 19.53139\nTotal prob of N-in: 0.94364\nPOSSIBLE N-term signal sequence\ninside 1 19\nTMhelix 20 39\noutside 40 325\nTMhelix 326 348\ninside 349 372

12179 GCF_001753525.1_ASM175352v1 Escherichia coli O157:H7 Proteobacteria; Gammaproteobacteria; Enterobacterales; Enterobacteriaceae; Escherichia; Escherichia coli
MNNEETFYQAMRRQGVTRRSFLKYCSLAATSLGLGAGMAPKIAWALENKPRIPVVWIHGL WP_001301613.1
hydrogenase [Escherichia coli] Length: 372\nNumber of predicted TMHs: 2\nExp number of AAs in TMHs: 37.46598\nExp number, first 60 AAs: 19.53139\nTotal prob of N-in: 0.94364\nPOSSIBLE N-term signal sequence\ninside 1 19\nTMhelix 20 39\noutside 40 325\nTMhelix 326 348\ninside 349 372

12180 GCF_001753565.1_ASM175356v1 Escherichia coli O157:H7 Proteobacteria; Gammaproteobacteria; Enterobacterales; Enterobacteriaceae; Escherichia; Escherichia coli
MNNEETFYQAMRRQGVTRRSFLKYCSLAATSLGLGAGMAPKIAWALENKPRIPVVWIHGL WP_001301613.1
hydrogenase [Escherichia coli] Length: 372\nNumber of predicted TMHs: 2\nExp number of AAs in TMHs: 37.46598\nExp number, first 60 AAs: 19.53139\nTotal prob of N-in: 0.94364\nPOSSIBLE N-term signal sequence\ninside 1 19\nTMhelix 20 39\noutside 40 325\nTMhelix 326 348\ninside 349 372

12181 GCF_001558995.2_ASM155899v2 Escherichia coli O157:H7 Proteobacteria; Gammaproteobacteria; Enterobacterales; Enterobacteriaceae; Escherichia; Escherichia coli
MNNEETFYQAMRRQGVTRRSFLKYCSLAATSLGLGAGMAPKIAWALENKPRIPVVWIHGL WP_001301613.1
hydrogenase [Escherichia coli] Length: 372\nNumber of predicted TMHs: 2\nExp number of AAs in TMHs: 37.46598\nExp number, first 60 AAs: 19.53139\nTotal prob of N-in: 0.94364\nPOSSIBLE N-term signal sequence\ninside 1 19\nTMhelix 20 39\noutside 40 325\nTMhelix 326 348\ninside 349 372

12182 GCF_001558995.2_ASM155899v2 Escherichia coli O157:H7 Proteobacteria; Gammaproteobacteria; Enterobacterales; Enterobacteriaceae; Escherichia; Escherichia coli
MTGDNTLIHSHGINRRDFMKLCAALAATMGLSSKAAAEMAESVTNPQRPPVIWIGAQECS WP_061069243.1
hydrogenase 2 small subunit [Escherichia coli] Length: 372\nNumber of predicted TMHs: 1\nExp number of AAs in TMHs: 23.72219\nExp number, first 60 AAs: 0.22034\nTotal prob of N-in: 0.12994\noutside 1 331\nTMhelix 332 354\ninside 355 372

12183 GCF_000978815.1_ASM97881v1 Escherichia coli O157:H7 Proteobacteria; Gammaproteobacteria; Enterobacterales; Enterobacteriaceae; Escherichia; Escherichia coli
MTGDNTLIHSHGINRRDFMKLCAALAATMGLSSKAAAEMAESVTNPQRPPVIWIGAQECT WP_000145410.1
MULTISPECIES: hydrogenase-2 small chain [Proteobacteria] Length: 372\nNumber of predicted TMHs: 1\nExp number of AAs in TMHs: 23.72226\nExp number, first 60 AAs: 0.22038\nTotal prob of N-in: 0.12994\noutside 1 331\nTMhelix 332 354\ninside 355 372

12184 GCF_000978815.1_ASM97881v1 Escherichia coli O157:H7 Proteobacteria; Gammaproteobacteria; Enterobacterales; Enterobacteriaceae; Escherichia; Escherichia coli
MNNEETFYQAMRRQGVTRRSFLKYCSLAATSLGLGAGMAPKIAWALENKPRIPVVWIHGL WP_001301613.1
hydrogenase [Escherichia coli] Length: 372\nNumber of predicted TMHs: 2\nExp number of AAs in TMHs: 37.46598\nExp number, first 60 AAs: 19.53139\nTotal prob of N-in: 0.94364\nPOSSIBLE N-term signal sequence\ninside 1 19\nTMhelix 20 39\noutside 40 325\nTMhelix 326 348\ninside 349 372

12185 GCF_001440045.1_ASM144004v1 Escherichia coli O157:H7 Proteobacteria; Gammaproteobacteria; Enterobacterales; Enterobacteriaceae; Escherichia; Escherichia coli

MNNEETFYQAMRRQGVTRRSFLKYCSLAATSLGLGAGMAPKIAWALENKPRIPVVWIHGL WP_001301613.1 hydrogenase [Escherichia coli] Length: 372\nNumber of predicted TMHs: 2\nExp number of AAs in TMHs: 37.46598\nExp number, first 60 AAs: 19.53139\nTotal prob of N-in: 0.94364\nPOSSIBLE N-term signal sequence\ninside 1 19\nTMhelix 20 39\noutside 40 325\nTMhelix 326 348\ninside 349 372

12186 GCF_001753465.1_ASM175346v1 Escherichia coli O157:H7 Proteobacteria; Gammaproteobacteria; Enterobacterales; Enterobacteriaceae; Escherichia; Escherichia coli MNNEETFYQAMRRQGVTRRSFLKYCSLAATSLGLGAGMAPKIAWALENKPRIPVVWIHGL WP_001301613.1 hydrogenase [Escherichia coli] Length: 372\nNumber of predicted TMHs: 2\nExp number of AAs in TMHs: 37.46598\nExp number, first 60 AAs: 19.53139\nTotal prob of N-in: 0.94364\nPOSSIBLE N-term signal sequence\ninside 1 19\nTMhelix 20 39\noutside 40 325\nTMhelix 326 348\ninside 349 372

12187 GCF_001651945.1_ASM165194v1 Escherichia coli O157:H7 Proteobacteria; Gammaproteobacteria; Enterobacterales; Enterobacteriaceae; Escherichia; Escherichia coli MNNEETFYQAMRRQGVTRRSFLKYCSLAATSLGLGAGMAPKIAWALENKPRIPVVWIHGL WP_001301613.1 hydrogenase [Escherichia coli] Length: 372\nNumber of predicted TMHs: 2\nExp number of AAs in TMHs: 37.46598\nExp number, first 60 AAs: 19.53139\nTotal prob of N-in: 0.94364\nPOSSIBLE N-term signal sequence\ninside 1 19\nTMhelix 20 39\noutside 40 325\nTMhelix 326 348\ninside 349 372

12188 GCF_001651965.1_ASM165196v1 Escherichia coli O157:H7 Proteobacteria; Gammaproteobacteria; Enterobacterales; Enterobacteriaceae; Escherichia; Escherichia coli MTGDNTLIHSHGINRRDFMKLCAALAATMGLSSKAAAEMAESVTNPQRPPVIWIGAQECT WP_000145410.1 MULTISPECIES: hydrogenase-2 small chain [Proteobacteria] Length: 372\nNumber of predicted TMHs: 1\nExp number of AAs in TMHs: 23.72226\nExp number, first 60 AAs: 0.22038\nTotal prob of N-in: 0.12994\noutside 1 331\nTMhelix 332 354\ninside 355 372

12189 GCF_001753465.1_ASM175346v1 Escherichia coli O157:H7 Proteobacteria; Gammaproteobacteria; Enterobacterales; Enterobacteriaceae; Escherichia; Escherichia coli MTGDNTLIHSHGINRRDFMKLCAALAATMGLSSKAAAEMAESVTNPQRPPVIWIGAQECT WP_000145410.1 MULTISPECIES: hydrogenase-2 small chain [Proteobacteria] Length: 372\nNumber of predicted TMHs: 1\nExp number of AAs in TMHs: 23.72226\nExp number, first 60 AAs: 0.22038\nTotal prob of N-in: 0.12994\noutside 1 331\nTMhelix 332 354\ninside 355 372

12190 GCF_001753505.1_ASM175350v1 Escherichia coli O157:H7 Proteobacteria; Gammaproteobacteria; Enterobacterales; Enterobacteriaceae; Escherichia; Escherichia coli MTGDNTLIHSHGINRRDFMKLCAALAATMGLSSKAAAEMAESVTNPQRPPVIWIGAQECT WP_000145410.1 MULTISPECIES: hydrogenase-2 small chain [Proteobacteria] Length: 372\nNumber of predicted TMHs: 1\nExp number of AAs in TMHs: 23.72226\nExp number, first 60 AAs: 0.22038\nTotal prob of N-in: 0.12994\noutside 1 331\nTMhelix 332 354\ninside 355 372

12191 GCF_001753525.1_ASM175352v1 Escherichia coli O157:H7 Proteobacteria; Gammaproteobacteria; Enterobacterales; Enterobacteriaceae; Escherichia; Escherichia coli MTGDNTLIHSHGINRRDFMKLCAALAATMGLSSKAAAEMAESVTNPQRPPVIWIGAQECT WP_000145410.1 MULTISPECIES: hydrogenase-2 small chain [Proteobacteria] Length: 372\nNumber of predicted TMHs: 1\nExp number of AAs in TMHs: 23.72226\nExp number, first 60 AAs: 0.22038\nTotal prob of N-in: 0.12994\noutside 1 331\nTMhelix 332 354\ninside 355 372

12192 GCF_001753565.1_ASM175356v1 Escherichia coli O157:H7 Proteobacteria; Gammaproteobacteria; Enterobacterales; Enterobacteriaceae; Escherichia; Escherichia coli MTGDNTLIHSHGINRRDFMKLCAALAATMGLSSKAAAEMAESVTNPQRPPVIWIGAQECT WP_000145410.1 MULTISPECIES: hydrogenase-2 small chain [Proteobacteria] Length: 372\nNumber of predicted TMHs: 1\nExp number of AAs in TMHs: 23.72226\nExp number, first 60 AAs: 0.22038\nTotal prob of N-in: 0.12994\noutside 1 331\nTMhelix 332 354\ninside 355 372

12193 GCF_000978845.1_ASM97884v1 Escherichia coli O157:H7 Proteobacteria; Gammaproteobacteria; Enterobacterales; Enterobacteriaceae; Escherichia; Escherichia coli MNNEETFYQAMRRQGVTRRSFLKYCSLAATSLGLGAGMAPKIAWALENKPRIPVVWIHGL WP_001301613.1 hydrogenase [Escherichia coli] Length: 372\nNumber of predicted TMHs: 2\nExp number of AAs in TMHs: 37.46598\nExp number, first 60 AAs: 19.53139\nTotal prob of N-in: 0.94364\nPOSSIBLE N-term signal sequence\ninside 1 19\nTMhelix 20 39\noutside 40 325\nTMhelix 326 348\ninside 349 372

12194 GCF_000978845.1_ASM97884v1 Escherichia coli O157:H7 Proteobacteria; Gammaproteobacteria; Enterobacterales; Enterobacteriaceae; Escherichia; Escherichia coli
MTGDNTLIHSHGINRRDFMKLCAALAATMGLSSKAAAEMAESVTNPQRPPVIWIGAQECT WP_000145410.1
MULTISPECIES: hydrogenase-2 small chain [Proteobacteria] Length: 372\nNumber of predicted TMHs: 1\nExp number of AAs in TMHs: 23.72226\nExp number, first 60 AAs: 0.22038\nTotal prob of N-in: 0.12994\noutside 1 331\nTMhelix 332 354\ninside 355 372

12195 GCF_001432175.2_WS4435_final_V2 Escherichia coli O157:H7 Proteobacteria; Gammaproteobacteria; Enterobacterales; Enterobacteriaceae; Escherichia; Escherichia coli
MNNEETFYQAMRRQGVTRRSFLKYCSLAATSLGLGAGMAPKIAWALENKPRIPVVWIHGL WP_001301613.1
hydrogenase [Escherichia coli] Length: 372\nNumber of predicted TMHs: 2\nExp number of AAs in TMHs: 37.46598\nExp number, first 60 AAs: 19.53139\nTotal prob of N-in: 0.94364\nPOSSIBLE N-term signal sequence\ninside 1 19\nTMhelix 20 39\noutside 40 325\nTMhelix 326 348\ninside 349 372

12196 GCF_001432175.2_WS4435_final_V2 Escherichia coli O157:H7 Proteobacteria; Gammaproteobacteria; Enterobacterales; Enterobacteriaceae; Escherichia; Escherichia coli
MTGDNTLIHSHGINRRDFMKLCAALAATMGLSSKAAAEMAESVTNPQRPPVIWIGAQECT WP_000145410.1
MULTISPECIES: hydrogenase-2 small chain [Proteobacteria] Length: 372\nNumber of predicted TMHs: 1\nExp number of AAs in TMHs: 23.72226\nExp number, first 60 AAs: 0.22038\nTotal prob of N-in: 0.12994\noutside 1 331\nTMhelix 332 354\ninside 355 372

12197 GCF_001307215.1_ASM130721v1 Escherichia coli O157:H7 Proteobacteria; Gammaproteobacteria; Enterobacterales; Enterobacteriaceae; Escherichia; Escherichia coli
MNNEETFYQAMRRQGVTRRSFLKYCSLAATSLGLGAGMAPKIAWALENKPRIPVVWIHGL WP_001301613.1
hydrogenase [Escherichia coli] Length: 372\nNumber of predicted TMHs: 2\nExp number of AAs in TMHs: 37.46598\nExp number, first 60 AAs: 19.53139\nTotal prob of N-in: 0.94364\nPOSSIBLE N-term signal sequence\ninside 1 19\nTMhelix 20 39\noutside 40 325\nTMhelix 326 348\ninside 349 372

12198 GCF_001307215.1_ASM130721v1 Escherichia coli O157:H7 Proteobacteria; Gammaproteobacteria; Enterobacterales; Enterobacteriaceae; Escherichia; Escherichia coli
MTGDNTLIHSHGINRRDFMKLCAALAATMGLSSKAAAEMAESVTNPQRPPVIWIGAQECT WP_000145410.1
MULTISPECIES: hydrogenase-2 small chain [Proteobacteria] Length: 372\nNumber of predicted TMHs: 1\nExp number of AAs in TMHs: 23.72226\nExp number, first 60 AAs: 0.22038\nTotal prob of N-in: 0.12994\noutside 1 331\nTMhelix 332 354\ninside 355 372

12199 GCF_001651965.1_ASM165196v1 Escherichia coli O157:H7 Proteobacteria; Gammaproteobacteria; Enterobacterales; Enterobacteriaceae; Escherichia; Escherichia coli
MNNEETFYQAMRRQGVTRRSFLKYCSLAATSLGLGAGMAPKIAWALENKPRIPVVWIHGL WP_001301613.1
hydrogenase [Escherichia coli] Length: 372\nNumber of predicted TMHs: 2\nExp number of AAs in TMHs: 37.46598\nExp number, first 60 AAs: 19.53139\nTotal prob of N-in: 0.94364\nPOSSIBLE N-term signal sequence\ninside 1 19\nTMhelix 20 39\noutside 40 325\nTMhelix 326 348\ninside 349 372

12200 GCF_001651945.1_ASM165194v1 Escherichia coli O157:H7 Proteobacteria; Gammaproteobacteria; Enterobacterales; Enterobacteriaceae; Escherichia; Escherichia coli
MTGDNTLIHSHGINRRDFMKLCAALAATMGLSSKAAAEMAESVTNPQRPPVIWIGAQECT WP_000145410.1
MULTISPECIES: hydrogenase-2 small chain [Proteobacteria] Length: 372\nNumber of predicted TMHs: 1\nExp number of AAs in TMHs: 23.72226\nExp number, first 60 AAs: 0.22038\nTotal prob of N-in: 0.12994\noutside 1 331\nTMhelix 332 354\ninside 355 372

12201 GCF_001695515.1_ASM169551v1 Escherichia coli O157:H7 Proteobacteria; Gammaproteobacteria; Enterobacterales; Enterobacteriaceae; Escherichia; Escherichia coli
MTGDNTLIHSHGINRRDFMKLCAALAATMGLSSKAAAEMAESVTNPQRPPVIWIGAQECT WP_000145410.1
MULTISPECIES: hydrogenase-2 small chain [Proteobacteria] Length: 372\nNumber of predicted TMHs: 1\nExp number of AAs in TMHs: 23.72226\nExp number, first 60 AAs: 0.22038\nTotal prob of N-in: 0.12994\noutside 1 331\nTMhelix 332 354\ninside 355 372

12202 GCF_001695515.1_ASM169551v1 Escherichia coli O157:H7 Proteobacteria; Gammaproteobacteria; Enterobacterales; Enterobacteriaceae; Escherichia; Escherichia coli
MNNEETFYQAMRRQGVTRRSFLKYCSLAATSLGLGAGMAPKIAWALENKPRIPVVWIHGL WP_001301613.1
hydrogenase [Escherichia coli] Length: 372\nNumber of predicted TMHs: 2\nExp number of AAs in

TMHs: 37.46598\nExp number, first 60 AAs: 19.53139\nTotal prob of N-in: 0.94364\nPOSSIBLE N-term signal sequence\ninside 1 19\nTMhelix 20 39\noutside 40 325\nTMhelix 326 348\ninside 349 372

12203 GCF_001753545.1_ASM175354v1 Escherichia coli O157:H7 Proteobacteria; Gammaproteobacteria; Enterobacterales; Enterobacteriaceae; Escherichia; Escherichia coli MNNEETFYQAMRRQGVTRRSFLKYCSLAATSLGLGAGMAPKIAWALENKPRIPVVWIHGL WP_001301613.1 hydrogenase [Escherichia coli] Length: 372\nNumber of predicted TMHs: 2\nExp number of AAs in TMHs: 37.46598\nExp number, first 60 AAs: 19.53139\nTotal prob of N-in: 0.94364\nPOSSIBLE N-term signal sequence\ninside 1 19\nTMhelix 20 39\noutside 40 325\nTMhelix 326 348\ninside 349 372

12204 GCF_001753485.1_ASM175348v1 Escherichia coli O157:H7 Proteobacteria; Gammaproteobacteria; Enterobacterales; Enterobacteriaceae; Escherichia; Escherichia coli MTGDNTLIHSHGINRRDFMKLCAALAATMGLSSKAAAEMAESVTNPQRPPVIWIGAQECT WP_000145410.1 MULTISPECIES: hydrogenase-2 small chain [Proteobacteria] Length: 372\nNumber of predicted TMHs: 1\nExp number of AAs in TMHs: 23.72226\nExp number, first 60 AAs: 0.22038\nTotal prob of N-in: 0.12994\noutside 1 331\nTMhelix 332 354\ninside 355 372

12205 GCF_001753545.1_ASM175354v1 Escherichia coli O157:H7 Proteobacteria; Gammaproteobacteria; Enterobacterales; Enterobacteriaceae; Escherichia; Escherichia coli MTGDNTLIHSHGINRRDFMKLCAALAATMGLSSKAAAEMAESVTNPQRPPVIWIGAQECT WP_000145410.1 MULTISPECIES: hydrogenase-2 small chain [Proteobacteria] Length: 372\nNumber of predicted TMHs: 1\nExp number of AAs in TMHs: 23.72226\nExp number, first 60 AAs: 0.22038\nTotal prob of N-in: 0.12994\noutside 1 331\nTMhelix 332 354\ninside 355 372

12206 GCF_002027645.1_ASM202764v1 Escherichia coli O157:H7 Proteobacteria; Gammaproteobacteria; Enterobacterales; Enterobacteriaceae; Escherichia; Escherichia coli MTGDNTLIHSHGINRRDFMKLCAALAATMGLSSKAAAEMAESVTNPQRPPVIWIGAQECT WP_000145410.1 MULTISPECIES: hydrogenase-2 small chain [Proteobacteria] Length: 372\nNumber of predicted TMHs: 1\nExp number of AAs in TMHs: 23.72226\nExp number, first 60 AAs: 0.22038\nTotal prob of N-in: 0.12994\noutside 1 331\nTMhelix 332 354\ninside 355 372

12207 GCF_002027685.1_ASM202768v1 Escherichia coli O157:H7 Proteobacteria; Gammaproteobacteria; Enterobacterales; Enterobacteriaceae; Escherichia; Escherichia coli MTGDNTLIHSHGINRRDFMKLCAALAATMGLSSKAAAEMAESVTNPQRPPVIWIGAQECT WP_000145410.1 MULTISPECIES: hydrogenase-2 small chain [Proteobacteria] Length: 372\nNumber of predicted TMHs: 1\nExp number of AAs in TMHs: 23.72226\nExp number, first 60 AAs: 0.22038\nTotal prob of N-in: 0.12994\noutside 1 331\nTMhelix 332 354\ninside 355 372

12208 GCF_001753485.1_ASM175348v1 Escherichia coli O157:H7 Proteobacteria; Gammaproteobacteria; Enterobacterales; Enterobacteriaceae; Escherichia; Escherichia coli MNNEETFYQAMRRQGVTRRSFLKYCSLAATSLGLGAGMAPKIAWALENKPRIPVVWIHGL WP_001301613.1 hydrogenase [Escherichia coli] Length: 372\nNumber of predicted TMHs: 2\nExp number of AAs in TMHs: 37.46598\nExp number, first 60 AAs: 19.53139\nTotal prob of N-in: 0.94364\nPOSSIBLE N-term signal sequence\ninside 1 19\nTMhelix 20 39\noutside 40 325\nTMhelix 326 348\ninside 349 372

12209 GCF_000647515.1_220_00_clc_bio_denovo Escherichia coli O174:H21 Proteobacteria; Gammaproteobacteria; Enterobacterales; Enterobacteriaceae; Escherichia; Escherichia coli MNNEETFYQAMRRQGVTRRSFLKYCSLAATSLGLGAGMAPKIAWALENKPRIPVVWIHGL WP_001058323.1 MULTISPECIES: hydrogenase-1 small chain [Proteobacteria] Length: 372\nNumber of predicted TMHs: 2\nExp number of AAs in TMHs: 37.2608499999999\nExp number, first 60 AAs: 19.52923\nTotal prob of N-in: 0.94351\nPOSSIBLE N-term signal sequence\ninside 1 19\nTMhelix 20 39\noutside 40 325\nTMhelix 326 348\ninside 349 372

12210 GCF_000647515.1_220_00_clc_bio_denovo Escherichia coli O174:H21 Proteobacteria; Gammaproteobacteria; Enterobacterales; Enterobacteriaceae; Escherichia; Escherichia coli MTGDNTLIHSHGINRRDFMKLCAALAATMGLSSKAAAEMAESVTNPQRPPVIWIGAQECT WP_000145410.1 MULTISPECIES: hydrogenase-2 small chain [Proteobacteria] Length: 372\nNumber of predicted TMHs: 1\nExp number of AAs in TMHs: 23.72226\nExp number, first 60 AAs: 0.22038\nTotal prob of N-in: 0.12994\noutside 1 331\nTMhelix 332 354\ninside 355 372

12211 GCF_001940375.1_ASM194037v1 Escherichia coli O25b:H4-ST131 Proteobacteria; Gammaproteobacteria; Enterobacterales; Enterobacteriaceae; Escherichia; Escherichia coli MTGDNTLIHSHGINRRDFMKLCAALAATMGLSSKAAAEMVESVTNPQRPPVIWIGAQECT WP_000145423.1 hydrogenase 2 small subunit [Escherichia coli] Length: 372\nNumber of predicted TMHs: 1\nExp number of AAs in TMHs: 23.78201\nExp number, first 60 AAs: 0.28355\nTotal prob of N-in: 0.13257\noutside 1 331\nTMhelix 332 354\ninside 355 372

12212 GCF_000285655.3_EC958.v1 Escherichia coli O25b:H4-ST131 Proteobacteria; Gammaproteobacteria; Enterobacterales; Enterobacteriaceae; Escherichia; Escherichia coli MTGDNTLIHSHGINRRDFMKLCAALAATMGLSSKAAAEMVESVTNPQRPPVIWIGAQECT WP_000145423.1 hydrogenase 2 small subunit [Escherichia coli] Length: 372\nNumber of predicted TMHs: 1\nExp number of AAs in TMHs: 23.78201\nExp number, first 60 AAs: 0.28355\nTotal prob of N-in: 0.13257\noutside 1 331\nTMhelix 332 354\ninside 355 372

12213 GCF_000285655.3_EC958.v1 Escherichia coli O25b:H4-ST131 Proteobacteria; Gammaproteobacteria; Enterobacterales; Enterobacteriaceae; Escherichia; Escherichia coli MNNEETFYQAMRRQGVTRRSFLKYCSLAATSLGLGAGMAPKIAWALENKPRIPVVWIHGL WP_001295940.1 MULTISPECIES: hydrogenase [Enterobacteriaceae] Length: 372\nNumber of predicted TMHs: 2\nExp number of AAs in TMHs: 37.2608899999999\nExp number, first 60 AAs: 19.52923\nTotal prob of N-in: 0.94351\nPOSSIBLE N-term signal sequence\ninside 1 19\nTMhelix 20 39\noutside 40 325\nTMhelix 326 348\ninside 349 372

12214 GCF_001940375.1_ASM194037v1 Escherichia coli O25b:H4-ST131 Proteobacteria; Gammaproteobacteria; Enterobacterales; Enterobacteriaceae; Escherichia; Escherichia coli MNNEETFYQAMRRQGVTRRSFLKYCSLAATSLGLGAGMAPKIAWALENKPRIPVVWIHGL WP_001295940.1 MULTISPECIES: hydrogenase [Enterobacteriaceae] Length: 372\nNumber of predicted TMHs: 2\nExp number of AAs in TMHs: 37.2608899999999\nExp number, first 60 AAs: 19.52923\nTotal prob of N-in: 0.94351\nPOSSIBLE N-term signal sequence\ninside 1 19\nTMhelix 20 39\noutside 40 325\nTMhelix 326 348\ninside 349 372

12215 GCF_900000205.1_Escherichia_coli_O26_H11_21765 Escherichia coli O26:H11 Proteobacteria; Gammaproteobacteria; Enterobacterales; Enterobacteriaceae; Escherichia; Escherichia coli MNNEETFYQAMRRQGVTRRSFLKYCSLAATSLGLGAGMAPKIAWALENKPRIPVVWIHGL WP_001058323.1 MULTISPECIES: hydrogenase-1 small chain [Proteobacteria] Length: 372\nNumber of predicted TMHs: 2\nExp number of AAs in TMHs: 37.2608499999999\nExp number, first 60 AAs: 19.52923\nTotal prob of N-in: 0.94351\nPOSSIBLE N-term signal sequence\ninside 1 19\nTMhelix 20 39\noutside 40 325\nTMhelix 326 348\ninside 349 372

12216 GCF_900000205.1_Escherichia_coli_O26_H11_21765 Escherichia coli O26:H11 Proteobacteria; Gammaproteobacteria; Enterobacterales; Enterobacteriaceae; Escherichia; Escherichia coli MTGDNTLIHSHGINRRDFMKLCAALAATMGLSSKAAAEMAESVTNPQRPPVIWIGAQECT WP_000145410.1 MULTISPECIES: hydrogenase-2 small chain [Proteobacteria] Length: 372\nNumber of predicted TMHs: 1\nExp number of AAs in TMHs: 23.72226\nExp number, first 60 AAs: 0.22038\nTotal prob of N-in: 0.12994\noutside 1 331\nTMhelix 332 354\ninside 355 372

12217 GCF_000403985.1_ASM40398v1 Escherichia coli O91 Proteobacteria; Gammaproteobacteria; Enterobacterales; Enterobacteriaceae; Escherichia; Escherichia coli MNNEETFYQAMRRQGVTRRSFLKYCSLAATSLGLGAGMAPKIAWALENKPRIPVVWIHGL WP_001058323.1 MULTISPECIES: hydrogenase-1 small chain [Proteobacteria] Length: 372\nNumber of predicted TMHs: 2\nExp number of AAs in TMHs: 37.2608499999999\nExp number, first 60 AAs: 19.52923\nTotal prob of N-in: 0.94351\nPOSSIBLE N-term signal sequence\ninside 1 19\nTMhelix 20 39\noutside 40 325\nTMhelix 326 348\ninside 349 372

12218 GCF_000403965.1_ASM40396v1 Escherichia coli O91 Proteobacteria; Gammaproteobacteria; Enterobacterales; Enterobacteriaceae; Escherichia; Escherichia coli MNNEETFYQAMRRQGVTRRSFLKYCSLAATSLGLGAGMAPKIAWALENKPRIPVVWIHGL WP_001058323.1 MULTISPECIES: hydrogenase-1 small chain [Proteobacteria] Length: 372\nNumber of predicted TMHs: 2\nExp number of AAs in TMHs: 37.2608499999999\nExp number, first 60 AAs: 19.52923\nTotal prob of N-in: 0.94351\nPOSSIBLE N-term signal sequence\ninside 1 19\nTMhelix 20 39\noutside 40 325\nTMhelix 326 348\ninside 349 372

12219 GCF_000403965.1_ASM40396v1 Escherichia coli O91 Proteobacteria; Gammaproteobacteria; Enterobacterales; Enterobacteriaceae; Escherichia; Escherichia coli
MTGDNTLIHSHGINRRDFMKLCAALAATMGLSSKAAAEMAESVTNPQRPPVIWIGAQECT WP_000145410.1
MULTISPECIES: hydrogenase-2 small chain [Proteobacteria] Length: 372\nNumber of predicted TMHs: 1\nExp number of AAs in TMHs: 23.72226\nExp number, first 60 AAs: 0.22038\nTotal prob of N-in: 0.12994\noutside 1 331\nTMhelix 332 354\ninside 355 372

12220 GCF_000403985.1_ASM40398v1 Escherichia coli O91 Proteobacteria; Gammaproteobacteria; Enterobacterales; Enterobacteriaceae; Escherichia; Escherichia coli
MTGDNTLIHSHGINRRDFMKLCAALAATMGLSSKAAAEMAESVTNPQRPPVIWIGAQECT WP_000145410.1
MULTISPECIES: hydrogenase-2 small chain [Proteobacteria] Length: 372\nNumber of predicted TMHs: 1\nExp number of AAs in TMHs: 23.72226\nExp number, first 60 AAs: 0.22038\nTotal prob of N-in: 0.12994\noutside 1 331\nTMhelix 332 354\ninside 355 372

12221 GCF_000188835.1_ASM18883v2 Escherichia coli OK1180 Proteobacteria; Gammaproteobacteria; Enterobacterales; Enterobacteriaceae; Escherichia; Escherichia coli
MNNEETFYQAMRRQGVTRRSFLKYCSLAATSLGLGAGMAPKIAWALENKPRIPVVWIHGL WP_001058323.1
MULTISPECIES: hydrogenase-1 small chain [Proteobacteria] Length: 372\nNumber of predicted TMHs: 2\nExp number of AAs in TMHs: 37.2608499999999\nExp number, first 60 AAs: 19.52923\nTotal prob of N-in: 0.94351\nPOSSIBLE N-term signal sequence\ninside 1 19\nTMhelix 20 39\noutside 40 325\nTMhelix 326 348\ninside 349 372

12222 GCF_000188835.1_ASM18883v2 Escherichia coli OK1180 Proteobacteria; Gammaproteobacteria; Enterobacterales; Enterobacteriaceae; Escherichia; Escherichia coli
MTGDNTLIHSHGINRRDFMKLCAALAATMGLSSKAAAEMAESVTNPQRPPVIWIGAQECT WP_000145410.1
MULTISPECIES: hydrogenase-2 small chain [Proteobacteria] Length: 372\nNumber of predicted TMHs: 1\nExp number of AAs in TMHs: 23.72226\nExp number, first 60 AAs: 0.22038\nTotal prob of N-in: 0.12994\noutside 1 331\nTMhelix 332 354\ninside 355 372

12223 GCF_000188855.1_ASM18885v2 Escherichia coli OK1357 Proteobacteria; Gammaproteobacteria; Enterobacterales; Enterobacteriaceae; Escherichia; Escherichia coli
MNNEETFYQAMRRQGVTRRSFLKYCSLAATSLGLGAGMAPKIAWALENKPRIPVVWIHGL WP_001295940.1
MULTISPECIES: hydrogenase [Enterobacteriaceae] Length: 372\nNumber of predicted TMHs: 2\nExp number of AAs in TMHs: 37.2608899999999\nExp number, first 60 AAs: 19.52923\nTotal prob of N-in: 0.94351\nPOSSIBLE N-term signal sequence\ninside 1 19\nTMhelix 20 39\noutside 40 325\nTMhelix 326 348\ninside 349 372

12224 GCF_000188855.1_ASM18885v2 Escherichia coli OK1357 Proteobacteria; Gammaproteobacteria; Enterobacterales; Enterobacteriaceae; Escherichia; Escherichia coli
MTGDNTLIHSHGINRRDFMKLCAALAATMGLSSKAAAEMAESVTNPQRPPVIWIGAQECT WP_000145410.1
MULTISPECIES: hydrogenase-2 small chain [Proteobacteria] Length: 372\nNumber of predicted TMHs: 1\nExp number of AAs in TMHs: 23.72226\nExp number, first 60 AAs: 0.22038\nTotal prob of N-in: 0.12994\noutside 1 331\nTMhelix 332 354\ninside 355 372

12225 GCF_000356165.2_ASM35616v2 Escherichia coli P0298942.1 Proteobacteria; Gammaproteobacteria; Enterobacterales; Enterobacteriaceae; Escherichia; Escherichia coli
MNNEETFYQAMRRQGVTRRSFLKYCSLAATSLGLGAGMAPKIAWALENKPRIPVVWIHGL WP_001058323.1
MULTISPECIES: hydrogenase-1 small chain [Proteobacteria] Length: 372\nNumber of predicted TMHs: 2\nExp number of AAs in TMHs: 37.2608499999999\nExp number, first 60 AAs: 19.52923\nTotal prob of N-in: 0.94351\nPOSSIBLE N-term signal sequence\ninside 1 19\nTMhelix 20 39\noutside 40 325\nTMhelix 326 348\ninside 349 372

12226 GCF_000356165.2_ASM35616v2 Escherichia coli P0298942.1 Proteobacteria; Gammaproteobacteria; Enterobacterales; Enterobacteriaceae; Escherichia; Escherichia coli
MTGDNTLIHSHGINRRDFMKLCAALAATMGLSSKAAAEMAESVTNPQRPPVIWIGAQECT WP_000145410.1
MULTISPECIES: hydrogenase-2 small chain [Proteobacteria] Length: 372\nNumber of predicted TMHs: 1\nExp number of AAs in TMHs: 23.72226\nExp number, first 60 AAs: 0.22038\nTotal prob of N-in: 0.12994\noutside 1 331\nTMhelix 332 354\ninside 355 372

12227 GCF_000356525.2_ASM35652v2 Escherichia coli P0298942.10 Proteobacteria; Gammaproteobacteria; Enterobacterales; Enterobacteriaceae; Escherichia; Escherichia coli

MNNEETFYQAMRRQGVTRRSFLKYCSLAATSLGLGAGMAPKIAWALENKPRIPVVWIHGL WP_001058323.1 MULTISPECIES: hydrogenase-1 small chain [Proteobacteria] Length: 372\nNumber of predicted TMHs: 2\nExp number of AAs in TMHs: 37.2608499999999\nExp number, first 60 AAs: 19.52923\nTotal prob of N-in: 0.94351\nPOSSIBLE N-term signal sequence\ninside 1 19\nTMhelix 20 39\noutside 40 325\nTMhelix 326 348\ninside 349 372

12228 GCF_000356525.2_ASM35652v2 Escherichia coli P0298942.10 Proteobacteria; Gammaproteobacteria; Enterobacterales; Enterobacteriaceae; Escherichia; Escherichia coli MTGDNTLIHSHGINRRDFMKLCAALAATMGLSSKAAAEMAESVTNPQRPPVIWIGAQECT WP_000145410.1 MULTISPECIES: hydrogenase-2 small chain [Proteobacteria] Length: 372\nNumber of predicted TMHs: 1\nExp number of AAs in TMHs: 23.72226\nExp number, first 60 AAs: 0.22038\nTotal prob of N-in: 0.12994\noutside 1 331\nTMhelix 332 354\ninside 355 372

12229 GCF_000357725.1_ASM35772v1 Escherichia coli P0298942.11 Proteobacteria; Gammaproteobacteria; Enterobacterales; Enterobacteriaceae; Escherichia; Escherichia coli MNNEETFYQAMRRQGVTRRSFLKYCSLAATSLGLGAGMAPKIAWALENKPRIPVVWIHGL WP_001425448.1 hydrogenase-1 small chain [Escherichia coli] Length: 372\nNumber of predicted TMHs: 2\nExp number of AAs in TMHs: 37.14002\nExp number, first 60 AAs: 19.53586\nTotal prob of N-in: 0.94339\nPOSSIBLE N-term signal sequence\ninside 1 19\nTMhelix 20 39\noutside 40 325\nTMhelix 326 348\ninside 349 372

12230 GCF_000357725.1_ASM35772v1 Escherichia coli P0298942.11 Proteobacteria; Gammaproteobacteria; Enterobacterales; Enterobacteriaceae; Escherichia; Escherichia coli MTGDNTLIHSHGINRRDFMKLCAALAATMGLSSKAAAEMAESVTNPQRPPVIWIGAQECT WP_000145410.1 MULTISPECIES: hydrogenase-2 small chain [Proteobacteria] Length: 372\nNumber of predicted TMHs: 1\nExp number of AAs in TMHs: 23.72226\nExp number, first 60 AAs: 0.22038\nTotal prob of N-in: 0.12994\noutside 1 331\nTMhelix 332 354\ninside 355 372

12231 GCF_000358915.1_ASM35891v1 Escherichia coli P0298942.12 Proteobacteria; Gammaproteobacteria; Enterobacterales; Enterobacteriaceae; Escherichia; Escherichia coli MTGDNTLIHSHGINRRDFMKLCAALAATMGLSSKAAAEMAESVTNPQRPPVIWIGAQECT WP_000145410.1 MULTISPECIES: hydrogenase-2 small chain [Proteobacteria] Length: 372\nNumber of predicted TMHs: 1\nExp number of AAs in TMHs: 23.72226\nExp number, first 60 AAs: 0.22038\nTotal prob of N-in: 0.12994\noutside 1 331\nTMhelix 332 354\ninside 355 372

12232 GCF_000358915.1_ASM35891v1 Escherichia coli P0298942.12 Proteobacteria; Gammaproteobacteria; Enterobacterales; Enterobacteriaceae; Escherichia; Escherichia coli MNNEETFYQAMRRQGVTRRSFLKYCSLAATSLGLGAGMAPKIAWALENKPRIPVVWIHGL WP_001427287.1 hydrogenase-1 small chain [Escherichia coli] Length: 372\nNumber of predicted TMHs: 2\nExp number of AAs in TMHs: 37.3304499999999\nExp number, first 60 AAs: 19.53932\nTotal prob of N-in: 0.94334\nPOSSIBLE N-term signal sequence\ninside 1 19\nTMhelix 20 39\noutside 40 325\nTMhelix 326 348\ninside 349 372

12233 GCF_000358815.1_ASM35881v1 Escherichia coli P0298942.14 Proteobacteria; Gammaproteobacteria; Enterobacterales; Enterobacteriaceae; Escherichia; Escherichia coli MNNEETFYQAMRRQGVTRRSFLKYCSLAATSLGLGAGMAPKIAWALENKPRIPVVWIHGL WP_001058323.1 MULTISPECIES: hydrogenase-1 small chain [Proteobacteria] Length: 372\nNumber of predicted TMHs: 2\nExp number of AAs in TMHs: 37.2608499999999\nExp number, first 60 AAs: 19.52923\nTotal prob of N-in: 0.94351\nPOSSIBLE N-term signal sequence\ninside 1 19\nTMhelix 20 39\noutside 40 325\nTMhelix 326 348\ninside 349 372

12234 GCF_000358815.1_ASM35881v1 Escherichia coli P0298942.14 Proteobacteria; Gammaproteobacteria; Enterobacterales; Enterobacteriaceae; Escherichia; Escherichia coli MTGDNTLIHSHGINRRDFMKLCAALAATMGLSSKAAAEMAESVTNPQRPPVIWIGAQECT WP_000145410.1 MULTISPECIES: hydrogenase-2 small chain [Proteobacteria] Length: 372\nNumber of predicted TMHs: 1\nExp number of AAs in TMHs: 23.72226\nExp number, first 60 AAs: 0.22038\nTotal prob of N-in: 0.12994\noutside 1 331\nTMhelix 332 354\ninside 355 372

12235 GCF_000357385.2_ASM35738v2 Escherichia coli P0298942.15 Proteobacteria; Gammaproteobacteria; Enterobacterales; Enterobacteriaceae; Escherichia; Escherichia coli MTGDNTLIHSHGINRRDFMKLCAALAATMGLSSKAAAEMAESVTNPQRPPVIWIGAQECT WP_000145410.1 MULTISPECIES: hydrogenase-2 small chain [Proteobacteria] Length: 372\nNumber of predicted TMHs:

1\nExp number of AAs in TMHs: 23.72226\nExp number, first 60 AAs: 0.22038\nTotal prob of N-in: 0.12994\noutside 1 331\nTMhelix 332 354\ninside 355 372

12236 GCF_000357385.2_ASM35738v2 Escherichia coli P0298942.15 Proteobacteria; Gammaproteobacteria; Enterobacterales; Enterobacteriaceae; Escherichia; Escherichia coli
MNNEETFYQAMRRQGVTRRSFLKYCSLAATSLGLGAGMAPKIAWALENKPRIPVVWIHGL WP_001058323.1
MULTISPECIES: hydrogenase-1 small chain [Proteobacteria] Length: 372\nNumber of predicted TMHs: 2\nExp number of AAs in TMHs: 37.2608499999999\nExp number, first 60 AAs: 19.52923\nTotal prob of N-in: 0.94351\nPOSSIBLE N-term signal sequence\ninside 1 19\nTMhelix 20 39\noutside 40 325\nTMhelix 326 348\ninside 349 372

12237 GCF_000357405.2_ASM35740v2 Escherichia coli P0298942.2 Proteobacteria; Gammaproteobacteria; Enterobacterales; Enterobacteriaceae; Escherichia; Escherichia coli
MNNEETFYQAMRRQGVTRRSFLKYCSLAATSLGLGAGMAPKIAWALENKPRIPVVWIHGL WP_001058323.1
MULTISPECIES: hydrogenase-1 small chain [Proteobacteria] Length: 372\nNumber of predicted TMHs: 2\nExp number of AAs in TMHs: 37.2608499999999\nExp number, first 60 AAs: 19.52923\nTotal prob of N-in: 0.94351\nPOSSIBLE N-term signal sequence\ninside 1 19\nTMhelix 20 39\noutside 40 325\nTMhelix 326 348\ninside 349 372

12238 GCF_000357405.2_ASM35740v2 Escherichia coli P0298942.2 Proteobacteria; Gammaproteobacteria; Enterobacterales; Enterobacteriaceae; Escherichia; Escherichia coli
MTGDNTLIHSHGINRRDFMKLCAALAATMGLSSKAAAEMAESVTNPQRPPVIWIGAQECT WP_000145410.1
MULTISPECIES: hydrogenase-2 small chain [Proteobacteria] Length: 372\nNumber of predicted TMHs: 1\nExp number of AAs in TMHs: 23.72226\nExp number, first 60 AAs: 0.22038\nTotal prob of N-in: 0.12994\noutside 1 331\nTMhelix 332 354\ninside 355 372

12239 GCF_000357445.2_ASM35744v2 Escherichia coli P0298942.3 Proteobacteria; Gammaproteobacteria; Enterobacterales; Enterobacteriaceae; Escherichia; Escherichia coli
MTGDNTLIHSHGINRRDFMKLCAALAATMGLSSKAAAEMAESVTNPQRPPVIWIGAQECT WP_000145410.1
MULTISPECIES: hydrogenase-2 small chain [Proteobacteria] Length: 372\nNumber of predicted TMHs: 1\nExp number of AAs in TMHs: 23.72226\nExp number, first 60 AAs: 0.22038\nTotal prob of N-in: 0.12994\noutside 1 331\nTMhelix 332 354\ninside 355 372

12240 GCF_000357445.2_ASM35744v2 Escherichia coli P0298942.3 Proteobacteria; Gammaproteobacteria; Enterobacterales; Enterobacteriaceae; Escherichia; Escherichia coli
MNNEETFYQAMRRQGVTRRSFLKYCSLAATSLGLGAGMAPKIAWALENKPRIPVVWIHGL WP_001058323.1
MULTISPECIES: hydrogenase-1 small chain [Proteobacteria] Length: 372\nNumber of predicted TMHs: 2\nExp number of AAs in TMHs: 37.2608499999999\nExp number, first 60 AAs: 19.52923\nTotal prob of N-in: 0.94351\nPOSSIBLE N-term signal sequence\ninside 1 19\nTMhelix 20 39\noutside 40 325\nTMhelix 326 348\ninside 349 372

12241 GCF_000357065.2_ASM35706v2 Escherichia coli P0298942.4 Proteobacteria; Gammaproteobacteria; Enterobacterales; Enterobacteriaceae; Escherichia; Escherichia coli
MNNEETFYQAMRRQGVTRRSFLKYCSLAATSLGLGAGMAPKIAWALENKPRIPVVWIHGL WP_001511681.1
hydrogenase-1 small chain [Escherichia coli] Length: 372\nNumber of predicted TMHs: 2\nExp number of AAs in TMHs: 37.3875\nExp number, first 60 AAs: 19.53624\nTotal prob of N-in: 0.94339\nPOSSIBLE N-term signal sequence\ninside 1 19\nTMhelix 20 39\noutside 40 325\nTMhelix 326 348\ninside 349 372

12242 GCF_000357065.2_ASM35706v2 Escherichia coli P0298942.4 Proteobacteria; Gammaproteobacteria; Enterobacterales; Enterobacteriaceae; Escherichia; Escherichia coli
MTGDNTLIHSHGINRRDFMKLCAALAATMGLSSKAAAEMAESVTNPQRPPVIWIGAQECT WP_000145410.1
MULTISPECIES: hydrogenase-2 small chain [Proteobacteria] Length: 372\nNumber of predicted TMHs: 1\nExp number of AAs in TMHs: 23.72226\nExp number, first 60 AAs: 0.22038\nTotal prob of N-in: 0.12994\noutside 1 331\nTMhelix 332 354\ninside 355 372

12243 GCF_000357425.2_ASM35742v2 Escherichia coli P0298942.6 Proteobacteria; Gammaproteobacteria; Enterobacterales; Enterobacteriaceae; Escherichia; Escherichia coli
MNNEETFYQAMRRQGVTRRSFLKYCSLAATSLGLGAGMAPKIAWALENKPRIPVVWIHGL WP_001058323.1
MULTISPECIES: hydrogenase-1 small chain [Proteobacteria] Length: 372\nNumber of predicted TMHs: 2\nExp number of AAs in TMHs: 37.2608499999999\nExp number, first 60 AAs: 19.52923\nTotal prob of N-in:

0.94351\nPOSSIBLE N-term signal sequence\ninside 1 19\nTMhelix 20 39\noutside 40 325\nTMhelix 326 348\ninside 349 372

12244 GCF_000357425.2_ASM35742v2 Escherichia coli P0298942.6 Proteobacteria; Gammaproteobacteria; Enterobacterales; Enterobacteriaceae; Escherichia; Escherichia coli MTGDNTLIHSHGINRRDFMKLCAALAATMGLSSKAAAEMAESVTNPQRPPVIWIGAQECT WP_000145410.1 MULTISPECIES: hydrogenase-2 small chain [Proteobacteria] Length: 372\nNumber of predicted TMHs: 1\nExp number of AAs in TMHs: 23.72226\nExp number, first 60 AAs: 0.22038\nTotal prob of N-in: 0.12994\noutside 1 331\nTMhelix 332 354\ninside 355 372

12245 GCF_000357545.2_ASM35754v2 Escherichia coli P0298942.7 Proteobacteria; Gammaproteobacteria; Enterobacterales; Enterobacteriaceae; Escherichia; Escherichia coli MTGDNTLIHSHGINRRDFMKLCAALAATMGLSSKAAAEMAESVTNPQRPPVIWIGAQECT WP_000145410.1 MULTISPECIES: hydrogenase-2 small chain [Proteobacteria] Length: 372\nNumber of predicted TMHs: 1\nExp number of AAs in TMHs: 23.72226\nExp number, first 60 AAs: 0.22038\nTotal prob of N-in: 0.12994\noutside 1 331\nTMhelix 332 354\ninside 355 372

12246 GCF_000357545.2_ASM35754v2 Escherichia coli P0298942.7 Proteobacteria; Gammaproteobacteria; Enterobacterales; Enterobacteriaceae; Escherichia; Escherichia coli MNNEETFYQAMRRQGVTRRSFLKYCSLAATSLGLGAGMAPKIAWALENKPRIPVVWIHGL WP_001456059.1 hydrogenase-1 small chain [Escherichia coli] Length: 372\nNumber of predicted TMHs: 2\nExp number of AAs in TMHs: 37.13915\nExp number, first 60 AAs: 19.53587\nTotal prob of N-in: 0.94339\nPOSSIBLE N-term signal sequence\ninside 1 19\nTMhelix 20 39\noutside 40 325\nTMhelix 326 348\ninside 349 372

12247 GCF_000357565.2_ASM35756v2 Escherichia coli P0298942.8 Proteobacteria; Gammaproteobacteria; Enterobacterales; Enterobacteriaceae; Escherichia; Escherichia coli MNNEETFYQAMRRQGVTRRSFLKYCSLAATSLGLGAGMAPKIAWALENKPRIPVVWIHGL WP_001058323.1 MULTISPECIES: hydrogenase-1 small chain [Proteobacteria] Length: 372\nNumber of predicted TMHs: 2\nExp number of AAs in TMHs: 37.2608499999999\nExp number, first 60 AAs: 19.52923\nTotal prob of N-in: 0.94351\nPOSSIBLE N-term signal sequence\ninside 1 19\nTMhelix 20 39\noutside 40 325\nTMhelix 326 348\ninside 349 372

12248 GCF_000357565.2_ASM35756v2 Escherichia coli P0298942.8 Proteobacteria; Gammaproteobacteria; Enterobacterales; Enterobacteriaceae; Escherichia; Escherichia coli MTGDNTLIHSHGINRRDFMKLCAALAATMGLSSKAAAEMAESVTNPQRPPVIWIGAQECT WP_000145410.1 MULTISPECIES: hydrogenase-2 small chain [Proteobacteria] Length: 372\nNumber of predicted TMHs: 1\nExp number of AAs in TMHs: 23.72226\nExp number, first 60 AAs: 0.22038\nTotal prob of N-in: 0.12994\noutside 1 331\nTMhelix 332 354\ninside 355 372

12249 GCF_000357745.1_ASM35774v1 Escherichia coli P0298942.9 Proteobacteria; Gammaproteobacteria; Enterobacterales; Enterobacteriaceae; Escherichia; Escherichia coli MTGDNTLIHSHGINRRDFMKLCAALAATMGLSSKAAAEMAESVTNPQRPPVIWIGAQECT WP_000145410.1 MULTISPECIES: hydrogenase-2 small chain [Proteobacteria] Length: 372\nNumber of predicted TMHs: 1\nExp number of AAs in TMHs: 23.72226\nExp number, first 60 AAs: 0.22038\nTotal prob of N-in: 0.12994\noutside 1 331\nTMhelix 332 354\ninside 355 372

12250 GCF_000357745.1_ASM35774v1 Escherichia coli P0298942.9 Proteobacteria; Gammaproteobacteria; Enterobacterales; Enterobacteriaceae; Escherichia; Escherichia coli MNNEETFYQAMRRQGVTRRSFLKYCSLAATSLGLGAGMAPKIAWALENKPRIPVVWIHGL WP_001058323.1 MULTISPECIES: hydrogenase-1 small chain [Proteobacteria] Length: 372\nNumber of predicted TMHs: 2\nExp number of AAs in TMHs: 37.2608499999999\nExp number, first 60 AAs: 19.52923\nTotal prob of N-in: 0.94351\nPOSSIBLE N-term signal sequence\ninside 1 19\nTMhelix 20 39\noutside 40 325\nTMhelix 326 348\ninside 349 372

12251 GCF_000357585.2_ASM35758v2 Escherichia coli P0299438.10 Proteobacteria; Gammaproteobacteria; Enterobacterales; Enterobacteriaceae; Escherichia; Escherichia coli MNNEETFYQAMRRQGVTRRSFLKYCSLAATSLGLGAGMAPKIAWALENKPRIPVVWIHGL WP_001058323.1 MULTISPECIES: hydrogenase-1 small chain [Proteobacteria] Length: 372\nNumber of predicted TMHs: 2\nExp number of AAs in TMHs: 37.2608499999999\nExp number, first 60 AAs: 19.52923\nTotal prob of N-in: 0.94351\nPOSSIBLE N-term signal sequence\ninside 1 19\nTMhelix 20 39\noutside 40 325\nTMhelix 326 348\ninside 349 372

12252 GCF_000357585.2_ASM35758v2 Escherichia coli P0299438.10 Proteobacteria; Gammaproteobacteria; Enterobacterales; Enterobacteriaceae; Escherichia; Escherichia coli
MTGDNTLIHSHGINRRDFMKLCAALAATMGLSSKAAAEMAESVTNPQRPPVIWIGAQECT WP_000145410.1
MULTISPECIES: hydrogenase-2 small chain [Proteobacteria] Length: 372\nNumber of predicted TMHs: 1\nExp number of AAs in TMHs: 23.72226\nExp number, first 60 AAs: 0.22038\nTotal prob of N-in: 0.12994\noutside 1 331\nTMhelix 332 354\ninside 355 372

12253 GCF_000356985.2_ASM35698v2 Escherichia coli P0299438.11 Proteobacteria; Gammaproteobacteria; Enterobacterales; Enterobacteriaceae; Escherichia; Escherichia coli
MNNEETFYQAMRRQGVTRRSFLKYCSLAATSLGLGAGMAPKIAWALENKPRIPVVWIHGL WP_001058323.1
MULTISPECIES: hydrogenase-1 small chain [Proteobacteria] Length: 372\nNumber of predicted TMHs: 2\nExp number of AAs in TMHs: 37.2608499999999\nExp number, first 60 AAs: 19.52923\nTotal prob of N-in: 0.94351\nPOSSIBLE N-term signal sequence\ninside 1 19\nTMhelix 20 39\noutside 40 325\nTMhelix 326 348\ninside 349 372

12254 GCF_000356985.2_ASM35698v2 Escherichia coli P0299438.11 Proteobacteria; Gammaproteobacteria; Enterobacterales; Enterobacteriaceae; Escherichia; Escherichia coli
MTGDNPLIHSHGINRRDFMKLCAALAATMGLSSKAAAEMAESVTNPQRPPVIWIGAQECT WP_001461571.1
hydrogenase-2 small chain [Escherichia coli] Length: 372\nNumber of predicted TMHs: 1\nExp number of AAs in TMHs: 22.60469\nExp number, first 60 AAs: 0.1106\nTotal prob of N-in: 0.06393\noutside 1 331\nTMhelix 332 354\ninside 355 372

12255 GCF_000356645.2_ASM35664v2 Escherichia coli P0299438.2 Proteobacteria; Gammaproteobacteria; Enterobacterales; Enterobacteriaceae; Escherichia; Escherichia coli
MTGDNTLIHSHGINRRDFMKLCAALAATMGLSSKAAAEMAESVTNPQRPPVIWIGAQECT WP_000145410.1
MULTISPECIES: hydrogenase-2 small chain [Proteobacteria] Length: 372\nNumber of predicted TMHs: 1\nExp number of AAs in TMHs: 23.72226\nExp number, first 60 AAs: 0.22038\nTotal prob of N-in: 0.12994\noutside 1 331\nTMhelix 332 354\ninside 355 372

12256 GCF_000356645.2_ASM35664v2 Escherichia coli P0299438.2 Proteobacteria; Gammaproteobacteria; Enterobacterales; Enterobacteriaceae; Escherichia; Escherichia coli
MNNEETFYQAMRRQGVTRRSFLKYCSLAATSLGLGAGMAPKIAWALENKPRIPVVWIHGL WP_001058323.1
MULTISPECIES: hydrogenase-1 small chain [Proteobacteria] Length: 372\nNumber of predicted TMHs: 2\nExp number of AAs in TMHs: 37.2608499999999\nExp number, first 60 AAs: 19.52923\nTotal prob of N-in: 0.94351\nPOSSIBLE N-term signal sequence\ninside 1 19\nTMhelix 20 39\noutside 40 325\nTMhelix 326 348\ninside 349 372

12257 GCF_000357005.2_ASM35700v2 Escherichia coli P0299438.3 Proteobacteria; Gammaproteobacteria; Enterobacterales; Enterobacteriaceae; Escherichia; Escherichia coli
MNNEETFYQAMRRQGVTRRSFLKYCSLAATSLGLGAGMAPKIAWALENKPRIPVVWIHGL WP_001058323.1
MULTISPECIES: hydrogenase-1 small chain [Proteobacteria] Length: 372\nNumber of predicted TMHs: 2\nExp number of AAs in TMHs: 37.2608499999999\nExp number, first 60 AAs: 19.52923\nTotal prob of N-in: 0.94351\nPOSSIBLE N-term signal sequence\ninside 1 19\nTMhelix 20 39\noutside 40 325\nTMhelix 326 348\ninside 349 372

12258 GCF_000357005.2_ASM35700v2 Escherichia coli P0299438.3 Proteobacteria; Gammaproteobacteria; Enterobacterales; Enterobacteriaceae; Escherichia; Escherichia coli
MTGDNTLIHSHGINRRDFMKLCAALAATMGLSSKAAAEMAESVTNPQRPPVIWIGAQECT WP_000145410.1
MULTISPECIES: hydrogenase-2 small chain [Proteobacteria] Length: 372\nNumber of predicted TMHs: 1\nExp number of AAs in TMHs: 23.72226\nExp number, first 60 AAs: 0.22038\nTotal prob of N-in: 0.12994\noutside 1 331\nTMhelix 332 354\ninside 355 372

12259 GCF_000357025.2_ASM35702v2 Escherichia coli P0299438.4 Proteobacteria; Gammaproteobacteria; Enterobacterales; Enterobacteriaceae; Escherichia; Escherichia coli
MTGDNTLIHSHGINRRDFMKLCAALAATMGLSSKAAAEMAESVTNPQRPPVIWIGAQECT WP_000145410.1
MULTISPECIES: hydrogenase-2 small chain [Proteobacteria] Length: 372\nNumber of predicted TMHs: 1\nExp number of AAs in TMHs: 23.72226\nExp number, first 60 AAs: 0.22038\nTotal prob of N-in: 0.12994\noutside 1 331\nTMhelix 332 354\ninside 355 372

12260 GCF_000357025.2_ASM35702v2 Escherichia coli P0299438.4 Proteobacteria; Gammaproteobacteria; Enterobacterales; Enterobacteriaceae; Escherichia; Escherichia coli

MNNEETFYQAMRRQGVTRRSFLKYCSLAATSLGLGAGMAPKIAWALENKPRIPVVWIHGL WP_001058323.1 MULTISPECIES: hydrogenase-1 small chain [Proteobacteria] Length: 372\nNumber of predicted TMHs: 2\nExp number of AAs in TMHs: 37.2608499999999\nExp number, first 60 AAs: 19.52923\nTotal prob of N-in: 0.94351\nPOSSIBLE N-term signal sequence\ninside 1 19\nTMhelix 20 39\noutside 40 325\nTMhelix 326 348\ninside 349 372

12261 GCF_000357045.2_ASM35704v2 Escherichia coli P0299438.5 Proteobacteria; Gammaproteobacteria; Enterobacterales; Enterobacteriaceae; Escherichia; Escherichia coli MNNEETFYQAMRRQGVTRRSFLKYCSLAATSLGLGAGMAPKIAWALENKPRIPVVWIHGL WP_001058323.1 MULTISPECIES: hydrogenase-1 small chain [Proteobacteria] Length: 372\nNumber of predicted TMHs: 2\nExp number of AAs in TMHs: 37.2608499999999\nExp number, first 60 AAs: 19.52923\nTotal prob of N-in: 0.94351\nPOSSIBLE N-term signal sequence\ninside 1 19\nTMhelix 20 39\noutside 40 325\nTMhelix 326 348\ninside 349 372

12262 GCF_000357045.2_ASM35704v2 Escherichia coli P0299438.5 Proteobacteria; Gammaproteobacteria; Enterobacterales; Enterobacteriaceae; Escherichia; Escherichia coli MTGDNTLIHSHGINRRDFMKLCAALAATMGLSSKAAAEMAESVTNPQRPPVIWIGAQECT WP_000145410.1 MULTISPECIES: hydrogenase-2 small chain [Proteobacteria] Length: 372\nNumber of predicted TMHs: 1\nExp number of AAs in TMHs: 23.72226\nExp number, first 60 AAs: 0.22038\nTotal prob of N-in: 0.12994\noutside 1 331\nTMhelix 332 354\ninside 355 372

12263 GCF_000357225.2_ASM35722v2 Escherichia coli P0299438.6 Proteobacteria; Gammaproteobacteria; Enterobacterales; Enterobacteriaceae; Escherichia; Escherichia coli MNNEETFYQAMRRQGVTRRSFLKYCSLAATSLGLGAGMAPKIAWALENKPRIPVVWIHGL WP_001058323.1 MULTISPECIES: hydrogenase-1 small chain [Proteobacteria] Length: 372\nNumber of predicted TMHs: 2\nExp number of AAs in TMHs: 37.2608499999999\nExp number, first 60 AAs: 19.52923\nTotal prob of N-in: 0.94351\nPOSSIBLE N-term signal sequence\ninside 1 19\nTMhelix 20 39\noutside 40 325\nTMhelix 326 348\ninside 349 372

12264 GCF_000357225.2_ASM35722v2 Escherichia coli P0299438.6 Proteobacteria; Gammaproteobacteria; Enterobacterales; Enterobacteriaceae; Escherichia; Escherichia coli MTGDNTLIHSHGINRRDFMKLCAALAATMGLSSKAAAEMAESVTNPQRPPVIWIGAQECT WP_000145410.1 MULTISPECIES: hydrogenase-2 small chain [Proteobacteria] Length: 372\nNumber of predicted TMHs: 1\nExp number of AAs in TMHs: 23.72226\nExp number, first 60 AAs: 0.22038\nTotal prob of N-in: 0.12994\noutside 1 331\nTMhelix 332 354\ninside 355 372

12265 GCF_000357245.1_ASM35724v1 Escherichia coli P0299438.7 Proteobacteria; Gammaproteobacteria; Enterobacterales; Enterobacteriaceae; Escherichia; Escherichia coli MNNEETFYQAMRRQGVTRRSFLKYCSLAATSLGLGAGMAPKIAWALENKPRIPVVWIHGL WP_001058323.1 MULTISPECIES: hydrogenase-1 small chain [Proteobacteria] Length: 372\nNumber of predicted TMHs: 2\nExp number of AAs in TMHs: 37.2608499999999\nExp number, first 60 AAs: 19.52923\nTotal prob of N-in: 0.94351\nPOSSIBLE N-term signal sequence\ninside 1 19\nTMhelix 20 39\noutside 40 325\nTMhelix 326 348\ninside 349 372

12266 GCF_000357245.1_ASM35724v1 Escherichia coli P0299438.7 Proteobacteria; Gammaproteobacteria; Enterobacterales; Enterobacteriaceae; Escherichia; Escherichia coli MTGDNTLIHSHGINRRDFMKLCAALAATMGLSSKAAAEMAESVTNPQRPPVIWIGAQECT WP_000145410.1 MULTISPECIES: hydrogenase-2 small chain [Proteobacteria] Length: 372\nNumber of predicted TMHs: 1\nExp number of AAs in TMHs: 23.72226\nExp number, first 60 AAs: 0.22038\nTotal prob of N-in: 0.12994\noutside 1 331\nTMhelix 332 354\ninside 355 372

12267 GCF_000357265.2_ASM35726v2 Escherichia coli P0299438.8 Proteobacteria; Gammaproteobacteria; Enterobacterales; Enterobacteriaceae; Escherichia; Escherichia coli MNNEETFYQAMRRQGVTRRSFLKYCSLAATSLGLGAGMAPKIAWALENKPRIPVVWIHGL WP_001058323.1 MULTISPECIES: hydrogenase-1 small chain [Proteobacteria] Length: 372\nNumber of predicted TMHs: 2\nExp number of AAs in TMHs: 37.2608499999999\nExp number, first 60 AAs: 19.52923\nTotal prob of N-in: 0.94351\nPOSSIBLE N-term signal sequence\ninside 1 19\nTMhelix 20 39\noutside 40 325\nTMhelix 326 348\ninside 349 372

12268 GCF_000357265.2_ASM35726v2 Escherichia coli P0299438.8 Proteobacteria; Gammaproteobacteria; Enterobacterales; Enterobacteriaceae; Escherichia; Escherichia coli

MTGDNTLIHSHGINRRDFMKLCAALAATMGLSSKAAAEMAESVTNPQRPPVIWIGAQECT WP_000145410.1 MULTISPECIES: hydrogenase-2 small chain [Proteobacteria] Length: 372\nNumber of predicted TMHs: 1\nExp number of AAs in TMHs: 23.72226\nExp number, first 60 AAs: 0.22038\nTotal prob of N-in: 0.12994\noutside 1 331\nTMhelix 332 354\ninside 355 372

12269 GCF_000357285.2_ASM35728v2 Escherichia coli P0299438.9 Proteobacteria; Gammaproteobacteria; Enterobacterales; Enterobacteriaceae; Escherichia; Escherichia coli MTGDNTLIHSHGINRRDFMKLCAALAATMGLSSKAAAEMAESVTNPQRPPVIWIGAQECT WP_000145410.1 MULTISPECIES: hydrogenase-2 small chain [Proteobacteria] Length: 372\nNumber of predicted TMHs: 1\nExp number of AAs in TMHs: 23.72226\nExp number, first 60 AAs: 0.22038\nTotal prob of N-in: 0.12994\noutside 1 331\nTMhelix 332 354\ninside 355 372

12270 GCF_000357285.2_ASM35728v2 Escherichia coli P0299438.9 Proteobacteria; Gammaproteobacteria; Enterobacterales; Enterobacteriaceae; Escherichia; Escherichia coli MNNEETFYQAMRRQGVTRRSFLKYCSLAATSLGLGAGMAPKIAWALENKPRIPVVWIHGL WP_001058323.1 MULTISPECIES: hydrogenase-1 small chain [Proteobacteria] Length: 372\nNumber of predicted TMHs: 2\nExp number of AAs in TMHs: 37.2608499999999\nExp number, first 60 AAs: 19.52923\nTotal prob of N-in: 0.94351\nPOSSIBLE N-term signal sequence\ninside 1 19\nTMhelix 20 39\noutside 40 325\nTMhelix 326 348\ninside 349 372

12271 GCF_000357465.2_ASM35746v2 Escherichia coli P0299483.1 Proteobacteria; Gammaproteobacteria; Enterobacterales; Enterobacteriaceae; Escherichia; Escherichia coli MNNEETFYQAMRRQGVTRRSFLKYCSLAATSLGLGAGMAPKIAWALENKPRIPVVWIHGL WP_001295940.1 MULTISPECIES: hydrogenase [Enterobacteriaceae] Length: 372\nNumber of predicted TMHs: 2\nExp number of AAs in TMHs: 37.2608899999999\nExp number, first 60 AAs: 19.52923\nTotal prob of N-in: 0.94351\nPOSSIBLE N-term signal sequence\ninside 1 19\nTMhelix 20 39\noutside 40 325\nTMhelix 326 348\ninside 349 372

12272 GCF_000357465.2_ASM35746v2 Escherichia coli P0299483.1 Proteobacteria; Gammaproteobacteria; Enterobacterales; Enterobacteriaceae; Escherichia; Escherichia coli MTGDNTLIHSHGINRRDFMKLCAALAATMGLSSKAAAEMAESVTNPQRPPVIWIGAQECT WP_000145410.1 MULTISPECIES: hydrogenase-2 small chain [Proteobacteria] Length: 372\nNumber of predicted TMHs: 1\nExp number of AAs in TMHs: 23.72226\nExp number, first 60 AAs: 0.22038\nTotal prob of N-in: 0.12994\noutside 1 331\nTMhelix 332 354\ninside 355 372

12273 GCF_000357105.1_ASM35710v1 Escherichia coli P0299483.2 Proteobacteria; Gammaproteobacteria; Enterobacterales; Enterobacteriaceae; Escherichia; Escherichia coli MNNEETFYQAMRRQGVTRRSFLKYCSLAATSLGLGAGMAPKIAWALENKPRIPVVWIHGL WP_001295940.1 MULTISPECIES: hydrogenase [Enterobacteriaceae] Length: 372\nNumber of predicted TMHs: 2\nExp number of AAs in TMHs: 37.2608899999999\nExp number, first 60 AAs: 19.52923\nTotal prob of N-in: 0.94351\nPOSSIBLE N-term signal sequence\ninside 1 19\nTMhelix 20 39\noutside 40 325\nTMhelix 326 348\ninside 349 372

12274 GCF_000357105.1_ASM35710v1 Escherichia coli P0299483.2 Proteobacteria; Gammaproteobacteria; Enterobacterales; Enterobacteriaceae; Escherichia; Escherichia coli MTGDNTLIHSHGINRRDFMKLCAALAATMGLSSKAAAEMAESVTNPQRPPVIWIGAQECT WP_000145410.1 MULTISPECIES: hydrogenase-2 small chain [Proteobacteria] Length: 372\nNumber of predicted TMHs: 1\nExp number of AAs in TMHs: 23.72226\nExp number, first 60 AAs: 0.22038\nTotal prob of N-in: 0.12994\noutside 1 331\nTMhelix 332 354\ninside 355 372

12275 GCF_000357125.1_ASM35712v1 Escherichia coli P0299483.3 Proteobacteria; Gammaproteobacteria; Enterobacterales; Enterobacteriaceae; Escherichia; Escherichia coli MTGDNTLIHSHGINRRDFMKLCAALAATMGLSSKAAAEMAESVTNPQRPPVIWIGAQECT WP_000145410.1 MULTISPECIES: hydrogenase-2 small chain [Proteobacteria] Length: 372\nNumber of predicted TMHs: 1\nExp number of AAs in TMHs: 23.72226\nExp number, first 60 AAs: 0.22038\nTotal prob of N-in: 0.12994\noutside 1 331\nTMhelix 332 354\ninside 355 372

12276 GCF_000357125.1_ASM35712v1 Escherichia coli P0299483.3 Proteobacteria; Gammaproteobacteria; Enterobacterales; Enterobacteriaceae; Escherichia; Escherichia coli MNNEETFYQAMRRQGVTRRSFLKYCSLAATSLGLGAGMAPKIAWALENKPRIPVVWIHGL WP_001295940.1 MULTISPECIES: hydrogenase [Enterobacteriaceae] Length: 372\nNumber of predicted TMHs: 2\nExp

number of AAs in TMHs: 37.2608899999999\nExp number, first 60 AAs: 19.52923\nTotal prob of N-in: 0.94351\nPOSSIBLE N-term signal sequence\ninside 1 19\nTMhelix 20 39\noutside 40 325\nTMhelix 326 348\ninside 349 372

12277 GCF_000357145.2_ASM35714v2 Escherichia coli P02997067.6 Proteobacteria; Gammaproteobacteria; Enterobacterales; Enterobacteriaceae; Escherichia; Escherichia coli MNNEETFYQAMRRQGVTRRSFLKYCSLAATSLGLGAGMAPKIAWALENKPRIPVVWIHGL WP_001295940.1 MULTISPECIES: hydrogenase [Enterobacteriaceae] Length: 372\nNumber of predicted TMHs: 2\nExp number of AAs in TMHs: 37.2608899999999\nExp number, first 60 AAs: 19.52923\nTotal prob of N-in: 0.94351\nPOSSIBLE N-term signal sequence\ninside 1 19\nTMhelix 20 39\noutside 40 325\nTMhelix 326 348\ninside 349 372

12278 GCF_000357145.2_ASM35714v2 Escherichia coli P02997067.6 Proteobacteria; Gammaproteobacteria; Enterobacterales; Enterobacteriaceae; Escherichia; Escherichia coli MTGDNPLIHSHGINRRDFMKLCAALAATMGLSSKAAAEMAESVTNPQRPPVIWIGAQECT WP_001472289.1 hydrogenase-2 small chain [Escherichia coli] Length: 372\nNumber of predicted TMHs: 1\nExp number of AAs in TMHs: 22.59996\nExp number, first 60 AAs: 0.1106\nTotal prob of N-in: 0.06389\noutside 1 331\nTMhelix 332 354\ninside 355 372

12279 GCF_000356665.2_ASM35666v2 Escherichia coli P0299917.1 Proteobacteria; Gammaproteobacteria; Enterobacterales; Enterobacteriaceae; Escherichia; Escherichia coli MNNEETFYQAMRRQGVTRRSFLKYCSLAATSLGLGAGMAPKIAWALENKPRIPVVWIHGL WP_001295940.1 MULTISPECIES: hydrogenase [Enterobacteriaceae] Length: 372\nNumber of predicted TMHs: 2\nExp number of AAs in TMHs: 37.2608899999999\nExp number, first 60 AAs: 19.52923\nTotal prob of N-in: 0.94351\nPOSSIBLE N-term signal sequence\ninside 1 19\nTMhelix 20 39\noutside 40 325\nTMhelix 326 348\ninside 349 372

12280 GCF_000356665.2_ASM35666v2 Escherichia coli P0299917.1 Proteobacteria; Gammaproteobacteria; Enterobacterales; Enterobacteriaceae; Escherichia; Escherichia coli MTGDNTLIHSHGINRRDFMKLCAALAATMGLSSKAAAEMAESVTNPQRPPVIWIGAQECT WP_001748757.1 hydrogenase-2 small chain [Escherichia coli] Length: 372\nNumber of predicted TMHs: 1\nExp number of AAs in TMHs: 23.71576\nExp number, first 60 AAs: 0.22033\nTotal prob of N-in: 0.12987\noutside 1 331\nTMhelix 332 354\ninside 355 372

12281 GCF_000357165.1_ASM35716v1 Escherichia coli P0299917.10 Proteobacteria; Gammaproteobacteria; Enterobacterales; Enterobacteriaceae; Escherichia; Escherichia coli MNNEETFYQAMRRQGVTRRSFLKYCSLAATSLGLGAGMAPKIAWALENKPRIPVVWIHGL WP_001058323.1 MULTISPECIES: hydrogenase-1 small chain [Proteobacteria] Length: 372\nNumber of predicted TMHs: 2\nExp number of AAs in TMHs: 37.2608499999999\nExp number, first 60 AAs: 19.52923\nTotal prob of N-in: 0.94351\nPOSSIBLE N-term signal sequence\ninside 1 19\nTMhelix 20 39\noutside 40 325\nTMhelix 326 348\ninside 349 372

12282 GCF_000357165.1_ASM35716v1 Escherichia coli P0299917.10 Proteobacteria; Gammaproteobacteria; Enterobacterales; Enterobacteriaceae; Escherichia; Escherichia coli MTGDNTLIHSHGINRRDFMKLCAALAATMGLSSKAAAEMAESVTNPQRPPVIWIGAQECT WP_001474635.1 hydrogenase-2 small chain [Escherichia coli] Length: 372\nNumber of predicted TMHs: 1\nExp number of AAs in TMHs: 23.72052\nExp number, first 60 AAs: 0.22038\nTotal prob of N-in: 0.12989\noutside 1 331\nTMhelix 332 354\ninside 355 372

12283 GCF_000357185.2_ASM35718v2 Escherichia coli P0299917.2 Proteobacteria; Gammaproteobacteria; Enterobacterales; Enterobacteriaceae; Escherichia; Escherichia coli MTGDNTLIHSHGINRRDFMKLCAALAATMGLSSKAAAEMAESVTNPQRPPVIWIGAQECT WP_000145410.1 MULTISPECIES: hydrogenase-2 small chain [Proteobacteria] Length: 372\nNumber of predicted TMHs: 1\nExp number of AAs in TMHs: 23.72226\nExp number, first 60 AAs: 0.22038\nTotal prob of N-in: 0.12994\noutside 1 331\nTMhelix 332 354\ninside 355 372

12284 GCF_000357185.2_ASM35718v2 Escherichia coli P0299917.2 Proteobacteria; Gammaproteobacteria; Enterobacterales; Enterobacteriaceae; Escherichia; Escherichia coli MNNEETFYQAMRRQGVTRRSFLKYCSLAATSLGLGAGMAPKIAWALENKPRIPVVWIHGL WP_001058323.1 MULTISPECIES: hydrogenase-1 small chain [Proteobacteria] Length: 372\nNumber of predicted TMHs: 2\nExp number of AAs in TMHs: 37.2608499999999\nExp number, first 60 AAs: 19.52923\nTotal prob of N-in:

0.94351\nPOSSIBLE N-term signal sequence\ninside 1 19\nTMhelix 20 39\noutside 40 325\nTMhelix 326 348\ninside 349 372

12285 GCF_000357205.2_ASM35720v2 Escherichia coli P0299917.3 Proteobacteria; Gammaproteobacteria; Enterobacterales; Enterobacteriaceae; Escherichia; Escherichia coli
MNNEETFYQAMRRQGVTRRSFLKYCSLAATSLGLGAGMAPKIAWALENKPRIPVVWIHGL WP_001058323.1
MULTISPECIES: hydrogenase-1 small chain [Proteobacteria] Length: 372\nNumber of predicted TMHs: 2\nExp number of AAs in TMHs: 37.2608499999999\nExp number, first 60 AAs: 19.52923\nTotal prob of N-in: 0.94351\nPOSSIBLE N-term signal sequence\ninside 1 19\nTMhelix 20 39\noutside 40 325\nTMhelix 326 348\ninside 349 372

12286 GCF_000357205.2_ASM35720v2 Escherichia coli P0299917.3 Proteobacteria; Gammaproteobacteria; Enterobacterales; Enterobacteriaceae; Escherichia; Escherichia coli
MTGDNTLIHSHGINRRDFMKLCAALAATMGLSSKAAAEMAESVTNPQRPPVIWIGAQECT WP_000145410.1
MULTISPECIES: hydrogenase-2 small chain [Proteobacteria] Length: 372\nNumber of predicted TMHs: 1\nExp number of AAs in TMHs: 23.72226\nExp number, first 60 AAs: 0.22038\nTotal prob of N-in: 0.12994\noutside 1 331\nTMhelix 332 354\ninside 355 372

12287 GCF_000357305.2_ASM35730v2 Escherichia coli P0299917.4 Proteobacteria; Gammaproteobacteria; Enterobacterales; Enterobacteriaceae; Escherichia; Escherichia coli
MNNEETFYQAMRRQGVTRRSFLKYCSLAATSLGLGAGMAPKIAWALENKPRIPVVWIHGL WP_001058323.1
MULTISPECIES: hydrogenase-1 small chain [Proteobacteria] Length: 372\nNumber of predicted TMHs: 2\nExp number of AAs in TMHs: 37.2608499999999\nExp number, first 60 AAs: 19.52923\nTotal prob of N-in: 0.94351\nPOSSIBLE N-term signal sequence\ninside 1 19\nTMhelix 20 39\noutside 40 325\nTMhelix 326 348\ninside 349 372

12288 GCF_000357305.2_ASM35730v2 Escherichia coli P0299917.4 Proteobacteria; Gammaproteobacteria; Enterobacterales; Enterobacteriaceae; Escherichia; Escherichia coli
MTGDNTLIHSHGINRRDFMKLCAALAATMGLSSKAAAEMAESVTNPQRPPVIWIGAQECT WP_000145410.1
MULTISPECIES: hydrogenase-2 small chain [Proteobacteria] Length: 372\nNumber of predicted TMHs: 1\nExp number of AAs in TMHs: 23.72226\nExp number, first 60 AAs: 0.22038\nTotal prob of N-in: 0.12994\noutside 1 331\nTMhelix 332 354\ninside 355 372

12289 GCF_000357325.1_ASM35732v1 Escherichia coli P0299917.5 Proteobacteria; Gammaproteobacteria; Enterobacterales; Enterobacteriaceae; Escherichia; Escherichia coli
MNNEETFYQAMRRQGVTRRSFLKYCSLAATSLGLGAGMAPKIAWALENKPRIPVVWIHGL WP_001058323.1
MULTISPECIES: hydrogenase-1 small chain [Proteobacteria] Length: 372\nNumber of predicted TMHs: 2\nExp number of AAs in TMHs: 37.2608499999999\nExp number, first 60 AAs: 19.52923\nTotal prob of N-in: 0.94351\nPOSSIBLE N-term signal sequence\ninside 1 19\nTMhelix 20 39\noutside 40 325\nTMhelix 326 348\ninside 349 372

12290 GCF_000357325.1_ASM35732v1 Escherichia coli P0299917.5 Proteobacteria; Gammaproteobacteria; Enterobacterales; Enterobacteriaceae; Escherichia; Escherichia coli
MTGDNTLIHSHGINRRDFMKLCAALAATMGLSSKAAAEMAESVTNPQRPPVIWIGAQECT WP_000145410.1
MULTISPECIES: hydrogenase-2 small chain [Proteobacteria] Length: 372\nNumber of predicted TMHs: 1\nExp number of AAs in TMHs: 23.72226\nExp number, first 60 AAs: 0.22038\nTotal prob of N-in: 0.12994\noutside 1 331\nTMhelix 332 354\ninside 355 372

12291 GCF_000357345.2_ASM35734v2 Escherichia coli P0299917.6 Proteobacteria; Gammaproteobacteria; Enterobacterales; Enterobacteriaceae; Escherichia; Escherichia coli
MNNEETFYQAMRRQGVTRRSFLKYCSLAATSLGLGAGMAPKIAWALENKPRIPVVWIHGL WP_001058323.1
MULTISPECIES: hydrogenase-1 small chain [Proteobacteria] Length: 372\nNumber of predicted TMHs: 2\nExp number of AAs in TMHs: 37.2608499999999\nExp number, first 60 AAs: 19.52923\nTotal prob of N-in: 0.94351\nPOSSIBLE N-term signal sequence\ninside 1 19\nTMhelix 20 39\noutside 40 325\nTMhelix 326 348\ninside 349 372

12292 GCF_000357345.2_ASM35734v2 Escherichia coli P0299917.6 Proteobacteria; Gammaproteobacteria; Enterobacterales; Enterobacteriaceae; Escherichia; Escherichia coli
MTGDNTLIHSHGINRRDFMKLCAALAATMGLSSKAAAEMAESVTNPQRPPVIWIGAQECT WP_000145410.1
MULTISPECIES: hydrogenase-2 small chain [Proteobacteria] Length: 372\nNumber of predicted TMHs:

1\nExp number of AAs in TMHs: 23.72226\nExp number, first 60 AAs: 0.22038\nTotal prob of N-in: 0.12994\noutside 1 331\nTMhelix 332 354\ninside 355 372

12293 GCF_000357365.2_ASM35736v2 Escherichia coli P0299917.7 Proteobacteria; Gammaproteobacteria; Enterobacterales; Enterobacteriaceae; Escherichia; Escherichia coli MNNEETFYQAMRRQGVTRRSFLKYCSLAATSLGLGAGMAPKIAWALENKPRIPVVWIHGL WP_001058323.1 MULTISPECIES: hydrogenase-1 small chain [Proteobacteria] Length: 372\nNumber of predicted TMHs: 2\nExp number of AAs in TMHs: 37.2608499999999\nExp number, first 60 AAs: 19.52923\nTotal prob of N-in: 0.94351\nPOSSIBLE N-term signal sequence\ninside 1 19\nTMhelix 20 39\noutside 40 325\nTMhelix 326 348\ninside 349 372

12294 GCF_000357365.2_ASM35736v2 Escherichia coli P0299917.7 Proteobacteria; Gammaproteobacteria; Enterobacterales; Enterobacteriaceae; Escherichia; Escherichia coli MTGDNTLIHSHGINRRDFMKLCAALAATMGLSSKAAAEMAESVTNPQRPPVIWIGAQECT WP_000145410.1 MULTISPECIES: hydrogenase-2 small chain [Proteobacteria] Length: 372\nNumber of predicted TMHs: 1\nExp number of AAs in TMHs: 23.72226\nExp number, first 60 AAs: 0.22038\nTotal prob of N-in: 0.12994\noutside 1 331\nTMhelix 332 354\ninside 355 372

12295 GCF_000358795.2_ASM35879v2 Escherichia coli P0299917.8 Proteobacteria; Gammaproteobacteria; Enterobacterales; Enterobacteriaceae; Escherichia; Escherichia coli MNNEETFYQAMRRQGVTRRSFLKYCSLAATSLGLGAGMAPKIAWALENKPRIPVVWIHGL WP_001058323.1 MULTISPECIES: hydrogenase-1 small chain [Proteobacteria] Length: 372\nNumber of predicted TMHs: 2\nExp number of AAs in TMHs: 37.2608499999999\nExp number, first 60 AAs: 19.52923\nTotal prob of N-in: 0.94351\nPOSSIBLE N-term signal sequence\ninside 1 19\nTMhelix 20 39\noutside 40 325\nTMhelix 326 348\ninside 349 372

12296 GCF_000358795.2_ASM35879v2 Escherichia coli P0299917.8 Proteobacteria; Gammaproteobacteria; Enterobacterales; Enterobacteriaceae; Escherichia; Escherichia coli MTGDNPLIHSHGINRRDFMKLCAALAATMGLSSKAAAEMAESVTNPQRPPVIWIGAQECT WP_001461571.1 hydrogenase-2 small chain [Escherichia coli] Length: 372\nNumber of predicted TMHs: 1\nExp number of AAs in TMHs: 22.60469\nExp number, first 60 AAs: 0.1106\nTotal prob of N-in: 0.06393\noutside 1 331\nTMhelix 332 354\ninside 355 372

12297 GCF_000357505.2_ASM35750v2 Escherichia coli P0299917.9 Proteobacteria; Gammaproteobacteria; Enterobacterales; Enterobacteriaceae; Escherichia; Escherichia coli MTGDNTLIHSHGINRRDFMKLCAALAATMGLSSKAAAEMAESVTNPQRPPVIWIGAQECT WP_000145410.1 MULTISPECIES: hydrogenase-2 small chain [Proteobacteria] Length: 372\nNumber of predicted TMHs: 1\nExp number of AAs in TMHs: 23.72226\nExp number, first 60 AAs: 0.22038\nTotal prob of N-in: 0.12994\noutside 1 331\nTMhelix 332 354\ninside 355 372

12298 GCF_000357505.2_ASM35750v2 Escherichia coli P0299917.9 Proteobacteria; Gammaproteobacteria; Enterobacterales; Enterobacteriaceae; Escherichia; Escherichia coli MNNEETFYQAMRRQGVTRRSFLKYCSLAATSLGLGAGMAPKIAWALENKPRIPVVWIHGL WP_001058323.1 MULTISPECIES: hydrogenase-1 small chain [Proteobacteria] Length: 372\nNumber of predicted TMHs: 2\nExp number of AAs in TMHs: 37.2608499999999\nExp number, first 60 AAs: 19.52923\nTotal prob of N-in: 0.94351\nPOSSIBLE N-term signal sequence\ninside 1 19\nTMhelix 20 39\noutside 40 325\nTMhelix 326 348\ninside 349 372

12299 GCF_000355315.2_ASM35531v2 Escherichia coli P0301867.1 Proteobacteria; Gammaproteobacteria; Enterobacterales; Enterobacteriaceae; Escherichia; Escherichia coli MTGDNTLIHSHGINRRDFMKLCAALAATMGLSSKAAAEMAESVTNPQRPPVIWIGAQECT WP_000145410.1 MULTISPECIES: hydrogenase-2 small chain [Proteobacteria] Length: 372\nNumber of predicted TMHs: 1\nExp number of AAs in TMHs: 23.72226\nExp number, first 60 AAs: 0.22038\nTotal prob of N-in: 0.12994\noutside 1 331\nTMhelix 332 354\ninside 355 372

12300 GCF_000355315.2_ASM35531v2 Escherichia coli P0301867.1 Proteobacteria; Gammaproteobacteria; Enterobacterales; Enterobacteriaceae; Escherichia; Escherichia coli MNNEETFYQAMRRQGVTRRSFLKYCSLAATSLGLGAGMAPKIAWALENKPRIPVVWIHGL WP_001058323.1 MULTISPECIES: hydrogenase-1 small chain [Proteobacteria] Length: 372\nNumber of predicted TMHs: 2\nExp number of AAs in TMHs: 37.2608499999999\nExp number, first 60 AAs: 19.52923\nTotal prob of N-in:

0.94351\nPOSSIBLE N-term signal sequence\ninside 1 19\nTMhelix 20 39\noutside 40 325\nTMhelix 326 348\ninside 349 372

12301 GCF_000357525.2_ASM35752v2 Escherichia coli P0301867.11 Proteobacteria; Gammaproteobacteria; Enterobacterales; Enterobacteriaceae; Escherichia; Escherichia coli MNNEETFYQAMRRQGVTRRSFLKYCSLAATSLGLGAGMAPKIAWALENKPRIPVVWIHGL WP_001058323.1 MULTISPECIES: hydrogenase-1 small chain [Proteobacteria] Length: 372\nNumber of predicted TMHs: 2\nExp number of AAs in TMHs: 37.2608499999999\nExp number, first 60 AAs: 19.52923\nTotal prob of N-in: 0.94351\nPOSSIBLE N-term signal sequence\ninside 1 19\nTMhelix 20 39\noutside 40 325\nTMhelix 326 348\ninside 349 372

12302 GCF_000357525.2_ASM35752v2 Escherichia coli P0301867.11 Proteobacteria; Gammaproteobacteria; Enterobacterales; Enterobacteriaceae; Escherichia; Escherichia coli MTGDNTLIHSHGINRRDFMKLCAALAATMGLSSKAAAEMAESVTNPQRPPVIWIGAQECT WP_000145410.1 MULTISPECIES: hydrogenase-2 small chain [Proteobacteria] Length: 372\nNumber of predicted TMHs: 1\nExp number of AAs in TMHs: 23.72226\nExp number, first 60 AAs: 0.22038\nTotal prob of N-in: 0.12994\noutside 1 331\nTMhelix 332 354\ninside 355 372

12303 GCF_000357625.2_ASM35762v2 Escherichia coli P0301867.13 Proteobacteria; Gammaproteobacteria; Enterobacterales; Enterobacteriaceae; Escherichia; Escherichia coli MNNEETFYQAMRRQGVTRRSFLKYCSLAATSLGLGAGMAPKIAWALENKPRIPVVWIHGL WP_001058323.1 MULTISPECIES: hydrogenase-1 small chain [Proteobacteria] Length: 372\nNumber of predicted TMHs: 2\nExp number of AAs in TMHs: 37.2608499999999\nExp number, first 60 AAs: 19.52923\nTotal prob of N-in: 0.94351\nPOSSIBLE N-term signal sequence\ninside 1 19\nTMhelix 20 39\noutside 40 325\nTMhelix 326 348\ninside 349 372

12304 GCF_000357625.2_ASM35762v2 Escherichia coli P0301867.13 Proteobacteria; Gammaproteobacteria; Enterobacterales; Enterobacteriaceae; Escherichia; Escherichia coli MTGDNTLIHSHGINRRDFMKLCAALAATMGLSSKAAAEMAESVTNPQRPPVIWIGAQECT WP_000145410.1 MULTISPECIES: hydrogenase-2 small chain [Proteobacteria] Length: 372\nNumber of predicted TMHs: 1\nExp number of AAs in TMHs: 23.72226\nExp number, first 60 AAs: 0.22038\nTotal prob of N-in: 0.12994\noutside 1 331\nTMhelix 332 354\ninside 355 372

12305 GCF_000356025.2_ASM35602v2 Escherichia coli P0301867.2 Proteobacteria; Gammaproteobacteria; Enterobacterales; Enterobacteriaceae; Escherichia; Escherichia coli MTGDNPLIHSHGINRRDFMKLCAALAATMGLSSKAAAEMAESVTNPQRPPVIWIGAQECT WP_001461571.1 hydrogenase-2 small chain [Escherichia coli] Length: 372\nNumber of predicted TMHs: 1\nExp number of AAs in TMHs: 22.60469\nExp number, first 60 AAs: 0.1106\nTotal prob of N-in: 0.06393\noutside 1 331\nTMhelix 332 354\ninside 355 372

12306 GCF_000356025.2_ASM35602v2 Escherichia coli P0301867.2 Proteobacteria; Gammaproteobacteria; Enterobacterales; Enterobacteriaceae; Escherichia; Escherichia coli MNNEETFYQAMRRQGVTRRSFLKYCSLAATSLGLGAGMAPKIAWALENKPRIPVVWIHGL WP_001058323.1 MULTISPECIES: hydrogenase-1 small chain [Proteobacteria] Length: 372\nNumber of predicted TMHs: 2\nExp number of AAs in TMHs: 37.2608499999999\nExp number, first 60 AAs: 19.52923\nTotal prob of N-in: 0.94351\nPOSSIBLE N-term signal sequence\ninside 1 19\nTMhelix 20 39\noutside 40 325\nTMhelix 326 348\ninside 349 372

12307 GCF_000358835.2_ASM35883v2 Escherichia coli P0301867.3 Proteobacteria; Gammaproteobacteria; Enterobacterales; Enterobacteriaceae; Escherichia; Escherichia coli MTGDNTLIHSHGINRRDFMKLCAALAATMGLSSKAAAEMAESVTNPQRPPVIWIGAQECT WP_000145410.1 MULTISPECIES: hydrogenase-2 small chain [Proteobacteria] Length: 372\nNumber of predicted TMHs: 1\nExp number of AAs in TMHs: 23.72226\nExp number, first 60 AAs: 0.22038\nTotal prob of N-in: 0.12994\noutside 1 331\nTMhelix 332 354\ninside 355 372

12308 GCF_000358835.2_ASM35883v2 Escherichia coli P0301867.3 Proteobacteria; Gammaproteobacteria; Enterobacterales; Enterobacteriaceae; Escherichia; Escherichia coli MNNEETFYQAMRRQGVTRRSFLKYCSLAATSLGLGAGMAPKIAWALENKPRIPVVWIHGL WP_001058323.1 MULTISPECIES: hydrogenase-1 small chain [Proteobacteria] Length: 372\nNumber of predicted TMHs: 2\nExp number of AAs in TMHs: 37.2608499999999\nExp number, first 60 AAs: 19.52923\nTotal prob of N-in:

0.94351\nPOSSIBLE N-term signal sequence\ninside 1 19\nTMhelix 20 39\noutside 40 325\nTMhelix 326 348\ninside 349 372

12309 GCF_000356005.2_ASM35600v2 Escherichia coli P0301867.4 Proteobacteria; Gammaproteobacteria; Enterobacterales; Enterobacteriaceae; Escherichia; Escherichia coli
MTGDNTLIHSHGINRRDFMKLCAALAATMGLSSKAAAEMAESVTNPQRPPVIWIGAQECT WP_000145410.1
MULTISPECIES: hydrogenase-2 small chain [Proteobacteria] Length: 372\nNumber of predicted TMHs: 1\nExp number of AAs in TMHs: 23.72226\nExp number, first 60 AAs: 0.22038\nTotal prob of N-in: 0.12994\noutside 1 331\nTMhelix 332 354\ninside 355 372

12310 GCF_000356005.2_ASM35600v2 Escherichia coli P0301867.4 Proteobacteria; Gammaproteobacteria; Enterobacterales; Enterobacteriaceae; Escherichia; Escherichia coli
MNNEETFYQAMRRQGVTRRSFLKYCSLAATSLGLGAGMAPKIAWALENKPRIPVVWIHGL WP_001058323.1
MULTISPECIES: hydrogenase-1 small chain [Proteobacteria] Length: 372\nNumber of predicted TMHs: 2\nExp number of AAs in TMHs: 37.2608499999999\nExp number, first 60 AAs: 19.52923\nTotal prob of N-in: 0.94351\nPOSSIBLE N-term signal sequence\ninside 1 19\nTMhelix 20 39\noutside 40 325\nTMhelix 326 348\ninside 349 372

12311 GCF_000358855.2_ASM35885v2 Escherichia coli P0301867.5 Proteobacteria; Gammaproteobacteria; Enterobacterales; Enterobacteriaceae; Escherichia; Escherichia coli
MTGDNPLIHSHGINRRDFMKLCAALAATMGLSSKAAAEMAESVTNPQRPPVIWIGAQECT WP_001461571.1
hydrogenase-2 small chain [Escherichia coli] Length: 372\nNumber of predicted TMHs: 1\nExp number of AAs in TMHs: 22.60469\nExp number, first 60 AAs: 0.1106\nTotal prob of N-in: 0.06393\noutside 1 331\nTMhelix 332 354\ninside 355 372

12312 GCF_000358855.2_ASM35885v2 Escherichia coli P0301867.5 Proteobacteria; Gammaproteobacteria; Enterobacterales; Enterobacteriaceae; Escherichia; Escherichia coli
MNNEETFYQAMRRQGVTRRSFLKYCSLAATSLGLGAGMAPKIAWALENKPRIPVVWIHGL WP_001488656.1
hydrogenase-1 small chain [Escherichia coli] Length: 372\nNumber of predicted TMHs: 2\nExp number of AAs in TMHs: 37.0420999999999\nExp number, first 60 AAs: 19.52951\nTotal prob of N-in: 0.94339\nPOSSIBLE N-term signal sequence\ninside 1 19\nTMhelix 20 39\noutside 40 325\nTMhelix 326 348\ninside 349 372

12313 GCF_000358775.2_ASM35877v2 Escherichia coli P0301867.7 Proteobacteria; Gammaproteobacteria; Enterobacterales; Enterobacteriaceae; Escherichia; Escherichia coli
MNNEETFYQAMRRQGVTRRSFLKYCSLAATSLGLGAGMAPKIAWALENKPRIPVVWIHGL WP_001058323.1
MULTISPECIES: hydrogenase-1 small chain [Proteobacteria] Length: 372\nNumber of predicted TMHs: 2\nExp number of AAs in TMHs: 37.2608499999999\nExp number, first 60 AAs: 19.52923\nTotal prob of N-in: 0.94351\nPOSSIBLE N-term signal sequence\ninside 1 19\nTMhelix 20 39\noutside 40 325\nTMhelix 326 348\ninside 349 372

12314 GCF_000358775.2_ASM35877v2 Escherichia coli P0301867.7 Proteobacteria; Gammaproteobacteria; Enterobacterales; Enterobacteriaceae; Escherichia; Escherichia coli
MTGDNTLIHSHGINRRDFMKLCAALAATMGLSSKAAAEMAESVTNPQRPPVIWIGAQECT WP_000145410.1
MULTISPECIES: hydrogenase-2 small chain [Proteobacteria] Length: 372\nNumber of predicted TMHs: 1\nExp number of AAs in TMHs: 23.72226\nExp number, first 60 AAs: 0.22038\nTotal prob of N-in: 0.12994\noutside 1 331\nTMhelix 332 354\ninside 355 372

12315 GCF_000357085.1_ASM35708v1 Escherichia coli P0301867.8 Proteobacteria; Gammaproteobacteria; Enterobacterales; Enterobacteriaceae; Escherichia; Escherichia coli
MNNEETFYQAMRRQGVTRRSFLKYCSLAATSLGLGAGMAPKIAWALENKPRIPVVWIHGL WP_001058323.1
MULTISPECIES: hydrogenase-1 small chain [Proteobacteria] Length: 372\nNumber of predicted TMHs: 2\nExp number of AAs in TMHs: 37.2608499999999\nExp number, first 60 AAs: 19.52923\nTotal prob of N-in: 0.94351\nPOSSIBLE N-term signal sequence\ninside 1 19\nTMhelix 20 39\noutside 40 325\nTMhelix 326 348\ninside 349 372

12316 GCF_000357085.1_ASM35708v1 Escherichia coli P0301867.8 Proteobacteria; Gammaproteobacteria; Enterobacterales; Enterobacteriaceae; Escherichia; Escherichia coli
MTGDNTLIHSHGINRRDFMKLCAALAATMGLSSKAAAEMAESVTNPQRPPVIWIGAQECT WP_000145410.1
MULTISPECIES: hydrogenase-2 small chain [Proteobacteria] Length: 372\nNumber of predicted TMHs: 1\nExp number of AAs in TMHs: 23.72226\nExp number, first 60 AAs: 0.22038\nTotal prob of N-in: 0.12994\noutside 1 331\nTMhelix 332 354\ninside 355 372

12317 GCF_000357645.1_ASM35764v1 Escherichia coli P0301904.3 Proteobacteria; Gammaproteobacteria; Enterobacterales; Enterobacteriaceae; Escherichia; Escherichia coli
MTGDNTLIHSHGINRRDFMKLCAALAATMGLSSKAAAEMAESVTNPQRPPVIWIGAQECT WP_000145410.1
MULTISPECIES: hydrogenase-2 small chain [Proteobacteria] Length: 372\nNumber of predicted TMHs: 1\nExp number of AAs in TMHs: 23.72226\nExp number, first 60 AAs: 0.22038\nTotal prob of N-in: 0.12994\noutside 1 331\nTMhelix 332 354\ninside 355 372

12318 GCF_000357645.1_ASM35764v1 Escherichia coli P0301904.3 Proteobacteria; Gammaproteobacteria; Enterobacterales; Enterobacteriaceae; Escherichia; Escherichia coli
MNNEETFYQAMRRQGVTRRSFLKYCSLAATSLGLGAGMAPKIAWALENKPRIPVVWIHGL WP_001058323.1
MULTISPECIES: hydrogenase-1 small chain [Proteobacteria] Length: 372\nNumber of predicted TMHs: 2\nExp number of AAs in TMHs: 37.2608499999999\nExp number, first 60 AAs: 19.52923\nTotal prob of N-in: 0.94351\nPOSSIBLE N-term signal sequence\ninside 1 19\nTMhelix 20 39\noutside 40 325\nTMhelix 326 348\ninside 349 372

12319 GCF_000356765.2_ASM35676v2 Escherichia coli P0302293.10 Proteobacteria; Gammaproteobacteria; Enterobacterales; Enterobacteriaceae; Escherichia; Escherichia coli
MTGDNTLIHSHGINRRDFMKLCAALAATMGLSSKAAAEMAESVTNPQRPPVIWIGAQECT WP_000145410.1
MULTISPECIES: hydrogenase-2 small chain [Proteobacteria] Length: 372\nNumber of predicted TMHs: 1\nExp number of AAs in TMHs: 23.72226\nExp number, first 60 AAs: 0.22038\nTotal prob of N-in: 0.12994\noutside 1 331\nTMhelix 332 354\ninside 355 372

12320 GCF_000356765.2_ASM35676v2 Escherichia coli P0302293.10 Proteobacteria; Gammaproteobacteria; Enterobacterales; Enterobacteriaceae; Escherichia; Escherichia coli
MNNEETFYQAMRRQGVTRRSFLKYCSLAATSLGLGAGMAPKIAWALENKPRIPVVWIHGL WP_001058323.1
MULTISPECIES: hydrogenase-1 small chain [Proteobacteria] Length: 372\nNumber of predicted TMHs: 2\nExp number of AAs in TMHs: 37.2608499999999\nExp number, first 60 AAs: 19.52923\nTotal prob of N-in: 0.94351\nPOSSIBLE N-term signal sequence\ninside 1 19\nTMhelix 20 39\noutside 40 325\nTMhelix 326 348\ninside 349 372

12321 GCF_000356545.1_ASM35654v1 Escherichia coli P0302293.2 Proteobacteria; Gammaproteobacteria; Enterobacterales; Enterobacteriaceae; Escherichia; Escherichia coli
MTGDNTLIHSHGINRRDFMKLCAALAATMGLSSKAAAEMAESVTNPQRPPVIWIGAQECT WP_000145410.1
MULTISPECIES: hydrogenase-2 small chain [Proteobacteria] Length: 372\nNumber of predicted TMHs: 1\nExp number of AAs in TMHs: 23.72226\nExp number, first 60 AAs: 0.22038\nTotal prob of N-in: 0.12994\noutside 1 331\nTMhelix 332 354\ninside 355 372

12322 GCF_000356545.1_ASM35654v1 Escherichia coli P0302293.2 Proteobacteria; Gammaproteobacteria; Enterobacterales; Enterobacteriaceae; Escherichia; Escherichia coli
MNNEETFYQAMRRQGVTRRSFLKYCSLAATSLGLGAGMAPKIAWALENKPRIPVVWIHGL WP_001058323.1
MULTISPECIES: hydrogenase-1 small chain [Proteobacteria] Length: 372\nNumber of predicted TMHs: 2\nExp number of AAs in TMHs: 37.2608499999999\nExp number, first 60 AAs: 19.52923\nTotal prob of N-in: 0.94351\nPOSSIBLE N-term signal sequence\ninside 1 19\nTMhelix 20 39\noutside 40 325\nTMhelix 326 348\ninside 349 372

12323 GCF_000356805.2_ASM35680v2 Escherichia coli P0302293.3 Proteobacteria; Gammaproteobacteria; Enterobacterales; Enterobacteriaceae; Escherichia; Escherichia coli
MNNEETFYQAMRRQGVTRRSFLKYCSLAATSLGLGAGMAPKIAWALENKPRIPGVWIHGL WP_001563861.1
hydrogenase-1 small chain [Escherichia coli] Length: 372\nNumber of predicted TMHs: 2\nExp number of AAs in TMHs: 37.27987\nExp number, first 60 AAs: 19.55568\nTotal prob of N-in: 0.94386\nPOSSIBLE N-term signal sequence\ninside 1 19\nTMhelix 20 39\noutside 40 325\nTMhelix 326 348\ninside 349 372

12324 GCF_000356805.2_ASM35680v2 Escherichia coli P0302293.3 Proteobacteria; Gammaproteobacteria; Enterobacterales; Enterobacteriaceae; Escherichia; Escherichia coli
MTGDNTLIHSHGINRRDFMKLCAALAATMGLSSKAAAEMAESVTNPQRPPVIWIGAQECT WP_000145410.1
MULTISPECIES: hydrogenase-2 small chain [Proteobacteria] Length: 372\nNumber of predicted TMHs: 1\nExp number of AAs in TMHs: 23.72226\nExp number, first 60 AAs: 0.22038\nTotal prob of N-in: 0.12994\noutside 1 331\nTMhelix 332 354\ninside 355 372

12325 GCF_000356785.2_ASM35678v2 Escherichia coli P0302293.4 Proteobacteria; Gammaproteobacteria; Enterobacterales; Enterobacteriaceae; Escherichia; Escherichia coli

MNNEETFYQAMRRQGVTRRSFLKYCSLAATSLGLGAGMAPKIAWALENKPRIPVVWIHGL WP_001058323.1 MULTISPECIES: hydrogenase-1 small chain [Proteobacteria] Length: 372\nNumber of predicted TMHs: 2\nExp number of AAs in TMHs: 37.2608499999999\nExp number, first 60 AAs: 19.52923\nTotal prob of N-in: 0.94351\nPOSSIBLE N-term signal sequence\ninside 1 19\nTMhelix 20 39\noutside 40 325\nTMhelix 326 348\ninside 349 372

12326 GCF_000356785.2_ASM35678v2 Escherichia coli P0302293.4 Proteobacteria; Gammaproteobacteria; Enterobacterales; Enterobacteriaceae; Escherichia; Escherichia coli MTGDNTLIHSHGINRRDFMKLCAALAATMGLSSKAAAEMAESVTNPQRPPVIWIGAQECT WP_000145410.1 MULTISPECIES: hydrogenase-2 small chain [Proteobacteria] Length: 372\nNumber of predicted TMHs: 1\nExp number of AAs in TMHs: 23.72226\nExp number, first 60 AAs: 0.22038\nTotal prob of N-in: 0.12994\noutside 1 331\nTMhelix 332 354\ninside 355 372

12327 GCF_000356945.1_ASM35694v1 Escherichia coli P0302293.6 Proteobacteria; Gammaproteobacteria; Enterobacterales; Enterobacteriaceae; Escherichia; Escherichia coli MTGDNTLIHSHGINRRDFMKLCAALAATMGLSSKAAAEMAESVTNPQRPPVIWIGAQECT WP_000145410.1 MULTISPECIES: hydrogenase-2 small chain [Proteobacteria] Length: 372\nNumber of predicted TMHs: 1\nExp number of AAs in TMHs: 23.72226\nExp number, first 60 AAs: 0.22038\nTotal prob of N-in: 0.12994\noutside 1 331\nTMhelix 332 354\ninside 355 372

12328 GCF_000356945.1_ASM35694v1 Escherichia coli P0302293.6 Proteobacteria; Gammaproteobacteria; Enterobacterales; Enterobacteriaceae; Escherichia; Escherichia coli MNNEETFYQAMRRQGVTRRSFLKYCSLAATSLGLGAGMAPKIAWALENKPRIPVVWIHGL WP_001058323.1 MULTISPECIES: hydrogenase-1 small chain [Proteobacteria] Length: 372\nNumber of predicted TMHs: 2\nExp number of AAs in TMHs: 37.2608499999999\nExp number, first 60 AAs: 19.52923\nTotal prob of N-in: 0.94351\nPOSSIBLE N-term signal sequence\ninside 1 19\nTMhelix 20 39\noutside 40 325\nTMhelix 326 348\ninside 349 372

12329 GCF_000357665.2_ASM35766v2 Escherichia coli P0302293.7 Proteobacteria; Gammaproteobacteria; Enterobacterales; Enterobacteriaceae; Escherichia; Escherichia coli MNNEETFYQAMRRQGVTRRSFLKYCSLAATSLGLGAGMAPKIAWALENKPRIPVVWIHGL WP_001058323.1 MULTISPECIES: hydrogenase-1 small chain [Proteobacteria] Length: 372\nNumber of predicted TMHs: 2\nExp number of AAs in TMHs: 37.2608499999999\nExp number, first 60 AAs: 19.52923\nTotal prob of N-in: 0.94351\nPOSSIBLE N-term signal sequence\ninside 1 19\nTMhelix 20 39\noutside 40 325\nTMhelix 326 348\ninside 349 372

12330 GCF_000357665.2_ASM35766v2 Escherichia coli P0302293.7 Proteobacteria; Gammaproteobacteria; Enterobacterales; Enterobacteriaceae; Escherichia; Escherichia coli MTGDNPLIHSHGINRRDFMKLCAALAATMGLSSKAAAEMAESVTNPQRPPVIWIGAQECT WP_001461571.1 hydrogenase-2 small chain [Escherichia coli] Length: 372\nNumber of predicted TMHs: 1\nExp number of AAs in TMHs: 22.60469\nExp number, first 60 AAs: 0.1106\nTotal prob of N-in: 0.06393\noutside 1 331\nTMhelix 332 354\ninside 355 372

12331 GCF_000356965.1_ASM35696v1 Escherichia coli P0302293.8 Proteobacteria; Gammaproteobacteria; Enterobacterales; Enterobacteriaceae; Escherichia; Escherichia coli MTGDNTLIHSHGINRRDFMKLCAALAATMGLSSKAAAEMAESVTNPQRPPVIWIGAQECT WP_000145410.1 MULTISPECIES: hydrogenase-2 small chain [Proteobacteria] Length: 372\nNumber of predicted TMHs: 1\nExp number of AAs in TMHs: 23.72226\nExp number, first 60 AAs: 0.22038\nTotal prob of N-in: 0.12994\noutside 1 331\nTMhelix 332 354\ninside 355 372

12332 GCF_000356965.1_ASM35696v1 Escherichia coli P0302293.8 Proteobacteria; Gammaproteobacteria; Enterobacterales; Enterobacteriaceae; Escherichia; Escherichia coli MNNEETFYQAMRRQGVTRRSFLKYCSLAATSLGLGAGMAPKIAWALENKPRIPVVWIHGL WP_001058323.1 MULTISPECIES: hydrogenase-1 small chain [Proteobacteria] Length: 372\nNumber of predicted TMHs: 2\nExp number of AAs in TMHs: 37.2608499999999\nExp number, first 60 AAs: 19.52923\nTotal prob of N-in: 0.94351\nPOSSIBLE N-term signal sequence\ninside 1 19\nTMhelix 20 39\noutside 40 325\nTMhelix 326 348\ninside 349 372

12333 GCF_000358165.2_ASM35816v2 Escherichia coli P0302293.9 Proteobacteria; Gammaproteobacteria; Enterobacterales; Enterobacteriaceae; Escherichia; Escherichia coli MTGDNTLIHSHGINRRDFMKLCAALAATMGLSSKAAAEMAESVTNPQRPPVIWIGAQECT WP_001474635.1

hydrogenase-2 small chain [Escherichia coli]	Length: 372\nNumber of predicted TMHs:  1\nExp number of AAs in TMHs: 23.72052\nExp number, first 60 AAs:  0.22038\nTotal prob of N-in:   0.12989\noutside   1   331\nTMhelix   332   354\ninside   355   372

12334	GCF_000358165.2_ASM35816v2	Escherichia coli P0302293.9	Proteobacteria; Gammaproteobacteria; Enterobacterales; Enterobacteriaceae; Escherichia; Escherichia coli
MNNEETFYQAMRRQGVTRRSFLKYCSLAATSLGLGAGMAPKIAWALENKPRIPVVWIHGL	WP_001058323.1
MULTISPECIES: hydrogenase-1 small chain [Proteobacteria]	Length: 372\nNumber of predicted TMHs: 2\nExp number of AAs in TMHs: 37.2608499999999\nExp number, first 60 AAs:  19.52923\nTotal prob of N-in: 0.94351\nPOSSIBLE N-term signal sequence\ninside   1   19\nTMhelix   20   39\noutside   40   325\nTMhelix   326 348\ninside   349   372

12335	GCF_000355075.2_ASM35507v2	Escherichia coli P0302308.1	Proteobacteria; Gammaproteobacteria; Enterobacterales; Enterobacteriaceae; Escherichia; Escherichia coli
MNNEETFYQAMRRQGVTRRSFLKYCSLAATSLGLGAGMAPKIAWALENKPRIPVVWIHGL	WP_001058323.1
MULTISPECIES: hydrogenase-1 small chain [Proteobacteria]	Length: 372\nNumber of predicted TMHs: 2\nExp number of AAs in TMHs: 37.2608499999999\nExp number, first 60 AAs:  19.52923\nTotal prob of N-in: 0.94351\nPOSSIBLE N-term signal sequence\ninside   1   19\nTMhelix   20   39\noutside   40   325\nTMhelix   326 348\ninside   349   372

12336	GCF_000355075.2_ASM35507v2	Escherichia coli P0302308.1	Proteobacteria; Gammaproteobacteria; Enterobacterales; Enterobacteriaceae; Escherichia; Escherichia coli
MTGDNTLIHSHGINRRDFMKLCAALAATMGLSSKAAAEMAESVTNPQRPPVIWIGAQECT	WP_000145410.1
MULTISPECIES: hydrogenase-2 small chain [Proteobacteria]	Length: 372\nNumber of predicted TMHs: 1\nExp number of AAs in TMHs: 23.72226\nExp number, first 60 AAs:  0.22038\nTotal prob of N-in:   0.12994\noutside   1 331\nTMhelix   332   354\ninside   355   372

12337	GCF_000358875.1_ASM35887v1	Escherichia coli P0302308.10 Proteobacteria; Gammaproteobacteria; Enterobacterales; Enterobacteriaceae; Escherichia; Escherichia coli
MTGDNTLIHSHGINRRDFMKLCAALAATMGLSSKAAAEMAESVTNPQRPPVIWIGAQECT	WP_000145410.1
MULTISPECIES: hydrogenase-2 small chain [Proteobacteria]	Length: 372\nNumber of predicted TMHs: 1\nExp number of AAs in TMHs: 23.72226\nExp number, first 60 AAs:  0.22038\nTotal prob of N-in:   0.12994\noutside   1 331\nTMhelix   332   354\ninside   355   372

12338	GCF_000358875.1_ASM35887v1	Escherichia coli P0302308.10 Proteobacteria; Gammaproteobacteria; Enterobacterales; Enterobacteriaceae; Escherichia; Escherichia coli
MNNEETFYQAMRRQGVTRRSFLKYCSLAATSLGLGAGMAPKIAWALENKPRIPVVWIHGL	WP_001058323.1
MULTISPECIES: hydrogenase-1 small chain [Proteobacteria]	Length: 372\nNumber of predicted TMHs: 2\nExp number of AAs in TMHs: 37.2608499999999\nExp number, first 60 AAs:  19.52923\nTotal prob of N-in: 0.94351\nPOSSIBLE N-term signal sequence\ninside   1   19\nTMhelix   20   39\noutside   40   325\nTMhelix   326 348\ninside   349   372

12339	GCF_000357605.2_ASM35760v2	Escherichia coli P0302308.11 Proteobacteria; Gammaproteobacteria; Enterobacterales; Enterobacteriaceae; Escherichia; Escherichia coli
MNNEETFYQAMRRQGVTRRSFLKYCSLAATSLGLGAGMAPKIAWALENKPRIPVVWIHGL	WP_001058323.1
MULTISPECIES: hydrogenase-1 small chain [Proteobacteria]	Length: 372\nNumber of predicted TMHs: 2\nExp number of AAs in TMHs: 37.2608499999999\nExp number, first 60 AAs:  19.52923\nTotal prob of N-in: 0.94351\nPOSSIBLE N-term signal sequence\ninside   1   19\nTMhelix   20   39\noutside   40   325\nTMhelix   326 348\ninside   349   372

12340	GCF_000357605.2_ASM35760v2	Escherichia coli P0302308.11 Proteobacteria; Gammaproteobacteria; Enterobacterales; Enterobacteriaceae; Escherichia; Escherichia coli
MTGDNTLIHSHGINRRDFMKLCAALAATMGLSSKAAAEMAESVTNPQRPPVIWIGAQECT	WP_000145410.1
MULTISPECIES: hydrogenase-2 small chain [Proteobacteria]	Length: 372\nNumber of predicted TMHs: 1\nExp number of AAs in TMHs: 23.72226\nExp number, first 60 AAs:  0.22038\nTotal prob of N-in:   0.12994\noutside   1 331\nTMhelix   332   354\ninside   355   372

12341	GCF_000358735.1_ASM35873v1	Escherichia coli P0302308.12 Proteobacteria; Gammaproteobacteria; Enterobacterales; Enterobacteriaceae; Escherichia; Escherichia coli
MNNEETFYQAMRRQGVTRRSFLKYCSLAATSLGLGAGMAPKIAWALENKPRIPVVWIHGL	WP_001058323.1
MULTISPECIES: hydrogenase-1 small chain [Proteobacteria]	Length: 372\nNumber of predicted TMHs:

2\nExp number of AAs in TMHs: 37.2608499999999\nExp number, first 60 AAs: 19.52923\nTotal prob of N-in: 0.94351\nPOSSIBLE N-term signal sequence\ninside 1 19\nTMhelix 20 39\noutside 40 325\nTMhelix 326 348\ninside 349 372

12342 GCF_000358735.1_ASM35873v1 Escherichia coli P0302308.12 Proteobacteria; Gammaproteobacteria; Enterobacterales; Enterobacteriaceae; Escherichia; Escherichia coli
MTGDNTLIHSHGINRRDFMKLCAALAATMGLSSKAAAEMAESVTNPQRPPVIWIGAQECT WP_000145410.1
MULTISPECIES: hydrogenase-2 small chain [Proteobacteria] Length: 372\nNumber of predicted TMHs: 1\nExp number of AAs in TMHs: 23.72226\nExp number, first 60 AAs: 0.22038\nTotal prob of N-in: 0.12994\noutside 1 331\nTMhelix 332 354\ninside 355 372

12343 GCF_000358975.2_ASM35897v2 Escherichia coli P0302308.13 Proteobacteria; Gammaproteobacteria; Enterobacterales; Enterobacteriaceae; Escherichia; Escherichia coli
MTGDNTLIHSHGINRRDFMKLCAALAATMGLSSKAAAEMAESVTNPQRPPVIWIGAQECT WP_000145410.1
MULTISPECIES: hydrogenase-2 small chain [Proteobacteria] Length: 372\nNumber of predicted TMHs: 1\nExp number of AAs in TMHs: 23.72226\nExp number, first 60 AAs: 0.22038\nTotal prob of N-in: 0.12994\noutside 1 331\nTMhelix 332 354\ninside 355 372

12344 GCF_000358975.2_ASM35897v2 Escherichia coli P0302308.13 Proteobacteria; Gammaproteobacteria; Enterobacterales; Enterobacteriaceae; Escherichia; Escherichia coli
MNNEETFYQAMRRQGVTRRSFLKYCSLAATSLGLGAGMAPKIAWALENKPRIPVVWIHGL WP_001058323.1
MULTISPECIES: hydrogenase-1 small chain [Proteobacteria] Length: 372\nNumber of predicted TMHs: 2\nExp number of AAs in TMHs: 37.2608499999999\nExp number, first 60 AAs: 19.52923\nTotal prob of N-in: 0.94351\nPOSSIBLE N-term signal sequence\ninside 1 19\nTMhelix 20 39\noutside 40 325\nTMhelix 326 348\ninside 349 372

12345 GCF_000358935.2_ASM35893v2 Escherichia coli P0302308.14 Proteobacteria; Gammaproteobacteria; Enterobacterales; Enterobacteriaceae; Escherichia; Escherichia coli
MNNEETFYQAMRRQGVTRRSFLKYCSLAATSLGLGAGMAPKIAWALENKPRIPVVWIHGL WP_001058323.1
MULTISPECIES: hydrogenase-1 small chain [Proteobacteria] Length: 372\nNumber of predicted TMHs: 2\nExp number of AAs in TMHs: 37.2608499999999\nExp number, first 60 AAs: 19.52923\nTotal prob of N-in: 0.94351\nPOSSIBLE N-term signal sequence\ninside 1 19\nTMhelix 20 39\noutside 40 325\nTMhelix 326 348\ninside 349 372

12346 GCF_000358935.2_ASM35893v2 Escherichia coli P0302308.14 Proteobacteria; Gammaproteobacteria; Enterobacterales; Enterobacteriaceae; Escherichia; Escherichia coli
MTGDNTLIHSHGINRRDFMKLCAALAATMGLSSKAAAEMAESVTNPQRPPVIWIGAQECT WP_000145410.1
MULTISPECIES: hydrogenase-2 small chain [Proteobacteria] Length: 372\nNumber of predicted TMHs: 1\nExp number of AAs in TMHs: 23.72226\nExp number, first 60 AAs: 0.22038\nTotal prob of N-in: 0.12994\noutside 1 331\nTMhelix 332 354\ninside 355 372

12347 GCF_000357765.2_ASM35776v2 Escherichia coli P0302308.2 Proteobacteria; Gammaproteobacteria; Enterobacterales; Enterobacteriaceae; Escherichia; Escherichia coli
MTGDNTLIHSHGINRRDFMKLCAALAATMGLSSKAAAEMAESVTNPQRPPVIWIGAQECT WP_000145410.1
MULTISPECIES: hydrogenase-2 small chain [Proteobacteria] Length: 372\nNumber of predicted TMHs: 1\nExp number of AAs in TMHs: 23.72226\nExp number, first 60 AAs: 0.22038\nTotal prob of N-in: 0.12994\noutside 1 331\nTMhelix 332 354\ninside 355 372

12348 GCF_000357765.2_ASM35776v2 Escherichia coli P0302308.2 Proteobacteria; Gammaproteobacteria; Enterobacterales; Enterobacteriaceae; Escherichia; Escherichia coli
MNNEETFYQAMRRQGVTRRSFLKYCSLAATSLGLGAGMAPKIAWALENKPRIPVVWIHGL WP_001058323.1
MULTISPECIES: hydrogenase-1 small chain [Proteobacteria] Length: 372\nNumber of predicted TMHs: 2\nExp number of AAs in TMHs: 37.2608499999999\nExp number, first 60 AAs: 19.52923\nTotal prob of N-in: 0.94351\nPOSSIBLE N-term signal sequence\ninside 1 19\nTMhelix 20 39\noutside 40 325\nTMhelix 326 348\ninside 349 372

12349 GCF_000357785.2_ASM35778v2 Escherichia coli P0302308.3 Proteobacteria; Gammaproteobacteria; Enterobacterales; Enterobacteriaceae; Escherichia; Escherichia coli
MTGDNTLIHSHGINRRDFMKLCAALAATMGLSSKAAAEMAESVTNPQRPPVIWIGAQECT WP_000145410.1
MULTISPECIES: hydrogenase-2 small chain [Proteobacteria] Length: 372\nNumber of predicted TMHs:

1\nExp number of AAs in TMHs: 23.72226\nExp number, first 60 AAs: 0.22038\nTotal prob of N-in: 0.12994\noutside 1 331\nTMhelix 332 354\ninside 355 372

12350 GCF_000357785.2_ASM35778v2 Escherichia coli P0302308.3 Proteobacteria; Gammaproteobacteria; Enterobacterales; Enterobacteriaceae; Escherichia; Escherichia coli
MNNEETFYQAMRRQGVTRRSFLKYCSLAATSLGLGAGMAPKIAWALENKPRIPVVWIHGL WP_001058323.1
MULTISPECIES: hydrogenase-1 small chain [Proteobacteria] Length: 372\nNumber of predicted TMHs: 2\nExp number of AAs in TMHs: 37.2608499999999\nExp number, first 60 AAs: 19.52923\nTotal prob of N-in: 0.94351\nPOSSIBLE N-term signal sequence\ninside 1 19\nTMhelix 20 39\noutside 40 325\nTMhelix 326 348\ninside 349 372

12351 GCF_000357805.1_ASM35780v1 Escherichia coli P0302308.4 Proteobacteria; Gammaproteobacteria; Enterobacterales; Enterobacteriaceae; Escherichia; Escherichia coli
MTGDNTLIHSHGINRRDFMKLCAALAATMGLSSKAAAEMAESVTNPQRPPVIWIGAQECT WP_000145410.1
MULTISPECIES: hydrogenase-2 small chain [Proteobacteria] Length: 372\nNumber of predicted TMHs: 1\nExp number of AAs in TMHs: 23.72226\nExp number, first 60 AAs: 0.22038\nTotal prob of N-in: 0.12994\noutside 1 331\nTMhelix 332 354\ninside 355 372

12352 GCF_000357805.1_ASM35780v1 Escherichia coli P0302308.4 Proteobacteria; Gammaproteobacteria; Enterobacterales; Enterobacteriaceae; Escherichia; Escherichia coli
MNNEETFYQAMRRQGVTRRSFLKYCSLAATSLGLGAGMAPKIAWALENKPRIPVVWIHGL WP_001058323.1
MULTISPECIES: hydrogenase-1 small chain [Proteobacteria] Length: 372\nNumber of predicted TMHs: 2\nExp number of AAs in TMHs: 37.2608499999999\nExp number, first 60 AAs: 19.52923\nTotal prob of N-in: 0.94351\nPOSSIBLE N-term signal sequence\ninside 1 19\nTMhelix 20 39\noutside 40 325\nTMhelix 326 348\ninside 349 372

12353 GCF_000357825.2_ASM35782v2 Escherichia coli P0302308.5 Proteobacteria; Gammaproteobacteria; Enterobacterales; Enterobacteriaceae; Escherichia; Escherichia coli
MTGDNTLIHSHGINRRDFMKLCAALAATMGLSSKAAAEMAESVTNPQRPPVIWIGAQECT WP_000145410.1
MULTISPECIES: hydrogenase-2 small chain [Proteobacteria] Length: 372\nNumber of predicted TMHs: 1\nExp number of AAs in TMHs: 23.72226\nExp number, first 60 AAs: 0.22038\nTotal prob of N-in: 0.12994\noutside 1 331\nTMhelix 332 354\ninside 355 372

12354 GCF_000357825.2_ASM35782v2 Escherichia coli P0302308.5 Proteobacteria; Gammaproteobacteria; Enterobacterales; Enterobacteriaceae; Escherichia; Escherichia coli
MNNEETFYQAMRRQGVTRRSFLKYCSLAATSLGLGAGMAPKIAWALENKPRIPVVWIHGL WP_001488656.1
hydrogenase-1 small chain [Escherichia coli] Length: 372\nNumber of predicted TMHs: 2\nExp number of AAs in TMHs: 37.0420999999999\nExp number, first 60 AAs: 19.52951\nTotal prob of N-in: 0.94339\nPOSSIBLE N-term signal sequence\ninside 1 19\nTMhelix 20 39\noutside 40 325\nTMhelix 326 348\ninside 349 372

12355 GCF_000355095.2_ASM35509v2 Escherichia coli P0304777.1 Proteobacteria; Gammaproteobacteria; Enterobacterales; Enterobacteriaceae; Escherichia; Escherichia coli
MTGDNTLIHSHGINRRDFMKLCAALAATMGLSSKAAAEMAESVTNPQRPPVIWIGAQECT WP_000145410.1
MULTISPECIES: hydrogenase-2 small chain [Proteobacteria] Length: 372\nNumber of predicted TMHs: 1\nExp number of AAs in TMHs: 23.72226\nExp number, first 60 AAs: 0.22038\nTotal prob of N-in: 0.12994\noutside 1 331\nTMhelix 332 354\ninside 355 372

12356 GCF_000355095.2_ASM35509v2 Escherichia coli P0304777.1 Proteobacteria; Gammaproteobacteria; Enterobacterales; Enterobacteriaceae; Escherichia; Escherichia coli
MNNEETFYQAMRRQGVTRRSFLKYCSLAATSLGLGAGMAPKIAWALENKPRIPVVWIHGL WP_001058320.1
hydrogenase [Escherichia coli] Length: 372\nNumber of predicted TMHs: 2\nExp number of AAs in TMHs: 37.26082\nExp number, first 60 AAs: 19.52923\nTotal prob of N-in: 0.94351\nPOSSIBLE N-term signal sequence\ninside 1 19\nTMhelix 20 39\noutside 40 325\nTMhelix 326 348\ninside 349 372

12357 GCF_000358185.2_ASM35818v2 Escherichia coli P0304777.10 Proteobacteria; Gammaproteobacteria; Enterobacterales; Enterobacteriaceae; Escherichia; Escherichia coli
MNNEETFYQAMRRQGVTRRSFLKYCSLAATSLGLGAGMAPKIAWALENKPRIPVVWIHGL WP_001058320.1
hydrogenase [Escherichia coli] Length: 372\nNumber of predicted TMHs: 2\nExp number of AAs in TMHs: 37.26082\nExp number, first 60 AAs: 19.52923\nTotal prob of N-in: 0.94351\nPOSSIBLE N-term signal sequence\ninside 1 19\nTMhelix 20 39\noutside 40 325\nTMhelix 326 348\ninside 349 372

12358 GCF_000358185.2_ASM35818v2 Escherichia coli P0304777.10 Proteobacteria; Gammaproteobacteria; Enterobacterales; Enterobacteriaceae; Escherichia; Escherichia coli
MTGDNTLIHSHGINRRDFMKLCAALAATMGLSSKAAAEMAESVTNPQRPPVIWIGAQEGT WP_003599695.1
hydrogenase-2 small chain [Escherichia coli] Length: 372\nNumber of predicted TMHs: 1\nExp number of AAs in TMHs: 23.71689\nExp number, first 60 AAs: 0.22035\nTotal prob of N-in: 0.12979\noutside 1 331\nTMhelix 332 354\ninside 355 372

12359 GCF_000358205.2_ASM35820v2 Escherichia coli P0304777.11 Proteobacteria; Gammaproteobacteria; Enterobacterales; Enterobacteriaceae; Escherichia; Escherichia coli
MTGDNTLIHSHGINRRDFMKLCAALAATMGLSSKAAAEMAESVTNPQRPPVIWIGAQECT WP_000145410.1
MULTISPECIES: hydrogenase-2 small chain [Proteobacteria] Length: 372\nNumber of predicted TMHs: 1\nExp number of AAs in TMHs: 23.72226\nExp number, first 60 AAs: 0.22038\nTotal prob of N-in: 0.12994\noutside 1 331\nTMhelix 332 354\ninside 355 372

12360 GCF_000358205.2_ASM35820v2 Escherichia coli P0304777.11 Proteobacteria; Gammaproteobacteria; Enterobacterales; Enterobacteriaceae; Escherichia; Escherichia coli
MNNEETFYQAMRRQGVTRRSFLKYCSLAATSLGLGAGMAPKIAWALENKPRIPVVWIHGL WP_001058320.1
hydrogenase [Escherichia coli] Length: 372\nNumber of predicted TMHs: 2\nExp number of AAs in TMHs: 37.26082\nExp number, first 60 AAs: 19.52923\nTotal prob of N-in: 0.94351\nPOSSIBLE N-term signal sequence\ninside 1 19\nTMhelix 20 39\noutside 40 325\nTMhelix 326 348\ninside 349 372

12361 GCF_000358225.2_ASM35822v2 Escherichia coli P0304777.12 Proteobacteria; Gammaproteobacteria; Enterobacterales; Enterobacteriaceae; Escherichia; Escherichia coli
MNNEETFYQAMRRQGVTRRSFLKYCSLAATSLGLGAGMAPKIAWALENKPRIPVVWIHGL WP_001058320.1
hydrogenase [Escherichia coli] Length: 372\nNumber of predicted TMHs: 2\nExp number of AAs in TMHs: 37.26082\nExp number, first 60 AAs: 19.52923\nTotal prob of N-in: 0.94351\nPOSSIBLE N-term signal sequence\ninside 1 19\nTMhelix 20 39\noutside 40 325\nTMhelix 326 348\ninside 349 372

12362 GCF_000358225.2_ASM35822v2 Escherichia coli P0304777.12 Proteobacteria; Gammaproteobacteria; Enterobacterales; Enterobacteriaceae; Escherichia; Escherichia coli
MTGDNPLIHSHGINRRDFMKLCAALAATMGLSSKAAAEMAESVTNPQRPPVIWIGAQECT WP_001461571.1
hydrogenase-2 small chain [Escherichia coli] Length: 372\nNumber of predicted TMHs: 1\nExp number of AAs in TMHs: 22.60469\nExp number, first 60 AAs: 0.1106\nTotal prob of N-in: 0.06393\noutside 1 331\nTMhelix 332 354\ninside 355 372

12363 GCF_000358245.2_ASM35824v2 Escherichia coli P0304777.13 Proteobacteria; Gammaproteobacteria; Enterobacterales; Enterobacteriaceae; Escherichia; Escherichia coli
MTGDNTLIHSHGINRRDFMKLCAALAATMGLSSKAAAEMAESVTNPQRPPVIWIGAQECT WP_000145410.1
MULTISPECIES: hydrogenase-2 small chain [Proteobacteria] Length: 372\nNumber of predicted TMHs: 1\nExp number of AAs in TMHs: 23.72226\nExp number, first 60 AAs: 0.22038\nTotal prob of N-in: 0.12994\noutside 1 331\nTMhelix 332 354\ninside 355 372

12364 GCF_000358245.2_ASM35824v2 Escherichia coli P0304777.13 Proteobacteria; Gammaproteobacteria; Enterobacterales; Enterobacteriaceae; Escherichia; Escherichia coli
MNNEETFYQAMRRQGVTRRSFLKYCSLAATSLGLGAGMAPKIAWALENKPRIPVVWIHGL WP_001058320.1
hydrogenase [Escherichia coli] Length: 372\nNumber of predicted TMHs: 2\nExp number of AAs in TMHs: 37.26082\nExp number, first 60 AAs: 19.52923\nTotal prob of N-in: 0.94351\nPOSSIBLE N-term signal sequence\ninside 1 19\nTMhelix 20 39\noutside 40 325\nTMhelix 326 348\ninside 349 372

12365 GCF_000358265.2_ASM35826v2 Escherichia coli P0304777.14 Proteobacteria; Gammaproteobacteria; Enterobacterales; Enterobacteriaceae; Escherichia; Escherichia coli
MNNEETFYQAMRRQGVTRRSFLKYCSLAATSLGLGAGMAPKIAWALENKPRIPVVWIHGL WP_001058320.1
hydrogenase [Escherichia coli] Length: 372\nNumber of predicted TMHs: 2\nExp number of AAs in TMHs: 37.26082\nExp number, first 60 AAs: 19.52923\nTotal prob of N-in: 0.94351\nPOSSIBLE N-term signal sequence\ninside 1 19\nTMhelix 20 39\noutside 40 325\nTMhelix 326 348\ninside 349 372

12366 GCF_000358265.2_ASM35826v2 Escherichia coli P0304777.14 Proteobacteria; Gammaproteobacteria; Enterobacterales; Enterobacteriaceae; Escherichia; Escherichia coli
MTGDNTLIHSHGINRRDFMKLCAALAATMGLSSKAAAEMAESVTNPQRPPVIWIGAQECT WP_000145410.1
MULTISPECIES: hydrogenase-2 small chain [Proteobacteria] Length: 372\nNumber of predicted TMHs:

1\nExp number of AAs in TMHs: 23.72226\nExp number, first 60 AAs:  0.22038\nTotal prob of N-in:   0.12994\noutside   1   331\nTMhelix   332   354\ninside   355   372

12367 GCF_000358285.2_ASM35828v2 Escherichia coli P0304777.15 Proteobacteria; Gammaproteobacteria; Enterobacterales; Enterobacteriaceae; Escherichia; Escherichia coli
MNNEETFYQAMRRQGVTRRSFLKYCSLAATSLGLGAGMAPKIAWALENKPRIPVVWIHGL WP_001058320.1
hydrogenase [Escherichia coli] Length: 372\nNumber of predicted TMHs:  2\nExp number of AAs in TMHs: 37.26082\nExp number, first 60 AAs:  19.52923\nTotal prob of N-in:   0.94351\nPOSSIBLE N-term signal sequence\ninside   1   19\nTMhelix   20   39\noutside   40   325\nTMhelix   326   348\ninside   349   372

12368 GCF_000358285.2_ASM35828v2 Escherichia coli P0304777.15 Proteobacteria; Gammaproteobacteria; Enterobacterales; Enterobacteriaceae; Escherichia; Escherichia coli
MTGDNTLIHSHGINRRDFMKLCAALAATMGLSSKAAAEMAESVTNPQRPPVIWIGAQECT WP_000145410.1
MULTISPECIES: hydrogenase-2 small chain [Proteobacteria] Length: 372\nNumber of predicted TMHs: 1\nExp number of AAs in TMHs: 23.72226\nExp number, first 60 AAs:  0.22038\nTotal prob of N-in:   0.12994\noutside   1   331\nTMhelix   332   354\ninside   355   372

12369 GCF_000400895.2_ASM40089v2 Escherichia coli P0304777.2 Proteobacteria; Gammaproteobacteria; Enterobacterales; Enterobacteriaceae; Escherichia; Escherichia coli
MNNEETFYQAMRRQGVTRRSFLKYCSLAATSLGLGAGMAPKIAWALENKPRIPVVWIHGL WP_001058320.1
hydrogenase [Escherichia coli] Length: 372\nNumber of predicted TMHs:  2\nExp number of AAs in TMHs: 37.26082\nExp number, first 60 AAs:  19.52923\nTotal prob of N-in:   0.94351\nPOSSIBLE N-term signal sequence\ninside   1   19\nTMhelix   20   39\noutside   40   325\nTMhelix   326   348\ninside   349   372

12370 GCF_000400895.2_ASM40089v2 Escherichia coli P0304777.2 Proteobacteria; Gammaproteobacteria; Enterobacterales; Enterobacteriaceae; Escherichia; Escherichia coli
MTGDNTLIHSHGINRRDFMKLCAALAATMGLSSKAAAEMAESVTNPQRPPVIWIGAQECT WP_000145410.1
MULTISPECIES: hydrogenase-2 small chain [Proteobacteria] Length: 372\nNumber of predicted TMHs: 1\nExp number of AAs in TMHs: 23.72226\nExp number, first 60 AAs:  0.22038\nTotal prob of N-in:   0.12994\noutside   1   331\nTMhelix   332   354\ninside   355   372

12371 GCF_000358315.1_ASM35831v1 Escherichia coli P0304777.3 Proteobacteria; Gammaproteobacteria; Enterobacterales; Enterobacteriaceae; Escherichia; Escherichia coli
MTGDNTLIHSHGINRRDFMKLCAALAATMGLSSKAAAEMAESVTNPQRPPVIWIGAQECT WP_000145410.1
MULTISPECIES: hydrogenase-2 small chain [Proteobacteria] Length: 372\nNumber of predicted TMHs: 1\nExp number of AAs in TMHs: 23.72226\nExp number, first 60 AAs:  0.22038\nTotal prob of N-in:   0.12994\noutside   1   331\nTMhelix   332   354\ninside   355   372

12372 GCF_000358315.1_ASM35831v1 Escherichia coli P0304777.3 Proteobacteria; Gammaproteobacteria; Enterobacterales; Enterobacteriaceae; Escherichia; Escherichia coli
MNNEETFYQAMRRQGVTRRSFLKYCSLAATSLGLGAGMAPKIAWALENKPRIPVVWIHGL WP_001058320.1
hydrogenase [Escherichia coli] Length: 372\nNumber of predicted TMHs:  2\nExp number of AAs in TMHs: 37.26082\nExp number, first 60 AAs:  19.52923\nTotal prob of N-in:   0.94351\nPOSSIBLE N-term signal sequence\ninside   1   19\nTMhelix   20   39\noutside   40   325\nTMhelix   326   348\ninside   349   372

12373 GCF_000358355.2_ASM35835v2 Escherichia coli P0304777.4 Proteobacteria; Gammaproteobacteria; Enterobacterales; Enterobacteriaceae; Escherichia; Escherichia coli
MNNEETFYQAMRRQGVTRRSFLKYCSLAATSLGLGAGMAPKIAWALENKPRIPVVWIHGL WP_001058320.1
hydrogenase [Escherichia coli] Length: 372\nNumber of predicted TMHs:  2\nExp number of AAs in TMHs: 37.26082\nExp number, first 60 AAs:  19.52923\nTotal prob of N-in:   0.94351\nPOSSIBLE N-term signal sequence\ninside   1   19\nTMhelix   20   39\noutside   40   325\nTMhelix   326   348\ninside   349   372

12374 GCF_000358355.2_ASM35835v2 Escherichia coli P0304777.4 Proteobacteria; Gammaproteobacteria; Enterobacterales; Enterobacteriaceae; Escherichia; Escherichia coli
MTGDNTLIHSHGINRRDFMKLCAALAATMGLSSKAAAEMAESVTNPQRPPVIWIGAQECT WP_000145410.1
MULTISPECIES: hydrogenase-2 small chain [Proteobacteria] Length: 372\nNumber of predicted TMHs: 1\nExp number of AAs in TMHs: 23.72226\nExp number, first 60 AAs:  0.22038\nTotal prob of N-in:   0.12994\noutside   1   331\nTMhelix   332   354\ninside   355   372

12375 GCF_000358335.2_ASM35833v2 Escherichia coli P0304777.5 Proteobacteria; Gammaproteobacteria; Enterobacterales; Enterobacteriaceae; Escherichia; Escherichia coli

MTGDNTLIHSHGINRRDFMKLCAALAATMGLSSKAAAEMAESVTNPQRPPVIWIGAQECT WP_000145410.1 MULTISPECIES: hydrogenase-2 small chain [Proteobacteria] Length: 372\nNumber of predicted TMHs: 1\nExp number of AAs in TMHs: 23.72226\nExp number, first 60 AAs: 0.22038\nTotal prob of N-in: 0.12994\noutside 1 331\nTMhelix 332 354\ninside 355 372

12376 GCF_000358335.2_ASM35833v2 Escherichia coli P0304777.5 Proteobacteria; Gammaproteobacteria; Enterobacterales; Enterobacteriaceae; Escherichia; Escherichia coli MNNEETFYQAMRRQGVTRRSFLKYCSLAATSLGLGAGMAPKIAWALENKPRIPVVWIHGL WP_001058320.1 hydrogenase [Escherichia coli] Length: 372\nNumber of predicted TMHs: 2\nExp number of AAs in TMHs: 37.26082\nExp number, first 60 AAs: 19.52923\nTotal prob of N-in: 0.94351\nPOSSIBLE N-term signal sequence\ninside 1 19\nTMhelix 20 39\noutside 40 325\nTMhelix 326 348\ninside 349 372

12377 GCF_000358375.1_ASM35837v1 Escherichia coli P0304777.7 Proteobacteria; Gammaproteobacteria; Enterobacterales; Enterobacteriaceae; Escherichia; Escherichia coli MNNEETFYQAMRRQGVTRRSFLKYCSLAATSLGLGAGMAPKIAWALENKPRIPVVWIHGL WP_001599290.1 hydrogenase-1 small chain [Escherichia coli] Length: 372\nNumber of predicted TMHs: 2\nExp number of AAs in TMHs: 37.2496299999999\nExp number, first 60 AAs: 19.52619\nTotal prob of N-in: 0.94349\nPOSSIBLE N-term signal sequence\ninside 1 19\nTMhelix 20 39\noutside 40 325\nTMhelix 326 348\ninside 349 372

12378 GCF_000358375.1_ASM35837v1 Escherichia coli P0304777.7 Proteobacteria; Gammaproteobacteria; Enterobacterales; Enterobacteriaceae; Escherichia; Escherichia coli MTGDNTLIHSHGINRRDFMKLCAALAATMGLSSKAAAEMAESVTNPQRPPVIWIGAQECT WP_000145410.1 MULTISPECIES: hydrogenase-2 small chain [Proteobacteria] Length: 372\nNumber of predicted TMHs: 1\nExp number of AAs in TMHs: 23.72226\nExp number, first 60 AAs: 0.22038\nTotal prob of N-in: 0.12994\noutside 1 331\nTMhelix 332 354\ninside 355 372

12379 GCF_000358395.2_ASM35839v2 Escherichia coli P0304777.8 Proteobacteria; Gammaproteobacteria; Enterobacterales; Enterobacteriaceae; Escherichia; Escherichia coli MNNEETFYQAMRRQGVTRRSFLKYCSLAATSLGLGAGMAPKIAWALENKPRIPVVWIHGL WP_001058320.1 hydrogenase [Escherichia coli] Length: 372\nNumber of predicted TMHs: 2\nExp number of AAs in TMHs: 37.26082\nExp number, first 60 AAs: 19.52923\nTotal prob of N-in: 0.94351\nPOSSIBLE N-term signal sequence\ninside 1 19\nTMhelix 20 39\noutside 40 325\nTMhelix 326 348\ninside 349 372

12380 GCF_000358395.2_ASM35839v2 Escherichia coli P0304777.8 Proteobacteria; Gammaproteobacteria; Enterobacterales; Enterobacteriaceae; Escherichia; Escherichia coli MTGDNTLIHSHGINRRDFMKLCAALAATMGLSSKAAAEMAESVTNPQRPPVIWIGAQECT WP_000145410.1 MULTISPECIES: hydrogenase-2 small chain [Proteobacteria] Length: 372\nNumber of predicted TMHs: 1\nExp number of AAs in TMHs: 23.72226\nExp number, first 60 AAs: 0.22038\nTotal prob of N-in: 0.12994\noutside 1 331\nTMhelix 332 354\ninside 355 372

12381 GCF_000358415.2_ASM35841v2 Escherichia coli P0304777.9 Proteobacteria; Gammaproteobacteria; Enterobacterales; Enterobacteriaceae; Escherichia; Escherichia coli MTGDNTLIHSHGINRRDFMKLCAALAATMGLSSKAAAEMAESVTNPQRPPVIWIGAQECT WP_000145410.1 MULTISPECIES: hydrogenase-2 small chain [Proteobacteria] Length: 372\nNumber of predicted TMHs: 1\nExp number of AAs in TMHs: 23.72226\nExp number, first 60 AAs: 0.22038\nTotal prob of N-in: 0.12994\noutside 1 331\nTMhelix 332 354\ninside 355 372

12382 GCF_000358415.2_ASM35841v2 Escherichia coli P0304777.9 Proteobacteria; Gammaproteobacteria; Enterobacterales; Enterobacteriaceae; Escherichia; Escherichia coli MNNEETFYQAMRRQGVTRRSFLKYCSLAATSLGLGAGMAPKIAWALENKPRIPVVWIHGL WP_001058320.1 hydrogenase [Escherichia coli] Length: 372\nNumber of predicted TMHs: 2\nExp number of AAs in TMHs: 37.26082\nExp number, first 60 AAs: 19.52923\nTotal prob of N-in: 0.94351\nPOSSIBLE N-term signal sequence\ninside 1 19\nTMhelix 20 39\noutside 40 325\nTMhelix 326 348\ninside 349 372

12383 GCF_000357685.2_ASM35768v2 Escherichia coli P0304799.3 Proteobacteria; Gammaproteobacteria; Enterobacterales; Enterobacteriaceae; Escherichia; Escherichia coli MNNEETFYQAMRRQGVTRRSFLKYCSLAATSLGLGAGMAPKIAWALENKPRIPVVWIHGL WP_001058323.1 MULTISPECIES: hydrogenase-1 small chain [Proteobacteria] Length: 372\nNumber of predicted TMHs: 2\nExp number of AAs in TMHs: 37.2608499999999\nExp number, first 60 AAs: 19.52923\nTotal prob of N-in: 0.94351\nPOSSIBLE N-term signal sequence\ninside 1 19\nTMhelix 20 39\noutside 40 325\nTMhelix 326 348\ninside 349 372

12384 GCF_000357685.2_ASM35768v2 Escherichia coli P0304799.3 Proteobacteria; Gammaproteobacteria; Enterobacterales; Enterobacteriaceae; Escherichia; Escherichia coli
MTGDNTLIHSHGINRRDFMKLCAALAATMGLSSKAAAEMAESVTNPQRPPVIWIGAQECT WP_000145410.1
MULTISPECIES: hydrogenase-2 small chain [Proteobacteria] Length: 372\nNumber of predicted TMHs: 1\nExp number of AAs in TMHs: 23.72226\nExp number, first 60 AAs: 0.22038\nTotal prob of N-in: 0.12994\noutside 1 331\nTMhelix 332 354\ninside 355 372

12385 GCF_000356565.1_ASM35656v1 Escherichia coli P0304816.1 Proteobacteria; Gammaproteobacteria; Enterobacterales; Enterobacteriaceae; Escherichia; Escherichia coli
MTGDNTLIHSHGINRRDFMKLCAALAATMGLSSKAAAEMAESVTNPQRPPVIWIGAQECT WP_000145410.1
MULTISPECIES: hydrogenase-2 small chain [Proteobacteria] Length: 372\nNumber of predicted TMHs: 1\nExp number of AAs in TMHs: 23.72226\nExp number, first 60 AAs: 0.22038\nTotal prob of N-in: 0.12994\noutside 1 331\nTMhelix 332 354\ninside 355 372

12386 GCF_000356565.1_ASM35656v1 Escherichia coli P0304816.1 Proteobacteria; Gammaproteobacteria; Enterobacterales; Enterobacteriaceae; Escherichia; Escherichia coli
MNNEETFYQAMRRQGVTRRSFLKYCSLAATSLGLGAGMAPKIAWALENKPRIPVVWIHGL WP_001295940.1
MULTISPECIES: hydrogenase [Enterobacteriaceae] Length: 372\nNumber of predicted TMHs: 2\nExp number of AAs in TMHs: 37.2608899999999\nExp number, first 60 AAs: 19.52923\nTotal prob of N-in: 0.94351\nPOSSIBLE N-term signal sequence\ninside 1 19\nTMhelix 20 39\noutside 40 325\nTMhelix 326 348\ninside 349 372

12387 GCF_000358435.2_ASM35843v2 Escherichia coli P0304816.10 Proteobacteria; Gammaproteobacteria; Enterobacterales; Enterobacteriaceae; Escherichia; Escherichia coli
MTGDNTLIHSHGINRRDFMKLCAALAATMGLSSKAAAEMAESVTNPQRPPVIWIGAQECT WP_000145410.1
MULTISPECIES: hydrogenase-2 small chain [Proteobacteria] Length: 372\nNumber of predicted TMHs: 1\nExp number of AAs in TMHs: 23.72226\nExp number, first 60 AAs: 0.22038\nTotal prob of N-in: 0.12994\noutside 1 331\nTMhelix 332 354\ninside 355 372

12388 GCF_000358435.2_ASM35843v2 Escherichia coli P0304816.10 Proteobacteria; Gammaproteobacteria; Enterobacterales; Enterobacteriaceae; Escherichia; Escherichia coli
MNNEETFYQAMRRQGVTRRSFLKYCSLAATSLGLGAGMAPKIAWALENKPRIPVVWIHGL WP_001295940.1
MULTISPECIES: hydrogenase [Enterobacteriaceae] Length: 372\nNumber of predicted TMHs: 2\nExp number of AAs in TMHs: 37.2608899999999\nExp number, first 60 AAs: 19.52923\nTotal prob of N-in: 0.94351\nPOSSIBLE N-term signal sequence\ninside 1 19\nTMhelix 20 39\noutside 40 325\nTMhelix 326 348\ninside 349 372

12389 GCF_000358455.2_ASM35845v2 Escherichia coli P0304816.11 Proteobacteria; Gammaproteobacteria; Enterobacterales; Enterobacteriaceae; Escherichia; Escherichia coli
MNNEETFYQAMRRQGVTRRSFLKYCSLAATSLGLGAGMAPKIAWALENKPRIPVVWIHGL WP_001295940.1
MULTISPECIES: hydrogenase [Enterobacteriaceae] Length: 372\nNumber of predicted TMHs: 2\nExp number of AAs in TMHs: 37.2608899999999\nExp number, first 60 AAs: 19.52923\nTotal prob of N-in: 0.94351\nPOSSIBLE N-term signal sequence\ninside 1 19\nTMhelix 20 39\noutside 40 325\nTMhelix 326 348\ninside 349 372

12390 GCF_000358455.2_ASM35845v2 Escherichia coli P0304816.11 Proteobacteria; Gammaproteobacteria; Enterobacterales; Enterobacteriaceae; Escherichia; Escherichia coli
MTGDNTLIHSHGINRRDFMKLCAALAATMGLSSKAAAEMAESVTNPQRPPVIWIGAQECT WP_000145410.1
MULTISPECIES: hydrogenase-2 small chain [Proteobacteria] Length: 372\nNumber of predicted TMHs: 1\nExp number of AAs in TMHs: 23.72226\nExp number, first 60 AAs: 0.22038\nTotal prob of N-in: 0.12994\noutside 1 331\nTMhelix 332 354\ninside 355 372

12391 GCF_000358495.2_ASM35849v2 Escherichia coli P0304816.12 Proteobacteria; Gammaproteobacteria; Enterobacterales; Enterobacteriaceae; Escherichia; Escherichia coli
MTGDNTLIHSHGINRRDFMKLCAALAATMGLSSKAAAEMAESVTNPQRPPVIWIGAQECT WP_000145410.1
MULTISPECIES: hydrogenase-2 small chain [Proteobacteria] Length: 372\nNumber of predicted TMHs: 1\nExp number of AAs in TMHs: 23.72226\nExp number, first 60 AAs: 0.22038\nTotal prob of N-in: 0.12994\noutside 1 331\nTMhelix 332 354\ninside 355 372

12392 GCF_000358495.2_ASM35849v2 Escherichia coli P0304816.12 Proteobacteria; Gammaproteobacteria; Enterobacterales; Enterobacteriaceae; Escherichia; Escherichia coli

MNNEETFYQAMRRQGVTRRSFLKYCSLAATSLGLGAGMAPKIAWALENKPRIPVVWIHGL WP_003878210.1 hydrogenase-1 small chain [Escherichia coli] Length: 372\nNumber of predicted TMHs: 2\nExp number of AAs in TMHs: 37.38752\nExp number, first 60 AAs: 19.53624\nTotal prob of N-in: 0.94339\nPOSSIBLE N-term signal sequence\ninside 1 19\nTMhelix 20 39\noutside 40 325\nTMhelix 326 348\ninside 349 372

12393 GCF_000358475.1_ASM35847v1 Escherichia coli P0304816.13 Proteobacteria; Gammaproteobacteria; Enterobacterales; Enterobacteriaceae; Escherichia; Escherichia coli MTGDNPLIHSHGINRRDFMKLCAALAATMGLSSKAAAEMAESVTNPQRPPVIWIGAQECT WP_001461571.1 hydrogenase-2 small chain [Escherichia coli] Length: 372\nNumber of predicted TMHs: 1\nExp number of AAs in TMHs: 22.60469\nExp number, first 60 AAs: 0.1106\nTotal prob of N-in: 0.06393\noutside 1 331\nTMhelix 332 354\ninside 355 372

12394 GCF_000358475.1_ASM35847v1 Escherichia coli P0304816.13 Proteobacteria; Gammaproteobacteria; Enterobacterales; Enterobacteriaceae; Escherichia; Escherichia coli MNNEETFYQAMRRQGVTRRSFLKYCSLAATSLGLGAGMAPKIAWALENKPRIPVVWIHGL WP_001295940.1 MULTISPECIES: hydrogenase [Enterobacteriaceae] Length: 372\nNumber of predicted TMHs: 2\nExp number of AAs in TMHs: 37.2608899999999\nExp number, first 60 AAs: 19.52923\nTotal prob of N-in: 0.94351\nPOSSIBLE N-term signal sequence\ninside 1 19\nTMhelix 20 39\noutside 40 325\nTMhelix 326 348\ninside 349 372

12395 GCF_000358995.1_ASM35899v1 Escherichia coli P0304816.14 Proteobacteria; Gammaproteobacteria; Enterobacterales; Enterobacteriaceae; Escherichia; Escherichia coli MTGDNPLIHSHGINRRDFMKLCAALAATMGLSSKAAAEMAESVTNPQRPPVIWIGAQECT WP_003905497.1 hydrogenase-2 small chain [Escherichia coli] Length: 372\nNumber of predicted TMHs: 1\nExp number of AAs in TMHs: 22.6059\nExp number, first 60 AAs: 0.1106\nTotal prob of N-in: 0.06395\noutside 1 331\nTMhelix 332 354\ninside 355 372

12396 GCF_000358995.1_ASM35899v1 Escherichia coli P0304816.14 Proteobacteria; Gammaproteobacteria; Enterobacterales; Enterobacteriaceae; Escherichia; Escherichia coli MNNEETFYQAMRRQGVTRRSFLKYCSLAATSLGLGAGMAPKIAWALENKPRIPVVWIHGL WP_001295940.1 MULTISPECIES: hydrogenase [Enterobacteriaceae] Length: 372\nNumber of predicted TMHs: 2\nExp number of AAs in TMHs: 37.2608899999999\nExp number, first 60 AAs: 19.52923\nTotal prob of N-in: 0.94351\nPOSSIBLE N-term signal sequence\ninside 1 19\nTMhelix 20 39\noutside 40 325\nTMhelix 326 348\ninside 349 372

12397 GCF_000358515.2_ASM35851v2 Escherichia coli P0304816.15 Proteobacteria; Gammaproteobacteria; Enterobacterales; Enterobacteriaceae; Escherichia; Escherichia coli MNNEETFYQAMRRQGVTRRSFLKYCSLAATSLGLGAGMAPKIAWALENKPRIPVVWIHGL WP_001295940.1 MULTISPECIES: hydrogenase [Enterobacteriaceae] Length: 372\nNumber of predicted TMHs: 2\nExp number of AAs in TMHs: 37.2608899999999\nExp number, first 60 AAs: 19.52923\nTotal prob of N-in: 0.94351\nPOSSIBLE N-term signal sequence\ninside 1 19\nTMhelix 20 39\noutside 40 325\nTMhelix 326 348\ninside 349 372

12398 GCF_000358515.2_ASM35851v2 Escherichia coli P0304816.15 Proteobacteria; Gammaproteobacteria; Enterobacterales; Enterobacteriaceae; Escherichia; Escherichia coli MTGDNTLIHSHGINRRDFMKLCAALAATMGLSSKAAAEMAESVTNPQRPPVIWIGAQECT WP_000145410.1 MULTISPECIES: hydrogenase-2 small chain [Proteobacteria] Length: 372\nNumber of predicted TMHs: 1\nExp number of AAs in TMHs: 23.72226\nExp number, first 60 AAs: 0.22038\nTotal prob of N-in: 0.12994\noutside 1 331\nTMhelix 332 354\ninside 355 372

12399 GCF_000358535.2_ASM35853v2 Escherichia coli P0304816.2 Proteobacteria; Gammaproteobacteria; Enterobacterales; Enterobacteriaceae; Escherichia; Escherichia coli MNNEETFYQAMRRQGVTRRSFLKYCSLAATSLGLGAGMAPKIAWALENKPRIPVVWIHGL WP_001295940.1 MULTISPECIES: hydrogenase [Enterobacteriaceae] Length: 372\nNumber of predicted TMHs: 2\nExp number of AAs in TMHs: 37.2608899999999\nExp number, first 60 AAs: 19.52923\nTotal prob of N-in: 0.94351\nPOSSIBLE N-term signal sequence\ninside 1 19\nTMhelix 20 39\noutside 40 325\nTMhelix 326 348\ninside 349 372

12400 GCF_000358535.2_ASM35853v2 Escherichia coli P0304816.2 Proteobacteria; Gammaproteobacteria; Enterobacterales; Enterobacteriaceae; Escherichia; Escherichia coli MTGDNTLIHSHGINRRDFMKLCAALAATMGLSSKAAAEMAESVTNPQRPPVIWIGAQECT WP_000145410.1

MULTISPECIES: hydrogenase-2 small chain [Proteobacteria] Length: 372\nNumber of predicted TMHs: 1\nExp number of AAs in TMHs: 23.72226\nExp number, first 60 AAs: 0.22038\nTotal prob of N-in: 0.12994\noutside 1 331\nTMhelix 332 354\ninside 355 372

12401 GCF_000359015.2_ASM35901v2 Escherichia coli P0304816.3 Proteobacteria; Gammaproteobacteria; Enterobacterales; Enterobacteriaceae; Escherichia; Escherichia coli
MTGDNTLIHSHGINRRDFMKLCAALAATMGLSSKAAAEMAESVTNPQRPPVIWIGAQECT WP_000145410.1
MULTISPECIES: hydrogenase-2 small chain [Proteobacteria] Length: 372\nNumber of predicted TMHs: 1\nExp number of AAs in TMHs: 23.72226\nExp number, first 60 AAs: 0.22038\nTotal prob of N-in: 0.12994\noutside 1 331\nTMhelix 332 354\ninside 355 372

12402 GCF_000359015.2_ASM35901v2 Escherichia coli P0304816.3 Proteobacteria; Gammaproteobacteria; Enterobacterales; Enterobacteriaceae; Escherichia; Escherichia coli
MNNEETFYQAMRRQGVTRRSFLKYCSLAATSLGLGAGMAPKIAWALENKPRIPVVWIHGL WP_001625367.1
hydrogenase-1 small chain [Escherichia coli] Length: 372\nNumber of predicted TMHs: 2\nExp number of AAs in TMHs: 37.2496399999999\nExp number, first 60 AAs: 19.52619\nTotal prob of N-in: 0.94349\nPOSSIBLE N-term signal sequence\ninside 1 19\nTMhelix 20 39\noutside 40 325\nTMhelix 326 348\ninside 349 372

12403 GCF_000359035.2_ASM35903v2 Escherichia coli P0304816.4 Proteobacteria; Gammaproteobacteria; Enterobacterales; Enterobacteriaceae; Escherichia; Escherichia coli
MTGDNTLIHSHGINRRDFMKLCAALAATMGLSSKAAAEMAESVTNPQRPPVIWIGAQECT WP_000145410.1
MULTISPECIES: hydrogenase-2 small chain [Proteobacteria] Length: 372\nNumber of predicted TMHs: 1\nExp number of AAs in TMHs: 23.72226\nExp number, first 60 AAs: 0.22038\nTotal prob of N-in: 0.12994\noutside 1 331\nTMhelix 332 354\ninside 355 372

12404 GCF_000359035.2_ASM35903v2 Escherichia coli P0304816.4 Proteobacteria; Gammaproteobacteria; Enterobacterales; Enterobacteriaceae; Escherichia; Escherichia coli
MNNEETFYQAMRRQGVTRRSFLKYCSLAATSLGLGAGMAPKIAWALENKPRIPVVWIHGL WP_001295940.1
MULTISPECIES: hydrogenase [Enterobacteriaceae] Length: 372\nNumber of predicted TMHs: 2\nExp number of AAs in TMHs: 37.2608899999999\nExp number, first 60 AAs: 19.52923\nTotal prob of N-in: 0.94351\nPOSSIBLE N-term signal sequence\ninside 1 19\nTMhelix 20 39\noutside 40 325\nTMhelix 326 348\ninside 349 372

12405 GCF_000358955.2_ASM35895v2 Escherichia coli P0304816.5 Proteobacteria; Gammaproteobacteria; Enterobacterales; Enterobacteriaceae; Escherichia; Escherichia coli
MNNEETFYQAMRRQGVTRRSFLKYCSLAATSLGLGAGMAPKIAWALENKPRIPVVWIHGL WP_001295940.1
MULTISPECIES: hydrogenase [Enterobacteriaceae] Length: 372\nNumber of predicted TMHs: 2\nExp number of AAs in TMHs: 37.2608899999999\nExp number, first 60 AAs: 19.52923\nTotal prob of N-in: 0.94351\nPOSSIBLE N-term signal sequence\ninside 1 19\nTMhelix 20 39\noutside 40 325\nTMhelix 326 348\ninside 349 372

12406 GCF_000358955.2_ASM35895v2 Escherichia coli P0304816.5 Proteobacteria; Gammaproteobacteria; Enterobacterales; Enterobacteriaceae; Escherichia; Escherichia coli
MTGDNTLIHSHGINRRDFMKLCAALAATMGLSSKAAAEMAESVTNPQRPPVIWIGAQECT WP_000145410.1
MULTISPECIES: hydrogenase-2 small chain [Proteobacteria] Length: 372\nNumber of predicted TMHs: 1\nExp number of AAs in TMHs: 23.72226\nExp number, first 60 AAs: 0.22038\nTotal prob of N-in: 0.12994\noutside 1 331\nTMhelix 332 354\ninside 355 372

12407 GCF_000357845.2_ASM35784v2 Escherichia coli P0304816.6 Proteobacteria; Gammaproteobacteria; Enterobacterales; Enterobacteriaceae; Escherichia; Escherichia coli
MTGDNTLIHSHGINRRDFMKLCAALAATMGLSSKAAAEMAESVTNPQRPPVIWIGAQECT WP_000145410.1
MULTISPECIES: hydrogenase-2 small chain [Proteobacteria] Length: 372\nNumber of predicted TMHs: 1\nExp number of AAs in TMHs: 23.72226\nExp number, first 60 AAs: 0.22038\nTotal prob of N-in: 0.12994\noutside 1 331\nTMhelix 332 354\ninside 355 372

12408 GCF_000357845.2_ASM35784v2 Escherichia coli P0304816.6 Proteobacteria; Gammaproteobacteria; Enterobacterales; Enterobacteriaceae; Escherichia; Escherichia coli
MNNEETFYQAMRRQGVTRRSFLKYCSLAATSLGLGAGMAPKIAWALENKPRIPVVWIHGL WP_001295940.1
MULTISPECIES: hydrogenase [Enterobacteriaceae] Length: 372\nNumber of predicted TMHs: 2\nExp number of AAs in TMHs: 37.2608899999999\nExp number, first 60 AAs: 19.52923\nTotal prob of N-in:

0.94351\nPOSSIBLE N-term signal sequence\ninside 1 19\nTMhelix 20 39\noutside 40 325\nTMhelix 326 348\ninside 349 372

12409 GCF_000357865.2_ASM35786v2 Escherichia coli P0304816.7 Proteobacteria; Gammaproteobacteria; Enterobacterales; Enterobacteriaceae; Escherichia; Escherichia coli MTGDNTLIHSHGINRRDFMKLCAALAATMGLSSKAAAEMAESVTNPQRPPVIWIGAQECT WP_000145410.1 MULTISPECIES: hydrogenase-2 small chain [Proteobacteria] Length: 372\nNumber of predicted TMHs: 1\nExp number of AAs in TMHs: 23.72226\nExp number, first 60 AAs: 0.22038\nTotal prob of N-in: 0.12994\noutside 1 331\nTMhelix 332 354\ninside 355 372

12410 GCF_000357865.2_ASM35786v2 Escherichia coli P0304816.7 Proteobacteria; Gammaproteobacteria; Enterobacterales; Enterobacteriaceae; Escherichia; Escherichia coli MNNEETFYQAMRRQGVTRRSFLKYCSLAATSLGLGAGMAPKIAWALENKPRIPVVWIHGL WP_001631259.1 hydrogenase-1 small chain [Escherichia coli] Length: 372\nNumber of predicted TMHs: 2\nExp number of AAs in TMHs: 37.14002\nExp number, first 60 AAs: 19.53586\nTotal prob of N-in: 0.94339\nPOSSIBLE N-term signal sequence\ninside 1 19\nTMhelix 20 39\noutside 40 325\nTMhelix 326 348\ninside 349 372

12411 GCF_000357885.2_ASM35788v2 Escherichia coli P0304816.8 Proteobacteria; Gammaproteobacteria; Enterobacterales; Enterobacteriaceae; Escherichia; Escherichia coli MTGDNTLIHSHGINRRDFMKLCAALAATMGLSSKAAAEMAESVTNPQRPPVIWIGAQECT WP_000145410.1 MULTISPECIES: hydrogenase-2 small chain [Proteobacteria] Length: 372\nNumber of predicted TMHs: 1\nExp number of AAs in TMHs: 23.72226\nExp number, first 60 AAs: 0.22038\nTotal prob of N-in: 0.12994\noutside 1 331\nTMhelix 332 354\ninside 355 372

12412 GCF_000357885.2_ASM35788v2 Escherichia coli P0304816.8 Proteobacteria; Gammaproteobacteria; Enterobacterales; Enterobacteriaceae; Escherichia; Escherichia coli MNNEETFYQAMRRQGVTRRSFLKYCSLAATSLGLGAGMAPKIAWALENKPRIPVVWIHGL WP_001632769.1 hydrogenase-1 small chain [Escherichia coli] Length: 372\nNumber of predicted TMHs: 2\nExp number of AAs in TMHs: 37.2608099999999\nExp number, first 60 AAs: 19.52923\nTotal prob of N-in: 0.94351\nPOSSIBLE N-term signal sequence\ninside 1 19\nTMhelix 20 39\noutside 40 325\nTMhelix 326 348\ninside 349 372

12413 GCF_000357905.2_ASM35790v2 Escherichia coli P0304816.9 Proteobacteria; Gammaproteobacteria; Enterobacterales; Enterobacteriaceae; Escherichia; Escherichia coli MNNEETFYQAMRRQGVTRRSFLKYCSLAATSLGLGAGMAPKIAWALENKPRIPVVWIHGL WP_001295940.1 MULTISPECIES: hydrogenase [Enterobacteriaceae] Length: 372\nNumber of predicted TMHs: 2\nExp number of AAs in TMHs: 37.2608899999999\nExp number, first 60 AAs: 19.52923\nTotal prob of N-in: 0.94351\nPOSSIBLE N-term signal sequence\ninside 1 19\nTMhelix 20 39\noutside 40 325\nTMhelix 326 348\ninside 349 372

12414 GCF_000357905.2_ASM35790v2 Escherichia coli P0304816.9 Proteobacteria; Gammaproteobacteria; Enterobacterales; Enterobacteriaceae; Escherichia; Escherichia coli MTGDNTLIHSHGINRRDFMKLCAALAATMGLSSKAAAEMAESVTNPQRPPVIWIGAQECT WP_000145410.1 MULTISPECIES: hydrogenase-2 small chain [Proteobacteria] Length: 372\nNumber of predicted TMHs: 1\nExp number of AAs in TMHs: 23.72226\nExp number, first 60 AAs: 0.22038\nTotal prob of N-in: 0.12994\noutside 1 331\nTMhelix 332 354\ninside 355 372

12415 GCF_000356585.2_ASM35658v2 Escherichia coli P0305260.1 Proteobacteria; Gammaproteobacteria; Enterobacterales; Enterobacteriaceae; Escherichia; Escherichia coli MNNEETFYQAMRRQGVTRRSFLKYCSLAATSLGLGAGMAPKIAWALENKPRIPVVWIHGL WP_001058323.1 MULTISPECIES: hydrogenase-1 small chain [Proteobacteria] Length: 372\nNumber of predicted TMHs: 2\nExp number of AAs in TMHs: 37.2608499999999\nExp number, first 60 AAs: 19.52923\nTotal prob of N-in: 0.94351\nPOSSIBLE N-term signal sequence\ninside 1 19\nTMhelix 20 39\noutside 40 325\nTMhelix 326 348\ninside 349 372

12416 GCF_000356585.2_ASM35658v2 Escherichia coli P0305260.1 Proteobacteria; Gammaproteobacteria; Enterobacterales; Enterobacteriaceae; Escherichia; Escherichia coli MTGDNTLIHSHGINRRDFMKLCAALAATMGLSSKAAAEMAESVTNPQRPPVIWIGAQECT WP_000145410.1 MULTISPECIES: hydrogenase-2 small chain [Proteobacteria] Length: 372\nNumber of predicted TMHs: 1\nExp number of AAs in TMHs: 23.72226\nExp number, first 60 AAs: 0.22038\nTotal prob of N-in: 0.12994\noutside 1 331\nTMhelix 332 354\ninside 355 372

12417 GCF_000357925.2_ASM35792v2 Escherichia coli P0305260.10 Proteobacteria; Gammaproteobacteria; Enterobacterales; Enterobacteriaceae; Escherichia; Escherichia coli
MTGDNTLIHSHGINRRDFMKLCAALAATMGLSSKAAAEMAESVTNPQRPPVIWIGAQECT WP_000145410.1
MULTISPECIES: hydrogenase-2 small chain [Proteobacteria] Length: 372\nNumber of predicted TMHs: 1\nExp number of AAs in TMHs: 23.72226\nExp number, first 60 AAs: 0.22038\nTotal prob of N-in: 0.12994\noutside 1 331\nTMhelix 332 354\ninside 355 372

12418 GCF_000357925.2_ASM35792v2 Escherichia coli P0305260.10 Proteobacteria; Gammaproteobacteria; Enterobacterales; Enterobacteriaceae; Escherichia; Escherichia coli
MNNEETFYQAMRRQGVTRRSFLKYCSLAATSLGLGAGMAPKIAWALENKPRIPVVWIHGL WP_001058323.1
MULTISPECIES: hydrogenase-1 small chain [Proteobacteria] Length: 372\nNumber of predicted TMHs: 2\nExp number of AAs in TMHs: 37.2608499999999\nExp number, first 60 AAs: 19.52923\nTotal prob of N-in: 0.94351\nPOSSIBLE N-term signal sequence\ninside 1 19\nTMhelix 20 39\noutside 40 325\nTMhelix 326 348\ninside 349 372

12419 GCF_000357945.2_ASM35794v2 Escherichia coli P0305260.11 Proteobacteria; Gammaproteobacteria; Enterobacterales; Enterobacteriaceae; Escherichia; Escherichia coli
MNNEETFYQAMRRQGVTRRSFLKYCSLAATSLGLGAGMAPKIAWALENKPRIPVVWIHGL WP_001058323.1
MULTISPECIES: hydrogenase-1 small chain [Proteobacteria] Length: 372\nNumber of predicted TMHs: 2\nExp number of AAs in TMHs: 37.2608499999999\nExp number, first 60 AAs: 19.52923\nTotal prob of N-in: 0.94351\nPOSSIBLE N-term signal sequence\ninside 1 19\nTMhelix 20 39\noutside 40 325\nTMhelix 326 348\ninside 349 372

12420 GCF_000357945.2_ASM35794v2 Escherichia coli P0305260.11 Proteobacteria; Gammaproteobacteria; Enterobacterales; Enterobacteriaceae; Escherichia; Escherichia coli
MTGDNTLIHSHGINRRDFMKLCAALAATMGLSSKAAAEMAESVTNPQRPPVIWIGAQECT WP_000145410.1
MULTISPECIES: hydrogenase-2 small chain [Proteobacteria] Length: 372\nNumber of predicted TMHs: 1\nExp number of AAs in TMHs: 23.72226\nExp number, first 60 AAs: 0.22038\nTotal prob of N-in: 0.12994\noutside 1 331\nTMhelix 332 354\ninside 355 372

12421 GCF_000357965.2_ASM35796v2 Escherichia coli P0305260.12 Proteobacteria; Gammaproteobacteria; Enterobacterales; Enterobacteriaceae; Escherichia; Escherichia coli
MNNEETFYQAMRRQGVTRRSFLKYCSLAATSLGLGAGMAPKIAWALENKPRIPVVWIHGL WP_001058323.1
MULTISPECIES: hydrogenase-1 small chain [Proteobacteria] Length: 372\nNumber of predicted TMHs: 2\nExp number of AAs in TMHs: 37.2608499999999\nExp number, first 60 AAs: 19.52923\nTotal prob of N-in: 0.94351\nPOSSIBLE N-term signal sequence\ninside 1 19\nTMhelix 20 39\noutside 40 325\nTMhelix 326 348\ninside 349 372

12422 GCF_000357965.2_ASM35796v2 Escherichia coli P0305260.12 Proteobacteria; Gammaproteobacteria; Enterobacterales; Enterobacteriaceae; Escherichia; Escherichia coli
MTGDNTLIHSHGINRRDFMKLCAALAATMGLSSKAAAEMAESVTNPQRPPVIWIGAQECT WP_000145410.1
MULTISPECIES: hydrogenase-2 small chain [Proteobacteria] Length: 372\nNumber of predicted TMHs: 1\nExp number of AAs in TMHs: 23.72226\nExp number, first 60 AAs: 0.22038\nTotal prob of N-in: 0.12994\noutside 1 331\nTMhelix 332 354\ninside 355 372

12423 GCF_000357985.2_ASM35798v2 Escherichia coli P0305260.13 Proteobacteria; Gammaproteobacteria; Enterobacterales; Enterobacteriaceae; Escherichia; Escherichia coli
MNNEETFYQAMRRQGVTRRSFLKYCSLAATSLGLGAGMAPKIAWALENKPRIPVVWIHGL WP_001058323.1
MULTISPECIES: hydrogenase-1 small chain [Proteobacteria] Length: 372\nNumber of predicted TMHs: 2\nExp number of AAs in TMHs: 37.2608499999999\nExp number, first 60 AAs: 19.52923\nTotal prob of N-in: 0.94351\nPOSSIBLE N-term signal sequence\ninside 1 19\nTMhelix 20 39\noutside 40 325\nTMhelix 326 348\ninside 349 372

12424 GCF_000357985.2_ASM35798v2 Escherichia coli P0305260.13 Proteobacteria; Gammaproteobacteria; Enterobacterales; Enterobacteriaceae; Escherichia; Escherichia coli
MTGDNTLIHSHGINRRDFMKLCAALAATMGLSSKAAAEMAESVTNPQRPPVIWIGAQECT WP_000145410.1
MULTISPECIES: hydrogenase-2 small chain [Proteobacteria] Length: 372\nNumber of predicted TMHs: 1\nExp number of AAs in TMHs: 23.72226\nExp number, first 60 AAs: 0.22038\nTotal prob of N-in: 0.12994\noutside 1 331\nTMhelix 332 354\ninside 355 372

12425 GCF_000358005.2_ASM35800v2 Escherichia coli P0305260.15 Proteobacteria; Gammaproteobacteria; Enterobacterales; Enterobacteriaceae; Escherichia; Escherichia coli
MTGDNPLIHSHGINRRDFMKLCAALAATMGLSSKAAAEMAESVTNPQRPPVIWIGAQECT WP_001461571.1
hydrogenase-2 small chain [Escherichia coli] Length: 372\nNumber of predicted TMHs: 1\nExp number of AAs in TMHs: 22.60469\nExp number, first 60 AAs: 0.1106\nTotal prob of N-in: 0.06393\noutside 1 331\nTMhelix 332 354\ninside 355 372

12426 GCF_000358005.2_ASM35800v2 Escherichia coli P0305260.15 Proteobacteria; Gammaproteobacteria; Enterobacterales; Enterobacteriaceae; Escherichia; Escherichia coli
MNNEETFYQAMRRQGVTRRSFLKYCSLAATSLGLGAGMAPKIAWALENKPRIPVVWIHGL WP_001058323.1
MULTISPECIES: hydrogenase-1 small chain [Proteobacteria] Length: 372\nNumber of predicted TMHs: 2\nExp number of AAs in TMHs: 37.2608499999999\nExp number, first 60 AAs: 19.52923\nTotal prob of N-in: 0.94351\nPOSSIBLE N-term signal sequence\ninside 1 19\nTMhelix 20 39\noutside 40 325\nTMhelix 326 348\ninside 349 372

12427 GCF_000357705.2_ASM35770v2 Escherichia coli P0305260.2 Proteobacteria; Gammaproteobacteria; Enterobacterales; Enterobacteriaceae; Escherichia; Escherichia coli
MNNEETFYQAMRRQGVTRRSFLKYCSLAATSLGLGAGMAPKIAWALENKPRIPVVWIHGL WP_001058323.1
MULTISPECIES: hydrogenase-1 small chain [Proteobacteria] Length: 372\nNumber of predicted TMHs: 2\nExp number of AAs in TMHs: 37.2608499999999\nExp number, first 60 AAs: 19.52923\nTotal prob of N-in: 0.94351\nPOSSIBLE N-term signal sequence\ninside 1 19\nTMhelix 20 39\noutside 40 325\nTMhelix 326 348\ninside 349 372

12428 GCF_000357705.2_ASM35770v2 Escherichia coli P0305260.2 Proteobacteria; Gammaproteobacteria; Enterobacterales; Enterobacteriaceae; Escherichia; Escherichia coli
MTGDNTLIHSHGINRRDFMKLCAALAATMGLSSKAAAEMAESVTNPQRPPVIWIGAQECT WP_000145410.1
MULTISPECIES: hydrogenase-2 small chain [Proteobacteria] Length: 372\nNumber of predicted TMHs: 1\nExp number of AAs in TMHs: 23.72226\nExp number, first 60 AAs: 0.22038\nTotal prob of N-in: 0.12994\noutside 1 331\nTMhelix 332 354\ninside 355 372

12429 GCF_000358025.2_ASM35802v2 Escherichia coli P0305260.3 Proteobacteria; Gammaproteobacteria; Enterobacterales; Enterobacteriaceae; Escherichia; Escherichia coli
MTGDNTLIHSHGINRRDFMKLCAALAATMGLSSKAAAEMAESVTNPQRPPVIWIGAQECT WP_000145410.1
MULTISPECIES: hydrogenase-2 small chain [Proteobacteria] Length: 372\nNumber of predicted TMHs: 1\nExp number of AAs in TMHs: 23.72226\nExp number, first 60 AAs: 0.22038\nTotal prob of N-in: 0.12994\noutside 1 331\nTMhelix 332 354\ninside 355 372

12430 GCF_000358025.2_ASM35802v2 Escherichia coli P0305260.3 Proteobacteria; Gammaproteobacteria; Enterobacterales; Enterobacteriaceae; Escherichia; Escherichia coli
MNNEETFYQAMRRQGVTRRSFLKYCSLAATSLGLGAGMAPKIAWALENKPRIPVVWIHGL WP_001058323.1
MULTISPECIES: hydrogenase-1 small chain [Proteobacteria] Length: 372\nNumber of predicted TMHs: 2\nExp number of AAs in TMHs: 37.2608499999999\nExp number, first 60 AAs: 19.52923\nTotal prob of N-in: 0.94351\nPOSSIBLE N-term signal sequence\ninside 1 19\nTMhelix 20 39\noutside 40 325\nTMhelix 326 348\ninside 349 372

12431 GCF_000358045.2_ASM35804v2 Escherichia coli P0305260.4 Proteobacteria; Gammaproteobacteria; Enterobacterales; Enterobacteriaceae; Escherichia; Escherichia coli
MNNEETFYQAMRRQGVTRRSFLKYCSLAATSLGLGAGMAPKIAWALENKPRIPVVWIHGL WP_001058323.1
MULTISPECIES: hydrogenase-1 small chain [Proteobacteria] Length: 372\nNumber of predicted TMHs: 2\nExp number of AAs in TMHs: 37.2608499999999\nExp number, first 60 AAs: 19.52923\nTotal prob of N-in: 0.94351\nPOSSIBLE N-term signal sequence\ninside 1 19\nTMhelix 20 39\noutside 40 325\nTMhelix 326 348\ninside 349 372

12432 GCF_000358045.2_ASM35804v2 Escherichia coli P0305260.4 Proteobacteria; Gammaproteobacteria; Enterobacterales; Enterobacteriaceae; Escherichia; Escherichia coli
MTGDNPLIHSHGINRRDFMKLCAALAATMGLSSKAAAEMAESVTNPQRPPVIWIGAQECT WP_001647491.1
hydrogenase-2 small chain [Escherichia coli] Length: 372\nNumber of predicted TMHs: 1\nExp number of AAs in TMHs: 22.60351\nExp number, first 60 AAs: 0.1106\nTotal prob of N-in: 0.06390\noutside 1 331\nTMhelix 332 354\ninside 355 372

12433 GCF_000358065.2_ASM35806v2 Escherichia coli P0305260.5 Proteobacteria; Gammaproteobacteria; Enterobacterales; Enterobacteriaceae; Escherichia; Escherichia coli
MTGDNTLIHSHGINRRDFMKLCAALAATMGLSSKAAAEMAESVTNPQRPPVIWIGAQECT WP_001649269.1
hydrogenase-2 small chain [Escherichia coli] Length: 372\nNumber of predicted TMHs: 1\nExp number of AAs in TMHs: 23.72398\nExp number, first 60 AAs: 0.22038\nTotal prob of N-in: 0.12999\noutside 1 331\nTMhelix 332 354\ninside 355 372

12434 GCF_000358065.2_ASM35806v2 Escherichia coli P0305260.5 Proteobacteria; Gammaproteobacteria; Enterobacterales; Enterobacteriaceae; Escherichia; Escherichia coli
MNNEETFYQAMRRQGVTRRSFLKYCSLAATSLGLGAGMAPKIAWALENKPRIPVVWIHGL WP_001058323.1
MULTISPECIES: hydrogenase-1 small chain [Proteobacteria] Length: 372\nNumber of predicted TMHs: 2\nExp number of AAs in TMHs: 37.2608499999999\nExp number, first 60 AAs: 19.52923\nTotal prob of N-in: 0.94351\nPOSSIBLE N-term signal sequence\ninside 1 19\nTMhelix 20 39\noutside 40 325\nTMhelix 326 348\ninside 349 372

12435 GCF_000358085.2_ASM35808v2 Escherichia coli P0305260.6 Proteobacteria; Gammaproteobacteria; Enterobacterales; Enterobacteriaceae; Escherichia; Escherichia coli
MNNEETFYQAMRRQGVTRRSFLKYCSLAATSLGLGAGMAPKIAWALENKPRIPVVWIHGL WP_001058323.1
MULTISPECIES: hydrogenase-1 small chain [Proteobacteria] Length: 372\nNumber of predicted TMHs: 2\nExp number of AAs in TMHs: 37.2608499999999\nExp number, first 60 AAs: 19.52923\nTotal prob of N-in: 0.94351\nPOSSIBLE N-term signal sequence\ninside 1 19\nTMhelix 20 39\noutside 40 325\nTMhelix 326 348\ninside 349 372

12436 GCF_000358085.2_ASM35808v2 Escherichia coli P0305260.6 Proteobacteria; Gammaproteobacteria; Enterobacterales; Enterobacteriaceae; Escherichia; Escherichia coli
MTGDNTLIHSHGINRRDFMKLCAALAATMGLSSKAAAEMAESVTNPQRPPVIWIGAQECT WP_000145410.1
MULTISPECIES: hydrogenase-2 small chain [Proteobacteria] Length: 372\nNumber of predicted TMHs: 1\nExp number of AAs in TMHs: 23.72226\nExp number, first 60 AAs: 0.22038\nTotal prob of N-in: 0.12994\noutside 1 331\nTMhelix 332 354\ninside 355 372

12437 GCF_000358105.2_ASM35810v2 Escherichia coli P0305260.7 Proteobacteria; Gammaproteobacteria; Enterobacterales; Enterobacteriaceae; Escherichia; Escherichia coli
MNNEETFYQAMRRQGVTRRSFLKYCSLAATSLGLGAGMAPKIAWALENKPRIPVVWIHGL WP_001058323.1
MULTISPECIES: hydrogenase-1 small chain [Proteobacteria] Length: 372\nNumber of predicted TMHs: 2\nExp number of AAs in TMHs: 37.2608499999999\nExp number, first 60 AAs: 19.52923\nTotal prob of N-in: 0.94351\nPOSSIBLE N-term signal sequence\ninside 1 19\nTMhelix 20 39\noutside 40 325\nTMhelix 326 348\ninside 349 372

12438 GCF_000358105.2_ASM35810v2 Escherichia coli P0305260.7 Proteobacteria; Gammaproteobacteria; Enterobacterales; Enterobacteriaceae; Escherichia; Escherichia coli
MTGDNTLIHSHGINRRDFMKLCAALAATMGLSSKAAAEMAESVTNPQRPPVIWIGAQECT WP_000145410.1
MULTISPECIES: hydrogenase-2 small chain [Proteobacteria] Length: 372\nNumber of predicted TMHs: 1\nExp number of AAs in TMHs: 23.72226\nExp number, first 60 AAs: 0.22038\nTotal prob of N-in: 0.12994\noutside 1 331\nTMhelix 332 354\ninside 355 372

12439 GCF_000358125.1_ASM35812v1 Escherichia coli P0305260.8 Proteobacteria; Gammaproteobacteria; Enterobacterales; Enterobacteriaceae; Escherichia; Escherichia coli
MTGDNTLIHSHGINRRDFMKLCAALAATMGLSSKAAAEMAESVTNPQRPPVIWIGAQECT WP_000145410.1
MULTISPECIES: hydrogenase-2 small chain [Proteobacteria] Length: 372\nNumber of predicted TMHs: 1\nExp number of AAs in TMHs: 23.72226\nExp number, first 60 AAs: 0.22038\nTotal prob of N-in: 0.12994\noutside 1 331\nTMhelix 332 354\ninside 355 372

12440 GCF_000358125.1_ASM35812v1 Escherichia coli P0305260.8 Proteobacteria; Gammaproteobacteria; Enterobacterales; Enterobacteriaceae; Escherichia; Escherichia coli
MNNEETFYQAMRRQGVTRRSFLKYCSLAATSLGLGAGMAPKIAWALENKPRIPVVWIHGL WP_001058323.1
MULTISPECIES: hydrogenase-1 small chain [Proteobacteria] Length: 372\nNumber of predicted TMHs: 2\nExp number of AAs in TMHs: 37.2608499999999\nExp number, first 60 AAs: 19.52923\nTotal prob of N-in: 0.94351\nPOSSIBLE N-term signal sequence\ninside 1 19\nTMhelix 20 39\noutside 40 325\nTMhelix 326 348\ninside 349 372

12441 GCF_000356825.2_ASM35682v2 Escherichia coli P0305260.9 Proteobacteria; Gammaproteobacteria; Enterobacterales; Enterobacteriaceae; Escherichia; Escherichia coli
MTGDNTLIHSHGINRRDFMKLCAALAATMGLSSKAAAEMAESVTNPQRPPVIWIGAQECT WP_001655824.1
hydrogenase-2 small chain [Escherichia coli] Length: 372\nNumber of predicted TMHs: 1\nExp number of AAs in TMHs: 23.72049\nExp number, first 60 AAs: 0.22038\nTotal prob of N-in: 0.12989\noutside 1 331\nTMhelix 332 354\ninside 355 372

12442 GCF_000356825.2_ASM35682v2 Escherichia coli P0305260.9 Proteobacteria; Gammaproteobacteria; Enterobacterales; Enterobacteriaceae; Escherichia; Escherichia coli
MNNEETFYQAMRRQGVTRRSFLKYCSLAATSLGLGAGMAPKIAWALENKPRIPVVWIHGL WP_001058323.1
MULTISPECIES: hydrogenase-1 small chain [Proteobacteria] Length: 372\nNumber of predicted TMHs: 2\nExp number of AAs in TMHs: 37.2608499999999\nExp number, first 60 AAs: 19.52923\nTotal prob of N-in: 0.94351\nPOSSIBLE N-term signal sequence\ninside 1 19\nTMhelix 20 39\noutside 40 325\nTMhelix 326 348\ninside 349 372

12443 GCF_000356605.2_ASM35660v2 Escherichia coli p0305293.1 Proteobacteria; Gammaproteobacteria; Enterobacterales; Enterobacteriaceae; Escherichia; Escherichia coli
MNNEETFYQAMRRQGVTRRSFLKYCSLAATSLGLGAGMAPKIVWALENKPRIPVVWIHGL WP_001398876.1
hydrogenase-1 small chain [Escherichia coli] Length: 372\nNumber of predicted TMHs: 2\nExp number of AAs in TMHs: 37.60268\nExp number, first 60 AAs: 19.84677\nTotal prob of N-in: 0.94621\nPOSSIBLE N-term signal sequence\ninside 1 20\nTMhelix 21 43\noutside 44 325\nTMhelix 326 348\ninside 349 372

12444 GCF_000356605.2_ASM35660v2 Escherichia coli p0305293.1 Proteobacteria; Gammaproteobacteria; Enterobacterales; Enterobacteriaceae; Escherichia; Escherichia coli
MTGDNTLIHSHGINRRDFMKLCAALAATMGLSSKAAAEMAESVTNPQRPPVIWIGAQECT WP_000145410.1
MULTISPECIES: hydrogenase-2 small chain [Proteobacteria] Length: 372\nNumber of predicted TMHs: 1\nExp number of AAs in TMHs: 23.72226\nExp number, first 60 AAs: 0.22038\nTotal prob of N-in: 0.12994\noutside 1 331\nTMhelix 332 354\ninside 355 372

12445 GCF_000358555.1_ASM35855v1 Escherichia coli p0305293.10 Proteobacteria; Gammaproteobacteria; Enterobacterales; Enterobacteriaceae; Escherichia; Escherichia coli
MNNEETFYQAMRRQGVTRRSFLKYCSLAATSLGLGAGMAPKIVWALENKPRIPVVWIHGL WP_001398876.1
hydrogenase-1 small chain [Escherichia coli] Length: 372\nNumber of predicted TMHs: 2\nExp number of AAs in TMHs: 37.60268\nExp number, first 60 AAs: 19.84677\nTotal prob of N-in: 0.94621\nPOSSIBLE N-term signal sequence\ninside 1 20\nTMhelix 21 43\noutside 44 325\nTMhelix 326 348\ninside 349 372

12446 GCF_000358555.1_ASM35855v1 Escherichia coli p0305293.10 Proteobacteria; Gammaproteobacteria; Enterobacterales; Enterobacteriaceae; Escherichia; Escherichia coli
MTGDNTLIHSHGINRRDFMKLCAALAATMGLSSKAAAEMAESVTNPQRPPVIWIGAQECT WP_000145410.1
MULTISPECIES: hydrogenase-2 small chain [Proteobacteria] Length: 372\nNumber of predicted TMHs: 1\nExp number of AAs in TMHs: 23.72226\nExp number, first 60 AAs: 0.22038\nTotal prob of N-in: 0.12994\noutside 1 331\nTMhelix 332 354\ninside 355 372

12447 GCF_000358575.2_ASM35857v2 Escherichia coli p0305293.11 Proteobacteria; Gammaproteobacteria; Enterobacterales; Enterobacteriaceae; Escherichia; Escherichia coli
MNNEETFYQAMRRQGVTRRSFLKYCSLAATSLGLGAGMAPKIVWALENKPRIPVVWIHGL WP_001398876.1
hydrogenase-1 small chain [Escherichia coli] Length: 372\nNumber of predicted TMHs: 2\nExp number of AAs in TMHs: 37.60268\nExp number, first 60 AAs: 19.84677\nTotal prob of N-in: 0.94621\nPOSSIBLE N-term signal sequence\ninside 1 20\nTMhelix 21 43\noutside 44 325\nTMhelix 326 348\ninside 349 372

12448 GCF_000358575.2_ASM35857v2 Escherichia coli p0305293.11 Proteobacteria; Gammaproteobacteria; Enterobacterales; Enterobacteriaceae; Escherichia; Escherichia coli
MTGDNTLIHSHGINRRDFMKLCAALAATMGLSSKAAAEMAESVTNPQRPPVIWIGAQECT WP_000145410.1
MULTISPECIES: hydrogenase-2 small chain [Proteobacteria] Length: 372\nNumber of predicted TMHs: 1\nExp number of AAs in TMHs: 23.72226\nExp number, first 60 AAs: 0.22038\nTotal prob of N-in: 0.12994\noutside 1 331\nTMhelix 332 354\ninside 355 372

12449 GCF_000358595.2_ASM35859v2 Escherichia coli p0305293.12 Proteobacteria; Gammaproteobacteria; Enterobacterales; Enterobacteriaceae; Escherichia; Escherichia coli
MNNEETFYQAMRRQGVTRRSFLKYCSLAATSLGLGAGMAPKIVWALENKPRIPVVWIHGL WP_001398876.1
hydrogenase-1 small chain [Escherichia coli] Length: 372\nNumber of predicted TMHs: 2\nExp number of

AAs in TMHs: 37.60268\nExp number, first 60 AAs: 19.84677\nTotal prob of N-in: 0.94621\nPOSSIBLE N-term signal sequence\ninside 1 20\nTMhelix 21 43\noutside 44 325\nTMhelix 326 348\ninside 349 372

12450 GCF_000358595.2_ASM35859v2 Escherichia coli p0305293.12 Proteobacteria; Gammaproteobacteria; Enterobacterales; Enterobacteriaceae; Escherichia; Escherichia coli
MTGDNTLIHSHGINRRDFMKLCAALAATMGLSSKAAAEMAESVTNPQRPPVIWIGAQECT WP_000145410.1
MULTISPECIES: hydrogenase-2 small chain [Proteobacteria] Length: 372\nNumber of predicted TMHs: 1\nExp number of AAs in TMHs: 23.72226\nExp number, first 60 AAs: 0.22038\nTotal prob of N-in: 0.12994\noutside 1 331\nTMhelix 332 354\ninside 355 372

12451 GCF_000356865.2_ASM35686v2 Escherichia coli p0305293.13 Proteobacteria; Gammaproteobacteria; Enterobacterales; Enterobacteriaceae; Escherichia; Escherichia coli
MNNEETFYQAMRRQGVTRRSFLKYCSLAATSLGLGAGMAPKIVWALENKPRIPVVWIHGL WP_001398876.1
hydrogenase-1 small chain [Escherichia coli] Length: 372\nNumber of predicted TMHs: 2\nExp number of AAs in TMHs: 37.60268\nExp number, first 60 AAs: 19.84677\nTotal prob of N-in: 0.94621\nPOSSIBLE N-term signal sequence\ninside 1 20\nTMhelix 21 43\noutside 44 325\nTMhelix 326 348\ninside 349 372

12452 GCF_000356865.2_ASM35686v2 Escherichia coli p0305293.13 Proteobacteria; Gammaproteobacteria; Enterobacterales; Enterobacteriaceae; Escherichia; Escherichia coli
MTGDNTLIHSHGINRRDFMKLCAALAATMGLSSKAAAEMAESVTNPQRPPVIWIGAQECT WP_000145410.1
MULTISPECIES: hydrogenase-2 small chain [Proteobacteria] Length: 372\nNumber of predicted TMHs: 1\nExp number of AAs in TMHs: 23.72226\nExp number, first 60 AAs: 0.22038\nTotal prob of N-in: 0.12994\noutside 1 331\nTMhelix 332 354\ninside 355 372

12453 GCF_000357485.2_ASM35748v2 Escherichia coli p0305293.14 Proteobacteria; Gammaproteobacteria; Enterobacterales; Enterobacteriaceae; Escherichia; Escherichia coli
MNNEETFYQAMRRQGVTRRSFLKYCSLAATSLGLGAGMAPKIVWALENKPRIPVVWIHGL WP_001398876.1
hydrogenase-1 small chain [Escherichia coli] Length: 372\nNumber of predicted TMHs: 2\nExp number of AAs in TMHs: 37.60268\nExp number, first 60 AAs: 19.84677\nTotal prob of N-in: 0.94621\nPOSSIBLE N-term signal sequence\ninside 1 20\nTMhelix 21 43\noutside 44 325\nTMhelix 326 348\ninside 349 372

12454 GCF_000357485.2_ASM35748v2 Escherichia coli p0305293.14 Proteobacteria; Gammaproteobacteria; Enterobacterales; Enterobacteriaceae; Escherichia; Escherichia coli
MTGDNTLIHSHGINRRDFMKLCAALAATMGLSSKAAAEMAESVTNPQRPPVIWIGAQECT WP_000145410.1
MULTISPECIES: hydrogenase-2 small chain [Proteobacteria] Length: 372\nNumber of predicted TMHs: 1\nExp number of AAs in TMHs: 23.72226\nExp number, first 60 AAs: 0.22038\nTotal prob of N-in: 0.12994\noutside 1 331\nTMhelix 332 354\ninside 355 372

12455 GCF_000358615.2_ASM35861v2 Escherichia coli p0305293.15 Proteobacteria; Gammaproteobacteria; Enterobacterales; Enterobacteriaceae; Escherichia; Escherichia coli
MTGDNTLIHSHGINRRDFMKLCAALAATMGLSSKAAAEMAESVTNPQRPPVIWIGAQECT WP_000145410.1
MULTISPECIES: hydrogenase-2 small chain [Proteobacteria] Length: 372\nNumber of predicted TMHs: 1\nExp number of AAs in TMHs: 23.72226\nExp number, first 60 AAs: 0.22038\nTotal prob of N-in: 0.12994\noutside 1 331\nTMhelix 332 354\ninside 355 372

12456 GCF_000358615.2_ASM35861v2 Escherichia coli p0305293.15 Proteobacteria; Gammaproteobacteria; Enterobacterales; Enterobacteriaceae; Escherichia; Escherichia coli
MNNEETFYQAMRRQGVTRRSFLKYCSLAATSLGLGAGMAPKIVWALENKPRIPVVWIHGL WP_001398876.1
hydrogenase-1 small chain [Escherichia coli] Length: 372\nNumber of predicted TMHs: 2\nExp number of AAs in TMHs: 37.60268\nExp number, first 60 AAs: 19.84677\nTotal prob of N-in: 0.94621\nPOSSIBLE N-term signal sequence\ninside 1 20\nTMhelix 21 43\noutside 44 325\nTMhelix 326 348\ninside 349 372

12457 GCF_000358635.2_ASM35863v2 Escherichia coli p0305293.2 Proteobacteria; Gammaproteobacteria; Enterobacterales; Enterobacteriaceae; Escherichia; Escherichia coli
MNNEETFYQAMRRQGVTRRSFLKYCSLAATSLGLGAGMAPKIVWALENKPRIPVVWIHGL WP_001398876.1
hydrogenase-1 small chain [Escherichia coli] Length: 372\nNumber of predicted TMHs: 2\nExp number of AAs in TMHs: 37.60268\nExp number, first 60 AAs: 19.84677\nTotal prob of N-in: 0.94621\nPOSSIBLE N-term signal sequence\ninside 1 20\nTMhelix 21 43\noutside 44 325\nTMhelix 326 348\ninside 349 372

12458 GCF_000358635.2_ASM35863v2 Escherichia coli p0305293.2 Proteobacteria; Gammaproteobacteria; Enterobacterales; Enterobacteriaceae; Escherichia; Escherichia coli

MTGDNTLIHSHGINRRDFMKLCAALAATMGLSSKAAAEMAESVTNPQRPPVIWIGAQECT WP_000145410.1 MULTISPECIES: hydrogenase-2 small chain [Proteobacteria] Length: 372\nNumber of predicted TMHs: 1\nExp number of AAs in TMHs: 23.72226\nExp number, first 60 AAs: 0.22038\nTotal prob of N-in: 0.12994\noutside 1 331\nTMhelix 332 354\ninside 355 372

12459 GCF_000358655.2_ASM35865v2 Escherichia coli p0305293.3 Proteobacteria; Gammaproteobacteria; Enterobacterales; Enterobacteriaceae; Escherichia; Escherichia coli MTGDNTLIHSHGINRRDFMKLCAALAATMGLSSKAAAEMAESVTNPQRPPVIWIGAQECT WP_000145410.1 MULTISPECIES: hydrogenase-2 small chain [Proteobacteria] Length: 372\nNumber of predicted TMHs: 1\nExp number of AAs in TMHs: 23.72226\nExp number, first 60 AAs: 0.22038\nTotal prob of N-in: 0.12994\noutside 1 331\nTMhelix 332 354\ninside 355 372

12460 GCF_000358655.2_ASM35865v2 Escherichia coli p0305293.3 Proteobacteria; Gammaproteobacteria; Enterobacterales; Enterobacteriaceae; Escherichia; Escherichia coli MNNEETFYQAMRRQGVTRRSFLKYCSLAATSLGLGAGMAPKIVWALENKPRIPVVWIHGL WP_001398876.1 hydrogenase-1 small chain [Escherichia coli] Length: 372\nNumber of predicted TMHs: 2\nExp number of AAs in TMHs: 37.60268\nExp number, first 60 AAs: 19.84677\nTotal prob of N-in: 0.94621\nPOSSIBLE N-term signal sequence\ninside 1 20\nTMhelix 21 43\noutside 44 325\nTMhelix 326 348\ninside 349 372

12461 GCF_000358675.2_ASM35867v2 Escherichia coli p0305293.4 Proteobacteria; Gammaproteobacteria; Enterobacterales; Enterobacteriaceae; Escherichia; Escherichia coli MNNEETFYQAMRRQGVTRRSFLKYCSLAATSLGLGAGMAPKIVWALENKPRIPVVWIHGL WP_001398876.1 hydrogenase-1 small chain [Escherichia coli] Length: 372\nNumber of predicted TMHs: 2\nExp number of AAs in TMHs: 37.60268\nExp number, first 60 AAs: 19.84677\nTotal prob of N-in: 0.94621\nPOSSIBLE N-term signal sequence\ninside 1 20\nTMhelix 21 43\noutside 44 325\nTMhelix 326 348\ninside 349 372

12462 GCF_000358675.2_ASM35867v2 Escherichia coli p0305293.4 Proteobacteria; Gammaproteobacteria; Enterobacterales; Enterobacteriaceae; Escherichia; Escherichia coli MTGDNTLIHSHGINRRDFMKLCAALAATMGLSSKAAAEMAESVTNPQRPPVIWIGAQECT WP_000145410.1 MULTISPECIES: hydrogenase-2 small chain [Proteobacteria] Length: 372\nNumber of predicted TMHs: 1\nExp number of AAs in TMHs: 23.72226\nExp number, first 60 AAs: 0.22038\nTotal prob of N-in: 0.12994\noutside 1 331\nTMhelix 332 354\ninside 355 372

12463 GCF_000359055.2_ASM35905v2 Escherichia coli p0305293.5 Proteobacteria; Gammaproteobacteria; Enterobacterales; Enterobacteriaceae; Escherichia; Escherichia coli MTGDNTLIHSHGINRRDFMKLCAALAATMGLSSKAAAEMAESVTNPQRPPVIWIGAQECT WP_001669906.1 hydrogenase-2 small chain [Escherichia coli] Length: 372\nNumber of predicted TMHs: 1\nExp number of AAs in TMHs: 23.71665\nExp number, first 60 AAs: 0.22037\nTotal prob of N-in: 0.12978\noutside 1 331\nTMhelix 332 354\ninside 355 372

12464 GCF_000359055.2_ASM35905v2 Escherichia coli p0305293.5 Proteobacteria; Gammaproteobacteria; Enterobacterales; Enterobacteriaceae; Escherichia; Escherichia coli MNNEETFYQAMRRQGVTRRSFLKYCSLAATSLGLGAGMAPKIVWALENKPRIPVVWIHGL WP_001398876.1 hydrogenase-1 small chain [Escherichia coli] Length: 372\nNumber of predicted TMHs: 2\nExp number of AAs in TMHs: 37.60268\nExp number, first 60 AAs: 19.84677\nTotal prob of N-in: 0.94621\nPOSSIBLE N-term signal sequence\ninside 1 20\nTMhelix 21 43\noutside 44 325\nTMhelix 326 348\ninside 349 372

12465 GCF_000359075.1_ASM35907v1 Escherichia coli p0305293.6 Proteobacteria; Gammaproteobacteria; Enterobacterales; Enterobacteriaceae; Escherichia; Escherichia coli MTGDNTLIHSHGINRRDFMKLCAALAATMGLSSKAAAEMAESVTNPQRPPVIWIGAQECT WP_000145410.1 MULTISPECIES: hydrogenase-2 small chain [Proteobacteria] Length: 372\nNumber of predicted TMHs: 1\nExp number of AAs in TMHs: 23.72226\nExp number, first 60 AAs: 0.22038\nTotal prob of N-in: 0.12994\noutside 1 331\nTMhelix 332 354\ninside 355 372

12466 GCF_000359075.1_ASM35907v1 Escherichia coli p0305293.6 Proteobacteria; Gammaproteobacteria; Enterobacterales; Enterobacteriaceae; Escherichia; Escherichia coli MNNEETFYQAMRRQGVTRRSFLKYCSLAATSLGLGAGMAPKIVWALENKPRIPVVWIHGL WP_001398876.1 hydrogenase-1 small chain [Escherichia coli] Length: 372\nNumber of predicted TMHs: 2\nExp number of AAs in TMHs: 37.60268\nExp number, first 60 AAs: 19.84677\nTotal prob of N-in: 0.94621\nPOSSIBLE N-term signal sequence\ninside 1 20\nTMhelix 21 43\noutside 44 325\nTMhelix 326 348\ninside 349 372

12467 GCF_000359095.2_ASM35909v2 Escherichia coli p0305293.7 Proteobacteria; Gammaproteobacteria; Enterobacterales; Enterobacteriaceae; Escherichia; Escherichia coli
MTGDNTLIHSHGINRRDFMKLCAALAATMGLSSKAAAEMAESVTNPQRPPVIWIGAQECT WP_000145410.1
MULTISPECIES: hydrogenase-2 small chain [Proteobacteria] Length: 372\nNumber of predicted TMHs: 1\nExp number of AAs in TMHs: 23.72226\nExp number, first 60 AAs: 0.22038\nTotal prob of N-in: 0.12994\noutside 1 331\nTMhelix 332 354\ninside 355 372

12468 GCF_000359095.2_ASM35909v2 Escherichia coli p0305293.7 Proteobacteria; Gammaproteobacteria; Enterobacterales; Enterobacteriaceae; Escherichia; Escherichia coli
MNNEETFYQAMRRQGVTRRSFLKYCSLAATSLGLGAGMAPKIVWALENKPRIPVVWIHGL WP_001398876.1
hydrogenase-1 small chain [Escherichia coli] Length: 372\nNumber of predicted TMHs: 2\nExp number of AAs in TMHs: 37.60268\nExp number, first 60 AAs: 19.84677\nTotal prob of N-in: 0.94621\nPOSSIBLE N-term signal sequence\ninside 1 20\nTMhelix 21 43\noutside 44 325\nTMhelix 326 348\ninside 349 372

12469 GCF_000358695.2_ASM35869v2 Escherichia coli p0305293.8 Proteobacteria; Gammaproteobacteria; Enterobacterales; Enterobacteriaceae; Escherichia; Escherichia coli
MNNEETFYQAMRRQGVTRRSFLKYCSLAATSLGLGAGMAPKIVWALENKPRIPVVWIHGL WP_001398876.1
hydrogenase-1 small chain [Escherichia coli] Length: 372\nNumber of predicted TMHs: 2\nExp number of AAs in TMHs: 37.60268\nExp number, first 60 AAs: 19.84677\nTotal prob of N-in: 0.94621\nPOSSIBLE N-term signal sequence\ninside 1 20\nTMhelix 21 43\noutside 44 325\nTMhelix 326 348\ninside 349 372

12470 GCF_000358695.2_ASM35869v2 Escherichia coli p0305293.8 Proteobacteria; Gammaproteobacteria; Enterobacterales; Enterobacteriaceae; Escherichia; Escherichia coli
MTGDNTLIHSHGINRRDFMKLCAALAATMGLSSKAAAEMAESVTNPQRPPVIWIGAQECT WP_000145410.1
MULTISPECIES: hydrogenase-2 small chain [Proteobacteria] Length: 372\nNumber of predicted TMHs: 1\nExp number of AAs in TMHs: 23.72226\nExp number, first 60 AAs: 0.22038\nTotal prob of N-in: 0.12994\noutside 1 331\nTMhelix 332 354\ninside 355 372

12471 GCF_000358715.2_ASM35871v2 Escherichia coli p0305293.9 Proteobacteria; Gammaproteobacteria; Enterobacterales; Enterobacteriaceae; Escherichia; Escherichia coli
MNNEETFYQAMRRQGVTRRSFLKYCSLAATSLGLGAGMAPKIVWALENKPRIPVVWIHGL WP_001398876.1
hydrogenase-1 small chain [Escherichia coli] Length: 372\nNumber of predicted TMHs: 2\nExp number of AAs in TMHs: 37.60268\nExp number, first 60 AAs: 19.84677\nTotal prob of N-in: 0.94621\nPOSSIBLE N-term signal sequence\ninside 1 20\nTMhelix 21 43\noutside 44 325\nTMhelix 326 348\ninside 349 372

12472 GCF_000358715.2_ASM35871v2 Escherichia coli p0305293.9 Proteobacteria; Gammaproteobacteria; Enterobacterales; Enterobacteriaceae; Escherichia; Escherichia coli
MTGDNTLIHSHGINRRDFMKLCAALAATMGLSSKAAAEMAESVTNPQRPPVIWIGAQECT WP_000145410.1
MULTISPECIES: hydrogenase-2 small chain [Proteobacteria] Length: 372\nNumber of predicted TMHs: 1\nExp number of AAs in TMHs: 23.72226\nExp number, first 60 AAs: 0.22038\nTotal prob of N-in: 0.12994\noutside 1 331\nTMhelix 332 354\ninside 355 372

12473 GCF_000257275.1_ASM25727v1 Escherichia coli P12b Proteobacteria; Gammaproteobacteria; Enterobacterales; Enterobacteriaceae; Escherichia; Escherichia coli
MTGDNTLIHSHGINRRDFMKLCAALAATMGLSSKAAAEMAESVTNPQRPPVIWIGAQECT WP_000145410.1
MULTISPECIES: hydrogenase-2 small chain [Proteobacteria] Length: 372\nNumber of predicted TMHs: 1\nExp number of AAs in TMHs: 23.72226\nExp number, first 60 AAs: 0.22038\nTotal prob of N-in: 0.12994\noutside 1 331\nTMhelix 332 354\ninside 355 372

12474 GCF_000257275.1_ASM25727v1 Escherichia coli P12b Proteobacteria; Gammaproteobacteria; Enterobacterales; Enterobacteriaceae; Escherichia; Escherichia coli
MNNEETFYQAMRRQGVTRRSFLKYCSLAATSLGLGAGMAPKIAWALENKPRIPVVWIHGL WP_001058323.1
MULTISPECIES: hydrogenase-1 small chain [Proteobacteria] Length: 372\nNumber of predicted TMHs: 2\nExp number of AAs in TMHs: 37.2608499999999\nExp number, first 60 AAs: 19.52923\nTotal prob of N-in: 0.94351\nPOSSIBLE N-term signal sequence\ninside 1 19\nTMhelix 20 39\noutside 40 325\nTMhelix 326 348\ninside 349 372

12475 GCF_000498255.1_ASM49825v1 Escherichia coli P4-96 Proteobacteria; Gammaproteobacteria; Enterobacterales; Enterobacteriaceae; Escherichia; Escherichia coli
MNNEETFYQAMRRQGVTRRSFLKYCSLAATSLGLGAGMAPKIAWALENKPRIPVVWIHGL WP_001295940.1
MULTISPECIES: hydrogenase [Enterobacteriaceae] Length: 372\nNumber of predicted TMHs: 2\nExp

number of AAs in TMHs: 37.2608899999999\nExp number, first 60 AAs: 19.52923\nTotal prob of N-in: 0.94351\nPOSSIBLE N-term signal sequence\ninside 1 19\nTMhelix 20 39\noutside 40 325\nTMhelix 326 348\ninside 349 372

12476 GCF_000498255.1_ASM49825v1 Escherichia coli P4-96 Proteobacteria; Gammaproteobacteria; Enterobacterales; Enterobacteriaceae; Escherichia; Escherichia coli MTGDNTLIHSHGINRRDFMKLCAALAATMGLSSKAAAEMAESVTNPQRPPVIWIGAQECT WP_000145410.1 MULTISPECIES: hydrogenase-2 small chain [Proteobacteria] Length: 372\nNumber of predicted TMHs: 1\nExp number of AAs in TMHs: 23.72226\nExp number, first 60 AAs: 0.22038\nTotal prob of N-in: 0.12994\noutside 1 331\nTMhelix 332 354\ninside 355 372

12477 GCF_000498275.1_ASM49827v1 Escherichia coli P4-NR Proteobacteria; Gammaproteobacteria; Enterobacterales; Enterobacteriaceae; Escherichia; Escherichia coli MTGDNTLIHSHGINRRDFMKLCAALAATMGLSSKAAAEMAESVTNPQRPPVIWIGAQECT WP_000145410.1 MULTISPECIES: hydrogenase-2 small chain [Proteobacteria] Length: 372\nNumber of predicted TMHs: 1\nExp number of AAs in TMHs: 23.72226\nExp number, first 60 AAs: 0.22038\nTotal prob of N-in: 0.12994\noutside 1 331\nTMhelix 332 354\ninside 355 372

12478 GCF_000498275.1_ASM49827v1 Escherichia coli P4-NR Proteobacteria; Gammaproteobacteria; Enterobacterales; Enterobacteriaceae; Escherichia; Escherichia coli MNNEETFYQAMRRQGVTRRSFLKYCSLAATSLGLGAGMAPKIAWALENKPRIPVVWIHGL WP_001295940.1 MULTISPECIES: hydrogenase [Enterobacteriaceae] Length: 372\nNumber of predicted TMHs: 2\nExp number of AAs in TMHs: 37.2608899999999\nExp number, first 60 AAs: 19.52923\nTotal prob of N-in: 0.94351\nPOSSIBLE N-term signal sequence\ninside 1 19\nTMhelix 20 39\noutside 40 325\nTMhelix 326 348\ninside 349 372

12479 GCF_000267865.2_ASM26786v2 Escherichia coli PA10 Proteobacteria; Gammaproteobacteria; Enterobacterales; Enterobacteriaceae; Escherichia; Escherichia coli MNNEETFYQAMRRQGVTRRSFLKYCSLAATSLGLGAGMAPKIAWALENKPRIPVVWIHGL WP_001301613.1 hydrogenase [Escherichia coli] Length: 372\nNumber of predicted TMHs: 2\nExp number of AAs in TMHs: 37.46598\nExp number, first 60 AAs: 19.53139\nTotal prob of N-in: 0.94364\nPOSSIBLE N-term signal sequence\ninside 1 19\nTMhelix 20 39\noutside 40 325\nTMhelix 326 348\ninside 349 372

12480 GCF_000267865.2_ASM26786v2 Escherichia coli PA10 Proteobacteria; Gammaproteobacteria; Enterobacterales; Enterobacteriaceae; Escherichia; Escherichia coli MTGDNTLIHSHGINRRDFMKLCAALAATMGLSSKAAAEMAESVTNPQRPPVIWIGAQECT WP_000145410.1 MULTISPECIES: hydrogenase-2 small chain [Proteobacteria] Length: 372\nNumber of predicted TMHs: 1\nExp number of AAs in TMHs: 23.72226\nExp number, first 60 AAs: 0.22038\nTotal prob of N-in: 0.12994\noutside 1 331\nTMhelix 332 354\ninside 355 372

12481 GCF_000267845.2_ASM26784v2 Escherichia coli PA14 Proteobacteria; Gammaproteobacteria; Enterobacterales; Enterobacteriaceae; Escherichia; Escherichia coli MNNEETFYQAMRRQGVTRRSFLKYCSLAATSLGLGAGMAPKIAWALENKPRIPVVWIHGL WP_001301613.1 hydrogenase [Escherichia coli] Length: 372\nNumber of predicted TMHs: 2\nExp number of AAs in TMHs: 37.46598\nExp number, first 60 AAs: 19.53139\nTotal prob of N-in: 0.94364\nPOSSIBLE N-term signal sequence\ninside 1 19\nTMhelix 20 39\noutside 40 325\nTMhelix 326 348\ninside 349 372

12482 GCF_000267845.2_ASM26784v2 Escherichia coli PA14 Proteobacteria; Gammaproteobacteria; Enterobacterales; Enterobacteriaceae; Escherichia; Escherichia coli MTGDNTLIHSHGINRRDFMKLCAALAATMGLSSKAAAEMAESVTNPQRPPVIWIGAQECT WP_000145410.1 MULTISPECIES: hydrogenase-2 small chain [Proteobacteria] Length: 372\nNumber of predicted TMHs: 1\nExp number of AAs in TMHs: 23.72226\nExp number, first 60 AAs: 0.22038\nTotal prob of N-in: 0.12994\noutside 1 331\nTMhelix 332 354\ninside 355 372

12483 GCF_000267825.2_ASM26782v2 Escherichia coli PA15 Proteobacteria; Gammaproteobacteria; Enterobacterales; Enterobacteriaceae; Escherichia; Escherichia coli MNNEETFYQAMRRQGVTRRSFLKYCSLAATSLGLGAGMAPKIAWALENKPRIPVVWIHGL WP_001301613.1 hydrogenase [Escherichia coli] Length: 372\nNumber of predicted TMHs: 2\nExp number of AAs in TMHs: 37.46598\nExp number, first 60 AAs: 19.53139\nTotal prob of N-in: 0.94364\nPOSSIBLE N-term signal sequence\ninside 1 19\nTMhelix 20 39\noutside 40 325\nTMhelix 326 348\ninside 349 372

12484 GCF_000267825.2_ASM26782v2 Escherichia coli PA15 Proteobacteria; Gammaproteobacteria; Enterobacterales; Enterobacteriaceae; Escherichia; Escherichia coli
MTGDNTLIHSHGINRRDFMKLCAALAATMGLSSKAAAEMAESVTNPQRPPVIWIGAQECT WP_000145410.1
MULTISPECIES: hydrogenase-2 small chain [Proteobacteria] Length: 372\nNumber of predicted TMHs: 1\nExp number of AAs in TMHs: 23.72226\nExp number, first 60 AAs: 0.22038\nTotal prob of N-in: 0.12994\noutside 1 331\nTMhelix 332 354\ninside 355 372

12485 GCF_000335355.2_ASM33535v2 Escherichia coli PA2 Proteobacteria; Gammaproteobacteria; Enterobacterales; Enterobacteriaceae; Escherichia; Escherichia coli
MTGDNTLIHSHGINRRDFMKLCAALAATMGLSSKAAAEMAESVTNPQRPPVIWIGAQECT WP_000145410.1
MULTISPECIES: hydrogenase-2 small chain [Proteobacteria] Length: 372\nNumber of predicted TMHs: 1\nExp number of AAs in TMHs: 23.72226\nExp number, first 60 AAs: 0.22038\nTotal prob of N-in: 0.12994\noutside 1 331\nTMhelix 332 354\ninside 355 372

12486 GCF_000335355.2_ASM33535v2 Escherichia coli PA2 Proteobacteria; Gammaproteobacteria; Enterobacterales; Enterobacteriaceae; Escherichia; Escherichia coli
MNNEETFYQAMRRQGVTRRSFLKYCSLAATSLGLGAGMAPKIAWALENKPRIPVVWIHGL WP_001301613.1
hydrogenase [Escherichia coli] Length: 372\nNumber of predicted TMHs: 2\nExp number of AAs in TMHs: 37.46598\nExp number, first 60 AAs: 19.53139\nTotal prob of N-in: 0.94364\nPOSSIBLE N-term signal sequence\ninside 1 19\nTMhelix 20 39\noutside 40 325\nTMhelix 326 348\ninside 349 372

12487 GCF_000267085.1_ASM26708v1 Escherichia coli PA22 Proteobacteria; Gammaproteobacteria; Enterobacterales; Enterobacteriaceae; Escherichia; Escherichia coli
MTGDNTLIHSHGINRRDFMKLCAALAATMGLSSKAAAEMAESVTNPQRPPVIWIGAQECT WP_000145410.1
MULTISPECIES: hydrogenase-2 small chain [Proteobacteria] Length: 372\nNumber of predicted TMHs: 1\nExp number of AAs in TMHs: 23.72226\nExp number, first 60 AAs: 0.22038\nTotal prob of N-in: 0.12994\noutside 1 331\nTMhelix 332 354\ninside 355 372

12488 GCF_000267085.1_ASM26708v1 Escherichia coli PA22 Proteobacteria; Gammaproteobacteria; Enterobacterales; Enterobacteriaceae; Escherichia; Escherichia coli
MNNEETFYQAMRRQGVTRRSFLKYCSLAATSLGLGAGMAPKIAWALENKPRIPVVWIHGL WP_001301613.1
hydrogenase [Escherichia coli] Length: 372\nNumber of predicted TMHs: 2\nExp number of AAs in TMHs: 37.46598\nExp number, first 60 AAs: 19.53139\nTotal prob of N-in: 0.94364\nPOSSIBLE N-term signal sequence\ninside 1 19\nTMhelix 20 39\noutside 40 325\nTMhelix 326 348\ninside 349 372

12489 GCF_000303895.2_ASM30389v2 Escherichia coli PA23 Proteobacteria; Gammaproteobacteria; Enterobacterales; Enterobacteriaceae; Escherichia; Escherichia coli
MTGDNTLIHSHGINRRDFMKLCAALAATMGLSSKAAAEMAESVTNPQRPPVIWIGAQECT WP_000145410.1
MULTISPECIES: hydrogenase-2 small chain [Proteobacteria] Length: 372\nNumber of predicted TMHs: 1\nExp number of AAs in TMHs: 23.72226\nExp number, first 60 AAs: 0.22038\nTotal prob of N-in: 0.12994\noutside 1 331\nTMhelix 332 354\ninside 355 372

12490 GCF_000303895.2_ASM30389v2 Escherichia coli PA23 Proteobacteria; Gammaproteobacteria; Enterobacterales; Enterobacteriaceae; Escherichia; Escherichia coli
MNNEETFYQAMRRQGVTRRSFLKYCSLAATSLGLGAGMAPKIAWALENKPRIPVVWIHGL WP_001301613.1
hydrogenase [Escherichia coli] Length: 372\nNumber of predicted TMHs: 2\nExp number of AAs in TMHs: 37.46598\nExp number, first 60 AAs: 19.53139\nTotal prob of N-in: 0.94364\nPOSSIBLE N-term signal sequence\ninside 1 19\nTMhelix 20 39\noutside 40 325\nTMhelix 326 348\ninside 349 372

12491 GCF_000267805.2_ASM26780v2 Escherichia coli PA24 Proteobacteria; Gammaproteobacteria; Enterobacterales; Enterobacteriaceae; Escherichia; Escherichia coli
MTGDNTLIHSHGINRRDFMKLCAALAATMGLSSKAAAEMAESVTNPQRPPVIWIGAQECT WP_000145410.1
MULTISPECIES: hydrogenase-2 small chain [Proteobacteria] Length: 372\nNumber of predicted TMHs: 1\nExp number of AAs in TMHs: 23.72226\nExp number, first 60 AAs: 0.22038\nTotal prob of N-in: 0.12994\noutside 1 331\nTMhelix 332 354\ninside 355 372

12492 GCF_000267805.2_ASM26780v2 Escherichia coli PA24 Proteobacteria; Gammaproteobacteria; Enterobacterales; Enterobacteriaceae; Escherichia; Escherichia coli
MNNEETFYQAMRRQGVTRRSFLKYCSLAATSLGLGAGMAPKIAWALENKPRIPVVWIHGL WP_001301613.1
hydrogenase [Escherichia coli] Length: 372\nNumber of predicted TMHs: 2\nExp number of AAs in

TMHs: 37.46598\nExp number, first 60 AAs: 19.53139\nTotal prob of N-in: 0.94364\nPOSSIBLE N-term signal sequence\ninside 1 19\nTMhelix 20 39\noutside 40 325\nTMhelix 326 348\ninside 349 372

12493 GCF_000267785.2_ASM26778v2 Escherichia coli PA25 Proteobacteria; Gammaproteobacteria; Enterobacterales; Enterobacteriaceae; Escherichia; Escherichia coli MNNEETFYQAMRRQGVTRRSFLKYCSLAATSLGLGAGMAPKIAWALENKPRIPVVWIHGL WP_001301613.1 hydrogenase [Escherichia coli] Length: 372\nNumber of predicted TMHs: 2\nExp number of AAs in TMHs: 37.46598\nExp number, first 60 AAs: 19.53139\nTotal prob of N-in: 0.94364\nPOSSIBLE N-term signal sequence\ninside 1 19\nTMhelix 20 39\noutside 40 325\nTMhelix 326 348\ninside 349 372

12494 GCF_000267785.2_ASM26778v2 Escherichia coli PA25 Proteobacteria; Gammaproteobacteria; Enterobacterales; Enterobacteriaceae; Escherichia; Escherichia coli MTGDNTLIHSHGINRRDFMKLCAALAATMGLSSKAAAEMAESVTNPQRPPVIWIGAQECT WP_000145410.1 MULTISPECIES: hydrogenase-2 small chain [Proteobacteria] Length: 372\nNumber of predicted TMHs: 1\nExp number of AAs in TMHs: 23.72226\nExp number, first 60 AAs: 0.22038\nTotal prob of N-in: 0.12994\noutside 1 331\nTMhelix 332 354\ninside 355 372

12495 GCF_000267765.2_ASM26776v2 Escherichia coli PA28 Proteobacteria; Gammaproteobacteria; Enterobacterales; Enterobacteriaceae; Escherichia; Escherichia coli MNNEETFYQAMRRQGVTRRSFLKYCSLAATSLGLGAGMAPKIAWALENKPRIPVVWIHGL WP_001301613.1 hydrogenase [Escherichia coli] Length: 372\nNumber of predicted TMHs: 2\nExp number of AAs in TMHs: 37.46598\nExp number, first 60 AAs: 19.53139\nTotal prob of N-in: 0.94364\nPOSSIBLE N-term signal sequence\ninside 1 19\nTMhelix 20 39\noutside 40 325\nTMhelix 326 348\ninside 349 372

12496 GCF_000267765.2_ASM26776v2 Escherichia coli PA28 Proteobacteria; Gammaproteobacteria; Enterobacterales; Enterobacteriaceae; Escherichia; Escherichia coli MTGDNTLIHSHGINRRDFMKLCAALAATMGLSSKAAAEMAESVTNPQRPPVIWIGAQECT WP_000145410.1 MULTISPECIES: hydrogenase-2 small chain [Proteobacteria] Length: 372\nNumber of predicted TMHs: 1\nExp number of AAs in TMHs: 23.72226\nExp number, first 60 AAs: 0.22038\nTotal prob of N-in: 0.12994\noutside 1 331\nTMhelix 332 354\ninside 355 372

12497 GCF_000267925.2_ASM26792v2 Escherichia coli PA3 Proteobacteria; Gammaproteobacteria; Enterobacterales; Enterobacteriaceae; Escherichia; Escherichia coli MTGDNTLIHSHGINRRDFMKLCAALAATMGLSSKAAAEMAESVTNPQRPPVIWIGAQECT WP_000145410.1 MULTISPECIES: hydrogenase-2 small chain [Proteobacteria] Length: 372\nNumber of predicted TMHs: 1\nExp number of AAs in TMHs: 23.72226\nExp number, first 60 AAs: 0.22038\nTotal prob of N-in: 0.12994\noutside 1 331\nTMhelix 332 354\ninside 355 372

12498 GCF_000267925.2_ASM26792v2 Escherichia coli PA3 Proteobacteria; Gammaproteobacteria; Enterobacterales; Enterobacteriaceae; Escherichia; Escherichia coli MNNEETFYQAMRRQGVTRRSFLKYCSLAATSLGLGAGMAPKIAWALENKPRIPVVWIHGL WP_001301613.1 hydrogenase [Escherichia coli] Length: 372\nNumber of predicted TMHs: 2\nExp number of AAs in TMHs: 37.46598\nExp number, first 60 AAs: 19.53139\nTotal prob of N-in: 0.94364\nPOSSIBLE N-term signal sequence\ninside 1 19\nTMhelix 20 39\noutside 40 325\nTMhelix 326 348\ninside 349 372

12499 GCF_000267745.2_ASM26774v2 Escherichia coli PA31 Proteobacteria; Gammaproteobacteria; Enterobacterales; Enterobacteriaceae; Escherichia; Escherichia coli MNNEETFYQAMRRQGVTRRSFLKYCSLAATSLGLGAGMAPKIAWALENKPRIPVVWIHGL WP_001301613.1 hydrogenase [Escherichia coli] Length: 372\nNumber of predicted TMHs: 2\nExp number of AAs in TMHs: 37.46598\nExp number, first 60 AAs: 19.53139\nTotal prob of N-in: 0.94364\nPOSSIBLE N-term signal sequence\ninside 1 19\nTMhelix 20 39\noutside 40 325\nTMhelix 326 348\ninside 349 372

12500 GCF_000267745.2_ASM26774v2 Escherichia coli PA31 Proteobacteria; Gammaproteobacteria; Enterobacterales; Enterobacteriaceae; Escherichia; Escherichia coli MTGDNTLIHSHGINRRDFMKLCAALAATMGLSSKAAAEMAESVTNPQRPPVIWIGAQECT WP_000145410.1 MULTISPECIES: hydrogenase-2 small chain [Proteobacteria] Length: 372\nNumber of predicted TMHs: 1\nExp number of AAs in TMHs: 23.72226\nExp number, first 60 AAs: 0.22038\nTotal prob of N-in: 0.12994\noutside 1 331\nTMhelix 332 354\ninside 355 372

12501 GCF_000267725.2_ASM26772v2 Escherichia coli PA32 Proteobacteria; Gammaproteobacteria; Enterobacterales; Enterobacteriaceae; Escherichia; Escherichia coli

MNNEETFYQAMRRQGVTRRSFLKYCSLAATSLGLGAGMAPKIAWALENKPRIPVVWIHGL WP_001301613.1 hydrogenase [Escherichia coli] Length: 372\nNumber of predicted TMHs: 2\nExp number of AAs in TMHs: 37.46598\nExp number, first 60 AAs: 19.53139\nTotal prob of N-in: 0.94364\nPOSSIBLE N-term signal sequence\ninside 1 19\nTMhelix 20 39\noutside 40 325\nTMhelix 326 348\ninside 349 372

12502 GCF_000267725.2_ASM26772v2 Escherichia coli PA32 Proteobacteria; Gammaproteobacteria; Enterobacterales; Enterobacteriaceae; Escherichia; Escherichia coli MTGDNTLIHSHGINRRDFMKLCAALAATMGLSSKAAAEMAESVTNPQRPPVIWIGAQECT WP_000145410.1 MULTISPECIES: hydrogenase-2 small chain [Proteobacteria] Length: 372\nNumber of predicted TMHs: 1\nExp number of AAs in TMHs: 23.72226\nExp number, first 60 AAs: 0.22038\nTotal prob of N-in: 0.12994\noutside 1 331\nTMhelix 332 354\ninside 355 372

12503 GCF_000267705.2_ASM26770v2 Escherichia coli PA33 Proteobacteria; Gammaproteobacteria; Enterobacterales; Enterobacteriaceae; Escherichia; Escherichia coli MNNEETFYQAMRRQGVTRRSFLKYCSLAATSLGLGAGMAPKIAWALENKPRIPVVWIHGL WP_001301613.1 hydrogenase [Escherichia coli] Length: 372\nNumber of predicted TMHs: 2\nExp number of AAs in TMHs: 37.46598\nExp number, first 60 AAs: 19.53139\nTotal prob of N-in: 0.94364\nPOSSIBLE N-term signal sequence\ninside 1 19\nTMhelix 20 39\noutside 40 325\nTMhelix 326 348\ninside 349 372

12504 GCF_000267705.2_ASM26770v2 Escherichia coli PA33 Proteobacteria; Gammaproteobacteria; Enterobacterales; Enterobacteriaceae; Escherichia; Escherichia coli MTGDNTLIHSHGINRRDFMKLCAALAATMGLSSKAAAEMAESVTNPQRPPVIWIGAQECT WP_000145410.1 MULTISPECIES: hydrogenase-2 small chain [Proteobacteria] Length: 372\nNumber of predicted TMHs: 1\nExp number of AAs in TMHs: 23.72226\nExp number, first 60 AAs: 0.22038\nTotal prob of N-in: 0.12994\noutside 1 331\nTMhelix 332 354\ninside 355 372

12505 GCF_000303695.2_ASM30369v2 Escherichia coli PA34 Proteobacteria; Gammaproteobacteria; Enterobacterales; Enterobacteriaceae; Escherichia; Escherichia coli MTGDNTLIHSHGINRRDFMKLCAALAATMGLSSKAAAEMAESVTNPQRPPVIWIGAQECT WP_000145410.1 MULTISPECIES: hydrogenase-2 small chain [Proteobacteria] Length: 372\nNumber of predicted TMHs: 1\nExp number of AAs in TMHs: 23.72226\nExp number, first 60 AAs: 0.22038\nTotal prob of N-in: 0.12994\noutside 1 331\nTMhelix 332 354\ninside 355 372

12506 GCF_000303695.2_ASM30369v2 Escherichia coli PA34 Proteobacteria; Gammaproteobacteria; Enterobacterales; Enterobacteriaceae; Escherichia; Escherichia coli MNNEETFYQAMRRQGVTRRSFLKYCSLAATSLGLGAGMAPKIAWALENKPRIPVVWIHGL WP_001301613.1 hydrogenase [Escherichia coli] Length: 372\nNumber of predicted TMHs: 2\nExp number of AAs in TMHs: 37.46598\nExp number, first 60 AAs: 19.53139\nTotal prob of N-in: 0.94364\nPOSSIBLE N-term signal sequence\ninside 1 19\nTMhelix 20 39\noutside 40 325\nTMhelix 326 348\ninside 349 372

12507 GCF_000303675.2_ASM30367v2 Escherichia coli PA38 Proteobacteria; Gammaproteobacteria; Enterobacterales; Enterobacteriaceae; Escherichia; Escherichia coli MTGDNTLIHSHGINRRDFMKLCAALAATMGLSSKAAAEMAESVTNPQRPPVIWIGAQECT WP_000145410.1 MULTISPECIES: hydrogenase-2 small chain [Proteobacteria] Length: 372\nNumber of predicted TMHs: 1\nExp number of AAs in TMHs: 23.72226\nExp number, first 60 AAs: 0.22038\nTotal prob of N-in: 0.12994\noutside 1 331\nTMhelix 332 354\ninside 355 372

12508 GCF_000303675.2_ASM30367v2 Escherichia coli PA38 Proteobacteria; Gammaproteobacteria; Enterobacterales; Enterobacteriaceae; Escherichia; Escherichia coli MNNEETFYQAMRRQGVTRRSFLKYCSLAATSLGLGAGMAPKIAWALENKPRIPVVWIHGL WP_001301613.1 hydrogenase [Escherichia coli] Length: 372\nNumber of predicted TMHs: 2\nExp number of AAs in TMHs: 37.46598\nExp number, first 60 AAs: 19.53139\nTotal prob of N-in: 0.94364\nPOSSIBLE N-term signal sequence\ninside 1 19\nTMhelix 20 39\noutside 40 325\nTMhelix 326 348\ninside 349 372

12509 GCF_000267685.2_ASM26768v2 Escherichia coli PA39 Proteobacteria; Gammaproteobacteria; Enterobacterales; Enterobacteriaceae; Escherichia; Escherichia coli MNNEETFYQAMRRQGVTRRSFLKYCSLAATSLGLGAGMAPKIAWALENKPRIPVVWIHGL WP_001301613.1 hydrogenase [Escherichia coli] Length: 372\nNumber of predicted TMHs: 2\nExp number of AAs in TMHs: 37.46598\nExp number, first 60 AAs: 19.53139\nTotal prob of N-in: 0.94364\nPOSSIBLE N-term signal sequence\ninside 1 19\nTMhelix 20 39\noutside 40 325\nTMhelix 326 348\ninside 349 372

12510	GCF_000267685.2_ASM26768v2	Escherichia coli PA39	Proteobacteria; Gammaproteobacteria; Enterobacterales; Enterobacteriaceae; Escherichia; Escherichia coli
MTGDNTLIHSHGINRRDFMKLCAALAATMGLSSKAAAEMAESVTNPQRPPVIWIGAQECT	WP_000145410.1
MULTISPECIES: hydrogenase-2 small chain [Proteobacteria]	Length: 372\nNumber of predicted TMHs: 1\nExp number of AAs in TMHs: 23.72226\nExp number, first 60 AAs: 0.22038\nTotal prob of N-in: 0.12994\noutside 1 331\nTMhelix 332 354\ninside 355 372

12511	GCF_000303875.2_ASM30387v2	Escherichia coli PA4	Proteobacteria; Gammaproteobacteria; Enterobacterales; Enterobacteriaceae; Escherichia; Escherichia coli
MTGDNTLIHSHGINRRDFMKLCAALAATMGLSSKAAAEMAESVTNPQRPPVIWIGAQECT	WP_000145410.1
MULTISPECIES: hydrogenase-2 small chain [Proteobacteria]	Length: 372\nNumber of predicted TMHs: 1\nExp number of AAs in TMHs: 23.72226\nExp number, first 60 AAs: 0.22038\nTotal prob of N-in: 0.12994\noutside 1 331\nTMhelix 332 354\ninside 355 372

12512	GCF_000303875.2_ASM30387v2	Escherichia coli PA4	Proteobacteria; Gammaproteobacteria; Enterobacterales; Enterobacteriaceae; Escherichia; Escherichia coli
MNNEETFYQAMRRQGVTRRSFLKYCSLAATSLGLGAGMAPKIAWALENKPRIPVVWIHGL	WP_001301613.1
hydrogenase [Escherichia coli]	Length: 372\nNumber of predicted TMHs: 2\nExp number of AAs in TMHs: 37.46598\nExp number, first 60 AAs: 19.53139\nTotal prob of N-in: 0.94364\nPOSSIBLE N-term signal sequence\ninside 1 19\nTMhelix 20 39\noutside 40 325\nTMhelix 326 348\ninside 349 372

12513	GCF_000267105.1_ASM26710v1	Escherichia coli PA40	Proteobacteria; Gammaproteobacteria; Enterobacterales; Enterobacteriaceae; Escherichia; Escherichia coli
MTGDNTLIHSHGINRRDFMKLCAALAATMGLSSKAAAEMAESVTNPQRPPVIWIGAQECT	WP_000145410.1
MULTISPECIES: hydrogenase-2 small chain [Proteobacteria]	Length: 372\nNumber of predicted TMHs: 1\nExp number of AAs in TMHs: 23.72226\nExp number, first 60 AAs: 0.22038\nTotal prob of N-in: 0.12994\noutside 1 331\nTMhelix 332 354\ninside 355 372

12514	GCF_000267105.1_ASM26710v1	Escherichia coli PA40	Proteobacteria; Gammaproteobacteria; Enterobacterales; Enterobacteriaceae; Escherichia; Escherichia coli
MNNEETFYQAMRRQGVTRRSFLKYCSLAATSLGLGAGMAPKIAWALENKPRIPVVWIHGL	WP_001301613.1
hydrogenase [Escherichia coli]	Length: 372\nNumber of predicted TMHs: 2\nExp number of AAs in TMHs: 37.46598\nExp number, first 60 AAs: 19.53139\nTotal prob of N-in: 0.94364\nPOSSIBLE N-term signal sequence\ninside 1 19\nTMhelix 20 39\noutside 40 325\nTMhelix 326 348\ninside 349 372

12515	GCF_000267665.2_ASM26766v2	Escherichia coli PA41	Proteobacteria; Gammaproteobacteria; Enterobacterales; Enterobacteriaceae; Escherichia; Escherichia coli
MTGDNTLIHSHGINRRDFMKLCAALAATMGLSSKAAAEMAESVTNPQRPPVIWIGAQECT	WP_000145410.1
MULTISPECIES: hydrogenase-2 small chain [Proteobacteria]	Length: 372\nNumber of predicted TMHs: 1\nExp number of AAs in TMHs: 23.72226\nExp number, first 60 AAs: 0.22038\nTotal prob of N-in: 0.12994\noutside 1 331\nTMhelix 332 354\ninside 355 372

12516	GCF_000267665.2_ASM26766v2	Escherichia coli PA41	Proteobacteria; Gammaproteobacteria; Enterobacterales; Enterobacteriaceae; Escherichia; Escherichia coli
MNNEETFYQAMRRQGVTRRSFLKYCSLAATSLGLGAGMAPKIAWALENKPRIPVVWIHGL	WP_001301613.1
hydrogenase [Escherichia coli]	Length: 372\nNumber of predicted TMHs: 2\nExp number of AAs in TMHs: 37.46598\nExp number, first 60 AAs: 19.53139\nTotal prob of N-in: 0.94364\nPOSSIBLE N-term signal sequence\ninside 1 19\nTMhelix 20 39\noutside 40 325\nTMhelix 326 348\ninside 349 372

12517	GCF_000267645.2_ASM26764v2	Escherichia coli PA42	Proteobacteria; Gammaproteobacteria; Enterobacterales; Enterobacteriaceae; Escherichia; Escherichia coli
MNNEETFYQAMRRQGVTRRSFLKYCSLAATSLGLGAGMAPKIAWALENKPRIPVVWIHGL	WP_001301613.1
hydrogenase [Escherichia coli]	Length: 372\nNumber of predicted TMHs: 2\nExp number of AAs in TMHs: 37.46598\nExp number, first 60 AAs: 19.53139\nTotal prob of N-in: 0.94364\nPOSSIBLE N-term signal sequence\ninside 1 19\nTMhelix 20 39\noutside 40 325\nTMhelix 326 348\ninside 349 372

12518	GCF_000267645.2_ASM26764v2	Escherichia coli PA42	Proteobacteria; Gammaproteobacteria; Enterobacterales; Enterobacteriaceae; Escherichia; Escherichia coli
MTGDNTLIHSHGINRRDFMKLCAALAATMGLSSKAAAEMAESVTNPQRPPVIWIGAQECT	WP_000145410.1
MULTISPECIES: hydrogenase-2 small chain [Proteobacteria]	Length: 372\nNumber of predicted TMHs:

1\nExp number of AAs in TMHs: 23.72226\nExp number, first 60 AAs: 0.22038\nTotal prob of N-in: 0.12994\noutside 1 331\nTMhelix 332 354\ninside 355 372

12519 GCF_000303935.2_ASM30393v2 Escherichia coli PA45 Proteobacteria; Gammaproteobacteria; Enterobacterales; Enterobacteriaceae; Escherichia; Escherichia coli MNNEETFYQAMRRQGVTRRSFLKYCSLAATSLGLGAGMAPKIAWALENKPRIPVVWIHGL WP_001301613.1 hydrogenase [Escherichia coli] Length: 372\nNumber of predicted TMHs: 2\nExp number of AAs in TMHs: 37.46598\nExp number, first 60 AAs: 19.53139\nTotal prob of N-in: 0.94364\nPOSSIBLE N-term signal sequence\ninside 1 19\nTMhelix 20 39\noutside 40 325\nTMhelix 326 348\ninside 349 372

12520 GCF_000303935.2_ASM30393v2 Escherichia coli PA45 Proteobacteria; Gammaproteobacteria; Enterobacterales; Enterobacteriaceae; Escherichia; Escherichia coli MTGDNTLIHSHGINRRDFMKLCAALAATMGLSSKAAAEMAESVTNPQRPPVIWIGAQECT WP_000145410.1 MULTISPECIES: hydrogenase-2 small chain [Proteobacteria] Length: 372\nNumber of predicted TMHs: 1\nExp number of AAs in TMHs: 23.72226\nExp number, first 60 AAs: 0.22038\nTotal prob of N-in: 0.12994\noutside 1 331\nTMhelix 332 354\ninside 355 372

12521 GCF_000335075.2_ASM33507v2 Escherichia coli PA47 Proteobacteria; Gammaproteobacteria; Enterobacterales; Enterobacteriaceae; Escherichia; Escherichia coli MNNEETFYQAMRRQGVTRRSFLKYCSLAATSLGLGAGMAPKIAWALENKPRIPVVWIHGL WP_001301613.1 hydrogenase [Escherichia coli] Length: 372\nNumber of predicted TMHs: 2\nExp number of AAs in TMHs: 37.46598\nExp number, first 60 AAs: 19.53139\nTotal prob of N-in: 0.94364\nPOSSIBLE N-term signal sequence\ninside 1 19\nTMhelix 20 39\noutside 40 325\nTMhelix 326 348\ninside 349 372

12522 GCF_000335075.2_ASM33507v2 Escherichia coli PA47 Proteobacteria; Gammaproteobacteria; Enterobacterales; Enterobacteriaceae; Escherichia; Escherichia coli MTGDNTLIHSHGINRRDFMKLCAALAATMGLSSKAAAEMAESVTNPQRPPVIWIGAQECT WP_000145410.1 MULTISPECIES: hydrogenase-2 small chain [Proteobacteria] Length: 372\nNumber of predicted TMHs: 1\nExp number of AAs in TMHs: 23.72226\nExp number, first 60 AAs: 0.22038\nTotal prob of N-in: 0.12994\noutside 1 331\nTMhelix 332 354\ninside 355 372

12523 GCF_000335095.2_ASM33509v2 Escherichia coli PA48 Proteobacteria; Gammaproteobacteria; Enterobacterales; Enterobacteriaceae; Escherichia; Escherichia coli MIGDNTLIHSHGINRRDFMKLCAALAATMGLSSKAAAEMAESVTNPQRPPVIWIGAQECT WP_000578724.1 hydrogenase 2 small subunit [Escherichia coli] Length: 372\nNumber of predicted TMHs: 1\nExp number of AAs in TMHs: 23.05601\nExp number, first 60 AAs: 0.15431\nTotal prob of N-in: 0.09116\noutside 1 331\nTMhelix 332 354\ninside 355 372

12524 GCF_000335095.2_ASM33509v2 Escherichia coli PA48 Proteobacteria; Gammaproteobacteria; Enterobacterales; Enterobacteriaceae; Escherichia; Escherichia coli MNNEETFYQAMRRQGVTRRSFLKYCSLAATSLGLGAGMAPKIAWALENKPRIPVVWIHGL WP_001301613.1 hydrogenase [Escherichia coli] Length: 372\nNumber of predicted TMHs: 2\nExp number of AAs in TMHs: 37.46598\nExp number, first 60 AAs: 19.53139\nTotal prob of N-in: 0.94364\nPOSSIBLE N-term signal sequence\ninside 1 19\nTMhelix 20 39\noutside 40 325\nTMhelix 326 348\ninside 349 372

12525 GCF_000303915.2_ASM30391v2 Escherichia coli PA49 Proteobacteria; Gammaproteobacteria; Enterobacterales; Enterobacteriaceae; Escherichia; Escherichia coli MTGDNTLIHSHGINRRDFMKLCAALAATMGLSSKAAAEMAESVTNPQRPPVIWIGAQECT WP_000145410.1 MULTISPECIES: hydrogenase-2 small chain [Proteobacteria] Length: 372\nNumber of predicted TMHs: 1\nExp number of AAs in TMHs: 23.72226\nExp number, first 60 AAs: 0.22038\nTotal prob of N-in: 0.12994\noutside 1 331\nTMhelix 332 354\ninside 355 372

12526 GCF_000303915.2_ASM30391v2 Escherichia coli PA49 Proteobacteria; Gammaproteobacteria; Enterobacterales; Enterobacteriaceae; Escherichia; Escherichia coli MNNEETFYQAMRRQGVTRRSFLKYCSLAATSLGLGAGMAPKIAWALENKPRIPVVWIHGL WP_001301613.1 hydrogenase [Escherichia coli] Length: 372\nNumber of predicted TMHs: 2\nExp number of AAs in TMHs: 37.46598\nExp number, first 60 AAs: 19.53139\nTotal prob of N-in: 0.94364\nPOSSIBLE N-term signal sequence\ninside 1 19\nTMhelix 20 39\noutside 40 325\nTMhelix 326 348\ninside 349 372

12527 GCF_000267905.2_ASM26790v2 Escherichia coli PA5 Proteobacteria; Gammaproteobacteria; Enterobacterales; Enterobacteriaceae; Escherichia; Escherichia coli

MNNEETFYQAMRRQGVTRRSFLKYCSLAATSLGLGAGMAPKIAWALENKPRIPVVWIHGL WP_001301613.1 hydrogenase [Escherichia coli] Length: 372\nNumber of predicted TMHs: 2\nExp number of AAs in TMHs: 37.46598\nExp number, first 60 AAs: 19.53139\nTotal prob of N-in: 0.94364\nPOSSIBLE N-term signal sequence\ninside 1 19\nTMhelix 20 39\noutside 40 325\nTMhelix 326 348\ninside 349 372

12528 GCF_000267905.2_ASM26790v2 Escherichia coli PA5 Proteobacteria; Gammaproteobacteria; Enterobacterales; Enterobacteriaceae; Escherichia; Escherichia coli MTGDNTLIHSHGINRRDFMKLCAALAATMGLSSKAAAEMAESVTNPQRPPVIWIGAQECT WP_000145410.1 MULTISPECIES: hydrogenase-2 small chain [Proteobacteria] Length: 372\nNumber of predicted TMHs: 1\nExp number of AAs in TMHs: 23.72226\nExp number, first 60 AAs: 0.22038\nTotal prob of N-in: 0.12994\noutside 1 331\nTMhelix 332 354\ninside 355 372

12529 GCF_000302735.1_ASM30273v1 Escherichia coli PA7 Proteobacteria; Gammaproteobacteria; Enterobacterales; Enterobacteriaceae; Escherichia; Escherichia coli MNNEETFYQAMRRQGVTRRSFLKYCSLAATSLGLGAGMAPKIAWALENKPRIPVVWIHGL WP_001301613.1 hydrogenase [Escherichia coli] Length: 372\nNumber of predicted TMHs: 2\nExp number of AAs in TMHs: 37.46598\nExp number, first 60 AAs: 19.53139\nTotal prob of N-in: 0.94364\nPOSSIBLE N-term signal sequence\ninside 1 19\nTMhelix 20 39\noutside 40 325\nTMhelix 326 348\ninside 349 372

12530 GCF_000302735.1_ASM30273v1 Escherichia coli PA7 Proteobacteria; Gammaproteobacteria; Enterobacterales; Enterobacteriaceae; Escherichia; Escherichia coli MTGDNTLIHSHGINRRDFMKLCAALAATMGLSSKAAAEMAESVTNPQRPPVIWIGAQECT WP_000145410.1 MULTISPECIES: hydrogenase-2 small chain [Proteobacteria] Length: 372\nNumber of predicted TMHs: 1\nExp number of AAs in TMHs: 23.72226\nExp number, first 60 AAs: 0.22038\nTotal prob of N-in: 0.12994\noutside 1 331\nTMhelix 332 354\ninside 355 372

12531 GCF_000335455.2_ASM33545v2 Escherichia coli PA8 Proteobacteria; Gammaproteobacteria; Enterobacterales; Enterobacteriaceae; Escherichia; Escherichia coli MNNEETFYQAMRRQGVTRRSFLKYCSLAATSLGLGAGMAPKIAWALENKPRIPVVWIHGL WP_001301613.1 hydrogenase [Escherichia coli] Length: 372\nNumber of predicted TMHs: 2\nExp number of AAs in TMHs: 37.46598\nExp number, first 60 AAs: 19.53139\nTotal prob of N-in: 0.94364\nPOSSIBLE N-term signal sequence\ninside 1 19\nTMhelix 20 39\noutside 40 325\nTMhelix 326 348\ninside 349 372

12532 GCF_000335455.2_ASM33545v2 Escherichia coli PA8 Proteobacteria; Gammaproteobacteria; Enterobacterales; Enterobacteriaceae; Escherichia; Escherichia coli MTGDNTLIHSHGINRRDFMKLCAALAATMGLSSKAAAEMAESVTNPQRPPVIWIGAQECT WP_000145410.1 MULTISPECIES: hydrogenase-2 small chain [Proteobacteria] Length: 372\nNumber of predicted TMHs: 1\nExp number of AAs in TMHs: 23.72226\nExp number, first 60 AAs: 0.22038\nTotal prob of N-in: 0.12994\noutside 1 331\nTMhelix 332 354\ninside 355 372

12533 GCF_000267885.2_ASM26788v2 Escherichia coli PA9 Proteobacteria; Gammaproteobacteria; Enterobacterales; Enterobacteriaceae; Escherichia; Escherichia coli MNNEETFYQAMRRQGVTRRSFLKYCSLAATSLGLGAGMAPKIAWALENKPRIPVVWIHGL WP_001301613.1 hydrogenase [Escherichia coli] Length: 372\nNumber of predicted TMHs: 2\nExp number of AAs in TMHs: 37.46598\nExp number, first 60 AAs: 19.53139\nTotal prob of N-in: 0.94364\nPOSSIBLE N-term signal sequence\ninside 1 19\nTMhelix 20 39\noutside 40 325\nTMhelix 326 348\ninside 349 372

12534 GCF_000267885.2_ASM26788v2 Escherichia coli PA9 Proteobacteria; Gammaproteobacteria; Enterobacterales; Enterobacteriaceae; Escherichia; Escherichia coli MTGDNTLIHSHGINRRDFMKLCAALAATMGLSSKAAAEMAESVTNPQRPPVIWIGAQECT WP_000145410.1 MULTISPECIES: hydrogenase-2 small chain [Proteobacteria] Length: 372\nNumber of predicted TMHs: 1\nExp number of AAs in TMHs: 23.72226\nExp number, first 60 AAs: 0.22038\nTotal prob of N-in: 0.12994\noutside 1 331\nTMhelix 332 354\ninside 355 372

12535 GCF_001642285.1_ASM164228v1 Escherichia coli PCN009 Proteobacteria; Gammaproteobacteria; Enterobacterales; Enterobacteriaceae; Escherichia; Escherichia coli MNNEETFYQAMRRQGVTRRSFLKYCSLAATSLGLGAGMAPKIAWALENKPRIPVVWIHGL WP_001058323.1 MULTISPECIES: hydrogenase-1 small chain [Proteobacteria] Length: 372\nNumber of predicted TMHs: 2\nExp number of AAs in TMHs: 37.2608499999999\nExp number, first 60 AAs: 19.52923\nTotal prob of N-in: 0.94351\nPOSSIBLE N-term signal sequence\ninside 1 19\nTMhelix 20 39\noutside 40 325\nTMhelix 326 348\ninside 349 372

12536 GCF_001642285.1_ASM164228v1 Escherichia coli PCN009 Proteobacteria; Gammaproteobacteria; Enterobacterales; Enterobacteriaceae; Escherichia; Escherichia coli
MTGDNTLIHSHGINRRDFMKLCAALAATMGLSSKAAAEMAESVTNPQRPPVIWIGAQECT WP_032298447.1
hydrogenase 2 small subunit [Escherichia coli] Length: 372\nNumber of predicted TMHs: 1\nExp number of AAs in TMHs: 23.69807\nExp number, first 60 AAs: 0.221\nTotal prob of N-in: 0.12853\noutside 1 331\nTMhelix 332 354\ninside 355 372

12537 GCF_000219515.2_ASM21951v3 Escherichia coli PCN033 Proteobacteria; Gammaproteobacteria; Enterobacterales; Enterobacteriaceae; Escherichia; Escherichia coli
MTGDNTLIHSHGINRRDFMKLCAALAATMGLSSKAAAEMAESVTNPQRPPVIWIGAQECT WP_000145410.1
MULTISPECIES: hydrogenase-2 small chain [Proteobacteria] Length: 372\nNumber of predicted TMHs: 1\nExp number of AAs in TMHs: 23.72226\nExp number, first 60 AAs: 0.22038\nTotal prob of N-in: 0.12994\noutside 1 331\nTMhelix 332 354\ninside 355 372

12538 GCF_000219515.2_ASM21951v3 Escherichia coli PCN033 Proteobacteria; Gammaproteobacteria; Enterobacterales; Enterobacteriaceae; Escherichia; Escherichia coli
MNNEETFYQAMRRQGVTRRSFLKYCSLAATSLGLGAGMAPKIAWALENKPRIPVVWIHGL WP_001058323.1
MULTISPECIES: hydrogenase-1 small chain [Proteobacteria] Length: 372\nNumber of predicted TMHs: 2\nExp number of AAs in TMHs: 37.2608499999999\nExp number, first 60 AAs: 19.52923\nTotal prob of N-in: 0.94351\nPOSSIBLE N-term signal sequence\ninside 1 19\nTMhelix 20 39\noutside 40 325\nTMhelix 326 348\ninside 349 372

12539 GCF_001029125.1_ASM102912v1 Escherichia coli PCN061 Proteobacteria; Gammaproteobacteria; Enterobacterales; Enterobacteriaceae; Escherichia; Escherichia coli
MTGDNTLIHSHGINRRDFMKLCAALAATMGLSSKAAAEMAESVTNPQRPPVIWIGAQECT WP_032298447.1
hydrogenase 2 small subunit [Escherichia coli] Length: 372\nNumber of predicted TMHs: 1\nExp number of AAs in TMHs: 23.69807\nExp number, first 60 AAs: 0.221\nTotal prob of N-in: 0.12853\noutside 1 331\nTMhelix 332 354\ninside 355 372

12540 GCF_001029125.1_ASM102912v1 Escherichia coli PCN061 Proteobacteria; Gammaproteobacteria; Enterobacterales; Enterobacteriaceae; Escherichia; Escherichia coli
MNNEETFYQAMRRQGVTRRSFLKYCSLAATSLGLGAGMAPKIAWALENKPRIPVVWIHGL WP_001058323.1
MULTISPECIES: hydrogenase-1 small chain [Proteobacteria] Length: 372\nNumber of predicted TMHs: 2\nExp number of AAs in TMHs: 37.2608499999999\nExp number, first 60 AAs: 19.52923\nTotal prob of N-in: 0.94351\nPOSSIBLE N-term signal sequence\ninside 1 19\nTMhelix 20 39\noutside 40 325\nTMhelix 326 348\ninside 349 372

12541 GCF_001642295.1_ASM164229v1 Escherichia coli PCN079 Proteobacteria; Gammaproteobacteria; Enterobacterales; Enterobacteriaceae; Escherichia; Escherichia coli
MTGDNTLIHSHGINRRDFMKLCAALAATMGLSSKAAAEMAESVTNPQRPPVIWIGAQECT WP_000145410.1
MULTISPECIES: hydrogenase-2 small chain [Proteobacteria] Length: 372\nNumber of predicted TMHs: 1\nExp number of AAs in TMHs: 23.72226\nExp number, first 60 AAs: 0.22038\nTotal prob of N-in: 0.12994\noutside 1 331\nTMhelix 332 354\ninside 355 372

12542 GCF_001642295.1_ASM164229v1 Escherichia coli PCN079 Proteobacteria; Gammaproteobacteria; Enterobacterales; Enterobacteriaceae; Escherichia; Escherichia coli
MNNEETFYQAMRRQGVTRRSFLKYCSLAATSLGLGAGMAPKIAWALENKPRIPVVWIHGL WP_001058323.1
MULTISPECIES: hydrogenase-1 small chain [Proteobacteria] Length: 372\nNumber of predicted TMHs: 2\nExp number of AAs in TMHs: 37.2608499999999\nExp number, first 60 AAs: 19.52923\nTotal prob of N-in: 0.94351\nPOSSIBLE N-term signal sequence\ninside 1 19\nTMhelix 20 39\noutside 40 325\nTMhelix 326 348\ninside 349 372

12543 GCF_000493595.2_EcoPMV1 Escherichia coli PMV-1 Proteobacteria; Gammaproteobacteria; Enterobacterales; Enterobacteriaceae; Escherichia; Escherichia coli
MTGDNTLIHSHGINRRDFMKLCAALAATMGLSSKAAAEMAESVTNPQRPPVIWIGAQECT WP_000145410.1
MULTISPECIES: hydrogenase-2 small chain [Proteobacteria] Length: 372\nNumber of predicted TMHs: 1\nExp number of AAs in TMHs: 23.72226\nExp number, first 60 AAs: 0.22038\nTotal prob of N-in: 0.12994\noutside 1 331\nTMhelix 332 354\ninside 355 372

12544 GCF_002110005.1_ASM211000v1 Escherichia coli R424 Proteobacteria; Gammaproteobacteria; Enterobacterales; Enterobacteriaceae; Escherichia; Escherichia coli

MNNEETFYQAMRRQGVTRRSFLKYCSLAATSLGLGAGMAPKIAWALENKPRIPVVWIHGL WP_001058323.1 MULTISPECIES: hydrogenase-1 small chain [Proteobacteria] Length: 372\nNumber of predicted TMHs: 2\nExp number of AAs in TMHs: 37.2608499999999\nExp number, first 60 AAs: 19.52923\nTotal prob of N-in: 0.94351\nPOSSIBLE N-term signal sequence\ninside 1 19\nTMhelix 20 39\noutside 40 325\nTMhelix 326 348\ninside 349 372

12545 GCF_002110005.1_ASM211000v1 Escherichia coli R424 Proteobacteria; Gammaproteobacteria; Enterobacterales; Enterobacteriaceae; Escherichia; Escherichia coli MTGDNTLIHSHGINRRDFMKLCAALAATMGLSSKAAAEMAESVTNPQRPPVIWIGAQECT WP_085452583.1 hydrogenase 2 small subunit [Escherichia coli] Length: 372\nNumber of predicted TMHs: 1\nExp number of AAs in TMHs: 24.09491\nExp number, first 60 AAs: 0.22003\nTotal prob of N-in: 0.12522\noutside 1 331\nTMhelix 332 354\ninside 355 372

12546 GCF_002109575.1_ASM210957v1 Escherichia coli R527 Proteobacteria; Gammaproteobacteria; Enterobacterales; Enterobacteriaceae; Escherichia; Escherichia coli MTGDNTLIHSHGINRRDFMKLCAALAATMGLSSKAAAEMAESVTNPQRPPVIWIGAQECT WP_000145410.1 MULTISPECIES: hydrogenase-2 small chain [Proteobacteria] Length: 372\nNumber of predicted TMHs: 1\nExp number of AAs in TMHs: 23.72226\nExp number, first 60 AAs: 0.22038\nTotal prob of N-in: 0.12994\noutside 1 331\nTMhelix 332 354\ninside 355 372

12547 GCF_000188875.1_ASM18887v2 Escherichia coli RN587/1 Proteobacteria; Gammaproteobacteria; Enterobacterales; Enterobacteriaceae; Escherichia; Escherichia coli MTGDNTLIHSHGINRRDFMKLCAALAATMGLSSKAAAEMAESVTNPQRPPVIWIGAQECT WP_000145410.1 MULTISPECIES: hydrogenase-2 small chain [Proteobacteria] Length: 372\nNumber of predicted TMHs: 1\nExp number of AAs in TMHs: 23.72226\nExp number, first 60 AAs: 0.22038\nTotal prob of N-in: 0.12994\noutside 1 331\nTMhelix 332 354\ninside 355 372

12548 GCF_000188875.1_ASM18887v2 Escherichia coli RN587/1 Proteobacteria; Gammaproteobacteria; Enterobacterales; Enterobacteriaceae; Escherichia; Escherichia coli MNNEETFYQAMRRQGVTRRSFLKYCSLAATSLGLGAGMAPKIAWALENKPRIPVVWIHGL WP_001058323.1 MULTISPECIES: hydrogenase-1 small chain [Proteobacteria] Length: 372\nNumber of predicted TMHs: 2\nExp number of AAs in TMHs: 37.2608499999999\nExp number, first 60 AAs: 19.52923\nTotal prob of N-in: 0.94351\nPOSSIBLE N-term signal sequence\ninside 1 19\nTMhelix 20 39\noutside 40 325\nTMhelix 326 348\ninside 349 372

12549 GCF_000800845.1_ASM80084v2 Escherichia coli RS218 Proteobacteria; Gammaproteobacteria; Enterobacterales; Enterobacteriaceae; Escherichia; Escherichia coli MTGDNTLIHSHGINRRDFMKLCAALAATMGLSSKAAAEMAESVTNPQRPPVIWIGAQECT WP_000145410.1 MULTISPECIES: hydrogenase-2 small chain [Proteobacteria] Length: 372\nNumber of predicted TMHs: 1\nExp number of AAs in TMHs: 23.72226\nExp number, first 60 AAs: 0.22038\nTotal prob of N-in: 0.12994\noutside 1 331\nTMhelix 332 354\ninside 355 372

12550 GCF_000817345.1_ASM81734v1 Escherichia coli RS218 Proteobacteria; Gammaproteobacteria; Enterobacterales; Enterobacteriaceae; Escherichia; Escherichia coli MTGDNTLIHSHGINRRDFMKLCAALAATMGLSSKAAAEMAESVTNPQRPPVIWIGAQECT WP_000145410.1 MULTISPECIES: hydrogenase-2 small chain [Proteobacteria] Length: 372\nNumber of predicted TMHs: 1\nExp number of AAs in TMHs: 23.72226\nExp number, first 60 AAs: 0.22038\nTotal prob of N-in: 0.12994\noutside 1 331\nTMhelix 332 354\ninside 355 372

12551 GCF_000340255.1_S17 Escherichia coli S17 Proteobacteria; Gammaproteobacteria; Enterobacterales; Enterobacteriaceae; Escherichia; Escherichia coli MTGDNTLIHSHGINRRDFMKLCAALAATMGLSSKAAAEMAESVTNPQRPPVIWIGAQECT WP_000145410.1 MULTISPECIES: hydrogenase-2 small chain [Proteobacteria] Length: 372\nNumber of predicted TMHs: 1\nExp number of AAs in TMHs: 23.72226\nExp number, first 60 AAs: 0.22038\nTotal prob of N-in: 0.12994\noutside 1 331\nTMhelix 332 354\ninside 355 372

12552 GCF_000340255.1_S17 Escherichia coli S17 Proteobacteria; Gammaproteobacteria; Enterobacterales; Enterobacteriaceae; Escherichia; Escherichia coli MNNEETFYQAMRRQGVTRRSFLKYCSLAATSLGLGAGMAPKIAWALENKPRIPVVWIHGL WP_001058323.1 MULTISPECIES: hydrogenase-1 small chain [Proteobacteria] Length: 372\nNumber of predicted TMHs: 2\nExp number of AAs in TMHs: 37.2608499999999\nExp number, first 60 AAs: 19.52923\nTotal prob of N-in:

0.94351\nPOSSIBLE N-term signal sequence\ninside 1 19\nTMhelix 20 39\noutside 40 325\nTMhelix 326 348\ninside 349 372

12553 GCF_000482045.1_Escherichia_coli_SCD1 Escherichia coli SCD1 Proteobacteria; Gammaproteobacteria; Enterobacterales; Enterobacteriaceae; Escherichia; Escherichia coli
MNNEETFYQAMRRQGVTRRSFLKYCSLAATSLGLGAGMAPKIAWALENKPRIPVVWIHGL WP_001058323.1
MULTISPECIES: hydrogenase-1 small chain [Proteobacteria] Length: 372\nNumber of predicted TMHs: 2\nExp number of AAs in TMHs: 37.2608499999999\nExp number, first 60 AAs: 19.52923\nTotal prob of N-in: 0.94351\nPOSSIBLE N-term signal sequence\ninside 1 19\nTMhelix 20 39\noutside 40 325\nTMhelix 326 348\ninside 349 372

12554 GCF_000482065.1_Escherichia_coli_SCD2 Escherichia coli SCD2 Proteobacteria; Gammaproteobacteria; Enterobacterales; Enterobacteriaceae; Escherichia; Escherichia coli
MTGDNTLIHSHGINRRDFMKLCAALAATMGLSSKAAAEMAESVTNPQRPPVIWIGAQECT WP_000145410.1
MULTISPECIES: hydrogenase-2 small chain [Proteobacteria] Length: 372\nNumber of predicted TMHs: 1\nExp number of AAs in TMHs: 23.72226\nExp number, first 60 AAs: 0.22038\nTotal prob of N-in: 0.12994\noutside 1 331\nTMhelix 332 354\ninside 355 372

12555 GCF_000482065.1_Escherichia_coli_SCD2 Escherichia coli SCD2 Proteobacteria; Gammaproteobacteria; Enterobacterales; Enterobacteriaceae; Escherichia; Escherichia coli
MNNEETFYQAMRRQGVTRRSFLKYCSLAATSLGLGAGMAPKIAWALENKPRIPVVWIHGL WP_001058323.1
MULTISPECIES: hydrogenase-1 small chain [Proteobacteria] Length: 372\nNumber of predicted TMHs: 2\nExp number of AAs in TMHs: 37.2608499999999\nExp number, first 60 AAs: 19.52923\nTotal prob of N-in: 0.94351\nPOSSIBLE N-term signal sequence\ninside 1 19\nTMhelix 20 39\noutside 40 325\nTMhelix 326 348\ninside 349 372

12556 GCF_000252805.1_ASM25280v2 Escherichia coli SCI-07 Proteobacteria; Gammaproteobacteria; Enterobacterales; Enterobacteriaceae; Escherichia; Escherichia coli
MNNEETFYQAMRRQGVTRRSFLKYCSLAATSLGLGAGMAPKIAWALENKPRIPVVWIHGL WP_001058323.1
MULTISPECIES: hydrogenase-1 small chain [Proteobacteria] Length: 372\nNumber of predicted TMHs: 2\nExp number of AAs in TMHs: 37.2608499999999\nExp number, first 60 AAs: 19.52923\nTotal prob of N-in: 0.94351\nPOSSIBLE N-term signal sequence\ninside 1 19\nTMhelix 20 39\noutside 40 325\nTMhelix 326 348\ninside 349 372

12557 GCF_000252805.1_ASM25280v2 Escherichia coli SCI-07 Proteobacteria; Gammaproteobacteria; Enterobacterales; Enterobacteriaceae; Escherichia; Escherichia coli
MTGDNTLIHSHGINRRDFMKLCAALAATMGLSSKAAAEMAESVTNPQRPPVIWIGAQECT WP_000145410.1
MULTISPECIES: hydrogenase-2 small chain [Proteobacteria] Length: 372\nNumber of predicted TMHs: 1\nExp number of AAs in TMHs: 23.72226\nExp number, first 60 AAs: 0.22038\nTotal prob of N-in: 0.12994\noutside 1 331\nTMhelix 332 354\ninside 355 372

12558 GCF_000010385.1_ASM1038v1 Escherichia coli SE11 Proteobacteria; Gammaproteobacteria; Enterobacterales; Enterobacteriaceae; Escherichia; Escherichia coli
MNNEETFYQAMRRQGVTRRSFLKYCSLAATSLGLGAGMAPKIAWALENKPRIPVVWIHGL WP_001295940.1
MULTISPECIES: hydrogenase [Enterobacteriaceae] Length: 372\nNumber of predicted TMHs: 2\nExp number of AAs in TMHs: 37.2608899999999\nExp number, first 60 AAs: 19.52923\nTotal prob of N-in: 0.94351\nPOSSIBLE N-term signal sequence\ninside 1 19\nTMhelix 20 39\noutside 40 325\nTMhelix 326 348\ninside 349 372

12559 GCF_000010385.1_ASM1038v1 Escherichia coli SE11 Proteobacteria; Gammaproteobacteria; Enterobacterales; Enterobacteriaceae; Escherichia; Escherichia coli
MTGDNTLIHSHGINRRDFMKLCAALAATMGLSSKAAAEMAESVTNPQRPPVIWIGAQECT WP_000145410.1
MULTISPECIES: hydrogenase-2 small chain [Proteobacteria] Length: 372\nNumber of predicted TMHs: 1\nExp number of AAs in TMHs: 23.72226\nExp number, first 60 AAs: 0.22038\nTotal prob of N-in: 0.12994\noutside 1 331\nTMhelix 332 354\ninside 355 372

12560 GCF_000010485.1_ASM1048v1 Escherichia coli SE15 Proteobacteria; Gammaproteobacteria; Enterobacterales; Enterobacteriaceae; Escherichia; Escherichia coli
MTGDNTLIHSHGINRRDFMKLCAALAATMGLSSKAAAEMAESVTNPQRPPVIWIGAQECT WP_000145410.1
MULTISPECIES: hydrogenase-2 small chain [Proteobacteria] Length: 372\nNumber of predicted TMHs:

1\nExp number of AAs in TMHs: 23.72226\nExp number, first 60 AAs: 0.22038\nTotal prob of N-in: 0.12994\noutside 1 331\nTMhelix 332 354\ninside 355 372

12561 GCF_000010485.1_ASM1048v1 Escherichia coli SE15 Proteobacteria; Gammaproteobacteria; Enterobacterales; Enterobacteriaceae; Escherichia; Escherichia coli
MNNEETFYQAMRRQGVTRRSFLKYCSLAATSLGLGAGMAPKIAWALENKPRIPVVWIHGL WP_001058323.1
MULTISPECIES: hydrogenase-1 small chain [Proteobacteria] Length: 372\nNumber of predicted TMHs: 2\nExp number of AAs in TMHs: 37.2608499999999\nExp number, first 60 AAs: 19.52923\nTotal prob of N-in: 0.94351\nPOSSIBLE N-term signal sequence\ninside 1 19\nTMhelix 20 39\noutside 40 325\nTMhelix 326 348\ninside 349 372

12562 GCF_000340275.1_SEPT362 Escherichia coli SEPT362 Proteobacteria; Gammaproteobacteria; Enterobacterales; Enterobacteriaceae; Escherichia; Escherichia coli
MNNEETFYQAMRRQGVTRRSFLKYCSLAATSLGLGAGMAPKIAWALENKPRIPVVWIHGL WP_001058323.1
MULTISPECIES: hydrogenase-1 small chain [Proteobacteria] Length: 372\nNumber of predicted TMHs: 2\nExp number of AAs in TMHs: 37.2608499999999\nExp number, first 60 AAs: 19.52923\nTotal prob of N-in: 0.94351\nPOSSIBLE N-term signal sequence\ninside 1 19\nTMhelix 20 39\noutside 40 325\nTMhelix 326 348\ninside 349 372

12563 GCF_000340275.1_SEPT362 Escherichia coli SEPT362 Proteobacteria; Gammaproteobacteria; Enterobacterales; Enterobacteriaceae; Escherichia; Escherichia coli
MTGDNTLIHSHGINRRDFMKLCAALAATMGLSSKAAAEMAESVTNPQRPPVIWIGAQECT WP_000145410.1
MULTISPECIES: hydrogenase-2 small chain [Proteobacteria] Length: 372\nNumber of predicted TMHs: 1\nExp number of AAs in TMHs: 23.72226\nExp number, first 60 AAs: 0.22038\nTotal prob of N-in: 0.12994\noutside 1 331\nTMhelix 332 354\ninside 355 372

12564 GCF_002109305.1_E_coli_SHECO001_v1 Escherichia coli SHECO001 Proteobacteria; Gammaproteobacteria; Enterobacterales; Enterobacteriaceae; Escherichia; Escherichia coli
MNNEETFYQAMRRQGVTRRSFLKYCSLAATSLGLGAGMAPKIAWALENKPRIPVVWIHGL WP_001058323.1
MULTISPECIES: hydrogenase-1 small chain [Proteobacteria] Length: 372\nNumber of predicted TMHs: 2\nExp number of AAs in TMHs: 37.2608499999999\nExp number, first 60 AAs: 19.52923\nTotal prob of N-in: 0.94351\nPOSSIBLE N-term signal sequence\ninside 1 19\nTMhelix 20 39\noutside 40 325\nTMhelix 326 348\ninside 349 372

12565 GCF_002109305.1_E_coli_SHECO001_v1 Escherichia coli SHECO001 Proteobacteria; Gammaproteobacteria; Enterobacterales; Enterobacteriaceae; Escherichia; Escherichia coli
MTGDNTLIHSHGINRRDFMKLCAALAATMGLSSKAAAEMAESVTNPQRPPVIWIGAQECT WP_000145410.1
MULTISPECIES: hydrogenase-2 small chain [Proteobacteria] Length: 372\nNumber of predicted TMHs: 1\nExp number of AAs in TMHs: 23.72226\nExp number, first 60 AAs: 0.22038\nTotal prob of N-in: 0.12994\noutside 1 331\nTMhelix 332 354\ninside 355 372

12566 GCF_002111485.1_E_coli_SHECO002_v1 Escherichia coli SHECO002 Proteobacteria; Gammaproteobacteria; Enterobacterales; Enterobacteriaceae; Escherichia; Escherichia coli
MTGDNTLIHSHGINRRDFMKLCAALAATMGLSSKAAAEMAESVTNPQRPPVIWIGAQECT WP_000145410.1
MULTISPECIES: hydrogenase-2 small chain [Proteobacteria] Length: 372\nNumber of predicted TMHs: 1\nExp number of AAs in TMHs: 23.72226\nExp number, first 60 AAs: 0.22038\nTotal prob of N-in: 0.12994\noutside 1 331\nTMhelix 332 354\ninside 355 372

12567 GCF_002111485.1_E_coli_SHECO002_v1 Escherichia coli SHECO002 Proteobacteria; Gammaproteobacteria; Enterobacterales; Enterobacteriaceae; Escherichia; Escherichia coli
MNNEETFYQAMRRQGVTRRSFLKYCSLAATSLGLGAGMAPKIAWALENKPRIPVVWIHGL WP_001301613.1
hydrogenase [Escherichia coli] Length: 372\nNumber of predicted TMHs: 2\nExp number of AAs in TMHs: 37.46598\nExp number, first 60 AAs: 19.53139\nTotal prob of N-in: 0.94364\nPOSSIBLE N-term signal sequence\ninside 1 19\nTMhelix 20 39\noutside 40 325\nTMhelix 326 348\ninside 349 372

12568 GCF_002111495.1_E_coli_SHECO003_v1 Escherichia coli SHECO003 Proteobacteria; Gammaproteobacteria; Enterobacterales; Enterobacteriaceae; Escherichia; Escherichia coli
MNNEETFYQAMRRQGVTRRSFLKYCSLAATSLGLGAGMAPKIAWALENKPRIPVVWIHGL WP_001301613.1
hydrogenase [Escherichia coli] Length: 372\nNumber of predicted TMHs: 2\nExp number of AAs in TMHs: 37.46598\nExp number, first 60 AAs: 19.53139\nTotal prob of N-in: 0.94364\nPOSSIBLE N-term signal sequence\ninside 1 19\nTMhelix 20 39\noutside 40 325\nTMhelix 326 348\ninside 349 372

12569 GCF_002111495.1_E_coli_SHECO003_v1 Escherichia coli SHECO003 Proteobacteria; Gammaproteobacteria; Enterobacterales; Enterobacteriaceae; Escherichia; Escherichia coli
MTGDNTLIHSHGINRRDFMKLCAALAATMGLSSKAAAEMAESVTNPQRPPVIWIGAQECT WP_000145410.1
MULTISPECIES: hydrogenase-2 small chain [Proteobacteria] Length: 372\nNumber of predicted TMHs: 1\nExp number of AAs in TMHs: 23.72226\nExp number, first 60 AAs: 0.22038\nTotal prob of N-in: 0.12994\noutside 1 331\nTMhelix 332 354\ninside 355 372

12570 GCF_000019645.1_ASM1964v1 Escherichia coli SMS-3-5 Proteobacteria; Gammaproteobacteria; Enterobacterales; Enterobacteriaceae; Escherichia; Escherichia coli
MTGDNTLIHSHGINRRDFMKLCAALAATMGLSSKAAAEMAESVTNPQRPPVIWIGAQECT WP_000145410.1
MULTISPECIES: hydrogenase-2 small chain [Proteobacteria] Length: 372\nNumber of predicted TMHs: 1\nExp number of AAs in TMHs: 23.72226\nExp number, first 60 AAs: 0.22038\nTotal prob of N-in: 0.12994\noutside 1 331\nTMhelix 332 354\ninside 355 372

12571 GCF_000019645.1_ASM1964v1 Escherichia coli SMS-3-5 Proteobacteria; Gammaproteobacteria; Enterobacterales; Enterobacteriaceae; Escherichia; Escherichia coli
MNNEETFYQAMRRQGVTRRSFLKYCSLAATSLGLGAGMAPKIAWALENKPRIPVVWIHGL WP_001058323.1
MULTISPECIES: hydrogenase-1 small chain [Proteobacteria] Length: 372\nNumber of predicted TMHs: 2\nExp number of AAs in TMHs: 37.2608499999999\nExp number, first 60 AAs: 19.52923\nTotal prob of N-in: 0.94351\nPOSSIBLE N-term signal sequence\ninside 1 19\nTMhelix 20 39\noutside 40 325\nTMhelix 326 348\ninside 349 372

12572 GCF_000194575.1_ASM19457v2 Escherichia coli STEC_7v Proteobacteria; Gammaproteobacteria; Enterobacterales; Enterobacteriaceae; Escherichia; Escherichia coli
MNNEETFYQAMRRQGVTRRSFLKYCSLAATSLGLGAGMAPKIAWALENKPRIPVVWIHGL WP_001353311.1
MULTISPECIES: hydrogenase [Escherichia] Length: 372\nNumber of predicted TMHs: 2\nExp number of AAs in TMHs: 37.2638699999999\nExp number, first 60 AAs: 19.52924\nTotal prob of N-in: 0.94351\nPOSSIBLE N-term signal sequence\ninside 1 19\nTMhelix 20 39\noutside 40 325\nTMhelix 326 348\ninside 349 372

12573 GCF_000194575.1_ASM19457v2 Escherichia coli STEC_7v Proteobacteria; Gammaproteobacteria; Enterobacterales; Enterobacteriaceae; Escherichia; Escherichia coli
MTGDNTLIHSHGINRRDFMKLCAALAATMGLSSKAAAEMAESVTNPQRPPVIWIGAQECT WP_000145410.1
MULTISPECIES: hydrogenase-2 small chain [Proteobacteria] Length: 372\nNumber of predicted TMHs: 1\nExp number of AAs in TMHs: 23.72226\nExp number, first 60 AAs: 0.22038\nTotal prob of N-in: 0.12994\noutside 1 331\nTMhelix 332 354\ninside 355 372

12574 GCF_000225025.1_ASM22502v2 Escherichia coli STEC_B2F1 Proteobacteria; Gammaproteobacteria; Enterobacterales; Enterobacteriaceae; Escherichia; Escherichia coli
MTGDNTLIHSHGINRRDFMKLCAALAATMGLSSKAAAEMAESVTNPQRPPVIWIGAQECT WP_000145410.1
MULTISPECIES: hydrogenase-2 small chain [Proteobacteria] Length: 372\nNumber of predicted TMHs: 1\nExp number of AAs in TMHs: 23.72226\nExp number, first 60 AAs: 0.22038\nTotal prob of N-in: 0.12994\noutside 1 331\nTMhelix 332 354\ninside 355 372

12575 GCF_000225025.1_ASM22502v2 Escherichia coli STEC_B2F1 Proteobacteria; Gammaproteobacteria; Enterobacterales; Enterobacteriaceae; Escherichia; Escherichia coli
MNNEETFYQAMRRQGVTRRSFLKYCSLAATSLGLGAGMAPKIAWALENKPRIPVVWIHGL WP_001295940.1
MULTISPECIES: hydrogenase [Enterobacteriaceae] Length: 372\nNumber of predicted TMHs: 2\nExp number of AAs in TMHs: 37.2608899999999\nExp number, first 60 AAs: 19.52923\nTotal prob of N-in: 0.94351\nPOSSIBLE N-term signal sequence\ninside 1 19\nTMhelix 20 39\noutside 40 325\nTMhelix 326 348\ninside 349 372

12576 GCF_000225045.1_ASM22504v2 Escherichia coli STEC_C165-02 Proteobacteria; Gammaproteobacteria; Enterobacterales; Enterobacteriaceae; Escherichia; Escherichia coli
MTGDNTLIHSHGINRRDFMKLCAALAATMGLSSKAAAEMAESVTNPQRPPVIWIGAQECT WP_000145410.1
MULTISPECIES: hydrogenase-2 small chain [Proteobacteria] Length: 372\nNumber of predicted TMHs: 1\nExp number of AAs in TMHs: 23.72226\nExp number, first 60 AAs: 0.22038\nTotal prob of N-in: 0.12994\noutside 1 331\nTMhelix 332 354\ninside 355 372

12577 GCF_000225045.1_ASM22504v2 Escherichia coli STEC_C165-02 Proteobacteria; Gammaproteobacteria; Enterobacterales; Enterobacteriaceae; Escherichia; Escherichia coli
MNNEETFYQAMRRQGVTRRSFLKYCSLAATSLGLGAGMAPKIAWALENKPRIPVVWIHGL WP_001058323.1

MULTISPECIES: hydrogenase-1 small chain [Proteobacteria] Length: 372\nNumber of predicted TMHs: 2\nExp number of AAs in TMHs: 37.2608499999999\nExp number, first 60 AAs: 19.52923\nTotal prob of N-in: 0.94351\nPOSSIBLE N-term signal sequence\ninside 1 19\nTMhelix 20 39\noutside 40 325\nTMhelix 326 348\ninside 349 372

12578 GCF_000225145.1_ASM22514v2 Escherichia coli STEC_EH250 Proteobacteria; Gammaproteobacteria; Enterobacterales; Enterobacteriaceae; Escherichia; Escherichia coli
MNNEETFYQAMRRQGVTRRSFLKYCSLAATSLGLGAGMAPKIAWALENKPRIPVVWIHGL WP_001058323.1
MULTISPECIES: hydrogenase-1 small chain [Proteobacteria] Length: 372\nNumber of predicted TMHs: 2\nExp number of AAs in TMHs: 37.2608499999999\nExp number, first 60 AAs: 19.52923\nTotal prob of N-in: 0.94351\nPOSSIBLE N-term signal sequence\ninside 1 19\nTMhelix 20 39\noutside 40 325\nTMhelix 326 348\ninside 349 372

12579 GCF_000225145.1_ASM22514v2 Escherichia coli STEC_EH250 Proteobacteria; Gammaproteobacteria; Enterobacterales; Enterobacteriaceae; Escherichia; Escherichia coli
MTGDNTLIHSHGINRRDFMKLCAALAATMGLSSKAAAEMAESVTNPQRPPVIWIGAQECT WP_000145410.1
MULTISPECIES: hydrogenase-2 small chain [Proteobacteria] Length: 372\nNumber of predicted TMHs: 1\nExp number of AAs in TMHs: 23.72226\nExp number, first 60 AAs: 0.22038\nTotal prob of N-in: 0.12994\noutside 1 331\nTMhelix 332 354\ninside 355 372

12580 GCF_000225205.1_ASM22520v2 Escherichia coli STEC_MHI813 Proteobacteria; Gammaproteobacteria; Enterobacterales; Enterobacteriaceae; Escherichia; Escherichia coli
MTGDNTLIHSHGINRRDFMKLCAALAATMGLSSKAAAEMAESVTNPQRPPVIWIGAQECT WP_000145410.1
MULTISPECIES: hydrogenase-2 small chain [Proteobacteria] Length: 372\nNumber of predicted TMHs: 1\nExp number of AAs in TMHs: 23.72226\nExp number, first 60 AAs: 0.22038\nTotal prob of N-in: 0.12994\noutside 1 331\nTMhelix 332 354\ninside 355 372

12581 GCF_000225205.1_ASM22520v2 Escherichia coli STEC_MHI813 Proteobacteria; Gammaproteobacteria; Enterobacterales; Enterobacteriaceae; Escherichia; Escherichia coli
MNNEETFYQAMRRQGVTRRSFLKYCSLAATSLGLGAGMAPKIAWALENKPRIPVVWIHGL WP_001058323.1
MULTISPECIES: hydrogenase-1 small chain [Proteobacteria] Length: 372\nNumber of predicted TMHs: 2\nExp number of AAs in TMHs: 37.2608499999999\nExp number, first 60 AAs: 19.52923\nTotal prob of N-in: 0.94351\nPOSSIBLE N-term signal sequence\ninside 1 19\nTMhelix 20 39\noutside 40 325\nTMhelix 326 348\ninside 349 372

12582 GCF_000281775.1_ASM28177v1 Escherichia coli STEC_O31 Proteobacteria; Gammaproteobacteria; Enterobacterales; Enterobacteriaceae; Escherichia; Escherichia coli
MNNEETFYQAMRRQGVTRRSFLKYCSLAATSLGLGAGMAPKIAWALENKPRIPVVWIHGL WP_001058323.1
MULTISPECIES: hydrogenase-1 small chain [Proteobacteria] Length: 372\nNumber of predicted TMHs: 2\nExp number of AAs in TMHs: 37.2608499999999\nExp number, first 60 AAs: 19.52923\nTotal prob of N-in: 0.94351\nPOSSIBLE N-term signal sequence\ninside 1 19\nTMhelix 20 39\noutside 40 325\nTMhelix 326 348\ninside 349 372

12583 GCF_000281775.1_ASM28177v1 Escherichia coli STEC_O31 Proteobacteria; Gammaproteobacteria; Enterobacterales; Enterobacteriaceae; Escherichia; Escherichia coli
MTGDNTLIHSHGINRRDFMKLCAALAATMGLSSKAAAEMAESVTNPQRPPVIWIGAQECT WP_000145410.1
MULTISPECIES: hydrogenase-2 small chain [Proteobacteria] Length: 372\nNumber of predicted TMHs: 1\nExp number of AAs in TMHs: 23.72226\nExp number, first 60 AAs: 0.22038\nTotal prob of N-in: 0.12994\noutside 1 331\nTMhelix 332 354\ninside 355 372

12584 GCF_000225225.1_ASM22522v2 Escherichia coli STEC_S1191 Proteobacteria; Gammaproteobacteria; Enterobacterales; Enterobacteriaceae; Escherichia; Escherichia coli
MNNEETFYQAMRRQGVTRRSFLKYCSLAATSLGLGAGMAPKIAWALENKPRIPVVWIHGL WP_001058323.1
MULTISPECIES: hydrogenase-1 small chain [Proteobacteria] Length: 372\nNumber of predicted TMHs: 2\nExp number of AAs in TMHs: 37.2608499999999\nExp number, first 60 AAs: 19.52923\nTotal prob of N-in: 0.94351\nPOSSIBLE N-term signal sequence\ninside 1 19\nTMhelix 20 39\noutside 40 325\nTMhelix 326 348\ninside 349 372

12585 GCF_000225225.1_ASM22522v2 Escherichia coli STEC_S1191 Proteobacteria; Gammaproteobacteria; Enterobacterales; Enterobacteriaceae; Escherichia; Escherichia coli
MTGDNTLIHSHGINRRDFMKLCAALAATMGLSSKAAAEMAESVTNPQRPPVIWIGAQECT WP_000145410.1

MULTISPECIES: hydrogenase-2 small chain [Proteobacteria] Length: 372\nNumber of predicted TMHs: 1\nExp number of AAs in TMHs: 23.72226\nExp number, first 60 AAs: 0.22038\nTotal prob of N-in: 0.12994\noutside 1 331\nTMhelix 332 354\ninside 355 372

12586 GCF_000233895.1_ASM23389v1 Escherichia coli str. clone Proteobacteria; Gammaproteobacteria; Enterobacterales; Enterobacteriaceae; Escherichia; Escherichia coli MNNEETFYQAMRRQGVTRRSFLKYCSLAATSLGLGAGMAPKIAWALENKPRIPVVWIHGL WP_001058323.1 MULTISPECIES: hydrogenase-1 small chain [Proteobacteria] Length: 372\nNumber of predicted TMHs: 2\nExp number of AAs in TMHs: 37.2608499999999\nExp number, first 60 AAs: 19.52923\nTotal prob of N-in: 0.94351\nPOSSIBLE N-term signal sequence\ninside 1 19\nTMhelix 20 39\noutside 40 325\nTMhelix 326 348\ninside 349 372

12587 GCF_000233895.1_ASM23389v1 Escherichia coli str. clone Proteobacteria; Gammaproteobacteria; Enterobacterales; Enterobacteriaceae; Escherichia; Escherichia coli MTGDNTLIHSHGINRRDFMKLCAALAATMGLSSKAAAEMAESVTNPQRPPVIWIGAQECT WP_000145410.1 MULTISPECIES: hydrogenase-2 small chain [Proteobacteria] Length: 372\nNumber of predicted TMHs: 1\nExp number of AAs in TMHs: 23.72226\nExp number, first 60 AAs: 0.22038\nTotal prob of N-in: 0.12994\noutside 1 331\nTMhelix 332 354\ninside 355 372

12588 GCF_000233875.1_ASM23387v1 Escherichia coli str. clone Proteobacteria; Gammaproteobacteria; Enterobacterales; Enterobacteriaceae; Escherichia; Escherichia coli MTGDNTLIHSHGINRRDFMKLCAALAATMGLSSKAAAEMAESVTNPQRPPVIWIGAQECT WP_000145410.1 MULTISPECIES: hydrogenase-2 small chain [Proteobacteria] Length: 372\nNumber of predicted TMHs: 1\nExp number of AAs in TMHs: 23.72226\nExp number, first 60 AAs: 0.22038\nTotal prob of N-in: 0.12994\noutside 1 331\nTMhelix 332 354\ninside 355 372

12589 GCF_000233875.1_ASM23387v1 Escherichia coli str. clone Proteobacteria; Gammaproteobacteria; Enterobacterales; Enterobacteriaceae; Escherichia; Escherichia coli MNNEETFYQAMRRQGVTRRSFLKYCSLAATSLGLGAGMAPKIAWALENKPRIPVVWIHGL WP_001058323.1 MULTISPECIES: hydrogenase-1 small chain [Proteobacteria] Length: 372\nNumber of predicted TMHs: 2\nExp number of AAs in TMHs: 37.2608499999999\nExp number, first 60 AAs: 19.52923\nTotal prob of N-in: 0.94351\nPOSSIBLE N-term signal sequence\ninside 1 19\nTMhelix 20 39\noutside 40 325\nTMhelix 326 348\ninside 349 372

12590 GCF_001447405.1_ASM144740v1 Escherichia coli str. Deng Proteobacteria; Gammaproteobacteria; Enterobacterales; Enterobacteriaceae; Escherichia; Escherichia coli MTGDNTLIHSHGINRRDFMKLCAALAATMGLSSKAAAEMAESVTNPQRPPVIWIGAQECT WP_000145410.1 MULTISPECIES: hydrogenase-2 small chain [Proteobacteria] Length: 372\nNumber of predicted TMHs: 1\nExp number of AAs in TMHs: 23.72226\nExp number, first 60 AAs: 0.22038\nTotal prob of N-in: 0.12994\noutside 1 331\nTMhelix 332 354\ninside 355 372

12591 GCF_001447405.1_ASM144740v1 Escherichia coli str. Deng Proteobacteria; Gammaproteobacteria; Enterobacterales; Enterobacteriaceae; Escherichia; Escherichia coli MNNEETFYQAMRRQGVTRRSFLKYCSLAATSLGLGAGMAPKIAWALENKPRIPVVWIHGL WP_058101158.1 hydrogenase [Escherichia coli] Length: 372\nNumber of predicted TMHs: 2\nExp number of AAs in TMHs: 36.5323399999999\nExp number, first 60 AAs: 19.52466\nTotal prob of N-in: 0.94357\nPOSSIBLE N-term signal sequence\ninside 1 19\nTMhelix 20 39\noutside 40 325\nTMhelix 326 348\ninside 349 372

12592 GCF_001610755.1_ASM161075v1 Escherichia coli str. Sanji Proteobacteria; Gammaproteobacteria; Enterobacterales; Enterobacteriaceae; Escherichia; Escherichia coli MNNEETFYQAMRRQGVTRRSFLKYCSLAATSLGLGAGMAPKIAWALENKPRIPVVWIHGL WP_001058323.1 MULTISPECIES: hydrogenase-1 small chain [Proteobacteria] Length: 372\nNumber of predicted TMHs: 2\nExp number of AAs in TMHs: 37.2608499999999\nExp number, first 60 AAs: 19.52923\nTotal prob of N-in: 0.94351\nPOSSIBLE N-term signal sequence\ninside 1 19\nTMhelix 20 39\noutside 40 325\nTMhelix 326 348\ninside 349 372

12593 GCF_001610755.1_ASM161075v1 Escherichia coli str. Sanji Proteobacteria; Gammaproteobacteria; Enterobacterales; Enterobacteriaceae; Escherichia; Escherichia coli MTGDNTLIHSHGINRRDFMKLCAALAATMGLSSKAAAEMAESVTNPQRPPVIWIGAQECT WP_000145410.1 MULTISPECIES: hydrogenase-2 small chain [Proteobacteria] Length: 372\nNumber of predicted TMHs:

1\nExp number of AAs in TMHs: 23.72226\nExp number, first 60 AAs: 0.22038\nTotal prob of N-in: 0.12994\noutside 1 331\nTMhelix 332 354\ninside 355 372

12594 GCF_000364305.1_Esch_coli_SWW33_V1 Escherichia coli SWW33 Proteobacteria; Gammaproteobacteria; Enterobacterales; Enterobacteriaceae; Escherichia; Escherichia coli
MTGDNTLIHSHGINRRDFMKLCAALAATMGLSSKAAAEMVESVTNPQRPPVIWIGAQECT WP_000145423.1
hydrogenase 2 small subunit [Escherichia coli] Length: 372\nNumber of predicted TMHs: 1\nExp number of AAs in TMHs: 23.78201\nExp number, first 60 AAs: 0.28355\nTotal prob of N-in: 0.13257\noutside 1 331\nTMhelix 332 354\ninside 355 372

12595 GCF_000364305.1_Esch_coli_SWW33_V1 Escherichia coli SWW33 Proteobacteria; Gammaproteobacteria; Enterobacterales; Enterobacteriaceae; Escherichia; Escherichia coli
MNNEETFYQAMRRQGVTRRSFLKYCSLAATSLGLGAGMAPKIAWALENKPRIPVVWIHGL WP_001058323.1
MULTISPECIES: hydrogenase-1 small chain [Proteobacteria] Length: 372\nNumber of predicted TMHs: 2\nExp number of AAs in TMHs: 37.2608499999999\nExp number, first 60 AAs: 19.52923\nTotal prob of N-in: 0.94351\nPOSSIBLE N-term signal sequence\ninside 1 19\nTMhelix 20 39\noutside 40 325\nTMhelix 326 348\ninside 349 372

12596 GCF_000462145.2_ASM46214v2 Escherichia coli T1282_01 Proteobacteria; Gammaproteobacteria; Enterobacterales; Enterobacteriaceae; Escherichia; Escherichia coli
MTGDNTLIHSHGINRRDFMKLCAALAATMGLSSKAAAEMAESVTNPQRPPVIWIGAQECT WP_000145410.1
MULTISPECIES: hydrogenase-2 small chain [Proteobacteria] Length: 372\nNumber of predicted TMHs: 1\nExp number of AAs in TMHs: 23.72226\nExp number, first 60 AAs: 0.22038\nTotal prob of N-in: 0.12994\noutside 1 331\nTMhelix 332 354\ninside 355 372

12597 GCF_000462145.2_ASM46214v2 Escherichia coli T1282_01 Proteobacteria; Gammaproteobacteria; Enterobacterales; Enterobacteriaceae; Escherichia; Escherichia coli
MNNEETFYQAMRRQGVTRRSFLKYCSLAATSLGLGAGMAPKIAWALENKPRIPVVWIHGL WP_001301613.1
hydrogenase [Escherichia coli] Length: 372\nNumber of predicted TMHs: 2\nExp number of AAs in TMHs: 37.46598\nExp number, first 60 AAs: 19.53139\nTotal prob of N-in: 0.94364\nPOSSIBLE N-term signal sequence\ninside 1 19\nTMhelix 20 39\noutside 40 325\nTMhelix 326 348\ninside 349 372

12598 GCF_000462265.2_ASM46226v2 Escherichia coli T1840_97 Proteobacteria; Gammaproteobacteria; Enterobacterales; Enterobacteriaceae; Escherichia; Escherichia coli
MNNEETFYQAMRRQGVTRRSFLKYCSLAATSLGLGAGMAPKIAWALENKPRIPVVWIHGL WP_001301613.1
hydrogenase [Escherichia coli] Length: 372\nNumber of predicted TMHs: 2\nExp number of AAs in TMHs: 37.46598\nExp number, first 60 AAs: 19.53139\nTotal prob of N-in: 0.94364\nPOSSIBLE N-term signal sequence\ninside 1 19\nTMhelix 20 39\noutside 40 325\nTMhelix 326 348\ninside 349 372

12599 GCF_000462265.2_ASM46226v2 Escherichia coli T1840_97 Proteobacteria; Gammaproteobacteria; Enterobacterales; Enterobacteriaceae; Escherichia; Escherichia coli
MTGDNTLIHSHGINRRDFMKLCAALAATMGLSSKAAAEMAESVTNPQRPPVIWIGAQECT WP_000145410.1
MULTISPECIES: hydrogenase-2 small chain [Proteobacteria] Length: 372\nNumber of predicted TMHs: 1\nExp number of AAs in TMHs: 23.72226\nExp number, first 60 AAs: 0.22038\nTotal prob of N-in: 0.12994\noutside 1 331\nTMhelix 332 354\ninside 355 372

12600 GCF_000462385.2_ASM46238v2 Escherichia coli T234_00 Proteobacteria; Gammaproteobacteria; Enterobacterales; Enterobacteriaceae; Escherichia; Escherichia coli
MTGDNTLIHSHGINRRDFMKLCAALAATMGLSSKAAAEMAESVTNPQRPPVIWIGAQECT WP_000145410.1
MULTISPECIES: hydrogenase-2 small chain [Proteobacteria] Length: 372\nNumber of predicted TMHs: 1\nExp number of AAs in TMHs: 23.72226\nExp number, first 60 AAs: 0.22038\nTotal prob of N-in: 0.12994\noutside 1 331\nTMhelix 332 354\ninside 355 372

12601 GCF_000462385.2_ASM46238v2 Escherichia coli T234_00 Proteobacteria; Gammaproteobacteria; Enterobacterales; Enterobacteriaceae; Escherichia; Escherichia coli
MNNEETFYQAMRRQGVTRRSFLKYCSLAATSLGLGAGMAPKIAWALENKPRIPVVWIHGL WP_001301613.1
hydrogenase [Escherichia coli] Length: 372\nNumber of predicted TMHs: 2\nExp number of AAs in TMHs: 37.46598\nExp number, first 60 AAs: 19.53139\nTotal prob of N-in: 0.94364\nPOSSIBLE N-term signal sequence\ninside 1 19\nTMhelix 20 39\noutside 40 325\nTMhelix 326 348\ninside 349 372

12602 GCF_002109945.1_ASM210994v1 Escherichia coli T426 Proteobacteria; Gammaproteobacteria; Enterobacterales; Enterobacteriaceae; Escherichia; Escherichia coli
MTGDNTLIHSHGINRRDFMKLCAALAATMGLSSKAAAEMAESVTNPQRPPVIWIGAQECT WP_000145410.1
MULTISPECIES: hydrogenase-2 small chain [Proteobacteria] Length: 372\nNumber of predicted TMHs: 1\nExp number of AAs in TMHs: 23.72226\nExp number, first 60 AAs: 0.22038\nTotal prob of N-in: 0.12994\noutside 1 331\nTMhelix 332 354\ninside 355 372

12603 GCF_002109945.1_ASM210994v1 Escherichia coli T426 Proteobacteria; Gammaproteobacteria; Enterobacterales; Enterobacteriaceae; Escherichia; Escherichia coli
MNNEETFYQAMRRQGVTRRSFLKYCSLAATSLGLGAGMAPKIAWALENKPRIPVVWIHGL WP_001295940.1
MULTISPECIES: hydrogenase [Enterobacteriaceae] Length: 372\nNumber of predicted TMHs: 2\nExp number of AAs in TMHs: 37.2608899999999\nExp number, first 60 AAs: 19.52923\nTotal prob of N-in: 0.94351\nPOSSIBLE N-term signal sequence\ninside 1 19\nTMhelix 20 39\noutside 40 325\nTMhelix 326 348\ninside 349 372

12604 GCF_000462165.2_ASM46216v2 Escherichia coli T924_01 Proteobacteria; Gammaproteobacteria; Enterobacterales; Enterobacteriaceae; Escherichia; Escherichia coli
MNNEETFYQAMRRQGVTRRSFLKYCSLAATSLGLGAGMAPKIAWALENKPRIPVVWIHGL WP_001301613.1
hydrogenase [Escherichia coli] Length: 372\nNumber of predicted TMHs: 2\nExp number of AAs in TMHs: 37.46598\nExp number, first 60 AAs: 19.53139\nTotal prob of N-in: 0.94364\nPOSSIBLE N-term signal sequence\ninside 1 19\nTMhelix 20 39\noutside 40 325\nTMhelix 326 348\ninside 349 372

12605 GCF_000462165.2_ASM46216v2 Escherichia coli T924_01 Proteobacteria; Gammaproteobacteria; Enterobacterales; Enterobacteriaceae; Escherichia; Escherichia coli
MTGDNTLIHSHGINRRDFMKLCAALAATMGLSSKAAAEMAESVTNPQRPPVIWIGAQECT WP_000145410.1
MULTISPECIES: hydrogenase-2 small chain [Proteobacteria] Length: 372\nNumber of predicted TMHs: 1\nExp number of AAs in TMHs: 23.72226\nExp number, first 60 AAs: 0.22038\nTotal prob of N-in: 0.12994\noutside 1 331\nTMhelix 332 354\ninside 355 372

12606 GCF_002109875.1_ASM210987v1 Escherichia coli TA008 Proteobacteria; Gammaproteobacteria; Enterobacterales; Enterobacteriaceae; Escherichia; Escherichia coli
MTGDNTLIHSHGINRRDFMKLCAALAATMGLSSKAAAEMAESVTNPQRPPVIWIGAQECT WP_000145410.1
MULTISPECIES: hydrogenase-2 small chain [Proteobacteria] Length: 372\nNumber of predicted TMHs: 1\nExp number of AAs in TMHs: 23.72226\nExp number, first 60 AAs: 0.22038\nTotal prob of N-in: 0.12994\noutside 1 331\nTMhelix 332 354\ninside 355 372

12607 GCF_002109875.1_ASM210987v1 Escherichia coli TA008 Proteobacteria; Gammaproteobacteria; Enterobacterales; Enterobacteriaceae; Escherichia; Escherichia coli
MNNEETFYQAMRRQGVTRRSFLKYCSLAATSLGLGAGMAPKIAWALENKPRIPVVWIHGL WP_001058323.1
MULTISPECIES: hydrogenase-1 small chain [Proteobacteria] Length: 372\nNumber of predicted TMHs: 2\nExp number of AAs in TMHs: 37.2608499999999\nExp number, first 60 AAs: 19.52923\nTotal prob of N-in: 0.94351\nPOSSIBLE N-term signal sequence\ninside 1 19\nTMhelix 20 39\noutside 40 325\nTMhelix 326 348\ninside 349 372

12608 GCF_002109905.1_ASM210990v1 Escherichia coli TA014 Proteobacteria; Gammaproteobacteria; Enterobacterales; Enterobacteriaceae; Escherichia; Escherichia coli
MTGDNTLIHSHGINRRDFMKLCAALAATMGLSSKAAAEMAESVTNPQRPPVIWIGAQECT WP_000145410.1
MULTISPECIES: hydrogenase-2 small chain [Proteobacteria] Length: 372\nNumber of predicted TMHs: 1\nExp number of AAs in TMHs: 23.72226\nExp number, first 60 AAs: 0.22038\nTotal prob of N-in: 0.12994\noutside 1 331\nTMhelix 332 354\ninside 355 372

12609 GCF_002109905.1_ASM210990v1 Escherichia coli TA014 Proteobacteria; Gammaproteobacteria; Enterobacterales; Enterobacteriaceae; Escherichia; Escherichia coli
MNNEETFYQAMRRQGVTRRSFLKYCSLAATSLGLGAGMAPKIAWALENKPRIPVVWIHGL WP_001058323.1
MULTISPECIES: hydrogenase-1 small chain [Proteobacteria] Length: 372\nNumber of predicted TMHs: 2\nExp number of AAs in TMHs: 37.2608499999999\nExp number, first 60 AAs: 19.52923\nTotal prob of N-in: 0.94351\nPOSSIBLE N-term signal sequence\ninside 1 19\nTMhelix 20 39\noutside 40 325\nTMhelix 326 348\ninside 349 372

12610 GCF_002110285.1_ASM211028v1 Escherichia coli TA054 Proteobacteria; Gammaproteobacteria; Enterobacterales; Enterobacteriaceae; Escherichia; Escherichia coli

MNNEETFYQAMRRQGVTRRSFLKYCSLAATSLGLGAGMAPKIAWALENKPRIPVVWIHGL WP_001058323.1 MULTISPECIES: hydrogenase-1 small chain [Proteobacteria] Length: 372\nNumber of predicted TMHs: 2\nExp number of AAs in TMHs: 37.2608499999999\nExp number, first 60 AAs: 19.52923\nTotal prob of N-in: 0.94351\nPOSSIBLE N-term signal sequence\ninside 1 19\nTMhelix 20 39\noutside 40 325\nTMhelix 326 348\ninside 349 372

12611 GCF_002110285.1_ASM211028v1 Escherichia coli TA054 Proteobacteria; Gammaproteobacteria; Enterobacterales; Enterobacteriaceae; Escherichia; Escherichia coli MTGDNTLIHSHGINRRDFMKLCAALAATMGLSSKAAAEMAESVTNPQRPPVIWIGAQECT WP_000145410.1 MULTISPECIES: hydrogenase-2 small chain [Proteobacteria] Length: 372\nNumber of predicted TMHs: 1\nExp number of AAs in TMHs: 23.72226\nExp number, first 60 AAs: 0.22038\nTotal prob of N-in: 0.12994\noutside 1 331\nTMhelix 332 354\ninside 355 372

12612 GCF_000242055.1_Esch_coli_TA124_V1 Escherichia coli TA124 Proteobacteria; Gammaproteobacteria; Enterobacterales; Enterobacteriaceae; Escherichia; Escherichia coli MTGDNTLIHSHGINRRDFMKLCAALAATMGLSSKAAAEMAESVTNPQRPPVIWIGAQECT WP_000145410.1 MULTISPECIES: hydrogenase-2 small chain [Proteobacteria] Length: 372\nNumber of predicted TMHs: 1\nExp number of AAs in TMHs: 23.72226\nExp number, first 60 AAs: 0.22038\nTotal prob of N-in: 0.12994\noutside 1 331\nTMhelix 332 354\ninside 355 372

12613 GCF_000242055.1_Esch_coli_TA124_V1 Escherichia coli TA124 Proteobacteria; Gammaproteobacteria; Enterobacterales; Enterobacteriaceae; Escherichia; Escherichia coli MNNEETFYQAMRRQGVTRRSFLKYCSLAATSLGLGAGMAPKIAWALENKPRIPVVWIHGL WP_001058323.1 MULTISPECIES: hydrogenase-1 small chain [Proteobacteria] Length: 372\nNumber of predicted TMHs: 2\nExp number of AAs in TMHs: 37.2608499999999\nExp number, first 60 AAs: 19.52923\nTotal prob of N-in: 0.94351\nPOSSIBLE N-term signal sequence\ninside 1 19\nTMhelix 20 39\noutside 40 325\nTMhelix 326 348\ninside 349 372

12614 GCF_000176615.2_ASM17661v2 Escherichia coli TA143 Proteobacteria; Gammaproteobacteria; Enterobacterales; Enterobacteriaceae; Escherichia; Escherichia coli MNNEETFYQAMRRQGVTRRSFLKYCSLAATSLGLGAGMAPKIAWALENKPRIPVVWIHGL WP_001058323.1 MULTISPECIES: hydrogenase-1 small chain [Proteobacteria] Length: 372\nNumber of predicted TMHs: 2\nExp number of AAs in TMHs: 37.2608499999999\nExp number, first 60 AAs: 19.52923\nTotal prob of N-in: 0.94351\nPOSSIBLE N-term signal sequence\ninside 1 19\nTMhelix 20 39\noutside 40 325\nTMhelix 326 348\ninside 349 372

12615 GCF_000176615.2_ASM17661v2 Escherichia coli TA143 Proteobacteria; Gammaproteobacteria; Enterobacterales; Enterobacteriaceae; Escherichia; Escherichia coli MTGDNTLIHSHGINRRDFMKLCAALAATMGLSSKAAAEMAESVTNPQRPPVIWIGAQECT WP_000145410.1 MULTISPECIES: hydrogenase-2 small chain [Proteobacteria] Length: 372\nNumber of predicted TMHs: 1\nExp number of AAs in TMHs: 23.72226\nExp number, first 60 AAs: 0.22038\nTotal prob of N-in: 0.12994\noutside 1 331\nTMhelix 332 354\ninside 355 372

12616 GCF_002110025.1_ASM211002v1 Escherichia coli TA144 Proteobacteria; Gammaproteobacteria; Enterobacterales; Enterobacteriaceae; Escherichia; Escherichia coli MTGDNTLIHSHGINRRDFMKLCAALAATMGLSSKAAAEMAESVTNPQRPPVIWIGAQECT WP_000145410.1 MULTISPECIES: hydrogenase-2 small chain [Proteobacteria] Length: 372\nNumber of predicted TMHs: 1\nExp number of AAs in TMHs: 23.72226\nExp number, first 60 AAs: 0.22038\nTotal prob of N-in: 0.12994\noutside 1 331\nTMhelix 332 354\ninside 355 372

12617 GCF_002110025.1_ASM211002v1 Escherichia coli TA144 Proteobacteria; Gammaproteobacteria; Enterobacterales; Enterobacteriaceae; Escherichia; Escherichia coli MNNEETFYQAMRRQGVTRRSFLKYCSLAATSLGLGAGMAPKIAWALENKPRIPVVWIHGL WP_001058323.1 MULTISPECIES: hydrogenase-1 small chain [Proteobacteria] Length: 372\nNumber of predicted TMHs: 2\nExp number of AAs in TMHs: 37.2608499999999\nExp number, first 60 AAs: 19.52923\nTotal prob of N-in: 0.94351\nPOSSIBLE N-term signal sequence\ninside 1 19\nTMhelix 20 39\noutside 40 325\nTMhelix 326 348\ninside 349 372

12618 GCF_000176595.2_ASM17659v2 Escherichia coli TA206 Proteobacteria; Gammaproteobacteria; Enterobacterales; Enterobacteriaceae; Escherichia; Escherichia coli MNNEETFYQAMRRQGVTRRSFLKYCSLAATSLGLGAGMAPKIAWALENKPRIPVVWIHGL WP_001058323.1

MULTISPECIES: hydrogenase-1 small chain [Proteobacteria] Length: 372\nNumber of predicted TMHs: 2\nExp number of AAs in TMHs: 37.2608499999999\nExp number, first 60 AAs: 19.52923\nTotal prob of N-in: 0.94351\nPOSSIBLE N-term signal sequence\ninside 1 19\nTMhelix 20 39\noutside 40 325\nTMhelix 326 348\ninside 349 372

12619 GCF_000176595.2_ASM17659v2 Escherichia coli TA206 Proteobacteria; Gammaproteobacteria; Enterobacterales; Enterobacteriaceae; Escherichia; Escherichia coli
MTGDNTLIHSHGINRRDFMKLCAALAATMGLSSKAAAEMVESVTNPQRPPVIWIGAQECT WP_000145423.1
hydrogenase 2 small subunit [Escherichia coli] Length: 372\nNumber of predicted TMHs: 1\nExp number of AAs in TMHs: 23.78201\nExp number, first 60 AAs: 0.28355\nTotal prob of N-in: 0.13257\noutside 1 331\nTMhelix 332 354\ninside 355 372

12620 GCF_002109925.1_ASM210992v1 Escherichia coli TA249 Proteobacteria; Gammaproteobacteria; Enterobacterales; Enterobacteriaceae; Escherichia; Escherichia coli
MTGDNTLIHSHGINRRDFMKLCAALAATMGLSSKAAAEMAESVTNPQRPPVIWIGAQECT WP_000145410.1
MULTISPECIES: hydrogenase-2 small chain [Proteobacteria] Length: 372\nNumber of predicted TMHs: 1\nExp number of AAs in TMHs: 23.72226\nExp number, first 60 AAs: 0.22038\nTotal prob of N-in: 0.12994\noutside 1 331\nTMhelix 332 354\ninside 355 372

12621 GCF_002109925.1_ASM210992v1 Escherichia coli TA249 Proteobacteria; Gammaproteobacteria; Enterobacterales; Enterobacteriaceae; Escherichia; Escherichia coli
MNNEETFYQAMRRQGVTRRSFLKYCSLAATSLGLGAGMAPKIAWALENKPRIPVVWIHGL WP_001058323.1
MULTISPECIES: hydrogenase-1 small chain [Proteobacteria] Length: 372\nNumber of predicted TMHs: 2\nExp number of AAs in TMHs: 37.2608499999999\nExp number, first 60 AAs: 19.52923\nTotal prob of N-in: 0.94351\nPOSSIBLE N-term signal sequence\ninside 1 19\nTMhelix 20 39\noutside 40 325\nTMhelix 326 348\ninside 349 372

12622 GCF_002110185.1_ASM211018v1 Escherichia coli TA255 Proteobacteria; Gammaproteobacteria; Enterobacterales; Enterobacteriaceae; Escherichia; Escherichia coli
MTGDNTLIHSHGINRRDFMKLCAALAATMGLSSKAAAEMAESVTNPQRPPVIWIGAQECT WP_000145410.1
MULTISPECIES: hydrogenase-2 small chain [Proteobacteria] Length: 372\nNumber of predicted TMHs: 1\nExp number of AAs in TMHs: 23.72226\nExp number, first 60 AAs: 0.22038\nTotal prob of N-in: 0.12994\noutside 1 331\nTMhelix 332 354\ninside 355 372

12623 GCF_002110185.1_ASM211018v1 Escherichia coli TA255 Proteobacteria; Gammaproteobacteria; Enterobacterales; Enterobacteriaceae; Escherichia; Escherichia coli
MNNEETFYQAMRRQGVTRRSFLKYCSLAATSLGLGAGMAPKIAWALENKPRIPVVWIHGL WP_001058323.1
MULTISPECIES: hydrogenase-1 small chain [Proteobacteria] Length: 372\nNumber of predicted TMHs: 2\nExp number of AAs in TMHs: 37.2608499999999\nExp number, first 60 AAs: 19.52923\nTotal prob of N-in: 0.94351\nPOSSIBLE N-term signal sequence\ninside 1 19\nTMhelix 20 39\noutside 40 325\nTMhelix 326 348\ninside 349 372

12624 GCF_000176655.2_ASM17665v2 Escherichia coli TA280 Proteobacteria; Gammaproteobacteria; Enterobacterales; Enterobacteriaceae; Escherichia; Escherichia coli
MTGDNTLIHSHGINRRDFMKLCAALAATMGLSSKAAAEMAESVTNPQRPPVIWIGAQECT WP_000145410.1
MULTISPECIES: hydrogenase-2 small chain [Proteobacteria] Length: 372\nNumber of predicted TMHs: 1\nExp number of AAs in TMHs: 23.72226\nExp number, first 60 AAs: 0.22038\nTotal prob of N-in: 0.12994\noutside 1 331\nTMhelix 332 354\ninside 355 372

12625 GCF_000176655.2_ASM17665v2 Escherichia coli TA280 Proteobacteria; Gammaproteobacteria; Enterobacterales; Enterobacteriaceae; Escherichia; Escherichia coli
MNNEETFYQAMRRQGVTRRSFLKYCSLAATSLGLGAGMAPKIAWALENKPRIPVVWIHGL WP_001058323.1
MULTISPECIES: hydrogenase-1 small chain [Proteobacteria] Length: 372\nNumber of predicted TMHs: 2\nExp number of AAs in TMHs: 37.2608499999999\nExp number, first 60 AAs: 19.52923\nTotal prob of N-in: 0.94351\nPOSSIBLE N-term signal sequence\ninside 1 19\nTMhelix 20 39\noutside 40 325\nTMhelix 326 348\ninside 349 372

12626 GCF_002110175.1_ASM211017v1 Escherichia coli TA447 Proteobacteria; Gammaproteobacteria; Enterobacterales; Enterobacteriaceae; Escherichia; Escherichia coli
MNNEETFYQAMRRQGVTRRSFLKYCSLAATSLGLGAGMAPKIAWALENKPRIPVVWIHGL WP_001058323.1
MULTISPECIES: hydrogenase-1 small chain [Proteobacteria] Length: 372\nNumber of predicted TMHs:

2\nExp number of AAs in TMHs: 37.2608499999999\nExp number, first 60 AAs: 19.52923\nTotal prob of N-in: 0.94351\nPOSSIBLE N-term signal sequence\ninside 1 19\nTMhelix 20 39\noutside 40 325\nTMhelix 326 348\ninside 349 372

12627 GCF_002110175.1_ASM211017v1 Escherichia coli TA447 Proteobacteria; Gammaproteobacteria; Enterobacterales; Enterobacteriaceae; Escherichia; Escherichia coli
MTGDNTLIHSHGINRRDFMKLCAALAATMGLSSKAAAEMAESVTNPQRPPVIWIGAQECT WP_000145410.1
MULTISPECIES: hydrogenase-2 small chain [Proteobacteria] Length: 372\nNumber of predicted TMHs: 1\nExp number of AAs in TMHs: 23.72226\nExp number, first 60 AAs: 0.22038\nTotal prob of N-in: 0.12994\noutside 1 331\nTMhelix 332 354\ninside 355 372

12628 GCF_002109865.1_ASM210986v1 Escherichia coli TA464 Proteobacteria; Gammaproteobacteria; Enterobacterales; Enterobacteriaceae; Escherichia; Escherichia coli
MTGDNTLIHSHGINRRDFMKLCAALAATMGLSSKAAAEMAESVTNPQRPPVIWIGAQECT WP_000145410.1
MULTISPECIES: hydrogenase-2 small chain [Proteobacteria] Length: 372\nNumber of predicted TMHs: 1\nExp number of AAs in TMHs: 23.72226\nExp number, first 60 AAs: 0.22038\nTotal prob of N-in: 0.12994\noutside 1 331\nTMhelix 332 354\ninside 355 372

12629 GCF_002109865.1_ASM210986v1 Escherichia coli TA464 Proteobacteria; Gammaproteobacteria; Enterobacterales; Enterobacteriaceae; Escherichia; Escherichia coli
MNNEETFYQAMRRQGVTRRSFLKYCSLAATSLGLGAGMAPKIAWALENKPRIPVVWIHGL WP_001058323.1
MULTISPECIES: hydrogenase-1 small chain [Proteobacteria] Length: 372\nNumber of predicted TMHs: 2\nExp number of AAs in TMHs: 37.2608499999999\nExp number, first 60 AAs: 19.52923\nTotal prob of N-in: 0.94351\nPOSSIBLE N-term signal sequence\ninside 1 19\nTMhelix 20 39\noutside 40 325\nTMhelix 326 348\ninside 349 372

12630 GCF_000355115.2_ASM35511v2 Escherichia coli ThroopD Proteobacteria; Gammaproteobacteria; Enterobacterales; Enterobacteriaceae; Escherichia; Escherichia coli
MNNEETFYQAMRRQGVTRRSFLKYCSLAATSLGLGAGMAPKIAWALENKPRIPVVWIHGL WP_001058320.1
hydrogenase [Escherichia coli] Length: 372\nNumber of predicted TMHs: 2\nExp number of AAs in TMHs: 37.26082\nExp number, first 60 AAs: 19.52923\nTotal prob of N-in: 0.94351\nPOSSIBLE N-term signal sequence\ninside 1 19\nTMhelix 20 39\noutside 40 325\nTMhelix 326 348\ninside 349 372

12631 GCF_000355115.2_ASM35511v2 Escherichia coli ThroopD Proteobacteria; Gammaproteobacteria; Enterobacterales; Enterobacteriaceae; Escherichia; Escherichia coli
MTGDNTLIHSHGINRRDFMKLCAALAATMGLSSKAAAEMAESVTNPQRPPVIWIGAQECT WP_000145410.1
MULTISPECIES: hydrogenase-2 small chain [Proteobacteria] Length: 372\nNumber of predicted TMHs: 1\nExp number of AAs in TMHs: 23.72226\nExp number, first 60 AAs: 0.22038\nTotal prob of N-in: 0.12994\noutside 1 331\nTMhelix 332 354\ninside 355 372

12632 GCF_000397725.1_Ec72_E1U1_TOP2386_MIRA_120724 Escherichia coli TOP2386 Proteobacteria; Gammaproteobacteria; Enterobacterales; Enterobacteriaceae; Escherichia; Escherichia coli
MNNEETFYQAMRRQGVTRRSFLKYCSLAATSLGLGAGMAPKIAWALENKPRIPVVWIHGL WP_001058323.1
MULTISPECIES: hydrogenase-1 small chain [Proteobacteria] Length: 372\nNumber of predicted TMHs: 2\nExp number of AAs in TMHs: 37.2608499999999\nExp number, first 60 AAs: 19.52923\nTotal prob of N-in: 0.94351\nPOSSIBLE N-term signal sequence\ninside 1 19\nTMhelix 20 39\noutside 40 325\nTMhelix 326 348\ninside 349 372

12633 GCF_000397725.1_Ec72_E1U1_TOP2386_MIRA_120724 Escherichia coli TOP2386 Proteobacteria; Gammaproteobacteria; Enterobacterales; Enterobacteriaceae; Escherichia; Escherichia coli
MTGDNTLIHSHGINRRDFMKLCAALAATMGLSSKAAAEMAESVTNPQRPPVIWIGAQECT WP_000145410.1
MULTISPECIES: hydrogenase-2 small chain [Proteobacteria] Length: 372\nNumber of predicted TMHs: 1\nExp number of AAs in TMHs: 23.72226\nExp number, first 60 AAs: 0.22038\nTotal prob of N-in: 0.12994\noutside 1 331\nTMhelix 332 354\ninside 355 372

12634 GCF_000397505.1_Ec72_E1U1_TOP2386_120714 Escherichia coli TOP2386 Proteobacteria; Gammaproteobacteria; Enterobacterales; Enterobacteriaceae; Escherichia; Escherichia coli
MTGDNTLIHSHGINRRDFMKLCAALAATMGLSSKAAAEMAESVTNPQRPPVIWIGAQECT WP_000145410.1
MULTISPECIES: hydrogenase-2 small chain [Proteobacteria] Length: 372\nNumber of predicted TMHs: 1\nExp number of AAs in TMHs: 23.72226\nExp number, first 60 AAs: 0.22038\nTotal prob of N-in: 0.12994\noutside 1 331\nTMhelix 332 354\ninside 355 372

12635 GCF_000397505.1_Ec72_E1U1_TOP2386_120714 Escherichia coli TOP2386 Proteobacteria; Gammaproteobacteria; Enterobacterales; Enterobacteriaceae; Escherichia; Escherichia coli MNNEETFYQAMRRQGVTRRSFLKYCSLAATSLGLGAGMAPKIAWALENKPRIPVVSIHGL WP_020239278.1 hydrogenase [Escherichia coli] Length: 372\nNumber of predicted TMHs: 2\nExp number of AAs in TMHs: 37.3787999999999\nExp number, first 60 AAs: 19.53168\nTotal prob of N-in: 0.94341\nPOSSIBLE N-term signal sequence\ninside 1 19\nTMhelix 20 39\noutside 40 325\nTMhelix 326 348\ninside 349 372

12636 GCF_000397525.1_Ec72_E1F1_TOP2396-1_120714 Escherichia coli TOP2396-1 Proteobacteria; Gammaproteobacteria; Enterobacterales; Enterobacteriaceae; Escherichia; Escherichia coli MNNEETFYQAMRRQGVTRRSFLKYCSLAATSLGLGAGMAPKIAWALENKPRIPVVWIHGL WP_001058323.1 MULTISPECIES: hydrogenase-1 small chain [Proteobacteria] Length: 372\nNumber of predicted TMHs: 2\nExp number of AAs in TMHs: 37.2608499999999\nExp number, first 60 AAs: 19.52923\nTotal prob of N-in: 0.94351\nPOSSIBLE N-term signal sequence\ninside 1 19\nTMhelix 20 39\noutside 40 325\nTMhelix 326 348\ninside 349 372

12637 GCF_000397525.1_Ec72_E1F1_TOP2396-1_120714 Escherichia coli TOP2396-1 Proteobacteria; Gammaproteobacteria; Enterobacterales; Enterobacteriaceae; Escherichia; Escherichia coli MTGDNTLIHSHGINRRDFMKLCAALAATMGLSSKAAAEMAESVTNPQRPPVIWIGAQECT WP_000145410.1 MULTISPECIES: hydrogenase-2 small chain [Proteobacteria] Length: 372\nNumber of predicted TMHs: 1\nExp number of AAs in TMHs: 23.72226\nExp number, first 60 AAs: 0.22038\nTotal prob of N-in: 0.12994\noutside 1 331\nTMhelix 332 354\ninside 355 372

12638 GCF_000397565.1_Ec72_E1F3_TOP2396-3_120714 Escherichia coli TOP2396-3 Proteobacteria; Gammaproteobacteria; Enterobacterales; Enterobacteriaceae; Escherichia; Escherichia coli MTGDNTLIHSHGINRRDFMKLCAALAATMGLSSKAAAEMAESVTNPQRPPVIWIGAQECT WP_000145410.1 MULTISPECIES: hydrogenase-2 small chain [Proteobacteria] Length: 372\nNumber of predicted TMHs: 1\nExp number of AAs in TMHs: 23.72226\nExp number, first 60 AAs: 0.22038\nTotal prob of N-in: 0.12994\noutside 1 331\nTMhelix 332 354\ninside 355 372

12639 GCF_000397605.1_Ec72_E2F1_TOP2522-1_120714 Escherichia coli TOP2522-1 Proteobacteria; Gammaproteobacteria; Enterobacterales; Enterobacteriaceae; Escherichia; Escherichia coli MNNEETFYQAMRRQGVTRRSFLKYCSLAATSLGLGAGMAPKIAWALENKPRIPVVWIHGL WP_001058323.1 MULTISPECIES: hydrogenase-1 small chain [Proteobacteria] Length: 372\nNumber of predicted TMHs: 2\nExp number of AAs in TMHs: 37.2608499999999\nExp number, first 60 AAs: 19.52923\nTotal prob of N-in: 0.94351\nPOSSIBLE N-term signal sequence\ninside 1 19\nTMhelix 20 39\noutside 40 325\nTMhelix 326 348\ninside 349 372

12640 GCF_000397605.1_Ec72_E2F1_TOP2522-1_120714 Escherichia coli TOP2522-1 Proteobacteria; Gammaproteobacteria; Enterobacterales; Enterobacteriaceae; Escherichia; Escherichia coli MTGDNTLIHSHGINRRDFMKLCAALAATMGLSSKAAAEMAESVTNPQRPPVIWIGAQECT WP_000145410.1 MULTISPECIES: hydrogenase-2 small chain [Proteobacteria] Length: 372\nNumber of predicted TMHs: 1\nExp number of AAs in TMHs: 23.72226\nExp number, first 60 AAs: 0.22038\nTotal prob of N-in: 0.12994\noutside 1 331\nTMhelix 332 354\ninside 355 372

12641 GCF_000397745.1_Ec72_E3U1_TOP2652_MIRA_120724 Escherichia coli TOP2652 Proteobacteria; Gammaproteobacteria; Enterobacterales; Enterobacteriaceae; Escherichia; Escherichia coli MTGDNTLIHSHGINRRDFMKLCAALAATMGLSSKAAAEMAESVTNPQRPPVIWIGAQECT WP_000145410.1 MULTISPECIES: hydrogenase-2 small chain [Proteobacteria] Length: 372\nNumber of predicted TMHs: 1\nExp number of AAs in TMHs: 23.72226\nExp number, first 60 AAs: 0.22038\nTotal prob of N-in: 0.12994\noutside 1 331\nTMhelix 332 354\ninside 355 372

12642 GCF_000397745.1_Ec72_E3U1_TOP2652_MIRA_120724 Escherichia coli TOP2652 Proteobacteria; Gammaproteobacteria; Enterobacterales; Enterobacteriaceae; Escherichia; Escherichia coli MNNEETFYQAMRRQGVTRRSFLKYCSLAATSLGLGAGMAPKIAWALENKPRIPVVWIHGL WP_001058323.1 MULTISPECIES: hydrogenase-1 small chain [Proteobacteria] Length: 372\nNumber of predicted TMHs: 2\nExp number of AAs in TMHs: 37.2608499999999\nExp number, first 60 AAs: 19.52923\nTotal prob of N-in: 0.94351\nPOSSIBLE N-term signal sequence\ninside 1 19\nTMhelix 20 39\noutside 40 325\nTMhelix 326 348\ninside 349 372

12643 GCF_000397625.1_Ec72_E3U1_TOP2652_120714 Escherichia coli TOP2652 Proteobacteria; Gammaproteobacteria; Enterobacterales; Enterobacteriaceae; Escherichia; Escherichia coli

MTGDNTLIHSHGINRRDFMKLCAALAATMGLSSKAAAEMAESVTNPQRPPVIWIGAQECT WP_000145410.1 MULTISPECIES: hydrogenase-2 small chain [Proteobacteria] Length: 372\nNumber of predicted TMHs: 1\nExp number of AAs in TMHs: 23.72226\nExp number, first 60 AAs: 0.22038\nTotal prob of N-in: 0.12994\noutside 1 331\nTMhelix 332 354\ninside 355 372

12644 GCF_000397625.1_Ec72_E3U1_TOP2652_120714 Escherichia coli TOP2652 Proteobacteria; Gammaproteobacteria; Enterobacterales; Enterobacteriaceae; Escherichia; Escherichia coli MNNEETFYQAMRRQGVTRRSFLKYCSLAATSLGLGAGMAPKIAWALENKPRIPVVWIHGL WP_001058323.1 MULTISPECIES: hydrogenase-1 small chain [Proteobacteria] Length: 372\nNumber of predicted TMHs: 2\nExp number of AAs in TMHs: 37.2608499999999\nExp number, first 60 AAs: 19.52923\nTotal prob of N-in: 0.94351\nPOSSIBLE N-term signal sequence\ninside 1 19\nTMhelix 20 39\noutside 40 325\nTMhelix 326 348\ninside 349 372

12645 GCF_000397645.1_Ec72_E3F1_TOP2662-1_120714 Escherichia coli TOP2662-1 Proteobacteria; Gammaproteobacteria; Enterobacterales; Enterobacteriaceae; Escherichia; Escherichia coli MTGDNTLIHSHGINRRDFMKLCAALAATMGLSSKAAAEMAESVTNPQRPPVIWIGAQECT WP_000145410.1 MULTISPECIES: hydrogenase-2 small chain [Proteobacteria] Length: 372\nNumber of predicted TMHs: 1\nExp number of AAs in TMHs: 23.72226\nExp number, first 60 AAs: 0.22038\nTotal prob of N-in: 0.12994\noutside 1 331\nTMhelix 332 354\ninside 355 372

12646 GCF_000397665.1_Ec72_E3F2_TOP2662-2_120714 Escherichia coli TOP2662-2 Proteobacteria; Gammaproteobacteria; Enterobacterales; Enterobacteriaceae; Escherichia; Escherichia coli MNNEETFYQAMRRQGVTRRSFLKYCSLAATSLGLGAGMAPKIAWALENKPRIPVVWIHGL WP_001058323.1 MULTISPECIES: hydrogenase-1 small chain [Proteobacteria] Length: 372\nNumber of predicted TMHs: 2\nExp number of AAs in TMHs: 37.2608499999999\nExp number, first 60 AAs: 19.52923\nTotal prob of N-in: 0.94351\nPOSSIBLE N-term signal sequence\ninside 1 19\nTMhelix 20 39\noutside 40 325\nTMhelix 326 348\ninside 349 372

12647 GCF_000397665.1_Ec72_E3F2_TOP2662-2_120714 Escherichia coli TOP2662-2 Proteobacteria; Gammaproteobacteria; Enterobacterales; Enterobacteriaceae; Escherichia; Escherichia coli MTGDNTLIHSHGINRRDFMKLCAALAATMGLSSKAAAEMAESVTNPQRPPVIWIGAQECT WP_000145410.1 MULTISPECIES: hydrogenase-2 small chain [Proteobacteria] Length: 372\nNumber of predicted TMHs: 1\nExp number of AAs in TMHs: 23.72226\nExp number, first 60 AAs: 0.22038\nTotal prob of N-in: 0.12994\noutside 1 331\nTMhelix 332 354\ninside 355 372

12648 GCF_000397685.1_Ec72_E3F3_TOP2662-3_120714 Escherichia coli TOP2662-3 Proteobacteria; Gammaproteobacteria; Enterobacterales; Enterobacteriaceae; Escherichia; Escherichia coli MNNEETFYQAMRRQGVTRRSFLKYCSLAATSLGLGAGMAPKIAWALENKPRIPVVWIHGL WP_001058323.1 MULTISPECIES: hydrogenase-1 small chain [Proteobacteria] Length: 372\nNumber of predicted TMHs: 2\nExp number of AAs in TMHs: 37.2608499999999\nExp number, first 60 AAs: 19.52923\nTotal prob of N-in: 0.94351\nPOSSIBLE N-term signal sequence\ninside 1 19\nTMhelix 20 39\noutside 40 325\nTMhelix 326 348\ninside 349 372

12649 GCF_000397685.1_Ec72_E3F3_TOP2662-3_120714 Escherichia coli TOP2662-3 Proteobacteria; Gammaproteobacteria; Enterobacterales; Enterobacteriaceae; Escherichia; Escherichia coli MTGDNTLIHSHGINRRDFMKLCAALAATMGLSSKAAAEMAESVTNPQRPPVIWIGAQECT WP_000145410.1 MULTISPECIES: hydrogenase-2 small chain [Proteobacteria] Length: 372\nNumber of predicted TMHs: 1\nExp number of AAs in TMHs: 23.72226\nExp number, first 60 AAs: 0.22038\nTotal prob of N-in: 0.12994\noutside 1 331\nTMhelix 332 354\ninside 355 372

12650 GCF_000397705.1_Ec72_E3F4_TOP2662-4_120714 Escherichia coli TOP2662-4 Proteobacteria; Gammaproteobacteria; Enterobacterales; Enterobacteriaceae; Escherichia; Escherichia coli MTGDNTLIHSHGINRRDFMKLCAALAATMGLSSKAAAEMAESVTNPQRPPVIWIGAQECT WP_000145410.1 MULTISPECIES: hydrogenase-2 small chain [Proteobacteria] Length: 372\nNumber of predicted TMHs: 1\nExp number of AAs in TMHs: 23.72226\nExp number, first 60 AAs: 0.22038\nTotal prob of N-in: 0.12994\noutside 1 331\nTMhelix 332 354\ninside 355 372

12651 GCF_000397705.1_Ec72_E3F4_TOP2662-4_120714 Escherichia coli TOP2662-4 Proteobacteria; Gammaproteobacteria; Enterobacterales; Enterobacteriaceae; Escherichia; Escherichia coli MNNEETFYQAMRRQGVTRRSFLKYCSLAATSLGLGAGMAPKIAWALENKPRIPVVWIHGL WP_020245785.1 hydrogenase [Escherichia coli] Length: 372\nNumber of predicted TMHs: 2\nExp number of AAs in

TMHs: 39.90377\nExp number, first 60 AAs: 19.5243\nTotal prob of N-in: 0.94368\nPOSSIBLE N-term signal sequence\ninside 1 19\nTMhelix 20 39\noutside 40 325\nTMhelix 326 348\ninside 349 372

12652 GCF_000397245.1_Ec13_E1F1_TOP382-1_120714 Escherichia coli TOP382-1 Proteobacteria; Gammaproteobacteria; Enterobacterales; Enterobacteriaceae; Escherichia; Escherichia coli MNNEETFYQAMRRQGVTRRSFLKYCSLAATSLGLGAGMAPKIAWALENKPRIPVVWIHGL WP_001058323.1 MULTISPECIES: hydrogenase-1 small chain [Proteobacteria] Length: 372\nNumber of predicted TMHs: 2\nExp number of AAs in TMHs: 37.2608499999999\nExp number, first 60 AAs: 19.52923\nTotal prob of N-in: 0.94351\nPOSSIBLE N-term signal sequence\ninside 1 19\nTMhelix 20 39\noutside 40 325\nTMhelix 326 348\ninside 349 372

12653 GCF_000397245.1_Ec13_E1F1_TOP382-1_120714 Escherichia coli TOP382-1 Proteobacteria; Gammaproteobacteria; Enterobacterales; Enterobacteriaceae; Escherichia; Escherichia coli MTGDNTLIHSHGINRRDFMKLCAALAATMGLSSKAAAEMAESVTNPQRPPVIWIGAQECT WP_000145410.1 MULTISPECIES: hydrogenase-2 small chain [Proteobacteria] Length: 372\nNumber of predicted TMHs: 1\nExp number of AAs in TMHs: 23.72226\nExp number, first 60 AAs: 0.22038\nTotal prob of N-in: 0.12994\noutside 1 331\nTMhelix 332 354\ninside 355 372

12654 GCF_000397265.1_Ec13_E1F2_TOP382-2_120714 Escherichia coli TOP382-2 Proteobacteria; Gammaproteobacteria; Enterobacterales; Enterobacteriaceae; Escherichia; Escherichia coli MTGDNTLIHSHGINRRDFMKLCAALAATMGLSSKAAAEMAESVTNPQRPPVIWIGAQECT WP_000145410.1 MULTISPECIES: hydrogenase-2 small chain [Proteobacteria] Length: 372\nNumber of predicted TMHs: 1\nExp number of AAs in TMHs: 23.72226\nExp number, first 60 AAs: 0.22038\nTotal prob of N-in: 0.12994\noutside 1 331\nTMhelix 332 354\ninside 355 372

12655 GCF_000397265.1_Ec13_E1F2_TOP382-2_120714 Escherichia coli TOP382-2 Proteobacteria; Gammaproteobacteria; Enterobacterales; Enterobacteriaceae; Escherichia; Escherichia coli MNNEETFYQAMRRQGVTRRSFLKYCSLAATSLGLGAGMAPKIAWALENKPRIPVVWIHGL WP_001058323.1 MULTISPECIES: hydrogenase-1 small chain [Proteobacteria] Length: 372\nNumber of predicted TMHs: 2\nExp number of AAs in TMHs: 37.2608499999999\nExp number, first 60 AAs: 19.52923\nTotal prob of N-in: 0.94351\nPOSSIBLE N-term signal sequence\ninside 1 19\nTMhelix 20 39\noutside 40 325\nTMhelix 326 348\ninside 349 372

12656 GCF_000397285.1_Ec13_E1F3_TOP382-3_120714 Escherichia coli TOP382-3 Proteobacteria; Gammaproteobacteria; Enterobacterales; Enterobacteriaceae; Escherichia; Escherichia coli MTGDNTLIHSHGINRRDFMKLCAALAATMGLSSKAAAEMAESVTNPQRPPVIWIGAQECT WP_000145410.1 MULTISPECIES: hydrogenase-2 small chain [Proteobacteria] Length: 372\nNumber of predicted TMHs: 1\nExp number of AAs in TMHs: 23.72226\nExp number, first 60 AAs: 0.22038\nTotal prob of N-in: 0.12994\noutside 1 331\nTMhelix 332 354\ninside 355 372

12657 GCF_000397285.1_Ec13_E1F3_TOP382-3_120714 Escherichia coli TOP382-3 Proteobacteria; Gammaproteobacteria; Enterobacterales; Enterobacteriaceae; Escherichia; Escherichia coli MNNEETFYQAMRRQGVTRRSFLKYCSLAATSLGLGAGMAPKIAWALENKPRIPVVWIHGL WP_020232254.1 hydrogenase [Escherichia coli] Length: 372\nNumber of predicted TMHs: 2\nExp number of AAs in TMHs: 37.2612899999999\nExp number, first 60 AAs: 19.52921\nTotal prob of N-in: 0.94351\nPOSSIBLE N-term signal sequence\ninside 1 19\nTMhelix 20 39\noutside 40 325\nTMhelix 326 348\ninside 349 372

12658 GCF_000397445.1_Ec13_E3F2_TOP550-2_120714 Escherichia coli TOP550-2 Proteobacteria; Gammaproteobacteria; Enterobacterales; Enterobacteriaceae; Escherichia; Escherichia coli MTGDNTLIHSHGINRRDFMKLCAALAATMGLSSKAAAEMAESVTNPQRPPVIWIGAQECT WP_000145410.1 MULTISPECIES: hydrogenase-2 small chain [Proteobacteria] Length: 372\nNumber of predicted TMHs: 1\nExp number of AAs in TMHs: 23.72226\nExp number, first 60 AAs: 0.22038\nTotal prob of N-in: 0.12994\noutside 1 331\nTMhelix 332 354\ninside 355 372

12659 GCF_000397445.1_Ec13_E3F2_TOP550-2_120714 Escherichia coli TOP550-2 Proteobacteria; Gammaproteobacteria; Enterobacterales; Enterobacteriaceae; Escherichia; Escherichia coli MNNEETFYQAMRRQGVTRRSFLKYCSLAATSLGLGAGMAPKIAWALENKPRIPVVWIHGL WP_001058323.1 MULTISPECIES: hydrogenase-1 small chain [Proteobacteria] Length: 372\nNumber of predicted TMHs: 2\nExp number of AAs in TMHs: 37.2608499999999\nExp number, first 60 AAs: 19.52923\nTotal prob of N-in: 0.94351\nPOSSIBLE N-term signal sequence\ninside 1 19\nTMhelix 20 39\noutside 40 325\nTMhelix 326 348\ninside 349 372

12660 GCF_000397465.1_Ec13_E3F3_TOP550-3_120714 Escherichia coli TOP550-3 Proteobacteria; Gammaproteobacteria; Enterobacterales; Enterobacteriaceae; Escherichia; Escherichia coli MTGDNTLIHSHGINRRDFMKLCAALAATMGLSSKAAAEMAESVTNPQRPPVIWIGAQECT WP_000145410.1 MULTISPECIES: hydrogenase-2 small chain [Proteobacteria] Length: 372\nNumber of predicted TMHs: 1\nExp number of AAs in TMHs: 23.72226\nExp number, first 60 AAs: 0.22038\nTotal prob of N-in: 0.12994\noutside 1 331\nTMhelix 332 354\ninside 355 372

12661 GCF_000397485.1_Ec13_E3F4_TOP550-4_120714 Escherichia coli TOP550-4 Proteobacteria; Gammaproteobacteria; Enterobacterales; Enterobacteriaceae; Escherichia; Escherichia coli MTGDNTLIHSHGINRRDFMKLCAALAATMGLSSKAAAEMAESVTNPQRPPVIWIGAQECT WP_000145410.1 MULTISPECIES: hydrogenase-2 small chain [Proteobacteria] Length: 372\nNumber of predicted TMHs: 1\nExp number of AAs in TMHs: 23.72226\nExp number, first 60 AAs: 0.22038\nTotal prob of N-in: 0.12994\noutside 1 331\nTMhelix 332 354\ninside 355 372

12662 GCF_000397485.1_Ec13_E3F4_TOP550-4_120714 Escherichia coli TOP550-4 Proteobacteria; Gammaproteobacteria; Enterobacterales; Enterobacteriaceae; Escherichia; Escherichia coli MNNEETFYQAMRRQGVTRRSFLKYCSLAATSLGLGAGMAPKIAWALENKPRIPVVWIHGL WP_001058323.1 MULTISPECIES: hydrogenase-1 small chain [Proteobacteria] Length: 372\nNumber of predicted TMHs: 2\nExp number of AAs in TMHs: 37.2608499999999\nExp number, first 60 AAs: 19.52923\nTotal prob of N-in: 0.94351\nPOSSIBLE N-term signal sequence\ninside 1 19\nTMhelix 20 39\noutside 40 325\nTMhelix 326 348\ninside 349 372

12663 GCF_000303955.2_ASM30395v2 Escherichia coli TT12B Proteobacteria; Gammaproteobacteria; Enterobacterales; Enterobacteriaceae; Escherichia; Escherichia coli MNNEETFYQAMRRQGVTRRSFLKYCSLAATSLGLGAGMAPKIAWALENKPRIPVVWIHGL WP_001301613.1 hydrogenase [Escherichia coli] Length: 372\nNumber of predicted TMHs: 2\nExp number of AAs in TMHs: 37.46598\nExp number, first 60 AAs: 19.53139\nTotal prob of N-in: 0.94364\nPOSSIBLE N-term signal sequence\ninside 1 19\nTMhelix 20 39\noutside 40 325\nTMhelix 326 348\ninside 349 372

12664 GCF_000303955.2_ASM30395v2 Escherichia coli TT12B Proteobacteria; Gammaproteobacteria; Enterobacterales; Enterobacteriaceae; Escherichia; Escherichia coli MTGDNTLIHSHGINRRDFMKLCAALAATMGLSSKAAAEMAESVTNPQRPPVIWIGAQECT WP_000145410.1 MULTISPECIES: hydrogenase-2 small chain [Proteobacteria] Length: 372\nNumber of predicted TMHs: 1\nExp number of AAs in TMHs: 23.72226\nExp number, first 60 AAs: 0.22038\nTotal prob of N-in: 0.12994\noutside 1 331\nTMhelix 332 354\ninside 355 372

12665 GCF_000304115.2_ASM30411v2 Escherichia coli TW00353 Proteobacteria; Gammaproteobacteria; Enterobacterales; Enterobacteriaceae; Escherichia; Escherichia coli MNNEETFYQAMRRQGVTRRSFLKYCSLAATSLGLGAGMAPKIAWALENKPRIPVVWIHGL WP_001058323.1 MULTISPECIES: hydrogenase-1 small chain [Proteobacteria] Length: 372\nNumber of predicted TMHs: 2\nExp number of AAs in TMHs: 37.2608499999999\nExp number, first 60 AAs: 19.52923\nTotal prob of N-in: 0.94351\nPOSSIBLE N-term signal sequence\ninside 1 19\nTMhelix 20 39\noutside 40 325\nTMhelix 326 348\ninside 349 372

12666 GCF_000304115.2_ASM30411v2 Escherichia coli TW00353 Proteobacteria; Gammaproteobacteria; Enterobacterales; Enterobacteriaceae; Escherichia; Escherichia coli MTGDNTLIHSHGINRRDFMKLCAALAATMGLSSKAAAEMAESVTNPQRPPVIWIGAQECT WP_000145410.1 MULTISPECIES: hydrogenase-2 small chain [Proteobacteria] Length: 372\nNumber of predicted TMHs: 1\nExp number of AAs in TMHs: 23.72226\nExp number, first 60 AAs: 0.22038\nTotal prob of N-in: 0.12994\noutside 1 331\nTMhelix 332 354\ninside 355 372

12667 GCF_000267125.1_ASM26712v1 Escherichia coli TW06591 Proteobacteria; Gammaproteobacteria; Enterobacterales; Enterobacteriaceae; Escherichia; Escherichia coli MNNEETFYQAMRRQGVTRRSFLKYCSLAATSLGLGAGMAPKIAWALENKPRIPVVWIHGL WP_001301613.1 hydrogenase [Escherichia coli] Length: 372\nNumber of predicted TMHs: 2\nExp number of AAs in TMHs: 37.46598\nExp number, first 60 AAs: 19.53139\nTotal prob of N-in: 0.94364\nPOSSIBLE N-term signal sequence\ninside 1 19\nTMhelix 20 39\noutside 40 325\nTMhelix 326 348\ninside 349 372

12668 GCF_000267125.1_ASM26712v1 Escherichia coli TW06591 Proteobacteria; Gammaproteobacteria; Enterobacterales; Enterobacteriaceae; Escherichia; Escherichia coli MTGDNTLIHSHGINRRDFMKLCAALAATMGLSSKAAAEMAESVTNPQRPPVIWIGAQECT WP_000145410.1

MULTISPECIES: hydrogenase-2 small chain [Proteobacteria] Length: 372\nNumber of predicted TMHs: 1\nExp number of AAs in TMHs: 23.72226\nExp number, first 60 AAs: 0.22038\nTotal prob of N-in: 0.12994\noutside 1 331\nTMhelix 332 354\ninside 355 372

12669 GCF_000462285.2_ASM46228v2 Escherichia coli TW07509 Proteobacteria; Gammaproteobacteria; Enterobacterales; Enterobacteriaceae; Escherichia; Escherichia coli
MTGDNTLIHSHGINRRDFMKLCAALAATMGLSSKAAAEMAESVTNPQRPPVIWIGAQECT WP_000145410.1
MULTISPECIES: hydrogenase-2 small chain [Proteobacteria] Length: 372\nNumber of predicted TMHs: 1\nExp number of AAs in TMHs: 23.72226\nExp number, first 60 AAs: 0.22038\nTotal prob of N-in: 0.12994\noutside 1 331\nTMhelix 332 354\ninside 355 372

12670 GCF_000462285.2_ASM46228v2 Escherichia coli TW07509 Proteobacteria; Gammaproteobacteria; Enterobacterales; Enterobacteriaceae; Escherichia; Escherichia coli
MNNEETFYQAMRRQGVTRRSFLKYCSLAATSLGLGAGMAPKIAWALENKPRIPVVWIHGL WP_001058323.1
MULTISPECIES: hydrogenase-1 small chain [Proteobacteria] Length: 372\nNumber of predicted TMHs: 2\nExp number of AAs in TMHs: 37.2608499999999\nExp number, first 60 AAs: 19.52923\nTotal prob of N-in: 0.94351\nPOSSIBLE N-term signal sequence\ninside 1 19\nTMhelix 20 39\noutside 40 325\nTMhelix 326 348\ninside 349 372

12671 GCF_000194685.1_ASM19468v2 Escherichia coli TW07793 Proteobacteria; Gammaproteobacteria; Enterobacterales; Enterobacteriaceae; Escherichia; Escherichia coli
MNNEETFYQAMRRQGVTRRSFLKYCSLAATSLGLGAGMAPKIAWALENKPRIPVVWIHGL WP_001374998.1
hydrogenase-1 small chain [Escherichia coli] Length: 372\nNumber of predicted TMHs: 2\nExp number of AAs in TMHs: 37.6595\nExp number, first 60 AAs: 19.52942\nTotal prob of N-in: 0.94352\nPOSSIBLE N-term signal sequence\ninside 1 19\nTMhelix 20 39\noutside 40 325\nTMhelix 326 348\ninside 349 372

12672 GCF_000194685.1_ASM19468v2 Escherichia coli TW07793 Proteobacteria; Gammaproteobacteria; Enterobacterales; Enterobacteriaceae; Escherichia; Escherichia coli
MTGDNTLIHSHGINRRDFMKLCAALAATMGLSSKAAAEMAESVTNPQRPPVIWIGAQECT WP_000145410.1
MULTISPECIES: hydrogenase-2 small chain [Proteobacteria] Length: 372\nNumber of predicted TMHs: 1\nExp number of AAs in TMHs: 23.72226\nExp number, first 60 AAs: 0.22038\nTotal prob of N-in: 0.12994\noutside 1 331\nTMhelix 332 354\ninside 355 372

12673 GCF_000267625.2_ASM26762v2 Escherichia coli TW07945 Proteobacteria; Gammaproteobacteria; Enterobacterales; Enterobacteriaceae; Escherichia; Escherichia coli
MNNEETFYQAMRRQGVTRRSFLKYCSLAATSLGLGAGMAPKIAWALENKPRIPVVWIHGL WP_001301613.1
hydrogenase [Escherichia coli] Length: 372\nNumber of predicted TMHs: 2\nExp number of AAs in TMHs: 37.46598\nExp number, first 60 AAs: 19.53139\nTotal prob of N-in: 0.94364\nPOSSIBLE N-term signal sequence\ninside 1 19\nTMhelix 20 39\noutside 40 325\nTMhelix 326 348\ninside 349 372

12674 GCF_000267625.2_ASM26762v2 Escherichia coli TW07945 Proteobacteria; Gammaproteobacteria; Enterobacterales; Enterobacteriaceae; Escherichia; Escherichia coli
MTGDNTLIHSHGINRRDFMKLCAALAATMGLSSKAAAEMAESVTNPQRPPVIWIGAQECT WP_000145410.1
MULTISPECIES: hydrogenase-2 small chain [Proteobacteria] Length: 372\nNumber of predicted TMHs: 1\nExp number of AAs in TMHs: 23.72226\nExp number, first 60 AAs: 0.22038\nTotal prob of N-in: 0.12994\noutside 1 331\nTMhelix 332 354\ninside 355 372

12675 GCF_000267605.2_ASM26760v2 Escherichia coli TW09098 Proteobacteria; Gammaproteobacteria; Enterobacterales; Enterobacteriaceae; Escherichia; Escherichia coli
MTGDNTLIHSHGINRRDFMKLCAALAATMGLSSKAAAEMAESVTNPQRPPVIWIGAQECT WP_000145410.1
MULTISPECIES: hydrogenase-2 small chain [Proteobacteria] Length: 372\nNumber of predicted TMHs: 1\nExp number of AAs in TMHs: 23.72226\nExp number, first 60 AAs: 0.22038\nTotal prob of N-in: 0.12994\noutside 1 331\nTMhelix 332 354\ninside 355 372

12676 GCF_000267605.2_ASM26760v2 Escherichia coli TW09098 Proteobacteria; Gammaproteobacteria; Enterobacterales; Enterobacteriaceae; Escherichia; Escherichia coli
MNNEETFYQAMRRQGVTRRSFLKYCSLAATSLGLGAGMAPKIAWALENKPRIPVVWIHGL WP_001301613.1
hydrogenase [Escherichia coli] Length: 372\nNumber of predicted TMHs: 2\nExp number of AAs in TMHs: 37.46598\nExp number, first 60 AAs: 19.53139\nTotal prob of N-in: 0.94364\nPOSSIBLE N-term signal sequence\ninside 1 19\nTMhelix 20 39\noutside 40 325\nTMhelix 326 348\ninside 349 372

12677 GCF_000267065.1_ASM26706v1 Escherichia coli TW09109 Proteobacteria; Gammaproteobacteria; Enterobacterales; Enterobacteriaceae; Escherichia; Escherichia coli
MNNEETFYQAMRRQGVTRRSFLKYCSLAATSLGLGAGMAPKIAWALENKPRIPVVWIHGL WP_001301613.1
hydrogenase [Escherichia coli] Length: 372\nNumber of predicted TMHs: 2\nExp number of AAs in TMHs: 37.46598\nExp number, first 60 AAs: 19.53139\nTotal prob of N-in: 0.94364\nPOSSIBLE N-term signal sequence\ninside 1 19\nTMhelix 20 39\noutside 40 325\nTMhelix 326 348\ninside 349 372

12678 GCF_000267065.1_ASM26706v1 Escherichia coli TW09109 Proteobacteria; Gammaproteobacteria; Enterobacterales; Enterobacteriaceae; Escherichia; Escherichia coli
MTGDNTLIHSHGINRRDFMKLCAALAATMGLSSKAAAEMAESVTNPQRPPVIWIGAQECT WP_000145410.1
MULTISPECIES: hydrogenase-2 small chain [Proteobacteria] Length: 372\nNumber of predicted TMHs: 1\nExp number of AAs in TMHs: 23.72226\nExp number, first 60 AAs: 0.22038\nTotal prob of N-in: 0.12994\noutside 1 331\nTMhelix 332 354\ninside 355 372

12679 GCF_000267585.2_ASM26758v2 Escherichia coli TW09195 Proteobacteria; Gammaproteobacteria; Enterobacterales; Enterobacteriaceae; Escherichia; Escherichia coli
MNNEETFYQAMRRQGVTRRSFLKYCSLAATSLGLGAGMAPKIAWALENKPRIPVVWIHGL WP_001301613.1
hydrogenase [Escherichia coli] Length: 372\nNumber of predicted TMHs: 2\nExp number of AAs in TMHs: 37.46598\nExp number, first 60 AAs: 19.53139\nTotal prob of N-in: 0.94364\nPOSSIBLE N-term signal sequence\ninside 1 19\nTMhelix 20 39\noutside 40 325\nTMhelix 326 348\ninside 349 372

12680 GCF_000267585.2_ASM26758v2 Escherichia coli TW09195 Proteobacteria; Gammaproteobacteria; Enterobacterales; Enterobacteriaceae; Escherichia; Escherichia coli
MTGDNTLIHSHGINRRDFMKLCAALAATMGLSSKAAAEMAESVTNPQRPPVIWIGAQECT WP_000145410.1
MULTISPECIES: hydrogenase-2 small chain [Proteobacteria] Length: 372\nNumber of predicted TMHs: 1\nExp number of AAs in TMHs: 23.72226\nExp number, first 60 AAs: 0.22038\nTotal prob of N-in: 0.12994\noutside 1 331\nTMhelix 332 354\ninside 355 372

12681 GCF_000267185.1_ASM26718v1 Escherichia coli TW10119 Proteobacteria; Gammaproteobacteria; Enterobacterales; Enterobacteriaceae; Escherichia; Escherichia coli
MNNEETFYQAMRRQGVTRRSFLKYCSLAATSLGLGAGMAPKIAWALENKPRIPVVWIHGL WP_001301613.1
hydrogenase [Escherichia coli] Length: 372\nNumber of predicted TMHs: 2\nExp number of AAs in TMHs: 37.46598\nExp number, first 60 AAs: 19.53139\nTotal prob of N-in: 0.94364\nPOSSIBLE N-term signal sequence\ninside 1 19\nTMhelix 20 39\noutside 40 325\nTMhelix 326 348\ninside 349 372

12682 GCF_000267185.1_ASM26718v1 Escherichia coli TW10119 Proteobacteria; Gammaproteobacteria; Enterobacterales; Enterobacteriaceae; Escherichia; Escherichia coli
MTGDNTLIHSHGINRRDFMKLCAALAATMGLSSKAAAEMAESVTNPQRPPVIWIGAQECT WP_000145410.1
MULTISPECIES: hydrogenase-2 small chain [Proteobacteria] Length: 372\nNumber of predicted TMHs: 1\nExp number of AAs in TMHs: 23.72226\nExp number, first 60 AAs: 0.22038\nTotal prob of N-in: 0.12994\noutside 1 331\nTMhelix 332 354\ninside 355 372

12683 GCF_000267145.1_ASM26714v1 Escherichia coli TW10246 Proteobacteria; Gammaproteobacteria; Enterobacterales; Enterobacteriaceae; Escherichia; Escherichia coli
MNNEETFYQAMRRQGVTRRSFLKYCSLAATSLGLGAGMAPKIAWALENKPRIPVVWIHGL WP_001301613.1
hydrogenase [Escherichia coli] Length: 372\nNumber of predicted TMHs: 2\nExp number of AAs in TMHs: 37.46598\nExp number, first 60 AAs: 19.53139\nTotal prob of N-in: 0.94364\nPOSSIBLE N-term signal sequence\ninside 1 19\nTMhelix 20 39\noutside 40 325\nTMhelix 326 348\ninside 349 372

12684 GCF_000267145.1_ASM26714v1 Escherichia coli TW10246 Proteobacteria; Gammaproteobacteria; Enterobacterales; Enterobacteriaceae; Escherichia; Escherichia coli
MTGDNTLIHSHGINRRDFMKLCAALAATMGLSSKAAAEMAESVTNPQRPPVIWIGAQECT WP_000145410.1
MULTISPECIES: hydrogenase-2 small chain [Proteobacteria] Length: 372\nNumber of predicted TMHs: 1\nExp number of AAs in TMHs: 23.72226\nExp number, first 60 AAs: 0.22038\nTotal prob of N-in: 0.12994\noutside 1 331\nTMhelix 332 354\ninside 355 372

12685 GCF_000190995.1_ASM19099v1 Escherichia coli TW10509 Proteobacteria; Gammaproteobacteria; Enterobacterales; Enterobacteriaceae; Escherichia; Escherichia coli
MNNEETFYQAMRRQGVTRRSFLKYCSLAATSLGLGAGMAPKIAWALENKPRIPVVWIHGL WP_001353311.1
MULTISPECIES: hydrogenase [Escherichia] Length: 372\nNumber of predicted TMHs: 2\nExp number of

AAs in TMHs: 37.2638699999999\nExp number, first 60 AAs: 19.52924\nTotal prob of N-in: 0.94351\nPOSSIBLE N-term signal sequence\ninside 1 19\nTMhelix 20 39\noutside 40 325\nTMhelix 326 348\ninside 349 372

12686 GCF_000190995.1_ASM19099v1 Escherichia coli TW10509 Proteobacteria; Gammaproteobacteria; Enterobacterales; Enterobacteriaceae; Escherichia; Escherichia coli MTGDNTLIHSHGINRRDFMKLCAALAATMGLSSKAAAEMAESVTNPQRPPVIWIGAQECT WP_000145410.1 MULTISPECIES: hydrogenase-2 small chain [Proteobacteria] Length: 372\nNumber of predicted TMHs: 1\nExp number of AAs in TMHs: 23.72226\nExp number, first 60 AAs: 0.22038\nTotal prob of N-in: 0.12994\noutside 1 331\nTMhelix 332 354\ninside 355 372

12687 GCF_000166535.2_ASM16653v3 Escherichia coli TW10598 Proteobacteria; Gammaproteobacteria; Enterobacterales; Enterobacteriaceae; Escherichia; Escherichia coli MTGDNTLIHSHGINRRDFMKLCAALAATMGLSSKAAAEMAESVTNPQRPPVIWIGAQECT WP_000145410.1 MULTISPECIES: hydrogenase-2 small chain [Proteobacteria] Length: 372\nNumber of predicted TMHs: 1\nExp number of AAs in TMHs: 23.72226\nExp number, first 60 AAs: 0.22038\nTotal prob of N-in: 0.12994\noutside 1 331\nTMhelix 332 354\ninside 355 372

12688 GCF_000166535.2_ASM16653v3 Escherichia coli TW10598 Proteobacteria; Gammaproteobacteria; Enterobacterales; Enterobacteriaceae; Escherichia; Escherichia coli MNNEETFYQAMRRQGVTRRSFLKYCSLAATSLGLGAGMAPKIAWALENKPRIPVVWIHGL WP_001058323.1 MULTISPECIES: hydrogenase-1 small chain [Proteobacteria] Length: 372\nNumber of predicted TMHs: 2\nExp number of AAs in TMHs: 37.2608499999999\nExp number, first 60 AAs: 19.52923\nTotal prob of N-in: 0.94351\nPOSSIBLE N-term signal sequence\ninside 1 19\nTMhelix 20 39\noutside 40 325\nTMhelix 326 348\ninside 349 372

12689 GCF_000166555.1_ASM16655v2 Escherichia coli TW10722 Proteobacteria; Gammaproteobacteria; Enterobacterales; Enterobacteriaceae; Escherichia; Escherichia coli MTGDNTLIHSHGINRRDFMKLCAALAATMGLSSKAAAEMAESVTNPQRPPVIWIGAQECT WP_000145410.1 MULTISPECIES: hydrogenase-2 small chain [Proteobacteria] Length: 372\nNumber of predicted TMHs: 1\nExp number of AAs in TMHs: 23.72226\nExp number, first 60 AAs: 0.22038\nTotal prob of N-in: 0.12994\noutside 1 331\nTMhelix 332 354\ninside 355 372

12690 GCF_000166555.1_ASM16655v2 Escherichia coli TW10722 Proteobacteria; Gammaproteobacteria; Enterobacterales; Enterobacteriaceae; Escherichia; Escherichia coli MNNEETFYQAMRRQGVTRRSFLKYCSLAATSLGLGAGMAPKIAWALENKPRIPVVWIHGL WP_001058323.1 MULTISPECIES: hydrogenase-1 small chain [Proteobacteria] Length: 372\nNumber of predicted TMHs: 2\nExp number of AAs in TMHs: 37.2608499999999\nExp number, first 60 AAs: 19.52923\nTotal prob of N-in: 0.94351\nPOSSIBLE N-term signal sequence\ninside 1 19\nTMhelix 20 39\noutside 40 325\nTMhelix 326 348\ninside 349 372

12691 GCF_000166575.1_ASM16657v2 Escherichia coli TW10828 Proteobacteria; Gammaproteobacteria; Enterobacterales; Enterobacteriaceae; Escherichia; Escherichia coli MTGDNTLIHSHGINRRDFMKLCVALAATMGLSSKAAAEMAESVTNPQRPPVIWIGAQECT WP_000145425.1 hydrogenase 2 small subunit [Escherichia coli] Length: 372\nNumber of predicted TMHs: 1\nExp number of AAs in TMHs: 23.94354\nExp number, first 60 AAs: 0.45405\nTotal prob of N-in: 0.13997\noutside 1 331\nTMhelix 332 354\ninside 355 372

12692 GCF_000166575.1_ASM16657v2 Escherichia coli TW10828 Proteobacteria; Gammaproteobacteria; Enterobacterales; Enterobacteriaceae; Escherichia; Escherichia coli MNNEETFYQAMRRQGVTRRSFLKYCSLAATSLGLGAGMAPKIAWALENKPRIPVVWIHGL WP_001295940.1 MULTISPECIES: hydrogenase [Enterobacteriaceae] Length: 372\nNumber of predicted TMHs: 2\nExp number of AAs in TMHs: 37.2608899999999\nExp number, first 60 AAs: 19.52923\nTotal prob of N-in: 0.94351\nPOSSIBLE N-term signal sequence\ninside 1 19\nTMhelix 20 39\noutside 40 325\nTMhelix 326 348\ninside 349 372

12693 GCF_000267165.1_ASM26716v1 Escherichia coli TW11039 Proteobacteria; Gammaproteobacteria; Enterobacterales; Enterobacteriaceae; Escherichia; Escherichia coli MNNEETFYQAMRRQGVTRRSFLKYCSLAATSLGLGAGMAPKIAWALENKPRIPVVWIHGL WP_001455430.1 hydrogenase [Escherichia coli] Length: 372\nNumber of predicted TMHs: 2\nExp number of AAs in TMHs: 37.13248\nExp number, first 60 AAs: 19.53082\nTotal prob of N-in: 0.94345\nPOSSIBLE N-term signal sequence\ninside 1 19\nTMhelix 20 39\noutside 40 325\nTMhelix 326 348\ninside 349 372

12694 GCF_000267165.1_ASM26716v1 Escherichia coli TW11039 Proteobacteria; Gammaproteobacteria; Enterobacterales; Enterobacteriaceae; Escherichia; Escherichia coli
MTGDNTLIHSHGINRRDFMKLCAALAATMGLSSKAAAEMAESVTNPQRPPVIWIGAQECT WP_000145410.1
MULTISPECIES: hydrogenase-2 small chain [Proteobacteria] Length: 372\nNumber of predicted TMHs: 1\nExp number of AAs in TMHs: 23.72226\nExp number, first 60 AAs: 0.22038\nTotal prob of N-in: 0.12994\noutside 1 331\nTMhelix 332 354\ninside 355 372

12695 GCF_000166595.1_ASM16659v2 Escherichia coli TW11681 Proteobacteria; Gammaproteobacteria; Enterobacterales; Enterobacteriaceae; Escherichia; Escherichia coli
MTGDNTLIHSHGINRRDFMKLCAALAATMGLSSKAAAEMAESVTNPQRPPVIWIGAQECT WP_000145410.1
MULTISPECIES: hydrogenase-2 small chain [Proteobacteria] Length: 372\nNumber of predicted TMHs: 1\nExp number of AAs in TMHs: 23.72226\nExp number, first 60 AAs: 0.22038\nTotal prob of N-in: 0.12994\noutside 1 331\nTMhelix 332 354\ninside 355 372

12696 GCF_000166595.1_ASM16659v2 Escherichia coli TW11681 Proteobacteria; Gammaproteobacteria; Enterobacterales; Enterobacteriaceae; Escherichia; Escherichia coli
MNNEETFYQAMRRQGVTRRSFLKYCSLAATSLGLGAGMAPKIAWALENKPRIPVVWIHGL WP_001058323.1
MULTISPECIES: hydrogenase-1 small chain [Proteobacteria] Length: 372\nNumber of predicted TMHs: 2\nExp number of AAs in TMHs: 37.2608499999999\nExp number, first 60 AAs: 19.52923\nTotal prob of N-in: 0.94351\nPOSSIBLE N-term signal sequence\ninside 1 19\nTMhelix 20 39\noutside 40 325\nTMhelix 326 348\ninside 349 372

12697 GCF_000267405.2_ASM26740v2 Escherichia coli TW14301 Proteobacteria; Gammaproteobacteria; Enterobacterales; Enterobacteriaceae; Escherichia; Escherichia coli
MNNEETFYQAMRRQGVTRRSFLKYCSLAATSLGLGAGMAPKIAWALENKPRIPVVWIHGL WP_001301613.1
hydrogenase [Escherichia coli] Length: 372\nNumber of predicted TMHs: 2\nExp number of AAs in TMHs: 37.46598\nExp number, first 60 AAs: 19.53139\nTotal prob of N-in: 0.94364\nPOSSIBLE N-term signal sequence\ninside 1 19\nTMhelix 20 39\noutside 40 325\nTMhelix 326 348\ninside 349 372

12698 GCF_000267405.2_ASM26740v2 Escherichia coli TW14301 Proteobacteria; Gammaproteobacteria; Enterobacterales; Enterobacteriaceae; Escherichia; Escherichia coli
MTGDNTLIHSHGINRRDFMKLCAALAATMGLSSKAAAEMAESVTNPQRPPVIWIGAQECT WP_000145410.1
MULTISPECIES: hydrogenase-2 small chain [Proteobacteria] Length: 372\nNumber of predicted TMHs: 1\nExp number of AAs in TMHs: 23.72226\nExp number, first 60 AAs: 0.22038\nTotal prob of N-in: 0.12994\noutside 1 331\nTMhelix 332 354\ninside 355 372

12699 GCF_000166615.1_ASM16661v2 Escherichia coli TW14425 Proteobacteria; Gammaproteobacteria; Enterobacterales; Enterobacteriaceae; Escherichia; Escherichia coli
MNNEETFYQAMRRQGVTRRSFLKYCSLAATSLGLGAGMAPKIAWALENKPRIPVVWIHGL WP_001058320.1
hydrogenase [Escherichia coli] Length: 372\nNumber of predicted TMHs: 2\nExp number of AAs in TMHs: 37.26082\nExp number, first 60 AAs: 19.52923\nTotal prob of N-in: 0.94351\nPOSSIBLE N-term signal sequence\ninside 1 19\nTMhelix 20 39\noutside 40 325\nTMhelix 326 348\ninside 349 372

12700 GCF_000166615.1_ASM16661v2 Escherichia coli TW14425 Proteobacteria; Gammaproteobacteria; Enterobacterales; Enterobacteriaceae; Escherichia; Escherichia coli
MTGDNTLIHSHGINRRDFMKLCAALAATMGLSSKAAAEMAESVTNPQRPPVIWIGAQECT WP_000145410.1
MULTISPECIES: hydrogenase-2 small chain [Proteobacteria] Length: 372\nNumber of predicted TMHs: 1\nExp number of AAs in TMHs: 23.72226\nExp number, first 60 AAs: 0.22038\nTotal prob of N-in: 0.12994\noutside 1 331\nTMhelix 332 354\ninside 355 372

12701 GCF_000304075.2_ASM30407v2 Escherichia coli TW15901 Proteobacteria; Gammaproteobacteria; Enterobacterales; Enterobacteriaceae; Escherichia; Escherichia coli
MTGDNTLIHSHGINRRDFMKLCAALAATMGLSSKAAAEMAESVTNPQRPPVIWIGAQECT WP_000145410.1
MULTISPECIES: hydrogenase-2 small chain [Proteobacteria] Length: 372\nNumber of predicted TMHs: 1\nExp number of AAs in TMHs: 23.72226\nExp number, first 60 AAs: 0.22038\nTotal prob of N-in: 0.12994\noutside 1 331\nTMhelix 332 354\ninside 355 372

12702 GCF_000304075.2_ASM30407v2 Escherichia coli TW15901 Proteobacteria; Gammaproteobacteria; Enterobacterales; Enterobacteriaceae; Escherichia; Escherichia coli
MNNEETFYQAMRRQGVTRRSFLKYCSLAATSLGLGAGMAPKIAWALENKPRIPVVWIHGL WP_001058323.1
MULTISPECIES: hydrogenase-1 small chain [Proteobacteria] Length: 372\nNumber of predicted TMHs:

2\nExp number of AAs in TMHs: 37.2608499999999\nExp number, first 60 AAs: 19.52923\nTotal prob of N-in: 0.94351\nPOSSIBLE N-term signal sequence\ninside 1 19\nTMhelix 20 39\noutside 40 325\nTMhelix 326 348\ninside 349 372

12703 GCF_000462885.2_ASM46288v2 Escherichia coli Tx1686 Proteobacteria; Gammaproteobacteria; Enterobacterales; Enterobacteriaceae; Escherichia; Escherichia coli MTGDNTLIHSHGINRRDFMKLCAALAATMGLSSKAAAEMAESVTNPQRPPVIWIGAQECT WP_000145410.1 MULTISPECIES: hydrogenase-2 small chain [Proteobacteria] Length: 372\nNumber of predicted TMHs: 1\nExp number of AAs in TMHs: 23.72226\nExp number, first 60 AAs: 0.22038\nTotal prob of N-in: 0.12994\noutside 1 331\nTMhelix 332 354\ninside 355 372

12704 GCF_000462885.2_ASM46288v2 Escherichia coli Tx1686 Proteobacteria; Gammaproteobacteria; Enterobacterales; Enterobacteriaceae; Escherichia; Escherichia coli MNNEETFYQAMRRQGVTRRSFLKYCSLAATSLGLGAGMAPKIAWALENKPRIPVVWIHGL WP_001301613.1 hydrogenase [Escherichia coli] Length: 372\nNumber of predicted TMHs: 2\nExp number of AAs in TMHs: 37.46598\nExp number, first 60 AAs: 19.53139\nTotal prob of N-in: 0.94364\nPOSSIBLE N-term signal sequence\ninside 1 19\nTMhelix 20 39\noutside 40 325\nTMhelix 326 348\ninside 349 372

12705 GCF_000462905.2_ASM46290v2 Escherichia coli Tx3800 Proteobacteria; Gammaproteobacteria; Enterobacterales; Enterobacteriaceae; Escherichia; Escherichia coli MTGDNTLIHSHGINRRDFMKLCAALAATMGLSSKAAAEMAESVTNPQRPPVIWIGAQECT WP_000145410.1 MULTISPECIES: hydrogenase-2 small chain [Proteobacteria] Length: 372\nNumber of predicted TMHs: 1\nExp number of AAs in TMHs: 23.72226\nExp number, first 60 AAs: 0.22038\nTotal prob of N-in: 0.12994\noutside 1 331\nTMhelix 332 354\ninside 355 372

12706 GCF_000462905.2_ASM46290v2 Escherichia coli Tx3800 Proteobacteria; Gammaproteobacteria; Enterobacterales; Enterobacteriaceae; Escherichia; Escherichia coli MNNEETFYQAMRRQGVTRRSFLKYCSLAATSLGLGAGMAPKIAWALENKPRIPVVWIHGL WP_001301613.1 hydrogenase [Escherichia coli] Length: 372\nNumber of predicted TMHs: 2\nExp number of AAs in TMHs: 37.46598\nExp number, first 60 AAs: 19.53139\nTotal prob of N-in: 0.94364\nPOSSIBLE N-term signal sequence\ninside 1 19\nTMhelix 20 39\noutside 40 325\nTMhelix 326 348\ninside 349 372

12707 GCF_000692795.1_Esch_coli_UCI_51_V1 Escherichia coli UCI 51 Proteobacteria; Gammaproteobacteria; Enterobacterales; Enterobacteriaceae; Escherichia; Escherichia coli MNNEETFYQAMRRQGVTRRSFLKYCSLAATSLGLGAGMAPKIAWALENKPRIPVVWIHGL WP_001295940.1 MULTISPECIES: hydrogenase [Enterobacteriaceae] Length: 372\nNumber of predicted TMHs: 2\nExp number of AAs in TMHs: 37.2608899999999\nExp number, first 60 AAs: 19.52923\nTotal prob of N-in: 0.94351\nPOSSIBLE N-term signal sequence\ninside 1 19\nTMhelix 20 39\noutside 40 325\nTMhelix 326 348\ninside 349 372

12708 GCF_000692795.1_Esch_coli_UCI_51_V1 Escherichia coli UCI 51 Proteobacteria; Gammaproteobacteria; Enterobacterales; Enterobacteriaceae; Escherichia; Escherichia coli MTGDNTLIHSHGINRRDFMKLCAALAATMGLSSKAAAEMVESVTNPQRPPVIWIGAQECT WP_000145423.1 hydrogenase 2 small subunit [Escherichia coli] Length: 372\nNumber of predicted TMHs: 1\nExp number of AAs in TMHs: 23.78201\nExp number, first 60 AAs: 0.28355\nTotal prob of N-in: 0.13257\noutside 1 331\nTMhelix 332 354\ninside 355 372

12709 GCF_000692815.1_Esch_coli_UCI_53_V1 Escherichia coli UCI 53 Proteobacteria; Gammaproteobacteria; Enterobacterales; Enterobacteriaceae; Escherichia; Escherichia coli MNNEETFYQAMRRQGVTRRSFLKYCSLAATSLGLGAGMAPKIAWALENKPRIPVVWIHGL WP_001058320.1 hydrogenase [Escherichia coli] Length: 372\nNumber of predicted TMHs: 2\nExp number of AAs in TMHs: 37.26082\nExp number, first 60 AAs: 19.52923\nTotal prob of N-in: 0.94351\nPOSSIBLE N-term signal sequence\ninside 1 19\nTMhelix 20 39\noutside 40 325\nTMhelix 326 348\ninside 349 372

12710 GCF_000692815.1_Esch_coli_UCI_53_V1 Escherichia coli UCI 53 Proteobacteria; Gammaproteobacteria; Enterobacterales; Enterobacteriaceae; Escherichia; Escherichia coli MTGDNTLIHSHGINRRDFMKLCAALAATMGLSSKAAAEMAESVTNPQRPPVIWIGAQECT WP_000145410.1 MULTISPECIES: hydrogenase-2 small chain [Proteobacteria] Length: 372\nNumber of predicted TMHs: 1\nExp number of AAs in TMHs: 23.72226\nExp number, first 60 AAs: 0.22038\nTotal prob of N-in: 0.12994\noutside 1 331\nTMhelix 332 354\ninside 355 372

12711 GCF_000692835.1_Esch_coli_UCI_57_V1 Escherichia coli UCI 57 Proteobacteria; Gammaproteobacteria; Enterobacterales; Enterobacteriaceae; Escherichia; Escherichia coli MTGDNTLIHSHGINRRDFMKLCAALAATMGLSSKAAAEMAESVTNPQRPPVIWIGAQECT WP_000145410.1 MULTISPECIES: hydrogenase-2 small chain [Proteobacteria] Length: 372\nNumber of predicted TMHs: 1\nExp number of AAs in TMHs: 23.72226\nExp number, first 60 AAs: 0.22038\nTotal prob of N-in: 0.12994\noutside 1 331\nTMhelix 332 354\ninside 355 372

12712 GCF_000692835.1_Esch_coli_UCI_57_V1 Escherichia coli UCI 57 Proteobacteria; Gammaproteobacteria; Enterobacterales; Enterobacteriaceae; Escherichia; Escherichia coli MNNEETFYQAMRRQGVTRRSFLKYCSLAATSLGLGAGMAPKIAWALENKPRIPVVWIHGL WP_001295940.1 MULTISPECIES: hydrogenase [Enterobacteriaceae] Length: 372\nNumber of predicted TMHs: 2\nExp number of AAs in TMHs: 37.2608899999999\nExp number, first 60 AAs: 19.52923\nTotal prob of N-in: 0.94351\nPOSSIBLE N-term signal sequence\ninside 1 19\nTMhelix 20 39\noutside 40 325\nTMhelix 326 348\ninside 349 372

12713 GCF_000692855.1_Esch_coli_UCI_58_V1 Escherichia coli UCI 58 Proteobacteria; Gammaproteobacteria; Enterobacterales; Enterobacteriaceae; Escherichia; Escherichia coli MNNEETFYQAMRRQGVTRRSFLKYCSLAATSLGLGAGMAPKIAWALENKPRIPVVWIHGL WP_001295940.1 MULTISPECIES: hydrogenase [Enterobacteriaceae] Length: 372\nNumber of predicted TMHs: 2\nExp number of AAs in TMHs: 37.2608899999999\nExp number, first 60 AAs: 19.52923\nTotal prob of N-in: 0.94351\nPOSSIBLE N-term signal sequence\ninside 1 19\nTMhelix 20 39\noutside 40 325\nTMhelix 326 348\ninside 349 372

12714 GCF_000692855.1_Esch_coli_UCI_58_V1 Escherichia coli UCI 58 Proteobacteria; Gammaproteobacteria; Enterobacterales; Enterobacteriaceae; Escherichia; Escherichia coli MTGDNTLIHSHGINRRDFMKLCAALAATMGLSSKAAAEMAESVTNPQRPPVIWIGAQECT WP_000145410.1 MULTISPECIES: hydrogenase-2 small chain [Proteobacteria] Length: 372\nNumber of predicted TMHs: 1\nExp number of AAs in TMHs: 23.72226\nExp number, first 60 AAs: 0.22038\nTotal prob of N-in: 0.12994\noutside 1 331\nTMhelix 332 354\ninside 355 372

12715 GCF_000692875.1_Esch_coli_UCI_65_V1 Escherichia coli UCI 65 Proteobacteria; Gammaproteobacteria; Enterobacterales; Enterobacteriaceae; Escherichia; Escherichia coli MTGDNTLIHSHGINRRDFMKLCAALAATMGLSSKAAAEMAESVTNPQRPPVIWIGAQECT WP_000145410.1 MULTISPECIES: hydrogenase-2 small chain [Proteobacteria] Length: 372\nNumber of predicted TMHs: 1\nExp number of AAs in TMHs: 23.72226\nExp number, first 60 AAs: 0.22038\nTotal prob of N-in: 0.12994\noutside 1 331\nTMhelix 332 354\ninside 355 372

12716 GCF_000692875.1_Esch_coli_UCI_65_V1 Escherichia coli UCI 65 Proteobacteria; Gammaproteobacteria; Enterobacterales; Enterobacteriaceae; Escherichia; Escherichia coli MNNEETFYQAMRRQGVTRRSFLKYCSLAATSLGLGAGMAPKIAWALENKPRIPVVWIHGL WP_001058323.1 MULTISPECIES: hydrogenase-1 small chain [Proteobacteria] Length: 372\nNumber of predicted TMHs: 2\nExp number of AAs in TMHs: 37.2608499999999\nExp number, first 60 AAs: 19.52923\nTotal prob of N-in: 0.94351\nPOSSIBLE N-term signal sequence\ninside 1 19\nTMhelix 20 39\noutside 40 325\nTMhelix 326 348\ninside 349 372

12717 GCF_000692895.1_Esch_coli_UCI_66_V1 Escherichia coli UCI 66 Proteobacteria; Gammaproteobacteria; Enterobacterales; Enterobacteriaceae; Escherichia; Escherichia coli MNNEETFYQAMRRQGVTRRSFLKYCSLAATSLGLGAGMAPKIAWALENKPRIPVVWIHGL WP_001058320.1 hydrogenase [Escherichia coli] Length: 372\nNumber of predicted TMHs: 2\nExp number of AAs in TMHs: 37.26082\nExp number, first 60 AAs: 19.52923\nTotal prob of N-in: 0.94351\nPOSSIBLE N-term signal sequence\ninside 1 19\nTMhelix 20 39\noutside 40 325\nTMhelix 326 348\ninside 349 372

12718 GCF_000692895.1_Esch_coli_UCI_66_V1 Escherichia coli UCI 66 Proteobacteria; Gammaproteobacteria; Enterobacterales; Enterobacteriaceae; Escherichia; Escherichia coli MTGDNTLIHSHGINRRDFMKLCAALAATMGLSSKAAAEMAESVTNPQRPPVIWIGAQECT WP_000145410.1 MULTISPECIES: hydrogenase-2 small chain [Proteobacteria] Length: 372\nNumber of predicted TMHs: 1\nExp number of AAs in TMHs: 23.72226\nExp number, first 60 AAs: 0.22038\nTotal prob of N-in: 0.12994\noutside 1 331\nTMhelix 332 354\ninside 355 372

12719 GCF_000148605.1_ASM14860v1 Escherichia coli UM146 Proteobacteria; Gammaproteobacteria; Enterobacterales; Enterobacteriaceae; Escherichia; Escherichia coli

MTGDNTLIHSHGINRRDFMKLCAALAATMGLSSKAAAEMAESVTNPQRPPVIWIGAQECT WP_000145410.1 MULTISPECIES: hydrogenase-2 small chain [Proteobacteria] Length: 372\nNumber of predicted TMHs: 1\nExp number of AAs in TMHs: 23.72226\nExp number, first 60 AAs: 0.22038\nTotal prob of N-in: 0.12994\noutside 1 331\nTMhelix 332 354\ninside 355 372

12720 GCF_000459955.1_Esch_coli_UMEA_3014-1_V1 Escherichia coli UMEA 3014-1 Proteobacteria; Gammaproteobacteria; Enterobacterales; Enterobacteriaceae; Escherichia; Escherichia coli MNNEETFYQAMRRQGVTRRSFLKYCSLAATSLGLGAGMAPKIAWALENKPRIPVVWIHGL WP_001058323.1 MULTISPECIES: hydrogenase-1 small chain [Proteobacteria] Length: 372\nNumber of predicted TMHs: 2\nExp number of AAs in TMHs: 37.2608499999999\nExp number, first 60 AAs: 19.52923\nTotal prob of N-in: 0.94351\nPOSSIBLE N-term signal sequence\ninside 1 19\nTMhelix 20 39\noutside 40 325\nTMhelix 326 348\ninside 349 372

12721 GCF_000459955.1_Esch_coli_UMEA_3014-1_V1 Escherichia coli UMEA 3014-1 Proteobacteria; Gammaproteobacteria; Enterobacterales; Enterobacteriaceae; Escherichia; Escherichia coli MTGDNTLIHSHGINRRDFMKLCAALAATMGLSSKAAAEMAESVTNPQRPPVIWIGAQECT WP_000145410.1 MULTISPECIES: hydrogenase-2 small chain [Proteobacteria] Length: 372\nNumber of predicted TMHs: 1\nExp number of AAs in TMHs: 23.72226\nExp number, first 60 AAs: 0.22038\nTotal prob of N-in: 0.12994\noutside 1 331\nTMhelix 332 354\ninside 355 372

12722 GCF_000459975.1_Esch_coli_UMEA_3022-1_V1 Escherichia coli UMEA 3022-1 Proteobacteria; Gammaproteobacteria; Enterobacterales; Enterobacteriaceae; Escherichia; Escherichia coli MNNEETFYQAMRRQGVTRRSFLKYCSLAATSLGLGAGMAPKIAWALENKPRIPVVWIHGL WP_001058323.1 MULTISPECIES: hydrogenase-1 small chain [Proteobacteria] Length: 372\nNumber of predicted TMHs: 2\nExp number of AAs in TMHs: 37.2608499999999\nExp number, first 60 AAs: 19.52923\nTotal prob of N-in: 0.94351\nPOSSIBLE N-term signal sequence\ninside 1 19\nTMhelix 20 39\noutside 40 325\nTMhelix 326 348\ninside 349 372

12723 GCF_000459975.1_Esch_coli_UMEA_3022-1_V1 Escherichia coli UMEA 3022-1 Proteobacteria; Gammaproteobacteria; Enterobacterales; Enterobacteriaceae; Escherichia; Escherichia coli MTGDNTLIHSHGINRRDFMKLCAALAATMGLSSKAAAEMAESVTNPQRPPVIWIGAQECT WP_000145410.1 MULTISPECIES: hydrogenase-2 small chain [Proteobacteria] Length: 372\nNumber of predicted TMHs: 1\nExp number of AAs in TMHs: 23.72226\nExp number, first 60 AAs: 0.22038\nTotal prob of N-in: 0.12994\noutside 1 331\nTMhelix 332 354\ninside 355 372

12724 GCF_000459995.1_Esch_coli_UMEA_3033-1_V1 Escherichia coli UMEA 3033-1 Proteobacteria; Gammaproteobacteria; Enterobacterales; Enterobacteriaceae; Escherichia; Escherichia coli MTGDNTLIHSHGINRRDFMKLCAALAATMGLSSKAAAEMAESVTNPQRPPVIWIGAQECT WP_000145410.1 MULTISPECIES: hydrogenase-2 small chain [Proteobacteria] Length: 372\nNumber of predicted TMHs: 1\nExp number of AAs in TMHs: 23.72226\nExp number, first 60 AAs: 0.22038\nTotal prob of N-in: 0.12994\noutside 1 331\nTMhelix 332 354\ninside 355 372

12725 GCF_000459995.1_Esch_coli_UMEA_3033-1_V1 Escherichia coli UMEA 3033-1 Proteobacteria; Gammaproteobacteria; Enterobacterales; Enterobacteriaceae; Escherichia; Escherichia coli MNNEETFYQAMRRQGVTRRSFLKYCSLAATSLGLGAGMAPKIAWALENKPRIPVVWIHGL WP_001058323.1 MULTISPECIES: hydrogenase-1 small chain [Proteobacteria] Length: 372\nNumber of predicted TMHs: 2\nExp number of AAs in TMHs: 37.2608499999999\nExp number, first 60 AAs: 19.52923\nTotal prob of N-in: 0.94351\nPOSSIBLE N-term signal sequence\ninside 1 19\nTMhelix 20 39\noutside 40 325\nTMhelix 326 348\ninside 349 372

12726 GCF_000460015.1_Esch_coli_UMEA_3041-1_V1 Escherichia coli UMEA 3041-1 Proteobacteria; Gammaproteobacteria; Enterobacterales; Enterobacteriaceae; Escherichia; Escherichia coli MTGDNTLIHSHGINRRDFMKLCAALAATMGLSSKAAAEMAESVTNPQRPPVIWIGAQECT WP_000145410.1 MULTISPECIES: hydrogenase-2 small chain [Proteobacteria] Length: 372\nNumber of predicted TMHs: 1\nExp number of AAs in TMHs: 23.72226\nExp number, first 60 AAs: 0.22038\nTotal prob of N-in: 0.12994\noutside 1 331\nTMhelix 332 354\ninside 355 372

12727 GCF_000460035.1_Esch_coli_UMEA_3052-1_V1 Escherichia coli UMEA 3052-1 Proteobacteria; Gammaproteobacteria; Enterobacterales; Enterobacteriaceae; Escherichia; Escherichia coli MTGDNTLIHSHGINRRDFMKLCAALAATMGLSSKAAAEMAESVTNPQRPPVIWIGAQECT WP_000145410.1 MULTISPECIES: hydrogenase-2 small chain [Proteobacteria] Length: 372\nNumber of predicted TMHs:

1\nExp number of AAs in TMHs: 23.72226\nExp number, first 60 AAs: 0.22038\nTotal prob of N-in: 0.12994\noutside 1 331\nTMhelix 332 354\ninside 355 372

12728 GCF_000460035.1_Esch_coli_UMEA_3052-1_V1 Escherichia coli UMEA 3052-1 Proteobacteria; Gammaproteobacteria; Enterobacterales; Enterobacteriaceae; Escherichia; Escherichia coli MNNEETFYQAMRRQGVTRRSFLKYCSLAATSLGLGAGMAPKIAWALENKPRIPVVWIHGL WP_001058323.1 MULTISPECIES: hydrogenase-1 small chain [Proteobacteria] Length: 372\nNumber of predicted TMHs: 2\nExp number of AAs in TMHs: 37.2608499999999\nExp number, first 60 AAs: 19.52923\nTotal prob of N-in: 0.94351\nPOSSIBLE N-term signal sequence\ninside 1 19\nTMhelix 20 39\noutside 40 325\nTMhelix 326 348\ninside 349 372

12729 GCF_000460055.1_Esch_coli_UMEA_3053-1_V1 Escherichia coli UMEA 3053-1 Proteobacteria; Gammaproteobacteria; Enterobacterales; Enterobacteriaceae; Escherichia; Escherichia coli MNNEETFYQAMRRQGVTRRSFLKYCSLAATSLGLGAGMAPKIAWALENKPRIPVVWIHGL WP_001538532.1 hydrogenase-1 small chain [Escherichia coli] Length: 372\nNumber of predicted TMHs: 2\nExp number of AAs in TMHs: 37.27041\nExp number, first 60 AAs: 19.52927\nTotal prob of N-in: 0.94352\nPOSSIBLE N-term signal sequence\ninside 1 19\nTMhelix 20 39\noutside 40 325\nTMhelix 326 348\ninside 349 372

12730 GCF_000460055.1_Esch_coli_UMEA_3053-1_V1 Escherichia coli UMEA 3053-1 Proteobacteria; Gammaproteobacteria; Enterobacterales; Enterobacteriaceae; Escherichia; Escherichia coli MTGDNTLIHSHGINRRDFMKLCAALAATMGLSSKAAAEMVESVTNPQRPPVIWIGAQECT WP_000145423.1 hydrogenase 2 small subunit [Escherichia coli] Length: 372\nNumber of predicted TMHs: 1\nExp number of AAs in TMHs: 23.78201\nExp number, first 60 AAs: 0.28355\nTotal prob of N-in: 0.13257\noutside 1 331\nTMhelix 332 354\ninside 355 372

12731 GCF_000460075.1_Esch_coli_UMEA_3065-1_V1 Escherichia coli UMEA 3065-1 Proteobacteria; Gammaproteobacteria; Enterobacterales; Enterobacteriaceae; Escherichia; Escherichia coli MTGDNTLIHSHGINRRDFMKLCAALAATMGLSSKAAAEMAESVTNPQRPPVIWIGAQECT WP_000145410.1 MULTISPECIES: hydrogenase-2 small chain [Proteobacteria] Length: 372\nNumber of predicted TMHs: 1\nExp number of AAs in TMHs: 23.72226\nExp number, first 60 AAs: 0.22038\nTotal prob of N-in: 0.12994\noutside 1 331\nTMhelix 332 354\ninside 355 372

12732 GCF_000460075.1_Esch_coli_UMEA_3065-1_V1 Escherichia coli UMEA 3065-1 Proteobacteria; Gammaproteobacteria; Enterobacterales; Enterobacteriaceae; Escherichia; Escherichia coli MNNEETFYQAMRRQGVTRRSFLKYCSLAATSLGLGAGMAPKIAWALENKPRIPVVWIHGL WP_001058320.1 hydrogenase [Escherichia coli] Length: 372\nNumber of predicted TMHs: 2\nExp number of AAs in TMHs: 37.26082\nExp number, first 60 AAs: 19.52923\nTotal prob of N-in: 0.94351\nPOSSIBLE N-term signal sequence\ninside 1 19\nTMhelix 20 39\noutside 40 325\nTMhelix 326 348\ninside 349 372

12733 GCF_000460095.1_Esch_coli_UMEA_3087-1_V1 Escherichia coli UMEA 3087-1 Proteobacteria; Gammaproteobacteria; Enterobacterales; Enterobacteriaceae; Escherichia; Escherichia coli MNNEETFYQAMRRQGVTRRSFLKYCSLAATSLGLGAGMAPKIAWALENKPRIPVVWIHGL WP_001058323.1 MULTISPECIES: hydrogenase-1 small chain [Proteobacteria] Length: 372\nNumber of predicted TMHs: 2\nExp number of AAs in TMHs: 37.2608499999999\nExp number, first 60 AAs: 19.52923\nTotal prob of N-in: 0.94351\nPOSSIBLE N-term signal sequence\ninside 1 19\nTMhelix 20 39\noutside 40 325\nTMhelix 326 348\ninside 349 372

12734 GCF_000460095.1_Esch_coli_UMEA_3087-1_V1 Escherichia coli UMEA 3087-1 Proteobacteria; Gammaproteobacteria; Enterobacterales; Enterobacteriaceae; Escherichia; Escherichia coli MTGDNTLIHSHGINRRDFMKLCAALAATMGLSSKAAAEMAESVTNPQRPPVIWIGAQECT WP_000145410.1 MULTISPECIES: hydrogenase-2 small chain [Proteobacteria] Length: 372\nNumber of predicted TMHs: 1\nExp number of AAs in TMHs: 23.72226\nExp number, first 60 AAs: 0.22038\nTotal prob of N-in: 0.12994\noutside 1 331\nTMhelix 332 354\ninside 355 372

12735 GCF_000460115.1_Esch_coli_UMEA_3088-1_V1 Escherichia coli UMEA 3088-1 Proteobacteria; Gammaproteobacteria; Enterobacterales; Enterobacteriaceae; Escherichia; Escherichia coli MNNEETFYQAMRRQGVTRRSFLKYCSLAATSLGLGAGMAPKIAWALENKPRIPVVWIHGL WP_001538532.1 hydrogenase-1 small chain [Escherichia coli] Length: 372\nNumber of predicted TMHs: 2\nExp number of AAs in TMHs: 37.27041\nExp number, first 60 AAs: 19.52927\nTotal prob of N-in: 0.94352\nPOSSIBLE N-term signal sequence\ninside 1 19\nTMhelix 20 39\noutside 40 325\nTMhelix 326 348\ninside 349 372

12736 GCF_000460115.1_Esch_coli_UMEA_3088-1_V1 Escherichia coli UMEA 3088-1 Proteobacteria; Gammaproteobacteria; Enterobacterales; Enterobacteriaceae; Escherichia; Escherichia coli MTGDNTLIHSHGINRRDFMKLCAALAATMGLSSKAAAEMVESVTNPQRPPVIWIGAQECT WP_000145423.1 hydrogenase 2 small subunit [Escherichia coli] Length: 372\nNumber of predicted TMHs: 1\nExp number of AAs in TMHs: 23.78201\nExp number, first 60 AAs: 0.28355\nTotal prob of N-in: 0.13257\noutside 1 331\nTMhelix 332 354\ninside 355 372

12737 GCF_000460135.1_Esch_coli_UMEA_3097-1_V1 Escherichia coli UMEA 3097-1 Proteobacteria; Gammaproteobacteria; Enterobacterales; Enterobacteriaceae; Escherichia; Escherichia coli MTGDNTLIHSHGINRRDFMKLCAALAATMGLSSKAAAEMAESVTNPQRPPVIWIGAQECT WP_000145410.1 MULTISPECIES: hydrogenase-2 small chain [Proteobacteria] Length: 372\nNumber of predicted TMHs: 1\nExp number of AAs in TMHs: 23.72226\nExp number, first 60 AAs: 0.22038\nTotal prob of N-in: 0.12994\noutside 1 331\nTMhelix 332 354\ninside 355 372

12738 GCF_000460135.1_Esch_coli_UMEA_3097-1_V1 Escherichia coli UMEA 3097-1 Proteobacteria; Gammaproteobacteria; Enterobacterales; Enterobacteriaceae; Escherichia; Escherichia coli MNNEETFYQAMRRQGVTRRSFLKYCSLAATSLGLGAGMAPKIAWALENKPRIPVVWIHGL WP_001058323.1 MULTISPECIES: hydrogenase-1 small chain [Proteobacteria] Length: 372\nNumber of predicted TMHs: 2\nExp number of AAs in TMHs: 37.2608499999999\nExp number, first 60 AAs: 19.52923\nTotal prob of N-in: 0.94351\nPOSSIBLE N-term signal sequence\ninside 1 19\nTMhelix 20 39\noutside 40 325\nTMhelix 326 348\ninside 349 372

12739 GCF_000460155.1_Esch_coli_UMEA_3108-1_V1 Escherichia coli UMEA 3108-1 Proteobacteria; Gammaproteobacteria; Enterobacterales; Enterobacteriaceae; Escherichia; Escherichia coli MTGDNTLIHSHGINRRDFMKLCAALAATMGLSSKAAAEMAESVTNPQRPPVIWIGAQECT WP_000145410.1 MULTISPECIES: hydrogenase-2 small chain [Proteobacteria] Length: 372\nNumber of predicted TMHs: 1\nExp number of AAs in TMHs: 23.72226\nExp number, first 60 AAs: 0.22038\nTotal prob of N-in: 0.12994\noutside 1 331\nTMhelix 332 354\ninside 355 372

12740 GCF_000460155.1_Esch_coli_UMEA_3108-1_V1 Escherichia coli UMEA 3108-1 Proteobacteria; Gammaproteobacteria; Enterobacterales; Enterobacteriaceae; Escherichia; Escherichia coli MNNEETFYQAMRRQGVTRRSFLKYCSLAATSLGLGAGMAPKIAWALENKPRIPVVWIHGL WP_001295940.1 MULTISPECIES: hydrogenase [Enterobacteriaceae] Length: 372\nNumber of predicted TMHs: 2\nExp number of AAs in TMHs: 37.2608899999999\nExp number, first 60 AAs: 19.52923\nTotal prob of N-in: 0.94351\nPOSSIBLE N-term signal sequence\ninside 1 19\nTMhelix 20 39\noutside 40 325\nTMhelix 326 348\ninside 349 372

12741 GCF_000460175.1_Esch_coli_UMEA_3113-1_V1 Escherichia coli UMEA 3113-1 Proteobacteria; Gammaproteobacteria; Enterobacterales; Enterobacteriaceae; Escherichia; Escherichia coli MTGDNTLIHSHGINRRDFMKLCAALAATMGLSSKAAAEMAESVTNPQRPPVIWIGAQECT WP_000145410.1 MULTISPECIES: hydrogenase-2 small chain [Proteobacteria] Length: 372\nNumber of predicted TMHs: 1\nExp number of AAs in TMHs: 23.72226\nExp number, first 60 AAs: 0.22038\nTotal prob of N-in: 0.12994\noutside 1 331\nTMhelix 332 354\ninside 355 372

12742 GCF_000460175.1_Esch_coli_UMEA_3113-1_V1 Escherichia coli UMEA 3113-1 Proteobacteria; Gammaproteobacteria; Enterobacterales; Enterobacteriaceae; Escherichia; Escherichia coli MNNEETFYQAMRRQGVTRRSFLKYCSLAATSLGLGAGMAPKIAWALENKPRIPVVWIHGL WP_001058323.1 MULTISPECIES: hydrogenase-1 small chain [Proteobacteria] Length: 372\nNumber of predicted TMHs: 2\nExp number of AAs in TMHs: 37.2608499999999\nExp number, first 60 AAs: 19.52923\nTotal prob of N-in: 0.94351\nPOSSIBLE N-term signal sequence\ninside 1 19\nTMhelix 20 39\noutside 40 325\nTMhelix 326 348\ninside 349 372

12743 GCF_000460195.1_Esch_coli_UMEA_3117-1_V1 Escherichia coli UMEA 3117-1 Proteobacteria; Gammaproteobacteria; Enterobacterales; Enterobacteriaceae; Escherichia; Escherichia coli MTGDNTLIHSHGINRRDFMKLCAALAATMGLSSKAAAEMAESVTNPQRPPVIWIGAQECT WP_000145410.1 MULTISPECIES: hydrogenase-2 small chain [Proteobacteria] Length: 372\nNumber of predicted TMHs: 1\nExp number of AAs in TMHs: 23.72226\nExp number, first 60 AAs: 0.22038\nTotal prob of N-in: 0.12994\noutside 1 331\nTMhelix 332 354\ninside 355 372

12744 GCF_000460195.1_Esch_coli_UMEA_3117-1_V1 Escherichia coli UMEA 3117-1 Proteobacteria; Gammaproteobacteria; Enterobacterales; Enterobacteriaceae; Escherichia; Escherichia coli

MNNEETFYQAMRRQGVTRRSFLKYCSLAATSLGLGAGMAPKIAWALENKPRIPVVWIHGL WP_001058323.1 MULTISPECIES: hydrogenase-1 small chain [Proteobacteria] Length: 372\nNumber of predicted TMHs: 2\nExp number of AAs in TMHs: 37.2608499999999\nExp number, first 60 AAs: 19.52923\nTotal prob of N-in: 0.94351\nPOSSIBLE N-term signal sequence\ninside 1 19\nTMhelix 20 39\noutside 40 325\nTMhelix 326 348\ninside 349 372

12745 GCF_000460215.1_Esch_coli_UMEA_3121-1_V1 Escherichia coli UMEA 3121-1 Proteobacteria; Gammaproteobacteria; Enterobacterales; Enterobacteriaceae; Escherichia; Escherichia coli MTGDNTLIHSHGINRRDFMKLCAALAATMGLSSKAAAEMAESVTNPQRPPVIWIGAQECT WP_000145410.1 MULTISPECIES: hydrogenase-2 small chain [Proteobacteria] Length: 372\nNumber of predicted TMHs: 1\nExp number of AAs in TMHs: 23.72226\nExp number, first 60 AAs: 0.22038\nTotal prob of N-in: 0.12994\noutside 1 331\nTMhelix 332 354\ninside 355 372

12746 GCF_000460215.1_Esch_coli_UMEA_3121-1_V1 Escherichia coli UMEA 3121-1 Proteobacteria; Gammaproteobacteria; Enterobacterales; Enterobacteriaceae; Escherichia; Escherichia coli MNNEETFYQAMRRQGVTRRSFLKYCSLAATSLGLGAGMAPKIAWALENKPRIPVVWIHGL WP_001058323.1 MULTISPECIES: hydrogenase-1 small chain [Proteobacteria] Length: 372\nNumber of predicted TMHs: 2\nExp number of AAs in TMHs: 37.2608499999999\nExp number, first 60 AAs: 19.52923\nTotal prob of N-in: 0.94351\nPOSSIBLE N-term signal sequence\ninside 1 19\nTMhelix 20 39\noutside 40 325\nTMhelix 326 348\ninside 349 372

12747 GCF_000460235.1_Esch_coli_UMEA_3122-1_V1 Escherichia coli UMEA 3122-1 Proteobacteria; Gammaproteobacteria; Enterobacterales; Enterobacteriaceae; Escherichia; Escherichia coli MTGDNTLIHSHGINRRDFMKLCAALAATMGLSSKAAAEMVESVTNPQRPPVIWIGAQECT WP_000145423.1 hydrogenase 2 small subunit [Escherichia coli] Length: 372\nNumber of predicted TMHs: 1\nExp number of AAs in TMHs: 23.78201\nExp number, first 60 AAs: 0.28355\nTotal prob of N-in: 0.13257\noutside 1 331\nTMhelix 332 354\ninside 355 372

12748 GCF_000460235.1_Esch_coli_UMEA_3122-1_V1 Escherichia coli UMEA 3122-1 Proteobacteria; Gammaproteobacteria; Enterobacterales; Enterobacteriaceae; Escherichia; Escherichia coli MNNEETFYQAMRRQGVTRRSFLKYCSLAATSLGLGAGMAPKIAWALENKPRIPVVWIHGL WP_001538532.1 hydrogenase-1 small chain [Escherichia coli] Length: 372\nNumber of predicted TMHs: 2\nExp number of AAs in TMHs: 37.27041\nExp number, first 60 AAs: 19.52927\nTotal prob of N-in: 0.94352\nPOSSIBLE N-term signal sequence\ninside 1 19\nTMhelix 20 39\noutside 40 325\nTMhelix 326 348\ninside 349 372

12749 GCF_000460255.1_Esch_coli_UMEA_3124-1_V1 Escherichia coli UMEA 3124-1 Proteobacteria; Gammaproteobacteria; Enterobacterales; Enterobacteriaceae; Escherichia; Escherichia coli MTGDNTLIHSHGINRRDFMKLCAALAATMGLSSKAAAEMAESVTNPQRPPVIWIGAQECT WP_021557215.1 hydrogenase-2 small chain [Escherichia coli] Length: 372\nNumber of predicted TMHs: 1\nExp number of AAs in TMHs: 23.85895\nExp number, first 60 AAs: 0.2195\nTotal prob of N-in: 0.14303\noutside 1 331\nTMhelix 332 354\ninside 355 372

12750 GCF_000460255.1_Esch_coli_UMEA_3124-1_V1 Escherichia coli UMEA 3124-1 Proteobacteria; Gammaproteobacteria; Enterobacterales; Enterobacteriaceae; Escherichia; Escherichia coli MNNEETFYQAMRRQGVTRRSFLKYCSLAATSLGLGAGMAPKIAWALENKPRIPVVWIHGL WP_001058323.1 MULTISPECIES: hydrogenase-1 small chain [Proteobacteria] Length: 372\nNumber of predicted TMHs: 2\nExp number of AAs in TMHs: 37.2608499999999\nExp number, first 60 AAs: 19.52923\nTotal prob of N-in: 0.94351\nPOSSIBLE N-term signal sequence\ninside 1 19\nTMhelix 20 39\noutside 40 325\nTMhelix 326 348\ninside 349 372

12751 GCF_000460275.1_Esch_coli_UMEA_3139-1_V1 Escherichia coli UMEA 3139-1 Proteobacteria; Gammaproteobacteria; Enterobacterales; Enterobacteriaceae; Escherichia; Escherichia coli MTGDNTLIHSHGINRRDFMKLCAALAATMGLSSKAAAEMAESVTNPQRPPVIWIGAQECT WP_000145410.1 MULTISPECIES: hydrogenase-2 small chain [Proteobacteria] Length: 372\nNumber of predicted TMHs: 1\nExp number of AAs in TMHs: 23.72226\nExp number, first 60 AAs: 0.22038\nTotal prob of N-in: 0.12994\noutside 1 331\nTMhelix 332 354\ninside 355 372

12752 GCF_000460275.1_Esch_coli_UMEA_3139-1_V1 Escherichia coli UMEA 3139-1 Proteobacteria; Gammaproteobacteria; Enterobacterales; Enterobacteriaceae; Escherichia; Escherichia coli MNNEETFYQAMRRQGVTRRSFLKYCSLAATSLGLGAGMAPKIAWALENKPRIPVVWIHGL WP_021557704.1 hydrogenase-1 small chain [Escherichia coli] Length: 372\nNumber of predicted TMHs: 2\nExp number of

AAs in TMHs: 37.2609699999999\nExp number, first 60 AAs: 19.52923\nTotal prob of N-in: 0.94351\nPOSSIBLE N-term signal sequence\ninside 1 19\nTMhelix 20 39\noutside 40 325\nTMhelix 326 348\ninside 349 372

12753 GCF_000460295.1_Esch_coli_UMEA_3140-1_V1 Escherichia coli UMEA 3140-1 Proteobacteria; Gammaproteobacteria; Enterobacterales; Enterobacteriaceae; Escherichia; Escherichia coli MTGDNTLIHSHGINRRDFMKLCAALAATMGLSSKAAAEMAESVTNPQRPPVIWIGAQECT WP_000145410.1 MULTISPECIES: hydrogenase-2 small chain [Proteobacteria] Length: 372\nNumber of predicted TMHs: 1\nExp number of AAs in TMHs: 23.72226\nExp number, first 60 AAs: 0.22038\nTotal prob of N-in: 0.12994\noutside 1 331\nTMhelix 332 354\ninside 355 372

12754 GCF_000460315.1_Esch_coli_UMEA_3144-1_V1 Escherichia coli UMEA 3144-1 Proteobacteria; Gammaproteobacteria; Enterobacterales; Enterobacteriaceae; Escherichia; Escherichia coli MNNEETFYQAMRRQGVTRRSFLKYCSLAATSLGLGAGMAPKIAWALENKPRIPVVWIHGL WP_001058323.1 MULTISPECIES: hydrogenase-1 small chain [Proteobacteria] Length: 372\nNumber of predicted TMHs: 2\nExp number of AAs in TMHs: 37.2608499999999\nExp number, first 60 AAs: 19.52923\nTotal prob of N-in: 0.94351\nPOSSIBLE N-term signal sequence\ninside 1 19\nTMhelix 20 39\noutside 40 325\nTMhelix 326 348\ninside 349 372

12755 GCF_000460315.1_Esch_coli_UMEA_3144-1_V1 Escherichia coli UMEA 3144-1 Proteobacteria; Gammaproteobacteria; Enterobacterales; Enterobacteriaceae; Escherichia; Escherichia coli MTGDNTLIHSHGINRRDFMKLCAALAATMGLSSKAAAEMAESVTNPQRPPVIWIGAQECT WP_000145410.1 MULTISPECIES: hydrogenase-2 small chain [Proteobacteria] Length: 372\nNumber of predicted TMHs: 1\nExp number of AAs in TMHs: 23.72226\nExp number, first 60 AAs: 0.22038\nTotal prob of N-in: 0.12994\noutside 1 331\nTMhelix 332 354\ninside 355 372

12756 GCF_000495075.1_Esch_coli_UMEA_3148-1_V1 Escherichia coli UMEA 3148-1 Proteobacteria; Gammaproteobacteria; Enterobacterales; Enterobacteriaceae; Escherichia; Escherichia coli MNNEETFYQAMRRQGVTRRSFLKYCSLAATSLGLGAGMAPKIAWALENKPRIPVVWIHGL WP_001058323.1 MULTISPECIES: hydrogenase-1 small chain [Proteobacteria] Length: 372\nNumber of predicted TMHs: 2\nExp number of AAs in TMHs: 37.2608499999999\nExp number, first 60 AAs: 19.52923\nTotal prob of N-in: 0.94351\nPOSSIBLE N-term signal sequence\ninside 1 19\nTMhelix 20 39\noutside 40 325\nTMhelix 326 348\ninside 349 372

12757 GCF_000495075.1_Esch_coli_UMEA_3148-1_V1 Escherichia coli UMEA 3148-1 Proteobacteria; Gammaproteobacteria; Enterobacterales; Enterobacteriaceae; Escherichia; Escherichia coli MTGDNTLIHSHGINRRDFMKLCAALAATMGLSSKAAAEMAESVTNPQRPPVIWIGAQECT WP_000145410.1 MULTISPECIES: hydrogenase-2 small chain [Proteobacteria] Length: 372\nNumber of predicted TMHs: 1\nExp number of AAs in TMHs: 23.72226\nExp number, first 60 AAs: 0.22038\nTotal prob of N-in: 0.12994\noutside 1 331\nTMhelix 332 354\ninside 355 372

12758 GCF_000460335.1_Esch_coli_UMEA_3150-1_V1 Escherichia coli UMEA 3150-1 Proteobacteria; Gammaproteobacteria; Enterobacterales; Enterobacteriaceae; Escherichia; Escherichia coli MNNEETFYQAMRRQGVTRRSFLKYCSLAATSLGLGAGMAPKIAWALENKPRIPVVWIHGL WP_001058323.1 MULTISPECIES: hydrogenase-1 small chain [Proteobacteria] Length: 372\nNumber of predicted TMHs: 2\nExp number of AAs in TMHs: 37.2608499999999\nExp number, first 60 AAs: 19.52923\nTotal prob of N-in: 0.94351\nPOSSIBLE N-term signal sequence\ninside 1 19\nTMhelix 20 39\noutside 40 325\nTMhelix 326 348\ninside 349 372

12759 GCF_000460335.1_Esch_coli_UMEA_3150-1_V1 Escherichia coli UMEA 3150-1 Proteobacteria; Gammaproteobacteria; Enterobacterales; Enterobacteriaceae; Escherichia; Escherichia coli MTGDNTLIHSHGINRRDFMKLCAALAATMGLSSKAAAEMAESVTNPQRPPVIWIGAQECT WP_000145410.1 MULTISPECIES: hydrogenase-2 small chain [Proteobacteria] Length: 372\nNumber of predicted TMHs: 1\nExp number of AAs in TMHs: 23.72226\nExp number, first 60 AAs: 0.22038\nTotal prob of N-in: 0.12994\noutside 1 331\nTMhelix 332 354\ninside 355 372

12760 GCF_000460355.1_Esch_coli_UMEA_3151-1_V1 Escherichia coli UMEA 3151-1 Proteobacteria; Gammaproteobacteria; Enterobacterales; Enterobacteriaceae; Escherichia; Escherichia coli MNNEETFYQAMRRQGVTRRSFLKYCSLAATSLGLGAGMAPKIAWALENKPRIPVVWIHGL WP_001058323.1 MULTISPECIES: hydrogenase-1 small chain [Proteobacteria] Length: 372\nNumber of predicted TMHs: 2\nExp number of AAs in TMHs: 37.2608499999999\nExp number, first 60 AAs: 19.52923\nTotal prob of N-in:

0.94351\nPOSSIBLE N-term signal sequence\ninside 1 19\nTMhelix 20 39\noutside 40 325\nTMhelix 326 348\ninside 349 372

12761 GCF_000460355.1_Esch_coli_UMEA_3151-1_V1 Escherichia coli UMEA 3151-1 Proteobacteria; Gammaproteobacteria; Enterobacterales; Enterobacteriaceae; Escherichia; Escherichia coli MTGDNTLIHSHGINRRDFMKLCAALAATMGLSSKAAAEMAESVTNPQRPPVIWIGAQECT WP_000145410.1 MULTISPECIES: hydrogenase-2 small chain [Proteobacteria] Length: 372\nNumber of predicted TMHs: 1\nExp number of AAs in TMHs: 23.72226\nExp number, first 60 AAs: 0.22038\nTotal prob of N-in: 0.12994\noutside 1 331\nTMhelix 332 354\ninside 355 372

12762 GCF_000460375.1_Esch_coli_UMEA_3152-1_V1 Escherichia coli UMEA 3152-1 Proteobacteria; Gammaproteobacteria; Enterobacterales; Enterobacteriaceae; Escherichia; Escherichia coli MNNEETFYQAMRRQGVTRRSFLKYCSLAATSLGLGAGMAPKIAWALENKPRIPVVWIHGL WP_001058323.1 MULTISPECIES: hydrogenase-1 small chain [Proteobacteria] Length: 372\nNumber of predicted TMHs: 2\nExp number of AAs in TMHs: 37.2608499999999\nExp number, first 60 AAs: 19.52923\nTotal prob of N-in: 0.94351\nPOSSIBLE N-term signal sequence\ninside 1 19\nTMhelix 20 39\noutside 40 325\nTMhelix 326 348\ninside 349 372

12763 GCF_000460375.1_Esch_coli_UMEA_3152-1_V1 Escherichia coli UMEA 3152-1 Proteobacteria; Gammaproteobacteria; Enterobacterales; Enterobacteriaceae; Escherichia; Escherichia coli MTGDNTLIHSHGINRRDFMKLCAALAATMGLSSKAAAEMAESVTNPQRPPVIWIGAQECT WP_000145410.1 MULTISPECIES: hydrogenase-2 small chain [Proteobacteria] Length: 372\nNumber of predicted TMHs: 1\nExp number of AAs in TMHs: 23.72226\nExp number, first 60 AAs: 0.22038\nTotal prob of N-in: 0.12994\noutside 1 331\nTMhelix 332 354\ninside 355 372

12764 GCF_000460395.1_Esch_coli_UMEA_3155-1_V1 Escherichia coli UMEA 3155-1 Proteobacteria; Gammaproteobacteria; Enterobacterales; Enterobacteriaceae; Escherichia; Escherichia coli MTGDNTLIHSHGINRRDFMKLCAALAATMGLSSKAAAEMAESVTNPQRPPVIWIGAQECT WP_000145410.1 MULTISPECIES: hydrogenase-2 small chain [Proteobacteria] Length: 372\nNumber of predicted TMHs: 1\nExp number of AAs in TMHs: 23.72226\nExp number, first 60 AAs: 0.22038\nTotal prob of N-in: 0.12994\noutside 1 331\nTMhelix 332 354\ninside 355 372

12765 GCF_000460395.1_Esch_coli_UMEA_3155-1_V1 Escherichia coli UMEA 3155-1 Proteobacteria; Gammaproteobacteria; Enterobacterales; Enterobacteriaceae; Escherichia; Escherichia coli MNNEETFYQAMRRQGVTRRSFLKYCSLAATSLGLGAGMAPKIAWALENKPRIPVVWIHGL WP_001295940.1 MULTISPECIES: hydrogenase [Enterobacteriaceae] Length: 372\nNumber of predicted TMHs: 2\nExp number of AAs in TMHs: 37.2608899999999\nExp number, first 60 AAs: 19.52923\nTotal prob of N-in: 0.94351\nPOSSIBLE N-term signal sequence\ninside 1 19\nTMhelix 20 39\noutside 40 325\nTMhelix 326 348\ninside 349 372

12766 GCF_000460415.1_Esch_coli_UMEA_3159-1_V1 Escherichia coli UMEA 3159-1 Proteobacteria; Gammaproteobacteria; Enterobacterales; Enterobacteriaceae; Escherichia; Escherichia coli MNNEETFYQAMRRQGVTRRSFLKYCSLAATSLGLGAGMAPKIAWALENKPRIPVVWIHGL WP_001058323.1 MULTISPECIES: hydrogenase-1 small chain [Proteobacteria] Length: 372\nNumber of predicted TMHs: 2\nExp number of AAs in TMHs: 37.2608499999999\nExp number, first 60 AAs: 19.52923\nTotal prob of N-in: 0.94351\nPOSSIBLE N-term signal sequence\ninside 1 19\nTMhelix 20 39\noutside 40 325\nTMhelix 326 348\ninside 349 372

12767 GCF_000460415.1_Esch_coli_UMEA_3159-1_V1 Escherichia coli UMEA 3159-1 Proteobacteria; Gammaproteobacteria; Enterobacterales; Enterobacteriaceae; Escherichia; Escherichia coli MTGDNTLIHSHGINRRDFMKLCAALAATMGLSSKAAAEMAESVTNPQRPPVIWIGAQECT WP_000145410.1 MULTISPECIES: hydrogenase-2 small chain [Proteobacteria] Length: 372\nNumber of predicted TMHs: 1\nExp number of AAs in TMHs: 23.72226\nExp number, first 60 AAs: 0.22038\nTotal prob of N-in: 0.12994\noutside 1 331\nTMhelix 332 354\ninside 355 372

12768 GCF_000460435.1_Esch_coli_UMEA_3160-1_V1 Escherichia coli UMEA 3160-1 Proteobacteria; Gammaproteobacteria; Enterobacterales; Enterobacteriaceae; Escherichia; Escherichia coli MNNEETFYQAMRRQGVTRRSFLKYCSLAATSLGLGAGMAPKIAWALENKPRIPVVWIHGL WP_021520630.1 hydrogenase-1 small chain [Escherichia coli] Length: 372\nNumber of predicted TMHs: 2\nExp number of AAs in TMHs: 37.8197599999999\nExp number, first 60 AAs: 19.53295\nTotal prob of N-in: 0.94396\nPOSSIBLE N-term signal sequence\ninside 1 19\nTMhelix 20 39\noutside 40 325\nTMhelix 326 348\ninside 349 372

12769 GCF_000460435.1_Esch_coli_UMEA_3160-1_V1 Escherichia coli UMEA 3160-1 Proteobacteria; Gammaproteobacteria; Enterobacterales; Enterobacteriaceae; Escherichia; Escherichia coli MTGDNTLIHSHGINRRDFMKLCAALAATMGLSSKAAAEMAESVTNPQRPPVIWIGAQECT WP_000145418.1 hydrogenase 2 small subunit [Escherichia coli] Length: 372\nNumber of predicted TMHs: 1\nExp number of AAs in TMHs: 23.72313\nExp number, first 60 AAs: 0.22034\nTotal prob of N-in: 0.12997\noutside 1 331\nTMhelix 332 354\ninside 355 372

12770 GCF_000460455.1_Esch_coli_UMEA_3161-1_V1 Escherichia coli UMEA 3161-1 Proteobacteria; Gammaproteobacteria; Enterobacterales; Enterobacteriaceae; Escherichia; Escherichia coli MTGDNTLIHSHGINRRDFMKLCAALAATMGLSSKAAAEMAESVTNPQRPPVIWIGAQECT WP_021519494.1 hydrogenase-2 small chain [Escherichia coli] Length: 372\nNumber of predicted TMHs: 1\nExp number of AAs in TMHs: 23.72331\nExp number, first 60 AAs: 0.22039\nTotal prob of N-in: 0.12995\noutside 1 331\nTMhelix 332 354\ninside 355 372

12771 GCF_000460455.1_Esch_coli_UMEA_3161-1_V1 Escherichia coli UMEA 3161-1 Proteobacteria; Gammaproteobacteria; Enterobacterales; Enterobacteriaceae; Escherichia; Escherichia coli MNNEETFYQAMRRQGVTRRSFLKYCSLAATSLGLGAGMAPKIAWALENKPRIPVVWIHGL WP_001058323.1 MULTISPECIES: hydrogenase-1 small chain [Proteobacteria] Length: 372\nNumber of predicted TMHs: 2\nExp number of AAs in TMHs: 37.2608499999999\nExp number, first 60 AAs: 19.52923\nTotal prob of N-in: 0.94351\nPOSSIBLE N-term signal sequence\ninside 1 19\nTMhelix 20 39\noutside 40 325\nTMhelix 326 348\ninside 349 372

12772 GCF_000460475.1_Esch_coli_UMEA_3162-1_V1 Escherichia coli UMEA 3162-1 Proteobacteria; Gammaproteobacteria; Enterobacterales; Enterobacteriaceae; Escherichia; Escherichia coli MTGDNTLIHSHGINRRDFMKLCAALAATMGLSSKAAAEMAESVTNPQRPPVIWIGAQECT WP_000145410.1 MULTISPECIES: hydrogenase-2 small chain [Proteobacteria] Length: 372\nNumber of predicted TMHs: 1\nExp number of AAs in TMHs: 23.72226\nExp number, first 60 AAs: 0.22038\nTotal prob of N-in: 0.12994\noutside 1 331\nTMhelix 332 354\ninside 355 372

12773 GCF_000460495.1_Esch_coli_UMEA_3163-1_V1 Escherichia coli UMEA 3163-1 Proteobacteria; Gammaproteobacteria; Enterobacterales; Enterobacteriaceae; Escherichia; Escherichia coli MNNEETFYQAMRRQGVTRRSFLKYCSLAATSLGLGAGMAPKIAWALENKPRIPVVWIHGL WP_001058323.1 MULTISPECIES: hydrogenase-1 small chain [Proteobacteria] Length: 372\nNumber of predicted TMHs: 2\nExp number of AAs in TMHs: 37.2608499999999\nExp number, first 60 AAs: 19.52923\nTotal prob of N-in: 0.94351\nPOSSIBLE N-term signal sequence\ninside 1 19\nTMhelix 20 39\noutside 40 325\nTMhelix 326 348\ninside 349 372

12774 GCF_000460495.1_Esch_coli_UMEA_3163-1_V1 Escherichia coli UMEA 3163-1 Proteobacteria; Gammaproteobacteria; Enterobacterales; Enterobacteriaceae; Escherichia; Escherichia coli MTGDNTLIHSHGINRRDFMKLCAALAATMGLSSKAAAEMAESVTNPQRPPVIWIGAQECT WP_000145410.1 MULTISPECIES: hydrogenase-2 small chain [Proteobacteria] Length: 372\nNumber of predicted TMHs: 1\nExp number of AAs in TMHs: 23.72226\nExp number, first 60 AAs: 0.22038\nTotal prob of N-in: 0.12994\noutside 1 331\nTMhelix 332 354\ninside 355 372

12775 GCF_000460515.1_Esch_coli_UMEA_3172-1_V1 Escherichia coli UMEA 3172-1 Proteobacteria; Gammaproteobacteria; Enterobacterales; Enterobacteriaceae; Escherichia; Escherichia coli MTGDNTLIHSHGINRRDFMKLCAALAATMGLSSKAAAEMAESVTNPQRPPVIWIGAQECT WP_000145410.1 MULTISPECIES: hydrogenase-2 small chain [Proteobacteria] Length: 372\nNumber of predicted TMHs: 1\nExp number of AAs in TMHs: 23.72226\nExp number, first 60 AAs: 0.22038\nTotal prob of N-in: 0.12994\noutside 1 331\nTMhelix 332 354\ninside 355 372

12776 GCF_000460515.1_Esch_coli_UMEA_3172-1_V1 Escherichia coli UMEA 3172-1 Proteobacteria; Gammaproteobacteria; Enterobacterales; Enterobacteriaceae; Escherichia; Escherichia coli MNNEETFYQAMRRQGVTRRSFLKYCSLAATSLGLGAGMAPKIAWALENKPRIPVVWIHGL WP_001058323.1 MULTISPECIES: hydrogenase-1 small chain [Proteobacteria] Length: 372\nNumber of predicted TMHs: 2\nExp number of AAs in TMHs: 37.2608499999999\nExp number, first 60 AAs: 19.52923\nTotal prob of N-in: 0.94351\nPOSSIBLE N-term signal sequence\ninside 1 19\nTMhelix 20 39\noutside 40 325\nTMhelix 326 348\ninside 349 372

12777 GCF_000460535.1_Esch_coli_UMEA_3173-1_V1 Escherichia coli UMEA 3173-1 Proteobacteria; Gammaproteobacteria; Enterobacterales; Enterobacteriaceae; Escherichia; Escherichia coli

MNNEETFYQAMRRQGVTRRSFLKYCSLAATSLGLGAGMAPKIAWALENKPRIPVVWIHGL WP_001058323.1 MULTISPECIES: hydrogenase-1 small chain [Proteobacteria] Length: 372\nNumber of predicted TMHs: 2\nExp number of AAs in TMHs: 37.2608499999999\nExp number, first 60 AAs: 19.52923\nTotal prob of N-in: 0.94351\nPOSSIBLE N-term signal sequence\ninside 1 19\nTMhelix 20 39\noutside 40 325\nTMhelix 326 348\ninside 349 372

12778 GCF_000460535.1_Esch_coli_UMEA_3173-1_V1 Escherichia coli UMEA 3173-1 Proteobacteria; Gammaproteobacteria; Enterobacterales; Enterobacteriaceae; Escherichia; Escherichia coli MTGDNTLIHSHGINRRDFMKLCAALAATMGLSSKAAAEMAESVTNPQRPPVIWIGAQECT WP_000145410.1 MULTISPECIES: hydrogenase-2 small chain [Proteobacteria] Length: 372\nNumber of predicted TMHs: 1\nExp number of AAs in TMHs: 23.72226\nExp number, first 60 AAs: 0.22038\nTotal prob of N-in: 0.12994\noutside 1 331\nTMhelix 332 354\ninside 355 372

12779 GCF_000460555.1_Esch_coli_UMEA_3174-1_V1 Escherichia coli UMEA 3174-1 Proteobacteria; Gammaproteobacteria; Enterobacterales; Enterobacteriaceae; Escherichia; Escherichia coli MTGDNTLIHSHGINRRDFMKLCAALAATMGLSSKAAAEMAESVTNPQRPPVIWIGAQECT WP_000145410.1 MULTISPECIES: hydrogenase-2 small chain [Proteobacteria] Length: 372\nNumber of predicted TMHs: 1\nExp number of AAs in TMHs: 23.72226\nExp number, first 60 AAs: 0.22038\nTotal prob of N-in: 0.12994\noutside 1 331\nTMhelix 332 354\ninside 355 372

12780 GCF_000460555.1_Esch_coli_UMEA_3174-1_V1 Escherichia coli UMEA 3174-1 Proteobacteria; Gammaproteobacteria; Enterobacterales; Enterobacteriaceae; Escherichia; Escherichia coli MNNEETFYQAMRRQGVTRRSFLKYCSLAATSLGLGAGMAPKIAWALENKPRIPVVWIHGL WP_001058323.1 MULTISPECIES: hydrogenase-1 small chain [Proteobacteria] Length: 372\nNumber of predicted TMHs: 2\nExp number of AAs in TMHs: 37.2608499999999\nExp number, first 60 AAs: 19.52923\nTotal prob of N-in: 0.94351\nPOSSIBLE N-term signal sequence\ninside 1 19\nTMhelix 20 39\noutside 40 325\nTMhelix 326 348\ninside 349 372

12781 GCF_000460575.1_Esch_coli_UMEA_3175-1_V1 Escherichia coli UMEA 3175-1 Proteobacteria; Gammaproteobacteria; Enterobacterales; Enterobacteriaceae; Escherichia; Escherichia coli MNNEETFYQAMRRQGVTRRSFLKYCSLAATSLGLGAGMAPKIAWALENKPRIPVVWIHGL WP_001058323.1 MULTISPECIES: hydrogenase-1 small chain [Proteobacteria] Length: 372\nNumber of predicted TMHs: 2\nExp number of AAs in TMHs: 37.2608499999999\nExp number, first 60 AAs: 19.52923\nTotal prob of N-in: 0.94351\nPOSSIBLE N-term signal sequence\ninside 1 19\nTMhelix 20 39\noutside 40 325\nTMhelix 326 348\ninside 349 372

12782 GCF_000460575.1_Esch_coli_UMEA_3175-1_V1 Escherichia coli UMEA 3175-1 Proteobacteria; Gammaproteobacteria; Enterobacterales; Enterobacteriaceae; Escherichia; Escherichia coli MTGDNTLIHSHGINRRDFMKLCAALAATMGLSSKAAAEMAESVTNPQRPPVIWIGAQECT WP_000145410.1 MULTISPECIES: hydrogenase-2 small chain [Proteobacteria] Length: 372\nNumber of predicted TMHs: 1\nExp number of AAs in TMHs: 23.72226\nExp number, first 60 AAs: 0.22038\nTotal prob of N-in: 0.12994\noutside 1 331\nTMhelix 332 354\ninside 355 372

12783 GCF_000460595.1_Esch_coli_UMEA_3176-1_V1 Escherichia coli UMEA 3176-1 Proteobacteria; Gammaproteobacteria; Enterobacterales; Enterobacteriaceae; Escherichia; Escherichia coli MNNEETFYQAMRRQGVTRRSFLKYCSLAATSLGLGAGMAPKIAWALENKPRIPVVWIHGL WP_001058323.1 MULTISPECIES: hydrogenase-1 small chain [Proteobacteria] Length: 372\nNumber of predicted TMHs: 2\nExp number of AAs in TMHs: 37.2608499999999\nExp number, first 60 AAs: 19.52923\nTotal prob of N-in: 0.94351\nPOSSIBLE N-term signal sequence\ninside 1 19\nTMhelix 20 39\noutside 40 325\nTMhelix 326 348\ninside 349 372

12784 GCF_000460595.1_Esch_coli_UMEA_3176-1_V1 Escherichia coli UMEA 3176-1 Proteobacteria; Gammaproteobacteria; Enterobacterales; Enterobacteriaceae; Escherichia; Escherichia coli MTGDNTLIHSHGINRRDFMKLCAALAATMGLSSKAAAEMAESVTNPQRPPVIWIGAQECT WP_000145410.1 MULTISPECIES: hydrogenase-2 small chain [Proteobacteria] Length: 372\nNumber of predicted TMHs: 1\nExp number of AAs in TMHs: 23.72226\nExp number, first 60 AAs: 0.22038\nTotal prob of N-in: 0.12994\noutside 1 331\nTMhelix 332 354\ninside 355 372

12785 GCF_000460615.1_Esch_coli_UMEA_3178-1_V1 Escherichia coli UMEA 3178-1 Proteobacteria; Gammaproteobacteria; Enterobacterales; Enterobacteriaceae; Escherichia; Escherichia coli MTGDNTLIHSHGINRRDFMKLCAALAATMGLSSKAAAEMAESVTNPQRPPVIWIGAQECT WP_000145410.1

MULTISPECIES: hydrogenase-2 small chain [Proteobacteria] Length: 372\nNumber of predicted TMHs: 1\nExp number of AAs in TMHs: 23.72226\nExp number, first 60 AAs: 0.22038\nTotal prob of N-in: 0.12994\noutside 1 331\nTMhelix 332 354\ninside 355 372

12786 GCF_000460615.1_Esch_coli_UMEA_3178-1_V1 Escherichia coli UMEA 3178-1 Proteobacteria; Gammaproteobacteria; Enterobacterales; Enterobacteriaceae; Escherichia; Escherichia coli MNNEETFYQAMRRQGVTRRSFLKYCSLAATSLGLGAGMAPKIAWALENKPRIPVVWIHGL WP_001058323.1 MULTISPECIES: hydrogenase-1 small chain [Proteobacteria] Length: 372\nNumber of predicted TMHs: 2\nExp number of AAs in TMHs: 37.2608499999999\nExp number, first 60 AAs: 19.52923\nTotal prob of N-in: 0.94351\nPOSSIBLE N-term signal sequence\ninside 1 19\nTMhelix 20 39\noutside 40 325\nTMhelix 326 348\ninside 349 372

12787 GCF_000460635.1_Esch_coli_UMEA_3180-1_V1 Escherichia coli UMEA 3180-1 Proteobacteria; Gammaproteobacteria; Enterobacterales; Enterobacteriaceae; Escherichia; Escherichia coli MTGDNTLIHSHGINRRDFMKLCAALAATMGLSSKAAAEMAESVTNPQRPPVIWIGAQECT WP_000145410.1 MULTISPECIES: hydrogenase-2 small chain [Proteobacteria] Length: 372\nNumber of predicted TMHs: 1\nExp number of AAs in TMHs: 23.72226\nExp number, first 60 AAs: 0.22038\nTotal prob of N-in: 0.12994\noutside 1 331\nTMhelix 332 354\ninside 355 372

12788 GCF_000460635.1_Esch_coli_UMEA_3180-1_V1 Escherichia coli UMEA 3180-1 Proteobacteria; Gammaproteobacteria; Enterobacterales; Enterobacteriaceae; Escherichia; Escherichia coli MNNEETFYQAMRRQGVTRRSFLKYCSLAATSLGLGAGMAPKIAWALENKPRIPVVWIHGL WP_001058323.1 MULTISPECIES: hydrogenase-1 small chain [Proteobacteria] Length: 372\nNumber of predicted TMHs: 2\nExp number of AAs in TMHs: 37.2608499999999\nExp number, first 60 AAs: 19.52923\nTotal prob of N-in: 0.94351\nPOSSIBLE N-term signal sequence\ninside 1 19\nTMhelix 20 39\noutside 40 325\nTMhelix 326 348\ninside 349 372

12789 GCF_000460655.1_Esch_coli_UMEA_3185-1_V1 Escherichia coli UMEA 3185-1 Proteobacteria; Gammaproteobacteria; Enterobacterales; Enterobacteriaceae; Escherichia; Escherichia coli MTGDNTLIHSHGINRRDFMKLCAALAATMGLSSKAAAEMAESVTNPQRPPVIWIGAQECT WP_000145410.1 MULTISPECIES: hydrogenase-2 small chain [Proteobacteria] Length: 372\nNumber of predicted TMHs: 1\nExp number of AAs in TMHs: 23.72226\nExp number, first 60 AAs: 0.22038\nTotal prob of N-in: 0.12994\noutside 1 331\nTMhelix 332 354\ninside 355 372

12790 GCF_000460655.1_Esch_coli_UMEA_3185-1_V1 Escherichia coli UMEA 3185-1 Proteobacteria; Gammaproteobacteria; Enterobacterales; Enterobacteriaceae; Escherichia; Escherichia coli MNNEETFYQAMRRQGVTRRSFLKYCSLAATSLGLGAGMAPKIAWALENKPRIPVVWIHGL WP_001058323.1 MULTISPECIES: hydrogenase-1 small chain [Proteobacteria] Length: 372\nNumber of predicted TMHs: 2\nExp number of AAs in TMHs: 37.2608499999999\nExp number, first 60 AAs: 19.52923\nTotal prob of N-in: 0.94351\nPOSSIBLE N-term signal sequence\ninside 1 19\nTMhelix 20 39\noutside 40 325\nTMhelix 326 348\ninside 349 372

12791 GCF_000460675.1_Esch_coli_UMEA_3190-1_V1 Escherichia coli UMEA 3190-1 Proteobacteria; Gammaproteobacteria; Enterobacterales; Enterobacteriaceae; Escherichia; Escherichia coli MTGDNTLIHSHGINRRDFMKLCAALAATMGLSSKAAAEMAESVTNPQRPPVIWIGAQECT WP_000145410.1 MULTISPECIES: hydrogenase-2 small chain [Proteobacteria] Length: 372\nNumber of predicted TMHs: 1\nExp number of AAs in TMHs: 23.72226\nExp number, first 60 AAs: 0.22038\nTotal prob of N-in: 0.12994\noutside 1 331\nTMhelix 332 354\ninside 355 372

12792 GCF_000460675.1_Esch_coli_UMEA_3190-1_V1 Escherichia coli UMEA 3190-1 Proteobacteria; Gammaproteobacteria; Enterobacterales; Enterobacteriaceae; Escherichia; Escherichia coli MNNEETFYQAMRRQGVTRRSFLKYCSLAATSLGLGAGMAPKIAWALENKPRIPVVWIHGL WP_001058323.1 MULTISPECIES: hydrogenase-1 small chain [Proteobacteria] Length: 372\nNumber of predicted TMHs: 2\nExp number of AAs in TMHs: 37.2608499999999\nExp number, first 60 AAs: 19.52923\nTotal prob of N-in: 0.94351\nPOSSIBLE N-term signal sequence\ninside 1 19\nTMhelix 20 39\noutside 40 325\nTMhelix 326 348\ninside 349 372

12793 GCF_000460695.1_Esch_coli_UMEA_3193-1_V1 Escherichia coli UMEA 3193-1 Proteobacteria; Gammaproteobacteria; Enterobacterales; Enterobacteriaceae; Escherichia; Escherichia coli MNNEETFYQAMRRQGVTRRSFLKYCSLAATSLGLGAGMAPKIAWALENKPRIPVVWIHGL WP_001058323.1 MULTISPECIES: hydrogenase-1 small chain [Proteobacteria] Length: 372\nNumber of predicted TMHs:

2\nExp number of AAs in TMHs: 37.2608499999999\nExp number, first 60 AAs: 19.52923\nTotal prob of N-in: 0.94351\nPOSSIBLE N-term signal sequence\ninside 1 19\nTMhelix 20 39\noutside 40 325\nTMhelix 326 348\ninside 349 372

12794 GCF_000460695.1_Esch_coli_UMEA_3193-1_V1 Escherichia coli UMEA 3193-1 Proteobacteria; Gammaproteobacteria; Enterobacterales; Enterobacteriaceae; Escherichia; Escherichia coli MTGDNTLIHSHGINRRDFMKLCAALAATMGLSSKAAAEMAESVTNPQRPPVIWIGAQECT WP_000145410.1 MULTISPECIES: hydrogenase-2 small chain [Proteobacteria] Length: 372\nNumber of predicted TMHs: 1\nExp number of AAs in TMHs: 23.72226\nExp number, first 60 AAs: 0.22038\nTotal prob of N-in: 0.12994\noutside 1 331\nTMhelix 332 354\ninside 355 372

12795 GCF_000460715.1_Esch_coli_UMEA_3199-1_V1 Escherichia coli UMEA 3199-1 Proteobacteria; Gammaproteobacteria; Enterobacterales; Enterobacteriaceae; Escherichia; Escherichia coli MTGDNTLIHSHGINRRDFMKLCAALAATMGLSSKAAAEMAESVTNPQRPPVIWIGAQECT WP_000145410.1 MULTISPECIES: hydrogenase-2 small chain [Proteobacteria] Length: 372\nNumber of predicted TMHs: 1\nExp number of AAs in TMHs: 23.72226\nExp number, first 60 AAs: 0.22038\nTotal prob of N-in: 0.12994\noutside 1 331\nTMhelix 332 354\ninside 355 372

12796 GCF_000460715.1_Esch_coli_UMEA_3199-1_V1 Escherichia coli UMEA 3199-1 Proteobacteria; Gammaproteobacteria; Enterobacterales; Enterobacteriaceae; Escherichia; Escherichia coli MNNEETFYQAMRRQGVTRRSFLKYCSLAATSLGLGAGMAPKIAWALENKPRIPVVWIHGL WP_001058323.1 MULTISPECIES: hydrogenase-1 small chain [Proteobacteria] Length: 372\nNumber of predicted TMHs: 2\nExp number of AAs in TMHs: 37.2608499999999\nExp number, first 60 AAs: 19.52923\nTotal prob of N-in: 0.94351\nPOSSIBLE N-term signal sequence\ninside 1 19\nTMhelix 20 39\noutside 40 325\nTMhelix 326 348\ninside 349 372

12797 GCF_000460735.1_Esch_coli_UMEA_3200-1_V1 Escherichia coli UMEA 3200-1 Proteobacteria; Gammaproteobacteria; Enterobacterales; Enterobacteriaceae; Escherichia; Escherichia coli MTGDNTLIHSHGINRRDFMKLCAALAATMGLSSKAAAEMAESVTNPQRPPVIWIGAQECT WP_000145410.1 MULTISPECIES: hydrogenase-2 small chain [Proteobacteria] Length: 372\nNumber of predicted TMHs: 1\nExp number of AAs in TMHs: 23.72226\nExp number, first 60 AAs: 0.22038\nTotal prob of N-in: 0.12994\noutside 1 331\nTMhelix 332 354\ninside 355 372

12798 GCF_000460735.1_Esch_coli_UMEA_3200-1_V1 Escherichia coli UMEA 3200-1 Proteobacteria; Gammaproteobacteria; Enterobacterales; Enterobacteriaceae; Escherichia; Escherichia coli MNNEETFYQAMRRQGVTRRSFLKYCSLAATSLGLGAGMAPKIAWALENKPRIPVVWIHGL WP_001058323.1 MULTISPECIES: hydrogenase-1 small chain [Proteobacteria] Length: 372\nNumber of predicted TMHs: 2\nExp number of AAs in TMHs: 37.2608499999999\nExp number, first 60 AAs: 19.52923\nTotal prob of N-in: 0.94351\nPOSSIBLE N-term signal sequence\ninside 1 19\nTMhelix 20 39\noutside 40 325\nTMhelix 326 348\ninside 349 372

12799 GCF_000460755.1_Esch_coli_UMEA_3201-1_V1 Escherichia coli UMEA 3201-1 Proteobacteria; Gammaproteobacteria; Enterobacterales; Enterobacteriaceae; Escherichia; Escherichia coli MNNEETFYQAMRRQGVTRRSFLKYCSLAATSLGLGAGMAPKIAWALENKPRIPVVWIHGL WP_001058323.1 MULTISPECIES: hydrogenase-1 small chain [Proteobacteria] Length: 372\nNumber of predicted TMHs: 2\nExp number of AAs in TMHs: 37.2608499999999\nExp number, first 60 AAs: 19.52923\nTotal prob of N-in: 0.94351\nPOSSIBLE N-term signal sequence\ninside 1 19\nTMhelix 20 39\noutside 40 325\nTMhelix 326 348\ninside 349 372

12800 GCF_000460755.1_Esch_coli_UMEA_3201-1_V1 Escherichia coli UMEA 3201-1 Proteobacteria; Gammaproteobacteria; Enterobacterales; Enterobacteriaceae; Escherichia; Escherichia coli MTGDNTLIHSHGINRRDFMKLCAALAATMGLSSKAAAEMAESVTNPQRPPVIWIGAQECT WP_021564866.1 hydrogenase-2 small chain [Escherichia coli] Length: 372\nNumber of predicted TMHs: 1\nExp number of AAs in TMHs: 23.72269\nExp number, first 60 AAs: 0.22038\nTotal prob of N-in: 0.12995\noutside 1 331\nTMhelix 332 354\ninside 355 372

12801 GCF_000460775.1_Esch_coli_UMEA_3203-1_V1 Escherichia coli UMEA 3203-1 Proteobacteria; Gammaproteobacteria; Enterobacterales; Enterobacteriaceae; Escherichia; Escherichia coli MTGDNTLIHSHGINRRDFMKLCAALAATMGLSSKAAAEMAESVTNPQRPPVIWIGAQECT WP_000145410.1 MULTISPECIES: hydrogenase-2 small chain [Proteobacteria] Length: 372\nNumber of predicted TMHs:

1\nExp number of AAs in TMHs: 23.72226\nExp number, first 60 AAs: 0.22038\nTotal prob of N-in: 0.12994\noutside 1 331\nTMhelix 332 354\ninside 355 372

12802 GCF_000460795.1_Esch_coli_UMEA_3206-1_V1 Escherichia coli UMEA 3206-1 Proteobacteria; Gammaproteobacteria; Enterobacterales; Enterobacteriaceae; Escherichia; Escherichia coli MTGDNTLIHSHGINRRDFMKLCAALAATMGLSSKAAAEMAESVTNPQRPPVIWIGAQECT WP_000145410.1 MULTISPECIES: hydrogenase-2 small chain [Proteobacteria] Length: 372\nNumber of predicted TMHs: 1\nExp number of AAs in TMHs: 23.72226\nExp number, first 60 AAs: 0.22038\nTotal prob of N-in: 0.12994\noutside 1 331\nTMhelix 332 354\ninside 355 372

12803 GCF_000460815.1_Esch_coli_UMEA_3208-1_V1 Escherichia coli UMEA 3208-1 Proteobacteria; Gammaproteobacteria; Enterobacterales; Enterobacteriaceae; Escherichia; Escherichia coli MTGDNTLIHSHGINRRDFMKLCAALAATMGLSSKAAAEMAESVTNPQRPPVIWIGAQECT WP_000145410.1 MULTISPECIES: hydrogenase-2 small chain [Proteobacteria] Length: 372\nNumber of predicted TMHs: 1\nExp number of AAs in TMHs: 23.72226\nExp number, first 60 AAs: 0.22038\nTotal prob of N-in: 0.12994\noutside 1 331\nTMhelix 332 354\ninside 355 372

12804 GCF_000460815.1_Esch_coli_UMEA_3208-1_V1 Escherichia coli UMEA 3208-1 Proteobacteria; Gammaproteobacteria; Enterobacterales; Enterobacteriaceae; Escherichia; Escherichia coli MNNEETFYQAMRRQGVTRRSFLKYCSLAATSLGLGAGMAPKIAWALENKPRIPVVWIHGL WP_001058323.1 MULTISPECIES: hydrogenase-1 small chain [Proteobacteria] Length: 372\nNumber of predicted TMHs: 2\nExp number of AAs in TMHs: 37.2608499999999\nExp number, first 60 AAs: 19.52923\nTotal prob of N-in: 0.94351\nPOSSIBLE N-term signal sequence\ninside 1 19\nTMhelix 20 39\noutside 40 325\nTMhelix 326 348\ninside 349 372

12805 GCF_000460835.1_Esch_coli_UMEA_3212-1_V1 Escherichia coli UMEA 3212-1 Proteobacteria; Gammaproteobacteria; Enterobacterales; Enterobacteriaceae; Escherichia; Escherichia coli MNNEETFYQAMRRQGVTRRSFLKYCSLAATSLGLGAGMAPKIAWALENKPRIPVVWIHGL WP_001058323.1 MULTISPECIES: hydrogenase-1 small chain [Proteobacteria] Length: 372\nNumber of predicted TMHs: 2\nExp number of AAs in TMHs: 37.2608499999999\nExp number, first 60 AAs: 19.52923\nTotal prob of N-in: 0.94351\nPOSSIBLE N-term signal sequence\ninside 1 19\nTMhelix 20 39\noutside 40 325\nTMhelix 326 348\ninside 349 372

12806 GCF_000460835.1_Esch_coli_UMEA_3212-1_V1 Escherichia coli UMEA 3212-1 Proteobacteria; Gammaproteobacteria; Enterobacterales; Enterobacteriaceae; Escherichia; Escherichia coli MTGDNTLIHSHGINRRDFMKLCAALAATMGLSSKAAAEMAESVTNPQRPPVIWIGAQECT WP_000145410.1 MULTISPECIES: hydrogenase-2 small chain [Proteobacteria] Length: 372\nNumber of predicted TMHs: 1\nExp number of AAs in TMHs: 23.72226\nExp number, first 60 AAs: 0.22038\nTotal prob of N-in: 0.12994\noutside 1 331\nTMhelix 332 354\ninside 355 372

12807 GCF_000460855.1_Esch_coli_UMEA_3215-1_V1 Escherichia coli UMEA 3215-1 Proteobacteria; Gammaproteobacteria; Enterobacterales; Enterobacteriaceae; Escherichia; Escherichia coli MNNEETFYQAMRRQGVTRRSFLKYCSLAATSLGLGAGMAPKIAWALENKPRIPVVWIHGL WP_001538532.1 hydrogenase-1 small chain [Escherichia coli] Length: 372\nNumber of predicted TMHs: 2\nExp number of AAs in TMHs: 37.27041\nExp number, first 60 AAs: 19.52927\nTotal prob of N-in: 0.94352\nPOSSIBLE N-term signal sequence\ninside 1 19\nTMhelix 20 39\noutside 40 325\nTMhelix 326 348\ninside 349 372

12808 GCF_000460855.1_Esch_coli_UMEA_3215-1_V1 Escherichia coli UMEA 3215-1 Proteobacteria; Gammaproteobacteria; Enterobacterales; Enterobacteriaceae; Escherichia; Escherichia coli MTGDNTLIHSHGINRRDFMKLCAALAATMGLSSKAAAEMVESVTNPQRPPVIWIGAQECT WP_000145423.1 hydrogenase 2 small subunit [Escherichia coli] Length: 372\nNumber of predicted TMHs: 1\nExp number of AAs in TMHs: 23.78201\nExp number, first 60 AAs: 0.28355\nTotal prob of N-in: 0.13257\noutside 1 331\nTMhelix 332 354\ninside 355 372

12809 GCF_000460875.1_Esch_coli_UMEA_3216-1_V1 Escherichia coli UMEA 3216-1 Proteobacteria; Gammaproteobacteria; Enterobacterales; Enterobacteriaceae; Escherichia; Escherichia coli MTGDNTLIHSHGINRRDFMKLCAALAATMGLSSKAAAEMAESVTNPQRPPVIWIGAQECT WP_000145410.1 MULTISPECIES: hydrogenase-2 small chain [Proteobacteria] Length: 372\nNumber of predicted TMHs: 1\nExp number of AAs in TMHs: 23.72226\nExp number, first 60 AAs: 0.22038\nTotal prob of N-in: 0.12994\noutside 1 331\nTMhelix 332 354\ninside 355 372

12810 GCF_000460875.1_Esch_coli_UMEA_3216-1_V1 Escherichia coli UMEA 3216-1 Proteobacteria; Gammaproteobacteria; Enterobacterales; Enterobacteriaceae; Escherichia; Escherichia coli MNNEETFYQAMRRQGVTRRSFLKYCSLVATSLGLGAGMAPKIAWALENKPRIPVVWIHGL WP_021566792.1 hydrogenase-1 small chain [Escherichia coli] Length: 372\nNumber of predicted TMHs: 2\nExp number of AAs in TMHs: 37.76586\nExp number, first 60 AAs: 20.0399\nTotal prob of N-in: 0.96673\nPOSSIBLE N-term signal sequence\ninside 1 19\nTMhelix 20 39\noutside 40 325\nTMhelix 326 348\ninside 349 372

12811 GCF_000460895.1_Esch_coli_UMEA_3217-1_V1 Escherichia coli UMEA 3217-1 Proteobacteria; Gammaproteobacteria; Enterobacterales; Enterobacteriaceae; Escherichia; Escherichia coli MTGDNTLIHSHGINRRDFMKLCAALAATMGLSSKAAAEMAESVTNPQRPPVIWIGAQECT WP_000145410.1 MULTISPECIES: hydrogenase-2 small chain [Proteobacteria] Length: 372\nNumber of predicted TMHs: 1\nExp number of AAs in TMHs: 23.72226\nExp number, first 60 AAs: 0.22038\nTotal prob of N-in: 0.12994\noutside 1 331\nTMhelix 332 354\ninside 355 372

12812 GCF_000460895.1_Esch_coli_UMEA_3217-1_V1 Escherichia coli UMEA 3217-1 Proteobacteria; Gammaproteobacteria; Enterobacterales; Enterobacteriaceae; Escherichia; Escherichia coli MNNEETFYQAMRRQGVTRRSFLKYCSLAATSLGLGAGMAPKIAWALENKPRIPVVWIHGL WP_021566870.1 hydrogenase-1 small chain [Escherichia coli] Length: 372\nNumber of predicted TMHs: 2\nExp number of AAs in TMHs: 37.2608199999999\nExp number, first 60 AAs: 19.52923\nTotal prob of N-in: 0.94351\nPOSSIBLE N-term signal sequence\ninside 1 19\nTMhelix 20 39\noutside 40 325\nTMhelix 326 348\ninside 349 372

12813 GCF_000460915.1_Esch_coli_UMEA_3220-1_V1 Escherichia coli UMEA 3220-1 Proteobacteria; Gammaproteobacteria; Enterobacterales; Enterobacteriaceae; Escherichia; Escherichia coli MTGDNTLIHSHGINRRDFMKLCAALAATMGLSSKAAAEMAESVTNPQRPPVIWIGAQECT WP_000145410.1 MULTISPECIES: hydrogenase-2 small chain [Proteobacteria] Length: 372\nNumber of predicted TMHs: 1\nExp number of AAs in TMHs: 23.72226\nExp number, first 60 AAs: 0.22038\nTotal prob of N-in: 0.12994\noutside 1 331\nTMhelix 332 354\ninside 355 372

12814 GCF_000460915.1_Esch_coli_UMEA_3220-1_V1 Escherichia coli UMEA 3220-1 Proteobacteria; Gammaproteobacteria; Enterobacterales; Enterobacteriaceae; Escherichia; Escherichia coli MNNEETFYQAMRRQGVTRRSFLKYCSLAATSLGLGAGMAPKIAWALENKPRIPVVWIHGL WP_001058323.1 MULTISPECIES: hydrogenase-1 small chain [Proteobacteria] Length: 372\nNumber of predicted TMHs: 2\nExp number of AAs in TMHs: 37.2608499999999\nExp number, first 60 AAs: 19.52923\nTotal prob of N-in: 0.94351\nPOSSIBLE N-term signal sequence\ninside 1 19\nTMhelix 20 39\noutside 40 325\nTMhelix 326 348\ninside 349 372

12815 GCF_000460935.1_Esch_coli_UMEA_3221-1_V1 Escherichia coli UMEA 3221-1 Proteobacteria; Gammaproteobacteria; Enterobacterales; Enterobacteriaceae; Escherichia; Escherichia coli MNNEETFYQAMRRQGVTRRSFLKYCSLAATSLGLGAGMAPKIAWALENKPRIPVVWIHGL WP_001058323.1 MULTISPECIES: hydrogenase-1 small chain [Proteobacteria] Length: 372\nNumber of predicted TMHs: 2\nExp number of AAs in TMHs: 37.2608499999999\nExp number, first 60 AAs: 19.52923\nTotal prob of N-in: 0.94351\nPOSSIBLE N-term signal sequence\ninside 1 19\nTMhelix 20 39\noutside 40 325\nTMhelix 326 348\ninside 349 372

12816 GCF_000460935.1_Esch_coli_UMEA_3221-1_V1 Escherichia coli UMEA 3221-1 Proteobacteria; Gammaproteobacteria; Enterobacterales; Enterobacteriaceae; Escherichia; Escherichia coli MTGDNTLIHSHGINRRDFMKLCAALAATMGLSSKAAAEMAESVTNPQRPPVIWIGAQECT WP_000145410.1 MULTISPECIES: hydrogenase-2 small chain [Proteobacteria] Length: 372\nNumber of predicted TMHs: 1\nExp number of AAs in TMHs: 23.72226\nExp number, first 60 AAs: 0.22038\nTotal prob of N-in: 0.12994\noutside 1 331\nTMhelix 332 354\ninside 355 372

12817 GCF_000460955.1_Esch_coli_UMEA_3222-1_V1 Escherichia coli UMEA 3222-1 Proteobacteria; Gammaproteobacteria; Enterobacterales; Enterobacteriaceae; Escherichia; Escherichia coli MTGDNTLIHSHGINRRDFMKLCAALAATMGLSSKAAAEMAESVTNPQRPPVIWIGAQECT WP_000145410.1 MULTISPECIES: hydrogenase-2 small chain [Proteobacteria] Length: 372\nNumber of predicted TMHs: 1\nExp number of AAs in TMHs: 23.72226\nExp number, first 60 AAs: 0.22038\nTotal prob of N-in: 0.12994\noutside 1 331\nTMhelix 332 354\ninside 355 372

12818 GCF_000460955.1_Esch_coli_UMEA_3222-1_V1 Escherichia coli UMEA 3222-1 Proteobacteria; Gammaproteobacteria; Enterobacterales; Enterobacteriaceae; Escherichia; Escherichia coli MNNEETFYQAMRRQGVTRRSFLKYCSLAATSLGLGAGMAPKIAWALENKPRIPVVWIHGL WP_001058323.1

MULTISPECIES: hydrogenase-1 small chain [Proteobacteria] Length: 372\nNumber of predicted TMHs: 2\nExp number of AAs in TMHs: 37.2608499999999\nExp number, first 60 AAs: 19.52923\nTotal prob of N-in: 0.94351\nPOSSIBLE N-term signal sequence\ninside 1 19\nTMhelix 20 39\noutside 40 325\nTMhelix 326 348\ninside 349 372

12819 GCF_000460975.1_Esch_coli_UMEA_3230-1_V1 Escherichia coli UMEA 3230-1 Proteobacteria; Gammaproteobacteria; Enterobacterales; Enterobacteriaceae; Escherichia; Escherichia coli MTGDNTLIHSHGINRRDFMKLCAALAATMGLSSKAAAEMAESVTNPQRPPVIWIGAQECT WP_000145410.1 MULTISPECIES: hydrogenase-2 small chain [Proteobacteria] Length: 372\nNumber of predicted TMHs: 1\nExp number of AAs in TMHs: 23.72226\nExp number, first 60 AAs: 0.22038\nTotal prob of N-in: 0.12994\noutside 1 331\nTMhelix 332 354\ninside 355 372

12820 GCF_000460975.1_Esch_coli_UMEA_3230-1_V1 Escherichia coli UMEA 3230-1 Proteobacteria; Gammaproteobacteria; Enterobacterales; Enterobacteriaceae; Escherichia; Escherichia coli MNNEETFYQAMRRQGVTRRSFLKYCSLAATSLGLGAGMAPKIAWALENKPRIPVVWIHGL WP_001058323.1 MULTISPECIES: hydrogenase-1 small chain [Proteobacteria] Length: 372\nNumber of predicted TMHs: 2\nExp number of AAs in TMHs: 37.2608499999999\nExp number, first 60 AAs: 19.52923\nTotal prob of N-in: 0.94351\nPOSSIBLE N-term signal sequence\ninside 1 19\nTMhelix 20 39\noutside 40 325\nTMhelix 326 348\ninside 349 372

12821 GCF_000460995.1_Esch_coli_UMEA_3233-1_V1 Escherichia coli UMEA 3233-1 Proteobacteria; Gammaproteobacteria; Enterobacterales; Enterobacteriaceae; Escherichia; Escherichia coli MNNEETFYQAMRRQGVTRRSFLKYCSLAATSLGLGAGMAPKIAWALENKPRIPVVWIHGL WP_001058323.1 MULTISPECIES: hydrogenase-1 small chain [Proteobacteria] Length: 372\nNumber of predicted TMHs: 2\nExp number of AAs in TMHs: 37.2608499999999\nExp number, first 60 AAs: 19.52923\nTotal prob of N-in: 0.94351\nPOSSIBLE N-term signal sequence\ninside 1 19\nTMhelix 20 39\noutside 40 325\nTMhelix 326 348\ninside 349 372

12822 GCF_000460995.1_Esch_coli_UMEA_3233-1_V1 Escherichia coli UMEA 3233-1 Proteobacteria; Gammaproteobacteria; Enterobacterales; Enterobacteriaceae; Escherichia; Escherichia coli MTGDNTLIHSHGINRRDFMKLCAALAATMGLSSKAAAEMAESVTNPQRPPVIWIGAQECT WP_000145410.1 MULTISPECIES: hydrogenase-2 small chain [Proteobacteria] Length: 372\nNumber of predicted TMHs: 1\nExp number of AAs in TMHs: 23.72226\nExp number, first 60 AAs: 0.22038\nTotal prob of N-in: 0.12994\noutside 1 331\nTMhelix 332 354\ninside 355 372

12823 GCF_000461015.1_Esch_coli_UMEA_3240-1_V1 Escherichia coli UMEA 3240-1 Proteobacteria; Gammaproteobacteria; Enterobacterales; Enterobacteriaceae; Escherichia; Escherichia coli MNNEETFYQAMRRQGVTRRSFLKYCSLAATSLGLGAGMAPKIAWALENKPRIPVVWIHGL WP_001058323.1 MULTISPECIES: hydrogenase-1 small chain [Proteobacteria] Length: 372\nNumber of predicted TMHs: 2\nExp number of AAs in TMHs: 37.2608499999999\nExp number, first 60 AAs: 19.52923\nTotal prob of N-in: 0.94351\nPOSSIBLE N-term signal sequence\ninside 1 19\nTMhelix 20 39\noutside 40 325\nTMhelix 326 348\ninside 349 372

12824 GCF_000461015.1_Esch_coli_UMEA_3240-1_V1 Escherichia coli UMEA 3240-1 Proteobacteria; Gammaproteobacteria; Enterobacterales; Enterobacteriaceae; Escherichia; Escherichia coli MTGDNTLIHSHGINRRDFMKLCAALAATMGLSSKAAAEMAESVTNPQRPPVIWIGAQECT WP_000145410.1 MULTISPECIES: hydrogenase-2 small chain [Proteobacteria] Length: 372\nNumber of predicted TMHs: 1\nExp number of AAs in TMHs: 23.72226\nExp number, first 60 AAs: 0.22038\nTotal prob of N-in: 0.12994\noutside 1 331\nTMhelix 332 354\ninside 355 372

12825 GCF_000461035.1_Esch_coli_UMEA_3244-1_V1 Escherichia coli UMEA 3244-1 Proteobacteria; Gammaproteobacteria; Enterobacterales; Enterobacteriaceae; Escherichia; Escherichia coli MNNEETFYQAMRRQGVTRRSFLKYCSLAATSLGLGAGMAPKIAWALENKPRIPVVWIHGL WP_021567794.1 hydrogenase-1 small chain [Escherichia coli] Length: 372\nNumber of predicted TMHs: 2\nExp number of AAs in TMHs: 37.26051\nExp number, first 60 AAs: 19.52922\nTotal prob of N-in: 0.94351\nPOSSIBLE N-term signal sequence\ninside 1 19\nTMhelix 20 39\noutside 40 325\nTMhelix 326 348\ninside 349 372

12826 GCF_000461035.1_Esch_coli_UMEA_3244-1_V1 Escherichia coli UMEA 3244-1 Proteobacteria; Gammaproteobacteria; Enterobacterales; Enterobacteriaceae; Escherichia; Escherichia coli MTGDNTLIHSHGINRRDFMKLCAALAATMGLSSKAAAEMAESVTNPQRPPVIWIGAQECT WP_000145410.1 MULTISPECIES: hydrogenase-2 small chain [Proteobacteria] Length: 372\nNumber of predicted TMHs:

1\nExp number of AAs in TMHs: 23.72226\nExp number, first 60 AAs: 0.22038\nTotal prob of N-in: 0.12994\noutside 1 331\nTMhelix 332 354\ninside 355 372

12827 GCF_000461055.1_Esch_coli_UMEA_3257-1_V1 Escherichia coli UMEA 3257-1 Proteobacteria; Gammaproteobacteria; Enterobacterales; Enterobacteriaceae; Escherichia; Escherichia coli MNNEETFYQAMRRQGVTRRSFLKYCSLAATSLGLGAGMAPKIAWALENKPRIPVVWIHGL WP_001058323.1 MULTISPECIES: hydrogenase-1 small chain [Proteobacteria] Length: 372\nNumber of predicted TMHs: 2\nExp number of AAs in TMHs: 37.2608499999999\nExp number, first 60 AAs: 19.52923\nTotal prob of N-in: 0.94351\nPOSSIBLE N-term signal sequence\ninside 1 19\nTMhelix 20 39\noutside 40 325\nTMhelix 326 348\ninside 349 372

12828 GCF_000461055.1_Esch_coli_UMEA_3257-1_V1 Escherichia coli UMEA 3257-1 Proteobacteria; Gammaproteobacteria; Enterobacterales; Enterobacteriaceae; Escherichia; Escherichia coli MTGDNTLIHSHGINRRDFMKLCAALAATMGLSSKAAAEMAESVTNPQRPPVIWIGAQECT WP_000145410.1 MULTISPECIES: hydrogenase-2 small chain [Proteobacteria] Length: 372\nNumber of predicted TMHs: 1\nExp number of AAs in TMHs: 23.72226\nExp number, first 60 AAs: 0.22038\nTotal prob of N-in: 0.12994\noutside 1 331\nTMhelix 332 354\ninside 355 372

12829 GCF_000461075.1_Esch_coli_UMEA_3264-1_V1 Escherichia coli UMEA 3264-1 Proteobacteria; Gammaproteobacteria; Enterobacterales; Enterobacteriaceae; Escherichia; Escherichia coli MTGDNTLIHSHGINRRDFMKLCAALAATMGLSSKAAAEMAESVTNPQRPPVIWIGAQECT WP_000145410.1 MULTISPECIES: hydrogenase-2 small chain [Proteobacteria] Length: 372\nNumber of predicted TMHs: 1\nExp number of AAs in TMHs: 23.72226\nExp number, first 60 AAs: 0.22038\nTotal prob of N-in: 0.12994\noutside 1 331\nTMhelix 332 354\ninside 355 372

12830 GCF_000461075.1_Esch_coli_UMEA_3264-1_V1 Escherichia coli UMEA 3264-1 Proteobacteria; Gammaproteobacteria; Enterobacterales; Enterobacteriaceae; Escherichia; Escherichia coli MNNEETFYQAMRRQGVTRRSFLKYCSLAATSLGLGAGMAPKIAWALENKPRIPVVWIHGL WP_001058323.1 MULTISPECIES: hydrogenase-1 small chain [Proteobacteria] Length: 372\nNumber of predicted TMHs: 2\nExp number of AAs in TMHs: 37.2608499999999\nExp number, first 60 AAs: 19.52923\nTotal prob of N-in: 0.94351\nPOSSIBLE N-term signal sequence\ninside 1 19\nTMhelix 20 39\noutside 40 325\nTMhelix 326 348\ninside 349 372

12831 GCF_000461095.1_Esch_coli_UMEA_3268-1_V1 Escherichia coli UMEA 3268-1 Proteobacteria; Gammaproteobacteria; Enterobacterales; Enterobacteriaceae; Escherichia; Escherichia coli MTGDNTLIHSHGINRRDFMKLCAALAATMGLSSKAAAEMAESVTNPQRPPVIWIGAQECT WP_000145410.1 MULTISPECIES: hydrogenase-2 small chain [Proteobacteria] Length: 372\nNumber of predicted TMHs: 1\nExp number of AAs in TMHs: 23.72226\nExp number, first 60 AAs: 0.22038\nTotal prob of N-in: 0.12994\noutside 1 331\nTMhelix 332 354\ninside 355 372

12832 GCF_000461095.1_Esch_coli_UMEA_3268-1_V1 Escherichia coli UMEA 3268-1 Proteobacteria; Gammaproteobacteria; Enterobacterales; Enterobacteriaceae; Escherichia; Escherichia coli MNNEETFYQAMRRQGVTRRSFLKYCSLAATSLGLGAGMAPKIAWALENKPRIPVVWIHGL WP_001058323.1 MULTISPECIES: hydrogenase-1 small chain [Proteobacteria] Length: 372\nNumber of predicted TMHs: 2\nExp number of AAs in TMHs: 37.2608499999999\nExp number, first 60 AAs: 19.52923\nTotal prob of N-in: 0.94351\nPOSSIBLE N-term signal sequence\ninside 1 19\nTMhelix 20 39\noutside 40 325\nTMhelix 326 348\ninside 349 372

12833 GCF_000461115.1_Esch_coli_UMEA_3271-1_V1 Escherichia coli UMEA 3271-1 Proteobacteria; Gammaproteobacteria; Enterobacterales; Enterobacteriaceae; Escherichia; Escherichia coli MTGDNTLIHSHGINRRDFMKLCAALAATMGLSSKAAAEMAESVTNPQRPPVIWIGAQECT WP_000145410.1 MULTISPECIES: hydrogenase-2 small chain [Proteobacteria] Length: 372\nNumber of predicted TMHs: 1\nExp number of AAs in TMHs: 23.72226\nExp number, first 60 AAs: 0.22038\nTotal prob of N-in: 0.12994\noutside 1 331\nTMhelix 332 354\ninside 355 372

12834 GCF_000461115.1_Esch_coli_UMEA_3271-1_V1 Escherichia coli UMEA 3271-1 Proteobacteria; Gammaproteobacteria; Enterobacterales; Enterobacteriaceae; Escherichia; Escherichia coli MNNEETFYQAMRRQGVTRRSFLKYCSLAATSLGLGAGMAPKIAWALENKPRIPVVWIHGL WP_001058323.1 MULTISPECIES: hydrogenase-1 small chain [Proteobacteria] Length: 372\nNumber of predicted TMHs: 2\nExp number of AAs in TMHs: 37.2608499999999\nExp number, first 60 AAs: 19.52923\nTotal prob of N-in:

0.94351\nPOSSIBLE N-term signal sequence\ninside 1 19\nTMhelix 20 39\noutside 40 325\nTMhelix 326 348\ninside 349 372

12835 GCF_000488095.1_Esch_coli_UMEA_3290-1_V1 Escherichia coli UMEA 3290-1 Proteobacteria; Gammaproteobacteria; Enterobacterales; Enterobacteriaceae; Escherichia; Escherichia coli MTGDNTLIHSHGINRRDFMKLCAALAATMGLSSKAAAEMAESVTNPQRPPVIWIGAQECT WP_000145410.1 MULTISPECIES: hydrogenase-2 small chain [Proteobacteria] Length: 372\nNumber of predicted TMHs: 1\nExp number of AAs in TMHs: 23.72226\nExp number, first 60 AAs: 0.22038\nTotal prob of N-in: 0.12994\noutside 1 331\nTMhelix 332 354\ninside 355 372

12836 GCF_000488095.1_Esch_coli_UMEA_3290-1_V1 Escherichia coli UMEA 3290-1 Proteobacteria; Gammaproteobacteria; Enterobacterales; Enterobacteriaceae; Escherichia; Escherichia coli MNNEETFYQAMRRQGVTRRSFLKYCSLAATSLGLGAGMAPKIAWALENKPRIPVVWIHGL WP_001058323.1 MULTISPECIES: hydrogenase-1 small chain [Proteobacteria] Length: 372\nNumber of predicted TMHs: 2\nExp number of AAs in TMHs: 37.2608499999999\nExp number, first 60 AAs: 19.52923\nTotal prob of N-in: 0.94351\nPOSSIBLE N-term signal sequence\ninside 1 19\nTMhelix 20 39\noutside 40 325\nTMhelix 326 348\ninside 349 372

12837 GCF_000461135.1_Esch_coli_UMEA_3292-1_V1 Escherichia coli UMEA 3292-1 Proteobacteria; Gammaproteobacteria; Enterobacterales; Enterobacteriaceae; Escherichia; Escherichia coli MTGDNTLIHSHGINRRDFMKLCAALAATMGLSSKAAAEMAESVTNPQRPPVIWIGAQECT WP_000145410.1 MULTISPECIES: hydrogenase-2 small chain [Proteobacteria] Length: 372\nNumber of predicted TMHs: 1\nExp number of AAs in TMHs: 23.72226\nExp number, first 60 AAs: 0.22038\nTotal prob of N-in: 0.12994\noutside 1 331\nTMhelix 332 354\ninside 355 372

12838 GCF_000461135.1_Esch_coli_UMEA_3292-1_V1 Escherichia coli UMEA 3292-1 Proteobacteria; Gammaproteobacteria; Enterobacterales; Enterobacteriaceae; Escherichia; Escherichia coli MNNEETFYQAMRRQGVTRRSFLKYCSLAATSLGLGAGMAPKIAWALENKPRIPVVWIHGL WP_001058323.1 MULTISPECIES: hydrogenase-1 small chain [Proteobacteria] Length: 372\nNumber of predicted TMHs: 2\nExp number of AAs in TMHs: 37.2608499999999\nExp number, first 60 AAs: 19.52923\nTotal prob of N-in: 0.94351\nPOSSIBLE N-term signal sequence\ninside 1 19\nTMhelix 20 39\noutside 40 325\nTMhelix 326 348\ninside 349 372

12839 GCF_000461155.1_Esch_coli_UMEA_3298-1_V1 Escherichia coli UMEA 3298-1 Proteobacteria; Gammaproteobacteria; Enterobacterales; Enterobacteriaceae; Escherichia; Escherichia coli MTGDNTLIHSHGINRRDFMKLCAALAATMGLSSKAAAEMAESVTNPQRPPVIWIGAQECT WP_000145410.1 MULTISPECIES: hydrogenase-2 small chain [Proteobacteria] Length: 372\nNumber of predicted TMHs: 1\nExp number of AAs in TMHs: 23.72226\nExp number, first 60 AAs: 0.22038\nTotal prob of N-in: 0.12994\noutside 1 331\nTMhelix 332 354\ninside 355 372

12840 GCF_000461175.1_Esch_coli_UMEA_3304-1_V1 Escherichia coli UMEA 3304-1 Proteobacteria; Gammaproteobacteria; Enterobacterales; Enterobacteriaceae; Escherichia; Escherichia coli MTGDNTLIHSHGINRRDFMKLCAALAATMGLSSKAAAEMAESVTNPQRPPVIWIGAQECT WP_000145410.1 MULTISPECIES: hydrogenase-2 small chain [Proteobacteria] Length: 372\nNumber of predicted TMHs: 1\nExp number of AAs in TMHs: 23.72226\nExp number, first 60 AAs: 0.22038\nTotal prob of N-in: 0.12994\noutside 1 331\nTMhelix 332 354\ninside 355 372

12841 GCF_000461175.1_Esch_coli_UMEA_3304-1_V1 Escherichia coli UMEA 3304-1 Proteobacteria; Gammaproteobacteria; Enterobacterales; Enterobacteriaceae; Escherichia; Escherichia coli MNNEETFYQAMRRQGVTRRSFLKYCSLAATSLGLGAGMAPKIAWALENKPRIPVVWIHGL WP_001058323.1 MULTISPECIES: hydrogenase-1 small chain [Proteobacteria] Length: 372\nNumber of predicted TMHs: 2\nExp number of AAs in TMHs: 37.2608499999999\nExp number, first 60 AAs: 19.52923\nTotal prob of N-in: 0.94351\nPOSSIBLE N-term signal sequence\ninside 1 19\nTMhelix 20 39\noutside 40 325\nTMhelix 326 348\ninside 349 372

12842 GCF_000461195.1_Esch_coli_UMEA_3314-1_V1 Escherichia coli UMEA 3314-1 Proteobacteria; Gammaproteobacteria; Enterobacterales; Enterobacteriaceae; Escherichia; Escherichia coli MNNEETFYQAMRRQGVTRRSFLKYCSLAATSLGLGAGMAPKIAWALENKPRIPVVWIHGL WP_001058323.1 MULTISPECIES: hydrogenase-1 small chain [Proteobacteria] Length: 372\nNumber of predicted TMHs: 2\nExp number of AAs in TMHs: 37.2608499999999\nExp number, first 60 AAs: 19.52923\nTotal prob of N-in:

0.94351\nPOSSIBLE N-term signal sequence\ninside 1 19\nTMhelix 20 39\noutside 40 325\nTMhelix 326 348\ninside 349 372

12843 GCF_000461195.1_Esch_coli_UMEA_3314-1_V1 Escherichia coli UMEA 3314-1 Proteobacteria; Gammaproteobacteria; Enterobacterales; Enterobacteriaceae; Escherichia; Escherichia coli MTGDNTLIHSHGINRRDFMKLCAALAATMGLSSKAAAEMAESVTNPQRPPVIWIGAQECT WP_000145410.1 MULTISPECIES: hydrogenase-2 small chain [Proteobacteria] Length: 372\nNumber of predicted TMHs: 1\nExp number of AAs in TMHs: 23.72226\nExp number, first 60 AAs: 0.22038\nTotal prob of N-in: 0.12994\noutside 1 331\nTMhelix 332 354\ninside 355 372

12844 GCF_000461215.1_Esch_coli_UMEA_3317-1_V1 Escherichia coli UMEA 3317-1 Proteobacteria; Gammaproteobacteria; Enterobacterales; Enterobacteriaceae; Escherichia; Escherichia coli MNNEETFYQAMRRQGVTRRSFLKYCSLAATSLGLGAGMAPKIAWALENKPRIPVVWIHGL WP_001058323.1 MULTISPECIES: hydrogenase-1 small chain [Proteobacteria] Length: 372\nNumber of predicted TMHs: 2\nExp number of AAs in TMHs: 37.2608499999999\nExp number, first 60 AAs: 19.52923\nTotal prob of N-in: 0.94351\nPOSSIBLE N-term signal sequence\ninside 1 19\nTMhelix 20 39\noutside 40 325\nTMhelix 326 348\ninside 349 372

12845 GCF_000461215.1_Esch_coli_UMEA_3317-1_V1 Escherichia coli UMEA 3317-1 Proteobacteria; Gammaproteobacteria; Enterobacterales; Enterobacteriaceae; Escherichia; Escherichia coli MTGDNTLIHSHGINRRDFMKLCAALAATMGLSSKAAAEMAESVTNPQRPPVIWIGAQECT WP_000145410.1 MULTISPECIES: hydrogenase-2 small chain [Proteobacteria] Length: 372\nNumber of predicted TMHs: 1\nExp number of AAs in TMHs: 23.72226\nExp number, first 60 AAs: 0.22038\nTotal prob of N-in: 0.12994\noutside 1 331\nTMhelix 332 354\ninside 355 372

12846 GCF_000461235.1_Esch_coli_UMEA_3318-1_V1 Escherichia coli UMEA 3318-1 Proteobacteria; Gammaproteobacteria; Enterobacterales; Enterobacteriaceae; Escherichia; Escherichia coli MTGDNTLIHSHGINRRDFMKLCAALAATMGLSSKAAAEMAESVTNPQRPPVIWIGAQECT WP_000145410.1 MULTISPECIES: hydrogenase-2 small chain [Proteobacteria] Length: 372\nNumber of predicted TMHs: 1\nExp number of AAs in TMHs: 23.72226\nExp number, first 60 AAs: 0.22038\nTotal prob of N-in: 0.12994\noutside 1 331\nTMhelix 332 354\ninside 355 372

12847 GCF_000461235.1_Esch_coli_UMEA_3318-1_V1 Escherichia coli UMEA 3318-1 Proteobacteria; Gammaproteobacteria; Enterobacterales; Enterobacteriaceae; Escherichia; Escherichia coli MNNEETFYQAMRRQGVTRRSFLKYCSLAATSLGLGAGMAPKIAWALENKPRIPVVWIHGL WP_001058323.1 MULTISPECIES: hydrogenase-1 small chain [Proteobacteria] Length: 372\nNumber of predicted TMHs: 2\nExp number of AAs in TMHs: 37.2608499999999\nExp number, first 60 AAs: 19.52923\nTotal prob of N-in: 0.94351\nPOSSIBLE N-term signal sequence\ninside 1 19\nTMhelix 20 39\noutside 40 325\nTMhelix 326 348\ninside 349 372

12848 GCF_000488135.1_Esch_coli_UMEA_3323-1_V1 Escherichia coli UMEA 3323-1 Proteobacteria; Gammaproteobacteria; Enterobacterales; Enterobacteriaceae; Escherichia; Escherichia coli MNNEETFYQAMRRQGVTRRSFLKYCSLAATSLGLGAGMAPKIAWALENKPRIPVVWIHGL WP_001058323.1 MULTISPECIES: hydrogenase-1 small chain [Proteobacteria] Length: 372\nNumber of predicted TMHs: 2\nExp number of AAs in TMHs: 37.2608499999999\nExp number, first 60 AAs: 19.52923\nTotal prob of N-in: 0.94351\nPOSSIBLE N-term signal sequence\ninside 1 19\nTMhelix 20 39\noutside 40 325\nTMhelix 326 348\ninside 349 372

12849 GCF_000488135.1_Esch_coli_UMEA_3323-1_V1 Escherichia coli UMEA 3323-1 Proteobacteria; Gammaproteobacteria; Enterobacterales; Enterobacteriaceae; Escherichia; Escherichia coli MTGDNTLIHSHGINRRDFMKLCAALAATMGLSSKAAAEMAESVTNPQRPPVIWIGAQECT WP_000145410.1 MULTISPECIES: hydrogenase-2 small chain [Proteobacteria] Length: 372\nNumber of predicted TMHs: 1\nExp number of AAs in TMHs: 23.72226\nExp number, first 60 AAs: 0.22038\nTotal prob of N-in: 0.12994\noutside 1 331\nTMhelix 332 354\ninside 355 372

12850 GCF_000461255.1_Esch_coli_UMEA_3329-1_V1 Escherichia coli UMEA 3329-1 Proteobacteria; Gammaproteobacteria; Enterobacterales; Enterobacteriaceae; Escherichia; Escherichia coli MTGDNTLIHSHGINRRDFMKLCAALAATMGLSSKAAAEMAESVTNPQRPPVIWIGAQECT WP_000145410.1 MULTISPECIES: hydrogenase-2 small chain [Proteobacteria] Length: 372\nNumber of predicted TMHs: 1\nExp number of AAs in TMHs: 23.72226\nExp number, first 60 AAs: 0.22038\nTotal prob of N-in: 0.12994\noutside 1 331\nTMhelix 332 354\ninside 355 372

12851 GCF_000461255.1_Esch_coli_UMEA_3329-1_V1 Escherichia coli UMEA 3329-1 Proteobacteria; Gammaproteobacteria; Enterobacterales; Enterobacteriaceae; Escherichia; Escherichia coli MNNEETFYQAMRRQGVTRRSFLKYCSLAATSLGLGAGMAPKIAWALENKPRIPVVWIHGL WP_001058323.1 MULTISPECIES: hydrogenase-1 small chain [Proteobacteria] Length: 372\nNumber of predicted TMHs: 2\nExp number of AAs in TMHs: 37.2608499999999\nExp number, first 60 AAs: 19.52923\nTotal prob of N-in: 0.94351\nPOSSIBLE N-term signal sequence\ninside 1 19\nTMhelix 20 39\noutside 40 325\nTMhelix 326 348\ninside 349 372

12852 GCF_000488055.1_Esch_coli_UMEA_3336-1_V1 Escherichia coli UMEA 3336-1 Proteobacteria; Gammaproteobacteria; Enterobacterales; Enterobacteriaceae; Escherichia; Escherichia coli MTGDNTLIHSHGINRRDFMKLCAALAATMGLSSKAAAEMAESVTNPQRPPVIWIGAQECT WP_000145410.1 MULTISPECIES: hydrogenase-2 small chain [Proteobacteria] Length: 372\nNumber of predicted TMHs: 1\nExp number of AAs in TMHs: 23.72226\nExp number, first 60 AAs: 0.22038\nTotal prob of N-in: 0.12994\noutside 1 331\nTMhelix 332 354\ninside 355 372

12853 GCF_000488055.1_Esch_coli_UMEA_3336-1_V1 Escherichia coli UMEA 3336-1 Proteobacteria; Gammaproteobacteria; Enterobacterales; Enterobacteriaceae; Escherichia; Escherichia coli MNNEETFYQAMRRQGVTRRSFLKYCSLAATSLGLGAGMAPKIAWALENKPRIPVVWIHGL WP_001295940.1 MULTISPECIES: hydrogenase [Enterobacteriaceae] Length: 372\nNumber of predicted TMHs: 2\nExp number of AAs in TMHs: 37.2608899999999\nExp number, first 60 AAs: 19.52923\nTotal prob of N-in: 0.94351\nPOSSIBLE N-term signal sequence\ninside 1 19\nTMhelix 20 39\noutside 40 325\nTMhelix 326 348\ninside 349 372

12854 GCF_000461275.1_Esch_coli_UMEA_3337-1_V1 Escherichia coli UMEA 3337-1 Proteobacteria; Gammaproteobacteria; Enterobacterales; Enterobacteriaceae; Escherichia; Escherichia coli MTGDNTLIHSHGINRRDFMKLCAALAATMGLSSKAAAEMAESVTNPQRPPVIWIGAQECT WP_000145410.1 MULTISPECIES: hydrogenase-2 small chain [Proteobacteria] Length: 372\nNumber of predicted TMHs: 1\nExp number of AAs in TMHs: 23.72226\nExp number, first 60 AAs: 0.22038\nTotal prob of N-in: 0.12994\noutside 1 331\nTMhelix 332 354\ninside 355 372

12855 GCF_000461275.1_Esch_coli_UMEA_3337-1_V1 Escherichia coli UMEA 3337-1 Proteobacteria; Gammaproteobacteria; Enterobacterales; Enterobacteriaceae; Escherichia; Escherichia coli MNNEETFYQAMRRQGVTRRSFLKYCSLAATSLGLGAGMAPKIAWALENKPRIPVVWIHGL WP_001058323.1 MULTISPECIES: hydrogenase-1 small chain [Proteobacteria] Length: 372\nNumber of predicted TMHs: 2\nExp number of AAs in TMHs: 37.2608499999999\nExp number, first 60 AAs: 19.52923\nTotal prob of N-in: 0.94351\nPOSSIBLE N-term signal sequence\ninside 1 19\nTMhelix 20 39\noutside 40 325\nTMhelix 326 348\ninside 349 372

12856 GCF_000461295.1_Esch_coli_UMEA_3341-1_V1 Escherichia coli UMEA 3341-1 Proteobacteria; Gammaproteobacteria; Enterobacterales; Enterobacteriaceae; Escherichia; Escherichia coli MTGDNTLIHSHGINRRDFMKLCAALAATMGLSSKAAAEMAESVTNPQRPPVIWIGAQECT WP_000145410.1 MULTISPECIES: hydrogenase-2 small chain [Proteobacteria] Length: 372\nNumber of predicted TMHs: 1\nExp number of AAs in TMHs: 23.72226\nExp number, first 60 AAs: 0.22038\nTotal prob of N-in: 0.12994\noutside 1 331\nTMhelix 332 354\ninside 355 372

12857 GCF_000461295.1_Esch_coli_UMEA_3341-1_V1 Escherichia coli UMEA 3341-1 Proteobacteria; Gammaproteobacteria; Enterobacterales; Enterobacteriaceae; Escherichia; Escherichia coli MNNEETFYQAMRRQGVTRRSFLKYCSLAATSLGLGAGMAPKIAWALENKPRIPVVWIHGL WP_001058323.1 MULTISPECIES: hydrogenase-1 small chain [Proteobacteria] Length: 372\nNumber of predicted TMHs: 2\nExp number of AAs in TMHs: 37.2608499999999\nExp number, first 60 AAs: 19.52923\nTotal prob of N-in: 0.94351\nPOSSIBLE N-term signal sequence\ninside 1 19\nTMhelix 20 39\noutside 40 325\nTMhelix 326 348\ninside 349 372

12858 GCF_000488155.1_Esch_coli_UMEA_3342-1_V1 Escherichia coli UMEA 3342-1 Proteobacteria; Gammaproteobacteria; Enterobacterales; Enterobacteriaceae; Escherichia; Escherichia coli MTGDNTLIHSHGINRRDFMKLCAALAATMGLSSKAAAEMAESVTNPQRPPVIWIGAQECT WP_000145410.1 MULTISPECIES: hydrogenase-2 small chain [Proteobacteria] Length: 372\nNumber of predicted TMHs: 1\nExp number of AAs in TMHs: 23.72226\nExp number, first 60 AAs: 0.22038\nTotal prob of N-in: 0.12994\noutside 1 331\nTMhelix 332 354\ninside 355 372

12859 GCF_000488155.1_Esch_coli_UMEA_3342-1_V1 Escherichia coli UMEA 3342-1 Proteobacteria; Gammaproteobacteria; Enterobacterales; Enterobacteriaceae; Escherichia; Escherichia coli MNNEETFYQAMRRQGVTRRSFLKYCSLAATSLGLGAGMAPKIAWALENKPRIPVVWIHGL WP_001058323.1 MULTISPECIES: hydrogenase-1 small chain [Proteobacteria] Length: 372\nNumber of predicted TMHs: 2\nExp number of AAs in TMHs: 37.2608499999999\nExp number, first 60 AAs: 19.52923\nTotal prob of N-in: 0.94351\nPOSSIBLE N-term signal sequence\ninside 1 19\nTMhelix 20 39\noutside 40 325\nTMhelix 326 348\ninside 349 372

12860 GCF_000461315.1_Esch_coli_UMEA_3355-1_V1 Escherichia coli UMEA 3355-1 Proteobacteria; Gammaproteobacteria; Enterobacterales; Enterobacteriaceae; Escherichia; Escherichia coli MTGDNTLIHSHGINRRDFMKLCAALAATMGLSSKAAAEMAESVTNPQRPPVIWIGAQECT WP_000145410.1 MULTISPECIES: hydrogenase-2 small chain [Proteobacteria] Length: 372\nNumber of predicted TMHs: 1\nExp number of AAs in TMHs: 23.72226\nExp number, first 60 AAs: 0.22038\nTotal prob of N-in: 0.12994\noutside 1 331\nTMhelix 332 354\ninside 355 372

12861 GCF_000461315.1_Esch_coli_UMEA_3355-1_V1 Escherichia coli UMEA 3355-1 Proteobacteria; Gammaproteobacteria; Enterobacterales; Enterobacteriaceae; Escherichia; Escherichia coli MNNEETFYQAMRRQGVTRRSFLKYCSLAATSLGLGAGMAPKIAWALENKPRIPVVWIHGL WP_001058320.1 hydrogenase [Escherichia coli] Length: 372\nNumber of predicted TMHs: 2\nExp number of AAs in TMHs: 37.26082\nExp number, first 60 AAs: 19.52923\nTotal prob of N-in: 0.94351\nPOSSIBLE N-term signal sequence\ninside 1 19\nTMhelix 20 39\noutside 40 325\nTMhelix 326 348\ninside 349 372

12862 GCF_000461335.1_Esch_coli_UMEA_3391-1_V1 Escherichia coli UMEA 3391-1 Proteobacteria; Gammaproteobacteria; Enterobacterales; Enterobacteriaceae; Escherichia; Escherichia coli MTGDNTLIHSHGINRRDFMKLCAALAATMGLSSKAAAEMVESVTNPQRPPVIWIGAQECT WP_000145423.1 hydrogenase 2 small subunit [Escherichia coli] Length: 372\nNumber of predicted TMHs: 1\nExp number of AAs in TMHs: 23.78201\nExp number, first 60 AAs: 0.28355\nTotal prob of N-in: 0.13257\noutside 1 331\nTMhelix 332 354\ninside 355 372

12863 GCF_000461335.1_Esch_coli_UMEA_3391-1_V1 Escherichia coli UMEA 3391-1 Proteobacteria; Gammaproteobacteria; Enterobacterales; Enterobacteriaceae; Escherichia; Escherichia coli MNNEETFYQAMRRQGVTRRSFLKYCSLAATSLGLGAGMAPKIAWALENKPRIPVVWIHGL WP_001058323.1 MULTISPECIES: hydrogenase-1 small chain [Proteobacteria] Length: 372\nNumber of predicted TMHs: 2\nExp number of AAs in TMHs: 37.2608499999999\nExp number, first 60 AAs: 19.52923\nTotal prob of N-in: 0.94351\nPOSSIBLE N-term signal sequence\ninside 1 19\nTMhelix 20 39\noutside 40 325\nTMhelix 326 348\ninside 349 372

12864 GCF_000488075.1_Esch_coli_UMEA_3426-1_V1 Escherichia coli UMEA 3426-1 Proteobacteria; Gammaproteobacteria; Enterobacterales; Enterobacteriaceae; Escherichia; Escherichia coli MTGDNTLIHSHGINRRDFMKLCAALAATMGLSSKAAAEMAESVTNPQRPPVIWIGAQECT WP_000145410.1 MULTISPECIES: hydrogenase-2 small chain [Proteobacteria] Length: 372\nNumber of predicted TMHs: 1\nExp number of AAs in TMHs: 23.72226\nExp number, first 60 AAs: 0.22038\nTotal prob of N-in: 0.12994\noutside 1 331\nTMhelix 332 354\ninside 355 372

12865 GCF_000488075.1_Esch_coli_UMEA_3426-1_V1 Escherichia coli UMEA 3426-1 Proteobacteria; Gammaproteobacteria; Enterobacterales; Enterobacteriaceae; Escherichia; Escherichia coli MNNEETFYQAMRRQGVTRRSFLKYCSLAATSLGLGAGMAPKIAWALENKPRIPVVWIHGL WP_001058323.1 MULTISPECIES: hydrogenase-1 small chain [Proteobacteria] Length: 372\nNumber of predicted TMHs: 2\nExp number of AAs in TMHs: 37.2608499999999\nExp number, first 60 AAs: 19.52923\nTotal prob of N-in: 0.94351\nPOSSIBLE N-term signal sequence\ninside 1 19\nTMhelix 20 39\noutside 40 325\nTMhelix 326 348\ninside 349 372

12866 GCF_000507685.1_Esch_coli_UMEA_3489-1_V1 Escherichia coli UMEA 3489-1 Proteobacteria; Gammaproteobacteria; Enterobacterales; Enterobacteriaceae; Escherichia; Escherichia coli MTGDNTLIHSHGINRRDFMKLCAALAATMGLSSKAAAEMAESVTNPQRPPVIWIGAQECT WP_000145410.1 MULTISPECIES: hydrogenase-2 small chain [Proteobacteria] Length: 372\nNumber of predicted TMHs: 1\nExp number of AAs in TMHs: 23.72226\nExp number, first 60 AAs: 0.22038\nTotal prob of N-in: 0.12994\noutside 1 331\nTMhelix 332 354\ninside 355 372

12867 GCF_000507685.1_Esch_coli_UMEA_3489-1_V1 Escherichia coli UMEA 3489-1 Proteobacteria; Gammaproteobacteria; Enterobacterales; Enterobacteriaceae; Escherichia; Escherichia coli

MNNEETFYQAMRRQGVTRRSFLKYCSLAATSLGLGAGMAPKIAWALENKPRIPVVWIHGL WP_001058323.1 MULTISPECIES: hydrogenase-1 small chain [Proteobacteria] Length: 372\nNumber of predicted TMHs: 2\nExp number of AAs in TMHs: 37.2608499999999\nExp number, first 60 AAs: 19.52923\nTotal prob of N-in: 0.94351\nPOSSIBLE N-term signal sequence\ninside 1 19\nTMhelix 20 39\noutside 40 325\nTMhelix 326 348\ninside 349 372

12868 GCF_000461355.1_Esch_coli_UMEA_3490-1_V1 Escherichia coli UMEA 3490-1 Proteobacteria; Gammaproteobacteria; Enterobacterales; Enterobacteriaceae; Escherichia; Escherichia coli MNNEETFYQAMRRQGVTRRSFLKYCSLAATSLGLGAGMAPKIAWALENKPRIPVVWIHGL WP_001058323.1 MULTISPECIES: hydrogenase-1 small chain [Proteobacteria] Length: 372\nNumber of predicted TMHs: 2\nExp number of AAs in TMHs: 37.2608499999999\nExp number, first 60 AAs: 19.52923\nTotal prob of N-in: 0.94351\nPOSSIBLE N-term signal sequence\ninside 1 19\nTMhelix 20 39\noutside 40 325\nTMhelix 326 348\ninside 349 372

12869 GCF_000461355.1_Esch_coli_UMEA_3490-1_V1 Escherichia coli UMEA 3490-1 Proteobacteria; Gammaproteobacteria; Enterobacterales; Enterobacteriaceae; Escherichia; Escherichia coli MTGDNTLIHSHGINRRDFMKLCAALAATMGLSSKAAAEMAESVTNPQRPPVIWIGAQECT WP_000145410.1 MULTISPECIES: hydrogenase-2 small chain [Proteobacteria] Length: 372\nNumber of predicted TMHs: 1\nExp number of AAs in TMHs: 23.72226\nExp number, first 60 AAs: 0.22038\nTotal prob of N-in: 0.12994\noutside 1 331\nTMhelix 332 354\ninside 355 372

12870 GCF_000461375.1_Esch_coli_UMEA_3585-1_V1 Escherichia coli UMEA 3585-1 Proteobacteria; Gammaproteobacteria; Enterobacterales; Enterobacteriaceae; Escherichia; Escherichia coli MTGDNTLIHSHGINRRDFMKLCAALAATMGLSSKAAAEMAESVTNPQRPPVIWIGAQECT WP_000145410.1 MULTISPECIES: hydrogenase-2 small chain [Proteobacteria] Length: 372\nNumber of predicted TMHs: 1\nExp number of AAs in TMHs: 23.72226\nExp number, first 60 AAs: 0.22038\nTotal prob of N-in: 0.12994\noutside 1 331\nTMhelix 332 354\ninside 355 372

12871 GCF_000461375.1_Esch_coli_UMEA_3585-1_V1 Escherichia coli UMEA 3585-1 Proteobacteria; Gammaproteobacteria; Enterobacterales; Enterobacteriaceae; Escherichia; Escherichia coli MNNEETFYQAMRRQGVTRRSFLKYCSLAATSLGLGAGMAPKIAWALENKPRIPVVWIHGL WP_001058323.1 MULTISPECIES: hydrogenase-1 small chain [Proteobacteria] Length: 372\nNumber of predicted TMHs: 2\nExp number of AAs in TMHs: 37.2608499999999\nExp number, first 60 AAs: 19.52923\nTotal prob of N-in: 0.94351\nPOSSIBLE N-term signal sequence\ninside 1 19\nTMhelix 20 39\noutside 40 325\nTMhelix 326 348\ninside 349 372

12872 GCF_000461395.1_Esch_coli_UMEA_3592-1_V1 Escherichia coli UMEA 3592-1 Proteobacteria; Gammaproteobacteria; Enterobacterales; Enterobacteriaceae; Escherichia; Escherichia coli MTGDNTLIHSHGINRRDFMKLCAALAATMGLSSKAAAEMAESVTNPQRPPVIWIGAQECT WP_000145410.1 MULTISPECIES: hydrogenase-2 small chain [Proteobacteria] Length: 372\nNumber of predicted TMHs: 1\nExp number of AAs in TMHs: 23.72226\nExp number, first 60 AAs: 0.22038\nTotal prob of N-in: 0.12994\noutside 1 331\nTMhelix 332 354\ninside 355 372

12873 GCF_000461395.1_Esch_coli_UMEA_3592-1_V1 Escherichia coli UMEA 3592-1 Proteobacteria; Gammaproteobacteria; Enterobacterales; Enterobacteriaceae; Escherichia; Escherichia coli MNNEETFYQAMRRQGVTRRSFLKYCSLAATSLGLGAGMAPKIAWALENKPRIPVVWIHGL WP_001058323.1 MULTISPECIES: hydrogenase-1 small chain [Proteobacteria] Length: 372\nNumber of predicted TMHs: 2\nExp number of AAs in TMHs: 37.2608499999999\nExp number, first 60 AAs: 19.52923\nTotal prob of N-in: 0.94351\nPOSSIBLE N-term signal sequence\ninside 1 19\nTMhelix 20 39\noutside 40 325\nTMhelix 326 348\ninside 349 372

12874 GCF_000461415.1_Esch_coli_UMEA_3609-1_V1 Escherichia coli UMEA 3609-1 Proteobacteria; Gammaproteobacteria; Enterobacterales; Enterobacteriaceae; Escherichia; Escherichia coli MNNEETFYQAMRRQGVTRRSFLKYCSLAATSLGLGAGMAPKIAWALENKPRIPVVWIHGL WP_001058323.1 MULTISPECIES: hydrogenase-1 small chain [Proteobacteria] Length: 372\nNumber of predicted TMHs: 2\nExp number of AAs in TMHs: 37.2608499999999\nExp number, first 60 AAs: 19.52923\nTotal prob of N-in: 0.94351\nPOSSIBLE N-term signal sequence\ninside 1 19\nTMhelix 20 39\noutside 40 325\nTMhelix 326 348\ninside 349 372

12875 GCF_000461415.1_Esch_coli_UMEA_3609-1_V1 Escherichia coli UMEA 3609-1 Proteobacteria; Gammaproteobacteria; Enterobacterales; Enterobacteriaceae; Escherichia; Escherichia coli

MTGDNTLIHSHGINRRDFMKLCAALAATMGLSSKAAAEMAESVTNPQRPPVIWIGAQECT WP_000145410.1 MULTISPECIES: hydrogenase-2 small chain [Proteobacteria] Length: 372\nNumber of predicted TMHs: 1\nExp number of AAs in TMHs: 23.72226\nExp number, first 60 AAs: 0.22038\nTotal prob of N-in: 0.12994\noutside 1 331\nTMhelix 332 354\ninside 355 372

12876 GCF_000461435.1_Esch_coli_UMEA_3617-1_V1 Escherichia coli UMEA 3617-1 Proteobacteria; Gammaproteobacteria; Enterobacterales; Enterobacteriaceae; Escherichia; Escherichia coli MNNEETFYQAMRRQGVTRRSFLKYCSLAATSLGLGAGMAPKIAWALENKPRIPVVWIHGL WP_001058323.1 MULTISPECIES: hydrogenase-1 small chain [Proteobacteria] Length: 372\nNumber of predicted TMHs: 2\nExp number of AAs in TMHs: 37.2608499999999\nExp number, first 60 AAs: 19.52923\nTotal prob of N-in: 0.94351\nPOSSIBLE N-term signal sequence\ninside 1 19\nTMhelix 20 39\noutside 40 325\nTMhelix 326 348\ninside 349 372

12877 GCF_000461435.1_Esch_coli_UMEA_3617-1_V1 Escherichia coli UMEA 3617-1 Proteobacteria; Gammaproteobacteria; Enterobacterales; Enterobacteriaceae; Escherichia; Escherichia coli MTGDNTLIHSHGINRRDFMKLCAALAATMGLSSKAAAEMAESVTNPQRPPVIWIGAQECT WP_000145410.1 MULTISPECIES: hydrogenase-2 small chain [Proteobacteria] Length: 372\nNumber of predicted TMHs: 1\nExp number of AAs in TMHs: 23.72226\nExp number, first 60 AAs: 0.22038\nTotal prob of N-in: 0.12994\noutside 1 331\nTMhelix 332 354\ninside 355 372

12878 GCF_000461455.1_Esch_coli_UMEA_3632-1_V1 Escherichia coli UMEA 3632-1 Proteobacteria; Gammaproteobacteria; Enterobacterales; Enterobacteriaceae; Escherichia; Escherichia coli MTGDNTLIHSHGINRRDFMKLCAALAATMGLSSKAAAEMAESVTNPQRPPVIWIGAQECT WP_000145410.1 MULTISPECIES: hydrogenase-2 small chain [Proteobacteria] Length: 372\nNumber of predicted TMHs: 1\nExp number of AAs in TMHs: 23.72226\nExp number, first 60 AAs: 0.22038\nTotal prob of N-in: 0.12994\noutside 1 331\nTMhelix 332 354\ninside 355 372

12879 GCF_000463605.1_Esch_coli_UMEA_3652-1_V1 Escherichia coli UMEA 3652-1 Proteobacteria; Gammaproteobacteria; Enterobacterales; Enterobacteriaceae; Escherichia; Escherichia coli MNNEETFYQAMRRQGVTRRSFLKYCSLAATSLGLGTGMAPKIAWALENKPRIPVVWIHGL WP_021527074.1 hydrogenase-1 small chain [Escherichia coli] Length: 372\nNumber of predicted TMHs: 2\nExp number of AAs in TMHs: 36.47672\nExp number, first 60 AAs: 18.73629\nTotal prob of N-in: 0.90684\nPOSSIBLE N-term signal sequence\ninside 1 19\nTMhelix 20 39\noutside 40 325\nTMhelix 326 348\ninside 349 372

12880 GCF_000463605.1_Esch_coli_UMEA_3652-1_V1 Escherichia coli UMEA 3652-1 Proteobacteria; Gammaproteobacteria; Enterobacterales; Enterobacteriaceae; Escherichia; Escherichia coli MTGDNTLIHSHGINRRDFMKLCAALAATMGLSSKAAAEMAESVTNPQRPPVIWIGAQECT WP_000145410.1 MULTISPECIES: hydrogenase-2 small chain [Proteobacteria] Length: 372\nNumber of predicted TMHs: 1\nExp number of AAs in TMHs: 23.72226\nExp number, first 60 AAs: 0.22038\nTotal prob of N-in: 0.12994\noutside 1 331\nTMhelix 332 354\ninside 355 372

12881 GCF_000461475.1_Esch_coli_UMEA_3656-1_V1 Escherichia coli UMEA 3656-1 Proteobacteria; Gammaproteobacteria; Enterobacterales; Enterobacteriaceae; Escherichia; Escherichia coli MNNEETFYQAMRRQGVTRRSFLKYCSLAATSLGLGAGMAPKIAWALENKPRIPVVWIHGL WP_001058323.1 MULTISPECIES: hydrogenase-1 small chain [Proteobacteria] Length: 372\nNumber of predicted TMHs: 2\nExp number of AAs in TMHs: 37.2608499999999\nExp number, first 60 AAs: 19.52923\nTotal prob of N-in: 0.94351\nPOSSIBLE N-term signal sequence\ninside 1 19\nTMhelix 20 39\noutside 40 325\nTMhelix 326 348\ninside 349 372

12882 GCF_000461475.1_Esch_coli_UMEA_3656-1_V1 Escherichia coli UMEA 3656-1 Proteobacteria; Gammaproteobacteria; Enterobacterales; Enterobacteriaceae; Escherichia; Escherichia coli MTGDNTLIHSHGINRRDFMKLCAALAATMGLSSKAAAEMVESVTNPQRPPVIWIGAQECT WP_000145423.1 hydrogenase 2 small subunit [Escherichia coli] Length: 372\nNumber of predicted TMHs: 1\nExp number of AAs in TMHs: 23.78201\nExp number, first 60 AAs: 0.28355\nTotal prob of N-in: 0.13257\noutside 1 331\nTMhelix 332 354\ninside 355 372

12883 GCF_000461495.1_Esch_coli_UMEA_3662-1_V1 Escherichia coli UMEA 3662-1 Proteobacteria; Gammaproteobacteria; Enterobacterales; Enterobacteriaceae; Escherichia; Escherichia coli MTGDNTLIHSHGINRRDFMKLCAALAATMGLSSKAAAEMAESVTNPQRPPVIWIGAQECT WP_000145410.1 MULTISPECIES: hydrogenase-2 small chain [Proteobacteria] Length: 372\nNumber of predicted TMHs:

1\nExp number of AAs in TMHs: 23.72226\nExp number, first 60 AAs: 0.22038\nTotal prob of N-in: 0.12994\noutside 1 331\nTMhelix 332 354\ninside 355 372

12884 GCF_000461515.1_Esch_coli_UMEA_3671-1_V1 Escherichia coli UMEA 3671-1 Proteobacteria; Gammaproteobacteria; Enterobacterales; Enterobacteriaceae; Escherichia; Escherichia coli MNNEETFYQAMRRQGVTRRSFLKYCSLAATSLGLGAGMAPKIAWALENKPRIPVVWIHGL WP_001058323.1 MULTISPECIES: hydrogenase-1 small chain [Proteobacteria] Length: 372\nNumber of predicted TMHs: 2\nExp number of AAs in TMHs: 37.2608499999999\nExp number, first 60 AAs: 19.52923\nTotal prob of N-in: 0.94351\nPOSSIBLE N-term signal sequence\ninside 1 19\nTMhelix 20 39\noutside 40 325\nTMhelix 326 348\ninside 349 372

12885 GCF_000461515.1_Esch_coli_UMEA_3671-1_V1 Escherichia coli UMEA 3671-1 Proteobacteria; Gammaproteobacteria; Enterobacterales; Enterobacteriaceae; Escherichia; Escherichia coli MTGDNTLIHSHGINRRDFMKLCAALAATMGLSSKAAAEMAESVTNPQRPPVIWIGAQECT WP_000145410.1 MULTISPECIES: hydrogenase-2 small chain [Proteobacteria] Length: 372\nNumber of predicted TMHs: 1\nExp number of AAs in TMHs: 23.72226\nExp number, first 60 AAs: 0.22038\nTotal prob of N-in: 0.12994\noutside 1 331\nTMhelix 332 354\ninside 355 372

12886 GCF_000461535.1_Esch_coli_UMEA_3682-1_V1 Escherichia coli UMEA 3682-1 Proteobacteria; Gammaproteobacteria; Enterobacterales; Enterobacteriaceae; Escherichia; Escherichia coli MNNEETFYQAMRRQGVTRRSFLKYCSLAATSLGLGAGMAPKIAWALENKPRIPVVWIHGL WP_001058323.1 MULTISPECIES: hydrogenase-1 small chain [Proteobacteria] Length: 372\nNumber of predicted TMHs: 2\nExp number of AAs in TMHs: 37.2608499999999\nExp number, first 60 AAs: 19.52923\nTotal prob of N-in: 0.94351\nPOSSIBLE N-term signal sequence\ninside 1 19\nTMhelix 20 39\noutside 40 325\nTMhelix 326 348\ninside 349 372

12887 GCF_000461535.1_Esch_coli_UMEA_3682-1_V1 Escherichia coli UMEA 3682-1 Proteobacteria; Gammaproteobacteria; Enterobacterales; Enterobacteriaceae; Escherichia; Escherichia coli MTGDNTLIHSHGINRRDFMKLCAALAATMGLSSKAAAEMAESVTNPQRPPVIWIGAQECT WP_000145410.1 MULTISPECIES: hydrogenase-2 small chain [Proteobacteria] Length: 372\nNumber of predicted TMHs: 1\nExp number of AAs in TMHs: 23.72226\nExp number, first 60 AAs: 0.22038\nTotal prob of N-in: 0.12994\noutside 1 331\nTMhelix 332 354\ninside 355 372

12888 GCF_000461555.1_Esch_coli_UMEA_3687-1_V1 Escherichia coli UMEA 3687-1 Proteobacteria; Gammaproteobacteria; Enterobacterales; Enterobacteriaceae; Escherichia; Escherichia coli MTGDNTLIHSHGINRRDFMKLCAALAATMGLSSKAAAEMAESVTNPQRPPVIWIGAQECT WP_021575602.1 hydrogenase-2 small chain [Escherichia coli] Length: 372\nNumber of predicted TMHs: 1\nExp number of AAs in TMHs: 23.69215\nExp number, first 60 AAs: 0.22177\nTotal prob of N-in: 0.12758\noutside 1 331\nTMhelix 332 354\ninside 355 372

12889 GCF_000461555.1_Esch_coli_UMEA_3687-1_V1 Escherichia coli UMEA 3687-1 Proteobacteria; Gammaproteobacteria; Enterobacterales; Enterobacteriaceae; Escherichia; Escherichia coli MNNEETFYQAMRRQGVTRRSFLKYCSLAATSLGLGAGMAPKIAWALENKPRIPVVWIHGL WP_001058323.1 MULTISPECIES: hydrogenase-1 small chain [Proteobacteria] Length: 372\nNumber of predicted TMHs: 2\nExp number of AAs in TMHs: 37.2608499999999\nExp number, first 60 AAs: 19.52923\nTotal prob of N-in: 0.94351\nPOSSIBLE N-term signal sequence\ninside 1 19\nTMhelix 20 39\noutside 40 325\nTMhelix 326 348\ninside 349 372

12890 GCF_000488115.1_Esch_coli_UMEA_3693-1_V1 Escherichia coli UMEA 3693-1 Proteobacteria; Gammaproteobacteria; Enterobacterales; Enterobacteriaceae; Escherichia; Escherichia coli MNNEETFYQAMRRQGVTRRSFLKYCSLAATSLGLGAGMAPKIAWALENKPRIPVVWIHGL WP_001058323.1 MULTISPECIES: hydrogenase-1 small chain [Proteobacteria] Length: 372\nNumber of predicted TMHs: 2\nExp number of AAs in TMHs: 37.2608499999999\nExp number, first 60 AAs: 19.52923\nTotal prob of N-in: 0.94351\nPOSSIBLE N-term signal sequence\ninside 1 19\nTMhelix 20 39\noutside 40 325\nTMhelix 326 348\ninside 349 372

12891 GCF_000488115.1_Esch_coli_UMEA_3693-1_V1 Escherichia coli UMEA 3693-1 Proteobacteria; Gammaproteobacteria; Enterobacterales; Enterobacteriaceae; Escherichia; Escherichia coli MTGDNTLIHSHGINRRDFMKLCAALAATMGLSSKAAAEMAESVTNPQRPPVIWIGAQECT WP_000145410.1 MULTISPECIES: hydrogenase-2 small chain [Proteobacteria] Length: 372\nNumber of predicted TMHs:

1\nExp number of AAs in TMHs: 23.72226\nExp number, first 60 AAs: 0.22038\nTotal prob of N-in: 0.12994\noutside 1 331\nTMhelix 332 354\ninside 355 372

12892 GCF_000461575.1_Esch_coli_UMEA_3694-1_V1 Escherichia coli UMEA 3694-1 Proteobacteria; Gammaproteobacteria; Enterobacterales; Enterobacteriaceae; Escherichia; Escherichia coli MTGDNTLIHSHGINRRDFMKLCAALAATMGLSSKAAAEMAESVTNPQRPPVIWIGAQECT WP_000145410.1 MULTISPECIES: hydrogenase-2 small chain [Proteobacteria] Length: 372\nNumber of predicted TMHs: 1\nExp number of AAs in TMHs: 23.72226\nExp number, first 60 AAs: 0.22038\nTotal prob of N-in: 0.12994\noutside 1 331\nTMhelix 332 354\ninside 355 372

12893 GCF_000461575.1_Esch_coli_UMEA_3694-1_V1 Escherichia coli UMEA 3694-1 Proteobacteria; Gammaproteobacteria; Enterobacterales; Enterobacteriaceae; Escherichia; Escherichia coli MNNEETFYQAMRRQGVTRRSFLKYCSLAATSLGLGAGMAPKIAWALENKPRIPVVWIHGL WP_001058323.1 MULTISPECIES: hydrogenase-1 small chain [Proteobacteria] Length: 372\nNumber of predicted TMHs: 2\nExp number of AAs in TMHs: 37.2608499999999\nExp number, first 60 AAs: 19.52923\nTotal prob of N-in: 0.94351\nPOSSIBLE N-term signal sequence\ninside 1 19\nTMhelix 20 39\noutside 40 325\nTMhelix 326 348\ninside 349 372

12894 GCF_000461595.1_Esch_coli_UMEA_3702-1_V1 Escherichia coli UMEA 3702-1 Proteobacteria; Gammaproteobacteria; Enterobacterales; Enterobacteriaceae; Escherichia; Escherichia coli MTGDNTLIHSHGINRRDFMKLCAALAATMGLSSKAAAEMAESVTNPQRPPVIWIGAQECT WP_000145410.1 MULTISPECIES: hydrogenase-2 small chain [Proteobacteria] Length: 372\nNumber of predicted TMHs: 1\nExp number of AAs in TMHs: 23.72226\nExp number, first 60 AAs: 0.22038\nTotal prob of N-in: 0.12994\noutside 1 331\nTMhelix 332 354\ninside 355 372

12895 GCF_000461615.1_Esch_coli_UMEA_3703-1_V1 Escherichia coli UMEA 3703-1 Proteobacteria; Gammaproteobacteria; Enterobacterales; Enterobacteriaceae; Escherichia; Escherichia coli MTGDNTLIHSHGINRRDFMKLCAALAATMGLSSKAAAEMAESVTNPQRPPVIWIGAQECT WP_000145410.1 MULTISPECIES: hydrogenase-2 small chain [Proteobacteria] Length: 372\nNumber of predicted TMHs: 1\nExp number of AAs in TMHs: 23.72226\nExp number, first 60 AAs: 0.22038\nTotal prob of N-in: 0.12994\noutside 1 331\nTMhelix 332 354\ninside 355 372

12896 GCF_000461615.1_Esch_coli_UMEA_3703-1_V1 Escherichia coli UMEA 3703-1 Proteobacteria; Gammaproteobacteria; Enterobacterales; Enterobacteriaceae; Escherichia; Escherichia coli MNNEETFYQAMRRQGVTRRSFLKYCSLAATSLGLGAGMAPKIAWALENKPRIPVVWIHGL WP_001058323.1 MULTISPECIES: hydrogenase-1 small chain [Proteobacteria] Length: 372\nNumber of predicted TMHs: 2\nExp number of AAs in TMHs: 37.2608499999999\nExp number, first 60 AAs: 19.52923\nTotal prob of N-in: 0.94351\nPOSSIBLE N-term signal sequence\ninside 1 19\nTMhelix 20 39\noutside 40 325\nTMhelix 326 348\ninside 349 372

12897 GCF_000461635.1_Esch_coli_UMEA_3705-1_V1 Escherichia coli UMEA 3705-1 Proteobacteria; Gammaproteobacteria; Enterobacterales; Enterobacteriaceae; Escherichia; Escherichia coli MNNEETFYQAMRRQGVTRRSFLKYCSLAATSLGLGAGMAPKIAWALENKPRIPVVWIHGL WP_001058323.1 MULTISPECIES: hydrogenase-1 small chain [Proteobacteria] Length: 372\nNumber of predicted TMHs: 2\nExp number of AAs in TMHs: 37.2608499999999\nExp number, first 60 AAs: 19.52923\nTotal prob of N-in: 0.94351\nPOSSIBLE N-term signal sequence\ninside 1 19\nTMhelix 20 39\noutside 40 325\nTMhelix 326 348\ninside 349 372

12898 GCF_000461635.1_Esch_coli_UMEA_3705-1_V1 Escherichia coli UMEA 3705-1 Proteobacteria; Gammaproteobacteria; Enterobacterales; Enterobacteriaceae; Escherichia; Escherichia coli MTGDNTLIHSHGINRRDFMKLCAALAATMGLSSKAAAEMAESVTNPQRPPVIWIGAQECT WP_000145410.1 MULTISPECIES: hydrogenase-2 small chain [Proteobacteria] Length: 372\nNumber of predicted TMHs: 1\nExp number of AAs in TMHs: 23.72226\nExp number, first 60 AAs: 0.22038\nTotal prob of N-in: 0.12994\noutside 1 331\nTMhelix 332 354\ninside 355 372

12899 GCF_000461655.1_Esch_coli_UMEA_3707-1_V1 Escherichia coli UMEA 3707-1 Proteobacteria; Gammaproteobacteria; Enterobacterales; Enterobacteriaceae; Escherichia; Escherichia coli MTGDNTLIHSHGINRRDFMKLCAALAATMGLSSKAAAEMAESVTNPQRPPVIWIGAQECT WP_000145410.1 MULTISPECIES: hydrogenase-2 small chain [Proteobacteria] Length: 372\nNumber of predicted TMHs: 1\nExp number of AAs in TMHs: 23.72226\nExp number, first 60 AAs: 0.22038\nTotal prob of N-in: 0.12994\noutside 1 331\nTMhelix 332 354\ninside 355 372

12900 GCF_000461655.1_Esch_coli_UMEA_3707-1_V1 Escherichia coli UMEA 3707-1 Proteobacteria; Gammaproteobacteria; Enterobacterales; Enterobacteriaceae; Escherichia; Escherichia coli
MNNEETFYQAMRRQGVTRRSFLKYCSLAATSLGLGAGMAPKIAWALENKPRIPVVWIHGL WP_001058323.1
MULTISPECIES: hydrogenase-1 small chain [Proteobacteria] Length: 372\nNumber of predicted TMHs: 2\nExp number of AAs in TMHs: 37.2608499999999\nExp number, first 60 AAs: 19.52923\nTotal prob of N-in: 0.94351\nPOSSIBLE N-term signal sequence\ninside 1 19\nTMhelix 20 39\noutside 40 325\nTMhelix 326 348\ninside 349 372

12901 GCF_000461675.1_Esch_coli_UMEA_3718-1_V1 Escherichia coli UMEA 3718-1 Proteobacteria; Gammaproteobacteria; Enterobacterales; Enterobacteriaceae; Escherichia; Escherichia coli
MTGDNTLIHSHGINRRDFMKLCAALAATMGLSSKAAAEMAESVTNPQRPPVIWIGAQECT WP_000145410.1
MULTISPECIES: hydrogenase-2 small chain [Proteobacteria] Length: 372\nNumber of predicted TMHs: 1\nExp number of AAs in TMHs: 23.72226\nExp number, first 60 AAs: 0.22038\nTotal prob of N-in: 0.12994\noutside 1 331\nTMhelix 332 354\ninside 355 372

12902 GCF_000461675.1_Esch_coli_UMEA_3718-1_V1 Escherichia coli UMEA 3718-1 Proteobacteria; Gammaproteobacteria; Enterobacterales; Enterobacteriaceae; Escherichia; Escherichia coli
MNNEETFYQAMRRQGVTRRSFLKYCSLAATSLGLGAGMAPKIAWALENKPRIPVVWIHGL WP_001058323.1
MULTISPECIES: hydrogenase-1 small chain [Proteobacteria] Length: 372\nNumber of predicted TMHs: 2\nExp number of AAs in TMHs: 37.2608499999999\nExp number, first 60 AAs: 19.52923\nTotal prob of N-in: 0.94351\nPOSSIBLE N-term signal sequence\ninside 1 19\nTMhelix 20 39\noutside 40 325\nTMhelix 326 348\ninside 349 372

12903 GCF_000461695.1_Esch_coli_UMEA_3805-1_V1 Escherichia coli UMEA 3805-1 Proteobacteria; Gammaproteobacteria; Enterobacterales; Enterobacteriaceae; Escherichia; Escherichia coli
MNNEETFYQAMRRQGVTRRSFLKYCSLAATSLGLGAGMAPKIAWALENKPRIPVVWIHGL WP_001058323.1
MULTISPECIES: hydrogenase-1 small chain [Proteobacteria] Length: 372\nNumber of predicted TMHs: 2\nExp number of AAs in TMHs: 37.2608499999999\nExp number, first 60 AAs: 19.52923\nTotal prob of N-in: 0.94351\nPOSSIBLE N-term signal sequence\ninside 1 19\nTMhelix 20 39\noutside 40 325\nTMhelix 326 348\ninside 349 372

12904 GCF_000461695.1_Esch_coli_UMEA_3805-1_V1 Escherichia coli UMEA 3805-1 Proteobacteria; Gammaproteobacteria; Enterobacterales; Enterobacteriaceae; Escherichia; Escherichia coli
MTGDNTLIHSHGINRRDFMKLCAALAATMGLSSKAAAEMAESVTNPQRPPVIWIGAQECT WP_000145410.1
MULTISPECIES: hydrogenase-2 small chain [Proteobacteria] Length: 372\nNumber of predicted TMHs: 1\nExp number of AAs in TMHs: 23.72226\nExp number, first 60 AAs: 0.22038\nTotal prob of N-in: 0.12994\noutside 1 331\nTMhelix 332 354\ninside 355 372

12905 GCF_000461715.1_Esch_coli_UMEA_3821-1_V1 Escherichia coli UMEA 3821-1 Proteobacteria; Gammaproteobacteria; Enterobacterales; Enterobacteriaceae; Escherichia; Escherichia coli
MTGDNTLIHSHGINRRDFMKLCAALAATMGLSSKAAAEMAESVTNPQRPPVIWIGAQECT WP_000145410.1
MULTISPECIES: hydrogenase-2 small chain [Proteobacteria] Length: 372\nNumber of predicted TMHs: 1\nExp number of AAs in TMHs: 23.72226\nExp number, first 60 AAs: 0.22038\nTotal prob of N-in: 0.12994\noutside 1 331\nTMhelix 332 354\ninside 355 372

12906 GCF_000461715.1_Esch_coli_UMEA_3821-1_V1 Escherichia coli UMEA 3821-1 Proteobacteria; Gammaproteobacteria; Enterobacterales; Enterobacteriaceae; Escherichia; Escherichia coli
MNNEETFYQAMRRQGVTRRSFLKYCSLAATSLGLGAGMAPKIAWALENKPRIPVVWIHGL WP_001058323.1
MULTISPECIES: hydrogenase-1 small chain [Proteobacteria] Length: 372\nNumber of predicted TMHs: 2\nExp number of AAs in TMHs: 37.2608499999999\nExp number, first 60 AAs: 19.52923\nTotal prob of N-in: 0.94351\nPOSSIBLE N-term signal sequence\ninside 1 19\nTMhelix 20 39\noutside 40 325\nTMhelix 326 348\ninside 349 372

12907 GCF_000461735.1_Esch_coli_UMEA_3834-1_V1 Escherichia coli UMEA 3834-1 Proteobacteria; Gammaproteobacteria; Enterobacterales; Enterobacteriaceae; Escherichia; Escherichia coli
MTGDNTLIHSHGINRRDFMKLCAALAATMGLSSKAAAEMAESVTNPQRPPVIWIGAQECT WP_000145410.1
MULTISPECIES: hydrogenase-2 small chain [Proteobacteria] Length: 372\nNumber of predicted TMHs: 1\nExp number of AAs in TMHs: 23.72226\nExp number, first 60 AAs: 0.22038\nTotal prob of N-in: 0.12994\noutside 1 331\nTMhelix 332 354\ninside 355 372

12908 GCF_000461755.1_Esch_coli_UMEA_3889-1_V1 Escherichia coli UMEA 3889-1 Proteobacteria; Gammaproteobacteria; Enterobacterales; Enterobacteriaceae; Escherichia; Escherichia coli MNNEETFYQAMRRQGVTRRSFLKYCSLAATSLGLGAGMAPKIAWALENKPRIPVVWIHGL WP_001058323.1 MULTISPECIES: hydrogenase-1 small chain [Proteobacteria] Length: 372\nNumber of predicted TMHs: 2\nExp number of AAs in TMHs: 37.2608499999999\nExp number, first 60 AAs: 19.52923\nTotal prob of N-in: 0.94351\nPOSSIBLE N-term signal sequence\ninside 1 19\nTMhelix 20 39\noutside 40 325\nTMhelix 326 348\ninside 349 372

12909 GCF_000461755.1_Esch_coli_UMEA_3889-1_V1 Escherichia coli UMEA 3889-1 Proteobacteria; Gammaproteobacteria; Enterobacterales; Enterobacteriaceae; Escherichia; Escherichia coli MTGDNTLIHSHGINRRDFMKLCAALAATMGLSSKAAAEMAESVTNPQRPPVIWIGAQECT WP_000145410.1 MULTISPECIES: hydrogenase-2 small chain [Proteobacteria] Length: 372\nNumber of predicted TMHs: 1\nExp number of AAs in TMHs: 23.72226\nExp number, first 60 AAs: 0.22038\nTotal prob of N-in: 0.12994\noutside 1 331\nTMhelix 332 354\ninside 355 372

12910 GCF_000461775.1_Esch_coli_UMEA_3893-1_V1 Escherichia coli UMEA 3893-1 Proteobacteria; Gammaproteobacteria; Enterobacterales; Enterobacteriaceae; Escherichia; Escherichia coli MTGDNTLIHSHGINRRDFMKLCAALAATMGLSSKAAAEMAESVTNPQRPPVIWIGAQECT WP_000145410.1 MULTISPECIES: hydrogenase-2 small chain [Proteobacteria] Length: 372\nNumber of predicted TMHs: 1\nExp number of AAs in TMHs: 23.72226\nExp number, first 60 AAs: 0.22038\nTotal prob of N-in: 0.12994\noutside 1 331\nTMhelix 332 354\ninside 355 372

12911 GCF_000461795.1_Esch_coli_UMEA_3899-1_V1 Escherichia coli UMEA 3899-1 Proteobacteria; Gammaproteobacteria; Enterobacterales; Enterobacteriaceae; Escherichia; Escherichia coli MNNEETFYQAMRRQGVTRRSFLKYCSLAATSLGLGAGMAPKIAWALENKPRIPVVWIHGL WP_001295940.1 MULTISPECIES: hydrogenase [Enterobacteriaceae] Length: 372\nNumber of predicted TMHs: 2\nExp number of AAs in TMHs: 37.2608899999999\nExp number, first 60 AAs: 19.52923\nTotal prob of N-in: 0.94351\nPOSSIBLE N-term signal sequence\ninside 1 19\nTMhelix 20 39\noutside 40 325\nTMhelix 326 348\ninside 349 372

12912 GCF_000461795.1_Esch_coli_UMEA_3899-1_V1 Escherichia coli UMEA 3899-1 Proteobacteria; Gammaproteobacteria; Enterobacterales; Enterobacteriaceae; Escherichia; Escherichia coli MTGDNTLIHSHGINRRDFMKLCAALAATMGLSSKAAAEMAESVTNPQRPPVIWIGAQECT WP_000145410.1 MULTISPECIES: hydrogenase-2 small chain [Proteobacteria] Length: 372\nNumber of predicted TMHs: 1\nExp number of AAs in TMHs: 23.72226\nExp number, first 60 AAs: 0.22038\nTotal prob of N-in: 0.12994\noutside 1 331\nTMhelix 332 354\ninside 355 372

12913 GCF_000461815.1_Esch_coli_UMEA_3955-1_V1 Escherichia coli UMEA 3955-1 Proteobacteria; Gammaproteobacteria; Enterobacterales; Enterobacteriaceae; Escherichia; Escherichia coli MNNEETFYQAMRRQGVTRRSFLKYCSLAATSLGLGAGMAPKIAWALENKPRIPVVWIHGL WP_001058323.1 MULTISPECIES: hydrogenase-1 small chain [Proteobacteria] Length: 372\nNumber of predicted TMHs: 2\nExp number of AAs in TMHs: 37.2608499999999\nExp number, first 60 AAs: 19.52923\nTotal prob of N-in: 0.94351\nPOSSIBLE N-term signal sequence\ninside 1 19\nTMhelix 20 39\noutside 40 325\nTMhelix 326 348\ninside 349 372

12914 GCF_000461815.1_Esch_coli_UMEA_3955-1_V1 Escherichia coli UMEA 3955-1 Proteobacteria; Gammaproteobacteria; Enterobacterales; Enterobacteriaceae; Escherichia; Escherichia coli MTGDNTLIHSHGINRRDFMKLCAALAATMGLSSKAAAEMAESVTNPQRPPVIWIGAQECT WP_000145410.1 MULTISPECIES: hydrogenase-2 small chain [Proteobacteria] Length: 372\nNumber of predicted TMHs: 1\nExp number of AAs in TMHs: 23.72226\nExp number, first 60 AAs: 0.22038\nTotal prob of N-in: 0.12994\noutside 1 331\nTMhelix 332 354\ninside 355 372

12915 GCF_000461835.1_Esch_coli_UMEA_4075-1_V1 Escherichia coli UMEA 4075-1 Proteobacteria; Gammaproteobacteria; Enterobacterales; Enterobacteriaceae; Escherichia; Escherichia coli MNNEETFYQAMRRQGVTRRSFLKYCSLAATSLGLGAGMAPKIAWALENKPRIPVVWIHGL WP_001058323.1 MULTISPECIES: hydrogenase-1 small chain [Proteobacteria] Length: 372\nNumber of predicted TMHs: 2\nExp number of AAs in TMHs: 37.2608499999999\nExp number, first 60 AAs: 19.52923\nTotal prob of N-in: 0.94351\nPOSSIBLE N-term signal sequence\ninside 1 19\nTMhelix 20 39\noutside 40 325\nTMhelix 326 348\ninside 349 372

12916 GCF_000461835.1_Esch_coli_UMEA_4075-1_V1 Escherichia coli UMEA 4075-1 Proteobacteria; Gammaproteobacteria; Enterobacterales; Enterobacteriaceae; Escherichia; Escherichia coli
MTGDNTLIHSHGINRRDFMKLCAALAATMGLSSKAAAEMAESVTNPQRPPVIWIGAQECT WP_000145410.1
MULTISPECIES: hydrogenase-2 small chain [Proteobacteria] Length: 372\nNumber of predicted TMHs: 1\nExp number of AAs in TMHs: 23.72226\nExp number, first 60 AAs: 0.22038\nTotal prob of N-in: 0.12994\noutside 1 331\nTMhelix 332 354\ninside 355 372

12917 GCF_000461855.1_Esch_coli_UMEA_4076-1_V1 Escherichia coli UMEA 4076-1 Proteobacteria; Gammaproteobacteria; Enterobacterales; Enterobacteriaceae; Escherichia; Escherichia coli
MNNEETFYQAMRRQGVTRRSFLKYCSLAATSLGLGAGMAPKIAWALENKPRIPVVWIHGL WP_001058323.1
MULTISPECIES: hydrogenase-1 small chain [Proteobacteria] Length: 372\nNumber of predicted TMHs: 2\nExp number of AAs in TMHs: 37.2608499999999\nExp number, first 60 AAs: 19.52923\nTotal prob of N-in: 0.94351\nPOSSIBLE N-term signal sequence\ninside 1 19\nTMhelix 20 39\noutside 40 325\nTMhelix 326 348\ninside 349 372

12918 GCF_000461855.1_Esch_coli_UMEA_4076-1_V1 Escherichia coli UMEA 4076-1 Proteobacteria; Gammaproteobacteria; Enterobacterales; Enterobacteriaceae; Escherichia; Escherichia coli
MTGDNTLIHSHGINRRDFMKLCAALAATMGLSSKAAAEMVESVTNPQRPPVIWIGAQECT WP_000145423.1
hydrogenase 2 small subunit [Escherichia coli] Length: 372\nNumber of predicted TMHs: 1\nExp number of AAs in TMHs: 23.78201\nExp number, first 60 AAs: 0.28355\nTotal prob of N-in: 0.13257\noutside 1 331\nTMhelix 332 354\ninside 355 372

12919 GCF_000461875.1_Esch_coli_UMEA_4207-1_V1 Escherichia coli UMEA 4207-1 Proteobacteria; Gammaproteobacteria; Enterobacterales; Enterobacteriaceae; Escherichia; Escherichia coli
MTGDNTLIHSHGINRRDFMKLCAALAATMGLSSKAAAEMAESVTNPQRPPVIWIGAQECT WP_000145410.1
MULTISPECIES: hydrogenase-2 small chain [Proteobacteria] Length: 372\nNumber of predicted TMHs: 1\nExp number of AAs in TMHs: 23.72226\nExp number, first 60 AAs: 0.22038\nTotal prob of N-in: 0.12994\noutside 1 331\nTMhelix 332 354\ninside 355 372

12920 GCF_000461875.1_Esch_coli_UMEA_4207-1_V1 Escherichia coli UMEA 4207-1 Proteobacteria; Gammaproteobacteria; Enterobacterales; Enterobacteriaceae; Escherichia; Escherichia coli
MNNEETFYQAMRRQGVTRRSFLKYCSLAATSLGLGAGMAPKIAWALENKPRIPVVWIHGL WP_001058323.1
MULTISPECIES: hydrogenase-1 small chain [Proteobacteria] Length: 372\nNumber of predicted TMHs: 2\nExp number of AAs in TMHs: 37.2608499999999\nExp number, first 60 AAs: 19.52923\nTotal prob of N-in: 0.94351\nPOSSIBLE N-term signal sequence\ninside 1 19\nTMhelix 20 39\noutside 40 325\nTMhelix 326 348\ninside 349 372

12921 GCF_000026325.1_ASM2632v1 Escherichia coli UMN026 Proteobacteria; Gammaproteobacteria; Enterobacterales; Enterobacteriaceae; Escherichia; Escherichia coli
MTGDNTLIHSHGINRRDFMKLCAALAATMGLSSKAAAEMAESVTNPQRPPVIWIGAQECT YP_002414143.1
hydrogenase 2 small subunit [Escherichia coli UMN026] Length: 372\nNumber of predicted TMHs: 1\nExp number of AAs in TMHs: 23.72226\nExp number, first 60 AAs: 0.22038\nTotal prob of N-in: 0.12994\noutside 1 331\nTMhelix 332 354\ninside 355 372

12922 GCF_000026325.1_ASM2632v1 Escherichia coli UMN026 Proteobacteria; Gammaproteobacteria; Enterobacterales; Enterobacteriaceae; Escherichia; Escherichia coli
MNNEETFYQAMRRQGVTRRSFLKYCSLAATSLGLGAGMAPKIAWALENKPRIPVVWIHGL YP_002411915.1
hydrogenase 1 small subunit [Escherichia coli UMN026] Length: 372\nNumber of predicted TMHs: 2\nExp number of AAs in TMHs: 37.2608499999999\nExp number, first 60 AAs: 19.52923\nTotal prob of N-in: 0.94351\nPOSSIBLE N-term signal sequence\ninside 1 19\nTMhelix 20 39\noutside 40 325\nTMhelix 326 348\ninside 349 372

12923 GCF_000220005.1_ASM22000v2 Escherichia coli UMNF18 Proteobacteria; Gammaproteobacteria; Enterobacterales; Enterobacteriaceae; Escherichia; Escherichia coli
MTGDNTLIHSHGINRRDFMKLCAALAATMGLSSKAAAEMAESVTNPQRPPVIWIGAQECT WP_000145410.1
MULTISPECIES: hydrogenase-2 small chain [Proteobacteria] Length: 372\nNumber of predicted TMHs: 1\nExp number of AAs in TMHs: 23.72226\nExp number, first 60 AAs: 0.22038\nTotal prob of N-in: 0.12994\noutside 1 331\nTMhelix 332 354\ninside 355 372

12924 GCF_000220005.1_ASM22000v2 Escherichia coli UMNF18 Proteobacteria; Gammaproteobacteria; Enterobacterales; Enterobacteriaceae; Escherichia; Escherichia coli

MNNEETFYQAMRRQGVTRRSFLKYCSLAATSLGLGAGMAPKIAWALENKPRIPVVWIHGL WP_001058323.1 MULTISPECIES: hydrogenase-1 small chain [Proteobacteria] Length: 372\nNumber of predicted TMHs: 2\nExp number of AAs in TMHs: 37.2608499999999\nExp number, first 60 AAs: 19.52923\nTotal prob of N-in: 0.94351\nPOSSIBLE N-term signal sequence\ninside 1 19\nTMhelix 20 39\noutside 40 325\nTMhelix 326 348\ninside 349 372

12925 GCF_000212715.2_ASM21271v2 Escherichia coli UMNK88 Proteobacteria; Gammaproteobacteria; Enterobacterales; Enterobacteriaceae; Escherichia; Escherichia coli MNNEETFYQAMRRQGVTRRSFLKYCSLAATSLGLGAGMAPKIAWALENKPRIPVVWIHGL WP_001058323.1 MULTISPECIES: hydrogenase-1 small chain [Proteobacteria] Length: 372\nNumber of predicted TMHs: 2\nExp number of AAs in TMHs: 37.2608499999999\nExp number, first 60 AAs: 19.52923\nTotal prob of N-in: 0.94351\nPOSSIBLE N-term signal sequence\ninside 1 19\nTMhelix 20 39\noutside 40 325\nTMhelix 326 348\ninside 349 372

12926 GCF_000212715.2_ASM21271v2 Escherichia coli UMNK88 Proteobacteria; Gammaproteobacteria; Enterobacterales; Enterobacteriaceae; Escherichia; Escherichia coli MTGDNTLIHSHGINRRDFMKLCAALAATMGLSSKAAAEMAESVTNPQRPPVIWIGAQECT WP_000145410.1 MULTISPECIES: hydrogenase-2 small chain [Proteobacteria] Length: 372\nNumber of predicted TMHs: 1\nExp number of AAs in TMHs: 23.72226\nExp number, first 60 AAs: 0.22038\nTotal prob of N-in: 0.12994\noutside 1 331\nTMhelix 332 354\ninside 355 372

12927 GCF_000013265.1_ASM1326v1 Escherichia coli UTI89 Proteobacteria; Gammaproteobacteria; Enterobacterales; Enterobacteriaceae; Escherichia; Escherichia coli MTGDNTLIHSHGINRRDFMKLCAALAATMGLSSKAAAEMAESVTNPQRPPVIWIGAQECT WP_000145410.1 MULTISPECIES: hydrogenase-2 small chain [Proteobacteria] Length: 372\nNumber of predicted TMHs: 1\nExp number of AAs in TMHs: 23.72226\nExp number, first 60 AAs: 0.22038\nTotal prob of N-in: 0.12994\noutside 1 331\nTMhelix 332 354\ninside 355 372

12928 GCF_001441325.1_ASM144132v1 Escherichia coli VL2732 Proteobacteria; Gammaproteobacteria; Enterobacterales; Enterobacteriaceae; Escherichia; Escherichia coli MNNEETFYQAMRRQGVTRRSFLKYCSLAATSLGLGAGMAPKIAWALENKPRIPVVWIHGL WP_001058323.1 MULTISPECIES: hydrogenase-1 small chain [Proteobacteria] Length: 372\nNumber of predicted TMHs: 2\nExp number of AAs in TMHs: 37.2608499999999\nExp number, first 60 AAs: 19.52923\nTotal prob of N-in: 0.94351\nPOSSIBLE N-term signal sequence\ninside 1 19\nTMhelix 20 39\noutside 40 325\nTMhelix 326 348\ninside 349 372

12929 GCF_001441325.1_ASM144132v1 Escherichia coli VL2732 Proteobacteria; Gammaproteobacteria; Enterobacterales; Enterobacteriaceae; Escherichia; Escherichia coli MTGDNTLIHSHGINRRDFMKLCAALAATMGLSSKAAAEMAESVTNPQRPPVIWIGAQECT WP_000145410.1 MULTISPECIES: hydrogenase-2 small chain [Proteobacteria] Length: 372\nNumber of predicted TMHs: 1\nExp number of AAs in TMHs: 23.72226\nExp number, first 60 AAs: 0.22038\nTotal prob of N-in: 0.12994\noutside 1 331\nTMhelix 332 354\ninside 355 372

12930 GCF_001441335.1_ASM144133v1 Escherichia coli VL2874 Proteobacteria; Gammaproteobacteria; Enterobacterales; Enterobacteriaceae; Escherichia; Escherichia coli MTGDNTLIHSHGINRRDFMKLCAALAATMGLSSKAAAEMAESVTNPQRPPVIWIGAQECT WP_000145410.1 MULTISPECIES: hydrogenase-2 small chain [Proteobacteria] Length: 372\nNumber of predicted TMHs: 1\nExp number of AAs in TMHs: 23.72226\nExp number, first 60 AAs: 0.22038\nTotal prob of N-in: 0.12994\noutside 1 331\nTMhelix 332 354\ninside 355 372

12931 GCF_001441335.1_ASM144133v1 Escherichia coli VL2874 Proteobacteria; Gammaproteobacteria; Enterobacterales; Enterobacteriaceae; Escherichia; Escherichia coli MNNEETFYQAMRRQGVTRRSFLKYCSLAATSLGLGAGMAPKIAWALENKPRIPVVWIHGL WP_001058323.1 MULTISPECIES: hydrogenase-1 small chain [Proteobacteria] Length: 372\nNumber of predicted TMHs: 2\nExp number of AAs in TMHs: 37.2608499999999\nExp number, first 60 AAs: 19.52923\nTotal prob of N-in: 0.94351\nPOSSIBLE N-term signal sequence\ninside 1 19\nTMhelix 20 39\noutside 40 325\nTMhelix 326 348\ninside 349 372

12932 GCF_000968515.1_ASM96851v1 Escherichia coli VR50 Proteobacteria; Gammaproteobacteria; Enterobacterales; Enterobacteriaceae; Escherichia; Escherichia coli MTGDNTLIHSHGINRRDFMKLCAALAATMGLSSKAAAEMAESVTNPQRPPVIWIGAQECT WP_000145410.1

MULTISPECIES: hydrogenase-2 small chain [Proteobacteria] Length: 372\nNumber of predicted TMHs: 1\nExp number of AAs in TMHs: 23.72226\nExp number, first 60 AAs: 0.22038\nTotal prob of N-in: 0.12994\noutside 1 331\nTMhelix 332 354\ninside 355 372

12933 GCF_000968515.1_ASM96851v1 Escherichia coli VR50 Proteobacteria; Gammaproteobacteria; Enterobacterales; Enterobacteriaceae; Escherichia; Escherichia coli MNNEETFYQAMRRQGVTRRSFLKYCSLAATSLGLGAGMAPKIAWALENKPRIPVVWIHGL WP_001058323.1 MULTISPECIES: hydrogenase-1 small chain [Proteobacteria] Length: 372\nNumber of predicted TMHs: 2\nExp number of AAs in TMHs: 37.2608499999999\nExp number, first 60 AAs: 19.52923\nTotal prob of N-in: 0.94351\nPOSSIBLE N-term signal sequence\ninside 1 19\nTMhelix 20 39\noutside 40 325\nTMhelix 326 348\ninside 349 372

12934 GCF_000184185.1_ASM18418v1 Escherichia coli W Proteobacteria; Gammaproteobacteria; Enterobacterales; Enterobacteriaceae; Escherichia; Escherichia coli MTGDNTLIHSHGINRRDFMKLCAALAATMGLSSKAAAEMAESVTNPQRPPVIWIGAQECT WP_000145410.1 MULTISPECIES: hydrogenase-2 small chain [Proteobacteria] Length: 372\nNumber of predicted TMHs: 1\nExp number of AAs in TMHs: 23.72226\nExp number, first 60 AAs: 0.22038\nTotal prob of N-in: 0.12994\noutside 1 331\nTMhelix 332 354\ninside 355 372

12935 GCF_000258145.1_ASM25814v1 Escherichia coli W Proteobacteria; Gammaproteobacteria; Enterobacterales; Enterobacteriaceae; Escherichia; Escherichia coli MNNEETFYQAMRRQGVTRRSFLKYCSLAATSLGLGAGMAPKIAWALENKPRIPVVWIHGL WP_001349947.1 hydrogenase [Escherichia coli] Length: 372\nNumber of predicted TMHs: 2\nExp number of AAs in TMHs: 37.2615299999999\nExp number, first 60 AAs: 19.52923\nTotal prob of N-in: 0.94351\nPOSSIBLE N-term signal sequence\ninside 1 19\nTMhelix 20 39\noutside 40 325\nTMhelix 326 348\ninside 349 372

12936 GCF_000258145.1_ASM25814v1 Escherichia coli W Proteobacteria; Gammaproteobacteria; Enterobacterales; Enterobacteriaceae; Escherichia; Escherichia coli MTGDNTLIHSHGINRRDFMKLCAALAATMGLSSKAAAEMAESVTNPQRPPVIWIGAQECT WP_000145410.1 MULTISPECIES: hydrogenase-2 small chain [Proteobacteria] Length: 372\nNumber of predicted TMHs: 1\nExp number of AAs in TMHs: 23.72226\nExp number, first 60 AAs: 0.22038\nTotal prob of N-in: 0.12994\noutside 1 331\nTMhelix 332 354\ninside 355 372

12937 GCF_000147755.1_ASM14775v1 Escherichia coli W Proteobacteria; Gammaproteobacteria; Enterobacterales; Enterobacteriaceae; Escherichia; Escherichia coli MNNEETFYQAMRRQGVTRRSFLKYCSLAATSLGLGAGMAPKIAWALENKPRIPVVWIHGL WP_001349947.1 hydrogenase [Escherichia coli] Length: 372\nNumber of predicted TMHs: 2\nExp number of AAs in TMHs: 37.2615299999999\nExp number, first 60 AAs: 19.52923\nTotal prob of N-in: 0.94351\nPOSSIBLE N-term signal sequence\ninside 1 19\nTMhelix 20 39\noutside 40 325\nTMhelix 326 348\ninside 349 372

12938 GCF_000147755.1_ASM14775v1 Escherichia coli W Proteobacteria; Gammaproteobacteria; Enterobacterales; Enterobacteriaceae; Escherichia; Escherichia coli MTGDNTLIHSHGINRRDFMKLCAALAATMGLSSKAAAEMAESVTNPQRPPVIWIGAQECT WP_000145410.1 MULTISPECIES: hydrogenase-2 small chain [Proteobacteria] Length: 372\nNumber of predicted TMHs: 1\nExp number of AAs in TMHs: 23.72226\nExp number, first 60 AAs: 0.22038\nTotal prob of N-in: 0.12994\noutside 1 331\nTMhelix 332 354\ninside 355 372

12939 GCF_000184185.1_ASM18418v1 Escherichia coli W Proteobacteria; Gammaproteobacteria; Enterobacterales; Enterobacteriaceae; Escherichia; Escherichia coli MNNEETFYQAMRRQGVTRRSFLKYCSLAATSLGLGAGMAPKIAWALENKPRIPVVWIHGL WP_001349947.1 hydrogenase [Escherichia coli] Length: 372\nNumber of predicted TMHs: 2\nExp number of AAs in TMHs: 37.2615299999999\nExp number, first 60 AAs: 19.52923\nTotal prob of N-in: 0.94351\nPOSSIBLE N-term signal sequence\ninside 1 19\nTMhelix 20 39\noutside 40 325\nTMhelix 326 348\ninside 349 372

12940 GCF_000258225.1_ASM25822v1 Escherichia coli W26 Proteobacteria; Gammaproteobacteria; Enterobacterales; Enterobacteriaceae; Escherichia; Escherichia coli MTGDNTLIHSHGINRRDFMKLCAALAATMGLSSKAAAEMAESVTNPQRPPVIWIGAQECT WP_000145410.1 MULTISPECIES: hydrogenase-2 small chain [Proteobacteria] Length: 372\nNumber of predicted TMHs: 1\nExp number of AAs in TMHs: 23.72226\nExp number, first 60 AAs: 0.22038\nTotal prob of N-in: 0.12994\noutside 1 331\nTMhelix 332 354\ninside 355 372

12941 GCF_000258225.1_ASM25822v1 Escherichia coli W26 Proteobacteria; Gammaproteobacteria; Enterobacterales; Enterobacteriaceae; Escherichia; Escherichia coli
MNNEETFYQAMRRQGVTRRSFLKYCSLAATSLGLGAGMAPKIAWALENKPRIPVVWIHGL WP_001058323.1
MULTISPECIES: hydrogenase-1 small chain [Proteobacteria] Length: 372\nNumber of predicted TMHs: 2\nExp number of AAs in TMHs: 37.2608499999999\nExp number, first 60 AAs: 19.52923\nTotal prob of N-in: 0.94351\nPOSSIBLE N-term signal sequence\ninside 1 19\nTMhelix 20 39\noutside 40 325\nTMhelix 326 348\ninside 349 372

12942 GCF_000471585.1_E_coliWa1-1.0 Escherichia coli Wa1 Proteobacteria; Gammaproteobacteria; Enterobacterales; Enterobacteriaceae; Escherichia; Escherichia coli
MTGDNTLIHSHGINRRDFMKLCAALAATMGLSSKAAAEMAESVTNPQRPPVIWIGAQECT WP_000145410.1
MULTISPECIES: hydrogenase-2 small chain [Proteobacteria] Length: 372\nNumber of predicted TMHs: 1\nExp number of AAs in TMHs: 23.72226\nExp number, first 60 AAs: 0.22038\nTotal prob of N-in: 0.12994\noutside 1 331\nTMhelix 332 354\ninside 355 372

12943 GCF_000471585.1_E_coliWa1-1.0 Escherichia coli Wa1 Proteobacteria; Gammaproteobacteria; Enterobacterales; Enterobacteriaceae; Escherichia; Escherichia coli
MNNEETFYQAMRRQGVTRRSFLKYCSLAATSLGLGAGMAPKIAWALENKPRIPVVWIHGL WP_001301613.1
hydrogenase [Escherichia coli] Length: 372\nNumber of predicted TMHs: 2\nExp number of AAs in TMHs: 37.46598\nExp number, first 60 AAs: 19.53139\nTotal prob of N-in: 0.94364\nPOSSIBLE N-term signal sequence\ninside 1 19\nTMhelix 20 39\noutside 40 325\nTMhelix 326 348\ninside 349 372

12944 GCF_000471605.1_E_coliWa2-1.0 Escherichia coli Wa2 Proteobacteria; Gammaproteobacteria; Enterobacterales; Enterobacteriaceae; Escherichia; Escherichia coli
MNNEETFYQAMRRQGVTRRSFLKYCSLAATSLGLGAGMAPKIAWALENKPRIPVVWIHGL WP_001301613.1
hydrogenase [Escherichia coli] Length: 372\nNumber of predicted TMHs: 2\nExp number of AAs in TMHs: 37.46598\nExp number, first 60 AAs: 19.53139\nTotal prob of N-in: 0.94364\nPOSSIBLE N-term signal sequence\ninside 1 19\nTMhelix 20 39\noutside 40 325\nTMhelix 326 348\ninside 349 372

12945 GCF_000471605.1_E_coliWa2-1.0 Escherichia coli Wa2 Proteobacteria; Gammaproteobacteria; Enterobacterales; Enterobacteriaceae; Escherichia; Escherichia coli
MTGDNTLIHSHGINRRDFMKLCAALAATMGLSSKAAAEMAESVTNPQRPPVIWIGAQECT WP_000145410.1
MULTISPECIES: hydrogenase-2 small chain [Proteobacteria] Length: 372\nNumber of predicted TMHs: 1\nExp number of AAs in TMHs: 23.72226\nExp number, first 60 AAs: 0.22038\nTotal prob of N-in: 0.12994\noutside 1 331\nTMhelix 332 354\ninside 355 372

12946 GCF_000471545.1_E_coliWC1-1.0 Escherichia coli WC1 Proteobacteria; Gammaproteobacteria; Enterobacterales; Enterobacteriaceae; Escherichia; Escherichia coli
MTGDNTLIHSHGINRRDFMKLCAALAATMGLSSKAAAEMAESVTNPQRPPVIWIGAQECT WP_000145410.1
MULTISPECIES: hydrogenase-2 small chain [Proteobacteria] Length: 372\nNumber of predicted TMHs: 1\nExp number of AAs in TMHs: 23.72226\nExp number, first 60 AAs: 0.22038\nTotal prob of N-in: 0.12994\noutside 1 331\nTMhelix 332 354\ninside 355 372

12947 GCF_000471545.1_E_coliWC1-1.0 Escherichia coli WC1 Proteobacteria; Gammaproteobacteria; Enterobacterales; Enterobacteriaceae; Escherichia; Escherichia coli
MNNEETFYQAMRRQGVTRRSFLKYCSLAATSLGLGAGMAPKIAWALENKPRIPVVWIHGL WP_001301613.1
hydrogenase [Escherichia coli] Length: 372\nNumber of predicted TMHs: 2\nExp number of AAs in TMHs: 37.46598\nExp number, first 60 AAs: 19.53139\nTotal prob of N-in: 0.94364\nPOSSIBLE N-term signal sequence\ninside 1 19\nTMhelix 20 39\noutside 40 325\nTMhelix 326 348\ninside 349 372

12948 GCF_000471565.1_E_coliWC2-1.0 Escherichia coli WC2 Proteobacteria; Gammaproteobacteria; Enterobacterales; Enterobacteriaceae; Escherichia; Escherichia coli
MNNEETFYQAMRRQGVTRRSFLKYCSLAATSLGLGAGMAPKIAWALENKPRIPVVWIHGL WP_024251880.1
hydrogenase [Escherichia coli] Length: 372\nNumber of predicted TMHs: 2\nExp number of AAs in TMHs: 37.17638\nExp number, first 60 AAs: 19.53124\nTotal prob of N-in: 0.94348\nPOSSIBLE N-term signal sequence\ninside 1 19\nTMhelix 20 39\noutside 40 325\nTMhelix 326 348\ninside 349 372

12949 GCF_000471565.1_E_coliWC2-1.0 Escherichia coli WC2 Proteobacteria; Gammaproteobacteria; Enterobacterales; Enterobacteriaceae; Escherichia; Escherichia coli
MTGDNTLIHSHGINRRDFMKLCAALAATMGLSSKAAAEMAESVTNPQRPPVIWIGAQECT WP_000145410.1
MULTISPECIES: hydrogenase-2 small chain [Proteobacteria] Length: 372\nNumber of predicted TMHs:

1\nExp number of AAs in TMHs: 23.72226\nExp number, first 60 AAs: 0.22038\nTotal prob of N-in: 0.12994\noutside 1 331\nTMhelix 332 354\ninside 355 372

12950 GCF_000193975.1_ASM19397v2 Escherichia coli WV_060327 Proteobacteria; Gammaproteobacteria; Enterobacterales; Enterobacteriaceae; Escherichia; Escherichia coli MNNEETFYQAMRRQGVTRRSFLKYCSLAATSLGLGAGMAPKIAWALENKPRIPVVWIHGL WP_001058323.1 MULTISPECIES: hydrogenase-1 small chain [Proteobacteria] Length: 372\nNumber of predicted TMHs: 2\nExp number of AAs in TMHs: 37.2608499999999\nExp number, first 60 AAs: 19.52923\nTotal prob of N-in: 0.94351\nPOSSIBLE N-term signal sequence\ninside 1 19\nTMhelix 20 39\noutside 40 325\nTMhelix 326 348\ninside 349 372

12951 GCF_000193975.1_ASM19397v2 Escherichia coli WV_060327 Proteobacteria; Gammaproteobacteria; Enterobacterales; Enterobacteriaceae; Escherichia; Escherichia coli MTGDNTLIHSHGINRRDFMKLCAALAATMGLSSKAAAEMAESVTNPQRPPVIWIGAQECT WP_000145410.1 MULTISPECIES: hydrogenase-2 small chain [Proteobacteria] Length: 372\nNumber of predicted TMHs: 1\nExp number of AAs in TMHs: 23.72226\nExp number, first 60 AAs: 0.22038\nTotal prob of N-in: 0.12994\noutside 1 331\nTMhelix 332 354\ninside 355 372

12952 GCF_000224395.1_ASM22439v2 Escherichia coli XH001 Proteobacteria; Gammaproteobacteria; Enterobacterales; Enterobacteriaceae; Escherichia; Escherichia coli MNNEETFYQAMRRQGVTRRSFLKYCSLAATSLGLGAGMAPKIAWALENKPRIPVVWIHGL WP_001058323.1 MULTISPECIES: hydrogenase-1 small chain [Proteobacteria] Length: 372\nNumber of predicted TMHs: 2\nExp number of AAs in TMHs: 37.2608499999999\nExp number, first 60 AAs: 19.52923\nTotal prob of N-in: 0.94351\nPOSSIBLE N-term signal sequence\ninside 1 19\nTMhelix 20 39\noutside 40 325\nTMhelix 326 348\ninside 349 372

12953 GCF_000224395.1_ASM22439v2 Escherichia coli XH001 Proteobacteria; Gammaproteobacteria; Enterobacterales; Enterobacteriaceae; Escherichia; Escherichia coli MTGDNTLIHSHGINRRDFMKLCAALAATMGLSSKAAAEMAESVTNPQRPPVIWIGAQECT WP_000145410.1 MULTISPECIES: hydrogenase-2 small chain [Proteobacteria] Length: 372\nNumber of predicted TMHs: 1\nExp number of AAs in TMHs: 23.72226\nExp number, first 60 AAs: 0.22038\nTotal prob of N-in: 0.12994\noutside 1 331\nTMhelix 332 354\ninside 355 372

12954 GCF_000226585.1_ASM22658v1 Escherichia coli XH140A Proteobacteria; Gammaproteobacteria; Enterobacterales; Enterobacteriaceae; Escherichia; Escherichia coli MTGDNTLIHSHGINRRDFMKLCAALAATMGLSSKAAAEMAESVTNPQRPPVIWIGAQECT WP_000145410.1 MULTISPECIES: hydrogenase-2 small chain [Proteobacteria] Length: 372\nNumber of predicted TMHs: 1\nExp number of AAs in TMHs: 23.72226\nExp number, first 60 AAs: 0.22038\nTotal prob of N-in: 0.12994\noutside 1 331\nTMhelix 332 354\ninside 355 372

12955 GCF_000226585.1_ASM22658v1 Escherichia coli XH140A Proteobacteria; Gammaproteobacteria; Enterobacterales; Enterobacteriaceae; Escherichia; Escherichia coli MNNEETFYQAMRRQGVTRRSFLKYCSLAATSLGLGAGMAPKIAWALENKPRIPVVWIHGL WP_001058323.1 MULTISPECIES: hydrogenase-1 small chain [Proteobacteria] Length: 372\nNumber of predicted TMHs: 2\nExp number of AAs in TMHs: 37.2608499999999\nExp number, first 60 AAs: 19.52923\nTotal prob of N-in: 0.94351\nPOSSIBLE N-term signal sequence\ninside 1 19\nTMhelix 20 39\noutside 40 325\nTMhelix 326 348\ninside 349 372

12956 GCF_000262125.1_ASM26212v1 Escherichia coli Xuzhou21 Proteobacteria; Gammaproteobacteria; Enterobacterales; Enterobacteriaceae; Escherichia; Escherichia coli MTGDNTLIHSHGINRRDFMKLCAALAATMGLSSKAAAEMAESVTNPQRPPVIWIGAQECT WP_000145410.1 MULTISPECIES: hydrogenase-2 small chain [Proteobacteria] Length: 372\nNumber of predicted TMHs: 1\nExp number of AAs in TMHs: 23.72226\nExp number, first 60 AAs: 0.22038\nTotal prob of N-in: 0.12994\noutside 1 331\nTMhelix 332 354\ninside 355 372

12957 GCF_000262125.1_ASM26212v1 Escherichia coli Xuzhou21 Proteobacteria; Gammaproteobacteria; Enterobacterales; Enterobacteriaceae; Escherichia; Escherichia coli MNNEETFYQAMRRQGVTRRSFLKYCSLAATSLGLGAGMAPKIAWALENKPRIPVVWIHGL WP_001301613.1 hydrogenase [Escherichia coli] Length: 372\nNumber of predicted TMHs: 2\nExp number of AAs in TMHs: 37.46598\nExp number, first 60 AAs: 19.53139\nTotal prob of N-in: 0.94364\nPOSSIBLE N-term signal sequence\ninside 1 19\nTMhelix 20 39\noutside 40 325\nTMhelix 326 348\ninside 349 372

12958 GCF_000017985.1_ASM1798v1 Escherichia coli B str. REL606 Proteobacteria; Gammaproteobacteria; Enterobacterales; Enterobacteriaceae; Escherichia; Escherichia coli; Escherichia coli B
MNNEETFYQAMRRQGVTRRSFLKYCSLAATSLGLGAGMAPKIAWALENKPRIPVVWIHGL WP_001058323.1
MULTISPECIES: hydrogenase-1 small chain [Proteobacteria] Length: 372\nNumber of predicted TMHs: 2\nExp number of AAs in TMHs: 37.2608499999999\nExp number, first 60 AAs: 19.52923\nTotal prob of N-in: 0.94351\nPOSSIBLE N-term signal sequence\ninside 1 19\nTMhelix 20 39\noutside 40 325\nTMhelix 326 348\ninside 349 372

12959 GCF_000019385.1_ASM1938v1 Escherichia coli ATCC 8739 Proteobacteria; Gammaproteobacteria; Enterobacterales; Enterobacteriaceae; Escherichia; Escherichia coli; Escherichia coli C
MTGDNTLIHSHGINRRDFMKLCAALAATMGLSSKAAAEMAESVTNPQRPPVIWIGAQECT WP_000145410.1
MULTISPECIES: hydrogenase-2 small chain [Proteobacteria] Length: 372\nNumber of predicted TMHs: 1\nExp number of AAs in TMHs: 23.72226\nExp number, first 60 AAs: 0.22038\nTotal prob of N-in: 0.12994\noutside 1 331\nTMhelix 332 354\ninside 355 372

12960 GCF_000019385.1_ASM1938v1 Escherichia coli ATCC 8739 Proteobacteria; Gammaproteobacteria; Enterobacterales; Enterobacteriaceae; Escherichia; Escherichia coli; Escherichia coli C
MNNEETFYQAMRRQGVTRRSFLKYCSLAATSLGLGAGMAPKIAWALENKPRIPVVWIHGL WP_001058323.1
MULTISPECIES: hydrogenase-1 small chain [Proteobacteria] Length: 372\nNumber of predicted TMHs: 2\nExp number of AAs in TMHs: 37.2608499999999\nExp number, first 60 AAs: 19.52923\nTotal prob of N-in: 0.94351\nPOSSIBLE N-term signal sequence\ninside 1 19\nTMhelix 20 39\noutside 40 325\nTMhelix 326 348\ninside 349 372

12961 GCF_000750555.1_ASM75055v1 Escherichia coli BW25113 Proteobacteria; Gammaproteobacteria; Enterobacterales; Enterobacteriaceae; Escherichia; Escherichia coli; Escherichia coli K-12
MTGDNTLIHSHGINRRDFMKLCAALAATMGLSSKAAAEMAESVTNPQRPPVIWIGAQECT WP_000145410.1
MULTISPECIES: hydrogenase-2 small chain [Proteobacteria] Length: 372\nNumber of predicted TMHs: 1\nExp number of AAs in TMHs: 23.72226\nExp number, first 60 AAs: 0.22038\nTotal prob of N-in: 0.12994\noutside 1 331\nTMhelix 332 354\ninside 355 372

12962 GCF_000750555.1_ASM75055v1 Escherichia coli BW25113 Proteobacteria; Gammaproteobacteria; Enterobacterales; Enterobacteriaceae; Escherichia; Escherichia coli; Escherichia coli K-12
MNNEETFYQAMRRQGVTRRSFLKYCSLAATSLGLGAGMAPKIAWALENKPRIPVVWIHGL WP_001058323.1
MULTISPECIES: hydrogenase-1 small chain [Proteobacteria] Length: 372\nNumber of predicted TMHs: 2\nExp number of AAs in TMHs: 37.2608499999999\nExp number, first 60 AAs: 19.52923\nTotal prob of N-in: 0.94351\nPOSSIBLE N-term signal sequence\ninside 1 19\nTMhelix 20 39\noutside 40 325\nTMhelix 326 348\ninside 349 372

12963 GCF_000022345.1_ASM2234v1 Escherichia coli BW2952 Proteobacteria; Gammaproteobacteria; Enterobacterales; Enterobacteriaceae; Escherichia; Escherichia coli; Escherichia coli K-12
MTGDNTLIHSHGINRRDFMKLCAALAATMGLSSKAAAEMAESVTNPQRPPVIWIGAQECT WP_000145410.1
MULTISPECIES: hydrogenase-2 small chain [Proteobacteria] Length: 372\nNumber of predicted TMHs: 1\nExp number of AAs in TMHs: 23.72226\nExp number, first 60 AAs: 0.22038\nTotal prob of N-in: 0.12994\noutside 1 331\nTMhelix 332 354\ninside 355 372

12964 GCF_000022345.1_ASM2234v1 Escherichia coli BW2952 Proteobacteria; Gammaproteobacteria; Enterobacterales; Enterobacteriaceae; Escherichia; Escherichia coli; Escherichia coli K-12
MNNEETFYQAMRRQGVTRRSFLKYCSLAATSLGLGAGMAPKIAWALENKPRIPVVWIHGL WP_001058323.1
MULTISPECIES: hydrogenase-1 small chain [Proteobacteria] Length: 372\nNumber of predicted TMHs: 2\nExp number of AAs in TMHs: 37.2608499999999\nExp number, first 60 AAs: 19.52923\nTotal prob of N-in: 0.94351\nPOSSIBLE N-term signal sequence\ninside 1 19\nTMhelix 20 39\noutside 40 325\nTMhelix 326 348\ninside 349 372

12965 GCF_000800215.1_ASM80021v1 Escherichia coli ER2796 Proteobacteria; Gammaproteobacteria; Enterobacterales; Enterobacteriaceae; Escherichia; Escherichia coli; Escherichia coli K-12
MNNEETFYQAMRRQGVTRRSFLKYCSLAATSLGLGAGMAPKIAWALENKPRIPVVWIHGL WP_001058323.1
MULTISPECIES: hydrogenase-1 small chain [Proteobacteria] Length: 372\nNumber of predicted TMHs: 2\nExp number of AAs in TMHs: 37.2608499999999\nExp number, first 60 AAs: 19.52923\nTotal prob of N-in: 0.94351\nPOSSIBLE N-term signal sequence\ninside 1 19\nTMhelix 20 39\noutside 40 325\nTMhelix 326 348\ninside 349 372

12966 GCF_000800215.1_ASM80021v1 Escherichia coli ER2796 Proteobacteria; Gammaproteobacteria; Enterobacterales; Enterobacteriaceae; Escherichia; Escherichia coli; Escherichia coli K-12
MTGDNTLIHSHGINRRDFMKLCAALAATMGLSSKAAAEMAESVTNPQRPPVIWIGAQECT WP_000145410.1
MULTISPECIES: hydrogenase-2 small chain [Proteobacteria] Length: 372\nNumber of predicted TMHs: 1\nExp number of AAs in TMHs: 23.72226\nExp number, first 60 AAs: 0.22038\nTotal prob of N-in: 0.12994\noutside 1 331\nTMhelix 332 354\ninside 355 372

12967 GCF_000019425.1_ASM1942v1 Escherichia coli str. K-12 substr. DH10B Proteobacteria; Gammaproteobacteria; Enterobacterales; Enterobacteriaceae; Escherichia; Escherichia coli; Escherichia coli K-12
MTGDNTLIHSHGINRRDFMKLCAALAATMGLSSKAAAEMAESVTNPQRPPVIWIGAQECT WP_000145410.1
MULTISPECIES: hydrogenase-2 small chain [Proteobacteria] Length: 372\nNumber of predicted TMHs: 1\nExp number of AAs in TMHs: 23.72226\nExp number, first 60 AAs: 0.22038\nTotal prob of N-in: 0.12994\noutside 1 331\nTMhelix 332 354\ninside 355 372

12968 GCF_000019425.1_ASM1942v1 Escherichia coli str. K-12 substr. DH10B Proteobacteria; Gammaproteobacteria; Enterobacterales; Enterobacteriaceae; Escherichia; Escherichia coli; Escherichia coli K-12
MNNEETFYQAMRRQGVTRRSFLKYCSLAATSLGLGAGMAPKIAWALENKPRIPVVWIHGL WP_001058323.1
MULTISPECIES: hydrogenase-1 small chain [Proteobacteria] Length: 372\nNumber of predicted TMHs: 2\nExp number of AAs in TMHs: 37.2608499999999\nExp number, first 60 AAs: 19.52923\nTotal prob of N-in: 0.94351\nPOSSIBLE N-term signal sequence\ninside 1 19\nTMhelix 20 39\noutside 40 325\nTMhelix 326 348\ninside 349 372

12969 GCF_000499485.1_MYMC4100 Escherichia coli str. K-12 substr. MC4100 Proteobacteria; Gammaproteobacteria; Enterobacterales; Enterobacteriaceae; Escherichia; Escherichia coli; Escherichia coli K-12
MTGDNTLIHSHGINRRDFMKLCAALAATMGLSSKAAAEMAESVTNPQRPPVIWIGAQECT WP_000145410.1
MULTISPECIES: hydrogenase-2 small chain [Proteobacteria] Length: 372\nNumber of predicted TMHs: 1\nExp number of AAs in TMHs: 23.72226\nExp number, first 60 AAs: 0.22038\nTotal prob of N-in: 0.12994\noutside 1 331\nTMhelix 332 354\ninside 355 372

12970 GCF_000499485.1_MYMC4100 Escherichia coli str. K-12 substr. MC4100 Proteobacteria; Gammaproteobacteria; Enterobacterales; Enterobacteriaceae; Escherichia; Escherichia coli; Escherichia coli K-12
MNNEETFYQAMRRQGVTRRSFLKYCSLAATSLGLGAGMAPKIAWALENKPRIPVVWIHGL WP_001058323.1
MULTISPECIES: hydrogenase-1 small chain [Proteobacteria] Length: 372\nNumber of predicted TMHs: 2\nExp number of AAs in TMHs: 37.2608499999999\nExp number, first 60 AAs: 19.52923\nTotal prob of N-in: 0.94351\nPOSSIBLE N-term signal sequence\ninside 1 19\nTMhelix 20 39\noutside 40 325\nTMhelix 326 348\ninside 349 372

12971 GCF_000350185.1_ASM35018v1 Escherichia coli str. K-12 substr. MDS42Proteobacteria; Gammaproteobacteria; Enterobacterales; Enterobacteriaceae; Escherichia; Escherichia coli; Escherichia coli K-12
MNNEETFYQAMRRQGVTRRSFLKYCSLAATSLGLGAGMAPKIAWALENKPRIPVVWIHGL WP_001058323.1
MULTISPECIES: hydrogenase-1 small chain [Proteobacteria] Length: 372\nNumber of predicted TMHs: 2\nExp number of AAs in TMHs: 37.2608499999999\nExp number, first 60 AAs: 19.52923\nTotal prob of N-in: 0.94351\nPOSSIBLE N-term signal sequence\ninside 1 19\nTMhelix 20 39\noutside 40 325\nTMhelix 326 348\ninside 349 372

12972 GCF_000350185.1_ASM35018v1 Escherichia coli str. K-12 substr. MDS42Proteobacteria; Gammaproteobacteria; Enterobacterales; Enterobacteriaceae; Escherichia; Escherichia coli; Escherichia coli K-12
MTGDNTLIHSHGINRRDFMKLCAALAATMGLSSKAAAEMAESVTNPQRPPVIWIGAQECT WP_000145410.1
MULTISPECIES: hydrogenase-2 small chain [Proteobacteria] Length: 372\nNumber of predicted TMHs: 1\nExp number of AAs in TMHs: 23.72226\nExp number, first 60 AAs: 0.22038\nTotal prob of N-in: 0.12994\noutside 1 331\nTMhelix 332 354\ninside 355 372

12973 GCF_000364365.1_ASM36436v1 Escherichia coli str. K-12 substr. MG1655 Proteobacteria; Gammaproteobacteria; Enterobacterales; Enterobacteriaceae; Escherichia; Escherichia coli; Escherichia coli K-12
MTGDNTLIHSHGINRRDFMKLCAALAATMGLSSKAAAEMAESVTNPQRPPVIWIGAQECT WP_000145410.1
MULTISPECIES: hydrogenase-2 small chain [Proteobacteria] Length: 372\nNumber of predicted TMHs: 1\nExp number of AAs in TMHs: 23.72226\nExp number, first 60 AAs: 0.22038\nTotal prob of N-in: 0.12994\noutside 1 331\nTMhelix 332 354\ninside 355 372

12974 GCF_000364365.1_ASM36436v1 Escherichia coli str. K-12 substr. MG1655 Proteobacteria; Gammaproteobacteria; Enterobacterales; Enterobacteriaceae; Escherichia; Escherichia coli; Escherichia coli K-12

MNNEETFYQAMRRQGVTRRSFLKYCSLAATSLGLGAGMAPKIAWALENKPRIPVVWIHGL WP_001058323.1 MULTISPECIES: hydrogenase-1 small chain [Proteobacteria] Length: 372\nNumber of predicted TMHs: 2\nExp number of AAs in TMHs: 37.2608499999999\nExp number, first 60 AAs: 19.52923\nTotal prob of N-in: 0.94351\nPOSSIBLE N-term signal sequence\ninside 1 19\nTMhelix 20 39\noutside 40 325\nTMhelix 326 348\ninside 349 372

12975 GCF_000005845.2_ASM584v2 Escherichia coli str. K-12 substr. MG1655 Proteobacteria; Gammaproteobacteria; Enterobacterales; Enterobacteriaceae; Escherichia; Escherichia coli; Escherichia coli K-12 MTGDNTLIHSHGINRRDFMKLCAALAATMGLSSKAAAEMAESVTNPQRPPVIWIGAQECT NP_417471.1 hydrogenase 2, small subunit [Escherichia coli str. K-12 substr. MG1655] Length: 372\nNumber of predicted TMHs: 1\nExp number of AAs in TMHs: 23.72226\nExp number, first 60 AAs: 0.22038\nTotal prob of N-in: 0.12994\noutside 1 331\nTMhelix 332 354\ninside 355 372

12976 GCF_000005845.2_ASM584v2 Escherichia coli str. K-12 substr. MG1655 Proteobacteria; Gammaproteobacteria; Enterobacterales; Enterobacteriaceae; Escherichia; Escherichia coli; Escherichia coli K-12 MNNEETFYQAMRRQGVTRRSFLKYCSLAATSLGLGAGMAPKIAWALENKPRIPVVWIHGL NP_415491.1 hydrogenase 1, small subunit [Escherichia coli str. K-12 substr. MG1655] Length: 372\nNumber of predicted TMHs: 2\nExp number of AAs in TMHs: 37.2608499999999\nExp number, first 60 AAs: 19.52923\nTotal prob of N-in: 0.94351\nPOSSIBLE N-term signal sequence\ninside 1 19\nTMhelix 20 39\noutside 40 325\nTMhelix 326 348\ninside 349 372

12977 GCF_000273425.1_Esch_coli_MG12655_V1 Escherichia coli str. K-12 substr. MG1655 Proteobacteria; Gammaproteobacteria; Enterobacterales; Enterobacteriaceae; Escherichia; Escherichia coli; Escherichia coli K-12 MNNEETFYQAMRRQGVTRRSFLKYCSLAATSLGLGAGMAPKIAWALENKPRIPVVWIHGL WP_001058323.1 MULTISPECIES: hydrogenase-1 small chain [Proteobacteria] Length: 372\nNumber of predicted TMHs: 2\nExp number of AAs in TMHs: 37.2608499999999\nExp number, first 60 AAs: 19.52923\nTotal prob of N-in: 0.94351\nPOSSIBLE N-term signal sequence\ninside 1 19\nTMhelix 20 39\noutside 40 325\nTMhelix 326 348\ninside 349 372

12978 GCF_000482265.1_EC_K12_MG1655_Broad_SNP Escherichia coli str. K-12 substr. MG1655 Proteobacteria; Gammaproteobacteria; Enterobacterales; Enterobacteriaceae; Escherichia; Escherichia coli; Escherichia coli K-12 MNNEETFYQAMRRQGVTRRSFLKYCSLAATSLGLGAGMAPKIAWALENKPRIPVVWIHGL WP_001058323.1 MULTISPECIES: hydrogenase-1 small chain [Proteobacteria] Length: 372\nNumber of predicted TMHs: 2\nExp number of AAs in TMHs: 37.2608499999999\nExp number, first 60 AAs: 19.52923\nTotal prob of N-in: 0.94351\nPOSSIBLE N-term signal sequence\ninside 1 19\nTMhelix 20 39\noutside 40 325\nTMhelix 326 348\ninside 349 372

12979 GCF_000517165.1_ASM51716v1 Escherichia coli str. K-12 substr. MG1655 Proteobacteria; Gammaproteobacteria; Enterobacterales; Enterobacteriaceae; Escherichia; Escherichia coli; Escherichia coli K-12 MNNEETFYQAMRRQGVTRRSFLKYCSLAATSLGLGAGMAPKIAWALENKPRIPVVWIHGL WP_001058323.1 MULTISPECIES: hydrogenase-1 small chain [Proteobacteria] Length: 372\nNumber of predicted TMHs: 2\nExp number of AAs in TMHs: 37.2608499999999\nExp number, first 60 AAs: 19.52923\nTotal prob of N-in: 0.94351\nPOSSIBLE N-term signal sequence\ninside 1 19\nTMhelix 20 39\noutside 40 325\nTMhelix 326 348\ninside 349 372

12980 GCF_000482265.1_EC_K12_MG1655_Broad_SNP Escherichia coli str. K-12 substr. MG1655 Proteobacteria; Gammaproteobacteria; Enterobacterales; Enterobacteriaceae; Escherichia; Escherichia coli; Escherichia coli K-12 MTGDNTLIHSHGINRRDFMKLCAALAATMGLSSKAAAEMAESVTNPQRPPVIWIGAQECT WP_000145410.1 MULTISPECIES: hydrogenase-2 small chain [Proteobacteria] Length: 372\nNumber of predicted TMHs: 1\nExp number of AAs in TMHs: 23.72226\nExp number, first 60 AAs: 0.22038\nTotal prob of N-in: 0.12994\noutside 1 331\nTMhelix 332 354\ninside 355 372

12981 GCF_000269645.1_PB_Esch_coli_MG12655_V1 Escherichia coli str. K-12 substr. MG1655 Proteobacteria; Gammaproteobacteria; Enterobacterales; Enterobacteriaceae; Escherichia; Escherichia coli; Escherichia coli K-12 MNNEETFYQAMRRQGVTRRSFLKYCSLAATSLGLGAGMAPKIAWALENKPRIPVVWIHGL WP_001058323.1 MULTISPECIES: hydrogenase-1 small chain [Proteobacteria] Length: 372\nNumber of predicted TMHs: 2\nExp number of AAs in TMHs: 37.2608499999999\nExp number, first 60 AAs: 19.52923\nTotal prob of N-in: 0.94351\nPOSSIBLE N-term signal sequence\ninside 1 19\nTMhelix 20 39\noutside 40 325\nTMhelix 326 348\ninside 349 372

12982 GCF_000517165.1_ASM51716v1 Escherichia coli str. K-12 substr. MG1655 Proteobacteria; Gammaproteobacteria; Enterobacterales; Enterobacteriaceae; Escherichia; Escherichia coli; Escherichia coli K-12 MTGDNTLIHSHGINRRDFMKLCAALAATMGLSSKAAAEMAESVTNPQRPPVIWIGAQECT WP_000145410.1 MULTISPECIES: hydrogenase-2 small chain [Proteobacteria] Length: 372\nNumber of predicted TMHs: 1\nExp number of AAs in TMHs: 23.72226\nExp number, first 60 AAs: 0.22038\nTotal prob of N-in: 0.12994\noutside 1 331\nTMhelix 332 354\ninside 355 372

12983 GCF_000269645.1_PB_Esch_coli_MG12655_V1 Escherichia coli str. K-12 substr. MG1655 Proteobacteria; Gammaproteobacteria; Enterobacterales; Enterobacteriaceae; Escherichia; Escherichia coli; Escherichia coli K-12 MTGDNTLIHSHGINRRDFMKLCAALAATMGLSSKAAAEMAESVTNPQRPPVIWIGAQECT WP_000145410.1 MULTISPECIES: hydrogenase-2 small chain [Proteobacteria] Length: 372\nNumber of predicted TMHs: 1\nExp number of AAs in TMHs: 23.72226\nExp number, first 60 AAs: 0.22038\nTotal prob of N-in: 0.12994\noutside 1 331\nTMhelix 332 354\ninside 355 372

12984 GCF_000273425.1_Esch_coli_MG12655_V1 Escherichia coli str. K-12 substr. MG1655 Proteobacteria; Gammaproteobacteria; Enterobacterales; Enterobacteriaceae; Escherichia; Escherichia coli; Escherichia coli K-12 MTGDNTLIHSHGINRRDFMKLCAALAATMGLSSKAAAEMAESVTNPQRPPVIWIGAQECT WP_000145410.1 MULTISPECIES: hydrogenase-2 small chain [Proteobacteria] Length: 372\nNumber of predicted TMHs: 1\nExp number of AAs in TMHs: 23.72226\nExp number, first 60 AAs: 0.22038\nTotal prob of N-in: 0.12994\noutside 1 331\nTMhelix 332 354\ninside 355 372

12985 GCF_001544635.1_ASM154463v1 Escherichia coli str. K-12 substr. MG1655 Proteobacteria; Gammaproteobacteria; Enterobacterales; Enterobacteriaceae; Escherichia; Escherichia coli; Escherichia coli K-12 MNNEETFYQAMRRQGVTRRSFLKYCSLAATSLGLGAGMAPKIAWALENKPRIPVVWIHGL WP_001058323.1 MULTISPECIES: hydrogenase-1 small chain [Proteobacteria] Length: 372\nNumber of predicted TMHs: 2\nExp number of AAs in TMHs: 37.2608499999999\nExp number, first 60 AAs: 19.52923\nTotal prob of N-in: 0.94351\nPOSSIBLE N-term signal sequence\ninside 1 19\nTMhelix 20 39\noutside 40 325\nTMhelix 326 348\ninside 349 372

12986 GCF_001566335.1_ASM156633v1 Escherichia coli str. K-12 substr. MG1655 Proteobacteria; Gammaproteobacteria; Enterobacterales; Enterobacteriaceae; Escherichia; Escherichia coli; Escherichia coli K-12 MNNEETFYQAMRRQGVTRRSFLKYCSLAATSLGLGAGMAPKIAWALENKPRIPVVWIHGL WP_001058323.1 MULTISPECIES: hydrogenase-1 small chain [Proteobacteria] Length: 372\nNumber of predicted TMHs: 2\nExp number of AAs in TMHs: 37.2608499999999\nExp number, first 60 AAs: 19.52923\nTotal prob of N-in: 0.94351\nPOSSIBLE N-term signal sequence\ninside 1 19\nTMhelix 20 39\noutside 40 325\nTMhelix 326 348\ninside 349 372

12987 GCF_001566335.1_ASM156633v1 Escherichia coli str. K-12 substr. MG1655 Proteobacteria; Gammaproteobacteria; Enterobacterales; Enterobacteriaceae; Escherichia; Escherichia coli; Escherichia coli K-12 MTGDNTLIHSHGINRRDFMKLCAALAATMGLSSKAAAEMAESVTNPQRPPVIWIGAQECT WP_000145410.1 MULTISPECIES: hydrogenase-2 small chain [Proteobacteria] Length: 372\nNumber of predicted TMHs: 1\nExp number of AAs in TMHs: 23.72226\nExp number, first 60 AAs: 0.22038\nTotal prob of N-in: 0.12994\noutside 1 331\nTMhelix 332 354\ninside 355 372

12988 GCF_001308065.1_ASM130806v1 Escherichia coli str. K-12 substr. MG1655 Proteobacteria; Gammaproteobacteria; Enterobacterales; Enterobacteriaceae; Escherichia; Escherichia coli; Escherichia coli K-12 MNNEETFYQAMRRQGVTRRSFLKYCSLAATSLGLGAGMAPKIAWALENKPRIPVVWIHGL WP_001058323.1 MULTISPECIES: hydrogenase-1 small chain [Proteobacteria] Length: 372\nNumber of predicted TMHs: 2\nExp number of AAs in TMHs: 37.2608499999999\nExp number, first 60 AAs: 19.52923\nTotal prob of N-in: 0.94351\nPOSSIBLE N-term signal sequence\ninside 1 19\nTMhelix 20 39\noutside 40 325\nTMhelix 326 348\ninside 349 372

12989 GCF_001544635.1_ASM154463v1 Escherichia coli str. K-12 substr. MG1655 Proteobacteria; Gammaproteobacteria; Enterobacterales; Enterobacteriaceae; Escherichia; Escherichia coli; Escherichia coli K-12 MTGDNTLIHSHGINRRDFMKLCAALAATMGLSSKAAAEMAESVTNPQRPPVIWIGAQECT WP_000145410.1 MULTISPECIES: hydrogenase-2 small chain [Proteobacteria] Length: 372\nNumber of predicted TMHs: 1\nExp number of AAs in TMHs: 23.72226\nExp number, first 60 AAs: 0.22038\nTotal prob of N-in: 0.12994\noutside 1 331\nTMhelix 332 354\ninside 355 372

12990 GCF_000801205.1_ASM80120v1 Escherichia coli str. K-12 substr. MG1655 Proteobacteria; Gammaproteobacteria; Enterobacterales; Enterobacteriaceae; Escherichia; Escherichia coli; Escherichia coli K-12

MNNEETFYQAMRRQGVTRRSFLKYCSLAATSLGLGAGMAPKIAWALENKPRIPVVWIHGL WP_001058323.1 MULTISPECIES: hydrogenase-1 small chain [Proteobacteria] Length: 372\nNumber of predicted TMHs: 2\nExp number of AAs in TMHs: 37.2608499999999\nExp number, first 60 AAs: 19.52923\nTotal prob of N-in: 0.94351\nPOSSIBLE N-term signal sequence\ninside 1 19\nTMhelix 20 39\noutside 40 325\nTMhelix 326 348\ninside 349 372

12991 GCF_001308065.1_ASM130806v1 Escherichia coli str. K-12 substr. MG1655 Proteobacteria; Gammaproteobacteria; Enterobacterales; Enterobacteriaceae; Escherichia; Escherichia coli; Escherichia coli K-12 MTGDNTLIHSHGINRRDFMKLCAALAATMGLSSKAAAEMAESVTNPQRPPVIWIGAQECT WP_000145410.1 MULTISPECIES: hydrogenase-2 small chain [Proteobacteria] Length: 372\nNumber of predicted TMHs: 1\nExp number of AAs in TMHs: 23.72226\nExp number, first 60 AAs: 0.22038\nTotal prob of N-in: 0.12994\noutside 1 331\nTMhelix 332 354\ninside 355 372

12992 GCF_000801205.1_ASM80120v1 Escherichia coli str. K-12 substr. MG1655 Proteobacteria; Gammaproteobacteria; Enterobacterales; Enterobacteriaceae; Escherichia; Escherichia coli; Escherichia coli K-12 MTGDNTLIHSHGINRRDFMKLCAALAATMGLSSKAAAEMAESVTNPQRPPVIWIGAQECT WP_000145410.1 MULTISPECIES: hydrogenase-2 small chain [Proteobacteria] Length: 372\nNumber of predicted TMHs: 1\nExp number of AAs in TMHs: 23.72226\nExp number, first 60 AAs: 0.22038\nTotal prob of N-in: 0.12994\noutside 1 331\nTMhelix 332 354\ninside 355 372

12993 GCF_001878695.1_ASM187869v1 Escherichia coli str. K-12 substr. W3110Proteobacteria; Gammaproteobacteria; Enterobacterales; Enterobacteriaceae; Escherichia; Escherichia coli; Escherichia coli K-12 MNNEETFYQAMRRQGVTRRSFLKYCSLAATSLGLGAGMAPKIAWALENKPRIPVVWIHGL WP_001058323.1 MULTISPECIES: hydrogenase-1 small chain [Proteobacteria] Length: 372\nNumber of predicted TMHs: 2\nExp number of AAs in TMHs: 37.2608499999999\nExp number, first 60 AAs: 19.52923\nTotal prob of N-in: 0.94351\nPOSSIBLE N-term signal sequence\ninside 1 19\nTMhelix 20 39\noutside 40 325\nTMhelix 326 348\ninside 349 372

12994 GCF_000010245.2_ASM1024v1 Escherichia coli str. K-12 substr. W3110Proteobacteria; Gammaproteobacteria; Enterobacterales; Enterobacteriaceae; Escherichia; Escherichia coli; Escherichia coli K-12 MTGDNTLIHSHGINRRDFMKLCAALAATMGLSSKAAAEMAESVTNPQRPPVIWIGAQECT WP_000145410.1 MULTISPECIES: hydrogenase-2 small chain [Proteobacteria] Length: 372\nNumber of predicted TMHs: 1\nExp number of AAs in TMHs: 23.72226\nExp number, first 60 AAs: 0.22038\nTotal prob of N-in: 0.12994\noutside 1 331\nTMhelix 332 354\ninside 355 372

12995 GCF_000010245.2_ASM1024v1 Escherichia coli str. K-12 substr. W3110Proteobacteria; Gammaproteobacteria; Enterobacterales; Enterobacteriaceae; Escherichia; Escherichia coli; Escherichia coli K-12 MNNEETFYQAMRRQGVTRRSFLKYCSLAATSLGLGAGMAPKIAWALENKPRIPVVWIHGL WP_001058323.1 MULTISPECIES: hydrogenase-1 small chain [Proteobacteria] Length: 372\nNumber of predicted TMHs: 2\nExp number of AAs in TMHs: 37.2608499999999\nExp number, first 60 AAs: 19.52923\nTotal prob of N-in: 0.94351\nPOSSIBLE N-term signal sequence\ninside 1 19\nTMhelix 20 39\noutside 40 325\nTMhelix 326 348\ninside 349 372

12996 GCF_000146735.1_ASM14673v1 Escherichia coli str. K-12 substr. MG1655star Proteobacteria; Gammaproteobacteria; Enterobacterales; Enterobacteriaceae; Escherichia; Escherichia coli; Escherichia coli K-12; Escherichia coli str. K-12 substr. MG1655 MTGDNTLIHSHGINRRDFMKLCAALAATMGLSSKAAAEMAESVTNPQRPPVIWIGAQECT WP_000145410.1 MULTISPECIES: hydrogenase-2 small chain [Proteobacteria] Length: 372\nNumber of predicted TMHs: 1\nExp number of AAs in TMHs: 23.72226\nExp number, first 60 AAs: 0.22038\nTotal prob of N-in: 0.12994\noutside 1 331\nTMhelix 332 354\ninside 355 372

12997 GCF_000146735.1_ASM14673v1 Escherichia coli str. K-12 substr. MG1655star Proteobacteria; Gammaproteobacteria; Enterobacterales; Enterobacteriaceae; Escherichia; Escherichia coli; Escherichia coli K-12; Escherichia coli str. K-12 substr. MG1655 MNNEETFYQAMRRQGVTRRSFLKYCSLAATSLGLGAGMAPKIAWALENKPRIPVVWIHGL WP_001058323.1 MULTISPECIES: hydrogenase-1 small chain [Proteobacteria] Length: 372\nNumber of predicted TMHs: 2\nExp number of AAs in TMHs: 37.2608499999999\nExp number, first 60 AAs: 19.52923\nTotal prob of N-in: 0.94351\nPOSSIBLE N-term signal sequence\ninside 1 19\nTMhelix 20 39\noutside 40 325\nTMhelix 326 348\ninside 349 372

12998 GCF_000333175.1_ASM33317v1 Escherichia coli O10:K5(L):H4 str. ATCC 23506 Proteobacteria; Gammaproteobacteria; Enterobacterales; Enterobacteriaceae; Escherichia; Escherichia coli; Escherichia coli O10
MNNEETFYQAMRRQGVTRRSFLKYCSLAATSLGLGAGMAPKIAWALENKPRIPVVWIHGL WP_001058323.1
MULTISPECIES: hydrogenase-1 small chain [Proteobacteria] Length: 372\nNumber of predicted TMHs: 2\nExp number of AAs in TMHs: 37.2608499999999\nExp number, first 60 AAs: 19.52923\nTotal prob of N-in: 0.94351\nPOSSIBLE N-term signal sequence\ninside 1 19\nTMhelix 20 39\noutside 40 325\nTMhelix 326 348\ninside 349 372

12999 GCF_000333175.1_ASM33317v1 Escherichia coli O10:K5(L):H4 str. ATCC 23506 Proteobacteria; Gammaproteobacteria; Enterobacterales; Enterobacteriaceae; Escherichia; Escherichia coli; Escherichia coli O10
MTGDNTLIHSHGINRRDFMKLCAALAATMGLSSKAAAEMAESVTNPQRPPVIWIGAQECT WP_000145410.1
MULTISPECIES: hydrogenase-2 small chain [Proteobacteria] Length: 372\nNumber of predicted TMHs: 1\nExp number of AAs in TMHs: 23.72226\nExp number, first 60 AAs: 0.22038\nTotal prob of N-in: 0.12994\noutside 1 331\nTMhelix 332 354\ninside 355 372

13000 GCF_000167855.1_ASM16785v1 Escherichia coli E22 Proteobacteria; Gammaproteobacteria; Enterobacterales; Enterobacteriaceae; Escherichia; Escherichia coli; Escherichia coli O103
MNNEETFYQAMRRQGVTRRSFLKYCSLAATSLGLGAGMAPKIAWALENKPRIPVVWIHGL WP_001295940.1
MULTISPECIES: hydrogenase [Enterobacteriaceae] Length: 372\nNumber of predicted TMHs: 2\nExp number of AAs in TMHs: 37.2608899999999\nExp number, first 60 AAs: 19.52923\nTotal prob of N-in: 0.94351\nPOSSIBLE N-term signal sequence\ninside 1 19\nTMhelix 20 39\noutside 40 325\nTMhelix 326 348\ninside 349 372

13001 GCF_000167855.1_ASM16785v1 Escherichia coli E22 Proteobacteria; Gammaproteobacteria; Enterobacterales; Enterobacteriaceae; Escherichia; Escherichia coli; Escherichia coli O103
MTGDNTLIHSHGINRRDFMKLCAALAATMGLSSKAAAEMAESVTNPQRPPVIWIGAQECT WP_000145410.1
MULTISPECIES: hydrogenase-2 small chain [Proteobacteria] Length: 372\nNumber of predicted TMHs: 1\nExp number of AAs in TMHs: 23.72226\nExp number, first 60 AAs: 0.22038\nTotal prob of N-in: 0.12994\noutside 1 331\nTMhelix 332 354\ninside 355 372

13002 GCF_000617045.2_Ec04-3023 Escherichia coli O103:H11 str. 04-3023 Proteobacteria; Gammaproteobacteria; Enterobacterales; Enterobacteriaceae; Escherichia; Escherichia coli; Escherichia coli O103
MTGDNTLIHSHGINRRDFMKLCAALAATMGLSSKAAAEMAESVTNPQRPPVIWIGAQECT WP_032319252.1
hydrogenase-2 small chain [Escherichia coli] Length: 372\nNumber of predicted TMHs: 1\nExp number of AAs in TMHs: 23.56891\nExp number, first 60 AAs: 0.21932\nTotal prob of N-in: 0.12574\noutside 1 331\nTMhelix 332 354\ninside 355 372

13003 GCF_000617045.2_Ec04-3023 Escherichia coli O103:H11 str. 04-3023 Proteobacteria; Gammaproteobacteria; Enterobacterales; Enterobacteriaceae; Escherichia; Escherichia coli; Escherichia coli O103
MNNEETFYQAMRRQGVTRRSFLKYCSLAATSLGLGAGMAPKIAWALENKPRIPVVWIHGL WP_001058323.1
MULTISPECIES: hydrogenase-1 small chain [Proteobacteria] Length: 372\nNumber of predicted TMHs: 2\nExp number of AAs in TMHs: 37.2608499999999\nExp number, first 60 AAs: 19.52923\nTotal prob of N-in: 0.94351\nPOSSIBLE N-term signal sequence\ninside 1 19\nTMhelix 20 39\noutside 40 325\nTMhelix 326 348\ninside 349 372

13004 GCF_000615055.1_Ec2010C-3214 Escherichia coli O103:H11 str. 2010C-3214 Proteobacteria; Gammaproteobacteria; Enterobacterales; Enterobacteriaceae; Escherichia; Escherichia coli; Escherichia coli O103
MNNEETFYQAMRRQGVTRRSFLKYCSLAATSLGLGAGMAPKIAWALENKPRIPVVWIHGL WP_001058323.1
MULTISPECIES: hydrogenase-1 small chain [Proteobacteria] Length: 372\nNumber of predicted TMHs: 2\nExp number of AAs in TMHs: 37.2608499999999\nExp number, first 60 AAs: 19.52923\nTotal prob of N-in: 0.94351\nPOSSIBLE N-term signal sequence\ninside 1 19\nTMhelix 20 39\noutside 40 325\nTMhelix 326 348\ninside 349 372

13005 GCF_000615055.1_Ec2010C-3214 Escherichia coli O103:H11 str. 2010C-3214 Proteobacteria; Gammaproteobacteria; Enterobacterales; Enterobacteriaceae; Escherichia; Escherichia coli; Escherichia coli O103
MTGDNTLIHSHGINRRDFMKLCAALAATMGLSSKAAAEMAESVTNPQRPPVIWIGAQECT WP_000145410.1
MULTISPECIES: hydrogenase-2 small chain [Proteobacteria] Length: 372\nNumber of predicted TMHs: 1\nExp number of AAs in TMHs: 23.72226\nExp number, first 60 AAs: 0.22038\nTotal prob of N-in: 0.12994\noutside 1 331\nTMhelix 332 354\ninside 355 372

13006 GCF_000617745.1_Ec2010C-4529 Escherichia coli O103:H25 str. 2010C-4529 Proteobacteria; Gammaproteobacteria; Enterobacterales; Enterobacteriaceae; Escherichia; Escherichia coli; Escherichia coli O103 MNNEETFYQAMRRQGVTRRSFLKYCSLAATSLGLGAGMAPKIAWALENKPRIPVVWIHGL WP_001058323.1 MULTISPECIES: hydrogenase-1 small chain [Proteobacteria] Length: 372\nNumber of predicted TMHs: 2\nExp number of AAs in TMHs: 37.2608499999999\nExp number, first 60 AAs: 19.52923\nTotal prob of N-in: 0.94351\nPOSSIBLE N-term signal sequence\ninside 1 19\nTMhelix 20 39\noutside 40 325\nTMhelix 326 348\ninside 349 372

13007 GCF_000617745.1_Ec2010C-4529 Escherichia coli O103:H25 str. 2010C-4529 Proteobacteria; Gammaproteobacteria; Enterobacterales; Enterobacteriaceae; Escherichia; Escherichia coli; Escherichia coli O103 MTGDNTLIHSHGINRRDFMKLCAALAATMGLSSKAAAEMAESVTNPQRPPVIWIGAQECT WP_000145410.1 MULTISPECIES: hydrogenase-2 small chain [Proteobacteria] Length: 372\nNumber of predicted TMHs: 1\nExp number of AAs in TMHs: 23.72226\nExp number, first 60 AAs: 0.22038\nTotal prob of N-in: 0.12994\noutside 1 331\nTMhelix 332 354\ninside 355 372

13008 GCF_000263895.1_ASM26389v1 Escherichia coli O103:H25 str. CVM9340 Proteobacteria; Gammaproteobacteria; Enterobacterales; Enterobacteriaceae; Escherichia; Escherichia coli; Escherichia coli O103 MNNEETFYQAMRRQGVTRRSFLKYCSLAATSLGLGAGMAPKIAWALENKPRIPVVWIHGL WP_001058323.1 MULTISPECIES: hydrogenase-1 small chain [Proteobacteria] Length: 372\nNumber of predicted TMHs: 2\nExp number of AAs in TMHs: 37.2608499999999\nExp number, first 60 AAs: 19.52923\nTotal prob of N-in: 0.94351\nPOSSIBLE N-term signal sequence\ninside 1 19\nTMhelix 20 39\noutside 40 325\nTMhelix 326 348\ninside 349 372

13009 GCF_000263895.1_ASM26389v1 Escherichia coli O103:H25 str. CVM9340 Proteobacteria; Gammaproteobacteria; Enterobacterales; Enterobacteriaceae; Escherichia; Escherichia coli; Escherichia coli O103 MTGDNTLIHSHGINRRDFMKLCAALAATMGLSSKAAAEMAESVTNPQRPPVIWIGAQECT WP_000145410.1 MULTISPECIES: hydrogenase-2 small chain [Proteobacteria] Length: 372\nNumber of predicted TMHs: 1\nExp number of AAs in TMHs: 23.72226\nExp number, first 60 AAs: 0.22038\nTotal prob of N-in: 0.12994\noutside 1 331\nTMhelix 332 354\ninside 355 372

13010 GCF_000234605.1_ASM23460v2 Escherichia coli O103:H25 str. NIPH-11060424 Proteobacteria; Gammaproteobacteria; Enterobacterales; Enterobacteriaceae; Escherichia; Escherichia coli; Escherichia coli O103 MNNEETFYQAMRRQGVTRRSFLKYCSLAATSLGLGAGMAPKIAWALENKPRIPVVWIHGL WP_001058323.1 MULTISPECIES: hydrogenase-1 small chain [Proteobacteria] Length: 372\nNumber of predicted TMHs: 2\nExp number of AAs in TMHs: 37.2608499999999\nExp number, first 60 AAs: 19.52923\nTotal prob of N-in: 0.94351\nPOSSIBLE N-term signal sequence\ninside 1 19\nTMhelix 20 39\noutside 40 325\nTMhelix 326 348\ninside 349 372

13011 GCF_000234605.1_ASM23460v2 Escherichia coli O103:H25 str. NIPH-11060424 Proteobacteria; Gammaproteobacteria; Enterobacterales; Enterobacteriaceae; Escherichia; Escherichia coli; Escherichia coli O103 MTGDNTLIHSHGINRRDFMKLCAALAATMGLSSKAAAEMAESVTNPQRPPVIWIGAQECT WP_000145410.1 MULTISPECIES: hydrogenase-2 small chain [Proteobacteria] Length: 372\nNumber of predicted TMHs: 1\nExp number of AAs in TMHs: 23.72226\nExp number, first 60 AAs: 0.22038\nTotal prob of N-in: 0.12994\noutside 1 331\nTMhelix 332 354\ninside 355 372

13012 GCF_000010745.1_ASM1074v1 Escherichia coli O103:H2 str. 12009 Proteobacteria; Gammaproteobacteria; Enterobacterales; Enterobacteriaceae; Escherichia; Escherichia coli; Escherichia coli O103 MNNEETFYQAMRRQGVTRRSFLKYCSLAATSLGLGAGMAPKIAWALENKPRIPVVWIHGL WP_001295940.1 MULTISPECIES: hydrogenase [Enterobacteriaceae] Length: 372\nNumber of predicted TMHs: 2\nExp number of AAs in TMHs: 37.2608899999999\nExp number, first 60 AAs: 19.52923\nTotal prob of N-in: 0.94351\nPOSSIBLE N-term signal sequence\ninside 1 19\nTMhelix 20 39\noutside 40 325\nTMhelix 326 348\ninside 349 372

13013 GCF_000010745.1_ASM1074v1 Escherichia coli O103:H2 str. 12009 Proteobacteria; Gammaproteobacteria; Enterobacterales; Enterobacteriaceae; Escherichia; Escherichia coli; Escherichia coli O103 MTGDNTLIHSHGINRRDFMKLCAALAATMGLSSKAAAEMAESVTNPQRPPVIWIGAQECT WP_000145410.1 MULTISPECIES: hydrogenase-2 small chain [Proteobacteria] Length: 372\nNumber of predicted TMHs: 1\nExp number of AAs in TMHs: 23.72226\nExp number, first 60 AAs: 0.22038\nTotal prob of N-in: 0.12994\noutside 1 331\nTMhelix 332 354\ninside 355 372

13014 GCF_000619765.2_Ec2009C-3279 Escherichia coli O103:H2 str. 2009C-3279 Proteobacteria; Gammaproteobacteria; Enterobacterales; Enterobacteriaceae; Escherichia; Escherichia coli; Escherichia coli O103 MNNEETFYQAMRRQGVTRRSFLKYCSLAATSLGLGAGMAPKIAWALENKPRIPVVWIHGL WP_001295940.1 MULTISPECIES: hydrogenase [Enterobacteriaceae] Length: 372\nNumber of predicted TMHs: 2\nExp number of AAs in TMHs: 37.2608899999999\nExp number, first 60 AAs: 19.52923\nTotal prob of N-in: 0.94351\nPOSSIBLE N-term signal sequence\ninside 1 19\nTMhelix 20 39\noutside 40 325\nTMhelix 326 348\ninside 349 372

13015 GCF_000619765.2_Ec2009C-3279 Escherichia coli O103:H2 str. 2009C-3279 Proteobacteria; Gammaproteobacteria; Enterobacterales; Enterobacteriaceae; Escherichia; Escherichia coli; Escherichia coli O103 MTGDNTLIHSHGINRRDFMKLCAALAATMGLSSKAAAEMAESVTNPQRPPVIWIGAQECT WP_000145410.1 MULTISPECIES: hydrogenase-2 small chain [Proteobacteria] Length: 372\nNumber of predicted TMHs: 1\nExp number of AAs in TMHs: 23.72226\nExp number, first 60 AAs: 0.22038\nTotal prob of N-in: 0.12994\noutside 1 331\nTMhelix 332 354\ninside 355 372

13016 GCF_000617685.1_Ec2010C-4433 Escherichia coli O103:H2 str. 2010C-4433 Proteobacteria; Gammaproteobacteria; Enterobacterales; Enterobacteriaceae; Escherichia; Escherichia coli; Escherichia coli O103 MTGDNTLIHSHGINRRDFMKLCAALAATMGLSSKAAAEMAESVTNPQRPPVIWIGAQECT WP_000145410.1 MULTISPECIES: hydrogenase-2 small chain [Proteobacteria] Length: 372\nNumber of predicted TMHs: 1\nExp number of AAs in TMHs: 23.72226\nExp number, first 60 AAs: 0.22038\nTotal prob of N-in: 0.12994\noutside 1 331\nTMhelix 332 354\ninside 355 372

13017 GCF_000617685.1_Ec2010C-4433 Escherichia coli O103:H2 str. 2010C-4433 Proteobacteria; Gammaproteobacteria; Enterobacterales; Enterobacteriaceae; Escherichia; Escherichia coli; Escherichia coli O103 MNNEETFYQAMRRQGVTRRSFLKYCSLAATSLGLGAGMAPKIAWALENKPRIPVVWIHGL WP_001295940.1 MULTISPECIES: hydrogenase [Enterobacteriaceae] Length: 372\nNumber of predicted TMHs: 2\nExp number of AAs in TMHs: 37.2608899999999\nExp number, first 60 AAs: 19.52923\nTotal prob of N-in: 0.94351\nPOSSIBLE N-term signal sequence\ninside 1 19\nTMhelix 20 39\noutside 40 325\nTMhelix 326 348\ninside 349 372

13018 GCF_000616345.2_Ec2011C-3750 Escherichia coli O103:H2 str. 2011C-3750 Proteobacteria; Gammaproteobacteria; Enterobacterales; Enterobacteriaceae; Escherichia; Escherichia coli; Escherichia coli O103 MNNEETFYQAMRRQGVTRRSFLKYCSLAATSLGLGAGMAPKIAWALENKPRIPVVWIHGL WP_001295940.1 MULTISPECIES: hydrogenase [Enterobacteriaceae] Length: 372\nNumber of predicted TMHs: 2\nExp number of AAs in TMHs: 37.2608899999999\nExp number, first 60 AAs: 19.52923\nTotal prob of N-in: 0.94351\nPOSSIBLE N-term signal sequence\ninside 1 19\nTMhelix 20 39\noutside 40 325\nTMhelix 326 348\ninside 349 372

13019 GCF_000616345.2_Ec2011C-3750 Escherichia coli O103:H2 str. 2011C-3750 Proteobacteria; Gammaproteobacteria; Enterobacterales; Enterobacteriaceae; Escherichia; Escherichia coli; Escherichia coli O103 MTGDNTLIHSHGINRRDFMKLCAALAATMGLSSKAAAEMAESVTNPQRPPVIWIGAQECT WP_000145410.1 MULTISPECIES: hydrogenase-2 small chain [Proteobacteria] Length: 372\nNumber of predicted TMHs: 1\nExp number of AAs in TMHs: 23.72226\nExp number, first 60 AAs: 0.22038\nTotal prob of N-in: 0.12994\noutside 1 331\nTMhelix 332 354\ninside 355 372

13020 GCF_000263915.1_ASM26391v1 Escherichia coli O103:H2 str. CVM9450 Proteobacteria; Gammaproteobacteria; Enterobacterales; Enterobacteriaceae; Escherichia; Escherichia coli; Escherichia coli O103 MTGDNTLIHSHGINRRDFMKLCAALAATMGLSSKAAAEMAESVTNPQRPPVIWIGAQECT WP_000145410.1 MULTISPECIES: hydrogenase-2 small chain [Proteobacteria] Length: 372\nNumber of predicted TMHs: 1\nExp number of AAs in TMHs: 23.72226\nExp number, first 60 AAs: 0.22038\nTotal prob of N-in: 0.12994\noutside 1 331\nTMhelix 332 354\ninside 355 372

13021 GCF_000263915.1_ASM26391v1 Escherichia coli O103:H2 str. CVM9450 Proteobacteria; Gammaproteobacteria; Enterobacterales; Enterobacteriaceae; Escherichia; Escherichia coli; Escherichia coli O103 MNNEETFYQAMRRQGVTRRSFLKYCSLAATSLGLGAGMAPKIAWALENKPRIPVVWIHGL WP_001295940.1 MULTISPECIES: hydrogenase [Enterobacteriaceae] Length: 372\nNumber of predicted TMHs: 2\nExp number of AAs in TMHs: 37.2608899999999\nExp number, first 60 AAs: 19.52923\nTotal prob of N-in: 0.94351\nPOSSIBLE N-term signal sequence\ninside 1 19\nTMhelix 20 39\noutside 40 325\nTMhelix 326 348\ninside 349 372

13022 GCF_000618965.1_Ec94-3025 Escherichia coli O104:H21 str. 94-3025 Proteobacteria; Gammaproteobacteria; Enterobacterales; Enterobacteriaceae; Escherichia; Escherichia coli; Escherichia coli O104
MTGDNTLIHSHGINRRDFMKLCAALAATMGLSSKAAAEMAESVTNPQRPPVIWIGAQECT WP_000145410.1
MULTISPECIES: hydrogenase-2 small chain [Proteobacteria] Length: 372\nNumber of predicted TMHs: 1\nExp number of AAs in TMHs: 23.72226\nExp number, first 60 AAs: 0.22038\nTotal prob of N-in: 0.12994\noutside 1 331\nTMhelix 332 354\ninside 355 372

13023 GCF_000618965.1_Ec94-3025 Escherichia coli O104:H21 str. 94-3025 Proteobacteria; Gammaproteobacteria; Enterobacterales; Enterobacteriaceae; Escherichia; Escherichia coli; Escherichia coli O104
MNNEETFYQAMRRQGVTRRSFLKYCSLAATSLGLGAGMAPKIAWALENKPRIPVVWIHGL WP_001295940.1
MULTISPECIES: hydrogenase [Enterobacteriaceae] Length: 372\nNumber of predicted TMHs: 2\nExp number of AAs in TMHs: 37.2608899999999\nExp number, first 60 AAs: 19.52923\nTotal prob of N-in: 0.94351\nPOSSIBLE N-term signal sequence\ninside 1 19\nTMhelix 20 39\noutside 40 325\nTMhelix 326 348\ninside 349 372

13024 GCF_000235245.1_Esch_coli_04-8351_V2 Escherichia coli O104:H4 str. 04-8351 Proteobacteria; Gammaproteobacteria; Enterobacterales; Enterobacteriaceae; Escherichia; Escherichia coli; Escherichia coli O104
MTGDNTLIHSHGINRRDFMKLCAALAATMGLSSKAAAEMAESVTNPQRPPVIWIGAQECT WP_000145410.1
MULTISPECIES: hydrogenase-2 small chain [Proteobacteria] Length: 372\nNumber of predicted TMHs: 1\nExp number of AAs in TMHs: 23.72226\nExp number, first 60 AAs: 0.22038\nTotal prob of N-in: 0.12994\noutside 1 331\nTMhelix 332 354\ninside 355 372

13025 GCF_000235245.1_Esch_coli_04-8351_V2 Escherichia coli O104:H4 str. 04-8351 Proteobacteria; Gammaproteobacteria; Enterobacterales; Enterobacteriaceae; Escherichia; Escherichia coli; Escherichia coli O104
MNNEETFYQAMRRQGVTRRSFLKYCSLAATSLGLGAGMAPKIAWALENKPRIPVVWIHGL WP_001058323.1
MULTISPECIES: hydrogenase-1 small chain [Proteobacteria] Length: 372\nNumber of predicted TMHs: 2\nExp number of AAs in TMHs: 37.2608499999999\nExp number, first 60 AAs: 19.52923\nTotal prob of N-in: 0.94351\nPOSSIBLE N-term signal sequence\ninside 1 19\nTMhelix 20 39\noutside 40 325\nTMhelix 326 348\ninside 349 372

13026 GCF_000235225.1_Esch_coli_09-7901_V2 Escherichia coli O104:H4 str. 09-7901 Proteobacteria; Gammaproteobacteria; Enterobacterales; Enterobacteriaceae; Escherichia; Escherichia coli; Escherichia coli O104
MTGDNTLIHSHGINRRDFMKLCAALAATMGLSSKAAAEMAESVTNPQRPPVIWIGAQECT WP_000145410.1
MULTISPECIES: hydrogenase-2 small chain [Proteobacteria] Length: 372\nNumber of predicted TMHs: 1\nExp number of AAs in TMHs: 23.72226\nExp number, first 60 AAs: 0.22038\nTotal prob of N-in: 0.12994\noutside 1 331\nTMhelix 332 354\ninside 355 372

13027 GCF_000235225.1_Esch_coli_09-7901_V2 Escherichia coli O104:H4 str. 09-7901 Proteobacteria; Gammaproteobacteria; Enterobacterales; Enterobacteriaceae; Escherichia; Escherichia coli; Escherichia coli O104
MNNEETFYQAMRRQGVTRRSFLKYCSLAATSLGLGAGMAPKIAWALENKPRIPVVWIHGL WP_001058323.1
MULTISPECIES: hydrogenase-1 small chain [Proteobacteria] Length: 372\nNumber of predicted TMHs: 2\nExp number of AAs in TMHs: 37.2608499999999\nExp number, first 60 AAs: 19.52923\nTotal prob of N-in: 0.94351\nPOSSIBLE N-term signal sequence\ninside 1 19\nTMhelix 20 39\noutside 40 325\nTMhelix 326 348\ninside 349 372

13028 GCF_000319995.1_Esch_coli_11-02030_V1 Escherichia coli O104:H4 str. 11-02030 Proteobacteria; Gammaproteobacteria; Enterobacterales; Enterobacteriaceae; Escherichia; Escherichia coli; Escherichia coli O104
MTGDNTLIHSHGINRRDFMKLCAALAATMGLSSKAAAEMAESVTNPQRPPVIWIGAQECT WP_000145410.1
MULTISPECIES: hydrogenase-2 small chain [Proteobacteria] Length: 372\nNumber of predicted TMHs: 1\nExp number of AAs in TMHs: 23.72226\nExp number, first 60 AAs: 0.22038\nTotal prob of N-in: 0.12994\noutside 1 331\nTMhelix 332 354\ninside 355 372

13029 GCF_000319995.1_Esch_coli_11-02030_V1 Escherichia coli O104:H4 str. 11-02030 Proteobacteria; Gammaproteobacteria; Enterobacterales; Enterobacteriaceae; Escherichia; Escherichia coli; Escherichia coli O104
MNNEETFYQAMRRQGVTRRSFLKYCSLAATSLGLGAGMAPKIAWALENKPRIPVVWIHGL WP_001058323.1
MULTISPECIES: hydrogenase-1 small chain [Proteobacteria] Length: 372\nNumber of predicted TMHs: 2\nExp number of AAs in TMHs: 37.2608499999999\nExp number, first 60 AAs: 19.52923\nTotal prob of N-in: 0.94351\nPOSSIBLE N-term signal sequence\ninside 1 19\nTMhelix 20 39\noutside 40 325\nTMhelix 326 348\ninside 349 372

13030 GCF_000320015.1_Esch_coli_11-02033-1_V1 Escherichia coli O104:H4 str. 11-02033-1 Proteobacteria; Gammaproteobacteria; Enterobacterales; Enterobacteriaceae; Escherichia; Escherichia coli; Escherichia coli O104 MTGDNTLIHSHGINRRDFMKLCAALAATMGLSSKAAAEMAESVTNPQRPPVIWIGAQECT WP_000145410.1 MULTISPECIES: hydrogenase-2 small chain [Proteobacteria] Length: 372\nNumber of predicted TMHs: 1\nExp number of AAs in TMHs: 23.72226\nExp number, first 60 AAs: 0.22038\nTotal prob of N-in: 0.12994\noutside 1 331\nTMhelix 332 354\ninside 355 372

13031 GCF_000320015.1_Esch_coli_11-02033-1_V1 Escherichia coli O104:H4 str. 11-02033-1 Proteobacteria; Gammaproteobacteria; Enterobacterales; Enterobacteriaceae; Escherichia; Escherichia coli; Escherichia coli O104 MNNEETFYQAMRRQGVTRRSFLKYCSLAATSLGLGAGMAPKIAWALENKPRIPVVWIHGL WP_001058323.1 MULTISPECIES: hydrogenase-1 small chain [Proteobacteria] Length: 372\nNumber of predicted TMHs: 2\nExp number of AAs in TMHs: 37.2608499999999\nExp number, first 60 AAs: 19.52923\nTotal prob of N-in: 0.94351\nPOSSIBLE N-term signal sequence\ninside 1 19\nTMhelix 20 39\noutside 40 325\nTMhelix 326 348\ninside 349 372

13032 GCF_000320035.1_Esch_coli_11-02092_V1 Escherichia coli O104:H4 str. 11-02092 Proteobacteria; Gammaproteobacteria; Enterobacterales; Enterobacteriaceae; Escherichia; Escherichia coli; Escherichia coli O104 MNNEETFYQAMRRQGVTRRSFLKYCSLAATSLGLGAGMAPKIAWALENKPRIPVVWIHGL WP_001058323.1 MULTISPECIES: hydrogenase-1 small chain [Proteobacteria] Length: 372\nNumber of predicted TMHs: 2\nExp number of AAs in TMHs: 37.2608499999999\nExp number, first 60 AAs: 19.52923\nTotal prob of N-in: 0.94351\nPOSSIBLE N-term signal sequence\ninside 1 19\nTMhelix 20 39\noutside 40 325\nTMhelix 326 348\ninside 349 372

13033 GCF_000320035.1_Esch_coli_11-02092_V1 Escherichia coli O104:H4 str. 11-02092 Proteobacteria; Gammaproteobacteria; Enterobacterales; Enterobacteriaceae; Escherichia; Escherichia coli; Escherichia coli O104 MTGDNTLIHSHGINRRDFMKLCAALAATMGLSSKAAAEMAESVTNPQRPPVIWIGAQECT WP_000145410.1 MULTISPECIES: hydrogenase-2 small chain [Proteobacteria] Length: 372\nNumber of predicted TMHs: 1\nExp number of AAs in TMHs: 23.72226\nExp number, first 60 AAs: 0.22038\nTotal prob of N-in: 0.12994\noutside 1 331\nTMhelix 332 354\ninside 355 372

13034 GCF_000320075.1_Esch_coli_11-02093_V1 Escherichia coli O104:H4 str. 11-02093 Proteobacteria; Gammaproteobacteria; Enterobacterales; Enterobacteriaceae; Escherichia; Escherichia coli; Escherichia coli O104 MNNEETFYQAMRRQGVTRRSFLKYCSLAATSLGLGAGMAPKIAWALENKPRIPVVWIHGL WP_001058323.1 MULTISPECIES: hydrogenase-1 small chain [Proteobacteria] Length: 372\nNumber of predicted TMHs: 2\nExp number of AAs in TMHs: 37.2608499999999\nExp number, first 60 AAs: 19.52923\nTotal prob of N-in: 0.94351\nPOSSIBLE N-term signal sequence\ninside 1 19\nTMhelix 20 39\noutside 40 325\nTMhelix 326 348\ninside 349 372

13035 GCF_000320075.1_Esch_coli_11-02093_V1 Escherichia coli O104:H4 str. 11-02093 Proteobacteria; Gammaproteobacteria; Enterobacterales; Enterobacteriaceae; Escherichia; Escherichia coli; Escherichia coli O104 MTGDNTLIHSHGINRRDFMKLCAALAATMGLSSKAAAEMAESVTNPQRPPVIWIGAQECT WP_000145410.1 MULTISPECIES: hydrogenase-2 small chain [Proteobacteria] Length: 372\nNumber of predicted TMHs: 1\nExp number of AAs in TMHs: 23.72226\nExp number, first 60 AAs: 0.22038\nTotal prob of N-in: 0.12994\noutside 1 331\nTMhelix 332 354\ninside 355 372

13036 GCF_000320095.1_Esch_coli_11-02281_V1 Escherichia coli O104:H4 str. 11-02281 Proteobacteria; Gammaproteobacteria; Enterobacterales; Enterobacteriaceae; Escherichia; Escherichia coli; Escherichia coli O104 MTGDNTLIHSHGINRRDFMKLCAALAATMGLSSKAAAEMAESVTNPQRPPVIWIGAQECT WP_000145410.1 MULTISPECIES: hydrogenase-2 small chain [Proteobacteria] Length: 372\nNumber of predicted TMHs: 1\nExp number of AAs in TMHs: 23.72226\nExp number, first 60 AAs: 0.22038\nTotal prob of N-in: 0.12994\noutside 1 331\nTMhelix 332 354\ninside 355 372

13037 GCF_000320095.1_Esch_coli_11-02281_V1 Escherichia coli O104:H4 str. 11-02281 Proteobacteria; Gammaproteobacteria; Enterobacterales; Enterobacteriaceae; Escherichia; Escherichia coli; Escherichia coli O104 MNNEETFYQAMRRQGVTRRSFLKYCSLAATSLGLGAGMAPKIAWALENKPRIPVVWIHGL WP_001058323.1 MULTISPECIES: hydrogenase-1 small chain [Proteobacteria] Length: 372\nNumber of predicted TMHs: 2\nExp number of AAs in TMHs: 37.2608499999999\nExp number, first 60 AAs: 19.52923\nTotal prob of N-in: 0.94351\nPOSSIBLE N-term signal sequence\ninside 1 19\nTMhelix 20 39\noutside 40 325\nTMhelix 326 348\ninside 349 372

13038 GCF_000319835.1_Esch_coli_11-02318_V1 Escherichia coli O104:H4 str. 11-02318 Proteobacteria; Gammaproteobacteria; Enterobacterales; Enterobacteriaceae; Escherichia; Escherichia coli; Escherichia coli O104
MTGDNTLIHSHGINRRDFMKLCAALAATMGLSSKAAAEMAESVTNPQRPPVIWIGAQECT WP_000145410.1
MULTISPECIES: hydrogenase-2 small chain [Proteobacteria] Length: 372\nNumber of predicted TMHs: 1\nExp number of AAs in TMHs: 23.72226\nExp number, first 60 AAs: 0.22038\nTotal prob of N-in: 0.12994\noutside 1 331\nTMhelix 332 354\ninside 355 372

13039 GCF_000319835.1_Esch_coli_11-02318_V1 Escherichia coli O104:H4 str. 11-02318 Proteobacteria; Gammaproteobacteria; Enterobacterales; Enterobacteriaceae; Escherichia; Escherichia coli; Escherichia coli O104
MNNEETFYQAMRRQGVTRRSFLKYCSLAATSLGLGAGMAPKIAWALENKPRIPVVWIHGL WP_001058323.1
MULTISPECIES: hydrogenase-1 small chain [Proteobacteria] Length: 372\nNumber of predicted TMHs: 2\nExp number of AAs in TMHs: 37.2608499999999\nExp number, first 60 AAs: 19.52923\nTotal prob of N-in: 0.94351\nPOSSIBLE N-term signal sequence\ninside 1 19\nTMhelix 20 39\noutside 40 325\nTMhelix 326 348\ninside 349 372

13040 GCF_000320115.1_Esch_coli_11-02913_V1 Escherichia coli O104:H4 str. 11-02913 Proteobacteria; Gammaproteobacteria; Enterobacterales; Enterobacteriaceae; Escherichia; Escherichia coli; Escherichia coli O104
MTGDNTLIHSHGINRRDFMKLCAALAATMGLSSKAAAEMAESVTNPQRPPVIWIGAQECT WP_000145410.1
MULTISPECIES: hydrogenase-2 small chain [Proteobacteria] Length: 372\nNumber of predicted TMHs: 1\nExp number of AAs in TMHs: 23.72226\nExp number, first 60 AAs: 0.22038\nTotal prob of N-in: 0.12994\noutside 1 331\nTMhelix 332 354\ninside 355 372

13041 GCF_000320115.1_Esch_coli_11-02913_V1 Escherichia coli O104:H4 str. 11-02913 Proteobacteria; Gammaproteobacteria; Enterobacterales; Enterobacteriaceae; Escherichia; Escherichia coli; Escherichia coli O104
MNNEETFYQAMRRQGVTRRSFLKYCSLAATSLGLGAGMAPKIAWALENKPRIPVVWIHGL WP_001058323.1
MULTISPECIES: hydrogenase-1 small chain [Proteobacteria] Length: 372\nNumber of predicted TMHs: 2\nExp number of AAs in TMHs: 37.2608499999999\nExp number, first 60 AAs: 19.52923\nTotal prob of N-in: 0.94351\nPOSSIBLE N-term signal sequence\ninside 1 19\nTMhelix 20 39\noutside 40 325\nTMhelix 326 348\ninside 349 372

13042 GCF_000320135.1_Esch_coli_11-03439_V1 Escherichia coli O104:H4 str. 11-03439 Proteobacteria; Gammaproteobacteria; Enterobacterales; Enterobacteriaceae; Escherichia; Escherichia coli; Escherichia coli O104
MTGDNTLIHSHGINRRDFMKLCAALAATMGLSSKAAAEMAESVTNPQRPPVIWIGAQECT WP_000145410.1
MULTISPECIES: hydrogenase-2 small chain [Proteobacteria] Length: 372\nNumber of predicted TMHs: 1\nExp number of AAs in TMHs: 23.72226\nExp number, first 60 AAs: 0.22038\nTotal prob of N-in: 0.12994\noutside 1 331\nTMhelix 332 354\ninside 355 372

13043 GCF_000320135.1_Esch_coli_11-03439_V1 Escherichia coli O104:H4 str. 11-03439 Proteobacteria; Gammaproteobacteria; Enterobacterales; Enterobacteriaceae; Escherichia; Escherichia coli; Escherichia coli O104
MNNEETFYQAMRRQGVTRRSFLKYCSLAATSLGLGAGMAPKIAWALENKPRIPVVWIHGL WP_001058323.1
MULTISPECIES: hydrogenase-1 small chain [Proteobacteria] Length: 372\nNumber of predicted TMHs: 2\nExp number of AAs in TMHs: 37.2608499999999\nExp number, first 60 AAs: 19.52923\nTotal prob of N-in: 0.94351\nPOSSIBLE N-term signal sequence\ninside 1 19\nTMhelix 20 39\noutside 40 325\nTMhelix 326 348\ninside 349 372

13044 GCF_000320155.1_Esch_coli_11-03943_V1 Escherichia coli O104:H4 str. 11-03943 Proteobacteria; Gammaproteobacteria; Enterobacterales; Enterobacteriaceae; Escherichia; Escherichia coli; Escherichia coli O104
MTGDNTLIHSHGINRRDFMKLCAALAATMGLSSKAAAEMAESVTNPQRPPVIWIGAQECT WP_000145410.1
MULTISPECIES: hydrogenase-2 small chain [Proteobacteria] Length: 372\nNumber of predicted TMHs: 1\nExp number of AAs in TMHs: 23.72226\nExp number, first 60 AAs: 0.22038\nTotal prob of N-in: 0.12994\noutside 1 331\nTMhelix 332 354\ninside 355 372

13045 GCF_000320155.1_Esch_coli_11-03943_V1 Escherichia coli O104:H4 str. 11-03943 Proteobacteria; Gammaproteobacteria; Enterobacterales; Enterobacteriaceae; Escherichia; Escherichia coli; Escherichia coli O104
MNNEETFYQAMRRQGVTRRSFLKYCSLAATSLGLGAGMAPKIAWALENKPRIPVVWIHGL WP_001058323.1
MULTISPECIES: hydrogenase-1 small chain [Proteobacteria] Length: 372\nNumber of predicted TMHs: 2\nExp number of AAs in TMHs: 37.2608499999999\nExp number, first 60 AAs: 19.52923\nTotal prob of N-in: 0.94351\nPOSSIBLE N-term signal sequence\ninside 1 19\nTMhelix 20 39\noutside 40 325\nTMhelix 326 348\ninside 349 372

13046 GCF_000320175.1_Esch_coli_11-04080_V1 Escherichia coli O104:H4 str. 11-04080 Proteobacteria; Gammaproteobacteria; Enterobacterales; Enterobacteriaceae; Escherichia; Escherichia coli; Escherichia coli O104 MTGDNTLIHSHGINRRDFMKLCAALAATMGLSSKAAAEMAESVTNPQRPPVIWIGAQECT WP_000145410.1 MULTISPECIES: hydrogenase-2 small chain [Proteobacteria] Length: 372\nNumber of predicted TMHs: 1\nExp number of AAs in TMHs: 23.72226\nExp number, first 60 AAs: 0.22038\nTotal prob of N-in: 0.12994\noutside 1 331\nTMhelix 332 354\ninside 355 372

13047 GCF_000320175.1_Esch_coli_11-04080_V1 Escherichia coli O104:H4 str. 11-04080 Proteobacteria; Gammaproteobacteria; Enterobacterales; Enterobacteriaceae; Escherichia; Escherichia coli; Escherichia coli O104 MNNEETFYQAMRRQGVTRRSFLKYCSLAATSLGLGAGMAPKIAWALENKPRIPVVWIHGL WP_001058323.1 MULTISPECIES: hydrogenase-1 small chain [Proteobacteria] Length: 372\nNumber of predicted TMHs: 2\nExp number of AAs in TMHs: 37.2608499999999\nExp number, first 60 AAs: 19.52923\nTotal prob of N-in: 0.94351\nPOSSIBLE N-term signal sequence\ninside 1 19\nTMhelix 20 39\noutside 40 325\nTMhelix 326 348\ninside 349 372

13048 GCF_000235205.1_Esch_coli_11-3677_V2 Escherichia coli O104:H4 str. 11-3677 Proteobacteria; Gammaproteobacteria; Enterobacterales; Enterobacteriaceae; Escherichia; Escherichia coli; Escherichia coli O104 MTGDNTLIHSHGINRRDFMKLCAALAATMGLSSKAAAEMAESVTNPQRPPVIWIGAQECT WP_000145410.1 MULTISPECIES: hydrogenase-2 small chain [Proteobacteria] Length: 372\nNumber of predicted TMHs: 1\nExp number of AAs in TMHs: 23.72226\nExp number, first 60 AAs: 0.22038\nTotal prob of N-in: 0.12994\noutside 1 331\nTMhelix 332 354\ninside 355 372

13049 GCF_000235205.1_Esch_coli_11-3677_V2 Escherichia coli O104:H4 str. 11-3677 Proteobacteria; Gammaproteobacteria; Enterobacterales; Enterobacteriaceae; Escherichia; Escherichia coli; Escherichia coli O104 MNNEETFYQAMRRQGVTRRSFLKYCSLAATSLGLGAGMAPKIAWALENKPRIPVVWIHGL WP_001058323.1 MULTISPECIES: hydrogenase-1 small chain [Proteobacteria] Length: 372\nNumber of predicted TMHs: 2\nExp number of AAs in TMHs: 37.2608499999999\nExp number, first 60 AAs: 19.52923\nTotal prob of N-in: 0.94351\nPOSSIBLE N-term signal sequence\ninside 1 19\nTMhelix 20 39\noutside 40 325\nTMhelix 326 348\ninside 349 372

13050 GCF_000235185.1_Esch_coli_11-4404_V2 Escherichia coli O104:H4 str. 11-4404 Proteobacteria; Gammaproteobacteria; Enterobacterales; Enterobacteriaceae; Escherichia; Escherichia coli; Escherichia coli O104 MNNEETFYQAMRRQGVTRRSFLKYCSLAATSLGLGAGMAPKIAWALENKPRIPVVWIHGL WP_001058323.1 MULTISPECIES: hydrogenase-1 small chain [Proteobacteria] Length: 372\nNumber of predicted TMHs: 2\nExp number of AAs in TMHs: 37.2608499999999\nExp number, first 60 AAs: 19.52923\nTotal prob of N-in: 0.94351\nPOSSIBLE N-term signal sequence\ninside 1 19\nTMhelix 20 39\noutside 40 325\nTMhelix 326 348\ninside 349 372

13051 GCF_000235185.1_Esch_coli_11-4404_V2 Escherichia coli O104:H4 str. 11-4404 Proteobacteria; Gammaproteobacteria; Enterobacterales; Enterobacteriaceae; Escherichia; Escherichia coli; Escherichia coli O104 MTGDNTLIHSHGINRRDFMKLCAALAATMGLSSKAAAEMAESVTNPQRPPVIWIGAQECT WP_000145410.1 MULTISPECIES: hydrogenase-2 small chain [Proteobacteria] Length: 372\nNumber of predicted TMHs: 1\nExp number of AAs in TMHs: 23.72226\nExp number, first 60 AAs: 0.22038\nTotal prob of N-in: 0.12994\noutside 1 331\nTMhelix 332 354\ninside 355 372

13052 GCF_000235165.1_Esch_coli_11-4522_V2 Escherichia coli O104:H4 str. 11-4522 Proteobacteria; Gammaproteobacteria; Enterobacterales; Enterobacteriaceae; Escherichia; Escherichia coli; Escherichia coli O104 MNNEETFYQAMRRQGVTRRSFLKYCSLAATSLGLGAGMAPKIAWALENKPRIPVVWIHGL WP_001058323.1 MULTISPECIES: hydrogenase-1 small chain [Proteobacteria] Length: 372\nNumber of predicted TMHs: 2\nExp number of AAs in TMHs: 37.2608499999999\nExp number, first 60 AAs: 19.52923\nTotal prob of N-in: 0.94351\nPOSSIBLE N-term signal sequence\ninside 1 19\nTMhelix 20 39\noutside 40 325\nTMhelix 326 348\ninside 349 372

13053 GCF_000235165.1_Esch_coli_11-4522_V2 Escherichia coli O104:H4 str. 11-4522 Proteobacteria; Gammaproteobacteria; Enterobacterales; Enterobacteriaceae; Escherichia; Escherichia coli; Escherichia coli O104 MTGDNTLIHSHGINRRDFMKLCAALAATMGLSSKAAAEMAESVTNPQRPPVIWIGAQECT WP_000145410.1 MULTISPECIES: hydrogenase-2 small chain [Proteobacteria] Length: 372\nNumber of predicted TMHs: 1\nExp number of AAs in TMHs: 23.72226\nExp number, first 60 AAs: 0.22038\nTotal prob of N-in: 0.12994\noutside 1 331\nTMhelix 332 354\ninside 355 372

13054 GCF_000235145.1_Esch_coli_11-4623_V2 Escherichia coli O104:H4 str. 11-4623 Proteobacteria; Gammaproteobacteria; Enterobacterales; Enterobacteriaceae; Escherichia; Escherichia coli; Escherichia coli O104 MNNEETFYQAMRRQGVTRRSFLKYCSLAATSLGLGAGMAPKIAWALENKPRIPVVWIHGL WP_001058323.1 MULTISPECIES: hydrogenase-1 small chain [Proteobacteria] Length: 372\nNumber of predicted TMHs: 2\nExp number of AAs in TMHs: 37.2608499999999\nExp number, first 60 AAs: 19.52923\nTotal prob of N-in: 0.94351\nPOSSIBLE N-term signal sequence\ninside 1 19\nTMhelix 20 39\noutside 40 325\nTMhelix 326 348\ninside 349 372

13055 GCF_000235145.1_Esch_coli_11-4623_V2 Escherichia coli O104:H4 str. 11-4623 Proteobacteria; Gammaproteobacteria; Enterobacterales; Enterobacteriaceae; Escherichia; Escherichia coli; Escherichia coli O104 MTGDNTLIHSHGINRRDFMKLCAALAATMGLSSKAAAEMAESVTNPQRPPVIWIGAQECT WP_000145410.1 MULTISPECIES: hydrogenase-2 small chain [Proteobacteria] Length: 372\nNumber of predicted TMHs: 1\nExp number of AAs in TMHs: 23.72226\nExp number, first 60 AAs: 0.22038\nTotal prob of N-in: 0.12994\noutside 1 331\nTMhelix 332 354\ninside 355 372

13056 GCF_000235125.1_Esch_coli_11-4632_C1_V2 Escherichia coli O104:H4 str. 11-4632 C1 Proteobacteria; Gammaproteobacteria; Enterobacterales; Enterobacteriaceae; Escherichia; Escherichia coli; Escherichia coli O104 MTGDNTLIHSHGINRRDFMKLCAALAATMGLSSKAAAEMAESVTNPQRPPVIWIGAQECT WP_000145410.1 MULTISPECIES: hydrogenase-2 small chain [Proteobacteria] Length: 372\nNumber of predicted TMHs: 1\nExp number of AAs in TMHs: 23.72226\nExp number, first 60 AAs: 0.22038\nTotal prob of N-in: 0.12994\noutside 1 331\nTMhelix 332 354\ninside 355 372

13057 GCF_000235125.1_Esch_coli_11-4632_C1_V2 Escherichia coli O104:H4 str. 11-4632 C1 Proteobacteria; Gammaproteobacteria; Enterobacterales; Enterobacteriaceae; Escherichia; Escherichia coli; Escherichia coli O104 MNNEETFYQAMRRQGVTRRSFLKYCSLAATSLGLGAGMAPKIAWALENKPRIPVVWIHGL WP_001058323.1 MULTISPECIES: hydrogenase-1 small chain [Proteobacteria] Length: 372\nNumber of predicted TMHs: 2\nExp number of AAs in TMHs: 37.2608499999999\nExp number, first 60 AAs: 19.52923\nTotal prob of N-in: 0.94351\nPOSSIBLE N-term signal sequence\ninside 1 19\nTMhelix 20 39\noutside 40 325\nTMhelix 326 348\ninside 349 372

13058 GCF_000235105.1_Esch_coli_11-4632_C2_V2 Escherichia coli O104:H4 str. 11-4632 C2 Proteobacteria; Gammaproteobacteria; Enterobacterales; Enterobacteriaceae; Escherichia; Escherichia coli; Escherichia coli O104 MTGDNTLIHSHGINRRDFMKLCAALAATMGLSSKAAAEMAESVTNPQRPPVIWIGAQECT WP_000145410.1 MULTISPECIES: hydrogenase-2 small chain [Proteobacteria] Length: 372\nNumber of predicted TMHs: 1\nExp number of AAs in TMHs: 23.72226\nExp number, first 60 AAs: 0.22038\nTotal prob of N-in: 0.12994\noutside 1 331\nTMhelix 332 354\ninside 355 372

13059 GCF_000235105.1_Esch_coli_11-4632_C2_V2 Escherichia coli O104:H4 str. 11-4632 C2 Proteobacteria; Gammaproteobacteria; Enterobacterales; Enterobacteriaceae; Escherichia; Escherichia coli; Escherichia coli O104 MNNEETFYQAMRRQGVTRRSFLKYCSLAATSLGLGAGMAPKIAWALENKPRIPVVWIHGL WP_001058323.1 MULTISPECIES: hydrogenase-1 small chain [Proteobacteria] Length: 372\nNumber of predicted TMHs: 2\nExp number of AAs in TMHs: 37.2608499999999\nExp number, first 60 AAs: 19.52923\nTotal prob of N-in: 0.94351\nPOSSIBLE N-term signal sequence\ninside 1 19\nTMhelix 20 39\noutside 40 325\nTMhelix 326 348\ninside 349 372

13060 GCF_000235085.1_Esch_coli_11-4632_C3_V2 Escherichia coli O104:H4 str. 11-4632 C3 Proteobacteria; Gammaproteobacteria; Enterobacterales; Enterobacteriaceae; Escherichia; Escherichia coli; Escherichia coli O104 MNNEETFYQAMRRQGVTRRSFLKYCSLAATSLGLGAGMAPKIAWALENKPRIPVVWIHGL WP_001058323.1 MULTISPECIES: hydrogenase-1 small chain [Proteobacteria] Length: 372\nNumber of predicted TMHs: 2\nExp number of AAs in TMHs: 37.2608499999999\nExp number, first 60 AAs: 19.52923\nTotal prob of N-in: 0.94351\nPOSSIBLE N-term signal sequence\ninside 1 19\nTMhelix 20 39\noutside 40 325\nTMhelix 326 348\ninside 349 372

13061 GCF_000235085.1_Esch_coli_11-4632_C3_V2 Escherichia coli O104:H4 str. 11-4632 C3 Proteobacteria; Gammaproteobacteria; Enterobacterales; Enterobacteriaceae; Escherichia; Escherichia coli; Escherichia coli O104 MTGDNTLIHSHGINRRDFMKLCAALAATMGLSSKAAAEMAESVTNPQRPPVIWIGAQECT WP_000145410.1 MULTISPECIES: hydrogenase-2 small chain [Proteobacteria] Length: 372\nNumber of predicted TMHs: 1\nExp number of AAs in TMHs: 23.72226\nExp number, first 60 AAs: 0.22038\nTotal prob of N-in: 0.12994\noutside 1 331\nTMhelix 332 354\ninside 355 372

13062 GCF_000235065.1_Esch_coli_11-4632_C4_V2 Escherichia coli O104:H4 str. 11-4632 C4 Proteobacteria; Gammaproteobacteria; Enterobacterales; Enterobacteriaceae; Escherichia; Escherichia coli; Escherichia coli O104 MTGDNTLIHSHGINRRDFMKLCAALAATMGLSSKAAAEMAESVTNPQRPPVIWIGAQECT WP_000145410.1 MULTISPECIES: hydrogenase-2 small chain [Proteobacteria] Length: 372\nNumber of predicted TMHs: 1\nExp number of AAs in TMHs: 23.72226\nExp number, first 60 AAs: 0.22038\nTotal prob of N-in: 0.12994\noutside 1 331\nTMhelix 332 354\ninside 355 372

13063 GCF_000235065.1_Esch_coli_11-4632_C4_V2 Escherichia coli O104:H4 str. 11-4632 C4 Proteobacteria; Gammaproteobacteria; Enterobacterales; Enterobacteriaceae; Escherichia; Escherichia coli; Escherichia coli O104 MNNEETFYQAMRRQGVTRRSFLKYCSLAATSLGLGAGMAPKIAWALENKPRIPVVWIHGL WP_001058323.1 MULTISPECIES: hydrogenase-1 small chain [Proteobacteria] Length: 372\nNumber of predicted TMHs: 2\nExp number of AAs in TMHs: 37.2608499999999\nExp number, first 60 AAs: 19.52923\nTotal prob of N-in: 0.94351\nPOSSIBLE N-term signal sequence\ninside 1 19\nTMhelix 20 39\noutside 40 325\nTMhelix 326 348\ninside 349 372

13064 GCF_000235045.1_Esch_coli_11-4632_C5_V2 Escherichia coli O104:H4 str. 11-4632 C5 Proteobacteria; Gammaproteobacteria; Enterobacterales; Enterobacteriaceae; Escherichia; Escherichia coli; Escherichia coli O104 MTGDNTLIHSHGINRRDFMKLCAALAATMGLSSKAAAEMAESVTNPQRPPVIWIGAQECT WP_000145410.1 MULTISPECIES: hydrogenase-2 small chain [Proteobacteria] Length: 372\nNumber of predicted TMHs: 1\nExp number of AAs in TMHs: 23.72226\nExp number, first 60 AAs: 0.22038\nTotal prob of N-in: 0.12994\noutside 1 331\nTMhelix 332 354\ninside 355 372

13065 GCF_000235045.1_Esch_coli_11-4632_C5_V2 Escherichia coli O104:H4 str. 11-4632 C5 Proteobacteria; Gammaproteobacteria; Enterobacterales; Enterobacteriaceae; Escherichia; Escherichia coli; Escherichia coli O104 MNNEETFYQAMRRQGVTRRSFLKYCSLAATSLGLGAGMAPKIAWALENKPRIPVVWIHGL WP_001058323.1 MULTISPECIES: hydrogenase-1 small chain [Proteobacteria] Length: 372\nNumber of predicted TMHs: 2\nExp number of AAs in TMHs: 37.2608499999999\nExp number, first 60 AAs: 19.52923\nTotal prob of N-in: 0.94351\nPOSSIBLE N-term signal sequence\ninside 1 19\nTMhelix 20 39\noutside 40 325\nTMhelix 326 348\ninside 349 372

13066 GCF_000299255.1_ASM29925v1 Escherichia coli O104:H4 str. 2009EL-2050 Proteobacteria; Gammaproteobacteria; Enterobacterales; Enterobacteriaceae; Escherichia; Escherichia coli; Escherichia coli O104 MNNEETFYQAMRRQGVTRRSFLKYCSLAATSLGLGAGMAPKIAWALENKPRIPVVWIHGL WP_001058323.1 MULTISPECIES: hydrogenase-1 small chain [Proteobacteria] Length: 372\nNumber of predicted TMHs: 2\nExp number of AAs in TMHs: 37.2608499999999\nExp number, first 60 AAs: 19.52923\nTotal prob of N-in: 0.94351\nPOSSIBLE N-term signal sequence\ninside 1 19\nTMhelix 20 39\noutside 40 325\nTMhelix 326 348\ninside 349 372

13067 GCF_000299255.1_ASM29925v1 Escherichia coli O104:H4 str. 2009EL-2050 Proteobacteria; Gammaproteobacteria; Enterobacterales; Enterobacteriaceae; Escherichia; Escherichia coli; Escherichia coli O104 MTGDNTLIHSHGINRRDFMKLCAALAATMGLSSKAAAEMAESVTNPQRPPVIWIGAQECT WP_000145410.1 MULTISPECIES: hydrogenase-2 small chain [Proteobacteria] Length: 372\nNumber of predicted TMHs: 1\nExp number of AAs in TMHs: 23.72226\nExp number, first 60 AAs: 0.22038\nTotal prob of N-in: 0.12994\noutside 1 331\nTMhelix 332 354\ninside 355 372

13068 GCF_000299475.1_ASM29947v1 Escherichia coli O104:H4 str. 2009EL-2071 Proteobacteria; Gammaproteobacteria; Enterobacterales; Enterobacteriaceae; Escherichia; Escherichia coli; Escherichia coli O104 MTGDNTLIHSHGINRRDFMKLCAALAATMGLSSKAAAEMAESVTNPQRPPVIWIGAQECT WP_000145410.1 MULTISPECIES: hydrogenase-2 small chain [Proteobacteria] Length: 372\nNumber of predicted TMHs: 1\nExp number of AAs in TMHs: 23.72226\nExp number, first 60 AAs: 0.22038\nTotal prob of N-in: 0.12994\noutside 1 331\nTMhelix 332 354\ninside 355 372

13069 GCF_000299475.1_ASM29947v1 Escherichia coli O104:H4 str. 2009EL-2071 Proteobacteria; Gammaproteobacteria; Enterobacterales; Enterobacteriaceae; Escherichia; Escherichia coli; Escherichia coli O104 MNNEETFYQAMRRQGVTRRSFLKYCSLAATSLGLGAGMAPKIAWALENKPRIPVVWIHGL WP_001058323.1 MULTISPECIES: hydrogenase-1 small chain [Proteobacteria] Length: 372\nNumber of predicted TMHs: 2\nExp number of AAs in TMHs: 37.2608499999999\nExp number, first 60 AAs: 19.52923\nTotal prob of N-in: 0.94351\nPOSSIBLE N-term signal sequence\ninside 1 19\nTMhelix 20 39\noutside 40 325\nTMhelix 326 348\ninside 349 372

13070 GCF_000299455.1_ASM29945v1 Escherichia coli O104:H4 str. 2011C-3493 Proteobacteria; Gammaproteobacteria; Enterobacterales; Enterobacteriaceae; Escherichia; Escherichia coli; Escherichia coli O104 MNNEETFYQAMRRQGVTRRSFLKYCSLAATSLGLGAGMAPKIAWALENKPRIPVVWIHGL YP_006779971.1 hydrogenase 1, small subunit [Escherichia coli O104:H4 str. 2011C-3493] Length: 372\nNumber of predicted TMHs: 2\nExp number of AAs in TMHs: 37.2608499999999\nExp number, first 60 AAs: 19.52923\nTotal prob of N-in: 0.94351\nPOSSIBLE N-term signal sequence\ninside 1 19\nTMhelix 20 39\noutside 40 325\nTMhelix 326 348\ninside 349 372

13071 GCF_000299455.1_ASM29945v1 Escherichia coli O104:H4 str. 2011C-3493 Proteobacteria; Gammaproteobacteria; Enterobacterales; Enterobacteriaceae; Escherichia; Escherichia coli; Escherichia coli O104 MTGDNTLIHSHGINRRDFMKLCAALAATMGLSSKAAAEMAESVTNPQRPPVIWIGAQECT YP_006777516.1 hydrogenase 2 small subunit [Escherichia coli O104:H4 str. 2011C-3493] Length: 372\nNumber of predicted TMHs: 1\nExp number of AAs in TMHs: 23.72226\nExp number, first 60 AAs: 0.22038\nTotal prob of N-in: 0.12994\noutside 1 331\nTMhelix 332 354\ninside 355 372

13072 GCF_000616385.2_Ec2011EL-1675A Escherichia coli O104:H4 str. 2011EL-1675A Proteobacteria; Gammaproteobacteria; Enterobacterales; Enterobacteriaceae; Escherichia; Escherichia coli; Escherichia coli O104 MTGDNTLIHSHGINRRDFMKLCAALAATMGLSSKAAAEMAESVTNPQRPPVIWIGAQECT WP_000145410.1 MULTISPECIES: hydrogenase-2 small chain [Proteobacteria] Length: 372\nNumber of predicted TMHs: 1\nExp number of AAs in TMHs: 23.72226\nExp number, first 60 AAs: 0.22038\nTotal prob of N-in: 0.12994\noutside 1 331\nTMhelix 332 354\ninside 355 372

13073 GCF_000616385.2_Ec2011EL-1675A Escherichia coli O104:H4 str. 2011EL-1675A Proteobacteria; Gammaproteobacteria; Enterobacterales; Enterobacteriaceae; Escherichia; Escherichia coli; Escherichia coli O104 MNNEETFYQAMRRQGVTRRSFLKYCSLAATSLGLGAGMAPKIAWALENKPRIPVVWIHGL WP_001058323.1 MULTISPECIES: hydrogenase-1 small chain [Proteobacteria] Length: 372\nNumber of predicted TMHs: 2\nExp number of AAs in TMHs: 37.2608499999999\nExp number, first 60 AAs: 19.52923\nTotal prob of N-in: 0.94351\nPOSSIBLE N-term signal sequence\ninside 1 19\nTMhelix 20 39\noutside 40 325\nTMhelix 326 348\ninside 349 372

13074 GCF_000235285.1_Esch_coli_C227-11_V2 Escherichia coli O104:H4 str. C227-11 Proteobacteria; Gammaproteobacteria; Enterobacterales; Enterobacteriaceae; Escherichia; Escherichia coli; Escherichia coli O104 MTGDNTLIHSHGINRRDFMKLCAALAATMGLSSKAAAEMAESVTNPQRPPVIWIGAQECT WP_000145410.1 MULTISPECIES: hydrogenase-2 small chain [Proteobacteria] Length: 372\nNumber of predicted TMHs: 1\nExp number of AAs in TMHs: 23.72226\nExp number, first 60 AAs: 0.22038\nTotal prob of N-in: 0.12994\noutside 1 331\nTMhelix 332 354\ninside 355 372

13075 GCF_000235285.1_Esch_coli_C227-11_V2 Escherichia coli O104:H4 str. C227-11 Proteobacteria; Gammaproteobacteria; Enterobacterales; Enterobacteriaceae; Escherichia; Escherichia coli; Escherichia coli O104 MNNEETFYQAMRRQGVTRRSFLKYCSLAATSLGLGAGMAPKIAWALENKPRIPVVWIHGL WP_001058323.1 MULTISPECIES: hydrogenase-1 small chain [Proteobacteria] Length: 372\nNumber of predicted TMHs: 2\nExp number of AAs in TMHs: 37.2608499999999\nExp number, first 60 AAs: 19.52923\nTotal prob of N-in: 0.94351\nPOSSIBLE N-term signal sequence\ninside 1 19\nTMhelix 20 39\noutside 40 325\nTMhelix 326 348\ninside 349 372

13076 GCF_000986765.1_ASM98676v1 Escherichia coli O104:H4 str. C227-11 Proteobacteria; Gammaproteobacteria; Enterobacterales; Enterobacteriaceae; Escherichia; Escherichia coli; Escherichia coli O104 MNNEETFYQAMRRQGVTRRSFLKYCSLAATSLGLGAGMAPKIAWALENKPRIPVVWIHGL WP_001058323.1 MULTISPECIES: hydrogenase-1 small chain [Proteobacteria] Length: 372\nNumber of predicted TMHs: 2\nExp number of AAs in TMHs: 37.2608499999999\nExp number, first 60 AAs: 19.52923\nTotal prob of N-in: 0.94351\nPOSSIBLE N-term signal sequence\ninside 1 19\nTMhelix 20 39\noutside 40 325\nTMhelix 326 348\ninside 349 372

13077 GCF_000986765.1_ASM98676v1 Escherichia coli O104:H4 str. C227-11 Proteobacteria; Gammaproteobacteria; Enterobacterales; Enterobacteriaceae; Escherichia; Escherichia coli; Escherichia coli O104 MTGDNTLIHSHGINRRDFMKLCAALAATMGLSSKAAAEMAESVTNPQRPPVIWIGAQECT WP_000145410.1 MULTISPECIES: hydrogenase-2 small chain [Proteobacteria] Length: 372\nNumber of predicted TMHs: 1\nExp number of AAs in TMHs: 23.72226\nExp number, first 60 AAs: 0.22038\nTotal prob of N-in: 0.12994\noutside 1 331\nTMhelix 332 354\ninside 355 372

13078 GCF_000235265.1_Esch_coli_C236-11_V2 Escherichia coli O104:H4 str. C236-11 Proteobacteria; Gammaproteobacteria; Enterobacterales; Enterobacteriaceae; Escherichia; Escherichia coli; Escherichia coli O104 MTGDNTLIHSHGINRRDFMKLCAALAATMGLSSKAAAEMAESVTNPQRPPVIWIGAQECT WP_000145410.1 MULTISPECIES: hydrogenase-2 small chain [Proteobacteria] Length: 372\nNumber of predicted TMHs: 1\nExp number of AAs in TMHs: 23.72226\nExp number, first 60 AAs: 0.22038\nTotal prob of N-in: 0.12994\noutside 1 331\nTMhelix 332 354\ninside 355 372

13079 GCF_000235265.1_Esch_coli_C236-11_V2 Escherichia coli O104:H4 str. C236-11 Proteobacteria; Gammaproteobacteria; Enterobacterales; Enterobacteriaceae; Escherichia; Escherichia coli; Escherichia coli O104 MNNEETFYQAMRRQGVTRRSFLKYCSLAATSLGLGAGMAPKIAWALENKPRIPVVWIHGL WP_001058323.1 MULTISPECIES: hydrogenase-1 small chain [Proteobacteria] Length: 372\nNumber of predicted TMHs: 2\nExp number of AAs in TMHs: 37.2608499999999\nExp number, first 60 AAs: 19.52923\nTotal prob of N-in: 0.94351\nPOSSIBLE N-term signal sequence\ninside 1 19\nTMhelix 20 39\noutside 40 325\nTMhelix 326 348\ninside 349 372

13080 GCF_000350005.1_E112_10NrN01 Escherichia coli O104:H4 str. E112/10 Proteobacteria; Gammaproteobacteria; Enterobacterales; Enterobacteriaceae; Escherichia; Escherichia coli; Escherichia coli O104 MNNEETFYQAMRRQGVTRRSFLKYCSLAATSLGLGAGMAPKIAWALENKPRIPVVWIHGL WP_001058323.1 MULTISPECIES: hydrogenase-1 small chain [Proteobacteria] Length: 372\nNumber of predicted TMHs: 2\nExp number of AAs in TMHs: 37.2608499999999\nExp number, first 60 AAs: 19.52923\nTotal prob of N-in: 0.94351\nPOSSIBLE N-term signal sequence\ninside 1 19\nTMhelix 20 39\noutside 40 325\nTMhelix 326 348\ninside 349 372

13081 GCF_000350005.1_E112_10NrN01 Escherichia coli O104:H4 str. E112/10 Proteobacteria; Gammaproteobacteria; Enterobacterales; Enterobacteriaceae; Escherichia; Escherichia coli; Escherichia coli O104 MTGDNTLIHSHGINRRDFMKLCAALAATMGLSSKAAAEMAESVTNPQRPPVIWIGAQECT WP_000145410.1 MULTISPECIES: hydrogenase-2 small chain [Proteobacteria] Length: 372\nNumber of predicted TMHs: 1\nExp number of AAs in TMHs: 23.72226\nExp number, first 60 AAs: 0.22038\nTotal prob of N-in: 0.12994\noutside 1 331\nTMhelix 332 354\ninside 355 372

13082 GCF_000353545.1_E92_11NrN01 Escherichia coli O104:H4 str. E92/11 Proteobacteria; Gammaproteobacteria; Enterobacterales; Enterobacteriaceae; Escherichia; Escherichia coli; Escherichia coli O104 MNNEETFYQAMRRQGVTRRSFLKYCSLAATSLGLGAGMAPKIAWALENKPRIPVVWIHGL WP_001058323.1 MULTISPECIES: hydrogenase-1 small chain [Proteobacteria] Length: 372\nNumber of predicted TMHs: 2\nExp number of AAs in TMHs: 37.2608499999999\nExp number, first 60 AAs: 19.52923\nTotal prob of N-in: 0.94351\nPOSSIBLE N-term signal sequence\ninside 1 19\nTMhelix 20 39\noutside 40 325\nTMhelix 326 348\ninside 349 372

13083 GCF_000353545.1_E92_11NrN01 Escherichia coli O104:H4 str. E92/11 Proteobacteria; Gammaproteobacteria; Enterobacterales; Enterobacteriaceae; Escherichia; Escherichia coli; Escherichia coli O104 MTGDNTLIHSHGINRRDFMKLCAALAATMGLSSKAAAEMAESVTNPQRPPVIWIGAQECT WP_000145410.1 MULTISPECIES: hydrogenase-2 small chain [Proteobacteria] Length: 372\nNumber of predicted TMHs: 1\nExp number of AAs in TMHs: 23.72226\nExp number, first 60 AAs: 0.22038\nTotal prob of N-in: 0.12994\noutside 1 331\nTMhelix 332 354\ninside 355 372

13084 GCF_000319915.1_Esch_coli_Ec11-4984_V1 Escherichia coli O104:H4 str. Ec11-4984 Proteobacteria; Gammaproteobacteria; Enterobacterales; Enterobacteriaceae; Escherichia; Escherichia coli; Escherichia coli O104 MNNEETFYQAMRRQGVTRRSFLKYCSLAATSLGLGAGMAPKIAWALENKPRIPVVWIHGL WP_001058323.1 MULTISPECIES: hydrogenase-1 small chain [Proteobacteria] Length: 372\nNumber of predicted TMHs: 2\nExp number of AAs in TMHs: 37.2608499999999\nExp number, first 60 AAs: 19.52923\nTotal prob of N-in: 0.94351\nPOSSIBLE N-term signal sequence\ninside 1 19\nTMhelix 20 39\noutside 40 325\nTMhelix 326 348\ninside 349 372

13085 GCF_000319915.1_Esch_coli_Ec11-4984_V1 Escherichia coli O104:H4 str. Ec11-4984 Proteobacteria; Gammaproteobacteria; Enterobacterales; Enterobacteriaceae; Escherichia; Escherichia coli; Escherichia coli O104 MTGDNTLIHSHGINRRDFMKLCAALAATMGLSSKAAAEMAESVTNPQRPPVIWIGAQECT WP_000145410.1 MULTISPECIES: hydrogenase-2 small chain [Proteobacteria] Length: 372\nNumber of predicted TMHs: 1\nExp number of AAs in TMHs: 23.72226\nExp number, first 60 AAs: 0.22038\nTotal prob of N-in: 0.12994\noutside 1 331\nTMhelix 332 354\ninside 355 372

13086 GCF_000319955.1_Esch_coli_Ec11-4986_V1 Escherichia coli O104:H4 str. Ec11-4986 Proteobacteria; Gammaproteobacteria; Enterobacterales; Enterobacteriaceae; Escherichia; Escherichia coli; Escherichia coli O104 MTGDNTLIHSHGINRRDFMKLCAALAATMGLSSKAAAEMAESVTNPQRPPVIWIGAQECT WP_000145410.1 MULTISPECIES: hydrogenase-2 small chain [Proteobacteria] Length: 372\nNumber of predicted TMHs: 1\nExp number of AAs in TMHs: 23.72226\nExp number, first 60 AAs: 0.22038\nTotal prob of N-in: 0.12994\noutside 1 331\nTMhelix 332 354\ninside 355 372

13087 GCF_000319955.1_Esch_coli_Ec11-4986_V1 Escherichia coli O104:H4 str. Ec11-4986 Proteobacteria; Gammaproteobacteria; Enterobacterales; Enterobacteriaceae; Escherichia; Escherichia coli; Escherichia coli O104 MNNEETFYQAMRRQGVTRRSFLKYCSLAATSLGLGAGMAPKIAWALENKPRIPVVWIHGL WP_001058323.1 MULTISPECIES: hydrogenase-1 small chain [Proteobacteria] Length: 372\nNumber of predicted TMHs: 2\nExp number of AAs in TMHs: 37.2608499999999\nExp number, first 60 AAs: 19.52923\nTotal prob of N-in: 0.94351\nPOSSIBLE N-term signal sequence\ninside 1 19\nTMhelix 20 39\noutside 40 325\nTMhelix 326 348\ninside 349 372

13088 GCF_000319975.1_Esch_coli_Ec11-4987_V1 Escherichia coli O104:H4 str. Ec11-4987 Proteobacteria; Gammaproteobacteria; Enterobacterales; Enterobacteriaceae; Escherichia; Escherichia coli; Escherichia coli O104 MTGDNTLIHSHGINRRDFMKLCAALAATMGLSSKAAAEMAESVTNPQRPPVIWIGAQECT WP_000145410.1 MULTISPECIES: hydrogenase-2 small chain [Proteobacteria] Length: 372\nNumber of predicted TMHs: 1\nExp number of AAs in TMHs: 23.72226\nExp number, first 60 AAs: 0.22038\nTotal prob of N-in: 0.12994\noutside 1 331\nTMhelix 332 354\ninside 355 372

13089 GCF_000319975.1_Esch_coli_Ec11-4987_V1 Escherichia coli O104:H4 str. Ec11-4987 Proteobacteria; Gammaproteobacteria; Enterobacterales; Enterobacteriaceae; Escherichia; Escherichia coli; Escherichia coli O104 MNNEETFYQAMRRQGVTRRSFLKYCSLAATSLGLGAGMAPKIAWALENKPRIPVVWIHGL WP_001058323.1 MULTISPECIES: hydrogenase-1 small chain [Proteobacteria] Length: 372\nNumber of predicted TMHs: 2\nExp number of AAs in TMHs: 37.2608499999999\nExp number, first 60 AAs: 19.52923\nTotal prob of N-in: 0.94351\nPOSSIBLE N-term signal sequence\ninside 1 19\nTMhelix 20 39\noutside 40 325\nTMhelix 326 348\ninside 349 372

13090 GCF_000320195.1_Esch_coli_Ec11-4988_V1 Escherichia coli O104:H4 str. Ec11-4988 Proteobacteria; Gammaproteobacteria; Enterobacterales; Enterobacteriaceae; Escherichia; Escherichia coli; Escherichia coli O104 MTGDNTLIHSHGINRRDFMKLCAALAATMGLSSKAAAEMAESVTNPQRPPVIWIGAQECT WP_000145410.1 MULTISPECIES: hydrogenase-2 small chain [Proteobacteria] Length: 372\nNumber of predicted TMHs: 1\nExp number of AAs in TMHs: 23.72226\nExp number, first 60 AAs: 0.22038\nTotal prob of N-in: 0.12994\noutside 1 331\nTMhelix 332 354\ninside 355 372

13091 GCF_000320195.1_Esch_coli_Ec11-4988_V1 Escherichia coli O104:H4 str. Ec11-4988 Proteobacteria; Gammaproteobacteria; Enterobacterales; Enterobacteriaceae; Escherichia; Escherichia coli; Escherichia coli O104 MNNEETFYQAMRRQGVTRRSFLKYCSLAATSLGLGAGMAPKIAWALENKPRIPVVWIHGL WP_001058323.1 MULTISPECIES: hydrogenase-1 small chain [Proteobacteria] Length: 372\nNumber of predicted TMHs: 2\nExp number of AAs in TMHs: 37.2608499999999\nExp number, first 60 AAs: 19.52923\nTotal prob of N-in: 0.94351\nPOSSIBLE N-term signal sequence\ninside 1 19\nTMhelix 20 39\noutside 40 325\nTMhelix 326 348\ninside 349 372

13092 GCF_000320215.1_Esch_coli_Ec11-5603_V1 Escherichia coli O104:H4 str. Ec11-5603 Proteobacteria; Gammaproteobacteria; Enterobacterales; Enterobacteriaceae; Escherichia; Escherichia coli; Escherichia coli O104 MNNEETFYQAMRRQGVTRRSFLKYCSLAATSLGLGAGMAPKIAWALENKPRIPVVWIHGL WP_001058323.1 MULTISPECIES: hydrogenase-1 small chain [Proteobacteria] Length: 372\nNumber of predicted TMHs: 2\nExp number of AAs in TMHs: 37.2608499999999\nExp number, first 60 AAs: 19.52923\nTotal prob of N-in: 0.94351\nPOSSIBLE N-term signal sequence\ninside 1 19\nTMhelix 20 39\noutside 40 325\nTMhelix 326 348\ninside 349 372

13093 GCF_000320215.1_Esch_coli_Ec11-5603_V1 Escherichia coli O104:H4 str. Ec11-5603 Proteobacteria; Gammaproteobacteria; Enterobacterales; Enterobacteriaceae; Escherichia; Escherichia coli; Escherichia coli O104 MTGDNTLIHSHGINRRDFMKLCAALAATMGLSSKAAAEMAESVTNPQRPPVIWIGAQECT WP_000145410.1 MULTISPECIES: hydrogenase-2 small chain [Proteobacteria] Length: 372\nNumber of predicted TMHs: 1\nExp number of AAs in TMHs: 23.72226\nExp number, first 60 AAs: 0.22038\nTotal prob of N-in: 0.12994\noutside 1 331\nTMhelix 332 354\ninside 355 372

13094 GCF_000319935.1_Esch_coli_Ec11-5604_V1 Escherichia coli O104:H4 str. Ec11-5604 Proteobacteria; Gammaproteobacteria; Enterobacterales; Enterobacteriaceae; Escherichia; Escherichia coli; Escherichia coli O104 MTGDNTLIHSHGINRRDFMKLCAALAATMGLSSKAAAEMAESVTNPQRPPVIWIGAQECT WP_000145410.1 MULTISPECIES: hydrogenase-2 small chain [Proteobacteria] Length: 372\nNumber of predicted TMHs: 1\nExp number of AAs in TMHs: 23.72226\nExp number, first 60 AAs: 0.22038\nTotal prob of N-in: 0.12994\noutside 1 331\nTMhelix 332 354\ninside 355 372

13095 GCF_000319935.1_Esch_coli_Ec11-5604_V1 Escherichia coli O104:H4 str. Ec11-5604 Proteobacteria; Gammaproteobacteria; Enterobacterales; Enterobacteriaceae; Escherichia; Escherichia coli; Escherichia coli O104 MNNEETFYQAMRRQGVTRRSFLKYCSLAATSLGLGAGMAPKIAWALENKPRIPVVWIHGL WP_001058323.1 MULTISPECIES: hydrogenase-1 small chain [Proteobacteria] Length: 372\nNumber of predicted TMHs: 2\nExp number of AAs in TMHs: 37.2608499999999\nExp number, first 60 AAs: 19.52923\nTotal prob of N-in: 0.94351\nPOSSIBLE N-term signal sequence\ninside 1 19\nTMhelix 20 39\noutside 40 325\nTMhelix 326 348\ninside 349 372

13096 GCF_000320235.1_Esch_coli_Ec11-6006_V1 Escherichia coli O104:H4 str. Ec11-6006 Proteobacteria; Gammaproteobacteria; Enterobacterales; Enterobacteriaceae; Escherichia; Escherichia coli; Escherichia coli O104 MNNEETFYQAMRRQGVTRRSFLKYCSLAATSLGLGAGMAPKIAWALENKPRIPVVWIHGL WP_001058323.1 MULTISPECIES: hydrogenase-1 small chain [Proteobacteria] Length: 372\nNumber of predicted TMHs: 2\nExp number of AAs in TMHs: 37.2608499999999\nExp number, first 60 AAs: 19.52923\nTotal prob of N-in: 0.94351\nPOSSIBLE N-term signal sequence\ninside 1 19\nTMhelix 20 39\noutside 40 325\nTMhelix 326 348\ninside 349 372

13097 GCF_000320235.1_Esch_coli_Ec11-6006_V1 Escherichia coli O104:H4 str. Ec11-6006 Proteobacteria; Gammaproteobacteria; Enterobacterales; Enterobacteriaceae; Escherichia; Escherichia coli; Escherichia coli O104 MTGDNTLIHSHGINRRDFMKLCAALAATMGLSSKAAAEMAESVTNPQRPPVIWIGAQECT WP_000145410.1 MULTISPECIES: hydrogenase-2 small chain [Proteobacteria] Length: 372\nNumber of predicted TMHs: 1\nExp number of AAs in TMHs: 23.72226\nExp number, first 60 AAs: 0.22038\nTotal prob of N-in: 0.12994\noutside 1 331\nTMhelix 332 354\ninside 355 372

13098 GCF_000319855.1_Esch_coli_Ec_11-9450_V2 Escherichia coli O104:H4 str. Ec11-9450 Proteobacteria; Gammaproteobacteria; Enterobacterales; Enterobacteriaceae; Escherichia; Escherichia coli; Escherichia coli O104 MTGDNTLIHSHGINRRDFMKLCAALAATMGLSSKAAAEMAESVTNPQRPPVIWIGAQECT WP_000145410.1 MULTISPECIES: hydrogenase-2 small chain [Proteobacteria] Length: 372\nNumber of predicted TMHs: 1\nExp number of AAs in TMHs: 23.72226\nExp number, first 60 AAs: 0.22038\nTotal prob of N-in: 0.12994\noutside 1 331\nTMhelix 332 354\ninside 355 372

13099 GCF_000319855.1_Esch_coli_Ec_11-9450_V2 Escherichia coli O104:H4 str. Ec11-9450 Proteobacteria; Gammaproteobacteria; Enterobacterales; Enterobacteriaceae; Escherichia; Escherichia coli; Escherichia coli O104 MNNEETFYQAMRRQGVTRRSFLKYCSLAATSLGLGAGMAPKIAWALENKPRIPVVWIHGL WP_001058323.1 MULTISPECIES: hydrogenase-1 small chain [Proteobacteria] Length: 372\nNumber of predicted TMHs: 2\nExp number of AAs in TMHs: 37.2608499999999\nExp number, first 60 AAs: 19.52923\nTotal prob of N-in: 0.94351\nPOSSIBLE N-term signal sequence\ninside 1 19\nTMhelix 20 39\noutside 40 325\nTMhelix 326 348\ninside 349 372

13100 GCF_000320045.1_Esch_coli_Ec_11-9941_V6 Escherichia coli O104:H4 str. Ec11-9941 Proteobacteria; Gammaproteobacteria; Enterobacterales; Enterobacteriaceae; Escherichia; Escherichia coli; Escherichia coli O104 MNNEETFYQAMRRQGVTRRSFLKYCSLAATSLGLGAGMAPKIAWALENKPRIPVVWIHGL WP_001058323.1 MULTISPECIES: hydrogenase-1 small chain [Proteobacteria] Length: 372\nNumber of predicted TMHs: 2\nExp number of AAs in TMHs: 37.2608499999999\nExp number, first 60 AAs: 19.52923\nTotal prob of N-in: 0.94351\nPOSSIBLE N-term signal sequence\ninside 1 19\nTMhelix 20 39\noutside 40 325\nTMhelix 326 348\ninside 349 372

13101 GCF_000320045.1_Esch_coli_Ec_11-9941_V6 Escherichia coli O104:H4 str. Ec11-9941 Proteobacteria; Gammaproteobacteria; Enterobacterales; Enterobacteriaceae; Escherichia; Escherichia coli; Escherichia coli O104 MTGDNTLIHSHGINRRDFMKLCAALAATMGLSSKAAAEMAESVTNPQRPPVIWIGAQECT WP_000145410.1 MULTISPECIES: hydrogenase-2 small chain [Proteobacteria] Length: 372\nNumber of predicted TMHs: 1\nExp number of AAs in TMHs: 23.72226\nExp number, first 60 AAs: 0.22038\nTotal prob of N-in: 0.12994\noutside 1 331\nTMhelix 332 354\ninside 355 372

13102 GCF_000319875.1_Esch_coli_Ec_11-9990_V2 Escherichia coli O104:H4 str. Ec11-9990 Proteobacteria; Gammaproteobacteria; Enterobacterales; Enterobacteriaceae; Escherichia; Escherichia coli; Escherichia coli O104 MTGDNTLIHSHGINRRDFMKLCAALAATMGLSSKAAAEMAESVTNPQRPPVIWIGAQECT WP_000145410.1 MULTISPECIES: hydrogenase-2 small chain [Proteobacteria] Length: 372\nNumber of predicted TMHs: 1\nExp number of AAs in TMHs: 23.72226\nExp number, first 60 AAs: 0.22038\nTotal prob of N-in: 0.12994\noutside 1 331\nTMhelix 332 354\ninside 355 372

13103 GCF_000319875.1_Esch_coli_Ec_11-9990_V2 Escherichia coli O104:H4 str. Ec11-9990 Proteobacteria; Gammaproteobacteria; Enterobacterales; Enterobacteriaceae; Escherichia; Escherichia coli; Escherichia coli O104 MNNEETFYQAMRRQGVTRRSFLKYCSLAATSLGLGAGMAPKIAWALENKPRIPVVWIHGL WP_001058323.1 MULTISPECIES: hydrogenase-1 small chain [Proteobacteria] Length: 372\nNumber of predicted TMHs: 2\nExp number of AAs in TMHs: 37.2608499999999\nExp number, first 60 AAs: 19.52923\nTotal prob of N-in: 0.94351\nPOSSIBLE N-term signal sequence\ninside 1 19\nTMhelix 20 39\noutside 40 325\nTMhelix 326 348\ninside 349 372

13104 GCF_000320255.1_Esch_coli_Ec12-0465_V1 Escherichia coli O104:H4 str. Ec12-0465 Proteobacteria; Gammaproteobacteria; Enterobacterales; Enterobacteriaceae; Escherichia; Escherichia coli; Escherichia coli O104 MNNEETFYQAMRRQGVTRRSFLKYCSLAATSLGLGAGMAPKIAWALENKPRIPVVWIHGL WP_001058323.1 MULTISPECIES: hydrogenase-1 small chain [Proteobacteria] Length: 372\nNumber of predicted TMHs: 2\nExp number of AAs in TMHs: 37.2608499999999\nExp number, first 60 AAs: 19.52923\nTotal prob of N-in: 0.94351\nPOSSIBLE N-term signal sequence\ninside 1 19\nTMhelix 20 39\noutside 40 325\nTMhelix 326 348\ninside 349 372

13105 GCF_000320255.1_Esch_coli_Ec12-0465_V1 Escherichia coli O104:H4 str. Ec12-0465 Proteobacteria; Gammaproteobacteria; Enterobacterales; Enterobacteriaceae; Escherichia; Escherichia coli; Escherichia coli O104 MTGDNTLIHSHGINRRDFMKLCAALAATMGLSSKAAAEMAESVTNPQRPPVIWIGAQECT WP_000145410.1 MULTISPECIES: hydrogenase-2 small chain [Proteobacteria] Length: 372\nNumber of predicted TMHs: 1\nExp number of AAs in TMHs: 23.72226\nExp number, first 60 AAs: 0.22038\nTotal prob of N-in: 0.12994\noutside 1 331\nTMhelix 332 354\ninside 355 372

13106 GCF_000320275.1_Esch_coli_Ec12-0466_V1 Escherichia coli O104:H4 str. Ec12-0466 Proteobacteria; Gammaproteobacteria; Enterobacterales; Enterobacteriaceae; Escherichia; Escherichia coli; Escherichia coli O104 MNNEETFYQAMRRQGVTRRSFLKYCSLAATSLGLGAGMAPKIAWALENKPRIPVVWIHGL WP_001058323.1 MULTISPECIES: hydrogenase-1 small chain [Proteobacteria] Length: 372\nNumber of predicted TMHs: 2\nExp number of AAs in TMHs: 37.2608499999999\nExp number, first 60 AAs: 19.52923\nTotal prob of N-in: 0.94351\nPOSSIBLE N-term signal sequence\ninside 1 19\nTMhelix 20 39\noutside 40 325\nTMhelix 326 348\ninside 349 372

13107 GCF_000320275.1_Esch_coli_Ec12-0466_V1 Escherichia coli O104:H4 str. Ec12-0466 Proteobacteria; Gammaproteobacteria; Enterobacterales; Enterobacteriaceae; Escherichia; Escherichia coli; Escherichia coli O104 MTGDNTLIHSHGINRRDFMKLCAALAATMGLSSKAAAEMAESVTNPQRPPVIWIGAQECT WP_000145410.1 MULTISPECIES: hydrogenase-2 small chain [Proteobacteria] Length: 372\nNumber of predicted TMHs: 1\nExp number of AAs in TMHs: 23.72226\nExp number, first 60 AAs: 0.22038\nTotal prob of N-in: 0.12994\noutside 1 331\nTMhelix 332 354\ninside 355 372

13108 GCF_000223015.1_ASM22301v2 Escherichia coli O104:H4 str. GOS1 Proteobacteria; Gammaproteobacteria; Enterobacterales; Enterobacteriaceae; Escherichia; Escherichia coli; Escherichia coli O104 MNNEETFYQAMRRQGVTRRSFLKYCSLAATSLGLGAGMAPKIAWALENKPRIPVVWIHGL WP_001058323.1 MULTISPECIES: hydrogenase-1 small chain [Proteobacteria] Length: 372\nNumber of predicted TMHs: 2\nExp number of AAs in TMHs: 37.2608499999999\nExp number, first 60 AAs: 19.52923\nTotal prob of N-in: 0.94351\nPOSSIBLE N-term signal sequence\ninside 1 19\nTMhelix 20 39\noutside 40 325\nTMhelix 326 348\ninside 349 372

13109 GCF_000223015.1_ASM22301v2 Escherichia coli O104:H4 str. GOS1 Proteobacteria; Gammaproteobacteria; Enterobacterales; Enterobacteriaceae; Escherichia; Escherichia coli; Escherichia coli O104 MTGDNTLIHSHGINRRDFMKLCAALAATMGLSSKAAAEMAESVTNPQRPPVIWIGAQECT WP_000145410.1 MULTISPECIES: hydrogenase-2 small chain [Proteobacteria] Length: 372\nNumber of predicted TMHs: 1\nExp number of AAs in TMHs: 23.72226\nExp number, first 60 AAs: 0.22038\nTotal prob of N-in: 0.12994\noutside 1 331\nTMhelix 332 354\ninside 355 372

13110 GCF_000223035.1_ASM22303v2 Escherichia coli O104:H4 str. GOS2 Proteobacteria; Gammaproteobacteria; Enterobacterales; Enterobacteriaceae; Escherichia; Escherichia coli; Escherichia coli O104 MNNEETFYQAMRRQGVTRRSFLKYCSLAATSLGLGAGMAPKIAWALENKPRIPVVWIHGL WP_001058323.1 MULTISPECIES: hydrogenase-1 small chain [Proteobacteria] Length: 372\nNumber of predicted TMHs: 2\nExp number of AAs in TMHs: 37.2608499999999\nExp number, first 60 AAs: 19.52923\nTotal prob of N-in: 0.94351\nPOSSIBLE N-term signal sequence\ninside 1 19\nTMhelix 20 39\noutside 40 325\nTMhelix 326 348\ninside 349 372

13111 GCF_000223035.1_ASM22303v2 Escherichia coli O104:H4 str. GOS2 Proteobacteria; Gammaproteobacteria; Enterobacterales; Enterobacteriaceae; Escherichia; Escherichia coli; Escherichia coli O104 MTGDNTLIHSHGINRRDFMKLCAALAATMGLSSKAAAEMAESVTNPQRPPVIWIGAQECT WP_000145410.1 MULTISPECIES: hydrogenase-2 small chain [Proteobacteria] Length: 372\nNumber of predicted TMHs: 1\nExp number of AAs in TMHs: 23.72226\nExp number, first 60 AAs: 0.22038\nTotal prob of N-in: 0.12994\noutside 1 331\nTMhelix 332 354\ninside 355 372

13112 GCF_000217975.1_ASM21797v2 Escherichia coli O104:H4 str. H112180280 Proteobacteria; Gammaproteobacteria; Enterobacterales; Enterobacteriaceae; Escherichia; Escherichia coli; Escherichia coli O104 MNNEETFYQAMRRQGVTRRSFLKYCSLAATSLGLGAGMAPKIAWALENKPRIPVVWIHGL WP_001058323.1 MULTISPECIES: hydrogenase-1 small chain [Proteobacteria] Length: 372\nNumber of predicted TMHs: 2\nExp number of AAs in TMHs: 37.2608499999999\nExp number, first 60 AAs: 19.52923\nTotal prob of N-in: 0.94351\nPOSSIBLE N-term signal sequence\ninside 1 19\nTMhelix 20 39\noutside 40 325\nTMhelix 326 348\ninside 349 372

13113 GCF_000217975.1_ASM21797v2 Escherichia coli O104:H4 str. H112180280 Proteobacteria; Gammaproteobacteria; Enterobacterales; Enterobacteriaceae; Escherichia; Escherichia coli; Escherichia coli O104 MTGDNTLIHSHGINRRDFMKLCAALAATMGLSSKAAAEMAESVTNPQRPPVIWIGAQECT WP_000145410.1 MULTISPECIES: hydrogenase-2 small chain [Proteobacteria] Length: 372\nNumber of predicted TMHs: 1\nExp number of AAs in TMHs: 23.72226\nExp number, first 60 AAs: 0.22038\nTotal prob of N-in: 0.12994\noutside 1 331\nTMhelix 332 354\ninside 355 372

13114 GCF_000222505.1_ASM22250v2 Escherichia coli O104:H4 str. H112180282 Proteobacteria; Gammaproteobacteria; Enterobacterales; Enterobacteriaceae; Escherichia; Escherichia coli; Escherichia coli O104 MTGDNTLIHSHGINRRDFMKLCAALAATMGLSSKAAAEMAESVTNPQRPPVIWIGAQECT WP_000145410.1 MULTISPECIES: hydrogenase-2 small chain [Proteobacteria] Length: 372\nNumber of predicted TMHs: 1\nExp number of AAs in TMHs: 23.72226\nExp number, first 60 AAs: 0.22038\nTotal prob of N-in: 0.12994\noutside 1 331\nTMhelix 332 354\ninside 355 372

13115 GCF_000222505.1_ASM22250v2 Escherichia coli O104:H4 str. H112180282 Proteobacteria; Gammaproteobacteria; Enterobacterales; Enterobacteriaceae; Escherichia; Escherichia coli; Escherichia coli O104 MNNEETFYQAMRRQGVTRRSFLKYCSLAATSLGLGAGMAPKIAWALENKPRIPVVWIHGL WP_001058323.1 MULTISPECIES: hydrogenase-1 small chain [Proteobacteria] Length: 372\nNumber of predicted TMHs: 2\nExp number of AAs in TMHs: 37.2608499999999\nExp number, first 60 AAs: 19.52923\nTotal prob of N-in: 0.94351\nPOSSIBLE N-term signal sequence\ninside 1 19\nTMhelix 20 39\noutside 40 325\nTMhelix 326 348\ninside 349 372

13116 GCF_000258635.1_ASM25863v1 Escherichia coli O104:H4 str. ON2010 Proteobacteria; Gammaproteobacteria; Enterobacterales; Enterobacteriaceae; Escherichia; Escherichia coli; Escherichia coli O104 MTGDNTLIHSHGINRRDFMKLCAALAATMGLSSKAAAEMAESVTNPQRPPVIWIGAQECT WP_000145410.1 MULTISPECIES: hydrogenase-2 small chain [Proteobacteria] Length: 372\nNumber of predicted TMHs: 1\nExp number of AAs in TMHs: 23.72226\nExp number, first 60 AAs: 0.22038\nTotal prob of N-in: 0.12994\noutside 1 331\nTMhelix 332 354\ninside 355 372

13117 GCF_000258635.1_ASM25863v1 Escherichia coli O104:H4 str. ON2010 Proteobacteria; Gammaproteobacteria; Enterobacterales; Enterobacteriaceae; Escherichia; Escherichia coli; Escherichia coli O104 MNNEETFYQAMRRQGVTRRSFLKYCSLAATSLGLGAGMAPKIAWALENKPRIPVVWIHGL WP_001058323.1 MULTISPECIES: hydrogenase-1 small chain [Proteobacteria] Length: 372\nNumber of predicted TMHs: 2\nExp number of AAs in TMHs: 37.2608499999999\nExp number, first 60 AAs: 19.52923\nTotal prob of N-in: 0.94351\nPOSSIBLE N-term signal sequence\ninside 1 19\nTMhelix 20 39\noutside 40 325\nTMhelix 326 348\ninside 349 372

13118 GCF_000258615.1_ASM25861v1 Escherichia coli O104:H4 str. ON2011 Proteobacteria; Gammaproteobacteria; Enterobacterales; Enterobacteriaceae; Escherichia; Escherichia coli; Escherichia coli O104 MNNEETFYQAMRRQGVTRRSFLKYCSLAATSLGLGAGMAPKIAWALENKPRIPVVWIHGL WP_001058323.1 MULTISPECIES: hydrogenase-1 small chain [Proteobacteria] Length: 372\nNumber of predicted TMHs: 2\nExp number of AAs in TMHs: 37.2608499999999\nExp number, first 60 AAs: 19.52923\nTotal prob of N-in: 0.94351\nPOSSIBLE N-term signal sequence\ninside 1 19\nTMhelix 20 39\noutside 40 325\nTMhelix 326 348\ninside 349 372

13119 GCF_000258615.1_ASM25861v1 Escherichia coli O104:H4 str. ON2011 Proteobacteria; Gammaproteobacteria; Enterobacterales; Enterobacteriaceae; Escherichia; Escherichia coli; Escherichia coli O104 MTGDNTLIHSHGINRRDFMKLCAALAATMGLSSKAAAEMAESVTNPQRPPVIWIGAQECT WP_000145410.1 MULTISPECIES: hydrogenase-2 small chain [Proteobacteria] Length: 372\nNumber of predicted TMHs: 1\nExp number of AAs in TMHs: 23.72226\nExp number, first 60 AAs: 0.22038\nTotal prob of N-in: 0.12994\noutside 1 331\nTMhelix 332 354\ninside 355 372

13120 GCF_000217695.1_ASM21769v2 Escherichia coli O104:H4 str. TY-2482 Proteobacteria; Gammaproteobacteria; Enterobacterales; Enterobacteriaceae; Escherichia; Escherichia coli; Escherichia coli O104 MNNEETFYQAMRRQGVTRRSFLKYCSLAATSLGLGAGMAPKIAWALENKPRIPVVWIHGL WP_001058323.1 MULTISPECIES: hydrogenase-1 small chain [Proteobacteria] Length: 372\nNumber of predicted TMHs: 2\nExp number of AAs in TMHs: 37.2608499999999\nExp number, first 60 AAs: 19.52923\nTotal prob of N-in: 0.94351\nPOSSIBLE N-term signal sequence\ninside 1 19\nTMhelix 20 39\noutside 40 325\nTMhelix 326 348\ninside 349 372

13121 GCF_000217695.1_ASM21769v2 Escherichia coli O104:H4 str. TY-2482 Proteobacteria; Gammaproteobacteria; Enterobacterales; Enterobacteriaceae; Escherichia; Escherichia coli; Escherichia coli O104 MTGDNTLIHSHGINRRDFMKLCAALAATMGLSSKAAAEMAESVTNPQRPPVIWIGAQECT WP_000145410.1 MULTISPECIES: hydrogenase-2 small chain [Proteobacteria] Length: 372\nNumber of predicted TMHs: 1\nExp number of AAs in TMHs: 23.72226\nExp number, first 60 AAs: 0.22038\nTotal prob of N-in: 0.12994\noutside 1 331\nTMhelix 332 354\ninside 355 372

13122 GCF_000221885.1_E.coli_0104_H4_Illumina_1.0 Escherichia coli O104:H4 str. TY-2482 Proteobacteria; Gammaproteobacteria; Enterobacterales; Enterobacteriaceae; Escherichia; Escherichia coli; Escherichia coli O104 MTGDNTLIHSHGINRRDFMKLCAALAATMGLSSKAAAEMAESVTNPQRPPVIWIGAQECT WP_000145410.1 MULTISPECIES: hydrogenase-2 small chain [Proteobacteria] Length: 372\nNumber of predicted TMHs: 1\nExp number of AAs in TMHs: 23.72226\nExp number, first 60 AAs: 0.22038\nTotal prob of N-in: 0.12994\noutside 1 331\nTMhelix 332 354\ninside 355 372

13123 GCF_000221885.1_E.coli_0104_H4_Illumina_1.0 Escherichia coli O104:H4 str. TY-2482 Proteobacteria; Gammaproteobacteria; Enterobacterales; Enterobacteriaceae; Escherichia; Escherichia coli; Escherichia coli O104 MNNEETFYQAMRRQGVTRRSFLKYCSLAATSLGLGAGMAPKIAWALENKPRIPVVWIHGL WP_001058323.1 MULTISPECIES: hydrogenase-1 small chain [Proteobacteria] Length: 372\nNumber of predicted TMHs: 2\nExp number of AAs in TMHs: 37.2608499999999\nExp number, first 60 AAs: 19.52923\nTotal prob of N-in: 0.94351\nPOSSIBLE N-term signal sequence\ninside 1 19\nTMhelix 20 39\noutside 40 325\nTMhelix 326 348\ninside 349 372

13124 GCF_000313425.1_CFSAN001630_1.0 Escherichia coli O111:H11 str. CFSAN001630 Proteobacteria; Gammaproteobacteria; Enterobacterales; Enterobacteriaceae; Escherichia; Escherichia coli; Escherichia coli O111 MNNEETFYQAMRRQGVTRRSFLKYCSLAATSLGLGAGMAPKIAWALENKPRIPVVWIHGL WP_001058323.1 MULTISPECIES: hydrogenase-1 small chain [Proteobacteria] Length: 372\nNumber of predicted TMHs: 2\nExp number of AAs in TMHs: 37.2608499999999\nExp number, first 60 AAs: 19.52923\nTotal prob of N-in: 0.94351\nPOSSIBLE N-term signal sequence\ninside 1 19\nTMhelix 20 39\noutside 40 325\nTMhelix 326 348\ninside 349 372

13125 GCF_000313425.1_CFSAN001630_1.0 Escherichia coli O111:H11 str. CFSAN001630 Proteobacteria; Gammaproteobacteria; Enterobacterales; Enterobacteriaceae; Escherichia; Escherichia coli; Escherichia coli O111 MTGDNTLIHSHGINRRDFMKLCAALAATMGLSSKAAAEMAESVTNPQRPPVIWIGAQECT WP_000145410.1 MULTISPECIES: hydrogenase-2 small chain [Proteobacteria] Length: 372\nNumber of predicted TMHs: 1\nExp number of AAs in TMHs: 23.72226\nExp number, first 60 AAs: 0.22038\nTotal prob of N-in: 0.12994\noutside 1 331\nTMhelix 332 354\ninside 355 372

13126 GCF_000263935.1_ASM26393v1 Escherichia coli O111:H11 str. CVM9534 Proteobacteria; Gammaproteobacteria; Enterobacterales; Enterobacteriaceae; Escherichia; Escherichia coli; Escherichia coli O111 MTGDNTLIHSHGINRRDFMKLCAALAATMGLSSKAAAEMAESVTNPQRPPVIWIGAQECT WP_000145410.1 MULTISPECIES: hydrogenase-2 small chain [Proteobacteria] Length: 372\nNumber of predicted TMHs: 1\nExp number of AAs in TMHs: 23.72226\nExp number, first 60 AAs: 0.22038\nTotal prob of N-in: 0.12994\noutside 1 331\nTMhelix 332 354\ninside 355 372

13127 GCF_000263935.1_ASM26393v1 Escherichia coli O111:H11 str. CVM9534 Proteobacteria; Gammaproteobacteria; Enterobacterales; Enterobacteriaceae; Escherichia; Escherichia coli; Escherichia coli O111 MNNEETFYQAMRRQGVTRRSFLKYCSLAATSLGLGAGMAPKIAWALENKPRIPVVWIHGL WP_001058323.1 MULTISPECIES: hydrogenase-1 small chain [Proteobacteria] Length: 372\nNumber of predicted TMHs: 2\nExp number of AAs in TMHs: 37.2608499999999\nExp number, first 60 AAs: 19.52923\nTotal prob of N-in: 0.94351\nPOSSIBLE N-term signal sequence\ninside 1 19\nTMhelix 20 39\noutside 40 325\nTMhelix 326 348\ninside 349 372

13128 GCF_000263955.1_ASM26395v1 Escherichia coli O111:H11 str. CVM9545 Proteobacteria; Gammaproteobacteria; Enterobacterales; Enterobacteriaceae; Escherichia; Escherichia coli; Escherichia coli O111 MNNEETFYQAMRRQGVTRRSFLKYCSLAATSLGLGAGMAPKIAWALENKPRIPVVWIHGL WP_001058323.1 MULTISPECIES: hydrogenase-1 small chain [Proteobacteria] Length: 372\nNumber of predicted TMHs: 2\nExp number of AAs in TMHs: 37.2608499999999\nExp number, first 60 AAs: 19.52923\nTotal prob of N-in: 0.94351\nPOSSIBLE N-term signal sequence\ninside 1 19\nTMhelix 20 39\noutside 40 325\nTMhelix 326 348\ninside 349 372

13129 GCF_000263955.1_ASM26395v1 Escherichia coli O111:H11 str. CVM9545 Proteobacteria; Gammaproteobacteria; Enterobacterales; Enterobacteriaceae; Escherichia; Escherichia coli; Escherichia coli O111 MTGDNTLIHSHGINRRDFMKLCAALAATMGLSSKAAAEMAESVTNPQRPPVIWIGAQECT WP_000145410.1 MULTISPECIES: hydrogenase-2 small chain [Proteobacteria] Length: 372\nNumber of predicted TMHs: 1\nExp number of AAs in TMHs: 23.72226\nExp number, first 60 AAs: 0.22038\nTotal prob of N-in: 0.12994\noutside 1 331\nTMhelix 332 354\ninside 355 372

13130 GCF_000276925.1_EC_O111_H11_CVM9553_1.0 Escherichia coli O111:H11 str. CVM9553 Proteobacteria; Gammaproteobacteria; Enterobacterales; Enterobacteriaceae; Escherichia; Escherichia coli; Escherichia coli O111 MNNEETFYQAMRRQGVTRRSFLKYCSLAATSLGLGAGMAPKIAWALENKPRIPVVWIHGL WP_001058323.1 MULTISPECIES: hydrogenase-1 small chain [Proteobacteria] Length: 372\nNumber of predicted TMHs: 2\nExp number of AAs in TMHs: 37.2608499999999\nExp number, first 60 AAs: 19.52923\nTotal prob of N-in: 0.94351\nPOSSIBLE N-term signal sequence\ninside 1 19\nTMhelix 20 39\noutside 40 325\nTMhelix 326 348\ninside 349 372

13131 GCF_000276925.1_EC_O111_H11_CVM9553_1.0 Escherichia coli O111:H11 str. CVM9553 Proteobacteria; Gammaproteobacteria; Enterobacterales; Enterobacteriaceae; Escherichia; Escherichia coli; Escherichia coli O111 MTGDNTLIHSHGINRRDFMKLCAALAATMGLSSKAAAEMAESVTNPQRPPVIWIGAQECT WP_000145410.1 MULTISPECIES: hydrogenase-2 small chain [Proteobacteria] Length: 372\nNumber of predicted TMHs: 1\nExp number of AAs in TMHs: 23.72226\nExp number, first 60 AAs: 0.22038\nTotal prob of N-in: 0.12994\noutside 1 331\nTMhelix 332 354\ninside 355 372

13132 GCF_000632635.1_Ec2009C-4126 Escherichia coli O111:H8 str. 2009C-4126 Proteobacteria; Gammaproteobacteria; Enterobacterales; Enterobacteriaceae; Escherichia; Escherichia coli; Escherichia coli O111 MNNEETFYQAMRRQGVTRRSFLKYCSLAATSLGLGAGMAPKIAWALENKPRIPVVWIHGL WP_001058323.1 MULTISPECIES: hydrogenase-1 small chain [Proteobacteria] Length: 372\nNumber of predicted TMHs: 2\nExp number of AAs in TMHs: 37.2608499999999\nExp number, first 60 AAs: 19.52923\nTotal prob of N-in: 0.94351\nPOSSIBLE N-term signal sequence\ninside 1 19\nTMhelix 20 39\noutside 40 325\nTMhelix 326 348\ninside 349 372

13133 GCF_000632635.1_Ec2009C-4126 Escherichia coli O111:H8 str. 2009C-4126 Proteobacteria; Gammaproteobacteria; Enterobacterales; Enterobacteriaceae; Escherichia; Escherichia coli; Escherichia coli O111 MTGDNTLIHSHGINRRDFMKLCAALAATMGLSSKAAAEMAESVTNPQRPPVIWIGAQECT WP_000145410.1 MULTISPECIES: hydrogenase-2 small chain [Proteobacteria] Length: 372\nNumber of predicted TMHs: 1\nExp number of AAs in TMHs: 23.72226\nExp number, first 60 AAs: 0.22038\nTotal prob of N-in: 0.12994\noutside 1 331\nTMhelix 332 354\ninside 355 372

13134 GCF_000632595.2_Ec2009EL-2169 Escherichia coli O111:H8 str. 2009EL-2169 Proteobacteria; Gammaproteobacteria; Enterobacterales; Enterobacteriaceae; Escherichia; Escherichia coli; Escherichia coli O111 MTGDNTLIHSHGINRRDFMKLCAALAATMGLSSKAAAEMAESVTNPQRPPVIWIGAQECT WP_000145410.1 MULTISPECIES: hydrogenase-2 small chain [Proteobacteria] Length: 372\nNumber of predicted TMHs: 1\nExp number of AAs in TMHs: 23.72226\nExp number, first 60 AAs: 0.22038\nTotal prob of N-in: 0.12994\noutside 1 331\nTMhelix 332 354\ninside 355 372

13135 GCF_000632595.2_Ec2009EL-2169 Escherichia coli O111:H8 str. 2009EL-2169 Proteobacteria; Gammaproteobacteria; Enterobacterales; Enterobacteriaceae; Escherichia; Escherichia coli; Escherichia coli O111 MNNEETFYQAMRRQGVTRRSFLKYCSLAATSLGLGAGMAPKIAWALENKPRIPVVWIHGL WP_001058323.1 MULTISPECIES: hydrogenase-1 small chain [Proteobacteria] Length: 372\nNumber of predicted TMHs: 2\nExp number of AAs in TMHs: 37.2608499999999\nExp number, first 60 AAs: 19.52923\nTotal prob of N-in: 0.94351\nPOSSIBLE N-term signal sequence\ninside 1 19\nTMhelix 20 39\noutside 40 325\nTMhelix 326 348\ninside 349 372

13136 GCF_000632675.2_Ec2011C-3453 Escherichia coli O111:H8 str. 2011C-3453 Proteobacteria; Gammaproteobacteria; Enterobacterales; Enterobacteriaceae; Escherichia; Escherichia coli; Escherichia coli O111 MNNEETFYQAMRRQGVTRRSFLKYCSLAATSLGLGAGMAPKIAWALENKPRIPVVWIHGL WP_001058323.1 MULTISPECIES: hydrogenase-1 small chain [Proteobacteria] Length: 372\nNumber of predicted TMHs: 2\nExp number of AAs in TMHs: 37.2608499999999\nExp number, first 60 AAs: 19.52923\nTotal prob of N-in: 0.94351\nPOSSIBLE N-term signal sequence\ninside 1 19\nTMhelix 20 39\noutside 40 325\nTMhelix 326 348\ninside 349 372

13137 GCF_000632675.2_Ec2011C-3453 Escherichia coli O111:H8 str. 2011C-3453 Proteobacteria; Gammaproteobacteria; Enterobacterales; Enterobacteriaceae; Escherichia; Escherichia coli; Escherichia coli O111 MTGDNTLIHSHGINRRDFMKLCAALAATMGLSSKAAAEMAESVTNPQRPPVIWIGAQECT WP_000145410.1 MULTISPECIES: hydrogenase-2 small chain [Proteobacteria] Length: 372\nNumber of predicted TMHs: 1\nExp number of AAs in TMHs: 23.72226\nExp number, first 60 AAs: 0.22038\nTotal prob of N-in: 0.12994\noutside 1 331\nTMhelix 332 354\ninside 355 372

13138 GCF_000313445.1_CFSAN001632_1.0 Escherichia coli O111:H8 str. CFSAN001632 Proteobacteria; Gammaproteobacteria; Enterobacterales; Enterobacteriaceae; Escherichia; Escherichia coli; Escherichia coli O111 MTGDNTLIHSHGINRRDFMKLCAALAATMGLSGKAAAEMAESVTNPQRPPVIWIGAQECT WP_009448969.1 hydrogenase 2 small subunit [Escherichia coli] Length: 372\nNumber of predicted TMHs: 1\nExp number of AAs in TMHs: 23.92168\nExp number, first 60 AAs: 0.43108\nTotal prob of N-in: 0.13898\noutside 1 331\nTMhelix 332 354\ninside 355 372

13139 GCF_000313445.1_CFSAN001632_1.0 Escherichia coli O111:H8 str. CFSAN001632 Proteobacteria; Gammaproteobacteria; Enterobacterales; Enterobacteriaceae; Escherichia; Escherichia coli; Escherichia coli O111 MNNEETFYQAMRRQGVTRRSFLKYCSLAATSLGLGAGMAPKIAWALENKPRIPVVWIHGL WP_001058323.1 MULTISPECIES: hydrogenase-1 small chain [Proteobacteria] Length: 372\nNumber of predicted TMHs: 2\nExp number of AAs in TMHs: 37.2608499999999\nExp number, first 60 AAs: 19.52923\nTotal prob of N-in: 0.94351\nPOSSIBLE N-term signal sequence\ninside 1 19\nTMhelix 20 39\noutside 40 325\nTMhelix 326 348\ninside 349 372

13140 GCF_000263975.1_ASM26397v1 Escherichia coli O111:H8 str. CVM9570 Proteobacteria; Gammaproteobacteria; Enterobacterales; Enterobacteriaceae; Escherichia; Escherichia coli; Escherichia coli O111 MNNEETFYQAMRRQGVTRRSFLKYCSLAATSLGLGAGMAPKIAWALENKPRIPVVWIHGL WP_001058323.1 MULTISPECIES: hydrogenase-1 small chain [Proteobacteria] Length: 372\nNumber of predicted TMHs: 2\nExp number of AAs in TMHs: 37.2608499999999\nExp number, first 60 AAs: 19.52923\nTotal prob of N-in: 0.94351\nPOSSIBLE N-term signal sequence\ninside 1 19\nTMhelix 20 39\noutside 40 325\nTMhelix 326 348\ninside 349 372

13141 GCF_000263975.1_ASM26397v1 Escherichia coli O111:H8 str. CVM9570 Proteobacteria; Gammaproteobacteria; Enterobacterales; Enterobacteriaceae; Escherichia; Escherichia coli; Escherichia coli O111 MTGDNTLIHSHGINRRDFMKLCAALAATMGLSSKAAAEMAESVTNPQRPPVIWIGAQECT WP_000145410.1 MULTISPECIES: hydrogenase-2 small chain [Proteobacteria] Length: 372\nNumber of predicted TMHs: 1\nExp number of AAs in TMHs: 23.72226\nExp number, first 60 AAs: 0.22038\nTotal prob of N-in: 0.12994\noutside 1 331\nTMhelix 332 354\ninside 355 372

13142 GCF_000263995.1_ASM26399v1 Escherichia coli O111:H8 str. CVM9574 Proteobacteria; Gammaproteobacteria; Enterobacterales; Enterobacteriaceae; Escherichia; Escherichia coli; Escherichia coli O111 MNNEETFYQAMRRQGVTRRSFLKYCSLAATSLGLGAGMAPKIAWALENKPRIPVVWIHGL WP_001058323.1 MULTISPECIES: hydrogenase-1 small chain [Proteobacteria] Length: 372\nNumber of predicted TMHs: 2\nExp number of AAs in TMHs: 37.2608499999999\nExp number, first 60 AAs: 19.52923\nTotal prob of N-in: 0.94351\nPOSSIBLE N-term signal sequence\ninside 1 19\nTMhelix 20 39\noutside 40 325\nTMhelix 326 348\ninside 349 372

13143 GCF_000263995.1_ASM26399v1 Escherichia coli O111:H8 str. CVM9574 Proteobacteria; Gammaproteobacteria; Enterobacterales; Enterobacteriaceae; Escherichia; Escherichia coli; Escherichia coli O111 MTGDNTLIHSHGINRRDFMKLCAALAATMGLSSKAAAEMAESVTNPQRPPVIWIGAQECT WP_000145410.1 MULTISPECIES: hydrogenase-2 small chain [Proteobacteria] Length: 372\nNumber of predicted TMHs: 1\nExp number of AAs in TMHs: 23.72226\nExp number, first 60 AAs: 0.22038\nTotal prob of N-in: 0.12994\noutside 1 331\nTMhelix 332 354\ninside 355 372

13144 GCF_000276745.1_EC_O111_H8_CVM9602_v01 Escherichia coli O111:H8 str. CVM9602 Proteobacteria; Gammaproteobacteria; Enterobacterales; Enterobacteriaceae; Escherichia; Escherichia coli; Escherichia coli O111 MNNEETFYQAMRRQGVTRRSFLKYCSLAATSLGLGAGMAPKIAWALENKPRIPVVWIHGL WP_001058323.1 MULTISPECIES: hydrogenase-1 small chain [Proteobacteria] Length: 372\nNumber of predicted TMHs: 2\nExp number of AAs in TMHs: 37.2608499999999\nExp number, first 60 AAs: 19.52923\nTotal prob of N-in: 0.94351\nPOSSIBLE N-term signal sequence\ninside 1 19\nTMhelix 20 39\noutside 40 325\nTMhelix 326 348\ninside 349 372

13145 GCF_000276745.1_EC_O111_H8_CVM9602_v01 Escherichia coli O111:H8 str. CVM9602 Proteobacteria; Gammaproteobacteria; Enterobacterales; Enterobacteriaceae; Escherichia; Escherichia coli; Escherichia coli O111 MTGDNTLIHSHGINRRDFMKLCAALAATMGLSSKAAAEMAESVTNPQRPPVIWIGAQECT WP_000145410.1 MULTISPECIES: hydrogenase-2 small chain [Proteobacteria] Length: 372\nNumber of predicted TMHs: 1\nExp number of AAs in TMHs: 23.72226\nExp number, first 60 AAs: 0.22038\nTotal prob of N-in: 0.12994\noutside 1 331\nTMhelix 332 354\ninside 355 372

13146 GCF_000276765.1_EC_O111_H8_CVM9634_1.0 Escherichia coli O111:H8 str. CVM9634 Proteobacteria; Gammaproteobacteria; Enterobacterales; Enterobacteriaceae; Escherichia; Escherichia coli; Escherichia coli O111 MTGDNTLIHSHGINRRDFMKLCAALAATMGLSSKAAAEMAESVTNPQRPPVIWIGAQECT WP_000145410.1 MULTISPECIES: hydrogenase-2 small chain [Proteobacteria] Length: 372\nNumber of predicted TMHs: 1\nExp number of AAs in TMHs: 23.72226\nExp number, first 60 AAs: 0.22038\nTotal prob of N-in: 0.12994\noutside 1 331\nTMhelix 332 354\ninside 355 372

13147 GCF_000276765.1_EC_O111_H8_CVM9634_1.0 Escherichia coli O111:H8 str. CVM9634 Proteobacteria; Gammaproteobacteria; Enterobacterales; Enterobacteriaceae; Escherichia; Escherichia coli; Escherichia coli O111 MNNEETFYQAMRRQGVTRRSFLKYCSLAATSLGLGAGMAPKIAWALENKPRIPVVWIHGL WP_001058323.1 MULTISPECIES: hydrogenase-1 small chain [Proteobacteria] Length: 372\nNumber of predicted TMHs: 2\nExp number of AAs in TMHs: 37.2608499999999\nExp number, first 60 AAs: 19.52923\nTotal prob of N-in: 0.94351\nPOSSIBLE N-term signal sequence\ninside 1 19\nTMhelix 20 39\noutside 40 325\nTMhelix 326 348\ninside 349 372

13148 GCF_000618125.1_EcF6627 Escherichia coli O111:H8 str. F6627 Proteobacteria; Gammaproteobacteria; Enterobacterales; Enterobacteriaceae; Escherichia; Escherichia coli; Escherichia coli O111 MTGDNTLIHSHGINRRDFMKLCAALAATMGLSSKAAAEMAESVTNPQRPPVIWIGAQECT WP_000145410.1 MULTISPECIES: hydrogenase-2 small chain [Proteobacteria] Length: 372\nNumber of predicted TMHs: 1\nExp number of AAs in TMHs: 23.72226\nExp number, first 60 AAs: 0.22038\nTotal prob of N-in: 0.12994\noutside 1 331\nTMhelix 332 354\ninside 355 372

13149 GCF_000618125.1_EcF6627 Escherichia coli O111:H8 str. F6627 Proteobacteria; Gammaproteobacteria; Enterobacterales; Enterobacteriaceae; Escherichia; Escherichia coli; Escherichia coli O111 MNNEETFYQAMRRQGVTRRSFLKYCSLAATSLGLGAGMAPKIAWALENKPRIPVVWIHGL WP_001058323.1 MULTISPECIES: hydrogenase-1 small chain [Proteobacteria] Length: 372\nNumber of predicted TMHs: 2\nExp number of AAs in TMHs: 37.2608499999999\nExp number, first 60 AAs: 19.52923\nTotal prob of N-in: 0.94351\nPOSSIBLE N-term signal sequence\ninside 1 19\nTMhelix 20 39\noutside 40 325\nTMhelix 326 348\ninside 349 372

13150	GCF_000010765.1_ASM1076v1	Escherichia coli O111:H- str. 11128	Proteobacteria; Gammaproteobacteria; Enterobacterales; Enterobacteriaceae; Escherichia; Escherichia coli; Escherichia coli O111	MNNEETFYQAMRRQGVTRRSFLKYCSLAATSLGLGAGMAPKIAWALENKPRIPVVWIHGL	WP_001058323.1	MULTISPECIES: hydrogenase-1 small chain [Proteobacteria]	Length: 372\nNumber of predicted TMHs: 2\nExp number of AAs in TMHs: 37.2608499999999\nExp number, first 60 AAs: 19.52923\nTotal prob of N-in: 0.94351\nPOSSIBLE N-term signal sequence\ninside 1 19\nTMhelix 20 39\noutside 40 325\nTMhelix 326 348\ninside 349 372

13151	GCF_000010765.1_ASM1076v1	Escherichia coli O111:H- str. 11128	Proteobacteria; Gammaproteobacteria; Enterobacterales; Enterobacteriaceae; Escherichia; Escherichia coli; Escherichia coli O111	MTGDNTLIHSHGINRRDFMKLCAALAATMGLSSKAAAEMAESVTNPQRPPVIWIGAQECT	WP_000145410.1	MULTISPECIES: hydrogenase-2 small chain [Proteobacteria]	Length: 372\nNumber of predicted TMHs: 1\nExp number of AAs in TMHs: 23.72226\nExp number, first 60 AAs: 0.22038\nTotal prob of N-in: 0.12994\noutside 1 331\nTMhelix 332 354\ninside 355 372

13152	GCF_000701125.2_ASM70112v2	Escherichia coli O111:NM str. 01-3076	Proteobacteria; Gammaproteobacteria; Enterobacterales; Enterobacteriaceae; Escherichia; Escherichia coli; Escherichia coli O111	MNNEETFYQAMRRQGVTRRSFLKYCSLAATSLGLGAGMAPKIAWALENKPRIPVVWIHGL	WP_001058323.1	MULTISPECIES: hydrogenase-1 small chain [Proteobacteria]	Length: 372\nNumber of predicted TMHs: 2\nExp number of AAs in TMHs: 37.2608499999999\nExp number, first 60 AAs: 19.52923\nTotal prob of N-in: 0.94351\nPOSSIBLE N-term signal sequence\ninside 1 19\nTMhelix 20 39\noutside 40 325\nTMhelix 326 348\ninside 349 372

13153	GCF_000701125.2_ASM70112v2	Escherichia coli O111:NM str. 01-3076	Proteobacteria; Gammaproteobacteria; Enterobacterales; Enterobacteriaceae; Escherichia; Escherichia coli; Escherichia coli O111	MTGDNTLIHSHGINRRDFMKLCAALAATMGLSSKAAAEMAESVTNPQRPPVIWIGAQECT	WP_000145410.1	MULTISPECIES: hydrogenase-2 small chain [Proteobacteria]	Length: 372\nNumber of predicted TMHs: 1\nExp number of AAs in TMHs: 23.72226\nExp number, first 60 AAs: 0.22038\nTotal prob of N-in: 0.12994\noutside 1 331\nTMhelix 332 354\ninside 355 372

13154	GCF_000617245.2_Ec03-3484	Escherichia coli O111:NM str. 03-3484	Proteobacteria; Gammaproteobacteria; Enterobacterales; Enterobacteriaceae; Escherichia; Escherichia coli; Escherichia coli O111	MTGDNTLIHSHGINRRDFMKLCAALAATMGLSSKAAAEMAESVTNPQRPPVIWIGAQECT	WP_000145410.1	MULTISPECIES: hydrogenase-2 small chain [Proteobacteria]	Length: 372\nNumber of predicted TMHs: 1\nExp number of AAs in TMHs: 23.72226\nExp number, first 60 AAs: 0.22038\nTotal prob of N-in: 0.12994\noutside 1 331\nTMhelix 332 354\ninside 355 372

13155	GCF_000617245.2_Ec03-3484	Escherichia coli O111:NM str. 03-3484	Proteobacteria; Gammaproteobacteria; Enterobacterales; Enterobacteriaceae; Escherichia; Escherichia coli; Escherichia coli O111	MNNEETFYQAMRRQGVTRRSFLKYCSLAATSLGLGAGMAPKIAWALENKPRIPVVWIHGL	WP_001058323.1	MULTISPECIES: hydrogenase-1 small chain [Proteobacteria]	Length: 372\nNumber of predicted TMHs: 2\nExp number of AAs in TMHs: 37.2608499999999\nExp number, first 60 AAs: 19.52923\nTotal prob of N-in: 0.94351\nPOSSIBLE N-term signal sequence\ninside 1 19\nTMhelix 20 39\noutside 40 325\nTMhelix 326 348\ninside 349 372

13156	GCF_000617265.1_Ec04-3211	Escherichia coli O111:NM str. 04-3211	Proteobacteria; Gammaproteobacteria; Enterobacterales; Enterobacteriaceae; Escherichia; Escherichia coli; Escherichia coli O111	MTGDNTLIHSHGINRRDFMKLCAALAATMGLSSKAAAEMAESVTNPQRPPVIWIGAQECT	WP_000145410.1	MULTISPECIES: hydrogenase-2 small chain [Proteobacteria]	Length: 372\nNumber of predicted TMHs: 1\nExp number of AAs in TMHs: 23.72226\nExp number, first 60 AAs: 0.22038\nTotal prob of N-in: 0.12994\noutside 1 331\nTMhelix 332 354\ninside 355 372

13157	GCF_000617265.1_Ec04-3211	Escherichia coli O111:NM str. 04-3211	Proteobacteria; Gammaproteobacteria; Enterobacterales; Enterobacteriaceae; Escherichia; Escherichia coli; Escherichia coli O111	MNNEETFYQAMRRQGVTRRSFLKYCSLAATSLGLGAGMAPKIAWALENKPRIPVVWIHGL	WP_001058323.1	MULTISPECIES: hydrogenase-1 small chain [Proteobacteria]	Length: 372\nNumber of predicted TMHs: 2\nExp number of AAs in TMHs: 37.2608499999999\nExp number, first 60 AAs: 19.52923\nTotal prob of N-in: 0.94351\nPOSSIBLE N-term signal sequence\ninside 1 19\nTMhelix 20 39\noutside 40 325\nTMhelix 326 348\ninside 349 372

13158 GCF_000616685.2_Ec08-4487 Escherichia coli O111:NM str. 08-4487 Proteobacteria; Gammaproteobacteria; Enterobacterales; Enterobacteriaceae; Escherichia; Escherichia coli; Escherichia coli O111 MTGDNTLIHSHGINRRDFMKLCAALAATMGLSSKAAAEMAESVTNPQRPPVIWIGAQECT WP_000145410.1 MULTISPECIES: hydrogenase-2 small chain [Proteobacteria] Length: 372\nNumber of predicted TMHs: 1\nExp number of AAs in TMHs: 23.72226\nExp number, first 60 AAs: 0.22038\nTotal prob of N-in: 0.12994\noutside 1 331\nTMhelix 332 354\ninside 355 372

13159 GCF_000616685.2_Ec08-4487 Escherichia coli O111:NM str. 08-4487 Proteobacteria; Gammaproteobacteria; Enterobacterales; Enterobacteriaceae; Escherichia; Escherichia coli; Escherichia coli O111 MNNEETFYQAMRRQGVTRRSFLKYCSLAATSLGLGAGMAPKIAWALENKPRIPVVWIHGL WP_001058323.1 MULTISPECIES: hydrogenase-1 small chain [Proteobacteria] Length: 372\nNumber of predicted TMHs: 2\nExp number of AAs in TMHs: 37.2608499999999\nExp number, first 60 AAs: 19.52923\nTotal prob of N-in: 0.94351\nPOSSIBLE N-term signal sequence\ninside 1 19\nTMhelix 20 39\noutside 40 325\nTMhelix 326 348\ninside 349 372

13160 GCF_000619225.2_Ec2009C-4006 Escherichia coli O111:NM str. 2009C-4006 Proteobacteria; Gammaproteobacteria; Enterobacterales; Enterobacteriaceae; Escherichia; Escherichia coli; Escherichia coli O111 MNNEETFYQAMRRQGVTRRSFLKYCSLAATSLGLGAGMAPKIAWALENKPRIPVVWIHGL WP_001058323.1 MULTISPECIES: hydrogenase-1 small chain [Proteobacteria] Length: 372\nNumber of predicted TMHs: 2\nExp number of AAs in TMHs: 37.2608499999999\nExp number, first 60 AAs: 19.52923\nTotal prob of N-in: 0.94351\nPOSSIBLE N-term signal sequence\ninside 1 19\nTMhelix 20 39\noutside 40 325\nTMhelix 326 348\ninside 349 372

13161 GCF_000619225.2_Ec2009C-4006 Escherichia coli O111:NM str. 2009C-4006 Proteobacteria; Gammaproteobacteria; Enterobacterales; Enterobacteriaceae; Escherichia; Escherichia coli; Escherichia coli O111 MTGDNTLIHSHGINRRDFMKLCAALAATMGLSSKAAAEMAESVTNPQRPPVIWIGAQECT WP_000145410.1 MULTISPECIES: hydrogenase-2 small chain [Proteobacteria] Length: 372\nNumber of predicted TMHs: 1\nExp number of AAs in TMHs: 23.72226\nExp number, first 60 AAs: 0.22038\nTotal prob of N-in: 0.12994\noutside 1 331\nTMhelix 332 354\ninside 355 372

13162 GCF_000619005.2_Ec2009C-4052 Escherichia coli O111:NM str. 2009C-4052 Proteobacteria; Gammaproteobacteria; Enterobacterales; Enterobacteriaceae; Escherichia; Escherichia coli; Escherichia coli O111 MNNEETFYQAMRRQGVTRRSFLKYCSLAATSLGLGAGMAPKIAWALENKPRIPVVWIHGL WP_001058323.1 MULTISPECIES: hydrogenase-1 small chain [Proteobacteria] Length: 372\nNumber of predicted TMHs: 2\nExp number of AAs in TMHs: 37.2608499999999\nExp number, first 60 AAs: 19.52923\nTotal prob of N-in: 0.94351\nPOSSIBLE N-term signal sequence\ninside 1 19\nTMhelix 20 39\noutside 40 325\nTMhelix 326 348\ninside 349 372

13163 GCF_000619005.2_Ec2009C-4052 Escherichia coli O111:NM str. 2009C-4052 Proteobacteria; Gammaproteobacteria; Enterobacterales; Enterobacteriaceae; Escherichia; Escherichia coli; Escherichia coli O111 MTGDNTLIHSHGINRRDFMKLCAALAATMGLSSKAAAEMAESVTNPQRPPVIWIGAQECT WP_000145410.1 MULTISPECIES: hydrogenase-2 small chain [Proteobacteria] Length: 372\nNumber of predicted TMHs: 1\nExp number of AAs in TMHs: 23.72226\nExp number, first 60 AAs: 0.22038\nTotal prob of N-in: 0.12994\noutside 1 331\nTMhelix 332 354\ninside 355 372

13164 GCF_000632615.2_Ec2010C-3053 Escherichia coli O111:NM str. 2010C-3053 Proteobacteria; Gammaproteobacteria; Enterobacterales; Enterobacteriaceae; Escherichia; Escherichia coli; Escherichia coli O111 MNNEETFYQAMRRQGVTRRSFLKYCSLAATSLGLGAGMAPKIAWALENKPRIPVVWIHGL WP_001058323.1 MULTISPECIES: hydrogenase-1 small chain [Proteobacteria] Length: 372\nNumber of predicted TMHs: 2\nExp number of AAs in TMHs: 37.2608499999999\nExp number, first 60 AAs: 19.52923\nTotal prob of N-in: 0.94351\nPOSSIBLE N-term signal sequence\ninside 1 19\nTMhelix 20 39\noutside 40 325\nTMhelix 326 348\ninside 349 372

13165 GCF_000632615.2_Ec2010C-3053 Escherichia coli O111:NM str. 2010C-3053 Proteobacteria; Gammaproteobacteria; Enterobacterales; Enterobacteriaceae; Escherichia; Escherichia coli; Escherichia coli O111 MTGDNTLIHSHGINRRDFMKLCAALAATMGLSSKAAAEMAESVTNPQRPPVIWIGAQECT WP_000145410.1 MULTISPECIES: hydrogenase-2 small chain [Proteobacteria] Length: 372\nNumber of predicted TMHs: 1\nExp number of AAs in TMHs: 23.72226\nExp number, first 60 AAs: 0.22038\nTotal prob of N-in: 0.12994\noutside 1 331\nTMhelix 332 354\ninside 355 372

13166 GCF_000615575.2_Ec2010C-3977 Escherichia coli O111:NM str. 2010C-3977 Proteobacteria; Gammaproteobacteria; Enterobacterales; Enterobacteriaceae; Escherichia; Escherichia coli; Escherichia coli O111 MTGDNTLIHSHGINRRDFMKLCAALAATMGLSSKAAAEMAESVTNPQRPPVIWIGAQECT WP_000145410.1 MULTISPECIES: hydrogenase-2 small chain [Proteobacteria] Length: 372\nNumber of predicted TMHs: 1\nExp number of AAs in TMHs: 23.72226\nExp number, first 60 AAs: 0.22038\nTotal prob of N-in: 0.12994\noutside 1 331\nTMhelix 332 354\ninside 355 372

13167 GCF_000615575.2_Ec2010C-3977 Escherichia coli O111:NM str. 2010C-3977 Proteobacteria; Gammaproteobacteria; Enterobacterales; Enterobacteriaceae; Escherichia; Escherichia coli; Escherichia coli O111 MNNEETFYQAMRRQGVTRRSFLKYCSLAATSLGLGAGMAPKIAWALENKPRIPVVWIHGL WP_001058323.1 MULTISPECIES: hydrogenase-1 small chain [Proteobacteria] Length: 372\nNumber of predicted TMHs: 2\nExp number of AAs in TMHs: 37.2608499999999\nExp number, first 60 AAs: 19.52923\nTotal prob of N-in: 0.94351\nPOSSIBLE N-term signal sequence\ninside 1 19\nTMhelix 20 39\noutside 40 325\nTMhelix 326 348\ninside 349 372

13168 GCF_000615605.2_Ec2010C-4086 Escherichia coli O111:NM str. 2010C-4086 Proteobacteria; Gammaproteobacteria; Enterobacterales; Enterobacteriaceae; Escherichia; Escherichia coli; Escherichia coli O111 MNNEETFYQAMRRQGVTRRSFLKYCSLAATSLGLGAGMAPKIAWALENKPRIPVVWIHGL WP_001058323.1 MULTISPECIES: hydrogenase-1 small chain [Proteobacteria] Length: 372\nNumber of predicted TMHs: 2\nExp number of AAs in TMHs: 37.2608499999999\nExp number, first 60 AAs: 19.52923\nTotal prob of N-in: 0.94351\nPOSSIBLE N-term signal sequence\ninside 1 19\nTMhelix 20 39\noutside 40 325\nTMhelix 326 348\ninside 349 372

13169 GCF_000615605.2_Ec2010C-4086 Escherichia coli O111:NM str. 2010C-4086 Proteobacteria; Gammaproteobacteria; Enterobacterales; Enterobacteriaceae; Escherichia; Escherichia coli; Escherichia coli O111 MTGDNTLIHSHGINRRDFMKLCAALAATMGLSSKAAAEMAESVTNPQRPPVIWIGAQECT WP_000145410.1 MULTISPECIES: hydrogenase-2 small chain [Proteobacteria] Length: 372\nNumber of predicted TMHs: 1\nExp number of AAs in TMHs: 23.72226\nExp number, first 60 AAs: 0.22038\nTotal prob of N-in: 0.12994\noutside 1 331\nTMhelix 332 354\ninside 355 372

13170 GCF_000615655.2_Ec2010C-4221 Escherichia coli O111:NM str. 2010C-4221 Proteobacteria; Gammaproteobacteria; Enterobacterales; Enterobacteriaceae; Escherichia; Escherichia coli; Escherichia coli O111 MTGDNTLIHSHGINRRDFMKLCAALAATMGLSSKAAAEMAESVTNPQRPPVIWIGAQECT WP_000145410.1 MULTISPECIES: hydrogenase-2 small chain [Proteobacteria] Length: 372\nNumber of predicted TMHs: 1\nExp number of AAs in TMHs: 23.72226\nExp number, first 60 AAs: 0.22038\nTotal prob of N-in: 0.12994\noutside 1 331\nTMhelix 332 354\ninside 355 372

13171 GCF_000615655.2_Ec2010C-4221 Escherichia coli O111:NM str. 2010C-4221 Proteobacteria; Gammaproteobacteria; Enterobacterales; Enterobacteriaceae; Escherichia; Escherichia coli; Escherichia coli O111 MNNEETFYQAMRRQGVTRRSFLKYCSLAATSLGLGAGMAPKIAWALENKPRIPVVWIHGL WP_001058323.1 MULTISPECIES: hydrogenase-1 small chain [Proteobacteria] Length: 372\nNumber of predicted TMHs: 2\nExp number of AAs in TMHs: 37.2608499999999\nExp number, first 60 AAs: 19.52923\nTotal prob of N-in: 0.94351\nPOSSIBLE N-term signal sequence\ninside 1 19\nTMhelix 20 39\noutside 40 325\nTMhelix 326 348\ninside 349 372

13172 GCF_000617805.2_Ec2010C-4592 Escherichia coli O111:NM str. 2010C-4592 Proteobacteria; Gammaproteobacteria; Enterobacterales; Enterobacteriaceae; Escherichia; Escherichia coli; Escherichia coli O111 MNNEETFYQAMRRQGVTRRSFLKYCSLAATSLGLGAGMAPKIAWALENKPRIPVVWIHGL WP_001058323.1 MULTISPECIES: hydrogenase-1 small chain [Proteobacteria] Length: 372\nNumber of predicted TMHs: 2\nExp number of AAs in TMHs: 37.2608499999999\nExp number, first 60 AAs: 19.52923\nTotal prob of N-in: 0.94351\nPOSSIBLE N-term signal sequence\ninside 1 19\nTMhelix 20 39\noutside 40 325\nTMhelix 326 348\ninside 349 372

13173 GCF_000617805.2_Ec2010C-4592 Escherichia coli O111:NM str. 2010C-4592 Proteobacteria; Gammaproteobacteria; Enterobacterales; Enterobacteriaceae; Escherichia; Escherichia coli; Escherichia coli O111 MTGDNTLIHSHGINRRDFMKLCAALAATMGLSSKAAAEMAESVTNPQRPPVIWIGAQECT WP_000145410.1 MULTISPECIES: hydrogenase-2 small chain [Proteobacteria] Length: 372\nNumber of predicted TMHs: 1\nExp number of AAs in TMHs: 23.72226\nExp number, first 60 AAs: 0.22038\nTotal prob of N-in: 0.12994\noutside 1 331\nTMhelix 332 354\ninside 355 372

13174 GCF_000617625.2_Ec2010C-4622 Escherichia coli O111:NM str. 2010C-4622 Proteobacteria; Gammaproteobacteria; Enterobacterales; Enterobacteriaceae; Escherichia; Escherichia coli; Escherichia coli O111 MNNEETFYQAMRRQGVTRRSFLKYCSLAATSLGLGAGMAPKIAWALENKPRIPVVWIHGL WP_001058323.1 MULTISPECIES: hydrogenase-1 small chain [Proteobacteria] Length: 372\nNumber of predicted TMHs: 2\nExp number of AAs in TMHs: 37.2608499999999\nExp number, first 60 AAs: 19.52923\nTotal prob of N-in: 0.94351\nPOSSIBLE N-term signal sequence\ninside 1 19\nTMhelix 20 39\noutside 40 325\nTMhelix 326 348\ninside 349 372

13175 GCF_000617625.2_Ec2010C-4622 Escherichia coli O111:NM str. 2010C-4622 Proteobacteria; Gammaproteobacteria; Enterobacterales; Enterobacteriaceae; Escherichia; Escherichia coli; Escherichia coli O111 MTGDNTLIHSHGINRRDFMKLCAALAATMGLSSKAAAEMAESVTNPQRPPVIWIGAQECT WP_000145410.1 MULTISPECIES: hydrogenase-2 small chain [Proteobacteria] Length: 372\nNumber of predicted TMHs: 1\nExp number of AAs in TMHs: 23.72226\nExp number, first 60 AAs: 0.22038\nTotal prob of N-in: 0.12994\noutside 1 331\nTMhelix 332 354\ninside 355 372

13176 GCF_000617025.2_Ec2010C-4715 Escherichia coli O111:NM str. 2010C-4715 Proteobacteria; Gammaproteobacteria; Enterobacterales; Enterobacteriaceae; Escherichia; Escherichia coli; Escherichia coli O111 MNNEETFYQAMRRQGVTRRSFLKYCSLAATSLGLGAGMAPKIAWALENKPRIPVVWIHGL WP_001058323.1 MULTISPECIES: hydrogenase-1 small chain [Proteobacteria] Length: 372\nNumber of predicted TMHs: 2\nExp number of AAs in TMHs: 37.2608499999999\nExp number, first 60 AAs: 19.52923\nTotal prob of N-in: 0.94351\nPOSSIBLE N-term signal sequence\ninside 1 19\nTMhelix 20 39\noutside 40 325\nTMhelix 326 348\ninside 349 372

13177 GCF_000617025.2_Ec2010C-4715 Escherichia coli O111:NM str. 2010C-4715 Proteobacteria; Gammaproteobacteria; Enterobacterales; Enterobacteriaceae; Escherichia; Escherichia coli; Escherichia coli O111 MTGDNTLIHSHGINRRDFMKLCAALAATMGLSSKAAAEMAESVTNPQRPPVIWIGAQECT WP_000145410.1 MULTISPECIES: hydrogenase-2 small chain [Proteobacteria] Length: 372\nNumber of predicted TMHs: 1\nExp number of AAs in TMHs: 23.72226\nExp number, first 60 AAs: 0.22038\nTotal prob of N-in: 0.12994\noutside 1 331\nTMhelix 332 354\ninside 355 372

13178 GCF_000617465.2_Ec2010C-4735 Escherichia coli O111:NM str. 2010C-4735 Proteobacteria; Gammaproteobacteria; Enterobacterales; Enterobacteriaceae; Escherichia; Escherichia coli; Escherichia coli O111 MTGDNTLIHSHGINRRDFMKLCAALAATMGLSSKAAAEMAESVTNPQRPPVIWIGAQECT WP_000145410.1 MULTISPECIES: hydrogenase-2 small chain [Proteobacteria] Length: 372\nNumber of predicted TMHs: 1\nExp number of AAs in TMHs: 23.72226\nExp number, first 60 AAs: 0.22038\nTotal prob of N-in: 0.12994\noutside 1 331\nTMhelix 332 354\ninside 355 372

13179 GCF_000617465.2_Ec2010C-4735 Escherichia coli O111:NM str. 2010C-4735 Proteobacteria; Gammaproteobacteria; Enterobacterales; Enterobacteriaceae; Escherichia; Escherichia coli; Escherichia coli O111 MNNEETFYQAMRRQGVTRRSFLKYCSLAATSLGLGAGMAPKIAWALENKPRIPVVWIHGL WP_001058323.1 MULTISPECIES: hydrogenase-1 small chain [Proteobacteria] Length: 372\nNumber of predicted TMHs: 2\nExp number of AAs in TMHs: 37.2608499999999\nExp number, first 60 AAs: 19.52923\nTotal prob of N-in: 0.94351\nPOSSIBLE N-term signal sequence\ninside 1 19\nTMhelix 20 39\noutside 40 325\nTMhelix 326 348\ninside 349 372

13180 GCF_000617485.2_Ec2010C-4746 Escherichia coli O111:NM str. 2010C-4746 Proteobacteria; Gammaproteobacteria; Enterobacterales; Enterobacteriaceae; Escherichia; Escherichia coli; Escherichia coli O111 MNNEETFYQAMRRQGVTRRSFLKYCSLAATSLGLGAGMAPKIAWALENKPRIPVVWIHGL WP_001058323.1 MULTISPECIES: hydrogenase-1 small chain [Proteobacteria] Length: 372\nNumber of predicted TMHs: 2\nExp number of AAs in TMHs: 37.2608499999999\nExp number, first 60 AAs: 19.52923\nTotal prob of N-in: 0.94351\nPOSSIBLE N-term signal sequence\ninside 1 19\nTMhelix 20 39\noutside 40 325\nTMhelix 326 348\ninside 349 372

13181 GCF_000617485.2_Ec2010C-4746 Escherichia coli O111:NM str. 2010C-4746 Proteobacteria; Gammaproteobacteria; Enterobacterales; Enterobacteriaceae; Escherichia; Escherichia coli; Escherichia coli O111 MTGDNTLIHSHGINRRDFMKLCAALAATMGLSSKAAAEMAESVTNPQRPPVIWIGAQECT WP_000145410.1 MULTISPECIES: hydrogenase-2 small chain [Proteobacteria] Length: 372\nNumber of predicted TMHs: 1\nExp number of AAs in TMHs: 23.72226\nExp number, first 60 AAs: 0.22038\nTotal prob of N-in: 0.12994\noutside 1 331\nTMhelix 332 354\ninside 355 372

13182 GCF_000617525.2_Ec2010C-4799 Escherichia coli O111:NM str. 2010C-4799 Proteobacteria; Gammaproteobacteria; Enterobacterales; Enterobacteriaceae; Escherichia; Escherichia coli; Escherichia coli O111 MNNEETFYQAMRRQGVTRRSFLKYCSLAATSLGLGAGMAPKIAWALENKPRIPVVWIHGL WP_001058323.1 MULTISPECIES: hydrogenase-1 small chain [Proteobacteria] Length: 372\nNumber of predicted TMHs: 2\nExp number of AAs in TMHs: 37.2608499999999\nExp number, first 60 AAs: 19.52923\nTotal prob of N-in: 0.94351\nPOSSIBLE N-term signal sequence\ninside 1 19\nTMhelix 20 39\noutside 40 325\nTMhelix 326 348\ninside 349 372

13183 GCF_000617525.2_Ec2010C-4799 Escherichia coli O111:NM str. 2010C-4799 Proteobacteria; Gammaproteobacteria; Enterobacterales; Enterobacteriaceae; Escherichia; Escherichia coli; Escherichia coli O111 MTGDNTLIHSHGINRRDFMKLCAALAATMGLSSKAAAEMAESVTNPQRPPVIWIGAQECT WP_000145410.1 MULTISPECIES: hydrogenase-2 small chain [Proteobacteria] Length: 372\nNumber of predicted TMHs: 1\nExp number of AAs in TMHs: 23.72226\nExp number, first 60 AAs: 0.22038\nTotal prob of N-in: 0.12994\noutside 1 331\nTMhelix 332 354\ninside 355 372

13184 GCF_000617545.2_Ec2010C-4818 Escherichia coli O111:NM str. 2010C-4818 Proteobacteria; Gammaproteobacteria; Enterobacterales; Enterobacteriaceae; Escherichia; Escherichia coli; Escherichia coli O111 MNNEETFYQAMRRQGVTRRSFLKYCSLAATSLGLGAGMAPKIAWALENKPRIPVVWIHGL WP_001058323.1 MULTISPECIES: hydrogenase-1 small chain [Proteobacteria] Length: 372\nNumber of predicted TMHs: 2\nExp number of AAs in TMHs: 37.2608499999999\nExp number, first 60 AAs: 19.52923\nTotal prob of N-in: 0.94351\nPOSSIBLE N-term signal sequence\ninside 1 19\nTMhelix 20 39\noutside 40 325\nTMhelix 326 348\ninside 349 372

13185 GCF_000617545.2_Ec2010C-4818 Escherichia coli O111:NM str. 2010C-4818 Proteobacteria; Gammaproteobacteria; Enterobacterales; Enterobacteriaceae; Escherichia; Escherichia coli; Escherichia coli O111 MTGDNTLIHSHGINRRDFMKLCAALAATMGLSSKAAAEMAESVTNPQRPPVIWIGAQECT WP_000145410.1 MULTISPECIES: hydrogenase-2 small chain [Proteobacteria] Length: 372\nNumber of predicted TMHs: 1\nExp number of AAs in TMHs: 23.72226\nExp number, first 60 AAs: 0.22038\nTotal prob of N-in: 0.12994\noutside 1 331\nTMhelix 332 354\ninside 355 372

13186 GCF_000614035.2_Ec2011C-3170 Escherichia coli O111:NM str. 2011C-3170 Proteobacteria; Gammaproteobacteria; Enterobacterales; Enterobacteriaceae; Escherichia; Escherichia coli; Escherichia coli O111 MTGDNTLIHSHGINRRDFMKLCAALAATMGLSSKAAAEMAESVTNPQRPPVIWIGAQECT WP_000145410.1 MULTISPECIES: hydrogenase-2 small chain [Proteobacteria] Length: 372\nNumber of predicted TMHs: 1\nExp number of AAs in TMHs: 23.72226\nExp number, first 60 AAs: 0.22038\nTotal prob of N-in: 0.12994\noutside 1 331\nTMhelix 332 354\ninside 355 372

13187 GCF_000614035.2_Ec2011C-3170 Escherichia coli O111:NM str. 2011C-3170 Proteobacteria; Gammaproteobacteria; Enterobacterales; Enterobacteriaceae; Escherichia; Escherichia coli; Escherichia coli O111 MNNEETFYQAMRRQGVTRRSFLKYCSLAATSLGLGAGMAPKIAWALENKPRIPVVWIHGL WP_001058323.1 MULTISPECIES: hydrogenase-1 small chain [Proteobacteria] Length: 372\nNumber of predicted TMHs: 2\nExp number of AAs in TMHs: 37.2608499999999\nExp number, first 60 AAs: 19.52923\nTotal prob of N-in: 0.94351\nPOSSIBLE N-term signal sequence\ninside 1 19\nTMhelix 20 39\noutside 40 325\nTMhelix 326 348\ninside 349 372

13188 GCF_000616195.2_Ec2011C-3362 Escherichia coli O111:NM str. 2011C-3362 Proteobacteria; Gammaproteobacteria; Enterobacterales; Enterobacteriaceae; Escherichia; Escherichia coli; Escherichia coli O111 MNNEETFYQAMRRQGVTRRSFLKYCSLAATSLGLGAGMAPKIAWALENKPRIPVVWIHGL WP_001058323.1 MULTISPECIES: hydrogenase-1 small chain [Proteobacteria] Length: 372\nNumber of predicted TMHs: 2\nExp number of AAs in TMHs: 37.2608499999999\nExp number, first 60 AAs: 19.52923\nTotal prob of N-in: 0.94351\nPOSSIBLE N-term signal sequence\ninside 1 19\nTMhelix 20 39\noutside 40 325\nTMhelix 326 348\ninside 349 372

13189 GCF_000616195.2_Ec2011C-3362 Escherichia coli O111:NM str. 2011C-3362 Proteobacteria; Gammaproteobacteria; Enterobacterales; Enterobacteriaceae; Escherichia; Escherichia coli; Escherichia coli O111 MTGDNTLIHSHGINRRDFMKLCAALAATMGLSSKAAAEMAESVTNPQRPPVIWIGAQECT WP_000145410.1 MULTISPECIES: hydrogenase-2 small chain [Proteobacteria] Length: 372\nNumber of predicted TMHs: 1\nExp number of AAs in TMHs: 23.72226\nExp number, first 60 AAs: 0.22038\nTotal prob of N-in: 0.12994\noutside 1 331\nTMhelix 332 354\ninside 355 372

13190 GCF_000616265.1_Ec2011C-3573 Escherichia coli O111:NM str. 2011C-3573 Proteobacteria; Gammaproteobacteria; Enterobacterales; Enterobacteriaceae; Escherichia; Escherichia coli; Escherichia coli O111 MTGDNTLIHSHGINRRDFMKLCAALAATMGLSSKAAAEMAESVTNPQRPPVIWIGAQECT WP_000145410.1 MULTISPECIES: hydrogenase-2 small chain [Proteobacteria] Length: 372\nNumber of predicted TMHs: 1\nExp number of AAs in TMHs: 23.72226\nExp number, first 60 AAs: 0.22038\nTotal prob of N-in: 0.12994\noutside 1 331\nTMhelix 332 354\ninside 355 372

13191 GCF_000616265.1_Ec2011C-3573 Escherichia coli O111:NM str. 2011C-3573 Proteobacteria; Gammaproteobacteria; Enterobacterales; Enterobacteriaceae; Escherichia; Escherichia coli; Escherichia coli O111 MNNEETFYQAMRRQGVTRRSFLKYCSLAATSLGLGAGMAPKIAWALENKPRIPVVWIHGL WP_001058323.1 MULTISPECIES: hydrogenase-1 small chain [Proteobacteria] Length: 372\nNumber of predicted TMHs: 2\nExp number of AAs in TMHs: 37.2608499999999\nExp number, first 60 AAs: 19.52923\nTotal prob of N-in: 0.94351\nPOSSIBLE N-term signal sequence\ninside 1 19\nTMhelix 20 39\noutside 40 325\nTMhelix 326 348\ninside 349 372

13192 GCF_000616305.2_Ec2011C-3632 Escherichia coli O111:NM str. 2011C-3632 Proteobacteria; Gammaproteobacteria; Enterobacterales; Enterobacteriaceae; Escherichia; Escherichia coli; Escherichia coli O111 MTGDNTLIHSHGINRRDFMKLCAALAATMGLSSKAAAEMAESVTNPQRPPVIWIGAQECT WP_000145410.1 MULTISPECIES: hydrogenase-2 small chain [Proteobacteria] Length: 372\nNumber of predicted TMHs: 1\nExp number of AAs in TMHs: 23.72226\nExp number, first 60 AAs: 0.22038\nTotal prob of N-in: 0.12994\noutside 1 331\nTMhelix 332 354\ninside 355 372

13193 GCF_000616305.2_Ec2011C-3632 Escherichia coli O111:NM str. 2011C-3632 Proteobacteria; Gammaproteobacteria; Enterobacterales; Enterobacteriaceae; Escherichia; Escherichia coli; Escherichia coli O111 MNNEETFYQAMRRQGVTRRSFLKYCSLAATSLGLGAGMAPKIAWALENKPRIPVVWIHGL WP_001058323.1 MULTISPECIES: hydrogenase-1 small chain [Proteobacteria] Length: 372\nNumber of predicted TMHs: 2\nExp number of AAs in TMHs: 37.2608499999999\nExp number, first 60 AAs: 19.52923\nTotal prob of N-in: 0.94351\nPOSSIBLE N-term signal sequence\ninside 1 19\nTMhelix 20 39\noutside 40 325\nTMhelix 326 348\ninside 349 372

13194 GCF_000616325.2_Ec2011C-3679 Escherichia coli O111:NM str. 2011C-3679 Proteobacteria; Gammaproteobacteria; Enterobacterales; Enterobacteriaceae; Escherichia; Escherichia coli; Escherichia coli O111 MTGDNTLIHSHGINRRDFMKLCAALAATMGLSSKAAAEMAESVTNPQRPPVIWIGAQECT WP_000145410.1 MULTISPECIES: hydrogenase-2 small chain [Proteobacteria] Length: 372\nNumber of predicted TMHs: 1\nExp number of AAs in TMHs: 23.72226\nExp number, first 60 AAs: 0.22038\nTotal prob of N-in: 0.12994\noutside 1 331\nTMhelix 332 354\ninside 355 372

13195 GCF_000616325.2_Ec2011C-3679 Escherichia coli O111:NM str. 2011C-3679 Proteobacteria; Gammaproteobacteria; Enterobacterales; Enterobacteriaceae; Escherichia; Escherichia coli; Escherichia coli O111 MNNEETFYQAMRRQGVTRRSFLKYCSLAATSLGLGAGMAPKIAWALENKPRIPVVWIHGL WP_001058323.1 MULTISPECIES: hydrogenase-1 small chain [Proteobacteria] Length: 372\nNumber of predicted TMHs: 2\nExp number of AAs in TMHs: 37.2608499999999\nExp number, first 60 AAs: 19.52923\nTotal prob of N-in: 0.94351\nPOSSIBLE N-term signal sequence\ninside 1 19\nTMhelix 20 39\noutside 40 325\nTMhelix 326 348\ninside 349 372

13196 GCF_000619465.2_EcK6722 Escherichia coli O111:NM str. K6722 Proteobacteria; Gammaproteobacteria; Enterobacterales; Enterobacteriaceae; Escherichia; Escherichia coli; Escherichia coli O111 MTGDNTLIHSHGINRRDFMKLCAALAATMGLSSKAAAEMAESVTNPQRPPVIWIGAQECT WP_000145410.1 MULTISPECIES: hydrogenase-2 small chain [Proteobacteria] Length: 372\nNumber of predicted TMHs: 1\nExp number of AAs in TMHs: 23.72226\nExp number, first 60 AAs: 0.22038\nTotal prob of N-in: 0.12994\noutside 1 331\nTMhelix 332 354\ninside 355 372

13197 GCF_000619465.2_EcK6722 Escherichia coli O111:NM str. K6722 Proteobacteria; Gammaproteobacteria; Enterobacterales; Enterobacteriaceae; Escherichia; Escherichia coli; Escherichia coli O111 MNNEETFYQAMRRQGVTRRSFLKYCSLAATSLGLGAGMAPKIAWALENKPRIPVVWIHGL WP_001058323.1 MULTISPECIES: hydrogenase-1 small chain [Proteobacteria] Length: 372\nNumber of predicted TMHs: 2\nExp number of AAs in TMHs: 37.2608499999999\nExp number, first 60 AAs: 19.52923\nTotal prob of N-in: 0.94351\nPOSSIBLE N-term signal sequence\ninside 1 19\nTMhelix 20 39\noutside 40 325\nTMhelix 326 348\ninside 349 372

13198 GCF_000619485.1_EcK6723 Escherichia coli O111:NM str. K6723 Proteobacteria; Gammaproteobacteria; Enterobacterales; Enterobacteriaceae; Escherichia; Escherichia coli; Escherichia coli O111 MTGDNTLIHSHGINRRDFMKLCAALAATMGLSSKAAAEMAESVTNPQRPPVIWIGAQECT WP_000145410.1 MULTISPECIES: hydrogenase-2 small chain [Proteobacteria] Length: 372\nNumber of predicted TMHs: 1\nExp number of AAs in TMHs: 23.72226\nExp number, first 60 AAs: 0.22038\nTotal prob of N-in: 0.12994\noutside 1 331\nTMhelix 332 354\ninside 355 372

13199 GCF_000619485.1_EcK6723 Escherichia coli O111:NM str. K6723 Proteobacteria; Gammaproteobacteria; Enterobacterales; Enterobacteriaceae; Escherichia; Escherichia coli; Escherichia coli O111 MNNEETFYQAMRRQGVTRRSFLKYCSLAATSLGLGAGMAPKIAWALENKPRIPVVWIHGL WP_001058323.1 MULTISPECIES: hydrogenase-1 small chain [Proteobacteria] Length: 372\nNumber of predicted TMHs: 2\nExp number of AAs in TMHs: 37.2608499999999\nExp number, first 60 AAs: 19.52923\nTotal prob of N-in: 0.94351\nPOSSIBLE N-term signal sequence\ninside 1 19\nTMhelix 20 39\noutside 40 325\nTMhelix 326 348\ninside 349 372

13200 GCF_000619505.1_EcK6728 Escherichia coli O111:NM str. K6728 Proteobacteria; Gammaproteobacteria; Enterobacterales; Enterobacteriaceae; Escherichia; Escherichia coli; Escherichia coli O111 MNNEETFYQAMRRQGVTRRSFLKYCSLAATSLGLGAGMAPKIAWALENKPRIPVVWIHGL WP_001058323.1 MULTISPECIES: hydrogenase-1 small chain [Proteobacteria] Length: 372\nNumber of predicted TMHs: 2\nExp number of AAs in TMHs: 37.2608499999999\nExp number, first 60 AAs: 19.52923\nTotal prob of N-in: 0.94351\nPOSSIBLE N-term signal sequence\ninside 1 19\nTMhelix 20 39\noutside 40 325\nTMhelix 326 348\ninside 349 372

13201 GCF_000619505.1_EcK6728 Escherichia coli O111:NM str. K6728 Proteobacteria; Gammaproteobacteria; Enterobacterales; Enterobacteriaceae; Escherichia; Escherichia coli; Escherichia coli O111 MTGDNTLIHSHGINRRDFMKLCAALAATMGLSSKAAAEMAESVTNPQRPPVIWIGAQECT WP_000145410.1 MULTISPECIES: hydrogenase-2 small chain [Proteobacteria] Length: 372\nNumber of predicted TMHs: 1\nExp number of AAs in TMHs: 23.72226\nExp number, first 60 AAs: 0.22038\nTotal prob of N-in: 0.12994\noutside 1 331\nTMhelix 332 354\ninside 355 372

13202 GCF_000619525.1_EcK6890 Escherichia coli O111:NM str. K6890 Proteobacteria; Gammaproteobacteria; Enterobacterales; Enterobacteriaceae; Escherichia; Escherichia coli; Escherichia coli O111 MNNEETFYQAMRRQGVTRRSFLKYCSLAATSLGLGAGMAPKIAWALENKPRIPVVWIHGL WP_001058323.1 MULTISPECIES: hydrogenase-1 small chain [Proteobacteria] Length: 372\nNumber of predicted TMHs: 2\nExp number of AAs in TMHs: 37.2608499999999\nExp number, first 60 AAs: 19.52923\nTotal prob of N-in: 0.94351\nPOSSIBLE N-term signal sequence\ninside 1 19\nTMhelix 20 39\noutside 40 325\nTMhelix 326 348\ninside 349 372

13203 GCF_000619525.1_EcK6890 Escherichia coli O111:NM str. K6890 Proteobacteria; Gammaproteobacteria; Enterobacterales; Enterobacteriaceae; Escherichia; Escherichia coli; Escherichia coli O111 MTGDNTLIHSHGINRRDFMKLCAALAATMGLSSKAAAEMAESVTNPQRPPVIWIGAQECT WP_000145410.1 MULTISPECIES: hydrogenase-2 small chain [Proteobacteria] Length: 372\nNumber of predicted TMHs: 1\nExp number of AAs in TMHs: 23.72226\nExp number, first 60 AAs: 0.22038\nTotal prob of N-in: 0.12994\noutside 1 331\nTMhelix 332 354\ninside 355 372

13204 GCF_000619545.2_EcK6895 Escherichia coli O111:NM str. K6895 Proteobacteria; Gammaproteobacteria; Enterobacterales; Enterobacteriaceae; Escherichia; Escherichia coli; Escherichia coli O111 MNNEETFYQAMRRQGVTRRSFLKYCSLAATSLGLGAGMAPKIAWALENKPRIPVVWIHGL WP_001058323.1 MULTISPECIES: hydrogenase-1 small chain [Proteobacteria] Length: 372\nNumber of predicted TMHs: 2\nExp number of AAs in TMHs: 37.2608499999999\nExp number, first 60 AAs: 19.52923\nTotal prob of N-in: 0.94351\nPOSSIBLE N-term signal sequence\ninside 1 19\nTMhelix 20 39\noutside 40 325\nTMhelix 326 348\ninside 349 372

13205 GCF_000619545.2_EcK6895 Escherichia coli O111:NM str. K6895 Proteobacteria; Gammaproteobacteria; Enterobacterales; Enterobacteriaceae; Escherichia; Escherichia coli; Escherichia coli O111 MTGDNTLIHSHGINRRDFMKLCAALAATMGLSSKAAAEMAESVTNPQRPPVIWIGAQECT WP_000145410.1 MULTISPECIES: hydrogenase-2 small chain [Proteobacteria] Length: 372\nNumber of predicted TMHs: 1\nExp number of AAs in TMHs: 23.72226\nExp number, first 60 AAs: 0.22038\nTotal prob of N-in: 0.12994\noutside 1 331\nTMhelix 332 354\ninside 355 372

13206 GCF_000619565.2_EcK6897 Escherichia coli O111:NM str. K6897 Proteobacteria; Gammaproteobacteria; Enterobacterales; Enterobacteriaceae; Escherichia; Escherichia coli; Escherichia coli O111
MTGDNTLIHSHGINRRDFMKLCAALAATMGLSSKAAAEMAESVTNPQRPPVIWIGAQECT WP_000145410.1
MULTISPECIES: hydrogenase-2 small chain [Proteobacteria] Length: 372\nNumber of predicted TMHs: 1\nExp number of AAs in TMHs: 23.72226\nExp number, first 60 AAs: 0.22038\nTotal prob of N-in: 0.12994\noutside 1 331\nTMhelix 332 354\ninside 355 372

13207 GCF_000619565.2_EcK6897 Escherichia coli O111:NM str. K6897 Proteobacteria; Gammaproteobacteria; Enterobacterales; Enterobacteriaceae; Escherichia; Escherichia coli; Escherichia coli O111
MNNEETFYQAMRRQGVTRRSFLKYCSLAATSLGLGAGMAPKIAWALENKPRIPVVWIHGL WP_001058323.1
MULTISPECIES: hydrogenase-1 small chain [Proteobacteria] Length: 372\nNumber of predicted TMHs: 2\nExp number of AAs in TMHs: 37.2608499999999\nExp number, first 60 AAs: 19.52923\nTotal prob of N-in: 0.94351\nPOSSIBLE N-term signal sequence\ninside 1 19\nTMhelix 20 39\noutside 40 325\nTMhelix 326 348\ninside 349 372

13208 GCF_000619585.2_EcK6898 Escherichia coli O111:NM str. K6898 Proteobacteria; Gammaproteobacteria; Enterobacterales; Enterobacteriaceae; Escherichia; Escherichia coli; Escherichia coli O111
MNNEETFYQAMRRQGVTRRSFLKYCSLAATSLGLGAGMAPKIAWALENKPRIPVVWIHGL WP_001058323.1
MULTISPECIES: hydrogenase-1 small chain [Proteobacteria] Length: 372\nNumber of predicted TMHs: 2\nExp number of AAs in TMHs: 37.2608499999999\nExp number, first 60 AAs: 19.52923\nTotal prob of N-in: 0.94351\nPOSSIBLE N-term signal sequence\ninside 1 19\nTMhelix 20 39\noutside 40 325\nTMhelix 326 348\ninside 349 372

13209 GCF_000619585.2_EcK6898 Escherichia coli O111:NM str. K6898 Proteobacteria; Gammaproteobacteria; Enterobacterales; Enterobacteriaceae; Escherichia; Escherichia coli; Escherichia coli O111
MTGDNTLIHSHGINRRDFMKLCAALAATMGLSSKAAAEMAESVTNPQRPPVIWIGAQECT WP_000145410.1
MULTISPECIES: hydrogenase-2 small chain [Proteobacteria] Length: 372\nNumber of predicted TMHs: 1\nExp number of AAs in TMHs: 23.72226\nExp number, first 60 AAs: 0.22038\nTotal prob of N-in: 0.12994\noutside 1 331\nTMhelix 332 354\ninside 355 372

13210 GCF_000619625.1_EcK6904 Escherichia coli O111:NM str. K6904 Proteobacteria; Gammaproteobacteria; Enterobacterales; Enterobacteriaceae; Escherichia; Escherichia coli; Escherichia coli O111
MTGDNTLIHSHGINRRDFMKLCAALAATMGLSSKAAAEMAESVTNPQRPPVIWIGAQECT WP_000145410.1
MULTISPECIES: hydrogenase-2 small chain [Proteobacteria] Length: 372\nNumber of predicted TMHs: 1\nExp number of AAs in TMHs: 23.72226\nExp number, first 60 AAs: 0.22038\nTotal prob of N-in: 0.12994\noutside 1 331\nTMhelix 332 354\ninside 355 372

13211 GCF_000619625.1_EcK6904 Escherichia coli O111:NM str. K6904 Proteobacteria; Gammaproteobacteria; Enterobacterales; Enterobacteriaceae; Escherichia; Escherichia coli; Escherichia coli O111
MNNEETFYQAMRRQGVTRRSFLKYCSLAATSLGLGAGMAPKIAWALENKPRIPVVWIHGL WP_001058323.1
MULTISPECIES: hydrogenase-1 small chain [Proteobacteria] Length: 372\nNumber of predicted TMHs: 2\nExp number of AAs in TMHs: 37.2608499999999\nExp number, first 60 AAs: 19.52923\nTotal prob of N-in: 0.94351\nPOSSIBLE N-term signal sequence\ninside 1 19\nTMhelix 20 39\noutside 40 325\nTMhelix 326 348\ninside 349 372

13212 GCF_000619605.1_EcK6908 Escherichia coli O111:NM str. K6908 Proteobacteria; Gammaproteobacteria; Enterobacterales; Enterobacteriaceae; Escherichia; Escherichia coli; Escherichia coli O111
MNNEETFYQAMRRQGVTRRSFLKYCSLAATSLGLGAGMAPKIAWALENKPRIPVVWIHGL WP_001058323.1
MULTISPECIES: hydrogenase-1 small chain [Proteobacteria] Length: 372\nNumber of predicted TMHs: 2\nExp number of AAs in TMHs: 37.2608499999999\nExp number, first 60 AAs: 19.52923\nTotal prob of N-in: 0.94351\nPOSSIBLE N-term signal sequence\ninside 1 19\nTMhelix 20 39\noutside 40 325\nTMhelix 326 348\ninside 349 372

13213 GCF_000619605.1_EcK6908 Escherichia coli O111:NM str. K6908 Proteobacteria; Gammaproteobacteria; Enterobacterales; Enterobacteriaceae; Escherichia; Escherichia coli; Escherichia coli O111
MTGDNTLIHSHGINRRDFMKLCAALAATMGLSSKAAAEMAESVTNPQRPPVIWIGAQECT WP_000145410.1
MULTISPECIES: hydrogenase-2 small chain [Proteobacteria] Length: 372\nNumber of predicted TMHs: 1\nExp number of AAs in TMHs: 23.72226\nExp number, first 60 AAs: 0.22038\nTotal prob of N-in: 0.12994\noutside 1 331\nTMhelix 332 354\ninside 355 372

13214 GCF_000619645.2_EcK6915 Escherichia coli O111:NM str. K6915 Proteobacteria; Gammaproteobacteria; Enterobacterales; Enterobacteriaceae; Escherichia; Escherichia coli; Escherichia coli O111
MNNEETFYQAMRRQGVTRRSFLKYCSLAATSLGLGAGMAPKIAWALENKPRIPVVWIHGL WP_001058323.1
MULTISPECIES: hydrogenase-1 small chain [Proteobacteria] Length: 372\nNumber of predicted TMHs: 2\nExp number of AAs in TMHs: 37.2608499999999\nExp number, first 60 AAs: 19.52923\nTotal prob of N-in: 0.94351\nPOSSIBLE N-term signal sequence\ninside 1 19\nTMhelix 20 39\noutside 40 325\nTMhelix 326 348\ninside 349 372

13215 GCF_000619645.2_EcK6915 Escherichia coli O111:NM str. K6915 Proteobacteria; Gammaproteobacteria; Enterobacterales; Enterobacteriaceae; Escherichia; Escherichia coli; Escherichia coli O111
MTGDNTLIHSHGINRRDFMKLCAALAATMGLSSKAAAEMAESVTNPQRPPVIWIGAQECT WP_000145410.1
MULTISPECIES: hydrogenase-2 small chain [Proteobacteria] Length: 372\nNumber of predicted TMHs: 1\nExp number of AAs in TMHs: 23.72226\nExp number, first 60 AAs: 0.22038\nTotal prob of N-in: 0.12994\noutside 1 331\nTMhelix 332 354\ninside 355 372

13216 GCF_000234235.1_ASM23423v2 Escherichia coli O113:H21 str. CL-3 Proteobacteria; Gammaproteobacteria; Enterobacterales; Enterobacteriaceae; Escherichia; Escherichia coli; Escherichia coli O113
MNNEETFYQAMRRQGVTRRSFLKYCSLAATSLGLGAGMAPKIAWALENKPRIPVVWIHGL WP_001058323.1
MULTISPECIES: hydrogenase-1 small chain [Proteobacteria] Length: 372\nNumber of predicted TMHs: 2\nExp number of AAs in TMHs: 37.2608499999999\nExp number, first 60 AAs: 19.52923\nTotal prob of N-in: 0.94351\nPOSSIBLE N-term signal sequence\ninside 1 19\nTMhelix 20 39\noutside 40 325\nTMhelix 326 348\ninside 349 372

13217 GCF_000234235.1_ASM23423v2 Escherichia coli O113:H21 str. CL-3 Proteobacteria; Gammaproteobacteria; Enterobacterales; Enterobacteriaceae; Escherichia; Escherichia coli; Escherichia coli O113
MTGDNTLIHSHGINRRDFMKLCAALAATMGLSSKAAAEMAESVTNPQRPPVIWIGAQECT WP_000145410.1
MULTISPECIES: hydrogenase-2 small chain [Proteobacteria] Length: 372\nNumber of predicted TMHs: 1\nExp number of AAs in TMHs: 23.72226\nExp number, first 60 AAs: 0.22038\nTotal prob of N-in: 0.12994\noutside 1 331\nTMhelix 332 354\ninside 355 372

13218 GCF_000617305.2_Ec06-3256 Escherichia coli O118:H16 str. 06-3256 Proteobacteria; Gammaproteobacteria; Enterobacterales; Enterobacteriaceae; Escherichia; Escherichia coli; Escherichia coli O118
MTGDNTLIHSHGINRRDFMKLCAALAATMGLSSKAAAEMAESVTNPQRPPVIWIGAQECT WP_000145410.1
MULTISPECIES: hydrogenase-2 small chain [Proteobacteria] Length: 372\nNumber of predicted TMHs: 1\nExp number of AAs in TMHs: 23.72226\nExp number, first 60 AAs: 0.22038\nTotal prob of N-in: 0.12994\noutside 1 331\nTMhelix 332 354\ninside 355 372

13219 GCF_000617305.2_Ec06-3256 Escherichia coli O118:H16 str. 06-3256 Proteobacteria; Gammaproteobacteria; Enterobacterales; Enterobacteriaceae; Escherichia; Escherichia coli; Escherichia coli O118
MNNEETFYQAMRRQGVTRRSFLKYCSLAATSLGLGAGMAPKIAWALENKPRIPVVWIHGL WP_001058323.1
MULTISPECIES: hydrogenase-1 small chain [Proteobacteria] Length: 372\nNumber of predicted TMHs: 2\nExp number of AAs in TMHs: 37.2608499999999\nExp number, first 60 AAs: 19.52923\nTotal prob of N-in: 0.94351\nPOSSIBLE N-term signal sequence\ninside 1 19\nTMhelix 20 39\noutside 40 325\nTMhelix 326 348\ninside 349 372

13220 GCF_000617405.2_Ec06-3612 Escherichia coli O118:H16 str. 06-3612 Proteobacteria; Gammaproteobacteria; Enterobacterales; Enterobacteriaceae; Escherichia; Escherichia coli; Escherichia coli O118
MTGDNTLIHSHGINRRDFMKLCAALAATMGLSSKAAAEMAESVTNPQRPPVIWIGAQECT WP_000145410.1
MULTISPECIES: hydrogenase-2 small chain [Proteobacteria] Length: 372\nNumber of predicted TMHs: 1\nExp number of AAs in TMHs: 23.72226\nExp number, first 60 AAs: 0.22038\nTotal prob of N-in: 0.12994\noutside 1 331\nTMhelix 332 354\ninside 355 372

13221 GCF_000617405.2_Ec06-3612 Escherichia coli O118:H16 str. 06-3612 Proteobacteria; Gammaproteobacteria; Enterobacterales; Enterobacteriaceae; Escherichia; Escherichia coli; Escherichia coli O118
MNNEETFYQAMRRQGVTRRSFLKYCSLAATSLGLGAGMAPKIAWALENKPRIPVVWIHGL WP_001058323.1
MULTISPECIES: hydrogenase-1 small chain [Proteobacteria] Length: 372\nNumber of predicted TMHs: 2\nExp number of AAs in TMHs: 37.2608499999999\nExp number, first 60 AAs: 19.52923\nTotal prob of N-in: 0.94351\nPOSSIBLE N-term signal sequence\ninside 1 19\nTMhelix 20 39\noutside 40 325\nTMhelix 326 348\ninside 349 372

13222 GCF_000703245.1_Ec07-4255 Escherichia coli O118:H16 str. 07-4255 Proteobacteria; Gammaproteobacteria; Enterobacterales; Enterobacteriaceae; Escherichia; Escherichia coli; Escherichia coli O118 MTGDNTLIHSHGINRRDFMKLCAALAATMGLSSKAAAEMAESVTNPQRPPVIWIGAQECT WP_000145410.1 MULTISPECIES: hydrogenase-2 small chain [Proteobacteria] Length: 372\nNumber of predicted TMHs: 1\nExp number of AAs in TMHs: 23.72226\nExp number, first 60 AAs: 0.22038\nTotal prob of N-in: 0.12994\noutside 1 331\nTMhelix 332 354\ninside 355 372

13223 GCF_000703245.1_Ec07-4255 Escherichia coli O118:H16 str. 07-4255 Proteobacteria; Gammaproteobacteria; Enterobacterales; Enterobacteriaceae; Escherichia; Escherichia coli; Escherichia coli O118 MNNEETFYQAMRRQGVTRRSFLKYCSLAATSLGLGAGMAPKIAWALENKPRIPVVWIHGL WP_001058323.1 MULTISPECIES: hydrogenase-1 small chain [Proteobacteria] Length: 372\nNumber of predicted TMHs: 2\nExp number of AAs in TMHs: 37.2608499999999\nExp number, first 60 AAs: 19.52923\nTotal prob of N-in: 0.94351\nPOSSIBLE N-term signal sequence\ninside 1 19\nTMhelix 20 39\noutside 40 325\nTMhelix 326 348\ninside 349 372

13224 GCF_000616625.1_Ec08-3651 Escherichia coli O118:H16 str. 08-3651 Proteobacteria; Gammaproteobacteria; Enterobacterales; Enterobacteriaceae; Escherichia; Escherichia coli; Escherichia coli O118 MTGDNTLIHSHGINRRDFMKLCAALAATMGLSSKAAAEMAESVTNPQRPPVIWIGAQECT WP_000145410.1 MULTISPECIES: hydrogenase-2 small chain [Proteobacteria] Length: 372\nNumber of predicted TMHs: 1\nExp number of AAs in TMHs: 23.72226\nExp number, first 60 AAs: 0.22038\nTotal prob of N-in: 0.12994\noutside 1 331\nTMhelix 332 354\ninside 355 372

13225 GCF_000616625.1_Ec08-3651 Escherichia coli O118:H16 str. 08-3651 Proteobacteria; Gammaproteobacteria; Enterobacterales; Enterobacteriaceae; Escherichia; Escherichia coli; Escherichia coli O118 MNNEETFYQAMRRQGVTRRSFLKYCSLAATSLGLGAGMAPKIAWALENKPRIPVVWIHGL WP_001058323.1 MULTISPECIES: hydrogenase-1 small chain [Proteobacteria] Length: 372\nNumber of predicted TMHs: 2\nExp number of AAs in TMHs: 37.2608499999999\nExp number, first 60 AAs: 19.52923\nTotal prob of N-in: 0.94351\nPOSSIBLE N-term signal sequence\ninside 1 19\nTMhelix 20 39\noutside 40 325\nTMhelix 326 348\ninside 349 372

13226 GCF_000619025.1_Ec2009C-4446 Escherichia coli O118:H16 str. 2009C-4446 Proteobacteria; Gammaproteobacteria; Enterobacterales; Enterobacteriaceae; Escherichia; Escherichia coli; Escherichia coli O118 MTGDNTLIHSHGINRRDFMKLCAALAATMGLSSKAAAEMAESVTNPQRPPVIWIGAQECT WP_000145410.1 MULTISPECIES: hydrogenase-2 small chain [Proteobacteria] Length: 372\nNumber of predicted TMHs: 1\nExp number of AAs in TMHs: 23.72226\nExp number, first 60 AAs: 0.22038\nTotal prob of N-in: 0.12994\noutside 1 331\nTMhelix 332 354\ninside 355 372

13227 GCF_000619025.1_Ec2009C-4446 Escherichia coli O118:H16 str. 2009C-4446 Proteobacteria; Gammaproteobacteria; Enterobacterales; Enterobacteriaceae; Escherichia; Escherichia coli; Escherichia coli O118 MNNEETFYQAMRRQGVTRRSFLKYCSLAATSLGLGAGMAPKIAWALENKPRIPVVWIHGL WP_001058323.1 MULTISPECIES: hydrogenase-1 small chain [Proteobacteria] Length: 372\nNumber of predicted TMHs: 2\nExp number of AAs in TMHs: 37.2608499999999\nExp number, first 60 AAs: 19.52923\nTotal prob of N-in: 0.94351\nPOSSIBLE N-term signal sequence\ninside 1 19\nTMhelix 20 39\noutside 40 325\nTMhelix 326 348\ninside 349 372

13228 GCF_000617225.1_Ec03-3458 Escherichia coli O119:H4 str. 03-3458 Proteobacteria; Gammaproteobacteria; Enterobacterales; Enterobacteriaceae; Escherichia; Escherichia coli; Escherichia coli O119 MNNEETFYQAMRRQGVTRRSFLKYCSLAATSLGLGAGMAPKIAWALENKPRIPVVWIHGL WP_001058323.1 MULTISPECIES: hydrogenase-1 small chain [Proteobacteria] Length: 372\nNumber of predicted TMHs: 2\nExp number of AAs in TMHs: 37.2608499999999\nExp number, first 60 AAs: 19.52923\nTotal prob of N-in: 0.94351\nPOSSIBLE N-term signal sequence\ninside 1 19\nTMhelix 20 39\noutside 40 325\nTMhelix 326 348\ninside 349 372

13229 GCF_000617225.1_Ec03-3458 Escherichia coli O119:H4 str. 03-3458 Proteobacteria; Gammaproteobacteria; Enterobacterales; Enterobacteriaceae; Escherichia; Escherichia coli; Escherichia coli O119 MTGDNTLIHSHGINRRDFMKLCAALAATMGLSSKAAAEMAESVTNPQRPPVIWIGAQECT WP_000145410.1 MULTISPECIES: hydrogenase-2 small chain [Proteobacteria] Length: 372\nNumber of predicted TMHs: 1\nExp number of AAs in TMHs: 23.72226\nExp number, first 60 AAs: 0.22038\nTotal prob of N-in: 0.12994\noutside 1 331\nTMhelix 332 354\ninside 355 372

13230 GCF_000617185.2_Ec03-3227 Escherichia coli O121:H19 str. 03-3227 Proteobacteria; Gammaproteobacteria; Enterobacterales; Enterobacteriaceae; Escherichia; Escherichia coli; Escherichia coli O121 MTGDNTLIHSHGINRRDFMKLCAALAATMGLSSKAAAEMAESVTNPQRPPVIWIGAQECT WP_000145410.1 MULTISPECIES: hydrogenase-2 small chain [Proteobacteria] Length: 372\nNumber of predicted TMHs: 1\nExp number of AAs in TMHs: 23.72226\nExp number, first 60 AAs: 0.22038\nTotal prob of N-in: 0.12994\noutside 1 331\nTMhelix 332 354\ninside 355 372

13231 GCF_000617285.2_Ec06-3003 Escherichia coli O121:H19 str. 06-3003 Proteobacteria; Gammaproteobacteria; Enterobacterales; Enterobacteriaceae; Escherichia; Escherichia coli; Escherichia coli O121 MTGDNTLIHSHGINRRDFMKLCAALAATMGLSSKAAAEMAESVTNPQRPPVIWIGAQECT WP_000145410.1 MULTISPECIES: hydrogenase-2 small chain [Proteobacteria] Length: 372\nNumber of predicted TMHs: 1\nExp number of AAs in TMHs: 23.72226\nExp number, first 60 AAs: 0.22038\nTotal prob of N-in: 0.12994\noutside 1 331\nTMhelix 332 354\ninside 355 372

13232 GCF_000617645.2_Ec06-3822 Escherichia coli O121:H19 str. 06-3822 Proteobacteria; Gammaproteobacteria; Enterobacterales; Enterobacteriaceae; Escherichia; Escherichia coli; Escherichia coli O121 MTGDNTLIHSHGINRRDFMKLCAALAATMGLSSKAAAEMAESVTNPQRPPVIWIGAQECT WP_000145410.1 MULTISPECIES: hydrogenase-2 small chain [Proteobacteria] Length: 372\nNumber of predicted TMHs: 1\nExp number of AAs in TMHs: 23.72226\nExp number, first 60 AAs: 0.22038\nTotal prob of N-in: 0.12994\noutside 1 331\nTMhelix 332 354\ninside 355 372

13233 GCF_000619745.2_Ec2009C-4050 Escherichia coli O121:H19 str. 2009C-4050 Proteobacteria; Gammaproteobacteria; Enterobacterales; Enterobacteriaceae; Escherichia; Escherichia coli; Escherichia coli O121 MTGDNTLIHSHGINRRDFMKLCAALAATMGLSSKAAAEMAESVTNPQRPPVIWIGAQECT WP_000145410.1 MULTISPECIES: hydrogenase-2 small chain [Proteobacteria] Length: 372\nNumber of predicted TMHs: 1\nExp number of AAs in TMHs: 23.72226\nExp number, first 60 AAs: 0.22038\nTotal prob of N-in: 0.12994\noutside 1 331\nTMhelix 332 354\ninside 355 372

13234 GCF_000614765.2_Ec2009C-4659 Escherichia coli O121:H19 str. 2009C-4659 Proteobacteria; Gammaproteobacteria; Enterobacterales; Enterobacteriaceae; Escherichia; Escherichia coli; Escherichia coli O121 MTGDNTLIHSHGINRRDFMKLCAALAATMGLSSKAAAEMAESVTNPQRPPVIWIGAQECT WP_000145410.1 MULTISPECIES: hydrogenase-2 small chain [Proteobacteria] Length: 372\nNumber of predicted TMHs: 1\nExp number of AAs in TMHs: 23.72226\nExp number, first 60 AAs: 0.22038\nTotal prob of N-in: 0.12994\noutside 1 331\nTMhelix 332 354\ninside 355 372

13235 GCF_000619085.2_Ec2009C-4750 Escherichia coli O121:H19 str. 2009C-4750 Proteobacteria; Gammaproteobacteria; Enterobacterales; Enterobacteriaceae; Escherichia; Escherichia coli; Escherichia coli O121 MTGDNTLIHSHGINRRDFMKLCAALAATMGLSSKAAAEMAESVTNPQRPPVIWIGAQECT WP_000145410.1 MULTISPECIES: hydrogenase-2 small chain [Proteobacteria] Length: 372\nNumber of predicted TMHs: 1\nExp number of AAs in TMHs: 23.72226\nExp number, first 60 AAs: 0.22038\nTotal prob of N-in: 0.12994\noutside 1 331\nTMhelix 332 354\ninside 355 372

13236 GCF_000619125.2_Ec2009EL1302 Escherichia coli O121:H19 str. 2009EL1302 Proteobacteria; Gammaproteobacteria; Enterobacterales; Enterobacteriaceae; Escherichia; Escherichia coli; Escherichia coli O121 MTGDNTLIHSHGINRRDFMKLCAALAATMGLSSKAAAEMAESVTNPQRPPVIWIGAQECT WP_000145410.1 MULTISPECIES: hydrogenase-2 small chain [Proteobacteria] Length: 372\nNumber of predicted TMHs: 1\nExp number of AAs in TMHs: 23.72226\nExp number, first 60 AAs: 0.22038\nTotal prob of N-in: 0.12994\noutside 1 331\nTMhelix 332 354\ninside 355 372

13237 GCF_000614925.2_Ec2009EL1412 Escherichia coli O121:H19 str. 2009EL1412 Proteobacteria; Gammaproteobacteria; Enterobacterales; Enterobacteriaceae; Escherichia; Escherichia coli; Escherichia coli O121 MTGDNTLIHSHGINRRDFMKLCAALAATMGLSSKAAAEMAESVTNPQRPPVIWIGAQECT WP_000145410.1 MULTISPECIES: hydrogenase-2 small chain [Proteobacteria] Length: 372\nNumber of predicted TMHs: 1\nExp number of AAs in TMHs: 23.72226\nExp number, first 60 AAs: 0.22038\nTotal prob of N-in: 0.12994\noutside 1 331\nTMhelix 332 354\ninside 355 372

13238 GCF_000615455.2_Ec2010C-3609 Escherichia coli O121:H19 str. 2010C-3609 Proteobacteria; Gammaproteobacteria; Enterobacterales; Enterobacteriaceae; Escherichia; Escherichia coli; Escherichia coli O121 MTGDNTLIHSHGINRRDFMKLCAALAATMGLSSKAAAEMAESVTNPQRPPVIWIGAQECT WP_000145410.1 MULTISPECIES: hydrogenase-2 small chain [Proteobacteria] Length: 372\nNumber of predicted TMHs:

1\nExp number of AAs in TMHs: 23.72226\nExp number, first 60 AAs: 0.22038\nTotal prob of N-in: 0.12994\noutside 1 331\nTMhelix 332 354\ninside 355 372

13239 GCF_000619205.2_Ec2010C-3794 Escherichia coli O121:H19 str. 2010C-3794 Proteobacteria; Gammaproteobacteria; Enterobacterales; Enterobacteriaceae; Escherichia; Escherichia coli; Escherichia coli O121 MTGDNTLIHSHGINRRDFMKLCAALAATMGLSSKAAAEMAESVTNPQRPPVIWIGAQECT WP_000145410.1 MULTISPECIES: hydrogenase-2 small chain [Proteobacteria] Length: 372\nNumber of predicted TMHs: 1\nExp number of AAs in TMHs: 23.72226\nExp number, first 60 AAs: 0.22038\nTotal prob of N-in: 0.12994\noutside 1 331\nTMhelix 332 354\ninside 355 372

13240 GCF_000615505.1_Ec2010C-3840 Escherichia coli O121:H19 str. 2010C-3840 Proteobacteria; Gammaproteobacteria; Enterobacterales; Enterobacteriaceae; Escherichia; Escherichia coli; Escherichia coli O121 MTGDNTLIHSHGINRRDFMKLCAALAATMGLSSKAAAEMAESVTNPQRPPVIWIGAQECT WP_000145410.1 MULTISPECIES: hydrogenase-2 small chain [Proteobacteria] Length: 372\nNumber of predicted TMHs: 1\nExp number of AAs in TMHs: 23.72226\nExp number, first 60 AAs: 0.22038\nTotal prob of N-in: 0.12994\noutside 1 331\nTMhelix 332 354\ninside 355 372

13241 GCF_000614725.2_Ec2010C-4254 Escherichia coli O121:H19 str. 2010C-4254 Proteobacteria; Gammaproteobacteria; Enterobacterales; Enterobacteriaceae; Escherichia; Escherichia coli; Escherichia coli O121 MTGDNTLIHSHGINRRDFMKLCAALAATMGLSSKAAAEMAESVTNPQRPPVIWIGAQECT WP_000145410.1 MULTISPECIES: hydrogenase-2 small chain [Proteobacteria] Length: 372\nNumber of predicted TMHs: 1\nExp number of AAs in TMHs: 23.72226\nExp number, first 60 AAs: 0.22038\nTotal prob of N-in: 0.12994\noutside 1 331\nTMhelix 332 354\ninside 355 372

13242 GCF_000617005.2_Ec2010C-4732 Escherichia coli O121:H19 str. 2010C-4732 Proteobacteria; Gammaproteobacteria; Enterobacterales; Enterobacteriaceae; Escherichia; Escherichia coli; Escherichia coli O121 MTGDNTLIHSHGINRRDFMKLCAALAATMGLSSKAAAEMAESVTNPQRPPVIWIGAQECT WP_000145410.1 MULTISPECIES: hydrogenase-2 small chain [Proteobacteria] Length: 372\nNumber of predicted TMHs: 1\nExp number of AAs in TMHs: 23.72226\nExp number, first 60 AAs: 0.22038\nTotal prob of N-in: 0.12994\noutside 1 331\nTMhelix 332 354\ninside 355 372

13243 GCF_000617565.1_Ec2010C-4824 Escherichia coli O121:H19 str. 2010C-4824 Proteobacteria; Gammaproteobacteria; Enterobacterales; Enterobacteriaceae; Escherichia; Escherichia coli; Escherichia coli O121 MTGDNTLIHSHGINRRDFMKLCAALAATMGLSSKAAAEMAESVTNPQRPPVIWIGAQECT WP_000145410.1 MULTISPECIES: hydrogenase-2 small chain [Proteobacteria] Length: 372\nNumber of predicted TMHs: 1\nExp number of AAs in TMHs: 23.72226\nExp number, first 60 AAs: 0.22038\nTotal prob of N-in: 0.12994\noutside 1 331\nTMhelix 332 354\ninside 355 372

13244 GCF_000617605.2_Ec2010C-4966 Escherichia coli O121:H19 str. 2010C-4966 Proteobacteria; Gammaproteobacteria; Enterobacterales; Enterobacteriaceae; Escherichia; Escherichia coli; Escherichia coli O121 MTGDNTLIHSHGINRRDFMKLCAALAATMGLSSKAAAEMAESVTNPQRPPVIWIGAQECT WP_000145410.1 MULTISPECIES: hydrogenase-2 small chain [Proteobacteria] Length: 372\nNumber of predicted TMHs: 1\nExp number of AAs in TMHs: 23.72226\nExp number, first 60 AAs: 0.22038\nTotal prob of N-in: 0.12994\noutside 1 331\nTMhelix 332 354\ninside 355 372

13245 GCF_000614515.1_Ec2010C-4989 Escherichia coli O121:H19 str. 2010C-4989 Proteobacteria; Gammaproteobacteria; Enterobacterales; Enterobacteriaceae; Escherichia; Escherichia coli; Escherichia coli O121 MTGDNTLIHSHGINRRDFMKLCAALAATMGLSSKAAAEMAESVTNPQRPPVIWIGAQECT WP_000145410.1 MULTISPECIES: hydrogenase-2 small chain [Proteobacteria] Length: 372\nNumber of predicted TMHs: 1\nExp number of AAs in TMHs: 23.72226\nExp number, first 60 AAs: 0.22038\nTotal prob of N-in: 0.12994\noutside 1 331\nTMhelix 332 354\ninside 355 372

13246 GCF_000614595.1_Ec2010EL1058 Escherichia coli O121:H19 str. 2010EL1058 Proteobacteria; Gammaproteobacteria; Enterobacterales; Enterobacteriaceae; Escherichia; Escherichia coli; Escherichia coli O121 MTGDNTLIHSHGINRRDFMKLCAALAATMGLSSKAAAEMAESVTNPQRPPVIWIGAQECT WP_000145410.1 MULTISPECIES: hydrogenase-2 small chain [Proteobacteria] Length: 372\nNumber of predicted TMHs: 1\nExp number of AAs in TMHs: 23.72226\nExp number, first 60 AAs: 0.22038\nTotal prob of N-in: 0.12994\noutside 1 331\nTMhelix 332 354\ninside 355 372

13247 GCF_000616725.2_Ec2011C-3072 Escherichia coli O121:H19 str. 2011C-3072 Proteobacteria; Gammaproteobacteria; Enterobacterales; Enterobacteriaceae; Escherichia; Escherichia coli; Escherichia coli O121

MTGDNTLIHSHGINRRDFMKLCAALAATMGLSSKAAAEMAESVTNPQRPPVIWIGAQECT WP_000145410.1 MULTISPECIES: hydrogenase-2 small chain [Proteobacteria] Length: 372\nNumber of predicted TMHs: 1\nExp number of AAs in TMHs: 23.72226\nExp number, first 60 AAs: 0.22038\nTotal prob of N-in: 0.12994\noutside 1 331\nTMhelix 332 354\ninside 355 372

13248 GCF_000614215.2_Ec2011C-3108 Escherichia coli O121:H19 str. 2011C-3108 Proteobacteria; Gammaproteobacteria; Enterobacterales; Enterobacteriaceae; Escherichia; Escherichia coli; Escherichia coli O121 MTGDNTLIHSHGINRRDFMKLCAALAATMGLSSKAAAEMAESVTNPQRPPVIWIGAQECT WP_000145410.1 MULTISPECIES: hydrogenase-2 small chain [Proteobacteria] Length: 372\nNumber of predicted TMHs: 1\nExp number of AAs in TMHs: 23.72226\nExp number, first 60 AAs: 0.22038\nTotal prob of N-in: 0.12994\noutside 1 331\nTMhelix 332 354\ninside 355 372

13249 GCF_000616135.2_Ec2011C-3216 Escherichia coli O121:H19 str. 2011C-3216 Proteobacteria; Gammaproteobacteria; Enterobacterales; Enterobacteriaceae; Escherichia; Escherichia coli; Escherichia coli O121 MTGDNTLIHSHGINRRDFMKLCAALAATMGLSSKAAAEMAESVTNPQRPPVIWIGAQECT WP_000145410.1 MULTISPECIES: hydrogenase-2 small chain [Proteobacteria] Length: 372\nNumber of predicted TMHs: 1\nExp number of AAs in TMHs: 23.72226\nExp number, first 60 AAs: 0.22038\nTotal prob of N-in: 0.12994\noutside 1 331\nTMhelix 332 354\ninside 355 372

13250 GCF_000616225.2_Ec2011C-3500 Escherichia coli O121:H19 str. 2011C-3500 Proteobacteria; Gammaproteobacteria; Enterobacterales; Enterobacteriaceae; Escherichia; Escherichia coli; Escherichia coli O121 MTGDNTLIHSHGINRRDFMKLCAALAATMGLSSKAAAEMAESVTNPQRPPVIWIGAQECT WP_000145410.1 MULTISPECIES: hydrogenase-2 small chain [Proteobacteria] Length: 372\nNumber of predicted TMHs: 1\nExp number of AAs in TMHs: 23.72226\nExp number, first 60 AAs: 0.22038\nTotal prob of N-in: 0.12994\noutside 1 331\nTMhelix 332 354\ninside 355 372

13251 GCF_000616245.1_Ec2011C-3537 Escherichia coli O121:H19 str. 2011C-3537 Proteobacteria; Gammaproteobacteria; Enterobacterales; Enterobacteriaceae; Escherichia; Escherichia coli; Escherichia coli O121 MTGDNTLIHSHGINRRDFMKLCAALAATMGLSSKAAAEMAESVTNPQRPPVIWIGAQECT WP_000145410.1 MULTISPECIES: hydrogenase-2 small chain [Proteobacteria] Length: 372\nNumber of predicted TMHs: 1\nExp number of AAs in TMHs: 23.72226\nExp number, first 60 AAs: 0.22038\nTotal prob of N-in: 0.12994\noutside 1 331\nTMhelix 332 354\ninside 355 372

13252 GCF_000703365.1_Ec2011C-3609 Escherichia coli O121:H19 str. 2011C-3609 Proteobacteria; Gammaproteobacteria; Enterobacterales; Enterobacteriaceae; Escherichia; Escherichia coli; Escherichia coli O121 MTGDNTLIHSHGINRRDFMKLCAALAATMGLSSKAAAEMAESVTNPQRPPVIWIGAQECT WP_000145410.1 MULTISPECIES: hydrogenase-2 small chain [Proteobacteria] Length: 372\nNumber of predicted TMHs: 1\nExp number of AAs in TMHs: 23.72226\nExp number, first 60 AAs: 0.22038\nTotal prob of N-in: 0.12994\noutside 1 331\nTMhelix 332 354\ninside 355 372

13253 GCF_000618165.2_EcF6714 Escherichia coli O121:H19 str. F6714 Proteobacteria; Gammaproteobacteria; Enterobacterales; Enterobacteriaceae; Escherichia; Escherichia coli; Escherichia coli O121 MTGDNTLIHSHGINRRDFMKLCAALAATMGLSSKAAAEMAESVTNPQRPPVIWIGAQECT WP_000145410.1 MULTISPECIES: hydrogenase-2 small chain [Proteobacteria] Length: 372\nNumber of predicted TMHs: 1\nExp number of AAs in TMHs: 23.72226\nExp number, first 60 AAs: 0.22038\nTotal prob of N-in: 0.12994\noutside 1 331\nTMhelix 332 354\ninside 355 372

13254 GCF_000618705.2_EcK5198 Escherichia coli O121:H19 str. K5198 Proteobacteria; Gammaproteobacteria; Enterobacterales; Enterobacteriaceae; Escherichia; Escherichia coli; Escherichia coli O121 MTGDNTLIHSHGINRRDFMKLCAALAATMGLSSKAAAEMAESVTNPQRPPVIWIGAQECT WP_000145410.1 MULTISPECIES: hydrogenase-2 small chain [Proteobacteria] Length: 372\nNumber of predicted TMHs: 1\nExp number of AAs in TMHs: 23.72226\nExp number, first 60 AAs: 0.22038\nTotal prob of N-in: 0.12994\noutside 1 331\nTMhelix 332 354\ninside 355 372

13255 GCF_000618725.2_EcK5269 Escherichia coli O121:H19 str. K5269 Proteobacteria; Gammaproteobacteria; Enterobacterales; Enterobacteriaceae; Escherichia; Escherichia coli; Escherichia coli O121 MTGDNTLIHSHGINRRDFMKLCAALAATMGLSSKAAAEMAESVTNPQRPPVIWIGAQECT WP_000145410.1 MULTISPECIES: hydrogenase-2 small chain [Proteobacteria] Length: 372\nNumber of predicted TMHs: 1\nExp number of AAs in TMHs: 23.72226\nExp number, first 60 AAs: 0.22038\nTotal prob of N-in: 0.12994\noutside 1 331\nTMhelix 332 354\ninside 355 372

13256 GCF_000234275.2_ASM23427v3 Escherichia coli O121:H19 str. MT#2 Proteobacteria; Gammaproteobacteria; Enterobacterales; Enterobacteriaceae; Escherichia; Escherichia coli; Escherichia coli O121 MTGDNTLIHSHGINRRDFMKLCAALAATMGLSSKAAAEMAESVTNPQRPPVIWIGAQECT WP_000145410.1 MULTISPECIES: hydrogenase-2 small chain [Proteobacteria] Length: 372\nNumber of predicted TMHs: 1\nExp number of AAs in TMHs: 23.72226\nExp number, first 60 AAs: 0.22038\nTotal prob of N-in: 0.12994\noutside 1 331\nTMhelix 332 354\ninside 355 372

13257 GCF_000619245.1_Ec2009C-3299 Escherichia coli O121:H7 str. 2009C-3299 Proteobacteria; Gammaproteobacteria; Enterobacterales; Enterobacteriaceae; Escherichia; Escherichia coli; Escherichia coli O121 MNNEETFYQAMRRQGVTRRSFLKYCSLAATSLGLGAGMAPKIAWALENKPRIPVVWIHGL WP_001058323.1 MULTISPECIES: hydrogenase-1 small chain [Proteobacteria] Length: 372\nNumber of predicted TMHs: 2\nExp number of AAs in TMHs: 37.2608499999999\nExp number, first 60 AAs: 19.52923\nTotal prob of N-in: 0.94351\nPOSSIBLE N-term signal sequence\ninside 1 19\nTMhelix 20 39\noutside 40 325\nTMhelix 326 348\ninside 349 372

13258 GCF_000619245.1_Ec2009C-3299 Escherichia coli O121:H7 str. 2009C-3299 Proteobacteria; Gammaproteobacteria; Enterobacterales; Enterobacteriaceae; Escherichia; Escherichia coli; Escherichia coli O121 MTGDNTLIHSHGINRRDFMKLCAALAATMGLSSKAAAEMAESVTNPQRPPVIWIGAQECT WP_000145410.1 MULTISPECIES: hydrogenase-2 small chain [Proteobacteria] Length: 372\nNumber of predicted TMHs: 1\nExp number of AAs in TMHs: 23.72226\nExp number, first 60 AAs: 0.22038\nTotal prob of N-in: 0.12994\noutside 1 331\nTMhelix 332 354\ninside 355 372

13259 GCF_000619325.2_Ec2009C-3307 Escherichia coli O123:H11 str. 2009C-3307 Proteobacteria; Gammaproteobacteria; Enterobacterales; Enterobacteriaceae; Escherichia; Escherichia coli; Escherichia coli O123 MNNEETFYQAMRRQGVTRRSFLKYCSLAATSLGLGAGMAPKIAWALENKPRIPVVWIHGL WP_001058323.1 MULTISPECIES: hydrogenase-1 small chain [Proteobacteria] Length: 372\nNumber of predicted TMHs: 2\nExp number of AAs in TMHs: 37.2608499999999\nExp number, first 60 AAs: 19.52923\nTotal prob of N-in: 0.94351\nPOSSIBLE N-term signal sequence\ninside 1 19\nTMhelix 20 39\noutside 40 325\nTMhelix 326 348\ninside 349 372

13260 GCF_000619325.2_Ec2009C-3307 Escherichia coli O123:H11 str. 2009C-3307 Proteobacteria; Gammaproteobacteria; Enterobacterales; Enterobacteriaceae; Escherichia; Escherichia coli; Escherichia coli O123 MTGDNTLIHSHGINRRDFMKLCAALAATMGLSSKAAAEMAESVTNPQRPPVIWIGAQECT WP_000145410.1 MULTISPECIES: hydrogenase-2 small chain [Proteobacteria] Length: 372\nNumber of predicted TMHs: 1\nExp number of AAs in TMHs: 23.72226\nExp number, first 60 AAs: 0.22038\nTotal prob of N-in: 0.12994\noutside 1 331\nTMhelix 332 354\ninside 355 372

13261 GCF_000350045.2_C43_90_IrSpV01 Escherichia coli O127:H27 str. C43/90 Proteobacteria; Gammaproteobacteria; Enterobacterales; Enterobacteriaceae; Escherichia; Escherichia coli; Escherichia coli O127 MTGDNTLIHSHGINRRDFMKLCAALAATMGLSSKAAAEMAESVTNPQRPPVIWIGAQECT WP_000145410.1 MULTISPECIES: hydrogenase-2 small chain [Proteobacteria] Length: 372\nNumber of predicted TMHs: 1\nExp number of AAs in TMHs: 23.72226\nExp number, first 60 AAs: 0.22038\nTotal prob of N-in: 0.12994\noutside 1 331\nTMhelix 332 354\ninside 355 372

13262 GCF_000350045.2_C43_90_IrSpV01 Escherichia coli O127:H27 str. C43/90 Proteobacteria; Gammaproteobacteria; Enterobacterales; Enterobacteriaceae; Escherichia; Escherichia coli; Escherichia coli O127 MNNEETFYQAMRRQGVTRRSFLKYCSLAATSLGLGAGMAPKIAWALENKPRIPVVWIHGL WP_001058323.1 MULTISPECIES: hydrogenase-1 small chain [Proteobacteria] Length: 372\nNumber of predicted TMHs: 2\nExp number of AAs in TMHs: 37.2608499999999\nExp number, first 60 AAs: 19.52923\nTotal prob of N-in: 0.94351\nPOSSIBLE N-term signal sequence\ninside 1 19\nTMhelix 20 39\noutside 40 325\nTMhelix 326 348\ninside 349 372

13263 GCF_000026545.1_ASM2654v1 Escherichia coli O127:H6 str. E2348/69 Proteobacteria; Gammaproteobacteria; Enterobacterales; Enterobacteriaceae; Escherichia; Escherichia coli; Escherichia coli O127 MTGDNTLIHSHGINRRDFMKLCAALAATMGLSSKAAAEMAESVTNPQRPPVIWIGAQECT WP_000145408.1 hydrogenase 2 small subunit [Escherichia coli] Length: 372\nNumber of predicted TMHs: 1\nExp number of AAs in TMHs: 23.72255\nExp number, first 60 AAs: 0.22038\nTotal prob of N-in: 0.12995\noutside 1 331\nTMhelix 332 354\ninside 355 372

13264 GCF_000026545.1_ASM2654v1 Escherichia coli O127:H6 str. E2348/69 Proteobacteria; Gammaproteobacteria; Enterobacterales; Enterobacteriaceae; Escherichia; Escherichia coli; Escherichia coli O127

MNNEETFYQAMRRQGVTRRSFLKYCSLAATSLGLGAGMAPKIAWALENKPRIPVVWIHGL WP_001058323.1 MULTISPECIES: hydrogenase-1 small chain [Proteobacteria] Length: 372\nNumber of predicted TMHs: 2\nExp number of AAs in TMHs: 37.2608499999999\nExp number, first 60 AAs: 19.52923\nTotal prob of N-in: 0.94351\nPOSSIBLE N-term signal sequence\ninside 1 19\nTMhelix 20 39\noutside 40 325\nTMhelix 326 348\ninside 349 372

13265 GCF_000442065.2_Escherichia_coli_O127_H6_str._E2348_69_substr._CVDNalr Escherichia coli O127:H6 str. E2348/69 substr. CVDNalr Proteobacteria; Gammaproteobacteria; Enterobacterales; Enterobacteriaceae; Escherichia; Escherichia coli; Escherichia coli O127; Escherichia coli O127 MNNEETFYQAMRRQGVTRRSFLKYCSLAATSLGLGAGMAPKIAWALENKPRIPVVWIHGL WP_001058323.1 MULTISPECIES: hydrogenase-1 small chain [Proteobacteria] Length: 372\nNumber of predicted TMHs: 2\nExp number of AAs in TMHs: 37.2608499999999\nExp number, first 60 AAs: 19.52923\nTotal prob of N-in: 0.94351\nPOSSIBLE N-term signal sequence\ninside 1 19\nTMhelix 20 39\noutside 40 325\nTMhelix 326 348\ninside 349 372

13266 GCF_000442065.2_Escherichia_coli_O127_H6_str._E2348_69_substr._CVDNalr Escherichia coli O127:H6 str. E2348/69 substr. CVDNalr Proteobacteria; Gammaproteobacteria; Enterobacterales; Enterobacteriaceae; Escherichia; Escherichia coli; Escherichia coli O127; Escherichia coli O127 MTGDNTLIHSHGINRRDFMKLCAALAATMGLSSKAAAEMAESVTNPQRPPVIWIGAQECT WP_000145408.1 hydrogenase 2 small subunit [Escherichia coli] Length: 372\nNumber of predicted TMHs: 1\nExp number of AAs in TMHs: 23.72255\nExp number, first 60 AAs: 0.22038\nTotal prob of N-in: 0.12995\noutside 1 331\nTMhelix 332 354\ninside 355 372

13267 GCF_000442085.2_Escherichia_coli_O127_H6_str._E2348_69_substr._UMD753 Escherichia coli O127:H6 str. E2348/69 substr. UMD753 Proteobacteria; Gammaproteobacteria; Enterobacterales; Enterobacteriaceae; Escherichia; Escherichia coli; Escherichia coli O127; Escherichia coli O127 MNNEETFYQAMRRQGVTRRSFLKYCSLAATSLGLGAGMAPKIAWALENKPRIPVVWIHGL WP_001058323.1 MULTISPECIES: hydrogenase-1 small chain [Proteobacteria] Length: 372\nNumber of predicted TMHs: 2\nExp number of AAs in TMHs: 37.2608499999999\nExp number, first 60 AAs: 19.52923\nTotal prob of N-in: 0.94351\nPOSSIBLE N-term signal sequence\ninside 1 19\nTMhelix 20 39\noutside 40 325\nTMhelix 326 348\ninside 349 372

13268 GCF_000442085.2_Escherichia_coli_O127_H6_str._E2348_69_substr._UMD753 Escherichia coli O127:H6 str. E2348/69 substr. UMD753 Proteobacteria; Gammaproteobacteria; Enterobacterales; Enterobacteriaceae; Escherichia; Escherichia coli; Escherichia coli O127; Escherichia coli O127 MTGDNTLIHSHGINRRDFMKLCAALAATMGLSSKAAAEMAESVTNPQRPPVIWIGAQECT WP_000145408.1 hydrogenase 2 small subunit [Escherichia coli] Length: 372\nNumber of predicted TMHs: 1\nExp number of AAs in TMHs: 23.72255\nExp number, first 60 AAs: 0.22038\nTotal prob of N-in: 0.12995\noutside 1 331\nTMhelix 332 354\ninside 355 372

13269 GCF_000703345.1_Ec2011C-3317 Escherichia coli O128:H2 str. 2011C-3317 Proteobacteria; Gammaproteobacteria; Enterobacterales; Enterobacteriaceae; Escherichia; Escherichia coli; Escherichia coli O128 MTGDNTLIHSHGINRRDFMKLCAALAATMGLSSKAAAEMAESVTNPQRPPVIWIGAQECT WP_000145410.1 MULTISPECIES: hydrogenase-2 small chain [Proteobacteria] Length: 372\nNumber of predicted TMHs: 1\nExp number of AAs in TMHs: 23.72226\nExp number, first 60 AAs: 0.22038\nTotal prob of N-in: 0.12994\noutside 1 331\nTMhelix 332 354\ninside 355 372

13270 GCF_000703345.1_Ec2011C-3317 Escherichia coli O128:H2 str. 2011C-3317 Proteobacteria; Gammaproteobacteria; Enterobacterales; Enterobacteriaceae; Escherichia; Escherichia coli; Escherichia coli O128 MNNEETFYQAMRRQGVTRRSFLKYCSLAATSLGLGAGMAPKIAWALENKPRIPVVWIHGL WP_001058323.1 MULTISPECIES: hydrogenase-1 small chain [Proteobacteria] Length: 372\nNumber of predicted TMHs: 2\nExp number of AAs in TMHs: 37.2608499999999\nExp number, first 60 AAs: 19.52923\nTotal prob of N-in: 0.94351\nPOSSIBLE N-term signal sequence\ninside 1 19\nTMhelix 20 39\noutside 40 325\nTMhelix 326 348\ninside 349 372

13271 GCF_000017745.1_ASM1774v1 Escherichia coli O139:H28 str. E24377A Proteobacteria; Gammaproteobacteria; Enterobacterales; Enterobacteriaceae; Escherichia; Escherichia coli; Escherichia coli O139 MTGDNTLIHSHGINRRDFMKLCAALAATMGLSSKAAAEMAESVTNPQRPPVIWIGAQECT WP_000145410.1 MULTISPECIES: hydrogenase-2 small chain [Proteobacteria] Length: 372\nNumber of predicted TMHs: 1\nExp number of AAs in TMHs: 23.72226\nExp number, first 60 AAs: 0.22038\nTotal prob of N-in: 0.12994\noutside 1 331\nTMhelix 332 354\ninside 355 372

13272 GCF_000017745.1_ASM1774v1 Escherichia coli O139:H28 str. E24377A Proteobacteria; Gammaproteobacteria; Enterobacterales; Enterobacteriaceae; Escherichia; Escherichia coli; Escherichia coli O139 MNNEETFYQVMRRQGVTRRSFLKYCSLAATSLGLGAGMAPKIAWALENKPRIPVVWIHGL WP_001058328.1 hydrogenase [Escherichia coli] Length: 372\nNumber of predicted TMHs: 2\nExp number of AAs in TMHs: 37.1484199999999\nExp number, first 60 AAs: 19.41882\nTotal prob of N-in: 0.93813\nPOSSIBLE N-term signal sequence\ninside 1 19\nTMhelix 20 39\noutside 40 325\nTMhelix 326 348\ninside 349 372

13273 GCF_000703225.1_Ec07-3858 Escherichia coli O145:H25 str. 07-3858 Proteobacteria; Gammaproteobacteria; Enterobacterales; Enterobacteriaceae; Escherichia; Escherichia coli; Escherichia coli O145 MTGDNTLIHSHGINRRDFMKLCAALAATMGLSSKAAAEMAESVTNPQRPPVIWIGAQECT WP_000145410.1 MULTISPECIES: hydrogenase-2 small chain [Proteobacteria] Length: 372\nNumber of predicted TMHs: 1\nExp number of AAs in TMHs: 23.72226\nExp number, first 60 AAs: 0.22038\nTotal prob of N-in: 0.12994\noutside 1 331\nTMhelix 332 354\ninside 355 372

13274 GCF_000703225.1_Ec07-3858 Escherichia coli O145:H25 str. 07-3858 Proteobacteria; Gammaproteobacteria; Enterobacterales; Enterobacteriaceae; Escherichia; Escherichia coli; Escherichia coli O145 MNNEETFYQAMRRQGVTRRSFLKYCSLAATSLGLGAGMAPKIAWALENKPRIPVVWIHGL WP_001058323.1 MULTISPECIES: hydrogenase-1 small chain [Proteobacteria] Length: 372\nNumber of predicted TMHs: 2\nExp number of AAs in TMHs: 37.2608499999999\nExp number, first 60 AAs: 19.52923\nTotal prob of N-in: 0.94351\nPOSSIBLE N-term signal sequence\ninside 1 19\nTMhelix 20 39\noutside 40 325\nTMhelix 326 348\ninside 349 372

13275 GCF_000619345.1_Ec2009C-3292 Escherichia coli O145:H28 str. 2009C-3292 Proteobacteria; Gammaproteobacteria; Enterobacterales; Enterobacteriaceae; Escherichia; Escherichia coli; Escherichia coli O145 MNNEETFYQAMRRQGVTRRSFLKYCSLAATSLGLGAGMAPKIAWALENKPRIPVVWIHGL WP_001058323.1 MULTISPECIES: hydrogenase-1 small chain [Proteobacteria] Length: 372\nNumber of predicted TMHs: 2\nExp number of AAs in TMHs: 37.2608499999999\nExp number, first 60 AAs: 19.52923\nTotal prob of N-in: 0.94351\nPOSSIBLE N-term signal sequence\ninside 1 19\nTMhelix 20 39\noutside 40 325\nTMhelix 326 348\ninside 349 372

13276 GCF_000619345.1_Ec2009C-3292 Escherichia coli O145:H28 str. 2009C-3292 Proteobacteria; Gammaproteobacteria; Enterobacterales; Enterobacteriaceae; Escherichia; Escherichia coli; Escherichia coli O145 MTGDNTLIHSHGINRRDFMKLCAALAATMGLSSKAAAEMAESVTNPQRPPVIWIGAQECT WP_000145410.1 MULTISPECIES: hydrogenase-2 small chain [Proteobacteria] Length: 372\nNumber of predicted TMHs: 1\nExp number of AAs in TMHs: 23.72226\nExp number, first 60 AAs: 0.22038\nTotal prob of N-in: 0.12994\noutside 1 331\nTMhelix 332 354\ninside 355 372

13277 GCF_000234315.1_ASM23431v2 Escherichia coli O145:H28 str. 4865/96 Proteobacteria; Gammaproteobacteria; Enterobacterales; Enterobacteriaceae; Escherichia; Escherichia coli; Escherichia coli O145 MTGDNTLIHSHGINRRDFMKLCAALAATMGLSSKAAAEMAESVTNPQRPPVIWIGAQECT WP_000145410.1 MULTISPECIES: hydrogenase-2 small chain [Proteobacteria] Length: 372\nNumber of predicted TMHs: 1\nExp number of AAs in TMHs: 23.72226\nExp number, first 60 AAs: 0.22038\nTotal prob of N-in: 0.12994\noutside 1 331\nTMhelix 332 354\ninside 355 372

13278 GCF_000234315.1_ASM23431v2 Escherichia coli O145:H28 str. 4865/96 Proteobacteria; Gammaproteobacteria; Enterobacterales; Enterobacteriaceae; Escherichia; Escherichia coli; Escherichia coli O145 MNNEETFYQAMRRQGVTRRSFLKYCSLAATSLGLGAGMAPKIAWALENKPRIPVVWIHGL WP_001058323.1 MULTISPECIES: hydrogenase-1 small chain [Proteobacteria] Length: 372\nNumber of predicted TMHs: 2\nExp number of AAs in TMHs: 37.2608499999999\nExp number, first 60 AAs: 19.52923\nTotal prob of N-in: 0.94351\nPOSSIBLE N-term signal sequence\ninside 1 19\nTMhelix 20 39\noutside 40 325\nTMhelix 326 348\ninside 349 372

13279 GCF_000695155.1_Ecoli4865_96v1 Escherichia coli O145:H28 str. 4865/96 Proteobacteria; Gammaproteobacteria; Enterobacterales; Enterobacteriaceae; Escherichia; Escherichia coli; Escherichia coli O145 MNNEETFYQAMRRQGVTRRSFLKYCSLAATSLGLGAGMAPKIAWALENKPRIPVVWIHGL WP_001058323.1 MULTISPECIES: hydrogenase-1 small chain [Proteobacteria] Length: 372\nNumber of predicted TMHs: 2\nExp number of AAs in TMHs: 37.2608499999999\nExp number, first 60 AAs: 19.52923\nTotal prob of N-in: 0.94351\nPOSSIBLE N-term signal sequence\ninside 1 19\nTMhelix 20 39\noutside 40 325\nTMhelix 326 348\ninside 349 372

13280 GCF_000695155.1_Ecoli4865_96v1 Escherichia coli O145:H28 str. 4865/96 Proteobacteria; Gammaproteobacteria; Enterobacterales; Enterobacteriaceae; Escherichia; Escherichia coli; Escherichia coli O145 MTGDNTLIHSHGINRRDFMKLCAALAATMGLSSKAAAEMAESVTNPQRPPVIWIGAQECT WP_000145410.1 MULTISPECIES: hydrogenase-2 small chain [Proteobacteria] Length: 372\nNumber of predicted TMHs: 1\nExp number of AAs in TMHs: 23.72226\nExp number, first 60 AAs: 0.22038\nTotal prob of N-in: 0.12994\noutside 1 331\nTMhelix 332 354\ninside 355 372

13281 GCF_000671295.1_ASM67129v1 Escherichia coli O145:H28 str. RM12581 Proteobacteria; Gammaproteobacteria; Enterobacterales; Enterobacteriaceae; Escherichia; Escherichia coli; Escherichia coli O145 MTGDNTLIHSHGINRRDFMKLCAALAATMGLSSKAAAEMAESVTNPQRPPVIWIGAQECT WP_000145410.1 MULTISPECIES: hydrogenase-2 small chain [Proteobacteria] Length: 372\nNumber of predicted TMHs: 1\nExp number of AAs in TMHs: 23.72226\nExp number, first 60 AAs: 0.22038\nTotal prob of N-in: 0.12994\noutside 1 331\nTMhelix 332 354\ninside 355 372

13282 GCF_000671295.1_ASM67129v1 Escherichia coli O145:H28 str. RM12581 Proteobacteria; Gammaproteobacteria; Enterobacterales; Enterobacteriaceae; Escherichia; Escherichia coli; Escherichia coli O145 MNNEETFYQAMRRQGVTRRSFLKYCSLAATSLGLGAGMAPKIAWALENKPRIPVVWIHGL WP_001058323.1 MULTISPECIES: hydrogenase-1 small chain [Proteobacteria] Length: 372\nNumber of predicted TMHs: 2\nExp number of AAs in TMHs: 37.2608499999999\nExp number, first 60 AAs: 19.52923\nTotal prob of N-in: 0.94351\nPOSSIBLE N-term signal sequence\ninside 1 19\nTMhelix 20 39\noutside 40 325\nTMhelix 326 348\ninside 349 372

13283 GCF_000662395.1_ASM66239v1 Escherichia coli O145:H28 str. RM12761 Proteobacteria; Gammaproteobacteria; Enterobacterales; Enterobacteriaceae; Escherichia; Escherichia coli; Escherichia coli O145 MNNEETFYQAMRRQGVTRRSFLKYCSLAATSLGLGAGMAPKIAWALENKPRIPVVWIHGL WP_001058323.1 MULTISPECIES: hydrogenase-1 small chain [Proteobacteria] Length: 372\nNumber of predicted TMHs: 2\nExp number of AAs in TMHs: 37.2608499999999\nExp number, first 60 AAs: 19.52923\nTotal prob of N-in: 0.94351\nPOSSIBLE N-term signal sequence\ninside 1 19\nTMhelix 20 39\noutside 40 325\nTMhelix 326 348\ninside 349 372

13284 GCF_000662395.1_ASM66239v1 Escherichia coli O145:H28 str. RM12761 Proteobacteria; Gammaproteobacteria; Enterobacterales; Enterobacteriaceae; Escherichia; Escherichia coli; Escherichia coli O145 MTGDNTLIHSHGINRRDFMKLCAALAATMGLSSKAAAEMAESVTNPQRPPVIWIGAQECT WP_000145410.1 MULTISPECIES: hydrogenase-2 small chain [Proteobacteria] Length: 372\nNumber of predicted TMHs: 1\nExp number of AAs in TMHs: 23.72226\nExp number, first 60 AAs: 0.22038\nTotal prob of N-in: 0.12994\noutside 1 331\nTMhelix 332 354\ninside 355 372

13285 GCF_000520035.1_ASM52003v1 Escherichia coli O145:H28 str. RM13514 Proteobacteria; Gammaproteobacteria; Enterobacterales; Enterobacteriaceae; Escherichia; Escherichia coli; Escherichia coli O145 MNNEETFYQAMRRQGVTRRSFLKYCSLAATSLGLGAGMAPKIAWALENKPRIPVVWIHGL WP_001058323.1 MULTISPECIES: hydrogenase-1 small chain [Proteobacteria] Length: 372\nNumber of predicted TMHs: 2\nExp number of AAs in TMHs: 37.2608499999999\nExp number, first 60 AAs: 19.52923\nTotal prob of N-in: 0.94351\nPOSSIBLE N-term signal sequence\ninside 1 19\nTMhelix 20 39\noutside 40 325\nTMhelix 326 348\ninside 349 372

13286 GCF_000520035.1_ASM52003v1 Escherichia coli O145:H28 str. RM13514 Proteobacteria; Gammaproteobacteria; Enterobacterales; Enterobacteriaceae; Escherichia; Escherichia coli; Escherichia coli O145 MTGDNTLIHSHGINRRDFMKLCAALAATMGLSSKAAAEMAESVTNPQRPPVIWIGAQECT WP_000145410.1 MULTISPECIES: hydrogenase-2 small chain [Proteobacteria] Length: 372\nNumber of predicted TMHs: 1\nExp number of AAs in TMHs: 23.72226\nExp number, first 60 AAs: 0.22038\nTotal prob of N-in: 0.12994\noutside 1 331\nTMhelix 332 354\ninside 355 372

13287 GCF_000520055.1_ASM52005v1 Escherichia coli O145:H28 str. RM13516 Proteobacteria; Gammaproteobacteria; Enterobacterales; Enterobacteriaceae; Escherichia; Escherichia coli; Escherichia coli O145 MTGDNTLIHSHGINRRDFMKLCAALAATMGLSSKAAAEMAESVTNPQRPPVIWIGAQECT WP_000145410.1 MULTISPECIES: hydrogenase-2 small chain [Proteobacteria] Length: 372\nNumber of predicted TMHs: 1\nExp number of AAs in TMHs: 23.72226\nExp number, first 60 AAs: 0.22038\nTotal prob of N-in: 0.12994\noutside 1 331\nTMhelix 332 354\ninside 355 372

13288 GCF_000520055.1_ASM52005v1 Escherichia coli O145:H28 str. RM13516 Proteobacteria; Gammaproteobacteria; Enterobacterales; Enterobacteriaceae; Escherichia; Escherichia coli; Escherichia coli O145

MNNEETFYQAMRRQGVTRRSFLKYCSLAATSLGLGAGMAPKIAWALENKPRIPVVWIHGL WP_001058323.1 MULTISPECIES: hydrogenase-1 small chain [Proteobacteria] Length: 372\nNumber of predicted TMHs: 2\nExp number of AAs in TMHs: 37.2608499999999\nExp number, first 60 AAs: 19.52923\nTotal prob of N-in: 0.94351\nPOSSIBLE N-term signal sequence\ninside 1 19\nTMhelix 20 39\noutside 40 325\nTMhelix 326 348\ninside 349 372

13289 GCF_000617345.1_Ec06-3484 Escherichia coli O145:NM str. 06-3484 Proteobacteria; Gammaproteobacteria; Enterobacterales; Enterobacteriaceae; Escherichia; Escherichia coli; Escherichia coli O145 MTGDNTLIHSHGINRRDFMKLCAALAATMGLSSKAAAEMAESVTNPQRPPVIWIGAQECT WP_000145410.1 MULTISPECIES: hydrogenase-2 small chain [Proteobacteria] Length: 372\nNumber of predicted TMHs: 1\nExp number of AAs in TMHs: 23.72226\nExp number, first 60 AAs: 0.22038\nTotal prob of N-in: 0.12994\noutside 1 331\nTMhelix 332 354\ninside 355 372

13290 GCF_000617345.1_Ec06-3484 Escherichia coli O145:NM str. 06-3484 Proteobacteria; Gammaproteobacteria; Enterobacterales; Enterobacteriaceae; Escherichia; Escherichia coli; Escherichia coli O145 MNNEETFYQAMRRQGVTRRSFLKYCSLAATSLGLGAGMAPKIAWALENKPRIPVVWIHGL WP_001058323.1 MULTISPECIES: hydrogenase-1 small chain [Proteobacteria] Length: 372\nNumber of predicted TMHs: 2\nExp number of AAs in TMHs: 37.2608499999999\nExp number, first 60 AAs: 19.52923\nTotal prob of N-in: 0.94351\nPOSSIBLE N-term signal sequence\ninside 1 19\nTMhelix 20 39\noutside 40 325\nTMhelix 326 348\ninside 349 372

13291 GCF_000616665.2_Ec08-4270 Escherichia coli O145:NM str. 08-4270 Proteobacteria; Gammaproteobacteria; Enterobacterales; Enterobacteriaceae; Escherichia; Escherichia coli; Escherichia coli O145 MTGDNTLIHSHGINRRDFMKLCAALAATMGLSSKAAAEMAESVTNPQRPPVIWIGAQECT WP_000145410.1 MULTISPECIES: hydrogenase-2 small chain [Proteobacteria] Length: 372\nNumber of predicted TMHs: 1\nExp number of AAs in TMHs: 23.72226\nExp number, first 60 AAs: 0.22038\nTotal prob of N-in: 0.12994\noutside 1 331\nTMhelix 332 354\ninside 355 372

13292 GCF_000616665.2_Ec08-4270 Escherichia coli O145:NM str. 08-4270 Proteobacteria; Gammaproteobacteria; Enterobacterales; Enterobacteriaceae; Escherichia; Escherichia coli; Escherichia coli O145 MNNEETFYQAMRRQGVTRRSFLKYCSLAATSLGLGAGMAPKIAWALENKPRIPVVWIHGL WP_001058323.1 MULTISPECIES: hydrogenase-1 small chain [Proteobacteria] Length: 372\nNumber of predicted TMHs: 2\nExp number of AAs in TMHs: 37.2608499999999\nExp number, first 60 AAs: 19.52923\nTotal prob of N-in: 0.94351\nPOSSIBLE N-term signal sequence\ninside 1 19\nTMhelix 20 39\noutside 40 325\nTMhelix 326 348\ninside 349 372

13293 GCF_000615085.1_Ec2010C-3507 Escherichia coli O145:NM str. 2010C-3507 Proteobacteria; Gammaproteobacteria; Enterobacterales; Enterobacteriaceae; Escherichia; Escherichia coli; Escherichia coli O145 MNNEETFYQAMRRQGVTRRSFLKYCSLAATSLGLGAGMAPKIAWALENKPRIPVVWIHGL WP_001058323.1 MULTISPECIES: hydrogenase-1 small chain [Proteobacteria] Length: 372\nNumber of predicted TMHs: 2\nExp number of AAs in TMHs: 37.2608499999999\nExp number, first 60 AAs: 19.52923\nTotal prob of N-in: 0.94351\nPOSSIBLE N-term signal sequence\ninside 1 19\nTMhelix 20 39\noutside 40 325\nTMhelix 326 348\ninside 349 372

13294 GCF_000615085.1_Ec2010C-3507 Escherichia coli O145:NM str. 2010C-3507 Proteobacteria; Gammaproteobacteria; Enterobacterales; Enterobacteriaceae; Escherichia; Escherichia coli; Escherichia coli O145 MTGDNTLIHSHGINRRDFMKLCAALAATMGLSSKAAAEMAESVTNPQRPPVIWIGAQECT WP_000145410.1 MULTISPECIES: hydrogenase-2 small chain [Proteobacteria] Length: 372\nNumber of predicted TMHs: 1\nExp number of AAs in TMHs: 23.72226\nExp number, first 60 AAs: 0.22038\nTotal prob of N-in: 0.12994\noutside 1 331\nTMhelix 332 354\ninside 355 372

13295 GCF_000619185.1_Ec2010C-3508 Escherichia coli O145:NM str. 2010C-3508 Proteobacteria; Gammaproteobacteria; Enterobacterales; Enterobacteriaceae; Escherichia; Escherichia coli; Escherichia coli O145 MNNEETFYQAMRRQGVTRRSFLKYCSLAATSLGLGAGMAPKIAWALENKPRIPVVWIHGL WP_001058323.1 MULTISPECIES: hydrogenase-1 small chain [Proteobacteria] Length: 372\nNumber of predicted TMHs: 2\nExp number of AAs in TMHs: 37.2608499999999\nExp number, first 60 AAs: 19.52923\nTotal prob of N-in: 0.94351\nPOSSIBLE N-term signal sequence\ninside 1 19\nTMhelix 20 39\noutside 40 325\nTMhelix 326 348\ninside 349 372

13296 GCF_000619185.1_Ec2010C-3508 Escherichia coli O145:NM str. 2010C-3508 Proteobacteria; Gammaproteobacteria; Enterobacterales; Enterobacteriaceae; Escherichia; Escherichia coli; Escherichia coli O145

MTGDNTLIHSHGINRRDFMKLCAALAATMGLSSKAAAEMAESVTNPQRPPVIWIGAQECT WP_000145410.1 MULTISPECIES: hydrogenase-2 small chain [Proteobacteria] Length: 372\nNumber of predicted TMHs: 1\nExp number of AAs in TMHs: 23.72226\nExp number, first 60 AAs: 0.22038\nTotal prob of N-in: 0.12994\noutside 1 331\nTMhelix 332 354\ninside 355 372

13297 GCF_000615145.1_Ec2010C-3509 Escherichia coli O145:NM str. 2010C-3509 Proteobacteria; Gammaproteobacteria; Enterobacterales; Enterobacteriaceae; Escherichia; Escherichia coli; Escherichia coli O145 MTGDNTLIHSHGINRRDFMKLCAALAATMGLSSKAAAEMAESVTNPQRPPVIWIGAQECT WP_000145410.1 MULTISPECIES: hydrogenase-2 small chain [Proteobacteria] Length: 372\nNumber of predicted TMHs: 1\nExp number of AAs in TMHs: 23.72226\nExp number, first 60 AAs: 0.22038\nTotal prob of N-in: 0.12994\noutside 1 331\nTMhelix 332 354\ninside 355 372

13298 GCF_000615145.1_Ec2010C-3509 Escherichia coli O145:NM str. 2010C-3509 Proteobacteria; Gammaproteobacteria; Enterobacterales; Enterobacteriaceae; Escherichia; Escherichia coli; Escherichia coli O145 MNNEETFYQAMRRQGVTRRSFLKYCSLAATSLGLGAGMAPKIAWALENKPRIPVVWIHGL WP_001058323.1 MULTISPECIES: hydrogenase-1 small chain [Proteobacteria] Length: 372\nNumber of predicted TMHs: 2\nExp number of AAs in TMHs: 37.2608499999999\nExp number, first 60 AAs: 19.52923\nTotal prob of N-in: 0.94351\nPOSSIBLE N-term signal sequence\ninside 1 19\nTMhelix 20 39\noutside 40 325\nTMhelix 326 348\ninside 349 372

13299 GCF_000615175.1_Ec2010C-3510 Escherichia coli O145:NM str. 2010C-3510 Proteobacteria; Gammaproteobacteria; Enterobacterales; Enterobacteriaceae; Escherichia; Escherichia coli; Escherichia coli O145 MTGDNTLIHSHGINRRDFMKLCAALAATMGLSSKAAAEMAESVTNPQRPPVIWIGAQECT WP_000145410.1 MULTISPECIES: hydrogenase-2 small chain [Proteobacteria] Length: 372\nNumber of predicted TMHs: 1\nExp number of AAs in TMHs: 23.72226\nExp number, first 60 AAs: 0.22038\nTotal prob of N-in: 0.12994\noutside 1 331\nTMhelix 332 354\ninside 355 372

13300 GCF_000615175.1_Ec2010C-3510 Escherichia coli O145:NM str. 2010C-3510 Proteobacteria; Gammaproteobacteria; Enterobacterales; Enterobacteriaceae; Escherichia; Escherichia coli; Escherichia coli O145 MNNEETFYQAMRRQGVTRRSFLKYCSLAATSLGLGAGMAPKIAWALENKPRIPVVWIHGL WP_001058323.1 MULTISPECIES: hydrogenase-1 small chain [Proteobacteria] Length: 372\nNumber of predicted TMHs: 2\nExp number of AAs in TMHs: 37.2608499999999\nExp number, first 60 AAs: 19.52923\nTotal prob of N-in: 0.94351\nPOSSIBLE N-term signal sequence\ninside 1 19\nTMhelix 20 39\noutside 40 325\nTMhelix 326 348\ninside 349 372

13301 GCF_000615225.2_Ec2010C-3511 Escherichia coli O145:NM str. 2010C-3511 Proteobacteria; Gammaproteobacteria; Enterobacterales; Enterobacteriaceae; Escherichia; Escherichia coli; Escherichia coli O145 MTGDNTLIHSHGINRRDFMKLCAALAATMGLSSKAAAEMAESVTNPQRPPVIWIGAQECT WP_000145410.1 MULTISPECIES: hydrogenase-2 small chain [Proteobacteria] Length: 372\nNumber of predicted TMHs: 1\nExp number of AAs in TMHs: 23.72226\nExp number, first 60 AAs: 0.22038\nTotal prob of N-in: 0.12994\noutside 1 331\nTMhelix 332 354\ninside 355 372

13302 GCF_000615225.2_Ec2010C-3511 Escherichia coli O145:NM str. 2010C-3511 Proteobacteria; Gammaproteobacteria; Enterobacterales; Enterobacteriaceae; Escherichia; Escherichia coli; Escherichia coli O145 MNNEETFYQAMRRQGVTRRSFLKYCSLAATSLGLGAGMAPKIAWALENKPRIPVVWIHGL WP_001058323.1 MULTISPECIES: hydrogenase-1 small chain [Proteobacteria] Length: 372\nNumber of predicted TMHs: 2\nExp number of AAs in TMHs: 37.2608499999999\nExp number, first 60 AAs: 19.52923\nTotal prob of N-in: 0.94351\nPOSSIBLE N-term signal sequence\ninside 1 19\nTMhelix 20 39\noutside 40 325\nTMhelix 326 348\ninside 349 372

13303 GCF_000615265.1_Ec2010C-3516 Escherichia coli O145:NM str. 2010C-3516 Proteobacteria; Gammaproteobacteria; Enterobacterales; Enterobacteriaceae; Escherichia; Escherichia coli; Escherichia coli O145 MNNEETFYQAMRRQGVTRRSFLKYCSLAATSLGLGAGMAPKIAWALENKPRIPVVWIHGL WP_001058323.1 MULTISPECIES: hydrogenase-1 small chain [Proteobacteria] Length: 372\nNumber of predicted TMHs: 2\nExp number of AAs in TMHs: 37.2608499999999\nExp number, first 60 AAs: 19.52923\nTotal prob of N-in: 0.94351\nPOSSIBLE N-term signal sequence\ninside 1 19\nTMhelix 20 39\noutside 40 325\nTMhelix 326 348\ninside 349 372

13304 GCF_000615265.1_Ec2010C-3516 Escherichia coli O145:NM str. 2010C-3516 Proteobacteria; Gammaproteobacteria; Enterobacterales; Enterobacteriaceae; Escherichia; Escherichia coli; Escherichia coli O145 MTGDNTLIHSHGINRRDFMKLCAALAATMGLSSKAAAEMAESVTNPQRPPVIWIGAQECT WP_000145410.1

MULTISPECIES: hydrogenase-2 small chain [Proteobacteria] Length: 372\nNumber of predicted TMHs: 1\nExp number of AAs in TMHs: 23.72226\nExp number, first 60 AAs: 0.22038\nTotal prob of N-in: 0.12994\noutside 1 331\nTMhelix 332 354\ninside 355 372

13305 GCF_000615305.1_Ec2010C-3517 Escherichia coli O145:NM str. 2010C-3517 Proteobacteria; Gammaproteobacteria; Enterobacterales; Enterobacteriaceae; Escherichia; Escherichia coli; Escherichia coli O145 MTGDNTLIHSHGINRRDFMKLCAALAATMGLSSKAAAEMAESVTNPQRPPVIWIGAQECT WP_000145410.1 MULTISPECIES: hydrogenase-2 small chain [Proteobacteria] Length: 372\nNumber of predicted TMHs: 1\nExp number of AAs in TMHs: 23.72226\nExp number, first 60 AAs: 0.22038\nTotal prob of N-in: 0.12994\noutside 1 331\nTMhelix 332 354\ninside 355 372

13306 GCF_000615305.1_Ec2010C-3517 Escherichia coli O145:NM str. 2010C-3517 Proteobacteria; Gammaproteobacteria; Enterobacterales; Enterobacteriaceae; Escherichia; Escherichia coli; Escherichia coli O145 MNNEETFYQAMRRQGVTRRSFLKYCSLAATSLGLGAGMAPKIAWALENKPRIPVVWIHGL WP_001058323.1 MULTISPECIES: hydrogenase-1 small chain [Proteobacteria] Length: 372\nNumber of predicted TMHs: 2\nExp number of AAs in TMHs: 37.2608499999999\nExp number, first 60 AAs: 19.52923\nTotal prob of N-in: 0.94351\nPOSSIBLE N-term signal sequence\ninside 1 19\nTMhelix 20 39\noutside 40 325\nTMhelix 326 348\ninside 349 372

13307 GCF_000615345.1_Ec2010C-3518 Escherichia coli O145:NM str. 2010C-3518 Proteobacteria; Gammaproteobacteria; Enterobacterales; Enterobacteriaceae; Escherichia; Escherichia coli; Escherichia coli O145 MTGDNTLIHSHGINRRDFMKLCAALAATMGLSSKAAAEMAESVTNPQRPPVIWIGAQECT WP_000145410.1 MULTISPECIES: hydrogenase-2 small chain [Proteobacteria] Length: 372\nNumber of predicted TMHs: 1\nExp number of AAs in TMHs: 23.72226\nExp number, first 60 AAs: 0.22038\nTotal prob of N-in: 0.12994\noutside 1 331\nTMhelix 332 354\ninside 355 372

13308 GCF_000615345.1_Ec2010C-3518 Escherichia coli O145:NM str. 2010C-3518 Proteobacteria; Gammaproteobacteria; Enterobacterales; Enterobacteriaceae; Escherichia; Escherichia coli; Escherichia coli O145 MNNEETFYQAMRRQGVTRRSFLKYCSLAATSLGLGAGMAPKIAWALENKPRIPVVWIHGL WP_001058323.1 MULTISPECIES: hydrogenase-1 small chain [Proteobacteria] Length: 372\nNumber of predicted TMHs: 2\nExp number of AAs in TMHs: 37.2608499999999\nExp number, first 60 AAs: 19.52923\nTotal prob of N-in: 0.94351\nPOSSIBLE N-term signal sequence\ninside 1 19\nTMhelix 20 39\noutside 40 325\nTMhelix 326 348\ninside 349 372

13309 GCF_000615385.1_Ec2010C-3521 Escherichia coli O145:NM str. 2010C-3521 Proteobacteria; Gammaproteobacteria; Enterobacterales; Enterobacteriaceae; Escherichia; Escherichia coli; Escherichia coli O145 MNNEETFYQAMRRQGVTRRSFLKYCSLAATSLGLGAGMAPKIAWALENKPRIPVVWIHGL WP_001058323.1 MULTISPECIES: hydrogenase-1 small chain [Proteobacteria] Length: 372\nNumber of predicted TMHs: 2\nExp number of AAs in TMHs: 37.2608499999999\nExp number, first 60 AAs: 19.52923\nTotal prob of N-in: 0.94351\nPOSSIBLE N-term signal sequence\ninside 1 19\nTMhelix 20 39\noutside 40 325\nTMhelix 326 348\ninside 349 372

13310 GCF_000615385.1_Ec2010C-3521 Escherichia coli O145:NM str. 2010C-3521 Proteobacteria; Gammaproteobacteria; Enterobacterales; Enterobacteriaceae; Escherichia; Escherichia coli; Escherichia coli O145 MTGDNTLIHSHGINRRDFMKLCAALAATMGLSSKAAAEMAESVTNPQRPPVIWIGAQECT WP_000145410.1 MULTISPECIES: hydrogenase-2 small chain [Proteobacteria] Length: 372\nNumber of predicted TMHs: 1\nExp number of AAs in TMHs: 23.72226\nExp number, first 60 AAs: 0.22038\nTotal prob of N-in: 0.12994\noutside 1 331\nTMhelix 332 354\ninside 355 372

13311 GCF_000615415.2_Ec2010C-3526 Escherichia coli O145:NM str. 2010C-3526 Proteobacteria; Gammaproteobacteria; Enterobacterales; Enterobacteriaceae; Escherichia; Escherichia coli; Escherichia coli O145 MTGDNTLIHSHGINRRDFMKLCAALAATMGLSSKAAAEMAESVTNPQRPPVIWIGAQECT WP_000145410.1 MULTISPECIES: hydrogenase-2 small chain [Proteobacteria] Length: 372\nNumber of predicted TMHs: 1\nExp number of AAs in TMHs: 23.72226\nExp number, first 60 AAs: 0.22038\nTotal prob of N-in: 0.12994\noutside 1 331\nTMhelix 332 354\ninside 355 372

13312 GCF_000615415.2_Ec2010C-3526 Escherichia coli O145:NM str. 2010C-3526 Proteobacteria; Gammaproteobacteria; Enterobacterales; Enterobacteriaceae; Escherichia; Escherichia coli; Escherichia coli O145 MNNEETFYQAMRRQGVTRRSFLKYCSLAATSLGLGAGMAPKIAWALENKPRIPVVWIHGL WP_001058323.1 MULTISPECIES: hydrogenase-1 small chain [Proteobacteria] Length: 372\nNumber of predicted TMHs: 2\nExp number of AAs in TMHs: 37.2608499999999\nExp number, first 60 AAs: 19.52923\nTotal prob of N-in:

0.94351\nPOSSIBLE N-term signal sequence\ninside 1 19\nTMhelix 20 39\noutside 40 325\nTMhelix 326 348\ninside 349 372

13313 GCF_000617765.2_Ec2010C-4557C2 Escherichia coli O145:NM str. 2010C-4557C2 Proteobacteria; Gammaproteobacteria; Enterobacterales; Enterobacteriaceae; Escherichia; Escherichia coli; Escherichia coli O145 MNNEETFYQAMRRQGVTRRSFLKYCSLAATSLGLGAGMAPKIAWALENKPRIPVVWIHGL WP_001058323.1 MULTISPECIES: hydrogenase-1 small chain [Proteobacteria] Length: 372\nNumber of predicted TMHs: 2\nExp number of AAs in TMHs: 37.2608499999999\nExp number, first 60 AAs: 19.52923\nTotal prob of N-in: 0.94351\nPOSSIBLE N-term signal sequence\ninside 1 19\nTMhelix 20 39\noutside 40 325\nTMhelix 326 348\ninside 349 372

13314 GCF_000617765.2_Ec2010C-4557C2 Escherichia coli O145:NM str. 2010C-4557C2 Proteobacteria; Gammaproteobacteria; Enterobacterales; Enterobacteriaceae; Escherichia; Escherichia coli; Escherichia coli O145 MTGDNTLIHSHGINRRDFMKLCAALAATMGLSSKAAAEMAESVTNPQRPPVIWIGAQECT WP_000145410.1 MULTISPECIES: hydrogenase-2 small chain [Proteobacteria] Length: 372\nNumber of predicted TMHs: 1\nExp number of AAs in TMHs: 23.72226\nExp number, first 60 AAs: 0.22038\nTotal prob of N-in: 0.12994\noutside 1 331\nTMhelix 332 354\ninside 355 372

13315 GCF_000703285.1_Ec2010C-3325 Escherichia coli O146:H21 str. 2010C-3325 Proteobacteria; Gammaproteobacteria; Enterobacterales; Enterobacteriaceae; Escherichia; Escherichia coli; Escherichia coli O146 MNNEETFYQAMRRQGVTRRSFLKYCSLAATSLGLGAGMAPKIAWALENKPRIPVVWIHGL WP_001295940.1 MULTISPECIES: hydrogenase [Enterobacteriaceae] Length: 372\nNumber of predicted TMHs: 2\nExp number of AAs in TMHs: 37.2608899999999\nExp number, first 60 AAs: 19.52923\nTotal prob of N-in: 0.94351\nPOSSIBLE N-term signal sequence\ninside 1 19\nTMhelix 20 39\noutside 40 325\nTMhelix 326 348\ninside 349 372

13316 GCF_000703285.1_Ec2010C-3325 Escherichia coli O146:H21 str. 2010C-3325 Proteobacteria; Gammaproteobacteria; Enterobacterales; Enterobacteriaceae; Escherichia; Escherichia coli; Escherichia coli O146 MTGDNTLIHSHGINRRDFMKLCAALAATMGLSSKAAAEMAESVTNPQRPPVIWIGAQECT WP_000145410.1 MULTISPECIES: hydrogenase-2 small chain [Proteobacteria] Length: 372\nNumber of predicted TMHs: 1\nExp number of AAs in TMHs: 23.72226\nExp number, first 60 AAs: 0.22038\nTotal prob of N-in: 0.12994\noutside 1 331\nTMhelix 332 354\ninside 355 372

13317 GCF_000617985.2_EcK1516 Escherichia coli O15:H18 str. K1516 Proteobacteria; Gammaproteobacteria; Enterobacterales; Enterobacteriaceae; Escherichia; Escherichia coli; Escherichia coli O15 MTGDNTLIHSHGINRRDFMKLCAALAATMGLSSKAAAEMAESVTNPQRPPVIWIGAQECT WP_000145410.1 MULTISPECIES: hydrogenase-2 small chain [Proteobacteria] Length: 372\nNumber of predicted TMHs: 1\nExp number of AAs in TMHs: 23.72226\nExp number, first 60 AAs: 0.22038\nTotal prob of N-in: 0.12994\noutside 1 331\nTMhelix 332 354\ninside 355 372

13318 GCF_000617985.2_EcK1516 Escherichia coli O15:H18 str. K1516 Proteobacteria; Gammaproteobacteria; Enterobacterales; Enterobacteriaceae; Escherichia; Escherichia coli; Escherichia coli O15 MNNEETFYQAMRRQGVTRRSFLKYCSLAATSLGLGAGMAPKIAWALENKPRIPVVWIHGL WP_001058323.1 MULTISPECIES: hydrogenase-1 small chain [Proteobacteria] Length: 372\nNumber of predicted TMHs: 2\nExp number of AAs in TMHs: 37.2608499999999\nExp number, first 60 AAs: 19.52923\nTotal prob of N-in: 0.94351\nPOSSIBLE N-term signal sequence\ninside 1 19\nTMhelix 20 39\noutside 40 325\nTMhelix 326 348\ninside 349 372

13319 GCF_000614565.2_Ec2010C-5034 Escherichia coli O153:H2 str. 2010C-5034 Proteobacteria; Gammaproteobacteria; Enterobacterales; Enterobacteriaceae; Escherichia; Escherichia coli; Escherichia coli O153 MTGDNTLIHSHGINRRDFMKLCAALAATMGLSSKAAAEMAESVTNPQRPPVIWIGAQECT WP_000145410.1 MULTISPECIES: hydrogenase-2 small chain [Proteobacteria] Length: 372\nNumber of predicted TMHs: 1\nExp number of AAs in TMHs: 23.72226\nExp number, first 60 AAs: 0.22038\nTotal prob of N-in: 0.12994\noutside 1 331\nTMhelix 332 354\ninside 355 372

13320 GCF_000614565.2_Ec2010C-5034 Escherichia coli O153:H2 str. 2010C-5034 Proteobacteria; Gammaproteobacteria; Enterobacterales; Enterobacteriaceae; Escherichia; Escherichia coli; Escherichia coli O153 MNNEETFYQAMRRQGVTRRSFLKYCSLAATSLGLGAGMAPKIAWALENKPRIPVVWIHGL WP_001295940.1 MULTISPECIES: hydrogenase [Enterobacteriaceae] Length: 372\nNumber of predicted TMHs: 2\nExp number of AAs in TMHs: 37.2608899999999\nExp number, first 60 AAs: 19.52923\nTotal prob of N-in:

0.94351\nPOSSIBLE N-term signal sequence\ninside 1 19\nTMhelix 20 39\noutside 40 325\nTMhelix 326 348\ninside 349 372

13321 GCF_000616285.1_Ec2011C-3602 Escherichia coli O156:H25 str. 2011C-3602 Proteobacteria; Gammaproteobacteria; Enterobacterales; Enterobacteriaceae; Escherichia; Escherichia coli; Escherichia coli O156 MNNEETFYQAMRRQGVTRRSFLKYCSLAATSLGLGAGMAPKIAWALENKPRIPVVWIHGL WP_001058323.1 MULTISPECIES: hydrogenase-1 small chain [Proteobacteria] Length: 372\nNumber of predicted TMHs: 2\nExp number of AAs in TMHs: 37.2608499999999\nExp number, first 60 AAs: 19.52923\nTotal prob of N-in: 0.94351\nPOSSIBLE N-term signal sequence\ninside 1 19\nTMhelix 20 39\noutside 40 325\nTMhelix 326 348\ninside 349 372

13322 GCF_000616285.1_Ec2011C-3602 Escherichia coli O156:H25 str. 2011C-3602 Proteobacteria; Gammaproteobacteria; Enterobacterales; Enterobacteriaceae; Escherichia; Escherichia coli; Escherichia coli O156 MTGDNTLIHSHGINRRDFMKLCAALAATMGLSSKAAAEMAESVTNPQRPPVIWIGAQECT WP_000145410.1 MULTISPECIES: hydrogenase-2 small chain [Proteobacteria] Length: 372\nNumber of predicted TMHs: 1\nExp number of AAs in TMHs: 23.72226\nExp number, first 60 AAs: 0.22038\nTotal prob of N-in: 0.12994\noutside 1 331\nTMhelix 332 354\ninside 355 372

13323 GCF_000335195.2_ASM33519v2 Escherichia coli 3.4880 Proteobacteria; Gammaproteobacteria; Enterobacterales; Enterobacteriaceae; Escherichia; Escherichia coli; Escherichia coli O157 MTGDNTLIHSHGINRRDFMKLCAALAATMGLSSKAAAEMAESVTNPQRPPVIWIGAQECT WP_000145410.1 MULTISPECIES: hydrogenase-2 small chain [Proteobacteria] Length: 372\nNumber of predicted TMHs: 1\nExp number of AAs in TMHs: 23.72226\nExp number, first 60 AAs: 0.22038\nTotal prob of N-in: 0.12994\noutside 1 331\nTMhelix 332 354\ninside 355 372

13324 GCF_000335195.2_ASM33519v2 Escherichia coli 3.4880 Proteobacteria; Gammaproteobacteria; Enterobacterales; Enterobacteriaceae; Escherichia; Escherichia coli; Escherichia coli O157 MNNEETFYQAMRRQGVTRRSFLKYCSLAATSLGLGAGMAPKIAWALENKPRIPVVWIHGL WP_001301613.1 hydrogenase [Escherichia coli] Length: 372\nNumber of predicted TMHs: 2\nExp number of AAs in TMHs: 37.46598\nExp number, first 60 AAs: 19.53139\nTotal prob of N-in: 0.94364\nPOSSIBLE N-term signal sequence\ninside 1 19\nTMhelix 20 39\noutside 40 325\nTMhelix 326 348\ninside 349 372

13325 GCF_000335115.1_ASM33511v1 Escherichia coli 7.1982 Proteobacteria; Gammaproteobacteria; Enterobacterales; Enterobacteriaceae; Escherichia; Escherichia coli; Escherichia coli O157 MTGDNTLIHSHGINRRDFMKLCAALAATMGLSSKAAAEMAESVTNPQRPPVIWIGAQECT WP_000145410.1 MULTISPECIES: hydrogenase-2 small chain [Proteobacteria] Length: 372\nNumber of predicted TMHs: 1\nExp number of AAs in TMHs: 23.72226\nExp number, first 60 AAs: 0.22038\nTotal prob of N-in: 0.12994\noutside 1 331\nTMhelix 332 354\ninside 355 372

13326 GCF_000335115.1_ASM33511v1 Escherichia coli 7.1982 Proteobacteria; Gammaproteobacteria; Enterobacterales; Enterobacteriaceae; Escherichia; Escherichia coli; Escherichia coli O157 MNNEETFYQAMRRQGVTRRSFLKYCSLAATSLGLGAGMAPKIAWALENKPRIPVVWIHGL WP_001301613.1 hydrogenase [Escherichia coli] Length: 372\nNumber of predicted TMHs: 2\nExp number of AAs in TMHs: 37.46598\nExp number, first 60 AAs: 19.53139\nTotal prob of N-in: 0.94364\nPOSSIBLE N-term signal sequence\ninside 1 19\nTMhelix 20 39\noutside 40 325\nTMhelix 326 348\ninside 349 372

13327 GCF_000335375.2_ASM33537v2 Escherichia coli 95.0083 Proteobacteria; Gammaproteobacteria; Enterobacterales; Enterobacteriaceae; Escherichia; Escherichia coli; Escherichia coli O157 MTGDNTLIHSHGINRRDFMKLCAALAATMGLSSKAAAEMAESVTNPQRPPVIWIGAQECT WP_000145410.1 MULTISPECIES: hydrogenase-2 small chain [Proteobacteria] Length: 372\nNumber of predicted TMHs: 1\nExp number of AAs in TMHs: 23.72226\nExp number, first 60 AAs: 0.22038\nTotal prob of N-in: 0.12994\noutside 1 331\nTMhelix 332 354\ninside 355 372

13328 GCF_000335055.2_ASM33505v2 Escherichia coli ATCC 700728 Proteobacteria; Gammaproteobacteria; Enterobacterales; Enterobacteriaceae; Escherichia; Escherichia coli; Escherichia coli O157 MTGDNTLIHSHGINRRDFMKLCAALAATMGLSSKAAAEMAESVTNPQRPPVIWIGAQECT WP_000145410.1 MULTISPECIES: hydrogenase-2 small chain [Proteobacteria] Length: 372\nNumber of predicted TMHs: 1\nExp number of AAs in TMHs: 23.72226\nExp number, first 60 AAs: 0.22038\nTotal prob of N-in: 0.12994\noutside 1 331\nTMhelix 332 354\ninside 355 372

13329 GCF_000335055.2_ASM33505v2 Escherichia coli ATCC 700728 Proteobacteria; Gammaproteobacteria; Enterobacterales; Enterobacteriaceae; Escherichia; Escherichia coli; Escherichia coli O157
MNNEETFYQAMRRQGVTRRSFLKYCSLAATSLGLGAGMAPKIAWALENKPRIPVVWIHGL WP_001301613.1
hydrogenase [Escherichia coli] Length: 372\nNumber of predicted TMHs: 2\nExp number of AAs in TMHs: 37.46598\nExp number, first 60 AAs: 19.53139\nTotal prob of N-in: 0.94364\nPOSSIBLE N-term signal sequence\ninside 1 19\nTMhelix 20 39\noutside 40 325\nTMhelix 326 348\ninside 349 372

13330 GCF_000618005.2_Ec98-3133 Escherichia coli O157:H16 str. 98-3133 Proteobacteria; Gammaproteobacteria; Enterobacterales; Enterobacteriaceae; Escherichia; Escherichia coli; Escherichia coli O157
MNNEETFYQAMRRQGVTRRSFLKYCSLAATSLGLGAGMAPKIAWALENKPRIPVVWIHGL WP_001058323.1
MULTISPECIES: hydrogenase-1 small chain [Proteobacteria] Length: 372\nNumber of predicted TMHs: 2\nExp number of AAs in TMHs: 37.2608499999999\nExp number, first 60 AAs: 19.52923\nTotal prob of N-in: 0.94351\nPOSSIBLE N-term signal sequence\ninside 1 19\nTMhelix 20 39\noutside 40 325\nTMhelix 326 348\ninside 349 372

13331 GCF_000618005.2_Ec98-3133 Escherichia coli O157:H16 str. 98-3133 Proteobacteria; Gammaproteobacteria; Enterobacterales; Enterobacteriaceae; Escherichia; Escherichia coli; Escherichia coli O157
MTGDNTLIHSHGINRRDFMKLCAALAATMGLSSKAAAEMAESVTNPQRPPVIWIGAQECT WP_000145410.1
MULTISPECIES: hydrogenase-2 small chain [Proteobacteria] Length: 372\nNumber of predicted TMHs: 1\nExp number of AAs in TMHs: 23.72226\nExp number, first 60 AAs: 0.22038\nTotal prob of N-in: 0.12994\noutside 1 331\nTMhelix 332 354\ninside 355 372

13332 GCF_000247665.3_EcoliT22_2.0 Escherichia coli O157:H43 str. T22 Proteobacteria; Gammaproteobacteria; Enterobacterales; Enterobacteriaceae; Escherichia; Escherichia coli; Escherichia coli O157
MNNEETFYQAMRRQGVTRRSFLKYCSLAATSLGLGAGMAPKIAWALENKPRIPVVWIHGL WP_001058323.1
MULTISPECIES: hydrogenase-1 small chain [Proteobacteria] Length: 372\nNumber of predicted TMHs: 2\nExp number of AAs in TMHs: 37.2608499999999\nExp number, first 60 AAs: 19.52923\nTotal prob of N-in: 0.94351\nPOSSIBLE N-term signal sequence\ninside 1 19\nTMhelix 20 39\noutside 40 325\nTMhelix 326 348\ninside 349 372

13333 GCF_000247665.3_EcoliT22_2.0 Escherichia coli O157:H43 str. T22 Proteobacteria; Gammaproteobacteria; Enterobacterales; Enterobacteriaceae; Escherichia; Escherichia coli; Escherichia coli O157
MTGDNTLIHSHGINRRDFMKLCAALAATMGLSSKAAAEMAESVTNPQRPPVIWIGAQECT WP_000145410.1
MULTISPECIES: hydrogenase-2 small chain [Proteobacteria] Length: 372\nNumber of predicted TMHs: 1\nExp number of AAs in TMHs: 23.72226\nExp number, first 60 AAs: 0.22038\nTotal prob of N-in: 0.12994\noutside 1 331\nTMhelix 332 354\ninside 355 372

13334 GCF_000617445.2_Ec06-3745 Escherichia coli O157:H7 str. 06-3745 Proteobacteria; Gammaproteobacteria; Enterobacterales; Enterobacteriaceae; Escherichia; Escherichia coli; Escherichia coli O157
MTGDNTLIHSHGINRRDFMKLCAALAATMGLSSKAAAEMAESVTNPQRPPVIWIGAQECT WP_000145410.1
MULTISPECIES: hydrogenase-2 small chain [Proteobacteria] Length: 372\nNumber of predicted TMHs: 1\nExp number of AAs in TMHs: 23.72226\nExp number, first 60 AAs: 0.22038\nTotal prob of N-in: 0.12994\noutside 1 331\nTMhelix 332 354\ninside 355 372

13335 GCF_000617445.2_Ec06-3745 Escherichia coli O157:H7 str. 06-3745 Proteobacteria; Gammaproteobacteria; Enterobacterales; Enterobacteriaceae; Escherichia; Escherichia coli; Escherichia coli O157
MNNEETFYQAMRRQGVTRRSFLKYCSLAATSLGLGAGMAPKIAWALENKPRIPVVWIHGL WP_001301613.1
hydrogenase [Escherichia coli] Length: 372\nNumber of predicted TMHs: 2\nExp number of AAs in TMHs: 37.46598\nExp number, first 60 AAs: 19.53139\nTotal prob of N-in: 0.94364\nPOSSIBLE N-term signal sequence\ninside 1 19\nTMhelix 20 39\noutside 40 325\nTMhelix 326 348\ninside 349 372

13336 GCF_000617705.2_Ec06-4039 Escherichia coli O157:H7 str. 06-4039 Proteobacteria; Gammaproteobacteria; Enterobacterales; Enterobacteriaceae; Escherichia; Escherichia coli; Escherichia coli O157
MTGDNTLIHSHGINRRDFMKLCAALAATMGLSSKAAAEMAESVTNPQRPPVIWIGAQECT WP_000145410.1
MULTISPECIES: hydrogenase-2 small chain [Proteobacteria] Length: 372\nNumber of predicted TMHs: 1\nExp number of AAs in TMHs: 23.72226\nExp number, first 60 AAs: 0.22038\nTotal prob of N-in: 0.12994\noutside 1 331\nTMhelix 332 354\ninside 355 372

13337 GCF_000617705.2_Ec06-4039 Escherichia coli O157:H7 str. 06-4039 Proteobacteria; Gammaproteobacteria; Enterobacterales; Enterobacteriaceae; Escherichia; Escherichia coli; Escherichia coli O157
MNNEETFYQAMRRQGVTRRSFLKYCSLAATSLGLGAGMAPKIAWALENKPRIPVVWIHGL WP_001301613.1

hydrogenase [Escherichia coli] Length: 372\nNumber of predicted TMHs: 2\nExp number of AAs in TMHs: 37.46598\nExp number, first 60 AAs: 19.53139\nTotal prob of N-in: 0.94364\nPOSSIBLE N-term signal sequence\ninside 1 19\nTMhelix 20 39\noutside 40 325\nTMhelix 326 348\ninside 349 372

13338 GCF_000617665.2_Ec07-3091 Escherichia coli O157:H7 str. 07-3091 Proteobacteria; Gammaproteobacteria; Enterobacterales; Enterobacteriaceae; Escherichia; Escherichia coli; Escherichia coli O157 MNNEETFYQAMRRQGVTRRSFLKYCSLAATSLGLGAGMAPKIAWALENKPRIPVVWIHGL WP_001301613.1 hydrogenase [Escherichia coli] Length: 372\nNumber of predicted TMHs: 2\nExp number of AAs in TMHs: 37.46598\nExp number, first 60 AAs: 19.53139\nTotal prob of N-in: 0.94364\nPOSSIBLE N-term signal sequence\ninside 1 19\nTMhelix 20 39\noutside 40 325\nTMhelix 326 348\ninside 349 372

13339 GCF_000617665.2_Ec07-3091 Escherichia coli O157:H7 str. 07-3091 Proteobacteria; Gammaproteobacteria; Enterobacterales; Enterobacteriaceae; Escherichia; Escherichia coli; Escherichia coli O157 MTGDNTLIHSHGINRRDFMKLCAALAATMGLSSKAAAEMAESVTNPQRPPVIWIGAQECT WP_000145410.1 MULTISPECIES: hydrogenase-2 small chain [Proteobacteria] Length: 372\nNumber of predicted TMHs: 1\nExp number of AAs in TMHs: 23.72226\nExp number, first 60 AAs: 0.22038\nTotal prob of N-in: 0.12994\noutside 1 331\nTMhelix 332 354\ninside 355 372

13340 GCF_000617725.2_Ec07-3391 Escherichia coli O157:H7 str. 07-3391 Proteobacteria; Gammaproteobacteria; Enterobacterales; Enterobacteriaceae; Escherichia; Escherichia coli; Escherichia coli O157 MTGDNTLIHSHGINRRDFMKLCAALAATMGLSSKAAAEMAESVTNPQRPPVIWIGAQECT WP_000145410.1 MULTISPECIES: hydrogenase-2 small chain [Proteobacteria] Length: 372\nNumber of predicted TMHs: 1\nExp number of AAs in TMHs: 23.72226\nExp number, first 60 AAs: 0.22038\nTotal prob of N-in: 0.12994\noutside 1 331\nTMhelix 332 354\ninside 355 372

13341 GCF_000617725.2_Ec07-3391 Escherichia coli O157:H7 str. 07-3391 Proteobacteria; Gammaproteobacteria; Enterobacterales; Enterobacteriaceae; Escherichia; Escherichia coli; Escherichia coli O157 MNNEETFYQAMRRQGVTRRSFLKYCSLAATSLGLGAGMAPKIAWALENKPRIPVVWIHGL WP_001301613.1 hydrogenase [Escherichia coli] Length: 372\nNumber of predicted TMHs: 2\nExp number of AAs in TMHs: 37.46598\nExp number, first 60 AAs: 19.53139\nTotal prob of N-in: 0.94364\nPOSSIBLE N-term signal sequence\ninside 1 19\nTMhelix 20 39\noutside 40 325\nTMhelix 326 348\ninside 349 372

13342 GCF_000616585.2_Ec08-3037 Escherichia coli O157:H7 str. 08-3037 Proteobacteria; Gammaproteobacteria; Enterobacterales; Enterobacteriaceae; Escherichia; Escherichia coli; Escherichia coli O157 MTGDNTLIHSHGINRRDFMKLCAALAATMGLSSKAAAEMAESVTNPQRPPVIWIGAQECT WP_000145410.1 MULTISPECIES: hydrogenase-2 small chain [Proteobacteria] Length: 372\nNumber of predicted TMHs: 1\nExp number of AAs in TMHs: 23.72226\nExp number, first 60 AAs: 0.22038\nTotal prob of N-in: 0.12994\noutside 1 331\nTMhelix 332 354\ninside 355 372

13343 GCF_000616585.2_Ec08-3037 Escherichia coli O157:H7 str. 08-3037 Proteobacteria; Gammaproteobacteria; Enterobacterales; Enterobacteriaceae; Escherichia; Escherichia coli; Escherichia coli O157 MNNEETFYQAMRRQGVTRRSFLKYCSLAATSLGLGAGMAPKIAWALENKPRIPVVWIHGL WP_001301613.1 hydrogenase [Escherichia coli] Length: 372\nNumber of predicted TMHs: 2\nExp number of AAs in TMHs: 37.46598\nExp number, first 60 AAs: 19.53139\nTotal prob of N-in: 0.94364\nPOSSIBLE N-term signal sequence\ninside 1 19\nTMhelix 20 39\noutside 40 325\nTMhelix 326 348\ninside 349 372

13344 GCF_000616605.2_Ec08-3527 Escherichia coli O157:H7 str. 08-3527 Proteobacteria; Gammaproteobacteria; Enterobacterales; Enterobacteriaceae; Escherichia; Escherichia coli; Escherichia coli O157 MTGDNTLIHSHGINRRDFMKLCAALAATMGLSSKAAAEMAESVTNPQRPPVIWIGAQECT WP_000145410.1 MULTISPECIES: hydrogenase-2 small chain [Proteobacteria] Length: 372\nNumber of predicted TMHs: 1\nExp number of AAs in TMHs: 23.72226\nExp number, first 60 AAs: 0.22038\nTotal prob of N-in: 0.12994\noutside 1 331\nTMhelix 332 354\ninside 355 372

13345 GCF_000616605.2_Ec08-3527 Escherichia coli O157:H7 str. 08-3527 Proteobacteria; Gammaproteobacteria; Enterobacterales; Enterobacteriaceae; Escherichia; Escherichia coli; Escherichia coli O157 MNNEETFYQAMRRQGVTRRSFLKYCSLAATSLGLGAGMAPKIAWALENKPRIPVVWIHGL WP_001301613.1 hydrogenase [Escherichia coli] Length: 372\nNumber of predicted TMHs: 2\nExp number of AAs in TMHs: 37.46598\nExp number, first 60 AAs: 19.53139\nTotal prob of N-in: 0.94364\nPOSSIBLE N-term signal sequence\ninside 1 19\nTMhelix 20 39\noutside 40 325\nTMhelix 326 348\ninside 349 372

13346 GCF_000616645.2_Ec08-4169 Escherichia coli O157:H7 str. 08-4169 Proteobacteria; Gammaproteobacteria; Enterobacterales; Enterobacteriaceae; Escherichia; Escherichia coli; Escherichia coli O157 MNNEETFYQAMRRQGVTRRSFLKYCSLAATSLGLGAGMAPKIAWALENKPRIPVVWIHGL WP_001301613.1 hydrogenase [Escherichia coli] Length: 372\nNumber of predicted TMHs: 2\nExp number of AAs in TMHs: 37.46598\nExp number, first 60 AAs: 19.53139\nTotal prob of N-in: 0.94364\nPOSSIBLE N-term signal sequence\ninside 1 19\nTMhelix 20 39\noutside 40 325\nTMhelix 326 348\ninside 349 372

13347 GCF_000616645.2_Ec08-4169 Escherichia coli O157:H7 str. 08-4169 Proteobacteria; Gammaproteobacteria; Enterobacterales; Enterobacteriaceae; Escherichia; Escherichia coli; Escherichia coli O157 MTGDNTLIHSHGINRRDFMKLCAALAATMGLSSKAAAEMAESVTNPQRPPVIWIGAQECT WP_000145410.1 MULTISPECIES: hydrogenase-2 small chain [Proteobacteria] Length: 372\nNumber of predicted TMHs: 1\nExp number of AAs in TMHs: 23.72226\nExp number, first 60 AAs: 0.22038\nTotal prob of N-in: 0.12994\noutside 1 331\nTMhelix 332 354\ninside 355 372

13348 GCF_000619665.1_Ec08-4529 Escherichia coli O157:H7 str. 08-4529 Proteobacteria; Gammaproteobacteria; Enterobacterales; Enterobacteriaceae; Escherichia; Escherichia coli; Escherichia coli O157 MNNEETFYQAMRRQGVTRRSFLKYCSLAATSLGLGAGMAPKIAWALENKPRIPVVWIHGL WP_001301613.1 hydrogenase [Escherichia coli] Length: 372\nNumber of predicted TMHs: 2\nExp number of AAs in TMHs: 37.46598\nExp number, first 60 AAs: 19.53139\nTotal prob of N-in: 0.94364\nPOSSIBLE N-term signal sequence\ninside 1 19\nTMhelix 20 39\noutside 40 325\nTMhelix 326 348\ninside 349 372

13349 GCF_000619665.1_Ec08-4529 Escherichia coli O157:H7 str. 08-4529 Proteobacteria; Gammaproteobacteria; Enterobacterales; Enterobacteriaceae; Escherichia; Escherichia coli; Escherichia coli O157 MTGDNTLIHSHGINRRDFMKLCAALAATMGLSSKAAAEMAESVTNPQRPPVIWIGAQECT WP_000145410.1 MULTISPECIES: hydrogenase-2 small chain [Proteobacteria] Length: 372\nNumber of predicted TMHs: 1\nExp number of AAs in TMHs: 23.72226\nExp number, first 60 AAs: 0.22038\nTotal prob of N-in: 0.12994\noutside 1 331\nTMhelix 332 354\ninside 355 372

13350 GCF_000747465.1_O157_08BKT61141_454 Escherichia coli O157:H7 str. 08BKT061141 Proteobacteria; Gammaproteobacteria; Enterobacterales; Enterobacteriaceae; Escherichia; Escherichia coli; Escherichia coli O157 MTGDNTLIHSHGINRRDFMKLCAALAATMGLSSKAAAEMAESVTNPQRPPVIWIGAQECT WP_000145410.1 MULTISPECIES: hydrogenase-2 small chain [Proteobacteria] Length: 372\nNumber of predicted TMHs: 1\nExp number of AAs in TMHs: 23.72226\nExp number, first 60 AAs: 0.22038\nTotal prob of N-in: 0.12994\noutside 1 331\nTMhelix 332 354\ninside 355 372

13351 GCF_000747445.1_O157_09BKT048303_454 Escherichia coli O157:H7 str. 09BKT048303 Proteobacteria; Gammaproteobacteria; Enterobacterales; Enterobacteriaceae; Escherichia; Escherichia coli; Escherichia coli O157 MNNEETFYQAMRRQGVTRRSFLKYCSLAATSLGLGAGMAPKIAWALENKPRIPVVWIHGL WP_001301613.1 hydrogenase [Escherichia coli] Length: 372\nNumber of predicted TMHs: 2\nExp number of AAs in TMHs: 37.46598\nExp number, first 60 AAs: 19.53139\nTotal prob of N-in: 0.94364\nPOSSIBLE N-term signal sequence\ninside 1 19\nTMhelix 20 39\noutside 40 325\nTMhelix 326 348\ninside 349 372

13352 GCF_000747445.1_O157_09BKT048303_454 Escherichia coli O157:H7 str. 09BKT048303 Proteobacteria; Gammaproteobacteria; Enterobacterales; Enterobacteriaceae; Escherichia; Escherichia coli; Escherichia coli O157 MTGDNTLIHSHGINRRDFMKLCAALAATMGLSSKAAAEMAESVTNPQRPPVIWIGAQECT WP_000145410.1 MULTISPECIES: hydrogenase-2 small chain [Proteobacteria] Length: 372\nNumber of predicted TMHs: 1\nExp number of AAs in TMHs: 23.72226\nExp number, first 60 AAs: 0.22038\nTotal prob of N-in: 0.12994\noutside 1 331\nTMhelix 332 354\ninside 355 372

13353 GCF_000192665.1_ASM19266v2 Escherichia coli O157:H7 str. 1044 Proteobacteria; Gammaproteobacteria; Enterobacterales; Enterobacteriaceae; Escherichia; Escherichia coli; Escherichia coli O157 MNNEETFYQAMRRQGVTRRSFLKYCSLAATSLGLGAGMAPKIAWALENKPRIPVVWIHGL WP_001301613.1 hydrogenase [Escherichia coli] Length: 372\nNumber of predicted TMHs: 2\nExp number of AAs in TMHs: 37.46598\nExp number, first 60 AAs: 19.53139\nTotal prob of N-in: 0.94364\nPOSSIBLE N-term signal sequence\ninside 1 19\nTMhelix 20 39\noutside 40 325\nTMhelix 326 348\ninside 349 372

13354 GCF_000192665.1_ASM19266v2 Escherichia coli O157:H7 str. 1044 Proteobacteria; Gammaproteobacteria; Enterobacterales; Enterobacteriaceae; Escherichia; Escherichia coli; Escherichia coli O157 MTGDNTLIHSHGINRRDFMKLCAALAATMGLSSKAAAEMAESVTNPQRPPVIWIGAQECT WP_000145410.1 MULTISPECIES: hydrogenase-2 small chain [Proteobacteria] Length: 372\nNumber of predicted TMHs:

1\nExp number of AAs in TMHs: 23.72226\nExp number, first 60 AAs: 0.22038\nTotal prob of N-in: 0.12994\noutside 1 331\nTMhelix 332 354\ninside 355 372

13355 GCF_000192685.1_ASM19268v2 Escherichia coli O157:H7 str. 1125 Proteobacteria; Gammaproteobacteria; Enterobacterales; Enterobacteriaceae; Escherichia; Escherichia coli; Escherichia coli O157 MTGDNTLIHSHGINRRDFMKLCAALAATMGLSSKAAAEMAESVTNPQRPPVIWIGAQECT WP_000145410.1 MULTISPECIES: hydrogenase-2 small chain [Proteobacteria] Length: 372\nNumber of predicted TMHs: 1\nExp number of AAs in TMHs: 23.72226\nExp number, first 60 AAs: 0.22038\nTotal prob of N-in: 0.12994\noutside 1 331\nTMhelix 332 354\ninside 355 372

13356 GCF_000192685.1_ASM19268v2 Escherichia coli O157:H7 str. 1125 Proteobacteria; Gammaproteobacteria; Enterobacterales; Enterobacteriaceae; Escherichia; Escherichia coli; Escherichia coli O157 MNNEETFYQAMRRQGVTRRSFLKYCSLAATSLGLGAGMAPKIAWALENKPRIPVVWIHGL WP_001301613.1 hydrogenase [Escherichia coli] Length: 372\nNumber of predicted TMHs: 2\nExp number of AAs in TMHs: 37.46598\nExp number, first 60 AAs: 19.53139\nTotal prob of N-in: 0.94364\nPOSSIBLE N-term signal sequence\ninside 1 19\nTMhelix 20 39\noutside 40 325\nTMhelix 326 348\ninside 349 372

13357 GCF_000619045.2_Ec2009C-4258 Escherichia coli O157:H7 str. 2009C-4258 Proteobacteria; Gammaproteobacteria; Enterobacterales; Enterobacteriaceae; Escherichia; Escherichia coli; Escherichia coli O157 MNNEETFYQAMRRQGVTRRSFLKYCSLAATSLGLGAGMAPKIAWALENKPRIPVVWIHGL WP_001301613.1 hydrogenase [Escherichia coli] Length: 372\nNumber of predicted TMHs: 2\nExp number of AAs in TMHs: 37.46598\nExp number, first 60 AAs: 19.53139\nTotal prob of N-in: 0.94364\nPOSSIBLE N-term signal sequence\ninside 1 19\nTMhelix 20 39\noutside 40 325\nTMhelix 326 348\ninside 349 372

13358 GCF_000619045.2_Ec2009C-4258 Escherichia coli O157:H7 str. 2009C-4258 Proteobacteria; Gammaproteobacteria; Enterobacterales; Enterobacteriaceae; Escherichia; Escherichia coli; Escherichia coli O157 MTGDNTLIHSHGINRRDFMKLCAALAATMGLSSKAAAEMAESVTNPQRPPVIWIGAQECT WP_000145410.1 MULTISPECIES: hydrogenase-2 small chain [Proteobacteria] Length: 372\nNumber of predicted TMHs: 1\nExp number of AAs in TMHs: 23.72226\nExp number, first 60 AAs: 0.22038\nTotal prob of N-in: 0.12994\noutside 1 331\nTMhelix 332 354\ninside 355 372

13359 GCF_000619145.1_Ec2009EL1449 Escherichia coli O157:H7 str. 2009EL1449 Proteobacteria; Gammaproteobacteria; Enterobacterales; Enterobacteriaceae; Escherichia; Escherichia coli; Escherichia coli O157 MNNEETFYQAMRRQGVTRRSFLKYCSLAATSLGLGAGMAPKIAWALENKPRIPVVWIHGL WP_001301613.1 hydrogenase [Escherichia coli] Length: 372\nNumber of predicted TMHs: 2\nExp number of AAs in TMHs: 37.46598\nExp number, first 60 AAs: 19.53139\nTotal prob of N-in: 0.94364\nPOSSIBLE N-term signal sequence\ninside 1 19\nTMhelix 20 39\noutside 40 325\nTMhelix 326 348\ninside 349 372

13360 GCF_000619145.1_Ec2009EL1449 Escherichia coli O157:H7 str. 2009EL1449 Proteobacteria; Gammaproteobacteria; Enterobacterales; Enterobacteriaceae; Escherichia; Escherichia coli; Escherichia coli O157 MTGDNTLIHSHGINRRDFMKLCAALAATMGLSSKAAAEMAESVTNPQRPPVIWIGAQECT WP_000145410.1 MULTISPECIES: hydrogenase-2 small chain [Proteobacteria] Length: 372\nNumber of predicted TMHs: 1\nExp number of AAs in TMHs: 23.72226\nExp number, first 60 AAs: 0.22038\nTotal prob of N-in: 0.12994\noutside 1 331\nTMhelix 332 354\ninside 355 372

13361 GCF_000614965.1_Ec2009EL1705 Escherichia coli O157:H7 str. 2009EL1705 Proteobacteria; Gammaproteobacteria; Enterobacterales; Enterobacteriaceae; Escherichia; Escherichia coli; Escherichia coli O157 MTGDNTLIHSHGINRRDFMKLCAALAATMGLSSKAAAEMAESVTNPQRPPVIWIGAQECT WP_000145410.1 MULTISPECIES: hydrogenase-2 small chain [Proteobacteria] Length: 372\nNumber of predicted TMHs: 1\nExp number of AAs in TMHs: 23.72226\nExp number, first 60 AAs: 0.22038\nTotal prob of N-in: 0.12994\noutside 1 331\nTMhelix 332 354\ninside 355 372

13362 GCF_000614965.1_Ec2009EL1705 Escherichia coli O157:H7 str. 2009EL1705 Proteobacteria; Gammaproteobacteria; Enterobacterales; Enterobacteriaceae; Escherichia; Escherichia coli; Escherichia coli O157 MNNEETFYQAMRRQGVTRRSFLKYCSLAATSLGLGAGMAPKIAWALENKPRIPVVWIHGL WP_001301613.1 hydrogenase [Escherichia coli] Length: 372\nNumber of predicted TMHs: 2\nExp number of AAs in TMHs: 37.46598\nExp number, first 60 AAs: 19.53139\nTotal prob of N-in: 0.94364\nPOSSIBLE N-term signal sequence\ninside 1 19\nTMhelix 20 39\noutside 40 325\nTMhelix 326 348\ninside 349 372

13363 GCF_000619165.1_Ec2009EL1913 Escherichia coli O157:H7 str. 2009EL1913 Proteobacteria; Gammaproteobacteria; Enterobacterales; Enterobacteriaceae; Escherichia; Escherichia coli; Escherichia coli O157

MNNEETFYQAMRRQGVTRRSFLKYCSLAATSLGLGAGMAPKIAWALENKPRIPVVWIHGL WP_001301613.1 hydrogenase [Escherichia coli] Length: 372\nNumber of predicted TMHs: 2\nExp number of AAs in TMHs: 37.46598\nExp number, first 60 AAs: 19.53139\nTotal prob of N-in: 0.94364\nPOSSIBLE N-term signal sequence\ninside 1 19\nTMhelix 20 39\noutside 40 325\nTMhelix 326 348\ninside 349 372

13364 GCF_000619165.1_Ec2009EL1913 Escherichia coli O157:H7 str. 2009EL1913 Proteobacteria; Gammaproteobacteria; Enterobacterales; Enterobacteriaceae; Escherichia; Escherichia coli; Escherichia coli O157 MTGDNTLIHSHGINRRDFMKLCAALAATMGLSSKAAAEMAESVTNPQRPPVIWIGAQECT WP_000145410.1 MULTISPECIES: hydrogenase-2 small chain [Proteobacteria] Length: 372\nNumber of predicted TMHs: 1\nExp number of AAs in TMHs: 23.72226\nExp number, first 60 AAs: 0.22038\nTotal prob of N-in: 0.12994\noutside 1 331\nTMhelix 332 354\ninside 355 372

13365 GCF_000615015.1_Ec2009EL2109 Escherichia coli O157:H7 str. 2009EL2109 Proteobacteria; Gammaproteobacteria; Enterobacterales; Enterobacteriaceae; Escherichia; Escherichia coli; Escherichia coli O157 MTGDNTLIHSHGINRRDFMKLCAALAATMGLSSKAAAEMAESVTNPQRPPVIWIGAQECT WP_000145410.1 MULTISPECIES: hydrogenase-2 small chain [Proteobacteria] Length: 372\nNumber of predicted TMHs: 1\nExp number of AAs in TMHs: 23.72226\nExp number, first 60 AAs: 0.22038\nTotal prob of N-in: 0.12994\noutside 1 331\nTMhelix 332 354\ninside 355 372

13366 GCF_000615015.1_Ec2009EL2109 Escherichia coli O157:H7 str. 2009EL2109 Proteobacteria; Gammaproteobacteria; Enterobacterales; Enterobacteriaceae; Escherichia; Escherichia coli; Escherichia coli O157 MNNEETFYQAMRRQGVTRRSFLKYCSLAATSLGLGAGMAPKIAWALENKPRIPVVWIHGL WP_001301613.1 hydrogenase [Escherichia coli] Length: 372\nNumber of predicted TMHs: 2\nExp number of AAs in TMHs: 37.46598\nExp number, first 60 AAs: 19.53139\nTotal prob of N-in: 0.94364\nPOSSIBLE N-term signal sequence\ninside 1 19\nTMhelix 20 39\noutside 40 325\nTMhelix 326 348\ninside 349 372

13367 GCF_000614885.2_Ec2010C-4979C1 Escherichia coli O157:H7 str. 2010C-4979C1 Proteobacteria; Gammaproteobacteria; Enterobacterales; Enterobacteriaceae; Escherichia; Escherichia coli; Escherichia coli O157 MIGDNTLIHSHGINRRDFMKLCAALAATMGLSSKAAAEMAESVTNPQRPPVIWIGAQECT WP_000578724.1 hydrogenase 2 small subunit [Escherichia coli] Length: 372\nNumber of predicted TMHs: 1\nExp number of AAs in TMHs: 23.05601\nExp number, first 60 AAs: 0.15431\nTotal prob of N-in: 0.09116\noutside 1 331\nTMhelix 332 354\ninside 355 372

13368 GCF_000614885.2_Ec2010C-4979C1 Escherichia coli O157:H7 str. 2010C-4979C1 Proteobacteria; Gammaproteobacteria; Enterobacterales; Enterobacteriaceae; Escherichia; Escherichia coli; Escherichia coli O157 MNNEETFYQAMRRQGVTRRSFLKYCSLAATSLGLGAGMAPKIAWALENKPRIPVVWIHGL WP_001301613.1 hydrogenase [Escherichia coli] Length: 372\nNumber of predicted TMHs: 2\nExp number of AAs in TMHs: 37.46598\nExp number, first 60 AAs: 19.53139\nTotal prob of N-in: 0.94364\nPOSSIBLE N-term signal sequence\ninside 1 19\nTMhelix 20 39\noutside 40 325\nTMhelix 326 348\ninside 349 372

13369 GCF_000616365.1_Ec2011EL-1107 Escherichia coli O157:H7 str. 2011EL-1107 Proteobacteria; Gammaproteobacteria; Enterobacterales; Enterobacteriaceae; Escherichia; Escherichia coli; Escherichia coli O157 MNNEETFYQAMRRQGVTRRSFLKYCSLAATSLGLGAGMAPKIAWALENKPRIPVVWIHGL WP_001301613.1 hydrogenase [Escherichia coli] Length: 372\nNumber of predicted TMHs: 2\nExp number of AAs in TMHs: 37.46598\nExp number, first 60 AAs: 19.53139\nTotal prob of N-in: 0.94364\nPOSSIBLE N-term signal sequence\ninside 1 19\nTMhelix 20 39\noutside 40 325\nTMhelix 326 348\ninside 349 372

13370 GCF_000616365.1_Ec2011EL-1107 Escherichia coli O157:H7 str. 2011EL-1107 Proteobacteria; Gammaproteobacteria; Enterobacterales; Enterobacteriaceae; Escherichia; Escherichia coli; Escherichia coli O157 MTGDNTLIHSHGINRRDFMKLCAALAATMGLSSKAAAEMAESVTNPQRPPVIWIGAQECT WP_000145410.1 MULTISPECIES: hydrogenase-2 small chain [Proteobacteria] Length: 372\nNumber of predicted TMHs: 1\nExp number of AAs in TMHs: 23.72226\nExp number, first 60 AAs: 0.22038\nTotal prob of N-in: 0.12994\noutside 1 331\nTMhelix 332 354\ninside 355 372

13371 GCF_000616405.2_Ec2011EL-2090 Escherichia coli O157:H7 str. 2011EL-2090 Proteobacteria; Gammaproteobacteria; Enterobacterales; Enterobacteriaceae; Escherichia; Escherichia coli; Escherichia coli O157 MTGDNTLIHSHGINRRDFMKLCAALAATMGLSSKAAAEMAESVTNPQRPPVIWIGAQECT WP_000145410.1 MULTISPECIES: hydrogenase-2 small chain [Proteobacteria] Length: 372\nNumber of predicted TMHs: 1\nExp number of AAs in TMHs: 23.72226\nExp number, first 60 AAs: 0.22038\nTotal prob of N-in: 0.12994\noutside 1 331\nTMhelix 332 354\ninside 355 372

13372 GCF_000616405.2_Ec2011EL-2090 Escherichia coli O157:H7 str. 2011EL-2090 Proteobacteria; Gammaproteobacteria; Enterobacterales; Enterobacteriaceae; Escherichia; Escherichia coli; Escherichia coli O157 MNNEETFYQAMRRQGVTRRSFLKYCSLAATSLGLGAGMAPKIAWALENKPRIPVVWIHGL WP_001301613.1 hydrogenase [Escherichia coli] Length: 372\nNumber of predicted TMHs: 2\nExp number of AAs in TMHs: 37.46598\nExp number, first 60 AAs: 19.53139\nTotal prob of N-in: 0.94364\nPOSSIBLE N-term signal sequence\ninside 1 19\nTMhelix 20 39\noutside 40 325\nTMhelix 326 348\ninside 349 372

13373 GCF_000616425.2_Ec2011EL-2091 Escherichia coli O157:H7 str. 2011EL-2091 Proteobacteria; Gammaproteobacteria; Enterobacterales; Enterobacteriaceae; Escherichia; Escherichia coli; Escherichia coli O157 MNNEETFYQAMRRQGVTRRSFLKYCSLAATSLGLGAGMAPKIAWALENKPRIPVVWIHGL WP_001301613.1 hydrogenase [Escherichia coli] Length: 372\nNumber of predicted TMHs: 2\nExp number of AAs in TMHs: 37.46598\nExp number, first 60 AAs: 19.53139\nTotal prob of N-in: 0.94364\nPOSSIBLE N-term signal sequence\ninside 1 19\nTMhelix 20 39\noutside 40 325\nTMhelix 326 348\ninside 349 372

13374 GCF_000616425.2_Ec2011EL-2091 Escherichia coli O157:H7 str. 2011EL-2091 Proteobacteria; Gammaproteobacteria; Enterobacterales; Enterobacteriaceae; Escherichia; Escherichia coli; Escherichia coli O157 MTGDNTLIHSHGINRRDFMKLCAALAATMGLSSKAAAEMAESVTNPQRPPVIWIGAQECT WP_000145410.1 MULTISPECIES: hydrogenase-2 small chain [Proteobacteria] Length: 372\nNumber of predicted TMHs: 1\nExp number of AAs in TMHs: 23.72226\nExp number, first 60 AAs: 0.22038\nTotal prob of N-in: 0.12994\noutside 1 331\nTMhelix 332 354\ninside 355 372

13375 GCF_000616445.2_Ec2011EL-2092 Escherichia coli O157:H7 str. 2011EL-2092 Proteobacteria; Gammaproteobacteria; Enterobacterales; Enterobacteriaceae; Escherichia; Escherichia coli; Escherichia coli O157 MNNEETFYQAMRRQGVTRRSFLKYCSLAATSLGLGAGMAPKIAWALENKPRIPVVWIHGL WP_001301613.1 hydrogenase [Escherichia coli] Length: 372\nNumber of predicted TMHs: 2\nExp number of AAs in TMHs: 37.46598\nExp number, first 60 AAs: 19.53139\nTotal prob of N-in: 0.94364\nPOSSIBLE N-term signal sequence\ninside 1 19\nTMhelix 20 39\noutside 40 325\nTMhelix 326 348\ninside 349 372

13376 GCF_000616445.2_Ec2011EL-2092 Escherichia coli O157:H7 str. 2011EL-2092 Proteobacteria; Gammaproteobacteria; Enterobacterales; Enterobacteriaceae; Escherichia; Escherichia coli; Escherichia coli O157 MTGDNTLIHSHGINRRDFMKLCAALAATMGLSSKAAAEMAESVTNPQRPPVIWIGAQECT WP_000145410.1 MULTISPECIES: hydrogenase-2 small chain [Proteobacteria] Length: 372\nNumber of predicted TMHs: 1\nExp number of AAs in TMHs: 23.72226\nExp number, first 60 AAs: 0.22038\nTotal prob of N-in: 0.12994\noutside 1 331\nTMhelix 332 354\ninside 355 372

13377 GCF_000616465.2_Ec2011EL-2093 Escherichia coli O157:H7 str. 2011EL-2093 Proteobacteria; Gammaproteobacteria; Enterobacterales; Enterobacteriaceae; Escherichia; Escherichia coli; Escherichia coli O157 MNNEETFYQAMRRQGVTRRSFLKYCSLAATSLGLGAGMAPKIAWALENKPRIPVVWIHGL WP_001301613.1 hydrogenase [Escherichia coli] Length: 372\nNumber of predicted TMHs: 2\nExp number of AAs in TMHs: 37.46598\nExp number, first 60 AAs: 19.53139\nTotal prob of N-in: 0.94364\nPOSSIBLE N-term signal sequence\ninside 1 19\nTMhelix 20 39\noutside 40 325\nTMhelix 326 348\ninside 349 372

13378 GCF_000616465.2_Ec2011EL-2093 Escherichia coli O157:H7 str. 2011EL-2093 Proteobacteria; Gammaproteobacteria; Enterobacterales; Enterobacteriaceae; Escherichia; Escherichia coli; Escherichia coli O157 MTGDNTLIHSHGINRRDFMKLCAALAATMGLSSKAAAEMAESVTNPQRPPVIWIGAQECT WP_000145410.1 MULTISPECIES: hydrogenase-2 small chain [Proteobacteria] Length: 372\nNumber of predicted TMHs: 1\nExp number of AAs in TMHs: 23.72226\nExp number, first 60 AAs: 0.22038\nTotal prob of N-in: 0.12994\noutside 1 331\nTMhelix 332 354\ninside 355 372

13379 GCF_000616485.2_Ec2011EL-2094 Escherichia coli O157:H7 str. 2011EL-2094 Proteobacteria; Gammaproteobacteria; Enterobacterales; Enterobacteriaceae; Escherichia; Escherichia coli; Escherichia coli O157 MNNEETFYQAMRRQGVTRRSFLKYCSLAATSLGLGAGMAPKIAWALENKPRIPVVWIHGL WP_001301613.1 hydrogenase [Escherichia coli] Length: 372\nNumber of predicted TMHs: 2\nExp number of AAs in TMHs: 37.46598\nExp number, first 60 AAs: 19.53139\nTotal prob of N-in: 0.94364\nPOSSIBLE N-term signal sequence\ninside 1 19\nTMhelix 20 39\noutside 40 325\nTMhelix 326 348\ninside 349 372

13380 GCF_000616485.2_Ec2011EL-2094 Escherichia coli O157:H7 str. 2011EL-2094 Proteobacteria; Gammaproteobacteria; Enterobacterales; Enterobacteriaceae; Escherichia; Escherichia coli; Escherichia coli O157 MTGDNTLIHSHGINRRDFMKLCAALAATMGLSSKAAAEMAESVTNPQRPPVIWIGAQECT WP_000145410.1 MULTISPECIES: hydrogenase-2 small chain [Proteobacteria] Length: 372\nNumber of predicted TMHs:

1\nExp number of AAs in TMHs: 23.72226\nExp number, first 60 AAs: 0.22038\nTotal prob of N-in: 0.12994\noutside 1 331\nTMhelix 332 354\ninside 355 372

13381 GCF_000616505.2_Ec2011EL-2096 Escherichia coli O157:H7 str. 2011EL-2096 Proteobacteria; Gammaproteobacteria; Enterobacterales; Enterobacteriaceae; Escherichia; Escherichia coli; Escherichia coli O157 MTGDNTLIHSHGINRRDFMKLCAALAATMGLSSKAAAEMAESVTNPQRPPVIWIGAQECT WP_000145410.1 MULTISPECIES: hydrogenase-2 small chain [Proteobacteria] Length: 372\nNumber of predicted TMHs: 1\nExp number of AAs in TMHs: 23.72226\nExp number, first 60 AAs: 0.22038\nTotal prob of N-in: 0.12994\noutside 1 331\nTMhelix 332 354\ninside 355 372

13382 GCF_000616505.2_Ec2011EL-2096 Escherichia coli O157:H7 str. 2011EL-2096 Proteobacteria; Gammaproteobacteria; Enterobacterales; Enterobacteriaceae; Escherichia; Escherichia coli; Escherichia coli O157 MNNEETFYQAMRRQGVTRRSFLKYCSLAATSLGLGAGMAPKIAWALENKPRIPVVWIHGL WP_001301613.1 hydrogenase [Escherichia coli] Length: 372\nNumber of predicted TMHs: 2\nExp number of AAs in TMHs: 37.46598\nExp number, first 60 AAs: 19.53139\nTotal prob of N-in: 0.94364\nPOSSIBLE N-term signal sequence\ninside 1 19\nTMhelix 20 39\noutside 40 325\nTMhelix 326 348\ninside 349 372

13383 GCF_000616525.2_Ec2011EL-2097 Escherichia coli O157:H7 str. 2011EL-2097 Proteobacteria; Gammaproteobacteria; Enterobacterales; Enterobacteriaceae; Escherichia; Escherichia coli; Escherichia coli O157 MNNEETFYQAMRRQGVTRRSFLKYCSLAATSLGLGAGMAPKIAWALENKPRIPVVWIHGL WP_001301613.1 hydrogenase [Escherichia coli] Length: 372\nNumber of predicted TMHs: 2\nExp number of AAs in TMHs: 37.46598\nExp number, first 60 AAs: 19.53139\nTotal prob of N-in: 0.94364\nPOSSIBLE N-term signal sequence\ninside 1 19\nTMhelix 20 39\noutside 40 325\nTMhelix 326 348\ninside 349 372

13384 GCF_000616525.2_Ec2011EL-2097 Escherichia coli O157:H7 str. 2011EL-2097 Proteobacteria; Gammaproteobacteria; Enterobacterales; Enterobacteriaceae; Escherichia; Escherichia coli; Escherichia coli O157 MTGDNTLIHSHGINRRDFMKLCAALAATMGLSSKAAAEMAESVTNPQRPPVIWIGAQECT WP_000145410.1 MULTISPECIES: hydrogenase-2 small chain [Proteobacteria] Length: 372\nNumber of predicted TMHs: 1\nExp number of AAs in TMHs: 23.72226\nExp number, first 60 AAs: 0.22038\nTotal prob of N-in: 0.12994\noutside 1 331\nTMhelix 332 354\ninside 355 372

13385 GCF_000616545.2_Ec2011EL-2098 Escherichia coli O157:H7 str. 2011EL-2098 Proteobacteria; Gammaproteobacteria; Enterobacterales; Enterobacteriaceae; Escherichia; Escherichia coli; Escherichia coli O157 MTGDNTLIHSHGINRRDFMKLCAALAATMGLSSKAAAEMAESVTNPQRPPVIWIGAQECT WP_000145410.1 MULTISPECIES: hydrogenase-2 small chain [Proteobacteria] Length: 372\nNumber of predicted TMHs: 1\nExp number of AAs in TMHs: 23.72226\nExp number, first 60 AAs: 0.22038\nTotal prob of N-in: 0.12994\noutside 1 331\nTMhelix 332 354\ninside 355 372

13386 GCF_000616545.2_Ec2011EL-2098 Escherichia coli O157:H7 str. 2011EL-2098 Proteobacteria; Gammaproteobacteria; Enterobacterales; Enterobacteriaceae; Escherichia; Escherichia coli; Escherichia coli O157 MNNEETFYQAMRRQGVTRRSFLKYCSLAATSLGLGAGMAPKIAWALENKPRIPVVWIHGL WP_001301613.1 hydrogenase [Escherichia coli] Length: 372\nNumber of predicted TMHs: 2\nExp number of AAs in TMHs: 37.46598\nExp number, first 60 AAs: 19.53139\nTotal prob of N-in: 0.94364\nPOSSIBLE N-term signal sequence\ninside 1 19\nTMhelix 20 39\noutside 40 325\nTMhelix 326 348\ninside 349 372

13387 GCF_000616705.2_Ec2011EL-2099 Escherichia coli O157:H7 str. 2011EL-2099 Proteobacteria; Gammaproteobacteria; Enterobacterales; Enterobacteriaceae; Escherichia; Escherichia coli; Escherichia coli O157 MNNEETFYQAMRRQGVTRRSFLKYCSLAATSLGLGAGMAPKIAWALENKPRIPVVWIHGL WP_001301613.1 hydrogenase [Escherichia coli] Length: 372\nNumber of predicted TMHs: 2\nExp number of AAs in TMHs: 37.46598\nExp number, first 60 AAs: 19.53139\nTotal prob of N-in: 0.94364\nPOSSIBLE N-term signal sequence\ninside 1 19\nTMhelix 20 39\noutside 40 325\nTMhelix 326 348\ninside 349 372

13388 GCF_000616705.2_Ec2011EL-2099 Escherichia coli O157:H7 str. 2011EL-2099 Proteobacteria; Gammaproteobacteria; Enterobacterales; Enterobacteriaceae; Escherichia; Escherichia coli; Escherichia coli O157 MTGDNTLIHSHGINRRDFMKLCAALAATMGLSSKAAAEMAESVTNPQRPPVIWIGAQECT WP_000145410.1 MULTISPECIES: hydrogenase-2 small chain [Proteobacteria] Length: 372\nNumber of predicted TMHs: 1\nExp number of AAs in TMHs: 23.72226\nExp number, first 60 AAs: 0.22038\nTotal prob of N-in: 0.12994\noutside 1 331\nTMhelix 332 354\ninside 355 372

13389 GCF_000615695.1_Ec2011EL-2101 Escherichia coli O157:H7 str. 2011EL-2101 Proteobacteria; Gammaproteobacteria; Enterobacterales; Enterobacteriaceae; Escherichia; Escherichia coli; Escherichia coli O157

MTGDNTLIHSHGINRRDFMKLCAALAATMGLSSKAAAEMAESVTNPQRPPVIWIGAQECT WP_000145410.1 MULTISPECIES: hydrogenase-2 small chain [Proteobacteria] Length: 372\nNumber of predicted TMHs: 1\nExp number of AAs in TMHs: 23.72226\nExp number, first 60 AAs: 0.22038\nTotal prob of N-in: 0.12994\noutside 1 331\nTMhelix 332 354\ninside 355 372

13390 GCF_000615695.1_Ec2011EL-2101 Escherichia coli O157:H7 str. 2011EL-2101 Proteobacteria; Gammaproteobacteria; Enterobacterales; Enterobacteriaceae; Escherichia; Escherichia coli; Escherichia coli O157 MNNEETFYQAMRRQGVTRRSFLKYCSLAATSLGLGAGMAPKIAWALENKPRIPVVWIHGL WP_001301613.1 hydrogenase [Escherichia coli] Length: 372\nNumber of predicted TMHs: 2\nExp number of AAs in TMHs: 37.46598\nExp number, first 60 AAs: 19.53139\nTotal prob of N-in: 0.94364\nPOSSIBLE N-term signal sequence\ninside 1 19\nTMhelix 20 39\noutside 40 325\nTMhelix 326 348\ninside 349 372

13391 GCF_000615745.2_Ec2011EL-2103 Escherichia coli O157:H7 str. 2011EL-2103 Proteobacteria; Gammaproteobacteria; Enterobacterales; Enterobacteriaceae; Escherichia; Escherichia coli; Escherichia coli O157 MTGDNTLIHSHGINRRDFMKLCAALAATMGLSSKAAAEMAESVTNPQRPPVIWIGAQECT WP_000145410.1 MULTISPECIES: hydrogenase-2 small chain [Proteobacteria] Length: 372\nNumber of predicted TMHs: 1\nExp number of AAs in TMHs: 23.72226\nExp number, first 60 AAs: 0.22038\nTotal prob of N-in: 0.12994\noutside 1 331\nTMhelix 332 354\ninside 355 372

13392 GCF_000615745.2_Ec2011EL-2103 Escherichia coli O157:H7 str. 2011EL-2103 Proteobacteria; Gammaproteobacteria; Enterobacterales; Enterobacteriaceae; Escherichia; Escherichia coli; Escherichia coli O157 MNNEETFYQAMRRQGVTRRSFLKYCSLAATSLGLGAGMAPKIAWALENKPRIPVVWIHGL WP_001301613.1 hydrogenase [Escherichia coli] Length: 372\nNumber of predicted TMHs: 2\nExp number of AAs in TMHs: 37.46598\nExp number, first 60 AAs: 19.53139\nTotal prob of N-in: 0.94364\nPOSSIBLE N-term signal sequence\ninside 1 19\nTMhelix 20 39\noutside 40 325\nTMhelix 326 348\ninside 349 372

13393 GCF_000615785.2_Ec2011EL-2104 Escherichia coli O157:H7 str. 2011EL-2104 Proteobacteria; Gammaproteobacteria; Enterobacterales; Enterobacteriaceae; Escherichia; Escherichia coli; Escherichia coli O157 MTGDNTLIHSHGINRRDFMKLCAALAATMGLSSKAAAEMAESVTNPQRPPVIWIGAQECT WP_000145410.1 MULTISPECIES: hydrogenase-2 small chain [Proteobacteria] Length: 372\nNumber of predicted TMHs: 1\nExp number of AAs in TMHs: 23.72226\nExp number, first 60 AAs: 0.22038\nTotal prob of N-in: 0.12994\noutside 1 331\nTMhelix 332 354\ninside 355 372

13394 GCF_000615785.2_Ec2011EL-2104 Escherichia coli O157:H7 str. 2011EL-2104 Proteobacteria; Gammaproteobacteria; Enterobacterales; Enterobacteriaceae; Escherichia; Escherichia coli; Escherichia coli O157 MNNEETFYQAMRRQGVTRRSFLKYCSLAATSLGLGAGMAPKIAWALENKPRIPVVWIHGL WP_001301613.1 hydrogenase [Escherichia coli] Length: 372\nNumber of predicted TMHs: 2\nExp number of AAs in TMHs: 37.46598\nExp number, first 60 AAs: 19.53139\nTotal prob of N-in: 0.94364\nPOSSIBLE N-term signal sequence\ninside 1 19\nTMhelix 20 39\noutside 40 325\nTMhelix 326 348\ninside 349 372

13395 GCF_000615815.2_Ec2011EL-2105 Escherichia coli O157:H7 str. 2011EL-2105 Proteobacteria; Gammaproteobacteria; Enterobacterales; Enterobacteriaceae; Escherichia; Escherichia coli; Escherichia coli O157 MNNEETFYQAMRRQGVTRRSFLKYCSLAATSLGLGAGMAPKIAWALENKPRIPVVWIHGL WP_001301613.1 hydrogenase [Escherichia coli] Length: 372\nNumber of predicted TMHs: 2\nExp number of AAs in TMHs: 37.46598\nExp number, first 60 AAs: 19.53139\nTotal prob of N-in: 0.94364\nPOSSIBLE N-term signal sequence\ninside 1 19\nTMhelix 20 39\noutside 40 325\nTMhelix 326 348\ninside 349 372

13396 GCF_000615815.2_Ec2011EL-2105 Escherichia coli O157:H7 str. 2011EL-2105 Proteobacteria; Gammaproteobacteria; Enterobacterales; Enterobacteriaceae; Escherichia; Escherichia coli; Escherichia coli O157 MTGDNTLIHSHGINRRDFMKLCAALAATMGLSSKAAAEMAESVTNPQRPPVIWIGAQECT WP_000145410.1 MULTISPECIES: hydrogenase-2 small chain [Proteobacteria] Length: 372\nNumber of predicted TMHs: 1\nExp number of AAs in TMHs: 23.72226\nExp number, first 60 AAs: 0.22038\nTotal prob of N-in: 0.12994\noutside 1 331\nTMhelix 332 354\ninside 355 372

13397 GCF_000615865.2_Ec2011EL-2106 Escherichia coli O157:H7 str. 2011EL-2106 Proteobacteria; Gammaproteobacteria; Enterobacterales; Enterobacteriaceae; Escherichia; Escherichia coli; Escherichia coli O157 MNNEETFYQAMRRQGVTRRSFLKYCSLAATSLGLGAGMAPKIAWALENKPRIPVVWIHGL WP_001301613.1 hydrogenase [Escherichia coli] Length: 372\nNumber of predicted TMHs: 2\nExp number of AAs in TMHs: 37.46598\nExp number, first 60 AAs: 19.53139\nTotal prob of N-in: 0.94364\nPOSSIBLE N-term signal sequence\ninside 1 19\nTMhelix 20 39\noutside 40 325\nTMhelix 326 348\ninside 349 372

13398 GCF_000615865.2_Ec2011EL-2106 Escherichia coli O157:H7 str. 2011EL-2106 Proteobacteria; Gammaproteobacteria; Enterobacterales; Enterobacteriaceae; Escherichia; Escherichia coli; Escherichia coli O157 MTGDNTLIHSHGINRRDFMKLCAALAATMGLSSKAAAEMAESVTNPQRPPVIWIGAQECT WP_000145410.1 MULTISPECIES: hydrogenase-2 small chain [Proteobacteria] Length: 372\nNumber of predicted TMHs: 1\nExp number of AAs in TMHs: 23.72226\nExp number, first 60 AAs: 0.22038\nTotal prob of N-in: 0.12994\noutside 1 331\nTMhelix 332 354\ninside 355 372

13399 GCF_000615925.2_Ec2011EL-2107 Escherichia coli O157:H7 str. 2011EL-2107 Proteobacteria; Gammaproteobacteria; Enterobacterales; Enterobacteriaceae; Escherichia; Escherichia coli; Escherichia coli O157 MTGDNTLIHSHGINRRDFMKLCAALAATMGLSSKAAAEMAESVTNPQRPPVIWIGAQECT WP_000145410.1 MULTISPECIES: hydrogenase-2 small chain [Proteobacteria] Length: 372\nNumber of predicted TMHs: 1\nExp number of AAs in TMHs: 23.72226\nExp number, first 60 AAs: 0.22038\nTotal prob of N-in: 0.12994\noutside 1 331\nTMhelix 332 354\ninside 355 372

13400 GCF_000615925.2_Ec2011EL-2107 Escherichia coli O157:H7 str. 2011EL-2107 Proteobacteria; Gammaproteobacteria; Enterobacterales; Enterobacteriaceae; Escherichia; Escherichia coli; Escherichia coli O157 MNNEETFYQAMRRQGVTRRSFLKYCSLAATSLGLGAGMAPKIAWALENKPRIPVVWIHGL WP_001301613.1 hydrogenase [Escherichia coli] Length: 372\nNumber of predicted TMHs: 2\nExp number of AAs in TMHs: 37.46598\nExp number, first 60 AAs: 19.53139\nTotal prob of N-in: 0.94364\nPOSSIBLE N-term signal sequence\ninside 1 19\nTMhelix 20 39\noutside 40 325\nTMhelix 326 348\ninside 349 372

13401 GCF_000615965.2_Ec2011EL-2108 Escherichia coli O157:H7 str. 2011EL-2108 Proteobacteria; Gammaproteobacteria; Enterobacterales; Enterobacteriaceae; Escherichia; Escherichia coli; Escherichia coli O157 MTGDNTLIHSHGINRRDFMKLCAALAATMGLSSKAAAEMAESVTNPQRPPVIWIGAQECT WP_000145410.1 MULTISPECIES: hydrogenase-2 small chain [Proteobacteria] Length: 372\nNumber of predicted TMHs: 1\nExp number of AAs in TMHs: 23.72226\nExp number, first 60 AAs: 0.22038\nTotal prob of N-in: 0.12994\noutside 1 331\nTMhelix 332 354\ninside 355 372

13402 GCF_000615965.2_Ec2011EL-2108 Escherichia coli O157:H7 str. 2011EL-2108 Proteobacteria; Gammaproteobacteria; Enterobacterales; Enterobacteriaceae; Escherichia; Escherichia coli; Escherichia coli O157 MNNEETFYQAMRRQGVTRRSFLKYCSLAATSLGLGAGMAPKIAWALENKPRIPVVWIHGL WP_001301613.1 hydrogenase [Escherichia coli] Length: 372\nNumber of predicted TMHs: 2\nExp number of AAs in TMHs: 37.46598\nExp number, first 60 AAs: 19.53139\nTotal prob of N-in: 0.94364\nPOSSIBLE N-term signal sequence\ninside 1 19\nTMhelix 20 39\noutside 40 325\nTMhelix 326 348\ninside 349 372

13403 GCF_000616005.2_Ec2011EL-2109 Escherichia coli O157:H7 str. 2011EL-2109 Proteobacteria; Gammaproteobacteria; Enterobacterales; Enterobacteriaceae; Escherichia; Escherichia coli; Escherichia coli O157 MTGDNTLIHSHGINRRDFMKLCAALAATMGLSSKAAAEMAESVTNPQRPPVIWIGAQECT WP_000145410.1 MULTISPECIES: hydrogenase-2 small chain [Proteobacteria] Length: 372\nNumber of predicted TMHs: 1\nExp number of AAs in TMHs: 23.72226\nExp number, first 60 AAs: 0.22038\nTotal prob of N-in: 0.12994\noutside 1 331\nTMhelix 332 354\ninside 355 372

13404 GCF_000616005.2_Ec2011EL-2109 Escherichia coli O157:H7 str. 2011EL-2109 Proteobacteria; Gammaproteobacteria; Enterobacterales; Enterobacteriaceae; Escherichia; Escherichia coli; Escherichia coli O157 MNNEETFYQAMRRQGVTRRSFLKYCSLAATSLGLGAGMAPKIAWALENKPRIPVVWIHGL WP_001301613.1 hydrogenase [Escherichia coli] Length: 372\nNumber of predicted TMHs: 2\nExp number of AAs in TMHs: 37.46598\nExp number, first 60 AAs: 19.53139\nTotal prob of N-in: 0.94364\nPOSSIBLE N-term signal sequence\ninside 1 19\nTMhelix 20 39\noutside 40 325\nTMhelix 326 348\ninside 349 372

13405 GCF_000616035.2_Ec2011EL-2111 Escherichia coli O157:H7 str. 2011EL-2111 Proteobacteria; Gammaproteobacteria; Enterobacterales; Enterobacteriaceae; Escherichia; Escherichia coli; Escherichia coli O157 MNNEETFYQAMRRQGVTRRSFLKYCSLAATSLGLGAGMAPKIAWALENKPRIPVVWIHGL WP_001301613.1 hydrogenase [Escherichia coli] Length: 372\nNumber of predicted TMHs: 2\nExp number of AAs in TMHs: 37.46598\nExp number, first 60 AAs: 19.53139\nTotal prob of N-in: 0.94364\nPOSSIBLE N-term signal sequence\ninside 1 19\nTMhelix 20 39\noutside 40 325\nTMhelix 326 348\ninside 349 372

13406 GCF_000616035.2_Ec2011EL-2111 Escherichia coli O157:H7 str. 2011EL-2111 Proteobacteria; Gammaproteobacteria; Enterobacterales; Enterobacteriaceae; Escherichia; Escherichia coli; Escherichia coli O157 MTGDNTLIHSHGINRRDFMKLCAALAATMGLSSKAAAEMAESVTNPQRPPVIWIGAQECT WP_000145410.1 MULTISPECIES: hydrogenase-2 small chain [Proteobacteria] Length: 372\nNumber of predicted TMHs:

1\nExp number of AAs in TMHs: 23.72226\nExp number, first 60 AAs: 0.22038\nTotal prob of N-in: 0.12994\noutside 1 331\nTMhelix 332 354\ninside 355 372

13407 GCF_000616075.2_Ec2011EL-2112 Escherichia coli O157:H7 str. 2011EL-2112 Proteobacteria; Gammaproteobacteria; Enterobacterales; Enterobacteriaceae; Escherichia; Escherichia coli; Escherichia coli O157 MNNEETFYQAMRRQGVTRRSFLKYCSLAATSLGLGAGMAPKIAWALENKPRIPVVWIHGL WP_001301613.1 hydrogenase [Escherichia coli] Length: 372\nNumber of predicted TMHs: 2\nExp number of AAs in TMHs: 37.46598\nExp number, first 60 AAs: 19.53139\nTotal prob of N-in: 0.94364\nPOSSIBLE N-term signal sequence\ninside 1 19\nTMhelix 20 39\noutside 40 325\nTMhelix 326 348\ninside 349 372

13408 GCF_000616075.2_Ec2011EL-2112 Escherichia coli O157:H7 str. 2011EL-2112 Proteobacteria; Gammaproteobacteria; Enterobacterales; Enterobacteriaceae; Escherichia; Escherichia coli; Escherichia coli O157 MTGDNTLIHSHGINRRDFMKLCAALAATMGLSSKAAAEMAESVTNPQRPPVIWIGAQECT WP_000145410.1 MULTISPECIES: hydrogenase-2 small chain [Proteobacteria] Length: 372\nNumber of predicted TMHs: 1\nExp number of AAs in TMHs: 23.72226\nExp number, first 60 AAs: 0.22038\nTotal prob of N-in: 0.12994\noutside 1 331\nTMhelix 332 354\ninside 355 372

13409 GCF_000614645.2_Ec2011EL-2113 Escherichia coli O157:H7 str. 2011EL-2113 Proteobacteria; Gammaproteobacteria; Enterobacterales; Enterobacteriaceae; Escherichia; Escherichia coli; Escherichia coli O157 MNNEETFYQAMRRQGVTRRSFLKYCSLAATSLGLGAGMAPKIAWALENKPRIPVVWIHGL WP_001301613.1 hydrogenase [Escherichia coli] Length: 372\nNumber of predicted TMHs: 2\nExp number of AAs in TMHs: 37.46598\nExp number, first 60 AAs: 19.53139\nTotal prob of N-in: 0.94364\nPOSSIBLE N-term signal sequence\ninside 1 19\nTMhelix 20 39\noutside 40 325\nTMhelix 326 348\ninside 349 372

13410 GCF_000614645.2_Ec2011EL-2113 Escherichia coli O157:H7 str. 2011EL-2113 Proteobacteria; Gammaproteobacteria; Enterobacterales; Enterobacteriaceae; Escherichia; Escherichia coli; Escherichia coli O157 MTGDNTLIHSHGINRRDFMKLCAALAATMGLSSKAAAEMAESVTNPQRPPVIWIGAQECT WP_000145410.1 MULTISPECIES: hydrogenase-2 small chain [Proteobacteria] Length: 372\nNumber of predicted TMHs: 1\nExp number of AAs in TMHs: 23.72226\nExp number, first 60 AAs: 0.22038\nTotal prob of N-in: 0.12994\noutside 1 331\nTMhelix 332 354\ninside 355 372

13411 GCF_000614255.1_Ec2011EL-2114 Escherichia coli O157:H7 str. 2011EL-2114 Proteobacteria; Gammaproteobacteria; Enterobacterales; Enterobacteriaceae; Escherichia; Escherichia coli; Escherichia coli O157 MNNEETFYQAMRRQGVTRRSFLKYCSLAATSLGLGAGMAPKIAWALENKPRIPVVWIHGL WP_001301613.1 hydrogenase [Escherichia coli] Length: 372\nNumber of predicted TMHs: 2\nExp number of AAs in TMHs: 37.46598\nExp number, first 60 AAs: 19.53139\nTotal prob of N-in: 0.94364\nPOSSIBLE N-term signal sequence\ninside 1 19\nTMhelix 20 39\noutside 40 325\nTMhelix 326 348\ninside 349 372

13412 GCF_000614255.1_Ec2011EL-2114 Escherichia coli O157:H7 str. 2011EL-2114 Proteobacteria; Gammaproteobacteria; Enterobacterales; Enterobacteriaceae; Escherichia; Escherichia coli; Escherichia coli O157 MTGDNTLIHSHGINRRDFMKLCAALAATMGLSSKAAAEMAESVTNPQRPPVIWIGAQECT WP_000145410.1 MULTISPECIES: hydrogenase-2 small chain [Proteobacteria] Length: 372\nNumber of predicted TMHs: 1\nExp number of AAs in TMHs: 23.72226\nExp number, first 60 AAs: 0.22038\nTotal prob of N-in: 0.12994\noutside 1 331\nTMhelix 332 354\ninside 355 372

13413 GCF_000614305.2_Ec2011EL-2286 Escherichia coli O157:H7 str. 2011EL-2286 Proteobacteria; Gammaproteobacteria; Enterobacterales; Enterobacteriaceae; Escherichia; Escherichia coli; Escherichia coli O157 MTGDNTLIHSHGINRRDFMKLCAALAATMGLSSKAAAEMAESVTNPQRPPVIWIGAQECT WP_000145410.1 MULTISPECIES: hydrogenase-2 small chain [Proteobacteria] Length: 372\nNumber of predicted TMHs: 1\nExp number of AAs in TMHs: 23.72226\nExp number, first 60 AAs: 0.22038\nTotal prob of N-in: 0.12994\noutside 1 331\nTMhelix 332 354\ninside 355 372

13414 GCF_000614305.2_Ec2011EL-2286 Escherichia coli O157:H7 str. 2011EL-2286 Proteobacteria; Gammaproteobacteria; Enterobacterales; Enterobacteriaceae; Escherichia; Escherichia coli; Escherichia coli O157 MNNEETFYQAMRRQGVTRRSFLKYCSLAATSLGLGAGMAPKIAWALENKPRIPVVWIHGL WP_001301613.1 hydrogenase [Escherichia coli] Length: 372\nNumber of predicted TMHs: 2\nExp number of AAs in TMHs: 37.46598\nExp number, first 60 AAs: 19.53139\nTotal prob of N-in: 0.94364\nPOSSIBLE N-term signal sequence\ninside 1 19\nTMhelix 20 39\noutside 40 325\nTMhelix 326 348\ninside 349 372

13415 GCF_000614405.2_Ec2011EL-2287 Escherichia coli O157:H7 str. 2011EL-2287 Proteobacteria; Gammaproteobacteria; Enterobacterales; Enterobacteriaceae; Escherichia; Escherichia coli; Escherichia coli O157

MTGDNTLIHSHGINRRDFMKLCAALAATMGLSSKAAAEMAESVTNPQRPPVIWIGAQECT WP_000145410.1 MULTISPECIES: hydrogenase-2 small chain [Proteobacteria] Length: 372\nNumber of predicted TMHs: 1\nExp number of AAs in TMHs: 23.72226\nExp number, first 60 AAs: 0.22038\nTotal prob of N-in: 0.12994\noutside 1 331\nTMhelix 332 354\ninside 355 372

13416 GCF_000614405.2_Ec2011EL-2287 Escherichia coli O157:H7 str. 2011EL-2287 Proteobacteria; Gammaproteobacteria; Enterobacterales; Enterobacteriaceae; Escherichia; Escherichia coli; Escherichia coli O157 MNNEETFYQAMRRQGVTRRSFLKYCSLAATSLGLGAGMAPKIAWALENKPRIPVVWIHGL WP_001301613.1 hydrogenase [Escherichia coli] Length: 372\nNumber of predicted TMHs: 2\nExp number of AAs in TMHs: 37.46598\nExp number, first 60 AAs: 19.53139\nTotal prob of N-in: 0.94364\nPOSSIBLE N-term signal sequence\ninside 1 19\nTMhelix 20 39\noutside 40 325\nTMhelix 326 348\ninside 349 372

13417 GCF_000614085.2_Ec2011EL-2288 Escherichia coli O157:H7 str. 2011EL-2288 Proteobacteria; Gammaproteobacteria; Enterobacterales; Enterobacteriaceae; Escherichia; Escherichia coli; Escherichia coli O157 MNNEETFYQAMRRQGVTRRSFLKYCSLAATSLGLGAGMAPKIAWALENKPRIPVVWIHGL WP_001301613.1 hydrogenase [Escherichia coli] Length: 372\nNumber of predicted TMHs: 2\nExp number of AAs in TMHs: 37.46598\nExp number, first 60 AAs: 19.53139\nTotal prob of N-in: 0.94364\nPOSSIBLE N-term signal sequence\ninside 1 19\nTMhelix 20 39\noutside 40 325\nTMhelix 326 348\ninside 349 372

13418 GCF_000614085.2_Ec2011EL-2288 Escherichia coli O157:H7 str. 2011EL-2288 Proteobacteria; Gammaproteobacteria; Enterobacterales; Enterobacteriaceae; Escherichia; Escherichia coli; Escherichia coli O157 MTGDNTLIHSHGINRRDFMKLCAALAATMGLSSKAAAEMAESVTNPQRPPVIWIGAQECT WP_000145410.1 MULTISPECIES: hydrogenase-2 small chain [Proteobacteria] Length: 372\nNumber of predicted TMHs: 1\nExp number of AAs in TMHs: 23.72226\nExp number, first 60 AAs: 0.22038\nTotal prob of N-in: 0.12994\noutside 1 331\nTMhelix 332 354\ninside 355 372

13419 GCF_000614685.2_Ec2011EL-2289 Escherichia coli O157:H7 str. 2011EL-2289 Proteobacteria; Gammaproteobacteria; Enterobacterales; Enterobacteriaceae; Escherichia; Escherichia coli; Escherichia coli O157 MTGDNTLIHSHGINRRDFMKLCAALAATMGLSSKAAAEMAESVTNPQRPPVIWIGAQECT WP_000145410.1 MULTISPECIES: hydrogenase-2 small chain [Proteobacteria] Length: 372\nNumber of predicted TMHs: 1\nExp number of AAs in TMHs: 23.72226\nExp number, first 60 AAs: 0.22038\nTotal prob of N-in: 0.12994\noutside 1 331\nTMhelix 332 354\ninside 355 372

13420 GCF_000614685.2_Ec2011EL-2289 Escherichia coli O157:H7 str. 2011EL-2289 Proteobacteria; Gammaproteobacteria; Enterobacterales; Enterobacteriaceae; Escherichia; Escherichia coli; Escherichia coli O157 MNNEETFYQAMRRQGVTRRSFLKYCSLAATSLGLGAGMAPKIAWALENKPRIPVVWIHGL WP_001301613.1 hydrogenase [Escherichia coli] Length: 372\nNumber of predicted TMHs: 2\nExp number of AAs in TMHs: 37.46598\nExp number, first 60 AAs: 19.53139\nTotal prob of N-in: 0.94364\nPOSSIBLE N-term signal sequence\ninside 1 19\nTMhelix 20 39\noutside 40 325\nTMhelix 326 348\ninside 349 372

13421 GCF_000618945.2_Ec2011EL-2290 Escherichia coli O157:H7 str. 2011EL-2290 Proteobacteria; Gammaproteobacteria; Enterobacterales; Enterobacteriaceae; Escherichia; Escherichia coli; Escherichia coli O157 MNNEETFYQAMRRQGVTRRSFLKYCSLAATSLGLGAGMAPKIAWALENKPRIPVVWIHGL WP_001301613.1 hydrogenase [Escherichia coli] Length: 372\nNumber of predicted TMHs: 2\nExp number of AAs in TMHs: 37.46598\nExp number, first 60 AAs: 19.53139\nTotal prob of N-in: 0.94364\nPOSSIBLE N-term signal sequence\ninside 1 19\nTMhelix 20 39\noutside 40 325\nTMhelix 326 348\ninside 349 372

13422 GCF_000618945.2_Ec2011EL-2290 Escherichia coli O157:H7 str. 2011EL-2290 Proteobacteria; Gammaproteobacteria; Enterobacterales; Enterobacteriaceae; Escherichia; Escherichia coli; Escherichia coli O157 MTGDNTLIHSHGINRRDFMKLCAALAATMGLSSKAAAEMAESVTNPQRPPVIWIGAQECT WP_000145410.1 MULTISPECIES: hydrogenase-2 small chain [Proteobacteria] Length: 372\nNumber of predicted TMHs: 1\nExp number of AAs in TMHs: 23.72226\nExp number, first 60 AAs: 0.22038\nTotal prob of N-in: 0.12994\noutside 1 331\nTMhelix 332 354\ninside 355 372

13423 GCF_000614345.2_Ec2011EL-2312 Escherichia coli O157:H7 str. 2011EL-2312 Proteobacteria; Gammaproteobacteria; Enterobacterales; Enterobacteriaceae; Escherichia; Escherichia coli; Escherichia coli O157 MNNEETFYQAMRRQGVTRRSFLKYCSLAATSLGLGAGMAPKIAWALENKPRIPVVWIHGL WP_001301613.1 hydrogenase [Escherichia coli] Length: 372\nNumber of predicted TMHs: 2\nExp number of AAs in TMHs: 37.46598\nExp number, first 60 AAs: 19.53139\nTotal prob of N-in: 0.94364\nPOSSIBLE N-term signal sequence\ninside 1 19\nTMhelix 20 39\noutside 40 325\nTMhelix 326 348\ninside 349 372

13424 GCF_000614345.2_Ec2011EL-2312 Escherichia coli O157:H7 str. 2011EL-2312 Proteobacteria; Gammaproteobacteria; Enterobacterales; Enterobacteriaceae; Escherichia; Escherichia coli; Escherichia coli O157 MTGDNTLIHSHGINRRDFMKLCAALAATMGLSSKAAAEMAESVTNPQRPPVIWIGAQECT WP_000145410.1 MULTISPECIES: hydrogenase-2 small chain [Proteobacteria] Length: 372\nNumber of predicted TMHs: 1\nExp number of AAs in TMHs: 23.72226\nExp number, first 60 AAs: 0.22038\nTotal prob of N-in: 0.12994\noutside 1 331\nTMhelix 332 354\ninside 355 372

13425 GCF_000618985.2_Ec2011EL-2313 Escherichia coli O157:H7 str. 2011EL-2313 Proteobacteria; Gammaproteobacteria; Enterobacterales; Enterobacteriaceae; Escherichia; Escherichia coli; Escherichia coli O157 MNNEETFYQAMRRQGVTRRSFLKYCSLAATSLGLGAGMAPKIAWALENKPRIPVVWIHGL WP_001301613.1 hydrogenase [Escherichia coli] Length: 372\nNumber of predicted TMHs: 2\nExp number of AAs in TMHs: 37.46598\nExp number, first 60 AAs: 19.53139\nTotal prob of N-in: 0.94364\nPOSSIBLE N-term signal sequence\ninside 1 19\nTMhelix 20 39\noutside 40 325\nTMhelix 326 348\ninside 349 372

13426 GCF_000618985.2_Ec2011EL-2313 Escherichia coli O157:H7 str. 2011EL-2313 Proteobacteria; Gammaproteobacteria; Enterobacterales; Enterobacteriaceae; Escherichia; Escherichia coli; Escherichia coli O157 MTGDNTLIHSHGINRRDFMKLCAALAATMGLSSKAAAEMAESVTNPQRPPVIWIGAQECT WP_000145410.1 MULTISPECIES: hydrogenase-2 small chain [Proteobacteria] Length: 372\nNumber of predicted TMHs: 1\nExp number of AAs in TMHs: 23.72226\nExp number, first 60 AAs: 0.22038\nTotal prob of N-in: 0.12994\noutside 1 331\nTMhelix 332 354\ninside 355 372

13427 GCF_000193955.1_ASM19395v2 Escherichia coli O157:H7 str. EC1212 Proteobacteria; Gammaproteobacteria; Enterobacterales; Enterobacteriaceae; Escherichia; Escherichia coli; Escherichia coli O157 MNNEETFYQAMRRQGVTRRSFLKYCSLAATSLGLGAGMAPKIAWALENKPRIPVVWIHGL WP_001301613.1 hydrogenase [Escherichia coli] Length: 372\nNumber of predicted TMHs: 2\nExp number of AAs in TMHs: 37.46598\nExp number, first 60 AAs: 19.53139\nTotal prob of N-in: 0.94364\nPOSSIBLE N-term signal sequence\ninside 1 19\nTMhelix 20 39\noutside 40 325\nTMhelix 326 348\ninside 349 372

13428 GCF_000193955.1_ASM19395v2 Escherichia coli O157:H7 str. EC1212 Proteobacteria; Gammaproteobacteria; Enterobacterales; Enterobacteriaceae; Escherichia; Escherichia coli; Escherichia coli O157 MTGDNTLIHSHGINRRDFMKLCAALAATMGLSSKAAAEMAESVTNPQRPPVIWIGAQECT WP_000145410.1 MULTISPECIES: hydrogenase-2 small chain [Proteobacteria] Length: 372\nNumber of predicted TMHs: 1\nExp number of AAs in TMHs: 23.72226\nExp number, first 60 AAs: 0.22038\nTotal prob of N-in: 0.12994\noutside 1 331\nTMhelix 332 354\ninside 355 372

13429 GCF_000178315.1_ASM17831v1 Escherichia coli O157:H7 str. EC4009 Proteobacteria; Gammaproteobacteria; Enterobacterales; Enterobacteriaceae; Escherichia; Escherichia coli; Escherichia coli O157 MNNEETFYQAMRRQGVTRRSFLKYCSLAATSLGLGAGMAPKIAWALENKPRIPVVWIHGL WP_001301613.1 hydrogenase [Escherichia coli] Length: 372\nNumber of predicted TMHs: 2\nExp number of AAs in TMHs: 37.46598\nExp number, first 60 AAs: 19.53139\nTotal prob of N-in: 0.94364\nPOSSIBLE N-term signal sequence\ninside 1 19\nTMhelix 20 39\noutside 40 325\nTMhelix 326 348\ninside 349 372

13430 GCF_000178315.1_ASM17831v1 Escherichia coli O157:H7 str. EC4009 Proteobacteria; Gammaproteobacteria; Enterobacterales; Enterobacteriaceae; Escherichia; Escherichia coli; Escherichia coli O157 MTGDNTLIHSHGINRRDFMKLCAALAATMGLSSKAAAEMAESVTNPQRPPVIWIGAQECT WP_000145410.1 MULTISPECIES: hydrogenase-2 small chain [Proteobacteria] Length: 372\nNumber of predicted TMHs: 1\nExp number of AAs in TMHs: 23.72226\nExp number, first 60 AAs: 0.22038\nTotal prob of N-in: 0.12994\noutside 1 331\nTMhelix 332 354\ninside 355 372

13431 GCF_000181775.1_ASM18177v1 Escherichia coli O157:H7 str. EC4042 Proteobacteria; Gammaproteobacteria; Enterobacterales; Enterobacteriaceae; Escherichia; Escherichia coli; Escherichia coli O157 MNNEETFYQAMRRQGVTRRSFLKYCSLAATSLGLGAGMAPKIAWALENKPRIPVVWIHGL WP_001301613.1 hydrogenase [Escherichia coli] Length: 372\nNumber of predicted TMHs: 2\nExp number of AAs in TMHs: 37.46598\nExp number, first 60 AAs: 19.53139\nTotal prob of N-in: 0.94364\nPOSSIBLE N-term signal sequence\ninside 1 19\nTMhelix 20 39\noutside 40 325\nTMhelix 326 348\ninside 349 372

13432 GCF_000181775.1_ASM18177v1 Escherichia coli O157:H7 str. EC4042 Proteobacteria; Gammaproteobacteria; Enterobacterales; Enterobacteriaceae; Escherichia; Escherichia coli; Escherichia coli O157 MTGDNTLIHSHGINRRDFMKLCAALAATMGLSSKAAAEMAESVTNPQRPPVIWIGAQECT WP_000145410.1 MULTISPECIES: hydrogenase-2 small chain [Proteobacteria] Length: 372\nNumber of predicted TMHs:

1\nExp number of AAs in TMHs: 23.72226\nExp number, first 60 AAs: 0.22038\nTotal prob of N-in: 0.12994\noutside 1 331\nTMhelix 332 354\ninside 355 372

13433 GCF_000181755.1_ASM18175v1 Escherichia coli O157:H7 str. EC4045 Proteobacteria; Gammaproteobacteria; Enterobacterales; Enterobacteriaceae; Escherichia; Escherichia coli; Escherichia coli O157 MNNEETFYQAMRRQGVTRRSFLKYCSLAATSLGLGAGMAPKIAWALENKPRIPVVWIHGL WP_001301613.1 hydrogenase [Escherichia coli] Length: 372\nNumber of predicted TMHs: 2\nExp number of AAs in TMHs: 37.46598\nExp number, first 60 AAs: 19.53139\nTotal prob of N-in: 0.94364\nPOSSIBLE N-term signal sequence\ninside 1 19\nTMhelix 20 39\noutside 40 325\nTMhelix 326 348\ninside 349 372

13434 GCF_000181755.1_ASM18175v1 Escherichia coli O157:H7 str. EC4045 Proteobacteria; Gammaproteobacteria; Enterobacterales; Enterobacteriaceae; Escherichia; Escherichia coli; Escherichia coli O157 MTGDNTLIHSHGINRRDFMKLCAALAATMGLSSKAAAEMAESVTNPQRPPVIWIGAQECT WP_000145410.1 MULTISPECIES: hydrogenase-2 small chain [Proteobacteria] Length: 372\nNumber of predicted TMHs: 1\nExp number of AAs in TMHs: 23.72226\nExp number, first 60 AAs: 0.22038\nTotal prob of N-in: 0.12994\noutside 1 331\nTMhelix 332 354\ninside 355 372

13435 GCF_000171955.1_ASM17195v1 Escherichia coli O157:H7 str. EC4076 Proteobacteria; Gammaproteobacteria; Enterobacterales; Enterobacteriaceae; Escherichia; Escherichia coli; Escherichia coli O157 MTGDNTLIHSHGINRRDFMKLCAALAATMGLSSKAAAEMAESVTNPQRPPVIWIGAQECT WP_000145410.1 MULTISPECIES: hydrogenase-2 small chain [Proteobacteria] Length: 372\nNumber of predicted TMHs: 1\nExp number of AAs in TMHs: 23.72226\nExp number, first 60 AAs: 0.22038\nTotal prob of N-in: 0.12994\noutside 1 331\nTMhelix 332 354\ninside 355 372

13436 GCF_000171955.1_ASM17195v1 Escherichia coli O157:H7 str. EC4076 Proteobacteria; Gammaproteobacteria; Enterobacterales; Enterobacteriaceae; Escherichia; Escherichia coli; Escherichia coli O157 MNNEETFYQAMRRQGVTRRSFLKYCSLAATSLGLGAGMAPKIAWALENKPRIPVVWIHGL WP_001301613.1 hydrogenase [Escherichia coli] Length: 372\nNumber of predicted TMHs: 2\nExp number of AAs in TMHs: 37.46598\nExp number, first 60 AAs: 19.53139\nTotal prob of N-in: 0.94364\nPOSSIBLE N-term signal sequence\ninside 1 19\nTMhelix 20 39\noutside 40 325\nTMhelix 326 348\ninside 349 372

13437 GCF_000171935.1_ASM17193v1 Escherichia coli O157:H7 str. EC4113 Proteobacteria; Gammaproteobacteria; Enterobacterales; Enterobacteriaceae; Escherichia; Escherichia coli; Escherichia coli O157 MNNEETFYQAMRRQGVTRRSFLKYCSLAATSLGLGAGMAPKIAWALENKPRIPVVWIHGL WP_001301613.1 hydrogenase [Escherichia coli] Length: 372\nNumber of predicted TMHs: 2\nExp number of AAs in TMHs: 37.46598\nExp number, first 60 AAs: 19.53139\nTotal prob of N-in: 0.94364\nPOSSIBLE N-term signal sequence\ninside 1 19\nTMhelix 20 39\noutside 40 325\nTMhelix 326 348\ninside 349 372

13438 GCF_000171935.1_ASM17193v1 Escherichia coli O157:H7 str. EC4113 Proteobacteria; Gammaproteobacteria; Enterobacterales; Enterobacteriaceae; Escherichia; Escherichia coli; Escherichia coli O157 MTGDNTLIHSHGINRRDFMKLCAALAATMGLSSKAAAEMAESVTNPQRPPVIWIGAQECT WP_000145410.1 MULTISPECIES: hydrogenase-2 small chain [Proteobacteria] Length: 372\nNumber of predicted TMHs: 1\nExp number of AAs in TMHs: 23.72226\nExp number, first 60 AAs: 0.22038\nTotal prob of N-in: 0.12994\noutside 1 331\nTMhelix 332 354\ninside 355 372

13439 GCF_000021125.1_ASM2112v1 Escherichia coli O157:H7 str. EC4115 Proteobacteria; Gammaproteobacteria; Enterobacterales; Enterobacteriaceae; Escherichia; Escherichia coli; Escherichia coli O157 MNNEETFYQAMRRQGVTRRSFLKYCSLAATSLGLGAGMAPKIAWALENKPRIPVVWIHGL WP_001301613.1 hydrogenase [Escherichia coli] Length: 372\nNumber of predicted TMHs: 2\nExp number of AAs in TMHs: 37.46598\nExp number, first 60 AAs: 19.53139\nTotal prob of N-in: 0.94364\nPOSSIBLE N-term signal sequence\ninside 1 19\nTMhelix 20 39\noutside 40 325\nTMhelix 326 348\ninside 349 372

13440 GCF_000021125.1_ASM2112v1 Escherichia coli O157:H7 str. EC4115 Proteobacteria; Gammaproteobacteria; Enterobacterales; Enterobacteriaceae; Escherichia; Escherichia coli; Escherichia coli O157 MTGDNTLIHSHGINRRDFMKLCAALAATMGLSSKAAAEMAESVTNPQRPPVIWIGAQECT WP_000145410.1 MULTISPECIES: hydrogenase-2 small chain [Proteobacteria] Length: 372\nNumber of predicted TMHs: 1\nExp number of AAs in TMHs: 23.72226\nExp number, first 60 AAs: 0.22038\nTotal prob of N-in: 0.12994\noutside 1 331\nTMhelix 332 354\ninside 355 372

13441 GCF_000178735.1_ASM17873v1 Escherichia coli O157:H7 str. EC4127 Proteobacteria; Gammaproteobacteria; Enterobacterales; Enterobacteriaceae; Escherichia; Escherichia coli; Escherichia coli O157

MNNEETFYQAMRRQGVTRRSFLKYCSLAATSLGLGAGMAPKIAWALENKPRIPVVWIHGL WP_001301613.1 hydrogenase [Escherichia coli] Length: 372\nNumber of predicted TMHs: 2\nExp number of AAs in TMHs: 37.46598\nExp number, first 60 AAs: 19.53139\nTotal prob of N-in: 0.94364\nPOSSIBLE N-term signal sequence\ninside 1 19\nTMhelix 20 39\noutside 40 325\nTMhelix 326 348\ninside 349 372

13442 GCF_000178735.1_ASM17873v1 Escherichia coli O157:H7 str. EC4127 Proteobacteria; Gammaproteobacteria; Enterobacterales; Enterobacteriaceae; Escherichia; Escherichia coli; Escherichia coli O157 MTGDNTLIHSHGINRRDFMKLCAALAATMGLSSKAAAEMAESVTNPQRPPVIWIGAQECT WP_000145410.1 MULTISPECIES: hydrogenase-2 small chain [Proteobacteria] Length: 372\nNumber of predicted TMHs: 1\nExp number of AAs in TMHs: 23.72226\nExp number, first 60 AAs: 0.22038\nTotal prob of N-in: 0.12994\noutside 1 331\nTMhelix 332 354\ninside 355 372

13443 GCF_000178755.1_ASM17875v1 Escherichia coli O157:H7 str. EC4191 Proteobacteria; Gammaproteobacteria; Enterobacterales; Enterobacteriaceae; Escherichia; Escherichia coli; Escherichia coli O157 MNNEETFYQAMRRQGVTRRSFLKYCSLAATSLGLGAGMAPKIAWALENKPRIPVVWIHGL WP_001301613.1 hydrogenase [Escherichia coli] Length: 372\nNumber of predicted TMHs: 2\nExp number of AAs in TMHs: 37.46598\nExp number, first 60 AAs: 19.53139\nTotal prob of N-in: 0.94364\nPOSSIBLE N-term signal sequence\ninside 1 19\nTMhelix 20 39\noutside 40 325\nTMhelix 326 348\ninside 349 372

13444 GCF_000178755.1_ASM17875v1 Escherichia coli O157:H7 str. EC4191 Proteobacteria; Gammaproteobacteria; Enterobacterales; Enterobacteriaceae; Escherichia; Escherichia coli; Escherichia coli O157 MTGDNTLIHSHGINRRDFMKLCAALAATMGLSSKAAAEMAESVTNPQRPPVIWIGAQECT WP_000145410.1 MULTISPECIES: hydrogenase-2 small chain [Proteobacteria] Length: 372\nNumber of predicted TMHs: 1\nExp number of AAs in TMHs: 23.72226\nExp number, first 60 AAs: 0.22038\nTotal prob of N-in: 0.12994\noutside 1 331\nTMhelix 332 354\ninside 355 372

13445 GCF_000178695.1_ASM17869v1 Escherichia coli O157:H7 str. EC4192 Proteobacteria; Gammaproteobacteria; Enterobacterales; Enterobacteriaceae; Escherichia; Escherichia coli; Escherichia coli O157 MNNEETFYQAMRRQGVTRRSFLKYCSLAATSLGLGAGMAPKIAWALENKPRIPVVWIHGL WP_001301613.1 hydrogenase [Escherichia coli] Length: 372\nNumber of predicted TMHs: 2\nExp number of AAs in TMHs: 37.46598\nExp number, first 60 AAs: 19.53139\nTotal prob of N-in: 0.94364\nPOSSIBLE N-term signal sequence\ninside 1 19\nTMhelix 20 39\noutside 40 325\nTMhelix 326 348\ninside 349 372

13446 GCF_000178695.1_ASM17869v1 Escherichia coli O157:H7 str. EC4192 Proteobacteria; Gammaproteobacteria; Enterobacterales; Enterobacteriaceae; Escherichia; Escherichia coli; Escherichia coli O157 MTGDNTLIHSHGINRRDFMKLCAALAATMGLSSKAAAEMAESVTNPQRPPVIWIGAQECT WP_000145410.1 MULTISPECIES: hydrogenase-2 small chain [Proteobacteria] Length: 372\nNumber of predicted TMHs: 1\nExp number of AAs in TMHs: 23.72226\nExp number, first 60 AAs: 0.22038\nTotal prob of N-in: 0.12994\noutside 1 331\nTMhelix 332 354\ninside 355 372

13447 GCF_000171915.1_ASM17191v1 Escherichia coli O157:H7 str. EC4196 Proteobacteria; Gammaproteobacteria; Enterobacterales; Enterobacteriaceae; Escherichia; Escherichia coli; Escherichia coli O157 MTGDNTLIHSHGINRRDFMKLCAALAATMGLSSKAAAEMAESVTNPQRPPVIWIGAQECT WP_000145410.1 MULTISPECIES: hydrogenase-2 small chain [Proteobacteria] Length: 372\nNumber of predicted TMHs: 1\nExp number of AAs in TMHs: 23.72226\nExp number, first 60 AAs: 0.22038\nTotal prob of N-in: 0.12994\noutside 1 331\nTMhelix 332 354\ninside 355 372

13448 GCF_000171915.1_ASM17191v1 Escherichia coli O157:H7 str. EC4196 Proteobacteria; Gammaproteobacteria; Enterobacterales; Enterobacteriaceae; Escherichia; Escherichia coli; Escherichia coli O157 MNNEETFYQAMRRQGVTRRSFLKYCSLAATSLGLGAGMAPKIAWALENKPRIPVVWIHGL WP_001301613.1 hydrogenase [Escherichia coli] Length: 372\nNumber of predicted TMHs: 2\nExp number of AAs in TMHs: 37.46598\nExp number, first 60 AAs: 19.53139\nTotal prob of N-in: 0.94364\nPOSSIBLE N-term signal sequence\ninside 1 19\nTMhelix 20 39\noutside 40 325\nTMhelix 326 348\ninside 349 372

13449 GCF_000181735.1_ASM18173v1 Escherichia coli O157:H7 str. EC4206 Proteobacteria; Gammaproteobacteria; Enterobacterales; Enterobacteriaceae; Escherichia; Escherichia coli; Escherichia coli O157 MTGDNTLIHSHGINRRDFMKLCAALAATMGLSSKAAAEMAESVTNPQRPPVIWIGAQECT WP_000145410.1 MULTISPECIES: hydrogenase-2 small chain [Proteobacteria] Length: 372\nNumber of predicted TMHs: 1\nExp number of AAs in TMHs: 23.72226\nExp number, first 60 AAs: 0.22038\nTotal prob of N-in: 0.12994\noutside 1 331\nTMhelix 332 354\ninside 355 372

13450 GCF_000181735.1_ASM18173v1 Escherichia coli O157:H7 str. EC4206 Proteobacteria; Gammaproteobacteria; Enterobacterales; Enterobacteriaceae; Escherichia; Escherichia coli; Escherichia coli O157
MNNEETFYQAMRRQGVTRRSFLKYCSLAATSLGLGAGMAPKIAWALENKPRIPVVWIHGL WP_001301613.1
hydrogenase [Escherichia coli] Length: 372\nNumber of predicted TMHs: 2\nExp number of AAs in TMHs: 37.46598\nExp number, first 60 AAs: 19.53139\nTotal prob of N-in: 0.94364\nPOSSIBLE N-term signal sequence\ninside 1 19\nTMhelix 20 39\noutside 40 325\nTMhelix 326 348\ninside 349 372

13451 GCF_000171975.1_ASM17197v1 Escherichia coli O157:H7 str. EC4401 Proteobacteria; Gammaproteobacteria; Enterobacterales; Enterobacteriaceae; Escherichia; Escherichia coli; Escherichia coli O157
MTGDNTLIHSHGINRRDFMKLCAALAATMGLSSKAAAEMAESVTNPQRPPVIWIGAQECT WP_000145410.1
MULTISPECIES: hydrogenase-2 small chain [Proteobacteria] Length: 372\nNumber of predicted TMHs: 1\nExp number of AAs in TMHs: 23.72226\nExp number, first 60 AAs: 0.22038\nTotal prob of N-in: 0.12994\noutside 1 331\nTMhelix 332 354\ninside 355 372

13452 GCF_000171975.1_ASM17197v1 Escherichia coli O157:H7 str. EC4401 Proteobacteria; Gammaproteobacteria; Enterobacterales; Enterobacteriaceae; Escherichia; Escherichia coli; Escherichia coli O157
MNNEETFYQAMRRQGVTRRSFLKYCSLAATSLGLGAGMAPKIAWALENKPRIPVVWIHGL WP_001301613.1
hydrogenase [Escherichia coli] Length: 372\nNumber of predicted TMHs: 2\nExp number of AAs in TMHs: 37.46598\nExp number, first 60 AAs: 19.53139\nTotal prob of N-in: 0.94364\nPOSSIBLE N-term signal sequence\ninside 1 19\nTMhelix 20 39\noutside 40 325\nTMhelix 326 348\ninside 349 372

13453 GCF_000171995.1_ASM17199v1 Escherichia coli O157:H7 str. EC4486 Proteobacteria; Gammaproteobacteria; Enterobacterales; Enterobacteriaceae; Escherichia; Escherichia coli; Escherichia coli O157
MTGDNTLIHSHGINRRDFMKLCAALAATMGLSSKAAAEMAESVTNPQRPPVIWIGAQECT WP_000145410.1
MULTISPECIES: hydrogenase-2 small chain [Proteobacteria] Length: 372\nNumber of predicted TMHs: 1\nExp number of AAs in TMHs: 23.72226\nExp number, first 60 AAs: 0.22038\nTotal prob of N-in: 0.12994\noutside 1 331\nTMhelix 332 354\ninside 355 372

13454 GCF_000171995.1_ASM17199v1 Escherichia coli O157:H7 str. EC4486 Proteobacteria; Gammaproteobacteria; Enterobacterales; Enterobacteriaceae; Escherichia; Escherichia coli; Escherichia coli O157
MNNEETFYQAMRRQGVTRRSFLKYCSLAATSLGLGAGMAPKIAWALENKPRIPVVWIHGL WP_001301613.1
hydrogenase [Escherichia coli] Length: 372\nNumber of predicted TMHs: 2\nExp number of AAs in TMHs: 37.46598\nExp number, first 60 AAs: 19.53139\nTotal prob of N-in: 0.94364\nPOSSIBLE N-term signal sequence\ninside 1 19\nTMhelix 20 39\noutside 40 325\nTMhelix 326 348\ninside 349 372

13455 GCF_000172015.1_ASM17201v1 Escherichia coli O157:H7 str. EC4501 Proteobacteria; Gammaproteobacteria; Enterobacterales; Enterobacteriaceae; Escherichia; Escherichia coli; Escherichia coli O157
MNNEETFYQAMRRQGVTRRSFLKYCSLAATSLGLGAGMAPKIAWALENKPRIPVVWIHGL WP_001301613.1
hydrogenase [Escherichia coli] Length: 372\nNumber of predicted TMHs: 2\nExp number of AAs in TMHs: 37.46598\nExp number, first 60 AAs: 19.53139\nTotal prob of N-in: 0.94364\nPOSSIBLE N-term signal sequence\ninside 1 19\nTMhelix 20 39\noutside 40 325\nTMhelix 326 348\ninside 349 372

13456 GCF_000172015.1_ASM17201v1 Escherichia coli O157:H7 str. EC4501 Proteobacteria; Gammaproteobacteria; Enterobacterales; Enterobacteriaceae; Escherichia; Escherichia coli; Escherichia coli O157
MTGDNTLIHSHGINRRDFMKLCAALAATMGLSSKAAAEMAESVTNPQRPPVIWIGAQECT WP_000145410.1
MULTISPECIES: hydrogenase-2 small chain [Proteobacteria] Length: 372\nNumber of predicted TMHs: 1\nExp number of AAs in TMHs: 23.72226\nExp number, first 60 AAs: 0.22038\nTotal prob of N-in: 0.12994\noutside 1 331\nTMhelix 332 354\ninside 355 372

13457 GCF_000172055.1_ASM17205v1 Escherichia coli O157:H7 str. EC508 Proteobacteria; Gammaproteobacteria; Enterobacterales; Enterobacteriaceae; Escherichia; Escherichia coli; Escherichia coli O157
MNNEETFYQAMRRQGVTRRSFLKYCSLAATSLGLGAGMAPKIAWALENKPRIPVVWIHGL WP_001301613.1
hydrogenase [Escherichia coli] Length: 372\nNumber of predicted TMHs: 2\nExp number of AAs in TMHs: 37.46598\nExp number, first 60 AAs: 19.53139\nTotal prob of N-in: 0.94364\nPOSSIBLE N-term signal sequence\ninside 1 19\nTMhelix 20 39\noutside 40 325\nTMhelix 326 348\ninside 349 372

13458 GCF_000172055.1_ASM17205v1 Escherichia coli O157:H7 str. EC508 Proteobacteria; Gammaproteobacteria; Enterobacterales; Enterobacteriaceae; Escherichia; Escherichia coli; Escherichia coli O157
MTGDNTLIHSHGINRRDFMKLCAALAATMGLSSKAAAEMAESVTNPQRPPVIWIGAQECT WP_000145410.1
MULTISPECIES: hydrogenase-2 small chain [Proteobacteria] Length: 372\nNumber of predicted TMHs:

1\nExp number of AAs in TMHs: 23.72226\nExp number, first 60 AAs: 0.22038\nTotal prob of N-in: 0.12994\noutside 1 331\nTMhelix 332 354\ninside 355 372

13459 GCF_000178795.1_ASM17879v1 Escherichia coli O157:H7 str. EC536 Proteobacteria; Gammaproteobacteria; Enterobacterales; Enterobacteriaceae; Escherichia; Escherichia coli; Escherichia coli O157 MTGDNTLIHSHGINRRDFMKLCAALAATMGLSSKAAAEMAESVTNPQRPPVIWIGAQECT WP_000145410.1 MULTISPECIES: hydrogenase-2 small chain [Proteobacteria] Length: 372\nNumber of predicted TMHs: 1\nExp number of AAs in TMHs: 23.72226\nExp number, first 60 AAs: 0.22038\nTotal prob of N-in: 0.12994\noutside 1 331\nTMhelix 332 354\ninside 355 372

13460 GCF_000178795.1_ASM17879v1 Escherichia coli O157:H7 str. EC536 Proteobacteria; Gammaproteobacteria; Enterobacterales; Enterobacteriaceae; Escherichia; Escherichia coli; Escherichia coli O157 MNNEETFYQAMRRQGVTRRSFLKYCSLAATSLGLGAGMAPKIAWALENKPRIPVVWIHGL WP_001301613.1 hydrogenase [Escherichia coli] Length: 372\nNumber of predicted TMHs: 2\nExp number of AAs in TMHs: 37.46598\nExp number, first 60 AAs: 19.53139\nTotal prob of N-in: 0.94364\nPOSSIBLE N-term signal sequence\ninside 1 19\nTMhelix 20 39\noutside 40 325\nTMhelix 326 348\ninside 349 372

13461 GCF_000172035.1_ASM17203v1 Escherichia coli O157:H7 str. EC869 Proteobacteria; Gammaproteobacteria; Enterobacterales; Enterobacteriaceae; Escherichia; Escherichia coli; Escherichia coli O157 MTGDNTLIHSHGINRRDFMKLCAALAATMGLSSKAAAEMAESVTNPQRPPVIWIGAQECT WP_000145410.1 MULTISPECIES: hydrogenase-2 small chain [Proteobacteria] Length: 372\nNumber of predicted TMHs: 1\nExp number of AAs in TMHs: 23.72226\nExp number, first 60 AAs: 0.22038\nTotal prob of N-in: 0.12994\noutside 1 331\nTMhelix 332 354\ninside 355 372

13462 GCF_000172035.1_ASM17203v1 Escherichia coli O157:H7 str. EC869 Proteobacteria; Gammaproteobacteria; Enterobacterales; Enterobacteriaceae; Escherichia; Escherichia coli; Escherichia coli O157 MNNEETFYQAMRRQGVTRRSFLKYCSLAATSLGLGAGMAPKIAWALENKPRIPVVWIHGL WP_001301613.1 hydrogenase [Escherichia coli] Length: 372\nNumber of predicted TMHs: 2\nExp number of AAs in TMHs: 37.46598\nExp number, first 60 AAs: 19.53139\nTotal prob of N-in: 0.94364\nPOSSIBLE N-term signal sequence\ninside 1 19\nTMhelix 20 39\noutside 40 325\nTMhelix 326 348\ninside 349 372

13463 GCF_000006665.1_ASM666v1 Escherichia coli O157:H7 str. EDL933 Proteobacteria; Gammaproteobacteria; Enterobacterales; Enterobacteriaceae; Escherichia; Escherichia coli; Escherichia coli O157 MNNEETFYQAMRRQGVTRRSFLKYCSLAATSLGLGAGMAPKIAWALENKPRIPVVWIHGL WP_001301613.1 hydrogenase [Escherichia coli] Length: 372\nNumber of predicted TMHs: 2\nExp number of AAs in TMHs: 37.46598\nExp number, first 60 AAs: 19.53139\nTotal prob of N-in: 0.94364\nPOSSIBLE N-term signal sequence\ninside 1 19\nTMhelix 20 39\noutside 40 325\nTMhelix 326 348\ninside 349 372

13464 GCF_000006665.1_ASM666v1 Escherichia coli O157:H7 str. EDL933 Proteobacteria; Gammaproteobacteria; Enterobacterales; Enterobacteriaceae; Escherichia; Escherichia coli; Escherichia coli O157 MTGDNTLIHSHGINRRDFMKLCAALAATMGLSSKAAAEMAESVTNPQRPPVIWIGAQECT WP_000145410.1 MULTISPECIES: hydrogenase-2 small chain [Proteobacteria] Length: 372\nNumber of predicted TMHs: 1\nExp number of AAs in TMHs: 23.72226\nExp number, first 60 AAs: 0.22038\nTotal prob of N-in: 0.12994\noutside 1 331\nTMhelix 332 354\ninside 355 372

13465 GCF_000948445.1_ASM94844v1 Escherichia coli O157:H7 str. EDL933 Proteobacteria; Gammaproteobacteria; Enterobacterales; Enterobacteriaceae; Escherichia; Escherichia coli; Escherichia coli O157 MTGDNTLIHSHGINRRDFMKLCAALAATMGLSSKAAAEMAESVTNPQRPPVIWIGAQECT WP_000145410.1 MULTISPECIES: hydrogenase-2 small chain [Proteobacteria] Length: 372\nNumber of predicted TMHs: 1\nExp number of AAs in TMHs: 23.72226\nExp number, first 60 AAs: 0.22038\nTotal prob of N-in: 0.12994\noutside 1 331\nTMhelix 332 354\ninside 355 372

13466 GCF_000948445.1_ASM94844v1 Escherichia coli O157:H7 str. EDL933 Proteobacteria; Gammaproteobacteria; Enterobacterales; Enterobacteriaceae; Escherichia; Escherichia coli; Escherichia coli O157 MNNEETFYQAMRRQGVTRRSFLKYCSLAATSLGLGAGMAPKIAWALENKPRIPVVWIHGL WP_001301613.1 hydrogenase [Escherichia coli] Length: 372\nNumber of predicted TMHs: 2\nExp number of AAs in TMHs: 37.46598\nExp number, first 60 AAs: 19.53139\nTotal prob of N-in: 0.94364\nPOSSIBLE N-term signal sequence\ninside 1 19\nTMhelix 20 39\noutside 40 325\nTMhelix 326 348\ninside 349 372

13467 GCF_000732965.1_ASM73296v1 Escherichia coli O157:H7 str. EDL933 Proteobacteria; Gammaproteobacteria; Enterobacterales; Enterobacteriaceae; Escherichia; Escherichia coli; Escherichia coli O157

MTGDNTLIHSHGINRRDFMKLCAALAATMGLSSKAAAEMAESVTNPQRPPVIWIGAQECT WP_000145410.1 MULTISPECIES: hydrogenase-2 small chain [Proteobacteria] Length: 372\nNumber of predicted TMHs: 1\nExp number of AAs in TMHs: 23.72226\nExp number, first 60 AAs: 0.22038\nTotal prob of N-in: 0.12994\noutside 1 331\nTMhelix 332 354\ninside 355 372

13468 GCF_000732965.1_ASM73296v1 Escherichia coli O157:H7 str. EDL933 Proteobacteria; Gammaproteobacteria; Enterobacterales; Enterobacteriaceae; Escherichia; Escherichia coli; Escherichia coli O157 MNNEETFYQAMRRQGVTRRSFLKYCSLAATSLGLGAGMAPKIAWALENKPRIPVVWIHGL WP_001301613.1 hydrogenase [Escherichia coli] Length: 372\nNumber of predicted TMHs: 2\nExp number of AAs in TMHs: 37.46598\nExp number, first 60 AAs: 19.53139\nTotal prob of N-in: 0.94364\nPOSSIBLE N-term signal sequence\ninside 1 19\nTMhelix 20 39\noutside 40 325\nTMhelix 326 348\ninside 349 372

13469 GCF_000618105.2_EcF6142 Escherichia coli O157:H7 str. F6142 Proteobacteria; Gammaproteobacteria; Enterobacterales; Enterobacteriaceae; Escherichia; Escherichia coli; Escherichia coli O157 MNNEETFYQAMRRQGVTRRSFLKYCSLAATSLGLGAGMAPKIAWALENKPRIPVVWIHGL WP_001301613.1 hydrogenase [Escherichia coli] Length: 372\nNumber of predicted TMHs: 2\nExp number of AAs in TMHs: 37.46598\nExp number, first 60 AAs: 19.53139\nTotal prob of N-in: 0.94364\nPOSSIBLE N-term signal sequence\ninside 1 19\nTMhelix 20 39\noutside 40 325\nTMhelix 326 348\ninside 349 372

13470 GCF_000618105.2_EcF6142 Escherichia coli O157:H7 str. F6142 Proteobacteria; Gammaproteobacteria; Enterobacterales; Enterobacteriaceae; Escherichia; Escherichia coli; Escherichia coli O157 MTGDNTLIHSHGINRRDFMKLCAALAATMGLSSKAAAEMAESVTNPQRPPVIWIGAQECT WP_000145410.1 MULTISPECIES: hydrogenase-2 small chain [Proteobacteria] Length: 372\nNumber of predicted TMHs: 1\nExp number of AAs in TMHs: 23.72226\nExp number, first 60 AAs: 0.22038\nTotal prob of N-in: 0.12994\noutside 1 331\nTMhelix 332 354\ninside 355 372

13471 GCF_000618045.1_EcF6749 Escherichia coli O157:H7 str. F6749 Proteobacteria; Gammaproteobacteria; Enterobacterales; Enterobacteriaceae; Escherichia; Escherichia coli; Escherichia coli O157 MTGDNTLIHSHGINRRDFMKLCAALAATMGLSSKAAAEMAESVTNPQRPPVIWIGAQECT WP_000145410.1 MULTISPECIES: hydrogenase-2 small chain [Proteobacteria] Length: 372\nNumber of predicted TMHs: 1\nExp number of AAs in TMHs: 23.72226\nExp number, first 60 AAs: 0.22038\nTotal prob of N-in: 0.12994\noutside 1 331\nTMhelix 332 354\ninside 355 372

13472 GCF_000618045.1_EcF6749 Escherichia coli O157:H7 str. F6749 Proteobacteria; Gammaproteobacteria; Enterobacterales; Enterobacteriaceae; Escherichia; Escherichia coli; Escherichia coli O157 MNNEETFYQAMRRQGVTRRSFLKYCSLAATSLGLGAGMAPKIAWALENKPRIPVVWIHGL WP_001301613.1 hydrogenase [Escherichia coli] Length: 372\nNumber of predicted TMHs: 2\nExp number of AAs in TMHs: 37.46598\nExp number, first 60 AAs: 19.53139\nTotal prob of N-in: 0.94364\nPOSSIBLE N-term signal sequence\ninside 1 19\nTMhelix 20 39\noutside 40 325\nTMhelix 326 348\ninside 349 372

13473 GCF_000618305.2_EcF6750 Escherichia coli O157:H7 str. F6750 Proteobacteria; Gammaproteobacteria; Enterobacterales; Enterobacteriaceae; Escherichia; Escherichia coli; Escherichia coli O157 MNNEETFYQAMRRQGVTRRSFLKYCSLAATSLGLGAGMAPKIAWALENKPRIPVVWIHGL WP_001301613.1 hydrogenase [Escherichia coli] Length: 372\nNumber of predicted TMHs: 2\nExp number of AAs in TMHs: 37.46598\nExp number, first 60 AAs: 19.53139\nTotal prob of N-in: 0.94364\nPOSSIBLE N-term signal sequence\ninside 1 19\nTMhelix 20 39\noutside 40 325\nTMhelix 326 348\ninside 349 372

13474 GCF_000618305.2_EcF6750 Escherichia coli O157:H7 str. F6750 Proteobacteria; Gammaproteobacteria; Enterobacterales; Enterobacteriaceae; Escherichia; Escherichia coli; Escherichia coli O157 MTGDNTLIHSHGINRRDFMKLCAALAATMGLSSKAAAEMAESVTNPQRPPVIWIGAQECT WP_000145410.1 MULTISPECIES: hydrogenase-2 small chain [Proteobacteria] Length: 372\nNumber of predicted TMHs: 1\nExp number of AAs in TMHs: 23.72226\nExp number, first 60 AAs: 0.22038\nTotal prob of N-in: 0.12994\noutside 1 331\nTMhelix 332 354\ninside 355 372

13475 GCF_000618265.2_EcF6751 Escherichia coli O157:H7 str. F6751 Proteobacteria; Gammaproteobacteria; Enterobacterales; Enterobacteriaceae; Escherichia; Escherichia coli; Escherichia coli O157 MTGDNTLIHSHGINRRDFMKLCAALAATMGLSSKAAAEMAESVTNPQRPPVIWIGAQECT WP_000145410.1 MULTISPECIES: hydrogenase-2 small chain [Proteobacteria] Length: 372\nNumber of predicted TMHs: 1\nExp number of AAs in TMHs: 23.72226\nExp number, first 60 AAs: 0.22038\nTotal prob of N-in: 0.12994\noutside 1 331\nTMhelix 332 354\ninside 355 372

13476 GCF_000618265.2_EcF6751 Escherichia coli O157:H7 str. F6751 Proteobacteria; Gammaproteobacteria; Enterobacterales; Enterobacteriaceae; Escherichia; Escherichia coli; Escherichia coli O157
MNNEETFYQAMRRQGVTRRSFLKYCSLAATSLGLGAGMAPKIAWALENKPRIPVVWIHGL WP_001301613.1
hydrogenase [Escherichia coli] Length: 372\nNumber of predicted TMHs: 2\nExp number of AAs in TMHs: 37.46598\nExp number, first 60 AAs: 19.53139\nTotal prob of N-in: 0.94364\nPOSSIBLE N-term signal sequence\ninside 1 19\nTMhelix 20 39\noutside 40 325\nTMhelix 326 348\ninside 349 372

13477 GCF_000614805.2_EcF7350 Escherichia coli O157:H7 str. F7350 Proteobacteria; Gammaproteobacteria; Enterobacterales; Enterobacteriaceae; Escherichia; Escherichia coli; Escherichia coli O157
MNNEETFYQAMRRQGVTRRSFLKYCSLAATSLGLGAGMAPKIAWALENKPRIPVVWIHGL WP_001301613.1
hydrogenase [Escherichia coli] Length: 372\nNumber of predicted TMHs: 2\nExp number of AAs in TMHs: 37.46598\nExp number, first 60 AAs: 19.53139\nTotal prob of N-in: 0.94364\nPOSSIBLE N-term signal sequence\ninside 1 19\nTMhelix 20 39\noutside 40 325\nTMhelix 326 348\ninside 349 372

13478 GCF_000614805.2_EcF7350 Escherichia coli O157:H7 str. F7350 Proteobacteria; Gammaproteobacteria; Enterobacterales; Enterobacteriaceae; Escherichia; Escherichia coli; Escherichia coli O157
MTGDNTLIHSHGINRRDFMKLCAALAATMGLSSKAAAEMAESVTNPQRPPVIWIGAQECT WP_000145410.1
MULTISPECIES: hydrogenase-2 small chain [Proteobacteria] Length: 372\nNumber of predicted TMHs: 1\nExp number of AAs in TMHs: 23.72226\nExp number, first 60 AAs: 0.22038\nTotal prob of N-in: 0.12994\noutside 1 331\nTMhelix 332 354\ninside 355 372

13479 GCF_000618225.2_EcF7377 Escherichia coli O157:H7 str. F7377 Proteobacteria; Gammaproteobacteria; Enterobacterales; Enterobacteriaceae; Escherichia; Escherichia coli; Escherichia coli O157
MNNEETFYQAMRRQGVTRRSFLKYCSLAATSLGLGAGMAPKIAWALENKPRIPVVWIHGL WP_001301613.1
hydrogenase [Escherichia coli] Length: 372\nNumber of predicted TMHs: 2\nExp number of AAs in TMHs: 37.46598\nExp number, first 60 AAs: 19.53139\nTotal prob of N-in: 0.94364\nPOSSIBLE N-term signal sequence\ninside 1 19\nTMhelix 20 39\noutside 40 325\nTMhelix 326 348\ninside 349 372

13480 GCF_000618225.2_EcF7377 Escherichia coli O157:H7 str. F7377 Proteobacteria; Gammaproteobacteria; Enterobacterales; Enterobacteriaceae; Escherichia; Escherichia coli; Escherichia coli O157
MTGDNTLIHSHGINRRDFMKLCAALAATMGLSSKAAAEMAESVTNPQRPPVIWIGAQECT WP_000145410.1
MULTISPECIES: hydrogenase-2 small chain [Proteobacteria] Length: 372\nNumber of predicted TMHs: 1\nExp number of AAs in TMHs: 23.72226\nExp number, first 60 AAs: 0.22038\nTotal prob of N-in: 0.12994\noutside 1 331\nTMhelix 332 354\ninside 355 372

13481 GCF_000618065.2_EcF7384 Escherichia coli O157:H7 str. F7384 Proteobacteria; Gammaproteobacteria; Enterobacterales; Enterobacteriaceae; Escherichia; Escherichia coli; Escherichia coli O157
MNNEETFYQAMRRQGVTRRSFLKYCSLAATSLGLGAGMAPKIAWALENKPRIPVVWIHGL WP_001301613.1
hydrogenase [Escherichia coli] Length: 372\nNumber of predicted TMHs: 2\nExp number of AAs in TMHs: 37.46598\nExp number, first 60 AAs: 19.53139\nTotal prob of N-in: 0.94364\nPOSSIBLE N-term signal sequence\ninside 1 19\nTMhelix 20 39\noutside 40 325\nTMhelix 326 348\ninside 349 372

13482 GCF_000618065.2_EcF7384 Escherichia coli O157:H7 str. F7384 Proteobacteria; Gammaproteobacteria; Enterobacterales; Enterobacteriaceae; Escherichia; Escherichia coli; Escherichia coli O157
MTGDNTLIHSHGINRRDFMKLCAALAATMGLSSKAAAEMAESVTNPQRPPVIWIGAQECT WP_000145410.1
MULTISPECIES: hydrogenase-2 small chain [Proteobacteria] Length: 372\nNumber of predicted TMHs: 1\nExp number of AAs in TMHs: 23.72226\nExp number, first 60 AAs: 0.22038\nTotal prob of N-in: 0.12994\noutside 1 331\nTMhelix 332 354\ninside 355 372

13483 GCF_000618145.2_EcF7410 Escherichia coli O157:H7 str. F7410 Proteobacteria; Gammaproteobacteria; Enterobacterales; Enterobacteriaceae; Escherichia; Escherichia coli; Escherichia coli O157
MTGDNTLIHSHGINRRDFMKLCAALAATMGLSSKAAAEMAESVTNPQRPPVIWIGAQECT WP_000145410.1
MULTISPECIES: hydrogenase-2 small chain [Proteobacteria] Length: 372\nNumber of predicted TMHs: 1\nExp number of AAs in TMHs: 23.72226\nExp number, first 60 AAs: 0.22038\nTotal prob of N-in: 0.12994\noutside 1 331\nTMhelix 332 354\ninside 355 372

13484 GCF_000618145.2_EcF7410 Escherichia coli O157:H7 str. F7410 Proteobacteria; Gammaproteobacteria; Enterobacterales; Enterobacteriaceae; Escherichia; Escherichia coli; Escherichia coli O157
MNNEETFYQAMRRQGVTRRSFLKYCSLAATSLGLGAGMAPKIAWALENKPRIPVVWIHGL WP_001301613.1
hydrogenase [Escherichia coli] Length: 372\nNumber of predicted TMHs: 2\nExp number of AAs in

TMHs: 37.46598\nExp number, first 60 AAs: 19.53139\nTotal prob of N-in: 0.94364\nPOSSIBLE N-term signal sequence\ninside 1 19\nTMhelix 20 39\noutside 40 325\nTMhelix 326 348\ninside 349 372

13485 GCF_000513035.1_E._coli_O157 Escherichia coli O157:H7 str. F8092B Proteobacteria; Gammaproteobacteria; Enterobacterales; Enterobacteriaceae; Escherichia; Escherichia coli; Escherichia coli O157 MNNEETFYQAMRRQGVTRRSFLKYCSLAATSLGLGAGMAPKIAWALENKPRIPVVWIHGL WP_001301613.1 hydrogenase [Escherichia coli] Length: 372\nNumber of predicted TMHs: 2\nExp number of AAs in TMHs: 37.46598\nExp number, first 60 AAs: 19.53139\nTotal prob of N-in: 0.94364\nPOSSIBLE N-term signal sequence\ninside 1 19\nTMhelix 20 39\noutside 40 325\nTMhelix 326 348\ninside 349 372

13486 GCF_000513035.1_E._coli_O157 Escherichia coli O157:H7 str. F8092B Proteobacteria; Gammaproteobacteria; Enterobacterales; Enterobacteriaceae; Escherichia; Escherichia coli; Escherichia coli O157 MTGDNTLIHSHGINRRDFMKLCAALAATMGLSSKAAAEMAESVTNPQRPPVIWIGAQECT WP_000145410.1 MULTISPECIES: hydrogenase-2 small chain [Proteobacteria] Length: 372\nNumber of predicted TMHs: 1\nExp number of AAs in TMHs: 23.72226\nExp number, first 60 AAs: 0.22038\nTotal prob of N-in: 0.12994\noutside 1 331\nTMhelix 332 354\ninside 355 372

13487 GCF_000175755.1_ASM17575v1 Escherichia coli O157:H7 str. FRIK2000 Proteobacteria; Gammaproteobacteria; Enterobacterales; Enterobacteriaceae; Escherichia; Escherichia coli; Escherichia coli O157 MNNEETFYQAMRRQGVTRRSFLKYCSLAATSLGLGAGMAPKIAWALENKPRIPVVWIHGL WP_001301613.1 hydrogenase [Escherichia coli] Length: 372\nNumber of predicted TMHs: 2\nExp number of AAs in TMHs: 37.46598\nExp number, first 60 AAs: 19.53139\nTotal prob of N-in: 0.94364\nPOSSIBLE N-term signal sequence\ninside 1 19\nTMhelix 20 39\noutside 40 325\nTMhelix 326 348\ninside 349 372

13488 GCF_000175755.1_ASM17575v1 Escherichia coli O157:H7 str. FRIK2000 Proteobacteria; Gammaproteobacteria; Enterobacterales; Enterobacteriaceae; Escherichia; Escherichia coli; Escherichia coli O157 MTGDNTLIHSHGINRRDFMKLCAALAATMGLSSKAAAEMAESVTNPQRPPVIWIGAQECT WP_000145410.1 MULTISPECIES: hydrogenase-2 small chain [Proteobacteria] Length: 372\nNumber of predicted TMHs: 1\nExp number of AAs in TMHs: 23.72226\nExp number, first 60 AAs: 0.22038\nTotal prob of N-in: 0.12994\noutside 1 331\nTMhelix 332 354\ninside 355 372

13489 GCF_000175735.1_ASM17573v1 Escherichia coli O157:H7 str. FRIK966 Proteobacteria; Gammaproteobacteria; Enterobacterales; Enterobacteriaceae; Escherichia; Escherichia coli; Escherichia coli O157 MNNEETFYQAMRRQGVTRRSFLKYCSLAATSLGLGAGMAPKIAWALENKPRIPVVWIHGL WP_001301613.1 hydrogenase [Escherichia coli] Length: 372\nNumber of predicted TMHs: 2\nExp number of AAs in TMHs: 37.46598\nExp number, first 60 AAs: 19.53139\nTotal prob of N-in: 0.94364\nPOSSIBLE N-term signal sequence\ninside 1 19\nTMhelix 20 39\noutside 40 325\nTMhelix 326 348\ninside 349 372

13490 GCF_000175735.1_ASM17573v1 Escherichia coli O157:H7 str. FRIK966 Proteobacteria; Gammaproteobacteria; Enterobacterales; Enterobacteriaceae; Escherichia; Escherichia coli; Escherichia coli O157 MTGDNTLIHSHGINRRDFMKLCAALAATMGLSSKAAAEMAESVTNPQRPPVIWIGAQECT WP_000145410.1 MULTISPECIES: hydrogenase-2 small chain [Proteobacteria] Length: 372\nNumber of predicted TMHs: 1\nExp number of AAs in TMHs: 23.72226\nExp number, first 60 AAs: 0.22038\nTotal prob of N-in: 0.12994\noutside 1 331\nTMhelix 332 354\ninside 355 372

13491 GCF_000187285.1_ASM18728v4 Escherichia coli O157:H7 str. G5101 Proteobacteria; Gammaproteobacteria; Enterobacterales; Enterobacteriaceae; Escherichia; Escherichia coli; Escherichia coli O157 MIGDNTLIHSHGINRRDFMKLCAALAATMGLSSKAAAEMAESVTNPQRPPVIWIGAQECT WP_000578724.1 hydrogenase 2 small subunit [Escherichia coli] Length: 372\nNumber of predicted TMHs: 1\nExp number of AAs in TMHs: 23.05601\nExp number, first 60 AAs: 0.15431\nTotal prob of N-in: 0.09116\noutside 1 331\nTMhelix 332 354\ninside 355 372

13492 GCF_000187285.1_ASM18728v4 Escherichia coli O157:H7 str. G5101 Proteobacteria; Gammaproteobacteria; Enterobacterales; Enterobacteriaceae; Escherichia; Escherichia coli; Escherichia coli O157 MNNEETFYQAMRRQGVTRRSFLKYCSLAATSLGLGAGMAPKIAWALENKPRIPVVWIHGL WP_001301613.1 hydrogenase [Escherichia coli] Length: 372\nNumber of predicted TMHs: 2\nExp number of AAs in TMHs: 37.46598\nExp number, first 60 AAs: 19.53139\nTotal prob of N-in: 0.94364\nPOSSIBLE N-term signal sequence\ninside 1 19\nTMhelix 20 39\noutside 40 325\nTMhelix 326 348\ninside 349 372

13493 GCF_000618185.1_EcG5303 Escherichia coli O157:H7 str. G5303 Proteobacteria; Gammaproteobacteria; Enterobacterales; Enterobacteriaceae; Escherichia; Escherichia coli; Escherichia coli O157

MNNEETFYQAMRRQGVTRRSFLKYCSLAATSLGLGAGMAPKIAWALENKPRIPVVWIHGL WP_001301613.1 hydrogenase [Escherichia coli] Length: 372\nNumber of predicted TMHs: 2\nExp number of AAs in TMHs: 37.46598\nExp number, first 60 AAs: 19.53139\nTotal prob of N-in: 0.94364\nPOSSIBLE N-term signal sequence\ninside 1 19\nTMhelix 20 39\noutside 40 325\nTMhelix 326 348\ninside 349 372

13494 GCF_000618185.1_EcG5303 Escherichia coli O157:H7 str. G5303 Proteobacteria; Gammaproteobacteria; Enterobacterales; Enterobacteriaceae; Escherichia; Escherichia coli; Escherichia coli O157 MTGDNTLIHSHGINRRDFMKLCAALAATMGLSSKAAAEMAESVTNPQRPPVIWIGAQECT WP_000145410.1 MULTISPECIES: hydrogenase-2 small chain [Proteobacteria] Length: 372\nNumber of predicted TMHs: 1\nExp number of AAs in TMHs: 23.72226\nExp number, first 60 AAs: 0.22038\nTotal prob of N-in: 0.12994\noutside 1 331\nTMhelix 332 354\ninside 355 372

13495 GCF_000321145.1_ASM32114v1 Escherichia coli O157:H7 str. H093800014 Proteobacteria; Gammaproteobacteria; Enterobacterales; Enterobacteriaceae; Escherichia; Escherichia coli; Escherichia coli O157 MNNEETFYQAMRRQGVTRRSFLKYCSLAATSLGLGAGMAPKIAWALENKPRIPVVWIHGL WP_001301613.1 hydrogenase [Escherichia coli] Length: 372\nNumber of predicted TMHs: 2\nExp number of AAs in TMHs: 37.46598\nExp number, first 60 AAs: 19.53139\nTotal prob of N-in: 0.94364\nPOSSIBLE N-term signal sequence\ninside 1 19\nTMhelix 20 39\noutside 40 325\nTMhelix 326 348\ninside 349 372

13496 GCF_000321145.1_ASM32114v1 Escherichia coli O157:H7 str. H093800014 Proteobacteria; Gammaproteobacteria; Enterobacterales; Enterobacteriaceae; Escherichia; Escherichia coli; Escherichia coli O157 MTGDNTLIHSHGINRRDFMKLCAALAATMGLSSKAAAEMAESVTNPQRPPVIWIGAQECT WP_000145410.1 MULTISPECIES: hydrogenase-2 small chain [Proteobacteria] Length: 372\nNumber of predicted TMHs: 1\nExp number of AAs in TMHs: 23.72226\nExp number, first 60 AAs: 0.22038\nTotal prob of N-in: 0.12994\noutside 1 331\nTMhelix 332 354\ninside 355 372

13497 GCF_000618245.1_EcH2495 Escherichia coli O157:H7 str. H2495 Proteobacteria; Gammaproteobacteria; Enterobacterales; Enterobacteriaceae; Escherichia; Escherichia coli; Escherichia coli O157 MTGDNTLIHSHGINRRDFMKLCAALAATMGLSSKAAAEMAESVTNPQRPPVIWIGAQECT WP_000145410.1 MULTISPECIES: hydrogenase-2 small chain [Proteobacteria] Length: 372\nNumber of predicted TMHs: 1\nExp number of AAs in TMHs: 23.72226\nExp number, first 60 AAs: 0.22038\nTotal prob of N-in: 0.12994\noutside 1 331\nTMhelix 332 354\ninside 355 372

13498 GCF_000618245.1_EcH2495 Escherichia coli O157:H7 str. H2495 Proteobacteria; Gammaproteobacteria; Enterobacterales; Enterobacteriaceae; Escherichia; Escherichia coli; Escherichia coli O157 MNNEETFYQAMRRQGVTRRSFLKYCSLAATSLGLGAGMAPKIAWALENKPRIPVVWIHGL WP_001301613.1 hydrogenase [Escherichia coli] Length: 372\nNumber of predicted TMHs: 2\nExp number of AAs in TMHs: 37.46598\nExp number, first 60 AAs: 19.53139\nTotal prob of N-in: 0.94364\nPOSSIBLE N-term signal sequence\ninside 1 19\nTMhelix 20 39\noutside 40 325\nTMhelix 326 348\ninside 349 372

13499 GCF_000618285.2_EcH2498 Escherichia coli O157:H7 str. H2498 Proteobacteria; Gammaproteobacteria; Enterobacterales; Enterobacteriaceae; Escherichia; Escherichia coli; Escherichia coli O157 MTGDNTLIHSHGINRRDFMKLCAALAATMGLSSKAAAEMAESVTNPQRPPVIWIGAQECT WP_000145410.1 MULTISPECIES: hydrogenase-2 small chain [Proteobacteria] Length: 372\nNumber of predicted TMHs: 1\nExp number of AAs in TMHs: 23.72226\nExp number, first 60 AAs: 0.22038\nTotal prob of N-in: 0.12994\noutside 1 331\nTMhelix 332 354\ninside 355 372

13500 GCF_000618285.2_EcH2498 Escherichia coli O157:H7 str. H2498 Proteobacteria; Gammaproteobacteria; Enterobacterales; Enterobacteriaceae; Escherichia; Escherichia coli; Escherichia coli O157 MNNEETFYQAMRRQGVTRRSFLKYCSLAATSLGLGAGMAPKIAWALENKPRIPVVWIHGL WP_001301613.1 hydrogenase [Escherichia coli] Length: 372\nNumber of predicted TMHs: 2\nExp number of AAs in TMHs: 37.46598\nExp number, first 60 AAs: 19.53139\nTotal prob of N-in: 0.94364\nPOSSIBLE N-term signal sequence\ninside 1 19\nTMhelix 20 39\noutside 40 325\nTMhelix 326 348\ninside 349 372

13501 GCF_000618025.1_EcK1420 Escherichia coli O157:H7 str. K1420 Proteobacteria; Gammaproteobacteria; Enterobacterales; Enterobacteriaceae; Escherichia; Escherichia coli; Escherichia coli O157 MTGDNTLIHSHGINRRDFMKLCAALAATMGLSSKAAAEMAESVTNPQRPPVIWIGAQECT WP_000145410.1 MULTISPECIES: hydrogenase-2 small chain [Proteobacteria] Length: 372\nNumber of predicted TMHs: 1\nExp number of AAs in TMHs: 23.72226\nExp number, first 60 AAs: 0.22038\nTotal prob of N-in: 0.12994\noutside 1 331\nTMhelix 332 354\ninside 355 372

13502 GCF_000618025.1_EcK1420 Escherichia coli O157:H7 str. K1420 Proteobacteria; Gammaproteobacteria; Enterobacterales; Enterobacteriaceae; Escherichia; Escherichia coli; Escherichia coli O157
MNNEETFYQAMRRQGVTRRSFLKYCSLAATSLGLGAGMAPKIAWALENKPRIPVVWIHGL WP_001301613.1
hydrogenase [Escherichia coli] Length: 372\nNumber of predicted TMHs: 2\nExp number of AAs in TMHs: 37.46598\nExp number, first 60 AAs: 19.53139\nTotal prob of N-in: 0.94364\nPOSSIBLE N-term signal sequence\ninside 1 19\nTMhelix 20 39\noutside 40 325\nTMhelix 326 348\ninside 349 372

13503 GCF_000618425.1_EcK1792 Escherichia coli O157:H7 str. K1792 Proteobacteria; Gammaproteobacteria; Enterobacterales; Enterobacteriaceae; Escherichia; Escherichia coli; Escherichia coli O157
MNNEETFYQAMRRQGVTRRSFLKYCSLAATSLGLGAGMAPKIAWALENKPRIPVVWIHGL WP_001301613.1
hydrogenase [Escherichia coli] Length: 372\nNumber of predicted TMHs: 2\nExp number of AAs in TMHs: 37.46598\nExp number, first 60 AAs: 19.53139\nTotal prob of N-in: 0.94364\nPOSSIBLE N-term signal sequence\ninside 1 19\nTMhelix 20 39\noutside 40 325\nTMhelix 326 348\ninside 349 372

13504 GCF_000618425.1_EcK1792 Escherichia coli O157:H7 str. K1792 Proteobacteria; Gammaproteobacteria; Enterobacterales; Enterobacteriaceae; Escherichia; Escherichia coli; Escherichia coli O157
MTGDNTLIHSHGINRRDFMKLCAALAATMGLSSKAAAEMAESVTNPQRPPVIWIGAQECT WP_000145410.1
MULTISPECIES: hydrogenase-2 small chain [Proteobacteria] Length: 372\nNumber of predicted TMHs: 1\nExp number of AAs in TMHs: 23.72226\nExp number, first 60 AAs: 0.22038\nTotal prob of N-in: 0.12994\noutside 1 331\nTMhelix 332 354\ninside 355 372

13505 GCF_000618365.1_EcK1793 Escherichia coli O157:H7 str. K1793 Proteobacteria; Gammaproteobacteria; Enterobacterales; Enterobacteriaceae; Escherichia; Escherichia coli; Escherichia coli O157
MTGDNTLIHSHGINRRDFMKLCAALAATMGLSSKAAAEMAESVTNPQRPPVIWIGAQECT WP_000145410.1
MULTISPECIES: hydrogenase-2 small chain [Proteobacteria] Length: 372\nNumber of predicted TMHs: 1\nExp number of AAs in TMHs: 23.72226\nExp number, first 60 AAs: 0.22038\nTotal prob of N-in: 0.12994\noutside 1 331\nTMhelix 332 354\ninside 355 372

13506 GCF_000618365.1_EcK1793 Escherichia coli O157:H7 str. K1793 Proteobacteria; Gammaproteobacteria; Enterobacterales; Enterobacteriaceae; Escherichia; Escherichia coli; Escherichia coli O157
MNNEETFYQAMRRQGVTRRSFLKYCSLAATSLGLGAGMAPKIAWALENKPRIPVVWIHGL WP_001301613.1
hydrogenase [Escherichia coli] Length: 372\nNumber of predicted TMHs: 2\nExp number of AAs in TMHs: 37.46598\nExp number, first 60 AAs: 19.53139\nTotal prob of N-in: 0.94364\nPOSSIBLE N-term signal sequence\ninside 1 19\nTMhelix 20 39\noutside 40 325\nTMhelix 326 348\ninside 349 372

13507 GCF_000618385.1_EcK1795 Escherichia coli O157:H7 str. K1795 Proteobacteria; Gammaproteobacteria; Enterobacterales; Enterobacteriaceae; Escherichia; Escherichia coli; Escherichia coli O157
MTGDNTLIHSHGINRRDFMKLCAALAATMGLSSKAAAEMAESVTNPQRPPVIWIGAQECT WP_000145410.1
MULTISPECIES: hydrogenase-2 small chain [Proteobacteria] Length: 372\nNumber of predicted TMHs: 1\nExp number of AAs in TMHs: 23.72226\nExp number, first 60 AAs: 0.22038\nTotal prob of N-in: 0.12994\noutside 1 331\nTMhelix 332 354\ninside 355 372

13508 GCF_000618385.1_EcK1795 Escherichia coli O157:H7 str. K1795 Proteobacteria; Gammaproteobacteria; Enterobacterales; Enterobacteriaceae; Escherichia; Escherichia coli; Escherichia coli O157
MNNEETFYQAMRRQGVTRRSFLKYCSLAATSLGLGAGMAPKIAWALENKPRIPVVWIHGL WP_001301613.1
hydrogenase [Escherichia coli] Length: 372\nNumber of predicted TMHs: 2\nExp number of AAs in TMHs: 37.46598\nExp number, first 60 AAs: 19.53139\nTotal prob of N-in: 0.94364\nPOSSIBLE N-term signal sequence\ninside 1 19\nTMhelix 20 39\noutside 40 325\nTMhelix 326 348\ninside 349 372

13509 GCF_000618325.1_EcK1796 Escherichia coli O157:H7 str. K1796 Proteobacteria; Gammaproteobacteria; Enterobacterales; Enterobacteriaceae; Escherichia; Escherichia coli; Escherichia coli O157
MNNEETFYQAMRRQGVTRRSFLKYCSLAATSLGLGAGMAPKIAWALENKPRIPVVWIHGL WP_001301613.1
hydrogenase [Escherichia coli] Length: 372\nNumber of predicted TMHs: 2\nExp number of AAs in TMHs: 37.46598\nExp number, first 60 AAs: 19.53139\nTotal prob of N-in: 0.94364\nPOSSIBLE N-term signal sequence\ninside 1 19\nTMhelix 20 39\noutside 40 325\nTMhelix 326 348\ninside 349 372

13510 GCF_000618325.1_EcK1796 Escherichia coli O157:H7 str. K1796 Proteobacteria; Gammaproteobacteria; Enterobacterales; Enterobacteriaceae; Escherichia; Escherichia coli; Escherichia coli O157
MTGDNTLIHSHGINRRDFMKLCAALAATMGLSSKAAAEMAESVTNPQRPPVIWIGAQECT WP_000145410.1
MULTISPECIES: hydrogenase-2 small chain [Proteobacteria] Length: 372\nNumber of predicted TMHs:

1\nExp number of AAs in TMHs: 23.72226\nExp number, first 60 AAs: 0.22038\nTotal prob of N-in: 0.12994\noutside 1 331\nTMhelix 332 354\ninside 355 372

13511 GCF_000618345.1_EcK1845 Escherichia coli O157:H7 str. K1845 Proteobacteria; Gammaproteobacteria; Enterobacterales; Enterobacteriaceae; Escherichia; Escherichia coli; Escherichia coli O157 MTGDNTLIHSHGINRRDFMKLCAALAATMGLSSKAAAEMAESVTNPQRPPVIWIGAQECT WP_000145410.1 MULTISPECIES: hydrogenase-2 small chain [Proteobacteria] Length: 372\nNumber of predicted TMHs: 1\nExp number of AAs in TMHs: 23.72226\nExp number, first 60 AAs: 0.22038\nTotal prob of N-in: 0.12994\noutside 1 331\nTMhelix 332 354\ninside 355 372

13512 GCF_000618345.1_EcK1845 Escherichia coli O157:H7 str. K1845 Proteobacteria; Gammaproteobacteria; Enterobacterales; Enterobacteriaceae; Escherichia; Escherichia coli; Escherichia coli O157 MNNEETFYQAMRRQGVTRRSFLKYCSLAATSLGLGAGMAPKIAWALENKPRIPVVWIHGL WP_001301613.1 hydrogenase [Escherichia coli] Length: 372\nNumber of predicted TMHs: 2\nExp number of AAs in TMHs: 37.46598\nExp number, first 60 AAs: 19.53139\nTotal prob of N-in: 0.94364\nPOSSIBLE N-term signal sequence\ninside 1 19\nTMhelix 20 39\noutside 40 325\nTMhelix 326 348\ninside 349 372

13513 GCF_000618445.1_EcK1921 Escherichia coli O157:H7 str. K1921 Proteobacteria; Gammaproteobacteria; Enterobacterales; Enterobacteriaceae; Escherichia; Escherichia coli; Escherichia coli O157 MNNEETFYQAMRRQGVTRRSFLKYCSLAATSLGLGAGMAPKIAWALENKPRIPVVWIHGL WP_001301613.1 hydrogenase [Escherichia coli] Length: 372\nNumber of predicted TMHs: 2\nExp number of AAs in TMHs: 37.46598\nExp number, first 60 AAs: 19.53139\nTotal prob of N-in: 0.94364\nPOSSIBLE N-term signal sequence\ninside 1 19\nTMhelix 20 39\noutside 40 325\nTMhelix 326 348\ninside 349 372

13514 GCF_000618445.1_EcK1921 Escherichia coli O157:H7 str. K1921 Proteobacteria; Gammaproteobacteria; Enterobacterales; Enterobacteriaceae; Escherichia; Escherichia coli; Escherichia coli O157 MTGDNTLIHSHGINRRDFMKLCAALAATMGLSSKAAAEMAESVTNPQRPPVIWIGAQECT WP_000145410.1 MULTISPECIES: hydrogenase-2 small chain [Proteobacteria] Length: 372\nNumber of predicted TMHs: 1\nExp number of AAs in TMHs: 23.72226\nExp number, first 60 AAs: 0.22038\nTotal prob of N-in: 0.12994\noutside 1 331\nTMhelix 332 354\ninside 355 372

13515 GCF_000618465.1_EcK1927 Escherichia coli O157:H7 str. K1927 Proteobacteria; Gammaproteobacteria; Enterobacterales; Enterobacteriaceae; Escherichia; Escherichia coli; Escherichia coli O157 MNNEETFYQAMRRQGVTRRSFLKYCSLAATSLGLGAGMAPKIAWALENKPRIPVVWIHGL WP_001301613.1 hydrogenase [Escherichia coli] Length: 372\nNumber of predicted TMHs: 2\nExp number of AAs in TMHs: 37.46598\nExp number, first 60 AAs: 19.53139\nTotal prob of N-in: 0.94364\nPOSSIBLE N-term signal sequence\ninside 1 19\nTMhelix 20 39\noutside 40 325\nTMhelix 326 348\ninside 349 372

13516 GCF_000618465.1_EcK1927 Escherichia coli O157:H7 str. K1927 Proteobacteria; Gammaproteobacteria; Enterobacterales; Enterobacteriaceae; Escherichia; Escherichia coli; Escherichia coli O157 MTGDNTLIHSHGINRRDFMKLCAALAATMGLSSKAAAEMAESVTNPQRPPVIWIGAQECT WP_000145410.1 MULTISPECIES: hydrogenase-2 small chain [Proteobacteria] Length: 372\nNumber of predicted TMHs: 1\nExp number of AAs in TMHs: 23.72226\nExp number, first 60 AAs: 0.22038\nTotal prob of N-in: 0.12994\noutside 1 331\nTMhelix 332 354\ninside 355 372

13517 GCF_000618485.1_EcK2188 Escherichia coli O157:H7 str. K2188 Proteobacteria; Gammaproteobacteria; Enterobacterales; Enterobacteriaceae; Escherichia; Escherichia coli; Escherichia coli O157 MNNEETFYQAMRRQGVTRRSFLKYCSLAATSLGLGAGMAPKIAWALENKPRIPVVWIHGL WP_001301613.1 hydrogenase [Escherichia coli] Length: 372\nNumber of predicted TMHs: 2\nExp number of AAs in TMHs: 37.46598\nExp number, first 60 AAs: 19.53139\nTotal prob of N-in: 0.94364\nPOSSIBLE N-term signal sequence\ninside 1 19\nTMhelix 20 39\noutside 40 325\nTMhelix 326 348\ninside 349 372

13518 GCF_000618485.1_EcK2188 Escherichia coli O157:H7 str. K2188 Proteobacteria; Gammaproteobacteria; Enterobacterales; Enterobacteriaceae; Escherichia; Escherichia coli; Escherichia coli O157 MTGDNTLIHSHGINRRDFMKLCAALAATMGLSSKAAAEMAESVTNPQRPPVIWIGAQECT WP_000145410.1 MULTISPECIES: hydrogenase-2 small chain [Proteobacteria] Length: 372\nNumber of predicted TMHs: 1\nExp number of AAs in TMHs: 23.72226\nExp number, first 60 AAs: 0.22038\nTotal prob of N-in: 0.12994\noutside 1 331\nTMhelix 332 354\ninside 355 372

13519 GCF_000618505.1_EcK2191 Escherichia coli O157:H7 str. K2191 Proteobacteria; Gammaproteobacteria; Enterobacterales; Enterobacteriaceae; Escherichia; Escherichia coli; Escherichia coli O157

MNNEETFYQAMRRQGVTRRSFLKYCSLAATSLGLGAGMAPKIAWALENKPRIPVVWIHGL WP_001301613.1 hydrogenase [Escherichia coli] Length: 372\nNumber of predicted TMHs: 2\nExp number of AAs in TMHs: 37.46598\nExp number, first 60 AAs: 19.53139\nTotal prob of N-in: 0.94364\nPOSSIBLE N-term signal sequence\ninside 1 19\nTMhelix 20 39\noutside 40 325\nTMhelix 326 348\ninside 349 372

13520 GCF_000618505.1_EcK2191 Escherichia coli O157:H7 str. K2191 Proteobacteria; Gammaproteobacteria; Enterobacterales; Enterobacteriaceae; Escherichia; Escherichia coli; Escherichia coli O157 MTGDNTLIHSHGINRRDFMKLCAALAATMGLSSKAAAEMAESVTNPQRPPVIWIGAQECT WP_000145410.1 MULTISPECIES: hydrogenase-2 small chain [Proteobacteria] Length: 372\nNumber of predicted TMHs: 1\nExp number of AAs in TMHs: 23.72226\nExp number, first 60 AAs: 0.22038\nTotal prob of N-in: 0.12994\noutside 1 331\nTMhelix 332 354\ninside 355 372

13521 GCF_000618645.1_EcK2192 Escherichia coli O157:H7 str. K2192 Proteobacteria; Gammaproteobacteria; Enterobacterales; Enterobacteriaceae; Escherichia; Escherichia coli; Escherichia coli O157 MTGDNTLIHSHGINRRDFMKLCAALAATMGLSSKAAAEMAESVTNPQRPPVIWIGAQECT WP_000145410.1 MULTISPECIES: hydrogenase-2 small chain [Proteobacteria] Length: 372\nNumber of predicted TMHs: 1\nExp number of AAs in TMHs: 23.72226\nExp number, first 60 AAs: 0.22038\nTotal prob of N-in: 0.12994\noutside 1 331\nTMhelix 332 354\ninside 355 372

13522 GCF_000618645.1_EcK2192 Escherichia coli O157:H7 str. K2192 Proteobacteria; Gammaproteobacteria; Enterobacterales; Enterobacteriaceae; Escherichia; Escherichia coli; Escherichia coli O157 MNNEETFYQAMRRQGVTRRSFLKYCSLAATSLGLGAGMAPKIAWALENKPRIPVVWIHGL WP_001301613.1 hydrogenase [Escherichia coli] Length: 372\nNumber of predicted TMHs: 2\nExp number of AAs in TMHs: 37.46598\nExp number, first 60 AAs: 19.53139\nTotal prob of N-in: 0.94364\nPOSSIBLE N-term signal sequence\ninside 1 19\nTMhelix 20 39\noutside 40 325\nTMhelix 326 348\ninside 349 372

13523 GCF_000618665.1_EcK2324 Escherichia coli O157:H7 str. K2324 Proteobacteria; Gammaproteobacteria; Enterobacterales; Enterobacteriaceae; Escherichia; Escherichia coli; Escherichia coli O157 MNNEETFYQAMRRQGVTRRSFLKYCSLAATSLGLGAGMAPKIAWALENKPRIPVVWIHGL WP_001301613.1 hydrogenase [Escherichia coli] Length: 372\nNumber of predicted TMHs: 2\nExp number of AAs in TMHs: 37.46598\nExp number, first 60 AAs: 19.53139\nTotal prob of N-in: 0.94364\nPOSSIBLE N-term signal sequence\ninside 1 19\nTMhelix 20 39\noutside 40 325\nTMhelix 326 348\ninside 349 372

13524 GCF_000618665.1_EcK2324 Escherichia coli O157:H7 str. K2324 Proteobacteria; Gammaproteobacteria; Enterobacterales; Enterobacteriaceae; Escherichia; Escherichia coli; Escherichia coli O157 MTGDNTLIHSHGINRRDFMKLCAALAATMGLSSKAAAEMAESVTNPQRPPVIWIGAQECT WP_000145410.1 MULTISPECIES: hydrogenase-2 small chain [Proteobacteria] Length: 372\nNumber of predicted TMHs: 1\nExp number of AAs in TMHs: 23.72226\nExp number, first 60 AAs: 0.22038\nTotal prob of N-in: 0.12994\noutside 1 331\nTMhelix 332 354\ninside 355 372

13525 GCF_000618525.1_EcK2581 Escherichia coli O157:H7 str. K2581 Proteobacteria; Gammaproteobacteria; Enterobacterales; Enterobacteriaceae; Escherichia; Escherichia coli; Escherichia coli O157 MNNEETFYQAMRRQGVTRRSFLKYCSLAATSLGLGAGMAPKIAWALENKPRIPVVWIHGL WP_001301613.1 hydrogenase [Escherichia coli] Length: 372\nNumber of predicted TMHs: 2\nExp number of AAs in TMHs: 37.46598\nExp number, first 60 AAs: 19.53139\nTotal prob of N-in: 0.94364\nPOSSIBLE N-term signal sequence\ninside 1 19\nTMhelix 20 39\noutside 40 325\nTMhelix 326 348\ninside 349 372

13526 GCF_000618525.1_EcK2581 Escherichia coli O157:H7 str. K2581 Proteobacteria; Gammaproteobacteria; Enterobacterales; Enterobacteriaceae; Escherichia; Escherichia coli; Escherichia coli O157 MTGDNTLIHSHGINRRDFMKLCAALAATMGLSSKAAAEMAESVTNPQRPPVIWIGAQECT WP_000145410.1 MULTISPECIES: hydrogenase-2 small chain [Proteobacteria] Length: 372\nNumber of predicted TMHs: 1\nExp number of AAs in TMHs: 23.72226\nExp number, first 60 AAs: 0.22038\nTotal prob of N-in: 0.12994\noutside 1 331\nTMhelix 332 354\ninside 355 372

13527 GCF_000618545.1_EcK2622 Escherichia coli O157:H7 str. K2622 Proteobacteria; Gammaproteobacteria; Enterobacterales; Enterobacteriaceae; Escherichia; Escherichia coli; Escherichia coli O157 MTGDNTLIHSHGINRRDFMKLCAALAATMGLSSKAAAEMAESVTNPQRPPVIWIGAQECT WP_000145410.1 MULTISPECIES: hydrogenase-2 small chain [Proteobacteria] Length: 372\nNumber of predicted TMHs: 1\nExp number of AAs in TMHs: 23.72226\nExp number, first 60 AAs: 0.22038\nTotal prob of N-in: 0.12994\noutside 1 331\nTMhelix 332 354\ninside 355 372

13528 GCF_000618545.1_EcK2622 Escherichia coli O157:H7 str. K2622 Proteobacteria; Gammaproteobacteria; Enterobacterales; Enterobacteriaceae; Escherichia; Escherichia coli; Escherichia coli O157
MNNEETFYQAMRRQGVTRRSFLKYCSLAATSLGLGAGMAPKIAWALENKPRIPVVWIHGL WP_001301613.1
hydrogenase [Escherichia coli] Length: 372\nNumber of predicted TMHs: 2\nExp number of AAs in TMHs: 37.46598\nExp number, first 60 AAs: 19.53139\nTotal prob of N-in: 0.94364\nPOSSIBLE N-term signal sequence\ninside 1 19\nTMhelix 20 39\noutside 40 325\nTMhelix 326 348\ninside 349 372

13529 GCF_000618565.1_EcK2845 Escherichia coli O157:H7 str. K2845 Proteobacteria; Gammaproteobacteria; Enterobacterales; Enterobacteriaceae; Escherichia; Escherichia coli; Escherichia coli O157
MTGDNTLIHSHGINRRDFMKLCAALAATMGLSSKAAAEMAESVTNPQRPPVIWIGAQECT WP_000145410.1
MULTISPECIES: hydrogenase-2 small chain [Proteobacteria] Length: 372\nNumber of predicted TMHs: 1\nExp number of AAs in TMHs: 23.72226\nExp number, first 60 AAs: 0.22038\nTotal prob of N-in: 0.12994\noutside 1 331\nTMhelix 332 354\ninside 355 372

13530 GCF_000618565.1_EcK2845 Escherichia coli O157:H7 str. K2845 Proteobacteria; Gammaproteobacteria; Enterobacterales; Enterobacteriaceae; Escherichia; Escherichia coli; Escherichia coli O157
MNNEETFYQAMRRQGVTRRSFLKYCSLAATSLGLGAGMAPKIAWALENKPRIPVVWIHGL WP_001301613.1
hydrogenase [Escherichia coli] Length: 372\nNumber of predicted TMHs: 2\nExp number of AAs in TMHs: 37.46598\nExp number, first 60 AAs: 19.53139\nTotal prob of N-in: 0.94364\nPOSSIBLE N-term signal sequence\ninside 1 19\nTMhelix 20 39\noutside 40 325\nTMhelix 326 348\ninside 349 372

13531 GCF_000618585.1_EcK2854 Escherichia coli O157:H7 str. K2854 Proteobacteria; Gammaproteobacteria; Enterobacterales; Enterobacteriaceae; Escherichia; Escherichia coli; Escherichia coli O157
MNNEETFYQAMRRQGVTRRSFLKYCSLAATSLGLGAGMAPKIAWALENKPRIPVVWIHGL WP_001301613.1
hydrogenase [Escherichia coli] Length: 372\nNumber of predicted TMHs: 2\nExp number of AAs in TMHs: 37.46598\nExp number, first 60 AAs: 19.53139\nTotal prob of N-in: 0.94364\nPOSSIBLE N-term signal sequence\ninside 1 19\nTMhelix 20 39\noutside 40 325\nTMhelix 326 348\ninside 349 372

13532 GCF_000618585.1_EcK2854 Escherichia coli O157:H7 str. K2854 Proteobacteria; Gammaproteobacteria; Enterobacterales; Enterobacteriaceae; Escherichia; Escherichia coli; Escherichia coli O157
MTGDNTLIHSHGINRRDFMKLCAALAATMGLSSKAAAEMAESVTNPQRPPVIWIGAQECT WP_000145410.1
MULTISPECIES: hydrogenase-2 small chain [Proteobacteria] Length: 372\nNumber of predicted TMHs: 1\nExp number of AAs in TMHs: 23.72226\nExp number, first 60 AAs: 0.22038\nTotal prob of N-in: 0.12994\noutside 1 331\nTMhelix 332 354\ninside 355 372

13533 GCF_000618605.1_EcK4396 Escherichia coli O157:H7 str. K4396 Proteobacteria; Gammaproteobacteria; Enterobacterales; Enterobacteriaceae; Escherichia; Escherichia coli; Escherichia coli O157
MNNEETFYQAMRRQGVTRRSFLKYCSLAATSLGLGAGMAPKIAWALENKPRIPVVWIHGL WP_001301613.1
hydrogenase [Escherichia coli] Length: 372\nNumber of predicted TMHs: 2\nExp number of AAs in TMHs: 37.46598\nExp number, first 60 AAs: 19.53139\nTotal prob of N-in: 0.94364\nPOSSIBLE N-term signal sequence\ninside 1 19\nTMhelix 20 39\noutside 40 325\nTMhelix 326 348\ninside 349 372

13534 GCF_000618605.1_EcK4396 Escherichia coli O157:H7 str. K4396 Proteobacteria; Gammaproteobacteria; Enterobacterales; Enterobacteriaceae; Escherichia; Escherichia coli; Escherichia coli O157
MTGDNTLIHSHGINRRDFMKLCAALAATMGLSSKAAAEMAESVTNPQRPPVIWIGAQECT WP_000145410.1
MULTISPECIES: hydrogenase-2 small chain [Proteobacteria] Length: 372\nNumber of predicted TMHs: 1\nExp number of AAs in TMHs: 23.72226\nExp number, first 60 AAs: 0.22038\nTotal prob of N-in: 0.12994\noutside 1 331\nTMhelix 332 354\ninside 355 372

13535 GCF_000617965.1_EcK4405 Escherichia coli O157:H7 str. K4405 Proteobacteria; Gammaproteobacteria; Enterobacterales; Enterobacteriaceae; Escherichia; Escherichia coli; Escherichia coli O157
MNNEETFYQAMRRQGVTRRSFLKYCSLAATSLGLGAGMAPKIAWALENKPRIPVVWIHGL WP_001301613.1
hydrogenase [Escherichia coli] Length: 372\nNumber of predicted TMHs: 2\nExp number of AAs in TMHs: 37.46598\nExp number, first 60 AAs: 19.53139\nTotal prob of N-in: 0.94364\nPOSSIBLE N-term signal sequence\ninside 1 19\nTMhelix 20 39\noutside 40 325\nTMhelix 326 348\ninside 349 372

13536 GCF_000617965.1_EcK4405 Escherichia coli O157:H7 str. K4405 Proteobacteria; Gammaproteobacteria; Enterobacterales; Enterobacteriaceae; Escherichia; Escherichia coli; Escherichia coli O157
MTGDNTLIHSHGINRRDFMKLCAALAATMGLSSKAAAEMAESVTNPQRPPVIWIGAQECT WP_000145410.1
MULTISPECIES: hydrogenase-2 small chain [Proteobacteria] Length: 372\nNumber of predicted TMHs:

1\nExp number of AAs in TMHs: 23.72226\nExp number, first 60 AAs: 0.22038\nTotal prob of N-in: 0.12994\noutside 1 331\nTMhelix 332 354\ninside 355 372

13537 GCF_000618625.1_EcK4406 Escherichia coli O157:H7 str. K4406 Proteobacteria; Gammaproteobacteria; Enterobacterales; Enterobacteriaceae; Escherichia; Escherichia coli; Escherichia coli O157 MTGDNTLIHSHGINRRDFMKLCAALAATMGLSSKAAAEMAESVTNPQRPPVIWIGAQECT WP_000145410.1 MULTISPECIES: hydrogenase-2 small chain [Proteobacteria] Length: 372\nNumber of predicted TMHs: 1\nExp number of AAs in TMHs: 23.72226\nExp number, first 60 AAs: 0.22038\nTotal prob of N-in: 0.12994\noutside 1 331\nTMhelix 332 354\ninside 355 372

13538 GCF_000618625.1_EcK4406 Escherichia coli O157:H7 str. K4406 Proteobacteria; Gammaproteobacteria; Enterobacterales; Enterobacteriaceae; Escherichia; Escherichia coli; Escherichia coli O157 MNNEETFYQAMRRQGVTRRSFLKYCSLAATSLGLGAGMAPKIAWALENKPRIPVVWIHGL WP_001301613.1 hydrogenase [Escherichia coli] Length: 372\nNumber of predicted TMHs: 2\nExp number of AAs in TMHs: 37.46598\nExp number, first 60 AAs: 19.53139\nTotal prob of N-in: 0.94364\nPOSSIBLE N-term signal sequence\ninside 1 19\nTMhelix 20 39\noutside 40 325\nTMhelix 326 348\ninside 349 372

13539 GCF_000618685.1_EcK4527 Escherichia coli O157:H7 str. K4527 Proteobacteria; Gammaproteobacteria; Enterobacterales; Enterobacteriaceae; Escherichia; Escherichia coli; Escherichia coli O157 MNNEETFYQAMRRQGVTRRSFLKYCSLAATSLGLGAGMAPKIAWALENKPRIPVVWIHGL WP_001301613.1 hydrogenase [Escherichia coli] Length: 372\nNumber of predicted TMHs: 2\nExp number of AAs in TMHs: 37.46598\nExp number, first 60 AAs: 19.53139\nTotal prob of N-in: 0.94364\nPOSSIBLE N-term signal sequence\ninside 1 19\nTMhelix 20 39\noutside 40 325\nTMhelix 326 348\ninside 349 372

13540 GCF_000618685.1_EcK4527 Escherichia coli O157:H7 str. K4527 Proteobacteria; Gammaproteobacteria; Enterobacterales; Enterobacteriaceae; Escherichia; Escherichia coli; Escherichia coli O157 MTGDNTLIHSHGINRRDFMKLCAALAATMGLSSKAAAEMAESVTNPQRPPVIWIGAQECT WP_000145410.1 MULTISPECIES: hydrogenase-2 small chain [Proteobacteria] Length: 372\nNumber of predicted TMHs: 1\nExp number of AAs in TMHs: 23.72226\nExp number, first 60 AAs: 0.22038\nTotal prob of N-in: 0.12994\noutside 1 331\nTMhelix 332 354\ninside 355 372

13541 GCF_000618745.1_EcK5418 Escherichia coli O157:H7 str. K5418 Proteobacteria; Gammaproteobacteria; Enterobacterales; Enterobacteriaceae; Escherichia; Escherichia coli; Escherichia coli O157 MTGDNTLIHSHGINRRDFMKLCAALAATMGLSSKAAAEMAESVTNPQRPPVIWIGAQECT WP_000145410.1 MULTISPECIES: hydrogenase-2 small chain [Proteobacteria] Length: 372\nNumber of predicted TMHs: 1\nExp number of AAs in TMHs: 23.72226\nExp number, first 60 AAs: 0.22038\nTotal prob of N-in: 0.12994\noutside 1 331\nTMhelix 332 354\ninside 355 372

13542 GCF_000618745.1_EcK5418 Escherichia coli O157:H7 str. K5418 Proteobacteria; Gammaproteobacteria; Enterobacterales; Enterobacteriaceae; Escherichia; Escherichia coli; Escherichia coli O157 MNNEETFYQAMRRQGVTRRSFLKYCSLAATSLGLGAGMAPKIAWALENKPRIPVVWIHGL WP_001301613.1 hydrogenase [Escherichia coli] Length: 372\nNumber of predicted TMHs: 2\nExp number of AAs in TMHs: 37.46598\nExp number, first 60 AAs: 19.53139\nTotal prob of N-in: 0.94364\nPOSSIBLE N-term signal sequence\ninside 1 19\nTMhelix 20 39\noutside 40 325\nTMhelix 326 348\ninside 349 372

13543 GCF_000618765.1_EcK5448 Escherichia coli O157:H7 str. K5448 Proteobacteria; Gammaproteobacteria; Enterobacterales; Enterobacteriaceae; Escherichia; Escherichia coli; Escherichia coli O157 MTGDNTLIHSHGINRRDFMKLCAALAATMGLSSKAAAEMAESVTNPQRPPVIWIGAQECT WP_000145410.1 MULTISPECIES: hydrogenase-2 small chain [Proteobacteria] Length: 372\nNumber of predicted TMHs: 1\nExp number of AAs in TMHs: 23.72226\nExp number, first 60 AAs: 0.22038\nTotal prob of N-in: 0.12994\noutside 1 331\nTMhelix 332 354\ninside 355 372

13544 GCF_000618765.1_EcK5448 Escherichia coli O157:H7 str. K5448 Proteobacteria; Gammaproteobacteria; Enterobacterales; Enterobacteriaceae; Escherichia; Escherichia coli; Escherichia coli O157 MNNEETFYQAMRRQGVTRRSFLKYCSLAATSLGLGAGMAPKIAWALENKPRIPVVWIHGL WP_032166558.1 hydrogenase [Escherichia coli] Length: 372\nNumber of predicted TMHs: 2\nExp number of AAs in TMHs: 37.2028\nExp number, first 60 AAs: 19.53107\nTotal prob of N-in: 0.94349\nPOSSIBLE N-term signal sequence\ninside 1 19\nTMhelix 20 39\noutside 40 325\nTMhelix 326 348\ninside 349 372

13545 GCF_000618785.1_EcK5449 Escherichia coli O157:H7 str. K5449 Proteobacteria; Gammaproteobacteria; Enterobacterales; Enterobacteriaceae; Escherichia; Escherichia coli; Escherichia coli O157

MTGDNTLIHSHGINRRDFMKLCAALAATMGLSSKAAAEMAESVTNPQRPPVIWIGAQECT WP_000145410.1 MULTISPECIES: hydrogenase-2 small chain [Proteobacteria] Length: 372\nNumber of predicted TMHs: 1\nExp number of AAs in TMHs: 23.72226\nExp number, first 60 AAs: 0.22038\nTotal prob of N-in: 0.12994\noutside 1 331\nTMhelix 332 354\ninside 355 372

13546 GCF_000618785.1_EcK5449 Escherichia coli O157:H7 str. K5449 Proteobacteria; Gammaproteobacteria; Enterobacterales; Enterobacteriaceae; Escherichia; Escherichia coli; Escherichia coli O157 MNNEETFYQAMRRQGVTRRSFLKYCSLAATSLGLGAGMAPKIAWALENKPRIPVVWIHGL WP_032166558.1 hydrogenase [Escherichia coli] Length: 372\nNumber of predicted TMHs: 2\nExp number of AAs in TMHs: 37.2028\nExp number, first 60 AAs: 19.53107\nTotal prob of N-in: 0.94349\nPOSSIBLE N-term signal sequence\ninside 1 19\nTMhelix 20 39\noutside 40 325\nTMhelix 326 348\ninside 349 372

13547 GCF_000618805.1_EcK5453 Escherichia coli O157:H7 str. K5453 Proteobacteria; Gammaproteobacteria; Enterobacterales; Enterobacteriaceae; Escherichia; Escherichia coli; Escherichia coli O157 MNNEETFYQAMRRQGVTRRSFLKYCSLAATSLGLGAGMAPKIAWALENKPRIPVVWIHGL WP_032166558.1 hydrogenase [Escherichia coli] Length: 372\nNumber of predicted TMHs: 2\nExp number of AAs in TMHs: 37.2028\nExp number, first 60 AAs: 19.53107\nTotal prob of N-in: 0.94349\nPOSSIBLE N-term signal sequence\ninside 1 19\nTMhelix 20 39\noutside 40 325\nTMhelix 326 348\ninside 349 372

13548 GCF_000618805.1_EcK5453 Escherichia coli O157:H7 str. K5453 Proteobacteria; Gammaproteobacteria; Enterobacterales; Enterobacteriaceae; Escherichia; Escherichia coli; Escherichia coli O157 MTGDNTLIHSHGINRRDFMKLCAALAATMGLSSKAAAEMAESVTNPQRPPVIWIGAQECT WP_000145410.1 MULTISPECIES: hydrogenase-2 small chain [Proteobacteria] Length: 372\nNumber of predicted TMHs: 1\nExp number of AAs in TMHs: 23.72226\nExp number, first 60 AAs: 0.22038\nTotal prob of N-in: 0.12994\noutside 1 331\nTMhelix 332 354\ninside 355 372

13549 GCF_000618885.1_EcK5460 Escherichia coli O157:H7 str. K5460 Proteobacteria; Gammaproteobacteria; Enterobacterales; Enterobacteriaceae; Escherichia; Escherichia coli; Escherichia coli O157 MNNEETFYQAMRRQGVTRRSFLKYCSLAATSLGLGAGMAPKIAWALENKPRIPVVWIHGL WP_032166558.1 hydrogenase [Escherichia coli] Length: 372\nNumber of predicted TMHs: 2\nExp number of AAs in TMHs: 37.2028\nExp number, first 60 AAs: 19.53107\nTotal prob of N-in: 0.94349\nPOSSIBLE N-term signal sequence\ninside 1 19\nTMhelix 20 39\noutside 40 325\nTMhelix 326 348\ninside 349 372

13550 GCF_000618885.1_EcK5460 Escherichia coli O157:H7 str. K5460 Proteobacteria; Gammaproteobacteria; Enterobacterales; Enterobacteriaceae; Escherichia; Escherichia coli; Escherichia coli O157 MTGDNTLIHSHGINRRDFMKLCAALAATMGLSSKAAAEMAESVTNPQRPPVIWIGAQECT WP_000145410.1 MULTISPECIES: hydrogenase-2 small chain [Proteobacteria] Length: 372\nNumber of predicted TMHs: 1\nExp number of AAs in TMHs: 23.72226\nExp number, first 60 AAs: 0.22038\nTotal prob of N-in: 0.12994\noutside 1 331\nTMhelix 332 354\ninside 355 372

13551 GCF_000618825.1_EcK5467 Escherichia coli O157:H7 str. K5467 Proteobacteria; Gammaproteobacteria; Enterobacterales; Enterobacteriaceae; Escherichia; Escherichia coli; Escherichia coli O157 MNNEETFYQAMRRQGVTRRSFLKYCSLAATSLGLGAGMAPKIAWALENKPRIPVVWIHGL WP_032166558.1 hydrogenase [Escherichia coli] Length: 372\nNumber of predicted TMHs: 2\nExp number of AAs in TMHs: 37.2028\nExp number, first 60 AAs: 19.53107\nTotal prob of N-in: 0.94349\nPOSSIBLE N-term signal sequence\ninside 1 19\nTMhelix 20 39\noutside 40 325\nTMhelix 326 348\ninside 349 372

13552 GCF_000618825.1_EcK5467 Escherichia coli O157:H7 str. K5467 Proteobacteria; Gammaproteobacteria; Enterobacterales; Enterobacteriaceae; Escherichia; Escherichia coli; Escherichia coli O157 MTGDNTLIHSHGINRRDFMKLCAALAATMGLSSKAAAEMAESVTNPQRPPVIWIGAQECT WP_000145410.1 MULTISPECIES: hydrogenase-2 small chain [Proteobacteria] Length: 372\nNumber of predicted TMHs: 1\nExp number of AAs in TMHs: 23.72226\nExp number, first 60 AAs: 0.22038\nTotal prob of N-in: 0.12994\noutside 1 331\nTMhelix 332 354\ninside 355 372

13553 GCF_000618845.1_EcK5602 Escherichia coli O157:H7 str. K5602 Proteobacteria; Gammaproteobacteria; Enterobacterales; Enterobacteriaceae; Escherichia; Escherichia coli; Escherichia coli O157 MTGDNTLIHSHGINRRDFMKLCAALAATMGLSSKAAAEMAESVTNPQRPPVIWIGAQECT WP_000145410.1 MULTISPECIES: hydrogenase-2 small chain [Proteobacteria] Length: 372\nNumber of predicted TMHs: 1\nExp number of AAs in TMHs: 23.72226\nExp number, first 60 AAs: 0.22038\nTotal prob of N-in: 0.12994\noutside 1 331\nTMhelix 332 354\ninside 355 372

13554 GCF_000618845.1_EcK5602 Escherichia coli O157:H7 str. K5602 Proteobacteria; Gammaproteobacteria; Enterobacterales; Enterobacteriaceae; Escherichia; Escherichia coli; Escherichia coli O157
MNNEETFYQAMRRQGVTRRSFLKYCSLAATSLGLGAGMAPKIAWALENKPRIPVVWIHGL WP_032166558.1
hydrogenase [Escherichia coli] Length: 372\nNumber of predicted TMHs: 2\nExp number of AAs in TMHs: 37.2028\nExp number, first 60 AAs: 19.53107\nTotal prob of N-in: 0.94349\nPOSSIBLE N-term signal sequence\ninside 1 19\nTMhelix 20 39\noutside 40 325\nTMhelix 326 348\ninside 349 372

13555 GCF_000618865.1_EcK5607 Escherichia coli O157:H7 str. K5607 Proteobacteria; Gammaproteobacteria; Enterobacterales; Enterobacteriaceae; Escherichia; Escherichia coli; Escherichia coli O157
MNNEETFYQAMRRQGVTRRSFLKYCSLAATSLGLGAGMAPKIAWALENKPRIPVVWIHGL WP_032166558.1
hydrogenase [Escherichia coli] Length: 372\nNumber of predicted TMHs: 2\nExp number of AAs in TMHs: 37.2028\nExp number, first 60 AAs: 19.53107\nTotal prob of N-in: 0.94349\nPOSSIBLE N-term signal sequence\ninside 1 19\nTMhelix 20 39\noutside 40 325\nTMhelix 326 348\ninside 349 372

13556 GCF_000618865.1_EcK5607 Escherichia coli O157:H7 str. K5607 Proteobacteria; Gammaproteobacteria; Enterobacterales; Enterobacteriaceae; Escherichia; Escherichia coli; Escherichia coli O157
MTGDNTLIHSHGINRRDFMKLCAALAATMGLSSKAAAEMAESVTNPQRPPVIWIGAQECT WP_000145410.1
MULTISPECIES: hydrogenase-2 small chain [Proteobacteria] Length: 372\nNumber of predicted TMHs: 1\nExp number of AAs in TMHs: 23.72226\nExp number, first 60 AAs: 0.22038\nTotal prob of N-in: 0.12994\noutside 1 331\nTMhelix 332 354\ninside 355 372

13557 GCF_000619445.1_EcK5609 Escherichia coli O157:H7 str. K5609 Proteobacteria; Gammaproteobacteria; Enterobacterales; Enterobacteriaceae; Escherichia; Escherichia coli; Escherichia coli O157
MNNEETFYQAMRRQGVTRRSFLKYCSLAATSLGLGAGMAPKIAWALENKPRIPVVWIHGL WP_032166558.1
hydrogenase [Escherichia coli] Length: 372\nNumber of predicted TMHs: 2\nExp number of AAs in TMHs: 37.2028\nExp number, first 60 AAs: 19.53107\nTotal prob of N-in: 0.94349\nPOSSIBLE N-term signal sequence\ninside 1 19\nTMhelix 20 39\noutside 40 325\nTMhelix 326 348\ninside 349 372

13558 GCF_000619445.1_EcK5609 Escherichia coli O157:H7 str. K5609 Proteobacteria; Gammaproteobacteria; Enterobacterales; Enterobacteriaceae; Escherichia; Escherichia coli; Escherichia coli O157
MTGDNTLIHSHGINRRDFMKLCAALAATMGLSSKAAAEMAESVTNPQRPPVIWIGAQECT WP_000145410.1
MULTISPECIES: hydrogenase-2 small chain [Proteobacteria] Length: 372\nNumber of predicted TMHs: 1\nExp number of AAs in TMHs: 23.72226\nExp number, first 60 AAs: 0.22038\nTotal prob of N-in: 0.12994\noutside 1 331\nTMhelix 332 354\ninside 355 372

13559 GCF_000616745.1_EcK5806 Escherichia coli O157:H7 str. K5806 Proteobacteria; Gammaproteobacteria; Enterobacterales; Enterobacteriaceae; Escherichia; Escherichia coli; Escherichia coli O157
MNNEETFYQAMRRQGVTRRSFLKYCSLAATSLGLGAGMAPKIAWALENKPRIPVVWIHGL WP_001301613.1
hydrogenase [Escherichia coli] Length: 372\nNumber of predicted TMHs: 2\nExp number of AAs in TMHs: 37.46598\nExp number, first 60 AAs: 19.53139\nTotal prob of N-in: 0.94364\nPOSSIBLE N-term signal sequence\ninside 1 19\nTMhelix 20 39\noutside 40 325\nTMhelix 326 348\ninside 349 372

13560 GCF_000616745.1_EcK5806 Escherichia coli O157:H7 str. K5806 Proteobacteria; Gammaproteobacteria; Enterobacterales; Enterobacteriaceae; Escherichia; Escherichia coli; Escherichia coli O157
MTGDNTLIHSHGINRRDFMKLCAALAATMGLSSKAAAEMAESVTNPQRPPVIWIGAQECT WP_000145410.1
MULTISPECIES: hydrogenase-2 small chain [Proteobacteria] Length: 372\nNumber of predicted TMHs: 1\nExp number of AAs in TMHs: 23.72226\nExp number, first 60 AAs: 0.22038\nTotal prob of N-in: 0.12994\noutside 1 331\nTMhelix 332 354\ninside 355 372

13561 GCF_000619365.1_EcK5852 Escherichia coli O157:H7 str. K5852 Proteobacteria; Gammaproteobacteria; Enterobacterales; Enterobacteriaceae; Escherichia; Escherichia coli; Escherichia coli O157
MTGDNTLIHSHGINRRDFMKLCAALAATMGLSSKAAAEMAESVTNPQRPPVIWIGAQECT WP_000145410.1
MULTISPECIES: hydrogenase-2 small chain [Proteobacteria] Length: 372\nNumber of predicted TMHs: 1\nExp number of AAs in TMHs: 23.72226\nExp number, first 60 AAs: 0.22038\nTotal prob of N-in: 0.12994\noutside 1 331\nTMhelix 332 354\ninside 355 372

13562 GCF_000619365.1_EcK5852 Escherichia coli O157:H7 str. K5852 Proteobacteria; Gammaproteobacteria; Enterobacterales; Enterobacteriaceae; Escherichia; Escherichia coli; Escherichia coli O157
MNNEETFYQAMRRQGVTRRSFLKYCSLAATSLGLGAGMAPKIAWALENKPRIPVVWIHGL WP_001301613.1
hydrogenase [Escherichia coli] Length: 372\nNumber of predicted TMHs: 2\nExp number of AAs in

TMHs: 37.46598\nExp number, first 60 AAs: 19.53139\nTotal prob of N-in: 0.94364\nPOSSIBLE N-term signal sequence\ninside 1 19\nTMhelix 20 39\noutside 40 325\nTMhelix 326 348\ninside 349 372

13563 GCF_000619385.1_EcK6590 Escherichia coli O157:H7 str. K6590 Proteobacteria; Gammaproteobacteria; Enterobacterales; Enterobacteriaceae; Escherichia; Escherichia coli; Escherichia coli O157
MTGDNTLIHSHGINRRDFMKLCAALAATMGLSSKAAAEMAESVTNPQRPPVIWIGAQECT WP_000145410.1
MULTISPECIES: hydrogenase-2 small chain [Proteobacteria] Length: 372\nNumber of predicted TMHs: 1\nExp number of AAs in TMHs: 23.72226\nExp number, first 60 AAs: 0.22038\nTotal prob of N-in: 0.12994\noutside 1 331\nTMhelix 332 354\ninside 355 372

13564 GCF_000619385.1_EcK6590 Escherichia coli O157:H7 str. K6590 Proteobacteria; Gammaproteobacteria; Enterobacterales; Enterobacteriaceae; Escherichia; Escherichia coli; Escherichia coli O157
MNNEETFYQAMRRQGVTRRSFLKYCSLAATSLGLGAGMAPKIAWALENKPRIPVVWIHGL WP_001301613.1
hydrogenase [Escherichia coli] Length: 372\nNumber of predicted TMHs: 2\nExp number of AAs in TMHs: 37.46598\nExp number, first 60 AAs: 19.53139\nTotal prob of N-in: 0.94364\nPOSSIBLE N-term signal sequence\ninside 1 19\nTMhelix 20 39\noutside 40 325\nTMhelix 326 348\ninside 349 372

13565 GCF_000619405.1_EcK6676 Escherichia coli O157:H7 str. K6676 Proteobacteria; Gammaproteobacteria; Enterobacterales; Enterobacteriaceae; Escherichia; Escherichia coli; Escherichia coli O157
MTGDNTLIHSHGINRRDFMKLCAALAATMGLSSKAAAEMAESVTNPQRPPVIWIGAQECT WP_000145410.1
MULTISPECIES: hydrogenase-2 small chain [Proteobacteria] Length: 372\nNumber of predicted TMHs: 1\nExp number of AAs in TMHs: 23.72226\nExp number, first 60 AAs: 0.22038\nTotal prob of N-in: 0.12994\noutside 1 331\nTMhelix 332 354\ninside 355 372

13566 GCF_000619405.1_EcK6676 Escherichia coli O157:H7 str. K6676 Proteobacteria; Gammaproteobacteria; Enterobacterales; Enterobacteriaceae; Escherichia; Escherichia coli; Escherichia coli O157
MNNEETFYQAMRRQGVTRRSFLKYCSLAATSLGLGAGMAPKIAWALENKPRIPVVWIHGL WP_001301613.1
hydrogenase [Escherichia coli] Length: 372\nNumber of predicted TMHs: 2\nExp number of AAs in TMHs: 37.46598\nExp number, first 60 AAs: 19.53139\nTotal prob of N-in: 0.94364\nPOSSIBLE N-term signal sequence\ninside 1 19\nTMhelix 20 39\noutside 40 325\nTMhelix 326 348\ninside 349 372

13567 GCF_000619425.1_EcK6687 Escherichia coli O157:H7 str. K6687 Proteobacteria; Gammaproteobacteria; Enterobacterales; Enterobacteriaceae; Escherichia; Escherichia coli; Escherichia coli O157
MNNEETFYQAMRRQGVTRRSFLKYCSLAATSLGLGAGMAPKIAWALENKPRIPVVWIHGL WP_001301613.1
hydrogenase [Escherichia coli] Length: 372\nNumber of predicted TMHs: 2\nExp number of AAs in TMHs: 37.46598\nExp number, first 60 AAs: 19.53139\nTotal prob of N-in: 0.94364\nPOSSIBLE N-term signal sequence\ninside 1 19\nTMhelix 20 39\noutside 40 325\nTMhelix 326 348\ninside 349 372

13568 GCF_000619425.1_EcK6687 Escherichia coli O157:H7 str. K6687 Proteobacteria; Gammaproteobacteria; Enterobacterales; Enterobacteriaceae; Escherichia; Escherichia coli; Escherichia coli O157
MTGDNTLIHSHGINRRDFMKLCAALAATMGLSSKAAAEMAESVTNPQRPPVIWIGAQECT WP_000145410.1
MULTISPECIES: hydrogenase-2 small chain [Proteobacteria] Length: 372\nNumber of predicted TMHs: 1\nExp number of AAs in TMHs: 23.72226\nExp number, first 60 AAs: 0.22038\nTotal prob of N-in: 0.12994\noutside 1 331\nTMhelix 332 354\ninside 355 372

13569 GCF_000618925.2_EcK7140 Escherichia coli O157:H7 str. K7140 Proteobacteria; Gammaproteobacteria; Enterobacterales; Enterobacteriaceae; Escherichia; Escherichia coli; Escherichia coli O157
MTGDNTLIHSHGINRRDFMKLCAALAATMGLSSKAAAEMAESVTNPQRPPVIWIGAQECT WP_000145410.1
MULTISPECIES: hydrogenase-2 small chain [Proteobacteria] Length: 372\nNumber of predicted TMHs: 1\nExp number of AAs in TMHs: 23.72226\nExp number, first 60 AAs: 0.22038\nTotal prob of N-in: 0.12994\noutside 1 331\nTMhelix 332 354\ninside 355 372

13570 GCF_000618925.2_EcK7140 Escherichia coli O157:H7 str. K7140 Proteobacteria; Gammaproteobacteria; Enterobacterales; Enterobacteriaceae; Escherichia; Escherichia coli; Escherichia coli O157
MNNEETFYQAMRRQGVTRRSFLKYCSLAATSLGLGAGMAPKIAWALENKPRIPVVWIHGL WP_001301613.1
hydrogenase [Escherichia coli] Length: 372\nNumber of predicted TMHs: 2\nExp number of AAs in TMHs: 37.46598\nExp number, first 60 AAs: 19.53139\nTotal prob of N-in: 0.94364\nPOSSIBLE N-term signal sequence\ninside 1 19\nTMhelix 20 39\noutside 40 325\nTMhelix 326 348\ninside 349 372

13571 GCF_000187385.1_ASM18738v2 Escherichia coli O157:H7 str. LSU-61 Proteobacteria; Gammaproteobacteria; Enterobacterales; Enterobacteriaceae; Escherichia; Escherichia coli; Escherichia coli O157

MTGDNTLIHSHGINRRDFMKLCAALAATMGLSSKAAAEMAESVTNPQRPPVIWIGAQECT WP_000145410.1 MULTISPECIES: hydrogenase-2 small chain [Proteobacteria] Length: 372\nNumber of predicted TMHs: 1\nExp number of AAs in TMHs: 23.72226\nExp number, first 60 AAs: 0.22038\nTotal prob of N-in: 0.12994\noutside 1 331\nTMhelix 332 354\ninside 355 372

13572 GCF_000187385.1_ASM18738v2 Escherichia coli O157:H7 str. LSU-61 Proteobacteria; Gammaproteobacteria; Enterobacterales; Enterobacteriaceae; Escherichia; Escherichia coli; Escherichia coli O157 MNNEETFYQAMRRQGVTRRSFLKYCSLAATSLGLGAGMAPKIAWALENKPRIPVVWIHGL WP_001301613.1 hydrogenase [Escherichia coli] Length: 372\nNumber of predicted TMHs: 2\nExp number of AAs in TMHs: 37.46598\nExp number, first 60 AAs: 19.53139\nTotal prob of N-in: 0.94364\nPOSSIBLE N-term signal sequence\ninside 1 19\nTMhelix 20 39\noutside 40 325\nTMhelix 326 348\ninside 349 372

13573 GCF_000008865.1_ASM886v1 Escherichia coli O157:H7 str. Sakai Proteobacteria; Gammaproteobacteria; Enterobacterales; Enterobacteriaceae; Escherichia; Escherichia coli; Escherichia coli O157 MNNEETFYQAMRRQGVTRRSFLKYCSLAATSLGLGAGMAPKIAWALENKPRIPVVWIHGL NP_309155.1 hydrogenase-1 small subunit [Escherichia coli O157:H7 str. Sakai] Length: 372\nNumber of predicted TMHs: 2\nExp number of AAs in TMHs: 37.46598\nExp number, first 60 AAs: 19.53139\nTotal prob of N-in: 0.94364\nPOSSIBLE N-term signal sequence\ninside 1 19\nTMhelix 20 39\noutside 40 325\nTMhelix 326 348\ninside 349 372

13574 GCF_000008865.1_ASM886v1 Escherichia coli O157:H7 str. Sakai Proteobacteria; Gammaproteobacteria; Enterobacterales; Enterobacteriaceae; Escherichia; Escherichia coli; Escherichia coli O157 MTGDNTLIHSHGINRRDFMKLCAALAATMGLSSKAAAEMAESVTNPQRPPVIWIGAQECT NP_311909.1 hydrogenase 2 small subunit [Escherichia coli O157:H7 str. Sakai] Length: 372\nNumber of predicted TMHs: 1\nExp number of AAs in TMHs: 23.72226\nExp number, first 60 AAs: 0.22038\nTotal prob of N-in: 0.12994\noutside 1 331\nTMhelix 332 354\ninside 355 372

13575 GCF_000730345.1_ASM73034v1 Escherichia coli O157:H7 str. SS17 Proteobacteria; Gammaproteobacteria; Enterobacterales; Enterobacteriaceae; Escherichia; Escherichia coli; Escherichia coli O157 MNNEETFYQAMRRQGVTRRSFLKYCSLAATSLGLGAGMAPKIAWALENKPRIPVVWIHGL WP_001301613.1 hydrogenase [Escherichia coli] Length: 372\nNumber of predicted TMHs: 2\nExp number of AAs in TMHs: 37.46598\nExp number, first 60 AAs: 19.53139\nTotal prob of N-in: 0.94364\nPOSSIBLE N-term signal sequence\ninside 1 19\nTMhelix 20 39\noutside 40 325\nTMhelix 326 348\ninside 349 372

13576 GCF_000730345.1_ASM73034v1 Escherichia coli O157:H7 str. SS17 Proteobacteria; Gammaproteobacteria; Enterobacterales; Enterobacteriaceae; Escherichia; Escherichia coli; Escherichia coli O157 MTGDNTLIHSHGINRRDFMKLCAALAATMGLSSKAAAEMAESVTNPQRPPVIWIGAQECT WP_000145410.1 MULTISPECIES: hydrogenase-2 small chain [Proteobacteria] Length: 372\nNumber of predicted TMHs: 1\nExp number of AAs in TMHs: 23.72226\nExp number, first 60 AAs: 0.22038\nTotal prob of N-in: 0.12994\noutside 1 331\nTMhelix 332 354\ninside 355 372

13577 GCF_000803705.1_ASM80370v1 Escherichia coli O157:H7 str. SS52 Proteobacteria; Gammaproteobacteria; Enterobacterales; Enterobacteriaceae; Escherichia; Escherichia coli; Escherichia coli O157 MTGDNTLIHSHGINRRDFMKLCAALAATMGLSSKAAAEMAESVTNPQRPPVIWIGAQECT WP_000145410.1 MULTISPECIES: hydrogenase-2 small chain [Proteobacteria] Length: 372\nNumber of predicted TMHs: 1\nExp number of AAs in TMHs: 23.72226\nExp number, first 60 AAs: 0.22038\nTotal prob of N-in: 0.12994\noutside 1 331\nTMhelix 332 354\ninside 355 372

13578 GCF_000803705.1_ASM80370v1 Escherichia coli O157:H7 str. SS52 Proteobacteria; Gammaproteobacteria; Enterobacterales; Enterobacteriaceae; Escherichia; Escherichia coli; Escherichia coli O157 MNNEETFYQAMRRQGVTRRSFLKYCSLAATSLGLGAGMAPKIAWALENKPRIPVVWIHGL WP_001301613.1 hydrogenase [Escherichia coli] Length: 372\nNumber of predicted TMHs: 2\nExp number of AAs in TMHs: 37.46598\nExp number, first 60 AAs: 19.53139\nTotal prob of N-in: 0.94364\nPOSSIBLE N-term signal sequence\ninside 1 19\nTMhelix 20 39\noutside 40 325\nTMhelix 326 348\ninside 349 372

13579 GCF_000747455.1_O157_T154306_454 Escherichia coli O157:H7 str. T1543_06 Proteobacteria; Gammaproteobacteria; Enterobacterales; Enterobacteriaceae; Escherichia; Escherichia coli; Escherichia coli O157 MNNEETFYQAMRRQGVTRRSFLKYCSLAATSLGLGAGMAPKIAWALENKPRIPVVWIHGL WP_001301613.1 hydrogenase [Escherichia coli] Length: 372\nNumber of predicted TMHs: 2\nExp number of AAs in TMHs: 37.46598\nExp number, first 60 AAs: 19.53139\nTotal prob of N-in: 0.94364\nPOSSIBLE N-term signal sequence\ninside 1 19\nTMhelix 20 39\noutside 40 325\nTMhelix 326 348\ninside 349 372

13580	GCF_000747455.1_O157_T154306_454	Escherichia coli O157:H7 str. T1543_06 Proteobacteria; Gammaproteobacteria; Enterobacterales; Enterobacteriaceae; Escherichia; Escherichia coli; Escherichia coli O157
MTGDNTLIHSHGINRRDFMKLCAALAATMGLSSKAAAEMAESVTNPQRPPVIWIGAQECT	WP_000145410.1
MULTISPECIES: hydrogenase-2 small chain [Proteobacteria]	Length: 372\nNumber of predicted TMHs: 1\nExp number of AAs in TMHs: 23.72226\nExp number, first 60 AAs: 0.22038\nTotal prob of N-in: 0.12994\noutside 1 331\nTMhelix 332 354\ninside 355 372

13581	GCF_000267425.2_ASM26742v2	Escherichia coli O157:H7 str. TW14313 Proteobacteria; Gammaproteobacteria; Enterobacterales; Enterobacteriaceae; Escherichia; Escherichia coli; Escherichia coli O157
MNNEETFYQAMRRQGVTRRSFLKYCSLAATSLGLGAGMAPKIAWALENKPRIPVVWIHGL	WP_001301613.1
hydrogenase [Escherichia coli]	Length: 372\nNumber of predicted TMHs: 2\nExp number of AAs in TMHs: 37.46598\nExp number, first 60 AAs: 19.53139\nTotal prob of N-in: 0.94364\nPOSSIBLE N-term signal sequence\ninside 1 19\nTMhelix 20 39\noutside 40 325\nTMhelix 326 348\ninside 349 372

13582	GCF_000267425.2_ASM26742v2	Escherichia coli O157:H7 str. TW14313 Proteobacteria; Gammaproteobacteria; Enterobacterales; Enterobacteriaceae; Escherichia; Escherichia coli; Escherichia coli O157
MTGDNTLIHSHGINRRDFMKLCAALAATMGLSSKAAAEMAESVTNPQRPPVIWIGAQECT	WP_000145410.1
MULTISPECIES: hydrogenase-2 small chain [Proteobacteria]	Length: 372\nNumber of predicted TMHs: 1\nExp number of AAs in TMHs: 23.72226\nExp number, first 60 AAs: 0.22038\nTotal prob of N-in: 0.12994\noutside 1 331\nTMhelix 332 354\ninside 355 372

13583	GCF_000022225.1_ASM2222v1	Escherichia coli O157:H7 str. TW14359 Proteobacteria; Gammaproteobacteria; Enterobacterales; Enterobacteriaceae; Escherichia; Escherichia coli; Escherichia coli O157
MNNEETFYQAMRRQGVTRRSFLKYCSLAATSLGLGAGMAPKIAWALENKPRIPVVWIHGL	WP_001301613.1
hydrogenase [Escherichia coli]	Length: 372\nNumber of predicted TMHs: 2\nExp number of AAs in TMHs: 37.46598\nExp number, first 60 AAs: 19.53139\nTotal prob of N-in: 0.94364\nPOSSIBLE N-term signal sequence\ninside 1 19\nTMhelix 20 39\noutside 40 325\nTMhelix 326 348\ninside 349 372

13584	GCF_000022225.1_ASM2222v1	Escherichia coli O157:H7 str. TW14359 Proteobacteria; Gammaproteobacteria; Enterobacterales; Enterobacteriaceae; Escherichia; Escherichia coli; Escherichia coli O157
MTGDNTLIHSHGINRRDFMKLCAALAATMGLSSKAAAEMAESVTNPQRPPVIWIGAQECT	WP_000145410.1
MULTISPECIES: hydrogenase-2 small chain [Proteobacteria]	Length: 372\nNumber of predicted TMHs: 1\nExp number of AAs in TMHs: 23.72226\nExp number, first 60 AAs: 0.22038\nTotal prob of N-in: 0.12994\noutside 1 331\nTMhelix 332 354\ninside 355 372

13585	GCF_000155125.1_ASM15512v1	Escherichia coli O157:H7 str. TW14588 Proteobacteria; Gammaproteobacteria; Enterobacterales; Enterobacteriaceae; Escherichia; Escherichia coli; Escherichia coli O157
MTGDNTLIHSHGINRRDFMKLCAALAATMGLSSKAAAEMAESVTNPQRPPVIWIGAQECT	WP_000145410.1
MULTISPECIES: hydrogenase-2 small chain [Proteobacteria]	Length: 372\nNumber of predicted TMHs: 1\nExp number of AAs in TMHs: 23.72226\nExp number, first 60 AAs: 0.22038\nTotal prob of N-in: 0.12994\noutside 1 331\nTMhelix 332 354\ninside 355 372

13586	GCF_000155125.1_ASM15512v1	Escherichia coli O157:H7 str. TW14588 Proteobacteria; Gammaproteobacteria; Enterobacterales; Enterobacteriaceae; Escherichia; Escherichia coli; Escherichia coli O157
MNNEETFYQAMRRQGVTRRSFLKYCSLAATSLGLGAGMAPKIAWALENKPRIPVVWIHGL	WP_001301613.1
hydrogenase [Escherichia coli]	Length: 372\nNumber of predicted TMHs: 2\nExp number of AAs in TMHs: 37.46598\nExp number, first 60 AAs: 19.53139\nTotal prob of N-in: 0.94364\nPOSSIBLE N-term signal sequence\ninside 1 19\nTMhelix 20 39\noutside 40 325\nTMhelix 326 348\ninside 349 372

13587	GCF_000187305.1_ASM18730v2	Escherichia coli O157:H- str. 493-89	Proteobacteria; Gammaproteobacteria; Enterobacterales; Enterobacteriaceae; Escherichia; Escherichia coli; Escherichia coli O157
MNNEETFYQAMRRQGVTRRSFLKYCSLAATSLGLGAGMAPKIAWALENKPRIPVVWIHGL	WP_001301613.1
hydrogenase [Escherichia coli]	Length: 372\nNumber of predicted TMHs: 2\nExp number of AAs in TMHs: 37.46598\nExp number, first 60 AAs: 19.53139\nTotal prob of N-in: 0.94364\nPOSSIBLE N-term signal sequence\ninside 1 19\nTMhelix 20 39\noutside 40 325\nTMhelix 326 348\ninside 349 372

13588	GCF_000187305.1_ASM18730v2	Escherichia coli O157:H- str. 493-89	Proteobacteria; Gammaproteobacteria; Enterobacterales; Enterobacteriaceae; Escherichia; Escherichia coli; Escherichia coli O157
MTGDNTLIHSHGINRRDFMKLCAALAATMGLSSKAAAEMAESVTNPQRPPVIWIGAQECT	WP_000145410.1
MULTISPECIES: hydrogenase-2 small chain [Proteobacteria]	Length: 372\nNumber of predicted TMHs:

1\nExp number of AAs in TMHs: 23.72226\nExp number, first 60 AAs: 0.22038\nTotal prob of N-in: 0.12994\noutside 1 331\nTMhelix 332 354\ninside 355 372

13589 GCF_000234215.1_ASM23421v2 Escherichia coli O157:H- str. 493-89 Proteobacteria; Gammaproteobacteria; Enterobacterales; Enterobacteriaceae; Escherichia; Escherichia coli; Escherichia coli O157 MTGDNTLIHSHGINRRDFMKLCAALAATMGLSSKAAAEMAESVTNPQRPPVIWIGAQECT WP_000145410.1 MULTISPECIES: hydrogenase-2 small chain [Proteobacteria] Length: 372\nNumber of predicted TMHs: 1\nExp number of AAs in TMHs: 23.72226\nExp number, first 60 AAs: 0.22038\nTotal prob of N-in: 0.12994\noutside 1 331\nTMhelix 332 354\ninside 355 372

13590 GCF_000234215.1_ASM23421v2 Escherichia coli O157:H- str. 493-89 Proteobacteria; Gammaproteobacteria; Enterobacterales; Enterobacteriaceae; Escherichia; Escherichia coli; Escherichia coli O157 MNNEETFYQAMRRQGVTRRSFLKYCSLAATSLGLGAGMAPKIAWALENKPRIPVVWIHGL WP_001301613.1 hydrogenase [Escherichia coli] Length: 372\nNumber of predicted TMHs: 2\nExp number of AAs in TMHs: 37.46598\nExp number, first 60 AAs: 19.53139\nTotal prob of N-in: 0.94364\nPOSSIBLE N-term signal sequence\ninside 1 19\nTMhelix 20 39\noutside 40 325\nTMhelix 326 348\ninside 349 372

13591 GCF_000187325.1_ASM18732v2 Escherichia coli O157:H- str. H 2687 Proteobacteria; Gammaproteobacteria; Enterobacterales; Enterobacteriaceae; Escherichia; Escherichia coli; Escherichia coli O157 MTGDNTLIHSHGINRRDFMKLCAALAATMGLSSKAAAEMAESVTNPQRPPVIWIGAQECT WP_000145410.1 MULTISPECIES: hydrogenase-2 small chain [Proteobacteria] Length: 372\nNumber of predicted TMHs: 1\nExp number of AAs in TMHs: 23.72226\nExp number, first 60 AAs: 0.22038\nTotal prob of N-in: 0.12994\noutside 1 331\nTMhelix 332 354\ninside 355 372

13592 GCF_000187325.1_ASM18732v2 Escherichia coli O157:H- str. H 2687 Proteobacteria; Gammaproteobacteria; Enterobacterales; Enterobacteriaceae; Escherichia; Escherichia coli; Escherichia coli O157 MNNEETFYQAMRRQGVTRRSFLKYCSLAATSLGLGAGMAPKIAWALENKPRIPVVWIHGL WP_001301613.1 hydrogenase [Escherichia coli] Length: 372\nNumber of predicted TMHs: 2\nExp number of AAs in TMHs: 37.46598\nExp number, first 60 AAs: 19.53139\nTotal prob of N-in: 0.94364\nPOSSIBLE N-term signal sequence\ninside 1 19\nTMhelix 20 39\noutside 40 325\nTMhelix 326 348\ninside 349 372

13593 GCF_000619685.2_Ec08-4540 Escherichia coli O157:NM str. 08-4540 Proteobacteria; Gammaproteobacteria; Enterobacterales; Enterobacteriaceae; Escherichia; Escherichia coli; Escherichia coli O157 MTGDNTLIHSHGINRRDFMKLCAALAATMGLSSKAAAEMAESVTNPQRPPVIWIGAQECT WP_000145410.1 MULTISPECIES: hydrogenase-2 small chain [Proteobacteria] Length: 372\nNumber of predicted TMHs: 1\nExp number of AAs in TMHs: 23.72226\nExp number, first 60 AAs: 0.22038\nTotal prob of N-in: 0.12994\noutside 1 331\nTMhelix 332 354\ninside 355 372

13594 GCF_000619685.2_Ec08-4540 Escherichia coli O157:NM str. 08-4540 Proteobacteria; Gammaproteobacteria; Enterobacterales; Enterobacteriaceae; Escherichia; Escherichia coli; Escherichia coli O157 MNNEETFYQAMRRQGVTRRSFLKYCSLAATSLGLGAGMAPKIAWALENKPRIPVVWIHGL WP_001301613.1 hydrogenase [Escherichia coli] Length: 372\nNumber of predicted TMHs: 2\nExp number of AAs in TMHs: 37.46598\nExp number, first 60 AAs: 19.53139\nTotal prob of N-in: 0.94364\nPOSSIBLE N-term signal sequence\ninside 1 19\nTMhelix 20 39\noutside 40 325\nTMhelix 326 348\ninside 349 372

13595 GCF_000632555.1_Ec2010EL-2044 Escherichia coli O157: str. 2010EL-2044 Proteobacteria; Gammaproteobacteria; Enterobacterales; Enterobacteriaceae; Escherichia; Escherichia coli; Escherichia coli O157 MNNEETFYQAMRRQGVTRRSFLKYCSLAATSLGLGAGMAPKIAWALENKPRIPVVWIHGL WP_001301613.1 hydrogenase [Escherichia coli] Length: 372\nNumber of predicted TMHs: 2\nExp number of AAs in TMHs: 37.46598\nExp number, first 60 AAs: 19.53139\nTotal prob of N-in: 0.94364\nPOSSIBLE N-term signal sequence\ninside 1 19\nTMhelix 20 39\noutside 40 325\nTMhelix 326 348\ninside 349 372

13596 GCF_000632555.1_Ec2010EL-2044 Escherichia coli O157: str. 2010EL-2044 Proteobacteria; Gammaproteobacteria; Enterobacterales; Enterobacteriaceae; Escherichia; Escherichia coli; Escherichia coli O157 MTGDNTLIHSHGINRRDFMKLCAALAATMGLSSKAAAEMAESVTNPQRPPVIWIGAQECT WP_000145410.1 MULTISPECIES: hydrogenase-2 small chain [Proteobacteria] Length: 372\nNumber of predicted TMHs: 1\nExp number of AAs in TMHs: 23.72226\nExp number, first 60 AAs: 0.22038\nTotal prob of N-in: 0.12994\noutside 1 331\nTMhelix 332 354\ninside 355 372

13597 GCF_000632655.1_Ec2010EL-2045 Escherichia coli O157: str. 2010EL-2045 Proteobacteria; Gammaproteobacteria; Enterobacterales; Enterobacteriaceae; Escherichia; Escherichia coli; Escherichia coli O157

MNNEETFYQAMRRQGVTRRSFLKYCSLAATSLGLGAGMAPKIAWALENKPRIPVVWIHGL WP_001301613.1 hydrogenase [Escherichia coli] Length: 372\nNumber of predicted TMHs: 2\nExp number of AAs in TMHs: 37.46598\nExp number, first 60 AAs: 19.53139\nTotal prob of N-in: 0.94364\nPOSSIBLE N-term signal sequence\ninside 1 19\nTMhelix 20 39\noutside 40 325\nTMhelix 326 348\ninside 349 372

13598 GCF_000632655.1_Ec2010EL-2045 Escherichia coli O157: str. 2010EL-2045 Proteobacteria; Gammaproteobacteria; Enterobacterales; Enterobacteriaceae; Escherichia; Escherichia coli; Escherichia coli O157 MTGDNTLIHSHGINRRDFMKLCAALAATMGLSSKAAAEMAESVTNPQRPPVIWIGAQECT WP_000145410.1 MULTISPECIES: hydrogenase-2 small chain [Proteobacteria] Length: 372\nNumber of predicted TMHs: 1\nExp number of AAs in TMHs: 23.72226\nExp number, first 60 AAs: 0.22038\nTotal prob of N-in: 0.12994\noutside 1 331\nTMhelix 332 354\ninside 355 372

13599 GCF_000392575.1_TG1203D3338 Escherichia coli O157 str. NCCP15738 Proteobacteria; Gammaproteobacteria; Enterobacterales; Enterobacteriaceae; Escherichia; Escherichia coli; Escherichia coli O157 MNNEETFYQAMRRQGVTRRSFLKYCSLAATSLGLGAGMAPKIAWALENKPRIPVVWIHGL WP_001058323.1 MULTISPECIES: hydrogenase-1 small chain [Proteobacteria] Length: 372\nNumber of predicted TMHs: 2\nExp number of AAs in TMHs: 37.2608499999999\nExp number, first 60 AAs: 19.52923\nTotal prob of N-in: 0.94351\nPOSSIBLE N-term signal sequence\ninside 1 19\nTMhelix 20 39\noutside 40 325\nTMhelix 326 348\ninside 349 372

13600 GCF_000392575.1_TG1203D3338 Escherichia coli O157 str. NCCP15738 Proteobacteria; Gammaproteobacteria; Enterobacterales; Enterobacteriaceae; Escherichia; Escherichia coli; Escherichia coli O157 MTGDNTLIHSHGINRRDFMKLCAALAATMGLSSKAAAEMAESVTNPQRPPVIWIGAQECT WP_020218970.1 hydrogenase-2 small chain [Escherichia coli] Length: 372\nNumber of predicted TMHs: 1\nExp number of AAs in TMHs: 23.7222\nExp number, first 60 AAs: 0.22035\nTotal prob of N-in: 0.12994\noutside 1 331\nTMhelix 332 354\ninside 355 372

13601 GCF_000392555.1_TG1203D3337 Escherichia coli O157 str. NCCP15739 Proteobacteria; Gammaproteobacteria; Enterobacterales; Enterobacteriaceae; Escherichia; Escherichia coli; Escherichia coli O157 MTGDNTLIHSHGINRRDFMKLCAALAATMGLSSKAAAEMAESVTNPQRPPVIWIGAQECT WP_000145410.1 MULTISPECIES: hydrogenase-2 small chain [Proteobacteria] Length: 372\nNumber of predicted TMHs: 1\nExp number of AAs in TMHs: 23.72226\nExp number, first 60 AAs: 0.22038\nTotal prob of N-in: 0.12994\noutside 1 331\nTMhelix 332 354\ninside 355 372

13602 GCF_000392555.1_TG1203D3337 Escherichia coli O157 str. NCCP15739 Proteobacteria; Gammaproteobacteria; Enterobacterales; Enterobacteriaceae; Escherichia; Escherichia coli; Escherichia coli O157 MNNEETFYQAMRRQGVTRRSFLKYCSLAATSLGLGAGMAPKIAWALENKPRIPVVWIHGL WP_001301613.1 hydrogenase [Escherichia coli] Length: 372\nNumber of predicted TMHs: 2\nExp number of AAs in TMHs: 37.46598\nExp number, first 60 AAs: 19.53139\nTotal prob of N-in: 0.94364\nPOSSIBLE N-term signal sequence\ninside 1 19\nTMhelix 20 39\noutside 40 325\nTMhelix 326 348\ninside 349 372

13603 GCF_000335415.2_ASM33541v2 Escherichia coli PA11 Proteobacteria; Gammaproteobacteria; Enterobacterales; Enterobacteriaceae; Escherichia; Escherichia coli; Escherichia coli O157 MNNEETFYQAMRRQGVTRRSFLKYCSLAATSLGLGAGMAPKIAWALENKPRIPVVWIHGL WP_001301613.1 hydrogenase [Escherichia coli] Length: 372\nNumber of predicted TMHs: 2\nExp number of AAs in TMHs: 37.46598\nExp number, first 60 AAs: 19.53139\nTotal prob of N-in: 0.94364\nPOSSIBLE N-term signal sequence\ninside 1 19\nTMhelix 20 39\noutside 40 325\nTMhelix 326 348\ninside 349 372

13604 GCF_000335415.2_ASM33541v2 Escherichia coli PA11 Proteobacteria; Gammaproteobacteria; Enterobacterales; Enterobacteriaceae; Escherichia; Escherichia coli; Escherichia coli O157 MTGDNTLIHSHGINRRDFMKLCAALAATMGLSSKAAAEMAESVTNPQRPPVIWIGAQECT WP_000145410.1 MULTISPECIES: hydrogenase-2 small chain [Proteobacteria] Length: 372\nNumber of predicted TMHs: 1\nExp number of AAs in TMHs: 23.72226\nExp number, first 60 AAs: 0.22038\nTotal prob of N-in: 0.12994\noutside 1 331\nTMhelix 332 354\ninside 355 372

13605 GCF_000335435.2_ASM33543v2 Escherichia coli PA13 Proteobacteria; Gammaproteobacteria; Enterobacterales; Enterobacteriaceae; Escherichia; Escherichia coli; Escherichia coli O157 MTGDNTLIHSHGINRRDFMKLCAALAATMGLSSKAAAEMAESVTNPQRPPVIWIGAQECT WP_000145410.1 MULTISPECIES: hydrogenase-2 small chain [Proteobacteria] Length: 372\nNumber of predicted TMHs: 1\nExp number of AAs in TMHs: 23.72226\nExp number, first 60 AAs: 0.22038\nTotal prob of N-in: 0.12994\noutside 1 331\nTMhelix 332 354\ninside 355 372

13606 GCF_000335435.2_ASM33543v2 Escherichia coli PA13 Proteobacteria; Gammaproteobacteria; Enterobacterales; Enterobacteriaceae; Escherichia; Escherichia coli; Escherichia coli O157
MNNEETFYQAMRRQGVTRRSFLKYCSLAATSLGLGAGMAPKIAWALENKPRIPVVWIHGL WP_001301613.1
hydrogenase [Escherichia coli] Length: 372\nNumber of predicted TMHs: 2\nExp number of AAs in TMHs: 37.46598\nExp number, first 60 AAs: 19.53139\nTotal prob of N-in: 0.94364\nPOSSIBLE N-term signal sequence\ninside 1 19\nTMhelix 20 39\noutside 40 325\nTMhelix 326 348\ninside 349 372

13607 GCF_000335335.2_ASM33533v2 Escherichia coli PA19 Proteobacteria; Gammaproteobacteria; Enterobacterales; Enterobacteriaceae; Escherichia; Escherichia coli; Escherichia coli O157
MNNEETFYQAMRRQGVTRRSFLKYCSLAATSLGLGAGMAPKIAWALENKPRIPVVWIHGL WP_001301613.1
hydrogenase [Escherichia coli] Length: 372\nNumber of predicted TMHs: 2\nExp number of AAs in TMHs: 37.46598\nExp number, first 60 AAs: 19.53139\nTotal prob of N-in: 0.94364\nPOSSIBLE N-term signal sequence\ninside 1 19\nTMhelix 20 39\noutside 40 325\nTMhelix 326 348\ninside 349 372

13608 GCF_000335335.2_ASM33533v2 Escherichia coli PA19 Proteobacteria; Gammaproteobacteria; Enterobacterales; Enterobacteriaceae; Escherichia; Escherichia coli; Escherichia coli O157
MTGDNTLIHSHGINRRDFMKLCAALAATMGLSSKAAAEMAESVTNPQRPPVIWIGAQECT WP_000145410.1
MULTISPECIES: hydrogenase-2 small chain [Proteobacteria] Length: 372\nNumber of predicted TMHs: 1\nExp number of AAs in TMHs: 23.72226\nExp number, first 60 AAs: 0.22038\nTotal prob of N-in: 0.12994\noutside 1 331\nTMhelix 332 354\ninside 355 372

13609 GCF_000335175.2_ASM33517v2 Escherichia coli PA35 Proteobacteria; Gammaproteobacteria; Enterobacterales; Enterobacteriaceae; Escherichia; Escherichia coli; Escherichia coli O157
MNNEETFYQAMRRQGVTRRSFLKYCSLAATSLGLGAGMAPKIAWALENKPRIPVVWIHGL WP_001301613.1
hydrogenase [Escherichia coli] Length: 372\nNumber of predicted TMHs: 2\nExp number of AAs in TMHs: 37.46598\nExp number, first 60 AAs: 19.53139\nTotal prob of N-in: 0.94364\nPOSSIBLE N-term signal sequence\ninside 1 19\nTMhelix 20 39\noutside 40 325\nTMhelix 326 348\ninside 349 372

13610 GCF_000335175.2_ASM33517v2 Escherichia coli PA35 Proteobacteria; Gammaproteobacteria; Enterobacterales; Enterobacteriaceae; Escherichia; Escherichia coli; Escherichia coli O157
MTGDNTLIHSHGINRRDFMKLCAALAATMGLSSKAAAEMAESVTNPQRPPVIWIGAQECT WP_000145410.1
MULTISPECIES: hydrogenase-2 small chain [Proteobacteria] Length: 372\nNumber of predicted TMHs: 1\nExp number of AAs in TMHs: 23.72226\nExp number, first 60 AAs: 0.22038\nTotal prob of N-in: 0.12994\noutside 1 331\nTMhelix 332 354\ninside 355 372

13611 GCF_000617585.1_Ec2010C-4874 Escherichia coli O165:H25 str. 2010C-4874 Proteobacteria; Gammaproteobacteria; Enterobacterales; Enterobacteriaceae; Escherichia; Escherichia coli; Escherichia coli O165
MNNEETFYQAMRRQGVTRRSFLKYCSLAATSLGLGAGMAPKIAWALENKPRIPVVWIHGL WP_001058323.1
MULTISPECIES: hydrogenase-1 small chain [Proteobacteria] Length: 372\nNumber of predicted TMHs: 2\nExp number of AAs in TMHs: 37.2608499999999\nExp number, first 60 AAs: 19.52923\nTotal prob of N-in: 0.94351\nPOSSIBLE N-term signal sequence\ninside 1 19\nTMhelix 20 39\noutside 40 325\nTMhelix 326 348\ninside 349 372

13612 GCF_000617585.1_Ec2010C-4874 Escherichia coli O165:H25 str. 2010C-4874 Proteobacteria; Gammaproteobacteria; Enterobacterales; Enterobacteriaceae; Escherichia; Escherichia coli; Escherichia coli O165
MTGDNTLIHSHGINRRDFMKLCAALAATMGLSSKAAAEMAESVTNPQRPPVIWIGAQECT WP_000145410.1
MULTISPECIES: hydrogenase-2 small chain [Proteobacteria] Length: 372\nNumber of predicted TMHs: 1\nExp number of AAs in TMHs: 23.72226\nExp number, first 60 AAs: 0.22038\nTotal prob of N-in: 0.12994\noutside 1 331\nTMhelix 332 354\ninside 355 372

13613 GCF_000618205.2_EcF9792 Escherichia coli O169:H41 str. F9792 Proteobacteria; Gammaproteobacteria; Enterobacterales; Enterobacteriaceae; Escherichia; Escherichia coli; Escherichia coli O169
MTGDNTLIHSHGINRRDFMKLCAALAATMGLSSKAAAEMAESVTNPQRPPVIWIGAQECT WP_000145410.1
MULTISPECIES: hydrogenase-2 small chain [Proteobacteria] Length: 372\nNumber of predicted TMHs: 1\nExp number of AAs in TMHs: 23.72226\nExp number, first 60 AAs: 0.22038\nTotal prob of N-in: 0.12994\noutside 1 331\nTMhelix 332 354\ninside 355 372

13614 GCF_000618205.2_EcF9792 Escherichia coli O169:H41 str. F9792 Proteobacteria; Gammaproteobacteria; Enterobacterales; Enterobacteriaceae; Escherichia; Escherichia coli; Escherichia coli O169
MNNEETFYQAMRRQGVTRRSFLKYCSLAATSLGLGAGMAPKIAWALENKPRIPVVWIHGL WP_001717971.1
hydrogenase-1 small chain [Escherichia coli] Length: 372\nNumber of predicted TMHs: 2\nExp number of

AAs in TMHs: 38.00708\nExp number, first 60 AAs:  19.5286\nTotal prob of N-in:   0.94354\nPOSSIBLE N-term signal sequence\ninside   1   19\nTMhelix   20   39\noutside   40   325\nTMhelix   326   348\ninside   349   372

13615 GCF_000617205.1_Ec03-3269 Escherichia coli O174:H21 str. 03-3269 Proteobacteria; Gammaproteobacteria; Enterobacterales; Enterobacteriaceae; Escherichia; Escherichia coli; Escherichia coli O174 MTGDNTLIHSHGINRRDFMKLCAALAATMGLSSKAAAEMAESVTNPQRPPVIWIGAQECT WP_000145410.1 MULTISPECIES: hydrogenase-2 small chain [Proteobacteria] Length: 372\nNumber of predicted TMHs: 1\nExp number of AAs in TMHs: 23.72226\nExp number, first 60 AAs:  0.22038\nTotal prob of N-in:   0.12994\noutside   1   331\nTMhelix   332   354\ninside   355   372

13616 GCF_000617205.1_Ec03-3269 Escherichia coli O174:H21 str. 03-3269 Proteobacteria; Gammaproteobacteria; Enterobacterales; Enterobacteriaceae; Escherichia; Escherichia coli; Escherichia coli O174 MNNEETFYQAMRRQGVTRRSFLKYCSLAATSLGLGAGMAPKIAWALENKPRIPVVWIHGL WP_001058323.1 MULTISPECIES: hydrogenase-1 small chain [Proteobacteria] Length: 372\nNumber of predicted TMHs: 2\nExp number of AAs in TMHs: 37.2608499999999\nExp number, first 60 AAs:  19.52923\nTotal prob of N-in: 0.94351\nPOSSIBLE N-term signal sequence\ninside   1   19\nTMhelix   20   39\noutside   40   325\nTMhelix   326 348\ninside   349   372

13617 GCF_000617085.1_Ec04-3038 Escherichia coli O174:H8 str. 04-3038 Proteobacteria; Gammaproteobacteria; Enterobacterales; Enterobacteriaceae; Escherichia; Escherichia coli; Escherichia coli O174 MTGDNTLIHSHGINRRDFMKLCAALAATMGLSSKAAAEMAESVTNPQRPPVIWIGAQECT WP_000145410.1 MULTISPECIES: hydrogenase-2 small chain [Proteobacteria] Length: 372\nNumber of predicted TMHs: 1\nExp number of AAs in TMHs: 23.72226\nExp number, first 60 AAs:  0.22038\nTotal prob of N-in:   0.12994\noutside   1 331\nTMhelix   332   354\ninside   355   372

13618 GCF_000617085.1_Ec04-3038 Escherichia coli O174:H8 str. 04-3038 Proteobacteria; Gammaproteobacteria; Enterobacterales; Enterobacteriaceae; Escherichia; Escherichia coli; Escherichia coli O174 MNNEETFYQAMRRQGVTRRSFLKYCSLAATSLGLGAGMAPKIAWALENKPRIPVVWIHGL WP_001058323.1 MULTISPECIES: hydrogenase-1 small chain [Proteobacteria] Length: 372\nNumber of predicted TMHs: 2\nExp number of AAs in TMHs: 37.2608499999999\nExp number, first 60 AAs:  19.52923\nTotal prob of N-in: 0.94351\nPOSSIBLE N-term signal sequence\ninside   1   19\nTMhelix   20   39\noutside   40   325\nTMhelix   326 348\ninside   349   372

13619 GCF_000647455.1_02-04446_clc_bio_denovo Escherichia coli STEC O174:H2 str. 02-04446 Proteobacteria; Gammaproteobacteria; Enterobacterales; Enterobacteriaceae; Escherichia; Escherichia coli; Escherichia coli O174 MTGDNTLIHSHGINRRDFMKLCAALAATMGLSSKAAAEMAESVTNPQRPPVIWIGAQECT WP_000145410.1 MULTISPECIES: hydrogenase-2 small chain [Proteobacteria] Length: 372\nNumber of predicted TMHs:  1\nExp number of AAs in TMHs: 23.72226\nExp number, first 60 AAs:  0.22038\nTotal prob of N-in: 0.12994\noutside   1   331\nTMhelix   332   354\ninside   355   372

13620 GCF_000647455.1_02-04446_clc_bio_denovo Escherichia coli STEC O174:H2 str. 02-04446 Proteobacteria; Gammaproteobacteria; Enterobacterales; Enterobacteriaceae; Escherichia; Escherichia coli; Escherichia coli O174 MNNEETFYQAMRRQGVTRRSFLKYCSLAATSLGLGAGMAPKIAWALENKPRIPVVWIHGL WP_001058323.1 MULTISPECIES: hydrogenase-1 small chain [Proteobacteria] Length: 372\nNumber of predicted TMHs:  2\nExp number of AAs in TMHs: 37.2608499999999\nExp number, first 60 AAs:  19.52923\nTotal prob of N-in:   0.94351\nPOSSIBLE N-term signal sequence\ninside   1   19\nTMhelix   20   39\noutside   40   325\nTMhelix   326 348\ninside   349   372

13621 GCF_000647475.1_I-151_clc_bio_denovo Escherichia coli STEC O174:H46 str. I-151 Proteobacteria; Gammaproteobacteria; Enterobacterales; Enterobacteriaceae; Escherichia; Escherichia coli; Escherichia coli O174 MNNEETFYQAMRRQGVTRRSFLKYCSLAATSLGLGAGMAPKIAWALENKPRIPVVWIHGL WP_001058323.1 MULTISPECIES: hydrogenase-1 small chain [Proteobacteria] Length: 372\nNumber of predicted TMHs:  2\nExp number of AAs in TMHs: 37.2608499999999\nExp number, first 60 AAs:  19.52923\nTotal prob of N-in:   0.94351\nPOSSIBLE N-term signal sequence\ninside   1   19\nTMhelix   20   39\noutside   40   325\nTMhelix   326 348\ninside   349   372

13622 GCF_000647475.1_I-151_clc_bio_denovo Escherichia coli STEC O174:H46 str. I-151 Proteobacteria; Gammaproteobacteria; Enterobacterales; Enterobacteriaceae; Escherichia; Escherichia coli; Escherichia coli O174 MTGDNTLIHSHGINRRDFMKLCAALAATMGLSSKAAAEMAESVTNPQRPPVIWIGAQECT WP_028131934.1 hydrogenase-2 small chain [Escherichia coli] Length: 372\nNumber of predicted TMHs:

1\nExp number of AAs in TMHs: 24.01565\nExp number, first 60 AAs: 0.22022\nTotal prob of N-in: 0.13096\noutside 1 331\nTMhelix 332 354\ninside 355 372

13623 GCF_000647495.1_02-07607_clc_bio_denovo Escherichia coli STEC O174:H8 str. 02-07607 Proteobacteria; Gammaproteobacteria; Enterobacterales; Enterobacteriaceae; Escherichia; Escherichia coli; Escherichia coli O174 MTGDNTLIHSHGINRRDFMKLCAALAATMGLSSKAAAEMAESVTNPQRPPVIWIGAQECT WP_000145410.1 MULTISPECIES: hydrogenase-2 small chain [Proteobacteria] Length: 372\nNumber of predicted TMHs: 1\nExp number of AAs in TMHs: 23.72226\nExp number, first 60 AAs: 0.22038\nTotal prob of N-in: 0.12994\noutside 1 331\nTMhelix 332 354\ninside 355 372

13624 GCF_000647495.1_02-07607_clc_bio_denovo Escherichia coli STEC O174:H8 str. 02-07607 Proteobacteria; Gammaproteobacteria; Enterobacterales; Enterobacteriaceae; Escherichia; Escherichia coli; Escherichia coli O174 MNNEETFYQAMRRQGVTRRSFLKYCSLAATSLGLGAGMAPKIAWALENKPRIPVVWIHGL WP_001058323.1 MULTISPECIES: hydrogenase-1 small chain [Proteobacteria] Length: 372\nNumber of predicted TMHs: 2\nExp number of AAs in TMHs: 37.2608499999999\nExp number, first 60 AAs: 19.52923\nTotal prob of N-in: 0.94351\nPOSSIBLE N-term signal sequence\ninside 1 19\nTMhelix 20 39\noutside 40 325\nTMhelix 326 348\ninside 349 372

13625 GCF_000617785.1_Ec2010C-4558 Escherichia coli O177:NM str. 2010C-4558 Proteobacteria; Gammaproteobacteria; Enterobacterales; Enterobacteriaceae; Escherichia; Escherichia coli; Escherichia coli O177 MTGDNTLIHSHGINRRDFMKLCAALAATMGLSSKAAAEMAESVTNPQRPPVIWIGAQECT WP_000145410.1 MULTISPECIES: hydrogenase-2 small chain [Proteobacteria] Length: 372\nNumber of predicted TMHs: 1\nExp number of AAs in TMHs: 23.72226\nExp number, first 60 AAs: 0.22038\nTotal prob of N-in: 0.12994\noutside 1 331\nTMhelix 332 354\ninside 355 372

13626 GCF_000617785.1_Ec2010C-4558 Escherichia coli O177:NM str. 2010C-4558 Proteobacteria; Gammaproteobacteria; Enterobacterales; Enterobacteriaceae; Escherichia; Escherichia coli; Escherichia coli O177 MNNEETFYQAMRRQGVTRRSFLKYCSLAATSLGLGAGMAPKIAWALENKPRIPVVWIHGL WP_032204866.1 hydrogenase [Escherichia coli] Length: 372\nNumber of predicted TMHs: 2\nExp number of AAs in TMHs: 37.2621299999999\nExp number, first 60 AAs: 19.52923\nTotal prob of N-in: 0.94351\nPOSSIBLE N-term signal sequence\ninside 1 19\nTMhelix 20 39\noutside 40 325\nTMhelix 326 348\ninside 349 372

13627 GCF_000618405.2_EcE2539C1 Escherichia coli O25:NM str. E2539C1 Proteobacteria; Gammaproteobacteria; Enterobacterales; Enterobacteriaceae; Escherichia; Escherichia coli; Escherichia coli O25 MNNEETFYQAMRRQGVTRRSFLKYCSLAATSLGLGAGMAPKIAWALENKPRIPVVWIHGL WP_001058323.1 MULTISPECIES: hydrogenase-1 small chain [Proteobacteria] Length: 372\nNumber of predicted TMHs: 2\nExp number of AAs in TMHs: 37.2608499999999\nExp number, first 60 AAs: 19.52923\nTotal prob of N-in: 0.94351\nPOSSIBLE N-term signal sequence\ninside 1 19\nTMhelix 20 39\noutside 40 325\nTMhelix 326 348\ninside 349 372

13628 GCF_000618405.2_EcE2539C1 Escherichia coli O25:NM str. E2539C1 Proteobacteria; Gammaproteobacteria; Enterobacterales; Enterobacteriaceae; Escherichia; Escherichia coli; Escherichia coli O25 MTGDNTLIHSHGINRRDFMKLCAALAATMGLSSKAAAEMAESVTNPQRPPVIWIGAQECT WP_000145410.1 MULTISPECIES: hydrogenase-2 small chain [Proteobacteria] Length: 372\nNumber of predicted TMHs: 1\nExp number of AAs in TMHs: 23.72226\nExp number, first 60 AAs: 0.22038\nTotal prob of N-in: 0.12994\noutside 1 331\nTMhelix 332 354\ninside 355 372

13629 GCF_000622445.2_Ec03-3500 Escherichia coli O26:H11 str. 03-3500 Proteobacteria; Gammaproteobacteria; Enterobacterales; Enterobacteriaceae; Escherichia; Escherichia coli; Escherichia coli O26 MTGDNTLIHSHGINRRDFMKLCAALAATMGLSSKAAAEMAESVTNPQRPPVIWIGAQECT WP_000145410.1 MULTISPECIES: hydrogenase-2 small chain [Proteobacteria] Length: 372\nNumber of predicted TMHs: 1\nExp number of AAs in TMHs: 23.72226\nExp number, first 60 AAs: 0.22038\nTotal prob of N-in: 0.12994\noutside 1 331\nTMhelix 332 354\ninside 355 372

13630 GCF_000622445.2_Ec03-3500 Escherichia coli O26:H11 str. 03-3500 Proteobacteria; Gammaproteobacteria; Enterobacterales; Enterobacteriaceae; Escherichia; Escherichia coli; Escherichia coli O26 MNNEETFYQAMRRQGVTRRSFLKYCSLAATSLGLGAGMAPKIAWALENKPRIPVVWIHGL WP_001058323.1 MULTISPECIES: hydrogenase-1 small chain [Proteobacteria] Length: 372\nNumber of predicted TMHs: 2\nExp number of AAs in TMHs: 37.2608499999999\nExp number, first 60 AAs: 19.52923\nTotal prob of N-in: 0.94351\nPOSSIBLE N-term signal sequence\ninside 1 19\nTMhelix 20 39\noutside 40 325\nTMhelix 326 348\ninside 349 372

13631 GCF_000617065.1_Ec05-3646 Escherichia coli O26:H11 str. 05-3646 Proteobacteria; Gammaproteobacteria; Enterobacterales; Enterobacteriaceae; Escherichia; Escherichia coli; Escherichia coli O26 MNNEETFYQAMRRQGVTRRSFLKYCSLAATSLGLGAGMAPKIAWALENKPRIPVVWIHGL WP_001058323.1 MULTISPECIES: hydrogenase-1 small chain [Proteobacteria] Length: 372\nNumber of predicted TMHs: 2\nExp number of AAs in TMHs: 37.2608499999999\nExp number, first 60 AAs: 19.52923\nTotal prob of N-in: 0.94351\nPOSSIBLE N-term signal sequence\ninside 1 19\nTMhelix 20 39\noutside 40 325\nTMhelix 326 348\ninside 349 372

13632 GCF_000617065.1_Ec05-3646 Escherichia coli O26:H11 str. 05-3646 Proteobacteria; Gammaproteobacteria; Enterobacterales; Enterobacteriaceae; Escherichia; Escherichia coli; Escherichia coli O26 MTGDNTLIHSHGINRRDFMKLCAALAATMGLSSKAAAEMAESVTNPQRPPVIWIGAQECT WP_000145410.1 MULTISPECIES: hydrogenase-2 small chain [Proteobacteria] Length: 372\nNumber of predicted TMHs: 1\nExp number of AAs in TMHs: 23.72226\nExp number, first 60 AAs: 0.22038\nTotal prob of N-in: 0.12994\noutside 1 331\nTMhelix 332 354\ninside 355 372

13633 GCF_000622465.2_Ec06-3464 Escherichia coli O26:H11 str. 06-3464 Proteobacteria; Gammaproteobacteria; Enterobacterales; Enterobacteriaceae; Escherichia; Escherichia coli; Escherichia coli O26 MNNEETFYQAMRRQGVTRRSFLKYCSLAATSLGLGAGMAPKIAWALENKPRIPVVWIHGL WP_001058323.1 MULTISPECIES: hydrogenase-1 small chain [Proteobacteria] Length: 372\nNumber of predicted TMHs: 2\nExp number of AAs in TMHs: 37.2608499999999\nExp number, first 60 AAs: 19.52923\nTotal prob of N-in: 0.94351\nPOSSIBLE N-term signal sequence\ninside 1 19\nTMhelix 20 39\noutside 40 325\nTMhelix 326 348\ninside 349 372

13634 GCF_000622465.2_Ec06-3464 Escherichia coli O26:H11 str. 06-3464 Proteobacteria; Gammaproteobacteria; Enterobacterales; Enterobacteriaceae; Escherichia; Escherichia coli; Escherichia coli O26 MTGDNTLIHSHGINRRDFMKLCAALAATMGLSSKAAAEMAESVTNPQRPPVIWIGAQECT WP_000145410.1 MULTISPECIES: hydrogenase-2 small chain [Proteobacteria] Length: 372\nNumber of predicted TMHs: 1\nExp number of AAs in TMHs: 23.72226\nExp number, first 60 AAs: 0.22038\nTotal prob of N-in: 0.12994\noutside 1 331\nTMhelix 332 354\ninside 355 372

13635 GCF_000091005.1_ASM9100v1 Escherichia coli O26:H11 str. 11368 Proteobacteria; Gammaproteobacteria; Enterobacterales; Enterobacteriaceae; Escherichia; Escherichia coli; Escherichia coli O26 MTGDNTLIHSHGINRRDFMKLCAALAATMGLSSKAAAEMAESVTNPQRPPVIWIGAQECT WP_000145410.1 MULTISPECIES: hydrogenase-2 small chain [Proteobacteria] Length: 372\nNumber of predicted TMHs: 1\nExp number of AAs in TMHs: 23.72226\nExp number, first 60 AAs: 0.22038\nTotal prob of N-in: 0.12994\noutside 1 331\nTMhelix 332 354\ninside 355 372

13636 GCF_000091005.1_ASM9100v1 Escherichia coli O26:H11 str. 11368 Proteobacteria; Gammaproteobacteria; Enterobacterales; Enterobacteriaceae; Escherichia; Escherichia coli; Escherichia coli O26 MNNEETFYQAMRRQGVTRRSFLKYCSLAATSLGLGAGMAPKIAWALENKPRIPVVWIHGL WP_001058323.1 MULTISPECIES: hydrogenase-1 small chain [Proteobacteria] Length: 372\nNumber of predicted TMHs: 2\nExp number of AAs in TMHs: 37.2608499999999\nExp number, first 60 AAs: 19.52923\nTotal prob of N-in: 0.94351\nPOSSIBLE N-term signal sequence\ninside 1 19\nTMhelix 20 39\noutside 40 325\nTMhelix 326 348\ninside 349 372

13637 GCF_000622875.2_Ec2009C-3612 Escherichia coli O26:H11 str. 2009C-3612 Proteobacteria; Gammaproteobacteria; Enterobacterales; Enterobacteriaceae; Escherichia; Escherichia coli; Escherichia coli O26 MTGDNTLIHSHGINRRDFMKLCAALAATMGLSSKAAAEMAESVTNPQRPPVIWIGAQECT WP_000145410.1 MULTISPECIES: hydrogenase-2 small chain [Proteobacteria] Length: 372\nNumber of predicted TMHs: 1\nExp number of AAs in TMHs: 23.72226\nExp number, first 60 AAs: 0.22038\nTotal prob of N-in: 0.12994\noutside 1 331\nTMhelix 332 354\ninside 355 372

13638 GCF_000622875.2_Ec2009C-3612 Escherichia coli O26:H11 str. 2009C-3612 Proteobacteria; Gammaproteobacteria; Enterobacterales; Enterobacteriaceae; Escherichia; Escherichia coli; Escherichia coli O26 MNNEETFYQAMRRQGVTRRSFLKYCSLAATSLGLGAGMAPKIAWALENKPRIPVVWIHGL WP_001058323.1 MULTISPECIES: hydrogenase-1 small chain [Proteobacteria] Length: 372\nNumber of predicted TMHs: 2\nExp number of AAs in TMHs: 37.2608499999999\nExp number, first 60 AAs: 19.52923\nTotal prob of N-in: 0.94351\nPOSSIBLE N-term signal sequence\ninside 1 19\nTMhelix 20 39\noutside 40 325\nTMhelix 326 348\ninside 349 372

13639 GCF_000622755.2_Ec2009C-3689 Escherichia coli O26:H11 str. 2009C-3689 Proteobacteria; Gammaproteobacteria; Enterobacterales; Enterobacteriaceae; Escherichia; Escherichia coli; Escherichia coli O26 MTGDNTLIHSHGINRRDFMKLCAALAATMGLSSKAAAEMAESVTNPQRPPVIWIGAQECT WP_000145410.1 MULTISPECIES: hydrogenase-2 small chain [Proteobacteria] Length: 372\nNumber of predicted TMHs: 1\nExp number of AAs in TMHs: 23.72226\nExp number, first 60 AAs: 0.22038\nTotal prob of N-in: 0.12994\noutside 1 331\nTMhelix 332 354\ninside 355 372

13640 GCF_000622755.2_Ec2009C-3689 Escherichia coli O26:H11 str. 2009C-3689 Proteobacteria; Gammaproteobacteria; Enterobacterales; Enterobacteriaceae; Escherichia; Escherichia coli; Escherichia coli O26 MNNEETFYQAMRRQGVTRRSFLKYCSLAATSLGLGAGMAPKIAWALENKPRIPVVWIHGL WP_001058323.1 MULTISPECIES: hydrogenase-1 small chain [Proteobacteria] Length: 372\nNumber of predicted TMHs: 2\nExp number of AAs in TMHs: 37.2608499999999\nExp number, first 60 AAs: 19.52923\nTotal prob of N-in: 0.94351\nPOSSIBLE N-term signal sequence\ninside 1 19\nTMhelix 20 39\noutside 40 325\nTMhelix 326 348\ninside 349 372

13641 GCF_000622595.2_Ec2009C-3996 Escherichia coli O26:H11 str. 2009C-3996 Proteobacteria; Gammaproteobacteria; Enterobacterales; Enterobacteriaceae; Escherichia; Escherichia coli; Escherichia coli O26 MNNEETFYQAMRRQGVTRRSFLKYCSLAATSLGLGAGMAPKIAWALENKPRIPVVWIHGL WP_001058323.1 MULTISPECIES: hydrogenase-1 small chain [Proteobacteria] Length: 372\nNumber of predicted TMHs: 2\nExp number of AAs in TMHs: 37.2608499999999\nExp number, first 60 AAs: 19.52923\nTotal prob of N-in: 0.94351\nPOSSIBLE N-term signal sequence\ninside 1 19\nTMhelix 20 39\noutside 40 325\nTMhelix 326 348\ninside 349 372

13642 GCF_000622595.2_Ec2009C-3996 Escherichia coli O26:H11 str. 2009C-3996 Proteobacteria; Gammaproteobacteria; Enterobacterales; Enterobacteriaceae; Escherichia; Escherichia coli; Escherichia coli O26 MTGDNTLIHSHGINRRDFMKLCAALAATMGLSSKAAAEMAESVTNPQRPPVIWIGAQECT WP_000145410.1 MULTISPECIES: hydrogenase-2 small chain [Proteobacteria] Length: 372\nNumber of predicted TMHs: 1\nExp number of AAs in TMHs: 23.72226\nExp number, first 60 AAs: 0.22038\nTotal prob of N-in: 0.12994\noutside 1 331\nTMhelix 332 354\ninside 355 372

13643 GCF_000622615.2_Ec2009C-4760 Escherichia coli O26:H11 str. 2009C-4760 Proteobacteria; Gammaproteobacteria; Enterobacterales; Enterobacteriaceae; Escherichia; Escherichia coli; Escherichia coli O26 MTGDNTLIHSHGINRRDFMKLCAALAATMGLSSKAAAEMAESVTNPQRPPVIWIGAQECT WP_000145410.1 MULTISPECIES: hydrogenase-2 small chain [Proteobacteria] Length: 372\nNumber of predicted TMHs: 1\nExp number of AAs in TMHs: 23.72226\nExp number, first 60 AAs: 0.22038\nTotal prob of N-in: 0.12994\noutside 1 331\nTMhelix 332 354\ninside 355 372

13644 GCF_000622615.2_Ec2009C-4760 Escherichia coli O26:H11 str. 2009C-4760 Proteobacteria; Gammaproteobacteria; Enterobacterales; Enterobacteriaceae; Escherichia; Escherichia coli; Escherichia coli O26 MNNEETFYQAMRRQGVTRRSFLKYCSLAATSLGLGAGMAPKIAWALENKPRIPVVWIHGL WP_001058323.1 MULTISPECIES: hydrogenase-1 small chain [Proteobacteria] Length: 372\nNumber of predicted TMHs: 2\nExp number of AAs in TMHs: 37.2608499999999\nExp number, first 60 AAs: 19.52923\nTotal prob of N-in: 0.94351\nPOSSIBLE N-term signal sequence\ninside 1 19\nTMhelix 20 39\noutside 40 325\nTMhelix 326 348\ninside 349 372

13645 GCF_000622635.2_Ec2009C-4826 Escherichia coli O26:H11 str. 2009C-4826 Proteobacteria; Gammaproteobacteria; Enterobacterales; Enterobacteriaceae; Escherichia; Escherichia coli; Escherichia coli O26 MNNEETFYQAMRRQGVTRRSFLKYCSLAATSLGLGAGMAPKIAWALENKPRIPVVWIHGL WP_001058323.1 MULTISPECIES: hydrogenase-1 small chain [Proteobacteria] Length: 372\nNumber of predicted TMHs: 2\nExp number of AAs in TMHs: 37.2608499999999\nExp number, first 60 AAs: 19.52923\nTotal prob of N-in: 0.94351\nPOSSIBLE N-term signal sequence\ninside 1 19\nTMhelix 20 39\noutside 40 325\nTMhelix 326 348\ninside 349 372

13646 GCF_000622635.2_Ec2009C-4826 Escherichia coli O26:H11 str. 2009C-4826 Proteobacteria; Gammaproteobacteria; Enterobacterales; Enterobacteriaceae; Escherichia; Escherichia coli; Escherichia coli O26 MTGDNTLIHSHGINRRDFMKLCAALAATMGLSSKAAAEMAESVTNPQRPPVIWIGAQECT WP_000145410.1 MULTISPECIES: hydrogenase-2 small chain [Proteobacteria] Length: 372\nNumber of predicted TMHs: 1\nExp number of AAs in TMHs: 23.72226\nExp number, first 60 AAs: 0.22038\nTotal prob of N-in: 0.12994\noutside 1 331\nTMhelix 332 354\ninside 355 372

13647 GCF_000622675.2_Ec2010C-3051 Escherichia coli O26:H11 str. 2010C-3051 Proteobacteria; Gammaproteobacteria; Enterobacterales; Enterobacteriaceae; Escherichia; Escherichia coli; Escherichia coli O26 MNNEETFYQAMRRQGVTRRSFLKYCSLAATSLGLGAGMAPKIAWALENKPRIPVVWIHGL WP_001058323.1 MULTISPECIES: hydrogenase-1 small chain [Proteobacteria] Length: 372\nNumber of predicted TMHs: 2\nExp number of AAs in TMHs: 37.2608499999999\nExp number, first 60 AAs: 19.52923\nTotal prob of N-in: 0.94351\nPOSSIBLE N-term signal sequence\ninside 1 19\nTMhelix 20 39\noutside 40 325\nTMhelix 326 348\ninside 349 372

13648 GCF_000622675.2_Ec2010C-3051 Escherichia coli O26:H11 str. 2010C-3051 Proteobacteria; Gammaproteobacteria; Enterobacterales; Enterobacteriaceae; Escherichia; Escherichia coli; Escherichia coli O26 MTGDNTLIHSHGINRRDFMKLCAALAATMGLSSKAAAEMAESVTNPQRPPVIWIGAQECT WP_000145410.1 MULTISPECIES: hydrogenase-2 small chain [Proteobacteria] Length: 372\nNumber of predicted TMHs: 1\nExp number of AAs in TMHs: 23.72226\nExp number, first 60 AAs: 0.22038\nTotal prob of N-in: 0.12994\noutside 1 331\nTMhelix 332 354\ninside 355 372

13649 GCF_000622695.2_Ec2010C-3472 Escherichia coli O26:H11 str. 2010C-3472 Proteobacteria; Gammaproteobacteria; Enterobacterales; Enterobacteriaceae; Escherichia; Escherichia coli; Escherichia coli O26 MNNEETFYQAMRRQGVTRRSFLKYCSLAATSLGLGAGMAPKIAWALENKPRIPVVWIHGL WP_001058323.1 MULTISPECIES: hydrogenase-1 small chain [Proteobacteria] Length: 372\nNumber of predicted TMHs: 2\nExp number of AAs in TMHs: 37.2608499999999\nExp number, first 60 AAs: 19.52923\nTotal prob of N-in: 0.94351\nPOSSIBLE N-term signal sequence\ninside 1 19\nTMhelix 20 39\noutside 40 325\nTMhelix 326 348\ninside 349 372

13650 GCF_000622695.2_Ec2010C-3472 Escherichia coli O26:H11 str. 2010C-3472 Proteobacteria; Gammaproteobacteria; Enterobacterales; Enterobacteriaceae; Escherichia; Escherichia coli; Escherichia coli O26 MTGDNTLIHSHGINRRDFMKLCAALAATMGLSSKAAAEMAESVTNPQRPPVIWIGAQECT WP_000145410.1 MULTISPECIES: hydrogenase-2 small chain [Proteobacteria] Length: 372\nNumber of predicted TMHs: 1\nExp number of AAs in TMHs: 23.72226\nExp number, first 60 AAs: 0.22038\nTotal prob of N-in: 0.12994\noutside 1 331\nTMhelix 332 354\ninside 355 372

13651 GCF_000622715.2_Ec2010C-3871 Escherichia coli O26:H11 str. 2010C-3871 Proteobacteria; Gammaproteobacteria; Enterobacterales; Enterobacteriaceae; Escherichia; Escherichia coli; Escherichia coli O26 MTGDNTLIHSHGINRRDFMKLCAALAATMGLSSKAAAEMAESVTNPQRPPVIWIGAQECT WP_000145410.1 MULTISPECIES: hydrogenase-2 small chain [Proteobacteria] Length: 372\nNumber of predicted TMHs: 1\nExp number of AAs in TMHs: 23.72226\nExp number, first 60 AAs: 0.22038\nTotal prob of N-in: 0.12994\noutside 1 331\nTMhelix 332 354\ninside 355 372

13652 GCF_000622715.2_Ec2010C-3871 Escherichia coli O26:H11 str. 2010C-3871 Proteobacteria; Gammaproteobacteria; Enterobacterales; Enterobacteriaceae; Escherichia; Escherichia coli; Escherichia coli O26 MNNEETFYQAMRRQGVTRRSFLKYCSLAATSLGLGAGMAPKIAWALENKPRIPVVWIHGL WP_001058323.1 MULTISPECIES: hydrogenase-1 small chain [Proteobacteria] Length: 372\nNumber of predicted TMHs: 2\nExp number of AAs in TMHs: 37.2608499999999\nExp number, first 60 AAs: 19.52923\nTotal prob of N-in: 0.94351\nPOSSIBLE N-term signal sequence\ninside 1 19\nTMhelix 20 39\noutside 40 325\nTMhelix 326 348\ninside 349 372

13653 GCF_000622735.2_Ec2010C-3902 Escherichia coli O26:H11 str. 2010C-3902 Proteobacteria; Gammaproteobacteria; Enterobacterales; Enterobacteriaceae; Escherichia; Escherichia coli; Escherichia coli O26 MTGDNTLIHSHGINRRDFMKLCAALAATMGLSSKAAAEMAESVTNPQRPPVIWIGAQECT WP_000145410.1 MULTISPECIES: hydrogenase-2 small chain [Proteobacteria] Length: 372\nNumber of predicted TMHs: 1\nExp number of AAs in TMHs: 23.72226\nExp number, first 60 AAs: 0.22038\nTotal prob of N-in: 0.12994\noutside 1 331\nTMhelix 332 354\ninside 355 372

13654 GCF_000622735.2_Ec2010C-3902 Escherichia coli O26:H11 str. 2010C-3902 Proteobacteria; Gammaproteobacteria; Enterobacterales; Enterobacteriaceae; Escherichia; Escherichia coli; Escherichia coli O26 MNNEETFYQAMRRQGVTRRSFLKYCSLAATSLGLGAGMAPKIAWALENKPRIPVVWIHGL WP_001058323.1 MULTISPECIES: hydrogenase-1 small chain [Proteobacteria] Length: 372\nNumber of predicted TMHs: 2\nExp number of AAs in TMHs: 37.2608499999999\nExp number, first 60 AAs: 19.52923\nTotal prob of N-in: 0.94351\nPOSSIBLE N-term signal sequence\ninside 1 19\nTMhelix 20 39\noutside 40 325\nTMhelix 326 348\ninside 349 372

13655 GCF_000622575.2_Ec2010C-4244 Escherichia coli O26:H11 str. 2010C-4244 Proteobacteria; Gammaproteobacteria; Enterobacterales; Enterobacteriaceae; Escherichia; Escherichia coli; Escherichia coli O26 MTGDNTLIHSHGINRRDFMKLCAALAATMGLSSKAAAEMAESVTNPQRPPVIWIGAQECT WP_000145410.1 MULTISPECIES: hydrogenase-2 small chain [Proteobacteria] Length: 372\nNumber of predicted TMHs: 1\nExp number of AAs in TMHs: 23.72226\nExp number, first 60 AAs: 0.22038\nTotal prob of N-in: 0.12994\noutside 1 331\nTMhelix 332 354\ninside 355 372

13656 GCF_000622575.2_Ec2010C-4244 Escherichia coli O26:H11 str. 2010C-4244 Proteobacteria; Gammaproteobacteria; Enterobacterales; Enterobacteriaceae; Escherichia; Escherichia coli; Escherichia coli O26 MNNEETFYQAMRRQGVTRRSFLKYCSLAATSLGLGAGMAPKIAWALENKPRIPVVWIHGL WP_001058323.1 MULTISPECIES: hydrogenase-1 small chain [Proteobacteria] Length: 372\nNumber of predicted TMHs: 2\nExp number of AAs in TMHs: 37.2608499999999\nExp number, first 60 AAs: 19.52923\nTotal prob of N-in: 0.94351\nPOSSIBLE N-term signal sequence\ninside 1 19\nTMhelix 20 39\noutside 40 325\nTMhelix 326 348\ninside 349 372

13657 GCF_000622505.2_Ec2010C-4430 Escherichia coli O26:H11 str. 2010C-4430 Proteobacteria; Gammaproteobacteria; Enterobacterales; Enterobacteriaceae; Escherichia; Escherichia coli; Escherichia coli O26 MNNEETFYQAMRRQGVTRRSFLKYCSLAATSLGLGAGMAPKIAWALENKPRIPVVWIHGL WP_001058323.1 MULTISPECIES: hydrogenase-1 small chain [Proteobacteria] Length: 372\nNumber of predicted TMHs: 2\nExp number of AAs in TMHs: 37.2608499999999\nExp number, first 60 AAs: 19.52923\nTotal prob of N-in: 0.94351\nPOSSIBLE N-term signal sequence\ninside 1 19\nTMhelix 20 39\noutside 40 325\nTMhelix 326 348\ninside 349 372

13658 GCF_000622505.2_Ec2010C-4430 Escherichia coli O26:H11 str. 2010C-4430 Proteobacteria; Gammaproteobacteria; Enterobacterales; Enterobacteriaceae; Escherichia; Escherichia coli; Escherichia coli O26 MTGDNTLIHSHGINRRDFMKLCAALAATMGLSSKAAAEMAESVTNPQRPPVIWIGAQECT WP_000145410.1 MULTISPECIES: hydrogenase-2 small chain [Proteobacteria] Length: 372\nNumber of predicted TMHs: 1\nExp number of AAs in TMHs: 23.72226\nExp number, first 60 AAs: 0.22038\nTotal prob of N-in: 0.12994\noutside 1 331\nTMhelix 332 354\ninside 355 372

13659 GCF_000622485.2_Ec2010C-4819 Escherichia coli O26:H11 str. 2010C-4819 Proteobacteria; Gammaproteobacteria; Enterobacterales; Enterobacteriaceae; Escherichia; Escherichia coli; Escherichia coli O26 MNNEETFYQAMRRQGVTRRSFLKYCSLAATSLGLGAGMAPKIAWALENKPRIPVVWIHGL WP_001058323.1 MULTISPECIES: hydrogenase-1 small chain [Proteobacteria] Length: 372\nNumber of predicted TMHs: 2\nExp number of AAs in TMHs: 37.2608499999999\nExp number, first 60 AAs: 19.52923\nTotal prob of N-in: 0.94351\nPOSSIBLE N-term signal sequence\ninside 1 19\nTMhelix 20 39\noutside 40 325\nTMhelix 326 348\ninside 349 372

13660 GCF_000622485.2_Ec2010C-4819 Escherichia coli O26:H11 str. 2010C-4819 Proteobacteria; Gammaproteobacteria; Enterobacterales; Enterobacteriaceae; Escherichia; Escherichia coli; Escherichia coli O26 MTGDNTLIHSHGINRRDFMKLCAALAATMGLSSKAAAEMAESVTNPQRPPVIWIGAQECT WP_000145410.1 MULTISPECIES: hydrogenase-2 small chain [Proteobacteria] Length: 372\nNumber of predicted TMHs: 1\nExp number of AAs in TMHs: 23.72226\nExp number, first 60 AAs: 0.22038\nTotal prob of N-in: 0.12994\noutside 1 331\nTMhelix 332 354\ninside 355 372

13661 GCF_000622655.2_Ec2010C-4834 Escherichia coli O26:H11 str. 2010C-4834 Proteobacteria; Gammaproteobacteria; Enterobacterales; Enterobacteriaceae; Escherichia; Escherichia coli; Escherichia coli O26 MNNEETFYQAMRRQGVTRRSFLKYCSLAATSLGLGAGMAPKIAWALENKPRIPVVWIHGL WP_001058323.1 MULTISPECIES: hydrogenase-1 small chain [Proteobacteria] Length: 372\nNumber of predicted TMHs: 2\nExp number of AAs in TMHs: 37.2608499999999\nExp number, first 60 AAs: 19.52923\nTotal prob of N-in: 0.94351\nPOSSIBLE N-term signal sequence\ninside 1 19\nTMhelix 20 39\noutside 40 325\nTMhelix 326 348\ninside 349 372

13662 GCF_000622655.2_Ec2010C-4834 Escherichia coli O26:H11 str. 2010C-4834 Proteobacteria; Gammaproteobacteria; Enterobacterales; Enterobacteriaceae; Escherichia; Escherichia coli; Escherichia coli O26 MTGDNTLIHSHGINRRDFMKLCAALAATMGLSSKAAAEMAESVTNPQRPPVIWIGAQECT WP_000145410.1 MULTISPECIES: hydrogenase-2 small chain [Proteobacteria] Length: 372\nNumber of predicted TMHs: 1\nExp number of AAs in TMHs: 23.72226\nExp number, first 60 AAs: 0.22038\nTotal prob of N-in: 0.12994\noutside 1 331\nTMhelix 332 354\ninside 355 372

13663	GCF_000622535.2_Ec2010C-5028	Escherichia coli O26:H11 str. 2010C-5028	Proteobacteria; Gammaproteobacteria; Enterobacterales; Enterobacteriaceae; Escherichia; Escherichia coli; Escherichia coli O26	MNNEETFYQAMRRQGVTRRSFLKYCSLAATSLGLGAGMAPKIAWALENKPRIPVVWIHGL	WP_001058323.1	MULTISPECIES: hydrogenase-1 small chain [Proteobacteria]	Length: 372\nNumber of predicted TMHs: 2\nExp number of AAs in TMHs: 37.2608499999999\nExp number, first 60 AAs: 19.52923\nTotal prob of N-in: 0.94351\nPOSSIBLE N-term signal sequence\ninside 1 19\nTMhelix 20 39\noutside 40 325\nTMhelix 326 348\ninside 349 372

13664	GCF_000622535.2_Ec2010C-5028	Escherichia coli O26:H11 str. 2010C-5028	Proteobacteria; Gammaproteobacteria; Enterobacterales; Enterobacteriaceae; Escherichia; Escherichia coli; Escherichia coli O26	MTGDNTLIHSHGINRRDFMKLCAALAATMGLSSKAAAEMAESVTNPQRPPVIWIGAQECT	WP_000145410.1	MULTISPECIES: hydrogenase-2 small chain [Proteobacteria]	Length: 372\nNumber of predicted TMHs: 1\nExp number of AAs in TMHs: 23.72226\nExp number, first 60 AAs: 0.22038\nTotal prob of N-in: 0.12994\noutside 1 331\nTMhelix 332 354\ninside 355 372

13665	GCF_000622555.1_Ec2010EL-1699	Escherichia coli O26:H11 str. 2010EL-1699	Proteobacteria; Gammaproteobacteria; Enterobacterales; Enterobacteriaceae; Escherichia; Escherichia coli; Escherichia coli O26	MTGDNTLIHSHGINRRDFMKLCAALAATMGLSSKAAAEMAESVTNPQRPPVIWIGAQECT	WP_000145410.1	MULTISPECIES: hydrogenase-2 small chain [Proteobacteria]	Length: 372\nNumber of predicted TMHs: 1\nExp number of AAs in TMHs: 23.72226\nExp number, first 60 AAs: 0.22038\nTotal prob of N-in: 0.12994\noutside 1 331\nTMhelix 332 354\ninside 355 372

13666	GCF_000622555.1_Ec2010EL-1699	Escherichia coli O26:H11 str. 2010EL-1699	Proteobacteria; Gammaproteobacteria; Enterobacterales; Enterobacteriaceae; Escherichia; Escherichia coli; Escherichia coli O26	MNNEETFYQAMRRQGVTRRSFLKYCSLAATSLGLGAGMAPKIAWALENKPRIPVVWIHGL	WP_001058323.1	MULTISPECIES: hydrogenase-1 small chain [Proteobacteria]	Length: 372\nNumber of predicted TMHs: 2\nExp number of AAs in TMHs: 37.2608499999999\nExp number, first 60 AAs: 19.52923\nTotal prob of N-in: 0.94351\nPOSSIBLE N-term signal sequence\ninside 1 19\nTMhelix 20 39\noutside 40 325\nTMhelix 326 348\ninside 349 372

13667	GCF_000622775.2_Ec2011C-3270	Escherichia coli O26:H11 str. 2011C-3270	Proteobacteria; Gammaproteobacteria; Enterobacterales; Enterobacteriaceae; Escherichia; Escherichia coli; Escherichia coli O26	MTGDNTLIHSHGINRRDFMKLCAALAATMGLSSKAAAEMAESVTNPQRPPVIWIGAQECT	WP_000145410.1	MULTISPECIES: hydrogenase-2 small chain [Proteobacteria]	Length: 372\nNumber of predicted TMHs: 1\nExp number of AAs in TMHs: 23.72226\nExp number, first 60 AAs: 0.22038\nTotal prob of N-in: 0.12994\noutside 1 331\nTMhelix 332 354\ninside 355 372

13668	GCF_000622775.2_Ec2011C-3270	Escherichia coli O26:H11 str. 2011C-3270	Proteobacteria; Gammaproteobacteria; Enterobacterales; Enterobacteriaceae; Escherichia; Escherichia coli; Escherichia coli O26	MNNEETFYQAMRRQGVTRRSFLKYCSLAATSLGLGAGMAPKIAWALENKPRIPVVWIHGL	WP_001058323.1	MULTISPECIES: hydrogenase-1 small chain [Proteobacteria]	Length: 372\nNumber of predicted TMHs: 2\nExp number of AAs in TMHs: 37.2608499999999\nExp number, first 60 AAs: 19.52923\nTotal prob of N-in: 0.94351\nPOSSIBLE N-term signal sequence\ninside 1 19\nTMhelix 20 39\noutside 40 325\nTMhelix 326 348\ninside 349 372

13669	GCF_000703325.1_Ec2011C-3274	Escherichia coli O26:H11 str. 2011C-3274	Proteobacteria; Gammaproteobacteria; Enterobacterales; Enterobacteriaceae; Escherichia; Escherichia coli; Escherichia coli O26	MTGDNTLIHSHGINRRDFMKLCAALAATMGLSSKAAAEMAESVTNPQRPPVIWIGAQECT	WP_000145410.1	MULTISPECIES: hydrogenase-2 small chain [Proteobacteria]	Length: 372\nNumber of predicted TMHs: 1\nExp number of AAs in TMHs: 23.72226\nExp number, first 60 AAs: 0.22038\nTotal prob of N-in: 0.12994\noutside 1 331\nTMhelix 332 354\ninside 355 372

13670	GCF_000703325.1_Ec2011C-3274	Escherichia coli O26:H11 str. 2011C-3274	Proteobacteria; Gammaproteobacteria; Enterobacterales; Enterobacteriaceae; Escherichia; Escherichia coli; Escherichia coli O26	MNNEETFYQAMRRQGVTRRSFLKYCSLAATSLGLGAGMAPKIAWALENKPRIPVVWIHGL	WP_001058323.1	MULTISPECIES: hydrogenase-1 small chain [Proteobacteria]	Length: 372\nNumber of predicted TMHs: 2\nExp number of AAs in TMHs: 37.2608499999999\nExp number, first 60 AAs: 19.52923\nTotal prob of N-in: 0.94351\nPOSSIBLE N-term signal sequence\ninside 1 19\nTMhelix 20 39\noutside 40 325\nTMhelix 326 348\ninside 349 372

13671 GCF_000622795.2_Ec2011C-3282 Escherichia coli O26:H11 str. 2011C-3282 Proteobacteria; Gammaproteobacteria; Enterobacterales; Enterobacteriaceae; Escherichia; Escherichia coli; Escherichia coli O26 MTGDNTLIHSHGINRRDFMKLCAALAATMGLSSKAAAEMAESVTNPQRPPVIWIGAQECT WP_000145410.1 MULTISPECIES: hydrogenase-2 small chain [Proteobacteria] Length: 372\nNumber of predicted TMHs: 1\nExp number of AAs in TMHs: 23.72226\nExp number, first 60 AAs: 0.22038\nTotal prob of N-in: 0.12994\noutside 1 331\nTMhelix 332 354\ninside 355 372

13672 GCF_000622795.2_Ec2011C-3282 Escherichia coli O26:H11 str. 2011C-3282 Proteobacteria; Gammaproteobacteria; Enterobacterales; Enterobacteriaceae; Escherichia; Escherichia coli; Escherichia coli O26 MNNEETFYQAMRRQGVTRRSFLKYCSLAATSLGLGAGMAPKIAWALENKPRIPVVWIHGL WP_001058323.1 MULTISPECIES: hydrogenase-1 small chain [Proteobacteria] Length: 372\nNumber of predicted TMHs: 2\nExp number of AAs in TMHs: 37.2608499999999\nExp number, first 60 AAs: 19.52923\nTotal prob of N-in: 0.94351\nPOSSIBLE N-term signal sequence\ninside 1 19\nTMhelix 20 39\noutside 40 325\nTMhelix 326 348\ninside 349 372

13673 GCF_000622815.1_Ec2011C-3387 Escherichia coli O26:H11 str. 2011C-3387 Proteobacteria; Gammaproteobacteria; Enterobacterales; Enterobacteriaceae; Escherichia; Escherichia coli; Escherichia coli O26 MTGDNTLIHSHGINRRDFMKLCAALAATMGLSSKAAAEMAESVTNPQRPPVIWIGAQECT WP_000145410.1 MULTISPECIES: hydrogenase-2 small chain [Proteobacteria] Length: 372\nNumber of predicted TMHs: 1\nExp number of AAs in TMHs: 23.72226\nExp number, first 60 AAs: 0.22038\nTotal prob of N-in: 0.12994\noutside 1 331\nTMhelix 332 354\ninside 355 372

13674 GCF_000622815.1_Ec2011C-3387 Escherichia coli O26:H11 str. 2011C-3387 Proteobacteria; Gammaproteobacteria; Enterobacterales; Enterobacteriaceae; Escherichia; Escherichia coli; Escherichia coli O26 MNNEETFYQAMRRQGVTRRSFLKYCSLAATSLGLGAGMAPKIAWALENKPRIPVVWIHGL WP_001058323.1 MULTISPECIES: hydrogenase-1 small chain [Proteobacteria] Length: 372\nNumber of predicted TMHs: 2\nExp number of AAs in TMHs: 37.2608499999999\nExp number, first 60 AAs: 19.52923\nTotal prob of N-in: 0.94351\nPOSSIBLE N-term signal sequence\ninside 1 19\nTMhelix 20 39\noutside 40 325\nTMhelix 326 348\ninside 349 372

13675 GCF_000622835.1_Ec2011C-3506 Escherichia coli O26:H11 str. 2011C-3506 Proteobacteria; Gammaproteobacteria; Enterobacterales; Enterobacteriaceae; Escherichia; Escherichia coli; Escherichia coli O26 MTGDNTLIHSHGINRRDFMKLCAALAATMGLSSKAAAEMAESVTNPQRPPVIWIGAQECT WP_000145410.1 MULTISPECIES: hydrogenase-2 small chain [Proteobacteria] Length: 372\nNumber of predicted TMHs: 1\nExp number of AAs in TMHs: 23.72226\nExp number, first 60 AAs: 0.22038\nTotal prob of N-in: 0.12994\noutside 1 331\nTMhelix 332 354\ninside 355 372

13676 GCF_000622835.1_Ec2011C-3506 Escherichia coli O26:H11 str. 2011C-3506 Proteobacteria; Gammaproteobacteria; Enterobacterales; Enterobacteriaceae; Escherichia; Escherichia coli; Escherichia coli O26 MNNEETFYQAMRRQGVTRRSFLKYCSLAATSLGLGAGMAPKIAWALENKPRIPVVWIHGL WP_001058323.1 MULTISPECIES: hydrogenase-1 small chain [Proteobacteria] Length: 372\nNumber of predicted TMHs: 2\nExp number of AAs in TMHs: 37.2608499999999\nExp number, first 60 AAs: 19.52923\nTotal prob of N-in: 0.94351\nPOSSIBLE N-term signal sequence\ninside 1 19\nTMhelix 20 39\noutside 40 325\nTMhelix 326 348\ninside 349 372

13677 GCF_000622855.2_Ec2011C-3655 Escherichia coli O26:H11 str. 2011C-3655 Proteobacteria; Gammaproteobacteria; Enterobacterales; Enterobacteriaceae; Escherichia; Escherichia coli; Escherichia coli O26 MNNEETFYQAMRRQGVTRRSFLKYCSLAATSLGLGAGMAPKIAWALENKPRIPVVWIHGL WP_001058323.1 MULTISPECIES: hydrogenase-1 small chain [Proteobacteria] Length: 372\nNumber of predicted TMHs: 2\nExp number of AAs in TMHs: 37.2608499999999\nExp number, first 60 AAs: 19.52923\nTotal prob of N-in: 0.94351\nPOSSIBLE N-term signal sequence\ninside 1 19\nTMhelix 20 39\noutside 40 325\nTMhelix 326 348\ninside 349 372

13678 GCF_000622855.2_Ec2011C-3655 Escherichia coli O26:H11 str. 2011C-3655 Proteobacteria; Gammaproteobacteria; Enterobacterales; Enterobacteriaceae; Escherichia; Escherichia coli; Escherichia coli O26 MTGDNTLIHSHGINRRDFMKLCAALAATMGLSSKAAAEMAESVTNPQRPPVIWIGAQECT WP_000145410.1 MULTISPECIES: hydrogenase-2 small chain [Proteobacteria] Length: 372\nNumber of predicted TMHs: 1\nExp number of AAs in TMHs: 23.72226\nExp number, first 60 AAs: 0.22038\nTotal prob of N-in: 0.12994\noutside 1 331\nTMhelix 332 354\ninside 355 372

13679 GCF_000313405.1_CFSAN001629_1.0 Escherichia coli O26:H11 str. CFSAN001629 Proteobacteria; Gammaproteobacteria; Enterobacterales; Enterobacteriaceae; Escherichia; Escherichia coli; Escherichia coli O26 MNNEETFYQAMRRQGVTRRSFLKYCSLAATSLGLGAGMAPKIAWALENKPRIPVVWIHGL WP_001058323.1 MULTISPECIES: hydrogenase-1 small chain [Proteobacteria] Length: 372\nNumber of predicted TMHs: 2\nExp number of AAs in TMHs: 37.2608499999999\nExp number, first 60 AAs: 19.52923\nTotal prob of N-in: 0.94351\nPOSSIBLE N-term signal sequence\ninside 1 19\nTMhelix 20 39\noutside 40 325\nTMhelix 326 348\ninside 349 372

13680 GCF_000313405.1_CFSAN001629_1.0 Escherichia coli O26:H11 str. CFSAN001629 Proteobacteria; Gammaproteobacteria; Enterobacterales; Enterobacteriaceae; Escherichia; Escherichia coli; Escherichia coli O26 MTGDNTLIHSHGINRRDFMKLCAALAATMGLSSKAAAEMAESVTNPQRPPVIWIGAQECT WP_000145410.1 MULTISPECIES: hydrogenase-2 small chain [Proteobacteria] Length: 372\nNumber of predicted TMHs: 1\nExp number of AAs in TMHs: 23.72226\nExp number, first 60 AAs: 0.22038\nTotal prob of N-in: 0.12994\noutside 1 331\nTMhelix 332 354\ninside 355 372

13681 GCF_000276945.1_EC_O26_H11_CVM10021_1.0 Escherichia coli O26:H11 str. CVM10021 Proteobacteria; Gammaproteobacteria; Enterobacterales; Enterobacteriaceae; Escherichia; Escherichia coli; Escherichia coli O26 MTGDNTLIHSHGINRRDFMKLCAALAATMGLSSKAAAEMAESVTNPQRPPVIWIGAQECT WP_000145410.1 MULTISPECIES: hydrogenase-2 small chain [Proteobacteria] Length: 372\nNumber of predicted TMHs: 1\nExp number of AAs in TMHs: 23.72226\nExp number, first 60 AAs: 0.22038\nTotal prob of N-in: 0.12994\noutside 1 331\nTMhelix 332 354\ninside 355 372

13682 GCF_000264035.1_ASM26403v1 Escherichia coli O26:H11 str. CVM10026 Proteobacteria; Gammaproteobacteria; Enterobacterales; Enterobacteriaceae; Escherichia; Escherichia coli; Escherichia coli O26 MTGDNTLIHSHGINRRDFMKLCAALAATMGLSSKAAAEMAESVTNPQRPPVIWIGAQECT WP_000145410.1 MULTISPECIES: hydrogenase-2 small chain [Proteobacteria] Length: 372\nNumber of predicted TMHs: 1\nExp number of AAs in TMHs: 23.72226\nExp number, first 60 AAs: 0.22038\nTotal prob of N-in: 0.12994\noutside 1 331\nTMhelix 332 354\ninside 355 372

13683 GCF_000264035.1_ASM26403v1 Escherichia coli O26:H11 str. CVM10026 Proteobacteria; Gammaproteobacteria; Enterobacterales; Enterobacteriaceae; Escherichia; Escherichia coli; Escherichia coli O26 MNNEETFYQAMRRQGVTRRSFLKYCSLAATSLGLGAGMAPKIAWALENKPRIPVVWIHGL WP_001058323.1 MULTISPECIES: hydrogenase-1 small chain [Proteobacteria] Length: 372\nNumber of predicted TMHs: 2\nExp number of AAs in TMHs: 37.2608499999999\nExp number, first 60 AAs: 19.52923\nTotal prob of N-in: 0.94351\nPOSSIBLE N-term signal sequence\ninside 1 19\nTMhelix 20 39\noutside 40 325\nTMhelix 326 348\ninside 349 372

13684 GCF_000276845.1_EC_O26_H11_CVM10030_1.0 Escherichia coli O26:H11 str. CVM10030 Proteobacteria; Gammaproteobacteria; Enterobacterales; Enterobacteriaceae; Escherichia; Escherichia coli; Escherichia coli O26 MNNEETFYQAMRRQGVTRRSFLKYCSLAATSLGLGAGMAPKIAWALENKPRIPVVWIHGL WP_001058323.1 MULTISPECIES: hydrogenase-1 small chain [Proteobacteria] Length: 372\nNumber of predicted TMHs: 2\nExp number of AAs in TMHs: 37.2608499999999\nExp number, first 60 AAs: 19.52923\nTotal prob of N-in: 0.94351\nPOSSIBLE N-term signal sequence\ninside 1 19\nTMhelix 20 39\noutside 40 325\nTMhelix 326 348\ninside 349 372

13685 GCF_000276845.1_EC_O26_H11_CVM10030_1.0 Escherichia coli O26:H11 str. CVM10030 Proteobacteria; Gammaproteobacteria; Enterobacterales; Enterobacteriaceae; Escherichia; Escherichia coli; Escherichia coli O26 MTGDNTLIHSHGINRRDFMKLCAALAATMGLSSKAAAEMAESVTNPQRPPVIWIGAQECT WP_000145410.1 MULTISPECIES: hydrogenase-2 small chain [Proteobacteria] Length: 372\nNumber of predicted TMHs: 1\nExp number of AAs in TMHs: 23.72226\nExp number, first 60 AAs: 0.22038\nTotal prob of N-in: 0.12994\noutside 1 331\nTMhelix 332 354\ninside 355 372

13686 GCF_000276785.1_EC_O26_H11_CVM10224_1.0 Escherichia coli O26:H11 str. CVM10224 Proteobacteria; Gammaproteobacteria; Enterobacterales; Enterobacteriaceae; Escherichia; Escherichia coli; Escherichia coli O26 MNNEETFYQAMRRQGVTRRSFLKYCSLAATSLGLGAGMAPKIAWALENKPRIPVVWIHGL WP_001058323.1 MULTISPECIES: hydrogenase-1 small chain [Proteobacteria] Length: 372\nNumber of predicted TMHs: 2\nExp number of AAs in TMHs: 37.2608499999999\nExp number, first 60 AAs: 19.52923\nTotal prob of N-in: 0.94351\nPOSSIBLE N-term signal sequence\ninside 1 19\nTMhelix 20 39\noutside 40 325\nTMhelix 326 348\ninside 349 372

13687 GCF_000276785.1_EC_O26_H11_CVM10224_1.0 Escherichia coli O26:H11 str. CVM10224 Proteobacteria; Gammaproteobacteria; Enterobacterales; Enterobacteriaceae; Escherichia; Escherichia coli; Escherichia coli O26 MTGDNTLIHSHGINRRDFMKLCAALAATMGLSSKAAAEMAESVTNPQRPPVIWIGAQECT WP_000145410.1 MULTISPECIES: hydrogenase-2 small chain [Proteobacteria] Length: 372\nNumber of predicted TMHs: 1\nExp number of AAs in TMHs: 23.72226\nExp number, first 60 AAs: 0.22038\nTotal prob of N-in: 0.12994\noutside 1 331\nTMhelix 332 354\ninside 355 372

13688 GCF_000264015.1_ASM26401v1 Escherichia coli O26:H11 str. CVM9942 Proteobacteria; Gammaproteobacteria; Enterobacterales; Enterobacteriaceae; Escherichia; Escherichia coli; Escherichia coli O26 MTGDNTLIHSHGINRRDFMKLCAALAATMGLSSKAAAEMAESVTNPQRPPVIWIGAQECT WP_000145410.1 MULTISPECIES: hydrogenase-2 small chain [Proteobacteria] Length: 372\nNumber of predicted TMHs: 1\nExp number of AAs in TMHs: 23.72226\nExp number, first 60 AAs: 0.22038\nTotal prob of N-in: 0.12994\noutside 1 331\nTMhelix 332 354\ninside 355 372

13689 GCF_000264015.1_ASM26401v1 Escherichia coli O26:H11 str. CVM9942 Proteobacteria; Gammaproteobacteria; Enterobacterales; Enterobacteriaceae; Escherichia; Escherichia coli; Escherichia coli O26 MNNEETFYQAMRRQGVTRRSFLKYCSLAATSLGLGAGMAPKIAWALENKPRIPVVWIHGL WP_001058323.1 MULTISPECIES: hydrogenase-1 small chain [Proteobacteria] Length: 372\nNumber of predicted TMHs: 2\nExp number of AAs in TMHs: 37.2608499999999\nExp number, first 60 AAs: 19.52923\nTotal prob of N-in: 0.94351\nPOSSIBLE N-term signal sequence\ninside 1 19\nTMhelix 20 39\noutside 40 325\nTMhelix 326 348\ninside 349 372

13690 GCF_000276885.1_EC_O26_H11_CVM9952_1.0 Escherichia coli O26:H11 str. CVM9952 Proteobacteria; Gammaproteobacteria; Enterobacterales; Enterobacteriaceae; Escherichia; Escherichia coli; Escherichia coli O26 MNNEETFYQAMRRQGVTRRSFLKYCSLAATSLGLGAGMAPKIAWALENKPRIPVVWIHGL WP_001058323.1 MULTISPECIES: hydrogenase-1 small chain [Proteobacteria] Length: 372\nNumber of predicted TMHs: 2\nExp number of AAs in TMHs: 37.2608499999999\nExp number, first 60 AAs: 19.52923\nTotal prob of N-in: 0.94351\nPOSSIBLE N-term signal sequence\ninside 1 19\nTMhelix 20 39\noutside 40 325\nTMhelix 326 348\ninside 349 372

13691 GCF_000276885.1_EC_O26_H11_CVM9952_1.0 Escherichia coli O26:H11 str. CVM9952 Proteobacteria; Gammaproteobacteria; Enterobacterales; Enterobacteriaceae; Escherichia; Escherichia coli; Escherichia coli O26 MTGDNTLIHSHGINRRDFMKLCAALAATMGLSSKAAAEMAESVTNPQRPPVIWIGAQECT WP_000145410.1 MULTISPECIES: hydrogenase-2 small chain [Proteobacteria] Length: 372\nNumber of predicted TMHs: 1\nExp number of AAs in TMHs: 23.72226\nExp number, first 60 AAs: 0.22038\nTotal prob of N-in: 0.12994\noutside 1 331\nTMhelix 332 354\ninside 355 372

13692 GCF_000632575.1_Ec2009C-4747 Escherichia coli O26:H1 str. 2009C-4747 Proteobacteria; Gammaproteobacteria; Enterobacterales; Enterobacteriaceae; Escherichia; Escherichia coli; Escherichia coli O26 MNNEETFYQAMRRQGVTRRSFLKYCSLAATSLGLGAGMAPKIAWALENKPRIPVVWIHGL WP_001058323.1 MULTISPECIES: hydrogenase-1 small chain [Proteobacteria] Length: 372\nNumber of predicted TMHs: 2\nExp number of AAs in TMHs: 37.2608499999999\nExp number, first 60 AAs: 19.52923\nTotal prob of N-in: 0.94351\nPOSSIBLE N-term signal sequence\ninside 1 19\nTMhelix 20 39\noutside 40 325\nTMhelix 326 348\ninside 349 372

13693 GCF_000632575.1_Ec2009C-4747 Escherichia coli O26:H1 str. 2009C-4747 Proteobacteria; Gammaproteobacteria; Enterobacterales; Enterobacteriaceae; Escherichia; Escherichia coli; Escherichia coli O26 MTGDNTLIHSHGINRRDFMKLCAALAATMGLSSKAAAEMAESVTNPQRPPVIWIGAQECT WP_000145410.1 MULTISPECIES: hydrogenase-2 small chain [Proteobacteria] Length: 372\nNumber of predicted TMHs: 1\nExp number of AAs in TMHs: 23.72226\nExp number, first 60 AAs: 0.22038\nTotal prob of N-in: 0.12994\noutside 1 331\nTMhelix 332 354\ninside 355 372

13694 GCF_000614845.2_Ec2010C-4347 Escherichia coli O26:NM str. 2010C-4347 Proteobacteria; Gammaproteobacteria; Enterobacterales; Enterobacteriaceae; Escherichia; Escherichia coli; Escherichia coli O26 MTGDNTLIHSHGINRRDFMKLCAALAATMGLSSKAAAEMAESVTNPQRPPVIWIGAQECT WP_000145410.1 MULTISPECIES: hydrogenase-2 small chain [Proteobacteria] Length: 372\nNumber of predicted TMHs: 1\nExp number of AAs in TMHs: 23.72226\nExp number, first 60 AAs: 0.22038\nTotal prob of N-in: 0.12994\noutside 1 331\nTMhelix 332 354\ninside 355 372

13695 GCF_000614845.2_Ec2010C-4347 Escherichia coli O26:NM str. 2010C-4347 Proteobacteria; Gammaproteobacteria; Enterobacterales; Enterobacteriaceae; Escherichia; Escherichia coli; Escherichia coli O26

MNNEETFYQAMRRQGVTRRSFLKYCSLAATSLGLGAGMAPKIAWALENKPRIPVVWIHGL WP_001058323.1 MULTISPECIES: hydrogenase-1 small chain [Proteobacteria] Length: 372\nNumber of predicted TMHs: 2\nExp number of AAs in TMHs: 37.2608499999999\nExp number, first 60 AAs: 19.52923\nTotal prob of N-in: 0.94351\nPOSSIBLE N-term signal sequence\ninside 1 19\nTMhelix 20 39\noutside 40 325\nTMhelix 326 348\ninside 349 372

13696 GCF_000617505.2_Ec2010C-4788 Escherichia coli O26:NM str. 2010C-4788 Proteobacteria; Gammaproteobacteria; Enterobacterales; Enterobacteriaceae; Escherichia; Escherichia coli; Escherichia coli O26 MTGDNTLIHSHGINRRDFMKLCAALAATMGLSSKAAAEMAESVTNPQRPPVIWIGAQECT WP_000145410.1 MULTISPECIES: hydrogenase-2 small chain [Proteobacteria] Length: 372\nNumber of predicted TMHs: 1\nExp number of AAs in TMHs: 23.72226\nExp number, first 60 AAs: 0.22038\nTotal prob of N-in: 0.12994\noutside 1 331\nTMhelix 332 354\ninside 355 372

13697 GCF_000617505.2_Ec2010C-4788 Escherichia coli O26:NM str. 2010C-4788 Proteobacteria; Gammaproteobacteria; Enterobacterales; Enterobacteriaceae; Escherichia; Escherichia coli; Escherichia coli O26 MNNEETFYQAMRRQGVTRRSFLKYCSLAATSLGLGAGMAPKIAWALENKPRIPVVWIHGL WP_001058323.1 MULTISPECIES: hydrogenase-1 small chain [Proteobacteria] Length: 372\nNumber of predicted TMHs: 2\nExp number of AAs in TMHs: 37.2608499999999\nExp number, first 60 AAs: 19.52923\nTotal prob of N-in: 0.94351\nPOSSIBLE N-term signal sequence\ninside 1 19\nTMhelix 20 39\noutside 40 325\nTMhelix 326 348\ninside 349 372

13698 GCF_000617165.2_Ec02-3404 Escherichia coli O28ac:NM str. 02-3404 Proteobacteria; Gammaproteobacteria; Enterobacterales; Enterobacteriaceae; Escherichia; Escherichia coli; Escherichia coli O28ac MTGDNTLIHSHGINRRDFMKLCAALAATMGLSSKAAAEMAESVTNPQRPPVIWIGAQECT WP_000145410.1 MULTISPECIES: hydrogenase-2 small chain [Proteobacteria] Length: 372\nNumber of predicted TMHs: 1\nExp number of AAs in TMHs: 23.72226\nExp number, first 60 AAs: 0.22038\nTotal prob of N-in: 0.12994\noutside 1 331\nTMhelix 332 354\ninside 355 372

13699 GCF_000617165.2_Ec02-3404 Escherichia coli O28ac:NM str. 02-3404 Proteobacteria; Gammaproteobacteria; Enterobacterales; Enterobacteriaceae; Escherichia; Escherichia coli; Escherichia coli O28ac MNNEETFYQAMRRQGVTRRSFLKYCSLAATSLGLGAGMAPKIAWALENKPRIPVVWIHGL WP_001058323.1 MULTISPECIES: hydrogenase-1 small chain [Proteobacteria] Length: 372\nNumber of predicted TMHs: 2\nExp number of AAs in TMHs: 37.2608499999999\nExp number, first 60 AAs: 19.52923\nTotal prob of N-in: 0.94351\nPOSSIBLE N-term signal sequence\ninside 1 19\nTMhelix 20 39\noutside 40 325\nTMhelix 326 348\ninside 349 372

13700 GCF_000259425.1_ASM25942v1 Escherichia coli O32:H37 str. P4 Proteobacteria; Gammaproteobacteria; Enterobacterales; Enterobacteriaceae; Escherichia; Escherichia coli; Escherichia coli O32 MNNEETFYQAMRRQGVTRRSFLKYCSLAATSLGLGAGMAPKIAWALENKPRIPVVWIHGL WP_001058323.1 MULTISPECIES: hydrogenase-1 small chain [Proteobacteria] Length: 372\nNumber of predicted TMHs: 2\nExp number of AAs in TMHs: 37.2608499999999\nExp number, first 60 AAs: 19.52923\nTotal prob of N-in: 0.94351\nPOSSIBLE N-term signal sequence\ninside 1 19\nTMhelix 20 39\noutside 40 325\nTMhelix 326 348\ninside 349 372

13701 GCF_000259425.1_ASM25942v1 Escherichia coli O32:H37 str. P4 Proteobacteria; Gammaproteobacteria; Enterobacterales; Enterobacteriaceae; Escherichia; Escherichia coli; Escherichia coli O32 MTGDNTLIHSHGINRRDFMKLCAALAATMGLSSKAAAEMAESVTNPQRPPVIWIGAQECT WP_000145410.1 MULTISPECIES: hydrogenase-2 small chain [Proteobacteria] Length: 372\nNumber of predicted TMHs: 1\nExp number of AAs in TMHs: 23.72226\nExp number, first 60 AAs: 0.22038\nTotal prob of N-in: 0.12994\noutside 1 331\nTMhelix 332 354\ninside 355 372

13702 GCF_000618905.2_EcF8704-2 Escherichia coli O39:NM str. F8704-2 Proteobacteria; Gammaproteobacteria; Enterobacterales; Enterobacteriaceae; Escherichia; Escherichia coli; Escherichia coli O39 MNNEETFYQAMRRQGVTRRSFLKYCSLAATSLGLGAGMAPKIAWALENKPRIPVVWIHGL WP_001058323.1 MULTISPECIES: hydrogenase-1 small chain [Proteobacteria] Length: 372\nNumber of predicted TMHs: 2\nExp number of AAs in TMHs: 37.2608499999999\nExp number, first 60 AAs: 19.52923\nTotal prob of N-in: 0.94351\nPOSSIBLE N-term signal sequence\ninside 1 19\nTMhelix 20 39\noutside 40 325\nTMhelix 326 348\ninside 349 372

13703 GCF_000618905.2_EcF8704-2 Escherichia coli O39:NM str. F8704-2 Proteobacteria; Gammaproteobacteria; Enterobacterales; Enterobacteriaceae; Escherichia; Escherichia coli; Escherichia coli O39

MTGDNTLIHSHGINRRDFMKLCAALAATMGLSSKAAAEMAESVTNPQRPPVIWIGAQECT WP_001685818.1 MULTISPECIES: hydrogenase-2 small chain [Enterobacteriaceae] Length: 372\nNumber of predicted TMHs: 1\nExp number of AAs in TMHs: 23.82505\nExp number, first 60 AAs: 0.21974\nTotal prob of N-in: 0.13379\noutside 1 331\nTMhelix 332 354\ninside 355 372

13704 GCF_000027125.1_ASM2712v1 Escherichia coli 042 Proteobacteria; Gammaproteobacteria; Enterobacterales; Enterobacteriaceae; Escherichia; Escherichia coli; Escherichia coli O44 MNNEETFYQAMRRQGVTRRSFLKYCSLAATSLGLGAGMAPKIAWALENKPRIPVVWIHGL WP_001058323.1 MULTISPECIES: hydrogenase-1 small chain [Proteobacteria] Length: 372\nNumber of predicted TMHs: 2\nExp number of AAs in TMHs: 37.2608499999999\nExp number, first 60 AAs: 19.52923\nTotal prob of N-in: 0.94351\nPOSSIBLE N-term signal sequence\ninside 1 19\nTMhelix 20 39\noutside 40 325\nTMhelix 326 348\ninside 349 372

13705 GCF_000027125.1_ASM2712v1 Escherichia coli 042 Proteobacteria; Gammaproteobacteria; Enterobacterales; Enterobacteriaceae; Escherichia; Escherichia coli; Escherichia coli O44 MTGDNTLIHSHGINRRDFMKLCAALAATMGLSSKAAAEMAESVTNPQRPPVIWIGAQECT WP_000145410.1 MULTISPECIES: hydrogenase-2 small chain [Proteobacteria] Length: 372\nNumber of predicted TMHs: 1\nExp number of AAs in TMHs: 23.72226\nExp number, first 60 AAs: 0.22038\nTotal prob of N-in: 0.12994\noutside 1 331\nTMhelix 332 354\ninside 355 372

13706 GCF_000617125.2_Ec01-3147 Escherichia coli O45:H2 str. 01-3147 Proteobacteria; Gammaproteobacteria; Enterobacterales; Enterobacteriaceae; Escherichia; Escherichia coli; Escherichia coli O45 MTGDNTLIHSHGINRRDFMKLCAALAATMGLSSKAAAEMAESVTNPQRPPVIWIGAQECT WP_000145410.1 MULTISPECIES: hydrogenase-2 small chain [Proteobacteria] Length: 372\nNumber of predicted TMHs: 1\nExp number of AAs in TMHs: 23.72226\nExp number, first 60 AAs: 0.22038\nTotal prob of N-in: 0.12994\noutside 1 331\nTMhelix 332 354\ninside 355 372

13707 GCF_000617125.2_Ec01-3147 Escherichia coli O45:H2 str. 01-3147 Proteobacteria; Gammaproteobacteria; Enterobacterales; Enterobacteriaceae; Escherichia; Escherichia coli; Escherichia coli O45 MNNEETFYQAMRRQGVTRRSFLKYCSLAATSLGLGAGIAPKIAWALENKPRIPVVWIHGL WP_023982790.1 hydrogenase [Escherichia coli] Length: 372\nNumber of predicted TMHs: 2\nExp number of AAs in TMHs: 37.8310099999999\nExp number, first 60 AAs: 20.15722\nTotal prob of N-in: 0.95896\nPOSSIBLE N-term signal sequence\ninside 1 20\nTMhelix 21 43\noutside 44 325\nTMhelix 326 348\ninside 349 372

13708 GCF_000234295.1_ASM23429v2 Escherichia coli O45:H2 str. 03-EN-705 Proteobacteria; Gammaproteobacteria; Enterobacterales; Enterobacteriaceae; Escherichia; Escherichia coli; Escherichia coli O45 MTGDNTLIHSHGINRRDFMKLCAALAATMGLSSKAAAEMAESVTNPQRPPVIWIGAQECT WP_000145410.1 MULTISPECIES: hydrogenase-2 small chain [Proteobacteria] Length: 372\nNumber of predicted TMHs: 1\nExp number of AAs in TMHs: 23.72226\nExp number, first 60 AAs: 0.22038\nTotal prob of N-in: 0.12994\noutside 1 331\nTMhelix 332 354\ninside 355 372

13709 GCF_000619265.2_Ec2009C-3686 Escherichia coli O45:H2 str. 2009C-3686 Proteobacteria; Gammaproteobacteria; Enterobacterales; Enterobacteriaceae; Escherichia; Escherichia coli; Escherichia coli O45 MTGDNTLIHSHGINRRDFMKLCAALAATMGLSSKAAAEMAESVTNPQRPPVIWIGAQECT WP_000145410.1 MULTISPECIES: hydrogenase-2 small chain [Proteobacteria] Length: 372\nNumber of predicted TMHs: 1\nExp number of AAs in TMHs: 23.72226\nExp number, first 60 AAs: 0.22038\nTotal prob of N-in: 0.12994\noutside 1 331\nTMhelix 332 354\ninside 355 372

13710 GCF_000619265.2_Ec2009C-3686 Escherichia coli O45:H2 str. 2009C-3686 Proteobacteria; Gammaproteobacteria; Enterobacterales; Enterobacteriaceae; Escherichia; Escherichia coli; Escherichia coli O45 MNNEETFYQAMRRQGVTRRSFLKYCSLAATSLGLGAGIAPKIAWALENKPRIPVVWIHGL WP_023982790.1 hydrogenase [Escherichia coli] Length: 372\nNumber of predicted TMHs: 2\nExp number of AAs in TMHs: 37.8310099999999\nExp number, first 60 AAs: 20.15722\nTotal prob of N-in: 0.95896\nPOSSIBLE N-term signal sequence\ninside 1 20\nTMhelix 21 43\noutside 44 325\nTMhelix 326 348\ninside 349 372

13711 GCF_000619105.1_Ec2009C-4780 Escherichia coli O45:H2 str. 2009C-4780 Proteobacteria; Gammaproteobacteria; Enterobacterales; Enterobacteriaceae; Escherichia; Escherichia coli; Escherichia coli O45 MTGDNTLIHSHGINRRDFMKLCAALAATMGLSSKAAAEMAESVTNPQRPPVIWIGAQECT WP_000145410.1 MULTISPECIES: hydrogenase-2 small chain [Proteobacteria] Length: 372\nNumber of predicted TMHs: 1\nExp number of AAs in TMHs: 23.72226\nExp number, first 60 AAs: 0.22038\nTotal prob of N-in: 0.12994\noutside 1 331\nTMhelix 332 354\ninside 355 372

13712 GCF_000619105.1_Ec2009C-4780 Escherichia coli O45:H2 str. 2009C-4780 Proteobacteria; Gammaproteobacteria; Enterobacterales; Enterobacteriaceae; Escherichia; Escherichia coli; Escherichia coli O45 MNNEETFYQAMRRQGVTRRSFLKYCSLAATSLGLGAGIAPKIAWALENKPRIPVVWIHGL WP_023982790.1 hydrogenase [Escherichia coli] Length: 372\nNumber of predicted TMHs: 2\nExp number of AAs in TMHs: 37.8310099999999\nExp number, first 60 AAs: 20.15722\nTotal prob of N-in: 0.95896\nPOSSIBLE N-term signal sequence\ninside 1 20\nTMhelix 21 43\noutside 44 325\nTMhelix 326 348\ninside 349 372

13713 GCF_000615535.1_Ec2010C-3876 Escherichia coli O45:H2 str. 2010C-3876 Proteobacteria; Gammaproteobacteria; Enterobacterales; Enterobacteriaceae; Escherichia; Escherichia coli; Escherichia coli O45 MNNEETFYQAMRRQGVTRRSFLKYCSLAATSLGLGAGIAPKIAWALENKPRIPVVWIHGL WP_023982790.1 hydrogenase [Escherichia coli] Length: 372\nNumber of predicted TMHs: 2\nExp number of AAs in TMHs: 37.8310099999999\nExp number, first 60 AAs: 20.15722\nTotal prob of N-in: 0.95896\nPOSSIBLE N-term signal sequence\ninside 1 20\nTMhelix 21 43\noutside 44 325\nTMhelix 326 348\ninside 349 372

13714 GCF_000615535.1_Ec2010C-3876 Escherichia coli O45:H2 str. 2010C-3876 Proteobacteria; Gammaproteobacteria; Enterobacterales; Enterobacteriaceae; Escherichia; Escherichia coli; Escherichia coli O45 MTGDNTLIHSHGINRRDFMKLCAALAATMGLSSKAAAEMAESVTNPQRPPVIWIGAQECT WP_000145410.1 MULTISPECIES: hydrogenase-2 small chain [Proteobacteria] Length: 372\nNumber of predicted TMHs: 1\nExp number of AAs in TMHs: 23.72226\nExp number, first 60 AAs: 0.22038\nTotal prob of N-in: 0.12994\noutside 1 331\nTMhelix 332 354\ninside 355 372

13715 GCF_000703305.1_Ec2010C-4211 Escherichia coli O45:H2 str. 2010C-4211 Proteobacteria; Gammaproteobacteria; Enterobacterales; Enterobacteriaceae; Escherichia; Escherichia coli; Escherichia coli O45 MTGDNTLIHSHGINRRDFMKLCAALAATMGLSSKAAAEMAESVTNPQRPPVIWIGAQECT WP_000145410.1 MULTISPECIES: hydrogenase-2 small chain [Proteobacteria] Length: 372\nNumber of predicted TMHs: 1\nExp number of AAs in TMHs: 23.72226\nExp number, first 60 AAs: 0.22038\nTotal prob of N-in: 0.12994\noutside 1 331\nTMhelix 332 354\ninside 355 372

13716 GCF_000703305.1_Ec2010C-4211 Escherichia coli O45:H2 str. 2010C-4211 Proteobacteria; Gammaproteobacteria; Enterobacterales; Enterobacteriaceae; Escherichia; Escherichia coli; Escherichia coli O45 MNNEETFYQAMRRQGVTRRSFLKYCSLAATSLGLGAGIAPKIAWALENKPRIPVVWIHGL WP_023982790.1 hydrogenase [Escherichia coli] Length: 372\nNumber of predicted TMHs: 2\nExp number of AAs in TMHs: 37.8310099999999\nExp number, first 60 AAs: 20.15722\nTotal prob of N-in: 0.95896\nPOSSIBLE N-term signal sequence\ninside 1 20\nTMhelix 21 43\noutside 44 325\nTMhelix 326 348\ninside 349 372

13717 GCF_000333195.1_ASM33319v1 Escherichia coli O5:K4(L):H4 str. ATCC 23502 Proteobacteria; Gammaproteobacteria; Enterobacterales; Enterobacteriaceae; Escherichia; Escherichia coli; Escherichia coli O5 MTGDNTLIHSHGINRRDFMKLCAALAATMGLSSKAAAEMAESVTNPQRPPVIWIGAQECT WP_000145410.1 MULTISPECIES: hydrogenase-2 small chain [Proteobacteria] Length: 372\nNumber of predicted TMHs: 1\nExp number of AAs in TMHs: 23.72226\nExp number, first 60 AAs: 0.22038\nTotal prob of N-in: 0.12994\noutside 1 331\nTMhelix 332 354\ninside 355 372

13718 GCF_000333195.1_ASM33319v1 Escherichia coli O5:K4(L):H4 str. ATCC 23502 Proteobacteria; Gammaproteobacteria; Enterobacterales; Enterobacteriaceae; Escherichia; Escherichia coli; Escherichia coli O5 MNNEETFYQAMRRQGVTRRSFLKYCSLAATSLGLGAGMAPKIAWALENKPRIPVVWIHGL WP_001058323.1 MULTISPECIES: hydrogenase-1 small chain [Proteobacteria] Length: 372\nNumber of predicted TMHs: 2\nExp number of AAs in TMHs: 37.2608499999999\nExp number, first 60 AAs: 19.52923\nTotal prob of N-in: 0.94351\nPOSSIBLE N-term signal sequence\ninside 1 19\nTMhelix 20 39\noutside 40 325\nTMhelix 326 348\ninside 349 372

13719 GCF_000617385.1_Ec06-3555 Escherichia coli O55:H7 str. 06-3555 Proteobacteria; Gammaproteobacteria; Enterobacterales; Enterobacteriaceae; Escherichia; Escherichia coli; Escherichia coli O55 MNNEETFYQAMRRQGVTRRSFLKYCSLAATSLGLGAGMAPKIAWALENKPRIPVVWIHGL WP_001359719.1 hydrogenase [Escherichia coli] Length: 372\nNumber of predicted TMHs: 2\nExp number of AAs in TMHs: 37.2176\nExp number, first 60 AAs: 19.53448\nTotal prob of N-in: 0.94360\nPOSSIBLE N-term signal sequence\ninside 1 19\nTMhelix 20 39\noutside 40 325\nTMhelix 326 348\ninside 349 372

13720 GCF_000617385.1_Ec06-3555 Escherichia coli O55:H7 str. 06-3555 Proteobacteria; Gammaproteobacteria; Enterobacterales; Enterobacteriaceae; Escherichia; Escherichia coli; Escherichia coli O55 MTGDNTLIHSHGINRRDFMKLCAALAATMGLSSKAAAEMAESVTNPQRPPVIWIGAQECT WP_000145410.1 MULTISPECIES: hydrogenase-2 small chain [Proteobacteria] Length: 372\nNumber of predicted TMHs:

1\nExp number of AAs in TMHs: 23.72226\nExp number, first 60 AAs: 0.22038\nTotal prob of N-in: 0.12994\noutside 1 331\nTMhelix 332 354\ninside 355 372

13721 GCF_000187345.1_ASM18734v2 Escherichia coli O55:H7 str. 3256-97 Proteobacteria; Gammaproteobacteria; Enterobacterales; Enterobacteriaceae; Escherichia; Escherichia coli; Escherichia coli O55 MTGDNTLIHSHGINRRDFMKLCAALAATMGLSSKAAAEMAESVTNPQRPPVIWIGAQECT WP_000145410.1 MULTISPECIES: hydrogenase-2 small chain [Proteobacteria] Length: 372\nNumber of predicted TMHs: 1\nExp number of AAs in TMHs: 23.72226\nExp number, first 60 AAs: 0.22038\nTotal prob of N-in: 0.12994\noutside 1 331\nTMhelix 332 354\ninside 355 372

13722 GCF_000187345.1_ASM18734v2 Escherichia coli O55:H7 str. 3256-97 Proteobacteria; Gammaproteobacteria; Enterobacterales; Enterobacteriaceae; Escherichia; Escherichia coli; Escherichia coli O55 MNNEETFYQAMRRQGVTRRSFLKYCSLAATSLGLGAGMAPKIAWALENKPRIPVVWIHGL WP_001301613.1 hydrogenase [Escherichia coli] Length: 372\nNumber of predicted TMHs: 2\nExp number of AAs in TMHs: 37.46598\nExp number, first 60 AAs: 19.53139\nTotal prob of N-in: 0.94364\nPOSSIBLE N-term signal sequence\ninside 1 19\nTMhelix 20 39\noutside 40 325\nTMhelix 326 348\ninside 349 372

13723 GCF_000025165.1_ASM2516v1 Escherichia coli O55:H7 str. CB9615 Proteobacteria; Gammaproteobacteria; Enterobacterales; Enterobacteriaceae; Escherichia; Escherichia coli; Escherichia coli O55 MNNEETFYQAMRRQGVTRRSFLKYCSLAATSLGLGAGMAPKIAWALENKPRIPVVWIHGL WP_001359719.1 hydrogenase [Escherichia coli] Length: 372\nNumber of predicted TMHs: 2\nExp number of AAs in TMHs: 37.2176\nExp number, first 60 AAs: 19.53448\nTotal prob of N-in: 0.94360\nPOSSIBLE N-term signal sequence\ninside 1 19\nTMhelix 20 39\noutside 40 325\nTMhelix 326 348\ninside 349 372

13724 GCF_000025165.1_ASM2516v1 Escherichia coli O55:H7 str. CB9615 Proteobacteria; Gammaproteobacteria; Enterobacterales; Enterobacteriaceae; Escherichia; Escherichia coli; Escherichia coli O55 MTGDNTLIHSHGINRRDFMKLCAALAATMGLSSKAAAEMAESVTNPQRPPVIWIGAQECT WP_000145410.1 MULTISPECIES: hydrogenase-2 small chain [Proteobacteria] Length: 372\nNumber of predicted TMHs: 1\nExp number of AAs in TMHs: 23.72226\nExp number, first 60 AAs: 0.22038\nTotal prob of N-in: 0.12994\noutside 1 331\nTMhelix 332 354\ninside 355 372

13725 GCF_000245515.1_ASM24551v1 Escherichia coli O55:H7 str. RM12579 Proteobacteria; Gammaproteobacteria; Enterobacterales; Enterobacteriaceae; Escherichia; Escherichia coli; Escherichia coli O55 MNNEETFYQAMRRQGVTRRSFLKYCSLAATSLGLGAGMAPKIAWALENKPRIPVVWIHGL WP_001359719.1 hydrogenase [Escherichia coli] Length: 372\nNumber of predicted TMHs: 2\nExp number of AAs in TMHs: 37.2176\nExp number, first 60 AAs: 19.53448\nTotal prob of N-in: 0.94360\nPOSSIBLE N-term signal sequence\ninside 1 19\nTMhelix 20 39\noutside 40 325\nTMhelix 326 348\ninside 349 372

13726 GCF_000245515.1_ASM24551v1 Escherichia coli O55:H7 str. RM12579 Proteobacteria; Gammaproteobacteria; Enterobacterales; Enterobacteriaceae; Escherichia; Escherichia coli; Escherichia coli O55 MTGDNTLIHSHGINRRDFMKLCAALAATMGLSSKAAAEMAESVTNPQRPPVIWIGAQECT WP_000145410.1 MULTISPECIES: hydrogenase-2 small chain [Proteobacteria] Length: 372\nNumber of predicted TMHs: 1\nExp number of AAs in TMHs: 23.72226\nExp number, first 60 AAs: 0.22038\nTotal prob of N-in: 0.12994\noutside 1 331\nTMhelix 332 354\ninside 355 372

13727 GCF_000187365.1_ASM18736v2 Escherichia coli O55:H7 str. USDA 5905 Proteobacteria; Gammaproteobacteria; Enterobacterales; Enterobacteriaceae; Escherichia; Escherichia coli; Escherichia coli O55 MNNEETFYQAMRRQGVTRRSFLKYCSLAATSLGLGAGMAPKIAWALENKPRIPVVWIHGL WP_001359719.1 hydrogenase [Escherichia coli] Length: 372\nNumber of predicted TMHs: 2\nExp number of AAs in TMHs: 37.2176\nExp number, first 60 AAs: 19.53448\nTotal prob of N-in: 0.94360\nPOSSIBLE N-term signal sequence\ninside 1 19\nTMhelix 20 39\noutside 40 325\nTMhelix 326 348\ninside 349 372

13728 GCF_000187365.1_ASM18736v2 Escherichia coli O55:H7 str. USDA 5905 Proteobacteria; Gammaproteobacteria; Enterobacterales; Enterobacteriaceae; Escherichia; Escherichia coli; Escherichia coli O55 MTGDNTLIHSHGINRRDFMKLCAALAATMGLSSKAAAEMAESVTNPQRPPVIWIGAQECT WP_000145410.1 MULTISPECIES: hydrogenase-2 small chain [Proteobacteria] Length: 372\nNumber of predicted TMHs: 1\nExp number of AAs in TMHs: 23.72226\nExp number, first 60 AAs: 0.22038\nTotal prob of N-in: 0.12994\noutside 1 331\nTMhelix 332 354\ninside 355 372

13729 GCF_000517245.1_ETEC_B2C Escherichia coli O6:H16:CFA/II str. B2C Proteobacteria; Gammaproteobacteria; Enterobacterales; Enterobacteriaceae; Escherichia; Escherichia coli; Escherichia coli O6

MTGDNTLIHSHGINRRDFMKLCAALAATMGLSSKAAAEMAESVTNPQRPPVIWIGAQECT WP_000145410.1 MULTISPECIES: hydrogenase-2 small chain [Proteobacteria] Length: 372\nNumber of predicted TMHs: 1\nExp number of AAs in TMHs: 23.72226\nExp number, first 60 AAs: 0.22038\nTotal prob of N-in: 0.12994\noutside 1 331\nTMhelix 332 354\ninside 355 372

13730 GCF_000517245.1_ETEC_B2C Escherichia coli O6:H16:CFA/II str. B2C Proteobacteria; Gammaproteobacteria; Enterobacterales; Enterobacteriaceae; Escherichia; Escherichia coli; Escherichia coli O6 MNNEETFYQAMRRQGVTRRSFLKYCSLAATSLGLGAGMAPKIAWALENKPRIPVVWIHGL WP_001058323.1 MULTISPECIES: hydrogenase-1 small chain [Proteobacteria] Length: 372\nNumber of predicted TMHs: 2\nExp number of AAs in TMHs: 37.2608499999999\nExp number, first 60 AAs: 19.52923\nTotal prob of N-in: 0.94351\nPOSSIBLE N-term signal sequence\ninside 1 19\nTMhelix 20 39\noutside 40 325\nTMhelix 326 348\ninside 349 372

13731 GCF_000614435.2_Ec99-3165 Escherichia coli O6:H16 str. 99-3165 Proteobacteria; Gammaproteobacteria; Enterobacterales; Enterobacteriaceae; Escherichia; Escherichia coli; Escherichia coli O6 MTGDNTLIHSHGINRRDFMKLCAALAATMGLSSKAAAEMAESVTNPQRPPVIWIGAQECT WP_000145410.1 MULTISPECIES: hydrogenase-2 small chain [Proteobacteria] Length: 372\nNumber of predicted TMHs: 1\nExp number of AAs in TMHs: 23.72226\nExp number, first 60 AAs: 0.22038\nTotal prob of N-in: 0.12994\noutside 1 331\nTMhelix 332 354\ninside 355 372

13732 GCF_000614435.2_Ec99-3165 Escherichia coli O6:H16 str. 99-3165 Proteobacteria; Gammaproteobacteria; Enterobacterales; Enterobacteriaceae; Escherichia; Escherichia coli; Escherichia coli O6 MNNEETFYQAMRRQGVTRRSFLKYCSLAATSLGLGAGMAPKIAWALENKPRIPVVWIHGL WP_001058323.1 MULTISPECIES: hydrogenase-1 small chain [Proteobacteria] Length: 372\nNumber of predicted TMHs: 2\nExp number of AAs in TMHs: 37.2608499999999\nExp number, first 60 AAs: 19.52923\nTotal prob of N-in: 0.94351\nPOSSIBLE N-term signal sequence\ninside 1 19\nTMhelix 20 39\noutside 40 325\nTMhelix 326 348\ninside 349 372

13733 GCF_000618085.1_EcF5656C1 Escherichia coli O6:H16 str. F5656C1 Proteobacteria; Gammaproteobacteria; Enterobacterales; Enterobacteriaceae; Escherichia; Escherichia coli; Escherichia coli O6 MNNEETFYQAMRRQGVTRRSFLKYCSLAATSLGLGAGMAPKIAWALENKPRIPVVWIHGL WP_001058323.1 MULTISPECIES: hydrogenase-1 small chain [Proteobacteria] Length: 372\nNumber of predicted TMHs: 2\nExp number of AAs in TMHs: 37.2608499999999\nExp number, first 60 AAs: 19.52923\nTotal prob of N-in: 0.94351\nPOSSIBLE N-term signal sequence\ninside 1 19\nTMhelix 20 39\noutside 40 325\nTMhelix 326 348\ninside 349 372

13734 GCF_000618085.1_EcF5656C1 Escherichia coli O6:H16 str. F5656C1 Proteobacteria; Gammaproteobacteria; Enterobacterales; Enterobacteriaceae; Escherichia; Escherichia coli; Escherichia coli O6 MTGDNTLIHSHGINRRDFMKLCAALAATMGLSSKAAAEMAESVTNPQRPPVIWIGAQECT WP_000145410.1 MULTISPECIES: hydrogenase-2 small chain [Proteobacteria] Length: 372\nNumber of predicted TMHs: 1\nExp number of AAs in TMHs: 23.72226\nExp number, first 60 AAs: 0.22038\nTotal prob of N-in: 0.12994\noutside 1 331\nTMhelix 332 354\ninside 355 372

13735 GCF_000617325.1_Ec06-3325 Escherichia coli O69:H11 str. 06-3325 Proteobacteria; Gammaproteobacteria; Enterobacterales; Enterobacteriaceae; Escherichia; Escherichia coli; Escherichia coli O69 MTGDNTLIHSHGINRRDFMKLCAALAATMGLSSKAAAEMAESVTNPQRPPVIWIGAQECT WP_000145410.1 MULTISPECIES: hydrogenase-2 small chain [Proteobacteria] Length: 372\nNumber of predicted TMHs: 1\nExp number of AAs in TMHs: 23.72226\nExp number, first 60 AAs: 0.22038\nTotal prob of N-in: 0.12994\noutside 1 331\nTMhelix 332 354\ninside 355 372

13736 GCF_000617325.1_Ec06-3325 Escherichia coli O69:H11 str. 06-3325 Proteobacteria; Gammaproteobacteria; Enterobacterales; Enterobacteriaceae; Escherichia; Escherichia coli; Escherichia coli O69 MNNEETFYQAMRRQGVTRRSFLKYCSLAATSLGLGAGMAPKIAWALENKPRIPVVWIHGL WP_001058323.1 MULTISPECIES: hydrogenase-1 small chain [Proteobacteria] Length: 372\nNumber of predicted TMHs: 2\nExp number of AAs in TMHs: 37.2608499999999\nExp number, first 60 AAs: 19.52923\nTotal prob of N-in: 0.94351\nPOSSIBLE N-term signal sequence\ninside 1 19\nTMhelix 20 39\noutside 40 325\nTMhelix 326 348\ninside 349 372

13737 GCF_000703205.1_Ec07-3763 Escherichia coli O69:H11 str. 07-3763 Proteobacteria; Gammaproteobacteria; Enterobacterales; Enterobacteriaceae; Escherichia; Escherichia coli; Escherichia coli O69 MNNEETFYQAMRRQGVTRRSFLKYCSLAATSLGLGAGMAPKIAWALENKPRIPVVWIHGL WP_001058323.1

MULTISPECIES: hydrogenase-1 small chain [Proteobacteria] Length: 372\nNumber of predicted TMHs: 2\nExp number of AAs in TMHs: 37.2608499999999\nExp number, first 60 AAs: 19.52923\nTotal prob of N-in: 0.94351\nPOSSIBLE N-term signal sequence\ninside 1 19\nTMhelix 20 39\noutside 40 325\nTMhelix 326 348\ninside 349 372

13738 GCF_000703205.1_Ec07-3763 Escherichia coli O69:H11 str. 07-3763 Proteobacteria; Gammaproteobacteria; Enterobacterales; Enterobacteriaceae; Escherichia; Escherichia coli; Escherichia coli O69 MTGDNTLIHSHGINRRDFMKLCAALAATMGLSSKAAAEMAESVTNPQRPPVIWIGAQECT WP_000145410.1 MULTISPECIES: hydrogenase-2 small chain [Proteobacteria] Length: 372\nNumber of predicted TMHs: 1\nExp number of AAs in TMHs: 23.72226\nExp number, first 60 AAs: 0.22038\nTotal prob of N-in: 0.12994\noutside 1 331\nTMhelix 332 354\ninside 355 372

13739 GCF_000616565.1_Ec07-4281 Escherichia coli O69:H11 str. 07-4281 Proteobacteria; Gammaproteobacteria; Enterobacterales; Enterobacteriaceae; Escherichia; Escherichia coli; Escherichia coli O69 MTGDNTLIHSHGINRRDFMKLCAALAATMGLSSKAAAEMAESVTNPQRPPVIWIGAQECT WP_000145410.1 MULTISPECIES: hydrogenase-2 small chain [Proteobacteria] Length: 372\nNumber of predicted TMHs: 1\nExp number of AAs in TMHs: 23.72226\nExp number, first 60 AAs: 0.22038\nTotal prob of N-in: 0.12994\noutside 1 331\nTMhelix 332 354\ninside 355 372

13740 GCF_000616565.1_Ec07-4281 Escherichia coli O69:H11 str. 07-4281 Proteobacteria; Gammaproteobacteria; Enterobacterales; Enterobacteriaceae; Escherichia; Escherichia coli; Escherichia coli O69 MNNEETFYQAMRRQGVTRRSFLKYCSLAATSLGLGAGMAPKIAWALENKPRIPVVWIHGL WP_001058323.1 MULTISPECIES: hydrogenase-1 small chain [Proteobacteria] Length: 372\nNumber of predicted TMHs: 2\nExp number of AAs in TMHs: 37.2608499999999\nExp number, first 60 AAs: 19.52923\nTotal prob of N-in: 0.94351\nPOSSIBLE N-term signal sequence\ninside 1 19\nTMhelix 20 39\noutside 40 325\nTMhelix 326 348\ninside 349 372

13741 GCF_000619705.2_Ec08-4661 Escherichia coli O69:H11 str. 08-4661 Proteobacteria; Gammaproteobacteria; Enterobacterales; Enterobacteriaceae; Escherichia; Escherichia coli; Escherichia coli O69 MTGDNTLIHSHGINRRDFMKLCAALAATMGLSSKAAAEMAESVTNPQRPPVIWIGAQECT WP_000145410.1 MULTISPECIES: hydrogenase-2 small chain [Proteobacteria] Length: 372\nNumber of predicted TMHs: 1\nExp number of AAs in TMHs: 23.72226\nExp number, first 60 AAs: 0.22038\nTotal prob of N-in: 0.12994\noutside 1 331\nTMhelix 332 354\ninside 355 372

13742 GCF_000619705.2_Ec08-4661 Escherichia coli O69:H11 str. 08-4661 Proteobacteria; Gammaproteobacteria; Enterobacterales; Enterobacteriaceae; Escherichia; Escherichia coli; Escherichia coli O69 MNNEETFYQAMRRQGVTRRSFLKYCSLAATSLGLGAGMAPKIAWALENKPRIPVVWIHGL WP_001058323.1 MULTISPECIES: hydrogenase-1 small chain [Proteobacteria] Length: 372\nNumber of predicted TMHs: 2\nExp number of AAs in TMHs: 37.2608499999999\nExp number, first 60 AAs: 19.52923\nTotal prob of N-in: 0.94351\nPOSSIBLE N-term signal sequence\ninside 1 19\nTMhelix 20 39\noutside 40 325\nTMhelix 326 348\ninside 349 372

13743 GCF_000619285.2_Ec2009C-3601 Escherichia coli O69:H11 str. 2009C-3601 Proteobacteria; Gammaproteobacteria; Enterobacterales; Enterobacteriaceae; Escherichia; Escherichia coli; Escherichia coli O69 MNNEETFYQAMRRQGVTRRSFLKYCSLAATSLGLGAGMAPKIAWALENKPRIPVVWIHGL WP_001058323.1 MULTISPECIES: hydrogenase-1 small chain [Proteobacteria] Length: 372\nNumber of predicted TMHs: 2\nExp number of AAs in TMHs: 37.2608499999999\nExp number, first 60 AAs: 19.52923\nTotal prob of N-in: 0.94351\nPOSSIBLE N-term signal sequence\ninside 1 19\nTMhelix 20 39\noutside 40 325\nTMhelix 326 348\ninside 349 372

13744 GCF_000619285.2_Ec2009C-3601 Escherichia coli O69:H11 str. 2009C-3601 Proteobacteria; Gammaproteobacteria; Enterobacterales; Enterobacteriaceae; Escherichia; Escherichia coli; Escherichia coli O69 MTGDNTLIHSHGINRRDFMKLCAALAATMGLSSKAAAEMAESVTNPQRPPVIWIGAQECT WP_000145410.1 MULTISPECIES: hydrogenase-2 small chain [Proteobacteria] Length: 372\nNumber of predicted TMHs: 1\nExp number of AAs in TMHs: 23.72226\nExp number, first 60 AAs: 0.22038\nTotal prob of N-in: 0.12994\noutside 1 331\nTMhelix 332 354\ninside 355 372

13745 GCF_000227625.1_ASM22762v1 Escherichia coli O7:K1 str. CE10 Proteobacteria; Gammaproteobacteria; Enterobacterales; Enterobacteriaceae; Escherichia; Escherichia coli; Escherichia coli O7 MTGDNTLIHSHGINRRDFMKLCAALAATMGLSSKAAAEMAESVTNPQRPPVIWIGAQECT WP_000145410.1 MULTISPECIES: hydrogenase-2 small chain [Proteobacteria] Length: 372\nNumber of predicted TMHs:

1\nExp number of AAs in TMHs: 23.72226\nExp number, first 60 AAs: 0.22038\nTotal prob of N-in: 0.12994\noutside 1 331\nTMhelix 332 354\ninside 355 372

13746 GCF_000227625.1_ASM22762v1 Escherichia coli O7:K1 str. CE10 Proteobacteria; Gammaproteobacteria; Enterobacterales; Enterobacteriaceae; Escherichia; Escherichia coli; Escherichia coli O7 MNNEETFYQAMRRQGVTRRSFLKYCSLAATSLGLGAGMAPKIAWALENKPRIPVVWIHGL WP_001295940.1 MULTISPECIES: hydrogenase [Enterobacteriaceae] Length: 372\nNumber of predicted TMHs: 2\nExp number of AAs in TMHs: 37.2608899999999\nExp number, first 60 AAs: 19.52923\nTotal prob of N-in: 0.94351\nPOSSIBLE N-term signal sequence\ninside 1 19\nTMhelix 20 39\noutside 40 325\nTMhelix 326 348\ninside 349 372

13747 GCF_000701105.2_ASM70110v2 Escherichia coli O78:H12 str. 00-3279 Proteobacteria; Gammaproteobacteria; Enterobacterales; Enterobacteriaceae; Escherichia; Escherichia coli; Escherichia coli O78 MNNEETFYQAMRRQGVTRRSFLKYCSLAATSLGLGAGMAPKIAWALENKPRIPVVWIHGL WP_001295940.1 MULTISPECIES: hydrogenase [Enterobacteriaceae] Length: 372\nNumber of predicted TMHs: 2\nExp number of AAs in TMHs: 37.2608899999999\nExp number, first 60 AAs: 19.52923\nTotal prob of N-in: 0.94351\nPOSSIBLE N-term signal sequence\ninside 1 19\nTMhelix 20 39\noutside 40 325\nTMhelix 326 348\ninside 349 372

13748 GCF_000701105.2_ASM70110v2 Escherichia coli O78:H12 str. 00-3279 Proteobacteria; Gammaproteobacteria; Enterobacterales; Enterobacteriaceae; Escherichia; Escherichia coli; Escherichia coli O78 MTGDNTLIHSHGINRRDFMKLCAALAATMGLSSKAAAEMAESVTNPQRPPVIWIGAQECT WP_000145410.1 MULTISPECIES: hydrogenase-2 small chain [Proteobacteria] Length: 372\nNumber of predicted TMHs: 1\nExp number of AAs in TMHs: 23.72226\nExp number, first 60 AAs: 0.22038\nTotal prob of N-in: 0.12994\noutside 1 331\nTMhelix 332 354\ninside 355 372

13749 GCF_000617365.2_Ec06-3501 Escherichia coli O79:H7 str. 06-3501 Proteobacteria; Gammaproteobacteria; Enterobacterales; Enterobacteriaceae; Escherichia; Escherichia coli; Escherichia coli O79 MNNEETFYQAMRRQGVTRRSFLKYCSLAATSLGLGAGMAPKIAWALENKPRIPVVWIHGL WP_001058323.1 MULTISPECIES: hydrogenase-1 small chain [Proteobacteria] Length: 372\nNumber of predicted TMHs: 2\nExp number of AAs in TMHs: 37.2608499999999\nExp number, first 60 AAs: 19.52923\nTotal prob of N-in: 0.94351\nPOSSIBLE N-term signal sequence\ninside 1 19\nTMhelix 20 39\noutside 40 325\nTMhelix 326 348\ninside 349 372

13750 GCF_000617365.2_Ec06-3501 Escherichia coli O79:H7 str. 06-3501 Proteobacteria; Gammaproteobacteria; Enterobacterales; Enterobacteriaceae; Escherichia; Escherichia coli; Escherichia coli O79 MTGDNTLIHSHGINRRDFMKLCAALAATMGLSSKAAAEMAESVTNPQRPPVIWIGAQECT WP_000145410.1 MULTISPECIES: hydrogenase-2 small chain [Proteobacteria] Length: 372\nNumber of predicted TMHs: 1\nExp number of AAs in TMHs: 23.72226\nExp number, first 60 AAs: 0.22038\nTotal prob of N-in: 0.12994\noutside 1 331\nTMhelix 332 354\ninside 355 372

13751 GCF_000617145.2_Ec02-3012 Escherichia coli O81:NM str. 02-3012 Proteobacteria; Gammaproteobacteria; Enterobacterales; Enterobacteriaceae; Escherichia; Escherichia coli; Escherichia coli O81 MNNEETFYQAMRRQGVTRRSFLKYCSLAATSLGLGAGMAPKIAWALENKPRIPVVWIHGL WP_032325754.1 hydrogenase [Escherichia coli] Length: 372\nNumber of predicted TMHs: 2\nExp number of AAs in TMHs: 38.85546\nExp number, first 60 AAs: 19.52672\nTotal prob of N-in: 0.94441\nPOSSIBLE N-term signal sequence\ninside 1 19\nTMhelix 20 39\noutside 40 325\nTMhelix 326 348\ninside 349 372

13752 GCF_000617145.2_Ec02-3012 Escherichia coli O81:NM str. 02-3012 Proteobacteria; Gammaproteobacteria; Enterobacterales; Enterobacteriaceae; Escherichia; Escherichia coli; Escherichia coli O81 MTGDNTLIHSHGINRRDFMKLCAALAATMGLSSKAAAEMAESVTNPQRPPVIWIGAQECT WP_000145418.1 hydrogenase 2 small subunit [Escherichia coli] Length: 372\nNumber of predicted TMHs: 1\nExp number of AAs in TMHs: 23.72313\nExp number, first 60 AAs: 0.22034\nTotal prob of N-in: 0.12997\noutside 1 331\nTMhelix 332 354\ninside 355 372

13753 GCF_000183345.1_ASM18334v1 Escherichia coli O83:H1 str. NRG 857C Proteobacteria; Gammaproteobacteria; Enterobacterales; Enterobacteriaceae; Escherichia; Escherichia coli; Escherichia coli O83 MNNEETFYQAMRRQGVTRRSFLKYCSLAATSLGLGAGMAPKIAWALENKPRIPVVWIHGL YP_006119245.1 hydrogenase-1 small subunit [Escherichia coli O83:H1 str. NRG 857C] Length: 372\nNumber of predicted TMHs: 2\nExp number of AAs in TMHs: 37.2608499999999\nExp number, first 60 AAs: 19.52923\nTotal prob of N-in:

0.94351\nPOSSIBLE N-term signal sequence\ninside 1 19\nTMhelix 20 39\noutside 40 325\nTMhelix 326 348\ninside 349 372

13754 GCF_000183345.1_ASM18334v1 Escherichia coli O83:H1 str. NRG 857C Proteobacteria; Gammaproteobacteria; Enterobacterales; Enterobacteriaceae; Escherichia; Escherichia coli; Escherichia coli O83 MTGDNTLIHSHGINRRDFMKLCAALAATMGLSSKAAAEMAESVTNPQRPPVIWIGAQECT YP_006121319.1 hydrogenase 2 small subunit [Escherichia coli O83:H1 str. NRG 857C] Length: 372\nNumber of predicted TMHs: 1\nExp number of AAs in TMHs: 23.72313\nExp number, first 60 AAs: 0.22034\nTotal prob of N-in: 0.12997\noutside 1 331\nTMhelix 332 354\ninside 355 372

13755 GCF_000614465.2_Ec99-3124 Escherichia coli O86:H34 str. 99-3124 Proteobacteria; Gammaproteobacteria; Enterobacterales; Enterobacteriaceae; Escherichia; Escherichia coli; Escherichia coli O86 MTGDNTLIHSHGINRRDFMKLCAALAATMGLSSKAAAEMAESVTNPQRPPVIWIGAQECT WP_000145410.1 MULTISPECIES: hydrogenase-2 small chain [Proteobacteria] Length: 372\nNumber of predicted TMHs: 1\nExp number of AAs in TMHs: 23.72226\nExp number, first 60 AAs: 0.22038\nTotal prob of N-in: 0.12994\noutside 1 331\nTMhelix 332 354\ninside 355 372

13756 GCF_000614465.2_Ec99-3124 Escherichia coli O86:H34 str. 99-3124 Proteobacteria; Gammaproteobacteria; Enterobacterales; Enterobacteriaceae; Escherichia; Escherichia coli; Escherichia coli O86 MNNEETFYQAMRRQGVTRRSFLKYCSLAATSLGLGAGMAPKIAWALENKPRIPVVWIHGL WP_001058323.1 MULTISPECIES: hydrogenase-1 small chain [Proteobacteria] Length: 372\nNumber of predicted TMHs: 2\nExp number of AAs in TMHs: 37.2608499999999\nExp number, first 60 AAs: 19.52923\nTotal prob of N-in: 0.94351\nPOSSIBLE N-term signal sequence\ninside 1 19\nTMhelix 20 39\noutside 40 325\nTMhelix 326 348\ninside 349 372

13757 GCF_000617425.1_Ec06-3691 Escherichia coli O91:H14 str. 06-3691 Proteobacteria; Gammaproteobacteria; Enterobacterales; Enterobacteriaceae; Escherichia; Escherichia coli; Escherichia coli O91 MNNEETFYQAMRRQGVTRRSFLKYCSLAATSLGLGAGMAPKIAWALENKPRIPVVWIHGL WP_001058323.1 MULTISPECIES: hydrogenase-1 small chain [Proteobacteria] Length: 372\nNumber of predicted TMHs: 2\nExp number of AAs in TMHs: 37.2608499999999\nExp number, first 60 AAs: 19.52923\nTotal prob of N-in: 0.94351\nPOSSIBLE N-term signal sequence\ninside 1 19\nTMhelix 20 39\noutside 40 325\nTMhelix 326 348\ninside 349 372

13758 GCF_000617425.1_Ec06-3691 Escherichia coli O91:H14 str. 06-3691 Proteobacteria; Gammaproteobacteria; Enterobacterales; Enterobacteriaceae; Escherichia; Escherichia coli; Escherichia coli O91 MTGDNTLIHSHGINRRDFMKLCAALAATMGLSSKAAAEMAESVTNPQRPPVIWIGAQECT WP_000145410.1 MULTISPECIES: hydrogenase-2 small chain [Proteobacteria] Length: 372\nNumber of predicted TMHs: 1\nExp number of AAs in TMHs: 23.72226\nExp number, first 60 AAs: 0.22038\nTotal prob of N-in: 0.12994\noutside 1 331\nTMhelix 332 354\ninside 355 372

13759 GCF_000619725.2_Ec2009C-3227 Escherichia coli O91:H14 str. 2009C-3227 Proteobacteria; Gammaproteobacteria; Enterobacterales; Enterobacteriaceae; Escherichia; Escherichia coli; Escherichia coli O91 MTGDNTLIHSHGINRRDFMKLCAALAATMGLSSKAAAEMAESVTNPQRPPVIWIGAQECT WP_000145410.1 MULTISPECIES: hydrogenase-2 small chain [Proteobacteria] Length: 372\nNumber of predicted TMHs: 1\nExp number of AAs in TMHs: 23.72226\nExp number, first 60 AAs: 0.22038\nTotal prob of N-in: 0.12994\noutside 1 331\nTMhelix 332 354\ninside 355 372

13760 GCF_000619725.2_Ec2009C-3227 Escherichia coli O91:H14 str. 2009C-3227 Proteobacteria; Gammaproteobacteria; Enterobacterales; Enterobacteriaceae; Escherichia; Escherichia coli; Escherichia coli O91 MNNEETFYQAMRRQGVTRRSFLKYCSLAATSLGLGAGMAPKIAWALENKPRIPVVWIHGL WP_001058323.1 MULTISPECIES: hydrogenase-1 small chain [Proteobacteria] Length: 372\nNumber of predicted TMHs: 2\nExp number of AAs in TMHs: 37.2608499999999\nExp number, first 60 AAs: 19.52923\nTotal prob of N-in: 0.94351\nPOSSIBLE N-term signal sequence\ninside 1 19\nTMhelix 20 39\noutside 40 325\nTMhelix 326 348\ninside 349 372

13761 GCF_000703265.1_Ec2009C-3740 Escherichia coli O91:H21 str. 2009C-3740 Proteobacteria; Gammaproteobacteria; Enterobacterales; Enterobacteriaceae; Escherichia; Escherichia coli; Escherichia coli O91 MNNEETFYQAMRRQGVTRRSFLKYCSLAATSLGLGAGMAPKIAWALENKPRIPVVWIHGL WP_001295940.1 MULTISPECIES: hydrogenase [Enterobacteriaceae] Length: 372\nNumber of predicted TMHs: 2\nExp number of AAs in TMHs: 37.2608899999999\nExp number, first 60 AAs: 19.52923\nTotal prob of N-in:

0.94351\nPOSSIBLE N-term signal sequence\ninside 1 19\nTMhelix 20 39\noutside 40 325\nTMhelix 326 348\ninside 349 372

13762 GCF_000703265.1_Ec2009C-3740 Escherichia coli O91:H21 str. 2009C-3740 Proteobacteria; Gammaproteobacteria; Enterobacterales; Enterobacteriaceae; Escherichia; Escherichia coli; Escherichia coli O91 MTGDNTLIHSHGINRRDFMKLCAALAATMGLSSKAAAEMAESVTNPQRPPVIWIGAQECT WP_000145410.1 MULTISPECIES: hydrogenase-2 small chain [Proteobacteria] Length: 372\nNumber of predicted TMHs: 1\nExp number of AAs in TMHs: 23.72226\nExp number, first 60 AAs: 0.22038\nTotal prob of N-in: 0.12994\noutside 1 331\nTMhelix 332 354\ninside 355 372

13763 GCF_000619065.2_Ec2009C-4646 Escherichia coli O91:H21 str. 2009C-4646 Proteobacteria; Gammaproteobacteria; Enterobacterales; Enterobacteriaceae; Escherichia; Escherichia coli; Escherichia coli O91 MTGDNTLIHSHGINRRDFMKLCAALAATMGLSSKAAAEMAESVTNPQRPPVIWIGAQECT WP_000145410.1 MULTISPECIES: hydrogenase-2 small chain [Proteobacteria] Length: 372\nNumber of predicted TMHs: 1\nExp number of AAs in TMHs: 23.72226\nExp number, first 60 AAs: 0.22038\nTotal prob of N-in: 0.12994\noutside 1 331\nTMhelix 332 354\ninside 355 372

13764 GCF_000619065.2_Ec2009C-4646 Escherichia coli O91:H21 str. 2009C-4646 Proteobacteria; Gammaproteobacteria; Enterobacterales; Enterobacteriaceae; Escherichia; Escherichia coli; Escherichia coli O91 MNNEETFYQAMRRQGVTRRSFLKYCSLAATSLGLGAGMAPKIAWALENKPRIPVVWIHGL WP_001295940.1 MULTISPECIES: hydrogenase [Enterobacteriaceae] Length: 372\nNumber of predicted TMHs: 2\nExp number of AAs in TMHs: 37.2608899999999\nExp number, first 60 AAs: 19.52923\nTotal prob of N-in: 0.94351\nPOSSIBLE N-term signal sequence\ninside 1 19\nTMhelix 20 39\noutside 40 325\nTMhelix 326 348\ninside 349 372

13765 GCF_000234255.2_ASM23425v3 Escherichia coli O91:H21 str. B2F1 Proteobacteria; Gammaproteobacteria; Enterobacterales; Enterobacteriaceae; Escherichia; Escherichia coli; Escherichia coli O91 MTGDNTLIHSHGINRRDFMKLCAALAATMGLSSKAAAEMAESVTNPQRPPVIWIGAQECT WP_000145410.1 MULTISPECIES: hydrogenase-2 small chain [Proteobacteria] Length: 372\nNumber of predicted TMHs: 1\nExp number of AAs in TMHs: 23.72226\nExp number, first 60 AAs: 0.22038\nTotal prob of N-in: 0.12994\noutside 1 331\nTMhelix 332 354\ninside 355 372

13766 GCF_000234255.2_ASM23425v3 Escherichia coli O91:H21 str. B2F1 Proteobacteria; Gammaproteobacteria; Enterobacterales; Enterobacteriaceae; Escherichia; Escherichia coli; Escherichia coli O91 MNNEETFYQAMRRQGVTRRSFLKYCSLAATSLGLGAGMAPKIAWALENKPRIPVVWIHGL WP_001295940.1 MULTISPECIES: hydrogenase [Enterobacteriaceae] Length: 372\nNumber of predicted TMHs: 2\nExp number of AAs in TMHs: 37.2608899999999\nExp number, first 60 AAs: 19.52923\nTotal prob of N-in: 0.94351\nPOSSIBLE N-term signal sequence\ninside 1 19\nTMhelix 20 39\noutside 40 325\nTMhelix 326 348\ninside 349 372

13767 GCF_000619305.1_Ec2009C-3745 Escherichia coli O91:NM str. 2009C-3745 Proteobacteria; Gammaproteobacteria; Enterobacterales; Enterobacteriaceae; Escherichia; Escherichia coli; Escherichia coli O91 MTGDNTLIHSHGINRRDFMKLCAALAATMGLSSKAAAEMAESVTNPQRPPVIWIGAQECT WP_000145410.1 MULTISPECIES: hydrogenase-2 small chain [Proteobacteria] Length: 372\nNumber of predicted TMHs: 1\nExp number of AAs in TMHs: 23.72226\nExp number, first 60 AAs: 0.22038\nTotal prob of N-in: 0.12994\noutside 1 331\nTMhelix 332 354\ninside 355 372

13768 GCF_000619305.1_Ec2009C-3745 Escherichia coli O91:NM str. 2009C-3745 Proteobacteria; Gammaproteobacteria; Enterobacterales; Enterobacteriaceae; Escherichia; Escherichia coli; Escherichia coli O91 MNNEETFYQAMRRQGVTRRSFLKYCSLAATSLGLGAGMAPKIAWALENKPRIPVVWIHGL WP_001058323.1 MULTISPECIES: hydrogenase-1 small chain [Proteobacteria] Length: 372\nNumber of predicted TMHs: 2\nExp number of AAs in TMHs: 37.2608499999999\nExp number, first 60 AAs: 19.52923\nTotal prob of N-in: 0.94351\nPOSSIBLE N-term signal sequence\ninside 1 19\nTMhelix 20 39\noutside 40 325\nTMhelix 326 348\ninside 349 372

13769 GCF_000350025.2_C48_93_IrSpV01 Escherichia coli ONT:H33 str. C48/93 Proteobacteria; Gammaproteobacteria; Enterobacterales; Enterobacteriaceae; Escherichia; Escherichia coli; Escherichia coli ONT MNNEETFYQAMRRQGVTRRSFLKYCSLAATSLGLGAGMAPKIAWALENKPRIPVVWIHGL WP_001058323.1 MULTISPECIES: hydrogenase-1 small chain [Proteobacteria] Length: 372\nNumber of predicted TMHs: 2\nExp number of AAs in TMHs: 37.2608499999999\nExp number, first 60 AAs: 19.52923\nTotal prob of N-in:

0.94351\nPOSSIBLE N-term signal sequence\ninside 1 19\nTMhelix 20 39\noutside 40 325\nTMhelix 326 348\ninside 349 372

13770 GCF_000350025.2_C48_93_IrSpV01 Escherichia coli ONT:H33 str. C48/93 Proteobacteria; Gammaproteobacteria; Enterobacterales; Enterobacteriaceae; Escherichia; Escherichia coli; Escherichia coli ONT MTGDNTLIHSHGINRRDFMKLCAALAATMGLSSKAAAEMAESVTNPQRPPVIWIGAQECT WP_000145410.1 MULTISPECIES: hydrogenase-2 small chain [Proteobacteria] Length: 372\nNumber of predicted TMHs: 1\nExp number of AAs in TMHs: 23.72226\nExp number, first 60 AAs: 0.22038\nTotal prob of N-in: 0.12994\noutside 1 331\nTMhelix 332 354\ninside 355 372

13771 GCF_000026225.1_ASM2622v1 Escherichia fergusonii ATCC 35469 Proteobacteria; Gammaproteobacteria; Enterobacterales; Enterobacteriaceae; Escherichia; Escherichia fergusonii MTGDNTLDNSHGINRRDFMKLCAALAATMGLSSKAAAEMAESVANPQRPPVIWIGAQECT WP_000145402.1 hydrogenase 2 small subunit [Escherichia fergusonii] Length: 372\nNumber of predicted TMHs: 1\nExp number of AAs in TMHs: 24.63122\nExp number, first 60 AAs: 0.32267\nTotal prob of N-in: 0.18195\noutside 1 331\nTMhelix 332 354\ninside 355 372

13772 GCF_000026225.1_ASM2622v1 Escherichia fergusonii ATCC 35469 Proteobacteria; Gammaproteobacteria; Enterobacterales; Enterobacteriaceae; Escherichia; Escherichia fergusonii MNNEETFYQAMRRQGVTRRSFLKYCSLAATSLGLGAGMAPKIAWALENKPRIPVVWIHGL WP_002431613.1 hydrogenase-1 small chain [Escherichia fergusonii] Length: 372\nNumber of predicted TMHs: 2\nExp number of AAs in TMHs: 37.25693\nExp number, first 60 AAs: 19.52926\nTotal prob of N-in: 0.94351\nPOSSIBLE N-term signal sequence\ninside 1 19\nTMhelix 20 39\noutside 40 325\nTMhelix 326 348\ninside 349 372

13773 GCF_000190495.1_ASM19049v1 Escherichia fergusonii B253 Proteobacteria; Gammaproteobacteria; Enterobacterales; Enterobacteriaceae; Escherichia; Escherichia fergusonii MTGDNTLDNSHGINRRDFMKLCAALAATMGLSSKAAAEMAESVANPQRPPVIWIGAQECT WP_000145402.1 hydrogenase 2 small subunit [Escherichia fergusonii] Length: 372\nNumber of predicted TMHs: 1\nExp number of AAs in TMHs: 24.63122\nExp number, first 60 AAs: 0.32267\nTotal prob of N-in: 0.18195\noutside 1 331\nTMhelix 332 354\ninside 355 372

13774 GCF_000190495.1_ASM19049v1 Escherichia fergusonii B253 Proteobacteria; Gammaproteobacteria; Enterobacterales; Enterobacteriaceae; Escherichia; Escherichia fergusonii MNNEETFYQAMRRQGVTRRSFLKYCSLAATSLGLGAGMAPKIAWALENKPRIPVVWIHGL WP_002431613.1 hydrogenase-1 small chain [Escherichia fergusonii] Length: 372\nNumber of predicted TMHs: 2\nExp number of AAs in TMHs: 37.25693\nExp number, first 60 AAs: 19.52926\nTotal prob of N-in: 0.94351\nPOSSIBLE N-term signal sequence\ninside 1 19\nTMhelix 20 39\noutside 40 325\nTMhelix 326 348\ninside 349 372

13775 GCF_000191665.1_ecmda7 Escherichia fergusonii ECD227 Proteobacteria; Gammaproteobacteria; Enterobacterales; Enterobacteriaceae; Escherichia; Escherichia fergusonii MNNEETFYQAMRRQGVTRRSFLKYCSLAATSLGLGAGMAPKIAWALENKPRIPVVWIHGL WP_002431613.1 hydrogenase-1 small chain [Escherichia fergusonii] Length: 372\nNumber of predicted TMHs: 2\nExp number of AAs in TMHs: 37.25693\nExp number, first 60 AAs: 19.52926\nTotal prob of N-in: 0.94351\nPOSSIBLE N-term signal sequence\ninside 1 19\nTMhelix 20 39\noutside 40 325\nTMhelix 326 348\ninside 349 372

13776 GCF_000191665.1_ecmda7 Escherichia fergusonii ECD227 Proteobacteria; Gammaproteobacteria; Enterobacterales; Enterobacteriaceae; Escherichia; Escherichia fergusonii MTGDNTLDNSHGINRRDFMKLCAALAATMGLSSKAAAEMAESVANPQRPPVIWIGAQECT WP_000145402.1 hydrogenase 2 small subunit [Escherichia fergusonii] Length: 372\nNumber of predicted TMHs: 1\nExp number of AAs in TMHs: 24.63122\nExp number, first 60 AAs: 0.32267\nTotal prob of N-in: 0.18195\noutside 1 331\nTMhelix 332 354\ninside 355 372

13777 GCF_000957705.1_ASM95770v1 Klebsiella aerogenes Proteobacteria; Gammaproteobacteria; Enterobacterales; Enterobacteriaceae; Klebsiella MHYLNREARREVIMQAAMQVALNGGFSAMTVRQIASEAGVAAGQLHHHFTSTGELKAQAF WP_045382429.1 TetR family transcriptional regulator [Klebsiella aerogenes] Length: 193\nNumber of predicted TMHs: 1\nExp number of AAs in TMHs: 14.61655\nExp number, first 60 AAs: 0.26037\nTotal prob of N-in: 0.75782\ninside 1 150\nTMhelix 151 173\noutside 174 193

13778 GCF_001058645.1_ASM105864v1 Klebsiella aerogenes Proteobacteria; Gammaproteobacteria; Enterobacterales; Enterobacteriaceae; Klebsiella

MHYLNREARREVIMQAAMQVALNGGFSAMTVRQIASEAGVAAGQLHHHFTSTGELKAQAF WP_049058166.1 TetR family transcriptional regulator [Klebsiella aerogenes] Length: 193\nNumber of predicted TMHs: 1\nExp number of AAs in TMHs: 14.6165\nExp number, first 60 AAs: 0.26033\nTotal prob of N-in: 0.75782\ninside 1 150\nTMhelix 151 173\noutside 174 193

13779 GCF_000878365.1_ASM87836v1 Klebsiella aerogenes Proteobacteria; Gammaproteobacteria; Enterobacterales; Enterobacteriaceae; Klebsiella
MSEGQNAFYHNGISRRDFMTLCAALSATMGLTGNASAEMVEALASPSRPPVVWIGAQECT WP_043866230.1 hydrogenase 2 small subunit [Klebsiella aerogenes] Length: 376\nNumber of predicted TMHs: 1\nExp number of AAs in TMHs: 21.84784\nExp number, first 60 AAs: 0.66622\nTotal prob of N-in: 0.11050\noutside 1 330\nTMhelix 331 353\ninside 354 376

13780 GCF_001011595.1_ASM101159v1 Klebsiella aerogenes Proteobacteria; Gammaproteobacteria; Enterobacterales; Enterobacteriaceae; Klebsiella
MHYLNREARREVIMQAAMQVALNGGFSAMTVRQIASEAGVAAGQLHHHFTSTGELKAQAF WP_045382429.1 TetR family transcriptional regulator [Klebsiella aerogenes] Length: 193\nNumber of predicted TMHs: 1\nExp number of AAs in TMHs: 14.61655\nExp number, first 60 AAs: 0.26037\nTotal prob of N-in: 0.75782\ninside 1 150\nTMhelix 151 173\noutside 174 193

13781 GCF_001631645.1_ASM163164v1 Klebsiella aerogenes Proteobacteria; Gammaproteobacteria; Enterobacterales; Enterobacteriaceae; Klebsiella
MKEVLEKCTVEDFNYISATLDSYVSFTNDKRRKTLLSAYQNNPALHAELISLIDTQIKYF WP_063444545.1 hypothetical protein [Klebsiella aerogenes] Length: 249\nNumber of predicted TMHs: 2\nExp number of AAs in TMHs: 39.22785\nExp number, first 60 AAs: 0.0234\nTotal prob of N-in: 0.11298\noutside 1 157\nTMhelix 158 180\ninside 181 200\nTMhelix 201 220\noutside 221 249

13782 GCF_001057945.1_ASM105794v1 Klebsiella aerogenes Proteobacteria; Gammaproteobacteria; Enterobacterales; Enterobacteriaceae; Klebsiella
MHYLNREARREVIMQAAMQVALNGGFSAMTVRQIASEAGVAAGQLHHHFTSTGELKAQAF WP_045382429.1 TetR family transcriptional regulator [Klebsiella aerogenes] Length: 193\nNumber of predicted TMHs: 1\nExp number of AAs in TMHs: 14.61655\nExp number, first 60 AAs: 0.26037\nTotal prob of N-in: 0.75782\ninside 1 150\nTMhelix 151 173\noutside 174 193

13783 GCF_900083925.1_12045_7_1 Klebsiella oxytoca Proteobacteria; Gammaproteobacteria; Enterobacterales; Enterobacteriaceae; Klebsiella
MNKRRTTLTVLALIASLGLSSAPALADKGGNGNGNGNGHGNSGNNGNHGNNGNHGNKGNK WP_049200367.1 MULTISPECIES: membrane protein [Enterobacterales] Length: 164\nNumber of predicted TMHs: 2\nExp number of AAs in TMHs: 30.51526\nExp number, first 60 AAs: 14.42673\nTotal prob of N-in: 0.94579\nPOSSIBLE N-term signal sequence\ninside 1 6\nTMhelix 7 26\noutside 27 144\nTMhelix 145 163\ninside 164 164

13784 GCF_900083935.1_11983_8_79 Klebsiella oxytoca Proteobacteria; Gammaproteobacteria; Enterobacterales; Enterobacteriaceae; Klebsiella
MNNEETFYQAMRRQGVTRRSFLKYCSLAATSLGLGAGMAPKIAWALENKPRIPVVWIHGL WP_001058323.1 MULTISPECIES: hydrogenase-1 small chain [Proteobacteria] Length: 372\nNumber of predicted TMHs: 2\nExp number of AAs in TMHs: 37.2608499999999\nExp number, first 60 AAs: 19.52923\nTotal prob of N-in: 0.94351\nPOSSIBLE N-term signal sequence\ninside 1 19\nTMhelix 20 39\noutside 40 325\nTMhelix 326 348\ninside 349 372

13785 GCF_900083935.1_11983_8_79 Klebsiella oxytoca Proteobacteria; Gammaproteobacteria; Enterobacterales; Enterobacteriaceae; Klebsiella
MTGDNTLIHSHGINRRDFMKLCAALAATMGLSSKAAAEMAESVTNPQRPPVIWIGAQECT WP_000145410.1 MULTISPECIES: hydrogenase-2 small chain [Proteobacteria] Length: 372\nNumber of predicted TMHs: 1\nExp number of AAs in TMHs: 23.72226\nExp number, first 60 AAs: 0.22038\nTotal prob of N-in: 0.12994\noutside 1 331\nTMhelix 332 354\ninside 355 372

13786 GCF_001682915.1_ASM168291v1 Kluyvera ascorbata Proteobacteria; Gammaproteobacteria; Enterobacterales; Enterobacteriaceae; Kluyvera
MKRLLPRTLFLTRKLRIIRQQGWSRVQRLARRGHLWQWMIVAPMALLIVYLAIFSQPRFV WP_065357034.1 capsule biosynthesis protein [Kluyvera ascorbata] Length: 389\nNumber of predicted TMHs: 2\nExp number of AAs in TMHs: 42.78902\nExp number, first 60 AAs: 20.84196\nTotal prob of N-in: 0.77775\nPOSSIBLE N-term signal sequence\ninside 1 34\nTMhelix 35 54\noutside 55 361\nTMhelix 362 384\ninside 385 389

13787 GCF_001022135.1_ASM102213v1 Kluyvera intermedia Proteobacteria; Gammaproteobacteria; Enterobacterales; Enterobacteriaceae; Kluyvera
MSFTSCGINRRDFMKLCTALAATMGLSSNAAAKMAQAVTQNKRPPVVWIGAQECTGCTES WP_047373483.1
hydrogenase 2 small subunit [Kluyvera intermedia] Length: 367\nNumber of predicted TMHs: 1\nExp number of AAs in TMHs: 25.29225\nExp number, first 60 AAs: 1.03612\nTotal prob of N-in: 0.20333\noutside 1 325\nTMhelix 326 348\ninside 349 367

13788 GCF_002092975.1_ASM209297v1 Kluyvera intermedia Proteobacteria; Gammaproteobacteria; Enterobacterales; Enterobacteriaceae; Kluyvera
MADEIPQLTSFTSCGINRRDFMKLCTALAATMGLSSNAAAKMAQAVTQNKRPPVVWIGAQ WP_085005770.1
hydrogenase 2 small subunit [Kluyvera intermedia] Length: 375\nNumber of predicted TMHs: 1\nExp number of AAs in TMHs: 23.27892\nExp number, first 60 AAs: 0.52471\nTotal prob of N-in: 0.08738\noutside 1 333\nTMhelix 334 356\ninside 357 375

13789 GCF_001856865.1_ASM185686v1 Kluyvera intestini Proteobacteria; Gammaproteobacteria; Enterobacterales; Enterobacteriaceae; Kluyvera
MADEIPQLTSFTSCGINRRDFMKLCTALAATMGLSSTAAAKMAQAVTQSKRPPVVWIGAQ WP_071194554.1
hydrogenase 2 small subunit [Kluyvera intestini] Length: 375\nNumber of predicted TMHs: 1\nExp number of AAs in TMHs: 24.47449\nExp number, first 60 AAs: 1.76766\nTotal prob of N-in: 0.14301\noutside 1 333\nTMhelix 334 356\ninside 357 375

13790 GCF_001654985.1_Kge51603_DRAFTv1 Kluyvera georgiana ATCC 51603 Proteobacteria; Gammaproteobacteria; Enterobacterales; Enterobacteriaceae; Kluyvera; Kluyvera georgiana
MKRLLPRTLFLTRKLRIIRQQGWSRVQRLARRGHLWQWMIVAPMALLIVYLAIFSQPRFV WP_064547219.1
hypothetical protein [Kluyvera georgiana] Length: 389\nNumber of predicted TMHs: 2\nExp number of AAs in TMHs: 42.55483\nExp number, first 60 AAs: 20.83832\nTotal prob of N-in: 0.76850\nPOSSIBLE N-term signal sequence\ninside 1 34\nTMhelix 35 54\noutside 55 361\nTMhelix 362 384\ninside 385 389

13791 GCF_900168185.1_IMG-taxon_2667527231_annotated_assembly Kosakonia oryzae Proteobacteria; Gammaproteobacteria; Enterobacterales; Enterobacteriaceae; Kosakonia
MAGDNLHLASQGVSRRDFMKLCAALAATMGLSGKAAAEMAEAISHPQRPPVVWIGAQECT WP_007370143.1
MULTISPECIES: hydrogenase 2 small subunit [Enterobacteriaceae] Length: 372\nNumber of predicted TMHs: 1\nExp number of AAs in TMHs: 24.18752\nExp number, first 60 AAs: 0.2398\nTotal prob of N-in: 0.11792\noutside 1 332\nTMhelix 333 355\ninside 356 372

13792 GCF_900112145.1_IMG-taxon_2675902965_annotated_assembly Kosakonia oryzae Proteobacteria; Gammaproteobacteria; Enterobacterales; Enterobacteriaceae; Kosakonia
MAGDNLHLASQGVSRRDFMKLCAALAATMGLSGKAAAEMAEAISHPQRPPVVWIGAQECT WP_007370143.1
MULTISPECIES: hydrogenase 2 small subunit [Enterobacteriaceae] Length: 372\nNumber of predicted TMHs: 1\nExp number of AAs in TMHs: 24.18752\nExp number, first 60 AAs: 0.2398\nTotal prob of N-in: 0.11792\noutside 1 332\nTMhelix 333 355\ninside 356 372

13793 GCF_000958895.1_ASM95889v1 Kosakonia oryzae Proteobacteria; Gammaproteobacteria; Enterobacterales; Enterobacteriaceae; Kosakonia
MAGENQQFASHGVSRRDFMKLCAALAATMGLSGKAAAEMAEAVSHPQRPPVIWIGAQECT WP_045512138.1
hydrogenase 2 small subunit [Kosakonia oryzae] Length: 372\nNumber of predicted TMHs: 1\nExp number of AAs in TMHs: 25.61475\nExp number, first 60 AAs: 0.31978\nTotal prob of N-in: 0.21983\noutside 1 330\nTMhelix 331 353\ninside 354 372

13794 GCF_001658025.1_ASM165802v1 Kosakonia oryzae Proteobacteria; Gammaproteobacteria; Enterobacterales; Enterobacteriaceae; Kosakonia
MAGDNLHLASQGVSRRDFMKLCAALAATMGLSGKAAAEMAEAISHPQRPPVVWIGAQECT WP_007370143.1
MULTISPECIES: hydrogenase 2 small subunit [Enterobacteriaceae] Length: 372\nNumber of predicted TMHs: 1\nExp number of AAs in TMHs: 24.18752\nExp number, first 60 AAs: 0.2398\nTotal prob of N-in: 0.11792\noutside 1 332\nTMhelix 333 355\ninside 356 372

13795 GCF_900185945.1_ASM90018594v1 Kosakonia oryzendophytica Proteobacteria; Gammaproteobacteria; Enterobacterales; Enterobacteriaceae; Kosakonia
MAGENQHFVSQGVSRRDFMKLCAALAATMGLSGKAAAEMAEAVSHPQRPPVIWIGAQECT WP_088238403.1
hydrogenase 2 small subunit [Kosakonia oryzendophytica] Length: 372\nNumber of predicted TMHs: 1\nExp

number of AAs in TMHs: 23.58996\nExp number, first 60 AAs: 0.32199\nTotal prob of N-in: 0.09013\noutside 1 330\nTMhelix 331 353\ninside 354 372

13796 GCF_900184035.1_Kosakonia_pseudosacchari_JM-387_T Kosakonia pseudosacchari Proteobacteria; Gammaproteobacteria; Enterobacterales; Enterobacteriaceae; Kosakonia
MAGENQQFASHGVSRRDFMKLCAALAATMGLSGKAAAEMAEAVSHPQRPPVIWIGAQECT WP_086873409.1
hydrogenase 2 small subunit [Kosakonia pseudosacchari] Length: 372\nNumber of predicted TMHs: 1\nExp number of AAs in TMHs: 25.56529\nExp number, first 60 AAs: 0.31976\nTotal prob of N-in: 0.21964\noutside 1 330\nTMhelix 331 353\ninside 354 372

13797 GCF_001887675.1_ASM188767v1 Kosakonia radicincitans Proteobacteria; Gammaproteobacteria; Enterobacterales; Enterobacteriaceae; Kosakonia
MAGDNLHLASQGVSRRDFMKLCAALAATMGLSGKAAAEMAEAISHPQRPPVVWIGAQECT WP_007370143.1
MULTISPECIES: hydrogenase 2 small subunit [Enterobacteriaceae] Length: 372\nNumber of predicted TMHs: 1\nExp number of AAs in TMHs: 24.18752\nExp number, first 60 AAs: 0.2398\nTotal prob of N-in: 0.11792\noutside 1 332\nTMhelix 333 355\ninside 356 372

13798 GCF_900109485.1_IMG-taxon_2623620455_annotated_assembly Kosakonia sacchari Proteobacteria; Gammaproteobacteria; Enterobacterales; Enterobacteriaceae; Kosakonia
MAGDNLHLASQGVSRRDFMKLCAALAATMGLSGKAAAEMAEAISHPQRPPVVWIGAQECT WP_007370143.1
MULTISPECIES: hydrogenase 2 small subunit [Enterobacteriaceae] Length: 372\nNumber of predicted TMHs: 1\nExp number of AAs in TMHs: 24.18752\nExp number, first 60 AAs: 0.2398\nTotal prob of N-in: 0.11792\noutside 1 332\nTMhelix 333 355\ninside 356 372

13799 GCF_900100995.1_IMG-taxon_2596583580_annotated_assembly Kosakonia sacchari Proteobacteria; Gammaproteobacteria; Enterobacterales; Enterobacteriaceae; Kosakonia
MAGENQQFASHGVSRRDFMKLCAALAATMGLSGKAAAEMAKAVSHPQRPPVIWIGAQECT WP_017457491.1
hydrogenase 2 small subunit [Kosakonia sacchari] Length: 372\nNumber of predicted TMHs: 1\nExp number of AAs in TMHs: 26.66562\nExp number, first 60 AAs: 0.34625\nTotal prob of N-in: 0.26285\noutside 1 330\nTMhelix 331 353\ninside 354 372

13800 GCF_001683395.1_ASM168339v1 Kosakonia sacchari Proteobacteria; Gammaproteobacteria; Enterobacterales; Enterobacteriaceae; Kosakonia
MAGENQQFASHGVSRRDFMKLCAALAATMGLSGKAAAEMAEAVSHPQRPPVIWIGAQECT WP_065370054.1
hydrogenase 2 small subunit [Kosakonia sacchari] Length: 372\nNumber of predicted TMHs: 1\nExp number of AAs in TMHs: 27.4582\nExp number, first 60 AAs: 0.30352\nTotal prob of N-in: 0.26440\noutside 1 330\nTMhelix 331 353\ninside 354 372

13801 GCF_000280495.2_ASM28049v2 Kosakonia radicincitans DSM 16656 Proteobacteria; Gammaproteobacteria; Enterobacterales; Enterobacteriaceae; Kosakonia; Kosakonia radicincitans
MAGDNLHLASQGVSRRDFMKLCAALAATMGLSGKAAAEMAEAISHPQRPPVVWIGAQECT WP_007370143.1
MULTISPECIES: hydrogenase 2 small subunit [Enterobacteriaceae] Length: 372\nNumber of predicted TMHs: 1\nExp number of AAs in TMHs: 24.18752\nExp number, first 60 AAs: 0.2398\nTotal prob of N-in: 0.11792\noutside 1 332\nTMhelix 333 355\ninside 356 372

13802 GCF_000691205.1_UMEnt01_12.1 Kosakonia radicincitans UMEnt01/12 Proteobacteria; Gammaproteobacteria; Enterobacterales; Enterobacteriaceae; Kosakonia; Kosakonia radicincitans
MAGDNLHLASQGVSRRDFMKLCAALAATMGLSGKAAAEMAEAISHPQRPPVVWIGAQECT WP_007370143.1
MULTISPECIES: hydrogenase 2 small subunit [Enterobacteriaceae] Length: 372\nNumber of predicted TMHs: 1\nExp number of AAs in TMHs: 24.18752\nExp number, first 60 AAs: 0.2398\nTotal prob of N-in: 0.11792\noutside 1 332\nTMhelix 333 355\ninside 356 372

13803 GCF_000877295.1_GFC_25 Kosakonia radicincitans YD4 Proteobacteria; Gammaproteobacteria; Enterobacterales; Enterobacteriaceae; Kosakonia; Kosakonia radicincitans
MAGDNLHLASQGVSRRDFMKLCAALAATMGLSGKAAAEMAEAISHPQRPPVVWIGAQECT WP_043956539.1
hydrogenase 2 small subunit [Kosakonia radicincitans] Length: 372\nNumber of predicted TMHs: 1\nExp number of AAs in TMHs: 24.16605\nExp number, first 60 AAs: 0.24134\nTotal prob of N-in: 0.11649\noutside 1 332\nTMhelix 333 355\ninside 356 372

13804 GCF_000300455.3_ASM30045v4 Kosakonia sacchari SP1 Proteobacteria; Gammaproteobacteria; Enterobacterales; Enterobacteriaceae; Kosakonia; Kosakonia sacchari

MAGENQQFASHGVSRRDFMKLCAALAATMGLSGKAAAEMAKAVSHPQRPPVIWIGAQECT WP_017457491.1 hydrogenase 2 small subunit [Kosakonia sacchari] Length: 372\nNumber of predicted TMHs: 1\nExp number of AAs in TMHs: 26.66562\nExp number, first 60 AAs: 0.34625\nTotal prob of N-in: 0.26285\noutside 1 330\nTMhelix 331 353\ninside 354 372

13805 GCF_001655675.1_ASM165567v1 Mangrovibacter phragmitis Proteobacteria; Gammaproteobacteria; Enterobacterales; Enterobacteriaceae; Mangrovibacter MKGVLEKCTVEDFNYISATLDSYVSFTNDKRRKTLLSAYQNNPALHAELISLIDTQIKYF WP_001515206.1 MULTISPECIES: hypothetical protein [Enterobacteriaceae] Length: 249\nNumber of predicted TMHs: 2\nExp number of AAs in TMHs: 39.68696\nExp number, first 60 AAs: 0.01961\nTotal prob of N-in: 0.09053\noutside 1 157\nTMhelix 158 180\ninside 181 200\nTMhelix 201 220\noutside 221 249

13806 GCF_002035475.1_ASM203547v1 Salmonella bongori Proteobacteria; Gammaproteobacteria; Enterobacterales; Enterobacteriaceae; Salmonella MTGDNTLITSHGINRRDFMKLCAALAATMGLSSKAAAEMAESVSNPQRPPVIWIGAQECT WP_000145426.1 hydrogenase 2 small subunit [Salmonella bongori] Length: 372\nNumber of predicted TMHs: 1\nExp number of AAs in TMHs: 23.21866\nExp number, first 60 AAs: 0.2769\nTotal prob of N-in: 0.08285\noutside 1 331\nTMhelix 332 354\ninside 355 372

13807 GCF_002035285.1_ASM203528v1 Salmonella bongori Proteobacteria; Gammaproteobacteria; Enterobacterales; Enterobacteriaceae; Salmonella MTGDNTLITSHGINRRDFMKLCAALAATMGLSSKAAAEMAESVSNPQRPPVIWIGAQECT WP_000145426.1 hydrogenase 2 small subunit [Salmonella bongori] Length: 372\nNumber of predicted TMHs: 1\nExp number of AAs in TMHs: 23.21866\nExp number, first 60 AAs: 0.2769\nTotal prob of N-in: 0.08285\noutside 1 331\nTMhelix 332 354\ninside 355 372

13808 GCF_002119365.1_ASM211936v1 Salmonella bongori Proteobacteria; Gammaproteobacteria; Enterobacterales; Enterobacteriaceae; Salmonella MTGDNTLITSHGINRRDFMKLCAALAATMGLSSKAAAEMAESVSNPQRPPVIWIGAQECT WP_000145426.1 hydrogenase 2 small subunit [Salmonella bongori] Length: 372\nNumber of predicted TMHs: 1\nExp number of AAs in TMHs: 23.21866\nExp number, first 60 AAs: 0.2769\nTotal prob of N-in: 0.08285\noutside 1 331\nTMhelix 332 354\ninside 355 372

13809 GCF_001474515.1_Salmonella_enterica_CVM_43785_v1.0Salmonella enterica Proteobacteria; Gammaproteobacteria; Enterobacterales; Enterobacteriaceae; Salmonella MTGDNTLITSHGINRRDFMKLCAALAATMGLSSKAAAEMAESVSNPQRPPVIWIGAQECT WP_000145429.1 hydrogenase 2 small subunit [Salmonella enterica] Length: 372\nNumber of predicted TMHs: 1\nExp number of AAs in TMHs: 24.51309\nExp number, first 60 AAs: 0.2682\nTotal prob of N-in: 0.14402\noutside 1 331\nTMhelix 332 354\ninside 355 372

13810 GCF_001473795.1_Salmonella_enterica_CVM_43823_v1.0Salmonella enterica Proteobacteria; Gammaproteobacteria; Enterobacterales; Enterobacteriaceae; Salmonella MNNEETFYQAMRRKGVTRRSFLKFCSLAATSLGLGAGMTPKIAWALENKPRIPVVWIHGL WP_001058312.1 hydrogenase [Salmonella enterica] Length: 372\nNumber of predicted TMHs: 2\nExp number of AAs in TMHs: 36.51738\nExp number, first 60 AAs: 19.72849\nTotal prob of N-in: 0.96170\nPOSSIBLE N-term signal sequence\ninside 1 19\nTMhelix 20 39\noutside 40 325\nTMhelix 326 348\ninside 349 372

13811 GCF_001473795.1_Salmonella_enterica_CVM_43823_v1.0Salmonella enterica Proteobacteria; Gammaproteobacteria; Enterobacterales; Enterobacteriaceae; Salmonella MTGDNTLITSHGINRRDFMKLCAALAATMGLSSKAAAEMAESVSNPQRPPVIWIGAQECT WP_000145429.1 hydrogenase 2 small subunit [Salmonella enterica] Length: 372\nNumber of predicted TMHs: 1\nExp number of AAs in TMHs: 24.51309\nExp number, first 60 AAs: 0.2682\nTotal prob of N-in: 0.14402\noutside 1 331\nTMhelix 332 354\ninside 355 372

13812 GCF_001835005.1_ASM183500v1 Salmonella enterica Proteobacteria; Gammaproteobacteria; Enterobacterales; Enterobacteriaceae; Salmonella MTGDNTLITSHGINRRDFMKLCAALAATMGLSSKAAAEMAESVSNPQRPPVIWIGAQECT WP_000145429.1 hydrogenase 2 small subunit [Salmonella enterica] Length: 372\nNumber of predicted TMHs: 1\nExp number of AAs in TMHs: 24.51309\nExp number, first 60 AAs: 0.2682\nTotal prob of N-in: 0.14402\noutside 1 331\nTMhelix 332 354\ninside 355 372

13813 GCF_001834845.1_ASM183484v1 Salmonella enterica Proteobacteria; Gammaproteobacteria; Enterobacterales; Enterobacteriaceae; Salmonella
MTGDNTLITSHGINRRDFMKLCAALAATMGLSSKAAAEMAESVSNPQRPPVIWIGAQECT WP_000145429.1
hydrogenase 2 small subunit [Salmonella enterica] Length: 372\nNumber of predicted TMHs: 1\nExp number of AAs in TMHs: 24.51309\nExp number, first 60 AAs: 0.2682\nTotal prob of N-in: 0.14402\noutside 1 331\nTMhelix 332 354\ninside 355 372

13814 GCF_001474825.1_Salmonella_enterica_CVM_43810_v1.0Salmonella enterica Proteobacteria; Gammaproteobacteria; Enterobacterales; Enterobacteriaceae; Salmonella
MTGDNTLITSHGINRRDFMKLCAALAATMGLSSKAAAEMAESVSNPQRPPVIWIGAQECT WP_000145429.1
hydrogenase 2 small subunit [Salmonella enterica] Length: 372\nNumber of predicted TMHs: 1\nExp number of AAs in TMHs: 24.51309\nExp number, first 60 AAs: 0.2682\nTotal prob of N-in: 0.14402\noutside 1 331\nTMhelix 332 354\ninside 355 372

13815 GCF_001569805.1_ASM156980v1 Salmonella enterica Proteobacteria; Gammaproteobacteria; Enterobacterales; Enterobacteriaceae; Salmonella
MTGDNTLITSHGINRRDFMKLCAALAATMGLSSKAAAEMAESVSNPQRPPVIWIGAQECT WP_000145429.1
hydrogenase 2 small subunit [Salmonella enterica] Length: 372\nNumber of predicted TMHs: 1\nExp number of AAs in TMHs: 24.51309\nExp number, first 60 AAs: 0.2682\nTotal prob of N-in: 0.14402\noutside 1 331\nTMhelix 332 354\ninside 355 372

13816 GCF_001078155.1_ASM107815v1 Salmonella enterica Proteobacteria; Gammaproteobacteria; Enterobacterales; Enterobacteriaceae; Salmonella
MTGDNTLITSHGINRRDFMKLCAALAATMGLSSKAAAEMAESVSNPQRPPVIWIGAQECT WP_000145429.1
hydrogenase 2 small subunit [Salmonella enterica] Length: 372\nNumber of predicted TMHs: 1\nExp number of AAs in TMHs: 24.51309\nExp number, first 60 AAs: 0.2682\nTotal prob of N-in: 0.14402\noutside 1 331\nTMhelix 332 354\ninside 355 372

13817 GCF_001473785.1_Salmonella_enterica_CVM_43822_v1.0Salmonella enterica Proteobacteria; Gammaproteobacteria; Enterobacterales; Enterobacteriaceae; Salmonella
MTGDNTLITSHGINRRDFMKLCAALAATMGLSSKAAAEMAESVSNPQRPPVIWIGAQECT WP_000145429.1
hydrogenase 2 small subunit [Salmonella enterica] Length: 372\nNumber of predicted TMHs: 1\nExp number of AAs in TMHs: 24.51309\nExp number, first 60 AAs: 0.2682\nTotal prob of N-in: 0.14402\noutside 1 331\nTMhelix 332 354\ninside 355 372

13818 GCF_001643825.1_CRJJGF_00061_v1.0Salmonella enterica Proteobacteria; Gammaproteobacteria; Enterobacterales; Enterobacteriaceae; Salmonella
MLKFRTIFLKKIFTLKNSILARRGAVIVIVSAVFTSIMFFAHSWASDKEVAMTLSVLNSN WP_023226644.1 hypothetical protein [Salmonella enterica] Length: 204\nNumber of predicted TMHs: 2\nExp number of AAs in TMHs: 44.99526\nExp number, first 60 AAs: 22.33786\nTotal prob of N-in: 0.87083\nPOSSIBLE N-term signal sequence\ninside 1 19\nTMhelix 20 42\noutside 43 171\nTMhelix 172 194\ninside 195 204

13819 GCF_001569765.1_ASM156976v1 Salmonella enterica Proteobacteria; Gammaproteobacteria; Enterobacterales; Enterobacteriaceae; Salmonella
MTGDNTLITSHGINRRDFMKLCAALAATMGLSSKAAAEMAESVSNPQRPPVIWIGAQECT WP_000145429.1
hydrogenase 2 small subunit [Salmonella enterica] Length: 372\nNumber of predicted TMHs: 1\nExp number of AAs in TMHs: 24.51309\nExp number, first 60 AAs: 0.2682\nTotal prob of N-in: 0.14402\noutside 1 331\nTMhelix 332 354\ninside 355 372

13820 GCF_002043285.1_ASM204328v1 Salmonella enterica Proteobacteria; Gammaproteobacteria; Enterobacterales; Enterobacteriaceae; Salmonella
MTGDNTLITSHGINRRDFMKLCAALAATMGLSSKAAAEMAESVSNPQRPPVIWIGAQECT WP_000145429.1
hydrogenase 2 small subunit [Salmonella enterica] Length: 372\nNumber of predicted TMHs: 1\nExp number of AAs in TMHs: 24.51309\nExp number, first 60 AAs: 0.2682\nTotal prob of N-in: 0.14402\noutside 1 331\nTMhelix 332 354\ninside 355 372

13821 GCF_002033515.1_ASM203351v1 Salmonella enterica Proteobacteria; Gammaproteobacteria; Enterobacterales; Enterobacteriaceae; Salmonella
MTGDNTLITSHGINRRDFMKLCAALAATMGLSSKAAAEMAESVSNPQRPPVIWIGAQECT WP_000145429.1
hydrogenase 2 small subunit [Salmonella enterica] Length: 372\nNumber of predicted TMHs: 1\nExp

number of AAs in TMHs: 24.51309\nExp number, first 60 AAs: 0.2682\nTotal prob of N-in: 0.14402\noutside 1 331\nTMhelix 332 354\ninside 355 372

13822 GCF_001473355.1_Salmonella_enterica_CVM_43753_v1.0Salmonella enterica Proteobacteria; Gammaproteobacteria; Enterobacterales; Enterobacteriaceae; Salmonella MTGDNTLITSHGINRRDFMKLCAALAATMGLSSKAAAEMAESVSNPQRPPVIWIGAQECT WP_000145429.1 hydrogenase 2 small subunit [Salmonella enterica] Length: 372\nNumber of predicted TMHs: 1\nExp number of AAs in TMHs: 24.51309\nExp number, first 60 AAs: 0.2682\nTotal prob of N-in: 0.14402\noutside 1 331\nTMhelix 332 354\ninside 355 372

13823 GCF_001834805.1_ASM183480v1 Salmonella enterica Proteobacteria; Gammaproteobacteria; Enterobacterales; Enterobacteriaceae; Salmonella MTGDNTLITSHGINRRDFMKLCAALAATMGLSSKAAAEMAESVSNPQRPPVIWIGAQECT WP_000145429.1 hydrogenase 2 small subunit [Salmonella enterica] Length: 372\nNumber of predicted TMHs: 1\nExp number of AAs in TMHs: 24.51309\nExp number, first 60 AAs: 0.2682\nTotal prob of N-in: 0.14402\noutside 1 331\nTMhelix 332 354\ninside 355 372

13824 GCF_001473305.1_Salmonella_enterica_CVM_43756_v1.0Salmonella enterica Proteobacteria; Gammaproteobacteria; Enterobacterales; Enterobacteriaceae; Salmonella MTGDNTLITSHGINRRDFMKLCAALAATMGLSSKAAAEMAESVSNPQRPPVIWIGAQECT WP_000145429.1 hydrogenase 2 small subunit [Salmonella enterica] Length: 372\nNumber of predicted TMHs: 1\nExp number of AAs in TMHs: 24.51309\nExp number, first 60 AAs: 0.2682\nTotal prob of N-in: 0.14402\noutside 1 331\nTMhelix 332 354\ninside 355 372

13825 GCF_001834965.1_ASM183496v1 Salmonella enterica Proteobacteria; Gammaproteobacteria; Enterobacterales; Enterobacteriaceae; Salmonella MTGDNTLITSHGINRRDFMKLCAALAATMGLSSKAAAEMAESVSNPQRPPVIWIGAQECT WP_000145429.1 hydrogenase 2 small subunit [Salmonella enterica] Length: 372\nNumber of predicted TMHs: 1\nExp number of AAs in TMHs: 24.51309\nExp number, first 60 AAs: 0.2682\nTotal prob of N-in: 0.14402\noutside 1 331\nTMhelix 332 354\ninside 355 372

13826 GCF_001833625.1_ASM183362v1 Salmonella enterica Proteobacteria; Gammaproteobacteria; Enterobacterales; Enterobacteriaceae; Salmonella MNNEETFYQAMRRKGVTRRSFLKFCSLAATSLGLGAGMTPKIAWALENKPRIPVVWIHGL WP_001058312.1 hydrogenase [Salmonella enterica] Length: 372\nNumber of predicted TMHs: 2\nExp number of AAs in TMHs: 36.51738\nExp number, first 60 AAs: 19.72849\nTotal prob of N-in: 0.96170\nPOSSIBLE N-term signal sequence\ninside 1 19\nTMhelix 20 39\noutside 40 325\nTMhelix 326 348\ninside 349 372

13827 GCF_001834885.1_ASM183488v1 Salmonella enterica Proteobacteria; Gammaproteobacteria; Enterobacterales; Enterobacteriaceae; Salmonella MNNEETFYQAMRRKGVTRRSFLKFCSLAATSLGLGAGMTPKIAWALENKPRIPVVWIHGL WP_001058306.1 hydrogenase [Salmonella enterica] Length: 372\nNumber of predicted TMHs: 2\nExp number of AAs in TMHs: 36.5540099999999\nExp number, first 60 AAs: 19.72732\nTotal prob of N-in: 0.96171\nPOSSIBLE N-term signal sequence\ninside 1 19\nTMhelix 20 39\noutside 40 325\nTMhelix 326 348\ninside 349 372

13828 GCF_001834815.1_ASM183481v1 Salmonella enterica Proteobacteria; Gammaproteobacteria; Enterobacterales; Enterobacteriaceae; Salmonella MTGDNTLITSHGINRRDFMKLCAALAATMGLSSKAAAEMAESVSNPQRPPVIWIGAQECT WP_000145429.1 hydrogenase 2 small subunit [Salmonella enterica] Length: 372\nNumber of predicted TMHs: 1\nExp number of AAs in TMHs: 24.51309\nExp number, first 60 AAs: 0.2682\nTotal prob of N-in: 0.14402\noutside 1 331\nTMhelix 332 354\ninside 355 372

13829 GCF_001834755.1_ASM183475v1 Salmonella enterica Proteobacteria; Gammaproteobacteria; Enterobacterales; Enterobacteriaceae; Salmonella MNNEETFYQAMRRKGVTRRSFLKFCSLAATSLGLGAGMTPKIAWALENKPRIPVVWIHGL WP_001058312.1 hydrogenase [Salmonella enterica] Length: 372\nNumber of predicted TMHs: 2\nExp number of AAs in TMHs: 36.51738\nExp number, first 60 AAs: 19.72849\nTotal prob of N-in: 0.96170\nPOSSIBLE N-term signal sequence\ninside 1 19\nTMhelix 20 39\noutside 40 325\nTMhelix 326 348\ninside 349 372

13830 GCF_001834785.1_ASM183478v1 Salmonella enterica Proteobacteria; Gammaproteobacteria; Enterobacterales; Enterobacteriaceae; Salmonella

MNNEETFYQAMRRKGVTRRSFLKFCSLAATSLGLGAGMTPKIAWALENKPRIPVVWIHGL WP_001058312.1 hydrogenase [Salmonella enterica] Length: 372\nNumber of predicted TMHs: 2\nExp number of AAs in TMHs: 36.51738\nExp number, first 60 AAs: 19.72849\nTotal prob of N-in: 0.96170\nPOSSIBLE N-term signal sequence\ninside 1 19\nTMhelix 20 39\noutside 40 325\nTMhelix 326 348\ninside 349 372

13831 GCF_001835045.1_ASM183504v1 Salmonella enterica Proteobacteria; Gammaproteobacteria; Enterobacterales; Enterobacteriaceae; Salmonella MNNEETFYQAMRRKGVTRRSFLKFCSLAATSLGLGAGMTPKIAWALENKPRIPVVWIHGL WP_001058312.1 hydrogenase [Salmonella enterica] Length: 372\nNumber of predicted TMHs: 2\nExp number of AAs in TMHs: 36.51738\nExp number, first 60 AAs: 19.72849\nTotal prob of N-in: 0.96170\nPOSSIBLE N-term signal sequence\ninside 1 19\nTMhelix 20 39\noutside 40 325\nTMhelix 326 348\ninside 349 372

13832 GCF_001835235.1_ASM183523v1 Salmonella enterica Proteobacteria; Gammaproteobacteria; Enterobacterales; Enterobacteriaceae; Salmonella MTGDNTLITSHGINRRDFMKLCAALAATMGLSSKAAAEMAESVSNPQRPPVIWIGAQECT WP_000145429.1 hydrogenase 2 small subunit [Salmonella enterica] Length: 372\nNumber of predicted TMHs: 1\nExp number of AAs in TMHs: 24.51309\nExp number, first 60 AAs: 0.2682\nTotal prob of N-in: 0.14402\noutside 1 331\nTMhelix 332 354\ninside 355 372

13833 GCF_001879775.1_ASM187977v1 Salmonella enterica Proteobacteria; Gammaproteobacteria; Enterobacterales; Enterobacteriaceae; Salmonella MTGDNTLITSHGINRRDFMKLCAALAATMGLSSKAAAEMAESVSNPQRPPVIWIGAQECT WP_000145431.1 hydrogenase 2 small subunit [Salmonella enterica] Length: 372\nNumber of predicted TMHs: 1\nExp number of AAs in TMHs: 24.51505\nExp number, first 60 AAs: 0.2682\nTotal prob of N-in: 0.14406\noutside 1 331\nTMhelix 332 354\ninside 355 372

13834 GCF_001952055.1_ASM195205v1 Salmonella enterica Proteobacteria; Gammaproteobacteria; Enterobacterales; Enterobacteriaceae; Salmonella MNNEETLYQAMRRKGVTRRSFLKFCSLAATSLGLGAGMTPKIAWALENKPRIPVVWIHGL WP_001752445.1 hydrogenase-1 small chain [Salmonella enterica] Length: 372\nNumber of predicted TMHs: 2\nExp number of AAs in TMHs: 36.5123\nExp number, first 60 AAs: 19.72341\nTotal prob of N-in: 0.96144\nPOSSIBLE N-term signal sequence\ninside 1 19\nTMhelix 20 39\noutside 40 325\nTMhelix 326 348\ninside 349 372

13835 GCF_001952055.1_ASM195205v1 Salmonella enterica Proteobacteria; Gammaproteobacteria; Enterobacterales; Enterobacteriaceae; Salmonella MTGDNTLITSHGINRRDFMKLCAALAATMGLSSKAAAEMAESVSNPQRPPVIWIGAQECT WP_000145428.1 hydrogenase 2 small subunit [Salmonella enterica] Length: 372\nNumber of predicted TMHs: 1\nExp number of AAs in TMHs: 24.50677\nExp number, first 60 AAs: 0.2682\nTotal prob of N-in: 0.14390\noutside 1 331\nTMhelix 332 354\ninside 355 372

13836 GCF_001951875.1_ASM195187v1 Salmonella enterica Proteobacteria; Gammaproteobacteria; Enterobacterales; Enterobacteriaceae; Salmonella MNNEETFYQAMRRKGVTRRSFLKFCSLAATSLGLGAGMTPKIAWALENKPRIPVVWIHGL WP_001058312.1 hydrogenase [Salmonella enterica] Length: 372\nNumber of predicted TMHs: 2\nExp number of AAs in TMHs: 36.51738\nExp number, first 60 AAs: 19.72849\nTotal prob of N-in: 0.96170\nPOSSIBLE N-term signal sequence\ninside 1 19\nTMhelix 20 39\noutside 40 325\nTMhelix 326 348\ninside 349 372

13837 GCF_002059645.1_ASM205964v1 Salmonella enterica Proteobacteria; Gammaproteobacteria; Enterobacterales; Enterobacteriaceae; Salmonella MNNEETFYQAMRRKGVTRRSFLKFCSLAATSLGLGAGMTPKIAWALENKPRIPVVWIHGL WP_001058310.1 hydrogenase [Salmonella enterica] Length: 372\nNumber of predicted TMHs: 2\nExp number of AAs in TMHs: 37.4567299999999\nExp number, first 60 AAs: 19.72694\nTotal prob of N-in: 0.96171\nPOSSIBLE N-term signal sequence\ninside 1 19\nTMhelix 20 39\noutside 40 325\nTMhelix 326 348\ninside 349 372

13838 GCF_002059635.1_ASM205963v1 Salmonella enterica Proteobacteria; Gammaproteobacteria; Enterobacterales; Enterobacteriaceae; Salmonella MTGDNTLITSHGINRRDFMKLCAALAATMGLSSKAAAEMAESVSNPQRPPVIWIGAQECT WP_000145429.1 hydrogenase 2 small subunit [Salmonella enterica] Length: 372\nNumber of predicted TMHs: 1\nExp number of AAs in TMHs: 24.51309\nExp number, first 60 AAs: 0.2682\nTotal prob of N-in: 0.14402\noutside 1 331\nTMhelix 332 354\ninside 355 372

13839 GCF_002059455.1_ASM205945v1 Salmonella enterica Proteobacteria; Gammaproteobacteria; Enterobacterales; Enterobacteriaceae; Salmonella
MNNEETFYQAMRRKGVTRRSFLKFCSLAATSLGLGAGMTPKIAWALENKPRIPVVWIHGL WP_001058312.1
hydrogenase [Salmonella enterica] Length: 372\nNumber of predicted TMHs: 2\nExp number of AAs in TMHs: 36.51738\nExp number, first 60 AAs: 19.72849\nTotal prob of N-in: 0.96170\nPOSSIBLE N-term signal sequence\ninside 1 19\nTMhelix 20 39\noutside 40 325\nTMhelix 326 348\ninside 349 372

13840 GCF_002059455.1_ASM205945v1 Salmonella enterica Proteobacteria; Gammaproteobacteria; Enterobacterales; Enterobacteriaceae; Salmonella
MTGDNTLITSHGINRRDFMKLCAALAATMGLSSKAAAEMAESVSNPQRPPVIWIGAQECT WP_000145429.1
hydrogenase 2 small subunit [Salmonella enterica] Length: 372\nNumber of predicted TMHs: 1\nExp number of AAs in TMHs: 24.51309\nExp number, first 60 AAs: 0.2682\nTotal prob of N-in: 0.14402\noutside 1 331\nTMhelix 332 354\ninside 355 372

13841 GCF_002059445.1_ASM205944v1 Salmonella enterica Proteobacteria; Gammaproteobacteria; Enterobacterales; Enterobacteriaceae; Salmonella
MTGDNTLITSHGINRRDFMKLCAALAATMGLSSKAAAEMAESVSNPQRPPVIWIGAQECT WP_000145429.1
hydrogenase 2 small subunit [Salmonella enterica] Length: 372\nNumber of predicted TMHs: 1\nExp number of AAs in TMHs: 24.51309\nExp number, first 60 AAs: 0.2682\nTotal prob of N-in: 0.14402\noutside 1 331\nTMhelix 332 354\ninside 355 372

13842 GCF_002059395.1_ASM205939v1 Salmonella enterica Proteobacteria; Gammaproteobacteria; Enterobacterales; Enterobacteriaceae; Salmonella
MTGDNTLITSHGINRRDFMKLCAALAATMGLSSKAAAEMAESVSNPQRPPVIWIGAQECT WP_000145429.1
hydrogenase 2 small subunit [Salmonella enterica] Length: 372\nNumber of predicted TMHs: 1\nExp number of AAs in TMHs: 24.51309\nExp number, first 60 AAs: 0.2682\nTotal prob of N-in: 0.14402\noutside 1 331\nTMhelix 332 354\ninside 355 372

13843 GCF_002059375.1_ASM205937v1 Salmonella enterica Proteobacteria; Gammaproteobacteria; Enterobacterales; Enterobacteriaceae; Salmonella
MTGDNTLITSHGINRRDFMKLCAALAATMGLSSKAAAEMAESVSNPQRPPVIWIGAQECT WP_000145429.1
hydrogenase 2 small subunit [Salmonella enterica] Length: 372\nNumber of predicted TMHs: 1\nExp number of AAs in TMHs: 24.51309\nExp number, first 60 AAs: 0.2682\nTotal prob of N-in: 0.14402\noutside 1 331\nTMhelix 332 354\ninside 355 372

13844 GCF_002058675.1_ASM205867v1 Salmonella enterica Proteobacteria; Gammaproteobacteria; Enterobacterales; Enterobacteriaceae; Salmonella
MTGDNTLITSHGINRRDFMKLCAALAATMGLSSKAAAEMAESVSNPQRPPVIWIGAQECT WP_000145429.1
hydrogenase 2 small subunit [Salmonella enterica] Length: 372\nNumber of predicted TMHs: 1\nExp number of AAs in TMHs: 24.51309\nExp number, first 60 AAs: 0.2682\nTotal prob of N-in: 0.14402\noutside 1 331\nTMhelix 332 354\ninside 355 372

13845 GCF_002057755.1_ASM205775v1 Salmonella enterica Proteobacteria; Gammaproteobacteria; Enterobacterales; Enterobacteriaceae; Salmonella
MNNEETFYQAMRRKGVTRRSFLKFCSLAATSLGLGAGMTPKIAWALENKPRIPVVWIHGL WP_001058307.1
hydrogenase [Salmonella enterica] Length: 372\nNumber of predicted TMHs: 2\nExp number of AAs in TMHs: 36.51738\nExp number, first 60 AAs: 19.72849\nTotal prob of N-in: 0.96170\nPOSSIBLE N-term signal sequence\ninside 1 19\nTMhelix 20 39\noutside 40 325\nTMhelix 326 348\ninside 349 372

13846 GCF_002057435.1_ASM205743v1 Salmonella enterica Proteobacteria; Gammaproteobacteria; Enterobacterales; Enterobacteriaceae; Salmonella
MNNEETFYQAMRRKGVTRRSFLKFCSLAATSLGLGAGMTPKIAWALENKPRIPVVWIHGL WP_001058312.1
hydrogenase [Salmonella enterica] Length: 372\nNumber of predicted TMHs: 2\nExp number of AAs in TMHs: 36.51738\nExp number, first 60 AAs: 19.72849\nTotal prob of N-in: 0.96170\nPOSSIBLE N-term signal sequence\ninside 1 19\nTMhelix 20 39\noutside 40 325\nTMhelix 326 348\ninside 349 372

13847 GCF_002057495.1_ASM205749v1 Salmonella enterica Proteobacteria; Gammaproteobacteria; Enterobacterales; Enterobacteriaceae; Salmonella
MNNEETFYQAMRRKGVTRRSFLKFCSLAATSLGLGAGMTPKIAWALENKPRIPVVWIHGL WP_001058312.1
hydrogenase [Salmonella enterica] Length: 372\nNumber of predicted TMHs: 2\nExp number of AAs in TMHs:

TMHs: 36.51738\nExp number, first 60 AAs: 19.72849\nTotal prob of N-in: 0.96170\nPOSSIBLE N-term signal sequence\ninside 1 19\nTMhelix 20 39\noutside 40 325\nTMhelix 326 348\ninside 349 372

13848 GCF_002057585.1_ASM205758v1 Salmonella enterica Proteobacteria; Gammaproteobacteria; Enterobacterales; Enterobacteriaceae; Salmonella MNNEETFYQAMRRKGVTRRSFLKFCSLAATSLGLGAGMTPKIAWALENKPRIPVVWIHGL WP_001058307.1 hydrogenase [Salmonella enterica] Length: 372\nNumber of predicted TMHs: 2\nExp number of AAs in TMHs: 36.51738\nExp number, first 60 AAs: 19.72849\nTotal prob of N-in: 0.96170\nPOSSIBLE N-term signal sequence\ninside 1 19\nTMhelix 20 39\noutside 40 325\nTMhelix 326 348\ninside 349 372

13849 GCF_002057585.1_ASM205758v1 Salmonella enterica Proteobacteria; Gammaproteobacteria; Enterobacterales; Enterobacteriaceae; Salmonella MTGDNTLITSHGINRRDFMKLCAALAATMGLSSKAAAEMAESVSNPQRPPVIWIGAQECT WP_000145429.1 hydrogenase 2 small subunit [Salmonella enterica] Length: 372\nNumber of predicted TMHs: 1\nExp number of AAs in TMHs: 24.51309\nExp number, first 60 AAs: 0.2682\nTotal prob of N-in: 0.14402\noutside 1 331\nTMhelix 332 354\ninside 355 372

13850 GCF_002057395.1_ASM205739v1 Salmonella enterica Proteobacteria; Gammaproteobacteria; Enterobacterales; Enterobacteriaceae; Salmonella MNNEETFYQAMRRKGVTRRSFLKFCSLAATSLGLGAGMTPKIAWALENKPRIPVVWIHGL WP_001058312.1 hydrogenase [Salmonella enterica] Length: 372\nNumber of predicted TMHs: 2\nExp number of AAs in TMHs: 36.51738\nExp number, first 60 AAs: 19.72849\nTotal prob of N-in: 0.96170\nPOSSIBLE N-term signal sequence\ninside 1 19\nTMhelix 20 39\noutside 40 325\nTMhelix 326 348\ninside 349 372

13851 GCF_002057575.1_ASM205757v1 Salmonella enterica Proteobacteria; Gammaproteobacteria; Enterobacterales; Enterobacteriaceae; Salmonella MNNEETFYQAMRRKGVTRRSFLKFCSLAATSLGLGAGMTPKIAWALENKPRIPVVWIHGL WP_001058312.1 hydrogenase [Salmonella enterica] Length: 372\nNumber of predicted TMHs: 2\nExp number of AAs in TMHs: 36.51738\nExp number, first 60 AAs: 19.72849\nTotal prob of N-in: 0.96170\nPOSSIBLE N-term signal sequence\ninside 1 19\nTMhelix 20 39\noutside 40 325\nTMhelix 326 348\ninside 349 372

13852 GCF_002057395.1_ASM205739v1 Salmonella enterica Proteobacteria; Gammaproteobacteria; Enterobacterales; Enterobacteriaceae; Salmonella MTGDNTLITSHGINRRDFMKLCAALAATMGLSSKAAAEMAESVSNPQRPPVIWIGAQECT WP_000145429.1 hydrogenase 2 small subunit [Salmonella enterica] Length: 372\nNumber of predicted TMHs: 1\nExp number of AAs in TMHs: 24.51309\nExp number, first 60 AAs: 0.2682\nTotal prob of N-in: 0.14402\noutside 1 331\nTMhelix 332 354\ninside 355 372

13853 GCF_002057295.1_ASM205729v1 Salmonella enterica Proteobacteria; Gammaproteobacteria; Enterobacterales; Enterobacteriaceae; Salmonella MTGDNTLITSHGINRRDFMKLCAALAATMGLSSKAAAEMAESVSNPQRPPVIWIGAQECT WP_000145429.1 hydrogenase 2 small subunit [Salmonella enterica] Length: 372\nNumber of predicted TMHs: 1\nExp number of AAs in TMHs: 24.51309\nExp number, first 60 AAs: 0.2682\nTotal prob of N-in: 0.14402\noutside 1 331\nTMhelix 332 354\ninside 355 372

13854 GCF_002057345.1_ASM205734v1 Salmonella enterica Proteobacteria; Gammaproteobacteria; Enterobacterales; Enterobacteriaceae; Salmonella MTGDNTLITSHGINRRDFMKLCAALAATMGLSSKAAAEMAESVSNPQRPPVIWIGAQECT WP_000145429.1 hydrogenase 2 small subunit [Salmonella enterica] Length: 372\nNumber of predicted TMHs: 1\nExp number of AAs in TMHs: 24.51309\nExp number, first 60 AAs: 0.2682\nTotal prob of N-in: 0.14402\noutside 1 331\nTMhelix 332 354\ninside 355 372

13855 GCF_002056955.1_ASM205695v1 Salmonella enterica Proteobacteria; Gammaproteobacteria; Enterobacterales; Enterobacteriaceae; Salmonella MNNEETLYQAMRRKGVTRRSFLKFCSLAATSLGLGAGMTPKIAWALENKPRIPVVWIHGL WP_001752445.1 hydrogenase-1 small chain [Salmonella enterica] Length: 372\nNumber of predicted TMHs: 2\nExp number of AAs in TMHs: 36.5123\nExp number, first 60 AAs: 19.72341\nTotal prob of N-in: 0.96144\nPOSSIBLE N-term signal sequence\ninside 1 19\nTMhelix 20 39\noutside 40 325\nTMhelix 326 348\ninside 349 372

13856 GCF_002057275.1_ASM205727v1 Salmonella enterica Proteobacteria; Gammaproteobacteria; Enterobacterales; Enterobacteriaceae; Salmonella

MNNEETFYQAMRRKGVTRRSFLKFCSLAATSLGLGAGMTPKIAWALENKPRIPVVWIHGL WP_001058310.1 hydrogenase [Salmonella enterica] Length: 372\nNumber of predicted TMHs: 2\nExp number of AAs in TMHs: 37.4567299999999\nExp number, first 60 AAs: 19.72694\nTotal prob of N-in: 0.96171\nPOSSIBLE N-term signal sequence\ninside 1 19\nTMhelix 20 39\noutside 40 325\nTMhelix 326 348\ninside 349 372

13857 GCF_002057275.1_ASM205727v1 Salmonella enterica Proteobacteria; Gammaproteobacteria; Enterobacterales; Enterobacteriaceae; Salmonella MTGDNTLITSHGINRRDFMKLCAALAATMGLSSKAAAEMAESVSNPQRPPVIWIGAQECT WP_000145429.1 hydrogenase 2 small subunit [Salmonella enterica] Length: 372\nNumber of predicted TMHs: 1\nExp number of AAs in TMHs: 24.51309\nExp number, first 60 AAs: 0.2682\nTotal prob of N-in: 0.14402\noutside 1 331\nTMhelix 332 354\ninside 355 372

13858 GCF_002056955.1_ASM205695v1 Salmonella enterica Proteobacteria; Gammaproteobacteria; Enterobacterales; Enterobacteriaceae; Salmonella MTGDNTLITSHGINRRDFMKLCAALAATMGLSSKAAAEMAESVSNPQRPPVIWIGAQECT WP_000145428.1 hydrogenase 2 small subunit [Salmonella enterica] Length: 372\nNumber of predicted TMHs: 1\nExp number of AAs in TMHs: 24.50677\nExp number, first 60 AAs: 0.2682\nTotal prob of N-in: 0.14390\noutside 1 331\nTMhelix 332 354\ninside 355 372

13859 GCF_002056485.1_ASM205648v1 Salmonella enterica Proteobacteria; Gammaproteobacteria; Enterobacterales; Enterobacteriaceae; Salmonella MTGDNTLITSHGINRRDFMKLCAALAATMGLSSKAAAEMAESVSNPQRPPVIWIGAQECT WP_079776618.1 MULTISPECIES: hydrogenase 2 small subunit [Salmonella] Length: 372\nNumber of predicted TMHs: 1\nExp number of AAs in TMHs: 24.65705\nExp number, first 60 AAs: 0.26805\nTotal prob of N-in: 0.13761\noutside 1 331\nTMhelix 332 354\ninside 355 372

13860 GCF_002056535.1_ASM205653v1 Salmonella enterica Proteobacteria; Gammaproteobacteria; Enterobacterales; Enterobacteriaceae; Salmonella MNNEETFYQAMRRKGVTRRSFLKFCSLAATSLGLGAGMTPKIAWALENKPRIPVVWIHGL WP_001058307.1 hydrogenase [Salmonella enterica] Length: 372\nNumber of predicted TMHs: 2\nExp number of AAs in TMHs: 36.51738\nExp number, first 60 AAs: 19.72849\nTotal prob of N-in: 0.96170\nPOSSIBLE N-term signal sequence\ninside 1 19\nTMhelix 20 39\noutside 40 325\nTMhelix 326 348\ninside 349 372

13861 GCF_002057495.1_ASM205749v1 Salmonella enterica Proteobacteria; Gammaproteobacteria; Enterobacterales; Enterobacteriaceae; Salmonella MTGDNTLITSHGINRRDFMKLCAALAATMGLSSKAAAEMAESVSNPQRPPVIWIGAQECT WP_000145429.1 hydrogenase 2 small subunit [Salmonella enterica] Length: 372\nNumber of predicted TMHs: 1\nExp number of AAs in TMHs: 24.51309\nExp number, first 60 AAs: 0.2682\nTotal prob of N-in: 0.14402\noutside 1 331\nTMhelix 332 354\ninside 355 372

13862 GCF_002056445.1_ASM205644v1 Salmonella enterica Proteobacteria; Gammaproteobacteria; Enterobacterales; Enterobacteriaceae; Salmonella MNNEETFYQAMRRKGVTRRSFLKFCSLAATSLGLGAGMTPKIAWALENKPRIPVVWIHGL WP_001058307.1 hydrogenase [Salmonella enterica] Length: 372\nNumber of predicted TMHs: 2\nExp number of AAs in TMHs: 36.51738\nExp number, first 60 AAs: 19.72849\nTotal prob of N-in: 0.96170\nPOSSIBLE N-term signal sequence\ninside 1 19\nTMhelix 20 39\noutside 40 325\nTMhelix 326 348\ninside 349 372

13863 GCF_002056625.1_ASM205662v1 Salmonella enterica Proteobacteria; Gammaproteobacteria; Enterobacterales; Enterobacteriaceae; Salmonella MTGDNTLITSHGINRRDFMKLCAALAATMGLSSKAAAEMAESVSNPQRPPVIWIGAQECT WP_000145429.1 hydrogenase 2 small subunit [Salmonella enterica] Length: 372\nNumber of predicted TMHs: 1\nExp number of AAs in TMHs: 24.51309\nExp number, first 60 AAs: 0.2682\nTotal prob of N-in: 0.14402\noutside 1 331\nTMhelix 332 354\ninside 355 372

13864 GCF_002056265.1_ASM205626v1 Salmonella enterica Proteobacteria; Gammaproteobacteria; Enterobacterales; Enterobacteriaceae; Salmonella MNNEETFYQAMRRKGVTRRSFLKFCSLAATSLGLGAGMTPKIAWALENKPRIPVVWIHGL WP_001058307.1 hydrogenase [Salmonella enterica] Length: 372\nNumber of predicted TMHs: 2\nExp number of AAs in TMHs: 36.51738\nExp number, first 60 AAs: 19.72849\nTotal prob of N-in: 0.96170\nPOSSIBLE N-term signal sequence\ninside 1 19\nTMhelix 20 39\noutside 40 325\nTMhelix 326 348\ninside 349 372

13865 GCF_002056395.1_ASM205639v1 Salmonella enterica Proteobacteria; Gammaproteobacteria; Enterobacterales; Enterobacteriaceae; Salmonella
MTGDNTLITSHGINRRDFMKLCAALAATMGLSSKAAAEMAESVSNPQRPPVIWIGAQECT WP_000145429.1
hydrogenase 2 small subunit [Salmonella enterica] Length: 372\nNumber of predicted TMHs: 1\nExp number of AAs in TMHs: 24.51309\nExp number, first 60 AAs: 0.2682\nTotal prob of N-in: 0.14402\noutside 1 331\nTMhelix 332 354\ninside 355 372

13866 GCF_002056335.1_ASM205633v1 Salmonella enterica Proteobacteria; Gammaproteobacteria; Enterobacterales; Enterobacteriaceae; Salmonella
MTGDNTLITSHGINRRDFMKLCAALAATMGLSSKAAAEMAESVSNPQRPPVIWIGAQECT WP_000145429.1
hydrogenase 2 small subunit [Salmonella enterica] Length: 372\nNumber of predicted TMHs: 1\nExp number of AAs in TMHs: 24.51309\nExp number, first 60 AAs: 0.2682\nTotal prob of N-in: 0.14402\noutside 1 331\nTMhelix 332 354\ninside 355 372

13867 GCF_002056265.1_ASM205626v1 Salmonella enterica Proteobacteria; Gammaproteobacteria; Enterobacterales; Enterobacteriaceae; Salmonella
MTGDNTLITSHGINRRDFMKLCAALAATMGLSSKAAAEMAESVSNPQRPPVIWIGAQECT WP_000145429.1
hydrogenase 2 small subunit [Salmonella enterica] Length: 372\nNumber of predicted TMHs: 1\nExp number of AAs in TMHs: 24.51309\nExp number, first 60 AAs: 0.2682\nTotal prob of N-in: 0.14402\noutside 1 331\nTMhelix 332 354\ninside 355 372

13868 GCF_002056155.1_ASM205615v1 Salmonella enterica Proteobacteria; Gammaproteobacteria; Enterobacterales; Enterobacteriaceae; Salmonella
MNNEETFYQAMRRKGVTRRSFLKFCSLAATSLGLGAGMTPKIAWALENKPRIPVVWIHGL WP_001058307.1
hydrogenase [Salmonella enterica] Length: 372\nNumber of predicted TMHs: 2\nExp number of AAs in TMHs: 36.51738\nExp number, first 60 AAs: 19.72849\nTotal prob of N-in: 0.96170\nPOSSIBLE N-term signal sequence\ninside 1 19\nTMhelix 20 39\noutside 40 325\nTMhelix 326 348\ninside 349 372

13869 GCF_002056235.1_ASM205623v1 Salmonella enterica Proteobacteria; Gammaproteobacteria; Enterobacterales; Enterobacteriaceae; Salmonella
MNNEETFYQAMRRKGVTRRSFLKFCSLAATSLGLGAGMTPKIAWALENKPRIPVVWIHGL WP_001058307.1
hydrogenase [Salmonella enterica] Length: 372\nNumber of predicted TMHs: 2\nExp number of AAs in TMHs: 36.51738\nExp number, first 60 AAs: 19.72849\nTotal prob of N-in: 0.96170\nPOSSIBLE N-term signal sequence\ninside 1 19\nTMhelix 20 39\noutside 40 325\nTMhelix 326 348\ninside 349 372

13870 GCF_002056235.1_ASM205623v1 Salmonella enterica Proteobacteria; Gammaproteobacteria; Enterobacterales; Enterobacteriaceae; Salmonella
MTGDNTLITSHGINRRDFMKLCAALAATMGLSSKAAAEMAESVSNPQRPPVIWIGAQECT WP_000145429.1
hydrogenase 2 small subunit [Salmonella enterica] Length: 372\nNumber of predicted TMHs: 1\nExp number of AAs in TMHs: 24.51309\nExp number, first 60 AAs: 0.2682\nTotal prob of N-in: 0.14402\noutside 1 331\nTMhelix 332 354\ninside 355 372

13871 GCF_002056055.1_ASM205605v1 Salmonella enterica Proteobacteria; Gammaproteobacteria; Enterobacterales; Enterobacteriaceae; Salmonella
MNNEETFYQAMRRKGVTRRSFLKFCSLAATSLGLGAGMTPKIAWALENKPRIPVVWIHGL WP_001058307.1
hydrogenase [Salmonella enterica] Length: 372\nNumber of predicted TMHs: 2\nExp number of AAs in TMHs: 36.51738\nExp number, first 60 AAs: 19.72849\nTotal prob of N-in: 0.96170\nPOSSIBLE N-term signal sequence\ninside 1 19\nTMhelix 20 39\noutside 40 325\nTMhelix 326 348\ninside 349 372

13872 GCF_002056055.1_ASM205605v1 Salmonella enterica Proteobacteria; Gammaproteobacteria; Enterobacterales; Enterobacteriaceae; Salmonella
MTGDNTLITSHGINRRDFMKLCAALAATMGLSSKAAAEMAESVSNPQRPPVIWIGAQECT WP_000145429.1
hydrogenase 2 small subunit [Salmonella enterica] Length: 372\nNumber of predicted TMHs: 1\nExp number of AAs in TMHs: 24.51309\nExp number, first 60 AAs: 0.2682\nTotal prob of N-in: 0.14402\noutside 1 331\nTMhelix 332 354\ninside 355 372

13873 GCF_002055555.1_ASM205555v1 Salmonella enterica Proteobacteria; Gammaproteobacteria; Enterobacterales; Enterobacteriaceae; Salmonella
MNNEETFYQAMRRKGVTRRSFLKFCSLAATSLGLGAGMTPKIAWALENKPRIPVVWIHGL WP_080107816.1
[Ni/Fe] hydrogenase small subunit [Salmonella enterica] Length: 372\nNumber of predicted TMHs: 2\nExp

number of AAs in TMHs: 36.51739\nExp number, first 60 AAs: 19.72849\nTotal prob of N-in: 0.96170\nPOSSIBLE N-term signal sequence\ninside 1 19\nTMhelix 20 39\noutside 40 325\nTMhelix 326 348\ninside 349 372

13874 GCF_002055905.1_ASM205590v1 Salmonella enterica Proteobacteria; Gammaproteobacteria; Enterobacterales; Enterobacteriaceae; Salmonella
MNNEETFYQAMRRKGVTRRSFLKFCSLAATSLGLGAGMTPKIAWALENKPRIPVVWIHGL WP_001058312.1
hydrogenase [Salmonella enterica] Length: 372\nNumber of predicted TMHs: 2\nExp number of AAs in TMHs: 36.51738\nExp number, first 60 AAs: 19.72849\nTotal prob of N-in: 0.96170\nPOSSIBLE N-term signal sequence\ninside 1 19\nTMhelix 20 39\noutside 40 325\nTMhelix 326 348\ninside 349 372

13875 GCF_002055775.1_ASM205577v1 Salmonella enterica Proteobacteria; Gammaproteobacteria; Enterobacterales; Enterobacteriaceae; Salmonella
MNNEETFYQAMRRKGVTRRSFLKFCSLAATSLGLGAGMTPKIAWALENKPRIPVVWIHGL WP_001058312.1
hydrogenase [Salmonella enterica] Length: 372\nNumber of predicted TMHs: 2\nExp number of AAs in TMHs: 36.51738\nExp number, first 60 AAs: 19.72849\nTotal prob of N-in: 0.96170\nPOSSIBLE N-term signal sequence\ninside 1 19\nTMhelix 20 39\noutside 40 325\nTMhelix 326 348\ninside 349 372

13876 GCF_002055805.1_ASM205580v1 Salmonella enterica Proteobacteria; Gammaproteobacteria; Enterobacterales; Enterobacteriaceae; Salmonella
MNNEETFYQAMRRKGVTRRSFLKFCSLAATSLGLGAGMTPKIAWALENKPRIPVVWIHGL WP_001058312.1
hydrogenase [Salmonella enterica] Length: 372\nNumber of predicted TMHs: 2\nExp number of AAs in TMHs: 36.51738\nExp number, first 60 AAs: 19.72849\nTotal prob of N-in: 0.96170\nPOSSIBLE N-term signal sequence\ninside 1 19\nTMhelix 20 39\noutside 40 325\nTMhelix 326 348\ninside 349 372

13877 GCF_002055715.1_ASM205571v1 Salmonella enterica Proteobacteria; Gammaproteobacteria; Enterobacterales; Enterobacteriaceae; Salmonella
MNNEETFYQAMRRKGVTRRSFLKFCSLAATSLGLGAGMTPKIAWALENKPRIPVVWIHGL WP_001058312.1
hydrogenase [Salmonella enterica] Length: 372\nNumber of predicted TMHs: 2\nExp number of AAs in TMHs: 36.51738\nExp number, first 60 AAs: 19.72849\nTotal prob of N-in: 0.96170\nPOSSIBLE N-term signal sequence\ninside 1 19\nTMhelix 20 39\noutside 40 325\nTMhelix 326 348\ninside 349 372

13878 GCF_002055805.1_ASM205580v1 Salmonella enterica Proteobacteria; Gammaproteobacteria; Enterobacterales; Enterobacteriaceae; Salmonella
MTGDNTLITSHGINRRDFMKLCAALAATMGLSSKAAAEMAESVSNPQRPPVIWIGAQECT WP_000145429.1
hydrogenase 2 small subunit [Salmonella enterica] Length: 372\nNumber of predicted TMHs: 1\nExp number of AAs in TMHs: 24.51309\nExp number, first 60 AAs: 0.2682\nTotal prob of N-in: 0.14402\noutside 1 331\nTMhelix 332 354\ninside 355 372

13879 GCF_002055775.1_ASM205577v1 Salmonella enterica Proteobacteria; Gammaproteobacteria; Enterobacterales; Enterobacteriaceae; Salmonella
MTGDNTLITSHGINRRDFMKLCAALAATMGLSSKAAAEMAESVSNPQRPPVIWIGAQECT WP_000145429.1
hydrogenase 2 small subunit [Salmonella enterica] Length: 372\nNumber of predicted TMHs: 1\nExp number of AAs in TMHs: 24.51309\nExp number, first 60 AAs: 0.2682\nTotal prob of N-in: 0.14402\noutside 1 331\nTMhelix 332 354\ninside 355 372

13880 GCF_002055715.1_ASM205571v1 Salmonella enterica Proteobacteria; Gammaproteobacteria; Enterobacterales; Enterobacteriaceae; Salmonella
MTGDNTLITSHGINRRDFMKLCAALAATMGLSSKAAAEMAESVSNPQRPPVIWIGAQECT WP_000145429.1
hydrogenase 2 small subunit [Salmonella enterica] Length: 372\nNumber of predicted TMHs: 1\nExp number of AAs in TMHs: 24.51309\nExp number, first 60 AAs: 0.2682\nTotal prob of N-in: 0.14402\noutside 1 331\nTMhelix 332 354\ninside 355 372

13881 GCF_002055645.1_ASM205564v1 Salmonella enterica Proteobacteria; Gammaproteobacteria; Enterobacterales; Enterobacteriaceae; Salmonella
MTGDNTLITSHGINRRDFMKLCAALAATMGLSSKAAAEMAESVSNPQRPPVIWIGAQECT WP_000145429.1
hydrogenase 2 small subunit [Salmonella enterica] Length: 372\nNumber of predicted TMHs: 1\nExp number of AAs in TMHs: 24.51309\nExp number, first 60 AAs: 0.2682\nTotal prob of N-in: 0.14402\noutside 1 331\nTMhelix 332 354\ninside 355 372

13882 GCF_002055555.1_ASM205555v1 Salmonella enterica Proteobacteria; Gammaproteobacteria; Enterobacterales; Enterobacteriaceae; Salmonella

MTGDNTLITSHGINRRDFMKLCAALAATMGLSSKAAAEMAESVSNPQRPPVIWIGAQECT WP_000145429.1 hydrogenase 2 small subunit [Salmonella enterica] Length: 372\nNumber of predicted TMHs: 1\nExp number of AAs in TMHs: 24.51309\nExp number, first 60 AAs: 0.2682\nTotal prob of N-in: 0.14402\noutside 1 331\nTMhelix 332 354\ninside 355 372

13883 GCF_001648855.1_ASM164885v1 Salmonella enterica Proteobacteria; Gammaproteobacteria; Enterobacterales; Enterobacteriaceae; Salmonella MLKFRTIFLKKIFTLKNSILARRGAVIVIVSAVFTSIMFFAHSWASDKEVAMTLSVLNSN WP_023226644.1 hypothetical protein [Salmonella enterica] Length: 204\nNumber of predicted TMHs: 2\nExp number of AAs in TMHs: 44.99526\nExp number, first 60 AAs: 22.33786\nTotal prob of N-in: 0.87083\nPOSSIBLE N-term signal sequence\ninside 1 19\nTMhelix 20 42\noutside 43 171\nTMhelix 172 194\ninside 195 204

13884 GCF_002055315.1_ASM205531v1 Salmonella enterica Proteobacteria; Gammaproteobacteria; Enterobacterales; Enterobacteriaceae; Salmonella MNNEETFYQAMRRKGVTRRSFLKFCSLAATSLGLGAGMTPKIAWALENKPRIPVVWIHGL WP_001058308.1 hydrogenase [Salmonella enterica] Length: 372\nNumber of predicted TMHs: 2\nExp number of AAs in TMHs: 39.08204\nExp number, first 60 AAs: 19.72474\nTotal prob of N-in: 0.96174\nPOSSIBLE N-term signal sequence\ninside 1 19\nTMhelix 20 39\noutside 40 325\nTMhelix 326 348\ninside 349 372

13885 GCF_002055495.1_ASM205549v1 Salmonella enterica Proteobacteria; Gammaproteobacteria; Enterobacterales; Enterobacteriaceae; Salmonella MNNEETFYQAMRRKGVTRRSFLKFCSLAATSLGLGAGMTPKIAWALENKPRIPVVWIHGL WP_001058312.1 hydrogenase [Salmonella enterica] Length: 372\nNumber of predicted TMHs: 2\nExp number of AAs in TMHs: 36.51738\nExp number, first 60 AAs: 19.72849\nTotal prob of N-in: 0.96170\nPOSSIBLE N-term signal sequence\ninside 1 19\nTMhelix 20 39\noutside 40 325\nTMhelix 326 348\ninside 349 372

13886 GCF_002055495.1_ASM205549v1 Salmonella enterica Proteobacteria; Gammaproteobacteria; Enterobacterales; Enterobacteriaceae; Salmonella MTGDNTLITSHGINRRDFMKLCAALAATMGLSSKAAAEMAESVSNPQRPPVIWIGAQECT WP_000145429.1 hydrogenase 2 small subunit [Salmonella enterica] Length: 372\nNumber of predicted TMHs: 1\nExp number of AAs in TMHs: 24.51309\nExp number, first 60 AAs: 0.2682\nTotal prob of N-in: 0.14402\noutside 1 331\nTMhelix 332 354\ninside 355 372

13887 GCF_002055395.1_ASM205539v1 Salmonella enterica Proteobacteria; Gammaproteobacteria; Enterobacterales; Enterobacteriaceae; Salmonella MNNEETFYQAMRRKGVTRRSFLKFCSLAATSLGLGAGMTPKIAWALENKPRIPVVWIHGL WP_001058312.1 hydrogenase [Salmonella enterica] Length: 372\nNumber of predicted TMHs: 2\nExp number of AAs in TMHs: 36.51738\nExp number, first 60 AAs: 19.72849\nTotal prob of N-in: 0.96170\nPOSSIBLE N-term signal sequence\ninside 1 19\nTMhelix 20 39\noutside 40 325\nTMhelix 326 348\ninside 349 372

13888 GCF_002055395.1_ASM205539v1 Salmonella enterica Proteobacteria; Gammaproteobacteria; Enterobacterales; Enterobacteriaceae; Salmonella MTGDNTLITSHGINRRDFMKLCAALAATMGLSSKAAAEMAESVSNPQRPPVIWIGAQECT WP_000145429.1 hydrogenase 2 small subunit [Salmonella enterica] Length: 372\nNumber of predicted TMHs: 1\nExp number of AAs in TMHs: 24.51309\nExp number, first 60 AAs: 0.2682\nTotal prob of N-in: 0.14402\noutside 1 331\nTMhelix 332 354\ninside 355 372

13889 GCF_002055345.1_ASM205534v1 Salmonella enterica Proteobacteria; Gammaproteobacteria; Enterobacterales; Enterobacteriaceae; Salmonella MTGDNTLITSHGINRRDFMKLCAALAATMGLSSKAAAEMAESVSNPQRPPVIWIGAQECT WP_000145429.1 hydrogenase 2 small subunit [Salmonella enterica] Length: 372\nNumber of predicted TMHs: 1\nExp number of AAs in TMHs: 24.51309\nExp number, first 60 AAs: 0.2682\nTotal prob of N-in: 0.14402\noutside 1 331\nTMhelix 332 354\ninside 355 372

13890 GCF_002055175.1_ASM205517v1 Salmonella enterica Proteobacteria; Gammaproteobacteria; Enterobacterales; Enterobacteriaceae; Salmonella MNNEETFYQAMRRKGVTRRSFLKFCSLAATSLGLGAGMTPKIAWALENKPRIPVVWIHGL WP_001058312.1 hydrogenase [Salmonella enterica] Length: 372\nNumber of predicted TMHs: 2\nExp number of AAs in TMHs: 36.51738\nExp number, first 60 AAs: 19.72849\nTotal prob of N-in: 0.96170\nPOSSIBLE N-term signal sequence\ninside 1 19\nTMhelix 20 39\noutside 40 325\nTMhelix 326 348\ninside 349 372

13891 GCF_002055255.1_ASM205525v1 Salmonella enterica Proteobacteria; Gammaproteobacteria; Enterobacterales; Enterobacteriaceae; Salmonella
MTGDNTLITSHGINRRDFMKLCAALAATMGLSSKAAAEMAESVSNPQRPPVIWIGAQECT WP_000145429.1
hydrogenase 2 small subunit [Salmonella enterica] Length: 372\nNumber of predicted TMHs: 1\nExp number of AAs in TMHs: 24.51309\nExp number, first 60 AAs: 0.2682\nTotal prob of N-in: 0.14402\noutside 1 331\nTMhelix 332 354\ninside 355 372

13892 GCF_002054815.1_ASM205481v1 Salmonella enterica Proteobacteria; Gammaproteobacteria; Enterobacterales; Enterobacteriaceae; Salmonella
MNNEETFYQAMRRKGVTRRSFLKFCSLAATSLGLGAGMTPKIAWALENKPRIPVVWIHGL WP_001058312.1
hydrogenase [Salmonella enterica] Length: 372\nNumber of predicted TMHs: 2\nExp number of AAs in TMHs: 36.51738\nExp number, first 60 AAs: 19.72849\nTotal prob of N-in: 0.96170\nPOSSIBLE N-term signal sequence\ninside 1 19\nTMhelix 20 39\noutside 40 325\nTMhelix 326 348\ninside 349 372

13893 GCF_002054905.1_ASM205490v1 Salmonella enterica Proteobacteria; Gammaproteobacteria; Enterobacterales; Enterobacteriaceae; Salmonella
MNNEETFYQAMRRKGVTRRSFLKFCSLAATSLGLGAGMTPKIAWALENKPRIPVVWIHGL WP_001058310.1
hydrogenase [Salmonella enterica] Length: 372\nNumber of predicted TMHs: 2\nExp number of AAs in TMHs: 37.4567299999999\nExp number, first 60 AAs: 19.72694\nTotal prob of N-in: 0.96171\nPOSSIBLE N-term signal sequence\ninside 1 19\nTMhelix 20 39\noutside 40 325\nTMhelix 326 348\ninside 349 372

13894 GCF_002055315.1_ASM205531v1 Salmonella enterica Proteobacteria; Gammaproteobacteria; Enterobacterales; Enterobacteriaceae; Salmonella
MTGDNTLITSHGINRRDFMKLCAALAATMGLSSKAAAEMAESVSNPQRPPVIWIGAQECT WP_000145429.1
hydrogenase 2 small subunit [Salmonella enterica] Length: 372\nNumber of predicted TMHs: 1\nExp number of AAs in TMHs: 24.51309\nExp number, first 60 AAs: 0.2682\nTotal prob of N-in: 0.14402\noutside 1 331\nTMhelix 332 354\ninside 355 372

13895 GCF_002054935.1_ASM205493v1 Salmonella enterica Proteobacteria; Gammaproteobacteria; Enterobacterales; Enterobacteriaceae; Salmonella
MNNEETFYQAMRRKGVTRRSFLKFCSLAATSLGLGAGMTPKIAWALENKPRIPVVWIHGL WP_001058307.1
hydrogenase [Salmonella enterica] Length: 372\nNumber of predicted TMHs: 2\nExp number of AAs in TMHs: 36.51738\nExp number, first 60 AAs: 19.72849\nTotal prob of N-in: 0.96170\nPOSSIBLE N-term signal sequence\ninside 1 19\nTMhelix 20 39\noutside 40 325\nTMhelix 326 348\ninside 349 372

13896 GCF_002054985.1_ASM205498v1 Salmonella enterica Proteobacteria; Gammaproteobacteria; Enterobacterales; Enterobacteriaceae; Salmonella
MTGDNTLITSHGINRRDFMKLCAALAATMGLSSKAAAEMAESVSNPQRPPVIWIGAQECT WP_000145429.1
hydrogenase 2 small subunit [Salmonella enterica] Length: 372\nNumber of predicted TMHs: 1\nExp number of AAs in TMHs: 24.51309\nExp number, first 60 AAs: 0.2682\nTotal prob of N-in: 0.14402\noutside 1 331\nTMhelix 332 354\ninside 355 372

13897 GCF_002054905.1_ASM205490v1 Salmonella enterica Proteobacteria; Gammaproteobacteria; Enterobacterales; Enterobacteriaceae; Salmonella
MTGDNTLITSHGINRRDFMKLCAALAATMGLSSKAAAEMAESVSNPQRPPVIWIGAQECT WP_000145429.1
hydrogenase 2 small subunit [Salmonella enterica] Length: 372\nNumber of predicted TMHs: 1\nExp number of AAs in TMHs: 24.51309\nExp number, first 60 AAs: 0.2682\nTotal prob of N-in: 0.14402\noutside 1 331\nTMhelix 332 354\ninside 355 372

13898 GCF_002054875.1_ASM205487v1 Salmonella enterica Proteobacteria; Gammaproteobacteria; Enterobacterales; Enterobacteriaceae; Salmonella
MTGDNTLITSHGINRRDFMKLCAALAATMGLSSKAAAEMAESVSNPQRPPVIWIGAQECT WP_000145429.1
hydrogenase 2 small subunit [Salmonella enterica] Length: 372\nNumber of predicted TMHs: 1\nExp number of AAs in TMHs: 24.51309\nExp number, first 60 AAs: 0.2682\nTotal prob of N-in: 0.14402\noutside 1 331\nTMhelix 332 354\ninside 355 372

13899 GCF_002054495.1_ASM205449v1 Salmonella enterica Proteobacteria; Gammaproteobacteria; Enterobacterales; Enterobacteriaceae; Salmonella
MNNEETFYQAMRRKGVTRRSFLKFCSLAATSLGLGAGMTPKIAWALENKPRIPVVWIHGL WP_001058311.1
hydrogenase [Salmonella enterica] Length: 372\nNumber of predicted TMHs: 2\nExp number of AAs in

TMHs: 37.4699899999999\nExp number, first 60 AAs: 19.72699\nTotal prob of N-in: 0.96171\nPOSSIBLE N-term signal sequence\ninside 1 19\nTMhelix 20 39\noutside 40 325\nTMhelix 326 348\ninside 349 372

13900 GCF_002054545.1_ASM205454v1 Salmonella enterica Proteobacteria; Gammaproteobacteria; Enterobacterales; Enterobacteriaceae; Salmonella
MTGDNTLITSHGINRRDFMKLCAALAATMGLSSKAAAEMAESVSNPQRPPVIWIGAQECT WP_000145429.1
hydrogenase 2 small subunit [Salmonella enterica] Length: 372\nNumber of predicted TMHs: 1\nExp number of AAs in TMHs: 24.51309\nExp number, first 60 AAs: 0.2682\nTotal prob of N-in: 0.14402\noutside 1 331\nTMhelix 332 354\ninside 355 372

13901 GCF_002054495.1_ASM205449v1 Salmonella enterica Proteobacteria; Gammaproteobacteria; Enterobacterales; Enterobacteriaceae; Salmonella
MTGDNTLITSHGINRRDFMKLCAALAATMGLSSKAAAEMAESVSNPQRPPVIWIGAQECT WP_000145429.1
hydrogenase 2 small subunit [Salmonella enterica] Length: 372\nNumber of predicted TMHs: 1\nExp number of AAs in TMHs: 24.51309\nExp number, first 60 AAs: 0.2682\nTotal prob of N-in: 0.14402\noutside 1 331\nTMhelix 332 354\ninside 355 372

13902 GCF_002054355.1_ASM205435v1 Salmonella enterica Proteobacteria; Gammaproteobacteria; Enterobacterales; Enterobacteriaceae; Salmonella
MNNEETFYQAMRRKGVTRRSFLKFCSLAATSLGLGAGMTPKIAWALENKPRIPVVWIHGL WP_001058310.1
hydrogenase [Salmonella enterica] Length: 372\nNumber of predicted TMHs: 2\nExp number of AAs in TMHs: 37.4567299999999\nExp number, first 60 AAs: 19.72694\nTotal prob of N-in: 0.96171\nPOSSIBLE N-term signal sequence\ninside 1 19\nTMhelix 20 39\noutside 40 325\nTMhelix 326 348\ninside 349 372

13903 GCF_002054455.1_ASM205445v1 Salmonella enterica Proteobacteria; Gammaproteobacteria; Enterobacterales; Enterobacteriaceae; Salmonella
MNNEETFYQAMRRKGVTRRSFLKFCSLAATSLGLGAGMTPKIAWALENKPRIPVVWIHGL WP_001058310.1
hydrogenase [Salmonella enterica] Length: 372\nNumber of predicted TMHs: 2\nExp number of AAs in TMHs: 37.4567299999999\nExp number, first 60 AAs: 19.72694\nTotal prob of N-in: 0.96171\nPOSSIBLE N-term signal sequence\ninside 1 19\nTMhelix 20 39\noutside 40 325\nTMhelix 326 348\ninside 349 372

13904 GCF_002054515.1_ASM205451v1 Salmonella enterica Proteobacteria; Gammaproteobacteria; Enterobacterales; Enterobacteriaceae; Salmonella
MNNEETFYQAMRRKGVTRRSFLKFCSLAATSLGLGAGMTPKIAWALENKPRIPVVWIHGL WP_001058312.1
hydrogenase [Salmonella enterica] Length: 372\nNumber of predicted TMHs: 2\nExp number of AAs in TMHs: 36.51738\nExp number, first 60 AAs: 19.72849\nTotal prob of N-in: 0.96170\nPOSSIBLE N-term signal sequence\ninside 1 19\nTMhelix 20 39\noutside 40 325\nTMhelix 326 348\ninside 349 372

13905 GCF_002054515.1_ASM205451v1 Salmonella enterica Proteobacteria; Gammaproteobacteria; Enterobacterales; Enterobacteriaceae; Salmonella
MTGDNTLITSHGINRRDFMKLCAALAATMGLSSKAAAEMAESVSNPQRPPVIWIGAQECT WP_000145429.1
hydrogenase 2 small subunit [Salmonella enterica] Length: 372\nNumber of predicted TMHs: 1\nExp number of AAs in TMHs: 24.51309\nExp number, first 60 AAs: 0.2682\nTotal prob of N-in: 0.14402\noutside 1 331\nTMhelix 332 354\ninside 355 372

13906 GCF_002054165.1_ASM205416v1 Salmonella enterica Proteobacteria; Gammaproteobacteria; Enterobacterales; Enterobacteriaceae; Salmonella
MTGDNTLITSHGINRRDFMKLCAALAATMGLSSKAAAEMAESVSNPQRPPVIWIGAQECT WP_000145429.1
hydrogenase 2 small subunit [Salmonella enterica] Length: 372\nNumber of predicted TMHs: 1\nExp number of AAs in TMHs: 24.51309\nExp number, first 60 AAs: 0.2682\nTotal prob of N-in: 0.14402\noutside 1 331\nTMhelix 332 354\ninside 355 372

13907 GCF_002053865.1_ASM205386v1 Salmonella enterica Proteobacteria; Gammaproteobacteria; Enterobacterales; Enterobacteriaceae; Salmonella
MNNEETFYQAMRRKGVTRRSFLKFCSLAATSLGLGAGMTPKIAWALENKPRIPVVWIHGL WP_001058312.1
hydrogenase [Salmonella enterica] Length: 372\nNumber of predicted TMHs: 2\nExp number of AAs in TMHs: 36.51738\nExp number, first 60 AAs: 19.72849\nTotal prob of N-in: 0.96170\nPOSSIBLE N-term signal sequence\ninside 1 19\nTMhelix 20 39\noutside 40 325\nTMhelix 326 348\ninside 349 372

13908 GCF_002054025.1_ASM205402v1 Salmonella enterica Proteobacteria; Gammaproteobacteria; Enterobacterales; Enterobacteriaceae; Salmonella

MNNEETFYQAMRRKGVTRRSFLKFCSLAATSLGLGAGMTPKIAWALENKPRIPVVWIHGL WP_001058308.1 hydrogenase [Salmonella enterica] Length: 372\nNumber of predicted TMHs: 2\nExp number of AAs in TMHs: 39.08204\nExp number, first 60 AAs: 19.72474\nTotal prob of N-in: 0.96174\nPOSSIBLE N-term signal sequence\ninside 1 19\nTMhelix 20 39\noutside 40 325\nTMhelix 326 348\ninside 349 372

13909 GCF_002054075.1_ASM205407v1 Salmonella enterica Proteobacteria; Gammaproteobacteria; Enterobacterales; Enterobacteriaceae; Salmonella MNNEETFYQAMRRKGVTRRSFLKFCSLAATSLGLGAGMTPKIAWALENKPRIPVVWIHGL WP_001058312.1 hydrogenase [Salmonella enterica] Length: 372\nNumber of predicted TMHs: 2\nExp number of AAs in TMHs: 36.51738\nExp number, first 60 AAs: 19.72849\nTotal prob of N-in: 0.96170\nPOSSIBLE N-term signal sequence\ninside 1 19\nTMhelix 20 39\noutside 40 325\nTMhelix 326 348\ninside 349 372

13910 GCF_002054075.1_ASM205407v1 Salmonella enterica Proteobacteria; Gammaproteobacteria; Enterobacterales; Enterobacteriaceae; Salmonella MTGDNTLITSHGINRRDFMKLCAALAATMGLSSKAAAEMAESVSNPQRPPVIWIGAQECT WP_000145429.1 hydrogenase 2 small subunit [Salmonella enterica] Length: 372\nNumber of predicted TMHs: 1\nExp number of AAs in TMHs: 24.51309\nExp number, first 60 AAs: 0.2682\nTotal prob of N-in: 0.14402\noutside 1 331\nTMhelix 332 354\ninside 355 372

13911 GCF_002054025.1_ASM205402v1 Salmonella enterica Proteobacteria; Gammaproteobacteria; Enterobacterales; Enterobacteriaceae; Salmonella MTGDNTLITSHGINRRDFMKLCAALAATMGLSSKAAAEMAESVSNPQRPPVIWIGAQECT WP_000145429.1 hydrogenase 2 small subunit [Salmonella enterica] Length: 372\nNumber of predicted TMHs: 1\nExp number of AAs in TMHs: 24.51309\nExp number, first 60 AAs: 0.2682\nTotal prob of N-in: 0.14402\noutside 1 331\nTMhelix 332 354\ninside 355 372

13912 GCF_002053955.1_ASM205395v1 Salmonella enterica Proteobacteria; Gammaproteobacteria; Enterobacterales; Enterobacteriaceae; Salmonella MNNEETFYQAMRRKGVTRRSFLKFCSLAATSLGLGAGMTPKIAWALENKPRIPVVWIHGL WP_001058312.1 hydrogenase [Salmonella enterica] Length: 372\nNumber of predicted TMHs: 2\nExp number of AAs in TMHs: 36.51738\nExp number, first 60 AAs: 19.72849\nTotal prob of N-in: 0.96170\nPOSSIBLE N-term signal sequence\ninside 1 19\nTMhelix 20 39\noutside 40 325\nTMhelix 326 348\ninside 349 372

13913 GCF_002053375.1_ASM205337v1 Salmonella enterica Proteobacteria; Gammaproteobacteria; Enterobacterales; Enterobacteriaceae; Salmonella MNNEETFYQAMRRKGVTRRSFLKFCSLAATSLGLGAGMTPKIAWALENKPRIPVVWIHGL WP_001058312.1 hydrogenase [Salmonella enterica] Length: 372\nNumber of predicted TMHs: 2\nExp number of AAs in TMHs: 36.51738\nExp number, first 60 AAs: 19.72849\nTotal prob of N-in: 0.96170\nPOSSIBLE N-term signal sequence\ninside 1 19\nTMhelix 20 39\noutside 40 325\nTMhelix 326 348\ninside 349 372

13914 GCF_002053405.1_ASM205340v1 Salmonella enterica Proteobacteria; Gammaproteobacteria; Enterobacterales; Enterobacteriaceae; Salmonella MNNEETFYQAMRRKGVTRRSFLKFCSLAATSLGLGAGMTPKIAWALENKPRIPVVWIHGL WP_001058312.1 hydrogenase [Salmonella enterica] Length: 372\nNumber of predicted TMHs: 2\nExp number of AAs in TMHs: 36.51738\nExp number, first 60 AAs: 19.72849\nTotal prob of N-in: 0.96170\nPOSSIBLE N-term signal sequence\ninside 1 19\nTMhelix 20 39\noutside 40 325\nTMhelix 326 348\ninside 349 372

13915 GCF_002053565.1_ASM205356v1 Salmonella enterica Proteobacteria; Gammaproteobacteria; Enterobacterales; Enterobacteriaceae; Salmonella MNNEETFYQAMRRKGVTRRSFLKFCSLAATSLGLGAGMTPKIAWALENKPRIPVVWIHGL WP_001058311.1 hydrogenase [Salmonella enterica] Length: 372\nNumber of predicted TMHs: 2\nExp number of AAs in TMHs: 37.4699899999999\nExp number, first 60 AAs: 19.72699\nTotal prob of N-in: 0.96171\nPOSSIBLE N-term signal sequence\ninside 1 19\nTMhelix 20 39\noutside 40 325\nTMhelix 326 348\ninside 349 372

13916 GCF_002053565.1_ASM205356v1 Salmonella enterica Proteobacteria; Gammaproteobacteria; Enterobacterales; Enterobacteriaceae; Salmonella MTGDNTLITSHGINRRDFMKLCAALAATMGLSSKAAAEMAESVSNPQRPPVIWIGAQECT WP_000145429.1 hydrogenase 2 small subunit [Salmonella enterica] Length: 372\nNumber of predicted TMHs: 1\nExp number of AAs in TMHs: 24.51309\nExp number, first 60 AAs: 0.2682\nTotal prob of N-in: 0.14402\noutside 1 331\nTMhelix 332 354\ninside 355 372

13917 GCF_002053505.1_ASM205350v1 Salmonella enterica Proteobacteria; Gammaproteobacteria; Enterobacterales; Enterobacteriaceae; Salmonella
MNNEETFYQAMRRKGVTRRSFLKFCSLAATSLGLGAGMTPKIAWALENKPRIPVVWIHGL WP_001058312.1
hydrogenase [Salmonella enterica] Length: 372\nNumber of predicted TMHs: 2\nExp number of AAs in TMHs: 36.51738\nExp number, first 60 AAs: 19.72849\nTotal prob of N-in: 0.96170\nPOSSIBLE N-term signal sequence\ninside 1 19\nTMhelix 20 39\noutside 40 325\nTMhelix 326 348\ninside 349 372

13918 GCF_002053505.1_ASM205350v1 Salmonella enterica Proteobacteria; Gammaproteobacteria; Enterobacterales; Enterobacteriaceae; Salmonella
MTGDNTLITSHGINRRDFMKLCAALAATMGLSSKAAAEMAESVSNPQRPPVIWIGAQECT WP_000145431.1
hydrogenase 2 small subunit [Salmonella enterica] Length: 372\nNumber of predicted TMHs: 1\nExp number of AAs in TMHs: 24.51505\nExp number, first 60 AAs: 0.2682\nTotal prob of N-in: 0.14406\noutside 1 331\nTMhelix 332 354\ninside 355 372

13919 GCF_002053225.1_ASM205322v1 Salmonella enterica Proteobacteria; Gammaproteobacteria; Enterobacterales; Enterobacteriaceae; Salmonella
MTGDNTLITSHGINRRDFMKLCAALAATMGLSSKAAAEMAESVSNPQRPPVIWIGAQECT WP_000145429.1
hydrogenase 2 small subunit [Salmonella enterica] Length: 372\nNumber of predicted TMHs: 1\nExp number of AAs in TMHs: 24.51309\nExp number, first 60 AAs: 0.2682\nTotal prob of N-in: 0.14402\noutside 1 331\nTMhelix 332 354\ninside 355 372

13920 GCF_002053075.1_ASM205307v1 Salmonella enterica Proteobacteria; Gammaproteobacteria; Enterobacterales; Enterobacteriaceae; Salmonella
MTGDNTLITSHGINRRDFMKLCAALAATMGLSSKAAAEMAESVSNPQRPPVIWIGAQECT WP_000145429.1
hydrogenase 2 small subunit [Salmonella enterica] Length: 372\nNumber of predicted TMHs: 1\nExp number of AAs in TMHs: 24.51309\nExp number, first 60 AAs: 0.2682\nTotal prob of N-in: 0.14402\noutside 1 331\nTMhelix 332 354\ninside 355 372

13921 GCF_002053175.1_ASM205317v1 Salmonella enterica Proteobacteria; Gammaproteobacteria; Enterobacterales; Enterobacteriaceae; Salmonella
MTGDNTLITSHGINRRDFMKLCAALAATMGLSSKAAAEMAESVSNPQRPPVIWIGAQECT WP_000145429.1
hydrogenase 2 small subunit [Salmonella enterica] Length: 372\nNumber of predicted TMHs: 1\nExp number of AAs in TMHs: 24.51309\nExp number, first 60 AAs: 0.2682\nTotal prob of N-in: 0.14402\noutside 1 331\nTMhelix 332 354\ninside 355 372

13922 GCF_002053085.1_ASM205308v1 Salmonella enterica Proteobacteria; Gammaproteobacteria; Enterobacterales; Enterobacteriaceae; Salmonella
MTGDNTLITSHGINRRDFMKLCAALAATMGLSSKAAAEMAESVSNPQRPPVIWIGAQECT WP_000145429.1
hydrogenase 2 small subunit [Salmonella enterica] Length: 372\nNumber of predicted TMHs: 1\nExp number of AAs in TMHs: 24.51309\nExp number, first 60 AAs: 0.2682\nTotal prob of N-in: 0.14402\noutside 1 331\nTMhelix 332 354\ninside 355 372

13923 GCF_002052805.1_ASM205280v1 Salmonella enterica Proteobacteria; Gammaproteobacteria; Enterobacterales; Enterobacteriaceae; Salmonella
MNNEETFYQAMRRKGVTRRSFLKFCSLAATSLGLGAGMTPKIAWALENKPRIPVVWIHGL WP_001058312.1
hydrogenase [Salmonella enterica] Length: 372\nNumber of predicted TMHs: 2\nExp number of AAs in TMHs: 36.51738\nExp number, first 60 AAs: 19.72849\nTotal prob of N-in: 0.96170\nPOSSIBLE N-term signal sequence\ninside 1 19\nTMhelix 20 39\noutside 40 325\nTMhelix 326 348\ninside 349 372

13924 GCF_002052995.1_ASM205299v1 Salmonella enterica Proteobacteria; Gammaproteobacteria; Enterobacterales; Enterobacteriaceae; Salmonella
MNNEETFYQAMRRKGVTRRSFLKFCSLAATSLGLGAGMTPKIAWALENKPRIPVVWIHGL WP_001058312.1
hydrogenase [Salmonella enterica] Length: 372\nNumber of predicted TMHs: 2\nExp number of AAs in TMHs: 36.51738\nExp number, first 60 AAs: 19.72849\nTotal prob of N-in: 0.96170\nPOSSIBLE N-term signal sequence\ninside 1 19\nTMhelix 20 39\noutside 40 325\nTMhelix 326 348\ninside 349 372

13925 GCF_002052375.1_ASM205237v1 Salmonella enterica Proteobacteria; Gammaproteobacteria; Enterobacterales; Enterobacteriaceae; Salmonella
MNNEETFYQAMRRKGVTRRSFLKFCSLAATSLGLGAGMTPKIAWALENKPRIPVVWIHGL WP_001058312.1
hydrogenase [Salmonella enterica] Length: 372\nNumber of predicted TMHs: 2\nExp number of AAs in TMHs: 36.51738\nExp number, first 60 AAs: 19.72849\nTotal prob of N-in: 0.96170\nPOSSIBLE N-term signal sequence\ninside 1 19\nTMhelix 20 39\noutside 40 325\nTMhelix 326 348\ninside 349 372

TMHs: 36.51738\nExp number, first 60 AAs: 19.72849\nTotal prob of N-in: 0.96170\nPOSSIBLE N-term signal sequence\ninside 1 19\nTMhelix 20 39\noutside 40 325\nTMhelix 326 348\ninside 349 372

13926 GCF_002052565.1_ASM205256v1 Salmonella enterica Proteobacteria; Gammaproteobacteria; Enterobacterales; Enterobacteriaceae; Salmonella
MNNEETFYQAMRRKGVTRRSFLKFCSLAATSLGLGAGMTPKIAWALENKPRIPVVWIHGL WP_001058312.1
hydrogenase [Salmonella enterica] Length: 372\nNumber of predicted TMHs: 2\nExp number of AAs in TMHs: 36.51738\nExp number, first 60 AAs: 19.72849\nTotal prob of N-in: 0.96170\nPOSSIBLE N-term signal sequence\ninside 1 19\nTMhelix 20 39\noutside 40 325\nTMhelix 326 348\ninside 349 372

13927 GCF_002052415.1_ASM205241v1 Salmonella enterica Proteobacteria; Gammaproteobacteria; Enterobacterales; Enterobacteriaceae; Salmonella
MNNEETFYQAMRRKGVTRRSFLKFCSLAATSLGLGAGMTPKIAWALENKPRIPVVWIHGL WP_001058311.1
hydrogenase [Salmonella enterica] Length: 372\nNumber of predicted TMHs: 2\nExp number of AAs in TMHs: 37.4699899999999\nExp number, first 60 AAs: 19.72699\nTotal prob of N-in: 0.96171\nPOSSIBLE N-term signal sequence\ninside 1 19\nTMhelix 20 39\noutside 40 325\nTMhelix 326 348\ninside 349 372

13928 GCF_002052565.1_ASM205256v1 Salmonella enterica Proteobacteria; Gammaproteobacteria; Enterobacterales; Enterobacteriaceae; Salmonella
MTGDNTLITSHGINRRDFMKLCAALAATMGLSSKAAAEMAESVSNPQRPPVIWIGAQECT WP_000145429.1
hydrogenase 2 small subunit [Salmonella enterica] Length: 372\nNumber of predicted TMHs: 1\nExp number of AAs in TMHs: 24.51309\nExp number, first 60 AAs: 0.2682\nTotal prob of N-in: 0.14402\noutside 1 331\nTMhelix 332 354\ninside 355 372

13929 GCF_002052375.1_ASM205237v1 Salmonella enterica Proteobacteria; Gammaproteobacteria; Enterobacterales; Enterobacteriaceae; Salmonella
MTGDNTLITSHGINRRDFMKLCAALAATMGLSSKAAAEMAESVSNPQRPPVIWIGAQECT WP_000145429.1
hydrogenase 2 small subunit [Salmonella enterica] Length: 372\nNumber of predicted TMHs: 1\nExp number of AAs in TMHs: 24.51309\nExp number, first 60 AAs: 0.2682\nTotal prob of N-in: 0.14402\noutside 1 331\nTMhelix 332 354\ninside 355 372

13930 GCF_002052415.1_ASM205241v1 Salmonella enterica Proteobacteria; Gammaproteobacteria; Enterobacterales; Enterobacteriaceae; Salmonella
MTGDNTLITSHGINRRDFMKLCAALAATMGLSSKAAAEMAESVSNPQRPPVIWIGAQECT WP_000145429.1
hydrogenase 2 small subunit [Salmonella enterica] Length: 372\nNumber of predicted TMHs: 1\nExp number of AAs in TMHs: 24.51309\nExp number, first 60 AAs: 0.2682\nTotal prob of N-in: 0.14402\noutside 1 331\nTMhelix 332 354\ninside 355 372

13931 GCF_002052175.1_ASM205217v1 Salmonella enterica Proteobacteria; Gammaproteobacteria; Enterobacterales; Enterobacteriaceae; Salmonella
MNNEETFYQAMRRKGVTRRSFLKFCSLAATSLGLGAGMTPKIAWALENKPRIPVVWIHGL WP_001058312.1
hydrogenase [Salmonella enterica] Length: 372\nNumber of predicted TMHs: 2\nExp number of AAs in TMHs: 36.51738\nExp number, first 60 AAs: 19.72849\nTotal prob of N-in: 0.96170\nPOSSIBLE N-term signal sequence\ninside 1 19\nTMhelix 20 39\noutside 40 325\nTMhelix 326 348\ninside 349 372

13932 GCF_002051965.1_ASM205196v1 Salmonella enterica Proteobacteria; Gammaproteobacteria; Enterobacterales; Enterobacteriaceae; Salmonella
MTGDNTLITSHGINRRDFMKLCAALAATMGLSSKAAAEMAESVSNPQRPPVIWIGAQECT WP_000145429.1
hydrogenase 2 small subunit [Salmonella enterica] Length: 372\nNumber of predicted TMHs: 1\nExp number of AAs in TMHs: 24.51309\nExp number, first 60 AAs: 0.2682\nTotal prob of N-in: 0.14402\noutside 1 331\nTMhelix 332 354\ninside 355 372

13933 GCF_002052075.1_ASM205207v1 Salmonella enterica Proteobacteria; Gammaproteobacteria; Enterobacterales; Enterobacteriaceae; Salmonella
MTGDNTLITSHGINRRDFMKLCAALAATMGLSSKAAAEMAESVSNPQRPPVIWIGAQECT WP_000145429.1
hydrogenase 2 small subunit [Salmonella enterica] Length: 372\nNumber of predicted TMHs: 1\nExp number of AAs in TMHs: 24.51309\nExp number, first 60 AAs: 0.2682\nTotal prob of N-in: 0.14402\noutside 1 331\nTMhelix 332 354\ninside 355 372

13934 GCF_002051825.1_ASM205182v1 Salmonella enterica Proteobacteria; Gammaproteobacteria; Enterobacterales; Enterobacteriaceae; Salmonella

MNNEETFYQAMRRKGVTRRSFLKFCSLAATSLGLGAGMTPKIAWALENKPRIPVVWIHGL WP_001058312.1 hydrogenase [Salmonella enterica] Length: 372\nNumber of predicted TMHs: 2\nExp number of AAs in TMHs: 36.51738\nExp number, first 60 AAs: 19.72849\nTotal prob of N-in: 0.96170\nPOSSIBLE N-term signal sequence\ninside 1 19\nTMhelix 20 39\noutside 40 325\nTMhelix 326 348\ninside 349 372

13935 GCF_002051735.1_ASM205173v1 Salmonella enterica Proteobacteria; Gammaproteobacteria; Enterobacterales; Enterobacteriaceae; Salmonella MNNEETFYQAMRRKGVTRRSFLKFCSLAATSLGLGAGMTPKIAWALENKPRIPVVWIHGL WP_001058312.1 hydrogenase [Salmonella enterica] Length: 372\nNumber of predicted TMHs: 2\nExp number of AAs in TMHs: 36.51738\nExp number, first 60 AAs: 19.72849\nTotal prob of N-in: 0.96170\nPOSSIBLE N-term signal sequence\ninside 1 19\nTMhelix 20 39\noutside 40 325\nTMhelix 326 348\ninside 349 372

13936 GCF_002051705.1_ASM205170v1 Salmonella enterica Proteobacteria; Gammaproteobacteria; Enterobacterales; Enterobacteriaceae; Salmonella MNNEETFYQAMRRKGVTRRSFLKFCSLAATSLGLGAGMTPKIAWALENKPRIPVVWIHGL WP_001058312.1 hydrogenase [Salmonella enterica] Length: 372\nNumber of predicted TMHs: 2\nExp number of AAs in TMHs: 36.51738\nExp number, first 60 AAs: 19.72849\nTotal prob of N-in: 0.96170\nPOSSIBLE N-term signal sequence\ninside 1 19\nTMhelix 20 39\noutside 40 325\nTMhelix 326 348\ninside 349 372

13937 GCF_002051825.1_ASM205182v1 Salmonella enterica Proteobacteria; Gammaproteobacteria; Enterobacterales; Enterobacteriaceae; Salmonella MTGDNTLITSHGINRRDFMKLCAALAATMGLSSKAAAEMAESVSNPQRPPVIWIGAQECT WP_000145429.1 hydrogenase 2 small subunit [Salmonella enterica] Length: 372\nNumber of predicted TMHs: 1\nExp number of AAs in TMHs: 24.51309\nExp number, first 60 AAs: 0.2682\nTotal prob of N-in: 0.14402\noutside 1 331\nTMhelix 332 354\ninside 355 372

13938 GCF_002051735.1_ASM205173v1 Salmonella enterica Proteobacteria; Gammaproteobacteria; Enterobacterales; Enterobacteriaceae; Salmonella MTGDNTLITSHGINRRDFMKLCAALAATMGLSSKAAAEMAESVSNPQRPPVIWIGAQECT WP_000145429.1 hydrogenase 2 small subunit [Salmonella enterica] Length: 372\nNumber of predicted TMHs: 1\nExp number of AAs in TMHs: 24.51309\nExp number, first 60 AAs: 0.2682\nTotal prob of N-in: 0.14402\noutside 1 331\nTMhelix 332 354\ninside 355 372

13939 GCF_002051485.1_ASM205148v1 Salmonella enterica Proteobacteria; Gammaproteobacteria; Enterobacterales; Enterobacteriaceae; Salmonella MTGDNTLITSHGINRRDFMKLCAALAATMGLSSKAAAEMAESVSNPQRPPVIWIGAQECT WP_000145429.1 hydrogenase 2 small subunit [Salmonella enterica] Length: 372\nNumber of predicted TMHs: 1\nExp number of AAs in TMHs: 24.51309\nExp number, first 60 AAs: 0.2682\nTotal prob of N-in: 0.14402\noutside 1 331\nTMhelix 332 354\ninside 355 372

13940 GCF_002051305.1_ASM205130v1 Salmonella enterica Proteobacteria; Gammaproteobacteria; Enterobacterales; Enterobacteriaceae; Salmonella MNNEETFYQAMRRKGVTRRSFLKFCSLAATSLGLGAGMTPKIAWALENKPRIPVVWIHGL WP_001058312.1 hydrogenase [Salmonella enterica] Length: 372\nNumber of predicted TMHs: 2\nExp number of AAs in TMHs: 36.51738\nExp number, first 60 AAs: 19.72849\nTotal prob of N-in: 0.96170\nPOSSIBLE N-term signal sequence\ninside 1 19\nTMhelix 20 39\noutside 40 325\nTMhelix 326 348\ninside 349 372

13941 GCF_002051405.1_ASM205140v1 Salmonella enterica Proteobacteria; Gammaproteobacteria; Enterobacterales; Enterobacteriaceae; Salmonella MTGDNTLITSHGINRRDFMKLCAALAATMGLSSKAAAEMAESVSNPQRPPVIWIGAQECT WP_000145429.1 hydrogenase 2 small subunit [Salmonella enterica] Length: 372\nNumber of predicted TMHs: 1\nExp number of AAs in TMHs: 24.51309\nExp number, first 60 AAs: 0.2682\nTotal prob of N-in: 0.14402\noutside 1 331\nTMhelix 332 354\ninside 355 372

13942 GCF_002051385.1_ASM205138v1 Salmonella enterica Proteobacteria; Gammaproteobacteria; Enterobacterales; Enterobacteriaceae; Salmonella MTGDNTLITSHGINRRDFMKLCAALAATMGLSSKAAAEMAESVSNPQRPPVIWIGAQECT WP_000145429.1 hydrogenase 2 small subunit [Salmonella enterica] Length: 372\nNumber of predicted TMHs: 1\nExp number of AAs in TMHs: 24.51309\nExp number, first 60 AAs: 0.2682\nTotal prob of N-in: 0.14402\noutside 1 331\nTMhelix 332 354\ninside 355 372

13943 GCF_002051445.1_ASM205144v1 Salmonella enterica Proteobacteria; Gammaproteobacteria; Enterobacterales; Enterobacteriaceae; Salmonella
MTGDNTLITSHGINRRDFMKLCAALAATMGLSSKAAAEMAESVSNPQRPPVIWIGAQECT WP_000145429.1
hydrogenase 2 small subunit [Salmonella enterica] Length: 372\nNumber of predicted TMHs: 1\nExp number of AAs in TMHs: 24.51309\nExp number, first 60 AAs: 0.2682\nTotal prob of N-in: 0.14402\noutside 1 331\nTMhelix 332 354\ninside 355 372

13944 GCF_002051355.1_ASM205135v1 Salmonella enterica Proteobacteria; Gammaproteobacteria; Enterobacterales; Enterobacteriaceae; Salmonella
MTGDNTLITSHGINRRDFMKLCAALAATMGLSSKAAAEMAESVSNPQRPPVIWIGAQECT WP_000145429.1
hydrogenase 2 small subunit [Salmonella enterica] Length: 372\nNumber of predicted TMHs: 1\nExp number of AAs in TMHs: 24.51309\nExp number, first 60 AAs: 0.2682\nTotal prob of N-in: 0.14402\noutside 1 331\nTMhelix 332 354\ninside 355 372

13945 GCF_002051275.1_ASM205127v1 Salmonella enterica Proteobacteria; Gammaproteobacteria; Enterobacterales; Enterobacteriaceae; Salmonella
MNNEETFYQAMRRKGVTRRSFLKFCSLAATSLGLGAGMTPKIAWALENKPRIPVVWIHGL WP_001058312.1
hydrogenase [Salmonella enterica] Length: 372\nNumber of predicted TMHs: 2\nExp number of AAs in TMHs: 36.51738\nExp number, first 60 AAs: 19.72849\nTotal prob of N-in: 0.96170\nPOSSIBLE N-term signal sequence\ninside 1 19\nTMhelix 20 39\noutside 40 325\nTMhelix 326 348\ninside 349 372

13946 GCF_002050775.1_ASM205077v1 Salmonella enterica Proteobacteria; Gammaproteobacteria; Enterobacterales; Enterobacteriaceae; Salmonella
MNNEETFYQAMRRKGVTRRSFLKFCSLAATSLGLGAGMTPKIAWALENKPRIPVVWIHGL WP_001058312.1
hydrogenase [Salmonella enterica] Length: 372\nNumber of predicted TMHs: 2\nExp number of AAs in TMHs: 36.51738\nExp number, first 60 AAs: 19.72849\nTotal prob of N-in: 0.96170\nPOSSIBLE N-term signal sequence\ninside 1 19\nTMhelix 20 39\noutside 40 325\nTMhelix 326 348\ninside 349 372

13947 GCF_002050795.1_ASM205079v1 Salmonella enterica Proteobacteria; Gammaproteobacteria; Enterobacterales; Enterobacteriaceae; Salmonella
MTGDNTLITSHGINRRDFMKLCAALAATMGLSSKAAAEMAESVSNPQRPPVIWIGAQECT WP_000145429.1
hydrogenase 2 small subunit [Salmonella enterica] Length: 372\nNumber of predicted TMHs: 1\nExp number of AAs in TMHs: 24.51309\nExp number, first 60 AAs: 0.2682\nTotal prob of N-in: 0.14402\noutside 1 331\nTMhelix 332 354\ninside 355 372

13948 GCF_002047865.1_ASM204786v1 Salmonella enterica Proteobacteria; Gammaproteobacteria; Enterobacterales; Enterobacteriaceae; Salmonella
MNNEETLYQAMRRKGVTRRSFLKFCSLAATSLGLGAGMTPKIAWALENKPRIPVVWIHGL WP_001752445.1
hydrogenase-1 small chain [Salmonella enterica] Length: 372\nNumber of predicted TMHs: 2\nExp number of AAs in TMHs: 36.5123\nExp number, first 60 AAs: 19.72341\nTotal prob of N-in: 0.96144\nPOSSIBLE N-term signal sequence\ninside 1 19\nTMhelix 20 39\noutside 40 325\nTMhelix 326 348\ninside 349 372

13949 GCF_002047865.1_ASM204786v1 Salmonella enterica Proteobacteria; Gammaproteobacteria; Enterobacterales; Enterobacteriaceae; Salmonella
MTGDNTLITSHGINRRDFMKLCAALAATMGLSSKAAAEMAESVSNPQRPPVIWIGAQECT WP_000145428.1
hydrogenase 2 small subunit [Salmonella enterica] Length: 372\nNumber of predicted TMHs: 1\nExp number of AAs in TMHs: 24.50677\nExp number, first 60 AAs: 0.2682\nTotal prob of N-in: 0.14390\noutside 1 331\nTMhelix 332 354\ninside 355 372

13950 GCF_002047635.1_ASM204763v1 Salmonella enterica Proteobacteria; Gammaproteobacteria; Enterobacterales; Enterobacteriaceae; Salmonella
MTGDNTLITSHGINRRDFMKLCAALAATMGLSSKAAAEMAESVSNPQRPPVIWIGAQECT WP_000145429.1
hydrogenase 2 small subunit [Salmonella enterica] Length: 372\nNumber of predicted TMHs: 1\nExp number of AAs in TMHs: 24.51309\nExp number, first 60 AAs: 0.2682\nTotal prob of N-in: 0.14402\noutside 1 331\nTMhelix 332 354\ninside 355 372

13951 GCF_002046415.1_ASM204641v1 Salmonella enterica Proteobacteria; Gammaproteobacteria; Enterobacterales; Enterobacteriaceae; Salmonella
MNNEETFYQAMRRKGVTRRSFLKFCSLAATSLGLGAGMTPKIAWALENKPRIPVVWIHGL WP_001058311.1
hydrogenase [Salmonella enterica] Length: 372\nNumber of predicted TMHs: 2\nExp number of AAs in

TMHs: 37.4699899999999\nExp number, first 60 AAs: 19.72699\nTotal prob of N-in: 0.96171\nPOSSIBLE N-term signal sequence\ninside 1 19\nTMhelix 20 39\noutside 40 325\nTMhelix 326 348\ninside 349 372

13952 GCF_002046265.1_ASM204626v1 Salmonella enterica Proteobacteria; Gammaproteobacteria; Enterobacterales; Enterobacteriaceae; Salmonella
MTGDNTLITSHGINRRDFMKLCAALAATMGLSSKAAAEMAESVSNPQRPPVIWIGAQECT WP_000145429.1
hydrogenase 2 small subunit [Salmonella enterica] Length: 372\nNumber of predicted TMHs: 1\nExp number of AAs in TMHs: 24.51309\nExp number, first 60 AAs: 0.2682\nTotal prob of N-in: 0.14402\noutside 1 331\nTMhelix 332 354\ninside 355 372

13953 GCF_002046165.1_ASM204616v1 Salmonella enterica Proteobacteria; Gammaproteobacteria; Enterobacterales; Enterobacteriaceae; Salmonella
MNNEETFYQAMRRKGVTRRSFLKFCSLAATSLGLGAGMTPKIAWALENKPRIPVVWIHGL WP_001058311.1
hydrogenase [Salmonella enterica] Length: 372\nNumber of predicted TMHs: 2\nExp number of AAs in TMHs: 37.4699899999999\nExp number, first 60 AAs: 19.72699\nTotal prob of N-in: 0.96171\nPOSSIBLE N-term signal sequence\ninside 1 19\nTMhelix 20 39\noutside 40 325\nTMhelix 326 348\ninside 349 372

13954 GCF_002046165.1_ASM204616v1 Salmonella enterica Proteobacteria; Gammaproteobacteria; Enterobacterales; Enterobacteriaceae; Salmonella
MTGDNTLITSHGINRRDFMKLCAALAATMGLSSKAAAEMAESVSNPQRPPVIWIGAQECT WP_000145429.1
hydrogenase 2 small subunit [Salmonella enterica] Length: 372\nNumber of predicted TMHs: 1\nExp number of AAs in TMHs: 24.51309\nExp number, first 60 AAs: 0.2682\nTotal prob of N-in: 0.14402\noutside 1 331\nTMhelix 332 354\ninside 355 372

13955 GCF_002045745.1_ASM204574v1 Salmonella enterica Proteobacteria; Gammaproteobacteria; Enterobacterales; Enterobacteriaceae; Salmonella
MTGDNTLITSHGINRRDFMKLCAALAATMGLSSKAAAEMAESVSNPQRPPVIWIGAQECT WP_000145429.1
hydrogenase 2 small subunit [Salmonella enterica] Length: 372\nNumber of predicted TMHs: 1\nExp number of AAs in TMHs: 24.51309\nExp number, first 60 AAs: 0.2682\nTotal prob of N-in: 0.14402\noutside 1 331\nTMhelix 332 354\ninside 355 372

13956 GCF_002045615.1_ASM204561v1 Salmonella enterica Proteobacteria; Gammaproteobacteria; Enterobacterales; Enterobacteriaceae; Salmonella
MNNEETFYQAMRRKGVTRRSFLKFCSLAATSLGLGAGMTPKIAWALENKPRIPVVWIHGL WP_001058311.1
hydrogenase [Salmonella enterica] Length: 372\nNumber of predicted TMHs: 2\nExp number of AAs in TMHs: 37.4699899999999\nExp number, first 60 AAs: 19.72699\nTotal prob of N-in: 0.96171\nPOSSIBLE N-term signal sequence\ninside 1 19\nTMhelix 20 39\noutside 40 325\nTMhelix 326 348\ninside 349 372

13957 GCF_002045735.1_ASM204573v1 Salmonella enterica Proteobacteria; Gammaproteobacteria; Enterobacterales; Enterobacteriaceae; Salmonella
MNNEETFYQAMRRKGVTRRSFLKFCSLAATSLGLGAGMTPKIAWALENKPRIPVVWIHGL WP_001058311.1
hydrogenase [Salmonella enterica] Length: 372\nNumber of predicted TMHs: 2\nExp number of AAs in TMHs: 37.4699899999999\nExp number, first 60 AAs: 19.72699\nTotal prob of N-in: 0.96171\nPOSSIBLE N-term signal sequence\ninside 1 19\nTMhelix 20 39\noutside 40 325\nTMhelix 326 348\ninside 349 372

13958 GCF_002045665.1_ASM204566v1 Salmonella enterica Proteobacteria; Gammaproteobacteria; Enterobacterales; Enterobacteriaceae; Salmonella
MNNEETFYQAMRRKGVTRRSFLKFCSLAATSLGLGAGMTPKIAWALENKPRIPVVWIHGL WP_001058312.1
hydrogenase [Salmonella enterica] Length: 372\nNumber of predicted TMHs: 2\nExp number of AAs in TMHs: 36.51738\nExp number, first 60 AAs: 19.72849\nTotal prob of N-in: 0.96170\nPOSSIBLE N-term signal sequence\ninside 1 19\nTMhelix 20 39\noutside 40 325\nTMhelix 326 348\ninside 349 372

13959 GCF_002045585.1_ASM204558v1 Salmonella enterica Proteobacteria; Gammaproteobacteria; Enterobacterales; Enterobacteriaceae; Salmonella
MNNEETFYQAMRRKGVTRRSFLKFCSLAATSLGLGAGMTPKIAWALENKPRIPVVWIHGL WP_001058311.1
hydrogenase [Salmonella enterica] Length: 372\nNumber of predicted TMHs: 2\nExp number of AAs in TMHs: 37.4699899999999\nExp number, first 60 AAs: 19.72699\nTotal prob of N-in: 0.96171\nPOSSIBLE N-term signal sequence\ninside 1 19\nTMhelix 20 39\noutside 40 325\nTMhelix 326 348\ninside 349 372

13960 GCF_002045085.1_ASM204508v1 Salmonella enterica Proteobacteria; Gammaproteobacteria; Enterobacterales; Enterobacteriaceae; Salmonella

MTGDNTLITSHGINRRDFMKLCAALAATMGLSSKAAAEMAESVSNPQRPPVIWIGAQECT WP_000145429.1 hydrogenase 2 small subunit [Salmonella enterica] Length: 372\nNumber of predicted TMHs: 1\nExp number of AAs in TMHs: 24.51309\nExp number, first 60 AAs: 0.2682\nTotal prob of N-in: 0.14402\noutside 1 331\nTMhelix 332 354\ninside 355 372

13961 GCF_002044705.1_ASM204470v1 Salmonella enterica Proteobacteria; Gammaproteobacteria; Enterobacterales; Enterobacteriaceae; Salmonella MTGDNTLITSHGINRRDFMKLCAALAATMGLSSKAAAEMAESVSNPQRPPVIWIGAQECT WP_000145429.1 hydrogenase 2 small subunit [Salmonella enterica] Length: 372\nNumber of predicted TMHs: 1\nExp number of AAs in TMHs: 24.51309\nExp number, first 60 AAs: 0.2682\nTotal prob of N-in: 0.14402\noutside 1 331\nTMhelix 332 354\ninside 355 372

13962 GCF_002044675.1_ASM204467v1 Salmonella enterica Proteobacteria; Gammaproteobacteria; Enterobacterales; Enterobacteriaceae; Salmonella MTGDNTLITSHGINRRDFMKLCAALAATMGLSSKAAAEMAESVSNPQRPPVIWIGAQECT WP_000145429.1 hydrogenase 2 small subunit [Salmonella enterica] Length: 372\nNumber of predicted TMHs: 1\nExp number of AAs in TMHs: 24.51309\nExp number, first 60 AAs: 0.2682\nTotal prob of N-in: 0.14402\noutside 1 331\nTMhelix 332 354\ninside 355 372

13963 GCF_002044625.1_ASM204462v1 Salmonella enterica Proteobacteria; Gammaproteobacteria; Enterobacterales; Enterobacteriaceae; Salmonella MTGDNTLITSHGINRRDFMKLCAALAATMGLSSKAAAEMAESVSNPQRPPVIWIGAQECT WP_000145429.1 hydrogenase 2 small subunit [Salmonella enterica] Length: 372\nNumber of predicted TMHs: 1\nExp number of AAs in TMHs: 24.51309\nExp number, first 60 AAs: 0.2682\nTotal prob of N-in: 0.14402\noutside 1 331\nTMhelix 332 354\ninside 355 372

13964 GCF_002044565.1_ASM204456v1 Salmonella enterica Proteobacteria; Gammaproteobacteria; Enterobacterales; Enterobacteriaceae; Salmonella MTGDNTLITSHGINRRDFMKLCAALAATMGLSSKAAAEMAESVSNPQRPPVIWIGAQECT WP_000145429.1 hydrogenase 2 small subunit [Salmonella enterica] Length: 372\nNumber of predicted TMHs: 1\nExp number of AAs in TMHs: 24.51309\nExp number, first 60 AAs: 0.2682\nTotal prob of N-in: 0.14402\noutside 1 331\nTMhelix 332 354\ninside 355 372

13965 GCF_002044505.1_ASM204450v1 Salmonella enterica Proteobacteria; Gammaproteobacteria; Enterobacterales; Enterobacteriaceae; Salmonella MNNEETFYQAMRRKGVTRRSFLKFCSLAATSLGLGAGMTPKIAWALENKPRIPVVWIHGL WP_001058312.1 hydrogenase [Salmonella enterica] Length: 372\nNumber of predicted TMHs: 2\nExp number of AAs in TMHs: 36.51738\nExp number, first 60 AAs: 19.72849\nTotal prob of N-in: 0.96170\nPOSSIBLE N-term signal sequence\ninside 1 19\nTMhelix 20 39\noutside 40 325\nTMhelix 326 348\ninside 349 372

13966 GCF_002044555.1_ASM204455v1 Salmonella enterica Proteobacteria; Gammaproteobacteria; Enterobacterales; Enterobacteriaceae; Salmonella MTGDNTLITSHGINRRDFMKLCAALAATMGLSSKAAAEMAESVSNPQRPPVIWIGAQECT WP_000145429.1 hydrogenase 2 small subunit [Salmonella enterica] Length: 372\nNumber of predicted TMHs: 1\nExp number of AAs in TMHs: 24.51309\nExp number, first 60 AAs: 0.2682\nTotal prob of N-in: 0.14402\noutside 1 331\nTMhelix 332 354\ninside 355 372

13967 GCF_002044505.1_ASM204450v1 Salmonella enterica Proteobacteria; Gammaproteobacteria; Enterobacterales; Enterobacteriaceae; Salmonella MTGDNTLITSHGINRRDFMKLCAALAATMGLSSKAAAEMAESVSNPQRPPVIWIGAQECT WP_000145429.1 hydrogenase 2 small subunit [Salmonella enterica] Length: 372\nNumber of predicted TMHs: 1\nExp number of AAs in TMHs: 24.51309\nExp number, first 60 AAs: 0.2682\nTotal prob of N-in: 0.14402\noutside 1 331\nTMhelix 332 354\ninside 355 372

13968 GCF_002043945.1_ASM204394v1 Salmonella enterica Proteobacteria; Gammaproteobacteria; Enterobacterales; Enterobacteriaceae; Salmonella MNNEETFYQAMRRKGVTRRSFLKFCSLAATSLGLGAGMTPKIAWALENKPRIPVVWIHGL WP_023233644.1 hydrogenase 1, small subunit [Salmonella enterica] Length: 372\nNumber of predicted TMHs: 2\nExp number of AAs in TMHs: 37.4208299999999\nExp number, first 60 AAs: 19.72732\nTotal prob of N-in: 0.96168\nPOSSIBLE N-term signal sequence\ninside 1 19\nTMhelix 20 39\noutside 40 325\nTMhelix 326 348\ninside 349 372

13969 GCF_002044225.1_ASM204422v1 Salmonella enterica Proteobacteria; Gammaproteobacteria; Enterobacterales; Enterobacteriaceae; Salmonella
MNNEETFYQAMRRKGVTRRSFLKFCSLAATSLGLGAGMTPKIAWALENKPRIPVVWIHGL WP_001058311.1
hydrogenase [Salmonella enterica] Length: 372\nNumber of predicted TMHs: 2\nExp number of AAs in TMHs: 37.4699899999999\nExp number, first 60 AAs: 19.72699\nTotal prob of N-in: 0.96171\nPOSSIBLE N-term signal sequence\ninside 1 19\nTMhelix 20 39\noutside 40 325\nTMhelix 326 348\ninside 349 372

13970 GCF_002044225.1_ASM204422v1 Salmonella enterica Proteobacteria; Gammaproteobacteria; Enterobacterales; Enterobacteriaceae; Salmonella
MTGDNTLITSHGINRRDFMKLCAALAATMGLSSKAAAEMAESVSNPQRPPVIWIGAQECT WP_000145429.1
hydrogenase 2 small subunit [Salmonella enterica] Length: 372\nNumber of predicted TMHs: 1\nExp number of AAs in TMHs: 24.51309\nExp number, first 60 AAs: 0.2682\nTotal prob of N-in: 0.14402\noutside 1 331\nTMhelix 332 354\ninside 355 372

13971 GCF_002044035.1_ASM204403v1 Salmonella enterica Proteobacteria; Gammaproteobacteria; Enterobacterales; Enterobacteriaceae; Salmonella
MNNEETFYQAMRRKGVTRRSFLKFCSLAATSLGLGAGMTPKIAWALENKPRIPVVWIHGL WP_001058312.1
hydrogenase [Salmonella enterica] Length: 372\nNumber of predicted TMHs: 2\nExp number of AAs in TMHs: 36.51738\nExp number, first 60 AAs: 19.72849\nTotal prob of N-in: 0.96170\nPOSSIBLE N-term signal sequence\ninside 1 19\nTMhelix 20 39\noutside 40 325\nTMhelix 326 348\ninside 349 372

13972 GCF_002044195.1_ASM204419v1 Salmonella enterica Proteobacteria; Gammaproteobacteria; Enterobacterales; Enterobacteriaceae; Salmonella
MNNEETFYQAMRRKGVTRRSFLKFCSLAATSLGLGAGMTPKIAWALENKPRIPVVWIHGL WP_001058312.1
hydrogenase [Salmonella enterica] Length: 372\nNumber of predicted TMHs: 2\nExp number of AAs in TMHs: 36.51738\nExp number, first 60 AAs: 19.72849\nTotal prob of N-in: 0.96170\nPOSSIBLE N-term signal sequence\ninside 1 19\nTMhelix 20 39\noutside 40 325\nTMhelix 326 348\ninside 349 372

13973 GCF_002044195.1_ASM204419v1 Salmonella enterica Proteobacteria; Gammaproteobacteria; Enterobacterales; Enterobacteriaceae; Salmonella
MTGDNTLITSHGINRRDFMKLCAALAATMGLSSKAAAEMAESVSNPQRPPVIWIGAQECT WP_000145429.1
hydrogenase 2 small subunit [Salmonella enterica] Length: 372\nNumber of predicted TMHs: 1\nExp number of AAs in TMHs: 24.51309\nExp number, first 60 AAs: 0.2682\nTotal prob of N-in: 0.14402\noutside 1 331\nTMhelix 332 354\ninside 355 372

13974 GCF_002044005.1_ASM204400v1 Salmonella enterica Proteobacteria; Gammaproteobacteria; Enterobacterales; Enterobacteriaceae; Salmonella
MNNEETFYQAMRRKGVTRRSFLKFCSLAATSLGLGAGMTPKIAWALENKPRIPVVWIHGL WP_001058311.1
hydrogenase [Salmonella enterica] Length: 372\nNumber of predicted TMHs: 2\nExp number of AAs in TMHs: 37.4699899999999\nExp number, first 60 AAs: 19.72699\nTotal prob of N-in: 0.96171\nPOSSIBLE N-term signal sequence\ninside 1 19\nTMhelix 20 39\noutside 40 325\nTMhelix 326 348\ninside 349 372

13975 GCF_002043725.1_ASM204372v1 Salmonella enterica Proteobacteria; Gammaproteobacteria; Enterobacterales; Enterobacteriaceae; Salmonella
MTGDNTLITSHGINRRDFMKLCAALAATMGLSSKAAAEMAESVSNPQRPPVIWIGAQECT WP_000145429.1
hydrogenase 2 small subunit [Salmonella enterica] Length: 372\nNumber of predicted TMHs: 1\nExp number of AAs in TMHs: 24.51309\nExp number, first 60 AAs: 0.2682\nTotal prob of N-in: 0.14402\noutside 1 331\nTMhelix 332 354\ninside 355 372

13976 GCF_002043785.1_ASM204378v1 Salmonella enterica Proteobacteria; Gammaproteobacteria; Enterobacterales; Enterobacteriaceae; Salmonella
MTGDNTLITSHGINRRDFMKLCAALAATMGLSSKAAAEMAESVSNPQRPPVIWIGAQECT WP_000145429.1
hydrogenase 2 small subunit [Salmonella enterica] Length: 372\nNumber of predicted TMHs: 1\nExp number of AAs in TMHs: 24.51309\nExp number, first 60 AAs: 0.2682\nTotal prob of N-in: 0.14402\noutside 1 331\nTMhelix 332 354\ninside 355 372

13977 GCF_002043665.1_ASM204366v1 Salmonella enterica Proteobacteria; Gammaproteobacteria; Enterobacterales; Enterobacteriaceae; Salmonella
MTGDNTLITSHGINRRDFMKLCAALAATMGLSSKAAAEMAESVSNPQRPPVIWIGAQECT WP_000145429.1
hydrogenase 2 small subunit [Salmonella enterica] Length: 372\nNumber of predicted TMHs: 1\nExp

number of AAs in TMHs: 24.51309\nExp number, first 60 AAs: 0.2682\nTotal prob of N-in: 0.14402\noutside 1 331\nTMhelix 332 354\ninside 355 372

13978 GCF_002043255.1_ASM204325v1 Salmonella enterica Proteobacteria; Gammaproteobacteria; Enterobacterales; Enterobacteriaceae; Salmonella
MNNEETFYQAMRRKGVTRRSFLKFCSLAATSLGLGAGMTPKIAWALENKPRIPVVWIHGL WP_023213673.1
hydrogenase 1, small subunit [Salmonella enterica] Length: 372\nNumber of predicted TMHs: 2\nExp number of AAs in TMHs: 36.51753\nExp number, first 60 AAs: 19.72849\nTotal prob of N-in: 0.96170\nPOSSIBLE N-term signal sequence\ninside 1 19\nTMhelix 20 39\noutside 40 325\nTMhelix 326 348\ninside 349 372

13979 GCF_002043605.1_ASM204360v1 Salmonella enterica Proteobacteria; Gammaproteobacteria; Enterobacterales; Enterobacteriaceae; Salmonella
MNNEETFYQAMRRKGVTRRSFLKFCSLAATSLGLGAGMTPKIAWALENKPRIPVVWIHGL WP_001058311.1
hydrogenase [Salmonella enterica] Length: 372\nNumber of predicted TMHs: 2\nExp number of AAs in TMHs: 37.4699899999999\nExp number, first 60 AAs: 19.72699\nTotal prob of N-in: 0.96171\nPOSSIBLE N-term signal sequence\ninside 1 19\nTMhelix 20 39\noutside 40 325\nTMhelix 326 348\ninside 349 372

13980 GCF_002043545.1_ASM204354v1 Salmonella enterica Proteobacteria; Gammaproteobacteria; Enterobacterales; Enterobacteriaceae; Salmonella
MNNEETFYQAMRRKGVTRRSFLKFCSLAATSLGLGAGMTPKIAWALENKPRIPVVWIHGL WP_001058312.1
hydrogenase [Salmonella enterica] Length: 372\nNumber of predicted TMHs: 2\nExp number of AAs in TMHs: 36.51738\nExp number, first 60 AAs: 19.72849\nTotal prob of N-in: 0.96170\nPOSSIBLE N-term signal sequence\ninside 1 19\nTMhelix 20 39\noutside 40 325\nTMhelix 326 348\ninside 349 372

13981 GCF_002043605.1_ASM204360v1 Salmonella enterica Proteobacteria; Gammaproteobacteria; Enterobacterales; Enterobacteriaceae; Salmonella
MTGDNTLITSHGINRRDFMKLCAALAATMGLSSKAAAEMAESVSNPQRPPVIWIGAQECT WP_000145429.1
hydrogenase 2 small subunit [Salmonella enterica] Length: 372\nNumber of predicted TMHs: 1\nExp number of AAs in TMHs: 24.51309\nExp number, first 60 AAs: 0.2682\nTotal prob of N-in: 0.14402\noutside 1 331\nTMhelix 332 354\ninside 355 372

13982 GCF_002043255.1_ASM204325v1 Salmonella enterica Proteobacteria; Gammaproteobacteria; Enterobacterales; Enterobacteriaceae; Salmonella
MTGDNTLITSHGINRRDFMKLCAALAATMGLSSKAAAEMAESVSNPQRPPVIWIGAQECT WP_000145429.1
hydrogenase 2 small subunit [Salmonella enterica] Length: 372\nNumber of predicted TMHs: 1\nExp number of AAs in TMHs: 24.51309\nExp number, first 60 AAs: 0.2682\nTotal prob of N-in: 0.14402\noutside 1 331\nTMhelix 332 354\ninside 355 372

13983 GCF_002043065.1_ASM204306v1 Salmonella enterica Proteobacteria; Gammaproteobacteria; Enterobacterales; Enterobacteriaceae; Salmonella
MNNEETFYQAMRRKGVTRRSFLKFCSLAATSLGLGAGMTPKIAWALENKPRIPVVWIHGL WP_001058312.1
hydrogenase [Salmonella enterica] Length: 372\nNumber of predicted TMHs: 2\nExp number of AAs in TMHs: 36.51738\nExp number, first 60 AAs: 19.72849\nTotal prob of N-in: 0.96170\nPOSSIBLE N-term signal sequence\ninside 1 19\nTMhelix 20 39\noutside 40 325\nTMhelix 326 348\ninside 349 372

13984 GCF_002043215.1_ASM204321v1 Salmonella enterica Proteobacteria; Gammaproteobacteria; Enterobacterales; Enterobacteriaceae; Salmonella
MNNEETLYQAMRRKGVTRRSFLKFCSLAATSLGLGAGMTPKIAWALENKPRIPVVWIHGL WP_001752445.1
hydrogenase-1 small chain [Salmonella enterica] Length: 372\nNumber of predicted TMHs: 2\nExp number of AAs in TMHs: 36.5123\nExp number, first 60 AAs: 19.72341\nTotal prob of N-in: 0.96144\nPOSSIBLE N-term signal sequence\ninside 1 19\nTMhelix 20 39\noutside 40 325\nTMhelix 326 348\ninside 349 372

13985 GCF_002043165.1_ASM204316v1 Salmonella enterica Proteobacteria; Gammaproteobacteria; Enterobacterales; Enterobacteriaceae; Salmonella
MNNEETFYQAMRRKGVTRRSFLKFCSLAATSLGLGAGMTPKIAWALENKPRIPVVWIHGL WP_001058311.1
hydrogenase [Salmonella enterica] Length: 372\nNumber of predicted TMHs: 2\nExp number of AAs in TMHs: 37.4699899999999\nExp number, first 60 AAs: 19.72699\nTotal prob of N-in: 0.96171\nPOSSIBLE N-term signal sequence\ninside 1 19\nTMhelix 20 39\noutside 40 325\nTMhelix 326 348\ninside 349 372

13986 GCF_002043065.1_ASM204306v1 Salmonella enterica Proteobacteria; Gammaproteobacteria; Enterobacterales; Enterobacteriaceae; Salmonella

MTGDNTLITSHGINRRDFMKLCAALAATMGLSSKAAAEMAESVSNPQRPPVIWIGAQECT WP_000145429.1 hydrogenase 2 small subunit [Salmonella enterica] Length: 372\nNumber of predicted TMHs: 1\nExp number of AAs in TMHs: 24.51309\nExp number, first 60 AAs: 0.2682\nTotal prob of N-in: 0.14402\noutside 1 331\nTMhelix 332 354\ninside 355 372

13987 GCF_002043075.1_ASM204307v1 Salmonella enterica Proteobacteria; Gammaproteobacteria; Enterobacterales; Enterobacteriaceae; Salmonella MTGDNTLITSHGINRRDFMKLCAALAATMGLSSKAAAEMAESVSNPQRPPVIWIGAQECT WP_000145428.1 hydrogenase 2 small subunit [Salmonella enterica] Length: 372\nNumber of predicted TMHs: 1\nExp number of AAs in TMHs: 24.50677\nExp number, first 60 AAs: 0.2682\nTotal prob of N-in: 0.14390\noutside 1 331\nTMhelix 332 354\ninside 355 372

13988 GCF_000783815.1_ASM78381v1 Salmonella enterica Proteobacteria; Gammaproteobacteria; Enterobacterales; Enterobacteriaceae; Salmonella MNNEETFYQAMRRKGVTRRSFLKFCSLAATSLGLGAGMTPKIAWALENKPRIPVVWIHGL WP_001058307.1 hydrogenase [Salmonella enterica] Length: 372\nNumber of predicted TMHs: 2\nExp number of AAs in TMHs: 36.51738\nExp number, first 60 AAs: 19.72849\nTotal prob of N-in: 0.96170\nPOSSIBLE N-term signal sequence\ninside 1 19\nTMhelix 20 39\noutside 40 325\nTMhelix 326 348\ninside 349 372

13989 GCF_002061945.1_ASM206194v1 Salmonella enterica Proteobacteria; Gammaproteobacteria; Enterobacterales; Enterobacteriaceae; Salmonella MTGDNTLITSHGINRRDFMKLCAALAATMGLSSKAAAEMAESVSNPQRPPVIWIGAQECT WP_000145429.1 hydrogenase 2 small subunit [Salmonella enterica] Length: 372\nNumber of predicted TMHs: 1\nExp number of AAs in TMHs: 24.51309\nExp number, first 60 AAs: 0.2682\nTotal prob of N-in: 0.14402\noutside 1 331\nTMhelix 332 354\ninside 355 372

13990 GCF_002037265.1_ASM203726v1 Salmonella enterica Proteobacteria; Gammaproteobacteria; Enterobacterales; Enterobacteriaceae; Salmonella MNNEETFYQAMRRKGVTRRSFLKFCSLAATSLGLGAGMTPKIAWALENKPRIPVVWIHGL WP_001058312.1 hydrogenase [Salmonella enterica] Length: 372\nNumber of predicted TMHs: 2\nExp number of AAs in TMHs: 36.51738\nExp number, first 60 AAs: 19.72849\nTotal prob of N-in: 0.96170\nPOSSIBLE N-term signal sequence\ninside 1 19\nTMhelix 20 39\noutside 40 325\nTMhelix 326 348\ninside 349 372

13991 GCF_002037245.1_ASM203724v1 Salmonella enterica Proteobacteria; Gammaproteobacteria; Enterobacterales; Enterobacteriaceae; Salmonella MNNEETFYQAMRRKGVTRRSFLKFCSLAATSLGLGAGMTPKIAWALENKPRIPVVWIHGL WP_001058312.1 hydrogenase [Salmonella enterica] Length: 372\nNumber of predicted TMHs: 2\nExp number of AAs in TMHs: 36.51738\nExp number, first 60 AAs: 19.72849\nTotal prob of N-in: 0.96170\nPOSSIBLE N-term signal sequence\ninside 1 19\nTMhelix 20 39\noutside 40 325\nTMhelix 326 348\ninside 349 372

13992 GCF_002061905.1_ASM206190v1 Salmonella enterica Proteobacteria; Gammaproteobacteria; Enterobacterales; Enterobacteriaceae; Salmonella MNNEETFYQAMRRKGVTRRSFLKFCSLAATSLGLGAGMTPKIAWALENKPRIPVVWIHGL WP_001058312.1 hydrogenase [Salmonella enterica] Length: 372\nNumber of predicted TMHs: 2\nExp number of AAs in TMHs: 36.51738\nExp number, first 60 AAs: 19.72849\nTotal prob of N-in: 0.96170\nPOSSIBLE N-term signal sequence\ninside 1 19\nTMhelix 20 39\noutside 40 325\nTMhelix 326 348\ninside 349 372

13993 GCF_002061865.1_ASM206186v1 Salmonella enterica Proteobacteria; Gammaproteobacteria; Enterobacterales; Enterobacteriaceae; Salmonella MTGDNTLITSHGINRRDFMKLCAALAATMGLSSKAAAEMAESVSNPQRPPVIWIGAQECT WP_000145429.1 hydrogenase 2 small subunit [Salmonella enterica] Length: 372\nNumber of predicted TMHs: 1\nExp number of AAs in TMHs: 24.51309\nExp number, first 60 AAs: 0.2682\nTotal prob of N-in: 0.14402\noutside 1 331\nTMhelix 332 354\ninside 355 372

13994 GCF_002061905.1_ASM206190v1 Salmonella enterica Proteobacteria; Gammaproteobacteria; Enterobacterales; Enterobacteriaceae; Salmonella MTGDNTLITSHGINRRDFMKLCAALAATMGLSSKAAAEMAESVSNPQRPPVIWIGAQECT WP_000145429.1 hydrogenase 2 small subunit [Salmonella enterica] Length: 372\nNumber of predicted TMHs: 1\nExp number of AAs in TMHs: 24.51309\nExp number, first 60 AAs: 0.2682\nTotal prob of N-in: 0.14402\noutside 1 331\nTMhelix 332 354\ninside 355 372

13995 GCF_002037265.1_ASM203726v1 Salmonella enterica Proteobacteria; Gammaproteobacteria; Enterobacterales; Enterobacteriaceae; Salmonella
MTGDNTLITSHGINRRDFMKLCAALAATMGLSSKAAAEMAESVSNPQRPPVIWIGAQECT WP_000145429.1
hydrogenase 2 small subunit [Salmonella enterica] Length: 372\nNumber of predicted TMHs: 1\nExp number of AAs in TMHs: 24.51309\nExp number, first 60 AAs: 0.2682\nTotal prob of N-in: 0.14402\noutside 1 331\nTMhelix 332 354\ninside 355 372

13996 GCF_002037245.1_ASM203724v1 Salmonella enterica Proteobacteria; Gammaproteobacteria; Enterobacterales; Enterobacteriaceae; Salmonella
MTGDNTLITSHGINRRDFMKLCAALAATMGLSSKAAAEMAESVSNPQRPPVIWIGAQECT WP_000145429.1
hydrogenase 2 small subunit [Salmonella enterica] Length: 372\nNumber of predicted TMHs: 1\nExp number of AAs in TMHs: 24.51309\nExp number, first 60 AAs: 0.2682\nTotal prob of N-in: 0.14402\noutside 1 331\nTMhelix 332 354\ninside 355 372

13997 GCF_002036945.1_ASM203694v1 Salmonella enterica Proteobacteria; Gammaproteobacteria; Enterobacterales; Enterobacteriaceae; Salmonella
MNNEETFYQAMRRKGVTRRSFLKFCSLAATSLGLGAGMTPKIAWALENKPRIPVVWIHGL WP_001058312.1
hydrogenase [Salmonella enterica] Length: 372\nNumber of predicted TMHs: 2\nExp number of AAs in TMHs: 36.51738\nExp number, first 60 AAs: 19.72849\nTotal prob of N-in: 0.96170\nPOSSIBLE N-term signal sequence\ninside 1 19\nTMhelix 20 39\noutside 40 325\nTMhelix 326 348\ninside 349 372

13998 GCF_002036995.1_ASM203699v1 Salmonella enterica Proteobacteria; Gammaproteobacteria; Enterobacterales; Enterobacteriaceae; Salmonella
MTGDNTLITSHGINRRDFMKLCAALAATMGLSSKAAAEMAESVSNPQRPPVIWIGAQECT WP_000145431.1
hydrogenase 2 small subunit [Salmonella enterica] Length: 372\nNumber of predicted TMHs: 1\nExp number of AAs in TMHs: 24.51505\nExp number, first 60 AAs: 0.2682\nTotal prob of N-in: 0.14406\noutside 1 331\nTMhelix 332 354\ninside 355 372

13999 GCF_002036965.1_ASM203696v1 Salmonella enterica Proteobacteria; Gammaproteobacteria; Enterobacterales; Enterobacteriaceae; Salmonella
MTGDNTLITSHGINRRDFMKLCAALAATMGLSSKAAAEMAESVSNPQRPPVIWIGAQECT WP_000145431.1
hydrogenase 2 small subunit [Salmonella enterica] Length: 372\nNumber of predicted TMHs: 1\nExp number of AAs in TMHs: 24.51505\nExp number, first 60 AAs: 0.2682\nTotal prob of N-in: 0.14406\noutside 1 331\nTMhelix 332 354\ninside 355 372

14000 GCF_002036945.1_ASM203694v1 Salmonella enterica Proteobacteria; Gammaproteobacteria; Enterobacterales; Enterobacteriaceae; Salmonella
MTGDNTLITSHGINRRDFMKLCAALAATMGLSSKAAAEMAESVSNPQRPPVIWIGAQECT WP_000145429.1
hydrogenase 2 small subunit [Salmonella enterica] Length: 372\nNumber of predicted TMHs: 1\nExp number of AAs in TMHs: 24.51309\nExp number, first 60 AAs: 0.2682\nTotal prob of N-in: 0.14402\noutside 1 331\nTMhelix 332 354\ninside 355 372

14001 GCF_002036795.1_ASM203679v1 Salmonella enterica Proteobacteria; Gammaproteobacteria; Enterobacterales; Enterobacteriaceae; Salmonella
MNNEETFYQAMRRKGVTRRSFLKFCSLAATSLGLGAGMTPKIAWALENKPRIPVVWIHGL WP_001058312.1
hydrogenase [Salmonella enterica] Length: 372\nNumber of predicted TMHs: 2\nExp number of AAs in TMHs: 36.51738\nExp number, first 60 AAs: 19.72849\nTotal prob of N-in: 0.96170\nPOSSIBLE N-term signal sequence\ninside 1 19\nTMhelix 20 39\noutside 40 325\nTMhelix 326 348\ninside 349 372

14002 GCF_002036895.1_ASM203689v1 Salmonella enterica Proteobacteria; Gammaproteobacteria; Enterobacterales; Enterobacteriaceae; Salmonella
MNNEETFYQAMRRKGVTRRSFLKFCSLAATSLGLGAGMTPKIAWALENKPRIPVVWIHGL WP_001058312.1
hydrogenase [Salmonella enterica] Length: 372\nNumber of predicted TMHs: 2\nExp number of AAs in TMHs: 36.51738\nExp number, first 60 AAs: 19.72849\nTotal prob of N-in: 0.96170\nPOSSIBLE N-term signal sequence\ninside 1 19\nTMhelix 20 39\noutside 40 325\nTMhelix 326 348\ninside 349 372

14003 GCF_002036895.1_ASM203689v1 Salmonella enterica Proteobacteria; Gammaproteobacteria; Enterobacterales; Enterobacteriaceae; Salmonella
MTGDNTLITSHGINRRDFMKLCAALAATMGLSSKAAAEMAESVSNPQRPPVIWIGAQECT WP_000145429.1
hydrogenase 2 small subunit [Salmonella enterica] Length: 372\nNumber of predicted TMHs: 1\nExp

number of AAs in TMHs: 24.51309\nExp number, first 60 AAs: 0.2682\nTotal prob of N-in: 0.14402\noutside 1 331\nTMhelix 332 354\ninside 355 372

14004 GCF_002036525.1_ASM203652v1 Salmonella enterica Proteobacteria; Gammaproteobacteria; Enterobacterales; Enterobacteriaceae; Salmonella
MNNEETFYQAMRRKGVTRRSFLKFCSLAATSLGLGAGMTPKIAWALENKPRIPVVWIHGL WP_001058307.1
hydrogenase [Salmonella enterica] Length: 372\nNumber of predicted TMHs: 2\nExp number of AAs in TMHs: 36.51738\nExp number, first 60 AAs: 19.72849\nTotal prob of N-in: 0.96170\nPOSSIBLE N-term signal sequence\ninside 1 19\nTMhelix 20 39\noutside 40 325\nTMhelix 326 348\ninside 349 372

14005 GCF_002036595.1_ASM203659v1 Salmonella enterica Proteobacteria; Gammaproteobacteria; Enterobacterales; Enterobacteriaceae; Salmonella
MNNEETFYQAMRRKGVTRRSFLKFCSLAATSLGLGAGMTPKIAWALENKPRIPVVWIHGL WP_001058307.1
hydrogenase [Salmonella enterica] Length: 372\nNumber of predicted TMHs: 2\nExp number of AAs in TMHs: 36.51738\nExp number, first 60 AAs: 19.72849\nTotal prob of N-in: 0.96170\nPOSSIBLE N-term signal sequence\ninside 1 19\nTMhelix 20 39\noutside 40 325\nTMhelix 326 348\ninside 349 372

14006 GCF_002036565.1_ASM203656v1 Salmonella enterica Proteobacteria; Gammaproteobacteria; Enterobacterales; Enterobacteriaceae; Salmonella
MNNEETFYQAMRRKGVTRRSFLKFCSLAATSLGLGAGMTPKIAWALENKPRIPVVWIHGL WP_001058307.1
hydrogenase [Salmonella enterica] Length: 372\nNumber of predicted TMHs: 2\nExp number of AAs in TMHs: 36.51738\nExp number, first 60 AAs: 19.72849\nTotal prob of N-in: 0.96170\nPOSSIBLE N-term signal sequence\ninside 1 19\nTMhelix 20 39\noutside 40 325\nTMhelix 326 348\ninside 349 372

14007 GCF_002036595.1_ASM203659v1 Salmonella enterica Proteobacteria; Gammaproteobacteria; Enterobacterales; Enterobacteriaceae; Salmonella
MTGDNTLITSHGINRRDFMKLCAALAATMGLSSKAAAEMAESVSNPQRPPVIWIGAQECT WP_000145429.1
hydrogenase 2 small subunit [Salmonella enterica] Length: 372\nNumber of predicted TMHs: 1\nExp number of AAs in TMHs: 24.51309\nExp number, first 60 AAs: 0.2682\nTotal prob of N-in: 0.14402\noutside 1 331\nTMhelix 332 354\ninside 355 372

14008 GCF_002036565.1_ASM203656v1 Salmonella enterica Proteobacteria; Gammaproteobacteria; Enterobacterales; Enterobacteriaceae; Salmonella
MTGDNTLITSHGINRRDFMKLCAALAATMGLSSKAAAEMAESVSNPQRPPVIWIGAQECT WP_000145429.1
hydrogenase 2 small subunit [Salmonella enterica] Length: 372\nNumber of predicted TMHs: 1\nExp number of AAs in TMHs: 24.51309\nExp number, first 60 AAs: 0.2682\nTotal prob of N-in: 0.14402\noutside 1 331\nTMhelix 332 354\ninside 355 372

14009 GCF_002036455.1_ASM203645v1 Salmonella enterica Proteobacteria; Gammaproteobacteria; Enterobacterales; Enterobacteriaceae; Salmonella
MNNEETFYQAMRRKGVTRRSFLKFCSLAATSLGLGAGMTPKIAWALENKPRIPVVWIHGL WP_001058307.1
hydrogenase [Salmonella enterica] Length: 372\nNumber of predicted TMHs: 2\nExp number of AAs in TMHs: 36.51738\nExp number, first 60 AAs: 19.72849\nTotal prob of N-in: 0.96170\nPOSSIBLE N-term signal sequence\ninside 1 19\nTMhelix 20 39\noutside 40 325\nTMhelix 326 348\ninside 349 372

14010 GCF_002036285.1_ASM203628v1 Salmonella enterica Proteobacteria; Gammaproteobacteria; Enterobacterales; Enterobacteriaceae; Salmonella
MTGDNTLITSHGINRRDFMKLCAALAATMGLSSKAAAEMAESVSNPQRPPVIWIGAQECT WP_000145429.1
hydrogenase 2 small subunit [Salmonella enterica] Length: 372\nNumber of predicted TMHs: 1\nExp number of AAs in TMHs: 24.51309\nExp number, first 60 AAs: 0.2682\nTotal prob of N-in: 0.14402\noutside 1 331\nTMhelix 332 354\ninside 355 372

14011 GCF_002035765.1_ASM203576v1 Salmonella enterica Proteobacteria; Gammaproteobacteria; Enterobacterales; Enterobacteriaceae; Salmonella
MTGDNTLITSHGINRRDFMKLCAALAATMGLSSKAAAEMAESVSNPQRPPVIWIGAQECT WP_000145429.1
hydrogenase 2 small subunit [Salmonella enterica] Length: 372\nNumber of predicted TMHs: 1\nExp number of AAs in TMHs: 24.51309\nExp number, first 60 AAs: 0.2682\nTotal prob of N-in: 0.14402\noutside 1 331\nTMhelix 332 354\ninside 355 372

14012 GCF_002035695.1_ASM203569v1 Salmonella enterica Proteobacteria; Gammaproteobacteria; Enterobacterales; Enterobacteriaceae; Salmonella

MTGDNTLITSHGINRRDFMKLCAALAATMGLSSKAAAEMAESVSNPQRPPVIWIGAQECT WP_023248548.1 hydrogenase 2 small subunit [Salmonella enterica] Length: 372\nNumber of predicted TMHs: 1\nExp number of AAs in TMHs: 24.51281\nExp number, first 60 AAs: 0.26822\nTotal prob of N-in: 0.14402\noutside 1 331\nTMhelix 332 354\ninside 355 372

14013 GCF_002035625.1_ASM203562v1 Salmonella enterica Proteobacteria; Gammaproteobacteria; Enterobacterales; Enterobacteriaceae; Salmonella MNNEETFYQAMRRKGVTRRSFLKFCSLAATSLGLGAGMTPKIAWALENKPRIPVVWIHGL WP_001058312.1 hydrogenase [Salmonella enterica] Length: 372\nNumber of predicted TMHs: 2\nExp number of AAs in TMHs: 36.51738\nExp number, first 60 AAs: 19.72849\nTotal prob of N-in: 0.96170\nPOSSIBLE N-term signal sequence\ninside 1 19\nTMhelix 20 39\noutside 40 325\nTMhelix 326 348\ninside 349 372

14014 GCF_002035495.1_ASM203549v1 Salmonella enterica Proteobacteria; Gammaproteobacteria; Enterobacterales; Enterobacteriaceae; Salmonella MNNEETFYQAMRRKGVTRRSFLKFCSLAATSLGLGAGMTPKIAWALENKPRIPVVWIHGL WP_001058310.1 hydrogenase [Salmonella enterica] Length: 372\nNumber of predicted TMHs: 2\nExp number of AAs in TMHs: 37.4567299999999\nExp number, first 60 AAs: 19.72694\nTotal prob of N-in: 0.96171\nPOSSIBLE N-term signal sequence\ninside 1 19\nTMhelix 20 39\noutside 40 325\nTMhelix 326 348\ninside 349 372

14015 GCF_002035495.1_ASM203549v1 Salmonella enterica Proteobacteria; Gammaproteobacteria; Enterobacterales; Enterobacteriaceae; Salmonella MTGDNTLITSHGINRRDFMKLCAALAATMGLSSKAAAEMAESVSNPQRPPVIWIGAQECT WP_000145429.1 hydrogenase 2 small subunit [Salmonella enterica] Length: 372\nNumber of predicted TMHs: 1\nExp number of AAs in TMHs: 24.51309\nExp number, first 60 AAs: 0.2682\nTotal prob of N-in: 0.14402\noutside 1 331\nTMhelix 332 354\ninside 355 372

14016 GCF_002034695.1_ASM203469v1 Salmonella enterica Proteobacteria; Gammaproteobacteria; Enterobacterales; Enterobacteriaceae; Salmonella MTGDNTLITSHGINRRDFMKLCAALAATMGLSSKAAAEMAESVSNPQRPPVIWIGAQECT WP_079832985.1 hydrogenase 2 small subunit [Salmonella enterica] Length: 372\nNumber of predicted TMHs: 1\nExp number of AAs in TMHs: 24.51448\nExp number, first 60 AAs: 0.26822\nTotal prob of N-in: 0.14406\noutside 1 331\nTMhelix 332 354\ninside 355 372

14017 GCF_002034695.1_ASM203469v1 Salmonella enterica Proteobacteria; Gammaproteobacteria; Enterobacterales; Enterobacteriaceae; Salmonella MNNEETFYQAMRRKGVTRRSFLKFCSLAATSLGLGAGMTPKIAWALENKPRIPVVWIHGL WP_001058306.1 hydrogenase [Salmonella enterica] Length: 372\nNumber of predicted TMHs: 2\nExp number of AAs in TMHs: 36.5540099999999\nExp number, first 60 AAs: 19.72732\nTotal prob of N-in: 0.96171\nPOSSIBLE N-term signal sequence\ninside 1 19\nTMhelix 20 39\noutside 40 325\nTMhelix 326 348\ninside 349 372

14018 GCF_002061815.1_ASM206181v1 Salmonella enterica Proteobacteria; Gammaproteobacteria; Enterobacterales; Enterobacteriaceae; Salmonella MNNEETFYQAMRRKGVTRRSFLKFCSLAATSLGLGAGMTPKIAWALENKPRIPVVWIHGL WP_001058312.1 hydrogenase [Salmonella enterica] Length: 372\nNumber of predicted TMHs: 2\nExp number of AAs in TMHs: 36.51738\nExp number, first 60 AAs: 19.72849\nTotal prob of N-in: 0.96170\nPOSSIBLE N-term signal sequence\ninside 1 19\nTMhelix 20 39\noutside 40 325\nTMhelix 326 348\ninside 349 372

14019 GCF_002034185.1_ASM203418v1 Salmonella enterica Proteobacteria; Gammaproteobacteria; Enterobacterales; Enterobacteriaceae; Salmonella MNNEETFYQAMRRKGVTRRSFLKFCSLAATSLGLGAGMTPKIAWALENKPRIPVVWIHGL WP_001058312.1 hydrogenase [Salmonella enterica] Length: 372\nNumber of predicted TMHs: 2\nExp number of AAs in TMHs: 36.51738\nExp number, first 60 AAs: 19.72849\nTotal prob of N-in: 0.96170\nPOSSIBLE N-term signal sequence\ninside 1 19\nTMhelix 20 39\noutside 40 325\nTMhelix 326 348\ninside 349 372

14020 GCF_002034245.1_ASM203424v1 Salmonella enterica Proteobacteria; Gammaproteobacteria; Enterobacterales; Enterobacteriaceae; Salmonella MNNEETFYQAMRRKGVTRRSFLKFCSLAATSLGLGAGMTPKIAWALENKPRIPVVWIHGL WP_001058310.1 hydrogenase [Salmonella enterica] Length: 372\nNumber of predicted TMHs: 2\nExp number of AAs in TMHs: 37.4567299999999\nExp number, first 60 AAs: 19.72694\nTotal prob of N-in: 0.96171\nPOSSIBLE N-term signal sequence\ninside 1 19\nTMhelix 20 39\noutside 40 325\nTMhelix 326 348\ninside 349 372

14021 GCF_002034185.1_ASM203418v1 Salmonella enterica Proteobacteria; Gammaproteobacteria; Enterobacterales; Enterobacteriaceae; Salmonella
MTGDNTLITSHGINRRDFMKLCAALAATMGLSSKAAAEMAESVSNPQRPPVIWIGAQECT WP_000145429.1
hydrogenase 2 small subunit [Salmonella enterica] Length: 372\nNumber of predicted TMHs: 1\nExp number of AAs in TMHs: 24.51309\nExp number, first 60 AAs: 0.2682\nTotal prob of N-in: 0.14402\noutside 1 331\nTMhelix 332 354\ninside 355 372

14022 GCF_002034245.1_ASM203424v1 Salmonella enterica Proteobacteria; Gammaproteobacteria; Enterobacterales; Enterobacteriaceae; Salmonella
MTGDNTLITSHGINRRDFMKLCAALAATMGLSSKAAAEMAESVSNPQRPPVIWIGAQECT WP_000145429.1
hydrogenase 2 small subunit [Salmonella enterica] Length: 372\nNumber of predicted TMHs: 1\nExp number of AAs in TMHs: 24.51309\nExp number, first 60 AAs: 0.2682\nTotal prob of N-in: 0.14402\noutside 1 331\nTMhelix 332 354\ninside 355 372

14023 GCF_002033995.1_ASM203399v1 Salmonella enterica Proteobacteria; Gammaproteobacteria; Enterobacterales; Enterobacteriaceae; Salmonella
MNNEETFYQAMRRKGVTRRSFLKFCSLAATSLGLGAGMTPKIAWALENKPRIPVVWIHGL WP_001058311.1
hydrogenase [Salmonella enterica] Length: 372\nNumber of predicted TMHs: 2\nExp number of AAs in TMHs: 37.4699899999999\nExp number, first 60 AAs: 19.72699\nTotal prob of N-in: 0.96171\nPOSSIBLE N-term signal sequence\ninside 1 19\nTMhelix 20 39\noutside 40 325\nTMhelix 326 348\ninside 349 372

14024 GCF_002033995.1_ASM203399v1 Salmonella enterica Proteobacteria; Gammaproteobacteria; Enterobacterales; Enterobacteriaceae; Salmonella
MTGDNTLITSHGINRRDFMKLCAALAATMGLSSKAAAEMAESVSNPQRPPVIWIGAQECT WP_000145429.1
hydrogenase 2 small subunit [Salmonella enterica] Length: 372\nNumber of predicted TMHs: 1\nExp number of AAs in TMHs: 24.51309\nExp number, first 60 AAs: 0.2682\nTotal prob of N-in: 0.14402\noutside 1 331\nTMhelix 332 354\ninside 355 372

14025 GCF_002033945.1_ASM203394v1 Salmonella enterica Proteobacteria; Gammaproteobacteria; Enterobacterales; Enterobacteriaceae; Salmonella
MTGDNTLITSHGINRRDFMKLCAALAATMGLSSKAAAEMAESVSNPQRPPVIWIGAQECT WP_000145429.1
hydrogenase 2 small subunit [Salmonella enterica] Length: 372\nNumber of predicted TMHs: 1\nExp number of AAs in TMHs: 24.51309\nExp number, first 60 AAs: 0.2682\nTotal prob of N-in: 0.14402\noutside 1 331\nTMhelix 332 354\ninside 355 372

14026 GCF_002033885.1_ASM203388v1 Salmonella enterica Proteobacteria; Gammaproteobacteria; Enterobacterales; Enterobacteriaceae; Salmonella
MTGDNTLITSHGINRRDFMKLCAALAATMGLSSKAAAEMAESVSNPQRPPVIWIGAQECT WP_000145430.1
hydrogenase 2 small subunit [Salmonella enterica] Length: 372\nNumber of predicted TMHs: 1\nExp number of AAs in TMHs: 24.51306\nExp number, first 60 AAs: 0.2682\nTotal prob of N-in: 0.14402\noutside 1 331\nTMhelix 332 354\ninside 355 372

14027 GCF_002033525.1_ASM203352v1 Salmonella enterica Proteobacteria; Gammaproteobacteria; Enterobacterales; Enterobacteriaceae; Salmonella
MTGDNTLITSHGINRRDFMKLCAALAATMGLSSKAAAEMAESVSNPQRPPVIWIGAQECT WP_000145429.1
hydrogenase 2 small subunit [Salmonella enterica] Length: 372\nNumber of predicted TMHs: 1\nExp number of AAs in TMHs: 24.51309\nExp number, first 60 AAs: 0.2682\nTotal prob of N-in: 0.14402\noutside 1 331\nTMhelix 332 354\ninside 355 372

14028 GCF_002033375.1_ASM203337v1 Salmonella enterica Proteobacteria; Gammaproteobacteria; Enterobacterales; Enterobacteriaceae; Salmonella
MNNEETFYQAMRRKGVTRRSFLKFCSLAATSLGLGAGMTPKIAWALENKPRIPVVWIHGL WP_001058312.1
hydrogenase [Salmonella enterica] Length: 372\nNumber of predicted TMHs: 2\nExp number of AAs in TMHs: 36.51738\nExp number, first 60 AAs: 19.72849\nTotal prob of N-in: 0.96170\nPOSSIBLE N-term signal sequence\ninside 1 19\nTMhelix 20 39\noutside 40 325\nTMhelix 326 348\ninside 349 372

14029 GCF_002033415.1_ASM203341v1 Salmonella enterica Proteobacteria; Gammaproteobacteria; Enterobacterales; Enterobacteriaceae; Salmonella
MNNEETFYQAMRRKGVTRRSFLKFCSLAATSLGLGAGMTPKIAWALENKPRIPVVWIHGL WP_001058311.1
hydrogenase [Salmonella enterica] Length: 372\nNumber of predicted TMHs: 2\nExp number of AAs in

TMHs: 37.4699899999999\nExp number, first 60 AAs: 19.72699\nTotal prob of N-in: 0.96171\nPOSSIBLE N-term signal sequence\ninside 1 19\nTMhelix 20 39\noutside 40 325\nTMhelix 326 348\ninside 349 372

14030 GCF_002033645.1_ASM203364v1 Salmonella enterica Proteobacteria; Gammaproteobacteria; Enterobacterales; Enterobacteriaceae; Salmonella MNNEETFYQAMRRKGVTRRSFLKFCSLAATSLGLGAGMTPKIAWALENKPRIPVVWIHGL WP_023233644.1 hydrogenase 1, small subunit [Salmonella enterica] Length: 372\nNumber of predicted TMHs: 2\nExp number of AAs in TMHs: 37.4208299999999\nExp number, first 60 AAs: 19.72732\nTotal prob of N-in: 0.96168\nPOSSIBLE N-term signal sequence\ninside 1 19\nTMhelix 20 39\noutside 40 325\nTMhelix 326 348\ninside 349 372

14031 GCF_002033645.1_ASM203364v1 Salmonella enterica Proteobacteria; Gammaproteobacteria; Enterobacterales; Enterobacteriaceae; Salmonella MTGDNTLITSHGINRRDFMKLCAALAATMGLSSKAAAEMAESVSNPQRPPVIWIGAQECT WP_000145429.1 hydrogenase 2 small subunit [Salmonella enterica] Length: 372\nNumber of predicted TMHs: 1\nExp number of AAs in TMHs: 24.51309\nExp number, first 60 AAs: 0.2682\nTotal prob of N-in: 0.14402\noutside 1 331\nTMhelix 332 354\ninside 355 372

14032 GCF_002033335.1_ASM203333v1 Salmonella enterica Proteobacteria; Gammaproteobacteria; Enterobacterales; Enterobacteriaceae; Salmonella MNNEETFYQAMRRKGVTRRSFLKFCSLAATSLGLGAGMTPKIAWALENKPRIPVVWIHGL WP_001058307.1 hydrogenase [Salmonella enterica] Length: 372\nNumber of predicted TMHs: 2\nExp number of AAs in TMHs: 36.51738\nExp number, first 60 AAs: 19.72849\nTotal prob of N-in: 0.96170\nPOSSIBLE N-term signal sequence\ninside 1 19\nTMhelix 20 39\noutside 40 325\nTMhelix 326 348\ninside 349 372

14033 GCF_002032855.1_ASM203285v1 Salmonella enterica Proteobacteria; Gammaproteobacteria; Enterobacterales; Enterobacteriaceae; Salmonella MNNEETFYQAMRRKGVTRRSFLKFCSLAATSLGLGAGMTPKIAWALENKPRIPVVWIHGL WP_001058307.1 hydrogenase [Salmonella enterica] Length: 372\nNumber of predicted TMHs: 2\nExp number of AAs in TMHs: 36.51738\nExp number, first 60 AAs: 19.72849\nTotal prob of N-in: 0.96170\nPOSSIBLE N-term signal sequence\ninside 1 19\nTMhelix 20 39\noutside 40 325\nTMhelix 326 348\ninside 349 372

14034 GCF_002032675.1_ASM203267v1 Salmonella enterica Proteobacteria; Gammaproteobacteria; Enterobacterales; Enterobacteriaceae; Salmonella MTGDNTLITSHGINRRDFMKLCAALAATMGLSSKAAAEMAESVSNPQRPPVIWIGAQECT WP_000145429.1 hydrogenase 2 small subunit [Salmonella enterica] Length: 372\nNumber of predicted TMHs: 1\nExp number of AAs in TMHs: 24.51309\nExp number, first 60 AAs: 0.2682\nTotal prob of N-in: 0.14402\noutside 1 331\nTMhelix 332 354\ninside 355 372

14035 GCF_002032615.1_ASM203261v1 Salmonella enterica Proteobacteria; Gammaproteobacteria; Enterobacterales; Enterobacteriaceae; Salmonella MTGDNTLITSHGINRRDFMKLCAALAATMGLSSKAAAEMAESVSNPQRPPVIWIGAQECT WP_000145429.1 hydrogenase 2 small subunit [Salmonella enterica] Length: 372\nNumber of predicted TMHs: 1\nExp number of AAs in TMHs: 24.51309\nExp number, first 60 AAs: 0.2682\nTotal prob of N-in: 0.14402\noutside 1 331\nTMhelix 332 354\ninside 355 372

14036 GCF_002032585.1_ASM203258v1 Salmonella enterica Proteobacteria; Gammaproteobacteria; Enterobacterales; Enterobacteriaceae; Salmonella MTGDNTLITSHGINRRDFMKLCAALAATMGLSSKAAAEMAESVSNPQRPPVIWIGAQECT WP_000145429.1 hydrogenase 2 small subunit [Salmonella enterica] Length: 372\nNumber of predicted TMHs: 1\nExp number of AAs in TMHs: 24.51309\nExp number, first 60 AAs: 0.2682\nTotal prob of N-in: 0.14402\noutside 1 331\nTMhelix 332 354\ninside 355 372

14037 GCF_002032565.1_ASM203256v1 Salmonella enterica Proteobacteria; Gammaproteobacteria; Enterobacterales; Enterobacteriaceae; Salmonella MNNEETFYQAMRRKGVTRRSFLKFCSLAATSLGLGAGMTPKIAWALENKPRIPVVWIHGL WP_001058312.1 hydrogenase [Salmonella enterica] Length: 372\nNumber of predicted TMHs: 2\nExp number of AAs in TMHs: 36.51738\nExp number, first 60 AAs: 19.72849\nTotal prob of N-in: 0.96170\nPOSSIBLE N-term signal sequence\ninside 1 19\nTMhelix 20 39\noutside 40 325\nTMhelix 326 348\ninside 349 372

14038 GCF_002032565.1_ASM203256v1 Salmonella enterica Proteobacteria; Gammaproteobacteria; Enterobacterales; Enterobacteriaceae; Salmonella
MTGDNTLITSHGINRRDFMKLCAALAATMGLSSKAAAEMAESVSNPQRPPVIWIGAQECT WP_000145429.1
hydrogenase 2 small subunit [Salmonella enterica] Length: 372\nNumber of predicted TMHs: 1\nExp number of AAs in TMHs: 24.51309\nExp number, first 60 AAs: 0.2682\nTotal prob of N-in: 0.14402\noutside 1 331\nTMhelix 332 354\ninside 355 372

14039 GCF_002032485.1_ASM203248v1 Salmonella enterica Proteobacteria; Gammaproteobacteria; Enterobacterales; Enterobacteriaceae; Salmonella
MNNEETFYQAMRRKGVTRRSFLKFCSLAATSLGLGAGMTPKIAWALENKPRIPVVWIHGL WP_001058312.1
hydrogenase [Salmonella enterica] Length: 372\nNumber of predicted TMHs: 2\nExp number of AAs in TMHs: 36.51738\nExp number, first 60 AAs: 19.72849\nTotal prob of N-in: 0.96170\nPOSSIBLE N-term signal sequence\ninside 1 19\nTMhelix 20 39\noutside 40 325\nTMhelix 326 348\ninside 349 372

14040 GCF_002030895.1_ASM203089v1 Salmonella enterica Proteobacteria; Gammaproteobacteria; Enterobacterales; Enterobacteriaceae; Salmonella
MTGDNTLITSHGINRRDFMKLCAALAATMGLSSKAAAEMAESVSNPQRPPVIWIGAQECT WP_000145429.1
hydrogenase 2 small subunit [Salmonella enterica] Length: 372\nNumber of predicted TMHs: 1\nExp number of AAs in TMHs: 24.51309\nExp number, first 60 AAs: 0.2682\nTotal prob of N-in: 0.14402\noutside 1 331\nTMhelix 332 354\ninside 355 372

14041 GCF_002030825.1_ASM203082v1 Salmonella enterica Proteobacteria; Gammaproteobacteria; Enterobacterales; Enterobacteriaceae; Salmonella
MTGDNTLITSHGINRRDFMKLCAALAATMGLSSKAAAEMAESVSNPQRPPVIWIGAQECT WP_000145429.1
hydrogenase 2 small subunit [Salmonella enterica] Length: 372\nNumber of predicted TMHs: 1\nExp number of AAs in TMHs: 24.51309\nExp number, first 60 AAs: 0.2682\nTotal prob of N-in: 0.14402\noutside 1 331\nTMhelix 332 354\ninside 355 372

14042 GCF_002061815.1_ASM206181v1 Salmonella enterica Proteobacteria; Gammaproteobacteria; Enterobacterales; Enterobacteriaceae; Salmonella
MTGDNTLITSHGINRRDFMKLCAALAATMGLSSKAAAEMAESVSNPQRPPVIWIGAQECT WP_000145429.1
hydrogenase 2 small subunit [Salmonella enterica] Length: 372\nNumber of predicted TMHs: 1\nExp number of AAs in TMHs: 24.51309\nExp number, first 60 AAs: 0.2682\nTotal prob of N-in: 0.14402\noutside 1 331\nTMhelix 332 354\ninside 355 372

14043 GCF_002030295.1_ASM203029v1 Salmonella enterica Proteobacteria; Gammaproteobacteria; Enterobacterales; Enterobacteriaceae; Salmonella
MNNEETFYQAMRRKGVTRRSFLKFCSLAATSLGLGAGMTPKIAWALENKPRIPVVWIHGL WP_001058311.1
hydrogenase [Salmonella enterica] Length: 372\nNumber of predicted TMHs: 2\nExp number of AAs in TMHs: 37.4699899999999\nExp number, first 60 AAs: 19.72699\nTotal prob of N-in: 0.96171\nPOSSIBLE N-term signal sequence\ninside 1 19\nTMhelix 20 39\noutside 40 325\nTMhelix 326 348\ninside 349 372

14044 GCF_001648855.1_ASM164885v1 Salmonella enterica Proteobacteria; Gammaproteobacteria; Enterobacterales; Enterobacteriaceae; Salmonella
MTGDNTLITSHGINRRDFMKLCAALAATMGLSSKAAAEMAESVSNPQRPPVIWIGAQECT WP_000145429.1
hydrogenase 2 small subunit [Salmonella enterica] Length: 372\nNumber of predicted TMHs: 1\nExp number of AAs in TMHs: 24.51309\nExp number, first 60 AAs: 0.2682\nTotal prob of N-in: 0.14402\noutside 1 331\nTMhelix 332 354\ninside 355 372

14045 GCF_001716745.1_ASM171674v1 Salmonella enterica Proteobacteria; Gammaproteobacteria; Enterobacterales; Enterobacteriaceae; Salmonella
MTGDNTLITSHGINRRDFMKLCAALAATMGLSSKAAAEMAESVSNPQRPPVIWIGAQECT WP_000145429.1
hydrogenase 2 small subunit [Salmonella enterica] Length: 372\nNumber of predicted TMHs: 1\nExp number of AAs in TMHs: 24.51309\nExp number, first 60 AAs: 0.2682\nTotal prob of N-in: 0.14402\noutside 1 331\nTMhelix 332 354\ninside 355 372

14046 GCF_001275185.1_ASM127518v1 Salmonella enterica Proteobacteria; Gammaproteobacteria; Enterobacterales; Enterobacteriaceae; Salmonella
MNNEETFYQAMRRKGVTRRSFLKFCSLAATSLGLGAGMTPKIAWALENKPRIPVVWIHGL WP_001058311.1
hydrogenase [Salmonella enterica] Length: 372\nNumber of predicted TMHs: 2\nExp number of AAs in

TMHs: 37.4699899999999\nExp number, first 60 AAs: 19.72699\nTotal prob of N-in: 0.96171\nPOSSIBLE N-term signal sequence\ninside 1 19\nTMhelix 20 39\noutside 40 325\nTMhelix 326 348\ninside 349 372

14047 GCF_002061105.1_ASM206110v1 Salmonella enterica Proteobacteria; Gammaproteobacteria; Enterobacterales; Enterobacteriaceae; Salmonella
MNNEETFYQAMRRKGVTRRSFLKFCSLAATSLGLGAGMTPKIAWALENKPRIPVVWIHGL WP_023234992.1
hydrogenase 1, small subunit [Salmonella enterica] Length: 372\nNumber of predicted TMHs: 2\nExp number of AAs in TMHs: 37.41309\nExp number, first 60 AAs: 19.72865\nTotal prob of N-in: 0.96169\nPOSSIBLE N-term signal sequence\ninside 1 19\nTMhelix 20 39\noutside 40 325\nTMhelix 326 348\ninside 349 372

14048 GCF_001475085.1_Salmonella_enterica_CVM_43840_v1.0Salmonella enterica Proteobacteria; Gammaproteobacteria; Enterobacterales; Enterobacteriaceae; Salmonella
MTGDNTLITSHGINRRDFMKLCAALAATMGLSSKAAAEMAESVSNPQRPPVIWIGAQECT WP_000145428.1
hydrogenase 2 small subunit [Salmonella enterica] Length: 372\nNumber of predicted TMHs: 1\nExp number of AAs in TMHs: 24.50677\nExp number, first 60 AAs: 0.2682\nTotal prob of N-in: 0.14390\noutside 1 331\nTMhelix 332 354\ninside 355 372

14049 GCF_001475165.1_Salmonella_enterica_CVM_43845_v1.0Salmonella enterica Proteobacteria; Gammaproteobacteria; Enterobacterales; Enterobacteriaceae; Salmonella
MTGDNTLITSHGINRRDFMKLCAALAATMGLSSKAAAEMAESVSNPQRPPVIWIGAQECT WP_000145429.1
hydrogenase 2 small subunit [Salmonella enterica] Length: 372\nNumber of predicted TMHs: 1\nExp number of AAs in TMHs: 24.51309\nExp number, first 60 AAs: 0.2682\nTotal prob of N-in: 0.14402\noutside 1 331\nTMhelix 332 354\ninside 355 372

14050 GCF_001475165.1_Salmonella_enterica_CVM_43845_v1.0Salmonella enterica Proteobacteria; Gammaproteobacteria; Enterobacterales; Enterobacteriaceae; Salmonella
MNNEETFYQAMRRKGVTRRSFLKFCSLAATSLGLGAGMTPKIAWALENKPRIPVVWIHGL WP_001058312.1
hydrogenase [Salmonella enterica] Length: 372\nNumber of predicted TMHs: 2\nExp number of AAs in TMHs: 36.51738\nExp number, first 60 AAs: 19.72849\nTotal prob of N-in: 0.96170\nPOSSIBLE N-term signal sequence\ninside 1 19\nTMhelix 20 39\noutside 40 325\nTMhelix 326 348\ninside 349 372

14051 GCF_001474995.1_Salmonella_enterica_CVM_43830_v1.0Salmonella enterica Proteobacteria; Gammaproteobacteria; Enterobacterales; Enterobacteriaceae; Salmonella
MNNEETFYQAMRRKGVTRRSFLKFCSLAATSLGLGAGMTPKIAWALENKPRIPVVWIHGL WP_001058312.1
hydrogenase [Salmonella enterica] Length: 372\nNumber of predicted TMHs: 2\nExp number of AAs in TMHs: 36.51738\nExp number, first 60 AAs: 19.72849\nTotal prob of N-in: 0.96170\nPOSSIBLE N-term signal sequence\ninside 1 19\nTMhelix 20 39\noutside 40 325\nTMhelix 326 348\ninside 349 372

14052 GCF_001475005.1_Salmonella_enterica_CVM_43831_v1.0Salmonella enterica Proteobacteria; Gammaproteobacteria; Enterobacterales; Enterobacteriaceae; Salmonella
MNNEETFYQAMRRKGVTRRSFLKFCSLAATSLGLGAGMTPKIAWALENKPRIPVVWIHGL WP_001058312.1
hydrogenase [Salmonella enterica] Length: 372\nNumber of predicted TMHs: 2\nExp number of AAs in TMHs: 36.51738\nExp number, first 60 AAs: 19.72849\nTotal prob of N-in: 0.96170\nPOSSIBLE N-term signal sequence\ninside 1 19\nTMhelix 20 39\noutside 40 325\nTMhelix 326 348\ninside 349 372

14053 GCF_001474965.1_Salmonella_enterica_CVM_43827_v1.0Salmonella enterica Proteobacteria; Gammaproteobacteria; Enterobacterales; Enterobacteriaceae; Salmonella
MNNEETFYQAMRRKGVTRRSFLKFCSLAATSLGLGAGMTPKIAWALENKPRIPVVWIHGL WP_001058307.1
hydrogenase [Salmonella enterica] Length: 372\nNumber of predicted TMHs: 2\nExp number of AAs in TMHs: 36.51738\nExp number, first 60 AAs: 19.72849\nTotal prob of N-in: 0.96170\nPOSSIBLE N-term signal sequence\ninside 1 19\nTMhelix 20 39\noutside 40 325\nTMhelix 326 348\ninside 349 372

14054 GCF_001475105.1_Salmonella_enterica_CVM_43841_v1.0Salmonella enterica Proteobacteria; Gammaproteobacteria; Enterobacterales; Enterobacteriaceae; Salmonella
MTGDNTLITSHGINRRDFMKLCAALAATMGLSSKAAAEMAESVSNPQRPPVIWIGAQECT WP_000145429.1
hydrogenase 2 small subunit [Salmonella enterica] Length: 372\nNumber of predicted TMHs: 1\nExp number of AAs in TMHs: 24.51309\nExp number, first 60 AAs: 0.2682\nTotal prob of N-in: 0.14402\noutside 1 331\nTMhelix 332 354\ninside 355 372

14055 GCF_001475105.1_Salmonella_enterica_CVM_43841_v1.0Salmonella enterica Proteobacteria; Gammaproteobacteria; Enterobacterales; Enterobacteriaceae; Salmonella

MNNEETFYQAMRRKGVTRRSFLKFCSLAATSLGLGAGMTPKIAWALENKPRIPVVWIHGL WP_001058308.1 hydrogenase [Salmonella enterica] Length: 372\nNumber of predicted TMHs: 2\nExp number of AAs in TMHs: 39.08204\nExp number, first 60 AAs: 19.72474\nTotal prob of N-in: 0.96174\nPOSSIBLE N-term signal sequence\ninside 1 19\nTMhelix 20 39\noutside 40 325\nTMhelix 326 348\ninside 349 372

14056 GCF_001474985.1_Salmonella_enterica_CVM_43829_v1.0Salmonella enterica Proteobacteria; Gammaproteobacteria; Enterobacterales; Enterobacteriaceae; Salmonella MNNEETFYQAMRRKGVTRRSFLKFCSLAATSLGLGAGMTPKIAWALENKPRIPVVWIHGL WP_001058312.1 hydrogenase [Salmonella enterica] Length: 372\nNumber of predicted TMHs: 2\nExp number of AAs in TMHs: 36.51738\nExp number, first 60 AAs: 19.72849\nTotal prob of N-in: 0.96170\nPOSSIBLE N-term signal sequence\ninside 1 19\nTMhelix 20 39\noutside 40 325\nTMhelix 326 348\ninside 349 372

14057 GCF_001475175.1_Salmonella_enterica_CVM_43846_v1.0Salmonella enterica Proteobacteria; Gammaproteobacteria; Enterobacterales; Enterobacteriaceae; Salmonella MTGDNTLITSHGINRRDFMKLCAALAATMGLSSKAAAEMAESVSNPQRPPVIWIGAQECT WP_000145429.1 hydrogenase 2 small subunit [Salmonella enterica] Length: 372\nNumber of predicted TMHs: 1\nExp number of AAs in TMHs: 24.51309\nExp number, first 60 AAs: 0.2682\nTotal prob of N-in: 0.14402\noutside 1 331\nTMhelix 332 354\ninside 355 372

14058 GCF_001475175.1_Salmonella_enterica_CVM_43846_v1.0Salmonella enterica Proteobacteria; Gammaproteobacteria; Enterobacterales; Enterobacteriaceae; Salmonella MNNEETFYQAMRRKGVTRRSFLKFCSLAATSLGLGAGMTPKIAWALENKPRIPVVWIHGL WP_001058312.1 hydrogenase [Salmonella enterica] Length: 372\nNumber of predicted TMHs: 2\nExp number of AAs in TMHs: 36.51738\nExp number, first 60 AAs: 19.72849\nTotal prob of N-in: 0.96170\nPOSSIBLE N-term signal sequence\ninside 1 19\nTMhelix 20 39\noutside 40 325\nTMhelix 326 348\ninside 349 372

14059 GCF_001475065.1_Salmonella_enterica_CVM_43838_v1.0Salmonella enterica Proteobacteria; Gammaproteobacteria; Enterobacterales; Enterobacteriaceae; Salmonella MNNEETFYQAMRRKGVTRRSFLKFCSLAATSLGLGAGMTPKIAWALENKPRIPVVWIHGL WP_001058312.1 hydrogenase [Salmonella enterica] Length: 372\nNumber of predicted TMHs: 2\nExp number of AAs in TMHs: 36.51738\nExp number, first 60 AAs: 19.72849\nTotal prob of N-in: 0.96170\nPOSSIBLE N-term signal sequence\ninside 1 19\nTMhelix 20 39\noutside 40 325\nTMhelix 326 348\ninside 349 372

14060 GCF_001475225.1_Salmonella_enterica_CVM_43820_v1.0Salmonella enterica Proteobacteria; Gammaproteobacteria; Enterobacterales; Enterobacteriaceae; Salmonella MTGDNTLITSHGINRRDFMKLCAALAATMGLSSKAAAEMAESVSNPQRPPVIWIGAQECT WP_000145429.1 hydrogenase 2 small subunit [Salmonella enterica] Length: 372\nNumber of predicted TMHs: 1\nExp number of AAs in TMHs: 24.51309\nExp number, first 60 AAs: 0.2682\nTotal prob of N-in: 0.14402\noutside 1 331\nTMhelix 332 354\ninside 355 372

14061 GCF_001475335.1_Salmonella_enterica_CVM_43815_v1.0Salmonella enterica Proteobacteria; Gammaproteobacteria; Enterobacterales; Enterobacteriaceae; Salmonella MNNEETFYQAMRRKGVTRRSFLKFCSLAATSLGLGAGMTPKIAWALENKPRIPVVWIHGL WP_001058312.1 hydrogenase [Salmonella enterica] Length: 372\nNumber of predicted TMHs: 2\nExp number of AAs in TMHs: 36.51738\nExp number, first 60 AAs: 19.72849\nTotal prob of N-in: 0.96170\nPOSSIBLE N-term signal sequence\ninside 1 19\nTMhelix 20 39\noutside 40 325\nTMhelix 326 348\ninside 349 372

14062 GCF_001831805.1_ASM183180v1 Salmonella enterica Proteobacteria; Gammaproteobacteria; Enterobacterales; Enterobacteriaceae; Salmonella MTGDNTLITSHGINRRDFMKLCAALAATMGLSSKAAAEMAESVSNPQRPPVIWIGAQECT WP_000145429.1 hydrogenase 2 small subunit [Salmonella enterica] Length: 372\nNumber of predicted TMHs: 1\nExp number of AAs in TMHs: 24.51309\nExp number, first 60 AAs: 0.2682\nTotal prob of N-in: 0.14402\noutside 1 331\nTMhelix 332 354\ninside 355 372

14063 GCF_001569585.1_ASM156958v1 Salmonella enterica Proteobacteria; Gammaproteobacteria; Enterobacterales; Enterobacteriaceae; Salmonella MTGDNTLITSHGINRRDFMKLCAALAATMGLSSKAAAEMAESVSNPQRPPVIWIGAQECT WP_000145429.1 hydrogenase 2 small subunit [Salmonella enterica] Length: 372\nNumber of predicted TMHs: 1\nExp number of AAs in TMHs: 24.51309\nExp number, first 60 AAs: 0.2682\nTotal prob of N-in: 0.14402\noutside 1 331\nTMhelix 332 354\ninside 355 372

14064 GCF_001569645.1_ASM156964v1 Salmonella enterica Proteobacteria; Gammaproteobacteria; Enterobacterales; Enterobacteriaceae; Salmonella
MTGDNTLITSHGINRRDFMKLCAALAATMGLSSKAAAEMAESVSNPQRPPVIWIGAQECT WP_000145429.1
hydrogenase 2 small subunit [Salmonella enterica] Length: 372\nNumber of predicted TMHs: 1\nExp number of AAs in TMHs: 24.51309\nExp number, first 60 AAs: 0.2682\nTotal prob of N-in: 0.14402\noutside 1 331\nTMhelix 332 354\ninside 355 372

14065 GCF_001569585.1_ASM156958v1 Salmonella enterica Proteobacteria; Gammaproteobacteria; Enterobacterales; Enterobacteriaceae; Salmonella
MNNEETFYQAMRRKGVTRRSFLKFCSLAATSLGLGAGMTPKIAWALENKPRIPVIWIHGL WP_061376537.1
hydrogenase [Salmonella enterica] Length: 372\nNumber of predicted TMHs: 2\nExp number of AAs in TMHs: 36.5556\nExp number, first 60 AAs: 19.76633\nTotal prob of N-in: 0.96197\nPOSSIBLE N-term signal sequence\ninside 1 19\nTMhelix 20 39\noutside 40 325\nTMhelix 326 348\ninside 349 372

14066 GCF_001569545.1_ASM156954v1 Salmonella enterica Proteobacteria; Gammaproteobacteria; Enterobacterales; Enterobacteriaceae; Salmonella
MNNEETFYQAMRRKGVTRRSFLKFCSLAATSLGLGAGMTPKIAWALENKPRIPVVWIHGL WP_001058307.1
hydrogenase [Salmonella enterica] Length: 372\nNumber of predicted TMHs: 2\nExp number of AAs in TMHs: 36.51738\nExp number, first 60 AAs: 19.72849\nTotal prob of N-in: 0.96170\nPOSSIBLE N-term signal sequence\ninside 1 19\nTMhelix 20 39\noutside 40 325\nTMhelix 326 348\ninside 349 372

14067 GCF_001569685.1_ASM156968v1 Salmonella enterica Proteobacteria; Gammaproteobacteria; Enterobacterales; Enterobacteriaceae; Salmonella
MTGDNTLITSHGINRRDFMKLCAALAATMGLSSKAAAEMAESVSNPQRPPVIWIGAQECT WP_000145429.1
hydrogenase 2 small subunit [Salmonella enterica] Length: 372\nNumber of predicted TMHs: 1\nExp number of AAs in TMHs: 24.51309\nExp number, first 60 AAs: 0.2682\nTotal prob of N-in: 0.14402\noutside 1 331\nTMhelix 332 354\ninside 355 372

14068 GCF_001569945.1_ASM156994v1 Salmonella enterica Proteobacteria; Gammaproteobacteria; Enterobacterales; Enterobacteriaceae; Salmonella
MNNEETFYQAMRRKGVTRRSFLKFCSLAATSLGLGAGMTPKIAWALENKPRIPVVWIHGL WP_001058307.1
hydrogenase [Salmonella enterica] Length: 372\nNumber of predicted TMHs: 2\nExp number of AAs in TMHs: 36.51738\nExp number, first 60 AAs: 19.72849\nTotal prob of N-in: 0.96170\nPOSSIBLE N-term signal sequence\ninside 1 19\nTMhelix 20 39\noutside 40 325\nTMhelix 326 348\ninside 349 372

14069 GCF_001642515.1_ASM164251v1 Salmonella enterica Proteobacteria; Gammaproteobacteria; Enterobacterales; Enterobacteriaceae; Salmonella
MTGDNTLITSHGINRRDFMKLCAALAATMGLSSKAAAEMAESVSNPQRPPVIWIGAQECT WP_000145429.1
hydrogenase 2 small subunit [Salmonella enterica] Length: 372\nNumber of predicted TMHs: 1\nExp number of AAs in TMHs: 24.51309\nExp number, first 60 AAs: 0.2682\nTotal prob of N-in: 0.14402\noutside 1 331\nTMhelix 332 354\ninside 355 372

14070 GCF_001569985.1_ASM156998v1 Salmonella enterica Proteobacteria; Gammaproteobacteria; Enterobacterales; Enterobacteriaceae; Salmonella
MNNEETFYQAMRRKGVTRRSFLKFCSLAATSLGLGAGMTPKIAWALENKPRIPVVWIHGL WP_001058307.1
hydrogenase [Salmonella enterica] Length: 372\nNumber of predicted TMHs: 2\nExp number of AAs in TMHs: 36.51738\nExp number, first 60 AAs: 19.72849\nTotal prob of N-in: 0.96170\nPOSSIBLE N-term signal sequence\ninside 1 19\nTMhelix 20 39\noutside 40 325\nTMhelix 326 348\ninside 349 372

14071 GCF_001570125.1_ASM157012v1 Salmonella enterica Proteobacteria; Gammaproteobacteria; Enterobacterales; Enterobacteriaceae; Salmonella
MTGDNTLITSHGINRRDFMKLCAALAATMGLSSKAAAEMAESVSNPQRPPVIWIGAQECT WP_000145429.1
hydrogenase 2 small subunit [Salmonella enterica] Length: 372\nNumber of predicted TMHs: 1\nExp number of AAs in TMHs: 24.51309\nExp number, first 60 AAs: 0.2682\nTotal prob of N-in: 0.14402\noutside 1 331\nTMhelix 332 354\ninside 355 372

14072 GCF_001570125.1_ASM157012v1 Salmonella enterica Proteobacteria; Gammaproteobacteria; Enterobacterales; Enterobacteriaceae; Salmonella
MNNEETFYQAMRRKGVTRRSFLKFCSLAATSLGLGAGMTPKIAWALENKPRIPVVWIHGL WP_001058307.1
hydrogenase [Salmonella enterica] Length: 372\nNumber of predicted TMHs: 2\nExp number of AAs in

TMHs: 36.51738\nExp number, first 60 AAs:  19.72849\nTotal prob of N-in:   0.96170\nPOSSIBLE N-term signal sequence\ninside   1   19\nTMhelix   20   39\noutside   40   325\nTMhelix   326   348\ninside   349   372

14073 GCF_001570085.1_ASM157008v1 Salmonella enterica Proteobacteria; Gammaproteobacteria; Enterobacterales; Enterobacteriaceae; Salmonella
MNNEETFYQAMRRKGVTRRSFLKFCSLAATSLGLGAGMTPKIAWALENKPRIPVVWIHGL WP_001058307.1
hydrogenase [Salmonella enterica] Length: 372\nNumber of predicted TMHs:  2\nExp number of AAs in TMHs: 36.51738\nExp number, first 60 AAs:  19.72849\nTotal prob of N-in:   0.96170\nPOSSIBLE N-term signal sequence\ninside   1   19\nTMhelix   20   39\noutside   40   325\nTMhelix   326   348\ninside   349   372

14074 GCF_001570045.1_ASM157004v1 Salmonella enterica Proteobacteria; Gammaproteobacteria; Enterobacterales; Enterobacteriaceae; Salmonella
MNNEETFYQAMRRKGVTRRSFLKFCSLAATSLGLGAGMTPKIAWALENKPRIPVVWIHGL WP_001058311.1
hydrogenase [Salmonella enterica] Length: 372\nNumber of predicted TMHs:  2\nExp number of AAs in TMHs: 37.4699899999999\nExp number, first 60 AAs:  19.72699\nTotal prob of N-in:   0.96171\nPOSSIBLE N-term signal sequence\ninside   1   19\nTMhelix   20   39\noutside   40   325\nTMhelix   326   348\ninside   349   372

14075 GCF_002061035.1_ASM206103v1 Salmonella enterica Proteobacteria; Gammaproteobacteria; Enterobacterales; Enterobacteriaceae; Salmonella
MNNEETFYQAMRRKGVTRRSFLKFCSLAATSLGLGAGMTPKIAWALENKPRIPVVWIHGL WP_001058311.1
hydrogenase [Salmonella enterica] Length: 372\nNumber of predicted TMHs:  2\nExp number of AAs in TMHs: 37.4699899999999\nExp number, first 60 AAs:  19.72699\nTotal prob of N-in:   0.96171\nPOSSIBLE N-term signal sequence\ninside   1   19\nTMhelix   20   39\noutside   40   325\nTMhelix   326   348\ninside   349   372

14076 GCF_002061855.1_ASM206185v1 Salmonella enterica Proteobacteria; Gammaproteobacteria; Enterobacterales; Enterobacteriaceae; Salmonella
MTGDNTLITSHGINRRDFMKLCAALAATMGLSSKAAAEMAESVSNPQRPPVIWIGAQECT WP_000145427.1
hydrogenase 2 small subunit [Salmonella enterica] Length: 372\nNumber of predicted TMHs:  1\nExp number of AAs in TMHs: 24.48576\nExp number, first 60 AAs:  0.2692\nTotal prob of N-in:   0.14251\noutside   1   331\nTMhelix   332   354\ninside   355   372

14077 GCF_002060935.1_ASM206093v1 Salmonella enterica Proteobacteria; Gammaproteobacteria; Enterobacterales; Enterobacteriaceae; Salmonella
MNNEETFYQAMRRKGVTRRSFLKFCSLAATSLGLGAGMTPKIAWALENKPRIPVVWIHGL WP_001058312.1
hydrogenase [Salmonella enterica] Length: 372\nNumber of predicted TMHs:  2\nExp number of AAs in TMHs: 36.51738\nExp number, first 60 AAs:  19.72849\nTotal prob of N-in:   0.96170\nPOSSIBLE N-term signal sequence\ninside   1   19\nTMhelix   20   39\noutside   40   325\nTMhelix   326   348\ninside   349   372

14078 GCF_002060935.1_ASM206093v1 Salmonella enterica Proteobacteria; Gammaproteobacteria; Enterobacterales; Enterobacteriaceae; Salmonella
MTGDNTLITSHGINRRDFMKLCAALAATMGLSSKAAAEMAESVSNPQRPPVIWIGAQECT WP_000145429.1
hydrogenase 2 small subunit [Salmonella enterica] Length: 372\nNumber of predicted TMHs:  1\nExp number of AAs in TMHs: 24.51309\nExp number, first 60 AAs:  0.2682\nTotal prob of N-in:   0.14402\noutside   1   331\nTMhelix   332   354\ninside   355   372

14079 GCF_002060875.1_ASM206087v1 Salmonella enterica Proteobacteria; Gammaproteobacteria; Enterobacterales; Enterobacteriaceae; Salmonella
MTGDNTLITSHGINRRDFMKLCAALAATMGLSSKAAAEMAESVSNPQRPPVIWIGAQECT WP_023248548.1
hydrogenase 2 small subunit [Salmonella enterica] Length: 372\nNumber of predicted TMHs:  1\nExp number of AAs in TMHs: 24.51281\nExp number, first 60 AAs:  0.26822\nTotal prob of N-in:   0.14402\noutside   1   331\nTMhelix   332   354\ninside   355   372

14080 GCF_002060855.1_ASM206085v1 Salmonella enterica Proteobacteria; Gammaproteobacteria; Enterobacterales; Enterobacteriaceae; Salmonella
MTGDNTLITSHGINRRDFMKLCAALAATMGLSSKAAAEMAESVSNPQRPPVIWIGAQECT WP_000145429.1
hydrogenase 2 small subunit [Salmonella enterica] Length: 372\nNumber of predicted TMHs:  1\nExp number of AAs in TMHs: 24.51309\nExp number, first 60 AAs:  0.2682\nTotal prob of N-in:   0.14402\noutside   1   331\nTMhelix   332   354\ninside   355   372

14081 GCF_002060785.1_ASM206078v1 Salmonella enterica Proteobacteria; Gammaproteobacteria; Enterobacterales; Enterobacteriaceae; Salmonella

MNNEETFYQAMRRKGVTRRSFLKFCSLAATSLGLGAGMTPKIAWALENKPRIPVVWIHGL WP_001058312.1 hydrogenase [Salmonella enterica] Length: 372\nNumber of predicted TMHs: 2\nExp number of AAs in TMHs: 36.51738\nExp number, first 60 AAs: 19.72849\nTotal prob of N-in: 0.96170\nPOSSIBLE N-term signal sequence\ninside 1 19\nTMhelix 20 39\noutside 40 325\nTMhelix 326 348\ninside 349 372

14082 GCF_002060875.1_ASM206087v1 Salmonella enterica Proteobacteria; Gammaproteobacteria; Enterobacterales; Enterobacteriaceae; Salmonella MNNEETFYQAMRRKGVTRRSFLKFCSLAATSLGLGAGMTPKIAWALENKPRIPVVWIHGL WP_001058306.1 hydrogenase [Salmonella enterica] Length: 372\nNumber of predicted TMHs: 2\nExp number of AAs in TMHs: 36.5540099999999\nExp number, first 60 AAs: 19.72732\nTotal prob of N-in: 0.96171\nPOSSIBLE N-term signal sequence\ninside 1 19\nTMhelix 20 39\noutside 40 325\nTMhelix 326 348\ninside 349 372

14083 GCF_002060695.1_ASM206069v1 Salmonella enterica Proteobacteria; Gammaproteobacteria; Enterobacterales; Enterobacteriaceae; Salmonella MTGDNTLITSHGINRRDFMKLCAALAATMGLSSKAAAEMAESVSNPQRPPVIWIGAQECT WP_000145429.1 hydrogenase 2 small subunit [Salmonella enterica] Length: 372\nNumber of predicted TMHs: 1\nExp number of AAs in TMHs: 24.51309\nExp number, first 60 AAs: 0.2682\nTotal prob of N-in: 0.14402\noutside 1 331\nTMhelix 332 354\ninside 355 372

14084 GCF_002060645.1_ASM206064v1 Salmonella enterica Proteobacteria; Gammaproteobacteria; Enterobacterales; Enterobacteriaceae; Salmonella MNNEETFYQAMRRKGVTRRSFLKFCSLAATSLGLGAGMTPKIAWALENKPRIPVVWIHGL WP_001058311.1 hydrogenase [Salmonella enterica] Length: 372\nNumber of predicted TMHs: 2\nExp number of AAs in TMHs: 37.4699899999999\nExp number, first 60 AAs: 19.72699\nTotal prob of N-in: 0.96171\nPOSSIBLE N-term signal sequence\ninside 1 19\nTMhelix 20 39\noutside 40 325\nTMhelix 326 348\ninside 349 372

14085 GCF_002060645.1_ASM206064v1 Salmonella enterica Proteobacteria; Gammaproteobacteria; Enterobacterales; Enterobacteriaceae; Salmonella MTGDNTLITSHGINRRDFMKLCAALAATMGLSSKAAAEMAESVSNPQRPPVIWIGAQECT WP_000145429.1 hydrogenase 2 small subunit [Salmonella enterica] Length: 372\nNumber of predicted TMHs: 1\nExp number of AAs in TMHs: 24.51309\nExp number, first 60 AAs: 0.2682\nTotal prob of N-in: 0.14402\noutside 1 331\nTMhelix 332 354\ninside 355 372

14086 GCF_002060415.1_ASM206041v1 Salmonella enterica Proteobacteria; Gammaproteobacteria; Enterobacterales; Enterobacteriaceae; Salmonella MNNEETFYQAMRRKGVTRRSFLKFCSLAATSLGLGAGMTPKIAWALENKPRIPVVWIHGL WP_001540183.1 hydrogenase [Salmonella enterica] Length: 372\nNumber of predicted TMHs: 2\nExp number of AAs in TMHs: 37.41804\nExp number, first 60 AAs: 19.72808\nTotal prob of N-in: 0.96169\nPOSSIBLE N-term signal sequence\ninside 1 19\nTMhelix 20 39\noutside 40 325\nTMhelix 326 348\ninside 349 372

14087 GCF_002060385.1_ASM206038v1 Salmonella enterica Proteobacteria; Gammaproteobacteria; Enterobacterales; Enterobacteriaceae; Salmonella MNNEETFYQAMRRKGVTRRSFLKFCSLAATSLGLGAGMTPKIAWALENKPRIPVVWIHGL WP_001058312.1 hydrogenase [Salmonella enterica] Length: 372\nNumber of predicted TMHs: 2\nExp number of AAs in TMHs: 36.51738\nExp number, first 60 AAs: 19.72849\nTotal prob of N-in: 0.96170\nPOSSIBLE N-term signal sequence\ninside 1 19\nTMhelix 20 39\noutside 40 325\nTMhelix 326 348\ninside 349 372

14088 GCF_002060465.1_ASM206046v1 Salmonella enterica Proteobacteria; Gammaproteobacteria; Enterobacterales; Enterobacteriaceae; Salmonella MTGDNTLITSHGINRRDFMKLCAALAATMGLSSKAAAEMAESVSNPQRPPVIWIGAQECT WP_000145428.1 hydrogenase 2 small subunit [Salmonella enterica] Length: 372\nNumber of predicted TMHs: 1\nExp number of AAs in TMHs: 24.50677\nExp number, first 60 AAs: 0.2682\nTotal prob of N-in: 0.14390\noutside 1 331\nTMhelix 332 354\ninside 355 372

14089 GCF_002060435.1_ASM206043v1 Salmonella enterica Proteobacteria; Gammaproteobacteria; Enterobacterales; Enterobacteriaceae; Salmonella MTGDNTLITSHGINRRDFMKLCAALAATMGLSSKAAAEMAESVSNPQRPPVIWIGAQECT WP_000145429.1 hydrogenase 2 small subunit [Salmonella enterica] Length: 372\nNumber of predicted TMHs: 1\nExp number of AAs in TMHs: 24.51309\nExp number, first 60 AAs: 0.2682\nTotal prob of N-in: 0.14402\noutside 1 331\nTMhelix 332 354\ninside 355 372

14090 GCF_001952495.1_ASM195249v1 Salmonella enterica Proteobacteria; Gammaproteobacteria; Enterobacterales; Enterobacteriaceae; Salmonella
MNNEETFYQAMRRKGVTRRSFLKFCSLAATSLGLGAGMTPKIAWALENKPRIPVVWIHGL WP_001058312.1
hydrogenase [Salmonella enterica] Length: 372\nNumber of predicted TMHs: 2\nExp number of AAs in TMHs: 36.51738\nExp number, first 60 AAs: 19.72849\nTotal prob of N-in: 0.96170\nPOSSIBLE N-term signal sequence\ninside 1 19\nTMhelix 20 39\noutside 40 325\nTMhelix 326 348\ninside 349 372

14091 GCF_001952415.1_ASM195241v1 Salmonella enterica Proteobacteria; Gammaproteobacteria; Enterobacterales; Enterobacteriaceae; Salmonella
MTGDNTLITSHGINRRDFMKLCAALAATMGLSSKAAAEMAESVSNPQRPPVIWIGAQECT WP_000145429.1
hydrogenase 2 small subunit [Salmonella enterica] Length: 372\nNumber of predicted TMHs: 1\nExp number of AAs in TMHs: 24.51309\nExp number, first 60 AAs: 0.2682\nTotal prob of N-in: 0.14402\noutside 1 331\nTMhelix 332 354\ninside 355 372

14092 GCF_001643915.1_CRJJGF_00159_v1.0Salmonella enterica Proteobacteria; Gammaproteobacteria; Enterobacterales; Enterobacteriaceae; Salmonella
MNNEETFYQAMRRKGVTRRSFLKFCSLAATSLGLGAGMTPKIAWALENKPRIPVVWIHGL WP_001058311.1
hydrogenase [Salmonella enterica] Length: 372\nNumber of predicted TMHs: 2\nExp number of AAs in TMHs: 37.4699899999999\nExp number, first 60 AAs: 19.72699\nTotal prob of N-in: 0.96171\nPOSSIBLE N-term signal sequence\ninside 1 19\nTMhelix 20 39\noutside 40 325\nTMhelix 326 348\ninside 349 372

14093 GCF_001642525.1_ASM164252v1 Salmonella enterica Proteobacteria; Gammaproteobacteria; Enterobacterales; Enterobacteriaceae; Salmonella
MNNEETFYQAMRRKGVTRRSFLKFCSLAATSLGLGAGMTPKIAWALENKPRIPVVWIHGL WP_023233644.1
hydrogenase 1, small subunit [Salmonella enterica] Length: 372\nNumber of predicted TMHs: 2\nExp number of AAs in TMHs: 37.4208299999999\nExp number, first 60 AAs: 19.72732\nTotal prob of N-in: 0.96168\nPOSSIBLE N-term signal sequence\ninside 1 19\nTMhelix 20 39\noutside 40 325\nTMhelix 326 348\ninside 349 372

14094 GCF_001952385.1_ASM195238v1 Salmonella enterica Proteobacteria; Gammaproteobacteria; Enterobacterales; Enterobacteriaceae; Salmonella
MTGDNTLITSHGINRRDFMKLCAALAATMGLSSKAAAEMAESVSNPQRPPVIWIGAQECT WP_000145429.1
hydrogenase 2 small subunit [Salmonella enterica] Length: 372\nNumber of predicted TMHs: 1\nExp number of AAs in TMHs: 24.51309\nExp number, first 60 AAs: 0.2682\nTotal prob of N-in: 0.14402\noutside 1 331\nTMhelix 332 354\ninside 355 372

14095 GCF_001643895.1_CRJJGF_00147_v1.0Salmonella enterica Proteobacteria; Gammaproteobacteria; Enterobacterales; Enterobacteriaceae; Salmonella
MNNEETFYQAMRRKGVTRRSFLKFCSLAATSLGLGAGMTPKIAWALENKPRIPVVWIHGL WP_001058310.1
hydrogenase [Salmonella enterica] Length: 372\nNumber of predicted TMHs: 2\nExp number of AAs in TMHs: 37.4567299999999\nExp number, first 60 AAs: 19.72694\nTotal prob of N-in: 0.96171\nPOSSIBLE N-term signal sequence\ninside 1 19\nTMhelix 20 39\noutside 40 325\nTMhelix 326 348\ninside 349 372

14096 GCF_001641105.1_ASM164110v1 Salmonella enterica Proteobacteria; Gammaproteobacteria; Enterobacterales; Enterobacteriaceae; Salmonella
MNNEETLYQAMRRKGVTRRSFLKFCSLAATSLGLGAGMTPKIAWALENKPRIPVVWIHGL WP_001752445.1
hydrogenase-1 small chain [Salmonella enterica] Length: 372\nNumber of predicted TMHs: 2\nExp number of AAs in TMHs: 36.5123\nExp number, first 60 AAs: 19.72341\nTotal prob of N-in: 0.96144\nPOSSIBLE N-term signal sequence\ninside 1 19\nTMhelix 20 39\noutside 40 325\nTMhelix 326 348\ninside 349 372

14097 GCF_001691895.1_ASM169189v1 Salmonella enterica Proteobacteria; Gammaproteobacteria; Enterobacterales; Enterobacteriaceae; Salmonella
MNNEETFYQAMRRKGVTRRSFLKFCSLAATSLGLGAGMTPKIAWALENKPRIPVVWIHGL WP_001058312.1
hydrogenase [Salmonella enterica] Length: 372\nNumber of predicted TMHs: 2\nExp number of AAs in TMHs: 36.51738\nExp number, first 60 AAs: 19.72849\nTotal prob of N-in: 0.96170\nPOSSIBLE N-term signal sequence\ninside 1 19\nTMhelix 20 39\noutside 40 325\nTMhelix 326 348\ninside 349 372

14098 GCF_002060415.1_ASM206041v1 Salmonella enterica Proteobacteria; Gammaproteobacteria; Enterobacterales; Enterobacteriaceae; Salmonella
MTGDNTLITSHGINRRDFMKLCAALAATMGLSSKAAAEMAESVSNPQRPPVIWIGAQECT WP_000145429.1
hydrogenase 2 small subunit [Salmonella enterica] Length: 372\nNumber of predicted TMHs: 1\nExp

number of AAs in TMHs: 24.51309\nExp number, first 60 AAs: 0.2682\nTotal prob of N-in: 0.14402\noutside 1 331\nTMhelix 332 354\ninside 355 372

14099 GCF_002060385.1_ASM206038v1 Salmonella enterica Proteobacteria; Gammaproteobacteria; Enterobacterales; Enterobacteriaceae; Salmonella
MTGDNTLITSHGINRRDFMKLCAALAATMGLSSKAAAEMAESVSNPQRPPVIWIGAQECT WP_000145429.1
hydrogenase 2 small subunit [Salmonella enterica] Length: 372\nNumber of predicted TMHs: 1\nExp number of AAs in TMHs: 24.51309\nExp number, first 60 AAs: 0.2682\nTotal prob of N-in: 0.14402\noutside 1 331\nTMhelix 332 354\ninside 355 372

14100 GCF_001716375.1_ASM171637v1 Salmonella enterica Proteobacteria; Gammaproteobacteria; Enterobacterales; Enterobacteriaceae; Salmonella
MTGDNTLITSHGINRRDFMKLCAALAATMGLSSKAAAEMAESVSNPQRPPVIWIGAQECT WP_000145429.1
hydrogenase 2 small subunit [Salmonella enterica] Length: 372\nNumber of predicted TMHs: 1\nExp number of AAs in TMHs: 24.51309\nExp number, first 60 AAs: 0.2682\nTotal prob of N-in: 0.14402\noutside 1 331\nTMhelix 332 354\ninside 355 372

14101 GCF_001716725.1_ASM171672v1 Salmonella enterica Proteobacteria; Gammaproteobacteria; Enterobacterales; Enterobacteriaceae; Salmonella
MTGDNTLITSHGINRRDFMKLCAALAATMGLSSKAAAEMAESVSNPQRPPVIWIGAQECT WP_000145429.1
hydrogenase 2 small subunit [Salmonella enterica] Length: 372\nNumber of predicted TMHs: 1\nExp number of AAs in TMHs: 24.51309\nExp number, first 60 AAs: 0.2682\nTotal prob of N-in: 0.14402\noutside 1 331\nTMhelix 332 354\ninside 355 372

14102 GCF_002060125.1_ASM206012v1 Salmonella enterica Proteobacteria; Gammaproteobacteria; Enterobacterales; Enterobacteriaceae; Salmonella
MNNEETFYQAMRRKGVTRRSFLKFCSLAATSLGLGAGMTPKIAWALENKPRIPVVWIHGL WP_001058312.1
hydrogenase [Salmonella enterica] Length: 372\nNumber of predicted TMHs: 2\nExp number of AAs in TMHs: 36.51738\nExp number, first 60 AAs: 19.72849\nTotal prob of N-in: 0.96170\nPOSSIBLE N-term signal sequence\ninside 1 19\nTMhelix 20 39\noutside 40 325\nTMhelix 326 348\ninside 349 372

14103 GCF_002060125.1_ASM206012v1 Salmonella enterica Proteobacteria; Gammaproteobacteria; Enterobacterales; Enterobacteriaceae; Salmonella
MTGDNTLITSHGINRRDFMKLCAALAATMGLSSKAAAEMAESVSNPQRPPVIWIGAQECT WP_000145429.1
hydrogenase 2 small subunit [Salmonella enterica] Length: 372\nNumber of predicted TMHs: 1\nExp number of AAs in TMHs: 24.51309\nExp number, first 60 AAs: 0.2682\nTotal prob of N-in: 0.14402\noutside 1 331\nTMhelix 332 354\ninside 355 372

14104 GCF_001831555.1_ASM183155v1 Salmonella enterica Proteobacteria; Gammaproteobacteria; Enterobacterales; Enterobacteriaceae; Salmonella
MNNEETFYQAMRRKGVTRRSFLKFCSLAATSLGLGAGMTPKIAWALENKPRIPVVWIHGL WP_001058306.1
hydrogenase [Salmonella enterica] Length: 372\nNumber of predicted TMHs: 2\nExp number of AAs in TMHs: 36.5540099999999\nExp number, first 60 AAs: 19.72732\nTotal prob of N-in: 0.96171\nPOSSIBLE N-term signal sequence\ninside 1 19\nTMhelix 20 39\noutside 40 325\nTMhelix 326 348\ninside 349 372

14105 GCF_001831625.1_ASM183162v1 Salmonella enterica Proteobacteria; Gammaproteobacteria; Enterobacterales; Enterobacteriaceae; Salmonella
MNNEETFYQAMRRKGVTRRSFLKFCSLAATSLGLGAGMTPKIAWALENKPRIPVVWIHGL WP_001058306.1
hydrogenase [Salmonella enterica] Length: 372\nNumber of predicted TMHs: 2\nExp number of AAs in TMHs: 36.5540099999999\nExp number, first 60 AAs: 19.72732\nTotal prob of N-in: 0.96171\nPOSSIBLE N-term signal sequence\ninside 1 19\nTMhelix 20 39\noutside 40 325\nTMhelix 326 348\ninside 349 372

14106 GCF_001831565.1_ASM183156v1 Salmonella enterica Proteobacteria; Gammaproteobacteria; Enterobacterales; Enterobacteriaceae; Salmonella
MTGDNTLITSHGINRRDFMKLCAALAATMGLSSKAAAEMAESVSNPQRPPVIWIGAQECT WP_000145429.1
hydrogenase 2 small subunit [Salmonella enterica] Length: 372\nNumber of predicted TMHs: 1\nExp number of AAs in TMHs: 24.51309\nExp number, first 60 AAs: 0.2682\nTotal prob of N-in: 0.14402\noutside 1 331\nTMhelix 332 354\ninside 355 372

14107 GCF_001831835.1_ASM183183v1 Salmonella enterica Proteobacteria; Gammaproteobacteria; Enterobacterales; Enterobacteriaceae; Salmonella

MTGDNTLITSHGINRRDFMKLCAALAATMGLSSKAAAEMAESVSNPQRPPVIWIGAQECT WP_000145429.1 hydrogenase 2 small subunit [Salmonella enterica] Length: 372\nNumber of predicted TMHs: 1\nExp number of AAs in TMHs: 24.51309\nExp number, first 60 AAs: 0.2682\nTotal prob of N-in: 0.14402\noutside 1 331\nTMhelix 332 354\ninside 355 372

14108 GCF_001831805.1_ASM183180v1 Salmonella enterica Proteobacteria; Gammaproteobacteria; Enterobacterales; Enterobacteriaceae; Salmonella MNNEETFYQAMRRKGVTRRSFLKFCSLAATSLGLGAGMTPKIAWALENKPRIPVVWIHGL WP_001058312.1 hydrogenase [Salmonella enterica] Length: 372\nNumber of predicted TMHs: 2\nExp number of AAs in TMHs: 36.51738\nExp number, first 60 AAs: 19.72849\nTotal prob of N-in: 0.96170\nPOSSIBLE N-term signal sequence\ninside 1 19\nTMhelix 20 39\noutside 40 325\nTMhelix 326 348\ninside 349 372

14109 GCF_001831945.1_ASM183194v1 Salmonella enterica Proteobacteria; Gammaproteobacteria; Enterobacterales; Enterobacteriaceae; Salmonella MTGDNTLITSHGINRRDFMKLCAALAATMGLSSKAAAEMAESVSNPQRPPVIWIGAQECT WP_000145429.1 hydrogenase 2 small subunit [Salmonella enterica] Length: 372\nNumber of predicted TMHs: 1\nExp number of AAs in TMHs: 24.51309\nExp number, first 60 AAs: 0.2682\nTotal prob of N-in: 0.14402\noutside 1 331\nTMhelix 332 354\ninside 355 372

14110 GCF_001831915.1_ASM183191v1 Salmonella enterica Proteobacteria; Gammaproteobacteria; Enterobacterales; Enterobacteriaceae; Salmonella MTGDNTLITSHGINRRDFMKLCAALAATMGLSSKAAAEMAESVSNPQRPPVIWIGAQECT WP_000145429.1 hydrogenase 2 small subunit [Salmonella enterica] Length: 372\nNumber of predicted TMHs: 1\nExp number of AAs in TMHs: 24.51309\nExp number, first 60 AAs: 0.2682\nTotal prob of N-in: 0.14402\noutside 1 331\nTMhelix 332 354\ninside 355 372

14111 GCF_001831985.1_ASM183198v1 Salmonella enterica Proteobacteria; Gammaproteobacteria; Enterobacterales; Enterobacteriaceae; Salmonella MTGDNTLITSHGINRRDFMKLCAALAATMGLSSKAAAEMAESVSNPQRPPVIWIGAQECT WP_000145429.1 hydrogenase 2 small subunit [Salmonella enterica] Length: 372\nNumber of predicted TMHs: 1\nExp number of AAs in TMHs: 24.51309\nExp number, first 60 AAs: 0.2682\nTotal prob of N-in: 0.14402\noutside 1 331\nTMhelix 332 354\ninside 355 372

14112 GCF_001831835.1_ASM183183v1 Salmonella enterica Proteobacteria; Gammaproteobacteria; Enterobacterales; Enterobacteriaceae; Salmonella MNNEETFYQAMRRKGVTRRSFLKFCSLAATSLGLGAGMTPKIAWALENKPRIPVVWIHGL WP_001058312.1 hydrogenase [Salmonella enterica] Length: 372\nNumber of predicted TMHs: 2\nExp number of AAs in TMHs: 36.51738\nExp number, first 60 AAs: 19.72849\nTotal prob of N-in: 0.96170\nPOSSIBLE N-term signal sequence\ninside 1 19\nTMhelix 20 39\noutside 40 325\nTMhelix 326 348\ninside 349 372

14113 GCF_001832025.1_ASM183202v1 Salmonella enterica Proteobacteria; Gammaproteobacteria; Enterobacterales; Enterobacteriaceae; Salmonella MTGDNTLITSHGINRRDFMKLCAALAATMGLSSKAAAEMAESVSNPQRPPVIWIGAQECT WP_000145429.1 hydrogenase 2 small subunit [Salmonella enterica] Length: 372\nNumber of predicted TMHs: 1\nExp number of AAs in TMHs: 24.51309\nExp number, first 60 AAs: 0.2682\nTotal prob of N-in: 0.14402\noutside 1 331\nTMhelix 332 354\ninside 355 372

14114 GCF_001832095.1_ASM183209v1 Salmonella enterica Proteobacteria; Gammaproteobacteria; Enterobacterales; Enterobacteriaceae; Salmonella MTGDNTLITSHGINRRDFMKLCAALAATMGLSSKAAAEMAESVSNPQRPPVIWIGAQECT WP_000145429.1 hydrogenase 2 small subunit [Salmonella enterica] Length: 372\nNumber of predicted TMHs: 1\nExp number of AAs in TMHs: 24.51309\nExp number, first 60 AAs: 0.2682\nTotal prob of N-in: 0.14402\noutside 1 331\nTMhelix 332 354\ninside 355 372

14115 GCF_001832145.1_ASM183214v1 Salmonella enterica Proteobacteria; Gammaproteobacteria; Enterobacterales; Enterobacteriaceae; Salmonella MNNEETFYQAMRRKGVTRRSFLKFCSLAATSLGLGAGMTPKIAWALENKPRIPVVWIHGL WP_001058312.1 hydrogenase [Salmonella enterica] Length: 372\nNumber of predicted TMHs: 2\nExp number of AAs in TMHs: 36.51738\nExp number, first 60 AAs: 19.72849\nTotal prob of N-in: 0.96170\nPOSSIBLE N-term signal sequence\ninside 1 19\nTMhelix 20 39\noutside 40 325\nTMhelix 326 348\ninside 349 372

14116 GCF_001832205.1_ASM183220v1 Salmonella enterica Proteobacteria; Gammaproteobacteria; Enterobacterales; Enterobacteriaceae; Salmonella
MTGDNTLITSHGINRRDFMKLCAALAATMGLSSKAAAEMAESVSNPQRPPVIWIGAQECT WP_000145429.1
hydrogenase 2 small subunit [Salmonella enterica] Length: 372\nNumber of predicted TMHs: 1\nExp number of AAs in TMHs: 24.51309\nExp number, first 60 AAs: 0.2682\nTotal prob of N-in: 0.14402\noutside 1 331\nTMhelix 332 354\ninside 355 372

14117 GCF_001832205.1_ASM183220v1 Salmonella enterica Proteobacteria; Gammaproteobacteria; Enterobacterales; Enterobacteriaceae; Salmonella
MNNEETFYQAMRRKGVTRRSFLKFCSLAATSLGLGAGMTPKIAWALENKPRIPVVWIHGL WP_001058312.1
hydrogenase [Salmonella enterica] Length: 372\nNumber of predicted TMHs: 2\nExp number of AAs in TMHs: 36.51738\nExp number, first 60 AAs: 19.72849\nTotal prob of N-in: 0.96170\nPOSSIBLE N-term signal sequence\ninside 1 19\nTMhelix 20 39\noutside 40 325\nTMhelix 326 348\ninside 349 372

14118 GCF_001832675.1_ASM183267v1 Salmonella enterica Proteobacteria; Gammaproteobacteria; Enterobacterales; Enterobacteriaceae; Salmonella
MTGDNTLITSHGINRRDFMKLCAALAATMGLSSKAAAEMAESVSNPQRPPVIWIGAQECT WP_000145429.1
hydrogenase 2 small subunit [Salmonella enterica] Length: 372\nNumber of predicted TMHs: 1\nExp number of AAs in TMHs: 24.51309\nExp number, first 60 AAs: 0.2682\nTotal prob of N-in: 0.14402\noutside 1 331\nTMhelix 332 354\ninside 355 372

14119 GCF_001832675.1_ASM183267v1 Salmonella enterica Proteobacteria; Gammaproteobacteria; Enterobacterales; Enterobacteriaceae; Salmonella
MNNEETFYQAMRRKGVTRRSFLKFCSLAATSLGLGAGMTPKIAWALENKPRIPVVWIHGL WP_001058312.1
hydrogenase [Salmonella enterica] Length: 372\nNumber of predicted TMHs: 2\nExp number of AAs in TMHs: 36.51738\nExp number, first 60 AAs: 19.72849\nTotal prob of N-in: 0.96170\nPOSSIBLE N-term signal sequence\ninside 1 19\nTMhelix 20 39\noutside 40 325\nTMhelix 326 348\ninside 349 372

14120 GCF_001833095.1_ASM183309v1 Salmonella enterica Proteobacteria; Gammaproteobacteria; Enterobacterales; Enterobacteriaceae; Salmonella
MNNEETFYQAMRRKGVTRRSFLKFCSLAATSLGLGAGMTPKIAWALENKPRIPVVWIHGL WP_001058311.1
hydrogenase [Salmonella enterica] Length: 372\nNumber of predicted TMHs: 2\nExp number of AAs in TMHs: 37.4699899999999\nExp number, first 60 AAs: 19.72699\nTotal prob of N-in: 0.96171\nPOSSIBLE N-term signal sequence\ninside 1 19\nTMhelix 20 39\noutside 40 325\nTMhelix 326 348\ninside 349 372

14121 GCF_001833165.1_ASM183316v1 Salmonella enterica Proteobacteria; Gammaproteobacteria; Enterobacterales; Enterobacteriaceae; Salmonella
MNNEETFYQAMRRKGVTRRSFLKFCSLAATSLGLGAGMTPKIAWALENKPRIPVVWIHGL WP_001058312.1
hydrogenase [Salmonella enterica] Length: 372\nNumber of predicted TMHs: 2\nExp number of AAs in TMHs: 36.51738\nExp number, first 60 AAs: 19.72849\nTotal prob of N-in: 0.96170\nPOSSIBLE N-term signal sequence\ninside 1 19\nTMhelix 20 39\noutside 40 325\nTMhelix 326 348\ninside 349 372

14122 GCF_001833585.1_ASM183358v1 Salmonella enterica Proteobacteria; Gammaproteobacteria; Enterobacterales; Enterobacteriaceae; Salmonella
MTGDNTLITSHGINRRDFMKLCAALAATMGLSSKAAAEMAESVSNPQRPPVIWIGAQECT WP_000145429.1
hydrogenase 2 small subunit [Salmonella enterica] Length: 372\nNumber of predicted TMHs: 1\nExp number of AAs in TMHs: 24.51309\nExp number, first 60 AAs: 0.2682\nTotal prob of N-in: 0.14402\noutside 1 331\nTMhelix 332 354\ninside 355 372

14123 GCF_001833565.1_ASM183356v1 Salmonella enterica Proteobacteria; Gammaproteobacteria; Enterobacterales; Enterobacteriaceae; Salmonella
MNNEETFYQAMRRKGVTRRSFLKFCSLAATSLGLGAGMTPKIAWALENKPRIPVVWIHGL WP_001058312.1
hydrogenase [Salmonella enterica] Length: 372\nNumber of predicted TMHs: 2\nExp number of AAs in TMHs: 36.51738\nExp number, first 60 AAs: 19.72849\nTotal prob of N-in: 0.96170\nPOSSIBLE N-term signal sequence\ninside 1 19\nTMhelix 20 39\noutside 40 325\nTMhelix 326 348\ninside 349 372

14124 GCF_001833585.1_ASM183358v1 Salmonella enterica Proteobacteria; Gammaproteobacteria; Enterobacterales; Enterobacteriaceae; Salmonella
MNNEETFYQAMRRKGVTRRSFLKFCSLAATSLGLGAGMTPKIAWALENKPRIPVVWIHGL WP_001058311.1
hydrogenase [Salmonella enterica] Length: 372\nNumber of predicted TMHs: 2\nExp number of AAs in

TMHs: 37.4699899999999\nExp number, first 60 AAs: 19.72699\nTotal prob of N-in: 0.96171\nPOSSIBLE N-term signal sequence\ninside 1 19\nTMhelix 20 39\noutside 40 325\nTMhelix 326 348\ninside 349 372

14125 GCF_001833495.1_ASM183349v1 Salmonella enterica Proteobacteria; Gammaproteobacteria; Enterobacterales; Enterobacteriaceae; Salmonella
MNNEETFYQAMRRKGVTRRSFLKFCSLAATSLGLGAGMTPKIAWALENKPRIPVVWIHGL WP_001058311.1
hydrogenase [Salmonella enterica] Length: 372\nNumber of predicted TMHs: 2\nExp number of AAs in TMHs: 37.4699899999999\nExp number, first 60 AAs: 19.72699\nTotal prob of N-in: 0.96171\nPOSSIBLE N-term signal sequence\ninside 1 19\nTMhelix 20 39\noutside 40 325\nTMhelix 326 348\ninside 349 372

14126 GCF_001833625.1_ASM183362v1 Salmonella enterica Proteobacteria; Gammaproteobacteria; Enterobacterales; Enterobacteriaceae; Salmonella
MTGDNTLITSHGINRRDFMKLCAALAATMGLSSKAAAEMAESVSNPQRPPVIWIGAQECT WP_000145429.1
hydrogenase 2 small subunit [Salmonella enterica] Length: 372\nNumber of predicted TMHs: 1\nExp number of AAs in TMHs: 24.51309\nExp number, first 60 AAs: 0.2682\nTotal prob of N-in: 0.14402\noutside 1 331\nTMhelix 332 354\ninside 355 372

14127 GCF_001833645.1_ASM183364v1 Salmonella enterica Proteobacteria; Gammaproteobacteria; Enterobacterales; Enterobacteriaceae; Salmonella
MTGDNTLITSHGINRRDFMKLCAALAATMGLSSKAAAEMAESVSNPQRPPVIWIGAQECT WP_000145429.1
hydrogenase 2 small subunit [Salmonella enterica] Length: 372\nNumber of predicted TMHs: 1\nExp number of AAs in TMHs: 24.51309\nExp number, first 60 AAs: 0.2682\nTotal prob of N-in: 0.14402\noutside 1 331\nTMhelix 332 354\ninside 355 372

14128 GCF_002056285.1_ASM205628v1 Salmonella enterica Proteobacteria; Gammaproteobacteria; Enterobacterales; Enterobacteriaceae; Salmonella
MTGDNTLITSHGINRRDFMKLCAALAATMGLSSKAAAEMAESVSNPQRPPVIWIGAQECT WP_000145429.1
hydrogenase 2 small subunit [Salmonella enterica] Length: 372\nNumber of predicted TMHs: 1\nExp number of AAs in TMHs: 24.51309\nExp number, first 60 AAs: 0.2682\nTotal prob of N-in: 0.14402\noutside 1 331\nTMhelix 332 354\ninside 355 372

14129 GCF_002056695.1_ASM205669v1 Salmonella enterica Proteobacteria; Gammaproteobacteria; Enterobacterales; Enterobacteriaceae; Salmonella
MNNEETFYQAMRRKGVTRRSFLKFCSLAATSLGLGAGMTPKIAWALENKPRIPVVWIHGL WP_001058312.1
hydrogenase [Salmonella enterica] Length: 372\nNumber of predicted TMHs: 2\nExp number of AAs in TMHs: 36.51738\nExp number, first 60 AAs: 19.72849\nTotal prob of N-in: 0.96170\nPOSSIBLE N-term signal sequence\ninside 1 19\nTMhelix 20 39\noutside 40 325\nTMhelix 326 348\ninside 349 372

14130 GCF_002056585.1_ASM205658v1 Salmonella enterica Proteobacteria; Gammaproteobacteria; Enterobacterales; Enterobacteriaceae; Salmonella
MNNEETFYQAMRRKGVTRRSFLKFCSLAATSLGLGAGMTPKIAWALENKPRIPVVWIHGL WP_001058311.1
hydrogenase [Salmonella enterica] Length: 372\nNumber of predicted TMHs: 2\nExp number of AAs in TMHs: 37.4699899999999\nExp number, first 60 AAs: 19.72699\nTotal prob of N-in: 0.96171\nPOSSIBLE N-term signal sequence\ninside 1 19\nTMhelix 20 39\noutside 40 325\nTMhelix 326 348\ninside 349 372

14131 GCF_002056285.1_ASM205628v1 Salmonella enterica Proteobacteria; Gammaproteobacteria; Enterobacterales; Enterobacteriaceae; Salmonella
MNNEETFYQAMRRKGVTRRSFLKFCSLAATSLGLGAGMTPKIAWALENKPRIPVVWIHGL WP_001058307.1
hydrogenase [Salmonella enterica] Length: 372\nNumber of predicted TMHs: 2\nExp number of AAs in TMHs: 36.51738\nExp number, first 60 AAs: 19.72849\nTotal prob of N-in: 0.96170\nPOSSIBLE N-term signal sequence\ninside 1 19\nTMhelix 20 39\noutside 40 325\nTMhelix 326 348\ninside 349 372

14132 GCF_002056345.1_ASM205634v1 Salmonella enterica Proteobacteria; Gammaproteobacteria; Enterobacterales; Enterobacteriaceae; Salmonella
MNNEETFYQAMRRKGVTRRSFLKFCSLAATSLGLGAGMTPKIAWALENKPRIPVVWIHGL WP_001058307.1
hydrogenase [Salmonella enterica] Length: 372\nNumber of predicted TMHs: 2\nExp number of AAs in TMHs: 36.51738\nExp number, first 60 AAs: 19.72849\nTotal prob of N-in: 0.96170\nPOSSIBLE N-term signal sequence\ninside 1 19\nTMhelix 20 39\noutside 40 325\nTMhelix 326 348\ninside 349 372

14133 GCF_002057675.1_ASM205767v1 Salmonella enterica Proteobacteria; Gammaproteobacteria; Enterobacterales; Enterobacteriaceae; Salmonella

MTGDNTLITSHGINRRDFMKLCAALAATMGLSSKAAAEMAESVSNPQRPPVIWIGAQECT WP_000145429.1 hydrogenase 2 small subunit [Salmonella enterica] Length: 372\nNumber of predicted TMHs: 1\nExp number of AAs in TMHs: 24.51309\nExp number, first 60 AAs: 0.2682\nTotal prob of N-in: 0.14402\noutside 1 331\nTMhelix 332 354\ninside 355 372

14134 GCF_002057855.1_ASM205785v1 Salmonella enterica Proteobacteria; Gammaproteobacteria; Enterobacterales; Enterobacteriaceae; Salmonella MTGDNTLITSHGINRRDFMKLCAALAATMGLSSKAAAEMAESVSNPQRPPVIWIGAQECT WP_000145428.1 hydrogenase 2 small subunit [Salmonella enterica] Length: 372\nNumber of predicted TMHs: 1\nExp number of AAs in TMHs: 24.50677\nExp number, first 60 AAs: 0.2682\nTotal prob of N-in: 0.14390\noutside 1 331\nTMhelix 332 354\ninside 355 372

14135 GCF_002056375.1_ASM205637v1 Salmonella enterica Proteobacteria; Gammaproteobacteria; Enterobacterales; Enterobacteriaceae; Salmonella MNNEETFYQAMRRKGVTRRSFLKFCSLAATSLGLGAGMTPKIAWALENKPRIPVVWIHGL WP_001058307.1 hydrogenase [Salmonella enterica] Length: 372\nNumber of predicted TMHs: 2\nExp number of AAs in TMHs: 36.51738\nExp number, first 60 AAs: 19.72849\nTotal prob of N-in: 0.96170\nPOSSIBLE N-term signal sequence\ninside 1 19\nTMhelix 20 39\noutside 40 325\nTMhelix 326 348\ninside 349 372

14136 GCF_002057005.1_ASM205700v1 Salmonella enterica Proteobacteria; Gammaproteobacteria; Enterobacterales; Enterobacteriaceae; Salmonella MTGDNTLITSHGINRRDFMKLCAALAATMGLSSKAAAEMAESVSNPQRPPVIWIGAQECT WP_000145429.1 hydrogenase 2 small subunit [Salmonella enterica] Length: 372\nNumber of predicted TMHs: 1\nExp number of AAs in TMHs: 24.51309\nExp number, first 60 AAs: 0.2682\nTotal prob of N-in: 0.14402\noutside 1 331\nTMhelix 332 354\ninside 355 372

14137 GCF_002054225.1_ASM205422v1 Salmonella enterica Proteobacteria; Gammaproteobacteria; Enterobacterales; Enterobacteriaceae; Salmonella MNNEETFYQAMRRKGVTRRSFLKFCSLAATSLGLGAGMTPKIAWALENKPRIPVVWIHGL WP_001058312.1 hydrogenase [Salmonella enterica] Length: 372\nNumber of predicted TMHs: 2\nExp number of AAs in TMHs: 36.51738\nExp number, first 60 AAs: 19.72849\nTotal prob of N-in: 0.96170\nPOSSIBLE N-term signal sequence\ninside 1 19\nTMhelix 20 39\noutside 40 325\nTMhelix 326 348\ninside 349 372

14138 GCF_002057155.1_ASM205715v1 Salmonella enterica Proteobacteria; Gammaproteobacteria; Enterobacterales; Enterobacteriaceae; Salmonella MTGDNTLITSHGINRRDFMKLCAALAATMGLSSKAAAEMAESVSNPQRPPVIWIGAQECT WP_000145429.1 hydrogenase 2 small subunit [Salmonella enterica] Length: 372\nNumber of predicted TMHs: 1\nExp number of AAs in TMHs: 24.51309\nExp number, first 60 AAs: 0.2682\nTotal prob of N-in: 0.14402\noutside 1 331\nTMhelix 332 354\ninside 355 372

14139 GCF_002054295.1_ASM205429v1 Salmonella enterica Proteobacteria; Gammaproteobacteria; Enterobacterales; Enterobacteriaceae; Salmonella MTGDNTLITSHGINRRDFMKLCAALAATMGLSSKAAAEMAESVSNPQRPPVIWIGAQECT WP_000145429.1 hydrogenase 2 small subunit [Salmonella enterica] Length: 372\nNumber of predicted TMHs: 1\nExp number of AAs in TMHs: 24.51309\nExp number, first 60 AAs: 0.2682\nTotal prob of N-in: 0.14402\noutside 1 331\nTMhelix 332 354\ninside 355 372

14140 GCF_002054225.1_ASM205422v1 Salmonella enterica Proteobacteria; Gammaproteobacteria; Enterobacterales; Enterobacteriaceae; Salmonella MTGDNTLITSHGINRRDFMKLCAALAATMGLSSKAAAEMAESVSNPQRPPVIWIGAQECT WP_000145429.1 hydrogenase 2 small subunit [Salmonella enterica] Length: 372\nNumber of predicted TMHs: 1\nExp number of AAs in TMHs: 24.51309\nExp number, first 60 AAs: 0.2682\nTotal prob of N-in: 0.14402\noutside 1 331\nTMhelix 332 354\ninside 355 372

14141 GCF_002057205.1_ASM205720v1 Salmonella enterica Proteobacteria; Gammaproteobacteria; Enterobacterales; Enterobacteriaceae; Salmonella MTGDNTLITSHGINRRDFMKLCAALAATMGLSSKAAAEMAESVSNPQRPPVIWIGAQECT WP_000145429.1 hydrogenase 2 small subunit [Salmonella enterica] Length: 372\nNumber of predicted TMHs: 1\nExp number of AAs in TMHs: 24.51309\nExp number, first 60 AAs: 0.2682\nTotal prob of N-in: 0.14402\noutside 1 331\nTMhelix 332 354\ninside 355 372

14142 GCF_002057155.1_ASM205715v1 Salmonella enterica Proteobacteria; Gammaproteobacteria; Enterobacterales; Enterobacteriaceae; Salmonella
MNNEETFYQAMRRKGVTRRSFLKFCSLAATSLGLGAGMTPKIAWALENKPRIPVVWIHGL WP_001058312.1
hydrogenase [Salmonella enterica] Length: 372\nNumber of predicted TMHs: 2\nExp number of AAs in TMHs: 36.51738\nExp number, first 60 AAs: 19.72849\nTotal prob of N-in: 0.96170\nPOSSIBLE N-term signal sequence\ninside 1 19\nTMhelix 20 39\noutside 40 325\nTMhelix 326 348\ninside 349 372

14143 GCF_002057565.1_ASM205756v1 Salmonella enterica Proteobacteria; Gammaproteobacteria; Enterobacterales; Enterobacteriaceae; Salmonella
MTGDNTLITSHGINRRDFMKLCAALAATMGLSSKAAAEMAESVSNPQRPPVIWIGAQECT WP_000145429.1
hydrogenase 2 small subunit [Salmonella enterica] Length: 372\nNumber of predicted TMHs: 1\nExp number of AAs in TMHs: 24.51309\nExp number, first 60 AAs: 0.2682\nTotal prob of N-in: 0.14402\noutside 1 331\nTMhelix 332 354\ninside 355 372

14144 GCF_002057505.1_ASM205750v1 Salmonella enterica Proteobacteria; Gammaproteobacteria; Enterobacterales; Enterobacteriaceae; Salmonella
MTGDNTLITSHGINRRDFMKLCAALAATMGLSSKAAAEMAESVSNPQRPPVIWIGAQECT WP_000145429.1
hydrogenase 2 small subunit [Salmonella enterica] Length: 372\nNumber of predicted TMHs: 1\nExp number of AAs in TMHs: 24.51309\nExp number, first 60 AAs: 0.2682\nTotal prob of N-in: 0.14402\noutside 1 331\nTMhelix 332 354\ninside 355 372

14145 GCF_002057565.1_ASM205756v1 Salmonella enterica Proteobacteria; Gammaproteobacteria; Enterobacterales; Enterobacteriaceae; Salmonella
MNNEETFYQAMRRKGVTRRSFLKFCSLAATSLGLGAGMTPKIAWALENKPRIPVVWIHGL WP_001058312.1
hydrogenase [Salmonella enterica] Length: 372\nNumber of predicted TMHs: 2\nExp number of AAs in TMHs: 36.51738\nExp number, first 60 AAs: 19.72849\nTotal prob of N-in: 0.96170\nPOSSIBLE N-term signal sequence\ninside 1 19\nTMhelix 20 39\noutside 40 325\nTMhelix 326 348\ninside 349 372

14146 GCF_002057645.1_ASM205764v1 Salmonella enterica Proteobacteria; Gammaproteobacteria; Enterobacterales; Enterobacteriaceae; Salmonella
MTGDNTLITSHGINRRDFMKLCAALAATMGLSSKAAAEMAESVSNPQRPPVIWIGAQECT WP_000145429.1
hydrogenase 2 small subunit [Salmonella enterica] Length: 372\nNumber of predicted TMHs: 1\nExp number of AAs in TMHs: 24.51309\nExp number, first 60 AAs: 0.2682\nTotal prob of N-in: 0.14402\noutside 1 331\nTMhelix 332 354\ninside 355 372

14147 GCF_002053965.1_ASM205396v1 Salmonella enterica Proteobacteria; Gammaproteobacteria; Enterobacterales; Enterobacteriaceae; Salmonella
MNNEETFYQAMRRKGVTRRSFLKFCSLAATSLGLGAGMTPKIAWALENKPRIPVVWIHGL WP_001058312.1
hydrogenase [Salmonella enterica] Length: 372\nNumber of predicted TMHs: 2\nExp number of AAs in TMHs: 36.51738\nExp number, first 60 AAs: 19.72849\nTotal prob of N-in: 0.96170\nPOSSIBLE N-term signal sequence\ninside 1 19\nTMhelix 20 39\noutside 40 325\nTMhelix 326 348\ninside 349 372

14148 GCF_002054055.1_ASM205405v1 Salmonella enterica Proteobacteria; Gammaproteobacteria; Enterobacterales; Enterobacteriaceae; Salmonella
MNNEETFYQAMRRKGVTRRSFLKFCSLAATSLGLGAGMTPKIAWALENKPRIPVVWIHGL WP_001058310.1
hydrogenase [Salmonella enterica] Length: 372\nNumber of predicted TMHs: 2\nExp number of AAs in TMHs: 37.4567299999999\nExp number, first 60 AAs: 19.72694\nTotal prob of N-in: 0.96171\nPOSSIBLE N-term signal sequence\ninside 1 19\nTMhelix 20 39\noutside 40 325\nTMhelix 326 348\ninside 349 372

14149 GCF_001705055.1_ASM170505v1 Salmonella enterica Proteobacteria; Gammaproteobacteria; Enterobacterales; Enterobacteriaceae; Salmonella
MNNEETFYQAMRRKGVTRRSFLKFCSLAATSLGLGAGMTPKIAWALENKPRIPVVWIHGL WP_001058307.1
hydrogenase [Salmonella enterica] Length: 372\nNumber of predicted TMHs: 2\nExp number of AAs in TMHs: 36.51738\nExp number, first 60 AAs: 19.72849\nTotal prob of N-in: 0.96170\nPOSSIBLE N-term signal sequence\ninside 1 19\nTMhelix 20 39\noutside 40 325\nTMhelix 326 348\ninside 349 372

14150 GCF_002053895.1_ASM205389v1 Salmonella enterica Proteobacteria; Gammaproteobacteria; Enterobacterales; Enterobacteriaceae; Salmonella
MNNEETFYQAMRRKGVTRRSFLKFCSLAATSLGLGAGMTPKIAWALENKPRIPVVWIHGL WP_001058312.1
hydrogenase [Salmonella enterica] Length: 372\nNumber of predicted TMHs: 2\nExp number of AAs in TMHs:

TMHs: 36.51738\nExp number, first 60 AAs:  19.72849\nTotal prob of N-in:   0.96170\nPOSSIBLE N-term signal sequence\ninside   1   19\nTMhelix   20   39\noutside   40   325\nTMhelix   326   348\ninside   349   372

14151 GCF_002053575.1_ASM205357v1 Salmonella enterica Proteobacteria; Gammaproteobacteria; Enterobacterales; Enterobacteriaceae; Salmonella MNNEETFYQAMRRKGVTRRSFLKFCSLAATSLGLGAGMTPKIAWALENKPRIPVVWIHGL WP_001058312.1 hydrogenase [Salmonella enterica] Length: 372\nNumber of predicted TMHs:  2\nExp number of AAs in TMHs: 36.51738\nExp number, first 60 AAs:  19.72849\nTotal prob of N-in:   0.96170\nPOSSIBLE N-term signal sequence\ninside   1   19\nTMhelix   20   39\noutside   40   325\nTMhelix   326   348\ninside   349   372

14152 GCF_002043715.1_ASM204371v1 Salmonella enterica Proteobacteria; Gammaproteobacteria; Enterobacterales; Enterobacteriaceae; Salmonella MNNEETFYQAMRRKGVTRRSFLKFCSLAATSLGLGAGMTPKIAWALENKPRIPVVWIHGL WP_001058311.1 hydrogenase [Salmonella enterica] Length: 372\nNumber of predicted TMHs:  2\nExp number of AAs in TMHs: 37.4699899999999\nExp number, first 60 AAs:  19.72699\nTotal prob of N-in:   0.96171\nPOSSIBLE N-term signal sequence\ninside   1   19\nTMhelix   20   39\noutside   40   325\nTMhelix   326   348\ninside   349   372

14153 GCF_002054085.1_ASM205408v1 Salmonella enterica Proteobacteria; Gammaproteobacteria; Enterobacterales; Enterobacteriaceae; Salmonella MTGDNTLITSHGINRRDFMKLCAALAATMGLSSKAAAEMAESVSNPQRPPVIWIGAQECT WP_000145429.1 hydrogenase 2 small subunit [Salmonella enterica] Length: 372\nNumber of predicted TMHs:  1\nExp number of AAs in TMHs: 24.51309\nExp number, first 60 AAs:  0.2682\nTotal prob of N-in:   0.14402\noutside   1   331\nTMhelix   332   354\ninside   355   372

14154 GCF_002053965.1_ASM205396v1 Salmonella enterica Proteobacteria; Gammaproteobacteria; Enterobacterales; Enterobacteriaceae; Salmonella MTGDNTLITSHGINRRDFMKLCAALAATMGLSSKAAAEMAESVSNPQRPPVIWIGAQECT WP_000145429.1 hydrogenase 2 small subunit [Salmonella enterica] Length: 372\nNumber of predicted TMHs:  1\nExp number of AAs in TMHs: 24.51309\nExp number, first 60 AAs:  0.2682\nTotal prob of N-in:   0.14402\noutside   1   331\nTMhelix   332   354\ninside   355   372

14155 GCF_002053575.1_ASM205357v1 Salmonella enterica Proteobacteria; Gammaproteobacteria; Enterobacterales; Enterobacteriaceae; Salmonella MTGDNTLITSHGINRRDFMKLCAALAATMGLSSKAAAEMAESVSNPQRPPVIWIGAQECT WP_000145429.1 hydrogenase 2 small subunit [Salmonella enterica] Length: 372\nNumber of predicted TMHs:  1\nExp number of AAs in TMHs: 24.51309\nExp number, first 60 AAs:  0.2682\nTotal prob of N-in:   0.14402\noutside   1   331\nTMhelix   332   354\ninside   355   372

14156 GCF_002053695.1_ASM205369v1 Salmonella enterica Proteobacteria; Gammaproteobacteria; Enterobacterales; Enterobacteriaceae; Salmonella MTGDNTLITSHGINRRDFMKLCAALAATMGLSSKAAAEMAESVSNPQRPPVIWIGAQECT WP_000145429.1 hydrogenase 2 small subunit [Salmonella enterica] Length: 372\nNumber of predicted TMHs:  1\nExp number of AAs in TMHs: 24.51309\nExp number, first 60 AAs:  0.2682\nTotal prob of N-in:   0.14402\noutside   1   331\nTMhelix   332   354\ninside   355   372

14157 GCF_001716225.1_ASM171622v1 Salmonella enterica Proteobacteria; Gammaproteobacteria; Enterobacterales; Enterobacteriaceae; Salmonella MNNEETFYQAMRRKGVTRRSFLKFCSLAATSLGLGAGMTPKIAWALENKPRIPVVWIHGL WP_001058311.1 hydrogenase [Salmonella enterica] Length: 372\nNumber of predicted TMHs:  2\nExp number of AAs in TMHs: 37.4699899999999\nExp number, first 60 AAs:  19.72699\nTotal prob of N-in:   0.96171\nPOSSIBLE N-term signal sequence\ninside   1   19\nTMhelix   20   39\noutside   40   325\nTMhelix   326   348\ninside   349   372

14158 GCF_002053625.1_ASM205362v1 Salmonella enterica Proteobacteria; Gammaproteobacteria; Enterobacterales; Enterobacteriaceae; Salmonella MTGDNTLITSHGINRRDFMKLCAALAATMGLSSKAAAEMAESVSNPQRPPVIWIGAQECT WP_000145429.1 hydrogenase 2 small subunit [Salmonella enterica] Length: 372\nNumber of predicted TMHs:  1\nExp number of AAs in TMHs: 24.51309\nExp number, first 60 AAs:  0.2682\nTotal prob of N-in:   0.14402\noutside   1   331\nTMhelix   332   354\ninside   355   372

14159 GCF_001742445.1_ASM174244v1 Salmonella enterica Proteobacteria; Gammaproteobacteria; Enterobacterales; Enterobacteriaceae; Salmonella

MTGDNTLITSHGINRRDFMKLCAALAATMGLSSKAAAEMAESVSNPQRPPVIWIGAQECT WP_000145431.1 hydrogenase 2 small subunit [Salmonella enterica] Length: 372\nNumber of predicted TMHs: 1\nExp number of AAs in TMHs: 24.51505\nExp number, first 60 AAs: 0.2682\nTotal prob of N-in: 0.14406\noutside 1 331\nTMhelix 332 354\ninside 355 372

14160 GCF_002052965.1_ASM205296v1 Salmonella enterica Proteobacteria; Gammaproteobacteria; Enterobacterales; Enterobacteriaceae; Salmonella MNNEETFYQAMRRKGVTRRSFLKFCSLAATSLGLGAGMTPKIAWALENKPRIPVVWIHGL WP_080085110.1 [Ni/Fe] hydrogenase small subunit [Salmonella enterica] Length: 372\nNumber of predicted TMHs: 2\nExp number of AAs in TMHs: 37.4594399999999\nExp number, first 60 AAs: 19.72734\nTotal prob of N-in: 0.96171\nPOSSIBLE N-term signal sequence\ninside 1 19\nTMhelix 20 39\noutside 40 325\nTMhelix 326 348\ninside 349 372

14161 GCF_002043375.1_ASM204337v1 Salmonella enterica Proteobacteria; Gammaproteobacteria; Enterobacterales; Enterobacteriaceae; Salmonella MNNEETFYQAMRRKGVTRRSFLKFCSLAATSLGLGAGMTPKIAWALENKPRIPVVWIHGL WP_001058311.1 hydrogenase [Salmonella enterica] Length: 372\nNumber of predicted TMHs: 2\nExp number of AAs in TMHs: 37.4699899999999\nExp number, first 60 AAs: 19.72699\nTotal prob of N-in: 0.96171\nPOSSIBLE N-term signal sequence\ninside 1 19\nTMhelix 20 39\noutside 40 325\nTMhelix 326 348\ninside 349 372

14162 GCF_002043145.1_ASM204314v1 Salmonella enterica Proteobacteria; Gammaproteobacteria; Enterobacterales; Enterobacteriaceae; Salmonella MNNEETLYQAMRRKGVTRRSFLKFCSLAATSLGLGAGMTPKIAWALENKPRIPVVWIHGL WP_001752445.1 hydrogenase-1 small chain [Salmonella enterica] Length: 372\nNumber of predicted TMHs: 2\nExp number of AAs in TMHs: 36.5123\nExp number, first 60 AAs: 19.72341\nTotal prob of N-in: 0.96144\nPOSSIBLE N-term signal sequence\ninside 1 19\nTMhelix 20 39\noutside 40 325\nTMhelix 326 348\ninside 349 372

14163 GCF_001742445.1_ASM174244v1 Salmonella enterica Proteobacteria; Gammaproteobacteria; Enterobacterales; Enterobacteriaceae; Salmonella MNNEETFYQAMRRKGVTRRSFLKFCSLAATSLGLGAGMTPKIAWALENKPRIPVVWIHGL WP_001662439.1 hydrogenase [Salmonella enterica] Length: 372\nNumber of predicted TMHs: 2\nExp number of AAs in TMHs: 36.5172\nExp number, first 60 AAs: 19.72849\nTotal prob of N-in: 0.96169\nPOSSIBLE N-term signal sequence\ninside 1 19\nTMhelix 20 39\noutside 40 325\nTMhelix 326 348\ninside 349 372

14164 GCF_002053495.1_ASM205349v1 Salmonella enterica Proteobacteria; Gammaproteobacteria; Enterobacterales; Enterobacteriaceae; Salmonella MNNEETFYQAMRRKGVTRRSFLKFCSLAATSLGLGAGMTPKIAWALENKPRIPVVWIHGL WP_001058312.1 hydrogenase [Salmonella enterica] Length: 372\nNumber of predicted TMHs: 2\nExp number of AAs in TMHs: 36.51738\nExp number, first 60 AAs: 19.72849\nTotal prob of N-in: 0.96170\nPOSSIBLE N-term signal sequence\ninside 1 19\nTMhelix 20 39\noutside 40 325\nTMhelix 326 348\ninside 349 372

14165 GCF_001831545.1_ASM183154v1 Salmonella enterica Proteobacteria; Gammaproteobacteria; Enterobacterales; Enterobacteriaceae; Salmonella MTGDNTLITSHGINRRDFMKLCAALAATMGLSSKAAAEMAESVSNPQRPPVIWIGAQECT WP_000145429.1 hydrogenase 2 small subunit [Salmonella enterica] Length: 372\nNumber of predicted TMHs: 1\nExp number of AAs in TMHs: 24.51309\nExp number, first 60 AAs: 0.2682\nTotal prob of N-in: 0.14402\noutside 1 331\nTMhelix 332 354\ninside 355 372

14166 GCF_001831575.1_ASM183157v1 Salmonella enterica Proteobacteria; Gammaproteobacteria; Enterobacterales; Enterobacteriaceae; Salmonella MTGDNTLITSHGINRRDFMKLCAALAATMGLSSKAAAEMAESVSNPQRPPVIWIGAQECT WP_000145429.1 hydrogenase 2 small subunit [Salmonella enterica] Length: 372\nNumber of predicted TMHs: 1\nExp number of AAs in TMHs: 24.51309\nExp number, first 60 AAs: 0.2682\nTotal prob of N-in: 0.14402\noutside 1 331\nTMhelix 332 354\ninside 355 372

14167 GCF_001831545.1_ASM183154v1 Salmonella enterica Proteobacteria; Gammaproteobacteria; Enterobacterales; Enterobacteriaceae; Salmonella MNNEETFYQAMRRKGVTRRSFLKFCSLAATSLGLGAGMTPKIAWALENKPRIPVVWIHGL WP_001058312.1 hydrogenase [Salmonella enterica] Length: 372\nNumber of predicted TMHs: 2\nExp number of AAs in TMHs: 36.51738\nExp number, first 60 AAs: 19.72849\nTotal prob of N-in: 0.96170\nPOSSIBLE N-term signal sequence\ninside 1 19\nTMhelix 20 39\noutside 40 325\nTMhelix 326 348\ninside 349 372

14168 GCF_001831895.1_ASM183189v1 Salmonella enterica Proteobacteria; Gammaproteobacteria; Enterobacterales; Enterobacteriaceae; Salmonella
MTGDNTLITSHGINRRDFMKLCAALAATMGLSSKAAAEMAESVSNPQRPPVIWIGAQECT WP_000145429.1
hydrogenase 2 small subunit [Salmonella enterica] Length: 372\nNumber of predicted TMHs: 1\nExp number of AAs in TMHs: 24.51309\nExp number, first 60 AAs: 0.2682\nTotal prob of N-in: 0.14402\noutside 1 331\nTMhelix 332 354\ninside 355 372

14169 GCF_001831575.1_ASM183157v1 Salmonella enterica Proteobacteria; Gammaproteobacteria; Enterobacterales; Enterobacteriaceae; Salmonella
MNNEETFYQAMRRKGVTRRSFLKFCSLAATSLGLGAGMTPKIAWALENKPRIPVVWIHGL WP_001058312.1
hydrogenase [Salmonella enterica] Length: 372\nNumber of predicted TMHs: 2\nExp number of AAs in TMHs: 36.51738\nExp number, first 60 AAs: 19.72849\nTotal prob of N-in: 0.96170\nPOSSIBLE N-term signal sequence\ninside 1 19\nTMhelix 20 39\noutside 40 325\nTMhelix 326 348\ninside 349 372

14170 GCF_001831895.1_ASM183189v1 Salmonella enterica Proteobacteria; Gammaproteobacteria; Enterobacterales; Enterobacteriaceae; Salmonella
MNNEETFYQAMRRKGVTRRSFLKFCSLAATSLGLGAGMTPKIAWALENKPRIPVVWIHGL WP_001058311.1
hydrogenase [Salmonella enterica] Length: 372\nNumber of predicted TMHs: 2\nExp number of AAs in TMHs: 37.4699899999999\nExp number, first 60 AAs: 19.72699\nTotal prob of N-in: 0.96171\nPOSSIBLE N-term signal sequence\ninside 1 19\nTMhelix 20 39\noutside 40 325\nTMhelix 326 348\ninside 349 372

14171 GCF_001832065.1_ASM183206v1 Salmonella enterica Proteobacteria; Gammaproteobacteria; Enterobacterales; Enterobacteriaceae; Salmonella
MTGDNTLITSHGINRRDFMKLCAALAATMGLSSKAAAEMAESVSNPQRPPVIWIGAQECT WP_000145429.1
hydrogenase 2 small subunit [Salmonella enterica] Length: 372\nNumber of predicted TMHs: 1\nExp number of AAs in TMHs: 24.51309\nExp number, first 60 AAs: 0.2682\nTotal prob of N-in: 0.14402\noutside 1 331\nTMhelix 332 354\ninside 355 372

14172 GCF_001831995.1_ASM183199v1 Salmonella enterica Proteobacteria; Gammaproteobacteria; Enterobacterales; Enterobacteriaceae; Salmonella
MTGDNTLITSHGINRRDFMKLCAALAATMGLSSKAAAEMAESVSNPQRPPVIWIGAQECT WP_000145429.1
hydrogenase 2 small subunit [Salmonella enterica] Length: 372\nNumber of predicted TMHs: 1\nExp number of AAs in TMHs: 24.51309\nExp number, first 60 AAs: 0.2682\nTotal prob of N-in: 0.14402\noutside 1 331\nTMhelix 332 354\ninside 355 372

14173 GCF_001832125.1_ASM183212v1 Salmonella enterica Proteobacteria; Gammaproteobacteria; Enterobacterales; Enterobacteriaceae; Salmonella
MTGDNTLITSHGINRRDFMKLCAALAATMGLSSKAAAEMAESVSNPQRPPVIWIGAQECT WP_000145429.1
hydrogenase 2 small subunit [Salmonella enterica] Length: 372\nNumber of predicted TMHs: 1\nExp number of AAs in TMHs: 24.51309\nExp number, first 60 AAs: 0.2682\nTotal prob of N-in: 0.14402\noutside 1 331\nTMhelix 332 354\ninside 355 372

14174 GCF_001832065.1_ASM183206v1 Salmonella enterica Proteobacteria; Gammaproteobacteria; Enterobacterales; Enterobacteriaceae; Salmonella
MNNEETFYQAMRRKGVTRRSFLKFCSLAATSLGLGAGMTPKIAWALENKPRIPVVWIHGL WP_001058312.1
hydrogenase [Salmonella enterica] Length: 372\nNumber of predicted TMHs: 2\nExp number of AAs in TMHs: 36.51738\nExp number, first 60 AAs: 19.72849\nTotal prob of N-in: 0.96170\nPOSSIBLE N-term signal sequence\ninside 1 19\nTMhelix 20 39\noutside 40 325\nTMhelix 326 348\ninside 349 372

14175 GCF_002043435.1_ASM204343v1 Salmonella enterica Proteobacteria; Gammaproteobacteria; Enterobacterales; Enterobacteriaceae; Salmonella
MNNEETFYQAMRRKGVTRRSFLKFCSLAATSLGLGAGMTPKIAWALENKPRIPVVWIHGL WP_001058312.1
hydrogenase [Salmonella enterica] Length: 372\nNumber of predicted TMHs: 2\nExp number of AAs in TMHs: 36.51738\nExp number, first 60 AAs: 19.72849\nTotal prob of N-in: 0.96170\nPOSSIBLE N-term signal sequence\ninside 1 19\nTMhelix 20 39\noutside 40 325\nTMhelix 326 348\ninside 349 372

14176 GCF_002043245.1_ASM204324v1 Salmonella enterica Proteobacteria; Gammaproteobacteria; Enterobacterales; Enterobacteriaceae; Salmonella
MNNEETFYQAMRRKGVTRRSFLKFCSLAATSLGLGAGMTPKIAWALENKPRIPVVWIHGL WP_001058312.1
hydrogenase [Salmonella enterica] Length: 372\nNumber of predicted TMHs: 2\nExp number of AAs in

TMHs: 36.51738\nExp number, first 60 AAs: 19.72849\nTotal prob of N-in: 0.96170\nPOSSIBLE N-term signal sequence\ninside 1 19\nTMhelix 20 39\noutside 40 325\nTMhelix 326 348\ninside 349 372

14177 GCF_002043485.1_ASM204348v1 Salmonella enterica Proteobacteria; Gammaproteobacteria; Enterobacterales; Enterobacteriaceae; Salmonella
MTGDNTLITSHGINRRDFMKLCAALAATMGLSSKAAAEMAESVSNPQRPPVIWIGAQECT WP_000145429.1
hydrogenase 2 small subunit [Salmonella enterica] Length: 372\nNumber of predicted TMHs: 1\nExp number of AAs in TMHs: 24.51309\nExp number, first 60 AAs: 0.2682\nTotal prob of N-in: 0.14402\noutside 1 331\nTMhelix 332 354\ninside 355 372

14178 GCF_001832125.1_ASM183212v1 Salmonella enterica Proteobacteria; Gammaproteobacteria; Enterobacterales; Enterobacteriaceae; Salmonella
MNNEETFYQAMRRKGVTRRSFLKFCSLAATSLGLGAGMTPKIAWALENKPRIPVVWIHGL WP_001058312.1
hydrogenase [Salmonella enterica] Length: 372\nNumber of predicted TMHs: 2\nExp number of AAs in TMHs: 36.51738\nExp number, first 60 AAs: 19.72849\nTotal prob of N-in: 0.96170\nPOSSIBLE N-term signal sequence\ninside 1 19\nTMhelix 20 39\noutside 40 325\nTMhelix 326 348\ninside 349 372

14179 GCF_002043755.1_ASM204375v1 Salmonella enterica Proteobacteria; Gammaproteobacteria; Enterobacterales; Enterobacteriaceae; Salmonella
MTGDNTLITSHGINRRDFMKLCAALAATMGLSSKAAAEMAESVSNPQRPPVIWIGAQECT WP_000145429.1
hydrogenase 2 small subunit [Salmonella enterica] Length: 372\nNumber of predicted TMHs: 1\nExp number of AAs in TMHs: 24.51309\nExp number, first 60 AAs: 0.2682\nTotal prob of N-in: 0.14402\noutside 1 331\nTMhelix 332 354\ninside 355 372

14180 GCF_001832165.1_ASM183216v1 Salmonella enterica Proteobacteria; Gammaproteobacteria; Enterobacterales; Enterobacteriaceae; Salmonella
MNNEETFYQAMRRKGVTRRSFLKFCSLAATSLGLGAGMTPKIAWALENKPRIPVVWIHGL WP_001058312.1
hydrogenase [Salmonella enterica] Length: 372\nNumber of predicted TMHs: 2\nExp number of AAs in TMHs: 36.51738\nExp number, first 60 AAs: 19.72849\nTotal prob of N-in: 0.96170\nPOSSIBLE N-term signal sequence\ninside 1 19\nTMhelix 20 39\noutside 40 325\nTMhelix 326 348\ninside 349 372

14181 GCF_002043755.1_ASM204375v1 Salmonella enterica Proteobacteria; Gammaproteobacteria; Enterobacterales; Enterobacteriaceae; Salmonella
MNNEETFYQAMRRKGVTRRSFLKFCSLAATSLGLGAGMTPKIAWALENKPRIPVVWIHGL WP_001058311.1
hydrogenase [Salmonella enterica] Length: 372\nNumber of predicted TMHs: 2\nExp number of AAs in TMHs: 37.4699899999999\nExp number, first 60 AAs: 19.72699\nTotal prob of N-in: 0.96171\nPOSSIBLE N-term signal sequence\ninside 1 19\nTMhelix 20 39\noutside 40 325\nTMhelix 326 348\ninside 349 372

14182 GCF_002043205.1_ASM204320v1 Salmonella enterica Proteobacteria; Gammaproteobacteria; Enterobacterales; Enterobacteriaceae; Salmonella
MNNEETLYQAMRRKGVTRRSFLKFCSLAATSLGLGAGMTPKIAWALENKPRIPVVWIHGL WP_001752445.1
hydrogenase-1 small chain [Salmonella enterica] Length: 372\nNumber of predicted TMHs: 2\nExp number of AAs in TMHs: 36.5123\nExp number, first 60 AAs: 19.72341\nTotal prob of N-in: 0.96144\nPOSSIBLE N-term signal sequence\ninside 1 19\nTMhelix 20 39\noutside 40 325\nTMhelix 326 348\ninside 349 372

14183 GCF_002043815.1_ASM204381v1 Salmonella enterica Proteobacteria; Gammaproteobacteria; Enterobacterales; Enterobacteriaceae; Salmonella
MTGDNTLITSHGINRRDFMKLCAALAATMGLSSKAAAEMAESVSNPQRPPVIWIGAQECT WP_000145428.1
hydrogenase 2 small subunit [Salmonella enterica] Length: 372\nNumber of predicted TMHs: 1\nExp number of AAs in TMHs: 24.50677\nExp number, first 60 AAs: 0.2682\nTotal prob of N-in: 0.14390\noutside 1 331\nTMhelix 332 354\ninside 355 372

14184 GCF_002043815.1_ASM204381v1 Salmonella enterica Proteobacteria; Gammaproteobacteria; Enterobacterales; Enterobacteriaceae; Salmonella
MNNEETLYQAMRRKGVTRRSFLKFCSLAATSLGLGAGMTPKIAWALENKPRIPVVWIHGL WP_001752445.1
hydrogenase-1 small chain [Salmonella enterica] Length: 372\nNumber of predicted TMHs: 2\nExp number of AAs in TMHs: 36.5123\nExp number, first 60 AAs: 19.72341\nTotal prob of N-in: 0.96144\nPOSSIBLE N-term signal sequence\ninside 1 19\nTMhelix 20 39\noutside 40 325\nTMhelix 326 348\ninside 349 372

14185 GCF_002037665.1_ASM203766v1 Salmonella enterica Proteobacteria; Gammaproteobacteria; Enterobacterales; Enterobacteriaceae; Salmonella

MNNEETFYQAMRRKGVTRRSFLKFCSLAATSLGLGAGMTPKIAWALENKPRIPVVWIHGL WP_001058311.1 hydrogenase [Salmonella enterica] Length: 372\nNumber of predicted TMHs: 2\nExp number of AAs in TMHs: 37.4699899999999\nExp number, first 60 AAs: 19.72699\nTotal prob of N-in: 0.96171\nPOSSIBLE N-term signal sequence\ninside 1 19\nTMhelix 20 39\noutside 40 325\nTMhelix 326 348\ninside 349 372

14186 GCF_002053395.1_ASM205339v1 Salmonella enterica Proteobacteria; Gammaproteobacteria; Enterobacterales; Enterobacteriaceae; Salmonella MTGDNTLITSHGINRRDFMKLCAALAATMGLSSKAAAEMAESVSNPQRPPVIWIGAQECT WP_000145429.1 hydrogenase 2 small subunit [Salmonella enterica] Length: 372\nNumber of predicted TMHs: 1\nExp number of AAs in TMHs: 24.51309\nExp number, first 60 AAs: 0.2682\nTotal prob of N-in: 0.14402\noutside 1 331\nTMhelix 332 354\ninside 355 372

14187 GCF_002043485.1_ASM204348v1 Salmonella enterica Proteobacteria; Gammaproteobacteria; Enterobacterales; Enterobacteriaceae; Salmonella MNNEETFYQAMRRKGVTRRSFLKFCSLAATSLGLGAGMTPKIAWALENKPRIPVVWIHGL WP_001058311.1 hydrogenase [Salmonella enterica] Length: 372\nNumber of predicted TMHs: 2\nExp number of AAs in TMHs: 37.4699899999999\nExp number, first 60 AAs: 19.72699\nTotal prob of N-in: 0.96171\nPOSSIBLE N-term signal sequence\ninside 1 19\nTMhelix 20 39\noutside 40 325\nTMhelix 326 348\ninside 349 372

14188 GCF_001832695.1_ASM183269v1 Salmonella enterica Proteobacteria; Gammaproteobacteria; Enterobacterales; Enterobacteriaceae; Salmonella MTGDNTLITSHGINRRDFMKLCAALAATMGLSSKAAAEMAESVSNPQRPPVIWIGAQECT WP_000145429.1 hydrogenase 2 small subunit [Salmonella enterica] Length: 372\nNumber of predicted TMHs: 1\nExp number of AAs in TMHs: 24.51309\nExp number, first 60 AAs: 0.2682\nTotal prob of N-in: 0.14402\noutside 1 331\nTMhelix 332 354\ninside 355 372

14189 GCF_002037665.1_ASM203766v1 Salmonella enterica Proteobacteria; Gammaproteobacteria; Enterobacterales; Enterobacteriaceae; Salmonella MTGDNTLITSHGINRRDFMKLCAALAATMGLSSKAAAEMAESVSNPQRPPVIWIGAQECT WP_000145429.1 hydrogenase 2 small subunit [Salmonella enterica] Length: 372\nNumber of predicted TMHs: 1\nExp number of AAs in TMHs: 24.51309\nExp number, first 60 AAs: 0.2682\nTotal prob of N-in: 0.14402\noutside 1 331\nTMhelix 332 354\ninside 355 372

14190 GCF_001832695.1_ASM183269v1 Salmonella enterica Proteobacteria; Gammaproteobacteria; Enterobacterales; Enterobacteriaceae; Salmonella MNNEETFYQAMRRKGVTRRSFLKFCSLAATSLGLGAGMTPKIAWALENKPRIPVVWIHGL WP_001058312.1 hydrogenase [Salmonella enterica] Length: 372\nNumber of predicted TMHs: 2\nExp number of AAs in TMHs: 36.51738\nExp number, first 60 AAs: 19.72849\nTotal prob of N-in: 0.96170\nPOSSIBLE N-term signal sequence\ninside 1 19\nTMhelix 20 39\noutside 40 325\nTMhelix 326 348\ninside 349 372

14191 GCF_001833185.1_ASM183318v1 Salmonella enterica Proteobacteria; Gammaproteobacteria; Enterobacterales; Enterobacteriaceae; Salmonella MTGDNTLITSHGINRRDFMKLCAALAATMGLSSKAAAEMAESVSNPQRPPVIWIGAQECT WP_000145429.1 hydrogenase 2 small subunit [Salmonella enterica] Length: 372\nNumber of predicted TMHs: 1\nExp number of AAs in TMHs: 24.51309\nExp number, first 60 AAs: 0.2682\nTotal prob of N-in: 0.14402\noutside 1 331\nTMhelix 332 354\ninside 355 372

14192 GCF_001833155.1_ASM183315v1 Salmonella enterica Proteobacteria; Gammaproteobacteria; Enterobacterales; Enterobacteriaceae; Salmonella MTGDNTLITSHGINRRDFMKLCAALAATMGLSSKAAAEMAESVSNPQRPPVIWIGAQECT WP_000145429.1 hydrogenase 2 small subunit [Salmonella enterica] Length: 372\nNumber of predicted TMHs: 1\nExp number of AAs in TMHs: 24.51309\nExp number, first 60 AAs: 0.2682\nTotal prob of N-in: 0.14402\noutside 1 331\nTMhelix 332 354\ninside 355 372

14193 GCF_001833185.1_ASM183318v1 Salmonella enterica Proteobacteria; Gammaproteobacteria; Enterobacterales; Enterobacteriaceae; Salmonella MNNEETFYQAMRRKGVTRRSFLKFCSLAATSLGLGAGMTPKIAWALENKPRIPVVWIHGL WP_001058312.1 hydrogenase [Salmonella enterica] Length: 372\nNumber of predicted TMHs: 2\nExp number of AAs in TMHs: 36.51738\nExp number, first 60 AAs: 19.72849\nTotal prob of N-in: 0.96170\nPOSSIBLE N-term signal sequence\ninside 1 19\nTMhelix 20 39\noutside 40 325\nTMhelix 326 348\ninside 349 372

14194 GCF_001833155.1_ASM183315v1 Salmonella enterica Proteobacteria; Gammaproteobacteria; Enterobacterales; Enterobacteriaceae; Salmonella
MNNEETFYQAMRRKGVTRRSFLKFCSLAATSLGLGAGMTPKIAWALENKPRIPVVWIHGL WP_001058312.1
hydrogenase [Salmonella enterica] Length: 372\nNumber of predicted TMHs: 2\nExp number of AAs in TMHs: 36.51738\nExp number, first 60 AAs: 19.72849\nTotal prob of N-in: 0.96170\nPOSSIBLE N-term signal sequence\ninside 1 19\nTMhelix 20 39\noutside 40 325\nTMhelix 326 348\ninside 349 372

14195 GCF_002043865.1_ASM204386v1 Salmonella enterica Proteobacteria; Gammaproteobacteria; Enterobacterales; Enterobacteriaceae; Salmonella
MTGDNTLITSHGINRRDFMKLCAALAATMGLSSKAAAEMAESVSNPQRPPVIWIGAQECT WP_000145429.1
hydrogenase 2 small subunit [Salmonella enterica] Length: 372\nNumber of predicted TMHs: 1\nExp number of AAs in TMHs: 24.51309\nExp number, first 60 AAs: 0.2682\nTotal prob of N-in: 0.14402\noutside 1 331\nTMhelix 332 354\ninside 355 372

14196 GCF_002066635.1_ASM206663v1 Salmonella enterica Proteobacteria; Gammaproteobacteria; Enterobacterales; Enterobacteriaceae; Salmonella
MTGDNTLITSHGINRRDFMKLCAALAATMGLSSKAAAEMAESVSNPQRPPVIWIGAQECT WP_000145429.1
hydrogenase 2 small subunit [Salmonella enterica] Length: 372\nNumber of predicted TMHs: 1\nExp number of AAs in TMHs: 24.51309\nExp number, first 60 AAs: 0.2682\nTotal prob of N-in: 0.14402\noutside 1 331\nTMhelix 332 354\ninside 355 372

14197 GCF_002043865.1_ASM204386v1 Salmonella enterica Proteobacteria; Gammaproteobacteria; Enterobacterales; Enterobacteriaceae; Salmonella
MNNEETFYQAMRRKGVTRRSFLKFCSLAATSLGLGAGMTPKIAWALENKPRIPVVWIHGL WP_001058311.1
hydrogenase [Salmonella enterica] Length: 372\nNumber of predicted TMHs: 2\nExp number of AAs in TMHs: 37.4699899999999\nExp number, first 60 AAs: 19.72699\nTotal prob of N-in: 0.96171\nPOSSIBLE N-term signal sequence\ninside 1 19\nTMhelix 20 39\noutside 40 325\nTMhelix 326 348\ninside 349 372

14198 GCF_002053335.1_ASM205333v1 Salmonella enterica Proteobacteria; Gammaproteobacteria; Enterobacterales; Enterobacteriaceae; Salmonella
MTGDNTLITSHGINRRDFMKLCAALAATMGLSSKAAAEMAESVSNPQRPPVIWIGAQECT WP_000145427.1
hydrogenase 2 small subunit [Salmonella enterica] Length: 372\nNumber of predicted TMHs: 1\nExp number of AAs in TMHs: 24.48576\nExp number, first 60 AAs: 0.2692\nTotal prob of N-in: 0.14251\noutside 1 331\nTMhelix 332 354\ninside 355 372

14199 GCF_002053215.1_ASM205321v1 Salmonella enterica Proteobacteria; Gammaproteobacteria; Enterobacterales; Enterobacteriaceae; Salmonella
MNNEETFYQAMRRKGVTRRSFLKFCSLAATSLGLGAGMTPKIAWALENKPRIPVVWIHGL WP_001058312.1
hydrogenase [Salmonella enterica] Length: 372\nNumber of predicted TMHs: 2\nExp number of AAs in TMHs: 36.51738\nExp number, first 60 AAs: 19.72849\nTotal prob of N-in: 0.96170\nPOSSIBLE N-term signal sequence\ninside 1 19\nTMhelix 20 39\noutside 40 325\nTMhelix 326 348\ninside 349 372

14200 GCF_001835055.1_ASM183505v1 Salmonella enterica Proteobacteria; Gammaproteobacteria; Enterobacterales; Enterobacteriaceae; Salmonella
MTGDNTLITSHGINRRDFMKLCAALAATMGLSSKAAAEMAESVSNPQRPPVIWIGAQECT WP_000145429.1
hydrogenase 2 small subunit [Salmonella enterica] Length: 372\nNumber of predicted TMHs: 1\nExp number of AAs in TMHs: 24.51309\nExp number, first 60 AAs: 0.2682\nTotal prob of N-in: 0.14402\noutside 1 331\nTMhelix 332 354\ninside 355 372

14201 GCF_001835055.1_ASM183505v1 Salmonella enterica Proteobacteria; Gammaproteobacteria; Enterobacterales; Enterobacteriaceae; Salmonella
MNNEETFYQAMRRKGVTRRSFLKFCSLAATSLGLGAGMTPKIAWALENKPRIPVVWIHGL WP_001058312.1
hydrogenase [Salmonella enterica] Length: 372\nNumber of predicted TMHs: 2\nExp number of AAs in TMHs: 36.51738\nExp number, first 60 AAs: 19.72849\nTotal prob of N-in: 0.96170\nPOSSIBLE N-term signal sequence\ninside 1 19\nTMhelix 20 39\noutside 40 325\nTMhelix 326 348\ninside 349 372

14202 GCF_002053305.1_ASM205330v1 Salmonella enterica Proteobacteria; Gammaproteobacteria; Enterobacterales; Enterobacteriaceae; Salmonella
MTGDNTLITSHGINRRDFMKLCAALAATMGLSSKAAAEMAESVSNPQRPPVIWIGAQECT WP_000145427.1
hydrogenase 2 small subunit [Salmonella enterica] Length: 372\nNumber of predicted TMHs: 1\nExp

number of AAs in TMHs: 24.48576\nExp number, first 60 AAs: 0.2692\nTotal prob of N-in: 0.14251\noutside 1 331\nTMhelix 332 354\ninside 355 372

14203 GCF_002052775.1_ASM205277v1 Salmonella enterica Proteobacteria; Gammaproteobacteria; Enterobacterales; Enterobacteriaceae; Salmonella MNNEETFYQAMRRKGVTRRSFLKFCSLAATSLGLGAGMTPKIAWALENKPRIPVVWIHGL WP_001058312.1 hydrogenase [Salmonella enterica] Length: 372\nNumber of predicted TMHs: 2\nExp number of AAs in TMHs: 36.51738\nExp number, first 60 AAs: 19.72849\nTotal prob of N-in: 0.96170\nPOSSIBLE N-term signal sequence\ninside 1 19\nTMhelix 20 39\noutside 40 325\nTMhelix 326 348\ninside 349 372

14204 GCF_002044105.1_ASM204410v1 Salmonella enterica Proteobacteria; Gammaproteobacteria; Enterobacterales; Enterobacteriaceae; Salmonella MNNEETFYQAMRRKGVTRRSFLKFCSLAATSLGLGAGMTPKIAWALENKPRIPVVWIHGL WP_001058311.1 hydrogenase [Salmonella enterica] Length: 372\nNumber of predicted TMHs: 2\nExp number of AAs in TMHs: 37.4699899999999\nExp number, first 60 AAs: 19.72699\nTotal prob of N-in: 0.96171\nPOSSIBLE N-term signal sequence\ninside 1 19\nTMhelix 20 39\noutside 40 325\nTMhelix 326 348\ninside 349 372

14205 GCF_002066465.1_ASM206646v1 Salmonella enterica Proteobacteria; Gammaproteobacteria; Enterobacterales; Enterobacteriaceae; Salmonella MNNEETFYQAMRRKGVTRRSFLKFCSLAATSLGLGAGMTPKIAWALENKPRIPVVWIHGL WP_001058306.1 hydrogenase [Salmonella enterica] Length: 372\nNumber of predicted TMHs: 2\nExp number of AAs in TMHs: 36.5540099999999\nExp number, first 60 AAs: 19.72732\nTotal prob of N-in: 0.96171\nPOSSIBLE N-term signal sequence\ninside 1 19\nTMhelix 20 39\noutside 40 325\nTMhelix 326 348\ninside 349 372

14206 GCF_002044385.1_ASM204438v1 Salmonella enterica Proteobacteria; Gammaproteobacteria; Enterobacterales; Enterobacteriaceae; Salmonella MNNEETFYQAMRRKGVTRRSFLKFCSLAATSLGLGAGMTPKIAWALENKPRIPVVWIHGL WP_001058312.1 hydrogenase [Salmonella enterica] Length: 372\nNumber of predicted TMHs: 2\nExp number of AAs in TMHs: 36.51738\nExp number, first 60 AAs: 19.72849\nTotal prob of N-in: 0.96170\nPOSSIBLE N-term signal sequence\ninside 1 19\nTMhelix 20 39\noutside 40 325\nTMhelix 326 348\ninside 349 372

14207 GCF_002053045.1_ASM205304v1 Salmonella enterica Proteobacteria; Gammaproteobacteria; Enterobacterales; Enterobacteriaceae; Salmonella MNNEETFYQAMRRKGVTRRSFLKFCSLAATSLGLGAGMTPKIAWALENKPRIPVVWIHGL WP_001058311.1 hydrogenase [Salmonella enterica] Length: 372\nNumber of predicted TMHs: 2\nExp number of AAs in TMHs: 37.4699899999999\nExp number, first 60 AAs: 19.72699\nTotal prob of N-in: 0.96171\nPOSSIBLE N-term signal sequence\ninside 1 19\nTMhelix 20 39\noutside 40 325\nTMhelix 326 348\ninside 349 372

14208 GCF_002044265.1_ASM204426v1 Salmonella enterica Proteobacteria; Gammaproteobacteria; Enterobacterales; Enterobacteriaceae; Salmonella MNNEETFYQAMRRKGVTRRSFLKFCSLAATSLGLGAGMTPKIAWALENKPRIPVVWIHGL WP_001058311.1 hydrogenase [Salmonella enterica] Length: 372\nNumber of predicted TMHs: 2\nExp number of AAs in TMHs: 37.4699899999999\nExp number, first 60 AAs: 19.72699\nTotal prob of N-in: 0.96171\nPOSSIBLE N-term signal sequence\ninside 1 19\nTMhelix 20 39\noutside 40 325\nTMhelix 326 348\ninside 349 372

14209 GCF_002066185.1_ASM206618v1 Salmonella enterica Proteobacteria; Gammaproteobacteria; Enterobacterales; Enterobacteriaceae; Salmonella MNNEETFYQAMRRKGVTRRSFLKFCSLAATSLGLGAGMTPKIAWALENKPRIPVVWIHGL WP_001058311.1 hydrogenase [Salmonella enterica] Length: 372\nNumber of predicted TMHs: 2\nExp number of AAs in TMHs: 37.4699899999999\nExp number, first 60 AAs: 19.72699\nTotal prob of N-in: 0.96171\nPOSSIBLE N-term signal sequence\ninside 1 19\nTMhelix 20 39\noutside 40 325\nTMhelix 326 348\ninside 349 372

14210 GCF_002036845.1_ASM203684v1 Salmonella enterica Proteobacteria; Gammaproteobacteria; Enterobacterales; Enterobacteriaceae; Salmonella MNNEETFYQAMRRKGVTRRSFLKFCSLAATSLGLGAGMTPKIAWALENKPRIPVVWIHGL WP_001058312.1 hydrogenase [Salmonella enterica] Length: 372\nNumber of predicted TMHs: 2\nExp number of AAs in TMHs: 36.51738\nExp number, first 60 AAs: 19.72849\nTotal prob of N-in: 0.96170\nPOSSIBLE N-term signal sequence\ninside 1 19\nTMhelix 20 39\noutside 40 325\nTMhelix 326 348\ninside 349 372

14211 GCF_002036985.1_ASM203698v1 Salmonella enterica Proteobacteria; Gammaproteobacteria; Enterobacterales; Enterobacteriaceae; Salmonella

MNNEETFYQAMRRKGVTRRSFLKFCSLAATSLGLGAGMTPKIAWALENKPRIPVVWIHGL WP_001058312.1 hydrogenase [Salmonella enterica] Length: 372\nNumber of predicted TMHs: 2\nExp number of AAs in TMHs: 36.51738\nExp number, first 60 AAs: 19.72849\nTotal prob of N-in: 0.96170\nPOSSIBLE N-term signal sequence\ninside 1 19\nTMhelix 20 39\noutside 40 325\nTMhelix 326 348\ninside 349 372

14212 GCF_002037115.1_ASM203711v1 Salmonella enterica Proteobacteria; Gammaproteobacteria; Enterobacterales; Enterobacteriaceae; Salmonella MNNEETFYQAMRRKGVTRRSFLKFCSLAATSLGLGAGMTPKIAWALENKPRIPVVWIHGL WP_001058311.1 hydrogenase [Salmonella enterica] Length: 372\nNumber of predicted TMHs: 2\nExp number of AAs in TMHs: 37.4699899999999\nExp number, first 60 AAs: 19.72699\nTotal prob of N-in: 0.96171\nPOSSIBLE N-term signal sequence\ninside 1 19\nTMhelix 20 39\noutside 40 325\nTMhelix 326 348\ninside 349 372

14213 GCF_002037115.1_ASM203711v1 Salmonella enterica Proteobacteria; Gammaproteobacteria; Enterobacterales; Enterobacteriaceae; Salmonella MTGDNTLITSHGINRRDFMKLCAALAATMGLSSKAAAEMAESVSNPQRPPVIWIGAQECT WP_000145429.1 hydrogenase 2 small subunit [Salmonella enterica] Length: 372\nNumber of predicted TMHs: 1\nExp number of AAs in TMHs: 24.51309\nExp number, first 60 AAs: 0.2682\nTotal prob of N-in: 0.14402\noutside 1 331\nTMhelix 332 354\ninside 355 372

14214 GCF_002054375.1_ASM205437v1 Salmonella enterica Proteobacteria; Gammaproteobacteria; Enterobacterales; Enterobacteriaceae; Salmonella MNNEETFYQAMRRKGVTRRSFLKFCSLAATSLGLGAGMTPKIAWALENKPRIPVVWIHGL WP_001058312.1 hydrogenase [Salmonella enterica] Length: 372\nNumber of predicted TMHs: 2\nExp number of AAs in TMHs: 36.51738\nExp number, first 60 AAs: 19.72849\nTotal prob of N-in: 0.96170\nPOSSIBLE N-term signal sequence\ninside 1 19\nTMhelix 20 39\noutside 40 325\nTMhelix 326 348\ninside 349 372

14215 GCF_002061975.1_ASM206197v1 Salmonella enterica Proteobacteria; Gammaproteobacteria; Enterobacterales; Enterobacteriaceae; Salmonella MTGDNTLITSHGINRRDFMKLCAALAATMGLSSKAAAEMAESVSNPQRPPVIWIGAQECT WP_000145429.1 hydrogenase 2 small subunit [Salmonella enterica] Length: 372\nNumber of predicted TMHs: 1\nExp number of AAs in TMHs: 24.51309\nExp number, first 60 AAs: 0.2682\nTotal prob of N-in: 0.14402\noutside 1 331\nTMhelix 332 354\ninside 355 372

14216 GCF_002062005.1_ASM206200v1 Salmonella enterica Proteobacteria; Gammaproteobacteria; Enterobacterales; Enterobacteriaceae; Salmonella MNNEETFYQAMRRKGVTRRSFLKFCSLAATSLGLGAGMTPKIAWALENKPRIPVVWIHGL WP_001058310.1 hydrogenase [Salmonella enterica] Length: 372\nNumber of predicted TMHs: 2\nExp number of AAs in TMHs: 37.4567299999999\nExp number, first 60 AAs: 19.72694\nTotal prob of N-in: 0.96171\nPOSSIBLE N-term signal sequence\ninside 1 19\nTMhelix 20 39\noutside 40 325\nTMhelix 326 348\ninside 349 372

14217 GCF_002066095.1_ASM206609v1 Salmonella enterica Proteobacteria; Gammaproteobacteria; Enterobacterales; Enterobacteriaceae; Salmonella MNNEETFYQAMRRKGVTRRSFLKFCSLAATSLGLGAGMTPKIAWALENKPRIPVVWIHGL WP_001058312.1 hydrogenase [Salmonella enterica] Length: 372\nNumber of predicted TMHs: 2\nExp number of AAs in TMHs: 36.51738\nExp number, first 60 AAs: 19.72849\nTotal prob of N-in: 0.96170\nPOSSIBLE N-term signal sequence\ninside 1 19\nTMhelix 20 39\noutside 40 325\nTMhelix 326 348\ninside 349 372

14218 GCF_002036745.1_ASM203674v1 Salmonella enterica Proteobacteria; Gammaproteobacteria; Enterobacterales; Enterobacteriaceae; Salmonella MTGDNTLITSHGINRRDFMKLCAALAATMGLSSKAAAEMAESVSNPQRPPVIWIGAQECT WP_000145429.1 hydrogenase 2 small subunit [Salmonella enterica] Length: 372\nNumber of predicted TMHs: 1\nExp number of AAs in TMHs: 24.51309\nExp number, first 60 AAs: 0.2682\nTotal prob of N-in: 0.14402\noutside 1 331\nTMhelix 332 354\ninside 355 372

14219 GCF_002066185.1_ASM206618v1 Salmonella enterica Proteobacteria; Gammaproteobacteria; Enterobacterales; Enterobacteriaceae; Salmonella MTGDNTLITSHGINRRDFMKLCAALAATMGLSSKAAAEMAESVSNPQRPPVIWIGAQECT WP_000145429.1 hydrogenase 2 small subunit [Salmonella enterica] Length: 372\nNumber of predicted TMHs: 1\nExp number of AAs in TMHs: 24.51309\nExp number, first 60 AAs: 0.2682\nTotal prob of N-in: 0.14402\noutside 1 331\nTMhelix 332 354\ninside 355 372

14220 GCF_002044435.1_ASM204443v1 Salmonella enterica Proteobacteria; Gammaproteobacteria; Enterobacterales; Enterobacteriaceae; Salmonella
MTGDNTLITSHGINRRDFMKLCAALAATMGLSSKAAAEMAESVSNPQRPPVIWIGAQECT WP_000145429.1
hydrogenase 2 small subunit [Salmonella enterica] Length: 372\nNumber of predicted TMHs: 1\nExp number of AAs in TMHs: 24.51309\nExp number, first 60 AAs: 0.2682\nTotal prob of N-in: 0.14402\noutside 1 331\nTMhelix 332 354\ninside 355 372

14221 GCF_002044435.1_ASM204443v1 Salmonella enterica Proteobacteria; Gammaproteobacteria; Enterobacterales; Enterobacteriaceae; Salmonella
MNNEETFYQAMRRKGVTRRSFLKFCSLAATSLGLGAGMTPKIAWALENKPRIPVVWIHGL WP_023198237.1
hydrogenase 1, small subunit [Salmonella enterica] Length: 372\nNumber of predicted TMHs: 2\nExp number of AAs in TMHs: 36.39423\nExp number, first 60 AAs: 19.72877\nTotal prob of N-in: 0.96162\nPOSSIBLE N-term signal sequence\ninside 1 19\nTMhelix 20 39\noutside 40 325\nTMhelix 326 348\ninside 349 372

14222 GCF_002044165.1_ASM204416v1 Salmonella enterica Proteobacteria; Gammaproteobacteria; Enterobacterales; Enterobacteriaceae; Salmonella
MNNEETFYQAMRRKGVTRRSFLKFCSLAATSLGLGAGMTPKIAWALENKPRIPVVWIHGL WP_001058311.1
hydrogenase [Salmonella enterica] Length: 372\nNumber of predicted TMHs: 2\nExp number of AAs in TMHs: 37.4699899999999\nExp number, first 60 AAs: 19.72699\nTotal prob of N-in: 0.96171\nPOSSIBLE N-term signal sequence\ninside 1 19\nTMhelix 20 39\noutside 40 325\nTMhelix 326 348\ninside 349 372

14223 GCF_002066095.1_ASM206609v1 Salmonella enterica Proteobacteria; Gammaproteobacteria; Enterobacterales; Enterobacteriaceae; Salmonella
MTGDNTLITSHGINRRDFMKLCAALAATMGLSSKAAAEMAESVSNPQRPPVIWIGAQECT WP_000145429.1
hydrogenase 2 small subunit [Salmonella enterica] Length: 372\nNumber of predicted TMHs: 1\nExp number of AAs in TMHs: 24.51309\nExp number, first 60 AAs: 0.2682\nTotal prob of N-in: 0.14402\noutside 1 331\nTMhelix 332 354\ninside 355 372

14224 GCF_002061895.1_ASM206189v1 Salmonella enterica Proteobacteria; Gammaproteobacteria; Enterobacterales; Enterobacteriaceae; Salmonella
MNNEETFYQAMRRKGVTRRSFLKFCSLAATSLGLGAGMTPKIAWALENKPRIPVVWIHGL WP_001058312.1
hydrogenase [Salmonella enterica] Length: 372\nNumber of predicted TMHs: 2\nExp number of AAs in TMHs: 36.51738\nExp number, first 60 AAs: 19.72849\nTotal prob of N-in: 0.96170\nPOSSIBLE N-term signal sequence\ninside 1 19\nTMhelix 20 39\noutside 40 325\nTMhelix 326 348\ninside 349 372

14225 GCF_002036505.1_ASM203650v1 Salmonella enterica Proteobacteria; Gammaproteobacteria; Enterobacterales; Enterobacteriaceae; Salmonella
MTGDNTLITSHGINRRDFMKLCAALAATMGLSSKAAAEMAESVSNPQRPPVIWIGAQECT WP_000145429.1
hydrogenase 2 small subunit [Salmonella enterica] Length: 372\nNumber of predicted TMHs: 1\nExp number of AAs in TMHs: 24.51309\nExp number, first 60 AAs: 0.2682\nTotal prob of N-in: 0.14402\noutside 1 331\nTMhelix 332 354\ninside 355 372

14226 GCF_002044245.1_ASM204424v1 Salmonella enterica Proteobacteria; Gammaproteobacteria; Enterobacterales; Enterobacteriaceae; Salmonella
MNNEETFYQAMRRKGVTRRSFLKFCSLAATSLGLGAGMTPKIAWALENKPRIPVVWIHGL WP_001058311.1
hydrogenase [Salmonella enterica] Length: 372\nNumber of predicted TMHs: 2\nExp number of AAs in TMHs: 37.4699899999999\nExp number, first 60 AAs: 19.72699\nTotal prob of N-in: 0.96171\nPOSSIBLE N-term signal sequence\ninside 1 19\nTMhelix 20 39\noutside 40 325\nTMhelix 326 348\ninside 349 372

14227 GCF_002052895.1_ASM205289v1 Salmonella enterica Proteobacteria; Gammaproteobacteria; Enterobacterales; Enterobacteriaceae; Salmonella
MNNEETFYQAMRRKGVTRRSFLKFCSLAATSLGLGAGMTPKIAWALENKPRIPVVWIHGL WP_001058311.1
hydrogenase [Salmonella enterica] Length: 372\nNumber of predicted TMHs: 2\nExp number of AAs in TMHs: 37.4699899999999\nExp number, first 60 AAs: 19.72699\nTotal prob of N-in: 0.96171\nPOSSIBLE N-term signal sequence\ninside 1 19\nTMhelix 20 39\noutside 40 325\nTMhelix 326 348\ninside 349 372

14228 GCF_002062045.1_ASM206204v1 Salmonella enterica Proteobacteria; Gammaproteobacteria; Enterobacterales; Enterobacteriaceae; Salmonella
MTGDNTLITSHGINRRDFMKLCAALAATMGLSSKAAAEMAESVSNPQRPPVIWIGAQECT WP_000145429.1
hydrogenase 2 small subunit [Salmonella enterica] Length: 372\nNumber of predicted TMHs: 1\nExp

number of AAs in TMHs: 24.51309\nExp number, first 60 AAs: 0.2682\nTotal prob of N-in: 0.14402\noutside 1 331\nTMhelix 332 354\ninside 355 372

14229 GCF_002066465.1_ASM206646v1 Salmonella enterica Proteobacteria; Gammaproteobacteria; Enterobacterales; Enterobacteriaceae; Salmonella
MTGDNTLITSHGINRRDFMKLCAALAATMGLSSKAAAEMAESVSNPQRPPVIWIGAQECT WP_023248548.1
hydrogenase 2 small subunit [Salmonella enterica] Length: 372\nNumber of predicted TMHs: 1\nExp number of AAs in TMHs: 24.51281\nExp number, first 60 AAs: 0.26822\nTotal prob of N-in: 0.14402\noutside 1 331\nTMhelix 332 354\ninside 355 372

14230 GCF_002062005.1_ASM206200v1 Salmonella enterica Proteobacteria; Gammaproteobacteria; Enterobacterales; Enterobacteriaceae; Salmonella
MTGDNTLITSHGINRRDFMKLCAALAATMGLSSKAAAEMAESVSNPQRPPVIWIGAQECT WP_000145429.1
hydrogenase 2 small subunit [Salmonella enterica] Length: 372\nNumber of predicted TMHs: 1\nExp number of AAs in TMHs: 24.51309\nExp number, first 60 AAs: 0.2682\nTotal prob of N-in: 0.14402\noutside 1 331\nTMhelix 332 354\ninside 355 372

14231 GCF_002061785.1_ASM206178v1 Salmonella enterica Proteobacteria; Gammaproteobacteria; Enterobacterales; Enterobacteriaceae; Salmonella
MNNEETFYQAMRRKGVTRRSFLKFCSLAATSLGLGAGMTPKIAWALENKPRIPVVWIHGL WP_001058311.1
hydrogenase [Salmonella enterica] Length: 372\nNumber of predicted TMHs: 2\nExp number of AAs in TMHs: 37.4699899999999\nExp number, first 60 AAs: 19.72699\nTotal prob of N-in: 0.96171\nPOSSIBLE N-term signal sequence\ninside 1 19\nTMhelix 20 39\noutside 40 325\nTMhelix 326 348\ninside 349 372

14232 GCF_002052895.1_ASM205289v1 Salmonella enterica Proteobacteria; Gammaproteobacteria; Enterobacterales; Enterobacteriaceae; Salmonella
MTGDNTLITSHGINRRDFMKLCAALAATMGLSSKAAAEMAESVSNPQRPPVIWIGAQECT WP_000145431.1
hydrogenase 2 small subunit [Salmonella enterica] Length: 372\nNumber of predicted TMHs: 1\nExp number of AAs in TMHs: 24.51505\nExp number, first 60 AAs: 0.2682\nTotal prob of N-in: 0.14406\noutside 1 331\nTMhelix 332 354\ninside 355 372

14233 GCF_002052775.1_ASM205277v1 Salmonella enterica Proteobacteria; Gammaproteobacteria; Enterobacterales; Enterobacteriaceae; Salmonella
MTGDNTLITSHGINRRDFMKLCAALAATMGLSSKAAAEMAESVSNPQRPPVIWIGAQECT WP_000145429.1
hydrogenase 2 small subunit [Salmonella enterica] Length: 372\nNumber of predicted TMHs: 1\nExp number of AAs in TMHs: 24.51309\nExp number, first 60 AAs: 0.2682\nTotal prob of N-in: 0.14402\noutside 1 331\nTMhelix 332 354\ninside 355 372

14234 GCF_002052675.1_ASM205267v1 Salmonella enterica Proteobacteria; Gammaproteobacteria; Enterobacterales; Enterobacteriaceae; Salmonella
MTGDNTLITSHGINRRDFMKLCAALAATMGLSSKAAAEMAESVSNPQRPPVIWIGAQECT WP_000145429.1
hydrogenase 2 small subunit [Salmonella enterica] Length: 372\nNumber of predicted TMHs: 1\nExp number of AAs in TMHs: 24.51309\nExp number, first 60 AAs: 0.2682\nTotal prob of N-in: 0.14402\noutside 1 331\nTMhelix 332 354\ninside 355 372

14235 GCF_001834845.1_ASM183484v1 Salmonella enterica Proteobacteria; Gammaproteobacteria; Enterobacterales; Enterobacteriaceae; Salmonella
MNNEETFYQAMRRKGVTRRSFLKFCSLAATSLGLGAGMTPKIAWALENKPRIPVVWIHGL WP_001058311.1
hydrogenase [Salmonella enterica] Length: 372\nNumber of predicted TMHs: 2\nExp number of AAs in TMHs: 37.4699899999999\nExp number, first 60 AAs: 19.72699\nTotal prob of N-in: 0.96171\nPOSSIBLE N-term signal sequence\ninside 1 19\nTMhelix 20 39\noutside 40 325\nTMhelix 326 348\ninside 349 372

14236 GCF_001835005.1_ASM183500v1 Salmonella enterica Proteobacteria; Gammaproteobacteria; Enterobacterales; Enterobacteriaceae; Salmonella
MNNEETFYQAMRRKGVTRRSFLKFCSLAATSLGLGAGMTPKIAWALENKPRIPVVWIHGL WP_001058311.1
hydrogenase [Salmonella enterica] Length: 372\nNumber of predicted TMHs: 2\nExp number of AAs in TMHs: 37.4699899999999\nExp number, first 60 AAs: 19.72699\nTotal prob of N-in: 0.96171\nPOSSIBLE N-term signal sequence\ninside 1 19\nTMhelix 20 39\noutside 40 325\nTMhelix 326 348\ninside 349 372

14237 GCF_001834965.1_ASM183496v1 Salmonella enterica Proteobacteria; Gammaproteobacteria; Enterobacterales; Enterobacteriaceae; Salmonella

MNNEETFYQAMRRKGVTRRSFLKFCSLAATSLGLGAGMTPKIAWALENKPRIPVVWIHGL WP_001058312.1 hydrogenase [Salmonella enterica] Length: 372\nNumber of predicted TMHs: 2\nExp number of AAs in TMHs: 36.51738\nExp number, first 60 AAs: 19.72849\nTotal prob of N-in: 0.96170\nPOSSIBLE N-term signal sequence\ninside 1 19\nTMhelix 20 39\noutside 40 325\nTMhelix 326 348\ninside 349 372

14238 GCF_002062045.1_ASM206204v1 Salmonella enterica Proteobacteria; Gammaproteobacteria; Enterobacterales; Enterobacteriaceae; Salmonella MNNEETFYQAMRRKGVTRRSFLKFCSLAATSLGLGAGMTPKIAWALENKPRIPVVWIHGL WP_001058307.1 hydrogenase [Salmonella enterica] Length: 372\nNumber of predicted TMHs: 2\nExp number of AAs in TMHs: 36.51738\nExp number, first 60 AAs: 19.72849\nTotal prob of N-in: 0.96170\nPOSSIBLE N-term signal sequence\ninside 1 19\nTMhelix 20 39\noutside 40 325\nTMhelix 326 348\ninside 349 372

14239 GCF_002036145.1_ASM203614v1 Salmonella enterica Proteobacteria; Gammaproteobacteria; Enterobacterales; Enterobacteriaceae; Salmonella MTGDNTLITSHGINRRDFMKLCAALAATMGLSSKAAAEMAESVSNPQRPPVIWIGAQECT WP_000145429.1 hydrogenase 2 small subunit [Salmonella enterica] Length: 372\nNumber of predicted TMHs: 1\nExp number of AAs in TMHs: 24.51309\nExp number, first 60 AAs: 0.2682\nTotal prob of N-in: 0.14402\noutside 1 331\nTMhelix 332 354\ninside 355 372

14240 GCF_002061935.1_ASM206193v1 Salmonella enterica Proteobacteria; Gammaproteobacteria; Enterobacterales; Enterobacteriaceae; Salmonella MTGDNTLITSHGINRRDFMKLCAALAATMGLSSKAAAEMAESVSNPQRPPVIWIGAQECT WP_023248548.1 hydrogenase 2 small subunit [Salmonella enterica] Length: 372\nNumber of predicted TMHs: 1\nExp number of AAs in TMHs: 24.51281\nExp number, first 60 AAs: 0.26822\nTotal prob of N-in: 0.14402\noutside 1 331\nTMhelix 332 354\ninside 355 372

14241 GCF_002052625.1_ASM205262v1 Salmonella enterica Proteobacteria; Gammaproteobacteria; Enterobacterales; Enterobacteriaceae; Salmonella MTGDNTLITSHGINRRDFMKLCAALAATMGLSSKAAAEMAESVSNPQRPPVIWIGAQECT WP_000145427.1 hydrogenase 2 small subunit [Salmonella enterica] Length: 372\nNumber of predicted TMHs: 1\nExp number of AAs in TMHs: 24.48576\nExp number, first 60 AAs: 0.2692\nTotal prob of N-in: 0.14251\noutside 1 331\nTMhelix 332 354\ninside 355 372

14242 GCF_002052145.1_ASM205214v1 Salmonella enterica Proteobacteria; Gammaproteobacteria; Enterobacterales; Enterobacteriaceae; Salmonella MNNEETFYQAMRRKGVTRRSFLKFCSLAATSLGLGAGMTPKIAWALENKPRIPVVWIHGL WP_001058312.1 hydrogenase [Salmonella enterica] Length: 372\nNumber of predicted TMHs: 2\nExp number of AAs in TMHs: 36.51738\nExp number, first 60 AAs: 19.72849\nTotal prob of N-in: 0.96170\nPOSSIBLE N-term signal sequence\ninside 1 19\nTMhelix 20 39\noutside 40 325\nTMhelix 326 348\ninside 349 372

14243 GCF_002052395.1_ASM205239v1 Salmonella enterica Proteobacteria; Gammaproteobacteria; Enterobacterales; Enterobacteriaceae; Salmonella MNNEETFYQAMRRKGVTRRSFLKFCSLAATSLGLGAGMTPKIAWALENKPRIPVVWIHGL WP_001058312.1 hydrogenase [Salmonella enterica] Length: 372\nNumber of predicted TMHs: 2\nExp number of AAs in TMHs: 36.51738\nExp number, first 60 AAs: 19.72849\nTotal prob of N-in: 0.96170\nPOSSIBLE N-term signal sequence\ninside 1 19\nTMhelix 20 39\noutside 40 325\nTMhelix 326 348\ninside 349 372

14244 GCF_002051725.1_ASM205172v1 Salmonella enterica Proteobacteria; Gammaproteobacteria; Enterobacterales; Enterobacteriaceae; Salmonella MNNEETFYQAMRRKGVTRRSFLKFCSLAATSLGLGAGMTPKIAWALENKPRIPVVWIHGL WP_001058312.1 hydrogenase [Salmonella enterica] Length: 372\nNumber of predicted TMHs: 2\nExp number of AAs in TMHs: 36.51738\nExp number, first 60 AAs: 19.72849\nTotal prob of N-in: 0.96170\nPOSSIBLE N-term signal sequence\ninside 1 19\nTMhelix 20 39\noutside 40 325\nTMhelix 326 348\ninside 349 372

14245 GCF_002051865.1_ASM205186v1 Salmonella enterica Proteobacteria; Gammaproteobacteria; Enterobacterales; Enterobacteriaceae; Salmonella MNNEETFYQAMRRKGVTRRSFLKFCSLAATSLGLGAGMTPKIAWALENKPRIPVVWIHGL WP_001058312.1 hydrogenase [Salmonella enterica] Length: 372\nNumber of predicted TMHs: 2\nExp number of AAs in TMHs: 36.51738\nExp number, first 60 AAs: 19.72849\nTotal prob of N-in: 0.96170\nPOSSIBLE N-term signal sequence\ninside 1 19\nTMhelix 20 39\noutside 40 325\nTMhelix 326 348\ninside 349 372

14246 GCF_002051865.1_ASM205186v1 Salmonella enterica Proteobacteria; Gammaproteobacteria; Enterobacterales; Enterobacteriaceae; Salmonella
MTGDNTLITSHGINRRDFMKLCAALAATMGLSSKAAAEMAESVSNPQRPPVIWIGAQECT WP_000145429.1
hydrogenase 2 small subunit [Salmonella enterica] Length: 372\nNumber of predicted TMHs: 1\nExp number of AAs in TMHs: 24.51309\nExp number, first 60 AAs: 0.2682\nTotal prob of N-in: 0.14402\noutside 1 331\nTMhelix 332 354\ninside 355 372

14247 GCF_002051645.1_ASM205164v1 Salmonella enterica Proteobacteria; Gammaproteobacteria; Enterobacterales; Enterobacteriaceae; Salmonella
MNNEETFYQAMRRKGVTRRSFLKFCSLAATSLGLGAGMTPKIAWALENKPRIPVVWIHGL WP_001058312.1
hydrogenase [Salmonella enterica] Length: 372\nNumber of predicted TMHs: 2\nExp number of AAs in TMHs: 36.51738\nExp number, first 60 AAs: 19.72849\nTotal prob of N-in: 0.96170\nPOSSIBLE N-term signal sequence\ninside 1 19\nTMhelix 20 39\noutside 40 325\nTMhelix 326 348\ninside 349 372

14248 GCF_002051945.1_ASM205194v1 Salmonella enterica Proteobacteria; Gammaproteobacteria; Enterobacterales; Enterobacteriaceae; Salmonella
MTGDNTLITSHGINRRDFMKLCAALAATMGLSSKAAAEMAESVSNPQRPPVIWIGAQECT WP_000145429.1
hydrogenase 2 small subunit [Salmonella enterica] Length: 372\nNumber of predicted TMHs: 1\nExp number of AAs in TMHs: 24.51309\nExp number, first 60 AAs: 0.2682\nTotal prob of N-in: 0.14402\noutside 1 331\nTMhelix 332 354\ninside 355 372

14249 GCF_002036055.1_ASM203605v1 Salmonella enterica Proteobacteria; Gammaproteobacteria; Enterobacterales; Enterobacteriaceae; Salmonella
MNNEETFYQAMRRKGVTRRSFLKFCSLAATSLGLGAGMTPKIAWALENKPRIPVVWIHGL WP_001058306.1
hydrogenase [Salmonella enterica] Length: 372\nNumber of predicted TMHs: 2\nExp number of AAs in TMHs: 36.5540099999999\nExp number, first 60 AAs: 19.72732\nTotal prob of N-in: 0.96171\nPOSSIBLE N-term signal sequence\ninside 1 19\nTMhelix 20 39\noutside 40 325\nTMhelix 326 348\ninside 349 372

14250 GCF_002051315.1_ASM205131v1 Salmonella enterica Proteobacteria; Gammaproteobacteria; Enterobacterales; Enterobacteriaceae; Salmonella
MTGDNTLITSHGINRRDFMKLCAALAATMGLSSKAAAEMAESVSNPQRPPVIWIGAQECT WP_015589585.1
hydrogenase 2 small subunit [Salmonella enterica] Length: 372\nNumber of predicted TMHs: 1\nExp number of AAs in TMHs: 24.51304\nExp number, first 60 AAs: 0.2682\nTotal prob of N-in: 0.14402\noutside 1 331\nTMhelix 332 354\ninside 355 372

14251 GCF_001951925.1_ASM195192v1 Salmonella enterica Proteobacteria; Gammaproteobacteria; Enterobacterales; Enterobacteriaceae; Salmonella
MTGDNTLITSHGINRRDFMKLCAALAATMGLSSKAAAEMAESVSNPQRPPVIWIGAQECT WP_000145429.1
hydrogenase 2 small subunit [Salmonella enterica] Length: 372\nNumber of predicted TMHs: 1\nExp number of AAs in TMHs: 24.51309\nExp number, first 60 AAs: 0.2682\nTotal prob of N-in: 0.14402\noutside 1 331\nTMhelix 332 354\ninside 355 372

14252 GCF_002051785.1_ASM205178v1 Salmonella enterica Proteobacteria; Gammaproteobacteria; Enterobacterales; Enterobacteriaceae; Salmonella
MNNEETFYQAMRRKGVTRRSFLKFCSLAATSLGLGAGMTPKIAWALENKPRIPVVWIHGL WP_001058312.1
hydrogenase [Salmonella enterica] Length: 372\nNumber of predicted TMHs: 2\nExp number of AAs in TMHs: 36.51738\nExp number, first 60 AAs: 19.72849\nTotal prob of N-in: 0.96170\nPOSSIBLE N-term signal sequence\ninside 1 19\nTMhelix 20 39\noutside 40 325\nTMhelix 326 348\ninside 349 372

14253 GCF_002051525.1_ASM205152v1 Salmonella enterica Proteobacteria; Gammaproteobacteria; Enterobacterales; Enterobacteriaceae; Salmonella
MTGDNTLITSHGINRRDFMKLCAALAATMGLSSKAAAEMAESVSNPQRPPVIWIGAQECT WP_000145429.1
hydrogenase 2 small subunit [Salmonella enterica] Length: 372\nNumber of predicted TMHs: 1\nExp number of AAs in TMHs: 24.51309\nExp number, first 60 AAs: 0.2682\nTotal prob of N-in: 0.14402\noutside 1 331\nTMhelix 332 354\ninside 355 372

14254 GCF_002058405.1_ASM205840v1 Salmonella enterica Proteobacteria; Gammaproteobacteria; Enterobacterales; Enterobacteriaceae; Salmonella
MTGDNTLITSHGINRRDFMKLCAALAATMGLSSKAAAEMAESVSNPQRPPVIWIGAQECT WP_000145429.1
hydrogenase 2 small subunit [Salmonella enterica] Length: 372\nNumber of predicted TMHs: 1\nExp

number of AAs in TMHs: 24.51309\nExp number, first 60 AAs: 0.2682\nTotal prob of N-in: 0.14402\noutside 1 331\nTMhelix 332 354\ninside 355 372

14255 GCF_002030705.1_ASM203070v1 Salmonella enterica Proteobacteria; Gammaproteobacteria; Enterobacterales; Enterobacteriaceae; Salmonella
MTGDNTLITSHGINRRDFMKLCAALAATMGLSSKAAAEMAESVSNPQRPPVIWIGAQECT WP_000145429.1
hydrogenase 2 small subunit [Salmonella enterica] Length: 372\nNumber of predicted TMHs: 1\nExp number of AAs in TMHs: 24.51309\nExp number, first 60 AAs: 0.2682\nTotal prob of N-in: 0.14402\noutside 1 331\nTMhelix 332 354\ninside 355 372

14256 GCF_002054325.1_ASM205432v1 Salmonella enterica Proteobacteria; Gammaproteobacteria; Enterobacterales; Enterobacteriaceae; Salmonella
MNNEETFYQAMRRKGVTRRSFLKFCSLAATSLGLGAGMTPKIAWALENKPRIPVVWIHGL WP_001058312.1
hydrogenase [Salmonella enterica] Length: 372\nNumber of predicted TMHs: 2\nExp number of AAs in TMHs: 36.51738\nExp number, first 60 AAs: 19.72849\nTotal prob of N-in: 0.96170\nPOSSIBLE N-term signal sequence\ninside 1 19\nTMhelix 20 39\noutside 40 325\nTMhelix 326 348\ninside 349 372

14257 GCF_002054725.1_ASM205472v1 Salmonella enterica Proteobacteria; Gammaproteobacteria; Enterobacterales; Enterobacteriaceae; Salmonella
MNNEETFYQAMRRKGVTRRSFLKFCSLAATSLGLGAGMTPKIAWALENKPRIPVVWIHGL WP_001058311.1
hydrogenase [Salmonella enterica] Length: 372\nNumber of predicted TMHs: 2\nExp number of AAs in TMHs: 37.4699899999999\nExp number, first 60 AAs: 19.72699\nTotal prob of N-in: 0.96171\nPOSSIBLE N-term signal sequence\ninside 1 19\nTMhelix 20 39\noutside 40 325\nTMhelix 326 348\ninside 349 372

14258 GCF_002054385.1_ASM205438v1 Salmonella enterica Proteobacteria; Gammaproteobacteria; Enterobacterales; Enterobacteriaceae; Salmonella
MNNEETFYQAMRRKGVTRRSFLKFCSLAATSLGLSAGMTPKIAWALENKPRIPVVWIHGL WP_001058316.1
hydrogenase [Salmonella enterica] Length: 372\nNumber of predicted TMHs: 2\nExp number of AAs in TMHs: 35.76397\nExp number, first 60 AAs: 18.97201\nTotal prob of N-in: 0.92785\nPOSSIBLE N-term signal sequence\ninside 1 19\nTMhelix 20 39\noutside 40 325\nTMhelix 326 348\ninside 349 372

14259 GCF_002054655.1_ASM205465v1 Salmonella enterica Proteobacteria; Gammaproteobacteria; Enterobacterales; Enterobacteriaceae; Salmonella
MNNEETFYQAMRRKGVTRRSFLKFCSLAATSLGLGAGMTPKIAWALENKPRIPVVWIHGL WP_001058312.1
hydrogenase [Salmonella enterica] Length: 372\nNumber of predicted TMHs: 2\nExp number of AAs in TMHs: 36.51738\nExp number, first 60 AAs: 19.72849\nTotal prob of N-in: 0.96170\nPOSSIBLE N-term signal sequence\ninside 1 19\nTMhelix 20 39\noutside 40 325\nTMhelix 326 348\ninside 349 372

14260 GCF_002054435.1_ASM205443v1 Salmonella enterica Proteobacteria; Gammaproteobacteria; Enterobacterales; Enterobacteriaceae; Salmonella
MNNEETFYQAMRRKGVTRRSFLKFCSLAATSLGLSAGMTPKIAWALENKPRIPVVWIHGL WP_001058316.1
hydrogenase [Salmonella enterica] Length: 372\nNumber of predicted TMHs: 2\nExp number of AAs in TMHs: 35.76397\nExp number, first 60 AAs: 18.97201\nTotal prob of N-in: 0.92785\nPOSSIBLE N-term signal sequence\ninside 1 19\nTMhelix 20 39\noutside 40 325\nTMhelix 326 348\ninside 349 372

14261 GCF_002054865.1_ASM205486v1 Salmonella enterica Proteobacteria; Gammaproteobacteria; Enterobacterales; Enterobacteriaceae; Salmonella
MNNEETFYQAMRRKGVTRRSFLKFCSLAATSLGLGAGMTPKIAWALENKPRIPVVWIHGL WP_001058311.1
hydrogenase [Salmonella enterica] Length: 372\nNumber of predicted TMHs: 2\nExp number of AAs in TMHs: 37.4699899999999\nExp number, first 60 AAs: 19.72699\nTotal prob of N-in: 0.96171\nPOSSIBLE N-term signal sequence\ninside 1 19\nTMhelix 20 39\noutside 40 325\nTMhelix 326 348\ninside 349 372

14262 GCF_002054965.1_ASM205496v1 Salmonella enterica Proteobacteria; Gammaproteobacteria; Enterobacterales; Enterobacteriaceae; Salmonella
MTGDNTLITSHGINRRDFMKLCAALAATMGLSSKAAAEMAESVSNPQRPPVIWIGAQECT WP_000145429.1
hydrogenase 2 small subunit [Salmonella enterica] Length: 372\nNumber of predicted TMHs: 1\nExp number of AAs in TMHs: 24.51309\nExp number, first 60 AAs: 0.2682\nTotal prob of N-in: 0.14402\noutside 1 331\nTMhelix 332 354\ninside 355 372

14263 GCF_002055075.1_ASM205507v1 Salmonella enterica Proteobacteria; Gammaproteobacteria; Enterobacterales; Enterobacteriaceae; Salmonella

MTGDNTLITSHGINRRDFMKLCAALAATMGLSSKAAAEMAESVSNPQRPPVIWIGAQECT WP_000145429.1 hydrogenase 2 small subunit [Salmonella enterica] Length: 372\nNumber of predicted TMHs: 1\nExp number of AAs in TMHs: 24.51309\nExp number, first 60 AAs: 0.2682\nTotal prob of N-in: 0.14402\noutside 1 331\nTMhelix 332 354\ninside 355 372

14264 GCF_002055035.1_ASM205503v1 Salmonella enterica Proteobacteria; Gammaproteobacteria; Enterobacterales; Enterobacteriaceae; Salmonella MTGDNTLITSHGINRRDFMKLCAALAATMGLSSKAAAEMAESVSNPQRPPVIWIGAQECT WP_000145429.1 hydrogenase 2 small subunit [Salmonella enterica] Length: 372\nNumber of predicted TMHs: 1\nExp number of AAs in TMHs: 24.51309\nExp number, first 60 AAs: 0.2682\nTotal prob of N-in: 0.14402\noutside 1 331\nTMhelix 332 354\ninside 355 372

14265 GCF_002054925.1_ASM205492v1 Salmonella enterica Proteobacteria; Gammaproteobacteria; Enterobacterales; Enterobacteriaceae; Salmonella MNNEETFYQAMRRKGVTRRSFLKFCSLAATSLGLGAGMTPKIAWALENKPRIPVVWIHGL WP_001058311.1 hydrogenase [Salmonella enterica] Length: 372\nNumber of predicted TMHs: 2\nExp number of AAs in TMHs: 37.4699899999999\nExp number, first 60 AAs: 19.72699\nTotal prob of N-in: 0.96171\nPOSSIBLE N-term signal sequence\ninside 1 19\nTMhelix 20 39\noutside 40 325\nTMhelix 326 348\ninside 349 372

14266 GCF_002055235.1_ASM205523v1 Salmonella enterica Proteobacteria; Gammaproteobacteria; Enterobacterales; Enterobacteriaceae; Salmonella MTGDNTLITSHGINRRDFMKLCAALAATMGLSSKAAAEMAESVSNPQRPPVIWIGAQECT WP_000145429.1 hydrogenase 2 small subunit [Salmonella enterica] Length: 372\nNumber of predicted TMHs: 1\nExp number of AAs in TMHs: 24.51309\nExp number, first 60 AAs: 0.2682\nTotal prob of N-in: 0.14402\noutside 1 331\nTMhelix 332 354\ninside 355 372

14267 GCF_002055195.1_ASM205519v1 Salmonella enterica Proteobacteria; Gammaproteobacteria; Enterobacterales; Enterobacteriaceae; Salmonella MTGDNTLITSHGINRRDFMKLCAALAATMGLSSKAAAEMAESVSNPQRPPVIWIGAQECT WP_000145429.1 hydrogenase 2 small subunit [Salmonella enterica] Length: 372\nNumber of predicted TMHs: 1\nExp number of AAs in TMHs: 24.51309\nExp number, first 60 AAs: 0.2682\nTotal prob of N-in: 0.14402\noutside 1 331\nTMhelix 332 354\ninside 355 372

14268 GCF_002055335.1_ASM205533v1 Salmonella enterica Proteobacteria; Gammaproteobacteria; Enterobacterales; Enterobacteriaceae; Salmonella MTGDNTLITSHGINRRDFMKLCAALAATMGLSSKAAAEMAESVSNPQRPPVIWIGAQECT WP_000145429.1 hydrogenase 2 small subunit [Salmonella enterica] Length: 372\nNumber of predicted TMHs: 1\nExp number of AAs in TMHs: 24.51309\nExp number, first 60 AAs: 0.2682\nTotal prob of N-in: 0.14402\noutside 1 331\nTMhelix 332 354\ninside 355 372

14269 GCF_002055285.1_ASM205528v1 Salmonella enterica Proteobacteria; Gammaproteobacteria; Enterobacterales; Enterobacteriaceae; Salmonella MTGDNTLITSHGINRRDFMKLCAALAATMGLSSKAAAEMAESVSNPQRPPVIWIGAQECT WP_000145429.1 hydrogenase 2 small subunit [Salmonella enterica] Length: 372\nNumber of predicted TMHs: 1\nExp number of AAs in TMHs: 24.51309\nExp number, first 60 AAs: 0.2682\nTotal prob of N-in: 0.14402\noutside 1 331\nTMhelix 332 354\ninside 355 372

14270 GCF_002055335.1_ASM205533v1 Salmonella enterica Proteobacteria; Gammaproteobacteria; Enterobacterales; Enterobacteriaceae; Salmonella MNNEETFYQAMRRKGVTRRSFLKFCSLAATSLGLGAGMTPKIAWALENKPRIPVVWIHGL WP_001058312.1 hydrogenase [Salmonella enterica] Length: 372\nNumber of predicted TMHs: 2\nExp number of AAs in TMHs: 36.51738\nExp number, first 60 AAs: 19.72849\nTotal prob of N-in: 0.96170\nPOSSIBLE N-term signal sequence\ninside 1 19\nTMhelix 20 39\noutside 40 325\nTMhelix 326 348\ninside 349 372

14271 GCF_002055195.1_ASM205519v1 Salmonella enterica Proteobacteria; Gammaproteobacteria; Enterobacterales; Enterobacteriaceae; Salmonella MNNEETFYQAMRRKGVTRRSFLKFCSLAATSLGLSAGMTPKIAWALENKPRIPVVWIHGL WP_001058316.1 hydrogenase [Salmonella enterica] Length: 372\nNumber of predicted TMHs: 2\nExp number of AAs in TMHs: 35.76397\nExp number, first 60 AAs: 18.97201\nTotal prob of N-in: 0.92785\nPOSSIBLE N-term signal sequence\ninside 1 19\nTMhelix 20 39\noutside 40 325\nTMhelix 326 348\ninside 349 372

14272 GCF_002055285.1_ASM205528v1 Salmonella enterica Proteobacteria; Gammaproteobacteria; Enterobacterales; Enterobacteriaceae; Salmonella
MNNEETFYQAMRRKGVTRRSFLKFCSLAATSLGLSAGMTPKIAWALENKPRIPVVWIHGL WP_001058316.1
hydrogenase [Salmonella enterica] Length: 372\nNumber of predicted TMHs: 2\nExp number of AAs in TMHs: 35.76397\nExp number, first 60 AAs: 18.97201\nTotal prob of N-in: 0.92785\nPOSSIBLE N-term signal sequence\ninside 1 19\nTMhelix 20 39\noutside 40 325\nTMhelix 326 348\ninside 349 372

14273 GCF_002054965.1_ASM205496v1 Salmonella enterica Proteobacteria; Gammaproteobacteria; Enterobacterales; Enterobacteriaceae; Salmonella
MNNEETFYQAMRRKGVTRRSFLKFCSLAATSLGLGAGMTPKIAWALENKPRIPVVWIHGL WP_023213673.1
hydrogenase 1, small subunit [Salmonella enterica] Length: 372\nNumber of predicted TMHs: 2\nExp number of AAs in TMHs: 36.51753\nExp number, first 60 AAs: 19.72849\nTotal prob of N-in: 0.96170\nPOSSIBLE N-term signal sequence\ninside 1 19\nTMhelix 20 39\noutside 40 325\nTMhelix 326 348\ninside 349 372

14274 GCF_002055375.1_ASM205537v1 Salmonella enterica Proteobacteria; Gammaproteobacteria; Enterobacterales; Enterobacteriaceae; Salmonella
MTGDNTLITSHGINRRDFMKLCAALAATMGLSSKAAAEMAESVSNPQRPPVIWIGAQECT WP_000145429.1
hydrogenase 2 small subunit [Salmonella enterica] Length: 372\nNumber of predicted TMHs: 1\nExp number of AAs in TMHs: 24.51309\nExp number, first 60 AAs: 0.2682\nTotal prob of N-in: 0.14402\noutside 1 331\nTMhelix 332 354\ninside 355 372

14275 GCF_002055755.1_ASM205575v1 Salmonella enterica Proteobacteria; Gammaproteobacteria; Enterobacterales; Enterobacteriaceae; Salmonella
MNNEETFYQAMRRKGVTRRSFLKFCSLAATSLGLGAGMTPKIAWALENKPRIPVVWIHGL WP_001058312.1
hydrogenase [Salmonella enterica] Length: 372\nNumber of predicted TMHs: 2\nExp number of AAs in TMHs: 36.51738\nExp number, first 60 AAs: 19.72849\nTotal prob of N-in: 0.96170\nPOSSIBLE N-term signal sequence\ninside 1 19\nTMhelix 20 39\noutside 40 325\nTMhelix 326 348\ninside 349 372

14276 GCF_002055595.1_ASM205559v1 Salmonella enterica Proteobacteria; Gammaproteobacteria; Enterobacterales; Enterobacteriaceae; Salmonella
MNNEETFYQAMRRKGVTRRSFLKFCSLAATSLGLGAGMTPKIAWALENKPRIPVVWIHGL WP_080107816.1
[Ni/Fe] hydrogenase small subunit [Salmonella enterica] Length: 372\nNumber of predicted TMHs: 2\nExp number of AAs in TMHs: 36.51739\nExp number, first 60 AAs: 19.72849\nTotal prob of N-in: 0.96170\nPOSSIBLE N-term signal sequence\ninside 1 19\nTMhelix 20 39\noutside 40 325\nTMhelix 326 348\ninside 349 372

14277 GCF_002055475.1_ASM205547v1 Salmonella enterica Proteobacteria; Gammaproteobacteria; Enterobacterales; Enterobacteriaceae; Salmonella
MNNEETFYQAMRRKGVTRRSFLKFCSLAATSLGLGAGMTPKIAWALENKPRIPVVWIHGL WP_001058312.1
hydrogenase [Salmonella enterica] Length: 372\nNumber of predicted TMHs: 2\nExp number of AAs in TMHs: 36.51738\nExp number, first 60 AAs: 19.72849\nTotal prob of N-in: 0.96170\nPOSSIBLE N-term signal sequence\ninside 1 19\nTMhelix 20 39\noutside 40 325\nTMhelix 326 348\ninside 349 372

14278 GCF_002055815.1_ASM205581v1 Salmonella enterica Proteobacteria; Gammaproteobacteria; Enterobacterales; Enterobacteriaceae; Salmonella
MTGDNTLITSHGINRRDFMKLCAALAATMGLSSKAAAEMAESVSNPQRPPVIWIGAQECT WP_000145429.1
hydrogenase 2 small subunit [Salmonella enterica] Length: 372\nNumber of predicted TMHs: 1\nExp number of AAs in TMHs: 24.51309\nExp number, first 60 AAs: 0.2682\nTotal prob of N-in: 0.14402\noutside 1 331\nTMhelix 332 354\ninside 355 372

14279 GCF_002055885.1_ASM205588v1 Salmonella enterica Proteobacteria; Gammaproteobacteria; Enterobacterales; Enterobacteriaceae; Salmonella
MTGDNTLITSHGINRRDFMKLCAALAATMGLSSKAAAEMAESVSNPQRPPVIWIGAQECT WP_000145429.1
hydrogenase 2 small subunit [Salmonella enterica] Length: 372\nNumber of predicted TMHs: 1\nExp number of AAs in TMHs: 24.51309\nExp number, first 60 AAs: 0.2682\nTotal prob of N-in: 0.14402\noutside 1 331\nTMhelix 332 354\ninside 355 372

14280 GCF_002055955.1_ASM205595v1 Salmonella enterica Proteobacteria; Gammaproteobacteria; Enterobacterales; Enterobacteriaceae; Salmonella
MTGDNTLITSHGINRRDFMKLCAALAATMGLSSKAAAEMAESVSNPQRPPVIWIGAQECT WP_000145429.1
hydrogenase 2 small subunit [Salmonella enterica] Length: 372\nNumber of predicted TMHs: 1\nExp

number of AAs in TMHs: 24.51309\nExp number, first 60 AAs: 0.2682\nTotal prob of N-in: 0.14402\noutside 1 331\nTMhelix 332 354\ninside 355 372

14281 GCF_002055815.1_ASM205581v1 Salmonella enterica Proteobacteria; Gammaproteobacteria; Enterobacterales; Enterobacteriaceae; Salmonella MNNEETFYQAMRRKGVTRRSFLKFCSLAATSLGLGAGMTPKIAWALENKPRIPVVWIHGL WP_001058312.1 hydrogenase [Salmonella enterica] Length: 372\nNumber of predicted TMHs: 2\nExp number of AAs in TMHs: 36.51738\nExp number, first 60 AAs: 19.72849\nTotal prob of N-in: 0.96170\nPOSSIBLE N-term signal sequence\ninside 1 19\nTMhelix 20 39\noutside 40 325\nTMhelix 326 348\ninside 349 372

14282 GCF_002056135.1_ASM205613v1 Salmonella enterica Proteobacteria; Gammaproteobacteria; Enterobacterales; Enterobacteriaceae; Salmonella MNNEETFYQAMRRKGVTRRSFLKFCSLAATSLGLGAGMTPKIAWALENKPRIPVVWIHGL WP_001058307.1 hydrogenase [Salmonella enterica] Length: 372\nNumber of predicted TMHs: 2\nExp number of AAs in TMHs: 36.51738\nExp number, first 60 AAs: 19.72849\nTotal prob of N-in: 0.96170\nPOSSIBLE N-term signal sequence\ninside 1 19\nTMhelix 20 39\noutside 40 325\nTMhelix 326 348\ninside 349 372

14283 GCF_002056095.1_ASM205609v1 Salmonella enterica Proteobacteria; Gammaproteobacteria; Enterobacterales; Enterobacteriaceae; Salmonella MNNEETFYQAMRRKGVTRRSFLKFCSLAATSLGLGAGMTPKIAWALENKPRIPVVWIHGL WP_001058307.1 hydrogenase [Salmonella enterica] Length: 372\nNumber of predicted TMHs: 2\nExp number of AAs in TMHs: 36.51738\nExp number, first 60 AAs: 19.72849\nTotal prob of N-in: 0.96170\nPOSSIBLE N-term signal sequence\ninside 1 19\nTMhelix 20 39\noutside 40 325\nTMhelix 326 348\ninside 349 372

14284 GCF_002056245.1_ASM205624v1 Salmonella enterica Proteobacteria; Gammaproteobacteria; Enterobacterales; Enterobacteriaceae; Salmonella MTGDNTLITSHGINRRDFMKLCAALAATMGLSSKAAAEMAESVSNPQRPPVIWIGAQECT WP_000145429.1 hydrogenase 2 small subunit [Salmonella enterica] Length: 372\nNumber of predicted TMHs: 1\nExp number of AAs in TMHs: 24.51309\nExp number, first 60 AAs: 0.2682\nTotal prob of N-in: 0.14402\noutside 1 331\nTMhelix 332 354\ninside 355 372

14285 GCF_002056245.1_ASM205624v1 Salmonella enterica Proteobacteria; Gammaproteobacteria; Enterobacterales; Enterobacteriaceae; Salmonella MNNEETFYQAMRRKGVTRRSFLKFCSLAATSLGLGAGMTPKIAWALENKPRIPVVWIHGL WP_001058307.1 hydrogenase [Salmonella enterica] Length: 372\nNumber of predicted TMHs: 2\nExp number of AAs in TMHs: 36.51738\nExp number, first 60 AAs: 19.72849\nTotal prob of N-in: 0.96170\nPOSSIBLE N-term signal sequence\ninside 1 19\nTMhelix 20 39\noutside 40 325\nTMhelix 326 348\ninside 349 372

14286 GCF_002056195.1_ASM205619v1 Salmonella enterica Proteobacteria; Gammaproteobacteria; Enterobacterales; Enterobacteriaceae; Salmonella MNNEETFYQAMRRKGVTRRSFLKFCSLAATSLGLGAGMTPKIAWALENKPRIPVVWIHGL WP_001058307.1 hydrogenase [Salmonella enterica] Length: 372\nNumber of predicted TMHs: 2\nExp number of AAs in TMHs: 36.51738\nExp number, first 60 AAs: 19.72849\nTotal prob of N-in: 0.96170\nPOSSIBLE N-term signal sequence\ninside 1 19\nTMhelix 20 39\noutside 40 325\nTMhelix 326 348\ninside 349 372

14287 GCF_002056345.1_ASM205634v1 Salmonella enterica Proteobacteria; Gammaproteobacteria; Enterobacterales; Enterobacteriaceae; Salmonella MTGDNTLITSHGINRRDFMKLCAALAATMGLSSKAAAEMAESVSNPQRPPVIWIGAQECT WP_000145429.1 hydrogenase 2 small subunit [Salmonella enterica] Length: 372\nNumber of predicted TMHs: 1\nExp number of AAs in TMHs: 24.51309\nExp number, first 60 AAs: 0.2682\nTotal prob of N-in: 0.14402\noutside 1 331\nTMhelix 332 354\ninside 355 372

14288 GCF_002033395.1_ASM203339v1 Salmonella enterica Proteobacteria; Gammaproteobacteria; Enterobacterales; Enterobacteriaceae; Salmonella MNNEETFYQAMRRKGVTRRSFLKFCSLAATSLGLGAGMTPKIAWALENKPRIPVVWIHGL WP_001058307.1 hydrogenase [Salmonella enterica] Length: 372\nNumber of predicted TMHs: 2\nExp number of AAs in TMHs: 36.51738\nExp number, first 60 AAs: 19.72849\nTotal prob of N-in: 0.96170\nPOSSIBLE N-term signal sequence\ninside 1 19\nTMhelix 20 39\noutside 40 325\nTMhelix 326 348\ninside 349 372

14289 GCF_002044945.1_ASM204494v1 Salmonella enterica Proteobacteria; Gammaproteobacteria; Enterobacterales; Enterobacteriaceae; Salmonella

MNNEETFYQAMRRKGVTRRSFLKFCSLAATSLGLGAGMTPKIAWALENKPRIPVVWIHGL WP_001058312.1 hydrogenase [Salmonella enterica] Length: 372\nNumber of predicted TMHs: 2\nExp number of AAs in TMHs: 36.51738\nExp number, first 60 AAs: 19.72849\nTotal prob of N-in: 0.96170\nPOSSIBLE N-term signal sequence\ninside 1 19\nTMhelix 20 39\noutside 40 325\nTMhelix 326 348\ninside 349 372

14290 GCF_002033175.1_ASM203317v1 Salmonella enterica Proteobacteria; Gammaproteobacteria; Enterobacterales; Enterobacteriaceae; Salmonella MTGDNTLITSHGINRRDFMKLCAALAATMGLSSKAAAEMAESVSNPQRPPVIWIGAQECT WP_000145429.1 hydrogenase 2 small subunit [Salmonella enterica] Length: 372\nNumber of predicted TMHs: 1\nExp number of AAs in TMHs: 24.51309\nExp number, first 60 AAs: 0.2682\nTotal prob of N-in: 0.14402\noutside 1 331\nTMhelix 332 354\ninside 355 372

14291 GCF_002061575.1_ASM206157v1 Salmonella enterica Proteobacteria; Gammaproteobacteria; Enterobacterales; Enterobacteriaceae; Salmonella MTGDNTLITSHGINRRDFMKLCAALAATMGLSSKAAAEMAESVSNPQRPPVIWIGAQECT WP_000145431.1 hydrogenase 2 small subunit [Salmonella enterica] Length: 372\nNumber of predicted TMHs: 1\nExp number of AAs in TMHs: 24.51505\nExp number, first 60 AAs: 0.2682\nTotal prob of N-in: 0.14406\noutside 1 331\nTMhelix 332 354\ninside 355 372

14292 GCF_002032505.1_ASM203250v1 Salmonella enterica Proteobacteria; Gammaproteobacteria; Enterobacterales; Enterobacteriaceae; Salmonella MNNEETFYQAMRRKGVTRRSFLKFCSLAATSLGLGAGMTPKIAWALENKPRIPVVWIHGL WP_001058311.1 hydrogenase [Salmonella enterica] Length: 372\nNumber of predicted TMHs: 2\nExp number of AAs in TMHs: 37.4699899999999\nExp number, first 60 AAs: 19.72699\nTotal prob of N-in: 0.96171\nPOSSIBLE N-term signal sequence\ninside 1 19\nTMhelix 20 39\noutside 40 325\nTMhelix 326 348\ninside 349 372

14293 GCF_002032605.1_ASM203260v1 Salmonella enterica Proteobacteria; Gammaproteobacteria; Enterobacterales; Enterobacteriaceae; Salmonella MNNEETFYQAMRRKGVTRRSFLKFCSLAATSLGLGAGMTPKIAWALENKPRIPVVWIHGL WP_001058312.1 hydrogenase [Salmonella enterica] Length: 372\nNumber of predicted TMHs: 2\nExp number of AAs in TMHs: 36.51738\nExp number, first 60 AAs: 19.72849\nTotal prob of N-in: 0.96170\nPOSSIBLE N-term signal sequence\ninside 1 19\nTMhelix 20 39\noutside 40 325\nTMhelix 326 348\ninside 349 372

14294 GCF_002052255.1_ASM205225v1 Salmonella enterica Proteobacteria; Gammaproteobacteria; Enterobacterales; Enterobacteriaceae; Salmonella MTGDNTLITSHGINRRDFMKLCAALAATMGLSSKAAAEMAESVSNPQRPPVIWIGAQECT WP_000145429.1 hydrogenase 2 small subunit [Salmonella enterica] Length: 372\nNumber of predicted TMHs: 1\nExp number of AAs in TMHs: 24.51309\nExp number, first 60 AAs: 0.2682\nTotal prob of N-in: 0.14402\noutside 1 331\nTMhelix 332 354\ninside 355 372

14295 GCF_002032605.1_ASM203260v1 Salmonella enterica Proteobacteria; Gammaproteobacteria; Enterobacterales; Enterobacteriaceae; Salmonella MTGDNTLITSHGINRRDFMKLCAALAATMGLSSKAAAEMAESVSNPQRPPVIWIGAQECT WP_000145429.1 hydrogenase 2 small subunit [Salmonella enterica] Length: 372\nNumber of predicted TMHs: 1\nExp number of AAs in TMHs: 24.51309\nExp number, first 60 AAs: 0.2682\nTotal prob of N-in: 0.14402\noutside 1 331\nTMhelix 332 354\ninside 355 372

14296 GCF_001643905.1_CRJJGF_00101_v1.0Salmonella enterica Proteobacteria; Gammaproteobacteria; Enterobacterales; Enterobacteriaceae; Salmonella MTGDNTLITSHGINRRDFMKLCAALAATMGLSSKAAAEMAESVSNPQRPPVIWIGAQECT WP_000145429.1 hydrogenase 2 small subunit [Salmonella enterica] Length: 372\nNumber of predicted TMHs: 1\nExp number of AAs in TMHs: 24.51309\nExp number, first 60 AAs: 0.2682\nTotal prob of N-in: 0.14402\noutside 1 331\nTMhelix 332 354\ninside 355 372

14297 GCF_001643795.1_CRJJGF_00099_v1.0Salmonella enterica Proteobacteria; Gammaproteobacteria; Enterobacterales; Enterobacteriaceae; Salmonella MTGDNTLITSHGINRRDFMKLCAALAATMGLSSKAAAEMAESVSNPQRPPVIWIGAQECT WP_000145429.1 hydrogenase 2 small subunit [Salmonella enterica] Length: 372\nNumber of predicted TMHs: 1\nExp number of AAs in TMHs: 24.51309\nExp number, first 60 AAs: 0.2682\nTotal prob of N-in: 0.14402\noutside 1 331\nTMhelix 332 354\ninside 355 372

14298 GCF_002032645.1_ASM203264v1 Salmonella enterica Proteobacteria; Gammaproteobacteria; Enterobacterales; Enterobacteriaceae; Salmonella
MTGDNTLITSHGINRRDFMKLCAALAATMGLSSKAAAEMAESVSNPQRPPVIWIGAQECT WP_000145429.1
hydrogenase 2 small subunit [Salmonella enterica] Length: 372\nNumber of predicted TMHs: 1\nExp number of AAs in TMHs: 24.51309\nExp number, first 60 AAs: 0.2682\nTotal prob of N-in: 0.14402\noutside 1 331\nTMhelix 332 354\ninside 355 372

14299 GCF_001643825.1_CRJJGF_00061_v1.0Salmonella enterica Proteobacteria; Gammaproteobacteria; Enterobacterales; Enterobacteriaceae; Salmonella
MTGDNTLITSHGINRRDFMKLCAALAATMGLSSKAAAEMAESVSNPQRPPVIWIGAQECT WP_000145429.1
hydrogenase 2 small subunit [Salmonella enterica] Length: 372\nNumber of predicted TMHs: 1\nExp number of AAs in TMHs: 24.51309\nExp number, first 60 AAs: 0.2682\nTotal prob of N-in: 0.14402\noutside 1 331\nTMhelix 332 354\ninside 355 372

14300 GCF_002032505.1_ASM203250v1 Salmonella enterica Proteobacteria; Gammaproteobacteria; Enterobacterales; Enterobacteriaceae; Salmonella
MTGDNTLITSHGINRRDFMKLCAALAATMGLSSKAAAEMAESVSNPQRPPVIWIGAQECT WP_000145429.1
hydrogenase 2 small subunit [Salmonella enterica] Length: 372\nNumber of predicted TMHs: 1\nExp number of AAs in TMHs: 24.51309\nExp number, first 60 AAs: 0.2682\nTotal prob of N-in: 0.14402\noutside 1 331\nTMhelix 332 354\ninside 355 372

14301 GCF_002052145.1_ASM205214v1 Salmonella enterica Proteobacteria; Gammaproteobacteria; Enterobacterales; Enterobacteriaceae; Salmonella
MTGDNTLITSHGINRRDFMKLCAALAATMGLSSKAAAEMAESVSNPQRPPVIWIGAQECT WP_000145429.1
hydrogenase 2 small subunit [Salmonella enterica] Length: 372\nNumber of predicted TMHs: 1\nExp number of AAs in TMHs: 24.51309\nExp number, first 60 AAs: 0.2682\nTotal prob of N-in: 0.14402\noutside 1 331\nTMhelix 332 354\ninside 355 372

14302 GCF_002060675.1_ASM206067v1 Salmonella enterica Proteobacteria; Gammaproteobacteria; Enterobacterales; Enterobacteriaceae; Salmonella
MTGDNTLITSHGINRRDFMKLCAALAATMGLSSKAAAEMAESVSNPQRPPVIWIGAQECT WP_000145431.1
hydrogenase 2 small subunit [Salmonella enterica] Length: 372\nNumber of predicted TMHs: 1\nExp number of AAs in TMHs: 24.51505\nExp number, first 60 AAs: 0.2682\nTotal prob of N-in: 0.14406\noutside 1 331\nTMhelix 332 354\ninside 355 372

14303 GCF_001643905.1_CRJJGF_00101_v1.0Salmonella enterica Proteobacteria; Gammaproteobacteria; Enterobacterales; Enterobacteriaceae; Salmonella
MNNEETFYQAMRRKGVTRRSFLKFCSLAATSLGLGAGMTPKIAWALENKPRIPVVWIHGL WP_001058312.1
hydrogenase [Salmonella enterica] Length: 372\nNumber of predicted TMHs: 2\nExp number of AAs in TMHs: 36.51738\nExp number, first 60 AAs: 19.72849\nTotal prob of N-in: 0.96170\nPOSSIBLE N-term signal sequence\ninside 1 19\nTMhelix 20 39\noutside 40 325\nTMhelix 326 348\ninside 349 372

14304 GCF_002060975.1_ASM206097v1 Salmonella enterica Proteobacteria; Gammaproteobacteria; Enterobacterales; Enterobacteriaceae; Salmonella
MNNEETFYQAMRRKGVTRRSFLKFCSLAATSLGLGAGMTPKIAWALENKPRIPVVWIHGL WP_001058312.1
hydrogenase [Salmonella enterica] Length: 372\nNumber of predicted TMHs: 2\nExp number of AAs in TMHs: 36.51738\nExp number, first 60 AAs: 19.72849\nTotal prob of N-in: 0.96170\nPOSSIBLE N-term signal sequence\ninside 1 19\nTMhelix 20 39\noutside 40 325\nTMhelix 326 348\ninside 349 372

14305 GCF_001643795.1_CRJJGF_00099_v1.0Salmonella enterica Proteobacteria; Gammaproteobacteria; Enterobacterales; Enterobacteriaceae; Salmonella
MNNEETFYQAMRRKGVTRRSFLKFCSLAATSLGLGAGMTPKIAWALENKPRIPVVWIHGL WP_001058312.1
hydrogenase [Salmonella enterica] Length: 372\nNumber of predicted TMHs: 2\nExp number of AAs in TMHs: 36.51738\nExp number, first 60 AAs: 19.72849\nTotal prob of N-in: 0.96170\nPOSSIBLE N-term signal sequence\ninside 1 19\nTMhelix 20 39\noutside 40 325\nTMhelix 326 348\ninside 349 372

14306 GCF_002045605.1_ASM204560v1 Salmonella enterica Proteobacteria; Gammaproteobacteria; Enterobacterales; Enterobacteriaceae; Salmonella
MTGDNTLITSHGINRRDFMKLCAALAATMGLSSKAAAEMAESVSNPQRPPVIWIGAQECT WP_000145429.1
hydrogenase 2 small subunit [Salmonella enterica] Length: 372\nNumber of predicted TMHs: 1\nExp

number of AAs in TMHs: 24.51309\nExp number, first 60 AAs: 0.2682\nTotal prob of N-in: 0.14402\noutside 1 331\nTMhelix 332 354\ninside 355 372

14307 GCF_002066885.1_ASM206688v1 Salmonella enterica Proteobacteria; Gammaproteobacteria; Enterobacterales; Enterobacteriaceae; Salmonella MNNEETFYQAMRRKGVTRRSFLKFCSLAATSLGLGAGMTPKIAWALENKPRIPVVWIHGL WP_001058307.1 hydrogenase [Salmonella enterica] Length: 372\nNumber of predicted TMHs: 2\nExp number of AAs in TMHs: 36.51738\nExp number, first 60 AAs: 19.72849\nTotal prob of N-in: 0.96170\nPOSSIBLE N-term signal sequence\ninside 1 19\nTMhelix 20 39\noutside 40 325\nTMhelix 326 348\ninside 349 372

14308 GCF_002045605.1_ASM204560v1 Salmonella enterica Proteobacteria; Gammaproteobacteria; Enterobacterales; Enterobacteriaceae; Salmonella MNNEETFYQAMRRKGVTRRSFLKFCSLAATSLGLGAGMTPKIAWALENKPRIPVVWIHGL WP_001058311.1 hydrogenase [Salmonella enterica] Length: 372\nNumber of predicted TMHs: 2\nExp number of AAs in TMHs: 37.4699899999999\nExp number, first 60 AAs: 19.72699\nTotal prob of N-in: 0.96171\nPOSSIBLE N-term signal sequence\ninside 1 19\nTMhelix 20 39\noutside 40 325\nTMhelix 326 348\ninside 349 372

14309 GCF_002053895.1_ASM205389v1 Salmonella enterica Proteobacteria; Gammaproteobacteria; Enterobacterales; Enterobacteriaceae; Salmonella MTGDNTLITSHGINRRDFMKLCAALAATMGLSSKAAAEMAESVSNPQRPPVIWIGAQECT WP_000145429.1 hydrogenase 2 small subunit [Salmonella enterica] Length: 372\nNumber of predicted TMHs: 1\nExp number of AAs in TMHs: 24.51309\nExp number, first 60 AAs: 0.2682\nTotal prob of N-in: 0.14402\noutside 1 331\nTMhelix 332 354\ninside 355 372

14310 GCF_002045565.1_ASM204556v1 Salmonella enterica Proteobacteria; Gammaproteobacteria; Enterobacterales; Enterobacteriaceae; Salmonella MTGDNTLITSHGINRRDFMKLCAALAATMGLSSKAAAEMAESVSNPQRPPVIWIGAQECT WP_000145429.1 hydrogenase 2 small subunit [Salmonella enterica] Length: 372\nNumber of predicted TMHs: 1\nExp number of AAs in TMHs: 24.51309\nExp number, first 60 AAs: 0.2682\nTotal prob of N-in: 0.14402\noutside 1 331\nTMhelix 332 354\ninside 355 372

14311 GCF_002144455.1_ASM214445v1 Salmonella enterica Proteobacteria; Gammaproteobacteria; Enterobacterales; Enterobacteriaceae; Salmonella MNNEETFYQAMRRKGVTRRSFLKFCSLAATSLGLGAGMTPKIAWALENKPRIPVVWIHGL WP_001058312.1 hydrogenase [Salmonella enterica] Length: 372\nNumber of predicted TMHs: 2\nExp number of AAs in TMHs: 36.51738\nExp number, first 60 AAs: 19.72849\nTotal prob of N-in: 0.96170\nPOSSIBLE N-term signal sequence\ninside 1 19\nTMhelix 20 39\noutside 40 325\nTMhelix 326 348\ninside 349 372

14312 GCF_002045645.1_ASM204564v1 Salmonella enterica Proteobacteria; Gammaproteobacteria; Enterobacterales; Enterobacteriaceae; Salmonella MTGDNTLITSHGINRRDFMKLCAALAATMGLSSKAAAEMAESVSNPQRPPVIWIGAQECT WP_000145429.1 hydrogenase 2 small subunit [Salmonella enterica] Length: 372\nNumber of predicted TMHs: 1\nExp number of AAs in TMHs: 24.51309\nExp number, first 60 AAs: 0.2682\nTotal prob of N-in: 0.14402\noutside 1 331\nTMhelix 332 354\ninside 355 372

14313 GCF_002144415.1_ASM214441v1 Salmonella enterica Proteobacteria; Gammaproteobacteria; Enterobacterales; Enterobacteriaceae; Salmonella MNNEETFYQAMRRKGVTRRSFLKFCSLAATSLGLGAGMTPKIAWALENKPRIPVVWIHGL WP_001058312.1 hydrogenase [Salmonella enterica] Length: 372\nNumber of predicted TMHs: 2\nExp number of AAs in TMHs: 36.51738\nExp number, first 60 AAs: 19.72849\nTotal prob of N-in: 0.96170\nPOSSIBLE N-term signal sequence\ninside 1 19\nTMhelix 20 39\noutside 40 325\nTMhelix 326 348\ninside 349 372

14314 GCF_002144355.1_ASM214435v1 Salmonella enterica Proteobacteria; Gammaproteobacteria; Enterobacterales; Enterobacteriaceae; Salmonella MNNEETFYQAMRRKGVTRRSFLKFCSLAATSLGLGAGMTPKIAWALENKPRIPVVWIHGL WP_001058312.1 hydrogenase [Salmonella enterica] Length: 372\nNumber of predicted TMHs: 2\nExp number of AAs in TMHs: 36.51738\nExp number, first 60 AAs: 19.72849\nTotal prob of N-in: 0.96170\nPOSSIBLE N-term signal sequence\ninside 1 19\nTMhelix 20 39\noutside 40 325\nTMhelix 326 348\ninside 349 372

14315 GCF_002034525.1_ASM203452v1 Salmonella enterica Proteobacteria; Gammaproteobacteria; Enterobacterales; Enterobacteriaceae; Salmonella

MTGDNTLITSHGINRRDFMKLCAALAATMGLSSKAAAEMAESVSNPQRPPVIWIGAQECT WP_000145429.1 hydrogenase 2 small subunit [Salmonella enterica] Length: 372\nNumber of predicted TMHs: 1\nExp number of AAs in TMHs: 24.51309\nExp number, first 60 AAs: 0.2682\nTotal prob of N-in: 0.14402\noutside 1 331\nTMhelix 332 354\ninside 355 372

14316 GCF_002060885.1_ASM206088v1 Salmonella enterica Proteobacteria; Gammaproteobacteria; Enterobacterales; Enterobacteriaceae; Salmonella MNNEETFYQAMRRKGVTRRSFLKFCSLAATSLGLGAGMTPKIAWALENKPRIPVVWIHGL WP_001058312.1 hydrogenase [Salmonella enterica] Length: 372\nNumber of predicted TMHs: 2\nExp number of AAs in TMHs: 36.51738\nExp number, first 60 AAs: 19.72849\nTotal prob of N-in: 0.96170\nPOSSIBLE N-term signal sequence\ninside 1 19\nTMhelix 20 39\noutside 40 325\nTMhelix 326 348\ninside 349 372

14317 GCF_002045675.1_ASM204567v1 Salmonella enterica Proteobacteria; Gammaproteobacteria; Enterobacterales; Enterobacteriaceae; Salmonella MTGDNTLITSHGINRRDFMKLCAALAATMGLSSKAAAEMAESVSNPQRPPVIWIGAQECT WP_000145429.1 hydrogenase 2 small subunit [Salmonella enterica] Length: 372\nNumber of predicted TMHs: 1\nExp number of AAs in TMHs: 24.51309\nExp number, first 60 AAs: 0.2682\nTotal prob of N-in: 0.14402\noutside 1 331\nTMhelix 332 354\ninside 355 372

14318 GCF_002066635.1_ASM206663v1 Salmonella enterica Proteobacteria; Gammaproteobacteria; Enterobacterales; Enterobacteriaceae; Salmonella MNNEETFYQAMRRKGVTRRSFLKFCSLAATSLGLGAGMTPKIAWALENKPRIPVVWIHGL WP_001058312.1 hydrogenase [Salmonella enterica] Length: 372\nNumber of predicted TMHs: 2\nExp number of AAs in TMHs: 36.51738\nExp number, first 60 AAs: 19.72849\nTotal prob of N-in: 0.96170\nPOSSIBLE N-term signal sequence\ninside 1 19\nTMhelix 20 39\noutside 40 325\nTMhelix 326 348\ninside 349 372

14319 GCF_002061075.1_ASM206107v1 Salmonella enterica Proteobacteria; Gammaproteobacteria; Enterobacterales; Enterobacteriaceae; Salmonella MTGDNTLITSHGINRRDFMKLCAALAATMGLSSKAAAEMAESVSNPQRPPVIWIGAQECT WP_000145429.1 hydrogenase 2 small subunit [Salmonella enterica] Length: 372\nNumber of predicted TMHs: 1\nExp number of AAs in TMHs: 24.51309\nExp number, first 60 AAs: 0.2682\nTotal prob of N-in: 0.14402\noutside 1 331\nTMhelix 332 354\ninside 355 372

14320 GCF_002144455.1_ASM214445v1 Salmonella enterica Proteobacteria; Gammaproteobacteria; Enterobacterales; Enterobacteriaceae; Salmonella MTGDNTLITSHGINRRDFMKLCAALAATMGLSSKAAAEMAESVSNPQRPPVIWIGAQECT WP_000145429.1 hydrogenase 2 small subunit [Salmonella enterica] Length: 372\nNumber of predicted TMHs: 1\nExp number of AAs in TMHs: 24.51309\nExp number, first 60 AAs: 0.2682\nTotal prob of N-in: 0.14402\noutside 1 331\nTMhelix 332 354\ninside 355 372

14321 GCF_002144355.1_ASM214435v1 Salmonella enterica Proteobacteria; Gammaproteobacteria; Enterobacterales; Enterobacteriaceae; Salmonella MTGDNTLITSHGINRRDFMKLCAALAATMGLSSKAAAEMAESVSNPQRPPVIWIGAQECT WP_000145429.1 hydrogenase 2 small subunit [Salmonella enterica] Length: 372\nNumber of predicted TMHs: 1\nExp number of AAs in TMHs: 24.51309\nExp number, first 60 AAs: 0.2682\nTotal prob of N-in: 0.14402\noutside 1 331\nTMhelix 332 354\ninside 355 372

14322 GCF_002030705.1_ASM203070v1 Salmonella enterica Proteobacteria; Gammaproteobacteria; Enterobacterales; Enterobacteriaceae; Salmonella MNNEETFYQAMRRKGVTRRSFLKFCSLAATSLGLSAGMTPKIAWALENKPRIPVVWIHGL WP_001058316.1 hydrogenase [Salmonella enterica] Length: 372\nNumber of predicted TMHs: 2\nExp number of AAs in TMHs: 35.76397\nExp number, first 60 AAs: 18.97201\nTotal prob of N-in: 0.92785\nPOSSIBLE N-term signal sequence\ninside 1 19\nTMhelix 20 39\noutside 40 325\nTMhelix 326 348\ninside 349 372

14323 GCF_002051475.1_ASM205147v1 Salmonella enterica Proteobacteria; Gammaproteobacteria; Enterobacterales; Enterobacteriaceae; Salmonella MTGDNTLITSHGINRRDFMKLCAALAATMGLSSKAAAEMAESVSNPQRPPVIWIGAQECT WP_000145429.1 hydrogenase 2 small subunit [Salmonella enterica] Length: 372\nNumber of predicted TMHs: 1\nExp number of AAs in TMHs: 24.51309\nExp number, first 60 AAs: 0.2682\nTotal prob of N-in: 0.14402\noutside 1 331\nTMhelix 332 354\ninside 355 372

14324 GCF_002051435.1_ASM205143v1 Salmonella enterica Proteobacteria; Gammaproteobacteria; Enterobacterales; Enterobacteriaceae; Salmonella
MTGDNTLITSHGINRRDFMKLCAALAATMGLSSKAAAEMAESVSNPQRPPVIWIGAQECT WP_000145429.1
hydrogenase 2 small subunit [Salmonella enterica] Length: 372\nNumber of predicted TMHs: 1\nExp number of AAs in TMHs: 24.51309\nExp number, first 60 AAs: 0.2682\nTotal prob of N-in: 0.14402\noutside 1 331\nTMhelix 332 354\ninside 355 372

14325 GCF_002051195.1_ASM205119v1 Salmonella enterica Proteobacteria; Gammaproteobacteria; Enterobacterales; Enterobacteriaceae; Salmonella
MNNEETFYQAMRRKGVTRRSFLKFCSLAATSLGLGAGMTPKIAWALENKPRIPVVWIHGL WP_001058312.1
hydrogenase [Salmonella enterica] Length: 372\nNumber of predicted TMHs: 2\nExp number of AAs in TMHs: 36.51738\nExp number, first 60 AAs: 19.72849\nTotal prob of N-in: 0.96170\nPOSSIBLE N-term signal sequence\ninside 1 19\nTMhelix 20 39\noutside 40 325\nTMhelix 326 348\ninside 349 372

14326 GCF_002051365.1_ASM205136v1 Salmonella enterica Proteobacteria; Gammaproteobacteria; Enterobacterales; Enterobacteriaceae; Salmonella
MNNEETFYQAMRRKGVTRRSFLKFCSLAATSLGLGAGMTPKIAWALENKPRIPVVWIHGL WP_001058312.1
hydrogenase [Salmonella enterica] Length: 372\nNumber of predicted TMHs: 2\nExp number of AAs in TMHs: 36.51738\nExp number, first 60 AAs: 19.72849\nTotal prob of N-in: 0.96170\nPOSSIBLE N-term signal sequence\ninside 1 19\nTMhelix 20 39\noutside 40 325\nTMhelix 326 348\ninside 349 372

14327 GCF_002051365.1_ASM205136v1 Salmonella enterica Proteobacteria; Gammaproteobacteria; Enterobacterales; Enterobacteriaceae; Salmonella
MTGDNTLITSHGINRRDFMKLCAALAATMGLSSKAAAEMAESVSNPQRPPVIWIGAQECT WP_000145429.1
hydrogenase 2 small subunit [Salmonella enterica] Length: 372\nNumber of predicted TMHs: 1\nExp number of AAs in TMHs: 24.51309\nExp number, first 60 AAs: 0.2682\nTotal prob of N-in: 0.14402\noutside 1 331\nTMhelix 332 354\ninside 355 372

14328 GCF_002051045.1_ASM205104v1 Salmonella enterica Proteobacteria; Gammaproteobacteria; Enterobacterales; Enterobacteriaceae; Salmonella
MNNEETFYQAMRRKGVTRRSFLKFCSLAATSLGLGAGMTPQIAWALENKPRIPVVWIHGL WP_023227615.1
hydrogenase 1, small subunit [Salmonella enterica] Length: 372\nNumber of predicted TMHs: 2\nExp number of AAs in TMHs: 39.8591699999999\nExp number, first 60 AAs: 22.13093\nTotal prob of N-in: 0.98639\nPOSSIBLE N-term signal sequence\ninside 1 20\nTMhelix 21 43\noutside 44 325\nTMhelix 326 348\ninside 349 372

14329 GCF_002051025.1_ASM205102v1 Salmonella enterica Proteobacteria; Gammaproteobacteria; Enterobacterales; Enterobacteriaceae; Salmonella
MLKFRTIFLKKIFTLKNSILARRGAVIVIVSAVFTSIMFFAHSWASDKEVAMTLSVLNSN WP_023226644.1 hypothetical protein [Salmonella enterica] Length: 204\nNumber of predicted TMHs: 2\nExp number of AAs in TMHs: 44.99526\nExp number, first 60 AAs: 22.33786\nTotal prob of N-in: 0.87083\nPOSSIBLE N-term signal sequence\ninside 1 19\nTMhelix 20 42\noutside 43 171\nTMhelix 172 194\ninside 195 204

14330 GCF_002051095.1_ASM205109v1 Salmonella enterica Proteobacteria; Gammaproteobacteria; Enterobacterales; Enterobacteriaceae; Salmonella
MTGDNTLITSHGINRRDFMKLCAALAATMGLSSKAAAEMAESVSNPQRPPVIWIGAQECT WP_000145429.1
hydrogenase 2 small subunit [Salmonella enterica] Length: 372\nNumber of predicted TMHs: 1\nExp number of AAs in TMHs: 24.51309\nExp number, first 60 AAs: 0.2682\nTotal prob of N-in: 0.14402\noutside 1 331\nTMhelix 332 354\ninside 355 372

14331 GCF_002044725.1_ASM204472v1 Salmonella enterica Proteobacteria; Gammaproteobacteria; Enterobacterales; Enterobacteriaceae; Salmonella
MNNEETFYQAMRRKGVTRRSFLKFCSLAATSLGLGAGMTPKIAWALENKPRIPVVWIHGL WP_001058312.1
hydrogenase [Salmonella enterica] Length: 372\nNumber of predicted TMHs: 2\nExp number of AAs in TMHs: 36.51738\nExp number, first 60 AAs: 19.72849\nTotal prob of N-in: 0.96170\nPOSSIBLE N-term signal sequence\ninside 1 19\nTMhelix 20 39\noutside 40 325\nTMhelix 326 348\ninside 349 372

14332 GCF_001951945.1_ASM195194v1 Salmonella enterica Proteobacteria; Gammaproteobacteria; Enterobacterales; Enterobacteriaceae; Salmonella
MTGDNTLITSHGINRRDFMKLCAALAATMGLSSKAAAEMAESVSNPQRPPVIWIGAQECT WP_000145429.1
hydrogenase 2 small subunit [Salmonella enterica] Length: 372\nNumber of predicted TMHs: 1\nExp

number of AAs in TMHs: 24.51309\nExp number, first 60 AAs: 0.2682\nTotal prob of N-in: 0.14402\noutside 1 331\nTMhelix 332 354\ninside 355 372

14333 GCF_002051045.1_ASM205104v1 Salmonella enterica Proteobacteria; Gammaproteobacteria; Enterobacterales; Enterobacteriaceae; Salmonella
MTGDNTLITSHGINRRDFMKLCAALAATMGLSSKAAAEMAESVSNPQRPPVIWIGAQECT WP_000145429.1
hydrogenase 2 small subunit [Salmonella enterica] Length: 372\nNumber of predicted TMHs: 1\nExp number of AAs in TMHs: 24.51309\nExp number, first 60 AAs: 0.2682\nTotal prob of N-in: 0.14402\noutside 1 331\nTMhelix 332 354\ninside 355 372

14334 GCF_001951945.1_ASM195194v1 Salmonella enterica Proteobacteria; Gammaproteobacteria; Enterobacterales; Enterobacteriaceae; Salmonella
MNNEETFYQAMRRKGVTRRSFLKFCSLAATSLGLGAGMTPKIAWALENKPRIPVVWIHGL WP_001058312.1
hydrogenase [Salmonella enterica] Length: 372\nNumber of predicted TMHs: 2\nExp number of AAs in TMHs: 36.51738\nExp number, first 60 AAs: 19.72849\nTotal prob of N-in: 0.96170\nPOSSIBLE N-term signal sequence\ninside 1 19\nTMhelix 20 39\noutside 40 325\nTMhelix 326 348\ninside 349 372

14335 GCF_002035805.1_ASM203580v1 Salmonella enterica Proteobacteria; Gammaproteobacteria; Enterobacterales; Enterobacteriaceae; Salmonella
MNNEETFYQAMRRKGVTRRSFLKFCSLAATSLGLGAGMTPKIAWALENKPRIPVVWIHGL WP_001058314.1
hydrogenase [Salmonella enterica] Length: 372\nNumber of predicted TMHs: 2\nExp number of AAs in TMHs: 36.3110299999999\nExp number, first 60 AAs: 19.72774\nTotal prob of N-in: 0.96169\nPOSSIBLE N-term signal sequence\ninside 1 19\nTMhelix 20 39\noutside 40 325\nTMhelix 326 348\ninside 349 372

14336 GCF_001831755.1_ASM183175v1 Salmonella enterica Proteobacteria; Gammaproteobacteria; Enterobacterales; Enterobacteriaceae; Salmonella
MTGDNTLITSHGINRRDFMKLCAALAATMGLSSKAAAEMAESVSNPQRPPVIWIGAQECT WP_000145429.1
hydrogenase 2 small subunit [Salmonella enterica] Length: 372\nNumber of predicted TMHs: 1\nExp number of AAs in TMHs: 24.51309\nExp number, first 60 AAs: 0.2682\nTotal prob of N-in: 0.14402\noutside 1 331\nTMhelix 332 354\ninside 355 372

14337 GCF_002035805.1_ASM203580v1 Salmonella enterica Proteobacteria; Gammaproteobacteria; Enterobacterales; Enterobacteriaceae; Salmonella
MTGDNTLITSHGINRRDFMKLCAALAATMGLSSKAAAEMAESVSNPQRPPVIWIGAQECT WP_000145432.1
hydrogenase 2 small subunit [Salmonella enterica] Length: 372\nNumber of predicted TMHs: 1\nExp number of AAs in TMHs: 24.5101\nExp number, first 60 AAs: 0.2682\nTotal prob of N-in: 0.14397\noutside 1 331\nTMhelix 332 354\ninside 355 372

14338 GCF_002043585.1_ASM204358v1 Salmonella enterica Proteobacteria; Gammaproteobacteria; Enterobacterales; Enterobacteriaceae; Salmonella
MTGDNTLITSHGINRRDFMKLCAALAATMGLSSKAAAEMAESVSNPQRPPVIWIGAQECT WP_000145429.1
hydrogenase 2 small subunit [Salmonella enterica] Length: 372\nNumber of predicted TMHs: 1\nExp number of AAs in TMHs: 24.51309\nExp number, first 60 AAs: 0.2682\nTotal prob of N-in: 0.14402\noutside 1 331\nTMhelix 332 354\ninside 355 372

14339 GCF_002044605.1_ASM204460v1 Salmonella enterica Proteobacteria; Gammaproteobacteria; Enterobacterales; Enterobacteriaceae; Salmonella
MNNEETFYQAMRRKGVTRRSFLKFCSLAATSLGLGAGMTPKIAWALENKPRIPVVWIHGL WP_001058311.1
hydrogenase [Salmonella enterica] Length: 372\nNumber of predicted TMHs: 2\nExp number of AAs in TMHs: 37.4699899999999\nExp number, first 60 AAs: 19.72699\nTotal prob of N-in: 0.96171\nPOSSIBLE N-term signal sequence\ninside 1 19\nTMhelix 20 39\noutside 40 325\nTMhelix 326 348\ninside 349 372

14340 GCF_002035185.1_ASM203518v1 Salmonella enterica Proteobacteria; Gammaproteobacteria; Enterobacterales; Enterobacteriaceae; Salmonella
MTGDNTLITSHGINRRDFMKLCAALAATMGLSSKAAAEMAESVSNPQRPPVIWIGAQECT WP_079776618.1
MULTISPECIES: hydrogenase 2 small subunit [Salmonella] Length: 372\nNumber of predicted TMHs: 1\nExp number of AAs in TMHs: 24.65705\nExp number, first 60 AAs: 0.26805\nTotal prob of N-in: 0.13761\noutside 1 331\nTMhelix 332 354\ninside 355 372

14341 GCF_002061895.1_ASM206189v1 Salmonella enterica Proteobacteria; Gammaproteobacteria; Enterobacterales; Enterobacteriaceae; Salmonella

MTGDNTLITSHGINRRDFMKLCAALAATMGLSSKAAAEMAESVSNPQRPPVIWIGAQECT WP_000145429.1
hydrogenase 2 small subunit [Salmonella enterica] Length: 372\nNumber of predicted TMHs: 1\nExp number of AAs in TMHs: 24.51309\nExp number, first 60 AAs: 0.2682\nTotal prob of N-in: 0.14402\noutside 1 331\nTMhelix 332 354\ninside 355 372

14342 GCF_002061975.1_ASM206197v1 Salmonella enterica Proteobacteria; Gammaproteobacteria; Enterobacterales; Enterobacteriaceae; Salmonella
MNNEETFYQAMRRKGVTRRSFLKFCSLAATSLGLGAGMTPKIAWALENKPRIPVVWIHGL WP_001058312.1
hydrogenase [Salmonella enterica] Length: 372\nNumber of predicted TMHs: 2\nExp number of AAs in TMHs: 36.51738\nExp number, first 60 AAs: 19.72849\nTotal prob of N-in: 0.96170\nPOSSIBLE N-term signal sequence\ninside 1 19\nTMhelix 20 39\noutside 40 325\nTMhelix 326 348\ninside 349 372

14343 GCF_002052065.1_ASM205206v1 Salmonella enterica Proteobacteria; Gammaproteobacteria; Enterobacterales; Enterobacteriaceae; Salmonella
MTGDNTLITSHGINRRDFMKLCAALAATMGLSSKAAAEMAESVSNPQRPPVIWIGAQECT WP_000145429.1
hydrogenase 2 small subunit [Salmonella enterica] Length: 372\nNumber of predicted TMHs: 1\nExp number of AAs in TMHs: 24.51309\nExp number, first 60 AAs: 0.2682\nTotal prob of N-in: 0.14402\noutside 1 331\nTMhelix 332 354\ninside 355 372

14344 GCF_002051025.1_ASM205102v1 Salmonella enterica Proteobacteria; Gammaproteobacteria; Enterobacterales; Enterobacteriaceae; Salmonella
MTGDNTLITSHGINRRDFMKLCAALAATMGLSSKAAAEMAESVSNPQRPPVIWIGAQECT WP_000145429.1
hydrogenase 2 small subunit [Salmonella enterica] Length: 372\nNumber of predicted TMHs: 1\nExp number of AAs in TMHs: 24.51309\nExp number, first 60 AAs: 0.2682\nTotal prob of N-in: 0.14402\noutside 1 331\nTMhelix 332 354\ninside 355 372

14345 GCF_002057775.1_ASM205777v1 Salmonella enterica Proteobacteria; Gammaproteobacteria; Enterobacterales; Enterobacteriaceae; Salmonella
MTGDNTLITSHGINRRDFMKLCAALAATMGLSSKAAAEMAESVSNPQRPPVIWIGAQECT WP_000145429.1
hydrogenase 2 small subunit [Salmonella enterica] Length: 372\nNumber of predicted TMHs: 1\nExp number of AAs in TMHs: 24.51309\nExp number, first 60 AAs: 0.2682\nTotal prob of N-in: 0.14402\noutside 1 331\nTMhelix 332 354\ninside 355 372

14346 GCF_001952395.1_ASM195239v1 Salmonella enterica Proteobacteria; Gammaproteobacteria; Enterobacterales; Enterobacteriaceae; Salmonella
MTGDNTLITSHGINRRDFMKLCAALAATMGLSSKAAAEMAESVSNPQRPPVIWIGAQECT WP_000145429.1
hydrogenase 2 small subunit [Salmonella enterica] Length: 372\nNumber of predicted TMHs: 1\nExp number of AAs in TMHs: 24.51309\nExp number, first 60 AAs: 0.2682\nTotal prob of N-in: 0.14402\noutside 1 331\nTMhelix 332 354\ninside 355 372

14347 GCF_002057645.1_ASM205764v1 Salmonella enterica Proteobacteria; Gammaproteobacteria; Enterobacterales; Enterobacteriaceae; Salmonella
MNNEETFYQAMRRKGVTRRSFLKFCSLAATSLGLGAGMTPKIAWALENKPRIPVVWIHGL WP_001058307.1
hydrogenase [Salmonella enterica] Length: 372\nNumber of predicted TMHs: 2\nExp number of AAs in TMHs: 36.51738\nExp number, first 60 AAs: 19.72849\nTotal prob of N-in: 0.96170\nPOSSIBLE N-term signal sequence\ninside 1 19\nTMhelix 20 39\noutside 40 325\nTMhelix 326 348\ninside 349 372

14348 GCF_002058155.1_ASM205815v1 Salmonella enterica Proteobacteria; Gammaproteobacteria; Enterobacterales; Enterobacteriaceae; Salmonella
MTGDNTLITSHGINRRDFMKLCAALAATMGLSSKAAAEMAESVSNPQRPPVIWIGAQECT WP_000145428.1
hydrogenase 2 small subunit [Salmonella enterica] Length: 372\nNumber of predicted TMHs: 1\nExp number of AAs in TMHs: 24.50677\nExp number, first 60 AAs: 0.2682\nTotal prob of N-in: 0.14390\noutside 1 331\nTMhelix 332 354\ninside 355 372

14349 GCF_001952545.1_ASM195254v1 Salmonella enterica Proteobacteria; Gammaproteobacteria; Enterobacterales; Enterobacteriaceae; Salmonella
MTGDNTLITSHGINRRDFMKLCAALAATMGLSSKAAAEMAESVSNPQRPPVIWIGAQECT WP_000145429.1
hydrogenase 2 small subunit [Salmonella enterica] Length: 372\nNumber of predicted TMHs: 1\nExp number of AAs in TMHs: 24.51309\nExp number, first 60 AAs: 0.2682\nTotal prob of N-in: 0.14402\noutside 1 331\nTMhelix 332 354\ninside 355 372

14350 GCF_002035185.1_ASM203518v1 Salmonella enterica Proteobacteria; Gammaproteobacteria; Enterobacterales; Enterobacteriaceae; Salmonella
MNNEETFYQAMRRKGVTRRSFLKFCSLAATSLGLGAGMTPKIAWALENKPRIPVVWIHGL WP_023185544.1
hydrogenase 1, small subunit [Salmonella enterica] Length: 372\nNumber of predicted TMHs: 2\nExp number of AAs in TMHs: 36.55018\nExp number, first 60 AAs: 19.72756\nTotal prob of N-in: 0.96171\nPOSSIBLE N-term signal sequence\ninside 1 19\nTMhelix 20 39\noutside 40 325\nTMhelix 326 348\ninside 349 372

14351 GCF_001952625.1_ASM195262v1 Salmonella enterica Proteobacteria; Gammaproteobacteria; Enterobacterales; Enterobacteriaceae; Salmonella
MTGDNTLITSHGINRRDFMKLCAALAATMGLSSKAAAEMAESVSNPQRPPVIWIGAQECT WP_000145429.1
hydrogenase 2 small subunit [Salmonella enterica] Length: 372\nNumber of predicted TMHs: 1\nExp number of AAs in TMHs: 24.51309\nExp number, first 60 AAs: 0.2682\nTotal prob of N-in: 0.14402\noutside 1 331\nTMhelix 332 354\ninside 355 372

14352 GCF_002035125.1_ASM203512v1 Salmonella enterica Proteobacteria; Gammaproteobacteria; Enterobacterales; Enterobacteriaceae; Salmonella
MNNEETFYQAMRRKGVTRRSFLKFCSLAATSLGLGAGMTPKIAWALENKPRIPVVWIHGL WP_001058312.1
hydrogenase [Salmonella enterica] Length: 372\nNumber of predicted TMHs: 2\nExp number of AAs in TMHs: 36.51738\nExp number, first 60 AAs: 19.72849\nTotal prob of N-in: 0.96170\nPOSSIBLE N-term signal sequence\ninside 1 19\nTMhelix 20 39\noutside 40 325\nTMhelix 326 348\ninside 349 372

14353 GCF_001952625.1_ASM195262v1 Salmonella enterica Proteobacteria; Gammaproteobacteria; Enterobacterales; Enterobacteriaceae; Salmonella
MNNEETFYQAMRRKGVTRRSFLKFCSLAATSLGLGAGMTPKIAWALENKPRIPVVWIHGL WP_001058312.1
hydrogenase [Salmonella enterica] Length: 372\nNumber of predicted TMHs: 2\nExp number of AAs in TMHs: 36.51738\nExp number, first 60 AAs: 19.72849\nTotal prob of N-in: 0.96170\nPOSSIBLE N-term signal sequence\ninside 1 19\nTMhelix 20 39\noutside 40 325\nTMhelix 326 348\ninside 349 372

14354 GCF_002061935.1_ASM206193v1 Salmonella enterica Proteobacteria; Gammaproteobacteria; Enterobacterales; Enterobacteriaceae; Salmonella
MNNEETFYQAMRRKGVTRRSFLKFCSLAATSLGLGAGMTPKIAWALENKPRIPVVWIHGL WP_001058306.1
hydrogenase [Salmonella enterica] Length: 372\nNumber of predicted TMHs: 2\nExp number of AAs in TMHs: 36.5540099999999\nExp number, first 60 AAs: 19.72732\nTotal prob of N-in: 0.96171\nPOSSIBLE N-term signal sequence\ninside 1 19\nTMhelix 20 39\noutside 40 325\nTMhelix 326 348\ninside 349 372

14355 GCF_002054555.1_ASM205455v1 Salmonella enterica Proteobacteria; Gammaproteobacteria; Enterobacterales; Enterobacteriaceae; Salmonella
MTGDNTLITSHGINRRDFMKLCAALAATMGLSSKAAAEMAESVSNPQRPPVIWIGAQECT WP_000145429.1
hydrogenase 2 small subunit [Salmonella enterica] Length: 372\nNumber of predicted TMHs: 1\nExp number of AAs in TMHs: 24.51309\nExp number, first 60 AAs: 0.2682\nTotal prob of N-in: 0.14402\noutside 1 331\nTMhelix 332 354\ninside 355 372

14356 GCF_002044645.1_ASM204464v1 Salmonella enterica Proteobacteria; Gammaproteobacteria; Enterobacterales; Enterobacteriaceae; Salmonella
MNNEETFYQAMRRKGVTRRSFLKFCSLAATSLGLGAGMTPKIAWALENKPRIPVVWIHGL WP_001058311.1
hydrogenase [Salmonella enterica] Length: 372\nNumber of predicted TMHs: 2\nExp number of AAs in TMHs: 37.4699899999999\nExp number, first 60 AAs: 19.72699\nTotal prob of N-in: 0.96171\nPOSSIBLE N-term signal sequence\ninside 1 19\nTMhelix 20 39\noutside 40 325\nTMhelix 326 348\ninside 349 372

14357 GCF_002050805.1_ASM205080v1 Salmonella enterica Proteobacteria; Gammaproteobacteria; Enterobacterales; Enterobacteriaceae; Salmonella
MNNEETFYQAMRRKGVTRRSFLKFCSLAATSLGLGAGMTPQIAWALENKPRIPVVWIHGL WP_023227615.1
hydrogenase 1, small subunit [Salmonella enterica] Length: 372\nNumber of predicted TMHs: 2\nExp number of AAs in TMHs: 39.8591699999999\nExp number, first 60 AAs: 22.13093\nTotal prob of N-in: 0.98639\nPOSSIBLE N-term signal sequence\ninside 1 20\nTMhelix 21 43\noutside 44 325\nTMhelix 326 348\ninside 349 372

14358 GCF_002050805.1_ASM205080v1 Salmonella enterica Proteobacteria; Gammaproteobacteria; Enterobacterales; Enterobacteriaceae; Salmonella
MLKFRTIFLKKIFTLKNSILARRGAVIVIVSAVFTSIMFFAHSWASDKEVAMTLSVLNSN WP_023226644.1 hypothetical protein [Salmonella enterica] Length: 204\nNumber of predicted TMHs: 2\nExp number of AAs in TMHs: 44.99526\nExp

number, first 60 AAs: 22.33786\nTotal prob of N-in: 0.87083\nPOSSIBLE N-term signal sequence\ninside 1 19\nTMhelix 20 42\noutside 43 171\nTMhelix 172 194\ninside 195 204

14359 GCF_002062405.1_ASM206240v1 Salmonella enterica Proteobacteria; Gammaproteobacteria; Enterobacterales; Enterobacteriaceae; Salmonella MTGDNTLITSHGINRRDFMKLCAALAATMGLSSKAAAEMAESVSNPQRPPVIWIGAQECT WP_000145429.1 hydrogenase 2 small subunit [Salmonella enterica] Length: 372\nNumber of predicted TMHs: 1\nExp number of AAs in TMHs: 24.51309\nExp number, first 60 AAs: 0.2682\nTotal prob of N-in: 0.14402\noutside 1 331\nTMhelix 332 354\ninside 355 372

14360 GCF_002050805.1_ASM205080v1 Salmonella enterica Proteobacteria; Gammaproteobacteria; Enterobacterales; Enterobacteriaceae; Salmonella MTGDNTLITSHGINRRDFMKLCAALAATMGLSSKAAAEMAESVSNPQRPPVIWIGAQECT WP_000145429.1 hydrogenase 2 small subunit [Salmonella enterica] Length: 372\nNumber of predicted TMHs: 1\nExp number of AAs in TMHs: 24.51309\nExp number, first 60 AAs: 0.2682\nTotal prob of N-in: 0.14402\noutside 1 331\nTMhelix 332 354\ninside 355 372

14361 GCF_002034625.1_ASM203462v1 Salmonella enterica Proteobacteria; Gammaproteobacteria; Enterobacterales; Enterobacteriaceae; Salmonella MNNEETFYQAMRRKGVTRRSFLKFCSLAATSLGLGAGMTPKIAWALENKPRIPVVWIHGL WP_001058312.1 hydrogenase [Salmonella enterica] Length: 372\nNumber of predicted TMHs: 2\nExp number of AAs in TMHs: 36.51738\nExp number, first 60 AAs: 19.72849\nTotal prob of N-in: 0.96170\nPOSSIBLE N-term signal sequence\ninside 1 19\nTMhelix 20 39\noutside 40 325\nTMhelix 326 348\ninside 349 372

14362 GCF_002062405.1_ASM206240v1 Salmonella enterica Proteobacteria; Gammaproteobacteria; Enterobacterales; Enterobacteriaceae; Salmonella MNNEETFYQAMRRKGVTRRSFLKFCSLAATSLGLGAGMTPKIAWALENKPRIPVVWIHGL WP_001058312.1 hydrogenase [Salmonella enterica] Length: 372\nNumber of predicted TMHs: 2\nExp number of AAs in TMHs: 36.51738\nExp number, first 60 AAs: 19.72849\nTotal prob of N-in: 0.96170\nPOSSIBLE N-term signal sequence\ninside 1 19\nTMhelix 20 39\noutside 40 325\nTMhelix 326 348\ninside 349 372

14363 GCF_002058155.1_ASM205815v1 Salmonella enterica Proteobacteria; Gammaproteobacteria; Enterobacterales; Enterobacteriaceae; Salmonella MNNEETLYQAMRRKGVTRRSFLKFCSLAATSLGLGAGMTPKIAWALENKPRIPVVWIHGL WP_001752445.1 hydrogenase-1 small chain [Salmonella enterica] Length: 372\nNumber of predicted TMHs: 2\nExp number of AAs in TMHs: 36.5123\nExp number, first 60 AAs: 19.72341\nTotal prob of N-in: 0.96144\nPOSSIBLE N-term signal sequence\ninside 1 19\nTMhelix 20 39\noutside 40 325\nTMhelix 326 348\ninside 349 372

14364 GCF_002057675.1_ASM205767v1 Salmonella enterica Proteobacteria; Gammaproteobacteria; Enterobacterales; Enterobacteriaceae; Salmonella MNNEETFYQAMRRKGVTRRSFLKFCSLAATSLGLGAGMTPKIAWALENKPRIPVVWIHGL WP_001058307.1 hydrogenase [Salmonella enterica] Length: 372\nNumber of predicted TMHs: 2\nExp number of AAs in TMHs: 36.51738\nExp number, first 60 AAs: 19.72849\nTotal prob of N-in: 0.96170\nPOSSIBLE N-term signal sequence\ninside 1 19\nTMhelix 20 39\noutside 40 325\nTMhelix 326 348\ninside 349 372

14365 GCF_002034625.1_ASM203462v1 Salmonella enterica Proteobacteria; Gammaproteobacteria; Enterobacterales; Enterobacteriaceae; Salmonella MTGDNTLITSHGINRRDFMKLCAALAATMGLSSKAAAEMAESVSNPQRPPVIWIGAQECT WP_000145429.1 hydrogenase 2 small subunit [Salmonella enterica] Length: 372\nNumber of predicted TMHs: 1\nExp number of AAs in TMHs: 24.51309\nExp number, first 60 AAs: 0.2682\nTotal prob of N-in: 0.14402\noutside 1 331\nTMhelix 332 354\ninside 355 372

14366 GCF_002144415.1_ASM214441v1 Salmonella enterica Proteobacteria; Gammaproteobacteria; Enterobacterales; Enterobacteriaceae; Salmonella MTGDNTLITSHGINRRDFMKLCAALAATMGLSSKAAAEMAESVSNPQRPPVIWIGAQECT WP_000145429.1 hydrogenase 2 small subunit [Salmonella enterica] Length: 372\nNumber of predicted TMHs: 1\nExp number of AAs in TMHs: 24.51309\nExp number, first 60 AAs: 0.2682\nTotal prob of N-in: 0.14402\noutside 1 331\nTMhelix 332 354\ninside 355 372

14367 GCF_001642435.1_ASM164243v1 Salmonella enterica Proteobacteria; Gammaproteobacteria; Enterobacterales; Enterobacteriaceae; Salmonella

MTGDNTLITSHGINRRDFMKLCAALAATMGLSSKAAAEMAESVSNPQRPPVIWIGAQECT WP_000145429.1 hydrogenase 2 small subunit [Salmonella enterica] Length: 372\nNumber of predicted TMHs: 1\nExp number of AAs in TMHs: 24.51309\nExp number, first 60 AAs: 0.2682\nTotal prob of N-in: 0.14402\noutside 1 331\nTMhelix 332 354\ninside 355 372

14368 GCF_001642485.1_ASM164248v1 Salmonella enterica Proteobacteria; Gammaproteobacteria; Enterobacterales; Enterobacteriaceae; Salmonella MTGDNTLITSHGINRRDFMKLCAALAATMGLSSKAAAEMAESVSNPQRPPVIWIGAQECT WP_000145429.1 hydrogenase 2 small subunit [Salmonella enterica] Length: 372\nNumber of predicted TMHs: 1\nExp number of AAs in TMHs: 24.51309\nExp number, first 60 AAs: 0.2682\nTotal prob of N-in: 0.14402\noutside 1 331\nTMhelix 332 354\ninside 355 372

14369 GCF_002066885.1_ASM206688v1 Salmonella enterica Proteobacteria; Gammaproteobacteria; Enterobacterales; Enterobacteriaceae; Salmonella MTGDNTLITSHGINRRDFMKLCAALAATMGLSSKAAAEMAESVSNPQRPPVIWIGAQECT WP_000145429.1 hydrogenase 2 small subunit [Salmonella enterica] Length: 372\nNumber of predicted TMHs: 1\nExp number of AAs in TMHs: 24.51309\nExp number, first 60 AAs: 0.2682\nTotal prob of N-in: 0.14402\noutside 1 331\nTMhelix 332 354\ninside 355 372

14370 GCF_002036585.1_ASM203658v1 Salmonella enterica Proteobacteria; Gammaproteobacteria; Enterobacterales; Enterobacteriaceae; Salmonella MNNEETFYQAMRRKGVTRRSFLKFCSLAATSLGLGAGMTPKIAWALENKPRIPVVWIHGL WP_001058312.1 hydrogenase [Salmonella enterica] Length: 372\nNumber of predicted TMHs: 2\nExp number of AAs in TMHs: 36.51738\nExp number, first 60 AAs: 19.72849\nTotal prob of N-in: 0.96170\nPOSSIBLE N-term signal sequence\ninside 1 19\nTMhelix 20 39\noutside 40 325\nTMhelix 326 348\ninside 349 372

14371 GCF_001642435.1_ASM164243v1 Salmonella enterica Proteobacteria; Gammaproteobacteria; Enterobacterales; Enterobacteriaceae; Salmonella MNNEETFYQAMRRKGVTRRSFLKFCSLAATSLGLGAGMTPKIAWALENKPRIPVVWIHGL WP_001058311.1 hydrogenase [Salmonella enterica] Length: 372\nNumber of predicted TMHs: 2\nExp number of AAs in TMHs: 37.4699899999999\nExp number, first 60 AAs: 19.72699\nTotal prob of N-in: 0.96171\nPOSSIBLE N-term signal sequence\ninside 1 19\nTMhelix 20 39\noutside 40 325\nTMhelix 326 348\ninside 349 372

14372 GCF_002061785.1_ASM206178v1 Salmonella enterica Proteobacteria; Gammaproteobacteria; Enterobacterales; Enterobacteriaceae; Salmonella MTGDNTLITSHGINRRDFMKLCAALAATMGLSSKAAAEMAESVSNPQRPPVIWIGAQECT WP_000145429.1 hydrogenase 2 small subunit [Salmonella enterica] Length: 372\nNumber of predicted TMHs: 1\nExp number of AAs in TMHs: 24.51309\nExp number, first 60 AAs: 0.2682\nTotal prob of N-in: 0.14402\noutside 1 331\nTMhelix 332 354\ninside 355 372

14373 GCF_002058415.1_ASM205841v1 Salmonella enterica Proteobacteria; Gammaproteobacteria; Enterobacterales; Enterobacteriaceae; Salmonella MNNEETFYQAMRRKGVTRRSFLKFCSLAATSLGLGAGMTPKIAWALENKPRIPVVWIHGL WP_001058311.1 hydrogenase [Salmonella enterica] Length: 372\nNumber of predicted TMHs: 2\nExp number of AAs in TMHs: 37.4699899999999\nExp number, first 60 AAs: 19.72699\nTotal prob of N-in: 0.96171\nPOSSIBLE N-term signal sequence\ninside 1 19\nTMhelix 20 39\noutside 40 325\nTMhelix 326 348\ninside 349 372

14374 GCF_002061575.1_ASM206157v1 Salmonella enterica Proteobacteria; Gammaproteobacteria; Enterobacterales; Enterobacteriaceae; Salmonella MNNEETFYQAMRRKGVTRRSFLKFCSLAATSLGLGAGMTPKIAWALENKPRIPVVWIHGL WP_001058312.1 hydrogenase [Salmonella enterica] Length: 372\nNumber of predicted TMHs: 2\nExp number of AAs in TMHs: 36.51738\nExp number, first 60 AAs: 19.72849\nTotal prob of N-in: 0.96170\nPOSSIBLE N-term signal sequence\ninside 1 19\nTMhelix 20 39\noutside 40 325\nTMhelix 326 348\ninside 349 372

14375 GCF_002058485.1_ASM205848v1 Salmonella enterica Proteobacteria; Gammaproteobacteria; Enterobacterales; Enterobacteriaceae; Salmonella MNNEETFYQAMRRKGVTRRSFLKFCSLAATSLGLGAGMTPKIAWALENKPRIPVVWIHGL WP_001058312.1 hydrogenase [Salmonella enterica] Length: 372\nNumber of predicted TMHs: 2\nExp number of AAs in TMHs: 36.51738\nExp number, first 60 AAs: 19.72849\nTotal prob of N-in: 0.96170\nPOSSIBLE N-term signal sequence\ninside 1 19\nTMhelix 20 39\noutside 40 325\nTMhelix 326 348\ninside 349 372

14376 GCF_002048925.1_ASM204892v1 Salmonella enterica Proteobacteria; Gammaproteobacteria; Enterobacterales; Enterobacteriaceae; Salmonella
MNNEETFYQAMRRKGVTRRSFLKFCSLAATSLGLGAGMTPKIAWALENKPRIPVVWIHGL WP_001058307.1
hydrogenase [Salmonella enterica] Length: 372\nNumber of predicted TMHs: 2\nExp number of AAs in TMHs: 36.51738\nExp number, first 60 AAs: 19.72849\nTotal prob of N-in: 0.96170\nPOSSIBLE N-term signal sequence\ninside 1 19\nTMhelix 20 39\noutside 40 325\nTMhelix 326 348\ninside 349 372

14377 GCF_002034175.1_ASM203417v1 Salmonella enterica Proteobacteria; Gammaproteobacteria; Enterobacterales; Enterobacteriaceae; Salmonella
MNNEETFYQAMRRKGVTRRSFLKFCSLAATSLGLGAGMTPKIAWALENKPRIPVVWIHGL WP_001058312.1
hydrogenase [Salmonella enterica] Length: 372\nNumber of predicted TMHs: 2\nExp number of AAs in TMHs: 36.51738\nExp number, first 60 AAs: 19.72849\nTotal prob of N-in: 0.96170\nPOSSIBLE N-term signal sequence\ninside 1 19\nTMhelix 20 39\noutside 40 325\nTMhelix 326 348\ninside 349 372

14378 GCF_002034175.1_ASM203417v1 Salmonella enterica Proteobacteria; Gammaproteobacteria; Enterobacterales; Enterobacteriaceae; Salmonella
MTGDNTLITSHGINRRDFMKLCAALAATMGLSSKAAAEMAESVSNPQRPPVIWIGAQECT WP_000145429.1
hydrogenase 2 small subunit [Salmonella enterica] Length: 372\nNumber of predicted TMHs: 1\nExp number of AAs in TMHs: 24.51309\nExp number, first 60 AAs: 0.2682\nTotal prob of N-in: 0.14402\noutside 1 331\nTMhelix 332 354\ninside 355 372

14379 GCF_002034075.1_ASM203407v1 Salmonella enterica Proteobacteria; Gammaproteobacteria; Enterobacterales; Enterobacteriaceae; Salmonella
MNNEETFYQAMRRKGVTRRSFLKFCSLAATSLGLGAGMTPKIAWALENKPRIPVVWIHGL WP_001058311.1
hydrogenase [Salmonella enterica] Length: 372\nNumber of predicted TMHs: 2\nExp number of AAs in TMHs: 37.4699899999999\nExp number, first 60 AAs: 19.72699\nTotal prob of N-in: 0.96171\nPOSSIBLE N-term signal sequence\ninside 1 19\nTMhelix 20 39\noutside 40 325\nTMhelix 326 348\ninside 349 372

14380 GCF_002054635.1_ASM205463v1 Salmonella enterica Proteobacteria; Gammaproteobacteria; Enterobacterales; Enterobacteriaceae; Salmonella
MTGDNTLITSHGINRRDFMKLCAALAATMGLSSKAAAEMAESVSNPQRPPVIWIGAQECT WP_000145429.1
hydrogenase 2 small subunit [Salmonella enterica] Length: 372\nNumber of predicted TMHs: 1\nExp number of AAs in TMHs: 24.51309\nExp number, first 60 AAs: 0.2682\nTotal prob of N-in: 0.14402\noutside 1 331\nTMhelix 332 354\ninside 355 372

14381 GCF_002048795.1_ASM204879v1 Salmonella enterica Proteobacteria; Gammaproteobacteria; Enterobacterales; Enterobacteriaceae; Salmonella
MNNEETFYQAMRRKGVTRRSFLKFCSLAATSLGLGAGMTPKIAWALENKPRIPVVWIHGL WP_001058310.1
hydrogenase [Salmonella enterica] Length: 372\nNumber of predicted TMHs: 2\nExp number of AAs in TMHs: 37.4567299999999\nExp number, first 60 AAs: 19.72694\nTotal prob of N-in: 0.96171\nPOSSIBLE N-term signal sequence\ninside 1 19\nTMhelix 20 39\noutside 40 325\nTMhelix 326 348\ninside 349 372

14382 GCF_002048795.1_ASM204879v1 Salmonella enterica Proteobacteria; Gammaproteobacteria; Enterobacterales; Enterobacteriaceae; Salmonella
MTGDNTLITSHGINRRDFMKLCAALAATMGLSSKAAAEMAESVSNPQRPPVIWIGAQECT WP_000145429.1
hydrogenase 2 small subunit [Salmonella enterica] Length: 372\nNumber of predicted TMHs: 1\nExp number of AAs in TMHs: 24.51309\nExp number, first 60 AAs: 0.2682\nTotal prob of N-in: 0.14402\noutside 1 331\nTMhelix 332 354\ninside 355 372

14383 GCF_002058405.1_ASM205840v1 Salmonella enterica Proteobacteria; Gammaproteobacteria; Enterobacterales; Enterobacteriaceae; Salmonella
MNNEETFYQAMRRKGVTRRSFLKFCSLAATSLGLGAGMTPKIAWALENKPRIPVVWIHGL WP_001058312.1
hydrogenase [Salmonella enterica] Length: 372\nNumber of predicted TMHs: 2\nExp number of AAs in TMHs: 36.51738\nExp number, first 60 AAs: 19.72849\nTotal prob of N-in: 0.96170\nPOSSIBLE N-term signal sequence\ninside 1 19\nTMhelix 20 39\noutside 40 325\nTMhelix 326 348\ninside 349 372

14384 GCF_002034075.1_ASM203407v1 Salmonella enterica Proteobacteria; Gammaproteobacteria; Enterobacterales; Enterobacteriaceae; Salmonella
MTGDNTLITSHGINRRDFMKLCAALAATMGLSSKAAAEMAESVSNPQRPPVIWIGAQECT WP_000145429.1
hydrogenase 2 small subunit [Salmonella enterica] Length: 372\nNumber of predicted TMHs: 1\nExp

number of AAs in TMHs: 24.51309\nExp number, first 60 AAs: 0.2682\nTotal prob of N-in: 0.14402\noutside 1 331\nTMhelix 332 354\ninside 355 372

14385 GCF_002033935.1_ASM203393v1 Salmonella enterica Proteobacteria; Gammaproteobacteria; Enterobacterales; Enterobacteriaceae; Salmonella
MTGDNTLITSHGINRRDFMKLCAALAATMGLSSKAAAEMAESVSNPQRPPVIWIGAQECT WP_000145429.1
hydrogenase 2 small subunit [Salmonella enterica] Length: 372\nNumber of predicted TMHs: 1\nExp number of AAs in TMHs: 24.51309\nExp number, first 60 AAs: 0.2682\nTotal prob of N-in: 0.14402\noutside 1 331\nTMhelix 332 354\ninside 355 372

14386 GCF_002033755.1_ASM203375v1 Salmonella enterica Proteobacteria; Gammaproteobacteria; Enterobacterales; Enterobacteriaceae; Salmonella
MNNEETFYQAMRRKGVTRRSFLKFCSLAATSLGLGAGMTPKIAWALENKPRIPVVWIHGL WP_001058307.1
hydrogenase [Salmonella enterica] Length: 372\nNumber of predicted TMHs: 2\nExp number of AAs in TMHs: 36.51738\nExp number, first 60 AAs: 19.72849\nTotal prob of N-in: 0.96170\nPOSSIBLE N-term signal sequence\ninside 1 19\nTMhelix 20 39\noutside 40 325\nTMhelix 326 348\ninside 349 372

14387 GCF_002033755.1_ASM203375v1 Salmonella enterica Proteobacteria; Gammaproteobacteria; Enterobacterales; Enterobacteriaceae; Salmonella
MTGDNTLITSHGINRRDFMKLCAALAATMGLSSKAAAEMAESVSNPQRPPVIWIGAQECT WP_000145429.1
hydrogenase 2 small subunit [Salmonella enterica] Length: 372\nNumber of predicted TMHs: 1\nExp number of AAs in TMHs: 24.51309\nExp number, first 60 AAs: 0.2682\nTotal prob of N-in: 0.14402\noutside 1 331\nTMhelix 332 354\ninside 355 372

14388 GCF_002033455.1_ASM203345v1 Salmonella enterica Proteobacteria; Gammaproteobacteria; Enterobacterales; Enterobacteriaceae; Salmonella
MNNEETFYQAMRRKGVTRRSFLKFCSLAATSLGLGAGMTPKIAWALENKPRIPVVWIHGL WP_001058311.1
hydrogenase [Salmonella enterica] Length: 372\nNumber of predicted TMHs: 2\nExp number of AAs in TMHs: 37.4699899999999\nExp number, first 60 AAs: 19.72699\nTotal prob of N-in: 0.96171\nPOSSIBLE N-term signal sequence\ninside 1 19\nTMhelix 20 39\noutside 40 325\nTMhelix 326 348\ninside 349 372

14389 GCF_002047705.1_ASM204770v1 Salmonella enterica Proteobacteria; Gammaproteobacteria; Enterobacterales; Enterobacteriaceae; Salmonella
MNNEETFYQAMRRKGVTRRSFLKFCSLAATSLGLGAGMTPKIAWALENKPRIPVVWIHGL WP_001058312.1
hydrogenase [Salmonella enterica] Length: 372\nNumber of predicted TMHs: 2\nExp number of AAs in TMHs: 36.51738\nExp number, first 60 AAs: 19.72849\nTotal prob of N-in: 0.96170\nPOSSIBLE N-term signal sequence\ninside 1 19\nTMhelix 20 39\noutside 40 325\nTMhelix 326 348\ninside 349 372

14390 GCF_002033515.1_ASM203351v1 Salmonella enterica Proteobacteria; Gammaproteobacteria; Enterobacterales; Enterobacteriaceae; Salmonella
MNNEETFYQAMRRKGVTRRSFLKFCSLAATSLGLGAGMTPKIAWALENKPRIPVVWIHGL WP_023233644.1
hydrogenase 1, small subunit [Salmonella enterica] Length: 372\nNumber of predicted TMHs: 2\nExp number of AAs in TMHs: 37.4208299999999\nExp number, first 60 AAs: 19.72732\nTotal prob of N-in: 0.96168\nPOSSIBLE N-term signal sequence\ninside 1 19\nTMhelix 20 39\noutside 40 325\nTMhelix 326 348\ninside 349 372

14391 GCF_002054465.1_ASM205446v1 Salmonella enterica Proteobacteria; Gammaproteobacteria; Enterobacterales; Enterobacteriaceae; Salmonella
MNNEETFYQAMRRKGVTRRSFLKFCSLAATSLGLGAGMTPKIAWALENKPRIPVVWIHGL WP_001058311.1
hydrogenase [Salmonella enterica] Length: 372\nNumber of predicted TMHs: 2\nExp number of AAs in TMHs: 37.4699899999999\nExp number, first 60 AAs: 19.72699\nTotal prob of N-in: 0.96171\nPOSSIBLE N-term signal sequence\ninside 1 19\nTMhelix 20 39\noutside 40 325\nTMhelix 326 348\ninside 349 372

14392 GCF_002033665.1_ASM203366v1 Salmonella enterica Proteobacteria; Gammaproteobacteria; Enterobacterales; Enterobacteriaceae; Salmonella
MNNEETFYQAMRRKGVTRRSFLKFCSLAATSLGLGAGMTPKIAWALENKPRIPVVWIHGL WP_001058312.1
hydrogenase [Salmonella enterica] Length: 372\nNumber of predicted TMHs: 2\nExp number of AAs in TMHs: 36.51738\nExp number, first 60 AAs: 19.72849\nTotal prob of N-in: 0.96170\nPOSSIBLE N-term signal sequence\ninside 1 19\nTMhelix 20 39\noutside 40 325\nTMhelix 326 348\ninside 349 372

14393 GCF_002033665.1_ASM203366v1 Salmonella enterica Proteobacteria; Gammaproteobacteria; Enterobacterales; Enterobacteriaceae; Salmonella
MTGDNTLITSHGINRRDFMKLCAALAATMGLSSKAAAEMAESVSNPQRPPVIWIGAQECT WP_000145429.1
hydrogenase 2 small subunit [Salmonella enterica] Length: 372\nNumber of predicted TMHs: 1\nExp number of AAs in TMHs: 24.51309\nExp number, first 60 AAs: 0.2682\nTotal prob of N-in: 0.14402\noutside 1 331\nTMhelix 332 354\ninside 355 372

14394 GCF_001569725.1_ASM156972v1 Salmonella enterica Proteobacteria; Gammaproteobacteria; Enterobacterales; Enterobacteriaceae; Salmonella
MTGDNTLITSHGINRRDFMKLCAALAATMGLSSKAAAEMAESVSNPQRPPVIWIGAQECT WP_000145429.1
hydrogenase 2 small subunit [Salmonella enterica] Length: 372\nNumber of predicted TMHs: 1\nExp number of AAs in TMHs: 24.51309\nExp number, first 60 AAs: 0.2682\nTotal prob of N-in: 0.14402\noutside 1 331\nTMhelix 332 354\ninside 355 372

14395 GCF_001833645.1_ASM183364v1 Salmonella enterica Proteobacteria; Gammaproteobacteria; Enterobacterales; Enterobacteriaceae; Salmonella
MNNEETFYQAMRRKGVTRRSFLKFCSLAATSLGLGAGMTPKIAWALENKPRIPVVWIHGL WP_001058311.1
hydrogenase [Salmonella enterica] Length: 372\nNumber of predicted TMHs: 2\nExp number of AAs in TMHs: 37.4699899999999\nExp number, first 60 AAs: 19.72699\nTotal prob of N-in: 0.96171\nPOSSIBLE N-term signal sequence\ninside 1 19\nTMhelix 20 39\noutside 40 325\nTMhelix 326 348\ninside 349 372

14396 GCF_001833495.1_ASM183349v1 Salmonella enterica Proteobacteria; Gammaproteobacteria; Enterobacterales; Enterobacteriaceae; Salmonella
MTGDNTLITSHGINRRDFMKLCAALAATMGLSSKAAAEMAESVSNPQRPPVIWIGAQECT WP_000145429.1
hydrogenase 2 small subunit [Salmonella enterica] Length: 372\nNumber of predicted TMHs: 1\nExp number of AAs in TMHs: 24.51309\nExp number, first 60 AAs: 0.2682\nTotal prob of N-in: 0.14402\noutside 1 331\nTMhelix 332 354\ninside 355 372

14397 GCF_001833325.1_ASM183332v1 Salmonella enterica Proteobacteria; Gammaproteobacteria; Enterobacterales; Enterobacteriaceae; Salmonella
MNNEETFYQAMRRKGVTRRSFLKFCSLAATSLGLGAGMTPKIAWALENKPRIPVVWIHGL WP_001058311.1
hydrogenase [Salmonella enterica] Length: 372\nNumber of predicted TMHs: 2\nExp number of AAs in TMHs: 37.4699899999999\nExp number, first 60 AAs: 19.72699\nTotal prob of N-in: 0.96171\nPOSSIBLE N-term signal sequence\ninside 1 19\nTMhelix 20 39\noutside 40 325\nTMhelix 326 348\ninside 349 372

14398 GCF_001833485.1_ASM183348v1 Salmonella enterica Proteobacteria; Gammaproteobacteria; Enterobacterales; Enterobacteriaceae; Salmonella
MTGDNTLITSHGINRRDFMKLCAALAATMGLSSKAAAEMAESVSNPQRPPVIWIGAQECT WP_000145429.1
hydrogenase 2 small subunit [Salmonella enterica] Length: 372\nNumber of predicted TMHs: 1\nExp number of AAs in TMHs: 24.51309\nExp number, first 60 AAs: 0.2682\nTotal prob of N-in: 0.14402\noutside 1 331\nTMhelix 332 354\ninside 355 372

14399 GCF_001833405.1_ASM183340v1 Salmonella enterica Proteobacteria; Gammaproteobacteria; Enterobacterales; Enterobacteriaceae; Salmonella
MTGDNTLITSHGINRRDFMKLCAALAATMGLSSKAAAEMAESVSNPQRPPVIWIGAQECT WP_000145429.1
hydrogenase 2 small subunit [Salmonella enterica] Length: 372\nNumber of predicted TMHs: 1\nExp number of AAs in TMHs: 24.51309\nExp number, first 60 AAs: 0.2682\nTotal prob of N-in: 0.14402\noutside 1 331\nTMhelix 332 354\ninside 355 372

14400 GCF_001833395.1_ASM183339v1 Salmonella enterica Proteobacteria; Gammaproteobacteria; Enterobacterales; Enterobacteriaceae; Salmonella
MTGDNTLITSHGINRRDFMKLCAALAATMGLSSKAAAEMAESVSNPQRPPVIWIGAQECT WP_000145429.1
hydrogenase 2 small subunit [Salmonella enterica] Length: 372\nNumber of predicted TMHs: 1\nExp number of AAs in TMHs: 24.51309\nExp number, first 60 AAs: 0.2682\nTotal prob of N-in: 0.14402\noutside 1 331\nTMhelix 332 354\ninside 355 372

14401 GCF_001833325.1_ASM183332v1 Salmonella enterica Proteobacteria; Gammaproteobacteria; Enterobacterales; Enterobacteriaceae; Salmonella
MTGDNTLITSHGINRRDFMKLCAALAATMGLSSKAAAEMAESVSNPQRPPVIWIGAQECT WP_000145429.1
hydrogenase 2 small subunit [Salmonella enterica] Length: 372\nNumber of predicted TMHs: 1\nExp

number of AAs in TMHs: 24.51309\nExp number, first 60 AAs: 0.2682\nTotal prob of N-in: 0.14402\noutside 1 331\nTMhelix 332 354\ninside 355 372

14402 GCF_001833095.1_ASM183309v1 Salmonella enterica Proteobacteria; Gammaproteobacteria; Enterobacterales; Enterobacteriaceae; Salmonella MTGDNTLITSHGINRRDFMKLCAALAATMGLSSKAAAEMAESVSNPQRPPVIWIGAQECT WP_070809778.1 hydrogenase 2 small subunit [Salmonella enterica] Length: 372\nNumber of predicted TMHs: 1\nExp number of AAs in TMHs: 24.31185\nExp number, first 60 AAs: 0.26349\nTotal prob of N-in: 0.13958\noutside 1 331\nTMhelix 332 354\ninside 355 372

14403 GCF_001834755.1_ASM183475v1 Salmonella enterica Proteobacteria; Gammaproteobacteria; Enterobacterales; Enterobacteriaceae; Salmonella MTGDNTLITSHGINRRDFMKLCAALAATMGLSSKAAAEMAESVSNPQRPPVIWIGAQECT WP_000145429.1 hydrogenase 2 small subunit [Salmonella enterica] Length: 372\nNumber of predicted TMHs: 1\nExp number of AAs in TMHs: 24.51309\nExp number, first 60 AAs: 0.2682\nTotal prob of N-in: 0.14402\noutside 1 331\nTMhelix 332 354\ninside 355 372

14404 GCF_001834785.1_ASM183478v1 Salmonella enterica Proteobacteria; Gammaproteobacteria; Enterobacterales; Enterobacteriaceae; Salmonella MTGDNTLITSHGINRRDFMKLCAALAATMGLSSKAAAEMAESVSNPQRPPVIWIGAQECT WP_000145429.1 hydrogenase 2 small subunit [Salmonella enterica] Length: 372\nNumber of predicted TMHs: 1\nExp number of AAs in TMHs: 24.51309\nExp number, first 60 AAs: 0.2682\nTotal prob of N-in: 0.14402\noutside 1 331\nTMhelix 332 354\ninside 355 372

14405 GCF_001834815.1_ASM183481v1 Salmonella enterica Proteobacteria; Gammaproteobacteria; Enterobacterales; Enterobacteriaceae; Salmonella MNNEETFYQAMRRKGVTRRSFLKFCSLAATSLGLGAGMTPKIAWALENKPRIPVVWIHGL WP_001058312.1 hydrogenase [Salmonella enterica] Length: 372\nNumber of predicted TMHs: 2\nExp number of AAs in TMHs: 36.51738\nExp number, first 60 AAs: 19.72849\nTotal prob of N-in: 0.96170\nPOSSIBLE N-term signal sequence\ninside 1 19\nTMhelix 20 39\noutside 40 325\nTMhelix 326 348\ninside 349 372

14406 GCF_001834975.1_ASM183497v1 Salmonella enterica Proteobacteria; Gammaproteobacteria; Enterobacterales; Enterobacteriaceae; Salmonella MTGDNTLITSHGINRRDFMKLCAALAATMGLSSKAAAEMAESVSNPQRPPVIWIGAQECT WP_000145429.1 hydrogenase 2 small subunit [Salmonella enterica] Length: 372\nNumber of predicted TMHs: 1\nExp number of AAs in TMHs: 24.51309\nExp number, first 60 AAs: 0.2682\nTotal prob of N-in: 0.14402\noutside 1 331\nTMhelix 332 354\ninside 355 372

14407 GCF_001835015.1_ASM183501v1 Salmonella enterica Proteobacteria; Gammaproteobacteria; Enterobacterales; Enterobacteriaceae; Salmonella MTGDNTLITSHGINRRDFMKLCAALAATMGLSSKAAAEMAESVSNPQRPPVIWIGAQECT WP_000145429.1 hydrogenase 2 small subunit [Salmonella enterica] Length: 372\nNumber of predicted TMHs: 1\nExp number of AAs in TMHs: 24.51309\nExp number, first 60 AAs: 0.2682\nTotal prob of N-in: 0.14402\noutside 1 331\nTMhelix 332 354\ninside 355 372

14408 GCF_001834975.1_ASM183497v1 Salmonella enterica Proteobacteria; Gammaproteobacteria; Enterobacterales; Enterobacteriaceae; Salmonella MNNEETFYQAMRRKGVTRRSFLKFCSLAATSLGLGAGMTPKIAWALENKPRIPVVWIHGL WP_001058312.1 hydrogenase [Salmonella enterica] Length: 372\nNumber of predicted TMHs: 2\nExp number of AAs in TMHs: 36.51738\nExp number, first 60 AAs: 19.72849\nTotal prob of N-in: 0.96170\nPOSSIBLE N-term signal sequence\ninside 1 19\nTMhelix 20 39\noutside 40 325\nTMhelix 326 348\ninside 349 372

14409 GCF_001835015.1_ASM183501v1 Salmonella enterica Proteobacteria; Gammaproteobacteria; Enterobacterales; Enterobacteriaceae; Salmonella MNNEETFYQAMRRKGVTRRSFLKFCSLAATSLGLGAGMTPKIAWALENKPRIPVVWIHGL WP_001058312.1 hydrogenase [Salmonella enterica] Length: 372\nNumber of predicted TMHs: 2\nExp number of AAs in TMHs: 36.51738\nExp number, first 60 AAs: 19.72849\nTotal prob of N-in: 0.96170\nPOSSIBLE N-term signal sequence\ninside 1 19\nTMhelix 20 39\noutside 40 325\nTMhelix 326 348\ninside 349 372

14410 GCF_001834885.1_ASM183488v1 Salmonella enterica Proteobacteria; Gammaproteobacteria; Enterobacterales; Enterobacteriaceae; Salmonella

MTGDNTLITSHGINRRDFMKLCAALAATMGLSSKAAAEMAESVSNPQRPPVIWIGAQECT	WP_023248548.1	hydrogenase 2 small subunit [Salmonella enterica]	Length: 372\nNumber of predicted TMHs:  1\nExp number of AAs in TMHs: 24.51281\nExp number, first 60 AAs:  0.26822\nTotal prob of N-in:   0.14402\noutside   1   331\nTMhelix   332   354\ninside   355   372

14411	GCF_001835045.1_ASM183504v1	Salmonella enterica	Proteobacteria; Gammaproteobacteria; Enterobacterales; Enterobacteriaceae; Salmonella	MTGDNTLITSHGINRRDFMKLCAALAATMGLSSKAAAEMAESVSNPQRPPVIWIGAQECT	WP_000145429.1	hydrogenase 2 small subunit [Salmonella enterica]	Length: 372\nNumber of predicted TMHs:  1\nExp number of AAs in TMHs: 24.51309\nExp number, first 60 AAs:  0.2682\nTotal prob of N-in:   0.14402\noutside   1   331\nTMhelix   332   354\ninside   355   372

14412	GCF_001833075.1_ASM183307v1	Salmonella enterica	Proteobacteria; Gammaproteobacteria; Enterobacterales; Enterobacteriaceae; Salmonella	MNNEETFYQAMRRKGVTRRSFLKFCSLAATSLGLGAGMTPKIAWALENKPRIPVVWIHGL	WP_001058312.1	hydrogenase [Salmonella enterica]	Length: 372\nNumber of predicted TMHs:  2\nExp number of AAs in TMHs: 36.51738\nExp number, first 60 AAs:  19.72849\nTotal prob of N-in:   0.96170\nPOSSIBLE N-term signal sequence\ninside   1   19\nTMhelix   20   39\noutside   40   325\nTMhelix   326   348\ninside   349   372

14413	GCF_001833075.1_ASM183307v1	Salmonella enterica	Proteobacteria; Gammaproteobacteria; Enterobacterales; Enterobacteriaceae; Salmonella	MTGDNTLITSHGINRRDFMKLCAALAATMGLSSKAAAEMAESVSNPQRPPVIWIGAQECT	WP_000145429.1	hydrogenase 2 small subunit [Salmonella enterica]	Length: 372\nNumber of predicted TMHs:  1\nExp number of AAs in TMHs: 24.51309\nExp number, first 60 AAs:  0.2682\nTotal prob of N-in:   0.14402\noutside   1   331\nTMhelix   332   354\ninside   355   372

14414	GCF_001951995.1_ASM195199v1	Salmonella enterica	Proteobacteria; Gammaproteobacteria; Enterobacterales; Enterobacteriaceae; Salmonella	MTGDNTLITSHGINRRDFMKLCAALAATMGLSSKAAAEMAESVSNPQRPPVIWIGAQECT	WP_000145429.1	hydrogenase 2 small subunit [Salmonella enterica]	Length: 372\nNumber of predicted TMHs:  1\nExp number of AAs in TMHs: 24.51309\nExp number, first 60 AAs:  0.2682\nTotal prob of N-in:   0.14402\noutside   1   331\nTMhelix   332   354\ninside   355   372

14415	GCF_001832325.1_ASM183232v1	Salmonella enterica	Proteobacteria; Gammaproteobacteria; Enterobacterales; Enterobacteriaceae; Salmonella	MNNEETFYQAMRRKGVTRRSFLKFCSLAATSLGLGAGMTPKIAWALENKPRIPVVWIHGL	WP_001058311.1	hydrogenase [Salmonella enterica]	Length: 372\nNumber of predicted TMHs:  2\nExp number of AAs in TMHs: 37.4699899999999\nExp number, first 60 AAs:  19.72699\nTotal prob of N-in:   0.96171\nPOSSIBLE N-term signal sequence\ninside   1   19\nTMhelix   20   39\noutside   40   325\nTMhelix   326   348\ninside   349   372

14416	GCF_001831985.1_ASM183198v1	Salmonella enterica	Proteobacteria; Gammaproteobacteria; Enterobacterales; Enterobacteriaceae; Salmonella	MNNEETFYQAMRRKGVTRRSFLKFCSLAATSLGLGAGMTPKIAWALENKPRIPVVWIHGL	WP_001058312.1	hydrogenase [Salmonella enterica]	Length: 372\nNumber of predicted TMHs:  2\nExp number of AAs in TMHs: 36.51738\nExp number, first 60 AAs:  19.72849\nTotal prob of N-in:   0.96170\nPOSSIBLE N-term signal sequence\ninside   1   19\nTMhelix   20   39\noutside   40   325\nTMhelix   326   348\ninside   349   372

14417	GCF_001831915.1_ASM183191v1	Salmonella enterica	Proteobacteria; Gammaproteobacteria; Enterobacterales; Enterobacteriaceae; Salmonella	MNNEETFYQAMRRKGVTRRSFLKFCSLAATSLGLGAGMTPKIAWALENKPRIPVVWIHGL	WP_001058311.1	hydrogenase [Salmonella enterica]	Length: 372\nNumber of predicted TMHs:  2\nExp number of AAs in TMHs: 37.4699899999999\nExp number, first 60 AAs:  19.72699\nTotal prob of N-in:   0.96171\nPOSSIBLE N-term signal sequence\ninside   1   19\nTMhelix   20   39\noutside   40   325\nTMhelix   326   348\ninside   349   372

14418	GCF_001831945.1_ASM183194v1	Salmonella enterica	Proteobacteria; Gammaproteobacteria; Enterobacterales; Enterobacteriaceae; Salmonella	MNNEETFYQAMRRKGVTRRSFLKFCSLAATSLGLGAGMTPKIAWALENKPRIPVVWIHGL	WP_001058312.1	hydrogenase [Salmonella enterica]	Length: 372\nNumber of predicted TMHs:  2\nExp number of AAs in TMHs: 36.51738\nExp number, first 60 AAs:  19.72849\nTotal prob of N-in:   0.96170\nPOSSIBLE N-term signal sequence\ninside   1   19\nTMhelix   20   39\noutside   40   325\nTMhelix   326   348\ninside   349   372

14419 GCF_001832045.1_ASM183204v1 Salmonella enterica Proteobacteria; Gammaproteobacteria; Enterobacterales; Enterobacteriaceae; Salmonella
MNNEETFYQAMRRKGVTRRSFLKFCSLAATSLGLGAGMTPKIAWALENKPRIPVVWIHGL WP_001058312.1
hydrogenase [Salmonella enterica] Length: 372\nNumber of predicted TMHs: 2\nExp number of AAs in TMHs: 36.51738\nExp number, first 60 AAs: 19.72849\nTotal prob of N-in: 0.96170\nPOSSIBLE N-term signal sequence\ninside 1 19\nTMhelix 20 39\noutside 40 325\nTMhelix 326 348\ninside 349 372

14420 GCF_001951845.1_ASM195184v1 Salmonella enterica Proteobacteria; Gammaproteobacteria; Enterobacterales; Enterobacteriaceae; Salmonella
MNNEETFYQAMRRKGVTRRSFLKFCSLAATSLGLGAGMTPKIAWALENKPRIPVVWIHGL WP_001058312.1
hydrogenase [Salmonella enterica] Length: 372\nNumber of predicted TMHs: 2\nExp number of AAs in TMHs: 36.51738\nExp number, first 60 AAs: 19.72849\nTotal prob of N-in: 0.96170\nPOSSIBLE N-term signal sequence\ninside 1 19\nTMhelix 20 39\noutside 40 325\nTMhelix 326 348\ninside 349 372

14421 GCF_001951875.1_ASM195187v1 Salmonella enterica Proteobacteria; Gammaproteobacteria; Enterobacterales; Enterobacteriaceae; Salmonella
MTGDNTLITSHGINRRDFMKLCAALAATMGLSSKAAAEMAESVSNPQRPPVIWIGAQECT WP_000145429.1
hydrogenase 2 small subunit [Salmonella enterica] Length: 372\nNumber of predicted TMHs: 1\nExp number of AAs in TMHs: 24.51309\nExp number, first 60 AAs: 0.2682\nTotal prob of N-in: 0.14402\noutside 1 331\nTMhelix 332 354\ninside 355 372

14422 GCF_001951915.1_ASM195191v1 Salmonella enterica Proteobacteria; Gammaproteobacteria; Enterobacterales; Enterobacteriaceae; Salmonella
MTGDNTLITSHGINRRDFMKLCAALAATMGLSSKAAAEMAESVSNPQRPPVIWIGAQECT WP_000145429.1
hydrogenase 2 small subunit [Salmonella enterica] Length: 372\nNumber of predicted TMHs: 1\nExp number of AAs in TMHs: 24.51309\nExp number, first 60 AAs: 0.2682\nTotal prob of N-in: 0.14402\noutside 1 331\nTMhelix 332 354\ninside 355 372

14423 GCF_001951935.1_ASM195193v1 Salmonella enterica Proteobacteria; Gammaproteobacteria; Enterobacterales; Enterobacteriaceae; Salmonella
MTGDNTLITSHGINRRDFMKLCAALAATMGLSSKAAAEMAESVSNPQRPPVIWIGAQECT WP_000145429.1
hydrogenase 2 small subunit [Salmonella enterica] Length: 372\nNumber of predicted TMHs: 1\nExp number of AAs in TMHs: 24.51309\nExp number, first 60 AAs: 0.2682\nTotal prob of N-in: 0.14402\noutside 1 331\nTMhelix 332 354\ninside 355 372

14424 GCF_001952485.1_ASM195248v1 Salmonella enterica Proteobacteria; Gammaproteobacteria; Enterobacterales; Enterobacteriaceae; Salmonella
MTGDNTLITSHGINRRDFMKLCAALAATMGLSSKAAAEMAESVSNPQRPPVIWIGAQECT WP_000145429.1
hydrogenase 2 small subunit [Salmonella enterica] Length: 372\nNumber of predicted TMHs: 1\nExp number of AAs in TMHs: 24.51309\nExp number, first 60 AAs: 0.2682\nTotal prob of N-in: 0.14402\noutside 1 331\nTMhelix 332 354\ninside 355 372

14425 GCF_001832045.1_ASM183204v1 Salmonella enterica Proteobacteria; Gammaproteobacteria; Enterobacterales; Enterobacteriaceae; Salmonella
MTGDNTLITSHGINRRDFMKLCAALAATMGLSSKAAAEMAESVSNPQRPPVIWIGAQECT WP_000145429.1
hydrogenase 2 small subunit [Salmonella enterica] Length: 372\nNumber of predicted TMHs: 1\nExp number of AAs in TMHs: 24.51309\nExp number, first 60 AAs: 0.2682\nTotal prob of N-in: 0.14402\noutside 1 331\nTMhelix 332 354\ninside 355 372

14426 GCF_002060465.1_ASM206046v1 Salmonella enterica Proteobacteria; Gammaproteobacteria; Enterobacterales; Enterobacteriaceae; Salmonella
MNNEETLYQAMRRKGVTRRSFLKFCSLAATSLGLGAGMTPKIAWALENKPRIPVVWIHGL WP_001752445.1
hydrogenase-1 small chain [Salmonella enterica] Length: 372\nNumber of predicted TMHs: 2\nExp number of AAs in TMHs: 36.5123\nExp number, first 60 AAs: 19.72341\nTotal prob of N-in: 0.96144\nPOSSIBLE N-term signal sequence\ninside 1 19\nTMhelix 20 39\noutside 40 325\nTMhelix 326 348\ninside 349 372

14427 GCF_002060695.1_ASM206069v1 Salmonella enterica Proteobacteria; Gammaproteobacteria; Enterobacterales; Enterobacteriaceae; Salmonella
MNNEETFYQAMRRKGVTRRSFLKFCSLAATSLGLGAGMTPKIAWALENKPRIPVVWIHGL WP_001058307.1
hydrogenase [Salmonella enterica] Length: 372\nNumber of predicted TMHs: 2\nExp number of AAs in

TMHs: 36.51738\nExp number, first 60 AAs: 19.72849\nTotal prob of N-in: 0.96170\nPOSSIBLE N-term signal sequence\ninside 1 19\nTMhelix 20 39\noutside 40 325\nTMhelix 326 348\ninside 349 372

14428 GCF_002060705.1_ASM206070v1 Salmonella enterica Proteobacteria; Gammaproteobacteria; Enterobacterales; Enterobacteriaceae; Salmonella MNNEETFYQAMRRKGVTRRSFLKFCSLAATSLGLGAGMTPKIAWALENKPRIPVVWIHGL WP_001058310.1 hydrogenase [Salmonella enterica] Length: 372\nNumber of predicted TMHs: 2\nExp number of AAs in TMHs: 37.4567299999999\nExp number, first 60 AAs: 19.72694\nTotal prob of N-in: 0.96171\nPOSSIBLE N-term signal sequence\ninside 1 19\nTMhelix 20 39\noutside 40 325\nTMhelix 326 348\ninside 349 372

14429 GCF_002060705.1_ASM206070v1 Salmonella enterica Proteobacteria; Gammaproteobacteria; Enterobacterales; Enterobacteriaceae; Salmonella MTGDNTLITSHGINRRDFMKLCAALAATMGLSSKAAAEMAESVSNPQRPPVIWIGAQECT WP_000145429.1 hydrogenase 2 small subunit [Salmonella enterica] Length: 372\nNumber of predicted TMHs: 1\nExp number of AAs in TMHs: 24.51309\nExp number, first 60 AAs: 0.2682\nTotal prob of N-in: 0.14402\noutside 1 331\nTMhelix 332 354\ninside 355 372

14430 GCF_002060755.1_ASM206075v1 Salmonella enterica Proteobacteria; Gammaproteobacteria; Enterobacterales; Enterobacteriaceae; Salmonella MTGDNTLITSHGINRRDFMKLCAALAATMGLSSKAAAEMAESVSNPQRPPVIWIGAQECT WP_000145429.1 hydrogenase 2 small subunit [Salmonella enterica] Length: 372\nNumber of predicted TMHs: 1\nExp number of AAs in TMHs: 24.51309\nExp number, first 60 AAs: 0.2682\nTotal prob of N-in: 0.14402\noutside 1 331\nTMhelix 332 354\ninside 355 372

14431 GCF_002060435.1_ASM206043v1 Salmonella enterica Proteobacteria; Gammaproteobacteria; Enterobacterales; Enterobacteriaceae; Salmonella MNNEETFYQAMRRKGVTRRSFLKFCSLAATSLGLGAGMTPKIAWALENKPRIPVVWIHGL WP_001058312.1 hydrogenase [Salmonella enterica] Length: 372\nNumber of predicted TMHs: 2\nExp number of AAs in TMHs: 36.51738\nExp number, first 60 AAs: 19.72849\nTotal prob of N-in: 0.96170\nPOSSIBLE N-term signal sequence\ninside 1 19\nTMhelix 20 39\noutside 40 325\nTMhelix 326 348\ninside 349 372

14432 GCF_002060535.1_ASM206053v1 Salmonella enterica Proteobacteria; Gammaproteobacteria; Enterobacterales; Enterobacteriaceae; Salmonella MNNEETFYQAMRRKGVTRRSFLKFCSLAATSLGLGAGMTPKIAWALENKPRIPVVWIHGL WP_001058312.1 hydrogenase [Salmonella enterica] Length: 372\nNumber of predicted TMHs: 2\nExp number of AAs in TMHs: 36.51738\nExp number, first 60 AAs: 19.72849\nTotal prob of N-in: 0.96170\nPOSSIBLE N-term signal sequence\ninside 1 19\nTMhelix 20 39\noutside 40 325\nTMhelix 326 348\ninside 349 372

14433 GCF_002060525.1_ASM206052v1 Salmonella enterica Proteobacteria; Gammaproteobacteria; Enterobacterales; Enterobacteriaceae; Salmonella MNNEETFYQAMRRKGVTRRSFLKFCSLAATSLGLGAGMTPKIAWALENKPRIPVVWIHGL WP_001058311.1 hydrogenase [Salmonella enterica] Length: 372\nNumber of predicted TMHs: 2\nExp number of AAs in TMHs: 37.4699899999999\nExp number, first 60 AAs: 19.72699\nTotal prob of N-in: 0.96171\nPOSSIBLE N-term signal sequence\ninside 1 19\nTMhelix 20 39\noutside 40 325\nTMhelix 326 348\ninside 349 372

14434 GCF_002060535.1_ASM206053v1 Salmonella enterica Proteobacteria; Gammaproteobacteria; Enterobacterales; Enterobacteriaceae; Salmonella MTGDNTLITSHGINRRDFMKLCAALAATMGLSSKAAAEMAESVSNPQRPPVIWIGAQECT WP_000145431.1 hydrogenase 2 small subunit [Salmonella enterica] Length: 372\nNumber of predicted TMHs: 1\nExp number of AAs in TMHs: 24.51505\nExp number, first 60 AAs: 0.2682\nTotal prob of N-in: 0.14406\noutside 1 331\nTMhelix 332 354\ninside 355 372

14435 GCF_002060525.1_ASM206052v1 Salmonella enterica Proteobacteria; Gammaproteobacteria; Enterobacterales; Enterobacteriaceae; Salmonella MTGDNTLITSHGINRRDFMKLCAALAATMGLSSKAAAEMAESVSNPQRPPVIWIGAQECT WP_000145429.1 hydrogenase 2 small subunit [Salmonella enterica] Length: 372\nNumber of predicted TMHs: 1\nExp number of AAs in TMHs: 24.51309\nExp number, first 60 AAs: 0.2682\nTotal prob of N-in: 0.14402\noutside 1 331\nTMhelix 332 354\ninside 355 372

14436 GCF_001831875.1_ASM183187v1 Salmonella enterica Proteobacteria; Gammaproteobacteria; Enterobacterales; Enterobacteriaceae; Salmonella

MNNEETFYQAMRRKGVTRRSFLKFCSLAATSLGLGAGMTPKIAWALENKPRIPVVWIHGL WP_001058311.1 hydrogenase [Salmonella enterica] Length: 372\nNumber of predicted TMHs: 2\nExp number of AAs in TMHs: 37.4699899999999\nExp number, first 60 AAs: 19.72699\nTotal prob of N-in: 0.96171\nPOSSIBLE N-term signal sequence\ninside 1 19\nTMhelix 20 39\noutside 40 325\nTMhelix 326 348\ninside 349 372

14437 GCF_002060335.1_ASM206033v1 Salmonella enterica Proteobacteria; Gammaproteobacteria; Enterobacterales; Enterobacteriaceae; Salmonella MNNEETFYQAMRRKGVTRRSFLKFCSLAATSLGLGAGMTPKIAWALENKPRIPVVWIHGL WP_001058312.1 hydrogenase [Salmonella enterica] Length: 372\nNumber of predicted TMHs: 2\nExp number of AAs in TMHs: 36.51738\nExp number, first 60 AAs: 19.72849\nTotal prob of N-in: 0.96170\nPOSSIBLE N-term signal sequence\ninside 1 19\nTMhelix 20 39\noutside 40 325\nTMhelix 326 348\ninside 349 372

14438 GCF_002060335.1_ASM206033v1 Salmonella enterica Proteobacteria; Gammaproteobacteria; Enterobacterales; Enterobacteriaceae; Salmonella MTGDNTLITSHGINRRDFMKLCAALAATMGLSSKAAAEMAESVSNPQRPPVIWIGAQECT WP_000145429.1 hydrogenase 2 small subunit [Salmonella enterica] Length: 372\nNumber of predicted TMHs: 1\nExp number of AAs in TMHs: 24.51309\nExp number, first 60 AAs: 0.2682\nTotal prob of N-in: 0.14402\noutside 1 331\nTMhelix 332 354\ninside 355 372

14439 GCF_002060285.1_ASM206028v1 Salmonella enterica Proteobacteria; Gammaproteobacteria; Enterobacterales; Enterobacteriaceae; Salmonella MNNEETFYQAMRRKGVTRRSFLKFCSLAATSLGLGAGMTPKIAWALENKPRIPVVWIHGL WP_001058312.1 hydrogenase [Salmonella enterica] Length: 372\nNumber of predicted TMHs: 2\nExp number of AAs in TMHs: 36.51738\nExp number, first 60 AAs: 19.72849\nTotal prob of N-in: 0.96170\nPOSSIBLE N-term signal sequence\ninside 1 19\nTMhelix 20 39\noutside 40 325\nTMhelix 326 348\ninside 349 372

14440 GCF_002060285.1_ASM206028v1 Salmonella enterica Proteobacteria; Gammaproteobacteria; Enterobacterales; Enterobacteriaceae; Salmonella MTGDNTLITSHGINRRDFMKLCAALAATMGLSSKAAAEMAESVSNPQRPPVIWIGAQECT WP_000145429.1 hydrogenase 2 small subunit [Salmonella enterica] Length: 372\nNumber of predicted TMHs: 1\nExp number of AAs in TMHs: 24.51309\nExp number, first 60 AAs: 0.2682\nTotal prob of N-in: 0.14402\noutside 1 331\nTMhelix 332 354\ninside 355 372

14441 GCF_001832025.1_ASM183202v1 Salmonella enterica Proteobacteria; Gammaproteobacteria; Enterobacterales; Enterobacteriaceae; Salmonella MNNEETFYQAMRRKGVTRRSFLKFCSLAATSLGLGAGMTPKIAWALENKPRIPVVWIHGL WP_001058312.1 hydrogenase [Salmonella enterica] Length: 372\nNumber of predicted TMHs: 2\nExp number of AAs in TMHs: 36.51738\nExp number, first 60 AAs: 19.72849\nTotal prob of N-in: 0.96170\nPOSSIBLE N-term signal sequence\ninside 1 19\nTMhelix 20 39\noutside 40 325\nTMhelix 326 348\ninside 349 372

14442 GCF_001831715.1_ASM183171v1 Salmonella enterica Proteobacteria; Gammaproteobacteria; Enterobacterales; Enterobacteriaceae; Salmonella MNNEETFYQAMRRKGVTRRSFLKFCSLAATSLGLGAGMTPKIAWALENKPRIPVVWIHGL WP_001058312.1 hydrogenase [Salmonella enterica] Length: 372\nNumber of predicted TMHs: 2\nExp number of AAs in TMHs: 36.51738\nExp number, first 60 AAs: 19.72849\nTotal prob of N-in: 0.96170\nPOSSIBLE N-term signal sequence\ninside 1 19\nTMhelix 20 39\noutside 40 325\nTMhelix 326 348\ninside 349 372

14443 GCF_001831715.1_ASM183171v1 Salmonella enterica Proteobacteria; Gammaproteobacteria; Enterobacterales; Enterobacteriaceae; Salmonella MTGDNTLITSHGINRRDFMKLCAALAATMGLSSKAAAEMAESVSNPQRPPVIWIGAQECT WP_000145429.1 hydrogenase 2 small subunit [Salmonella enterica] Length: 372\nNumber of predicted TMHs: 1\nExp number of AAs in TMHs: 24.51309\nExp number, first 60 AAs: 0.2682\nTotal prob of N-in: 0.14402\noutside 1 331\nTMhelix 332 354\ninside 355 372

14444 GCF_002060025.1_ASM206002v1 Salmonella enterica Proteobacteria; Gammaproteobacteria; Enterobacterales; Enterobacteriaceae; Salmonella MNNEETFYQAMRRKGVTRRSFLKFCSLAATSLGLGAGMTPKIAWALENKPRIPVVWIHGL WP_001058312.1 hydrogenase [Salmonella enterica] Length: 372\nNumber of predicted TMHs: 2\nExp number of AAs in TMHs: 36.51738\nExp number, first 60 AAs: 19.72849\nTotal prob of N-in: 0.96170\nPOSSIBLE N-term signal sequence\ninside 1 19\nTMhelix 20 39\noutside 40 325\nTMhelix 326 348\ninside 349 372

14445 GCF_001831565.1_ASM183156v1 Salmonella enterica Proteobacteria; Gammaproteobacteria; Enterobacterales; Enterobacteriaceae; Salmonella
MNNEETFYQAMRRKGVTRRSFLKFCSLAATSLGLGAGMTPKIAWALENKPRIPVVWIHGL WP_001058312.1
hydrogenase [Salmonella enterica] Length: 372\nNumber of predicted TMHs: 2\nExp number of AAs in TMHs: 36.51738\nExp number, first 60 AAs: 19.72849\nTotal prob of N-in: 0.96170\nPOSSIBLE N-term signal sequence\ninside 1 19\nTMhelix 20 39\noutside 40 325\nTMhelix 326 348\ninside 349 372

14446 GCF_002060025.1_ASM206002v1 Salmonella enterica Proteobacteria; Gammaproteobacteria; Enterobacterales; Enterobacteriaceae; Salmonella
MTGDNTLITSHGINRRDFMKLCAALAATMGLSSKAAAEMAESVSNPQRPPVIWIGAQECT WP_000145429.1
hydrogenase 2 small subunit [Salmonella enterica] Length: 372\nNumber of predicted TMHs: 1\nExp number of AAs in TMHs: 24.51309\nExp number, first 60 AAs: 0.2682\nTotal prob of N-in: 0.14402\noutside 1 331\nTMhelix 332 354\ninside 355 372

14447 GCF_002059635.1_ASM205963v1 Salmonella enterica Proteobacteria; Gammaproteobacteria; Enterobacterales; Enterobacteriaceae; Salmonella
MNNEETFYQAMRRKGVTRRSFLKFCSLAATSLGLGAGMTPKIAWALENKPRIPVVWIHGL WP_001058312.1
hydrogenase [Salmonella enterica] Length: 372\nNumber of predicted TMHs: 2\nExp number of AAs in TMHs: 36.51738\nExp number, first 60 AAs: 19.72849\nTotal prob of N-in: 0.96170\nPOSSIBLE N-term signal sequence\ninside 1 19\nTMhelix 20 39\noutside 40 325\nTMhelix 326 348\ninside 349 372

14448 GCF_001831655.1_ASM183165v1 Salmonella enterica Proteobacteria; Gammaproteobacteria; Enterobacterales; Enterobacteriaceae; Salmonella
MNNEETFYQAMRRKGVTRRSFLKFCSLAATSLGLGAGMTPKIAWALENKPRIPVVWIHGL WP_001058312.1
hydrogenase [Salmonella enterica] Length: 372\nNumber of predicted TMHs: 2\nExp number of AAs in TMHs: 36.51738\nExp number, first 60 AAs: 19.72849\nTotal prob of N-in: 0.96170\nPOSSIBLE N-term signal sequence\ninside 1 19\nTMhelix 20 39\noutside 40 325\nTMhelix 326 348\ninside 349 372

14449 GCF_001831655.1_ASM183165v1 Salmonella enterica Proteobacteria; Gammaproteobacteria; Enterobacterales; Enterobacteriaceae; Salmonella
MTGDNTLITSHGINRRDFMKLCAALAATMGLSSKAAAEMAESVSNPQRPPVIWIGAQECT WP_000145429.1
hydrogenase 2 small subunit [Salmonella enterica] Length: 372\nNumber of predicted TMHs: 1\nExp number of AAs in TMHs: 24.51309\nExp number, first 60 AAs: 0.2682\nTotal prob of N-in: 0.14402\noutside 1 331\nTMhelix 332 354\ninside 355 372

14450 GCF_002060785.1_ASM206078v1 Salmonella enterica Proteobacteria; Gammaproteobacteria; Enterobacterales; Enterobacteriaceae; Salmonella
MTGDNTLITSHGINRRDFMKLCAALAATMGLSSKAAAEMAESVSNPQRPPVIWIGAQECT WP_000145429.1
hydrogenase 2 small subunit [Salmonella enterica] Length: 372\nNumber of predicted TMHs: 1\nExp number of AAs in TMHs: 24.51309\nExp number, first 60 AAs: 0.2682\nTotal prob of N-in: 0.14402\noutside 1 331\nTMhelix 332 354\ninside 355 372

14451 GCF_002060805.1_ASM206080v1 Salmonella enterica Proteobacteria; Gammaproteobacteria; Enterobacterales; Enterobacteriaceae; Salmonella
MTGDNTLITSHGINRRDFMKLCAALAATMGLSSKAAAEMAESVSNPQRPPVIWIGAQECT WP_000145429.1
hydrogenase 2 small subunit [Salmonella enterica] Length: 372\nNumber of predicted TMHs: 1\nExp number of AAs in TMHs: 24.51309\nExp number, first 60 AAs: 0.2682\nTotal prob of N-in: 0.14402\noutside 1 331\nTMhelix 332 354\ninside 355 372

14452 GCF_002058675.1_ASM205867v1 Salmonella enterica Proteobacteria; Gammaproteobacteria; Enterobacterales; Enterobacteriaceae; Salmonella
MNNEETFYQAMRRKGVTRRSFLKFCSLAATSLGLGAGMTPKIAWALENKPRIPVVWIHGL WP_001540183.1
hydrogenase [Salmonella enterica] Length: 372\nNumber of predicted TMHs: 2\nExp number of AAs in TMHs: 37.41804\nExp number, first 60 AAs: 19.72808\nTotal prob of N-in: 0.96169\nPOSSIBLE N-term signal sequence\ninside 1 19\nTMhelix 20 39\noutside 40 325\nTMhelix 326 348\ninside 349 372

14453 GCF_002059375.1_ASM205937v1 Salmonella enterica Proteobacteria; Gammaproteobacteria; Enterobacterales; Enterobacteriaceae; Salmonella
MNNEETFYQAMRRKGVTRRSFLKFCSLAATSLGLGAGMTPKIAWALENKPRIPVVWIHGL WP_001058311.1
hydrogenase [Salmonella enterica] Length: 372\nNumber of predicted TMHs: 2\nExp number of AAs in TMHs:

TMHs: 37.4699899999999\nExp number, first 60 AAs: 19.72699\nTotal prob of N-in: 0.96171\nPOSSIBLE N-term signal sequence\ninside 1 19\nTMhelix 20 39\noutside 40 325\nTMhelix 326 348\ninside 349 372

14454 GCF_002059265.1_ASM205926v1 Salmonella enterica Proteobacteria; Gammaproteobacteria; Enterobacterales; Enterobacteriaceae; Salmonella MNNEETFYQAMRRKGVTRRSFLKFCSLAATSLGLGAGMTPKIAWALENKPRIPVVWIHGL WP_001058312.1 hydrogenase [Salmonella enterica] Length: 372\nNumber of predicted TMHs: 2\nExp number of AAs in TMHs: 36.51738\nExp number, first 60 AAs: 19.72849\nTotal prob of N-in: 0.96170\nPOSSIBLE N-term signal sequence\ninside 1 19\nTMhelix 20 39\noutside 40 325\nTMhelix 326 348\ninside 349 372

14455 GCF_002060805.1_ASM206080v1 Salmonella enterica Proteobacteria; Gammaproteobacteria; Enterobacterales; Enterobacteriaceae; Salmonella MNNEETFYQAMRRKGVTRRSFLKFCSLAATSLGLGAGMTPKIAWALENKPRIPVVWIHGL WP_023234992.1 hydrogenase 1, small subunit [Salmonella enterica] Length: 372\nNumber of predicted TMHs: 2\nExp number of AAs in TMHs: 37.41309\nExp number, first 60 AAs: 19.72865\nTotal prob of N-in: 0.96169\nPOSSIBLE N-term signal sequence\ninside 1 19\nTMhelix 20 39\noutside 40 325\nTMhelix 326 348\ninside 349 372

14456 GCF_002059445.1_ASM205944v1 Salmonella enterica Proteobacteria; Gammaproteobacteria; Enterobacterales; Enterobacteriaceae; Salmonella MNNEETFYQAMRRKGVTRRSFLKFCSLAATSLGLGAGMTPKIAWALENKPRIPVVWIHGL WP_001058312.1 hydrogenase [Salmonella enterica] Length: 372\nNumber of predicted TMHs: 2\nExp number of AAs in TMHs: 36.51738\nExp number, first 60 AAs: 19.72849\nTotal prob of N-in: 0.96170\nPOSSIBLE N-term signal sequence\ninside 1 19\nTMhelix 20 39\noutside 40 325\nTMhelix 326 348\ninside 349 372

14457 GCF_002059395.1_ASM205939v1 Salmonella enterica Proteobacteria; Gammaproteobacteria; Enterobacterales; Enterobacteriaceae; Salmonella MNNEETFYQAMRRKGVTRRSFLKFCSLAATSLGLGAGMTPKIAWALENKPRIPVVWIHGL WP_001058310.1 hydrogenase [Salmonella enterica] Length: 372\nNumber of predicted TMHs: 2\nExp number of AAs in TMHs: 37.4567299999999\nExp number, first 60 AAs: 19.72694\nTotal prob of N-in: 0.96171\nPOSSIBLE N-term signal sequence\ninside 1 19\nTMhelix 20 39\noutside 40 325\nTMhelix 326 348\ninside 349 372

14458 GCF_002059265.1_ASM205926v1 Salmonella enterica Proteobacteria; Gammaproteobacteria; Enterobacterales; Enterobacteriaceae; Salmonella MTGDNTLITSHGINRRDFMKLCAALAATMGLSSKAAAEMAESVSNPQRPPVIWIGAQECT WP_000145429.1 hydrogenase 2 small subunit [Salmonella enterica] Length: 372\nNumber of predicted TMHs: 1\nExp number of AAs in TMHs: 24.51309\nExp number, first 60 AAs: 0.2682\nTotal prob of N-in: 0.14402\noutside 1 331\nTMhelix 332 354\ninside 355 372

14459 GCF_002060945.1_ASM206094v1 Salmonella enterica Proteobacteria; Gammaproteobacteria; Enterobacterales; Enterobacteriaceae; Salmonella MTGDNTLITSHGINRRDFMKLCAALAATMGLSSKAAAEMAESVSNPQRPPVIWIGAQECT WP_000145429.1 hydrogenase 2 small subunit [Salmonella enterica] Length: 372\nNumber of predicted TMHs: 1\nExp number of AAs in TMHs: 24.51309\nExp number, first 60 AAs: 0.2682\nTotal prob of N-in: 0.14402\noutside 1 331\nTMhelix 332 354\ninside 355 372

14460 GCF_002060855.1_ASM206085v1 Salmonella enterica Proteobacteria; Gammaproteobacteria; Enterobacterales; Enterobacteriaceae; Salmonella MNNEETFYQAMRRKGVTRRSFLKFCSLAATSLGLGAGMTPKIAWALENKPRIPVVWIHGL WP_001058312.1 hydrogenase [Salmonella enterica] Length: 372\nNumber of predicted TMHs: 2\nExp number of AAs in TMHs: 36.51738\nExp number, first 60 AAs: 19.72849\nTotal prob of N-in: 0.96170\nPOSSIBLE N-term signal sequence\ninside 1 19\nTMhelix 20 39\noutside 40 325\nTMhelix 326 348\ninside 349 372

14461 GCF_002060995.1_ASM206099v1 Salmonella enterica Proteobacteria; Gammaproteobacteria; Enterobacterales; Enterobacteriaceae; Salmonella MTGDNTLITSHGINRRDFMKLCAALAATMGLSSKAAAEMAESVSNPQRPPVIWIGAQECT WP_000145429.1 hydrogenase 2 small subunit [Salmonella enterica] Length: 372\nNumber of predicted TMHs: 1\nExp number of AAs in TMHs: 24.51309\nExp number, first 60 AAs: 0.2682\nTotal prob of N-in: 0.14402\noutside 1 331\nTMhelix 332 354\ninside 355 372

14462 GCF_002057815.1_ASM205781v1 Salmonella enterica Proteobacteria; Gammaproteobacteria; Enterobacterales; Enterobacteriaceae; Salmonella

MNNEETFYQAMRRKGVTRRSFLKFCSLAATSLGLGAGMTPKIAWALENKPRIPVVWIHGL WP_001058312.1 hydrogenase [Salmonella enterica] Length: 372\nNumber of predicted TMHs: 2\nExp number of AAs in TMHs: 36.51738\nExp number, first 60 AAs: 19.72849\nTotal prob of N-in: 0.96170\nPOSSIBLE N-term signal sequence\ninside 1 19\nTMhelix 20 39\noutside 40 325\nTMhelix 326 348\ninside 349 372

14463 GCF_002059645.1_ASM205964v1 Salmonella enterica Proteobacteria; Gammaproteobacteria; Enterobacterales; Enterobacteriaceae; Salmonella MTGDNTLITSHGINRRDFMKLCAALAATMGLSSKAAAEMAESVSNPQRPPVIWIGAQECT WP_000145429.1 hydrogenase 2 small subunit [Salmonella enterica] Length: 372\nNumber of predicted TMHs: 1\nExp number of AAs in TMHs: 24.51309\nExp number, first 60 AAs: 0.2682\nTotal prob of N-in: 0.14402\noutside 1 331\nTMhelix 332 354\ninside 355 372

14464 GCF_002057725.1_ASM205772v1 Salmonella enterica Proteobacteria; Gammaproteobacteria; Enterobacterales; Enterobacteriaceae; Salmonella MNNEETFYQAMRRKGVTRRSFLKFCSLAATSLGLGAGMTPKIAWALENKPRIPVVWIHGL WP_001058307.1 hydrogenase [Salmonella enterica] Length: 372\nNumber of predicted TMHs: 2\nExp number of AAs in TMHs: 36.51738\nExp number, first 60 AAs: 19.72849\nTotal prob of N-in: 0.96170\nPOSSIBLE N-term signal sequence\ninside 1 19\nTMhelix 20 39\noutside 40 325\nTMhelix 326 348\ninside 349 372

14465 GCF_002057815.1_ASM205781v1 Salmonella enterica Proteobacteria; Gammaproteobacteria; Enterobacterales; Enterobacteriaceae; Salmonella MTGDNTLITSHGINRRDFMKLCAALAATMGLSSKAAAEMAESVSNPQRPPVIWIGAQECT WP_000145429.1 hydrogenase 2 small subunit [Salmonella enterica] Length: 372\nNumber of predicted TMHs: 1\nExp number of AAs in TMHs: 24.51309\nExp number, first 60 AAs: 0.2682\nTotal prob of N-in: 0.14402\noutside 1 331\nTMhelix 332 354\ninside 355 372

14466 GCF_002057665.1_ASM205766v1 Salmonella enterica Proteobacteria; Gammaproteobacteria; Enterobacterales; Enterobacteriaceae; Salmonella MNNEETFYQAMRRKGVTRRSFLKFCSLAATSLGLGAGMTPKIAWALENKPRIPVVWIHGL WP_001058307.1 hydrogenase [Salmonella enterica] Length: 372\nNumber of predicted TMHs: 2\nExp number of AAs in TMHs: 36.51738\nExp number, first 60 AAs: 19.72849\nTotal prob of N-in: 0.96170\nPOSSIBLE N-term signal sequence\ninside 1 19\nTMhelix 20 39\noutside 40 325\nTMhelix 326 348\ninside 349 372

14467 GCF_002057755.1_ASM205775v1 Salmonella enterica Proteobacteria; Gammaproteobacteria; Enterobacterales; Enterobacteriaceae; Salmonella MTGDNTLITSHGINRRDFMKLCAALAATMGLSSKAAAEMAESVSNPQRPPVIWIGAQECT WP_000145429.1 hydrogenase 2 small subunit [Salmonella enterica] Length: 372\nNumber of predicted TMHs: 1\nExp number of AAs in TMHs: 24.51309\nExp number, first 60 AAs: 0.2682\nTotal prob of N-in: 0.14402\noutside 1 331\nTMhelix 332 354\ninside 355 372

14468 GCF_002057725.1_ASM205772v1 Salmonella enterica Proteobacteria; Gammaproteobacteria; Enterobacterales; Enterobacteriaceae; Salmonella MTGDNTLITSHGINRRDFMKLCAALAATMGLSSKAAAEMAESVSNPQRPPVIWIGAQECT WP_000145429.1 hydrogenase 2 small subunit [Salmonella enterica] Length: 372\nNumber of predicted TMHs: 1\nExp number of AAs in TMHs: 24.51309\nExp number, first 60 AAs: 0.2682\nTotal prob of N-in: 0.14402\noutside 1 331\nTMhelix 332 354\ninside 355 372

14469 GCF_002057665.1_ASM205766v1 Salmonella enterica Proteobacteria; Gammaproteobacteria; Enterobacterales; Enterobacteriaceae; Salmonella MTGDNTLITSHGINRRDFMKLCAALAATMGLSSKAAAEMAESVSNPQRPPVIWIGAQECT WP_000145429.1 hydrogenase 2 small subunit [Salmonella enterica] Length: 372\nNumber of predicted TMHs: 1\nExp number of AAs in TMHs: 24.51309\nExp number, first 60 AAs: 0.2682\nTotal prob of N-in: 0.14402\noutside 1 331\nTMhelix 332 354\ninside 355 372

14470 GCF_002057555.1_ASM205755v1 Salmonella enterica Proteobacteria; Gammaproteobacteria; Enterobacterales; Enterobacteriaceae; Salmonella MNNEETFYQAMRRKGVTRRSFLKFCSLAATSLGLGAGMTPKIAWALENKPRIPVVWIHGL WP_001058307.1 hydrogenase [Salmonella enterica] Length: 372\nNumber of predicted TMHs: 2\nExp number of AAs in TMHs: 36.51738\nExp number, first 60 AAs: 19.72849\nTotal prob of N-in: 0.96170\nPOSSIBLE N-term signal sequence\ninside 1 19\nTMhelix 20 39\noutside 40 325\nTMhelix 326 348\ninside 349 372

14471 GCF_002057555.1_ASM205755v1 Salmonella enterica Proteobacteria; Gammaproteobacteria; Enterobacterales; Enterobacteriaceae; Salmonella
MTGDNTLITSHGINRRDFMKLCAALAATMGLSSKAAAEMAESVSNPQRPPVIWIGAQECT WP_000145429.1
hydrogenase 2 small subunit [Salmonella enterica] Length: 372\nNumber of predicted TMHs: 1\nExp number of AAs in TMHs: 24.51309\nExp number, first 60 AAs: 0.2682\nTotal prob of N-in: 0.14402\noutside 1 331\nTMhelix 332 354\ninside 355 372

14472 GCF_002057575.1_ASM205757v1 Salmonella enterica Proteobacteria; Gammaproteobacteria; Enterobacterales; Enterobacteriaceae; Salmonella
MTGDNTLITSHGINRRDFMKLCAALAATMGLSSKAAAEMAESVSNPQRPPVIWIGAQECT WP_000145429.1
hydrogenase 2 small subunit [Salmonella enterica] Length: 372\nNumber of predicted TMHs: 1\nExp number of AAs in TMHs: 24.51309\nExp number, first 60 AAs: 0.2682\nTotal prob of N-in: 0.14402\noutside 1 331\nTMhelix 332 354\ninside 355 372

14473 GCF_002057295.1_ASM205729v1 Salmonella enterica Proteobacteria; Gammaproteobacteria; Enterobacterales; Enterobacteriaceae; Salmonella
MNNEETFYQAMRRKGVTRRSFLKFCSLAATSLGLGAGMTPKIAWALENKPRIPVVWIHGL WP_001058312.1
hydrogenase [Salmonella enterica] Length: 372\nNumber of predicted TMHs: 2\nExp number of AAs in TMHs: 36.51738\nExp number, first 60 AAs: 19.72849\nTotal prob of N-in: 0.96170\nPOSSIBLE N-term signal sequence\ninside 1 19\nTMhelix 20 39\noutside 40 325\nTMhelix 326 348\ninside 349 372

14474 GCF_002057345.1_ASM205734v1 Salmonella enterica Proteobacteria; Gammaproteobacteria; Enterobacterales; Enterobacteriaceae; Salmonella
MNNEETFYQAMRRKGVTRRSFLKFCSLAATSLGLGAGMTPKIAWALENKPRIPVVWIHGL WP_001058312.1
hydrogenase [Salmonella enterica] Length: 372\nNumber of predicted TMHs: 2\nExp number of AAs in TMHs: 36.51738\nExp number, first 60 AAs: 19.72849\nTotal prob of N-in: 0.96170\nPOSSIBLE N-term signal sequence\ninside 1 19\nTMhelix 20 39\noutside 40 325\nTMhelix 326 348\ninside 349 372

14475 GCF_002057435.1_ASM205743v1 Salmonella enterica Proteobacteria; Gammaproteobacteria; Enterobacterales; Enterobacteriaceae; Salmonella
MTGDNTLITSHGINRRDFMKLCAALAATMGLSSKAAAEMAESVSNPQRPPVIWIGAQECT WP_000145429.1
hydrogenase 2 small subunit [Salmonella enterica] Length: 372\nNumber of predicted TMHs: 1\nExp number of AAs in TMHs: 24.51309\nExp number, first 60 AAs: 0.2682\nTotal prob of N-in: 0.14402\noutside 1 331\nTMhelix 332 354\ninside 355 372

14476 GCF_002057195.1_ASM205719v1 Salmonella enterica Proteobacteria; Gammaproteobacteria; Enterobacterales; Enterobacteriaceae; Salmonella
MNNEETFYQAMRRKGVTRRSFLKFCSLAATSLGLGAGMTPKIAWALENKPRIPVVWIHGL WP_001058312.1
hydrogenase [Salmonella enterica] Length: 372\nNumber of predicted TMHs: 2\nExp number of AAs in TMHs: 36.51738\nExp number, first 60 AAs: 19.72849\nTotal prob of N-in: 0.96170\nPOSSIBLE N-term signal sequence\ninside 1 19\nTMhelix 20 39\noutside 40 325\nTMhelix 326 348\ninside 349 372

14477 GCF_002057185.1_ASM205718v1 Salmonella enterica Proteobacteria; Gammaproteobacteria; Enterobacterales; Enterobacteriaceae; Salmonella
MNNEETFYQAMRRKGVTRRSFLKFCSLAATSLGLGAGMTPKIAWALENKPRIPVVWIHGL WP_001058310.1
hydrogenase [Salmonella enterica] Length: 372\nNumber of predicted TMHs: 2\nExp number of AAs in TMHs: 37.4567299999999\nExp number, first 60 AAs: 19.72694\nTotal prob of N-in: 0.96171\nPOSSIBLE N-term signal sequence\ninside 1 19\nTMhelix 20 39\noutside 40 325\nTMhelix 326 348\ninside 349 372

14478 GCF_002057195.1_ASM205719v1 Salmonella enterica Proteobacteria; Gammaproteobacteria; Enterobacterales; Enterobacteriaceae; Salmonella
MTGDNTLITSHGINRRDFMKLCAALAATMGLSSKAAAEMAESVSNPQRPPVIWIGAQECT WP_000145431.1
hydrogenase 2 small subunit [Salmonella enterica] Length: 372\nNumber of predicted TMHs: 1\nExp number of AAs in TMHs: 24.51505\nExp number, first 60 AAs: 0.2682\nTotal prob of N-in: 0.14406\noutside 1 331\nTMhelix 332 354\ninside 355 372

14479 GCF_002057185.1_ASM205718v1 Salmonella enterica Proteobacteria; Gammaproteobacteria; Enterobacterales; Enterobacteriaceae; Salmonella
MTGDNTLITSHGINRRDFMKLCAALAATMGLSSKAAAEMAESVSNPQRPPVIWIGAQECT WP_000145429.1
hydrogenase 2 small subunit [Salmonella enterica] Length: 372\nNumber of predicted TMHs: 1\nExp

number of AAs in TMHs: 24.51309\nExp number, first 60 AAs: 0.2682\nTotal prob of N-in: 0.14402\noutside 1 331\nTMhelix 332 354\ninside 355 372

14480 GCF_002056625.1_ASM205662v1 Salmonella enterica Proteobacteria; Gammaproteobacteria; Enterobacterales; Enterobacteriaceae; Salmonella MNNEETFYQAMRRKGVTRRSFLKFCSLAATSLGLGAGMTPKIAWALENKPRIPVVWIHGL WP_001058312.1 hydrogenase [Salmonella enterica] Length: 372\nNumber of predicted TMHs: 2\nExp number of AAs in TMHs: 36.51738\nExp number, first 60 AAs: 19.72849\nTotal prob of N-in: 0.96170\nPOSSIBLE N-term signal sequence\ninside 1 19\nTMhelix 20 39\noutside 40 325\nTMhelix 326 348\ninside 349 372

14481 GCF_002056615.1_ASM205661v1 Salmonella enterica Proteobacteria; Gammaproteobacteria; Enterobacterales; Enterobacteriaceae; Salmonella MNNEETFYQAMRRKGVTRRSFLKFCSLAATSLGLGAGMTPKIAWALENKPRIPVVWIHGL WP_001058307.1 hydrogenase [Salmonella enterica] Length: 372\nNumber of predicted TMHs: 2\nExp number of AAs in TMHs: 36.51738\nExp number, first 60 AAs: 19.72849\nTotal prob of N-in: 0.96170\nPOSSIBLE N-term signal sequence\ninside 1 19\nTMhelix 20 39\noutside 40 325\nTMhelix 326 348\ninside 349 372

14482 GCF_002056615.1_ASM205661v1 Salmonella enterica Proteobacteria; Gammaproteobacteria; Enterobacterales; Enterobacteriaceae; Salmonella MTGDNTLITSHGINRRDFMKLCAALAATMGLSSKAAAEMAESVSNPQRPPVIWIGAQECT WP_000145429.1 hydrogenase 2 small subunit [Salmonella enterica] Length: 372\nNumber of predicted TMHs: 1\nExp number of AAs in TMHs: 24.51309\nExp number, first 60 AAs: 0.2682\nTotal prob of N-in: 0.14402\noutside 1 331\nTMhelix 332 354\ninside 355 372

14483 GCF_002056535.1_ASM205653v1 Salmonella enterica Proteobacteria; Gammaproteobacteria; Enterobacterales; Enterobacteriaceae; Salmonella MTGDNTLITSHGINRRDFMKLCAALAATMGLSSKAAAEMAESVSNPQRPPVIWIGAQECT WP_000145429.1 hydrogenase 2 small subunit [Salmonella enterica] Length: 372\nNumber of predicted TMHs: 1\nExp number of AAs in TMHs: 24.51309\nExp number, first 60 AAs: 0.2682\nTotal prob of N-in: 0.14402\noutside 1 331\nTMhelix 332 354\ninside 355 372

14484 GCF_002056575.1_ASM205657v1 Salmonella enterica Proteobacteria; Gammaproteobacteria; Enterobacterales; Enterobacteriaceae; Salmonella MTGDNTLITSHGINRRDFMKLCAALAATMGLSSKAAAEMAESVSNPQRPPVIWIGAQECT WP_000145429.1 hydrogenase 2 small subunit [Salmonella enterica] Length: 372\nNumber of predicted TMHs: 1\nExp number of AAs in TMHs: 24.51309\nExp number, first 60 AAs: 0.2682\nTotal prob of N-in: 0.14402\noutside 1 331\nTMhelix 332 354\ninside 355 372

14485 GCF_002056445.1_ASM205644v1 Salmonella enterica Proteobacteria; Gammaproteobacteria; Enterobacterales; Enterobacteriaceae; Salmonella MTGDNTLITSHGINRRDFMKLCAALAATMGLSSKAAAEMAESVSNPQRPPVIWIGAQECT WP_000145429.1 hydrogenase 2 small subunit [Salmonella enterica] Length: 372\nNumber of predicted TMHs: 1\nExp number of AAs in TMHs: 24.51309\nExp number, first 60 AAs: 0.2682\nTotal prob of N-in: 0.14402\noutside 1 331\nTMhelix 332 354\ninside 355 372

14486 GCF_002056205.1_ASM205620v1 Salmonella enterica Proteobacteria; Gammaproteobacteria; Enterobacterales; Enterobacteriaceae; Salmonella MNNEETFYQAMRRKGVTRRSFLKFCSLAATSLGLGAGMTPKIAWALENKPRIPVVWIHGL WP_001058307.1 hydrogenase [Salmonella enterica] Length: 372\nNumber of predicted TMHs: 2\nExp number of AAs in TMHs: 36.51738\nExp number, first 60 AAs: 19.72849\nTotal prob of N-in: 0.96170\nPOSSIBLE N-term signal sequence\ninside 1 19\nTMhelix 20 39\noutside 40 325\nTMhelix 326 348\ninside 349 372

14487 GCF_002056395.1_ASM205639v1 Salmonella enterica Proteobacteria; Gammaproteobacteria; Enterobacterales; Enterobacteriaceae; Salmonella MNNEETFYQAMRRKGVTRRSFLKFCSLAATSLGLGAGMTPKIAWALENKPRIPVVWIHGL WP_001058307.1 hydrogenase [Salmonella enterica] Length: 372\nNumber of predicted TMHs: 2\nExp number of AAs in TMHs: 36.51738\nExp number, first 60 AAs: 19.72849\nTotal prob of N-in: 0.96170\nPOSSIBLE N-term signal sequence\ninside 1 19\nTMhelix 20 39\noutside 40 325\nTMhelix 326 348\ninside 349 372

14488 GCF_002056335.1_ASM205633v1 Salmonella enterica Proteobacteria; Gammaproteobacteria; Enterobacterales; Enterobacteriaceae; Salmonella

MNNEETFYQAMRRKGVTRRSFLKFCSLAATSLGLGAGMTPKIAWALENKPRIPVVWIHGL WP_001058307.1 hydrogenase [Salmonella enterica] Length: 372\nNumber of predicted TMHs: 2\nExp number of AAs in TMHs: 36.51738\nExp number, first 60 AAs: 19.72849\nTotal prob of N-in: 0.96170\nPOSSIBLE N-term signal sequence\ninside 1 19\nTMhelix 20 39\noutside 40 325\nTMhelix 326 348\ninside 349 372

14489 GCF_002056105.1_ASM205610v1 Salmonella enterica Proteobacteria; Gammaproteobacteria; Enterobacterales; Enterobacteriaceae; Salmonella MNNEETFYQAMRRKGVTRRSFLKFCSLAATSLGLGAGMTPKIAWALENKPRIPVVWIHGL WP_001058307.1 hydrogenase [Salmonella enterica] Length: 372\nNumber of predicted TMHs: 2\nExp number of AAs in TMHs: 36.51738\nExp number, first 60 AAs: 19.72849\nTotal prob of N-in: 0.96170\nPOSSIBLE N-term signal sequence\ninside 1 19\nTMhelix 20 39\noutside 40 325\nTMhelix 326 348\ninside 349 372

14490 GCF_002056205.1_ASM205620v1 Salmonella enterica Proteobacteria; Gammaproteobacteria; Enterobacterales; Enterobacteriaceae; Salmonella MTGDNTLITSHGINRRDFMKLCAALAATMGLSSKAAAEMAESVSNPQRPPVIWIGAQECT WP_000145429.1 hydrogenase 2 small subunit [Salmonella enterica] Length: 372\nNumber of predicted TMHs: 1\nExp number of AAs in TMHs: 24.51309\nExp number, first 60 AAs: 0.2682\nTotal prob of N-in: 0.14402\noutside 1 331\nTMhelix 332 354\ninside 355 372

14491 GCF_002056155.1_ASM205615v1 Salmonella enterica Proteobacteria; Gammaproteobacteria; Enterobacterales; Enterobacteriaceae; Salmonella MTGDNTLITSHGINRRDFMKLCAALAATMGLSSKAAAEMAESVSNPQRPPVIWIGAQECT WP_000145429.1 hydrogenase 2 small subunit [Salmonella enterica] Length: 372\nNumber of predicted TMHs: 1\nExp number of AAs in TMHs: 24.51309\nExp number, first 60 AAs: 0.2682\nTotal prob of N-in: 0.14402\noutside 1 331\nTMhelix 332 354\ninside 355 372

14492 GCF_002056015.1_ASM205601v1 Salmonella enterica Proteobacteria; Gammaproteobacteria; Enterobacterales; Enterobacteriaceae; Salmonella MNNEETFYQAMRRKGVTRRSFLKFCSLAATSLGLGAGMTPKIAWALENKPRIPVVWIHGL WP_001058307.1 hydrogenase [Salmonella enterica] Length: 372\nNumber of predicted TMHs: 2\nExp number of AAs in TMHs: 36.51738\nExp number, first 60 AAs: 19.72849\nTotal prob of N-in: 0.96170\nPOSSIBLE N-term signal sequence\ninside 1 19\nTMhelix 20 39\noutside 40 325\nTMhelix 326 348\ninside 349 372

14493 GCF_002056105.1_ASM205610v1 Salmonella enterica Proteobacteria; Gammaproteobacteria; Enterobacterales; Enterobacteriaceae; Salmonella MTGDNTLITSHGINRRDFMKLCAALAATMGLSSKAAAEMAESVSNPQRPPVIWIGAQECT WP_000145429.1 hydrogenase 2 small subunit [Salmonella enterica] Length: 372\nNumber of predicted TMHs: 1\nExp number of AAs in TMHs: 24.51309\nExp number, first 60 AAs: 0.2682\nTotal prob of N-in: 0.14402\noutside 1 331\nTMhelix 332 354\ninside 355 372

14494 GCF_002056015.1_ASM205601v1 Salmonella enterica Proteobacteria; Gammaproteobacteria; Enterobacterales; Enterobacteriaceae; Salmonella MTGDNTLITSHGINRRDFMKLCAALAATMGLSSKAAAEMAESVSNPQRPPVIWIGAQECT WP_000145429.1 hydrogenase 2 small subunit [Salmonella enterica] Length: 372\nNumber of predicted TMHs: 1\nExp number of AAs in TMHs: 24.51309\nExp number, first 60 AAs: 0.2682\nTotal prob of N-in: 0.14402\noutside 1 331\nTMhelix 332 354\ninside 355 372

14495 GCF_002055865.1_ASM205586v1 Salmonella enterica Proteobacteria; Gammaproteobacteria; Enterobacterales; Enterobacteriaceae; Salmonella MNNEETFYQAMRRKGVTRRSFLKFCSLAATSLGLGAGMTPKIAWALENKPRIPVVWIHGL WP_001058312.1 hydrogenase [Salmonella enterica] Length: 372\nNumber of predicted TMHs: 2\nExp number of AAs in TMHs: 36.51738\nExp number, first 60 AAs: 19.72849\nTotal prob of N-in: 0.96170\nPOSSIBLE N-term signal sequence\ninside 1 19\nTMhelix 20 39\noutside 40 325\nTMhelix 326 348\ninside 349 372

14496 GCF_002055645.1_ASM205564v1 Salmonella enterica Proteobacteria; Gammaproteobacteria; Enterobacterales; Enterobacteriaceae; Salmonella MNNEETFYQAMRRKGVTRRSFLKFCSLAATSLGLGAGMTPKIAWALENKPRIPVVWIHGL WP_001058312.1 hydrogenase [Salmonella enterica] Length: 372\nNumber of predicted TMHs: 2\nExp number of AAs in TMHs: 36.51738\nExp number, first 60 AAs: 19.72849\nTotal prob of N-in: 0.96170\nPOSSIBLE N-term signal sequence\ninside 1 19\nTMhelix 20 39\noutside 40 325\nTMhelix 326 348\ninside 349 372

14497	GCF_002055935.1_ASM205593v1	Salmonella enterica	Proteobacteria; Gammaproteobacteria; Enterobacterales; Enterobacteriaceae; Salmonella
MNNEETFYQAMRRKGVTRRSFLKFCSLAATSLGLGAGMTPKIAWALENKPRIPVVWIHGL	WP_001058312.1
hydrogenase [Salmonella enterica]	Length: 372\nNumber of predicted TMHs:  2\nExp number of AAs in TMHs: 36.51738\nExp number, first 60 AAs:  19.72849\nTotal prob of N-in:   0.96170\nPOSSIBLE N-term signal sequence\ninside   1   19\nTMhelix   20   39\noutside   40   325\nTMhelix   326   348\ninside   349   372

14498	GCF_002055995.1_ASM205599v1	Salmonella enterica	Proteobacteria; Gammaproteobacteria; Enterobacterales; Enterobacteriaceae; Salmonella
MTGDNTLITSHGINRRDFMKLCAALAATMGLSSKAAAEMAESVSNPQRPPVIWIGAQECT	WP_023248548.1
hydrogenase 2 small subunit [Salmonella enterica]	Length: 372\nNumber of predicted TMHs:  1\nExp number of AAs in TMHs: 24.51281\nExp number, first 60 AAs:  0.26822\nTotal prob of N-in:   0.14402\noutside   1   331\nTMhelix   332   354\ninside   355   372

14499	GCF_002055995.1_ASM205599v1	Salmonella enterica	Proteobacteria; Gammaproteobacteria; Enterobacterales; Enterobacteriaceae; Salmonella
MNNEETFYQAMRRKGVTRRSFLKFCSLAATSLGLGAGMTPKIAWALENKPRIPVVWIHGL	WP_001058306.1
hydrogenase [Salmonella enterica]	Length: 372\nNumber of predicted TMHs:  2\nExp number of AAs in TMHs: 36.5540099999999\nExp number, first 60 AAs:  19.72732\nTotal prob of N-in:   0.96171\nPOSSIBLE N-term signal sequence\ninside   1   19\nTMhelix   20   39\noutside   40   325\nTMhelix   326   348\ninside   349   372

14500	GCF_002055935.1_ASM205593v1	Salmonella enterica	Proteobacteria; Gammaproteobacteria; Enterobacterales; Enterobacteriaceae; Salmonella
MTGDNTLITSHGINRRDFMKLCAALAATMGLSSKAAAEMAESVSNPQRPPVIWIGAQECT	WP_000145429.1
hydrogenase 2 small subunit [Salmonella enterica]	Length: 372\nNumber of predicted TMHs:  1\nExp number of AAs in TMHs: 24.51309\nExp number, first 60 AAs:  0.2682\nTotal prob of N-in:   0.14402\noutside   1   331\nTMhelix   332   354\ninside   355   372

14501	GCF_002055905.1_ASM205590v1	Salmonella enterica	Proteobacteria; Gammaproteobacteria; Enterobacterales; Enterobacteriaceae; Salmonella
MTGDNTLITSHGINRRDFMKLCAALAATMGLSSKAAAEMAESVSNPQRPPVIWIGAQECT	WP_000145429.1
hydrogenase 2 small subunit [Salmonella enterica]	Length: 372\nNumber of predicted TMHs:  1\nExp number of AAs in TMHs: 24.51309\nExp number, first 60 AAs:  0.2682\nTotal prob of N-in:   0.14402\noutside   1   331\nTMhelix   332   354\ninside   355   372

14502	GCF_002055865.1_ASM205586v1	Salmonella enterica	Proteobacteria; Gammaproteobacteria; Enterobacterales; Enterobacteriaceae; Salmonella
MTGDNTLITSHGINRRDFMKLCAALAATMGLSSKAAAEMAESVSNPQRPPVIWIGAQECT	WP_000145429.1
hydrogenase 2 small subunit [Salmonella enterica]	Length: 372\nNumber of predicted TMHs:  1\nExp number of AAs in TMHs: 24.51309\nExp number, first 60 AAs:  0.2682\nTotal prob of N-in:   0.14402\noutside   1   331\nTMhelix   332   354\ninside   355   372

14503	GCF_002055255.1_ASM205525v1	Salmonella enterica	Proteobacteria; Gammaproteobacteria; Enterobacterales; Enterobacteriaceae; Salmonella
MNNEETFYQAMRRKGVTRRSFLKFCSLAATSLGLSAGMTPKIAWALENKPRIPVVWIHGL	WP_001058316.1
hydrogenase [Salmonella enterica]	Length: 372\nNumber of predicted TMHs:  2\nExp number of AAs in TMHs: 35.76397\nExp number, first 60 AAs:  18.97201\nTotal prob of N-in:   0.92785\nPOSSIBLE N-term signal sequence\ninside   1   19\nTMhelix   20   39\noutside   40   325\nTMhelix   326   348\ninside   349   372

14504	GCF_002055345.1_ASM205534v1	Salmonella enterica	Proteobacteria; Gammaproteobacteria; Enterobacterales; Enterobacteriaceae; Salmonella
MNNEETFYQAMRRKGVTRRSFLKFCSLAATSLGLGAGMTPKIAWALENKPRIPVVWIHGL	WP_001058311.1
hydrogenase [Salmonella enterica]	Length: 372\nNumber of predicted TMHs:  2\nExp number of AAs in TMHs: 37.4699899999999\nExp number, first 60 AAs:  19.72699\nTotal prob of N-in:   0.96171\nPOSSIBLE N-term signal sequence\ninside   1   19\nTMhelix   20   39\noutside   40   325\nTMhelix   326   348\ninside   349   372

14505	GCF_002055425.1_ASM205542v1	Salmonella enterica	Proteobacteria; Gammaproteobacteria; Enterobacterales; Enterobacteriaceae; Salmonella
MNNEETFYQAMRRKGVTRRSFLKFCSLAATSLGLGAGMTPKIAWALENKPRIPVVWIHGL	WP_001058312.1
hydrogenase [Salmonella enterica]	Length: 372\nNumber of predicted TMHs:  2\nExp number of AAs in

TMHs: 36.51738\nExp number, first 60 AAs: 19.72849\nTotal prob of N-in: 0.96170\nPOSSIBLE N-term signal sequence\ninside 1 19\nTMhelix 20 39\noutside 40 325\nTMhelix 326 348\ninside 349 372

14506 GCF_002055425.1_ASM205542v1 Salmonella enterica Proteobacteria; Gammaproteobacteria; Enterobacterales; Enterobacteriaceae; Salmonella
MTGDNTLITSHGINRRDFMKLCAALAATMGLSSKAAAEMAESVSNPQRPPVIWIGAQECT WP_000145429.1
hydrogenase 2 small subunit [Salmonella enterica] Length: 372\nNumber of predicted TMHs: 1\nExp number of AAs in TMHs: 24.51309\nExp number, first 60 AAs: 0.2682\nTotal prob of N-in: 0.14402\noutside 1 331\nTMhelix 332 354\ninside 355 372

14507 GCF_002055205.1_ASM205520v1 Salmonella enterica Proteobacteria; Gammaproteobacteria; Enterobacterales; Enterobacteriaceae; Salmonella
MNNEETFYQAMRRKGVTRRSFLKFCSLAATSLGLGAGMTPKIAWALENKPRIPVVWIHGL WP_001058312.1
hydrogenase [Salmonella enterica] Length: 372\nNumber of predicted TMHs: 2\nExp number of AAs in TMHs: 36.51738\nExp number, first 60 AAs: 19.72849\nTotal prob of N-in: 0.96170\nPOSSIBLE N-term signal sequence\ninside 1 19\nTMhelix 20 39\noutside 40 325\nTMhelix 326 348\ninside 349 372

14508 GCF_002055405.1_ASM205540v1 Salmonella enterica Proteobacteria; Gammaproteobacteria; Enterobacterales; Enterobacteriaceae; Salmonella
MNNEETFYQAMRRKGVTRRSFLKFCSLAATSLGLGAGMTPKIAWALENKPRIPVVWIHGL WP_001058312.1
hydrogenase [Salmonella enterica] Length: 372\nNumber of predicted TMHs: 2\nExp number of AAs in TMHs: 36.51738\nExp number, first 60 AAs: 19.72849\nTotal prob of N-in: 0.96170\nPOSSIBLE N-term signal sequence\ninside 1 19\nTMhelix 20 39\noutside 40 325\nTMhelix 326 348\ninside 349 372

14509 GCF_002055405.1_ASM205540v1 Salmonella enterica Proteobacteria; Gammaproteobacteria; Enterobacterales; Enterobacteriaceae; Salmonella
MTGDNTLITSHGINRRDFMKLCAALAATMGLSSKAAAEMAESVSNPQRPPVIWIGAQECT WP_000145429.1
hydrogenase 2 small subunit [Salmonella enterica] Length: 372\nNumber of predicted TMHs: 1\nExp number of AAs in TMHs: 24.51309\nExp number, first 60 AAs: 0.2682\nTotal prob of N-in: 0.14402\noutside 1 331\nTMhelix 332 354\ninside 355 372

14510 GCF_002055205.1_ASM205520v1 Salmonella enterica Proteobacteria; Gammaproteobacteria; Enterobacterales; Enterobacteriaceae; Salmonella
MTGDNTLITSHGINRRDFMKLCAALAATMGLSSKAAAEMAESVSNPQRPPVIWIGAQECT WP_000145429.1
hydrogenase 2 small subunit [Salmonella enterica] Length: 372\nNumber of predicted TMHs: 1\nExp number of AAs in TMHs: 24.51309\nExp number, first 60 AAs: 0.2682\nTotal prob of N-in: 0.14402\noutside 1 331\nTMhelix 332 354\ninside 355 372

14511 GCF_002055175.1_ASM205517v1 Salmonella enterica Proteobacteria; Gammaproteobacteria; Enterobacterales; Enterobacteriaceae; Salmonella
MTGDNTLITSHGINRRDFMKLCAALAATMGLSSKAAAEMAESVSNPQRPPVIWIGAQECT WP_000145429.1
hydrogenase 2 small subunit [Salmonella enterica] Length: 372\nNumber of predicted TMHs: 1\nExp number of AAs in TMHs: 24.51309\nExp number, first 60 AAs: 0.2682\nTotal prob of N-in: 0.14402\noutside 1 331\nTMhelix 332 354\ninside 355 372

14512 GCF_002054985.1_ASM205498v1 Salmonella enterica Proteobacteria; Gammaproteobacteria; Enterobacterales; Enterobacteriaceae; Salmonella
MNNEETFYQAMRRKGVTRRSFLKFCSLAATSLGLGAGMTPKIAWALENKPRIPVVWIHGL WP_023213673.1
hydrogenase 1, small subunit [Salmonella enterica] Length: 372\nNumber of predicted TMHs: 2\nExp number of AAs in TMHs: 36.51753\nExp number, first 60 AAs: 19.72849\nTotal prob of N-in: 0.96170\nPOSSIBLE N-term signal sequence\ninside 1 19\nTMhelix 20 39\noutside 40 325\nTMhelix 326 348\ninside 349 372

14513 GCF_002055155.1_ASM205515v1 Salmonella enterica Proteobacteria; Gammaproteobacteria; Enterobacterales; Enterobacteriaceae; Salmonella
MNNEETFYQAMRRKGVTRRSFLKFCSLAATSLGLGAGMTPKIAWALENKPRIPVVWIHGL WP_001058312.1
hydrogenase [Salmonella enterica] Length: 372\nNumber of predicted TMHs: 2\nExp number of AAs in TMHs: 36.51738\nExp number, first 60 AAs: 19.72849\nTotal prob of N-in: 0.96170\nPOSSIBLE N-term signal sequence\ninside 1 19\nTMhelix 20 39\noutside 40 325\nTMhelix 326 348\ninside 349 372

14514 GCF_002055155.1_ASM205515v1 Salmonella enterica Proteobacteria; Gammaproteobacteria; Enterobacterales; Enterobacteriaceae; Salmonella

MTGDNTLITSHGINRRDFMKLCAALAATMGLSSKAAAEMAESVSNPQRPPVIWIGAQECT WP_000145429.1 hydrogenase 2 small subunit [Salmonella enterica] Length: 372\nNumber of predicted TMHs: 1\nExp number of AAs in TMHs: 24.51309\nExp number, first 60 AAs: 0.2682\nTotal prob of N-in: 0.14402\noutside 1 331\nTMhelix 332 354\ninside 355 372

14515 GCF_002054875.1_ASM205487v1 Salmonella enterica Proteobacteria; Gammaproteobacteria; Enterobacterales; Enterobacteriaceae; Salmonella MNNEETFYQAMRRKGVTRRSFLKFCSLAATSLGLGAGMTPKIAWALENKPRIPVVWIHGL WP_023243720.1 hydrogenase 1, small subunit [Salmonella enterica] Length: 372\nNumber of predicted TMHs: 2\nExp number of AAs in TMHs: 36.52147\nExp number, first 60 AAs: 19.72838\nTotal prob of N-in: 0.96170\nPOSSIBLE N-term signal sequence\ninside 1 19\nTMhelix 20 39\noutside 40 325\nTMhelix 326 348\ninside 349 372

14516 GCF_002054935.1_ASM205493v1 Salmonella enterica Proteobacteria; Gammaproteobacteria; Enterobacterales; Enterobacteriaceae; Salmonella MTGDNTLITSHGINRRDFMKLCAALAATMGLSSKAAAEMAESVSNPQRPPVIWIGAQECT WP_000145429.1 hydrogenase 2 small subunit [Salmonella enterica] Length: 372\nNumber of predicted TMHs: 1\nExp number of AAs in TMHs: 24.51309\nExp number, first 60 AAs: 0.2682\nTotal prob of N-in: 0.14402\noutside 1 331\nTMhelix 332 354\ninside 355 372

14517 GCF_002061035.1_ASM206103v1 Salmonella enterica Proteobacteria; Gammaproteobacteria; Enterobacterales; Enterobacteriaceae; Salmonella MLRFRTIFLKKIFTLKNSILARRGAVIVIVSAVFTSIMFFAHSWASDKEVAMTLSVLNSN WP_080151465.1 hypothetical protein [Salmonella enterica] Length: 204\nNumber of predicted TMHs: 2\nExp number of AAs in TMHs: 44.99538\nExp number, first 60 AAs: 22.34317\nTotal prob of N-in: 0.89744\nPOSSIBLE N-term signal sequence\ninside 1 19\nTMhelix 20 42\noutside 43 171\nTMhelix 172 194\ninside 195 204

14518 GCF_002054815.1_ASM205481v1 Salmonella enterica Proteobacteria; Gammaproteobacteria; Enterobacterales; Enterobacteriaceae; Salmonella MTGDNTLITSHGINRRDFMKLCAALAATMGLSSKAAAEMAESVSNPQRPPVIWIGAQECT WP_000145429.1 hydrogenase 2 small subunit [Salmonella enterica] Length: 372\nNumber of predicted TMHs: 1\nExp number of AAs in TMHs: 24.51309\nExp number, first 60 AAs: 0.2682\nTotal prob of N-in: 0.14402\noutside 1 331\nTMhelix 332 354\ninside 355 372

14519 GCF_002054615.1_ASM205461v1 Salmonella enterica Proteobacteria; Gammaproteobacteria; Enterobacterales; Enterobacteriaceae; Salmonella MNNEETFYQAMRRKGVTRRSFLKFCSLAATSLGLGAGMTPKIAWALENKPRIPVVWIHGL WP_001058312.1 hydrogenase [Salmonella enterica] Length: 372\nNumber of predicted TMHs: 2\nExp number of AAs in TMHs: 36.51738\nExp number, first 60 AAs: 19.72849\nTotal prob of N-in: 0.96170\nPOSSIBLE N-term signal sequence\ninside 1 19\nTMhelix 20 39\noutside 40 325\nTMhelix 326 348\ninside 349 372

14520 GCF_002054705.1_ASM205470v1 Salmonella enterica Proteobacteria; Gammaproteobacteria; Enterobacterales; Enterobacteriaceae; Salmonella MNNEETFYQAMRRKGVTRRSFLKFCSLAATSLGLGAGMTPKIAWALENKPRIPVVWIHGL WP_001058312.1 hydrogenase [Salmonella enterica] Length: 372\nNumber of predicted TMHs: 2\nExp number of AAs in TMHs: 36.51738\nExp number, first 60 AAs: 19.72849\nTotal prob of N-in: 0.96170\nPOSSIBLE N-term signal sequence\ninside 1 19\nTMhelix 20 39\noutside 40 325\nTMhelix 326 348\ninside 349 372

14521 GCF_002054745.1_ASM205474v1 Salmonella enterica Proteobacteria; Gammaproteobacteria; Enterobacterales; Enterobacteriaceae; Salmonella MNNEETFYQAMRRKGVTRRSFLKFCSLAATSLGLGAGMTPKIAWALENKPRIPVVWIHGL WP_001058312.1 hydrogenase [Salmonella enterica] Length: 372\nNumber of predicted TMHs: 2\nExp number of AAs in TMHs: 36.51738\nExp number, first 60 AAs: 19.72849\nTotal prob of N-in: 0.96170\nPOSSIBLE N-term signal sequence\ninside 1 19\nTMhelix 20 39\noutside 40 325\nTMhelix 326 348\ninside 349 372

14522 GCF_002054645.1_ASM205464v1 Salmonella enterica Proteobacteria; Gammaproteobacteria; Enterobacterales; Enterobacteriaceae; Salmonella MNNEETFYQAMRRKGVTRRSFLKFCSLAATSLGLGAGMTPKIAWALENKPRIPVVWIHGL WP_001058312.1 hydrogenase [Salmonella enterica] Length: 372\nNumber of predicted TMHs: 2\nExp number of AAs in TMHs: 36.51738\nExp number, first 60 AAs: 19.72849\nTotal prob of N-in: 0.96170\nPOSSIBLE N-term signal sequence\ninside 1 19\nTMhelix 20 39\noutside 40 325\nTMhelix 326 348\ninside 349 372

14523 GCF_002054255.1_ASM205425v1 Salmonella enterica Proteobacteria; Gammaproteobacteria; Enterobacterales; Enterobacteriaceae; Salmonella
MNNEETFYQAMRRKGVTRRSFLKFCSLAATSLGLGAGMTPKIAWALENKPRIPVVWIHGL WP_001058312.1
hydrogenase [Salmonella enterica] Length: 372\nNumber of predicted TMHs: 2\nExp number of AAs in TMHs: 36.51738\nExp number, first 60 AAs: 19.72849\nTotal prob of N-in: 0.96170\nPOSSIBLE N-term signal sequence\ninside 1 19\nTMhelix 20 39\noutside 40 325\nTMhelix 326 348\ninside 349 372

14524 GCF_002054395.1_ASM205439v1 Salmonella enterica Proteobacteria; Gammaproteobacteria; Enterobacterales; Enterobacteriaceae; Salmonella
MNNEETFYQAMRRKGVTRRSFLKFCSLAATSLGLGAGMTPKIAWALENKPRIPVVWIHGL WP_001058310.1
hydrogenase [Salmonella enterica] Length: 372\nNumber of predicted TMHs: 2\nExp number of AAs in TMHs: 37.4567299999999\nExp number, first 60 AAs: 19.72694\nTotal prob of N-in: 0.96171\nPOSSIBLE N-term signal sequence\ninside 1 19\nTMhelix 20 39\noutside 40 325\nTMhelix 326 348\ninside 349 372

14525 GCF_002054305.1_ASM205430v1 Salmonella enterica Proteobacteria; Gammaproteobacteria; Enterobacterales; Enterobacteriaceae; Salmonella
MNNEETFYQAMRRKGVTRRSFLKFCSLAATSLGLGAGMTPKIAWALENKPRIPVVWIHGL WP_001058308.1
hydrogenase [Salmonella enterica] Length: 372\nNumber of predicted TMHs: 2\nExp number of AAs in TMHs: 39.08204\nExp number, first 60 AAs: 19.72474\nTotal prob of N-in: 0.96174\nPOSSIBLE N-term signal sequence\ninside 1 19\nTMhelix 20 39\noutside 40 325\nTMhelix 326 348\ninside 349 372

14526 GCF_002054395.1_ASM205439v1 Salmonella enterica Proteobacteria; Gammaproteobacteria; Enterobacterales; Enterobacteriaceae; Salmonella
MTGDNTLITSHGINRRDFMKLCAALAATMGLSSKAAAEMAESVSNPQRPPVIWIGAQECT WP_000145429.1
hydrogenase 2 small subunit [Salmonella enterica] Length: 372\nNumber of predicted TMHs: 1\nExp number of AAs in TMHs: 24.51309\nExp number, first 60 AAs: 0.2682\nTotal prob of N-in: 0.14402\noutside 1 331\nTMhelix 332 354\ninside 355 372

14527 GCF_002054305.1_ASM205430v1 Salmonella enterica Proteobacteria; Gammaproteobacteria; Enterobacterales; Enterobacteriaceae; Salmonella
MTGDNTLITSHGINRRDFMKLCAALAATMGLSSKAAAEMAESVSNPQRPPVIWIGAQECT WP_000145429.1
hydrogenase 2 small subunit [Salmonella enterica] Length: 372\nNumber of predicted TMHs: 1\nExp number of AAs in TMHs: 24.51309\nExp number, first 60 AAs: 0.2682\nTotal prob of N-in: 0.14402\noutside 1 331\nTMhelix 332 354\ninside 355 372

14528 GCF_002054205.1_ASM205420v1 Salmonella enterica Proteobacteria; Gammaproteobacteria; Enterobacterales; Enterobacteriaceae; Salmonella
MNNEETFYQAMRRKGVTRRSFLKFCSLAATSLGLGAGMTPKIAWALENKPRIPVVWIHGL WP_001058308.1
hydrogenase [Salmonella enterica] Length: 372\nNumber of predicted TMHs: 2\nExp number of AAs in TMHs: 39.08204\nExp number, first 60 AAs: 19.72474\nTotal prob of N-in: 0.96174\nPOSSIBLE N-term signal sequence\ninside 1 19\nTMhelix 20 39\noutside 40 325\nTMhelix 326 348\ninside 349 372

14529 GCF_002053655.1_ASM205365v1 Salmonella enterica Proteobacteria; Gammaproteobacteria; Enterobacterales; Enterobacteriaceae; Salmonella
MNNEETFYQAMRRKGVTRRSFLKFCSLAATSLGLGAGMTPKIAWALENKPRIPVVWIHGL WP_001058312.1
hydrogenase [Salmonella enterica] Length: 372\nNumber of predicted TMHs: 2\nExp number of AAs in TMHs: 36.51738\nExp number, first 60 AAs: 19.72849\nTotal prob of N-in: 0.96170\nPOSSIBLE N-term signal sequence\ninside 1 19\nTMhelix 20 39\noutside 40 325\nTMhelix 326 348\ninside 349 372

14530 GCF_002054355.1_ASM205435v1 Salmonella enterica Proteobacteria; Gammaproteobacteria; Enterobacterales; Enterobacteriaceae; Salmonella
MTGDNTLITSHGINRRDFMKLCAALAATMGLSSKAAAEMAESVSNPQRPPVIWIGAQECT WP_000145429.1
hydrogenase 2 small subunit [Salmonella enterica] Length: 372\nNumber of predicted TMHs: 1\nExp number of AAs in TMHs: 24.51309\nExp number, first 60 AAs: 0.2682\nTotal prob of N-in: 0.14402\noutside 1 331\nTMhelix 332 354\ninside 355 372

14531 GCF_002053865.1_ASM205386v1 Salmonella enterica Proteobacteria; Gammaproteobacteria; Enterobacterales; Enterobacteriaceae; Salmonella
MTGDNTLITSHGINRRDFMKLCAALAATMGLSSKAAAEMAESVSNPQRPPVIWIGAQECT WP_000145429.1
hydrogenase 2 small subunit [Salmonella enterica] Length: 372\nNumber of predicted TMHs: 1\nExp

number of AAs in TMHs: 24.51309\nExp number, first 60 AAs: 0.2682\nTotal prob of N-in: 0.14402\noutside 1 331\nTMhelix 332 354\ninside 355 372

14532 GCF_002053955.1_ASM205395v1 Salmonella enterica Proteobacteria; Gammaproteobacteria; Enterobacterales; Enterobacteriaceae; Salmonella
MTGDNTLITSHGINRRDFMKLCAALAATMGLSSKAAAEMAESVSNPQRPPVIWIGAQECT WP_000145429.1
hydrogenase 2 small subunit [Salmonella enterica] Length: 372\nNumber of predicted TMHs: 1\nExp number of AAs in TMHs: 24.51309\nExp number, first 60 AAs: 0.2682\nTotal prob of N-in: 0.14402\noutside 1 331\nTMhelix 332 354\ninside 355 372

14533 GCF_002053825.1_ASM205382v1 Salmonella enterica Proteobacteria; Gammaproteobacteria; Enterobacterales; Enterobacteriaceae; Salmonella
MNNEETFYQAMRRKGVTRRSFLKFCSLAATSLGLGAGMTPKIAWALENKPRIPVVWIHGL WP_001058307.1
hydrogenase [Salmonella enterica] Length: 372\nNumber of predicted TMHs: 2\nExp number of AAs in TMHs: 36.51738\nExp number, first 60 AAs: 19.72849\nTotal prob of N-in: 0.96170\nPOSSIBLE N-term signal sequence\ninside 1 19\nTMhelix 20 39\noutside 40 325\nTMhelix 326 348\ninside 349 372

14534 GCF_002053825.1_ASM205382v1 Salmonella enterica Proteobacteria; Gammaproteobacteria; Enterobacterales; Enterobacteriaceae; Salmonella
MTGDNTLITSHGINRRDFMKLCAALAATMGLSSKAAAEMAESVSNPQRPPVIWIGAQECT WP_000145429.1
hydrogenase 2 small subunit [Salmonella enterica] Length: 372\nNumber of predicted TMHs: 1\nExp number of AAs in TMHs: 24.51309\nExp number, first 60 AAs: 0.2682\nTotal prob of N-in: 0.14402\noutside 1 331\nTMhelix 332 354\ninside 355 372

14535 GCF_002053655.1_ASM205365v1 Salmonella enterica Proteobacteria; Gammaproteobacteria; Enterobacterales; Enterobacteriaceae; Salmonella
MTGDNTLITSHGINRRDFMKLCAALAATMGLSSKAAAEMAESVSNPQRPPVIWIGAQECT WP_000145429.1
hydrogenase 2 small subunit [Salmonella enterica] Length: 372\nNumber of predicted TMHs: 1\nExp number of AAs in TMHs: 24.51309\nExp number, first 60 AAs: 0.2682\nTotal prob of N-in: 0.14402\noutside 1 331\nTMhelix 332 354\ninside 355 372

14536 GCF_002053585.1_ASM205358v1 Salmonella enterica Proteobacteria; Gammaproteobacteria; Enterobacterales; Enterobacteriaceae; Salmonella
MTGDNTLITSHGINRRDFMKLCAALAATMGLSSKAAAEMAESVSNPQRPPVIWIGAQECT WP_000145429.1
hydrogenase 2 small subunit [Salmonella enterica] Length: 372\nNumber of predicted TMHs: 1\nExp number of AAs in TMHs: 24.51309\nExp number, first 60 AAs: 0.2682\nTotal prob of N-in: 0.14402\noutside 1 331\nTMhelix 332 354\ninside 355 372

14537 GCF_002053405.1_ASM205340v1 Salmonella enterica Proteobacteria; Gammaproteobacteria; Enterobacterales; Enterobacteriaceae; Salmonella
MTGDNTLITSHGINRRDFMKLCAALAATMGLSSKAAAEMAESVSNPQRPPVIWIGAQECT WP_000145431.1
hydrogenase 2 small subunit [Salmonella enterica] Length: 372\nNumber of predicted TMHs: 1\nExp number of AAs in TMHs: 24.51505\nExp number, first 60 AAs: 0.2682\nTotal prob of N-in: 0.14406\noutside 1 331\nTMhelix 332 354\ninside 355 372

14538 GCF_002053415.1_ASM205341v1 Salmonella enterica Proteobacteria; Gammaproteobacteria; Enterobacterales; Enterobacteriaceae; Salmonella
MTGDNTLITSHGINRRDFMKLCAALAATMGLSSKAAAEMAESVSNPQRPPVIWIGAQECT WP_000145429.1
hydrogenase 2 small subunit [Salmonella enterica] Length: 372\nNumber of predicted TMHs: 1\nExp number of AAs in TMHs: 24.51309\nExp number, first 60 AAs: 0.2682\nTotal prob of N-in: 0.14402\noutside 1 331\nTMhelix 332 354\ninside 355 372

14539 GCF_002053175.1_ASM205317v1 Salmonella enterica Proteobacteria; Gammaproteobacteria; Enterobacterales; Enterobacteriaceae; Salmonella
MNNEETFYQAMRRKGVTRRSFLKFCSLAATSLGLGAGMTPKIAWALENKPRIPVVWIHGL WP_001058312.1
hydrogenase [Salmonella enterica] Length: 372\nNumber of predicted TMHs: 2\nExp number of AAs in TMHs: 36.51738\nExp number, first 60 AAs: 19.72849\nTotal prob of N-in: 0.96170\nPOSSIBLE N-term signal sequence\ninside 1 19\nTMhelix 20 39\noutside 40 325\nTMhelix 326 348\ninside 349 372

14540 GCF_002053325.1_ASM205332v1 Salmonella enterica Proteobacteria; Gammaproteobacteria; Enterobacterales; Enterobacteriaceae; Salmonella

MNNEETFYQAMRRKGVTRRSFLKFCSLAATSLGLGAGMTPKIAWALENKPRIPVVWIHGL WP_001058312.1 hydrogenase [Salmonella enterica] Length: 372\nNumber of predicted TMHs: 2\nExp number of AAs in TMHs: 36.51738\nExp number, first 60 AAs: 19.72849\nTotal prob of N-in: 0.96170\nPOSSIBLE N-term signal sequence\ninside 1 19\nTMhelix 20 39\noutside 40 325\nTMhelix 326 348\ninside 349 372

14541 GCF_002053325.1_ASM205332v1 Salmonella enterica Proteobacteria; Gammaproteobacteria; Enterobacterales; Enterobacteriaceae; Salmonella MTGDNTLITSHGINRRDFMKLCAALAATMGLSSKAAAEMAESVSNPQRPPVIWIGAQECT WP_000145429.1 hydrogenase 2 small subunit [Salmonella enterica] Length: 372\nNumber of predicted TMHs: 1\nExp number of AAs in TMHs: 24.51309\nExp number, first 60 AAs: 0.2682\nTotal prob of N-in: 0.14402\noutside 1 331\nTMhelix 332 354\ninside 355 372

14542 GCF_002053085.1_ASM205308v1 Salmonella enterica Proteobacteria; Gammaproteobacteria; Enterobacterales; Enterobacteriaceae; Salmonella MNNEETFYQAMRRKGVTRRSFLKFCSLAATSLGLGAGMTPKIAWALENKPRIPVVWIHGL WP_001058311.1 hydrogenase [Salmonella enterica] Length: 372\nNumber of predicted TMHs: 2\nExp number of AAs in TMHs: 37.4699899999999\nExp number, first 60 AAs: 19.72699\nTotal prob of N-in: 0.96171\nPOSSIBLE N-term signal sequence\ninside 1 19\nTMhelix 20 39\noutside 40 325\nTMhelix 326 348\ninside 349 372

14543 GCF_002052805.1_ASM205280v1 Salmonella enterica Proteobacteria; Gammaproteobacteria; Enterobacterales; Enterobacteriaceae; Salmonella MTGDNTLITSHGINRRDFMKLCAALAATMGLSSKAAAEMAESVSNPQRPPVIWIGAQECT WP_000145429.1 hydrogenase 2 small subunit [Salmonella enterica] Length: 372\nNumber of predicted TMHs: 1\nExp number of AAs in TMHs: 24.51309\nExp number, first 60 AAs: 0.2682\nTotal prob of N-in: 0.14402\noutside 1 331\nTMhelix 332 354\ninside 355 372

14544 GCF_002052905.1_ASM205290v1 Salmonella enterica Proteobacteria; Gammaproteobacteria; Enterobacterales; Enterobacteriaceae; Salmonella MTGDNTLITSHGINRRDFMKLCAALAATMGLSSKAAAEMAESVSNPQRPPVIWIGAQECT WP_000145429.1 hydrogenase 2 small subunit [Salmonella enterica] Length: 372\nNumber of predicted TMHs: 1\nExp number of AAs in TMHs: 24.51309\nExp number, first 60 AAs: 0.2682\nTotal prob of N-in: 0.14402\noutside 1 331\nTMhelix 332 354\ninside 355 372

14545 GCF_002052745.1_ASM205274v1 Salmonella enterica Proteobacteria; Gammaproteobacteria; Enterobacterales; Enterobacteriaceae; Salmonella MTGDNTLITSHGINRRDFMKLCAALAATMGLSSKAAAEMAESVSNPQRPPVIWIGAQECT WP_000145429.1 hydrogenase 2 small subunit [Salmonella enterica] Length: 372\nNumber of predicted TMHs: 1\nExp number of AAs in TMHs: 24.51309\nExp number, first 60 AAs: 0.2682\nTotal prob of N-in: 0.14402\noutside 1 331\nTMhelix 332 354\ninside 355 372

14546 GCF_002052685.1_ASM205268v1 Salmonella enterica Proteobacteria; Gammaproteobacteria; Enterobacterales; Enterobacteriaceae; Salmonella MTGDNTLITSHGINRRDFMKLCAALAATMGLSSKAAAEMAESVSNPQRPPVIWIGAQECT WP_000145429.1 hydrogenase 2 small subunit [Salmonella enterica] Length: 372\nNumber of predicted TMHs: 1\nExp number of AAs in TMHs: 24.51309\nExp number, first 60 AAs: 0.2682\nTotal prob of N-in: 0.14402\noutside 1 331\nTMhelix 332 354\ninside 355 372

14547 GCF_002052635.1_ASM205263v1 Salmonella enterica Proteobacteria; Gammaproteobacteria; Enterobacterales; Enterobacteriaceae; Salmonella MTGDNTLITSHGINRRDFMKLCAALAATMGLSSKAAAEMAESVSNPQRPPVIWIGAQECT WP_000145429.1 hydrogenase 2 small subunit [Salmonella enterica] Length: 372\nNumber of predicted TMHs: 1\nExp number of AAs in TMHs: 24.51309\nExp number, first 60 AAs: 0.2682\nTotal prob of N-in: 0.14402\noutside 1 331\nTMhelix 332 354\ninside 355 372

14548 GCF_002052615.1_ASM205261v1 Salmonella enterica Proteobacteria; Gammaproteobacteria; Enterobacterales; Enterobacteriaceae; Salmonella MNNEETFYQAMRRKGVTRRSFLKFCSLAATSLGLGAGMTPKIAWALENKPRIPVVWIHGL WP_001058311.1 hydrogenase [Salmonella enterica] Length: 372\nNumber of predicted TMHs: 2\nExp number of AAs in TMHs: 37.4699899999999\nExp number, first 60 AAs: 19.72699\nTotal prob of N-in: 0.96171\nPOSSIBLE N-term signal sequence\ninside 1 19\nTMhelix 20 39\noutside 40 325\nTMhelix 326 348\ninside 349 372

14549 GCF_002052615.1_ASM205261v1 Salmonella enterica Proteobacteria; Gammaproteobacteria; Enterobacterales; Enterobacteriaceae; Salmonella
MTGDNTLITSHGINRRDFMKLCAALAATMGLSSKAAAEMAESVSNPQRPPVIWIGAQECT WP_000145429.1
hydrogenase 2 small subunit [Salmonella enterica] Length: 372\nNumber of predicted TMHs: 1\nExp number of AAs in TMHs: 24.51309\nExp number, first 60 AAs: 0.2682\nTotal prob of N-in: 0.14402\noutside 1 331\nTMhelix 332 354\ninside 355 372

14550 GCF_002052245.1_ASM205224v1 Salmonella enterica Proteobacteria; Gammaproteobacteria; Enterobacterales; Enterobacteriaceae; Salmonella
MNNEETFYQAMRRKGVTRRSFLKFCSLAATSLGLGAGMTPKIAWALENKPRIPVVWIHGL WP_001058312.1
hydrogenase [Salmonella enterica] Length: 372\nNumber of predicted TMHs: 2\nExp number of AAs in TMHs: 36.51738\nExp number, first 60 AAs: 19.72849\nTotal prob of N-in: 0.96170\nPOSSIBLE N-term signal sequence\ninside 1 19\nTMhelix 20 39\noutside 40 325\nTMhelix 326 348\ninside 349 372

14551 GCF_002052855.1_ASM205285v1 Salmonella enterica Proteobacteria; Gammaproteobacteria; Enterobacterales; Enterobacteriaceae; Salmonella
MTGDNTLITSHGINRRDFMKLCAALAATMGLSSKAAAEMAESVSNPQRPPVIWIGAQECT WP_000145429.1
hydrogenase 2 small subunit [Salmonella enterica] Length: 372\nNumber of predicted TMHs: 1\nExp number of AAs in TMHs: 24.51309\nExp number, first 60 AAs: 0.2682\nTotal prob of N-in: 0.14402\noutside 1 331\nTMhelix 332 354\ninside 355 372

14552 GCF_002052185.1_ASM205218v1 Salmonella enterica Proteobacteria; Gammaproteobacteria; Enterobacterales; Enterobacteriaceae; Salmonella
MTGDNTLITSHGINRRDFMKLCAALAATMGLSSKAAAEMAESVSNPQRPPVIWIGAQECT WP_000145429.1
hydrogenase 2 small subunit [Salmonella enterica] Length: 372\nNumber of predicted TMHs: 1\nExp number of AAs in TMHs: 24.51309\nExp number, first 60 AAs: 0.2682\nTotal prob of N-in: 0.14402\noutside 1 331\nTMhelix 332 354\ninside 355 372

14553 GCF_002052245.1_ASM205224v1 Salmonella enterica Proteobacteria; Gammaproteobacteria; Enterobacterales; Enterobacteriaceae; Salmonella
MTGDNTLITSHGINRRDFMKLCAALAATMGLSSKAAAEMAESVSNPQRPPVIWIGAQECT WP_000145429.1
hydrogenase 2 small subunit [Salmonella enterica] Length: 372\nNumber of predicted TMHs: 1\nExp number of AAs in TMHs: 24.51309\nExp number, first 60 AAs: 0.2682\nTotal prob of N-in: 0.14402\noutside 1 331\nTMhelix 332 354\ninside 355 372

14554 GCF_002051905.1_ASM205190v1 Salmonella enterica Proteobacteria; Gammaproteobacteria; Enterobacterales; Enterobacteriaceae; Salmonella
MNNEETFYQAMRRKGVTRRSFLKFCSLAATSLGLGAGMTPKIAWALENKPRIPVVWIHGL WP_001058307.1
hydrogenase [Salmonella enterica] Length: 372\nNumber of predicted TMHs: 2\nExp number of AAs in TMHs: 36.51738\nExp number, first 60 AAs: 19.72849\nTotal prob of N-in: 0.96170\nPOSSIBLE N-term signal sequence\ninside 1 19\nTMhelix 20 39\noutside 40 325\nTMhelix 326 348\ninside 349 372

14555 GCF_002052175.1_ASM205217v1 Salmonella enterica Proteobacteria; Gammaproteobacteria; Enterobacterales; Enterobacteriaceae; Salmonella
MTGDNTLITSHGINRRDFMKLCAALAATMGLSSKAAAEMAESVSNPQRPPVIWIGAQECT WP_000145429.1
hydrogenase 2 small subunit [Salmonella enterica] Length: 372\nNumber of predicted TMHs: 1\nExp number of AAs in TMHs: 24.51309\nExp number, first 60 AAs: 0.2682\nTotal prob of N-in: 0.14402\noutside 1 331\nTMhelix 332 354\ninside 355 372

14556 GCF_002052135.1_ASM205213v1 Salmonella enterica Proteobacteria; Gammaproteobacteria; Enterobacterales; Enterobacteriaceae; Salmonella
MTGDNTLITSHGINRRDFMKLCAALAATMGLSSKAAAEMAESVSNPQRPPVIWIGAQECT WP_000145429.1
hydrogenase 2 small subunit [Salmonella enterica] Length: 372\nNumber of predicted TMHs: 1\nExp number of AAs in TMHs: 24.51309\nExp number, first 60 AAs: 0.2682\nTotal prob of N-in: 0.14402\noutside 1 331\nTMhelix 332 354\ninside 355 372

14557 GCF_002052035.1_ASM205203v1 Salmonella enterica Proteobacteria; Gammaproteobacteria; Enterobacterales; Enterobacteriaceae; Salmonella
MTGDNTLITSHGINRRDFMKLCAALAATMGLSSKAAAEMAESVSNPQRPPVIWIGAQECT WP_000145429.1
hydrogenase 2 small subunit [Salmonella enterica] Length: 372\nNumber of predicted TMHs: 1\nExp

number of AAs in TMHs: 24.51309\nExp number, first 60 AAs: 0.2682\nTotal prob of N-in: 0.14402\noutside 1 331\nTMhelix 332 354\ninside 355 372

14558 GCF_002051895.1_ASM205189v1 Salmonella enterica Proteobacteria; Gammaproteobacteria; Enterobacterales; Enterobacteriaceae; Salmonella MNNEETFYQAMRRKGVTRRSFLKFCSLAATSLGLGAGMTPKIAWALENKPRIPVVWIHGL WP_001058307.1 hydrogenase [Salmonella enterica] Length: 372\nNumber of predicted TMHs: 2\nExp number of AAs in TMHs: 36.51738\nExp number, first 60 AAs: 19.72849\nTotal prob of N-in: 0.96170\nPOSSIBLE N-term signal sequence\ninside 1 19\nTMhelix 20 39\noutside 40 325\nTMhelix 326 348\ninside 349 372

14559 GCF_002051655.1_ASM205165v1 Salmonella enterica Proteobacteria; Gammaproteobacteria; Enterobacterales; Enterobacteriaceae; Salmonella MTGDNTLITSHGINRRDFMKLCAALAATMGLSSKAAAEMAESVSNPQRPPVIWIGAQECT WP_000145429.1 hydrogenase 2 small subunit [Salmonella enterica] Length: 372\nNumber of predicted TMHs: 1\nExp number of AAs in TMHs: 24.51309\nExp number, first 60 AAs: 0.2682\nTotal prob of N-in: 0.14402\noutside 1 331\nTMhelix 332 354\ninside 355 372

14560 GCF_002051445.1_ASM205144v1 Salmonella enterica Proteobacteria; Gammaproteobacteria; Enterobacterales; Enterobacteriaceae; Salmonella MNNEETFYQAMRRKGVTRRSFLKFCSLAATSLGLGAGMTPKIAWALENKPRIPVVWIHGL WP_001058312.1 hydrogenase [Salmonella enterica] Length: 372\nNumber of predicted TMHs: 2\nExp number of AAs in TMHs: 36.51738\nExp number, first 60 AAs: 19.72849\nTotal prob of N-in: 0.96170\nPOSSIBLE N-term signal sequence\ninside 1 19\nTMhelix 20 39\noutside 40 325\nTMhelix 326 348\ninside 349 372

14561 GCF_002051515.1_ASM205151v1 Salmonella enterica Proteobacteria; Gammaproteobacteria; Enterobacterales; Enterobacteriaceae; Salmonella MNNEETFYQAMRRKGVTRRSFLKFCSLAATSLGLGAGMTPKIAWALENKPRIPVVWIHGL WP_001058312.1 hydrogenase [Salmonella enterica] Length: 372\nNumber of predicted TMHs: 2\nExp number of AAs in TMHs: 36.51738\nExp number, first 60 AAs: 19.72849\nTotal prob of N-in: 0.96170\nPOSSIBLE N-term signal sequence\ninside 1 19\nTMhelix 20 39\noutside 40 325\nTMhelix 326 348\ninside 349 372

14562 GCF_002051515.1_ASM205151v1 Salmonella enterica Proteobacteria; Gammaproteobacteria; Enterobacterales; Enterobacteriaceae; Salmonella MTGDNTLITSHGINRRDFMKLCAALAATMGLSSKAAAEMAESVSNPQRPPVIWIGAQECT WP_000145429.1 hydrogenase 2 small subunit [Salmonella enterica] Length: 372\nNumber of predicted TMHs: 1\nExp number of AAs in TMHs: 24.51309\nExp number, first 60 AAs: 0.2682\nTotal prob of N-in: 0.14402\noutside 1 331\nTMhelix 332 354\ninside 355 372

14563 GCF_002051355.1_ASM205135v1 Salmonella enterica Proteobacteria; Gammaproteobacteria; Enterobacterales; Enterobacteriaceae; Salmonella MNNEETFYQAMRRKGVTRRSFLKFCSLAATSLGLGAGMTPKIAWALENKPRIPVVWIHGL WP_001058312.1 hydrogenase [Salmonella enterica] Length: 372\nNumber of predicted TMHs: 2\nExp number of AAs in TMHs: 36.51738\nExp number, first 60 AAs: 19.72849\nTotal prob of N-in: 0.96170\nPOSSIBLE N-term signal sequence\ninside 1 19\nTMhelix 20 39\noutside 40 325\nTMhelix 326 348\ninside 349 372

14564 GCF_002051275.1_ASM205127v1 Salmonella enterica Proteobacteria; Gammaproteobacteria; Enterobacterales; Enterobacteriaceae; Salmonella MTGDNTLITSHGINRRDFMKLCAALAATMGLSSKAAAEMAESVSNPQRPPVIWIGAQECT WP_015589585.1 hydrogenase 2 small subunit [Salmonella enterica] Length: 372\nNumber of predicted TMHs: 1\nExp number of AAs in TMHs: 24.51304\nExp number, first 60 AAs: 0.2682\nTotal prob of N-in: 0.14402\noutside 1 331\nTMhelix 332 354\ninside 355 372

14565 GCF_002051485.1_ASM205148v1 Salmonella enterica Proteobacteria; Gammaproteobacteria; Enterobacterales; Enterobacteriaceae; Salmonella MNNEETFYQAMRRKGVTRRSFLKFCSLAATSLGLGAGMTPKIAWALENKPRIPVVWIHGL WP_001058312.1 hydrogenase [Salmonella enterica] Length: 372\nNumber of predicted TMHs: 2\nExp number of AAs in TMHs: 36.51738\nExp number, first 60 AAs: 19.72849\nTotal prob of N-in: 0.96170\nPOSSIBLE N-term signal sequence\ninside 1 19\nTMhelix 20 39\noutside 40 325\nTMhelix 326 348\ninside 349 372

14566 GCF_002051055.1_ASM205105v1 Salmonella enterica Proteobacteria; Gammaproteobacteria; Enterobacterales; Enterobacteriaceae; Salmonella

MNNEETFYQAMRRKGVTRRSFLKFCSLAATSLGLGAGMTPKIAWALENKPRIPVVWIHGL WP_001058312.1 hydrogenase [Salmonella enterica] Length: 372\nNumber of predicted TMHs: 2\nExp number of AAs in TMHs: 36.51738\nExp number, first 60 AAs: 19.72849\nTotal prob of N-in: 0.96170\nPOSSIBLE N-term signal sequence\ninside 1 19\nTMhelix 20 39\noutside 40 325\nTMhelix 326 348\ninside 349 372

14567 GCF_002051005.1_ASM205100v1 Salmonella enterica Proteobacteria; Gammaproteobacteria; Enterobacterales; Enterobacteriaceae; Salmonella MNNEETFYQAMRRKGVTRRSFLKFCSLAATSLGLGAGMTPKIAWALENKPRIPVVWIHGL WP_001058312.1 hydrogenase [Salmonella enterica] Length: 372\nNumber of predicted TMHs: 2\nExp number of AAs in TMHs: 36.51738\nExp number, first 60 AAs: 19.72849\nTotal prob of N-in: 0.96170\nPOSSIBLE N-term signal sequence\ninside 1 19\nTMhelix 20 39\noutside 40 325\nTMhelix 326 348\ninside 349 372

14568 GCF_002051165.1_ASM205116v1 Salmonella enterica Proteobacteria; Gammaproteobacteria; Enterobacterales; Enterobacteriaceae; Salmonella MTGDNTLITSHGINRRDFMKLCAALAATMGLSSKAAAEMAESVSNPQRPPVIWIGAQECT WP_000145429.1 hydrogenase 2 small subunit [Salmonella enterica] Length: 372\nNumber of predicted TMHs: 1\nExp number of AAs in TMHs: 24.51309\nExp number, first 60 AAs: 0.2682\nTotal prob of N-in: 0.14402\noutside 1 331\nTMhelix 332 354\ninside 355 372

14569 GCF_002051135.1_ASM205113v1 Salmonella enterica Proteobacteria; Gammaproteobacteria; Enterobacterales; Enterobacteriaceae; Salmonella MTGDNTLITSHGINRRDFMKLCAALAATMGLSSKAAAEMAESVSNPQRPPVIWIGAQECT WP_000145429.1 hydrogenase 2 small subunit [Salmonella enterica] Length: 372\nNumber of predicted TMHs: 1\nExp number of AAs in TMHs: 24.51309\nExp number, first 60 AAs: 0.2682\nTotal prob of N-in: 0.14402\noutside 1 331\nTMhelix 332 354\ninside 355 372

14570 GCF_002051115.1_ASM205111v1 Salmonella enterica Proteobacteria; Gammaproteobacteria; Enterobacterales; Enterobacteriaceae; Salmonella MTGDNTLITSHGINRRDFMKLCAALAATMGLSSKAAAEMAESVSNPQRPPVIWIGAQECT WP_000145429.1 hydrogenase 2 small subunit [Salmonella enterica] Length: 372\nNumber of predicted TMHs: 1\nExp number of AAs in TMHs: 24.51309\nExp number, first 60 AAs: 0.2682\nTotal prob of N-in: 0.14402\noutside 1 331\nTMhelix 332 354\ninside 355 372

14571 GCF_002051055.1_ASM205105v1 Salmonella enterica Proteobacteria; Gammaproteobacteria; Enterobacterales; Enterobacteriaceae; Salmonella MTGDNTLITSHGINRRDFMKLCAALAATMGLSSKAAAEMAESVSNPQRPPVIWIGAQECT WP_000145429.1 hydrogenase 2 small subunit [Salmonella enterica] Length: 372\nNumber of predicted TMHs: 1\nExp number of AAs in TMHs: 24.51309\nExp number, first 60 AAs: 0.2682\nTotal prob of N-in: 0.14402\noutside 1 331\nTMhelix 332 354\ninside 355 372

14572 GCF_002051005.1_ASM205100v1 Salmonella enterica Proteobacteria; Gammaproteobacteria; Enterobacterales; Enterobacteriaceae; Salmonella MTGDNTLITSHGINRRDFMKLCAALAATMGLSSKAAAEMAESVSNPQRPPVIWIGAQECT WP_000145429.1 hydrogenase 2 small subunit [Salmonella enterica] Length: 372\nNumber of predicted TMHs: 1\nExp number of AAs in TMHs: 24.51309\nExp number, first 60 AAs: 0.2682\nTotal prob of N-in: 0.14402\noutside 1 331\nTMhelix 332 354\ninside 355 372

14573 GCF_002050775.1_ASM205077v1 Salmonella enterica Proteobacteria; Gammaproteobacteria; Enterobacterales; Enterobacteriaceae; Salmonella MTGDNTLITSHGINRRDFMKLCAALAATMGLSSKAAAEMAESVSNPQRPPVIWIGAQECT WP_000145429.1 hydrogenase 2 small subunit [Salmonella enterica] Length: 372\nNumber of predicted TMHs: 1\nExp number of AAs in TMHs: 24.51309\nExp number, first 60 AAs: 0.2682\nTotal prob of N-in: 0.14402\noutside 1 331\nTMhelix 332 354\ninside 355 372

14574 GCF_002050755.1_ASM205075v1 Salmonella enterica Proteobacteria; Gammaproteobacteria; Enterobacterales; Enterobacteriaceae; Salmonella MNNEETFYQAMRRKGVTRRSFLKFCSLAATSLGLGAGMTPKIAWALENKPRIPVVWIHGL WP_001058311.1 hydrogenase [Salmonella enterica] Length: 372\nNumber of predicted TMHs: 2\nExp number of AAs in TMHs: 37.4699899999999\nExp number, first 60 AAs: 19.72699\nTotal prob of N-in: 0.96171\nPOSSIBLE N-term signal sequence\ninside 1 19\nTMhelix 20 39\noutside 40 325\nTMhelix 326 348\ninside 349 372

14575 GCF_002050755.1_ASM205075v1 Salmonella enterica Proteobacteria; Gammaproteobacteria; Enterobacterales; Enterobacteriaceae; Salmonella
MTGDNTLITSHGINRRDFMKLCAALAATMGLSSKAAAEMAESVSNPQRPPVIWIGAQECT WP_000145429.1
hydrogenase 2 small subunit [Salmonella enterica] Length: 372\nNumber of predicted TMHs: 1\nExp number of AAs in TMHs: 24.51309\nExp number, first 60 AAs: 0.2682\nTotal prob of N-in: 0.14402\noutside 1 331\nTMhelix 332 354\ninside 355 372

14576 GCF_002049155.1_ASM204915v1 Salmonella enterica Proteobacteria; Gammaproteobacteria; Enterobacterales; Enterobacteriaceae; Salmonella
MNNEETFYQAMRRKGVTRRSFLKFCSLAATSLGLGAGMTPKIAWALENKPRIPVVWIHGL WP_001058311.1
hydrogenase [Salmonella enterica] Length: 372\nNumber of predicted TMHs: 2\nExp number of AAs in TMHs: 37.4699899999999\nExp number, first 60 AAs: 19.72699\nTotal prob of N-in: 0.96171\nPOSSIBLE N-term signal sequence\ninside 1 19\nTMhelix 20 39\noutside 40 325\nTMhelix 326 348\ninside 349 372

14577 GCF_002049155.1_ASM204915v1 Salmonella enterica Proteobacteria; Gammaproteobacteria; Enterobacterales; Enterobacteriaceae; Salmonella
MTGDNTLITSHGINRRDFMKLCAALAATMGLSSKAAAEMAESVSNPQRPPVIWIGAQECT WP_000145429.1
hydrogenase 2 small subunit [Salmonella enterica] Length: 372\nNumber of predicted TMHs: 1\nExp number of AAs in TMHs: 24.51309\nExp number, first 60 AAs: 0.2682\nTotal prob of N-in: 0.14402\noutside 1 331\nTMhelix 332 354\ninside 355 372

14578 GCF_002047635.1_ASM204763v1 Salmonella enterica Proteobacteria; Gammaproteobacteria; Enterobacterales; Enterobacteriaceae; Salmonella
MNNEETFYQAMRRKGVTRRSFLKFCSLAATSLGLGAGMTPKIAWALENKPRIPVVWIHGL WP_001058312.1
hydrogenase [Salmonella enterica] Length: 372\nNumber of predicted TMHs: 2\nExp number of AAs in TMHs: 36.51738\nExp number, first 60 AAs: 19.72849\nTotal prob of N-in: 0.96170\nPOSSIBLE N-term signal sequence\ninside 1 19\nTMhelix 20 39\noutside 40 325\nTMhelix 326 348\ninside 349 372

14579 GCF_002047185.1_ASM204718v1 Salmonella enterica Proteobacteria; Gammaproteobacteria; Enterobacterales; Enterobacteriaceae; Salmonella
MNNEETFYQAMRRKGVTRRSFLKFCSLAATSLGLGAGMTPKIAWALENKPRIPVVWIHGL WP_001058307.1
hydrogenase [Salmonella enterica] Length: 372\nNumber of predicted TMHs: 2\nExp number of AAs in TMHs: 36.51738\nExp number, first 60 AAs: 19.72849\nTotal prob of N-in: 0.96170\nPOSSIBLE N-term signal sequence\ninside 1 19\nTMhelix 20 39\noutside 40 325\nTMhelix 326 348\ninside 349 372

14580 GCF_002046805.1_ASM204680v1 Salmonella enterica Proteobacteria; Gammaproteobacteria; Enterobacterales; Enterobacteriaceae; Salmonella
MNNEETFYQAMRRKGVTRRSFLKFCSLAATSLGLGAGMTPKIAWALENKPRIPVVWIHGL WP_001058312.1
hydrogenase [Salmonella enterica] Length: 372\nNumber of predicted TMHs: 2\nExp number of AAs in TMHs: 36.51738\nExp number, first 60 AAs: 19.72849\nTotal prob of N-in: 0.96170\nPOSSIBLE N-term signal sequence\ninside 1 19\nTMhelix 20 39\noutside 40 325\nTMhelix 326 348\ninside 349 372

14581 GCF_002046875.1_ASM204687v1 Salmonella enterica Proteobacteria; Gammaproteobacteria; Enterobacterales; Enterobacteriaceae; Salmonella
MNNEETFYQAMRRKGVTRRSFLKFCSLAATSLGLGAGMTPKIAWALENKPRIPVVWIHGL WP_001058312.1
hydrogenase [Salmonella enterica] Length: 372\nNumber of predicted TMHs: 2\nExp number of AAs in TMHs: 36.51738\nExp number, first 60 AAs: 19.72849\nTotal prob of N-in: 0.96170\nPOSSIBLE N-term signal sequence\ninside 1 19\nTMhelix 20 39\noutside 40 325\nTMhelix 326 348\ninside 349 372

14582 GCF_002046875.1_ASM204687v1 Salmonella enterica Proteobacteria; Gammaproteobacteria; Enterobacterales; Enterobacteriaceae; Salmonella
MTGDNTLITSHGINRRDFMKLCAALAATMGLSSKAAAEMAESVSNPQRPPVIWIGAQECT WP_000145429.1
hydrogenase 2 small subunit [Salmonella enterica] Length: 372\nNumber of predicted TMHs: 1\nExp number of AAs in TMHs: 24.51309\nExp number, first 60 AAs: 0.2682\nTotal prob of N-in: 0.14402\noutside 1 331\nTMhelix 332 354\ninside 355 372

14583 GCF_002045735.1_ASM204573v1 Salmonella enterica Proteobacteria; Gammaproteobacteria; Enterobacterales; Enterobacteriaceae; Salmonella
MTGDNTLITSHGINRRDFMKLCAALAATMGLSSKAAAEMAESVSNPQRPPVIWIGAQECT WP_000145429.1
hydrogenase 2 small subunit [Salmonella enterica] Length: 372\nNumber of predicted TMHs: 1\nExp

number of AAs in TMHs: 24.51309\nExp number, first 60 AAs:  0.2682\nTotal prob of N-in:   0.14402\noutside   1 331\nTMhelix   332   354\ninside   355   372

14584 GCF_002045665.1_ASM204566v1 Salmonella enterica Proteobacteria; Gammaproteobacteria; Enterobacterales; Enterobacteriaceae; Salmonella MTGDNTLITSHGINRRDFMKLCAALAATMGLSSKAAAEMAESVSNPQRPPVIWIGAQECT WP_000145429.1 hydrogenase 2 small subunit [Salmonella enterica] Length: 372\nNumber of predicted TMHs:  1\nExp number of AAs in TMHs: 24.51309\nExp number, first 60 AAs:  0.2682\nTotal prob of N-in:   0.14402\noutside   1 331\nTMhelix   332   354\ninside   355   372

14585 GCF_002045495.1_ASM204549v1 Salmonella enterica Proteobacteria; Gammaproteobacteria; Enterobacterales; Enterobacteriaceae; Salmonella MNNEETFYQAMRRKGVTRRSFLKFCSLAATSLGLGAGMTPKIAWALENKPRIPVVWIHGL WP_001058311.1 hydrogenase [Salmonella enterica] Length: 372\nNumber of predicted TMHs:  2\nExp number of AAs in TMHs: 37.4699899999999\nExp number, first 60 AAs:  19.72699\nTotal prob of N-in:   0.96171\nPOSSIBLE N-term signal sequence\ninside   1   19\nTMhelix   20   39\noutside   40   325\nTMhelix   326   348\ninside   349   372

14586 GCF_002045535.1_ASM204553v1 Salmonella enterica Proteobacteria; Gammaproteobacteria; Enterobacterales; Enterobacteriaceae; Salmonella MNNEETFYQAMRRKGVTRRSFLKFCSLAATSLGLGAGMTPKIAWALENKPRIPVVWIHGL WP_001058311.1 hydrogenase [Salmonella enterica] Length: 372\nNumber of predicted TMHs:  2\nExp number of AAs in TMHs: 37.4699899999999\nExp number, first 60 AAs:  19.72699\nTotal prob of N-in:   0.96171\nPOSSIBLE N-term signal sequence\ninside   1   19\nTMhelix   20   39\noutside   40   325\nTMhelix   326   348\ninside   349   372

14587 GCF_002045615.1_ASM204561v1 Salmonella enterica Proteobacteria; Gammaproteobacteria; Enterobacterales; Enterobacteriaceae; Salmonella MTGDNTLITSHGINRRDFMKLCAALAATMGLSSKAAAEMAESVSNPQRPPVIWIGAQECT WP_000145429.1 hydrogenase 2 small subunit [Salmonella enterica] Length: 372\nNumber of predicted TMHs:  1\nExp number of AAs in TMHs: 24.51309\nExp number, first 60 AAs:  0.2682\nTotal prob of N-in:   0.14402\noutside   1 331\nTMhelix   332   354\ninside   355   372

14588 GCF_002045085.1_ASM204508v1 Salmonella enterica Proteobacteria; Gammaproteobacteria; Enterobacterales; Enterobacteriaceae; Salmonella MNNEETFYQAMRRKGVTRRSFLKFCSLAATSLGLGAGMTPKIAWALENKPRIPVVWIHGL WP_064000476.1 hydrogenase [Salmonella enterica] Length: 372\nNumber of predicted TMHs:  2\nExp number of AAs in TMHs: 37.4700099999999\nExp number, first 60 AAs:  19.72698\nTotal prob of N-in:   0.96171\nPOSSIBLE N-term signal sequence\ninside   1   19\nTMhelix   20   39\noutside   40   325\nTMhelix   326   348\ninside   349   372

14589 GCF_002044425.1_ASM204442v1 Salmonella enterica Proteobacteria; Gammaproteobacteria; Enterobacterales; Enterobacteriaceae; Salmonella MNNEETFYQAMRRKGVTRRSFLKFCSLAATSLGLGAGMTPKIAWALENKPRIPVVWIHGL WP_079953656.1 [Ni/Fe] hydrogenase small subunit [Salmonella enterica] Length: 372\nNumber of predicted TMHs:  2\nExp number of AAs in TMHs: 36.5655299999999\nExp number, first 60 AAs:  19.72713\nTotal prob of N-in: 0.96172\nPOSSIBLE N-term signal sequence\ninside   1   19\nTMhelix   20   39\noutside   40   325\nTMhelix   326 348\ninside   349   372

14590 GCF_002044565.1_ASM204456v1 Salmonella enterica Proteobacteria; Gammaproteobacteria; Enterobacterales; Enterobacteriaceae; Salmonella MNNEETFYQAMRRKGVTRRSFLKFCSLAATSLGLGAGMTPKIAWALENKPRIPVVWIHGL WP_001058312.1 hydrogenase [Salmonella enterica] Length: 372\nNumber of predicted TMHs:  2\nExp number of AAs in TMHs: 36.51738\nExp number, first 60 AAs:  19.72849\nTotal prob of N-in:   0.96170\nPOSSIBLE N-term signal sequence\ninside   1   19\nTMhelix   20   39\noutside   40   325\nTMhelix   326   348\ninside   349   372

14591 GCF_002044625.1_ASM204462v1 Salmonella enterica Proteobacteria; Gammaproteobacteria; Enterobacterales; Enterobacteriaceae; Salmonella MNNEETFYQAMRRKGVTRRSFLKFCSLAATSLGLGAGMTPKIAWALENKPRIPVVWIHGL WP_001058311.1 hydrogenase [Salmonella enterica] Length: 372\nNumber of predicted TMHs:  2\nExp number of AAs in TMHs: 37.4699899999999\nExp number, first 60 AAs:  19.72699\nTotal prob of N-in:   0.96171\nPOSSIBLE N-term signal sequence\ninside   1   19\nTMhelix   20   39\noutside   40   325\nTMhelix   326   348\ninside   349   372

14592 GCF_002044675.1_ASM204467v1 Salmonella enterica Proteobacteria; Gammaproteobacteria; Enterobacterales; Enterobacteriaceae; Salmonella
MNNEETFYQAMRRKGVTRRSFLKFCSLAATSLGLGAGMTPKIAWALENKPRIPVVWIHGL WP_001058311.1
hydrogenase [Salmonella enterica] Length: 372\nNumber of predicted TMHs: 2\nExp number of AAs in TMHs: 37.4699899999999\nExp number, first 60 AAs: 19.72699\nTotal prob of N-in: 0.96171\nPOSSIBLE N-term signal sequence\ninside 1 19\nTMhelix 20 39\noutside 40 325\nTMhelix 326 348\ninside 349 372

14593 GCF_002043915.1_ASM204391v1 Salmonella enterica Proteobacteria; Gammaproteobacteria; Enterobacterales; Enterobacteriaceae; Salmonella
MNNEETLYQAMRRKGVTRRSFLKFCSLAATSLGLGAGMTPKIAWALENKPRIPVVWIHGL WP_001752445.1
hydrogenase-1 small chain [Salmonella enterica] Length: 372\nNumber of predicted TMHs: 2\nExp number of AAs in TMHs: 36.5123\nExp number, first 60 AAs: 19.72341\nTotal prob of N-in: 0.96144\nPOSSIBLE N-term signal sequence\ninside 1 19\nTMhelix 20 39\noutside 40 325\nTMhelix 326 348\ninside 349 372

14594 GCF_002044305.1_ASM204430v1 Salmonella enterica Proteobacteria; Gammaproteobacteria; Enterobacterales; Enterobacteriaceae; Salmonella
MNNEETFYQAMRRKGVTRRSFLKFCSLAATSLGLGAGMTPKIAWALENKPRIPVVWIHGL WP_001058312.1
hydrogenase [Salmonella enterica] Length: 372\nNumber of predicted TMHs: 2\nExp number of AAs in TMHs: 36.51738\nExp number, first 60 AAs: 19.72849\nTotal prob of N-in: 0.96170\nPOSSIBLE N-term signal sequence\ninside 1 19\nTMhelix 20 39\noutside 40 325\nTMhelix 326 348\ninside 349 372

14595 GCF_002044305.1_ASM204430v1 Salmonella enterica Proteobacteria; Gammaproteobacteria; Enterobacterales; Enterobacteriaceae; Salmonella
MTGDNTLITSHGINRRDFMKLCAALAATMGLSSKAAAEMAESVSNPQRPPVIWIGAQECT WP_000145429.1
hydrogenase 2 small subunit [Salmonella enterica] Length: 372\nNumber of predicted TMHs: 1\nExp number of AAs in TMHs: 24.51309\nExp number, first 60 AAs: 0.2682\nTotal prob of N-in: 0.14402\noutside 1 331\nTMhelix 332 354\ninside 355 372

14596 GCF_002044125.1_ASM204412v1 Salmonella enterica Proteobacteria; Gammaproteobacteria; Enterobacterales; Enterobacteriaceae; Salmonella
MNNEETFYQAMRRKGVTRRSFLKFCSLAATSLGLGAGMTPKIAWALENKPRIPVVWIHGL WP_001058311.1
hydrogenase [Salmonella enterica] Length: 372\nNumber of predicted TMHs: 2\nExp number of AAs in TMHs: 37.4699899999999\nExp number, first 60 AAs: 19.72699\nTotal prob of N-in: 0.96171\nPOSSIBLE N-term signal sequence\ninside 1 19\nTMhelix 20 39\noutside 40 325\nTMhelix 326 348\ninside 349 372

14597 GCF_002044075.1_ASM204407v1 Salmonella enterica Proteobacteria; Gammaproteobacteria; Enterobacterales; Enterobacteriaceae; Salmonella
MNNEETFYQAMRRKGVTRRSFLKFCSLAATSLGLGAGMTPKIAWALENKPRIPVVWIHGL WP_001058311.1
hydrogenase [Salmonella enterica] Length: 372\nNumber of predicted TMHs: 2\nExp number of AAs in TMHs: 37.4699899999999\nExp number, first 60 AAs: 19.72699\nTotal prob of N-in: 0.96171\nPOSSIBLE N-term signal sequence\ninside 1 19\nTMhelix 20 39\noutside 40 325\nTMhelix 326 348\ninside 349 372

14598 GCF_002044255.1_ASM204425v1 Salmonella enterica Proteobacteria; Gammaproteobacteria; Enterobacterales; Enterobacteriaceae; Salmonella
MNNEETFYQAMRRKGVTRRSFLKFCSLAATSLGLGAGMTPKIAWALENKPRIPVVWIHGL WP_001058311.1
hydrogenase [Salmonella enterica] Length: 372\nNumber of predicted TMHs: 2\nExp number of AAs in TMHs: 37.4699899999999\nExp number, first 60 AAs: 19.72699\nTotal prob of N-in: 0.96171\nPOSSIBLE N-term signal sequence\ninside 1 19\nTMhelix 20 39\noutside 40 325\nTMhelix 326 348\ninside 349 372

14599 GCF_002044255.1_ASM204425v1 Salmonella enterica Proteobacteria; Gammaproteobacteria; Enterobacterales; Enterobacteriaceae; Salmonella
MTGDNTLITSHGINRRDFMKLCAALAATMGLSSKAAAEMAESVSNPQRPPVIWIGAQECT WP_000145429.1
hydrogenase 2 small subunit [Salmonella enterica] Length: 372\nNumber of predicted TMHs: 1\nExp number of AAs in TMHs: 24.51309\nExp number, first 60 AAs: 0.2682\nTotal prob of N-in: 0.14402\noutside 1 331\nTMhelix 332 354\ninside 355 372

14600 GCF_002043845.1_ASM204384v1 Salmonella enterica Proteobacteria; Gammaproteobacteria; Enterobacterales; Enterobacteriaceae; Salmonella
MTGDNTLITSHGINRRDFMKLCAALAATMGLSSKAAAEMAESVSNPQRPPVIWIGAQECT WP_000145429.1
hydrogenase 2 small subunit [Salmonella enterica] Length: 372\nNumber of predicted TMHs: 1\nExp

number of AAs in TMHs: 24.51309\nExp number, first 60 AAs: 0.2682\nTotal prob of N-in: 0.14402\noutside 1 331\nTMhelix 332 354\ninside 355 372

14601 GCF_002043885.1_ASM204388v1 Salmonella enterica Proteobacteria; Gammaproteobacteria; Enterobacterales; Enterobacteriaceae; Salmonella MTGDNTLITSHGINRRDFMKLCAALAATMGLSSKAAAEMAESVSNPQRPPVIWIGAQECT WP_000145429.1 hydrogenase 2 small subunit [Salmonella enterica] Length: 372\nNumber of predicted TMHs: 1\nExp number of AAs in TMHs: 24.51309\nExp number, first 60 AAs: 0.2682\nTotal prob of N-in: 0.14402\noutside 1 331\nTMhelix 332 354\ninside 355 372

14602 GCF_002043725.1_ASM204372v1 Salmonella enterica Proteobacteria; Gammaproteobacteria; Enterobacterales; Enterobacteriaceae; Salmonella MNNEETFYQAMRRKGVTRRSFLKFCSLAATSLGLGAGMTPKIAWALENKPRIPVVWIHGL WP_001058311.1 hydrogenase [Salmonella enterica] Length: 372\nNumber of predicted TMHs: 2\nExp number of AAs in TMHs: 37.4699899999999\nExp number, first 60 AAs: 19.72699\nTotal prob of N-in: 0.96171\nPOSSIBLE N-term signal sequence\ninside 1 19\nTMhelix 20 39\noutside 40 325\nTMhelix 326 348\ninside 349 372

14603 GCF_002043785.1_ASM204378v1 Salmonella enterica Proteobacteria; Gammaproteobacteria; Enterobacterales; Enterobacteriaceae; Salmonella MNNEETFYQAMRRKGVTRRSFLKFCSLAATSLGLGAGMTPKIAWALENKPRIPVVWIHGL WP_001058311.1 hydrogenase [Salmonella enterica] Length: 372\nNumber of predicted TMHs: 2\nExp number of AAs in TMHs: 37.4699899999999\nExp number, first 60 AAs: 19.72699\nTotal prob of N-in: 0.96171\nPOSSIBLE N-term signal sequence\ninside 1 19\nTMhelix 20 39\noutside 40 325\nTMhelix 326 348\ninside 349 372

14604 GCF_002043795.1_ASM204379v1 Salmonella enterica Proteobacteria; Gammaproteobacteria; Enterobacterales; Enterobacteriaceae; Salmonella MTGDNTLITSHGINRRDFMKLCAALAATMGLSSKAAAEMAESVSNPQRPPVIWIGAQECT WP_000145429.1 hydrogenase 2 small subunit [Salmonella enterica] Length: 372\nNumber of predicted TMHs: 1\nExp number of AAs in TMHs: 24.51309\nExp number, first 60 AAs: 0.2682\nTotal prob of N-in: 0.14402\noutside 1 331\nTMhelix 332 354\ninside 355 372

14605 GCF_002043665.1_ASM204366v1 Salmonella enterica Proteobacteria; Gammaproteobacteria; Enterobacterales; Enterobacteriaceae; Salmonella MNNEETFYQAMRRKGVTRRSFLKFCSLAATSLGLGAGMTPKIAWALENKPRIPVVWIHGL WP_001058311.1 hydrogenase [Salmonella enterica] Length: 372\nNumber of predicted TMHs: 2\nExp number of AAs in TMHs: 37.4699899999999\nExp number, first 60 AAs: 19.72699\nTotal prob of N-in: 0.96171\nPOSSIBLE N-term signal sequence\ninside 1 19\nTMhelix 20 39\noutside 40 325\nTMhelix 326 348\ninside 349 372

14606 GCF_002043545.1_ASM204354v1 Salmonella enterica Proteobacteria; Gammaproteobacteria; Enterobacterales; Enterobacteriaceae; Salmonella MTGDNTLITSHGINRRDFMKLCAALAATMGLSSKAAAEMAESVSNPQRPPVIWIGAQECT WP_000145429.1 hydrogenase 2 small subunit [Salmonella enterica] Length: 372\nNumber of predicted TMHs: 1\nExp number of AAs in TMHs: 24.51309\nExp number, first 60 AAs: 0.2682\nTotal prob of N-in: 0.14402\noutside 1 331\nTMhelix 332 354\ninside 355 372

14607 GCF_002043155.1_ASM204315v1 Salmonella enterica Proteobacteria; Gammaproteobacteria; Enterobacterales; Enterobacteriaceae; Salmonella MNNEETLYQAMRRKGVTRRSFLKFCSLAATSLGLGAGMTPKIAWALENKPRIPVVWIHGL WP_001752445.1 hydrogenase-1 small chain [Salmonella enterica] Length: 372\nNumber of predicted TMHs: 2\nExp number of AAs in TMHs: 36.5123\nExp number, first 60 AAs: 19.72341\nTotal prob of N-in: 0.96144\nPOSSIBLE N-term signal sequence\ninside 1 19\nTMhelix 20 39\noutside 40 325\nTMhelix 326 348\ninside 349 372

14608 GCF_002043425.1_ASM204342v1 Salmonella enterica Proteobacteria; Gammaproteobacteria; Enterobacterales; Enterobacteriaceae; Salmonella MNNEETFYQAMRRKGVTRRSFLKFCSLAATSLGLGAGMTPKIAWALENKPRIPVVWIHGL WP_001058312.1 hydrogenase [Salmonella enterica] Length: 372\nNumber of predicted TMHs: 2\nExp number of AAs in TMHs: 36.51738\nExp number, first 60 AAs: 19.72849\nTotal prob of N-in: 0.96170\nPOSSIBLE N-term signal sequence\ninside 1 19\nTMhelix 20 39\noutside 40 325\nTMhelix 326 348\ninside 349 372

14609 GCF_002043345.1_ASM204334v1 Salmonella enterica Proteobacteria; Gammaproteobacteria; Enterobacterales; Enterobacteriaceae; Salmonella

MNNEETFYQAMRRKGVTRRSFLKFCSLAATSLGLGAGMTPKIAWALENKPRIPVVWIHGL WP_001058311.1 hydrogenase [Salmonella enterica] Length: 372\nNumber of predicted TMHs: 2\nExp number of AAs in TMHs: 37.4699899999999\nExp number, first 60 AAs: 19.72699\nTotal prob of N-in: 0.96171\nPOSSIBLE N-term signal sequence\ninside 1 19\nTMhelix 20 39\noutside 40 325\nTMhelix 326 348\ninside 349 372

14610 GCF_002043425.1_ASM204342v1 Salmonella enterica Proteobacteria; Gammaproteobacteria; Enterobacterales; Enterobacteriaceae; Salmonella MTGDNTLITSHGINRRDFMKLCAALAATMGLSSKAAAEMAESVSNPQRPPVIWIGAQECT WP_000145429.1 hydrogenase 2 small subunit [Salmonella enterica] Length: 372\nNumber of predicted TMHs: 1\nExp number of AAs in TMHs: 24.51309\nExp number, first 60 AAs: 0.2682\nTotal prob of N-in: 0.14402\noutside 1 331\nTMhelix 332 354\ninside 355 372

14611 GCF_002043365.1_ASM204336v1 Salmonella enterica Proteobacteria; Gammaproteobacteria; Enterobacterales; Enterobacteriaceae; Salmonella MTGDNTLITSHGINRRDFMKLCAALAATMGLSSKAAAEMAESVSNPQRPPVIWIGAQECT WP_000145429.1 hydrogenase 2 small subunit [Salmonella enterica] Length: 372\nNumber of predicted TMHs: 1\nExp number of AAs in TMHs: 24.51309\nExp number, first 60 AAs: 0.2682\nTotal prob of N-in: 0.14402\noutside 1 331\nTMhelix 332 354\ninside 355 372

14612 GCF_002043345.1_ASM204334v1 Salmonella enterica Proteobacteria; Gammaproteobacteria; Enterobacterales; Enterobacteriaceae; Salmonella MTGDNTLITSHGINRRDFMKLCAALAATMGLSSKAAAEMAESVSNPQRPPVIWIGAQECT WP_000145429.1 hydrogenase 2 small subunit [Salmonella enterica] Length: 372\nNumber of predicted TMHs: 1\nExp number of AAs in TMHs: 24.51309\nExp number, first 60 AAs: 0.2682\nTotal prob of N-in: 0.14402\noutside 1 331\nTMhelix 332 354\ninside 355 372

14613 GCF_001570145.1_ASM157014v1 Salmonella enterica Proteobacteria; Gammaproteobacteria; Enterobacterales; Enterobacteriaceae; Salmonella MTGDNTLITSHGINRRDFMKLCAALAATMGLSSKAAAEMAESVSNPQRPPVIWIGAQECT WP_000145429.1 hydrogenase 2 small subunit [Salmonella enterica] Length: 372\nNumber of predicted TMHs: 1\nExp number of AAs in TMHs: 24.51309\nExp number, first 60 AAs: 0.2682\nTotal prob of N-in: 0.14402\noutside 1 331\nTMhelix 332 354\ninside 355 372

14614 GCF_002037615.1_ASM203761v1 Salmonella enterica Proteobacteria; Gammaproteobacteria; Enterobacterales; Enterobacteriaceae; Salmonella MNNEETFYQAMRRKGVTRRSFLKFCSLAATSLGLGAGMTPKIAWALENKPRIPVVWIHGL WP_079805161.1 [Ni/Fe] hydrogenase small subunit [Salmonella enterica] Length: 372\nNumber of predicted TMHs: 2\nExp number of AAs in TMHs: 36.9928999999999\nExp number, first 60 AAs: 19.72755\nTotal prob of N-in: 0.96172\nPOSSIBLE N-term signal sequence\ninside 1 19\nTMhelix 20 39\noutside 40 325\nTMhelix 326 348\ninside 349 372

14615 GCF_002037595.1_ASM203759v1 Salmonella enterica Proteobacteria; Gammaproteobacteria; Enterobacterales; Enterobacteriaceae; Salmonella MNNEETFYQAMRRKGVTRRSFLKFCSLAATSLGLGAGMTPKIAWALENKPRIPVVWIHGL WP_023187532.1 hydrogenase 1, small subunit [Salmonella enterica] Length: 372\nNumber of predicted TMHs: 2\nExp number of AAs in TMHs: 36.9997\nExp number, first 60 AAs: 19.72759\nTotal prob of N-in: 0.96171\nPOSSIBLE N-term signal sequence\ninside 1 19\nTMhelix 20 39\noutside 40 325\nTMhelix 326 348\ninside 349 372

14616 GCF_002037595.1_ASM203759v1 Salmonella enterica Proteobacteria; Gammaproteobacteria; Enterobacterales; Enterobacteriaceae; Salmonella MTGDNTLITSHGINRRDFMKLCAALAATMGLSSKAAAEMAESVSNPQRPPVIWIGAQECT WP_000145429.1 hydrogenase 2 small subunit [Salmonella enterica] Length: 372\nNumber of predicted TMHs: 1\nExp number of AAs in TMHs: 24.51309\nExp number, first 60 AAs: 0.2682\nTotal prob of N-in: 0.14402\noutside 1 331\nTMhelix 332 354\ninside 355 372

14617 GCF_002037615.1_ASM203761v1 Salmonella enterica Proteobacteria; Gammaproteobacteria; Enterobacterales; Enterobacteriaceae; Salmonella MTGDNTLITSHGINRRDFMKLCAALAATMGLSSKAAAEMAESVSNPQRPPVIWIGAQECT WP_000145429.1 hydrogenase 2 small subunit [Salmonella enterica] Length: 372\nNumber of predicted TMHs: 1\nExp number of AAs in TMHs: 24.51309\nExp number, first 60 AAs: 0.2682\nTotal prob of N-in: 0.14402\noutside 1 331\nTMhelix 332 354\ninside 355 372

14618 GCF_002036995.1_ASM203699v1 Salmonella enterica Proteobacteria; Gammaproteobacteria; Enterobacterales; Enterobacteriaceae; Salmonella
MNNEETFYQAMRRKGVTRRSFLKFCSLAATSLGLGAGMTPKIAWALENKPRIPVVWIHGL WP_001058312.1 hydrogenase [Salmonella enterica] Length: 372\nNumber of predicted TMHs: 2\nExp number of AAs in TMHs: 36.51738\nExp number, first 60 AAs: 19.72849\nTotal prob of N-in: 0.96170\nPOSSIBLE N-term signal sequence\ninside 1 19\nTMhelix 20 39\noutside 40 325\nTMhelix 326 348\ninside 349 372

14619 GCF_002037135.1_ASM203713v1 Salmonella enterica Proteobacteria; Gammaproteobacteria; Enterobacterales; Enterobacteriaceae; Salmonella
MNNEETFYQAMRRKGVTRRSFLKFCSLAATSLGLGAGMTPKIAWALENKPRIPVVWIHGL WP_001058311.1 hydrogenase [Salmonella enterica] Length: 372\nNumber of predicted TMHs: 2\nExp number of AAs in TMHs: 37.4699899999999\nExp number, first 60 AAs: 19.72699\nTotal prob of N-in: 0.96171\nPOSSIBLE N-term signal sequence\ninside 1 19\nTMhelix 20 39\noutside 40 325\nTMhelix 326 348\ninside 349 372

14620 GCF_002037135.1_ASM203713v1 Salmonella enterica Proteobacteria; Gammaproteobacteria; Enterobacterales; Enterobacteriaceae; Salmonella
MTGDNTLITSHGINRRDFMKLCAALAATMGLSSKAAAEMAESVSNPQRPPVIWIGAQECT WP_000145429.1 hydrogenase 2 small subunit [Salmonella enterica] Length: 372\nNumber of predicted TMHs: 1\nExp number of AAs in TMHs: 24.51309\nExp number, first 60 AAs: 0.2682\nTotal prob of N-in: 0.14402\noutside 1 331\nTMhelix 332 354\ninside 355 372

14621 GCF_001691905.1_ASM169190v1 Salmonella enterica Proteobacteria; Gammaproteobacteria; Enterobacterales; Enterobacteriaceae; Salmonella
MNNEETFYQAMRRKGVTRRSFLKFCSLAATSLGLGAGMTPKIAWALENKPRIPVVWIHGL WP_001058312.1 hydrogenase [Salmonella enterica] Length: 372\nNumber of predicted TMHs: 2\nExp number of AAs in TMHs: 36.51738\nExp number, first 60 AAs: 19.72849\nTotal prob of N-in: 0.96170\nPOSSIBLE N-term signal sequence\ninside 1 19\nTMhelix 20 39\noutside 40 325\nTMhelix 326 348\ninside 349 372

14622 GCF_002061035.1_ASM206103v1 Salmonella enterica Proteobacteria; Gammaproteobacteria; Enterobacterales; Enterobacteriaceae; Salmonella
MTGDNTLITSHGINRRDFMKLCAALAATMGLSSKAAAEMAESVSNPQRPPVIWIGAQECT WP_000145429.1 hydrogenase 2 small subunit [Salmonella enterica] Length: 372\nNumber of predicted TMHs: 1\nExp number of AAs in TMHs: 24.51309\nExp number, first 60 AAs: 0.2682\nTotal prob of N-in: 0.14402\noutside 1 331\nTMhelix 332 354\ninside 355 372

14623 GCF_002061055.1_ASM206105v1 Salmonella enterica Proteobacteria; Gammaproteobacteria; Enterobacterales; Enterobacteriaceae; Salmonella
MTGDNTLITSHGINRRDFMKLCAALAATMGLSSKAAAEMAESVSNPQRPPVIWIGAQECT WP_000145429.1 hydrogenase 2 small subunit [Salmonella enterica] Length: 372\nNumber of predicted TMHs: 1\nExp number of AAs in TMHs: 24.51309\nExp number, first 60 AAs: 0.2682\nTotal prob of N-in: 0.14402\noutside 1 331\nTMhelix 332 354\ninside 355 372

14624 GCF_001647755.1_ASM164775v1 Salmonella enterica Proteobacteria; Gammaproteobacteria; Enterobacterales; Enterobacteriaceae; Salmonella
MNNEETFYQAMRRKGVTRRSFLKFCSLAATSLGLGAGMTPQIAWALENKPRIPVVWIHGL WP_023227615.1 hydrogenase 1, small subunit [Salmonella enterica] Length: 372\nNumber of predicted TMHs: 2\nExp number of AAs in TMHs: 39.8591699999999\nExp number, first 60 AAs: 22.13093\nTotal prob of N-in: 0.98639\nPOSSIBLE N-term signal sequence\ninside 1 20\nTMhelix 21 43\noutside 44 325\nTMhelix 326 348\ninside 349 372

14625 GCF_001643895.1_CRJJGF_00147_v1.0Salmonella enterica Proteobacteria; Gammaproteobacteria; Enterobacterales; Enterobacteriaceae; Salmonella
MTGDNTLITSHGINRRDFMKLCAALAATMGLSSKAAAEMAESVSNPQRPPVIWIGAQECT WP_000145429.1 hydrogenase 2 small subunit [Salmonella enterica] Length: 372\nNumber of predicted TMHs: 1\nExp number of AAs in TMHs: 24.51309\nExp number, first 60 AAs: 0.2682\nTotal prob of N-in: 0.14402\noutside 1 331\nTMhelix 332 354\ninside 355 372

14626 GCF_001643875.1_CRJJGF_00165_v1.0Salmonella enterica Proteobacteria; Gammaproteobacteria; Enterobacterales; Enterobacteriaceae; Salmonella
MNNEETFYQAMRRKGVTRRSFLKFCSLAATSLGLGAGMTPKIAWALENKPRIPVVWIHGL WP_001058306.1 hydrogenase [Salmonella enterica] Length: 372\nNumber of predicted TMHs: 2\nExp number of AAs in

TMHs: 36.5540099999999\nExp number, first 60 AAs: 19.72732\nTotal prob of N-in: 0.96171\nPOSSIBLE N-term signal sequence\ninside 1 19\nTMhelix 20 39\noutside 40 325\nTMhelix 326 348\ninside 349 372

14627 GCF_001643915.1_CRJJGF_00159_v1.0Salmonella enterica Proteobacteria; Gammaproteobacteria; Enterobacterales; Enterobacteriaceae; Salmonella
MTGDNTLITSHGINRRDFMKLCAALAATMGLSSKAAAEMAESVSNPQRPPVIWIGAQECT WP_000145429.1
hydrogenase 2 small subunit [Salmonella enterica] Length: 372\nNumber of predicted TMHs: 1\nExp number of AAs in TMHs: 24.51309\nExp number, first 60 AAs: 0.2682\nTotal prob of N-in: 0.14402\noutside 1 331\nTMhelix 332 354\ninside 355 372

14628 GCF_001642515.1_ASM164251v1 Salmonella enterica Proteobacteria; Gammaproteobacteria; Enterobacterales; Enterobacteriaceae; Salmonella
MNNEETFYQAMRRKGVTRRSFLKFCSLAATSLGLGAGMTPKIAWALENKPRIPVVWIHGL WP_001058311.1
hydrogenase [Salmonella enterica] Length: 372\nNumber of predicted TMHs: 2\nExp number of AAs in TMHs: 37.4699899999999\nExp number, first 60 AAs: 19.72699\nTotal prob of N-in: 0.96171\nPOSSIBLE N-term signal sequence\ninside 1 19\nTMhelix 20 39\noutside 40 325\nTMhelix 326 348\ninside 349 372

14629 GCF_001642445.1_ASM164244v1 Salmonella enterica Proteobacteria; Gammaproteobacteria; Enterobacterales; Enterobacteriaceae; Salmonella
MNNEETFYQAMRRKGVTRRSFLKFCSLAATSLGLGAGMTPKIAWALENKPRIPVVWIHGL WP_001058311.1
hydrogenase [Salmonella enterica] Length: 372\nNumber of predicted TMHs: 2\nExp number of AAs in TMHs: 37.4699899999999\nExp number, first 60 AAs: 19.72699\nTotal prob of N-in: 0.96171\nPOSSIBLE N-term signal sequence\ninside 1 19\nTMhelix 20 39\noutside 40 325\nTMhelix 326 348\ninside 349 372

14630 GCF_001647755.1_ASM164775v1 Salmonella enterica Proteobacteria; Gammaproteobacteria; Enterobacterales; Enterobacteriaceae; Salmonella
MTGDNTLITSHGINRRDFMKLCAALAATMGLSSKAAAEMAESVSNPQRPPVIWIGAQECT WP_000145429.1
hydrogenase 2 small subunit [Salmonella enterica] Length: 372\nNumber of predicted TMHs: 1\nExp number of AAs in TMHs: 24.51309\nExp number, first 60 AAs: 0.2682\nTotal prob of N-in: 0.14402\noutside 1 331\nTMhelix 332 354\ninside 355 372

14631 GCF_002036965.1_ASM203696v1 Salmonella enterica Proteobacteria; Gammaproteobacteria; Enterobacterales; Enterobacteriaceae; Salmonella
MNNEETFYQAMRRKGVTRRSFLKFCSLAATSLGLGAGMTPKIAWALENKPRIPVVWIHGL WP_001058312.1
hydrogenase [Salmonella enterica] Length: 372\nNumber of predicted TMHs: 2\nExp number of AAs in TMHs: 36.51738\nExp number, first 60 AAs: 19.72849\nTotal prob of N-in: 0.96170\nPOSSIBLE N-term signal sequence\ninside 1 19\nTMhelix 20 39\noutside 40 325\nTMhelix 326 348\ninside 349 372

14632 GCF_002037065.1_ASM203706v1 Salmonella enterica Proteobacteria; Gammaproteobacteria; Enterobacterales; Enterobacteriaceae; Salmonella
MTGDNTLITSHGINRRDFMKLCAALAATMGLSSKAAAEMAESVSNPQRPPVIWIGAQECT WP_000145429.1
hydrogenase 2 small subunit [Salmonella enterica] Length: 372\nNumber of predicted TMHs: 1\nExp number of AAs in TMHs: 24.51309\nExp number, first 60 AAs: 0.2682\nTotal prob of N-in: 0.14402\noutside 1 331\nTMhelix 332 354\ninside 355 372

14633 GCF_002037105.1_ASM203710v1 Salmonella enterica Proteobacteria; Gammaproteobacteria; Enterobacterales; Enterobacteriaceae; Salmonella
MTGDNTLITSHGINRRDFMKLCAALAATMGLSSKAAAEMAESVSNPQRPPVIWIGAQECT WP_000145429.1
hydrogenase 2 small subunit [Salmonella enterica] Length: 372\nNumber of predicted TMHs: 1\nExp number of AAs in TMHs: 24.51309\nExp number, first 60 AAs: 0.2682\nTotal prob of N-in: 0.14402\noutside 1 331\nTMhelix 332 354\ninside 355 372

14634 GCF_002037035.1_ASM203703v1 Salmonella enterica Proteobacteria; Gammaproteobacteria; Enterobacterales; Enterobacteriaceae; Salmonella
MTGDNTLITSHGINRRDFMKLCAALAATMGLSSKAAAEMAESVSNPQRPPVIWIGAQECT WP_000145431.1
hydrogenase 2 small subunit [Salmonella enterica] Length: 372\nNumber of predicted TMHs: 1\nExp number of AAs in TMHs: 24.51505\nExp number, first 60 AAs: 0.2682\nTotal prob of N-in: 0.14406\noutside 1 331\nTMhelix 332 354\ninside 355 372

14635 GCF_002036785.1_ASM203678v1 Salmonella enterica Proteobacteria; Gammaproteobacteria; Enterobacterales; Enterobacteriaceae; Salmonella

MNNEETFYQAMRRKGVTRRSFLKFCSLAATSLGLGAGMTPKIAWALENKPRIPVVWIHGL WP_001058312.1 hydrogenase [Salmonella enterica] Length: 372\nNumber of predicted TMHs: 2\nExp number of AAs in TMHs: 36.51738\nExp number, first 60 AAs: 19.72849\nTotal prob of N-in: 0.96170\nPOSSIBLE N-term signal sequence\ninside 1 19\nTMhelix 20 39\noutside 40 325\nTMhelix 326 348\ninside 349 372

14636 GCF_002061105.1_ASM206110v1 Salmonella enterica Proteobacteria; Gammaproteobacteria; Enterobacterales; Enterobacteriaceae; Salmonella MTGDNTLITSHGINRRDFMKLCAALAATMGLSSKAAAEMAESVSNPQRPPVIWIGAQECT WP_000145429.1 hydrogenase 2 small subunit [Salmonella enterica] Length: 372\nNumber of predicted TMHs: 1\nExp number of AAs in TMHs: 24.51309\nExp number, first 60 AAs: 0.2682\nTotal prob of N-in: 0.14402\noutside 1 331\nTMhelix 332 354\ninside 355 372

14637 GCF_002061055.1_ASM206105v1 Salmonella enterica Proteobacteria; Gammaproteobacteria; Enterobacterales; Enterobacteriaceae; Salmonella MNNEETFYQVMRRKGVTRRSFLKFCSLAATSLGLGAGMTPKIAWALENKPRIPVVWIHGL WP_080177115.1 [Ni/Fe] hydrogenase small subunit [Salmonella enterica] Length: 372\nNumber of predicted TMHs: 2\nExp number of AAs in TMHs: 36.44113\nExp number, first 60 AAs: 19.65261\nTotal prob of N-in: 0.95797\nPOSSIBLE N-term signal sequence\ninside 1 19\nTMhelix 20 39\noutside 40 325\nTMhelix 326 348\ninside 349 372

14638 GCF_002061415.1_ASM206141v1 Salmonella enterica Proteobacteria; Gammaproteobacteria; Enterobacterales; Enterobacteriaceae; Salmonella MTGDNTLITSHGINRRDFMKLCAALAATMGLSSKAAAEMAESVSNPQRPPVIWIGAQECT WP_000145429.1 hydrogenase 2 small subunit [Salmonella enterica] Length: 372\nNumber of predicted TMHs: 1\nExp number of AAs in TMHs: 24.51309\nExp number, first 60 AAs: 0.2682\nTotal prob of N-in: 0.14402\noutside 1 331\nTMhelix 332 354\ninside 355 372

14639 GCF_002061425.1_ASM206142v1 Salmonella enterica Proteobacteria; Gammaproteobacteria; Enterobacterales; Enterobacteriaceae; Salmonella MTGDNTLITSHGINRRDFMKLCAALAATMGLSSKAAAEMAESVSNPQRPPVIWIGAQECT WP_000145429.1 hydrogenase 2 small subunit [Salmonella enterica] Length: 372\nNumber of predicted TMHs: 1\nExp number of AAs in TMHs: 24.51309\nExp number, first 60 AAs: 0.2682\nTotal prob of N-in: 0.14402\noutside 1 331\nTMhelix 332 354\ninside 355 372

14640 GCF_002061415.1_ASM206141v1 Salmonella enterica Proteobacteria; Gammaproteobacteria; Enterobacterales; Enterobacteriaceae; Salmonella MNNEETFYQAMRRKGVTRRSFLKFCSLAATSLGLGAGMTPKIAWALENKPRIPVVWIHGL WP_001058312.1 hydrogenase [Salmonella enterica] Length: 372\nNumber of predicted TMHs: 2\nExp number of AAs in TMHs: 36.51738\nExp number, first 60 AAs: 19.72849\nTotal prob of N-in: 0.96170\nPOSSIBLE N-term signal sequence\ninside 1 19\nTMhelix 20 39\noutside 40 325\nTMhelix 326 348\ninside 349 372

14641 GCF_002061425.1_ASM206142v1 Salmonella enterica Proteobacteria; Gammaproteobacteria; Enterobacterales; Enterobacteriaceae; Salmonella MNNEETFYQAMRRKGVTRRSFLKFCSLAATSLGLGAGMTPKIAWALENKPRIPVVWIHGL WP_001058312.1 hydrogenase [Salmonella enterica] Length: 372\nNumber of predicted TMHs: 2\nExp number of AAs in TMHs: 36.51738\nExp number, first 60 AAs: 19.72849\nTotal prob of N-in: 0.96170\nPOSSIBLE N-term signal sequence\ninside 1 19\nTMhelix 20 39\noutside 40 325\nTMhelix 326 348\ninside 349 372

14642 GCF_002061585.1_ASM206158v1 Salmonella enterica Proteobacteria; Gammaproteobacteria; Enterobacterales; Enterobacteriaceae; Salmonella MTGDNTLITSHGINRRDFMKLCAALAATMGLSSKAAAEMAESVSNPQRPPVIWIGAQECT WP_000145429.1 hydrogenase 2 small subunit [Salmonella enterica] Length: 372\nNumber of predicted TMHs: 1\nExp number of AAs in TMHs: 24.51309\nExp number, first 60 AAs: 0.2682\nTotal prob of N-in: 0.14402\noutside 1 331\nTMhelix 332 354\ninside 355 372

14643 GCF_002061725.1_ASM206172v1 Salmonella enterica Proteobacteria; Gammaproteobacteria; Enterobacterales; Enterobacteriaceae; Salmonella MTGDNTLITSHGINRRDFMKLCAALAATMGLSSKAAAEMAESVSNPQRPPVIWIGAQECT WP_000145429.1 hydrogenase 2 small subunit [Salmonella enterica] Length: 372\nNumber of predicted TMHs: 1\nExp number of AAs in TMHs: 24.51309\nExp number, first 60 AAs: 0.2682\nTotal prob of N-in: 0.14402\noutside 1 331\nTMhelix 332 354\ninside 355 372

14644	GCF_002061725.1_ASM206172v1	Salmonella enterica	Proteobacteria; Gammaproteobacteria; Enterobacterales; Enterobacteriaceae; Salmonella
MNNEETFYQAMRRKGVTRRSFLKFCSLAATSLGLGAGMTPKIAWALENKPRIPVVWIHGL	WP_001058311.1
hydrogenase [Salmonella enterica]	Length: 372\nNumber of predicted TMHs: 2\nExp number of AAs in TMHs: 37.4699899999999\nExp number, first 60 AAs: 19.72699\nTotal prob of N-in: 0.96171\nPOSSIBLE N-term signal sequence\ninside 1 19\nTMhelix 20 39\noutside 40 325\nTMhelix 326 348\ninside 349 372

14645	GCF_002061585.1_ASM206158v1	Salmonella enterica	Proteobacteria; Gammaproteobacteria; Enterobacterales; Enterobacteriaceae; Salmonella
MNNEETFYQAMRRKGVTRRSFLKFCSLAATSLGLGAGMTPKIAWALENKPRIPVVWIHGL	WP_023187532.1
hydrogenase 1, small subunit [Salmonella enterica]	Length: 372\nNumber of predicted TMHs: 2\nExp number of AAs in TMHs: 36.9997\nExp number, first 60 AAs: 19.72759\nTotal prob of N-in: 0.96171\nPOSSIBLE N-term signal sequence\ninside 1 19\nTMhelix 20 39\noutside 40 325\nTMhelix 326 348\ninside 349 372

14646	GCF_002061865.1_ASM206186v1	Salmonella enterica	Proteobacteria; Gammaproteobacteria; Enterobacterales; Enterobacteriaceae; Salmonella
MNNEETFYQAMRRKGVTRRSFLKFCSLAATSLGLGAGMTPKIAWALENKPRIPVVWIHGL	WP_001058307.1
hydrogenase [Salmonella enterica]	Length: 372\nNumber of predicted TMHs: 2\nExp number of AAs in TMHs: 36.51738\nExp number, first 60 AAs: 19.72849\nTotal prob of N-in: 0.96170\nPOSSIBLE N-term signal sequence\ninside 1 19\nTMhelix 20 39\noutside 40 325\nTMhelix 326 348\ninside 349 372

14647	GCF_002061995.1_ASM206199v1	Salmonella enterica	Proteobacteria; Gammaproteobacteria; Enterobacterales; Enterobacteriaceae; Salmonella
MTGDNTLITSHGINRRDFMKLCAALAATMGLSSKAAAEMAESVSNPQRPPVIWIGAQECT	WP_000145429.1
hydrogenase 2 small subunit [Salmonella enterica]	Length: 372\nNumber of predicted TMHs: 1\nExp number of AAs in TMHs: 24.51309\nExp number, first 60 AAs: 0.2682\nTotal prob of N-in: 0.14402\noutside 1 331\nTMhelix 332 354\ninside 355 372

14648	GCF_001831875.1_ASM183187v1	Salmonella enterica	Proteobacteria; Gammaproteobacteria; Enterobacterales; Enterobacteriaceae; Salmonella
MTGDNTLITSHGINRRDFMKLCAALAATMGLSSKAAAEMAESVSNPQRPPVIWIGAQECT	WP_000145429.1
hydrogenase 2 small subunit [Salmonella enterica]	Length: 372\nNumber of predicted TMHs: 1\nExp number of AAs in TMHs: 24.51309\nExp number, first 60 AAs: 0.2682\nTotal prob of N-in: 0.14402\noutside 1 331\nTMhelix 332 354\ninside 355 372

14649	GCF_002061995.1_ASM206199v1	Salmonella enterica	Proteobacteria; Gammaproteobacteria; Enterobacterales; Enterobacteriaceae; Salmonella
MNNEETFYQAMRRKGVTRRSFLKFCSLAATSLGLGAGMTPKIAWALENKPRIPVVWIHGL	WP_001058312.1
hydrogenase [Salmonella enterica]	Length: 372\nNumber of predicted TMHs: 2\nExp number of AAs in TMHs: 36.51738\nExp number, first 60 AAs: 19.72849\nTotal prob of N-in: 0.96170\nPOSSIBLE N-term signal sequence\ninside 1 19\nTMhelix 20 39\noutside 40 325\nTMhelix 326 348\ninside 349 372

14650	GCF_002061945.1_ASM206194v1	Salmonella enterica	Proteobacteria; Gammaproteobacteria; Enterobacterales; Enterobacteriaceae; Salmonella
MNNEETFYQAMRRKGVTRRSFLKFCSLAATSLGLGAGMTPKIAWALENKPRIPVVWIHGL	WP_001058312.1
hydrogenase [Salmonella enterica]	Length: 372\nNumber of predicted TMHs: 2\nExp number of AAs in TMHs: 36.51738\nExp number, first 60 AAs: 19.72849\nTotal prob of N-in: 0.96170\nPOSSIBLE N-term signal sequence\ninside 1 19\nTMhelix 20 39\noutside 40 325\nTMhelix 326 348\ninside 349 372

14651	GCF_001578485.1_ASM157848v1	Salmonella enterica	Proteobacteria; Gammaproteobacteria; Enterobacterales; Enterobacteriaceae; Salmonella
MTGDNTLITSHGINRRDFMKLCAALAATMGLSSKAAAEMAESVSNPQRPPVIWIGAQECT	WP_000145429.1
hydrogenase 2 small subunit [Salmonella enterica]	Length: 372\nNumber of predicted TMHs: 1\nExp number of AAs in TMHs: 24.51309\nExp number, first 60 AAs: 0.2682\nTotal prob of N-in: 0.14402\noutside 1 331\nTMhelix 332 354\ninside 355 372

14652	GCF_001570305.1_ASM157030v1	Salmonella enterica	Proteobacteria; Gammaproteobacteria; Enterobacterales; Enterobacteriaceae; Salmonella
MNNEETFYQAMRRKGVTRRSFLKFCSLAATSLGLGAGMTPKIAWALENKPRIPVVWIHGL	WP_001058307.1
hydrogenase [Salmonella enterica]	Length: 372\nNumber of predicted TMHs: 2\nExp number of AAs in

TMHs: 36.51738\nExp number, first 60 AAs: 19.72849\nTotal prob of N-in: 0.96170\nPOSSIBLE N-term signal sequence\ninside 1 19\nTMhelix 20 39\noutside 40 325\nTMhelix 326 348\ninside 349 372

14653 GCF_001570265.1_ASM157026v1 Salmonella enterica Proteobacteria; Gammaproteobacteria; Enterobacterales; Enterobacteriaceae; Salmonella
MNNEETFYQAMRRKGVTRRSFLKFCSLAATSLGLGAGMTPKIAWALENKPRIPVVWIHGL WP_001058307.1
hydrogenase [Salmonella enterica] Length: 372\nNumber of predicted TMHs: 2\nExp number of AAs in TMHs: 36.51738\nExp number, first 60 AAs: 19.72849\nTotal prob of N-in: 0.96170\nPOSSIBLE N-term signal sequence\ninside 1 19\nTMhelix 20 39\noutside 40 325\nTMhelix 326 348\ninside 349 372

14654 GCF_001570325.1_ASM157032v1 Salmonella enterica Proteobacteria; Gammaproteobacteria; Enterobacterales; Enterobacteriaceae; Salmonella
MTGDNSLINSSGINRRDFMKLCAALAATMGLSSKAAAEMAESVSRPQRPPVIWIGAQECT WP_003024486.1
MULTISPECIES: hydrogenase 2 small subunit [Enterobacteriaceae] Length: 372\nNumber of predicted TMHs: 1\nExp number of AAs in TMHs: 27.14621\nExp number, first 60 AAs: 0.22683\nTotal prob of N-in: 0.23982\noutside 1 331\nTMhelix 332 354\ninside 355 372

14655 GCF_001570305.1_ASM157030v1 Salmonella enterica Proteobacteria; Gammaproteobacteria; Enterobacterales; Enterobacteriaceae; Salmonella
MTGDNTLITSHGINRRDFMKLCAALAATMGLSSKAAAEMAESVSNPQRPPVIWIGAQECT WP_000145429.1
hydrogenase 2 small subunit [Salmonella enterica] Length: 372\nNumber of predicted TMHs: 1\nExp number of AAs in TMHs: 24.51309\nExp number, first 60 AAs: 0.2682\nTotal prob of N-in: 0.14402\noutside 1 331\nTMhelix 332 354\ninside 355 372

14656 GCF_001570265.1_ASM157026v1 Salmonella enterica Proteobacteria; Gammaproteobacteria; Enterobacterales; Enterobacteriaceae; Salmonella
MTGDNTLITSHGINRRDFMKLCAALAATMGLSSKAAAEMAESVSNPQRPPVIWIGAQECT WP_000145429.1
hydrogenase 2 small subunit [Salmonella enterica] Length: 372\nNumber of predicted TMHs: 1\nExp number of AAs in TMHs: 24.51309\nExp number, first 60 AAs: 0.2682\nTotal prob of N-in: 0.14402\noutside 1 331\nTMhelix 332 354\ninside 355 372

14657 GCF_001569785.1_ASM156978v1 Salmonella enterica Proteobacteria; Gammaproteobacteria; Enterobacterales; Enterobacteriaceae; Salmonella
MNNEETFYQAMRRKGVTRRSFLKFCSLAATSLGLGAGMTPKIAWALENKPRIPVVWIHGL WP_001058311.1
hydrogenase [Salmonella enterica] Length: 372\nNumber of predicted TMHs: 2\nExp number of AAs in TMHs: 37.4699899999999\nExp number, first 60 AAs: 19.72699\nTotal prob of N-in: 0.96171\nPOSSIBLE N-term signal sequence\ninside 1 19\nTMhelix 20 39\noutside 40 325\nTMhelix 326 348\ninside 349 372

14658 GCF_001569845.1_ASM156984v1 Salmonella enterica Proteobacteria; Gammaproteobacteria; Enterobacterales; Enterobacteriaceae; Salmonella
MTGDNTLITSHGINRRDFMKLCAALAATMGLSSKAAAEMAESVSNPQRPPVIWIGAQECT WP_000145429.1
hydrogenase 2 small subunit [Salmonella enterica] Length: 372\nNumber of predicted TMHs: 1\nExp number of AAs in TMHs: 24.51309\nExp number, first 60 AAs: 0.2682\nTotal prob of N-in: 0.14402\noutside 1 331\nTMhelix 332 354\ninside 355 372

14659 GCF_001569825.1_ASM156982v1 Salmonella enterica Proteobacteria; Gammaproteobacteria; Enterobacterales; Enterobacteriaceae; Salmonella
MTGDNTLITSHGINRRDFMKLCAALAATMGLSSKAAAEMAESVSNPQRPPVIWIGAQECT WP_000145429.1
hydrogenase 2 small subunit [Salmonella enterica] Length: 372\nNumber of predicted TMHs: 1\nExp number of AAs in TMHs: 24.51309\nExp number, first 60 AAs: 0.2682\nTotal prob of N-in: 0.14402\noutside 1 331\nTMhelix 332 354\ninside 355 372

14660 GCF_001569785.1_ASM156978v1 Salmonella enterica Proteobacteria; Gammaproteobacteria; Enterobacterales; Enterobacteriaceae; Salmonella
MTGDNTLITSHGINRRDFMKLCAALAATMGLSSKAAAEMAESVSNPQRPPVIWIGAQECT WP_000145429.1
hydrogenase 2 small subunit [Salmonella enterica] Length: 372\nNumber of predicted TMHs: 1\nExp number of AAs in TMHs: 24.51309\nExp number, first 60 AAs: 0.2682\nTotal prob of N-in: 0.14402\noutside 1 331\nTMhelix 332 354\ninside 355 372

14661 GCF_001569745.1_ASM156974v1 Salmonella enterica Proteobacteria; Gammaproteobacteria; Enterobacterales; Enterobacteriaceae; Salmonella

MTGDNTLITSHGINRRDFMKLCAALAATMGLSSKAAAEMAESVSNPQRPPVIWIGAQECT WP_000145429.1 hydrogenase 2 small subunit [Salmonella enterica] Length: 372\nNumber of predicted TMHs: 1\nExp number of AAs in TMHs: 24.51309\nExp number, first 60 AAs: 0.2682\nTotal prob of N-in: 0.14402\noutside 1 331\nTMhelix 332 354\ninside 355 372

14662 GCF_001569725.1_ASM156972v1 Salmonella enterica Proteobacteria; Gammaproteobacteria; Enterobacterales; Enterobacteriaceae; Salmonella MNNEETFYQAMRRKGVTRRSFLKFCSLAATSLGLGAGMTPKIAWALENKPRIPVIWIHGL WP_061376537.1 hydrogenase [Salmonella enterica] Length: 372\nNumber of predicted TMHs: 2\nExp number of AAs in TMHs: 36.5556\nExp number, first 60 AAs: 19.76633\nTotal prob of N-in: 0.96197\nPOSSIBLE N-term signal sequence\ninside 1 19\nTMhelix 20 39\noutside 40 325\nTMhelix 326 348\ninside 349 372

14663 GCF_002036815.1_ASM203681v1 Salmonella enterica Proteobacteria; Gammaproteobacteria; Enterobacterales; Enterobacteriaceae; Salmonella MNNEETFYQAMRRKGVTRRSFLKFCSLAATSLGLGAGMTPKIAWALENKPRIPVVWIHGL WP_001058312.1 hydrogenase [Salmonella enterica] Length: 372\nNumber of predicted TMHs: 2\nExp number of AAs in TMHs: 36.51738\nExp number, first 60 AAs: 19.72849\nTotal prob of N-in: 0.96170\nPOSSIBLE N-term signal sequence\ninside 1 19\nTMhelix 20 39\noutside 40 325\nTMhelix 326 348\ninside 349 372

14664 GCF_001558355.1_ASM155835v1 Salmonella enterica Proteobacteria; Gammaproteobacteria; Enterobacterales; Enterobacteriaceae; Salmonella MNNEETFYQAMRRKGVTRRSFLKFCSLAATSLGLGAGMTPKIAWALENKPRIPVVWIHGL WP_001058312.1 hydrogenase [Salmonella enterica] Length: 372\nNumber of predicted TMHs: 2\nExp number of AAs in TMHs: 36.51738\nExp number, first 60 AAs: 19.72849\nTotal prob of N-in: 0.96170\nPOSSIBLE N-term signal sequence\ninside 1 19\nTMhelix 20 39\noutside 40 325\nTMhelix 326 348\ninside 349 372

14665 GCF_001480915.1_Salmonella_enterica_CVM_N40366-SQ_v1.0 Salmonella enterica Proteobacteria; Gammaproteobacteria; Enterobacterales; Enterobacteriaceae; Salmonella MNNEETFYQAMRRKGVTRRSFLKFCSLAATSLGLGAGMTPKIAWALENKPRIPVVWIHGL WP_001058312.1 hydrogenase [Salmonella enterica] Length: 372\nNumber of predicted TMHs: 2\nExp number of AAs in TMHs: 36.51738\nExp number, first 60 AAs: 19.72849\nTotal prob of N-in: 0.96170\nPOSSIBLE N-term signal sequence\ninside 1 19\nTMhelix 20 39\noutside 40 325\nTMhelix 326 348\ninside 349 372

14666 GCF_001475375.1_Salmonella_enterica_CVM_43842_v1.0Salmonella enterica Proteobacteria; Gammaproteobacteria; Enterobacterales; Enterobacteriaceae; Salmonella MNNEETFYQAMRRKGVTRRSFLKFCSLAATSLGLGAGMTPKIAWALENKPRIPVVWIHGL WP_001058312.1 hydrogenase [Salmonella enterica] Length: 372\nNumber of predicted TMHs: 2\nExp number of AAs in TMHs: 36.51738\nExp number, first 60 AAs: 19.72849\nTotal prob of N-in: 0.96170\nPOSSIBLE N-term signal sequence\ninside 1 19\nTMhelix 20 39\noutside 40 325\nTMhelix 326 348\ninside 349 372

14667 GCF_001475335.1_Salmonella_enterica_CVM_43815_v1.0Salmonella enterica Proteobacteria; Gammaproteobacteria; Enterobacterales; Enterobacteriaceae; Salmonella MTGDNTLITSHGINRRDFMKLCAALAATMGLSSKAAAEMAESVSNPQRPPVIWIGAQECT WP_000145429.1 hydrogenase 2 small subunit [Salmonella enterica] Length: 372\nNumber of predicted TMHs: 1\nExp number of AAs in TMHs: 24.51309\nExp number, first 60 AAs: 0.2682\nTotal prob of N-in: 0.14402\noutside 1 331\nTMhelix 332 354\ninside 355 372

14668 GCF_002036865.1_ASM203686v1 Salmonella enterica Proteobacteria; Gammaproteobacteria; Enterobacterales; Enterobacteriaceae; Salmonella MNNEETFYQAMRRKGVTRRSFLKFCSLAATSLGLGAGMTPKIAWALENKPRIPVVWIHGL WP_001058312.1 hydrogenase [Salmonella enterica] Length: 372\nNumber of predicted TMHs: 2\nExp number of AAs in TMHs: 36.51738\nExp number, first 60 AAs: 19.72849\nTotal prob of N-in: 0.96170\nPOSSIBLE N-term signal sequence\ninside 1 19\nTMhelix 20 39\noutside 40 325\nTMhelix 326 348\ninside 349 372

14669 GCF_002036815.1_ASM203681v1 Salmonella enterica Proteobacteria; Gammaproteobacteria; Enterobacterales; Enterobacteriaceae; Salmonella MTGDNTLITSHGINRRDFMKLCAALAATMGLSSKAAAEMAESVSNPQRPPVIWIGAQECT WP_000145429.1 hydrogenase 2 small subunit [Salmonella enterica] Length: 372\nNumber of predicted TMHs: 1\nExp number of AAs in TMHs: 24.51309\nExp number, first 60 AAs: 0.2682\nTotal prob of N-in: 0.14402\noutside 1 331\nTMhelix 332 354\ninside 355 372

14670 GCF_002036865.1_ASM203686v1 Salmonella enterica Proteobacteria; Gammaproteobacteria; Enterobacterales; Enterobacteriaceae; Salmonella
MTGDNTLITSHGINRRDFMKLCAALAATMGLSSKAAAEMAESVSNPQRPPVIWIGAQECT WP_000145429.1
hydrogenase 2 small subunit [Salmonella enterica] Length: 372\nNumber of predicted TMHs: 1\nExp number of AAs in TMHs: 24.51309\nExp number, first 60 AAs: 0.2682\nTotal prob of N-in: 0.14402\noutside 1 331\nTMhelix 332 354\ninside 355 372

14671 GCF_002036715.1_ASM203671v1 Salmonella enterica Proteobacteria; Gammaproteobacteria; Enterobacterales; Enterobacteriaceae; Salmonella
MNNEETFYQAMRRKGVTRRSFLKFCSLAATSLGLGAGMTPKIAWALENKPRIPVVWIHGL WP_001058312.1
hydrogenase [Salmonella enterica] Length: 372\nNumber of predicted TMHs: 2\nExp number of AAs in TMHs: 36.51738\nExp number, first 60 AAs: 19.72849\nTotal prob of N-in: 0.96170\nPOSSIBLE N-term signal sequence\ninside 1 19\nTMhelix 20 39\noutside 40 325\nTMhelix 326 348\ninside 349 372

14672 GCF_002036675.1_ASM203667v1 Salmonella enterica Proteobacteria; Gammaproteobacteria; Enterobacterales; Enterobacteriaceae; Salmonella
MNNEETFYQAMRRKGVTRRSFLKFCSLAATSLGLGAGMTPKIAWALENKPRIPVVWIHGL WP_001058308.1
hydrogenase [Salmonella enterica] Length: 372\nNumber of predicted TMHs: 2\nExp number of AAs in TMHs: 39.08204\nExp number, first 60 AAs: 19.72474\nTotal prob of N-in: 0.96174\nPOSSIBLE N-term signal sequence\ninside 1 19\nTMhelix 20 39\noutside 40 325\nTMhelix 326 348\ninside 349 372

14673 GCF_002036385.1_ASM203638v1 Salmonella enterica Proteobacteria; Gammaproteobacteria; Enterobacterales; Enterobacteriaceae; Salmonella
MNNEETFYQAMRRKGVTRRSFLKFCSLAATSLGLGAGMTPKIAWALENKPRIPVVWIHGL WP_001058307.1
hydrogenase [Salmonella enterica] Length: 372\nNumber of predicted TMHs: 2\nExp number of AAs in TMHs: 36.51738\nExp number, first 60 AAs: 19.72849\nTotal prob of N-in: 0.96170\nPOSSIBLE N-term signal sequence\ninside 1 19\nTMhelix 20 39\noutside 40 325\nTMhelix 326 348\ninside 349 372

14674 GCF_002036385.1_ASM203638v1 Salmonella enterica Proteobacteria; Gammaproteobacteria; Enterobacterales; Enterobacteriaceae; Salmonella
MTGDNTLITSHGINRRDFMKLCAALAATMGLSSKAAAEMAESVSNPQRPPVIWIGAQECT WP_000145429.1
hydrogenase 2 small subunit [Salmonella enterica] Length: 372\nNumber of predicted TMHs: 1\nExp number of AAs in TMHs: 24.51309\nExp number, first 60 AAs: 0.2682\nTotal prob of N-in: 0.14402\noutside 1 331\nTMhelix 332 354\ninside 355 372

14675 GCF_002036425.1_ASM203642v1 Salmonella enterica Proteobacteria; Gammaproteobacteria; Enterobacterales; Enterobacteriaceae; Salmonella
MNNEETFYQAMRRKGVTRRSFLKFCSLAATSLGLGAGMTPKIAWALENKPRIPVVWIHGL WP_079786335.1
[Ni/Fe] hydrogenase small subunit [Salmonella enterica] Length: 372\nNumber of predicted TMHs: 2\nExp number of AAs in TMHs: 36.51716\nExp number, first 60 AAs: 19.7285\nTotal prob of N-in: 0.96169\nPOSSIBLE N-term signal sequence\ninside 1 19\nTMhelix 20 39\noutside 40 325\nTMhelix 326 348\ninside 349 372

14676 GCF_002036425.1_ASM203642v1 Salmonella enterica Proteobacteria; Gammaproteobacteria; Enterobacterales; Enterobacteriaceae; Salmonella
MTGDNTLITSHGINRRDFMKLCAALAATMGLSSKAAAEMAESVSNPQRPPVIWIGAQECT WP_000145429.1
hydrogenase 2 small subunit [Salmonella enterica] Length: 372\nNumber of predicted TMHs: 1\nExp number of AAs in TMHs: 24.51309\nExp number, first 60 AAs: 0.2682\nTotal prob of N-in: 0.14402\noutside 1 331\nTMhelix 332 354\ninside 355 372

14677 GCF_002035765.1_ASM203576v1 Salmonella enterica Proteobacteria; Gammaproteobacteria; Enterobacterales; Enterobacteriaceae; Salmonella
MNNEETFYQAMRRKGVTRRSFLKFCSLAATSLGLGAGMTPKIAWALENKPRIPVVWIHGL WP_076937197.1
[Ni/Fe] hydrogenase small subunit [Salmonella enterica] Length: 372\nNumber of predicted TMHs: 2\nExp number of AAs in TMHs: 37.4699899999999\nExp number, first 60 AAs: 19.72699\nTotal prob of N-in: 0.96171\nPOSSIBLE N-term signal sequence\ninside 1 19\nTMhelix 20 39\noutside 40 325\nTMhelix 326 348\ninside 349 372

14678 GCF_002036045.1_ASM203604v1 Salmonella enterica Proteobacteria; Gammaproteobacteria; Enterobacterales; Enterobacteriaceae; Salmonella
MNNEETFYQAMRRKGVTRRSFLKFCSLAATSLGLGAGMTPKIAWALENKPRIPVVWIHGL WP_001058306.1
hydrogenase [Salmonella enterica] Length: 372\nNumber of predicted TMHs: 2\nExp number of AAs in

TMHs: 36.5540099999999\nExp number, first 60 AAs: 19.72732\nTotal prob of N-in: 0.96171\nPOSSIBLE N-term signal sequence\ninside 1 19\nTMhelix 20 39\noutside 40 325\nTMhelix 326 348\ninside 349 372

14679 GCF_002036045.1_ASM203604v1 Salmonella enterica Proteobacteria; Gammaproteobacteria; Enterobacterales; Enterobacteriaceae; Salmonella
MTGDNTLITSHGINRRDFMKLCAALAATMGLSSKAAAEMAESVSNPQRPPVIWIGAQECT WP_000145429.1
hydrogenase 2 small subunit [Salmonella enterica] Length: 372\nNumber of predicted TMHs: 1\nExp number of AAs in TMHs: 24.51309\nExp number, first 60 AAs: 0.2682\nTotal prob of N-in: 0.14402\noutside 1 331\nTMhelix 332 354\ninside 355 372

14680 GCF_002035695.1_ASM203569v1 Salmonella enterica Proteobacteria; Gammaproteobacteria; Enterobacterales; Enterobacteriaceae; Salmonella
MNNEETFYQAMRRKGVTRRSFLKFCSLAATSLGLGAGMTPKIAWALENKPRIPVVWIHGL WP_001058306.1
hydrogenase [Salmonella enterica] Length: 372\nNumber of predicted TMHs: 2\nExp number of AAs in TMHs: 36.5540099999999\nExp number, first 60 AAs: 19.72732\nTotal prob of N-in: 0.96171\nPOSSIBLE N-term signal sequence\ninside 1 19\nTMhelix 20 39\noutside 40 325\nTMhelix 326 348\ninside 349 372

14681 GCF_001570185.1_ASM157018v1 Salmonella enterica Proteobacteria; Gammaproteobacteria; Enterobacterales; Enterobacteriaceae; Salmonella
MTGDNTLITSHGINRRDFMKLCAALAATMGLSSKAAAEMAESVSNPQRPPVIWIGAQECT WP_000145429.1
hydrogenase 2 small subunit [Salmonella enterica] Length: 372\nNumber of predicted TMHs: 1\nExp number of AAs in TMHs: 24.51309\nExp number, first 60 AAs: 0.2682\nTotal prob of N-in: 0.14402\noutside 1 331\nTMhelix 332 354\ninside 355 372

14682 GCF_002035465.1_ASM203546v1 Salmonella enterica Proteobacteria; Gammaproteobacteria; Enterobacterales; Enterobacteriaceae; Salmonella
MTGDNTLITSHGINRRDFMKLCAALAATMGLSSKAAAEMAESVSNPQRPPVIWIGAQECT WP_079776618.1
MULTISPECIES: hydrogenase 2 small subunit [Salmonella] Length: 372\nNumber of predicted TMHs: 1\nExp number of AAs in TMHs: 24.65705\nExp number, first 60 AAs: 0.26805\nTotal prob of N-in: 0.13761\noutside 1 331\nTMhelix 332 354\ninside 355 372

14683 GCF_002035625.1_ASM203562v1 Salmonella enterica Proteobacteria; Gammaproteobacteria; Enterobacterales; Enterobacteriaceae; Salmonella
MTGDNTLITSHGINRRDFMKLCAALAATMGLSSKAAAEMAESVSNPQRPPVIWIGAQECT WP_000145429.1
hydrogenase 2 small subunit [Salmonella enterica] Length: 372\nNumber of predicted TMHs: 1\nExp number of AAs in TMHs: 24.51309\nExp number, first 60 AAs: 0.2682\nTotal prob of N-in: 0.14402\noutside 1 331\nTMhelix 332 354\ninside 355 372

14684 GCF_002035555.1_ASM203555v1 Salmonella enterica Proteobacteria; Gammaproteobacteria; Enterobacterales; Enterobacteriaceae; Salmonella
MNNEETFYQAMRRKGVTRRSFLKFCSLAATSLGLGAGMTPKIAWALENKPRIPVVWIHGL WP_001058311.1
hydrogenase [Salmonella enterica] Length: 372\nNumber of predicted TMHs: 2\nExp number of AAs in TMHs: 37.4699899999999\nExp number, first 60 AAs: 19.72699\nTotal prob of N-in: 0.96171\nPOSSIBLE N-term signal sequence\ninside 1 19\nTMhelix 20 39\noutside 40 325\nTMhelix 326 348\ninside 349 372

14685 GCF_002035205.1_ASM203520v1 Salmonella enterica Proteobacteria; Gammaproteobacteria; Enterobacterales; Enterobacteriaceae; Salmonella
MNNEETFYQAMRRKGVTRRSFLKFCSLAATSLGLGAGMTPKIAWALENKPRIPVVWIHGL WP_001058312.1
hydrogenase [Salmonella enterica] Length: 372\nNumber of predicted TMHs: 2\nExp number of AAs in TMHs: 36.51738\nExp number, first 60 AAs: 19.72849\nTotal prob of N-in: 0.96170\nPOSSIBLE N-term signal sequence\ninside 1 19\nTMhelix 20 39\noutside 40 325\nTMhelix 326 348\ninside 349 372

14686 GCF_002034895.1_ASM203489v1 Salmonella enterica Proteobacteria; Gammaproteobacteria; Enterobacterales; Enterobacteriaceae; Salmonella
MNNEETFYQAMRRKGVTRRSFLKFCSLAATSLGLGAGMTPKIAWALENKPRIPVVWIHGL WP_001058308.1
hydrogenase [Salmonella enterica] Length: 372\nNumber of predicted TMHs: 2\nExp number of AAs in TMHs: 39.08204\nExp number, first 60 AAs: 19.72474\nTotal prob of N-in: 0.96174\nPOSSIBLE N-term signal sequence\ninside 1 19\nTMhelix 20 39\noutside 40 325\nTMhelix 326 348\ninside 349 372

14687 GCF_002034815.1_ASM203481v1 Salmonella enterica Proteobacteria; Gammaproteobacteria; Enterobacterales; Enterobacteriaceae; Salmonella

MNNEETFYQAMRRKGVTRRSFLKFCSLAATSLGLGAGMTPKIAWALENKPRIPVVWIHGL WP_001058312.1 hydrogenase [Salmonella enterica] Length: 372\nNumber of predicted TMHs: 2\nExp number of AAs in TMHs: 36.51738\nExp number, first 60 AAs: 19.72849\nTotal prob of N-in: 0.96170\nPOSSIBLE N-term signal sequence\ninside 1 19\nTMhelix 20 39\noutside 40 325\nTMhelix 326 348\ninside 349 372

14688 GCF_002034455.1_ASM203445v1 Salmonella enterica Proteobacteria; Gammaproteobacteria; Enterobacterales; Enterobacteriaceae; Salmonella MNNEETFYQAMRRKGVTRRSFLKFCSLAATSLGLGAGMTPKIAWALENKPRIPVVWIHGL WP_001058307.1 hydrogenase [Salmonella enterica] Length: 372\nNumber of predicted TMHs: 2\nExp number of AAs in TMHs: 36.51738\nExp number, first 60 AAs: 19.72849\nTotal prob of N-in: 0.96170\nPOSSIBLE N-term signal sequence\ninside 1 19\nTMhelix 20 39\noutside 40 325\nTMhelix 326 348\ninside 349 372

14689 GCF_002034505.1_ASM203450v1 Salmonella enterica Proteobacteria; Gammaproteobacteria; Enterobacterales; Enterobacteriaceae; Salmonella MNNEETFYQAMRRKGVTRRSFLKFCSLAATSLGLGAGMTPKIAWALENKPRIPVVWIHGL WP_023243720.1 hydrogenase 1, small subunit [Salmonella enterica] Length: 372\nNumber of predicted TMHs: 2\nExp number of AAs in TMHs: 36.52147\nExp number, first 60 AAs: 19.72838\nTotal prob of N-in: 0.96170\nPOSSIBLE N-term signal sequence\ninside 1 19\nTMhelix 20 39\noutside 40 325\nTMhelix 326 348\ninside 349 372

14690 GCF_002034595.1_ASM203459v1 Salmonella enterica Proteobacteria; Gammaproteobacteria; Enterobacterales; Enterobacteriaceae; Salmonella MNNEETFYQAMRRKGVTRRSFLKFCSLAATSLGLGAGMTPKIAWALENKPRIPVVWIHGL WP_001058314.1 hydrogenase [Salmonella enterica] Length: 372\nNumber of predicted TMHs: 2\nExp number of AAs in TMHs: 36.3110299999999\nExp number, first 60 AAs: 19.72774\nTotal prob of N-in: 0.96169\nPOSSIBLE N-term signal sequence\ninside 1 19\nTMhelix 20 39\noutside 40 325\nTMhelix 326 348\ninside 349 372

14691 GCF_002034615.1_ASM203461v1 Salmonella enterica Proteobacteria; Gammaproteobacteria; Enterobacterales; Enterobacteriaceae; Salmonella MTGDNTLITSHGINRRDFMKLCAALAATMGLSSKAAAEMAESVSNPQRPPVIWIGAQECT WP_000145429.1 hydrogenase 2 small subunit [Salmonella enterica] Length: 372\nNumber of predicted TMHs: 1\nExp number of AAs in TMHs: 24.51309\nExp number, first 60 AAs: 0.2682\nTotal prob of N-in: 0.14402\noutside 1 331\nTMhelix 332 354\ninside 355 372

14692 GCF_002034595.1_ASM203459v1 Salmonella enterica Proteobacteria; Gammaproteobacteria; Enterobacterales; Enterobacteriaceae; Salmonella MTGDNTLITSHGINRRDFMKLCAALAATMGLSSKAAAEMAESVSNPQRPPVIWIGAQECT WP_000145432.1 hydrogenase 2 small subunit [Salmonella enterica] Length: 372\nNumber of predicted TMHs: 1\nExp number of AAs in TMHs: 24.5101\nExp number, first 60 AAs: 0.2682\nTotal prob of N-in: 0.14397\noutside 1 331\nTMhelix 332 354\ninside 355 372

14693 GCF_002034505.1_ASM203450v1 Salmonella enterica Proteobacteria; Gammaproteobacteria; Enterobacterales; Enterobacteriaceae; Salmonella MTGDNTLITSHGINRRDFMKLCAALAATMGLSSKAAAEMAESVSNPQRPPVIWIGAQECT WP_000145429.1 hydrogenase 2 small subunit [Salmonella enterica] Length: 372\nNumber of predicted TMHs: 1\nExp number of AAs in TMHs: 24.51309\nExp number, first 60 AAs: 0.2682\nTotal prob of N-in: 0.14402\noutside 1 331\nTMhelix 332 354\ninside 355 372

14694 GCF_002034165.1_ASM203416v1 Salmonella enterica Proteobacteria; Gammaproteobacteria; Enterobacterales; Enterobacteriaceae; Salmonella MNNEETFYQAMRRKGVTRRSFLKFCSLAATSLGLGAGMTPKIAWALENKPRIPVVWIHGL WP_001058312.1 hydrogenase [Salmonella enterica] Length: 372\nNumber of predicted TMHs: 2\nExp number of AAs in TMHs: 36.51738\nExp number, first 60 AAs: 19.72849\nTotal prob of N-in: 0.96170\nPOSSIBLE N-term signal sequence\ninside 1 19\nTMhelix 20 39\noutside 40 325\nTMhelix 326 348\ninside 349 372

14695 GCF_002034095.1_ASM203409v1 Salmonella enterica Proteobacteria; Gammaproteobacteria; Enterobacterales; Enterobacteriaceae; Salmonella MNNEETFYQAMRRKGVTRRSFLKFCSLAATSLGLGAGMTPKIAWALENKPRIPVVWIHGL WP_001058311.1 hydrogenase [Salmonella enterica] Length: 372\nNumber of predicted TMHs: 2\nExp number of AAs in TMHs: 37.4699899999999\nExp number, first 60 AAs: 19.72699\nTotal prob of N-in: 0.96171\nPOSSIBLE N-term signal sequence\ninside 1 19\nTMhelix 20 39\noutside 40 325\nTMhelix 326 348\ninside 349 372

14696 GCF_002034165.1_ASM203416v1 Salmonella enterica Proteobacteria; Gammaproteobacteria; Enterobacterales; Enterobacteriaceae; Salmonella
MTGDNTLITSHGINRRDFMKLCAALAATMGLSSKAAAEMAESVSNPQRPPVIWIGAQECT WP_000145429.1
hydrogenase 2 small subunit [Salmonella enterica] Length: 372\nNumber of predicted TMHs: 1\nExp number of AAs in TMHs: 24.51309\nExp number, first 60 AAs: 0.2682\nTotal prob of N-in: 0.14402\noutside 1 331\nTMhelix 332 354\ninside 355 372

14697 GCF_002033565.1_ASM203356v1 Salmonella enterica Proteobacteria; Gammaproteobacteria; Enterobacterales; Enterobacteriaceae; Salmonella
MNNEETFYQAMRRKGVTRRSFLKFCSLAATSLGLGAGMTPKIAWALENKPRIPVVWIHGL WP_001058307.1
hydrogenase [Salmonella enterica] Length: 372\nNumber of predicted TMHs: 2\nExp number of AAs in TMHs: 36.51738\nExp number, first 60 AAs: 19.72849\nTotal prob of N-in: 0.96170\nPOSSIBLE N-term signal sequence\ninside 1 19\nTMhelix 20 39\noutside 40 325\nTMhelix 326 348\ninside 349 372

14698 GCF_002033685.1_ASM203368v1 Salmonella enterica Proteobacteria; Gammaproteobacteria; Enterobacterales; Enterobacteriaceae; Salmonella
MTGDNTLITSHGINRRDFMKLCAALAATMGLSSKAAAEMAESVSNPQRPPVIWIGAQECT WP_000145428.1
hydrogenase 2 small subunit [Salmonella enterica] Length: 372\nNumber of predicted TMHs: 1\nExp number of AAs in TMHs: 24.50677\nExp number, first 60 AAs: 0.2682\nTotal prob of N-in: 0.14390\noutside 1 331\nTMhelix 332 354\ninside 355 372

14699 GCF_002033655.1_ASM203365v1 Salmonella enterica Proteobacteria; Gammaproteobacteria; Enterobacterales; Enterobacteriaceae; Salmonella
MTGDNTLITSHGINRRDFMKLCAALAATMGLSSKAAAEMAESVSNPQRPPVIWIGAQECT WP_000145429.1
hydrogenase 2 small subunit [Salmonella enterica] Length: 372\nNumber of predicted TMHs: 1\nExp number of AAs in TMHs: 24.51309\nExp number, first 60 AAs: 0.2682\nTotal prob of N-in: 0.14402\noutside 1 331\nTMhelix 332 354\ninside 355 372

14700 GCF_002033595.1_ASM203359v1 Salmonella enterica Proteobacteria; Gammaproteobacteria; Enterobacterales; Enterobacteriaceae; Salmonella
MTGDNTLITSHGINRRDFMKLCAALAATMGLSSKAAAEMAESVSNPQRPPVIWIGAQECT WP_000145429.1
hydrogenase 2 small subunit [Salmonella enterica] Length: 372\nNumber of predicted TMHs: 1\nExp number of AAs in TMHs: 24.51309\nExp number, first 60 AAs: 0.2682\nTotal prob of N-in: 0.14402\noutside 1 331\nTMhelix 332 354\ninside 355 372

14701 GCF_002033565.1_ASM203356v1 Salmonella enterica Proteobacteria; Gammaproteobacteria; Enterobacterales; Enterobacteriaceae; Salmonella
MTGDNTLITSHGINRRDFMKLCAALAATMGLSSKAAAEMAESVSNPQRPPVIWIGAQECT WP_000145429.1
hydrogenase 2 small subunit [Salmonella enterica] Length: 372\nNumber of predicted TMHs: 1\nExp number of AAs in TMHs: 24.51309\nExp number, first 60 AAs: 0.2682\nTotal prob of N-in: 0.14402\noutside 1 331\nTMhelix 332 354\ninside 355 372

14702 GCF_002033445.1_ASM203344v1 Salmonella enterica Proteobacteria; Gammaproteobacteria; Enterobacterales; Enterobacteriaceae; Salmonella
MNNEETFYQAMRRKGVTRRSFLKFCSLAATSLGLGAGMTPKIAWALENKPRIPVVWIHGL WP_001058307.1
hydrogenase [Salmonella enterica] Length: 372\nNumber of predicted TMHs: 2\nExp number of AAs in TMHs: 36.51738\nExp number, first 60 AAs: 19.72849\nTotal prob of N-in: 0.96170\nPOSSIBLE N-term signal sequence\ninside 1 19\nTMhelix 20 39\noutside 40 325\nTMhelix 326 348\ninside 349 372

14703 GCF_002033525.1_ASM203352v1 Salmonella enterica Proteobacteria; Gammaproteobacteria; Enterobacterales; Enterobacteriaceae; Salmonella
MNNEETFYQAMRRKGVTRRSFLKFCSLAATSLGLGAGMTPKIAWALENKPRIPVVWIHGL WP_001058307.1
hydrogenase [Salmonella enterica] Length: 372\nNumber of predicted TMHs: 2\nExp number of AAs in TMHs: 36.51738\nExp number, first 60 AAs: 19.72849\nTotal prob of N-in: 0.96170\nPOSSIBLE N-term signal sequence\ninside 1 19\nTMhelix 20 39\noutside 40 325\nTMhelix 326 348\ninside 349 372

14704 GCF_002033415.1_ASM203341v1 Salmonella enterica Proteobacteria; Gammaproteobacteria; Enterobacterales; Enterobacteriaceae; Salmonella
MTGDNTLITSHGINRRDFMKLCAALAATMGLSSKAAAEMAESVSNPQRPPVIWIGAQECT WP_000145429.1
hydrogenase 2 small subunit [Salmonella enterica] Length: 372\nNumber of predicted TMHs: 1\nExp

number of AAs in TMHs: 24.51309\nExp number, first 60 AAs: 0.2682\nTotal prob of N-in: 0.14402\noutside 1 331\nTMhelix 332 354\ninside 355 372

14705 GCF_002033335.1_ASM203333v1 Salmonella enterica Proteobacteria; Gammaproteobacteria; Enterobacterales; Enterobacteriaceae; Salmonella MTGDNTLITSHGINRRDFMKLCAALAATMGLSSKAAAEMAESVSNPQRPPVIWIGAQECT WP_000145429.1 hydrogenase 2 small subunit [Salmonella enterica] Length: 372\nNumber of predicted TMHs: 1\nExp number of AAs in TMHs: 24.51309\nExp number, first 60 AAs: 0.2682\nTotal prob of N-in: 0.14402\noutside 1 331\nTMhelix 332 354\ninside 355 372

14706 GCF_002033375.1_ASM203337v1 Salmonella enterica Proteobacteria; Gammaproteobacteria; Enterobacterales; Enterobacteriaceae; Salmonella MTGDNTLITSHGINRRDFMKLCAALAATMGLSSKAAAEMAESVSNPQRPPVIWIGAQECT WP_000145429.1 hydrogenase 2 small subunit [Salmonella enterica] Length: 372\nNumber of predicted TMHs: 1\nExp number of AAs in TMHs: 24.51309\nExp number, first 60 AAs: 0.2682\nTotal prob of N-in: 0.14402\noutside 1 331\nTMhelix 332 354\ninside 355 372

14707 GCF_002033325.1_ASM203332v1 Salmonella enterica Proteobacteria; Gammaproteobacteria; Enterobacterales; Enterobacteriaceae; Salmonella MNNEETFYQAMRRKGVTRRSFLKFCSLAATSLGLGAGMTPKIAWALENKPRIPVVWIHGL WP_001058312.1 hydrogenase [Salmonella enterica] Length: 372\nNumber of predicted TMHs: 2\nExp number of AAs in TMHs: 36.51738\nExp number, first 60 AAs: 19.72849\nTotal prob of N-in: 0.96170\nPOSSIBLE N-term signal sequence\ninside 1 19\nTMhelix 20 39\noutside 40 325\nTMhelix 326 348\ninside 349 372

14708 GCF_002033285.1_ASM203328v1 Salmonella enterica Proteobacteria; Gammaproteobacteria; Enterobacterales; Enterobacteriaceae; Salmonella MNNEETFYQAMRRKGVTRRSFLKFCSLAATSLGLGAGMTPKIAWALENKPRIPVVWIHGL WP_001058307.1 hydrogenase [Salmonella enterica] Length: 372\nNumber of predicted TMHs: 2\nExp number of AAs in TMHs: 36.51738\nExp number, first 60 AAs: 19.72849\nTotal prob of N-in: 0.96170\nPOSSIBLE N-term signal sequence\ninside 1 19\nTMhelix 20 39\noutside 40 325\nTMhelix 326 348\ninside 349 372

14709 GCF_002032855.1_ASM203285v1 Salmonella enterica Proteobacteria; Gammaproteobacteria; Enterobacterales; Enterobacteriaceae; Salmonella MTGDNTLITSHGINRRDFMKLCAALAATMGLSSKAAAEMAESVSNPQRPPVIWIGAQECT WP_000145429.1 hydrogenase 2 small subunit [Salmonella enterica] Length: 372\nNumber of predicted TMHs: 1\nExp number of AAs in TMHs: 24.51309\nExp number, first 60 AAs: 0.2682\nTotal prob of N-in: 0.14402\noutside 1 331\nTMhelix 332 354\ninside 355 372

14710 GCF_002032515.1_ASM203251v1 Salmonella enterica Proteobacteria; Gammaproteobacteria; Enterobacterales; Enterobacteriaceae; Salmonella MNNEETFYQAMRRKGVTRRSFLKFCSLAATSLGLGAGMTPKIAWALENKPRIPVVWIHGL WP_001058307.1 hydrogenase [Salmonella enterica] Length: 372\nNumber of predicted TMHs: 2\nExp number of AAs in TMHs: 36.51738\nExp number, first 60 AAs: 19.72849\nTotal prob of N-in: 0.96170\nPOSSIBLE N-term signal sequence\ninside 1 19\nTMhelix 20 39\noutside 40 325\nTMhelix 326 348\ninside 349 372

14711 GCF_002032515.1_ASM203251v1 Salmonella enterica Proteobacteria; Gammaproteobacteria; Enterobacterales; Enterobacteriaceae; Salmonella MTGDNTLITSHGINRRDFMKLCAALAATMGLSSKAAAEMAESVSNPQRPPVIWIGAQECT WP_000145429.1 hydrogenase 2 small subunit [Salmonella enterica] Length: 372\nNumber of predicted TMHs: 1\nExp number of AAs in TMHs: 24.51309\nExp number, first 60 AAs: 0.2682\nTotal prob of N-in: 0.14402\noutside 1 331\nTMhelix 332 354\ninside 355 372

14712 GCF_002032485.1_ASM203248v1 Salmonella enterica Proteobacteria; Gammaproteobacteria; Enterobacterales; Enterobacteriaceae; Salmonella MTGDNTLITSHGINRRDFMKLCAALAATMGLSSKAAAEMAESVSNPQRPPVIWIGAQECT WP_000145429.1 hydrogenase 2 small subunit [Salmonella enterica] Length: 372\nNumber of predicted TMHs: 1\nExp number of AAs in TMHs: 24.51309\nExp number, first 60 AAs: 0.2682\nTotal prob of N-in: 0.14402\noutside 1 331\nTMhelix 332 354\ninside 355 372

14713 GCF_002033445.1_ASM203344v1 Salmonella enterica Proteobacteria; Gammaproteobacteria; Enterobacterales; Enterobacteriaceae; Salmonella

MTGDNTLITSHGINRRDFMKLCAALAATMGLSSKAAAEMAESVSNPQRPPVIWIGAQECT WP_000145429.1 hydrogenase 2 small subunit [Salmonella enterica] Length: 372\nNumber of predicted TMHs: 1\nExp number of AAs in TMHs: 24.51309\nExp number, first 60 AAs: 0.2682\nTotal prob of N-in: 0.14402\noutside 1 331\nTMhelix 332 354\ninside 355 372

14714 GCF_002030895.1_ASM203089v1 Salmonella enterica Proteobacteria; Gammaproteobacteria; Enterobacterales; Enterobacteriaceae; Salmonella MNNEETFYQAMRRKGVTRRSFLKFCSLAATSLGLGAGMTPKIAWALENKPRIPVVWIHGL WP_001058308.1 hydrogenase [Salmonella enterica] Length: 372\nNumber of predicted TMHs: 2\nExp number of AAs in TMHs: 39.08204\nExp number, first 60 AAs: 19.72474\nTotal prob of N-in: 0.96174\nPOSSIBLE N-term signal sequence\ninside 1 19\nTMhelix 20 39\noutside 40 325\nTMhelix 326 348\ninside 349 372

14715 GCF_002030295.1_ASM203029v1 Salmonella enterica Proteobacteria; Gammaproteobacteria; Enterobacterales; Enterobacteriaceae; Salmonella MTGDNTLITSHGINRRDFMKLCAALAATMGLSSKAAAEMAESVSNPQRPPVIWIGAQECT WP_000145429.1 hydrogenase 2 small subunit [Salmonella enterica] Length: 372\nNumber of predicted TMHs: 1\nExp number of AAs in TMHs: 24.51309\nExp number, first 60 AAs: 0.2682\nTotal prob of N-in: 0.14402\noutside 1 331\nTMhelix 332 354\ninside 355 372

14716 GCF_002030285.1_ASM203028v1 Salmonella enterica Proteobacteria; Gammaproteobacteria; Enterobacterales; Enterobacteriaceae; Salmonella MTGDNTLITSHGINRRDFMKLCAALAATMGLSSKAAAEMAESVSNPQRPPVIWIGAQECT WP_000145429.1 hydrogenase 2 small subunit [Salmonella enterica] Length: 372\nNumber of predicted TMHs: 1\nExp number of AAs in TMHs: 24.51309\nExp number, first 60 AAs: 0.2682\nTotal prob of N-in: 0.14402\noutside 1 331\nTMhelix 332 354\ninside 355 372

14717 GCF_001275185.1_ASM127518v1 Salmonella enterica Proteobacteria; Gammaproteobacteria; Enterobacterales; Enterobacteriaceae; Salmonella MTGDNTLITSHGINRRDFMKLCAALAATMGLSSKAAAEMAESVSNPQRPPVIWIGAQECT WP_000145429.1 hydrogenase 2 small subunit [Salmonella enterica] Length: 372\nNumber of predicted TMHs: 1\nExp number of AAs in TMHs: 24.51309\nExp number, first 60 AAs: 0.2682\nTotal prob of N-in: 0.14402\noutside 1 331\nTMhelix 332 354\ninside 355 372

14718 GCF_001275215.1_ASM127521v1 Salmonella enterica Proteobacteria; Gammaproteobacteria; Enterobacterales; Enterobacteriaceae; Salmonella MNNEETFYQAMRRKGVTRRSFLKFCSLAATSLGLGAGMTPKIAWALENKPRIPVVWIHGL WP_001058312.1 hydrogenase [Salmonella enterica] Length: 372\nNumber of predicted TMHs: 2\nExp number of AAs in TMHs: 36.51738\nExp number, first 60 AAs: 19.72849\nTotal prob of N-in: 0.96170\nPOSSIBLE N-term signal sequence\ninside 1 19\nTMhelix 20 39\noutside 40 325\nTMhelix 326 348\ninside 349 372

14719 GCF_001473845.1_Salmonella_enterica_CVM_43828_v1.0Salmonella enterica Proteobacteria; Gammaproteobacteria; Enterobacterales; Enterobacteriaceae; Salmonella MTGDNTLITSHGINRRDFMKLCAALAATMGLSSKAAAEMAESVSNPQRPPVIWIGAQECT WP_000145429.1 hydrogenase 2 small subunit [Salmonella enterica] Length: 372\nNumber of predicted TMHs: 1\nExp number of AAs in TMHs: 24.51309\nExp number, first 60 AAs: 0.2682\nTotal prob of N-in: 0.14402\noutside 1 331\nTMhelix 332 354\ninside 355 372

14720 GCF_001473545.1_Salmonella_enterica_CVM_43783_v1.0Salmonella enterica Proteobacteria; Gammaproteobacteria; Enterobacterales; Enterobacteriaceae; Salmonella MTGDNTLITSHGINRRDFMKLCAALAATMGLSSKAAAEMAESVSNPQRPPVIWIGAQECT WP_000145429.1 hydrogenase 2 small subunit [Salmonella enterica] Length: 372\nNumber of predicted TMHs: 1\nExp number of AAs in TMHs: 24.51309\nExp number, first 60 AAs: 0.2682\nTotal prob of N-in: 0.14402\noutside 1 331\nTMhelix 332 354\ninside 355 372

14721 GCF_001275205.1_ASM127520v1 Salmonella enterica Proteobacteria; Gammaproteobacteria; Enterobacterales; Enterobacteriaceae; Salmonella MTGDNTLITSHGINRRDFMKLCAALAATMGLSSKAAAEMAESVSNPQRPPVIWIGAQECT WP_000145429.1 hydrogenase 2 small subunit [Salmonella enterica] Length: 372\nNumber of predicted TMHs: 1\nExp number of AAs in TMHs: 24.51309\nExp number, first 60 AAs: 0.2682\nTotal prob of N-in: 0.14402\noutside 1 331\nTMhelix 332 354\ninside 355 372

14722 GCF_001275205.1_ASM127520v1 Salmonella enterica Proteobacteria; Gammaproteobacteria; Enterobacterales; Enterobacteriaceae; Salmonella
MNNEETFYQAMRRKGVTRRSFLKFCSLAATSLGLGAGMTPKIAWALENKPRIPVVWIHGL WP_001058311.1
hydrogenase [Salmonella enterica] Length: 372\nNumber of predicted TMHs: 2\nExp number of AAs in TMHs: 37.4699899999999\nExp number, first 60 AAs: 19.72699\nTotal prob of N-in: 0.96171\nPOSSIBLE N-term signal sequence\ninside 1 19\nTMhelix 20 39\noutside 40 325\nTMhelix 326 348\ninside 349 372

14723 GCF_001473625.1_Salmonella_enterica_CVM_43790_v1.0Salmonella enterica Proteobacteria; Gammaproteobacteria; Enterobacterales; Enterobacteriaceae; Salmonella
MTGDNTLITSHGINRRDFMKLCAALAATMGLSSKAAAEMAESVSNPQRPPVIWIGAQECT WP_000145429.1
hydrogenase 2 small subunit [Salmonella enterica] Length: 372\nNumber of predicted TMHs: 1\nExp number of AAs in TMHs: 24.51309\nExp number, first 60 AAs: 0.2682\nTotal prob of N-in: 0.14402\noutside 1 331\nTMhelix 332 354\ninside 355 372

14724 GCF_001473815.1_Salmonella_enterica_CVM_43826_v1.0Salmonella enterica Proteobacteria; Gammaproteobacteria; Enterobacterales; Enterobacteriaceae; Salmonella
MNNEETFYQAMRRKGVTRRSFLKFCSLAATSLGLGAGMTPKIAWALENKPRIPVVWIHGL WP_001058311.1
hydrogenase [Salmonella enterica] Length: 372\nNumber of predicted TMHs: 2\nExp number of AAs in TMHs: 37.4699899999999\nExp number, first 60 AAs: 19.72699\nTotal prob of N-in: 0.96171\nPOSSIBLE N-term signal sequence\ninside 1 19\nTMhelix 20 39\noutside 40 325\nTMhelix 326 348\ninside 349 372

14725 GCF_001569965.1_ASM156996v1 Salmonella enterica Proteobacteria; Gammaproteobacteria; Enterobacterales; Enterobacteriaceae; Salmonella
MTGDNTLITSHGINRRDFMKLCAALAATMGLSSKAAAEMAESVSNPQRPPVIWIGAQECT WP_000145429.1
hydrogenase 2 small subunit [Salmonella enterica] Length: 372\nNumber of predicted TMHs: 1\nExp number of AAs in TMHs: 24.51309\nExp number, first 60 AAs: 0.2682\nTotal prob of N-in: 0.14402\noutside 1 331\nTMhelix 332 354\ninside 355 372

14726 GCF_001078155.1_ASM107815v1 Salmonella enterica Proteobacteria; Gammaproteobacteria; Enterobacterales; Enterobacteriaceae; Salmonella
MNNEETFYQAMRRKGVTRRSFLKFCSLAATSLGLGAGMTPKIAWALENKPRIPVVWIHGL WP_001058312.1
hydrogenase [Salmonella enterica] Length: 372\nNumber of predicted TMHs: 2\nExp number of AAs in TMHs: 36.51738\nExp number, first 60 AAs: 19.72849\nTotal prob of N-in: 0.96170\nPOSSIBLE N-term signal sequence\ninside 1 19\nTMhelix 20 39\noutside 40 325\nTMhelix 326 348\ninside 349 372

14727 GCF_001569965.1_ASM156996v1 Salmonella enterica Proteobacteria; Gammaproteobacteria; Enterobacterales; Enterobacteriaceae; Salmonella
MNNEETFYQAMRRKGVTRRSFLKFCSLAATSLGLGAGMTPKIAWALENKPRIPVIWIHGL WP_061376537.1
hydrogenase [Salmonella enterica] Length: 372\nNumber of predicted TMHs: 2\nExp number of AAs in TMHs: 36.5556\nExp number, first 60 AAs: 19.76633\nTotal prob of N-in: 0.96197\nPOSSIBLE N-term signal sequence\ninside 1 19\nTMhelix 20 39\noutside 40 325\nTMhelix 326 348\ninside 349 372

14728 GCF_001831645.1_ASM183164v1 Salmonella enterica Proteobacteria; Gammaproteobacteria; Enterobacterales; Enterobacteriaceae; Salmonella
MTGDNTLITSHGINRRDFMKLCAALAATMGLSSKAAAEMAESVSNPQRPPVIWIGAQECT WP_000145429.1
hydrogenase 2 small subunit [Salmonella enterica] Length: 372\nNumber of predicted TMHs: 1\nExp number of AAs in TMHs: 24.51309\nExp number, first 60 AAs: 0.2682\nTotal prob of N-in: 0.14402\noutside 1 331\nTMhelix 332 354\ninside 355 372

14729 GCF_001831705.1_ASM183170v1 Salmonella enterica Proteobacteria; Gammaproteobacteria; Enterobacterales; Enterobacteriaceae; Salmonella
MTGDNTLITSHGINRRDFMKLCAALAATMGLSSKAAAEMAESVSNPQRPPVIWIGAQECT WP_000145429.1
hydrogenase 2 small subunit [Salmonella enterica] Length: 372\nNumber of predicted TMHs: 1\nExp number of AAs in TMHs: 24.51309\nExp number, first 60 AAs: 0.2682\nTotal prob of N-in: 0.14402\noutside 1 331\nTMhelix 332 354\ninside 355 372

14730 GCF_001473465.1_Salmonella_enterica_CVM_43772_v1.0Salmonella enterica Proteobacteria; Gammaproteobacteria; Enterobacterales; Enterobacteriaceae; Salmonella
MTGDNTLITSHGINRRDFMKLCAALAATMGLSSKAAAEMAESVSNPQRPPVIWIGAQECT WP_000145429.1
hydrogenase 2 small subunit [Salmonella enterica] Length: 372\nNumber of predicted TMHs: 1\nExp

number of AAs in TMHs: 24.51309\nExp number, first 60 AAs: 0.2682\nTotal prob of N-in: 0.14402\noutside 1 331\nTMhelix 332 354\ninside 355 372

14731 GCF_001473485.1_Salmonella_enterica_CVM_43775_v1.0Salmonella enterica Proteobacteria; Gammaproteobacteria; Enterobacterales; Enterobacteriaceae; Salmonella MTGDNTLITSHGINRRDFMKLCAALAATMGLSSKAAAEMAESVSNPQRPPVIWIGAQECT WP_000145429.1 hydrogenase 2 small subunit [Salmonella enterica] Length: 372\nNumber of predicted TMHs: 1\nExp number of AAs in TMHs: 24.51309\nExp number, first 60 AAs: 0.2682\nTotal prob of N-in: 0.14402\noutside 1 331\nTMhelix 332 354\ninside 355 372

14732 GCF_001473725.1_Salmonella_enterica_CVM_43814_v1.0Salmonella enterica Proteobacteria; Gammaproteobacteria; Enterobacterales; Enterobacteriaceae; Salmonella MTGDNTLITSHGINRRDFMKLCAALAATMGLSSKAAAEMAESVSNPQRPPVIWIGAQECT WP_000145429.1 hydrogenase 2 small subunit [Salmonella enterica] Length: 372\nNumber of predicted TMHs: 1\nExp number of AAs in TMHs: 24.51309\nExp number, first 60 AAs: 0.2682\nTotal prob of N-in: 0.14402\noutside 1 331\nTMhelix 332 354\ninside 355 372

14733 GCF_001473665.1_Salmonella_enterica_CVM_43799_v1.0Salmonella enterica Proteobacteria; Gammaproteobacteria; Enterobacterales; Enterobacteriaceae; Salmonella MNNEETFYQAMRRKGVTRRSFLKFCSLAATSLGLGAGMTPKIAWALENKPRIPVVWIHGL WP_001058312.1 hydrogenase [Salmonella enterica] Length: 372\nNumber of predicted TMHs: 2\nExp number of AAs in TMHs: 36.51738\nExp number, first 60 AAs: 19.72849\nTotal prob of N-in: 0.96170\nPOSSIBLE N-term signal sequence\ninside 1 19\nTMhelix 20 39\noutside 40 325\nTMhelix 326 348\ninside 349 372

14734 GCF_001473665.1_Salmonella_enterica_CVM_43799_v1.0Salmonella enterica Proteobacteria; Gammaproteobacteria; Enterobacterales; Enterobacteriaceae; Salmonella MTGDNTLITSHGINRRDFMKLCAALAATMGLSSKAAAEMAESVSNPQRPPVIWIGAQECT WP_000145429.1 hydrogenase 2 small subunit [Salmonella enterica] Length: 372\nNumber of predicted TMHs: 1\nExp number of AAs in TMHs: 24.51309\nExp number, first 60 AAs: 0.2682\nTotal prob of N-in: 0.14402\noutside 1 331\nTMhelix 332 354\ninside 355 372

14735 GCF_001473845.1_Salmonella_enterica_CVM_43828_v1.0Salmonella enterica Proteobacteria; Gammaproteobacteria; Enterobacterales; Enterobacteriaceae; Salmonella MNNEETFYQAMRRKGVTRRSFLKFCSLAATSLGLGAGMTPKIAWALENKPRIPVVWIHGL WP_001058312.1 hydrogenase [Salmonella enterica] Length: 372\nNumber of predicted TMHs: 2\nExp number of AAs in TMHs: 36.51738\nExp number, first 60 AAs: 19.72849\nTotal prob of N-in: 0.96170\nPOSSIBLE N-term signal sequence\ninside 1 19\nTMhelix 20 39\noutside 40 325\nTMhelix 326 348\ninside 349 372

14736 GCF_001473885.1_Salmonella_enterica_CVM_43836_v1.0Salmonella enterica Proteobacteria; Gammaproteobacteria; Enterobacterales; Enterobacteriaceae; Salmonella MTGDNTLITSHGINRRDFMKLCAALAATMGLSSKAAAEMAESVSNPQRPPVIWIGAQECT WP_000145429.1 hydrogenase 2 small subunit [Salmonella enterica] Length: 372\nNumber of predicted TMHs: 1\nExp number of AAs in TMHs: 24.51309\nExp number, first 60 AAs: 0.2682\nTotal prob of N-in: 0.14402\noutside 1 331\nTMhelix 332 354\ninside 355 372

14737 GCF_002033395.1_ASM203339v1 Salmonella enterica Proteobacteria; Gammaproteobacteria; Enterobacterales; Enterobacteriaceae; Salmonella MTGDNTLITSHGINRRDFMKLCAALAATMGLSSKAAAEMAESVSNPQRPPVIWIGAQECT WP_000145429.1 hydrogenase 2 small subunit [Salmonella enterica] Length: 372\nNumber of predicted TMHs: 1\nExp number of AAs in TMHs: 24.51309\nExp number, first 60 AAs: 0.2682\nTotal prob of N-in: 0.14402\noutside 1 331\nTMhelix 332 354\ninside 355 372

14738 GCF_002033585.1_ASM203358v1 Salmonella enterica Proteobacteria; Gammaproteobacteria; Enterobacterales; Enterobacteriaceae; Salmonella MTGDNTLITSHGINRRDFMKLCAALAATMGLSSKAAAEMAESVSNPQRPPVIWIGAQECT WP_000145429.1 hydrogenase 2 small subunit [Salmonella enterica] Length: 372\nNumber of predicted TMHs: 1\nExp number of AAs in TMHs: 24.51309\nExp number, first 60 AAs: 0.2682\nTotal prob of N-in: 0.14402\noutside 1 331\nTMhelix 332 354\ninside 355 372

14739 GCF_001473435.1_Salmonella_enterica_CVM_43762_v1.0Salmonella enterica Proteobacteria; Gammaproteobacteria; Enterobacterales; Enterobacteriaceae; Salmonella

MTGDNTLITSHGINRRDFMKLCAALAATMGLSSKAAAEMAESVSNPQRPPVIWIGAQECT WP_000145429.1 hydrogenase 2 small subunit [Salmonella enterica] Length: 372\nNumber of predicted TMHs: 1\nExp number of AAs in TMHs: 24.51309\nExp number, first 60 AAs: 0.2682\nTotal prob of N-in: 0.14402\noutside 1 331\nTMhelix 332 354\ninside 355 372

14740 GCF_001473885.1_Salmonella_enterica_CVM_43836_v1.0Salmonella enterica Proteobacteria; Gammaproteobacteria; Enterobacterales; Enterobacteriaceae; Salmonella MNNEETFYQAMRRKGVTRRSFLKFCSLAATSLGLGAGMTPKIAWALENKPRIPVVWIHGL WP_001058312.1 hydrogenase [Salmonella enterica] Length: 372\nNumber of predicted TMHs: 2\nExp number of AAs in TMHs: 36.51738\nExp number, first 60 AAs: 19.72849\nTotal prob of N-in: 0.96170\nPOSSIBLE N-term signal sequence\ninside 1 19\nTMhelix 20 39\noutside 40 325\nTMhelix 326 348\ninside 349 372

14741 GCF_002033455.1_ASM203345v1 Salmonella enterica Proteobacteria; Gammaproteobacteria; Enterobacterales; Enterobacteriaceae; Salmonella MTGDNTLITSHGINRRDFMKLCAALAATMGLSSKAAAEMAESVSNPQRPPVIWIGAQECT WP_000145429.1 hydrogenase 2 small subunit [Salmonella enterica] Length: 372\nNumber of predicted TMHs: 1\nExp number of AAs in TMHs: 24.51309\nExp number, first 60 AAs: 0.2682\nTotal prob of N-in: 0.14402\noutside 1 331\nTMhelix 332 354\ninside 355 372

14742 GCF_001473815.1_Salmonella_enterica_CVM_43826_v1.0Salmonella enterica Proteobacteria; Gammaproteobacteria; Enterobacterales; Enterobacteriaceae; Salmonella MTGDNTLITSHGINRRDFMKLCAALAATMGLSSKAAAEMAESVSNPQRPPVIWIGAQECT WP_000145429.1 hydrogenase 2 small subunit [Salmonella enterica] Length: 372\nNumber of predicted TMHs: 1\nExp number of AAs in TMHs: 24.51309\nExp number, first 60 AAs: 0.2682\nTotal prob of N-in: 0.14402\noutside 1 331\nTMhelix 332 354\ninside 355 372

14743 GCF_001473595.1_Salmonella_enterica_CVM_43787_v1.0Salmonella enterica Proteobacteria; Gammaproteobacteria; Enterobacterales; Enterobacteriaceae; Salmonella MNNEETFYQAMRRKGVTRRSFLKFCSLAATSLGLGAGMTPKIAWALENKPRIPVVWIHGL WP_001058312.1 hydrogenase [Salmonella enterica] Length: 372\nNumber of predicted TMHs: 2\nExp number of AAs in TMHs: 36.51738\nExp number, first 60 AAs: 19.72849\nTotal prob of N-in: 0.96170\nPOSSIBLE N-term signal sequence\ninside 1 19\nTMhelix 20 39\noutside 40 325\nTMhelix 326 348\ninside 349 372

14744 GCF_001473635.1_Salmonella_enterica_CVM_43793_v1.0Salmonella enterica Proteobacteria; Gammaproteobacteria; Enterobacterales; Enterobacteriaceae; Salmonella MTGDNTLITSHGINRRDFMKLCAALAATMGLSSKAAAEMAESVSNPQRPPVIWIGAQECT WP_000145429.1 hydrogenase 2 small subunit [Salmonella enterica] Length: 372\nNumber of predicted TMHs: 1\nExp number of AAs in TMHs: 24.51309\nExp number, first 60 AAs: 0.2682\nTotal prob of N-in: 0.14402\noutside 1 331\nTMhelix 332 354\ninside 355 372

14745 GCF_001473905.1_Salmonella_enterica_CVM_43837_v1.0Salmonella enterica Proteobacteria; Gammaproteobacteria; Enterobacterales; Enterobacteriaceae; Salmonella MNNEETFYQAMRRKGVTRRSFLKFCSLAATSLGLGAGMTPKIAWALENKPRIPVVWIHGL WP_001058307.1 hydrogenase [Salmonella enterica] Length: 372\nNumber of predicted TMHs: 2\nExp number of AAs in TMHs: 36.51738\nExp number, first 60 AAs: 19.72849\nTotal prob of N-in: 0.96170\nPOSSIBLE N-term signal sequence\ninside 1 19\nTMhelix 20 39\noutside 40 325\nTMhelix 326 348\ninside 349 372

14746 GCF_002033625.1_ASM203362v1 Salmonella enterica Proteobacteria; Gammaproteobacteria; Enterobacterales; Enterobacteriaceae; Salmonella MTGDNTLITSHGINRRDFMKLCAALAATMGLSSKAAAEMAESVSNPQRPPVIWIGAQECT WP_000145429.1 hydrogenase 2 small subunit [Salmonella enterica] Length: 372\nNumber of predicted TMHs: 1\nExp number of AAs in TMHs: 24.51309\nExp number, first 60 AAs: 0.2682\nTotal prob of N-in: 0.14402\noutside 1 331\nTMhelix 332 354\ninside 355 372

14747 GCF_001570345.1_ASM157034v1 Salmonella enterica Proteobacteria; Gammaproteobacteria; Enterobacterales; Enterobacteriaceae; Salmonella MTGDNSLINSSGINRRDFMKLCAALAATMGLSSKAAAEMAESVSRPQRPPVIWIGAQECT WP_003024486.1 MULTISPECIES: hydrogenase 2 small subunit [Enterobacteriaceae] Length: 372\nNumber of predicted TMHs: 1\nExp number of AAs in TMHs: 27.14621\nExp number, first 60 AAs: 0.22683\nTotal prob of N-in: 0.23982\noutside 1 331\nTMhelix 332 354\ninside 355 372

14748 GCF_001473685.1_Salmonella_enterica_CVM_43804_v1.0Salmonella enterica Proteobacteria; Gammaproteobacteria; Enterobacterales; Enterobacteriaceae; Salmonella
MTGDNTLITSHGINRRDFMKLCAALAATMGLSSKAAAEMAESVSNPQRPPVIWIGAQECT WP_000145429.1
hydrogenase 2 small subunit [Salmonella enterica] Length: 372\nNumber of predicted TMHs: 1\nExp number of AAs in TMHs: 24.51309\nExp number, first 60 AAs: 0.2682\nTotal prob of N-in: 0.14402\noutside 1 331\nTMhelix 332 354\ninside 355 372

14749 GCF_001475145.1_Salmonella_enterica_CVM_43844_v1.0Salmonella enterica Proteobacteria; Gammaproteobacteria; Enterobacterales; Enterobacteriaceae; Salmonella
MTGDNTLITSHGINRRDFMKLCAALAATMGLSSKAAAEMAESVSNPQRPPVIWIGAQECT WP_000145429.1
hydrogenase 2 small subunit [Salmonella enterica] Length: 372\nNumber of predicted TMHs: 1\nExp number of AAs in TMHs: 24.51309\nExp number, first 60 AAs: 0.2682\nTotal prob of N-in: 0.14402\noutside 1 331\nTMhelix 332 354\ninside 355 372

14750 GCF_001473685.1_Salmonella_enterica_CVM_43804_v1.0Salmonella enterica Proteobacteria; Gammaproteobacteria; Enterobacterales; Enterobacteriaceae; Salmonella
MNNEETFYQAMRRKGVTRRSFLKFCSLAATSLGLGAGMTPKIAWALENKPRIPVVWIHGL WP_001058312.1
hydrogenase [Salmonella enterica] Length: 372\nNumber of predicted TMHs: 2\nExp number of AAs in TMHs: 36.51738\nExp number, first 60 AAs: 19.72849\nTotal prob of N-in: 0.96170\nPOSSIBLE N-term signal sequence\ninside 1 19\nTMhelix 20 39\noutside 40 325\nTMhelix 326 348\ninside 349 372

14751 GCF_001473905.1_Salmonella_enterica_CVM_43837_v1.0Salmonella enterica Proteobacteria; Gammaproteobacteria; Enterobacterales; Enterobacteriaceae; Salmonella
MTGDNTLITSHGINRRDFMKLCAALAATMGLSSKAAAEMAESVSNPQRPPVIWIGAQECT WP_000145429.1
hydrogenase 2 small subunit [Salmonella enterica] Length: 372\nNumber of predicted TMHs: 1\nExp number of AAs in TMHs: 24.51309\nExp number, first 60 AAs: 0.2682\nTotal prob of N-in: 0.14402\noutside 1 331\nTMhelix 332 354\ninside 355 372

14752 GCF_001474165.1_Salmonella_enterica_CVM_43758_v1.0Salmonella enterica Proteobacteria; Gammaproteobacteria; Enterobacterales; Enterobacteriaceae; Salmonella
MTGDNTLITSHGINRRDFMKLCAALAATMGLSSKAAAEMAESVSNPQRPPVIWIGAQECT WP_000145429.1
hydrogenase 2 small subunit [Salmonella enterica] Length: 372\nNumber of predicted TMHs: 1\nExp number of AAs in TMHs: 24.51309\nExp number, first 60 AAs: 0.2682\nTotal prob of N-in: 0.14402\noutside 1 331\nTMhelix 332 354\ninside 355 372

14753 GCF_001473915.1_Salmonella_enterica_CVM_43839_v1.0Salmonella enterica Proteobacteria; Gammaproteobacteria; Enterobacterales; Enterobacteriaceae; Salmonella
MNNEETFYQAMRRKGVTRRSFLKFCSLAATSLGLGAGMTPKIAWALENKPRIPVVWIHGL WP_001058312.1
hydrogenase [Salmonella enterica] Length: 372\nNumber of predicted TMHs: 2\nExp number of AAs in TMHs: 36.51738\nExp number, first 60 AAs: 19.72849\nTotal prob of N-in: 0.96170\nPOSSIBLE N-term signal sequence\ninside 1 19\nTMhelix 20 39\noutside 40 325\nTMhelix 326 348\ninside 349 372

14754 GCF_001473865.1_Salmonella_enterica_CVM_43832_v1.0Salmonella enterica Proteobacteria; Gammaproteobacteria; Enterobacterales; Enterobacteriaceae; Salmonella
MNNEETFYQAMRRKGVTRRSFLKFCSLAATSLGLGAGMTPKIAWALENKPRIPVVWIHGL WP_001058312.1
hydrogenase [Salmonella enterica] Length: 372\nNumber of predicted TMHs: 2\nExp number of AAs in TMHs: 36.51738\nExp number, first 60 AAs: 19.72849\nTotal prob of N-in: 0.96170\nPOSSIBLE N-term signal sequence\ninside 1 19\nTMhelix 20 39\noutside 40 325\nTMhelix 326 348\ninside 349 372

14755 GCF_002033725.1_ASM203372v1 Salmonella enterica Proteobacteria; Gammaproteobacteria; Enterobacterales; Enterobacteriaceae; Salmonella
MTGDNTLITSHGINRRDFMKLCAALAATMGLSSKAAAEMAESVSNPQRPPVIWIGAQECT WP_000145429.1
hydrogenase 2 small subunit [Salmonella enterica] Length: 372\nNumber of predicted TMHs: 1\nExp number of AAs in TMHs: 24.51309\nExp number, first 60 AAs: 0.2682\nTotal prob of N-in: 0.14402\noutside 1 331\nTMhelix 332 354\ninside 355 372

14756 GCF_001474775.1_Salmonella_enterica_CVM_43808_v1.0Salmonella enterica Proteobacteria; Gammaproteobacteria; Enterobacterales; Enterobacteriaceae; Salmonella
MTGDNTLITSHGINRRDFMKLCAALAATMGLSSKAAAEMAESVSNPQRPPVIWIGAQECT WP_000145429.1
hydrogenase 2 small subunit [Salmonella enterica] Length: 372\nNumber of predicted TMHs: 1\nExp

number of AAs in TMHs: 24.51309\nExp number, first 60 AAs: 0.2682\nTotal prob of N-in: 0.14402\noutside 1 331\nTMhelix 332 354\ninside 355 372

14757 GCF_001473705.1_Salmonella_enterica_CVM_43807_v1.0Salmonella enterica Proteobacteria; Gammaproteobacteria; Enterobacterales; Enterobacteriaceae; Salmonella MNNEETFYQAMRRKGVTRRSFLKFCSLAATSLGLGAGMTPKIAWALENKPRIPVVWIHGL WP_001058312.1 hydrogenase [Salmonella enterica] Length: 372\nNumber of predicted TMHs: 2\nExp number of AAs in TMHs: 36.51738\nExp number, first 60 AAs: 19.72849\nTotal prob of N-in: 0.96170\nPOSSIBLE N-term signal sequence\ninside 1 19\nTMhelix 20 39\noutside 40 325\nTMhelix 326 348\ninside 349 372

14758 GCF_001475075.1_Salmonella_enterica_CVM_43834_v1.0Salmonella enterica Proteobacteria; Gammaproteobacteria; Enterobacterales; Enterobacteriaceae; Salmonella MTGDNTLITSHGINRRDFMKLCAALAATMGLSSKAAAEMAESVSNPQRPPVIWIGAQECT WP_000145429.1 hydrogenase 2 small subunit [Salmonella enterica] Length: 372\nNumber of predicted TMHs: 1\nExp number of AAs in TMHs: 24.51309\nExp number, first 60 AAs: 0.2682\nTotal prob of N-in: 0.14402\noutside 1 331\nTMhelix 332 354\ninside 355 372

14759 GCF_001473705.1_Salmonella_enterica_CVM_43807_v1.0Salmonella enterica Proteobacteria; Gammaproteobacteria; Enterobacterales; Enterobacteriaceae; Salmonella MTGDNTLITSHGINRRDFMKLCAALAATMGLSSKAAAEMAESVSNPQRPPVIWIGAQECT WP_000145429.1 hydrogenase 2 small subunit [Salmonella enterica] Length: 372\nNumber of predicted TMHs: 1\nExp number of AAs in TMHs: 24.51309\nExp number, first 60 AAs: 0.2682\nTotal prob of N-in: 0.14402\noutside 1 331\nTMhelix 332 354\ninside 355 372

14760 GCF_001475045.1_Salmonella_enterica_CVM_43833_v1.0Salmonella enterica Proteobacteria; Gammaproteobacteria; Enterobacterales; Enterobacteriaceae; Salmonella MTGDNTLITSHGINRRDFMKLCAALAATMGLSSKAAAEMAESVSNPQRPPVIWIGAQECT WP_000145429.1 hydrogenase 2 small subunit [Salmonella enterica] Length: 372\nNumber of predicted TMHs: 1\nExp number of AAs in TMHs: 24.51309\nExp number, first 60 AAs: 0.2682\nTotal prob of N-in: 0.14402\noutside 1 331\nTMhelix 332 354\ninside 355 372

14761 GCF_001473635.1_Salmonella_enterica_CVM_43793_v1.0Salmonella enterica Proteobacteria; Gammaproteobacteria; Enterobacterales; Enterobacteriaceae; Salmonella MNNEETFYQAMRRKGVTRRSFLKFCSLAATSLGLGAGMTPKIAWALENKPRIPVVWIHGL WP_001058312.1 hydrogenase [Salmonella enterica] Length: 372\nNumber of predicted TMHs: 2\nExp number of AAs in TMHs: 36.51738\nExp number, first 60 AAs: 19.72849\nTotal prob of N-in: 0.96170\nPOSSIBLE N-term signal sequence\ninside 1 19\nTMhelix 20 39\noutside 40 325\nTMhelix 326 348\ninside 349 372

14762 GCF_001473915.1_Salmonella_enterica_CVM_43839_v1.0Salmonella enterica Proteobacteria; Gammaproteobacteria; Enterobacterales; Enterobacteriaceae; Salmonella MTGDNTLITSHGINRRDFMKLCAALAATMGLSSKAAAEMAESVSNPQRPPVIWIGAQECT WP_000145429.1 hydrogenase 2 small subunit [Salmonella enterica] Length: 372\nNumber of predicted TMHs: 1\nExp number of AAs in TMHs: 24.51309\nExp number, first 60 AAs: 0.2682\nTotal prob of N-in: 0.14402\noutside 1 331\nTMhelix 332 354\ninside 355 372

14763 GCF_001473625.1_Salmonella_enterica_CVM_43790_v1.0Salmonella enterica Proteobacteria; Gammaproteobacteria; Enterobacterales; Enterobacteriaceae; Salmonella MNNEETFYQAMRRKGVTRRSFLKFCSLAATSLGLGAGMTPKIAWALENKPRIPVVWIHGL WP_001058312.1 hydrogenase [Salmonella enterica] Length: 372\nNumber of predicted TMHs: 2\nExp number of AAs in TMHs: 36.51738\nExp number, first 60 AAs: 19.72849\nTotal prob of N-in: 0.96170\nPOSSIBLE N-term signal sequence\ninside 1 19\nTMhelix 20 39\noutside 40 325\nTMhelix 326 348\ninside 349 372

14764 GCF_001474505.1_Salmonella_enterica_CVM_43782_v1.0Salmonella enterica Proteobacteria; Gammaproteobacteria; Enterobacterales; Enterobacteriaceae; Salmonella MTGDNTLITSHGINRRDFMKLCAALAATMGLSSKAAAEMAESVSNPQRPPVIWIGAQECT WP_000145429.1 hydrogenase 2 small subunit [Salmonella enterica] Length: 372\nNumber of predicted TMHs: 1\nExp number of AAs in TMHs: 24.51309\nExp number, first 60 AAs: 0.2682\nTotal prob of N-in: 0.14402\noutside 1 331\nTMhelix 332 354\ninside 355 372

14765 GCF_001473725.1_Salmonella_enterica_CVM_43814_v1.0Salmonella enterica Proteobacteria; Gammaproteobacteria; Enterobacterales; Enterobacteriaceae; Salmonella

MNNEETFYQAMRRKGVTRRSFLKFCSLAATSLGLSAGMTPKIAWALENKPRIPVVWIHGL WP_001058316.1 hydrogenase [Salmonella enterica] Length: 372\nNumber of predicted TMHs: 2\nExp number of AAs in TMHs: 35.76397\nExp number, first 60 AAs: 18.97201\nTotal prob of N-in: 0.92785\nPOSSIBLE N-term signal sequence\ninside 1 19\nTMhelix 20 39\noutside 40 325\nTMhelix 326 348\ninside 349 372

14766 GCF_001474185.1_Salmonella_enterica_CVM_43760_v1.0Salmonella enterica Proteobacteria; Gammaproteobacteria; Enterobacterales; Enterobacteriaceae; Salmonella MTGDNTLITSHGINRRDFMKLCAALAATMGLSSKAAAEMAESVSNPQRPPVIWIGAQECT WP_000145429.1 hydrogenase 2 small subunit [Salmonella enterica] Length: 372\nNumber of predicted TMHs: 1\nExp number of AAs in TMHs: 24.51309\nExp number, first 60 AAs: 0.2682\nTotal prob of N-in: 0.14402\noutside 1 331\nTMhelix 332 354\ninside 355 372

14767 GCF_001474805.1_Salmonella_enterica_CVM_43809_v1.0Salmonella enterica Proteobacteria; Gammaproteobacteria; Enterobacterales; Enterobacteriaceae; Salmonella MTGDNTLITSHGINRRDFMKLCAALAATMGLSSKAAAEMAESVSNPQRPPVIWIGAQECT WP_000145429.1 hydrogenase 2 small subunit [Salmonella enterica] Length: 372\nNumber of predicted TMHs: 1\nExp number of AAs in TMHs: 24.51309\nExp number, first 60 AAs: 0.2682\nTotal prob of N-in: 0.14402\noutside 1 331\nTMhelix 332 354\ninside 355 372

14768 GCF_001474565.1_Salmonella_enterica_CVM_43789_v1.0Salmonella enterica Proteobacteria; Gammaproteobacteria; Enterobacterales; Enterobacteriaceae; Salmonella MTGDNTLITSHGINRRDFMKLCAALAATMGLSSKAAAEMAESVSNPQRPPVIWIGAQECT WP_000145429.1 hydrogenase 2 small subunit [Salmonella enterica] Length: 372\nNumber of predicted TMHs: 1\nExp number of AAs in TMHs: 24.51309\nExp number, first 60 AAs: 0.2682\nTotal prob of N-in: 0.14402\noutside 1 331\nTMhelix 332 354\ninside 355 372

14769 GCF_002033485.1_ASM203348v1 Salmonella enterica Proteobacteria; Gammaproteobacteria; Enterobacterales; Enterobacteriaceae; Salmonella MTGDNTLITSHGINRRDFMKLCAALAATMGLSSKAAAEMAESVSNPQRPPVIWIGAQECT WP_000145429.1 hydrogenase 2 small subunit [Salmonella enterica] Length: 372\nNumber of predicted TMHs: 1\nExp number of AAs in TMHs: 24.51309\nExp number, first 60 AAs: 0.2682\nTotal prob of N-in: 0.14402\noutside 1 331\nTMhelix 332 354\ninside 355 372

14770 GCF_001473865.1_Salmonella_enterica_CVM_43832_v1.0Salmonella enterica Proteobacteria; Gammaproteobacteria; Enterobacterales; Enterobacteriaceae; Salmonella MTGDNTLITSHGINRRDFMKLCAALAATMGLSSKAAAEMAESVSNPQRPPVIWIGAQECT WP_000145429.1 hydrogenase 2 small subunit [Salmonella enterica] Length: 372\nNumber of predicted TMHs: 1\nExp number of AAs in TMHs: 24.51309\nExp number, first 60 AAs: 0.2682\nTotal prob of N-in: 0.14402\noutside 1 331\nTMhelix 332 354\ninside 355 372

14771 GCF_002144435.1_ASM214443v1 Salmonella enterica Proteobacteria; Gammaproteobacteria; Enterobacterales; Enterobacteriaceae; Salmonella MTGDNTLITSHGINRRDFMKLCAALAATMGLSSKAAAEMAESVSNPQRPPVIWIGAQECT WP_000145429.1 hydrogenase 2 small subunit [Salmonella enterica] Length: 372\nNumber of predicted TMHs: 1\nExp number of AAs in TMHs: 24.51309\nExp number, first 60 AAs: 0.2682\nTotal prob of N-in: 0.14402\noutside 1 331\nTMhelix 332 354\ninside 355 372

14772 GCF_002066725.1_ASM206672v1 Salmonella enterica Proteobacteria; Gammaproteobacteria; Enterobacterales; Enterobacteriaceae; Salmonella MNNEETFYQAMRRKGVTRRSFLKFCSLAATSLGLGAGMTPKIAWALENKPRIPVVWIHGL WP_001058312.1 hydrogenase [Salmonella enterica] Length: 372\nNumber of predicted TMHs: 2\nExp number of AAs in TMHs: 36.51738\nExp number, first 60 AAs: 19.72849\nTotal prob of N-in: 0.96170\nPOSSIBLE N-term signal sequence\ninside 1 19\nTMhelix 20 39\noutside 40 325\nTMhelix 326 348\ninside 349 372

14773 GCF_002066685.1_ASM206668v1 Salmonella enterica Proteobacteria; Gammaproteobacteria; Enterobacterales; Enterobacteriaceae; Salmonella MNNEETFYQAMRRKGVTRRSFLKFCSLAATSLGLSAGMTPKIAWALENKPRIPVVWIHGL WP_001058316.1 hydrogenase [Salmonella enterica] Length: 372\nNumber of predicted TMHs: 2\nExp number of AAs in TMHs: 35.76397\nExp number, first 60 AAs: 18.97201\nTotal prob of N-in: 0.92785\nPOSSIBLE N-term signal sequence\ninside 1 19\nTMhelix 20 39\noutside 40 325\nTMhelix 326 348\ninside 349 372

14774 GCF_002066925.1_ASM206692v1 Salmonella enterica Proteobacteria; Gammaproteobacteria; Enterobacterales; Enterobacteriaceae; Salmonella
MTGDNTLITSHGINRRDFMKLCAALAATMGLSSKAAAEMAESVSNPQRPPVIWIGAQECT WP_080160933.1
hydrogenase 2 small subunit [Salmonella enterica] Length: 372\nNumber of predicted TMHs: 1\nExp number of AAs in TMHs: 24.51279\nExp number, first 60 AAs: 0.26821\nTotal prob of N-in: 0.14402\noutside 1 331\nTMhelix 332 354\ninside 355 372

14775 GCF_002066725.1_ASM206672v1 Salmonella enterica Proteobacteria; Gammaproteobacteria; Enterobacterales; Enterobacteriaceae; Salmonella
MTGDNTLITSHGINRRDFMKLCAALAATMGLSSKAAAEMAESVSNPQRPPVIWIGAQECT WP_000145429.1
hydrogenase 2 small subunit [Salmonella enterica] Length: 372\nNumber of predicted TMHs: 1\nExp number of AAs in TMHs: 24.51309\nExp number, first 60 AAs: 0.2682\nTotal prob of N-in: 0.14402\noutside 1 331\nTMhelix 332 354\ninside 355 372

14776 GCF_002066685.1_ASM206668v1 Salmonella enterica Proteobacteria; Gammaproteobacteria; Enterobacterales; Enterobacteriaceae; Salmonella
MTGDNTLITSHGINRRDFMKLCAALAATMGLSSKAAAEMAESVSNPQRPPVIWIGAQECT WP_000145429.1
hydrogenase 2 small subunit [Salmonella enterica] Length: 372\nNumber of predicted TMHs: 1\nExp number of AAs in TMHs: 24.51309\nExp number, first 60 AAs: 0.2682\nTotal prob of N-in: 0.14402\noutside 1 331\nTMhelix 332 354\ninside 355 372

14777 GCF_002144395.1_ASM214439v1 Salmonella enterica Proteobacteria; Gammaproteobacteria; Enterobacterales; Enterobacteriaceae; Salmonella
MTGDNTLITSHGINRRDFMKLCAALAATMGLSSKAAAEMAESVSNPQRPPVIWIGAQECT WP_000145429.1
hydrogenase 2 small subunit [Salmonella enterica] Length: 372\nNumber of predicted TMHs: 1\nExp number of AAs in TMHs: 24.51309\nExp number, first 60 AAs: 0.2682\nTotal prob of N-in: 0.14402\noutside 1 331\nTMhelix 332 354\ninside 355 372

14778 GCF_002066455.1_ASM206645v1 Salmonella enterica Proteobacteria; Gammaproteobacteria; Enterobacterales; Enterobacteriaceae; Salmonella
MNNEETFYQAMRRKGVTRRSFLKFCSLAATSLGLGAGMTPKIAWALENKPRIPVVWIHGL WP_001058307.1
hydrogenase [Salmonella enterica] Length: 372\nNumber of predicted TMHs: 2\nExp number of AAs in TMHs: 36.51738\nExp number, first 60 AAs: 19.72849\nTotal prob of N-in: 0.96170\nPOSSIBLE N-term signal sequence\ninside 1 19\nTMhelix 20 39\noutside 40 325\nTMhelix 326 348\ninside 349 372

14779 GCF_001569905.1_ASM156990v1 Salmonella enterica Proteobacteria; Gammaproteobacteria; Enterobacterales; Enterobacteriaceae; Salmonella
MTGDNTLITSHGINRRDFMKLCAALAATMGLSSKAAAEMAESVSNPQRPPVIWIGAQECT WP_000145429.1
hydrogenase 2 small subunit [Salmonella enterica] Length: 372\nNumber of predicted TMHs: 1\nExp number of AAs in TMHs: 24.51309\nExp number, first 60 AAs: 0.2682\nTotal prob of N-in: 0.14402\noutside 1 331\nTMhelix 332 354\ninside 355 372

14780 GCF_002066455.1_ASM206645v1 Salmonella enterica Proteobacteria; Gammaproteobacteria; Enterobacterales; Enterobacteriaceae; Salmonella
MTGDNTLITSHGINRRDFMKLCAALAATMGLSSKAAAEMAESVSNPQRPPVIWIGAQECT WP_000145429.1
hydrogenase 2 small subunit [Salmonella enterica] Length: 372\nNumber of predicted TMHs: 1\nExp number of AAs in TMHs: 24.51309\nExp number, first 60 AAs: 0.2682\nTotal prob of N-in: 0.14402\noutside 1 331\nTMhelix 332 354\ninside 355 372

14781 GCF_002066195.1_ASM206619v1 Salmonella enterica Proteobacteria; Gammaproteobacteria; Enterobacterales; Enterobacteriaceae; Salmonella
MNNEETFYQAMRRKGVTRRSFLKFCSLAATSLGLGAGMTPKIAWALENKPRIPVVWIHGL WP_023243720.1
hydrogenase 1, small subunit [Salmonella enterica] Length: 372\nNumber of predicted TMHs: 2\nExp number of AAs in TMHs: 36.52147\nExp number, first 60 AAs: 19.72838\nTotal prob of N-in: 0.96170\nPOSSIBLE N-term signal sequence\ninside 1 19\nTMhelix 20 39\noutside 40 325\nTMhelix 326 348\ninside 349 372

14782 GCF_002066395.1_ASM206639v1 Salmonella enterica Proteobacteria; Gammaproteobacteria; Enterobacterales; Enterobacteriaceae; Salmonella
MNNEETFYQAMRRKGVTRRSFLKFCSLAATSLGLGAGMTPKIAWALENKPRIPVVWIHGL WP_001058307.1
hydrogenase [Salmonella enterica] Length: 372\nNumber of predicted TMHs: 2\nExp number of AAs in

TMHs: 36.51738\nExp number, first 60 AAs: 19.72849\nTotal prob of N-in: 0.96170\nPOSSIBLE N-term signal sequence\ninside 1 19\nTMhelix 20 39\noutside 40 325\nTMhelix 326 348\ninside 349 372

14783 GCF_002066375.1_ASM206637v1 Salmonella enterica Proteobacteria; Gammaproteobacteria; Enterobacterales; Enterobacteriaceae; Salmonella
MNNEETFYQAMRRKGVTRRSFLKFCSLAATSLGLGAGMTPKIAWALENKPRIPVVWIHGL WP_001058307.1
hydrogenase [Salmonella enterica] Length: 372\nNumber of predicted TMHs: 2\nExp number of AAs in TMHs: 36.51738\nExp number, first 60 AAs: 19.72849\nTotal prob of N-in: 0.96170\nPOSSIBLE N-term signal sequence\ninside 1 19\nTMhelix 20 39\noutside 40 325\nTMhelix 326 348\ninside 349 372

14784 GCF_002066335.1_ASM206633v1 Salmonella enterica Proteobacteria; Gammaproteobacteria; Enterobacterales; Enterobacteriaceae; Salmonella
MNNEETFYQAMRRKGVTRRSFLKFCSLAATSLGLGAGMTPKIAWALENKPRIPVVWIHGL WP_001058307.1
hydrogenase [Salmonella enterica] Length: 372\nNumber of predicted TMHs: 2\nExp number of AAs in TMHs: 36.51738\nExp number, first 60 AAs: 19.72849\nTotal prob of N-in: 0.96170\nPOSSIBLE N-term signal sequence\ninside 1 19\nTMhelix 20 39\noutside 40 325\nTMhelix 326 348\ninside 349 372

14785 GCF_002066395.1_ASM206639v1 Salmonella enterica Proteobacteria; Gammaproteobacteria; Enterobacterales; Enterobacteriaceae; Salmonella
MTGDNTLITSHGINRRDFMKLCAALAATMGLSSKAAAEMAESVSNPQRPPVIWIGAQECT WP_000145429.1
hydrogenase 2 small subunit [Salmonella enterica] Length: 372\nNumber of predicted TMHs: 1\nExp number of AAs in TMHs: 24.51309\nExp number, first 60 AAs: 0.2682\nTotal prob of N-in: 0.14402\noutside 1 331\nTMhelix 332 354\ninside 355 372

14786 GCF_002066375.1_ASM206637v1 Salmonella enterica Proteobacteria; Gammaproteobacteria; Enterobacterales; Enterobacteriaceae; Salmonella
MTGDNTLITSHGINRRDFMKLCAALAATMGLSSKAAAEMAESVSNPQRPPVIWIGAQECT WP_000145429.1
hydrogenase 2 small subunit [Salmonella enterica] Length: 372\nNumber of predicted TMHs: 1\nExp number of AAs in TMHs: 24.51309\nExp number, first 60 AAs: 0.2682\nTotal prob of N-in: 0.14402\noutside 1 331\nTMhelix 332 354\ninside 355 372

14787 GCF_002066335.1_ASM206633v1 Salmonella enterica Proteobacteria; Gammaproteobacteria; Enterobacterales; Enterobacteriaceae; Salmonella
MTGDNTLITSHGINRRDFMKLCAALAATMGLSSKAAAEMAESVSNPQRPPVIWIGAQECT WP_000145429.1
hydrogenase 2 small subunit [Salmonella enterica] Length: 372\nNumber of predicted TMHs: 1\nExp number of AAs in TMHs: 24.51309\nExp number, first 60 AAs: 0.2682\nTotal prob of N-in: 0.14402\noutside 1 331\nTMhelix 332 354\ninside 355 372

14788 GCF_002065955.1_ASM206595v1 Salmonella enterica Proteobacteria; Gammaproteobacteria; Enterobacterales; Enterobacteriaceae; Salmonella
MNNEETFYQAMRRKGVTRRSFLKFCSLAATSLGLGAGMTPKIAWALENKPRIPVVWIHGL WP_001540183.1
hydrogenase [Salmonella enterica] Length: 372\nNumber of predicted TMHs: 2\nExp number of AAs in TMHs: 37.41804\nExp number, first 60 AAs: 19.72808\nTotal prob of N-in: 0.96169\nPOSSIBLE N-term signal sequence\ninside 1 19\nTMhelix 20 39\noutside 40 325\nTMhelix 326 348\ninside 349 372

14789 GCF_002066105.1_ASM206610v1 Salmonella enterica Proteobacteria; Gammaproteobacteria; Enterobacterales; Enterobacteriaceae; Salmonella
MNNEETFYQAMRRKGVTRRSFLKFCSLAATSLGLGAGMTPKIAWALENKPRIPVVWIHGL WP_001058310.1
hydrogenase [Salmonella enterica] Length: 372\nNumber of predicted TMHs: 2\nExp number of AAs in TMHs: 37.4567299999999\nExp number, first 60 AAs: 19.72694\nTotal prob of N-in: 0.96171\nPOSSIBLE N-term signal sequence\ninside 1 19\nTMhelix 20 39\noutside 40 325\nTMhelix 326 348\ninside 349 372

14790 GCF_002065945.1_ASM206594v1 Salmonella enterica Proteobacteria; Gammaproteobacteria; Enterobacterales; Enterobacteriaceae; Salmonella
MTGDNTLITSHCINRRDFMKLCAALAATMGLSSKAAAEMAESVSNPQRPPVIWIGAQECT WP_079970246.1
hydrogenase 2 small subunit [Salmonella enterica] Length: 372\nNumber of predicted TMHs: 1\nExp number of AAs in TMHs: 24.58192\nExp number, first 60 AAs: 0.34172\nTotal prob of N-in: 0.14721\noutside 1 331\nTMhelix 332 354\ninside 355 372

14791 GCF_002066195.1_ASM206619v1 Salmonella enterica Proteobacteria; Gammaproteobacteria; Enterobacterales; Enterobacteriaceae; Salmonella

MTGDNTLITSHGINRRDFMKLCAALAATMGLSSKAAAEMAESVSNPQRPPVIWIGAQECT WP_000145429.1 hydrogenase 2 small subunit [Salmonella enterica] Length: 372\nNumber of predicted TMHs: 1\nExp number of AAs in TMHs: 24.51309\nExp number, first 60 AAs: 0.2682\nTotal prob of N-in: 0.14402\noutside 1 331\nTMhelix 332 354\ninside 355 372

14792 GCF_002066045.1_ASM206604v1 Salmonella enterica Proteobacteria; Gammaproteobacteria; Enterobacterales; Enterobacteriaceae; Salmonella MNNEETFYQAMRRKGVTRRSFLKFCSLAATSLGLGAGMTPKIAWALENKPRIPVVWIHGL WP_001058312.1 hydrogenase [Salmonella enterica] Length: 372\nNumber of predicted TMHs: 2\nExp number of AAs in TMHs: 36.51738\nExp number, first 60 AAs: 19.72849\nTotal prob of N-in: 0.96170\nPOSSIBLE N-term signal sequence\ninside 1 19\nTMhelix 20 39\noutside 40 325\nTMhelix 326 348\ninside 349 372

14793 GCF_002065945.1_ASM206594v1 Salmonella enterica Proteobacteria; Gammaproteobacteria; Enterobacterales; Enterobacteriaceae; Salmonella MNNEETFYQAMRRKGVTRRSFLKFCSLAATSLGLGAGMTPKIAWALENKPRIPVVWIHGL WP_023234992.1 hydrogenase 1, small subunit [Salmonella enterica] Length: 372\nNumber of predicted TMHs: 2\nExp number of AAs in TMHs: 37.41309\nExp number, first 60 AAs: 19.72865\nTotal prob of N-in: 0.96169\nPOSSIBLE N-term signal sequence\ninside 1 19\nTMhelix 20 39\noutside 40 325\nTMhelix 326 348\ninside 349 372

14794 GCF_002066105.1_ASM206610v1 Salmonella enterica Proteobacteria; Gammaproteobacteria; Enterobacterales; Enterobacteriaceae; Salmonella MTGDNTLITSHGINRRDFMKLCAALAATMGLSSKAAAEMAESVSNPQRPPVIWIGAQECT WP_000145429.1 hydrogenase 2 small subunit [Salmonella enterica] Length: 372\nNumber of predicted TMHs: 1\nExp number of AAs in TMHs: 24.51309\nExp number, first 60 AAs: 0.2682\nTotal prob of N-in: 0.14402\noutside 1 331\nTMhelix 332 354\ninside 355 372

14795 GCF_002066045.1_ASM206604v1 Salmonella enterica Proteobacteria; Gammaproteobacteria; Enterobacterales; Enterobacteriaceae; Salmonella MTGDNTLITSHGINRRDFMKLCAALAATMGLSSKAAAEMAESVSNPQRPPVIWIGAQECT WP_000145429.1 hydrogenase 2 small subunit [Salmonella enterica] Length: 372\nNumber of predicted TMHs: 1\nExp number of AAs in TMHs: 24.51309\nExp number, first 60 AAs: 0.2682\nTotal prob of N-in: 0.14402\noutside 1 331\nTMhelix 332 354\ninside 355 372

14796 GCF_002065955.1_ASM206595v1 Salmonella enterica Proteobacteria; Gammaproteobacteria; Enterobacterales; Enterobacteriaceae; Salmonella MTGDNTLITSHGINRRDFMKLCAALAATMGLSSKAAAEMAESVSNPQRPPVIWIGAQECT WP_000145429.1 hydrogenase 2 small subunit [Salmonella enterica] Length: 372\nNumber of predicted TMHs: 1\nExp number of AAs in TMHs: 24.51309\nExp number, first 60 AAs: 0.2682\nTotal prob of N-in: 0.14402\noutside 1 331\nTMhelix 332 354\ninside 355 372

14797 GCF_002144365.1_ASM214436v1 Salmonella enterica Proteobacteria; Gammaproteobacteria; Enterobacterales; Enterobacteriaceae; Salmonella MNNEETFYQAMRRKGVTRRSFLKFCSLAATSLGLGAGMTPKIAWALENKPRIPVVWIHGL WP_001058312.1 hydrogenase [Salmonella enterica] Length: 372\nNumber of predicted TMHs: 2\nExp number of AAs in TMHs: 36.51738\nExp number, first 60 AAs: 19.72849\nTotal prob of N-in: 0.96170\nPOSSIBLE N-term signal sequence\ninside 1 19\nTMhelix 20 39\noutside 40 325\nTMhelix 326 348\ninside 349 372

14798 GCF_002066925.1_ASM206692v1 Salmonella enterica Proteobacteria; Gammaproteobacteria; Enterobacterales; Enterobacteriaceae; Salmonella MNNEETFYQAMRRKGVTRRSFLKFCSLAATSLGLGAGMTPKIAWALENKPRIPVVWIHGL WP_001058306.1 hydrogenase [Salmonella enterica] Length: 372\nNumber of predicted TMHs: 2\nExp number of AAs in TMHs: 36.5540099999999\nExp number, first 60 AAs: 19.72732\nTotal prob of N-in: 0.96171\nPOSSIBLE N-term signal sequence\ninside 1 19\nTMhelix 20 39\noutside 40 325\nTMhelix 326 348\ninside 349 372

14799 GCF_002209205.1_ASM220920v1 Salmonella enterica Proteobacteria; Gammaproteobacteria; Enterobacterales; Enterobacteriaceae; Salmonella MTGDNTLITSHGINRRDFMKLCAALAATMGLSSKAAAEMAESVSNPQRPPVIWIGAQECT WP_000145429.1 hydrogenase 2 small subunit [Salmonella enterica] Length: 372\nNumber of predicted TMHs: 1\nExp number of AAs in TMHs: 24.51309\nExp number, first 60 AAs: 0.2682\nTotal prob of N-in: 0.14402\noutside 1 331\nTMhelix 332 354\ninside 355 372

14800	GCF_002144475.1_ASM214447v1	Salmonella enterica	Proteobacteria; Gammaproteobacteria; Enterobacterales; Enterobacteriaceae; Salmonella
	MNNEETFYQAMRRKGVTRRSFLKFCSLAATSLGLGAGMTPKIAWALENKPRIPVVWIHGL	WP_001058312.1
	hydrogenase [Salmonella enterica]	Length: 372\nNumber of predicted TMHs: 2\nExp number of AAs in TMHs: 36.51738\nExp number, first 60 AAs: 19.72849\nTotal prob of N-in: 0.96170\nPOSSIBLE N-term signal sequence\ninside 1 19\nTMhelix 20 39\noutside 40 325\nTMhelix 326 348\ninside 349 372

14801	GCF_002144485.1_ASM214448v1	Salmonella enterica	Proteobacteria; Gammaproteobacteria; Enterobacterales; Enterobacteriaceae; Salmonella
	MNNEETFYQAMRRKGVTRRSFLKFCSLAATSLGLGAGMTPKIAWALENKPRIPVVWIHGL	WP_001058312.1
	hydrogenase [Salmonella enterica]	Length: 372\nNumber of predicted TMHs: 2\nExp number of AAs in TMHs: 36.51738\nExp number, first 60 AAs: 19.72849\nTotal prob of N-in: 0.96170\nPOSSIBLE N-term signal sequence\ninside 1 19\nTMhelix 20 39\noutside 40 325\nTMhelix 326 348\ninside 349 372

14802	GCF_002209205.1_ASM220920v1	Salmonella enterica	Proteobacteria; Gammaproteobacteria; Enterobacterales; Enterobacteriaceae; Salmonella
	MNNEETFYQAMRRKGVTRRSFLKFCSLAATSLGLGAGMTPKIAWALENKPRIPVVWIHGL	WP_001058308.1
	hydrogenase [Salmonella enterica]	Length: 372\nNumber of predicted TMHs: 2\nExp number of AAs in TMHs: 39.08204\nExp number, first 60 AAs: 19.72474\nTotal prob of N-in: 0.96174\nPOSSIBLE N-term signal sequence\ninside 1 19\nTMhelix 20 39\noutside 40 325\nTMhelix 326 348\ninside 349 372

14803	GCF_001473765.1_Salmonella_enterica_CVM_43819_v1.0Salmonella enterica	Proteobacteria; Gammaproteobacteria; Enterobacterales; Enterobacteriaceae; Salmonella
	MNNEETFYQAMRRKGVTRRSFLKFCSLAATSLGLGAGMTPKIAWALENKPRIPVVWIHGL	WP_001058307.1
	hydrogenase [Salmonella enterica]	Length: 372\nNumber of predicted TMHs: 2\nExp number of AAs in TMHs: 36.51738\nExp number, first 60 AAs: 19.72849\nTotal prob of N-in: 0.96170\nPOSSIBLE N-term signal sequence\ninside 1 19\nTMhelix 20 39\noutside 40 325\nTMhelix 326 348\ninside 349 372

14804	GCF_002064005.1_ASM206400v1	Salmonella enterica	Proteobacteria; Gammaproteobacteria; Enterobacterales; Enterobacteriaceae; Salmonella
	MTGDNTLITSHGINRRDFMKLCAALAATMGLSSKAAAEMAESVSNPQRPPVIWIGAQECT	WP_000145429.1
	hydrogenase 2 small subunit [Salmonella enterica]	Length: 372\nNumber of predicted TMHs: 1\nExp number of AAs in TMHs: 24.51309\nExp number, first 60 AAs: 0.2682\nTotal prob of N-in: 0.14402\noutside 1 331\nTMhelix 332 354\ninside 355 372

14805	GCF_002064005.1_ASM206400v1	Salmonella enterica	Proteobacteria; Gammaproteobacteria; Enterobacterales; Enterobacteriaceae; Salmonella
	MNNEETFYQAMRRKGVTRRSFLKFCSLAATSLGLGAGMTPKIAWALENKPRIPVVWIHGL	WP_001058311.1
	hydrogenase [Salmonella enterica]	Length: 372\nNumber of predicted TMHs: 2\nExp number of AAs in TMHs: 37.4699899999999\nExp number, first 60 AAs: 19.72699\nTotal prob of N-in: 0.96171\nPOSSIBLE N-term signal sequence\ninside 1 19\nTMhelix 20 39\noutside 40 325\nTMhelix 326 348\ninside 349 372

14806	GCF_001473765.1_Salmonella_enterica_CVM_43819_v1.0Salmonella enterica	Proteobacteria; Gammaproteobacteria; Enterobacterales; Enterobacteriaceae; Salmonella
	MTGDNTLITSHGINRRDFMKLCAALAATMGLSSKAAAEMAESVSNPQRPPVIWIGAQECT	WP_000145429.1
	hydrogenase 2 small subunit [Salmonella enterica]	Length: 372\nNumber of predicted TMHs: 1\nExp number of AAs in TMHs: 24.51309\nExp number, first 60 AAs: 0.2682\nTotal prob of N-in: 0.14402\noutside 1 331\nTMhelix 332 354\ninside 355 372

14807	GCF_002061825.1_ASM206182v1	Salmonella enterica	Proteobacteria; Gammaproteobacteria; Enterobacterales; Enterobacteriaceae; Salmonella
	MNNEETFYQAMRRKGVTRRSFLKFCSLAATSLGLGAGMTPKIAWALENKPRIPVVWIHGL	WP_001058311.1
	hydrogenase [Salmonella enterica]	Length: 372\nNumber of predicted TMHs: 2\nExp number of AAs in TMHs: 37.4699899999999\nExp number, first 60 AAs: 19.72699\nTotal prob of N-in: 0.96171\nPOSSIBLE N-term signal sequence\ninside 1 19\nTMhelix 20 39\noutside 40 325\nTMhelix 326 348\ninside 349 372

14808	GCF_002061825.1_ASM206182v1	Salmonella enterica	Proteobacteria; Gammaproteobacteria; Enterobacterales; Enterobacteriaceae; Salmonella
	MTGDNTLITSHGINRRDFMKLCAALAATMGLSSKAAAEMAESVSNPQRPPVIWIGAQECT	WP_000145429.1
	hydrogenase 2 small subunit [Salmonella enterica]	Length: 372\nNumber of predicted TMHs: 1\nExp

number of AAs in TMHs: 24.51309\nExp number, first 60 AAs: 0.2682\nTotal prob of N-in: 0.14402\noutside 1 331\nTMhelix 332 354\ninside 355 372

14809 GCF_002043435.1_ASM204343v1 Salmonella enterica Proteobacteria; Gammaproteobacteria; Enterobacterales; Enterobacteriaceae; Salmonella
MTGDNTLITSHGINRRDFMKLCAALAATMGLSSKAAAEMAESVSNPQRPPVIWIGAQECT WP_000145429.1
hydrogenase 2 small subunit [Salmonella enterica] Length: 372\nNumber of predicted TMHs: 1\nExp number of AAs in TMHs: 24.51309\nExp number, first 60 AAs: 0.2682\nTotal prob of N-in: 0.14402\noutside 1 331\nTMhelix 332 354\ninside 355 372

14810 GCF_001569565.1_ASM156956v1 Salmonella enterica Proteobacteria; Gammaproteobacteria; Enterobacterales; Enterobacteriaceae; Salmonella
MTGDNTLITSHGINRRDFMKLCAALAATMGLSSKAAAEMAESVSNPQRPPVIWIGAQECT WP_000145429.1
hydrogenase 2 small subunit [Salmonella enterica] Length: 372\nNumber of predicted TMHs: 1\nExp number of AAs in TMHs: 24.51309\nExp number, first 60 AAs: 0.2682\nTotal prob of N-in: 0.14402\noutside 1 331\nTMhelix 332 354\ninside 355 372

14811 GCF_002036605.1_ASM203660v1 Salmonella enterica Proteobacteria; Gammaproteobacteria; Enterobacterales; Enterobacteriaceae; Salmonella
MTGDNTLITSHGINRRDFMKLCAALAATMGLSSKAAAEMAESVSNPQRPPVIWIGAQECT WP_000145429.1
hydrogenase 2 small subunit [Salmonella enterica] Length: 372\nNumber of predicted TMHs: 1\nExp number of AAs in TMHs: 24.51309\nExp number, first 60 AAs: 0.2682\nTotal prob of N-in: 0.14402\noutside 1 331\nTMhelix 332 354\ninside 355 372

14812 GCF_002054925.1_ASM205492v1 Salmonella enterica Proteobacteria; Gammaproteobacteria; Enterobacterales; Enterobacteriaceae; Salmonella
MTGDNTLITSHGINRRDFMKLCAALAATMGLSSKAAAEMAESVSNPQRPPVIWIGAQECT WP_000145429.1
hydrogenase 2 small subunit [Salmonella enterica] Length: 372\nNumber of predicted TMHs: 1\nExp number of AAs in TMHs: 24.51309\nExp number, first 60 AAs: 0.2682\nTotal prob of N-in: 0.14402\noutside 1 331\nTMhelix 332 354\ninside 355 372

14813 GCF_001443185.1_ASM144318v1 Salmonella enterica Proteobacteria; Gammaproteobacteria; Enterobacterales; Enterobacteriaceae; Salmonella
MNNEETFYQAMRRKGVTRRSFLKFCSLAATSLGLGAGMTPKIAWALENKPRIPVVWIHGL WP_001058311.1
hydrogenase [Salmonella enterica] Length: 372\nNumber of predicted TMHs: 2\nExp number of AAs in TMHs: 37.4699899999999\nExp number, first 60 AAs: 19.72699\nTotal prob of N-in: 0.96171\nPOSSIBLE N-term signal sequence\ninside 1 19\nTMhelix 20 39\noutside 40 325\nTMhelix 326 348\ninside 349 372

14814 GCF_002047195.1_ASM204719v1 Salmonella enterica Proteobacteria; Gammaproteobacteria; Enterobacterales; Enterobacteriaceae; Salmonella
MTGDNTLITSHGINRRDFMKLCAALAATMGLSSKAAAEMAESVSNPQRPPVIWIGAQECT WP_000145429.1
hydrogenase 2 small subunit [Salmonella enterica] Length: 372\nNumber of predicted TMHs: 1\nExp number of AAs in TMHs: 24.51309\nExp number, first 60 AAs: 0.2682\nTotal prob of N-in: 0.14402\noutside 1 331\nTMhelix 332 354\ninside 355 372

14815 GCF_002055075.1_ASM205507v1 Salmonella enterica Proteobacteria; Gammaproteobacteria; Enterobacterales; Enterobacteriaceae; Salmonella
MNNEETFYQAMRRKGVTRRSFLKFCSLAATSLGLGAGMTPKIAWALENKPRIPVVWIHGL WP_001058310.1
hydrogenase [Salmonella enterica] Length: 372\nNumber of predicted TMHs: 2\nExp number of AAs in TMHs: 37.4567299999999\nExp number, first 60 AAs: 19.72694\nTotal prob of N-in: 0.96171\nPOSSIBLE N-term signal sequence\ninside 1 19\nTMhelix 20 39\noutside 40 325\nTMhelix 326 348\ninside 349 372

14816 GCF_002055135.1_ASM205513v1 Salmonella enterica Proteobacteria; Gammaproteobacteria; Enterobacterales; Enterobacteriaceae; Salmonella
MNNEETFYQAMRRKGVTRRSFLKFCSLAATSLGLGAGMTPKIAWALENKPRIPVVWIHGL WP_001058308.1
hydrogenase [Salmonella enterica] Length: 372\nNumber of predicted TMHs: 2\nExp number of AAs in TMHs: 39.08204\nExp number, first 60 AAs: 19.72474\nTotal prob of N-in: 0.96174\nPOSSIBLE N-term signal sequence\ninside 1 19\nTMhelix 20 39\noutside 40 325\nTMhelix 326 348\ninside 349 372

14817 GCF_002055275.1_ASM205527v1 Salmonella enterica Proteobacteria; Gammaproteobacteria; Enterobacterales; Enterobacteriaceae; Salmonella

MNNEETFYQAMRRKGVTRRSFLKFCSLAATSLGLGAGMTPKIAWALENKPRIPVVWIHGL WP_001058311.1 hydrogenase [Salmonella enterica] Length: 372\nNumber of predicted TMHs: 2\nExp number of AAs in TMHs: 37.4699899999999\nExp number, first 60 AAs: 19.72699\nTotal prob of N-in: 0.96171\nPOSSIBLE N-term signal sequence\ninside 1 19\nTMhelix 20 39\noutside 40 325\nTMhelix 326 348\ninside 349 372

14818 GCF_002055135.1_ASM205513v1 Salmonella enterica Proteobacteria; Gammaproteobacteria; Enterobacterales; Enterobacteriaceae; Salmonella MTGDNTLITSHGINRRDFMKLCAALAATMGLSSKAAAEMAESVSNPQRPPVIWIGAQECT WP_000145429.1 hydrogenase 2 small subunit [Salmonella enterica] Length: 372\nNumber of predicted TMHs: 1\nExp number of AAs in TMHs: 24.51309\nExp number, first 60 AAs: 0.2682\nTotal prob of N-in: 0.14402\noutside 1 331\nTMhelix 332 354\ninside 355 372

14819 GCF_002055275.1_ASM205527v1 Salmonella enterica Proteobacteria; Gammaproteobacteria; Enterobacterales; Enterobacteriaceae; Salmonella MTGDNTLITSHGINRRDFMKLCAALAATMGLSSKAAAEMAESVSNPQRPPVIWIGAQECT WP_000145429.1 hydrogenase 2 small subunit [Salmonella enterica] Length: 372\nNumber of predicted TMHs: 1\nExp number of AAs in TMHs: 24.51309\nExp number, first 60 AAs: 0.2682\nTotal prob of N-in: 0.14402\noutside 1 331\nTMhelix 332 354\ninside 355 372

14820 GCF_002055035.1_ASM205503v1 Salmonella enterica Proteobacteria; Gammaproteobacteria; Enterobacterales; Enterobacteriaceae; Salmonella MNNEETFYQAMRRKGVTRRSFLKFCSLAATSLGLGAGMTPKIAWALENKPRIPVVWIHGL WP_001058312.1 hydrogenase [Salmonella enterica] Length: 372\nNumber of predicted TMHs: 2\nExp number of AAs in TMHs: 36.51738\nExp number, first 60 AAs: 19.72849\nTotal prob of N-in: 0.96170\nPOSSIBLE N-term signal sequence\ninside 1 19\nTMhelix 20 39\noutside 40 325\nTMhelix 326 348\ninside 349 372

14821 GCF_002055235.1_ASM205523v1 Salmonella enterica Proteobacteria; Gammaproteobacteria; Enterobacterales; Enterobacteriaceae; Salmonella MNNEETFYQAMRRKGVTRRSFLKFCSLAATSLGLGAGMTPKIAWALENKPRIPVVWIHGL WP_001058312.1 hydrogenase [Salmonella enterica] Length: 372\nNumber of predicted TMHs: 2\nExp number of AAs in TMHs: 36.51738\nExp number, first 60 AAs: 19.72849\nTotal prob of N-in: 0.96170\nPOSSIBLE N-term signal sequence\ninside 1 19\nTMhelix 20 39\noutside 40 325\nTMhelix 326 348\ninside 349 372

14822 GCF_002054865.1_ASM205486v1 Salmonella enterica Proteobacteria; Gammaproteobacteria; Enterobacterales; Enterobacteriaceae; Salmonella MTGDNTLITSHGINRRDFMKLCAALAATMGLSSKAAAEMAESVSNPQRPPVIWIGAQECT WP_000145429.1 hydrogenase 2 small subunit [Salmonella enterica] Length: 372\nNumber of predicted TMHs: 1\nExp number of AAs in TMHs: 24.51309\nExp number, first 60 AAs: 0.2682\nTotal prob of N-in: 0.14402\noutside 1 331\nTMhelix 332 354\ninside 355 372

14823 GCF_002054555.1_ASM205455v1 Salmonella enterica Proteobacteria; Gammaproteobacteria; Enterobacterales; Enterobacteriaceae; Salmonella MNNEETFYQAMRRKGVTRRSFLKFCSLAATSLGLGAGMTPKIAWALENKPRIPVVWIHGL WP_001058308.1 hydrogenase [Salmonella enterica] Length: 372\nNumber of predicted TMHs: 2\nExp number of AAs in TMHs: 39.08204\nExp number, first 60 AAs: 19.72474\nTotal prob of N-in: 0.96174\nPOSSIBLE N-term signal sequence\ninside 1 19\nTMhelix 20 39\noutside 40 325\nTMhelix 326 348\ninside 349 372

14824 GCF_002054805.1_ASM205480v1 Salmonella enterica Proteobacteria; Gammaproteobacteria; Enterobacterales; Enterobacteriaceae; Salmonella MNNEETFYQAMRRKGVTRRSFLKFCSLAATSLGLGAGMTPKIAWALENKPRIPVVWIHGL WP_001058312.1 hydrogenase [Salmonella enterica] Length: 372\nNumber of predicted TMHs: 2\nExp number of AAs in TMHs: 36.51738\nExp number, first 60 AAs: 19.72849\nTotal prob of N-in: 0.96170\nPOSSIBLE N-term signal sequence\ninside 1 19\nTMhelix 20 39\noutside 40 325\nTMhelix 326 348\ninside 349 372

14825 GCF_002054805.1_ASM205480v1 Salmonella enterica Proteobacteria; Gammaproteobacteria; Enterobacterales; Enterobacteriaceae; Salmonella MTGDNTLITSHGINRRDFMKLCAALAATMGLSSKAAAEMAESVSNPQRPPVIWIGAQECT WP_000145429.1 hydrogenase 2 small subunit [Salmonella enterica] Length: 372\nNumber of predicted TMHs: 1\nExp number of AAs in TMHs: 24.51309\nExp number, first 60 AAs: 0.2682\nTotal prob of N-in: 0.14402\noutside 1 331\nTMhelix 332 354\ninside 355 372

14826 GCF_002054535.1_ASM205453v1 Salmonella enterica Proteobacteria; Gammaproteobacteria; Enterobacterales; Enterobacteriaceae; Salmonella
MNNEETFYQAMRRKGVTRRSFLKFCSLAATSLGLGAGMTPKIAWALENKPRIPVVWIHGL WP_001058308.1
hydrogenase [Salmonella enterica] Length: 372\nNumber of predicted TMHs: 2\nExp number of AAs in TMHs: 39.08204\nExp number, first 60 AAs: 19.72474\nTotal prob of N-in: 0.96174\nPOSSIBLE N-term signal sequence\ninside 1 19\nTMhelix 20 39\noutside 40 325\nTMhelix 326 348\ninside 349 372

14827 GCF_002054635.1_ASM205463v1 Salmonella enterica Proteobacteria; Gammaproteobacteria; Enterobacterales; Enterobacteriaceae; Salmonella
MNNEETFYQAMRRKGVTRRSFLKFCSLAATSLGLGAGMTPKIAWALENKPRIPVVWIHGL WP_001058312.1
hydrogenase [Salmonella enterica] Length: 372\nNumber of predicted TMHs: 2\nExp number of AAs in TMHs: 36.51738\nExp number, first 60 AAs: 19.72849\nTotal prob of N-in: 0.96170\nPOSSIBLE N-term signal sequence\ninside 1 19\nTMhelix 20 39\noutside 40 325\nTMhelix 326 348\ninside 349 372

14828 GCF_002054535.1_ASM205453v1 Salmonella enterica Proteobacteria; Gammaproteobacteria; Enterobacterales; Enterobacteriaceae; Salmonella
MTGDNTLITSHGINRRDFMKLCAALAATMGLSSKAAAEMAESVSNPQRPPVIWIGAQECT WP_000145429.1
hydrogenase 2 small subunit [Salmonella enterica] Length: 372\nNumber of predicted TMHs: 1\nExp number of AAs in TMHs: 24.51309\nExp number, first 60 AAs: 0.2682\nTotal prob of N-in: 0.14402\noutside 1 331\nTMhelix 332 354\ninside 355 372

14829 GCF_002054435.1_ASM205443v1 Salmonella enterica Proteobacteria; Gammaproteobacteria; Enterobacterales; Enterobacteriaceae; Salmonella
MTGDNTLITSHGINRRDFMKLCAALAATMGLSSKAAAEMAESVSNPQRPPVIWIGAQECT WP_000145429.1
hydrogenase 2 small subunit [Salmonella enterica] Length: 372\nNumber of predicted TMHs: 1\nExp number of AAs in TMHs: 24.51309\nExp number, first 60 AAs: 0.2682\nTotal prob of N-in: 0.14402\noutside 1 331\nTMhelix 332 354\ninside 355 372

14830 GCF_002054465.1_ASM205446v1 Salmonella enterica Proteobacteria; Gammaproteobacteria; Enterobacterales; Enterobacteriaceae; Salmonella
MTGDNTLITSHGINRRDFMKLCAALAATMGLSSKAAAEMAESVSNPQRPPVIWIGAQECT WP_000145429.1
hydrogenase 2 small subunit [Salmonella enterica] Length: 372\nNumber of predicted TMHs: 1\nExp number of AAs in TMHs: 24.51309\nExp number, first 60 AAs: 0.2682\nTotal prob of N-in: 0.14402\noutside 1 331\nTMhelix 332 354\ninside 355 372

14831 GCF_002054385.1_ASM205438v1 Salmonella enterica Proteobacteria; Gammaproteobacteria; Enterobacterales; Enterobacteriaceae; Salmonella
MTGDNTLITSHGINRRDFMKLCAALAATMGLSSKAAAEMAESVSNPQRPPVIWIGAQECT WP_000145429.1
hydrogenase 2 small subunit [Salmonella enterica] Length: 372\nNumber of predicted TMHs: 1\nExp number of AAs in TMHs: 24.51309\nExp number, first 60 AAs: 0.2682\nTotal prob of N-in: 0.14402\noutside 1 331\nTMhelix 332 354\ninside 355 372

14832 GCF_002054325.1_ASM205432v1 Salmonella enterica Proteobacteria; Gammaproteobacteria; Enterobacterales; Enterobacteriaceae; Salmonella
MTGDNTLITSHGINRRDFMKLCAALAATMGLSSKAAAEMAESVSNPQRPPVIWIGAQECT WP_000145429.1
hydrogenase 2 small subunit [Salmonella enterica] Length: 372\nNumber of predicted TMHs: 1\nExp number of AAs in TMHs: 24.51309\nExp number, first 60 AAs: 0.2682\nTotal prob of N-in: 0.14402\noutside 1 331\nTMhelix 332 354\ninside 355 372

14833 GCF_002054375.1_ASM205437v1 Salmonella enterica Proteobacteria; Gammaproteobacteria; Enterobacterales; Enterobacteriaceae; Salmonella
MTGDNTLITSHGINRRDFMKLCAALAATMGLSSKAAAEMAESVSNPQRPPVIWIGAQECT WP_000145429.1
hydrogenase 2 small subunit [Salmonella enterica] Length: 372\nNumber of predicted TMHs: 1\nExp number of AAs in TMHs: 24.51309\nExp number, first 60 AAs: 0.2682\nTotal prob of N-in: 0.14402\noutside 1 331\nTMhelix 332 354\ninside 355 372

14834 GCF_002054295.1_ASM205429v1 Salmonella enterica Proteobacteria; Gammaproteobacteria; Enterobacterales; Enterobacteriaceae; Salmonella
MNNEETFYQAMRRKGVTRRSFLKFCSLAATSLGLGAGMTPKIAWALENKPRIPVVWIHGL WP_001058311.1
hydrogenase [Salmonella enterica] Length: 372\nNumber of predicted TMHs: 2\nExp number of AAs in

TMHs: 37.4699899999999\nExp number, first 60 AAs: 19.72699\nTotal prob of N-in: 0.96171\nPOSSIBLE N-term signal sequence\ninside 1 19\nTMhelix 20 39\noutside 40 325\nTMhelix 326 348\ninside 349 372

14835 GCF_002054085.1_ASM205408v1 Salmonella enterica Proteobacteria; Gammaproteobacteria; Enterobacterales; Enterobacteriaceae; Salmonella MNNEETFYQAMRRKGVTRRSFLKFCSLAATSLGLGAGMTPKIAWALENKPRIPVVWIHGL WP_001058312.1 hydrogenase [Salmonella enterica] Length: 372\nNumber of predicted TMHs: 2\nExp number of AAs in TMHs: 36.51738\nExp number, first 60 AAs: 19.72849\nTotal prob of N-in: 0.96170\nPOSSIBLE N-term signal sequence\ninside 1 19\nTMhelix 20 39\noutside 40 325\nTMhelix 326 348\ninside 349 372

14836 GCF_002053695.1_ASM205369v1 Salmonella enterica Proteobacteria; Gammaproteobacteria; Enterobacterales; Enterobacteriaceae; Salmonella MNNEETFYQAMRRKGVTRRSFLKFCSLAATSLGLGAGMTPKIAWALENKPRIPVVWIHGL WP_001058312.1 hydrogenase [Salmonella enterica] Length: 372\nNumber of predicted TMHs: 2\nExp number of AAs in TMHs: 36.51738\nExp number, first 60 AAs: 19.72849\nTotal prob of N-in: 0.96170\nPOSSIBLE N-term signal sequence\ninside 1 19\nTMhelix 20 39\noutside 40 325\nTMhelix 326 348\ninside 349 372

14837 GCF_002054145.1_ASM205414v1 Salmonella enterica Proteobacteria; Gammaproteobacteria; Enterobacterales; Enterobacteriaceae; Salmonella MNNEETFYQAMRRKGVTRRSFLKFCSLAATSLGLSAGMTPKIAWALENKPRIPVVWIHGL WP_001058316.1 hydrogenase [Salmonella enterica] Length: 372\nNumber of predicted TMHs: 2\nExp number of AAs in TMHs: 35.76397\nExp number, first 60 AAs: 18.97201\nTotal prob of N-in: 0.92785\nPOSSIBLE N-term signal sequence\ninside 1 19\nTMhelix 20 39\noutside 40 325\nTMhelix 326 348\ninside 349 372

14838 GCF_002054195.1_ASM205419v1 Salmonella enterica Proteobacteria; Gammaproteobacteria; Enterobacterales; Enterobacteriaceae; Salmonella MNNEETFYQAMRRKGVTRRSFLKFCSLAATSLGLGAGMTPKIAWALENKPRIPVVWIHGL WP_001058312.1 hydrogenase [Salmonella enterica] Length: 372\nNumber of predicted TMHs: 2\nExp number of AAs in TMHs: 36.51738\nExp number, first 60 AAs: 19.72849\nTotal prob of N-in: 0.96170\nPOSSIBLE N-term signal sequence\ninside 1 19\nTMhelix 20 39\noutside 40 325\nTMhelix 326 348\ninside 349 372

14839 GCF_002054145.1_ASM205414v1 Salmonella enterica Proteobacteria; Gammaproteobacteria; Enterobacterales; Enterobacteriaceae; Salmonella MTGDNTLITSHGINRRDFMKLCAALAATMGLSSKAAAEMAESVSNPQRPPVIWIGAQECT WP_000145429.1 hydrogenase 2 small subunit [Salmonella enterica] Length: 372\nNumber of predicted TMHs: 1\nExp number of AAs in TMHs: 24.51309\nExp number, first 60 AAs: 0.2682\nTotal prob of N-in: 0.14402\noutside 1 331\nTMhelix 332 354\ninside 355 372

14840 GCF_002054195.1_ASM205419v1 Salmonella enterica Proteobacteria; Gammaproteobacteria; Enterobacterales; Enterobacteriaceae; Salmonella MTGDNTLITSHGINRRDFMKLCAALAATMGLSSKAAAEMAESVSNPQRPPVIWIGAQECT WP_000145429.1 hydrogenase 2 small subunit [Salmonella enterica] Length: 372\nNumber of predicted TMHs: 1\nExp number of AAs in TMHs: 24.51309\nExp number, first 60 AAs: 0.2682\nTotal prob of N-in: 0.14402\noutside 1 331\nTMhelix 332 354\ninside 355 372

14841 GCF_002053625.1_ASM205362v1 Salmonella enterica Proteobacteria; Gammaproteobacteria; Enterobacterales; Enterobacteriaceae; Salmonella MNNEETFYQAMRRKGVTRRSFLKFCSLAATSLGLGAGMTPKIAWALENKPRIPVVWIHGL WP_001058312.1 hydrogenase [Salmonella enterica] Length: 372\nNumber of predicted TMHs: 2\nExp number of AAs in TMHs: 36.51738\nExp number, first 60 AAs: 19.72849\nTotal prob of N-in: 0.96170\nPOSSIBLE N-term signal sequence\ninside 1 19\nTMhelix 20 39\noutside 40 325\nTMhelix 326 348\ninside 349 372

14842 GCF_002053475.1_ASM205347v1 Salmonella enterica Proteobacteria; Gammaproteobacteria; Enterobacterales; Enterobacteriaceae; Salmonella MNNEETFYQAMRRKGVTRRSFLKFCSLAATSLGLGAGMTPKIAWALENKPRIPVVWIHGL WP_001058312.1 hydrogenase [Salmonella enterica] Length: 372\nNumber of predicted TMHs: 2\nExp number of AAs in TMHs: 36.51738\nExp number, first 60 AAs: 19.72849\nTotal prob of N-in: 0.96170\nPOSSIBLE N-term signal sequence\ninside 1 19\nTMhelix 20 39\noutside 40 325\nTMhelix 326 348\ninside 349 372

14843 GCF_002053495.1_ASM205349v1 Salmonella enterica Proteobacteria; Gammaproteobacteria; Enterobacterales; Enterobacteriaceae; Salmonella

MTGDNTLITSHGINRRDFMKLCAALAATMGLSSKAAAEMAESVSNPQRPPVIWIGAQECT WP_000145429.1 hydrogenase 2 small subunit [Salmonella enterica] Length: 372\nNumber of predicted TMHs: 1\nExp number of AAs in TMHs: 24.51309\nExp number, first 60 AAs: 0.2682\nTotal prob of N-in: 0.14402\noutside 1 331\nTMhelix 332 354\ninside 355 372

14844 GCF_002035125.1_ASM203512v1 Salmonella enterica Proteobacteria; Gammaproteobacteria; Enterobacterales; Enterobacteriaceae; Salmonella MTGDNTLITSHGINRRDFMKLCAALAATMGLSSKAAAEMAESVSNPQRPPVIWIGAQECT WP_000145429.1 hydrogenase 2 small subunit [Salmonella enterica] Length: 372\nNumber of predicted TMHs: 1\nExp number of AAs in TMHs: 24.51309\nExp number, first 60 AAs: 0.2682\nTotal prob of N-in: 0.14402\noutside 1 331\nTMhelix 332 354\ninside 355 372

14845 GCF_002053475.1_ASM205347v1 Salmonella enterica Proteobacteria; Gammaproteobacteria; Enterobacterales; Enterobacteriaceae; Salmonella MTGDNTLITSHGINRRDFMKLCAALAATMGLSSKAAAEMAESVSNPQRPPVIWIGAQECT WP_000145429.1 hydrogenase 2 small subunit [Salmonella enterica] Length: 372\nNumber of predicted TMHs: 1\nExp number of AAs in TMHs: 24.51309\nExp number, first 60 AAs: 0.2682\nTotal prob of N-in: 0.14402\noutside 1 331\nTMhelix 332 354\ninside 355 372

14846 GCF_001570165.1_ASM157016v1 Salmonella enterica Proteobacteria; Gammaproteobacteria; Enterobacterales; Enterobacteriaceae; Salmonella MTGDNTLITSHGINRRDFMKLCAALAATMGLSSKAAAEMAESVSNPQRPPVIWIGAQECT WP_000145429.1 hydrogenase 2 small subunit [Salmonella enterica] Length: 372\nNumber of predicted TMHs: 1\nExp number of AAs in TMHs: 24.51309\nExp number, first 60 AAs: 0.2682\nTotal prob of N-in: 0.14402\noutside 1 331\nTMhelix 332 354\ninside 355 372

14847 GCF_002052825.1_ASM205282v1 Salmonella enterica Proteobacteria; Gammaproteobacteria; Enterobacterales; Enterobacteriaceae; Salmonella MNNEETFYQAMRRKGVTRRSFLKFCSLAATSLGLGAGMTPKIAWALENKPRIPVVWIHGL WP_080076830.1 [Ni/Fe] hydrogenase small subunit [Salmonella enterica] Length: 372\nNumber of predicted TMHs: 2\nExp number of AAs in TMHs: 36.51738\nExp number, first 60 AAs: 19.72849\nTotal prob of N-in: 0.96170\nPOSSIBLE N-term signal sequence\ninside 1 19\nTMhelix 20 39\noutside 40 325\nTMhelix 326 348\ninside 349 372

14848 GCF_002052825.1_ASM205282v1 Salmonella enterica Proteobacteria; Gammaproteobacteria; Enterobacterales; Enterobacteriaceae; Salmonella MTGDNTLITSHGINRRDFMKLCAALAATMGLSSKAAAEMAESVSNPQRPPVIWIGAQECT WP_022562441.1 hydrogenase-2 small chain protein [Salmonella enterica] Length: 372\nNumber of predicted TMHs: 1\nExp number of AAs in TMHs: 24.51309\nExp number, first 60 AAs: 0.2682\nTotal prob of N-in: 0.14402\noutside 1 331\nTMhelix 332 354\ninside 355 372

14849 GCF_002053095.1_ASM205309v1 Salmonella enterica Proteobacteria; Gammaproteobacteria; Enterobacterales; Enterobacteriaceae; Salmonella MNNEETFYQAMRRKGVTRRSFLKFCSLAATSLGLGAGMTPKIAWALENKPRIPVVWIHGL WP_001058312.1 hydrogenase [Salmonella enterica] Length: 372\nNumber of predicted TMHs: 2\nExp number of AAs in TMHs: 36.51738\nExp number, first 60 AAs: 19.72849\nTotal prob of N-in: 0.96170\nPOSSIBLE N-term signal sequence\ninside 1 19\nTMhelix 20 39\noutside 40 325\nTMhelix 326 348\ninside 349 372

14850 GCF_001570165.1_ASM157016v1 Salmonella enterica Proteobacteria; Gammaproteobacteria; Enterobacterales; Enterobacteriaceae; Salmonella MNNEETFYQAMRRKGVTRRSFLKFCSLAATSLGLGAGMTPKIAWALENKPRIPVVWIHGL WP_001058307.1 hydrogenase [Salmonella enterica] Length: 372\nNumber of predicted TMHs: 2\nExp number of AAs in TMHs: 36.51738\nExp number, first 60 AAs: 19.72849\nTotal prob of N-in: 0.96170\nPOSSIBLE N-term signal sequence\ninside 1 19\nTMhelix 20 39\noutside 40 325\nTMhelix 326 348\ninside 349 372

14851 GCF_002053395.1_ASM205339v1 Salmonella enterica Proteobacteria; Gammaproteobacteria; Enterobacterales; Enterobacteriaceae; Salmonella MNNEETFYQAMRRKGVTRRSFLKFCSLAATSLGLGAGMMPKIAWALENKPRIPVVWIHGL WP_057393961.1 hydrogenase [Salmonella enterica] Length: 372\nNumber of predicted TMHs: 2\nExp number of AAs in TMHs: 37.03037\nExp number, first 60 AAs: 20.25216\nTotal prob of N-in: 0.98577\nPOSSIBLE N-term signal sequence\ninside 1 19\nTMhelix 20 39\noutside 40 325\nTMhelix 326 348\ninside 349 372

14852 GCF_002053295.1_ASM205329v1 Salmonella enterica Proteobacteria; Gammaproteobacteria; Enterobacterales; Enterobacteriaceae; Salmonella
MTGDNTLITSHGINRRDFMKLCAALAATMGLSSKAAAEMAESVSNPQRPPVIWIGAQECT WP_000145429.1
hydrogenase 2 small subunit [Salmonella enterica] Length: 372\nNumber of predicted TMHs: 1\nExp number of AAs in TMHs: 24.51309\nExp number, first 60 AAs: 0.2682\nTotal prob of N-in: 0.14402\noutside 1 331\nTMhelix 332 354\ninside 355 372

14853 GCF_002053095.1_ASM205309v1 Salmonella enterica Proteobacteria; Gammaproteobacteria; Enterobacterales; Enterobacteriaceae; Salmonella
MTGDNTLITSHGINRRDFMKLCAALAATMGLSSKAAAEMAESVSNPQRPPVIWIGAQECT WP_000145429.1
hydrogenase 2 small subunit [Salmonella enterica] Length: 372\nNumber of predicted TMHs: 1\nExp number of AAs in TMHs: 24.51309\nExp number, first 60 AAs: 0.2682\nTotal prob of N-in: 0.14402\noutside 1 331\nTMhelix 332 354\ninside 355 372

14854 GCF_002053045.1_ASM205304v1 Salmonella enterica Proteobacteria; Gammaproteobacteria; Enterobacterales; Enterobacteriaceae; Salmonella
MTGDNTLITSHGINRRDFMKLCAALAATMGLSSKAAAEMAESVSNPQRPPVIWIGAQECT WP_000145429.1
hydrogenase 2 small subunit [Salmonella enterica] Length: 372\nNumber of predicted TMHs: 1\nExp number of AAs in TMHs: 24.51309\nExp number, first 60 AAs: 0.2682\nTotal prob of N-in: 0.14402\noutside 1 331\nTMhelix 332 354\ninside 355 372

14855 GCF_002053215.1_ASM205321v1 Salmonella enterica Proteobacteria; Gammaproteobacteria; Enterobacterales; Enterobacteriaceae; Salmonella
MTGDNTLITSHGINRRDFMKLCAALAATMGLSSKAAAEMAESVSNPQRPPVIWIGAQECT WP_000145429.1
hydrogenase 2 small subunit [Salmonella enterica] Length: 372\nNumber of predicted TMHs: 1\nExp number of AAs in TMHs: 24.51309\nExp number, first 60 AAs: 0.2682\nTotal prob of N-in: 0.14402\noutside 1 331\nTMhelix 332 354\ninside 355 372

14856 GCF_002052965.1_ASM205296v1 Salmonella enterica Proteobacteria; Gammaproteobacteria; Enterobacterales; Enterobacteriaceae; Salmonella
MTGDNTLITSHGINRRDFMKLCAALAATMGLSSKAAAEMAESVSNPQRPPVIWIGAQECT WP_000145429.1
hydrogenase 2 small subunit [Salmonella enterica] Length: 372\nNumber of predicted TMHs: 1\nExp number of AAs in TMHs: 24.51309\nExp number, first 60 AAs: 0.2682\nTotal prob of N-in: 0.14402\noutside 1 331\nTMhelix 332 354\ninside 355 372

14857 GCF_002052225.1_ASM205222v1 Salmonella enterica Proteobacteria; Gammaproteobacteria; Enterobacterales; Enterobacteriaceae; Salmonella
MNNEETFYQAMRRKGVTRRSFLKFCSLAATSLGLGAGMTPKIAWALENKPRIPVVWIHGL WP_080080011.1
[Ni/Fe] hydrogenase small subunit [Salmonella enterica] Length: 372\nNumber of predicted TMHs: 2\nExp number of AAs in TMHs: 36.51717\nExp number, first 60 AAs: 19.72849\nTotal prob of N-in: 0.96169\nPOSSIBLE N-term signal sequence\ninside 1 19\nTMhelix 20 39\noutside 40 325\nTMhelix 326 348\ninside 349 372

14858 GCF_002052675.1_ASM205267v1 Salmonella enterica Proteobacteria; Gammaproteobacteria; Enterobacterales; Enterobacteriaceae; Salmonella
MNNEETFYQAMRRKGVTRRSFLKFCSLAATSLGLGAGMTPKIAWALENKPRIPVVWIHGL WP_001058312.1
hydrogenase [Salmonella enterica] Length: 372\nNumber of predicted TMHs: 2\nExp number of AAs in TMHs: 36.51738\nExp number, first 60 AAs: 19.72849\nTotal prob of N-in: 0.96170\nPOSSIBLE N-term signal sequence\ninside 1 19\nTMhelix 20 39\noutside 40 325\nTMhelix 326 348\ninside 349 372

14859 GCF_002052255.1_ASM205225v1 Salmonella enterica Proteobacteria; Gammaproteobacteria; Enterobacterales; Enterobacteriaceae; Salmonella
MNNEETFYQAMRRKGVTRRSFLKFCSLAATSLGLGAGMTPKIAWALENKPRIPVVWIHGL WP_001058311.1
hydrogenase [Salmonella enterica] Length: 372\nNumber of predicted TMHs: 2\nExp number of AAs in TMHs: 37.4699899999999\nExp number, first 60 AAs: 19.72699\nTotal prob of N-in: 0.96171\nPOSSIBLE N-term signal sequence\ninside 1 19\nTMhelix 20 39\noutside 40 325\nTMhelix 326 348\ninside 349 372

14860 GCF_002052315.1_ASM205231v1 Salmonella enterica Proteobacteria; Gammaproteobacteria; Enterobacterales; Enterobacteriaceae; Salmonella
MNNEETFYQAMRRKGVTRRSFLKFCSLAATSLGLGAGMTPKIAWALENKPRIPVVWIHGL WP_001058312.1
hydrogenase [Salmonella enterica] Length: 372\nNumber of predicted TMHs: 2\nExp number of AAs in

TMHs: 36.51738\nExp number, first 60 AAs: 19.72849\nTotal prob of N-in: 0.96170\nPOSSIBLE N-term signal sequence\ninside 1 19\nTMhelix 20 39\noutside 40 325\nTMhelix 326 348\ninside 349 372

14861 GCF_002052505.1_ASM205250v1 Salmonella enterica Proteobacteria; Gammaproteobacteria; Enterobacterales; Enterobacteriaceae; Salmonella MNNEETFYQAMRRKGVTRRSFLKFCSLAATSLGLGAGMTPKIAWALENKPRIPVVWIHGL WP_001058312.1 hydrogenase [Salmonella enterica] Length: 372\nNumber of predicted TMHs: 2\nExp number of AAs in TMHs: 36.51738\nExp number, first 60 AAs: 19.72849\nTotal prob of N-in: 0.96170\nPOSSIBLE N-term signal sequence\ninside 1 19\nTMhelix 20 39\noutside 40 325\nTMhelix 326 348\ninside 349 372

14862 GCF_002052315.1_ASM205231v1 Salmonella enterica Proteobacteria; Gammaproteobacteria; Enterobacterales; Enterobacteriaceae; Salmonella MTGDNTLITSHGINRRDFMKLCAALAATMGLSSKAAAEMAESVSNPQRPPVIWIGAQECT WP_000145429.1 hydrogenase 2 small subunit [Salmonella enterica] Length: 372\nNumber of predicted TMHs: 1\nExp number of AAs in TMHs: 24.51309\nExp number, first 60 AAs: 0.2682\nTotal prob of N-in: 0.14402\noutside 1 331\nTMhelix 332 354\ninside 355 372

14863 GCF_002052505.1_ASM205250v1 Salmonella enterica Proteobacteria; Gammaproteobacteria; Enterobacterales; Enterobacteriaceae; Salmonella MTGDNTLITSHGINRRDFMKLCAALAATMGLSSKAAAEMAESVSNPQRPPVIWIGAQECT WP_000145431.1 hydrogenase 2 small subunit [Salmonella enterica] Length: 372\nNumber of predicted TMHs: 1\nExp number of AAs in TMHs: 24.51505\nExp number, first 60 AAs: 0.2682\nTotal prob of N-in: 0.14406\noutside 1 331\nTMhelix 332 354\ninside 355 372

14864 GCF_002052005.1_ASM205200v1 Salmonella enterica Proteobacteria; Gammaproteobacteria; Enterobacterales; Enterobacteriaceae; Salmonella MNNEETFYQAMRRKGVTRRSFLKFCSLAATSLGLGAGMTPKIAWALENKPRIPVVWIHGL WP_001058312.1 hydrogenase [Salmonella enterica] Length: 372\nNumber of predicted TMHs: 2\nExp number of AAs in TMHs: 36.51738\nExp number, first 60 AAs: 19.72849\nTotal prob of N-in: 0.96170\nPOSSIBLE N-term signal sequence\ninside 1 19\nTMhelix 20 39\noutside 40 325\nTMhelix 326 348\ninside 349 372

14865 GCF_002052065.1_ASM205206v1 Salmonella enterica Proteobacteria; Gammaproteobacteria; Enterobacterales; Enterobacteriaceae; Salmonella MNNEETFYQAMRRKGVTRRSFLKFCSLAATSLGLGAGMTPKIAWALENKPRIPVVWIHGL WP_001058312.1 hydrogenase [Salmonella enterica] Length: 372\nNumber of predicted TMHs: 2\nExp number of AAs in TMHs: 36.51738\nExp number, first 60 AAs: 19.72849\nTotal prob of N-in: 0.96170\nPOSSIBLE N-term signal sequence\ninside 1 19\nTMhelix 20 39\noutside 40 325\nTMhelix 326 348\ninside 349 372

14866 GCF_002051785.1_ASM205178v1 Salmonella enterica Proteobacteria; Gammaproteobacteria; Enterobacterales; Enterobacteriaceae; Salmonella MTGDNTLITSHGINRRDFMKLCAALAATMGLSSKAAAEMAESVSNPQRPPVIWIGAQECT WP_000145429.1 hydrogenase 2 small subunit [Salmonella enterica] Length: 372\nNumber of predicted TMHs: 1\nExp number of AAs in TMHs: 24.51309\nExp number, first 60 AAs: 0.2682\nTotal prob of N-in: 0.14402\noutside 1 331\nTMhelix 332 354\ninside 355 372

14867 GCF_002051525.1_ASM205152v1 Salmonella enterica Proteobacteria; Gammaproteobacteria; Enterobacterales; Enterobacteriaceae; Salmonella MNNEETFYQAMRRKGVTRRSFLKFCSLAATSLGLGAGMTPKIAWALENKPRIPVVWIHGL WP_001058312.1 hydrogenase [Salmonella enterica] Length: 372\nNumber of predicted TMHs: 2\nExp number of AAs in TMHs: 36.51738\nExp number, first 60 AAs: 19.72849\nTotal prob of N-in: 0.96170\nPOSSIBLE N-term signal sequence\ninside 1 19\nTMhelix 20 39\noutside 40 325\nTMhelix 326 348\ninside 349 372

14868 GCF_002051475.1_ASM205147v1 Salmonella enterica Proteobacteria; Gammaproteobacteria; Enterobacterales; Enterobacteriaceae; Salmonella MNNEETFYQAMRRKGVTRRSFLKFCSLAATSLGLGAGMTPKIAWALENKPRIPVVWIHGL WP_001058312.1 hydrogenase [Salmonella enterica] Length: 372\nNumber of predicted TMHs: 2\nExp number of AAs in TMHs: 36.51738\nExp number, first 60 AAs: 19.72849\nTotal prob of N-in: 0.96170\nPOSSIBLE N-term signal sequence\ninside 1 19\nTMhelix 20 39\noutside 40 325\nTMhelix 326 348\ninside 349 372

14869 GCF_002051435.1_ASM205143v1 Salmonella enterica Proteobacteria; Gammaproteobacteria; Enterobacterales; Enterobacteriaceae; Salmonella

MNNEETFYQAMRRKGVTRRSFLKFCSLAATSLGLGAGMTPKIAWALENKPRIPVVWIHGL WP_001058312.1 hydrogenase [Salmonella enterica] Length: 372\nNumber of predicted TMHs: 2\nExp number of AAs in TMHs: 36.51738\nExp number, first 60 AAs: 19.72849\nTotal prob of N-in: 0.96170\nPOSSIBLE N-term signal sequence\ninside 1 19\nTMhelix 20 39\noutside 40 325\nTMhelix 326 348\ninside 349 372

14870 GCF_002051645.1_ASM205164v1 Salmonella enterica Proteobacteria; Gammaproteobacteria; Enterobacterales; Enterobacteriaceae; Salmonella MTGDNTLITSHGINRRDFMKLCAALAATMGLSSKAAAEMAESVSNPQRPPVIWIGAQECT WP_000145429.1 hydrogenase 2 small subunit [Salmonella enterica] Length: 372\nNumber of predicted TMHs: 1\nExp number of AAs in TMHs: 24.51309\nExp number, first 60 AAs: 0.2682\nTotal prob of N-in: 0.14402\noutside 1 331\nTMhelix 332 354\ninside 355 372

14871 GCF_002051315.1_ASM205131v1 Salmonella enterica Proteobacteria; Gammaproteobacteria; Enterobacterales; Enterobacteriaceae; Salmonella MNNEETFYQAMRRKGVTRRSFLKFCSLAATSLGLGAGMTPKIAWALENKPRIPVVWIHGL WP_001058312.1 hydrogenase [Salmonella enterica] Length: 372\nNumber of predicted TMHs: 2\nExp number of AAs in TMHs: 36.51738\nExp number, first 60 AAs: 19.72849\nTotal prob of N-in: 0.96170\nPOSSIBLE N-term signal sequence\ninside 1 19\nTMhelix 20 39\noutside 40 325\nTMhelix 326 348\ninside 349 372

14872 GCF_002051285.1_ASM205128v1 Salmonella enterica Proteobacteria; Gammaproteobacteria; Enterobacterales; Enterobacteriaceae; Salmonella MNNEETFYQAMRRKGVTRRSFLKFCSLAATSLGLGAGMTPKIAWALENKPRIPVVWIHGL WP_001058312.1 hydrogenase [Salmonella enterica] Length: 372\nNumber of predicted TMHs: 2\nExp number of AAs in TMHs: 36.51738\nExp number, first 60 AAs: 19.72849\nTotal prob of N-in: 0.96170\nPOSSIBLE N-term signal sequence\ninside 1 19\nTMhelix 20 39\noutside 40 325\nTMhelix 326 348\ninside 349 372

14873 GCF_002051225.1_ASM205122v1 Salmonella enterica Proteobacteria; Gammaproteobacteria; Enterobacterales; Enterobacteriaceae; Salmonella MNNEETFYQAMRRKGVTRRSFLKFCSLAATSLGLGAGMTPKIAWALENKPRIPVVWIHGL WP_001058312.1 hydrogenase [Salmonella enterica] Length: 372\nNumber of predicted TMHs: 2\nExp number of AAs in TMHs: 36.51738\nExp number, first 60 AAs: 19.72849\nTotal prob of N-in: 0.96170\nPOSSIBLE N-term signal sequence\ninside 1 19\nTMhelix 20 39\noutside 40 325\nTMhelix 326 348\ninside 349 372

14874 GCF_002051125.1_ASM205112v1 Salmonella enterica Proteobacteria; Gammaproteobacteria; Enterobacterales; Enterobacteriaceae; Salmonella MNNEETFYQAMRRKGVTRRSFLKFCSLAATSLGLGAGMTPKIAWALENKPRIPVVWIHGL WP_001058312.1 hydrogenase [Salmonella enterica] Length: 372\nNumber of predicted TMHs: 2\nExp number of AAs in TMHs: 36.51738\nExp number, first 60 AAs: 19.72849\nTotal prob of N-in: 0.96170\nPOSSIBLE N-term signal sequence\ninside 1 19\nTMhelix 20 39\noutside 40 325\nTMhelix 326 348\ninside 349 372

14875 GCF_002051285.1_ASM205128v1 Salmonella enterica Proteobacteria; Gammaproteobacteria; Enterobacterales; Enterobacteriaceae; Salmonella MTGDNTLITSHGINRRDFMKLCAALAATMGLSSKAAAEMAESVSNPQRPPVIWIGAQECT WP_000145429.1 hydrogenase 2 small subunit [Salmonella enterica] Length: 372\nNumber of predicted TMHs: 1\nExp number of AAs in TMHs: 24.51309\nExp number, first 60 AAs: 0.2682\nTotal prob of N-in: 0.14402\noutside 1 331\nTMhelix 332 354\ninside 355 372

14876 GCF_002051225.1_ASM205122v1 Salmonella enterica Proteobacteria; Gammaproteobacteria; Enterobacterales; Enterobacteriaceae; Salmonella MTGDNTLITSHGINRRDFMKLCAALAATMGLSSKAAAEMAESVSNPQRPPVIWIGAQECT WP_000145429.1 hydrogenase 2 small subunit [Salmonella enterica] Length: 372\nNumber of predicted TMHs: 1\nExp number of AAs in TMHs: 24.51309\nExp number, first 60 AAs: 0.2682\nTotal prob of N-in: 0.14402\noutside 1 331\nTMhelix 332 354\ninside 355 372

14877 GCF_002051095.1_ASM205109v1 Salmonella enterica Proteobacteria; Gammaproteobacteria; Enterobacterales; Enterobacteriaceae; Salmonella MNNEETFYQAMRRKGVTRRSFLKFCSLAATSLGLGAGMTPKIAWALENKPRIPVVWIHGL WP_001058312.1 hydrogenase [Salmonella enterica] Length: 372\nNumber of predicted TMHs: 2\nExp number of AAs in TMHs: 36.51738\nExp number, first 60 AAs: 19.72849\nTotal prob of N-in: 0.96170\nPOSSIBLE N-term signal sequence\ninside 1 19\nTMhelix 20 39\noutside 40 325\nTMhelix 326 348\ninside 349 372

14878 GCF_002051025.1_ASM205102v1 Salmonella enterica Proteobacteria; Gammaproteobacteria; Enterobacterales; Enterobacteriaceae; Salmonella
MNNEETFYQAMRRKGVTRRSFLKFCSLAATSLGLGAGMTPQIAWALENKPRIPVVWIHGL WP_023227615.1 hydrogenase 1, small subunit [Salmonella enterica] Length: 372\nNumber of predicted TMHs: 2\nExp number of AAs in TMHs: 39.8591699999999\nExp number, first 60 AAs: 22.13093\nTotal prob of N-in: 0.98639\nPOSSIBLE N-term signal sequence\ninside 1 20\nTMhelix 21 43\noutside 44 325\nTMhelix 326 348\ninside 349 372

14879 GCF_001833085.1_ASM183308v1 Salmonella enterica Proteobacteria; Gammaproteobacteria; Enterobacterales; Enterobacteriaceae; Salmonella
MTGDNTLITSHGINRRDFMKLCAALAATMGLSSKAAAEMAESVSNPQRPPVIWIGAQECT WP_000145429.1 hydrogenase 2 small subunit [Salmonella enterica] Length: 372\nNumber of predicted TMHs: 1\nExp number of AAs in TMHs: 24.51309\nExp number, first 60 AAs: 0.2682\nTotal prob of N-in: 0.14402\noutside 1 331\nTMhelix 332 354\ninside 355 372

14880 GCF_001473305.1_Salmonella_enterica_CVM_43756_v1.0Salmonella enterica Proteobacteria; Gammaproteobacteria; Enterobacterales; Enterobacteriaceae; Salmonella
MNNEETFYQAMRRKGVTRRSFLKFCSLAATSLGLGAGMTPKIAWALENKPRIPVVWIHGL WP_001058307.1 hydrogenase [Salmonella enterica] Length: 372\nNumber of predicted TMHs: 2\nExp number of AAs in TMHs: 36.51738\nExp number, first 60 AAs: 19.72849\nTotal prob of N-in: 0.96170\nPOSSIBLE N-term signal sequence\ninside 1 19\nTMhelix 20 39\noutside 40 325\nTMhelix 326 348\ninside 349 372

14881 GCF_001570205.1_ASM157020v1 Salmonella enterica Proteobacteria; Gammaproteobacteria; Enterobacterales; Enterobacteriaceae; Salmonella
MTGDNTLITSHGINRRDFMKLCAALAATMGLSSKAAAEMAESVSNPQRPPVIWIGAQECT WP_000145429.1 hydrogenase 2 small subunit [Salmonella enterica] Length: 372\nNumber of predicted TMHs: 1\nExp number of AAs in TMHs: 24.51309\nExp number, first 60 AAs: 0.2682\nTotal prob of N-in: 0.14402\noutside 1 331\nTMhelix 332 354\ninside 355 372

14882 GCF_001473345.1_Salmonella_enterica_CVM_43745_v1.0Salmonella enterica Proteobacteria; Gammaproteobacteria; Enterobacterales; Enterobacteriaceae; Salmonella
MTGDNTLITSHGINRRDFMKLCAALAATMGLSSKAAAEMAESVSNPQRPPVIWIGAQECT WP_000145429.1 hydrogenase 2 small subunit [Salmonella enterica] Length: 372\nNumber of predicted TMHs: 1\nExp number of AAs in TMHs: 24.51309\nExp number, first 60 AAs: 0.2682\nTotal prob of N-in: 0.14402\noutside 1 331\nTMhelix 332 354\ninside 355 372

14883 GCF_001569605.1_ASM156960v1 Salmonella enterica Proteobacteria; Gammaproteobacteria; Enterobacterales; Enterobacteriaceae; Salmonella
MNNEETFYQAMRRKGVTRRSFLKFCSLAATSLGLGAGMTPKIAWALENKPRIPVVIWIHGL WP_061376537.1 hydrogenase [Salmonella enterica] Length: 372\nNumber of predicted TMHs: 2\nExp number of AAs in TMHs: 36.5556\nExp number, first 60 AAs: 19.76633\nTotal prob of N-in: 0.96197\nPOSSIBLE N-term signal sequence\ninside 1 19\nTMhelix 20 39\noutside 40 325\nTMhelix 326 348\ninside 349 372

14884 GCF_001473315.1_Salmonella_enterica_CVM_43747_v1.0Salmonella enterica Proteobacteria; Gammaproteobacteria; Enterobacterales; Enterobacteriaceae; Salmonella
MTGDNTLITSHGINRRDFMKLCAALAATMGLSSKAAAEMAESVSNPQRPPVIWIGAQECT WP_000145429.1 hydrogenase 2 small subunit [Salmonella enterica] Length: 372\nNumber of predicted TMHs: 1\nExp number of AAs in TMHs: 24.51309\nExp number, first 60 AAs: 0.2682\nTotal prob of N-in: 0.14402\noutside 1 331\nTMhelix 332 354\ninside 355 372

14885 GCF_002051045.1_ASM205104v1 Salmonella enterica Proteobacteria; Gammaproteobacteria; Enterobacterales; Enterobacteriaceae; Salmonella
MLKFRTIFLKKIFTLKNSILARRGAVIVIVSAVFTSIMFFAHSWASDKEVAMTLSVLNSN WP_023226644.1 hypothetical protein [Salmonella enterica] Length: 204\nNumber of predicted TMHs: 2\nExp number of AAs in TMHs: 44.99526\nExp number, first 60 AAs: 22.33786\nTotal prob of N-in: 0.87083\nPOSSIBLE N-term signal sequence\ninside 1 19\nTMhelix 20 42\noutside 43 171\nTMhelix 172 194\ninside 195 204

14886 GCF_002036605.1_ASM203660v1 Salmonella enterica Proteobacteria; Gammaproteobacteria; Enterobacterales; Enterobacteriaceae; Salmonella
MNNEETFYQAMRRKGVTRRSFLKFCSLAATSLGLGAGMTPKIAWALENKPRIPVVWIHGL WP_001058312.1 hydrogenase [Salmonella enterica] Length: 372\nNumber of predicted TMHs: 2\nExp number of AAs in

TMHs: 36.51738\nExp number, first 60 AAs: 19.72849\nTotal prob of N-in: 0.96170\nPOSSIBLE N-term signal sequence\ninside 1 19\nTMhelix 20 39\noutside 40 325\nTMhelix 326 348\ninside 349 372

14887 GCF_002048925.1_ASM204892v1 Salmonella enterica Proteobacteria; Gammaproteobacteria; Enterobacterales; Enterobacteriaceae; Salmonella
MTGDNTLITSHGINRRDFMKLCAALAATMGLSSKAAAEMAESVSNPQRPPVIWIGAQECT WP_000145429.1
hydrogenase 2 small subunit [Salmonella enterica] Length: 372\nNumber of predicted TMHs: 1\nExp number of AAs in TMHs: 24.51309\nExp number, first 60 AAs: 0.2682\nTotal prob of N-in: 0.14402\noutside 1 331\nTMhelix 332 354\ninside 355 372

14888 GCF_002043145.1_ASM204314v1 Salmonella enterica Proteobacteria; Gammaproteobacteria; Enterobacterales; Enterobacteriaceae; Salmonella
MTGDNTLITSHGINRRDFMKLCAALAATMGLSSKAAAEMAESVSNPQRPPVIWIGAQECT WP_000145428.1
hydrogenase 2 small subunit [Salmonella enterica] Length: 372\nNumber of predicted TMHs: 1\nExp number of AAs in TMHs: 24.50677\nExp number, first 60 AAs: 0.2682\nTotal prob of N-in: 0.14390\noutside 1 331\nTMhelix 332 354\ninside 355 372

14889 GCF_001831995.1_ASM183199v1 Salmonella enterica Proteobacteria; Gammaproteobacteria; Enterobacterales; Enterobacteriaceae; Salmonella
MNNEETFYQAMRRKGVTRRSFLKFCSLAATSLGLGAGMTPKIAWALENKPRIPVVWIHGL WP_001058312.1
hydrogenase [Salmonella enterica] Length: 372\nNumber of predicted TMHs: 2\nExp number of AAs in TMHs: 36.51738\nExp number, first 60 AAs: 19.72849\nTotal prob of N-in: 0.96170\nPOSSIBLE N-term signal sequence\ninside 1 19\nTMhelix 20 39\noutside 40 325\nTMhelix 326 348\ninside 349 372

14890 GCF_001716225.1_ASM171622v1 Salmonella enterica Proteobacteria; Gammaproteobacteria; Enterobacterales; Enterobacteriaceae; Salmonella
MTGDNTLITSHGINRRDFMKLCAALAATMGLSSKAAAEMAESVSNPQRPPVIWIGAQECT WP_000145429.1
hydrogenase 2 small subunit [Salmonella enterica] Length: 372\nNumber of predicted TMHs: 1\nExp number of AAs in TMHs: 24.51309\nExp number, first 60 AAs: 0.2682\nTotal prob of N-in: 0.14402\noutside 1 331\nTMhelix 332 354\ninside 355 372

14891 GCF_001831755.1_ASM183175v1 Salmonella enterica Proteobacteria; Gammaproteobacteria; Enterobacterales; Enterobacteriaceae; Salmonella
MNNEETFYQAMRRKGVTRRSFLKFCSLAATSLGLGAGMTPKIAWALENKPRIPVVWIHGL WP_001058312.1
hydrogenase [Salmonella enterica] Length: 372\nNumber of predicted TMHs: 2\nExp number of AAs in TMHs: 36.51738\nExp number, first 60 AAs: 19.72849\nTotal prob of N-in: 0.96170\nPOSSIBLE N-term signal sequence\ninside 1 19\nTMhelix 20 39\noutside 40 325\nTMhelix 326 348\ninside 349 372

14892 GCF_001473785.1_Salmonella_enterica_CVM_43822_v1.0Salmonella enterica Proteobacteria; Gammaproteobacteria; Enterobacterales; Enterobacteriaceae; Salmonella
MNNEETFYQAMRRKGVTRRSFLKFCSLAATSLGLGAGMTPKIAWALENKPRIPVVWIHGL WP_001058312.1
hydrogenase [Salmonella enterica] Length: 372\nNumber of predicted TMHs: 2\nExp number of AAs in TMHs: 36.51738\nExp number, first 60 AAs: 19.72849\nTotal prob of N-in: 0.96170\nPOSSIBLE N-term signal sequence\ninside 1 19\nTMhelix 20 39\noutside 40 325\nTMhelix 326 348\ninside 349 372

14893 GCF_001570225.1_ASM157022v1 Salmonella enterica Proteobacteria; Gammaproteobacteria; Enterobacterales; Enterobacteriaceae; Salmonella
MTGDNTLITSHGINRRDFMKLCAALAATMGLSSKAAAEMAESVSNPQRPPVIWIGAQECT WP_000145429.1
hydrogenase 2 small subunit [Salmonella enterica] Length: 372\nNumber of predicted TMHs: 1\nExp number of AAs in TMHs: 24.51309\nExp number, first 60 AAs: 0.2682\nTotal prob of N-in: 0.14402\noutside 1 331\nTMhelix 332 354\ninside 355 372

14894 GCF_001831965.1_ASM183196v1 Salmonella enterica Proteobacteria; Gammaproteobacteria; Enterobacterales; Enterobacteriaceae; Salmonella
MNNEETFYQAMRRKGVTRRSFLKFCSLAATSLGLGAGMTPKIAWALENKPRIPVVWIHGL WP_001058312.1
hydrogenase [Salmonella enterica] Length: 372\nNumber of predicted TMHs: 2\nExp number of AAs in TMHs: 36.51738\nExp number, first 60 AAs: 19.72849\nTotal prob of N-in: 0.96170\nPOSSIBLE N-term signal sequence\ninside 1 19\nTMhelix 20 39\noutside 40 325\nTMhelix 326 348\ninside 349 372

14895 GCF_001831965.1_ASM183196v1 Salmonella enterica Proteobacteria; Gammaproteobacteria; Enterobacterales; Enterobacteriaceae; Salmonella

MTGDNTLITSHGINRRDFMKLCAALAATMGLSSKAAAEMAESVSNPQRPPVIWIGAQECT WP_000145429.1 hydrogenase 2 small subunit [Salmonella enterica] Length: 372\nNumber of predicted TMHs: 1\nExp number of AAs in TMHs: 24.51309\nExp number, first 60 AAs: 0.2682\nTotal prob of N-in: 0.14402\noutside 1 331\nTMhelix 332 354\ninside 355 372

14896 GCF_001570245.1_ASM157024v1 Salmonella enterica Proteobacteria; Gammaproteobacteria; Enterobacterales; Enterobacteriaceae; Salmonella MTGDNTLITSHGINRRDFMKLCAALAATMGLSSKAAAEMAESVSNPQRPPVIWIGAQECT WP_000145429.1 hydrogenase 2 small subunit [Salmonella enterica] Length: 372\nNumber of predicted TMHs: 1\nExp number of AAs in TMHs: 24.51309\nExp number, first 60 AAs: 0.2682\nTotal prob of N-in: 0.14402\noutside 1 331\nTMhelix 332 354\ninside 355 372

14897 GCF_001831705.1_ASM183170v1 Salmonella enterica Proteobacteria; Gammaproteobacteria; Enterobacterales; Enterobacteriaceae; Salmonella MNNEETFYQAMRRKGVTRRSFLKFCSLAATSLGLGAGMTPKIAWALENKPRIPVVWIHGL WP_001058312.1 hydrogenase [Salmonella enterica] Length: 372\nNumber of predicted TMHs: 2\nExp number of AAs in TMHs: 36.51738\nExp number, first 60 AAs: 19.72849\nTotal prob of N-in: 0.96170\nPOSSIBLE N-term signal sequence\ninside 1 19\nTMhelix 20 39\noutside 40 325\nTMhelix 326 348\ninside 349 372

14898 GCF_001831645.1_ASM183164v1 Salmonella enterica Proteobacteria; Gammaproteobacteria; Enterobacterales; Enterobacteriaceae; Salmonella MNNEETFYQAMRRKGVTRRSFLKFCSLAATSLGLGAGMTPKIAWALENKPRIPVVWIHGL WP_001058312.1 hydrogenase [Salmonella enterica] Length: 372\nNumber of predicted TMHs: 2\nExp number of AAs in TMHs: 36.51738\nExp number, first 60 AAs: 19.72849\nTotal prob of N-in: 0.96170\nPOSSIBLE N-term signal sequence\ninside 1 19\nTMhelix 20 39\noutside 40 325\nTMhelix 326 348\ninside 349 372

14899 GCF_002036885.1_ASM203688v1 Salmonella enterica Proteobacteria; Gammaproteobacteria; Enterobacterales; Enterobacteriaceae; Salmonella MTGDNTLITSHGINRRDFMKLCAALAATMGLSSKAAAEMAESVSNPQRPPVIWIGAQECT WP_000145429.1 hydrogenase 2 small subunit [Salmonella enterica] Length: 372\nNumber of predicted TMHs: 1\nExp number of AAs in TMHs: 24.51309\nExp number, first 60 AAs: 0.2682\nTotal prob of N-in: 0.14402\noutside 1 331\nTMhelix 332 354\ninside 355 372

14900 GCF_001831865.1_ASM183186v1 Salmonella enterica Proteobacteria; Gammaproteobacteria; Enterobacterales; Enterobacteriaceae; Salmonella MNNEETFYQAMRRKGVTRRSFLKFCSLAATSLGLGAGMTPKIAWALENKPRIPVVWIHGL WP_001058312.1 hydrogenase [Salmonella enterica] Length: 372\nNumber of predicted TMHs: 2\nExp number of AAs in TMHs: 36.51738\nExp number, first 60 AAs: 19.72849\nTotal prob of N-in: 0.96170\nPOSSIBLE N-term signal sequence\ninside 1 19\nTMhelix 20 39\noutside 40 325\nTMhelix 326 348\ninside 349 372

14901 GCF_002047705.1_ASM204770v1 Salmonella enterica Proteobacteria; Gammaproteobacteria; Enterobacterales; Enterobacteriaceae; Salmonella MTGDNTLITSHGINRRDFMKLCAALAATMGLSSKAAAEMAESVSNPQRPPVIWIGAQECT WP_000145429.1 hydrogenase 2 small subunit [Salmonella enterica] Length: 372\nNumber of predicted TMHs: 1\nExp number of AAs in TMHs: 24.51309\nExp number, first 60 AAs: 0.2682\nTotal prob of N-in: 0.14402\noutside 1 331\nTMhelix 332 354\ninside 355 372

14902 GCF_001831795.1_ASM183179v1 Salmonella enterica Proteobacteria; Gammaproteobacteria; Enterobacterales; Enterobacteriaceae; Salmonella MNNEETFYQAMRRKGVTRRSFLKFCSLAATSLGLGAGMTPKIAWALENKPRIPVVWIHGL WP_001058312.1 hydrogenase [Salmonella enterica] Length: 372\nNumber of predicted TMHs: 2\nExp number of AAs in TMHs: 36.51738\nExp number, first 60 AAs: 19.72849\nTotal prob of N-in: 0.96170\nPOSSIBLE N-term signal sequence\ninside 1 19\nTMhelix 20 39\noutside 40 325\nTMhelix 326 348\ninside 349 372

14903 GCF_001570225.1_ASM157022v1 Salmonella enterica Proteobacteria; Gammaproteobacteria; Enterobacterales; Enterobacteriaceae; Salmonella MNNEETFYQAMRRKGVTRRSFLKFCSLAATSLGLGAGMTPKIAWALENKPRIPVIWIHGL WP_061376537.1 hydrogenase [Salmonella enterica] Length: 372\nNumber of predicted TMHs: 2\nExp number of AAs in TMHs: 36.5556\nExp number, first 60 AAs: 19.76633\nTotal prob of N-in: 0.96197\nPOSSIBLE N-term signal sequence\ninside 1 19\nTMhelix 20 39\noutside 40 325\nTMhelix 326 348\ninside 349 372

14904 GCF_002046325.1_ASM204632v1 Salmonella enterica Proteobacteria; Gammaproteobacteria; Enterobacterales; Enterobacteriaceae; Salmonella
MTGDNTLITSHGINRRDFMKLCAALAATMGLSSKAAAEMAESVSNPQRPPVIWIGAQECT WP_000145429.1
hydrogenase 2 small subunit [Salmonella enterica] Length: 372\nNumber of predicted TMHs: 1\nExp number of AAs in TMHs: 24.51309\nExp number, first 60 AAs: 0.2682\nTotal prob of N-in: 0.14402\noutside 1 331\nTMhelix 332 354\ninside 355 372

14905 GCF_002046325.1_ASM204632v1 Salmonella enterica Proteobacteria; Gammaproteobacteria; Enterobacterales; Enterobacteriaceae; Salmonella
MNNEETFYQAMRRKGVTRRSFLKFCSLAATSLGLGAGMTPKIAWALENKPRIPVVWIHGL WP_001058312.1
hydrogenase [Salmonella enterica] Length: 372\nNumber of predicted TMHs: 2\nExp number of AAs in TMHs: 36.51738\nExp number, first 60 AAs: 19.72849\nTotal prob of N-in: 0.96170\nPOSSIBLE N-term signal sequence\ninside 1 19\nTMhelix 20 39\noutside 40 325\nTMhelix 326 348\ninside 349 372

14906 GCF_001443185.1_ASM144318v1 Salmonella enterica Proteobacteria; Gammaproteobacteria; Enterobacterales; Enterobacteriaceae; Salmonella
MTGDNTLITSHGINRRDFMKLCAALAATMGLSSKAAAEMAESVSNPQRPPVIWIGAQECT WP_000145429.1
hydrogenase 2 small subunit [Salmonella enterica] Length: 372\nNumber of predicted TMHs: 1\nExp number of AAs in TMHs: 24.51309\nExp number, first 60 AAs: 0.2682\nTotal prob of N-in: 0.14402\noutside 1 331\nTMhelix 332 354\ninside 355 372

14907 GCF_002047195.1_ASM204719v1 Salmonella enterica Proteobacteria; Gammaproteobacteria; Enterobacterales; Enterobacteriaceae; Salmonella
MNNEETFYQAMRRKGVTRRSFLKFCSLAATSLGLGAGMTPKIAWALENKPRIPVVWIHGL WP_023198237.1
hydrogenase 1, small subunit [Salmonella enterica] Length: 372\nNumber of predicted TMHs: 2\nExp number of AAs in TMHs: 36.39423\nExp number, first 60 AAs: 19.72877\nTotal prob of N-in: 0.96162\nPOSSIBLE N-term signal sequence\ninside 1 19\nTMhelix 20 39\noutside 40 325\nTMhelix 326 348\ninside 349 372

14908 GCF_002044165.1_ASM204416v1 Salmonella enterica Proteobacteria; Gammaproteobacteria; Enterobacterales; Enterobacteriaceae; Salmonella
MTGDNTLITSHGINRRDFMKLCAALAATMGLSSKAAAEMAESVSNPQRPPVIWIGAQECT WP_000145429.1
hydrogenase 2 small subunit [Salmonella enterica] Length: 372\nNumber of predicted TMHs: 1\nExp number of AAs in TMHs: 24.51309\nExp number, first 60 AAs: 0.2682\nTotal prob of N-in: 0.14402\noutside 1 331\nTMhelix 332 354\ninside 355 372

14909 GCF_001569565.1_ASM156956v1 Salmonella enterica Proteobacteria; Gammaproteobacteria; Enterobacterales; Enterobacteriaceae; Salmonella
MNNEETFYQAMRRKGVTRRSFLKFCSLAATSLGLGAGMTPKIAWALENKPRIPVVWIHGL WP_001058307.1
hydrogenase [Salmonella enterica] Length: 372\nNumber of predicted TMHs: 2\nExp number of AAs in TMHs: 36.51738\nExp number, first 60 AAs: 19.72849\nTotal prob of N-in: 0.96170\nPOSSIBLE N-term signal sequence\ninside 1 19\nTMhelix 20 39\noutside 40 325\nTMhelix 326 348\ninside 349 372

14910 GCF_001569605.1_ASM156960v1 Salmonella enterica Proteobacteria; Gammaproteobacteria; Enterobacterales; Enterobacteriaceae; Salmonella
MTGDNTLITSHGINRRDFMKLCAALAATMGLSSKAAAEMAESVSNPQRPPVIWIGAQECT WP_000145429.1
hydrogenase 2 small subunit [Salmonella enterica] Length: 372\nNumber of predicted TMHs: 1\nExp number of AAs in TMHs: 24.51309\nExp number, first 60 AAs: 0.2682\nTotal prob of N-in: 0.14402\noutside 1 331\nTMhelix 332 354\ninside 355 372

14911 GCF_002043205.1_ASM204320v1 Salmonella enterica Proteobacteria; Gammaproteobacteria; Enterobacterales; Enterobacteriaceae; Salmonella
MTGDNTLITSHGINRRDFMKLCAALAATMGLSSKAAAEMAESVSNPQRPPVIWIGAQECT WP_000145428.1
hydrogenase 2 small subunit [Salmonella enterica] Length: 372\nNumber of predicted TMHs: 1\nExp number of AAs in TMHs: 24.50677\nExp number, first 60 AAs: 0.2682\nTotal prob of N-in: 0.14390\noutside 1 331\nTMhelix 332 354\ninside 355 372

14912 GCF_002043715.1_ASM204371v1 Salmonella enterica Proteobacteria; Gammaproteobacteria; Enterobacterales; Enterobacteriaceae; Salmonella
MTGDNTLITSHGINRRDFMKLCAALAATMGLSSKAAAEMAESVSNPQRPPVIWIGAQECT WP_000145429.1
hydrogenase 2 small subunit [Salmonella enterica] Length: 372\nNumber of predicted TMHs: 1\nExp

number of AAs in TMHs: 24.51309\nExp number, first 60 AAs: 0.2682\nTotal prob of N-in: 0.14402\noutside 1 331\nTMhelix 332 354\ninside 355 372

14913 GCF_001569665.1_ASM156966v1 Salmonella enterica Proteobacteria; Gammaproteobacteria; Enterobacterales; Enterobacteriaceae; Salmonella
MTGDNTLITSHGINRRDFMKLCAALAATMGLSSKAAAEMAESVSNPQRPPVIWIGAQECT WP_000145429.1
hydrogenase 2 small subunit [Salmonella enterica] Length: 372\nNumber of predicted TMHs: 1\nExp number of AAs in TMHs: 24.51309\nExp number, first 60 AAs: 0.2682\nTotal prob of N-in: 0.14402\noutside 1 331\nTMhelix 332 354\ninside 355 372

14914 GCF_002054655.1_ASM205465v1 Salmonella enterica Proteobacteria; Gammaproteobacteria; Enterobacterales; Enterobacteriaceae; Salmonella
MTGDNTLITSHGINRRDFMKLCAALAATMGLSSKAAAEMAESVSNPQRPPVIWIGAQECT WP_000145429.1
hydrogenase 2 small subunit [Salmonella enterica] Length: 372\nNumber of predicted TMHs: 1\nExp number of AAs in TMHs: 24.51309\nExp number, first 60 AAs: 0.2682\nTotal prob of N-in: 0.14402\noutside 1 331\nTMhelix 332 354\ninside 355 372

14915 GCF_001569665.1_ASM156966v1 Salmonella enterica Proteobacteria; Gammaproteobacteria; Enterobacterales; Enterobacteriaceae; Salmonella
MNNEETFYQAMRRKGVTRRSFLKFCSLAATSLGLGAGMTPKIAWALENKPRIPVVWIHGL WP_001058307.1
hydrogenase [Salmonella enterica] Length: 372\nNumber of predicted TMHs: 2\nExp number of AAs in TMHs: 36.51738\nExp number, first 60 AAs: 19.72849\nTotal prob of N-in: 0.96170\nPOSSIBLE N-term signal sequence\ninside 1 19\nTMhelix 20 39\noutside 40 325\nTMhelix 326 348\ninside 349 372

14916 GCF_001569865.1_ASM156986v1 Salmonella enterica Proteobacteria; Gammaproteobacteria; Enterobacterales; Enterobacteriaceae; Salmonella
MTGDNTLITSHGINRRDFMKLCAALAATMGLSSKAAAEMAESVSNPQRPPVIWIGAQECT WP_000145429.1
hydrogenase 2 small subunit [Salmonella enterica] Length: 372\nNumber of predicted TMHs: 1\nExp number of AAs in TMHs: 24.51309\nExp number, first 60 AAs: 0.2682\nTotal prob of N-in: 0.14402\noutside 1 331\nTMhelix 332 354\ninside 355 372

14917 GCF_001569705.1_ASM156970v1 Salmonella enterica Proteobacteria; Gammaproteobacteria; Enterobacterales; Enterobacteriaceae; Salmonella
MNNEETFYQAMRRKGVTRRSFLKFCSLAATSLGLGAGMTPKIAWALENKPRIPVIWIHGL WP_061376537.1
hydrogenase [Salmonella enterica] Length: 372\nNumber of predicted TMHs: 2\nExp number of AAs in TMHs: 36.5556\nExp number, first 60 AAs: 19.76633\nTotal prob of N-in: 0.96197\nPOSSIBLE N-term signal sequence\ninside 1 19\nTMhelix 20 39\noutside 40 325\nTMhelix 326 348\ninside 349 372

14918 GCF_002054055.1_ASM205405v1 Salmonella enterica Proteobacteria; Gammaproteobacteria; Enterobacterales; Enterobacteriaceae; Salmonella
MTGDNTLITSHGINRRDFMKLCAALAATMGLSSKAAAEMAESVSNPQRPPVIWIGAQECT WP_000145429.1
hydrogenase 2 small subunit [Salmonella enterica] Length: 372\nNumber of predicted TMHs: 1\nExp number of AAs in TMHs: 24.51309\nExp number, first 60 AAs: 0.2682\nTotal prob of N-in: 0.14402\noutside 1 331\nTMhelix 332 354\ninside 355 372

14919 GCF_002043285.1_ASM204328v1 Salmonella enterica Proteobacteria; Gammaproteobacteria; Enterobacterales; Enterobacteriaceae; Salmonella
MNNEETFYQAMRRKGVTRRSFLKFCSLAATSLGLGAGMTPKIAWALENKPRIPVVWIHGL WP_023243720.1
hydrogenase 1, small subunit [Salmonella enterica] Length: 372\nNumber of predicted TMHs: 2\nExp number of AAs in TMHs: 36.52147\nExp number, first 60 AAs: 19.72838\nTotal prob of N-in: 0.96170\nPOSSIBLE N-term signal sequence\ninside 1 19\nTMhelix 20 39\noutside 40 325\nTMhelix 326 348\ninside 349 372

14920 GCF_002043955.1_ASM204395v1 Salmonella enterica Proteobacteria; Gammaproteobacteria; Enterobacterales; Enterobacteriaceae; Salmonella
MTGDNTLITSHGINRRDFMKLCAALAATMGLSSKAAAEMAESVSNPQRPPVIWIGAQECT WP_000145429.1
hydrogenase 2 small subunit [Salmonella enterica] Length: 372\nNumber of predicted TMHs: 1\nExp number of AAs in TMHs: 24.51309\nExp number, first 60 AAs: 0.2682\nTotal prob of N-in: 0.14402\noutside 1 331\nTMhelix 332 354\ninside 355 372

14921 GCF_002032545.1_ASM203254v1 Salmonella enterica Proteobacteria; Gammaproteobacteria; Enterobacterales; Enterobacteriaceae; Salmonella

MTGDNTLITSHGINRRDFMKLCAALAATMGLSSKAAAEMAESVSNPQRPPVIWIGAQECT WP_000145429.1 hydrogenase 2 small subunit [Salmonella enterica] Length: 372\nNumber of predicted TMHs: 1\nExp number of AAs in TMHs: 24.51309\nExp number, first 60 AAs: 0.2682\nTotal prob of N-in: 0.14402\noutside 1 331\nTMhelix 332 354\ninside 355 372

14922 GCF_002044025.1_ASM204402v1 Salmonella enterica Proteobacteria; Gammaproteobacteria; Enterobacterales; Enterobacteriaceae; Salmonella MTGDNTLITSHGINRRDFMKLCAALAATMGLSSKAAAEMAESVSNPQRPPVIWIGAQECT WP_000145429.1 hydrogenase 2 small subunit [Salmonella enterica] Length: 372\nNumber of predicted TMHs: 1\nExp number of AAs in TMHs: 24.51309\nExp number, first 60 AAs: 0.2682\nTotal prob of N-in: 0.14402\noutside 1 331\nTMhelix 332 354\ninside 355 372

14923 GCF_002032645.1_ASM203264v1 Salmonella enterica Proteobacteria; Gammaproteobacteria; Enterobacterales; Enterobacteriaceae; Salmonella MNNEETFYQAMRRKGVTRRSFLKFCSLAATSLGLGAGMTPKIAWALENKPRIPVVWIHGL WP_023233644.1 hydrogenase 1, small subunit [Salmonella enterica] Length: 372\nNumber of predicted TMHs: 2\nExp number of AAs in TMHs: 37.4208299999999\nExp number, first 60 AAs: 19.72732\nTotal prob of N-in: 0.96168\nPOSSIBLE N-term signal sequence\ninside 1 19\nTMhelix 20 39\noutside 40 325\nTMhelix 326 348\ninside 349 372

14924 GCF_002032545.1_ASM203254v1 Salmonella enterica Proteobacteria; Gammaproteobacteria; Enterobacterales; Enterobacteriaceae; Salmonella MNNEETFYQAMRRKGVTRRSFLKFCSLAATSLGLGAGMTPKIAWALENKPRIPVVWIHGL WP_023243720.1 hydrogenase 1, small subunit [Salmonella enterica] Length: 372\nNumber of predicted TMHs: 2\nExp number of AAs in TMHs: 36.52147\nExp number, first 60 AAs: 19.72838\nTotal prob of N-in: 0.96170\nPOSSIBLE N-term signal sequence\ninside 1 19\nTMhelix 20 39\noutside 40 325\nTMhelix 326 348\ninside 349 372

14925 GCF_002052395.1_ASM205239v1 Salmonella enterica Proteobacteria; Gammaproteobacteria; Enterobacterales; Enterobacteriaceae; Salmonella MTGDNTLITSHGINRRDFMKLCAALAATMGLSSKAAAEMAESVSNPQRPPVIWIGAQECT WP_000145429.1 hydrogenase 2 small subunit [Salmonella enterica] Length: 372\nNumber of predicted TMHs: 1\nExp number of AAs in TMHs: 24.51309\nExp number, first 60 AAs: 0.2682\nTotal prob of N-in: 0.14402\noutside 1 331\nTMhelix 332 354\ninside 355 372

14926 GCF_002043895.1_ASM204389v1 Salmonella enterica Proteobacteria; Gammaproteobacteria; Enterobacterales; Enterobacteriaceae; Salmonella MTGDNTLITSHGINRRDFMKLCAALAATMGLSSKAAAEMAESVSNPQRPPVIWIGAQECT WP_000145429.1 hydrogenase 2 small subunit [Salmonella enterica] Length: 372\nNumber of predicted TMHs: 1\nExp number of AAs in TMHs: 24.51309\nExp number, first 60 AAs: 0.2682\nTotal prob of N-in: 0.14402\noutside 1 331\nTMhelix 332 354\ninside 355 372

14927 GCF_002033175.1_ASM203317v1 Salmonella enterica Proteobacteria; Gammaproteobacteria; Enterobacterales; Enterobacteriaceae; Salmonella MNNEETFYQAMRRKGVTRRSFLKFCSLAATSLGLGAGMTPKIAWALENKPRIPVVWIHGL WP_001058312.1 hydrogenase [Salmonella enterica] Length: 372\nNumber of predicted TMHs: 2\nExp number of AAs in TMHs: 36.51738\nExp number, first 60 AAs: 19.72849\nTotal prob of N-in: 0.96170\nPOSSIBLE N-term signal sequence\ninside 1 19\nTMhelix 20 39\noutside 40 325\nTMhelix 326 348\ninside 349 372

14928 GCF_002033365.1_ASM203336v1 Salmonella enterica Proteobacteria; Gammaproteobacteria; Enterobacterales; Enterobacteriaceae; Salmonella MTGDNTLITSHGINRRDFMKLCAALAATMGLSSKAAAEMAESVSNPQRPPVIWIGAQECT WP_000145429.1 hydrogenase 2 small subunit [Salmonella enterica] Length: 372\nNumber of predicted TMHs: 1\nExp number of AAs in TMHs: 24.51309\nExp number, first 60 AAs: 0.2682\nTotal prob of N-in: 0.14402\noutside 1 331\nTMhelix 332 354\ninside 355 372

14929 GCF_002033365.1_ASM203336v1 Salmonella enterica Proteobacteria; Gammaproteobacteria; Enterobacterales; Enterobacteriaceae; Salmonella MNNEETFYQAMRRKGVTRRSFLKFCSLAATSLGLGAGMTPKIAWALENKPRIPVVWIHGL WP_001058312.1 hydrogenase [Salmonella enterica] Length: 372\nNumber of predicted TMHs: 2\nExp number of AAs in TMHs: 36.51738\nExp number, first 60 AAs: 19.72849\nTotal prob of N-in: 0.96170\nPOSSIBLE N-term signal sequence\ninside 1 19\nTMhelix 20 39\noutside 40 325\nTMhelix 326 348\ninside 349 372

14930 GCF_002033295.1_ASM203329v1 Salmonella enterica Proteobacteria; Gammaproteobacteria; Enterobacterales; Enterobacteriaceae; Salmonella
MTGDNTLITSHGINRRDFMKLCAALAATMGLSSKAAAEMAESVSNPQRPPVIWIGAQECT WP_000145429.1
hydrogenase 2 small subunit [Salmonella enterica] Length: 372\nNumber of predicted TMHs: 1\nExp number of AAs in TMHs: 24.51309\nExp number, first 60 AAs: 0.2682\nTotal prob of N-in: 0.14402\noutside 1 331\nTMhelix 332 354\ninside 355 372

14931 GCF_002033295.1_ASM203329v1 Salmonella enterica Proteobacteria; Gammaproteobacteria; Enterobacterales; Enterobacteriaceae; Salmonella
MNNEETFYQAMRRKGVTRRSFLKFCSLAATSLGLGAGMTPKIAWALENKPRIPVVWIHGL WP_001058307.1
hydrogenase [Salmonella enterica] Length: 372\nNumber of predicted TMHs: 2\nExp number of AAs in TMHs: 36.51738\nExp number, first 60 AAs: 19.72849\nTotal prob of N-in: 0.96170\nPOSSIBLE N-term signal sequence\ninside 1 19\nTMhelix 20 39\noutside 40 325\nTMhelix 326 348\ninside 349 372

14932 GCF_002033255.1_ASM203325v1 Salmonella enterica Proteobacteria; Gammaproteobacteria; Enterobacterales; Enterobacteriaceae; Salmonella
MNNEETFYQAMRRKGVTRRSFLKFCSLAATSLGLGAGMTPKIAWALENKPRIPVVWIHGL WP_001058307.1
hydrogenase [Salmonella enterica] Length: 372\nNumber of predicted TMHs: 2\nExp number of AAs in TMHs: 36.51738\nExp number, first 60 AAs: 19.72849\nTotal prob of N-in: 0.96170\nPOSSIBLE N-term signal sequence\ninside 1 19\nTMhelix 20 39\noutside 40 325\nTMhelix 326 348\ninside 349 372

14933 GCF_002033725.1_ASM203372v1 Salmonella enterica Proteobacteria; Gammaproteobacteria; Enterobacterales; Enterobacteriaceae; Salmonella
MNNEETFYQAMRRKGVTRRSFLKFCSLAATSLGLGAGMTPKIAWALENKPRIPVVWIHGL WP_001058311.1
hydrogenase [Salmonella enterica] Length: 372\nNumber of predicted TMHs: 2\nExp number of AAs in TMHs: 37.4699899999999\nExp number, first 60 AAs: 19.72699\nTotal prob of N-in: 0.96171\nPOSSIBLE N-term signal sequence\ninside 1 19\nTMhelix 20 39\noutside 40 325\nTMhelix 326 348\ninside 349 372

14934 GCF_002033585.1_ASM203358v1 Salmonella enterica Proteobacteria; Gammaproteobacteria; Enterobacterales; Enterobacteriaceae; Salmonella
MNNEETFYQAMRRKGVTRRSFLKFCSLAATSLGLGAGMTPKIAWALENKPRIPVVWIHGL WP_001058307.1
hydrogenase [Salmonella enterica] Length: 372\nNumber of predicted TMHs: 2\nExp number of AAs in TMHs: 36.51738\nExp number, first 60 AAs: 19.72849\nTotal prob of N-in: 0.96170\nPOSSIBLE N-term signal sequence\ninside 1 19\nTMhelix 20 39\noutside 40 325\nTMhelix 326 348\ninside 349 372

14935 GCF_002033625.1_ASM203362v1 Salmonella enterica Proteobacteria; Gammaproteobacteria; Enterobacterales; Enterobacteriaceae; Salmonella
MNNEETFYQAMRRKGVTRRSFLKFCSLAATSLGLGAGMTPKIAWALENKPRIPVVWIHGL WP_001058307.1
hydrogenase [Salmonella enterica] Length: 372\nNumber of predicted TMHs: 2\nExp number of AAs in TMHs: 36.51738\nExp number, first 60 AAs: 19.72849\nTotal prob of N-in: 0.96170\nPOSSIBLE N-term signal sequence\ninside 1 19\nTMhelix 20 39\noutside 40 325\nTMhelix 326 348\ninside 349 372

14936 GCF_002033485.1_ASM203348v1 Salmonella enterica Proteobacteria; Gammaproteobacteria; Enterobacterales; Enterobacteriaceae; Salmonella
MNNEETFYQAMRRKGVTRRSFLKFCSLAATSLGLGAGMTPKIAWALENKPRIPVVWIHGL WP_079786335.1
[Ni/Fe] hydrogenase small subunit [Salmonella enterica] Length: 372\nNumber of predicted TMHs: 2\nExp number of AAs in TMHs: 36.51716\nExp number, first 60 AAs: 19.7285\nTotal prob of N-in: 0.96169\nPOSSIBLE N-term signal sequence\ninside 1 19\nTMhelix 20 39\noutside 40 325\nTMhelix 326 348\ninside 349 372

14937 GCF_002043955.1_ASM204395v1 Salmonella enterica Proteobacteria; Gammaproteobacteria; Enterobacterales; Enterobacteriaceae; Salmonella
MNNEETFYQAMRRKGVTRRSFLKFCSLAATSLGLGAGMTPKIAWALENKPRIPVVWIHGL WP_001058311.1
hydrogenase [Salmonella enterica] Length: 372\nNumber of predicted TMHs: 2\nExp number of AAs in TMHs: 37.4699899999999\nExp number, first 60 AAs: 19.72699\nTotal prob of N-in: 0.96171\nPOSSIBLE N-term signal sequence\ninside 1 19\nTMhelix 20 39\noutside 40 325\nTMhelix 326 348\ninside 349 372

14938 GCF_002037185.1_ASM203718v1 Salmonella enterica Proteobacteria; Gammaproteobacteria; Enterobacterales; Enterobacteriaceae; Salmonella
MTGDNTLITSHGINRRDFMKLCAALAATMGLSSKAAAEMAESVSNPQRPPVIWIGAQECT WP_000145429.1
hydrogenase 2 small subunit [Salmonella enterica] Length: 372\nNumber of predicted TMHs: 1\nExp

number of AAs in TMHs: 24.51309\nExp number, first 60 AAs:  0.2682\nTotal prob of N-in:   0.14402\noutside   1   331\nTMhelix   332   354\ninside   355   372

14939 GCF_002044065.1_ASM204406v1 Salmonella enterica Proteobacteria; Gammaproteobacteria; Enterobacterales; Enterobacteriaceae; Salmonella MTGDNTLITSHGINRRDFMKLCAALAATMGLSSKAAAEMAESVSNPQRPPVIWIGAQECT WP_000145429.1 hydrogenase 2 small subunit [Salmonella enterica] Length: 372\nNumber of predicted TMHs:  1\nExp number of AAs in TMHs: 24.51309\nExp number, first 60 AAs:  0.2682\nTotal prob of N-in:   0.14402\noutside   1   331\nTMhelix   332   354\ninside   355   372

14940 GCF_002043635.1_ASM204363v1 Salmonella enterica Proteobacteria; Gammaproteobacteria; Enterobacterales; Enterobacteriaceae; Salmonella MTGDNTLITSHGINRRDFMKLCAALAATMGLSSKAAAEMAESVSNPQRPPVIWIGAQECT WP_000145429.1 hydrogenase 2 small subunit [Salmonella enterica] Length: 372\nNumber of predicted TMHs:  1\nExp number of AAs in TMHs: 24.51309\nExp number, first 60 AAs:  0.2682\nTotal prob of N-in:   0.14402\noutside   1   331\nTMhelix   332   354\ninside   355   372

14941 GCF_002034235.1_ASM203423v1 Salmonella enterica Proteobacteria; Gammaproteobacteria; Enterobacterales; Enterobacteriaceae; Salmonella MTGDNTLITSHGINRRDFMKLCAALAATMGLSSKAAAEMAESVSNPQRPPVIWIGAQECT WP_000145429.1 hydrogenase 2 small subunit [Salmonella enterica] Length: 372\nNumber of predicted TMHs:  1\nExp number of AAs in TMHs: 24.51309\nExp number, first 60 AAs:  0.2682\nTotal prob of N-in:   0.14402\noutside   1   331\nTMhelix   332   354\ninside   355   372

14942 GCF_002034235.1_ASM203423v1 Salmonella enterica Proteobacteria; Gammaproteobacteria; Enterobacterales; Enterobacteriaceae; Salmonella MNNEETFYQAMRRKGVTRRSFLKFCSLAATSLGLGAGMTPKIAWALENKPRIPVVWIHGL WP_001058310.1 hydrogenase [Salmonella enterica] Length: 372\nNumber of predicted TMHs:  2\nExp number of AAs in TMHs: 37.4567299999999\nExp number, first 60 AAs:  19.72694\nTotal prob of N-in:   0.96171\nPOSSIBLE N-term signal sequence\ninside   1   19\nTMhelix   20   39\noutside   40   325\nTMhelix   326   348\ninside   349   372

14943 GCF_002033935.1_ASM203393v1 Salmonella enterica Proteobacteria; Gammaproteobacteria; Enterobacterales; Enterobacteriaceae; Salmonella MNNEETFYQAMRRKGVTRRSFLKFCSLAATSLGLGAGMTPKIAWALENKPRIPVVWIHGL WP_001058312.1 hydrogenase [Salmonella enterica] Length: 372\nNumber of predicted TMHs:  2\nExp number of AAs in TMHs: 36.51738\nExp number, first 60 AAs:  19.72849\nTotal prob of N-in:   0.96170\nPOSSIBLE N-term signal sequence\ninside   1   19\nTMhelix   20   39\noutside   40   325\nTMhelix   326   348\ninside   349   372

14944 GCF_002044265.1_ASM204426v1 Salmonella enterica Proteobacteria; Gammaproteobacteria; Enterobacterales; Enterobacteriaceae; Salmonella MTGDNTLITSHGINRRDFMKLCAALAATMGLSSKAAAEMAESVSNPQRPPVIWIGAQECT WP_000145429.1 hydrogenase 2 small subunit [Salmonella enterica] Length: 372\nNumber of predicted TMHs:  1\nExp number of AAs in TMHs: 24.51309\nExp number, first 60 AAs:  0.2682\nTotal prob of N-in:   0.14402\noutside   1   331\nTMhelix   332   354\ninside   355   372

14945 GCF_002044245.1_ASM204424v1 Salmonella enterica Proteobacteria; Gammaproteobacteria; Enterobacterales; Enterobacteriaceae; Salmonella MTGDNTLITSHGINRRDFMKLCAALAATMGLSSKAAAEMAESVSNPQRPPVIWIGAQECT WP_000145429.1 hydrogenase 2 small subunit [Salmonella enterica] Length: 372\nNumber of predicted TMHs:  1\nExp number of AAs in TMHs: 24.51309\nExp number, first 60 AAs:  0.2682\nTotal prob of N-in:   0.14402\noutside   1   331\nTMhelix   332   354\ninside   355   372

14946 GCF_002052225.1_ASM205222v1 Salmonella enterica Proteobacteria; Gammaproteobacteria; Enterobacterales; Enterobacteriaceae; Salmonella MTGDNTLITSHGINRRDFMKLCAALAATMGLSSKAAAEMAESVSNPQRPPVIWIGAQECT WP_000145429.1 hydrogenase 2 small subunit [Salmonella enterica] Length: 372\nNumber of predicted TMHs:  1\nExp number of AAs in TMHs: 24.51309\nExp number, first 60 AAs:  0.2682\nTotal prob of N-in:   0.14402\noutside   1   331\nTMhelix   332   354\ninside   355   372

14947 GCF_002037025.1_ASM203702v1 Salmonella enterica Proteobacteria; Gammaproteobacteria; Enterobacterales; Enterobacteriaceae; Salmonella

MTGDNTLITSHGINRRDFMKLCAALAATMGLSSKAAAEMAESVSNPQRPPVIWIGAQECT WP_000145431.1 hydrogenase 2 small subunit [Salmonella enterica] Length: 372\nNumber of predicted TMHs: 1\nExp number of AAs in TMHs: 24.51505\nExp number, first 60 AAs: 0.2682\nTotal prob of N-in: 0.14406\noutside 1 331\nTMhelix 332 354\ninside 355 372

14948 GCF_002044025.1_ASM204402v1 Salmonella enterica Proteobacteria; Gammaproteobacteria; Enterobacterales; Enterobacteriaceae; Salmonella MNNEETFYQAMRRKGVTRRSFLKFCSLAATSLGLGAGMTPKIAWALENKPRIPVVWIHGL WP_001058312.1 hydrogenase [Salmonella enterica] Length: 372\nNumber of predicted TMHs: 2\nExp number of AAs in TMHs: 36.51738\nExp number, first 60 AAs: 19.72849\nTotal prob of N-in: 0.96170\nPOSSIBLE N-term signal sequence\ninside 1 19\nTMhelix 20 39\noutside 40 325\nTMhelix 326 348\ninside 349 372

14949 GCF_002044345.1_ASM204434v1 Salmonella enterica Proteobacteria; Gammaproteobacteria; Enterobacterales; Enterobacteriaceae; Salmonella MNNEETFYQAMRRKGVTRRSFLKFCSLAATSLGLGAGMTPKIAWALENKPRIPVVWIHGL WP_001058311.1 hydrogenase [Salmonella enterica] Length: 372\nNumber of predicted TMHs: 2\nExp number of AAs in TMHs: 37.4699899999999\nExp number, first 60 AAs: 19.72699\nTotal prob of N-in: 0.96171\nPOSSIBLE N-term signal sequence\ninside 1 19\nTMhelix 20 39\noutside 40 325\nTMhelix 326 348\ninside 349 372

14950 GCF_002044525.1_ASM204452v1 Salmonella enterica Proteobacteria; Gammaproteobacteria; Enterobacterales; Enterobacteriaceae; Salmonella MTGDNTLITSHGINRRDFMKLCAALAATMGLSSKAAAEMAESVSNPQRPPVIWIGAQECT WP_000145429.1 hydrogenase 2 small subunit [Salmonella enterica] Length: 372\nNumber of predicted TMHs: 1\nExp number of AAs in TMHs: 24.51309\nExp number, first 60 AAs: 0.2682\nTotal prob of N-in: 0.14402\noutside 1 331\nTMhelix 332 354\ninside 355 372

14951 GCF_002033255.1_ASM203325v1 Salmonella enterica Proteobacteria; Gammaproteobacteria; Enterobacterales; Enterobacteriaceae; Salmonella MTGDNTLITSHGINRRDFMKLCAALAATMGLSSKAAAEMAESVSNPQRPPVIWIGAQECT WP_000145429.1 hydrogenase 2 small subunit [Salmonella enterica] Length: 372\nNumber of predicted TMHs: 1\nExp number of AAs in TMHs: 24.51309\nExp number, first 60 AAs: 0.2682\nTotal prob of N-in: 0.14402\noutside 1 331\nTMhelix 332 354\ninside 355 372

14952 GCF_002044455.1_ASM204445v1 Salmonella enterica Proteobacteria; Gammaproteobacteria; Enterobacterales; Enterobacteriaceae; Salmonella MTGDNTLITSHGINRRDFMKLCAALAATMGLSSKAAAEMAESVSNPQRPPVIWIGAQECT WP_000145429.1 hydrogenase 2 small subunit [Salmonella enterica] Length: 372\nNumber of predicted TMHs: 1\nExp number of AAs in TMHs: 24.51309\nExp number, first 60 AAs: 0.2682\nTotal prob of N-in: 0.14402\noutside 1 331\nTMhelix 332 354\ninside 355 372

14953 GCF_002044105.1_ASM204410v1 Salmonella enterica Proteobacteria; Gammaproteobacteria; Enterobacterales; Enterobacteriaceae; Salmonella MTGDNTLITSHGINRRDFMKLCAALAATMGLSSKAAAEMAESVSNPQRPPVIWIGAQECT WP_000145429.1 hydrogenase 2 small subunit [Salmonella enterica] Length: 372\nNumber of predicted TMHs: 1\nExp number of AAs in TMHs: 24.51309\nExp number, first 60 AAs: 0.2682\nTotal prob of N-in: 0.14402\noutside 1 331\nTMhelix 332 354\ninside 355 372

14954 GCF_002044605.1_ASM204460v1 Salmonella enterica Proteobacteria; Gammaproteobacteria; Enterobacterales; Enterobacteriaceae; Salmonella MTGDNTLITSHGINRRDFMKLCAALAATMGLSSKAAAEMAESVSNPQRPPVIWIGAQECT WP_000145429.1 hydrogenase 2 small subunit [Salmonella enterica] Length: 372\nNumber of predicted TMHs: 1\nExp number of AAs in TMHs: 24.51309\nExp number, first 60 AAs: 0.2682\nTotal prob of N-in: 0.14402\noutside 1 331\nTMhelix 332 354\ninside 355 372

14955 GCF_002034525.1_ASM203452v1 Salmonella enterica Proteobacteria; Gammaproteobacteria; Enterobacterales; Enterobacteriaceae; Salmonella MNNEETFYQAMRRKGVTRRSFLKFCSLAATSLGLGAGMTPKIAWALENKPRIPVVWIHGL WP_001058312.1 hydrogenase [Salmonella enterica] Length: 372\nNumber of predicted TMHs: 2\nExp number of AAs in TMHs: 36.51738\nExp number, first 60 AAs: 19.72849\nTotal prob of N-in: 0.96170\nPOSSIBLE N-term signal sequence\ninside 1 19\nTMhelix 20 39\noutside 40 325\nTMhelix 326 348\ninside 349 372

14956	GCF_002044645.1_ASM204464v1	Salmonella enterica	Proteobacteria; Gammaproteobacteria; Enterobacterales; Enterobacteriaceae; Salmonella
MTGDNTLITSHGINRRDFMKLCAALAATMGLSSKAAAEMAESVSNPQRPPVIWIGAQECT	WP_000145429.1
hydrogenase 2 small subunit [Salmonella enterica]	Length: 372\nNumber of predicted TMHs:  1\nExp number of AAs in TMHs: 24.51309\nExp number, first 60 AAs:  0.2682\nTotal prob of N-in:   0.14402\noutside   1   331\nTMhelix   332   354\ninside   355   372

14957	GCF_002044725.1_ASM204472v1	Salmonella enterica	Proteobacteria; Gammaproteobacteria; Enterobacterales; Enterobacteriaceae; Salmonella
MTGDNTLITSHGINRRDFMKLCAALAATMGLSSKAAAEMAESVSNPQRPPVIWIGAQECT	WP_000145429.1
hydrogenase 2 small subunit [Salmonella enterica]	Length: 372\nNumber of predicted TMHs:  1\nExp number of AAs in TMHs: 24.51309\nExp number, first 60 AAs:  0.2682\nTotal prob of N-in:   0.14402\noutside   1   331\nTMhelix   332   354\ninside   355   372

14958	GCF_002044525.1_ASM204452v1	Salmonella enterica	Proteobacteria; Gammaproteobacteria; Enterobacterales; Enterobacteriaceae; Salmonella
MNNEETFYQAMRRKGVTRRSFLKFCSLAATSLGLGAGMTPKIAWALENKPRIPVVWIHGL	WP_001058311.1
hydrogenase [Salmonella enterica]	Length: 372\nNumber of predicted TMHs:  2\nExp number of AAs in TMHs: 37.4699899999999\nExp number, first 60 AAs:  19.72699\nTotal prob of N-in:   0.96171\nPOSSIBLE N-term signal sequence\ninside   1   19\nTMhelix   20   39\noutside   40   325\nTMhelix   326   348\ninside   349   372

14959	GCF_002043635.1_ASM204363v1	Salmonella enterica	Proteobacteria; Gammaproteobacteria; Enterobacterales; Enterobacteriaceae; Salmonella
MNNEETFYQAMRRKGVTRRSFLKFCSLAATSLGLGAGMTPKIAWALENKPRIPVVWIHGL	WP_001058311.1
hydrogenase [Salmonella enterica]	Length: 372\nNumber of predicted TMHs:  2\nExp number of AAs in TMHs: 37.4699899999999\nExp number, first 60 AAs:  19.72699\nTotal prob of N-in:   0.96171\nPOSSIBLE N-term signal sequence\ninside   1   19\nTMhelix   20   39\noutside   40   325\nTMhelix   326   348\ninside   349   372

14960	GCF_002044455.1_ASM204445v1	Salmonella enterica	Proteobacteria; Gammaproteobacteria; Enterobacterales; Enterobacteriaceae; Salmonella
MNNEETFYQAMRRKGVTRRSFLKFCSLAATSLGLGAGMTPKIAWALENKPRIPVVWIHGL	WP_023198237.1
hydrogenase 1, small subunit [Salmonella enterica]	Length: 372\nNumber of predicted TMHs:  2\nExp number of AAs in TMHs: 36.39423\nExp number, first 60 AAs:  19.72877\nTotal prob of N-in:   0.96162\nPOSSIBLE N-term signal sequence\ninside   1   19\nTMhelix   20   39\noutside   40   325\nTMhelix   326   348\ninside   349   372

14961	GCF_002044945.1_ASM204494v1	Salmonella enterica	Proteobacteria; Gammaproteobacteria; Enterobacterales; Enterobacteriaceae; Salmonella
MTGDNTLITSHGINRRDFMKLCAALAATMGLSSKAAAEMAESVSNPQRPPVIWIGAQECT	WP_000145429.1
hydrogenase 2 small subunit [Salmonella enterica]	Length: 372\nNumber of predicted TMHs:  1\nExp number of AAs in TMHs: 24.51309\nExp number, first 60 AAs:  0.2682\nTotal prob of N-in:   0.14402\noutside   1   331\nTMhelix   332   354\ninside   355   372

14962	GCF_002045175.1_ASM204517v1	Salmonella enterica	Proteobacteria; Gammaproteobacteria; Enterobacterales; Enterobacteriaceae; Salmonella
MTGDNTLITSHGINRRDFMKLCAALAATMGLSSKAAAEMAESVSNPQRPPVIWIGAQECT	WP_000145429.1
hydrogenase 2 small subunit [Salmonella enterica]	Length: 372\nNumber of predicted TMHs:  1\nExp number of AAs in TMHs: 24.51309\nExp number, first 60 AAs:  0.2682\nTotal prob of N-in:   0.14402\noutside   1   331\nTMhelix   332   354\ninside   355   372

14963	GCF_002045175.1_ASM204517v1	Salmonella enterica	Proteobacteria; Gammaproteobacteria; Enterobacterales; Enterobacteriaceae; Salmonella
MNNEETFYQAMRRKGVTRRSFLKFCSLAATSLGLSAGMTPKIAWALENKPRIPVVWIHGL	WP_001058316.1
hydrogenase [Salmonella enterica]	Length: 372\nNumber of predicted TMHs:  2\nExp number of AAs in TMHs: 35.76397\nExp number, first 60 AAs:  18.97201\nTotal prob of N-in:   0.92785\nPOSSIBLE N-term signal sequence\ninside   1   19\nTMhelix   20   39\noutside   40   325\nTMhelix   326   348\ninside   349   372

14964	GCF_002044345.1_ASM204434v1	Salmonella enterica	Proteobacteria; Gammaproteobacteria; Enterobacterales; Enterobacteriaceae; Salmonella
MTGDNTLITSHGINRRDFMKLCAALAATMGLSSKAAAEMAESVSNPQRPPVIWIGAQECT	WP_000145429.1
hydrogenase 2 small subunit [Salmonella enterica]	Length: 372\nNumber of predicted TMHs:  1\nExp

number of AAs in TMHs: 24.51309\nExp number, first 60 AAs: 0.2682\nTotal prob of N-in: 0.14402\noutside 1 331\nTMhelix 332 354\ninside 355 372

14965 GCF_002035385.1_ASM203538v1 Salmonella enterica Proteobacteria; Gammaproteobacteria; Enterobacterales; Enterobacteriaceae; Salmonella
MTGDNTLITSHGINRRDFMKLCAALAATMGLSSKAAAEMAESVSNPQRPPVIWIGAQECT WP_079776618.1
MULTISPECIES: hydrogenase 2 small subunit [Salmonella] Length: 372\nNumber of predicted TMHs: 1\nExp number of AAs in TMHs: 24.65705\nExp number, first 60 AAs: 0.26805\nTotal prob of N-in: 0.13761\noutside 1 331\nTMhelix 332 354\ninside 355 372

14966 GCF_002035215.1_ASM203521v1 Salmonella enterica Proteobacteria; Gammaproteobacteria; Enterobacterales; Enterobacteriaceae; Salmonella
MTGDNTLITSHGINRRDFMKLCAALAATMGLSSKAAAEMAESVSNPQRPPVIWIGAQECT WP_079776618.1
MULTISPECIES: hydrogenase 2 small subunit [Salmonella] Length: 372\nNumber of predicted TMHs: 1\nExp number of AAs in TMHs: 24.65705\nExp number, first 60 AAs: 0.26805\nTotal prob of N-in: 0.13761\noutside 1 331\nTMhelix 332 354\ninside 355 372

14967 GCF_001831865.1_ASM183186v1 Salmonella enterica Proteobacteria; Gammaproteobacteria; Enterobacterales; Enterobacteriaceae; Salmonella
MTGDNTLITSHGINRRDFMKLCAALAATMGLSSKAAAEMAESVSNPQRPPVIWIGAQECT WP_000145429.1
hydrogenase 2 small subunit [Salmonella enterica] Length: 372\nNumber of predicted TMHs: 1\nExp number of AAs in TMHs: 24.51309\nExp number, first 60 AAs: 0.2682\nTotal prob of N-in: 0.14402\noutside 1 331\nTMhelix 332 354\ninside 355 372

14968 GCF_002045525.1_ASM204552v1 Salmonella enterica Proteobacteria; Gammaproteobacteria; Enterobacterales; Enterobacteriaceae; Salmonella
MTGDNTLITSHGINRRDFMKLCAALAATMGLSSKAAAEMAESVSNPQRPPVIWIGAQECT WP_000145429.1
hydrogenase 2 small subunit [Salmonella enterica] Length: 372\nNumber of predicted TMHs: 1\nExp number of AAs in TMHs: 24.51309\nExp number, first 60 AAs: 0.2682\nTotal prob of N-in: 0.14402\noutside 1 331\nTMhelix 332 354\ninside 355 372

14969 GCF_001831795.1_ASM183179v1 Salmonella enterica Proteobacteria; Gammaproteobacteria; Enterobacterales; Enterobacteriaceae; Salmonella
MTGDNTLITSHGINRRDFMKLCAALAATMGLSSKAAAEMAESVSNPQRPPVIWIGAQECT WP_000145429.1
hydrogenase 2 small subunit [Salmonella enterica] Length: 372\nNumber of predicted TMHs: 1\nExp number of AAs in TMHs: 24.51309\nExp number, first 60 AAs: 0.2682\nTotal prob of N-in: 0.14402\noutside 1 331\nTMhelix 332 354\ninside 355 372

14970 GCF_002043585.1_ASM204358v1 Salmonella enterica Proteobacteria; Gammaproteobacteria; Enterobacterales; Enterobacteriaceae; Salmonella
MNNEETFYQAMRRKGVTRRSFLKFCSLAATSLGLGAGMTPKIAWALENKPRIPVVWIHGL WP_001058312.1
hydrogenase [Salmonella enterica] Length: 372\nNumber of predicted TMHs: 2\nExp number of AAs in TMHs: 36.51738\nExp number, first 60 AAs: 19.72849\nTotal prob of N-in: 0.96170\nPOSSIBLE N-term signal sequence\ninside 1 19\nTMhelix 20 39\noutside 40 325\nTMhelix 326 348\ninside 349 372

14971 GCF_002044385.1_ASM204438v1 Salmonella enterica Proteobacteria; Gammaproteobacteria; Enterobacterales; Enterobacteriaceae; Salmonella
MTGDNTLITSHGINRRDFMKLCAALAATMGLSSKAAAEMAESVSNPQRPPVIWIGAQECT WP_000145429.1
hydrogenase 2 small subunit [Salmonella enterica] Length: 372\nNumber of predicted TMHs: 1\nExp number of AAs in TMHs: 24.51309\nExp number, first 60 AAs: 0.2682\nTotal prob of N-in: 0.14402\noutside 1 331\nTMhelix 332 354\ninside 355 372

14972 GCF_002035845.1_ASM203584v1 Salmonella enterica Proteobacteria; Gammaproteobacteria; Enterobacterales; Enterobacteriaceae; Salmonella
MTGDNTLITSHGINRRDFMKLCAALAATMGLSSKAAAEMAESVSNPQRPPVIWIGAQECT WP_000145429.1
hydrogenase 2 small subunit [Salmonella enterica] Length: 372\nNumber of predicted TMHs: 1\nExp number of AAs in TMHs: 24.51309\nExp number, first 60 AAs: 0.2682\nTotal prob of N-in: 0.14402\noutside 1 331\nTMhelix 332 354\ninside 355 372

14973 GCF_002043375.1_ASM204337v1 Salmonella enterica Proteobacteria; Gammaproteobacteria; Enterobacterales; Enterobacteriaceae; Salmonella

MTGDNTLITSHGINRRDFMKLCAALAATMGLSSKAAAEMAESVSNPQRPPVIWIGAQECT WP_000145429.1 hydrogenase 2 small subunit [Salmonella enterica] Length: 372\nNumber of predicted TMHs: 1\nExp number of AAs in TMHs: 24.51309\nExp number, first 60 AAs: 0.2682\nTotal prob of N-in: 0.14402\noutside 1 331\nTMhelix 332 354\ninside 355 372

14974 GCF_002035845.1_ASM203584v1 Salmonella enterica Proteobacteria; Gammaproteobacteria; Enterobacterales; Enterobacteriaceae; Salmonella MNNEETFYQAMRRKGVTRRSFLKFCSLAATSLGLGAGMTPKIAWALENKPRIPVVWIHGL WP_001058312.1 hydrogenase [Salmonella enterica] Length: 372\nNumber of predicted TMHs: 2\nExp number of AAs in TMHs: 36.51738\nExp number, first 60 AAs: 19.72849\nTotal prob of N-in: 0.96170\nPOSSIBLE N-term signal sequence\ninside 1 19\nTMhelix 20 39\noutside 40 325\nTMhelix 326 348\ninside 349 372

14975 GCF_001642485.1_ASM164248v1 Salmonella enterica Proteobacteria; Gammaproteobacteria; Enterobacterales; Enterobacteriaceae; Salmonella MNNEETFYQAMRRKGVTRRSFLKFCSLAATSLGLGAGMTPKIAWALENKPRIPVVWIHGL WP_001058312.1 hydrogenase [Salmonella enterica] Length: 372\nNumber of predicted TMHs: 2\nExp number of AAs in TMHs: 36.51738\nExp number, first 60 AAs: 19.72849\nTotal prob of N-in: 0.96170\nPOSSIBLE N-term signal sequence\ninside 1 19\nTMhelix 20 39\noutside 40 325\nTMhelix 326 348\ninside 349 372

14976 GCF_002043245.1_ASM204324v1 Salmonella enterica Proteobacteria; Gammaproteobacteria; Enterobacterales; Enterobacteriaceae; Salmonella MTGDNTLITSHGINRRDFMKLCAALAATMGLSSKAAAEMAESVSNPQRPPVIWIGAQECT WP_000145429.1 hydrogenase 2 small subunit [Salmonella enterica] Length: 372\nNumber of predicted TMHs: 1\nExp number of AAs in TMHs: 24.51309\nExp number, first 60 AAs: 0.2682\nTotal prob of N-in: 0.14402\noutside 1 331\nTMhelix 332 354\ninside 355 372

14977 GCF_001643805.1_CRJJGF_00093_v1.0Salmonella enterica Proteobacteria; Gammaproteobacteria; Enterobacterales; Enterobacteriaceae; Salmonella MTGDNTLITSHGINRRDFMKLCAALAATMGLSSKAAAEMAESVSNPQRPPVIWIGAQECT WP_000145429.1 hydrogenase 2 small subunit [Salmonella enterica] Length: 372\nNumber of predicted TMHs: 1\nExp number of AAs in TMHs: 24.51309\nExp number, first 60 AAs: 0.2682\nTotal prob of N-in: 0.14402\noutside 1 331\nTMhelix 332 354\ninside 355 372

14978 GCF_001643805.1_CRJJGF_00093_v1.0Salmonella enterica Proteobacteria; Gammaproteobacteria; Enterobacterales; Enterobacteriaceae; Salmonella MNNEETFYQAMRRKGVTRRSFLKFCSLAATSLGLGAGMTPKIAWALENKPRIPVVWIHGL WP_001058312.1 hydrogenase [Salmonella enterica] Length: 372\nNumber of predicted TMHs: 2\nExp number of AAs in TMHs: 36.51738\nExp number, first 60 AAs: 19.72849\nTotal prob of N-in: 0.96170\nPOSSIBLE N-term signal sequence\ninside 1 19\nTMhelix 20 39\noutside 40 325\nTMhelix 326 348\ninside 349 372

14979 GCF_002036055.1_ASM203605v1 Salmonella enterica Proteobacteria; Gammaproteobacteria; Enterobacterales; Enterobacteriaceae; Salmonella MTGDNTLITSHGINRRDFMKLCAALAATMGLSSKAAAEMAESVSNPQRPPVIWIGAQECT WP_000145429.1 hydrogenase 2 small subunit [Salmonella enterica] Length: 372\nNumber of predicted TMHs: 1\nExp number of AAs in TMHs: 24.51309\nExp number, first 60 AAs: 0.2682\nTotal prob of N-in: 0.14402\noutside 1 331\nTMhelix 332 354\ninside 355 372

14980 GCF_001643825.1_CRJJGF_00061_v1.0Salmonella enterica Proteobacteria; Gammaproteobacteria; Enterobacterales; Enterobacteriaceae; Salmonella MNNEETFYQAMRRKGVTRRSFLKFCSLAATSLGLGAGMTPKIAWALENKPRIPVVWIHGL WP_064000476.1 hydrogenase [Salmonella enterica] Length: 372\nNumber of predicted TMHs: 2\nExp number of AAs in TMHs: 37.4700099999999\nExp number, first 60 AAs: 19.72698\nTotal prob of N-in: 0.96171\nPOSSIBLE N-term signal sequence\ninside 1 19\nTMhelix 20 39\noutside 40 325\nTMhelix 326 348\ninside 349 372

14981 GCF_002045525.1_ASM204552v1 Salmonella enterica Proteobacteria; Gammaproteobacteria; Enterobacterales; Enterobacteriaceae; Salmonella MNNEETFYQAMRRKGVTRRSFLKFCSLAATSLGLGAGMTPKIAWALENKPRIPVVWIHGL WP_079953656.1 [Ni/Fe] hydrogenase small subunit [Salmonella enterica] Length: 372\nNumber of predicted TMHs: 2\nExp number of AAs in TMHs: 36.5655299999999\nExp number, first 60 AAs: 19.72713\nTotal prob of N-in: 0.96172\nPOSSIBLE N-term signal sequence\ninside 1 19\nTMhelix 20 39\noutside 40 325\nTMhelix 326 348\ninside 349 372

14982 GCF_002045565.1_ASM204556v1 Salmonella enterica Proteobacteria; Gammaproteobacteria; Enterobacterales; Enterobacteriaceae; Salmonella
MNNEETFYQAMRRKGVTRRSFLKFCSLAATSLGLGAGMTPKIAWALENKPRIPVVWIHGL WP_001058311.1
hydrogenase [Salmonella enterica] Length: 372\nNumber of predicted TMHs: 2\nExp number of AAs in TMHs: 37.4699899999999\nExp number, first 60 AAs: 19.72699\nTotal prob of N-in: 0.96171\nPOSSIBLE N-term signal sequence\ninside 1 19\nTMhelix 20 39\noutside 40 325\nTMhelix 326 348\ninside 349 372

14983 GCF_002036145.1_ASM203614v1 Salmonella enterica Proteobacteria; Gammaproteobacteria; Enterobacterales; Enterobacteriaceae; Salmonella
MNNEETFYQAMRRKGVTRRSFLKFCSLAATSLGLGAGMTPKIAWALENKPRIPVVWIHGL WP_001058310.1
hydrogenase [Salmonella enterica] Length: 372\nNumber of predicted TMHs: 2\nExp number of AAs in TMHs: 37.4567299999999\nExp number, first 60 AAs: 19.72694\nTotal prob of N-in: 0.96171\nPOSSIBLE N-term signal sequence\ninside 1 19\nTMhelix 20 39\noutside 40 325\nTMhelix 326 348\ninside 349 372

14984 GCF_002045645.1_ASM204564v1 Salmonella enterica Proteobacteria; Gammaproteobacteria; Enterobacterales; Enterobacteriaceae; Salmonella
MNNEETFYQAMRRKGVTRRSFLKFCSLAATSLGLGAGMTPKIAWALENKPRIPVVWIHGL WP_001058312.1
hydrogenase [Salmonella enterica] Length: 372\nNumber of predicted TMHs: 2\nExp number of AAs in TMHs: 36.51738\nExp number, first 60 AAs: 19.72849\nTotal prob of N-in: 0.96170\nPOSSIBLE N-term signal sequence\ninside 1 19\nTMhelix 20 39\noutside 40 325\nTMhelix 326 348\ninside 349 372

14985 GCF_002051195.1_ASM205119v1 Salmonella enterica Proteobacteria; Gammaproteobacteria; Enterobacterales; Enterobacteriaceae; Salmonella
MTGDNTLITSHGINRRDFMKLCAALAATMGLSSKAAAEMAESVSNPQRPPVIWIGAQECT WP_000145429.1
hydrogenase 2 small subunit [Salmonella enterica] Length: 372\nNumber of predicted TMHs: 1\nExp number of AAs in TMHs: 24.51309\nExp number, first 60 AAs: 0.2682\nTotal prob of N-in: 0.14402\noutside 1 331\nTMhelix 332 354\ninside 355 372

14986 GCF_002036745.1_ASM203674v1 Salmonella enterica Proteobacteria; Gammaproteobacteria; Enterobacterales; Enterobacteriaceae; Salmonella
MNNEETFYQAMRRKGVTRRSFLKFCSLAATSLGLGAGMTPKIAWALENKPRIPVVWIHGL WP_001058312.1
hydrogenase [Salmonella enterica] Length: 372\nNumber of predicted TMHs: 2\nExp number of AAs in TMHs: 36.51738\nExp number, first 60 AAs: 19.72849\nTotal prob of N-in: 0.96170\nPOSSIBLE N-term signal sequence\ninside 1 19\nTMhelix 20 39\noutside 40 325\nTMhelix 326 348\ninside 349 372

14987 GCF_002036505.1_ASM203650v1 Salmonella enterica Proteobacteria; Gammaproteobacteria; Enterobacterales; Enterobacteriaceae; Salmonella
MNNEETFYQAMRRKGVTRRSFLKFCSLAATSLGLGAGMTPKIAWALENKPRIPVVWIHGL WP_001058307.1
hydrogenase [Salmonella enterica] Length: 372\nNumber of predicted TMHs: 2\nExp number of AAs in TMHs: 36.51738\nExp number, first 60 AAs: 19.72849\nTotal prob of N-in: 0.96170\nPOSSIBLE N-term signal sequence\ninside 1 19\nTMhelix 20 39\noutside 40 325\nTMhelix 326 348\ninside 349 372

14988 GCF_002045675.1_ASM204567v1 Salmonella enterica Proteobacteria; Gammaproteobacteria; Enterobacterales; Enterobacteriaceae; Salmonella
MNNEETFYQAMRRKGVTRRSFLKFCSLAATSLGLGAGMTPKIAWALENKPRIPVVWIHGL WP_001058312.1
hydrogenase [Salmonella enterica] Length: 372\nNumber of predicted TMHs: 2\nExp number of AAs in TMHs: 36.51738\nExp number, first 60 AAs: 19.72849\nTotal prob of N-in: 0.96170\nPOSSIBLE N-term signal sequence\ninside 1 19\nTMhelix 20 39\noutside 40 325\nTMhelix 326 348\ninside 349 372

14989 GCF_002036845.1_ASM203684v1 Salmonella enterica Proteobacteria; Gammaproteobacteria; Enterobacterales; Enterobacteriaceae; Salmonella
MTGDNTLITSHGINRRDFMKLCAALAATMGLSSKAAAEMAESVSNPQRPPVIWIGAQECT WP_000145429.1
hydrogenase 2 small subunit [Salmonella enterica] Length: 372\nNumber of predicted TMHs: 1\nExp number of AAs in TMHs: 24.51309\nExp number, first 60 AAs: 0.2682\nTotal prob of N-in: 0.14402\noutside 1 331\nTMhelix 332 354\ninside 355 372

14990 GCF_002036985.1_ASM203698v1 Salmonella enterica Proteobacteria; Gammaproteobacteria; Enterobacterales; Enterobacteriaceae; Salmonella
MTGDNTLITSHGINRRDFMKLCAALAATMGLSSKAAAEMAESVSNPQRPPVIWIGAQECT WP_000145429.1
hydrogenase 2 small subunit [Salmonella enterica] Length: 372\nNumber of predicted TMHs: 1\nExp

number of AAs in TMHs: 24.51309\nExp number, first 60 AAs:  0.2682\nTotal prob of N-in:   0.14402\noutside   1 331\nTMhelix   332   354\ninside   355   372

14991 GCF_002036905.1_ASM203690v1 Salmonella enterica Proteobacteria; Gammaproteobacteria; Enterobacterales; Enterobacteriaceae; Salmonella MTGDNTLITSHGINRRDFMKLCAALAATMGLSSKAAAEMAESVSNPQRPPVIWIGAQECT WP_000145429.1 hydrogenase 2 small subunit [Salmonella enterica] Length: 372\nNumber of predicted TMHs:  1\nExp number of AAs in TMHs: 24.51309\nExp number, first 60 AAs:  0.2682\nTotal prob of N-in:   0.14402\noutside   1 331\nTMhelix   332   354\ninside   355   372

14992 GCF_002037185.1_ASM203718v1 Salmonella enterica Proteobacteria; Gammaproteobacteria; Enterobacterales; Enterobacteriaceae; Salmonella MNNEETFYQAMRRKGVTRRSFLKFCSLAATSLGLGAGMTPKIAWALENKPRIPVVWIHGL WP_001058311.1 hydrogenase [Salmonella enterica] Length: 372\nNumber of predicted TMHs:  2\nExp number of AAs in TMHs: 37.4699899999999\nExp number, first 60 AAs:  19.72699\nTotal prob of N-in:   0.96171\nPOSSIBLE N-term signal sequence\ninside   1   19\nTMhelix   20   39\noutside   40   325\nTMhelix   326   348\ninside   349   372

14993 GCF_002037025.1_ASM203702v1 Salmonella enterica Proteobacteria; Gammaproteobacteria; Enterobacterales; Enterobacteriaceae; Salmonella MNNEETFYQAMRRKGVTRRSFLKFCSLAATSLGLGAGMTPKIAWALENKPRIPVVWIHGL WP_001058312.1 hydrogenase [Salmonella enterica] Length: 372\nNumber of predicted TMHs:  2\nExp number of AAs in TMHs: 36.51738\nExp number, first 60 AAs:  19.72849\nTotal prob of N-in:   0.96170\nPOSSIBLE N-term signal sequence\ninside   1   19\nTMhelix   20   39\noutside   40   325\nTMhelix   326   348\ninside   349   372

14994 GCF_002036905.1_ASM203690v1 Salmonella enterica Proteobacteria; Gammaproteobacteria; Enterobacterales; Enterobacteriaceae; Salmonella MNNEETFYQAMRRKGVTRRSFLKFCSLAATSLGLGAGMTPKIAWALENKPRIPVVWIHGL WP_001058312.1 hydrogenase [Salmonella enterica] Length: 372\nNumber of predicted TMHs:  2\nExp number of AAs in TMHs: 36.51738\nExp number, first 60 AAs:  19.72849\nTotal prob of N-in:   0.96170\nPOSSIBLE N-term signal sequence\ninside   1   19\nTMhelix   20   39\noutside   40   325\nTMhelix   326   348\ninside   349   372

14995 GCF_002036885.1_ASM203688v1 Salmonella enterica Proteobacteria; Gammaproteobacteria; Enterobacterales; Enterobacteriaceae; Salmonella MNNEETFYQAMRRKGVTRRSFLKFCSLAATSLGLGAGMTPKIAWALENKPRIPVVWIHGL WP_001058312.1 hydrogenase [Salmonella enterica] Length: 372\nNumber of predicted TMHs:  2\nExp number of AAs in TMHs: 36.51738\nExp number, first 60 AAs:  19.72849\nTotal prob of N-in:   0.96170\nPOSSIBLE N-term signal sequence\ninside   1   19\nTMhelix   20   39\noutside   40   325\nTMhelix   326   348\ninside   349   372

14996 GCF_002037225.1_ASM203722v1 Salmonella enterica Proteobacteria; Gammaproteobacteria; Enterobacterales; Enterobacteriaceae; Salmonella MTGDNTLITSHGINRRDFMKLCAALAATMGLSSKAAAEMAESVSNPQRPPVIWIGAQECT WP_000145429.1 hydrogenase 2 small subunit [Salmonella enterica] Length: 372\nNumber of predicted TMHs:  1\nExp number of AAs in TMHs: 24.51309\nExp number, first 60 AAs:  0.2682\nTotal prob of N-in:   0.14402\noutside   1 331\nTMhelix   332   354\ninside   355   372

14997 GCF_002037225.1_ASM203722v1 Salmonella enterica Proteobacteria; Gammaproteobacteria; Enterobacterales; Enterobacteriaceae; Salmonella MNNEETFYQAMRRKGVTRRSFLKFCSLAATSLGLGAGMTPKIAWALENKPRIPVVWIHGL WP_001058311.1 hydrogenase [Salmonella enterica] Length: 372\nNumber of predicted TMHs:  2\nExp number of AAs in TMHs: 37.4699899999999\nExp number, first 60 AAs:  19.72699\nTotal prob of N-in:   0.96171\nPOSSIBLE N-term signal sequence\ninside   1   19\nTMhelix   20   39\noutside   40   325\nTMhelix   326   348\ninside   349   372

14998 GCF_001705055.1_ASM170505v1 Salmonella enterica Proteobacteria; Gammaproteobacteria; Enterobacterales; Enterobacteriaceae; Salmonella MTGDNTLITSHGINRRDFMKLCAALAATMGLSSKAAAEMAESVSNPQRPPVIWIGAQECT WP_000145429.1 hydrogenase 2 small subunit [Salmonella enterica] Length: 372\nNumber of predicted TMHs:  1\nExp number of AAs in TMHs: 24.51309\nExp number, first 60 AAs:  0.2682\nTotal prob of N-in:   0.14402\noutside   1 331\nTMhelix   332   354\ninside   355   372

14999 GCF_001475145.1_Salmonella_enterica_CVM_43844_v1.0Salmonella enterica Proteobacteria; Gammaproteobacteria; Enterobacterales; Enterobacteriaceae; Salmonella

MNNEETFYQAMRRKGVTRRSFLKFCSLAATSLGLSAGMTPKIAWALENKPRIPVVWIHGL WP_001058316.1 hydrogenase [Salmonella enterica] Length: 372\nNumber of predicted TMHs: 2\nExp number of AAs in TMHs: 35.76397\nExp number, first 60 AAs: 18.97201\nTotal prob of N-in: 0.92785\nPOSSIBLE N-term signal sequence\ninside 1 19\nTMhelix 20 39\noutside 40 325\nTMhelix 326 348\ninside 349 372

15000 GCF_001475235.1_Salmonella_enterica_CVM_43835_v1.0Salmonella enterica Proteobacteria; Gammaproteobacteria; Enterobacterales; Enterobacteriaceae; Salmonella MNNEETFYQAMRRKGVTRRSFLKFCSLAATSLGLGAGMTPKIAWALENKPRIPVVWIHGL WP_001058312.1 hydrogenase [Salmonella enterica] Length: 372\nNumber of predicted TMHs: 2\nExp number of AAs in TMHs: 36.51738\nExp number, first 60 AAs: 19.72849\nTotal prob of N-in: 0.96170\nPOSSIBLE N-term signal sequence\ninside 1 19\nTMhelix 20 39\noutside 40 325\nTMhelix 326 348\ninside 349 372

15001 GCF_001475305.1_Salmonella_enterica_CVM_43805_v1.0Salmonella enterica Proteobacteria; Gammaproteobacteria; Enterobacterales; Enterobacteriaceae; Salmonella MNNEETFYQAMRRKGVTRRSFLKFCSLAATSLGLGAGMTPKIAWALENKPRIPVVWIHGL WP_001058312.1 hydrogenase [Salmonella enterica] Length: 372\nNumber of predicted TMHs: 2\nExp number of AAs in TMHs: 36.51738\nExp number, first 60 AAs: 19.72849\nTotal prob of N-in: 0.96170\nPOSSIBLE N-term signal sequence\ninside 1 19\nTMhelix 20 39\noutside 40 325\nTMhelix 326 348\ninside 349 372

15002 GCF_001475305.1_Salmonella_enterica_CVM_43805_v1.0Salmonella enterica Proteobacteria; Gammaproteobacteria; Enterobacterales; Enterobacteriaceae; Salmonella MTGDNTLITSHGINRRDFMKLCAALAATMGLSSKAAAEMAESVSNPQRPPVIWIGAQECT WP_000145429.1 hydrogenase 2 small subunit [Salmonella enterica] Length: 372\nNumber of predicted TMHs: 1\nExp number of AAs in TMHs: 24.51309\nExp number, first 60 AAs: 0.2682\nTotal prob of N-in: 0.14402\noutside 1 331\nTMhelix 332 354\ninside 355 372

15003 GCF_001475075.1_Salmonella_enterica_CVM_43834_v1.0Salmonella enterica Proteobacteria; Gammaproteobacteria; Enterobacterales; Enterobacteriaceae; Salmonella MNNEETFYQAMRRKGVTRRSFLKFCSLAATSLGLGAGMTPKIAWALENKPRIPVVWIHGL WP_001058311.1 hydrogenase [Salmonella enterica] Length: 372\nNumber of predicted TMHs: 2\nExp number of AAs in TMHs: 37.4699899999999\nExp number, first 60 AAs: 19.72699\nTotal prob of N-in: 0.96171\nPOSSIBLE N-term signal sequence\ninside 1 19\nTMhelix 20 39\noutside 40 325\nTMhelix 326 348\ninside 349 372

15004 GCF_001475215.1_Salmonella_enterica_CVM_43811_v1.0Salmonella enterica Proteobacteria; Gammaproteobacteria; Enterobacterales; Enterobacteriaceae; Salmonella MNNEETFYQAMRRKGVTRRSFLKFCSLAATSLGLGAGMTPKIAWALENKPRIPVVWIHGL WP_001058312.1 hydrogenase [Salmonella enterica] Length: 372\nNumber of predicted TMHs: 2\nExp number of AAs in TMHs: 36.51738\nExp number, first 60 AAs: 19.72849\nTotal prob of N-in: 0.96170\nPOSSIBLE N-term signal sequence\ninside 1 19\nTMhelix 20 39\noutside 40 325\nTMhelix 326 348\ninside 349 372

15005 GCF_001475345.1_Salmonella_enterica_CVM_43812_v1.0Salmonella enterica Proteobacteria; Gammaproteobacteria; Enterobacterales; Enterobacteriaceae; Salmonella MNNEETFYQAMRRKGVTRRSFLKFCSLAATSLGLGAGMTPKIAWALENKPRIPVVWIHGL WP_001058312.1 hydrogenase [Salmonella enterica] Length: 372\nNumber of predicted TMHs: 2\nExp number of AAs in TMHs: 36.51738\nExp number, first 60 AAs: 19.72849\nTotal prob of N-in: 0.96170\nPOSSIBLE N-term signal sequence\ninside 1 19\nTMhelix 20 39\noutside 40 325\nTMhelix 326 348\ninside 349 372

15006 GCF_001475345.1_Salmonella_enterica_CVM_43812_v1.0Salmonella enterica Proteobacteria; Gammaproteobacteria; Enterobacterales; Enterobacteriaceae; Salmonella MTGDNTLITSHGINRRDFMKLCAALAATMGLSSKAAAEMAESVSNPQRPPVIWIGAQECT WP_000145429.1 hydrogenase 2 small subunit [Salmonella enterica] Length: 372\nNumber of predicted TMHs: 1\nExp number of AAs in TMHs: 24.51309\nExp number, first 60 AAs: 0.2682\nTotal prob of N-in: 0.14402\noutside 1 331\nTMhelix 332 354\ninside 355 372

15007 GCF_001475215.1_Salmonella_enterica_CVM_43811_v1.0Salmonella enterica Proteobacteria; Gammaproteobacteria; Enterobacterales; Enterobacteriaceae; Salmonella MTGDNTLITSHGINRRDFMKLCAALAATMGLSSKAAAEMAESVSNPQRPPVIWIGAQECT WP_000145429.1 hydrogenase 2 small subunit [Salmonella enterica] Length: 372\nNumber of predicted TMHs: 1\nExp number of AAs in TMHs: 24.51309\nExp number, first 60 AAs: 0.2682\nTotal prob of N-in: 0.14402\noutside 1 331\nTMhelix 332 354\ninside 355 372

15008 GCF_001475045.1_Salmonella_enterica_CVM_43833_v1.0Salmonella enterica Proteobacteria; Gammaproteobacteria; Enterobacterales; Enterobacteriaceae; Salmonella
MNNEETFYQAMRRKGVTRRSFLKFCSLAATSLGLGAGMTPKIAWALENKPRIPVVWIHGL WP_001058312.1
hydrogenase [Salmonella enterica] Length: 372\nNumber of predicted TMHs: 2\nExp number of AAs in TMHs: 36.51738\nExp number, first 60 AAs: 19.72849\nTotal prob of N-in: 0.96170\nPOSSIBLE N-term signal sequence\ninside 1 19\nTMhelix 20 39\noutside 40 325\nTMhelix 326 348\ninside 349 372

15009 GCF_001475235.1_Salmonella_enterica_CVM_43835_v1.0Salmonella enterica Proteobacteria; Gammaproteobacteria; Enterobacterales; Enterobacteriaceae; Salmonella
MTGDNTLITSHGINRRDFMKLCAALAATMGLSSKAAAEMAESVSNPQRPPVIWIGAQECT WP_000145429.1
hydrogenase 2 small subunit [Salmonella enterica] Length: 372\nNumber of predicted TMHs: 1\nExp number of AAs in TMHs: 24.51309\nExp number, first 60 AAs: 0.2682\nTotal prob of N-in: 0.14402\noutside 1 331\nTMhelix 332 354\ninside 355 372

15010 GCF_001474925.1_Salmonella_enterica_CVM_43825_v1.0Salmonella enterica Proteobacteria; Gammaproteobacteria; Enterobacterales; Enterobacteriaceae; Salmonella
MNNEETFYQAMRRKGVTRRSFLKFCSLAATSLGLGAGMTPKIAWALENKPRIPVVWIHGL WP_001058312.1
hydrogenase [Salmonella enterica] Length: 372\nNumber of predicted TMHs: 2\nExp number of AAs in TMHs: 36.51738\nExp number, first 60 AAs: 19.72849\nTotal prob of N-in: 0.96170\nPOSSIBLE N-term signal sequence\ninside 1 19\nTMhelix 20 39\noutside 40 325\nTMhelix 326 348\ninside 349 372

15011 GCF_001474925.1_Salmonella_enterica_CVM_43825_v1.0Salmonella enterica Proteobacteria; Gammaproteobacteria; Enterobacterales; Enterobacteriaceae; Salmonella
MTGDNTLITSHGINRRDFMKLCAALAATMGLSSKAAAEMAESVSNPQRPPVIWIGAQECT WP_000145429.1
hydrogenase 2 small subunit [Salmonella enterica] Length: 372\nNumber of predicted TMHs: 1\nExp number of AAs in TMHs: 24.51309\nExp number, first 60 AAs: 0.2682\nTotal prob of N-in: 0.14402\noutside 1 331\nTMhelix 332 354\ninside 355 372

15012 GCF_001474825.1_Salmonella_enterica_CVM_43810_v1.0Salmonella enterica Proteobacteria; Gammaproteobacteria; Enterobacterales; Enterobacteriaceae; Salmonella
MNNEETFYQAMRRKGVTRRSFLKFCSLAATSLGLGAGMTPKIAWALENKPRIPVVWIHGL WP_001058312.1
hydrogenase [Salmonella enterica] Length: 372\nNumber of predicted TMHs: 2\nExp number of AAs in TMHs: 36.51738\nExp number, first 60 AAs: 19.72849\nTotal prob of N-in: 0.96170\nPOSSIBLE N-term signal sequence\ninside 1 19\nTMhelix 20 39\noutside 40 325\nTMhelix 326 348\ninside 349 372

15013 GCF_001474915.1_Salmonella_enterica_CVM_43821_v1.0Salmonella enterica Proteobacteria; Gammaproteobacteria; Enterobacterales; Enterobacteriaceae; Salmonella
MNNEETFYQAMRRKGVTRRSFLKFCSLAATSLGLGAGMTPKIAWALENKPRIPVVWIHGL WP_001058312.1
hydrogenase [Salmonella enterica] Length: 372\nNumber of predicted TMHs: 2\nExp number of AAs in TMHs: 36.51738\nExp number, first 60 AAs: 19.72849\nTotal prob of N-in: 0.96170\nPOSSIBLE N-term signal sequence\ninside 1 19\nTMhelix 20 39\noutside 40 325\nTMhelix 326 348\ninside 349 372

15014 GCF_001474885.1_Salmonella_enterica_CVM_43818_v1.0Salmonella enterica Proteobacteria; Gammaproteobacteria; Enterobacterales; Enterobacteriaceae; Salmonella
MNNEETFYQAMRRKGVTRRSFLKFCSLAATSLGLGAGMTPKIAWALENKPRIPVVWIHGL WP_001058312.1
hydrogenase [Salmonella enterica] Length: 372\nNumber of predicted TMHs: 2\nExp number of AAs in TMHs: 36.51738\nExp number, first 60 AAs: 19.72849\nTotal prob of N-in: 0.96170\nPOSSIBLE N-term signal sequence\ninside 1 19\nTMhelix 20 39\noutside 40 325\nTMhelix 326 348\ninside 349 372

15015 GCF_001474915.1_Salmonella_enterica_CVM_43821_v1.0Salmonella enterica Proteobacteria; Gammaproteobacteria; Enterobacterales; Enterobacteriaceae; Salmonella
MTGDNTLITSHGINRRDFMKLCAALAATMGLSSKAAAEMAESVSNPQRPPVIWIGAQECT WP_000145429.1
hydrogenase 2 small subunit [Salmonella enterica] Length: 372\nNumber of predicted TMHs: 1\nExp number of AAs in TMHs: 24.51309\nExp number, first 60 AAs: 0.2682\nTotal prob of N-in: 0.14402\noutside 1 331\nTMhelix 332 354\ninside 355 372

15016 GCF_001474885.1_Salmonella_enterica_CVM_43818_v1.0Salmonella enterica Proteobacteria; Gammaproteobacteria; Enterobacterales; Enterobacteriaceae; Salmonella
MTGDNTLITSHGINRRDFMKLCAALAATMGLSSKAAAEMAESVSNPQRPPVIWIGAQECT WP_000145429.1
hydrogenase 2 small subunit [Salmonella enterica] Length: 372\nNumber of predicted TMHs: 1\nExp

number of AAs in TMHs: 24.51309\nExp number, first 60 AAs:  0.2682\nTotal prob of N-in:   0.14402\noutside   1   331\nTMhelix   332   354\ninside   355   372

15017 GCF_001474835.1_Salmonella_enterica_CVM_43813_v1.0Salmonella enterica Proteobacteria; Gammaproteobacteria; Enterobacterales; Enterobacteriaceae; Salmonella
MNNEETFYQAMRRKGVTRRSFLKFCSLAATSLGLGAGMTPKIAWALENKPRIPVVWIHGL WP_001058312.1
hydrogenase [Salmonella enterica] Length: 372\nNumber of predicted TMHs:  2\nExp number of AAs in TMHs: 36.51738\nExp number, first 60 AAs:  19.72849\nTotal prob of N-in:   0.96170\nPOSSIBLE N-term signal sequence\ninside   1   19\nTMhelix   20   39\noutside   40   325\nTMhelix   326   348\ninside   349   372

15018 GCF_001474845.1_Salmonella_enterica_CVM_43817_v1.0Salmonella enterica Proteobacteria; Gammaproteobacteria; Enterobacterales; Enterobacteriaceae; Salmonella
MNNEETFYQAMRRKGVTRRSFLKFCSLAATSLGLGAGMTPKIAWALENKPRIPVVWIHGL WP_001058312.1
hydrogenase [Salmonella enterica] Length: 372\nNumber of predicted TMHs:  2\nExp number of AAs in TMHs: 36.51738\nExp number, first 60 AAs:  19.72849\nTotal prob of N-in:   0.96170\nPOSSIBLE N-term signal sequence\ninside   1   19\nTMhelix   20   39\noutside   40   325\nTMhelix   326   348\ninside   349   372

15019 GCF_001474905.1_Salmonella_enterica_CVM_43824_v1.0Salmonella enterica Proteobacteria; Gammaproteobacteria; Enterobacterales; Enterobacteriaceae; Salmonella
MTGDNTLITSHGINRRDFMKLCAALAATMGLSSKAAAEMAESVSNPQRPPVIWIGAQECT WP_000145429.1
hydrogenase 2 small subunit [Salmonella enterica] Length: 372\nNumber of predicted TMHs:  1\nExp number of AAs in TMHs: 24.51309\nExp number, first 60 AAs:  0.2682\nTotal prob of N-in:   0.14402\noutside   1   331\nTMhelix   332   354\ninside   355   372

15020 GCF_001474805.1_Salmonella_enterica_CVM_43809_v1.0Salmonella enterica Proteobacteria; Gammaproteobacteria; Enterobacterales; Enterobacteriaceae; Salmonella
MNNEETFYQAMRRKGVTRRSFLKFCSLAATSLGLGAGMTPKIAWALENKPRIPVVWIHGL WP_001058312.1
hydrogenase [Salmonella enterica] Length: 372\nNumber of predicted TMHs:  2\nExp number of AAs in TMHs: 36.51738\nExp number, first 60 AAs:  19.72849\nTotal prob of N-in:   0.96170\nPOSSIBLE N-term signal sequence\ninside   1   19\nTMhelix   20   39\noutside   40   325\nTMhelix   326   348\ninside   349   372

15021 GCF_001474845.1_Salmonella_enterica_CVM_43817_v1.0Salmonella enterica Proteobacteria; Gammaproteobacteria; Enterobacterales; Enterobacteriaceae; Salmonella
MTGDNTLITSHGINRRDFMKLCAALAATMGLSSKAAAEMAESVSNPQRPPVIWIGAQECT WP_000145429.1
hydrogenase 2 small subunit [Salmonella enterica] Length: 372\nNumber of predicted TMHs:  1\nExp number of AAs in TMHs: 24.51309\nExp number, first 60 AAs:  0.2682\nTotal prob of N-in:   0.14402\noutside   1   331\nTMhelix   332   354\ninside   355   372

15022 GCF_001474835.1_Salmonella_enterica_CVM_43813_v1.0Salmonella enterica Proteobacteria; Gammaproteobacteria; Enterobacterales; Enterobacteriaceae; Salmonella
MTGDNTLITSHGINRRDFMKLCAALAATMGLSSKAAAEMAESVSNPQRPPVIWIGAQECT WP_000145429.1
hydrogenase 2 small subunit [Salmonella enterica] Length: 372\nNumber of predicted TMHs:  1\nExp number of AAs in TMHs: 24.51309\nExp number, first 60 AAs:  0.2682\nTotal prob of N-in:   0.14402\noutside   1   331\nTMhelix   332   354\ninside   355   372

15023 GCF_001474775.1_Salmonella_enterica_CVM_43808_v1.0Salmonella enterica Proteobacteria; Gammaproteobacteria; Enterobacterales; Enterobacteriaceae; Salmonella
MNNEETFYQAMRRKGVTRRSFLKFCSLAATSLGLGAGMTPKIAWALENKPRIPVVWIHGL WP_001058311.1
hydrogenase [Salmonella enterica] Length: 372\nNumber of predicted TMHs:  2\nExp number of AAs in TMHs: 37.4699899999999\nExp number, first 60 AAs:  19.72699\nTotal prob of N-in:   0.96171\nPOSSIBLE N-term signal sequence\ninside   1   19\nTMhelix   20   39\noutside   40   325\nTMhelix   326   348\ninside   349   372

15024 GCF_001474655.1_Salmonella_enterica_CVM_43796_v1.0Salmonella enterica Proteobacteria; Gammaproteobacteria; Enterobacterales; Enterobacteriaceae; Salmonella
MNNEETFYQAMRRKGVTRRSFLKFCSLAATSLGLSAGMTPKIAWALENKPRIPVVWIHGL WP_001058316.1
hydrogenase [Salmonella enterica] Length: 372\nNumber of predicted TMHs:  2\nExp number of AAs in TMHs: 35.76397\nExp number, first 60 AAs:  18.97201\nTotal prob of N-in:   0.92785\nPOSSIBLE N-term signal sequence\ninside   1   19\nTMhelix   20   39\noutside   40   325\nTMhelix   326   348\ninside   349   372

15025 GCF_001474685.1_Salmonella_enterica_CVM_43797_v1.0Salmonella enterica Proteobacteria; Gammaproteobacteria; Enterobacterales; Enterobacteriaceae; Salmonella

MNNEETFYQAMRRKGVTRRSFLKFCSLAATSLGLGAGMTPKIAWALENKPRIPVVWIHGL WP_001058307.1 hydrogenase [Salmonella enterica] Length: 372\nNumber of predicted TMHs: 2\nExp number of AAs in TMHs: 36.51738\nExp number, first 60 AAs: 19.72849\nTotal prob of N-in: 0.96170\nPOSSIBLE N-term signal sequence\ninside 1 19\nTMhelix 20 39\noutside 40 325\nTMhelix 326 348\ninside 349 372

15026 GCF_001474745.1_Salmonella_enterica_CVM_43803_v1.0Salmonella enterica Proteobacteria; Gammaproteobacteria; Enterobacterales; Enterobacteriaceae; Salmonella MNNEETFYQAMRRKGVTRRSFLKFCSLAATSLGLGAGMTPKIAWALENKPRIPVVWIHGL WP_001058312.1 hydrogenase [Salmonella enterica] Length: 372\nNumber of predicted TMHs: 2\nExp number of AAs in TMHs: 36.51738\nExp number, first 60 AAs: 19.72849\nTotal prob of N-in: 0.96170\nPOSSIBLE N-term signal sequence\ninside 1 19\nTMhelix 20 39\noutside 40 325\nTMhelix 326 348\ninside 349 372

15027 GCF_001474755.1_Salmonella_enterica_CVM_43806_v1.0Salmonella enterica Proteobacteria; Gammaproteobacteria; Enterobacterales; Enterobacteriaceae; Salmonella MNNEETFYQAMRRKGVTRRSFLKFCSLAATSLGLGAGMTPKIAWALENKPRIPVVWIHGL WP_001058312.1 hydrogenase [Salmonella enterica] Length: 372\nNumber of predicted TMHs: 2\nExp number of AAs in TMHs: 36.51738\nExp number, first 60 AAs: 19.72849\nTotal prob of N-in: 0.96170\nPOSSIBLE N-term signal sequence\ninside 1 19\nTMhelix 20 39\noutside 40 325\nTMhelix 326 348\ninside 349 372

15028 GCF_001474745.1_Salmonella_enterica_CVM_43803_v1.0Salmonella enterica Proteobacteria; Gammaproteobacteria; Enterobacterales; Enterobacteriaceae; Salmonella MTGDNTLITSHGINRRDFMKLCAALAATMGLSSKAAAEMAESVSNPQRPPVIWIGAQECT WP_000145429.1 hydrogenase 2 small subunit [Salmonella enterica] Length: 372\nNumber of predicted TMHs: 1\nExp number of AAs in TMHs: 24.51309\nExp number, first 60 AAs: 0.2682\nTotal prob of N-in: 0.14402\noutside 1 331\nTMhelix 332 354\ninside 355 372

15029 GCF_002054725.1_ASM205472v1 Salmonella enterica Proteobacteria; Gammaproteobacteria; Enterobacterales; Enterobacteriaceae; Salmonella MTGDNTLITSHGINRRDFMKLCAALAATMGLSSKAAAEMAESVSNPQRPPVIWIGAQECT WP_000145429.1 hydrogenase 2 small subunit [Salmonella enterica] Length: 372\nNumber of predicted TMHs: 1\nExp number of AAs in TMHs: 24.51309\nExp number, first 60 AAs: 0.2682\nTotal prob of N-in: 0.14402\noutside 1 331\nTMhelix 332 354\ninside 355 372

15030 GCF_001474755.1_Salmonella_enterica_CVM_43806_v1.0Salmonella enterica Proteobacteria; Gammaproteobacteria; Enterobacterales; Enterobacteriaceae; Salmonella MTGDNTLITSHGINRRDFMKLCAALAATMGLSSKAAAEMAESVSNPQRPPVIWIGAQECT WP_000145429.1 hydrogenase 2 small subunit [Salmonella enterica] Length: 372\nNumber of predicted TMHs: 1\nExp number of AAs in TMHs: 24.51309\nExp number, first 60 AAs: 0.2682\nTotal prob of N-in: 0.14402\noutside 1 331\nTMhelix 332 354\ninside 355 372

15031 GCF_001474705.1_Salmonella_enterica_CVM_43798_v1.0Salmonella enterica Proteobacteria; Gammaproteobacteria; Enterobacterales; Enterobacteriaceae; Salmonella MNNEETFYQAMRRKGVTRRSFLKFCSLAATSLGLGAGMTPKIAWALENKPRIPVVWIHGL WP_001058312.1 hydrogenase [Salmonella enterica] Length: 372\nNumber of predicted TMHs: 2\nExp number of AAs in TMHs: 36.51738\nExp number, first 60 AAs: 19.72849\nTotal prob of N-in: 0.96170\nPOSSIBLE N-term signal sequence\ninside 1 19\nTMhelix 20 39\noutside 40 325\nTMhelix 326 348\ninside 349 372

15032 GCF_001474725.1_Salmonella_enterica_CVM_43801_v1.0Salmonella enterica Proteobacteria; Gammaproteobacteria; Enterobacterales; Enterobacteriaceae; Salmonella MNNEETFYQAMRRKGVTRRSFLKFCSLAATSLGLGAGMTPKIAWALENKPRIPVVWIHGL WP_001058308.1 hydrogenase [Salmonella enterica] Length: 372\nNumber of predicted TMHs: 2\nExp number of AAs in TMHs: 39.08204\nExp number, first 60 AAs: 19.72474\nTotal prob of N-in: 0.96174\nPOSSIBLE N-term signal sequence\ninside 1 19\nTMhelix 20 39\noutside 40 325\nTMhelix 326 348\ninside 349 372

15033 GCF_001474725.1_Salmonella_enterica_CVM_43801_v1.0Salmonella enterica Proteobacteria; Gammaproteobacteria; Enterobacterales; Enterobacteriaceae; Salmonella MTGDNTLITSHGINRRDFMKLCAALAATMGLSSKAAAEMAESVSNPQRPPVIWIGAQECT WP_000145429.1 hydrogenase 2 small subunit [Salmonella enterica] Length: 372\nNumber of predicted TMHs: 1\nExp number of AAs in TMHs: 24.51309\nExp number, first 60 AAs: 0.2682\nTotal prob of N-in: 0.14402\noutside 1 331\nTMhelix 332 354\ninside 355 372

15034 GCF_001474605.1_Salmonella_enterica_CVM_43792_v1.0Salmonella enterica Proteobacteria; Gammaproteobacteria; Enterobacterales; Enterobacteriaceae; Salmonella
MNNEETFYQAMRRKGVTRRSFLKFCSLAATSLGLGAGMTPKIAWALENKPRIPVVWIHGL WP_001058312.1
hydrogenase [Salmonella enterica] Length: 372\nNumber of predicted TMHs: 2\nExp number of AAs in TMHs: 36.51738\nExp number, first 60 AAs: 19.72849\nTotal prob of N-in: 0.96170\nPOSSIBLE N-term signal sequence\ninside 1 19\nTMhelix 20 39\noutside 40 325\nTMhelix 326 348\ninside 349 372

15035 GCF_001474645.1_Salmonella_enterica_CVM_43795_v1.0Salmonella enterica Proteobacteria; Gammaproteobacteria; Enterobacterales; Enterobacteriaceae; Salmonella
MNNEETFYQAMRRKGVTRRSFLKFCSLAATSLGLGAGMTPKIAWALENKPRIPVVWIHGL WP_001058312.1
hydrogenase [Salmonella enterica] Length: 372\nNumber of predicted TMHs: 2\nExp number of AAs in TMHs: 36.51738\nExp number, first 60 AAs: 19.72849\nTotal prob of N-in: 0.96170\nPOSSIBLE N-term signal sequence\ninside 1 19\nTMhelix 20 39\noutside 40 325\nTMhelix 326 348\ninside 349 372

15036 GCF_001474685.1_Salmonella_enterica_CVM_43797_v1.0Salmonella enterica Proteobacteria; Gammaproteobacteria; Enterobacterales; Enterobacteriaceae; Salmonella
MTGDNTLITSHGINRRDFMKLCAALAATMGLSSKAAAEMAESVSNPQRPPVIWIGAQECT WP_000145429.1
hydrogenase 2 small subunit [Salmonella enterica] Length: 372\nNumber of predicted TMHs: 1\nExp number of AAs in TMHs: 24.51309\nExp number, first 60 AAs: 0.2682\nTotal prob of N-in: 0.14402\noutside 1 331\nTMhelix 332 354\ninside 355 372

15037 GCF_001474625.1_Salmonella_enterica_CVM_43794_v1.0Salmonella enterica Proteobacteria; Gammaproteobacteria; Enterobacterales; Enterobacteriaceae; Salmonella
MNNEETFYQAMRRKGVTRRSFLKFCSLAATSLGLGAGMTPKIAWALENKPRIPVVWIHGL WP_001058312.1
hydrogenase [Salmonella enterica] Length: 372\nNumber of predicted TMHs: 2\nExp number of AAs in TMHs: 36.51738\nExp number, first 60 AAs: 19.72849\nTotal prob of N-in: 0.96170\nPOSSIBLE N-term signal sequence\ninside 1 19\nTMhelix 20 39\noutside 40 325\nTMhelix 326 348\ninside 349 372

15038 GCF_001474705.1_Salmonella_enterica_CVM_43798_v1.0Salmonella enterica Proteobacteria; Gammaproteobacteria; Enterobacterales; Enterobacteriaceae; Salmonella
MTGDNTLITSHGINRRDFMKLCAALAATMGLSSKAAAEMAESVSNPQRPPVIWIGAQECT WP_000145429.1
hydrogenase 2 small subunit [Salmonella enterica] Length: 372\nNumber of predicted TMHs: 1\nExp number of AAs in TMHs: 24.51309\nExp number, first 60 AAs: 0.2682\nTotal prob of N-in: 0.14402\noutside 1 331\nTMhelix 332 354\ninside 355 372

15039 GCF_001474645.1_Salmonella_enterica_CVM_43795_v1.0Salmonella enterica Proteobacteria; Gammaproteobacteria; Enterobacterales; Enterobacteriaceae; Salmonella
MTGDNTLITSHGINRRDFMKLCAALAATMGLSSKAAAEMAESVSNPQRPPVIWIGAQECT WP_000145429.1
hydrogenase 2 small subunit [Salmonella enterica] Length: 372\nNumber of predicted TMHs: 1\nExp number of AAs in TMHs: 24.51309\nExp number, first 60 AAs: 0.2682\nTotal prob of N-in: 0.14402\noutside 1 331\nTMhelix 332 354\ninside 355 372

15040 GCF_001474605.1_Salmonella_enterica_CVM_43792_v1.0Salmonella enterica Proteobacteria; Gammaproteobacteria; Enterobacterales; Enterobacteriaceae; Salmonella
MTGDNTLITSHGINRRDFMKLCAALAATMGLSSKAAAEMAESVSNPQRPPVIWIGAQECT WP_000145429.1
hydrogenase 2 small subunit [Salmonella enterica] Length: 372\nNumber of predicted TMHs: 1\nExp number of AAs in TMHs: 24.51309\nExp number, first 60 AAs: 0.2682\nTotal prob of N-in: 0.14402\noutside 1 331\nTMhelix 332 354\ninside 355 372

15041 GCF_001474625.1_Salmonella_enterica_CVM_43794_v1.0Salmonella enterica Proteobacteria; Gammaproteobacteria; Enterobacterales; Enterobacteriaceae; Salmonella
MTGDNTLITSHGINRRDFMKLCAALAATMGLSSKAAAEMAESVSNPQRPPVIWIGAQECT WP_000145429.1
hydrogenase 2 small subunit [Salmonella enterica] Length: 372\nNumber of predicted TMHs: 1\nExp number of AAs in TMHs: 24.51309\nExp number, first 60 AAs: 0.2682\nTotal prob of N-in: 0.14402\noutside 1 331\nTMhelix 332 354\ninside 355 372

15042 GCF_001474505.1_Salmonella_enterica_CVM_43782_v1.0Salmonella enterica Proteobacteria; Gammaproteobacteria; Enterobacterales; Enterobacteriaceae; Salmonella
MNNEETFYQAMRRKGVTRRSFLKFCSLAATSLGLGAGMTPKIAWALENKPRIPVVWIHGL WP_023198237.1
hydrogenase 1, small subunit [Salmonella enterica] Length: 372\nNumber of predicted TMHs: 2\nExp

number of AAs in TMHs: 36.39423\nExp number, first 60 AAs: 19.72877\nTotal prob of N-in: 0.96162\nPOSSIBLE N-term signal sequence\ninside 1 19\nTMhelix 20 39\noutside 40 325\nTMhelix 326 348\ninside 349 372

15043 GCF_001474575.1_Salmonella_enterica_CVM_43791_v1.0Salmonella enterica Proteobacteria; Gammaproteobacteria; Enterobacterales; Enterobacteriaceae; Salmonella MNNEETLYQAMRRKGVTRRSFLKFCSLAATSLGLGAGMTPKIAWALENKPRIPVVWIHGL WP_001752445.1 hydrogenase-1 small chain [Salmonella enterica] Length: 372\nNumber of predicted TMHs: 2\nExp number of AAs in TMHs: 36.5123\nExp number, first 60 AAs: 19.72341\nTotal prob of N-in: 0.96144\nPOSSIBLE N-term signal sequence\ninside 1 19\nTMhelix 20 39\noutside 40 325\nTMhelix 326 348\ninside 349 372

15044 GCF_001474565.1_Salmonella_enterica_CVM_43789_v1.0Salmonella enterica Proteobacteria; Gammaproteobacteria; Enterobacterales; Enterobacteriaceae; Salmonella MNNEETFYQAMRRKGVTRRSFLKFCSLAATSLGLGAGMTPKIAWALENKPRIPVVWIHGL WP_001058312.1 hydrogenase [Salmonella enterica] Length: 372\nNumber of predicted TMHs: 2\nExp number of AAs in TMHs: 36.51738\nExp number, first 60 AAs: 19.72849\nTotal prob of N-in: 0.96170\nPOSSIBLE N-term signal sequence\ninside 1 19\nTMhelix 20 39\noutside 40 325\nTMhelix 326 348\ninside 349 372

15045 GCF_001474575.1_Salmonella_enterica_CVM_43791_v1.0Salmonella enterica Proteobacteria; Gammaproteobacteria; Enterobacterales; Enterobacteriaceae; Salmonella MTGDNTLITSHGINRRDFMKLCAALAATMGLSSKAAAEMAESVSNPQRPPVIWIGAQECT WP_000145428.1 hydrogenase 2 small subunit [Salmonella enterica] Length: 372\nNumber of predicted TMHs: 1\nExp number of AAs in TMHs: 24.50677\nExp number, first 60 AAs: 0.2682\nTotal prob of N-in: 0.14390\noutside 1 331\nTMhelix 332 354\ninside 355 372

15046 GCF_001474515.1_Salmonella_enterica_CVM_43785_v1.0Salmonella enterica Proteobacteria; Gammaproteobacteria; Enterobacterales; Enterobacteriaceae; Salmonella MNNEETFYQAMRRKGVTRRSFLKFCSLAATSLGLGAGMTPKIAWALENKPRIPVVWIHGL WP_001058312.1 hydrogenase [Salmonella enterica] Length: 372\nNumber of predicted TMHs: 2\nExp number of AAs in TMHs: 36.51738\nExp number, first 60 AAs: 19.72849\nTotal prob of N-in: 0.96170\nPOSSIBLE N-term signal sequence\ninside 1 19\nTMhelix 20 39\noutside 40 325\nTMhelix 326 348\ninside 349 372

15047 GCF_001474545.1_Salmonella_enterica_CVM_43788_v1.0Salmonella enterica Proteobacteria; Gammaproteobacteria; Enterobacterales; Enterobacteriaceae; Salmonella MNNEETFYQAMRRKGVTRRSFLKFCSLAATSLGLGAGMTPKIAWALENKPRIPVVWIHGL WP_001058312.1 hydrogenase [Salmonella enterica] Length: 372\nNumber of predicted TMHs: 2\nExp number of AAs in TMHs: 36.51738\nExp number, first 60 AAs: 19.72849\nTotal prob of N-in: 0.96170\nPOSSIBLE N-term signal sequence\ninside 1 19\nTMhelix 20 39\noutside 40 325\nTMhelix 326 348\ninside 349 372

15048 GCF_001474545.1_Salmonella_enterica_CVM_43788_v1.0Salmonella enterica Proteobacteria; Gammaproteobacteria; Enterobacterales; Enterobacteriaceae; Salmonella MTGDNTLITSHGINRRDFMKLCAALAATMGLSSKAAAEMAESVSNPQRPPVIWIGAQECT WP_000145429.1 hydrogenase 2 small subunit [Salmonella enterica] Length: 372\nNumber of predicted TMHs: 1\nExp number of AAs in TMHs: 24.51309\nExp number, first 60 AAs: 0.2682\nTotal prob of N-in: 0.14402\noutside 1 331\nTMhelix 332 354\ninside 355 372

15049 GCF_001474415.1_Salmonella_enterica_CVM_43776_v1.0Salmonella enterica Proteobacteria; Gammaproteobacteria; Enterobacterales; Enterobacteriaceae; Salmonella MNNEETFYQAMRRKGVTRRSFLKFCSLAATSLGLSAGMTPKIAWALENKPRIPVVWIHGL WP_001058316.1 hydrogenase [Salmonella enterica] Length: 372\nNumber of predicted TMHs: 2\nExp number of AAs in TMHs: 35.76397\nExp number, first 60 AAs: 18.97201\nTotal prob of N-in: 0.92785\nPOSSIBLE N-term signal sequence\ninside 1 19\nTMhelix 20 39\noutside 40 325\nTMhelix 326 348\ninside 349 372

15050 GCF_001474445.1_Salmonella_enterica_CVM_43777_v1.0Salmonella enterica Proteobacteria; Gammaproteobacteria; Enterobacterales; Enterobacteriaceae; Salmonella MNNEETFYQAMRRKGVTRRSFLKFCSLAATSLGLGAGMTPKIAWALENKPRIPVVWIHGL WP_001058312.1 hydrogenase [Salmonella enterica] Length: 372\nNumber of predicted TMHs: 2\nExp number of AAs in TMHs: 36.51738\nExp number, first 60 AAs: 19.72849\nTotal prob of N-in: 0.96170\nPOSSIBLE N-term signal sequence\ninside 1 19\nTMhelix 20 39\noutside 40 325\nTMhelix 326 348\ninside 349 372

15051 GCF_001474465.1_Salmonella_enterica_CVM_43779_v1.0Salmonella enterica Proteobacteria; Gammaproteobacteria; Enterobacterales; Enterobacteriaceae; Salmonella

MNNEETFYQAMRRKGVTRRSFLKFCSLAATSLGLGAGMTPKIAWALENKPRIPVVWIHGL WP_001058312.1 hydrogenase [Salmonella enterica] Length: 372\nNumber of predicted TMHs: 2\nExp number of AAs in TMHs: 36.51738\nExp number, first 60 AAs: 19.72849\nTotal prob of N-in: 0.96170\nPOSSIBLE N-term signal sequence\ninside 1 19\nTMhelix 20 39\noutside 40 325\nTMhelix 326 348\ninside 349 372

15052 GCF_001474485.1_Salmonella_enterica_CVM_43780_v1.0Salmonella enterica Proteobacteria; Gammaproteobacteria; Enterobacterales; Enterobacteriaceae; Salmonella MNNEETFYQAMRRKGVTRRSFLKFCSLAATSLGLGAGMTPKIAWALENKPRIPVVWIHGL WP_001058307.1 hydrogenase [Salmonella enterica] Length: 372\nNumber of predicted TMHs: 2\nExp number of AAs in TMHs: 36.51738\nExp number, first 60 AAs: 19.72849\nTotal prob of N-in: 0.96170\nPOSSIBLE N-term signal sequence\ninside 1 19\nTMhelix 20 39\noutside 40 325\nTMhelix 326 348\ninside 349 372

15053 GCF_001474415.1_Salmonella_enterica_CVM_43776_v1.0Salmonella enterica Proteobacteria; Gammaproteobacteria; Enterobacterales; Enterobacteriaceae; Salmonella MTGDNTLITSHGINRRDFMKLCAALAATMGLSSKAAAEMAESVSNPQRPPVIWIGAQECT WP_000145429.1 hydrogenase 2 small subunit [Salmonella enterica] Length: 372\nNumber of predicted TMHs: 1\nExp number of AAs in TMHs: 24.51309\nExp number, first 60 AAs: 0.2682\nTotal prob of N-in: 0.14402\noutside 1 331\nTMhelix 332 354\ninside 355 372

15054 GCF_001474445.1_Salmonella_enterica_CVM_43777_v1.0Salmonella enterica Proteobacteria; Gammaproteobacteria; Enterobacterales; Enterobacteriaceae; Salmonella MTGDNTLITSHGINRRDFMKLCAALAATMGLSSKAAAEMAESVSNPQRPPVIWIGAQECT WP_000145429.1 hydrogenase 2 small subunit [Salmonella enterica] Length: 372\nNumber of predicted TMHs: 1\nExp number of AAs in TMHs: 24.51309\nExp number, first 60 AAs: 0.2682\nTotal prob of N-in: 0.14402\noutside 1 331\nTMhelix 332 354\ninside 355 372

15055 GCF_001474485.1_Salmonella_enterica_CVM_43780_v1.0Salmonella enterica Proteobacteria; Gammaproteobacteria; Enterobacterales; Enterobacteriaceae; Salmonella MTGDNTLITSHGINRRDFMKLCAALAATMGLSSKAAAEMAESVSNPQRPPVIWIGAQECT WP_000145429.1 hydrogenase 2 small subunit [Salmonella enterica] Length: 372\nNumber of predicted TMHs: 1\nExp number of AAs in TMHs: 24.51309\nExp number, first 60 AAs: 0.2682\nTotal prob of N-in: 0.14402\noutside 1 331\nTMhelix 332 354\ninside 355 372

15056 GCF_001474465.1_Salmonella_enterica_CVM_43779_v1.0Salmonella enterica Proteobacteria; Gammaproteobacteria; Enterobacterales; Enterobacteriaceae; Salmonella MTGDNTLITSHGINRRDFMKLCAALAATMGLSSKAAAEMAESVSNPQRPPVIWIGAQECT WP_000145429.1 hydrogenase 2 small subunit [Salmonella enterica] Length: 372\nNumber of predicted TMHs: 1\nExp number of AAs in TMHs: 24.51309\nExp number, first 60 AAs: 0.2682\nTotal prob of N-in: 0.14402\noutside 1 331\nTMhelix 332 354\ninside 355 372

15057 GCF_001474355.1_Salmonella_enterica_CVM_43771_v1.0Salmonella enterica Proteobacteria; Gammaproteobacteria; Enterobacterales; Enterobacteriaceae; Salmonella MNNEETFYQAMRRKGVTRRSFLKFCSLAATSLGLGAGMTPKIAWALENKPRIPVVWIHGL WP_001058312.1 hydrogenase [Salmonella enterica] Length: 372\nNumber of predicted TMHs: 2\nExp number of AAs in TMHs: 36.51738\nExp number, first 60 AAs: 19.72849\nTotal prob of N-in: 0.96170\nPOSSIBLE N-term signal sequence\ninside 1 19\nTMhelix 20 39\noutside 40 325\nTMhelix 326 348\ninside 349 372

15058 GCF_001474325.1_Salmonella_enterica_CVM_43769_v1.0Salmonella enterica Proteobacteria; Gammaproteobacteria; Enterobacterales; Enterobacteriaceae; Salmonella MNNEETFYQAMRRKGVTRRSFLKFCSLAATSLGLGAGMTPKIAWALENKPRIPVVWIHGL WP_001058312.1 hydrogenase [Salmonella enterica] Length: 372\nNumber of predicted TMHs: 2\nExp number of AAs in TMHs: 36.51738\nExp number, first 60 AAs: 19.72849\nTotal prob of N-in: 0.96170\nPOSSIBLE N-term signal sequence\ninside 1 19\nTMhelix 20 39\noutside 40 325\nTMhelix 326 348\ninside 349 372

15059 GCF_001474405.1_Salmonella_enterica_CVM_43774_v1.0Salmonella enterica Proteobacteria; Gammaproteobacteria; Enterobacterales; Enterobacteriaceae; Salmonella MNNEETFYQAMRRKGVTRRSFLKFCSLAATSLGLGAGMTPKIAWALENKPRIPVVWIHGL WP_001058311.1 hydrogenase [Salmonella enterica] Length: 372\nNumber of predicted TMHs: 2\nExp number of AAs in TMHs: 37.4699899999999\nExp number, first 60 AAs: 19.72699\nTotal prob of N-in: 0.96171\nPOSSIBLE N-term signal sequence\ninside 1 19\nTMhelix 20 39\noutside 40 325\nTMhelix 326 348\ninside 349 372

15060	GCF_001474405.1_Salmonella_enterica_CVM_43774_v1.0Salmonella enterica	Proteobacteria; Gammaproteobacteria; Enterobacterales; Enterobacteriaceae; Salmonella
MTGDNTLITSHGINRRDFMKLCAALAATMGLSSKAAAEMAESVSNPQRPPVIWIGAQECT	WP_000145429.1
hydrogenase 2 small subunit [Salmonella enterica]	Length: 372\nNumber of predicted TMHs: 1\nExp number of AAs in TMHs: 24.51309\nExp number, first 60 AAs: 0.2682\nTotal prob of N-in: 0.14402\noutside 1 331\nTMhelix 332 354\ninside 355 372

15061	GCF_001570285.1_ASM157028v1	Salmonella enterica	Proteobacteria; Gammaproteobacteria; Enterobacterales; Enterobacteriaceae; Salmonella
MNNEETFYQAMRRKGVTRRSFLKFCSLAATSLGLGAGMTPKIAWALENKPRIPVVWIHGL	WP_001058310.1
hydrogenase [Salmonella enterica]	Length: 372\nNumber of predicted TMHs: 2\nExp number of AAs in TMHs: 37.4567299999999\nExp number, first 60 AAs: 19.72694\nTotal prob of N-in: 0.96171\nPOSSIBLE N-term signal sequence\ninside 1 19\nTMhelix 20 39\noutside 40 325\nTMhelix 326 348\ninside 349 372

15062	GCF_001474335.1_Salmonella_enterica_CVM_43770_v1.0Salmonella enterica	Proteobacteria; Gammaproteobacteria; Enterobacterales; Enterobacteriaceae; Salmonella
MNNEETFYQAMRRKGVTRRSFLKFCSLAATSLGLGAGMTPKIAWALENKPRIPVVWIHGL	WP_001058310.1
hydrogenase [Salmonella enterica]	Length: 372\nNumber of predicted TMHs: 2\nExp number of AAs in TMHs: 37.4567299999999\nExp number, first 60 AAs: 19.72694\nTotal prob of N-in: 0.96171\nPOSSIBLE N-term signal sequence\ninside 1 19\nTMhelix 20 39\noutside 40 325\nTMhelix 326 348\ninside 349 372

15063	GCF_001474385.1_Salmonella_enterica_CVM_43773_v1.0Salmonella enterica	Proteobacteria; Gammaproteobacteria; Enterobacterales; Enterobacteriaceae; Salmonella
MNNEETFYQAMRRKGVTRRSFLKFCSLAATSLGLGAGMTPKIAWALENKPRIPVVWIHGL	WP_001058311.1
hydrogenase [Salmonella enterica]	Length: 372\nNumber of predicted TMHs: 2\nExp number of AAs in TMHs: 37.4699899999999\nExp number, first 60 AAs: 19.72699\nTotal prob of N-in: 0.96171\nPOSSIBLE N-term signal sequence\ninside 1 19\nTMhelix 20 39\noutside 40 325\nTMhelix 326 348\ninside 349 372

15064	GCF_001474385.1_Salmonella_enterica_CVM_43773_v1.0Salmonella enterica	Proteobacteria; Gammaproteobacteria; Enterobacterales; Enterobacteriaceae; Salmonella
MTGDNTLITSHGINRRDFMKLCAALAATMGLSSKAAAEMAESVSNPQRPPVIWIGAQECT	WP_000145429.1
hydrogenase 2 small subunit [Salmonella enterica]	Length: 372\nNumber of predicted TMHs: 1\nExp number of AAs in TMHs: 24.51309\nExp number, first 60 AAs: 0.2682\nTotal prob of N-in: 0.14402\noutside 1 331\nTMhelix 332 354\ninside 355 372

15065	GCF_001474275.1_Salmonella_enterica_CVM_43767_v1.0Salmonella enterica	Proteobacteria; Gammaproteobacteria; Enterobacterales; Enterobacteriaceae; Salmonella
MNNEETFYQAMRRKGVTRRSFLKFCSLAATSLGLGAGMTPKIAWALENKPRIPVVWIHGL	WP_001058310.1
hydrogenase [Salmonella enterica]	Length: 372\nNumber of predicted TMHs: 2\nExp number of AAs in TMHs: 37.4567299999999\nExp number, first 60 AAs: 19.72694\nTotal prob of N-in: 0.96171\nPOSSIBLE N-term signal sequence\ninside 1 19\nTMhelix 20 39\noutside 40 325\nTMhelix 326 348\ninside 349 372

15066	GCF_001570285.1_ASM157028v1	Salmonella enterica	Proteobacteria; Gammaproteobacteria; Enterobacterales; Enterobacteriaceae; Salmonella
MTGDNTLITSHGINRRDFMKLCAALAATMGLSSKAAAEMAESVSNPQRPPVIWIGAQECT	WP_000145429.1
hydrogenase 2 small subunit [Salmonella enterica]	Length: 372\nNumber of predicted TMHs: 1\nExp number of AAs in TMHs: 24.51309\nExp number, first 60 AAs: 0.2682\nTotal prob of N-in: 0.14402\noutside 1 331\nTMhelix 332 354\ninside 355 372

15067	GCF_001474305.1_Salmonella_enterica_CVM_43768_v1.0Salmonella enterica	Proteobacteria; Gammaproteobacteria; Enterobacterales; Enterobacteriaceae; Salmonella
MNNEETFYQAMRRKGVTRRSFLKFCSLAATSLGLGAGMTPKIAWALENKPRIPVVWIHGL	WP_001058311.1
hydrogenase [Salmonella enterica]	Length: 372\nNumber of predicted TMHs: 2\nExp number of AAs in TMHs: 37.4699899999999\nExp number, first 60 AAs: 19.72699\nTotal prob of N-in: 0.96171\nPOSSIBLE N-term signal sequence\ninside 1 19\nTMhelix 20 39\noutside 40 325\nTMhelix 326 348\ninside 349 372

15068	GCF_001474325.1_Salmonella_enterica_CVM_43769_v1.0Salmonella enterica	Proteobacteria; Gammaproteobacteria; Enterobacterales; Enterobacteriaceae; Salmonella
MTGDNTLITSHGINRRDFMKLCAALAATMGLSSKAAAEMAESVSNPQRPPVIWIGAQECT	WP_000145429.1
hydrogenase 2 small subunit [Salmonella enterica]	Length: 372\nNumber of predicted TMHs: 1\nExp

number of AAs in TMHs: 24.51309\nExp number, first 60 AAs:  0.2682\nTotal prob of N-in:   0.14402\noutside   1   331\nTMhelix   332   354\ninside   355   372

15069 GCF_001474335.1_Salmonella_enterica_CVM_43770_v1.0Salmonella enterica Proteobacteria; Gammaproteobacteria; Enterobacterales; Enterobacteriaceae; Salmonella MTGDNTLITSHGINRRDFMKLCAALAATMGLSSKAAAEMAESVSNPQRPPVIWIGAQECT WP_000145429.1 hydrogenase 2 small subunit [Salmonella enterica] Length: 372\nNumber of predicted TMHs:  1\nExp number of AAs in TMHs: 24.51309\nExp number, first 60 AAs:  0.2682\nTotal prob of N-in:   0.14402\noutside   1   331\nTMhelix   332   354\ninside   355   372

15070 GCF_001474275.1_Salmonella_enterica_CVM_43767_v1.0Salmonella enterica Proteobacteria; Gammaproteobacteria; Enterobacterales; Enterobacteriaceae; Salmonella MTGDNTLITSHGINRRDFMKLCAALAATMGLSSKAAAEMAESVSNPQRPPVIWIGAQECT WP_000145429.1 hydrogenase 2 small subunit [Salmonella enterica] Length: 372\nNumber of predicted TMHs:  1\nExp number of AAs in TMHs: 24.51309\nExp number, first 60 AAs:  0.2682\nTotal prob of N-in:   0.14402\noutside   1   331\nTMhelix   332   354\ninside   355   372

15071 GCF_001474305.1_Salmonella_enterica_CVM_43768_v1.0Salmonella enterica Proteobacteria; Gammaproteobacteria; Enterobacterales; Enterobacteriaceae; Salmonella MTGDNTLITSHGINRRDFMKLCAALAATMGLSSKAAAEMAESVSNPQRPPVIWIGAQECT WP_000145429.1 hydrogenase 2 small subunit [Salmonella enterica] Length: 372\nNumber of predicted TMHs:  1\nExp number of AAs in TMHs: 24.51309\nExp number, first 60 AAs:  0.2682\nTotal prob of N-in:   0.14402\noutside   1   331\nTMhelix   332   354\ninside   355   372

15072 GCF_001474165.1_Salmonella_enterica_CVM_43758_v1.0Salmonella enterica Proteobacteria; Gammaproteobacteria; Enterobacterales; Enterobacteriaceae; Salmonella MNNEETFYQAMRRKGVTRRSFLKFCSLAATSLGLSAGMTPKIAWALENKPRIPVVWIHGL WP_001058316.1 hydrogenase [Salmonella enterica] Length: 372\nNumber of predicted TMHs:  2\nExp number of AAs in TMHs: 35.76397\nExp number, first 60 AAs:  18.97201\nTotal prob of N-in:   0.92785\nPOSSIBLE N-term signal sequence\ninside   1   19\nTMhelix   20   39\noutside   40   325\nTMhelix   326   348\ninside   349   372

15073 GCF_001474255.1_Salmonella_enterica_CVM_43766_v1.0Salmonella enterica Proteobacteria; Gammaproteobacteria; Enterobacterales; Enterobacteriaceae; Salmonella MNNEETLYQAMRRKGVTRRSFLKFCSLAATSLGLGAGMTPKIAWALENKPRIPVVWIHGL WP_001752445.1 hydrogenase-1 small chain [Salmonella enterica] Length: 372\nNumber of predicted TMHs:  2\nExp number of AAs in TMHs: 36.5123\nExp number, first 60 AAs:  19.72341\nTotal prob of N-in:   0.96144\nPOSSIBLE N-term signal sequence\ninside   1   19\nTMhelix   20   39\noutside   40   325\nTMhelix   326   348\ninside   349   372

15074 GCF_001474225.1_Salmonella_enterica_CVM_43763_v1.0Salmonella enterica Proteobacteria; Gammaproteobacteria; Enterobacterales; Enterobacteriaceae; Salmonella MNNEETFYQAMRRKGVTRRSFLKFCSLAATSLGLGAGMTPKIAWALENKPRIPVVWIHGL WP_001058312.1 hydrogenase [Salmonella enterica] Length: 372\nNumber of predicted TMHs:  2\nExp number of AAs in TMHs: 36.51738\nExp number, first 60 AAs:  19.72849\nTotal prob of N-in:   0.96170\nPOSSIBLE N-term signal sequence\ninside   1   19\nTMhelix   20   39\noutside   40   325\nTMhelix   326   348\ninside   349   372

15075 GCF_001474255.1_Salmonella_enterica_CVM_43766_v1.0Salmonella enterica Proteobacteria; Gammaproteobacteria; Enterobacterales; Enterobacteriaceae; Salmonella MTGDNTLITSHGINRRDFMKLCAALAATMGLSSKAAAEMAESVSNPQRPPVIWIGAQECT WP_000145428.1 hydrogenase 2 small subunit [Salmonella enterica] Length: 372\nNumber of predicted TMHs:  1\nExp number of AAs in TMHs: 24.50677\nExp number, first 60 AAs:  0.2682\nTotal prob of N-in:   0.14390\noutside   1   331\nTMhelix   332   354\ninside   355   372

15076 GCF_002036535.1_ASM203653v1 Salmonella enterica Proteobacteria; Gammaproteobacteria; Enterobacterales; Enterobacteriaceae; Salmonella MNNEETFYQAMRRKGVTRRSFLKFCSLAATSLGLGAGMTPKIAWALENKPRIPVVWIHGL WP_001058307.1 hydrogenase [Salmonella enterica] Length: 372\nNumber of predicted TMHs:  2\nExp number of AAs in TMHs: 36.51738\nExp number, first 60 AAs:  19.72849\nTotal prob of N-in:   0.96170\nPOSSIBLE N-term signal sequence\ninside   1   19\nTMhelix   20   39\noutside   40   325\nTMhelix   326   348\ninside   349   372

15077 GCF_002036535.1_ASM203653v1 Salmonella enterica Proteobacteria; Gammaproteobacteria; Enterobacterales; Enterobacteriaceae; Salmonella

MTGDNTLITSHGINRRDFMKLCAALAATMGLSSKAAAEMAESVSNPQRPPVIWIGAQECT WP_000145429.1 hydrogenase 2 small subunit [Salmonella enterica] Length: 372\nNumber of predicted TMHs: 1\nExp number of AAs in TMHs: 24.51309\nExp number, first 60 AAs: 0.2682\nTotal prob of N-in: 0.14402\noutside 1 331\nTMhelix 332 354\ninside 355 372

15078 GCF_002052105.1_ASM205210v1 Salmonella enterica Proteobacteria; Gammaproteobacteria; Enterobacterales; Enterobacteriaceae; Salmonella MTGDNTLITSHGINRRDFMKLCAALAATMGLSSKAAAEMAESVSNPQRPPVIWIGAQECT WP_000145429.1 hydrogenase 2 small subunit [Salmonella enterica] Length: 372\nNumber of predicted TMHs: 1\nExp number of AAs in TMHs: 24.51309\nExp number, first 60 AAs: 0.2682\nTotal prob of N-in: 0.14402\noutside 1 331\nTMhelix 332 354\ninside 355 372

15079 GCF_001474225.1_Salmonella_enterica_CVM_43763_v1.0Salmonella enterica Proteobacteria; Gammaproteobacteria; Enterobacterales; Enterobacteriaceae; Salmonella MTGDNTLITSHGINRRDFMKLCAALAATMGLSSKAAAEMAESVSNPQRPPVIWIGAQECT WP_000145429.1 hydrogenase 2 small subunit [Salmonella enterica] Length: 372\nNumber of predicted TMHs: 1\nExp number of AAs in TMHs: 24.51309\nExp number, first 60 AAs: 0.2682\nTotal prob of N-in: 0.14402\noutside 1 331\nTMhelix 332 354\ninside 355 372

15080 GCF_001474185.1_Salmonella_enterica_CVM_43760_v1.0Salmonella enterica Proteobacteria; Gammaproteobacteria; Enterobacterales; Enterobacteriaceae; Salmonella MNNEETFYQAMRRKGVTRRSFLKFCSLAATSLGLGAGMTPKIAWALENKPRIPVVWIHGL WP_001058311.1 hydrogenase [Salmonella enterica] Length: 372\nNumber of predicted TMHs: 2\nExp number of AAs in TMHs: 37.4699899999999\nExp number, first 60 AAs: 19.72699\nTotal prob of N-in: 0.96171\nPOSSIBLE N-term signal sequence\ninside 1 19\nTMhelix 20 39\noutside 40 325\nTMhelix 326 348\ninside 349 372

15081 GCF_002052005.1_ASM205200v1 Salmonella enterica Proteobacteria; Gammaproteobacteria; Enterobacterales; Enterobacteriaceae; Salmonella MTGDNTLITSHGINRRDFMKLCAALAATMGLSSKAAAEMAESVSNPQRPPVIWIGAQECT WP_000145429.1 hydrogenase 2 small subunit [Salmonella enterica] Length: 372\nNumber of predicted TMHs: 1\nExp number of AAs in TMHs: 24.51309\nExp number, first 60 AAs: 0.2682\nTotal prob of N-in: 0.14402\noutside 1 331\nTMhelix 332 354\ninside 355 372

15082 GCF_001474205.1_Salmonella_enterica_CVM_43761_v1.0Salmonella enterica Proteobacteria; Gammaproteobacteria; Enterobacterales; Enterobacteriaceae; Salmonella MNNEETFYQAMRRKGVTRRSFLKFCSLAATSLGLGAGMTPKIAWALENKPRIPVVWIHGL WP_001058312.1 hydrogenase [Salmonella enterica] Length: 372\nNumber of predicted TMHs: 2\nExp number of AAs in TMHs: 36.51738\nExp number, first 60 AAs: 19.72849\nTotal prob of N-in: 0.96170\nPOSSIBLE N-term signal sequence\ninside 1 19\nTMhelix 20 39\noutside 40 325\nTMhelix 326 348\ninside 349 372

15083 GCF_001474245.1_Salmonella_enterica_CVM_43764_v1.0Salmonella enterica Proteobacteria; Gammaproteobacteria; Enterobacterales; Enterobacteriaceae; Salmonella MNNEETFYQAMRRKGVTRRSFLKFCSLAATSLGLGAGMTPKIAWALENKPRIPVVWIHGL WP_001058311.1 hydrogenase [Salmonella enterica] Length: 372\nNumber of predicted TMHs: 2\nExp number of AAs in TMHs: 37.4699899999999\nExp number, first 60 AAs: 19.72699\nTotal prob of N-in: 0.96171\nPOSSIBLE N-term signal sequence\ninside 1 19\nTMhelix 20 39\noutside 40 325\nTMhelix 326 348\ninside 349 372

15084 GCF_001474245.1_Salmonella_enterica_CVM_43764_v1.0Salmonella enterica Proteobacteria; Gammaproteobacteria; Enterobacterales; Enterobacteriaceae; Salmonella MTGDNTLITSHGINRRDFMKLCAALAATMGLSSKAAAEMAESVSNPQRPPVIWIGAQECT WP_000145429.1 hydrogenase 2 small subunit [Salmonella enterica] Length: 372\nNumber of predicted TMHs: 1\nExp number of AAs in TMHs: 24.51309\nExp number, first 60 AAs: 0.2682\nTotal prob of N-in: 0.14402\noutside 1 331\nTMhelix 332 354\ninside 355 372

15085 GCF_001952395.1_ASM195239v1 Salmonella enterica Proteobacteria; Gammaproteobacteria; Enterobacterales; Enterobacteriaceae; Salmonella MNNEETFYQAMRRKGVTRRSFLKFCSLAATSLGLGAGMTPKIAWALENKPRIPVVWIHGL WP_001058312.1 hydrogenase [Salmonella enterica] Length: 372\nNumber of predicted TMHs: 2\nExp number of AAs in TMHs: 36.51738\nExp number, first 60 AAs: 19.72849\nTotal prob of N-in: 0.96170\nPOSSIBLE N-term signal sequence\ninside 1 19\nTMhelix 20 39\noutside 40 325\nTMhelix 326 348\ninside 349 372

15086	GCF_001474205.1_Salmonella_enterica_CVM_43761_v1.0Salmonella enterica	Proteobacteria; Gammaproteobacteria; Enterobacterales; Enterobacteriaceae; Salmonella
MTGDNTLITSHGINRRDFMKLCAALAATMGLSSKAAAEMAESVSNPQRPPVIWIGAQECT	WP_000145429.1
hydrogenase 2 small subunit [Salmonella enterica]	Length: 372\nNumber of predicted TMHs: 1\nExp number of AAs in TMHs: 24.51309\nExp number, first 60 AAs: 0.2682\nTotal prob of N-in: 0.14402\noutside 1 331\nTMhelix 332 354\ninside 355 372

15087	GCF_001952285.1_ASM195228v1	Salmonella enterica	Proteobacteria; Gammaproteobacteria; Enterobacterales; Enterobacteriaceae; Salmonella
MNNEETFYQAMRRKGVTRRSFLKFCSLAATSLGLGAGMTPKIAWALENKPRIPVVWIHGL	WP_001058312.1
hydrogenase [Salmonella enterica]	Length: 372\nNumber of predicted TMHs: 2\nExp number of AAs in TMHs: 36.51738\nExp number, first 60 AAs: 19.72849\nTotal prob of N-in: 0.96170\nPOSSIBLE N-term signal sequence\ninside 1 19\nTMhelix 20 39\noutside 40 325\nTMhelix 326 348\ninside 349 372

15088	GCF_001474135.1_Salmonella_enterica_CVM_43755_v1.0Salmonella enterica	Proteobacteria; Gammaproteobacteria; Enterobacterales; Enterobacteriaceae; Salmonella
MNNEETFYQAMRRKGVTRRSFLKFCSLAATSLGLGAGMTPKIAWALENKPRIPVVWIHGL	WP_001058307.1
hydrogenase [Salmonella enterica]	Length: 372\nNumber of predicted TMHs: 2\nExp number of AAs in TMHs: 36.51738\nExp number, first 60 AAs: 19.72849\nTotal prob of N-in: 0.96170\nPOSSIBLE N-term signal sequence\ninside 1 19\nTMhelix 20 39\noutside 40 325\nTMhelix 326 348\ninside 349 372

15089	GCF_001952545.1_ASM195254v1	Salmonella enterica	Proteobacteria; Gammaproteobacteria; Enterobacterales; Enterobacteriaceae; Salmonella
MNNEETFYQAMRRKGVTRRSFLKFCSLAATSLGLGAGMTPKIAWALENKPRIPVVWIHGL	WP_001058312.1
hydrogenase [Salmonella enterica]	Length: 372\nNumber of predicted TMHs: 2\nExp number of AAs in TMHs: 36.51738\nExp number, first 60 AAs: 19.72849\nTotal prob of N-in: 0.96170\nPOSSIBLE N-term signal sequence\ninside 1 19\nTMhelix 20 39\noutside 40 325\nTMhelix 326 348\ninside 349 372

15090	GCF_001474125.1_Salmonella_enterica_CVM_43754_v1.0Salmonella enterica	Proteobacteria; Gammaproteobacteria; Enterobacterales; Enterobacteriaceae; Salmonella
MNNEETFYQAMRRKGVTRRSFLKFCSLAATSLGLGAGMTPKIAWALENKPRIPVVWIHGL	WP_001058312.1
hydrogenase [Salmonella enterica]	Length: 372\nNumber of predicted TMHs: 2\nExp number of AAs in TMHs: 36.51738\nExp number, first 60 AAs: 19.72849\nTotal prob of N-in: 0.96170\nPOSSIBLE N-term signal sequence\ninside 1 19\nTMhelix 20 39\noutside 40 325\nTMhelix 326 348\ninside 349 372

15091	GCF_001474095.1_Salmonella_enterica_CVM_43752_v1.0Salmonella enterica	Proteobacteria; Gammaproteobacteria; Enterobacterales; Enterobacteriaceae; Salmonella
MNNEETFYQAMRRKGVTRRSFLKFCSLAATSLGLGAGMTPKIAWALENKPRIPVVWIHGL	WP_001058312.1
hydrogenase [Salmonella enterica]	Length: 372\nNumber of predicted TMHs: 2\nExp number of AAs in TMHs: 36.51738\nExp number, first 60 AAs: 19.72849\nTotal prob of N-in: 0.96170\nPOSSIBLE N-term signal sequence\ninside 1 19\nTMhelix 20 39\noutside 40 325\nTMhelix 326 348\ninside 349 372

15092	GCF_001952285.1_ASM195228v1	Salmonella enterica	Proteobacteria; Gammaproteobacteria; Enterobacterales; Enterobacteriaceae; Salmonella
MTGDNTLITSHGINRRDFMKLCAALAATMGLSSKAAAEMAESVSNPQRPPVIWIGAQECT	WP_000145429.1
hydrogenase 2 small subunit [Salmonella enterica]	Length: 372\nNumber of predicted TMHs: 1\nExp number of AAs in TMHs: 24.51309\nExp number, first 60 AAs: 0.2682\nTotal prob of N-in: 0.14402\noutside 1 331\nTMhelix 332 354\ninside 355 372

15093	GCF_001474135.1_Salmonella_enterica_CVM_43755_v1.0Salmonella enterica	Proteobacteria; Gammaproteobacteria; Enterobacterales; Enterobacteriaceae; Salmonella
MTGDNTLITSHGINRRDFMKLCAALAATMGLSSKAAAEMAESVSNPQRPPVIWIGAQECT	WP_000145429.1
hydrogenase 2 small subunit [Salmonella enterica]	Length: 372\nNumber of predicted TMHs: 1\nExp number of AAs in TMHs: 24.51309\nExp number, first 60 AAs: 0.2682\nTotal prob of N-in: 0.14402\noutside 1 331\nTMhelix 332 354\ninside 355 372

15094	GCF_001474125.1_Salmonella_enterica_CVM_43754_v1.0Salmonella enterica	Proteobacteria; Gammaproteobacteria; Enterobacterales; Enterobacteriaceae; Salmonella
MTGDNTLITSHGINRRDFMKLCAALAATMGLSSKAAAEMAESVSNPQRPPVIWIGAQECT	WP_000145429.1
hydrogenase 2 small subunit [Salmonella enterica]	Length: 372\nNumber of predicted TMHs: 1\nExp

number of AAs in TMHs: 24.51309\nExp number, first 60 AAs:  0.2682\nTotal prob of N-in:   0.14402\noutside   1   331\nTMhelix   332   354\ninside   355   372

15095 GCF_001474095.1_Salmonella_enterica_CVM_43752_v1.0Salmonella enterica Proteobacteria; Gammaproteobacteria; Enterobacterales; Enterobacteriaceae; Salmonella MTGDNTLITSHGINRRDFMKLCAALAATMGLSSKAAAEMAESVSNPQRPPVIWIGAQECT WP_000145429.1 hydrogenase 2 small subunit [Salmonella enterica] Length: 372\nNumber of predicted TMHs:  1\nExp number of AAs in TMHs: 24.51309\nExp number, first 60 AAs:  0.2682\nTotal prob of N-in:   0.14402\noutside   1   331\nTMhelix   332   354\ninside   355   372

15096 GCF_001473985.1_Salmonella_enterica_CVM_43744_v1.0Salmonella enterica Proteobacteria; Gammaproteobacteria; Enterobacterales; Enterobacteriaceae; Salmonella MNNEETFYQAMRRKGVTRRSFLKFCSLAATSLGLGAGMTPKIAWALENKPRIPVVWIHGL WP_001058307.1 hydrogenase [Salmonella enterica] Length: 372\nNumber of predicted TMHs:  2\nExp number of AAs in TMHs: 36.51738\nExp number, first 60 AAs:  19.72849\nTotal prob of N-in:   0.96170\nPOSSIBLE N-term signal sequence\ninside   1   19\nTMhelix   20   39\noutside   40   325\nTMhelix   326   348\ninside   349   372

15097 GCF_002051125.1_ASM205112v1 Salmonella enterica Proteobacteria; Gammaproteobacteria; Enterobacterales; Enterobacteriaceae; Salmonella MTGDNTLITSHGINRRDFMKLCAALAATMGLSSKAAAEMAESVSNPQRPPVIWIGAQECT WP_015589585.1 hydrogenase 2 small subunit [Salmonella enterica] Length: 372\nNumber of predicted TMHs:  1\nExp number of AAs in TMHs: 24.51304\nExp number, first 60 AAs:  0.2682\nTotal prob of N-in:   0.14402\noutside   1   331\nTMhelix   332   354\ninside   355   372

15098 GCF_001951865.1_ASM195186v1 Salmonella enterica Proteobacteria; Gammaproteobacteria; Enterobacterales; Enterobacteriaceae; Salmonella MNNEETFYQAMRRKGVTRRSFLKFCSLAATSLGLGAGMTPKIAWALENKPRIPVVWIHGL WP_001058312.1 hydrogenase [Salmonella enterica] Length: 372\nNumber of predicted TMHs:  2\nExp number of AAs in TMHs: 36.51738\nExp number, first 60 AAs:  19.72849\nTotal prob of N-in:   0.96170\nPOSSIBLE N-term signal sequence\ninside   1   19\nTMhelix   20   39\noutside   40   325\nTMhelix   326   348\ninside   349   372

15099 GCF_001951865.1_ASM195186v1 Salmonella enterica Proteobacteria; Gammaproteobacteria; Enterobacterales; Enterobacteriaceae; Salmonella MTGDNTLITSHGINRRDFMKLCAALAATMGLSSKAAAEMAESVSNPQRPPVIWIGAQECT WP_000145429.1 hydrogenase 2 small subunit [Salmonella enterica] Length: 372\nNumber of predicted TMHs:  1\nExp number of AAs in TMHs: 24.51309\nExp number, first 60 AAs:  0.2682\nTotal prob of N-in:   0.14402\noutside   1   331\nTMhelix   332   354\ninside   355   372

15100 GCF_001474075.1_Salmonella_enterica_CVM_43751_v1.0Salmonella enterica Proteobacteria; Gammaproteobacteria; Enterobacterales; Enterobacteriaceae; Salmonella MNNEETFYQAMRRKGVTRRSFLKFCSLAATSLGLGAGMTPKIAWALENKPRIPVVWIHGL WP_001058312.1 hydrogenase [Salmonella enterica] Length: 372\nNumber of predicted TMHs:  2\nExp number of AAs in TMHs: 36.51738\nExp number, first 60 AAs:  19.72849\nTotal prob of N-in:   0.96170\nPOSSIBLE N-term signal sequence\ninside   1   19\nTMhelix   20   39\noutside   40   325\nTMhelix   326   348\ninside   349   372

15101 GCF_001474075.1_Salmonella_enterica_CVM_43751_v1.0Salmonella enterica Proteobacteria; Gammaproteobacteria; Enterobacterales; Enterobacteriaceae; Salmonella MTGDNTLITSHGINRRDFMKLCAALAATMGLSSKAAAEMAESVSNPQRPPVIWIGAQECT WP_000145428.1 hydrogenase 2 small subunit [Salmonella enterica] Length: 372\nNumber of predicted TMHs:  1\nExp number of AAs in TMHs: 24.50677\nExp number, first 60 AAs:  0.2682\nTotal prob of N-in:   0.14390\noutside   1   331\nTMhelix   332   354\ninside   355   372

15102 GCF_001474025.1_Salmonella_enterica_CVM_43748_v1.0Salmonella enterica Proteobacteria; Gammaproteobacteria; Enterobacterales; Enterobacteriaceae; Salmonella MNNEETFYQAMRRKGVTRRSFLKFCSLAATSLGLGAGMTPKIAWALENKPRIPVVWIHGL WP_001058312.1 hydrogenase [Salmonella enterica] Length: 372\nNumber of predicted TMHs:  2\nExp number of AAs in TMHs: 36.51738\nExp number, first 60 AAs:  19.72849\nTotal prob of N-in:   0.96170\nPOSSIBLE N-term signal sequence\ninside   1   19\nTMhelix   20   39\noutside   40   325\nTMhelix   326   348\ninside   349   372

15103 GCF_001474045.1_Salmonella_enterica_CVM_43749_v1.0Salmonella enterica Proteobacteria; Gammaproteobacteria; Enterobacterales; Enterobacteriaceae; Salmonella

MNNEETFYQAMRRKGVTRRSFLKFCSLAATSLGLGAGMTPKIAWALENKPRIPVVWIHGL WP_001058307.1 hydrogenase [Salmonella enterica] Length: 372\nNumber of predicted TMHs: 2\nExp number of AAs in TMHs: 36.51738\nExp number, first 60 AAs: 19.72849\nTotal prob of N-in: 0.96170\nPOSSIBLE N-term signal sequence\ninside 1 19\nTMhelix 20 39\noutside 40 325\nTMhelix 326 348\ninside 349 372

15104 GCF_001474065.1_Salmonella_enterica_CVM_43750_v1.0Salmonella enterica Proteobacteria; Gammaproteobacteria; Enterobacterales; Enterobacteriaceae; Salmonella MNNEETFYQAMRRKGVTRRSFLKFCSLAATSLGLGAGMTPKIAWALENKPRIPVVWIHGL WP_001058312.1 hydrogenase [Salmonella enterica] Length: 372\nNumber of predicted TMHs: 2\nExp number of AAs in TMHs: 36.51738\nExp number, first 60 AAs: 19.72849\nTotal prob of N-in: 0.96170\nPOSSIBLE N-term signal sequence\ninside 1 19\nTMhelix 20 39\noutside 40 325\nTMhelix 326 348\ninside 349 372

15105 GCF_001474065.1_Salmonella_enterica_CVM_43750_v1.0Salmonella enterica Proteobacteria; Gammaproteobacteria; Enterobacterales; Enterobacteriaceae; Salmonella MTGDNTLITSHGINRRDFMKLCAALAATMGLSSKAAAEMAESVSNPQRPPVIWIGAQECT WP_000145429.1 hydrogenase 2 small subunit [Salmonella enterica] Length: 372\nNumber of predicted TMHs: 1\nExp number of AAs in TMHs: 24.51309\nExp number, first 60 AAs: 0.2682\nTotal prob of N-in: 0.14402\noutside 1 331\nTMhelix 332 354\ninside 355 372

15106 GCF_001474045.1_Salmonella_enterica_CVM_43749_v1.0Salmonella enterica Proteobacteria; Gammaproteobacteria; Enterobacterales; Enterobacteriaceae; Salmonella MTGDNTLITSHGINRRDFMKLCAALAATMGLSSKAAAEMAESVSNPQRPPVIWIGAQECT WP_000145429.1 hydrogenase 2 small subunit [Salmonella enterica] Length: 372\nNumber of predicted TMHs: 1\nExp number of AAs in TMHs: 24.51309\nExp number, first 60 AAs: 0.2682\nTotal prob of N-in: 0.14402\noutside 1 331\nTMhelix 332 354\ninside 355 372

15107 GCF_001473945.1_Salmonella_enterica_CVM_43843_v1.0Salmonella enterica Proteobacteria; Gammaproteobacteria; Enterobacterales; Enterobacteriaceae; Salmonella MNNEETFYQAMRRKGVTRRSFLKFCSLAATSLGLGAGMTPKIAWALENKPRIPVVWIHGL WP_001058312.1 hydrogenase [Salmonella enterica] Length: 372\nNumber of predicted TMHs: 2\nExp number of AAs in TMHs: 36.51738\nExp number, first 60 AAs: 19.72849\nTotal prob of N-in: 0.96170\nPOSSIBLE N-term signal sequence\ninside 1 19\nTMhelix 20 39\noutside 40 325\nTMhelix 326 348\ninside 349 372

15108 GCF_002061075.1_ASM206107v1 Salmonella enterica Proteobacteria; Gammaproteobacteria; Enterobacterales; Enterobacteriaceae; Salmonella MNNEETFYQAMRRKGVTRRSFLKFCSLAATSLGLGAGMTPKIAWALENKPRIPVVWIHGL WP_001058307.1 hydrogenase [Salmonella enterica] Length: 372\nNumber of predicted TMHs: 2\nExp number of AAs in TMHs: 36.51738\nExp number, first 60 AAs: 19.72849\nTotal prob of N-in: 0.96170\nPOSSIBLE N-term signal sequence\ninside 1 19\nTMhelix 20 39\noutside 40 325\nTMhelix 326 348\ninside 349 372

15109 GCF_002060865.1_ASM206086v1 Salmonella enterica Proteobacteria; Gammaproteobacteria; Enterobacterales; Enterobacteriaceae; Salmonella MNNEETFYQAMRRKGVTRRSFLKFCSLAATSLGLGAGMTPKIAWALENKPRIPVVWIHGL WP_001058312.1 hydrogenase [Salmonella enterica] Length: 372\nNumber of predicted TMHs: 2\nExp number of AAs in TMHs: 36.51738\nExp number, first 60 AAs: 19.72849\nTotal prob of N-in: 0.96170\nPOSSIBLE N-term signal sequence\ninside 1 19\nTMhelix 20 39\noutside 40 325\nTMhelix 326 348\ninside 349 372

15110 GCF_002060795.1_ASM206079v1 Salmonella enterica Proteobacteria; Gammaproteobacteria; Enterobacterales; Enterobacteriaceae; Salmonella MNNEETFYQAMRRKGVTRRSFLKFCSLAATSLGLGAGMTPKIAWALENKPRIPVVWIHGL WP_001058308.1 hydrogenase [Salmonella enterica] Length: 372\nNumber of predicted TMHs: 2\nExp number of AAs in TMHs: 39.08204\nExp number, first 60 AAs: 19.72474\nTotal prob of N-in: 0.96174\nPOSSIBLE N-term signal sequence\ninside 1 19\nTMhelix 20 39\noutside 40 325\nTMhelix 326 348\ninside 349 372

15111 GCF_002060885.1_ASM206088v1 Salmonella enterica Proteobacteria; Gammaproteobacteria; Enterobacterales; Enterobacteriaceae; Salmonella MTGDNTLITSHGINRRDFMKLCAALAATMGLSSKAAAEMAESVSNPQRPPVIWIGAQECT WP_000145429.1 hydrogenase 2 small subunit [Salmonella enterica] Length: 372\nNumber of predicted TMHs: 1\nExp number of AAs in TMHs: 24.51309\nExp number, first 60 AAs: 0.2682\nTotal prob of N-in: 0.14402\noutside 1 331\nTMhelix 332 354\ninside 355 372

15112 GCF_002060975.1_ASM206097v1 Salmonella enterica Proteobacteria; Gammaproteobacteria; Enterobacterales; Enterobacteriaceae; Salmonella
MTGDNTLITSHGINRRDFMKLCAALAATMGLSSKAAAEMAESVSNPQRPPVIWIGAQECT WP_000145429.1
hydrogenase 2 small subunit [Salmonella enterica] Length: 372\nNumber of predicted TMHs: 1\nExp number of AAs in TMHs: 24.51309\nExp number, first 60 AAs: 0.2682\nTotal prob of N-in: 0.14402\noutside 1 331\nTMhelix 332 354\ninside 355 372

15113 GCF_002060865.1_ASM206086v1 Salmonella enterica Proteobacteria; Gammaproteobacteria; Enterobacterales; Enterobacteriaceae; Salmonella
MTGDNTLITSHGINRRDFMKLCAALAATMGLSSKAAAEMAESVSNPQRPPVIWIGAQECT WP_000145429.1
hydrogenase 2 small subunit [Salmonella enterica] Length: 372\nNumber of predicted TMHs: 1\nExp number of AAs in TMHs: 24.51309\nExp number, first 60 AAs: 0.2682\nTotal prob of N-in: 0.14402\noutside 1 331\nTMhelix 332 354\ninside 355 372

15114 GCF_002060795.1_ASM206079v1 Salmonella enterica Proteobacteria; Gammaproteobacteria; Enterobacterales; Enterobacteriaceae; Salmonella
MTGDNTLITSHGINRRDFMKLCAALAATMGLSSKAAAEMAESVSNPQRPPVIWIGAQECT WP_000145429.1
hydrogenase 2 small subunit [Salmonella enterica] Length: 372\nNumber of predicted TMHs: 1\nExp number of AAs in TMHs: 24.51309\nExp number, first 60 AAs: 0.2682\nTotal prob of N-in: 0.14402\noutside 1 331\nTMhelix 332 354\ninside 355 372

15115 GCF_001473735.1_Salmonella_enterica_CVM_43816_v1.0Salmonella enterica Proteobacteria; Gammaproteobacteria; Enterobacterales; Enterobacteriaceae; Salmonella
MTGDNTLITSHGINRRDFMKLCAALAATMGLSSKAAAEMAESVSNPQRPPVIWIGAQECT WP_000145429.1
hydrogenase 2 small subunit [Salmonella enterica] Length: 372\nNumber of predicted TMHs: 1\nExp number of AAs in TMHs: 24.51309\nExp number, first 60 AAs: 0.2682\nTotal prob of N-in: 0.14402\noutside 1 331\nTMhelix 332 354\ninside 355 372

15116 GCF_001473735.1_Salmonella_enterica_CVM_43816_v1.0Salmonella enterica Proteobacteria; Gammaproteobacteria; Enterobacterales; Enterobacteriaceae; Salmonella
MNNEETFYQAMRRKGVTRRSFLKFCSLAATSLGLGAGMTPKIAWALENKPRIPVVWIHGL WP_001058312.1
hydrogenase [Salmonella enterica] Length: 372\nNumber of predicted TMHs: 2\nExp number of AAs in TMHs: 36.51738\nExp number, first 60 AAs: 19.72849\nTotal prob of N-in: 0.96170\nPOSSIBLE N-term signal sequence\ninside 1 19\nTMhelix 20 39\noutside 40 325\nTMhelix 326 348\ninside 349 372

15117 GCF_002060675.1_ASM206067v1 Salmonella enterica Proteobacteria; Gammaproteobacteria; Enterobacterales; Enterobacteriaceae; Salmonella
MNNEETFYQAMRRKGVTRRSFLKFCSLAATSLGLGAGMTPKIAWALENKPRIPVVWIHGL WP_001058312.1
hydrogenase [Salmonella enterica] Length: 372\nNumber of predicted TMHs: 2\nExp number of AAs in TMHs: 36.51738\nExp number, first 60 AAs: 19.72849\nTotal prob of N-in: 0.96170\nPOSSIBLE N-term signal sequence\ninside 1 19\nTMhelix 20 39\noutside 40 325\nTMhelix 326 348\ninside 349 372

15118 GCF_002060735.1_ASM206073v1 Salmonella enterica Proteobacteria; Gammaproteobacteria; Enterobacterales; Enterobacteriaceae; Salmonella
MNNEETFYQAMRRKGVTRRSFLKFCSLAATSLGLGAGMTPKIAWALENKPRIPVVWIHGL WP_001058312.1
hydrogenase [Salmonella enterica] Length: 372\nNumber of predicted TMHs: 2\nExp number of AAs in TMHs: 36.51738\nExp number, first 60 AAs: 19.72849\nTotal prob of N-in: 0.96170\nPOSSIBLE N-term signal sequence\ninside 1 19\nTMhelix 20 39\noutside 40 325\nTMhelix 326 348\ninside 349 372

15119 GCF_002060775.1_ASM206077v1 Salmonella enterica Proteobacteria; Gammaproteobacteria; Enterobacterales; Enterobacteriaceae; Salmonella
MNNEETFYQAMRRKGVTRRSFLKFCSLAATSLGLGAGMTPKIAWALENKPRIPVVWIHGL WP_023243720.1
hydrogenase 1, small subunit [Salmonella enterica] Length: 372\nNumber of predicted TMHs: 2\nExp number of AAs in TMHs: 36.52147\nExp number, first 60 AAs: 19.72838\nTotal prob of N-in: 0.96170\nPOSSIBLE N-term signal sequence\ninside 1 19\nTMhelix 20 39\noutside 40 325\nTMhelix 326 348\ninside 349 372

15120 GCF_002060775.1_ASM206077v1 Salmonella enterica Proteobacteria; Gammaproteobacteria; Enterobacterales; Enterobacteriaceae; Salmonella
MTGDNTLITSHGINRRDFMKLCAALAATMGLSSKAAAEMAESVSNPQRPPVIWIGAQECT WP_000145429.1
hydrogenase 2 small subunit [Salmonella enterica] Length: 372\nNumber of predicted TMHs: 1\nExp

number of AAs in TMHs: 24.51309\nExp number, first 60 AAs: 0.2682\nTotal prob of N-in: 0.14402\noutside 1 331\nTMhelix 332 354\ninside 355 372

15121 GCF_001473965.1_Salmonella_enterica_CVM_43743_v1.0Salmonella enterica Proteobacteria; Gammaproteobacteria; Enterobacterales; Enterobacteriaceae; Salmonella
MNNEETFYQAMRRKGVTRRSFLKFCSLAATSLGLGAGMTPKIAWALENKPRIPVVWIHGL WP_001058312.1
hydrogenase [Salmonella enterica] Length: 372\nNumber of predicted TMHs: 2\nExp number of AAs in TMHs: 36.51738\nExp number, first 60 AAs: 19.72849\nTotal prob of N-in: 0.96170\nPOSSIBLE N-term signal sequence\ninside 1 19\nTMhelix 20 39\noutside 40 325\nTMhelix 326 348\ninside 349 372

15122 GCF_001474005.1_Salmonella_enterica_CVM_43746_v1.0Salmonella enterica Proteobacteria; Gammaproteobacteria; Enterobacterales; Enterobacteriaceae; Salmonella
MNNEETFYQAMRRKGVTRRSFLKFCSLAATSLGLGAGMTPKIAWALENKPRIPVVWIHGL WP_001058312.1
hydrogenase [Salmonella enterica] Length: 372\nNumber of predicted TMHs: 2\nExp number of AAs in TMHs: 36.51738\nExp number, first 60 AAs: 19.72849\nTotal prob of N-in: 0.96170\nPOSSIBLE N-term signal sequence\ninside 1 19\nTMhelix 20 39\noutside 40 325\nTMhelix 326 348\ninside 349 372

15123 GCF_002060735.1_ASM206073v1 Salmonella enterica Proteobacteria; Gammaproteobacteria; Enterobacterales; Enterobacteriaceae; Salmonella
MTGDNTLITSHGINRRDFMKLCAALAATMGLSSKAAAEMAESVSNPQRPPVIWIGAQECT WP_000145429.1
hydrogenase 2 small subunit [Salmonella enterica] Length: 372\nNumber of predicted TMHs: 1\nExp number of AAs in TMHs: 24.51309\nExp number, first 60 AAs: 0.2682\nTotal prob of N-in: 0.14402\noutside 1 331\nTMhelix 332 354\ninside 355 372

15124 GCF_002060455.1_ASM206045v1 Salmonella enterica Proteobacteria; Gammaproteobacteria; Enterobacterales; Enterobacteriaceae; Salmonella
MNNEETFYQAMRRKGVTRRSFLKFCSLAATSLGLGAGMTPKIAWALENKPRIPVVWIHGL WP_001058311.1
hydrogenase [Salmonella enterica] Length: 372\nNumber of predicted TMHs: 2\nExp number of AAs in TMHs: 37.4699899999999\nExp number, first 60 AAs: 19.72699\nTotal prob of N-in: 0.96171\nPOSSIBLE N-term signal sequence\ninside 1 19\nTMhelix 20 39\noutside 40 325\nTMhelix 326 348\ninside 349 372

15125 GCF_001474025.1_Salmonella_enterica_CVM_43748_v1.0Salmonella enterica Proteobacteria; Gammaproteobacteria; Enterobacterales; Enterobacteriaceae; Salmonella
MTGDNTLITSHGINRRDFMKLCAALAATMGLSSKAAAEMAESVSNPQRPPVIWIGAQECT WP_000145431.1
hydrogenase 2 small subunit [Salmonella enterica] Length: 372\nNumber of predicted TMHs: 1\nExp number of AAs in TMHs: 24.51505\nExp number, first 60 AAs: 0.2682\nTotal prob of N-in: 0.14406\noutside 1 331\nTMhelix 332 354\ninside 355 372

15126 GCF_001473985.1_Salmonella_enterica_CVM_43744_v1.0Salmonella enterica Proteobacteria; Gammaproteobacteria; Enterobacterales; Enterobacteriaceae; Salmonella
MTGDNTLITSHGINRRDFMKLCAALAATMGLSSKAAAEMAESVSNPQRPPVIWIGAQECT WP_000145429.1
hydrogenase 2 small subunit [Salmonella enterica] Length: 372\nNumber of predicted TMHs: 1\nExp number of AAs in TMHs: 24.51309\nExp number, first 60 AAs: 0.2682\nTotal prob of N-in: 0.14402\noutside 1 331\nTMhelix 332 354\ninside 355 372

15127 GCF_002060455.1_ASM206045v1 Salmonella enterica Proteobacteria; Gammaproteobacteria; Enterobacterales; Enterobacteriaceae; Salmonella
MTGDNTLITSHGINRRDFMKLCAALAATMGLSSKAAAEMAESVSNPQRPPVIWIGAQECT WP_000145429.1
hydrogenase 2 small subunit [Salmonella enterica] Length: 372\nNumber of predicted TMHs: 1\nExp number of AAs in TMHs: 24.51309\nExp number, first 60 AAs: 0.2682\nTotal prob of N-in: 0.14402\noutside 1 331\nTMhelix 332 354\ninside 355 372

15128 GCF_001474005.1_Salmonella_enterica_CVM_43746_v1.0Salmonella enterica Proteobacteria; Gammaproteobacteria; Enterobacterales; Enterobacteriaceae; Salmonella
MTGDNTLITSHGINRRDFMKLCAALAATMGLSSKAAAEMAESVSNPQRPPVIWIGAQECT WP_000145429.1
hydrogenase 2 small subunit [Salmonella enterica] Length: 372\nNumber of predicted TMHs: 1\nExp number of AAs in TMHs: 24.51309\nExp number, first 60 AAs: 0.2682\nTotal prob of N-in: 0.14402\noutside 1 331\nTMhelix 332 354\ninside 355 372

15129 GCF_001473965.1_Salmonella_enterica_CVM_43743_v1.0Salmonella enterica Proteobacteria; Gammaproteobacteria; Enterobacterales; Enterobacteriaceae; Salmonella

MTGDNTLITSHGINRRDFMKLCAALAATMGLSSKAAAEMAESVSNPQRPPVIWIGAQECT WP_000145429.1 hydrogenase 2 small subunit [Salmonella enterica] Length: 372\nNumber of predicted TMHs: 1\nExp number of AAs in TMHs: 24.51309\nExp number, first 60 AAs: 0.2682\nTotal prob of N-in: 0.14402\noutside 1 331\nTMhelix 332 354\ninside 355 372

15130 GCF_001473555.1_Salmonella_enterica_CVM_43784_v1.0Salmonella enterica Proteobacteria; Gammaproteobacteria; Enterobacterales; Enterobacteriaceae; Salmonella MNNEETFYQAMRRKGVTRRSFLKFCSLAATSLGLGAGMTPKIAWALENKPRIPVVWIHGL WP_001058307.1 hydrogenase [Salmonella enterica] Length: 372\nNumber of predicted TMHs: 2\nExp number of AAs in TMHs: 36.51738\nExp number, first 60 AAs: 19.72849\nTotal prob of N-in: 0.96170\nPOSSIBLE N-term signal sequence\ninside 1 19\nTMhelix 20 39\noutside 40 325\nTMhelix 326 348\ninside 349 372

15131 GCF_002051725.1_ASM205172v1 Salmonella enterica Proteobacteria; Gammaproteobacteria; Enterobacterales; Enterobacteriaceae; Salmonella MTGDNTLITSHGINRRDFMKLCAALAATMGLSSKAAAEMAESVSNPQRPPVIWIGAQECT WP_000145429.1 hydrogenase 2 small subunit [Salmonella enterica] Length: 372\nNumber of predicted TMHs: 1\nExp number of AAs in TMHs: 24.51309\nExp number, first 60 AAs: 0.2682\nTotal prob of N-in: 0.14402\noutside 1 331\nTMhelix 332 354\ninside 355 372

15132 GCF_001951925.1_ASM195192v1 Salmonella enterica Proteobacteria; Gammaproteobacteria; Enterobacterales; Enterobacteriaceae; Salmonella MNNEETFYQAMRRKGVTRRSFLKFCSLAATSLGLGAGMTPKIAWALENKPRIPVVWIHGL WP_001058312.1 hydrogenase [Salmonella enterica] Length: 372\nNumber of predicted TMHs: 2\nExp number of AAs in TMHs: 36.51738\nExp number, first 60 AAs: 19.72849\nTotal prob of N-in: 0.96170\nPOSSIBLE N-term signal sequence\ninside 1 19\nTMhelix 20 39\noutside 40 325\nTMhelix 326 348\ninside 349 372

15133 GCF_001473585.1_Salmonella_enterica_CVM_43786_v1.0Salmonella enterica Proteobacteria; Gammaproteobacteria; Enterobacterales; Enterobacteriaceae; Salmonella MNNEETFYQAMRRKGVTRRSFLKFCSLAATSLGLGAGMTPKIAWALENKPRIPVVWIHGL WP_001058312.1 hydrogenase [Salmonella enterica] Length: 372\nNumber of predicted TMHs: 2\nExp number of AAs in TMHs: 36.51738\nExp number, first 60 AAs: 19.72849\nTotal prob of N-in: 0.96170\nPOSSIBLE N-term signal sequence\ninside 1 19\nTMhelix 20 39\noutside 40 325\nTMhelix 326 348\ninside 349 372

15134 GCF_001473525.1_Salmonella_enterica_CVM_43781_v1.0Salmonella enterica Proteobacteria; Gammaproteobacteria; Enterobacterales; Enterobacteriaceae; Salmonella MNNEETFYQAMRRKGVTRRSFLKFCSLAATSLGLGAGMTPKIAWALENKPRIPVVWIHGL WP_001058307.1 hydrogenase [Salmonella enterica] Length: 372\nNumber of predicted TMHs: 2\nExp number of AAs in TMHs: 36.51738\nExp number, first 60 AAs: 19.72849\nTotal prob of N-in: 0.96170\nPOSSIBLE N-term signal sequence\ninside 1 19\nTMhelix 20 39\noutside 40 325\nTMhelix 326 348\ninside 349 372

15135 GCF_001834805.1_ASM183480v1 Salmonella enterica Proteobacteria; Gammaproteobacteria; Enterobacterales; Enterobacteriaceae; Salmonella MNNEETFYQAMRRKGVTRRSFLKFCSLAATSLGLGAGMTPKIAWALENKPRIPVVWIHGL WP_001058311.1 hydrogenase [Salmonella enterica] Length: 372\nNumber of predicted TMHs: 2\nExp number of AAs in TMHs: 37.4699899999999\nExp number, first 60 AAs: 19.72699\nTotal prob of N-in: 0.96171\nPOSSIBLE N-term signal sequence\ninside 1 19\nTMhelix 20 39\noutside 40 325\nTMhelix 326 348\ninside 349 372

15136 GCF_001834895.1_ASM183489v1 Salmonella enterica Proteobacteria; Gammaproteobacteria; Enterobacterales; Enterobacteriaceae; Salmonella MNNEETFYQAMRRKGVTRRSFLKFCSLAATSLGLGAGMTPKIAWALENKPRIPVVWIHGL WP_001058312.1 hydrogenase [Salmonella enterica] Length: 372\nNumber of predicted TMHs: 2\nExp number of AAs in TMHs: 36.51738\nExp number, first 60 AAs: 19.72849\nTotal prob of N-in: 0.96170\nPOSSIBLE N-term signal sequence\ninside 1 19\nTMhelix 20 39\noutside 40 325\nTMhelix 326 348\ninside 349 372

15137 GCF_001834895.1_ASM183489v1 Salmonella enterica Proteobacteria; Gammaproteobacteria; Enterobacterales; Enterobacteriaceae; Salmonella MTGDNTLITSHGINRRDFMKLCAALAATMGLSSKAAAEMAESVSNPQRPPVIWIGAQECT WP_000145429.1 hydrogenase 2 small subunit [Salmonella enterica] Length: 372\nNumber of predicted TMHs: 1\nExp number of AAs in TMHs: 24.51309\nExp number, first 60 AAs: 0.2682\nTotal prob of N-in: 0.14402\noutside 1 331\nTMhelix 332 354\ninside 355 372

15138 GCF_001473945.1_Salmonella_enterica_CVM_43843_v1.0Salmonella enterica Proteobacteria; Gammaproteobacteria; Enterobacterales; Enterobacteriaceae; Salmonella MTGDNTLITSHGINRRDFMKLCAALAATMGLSSKAAAEMAESVSNPQRPPVIWIGAQECT WP_000145429.1 hydrogenase 2 small subunit [Salmonella enterica] Length: 372\nNumber of predicted TMHs: 1\nExp number of AAs in TMHs: 24.51309\nExp number, first 60 AAs: 0.2682\nTotal prob of N-in: 0.14402\noutside 1 331\nTMhelix 332 354\ninside 355 372

15139 GCF_001473555.1_Salmonella_enterica_CVM_43784_v1.0Salmonella enterica Proteobacteria; Gammaproteobacteria; Enterobacterales; Enterobacteriaceae; Salmonella MTGDNTLITSHGINRRDFMKLCAALAATMGLSSKAAAEMAESVSNPQRPPVIWIGAQECT WP_000145429.1 hydrogenase 2 small subunit [Salmonella enterica] Length: 372\nNumber of predicted TMHs: 1\nExp number of AAs in TMHs: 24.51309\nExp number, first 60 AAs: 0.2682\nTotal prob of N-in: 0.14402\noutside 1 331\nTMhelix 332 354\ninside 355 372

15140 GCF_001473585.1_Salmonella_enterica_CVM_43786_v1.0Salmonella enterica Proteobacteria; Gammaproteobacteria; Enterobacterales; Enterobacteriaceae; Salmonella MTGDNTLITSHGINRRDFMKLCAALAATMGLSSKAAAEMAESVSNPQRPPVIWIGAQECT WP_000145429.1 hydrogenase 2 small subunit [Salmonella enterica] Length: 372\nNumber of predicted TMHs: 1\nExp number of AAs in TMHs: 24.51309\nExp number, first 60 AAs: 0.2682\nTotal prob of N-in: 0.14402\noutside 1 331\nTMhelix 332 354\ninside 355 372

15141 GCF_001473525.1_Salmonella_enterica_CVM_43781_v1.0Salmonella enterica Proteobacteria; Gammaproteobacteria; Enterobacterales; Enterobacteriaceae; Salmonella MTGDNTLITSHGINRRDFMKLCAALAATMGLSSKAAAEMAESVSNPQRPPVIWIGAQECT WP_000145429.1 hydrogenase 2 small subunit [Salmonella enterica] Length: 372\nNumber of predicted TMHs: 1\nExp number of AAs in TMHs: 24.51309\nExp number, first 60 AAs: 0.2682\nTotal prob of N-in: 0.14402\noutside 1 331\nTMhelix 332 354\ninside 355 372

15142 GCF_002059765.1_ASM205976v1 Salmonella enterica Proteobacteria; Gammaproteobacteria; Enterobacterales; Enterobacteriaceae; Salmonella MNNEETFYQAMRRKGVTRRSFLKFCSLAATSLGLGAGMTPKIAWALENKPRIPVVWIHGL WP_001058312.1 hydrogenase [Salmonella enterica] Length: 372\nNumber of predicted TMHs: 2\nExp number of AAs in TMHs: 36.51738\nExp number, first 60 AAs: 19.72849\nTotal prob of N-in: 0.96170\nPOSSIBLE N-term signal sequence\ninside 1 19\nTMhelix 20 39\noutside 40 325\nTMhelix 326 348\ninside 349 372

15143 GCF_002059765.1_ASM205976v1 Salmonella enterica Proteobacteria; Gammaproteobacteria; Enterobacterales; Enterobacteriaceae; Salmonella MTGDNTLITSHGINRRDFMKLCAALAATMGLSSKAAAEMAESVSNPQRPPVIWIGAQECT WP_000145429.1 hydrogenase 2 small subunit [Salmonella enterica] Length: 372\nNumber of predicted TMHs: 1\nExp number of AAs in TMHs: 24.51309\nExp number, first 60 AAs: 0.2682\nTotal prob of N-in: 0.14402\noutside 1 331\nTMhelix 332 354\ninside 355 372

15144 GCF_001473485.1_Salmonella_enterica_CVM_43775_v1.0Salmonella enterica Proteobacteria; Gammaproteobacteria; Enterobacterales; Enterobacteriaceae; Salmonella MNNEETFYQAMRRKGVTRRSFLKFCSLAATSLGLGAGMTPKIAWALENKPRIPVVWIHGL WP_001058312.1 hydrogenase [Salmonella enterica] Length: 372\nNumber of predicted TMHs: 2\nExp number of AAs in TMHs: 36.51738\nExp number, first 60 AAs: 19.72849\nTotal prob of N-in: 0.96170\nPOSSIBLE N-term signal sequence\ninside 1 19\nTMhelix 20 39\noutside 40 325\nTMhelix 326 348\ninside 349 372

15145 GCF_001570005.1_ASM157000v1 Salmonella enterica Proteobacteria; Gammaproteobacteria; Enterobacterales; Enterobacteriaceae; Salmonella MTGDNTLITSHGINRRDFMKLCAALAATMGLSSKAAAEMAESVSNPQRPPVIWIGAQECT WP_000145429.1 hydrogenase 2 small subunit [Salmonella enterica] Length: 372\nNumber of predicted TMHs: 1\nExp number of AAs in TMHs: 24.51309\nExp number, first 60 AAs: 0.2682\nTotal prob of N-in: 0.14402\noutside 1 331\nTMhelix 332 354\ninside 355 372

15146 GCF_001833385.1_ASM183338v1 Salmonella enterica Proteobacteria; Gammaproteobacteria; Enterobacterales; Enterobacteriaceae; Salmonella MNNEETFYQAMRRKGVTRRSFLKFCSLAATSLGLGAGMTPKIAWALENKPRIPVVWIHGL WP_001058311.1 hydrogenase [Salmonella enterica] Length: 372\nNumber of predicted TMHs: 2\nExp number of AAs in

TMHs: 37.4699899999999\nExp number, first 60 AAs:  19.72699\nTotal prob of N-in:   0.96171\nPOSSIBLE N-term signal sequence\ninside   1   19\nTMhelix   20   39\noutside   40   325\nTMhelix   326   348\ninside   349   372

15147	GCF_001473545.1_Salmonella_enterica_CVM_43783_v1.0Salmonella enterica	Proteobacteria; Gammaproteobacteria; Enterobacterales; Enterobacteriaceae; Salmonella
	MNNEETFYQAMRRKGVTRRSFLKFCSLAATSLGLGAGMTPKIAWALENKPRIPVVWIHGL	WP_001058311.1
	hydrogenase [Salmonella enterica]	Length: 372\nNumber of predicted TMHs:  2\nExp number of AAs in TMHs: 37.4699899999999\nExp number, first 60 AAs:  19.72699\nTotal prob of N-in:   0.96171\nPOSSIBLE N-term signal sequence\ninside   1   19\nTMhelix   20   39\noutside   40   325\nTMhelix   326   348\ninside   349   372

15148	GCF_001473505.1_Salmonella_enterica_CVM_43778_v1.0Salmonella enterica	Proteobacteria; Gammaproteobacteria; Enterobacterales; Enterobacteriaceae; Salmonella
	MNNEETFYQAMRRKGVTRRSFLKFCSLAATSLGLGAGMTPKIAWALENKPRIPVVWIHGL	WP_001058307.1
	hydrogenase [Salmonella enterica]	Length: 372\nNumber of predicted TMHs:  2\nExp number of AAs in TMHs: 36.51738\nExp number, first 60 AAs:  19.72849\nTotal prob of N-in:   0.96170\nPOSSIBLE N-term signal sequence\ninside   1   19\nTMhelix   20   39\noutside   40   325\nTMhelix   326   348\ninside   349   372

15149	GCF_001833545.1_ASM183354v1	Salmonella enterica	Proteobacteria; Gammaproteobacteria; Enterobacterales; Enterobacteriaceae; Salmonella
	MNNEETFYQAMRRKGVTRRSFLKFCSLAATSLGLGAGMTPKIAWALENKPRIPVVWIHGL	WP_001058312.1
	hydrogenase [Salmonella enterica]	Length: 372\nNumber of predicted TMHs:  2\nExp number of AAs in TMHs: 36.51738\nExp number, first 60 AAs:  19.72849\nTotal prob of N-in:   0.96170\nPOSSIBLE N-term signal sequence\ninside   1   19\nTMhelix   20   39\noutside   40   325\nTMhelix   326   348\ninside   349   372

15150	GCF_001833545.1_ASM183354v1	Salmonella enterica	Proteobacteria; Gammaproteobacteria; Enterobacterales; Enterobacteriaceae; Salmonella
	MTGDNTLITSHGINRRDFMKLCAALAATMGLSSKAAAEMAESVSNPQRPPVIWIGAQECT	WP_000145429.1
	hydrogenase 2 small subunit [Salmonella enterica]	Length: 372\nNumber of predicted TMHs:  1\nExp number of AAs in TMHs: 24.51309\nExp number, first 60 AAs:  0.2682\nTotal prob of N-in:   0.14402\noutside   1   331\nTMhelix   332   354\ninside   355   372

15151	GCF_001473505.1_Salmonella_enterica_CVM_43778_v1.0Salmonella enterica	Proteobacteria; Gammaproteobacteria; Enterobacterales; Enterobacteriaceae; Salmonella
	MTGDNTLITSHGINRRDFMKLCAALAATMGLSSKAAAEMAESVSNPQRPPVIWIGAQECT	WP_000145429.1
	hydrogenase 2 small subunit [Salmonella enterica]	Length: 372\nNumber of predicted TMHs:  1\nExp number of AAs in TMHs: 24.51309\nExp number, first 60 AAs:  0.2682\nTotal prob of N-in:   0.14402\noutside   1   331\nTMhelix   332   354\ninside   355   372

15152	GCF_001833305.1_ASM183330v1	Salmonella enterica	Proteobacteria; Gammaproteobacteria; Enterobacterales; Enterobacteriaceae; Salmonella
	MNNEETFYQAMRRKGVTRRSFLKFCSLAATSLGLGAGMTPKIAWALENKPRIPVVWIHGL	WP_001058311.1
	hydrogenase [Salmonella enterica]	Length: 372\nNumber of predicted TMHs:  2\nExp number of AAs in TMHs: 37.4699899999999\nExp number, first 60 AAs:  19.72699\nTotal prob of N-in:   0.96171\nPOSSIBLE N-term signal sequence\ninside   1   19\nTMhelix   20   39\noutside   40   325\nTMhelix   326   348\ninside   349   372

15153	GCF_001473465.1_Salmonella_enterica_CVM_43772_v1.0Salmonella enterica	Proteobacteria; Gammaproteobacteria; Enterobacterales; Enterobacteriaceae; Salmonella
	MNNEETFYQAMRRKGVTRRSFLKFCSLAATSLGLGAGMTPKIAWALENKPRIPVVWIHGL	WP_001058312.1
	hydrogenase [Salmonella enterica]	Length: 372\nNumber of predicted TMHs:  2\nExp number of AAs in TMHs: 36.51738\nExp number, first 60 AAs:  19.72849\nTotal prob of N-in:   0.96170\nPOSSIBLE N-term signal sequence\ninside   1   19\nTMhelix   20   39\noutside   40   325\nTMhelix   326   348\ninside   349   372

15154	GCF_001833305.1_ASM183330v1	Salmonella enterica	Proteobacteria; Gammaproteobacteria; Enterobacterales; Enterobacteriaceae; Salmonella
	MTGDNTLITSHGINRRDFMKLCAALAATMGLSSKAAAEMAESVSNPQRPPVIWIGAQECT	WP_000145429.1
	hydrogenase 2 small subunit [Salmonella enterica]	Length: 372\nNumber of predicted TMHs:  1\nExp number of AAs in TMHs: 24.51309\nExp number, first 60 AAs:  0.2682\nTotal prob of N-in:   0.14402\noutside   1   331\nTMhelix   332   354\ninside   355   372

15155	GCF_001833275.1_ASM183327v1	Salmonella enterica	Proteobacteria; Gammaproteobacteria; Enterobacterales; Enterobacteriaceae; Salmonella

MNNEETFYQAMRRKGVTRRSFLKFCSLAATSLGLGAGMTPKIAWALENKPRIPVVWIHGL WP_001058312.1 hydrogenase [Salmonella enterica] Length: 372\nNumber of predicted TMHs: 2\nExp number of AAs in TMHs: 36.51738\nExp number, first 60 AAs: 19.72849\nTotal prob of N-in: 0.96170\nPOSSIBLE N-term signal sequence\ninside 1 19\nTMhelix 20 39\noutside 40 325\nTMhelix 326 348\ninside 349 372

15156 GCF_002058315.1_ASM205831v1 Salmonella enterica Proteobacteria; Gammaproteobacteria; Enterobacterales; Enterobacteriaceae; Salmonella MNNEETFYQAMRRKGVTRRSFLKFCSLAATSLGLSAGMTPKIAWALENKPRIPVVWIHGL WP_001058316.1 hydrogenase [Salmonella enterica] Length: 372\nNumber of predicted TMHs: 2\nExp number of AAs in TMHs: 35.76397\nExp number, first 60 AAs: 18.97201\nTotal prob of N-in: 0.92785\nPOSSIBLE N-term signal sequence\ninside 1 19\nTMhelix 20 39\noutside 40 325\nTMhelix 326 348\ninside 349 372

15157 GCF_001473435.1_Salmonella_enterica_CVM_43762_v1.0Salmonella enterica Proteobacteria; Gammaproteobacteria; Enterobacterales; Enterobacteriaceae; Salmonella MNNEETFYQAMRRKGVTRRSFLKFCSLAATSLGLGAGMTPKIAWALENKPRIPVVWIHGL WP_001058312.1 hydrogenase [Salmonella enterica] Length: 372\nNumber of predicted TMHs: 2\nExp number of AAs in TMHs: 36.51738\nExp number, first 60 AAs: 19.72849\nTotal prob of N-in: 0.96170\nPOSSIBLE N-term signal sequence\ninside 1 19\nTMhelix 20 39\noutside 40 325\nTMhelix 326 348\ninside 349 372

15158 GCF_002058485.1_ASM205848v1 Salmonella enterica Proteobacteria; Gammaproteobacteria; Enterobacterales; Enterobacteriaceae; Salmonella MTGDNTLITSHGINRRDFMKLCAALAATMGLSSKAAAEMAESVSNPQRPPVIWIGAQECT WP_000145429.1 hydrogenase 2 small subunit [Salmonella enterica] Length: 372\nNumber of predicted TMHs: 1\nExp number of AAs in TMHs: 24.51309\nExp number, first 60 AAs: 0.2682\nTotal prob of N-in: 0.14402\noutside 1 331\nTMhelix 332 354\ninside 355 372

15159 GCF_002058415.1_ASM205841v1 Salmonella enterica Proteobacteria; Gammaproteobacteria; Enterobacterales; Enterobacteriaceae; Salmonella MTGDNTLITSHGINRRDFMKLCAALAATMGLSSKAAAEMAESVSNPQRPPVIWIGAQECT WP_000145429.1 hydrogenase 2 small subunit [Salmonella enterica] Length: 372\nNumber of predicted TMHs: 1\nExp number of AAs in TMHs: 24.51309\nExp number, first 60 AAs: 0.2682\nTotal prob of N-in: 0.14402\noutside 1 331\nTMhelix 332 354\ninside 355 372

15160 GCF_001570005.1_ASM157000v1 Salmonella enterica Proteobacteria; Gammaproteobacteria; Enterobacterales; Enterobacteriaceae; Salmonella MNNEETFYQAMRRKGVTRRSFLKFCSLAATSLGLGAGMTPKIAWALENKPRIPVIWIHGL WP_061376537.1 hydrogenase [Salmonella enterica] Length: 372\nNumber of predicted TMHs: 2\nExp number of AAs in TMHs: 36.5556\nExp number, first 60 AAs: 19.76633\nTotal prob of N-in: 0.96197\nPOSSIBLE N-term signal sequence\ninside 1 19\nTMhelix 20 39\noutside 40 325\nTMhelix 326 348\ninside 349 372

15161 GCF_002058315.1_ASM205831v1 Salmonella enterica Proteobacteria; Gammaproteobacteria; Enterobacterales; Enterobacteriaceae; Salmonella MTGDNTLITSHGINRRDFMKLCAALAATMGLSSKAAAEMAESVSNPQRPPVIWIGAQECT WP_000145429.1 hydrogenase 2 small subunit [Salmonella enterica] Length: 372\nNumber of predicted TMHs: 1\nExp number of AAs in TMHs: 24.51309\nExp number, first 60 AAs: 0.2682\nTotal prob of N-in: 0.14402\noutside 1 331\nTMhelix 332 354\ninside 355 372

15162 GCF_001833275.1_ASM183327v1 Salmonella enterica Proteobacteria; Gammaproteobacteria; Enterobacterales; Enterobacteriaceae; Salmonella MTGDNTLITSHGINRRDFMKLCAALAATMGLSSKAAAEMAESVSNPQRPPVIWIGAQECT WP_000145429.1 hydrogenase 2 small subunit [Salmonella enterica] Length: 372\nNumber of predicted TMHs: 1\nExp number of AAs in TMHs: 24.51309\nExp number, first 60 AAs: 0.2682\nTotal prob of N-in: 0.14402\noutside 1 331\nTMhelix 332 354\ninside 355 372

15163 GCF_002057855.1_ASM205785v1 Salmonella enterica Proteobacteria; Gammaproteobacteria; Enterobacterales; Enterobacteriaceae; Salmonella MNNEETLYQAMRRKGVTRRSFLKFCSLAATSLGLGAGMTPKIAWALENKPRIPVVWIHGL WP_001752445.1 hydrogenase-1 small chain [Salmonella enterica] Length: 372\nNumber of predicted TMHs: 2\nExp number of AAs in TMHs: 36.5123\nExp number, first 60 AAs: 19.72341\nTotal prob of N-in: 0.96144\nPOSSIBLE N-term signal sequence\ninside 1 19\nTMhelix 20 39\noutside 40 325\nTMhelix 326 348\ninside 349 372

15164 GCF_002057915.1_ASM205791v1 Salmonella enterica Proteobacteria; Gammaproteobacteria; Enterobacterales; Enterobacteriaceae; Salmonella
MNNEETFYQAMRRKGVTRRSFLKFCSLAATSLGLGAGMTPKIAWALENKPRIPVVWIHGL WP_001058312.1
hydrogenase [Salmonella enterica] Length: 372\nNumber of predicted TMHs: 2\nExp number of AAs in TMHs: 36.51738\nExp number, first 60 AAs: 19.72849\nTotal prob of N-in: 0.96170\nPOSSIBLE N-term signal sequence\ninside 1 19\nTMhelix 20 39\noutside 40 325\nTMhelix 326 348\ninside 349 372

15165 GCF_002057915.1_ASM205791v1 Salmonella enterica Proteobacteria; Gammaproteobacteria; Enterobacterales; Enterobacteriaceae; Salmonella
MTGDNTLITSHGINRRDFMKLCAALAATMGLSSKAAAEMAESVSNPQRPPVIWIGAQECT WP_000145429.1
hydrogenase 2 small subunit [Salmonella enterica] Length: 372\nNumber of predicted TMHs: 1\nExp number of AAs in TMHs: 24.51309\nExp number, first 60 AAs: 0.2682\nTotal prob of N-in: 0.14402\noutside 1 331\nTMhelix 332 354\ninside 355 372

15166 GCF_002057505.1_ASM205750v1 Salmonella enterica Proteobacteria; Gammaproteobacteria; Enterobacterales; Enterobacteriaceae; Salmonella
MNNEETFYQAMRRKGVTRRSFLKFCSLAATSLGLGAGMTPKIAWALENKPRIPVVWIHGL WP_001058312.1
hydrogenase [Salmonella enterica] Length: 372\nNumber of predicted TMHs: 2\nExp number of AAs in TMHs: 36.51738\nExp number, first 60 AAs: 19.72849\nTotal prob of N-in: 0.96170\nPOSSIBLE N-term signal sequence\ninside 1 19\nTMhelix 20 39\noutside 40 325\nTMhelix 326 348\ninside 349 372

15167 GCF_002057775.1_ASM205777v1 Salmonella enterica Proteobacteria; Gammaproteobacteria; Enterobacterales; Enterobacteriaceae; Salmonella
MNNEETFYQAMRRKGVTRRSFLKFCSLAATSLGLGAGMTPKIAWALENKPRIPVVWIHGL WP_001058307.1
hydrogenase [Salmonella enterica] Length: 372\nNumber of predicted TMHs: 2\nExp number of AAs in TMHs: 36.51738\nExp number, first 60 AAs: 19.72849\nTotal prob of N-in: 0.96170\nPOSSIBLE N-term signal sequence\ninside 1 19\nTMhelix 20 39\noutside 40 325\nTMhelix 326 348\ninside 349 372

15168 GCF_001473405.1_Salmonella_enterica_CVM_43759_v1.0Salmonella enterica Proteobacteria; Gammaproteobacteria; Enterobacterales; Enterobacteriaceae; Salmonella
MNNEETFYQAMRRKGVTRRSFLKFCSLAATSLGLGAGMTPKIAWALENKPRIPVVWIHGL WP_001058312.1
hydrogenase [Salmonella enterica] Length: 372\nNumber of predicted TMHs: 2\nExp number of AAs in TMHs: 36.51738\nExp number, first 60 AAs: 19.72849\nTotal prob of N-in: 0.96170\nPOSSIBLE N-term signal sequence\ninside 1 19\nTMhelix 20 39\noutside 40 325\nTMhelix 326 348\ninside 349 372

15169 GCF_002057445.1_ASM205744v1 Salmonella enterica Proteobacteria; Gammaproteobacteria; Enterobacterales; Enterobacteriaceae; Salmonella
MNNEETFYQAMRRKGVTRRSFLKFCSLAATSLGLGAGMTPKIAWALENKPRIPVVWIHGL WP_001058312.1
hydrogenase [Salmonella enterica] Length: 372\nNumber of predicted TMHs: 2\nExp number of AAs in TMHs: 36.51738\nExp number, first 60 AAs: 19.72849\nTotal prob of N-in: 0.96170\nPOSSIBLE N-term signal sequence\ninside 1 19\nTMhelix 20 39\noutside 40 325\nTMhelix 326 348\ninside 349 372

15170 GCF_002057365.1_ASM205736v1 Salmonella enterica Proteobacteria; Gammaproteobacteria; Enterobacterales; Enterobacteriaceae; Salmonella
MNNEETFYQAMRRKGVTRRSFLKFCSLAATSLGLGAGMTPKIAWALENKPRIPVVWIHGL WP_001058312.1
hydrogenase [Salmonella enterica] Length: 372\nNumber of predicted TMHs: 2\nExp number of AAs in TMHs: 36.51738\nExp number, first 60 AAs: 19.72849\nTotal prob of N-in: 0.96170\nPOSSIBLE N-term signal sequence\ninside 1 19\nTMhelix 20 39\noutside 40 325\nTMhelix 326 348\ninside 349 372

15171 GCF_002057335.1_ASM205733v1 Salmonella enterica Proteobacteria; Gammaproteobacteria; Enterobacterales; Enterobacteriaceae; Salmonella
MNNEETFYQAMRRKGVTRRSFLKFCSLAATSLGLGAGMTPKIAWALENKPRIPVVWIHGL WP_001058312.1
hydrogenase [Salmonella enterica] Length: 372\nNumber of predicted TMHs: 2\nExp number of AAs in TMHs: 36.51738\nExp number, first 60 AAs: 19.72849\nTotal prob of N-in: 0.96170\nPOSSIBLE N-term signal sequence\ninside 1 19\nTMhelix 20 39\noutside 40 325\nTMhelix 326 348\ninside 349 372

15172 GCF_002057445.1_ASM205744v1 Salmonella enterica Proteobacteria; Gammaproteobacteria; Enterobacterales; Enterobacteriaceae; Salmonella
MTGDNTLITSHGINRRDFMKLCAALAATMGLSSKAAAEMAESVSNPQRPPVIWIGAQECT WP_000145429.1
hydrogenase 2 small subunit [Salmonella enterica] Length: 372\nNumber of predicted TMHs: 1\nExp

number of AAs in TMHs: 24.51309\nExp number, first 60 AAs:  0.2682\nTotal prob of N-in:   0.14402\noutside   1   331\nTMhelix   332   354\ninside   355   372

15173 GCF_002057365.1_ASM205736v1 Salmonella enterica Proteobacteria; Gammaproteobacteria; Enterobacterales; Enterobacteriaceae; Salmonella MTGDNTLITSHGINRRDFMKLCAALAATMGLSSKAAAEMAESVSNPQRPPVIWIGAQECT WP_000145429.1 hydrogenase 2 small subunit [Salmonella enterica] Length: 372\nNumber of predicted TMHs:  1\nExp number of AAs in TMHs: 24.51309\nExp number, first 60 AAs:  0.2682\nTotal prob of N-in:   0.14402\noutside   1   331\nTMhelix   332   354\ninside   355   372

15174 GCF_002057335.1_ASM205733v1 Salmonella enterica Proteobacteria; Gammaproteobacteria; Enterobacterales; Enterobacteriaceae; Salmonella MTGDNTLITSHGINRRDFMKLCAALAATMGLSSKAAAEMAESVSNPQRPPVIWIGAQECT WP_000145429.1 hydrogenase 2 small subunit [Salmonella enterica] Length: 372\nNumber of predicted TMHs:  1\nExp number of AAs in TMHs: 24.51309\nExp number, first 60 AAs:  0.2682\nTotal prob of N-in:   0.14402\noutside   1   331\nTMhelix   332   354\ninside   355   372

15175 GCF_002057005.1_ASM205700v1 Salmonella enterica Proteobacteria; Gammaproteobacteria; Enterobacterales; Enterobacteriaceae; Salmonella MNNEETFYQAMRRKGVTRRSFLKFCSLAATSLGLGAGMTPKIAWALENKPRIPVVWIHGL WP_001058312.1 hydrogenase [Salmonella enterica] Length: 372\nNumber of predicted TMHs:  2\nExp number of AAs in TMHs: 36.51738\nExp number, first 60 AAs:  19.72849\nTotal prob of N-in:   0.96170\nPOSSIBLE N-term signal sequence\ninside   1   19\nTMhelix   20   39\noutside   40   325\nTMhelix   326   348\ninside   349   372

15176 GCF_001570065.1_ASM157006v1 Salmonella enterica Proteobacteria; Gammaproteobacteria; Enterobacterales; Enterobacteriaceae; Salmonella MTGDNTLITSHGINRRDFMKLCAALAATMGLSSKAAAEMAESVSNPQRPPVIWIGAQECT WP_000145429.1 hydrogenase 2 small subunit [Salmonella enterica] Length: 372\nNumber of predicted TMHs:  1\nExp number of AAs in TMHs: 24.51309\nExp number, first 60 AAs:  0.2682\nTotal prob of N-in:   0.14402\noutside   1   331\nTMhelix   332   354\ninside   355   372

15177 GCF_002057285.1_ASM205728v1 Salmonella enterica Proteobacteria; Gammaproteobacteria; Enterobacterales; Enterobacteriaceae; Salmonella MNNEETFYQAMRRKGVTRRSFLKFCSLAATSLGLSAGMTPKIAWALENKPRIPVVWIHGL WP_001058316.1 hydrogenase [Salmonella enterica] Length: 372\nNumber of predicted TMHs:  2\nExp number of AAs in TMHs: 35.76397\nExp number, first 60 AAs:  18.97201\nTotal prob of N-in:   0.92785\nPOSSIBLE N-term signal sequence\ninside   1   19\nTMhelix   20   39\noutside   40   325\nTMhelix   326   348\ninside   349   372

15178 GCF_002057285.1_ASM205728v1 Salmonella enterica Proteobacteria; Gammaproteobacteria; Enterobacterales; Enterobacteriaceae; Salmonella MTGDNTLITSHGINRRDFMKLCAALAATMGLSSKAAAEMAESVSNPQRPPVIWIGAQECT WP_000145429.1 hydrogenase 2 small subunit [Salmonella enterica] Length: 372\nNumber of predicted TMHs:  1\nExp number of AAs in TMHs: 24.51309\nExp number, first 60 AAs:  0.2682\nTotal prob of N-in:   0.14402\noutside   1   331\nTMhelix   332   354\ninside   355   372

15179 GCF_001473385.1_Salmonella_enterica_CVM_43757_v1.0Salmonella enterica Proteobacteria; Gammaproteobacteria; Enterobacterales; Enterobacteriaceae; Salmonella MNNEETFYQAMRRKGVTRRSFLKFCSLAATSLGLGAGMTPKIAWALENKPRIPVVWIHGL WP_001058312.1 hydrogenase [Salmonella enterica] Length: 372\nNumber of predicted TMHs:  2\nExp number of AAs in TMHs: 36.51738\nExp number, first 60 AAs:  19.72849\nTotal prob of N-in:   0.96170\nPOSSIBLE N-term signal sequence\ninside   1   19\nTMhelix   20   39\noutside   40   325\nTMhelix   326   348\ninside   349   372

15180 GCF_001473355.1_Salmonella_enterica_CVM_43753_v1.0Salmonella enterica Proteobacteria; Gammaproteobacteria; Enterobacterales; Enterobacteriaceae; Salmonella MNNEETFYQAMRRKGVTRRSFLKFCSLAATSLGLGAGMTPKIAWALENKPRIPVVWIHGL WP_023198237.1 hydrogenase 1, small subunit [Salmonella enterica] Length: 372\nNumber of predicted TMHs:  2\nExp number of AAs in TMHs: 36.39423\nExp number, first 60 AAs:  19.72877\nTotal prob of N-in:   0.96162\nPOSSIBLE N-term signal sequence\ninside   1   19\nTMhelix   20   39\noutside   40   325\nTMhelix   326   348\ninside   349   372

15181 GCF_002057205.1_ASM205720v1 Salmonella enterica Proteobacteria; Gammaproteobacteria; Enterobacterales; Enterobacteriaceae; Salmonella

MNNEETFYQAMRRKGVTRRSFLKFCSLAATSLGLGAGMTPKIAWALENKPRIPVVWIHGL WP_001058310.1 hydrogenase [Salmonella enterica] Length: 372\nNumber of predicted TMHs: 2\nExp number of AAs in TMHs: 37.4567299999999\nExp number, first 60 AAs: 19.72694\nTotal prob of N-in: 0.96171\nPOSSIBLE N-term signal sequence\ninside 1 19\nTMhelix 20 39\noutside 40 325\nTMhelix 326 348\ninside 349 372

15182 GCF_001473425.1_Salmonella_enterica_CVM_43765_v1.0Salmonella enterica Proteobacteria; Gammaproteobacteria; Enterobacterales; Enterobacteriaceae; Salmonella MNNEETFYQAMRRKGVTRRSFLKFCSLAATSLGLGAGMTPKIAWALENKPRIPVVWIHGL WP_001058311.1 hydrogenase [Salmonella enterica] Length: 372\nNumber of predicted TMHs: 2\nExp number of AAs in TMHs: 37.4699899999999\nExp number, first 60 AAs: 19.72699\nTotal prob of N-in: 0.96171\nPOSSIBLE N-term signal sequence\ninside 1 19\nTMhelix 20 39\noutside 40 325\nTMhelix 326 348\ninside 349 372

15183 GCF_001473385.1_Salmonella_enterica_CVM_43757_v1.0Salmonella enterica Proteobacteria; Gammaproteobacteria; Enterobacterales; Enterobacteriaceae; Salmonella MTGDNTLITSHGINRRDFMKLCAALAATMGLSSKAAAEMAESVSNPQRPPVIWIGAQECT WP_000145429.1 hydrogenase 2 small subunit [Salmonella enterica] Length: 372\nNumber of predicted TMHs: 1\nExp number of AAs in TMHs: 24.51309\nExp number, first 60 AAs: 0.2682\nTotal prob of N-in: 0.14402\noutside 1 331\nTMhelix 332 354\ninside 355 372

15184 GCF_001473345.1_Salmonella_enterica_CVM_43745_v1.0Salmonella enterica Proteobacteria; Gammaproteobacteria; Enterobacterales; Enterobacteriaceae; Salmonella MNNEETFYQAMRRKGVTRRSFLKFCSLAATSLGLGAGMTPKIAWALENKPRIPVVWIHGL WP_001058312.1 hydrogenase [Salmonella enterica] Length: 372\nNumber of predicted TMHs: 2\nExp number of AAs in TMHs: 36.51738\nExp number, first 60 AAs: 19.72849\nTotal prob of N-in: 0.96170\nPOSSIBLE N-term signal sequence\ninside 1 19\nTMhelix 20 39\noutside 40 325\nTMhelix 326 348\ninside 349 372

15185 GCF_001833085.1_ASM183308v1 Salmonella enterica Proteobacteria; Gammaproteobacteria; Enterobacterales; Enterobacteriaceae; Salmonella MNNEETFYQAMRRKGVTRRSFLKFCSLAATSLGLGAGMTPKIAWALENKPRIPVVWIHGL WP_001058311.1 hydrogenase [Salmonella enterica] Length: 372\nNumber of predicted TMHs: 2\nExp number of AAs in TMHs: 37.4699899999999\nExp number, first 60 AAs: 19.72699\nTotal prob of N-in: 0.96171\nPOSSIBLE N-term signal sequence\ninside 1 19\nTMhelix 20 39\noutside 40 325\nTMhelix 326 348\ninside 349 372

15186 GCF_002056435.1_ASM205643v1 Salmonella enterica Proteobacteria; Gammaproteobacteria; Enterobacterales; Enterobacteriaceae; Salmonella MNNEETFYQAMRRKGVTRRSFLKFCSLAATSLGLGAGMTPKIAWALENKPRIPVVWIHGL WP_001058307.1 hydrogenase [Salmonella enterica] Length: 372\nNumber of predicted TMHs: 2\nExp number of AAs in TMHs: 36.51738\nExp number, first 60 AAs: 19.72849\nTotal prob of N-in: 0.96170\nPOSSIBLE N-term signal sequence\ninside 1 19\nTMhelix 20 39\noutside 40 325\nTMhelix 326 348\ninside 349 372

15187 GCF_002056505.1_ASM205650v1 Salmonella enterica Proteobacteria; Gammaproteobacteria; Enterobacterales; Enterobacteriaceae; Salmonella MNNEETFYQAMRRKGVTRRSFLKFCSLAATSLGLGAGMTPKIAWALENKPRIPVVWIHGL WP_001058307.1 hydrogenase [Salmonella enterica] Length: 372\nNumber of predicted TMHs: 2\nExp number of AAs in TMHs: 36.51738\nExp number, first 60 AAs: 19.72849\nTotal prob of N-in: 0.96170\nPOSSIBLE N-term signal sequence\ninside 1 19\nTMhelix 20 39\noutside 40 325\nTMhelix 326 348\ninside 349 372

15188 GCF_002056555.1_ASM205655v1 Salmonella enterica Proteobacteria; Gammaproteobacteria; Enterobacterales; Enterobacteriaceae; Salmonella MNNEETFYQAMRRKGVTRRSFLKFCSLAATSLGLGAGMTPKIAWALENKPRIPVVWIHGL WP_001058307.1 hydrogenase [Salmonella enterica] Length: 372\nNumber of predicted TMHs: 2\nExp number of AAs in TMHs: 36.51738\nExp number, first 60 AAs: 19.72849\nTotal prob of N-in: 0.96170\nPOSSIBLE N-term signal sequence\ninside 1 19\nTMhelix 20 39\noutside 40 325\nTMhelix 326 348\ninside 349 372

15189 GCF_002056555.1_ASM205655v1 Salmonella enterica Proteobacteria; Gammaproteobacteria; Enterobacterales; Enterobacteriaceae; Salmonella MTGDNTLITSHGINRRDFMKLCAALAATMGLSSKAAAEMAESVSNPQRPPVIWIGAQECT WP_000145429.1 hydrogenase 2 small subunit [Salmonella enterica] Length: 372\nNumber of predicted TMHs: 1\nExp number of AAs in TMHs: 24.51309\nExp number, first 60 AAs: 0.2682\nTotal prob of N-in: 0.14402\noutside 1 331\nTMhelix 332 354\ninside 355 372

15190 GCF_002056695.1_ASM205669v1 Salmonella enterica Proteobacteria; Gammaproteobacteria; Enterobacterales; Enterobacteriaceae; Salmonella
MTGDNTLITSHGINRRDFMKLCAALAATMGLSSKAAAEMAESVSNPQRPPVIWIGAQECT WP_000145429.1
hydrogenase 2 small subunit [Salmonella enterica] Length: 372\nNumber of predicted TMHs: 1\nExp number of AAs in TMHs: 24.51309\nExp number, first 60 AAs: 0.2682\nTotal prob of N-in: 0.14402\noutside 1 331\nTMhelix 332 354\ninside 355 372

15191 GCF_001473315.1_Salmonella_enterica_CVM_43747_v1.0Salmonella enterica Proteobacteria; Gammaproteobacteria; Enterobacterales; Enterobacteriaceae; Salmonella
MNNEETFYQAMRRKGVTRRSFLKFCSLAATSLGLGAGMTPKIAWALENKPRIPVVWIHGL WP_001058312.1
hydrogenase [Salmonella enterica] Length: 372\nNumber of predicted TMHs: 2\nExp number of AAs in TMHs: 36.51738\nExp number, first 60 AAs: 19.72849\nTotal prob of N-in: 0.96170\nPOSSIBLE N-term signal sequence\ninside 1 19\nTMhelix 20 39\noutside 40 325\nTMhelix 326 348\ninside 349 372

15192 GCF_001570105.1_ASM157010v1 Salmonella enterica Proteobacteria; Gammaproteobacteria; Enterobacterales; Enterobacteriaceae; Salmonella
MTGDNTLITSHGINRRDFMKLCAALAATMGLSSKAAAEMAESVSNPQRPPVIWIGAQECT WP_000145429.1
hydrogenase 2 small subunit [Salmonella enterica] Length: 372\nNumber of predicted TMHs: 1\nExp number of AAs in TMHs: 24.51309\nExp number, first 60 AAs: 0.2682\nTotal prob of N-in: 0.14402\noutside 1 331\nTMhelix 332 354\ninside 355 372

15193 GCF_002056505.1_ASM205650v1 Salmonella enterica Proteobacteria; Gammaproteobacteria; Enterobacterales; Enterobacteriaceae; Salmonella
MTGDNTLITSHGINRRDFMKLCAALAATMGLSSKAAAEMAESVSNPQRPPVIWIGAQECT WP_000145429.1
hydrogenase 2 small subunit [Salmonella enterica] Length: 372\nNumber of predicted TMHs: 1\nExp number of AAs in TMHs: 24.51309\nExp number, first 60 AAs: 0.2682\nTotal prob of N-in: 0.14402\noutside 1 331\nTMhelix 332 354\ninside 355 372

15194 GCF_002056585.1_ASM205658v1 Salmonella enterica Proteobacteria; Gammaproteobacteria; Enterobacterales; Enterobacteriaceae; Salmonella
MTGDNTLITSHGINRRDFMKLCAALAATMGLSSKAAAEMAESVSNPQRPPVIWIGAQECT WP_000145429.1
hydrogenase 2 small subunit [Salmonella enterica] Length: 372\nNumber of predicted TMHs: 1\nExp number of AAs in TMHs: 24.51309\nExp number, first 60 AAs: 0.2682\nTotal prob of N-in: 0.14402\noutside 1 331\nTMhelix 332 354\ninside 355 372

15195 GCF_002056435.1_ASM205643v1 Salmonella enterica Proteobacteria; Gammaproteobacteria; Enterobacterales; Enterobacteriaceae; Salmonella
MTGDNTLITSHGINRRDFMKLCAALAATMGLSSKAAAEMAESVSNPQRPPVIWIGAQECT WP_000145429.1
hydrogenase 2 small subunit [Salmonella enterica] Length: 372\nNumber of predicted TMHs: 1\nExp number of AAs in TMHs: 24.51309\nExp number, first 60 AAs: 0.2682\nTotal prob of N-in: 0.14402\noutside 1 331\nTMhelix 332 354\ninside 355 372

15196 GCF_002056375.1_ASM205637v1 Salmonella enterica Proteobacteria; Gammaproteobacteria; Enterobacterales; Enterobacteriaceae; Salmonella
MTGDNTLITSHGINRRDFMKLCAALAATMGLSSKAAAEMAESVSNPQRPPVIWIGAQECT WP_000145429.1
hydrogenase 2 small subunit [Salmonella enterica] Length: 372\nNumber of predicted TMHs: 1\nExp number of AAs in TMHs: 24.51309\nExp number, first 60 AAs: 0.2682\nTotal prob of N-in: 0.14402\noutside 1 331\nTMhelix 332 354\ninside 355 372

15197 GCF_002056195.1_ASM205619v1 Salmonella enterica Proteobacteria; Gammaproteobacteria; Enterobacterales; Enterobacteriaceae; Salmonella
MTGDNTLITSHGINRRDFMKLCAALAATMGLSSKAAAEMAESVSNPQRPPVIWIGAQECT WP_000145429.1
hydrogenase 2 small subunit [Salmonella enterica] Length: 372\nNumber of predicted TMHs: 1\nExp number of AAs in TMHs: 24.51309\nExp number, first 60 AAs: 0.2682\nTotal prob of N-in: 0.14402\noutside 1 331\nTMhelix 332 354\ninside 355 372

15198 GCF_002056005.1_ASM205600v1 Salmonella enterica Proteobacteria; Gammaproteobacteria; Enterobacterales; Enterobacteriaceae; Salmonella
MNNEETFYQAMRRKGVTRRSFLKFCSLAATSLGLGAGMTPKIAWALENKPRIPVVWIHGL WP_001058307.1
hydrogenase [Salmonella enterica] Length: 372\nNumber of predicted TMHs: 2\nExp number of AAs in

TMHs: 36.51738\nExp number, first 60 AAs: 19.72849\nTotal prob of N-in: 0.96170\nPOSSIBLE N-term signal sequence\ninside 1 19\nTMhelix 20 39\noutside 40 325\nTMhelix 326 348\ninside 349 372

15199 GCF_002056095.1_ASM205609v1 Salmonella enterica Proteobacteria; Gammaproteobacteria; Enterobacterales; Enterobacteriaceae; Salmonella
MTGDNTLITSHGINRRDFMKLCAALAATMGLSSKAAAEMAESVSNPQRPPVIWIGAQECT WP_000145429.1
hydrogenase 2 small subunit [Salmonella enterica] Length: 372\nNumber of predicted TMHs: 1\nExp number of AAs in TMHs: 24.51309\nExp number, first 60 AAs: 0.2682\nTotal prob of N-in: 0.14402\noutside 1 331\nTMhelix 332 354\ninside 355 372

15200 GCF_002056135.1_ASM205613v1 Salmonella enterica Proteobacteria; Gammaproteobacteria; Enterobacterales; Enterobacteriaceae; Salmonella
MTGDNTLITSHGINRRDFMKLCAALAATMGLSSKAAAEMAESVSNPQRPPVIWIGAQECT WP_000145429.1
hydrogenase 2 small subunit [Salmonella enterica] Length: 372\nNumber of predicted TMHs: 1\nExp number of AAs in TMHs: 24.51309\nExp number, first 60 AAs: 0.2682\nTotal prob of N-in: 0.14402\noutside 1 331\nTMhelix 332 354\ninside 355 372

15201 GCF_002055675.1_ASM205567v1 Salmonella enterica Proteobacteria; Gammaproteobacteria; Enterobacterales; Enterobacteriaceae; Salmonella
MNNEETFYQAMRRKGVTRRSFLKFCSLAATSLGLGAGMTPKIAWALENKPRIPVVWIHGL WP_001058312.1
hydrogenase [Salmonella enterica] Length: 372\nNumber of predicted TMHs: 2\nExp number of AAs in TMHs: 36.51738\nExp number, first 60 AAs: 19.72849\nTotal prob of N-in: 0.96170\nPOSSIBLE N-term signal sequence\ninside 1 19\nTMhelix 20 39\noutside 40 325\nTMhelix 326 348\ninside 349 372

15202 GCF_002055885.1_ASM205588v1 Salmonella enterica Proteobacteria; Gammaproteobacteria; Enterobacterales; Enterobacteriaceae; Salmonella
MNNEETFYQAMRRKGVTRRSFLKFCSLAATSLGLGAGMTPKIAWALENKPRIPVVWIHGL WP_001058312.1
hydrogenase [Salmonella enterica] Length: 372\nNumber of predicted TMHs: 2\nExp number of AAs in TMHs: 36.51738\nExp number, first 60 AAs: 19.72849\nTotal prob of N-in: 0.96170\nPOSSIBLE N-term signal sequence\ninside 1 19\nTMhelix 20 39\noutside 40 325\nTMhelix 326 348\ninside 349 372

15203 GCF_002055955.1_ASM205595v1 Salmonella enterica Proteobacteria; Gammaproteobacteria; Enterobacterales; Enterobacteriaceae; Salmonella
MNNEETFYQAMRRKGVTRRSFLKFCSLAATSLGLGAGMTPKIAWALENKPRIPVVWIHGL WP_001058312.1
hydrogenase [Salmonella enterica] Length: 372\nNumber of predicted TMHs: 2\nExp number of AAs in TMHs: 36.51738\nExp number, first 60 AAs: 19.72849\nTotal prob of N-in: 0.96170\nPOSSIBLE N-term signal sequence\ninside 1 19\nTMhelix 20 39\noutside 40 325\nTMhelix 326 348\ninside 349 372

15204 GCF_002055755.1_ASM205575v1 Salmonella enterica Proteobacteria; Gammaproteobacteria; Enterobacterales; Enterobacteriaceae; Salmonella
MTGDNTLITSHGINRRDFMKLCAALAATMGLSSKAAAEMAESVSNPQRPPVIWIGAQECT WP_000145429.1
hydrogenase 2 small subunit [Salmonella enterica] Length: 372\nNumber of predicted TMHs: 1\nExp number of AAs in TMHs: 24.51309\nExp number, first 60 AAs: 0.2682\nTotal prob of N-in: 0.14402\noutside 1 331\nTMhelix 332 354\ninside 355 372

15205 GCF_002055595.1_ASM205559v1 Salmonella enterica Proteobacteria; Gammaproteobacteria; Enterobacterales; Enterobacteriaceae; Salmonella
MTGDNTLITSHGINRRDFMKLCAALAATMGLSSKAAAEMAESVSNPQRPPVIWIGAQECT WP_000145429.1
hydrogenase 2 small subunit [Salmonella enterica] Length: 372\nNumber of predicted TMHs: 1\nExp number of AAs in TMHs: 24.51309\nExp number, first 60 AAs: 0.2682\nTotal prob of N-in: 0.14402\noutside 1 331\nTMhelix 332 354\ninside 355 372

15206 GCF_002055675.1_ASM205567v1 Salmonella enterica Proteobacteria; Gammaproteobacteria; Enterobacterales; Enterobacteriaceae; Salmonella
MTGDNTLITSHGINRRDFMKLCAALAATMGLSSKAAAEMAESVSNPQRPPVIWIGAQECT WP_000145429.1
hydrogenase 2 small subunit [Salmonella enterica] Length: 372\nNumber of predicted TMHs: 1\nExp number of AAs in TMHs: 24.51309\nExp number, first 60 AAs: 0.2682\nTotal prob of N-in: 0.14402\noutside 1 331\nTMhelix 332 354\ninside 355 372

15207 GCF_002055475.1_ASM205547v1 Salmonella enterica Proteobacteria; Gammaproteobacteria; Enterobacterales; Enterobacteriaceae; Salmonella

MTGDNTLITSHGINRRDFMKLCAALAATMGLSSKAAAEMAESVSNPQRPPVIWIGAQECT WP_000145429.1 hydrogenase 2 small subunit [Salmonella enterica] Length: 372\nNumber of predicted TMHs: 1\nExp number of AAs in TMHs: 24.51309\nExp number, first 60 AAs: 0.2682\nTotal prob of N-in: 0.14402\noutside 1 331\nTMhelix 332 354\ninside 355 372

15208 GCF_002055415.1_ASM205541v1 Salmonella enterica Proteobacteria; Gammaproteobacteria; Enterobacterales; Enterobacteriaceae; Salmonella MNNEETFYQAMRRKGVTRRSFLKFCSLAATSLGLGAGMTPKIAWALENKPRIPVVWIHGL WP_001058312.1 hydrogenase [Salmonella enterica] Length: 372\nNumber of predicted TMHs: 2\nExp number of AAs in TMHs: 36.51738\nExp number, first 60 AAs: 19.72849\nTotal prob of N-in: 0.96170\nPOSSIBLE N-term signal sequence\ninside 1 19\nTMhelix 20 39\noutside 40 325\nTMhelix 326 348\ninside 349 372

15209 GCF_002055375.1_ASM205537v1 Salmonella enterica Proteobacteria; Gammaproteobacteria; Enterobacterales; Enterobacteriaceae; Salmonella MNNEETFYQAMRRKGVTRRSFLKFCSLAATSLGLGAGMTPKIAWALENKPRIPVVWIHGL WP_001058311.1 hydrogenase [Salmonella enterica] Length: 372\nNumber of predicted TMHs: 2\nExp number of AAs in TMHs: 37.4699899999999\nExp number, first 60 AAs: 19.72699\nTotal prob of N-in: 0.96171\nPOSSIBLE N-term signal sequence\ninside 1 19\nTMhelix 20 39\noutside 40 325\nTMhelix 326 348\ninside 349 372

15210 GCF_002055415.1_ASM205541v1 Salmonella enterica Proteobacteria; Gammaproteobacteria; Enterobacterales; Enterobacteriaceae; Salmonella MTGDNTLITSHGINRRDFMKLCAALAATMGLSSKAAAEMAESVSNPQRPPVIWIGAQECT WP_000145429.1 hydrogenase 2 small subunit [Salmonella enterica] Length: 372\nNumber of predicted TMHs: 1\nExp number of AAs in TMHs: 24.51309\nExp number, first 60 AAs: 0.2682\nTotal prob of N-in: 0.14402\noutside 1 331\nTMhelix 332 354\ninside 355 372

15211 GCF_002055055.1_ASM205505v1 Salmonella enterica Proteobacteria; Gammaproteobacteria; Enterobacterales; Enterobacteriaceae; Salmonella MNNEETFYQAMRRKGVTRRSFLKFCSLAATSLGLGAGMTPKIAWALENKPRIPVVWIHGL WP_001058311.1 hydrogenase [Salmonella enterica] Length: 372\nNumber of predicted TMHs: 2\nExp number of AAs in TMHs: 37.4699899999999\nExp number, first 60 AAs: 19.72699\nTotal prob of N-in: 0.96171\nPOSSIBLE N-term signal sequence\ninside 1 19\nTMhelix 20 39\noutside 40 325\nTMhelix 326 348\ninside 349 372

15212 GCF_002055025.1_ASM205502v1 Salmonella enterica Proteobacteria; Gammaproteobacteria; Enterobacterales; Enterobacteriaceae; Salmonella MTGDNTLITSHGINRRDFMKLCAALAATMGLSSKAAAEMAESVSNPQRPPVIWIGAQECT WP_080069096.1 hydrogenase 2 small subunit [Salmonella enterica] Length: 372\nNumber of predicted TMHs: 1\nExp number of AAs in TMHs: 24.51237\nExp number, first 60 AAs: 0.2682\nTotal prob of N-in: 0.14401\noutside 1 331\nTMhelix 332 354\ninside 355 372

15213 GCF_002054775.1_ASM205477v1 Salmonella enterica Proteobacteria; Gammaproteobacteria; Enterobacterales; Enterobacteriaceae; Salmonella MNNEETFYQAMRRKGVTRRSFLKFCSLAATSLGLGAGMMPKIAWALENKPRIPVVWIHGL WP_057393961.1 hydrogenase [Salmonella enterica] Length: 372\nNumber of predicted TMHs: 2\nExp number of AAs in TMHs: 37.03037\nExp number, first 60 AAs: 20.25216\nTotal prob of N-in: 0.98577\nPOSSIBLE N-term signal sequence\ninside 1 19\nTMhelix 20 39\noutside 40 325\nTMhelix 326 348\ninside 349 372

15214 GCF_002055115.1_ASM205511v1 Salmonella enterica Proteobacteria; Gammaproteobacteria; Enterobacterales; Enterobacteriaceae; Salmonella MTGDNTLITSHGINRRDFMKLCAALAATMGLSSKAAAEMAESVSNPQRPPVIWIGAQECT WP_000145429.1 hydrogenase 2 small subunit [Salmonella enterica] Length: 372\nNumber of predicted TMHs: 1\nExp number of AAs in TMHs: 24.51309\nExp number, first 60 AAs: 0.2682\nTotal prob of N-in: 0.14402\noutside 1 331\nTMhelix 332 354\ninside 355 372

15215 GCF_002055115.1_ASM205511v1 Salmonella enterica Proteobacteria; Gammaproteobacteria; Enterobacterales; Enterobacteriaceae; Salmonella MNNEETFYQAMRRKGVTRRSFLKFCSLAATSLGLGAGMTPKIAWALENKPRIPVVWIHGL WP_001058310.1 hydrogenase [Salmonella enterica] Length: 372\nNumber of predicted TMHs: 2\nExp number of AAs in TMHs: 37.4567299999999\nExp number, first 60 AAs: 19.72694\nTotal prob of N-in: 0.96171\nPOSSIBLE N-term signal sequence\ninside 1 19\nTMhelix 20 39\noutside 40 325\nTMhelix 326 348\ninside 349 372

15216	GCF_002144365.1_ASM214436v1	Salmonella enterica	Proteobacteria; Gammaproteobacteria; Enterobacterales; Enterobacteriaceae; Salmonella
MTGDNTLITSHGINRRDFMKLCAALAATMGLSSKAAAEMAESVSNPQRPPVIWIGAQECT	WP_000145429.1
hydrogenase 2 small subunit [Salmonella enterica]	Length: 372\nNumber of predicted TMHs: 1\nExp number of AAs in TMHs: 24.51309\nExp number, first 60 AAs: 0.2682\nTotal prob of N-in: 0.14402\noutside 1 331\nTMhelix 332 354\ninside 355 372

15217	GCF_002144435.1_ASM214443v1	Salmonella enterica	Proteobacteria; Gammaproteobacteria; Enterobacterales; Enterobacteriaceae; Salmonella
MNNEETFYQAMRRKGVTRRSFLKFCSLAATSLGLGAGMTPKIAWALENKPRIPVVWIHGL	WP_001058312.1
hydrogenase [Salmonella enterica]	Length: 372\nNumber of predicted TMHs: 2\nExp number of AAs in TMHs: 36.51738\nExp number, first 60 AAs: 19.72849\nTotal prob of N-in: 0.96170\nPOSSIBLE N-term signal sequence\ninside 1 19\nTMhelix 20 39\noutside 40 325\nTMhelix 326 348\ninside 349 372

15218	GCF_002144395.1_ASM214439v1	Salmonella enterica	Proteobacteria; Gammaproteobacteria; Enterobacterales; Enterobacteriaceae; Salmonella
MNNEETFYQAMRRKGVTRRSFLKFCSLAATSLGLGAGMTPKIAWALENKPRIPVVWIHGL	WP_001058312.1
hydrogenase [Salmonella enterica]	Length: 372\nNumber of predicted TMHs: 2\nExp number of AAs in TMHs: 36.51738\nExp number, first 60 AAs: 19.72849\nTotal prob of N-in: 0.96170\nPOSSIBLE N-term signal sequence\ninside 1 19\nTMhelix 20 39\noutside 40 325\nTMhelix 326 348\ninside 349 372

15219	GCF_002144475.1_ASM214447v1	Salmonella enterica	Proteobacteria; Gammaproteobacteria; Enterobacterales; Enterobacteriaceae; Salmonella
MTGDNTLITSHGINRRDFMKLCAALAATMGLSSKAAAEMAESVSNPQRPPVIWIGAQECT	WP_000145429.1
hydrogenase 2 small subunit [Salmonella enterica]	Length: 372\nNumber of predicted TMHs: 1\nExp number of AAs in TMHs: 24.51309\nExp number, first 60 AAs: 0.2682\nTotal prob of N-in: 0.14402\noutside 1 331\nTMhelix 332 354\ninside 355 372

15220	GCF_002144485.1_ASM214448v1	Salmonella enterica	Proteobacteria; Gammaproteobacteria; Enterobacterales; Enterobacteriaceae; Salmonella
MTGDNTLITSHGINRRDFMKLCAALAATMGLSSKAAAEMAESVSNPQRPPVIWIGAQECT	WP_000145429.1
hydrogenase 2 small subunit [Salmonella enterica]	Length: 372\nNumber of predicted TMHs: 1\nExp number of AAs in TMHs: 24.51309\nExp number, first 60 AAs: 0.2682\nTotal prob of N-in: 0.14402\noutside 1 331\nTMhelix 332 354\ninside 355 372

15221	GCF_001569705.1_ASM156970v1	Salmonella enterica	Proteobacteria; Gammaproteobacteria; Enterobacterales; Enterobacteriaceae; Salmonella
MTGDNTLITSHGINRRDFMKLCAALAATMGLSSKAAAEMAESVSNPQRPPVIWIGAQECT	WP_000145429.1
hydrogenase 2 small subunit [Salmonella enterica]	Length: 372\nNumber of predicted TMHs: 1\nExp number of AAs in TMHs: 24.51309\nExp number, first 60 AAs: 0.2682\nTotal prob of N-in: 0.14402\noutside 1 331\nTMhelix 332 354\ninside 355 372

15222	GCF_001473595.1_Salmonella_enterica_CVM_43787_v1.0Salmonella enterica	Proteobacteria; Gammaproteobacteria; Enterobacterales; Enterobacteriaceae; Salmonella
MTGDNTLITSHGINRRDFMKLCAALAATMGLSSKAAAEMAESVSNPQRPPVIWIGAQECT	WP_000145429.1
hydrogenase 2 small subunit [Salmonella enterica]	Length: 372\nNumber of predicted TMHs: 1\nExp number of AAs in TMHs: 24.51309\nExp number, first 60 AAs: 0.2682\nTotal prob of N-in: 0.14402\noutside 1 331\nTMhelix 332 354\ninside 355 372

15223	GCF_001474355.1_Salmonella_enterica_CVM_43771_v1.0Salmonella enterica	Proteobacteria; Gammaproteobacteria; Enterobacterales; Enterobacteriaceae; Salmonella
MTGDNTLITSHGINRRDFMKLCAALAATMGLSSKAAAEMAESVSNPQRPPVIWIGAQECT	WP_000145429.1
hydrogenase 2 small subunit [Salmonella enterica]	Length: 372\nNumber of predicted TMHs: 1\nExp number of AAs in TMHs: 24.51309\nExp number, first 60 AAs: 0.2682\nTotal prob of N-in: 0.14402\noutside 1 331\nTMhelix 332 354\ninside 355 372

15224	GCF_001473425.1_Salmonella_enterica_CVM_43765_v1.0Salmonella enterica	Proteobacteria; Gammaproteobacteria; Enterobacterales; Enterobacteriaceae; Salmonella
MTGDNTLITSHGINRRDFMKLCAALAATMGLSSKAAAEMAESVSNPQRPPVIWIGAQECT	WP_000145429.1
hydrogenase 2 small subunit [Salmonella enterica]	Length: 372\nNumber of predicted TMHs: 1\nExp

number of AAs in TMHs: 24.51309\nExp number, first 60 AAs: 0.2682\nTotal prob of N-in: 0.14402\noutside 1 331\nTMhelix 332 354\ninside 355 372

15225 GCF_001473405.1_Salmonella_enterica_CVM_43759_v1.0Salmonella enterica Proteobacteria; Gammaproteobacteria; Enterobacterales; Enterobacteriaceae; Salmonella
MTGDNTLITSHGINRRDFMKLCAALAATMGLSSKAAAEMAESVSNPQRPPVIWIGAQECT WP_000145429.1
hydrogenase 2 small subunit [Salmonella enterica] Length: 372\nNumber of predicted TMHs: 1\nExp number of AAs in TMHs: 24.51309\nExp number, first 60 AAs: 0.2682\nTotal prob of N-in: 0.14402\noutside 1 331\nTMhelix 332 354\ninside 355 372

15226 GCF_001832165.1_ASM183216v1 Salmonella enterica Proteobacteria; Gammaproteobacteria; Enterobacterales; Enterobacteriaceae; Salmonella
MTGDNTLITSHGINRRDFMKLCAALAATMGLSSKAAAEMAESVSNPQRPPVIWIGAQECT WP_000145429.1
hydrogenase 2 small subunit [Salmonella enterica] Length: 372\nNumber of predicted TMHs: 1\nExp number of AAs in TMHs: 24.51309\nExp number, first 60 AAs: 0.2682\nTotal prob of N-in: 0.14402\noutside 1 331\nTMhelix 332 354\ninside 355 372

15227 GCF_001474655.1_Salmonella_enterica_CVM_43796_v1.0Salmonella enterica Proteobacteria; Gammaproteobacteria; Enterobacterales; Enterobacteriaceae; Salmonella
MTGDNTLITSHGINRRDFMKLCAALAATMGLSSKAAAEMAESVSNPQRPPVIWIGAQECT WP_000145429.1
hydrogenase 2 small subunit [Salmonella enterica] Length: 372\nNumber of predicted TMHs: 1\nExp number of AAs in TMHs: 24.51309\nExp number, first 60 AAs: 0.2682\nTotal prob of N-in: 0.14402\noutside 1 331\nTMhelix 332 354\ninside 355 372

15228 GCF_001833385.1_ASM183338v1 Salmonella enterica Proteobacteria; Gammaproteobacteria; Enterobacterales; Enterobacteriaceae; Salmonella
MTGDNTLITSHGINRRDFMKLCAALAATMGLSSKAAAEMAESVSNPQRPPVIWIGAQECT WP_000145429.1
hydrogenase 2 small subunit [Salmonella enterica] Length: 372\nNumber of predicted TMHs: 1\nExp number of AAs in TMHs: 24.51309\nExp number, first 60 AAs: 0.2682\nTotal prob of N-in: 0.14402\noutside 1 331\nTMhelix 332 354\ninside 355 372

15229 GCF_002036585.1_ASM203658v1 Salmonella enterica Proteobacteria; Gammaproteobacteria; Enterobacterales; Enterobacteriaceae; Salmonella
MTGDNTLITSHGINRRDFMKLCAALAATMGLSSKAAAEMAESVSNPQRPPVIWIGAQECT WP_000145429.1
hydrogenase 2 small subunit [Salmonella enterica] Length: 372\nNumber of predicted TMHs: 1\nExp number of AAs in TMHs: 24.51309\nExp number, first 60 AAs: 0.2682\nTotal prob of N-in: 0.14402\noutside 1 331\nTMhelix 332 354\ninside 355 372

15230 GCF_002055025.1_ASM205502v1 Salmonella enterica Proteobacteria; Gammaproteobacteria; Enterobacterales; Enterobacteriaceae; Salmonella
MNNEETFYQAMRRKGVTRRSFLKFCSLAATSLGLGAGMTPKIAWALENKPRIPVVWIHGL WP_001058312.1
hydrogenase [Salmonella enterica] Length: 372\nNumber of predicted TMHs: 2\nExp number of AAs in TMHs: 36.51738\nExp number, first 60 AAs: 19.72849\nTotal prob of N-in: 0.96170\nPOSSIBLE N-term signal sequence\ninside 1 19\nTMhelix 20 39\noutside 40 325\nTMhelix 326 348\ninside 349 372

15231 GCF_001569885.1_ASM156988v1 Salmonella enterica Proteobacteria; Gammaproteobacteria; Enterobacterales; Enterobacteriaceae; Salmonella
MTGDNTLITSHGINRRDFMKLCAALAATMGLSSKAAAEMAESVSNPQRPPVIWIGAQECT WP_000145429.1
hydrogenase 2 small subunit [Salmonella enterica] Length: 372\nNumber of predicted TMHs: 1\nExp number of AAs in TMHs: 24.51309\nExp number, first 60 AAs: 0.2682\nTotal prob of N-in: 0.14402\noutside 1 331\nTMhelix 332 354\ninside 355 372

15232 GCF_001570045.1_ASM157004v1 Salmonella enterica Proteobacteria; Gammaproteobacteria; Enterobacterales; Enterobacteriaceae; Salmonella
MTGDNTLITSHGINRRDFMKLCAALAATMGLSSKAAAEMAESVSNPQRPPVIWIGAQECT WP_000145429.1
hydrogenase 2 small subunit [Salmonella enterica] Length: 372\nNumber of predicted TMHs: 1\nExp number of AAs in TMHs: 24.51309\nExp number, first 60 AAs: 0.2682\nTotal prob of N-in: 0.14402\noutside 1 331\nTMhelix 332 354\ninside 355 372

15233 GCF_001570085.1_ASM157008v1 Salmonella enterica Proteobacteria; Gammaproteobacteria; Enterobacterales; Enterobacteriaceae; Salmonella

MTGDNTLITSHGINRRDFMKLCAALAATMGLSSKAAAEMAESVSNPQRPPVIWIGAQECT WP_000145429.1 hydrogenase 2 small subunit [Salmonella enterica] Length: 372\nNumber of predicted TMHs: 1\nExp number of AAs in TMHs: 24.51309\nExp number, first 60 AAs: 0.2682\nTotal prob of N-in: 0.14402\noutside 1 331\nTMhelix 332 354\ninside 355 372

15234 GCF_000783815.1_ASM78381v1 Salmonella enterica Proteobacteria; Gammaproteobacteria; Enterobacterales; Enterobacteriaceae; Salmonella MTGDNTLITSHGINRRDFMKLCAALAATMGLSSKAAAEMAESVSNPQRPPVIWIGAQECT WP_000145429.1 hydrogenase 2 small subunit [Salmonella enterica] Length: 372\nNumber of predicted TMHs: 1\nExp number of AAs in TMHs: 24.51309\nExp number, first 60 AAs: 0.2682\nTotal prob of N-in: 0.14402\noutside 1 331\nTMhelix 332 354\ninside 355 372

15235 GCF_001575015.1_ASM157501v1 Salmonella enterica Proteobacteria; Gammaproteobacteria; Enterobacterales; Enterobacteriaceae; Salmonella MTGDNTLITSHGINRRDFMKLCAALAATMGLSSKAAAEMAESVSNPQRPPVIWIGAQECT WP_000145429.1 hydrogenase 2 small subunit [Salmonella enterica] Length: 372\nNumber of predicted TMHs: 1\nExp number of AAs in TMHs: 24.51309\nExp number, first 60 AAs: 0.2682\nTotal prob of N-in: 0.14402\noutside 1 331\nTMhelix 332 354\ninside 355 372

15236 GCF_001569625.1_ASM156962v1 Salmonella enterica Proteobacteria; Gammaproteobacteria; Enterobacterales; Enterobacteriaceae; Salmonella MTGDNTLITSHGINRRDFMKLCAALAATMGLSSKAAAEMAESVSNPQRPPVIWIGAQECT WP_000145429.1 hydrogenase 2 small subunit [Salmonella enterica] Length: 372\nNumber of predicted TMHs: 1\nExp number of AAs in TMHs: 24.51309\nExp number, first 60 AAs: 0.2682\nTotal prob of N-in: 0.14402\noutside 1 331\nTMhelix 332 354\ninside 355 372

15237 GCF_001648865.1_ASM164886v1 Salmonella enterica Proteobacteria; Gammaproteobacteria; Enterobacterales; Enterobacteriaceae; Salmonella MTGDNTLITSHGINRRDFMKLCAALAATMGLSSKAAAEMAESVSNPQRPPVIWIGAQECT WP_000145429.1 hydrogenase 2 small subunit [Salmonella enterica] Length: 372\nNumber of predicted TMHs: 1\nExp number of AAs in TMHs: 24.51309\nExp number, first 60 AAs: 0.2682\nTotal prob of N-in: 0.14402\noutside 1 331\nTMhelix 332 354\ninside 355 372

15238 GCF_001691905.1_ASM169190v1 Salmonella enterica Proteobacteria; Gammaproteobacteria; Enterobacterales; Enterobacteriaceae; Salmonella MTGDNTLITSHGINRRDFMKLCAALAATMGLSSKAAAEMAESVSNPQRPPVIWIGAQECT WP_000145429.1 hydrogenase 2 small subunit [Salmonella enterica] Length: 372\nNumber of predicted TMHs: 1\nExp number of AAs in TMHs: 24.51309\nExp number, first 60 AAs: 0.2682\nTotal prob of N-in: 0.14402\noutside 1 331\nTMhelix 332 354\ninside 355 372

15239 GCF_001831635.1_ASM183163v1 Salmonella enterica Proteobacteria; Gammaproteobacteria; Enterobacterales; Enterobacteriaceae; Salmonella MTGDNTLITSHGINRRDFMKLCAALAATMGLSSKAAAEMAESVSNPQRPPVIWIGAQECT WP_000145429.1 hydrogenase 2 small subunit [Salmonella enterica] Length: 372\nNumber of predicted TMHs: 1\nExp number of AAs in TMHs: 24.51309\nExp number, first 60 AAs: 0.2682\nTotal prob of N-in: 0.14402\noutside 1 331\nTMhelix 332 354\ninside 355 372

15240 GCF_001275195.1_ASM127519v1 Salmonella enterica Proteobacteria; Gammaproteobacteria; Enterobacterales; Enterobacteriaceae; Salmonella MTGDNTLITSHGINRRDFMKLCAALAATMGLSSKAAAEMAESVSNPQRPPVIWIGAQECT WP_000145429.1 hydrogenase 2 small subunit [Salmonella enterica] Length: 372\nNumber of predicted TMHs: 1\nExp number of AAs in TMHs: 24.51309\nExp number, first 60 AAs: 0.2682\nTotal prob of N-in: 0.14402\noutside 1 331\nTMhelix 332 354\ninside 355 372

15241 GCF_001275215.1_ASM127521v1 Salmonella enterica Proteobacteria; Gammaproteobacteria; Enterobacterales; Enterobacteriaceae; Salmonella MTGDNTLITSHGINRRDFMKLCAALAATMGLSSKAAAEMAESVSNPQRPPVIWIGAQECT WP_000145429.1 hydrogenase 2 small subunit [Salmonella enterica] Length: 372\nNumber of predicted TMHs: 1\nExp number of AAs in TMHs: 24.51309\nExp number, first 60 AAs: 0.2682\nTotal prob of N-in: 0.14402\noutside 1 331\nTMhelix 332 354\ninside 355 372

15242 GCF_001275195.1_ASM127519v1 Salmonella enterica Proteobacteria; Gammaproteobacteria; Enterobacterales; Enterobacteriaceae; Salmonella
MNNEETFYQAMRRKGVTRRSFLKFCSLAATSLGLGAGMTPKIAWALENKPRIPVVWIHGL WP_001058312.1
hydrogenase [Salmonella enterica] Length: 372\nNumber of predicted TMHs: 2\nExp number of AAs in TMHs: 36.51738\nExp number, first 60 AAs: 19.72849\nTotal prob of N-in: 0.96170\nPOSSIBLE N-term signal sequence\ninside 1 19\nTMhelix 20 39\noutside 40 325\nTMhelix 326 348\ninside 349 372

15243 GCF_001474965.1_Salmonella_enterica_CVM_43827_v1.0Salmonella enterica Proteobacteria; Gammaproteobacteria; Enterobacterales; Enterobacteriaceae; Salmonella
MTGDNTLITSHGINRRDFMKLCAALAATMGLSSKAAAEMAESVSNPQRPPVIWIGAQECT WP_000145429.1
hydrogenase 2 small subunit [Salmonella enterica] Length: 372\nNumber of predicted TMHs: 1\nExp number of AAs in TMHs: 24.51309\nExp number, first 60 AAs: 0.2682\nTotal prob of N-in: 0.14402\noutside 1 331\nTMhelix 332 354\ninside 355 372

15244 GCF_001474995.1_Salmonella_enterica_CVM_43830_v1.0Salmonella enterica Proteobacteria; Gammaproteobacteria; Enterobacterales; Enterobacteriaceae; Salmonella
MTGDNTLITSHGINRRDFMKLCAALAATMGLSSKAAAEMAESVSNPQRPPVIWIGAQECT WP_000145429.1
hydrogenase 2 small subunit [Salmonella enterica] Length: 372\nNumber of predicted TMHs: 1\nExp number of AAs in TMHs: 24.51309\nExp number, first 60 AAs: 0.2682\nTotal prob of N-in: 0.14402\noutside 1 331\nTMhelix 332 354\ninside 355 372

15245 GCF_001475005.1_Salmonella_enterica_CVM_43831_v1.0Salmonella enterica Proteobacteria; Gammaproteobacteria; Enterobacterales; Enterobacteriaceae; Salmonella
MTGDNTLITSHGINRRDFMKLCAALAATMGLSSKAAAEMAESVSNPQRPPVIWIGAQECT WP_000145429.1
hydrogenase 2 small subunit [Salmonella enterica] Length: 372\nNumber of predicted TMHs: 1\nExp number of AAs in TMHs: 24.51309\nExp number, first 60 AAs: 0.2682\nTotal prob of N-in: 0.14402\noutside 1 331\nTMhelix 332 354\ninside 355 372

15246 GCF_001474985.1_Salmonella_enterica_CVM_43829_v1.0Salmonella enterica Proteobacteria; Gammaproteobacteria; Enterobacterales; Enterobacteriaceae; Salmonella
MTGDNTLITSHGINRRDFMKLCAALAATMGLSSKAAAEMAESVSNPQRPPVIWIGAQECT WP_000145429.1
hydrogenase 2 small subunit [Salmonella enterica] Length: 372\nNumber of predicted TMHs: 1\nExp number of AAs in TMHs: 24.51309\nExp number, first 60 AAs: 0.2682\nTotal prob of N-in: 0.14402\noutside 1 331\nTMhelix 332 354\ninside 355 372

15247 GCF_001475065.1_Salmonella_enterica_CVM_43838_v1.0Salmonella enterica Proteobacteria; Gammaproteobacteria; Enterobacterales; Enterobacteriaceae; Salmonella
MTGDNTLITSHGINRRDFMKLCAALAATMGLSSKAAAEMAESVSNPQRPPVIWIGAQECT WP_000145429.1
hydrogenase 2 small subunit [Salmonella enterica] Length: 372\nNumber of predicted TMHs: 1\nExp number of AAs in TMHs: 24.51309\nExp number, first 60 AAs: 0.2682\nTotal prob of N-in: 0.14402\noutside 1 331\nTMhelix 332 354\ninside 355 372

15248 GCF_001475295.1_Salmonella_enterica_CVM_43800_v1.0Salmonella enterica Proteobacteria; Gammaproteobacteria; Enterobacterales; Enterobacteriaceae; Salmonella
MTGDNTLITSHGINRRDFMKLCAALAATMGLSSKAAAEMAESVSNPQRPPVIWIGAQECT WP_000145429.1
hydrogenase 2 small subunit [Salmonella enterica] Length: 372\nNumber of predicted TMHs: 1\nExp number of AAs in TMHs: 24.51309\nExp number, first 60 AAs: 0.2682\nTotal prob of N-in: 0.14402\noutside 1 331\nTMhelix 332 354\ninside 355 372

15249 GCF_001475245.1_Salmonella_enterica_CVM_43802_v1.0Salmonella enterica Proteobacteria; Gammaproteobacteria; Enterobacterales; Enterobacteriaceae; Salmonella
MTGDNTLITSHGINRRDFMKLCAALAATMGLSSKAAAEMAESVSNPQRPPVIWIGAQECT WP_000145429.1
hydrogenase 2 small subunit [Salmonella enterica] Length: 372\nNumber of predicted TMHs: 1\nExp number of AAs in TMHs: 24.51309\nExp number, first 60 AAs: 0.2682\nTotal prob of N-in: 0.14402\noutside 1 331\nTMhelix 332 354\ninside 355 372

15250 GCF_001475225.1_Salmonella_enterica_CVM_43820_v1.0Salmonella enterica Proteobacteria; Gammaproteobacteria; Enterobacterales; Enterobacteriaceae; Salmonella
MNNEETFYQAMRRKGVTRRSFLKFCSLAATSLGLGAGMTPKIAWALENKPRIPVVWIHGL WP_001058312.1
hydrogenase [Salmonella enterica] Length: 372\nNumber of predicted TMHs: 2\nExp number of AAs in

TMHs: 36.51738\nExp number, first 60 AAs: 19.72849\nTotal prob of N-in: 0.96170\nPOSSIBLE N-term signal sequence\ninside 1 19\nTMhelix 20 39\noutside 40 325\nTMhelix 326 348\ninside 349 372

15251 GCF_001475295.1_Salmonella_enterica_CVM_43800_v1.0Salmonella enterica Proteobacteria; Gammaproteobacteria; Enterobacterales; Enterobacteriaceae; Salmonella
MNNEETFYQAMRRKGVTRRSFLKFCSLAATSLGLGAGMTPKIAWALENKPRIPVVWIHGL WP_001058312.1
hydrogenase [Salmonella enterica] Length: 372\nNumber of predicted TMHs: 2\nExp number of AAs in TMHs: 36.51738\nExp number, first 60 AAs: 19.72849\nTotal prob of N-in: 0.96170\nPOSSIBLE N-term signal sequence\ninside 1 19\nTMhelix 20 39\noutside 40 325\nTMhelix 326 348\ninside 349 372

15252 GCF_001475245.1_Salmonella_enterica_CVM_43802_v1.0Salmonella enterica Proteobacteria; Gammaproteobacteria; Enterobacterales; Enterobacteriaceae; Salmonella
MNNEETFYQAMRRKGVTRRSFLKFCSLAATSLGLGAGMTPKIAWALENKPRIPVVWIHGL WP_001058312.1
hydrogenase [Salmonella enterica] Length: 372\nNumber of predicted TMHs: 2\nExp number of AAs in TMHs: 36.51738\nExp number, first 60 AAs: 19.72849\nTotal prob of N-in: 0.96170\nPOSSIBLE N-term signal sequence\ninside 1 19\nTMhelix 20 39\noutside 40 325\nTMhelix 326 348\ninside 349 372

15253 GCF_001475085.1_Salmonella_enterica_CVM_43840_v1.0Salmonella enterica Proteobacteria; Gammaproteobacteria; Enterobacterales; Enterobacteriaceae; Salmonella
MNNEETLYQAMRRKGVTRRSFLKFCSLAATSLGLGAGMTPKIAWALENKPRIPVVWIHGL WP_001752445.1
hydrogenase-1 small chain [Salmonella enterica] Length: 372\nNumber of predicted TMHs: 2\nExp number of AAs in TMHs: 36.5123\nExp number, first 60 AAs: 19.72341\nTotal prob of N-in: 0.96144\nPOSSIBLE N-term signal sequence\ninside 1 19\nTMhelix 20 39\noutside 40 325\nTMhelix 326 348\ninside 349 372

15254 GCF_001475375.1_Salmonella_enterica_CVM_43842_v1.0Salmonella enterica Proteobacteria; Gammaproteobacteria; Enterobacterales; Enterobacteriaceae; Salmonella
MTGDNTLITSHGINRRDFMKLCAALAATMGLSSKAAAEMAESVSNPQRPPVIWIGAQECT WP_000145431.1
hydrogenase 2 small subunit [Salmonella enterica] Length: 372\nNumber of predicted TMHs: 1\nExp number of AAs in TMHs: 24.51505\nExp number, first 60 AAs: 0.2682\nTotal prob of N-in: 0.14406\noutside 1 331\nTMhelix 332 354\ninside 355 372

15255 GCF_001480915.1_Salmonella_enterica_CVM_N40366-SQ_v1.0 Salmonella enterica Proteobacteria; Gammaproteobacteria; Enterobacterales; Enterobacteriaceae; Salmonella
MTGDNTLITSHGINRRDFMKLCAALAATMGLSSKAAAEMAESVSNPQRPPVIWIGAQECT WP_000145429.1
hydrogenase 2 small subunit [Salmonella enterica] Length: 372\nNumber of predicted TMHs: 1\nExp number of AAs in TMHs: 24.51309\nExp number, first 60 AAs: 0.2682\nTotal prob of N-in: 0.14402\noutside 1 331\nTMhelix 332 354\ninside 355 372

15256 GCF_001558355.1_ASM155835v1 Salmonella enterica Proteobacteria; Gammaproteobacteria; Enterobacterales; Enterobacteriaceae; Salmonella
MTGDNTLITSHGINRRDFMKLCAALAATMGLSSKAAAEMAESVSNPQRPPVIWIGAQECT WP_000145429.1
hydrogenase 2 small subunit [Salmonella enterica] Length: 372\nNumber of predicted TMHs: 1\nExp number of AAs in TMHs: 24.51309\nExp number, first 60 AAs: 0.2682\nTotal prob of N-in: 0.14402\noutside 1 331\nTMhelix 332 354\ninside 355 372

15257 GCF_001569545.1_ASM156954v1 Salmonella enterica Proteobacteria; Gammaproteobacteria; Enterobacterales; Enterobacteriaceae; Salmonella
MTGDNTLITSHGINRRDFMKLCAALAATMGLSSKAAAEMAESVSNPQRPPVIWIGAQECT WP_000145429.1
hydrogenase 2 small subunit [Salmonella enterica] Length: 372\nNumber of predicted TMHs: 1\nExp number of AAs in TMHs: 24.51309\nExp number, first 60 AAs: 0.2682\nTotal prob of N-in: 0.14402\noutside 1 331\nTMhelix 332 354\ninside 355 372

15258 GCF_001569845.1_ASM156984v1 Salmonella enterica Proteobacteria; Gammaproteobacteria; Enterobacterales; Enterobacteriaceae; Salmonella
MNNEETFYQAMRRKGVTRRSFLKFCSLAATSLGLGAGMTPKIAWALENKPRIPVVWIHGL WP_001058307.1
hydrogenase [Salmonella enterica] Length: 372\nNumber of predicted TMHs: 2\nExp number of AAs in TMHs: 36.51738\nExp number, first 60 AAs: 19.72849\nTotal prob of N-in: 0.96170\nPOSSIBLE N-term signal sequence\ninside 1 19\nTMhelix 20 39\noutside 40 325\nTMhelix 326 348\ninside 349 372

15259 GCF_001569925.1_ASM156992v1 Salmonella enterica Proteobacteria; Gammaproteobacteria; Enterobacterales; Enterobacteriaceae; Salmonella

MTGDNTLITSHGINRRDFMKLCAALAATMGLSSKAAAEMAESVSNPQRPPVIWIGAQECT WP_000145429.1 hydrogenase 2 small subunit [Salmonella enterica] Length: 372\nNumber of predicted TMHs: 1\nExp number of AAs in TMHs: 24.51309\nExp number, first 60 AAs: 0.2682\nTotal prob of N-in: 0.14402\noutside 1 331\nTMhelix 332 354\ninside 355 372

15260 GCF_001569925.1_ASM156992v1 Salmonella enterica Proteobacteria; Gammaproteobacteria; Enterobacterales; Enterobacteriaceae; Salmonella MNNEETFYQAMRRKGVTRRSFLKFCSLAATSLGLGAGMTPKIAWALENKPRIPVVWIHGL WP_001058307.1 hydrogenase [Salmonella enterica] Length: 372\nNumber of predicted TMHs: 2\nExp number of AAs in TMHs: 36.51738\nExp number, first 60 AAs: 19.72849\nTotal prob of N-in: 0.96170\nPOSSIBLE N-term signal sequence\ninside 1 19\nTMhelix 20 39\noutside 40 325\nTMhelix 326 348\ninside 349 372

15261 GCF_001569885.1_ASM156988v1 Salmonella enterica Proteobacteria; Gammaproteobacteria; Enterobacterales; Enterobacteriaceae; Salmonella MNNEETFYQAMRRKGVTRRSFLKFCSLAATSLGLGAGMTPKIAWALENKPRIPVVWIHGL WP_001058307.1 hydrogenase [Salmonella enterica] Length: 372\nNumber of predicted TMHs: 2\nExp number of AAs in TMHs: 36.51738\nExp number, first 60 AAs: 19.72849\nTotal prob of N-in: 0.96170\nPOSSIBLE N-term signal sequence\ninside 1 19\nTMhelix 20 39\noutside 40 325\nTMhelix 326 348\ninside 349 372

15262 GCF_001569945.1_ASM156994v1 Salmonella enterica Proteobacteria; Gammaproteobacteria; Enterobacterales; Enterobacteriaceae; Salmonella MTGDNTLITSHGINRRDFMKLCAALAATMGLSSKAAAEMAESVSNPQRPPVIWIGAQECT WP_000145429.1 hydrogenase 2 small subunit [Salmonella enterica] Length: 372\nNumber of predicted TMHs: 1\nExp number of AAs in TMHs: 24.51309\nExp number, first 60 AAs: 0.2682\nTotal prob of N-in: 0.14402\noutside 1 331\nTMhelix 332 354\ninside 355 372

15263 GCF_001569985.1_ASM156998v1 Salmonella enterica Proteobacteria; Gammaproteobacteria; Enterobacterales; Enterobacteriaceae; Salmonella MTGDNTLITSHGINRRDFMKLCAALAATMGLSSKAAAEMAESVSNPQRPPVIWIGAQECT WP_000145429.1 hydrogenase 2 small subunit [Salmonella enterica] Length: 372\nNumber of predicted TMHs: 1\nExp number of AAs in TMHs: 24.51309\nExp number, first 60 AAs: 0.2682\nTotal prob of N-in: 0.14402\noutside 1 331\nTMhelix 332 354\ninside 355 372

15264 GCF_001570025.1_ASM157002v1 Salmonella enterica Proteobacteria; Gammaproteobacteria; Enterobacterales; Enterobacteriaceae; Salmonella MTGDNTLITSHGINRRDFMKLCAALAATMGLSSKAAAEMAESVSNPQRPPVIWIGAQECT WP_000145429.1 hydrogenase 2 small subunit [Salmonella enterica] Length: 372\nNumber of predicted TMHs: 1\nExp number of AAs in TMHs: 24.51309\nExp number, first 60 AAs: 0.2682\nTotal prob of N-in: 0.14402\noutside 1 331\nTMhelix 332 354\ninside 355 372

15265 GCF_001575015.1_ASM157501v1 Salmonella enterica Proteobacteria; Gammaproteobacteria; Enterobacterales; Enterobacteriaceae; Salmonella MNNEETFYQAMRRKGVTRRSFLKFCSLAATSLGLGAGMTPKIAWALENKPRIPVVWIHGL WP_001058311.1 hydrogenase [Salmonella enterica] Length: 372\nNumber of predicted TMHs: 2\nExp number of AAs in TMHs: 37.4699899999999\nExp number, first 60 AAs: 19.72699\nTotal prob of N-in: 0.96171\nPOSSIBLE N-term signal sequence\ninside 1 19\nTMhelix 20 39\noutside 40 325\nTMhelix 326 348\ninside 349 372

15266 GCF_001643875.1_CRJJGF_00165_v1.0Salmonella enterica Proteobacteria; Gammaproteobacteria; Enterobacterales; Enterobacteriaceae; Salmonella MTGDNTLITSHGINRRDFMKLCAALAATMGLSSKAAAEMAESVSNPQRPPVIWIGAQECT WP_023248548.1 hydrogenase 2 small subunit [Salmonella enterica] Length: 372\nNumber of predicted TMHs: 1\nExp number of AAs in TMHs: 24.51281\nExp number, first 60 AAs: 0.26822\nTotal prob of N-in: 0.14402\noutside 1 331\nTMhelix 332 354\ninside 355 372

15267 GCF_001641105.1_ASM164110v1 Salmonella enterica Proteobacteria; Gammaproteobacteria; Enterobacterales; Enterobacteriaceae; Salmonella MTGDNTLITSHGINRRDFMKLCAALAATMGLSSKAAAEMAESVSNPQRPPVIWIGAQECT WP_000145428.1 hydrogenase 2 small subunit [Salmonella enterica] Length: 372\nNumber of predicted TMHs: 1\nExp number of AAs in TMHs: 24.50677\nExp number, first 60 AAs: 0.2682\nTotal prob of N-in: 0.14390\noutside 1 331\nTMhelix 332 354\ninside 355 372

15268 GCF_001642525.1_ASM164252v1 Salmonella enterica Proteobacteria; Gammaproteobacteria; Enterobacterales; Enterobacteriaceae; Salmonella
MTGDNTLITSHGINRRDFMKLCAALAATMGLSSKAAAEMAESVSNPQRPPVIWIGAQECT WP_000145429.1
hydrogenase 2 small subunit [Salmonella enterica] Length: 372\nNumber of predicted TMHs: 1\nExp number of AAs in TMHs: 24.51309\nExp number, first 60 AAs: 0.2682\nTotal prob of N-in: 0.14402\noutside 1 331\nTMhelix 332 354\ninside 355 372

15269 GCF_001642445.1_ASM164244v1 Salmonella enterica Proteobacteria; Gammaproteobacteria; Enterobacterales; Enterobacteriaceae; Salmonella
MTGDNTLITSHGINRRDFMKLCAALAATMGLSSKAAAEMAESVSNPQRPPVIWIGAQECT WP_000145429.1
hydrogenase 2 small subunit [Salmonella enterica] Length: 372\nNumber of predicted TMHs: 1\nExp number of AAs in TMHs: 24.51309\nExp number, first 60 AAs: 0.2682\nTotal prob of N-in: 0.14402\noutside 1 331\nTMhelix 332 354\ninside 355 372

15270 GCF_001647755.1_ASM164775v1 Salmonella enterica Proteobacteria; Gammaproteobacteria; Enterobacterales; Enterobacteriaceae; Salmonella
MLKFRTIFLKKIFTLKNSILARRGAVIVIVSAVFTSIMFFAHSWASDKEVAMTLSVLNSN WP_023226644.1 hypothetical protein [Salmonella enterica] Length: 204\nNumber of predicted TMHs: 2\nExp number of AAs in TMHs: 44.99526\nExp number, first 60 AAs: 22.33786\nTotal prob of N-in: 0.87083\nPOSSIBLE N-term signal sequence\ninside 1 19\nTMhelix 20 42\noutside 43 171\nTMhelix 172 194\ninside 195 204

15271 GCF_001652385.1_CFSAN033950_02.0Salmonella enterica Proteobacteria; Gammaproteobacteria; Enterobacterales; Enterobacteriaceae; Salmonella
MTGDNTLITSHGINRRDFMKLCAALAATMGLSSKAAAEMAESVSNPQRPPVIWIGAQECT WP_000145429.1
hydrogenase 2 small subunit [Salmonella enterica] Length: 372\nNumber of predicted TMHs: 1\nExp number of AAs in TMHs: 24.51309\nExp number, first 60 AAs: 0.2682\nTotal prob of N-in: 0.14402\noutside 1 331\nTMhelix 332 354\ninside 355 372

15272 GCF_001648865.1_ASM164886v1 Salmonella enterica Proteobacteria; Gammaproteobacteria; Enterobacterales; Enterobacteriaceae; Salmonella
MNNEETFYQAMRRKGVTRRSFLKFCSLAATSLGLGAGMTPKIAWALENKPRIPVVWIHGL WP_001058311.1
hydrogenase [Salmonella enterica] Length: 372\nNumber of predicted TMHs: 2\nExp number of AAs in TMHs: 37.4699899999999\nExp number, first 60 AAs: 19.72699\nTotal prob of N-in: 0.96171\nPOSSIBLE N-term signal sequence\ninside 1 19\nTMhelix 20 39\noutside 40 325\nTMhelix 326 348\ninside 349 372

15273 GCF_001652385.1_CFSAN033950_02.0Salmonella enterica Proteobacteria; Gammaproteobacteria; Enterobacterales; Enterobacteriaceae; Salmonella
MNNEETFYQAMRRKGVTRRSFLKFCSLAATSLGLGAGMTPKIAWALENKPRIPVVWIHGL WP_001058312.1
hydrogenase [Salmonella enterica] Length: 372\nNumber of predicted TMHs: 2\nExp number of AAs in TMHs: 36.51738\nExp number, first 60 AAs: 19.72849\nTotal prob of N-in: 0.96170\nPOSSIBLE N-term signal sequence\ninside 1 19\nTMhelix 20 39\noutside 40 325\nTMhelix 326 348\ninside 349 372

15274 GCF_001648855.1_ASM164885v1 Salmonella enterica Proteobacteria; Gammaproteobacteria; Enterobacterales; Enterobacteriaceae; Salmonella
MNNEETFYQAMRRKGVTRRSFLKFCSLAATSLGLGAGMTPQIAWALENKPRIPVVWIHGL WP_023227615.1
hydrogenase 1, small subunit [Salmonella enterica] Length: 372\nNumber of predicted TMHs: 2\nExp number of AAs in TMHs: 39.8591699999999\nExp number, first 60 AAs: 22.13093\nTotal prob of N-in: 0.98639\nPOSSIBLE N-term signal sequence\ninside 1 20\nTMhelix 21 43\noutside 44 325\nTMhelix 326 348\ninside 349 372

15275 GCF_001691895.1_ASM169189v1 Salmonella enterica Proteobacteria; Gammaproteobacteria; Enterobacterales; Enterobacteriaceae; Salmonella
MTGDNTLITSHGINRRDFMKLCAALAATMGLSSKAAAEMAESVSNPQRPPVIWIGAQECT WP_000145429.1
hydrogenase 2 small subunit [Salmonella enterica] Length: 372\nNumber of predicted TMHs: 1\nExp number of AAs in TMHs: 24.51309\nExp number, first 60 AAs: 0.2682\nTotal prob of N-in: 0.14402\noutside 1 331\nTMhelix 332 354\ninside 355 372

15276 GCF_001713515.1_ASM171351v1 Salmonella enterica Proteobacteria; Gammaproteobacteria; Enterobacterales; Enterobacteriaceae; Salmonella
MTGDNTLITSHGINRRDFMKLCAALAATMGLSSKAAAEMAESVSNPQRPPVIWIGAQECT WP_000145429.1
hydrogenase 2 small subunit [Salmonella enterica] Length: 372\nNumber of predicted TMHs: 1\nExp

number of AAs in TMHs: 24.51309\nExp number, first 60 AAs: 0.2682\nTotal prob of N-in: 0.14402\noutside 1 331\nTMhelix 332 354\ninside 355 372

15277 GCF_001713515.1_ASM171351v1 Salmonella enterica Proteobacteria; Gammaproteobacteria; Enterobacterales; Enterobacteriaceae; Salmonella
MNNEETFYQAMRRKGVTRRSFLKFCSLAATSLGLGAGMTPKIAWALENKPRIPVVWIHGL WP_001058307.1
hydrogenase [Salmonella enterica] Length: 372\nNumber of predicted TMHs: 2\nExp number of AAs in TMHs: 36.51738\nExp number, first 60 AAs: 19.72849\nTotal prob of N-in: 0.96170\nPOSSIBLE N-term signal sequence\ninside 1 19\nTMhelix 20 39\noutside 40 325\nTMhelix 326 348\ninside 349 372

15278 GCF_001716375.1_ASM171637v1 Salmonella enterica Proteobacteria; Gammaproteobacteria; Enterobacterales; Enterobacteriaceae; Salmonella
MNNEETFYQAMRRKGVTRRSFLKFCSLAATSLGLGAGMTPKIAWALENKPRIPVVWIHGL WP_001058307.1
hydrogenase [Salmonella enterica] Length: 372\nNumber of predicted TMHs: 2\nExp number of AAs in TMHs: 36.51738\nExp number, first 60 AAs: 19.72849\nTotal prob of N-in: 0.96170\nPOSSIBLE N-term signal sequence\ninside 1 19\nTMhelix 20 39\noutside 40 325\nTMhelix 326 348\ninside 349 372

15279 GCF_001716745.1_ASM171674v1 Salmonella enterica Proteobacteria; Gammaproteobacteria; Enterobacterales; Enterobacteriaceae; Salmonella
MNNEETFYQAMRRKGVTRRSFLKFCSLAATSLGLGAGMTPKIAWALENKPRIPVVWIHGL WP_001058307.1
hydrogenase [Salmonella enterica] Length: 372\nNumber of predicted TMHs: 2\nExp number of AAs in TMHs: 36.51738\nExp number, first 60 AAs: 19.72849\nTotal prob of N-in: 0.96170\nPOSSIBLE N-term signal sequence\ninside 1 19\nTMhelix 20 39\noutside 40 325\nTMhelix 326 348\ninside 349 372

15280 GCF_001716725.1_ASM171672v1 Salmonella enterica Proteobacteria; Gammaproteobacteria; Enterobacterales; Enterobacteriaceae; Salmonella
MNNEETFYQAMRRKGVTRRSFLKFCSLAATSLGLGAGMTPKIAWALENKPRIPVVWIHGL WP_001058307.1
hydrogenase [Salmonella enterica] Length: 372\nNumber of predicted TMHs: 2\nExp number of AAs in TMHs: 36.51738\nExp number, first 60 AAs: 19.72849\nTotal prob of N-in: 0.96170\nPOSSIBLE N-term signal sequence\ninside 1 19\nTMhelix 20 39\noutside 40 325\nTMhelix 326 348\ninside 349 372

15281 GCF_001831635.1_ASM183163v1 Salmonella enterica Proteobacteria; Gammaproteobacteria; Enterobacterales; Enterobacteriaceae; Salmonella
MNNEETFYQAMRRKGVTRRSFLKFCSLAATSLGLGAGMTPKIAWALENKPRIPVVWIHGL WP_001058312.1
hydrogenase [Salmonella enterica] Length: 372\nNumber of predicted TMHs: 2\nExp number of AAs in TMHs: 36.51738\nExp number, first 60 AAs: 19.72849\nTotal prob of N-in: 0.96170\nPOSSIBLE N-term signal sequence\ninside 1 19\nTMhelix 20 39\noutside 40 325\nTMhelix 326 348\ninside 349 372

15282 GCF_001831725.1_ASM183172v1 Salmonella enterica Proteobacteria; Gammaproteobacteria; Enterobacterales; Enterobacteriaceae; Salmonella
MTGDNTLITSHGINRRDFMKLCAALAATMGLSSKAAAEMAESVSNPQRPPVIWIGAQECT WP_000145429.1
hydrogenase 2 small subunit [Salmonella enterica] Length: 372\nNumber of predicted TMHs: 1\nExp number of AAs in TMHs: 24.51309\nExp number, first 60 AAs: 0.2682\nTotal prob of N-in: 0.14402\noutside 1 331\nTMhelix 332 354\ninside 355 372

15283 GCF_001831725.1_ASM183172v1 Salmonella enterica Proteobacteria; Gammaproteobacteria; Enterobacterales; Enterobacteriaceae; Salmonella
MNNEETFYQAMRRKGVTRRSFLKFCSLAATSLGLGAGMTPKIAWALENKPRIPVVWIHGL WP_001058312.1
hydrogenase [Salmonella enterica] Length: 372\nNumber of predicted TMHs: 2\nExp number of AAs in TMHs: 36.51738\nExp number, first 60 AAs: 19.72849\nTotal prob of N-in: 0.96170\nPOSSIBLE N-term signal sequence\ninside 1 19\nTMhelix 20 39\noutside 40 325\nTMhelix 326 348\ninside 349 372

15284 GCF_001831555.1_ASM183155v1 Salmonella enterica Proteobacteria; Gammaproteobacteria; Enterobacterales; Enterobacteriaceae; Salmonella
MTGDNTLITSHGINRRDFMKLCAALAATMGLSSKAAAEMAESVSNPQRPPVIWIGAQECT WP_023248548.1
hydrogenase 2 small subunit [Salmonella enterica] Length: 372\nNumber of predicted TMHs: 1\nExp number of AAs in TMHs: 24.51281\nExp number, first 60 AAs: 0.26822\nTotal prob of N-in: 0.14402\noutside 1 331\nTMhelix 332 354\ninside 355 372

15285 GCF_001831785.1_ASM183178v1 Salmonella enterica Proteobacteria; Gammaproteobacteria; Enterobacterales; Enterobacteriaceae; Salmonella

MTGDNTLITSHGINRRDFMKLCAALAATMGLSSKAAAEMAESVSNPQRPPVIWIGAQECT WP_000145429.1 hydrogenase 2 small subunit [Salmonella enterica] Length: 372\nNumber of predicted TMHs: 1\nExp number of AAs in TMHs: 24.51309\nExp number, first 60 AAs: 0.2682\nTotal prob of N-in: 0.14402\noutside 1 331\nTMhelix 332 354\ninside 355 372

15286 GCF_001831785.1_ASM183178v1 Salmonella enterica Proteobacteria; Gammaproteobacteria; Enterobacterales; Enterobacteriaceae; Salmonella MNNEETFYQAMRRKGVTRRSFLKFCSLAATSLGLGAGMTPKIAWALENKPRIPVVWIHGL WP_001058312.1 hydrogenase [Salmonella enterica] Length: 372\nNumber of predicted TMHs: 2\nExp number of AAs in TMHs: 36.51738\nExp number, first 60 AAs: 19.72849\nTotal prob of N-in: 0.96170\nPOSSIBLE N-term signal sequence\ninside 1 19\nTMhelix 20 39\noutside 40 325\nTMhelix 326 348\ninside 349 372

15287 GCF_001831625.1_ASM183162v1 Salmonella enterica Proteobacteria; Gammaproteobacteria; Enterobacterales; Enterobacteriaceae; Salmonella MTGDNTLITSHGINRRDFMKLCAALAATMGLSSKAAAEMAESVSNPQRPPVIWIGAQECT WP_023248548.1 hydrogenase 2 small subunit [Salmonella enterica] Length: 372\nNumber of predicted TMHs: 1\nExp number of AAs in TMHs: 24.51281\nExp number, first 60 AAs: 0.26822\nTotal prob of N-in: 0.14402\noutside 1 331\nTMhelix 332 354\ninside 355 372

15288 GCF_001832075.1_ASM183207v1 Salmonella enterica Proteobacteria; Gammaproteobacteria; Enterobacterales; Enterobacteriaceae; Salmonella MTGDNTLITSHGINRRDFMKLCAALAATMGLSSKAAAEMAESVSNPQRPPVIWIGAQECT WP_000145429.1 hydrogenase 2 small subunit [Salmonella enterica] Length: 372\nNumber of predicted TMHs: 1\nExp number of AAs in TMHs: 24.51309\nExp number, first 60 AAs: 0.2682\nTotal prob of N-in: 0.14402\noutside 1 331\nTMhelix 332 354\ninside 355 372

15289 GCF_001832145.1_ASM183214v1 Salmonella enterica Proteobacteria; Gammaproteobacteria; Enterobacterales; Enterobacteriaceae; Salmonella MTGDNTLITSHGINRRDFMKLCAALAATMGLSSKAAAEMAESVSNPQRPPVIWIGAQECT WP_000145429.1 hydrogenase 2 small subunit [Salmonella enterica] Length: 372\nNumber of predicted TMHs: 1\nExp number of AAs in TMHs: 24.51309\nExp number, first 60 AAs: 0.2682\nTotal prob of N-in: 0.14402\noutside 1 331\nTMhelix 332 354\ninside 355 372

15290 GCF_001832075.1_ASM183207v1 Salmonella enterica Proteobacteria; Gammaproteobacteria; Enterobacterales; Enterobacteriaceae; Salmonella MNNEETFYQAMRRKGVTRRSFLKFCSLAATSLGLGAGMTPKIAWALENKPRIPVVWIHGL WP_001058312.1 hydrogenase [Salmonella enterica] Length: 372\nNumber of predicted TMHs: 2\nExp number of AAs in TMHs: 36.51738\nExp number, first 60 AAs: 19.72849\nTotal prob of N-in: 0.96170\nPOSSIBLE N-term signal sequence\ninside 1 19\nTMhelix 20 39\noutside 40 325\nTMhelix 326 348\ninside 349 372

15291 GCF_001832095.1_ASM183209v1 Salmonella enterica Proteobacteria; Gammaproteobacteria; Enterobacterales; Enterobacteriaceae; Salmonella MNNEETFYQAMRRKGVTRRSFLKFCSLAATSLGLGAGMTPKIAWALENKPRIPVVWIHGL WP_001058312.1 hydrogenase [Salmonella enterica] Length: 372\nNumber of predicted TMHs: 2\nExp number of AAs in TMHs: 36.51738\nExp number, first 60 AAs: 19.72849\nTotal prob of N-in: 0.96170\nPOSSIBLE N-term signal sequence\ninside 1 19\nTMhelix 20 39\noutside 40 325\nTMhelix 326 348\ninside 349 372

15292 GCF_001832175.1_ASM183217v1 Salmonella enterica Proteobacteria; Gammaproteobacteria; Enterobacterales; Enterobacteriaceae; Salmonella MTGDNTLITSHGINRRDFMKLCAALAATMGLSSKAAAEMAESVSNPQRPPVIWIGAQECT WP_000145429.1 hydrogenase 2 small subunit [Salmonella enterica] Length: 372\nNumber of predicted TMHs: 1\nExp number of AAs in TMHs: 24.51309\nExp number, first 60 AAs: 0.2682\nTotal prob of N-in: 0.14402\noutside 1 331\nTMhelix 332 354\ninside 355 372

15293 GCF_001832175.1_ASM183217v1 Salmonella enterica Proteobacteria; Gammaproteobacteria; Enterobacterales; Enterobacteriaceae; Salmonella MNNEETFYQAMRRKGVTRRSFLKFCSLAATSLGLGAGMTPKIAWALENKPRIPVVWIHGL WP_001058312.1 hydrogenase [Salmonella enterica] Length: 372\nNumber of predicted TMHs: 2\nExp number of AAs in TMHs: 36.51738\nExp number, first 60 AAs: 19.72849\nTotal prob of N-in: 0.96170\nPOSSIBLE N-term signal sequence\ninside 1 19\nTMhelix 20 39\noutside 40 325\nTMhelix 326 348\ninside 349 372

15294 GCF_001832325.1_ASM183232v1 Salmonella enterica Proteobacteria; Gammaproteobacteria; Enterobacterales; Enterobacteriaceae; Salmonella
MTGDNTLITSHGINRRDFMKLCAALAATMGLSSKAAAEMAESVSNPQRPPVIWIGAQECT WP_000145429.1
hydrogenase 2 small subunit [Salmonella enterica] Length: 372\nNumber of predicted TMHs: 1\nExp number of AAs in TMHs: 24.51309\nExp number, first 60 AAs: 0.2682\nTotal prob of N-in: 0.14402\noutside 1 331\nTMhelix 332 354\ninside 355 372

15295 GCF_001833145.1_ASM183314v1 Salmonella enterica Proteobacteria; Gammaproteobacteria; Enterobacterales; Enterobacteriaceae; Salmonella
MTGDNTLITSHGINRRDFMKLCAALAATMGLSSKAAAEMAESVSNPQRPPVIWIGAQECT WP_000145429.1
hydrogenase 2 small subunit [Salmonella enterica] Length: 372\nNumber of predicted TMHs: 1\nExp number of AAs in TMHs: 24.51309\nExp number, first 60 AAs: 0.2682\nTotal prob of N-in: 0.14402\noutside 1 331\nTMhelix 332 354\ninside 355 372

15296 GCF_001833065.1_ASM183306v1 Salmonella enterica Proteobacteria; Gammaproteobacteria; Enterobacterales; Enterobacteriaceae; Salmonella
MNNEETFYQAMRRKGVTRRSFLKFCSLAATSLGLGAGMTPKIAWALENKPRIPVVWIHGL WP_001058311.1
hydrogenase [Salmonella enterica] Length: 372\nNumber of predicted TMHs: 2\nExp number of AAs in TMHs: 37.4699899999999\nExp number, first 60 AAs: 19.72699\nTotal prob of N-in: 0.96171\nPOSSIBLE N-term signal sequence\ninside 1 19\nTMhelix 20 39\noutside 40 325\nTMhelix 326 348\ninside 349 372

15297 GCF_001833145.1_ASM183314v1 Salmonella enterica Proteobacteria; Gammaproteobacteria; Enterobacterales; Enterobacteriaceae; Salmonella
MNNEETFYQAMRRKGVTRRSFLKFCSLAATSLGLGAGMTPKIAWALENKPRIPVVWIHGL WP_001058312.1
hydrogenase [Salmonella enterica] Length: 372\nNumber of predicted TMHs: 2\nExp number of AAs in TMHs: 36.51738\nExp number, first 60 AAs: 19.72849\nTotal prob of N-in: 0.96170\nPOSSIBLE N-term signal sequence\ninside 1 19\nTMhelix 20 39\noutside 40 325\nTMhelix 326 348\ninside 349 372

15298 GCF_001833065.1_ASM183306v1 Salmonella enterica Proteobacteria; Gammaproteobacteria; Enterobacterales; Enterobacteriaceae; Salmonella
MTGDNTLITSHGINRRDFMKLCAALAATMGLSSKAAAEMAESVSNPQRPPVIWIGAQECT WP_070810661.1
hydrogenase 2 small subunit [Salmonella enterica] Length: 372\nNumber of predicted TMHs: 1\nExp number of AAs in TMHs: 22.61963\nExp number, first 60 AAs: 0.27879\nTotal prob of N-in: 0.06746\noutside 1 331\nTMhelix 332 354\ninside 355 372

15299 GCF_001833165.1_ASM183316v1 Salmonella enterica Proteobacteria; Gammaproteobacteria; Enterobacterales; Enterobacteriaceae; Salmonella
MTGDNTLITSHGINRRDFMKLCAALAATMGLSSKAAAEMAESVSNPQRPPVIWIGAQECT WP_000145429.1
hydrogenase 2 small subunit [Salmonella enterica] Length: 372\nNumber of predicted TMHs: 1\nExp number of AAs in TMHs: 24.51309\nExp number, first 60 AAs: 0.2682\nTotal prob of N-in: 0.14402\noutside 1 331\nTMhelix 332 354\ninside 355 372

15300 GCF_001833505.1_ASM183350v1 Salmonella enterica Proteobacteria; Gammaproteobacteria; Enterobacterales; Enterobacteriaceae; Salmonella
MTGDNTLITSHGINRRDFMKLCAALAATMGLSSKAAAEMAESVSNPQRPPVIWIGAQECT WP_000145429.1
hydrogenase 2 small subunit [Salmonella enterica] Length: 372\nNumber of predicted TMHs: 1\nExp number of AAs in TMHs: 24.51309\nExp number, first 60 AAs: 0.2682\nTotal prob of N-in: 0.14402\noutside 1 331\nTMhelix 332 354\ninside 355 372

15301 GCF_001833505.1_ASM183350v1 Salmonella enterica Proteobacteria; Gammaproteobacteria; Enterobacterales; Enterobacteriaceae; Salmonella
MNNEETFYQAMRRKGVTRRSFLKFCSLAATSLGLGAGMTPKIAWALENKPRIPVVWIHGL WP_001058311.1
hydrogenase [Salmonella enterica] Length: 372\nNumber of predicted TMHs: 2\nExp number of AAs in TMHs: 37.4699899999999\nExp number, first 60 AAs: 19.72699\nTotal prob of N-in: 0.96171\nPOSSIBLE N-term signal sequence\ninside 1 19\nTMhelix 20 39\noutside 40 325\nTMhelix 326 348\ninside 349 372

15302 GCF_001833395.1_ASM183339v1 Salmonella enterica Proteobacteria; Gammaproteobacteria; Enterobacterales; Enterobacteriaceae; Salmonella
MNNEETFYQAMRRKGVTRRSFLKFCSLAATSLGLGAGMTPKIAWALENKPRIPVVWIHGL WP_001058311.1
hydrogenase [Salmonella enterica] Length: 372\nNumber of predicted TMHs: 2\nExp number of AAs in

TMHs: 37.4699899999999\nExp number, first 60 AAs: 19.72699\nTotal prob of N-in: 0.96171\nPOSSIBLE N-term signal sequence\ninside 1 19\nTMhelix 20 39\noutside 40 325\nTMhelix 326 348\ninside 349 372

15303 GCF_001833405.1_ASM183340v1 Salmonella enterica Proteobacteria; Gammaproteobacteria; Enterobacterales; Enterobacteriaceae; Salmonella
MNNEETFYQAMRRKGVTRRSFLKFCSLAATSLGLGAGMTPKIAWALENKPRIPVVWIHGL WP_001058311.1
hydrogenase [Salmonella enterica] Length: 372\nNumber of predicted TMHs: 2\nExp number of AAs in TMHs: 37.4699899999999\nExp number, first 60 AAs: 19.72699\nTotal prob of N-in: 0.96171\nPOSSIBLE N-term signal sequence\ninside 1 19\nTMhelix 20 39\noutside 40 325\nTMhelix 326 348\ninside 349 372

15304 GCF_001833485.1_ASM183348v1 Salmonella enterica Proteobacteria; Gammaproteobacteria; Enterobacterales; Enterobacteriaceae; Salmonella
MNNEETFYQAMRRKGVTRRSFLKFCSLAATSLGLGAGMTPKIAWALENKPRIPVVWIHGL WP_001058311.1
hydrogenase [Salmonella enterica] Length: 372\nNumber of predicted TMHs: 2\nExp number of AAs in TMHs: 37.4699899999999\nExp number, first 60 AAs: 19.72699\nTotal prob of N-in: 0.96171\nPOSSIBLE N-term signal sequence\ninside 1 19\nTMhelix 20 39\noutside 40 325\nTMhelix 326 348\ninside 349 372

15305 GCF_001833565.1_ASM183356v1 Salmonella enterica Proteobacteria; Gammaproteobacteria; Enterobacterales; Enterobacteriaceae; Salmonella
MTGDNTLITSHGINRRDFMKLCAALAATMGLSSKAAAEMAESVSNPQRPPVIWIGAQECT WP_000145429.1
hydrogenase 2 small subunit [Salmonella enterica] Length: 372\nNumber of predicted TMHs: 1\nExp number of AAs in TMHs: 24.51309\nExp number, first 60 AAs: 0.2682\nTotal prob of N-in: 0.14402\noutside 1 331\nTMhelix 332 354\ninside 355 372

15306 GCF_001834335.1_ASM183433v1 Salmonella enterica Proteobacteria; Gammaproteobacteria; Enterobacterales; Enterobacteriaceae; Salmonella
MNNEETFYQAMRRKGVTRRSFLKFCSLAATSLGLGAGMTPKIAWALENKPRIPVVWIHGL WP_001058306.1
hydrogenase [Salmonella enterica] Length: 372\nNumber of predicted TMHs: 2\nExp number of AAs in TMHs: 36.5540099999999\nExp number, first 60 AAs: 19.72732\nTotal prob of N-in: 0.96171\nPOSSIBLE N-term signal sequence\ninside 1 19\nTMhelix 20 39\noutside 40 325\nTMhelix 326 348\ninside 349 372

15307 GCF_001834335.1_ASM183433v1 Salmonella enterica Proteobacteria; Gammaproteobacteria; Enterobacterales; Enterobacteriaceae; Salmonella
MTGDNTLITSHGINRRDFMKLCAALAATMGLSSKAAAEMAESVSNPQRPPVIWIGAQECT WP_023248548.1
hydrogenase 2 small subunit [Salmonella enterica] Length: 372\nNumber of predicted TMHs: 1\nExp number of AAs in TMHs: 24.51281\nExp number, first 60 AAs: 0.26822\nTotal prob of N-in: 0.14402\noutside 1 331\nTMhelix 332 354\ninside 355 372

15308 GCF_001834825.1_ASM183482v1 Salmonella enterica Proteobacteria; Gammaproteobacteria; Enterobacterales; Enterobacteriaceae; Salmonella
MTGDNTLITSHGINRRDFMKLCAALAATMGLSSKAAAEMAESVSNPQRPPVIWIGAQECT WP_023248548.1
hydrogenase 2 small subunit [Salmonella enterica] Length: 372\nNumber of predicted TMHs: 1\nExp number of AAs in TMHs: 24.51281\nExp number, first 60 AAs: 0.26822\nTotal prob of N-in: 0.14402\noutside 1 331\nTMhelix 332 354\ninside 355 372

15309 GCF_001834825.1_ASM183482v1 Salmonella enterica Proteobacteria; Gammaproteobacteria; Enterobacterales; Enterobacteriaceae; Salmonella
MNNEETFYQAMRRKGVTRRSFLKFCSLAATSLGLGAGMTPKIAWALENKPRIPVVWIHGL WP_070801304.1
hydrogenase [Salmonella enterica] Length: 372\nNumber of predicted TMHs: 2\nExp number of AAs in TMHs: 36.3895799999999\nExp number, first 60 AAs: 19.727\nTotal prob of N-in: 0.96170\nPOSSIBLE N-term signal sequence\ninside 1 19\nTMhelix 20 39\noutside 40 325\nTMhelix 326 348\ninside 349 372

15310 GCF_001834905.1_ASM183490v1 Salmonella enterica Proteobacteria; Gammaproteobacteria; Enterobacterales; Enterobacteriaceae; Salmonella
MTGDNTLITSHGINRRDFMKLCAALAATMGLSSKAAAEMAESVSNPQRPPVIWIGAQECT WP_000145429.1
hydrogenase 2 small subunit [Salmonella enterica] Length: 372\nNumber of predicted TMHs: 1\nExp number of AAs in TMHs: 24.51309\nExp number, first 60 AAs: 0.2682\nTotal prob of N-in: 0.14402\noutside 1 331\nTMhelix 332 354\ninside 355 372

15311 GCF_001834915.1_ASM183491v1 Salmonella enterica Proteobacteria; Gammaproteobacteria; Enterobacterales; Enterobacteriaceae; Salmonella

MTGDNTLITSHGINRRDFMKLCAALAATMGLSSKAAAEMAESVSNPQRPPVIWIGAQECT WP_000145429.1 hydrogenase 2 small subunit [Salmonella enterica] Length: 372\nNumber of predicted TMHs: 1\nExp number of AAs in TMHs: 24.51309\nExp number, first 60 AAs: 0.2682\nTotal prob of N-in: 0.14402\noutside 1 331\nTMhelix 332 354\ninside 355 372

15312 GCF_001834905.1_ASM183490v1 Salmonella enterica Proteobacteria; Gammaproteobacteria; Enterobacterales; Enterobacteriaceae; Salmonella MNNEETFYQAMRRKGVTRRSFLKFCSLAATSLGLGAGMTPKIAWALENKPRIPVVWIHGL WP_001058311.1 hydrogenase [Salmonella enterica] Length: 372\nNumber of predicted TMHs: 2\nExp number of AAs in TMHs: 37.4699899999999\nExp number, first 60 AAs: 19.72699\nTotal prob of N-in: 0.96171\nPOSSIBLE N-term signal sequence\ninside 1 19\nTMhelix 20 39\noutside 40 325\nTMhelix 326 348\ninside 349 372

15313 GCF_001834915.1_ASM183491v1 Salmonella enterica Proteobacteria; Gammaproteobacteria; Enterobacterales; Enterobacteriaceae; Salmonella MNNEETFYQAMRRKGVTRRSFLKFCSLAATSLGLGAGMTPKIAWALENKPRIPVVWIHGL WP_001058311.1 hydrogenase [Salmonella enterica] Length: 372\nNumber of predicted TMHs: 2\nExp number of AAs in TMHs: 37.4699899999999\nExp number, first 60 AAs: 19.72699\nTotal prob of N-in: 0.96171\nPOSSIBLE N-term signal sequence\ninside 1 19\nTMhelix 20 39\noutside 40 325\nTMhelix 326 348\ninside 349 372

15314 GCF_001835235.1_ASM183523v1 Salmonella enterica Proteobacteria; Gammaproteobacteria; Enterobacterales; Enterobacteriaceae; Salmonella MNNEETFYQAMRRKGVTRRSFLKFCSLAATSLGLGAGMTPKIAWALENKPRIPVVWIHGL WP_001058312.1 hydrogenase [Salmonella enterica] Length: 372\nNumber of predicted TMHs: 2\nExp number of AAs in TMHs: 36.51738\nExp number, first 60 AAs: 19.72849\nTotal prob of N-in: 0.96170\nPOSSIBLE N-term signal sequence\ninside 1 19\nTMhelix 20 39\noutside 40 325\nTMhelix 326 348\ninside 349 372

15315 GCF_001879775.1_ASM187977v1 Salmonella enterica Proteobacteria; Gammaproteobacteria; Enterobacterales; Enterobacteriaceae; Salmonella MNNEETFYQAMRRKGVTRRSFLKFCSLAATSLGLGAGMTPKIAWALENKPRIPVVWIHGL WP_001058312.1 hydrogenase [Salmonella enterica] Length: 372\nNumber of predicted TMHs: 2\nExp number of AAs in TMHs: 36.51738\nExp number, first 60 AAs: 19.72849\nTotal prob of N-in: 0.96170\nPOSSIBLE N-term signal sequence\ninside 1 19\nTMhelix 20 39\noutside 40 325\nTMhelix 326 348\ninside 349 372

15316 GCF_001952495.1_ASM195249v1 Salmonella enterica Proteobacteria; Gammaproteobacteria; Enterobacterales; Enterobacteriaceae; Salmonella MTGDNTLITSHGINRRDFMKLCAALAATMGLSSKAAAEMAESVSNPQRPPVIWIGAQECT WP_000145429.1 hydrogenase 2 small subunit [Salmonella enterica] Length: 372\nNumber of predicted TMHs: 1\nExp number of AAs in TMHs: 24.51309\nExp number, first 60 AAs: 0.2682\nTotal prob of N-in: 0.14402\noutside 1 331\nTMhelix 332 354\ninside 355 372

15317 GCF_001951845.1_ASM195184v1 Salmonella enterica Proteobacteria; Gammaproteobacteria; Enterobacterales; Enterobacteriaceae; Salmonella MTGDNTLITSHGINRRDFMKLCAALAATMGLSSKAAAEMAESVSNPQRPPVIWIGAQECT WP_000145429.1 hydrogenase 2 small subunit [Salmonella enterica] Length: 372\nNumber of predicted TMHs: 1\nExp number of AAs in TMHs: 24.51309\nExp number, first 60 AAs: 0.2682\nTotal prob of N-in: 0.14402\noutside 1 331\nTMhelix 332 354\ninside 355 372

15318 GCF_001951915.1_ASM195191v1 Salmonella enterica Proteobacteria; Gammaproteobacteria; Enterobacterales; Enterobacteriaceae; Salmonella MNNEETFYQAMRRKGVTRRSFLKFCSLAATSLGLGAGMTPKIAWALENKPRIPVVWIHGL WP_001058312.1 hydrogenase [Salmonella enterica] Length: 372\nNumber of predicted TMHs: 2\nExp number of AAs in TMHs: 36.51738\nExp number, first 60 AAs: 19.72849\nTotal prob of N-in: 0.96170\nPOSSIBLE N-term signal sequence\ninside 1 19\nTMhelix 20 39\noutside 40 325\nTMhelix 326 348\ninside 349 372

15319 GCF_001952235.1_ASM195223v1 Salmonella enterica Proteobacteria; Gammaproteobacteria; Enterobacterales; Enterobacteriaceae; Salmonella MTGDNTLITSHGINRRDFMKLCAALAATMGLSSKAAAEMAESVSNPQRPPVIWIGAQECT WP_000145429.1 hydrogenase 2 small subunit [Salmonella enterica] Length: 372\nNumber of predicted TMHs: 1\nExp number of AAs in TMHs: 24.51309\nExp number, first 60 AAs: 0.2682\nTotal prob of N-in: 0.14402\noutside 1 331\nTMhelix 332 354\ninside 355 372

15320 GCF_001952295.1_ASM195229v1 Salmonella enterica Proteobacteria; Gammaproteobacteria; Enterobacterales; Enterobacteriaceae; Salmonella
MTGDNTLITSHGINRRDFMKLCAALAATMGLSSKAAAEMAESVSNPQRPPVIWIGAQECT WP_000145429.1
hydrogenase 2 small subunit [Salmonella enterica] Length: 372\nNumber of predicted TMHs: 1\nExp number of AAs in TMHs: 24.51309\nExp number, first 60 AAs: 0.2682\nTotal prob of N-in: 0.14402\noutside 1 331\nTMhelix 332 354\ninside 355 372

15321 GCF_001952235.1_ASM195223v1 Salmonella enterica Proteobacteria; Gammaproteobacteria; Enterobacterales; Enterobacteriaceae; Salmonella
MNNEETFYQAMRRKGVTRRSFLKFCSLAATSLGLGAGMTPKIAWALENKPRIPVVWIHGL WP_001058312.1
hydrogenase [Salmonella enterica] Length: 372\nNumber of predicted TMHs: 2\nExp number of AAs in TMHs: 36.51738\nExp number, first 60 AAs: 19.72849\nTotal prob of N-in: 0.96170\nPOSSIBLE N-term signal sequence\ninside 1 19\nTMhelix 20 39\noutside 40 325\nTMhelix 326 348\ninside 349 372

15322 GCF_001952295.1_ASM195229v1 Salmonella enterica Proteobacteria; Gammaproteobacteria; Enterobacterales; Enterobacteriaceae; Salmonella
MNNEETFYQAMRRKGVTRRSFLKFCSLAATSLGLGAGMTPKIAWALENKPRIPVVWIHGL WP_001058312.1
hydrogenase [Salmonella enterica] Length: 372\nNumber of predicted TMHs: 2\nExp number of AAs in TMHs: 36.51738\nExp number, first 60 AAs: 19.72849\nTotal prob of N-in: 0.96170\nPOSSIBLE N-term signal sequence\ninside 1 19\nTMhelix 20 39\noutside 40 325\nTMhelix 326 348\ninside 349 372

15323 GCF_001951935.1_ASM195193v1 Salmonella enterica Proteobacteria; Gammaproteobacteria; Enterobacterales; Enterobacteriaceae; Salmonella
MNNEETFYQAMRRKGVTRRSFLKFCSLAATSLGLGAGMTPKIAWALENKPRIPVVWIHGL WP_001058312.1
hydrogenase [Salmonella enterica] Length: 372\nNumber of predicted TMHs: 2\nExp number of AAs in TMHs: 36.51738\nExp number, first 60 AAs: 19.72849\nTotal prob of N-in: 0.96170\nPOSSIBLE N-term signal sequence\ninside 1 19\nTMhelix 20 39\noutside 40 325\nTMhelix 326 348\ninside 349 372

15324 GCF_001951995.1_ASM195199v1 Salmonella enterica Proteobacteria; Gammaproteobacteria; Enterobacterales; Enterobacteriaceae; Salmonella
MNNEETFYQAMRRKGVTRRSFLKFCSLAATSLGLGAGMTPKIAWALENKPRIPVVWIHGL WP_001058312.1
hydrogenase [Salmonella enterica] Length: 372\nNumber of predicted TMHs: 2\nExp number of AAs in TMHs: 36.51738\nExp number, first 60 AAs: 19.72849\nTotal prob of N-in: 0.96170\nPOSSIBLE N-term signal sequence\ninside 1 19\nTMhelix 20 39\noutside 40 325\nTMhelix 326 348\ninside 349 372

15325 GCF_001952415.1_ASM195241v1 Salmonella enterica Proteobacteria; Gammaproteobacteria; Enterobacterales; Enterobacteriaceae; Salmonella
MNNEETFYQAMRRKGVTRRSFLKFCSLAATSLGLGAGMTPKIAWALENKPRIPVVWIHGL WP_001058312.1
hydrogenase [Salmonella enterica] Length: 372\nNumber of predicted TMHs: 2\nExp number of AAs in TMHs: 36.51738\nExp number, first 60 AAs: 19.72849\nTotal prob of N-in: 0.96170\nPOSSIBLE N-term signal sequence\ninside 1 19\nTMhelix 20 39\noutside 40 325\nTMhelix 326 348\ninside 349 372

15326 GCF_001952485.1_ASM195248v1 Salmonella enterica Proteobacteria; Gammaproteobacteria; Enterobacterales; Enterobacteriaceae; Salmonella
MNNEETFYQAMRRKGVTRRSFLKFCSLAATSLGLGAGMTPKIAWALENKPRIPVVWIHGL WP_001058312.1
hydrogenase [Salmonella enterica] Length: 372\nNumber of predicted TMHs: 2\nExp number of AAs in TMHs: 36.51738\nExp number, first 60 AAs: 19.72849\nTotal prob of N-in: 0.96170\nPOSSIBLE N-term signal sequence\ninside 1 19\nTMhelix 20 39\noutside 40 325\nTMhelix 326 348\ninside 349 372

15327 GCF_002033505.1_ASM203350v1 Salmonella enterica Proteobacteria; Gammaproteobacteria; Enterobacterales; Enterobacteriaceae; Salmonella
MTGDNTLITSHGINRRDFMKLCAALAATMGLSSKAAAEMAESVSNPQRPPVIWIGAQECT WP_000145429.1
hydrogenase 2 small subunit [Salmonella enterica] Length: 372\nNumber of predicted TMHs: 1\nExp number of AAs in TMHs: 24.51309\nExp number, first 60 AAs: 0.2682\nTotal prob of N-in: 0.14402\noutside 1 331\nTMhelix 332 354\ninside 355 372

15328 GCF_002030105.1_ASM203010v1 Salmonella enterica Proteobacteria; Gammaproteobacteria; Enterobacterales; Enterobacteriaceae; Salmonella
MTGDNTLITSHGINRRDFMKLCAALAATMGLSSKAAAEMAESVSNPQRPPVIWIGAQECT WP_000145429.1
hydrogenase 2 small subunit [Salmonella enterica] Length: 372\nNumber of predicted TMHs: 1\nExp

number of AAs in TMHs: 24.51309\nExp number, first 60 AAs: 0.2682\nTotal prob of N-in: 0.14402\noutside 1 331\nTMhelix 332 354\ninside 355 372

15329 GCF_002030095.1_ASM203009v1 Salmonella enterica Proteobacteria; Gammaproteobacteria; Enterobacterales; Enterobacteriaceae; Salmonella
MTGDNTLITSHGINRRDFMKLCAALAATMGLSSKAAAEMAESVSNPQRPPVIWIGAQECT WP_000145429.1
hydrogenase 2 small subunit [Salmonella enterica] Length: 372\nNumber of predicted TMHs: 1\nExp number of AAs in TMHs: 24.51309\nExp number, first 60 AAs: 0.2682\nTotal prob of N-in: 0.14402\noutside 1 331\nTMhelix 332 354\ninside 355 372

15330 GCF_002030105.1_ASM203010v1 Salmonella enterica Proteobacteria; Gammaproteobacteria; Enterobacterales; Enterobacteriaceae; Salmonella
MNNEETFYQAMRRKGVTRRSFLKFCSLAATSLGLGAGMTPKIAWALENKPRIPVVWIHGL WP_001058312.1
hydrogenase [Salmonella enterica] Length: 372\nNumber of predicted TMHs: 2\nExp number of AAs in TMHs: 36.51738\nExp number, first 60 AAs: 19.72849\nTotal prob of N-in: 0.96170\nPOSSIBLE N-term signal sequence\ninside 1 19\nTMhelix 20 39\noutside 40 325\nTMhelix 326 348\ninside 349 372

15331 GCF_002030095.1_ASM203009v1 Salmonella enterica Proteobacteria; Gammaproteobacteria; Enterobacterales; Enterobacteriaceae; Salmonella
MNNEETFYQAMRRKGVTRRSFLKFCSLAATSLGLGAGMTPKIAWALENKPRIPVVWIHGL WP_001058312.1
hydrogenase [Salmonella enterica] Length: 372\nNumber of predicted TMHs: 2\nExp number of AAs in TMHs: 36.51738\nExp number, first 60 AAs: 19.72849\nTotal prob of N-in: 0.96170\nPOSSIBLE N-term signal sequence\ninside 1 19\nTMhelix 20 39\noutside 40 325\nTMhelix 326 348\ninside 349 372

15332 GCF_002034095.1_ASM203409v1 Salmonella enterica Proteobacteria; Gammaproteobacteria; Enterobacterales; Enterobacteriaceae; Salmonella
MTGDNTLITSHGINRRDFMKLCAALAATMGLSSKAAAEMAESVSNPQRPPVIWIGAQECT WP_000145429.1
hydrogenase 2 small subunit [Salmonella enterica] Length: 372\nNumber of predicted TMHs: 1\nExp number of AAs in TMHs: 24.51309\nExp number, first 60 AAs: 0.2682\nTotal prob of N-in: 0.14402\noutside 1 331\nTMhelix 332 354\ninside 355 372

15333 GCF_002030825.1_ASM203082v1 Salmonella enterica Proteobacteria; Gammaproteobacteria; Enterobacterales; Enterobacteriaceae; Salmonella
MNNEETFYQAMRRKGVTRRSFLKFCSLAATSLGLGAGMTPKIAWALENKPRIPVVWIHGL WP_001058312.1
hydrogenase [Salmonella enterica] Length: 372\nNumber of predicted TMHs: 2\nExp number of AAs in TMHs: 36.51738\nExp number, first 60 AAs: 19.72849\nTotal prob of N-in: 0.96170\nPOSSIBLE N-term signal sequence\ninside 1 19\nTMhelix 20 39\noutside 40 325\nTMhelix 326 348\ninside 349 372

15334 GCF_002032585.1_ASM203258v1 Salmonella enterica Proteobacteria; Gammaproteobacteria; Enterobacterales; Enterobacteriaceae; Salmonella
MNNEETFYQAMRRKGVTRRSFLKFCSLAATSLGLGAGMTPKIAWALENKPRIPVVWIHGL WP_001058312.1
hydrogenase [Salmonella enterica] Length: 372\nNumber of predicted TMHs: 2\nExp number of AAs in TMHs: 36.51738\nExp number, first 60 AAs: 19.72849\nTotal prob of N-in: 0.96170\nPOSSIBLE N-term signal sequence\ninside 1 19\nTMhelix 20 39\noutside 40 325\nTMhelix 326 348\ninside 349 372

15335 GCF_002032615.1_ASM203261v1 Salmonella enterica Proteobacteria; Gammaproteobacteria; Enterobacterales; Enterobacteriaceae; Salmonella
MNNEETFYQAMRRKGVTRRSFLKFCSLAATSLGLGAGMTPKIAWALENKPRIPVVWIHGL WP_001058312.1
hydrogenase [Salmonella enterica] Length: 372\nNumber of predicted TMHs: 2\nExp number of AAs in TMHs: 36.51738\nExp number, first 60 AAs: 19.72849\nTotal prob of N-in: 0.96170\nPOSSIBLE N-term signal sequence\ninside 1 19\nTMhelix 20 39\noutside 40 325\nTMhelix 326 348\ninside 349 372

15336 GCF_002032675.1_ASM203267v1 Salmonella enterica Proteobacteria; Gammaproteobacteria; Enterobacterales; Enterobacteriaceae; Salmonella
MNNEETFYQAMRRKGVTRRSFLKFCSLAATSLGLGAGMTPKIAWALENKPRIPVVWIHGL WP_001058312.1
hydrogenase [Salmonella enterica] Length: 372\nNumber of predicted TMHs: 2\nExp number of AAs in TMHs: 36.51738\nExp number, first 60 AAs: 19.72849\nTotal prob of N-in: 0.96170\nPOSSIBLE N-term signal sequence\ninside 1 19\nTMhelix 20 39\noutside 40 325\nTMhelix 326 348\ninside 349 372

15337 GCF_002033285.1_ASM203328v1 Salmonella enterica Proteobacteria; Gammaproteobacteria; Enterobacterales; Enterobacteriaceae; Salmonella

MTGDNTLITSHGINRRDFMKLCAALAATMGLSSKAAAEMAESVSNPQRPPVIWIGAQECT WP_000145429.1 hydrogenase 2 small subunit [Salmonella enterica] Length: 372\nNumber of predicted TMHs: 1\nExp number of AAs in TMHs: 24.51309\nExp number, first 60 AAs: 0.2682\nTotal prob of N-in: 0.14402\noutside 1 331\nTMhelix 332 354\ninside 355 372

15338 GCF_002033325.1_ASM203332v1 Salmonella enterica Proteobacteria; Gammaproteobacteria; Enterobacterales; Enterobacteriaceae; Salmonella MTGDNTLITSHGINRRDFMKLCAALAATMGLSSKAAAEMAESVSNPQRPPVIWIGAQECT WP_000145429.1 hydrogenase 2 small subunit [Salmonella enterica] Length: 372\nNumber of predicted TMHs: 1\nExp number of AAs in TMHs: 24.51309\nExp number, first 60 AAs: 0.2682\nTotal prob of N-in: 0.14402\noutside 1 331\nTMhelix 332 354\ninside 355 372

15339 GCF_002033595.1_ASM203359v1 Salmonella enterica Proteobacteria; Gammaproteobacteria; Enterobacterales; Enterobacteriaceae; Salmonella MNNEETFYQAMRRKGVTRRSFLKFCSLAATSLGLGAGMTPKIAWALENKPRIPVVWIHGL WP_001058307.1 hydrogenase [Salmonella enterica] Length: 372\nNumber of predicted TMHs: 2\nExp number of AAs in TMHs: 36.51738\nExp number, first 60 AAs: 19.72849\nTotal prob of N-in: 0.96170\nPOSSIBLE N-term signal sequence\ninside 1 19\nTMhelix 20 39\noutside 40 325\nTMhelix 326 348\ninside 349 372

15340 GCF_002033505.1_ASM203350v1 Salmonella enterica Proteobacteria; Gammaproteobacteria; Enterobacterales; Enterobacteriaceae; Salmonella MNNEETFYQAMRRKGVTRRSFLKFCSLAATSLGLGAGMTPKIAWALENKPRIPVVWIHGL WP_079786335.1 [Ni/Fe] hydrogenase small subunit [Salmonella enterica] Length: 372\nNumber of predicted TMHs: 2\nExp number of AAs in TMHs: 36.51716\nExp number, first 60 AAs: 19.7285\nTotal prob of N-in: 0.96169\nPOSSIBLE N-term signal sequence\ninside 1 19\nTMhelix 20 39\noutside 40 325\nTMhelix 326 348\ninside 349 372

15341 GCF_002033655.1_ASM203365v1 Salmonella enterica Proteobacteria; Gammaproteobacteria; Enterobacterales; Enterobacteriaceae; Salmonella MNNEETFYQAMRRKGVTRRSFLKFCSLAATSLGLGAGMTPKIAWALENKPRIPVVWIHGL WP_001058310.1 hydrogenase [Salmonella enterica] Length: 372\nNumber of predicted TMHs: 2\nExp number of AAs in TMHs: 37.4567299999999\nExp number, first 60 AAs: 19.72694\nTotal prob of N-in: 0.96171\nPOSSIBLE N-term signal sequence\ninside 1 19\nTMhelix 20 39\noutside 40 325\nTMhelix 326 348\ninside 349 372

15342 GCF_002033885.1_ASM203388v1 Salmonella enterica Proteobacteria; Gammaproteobacteria; Enterobacterales; Enterobacteriaceae; Salmonella MNNEETFYQAMRRKGVTRRSFLKFCSLAATSLGLGAGMTPKIAWALENKPRIPVVWIHGL WP_001058309.1 hydrogenase [Salmonella enterica] Length: 372\nNumber of predicted TMHs: 2\nExp number of AAs in TMHs: 36.51725\nExp number, first 60 AAs: 19.7285\nTotal prob of N-in: 0.96170\nPOSSIBLE N-term signal sequence\ninside 1 19\nTMhelix 20 39\noutside 40 325\nTMhelix 326 348\ninside 349 372

15343 GCF_002033685.1_ASM203368v1 Salmonella enterica Proteobacteria; Gammaproteobacteria; Enterobacterales; Enterobacteriaceae; Salmonella MNNEETLYQAMRRKGVTRRSFLKFCSLAATSLGLGAGMTPKIAWALENKPRIPVVWIHGL WP_001752445.1 hydrogenase-1 small chain [Salmonella enterica] Length: 372\nNumber of predicted TMHs: 2\nExp number of AAs in TMHs: 36.5123\nExp number, first 60 AAs: 19.72341\nTotal prob of N-in: 0.96144\nPOSSIBLE N-term signal sequence\ninside 1 19\nTMhelix 20 39\noutside 40 325\nTMhelix 326 348\ninside 349 372

15344 GCF_002033945.1_ASM203394v1 Salmonella enterica Proteobacteria; Gammaproteobacteria; Enterobacterales; Enterobacteriaceae; Salmonella MNNEETFYQAMRRKGVTRRSFLKFCSLAATSLGLGAGMTPKIAWALENKPRIPVVWIHGL WP_001058311.1 hydrogenase [Salmonella enterica] Length: 372\nNumber of predicted TMHs: 2\nExp number of AAs in TMHs: 37.4699899999999\nExp number, first 60 AAs: 19.72699\nTotal prob of N-in: 0.96171\nPOSSIBLE N-term signal sequence\ninside 1 19\nTMhelix 20 39\noutside 40 325\nTMhelix 326 348\ninside 349 372

15345 GCF_002034025.1_ASM203402v1 Salmonella enterica Proteobacteria; Gammaproteobacteria; Enterobacterales; Enterobacteriaceae; Salmonella MTGDNTLITSHGINRRDFMKLCAALAATMGLSSKAAAEMAESVSNPQRPPVIWIGAQECT WP_000145429.1 hydrogenase 2 small subunit [Salmonella enterica] Length: 372\nNumber of predicted TMHs: 1\nExp number of AAs in TMHs: 24.51309\nExp number, first 60 AAs: 0.2682\nTotal prob of N-in: 0.14402\noutside 1 331\nTMhelix 332 354\ninside 355 372

15346 GCF_002034025.1_ASM203402v1 Salmonella enterica Proteobacteria; Gammaproteobacteria; Enterobacterales; Enterobacteriaceae; Salmonella
MNNEETFYQAMRRKGVTRRSFLKFCSLAATSLGLGAGMTPKIAWALENKPRIPVIWIHGL WP_079787412.1
[Ni/Fe] hydrogenase small subunit [Salmonella enterica] Length: 372\nNumber of predicted TMHs: 2\nExp number of AAs in TMHs: 37.49523\nExp number, first 60 AAs: 19.76485\nTotal prob of N-in: 0.96198\nPOSSIBLE N-term signal sequence\ninside 1 19\nTMhelix 20 39\noutside 40 325\nTMhelix 326 348\ninside 349 372

15347 GCF_002034465.1_ASM203446v1 Salmonella enterica Proteobacteria; Gammaproteobacteria; Enterobacterales; Enterobacteriaceae; Salmonella
MTGDNTLITSHGINRRDFMKLCAALAATMGLSSKAAAEMAESVSNPQRPPVIWIGAQECT WP_000145428.1
hydrogenase 2 small subunit [Salmonella enterica] Length: 372\nNumber of predicted TMHs: 1\nExp number of AAs in TMHs: 24.50677\nExp number, first 60 AAs: 0.2682\nTotal prob of N-in: 0.14390\noutside 1 331\nTMhelix 332 354\ninside 355 372

15348 GCF_002034455.1_ASM203445v1 Salmonella enterica Proteobacteria; Gammaproteobacteria; Enterobacterales; Enterobacteriaceae; Salmonella
MTGDNTLITSHGINRRDFMKLCAALAATMGLSSKAAAEMAESVSNPQRPPVIWIGAQECT WP_000145429.1
hydrogenase 2 small subunit [Salmonella enterica] Length: 372\nNumber of predicted TMHs: 1\nExp number of AAs in TMHs: 24.51309\nExp number, first 60 AAs: 0.2682\nTotal prob of N-in: 0.14402\noutside 1 331\nTMhelix 332 354\ninside 355 372

15349 GCF_002034465.1_ASM203446v1 Salmonella enterica Proteobacteria; Gammaproteobacteria; Enterobacterales; Enterobacteriaceae; Salmonella
MNNEETLYQAMRRKGVTRRSFLKFCSLAATSLGLGAGMTPKIAWALENKPRIPVVWIHGL WP_001752445.1
hydrogenase-1 small chain [Salmonella enterica] Length: 372\nNumber of predicted TMHs: 2\nExp number of AAs in TMHs: 36.5123\nExp number, first 60 AAs: 19.72341\nTotal prob of N-in: 0.96144\nPOSSIBLE N-term signal sequence\ninside 1 19\nTMhelix 20 39\noutside 40 325\nTMhelix 326 348\ninside 349 372

15350 GCF_002034815.1_ASM203481v1 Salmonella enterica Proteobacteria; Gammaproteobacteria; Enterobacterales; Enterobacteriaceae; Salmonella
MTGDNTLITSHGINRRDFMKLCAALAATMGLSSKAAAEMAESVSNPQRPPVIWIGAQECT WP_000145429.1
hydrogenase 2 small subunit [Salmonella enterica] Length: 372\nNumber of predicted TMHs: 1\nExp number of AAs in TMHs: 24.51309\nExp number, first 60 AAs: 0.2682\nTotal prob of N-in: 0.14402\noutside 1 331\nTMhelix 332 354\ninside 355 372

15351 GCF_002034895.1_ASM203489v1 Salmonella enterica Proteobacteria; Gammaproteobacteria; Enterobacterales; Enterobacteriaceae; Salmonella
MTGDNTLITSHGINRRDFMKLCAALAATMGLSSKAAAEMAESVSNPQRPPVIWIGAQECT WP_000145429.1
hydrogenase 2 small subunit [Salmonella enterica] Length: 372\nNumber of predicted TMHs: 1\nExp number of AAs in TMHs: 24.51309\nExp number, first 60 AAs: 0.2682\nTotal prob of N-in: 0.14402\noutside 1 331\nTMhelix 332 354\ninside 355 372

15352 GCF_002035205.1_ASM203520v1 Salmonella enterica Proteobacteria; Gammaproteobacteria; Enterobacterales; Enterobacteriaceae; Salmonella
MTGDNTLITSHGINRRDFMKLCAALAATMGLSSKAAAEMAESVSNPQRPPVIWIGAQECT WP_000145429.1
hydrogenase 2 small subunit [Salmonella enterica] Length: 372\nNumber of predicted TMHs: 1\nExp number of AAs in TMHs: 24.51309\nExp number, first 60 AAs: 0.2682\nTotal prob of N-in: 0.14402\noutside 1 331\nTMhelix 332 354\ninside 355 372

15353 GCF_002035405.1_ASM203540v1 Salmonella enterica Proteobacteria; Gammaproteobacteria; Enterobacterales; Enterobacteriaceae; Salmonella
MTGDNTLITSHGINRRDFMKLCAALAATMGLSSKAAAEMAESVSNPQRPPVIWIGAQECT WP_000145429.1
hydrogenase 2 small subunit [Salmonella enterica] Length: 372\nNumber of predicted TMHs: 1\nExp number of AAs in TMHs: 24.51309\nExp number, first 60 AAs: 0.2682\nTotal prob of N-in: 0.14402\noutside 1 331\nTMhelix 332 354\ninside 355 372

15354 GCF_002035405.1_ASM203540v1 Salmonella enterica Proteobacteria; Gammaproteobacteria; Enterobacterales; Enterobacteriaceae; Salmonella
MNNEETFYQAMRRKGVTRRSFLKFCSLAATSLGLGAGMTPKIAWALENKPRIPVVWIHGL WP_023187532.1
hydrogenase 1, small subunit [Salmonella enterica] Length: 372\nNumber of predicted TMHs: 2\nExp

number of AAs in TMHs: 36.9997\nExp number, first 60 AAs: 19.72759\nTotal prob of N-in: 0.96171\nPOSSIBLE N-term signal sequence\ninside 1 19\nTMhelix 20 39\noutside 40 325\nTMhelix 326 348\ninside 349 372

15355 GCF_002035865.1_ASM203586v1 Salmonella enterica Proteobacteria; Gammaproteobacteria; Enterobacterales; Enterobacteriaceae; Salmonella MTGDNTLITSHGINRRDFMKLCAALAATMGLSSKAAAEMAESVSNPQRPPVIWIGAQECT WP_000145432.1 hydrogenase 2 small subunit [Salmonella enterica] Length: 372\nNumber of predicted TMHs: 1\nExp number of AAs in TMHs: 24.5101\nExp number, first 60 AAs: 0.2682\nTotal prob of N-in: 0.14397\noutside 1 331\nTMhelix 332 354\ninside 355 372

15356 GCF_002035865.1_ASM203586v1 Salmonella enterica Proteobacteria; Gammaproteobacteria; Enterobacterales; Enterobacteriaceae; Salmonella MNNEETFYQAMRRKGVTRRSFLKFCSLAATSLGLGAGMTPKIAWALENKPRIPVVWIHGL WP_001058314.1 hydrogenase [Salmonella enterica] Length: 372\nNumber of predicted TMHs: 2\nExp number of AAs in TMHs: 36.3110299999999\nExp number, first 60 AAs: 19.72774\nTotal prob of N-in: 0.96169\nPOSSIBLE N-term signal sequence\ninside 1 19\nTMhelix 20 39\noutside 40 325\nTMhelix 326 348\ninside 349 372

15357 GCF_002036285.1_ASM203628v1 Salmonella enterica Proteobacteria; Gammaproteobacteria; Enterobacterales; Enterobacteriaceae; Salmonella MNNEETFYQAMRRKGVTRRSFLKFCSLAATSLGLGAGMTPKIAWALENKPRIPVVWIHGL WP_001058312.1 hydrogenase [Salmonella enterica] Length: 372\nNumber of predicted TMHs: 2\nExp number of AAs in TMHs: 36.51738\nExp number, first 60 AAs: 19.72849\nTotal prob of N-in: 0.96170\nPOSSIBLE N-term signal sequence\ninside 1 19\nTMhelix 20 39\noutside 40 325\nTMhelix 326 348\ninside 349 372

15358 GCF_002036465.1_ASM203646v1 Salmonella enterica Proteobacteria; Gammaproteobacteria; Enterobacterales; Enterobacteriaceae; Salmonella MTGDNTLITSHGINRRDFMKLCAALAATMGLSSKAAAEMAESVSNPQRPPVIWIGAQECT WP_000145429.1 hydrogenase 2 small subunit [Salmonella enterica] Length: 372\nNumber of predicted TMHs: 1\nExp number of AAs in TMHs: 24.51309\nExp number, first 60 AAs: 0.2682\nTotal prob of N-in: 0.14402\noutside 1 331\nTMhelix 332 354\ninside 355 372

15359 GCF_002036455.1_ASM203645v1 Salmonella enterica Proteobacteria; Gammaproteobacteria; Enterobacterales; Enterobacteriaceae; Salmonella MTGDNTLITSHGINRRDFMKLCAALAATMGLSSKAAAEMAESVSNPQRPPVIWIGAQECT WP_000145429.1 hydrogenase 2 small subunit [Salmonella enterica] Length: 372\nNumber of predicted TMHs: 1\nExp number of AAs in TMHs: 24.51309\nExp number, first 60 AAs: 0.2682\nTotal prob of N-in: 0.14402\noutside 1 331\nTMhelix 332 354\ninside 355 372

15360 GCF_002036525.1_ASM203652v1 Salmonella enterica Proteobacteria; Gammaproteobacteria; Enterobacterales; Enterobacteriaceae; Salmonella MTGDNTLITSHGINRRDFMKLCAALAATMGLSSKAAAEMAESVSNPQRPPVIWIGAQECT WP_000145429.1 hydrogenase 2 small subunit [Salmonella enterica] Length: 372\nNumber of predicted TMHs: 1\nExp number of AAs in TMHs: 24.51309\nExp number, first 60 AAs: 0.2682\nTotal prob of N-in: 0.14402\noutside 1 331\nTMhelix 332 354\ninside 355 372

15361 GCF_002036465.1_ASM203646v1 Salmonella enterica Proteobacteria; Gammaproteobacteria; Enterobacterales; Enterobacteriaceae; Salmonella MNNEETFYQAMRRKGVTRRSFLKFCSLAATSLGLGAGMTPKIAWALENKPRIPVVWIHGL WP_001058307.1 hydrogenase [Salmonella enterica] Length: 372\nNumber of predicted TMHs: 2\nExp number of AAs in TMHs: 36.51738\nExp number, first 60 AAs: 19.72849\nTotal prob of N-in: 0.96170\nPOSSIBLE N-term signal sequence\ninside 1 19\nTMhelix 20 39\noutside 40 325\nTMhelix 326 348\ninside 349 372

15362 GCF_002036645.1_ASM203664v1 Salmonella enterica Proteobacteria; Gammaproteobacteria; Enterobacterales; Enterobacteriaceae; Salmonella MTGDNTLITSHGINRRDFMKLCAALAATMGLSSKAAAEMAESVSNPQRPPVIWIGAQECT WP_000145429.1 hydrogenase 2 small subunit [Salmonella enterica] Length: 372\nNumber of predicted TMHs: 1\nExp number of AAs in TMHs: 24.51309\nExp number, first 60 AAs: 0.2682\nTotal prob of N-in: 0.14402\noutside 1 331\nTMhelix 332 354\ninside 355 372

15363 GCF_002036675.1_ASM203667v1 Salmonella enterica Proteobacteria; Gammaproteobacteria; Enterobacterales; Enterobacteriaceae; Salmonella

MTGDNTLITSHGINRRDFMKLCAALAATMGLSSKAAAEMAESVSNPQRPPVIWIGAQECT WP_000145429.1 hydrogenase 2 small subunit [Salmonella enterica] Length: 372\nNumber of predicted TMHs: 1\nExp number of AAs in TMHs: 24.51309\nExp number, first 60 AAs: 0.2682\nTotal prob of N-in: 0.14402\noutside 1 331\nTMhelix 332 354\ninside 355 372

15364 GCF_002036705.1_ASM203670v1 Salmonella enterica Proteobacteria; Gammaproteobacteria; Enterobacterales; Enterobacteriaceae; Salmonella MTGDNTLITSHGINRRDFMKLCAALAATMGLSSKAAAEMAESVSNPQRPPVIWIGAQECT WP_000145429.1 hydrogenase 2 small subunit [Salmonella enterica] Length: 372\nNumber of predicted TMHs: 1\nExp number of AAs in TMHs: 24.51309\nExp number, first 60 AAs: 0.2682\nTotal prob of N-in: 0.14402\noutside 1 331\nTMhelix 332 354\ninside 355 372

15365 GCF_002036715.1_ASM203671v1 Salmonella enterica Proteobacteria; Gammaproteobacteria; Enterobacterales; Enterobacteriaceae; Salmonella MTGDNTLITSHGINRRDFMKLCAALAATMGLSSKAAAEMAESVSNPQRPPVIWIGAQECT WP_000145429.1 hydrogenase 2 small subunit [Salmonella enterica] Length: 372\nNumber of predicted TMHs: 1\nExp number of AAs in TMHs: 24.51309\nExp number, first 60 AAs: 0.2682\nTotal prob of N-in: 0.14402\noutside 1 331\nTMhelix 332 354\ninside 355 372

15366 GCF_002036785.1_ASM203678v1 Salmonella enterica Proteobacteria; Gammaproteobacteria; Enterobacterales; Enterobacteriaceae; Salmonella MTGDNTLITSHGINRRDFMKLCAALAATMGLSSKAAAEMAESVSNPQRPPVIWIGAQECT WP_000145429.1 hydrogenase 2 small subunit [Salmonella enterica] Length: 372\nNumber of predicted TMHs: 1\nExp number of AAs in TMHs: 24.51309\nExp number, first 60 AAs: 0.2682\nTotal prob of N-in: 0.14402\noutside 1 331\nTMhelix 332 354\ninside 355 372

15367 GCF_002036795.1_ASM203679v1 Salmonella enterica Proteobacteria; Gammaproteobacteria; Enterobacterales; Enterobacteriaceae; Salmonella MTGDNTLITSHGINRRDFMKLCAALAATMGLSSKAAAEMAESVSNPQRPPVIWIGAQECT WP_000145429.1 hydrogenase 2 small subunit [Salmonella enterica] Length: 372\nNumber of predicted TMHs: 1\nExp number of AAs in TMHs: 24.51309\nExp number, first 60 AAs: 0.2682\nTotal prob of N-in: 0.14402\noutside 1 331\nTMhelix 332 354\ninside 355 372

15368 GCF_002036645.1_ASM203664v1 Salmonella enterica Proteobacteria; Gammaproteobacteria; Enterobacterales; Enterobacteriaceae; Salmonella MNNEETFYQAMRRKGVTRRSFLKFCSLAATSLGLGAGMTPKIAWALENKPRIPVVWIHGL WP_001058312.1 hydrogenase [Salmonella enterica] Length: 372\nNumber of predicted TMHs: 2\nExp number of AAs in TMHs: 36.51738\nExp number, first 60 AAs: 19.72849\nTotal prob of N-in: 0.96170\nPOSSIBLE N-term signal sequence\ninside 1 19\nTMhelix 20 39\noutside 40 325\nTMhelix 326 348\ninside 349 372

15369 GCF_002036705.1_ASM203670v1 Salmonella enterica Proteobacteria; Gammaproteobacteria; Enterobacterales; Enterobacteriaceae; Salmonella MNNEETFYQAMRRKGVTRRSFLKFCSLAATSLGLGAGMTPKIAWALENKPRIPVVWIHGL WP_001058312.1 hydrogenase [Salmonella enterica] Length: 372\nNumber of predicted TMHs: 2\nExp number of AAs in TMHs: 36.51738\nExp number, first 60 AAs: 19.72849\nTotal prob of N-in: 0.96170\nPOSSIBLE N-term signal sequence\ninside 1 19\nTMhelix 20 39\noutside 40 325\nTMhelix 326 348\ninside 349 372

15370 GCF_002037155.1_ASM203715v1 Salmonella enterica Proteobacteria; Gammaproteobacteria; Enterobacterales; Enterobacteriaceae; Salmonella MTGDNTLITSHGINRRDFMKLCAALAATMGLSSKAAAEMAESVSNPQRPPVIWIGAQECT WP_000145429.1 hydrogenase 2 small subunit [Salmonella enterica] Length: 372\nNumber of predicted TMHs: 1\nExp number of AAs in TMHs: 24.51309\nExp number, first 60 AAs: 0.2682\nTotal prob of N-in: 0.14402\noutside 1 331\nTMhelix 332 354\ninside 355 372

15371 GCF_002037195.1_ASM203719v1 Salmonella enterica Proteobacteria; Gammaproteobacteria; Enterobacterales; Enterobacteriaceae; Salmonella MTGDNTLITSHGINRRDFMKLCAALAATMGLSSKAAAEMAESVSNPQRPPVIWIGAQECT WP_000145429.1 hydrogenase 2 small subunit [Salmonella enterica] Length: 372\nNumber of predicted TMHs: 1\nExp number of AAs in TMHs: 24.51309\nExp number, first 60 AAs: 0.2682\nTotal prob of N-in: 0.14402\noutside 1 331\nTMhelix 332 354\ninside 355 372

15372 GCF_002037155.1_ASM203715v1 Salmonella enterica Proteobacteria; Gammaproteobacteria; Enterobacterales; Enterobacteriaceae; Salmonella
MNNEETFYQAMRRKGVTRRSFLKFCSLAATSLGLGAGMTPKIAWALENKPRIPVVWIHGL WP_001058311.1
hydrogenase [Salmonella enterica] Length: 372\nNumber of predicted TMHs: 2\nExp number of AAs in TMHs: 37.4699899999999\nExp number, first 60 AAs: 19.72699\nTotal prob of N-in: 0.96171\nPOSSIBLE N-term signal sequence\ninside 1 19\nTMhelix 20 39\noutside 40 325\nTMhelix 326 348\ninside 349 372

15373 GCF_002037195.1_ASM203719v1 Salmonella enterica Proteobacteria; Gammaproteobacteria; Enterobacterales; Enterobacteriaceae; Salmonella
MNNEETFYQAMRRKGVTRRSFLKFCSLAATSLGLGAGMTPKIAWALENKPRIPVVWIHGL WP_001058311.1
hydrogenase [Salmonella enterica] Length: 372\nNumber of predicted TMHs: 2\nExp number of AAs in TMHs: 37.4699899999999\nExp number, first 60 AAs: 19.72699\nTotal prob of N-in: 0.96171\nPOSSIBLE N-term signal sequence\ninside 1 19\nTMhelix 20 39\noutside 40 325\nTMhelix 326 348\ninside 349 372

15374 GCF_002037035.1_ASM203703v1 Salmonella enterica Proteobacteria; Gammaproteobacteria; Enterobacterales; Enterobacteriaceae; Salmonella
MNNEETFYQAMRRKGVTRRSFLKFCSLAATSLGLGAGMTPKIAWALENKPRIPVVWIHGL WP_001058312.1
hydrogenase [Salmonella enterica] Length: 372\nNumber of predicted TMHs: 2\nExp number of AAs in TMHs: 36.51738\nExp number, first 60 AAs: 19.72849\nTotal prob of N-in: 0.96170\nPOSSIBLE N-term signal sequence\ninside 1 19\nTMhelix 20 39\noutside 40 325\nTMhelix 326 348\ninside 349 372

15375 GCF_002037105.1_ASM203710v1 Salmonella enterica Proteobacteria; Gammaproteobacteria; Enterobacterales; Enterobacteriaceae; Salmonella
MNNEETFYQAMRRKGVTRRSFLKFCSLAATSLGLGAGMTPKIAWALENKPRIPVVWIHGL WP_001058311.1
hydrogenase [Salmonella enterica] Length: 372\nNumber of predicted TMHs: 2\nExp number of AAs in TMHs: 37.4699899999999\nExp number, first 60 AAs: 19.72699\nTotal prob of N-in: 0.96171\nPOSSIBLE N-term signal sequence\ninside 1 19\nTMhelix 20 39\noutside 40 325\nTMhelix 326 348\ninside 349 372

15376 GCF_002037065.1_ASM203706v1 Salmonella enterica Proteobacteria; Gammaproteobacteria; Enterobacterales; Enterobacteriaceae; Salmonella
MNNEETFYQAMRRKGVTRRSFLKFCSLAATSLGLGAGMTPKIAWALENKPRIPVVWIHGL WP_001058312.1
hydrogenase [Salmonella enterica] Length: 372\nNumber of predicted TMHs: 2\nExp number of AAs in TMHs: 36.51738\nExp number, first 60 AAs: 19.72849\nTotal prob of N-in: 0.96170\nPOSSIBLE N-term signal sequence\ninside 1 19\nTMhelix 20 39\noutside 40 325\nTMhelix 326 348\ninside 349 372

15377 GCF_002037525.1_ASM203752v1 Salmonella enterica Proteobacteria; Gammaproteobacteria; Enterobacterales; Enterobacteriaceae; Salmonella
MTGDNTLITSHGINRRDFMKLCAALAATMGLSSKAAAEMAESVSNPQRPPVIWIGAQECT WP_000145429.1
hydrogenase 2 small subunit [Salmonella enterica] Length: 372\nNumber of predicted TMHs: 1\nExp number of AAs in TMHs: 24.51309\nExp number, first 60 AAs: 0.2682\nTotal prob of N-in: 0.14402\noutside 1 331\nTMhelix 332 354\ninside 355 372

15378 GCF_002037525.1_ASM203752v1 Salmonella enterica Proteobacteria; Gammaproteobacteria; Enterobacterales; Enterobacteriaceae; Salmonella
MNNEETFYQAMRRKGVTRRSFLKFCSLAATSLGLGAGMTPKIAWALENKPRIPVVWIHGL WP_001058307.1
hydrogenase [Salmonella enterica] Length: 372\nNumber of predicted TMHs: 2\nExp number of AAs in TMHs: 36.51738\nExp number, first 60 AAs: 19.72849\nTotal prob of N-in: 0.96170\nPOSSIBLE N-term signal sequence\ninside 1 19\nTMhelix 20 39\noutside 40 325\nTMhelix 326 348\ninside 349 372

15379 GCF_002037545.1_ASM203754v1 Salmonella enterica Proteobacteria; Gammaproteobacteria; Enterobacterales; Enterobacteriaceae; Salmonella
MTGDNTLITSHGINRRDFMKLCAALAATMGLSSKAAAEMAESVSNPQRPPVIWIGAQECT WP_000145429.1
hydrogenase 2 small subunit [Salmonella enterica] Length: 372\nNumber of predicted TMHs: 1\nExp number of AAs in TMHs: 24.51309\nExp number, first 60 AAs: 0.2682\nTotal prob of N-in: 0.14402\noutside 1 331\nTMhelix 332 354\ninside 355 372

15380 GCF_002037545.1_ASM203754v1 Salmonella enterica Proteobacteria; Gammaproteobacteria; Enterobacterales; Enterobacteriaceae; Salmonella
MNNEETFYQAMRRKGVTRRSFLKFCSLAATSLGLGAGMTPKIAWALENKPRIPVVWIHGL WP_001058312.1
hydrogenase [Salmonella enterica] Length: 372\nNumber of predicted TMHs: 2\nExp number of AAs in

TMHs: 36.51738\nExp number, first 60 AAs: 19.72849\nTotal prob of N-in: 0.96170\nPOSSIBLE N-term signal sequence\ninside 1 19\nTMhelix 20 39\noutside 40 325\nTMhelix 326 348\ninside 349 372

15381 GCF_002043155.1_ASM204315v1 Salmonella enterica Proteobacteria; Gammaproteobacteria; Enterobacterales; Enterobacteriaceae; Salmonella
MTGDNTLITSHGINRRDFMKLCAALAATMGLSSKAAAEMAESVSNPQRPPVIWIGAQECT WP_000145428.1
hydrogenase 2 small subunit [Salmonella enterica] Length: 372\nNumber of predicted TMHs: 1\nExp number of AAs in TMHs: 24.50677\nExp number, first 60 AAs: 0.2682\nTotal prob of N-in: 0.14390\noutside 1 331\nTMhelix 332 354\ninside 355 372

15382 GCF_002043165.1_ASM204316v1 Salmonella enterica Proteobacteria; Gammaproteobacteria; Enterobacterales; Enterobacteriaceae; Salmonella
MTGDNTLITSHGINRRDFMKLCAALAATMGLSSKAAAEMAESVSNPQRPPVIWIGAQECT WP_000145429.1
hydrogenase 2 small subunit [Salmonella enterica] Length: 372\nNumber of predicted TMHs: 1\nExp number of AAs in TMHs: 24.51309\nExp number, first 60 AAs: 0.2682\nTotal prob of N-in: 0.14402\noutside 1 331\nTMhelix 332 354\ninside 355 372

15383 GCF_002043215.1_ASM204321v1 Salmonella enterica Proteobacteria; Gammaproteobacteria; Enterobacterales; Enterobacteriaceae; Salmonella
MTGDNTLITSHGINRRDFMKLCAALAATMGLSSKAAAEMAESVSNPQRPPVIWIGAQECT WP_000145428.1
hydrogenase 2 small subunit [Salmonella enterica] Length: 372\nNumber of predicted TMHs: 1\nExp number of AAs in TMHs: 24.50677\nExp number, first 60 AAs: 0.2682\nTotal prob of N-in: 0.14390\noutside 1 331\nTMhelix 332 354\ninside 355 372

15384 GCF_002043075.1_ASM204307v1 Salmonella enterica Proteobacteria; Gammaproteobacteria; Enterobacterales; Enterobacteriaceae; Salmonella
MNNEETLYQAMRRKGVTRRSFLKFCSLAATSLGLGAGMTPKIAWALENKPRIPVVWIHGL WP_001752445.1
hydrogenase-1 small chain [Salmonella enterica] Length: 372\nNumber of predicted TMHs: 2\nExp number of AAs in TMHs: 36.5123\nExp number, first 60 AAs: 19.72341\nTotal prob of N-in: 0.96144\nPOSSIBLE N-term signal sequence\ninside 1 19\nTMhelix 20 39\noutside 40 325\nTMhelix 326 348\ninside 349 372

15385 GCF_002043475.1_ASM204347v1 Salmonella enterica Proteobacteria; Gammaproteobacteria; Enterobacterales; Enterobacteriaceae; Salmonella
MTGDNTLITSHGINRRDFMKLCAALAATMGLSSKAAAEMAESVSNPQRPPVIWIGAQECT WP_000145429.1
hydrogenase 2 small subunit [Salmonella enterica] Length: 372\nNumber of predicted TMHs: 1\nExp number of AAs in TMHs: 24.51309\nExp number, first 60 AAs: 0.2682\nTotal prob of N-in: 0.14402\noutside 1 331\nTMhelix 332 354\ninside 355 372

15386 GCF_002043465.1_ASM204346v1 Salmonella enterica Proteobacteria; Gammaproteobacteria; Enterobacterales; Enterobacteriaceae; Salmonella
MTGDNTLITSHGINRRDFMKLCAALAATMGLSSKAAAEMAESVSNPQRPPVIWIGAQECT WP_000145429.1
hydrogenase 2 small subunit [Salmonella enterica] Length: 372\nNumber of predicted TMHs: 1\nExp number of AAs in TMHs: 24.51309\nExp number, first 60 AAs: 0.2682\nTotal prob of N-in: 0.14402\noutside 1 331\nTMhelix 332 354\ninside 355 372

15387 GCF_002043475.1_ASM204347v1 Salmonella enterica Proteobacteria; Gammaproteobacteria; Enterobacterales; Enterobacteriaceae; Salmonella
MNNEETFYQAMRRKGVTRRSFLKFCSLAATSLGLGAGMTPKIAWALENKPRIPVVWIHGL WP_001058312.1
hydrogenase [Salmonella enterica] Length: 372\nNumber of predicted TMHs: 2\nExp number of AAs in TMHs: 36.51738\nExp number, first 60 AAs: 19.72849\nTotal prob of N-in: 0.96170\nPOSSIBLE N-term signal sequence\ninside 1 19\nTMhelix 20 39\noutside 40 325\nTMhelix 326 348\ninside 349 372

15388 GCF_002043465.1_ASM204346v1 Salmonella enterica Proteobacteria; Gammaproteobacteria; Enterobacterales; Enterobacteriaceae; Salmonella
MNNEETFYQAMRRKGVTRRSFLKFCSLAATSLGLSAGMTPKIAWALENKPRIPVVWIHGL WP_001058316.1
hydrogenase [Salmonella enterica] Length: 372\nNumber of predicted TMHs: 2\nExp number of AAs in TMHs: 35.76397\nExp number, first 60 AAs: 18.97201\nTotal prob of N-in: 0.92785\nPOSSIBLE N-term signal sequence\ninside 1 19\nTMhelix 20 39\noutside 40 325\nTMhelix 326 348\ninside 349 372

15389 GCF_002043365.1_ASM204336v1 Salmonella enterica Proteobacteria; Gammaproteobacteria; Enterobacterales; Enterobacteriaceae; Salmonella

MNNEETFYQAMRRKGVTRRSFLKFCSLAATSLGLGAGMTPKIAWALENKPRIPVVWIHGL WP_001058310.1 hydrogenase [Salmonella enterica] Length: 372\nNumber of predicted TMHs: 2\nExp number of AAs in TMHs: 37.4567299999999\nExp number, first 60 AAs: 19.72694\nTotal prob of N-in: 0.96171\nPOSSIBLE N-term signal sequence\ninside 1 19\nTMhelix 20 39\noutside 40 325\nTMhelix 326 348\ninside 349 372

15390 GCF_002043685.1_ASM204368v1 Salmonella enterica Proteobacteria; Gammaproteobacteria; Enterobacterales; Enterobacteriaceae; Salmonella MTGDNTLITSHGINRRDFMKLCAALAATMGLSSKAAAEMAESVSNPQRPPVIWIGAQECT WP_000145429.1 hydrogenase 2 small subunit [Salmonella enterica] Length: 372\nNumber of predicted TMHs: 1\nExp number of AAs in TMHs: 24.51309\nExp number, first 60 AAs: 0.2682\nTotal prob of N-in: 0.14402\noutside 1 331\nTMhelix 332 354\ninside 355 372

15391 GCF_002043845.1_ASM204384v1 Salmonella enterica Proteobacteria; Gammaproteobacteria; Enterobacterales; Enterobacteriaceae; Salmonella MNNEETFYQAMRRKGVTRRSFLKFCSLAATSLGLGAGMTPKIAWALENKPRIPVVWIHGL WP_001058311.1 hydrogenase [Salmonella enterica] Length: 372\nNumber of predicted TMHs: 2\nExp number of AAs in TMHs: 37.4699899999999\nExp number, first 60 AAs: 19.72699\nTotal prob of N-in: 0.96171\nPOSSIBLE N-term signal sequence\ninside 1 19\nTMhelix 20 39\noutside 40 325\nTMhelix 326 348\ninside 349 372

15392 GCF_002043795.1_ASM204379v1 Salmonella enterica Proteobacteria; Gammaproteobacteria; Enterobacterales; Enterobacteriaceae; Salmonella MNNEETFYQAMRRKGVTRRSFLKFCSLAATSLGLGAGMTPKIAWALENKPRIPVVWIHGL WP_001058312.1 hydrogenase [Salmonella enterica] Length: 372\nNumber of predicted TMHs: 2\nExp number of AAs in TMHs: 36.51738\nExp number, first 60 AAs: 19.72849\nTotal prob of N-in: 0.96170\nPOSSIBLE N-term signal sequence\ninside 1 19\nTMhelix 20 39\noutside 40 325\nTMhelix 326 348\ninside 349 372

15393 GCF_002043945.1_ASM204394v1 Salmonella enterica Proteobacteria; Gammaproteobacteria; Enterobacterales; Enterobacteriaceae; Salmonella MTGDNTLITSHGINRRDFMKLCAALAATMGLSSKAAAEMAESVSNPQRPPVIWIGAQECT WP_000145429.1 hydrogenase 2 small subunit [Salmonella enterica] Length: 372\nNumber of predicted TMHs: 1\nExp number of AAs in TMHs: 24.51309\nExp number, first 60 AAs: 0.2682\nTotal prob of N-in: 0.14402\noutside 1 331\nTMhelix 332 354\ninside 355 372

15394 GCF_002043915.1_ASM204391v1 Salmonella enterica Proteobacteria; Gammaproteobacteria; Enterobacterales; Enterobacteriaceae; Salmonella MTGDNTLITSHGINRRDFMKLCAALAATMGLSSKAAAEMAESVSNPQRPPVIWIGAQECT WP_000145428.1 hydrogenase 2 small subunit [Salmonella enterica] Length: 372\nNumber of predicted TMHs: 1\nExp number of AAs in TMHs: 24.50677\nExp number, first 60 AAs: 0.2682\nTotal prob of N-in: 0.14390\noutside 1 331\nTMhelix 332 354\ninside 355 372

15395 GCF_002044005.1_ASM204400v1 Salmonella enterica Proteobacteria; Gammaproteobacteria; Enterobacterales; Enterobacteriaceae; Salmonella MTGDNTLITSHGINRRDFMKLCAALAATMGLSSKAAAEMAESVSNPQRPPVIWIGAQECT WP_000145429.1 hydrogenase 2 small subunit [Salmonella enterica] Length: 372\nNumber of predicted TMHs: 1\nExp number of AAs in TMHs: 24.51309\nExp number, first 60 AAs: 0.2682\nTotal prob of N-in: 0.14402\noutside 1 331\nTMhelix 332 354\ninside 355 372

15396 GCF_002044035.1_ASM204403v1 Salmonella enterica Proteobacteria; Gammaproteobacteria; Enterobacterales; Enterobacteriaceae; Salmonella MTGDNTLITSHGINRRDFMKLCAALAATMGLSSKAAAEMAESVSNPQRPPVIWIGAQECT WP_000145429.1 hydrogenase 2 small subunit [Salmonella enterica] Length: 372\nNumber of predicted TMHs: 1\nExp number of AAs in TMHs: 24.51309\nExp number, first 60 AAs: 0.2682\nTotal prob of N-in: 0.14402\noutside 1 331\nTMhelix 332 354\ninside 355 372

15397 GCF_002044075.1_ASM204407v1 Salmonella enterica Proteobacteria; Gammaproteobacteria; Enterobacterales; Enterobacteriaceae; Salmonella MTGDNTLITSHGINRRDFMKLCAALAATMGLSSKAAAEMAESVSNPQRPPVIWIGAQECT WP_000145429.1 hydrogenase 2 small subunit [Salmonella enterica] Length: 372\nNumber of predicted TMHs: 1\nExp number of AAs in TMHs: 24.51309\nExp number, first 60 AAs: 0.2682\nTotal prob of N-in: 0.14402\noutside 1 331\nTMhelix 332 354\ninside 355 372

15398 GCF_002044125.1_ASM204412v1 Salmonella enterica Proteobacteria; Gammaproteobacteria; Enterobacterales; Enterobacteriaceae; Salmonella
MTGDNTLITSHGINRRDFMKLCAALAATMGLSSKAAAEMAESVSNPQRPPVIWIGAQECT WP_000145429.1
hydrogenase 2 small subunit [Salmonella enterica] Length: 372\nNumber of predicted TMHs: 1\nExp number of AAs in TMHs: 24.51309\nExp number, first 60 AAs: 0.2682\nTotal prob of N-in: 0.14402\noutside 1 331\nTMhelix 332 354\ninside 355 372

15399 GCF_002044325.1_ASM204432v1 Salmonella enterica Proteobacteria; Gammaproteobacteria; Enterobacterales; Enterobacteriaceae; Salmonella
MTGDNTLITSHGINRRDFMKLCAALAATMGLSSKAAAEMAESVSNPQRPPVIWIGAQECT WP_000145429.1
hydrogenase 2 small subunit [Salmonella enterica] Length: 372\nNumber of predicted TMHs: 1\nExp number of AAs in TMHs: 24.51309\nExp number, first 60 AAs: 0.2682\nTotal prob of N-in: 0.14402\noutside 1 331\nTMhelix 332 354\ninside 355 372

15400 GCF_002044355.1_ASM204435v1 Salmonella enterica Proteobacteria; Gammaproteobacteria; Enterobacterales; Enterobacteriaceae; Salmonella
MTGDNTLITSHGINRRDFMKLCAALAATMGLSSKAAAEMAESVSNPQRPPVIWIGAQECT WP_000145429.1
hydrogenase 2 small subunit [Salmonella enterica] Length: 372\nNumber of predicted TMHs: 1\nExp number of AAs in TMHs: 24.51309\nExp number, first 60 AAs: 0.2682\nTotal prob of N-in: 0.14402\noutside 1 331\nTMhelix 332 354\ninside 355 372

15401 GCF_002044405.1_ASM204440v1 Salmonella enterica Proteobacteria; Gammaproteobacteria; Enterobacterales; Enterobacteriaceae; Salmonella
MTGDNTLITSHGINRRDFMKLCAALAATMGLSSKAAAEMAESVSNPQRPPVIWIGAQECT WP_000145429.1
hydrogenase 2 small subunit [Salmonella enterica] Length: 372\nNumber of predicted TMHs: 1\nExp number of AAs in TMHs: 24.51309\nExp number, first 60 AAs: 0.2682\nTotal prob of N-in: 0.14402\noutside 1 331\nTMhelix 332 354\ninside 355 372

15402 GCF_002044325.1_ASM204432v1 Salmonella enterica Proteobacteria; Gammaproteobacteria; Enterobacterales; Enterobacteriaceae; Salmonella
MNNEETFYQAMRRKGVTRRSFLKFCSLAATSLGLGAGMTPKIAWALENKPRIPVVWIHGL WP_001058311.1
hydrogenase [Salmonella enterica] Length: 372\nNumber of predicted TMHs: 2\nExp number of AAs in TMHs: 37.4699899999999\nExp number, first 60 AAs: 19.72699\nTotal prob of N-in: 0.96171\nPOSSIBLE N-term signal sequence\ninside 1 19\nTMhelix 20 39\noutside 40 325\nTMhelix 326 348\ninside 349 372

15403 GCF_002044425.1_ASM204442v1 Salmonella enterica Proteobacteria; Gammaproteobacteria; Enterobacterales; Enterobacteriaceae; Salmonella
MTGDNTLITSHGINRRDFMKLCAALAATMGLSSKAAAEMAESVSNPQRPPVIWIGAQECT WP_000145429.1
hydrogenase 2 small subunit [Salmonella enterica] Length: 372\nNumber of predicted TMHs: 1\nExp number of AAs in TMHs: 24.51309\nExp number, first 60 AAs: 0.2682\nTotal prob of N-in: 0.14402\noutside 1 331\nTMhelix 332 354\ninside 355 372

15404 GCF_002044355.1_ASM204435v1 Salmonella enterica Proteobacteria; Gammaproteobacteria; Enterobacterales; Enterobacteriaceae; Salmonella
MNNEETFYQAMRRKGVTRRSFLKFCSLAATSLGLGAGMTPKIAWALENKPRIPVVWIHGL WP_001058312.1
hydrogenase [Salmonella enterica] Length: 372\nNumber of predicted TMHs: 2\nExp number of AAs in TMHs: 36.51738\nExp number, first 60 AAs: 19.72849\nTotal prob of N-in: 0.96170\nPOSSIBLE N-term signal sequence\ninside 1 19\nTMhelix 20 39\noutside 40 325\nTMhelix 326 348\ninside 349 372

15405 GCF_002044445.1_ASM204444v1 Salmonella enterica Proteobacteria; Gammaproteobacteria; Enterobacterales; Enterobacteriaceae; Salmonella
MTGDNTLITSHGINRRDFMKLCAALAATMGLSSKAAAEMAESVSNPQRPPVIWIGAQECT WP_000145429.1
hydrogenase 2 small subunit [Salmonella enterica] Length: 372\nNumber of predicted TMHs: 1\nExp number of AAs in TMHs: 24.51309\nExp number, first 60 AAs: 0.2682\nTotal prob of N-in: 0.14402\noutside 1 331\nTMhelix 332 354\ninside 355 372

15406 GCF_002044445.1_ASM204444v1 Salmonella enterica Proteobacteria; Gammaproteobacteria; Enterobacterales; Enterobacteriaceae; Salmonella
MNNEETFYQAMRRKGVTRRSFLKFCSLAATSLGLGAGMTPKIAWALENKPRIPVVWIHGL WP_023198237.1
hydrogenase 1, small subunit [Salmonella enterica] Length: 372\nNumber of predicted TMHs: 2\nExp

number of AAs in TMHs: 36.39423\nExp number, first 60 AAs: 19.72877\nTotal prob of N-in: 0.96162\nPOSSIBLE N-term signal sequence\ninside 1 19\nTMhelix 20 39\noutside 40 325\nTMhelix 326 348\ninside 349 372

15407 GCF_002044755.1_ASM204475v1 Salmonella enterica Proteobacteria; Gammaproteobacteria; Enterobacterales; Enterobacteriaceae; Salmonella MTGDNTLITSHGINRRDFMKLCAALAATMGLSSKAAAEMAESVSNPQRPPVIWIGAQECT WP_000145429.1 hydrogenase 2 small subunit [Salmonella enterica] Length: 372\nNumber of predicted TMHs: 1\nExp number of AAs in TMHs: 24.51309\nExp number, first 60 AAs: 0.2682\nTotal prob of N-in: 0.14402\noutside 1 331\nTMhelix 332 354\ninside 355 372

15408 GCF_002044755.1_ASM204475v1 Salmonella enterica Proteobacteria; Gammaproteobacteria; Enterobacterales; Enterobacteriaceae; Salmonella MNNEETFYQAMRRKGVTRRSFLKFCSLAATSLGLGAGMTPKIAWALENKPRIPVVWIHGL WP_001058312.1 hydrogenase [Salmonella enterica] Length: 372\nNumber of predicted TMHs: 2\nExp number of AAs in TMHs: 36.51738\nExp number, first 60 AAs: 19.72849\nTotal prob of N-in: 0.96170\nPOSSIBLE N-term signal sequence\ninside 1 19\nTMhelix 20 39\noutside 40 325\nTMhelix 326 348\ninside 349 372

15409 GCF_002044555.1_ASM204455v1 Salmonella enterica Proteobacteria; Gammaproteobacteria; Enterobacterales; Enterobacteriaceae; Salmonella MNNEETFYQAMRRKGVTRRSFLKFCSLAATSLGLGAGMTPKIAWALENKPRIPVVWIHGL WP_001058312.1 hydrogenase [Salmonella enterica] Length: 372\nNumber of predicted TMHs: 2\nExp number of AAs in TMHs: 36.51738\nExp number, first 60 AAs: 19.72849\nTotal prob of N-in: 0.96170\nPOSSIBLE N-term signal sequence\ninside 1 19\nTMhelix 20 39\noutside 40 325\nTMhelix 326 348\ninside 349 372

15410 GCF_002044705.1_ASM204470v1 Salmonella enterica Proteobacteria; Gammaproteobacteria; Enterobacterales; Enterobacteriaceae; Salmonella MNNEETFYQAMRRKGVTRRSFLKFCSLAATSLGLGAGMTPKIAWALENKPRIPVVWIHGL WP_001058312.1 hydrogenase [Salmonella enterica] Length: 372\nNumber of predicted TMHs: 2\nExp number of AAs in TMHs: 36.51738\nExp number, first 60 AAs: 19.72849\nTotal prob of N-in: 0.96170\nPOSSIBLE N-term signal sequence\ninside 1 19\nTMhelix 20 39\noutside 40 325\nTMhelix 326 348\ninside 349 372

15411 GCF_002045085.1_ASM204508v1 Salmonella enterica Proteobacteria; Gammaproteobacteria; Enterobacterales; Enterobacteriaceae; Salmonella MLKFRTIFLKKIFTLKNSILARRGAVIVIVSAVFTSIMFFAHSWASDKEVAMTLSVLNSN WP_023226644.1 hypothetical protein [Salmonella enterica] Length: 204\nNumber of predicted TMHs: 2\nExp number of AAs in TMHs: 44.99526\nExp number, first 60 AAs: 22.33786\nTotal prob of N-in: 0.87083\nPOSSIBLE N-term signal sequence\ninside 1 19\nTMhelix 20 42\noutside 43 171\nTMhelix 172 194\ninside 195 204

15412 GCF_002045585.1_ASM204558v1 Salmonella enterica Proteobacteria; Gammaproteobacteria; Enterobacterales; Enterobacteriaceae; Salmonella MTGDNTLITSHGINRRDFMKLCAALAATMGLSSKAAAEMAESVSNPQRPPVIWIGAQECT WP_000145429.1 hydrogenase 2 small subunit [Salmonella enterica] Length: 372\nNumber of predicted TMHs: 1\nExp number of AAs in TMHs: 24.51309\nExp number, first 60 AAs: 0.2682\nTotal prob of N-in: 0.14402\noutside 1 331\nTMhelix 332 354\ninside 355 372

15413 GCF_002045475.1_ASM204547v1 Salmonella enterica Proteobacteria; Gammaproteobacteria; Enterobacterales; Enterobacteriaceae; Salmonella MTGDNTLITSHGINRRDFMKLCAALAATMGLSSKAAAEMAESVSNPQRPPVIWIGAQECT WP_000145429.1 hydrogenase 2 small subunit [Salmonella enterica] Length: 372\nNumber of predicted TMHs: 1\nExp number of AAs in TMHs: 24.51309\nExp number, first 60 AAs: 0.2682\nTotal prob of N-in: 0.14402\noutside 1 331\nTMhelix 332 354\ninside 355 372

15414 GCF_002045535.1_ASM204553v1 Salmonella enterica Proteobacteria; Gammaproteobacteria; Enterobacterales; Enterobacteriaceae; Salmonella MTGDNTLITSHGINRRDFMKLCAALAATMGLSSKAAAEMAESVSNPQRPPVIWIGAQECT WP_000145429.1 hydrogenase 2 small subunit [Salmonella enterica] Length: 372\nNumber of predicted TMHs: 1\nExp number of AAs in TMHs: 24.51309\nExp number, first 60 AAs: 0.2682\nTotal prob of N-in: 0.14402\noutside 1 331\nTMhelix 332 354\ninside 355 372

15415 GCF_002045495.1_ASM204549v1 Salmonella enterica Proteobacteria; Gammaproteobacteria; Enterobacterales; Enterobacteriaceae; Salmonella

MTGDNTLITSHGINRRDFMKLCAALAATMGLSSKAAAEMAESVSNPQRPPVIWIGAQECT WP_000145429.1 hydrogenase 2 small subunit [Salmonella enterica] Length: 372\nNumber of predicted TMHs: 1\nExp number of AAs in TMHs: 24.51309\nExp number, first 60 AAs: 0.2682\nTotal prob of N-in: 0.14402\noutside 1 331\nTMhelix 332 354\ninside 355 372

15416 GCF_002045745.1_ASM204574v1 Salmonella enterica Proteobacteria; Gammaproteobacteria; Enterobacterales; Enterobacteriaceae; Salmonella MNNEETFYQAMRRKGVTRRSFLKFCSLAATSLGLGAGMTPKIAWALENKPRIPVVWIHGL WP_001058311.1 hydrogenase [Salmonella enterica] Length: 372\nNumber of predicted TMHs: 2\nExp number of AAs in TMHs: 37.4699899999999\nExp number, first 60 AAs: 19.72699\nTotal prob of N-in: 0.96171\nPOSSIBLE N-term signal sequence\ninside 1 19\nTMhelix 20 39\noutside 40 325\nTMhelix 326 348\ninside 349 372

15417 GCF_002046265.1_ASM204626v1 Salmonella enterica Proteobacteria; Gammaproteobacteria; Enterobacterales; Enterobacteriaceae; Salmonella MNNEETFYQAMRRKGVTRRSFLKFCSLAATSLGLGAGMTPKIAWALENKPRIPVVWIHGL WP_001058310.1 hydrogenase [Salmonella enterica] Length: 372\nNumber of predicted TMHs: 2\nExp number of AAs in TMHs: 37.4567299999999\nExp number, first 60 AAs: 19.72694\nTotal prob of N-in: 0.96171\nPOSSIBLE N-term signal sequence\ninside 1 19\nTMhelix 20 39\noutside 40 325\nTMhelix 326 348\ninside 349 372

15418 GCF_002046415.1_ASM204641v1 Salmonella enterica Proteobacteria; Gammaproteobacteria; Enterobacterales; Enterobacteriaceae; Salmonella MTGDNTLITSHGINRRDFMKLCAALAATMGLSSKAAAEMAESVSNPQRPPVIWIGAQECT WP_000145429.1 hydrogenase 2 small subunit [Salmonella enterica] Length: 372\nNumber of predicted TMHs: 1\nExp number of AAs in TMHs: 24.51309\nExp number, first 60 AAs: 0.2682\nTotal prob of N-in: 0.14402\noutside 1 331\nTMhelix 332 354\ninside 355 372

15419 GCF_002046805.1_ASM204680v1 Salmonella enterica Proteobacteria; Gammaproteobacteria; Enterobacterales; Enterobacteriaceae; Salmonella MTGDNTLITSHGINRRDFMKLCAALAATMGLSSKAAAEMAESVSNPQRPPVIWIGAQECT WP_000145429.1 hydrogenase 2 small subunit [Salmonella enterica] Length: 372\nNumber of predicted TMHs: 1\nExp number of AAs in TMHs: 24.51309\nExp number, first 60 AAs: 0.2682\nTotal prob of N-in: 0.14402\noutside 1 331\nTMhelix 332 354\ninside 355 372

15420 GCF_002047185.1_ASM204718v1 Salmonella enterica Proteobacteria; Gammaproteobacteria; Enterobacterales; Enterobacteriaceae; Salmonella MTGDNTLITSHGINRRDFMKLCAALAATMGLSSKAAAEMAESVSNPQRPPVIWIGAQECT WP_000145429.1 hydrogenase 2 small subunit [Salmonella enterica] Length: 372\nNumber of predicted TMHs: 1\nExp number of AAs in TMHs: 24.51309\nExp number, first 60 AAs: 0.2682\nTotal prob of N-in: 0.14402\noutside 1 331\nTMhelix 332 354\ninside 355 372

15421 GCF_002048755.1_ASM204875v1 Salmonella enterica Proteobacteria; Gammaproteobacteria; Enterobacterales; Enterobacteriaceae; Salmonella MTGDNTLITSHGINRRDFMKLCAALAATMGLSSKAAAEMAESVSNPQRPPVIWIGAQECT WP_000145431.1 hydrogenase 2 small subunit [Salmonella enterica] Length: 372\nNumber of predicted TMHs: 1\nExp number of AAs in TMHs: 24.51505\nExp number, first 60 AAs: 0.2682\nTotal prob of N-in: 0.14406\noutside 1 331\nTMhelix 332 354\ninside 355 372

15422 GCF_002050795.1_ASM205079v1 Salmonella enterica Proteobacteria; Gammaproteobacteria; Enterobacterales; Enterobacteriaceae; Salmonella MNNEETFYQAMRRKGVTRRSFLKFCSLAATSLGLGAGMTPKIAWALENKPRIPVVWIHGL WP_001058307.1 hydrogenase [Salmonella enterica] Length: 372\nNumber of predicted TMHs: 2\nExp number of AAs in TMHs: 36.51738\nExp number, first 60 AAs: 19.72849\nTotal prob of N-in: 0.96170\nPOSSIBLE N-term signal sequence\ninside 1 19\nTMhelix 20 39\noutside 40 325\nTMhelix 326 348\ninside 349 372

15423 GCF_002051115.1_ASM205111v1 Salmonella enterica Proteobacteria; Gammaproteobacteria; Enterobacterales; Enterobacteriaceae; Salmonella MNNEETFYQAMRRKGVTRRSFLKFCSLAATSLGLGAGMTPKIAWALENKPRIPVVWIHGL WP_001058312.1 hydrogenase [Salmonella enterica] Length: 372\nNumber of predicted TMHs: 2\nExp number of AAs in TMHs: 36.51738\nExp number, first 60 AAs: 19.72849\nTotal prob of N-in: 0.96170\nPOSSIBLE N-term signal sequence\ninside 1 19\nTMhelix 20 39\noutside 40 325\nTMhelix 326 348\ninside 349 372

15424	GCF_002051215.1_ASM205121v1	Salmonella enterica	Proteobacteria; Gammaproteobacteria; Enterobacterales; Enterobacteriaceae; Salmonella
MTGDNTLITSHGINRRDFMKLCAALAATMGLSSKAAAEMAESVSNPQRPPVIWIGAQECT	WP_000145429.1
hydrogenase 2 small subunit [Salmonella enterica]	Length: 372\nNumber of predicted TMHs: 1\nExp number of AAs in TMHs: 24.51309\nExp number, first 60 AAs: 0.2682\nTotal prob of N-in: 0.14402\noutside 1 331\nTMhelix 332 354\ninside 355 372

15425	GCF_002051235.1_ASM205123v1	Salmonella enterica	Proteobacteria; Gammaproteobacteria; Enterobacterales; Enterobacteriaceae; Salmonella
MTGDNTLITSHGINRRDFMKLCAALAATMGLSSKAAAEMAESVSNPQRPPVIWIGAQECT	WP_000145429.1
hydrogenase 2 small subunit [Salmonella enterica]	Length: 372\nNumber of predicted TMHs: 1\nExp number of AAs in TMHs: 24.51309\nExp number, first 60 AAs: 0.2682\nTotal prob of N-in: 0.14402\noutside 1 331\nTMhelix 332 354\ninside 355 372

15426	GCF_002051215.1_ASM205121v1	Salmonella enterica	Proteobacteria; Gammaproteobacteria; Enterobacterales; Enterobacteriaceae; Salmonella
MNNEETFYQAMRRKGVTRRSFLKFCSLAATSLGLGAGMTPKIAWALENKPRIPVVWIHGL	WP_001058312.1
hydrogenase [Salmonella enterica]	Length: 372\nNumber of predicted TMHs: 2\nExp number of AAs in TMHs: 36.51738\nExp number, first 60 AAs: 19.72849\nTotal prob of N-in: 0.96170\nPOSSIBLE N-term signal sequence\ninside 1 19\nTMhelix 20 39\noutside 40 325\nTMhelix 326 348\ninside 349 372

15427	GCF_002051235.1_ASM205123v1	Salmonella enterica	Proteobacteria; Gammaproteobacteria; Enterobacterales; Enterobacteriaceae; Salmonella
MNNEETFYQAMRRKGVTRRSFLKFCSLAATSLGLGAGMTPKIAWALENKPRIPVVWIHGL	WP_001058312.1
hydrogenase [Salmonella enterica]	Length: 372\nNumber of predicted TMHs: 2\nExp number of AAs in TMHs: 36.51738\nExp number, first 60 AAs: 19.72849\nTotal prob of N-in: 0.96170\nPOSSIBLE N-term signal sequence\ninside 1 19\nTMhelix 20 39\noutside 40 325\nTMhelix 326 348\ninside 349 372

15428	GCF_002051165.1_ASM205116v1	Salmonella enterica	Proteobacteria; Gammaproteobacteria; Enterobacterales; Enterobacteriaceae; Salmonella
MNNEETFYQAMRRKGVTRRSFLKFCSLAATSLGLGAGMTPKIAWALENKPRIPVVWIHGL	WP_001058312.1
hydrogenase [Salmonella enterica]	Length: 372\nNumber of predicted TMHs: 2\nExp number of AAs in TMHs: 36.51738\nExp number, first 60 AAs: 19.72849\nTotal prob of N-in: 0.96170\nPOSSIBLE N-term signal sequence\ninside 1 19\nTMhelix 20 39\noutside 40 325\nTMhelix 326 348\ninside 349 372

15429	GCF_002051135.1_ASM205113v1	Salmonella enterica	Proteobacteria; Gammaproteobacteria; Enterobacterales; Enterobacteriaceae; Salmonella
MNNEETFYQAMRRKGVTRRSFLKFCSLAATSLGLGAGMTPKIAWALENKPRIPVVWIHGL	WP_001058307.1
hydrogenase [Salmonella enterica]	Length: 372\nNumber of predicted TMHs: 2\nExp number of AAs in TMHs: 36.51738\nExp number, first 60 AAs: 19.72849\nTotal prob of N-in: 0.96170\nPOSSIBLE N-term signal sequence\ninside 1 19\nTMhelix 20 39\noutside 40 325\nTMhelix 326 348\ninside 349 372

15430	GCF_002051655.1_ASM205165v1	Salmonella enterica	Proteobacteria; Gammaproteobacteria; Enterobacterales; Enterobacteriaceae; Salmonella
MNNEETFYQAMRRKGVTRRSFLKFCSLAATSLGLGAGMTPKIAWALENKPRIPVVWIHGL	WP_001058312.1
hydrogenase [Salmonella enterica]	Length: 372\nNumber of predicted TMHs: 2\nExp number of AAs in TMHs: 36.51738\nExp number, first 60 AAs: 19.72849\nTotal prob of N-in: 0.96170\nPOSSIBLE N-term signal sequence\ninside 1 19\nTMhelix 20 39\noutside 40 325\nTMhelix 326 348\ninside 349 372

15431	GCF_002051385.1_ASM205138v1	Salmonella enterica	Proteobacteria; Gammaproteobacteria; Enterobacterales; Enterobacteriaceae; Salmonella
MNNEETFYQAMRRKGVTRRSFLKFCSLAATSLGLGAGMTPKIAWALENKPRIPVVWIHGL	WP_001058312.1
hydrogenase [Salmonella enterica]	Length: 372\nNumber of predicted TMHs: 2\nExp number of AAs in TMHs: 36.51738\nExp number, first 60 AAs: 19.72849\nTotal prob of N-in: 0.96170\nPOSSIBLE N-term signal sequence\ninside 1 19\nTMhelix 20 39\noutside 40 325\nTMhelix 326 348\ninside 349 372

15432	GCF_002051625.1_ASM205162v1	Salmonella enterica	Proteobacteria; Gammaproteobacteria; Enterobacterales; Enterobacteriaceae; Salmonella
MTGDNTLITSHGINRRDFMKLCAALAATMGLSSKAAAEMAESVSNPQRPPVIWIGAQECT	WP_000145429.1
hydrogenase 2 small subunit [Salmonella enterica]	Length: 372\nNumber of predicted TMHs: 1\nExp

number of AAs in TMHs: 24.51309\nExp number, first 60 AAs: 0.2682\nTotal prob of N-in: 0.14402\noutside 1 331\nTMhelix 332 354\ninside 355 372

15433 GCF_002051625.1_ASM205162v1 Salmonella enterica Proteobacteria; Gammaproteobacteria; Enterobacterales; Enterobacteriaceae; Salmonella
MNNEETFYQAMRRKGVTRRSFLKFCSLAATSLGLGAGMTPKIAWALENKPRIPVVWIHGL WP_001058312.1
hydrogenase [Salmonella enterica] Length: 372\nNumber of predicted TMHs: 2\nExp number of AAs in TMHs: 36.51738\nExp number, first 60 AAs: 19.72849\nTotal prob of N-in: 0.96170\nPOSSIBLE N-term signal sequence\ninside 1 19\nTMhelix 20 39\noutside 40 325\nTMhelix 326 348\ninside 349 372

15434 GCF_002051305.1_ASM205130v1 Salmonella enterica Proteobacteria; Gammaproteobacteria; Enterobacterales; Enterobacteriaceae; Salmonella
MTGDNTLITSHGINRRDFMKLCAALAATMGLSSKAAAEMAESVSNPQRPPVIWIGAQECT WP_015589585.1
hydrogenase 2 small subunit [Salmonella enterica] Length: 372\nNumber of predicted TMHs: 1\nExp number of AAs in TMHs: 24.51304\nExp number, first 60 AAs: 0.2682\nTotal prob of N-in: 0.14402\noutside 1 331\nTMhelix 332 354\ninside 355 372

15435 GCF_002051405.1_ASM205140v1 Salmonella enterica Proteobacteria; Gammaproteobacteria; Enterobacterales; Enterobacteriaceae; Salmonella
MNNEETFYQAMRRKGVTRRSFLKFCSLAATSLGLGAGMTPKIAWALENKPRIPVVWIHGL WP_001058312.1
hydrogenase [Salmonella enterica] Length: 372\nNumber of predicted TMHs: 2\nExp number of AAs in TMHs: 36.51738\nExp number, first 60 AAs: 19.72849\nTotal prob of N-in: 0.96170\nPOSSIBLE N-term signal sequence\ninside 1 19\nTMhelix 20 39\noutside 40 325\nTMhelix 326 348\ninside 349 372

15436 GCF_002051705.1_ASM205170v1 Salmonella enterica Proteobacteria; Gammaproteobacteria; Enterobacterales; Enterobacteriaceae; Salmonella
MTGDNTLITSHGINRRDFMKLCAALAATMGLSSKAAAEMAESVSNPQRPPVIWIGAQECT WP_000145429.1
hydrogenase 2 small subunit [Salmonella enterica] Length: 372\nNumber of predicted TMHs: 1\nExp number of AAs in TMHs: 24.51309\nExp number, first 60 AAs: 0.2682\nTotal prob of N-in: 0.14402\noutside 1 331\nTMhelix 332 354\ninside 355 372

15437 GCF_002052185.1_ASM205218v1 Salmonella enterica Proteobacteria; Gammaproteobacteria; Enterobacterales; Enterobacteriaceae; Salmonella
MNNEETFYQAMRRKGVTRRSFLKFCSLAATSLGLGAGMTPKIAWALENKPRIPVVWIHGL WP_001058312.1
hydrogenase [Salmonella enterica] Length: 372\nNumber of predicted TMHs: 2\nExp number of AAs in TMHs: 36.51738\nExp number, first 60 AAs: 19.72849\nTotal prob of N-in: 0.96170\nPOSSIBLE N-term signal sequence\ninside 1 19\nTMhelix 20 39\noutside 40 325\nTMhelix 326 348\ninside 349 372

15438 GCF_002051965.1_ASM205196v1 Salmonella enterica Proteobacteria; Gammaproteobacteria; Enterobacterales; Enterobacteriaceae; Salmonella
MNNEETFYQAMRRKGVTRRSFLKFCSLAATSLGLGAGMTPKIAWALENKPRIPVVWIHGL WP_001058312.1
hydrogenase [Salmonella enterica] Length: 372\nNumber of predicted TMHs: 2\nExp number of AAs in TMHs: 36.51738\nExp number, first 60 AAs: 19.72849\nTotal prob of N-in: 0.96170\nPOSSIBLE N-term signal sequence\ninside 1 19\nTMhelix 20 39\noutside 40 325\nTMhelix 326 348\ninside 349 372

15439 GCF_002052135.1_ASM205213v1 Salmonella enterica Proteobacteria; Gammaproteobacteria; Enterobacterales; Enterobacteriaceae; Salmonella
MNNEETFYQAMRRKGVTRRSFLKFCSLAATSLGLGAGMTPKIAWALENKPRIPVVWIHGL WP_001058312.1
hydrogenase [Salmonella enterica] Length: 372\nNumber of predicted TMHs: 2\nExp number of AAs in TMHs: 36.51738\nExp number, first 60 AAs: 19.72849\nTotal prob of N-in: 0.96170\nPOSSIBLE N-term signal sequence\ninside 1 19\nTMhelix 20 39\noutside 40 325\nTMhelix 326 348\ninside 349 372

15440 GCF_002052035.1_ASM205203v1 Salmonella enterica Proteobacteria; Gammaproteobacteria; Enterobacterales; Enterobacteriaceae; Salmonella
MNNEETFYQAMRRKGVTRRSFLKFCSLAATSLGLGAGMTPKIAWALENKPRIPVVWIHGL WP_001058312.1
hydrogenase [Salmonella enterica] Length: 372\nNumber of predicted TMHs: 2\nExp number of AAs in TMHs: 36.51738\nExp number, first 60 AAs: 19.72849\nTotal prob of N-in: 0.96170\nPOSSIBLE N-term signal sequence\ninside 1 19\nTMhelix 20 39\noutside 40 325\nTMhelix 326 348\ninside 349 372

15441 GCF_002052305.1_ASM205230v1 Salmonella enterica Proteobacteria; Gammaproteobacteria; Enterobacterales; Enterobacteriaceae; Salmonella

MTGDNTLITSHGINRRDFMKLCAALAATMGLSSKAAAEMAESVSNPQRPPVIWIGAQECT WP_000145429.1 hydrogenase 2 small subunit [Salmonella enterica] Length: 372\nNumber of predicted TMHs: 1\nExp number of AAs in TMHs: 24.51309\nExp number, first 60 AAs: 0.2682\nTotal prob of N-in: 0.14402\noutside 1 331\nTMhelix 332 354\ninside 355 372

15442 GCF_002052355.1_ASM205235v1 Salmonella enterica Proteobacteria; Gammaproteobacteria; Enterobacterales; Enterobacteriaceae; Salmonella MTGDNTLITSHGINRRDFMKLCAALAATMGLSSKAAAEMAESVSNPQRPPVIWIGAQECT WP_000145429.1 hydrogenase 2 small subunit [Salmonella enterica] Length: 372\nNumber of predicted TMHs: 1\nExp number of AAs in TMHs: 24.51309\nExp number, first 60 AAs: 0.2682\nTotal prob of N-in: 0.14402\noutside 1 331\nTMhelix 332 354\ninside 355 372

15443 GCF_002052355.1_ASM205235v1 Salmonella enterica Proteobacteria; Gammaproteobacteria; Enterobacterales; Enterobacteriaceae; Salmonella MNNEETFYQAMRRKGVTRRSFLKFCSLAATSLGLGAGMTPKIAWALENKPRIPVVWIHGL WP_001058312.1 hydrogenase [Salmonella enterica] Length: 372\nNumber of predicted TMHs: 2\nExp number of AAs in TMHs: 36.51738\nExp number, first 60 AAs: 19.72849\nTotal prob of N-in: 0.96170\nPOSSIBLE N-term signal sequence\ninside 1 19\nTMhelix 20 39\noutside 40 325\nTMhelix 326 348\ninside 349 372

15444 GCF_002052855.1_ASM205285v1 Salmonella enterica Proteobacteria; Gammaproteobacteria; Enterobacterales; Enterobacteriaceae; Salmonella MNNEETFYQAMRRKGVTRRSFLKFCSLAATSLGLGAGMTPKIAWALENKPRIPVVWIHGL WP_001058312.1 hydrogenase [Salmonella enterica] Length: 372\nNumber of predicted TMHs: 2\nExp number of AAs in TMHs: 36.51738\nExp number, first 60 AAs: 19.72849\nTotal prob of N-in: 0.96170\nPOSSIBLE N-term signal sequence\ninside 1 19\nTMhelix 20 39\noutside 40 325\nTMhelix 326 348\ninside 349 372

15445 GCF_002052635.1_ASM205263v1 Salmonella enterica Proteobacteria; Gammaproteobacteria; Enterobacterales; Enterobacteriaceae; Salmonella MNNEETFYQAMRRKGVTRRSFLKFCSLAATSLGLGAGMTPKIAWALENKPRIPVVWIHGL WP_001058312.1 hydrogenase [Salmonella enterica] Length: 372\nNumber of predicted TMHs: 2\nExp number of AAs in TMHs: 36.51738\nExp number, first 60 AAs: 19.72849\nTotal prob of N-in: 0.96170\nPOSSIBLE N-term signal sequence\ninside 1 19\nTMhelix 20 39\noutside 40 325\nTMhelix 326 348\ninside 349 372

15446 GCF_002052925.1_ASM205292v1 Salmonella enterica Proteobacteria; Gammaproteobacteria; Enterobacterales; Enterobacteriaceae; Salmonella MTGDNTLITSHGINRRDFMKLCAALAATMGLSSKAAAEMAESVSNPQRPPVIWIGAQECT WP_000145429.1 hydrogenase 2 small subunit [Salmonella enterica] Length: 372\nNumber of predicted TMHs: 1\nExp number of AAs in TMHs: 24.51309\nExp number, first 60 AAs: 0.2682\nTotal prob of N-in: 0.14402\noutside 1 331\nTMhelix 332 354\ninside 355 372

15447 GCF_002052925.1_ASM205292v1 Salmonella enterica Proteobacteria; Gammaproteobacteria; Enterobacterales; Enterobacteriaceae; Salmonella MNNEETFYQAMRRKGVTRRSFLKFCSLAATSLGLGAGMTPKIAWALENKPRIPVVWIHGL WP_080084014.1 [Ni/Fe] hydrogenase small subunit, partial [Salmonella enterica] Length: 370\nNumber of predicted TMHs: 2\nExp number of AAs in TMHs: 37.4664\nExp number, first 60 AAs: 19.72616\nTotal prob of N-in: 0.96171\nPOSSIBLE N-term signal sequence\ninside 1 19\nTMhelix 20 39\noutside 40 325\nTMhelix 326 348\ninside 349 370

15448 GCF_002052685.1_ASM205268v1 Salmonella enterica Proteobacteria; Gammaproteobacteria; Enterobacterales; Enterobacteriaceae; Salmonella MNNEETFYQAMRRKGVTRRSFLKFCSLAATSLGLGAGMTPKIAWALENKPRIPVVWIHGL WP_001058312.1 hydrogenase [Salmonella enterica] Length: 372\nNumber of predicted TMHs: 2\nExp number of AAs in TMHs: 36.51738\nExp number, first 60 AAs: 19.72849\nTotal prob of N-in: 0.96170\nPOSSIBLE N-term signal sequence\ninside 1 19\nTMhelix 20 39\noutside 40 325\nTMhelix 326 348\ninside 349 372

15449 GCF_002052745.1_ASM205274v1 Salmonella enterica Proteobacteria; Gammaproteobacteria; Enterobacterales; Enterobacteriaceae; Salmonella MNNEETFYQAMRRKGVTRRSFLKFCSLAATSLGLGAGMTPKIAWALENKPRIPVVWIHGL WP_001058312.1 hydrogenase [Salmonella enterica] Length: 372\nNumber of predicted TMHs: 2\nExp number of AAs in TMHs: 36.51738\nExp number, first 60 AAs: 19.72849\nTotal prob of N-in: 0.96170\nPOSSIBLE N-term signal sequence\ninside 1 19\nTMhelix 20 39\noutside 40 325\nTMhelix 326 348\ninside 349 372

15450 GCF_002052995.1_ASM205299v1 Salmonella enterica Proteobacteria; Gammaproteobacteria; Enterobacterales; Enterobacteriaceae; Salmonella
MTGDNTLITSHGINRRDFMKLCAALAATMGLSSKAAAEMAESVSNPQRPPVIWIGAQECT WP_000145431.1
hydrogenase 2 small subunit [Salmonella enterica] Length: 372\nNumber of predicted TMHs: 1\nExp number of AAs in TMHs: 24.51505\nExp number, first 60 AAs: 0.2682\nTotal prob of N-in: 0.14406\noutside 1 331\nTMhelix 332 354\ninside 355 372

15451 GCF_002053375.1_ASM205337v1 Salmonella enterica Proteobacteria; Gammaproteobacteria; Enterobacterales; Enterobacteriaceae; Salmonella
MTGDNTLITSHGINRRDFMKLCAALAATMGLSSKAAAEMAESVSNPQRPPVIWIGAQECT WP_000145429.1
hydrogenase 2 small subunit [Salmonella enterica] Length: 372\nNumber of predicted TMHs: 1\nExp number of AAs in TMHs: 24.51309\nExp number, first 60 AAs: 0.2682\nTotal prob of N-in: 0.14402\noutside 1 331\nTMhelix 332 354\ninside 355 372

15452 GCF_002053485.1_ASM205348v1 Salmonella enterica Proteobacteria; Gammaproteobacteria; Enterobacterales; Enterobacteriaceae; Salmonella
MTGDNTLITSHGINRRDFMKLCAALAATMGLSSKAAAEMAESVSNPQRPPVIWIGAQECT WP_000145429.1
hydrogenase 2 small subunit [Salmonella enterica] Length: 372\nNumber of predicted TMHs: 1\nExp number of AAs in TMHs: 24.51309\nExp number, first 60 AAs: 0.2682\nTotal prob of N-in: 0.14402\noutside 1 331\nTMhelix 332 354\ninside 355 372

15453 GCF_002053415.1_ASM205341v1 Salmonella enterica Proteobacteria; Gammaproteobacteria; Enterobacterales; Enterobacteriaceae; Salmonella
MNNEETFYQAMRRKGVTRRSFLKFCSLAATSLGLGAGMTPKIAWALENKPRIPVVWIHGL WP_001058312.1
hydrogenase [Salmonella enterica] Length: 372\nNumber of predicted TMHs: 2\nExp number of AAs in TMHs: 36.51738\nExp number, first 60 AAs: 19.72849\nTotal prob of N-in: 0.96170\nPOSSIBLE N-term signal sequence\ninside 1 19\nTMhelix 20 39\noutside 40 325\nTMhelix 326 348\ninside 349 372

15454 GCF_002053485.1_ASM205348v1 Salmonella enterica Proteobacteria; Gammaproteobacteria; Enterobacterales; Enterobacteriaceae; Salmonella
MNNEETFYQAMRRKGVTRRSFLKFCSLAATSLGLGAGMTPKIAWALENKPRIPVVWIHGL WP_001058312.1
hydrogenase [Salmonella enterica] Length: 372\nNumber of predicted TMHs: 2\nExp number of AAs in TMHs: 36.51738\nExp number, first 60 AAs: 19.72849\nTotal prob of N-in: 0.96170\nPOSSIBLE N-term signal sequence\ninside 1 19\nTMhelix 20 39\noutside 40 325\nTMhelix 326 348\ninside 349 372

15455 GCF_002053075.1_ASM205307v1 Salmonella enterica Proteobacteria; Gammaproteobacteria; Enterobacterales; Enterobacteriaceae; Salmonella
MNNEETFYQAMRRKGVTRRSFLKFCSLAATSLGLGAGMTPKIAWALENKPRIPVVWIHGL WP_023243720.1
hydrogenase 1, small subunit [Salmonella enterica] Length: 372\nNumber of predicted TMHs: 2\nExp number of AAs in TMHs: 36.52147\nExp number, first 60 AAs: 19.72838\nTotal prob of N-in: 0.96170\nPOSSIBLE N-term signal sequence\ninside 1 19\nTMhelix 20 39\noutside 40 325\nTMhelix 326 348\ninside 349 372

15456 GCF_002053225.1_ASM205322v1 Salmonella enterica Proteobacteria; Gammaproteobacteria; Enterobacterales; Enterobacteriaceae; Salmonella
MNNEETFYQAMRRKGVTRRSFLKFCSLAATSLGLGAGMTPKIAWALENKPRIPVVWIHGL WP_001058312.1
hydrogenase [Salmonella enterica] Length: 372\nNumber of predicted TMHs: 2\nExp number of AAs in TMHs: 36.51738\nExp number, first 60 AAs: 19.72849\nTotal prob of N-in: 0.96170\nPOSSIBLE N-term signal sequence\ninside 1 19\nTMhelix 20 39\noutside 40 325\nTMhelix 326 348\ninside 349 372

15457 GCF_002053585.1_ASM205358v1 Salmonella enterica Proteobacteria; Gammaproteobacteria; Enterobacterales; Enterobacteriaceae; Salmonella
MNNEETFYQAMRRKGVTRRSFLKFCSLAATSLGLGAGMTPKIAWALENKPRIPVVWIHGL WP_001058312.1
hydrogenase [Salmonella enterica] Length: 372\nNumber of predicted TMHs: 2\nExp number of AAs in TMHs: 36.51738\nExp number, first 60 AAs: 19.72849\nTotal prob of N-in: 0.96170\nPOSSIBLE N-term signal sequence\ninside 1 19\nTMhelix 20 39\noutside 40 325\nTMhelix 326 348\ninside 349 372

15458 GCF_002054115.1_ASM205411v1 Salmonella enterica Proteobacteria; Gammaproteobacteria; Enterobacterales; Enterobacteriaceae; Salmonella
MTGDNTLITSHGINRRDFMKLCAALAATMGLSSKAAAEMAESVSNPQRPPVIWIGAQECT WP_000145429.1
hydrogenase 2 small subunit [Salmonella enterica] Length: 372\nNumber of predicted TMHs: 1\nExp

number of AAs in TMHs: 24.51309\nExp number, first 60 AAs: 0.2682\nTotal prob of N-in: 0.14402\noutside 1 331\nTMhelix 332 354\ninside 355 372

15459 GCF_002054005.1_ASM205400v1 Salmonella enterica Proteobacteria; Gammaproteobacteria; Enterobacterales; Enterobacteriaceae; Salmonella
MTGDNTLITSHGINRRDFMKLCAALAATMGLSSKAAAEMAESVSNPQRPPVIWIGAQECT WP_000145429.1
hydrogenase 2 small subunit [Salmonella enterica] Length: 372\nNumber of predicted TMHs: 1\nExp number of AAs in TMHs: 24.51309\nExp number, first 60 AAs: 0.2682\nTotal prob of N-in: 0.14402\noutside 1 331\nTMhelix 332 354\ninside 355 372

15460 GCF_002054125.1_ASM205412v1 Salmonella enterica Proteobacteria; Gammaproteobacteria; Enterobacterales; Enterobacteriaceae; Salmonella
MTGDNTLITSHGINRRDFMKLCAALAATMGLSSKAAAEMAESVSNPQRPPVIWIGAQECT WP_000145429.1
hydrogenase 2 small subunit [Salmonella enterica] Length: 372\nNumber of predicted TMHs: 1\nExp number of AAs in TMHs: 24.51309\nExp number, first 60 AAs: 0.2682\nTotal prob of N-in: 0.14402\noutside 1 331\nTMhelix 332 354\ninside 355 372

15461 GCF_002054115.1_ASM205411v1 Salmonella enterica Proteobacteria; Gammaproteobacteria; Enterobacterales; Enterobacteriaceae; Salmonella
MNNEETFYQAMRRKGVTRRSFLKFCSLAATSLGLGAGMTPKIAWALENKPRIPVVWIHGL WP_001058312.1
hydrogenase [Salmonella enterica] Length: 372\nNumber of predicted TMHs: 2\nExp number of AAs in TMHs: 36.51738\nExp number, first 60 AAs: 19.72849\nTotal prob of N-in: 0.96170\nPOSSIBLE N-term signal sequence\ninside 1 19\nTMhelix 20 39\noutside 40 325\nTMhelix 326 348\ninside 349 372

15462 GCF_002054125.1_ASM205412v1 Salmonella enterica Proteobacteria; Gammaproteobacteria; Enterobacterales; Enterobacteriaceae; Salmonella
MNNEETFYQAMRRKGVTRRSFLKFCSLAATSLGLGAGMTPKIAWALENKPRIPVVWIHGL WP_001058312.1
hydrogenase [Salmonella enterica] Length: 372\nNumber of predicted TMHs: 2\nExp number of AAs in TMHs: 36.51738\nExp number, first 60 AAs: 19.72849\nTotal prob of N-in: 0.96170\nPOSSIBLE N-term signal sequence\ninside 1 19\nTMhelix 20 39\noutside 40 325\nTMhelix 326 348\ninside 349 372

15463 GCF_002054005.1_ASM205400v1 Salmonella enterica Proteobacteria; Gammaproteobacteria; Enterobacterales; Enterobacteriaceae; Salmonella
MNNEETFYQAMRRKGVTRRSFLKFCSLAATSLGLGAGMTPKIAWALENKPRIPVVWIHGL WP_001058308.1
hydrogenase [Salmonella enterica] Length: 372\nNumber of predicted TMHs: 2\nExp number of AAs in TMHs: 39.08204\nExp number, first 60 AAs: 19.72474\nTotal prob of N-in: 0.96174\nPOSSIBLE N-term signal sequence\ninside 1 19\nTMhelix 20 39\noutside 40 325\nTMhelix 326 348\ninside 349 372

15464 GCF_002054205.1_ASM205420v1 Salmonella enterica Proteobacteria; Gammaproteobacteria; Enterobacterales; Enterobacteriaceae; Salmonella
MTGDNTLITSHGINRRDFMKLCAALAATMGLSSKAAAEMAESVSNPQRPPVIWIGAQECT WP_000145429.1
hydrogenase 2 small subunit [Salmonella enterica] Length: 372\nNumber of predicted TMHs: 1\nExp number of AAs in TMHs: 24.51309\nExp number, first 60 AAs: 0.2682\nTotal prob of N-in: 0.14402\noutside 1 331\nTMhelix 332 354\ninside 355 372

15465 GCF_002054265.1_ASM205426v1 Salmonella enterica Proteobacteria; Gammaproteobacteria; Enterobacterales; Enterobacteriaceae; Salmonella
MTGDNTLITSHGINRRDFMKLCAALAATMGLSSKAAAEMAESVSNPQRPPVIWIGAQECT WP_000145429.1
hydrogenase 2 small subunit [Salmonella enterica] Length: 372\nNumber of predicted TMHs: 1\nExp number of AAs in TMHs: 24.51309\nExp number, first 60 AAs: 0.2682\nTotal prob of N-in: 0.14402\noutside 1 331\nTMhelix 332 354\ninside 355 372

15466 GCF_002054255.1_ASM205425v1 Salmonella enterica Proteobacteria; Gammaproteobacteria; Enterobacterales; Enterobacteriaceae; Salmonella
MTGDNTLITSHGINRRDFMKLCAALAATMGLSSKAAAEMAESVSNPQRPPVIWIGAQECT WP_000145429.1
hydrogenase 2 small subunit [Salmonella enterica] Length: 372\nNumber of predicted TMHs: 1\nExp number of AAs in TMHs: 24.51309\nExp number, first 60 AAs: 0.2682\nTotal prob of N-in: 0.14402\noutside 1 331\nTMhelix 332 354\ninside 355 372

15467 GCF_002054165.1_ASM205416v1 Salmonella enterica Proteobacteria; Gammaproteobacteria; Enterobacterales; Enterobacteriaceae; Salmonella

MNNEETFYQAMRRKGVTRRSFLKFCSLAATSLGLSAGMTPKIAWALENKPRIPVVWIHGL WP_001058316.1 hydrogenase [Salmonella enterica] Length: 372\nNumber of predicted TMHs: 2\nExp number of AAs in TMHs: 35.76397\nExp number, first 60 AAs: 18.97201\nTotal prob of N-in: 0.92785\nPOSSIBLE N-term signal sequence\ninside 1 19\nTMhelix 20 39\noutside 40 325\nTMhelix 326 348\ninside 349 372

15468 GCF_002054265.1_ASM205426v1 Salmonella enterica Proteobacteria; Gammaproteobacteria; Enterobacterales; Enterobacteriaceae; Salmonella MNNEETFYQAMRRKGVTRRSFLKFCSLAATSLGLGAGMTPKIAWALENKPRIPVVWIHGL WP_001058312.1 hydrogenase [Salmonella enterica] Length: 372\nNumber of predicted TMHs: 2\nExp number of AAs in TMHs: 36.51738\nExp number, first 60 AAs: 19.72849\nTotal prob of N-in: 0.96170\nPOSSIBLE N-term signal sequence\ninside 1 19\nTMhelix 20 39\noutside 40 325\nTMhelix 326 348\ninside 349 372

15469 GCF_002054455.1_ASM205445v1 Salmonella enterica Proteobacteria; Gammaproteobacteria; Enterobacterales; Enterobacteriaceae; Salmonella MTGDNTLITSHGINRRDFMKLCAALAATMGLSSKAAAEMAESVSNPQRPPVIWIGAQECT WP_000145429.1 hydrogenase 2 small subunit [Salmonella enterica] Length: 372\nNumber of predicted TMHs: 1\nExp number of AAs in TMHs: 24.51309\nExp number, first 60 AAs: 0.2682\nTotal prob of N-in: 0.14402\noutside 1 331\nTMhelix 332 354\ninside 355 372

15470 GCF_002054595.1_ASM205459v1 Salmonella enterica Proteobacteria; Gammaproteobacteria; Enterobacterales; Enterobacteriaceae; Salmonella MTGDNTLITSHGINRRDFMKLCAALAATMGLSSKAAAEMAESVSNPQRPPVIWIGAQECT WP_000145429.1 hydrogenase 2 small subunit [Salmonella enterica] Length: 372\nNumber of predicted TMHs: 1\nExp number of AAs in TMHs: 24.51309\nExp number, first 60 AAs: 0.2682\nTotal prob of N-in: 0.14402\noutside 1 331\nTMhelix 332 354\ninside 355 372

15471 GCF_002054615.1_ASM205461v1 Salmonella enterica Proteobacteria; Gammaproteobacteria; Enterobacterales; Enterobacteriaceae; Salmonella MTGDNTLITSHGINRRDFMKLCAALAATMGLSSKAAAEMAESVSNPQRPPVIWIGAQECT WP_000145429.1 hydrogenase 2 small subunit [Salmonella enterica] Length: 372\nNumber of predicted TMHs: 1\nExp number of AAs in TMHs: 24.51309\nExp number, first 60 AAs: 0.2682\nTotal prob of N-in: 0.14402\noutside 1 331\nTMhelix 332 354\ninside 355 372

15472 GCF_002054545.1_ASM205454v1 Salmonella enterica Proteobacteria; Gammaproteobacteria; Enterobacterales; Enterobacteriaceae; Salmonella MNNEETFYQAMRRKGVTRRSFLKFCSLAATSLGLGAGMTPKIAWALENKPRIPVVWIHGL WP_001058312.1 hydrogenase [Salmonella enterica] Length: 372\nNumber of predicted TMHs: 2\nExp number of AAs in TMHs: 36.51738\nExp number, first 60 AAs: 19.72849\nTotal prob of N-in: 0.96170\nPOSSIBLE N-term signal sequence\ninside 1 19\nTMhelix 20 39\noutside 40 325\nTMhelix 326 348\ninside 349 372

15473 GCF_002054645.1_ASM205464v1 Salmonella enterica Proteobacteria; Gammaproteobacteria; Enterobacterales; Enterobacteriaceae; Salmonella MTGDNTLITSHGINRRDFMKLCAALAATMGLSSKAAAEMAESVSNPQRPPVIWIGAQECT WP_000145429.1 hydrogenase 2 small subunit [Salmonella enterica] Length: 372\nNumber of predicted TMHs: 1\nExp number of AAs in TMHs: 24.51309\nExp number, first 60 AAs: 0.2682\nTotal prob of N-in: 0.14402\noutside 1 331\nTMhelix 332 354\ninside 355 372

15474 GCF_002054745.1_ASM205474v1 Salmonella enterica Proteobacteria; Gammaproteobacteria; Enterobacterales; Enterobacteriaceae; Salmonella MTGDNTLITSHGINRRDFMKLCAALAATMGLSSKAAAEMAESVSNPQRPPVIWIGAQECT WP_000145429.1 hydrogenase 2 small subunit [Salmonella enterica] Length: 372\nNumber of predicted TMHs: 1\nExp number of AAs in TMHs: 24.51309\nExp number, first 60 AAs: 0.2682\nTotal prob of N-in: 0.14402\noutside 1 331\nTMhelix 332 354\ninside 355 372

15475 GCF_002054775.1_ASM205477v1 Salmonella enterica Proteobacteria; Gammaproteobacteria; Enterobacterales; Enterobacteriaceae; Salmonella MTGDNTLITSHGINRRDFMKLCAALAATMGLSSKAAAEMAESVSNPQRPPVIWIGAQECT WP_000145429.1 hydrogenase 2 small subunit [Salmonella enterica] Length: 372\nNumber of predicted TMHs: 1\nExp number of AAs in TMHs: 24.51309\nExp number, first 60 AAs: 0.2682\nTotal prob of N-in: 0.14402\noutside 1 331\nTMhelix 332 354\ninside 355 372

15476 GCF_002054705.1_ASM205470v1 Salmonella enterica Proteobacteria; Gammaproteobacteria; Enterobacterales; Enterobacteriaceae; Salmonella
MTGDNTLITSHGINRRDFMKLCAALAATMGLSSKAAAEMAESVSNPQRPPVIWIGAQECT WP_000145431.1
hydrogenase 2 small subunit [Salmonella enterica] Length: 372\nNumber of predicted TMHs: 1\nExp number of AAs in TMHs: 24.51505\nExp number, first 60 AAs: 0.2682\nTotal prob of N-in: 0.14406\noutside 1 331\nTMhelix 332 354\ninside 355 372

15477 GCF_002055055.1_ASM205505v1 Salmonella enterica Proteobacteria; Gammaproteobacteria; Enterobacterales; Enterobacteriaceae; Salmonella
MTGDNTLITSHGINRRDFMKLCAALAATMGLSSKAAAEMAESVSNPQRPPVIWIGAQECT WP_000145429.1
hydrogenase 2 small subunit [Salmonella enterica] Length: 372\nNumber of predicted TMHs: 1\nExp number of AAs in TMHs: 24.51309\nExp number, first 60 AAs: 0.2682\nTotal prob of N-in: 0.14402\noutside 1 331\nTMhelix 332 354\ninside 355 372

15478 GCF_001808015.1_ASM180801v1 Salmonella sp. HMSC13B08 Proteobacteria; Gammaproteobacteria; Enterobacterales; Enterobacteriaceae; Salmonella
MTGDNTLITSHGVNRRDFMKLCAALAATMGLSSKAAAEMAESVSRPQRPPVVWIGAQECT WP_012135130.1
MULTISPECIES: hydrogenase 2 small subunit [Enterobacteriaceae] Length: 372\nNumber of predicted TMHs: 1\nExp number of AAs in TMHs: 24.24593\nExp number, first 60 AAs: 0.31574\nTotal prob of N-in: 0.12382\noutside 1 331\nTMhelix 332 354\ninside 355 372

15479 GCF_000439255.1_ASM43925v1 Salmonella bongori N268-08 Proteobacteria; Gammaproteobacteria; Enterobacterales; Enterobacteriaceae; Salmonella; Salmonella bongori
MTGDNTLITSHGINRRDFMKLCAALAATMGLSSKAAAEMAESVSNPQRPPVIWIGAQECT WP_000145426.1
hydrogenase 2 small subunit [Salmonella bongori] Length: 372\nNumber of predicted TMHs: 1\nExp number of AAs in TMHs: 23.21866\nExp number, first 60 AAs: 0.2769\nTotal prob of N-in: 0.08285\noutside 1 331\nTMhelix 332 354\ninside 355 372

15480 GCF_000252995.1_ASM25299v1 Salmonella bongori NCTC 12419 Proteobacteria; Gammaproteobacteria; Enterobacterales; Enterobacteriaceae; Salmonella; Salmonella bongori
MTGDNTLITSHGINRRDFMKLCAALAATMGLSSKAAAEMAESVSNPQRPPVIWIGAQECT WP_000145426.1
hydrogenase 2 small subunit [Salmonella bongori] Length: 372\nNumber of predicted TMHs: 1\nExp number of AAs in TMHs: 23.21866\nExp number, first 60 AAs: 0.2769\nTotal prob of N-in: 0.08285\noutside 1 331\nTMhelix 332 354\ninside 355 372

15481 GCF_002035305.1_ASM203530v1 Salmonella bongori serovar 40:z35:- Proteobacteria; Gammaproteobacteria; Enterobacterales; Enterobacteriaceae; Salmonella; Salmonella bongori
MTGDNTLITSHGINRRDFMKLCAALAATMGLSSKAAAEMAESVSNPQRPPVIWIGAQECT WP_079776618.1
MULTISPECIES: hydrogenase 2 small subunit [Salmonella] Length: 372\nNumber of predicted TMHs: 1\nExp number of AAs in TMHs: 24.65705\nExp number, first 60 AAs: 0.26805\nTotal prob of N-in: 0.13761\noutside 1 331\nTMhelix 332 354\ninside 355 372

15482 GCF_002037625.1_ASM203762v1 Salmonella bongori serovar 44:z39:- Proteobacteria; Gammaproteobacteria; Enterobacterales; Enterobacteriaceae; Salmonella; Salmonella bongori
MTGDNTLITSHGINRRDFMKLCAALAATMGLSSKAAAEMAESVSNPQRPPVIWIGAQECT WP_000145426.1
hydrogenase 2 small subunit [Salmonella bongori] Length: 372\nNumber of predicted TMHs: 1\nExp number of AAs in TMHs: 23.21866\nExp number, first 60 AAs: 0.2769\nTotal prob of N-in: 0.08285\noutside 1 331\nTMhelix 332 354\ninside 355 372

15483 GCF_002033765.1_ASM203376v1 Salmonella bongori serovar 48:a:- Proteobacteria; Gammaproteobacteria; Enterobacterales; Enterobacteriaceae; Salmonella; Salmonella bongori
MTGDNTLITSHGINRRDFMKLCAALAATMGLSSKAAAEMAESVSNPQRPPVIWIGAQECT WP_000145426.1
hydrogenase 2 small subunit [Salmonella bongori] Length: 372\nNumber of predicted TMHs: 1\nExp number of AAs in TMHs: 23.21866\nExp number, first 60 AAs: 0.2769\nTotal prob of N-in: 0.08285\noutside 1 331\nTMhelix 332 354\ninside 355 372

15484 GCF_002035265.1_ASM203526v1 Salmonella bongori serovar 48:z35:- Proteobacteria; Gammaproteobacteria; Enterobacterales; Enterobacteriaceae; Salmonella; Salmonella bongori
MTGDNTLITSHGINRRDFMKLCAALAATMGLSSKAAAEMAESVSNPQRPPVIWIGAQECT WP_000145426.1
hydrogenase 2 small subunit [Salmonella bongori] Length: 372\nNumber of predicted TMHs: 1\nExp

number of AAs in TMHs: 23.21866\nExp number, first 60 AAs: 0.2769\nTotal prob of N-in: 0.08285\noutside 1 331\nTMhelix 332 354\ninside 355 372

15485 GCF_002035315.1_ASM203531v1 Salmonella bongori serovar 48:i:- Proteobacteria; Gammaproteobacteria; Enterobacterales; Enterobacteriaceae; Salmonella; Salmonella bongori; Salmonella bongori serovar 40 MTGDNTLITSHGINRRDFMKLCAALAATMGLSSKAAAEMAESVSNPQRPPVIWIGAQECT WP_000145426.1 hydrogenase 2 small subunit [Salmonella bongori] Length: 372\nNumber of predicted TMHs: 1\nExp number of AAs in TMHs: 23.21866\nExp number, first 60 AAs: 0.2769\nTotal prob of N-in: 0.08285\noutside 1 331\nTMhelix 332 354\ninside 355 372

15486 GCF_000709535.1_ASM70953v1 Salmonella bongori serovar 48:z41:-- str. RKS3044 Proteobacteria; Gammaproteobacteria; Enterobacterales; Enterobacteriaceae; Salmonella; Salmonella bongori; Salmonella bongori serovar 48 MTGDNTLITSHGINRRDFMKLCAALAATMGLSSKAAAEMAESVSNPQRPPVIWIGAQECT WP_000145426.1 hydrogenase 2 small subunit [Salmonella bongori] Length: 372\nNumber of predicted TMHs: 1\nExp number of AAs in TMHs: 23.21866\nExp number, first 60 AAs: 0.2769\nTotal prob of N-in: 0.08285\noutside 1 331\nTMhelix 332 354\ninside 355 372

15487 GCF_001276875.1_ASM127687v1 Salmonella enterica subsp. diarizonae Proteobacteria; Gammaproteobacteria; Enterobacterales; Enterobacteriaceae; Salmonella; Salmonella enterica MTGDNTLITSHGINRRDFMKLCAALAATMGLSSKAAAEMAESVSNPQRPPVIWIGAQECT WP_023248548.1 hydrogenase 2 small subunit [Salmonella enterica] Length: 372\nNumber of predicted TMHs: 1\nExp number of AAs in TMHs: 24.51281\nExp number, first 60 AAs: 0.26822\nTotal prob of N-in: 0.14402\noutside 1 331\nTMhelix 332 354\ninside 355 372

15488 GCF_001276795.1_ASM127679v1 Salmonella enterica subsp. diarizonae Proteobacteria; Gammaproteobacteria; Enterobacterales; Enterobacteriaceae; Salmonella; Salmonella enterica MNNEETFYQAMRRKGVTRRSFLKFCSLAATSLGLGAGMTPKIAWALENKPRIPVVWIHGL WP_001058306.1 hydrogenase [Salmonella enterica] Length: 372\nNumber of predicted TMHs: 2\nExp number of AAs in TMHs: 36.5540099999999\nExp number, first 60 AAs: 19.72732\nTotal prob of N-in: 0.96171\nPOSSIBLE N-term signal sequence\ninside 1 19\nTMhelix 20 39\noutside 40 325\nTMhelix 326 348\ninside 349 372

15489 GCF_001276795.1_ASM127679v1 Salmonella enterica subsp. diarizonae Proteobacteria; Gammaproteobacteria; Enterobacterales; Enterobacteriaceae; Salmonella; Salmonella enterica MTGDNTLITSHGINRRDFMKLCAALAATMGLSSKAAAEMAESVSNPQRPPVIWIGAQECT WP_023248548.1 hydrogenase 2 small subunit [Salmonella enterica] Length: 372\nNumber of predicted TMHs: 1\nExp number of AAs in TMHs: 24.51281\nExp number, first 60 AAs: 0.26822\nTotal prob of N-in: 0.14402\noutside 1 331\nTMhelix 332 354\ninside 355 372

15490 GCF_001629755.1_ASM162975v1 Salmonella enterica subsp. diarizonae Proteobacteria; Gammaproteobacteria; Enterobacterales; Enterobacteriaceae; Salmonella; Salmonella enterica MTGDNTLITSHGINRRDFMKLCAALAATMGLSSKAAAEMAESVSNPQRPPVIWIGAQECT WP_023248548.1 hydrogenase 2 small subunit [Salmonella enterica] Length: 372\nNumber of predicted TMHs: 1\nExp number of AAs in TMHs: 24.51281\nExp number, first 60 AAs: 0.26822\nTotal prob of N-in: 0.14402\noutside 1 331\nTMhelix 332 354\ninside 355 372

15491 GCF_001628755.1_ASM162875v1 Salmonella enterica subsp. diarizonae Proteobacteria; Gammaproteobacteria; Enterobacterales; Enterobacteriaceae; Salmonella; Salmonella enterica MTGDNTLITSHGINRRDFMKLCAALAATMGLSSKAAAEMAESVSNPQRPPVIWIGAQECT WP_023248548.1 hydrogenase 2 small subunit [Salmonella enterica] Length: 372\nNumber of predicted TMHs: 1\nExp number of AAs in TMHs: 24.51281\nExp number, first 60 AAs: 0.26822\nTotal prob of N-in: 0.14402\noutside 1 331\nTMhelix 332 354\ninside 355 372

15492 GCF_001628755.1_ASM162875v1 Salmonella enterica subsp. diarizonae Proteobacteria; Gammaproteobacteria; Enterobacterales; Enterobacteriaceae; Salmonella; Salmonella enterica MNNEETFYQAMRRKGVTRRSFLKFCSLAATSLGLGAGMTPKIAWALENKPRIPVVWIHGL WP_001058306.1 hydrogenase [Salmonella enterica] Length: 372\nNumber of predicted TMHs: 2\nExp number of AAs in TMHs: 36.5540099999999\nExp number, first 60 AAs: 19.72732\nTotal prob of N-in: 0.96171\nPOSSIBLE N-term signal sequence\ninside 1 19\nTMhelix 20 39\noutside 40 325\nTMhelix 326 348\ninside 349 372

15493 GCF_001629775.1_ASM162977v1 Salmonella enterica subsp. diarizonae Proteobacteria; Gammaproteobacteria; Enterobacterales; Enterobacteriaceae; Salmonella; Salmonella enterica

MNNEETFYQAMRRKGVTRRSFLKFCSLAATSLGLGAGMTPKIAWALENKPRIPVVWIHGL	WP_001058306.1	hydrogenase [Salmonella enterica]	Length: 372\nNumber of predicted TMHs:  2\nExp number of AAs in TMHs: 36.5540099999999\nExp number, first 60 AAs:  19.72732\nTotal prob of N-in:   0.96171\nPOSSIBLE N-term signal sequence\ninside   1   19\nTMhelix   20   39\noutside   40   325\nTMhelix   326   348\ninside   349   372

15494	GCF_001629775.1_ASM162977v1	Salmonella enterica subsp. diarizonae	Proteobacteria; Gammaproteobacteria; Enterobacterales; Enterobacteriaceae; Salmonella; Salmonella enterica	MTGDNTLITSHGINRRDFMKLCAALAATMGLSSKAAAEMAESVSNPQRPPVIWIGAQECT	WP_023248548.1	hydrogenase 2 small subunit [Salmonella enterica]	Length: 372\nNumber of predicted TMHs:  1\nExp number of AAs in TMHs: 24.51281\nExp number, first 60 AAs:  0.26822\nTotal prob of N-in:   0.14402\noutside   1   331\nTMhelix   332   354\ninside   355   372

15495	GCF_001276875.1_ASM127687v1	Salmonella enterica subsp. diarizonae	Proteobacteria; Gammaproteobacteria; Enterobacterales; Enterobacteriaceae; Salmonella; Salmonella enterica	MNNEETFYQAMRRKGVTRRSFLKFCSLAATSLGLGAGMTPKIAWALENKPRIPVVWIHGL	WP_001058306.1	hydrogenase [Salmonella enterica]	Length: 372\nNumber of predicted TMHs:  2\nExp number of AAs in TMHs: 36.5540099999999\nExp number, first 60 AAs:  19.72732\nTotal prob of N-in:   0.96171\nPOSSIBLE N-term signal sequence\ninside   1   19\nTMhelix   20   39\noutside   40   325\nTMhelix   326   348\ninside   349   372

15496	GCF_001629755.1_ASM162975v1	Salmonella enterica subsp. diarizonae	Proteobacteria; Gammaproteobacteria; Enterobacterales; Enterobacteriaceae; Salmonella; Salmonella enterica	MNNEETFYQAMRRKGVTRRSFLKFCSLAATSLGLGAGMTPKIAWALENKPRIPVVWIHGL	WP_001058306.1	hydrogenase [Salmonella enterica]	Length: 372\nNumber of predicted TMHs:  2\nExp number of AAs in TMHs: 36.5540099999999\nExp number, first 60 AAs:  19.72732\nTotal prob of N-in:   0.96171\nPOSSIBLE N-term signal sequence\ninside   1   19\nTMhelix   20   39\noutside   40   325\nTMhelix   326   348\ninside   349   372

15497	GCF_001716405.1_ASM171640v1	Salmonella enterica subsp. enterica	Proteobacteria; Gammaproteobacteria; Enterobacterales; Enterobacteriaceae; Salmonella; Salmonella enterica	MTGDNTLITSHGINRRDFMKLCAALAATMGLSSKAAAEMAESVSNPQRPPVIWIGAQECT	WP_000145429.1	hydrogenase 2 small subunit [Salmonella enterica]	Length: 372\nNumber of predicted TMHs:  1\nExp number of AAs in TMHs: 24.51309\nExp number, first 60 AAs:  0.2682\nTotal prob of N-in:   0.14402\noutside   1   331\nTMhelix   332   354\ninside   355   372

15498	GCF_001716585.1_ASM171658v1	Salmonella enterica subsp. enterica	Proteobacteria; Gammaproteobacteria; Enterobacterales; Enterobacteriaceae; Salmonella; Salmonella enterica	MNNEETFYQAMRRKGVTRRSFLKFCSLAATSLGLGAGMTPKIAWALENKPRIPVVWIHGL	WP_001058311.1	hydrogenase [Salmonella enterica]	Length: 372\nNumber of predicted TMHs:  2\nExp number of AAs in TMHs: 37.4699899999999\nExp number, first 60 AAs:  19.72699\nTotal prob of N-in:   0.96171\nPOSSIBLE N-term signal sequence\ninside   1   19\nTMhelix   20   39\noutside   40   325\nTMhelix   326   348\ninside   349   372

15499	GCF_001716455.1_ASM171645v1	Salmonella enterica subsp. enterica	Proteobacteria; Gammaproteobacteria; Enterobacterales; Enterobacteriaceae; Salmonella; Salmonella enterica	MTGDNTLITSHGINRRDFMKLCAALAATMGLSSKAAAEMAESVSNPQRPPVIWIGAQECT	WP_000145429.1	hydrogenase 2 small subunit [Salmonella enterica]	Length: 372\nNumber of predicted TMHs:  1\nExp number of AAs in TMHs: 24.51309\nExp number, first 60 AAs:  0.2682\nTotal prob of N-in:   0.14402\noutside   1   331\nTMhelix   332   354\ninside   355   372

15500	GCF_001586445.1_ASM158644v1	Salmonella enterica subsp. enterica	Proteobacteria; Gammaproteobacteria; Enterobacterales; Enterobacteriaceae; Salmonella; Salmonella enterica	MTGDNTLITSHGINRRDFMKLCAALAATMGLSSKAAAEMAESVSNPQRPPVIWIGAQECT	WP_000145429.1	hydrogenase 2 small subunit [Salmonella enterica]	Length: 372\nNumber of predicted TMHs:  1\nExp number of AAs in TMHs: 24.51309\nExp number, first 60 AAs:  0.2682\nTotal prob of N-in:   0.14402\noutside   1   331\nTMhelix   332   354\ninside   355   372

15501	GCF_001586515.1_ASM158651v1	Salmonella enterica subsp. enterica	Proteobacteria; Gammaproteobacteria; Enterobacterales; Enterobacteriaceae; Salmonella; Salmonella enterica	MTGDNTLITSHGINRRDFMKLCAALAATMGLSSKAAAEMAESVSNPQRPPVIWIGAQECT	WP_000145429.1	hydrogenase 2 small subunit [Salmonella enterica]	Length: 372\nNumber of predicted TMHs:  1\nExp number of AAs in TMHs: 24.51309\nExp number, first 60 AAs:  0.2682\nTotal prob of N-in:   0.14402\noutside   1   331\nTMhelix   332   354\ninside   355   372

15502 GCF_001716035.1_ASM171603v1 Salmonella enterica subsp. enterica Proteobacteria; Gammaproteobacteria; Enterobacterales; Enterobacteriaceae; Salmonella; Salmonella enterica
MNNEETFYQAMRRKGVTRRSFLKFCSLAATSLGLGAGMTPKIAWALENKPRIPVVWIHGL WP_001058311.1
hydrogenase [Salmonella enterica] Length: 372\nNumber of predicted TMHs: 2\nExp number of AAs in TMHs: 37.4699899999999\nExp number, first 60 AAs: 19.72699\nTotal prob of N-in: 0.96171\nPOSSIBLE N-term signal sequence\ninside 1 19\nTMhelix 20 39\noutside 40 325\nTMhelix 326 348\ninside 349 372

15503 GCF_001716485.1_ASM171648v1 Salmonella enterica subsp. enterica Proteobacteria; Gammaproteobacteria; Enterobacterales; Enterobacteriaceae; Salmonella; Salmonella enterica
MTGDNTLITSHGINRRDFMKLCAALAATMGLSSKAAAEMAESVSNPQRPPVIWIGAQECT WP_000145429.1
hydrogenase 2 small subunit [Salmonella enterica] Length: 372\nNumber of predicted TMHs: 1\nExp number of AAs in TMHs: 24.51309\nExp number, first 60 AAs: 0.2682\nTotal prob of N-in: 0.14402\noutside 1 331\nTMhelix 332 354\ninside 355 372

15504 GCF_001586535.1_ASM158653v1 Salmonella enterica subsp. enterica Proteobacteria; Gammaproteobacteria; Enterobacterales; Enterobacteriaceae; Salmonella; Salmonella enterica
MTGDNTLITSHGINRRDFMKLCAALAATMGLSSKAAAEMAESVSNPQRPPVIWIGAQECT WP_000145429.1
hydrogenase 2 small subunit [Salmonella enterica] Length: 372\nNumber of predicted TMHs: 1\nExp number of AAs in TMHs: 24.51309\nExp number, first 60 AAs: 0.2682\nTotal prob of N-in: 0.14402\noutside 1 331\nTMhelix 332 354\ninside 355 372

15505 GCF_001541975.1_ASM154197v1 Salmonella enterica subsp. enterica Proteobacteria; Gammaproteobacteria; Enterobacterales; Enterobacteriaceae; Salmonella; Salmonella enterica
MTGDNTLITSHGINRRDFMKLCAALAATMGLSSKAAAEMAESVSNPQRPPVIWIGAQECT WP_000145429.1
hydrogenase 2 small subunit [Salmonella enterica] Length: 372\nNumber of predicted TMHs: 1\nExp number of AAs in TMHs: 24.51309\nExp number, first 60 AAs: 0.2682\nTotal prob of N-in: 0.14402\noutside 1 331\nTMhelix 332 354\ninside 355 372

15506 GCF_001541975.1_ASM154197v1 Salmonella enterica subsp. enterica Proteobacteria; Gammaproteobacteria; Enterobacterales; Enterobacteriaceae; Salmonella; Salmonella enterica
MNNEETFYQAMRRKGVTRRSFLKFCSLAATSLGLGAGMTPKIAWALENKPRIPVVWIHGL WP_001058312.1
hydrogenase [Salmonella enterica] Length: 372\nNumber of predicted TMHs: 2\nExp number of AAs in TMHs: 36.51738\nExp number, first 60 AAs: 19.72849\nTotal prob of N-in: 0.96170\nPOSSIBLE N-term signal sequence\ninside 1 19\nTMhelix 20 39\noutside 40 325\nTMhelix 326 348\ninside 349 372

15507 GCF_001586445.1_ASM158644v1 Salmonella enterica subsp. enterica Proteobacteria; Gammaproteobacteria; Enterobacterales; Enterobacteriaceae; Salmonella; Salmonella enterica
MNNEETFYQAMRRKGVTRRSFLKFCSLAATSLGLGAGMTPKIAWALENKPRIPVVWIHGL WP_001058311.1
hydrogenase [Salmonella enterica] Length: 372\nNumber of predicted TMHs: 2\nExp number of AAs in TMHs: 37.4699899999999\nExp number, first 60 AAs: 19.72699\nTotal prob of N-in: 0.96171\nPOSSIBLE N-term signal sequence\ninside 1 19\nTMhelix 20 39\noutside 40 325\nTMhelix 326 348\ninside 349 372

15508 GCF_001586595.1_ASM158659v1 Salmonella enterica subsp. enterica Proteobacteria; Gammaproteobacteria; Enterobacterales; Enterobacteriaceae; Salmonella; Salmonella enterica
MTGDNTLITSHGINRRDFMKLCAALAATMGLSSKAAAEMAESVSNPQRPPVIWIGAQECT WP_000145429.1
hydrogenase 2 small subunit [Salmonella enterica] Length: 372\nNumber of predicted TMHs: 1\nExp number of AAs in TMHs: 24.51309\nExp number, first 60 AAs: 0.2682\nTotal prob of N-in: 0.14402\noutside 1 331\nTMhelix 332 354\ninside 355 372

15509 GCF_001586515.1_ASM158651v1 Salmonella enterica subsp. enterica Proteobacteria; Gammaproteobacteria; Enterobacterales; Enterobacteriaceae; Salmonella; Salmonella enterica
MNNEETFYQAMRRKGVTRRSFLKFCSLAATSLGLGAGMTPKIAWALENKPRIPVVWIHGL WP_001058311.1
hydrogenase [Salmonella enterica] Length: 372\nNumber of predicted TMHs: 2\nExp number of AAs in TMHs: 37.4699899999999\nExp number, first 60 AAs: 19.72699\nTotal prob of N-in: 0.96171\nPOSSIBLE N-term signal sequence\ninside 1 19\nTMhelix 20 39\noutside 40 325\nTMhelix 326 348\ninside 349 372

15510 GCF_001720445.1_ASM172044v1 Salmonella enterica subsp. enterica Proteobacteria; Gammaproteobacteria; Enterobacterales; Enterobacteriaceae; Salmonella; Salmonella enterica
MNNEETFYQAMRRKGVTRRSFLKFCSLAATSLGLGAGMTPKIAWALENKPRIPVVWIHGL WP_001058311.1
hydrogenase [Salmonella enterica] Length: 372\nNumber of predicted TMHs: 2\nExp number of AAs in TMHs:

TMHs: 37.4699899999999\nExp number, first 60 AAs:  19.72699\nTotal prob of N-in:   0.96171\nPOSSIBLE N-term signal sequence\ninside   1   19\nTMhelix   20   39\noutside   40   325\nTMhelix   326   348\ninside   349   372

15511 GCF_001586685.1_ASM158668v1 Salmonella enterica subsp. enterica Proteobacteria; Gammaproteobacteria; Enterobacterales; Enterobacteriaceae; Salmonella; Salmonella enterica MTGDNTLITSHGINRRDFMKLCAALAATMGLSSKAAAEMAESVSNPQRPPVIWIGAQECT WP_000145429.1 hydrogenase 2 small subunit [Salmonella enterica] Length: 372\nNumber of predicted TMHs:  1\nExp number of AAs in TMHs: 24.51309\nExp number, first 60 AAs:  0.2682\nTotal prob of N-in:   0.14402\noutside   1   331\nTMhelix   332   354\ninside   355   372

15512 GCF_001586685.1_ASM158668v1 Salmonella enterica subsp. enterica Proteobacteria; Gammaproteobacteria; Enterobacterales; Enterobacteriaceae; Salmonella; Salmonella enterica MNNEETFYQAMRRKGVTRRSFLKFCSLAATSLGLGAGMTPKIAWALENKPRIPVVWIHGL WP_001058311.1 hydrogenase [Salmonella enterica] Length: 372\nNumber of predicted TMHs:  2\nExp number of AAs in TMHs: 37.4699899999999\nExp number, first 60 AAs:  19.72699\nTotal prob of N-in:   0.96171\nPOSSIBLE N-term signal sequence\ninside   1   19\nTMhelix   20   39\noutside   40   325\nTMhelix   326   348\ninside   349   372

15513 GCF_001586535.1_ASM158653v1 Salmonella enterica subsp. enterica Proteobacteria; Gammaproteobacteria; Enterobacterales; Enterobacteriaceae; Salmonella; Salmonella enterica MNNEETFYQAMRRKGVTRRSFLKFCSLAATSLGLGAGMTPKIAWALENKPRIPVVWIHGL WP_001058311.1 hydrogenase [Salmonella enterica] Length: 372\nNumber of predicted TMHs:  2\nExp number of AAs in TMHs: 37.4699899999999\nExp number, first 60 AAs:  19.72699\nTotal prob of N-in:   0.96171\nPOSSIBLE N-term signal sequence\ninside   1   19\nTMhelix   20   39\noutside   40   325\nTMhelix   326   348\ninside   349   372

15514 GCF_001586595.1_ASM158659v1 Salmonella enterica subsp. enterica Proteobacteria; Gammaproteobacteria; Enterobacterales; Enterobacteriaceae; Salmonella; Salmonella enterica MNNEETFYQAMRRKGVTRRSFLKFCSLAATSLGLGAGMTPKIAWALENKPRIPVVWIHGL WP_001058311.1 hydrogenase [Salmonella enterica] Length: 372\nNumber of predicted TMHs:  2\nExp number of AAs in TMHs: 37.4699899999999\nExp number, first 60 AAs:  19.72699\nTotal prob of N-in:   0.96171\nPOSSIBLE N-term signal sequence\ninside   1   19\nTMhelix   20   39\noutside   40   325\nTMhelix   326   348\ninside   349   372

15515 GCF_001716485.1_ASM171648v1 Salmonella enterica subsp. enterica Proteobacteria; Gammaproteobacteria; Enterobacterales; Enterobacteriaceae; Salmonella; Salmonella enterica MNNEETFYQAMRRKGVTRRSFLKFCSLAATSLGLGAGMTPKIAWALENKPRIPVVWIHGL WP_001058311.1 hydrogenase [Salmonella enterica] Length: 372\nNumber of predicted TMHs:  2\nExp number of AAs in TMHs: 37.4699899999999\nExp number, first 60 AAs:  19.72699\nTotal prob of N-in:   0.96171\nPOSSIBLE N-term signal sequence\ninside   1   19\nTMhelix   20   39\noutside   40   325\nTMhelix   326   348\ninside   349   372

15516 GCF_001714305.1_ASM171430v1 Salmonella enterica subsp. enterica Proteobacteria; Gammaproteobacteria; Enterobacterales; Enterobacteriaceae; Salmonella; Salmonella enterica MNNEETFYQAMRRKGVTRRSFLKFCSLAATSLGLGAGMTPKIAWALENKPRIPVVWIHGL WP_001058307.1 hydrogenase [Salmonella enterica] Length: 372\nNumber of predicted TMHs:  2\nExp number of AAs in TMHs: 36.51738\nExp number, first 60 AAs:  19.72849\nTotal prob of N-in:   0.96170\nPOSSIBLE N-term signal sequence\ninside   1   19\nTMhelix   20   39\noutside   40   325\nTMhelix   326   348\ninside   349   372

15517 GCF_001716555.1_ASM171655v1 Salmonella enterica subsp. enterica Proteobacteria; Gammaproteobacteria; Enterobacterales; Enterobacteriaceae; Salmonella; Salmonella enterica MTGDNTLITSHGINRRDFMKLCAALAATMGLSSKAAAEMAESVSNPQRPPVIWIGAQECT WP_000145429.1 hydrogenase 2 small subunit [Salmonella enterica] Length: 372\nNumber of predicted TMHs:  1\nExp number of AAs in TMHs: 24.51309\nExp number, first 60 AAs:  0.2682\nTotal prob of N-in:   0.14402\noutside   1   331\nTMhelix   332   354\ninside   355   372

15518 GCF_001586655.1_ASM158665v1 Salmonella enterica subsp. enterica Proteobacteria; Gammaproteobacteria; Enterobacterales; Enterobacteriaceae; Salmonella; Salmonella enterica MTGDNTLITSHGINRRDFMKLCAALAATMGLSSKAAAEMAESVSNPQRPPVIWIGAQECT WP_000145429.1 hydrogenase 2 small subunit [Salmonella enterica] Length: 372\nNumber of predicted TMHs:  1\nExp number of AAs in TMHs: 24.51309\nExp number, first 60 AAs:  0.2682\nTotal prob of N-in:   0.14402\noutside   1   331\nTMhelix   332   354\ninside   355   372

15519 GCF_001586655.1_ASM158665v1 Salmonella enterica subsp. enterica Proteobacteria; Gammaproteobacteria; Enterobacterales; Enterobacteriaceae; Salmonella; Salmonella enterica

MNNEETFYQAMRRKGVTRRSFLKFCSLAATSLGLSAGMTPKIAWALENKPRIPVVWIHGL WP_001058316.1 hydrogenase [Salmonella enterica] Length: 372\nNumber of predicted TMHs: 2\nExp number of AAs in TMHs: 35.76397\nExp number, first 60 AAs: 18.97201\nTotal prob of N-in: 0.92785\nPOSSIBLE N-term signal sequence\ninside 1 19\nTMhelix 20 39\noutside 40 325\nTMhelix 326 348\ninside 349 372

15520 GCF_001716775.1_ASM171677v1 Salmonella enterica subsp. enterica Proteobacteria; Gammaproteobacteria; Enterobacterales; Enterobacteriaceae; Salmonella; Salmonella enterica MNNEETFYQAMRRKGVTRRSFLKFCSLAATSLGLGAGMTPKIAWALENKPRIPVVWIHGL WP_001058311.1 hydrogenase [Salmonella enterica] Length: 372\nNumber of predicted TMHs: 2\nExp number of AAs in TMHs: 37.4699899999999\nExp number, first 60 AAs: 19.72699\nTotal prob of N-in: 0.96171\nPOSSIBLE N-term signal sequence\ninside 1 19\nTMhelix 20 39\noutside 40 325\nTMhelix 326 348\ninside 349 372

15521 GCF_001714255.1_ASM171425v1 Salmonella enterica subsp. enterica Proteobacteria; Gammaproteobacteria; Enterobacterales; Enterobacteriaceae; Salmonella; Salmonella enterica MTGDNTLITSHGINRRDFMKLCAALAATMGLSSKAAAEMAESVSNPQRPPVIWIGAQECT WP_000145429.1 hydrogenase 2 small subunit [Salmonella enterica] Length: 372\nNumber of predicted TMHs: 1\nExp number of AAs in TMHs: 24.51309\nExp number, first 60 AAs: 0.2682\nTotal prob of N-in: 0.14402\noutside 1 331\nTMhelix 332 354\ninside 355 372

15522 GCF_001700365.1_ASM170036v1 Salmonella enterica subsp. enterica Proteobacteria; Gammaproteobacteria; Enterobacterales; Enterobacteriaceae; Salmonella; Salmonella enterica MTGDNTLITSHGINRRDFMKLCAALAATMGLSSKAAAEMAESVSNPQRPPVIWIGAQECT WP_000145429.1 hydrogenase 2 small subunit [Salmonella enterica] Length: 372\nNumber of predicted TMHs: 1\nExp number of AAs in TMHs: 24.51309\nExp number, first 60 AAs: 0.2682\nTotal prob of N-in: 0.14402\noutside 1 331\nTMhelix 332 354\ninside 355 372

15523 GCF_001700365.1_ASM170036v1 Salmonella enterica subsp. enterica Proteobacteria; Gammaproteobacteria; Enterobacterales; Enterobacteriaceae; Salmonella; Salmonella enterica MNNEETFYQAMRRKGVTRRSFLKFCSLAATSLGLGAGMTPKIAWALENKPRIPVVWIHGL WP_001058312.1 hydrogenase [Salmonella enterica] Length: 372\nNumber of predicted TMHs: 2\nExp number of AAs in TMHs: 36.51738\nExp number, first 60 AAs: 19.72849\nTotal prob of N-in: 0.96170\nPOSSIBLE N-term signal sequence\ninside 1 19\nTMhelix 20 39\noutside 40 325\nTMhelix 326 348\ninside 349 372

15524 GCF_001716195.1_ASM171619v1 Salmonella enterica subsp. enterica Proteobacteria; Gammaproteobacteria; Enterobacterales; Enterobacteriaceae; Salmonella; Salmonella enterica MTGDNTLITSHGINRRDFMKLCAALAATMGLSSKAAAEMAESVSNPQRPPVIWIGAQECT WP_000145429.1 hydrogenase 2 small subunit [Salmonella enterica] Length: 372\nNumber of predicted TMHs: 1\nExp number of AAs in TMHs: 24.51309\nExp number, first 60 AAs: 0.2682\nTotal prob of N-in: 0.14402\noutside 1 331\nTMhelix 332 354\ninside 355 372

15525 GCF_001714125.1_ASM171412v1 Salmonella enterica subsp. enterica Proteobacteria; Gammaproteobacteria; Enterobacterales; Enterobacteriaceae; Salmonella; Salmonella enterica MNNEETFYQAMRRKGVTRRSFLKFCSLAATSLGLGAGMTPKIAWALENKPRIPVVWIHGL WP_001058307.1 hydrogenase [Salmonella enterica] Length: 372\nNumber of predicted TMHs: 2\nExp number of AAs in TMHs: 36.51738\nExp number, first 60 AAs: 19.72849\nTotal prob of N-in: 0.96170\nPOSSIBLE N-term signal sequence\ninside 1 19\nTMhelix 20 39\noutside 40 325\nTMhelix 326 348\ninside 349 372

15526 GCF_001716065.1_ASM171606v1 Salmonella enterica subsp. enterica Proteobacteria; Gammaproteobacteria; Enterobacterales; Enterobacteriaceae; Salmonella; Salmonella enterica MNNEETFYQAMRRKGVTRRSFLKFCSLAATSLGLGAGMTPKIAWALENKPRIPVVWIHGL WP_001058311.1 hydrogenase [Salmonella enterica] Length: 372\nNumber of predicted TMHs: 2\nExp number of AAs in TMHs: 37.4699899999999\nExp number, first 60 AAs: 19.72699\nTotal prob of N-in: 0.96171\nPOSSIBLE N-term signal sequence\ninside 1 19\nTMhelix 20 39\noutside 40 325\nTMhelix 326 348\ninside 349 372

15527 GCF_001716155.1_ASM171615v1 Salmonella enterica subsp. enterica Proteobacteria; Gammaproteobacteria; Enterobacterales; Enterobacteriaceae; Salmonella; Salmonella enterica MTGDNTLITSHGINRRDFMKLCAALAATMGLSSKAAAEMAESVSNPQRPPVIWIGAQECT WP_000145429.1 hydrogenase 2 small subunit [Salmonella enterica] Length: 372\nNumber of predicted TMHs: 1\nExp number of AAs in TMHs: 24.51309\nExp number, first 60 AAs: 0.2682\nTotal prob of N-in: 0.14402\noutside 1 331\nTMhelix 332 354\ninside 355 372

15528 GCF_001716555.1_ASM171655v1 Salmonella enterica subsp. enterica Proteobacteria; Gammaproteobacteria; Enterobacterales; Enterobacteriaceae; Salmonella; Salmonella enterica
MNNEETFYQAMRRKGVTRRSFLKFCSLAATSLGLGAGMTPKIAWALENKPRIPVVWIHGL WP_001058311.1
hydrogenase [Salmonella enterica] Length: 372\nNumber of predicted TMHs: 2\nExp number of AAs in TMHs: 37.4699899999999\nExp number, first 60 AAs: 19.72699\nTotal prob of N-in: 0.96171\nPOSSIBLE N-term signal sequence\ninside 1 19\nTMhelix 20 39\noutside 40 325\nTMhelix 326 348\ninside 349 372

15529 GCF_001716805.1_ASM171680v1 Salmonella enterica subsp. enterica Proteobacteria; Gammaproteobacteria; Enterobacterales; Enterobacteriaceae; Salmonella; Salmonella enterica
MNNEETFYQAMRRKGVTRRSFLKFCSLAATSLGLGAGMTPKIAWALENKPRIPVVWIHGL WP_001058311.1
hydrogenase [Salmonella enterica] Length: 372\nNumber of predicted TMHs: 2\nExp number of AAs in TMHs: 37.4699899999999\nExp number, first 60 AAs: 19.72699\nTotal prob of N-in: 0.96171\nPOSSIBLE N-term signal sequence\ninside 1 19\nTMhelix 20 39\noutside 40 325\nTMhelix 326 348\ninside 349 372

15530 GCF_001720445.1_ASM172044v1 Salmonella enterica subsp. enterica Proteobacteria; Gammaproteobacteria; Enterobacterales; Enterobacteriaceae; Salmonella; Salmonella enterica
MTGDNTLITSHGINRRDFMKLCAALAATMGLSSKAAAEMAESVSNPQRPPVIWIGAQECT WP_000145429.1
hydrogenase 2 small subunit [Salmonella enterica] Length: 372\nNumber of predicted TMHs: 1\nExp number of AAs in TMHs: 24.51309\nExp number, first 60 AAs: 0.2682\nTotal prob of N-in: 0.14402\noutside 1 331\nTMhelix 332 354\ninside 355 372

15531 GCF_001647955.1_ASM164795v1 Salmonella enterica subsp. enterica Proteobacteria; Gammaproteobacteria; Enterobacterales; Enterobacteriaceae; Salmonella; Salmonella enterica
MTGDNTLITSHGINRRDFMKLCAALAATMGLSSKAAAEMAESVSNPQRPPVIWIGAQECT WP_000145429.1
hydrogenase 2 small subunit [Salmonella enterica] Length: 372\nNumber of predicted TMHs: 1\nExp number of AAs in TMHs: 24.51309\nExp number, first 60 AAs: 0.2682\nTotal prob of N-in: 0.14402\noutside 1 331\nTMhelix 332 354\ninside 355 372

15532 GCF_001716275.1_ASM171627v1 Salmonella enterica subsp. enterica Proteobacteria; Gammaproteobacteria; Enterobacterales; Enterobacteriaceae; Salmonella; Salmonella enterica
MTGDNTLITSHGINRRDFMKLCAALAATMGLSSKAAAEMAESVSNPQRPPVIWIGAQECT WP_000145429.1
hydrogenase 2 small subunit [Salmonella enterica] Length: 372\nNumber of predicted TMHs: 1\nExp number of AAs in TMHs: 24.51309\nExp number, first 60 AAs: 0.2682\nTotal prob of N-in: 0.14402\noutside 1 331\nTMhelix 332 354\ninside 355 372

15533 GCF_001716155.1_ASM171615v1 Salmonella enterica subsp. enterica Proteobacteria; Gammaproteobacteria; Enterobacterales; Enterobacteriaceae; Salmonella; Salmonella enterica
MNNEETFYQAMRRKGVTRRSFLKFCSLAATSLGLGAGMTPKIAWALENKPRIPVVWIHGL WP_001058311.1
hydrogenase [Salmonella enterica] Length: 372\nNumber of predicted TMHs: 2\nExp number of AAs in TMHs: 37.4699899999999\nExp number, first 60 AAs: 19.72699\nTotal prob of N-in: 0.96171\nPOSSIBLE N-term signal sequence\ninside 1 19\nTMhelix 20 39\noutside 40 325\nTMhelix 326 348\ninside 349 372

15534 GCF_001714125.1_ASM171412v1 Salmonella enterica subsp. enterica Proteobacteria; Gammaproteobacteria; Enterobacterales; Enterobacteriaceae; Salmonella; Salmonella enterica
MTGDNTLITSHGINRRDFMKLCAALAATMGLSSKAAAEMAESVSNPQRPPVIWIGAQECT WP_000145429.1
hydrogenase 2 small subunit [Salmonella enterica] Length: 372\nNumber of predicted TMHs: 1\nExp number of AAs in TMHs: 24.51309\nExp number, first 60 AAs: 0.2682\nTotal prob of N-in: 0.14402\noutside 1 331\nTMhelix 332 354\ninside 355 372

15535 GCF_001716135.1_ASM171613v1 Salmonella enterica subsp. enterica Proteobacteria; Gammaproteobacteria; Enterobacterales; Enterobacteriaceae; Salmonella; Salmonella enterica
MNNEETFYQAMRRKGVTRRSFLKFCSLAATSLGLGAGMTPKIAWALENKPRIPVVWIHGL WP_001058311.1
hydrogenase [Salmonella enterica] Length: 372\nNumber of predicted TMHs: 2\nExp number of AAs in TMHs: 37.4699899999999\nExp number, first 60 AAs: 19.72699\nTotal prob of N-in: 0.96171\nPOSSIBLE N-term signal sequence\ninside 1 19\nTMhelix 20 39\noutside 40 325\nTMhelix 326 348\ninside 349 372

15536 GCF_001714185.1_ASM171418v1 Salmonella enterica subsp. enterica Proteobacteria; Gammaproteobacteria; Enterobacterales; Enterobacteriaceae; Salmonella; Salmonella enterica
MTGDNTLITSHGINRRDFMKLCAALAATMGLSSKAAAEMAESVSNPQRPPVIWIGAQECT WP_000145429.1
hydrogenase 2 small subunit [Salmonella enterica] Length: 372\nNumber of predicted TMHs: 1\nExp

number of AAs in TMHs: 24.51309\nExp number, first 60 AAs: 0.2682\nTotal prob of N-in: 0.14402\noutside 1 331\nTMhelix 332 354\ninside 355 372

15537 GCF_001716195.1_ASM171619v1 Salmonella enterica subsp. enterica Proteobacteria; Gammaproteobacteria; Enterobacterales; Enterobacteriaceae; Salmonella; Salmonella enterica MNNEETFYQAMRRKGVTRRSFLKFCSLAATSLGLGAGMTPKIAWALENKPRIPVVWIHGL WP_001058311.1 hydrogenase [Salmonella enterica] Length: 372\nNumber of predicted TMHs: 2\nExp number of AAs in TMHs: 37.4699899999999\nExp number, first 60 AAs: 19.72699\nTotal prob of N-in: 0.96171\nPOSSIBLE N-term signal sequence\ninside 1 19\nTMhelix 20 39\noutside 40 325\nTMhelix 326 348\ninside 349 372

15538 GCF_001716585.1_ASM171658v1 Salmonella enterica subsp. enterica Proteobacteria; Gammaproteobacteria; Enterobacterales; Enterobacteriaceae; Salmonella; Salmonella enterica MTGDNTLITSHGINRRDFMKLCAALAATMGLSSKAAAEMAESVSNPQRPPVIWIGAQECT WP_000145429.1 hydrogenase 2 small subunit [Salmonella enterica] Length: 372\nNumber of predicted TMHs: 1\nExp number of AAs in TMHs: 24.51309\nExp number, first 60 AAs: 0.2682\nTotal prob of N-in: 0.14402\noutside 1 331\nTMhelix 332 354\ninside 355 372

15539 GCF_001716695.1_ASM171669v1 Salmonella enterica subsp. enterica Proteobacteria; Gammaproteobacteria; Enterobacterales; Enterobacteriaceae; Salmonella; Salmonella enterica MNNEETFYQAMRRKGVTRRSFLKFCSLAATSLGLGAGMTPKIAWALENKPRIPVVWIHGL WP_001058311.1 hydrogenase [Salmonella enterica] Length: 372\nNumber of predicted TMHs: 2\nExp number of AAs in TMHs: 37.4699899999999\nExp number, first 60 AAs: 19.72699\nTotal prob of N-in: 0.96171\nPOSSIBLE N-term signal sequence\ninside 1 19\nTMhelix 20 39\noutside 40 325\nTMhelix 326 348\ninside 349 372

15540 GCF_001716655.1_ASM171665v1 Salmonella enterica subsp. enterica Proteobacteria; Gammaproteobacteria; Enterobacterales; Enterobacteriaceae; Salmonella; Salmonella enterica MNNEETFYQAMRRKGVTRRSFLKFCSLAATSLGLGAGMTPKIAWALENKPRIPVVWIHGL WP_001058311.1 hydrogenase [Salmonella enterica] Length: 372\nNumber of predicted TMHs: 2\nExp number of AAs in TMHs: 37.4699899999999\nExp number, first 60 AAs: 19.72699\nTotal prob of N-in: 0.96171\nPOSSIBLE N-term signal sequence\ninside 1 19\nTMhelix 20 39\noutside 40 325\nTMhelix 326 348\ninside 349 372

15541 GCF_000806865.1_CFSAN024723_01.0Salmonella enterica subsp. enterica Proteobacteria; Gammaproteobacteria; Enterobacterales; Enterobacteriaceae; Salmonella; Salmonella enterica MTGDNTLITSHGINRRDFMKLCAALAATMGLSSKAAAEMAESVSNPQRPPVIWIGAQECT WP_000145429.1 hydrogenase 2 small subunit [Salmonella enterica] Length: 372\nNumber of predicted TMHs: 1\nExp number of AAs in TMHs: 24.51309\nExp number, first 60 AAs: 0.2682\nTotal prob of N-in: 0.14402\noutside 1 331\nTMhelix 332 354\ninside 355 372

15542 GCF_001716695.1_ASM171669v1 Salmonella enterica subsp. enterica Proteobacteria; Gammaproteobacteria; Enterobacterales; Enterobacteriaceae; Salmonella; Salmonella enterica MTGDNTLITSHGINRRDFMKLCAALAATMGLSSKAAAEMAESVSNPQRPPVIWIGAQECT WP_000145429.1 hydrogenase 2 small subunit [Salmonella enterica] Length: 372\nNumber of predicted TMHs: 1\nExp number of AAs in TMHs: 24.51309\nExp number, first 60 AAs: 0.2682\nTotal prob of N-in: 0.14402\noutside 1 331\nTMhelix 332 354\ninside 355 372

15543 GCF_001716385.1_ASM171638v1 Salmonella enterica subsp. enterica Proteobacteria; Gammaproteobacteria; Enterobacterales; Enterobacteriaceae; Salmonella; Salmonella enterica MTGDNTLITSHGINRRDFMKLCAALAATMGLSSKAAAEMAESVSNPQRPPVIWIGAQECT WP_000145429.1 hydrogenase 2 small subunit [Salmonella enterica] Length: 372\nNumber of predicted TMHs: 1\nExp number of AAs in TMHs: 24.51309\nExp number, first 60 AAs: 0.2682\nTotal prob of N-in: 0.14402\noutside 1 331\nTMhelix 332 354\ninside 355 372

15544 GCF_001716425.1_ASM171642v1 Salmonella enterica subsp. enterica Proteobacteria; Gammaproteobacteria; Enterobacterales; Enterobacteriaceae; Salmonella; Salmonella enterica MTGDNTLITSHGINRRDFMKLCAALAATMGLSSKAAAEMAESVSNPQRPPVIWIGAQECT WP_000145429.1 hydrogenase 2 small subunit [Salmonella enterica] Length: 372\nNumber of predicted TMHs: 1\nExp number of AAs in TMHs: 24.51309\nExp number, first 60 AAs: 0.2682\nTotal prob of N-in: 0.14402\noutside 1 331\nTMhelix 332 354\ninside 355 372

15545 GCF_001716145.1_ASM171614v1 Salmonella enterica subsp. enterica Proteobacteria; Gammaproteobacteria; Enterobacterales; Enterobacteriaceae; Salmonella; Salmonella enterica

MNNEETFYQAMRRKGVTRRSFLKFCSLAATSLGLGAGMTPKIAWALENKPRIPVVWIHGL WP_001058311.1 hydrogenase [Salmonella enterica] Length: 372\nNumber of predicted TMHs: 2\nExp number of AAs in TMHs: 37.4699899999999\nExp number, first 60 AAs: 19.72699\nTotal prob of N-in: 0.96171\nPOSSIBLE N-term signal sequence\ninside 1 19\nTMhelix 20 39\noutside 40 325\nTMhelix 326 348\ninside 349 372

15546 GCF_001716055.1_ASM171605v1 Salmonella enterica subsp. enterica Proteobacteria; Gammaproteobacteria; Enterobacterales; Enterobacteriaceae; Salmonella; Salmonella enterica MNNEETFYQAMRRKGVTRRSFLKFCSLAATSLGLGAGMTPKIAWALENKPRIPVVWIHGL WP_001058311.1 hydrogenase [Salmonella enterica] Length: 372\nNumber of predicted TMHs: 2\nExp number of AAs in TMHs: 37.4699899999999\nExp number, first 60 AAs: 19.72699\nTotal prob of N-in: 0.96171\nPOSSIBLE N-term signal sequence\ninside 1 19\nTMhelix 20 39\noutside 40 325\nTMhelix 326 348\ninside 349 372

15547 GCF_001716115.1_ASM171611v1 Salmonella enterica subsp. enterica Proteobacteria; Gammaproteobacteria; Enterobacterales; Enterobacteriaceae; Salmonella; Salmonella enterica MNNEETFYQAMRRKGVTRRSFLKFCSLAATSLGLGAGMTPKIAWALENKPRIPVVWIHGL WP_001058311.1 hydrogenase [Salmonella enterica] Length: 372\nNumber of predicted TMHs: 2\nExp number of AAs in TMHs: 37.4699899999999\nExp number, first 60 AAs: 19.72699\nTotal prob of N-in: 0.96171\nPOSSIBLE N-term signal sequence\ninside 1 19\nTMhelix 20 39\noutside 40 325\nTMhelix 326 348\ninside 349 372

15548 GCF_000806705.1_CFSAN024770_02.0Salmonella enterica subsp. enterica Proteobacteria; Gammaproteobacteria; Enterobacterales; Enterobacteriaceae; Salmonella; Salmonella enterica MNNEETFYQAMRRKGVTRRSFLKFCSLAATSLGLGAGMTPKIAWALENKPRIPVVWIHGL WP_001058311.1 hydrogenase [Salmonella enterica] Length: 372\nNumber of predicted TMHs: 2\nExp number of AAs in TMHs: 37.4699899999999\nExp number, first 60 AAs: 19.72699\nTotal prob of N-in: 0.96171\nPOSSIBLE N-term signal sequence\ninside 1 19\nTMhelix 20 39\noutside 40 325\nTMhelix 326 348\ninside 349 372

15549 GCF_001714095.1_ASM171409v1 Salmonella enterica subsp. enterica Proteobacteria; Gammaproteobacteria; Enterobacterales; Enterobacteriaceae; Salmonella; Salmonella enterica MNNEETFYQAMRRKGVTRRSFLKFCSLAATSLGLGAGMTPKIAWALENKPRIPVVWIHGL WP_001058307.1 hydrogenase [Salmonella enterica] Length: 372\nNumber of predicted TMHs: 2\nExp number of AAs in TMHs: 36.51738\nExp number, first 60 AAs: 19.72849\nTotal prob of N-in: 0.96170\nPOSSIBLE N-term signal sequence\ninside 1 19\nTMhelix 20 39\noutside 40 325\nTMhelix 326 348\ninside 349 372

15550 GCF_000806825.1_CFSAN024779_01.0Salmonella enterica subsp. enterica Proteobacteria; Gammaproteobacteria; Enterobacterales; Enterobacteriaceae; Salmonella; Salmonella enterica MNNEETFYQAMRRKGVTRRSFLKFCSLAATSLGLGAGMTPKIAWALENKPRIPVVWIHGL WP_001058311.1 hydrogenase [Salmonella enterica] Length: 372\nNumber of predicted TMHs: 2\nExp number of AAs in TMHs: 37.4699899999999\nExp number, first 60 AAs: 19.72699\nTotal prob of N-in: 0.96171\nPOSSIBLE N-term signal sequence\ninside 1 19\nTMhelix 20 39\noutside 40 325\nTMhelix 326 348\ninside 349 372

15551 GCF_001714325.1_ASM171432v1 Salmonella enterica subsp. enterica Proteobacteria; Gammaproteobacteria; Enterobacterales; Enterobacteriaceae; Salmonella; Salmonella enterica MNNEETFYQAMRRKGVTRRSFLKFCSLAATSLGLGAGMTPKIAWALENKPRIPVVWIHGL WP_001058307.1 hydrogenase [Salmonella enterica] Length: 372\nNumber of predicted TMHs: 2\nExp number of AAs in TMHs: 36.51738\nExp number, first 60 AAs: 19.72849\nTotal prob of N-in: 0.96170\nPOSSIBLE N-term signal sequence\ninside 1 19\nTMhelix 20 39\noutside 40 325\nTMhelix 326 348\ninside 349 372

15552 GCF_000807105.1_CFSAN024716_01.0Salmonella enterica subsp. enterica Proteobacteria; Gammaproteobacteria; Enterobacterales; Enterobacteriaceae; Salmonella; Salmonella enterica MTGDNTLITSHGINRRDFMKLCAALAATMGLSSKAAAEMAESVSNPQRPPVIWIGAQECT WP_000145429.1 hydrogenase 2 small subunit [Salmonella enterica] Length: 372\nNumber of predicted TMHs: 1\nExp number of AAs in TMHs: 24.51309\nExp number, first 60 AAs: 0.2682\nTotal prob of N-in: 0.14402\noutside 1 331\nTMhelix 332 354\ninside 355 372

15553 GCF_001707695.1_ASM170769v1 Salmonella enterica subsp. enterica Proteobacteria; Gammaproteobacteria; Enterobacterales; Enterobacteriaceae; Salmonella; Salmonella enterica MNNEETFYQAMRRKGVTRRSFLKFCSLAATSLGLGAGMTPKIAWALENKPRIPVVWIHGL WP_001058312.1 hydrogenase [Salmonella enterica] Length: 372\nNumber of predicted TMHs: 2\nExp number of AAs in TMHs: 36.51738\nExp number, first 60 AAs: 19.72849\nTotal prob of N-in: 0.96170\nPOSSIBLE N-term signal sequence\ninside 1 19\nTMhelix 20 39\noutside 40 325\nTMhelix 326 348\ninside 349 372

15554 GCF_000878845.1_10TTU468_a5 Salmonella enterica subsp. enterica Proteobacteria; Gammaproteobacteria; Enterobacterales; Enterobacteriaceae; Salmonella; Salmonella enterica MTGDNTLITSHGINRRDFMKLCAALAATMGLSSKAAAEMAESVSNPQRPPVIWIGAQECT WP_000145429.1 hydrogenase 2 small subunit [Salmonella enterica] Length: 372\nNumber of predicted TMHs: 1\nExp number of AAs in TMHs: 24.51309\nExp number, first 60 AAs: 0.2682\nTotal prob of N-in: 0.14402\noutside 1 331\nTMhelix 332 354\ninside 355 372

15555 GCF_000806965.1_CFSAN024777_01.0Salmonella enterica subsp. enterica Proteobacteria; Gammaproteobacteria; Enterobacterales; Enterobacteriaceae; Salmonella; Salmonella enterica MTGDNTLITSHGINRRDFMKLCAALAATMGLSSKAAAEMAESVSNPQRPPVIWIGAQECT WP_039509062.1 hydrogenase 2 small subunit [Salmonella enterica] Length: 372\nNumber of predicted TMHs: 1\nExp number of AAs in TMHs: 24.51305\nExp number, first 60 AAs: 0.26819\nTotal prob of N-in: 0.14402\noutside 1 331\nTMhelix 332 354\ninside 355 372

15556 GCF_000807025.1_CFSAN024720_01.0Salmonella enterica subsp. enterica Proteobacteria; Gammaproteobacteria; Enterobacterales; Enterobacteriaceae; Salmonella; Salmonella enterica MNNEETFYQAMRRKGVTRRSFLKFCSLAATSLGLGAGMTPKIAWALENKPRIPVVWIHGL WP_001058312.1 hydrogenase [Salmonella enterica] Length: 372\nNumber of predicted TMHs: 2\nExp number of AAs in TMHs: 36.51738\nExp number, first 60 AAs: 19.72849\nTotal prob of N-in: 0.96170\nPOSSIBLE N-term signal sequence\ninside 1 19\nTMhelix 20 39\noutside 40 325\nTMhelix 326 348\ninside 349 372

15557 GCF_001707695.1_ASM170769v1 Salmonella enterica subsp. enterica Proteobacteria; Gammaproteobacteria; Enterobacterales; Enterobacteriaceae; Salmonella; Salmonella enterica MTGDNTLITSHGINRRDFMKLCAALAATMGLSSKAAAEMAESVSNPQRPPVIWIGAQECT WP_000145429.1 hydrogenase 2 small subunit [Salmonella enterica] Length: 372\nNumber of predicted TMHs: 1\nExp number of AAs in TMHs: 24.51309\nExp number, first 60 AAs: 0.2682\nTotal prob of N-in: 0.14402\noutside 1 331\nTMhelix 332 354\ninside 355 372

15558 GCF_001714325.1_ASM171432v1 Salmonella enterica subsp. enterica Proteobacteria; Gammaproteobacteria; Enterobacterales; Enterobacteriaceae; Salmonella; Salmonella enterica MTGDNTLITSHGINRRDFMKLCAALAATMGLSSKAAAEMAESVSNPQRPPVIWIGAQECT WP_000145429.1 hydrogenase 2 small subunit [Salmonella enterica] Length: 372\nNumber of predicted TMHs: 1\nExp number of AAs in TMHs: 24.51309\nExp number, first 60 AAs: 0.2682\nTotal prob of N-in: 0.14402\noutside 1 331\nTMhelix 332 354\ninside 355 372

15559 GCF_000806605.1_CFSAN024761_02.0Salmonella enterica subsp. enterica Proteobacteria; Gammaproteobacteria; Enterobacterales; Enterobacteriaceae; Salmonella; Salmonella enterica MNNEETFYQAMRRKGVTRRSFLKFCSLAATSLGLSAGMTPKIAWALENKPRIPVVWIHGL WP_001058316.1 hydrogenase [Salmonella enterica] Length: 372\nNumber of predicted TMHs: 2\nExp number of AAs in TMHs: 35.76397\nExp number, first 60 AAs: 18.97201\nTotal prob of N-in: 0.92785\nPOSSIBLE N-term signal sequence\ninside 1 19\nTMhelix 20 39\noutside 40 325\nTMhelix 326 348\ninside 349 372

15560 GCF_001716785.1_ASM171678v1 Salmonella enterica subsp. enterica Proteobacteria; Gammaproteobacteria; Enterobacterales; Enterobacteriaceae; Salmonella; Salmonella enterica MTGDNTLITSHGINRRDFMKLCAALAATMGLSSKAAAEMAESVSNPQRPPVIWIGAQECT WP_000145429.1 hydrogenase 2 small subunit [Salmonella enterica] Length: 372\nNumber of predicted TMHs: 1\nExp number of AAs in TMHs: 24.51309\nExp number, first 60 AAs: 0.2682\nTotal prob of N-in: 0.14402\noutside 1 331\nTMhelix 332 354\ninside 355 372

15561 GCF_001716495.1_ASM171649v1 Salmonella enterica subsp. enterica Proteobacteria; Gammaproteobacteria; Enterobacterales; Enterobacteriaceae; Salmonella; Salmonella enterica MNNEETFYQAMRRKGVTRRSFLKFCSLAATSLGLGAGMTPKIAWALENKPRIPVVWIHGL WP_001058311.1 hydrogenase [Salmonella enterica] Length: 372\nNumber of predicted TMHs: 2\nExp number of AAs in TMHs: 37.4699899999999\nExp number, first 60 AAs: 19.72699\nTotal prob of N-in: 0.96171\nPOSSIBLE N-term signal sequence\ninside 1 19\nTMhelix 20 39\noutside 40 325\nTMhelix 326 348\ninside 349 372

15562 GCF_000806745.1_CFSAN024773_01.0Salmonella enterica subsp. enterica Proteobacteria; Gammaproteobacteria; Enterobacterales; Enterobacteriaceae; Salmonella; Salmonella enterica MTGDNTLITSHGINRRDFMKLCAALAATMGLSSKAAAEMAESVSNPQRPPVIWIGAQECT WP_000145429.1 hydrogenase 2 small subunit [Salmonella enterica] Length: 372\nNumber of predicted TMHs: 1\nExp

number of AAs in TMHs: 24.51309\nExp number, first 60 AAs: 0.2682\nTotal prob of N-in: 0.14402\noutside 1 331\nTMhelix 332 354\ninside 355 372

15563 GCF_000772635.1_PDT000034041.1 Salmonella enterica subsp. enterica Proteobacteria; Gammaproteobacteria; Enterobacterales; Enterobacteriaceae; Salmonella; Salmonella enterica
MTGDNTLITSHGINRRDFMKLCAALAATMGLSSKAAAEMAESVSNPQRPPVIWIGAQECT WP_000145429.1
hydrogenase 2 small subunit [Salmonella enterica] Length: 372\nNumber of predicted TMHs: 1\nExp number of AAs in TMHs: 24.51309\nExp number, first 60 AAs: 0.2682\nTotal prob of N-in: 0.14402\noutside 1 331\nTMhelix 332 354\ninside 355 372

15564 GCF_000772635.1_PDT000034041.1 Salmonella enterica subsp. enterica Proteobacteria; Gammaproteobacteria; Enterobacterales; Enterobacteriaceae; Salmonella; Salmonella enterica
MNNEETFYQAMRRKGVTRRSFLKFCSLAATSLGLGAGMTPKIAWALENKPRIPVVWIHGL WP_001058312.1
hydrogenase [Salmonella enterica] Length: 372\nNumber of predicted TMHs: 2\nExp number of AAs in TMHs: 36.51738\nExp number, first 60 AAs: 19.72849\nTotal prob of N-in: 0.96170\nPOSSIBLE N-term signal sequence\ninside 1 19\nTMhelix 20 39\noutside 40 325\nTMhelix 326 348\ninside 349 372

15565 GCF_000807025.1_CFSAN024720_01.0Salmonella enterica subsp. enterica Proteobacteria; Gammaproteobacteria; Enterobacterales; Enterobacteriaceae; Salmonella; Salmonella enterica
MTGDNTLITSHGINRRDFMKLCAALAATMGLSSKAAAEMAESVSNPQRPPVIWIGAQECT WP_000145429.1
hydrogenase 2 small subunit [Salmonella enterica] Length: 372\nNumber of predicted TMHs: 1\nExp number of AAs in TMHs: 24.51309\nExp number, first 60 AAs: 0.2682\nTotal prob of N-in: 0.14402\noutside 1 331\nTMhelix 332 354\ninside 355 372

15566 GCF_001647935.1_ASM164793v1 Salmonella enterica subsp. enterica Proteobacteria; Gammaproteobacteria; Enterobacterales; Enterobacteriaceae; Salmonella; Salmonella enterica
MTGDNTLITSHGINRRDFMKLCAALAATMGLSSKAAAEMAESVSNPQRPPVIWIGAQECT WP_000145429.1
hydrogenase 2 small subunit [Salmonella enterica] Length: 372\nNumber of predicted TMHs: 1\nExp number of AAs in TMHs: 24.51309\nExp number, first 60 AAs: 0.2682\nTotal prob of N-in: 0.14402\noutside 1 331\nTMhelix 332 354\ninside 355 372

15567 GCF_000806825.1_CFSAN024779_01.0Salmonella enterica subsp. enterica Proteobacteria; Gammaproteobacteria; Enterobacterales; Enterobacteriaceae; Salmonella; Salmonella enterica
MTGDNTLITSHGINRRDFMKLCAALAATMGLSSKAAAEMAESVSNPQRPPVIWIGAQECT WP_000145429.1
hydrogenase 2 small subunit [Salmonella enterica] Length: 372\nNumber of predicted TMHs: 1\nExp number of AAs in TMHs: 24.51309\nExp number, first 60 AAs: 0.2682\nTotal prob of N-in: 0.14402\noutside 1 331\nTMhelix 332 354\ninside 355 372

15568 GCF_001592925.1_ASM159292v1 Salmonella enterica subsp. enterica Proteobacteria; Gammaproteobacteria; Enterobacterales; Enterobacteriaceae; Salmonella; Salmonella enterica
MTGDNTLITSHGINRRDFMKLCAALAATMGLSSKAAAEMAESVSNPQRPPVIWIGAQECT WP_000145429.1
hydrogenase 2 small subunit [Salmonella enterica] Length: 372\nNumber of predicted TMHs: 1\nExp number of AAs in TMHs: 24.51309\nExp number, first 60 AAs: 0.2682\nTotal prob of N-in: 0.14402\noutside 1 331\nTMhelix 332 354\ninside 355 372

15569 GCF_001586435.1_ASM158643v1 Salmonella enterica subsp. enterica Proteobacteria; Gammaproteobacteria; Enterobacterales; Enterobacteriaceae; Salmonella; Salmonella enterica
MNNEETFYQAMRRKGVTRRSFLKFCSLAATSLGLGAGMTPKIAWALENKPRIPVVWIHGL WP_001058311.1
hydrogenase [Salmonella enterica] Length: 372\nNumber of predicted TMHs: 2\nExp number of AAs in TMHs: 37.4699899999999\nExp number, first 60 AAs: 19.72699\nTotal prob of N-in: 0.96171\nPOSSIBLE N-term signal sequence\ninside 1 19\nTMhelix 20 39\noutside 40 325\nTMhelix 326 348\ninside 349 372

15570 GCF_001586525.1_ASM158652v1 Salmonella enterica subsp. enterica Proteobacteria; Gammaproteobacteria; Enterobacterales; Enterobacteriaceae; Salmonella; Salmonella enterica
MTGDNTLITSHGINRRDFMKLCAALAATMGLSSKAAAEMAESVSNPQRPPVIWIGAQECT WP_000145429.1
hydrogenase 2 small subunit [Salmonella enterica] Length: 372\nNumber of predicted TMHs: 1\nExp number of AAs in TMHs: 24.51309\nExp number, first 60 AAs: 0.2682\nTotal prob of N-in: 0.14402\noutside 1 331\nTMhelix 332 354\ninside 355 372

15571 GCF_001586465.1_ASM158646v1 Salmonella enterica subsp. enterica Proteobacteria; Gammaproteobacteria; Enterobacterales; Enterobacteriaceae; Salmonella; Salmonella enterica

MTGDNTLITSHGINRRDFMKLCAALAATMGLSSKAAAEMAESVSNPQRPPVIWIGAQECT WP_000145429.1 hydrogenase 2 small subunit [Salmonella enterica] Length: 372\nNumber of predicted TMHs: 1\nExp number of AAs in TMHs: 24.51309\nExp number, first 60 AAs: 0.2682\nTotal prob of N-in: 0.14402\noutside 1 331\nTMhelix 332 354\ninside 355 372

15572 GCF_001586435.1_ASM158643v1 Salmonella enterica subsp. enterica Proteobacteria; Gammaproteobacteria; Enterobacterales; Enterobacteriaceae; Salmonella; Salmonella enterica MTGDNTLITSHGINRRDFMKLCAALAATMGLSSKAAAEMAESVSNPQRPPVIWIGAQECT WP_000145429.1 hydrogenase 2 small subunit [Salmonella enterica] Length: 372\nNumber of predicted TMHs: 1\nExp number of AAs in TMHs: 24.51309\nExp number, first 60 AAs: 0.2682\nTotal prob of N-in: 0.14402\noutside 1 331\nTMhelix 332 354\ninside 355 372

15573 GCF_002106315.1_ASM210631v1 Salmonella enterica subsp. enterica Proteobacteria; Gammaproteobacteria; Enterobacterales; Enterobacteriaceae; Salmonella; Salmonella enterica MNNEETFYQAMRRKGVTRRSFLKFCSLAATSLGLGAGMTPKIAWALENKPRIPVVWIHGL WP_042847606.1 hydrogenase [Salmonella enterica] Length: 372\nNumber of predicted TMHs: 2\nExp number of AAs in TMHs: 37.4207199999999\nExp number, first 60 AAs: 19.72732\nTotal prob of N-in: 0.96168\nPOSSIBLE N-term signal sequence\ninside 1 19\nTMhelix 20 39\noutside 40 325\nTMhelix 326 348\ninside 349 372

15574 GCF_002106225.1_ASM210622v1 Salmonella enterica subsp. enterica Proteobacteria; Gammaproteobacteria; Enterobacterales; Enterobacteriaceae; Salmonella; Salmonella enterica MNNEETFYQAMRRKGVTRRSFLKFCSLAATSLGLGAGMTPKIAWALENKPRIPVVWIHGL WP_042847606.1 hydrogenase [Salmonella enterica] Length: 372\nNumber of predicted TMHs: 2\nExp number of AAs in TMHs: 37.4207199999999\nExp number, first 60 AAs: 19.72732\nTotal prob of N-in: 0.96168\nPOSSIBLE N-term signal sequence\ninside 1 19\nTMhelix 20 39\noutside 40 325\nTMhelix 326 348\ninside 349 372

15575 GCF_002106225.1_ASM210622v1 Salmonella enterica subsp. enterica Proteobacteria; Gammaproteobacteria; Enterobacterales; Enterobacteriaceae; Salmonella; Salmonella enterica MTGDNTLITSHGINRRDFMKLCAALAATMGLSSKAAAEMAESVSNPQRPPVIWIGAQECT WP_000145429.1 hydrogenase 2 small subunit [Salmonella enterica] Length: 372\nNumber of predicted TMHs: 1\nExp number of AAs in TMHs: 24.51309\nExp number, first 60 AAs: 0.2682\nTotal prob of N-in: 0.14402\noutside 1 331\nTMhelix 332 354\ninside 355 372

15576 GCF_000807055.1_CFSAN024717_01.0Salmonella enterica subsp. enterica Proteobacteria; Gammaproteobacteria; Enterobacterales; Enterobacteriaceae; Salmonella; Salmonella enterica MTGDNTLITSHGINRRDFMKLCAALAATMGLSSKAAAEMAESVSNPQRPPVIWIGAQECT WP_000145429.1 hydrogenase 2 small subunit [Salmonella enterica] Length: 372\nNumber of predicted TMHs: 1\nExp number of AAs in TMHs: 24.51309\nExp number, first 60 AAs: 0.2682\nTotal prob of N-in: 0.14402\noutside 1 331\nTMhelix 332 354\ninside 355 372

15577 GCF_002107425.1_ASM210742v1 Salmonella enterica subsp. enterica Proteobacteria; Gammaproteobacteria; Enterobacterales; Enterobacteriaceae; Salmonella; Salmonella enterica MTGDNTLITSHGINRRDFMKLCAALAATMGLSSKAAAEMAESVSNPQRPPVIWIGAQECT WP_000145429.1 hydrogenase 2 small subunit [Salmonella enterica] Length: 372\nNumber of predicted TMHs: 1\nExp number of AAs in TMHs: 24.51309\nExp number, first 60 AAs: 0.2682\nTotal prob of N-in: 0.14402\noutside 1 331\nTMhelix 332 354\ninside 355 372

15578 GCF_002105805.1_ASM210580v1 Salmonella enterica subsp. enterica Proteobacteria; Gammaproteobacteria; Enterobacterales; Enterobacteriaceae; Salmonella; Salmonella enterica MNNEETFYQAMRRKGVTRRSFLKFCSLAATSLGLGAGMTPKIAWALENKPRIPVVWIHGL WP_042847606.1 hydrogenase [Salmonella enterica] Length: 372\nNumber of predicted TMHs: 2\nExp number of AAs in TMHs: 37.4207199999999\nExp number, first 60 AAs: 19.72732\nTotal prob of N-in: 0.96168\nPOSSIBLE N-term signal sequence\ninside 1 19\nTMhelix 20 39\noutside 40 325\nTMhelix 326 348\ninside 349 372

15579 GCF_002106315.1_ASM210631v1 Salmonella enterica subsp. enterica Proteobacteria; Gammaproteobacteria; Enterobacterales; Enterobacteriaceae; Salmonella; Salmonella enterica MTGDNTLITSHGINRRDFMKLCAALAATMGLSSKAAAEMAESVSNPQRPPVIWIGAQECT WP_000145429.1 hydrogenase 2 small subunit [Salmonella enterica] Length: 372\nNumber of predicted TMHs: 1\nExp number of AAs in TMHs: 24.51309\nExp number, first 60 AAs: 0.2682\nTotal prob of N-in: 0.14402\noutside 1 331\nTMhelix 332 354\ninside 355 372

15580 GCF_002105805.1_ASM210580v1 Salmonella enterica subsp. enterica Proteobacteria; Gammaproteobacteria; Enterobacterales; Enterobacteriaceae; Salmonella; Salmonella enterica MTGDNTLITSHGINRRDFMKLCAALAATMGLSSKAAAEMAESVSNPQRPPVIWIGAQECT WP_000145429.1 hydrogenase 2 small subunit [Salmonella enterica] Length: 372\nNumber of predicted TMHs: 1\nExp number of AAs in TMHs: 24.51309\nExp number, first 60 AAs: 0.2682\nTotal prob of N-in: 0.14402\noutside 1 331\nTMhelix 332 354\ninside 355 372

15581 GCF_002163995.1_ASM216399v1 Salmonella enterica subsp. enterica Proteobacteria; Gammaproteobacteria; Enterobacterales; Enterobacteriaceae; Salmonella; Salmonella enterica MTGDNTLITSHGINRRDFMKLCAALAATMGLSSKAAAEMAESVSNPQRPPVIWIGAQECT WP_000145429.1 hydrogenase 2 small subunit [Salmonella enterica] Length: 372\nNumber of predicted TMHs: 1\nExp number of AAs in TMHs: 24.51309\nExp number, first 60 AAs: 0.2682\nTotal prob of N-in: 0.14402\noutside 1 331\nTMhelix 332 354\ninside 355 372

15582 GCF_002163995.1_ASM216399v1 Salmonella enterica subsp. enterica Proteobacteria; Gammaproteobacteria; Enterobacterales; Enterobacteriaceae; Salmonella; Salmonella enterica MNNEETFYQAMRRKGVTRRSFLKFCSLAATSLGLGAGMTPKIAWALENKPRIPVVWIHGL WP_001058312.1 hydrogenase [Salmonella enterica] Length: 372\nNumber of predicted TMHs: 2\nExp number of AAs in TMHs: 36.51738\nExp number, first 60 AAs: 19.72849\nTotal prob of N-in: 0.96170\nPOSSIBLE N-term signal sequence\ninside 1 19\nTMhelix 20 39\noutside 40 325\nTMhelix 326 348\ninside 349 372

15583 GCF_002128385.1_ASM212838v1 Salmonella enterica subsp. enterica Proteobacteria; Gammaproteobacteria; Enterobacterales; Enterobacteriaceae; Salmonella; Salmonella enterica MTGDNTLITSHGINRRDFMKLCAALAATMGLSSKAAAEMAESVSNPQRPPVIWIGAQECT WP_000145429.1 hydrogenase 2 small subunit [Salmonella enterica] Length: 372\nNumber of predicted TMHs: 1\nExp number of AAs in TMHs: 24.51309\nExp number, first 60 AAs: 0.2682\nTotal prob of N-in: 0.14402\noutside 1 331\nTMhelix 332 354\ninside 355 372

15584 GCF_002128365.1_ASM212836v1 Salmonella enterica subsp. enterica Proteobacteria; Gammaproteobacteria; Enterobacterales; Enterobacteriaceae; Salmonella; Salmonella enterica MTGDNTLITSHGINRRDFMKLCAALAATMGLSSKAAAEMAESVSNPQRPPVIWIGAQECT WP_000145429.1 hydrogenase 2 small subunit [Salmonella enterica] Length: 372\nNumber of predicted TMHs: 1\nExp number of AAs in TMHs: 24.51309\nExp number, first 60 AAs: 0.2682\nTotal prob of N-in: 0.14402\noutside 1 331\nTMhelix 332 354\ninside 355 372

15585 GCF_000806645.1_CFSAN024765_02.0Salmonella enterica subsp. enterica Proteobacteria; Gammaproteobacteria; Enterobacterales; Enterobacteriaceae; Salmonella; Salmonella enterica MNNEETFYQAMRRKGVTRRSFLKFCSLAATSLGLGAGMTPKIAWALENKPRIPVVWIHGL WP_001058312.1 hydrogenase [Salmonella enterica] Length: 372\nNumber of predicted TMHs: 2\nExp number of AAs in TMHs: 36.51738\nExp number, first 60 AAs: 19.72849\nTotal prob of N-in: 0.96170\nPOSSIBLE N-term signal sequence\ninside 1 19\nTMhelix 20 39\noutside 40 325\nTMhelix 326 348\ninside 349 372

15586 GCF_000806945.1_CFSAN024767_02.0Salmonella enterica subsp. enterica Proteobacteria; Gammaproteobacteria; Enterobacterales; Enterobacteriaceae; Salmonella; Salmonella enterica MNNEETLYQAMRRKGVTRRSFLKFCSLAATSLGLGAGMTPKIAWALENKPRIPVVWIHGL WP_001752445.1 hydrogenase-1 small chain [Salmonella enterica] Length: 372\nNumber of predicted TMHs: 2\nExp number of AAs in TMHs: 36.5123\nExp number, first 60 AAs: 19.72341\nTotal prob of N-in: 0.96144\nPOSSIBLE N-term signal sequence\ninside 1 19\nTMhelix 20 39\noutside 40 325\nTMhelix 326 348\ninside 349 372

15587 GCF_000806625.1_CFSAN024764_02.0Salmonella enterica subsp. enterica Proteobacteria; Gammaproteobacteria; Enterobacterales; Enterobacteriaceae; Salmonella; Salmonella enterica MNNEETFYQAMRRKGVTRRSFLKFCSLAATSLGLGAGMTPKIAWALENKPRIPVVWIHGL WP_023243720.1 hydrogenase 1, small subunit [Salmonella enterica] Length: 372\nNumber of predicted TMHs: 2\nExp number of AAs in TMHs: 36.52147\nExp number, first 60 AAs: 19.72838\nTotal prob of N-in: 0.96170\nPOSSIBLE N-term signal sequence\ninside 1 19\nTMhelix 20 39\noutside 40 325\nTMhelix 326 348\ninside 349 372

15588 GCF_000974975.1_CFSAN003382_01.0Salmonella enterica subsp. enterica Proteobacteria; Gammaproteobacteria; Enterobacterales; Enterobacteriaceae; Salmonella; Salmonella enterica MNNEETFYQAMRRKGVTRRSFLKFCSLAATSLGLGAGMTPKIAWALENKPRIPVVWIHGL WP_001058312.1 hydrogenase [Salmonella enterica] Length: 372\nNumber of predicted TMHs: 2\nExp number of AAs in

TMHs: 36.51738\nExp number, first 60 AAs: 19.72849\nTotal prob of N-in: 0.96170\nPOSSIBLE N-term signal sequence\ninside 1 19\nTMhelix 20 39\noutside 40 325\nTMhelix 326 348\ninside 349 372

15589 GCF_000974975.1_CFSAN003382_01.0Salmonella enterica subsp. enterica Proteobacteria; Gammaproteobacteria; Enterobacterales; Enterobacteriaceae; Salmonella; Salmonella enterica MTGDNTLITSHGINRRDFMKLCAALAATMGLSSKAAAEMAESVSNPQRPPVIWIGAQECT WP_000145431.1 hydrogenase 2 small subunit [Salmonella enterica] Length: 372\nNumber of predicted TMHs: 1\nExp number of AAs in TMHs: 24.51505\nExp number, first 60 AAs: 0.2682\nTotal prob of N-in: 0.14406\noutside 1 331\nTMhelix 332 354\ninside 355 372

15590 GCF_000806845.1_CFSAN024722_01.0Salmonella enterica subsp. enterica Proteobacteria; Gammaproteobacteria; Enterobacterales; Enterobacteriaceae; Salmonella; Salmonella enterica MTGDNTLITSHGINRRDFMKLCAALAATMGLSSKAAAEMAESVSNPQRPPVIWIGAQECT WP_000145429.1 hydrogenase 2 small subunit [Salmonella enterica] Length: 372\nNumber of predicted TMHs: 1\nExp number of AAs in TMHs: 24.51309\nExp number, first 60 AAs: 0.2682\nTotal prob of N-in: 0.14402\noutside 1 331\nTMhelix 332 354\ninside 355 372

15591 GCF_000806805.1_CFSAN024778_01.0Salmonella enterica subsp. enterica Proteobacteria; Gammaproteobacteria; Enterobacterales; Enterobacteriaceae; Salmonella; Salmonella enterica MTGDNTLITSHGINRRDFMKLCAALAATMGLSSKAAAEMAESVSNPQRPPVIWIGAQECT WP_000145429.1 hydrogenase 2 small subunit [Salmonella enterica] Length: 372\nNumber of predicted TMHs: 1\nExp number of AAs in TMHs: 24.51309\nExp number, first 60 AAs: 0.2682\nTotal prob of N-in: 0.14402\noutside 1 331\nTMhelix 332 354\ninside 355 372

15592 GCF_000806765.1_CFSAN024774_01.0Salmonella enterica subsp. enterica Proteobacteria; Gammaproteobacteria; Enterobacterales; Enterobacteriaceae; Salmonella; Salmonella enterica MTGDNTLITSHGINRRDFMKLCAALAATMGLSSKAAAEMAESVSNPQRPPVIWIGAQECT WP_000145429.1 hydrogenase 2 small subunit [Salmonella enterica] Length: 372\nNumber of predicted TMHs: 1\nExp number of AAs in TMHs: 24.51309\nExp number, first 60 AAs: 0.2682\nTotal prob of N-in: 0.14402\noutside 1 331\nTMhelix 332 354\ninside 355 372

15593 GCF_000806905.1_CFSAN024759_02.0Salmonella enterica subsp. enterica Proteobacteria; Gammaproteobacteria; Enterobacterales; Enterobacteriaceae; Salmonella; Salmonella enterica MTGDNTLITSHGINRRDFMKLCAALAATMGLSSKAAAEMAESVSNPQRPPVIWIGAQECT WP_000145429.1 hydrogenase 2 small subunit [Salmonella enterica] Length: 372\nNumber of predicted TMHs: 1\nExp number of AAs in TMHs: 24.51309\nExp number, first 60 AAs: 0.2682\nTotal prob of N-in: 0.14402\noutside 1 331\nTMhelix 332 354\ninside 355 372

15594 GCF_000806845.1_CFSAN024722_01.0Salmonella enterica subsp. enterica Proteobacteria; Gammaproteobacteria; Enterobacterales; Enterobacteriaceae; Salmonella; Salmonella enterica MNNEETFYQAMRRKGVTRRSFLKFCSLAATSLGLGAGMTPKIAWALENKPRIPVVWIHGL WP_001058312.1 hydrogenase [Salmonella enterica] Length: 372\nNumber of predicted TMHs: 2\nExp number of AAs in TMHs: 36.51738\nExp number, first 60 AAs: 19.72849\nTotal prob of N-in: 0.96170\nPOSSIBLE N-term signal sequence\ninside 1 19\nTMhelix 20 39\noutside 40 325\nTMhelix 326 348\ninside 349 372

15595 GCF_000806905.1_CFSAN024759_02.0Salmonella enterica subsp. enterica Proteobacteria; Gammaproteobacteria; Enterobacterales; Enterobacteriaceae; Salmonella; Salmonella enterica MNNEETFYQAMRRKGVTRRSFLKFCSLAATSLGLSAGMTPKIAWALENKPRIPVVWIHGL WP_001058316.1 hydrogenase [Salmonella enterica] Length: 372\nNumber of predicted TMHs: 2\nExp number of AAs in TMHs: 35.76397\nExp number, first 60 AAs: 18.97201\nTotal prob of N-in: 0.92785\nPOSSIBLE N-term signal sequence\ninside 1 19\nTMhelix 20 39\noutside 40 325\nTMhelix 326 348\ninside 349 372

15596 GCF_000807035.1_CFSAN024719_01.0Salmonella enterica subsp. enterica Proteobacteria; Gammaproteobacteria; Enterobacterales; Enterobacteriaceae; Salmonella; Salmonella enterica MTGDNTLITSHGINRRDFMKLCAALAATMGLSSKAAAEMAESVSNPQRPPVIWIGAQECT WP_000145429.1 hydrogenase 2 small subunit [Salmonella enterica] Length: 372\nNumber of predicted TMHs: 1\nExp number of AAs in TMHs: 24.51309\nExp number, first 60 AAs: 0.2682\nTotal prob of N-in: 0.14402\noutside 1 331\nTMhelix 332 354\ninside 355 372

15597 GCF_000807035.1_CFSAN024719_01.0Salmonella enterica subsp. enterica Proteobacteria; Gammaproteobacteria; Enterobacterales; Enterobacteriaceae; Salmonella; Salmonella enterica

MNNEETFYQAMRRKGVTRRSFLKFCSLAATSLGLGAGMTPKIAWALENKPRIPVVWIHGL WP_001058312.1 hydrogenase [Salmonella enterica] Length: 372\nNumber of predicted TMHs: 2\nExp number of AAs in TMHs: 36.51738\nExp number, first 60 AAs: 19.72849\nTotal prob of N-in: 0.96170\nPOSSIBLE N-term signal sequence\ninside 1 19\nTMhelix 20 39\noutside 40 325\nTMhelix 326 348\ninside 349 372

15598 GCF_000806985.1_CFSAN024780_01.0Salmonella enterica subsp. enterica Proteobacteria; Gammaproteobacteria; Enterobacterales; Enterobacteriaceae; Salmonella; Salmonella enterica MNNEETFYQAMRRKGVTRRSFLKFCSLAATSLGLGAGMTPKIAWALENKPRIPVVWIHGL WP_001058311.1 hydrogenase [Salmonella enterica] Length: 372\nNumber of predicted TMHs: 2\nExp number of AAs in TMHs: 37.4699899999999\nExp number, first 60 AAs: 19.72699\nTotal prob of N-in: 0.96171\nPOSSIBLE N-term signal sequence\ninside 1 19\nTMhelix 20 39\noutside 40 325\nTMhelix 326 348\ninside 349 372

15599 GCF_000807055.1_CFSAN024717_01.0Salmonella enterica subsp. enterica Proteobacteria; Gammaproteobacteria; Enterobacterales; Enterobacteriaceae; Salmonella; Salmonella enterica MNNEETFYQAMRRKGVTRRSFLKFCSLAATSLGLGAGMTPKIAWALENKPRIPVVWIHGL WP_001058311.1 hydrogenase [Salmonella enterica] Length: 372\nNumber of predicted TMHs: 2\nExp number of AAs in TMHs: 37.4699899999999\nExp number, first 60 AAs: 19.72699\nTotal prob of N-in: 0.96171\nPOSSIBLE N-term signal sequence\ninside 1 19\nTMhelix 20 39\noutside 40 325\nTMhelix 326 348\ninside 349 372

15600 GCF_000878855.1_11TTU1590_a5 Salmonella enterica subsp. enterica Proteobacteria; Gammaproteobacteria; Enterobacterales; Enterobacteriaceae; Salmonella; Salmonella enterica MTGDNTLITSHGINRRDFMKLCAALAATMGLSSKAAAEMAESVSNPQRPPVIWIGAQECT WP_000145429.1 hydrogenase 2 small subunit [Salmonella enterica] Length: 372\nNumber of predicted TMHs: 1\nExp number of AAs in TMHs: 24.51309\nExp number, first 60 AAs: 0.2682\nTotal prob of N-in: 0.14402\noutside 1 331\nTMhelix 332 354\ninside 355 372

15601 GCF_000807125.1_CFSAN024718_01.0Salmonella enterica subsp. enterica Proteobacteria; Gammaproteobacteria; Enterobacterales; Enterobacteriaceae; Salmonella; Salmonella enterica MNNEETFYQAMRRKGVTRRSFLKFCSLAATSLGLGAGMTPKIAWALENKPRIPVVWIHGL WP_001058312.1 hydrogenase [Salmonella enterica] Length: 372\nNumber of predicted TMHs: 2\nExp number of AAs in TMHs: 36.51738\nExp number, first 60 AAs: 19.72849\nTotal prob of N-in: 0.96170\nPOSSIBLE N-term signal sequence\ninside 1 19\nTMhelix 20 39\noutside 40 325\nTMhelix 326 348\ninside 349 372

15602 GCF_000807125.1_CFSAN024718_01.0Salmonella enterica subsp. enterica Proteobacteria; Gammaproteobacteria; Enterobacterales; Enterobacteriaceae; Salmonella; Salmonella enterica MTGDNTLITSHGINRRDFMKLCAALAATMGLSSKAAAEMAESVSNPQRPPVIWIGAQECT WP_000145429.1 hydrogenase 2 small subunit [Salmonella enterica] Length: 372\nNumber of predicted TMHs: 1\nExp number of AAs in TMHs: 24.51309\nExp number, first 60 AAs: 0.2682\nTotal prob of N-in: 0.14402\noutside 1 331\nTMhelix 332 354\ninside 355 372

15603 GCF_000878855.1_11TTU1590_a5 Salmonella enterica subsp. enterica Proteobacteria; Gammaproteobacteria; Enterobacterales; Enterobacteriaceae; Salmonella; Salmonella enterica MNNEETFYQAMRRKGVTRRSFLKFCSLAATSLGLGAGMTPKIAWALENKPRIPVVWIHGL WP_001058311.1 hydrogenase [Salmonella enterica] Length: 372\nNumber of predicted TMHs: 2\nExp number of AAs in TMHs: 37.4699899999999\nExp number, first 60 AAs: 19.72699\nTotal prob of N-in: 0.96171\nPOSSIBLE N-term signal sequence\ninside 1 19\nTMhelix 20 39\noutside 40 325\nTMhelix 326 348\ninside 349 372

15604 GCF_000806805.1_CFSAN024778_01.0Salmonella enterica subsp. enterica Proteobacteria; Gammaproteobacteria; Enterobacterales; Enterobacteriaceae; Salmonella; Salmonella enterica MNNEETFYQAMRRKGVTRRSFLKFCSLAATSLGLGAGMTPKIAWALENKPRIPVVWIHGL WP_001058311.1 hydrogenase [Salmonella enterica] Length: 372\nNumber of predicted TMHs: 2\nExp number of AAs in TMHs: 37.4699899999999\nExp number, first 60 AAs: 19.72699\nTotal prob of N-in: 0.96171\nPOSSIBLE N-term signal sequence\ninside 1 19\nTMhelix 20 39\noutside 40 325\nTMhelix 326 348\ninside 349 372

15605 GCF_001714545.1_ASM171454v1 Salmonella enterica subsp. enterica Proteobacteria; Gammaproteobacteria; Enterobacterales; Enterobacteriaceae; Salmonella; Salmonella enterica MTGDNTLITSHGINRRDFMKLCAALAATMGLSSKAAAEMAESVSNPQRPPVIWIGAQECT WP_000145429.1 hydrogenase 2 small subunit [Salmonella enterica] Length: 372\nNumber of predicted TMHs: 1\nExp number of AAs in TMHs: 24.51309\nExp number, first 60 AAs: 0.2682\nTotal prob of N-in: 0.14402\noutside 1 331\nTMhelix 332 354\ninside 355 372

15606 GCF_000806525.1_CFSAN024725_01.0Salmonella enterica subsp. enterica Proteobacteria; Gammaproteobacteria; Enterobacterales; Enterobacteriaceae; Salmonella; Salmonella enterica MTGDNTLITSHGINRRDFMKLCAALAATMGLSSKAAAEMAESVSNPQRPPVIWIGAQECT WP_000145429.1 hydrogenase 2 small subunit [Salmonella enterica] Length: 372\nNumber of predicted TMHs: 1\nExp number of AAs in TMHs: 24.51309\nExp number, first 60 AAs: 0.2682\nTotal prob of N-in: 0.14402\noutside 1 331\nTMhelix 332 354\ninside 355 372

15607 GCF_001714165.1_ASM171416v1 Salmonella enterica subsp. enterica Proteobacteria; Gammaproteobacteria; Enterobacterales; Enterobacteriaceae; Salmonella; Salmonella enterica MTGDNTLITSHGINRRDFMKLCAALAATMGLSSKAAAEMAESVSNPQRPPVIWIGAQECT WP_000145429.1 hydrogenase 2 small subunit [Salmonella enterica] Length: 372\nNumber of predicted TMHs: 1\nExp number of AAs in TMHs: 24.51309\nExp number, first 60 AAs: 0.2682\nTotal prob of N-in: 0.14402\noutside 1 331\nTMhelix 332 354\ninside 355 372

15608 GCF_000806625.1_CFSAN024764_02.0Salmonella enterica subsp. enterica Proteobacteria; Gammaproteobacteria; Enterobacterales; Enterobacteriaceae; Salmonella; Salmonella enterica MTGDNTLITSHGINRRDFMKLCAALAATMGLSSKAAAEMAESVSNPQRPPVIWIGAQECT WP_000145429.1 hydrogenase 2 small subunit [Salmonella enterica] Length: 372\nNumber of predicted TMHs: 1\nExp number of AAs in TMHs: 24.51309\nExp number, first 60 AAs: 0.2682\nTotal prob of N-in: 0.14402\noutside 1 331\nTMhelix 332 354\ninside 355 372

15609 GCF_001716625.1_ASM171662v1 Salmonella enterica subsp. enterica Proteobacteria; Gammaproteobacteria; Enterobacterales; Enterobacteriaceae; Salmonella; Salmonella enterica MTGDNTLITSHGINRRDFMKLCAALAATMGLSSKAAAEMAESVSNPQRPPVIWIGAQECT WP_000145429.1 hydrogenase 2 small subunit [Salmonella enterica] Length: 372\nNumber of predicted TMHs: 1\nExp number of AAs in TMHs: 24.51309\nExp number, first 60 AAs: 0.2682\nTotal prob of N-in: 0.14402\noutside 1 331\nTMhelix 332 354\ninside 355 372

15610 GCF_000806585.1_CFSAN024760_02.0Salmonella enterica subsp. enterica Proteobacteria; Gammaproteobacteria; Enterobacterales; Enterobacteriaceae; Salmonella; Salmonella enterica MTGDNTLITSHGINRRDFMKLCAALAATMGLSSKAAAEMAESVSNPQRPPVIWIGAQECT WP_000145429.1 hydrogenase 2 small subunit [Salmonella enterica] Length: 372\nNumber of predicted TMHs: 1\nExp number of AAs in TMHs: 24.51309\nExp number, first 60 AAs: 0.2682\nTotal prob of N-in: 0.14402\noutside 1 331\nTMhelix 332 354\ninside 355 372

15611 GCF_000806645.1_CFSAN024765_02.0Salmonella enterica subsp. enterica Proteobacteria; Gammaproteobacteria; Enterobacterales; Enterobacteriaceae; Salmonella; Salmonella enterica MTGDNTLITSHGINRRDFMKLCAALAATMGLSSKAAAEMAESVSNPQRPPVIWIGAQECT WP_000145429.1 hydrogenase 2 small subunit [Salmonella enterica] Length: 372\nNumber of predicted TMHs: 1\nExp number of AAs in TMHs: 24.51309\nExp number, first 60 AAs: 0.2682\nTotal prob of N-in: 0.14402\noutside 1 331\nTMhelix 332 354\ninside 355 372

15612 GCF_000806525.1_CFSAN024725_01.0Salmonella enterica subsp. enterica Proteobacteria; Gammaproteobacteria; Enterobacterales; Enterobacteriaceae; Salmonella; Salmonella enterica MNNEETFYQAMRRKGVTRRSFLKFCSLAATSLGLGAGMTPKIAWALENKPRIPVVWIHGL WP_001058312.1 hydrogenase [Salmonella enterica] Length: 372\nNumber of predicted TMHs: 2\nExp number of AAs in TMHs: 36.51738\nExp number, first 60 AAs: 19.72849\nTotal prob of N-in: 0.96170\nPOSSIBLE N-term signal sequence\ninside 1 19\nTMhelix 20 39\noutside 40 325\nTMhelix 326 348\ninside 349 372

15613 GCF_000806665.1_CFSAN024768_02.0Salmonella enterica subsp. enterica Proteobacteria; Gammaproteobacteria; Enterobacterales; Enterobacteriaceae; Salmonella; Salmonella enterica MNNEETLYQAMRRKGVTRRSFLKFCSLAATSLGLGAGMTPKIAWALENKPRIPVVWIHGL WP_001752445.1 hydrogenase-1 small chain [Salmonella enterica] Length: 372\nNumber of predicted TMHs: 2\nExp number of AAs in TMHs: 36.5123\nExp number, first 60 AAs: 19.72341\nTotal prob of N-in: 0.96144\nPOSSIBLE N-term signal sequence\ninside 1 19\nTMhelix 20 39\noutside 40 325\nTMhelix 326 348\ninside 349 372

15614 GCF_000806725.1_CFSAN024771_01.0Salmonella enterica subsp. enterica Proteobacteria; Gammaproteobacteria; Enterobacterales; Enterobacteriaceae; Salmonella; Salmonella enterica MTGDNTLITSHGINRRDFMKLCAALAATMGLSSKAAAEMAESVSNPQRPPVIWIGAQECT WP_000145429.1 hydrogenase 2 small subunit [Salmonella enterica] Length: 372\nNumber of predicted TMHs: 1\nExp

number of AAs in TMHs: 24.51309\nExp number, first 60 AAs: 0.2682\nTotal prob of N-in: 0.14402\noutside 1 331\nTMhelix 332 354\ninside 355 372

15615 GCF_001716665.1_ASM171666v1 Salmonella enterica subsp. enterica Proteobacteria; Gammaproteobacteria; Enterobacterales; Enterobacteriaceae; Salmonella; Salmonella enterica MTGDNTLITSHGINRRDFMKLCAALAATMGLSSKAAAEMAESVSNPQRPPVIWIGAQECT WP_000145429.1 hydrogenase 2 small subunit [Salmonella enterica] Length: 372\nNumber of predicted TMHs: 1\nExp number of AAs in TMHs: 24.51309\nExp number, first 60 AAs: 0.2682\nTotal prob of N-in: 0.14402\noutside 1 331\nTMhelix 332 354\ninside 355 372

15616 GCF_000806725.1_CFSAN024771_01.0Salmonella enterica subsp. enterica Proteobacteria; Gammaproteobacteria; Enterobacterales; Enterobacteriaceae; Salmonella; Salmonella enterica MNNEETFYQAMRRKGVTRRSFLKFCSLAATSLGLGAGMTPKIAWALENKPRIPVVWIHGL WP_001058311.1 hydrogenase [Salmonella enterica] Length: 372\nNumber of predicted TMHs: 2\nExp number of AAs in TMHs: 37.4699899999999\nExp number, first 60 AAs: 19.72699\nTotal prob of N-in: 0.96171\nPOSSIBLE N-term signal sequence\ninside 1 19\nTMhelix 20 39\noutside 40 325\nTMhelix 326 348\ninside 349 372

15617 GCF_001188125.1_ASM118812v1 Salmonella enterica subsp. enterica Proteobacteria; Gammaproteobacteria; Enterobacterales; Enterobacteriaceae; Salmonella; Salmonella enterica MNNEETFYQAMRRKGVTRRSFLKFCSLAATSLGLGAGMTPKIAWALENKPRIPVVWIHGL WP_001058312.1 hydrogenase [Salmonella enterica] Length: 372\nNumber of predicted TMHs: 2\nExp number of AAs in TMHs: 36.51738\nExp number, first 60 AAs: 19.72849\nTotal prob of N-in: 0.96170\nPOSSIBLE N-term signal sequence\ninside 1 19\nTMhelix 20 39\noutside 40 325\nTMhelix 326 348\ninside 349 372

15618 GCF_000806765.1_CFSAN024774_01.0Salmonella enterica subsp. enterica Proteobacteria; Gammaproteobacteria; Enterobacterales; Enterobacteriaceae; Salmonella; Salmonella enterica MNNEETFYQAMRRKGVTRRSFLKFCSLAATSLGLGAGMTPKIAWALENKPRIPVVWIHGL WP_001058312.1 hydrogenase [Salmonella enterica] Length: 372\nNumber of predicted TMHs: 2\nExp number of AAs in TMHs: 36.51738\nExp number, first 60 AAs: 19.72849\nTotal prob of N-in: 0.96170\nPOSSIBLE N-term signal sequence\ninside 1 19\nTMhelix 20 39\noutside 40 325\nTMhelix 326 348\ninside 349 372

15619 GCF_000806585.1_CFSAN024760_02.0Salmonella enterica subsp. enterica Proteobacteria; Gammaproteobacteria; Enterobacterales; Enterobacteriaceae; Salmonella; Salmonella enterica MNNEETFYQAMRRKGVTRRSFLKFCSLAATSLGLSAGMTPKIAWALENKPRIPVVWIHGL WP_001058316.1 hydrogenase [Salmonella enterica] Length: 372\nNumber of predicted TMHs: 2\nExp number of AAs in TMHs: 35.76397\nExp number, first 60 AAs: 18.97201\nTotal prob of N-in: 0.92785\nPOSSIBLE N-term signal sequence\ninside 1 19\nTMhelix 20 39\noutside 40 325\nTMhelix 326 348\ninside 349 372

15620 GCF_000806945.1_CFSAN024767_02.0Salmonella enterica subsp. enterica Proteobacteria; Gammaproteobacteria; Enterobacterales; Enterobacteriaceae; Salmonella; Salmonella enterica MTGDNTLITSHGINRRDFMKLCAALAATMGLSSKAAAEMAESVSNPQRPPVIWIGAQECT WP_000145428.1 hydrogenase 2 small subunit [Salmonella enterica] Length: 372\nNumber of predicted TMHs: 1\nExp number of AAs in TMHs: 24.50677\nExp number, first 60 AAs: 0.2682\nTotal prob of N-in: 0.14390\noutside 1 331\nTMhelix 332 354\ninside 355 372

15621 GCF_002128365.1_ASM212836v1 Salmonella enterica subsp. enterica Proteobacteria; Gammaproteobacteria; Enterobacterales; Enterobacteriaceae; Salmonella; Salmonella enterica MNNEETFYQAMRRKGVTRRSFLKFCSLAATSLGLGAGMTPKIAWALENKPRIPVVWIHGL WP_001058307.1 hydrogenase [Salmonella enterica] Length: 372\nNumber of predicted TMHs: 2\nExp number of AAs in TMHs: 36.51738\nExp number, first 60 AAs: 19.72849\nTotal prob of N-in: 0.96170\nPOSSIBLE N-term signal sequence\ninside 1 19\nTMhelix 20 39\noutside 40 325\nTMhelix 326 348\ninside 349 372

15622 GCF_001188125.1_ASM118812v1 Salmonella enterica subsp. enterica Proteobacteria; Gammaproteobacteria; Enterobacterales; Enterobacteriaceae; Salmonella; Salmonella enterica MTGDNTLITSHGINRRDFMKLCAALAATMGLSSKAAAEMAESVSNPQRPPVIWIGAQECT WP_000145429.1 hydrogenase 2 small subunit [Salmonella enterica] Length: 372\nNumber of predicted TMHs: 1\nExp number of AAs in TMHs: 24.51309\nExp number, first 60 AAs: 0.2682\nTotal prob of N-in: 0.14402\noutside 1 331\nTMhelix 332 354\ninside 355 372

15623 GCF_002128385.1_ASM212838v1 Salmonella enterica subsp. enterica Proteobacteria; Gammaproteobacteria; Enterobacterales; Enterobacteriaceae; Salmonella; Salmonella enterica

MNNEETFYQAMRRKGVTRRSFLKFCSLAATSLGLGAGMTPKIAWALENKPRIPVVWIHGL WP_001058307.1 hydrogenase [Salmonella enterica] Length: 372\nNumber of predicted TMHs: 2\nExp number of AAs in TMHs: 36.51738\nExp number, first 60 AAs: 19.72849\nTotal prob of N-in: 0.96170\nPOSSIBLE N-term signal sequence\ninside 1 19\nTMhelix 20 39\noutside 40 325\nTMhelix 326 348\ninside 349 372

15624 GCF_002176715.1_ASM217671v1 Salmonella enterica subsp. enterica Proteobacteria; Gammaproteobacteria; Enterobacterales; Enterobacteriaceae; Salmonella; Salmonella enterica MNNEETFYQAMRRKGVTRRSFLKFCSLAATSLGLGAGMTPKIAWALENKPRIPVVWIHGL WP_001058307.1 hydrogenase [Salmonella enterica] Length: 372\nNumber of predicted TMHs: 2\nExp number of AAs in TMHs: 36.51738\nExp number, first 60 AAs: 19.72849\nTotal prob of N-in: 0.96170\nPOSSIBLE N-term signal sequence\ninside 1 19\nTMhelix 20 39\noutside 40 325\nTMhelix 326 348\ninside 349 372

15625 GCF_002105275.1_ASM210527v1 Salmonella enterica subsp. enterica Proteobacteria; Gammaproteobacteria; Enterobacterales; Enterobacteriaceae; Salmonella; Salmonella enterica MNNEETFYQAMRRKGVTRRSFLKFCSLAATSLGLGAGMTPKIAWALENKPRIPVVWIHGL WP_001058307.1 hydrogenase [Salmonella enterica] Length: 372\nNumber of predicted TMHs: 2\nExp number of AAs in TMHs: 36.51738\nExp number, first 60 AAs: 19.72849\nTotal prob of N-in: 0.96170\nPOSSIBLE N-term signal sequence\ninside 1 19\nTMhelix 20 39\noutside 40 325\nTMhelix 326 348\ninside 349 372

15626 GCF_002105275.1_ASM210527v1 Salmonella enterica subsp. enterica Proteobacteria; Gammaproteobacteria; Enterobacterales; Enterobacteriaceae; Salmonella; Salmonella enterica MTGDNTLITSHGINRRDFMKLCAALAATMGLSSKAAAEMAESVSNPQRPPVIWIGAQECT WP_000145429.1 hydrogenase 2 small subunit [Salmonella enterica] Length: 372\nNumber of predicted TMHs: 1\nExp number of AAs in TMHs: 24.51309\nExp number, first 60 AAs: 0.2682\nTotal prob of N-in: 0.14402\noutside 1 331\nTMhelix 332 354\ninside 355 372

15627 GCF_002107425.1_ASM210742v1 Salmonella enterica subsp. enterica Proteobacteria; Gammaproteobacteria; Enterobacterales; Enterobacteriaceae; Salmonella; Salmonella enterica MNNEETFYQAMRRKGVTRRSFLKFCSLAATSLGLGAGMTPKIAWALENKPRIPVVWIHGL WP_001058312.1 hydrogenase [Salmonella enterica] Length: 372\nNumber of predicted TMHs: 2\nExp number of AAs in TMHs: 36.51738\nExp number, first 60 AAs: 19.72849\nTotal prob of N-in: 0.96170\nPOSSIBLE N-term signal sequence\ninside 1 19\nTMhelix 20 39\noutside 40 325\nTMhelix 326 348\ninside 349 372

15628 GCF_002176715.1_ASM217671v1 Salmonella enterica subsp. enterica Proteobacteria; Gammaproteobacteria; Enterobacterales; Enterobacteriaceae; Salmonella; Salmonella enterica MTGDNTLITSHGINRRDFMKLCAALAATMGLSSKAAAEMAESVSNPQRPPVIWIGAQECT WP_000145429.1 hydrogenase 2 small subunit [Salmonella enterica] Length: 372\nNumber of predicted TMHs: 1\nExp number of AAs in TMHs: 24.51309\nExp number, first 60 AAs: 0.2682\nTotal prob of N-in: 0.14402\noutside 1 331\nTMhelix 332 354\ninside 355 372

15629 GCF_002066745.1_ASM206674v1 Salmonella enterica subsp. enterica Proteobacteria; Gammaproteobacteria; Enterobacterales; Enterobacteriaceae; Salmonella; Salmonella enterica MTGDNTLITSHGINRRDFMKLCAALAATMGLSSKAAAEMAESVSNPQRPPVIWIGAQECT WP_000145429.1 hydrogenase 2 small subunit [Salmonella enterica] Length: 372\nNumber of predicted TMHs: 1\nExp number of AAs in TMHs: 24.51309\nExp number, first 60 AAs: 0.2682\nTotal prob of N-in: 0.14402\noutside 1 331\nTMhelix 332 354\ninside 355 372

15630 GCF_002066745.1_ASM206674v1 Salmonella enterica subsp. enterica Proteobacteria; Gammaproteobacteria; Enterobacterales; Enterobacteriaceae; Salmonella; Salmonella enterica MNNEETFYQAMRRKGVTRRSFLKFCSLAATSLGLGAGMTPKIAWALENKPRIPVVWIHGL WP_001540183.1 hydrogenase [Salmonella enterica] Length: 372\nNumber of predicted TMHs: 2\nExp number of AAs in TMHs: 37.41804\nExp number, first 60 AAs: 19.72808\nTotal prob of N-in: 0.96169\nPOSSIBLE N-term signal sequence\ninside 1 19\nTMhelix 20 39\noutside 40 325\nTMhelix 326 348\ninside 349 372

15631 GCF_002058255.1_ASM205825v1 Salmonella enterica subsp. enterica Proteobacteria; Gammaproteobacteria; Enterobacterales; Enterobacteriaceae; Salmonella; Salmonella enterica MTGDNTLITSHGINRRDFMKLCAALAATMGLSSKAAAEMAESVSNPQRPPVIWIGAQECT WP_000145429.1 hydrogenase 2 small subunit [Salmonella enterica] Length: 372\nNumber of predicted TMHs: 1\nExp number of AAs in TMHs: 24.51309\nExp number, first 60 AAs: 0.2682\nTotal prob of N-in: 0.14402\noutside 1 331\nTMhelix 332 354\ninside 355 372

15632 GCF_000806545.1_CFSAN024757_02.0Salmonella enterica subsp. enterica Proteobacteria; Gammaproteobacteria; Enterobacterales; Enterobacteriaceae; Salmonella; Salmonella enterica
MTGDNTLITSHGINRRDFMKLCAALAATMGLSSKAAAEMAESVSNPQRPPVIWIGAQECT WP_000145429.1
hydrogenase 2 small subunit [Salmonella enterica] Length: 372\nNumber of predicted TMHs: 1\nExp number of AAs in TMHs: 24.51309\nExp number, first 60 AAs: 0.2682\nTotal prob of N-in: 0.14402\noutside 1 331\nTMhelix 332 354\ninside 355 372

15633 GCF_001716825.1_ASM171682v1 Salmonella enterica subsp. enterica Proteobacteria; Gammaproteobacteria; Enterobacterales; Enterobacteriaceae; Salmonella; Salmonella enterica
MNNEETFYQAMRRKGVTRRSFLKFCSLAATSLGLGAGMTPKIAWALENKPRIPVVWIHGL WP_001058311.1
hydrogenase [Salmonella enterica] Length: 372\nNumber of predicted TMHs: 2\nExp number of AAs in TMHs: 37.4699899999999\nExp number, first 60 AAs: 19.72699\nTotal prob of N-in: 0.96171\nPOSSIBLE N-term signal sequence\ninside 1 19\nTMhelix 20 39\noutside 40 325\nTMhelix 326 348\ninside 349 372

15634 GCF_001707605.1_ASM170760v1 Salmonella enterica subsp. enterica Proteobacteria; Gammaproteobacteria; Enterobacterales; Enterobacteriaceae; Salmonella; Salmonella enterica
MNNEETFYQAMRRKGVTRRSFLKFCSLAATSLGLGAGMTPKIAWALENKPRIPVVWIHGL WP_023243720.1
hydrogenase 1, small subunit [Salmonella enterica] Length: 372\nNumber of predicted TMHs: 2\nExp number of AAs in TMHs: 36.52147\nExp number, first 60 AAs: 19.72838\nTotal prob of N-in: 0.96170\nPOSSIBLE N-term signal sequence\ninside 1 19\nTMhelix 20 39\noutside 40 325\nTMhelix 326 348\ninside 349 372

15635 GCF_001716575.1_ASM171657v1 Salmonella enterica subsp. enterica Proteobacteria; Gammaproteobacteria; Enterobacterales; Enterobacteriaceae; Salmonella; Salmonella enterica
MNNEETFYQAMRRKGVTRRSFLKFCSLAATSLGLGAGMTPKIAWALENKPRIPVVWIHGL WP_001058311.1
hydrogenase [Salmonella enterica] Length: 372\nNumber of predicted TMHs: 2\nExp number of AAs in TMHs: 37.4699899999999\nExp number, first 60 AAs: 19.72699\nTotal prob of N-in: 0.96171\nPOSSIBLE N-term signal sequence\ninside 1 19\nTMhelix 20 39\noutside 40 325\nTMhelix 326 348\ninside 349 372

15636 GCF_001592925.1_ASM159292v1 Salmonella enterica subsp. enterica Proteobacteria; Gammaproteobacteria; Enterobacterales; Enterobacteriaceae; Salmonella; Salmonella enterica
MNNEETFYQAMRRKGVTRRSFLKFCSLAATSLGLGAGMTPKIAWALENKPRIPVVWIHGL WP_001058307.1
hydrogenase [Salmonella enterica] Length: 372\nNumber of predicted TMHs: 2\nExp number of AAs in TMHs: 36.51738\nExp number, first 60 AAs: 19.72849\nTotal prob of N-in: 0.96170\nPOSSIBLE N-term signal sequence\ninside 1 19\nTMhelix 20 39\noutside 40 325\nTMhelix 326 348\ninside 349 372

15637 GCF_002037475.1_ASM203747v1 Salmonella enterica subsp. enterica Proteobacteria; Gammaproteobacteria; Enterobacterales; Enterobacteriaceae; Salmonella; Salmonella enterica
MNNEETFYQAMRRKGVTRRSFLKFCSLAATSLGLGAGMTPKIAWALENKPRIPVVWIHGL WP_001058312.1
hydrogenase [Salmonella enterica] Length: 372\nNumber of predicted TMHs: 2\nExp number of AAs in TMHs: 36.51738\nExp number, first 60 AAs: 19.72849\nTotal prob of N-in: 0.96170\nPOSSIBLE N-term signal sequence\ninside 1 19\nTMhelix 20 39\noutside 40 325\nTMhelix 326 348\ninside 349 372

15638 GCF_002058255.1_ASM205825v1 Salmonella enterica subsp. enterica Proteobacteria; Gammaproteobacteria; Enterobacterales; Enterobacteriaceae; Salmonella; Salmonella enterica
MNNEETFYQAMRRKGVTRRSFLKFCSLAATSLGLGAGMTPKIAWALENKPRIPVVWIHGL WP_001058312.1
hydrogenase [Salmonella enterica] Length: 372\nNumber of predicted TMHs: 2\nExp number of AAs in TMHs: 36.51738\nExp number, first 60 AAs: 19.72849\nTotal prob of N-in: 0.96170\nPOSSIBLE N-term signal sequence\ninside 1 19\nTMhelix 20 39\noutside 40 325\nTMhelix 326 348\ninside 349 372

15639 GCF_001720385.1_ASM172038v1 Salmonella enterica subsp. enterica Proteobacteria; Gammaproteobacteria; Enterobacterales; Enterobacteriaceae; Salmonella; Salmonella enterica
MTGDNTLITSHGINRRDFMKLCAALAATMGLSSKAAAEMAESVSNPQRPPVIWIGAQECT WP_000145429.1
hydrogenase 2 small subunit [Salmonella enterica] Length: 372\nNumber of predicted TMHs: 1\nExp number of AAs in TMHs: 24.51309\nExp number, first 60 AAs: 0.2682\nTotal prob of N-in: 0.14402\noutside 1 331\nTMhelix 332 354\ninside 355 372

15640 GCF_002037475.1_ASM203747v1 Salmonella enterica subsp. enterica Proteobacteria; Gammaproteobacteria; Enterobacterales; Enterobacteriaceae; Salmonella; Salmonella enterica
MTGDNTLITSHGINRRDFMKLCAALAATMGLSSKAAAEMAESVSNPQRPPVIWIGAQECT WP_000145429.1
hydrogenase 2 small subunit [Salmonella enterica] Length: 372\nNumber of predicted TMHs: 1\nExp

number of AAs in TMHs: 24.51309\nExp number, first 60 AAs: 0.2682\nTotal prob of N-in: 0.14402\noutside 1 331\nTMhelix 332 354\ninside 355 372

15641 GCF_001716615.1_ASM171661v1 Salmonella enterica subsp. enterica Proteobacteria; Gammaproteobacteria; Enterobacterales; Enterobacteriaceae; Salmonella; Salmonella enterica MNNEETFYQAMRRKGVTRRSFLKFCSLAATSLGLGAGMTPKIAWALENKPRIPVVWIHGL WP_001058311.1 hydrogenase [Salmonella enterica] Length: 372\nNumber of predicted TMHs: 2\nExp number of AAs in TMHs: 37.4699899999999\nExp number, first 60 AAs: 19.72699\nTotal prob of N-in: 0.96171\nPOSSIBLE N-term signal sequence\ninside 1 19\nTMhelix 20 39\noutside 40 325\nTMhelix 326 348\ninside 349 372

15642 GCF_001714095.1_ASM171409v1 Salmonella enterica subsp. enterica Proteobacteria; Gammaproteobacteria; Enterobacterales; Enterobacteriaceae; Salmonella; Salmonella enterica MTGDNTLITSHGINRRDFMKLCAALAATMGLSSKAAAEMAESVSNPQRPPVIWIGAQECT WP_000145429.1 hydrogenase 2 small subunit [Salmonella enterica] Length: 372\nNumber of predicted TMHs: 1\nExp number of AAs in TMHs: 24.51309\nExp number, first 60 AAs: 0.2682\nTotal prob of N-in: 0.14402\noutside 1 331\nTMhelix 332 354\ninside 355 372

15643 GCF_000807005.1_CFSAN024781_01.0Salmonella enterica subsp. enterica Proteobacteria; Gammaproteobacteria; Enterobacterales; Enterobacteriaceae; Salmonella; Salmonella enterica MNNEETFYQAMRRKGVTRRSFLKFCSLAATSLGLGAGMTPKIAWALENKPRIPVVWIHGL WP_001058311.1 hydrogenase [Salmonella enterica] Length: 372\nNumber of predicted TMHs: 2\nExp number of AAs in TMHs: 37.4699899999999\nExp number, first 60 AAs: 19.72699\nTotal prob of N-in: 0.96171\nPOSSIBLE N-term signal sequence\ninside 1 19\nTMhelix 20 39\noutside 40 325\nTMhelix 326 348\ninside 349 372

15644 GCF_000806565.1_CFSAN024758_02.0Salmonella enterica subsp. enterica Proteobacteria; Gammaproteobacteria; Enterobacterales; Enterobacteriaceae; Salmonella; Salmonella enterica MNNEETFYQAMRRKGVTRRSFLKFCSLAATSLGLSAGMTPKIAWALENKPRIPVVWIHGL WP_001058316.1 hydrogenase [Salmonella enterica] Length: 372\nNumber of predicted TMHs: 2\nExp number of AAs in TMHs: 35.76397\nExp number, first 60 AAs: 18.97201\nTotal prob of N-in: 0.92785\nPOSSIBLE N-term signal sequence\ninside 1 19\nTMhelix 20 39\noutside 40 325\nTMhelix 326 348\ninside 349 372

15645 GCF_001714145.1_ASM171414v1 Salmonella enterica subsp. enterica Proteobacteria; Gammaproteobacteria; Enterobacterales; Enterobacteriaceae; Salmonella; Salmonella enterica MTGDNTLITSHGINRRDFMKLCAALAATMGLSSKAAAEMAESVSNPQRPPVIWIGAQECT WP_000145429.1 hydrogenase 2 small subunit [Salmonella enterica] Length: 372\nNumber of predicted TMHs: 1\nExp number of AAs in TMHs: 24.51309\nExp number, first 60 AAs: 0.2682\nTotal prob of N-in: 0.14402\noutside 1 331\nTMhelix 332 354\ninside 355 372

15646 GCF_000806545.1_CFSAN024757_02.0Salmonella enterica subsp. enterica Proteobacteria; Gammaproteobacteria; Enterobacterales; Enterobacteriaceae; Salmonella; Salmonella enterica MNNEETFYQAMRRKGVTRRSFLKFCSLAATSLGLSAGMTPKIAWALENKPRIPVVWIHGL WP_001058316.1 hydrogenase [Salmonella enterica] Length: 372\nNumber of predicted TMHs: 2\nExp number of AAs in TMHs: 35.76397\nExp number, first 60 AAs: 18.97201\nTotal prob of N-in: 0.92785\nPOSSIBLE N-term signal sequence\ninside 1 19\nTMhelix 20 39\noutside 40 325\nTMhelix 326 348\ninside 349 372

15647 GCF_001714145.1_ASM171414v1 Salmonella enterica subsp. enterica Proteobacteria; Gammaproteobacteria; Enterobacterales; Enterobacteriaceae; Salmonella; Salmonella enterica MNNEETFYQAMRRKGVTRRSFLKFCSLAATSLGLGAGMTPKIAWALENKPRIPVVWIHGL WP_001058307.1 hydrogenase [Salmonella enterica] Length: 372\nNumber of predicted TMHs: 2\nExp number of AAs in TMHs: 36.51738\nExp number, first 60 AAs: 19.72849\nTotal prob of N-in: 0.96170\nPOSSIBLE N-term signal sequence\ninside 1 19\nTMhelix 20 39\noutside 40 325\nTMhelix 326 348\ninside 349 372

15648 GCF_001714245.1_ASM171424v1 Salmonella enterica subsp. enterica Proteobacteria; Gammaproteobacteria; Enterobacterales; Enterobacteriaceae; Salmonella; Salmonella enterica MNNEETFYQAMRRKGVTRRSFLKFCSLAATSLGLGAGMTPKIAWALENKPRIPVVWIHGL WP_001058307.1 hydrogenase [Salmonella enterica] Length: 372\nNumber of predicted TMHs: 2\nExp number of AAs in TMHs: 36.51738\nExp number, first 60 AAs: 19.72849\nTotal prob of N-in: 0.96170\nPOSSIBLE N-term signal sequence\ninside 1 19\nTMhelix 20 39\noutside 40 325\nTMhelix 326 348\ninside 349 372

15649 GCF_001714265.1_ASM171426v1 Salmonella enterica subsp. enterica Proteobacteria; Gammaproteobacteria; Enterobacterales; Enterobacteriaceae; Salmonella; Salmonella enterica

MNNEETFYQAMRRKGVTRRSFLKFCSLAATSLGLGAGMTPKIAWALENKPRIPVVWIHGL WP_001058307.1 hydrogenase [Salmonella enterica] Length: 372\nNumber of predicted TMHs: 2\nExp number of AAs in TMHs: 36.51738\nExp number, first 60 AAs: 19.72849\nTotal prob of N-in: 0.96170\nPOSSIBLE N-term signal sequence\ninside 1 19\nTMhelix 20 39\noutside 40 325\nTMhelix 326 348\ninside 349 372

15650 GCF_001716045.1_ASM171604v1 Salmonella enterica subsp. enterica Proteobacteria; Gammaproteobacteria; Enterobacterales; Enterobacteriaceae; Salmonella; Salmonella enterica MTGDNTLITSHGINRRDFMKLCAALAATMGLSSKAAAEMAESVSNPQRPPVIWIGAQECT WP_000145429.1 hydrogenase 2 small subunit [Salmonella enterica] Length: 372\nNumber of predicted TMHs: 1\nExp number of AAs in TMHs: 24.51309\nExp number, first 60 AAs: 0.2682\nTotal prob of N-in: 0.14402\noutside 1 331\nTMhelix 332 354\ninside 355 372

15651 GCF_000807045.1_CFSAN024715_01.0Salmonella enterica subsp. enterica Proteobacteria; Gammaproteobacteria; Enterobacterales; Enterobacteriaceae; Salmonella; Salmonella enterica MNNEETFYQAMRRKGVTRRSFLKFCSLAATSLGLGAGMTPKIAWALENKPRIPVVWIHGL WP_001058311.1 hydrogenase [Salmonella enterica] Length: 372\nNumber of predicted TMHs: 2\nExp number of AAs in TMHs: 37.4699899999999\nExp number, first 60 AAs: 19.72699\nTotal prob of N-in: 0.96171\nPOSSIBLE N-term signal sequence\ninside 1 19\nTMhelix 20 39\noutside 40 325\nTMhelix 326 348\ninside 349 372

15652 GCF_000806885.1_CFSAN024756_02.0Salmonella enterica subsp. enterica Proteobacteria; Gammaproteobacteria; Enterobacterales; Enterobacteriaceae; Salmonella; Salmonella enterica MNNEETFYQAMRRKGVTRRSFLKFCSLAATSLGLSAGMTPKIAWALENKPRIPVVWIHGL WP_001058316.1 hydrogenase [Salmonella enterica] Length: 372\nNumber of predicted TMHs: 2\nExp number of AAs in TMHs: 35.76397\nExp number, first 60 AAs: 18.97201\nTotal prob of N-in: 0.92785\nPOSSIBLE N-term signal sequence\ninside 1 19\nTMhelix 20 39\noutside 40 325\nTMhelix 326 348\ninside 349 372

15653 GCF_000806505.1_CFSAN024724_01.0Salmonella enterica subsp. enterica Proteobacteria; Gammaproteobacteria; Enterobacterales; Enterobacteriaceae; Salmonella; Salmonella enterica MNNEETFYQAMRRKGVTRRSFLKFCSLAATSLGLGAGMTPKIAWALENKPRIPVVWIHGL WP_001058312.1 hydrogenase [Salmonella enterica] Length: 372\nNumber of predicted TMHs: 2\nExp number of AAs in TMHs: 36.51738\nExp number, first 60 AAs: 19.72849\nTotal prob of N-in: 0.96170\nPOSSIBLE N-term signal sequence\ninside 1 19\nTMhelix 20 39\noutside 40 325\nTMhelix 326 348\ninside 349 372

15654 GCF_000806685.1_CFSAN024769_02.0Salmonella enterica subsp. enterica Proteobacteria; Gammaproteobacteria; Enterobacterales; Enterobacteriaceae; Salmonella; Salmonella enterica MTGDNTLITSHGINRRDFMKLCAALAATMGLSSKAAAEMAESVSNPQRPPVIWIGAQECT WP_000145428.1 hydrogenase 2 small subunit [Salmonella enterica] Length: 372\nNumber of predicted TMHs: 1\nExp number of AAs in TMHs: 24.50677\nExp number, first 60 AAs: 0.2682\nTotal prob of N-in: 0.14390\noutside 1 331\nTMhelix 332 354\ninside 355 372

15655 GCF_000806885.1_CFSAN024756_02.0Salmonella enterica subsp. enterica Proteobacteria; Gammaproteobacteria; Enterobacterales; Enterobacteriaceae; Salmonella; Salmonella enterica MTGDNTLITSHGINRRDFMKLCAALAATMGLSSKAAAEMAESVSNPQRPPVIWIGAQECT WP_000145429.1 hydrogenase 2 small subunit [Salmonella enterica] Length: 372\nNumber of predicted TMHs: 1\nExp number of AAs in TMHs: 24.51309\nExp number, first 60 AAs: 0.2682\nTotal prob of N-in: 0.14402\noutside 1 331\nTMhelix 332 354\ninside 355 372

15656 GCF_001716115.1_ASM171611v1 Salmonella enterica subsp. enterica Proteobacteria; Gammaproteobacteria; Enterobacterales; Enterobacteriaceae; Salmonella; Salmonella enterica MTGDNTLITSHGINRRDFMKLCAALAATMGLSSKAAAEMAESVSNPQRPPVIWIGAQECT WP_000145429.1 hydrogenase 2 small subunit [Salmonella enterica] Length: 372\nNumber of predicted TMHs: 1\nExp number of AAs in TMHs: 24.51309\nExp number, first 60 AAs: 0.2682\nTotal prob of N-in: 0.14402\noutside 1 331\nTMhelix 332 354\ninside 355 372

15657 GCF_000806705.1_CFSAN024770_02.0Salmonella enterica subsp. enterica Proteobacteria; Gammaproteobacteria; Enterobacterales; Enterobacteriaceae; Salmonella; Salmonella enterica MTGDNTLITSHGINRRDFMKLCAALAATMGLSSKAAAEMAESVSNPQRPPVIWIGAQECT WP_000145429.1 hydrogenase 2 small subunit [Salmonella enterica] Length: 372\nNumber of predicted TMHs: 1\nExp number of AAs in TMHs: 24.51309\nExp number, first 60 AAs: 0.2682\nTotal prob of N-in: 0.14402\noutside 1 331\nTMhelix 332 354\ninside 355 372

15658	GCF_001716055.1_ASM171605v1	Salmonella enterica subsp. enterica	Proteobacteria; Gammaproteobacteria; Enterobacterales; Enterobacteriaceae; Salmonella; Salmonella enterica	MTGDNTLITSHGINRRDFMKLCAALAATMGLSSKAAAEMAESVSNPQRPPVIWIGAQECT	WP_000145429.1	hydrogenase 2 small subunit [Salmonella enterica]	Length: 372\nNumber of predicted TMHs:  1\nExp number of AAs in TMHs: 24.51309\nExp number, first 60 AAs:  0.2682\nTotal prob of N-in:   0.14402\noutside   1   331\nTMhelix   332   354\ninside   355   372

15659	GCF_000807045.1_CFSAN024715_01.0Salmonella enterica subsp. enterica	Proteobacteria; Gammaproteobacteria; Enterobacterales; Enterobacteriaceae; Salmonella; Salmonella enterica	MTGDNTLITSHGINRRDFMKLCAALAATMGLSSKAAAEMAESVSNPQRPPVIWIGAQECT	WP_000145429.1	hydrogenase 2 small subunit [Salmonella enterica]	Length: 372\nNumber of predicted TMHs:  1\nExp number of AAs in TMHs: 24.51309\nExp number, first 60 AAs:  0.2682\nTotal prob of N-in:   0.14402\noutside   1   331\nTMhelix   332   354\ninside   355   372

15660	GCF_001716145.1_ASM171614v1	Salmonella enterica subsp. enterica	Proteobacteria; Gammaproteobacteria; Enterobacterales; Enterobacteriaceae; Salmonella; Salmonella enterica	MTGDNTLITSHGINRRDFMKLCAALAATMGLSSKAAAEMAESVSNPQRPPVIWIGAQECT	WP_000145429.1	hydrogenase 2 small subunit [Salmonella enterica]	Length: 372\nNumber of predicted TMHs:  1\nExp number of AAs in TMHs: 24.51309\nExp number, first 60 AAs:  0.2682\nTotal prob of N-in:   0.14402\noutside   1   331\nTMhelix   332   354\ninside   355   372

15661	GCF_001716425.1_ASM171642v1	Salmonella enterica subsp. enterica	Proteobacteria; Gammaproteobacteria; Enterobacterales; Enterobacteriaceae; Salmonella; Salmonella enterica	MNNEETFYQAMRRKGVTRRSFLKFCSLAATSLGLGAGMTPKIAWALENKPRIPVVWIHGL	WP_001058311.1	hydrogenase [Salmonella enterica]	Length: 372\nNumber of predicted TMHs:  2\nExp number of AAs in TMHs: 37.4699899999999\nExp number, first 60 AAs:  19.72699\nTotal prob of N-in:   0.96171\nPOSSIBLE N-term signal sequence\ninside   1   19\nTMhelix   20   39\noutside   40   325\nTMhelix   326   348\ninside   349   372

15662	GCF_001716385.1_ASM171638v1	Salmonella enterica subsp. enterica	Proteobacteria; Gammaproteobacteria; Enterobacterales; Enterobacteriaceae; Salmonella; Salmonella enterica	MNNEETFYQAMRRKGVTRRSFLKFCSLAATSLGLGAGMTPKIAWALENKPRIPVVWIHGL	WP_001058311.1	hydrogenase [Salmonella enterica]	Length: 372\nNumber of predicted TMHs:  2\nExp number of AAs in TMHs: 37.4699899999999\nExp number, first 60 AAs:  19.72699\nTotal prob of N-in:   0.96171\nPOSSIBLE N-term signal sequence\ninside   1   19\nTMhelix   20   39\noutside   40   325\nTMhelix   326   348\ninside   349   372

15663	GCF_001716215.1_ASM171621v1	Salmonella enterica subsp. enterica	Proteobacteria; Gammaproteobacteria; Enterobacterales; Enterobacteriaceae; Salmonella; Salmonella enterica	MNNEETFYQAMRRKGVTRRSFLKFCSLAATSLGLGAGMTPKIAWALENKPRIPVVWIHGL	WP_001058311.1	hydrogenase [Salmonella enterica]	Length: 372\nNumber of predicted TMHs:  2\nExp number of AAs in TMHs: 37.4699899999999\nExp number, first 60 AAs:  19.72699\nTotal prob of N-in:   0.96171\nPOSSIBLE N-term signal sequence\ninside   1   19\nTMhelix   20   39\noutside   40   325\nTMhelix   326   348\ninside   349   372

15664	GCF_001716235.1_ASM171623v1	Salmonella enterica subsp. enterica	Proteobacteria; Gammaproteobacteria; Enterobacterales; Enterobacteriaceae; Salmonella; Salmonella enterica	MNNEETFYQAMRRKGVTRRSFLKFCSLAATSLGLGAGMTPKIAWALENKPRIPVVWIHGL	WP_001058311.1	hydrogenase [Salmonella enterica]	Length: 372\nNumber of predicted TMHs:  2\nExp number of AAs in TMHs: 37.4699899999999\nExp number, first 60 AAs:  19.72699\nTotal prob of N-in:   0.96171\nPOSSIBLE N-term signal sequence\ninside   1   19\nTMhelix   20   39\noutside   40   325\nTMhelix   326   348\ninside   349   372

15665	GCF_001586525.1_ASM158652v1	Salmonella enterica subsp. enterica	Proteobacteria; Gammaproteobacteria; Enterobacterales; Enterobacteriaceae; Salmonella; Salmonella enterica	MNNEETFYQAMRRKGVTRRSFLKFCSLAATSLGLGAGMTPKIAWALENKPRIPVVWIHGL	WP_001058311.1	hydrogenase [Salmonella enterica]	Length: 372\nNumber of predicted TMHs:  2\nExp number of AAs in TMHs: 37.4699899999999\nExp number, first 60 AAs:  19.72699\nTotal prob of N-in:   0.96171\nPOSSIBLE N-term signal sequence\ninside   1   19\nTMhelix   20   39\noutside   40   325\nTMhelix   326   348\ninside   349   372

15666	GCF_001586675.1_ASM158667v1	Salmonella enterica subsp. enterica	Proteobacteria; Gammaproteobacteria; Enterobacterales; Enterobacteriaceae; Salmonella; Salmonella enterica	MTGDNTLITSHGINRRDFMKLCAALAATMGLSSKAAAEMAESVSNPQRPPVIWIGAQECT	WP_000145429.1	hydrogenase 2 small subunit [Salmonella enterica]	Length: 372\nNumber of predicted TMHs:  1\nExp

number of AAs in TMHs: 24.51309\nExp number, first 60 AAs:  0.2682\nTotal prob of N-in:   0.14402\noutside   1   331\nTMhelix   332   354\ninside   355   372

15667 GCF_001586695.1_ASM158669v1 Salmonella enterica subsp. enterica Proteobacteria; Gammaproteobacteria; Enterobacterales; Enterobacteriaceae; Salmonella; Salmonella enterica MTGDNTLITSHGINRRDFMKLCAALAATMGLSSKAAAEMAESVSNPQRPPVIWIGAQECT WP_000145429.1 hydrogenase 2 small subunit [Salmonella enterica] Length: 372\nNumber of predicted TMHs:  1\nExp number of AAs in TMHs: 24.51309\nExp number, first 60 AAs:  0.2682\nTotal prob of N-in:   0.14402\noutside   1   331\nTMhelix   332   354\ninside   355   372

15668 GCF_001716295.1_ASM171629v1 Salmonella enterica subsp. enterica Proteobacteria; Gammaproteobacteria; Enterobacterales; Enterobacteriaceae; Salmonella; Salmonella enterica MNNEETFYQAMRRKGVTRRSFLKFCSLAATSLGLGAGMTPKIAWALENKPRIPVVWIHGL WP_001058311.1 hydrogenase [Salmonella enterica] Length: 372\nNumber of predicted TMHs:  2\nExp number of AAs in TMHs: 37.4699899999999\nExp number, first 60 AAs:  19.72699\nTotal prob of N-in:   0.96171\nPOSSIBLE N-term signal sequence\ninside   1   19\nTMhelix   20   39\noutside   40   325\nTMhelix   326   348\ninside   349   372

15669 GCF_001586675.1_ASM158667v1 Salmonella enterica subsp. enterica Proteobacteria; Gammaproteobacteria; Enterobacterales; Enterobacteriaceae; Salmonella; Salmonella enterica MNNEETFYQAMRRKGVTRRSFLKFCSLAATSLGLGAGMTPKIAWALENKPRIPVVWIHGL WP_001058311.1 hydrogenase [Salmonella enterica] Length: 372\nNumber of predicted TMHs:  2\nExp number of AAs in TMHs: 37.4699899999999\nExp number, first 60 AAs:  19.72699\nTotal prob of N-in:   0.96171\nPOSSIBLE N-term signal sequence\ninside   1   19\nTMhelix   20   39\noutside   40   325\nTMhelix   326   348\ninside   349   372

15670 GCF_001586695.1_ASM158669v1 Salmonella enterica subsp. enterica Proteobacteria; Gammaproteobacteria; Enterobacterales; Enterobacteriaceae; Salmonella; Salmonella enterica MNNEETFYQAMRRKGVTRRSFLKFCSLAATSLGLGAGMTPKIAWALENKPRIPVVWIHGL WP_001058311.1 hydrogenase [Salmonella enterica] Length: 372\nNumber of predicted TMHs:  2\nExp number of AAs in TMHs: 37.4699899999999\nExp number, first 60 AAs:  19.72699\nTotal prob of N-in:   0.96171\nPOSSIBLE N-term signal sequence\ninside   1   19\nTMhelix   20   39\noutside   40   325\nTMhelix   326   348\ninside   349   372

15671 GCF_001716335.1_ASM171633v1 Salmonella enterica subsp. enterica Proteobacteria; Gammaproteobacteria; Enterobacterales; Enterobacteriaceae; Salmonella; Salmonella enterica MNNEETFYQAMRRKGVTRRSFLKFCSLAATSLGLGAGMTPKIAWALENKPRIPVVWIHGL WP_001058311.1 hydrogenase [Salmonella enterica] Length: 372\nNumber of predicted TMHs:  2\nExp number of AAs in TMHs: 37.4699899999999\nExp number, first 60 AAs:  19.72699\nTotal prob of N-in:   0.96171\nPOSSIBLE N-term signal sequence\ninside   1   19\nTMhelix   20   39\noutside   40   325\nTMhelix   326   348\ninside   349   372

15672 GCF_001716475.1_ASM171647v1 Salmonella enterica subsp. enterica Proteobacteria; Gammaproteobacteria; Enterobacterales; Enterobacteriaceae; Salmonella; Salmonella enterica MTGDNTLITSHGINRRDFMKLCAALAATMGLSSKAAAEMAESVSNPQRPPVIWIGAQECT WP_000145429.1 hydrogenase 2 small subunit [Salmonella enterica] Length: 372\nNumber of predicted TMHs:  1\nExp number of AAs in TMHs: 24.51309\nExp number, first 60 AAs:  0.2682\nTotal prob of N-in:   0.14402\noutside   1   331\nTMhelix   332   354\ninside   355   372

15673 GCF_000807105.1_CFSAN024716_01.0Salmonella enterica subsp. enterica Proteobacteria; Gammaproteobacteria; Enterobacterales; Enterobacteriaceae; Salmonella; Salmonella enterica MNNEETFYQAMRRKGVTRRSFLKFCSLAATSLGLGAGMTPKIAWALENKPRIPVVWIHGL WP_001058311.1 hydrogenase [Salmonella enterica] Length: 372\nNumber of predicted TMHs:  2\nExp number of AAs in TMHs: 37.4699899999999\nExp number, first 60 AAs:  19.72699\nTotal prob of N-in:   0.96171\nPOSSIBLE N-term signal sequence\ninside   1   19\nTMhelix   20   39\noutside   40   325\nTMhelix   326   348\ninside   349   372

15674 GCF_001716785.1_ASM171678v1 Salmonella enterica subsp. enterica Proteobacteria; Gammaproteobacteria; Enterobacterales; Enterobacteriaceae; Salmonella; Salmonella enterica MNNEETFYQAMRRKGVTRRSFLKFCSLAATSLGLGAGMTPKIAWALENKPRIPVVWIHGL WP_001058311.1 hydrogenase [Salmonella enterica] Length: 372\nNumber of predicted TMHs:  2\nExp number of AAs in TMHs: 37.4699899999999\nExp number, first 60 AAs:  19.72699\nTotal prob of N-in:   0.96171\nPOSSIBLE N-term signal sequence\ninside   1   19\nTMhelix   20   39\noutside   40   325\nTMhelix   326   348\ninside   349   372

15675 GCF_000806605.1_CFSAN024761_02.0Salmonella enterica subsp. enterica Proteobacteria; Gammaproteobacteria; Enterobacterales; Enterobacteriaceae; Salmonella; Salmonella enterica

MTGDNTLITSHGINRRDFMKLCAALAATMGLSSKAAAEMAESVSNPQRPPVIWIGAQECT WP_000145429.1 hydrogenase 2 small subunit [Salmonella enterica] Length: 372\nNumber of predicted TMHs: 1\nExp number of AAs in TMHs: 24.51309\nExp number, first 60 AAs: 0.2682\nTotal prob of N-in: 0.14402\noutside 1 331\nTMhelix 332 354\ninside 355 372

15676 GCF_001647935.1_ASM164793v1 Salmonella enterica subsp. enterica Proteobacteria; Gammaproteobacteria; Enterobacterales; Enterobacteriaceae; Salmonella; Salmonella enterica MNNEETFYQAMRRKGVTRRSFLKFCSLAATSLGLGAGMTPKIAWALENKPRIPVVWIHGL WP_001058312.1 hydrogenase [Salmonella enterica] Length: 372\nNumber of predicted TMHs: 2\nExp number of AAs in TMHs: 36.51738\nExp number, first 60 AAs: 19.72849\nTotal prob of N-in: 0.96170\nPOSSIBLE N-term signal sequence\ninside 1 19\nTMhelix 20 39\noutside 40 325\nTMhelix 326 348\ninside 349 372

15677 GCF_001716535.1_ASM171653v1 Salmonella enterica subsp. enterica Proteobacteria; Gammaproteobacteria; Enterobacterales; Enterobacteriaceae; Salmonella; Salmonella enterica MNNEETFYQAMRRKGVTRRSFLKFCSLAATSLGLGAGMTPKIAWALENKPRIPVVWIHGL WP_001058311.1 hydrogenase [Salmonella enterica] Length: 372\nNumber of predicted TMHs: 2\nExp number of AAs in TMHs: 37.4699899999999\nExp number, first 60 AAs: 19.72699\nTotal prob of N-in: 0.96171\nPOSSIBLE N-term signal sequence\ninside 1 19\nTMhelix 20 39\noutside 40 325\nTMhelix 326 348\ninside 349 372

15678 GCF_001716825.1_ASM171682v1 Salmonella enterica subsp. enterica Proteobacteria; Gammaproteobacteria; Enterobacterales; Enterobacteriaceae; Salmonella; Salmonella enterica MTGDNTLITSHGINRRDFMKLCAALAATMGLSSKAAAEMAESVSNPQRPPVIWIGAQECT WP_000145429.1 hydrogenase 2 small subunit [Salmonella enterica] Length: 372\nNumber of predicted TMHs: 1\nExp number of AAs in TMHs: 24.51309\nExp number, first 60 AAs: 0.2682\nTotal prob of N-in: 0.14402\noutside 1 331\nTMhelix 332 354\ninside 355 372

15679 GCF_000806965.1_CFSAN024777_01.0Salmonella enterica subsp. enterica Proteobacteria; Gammaproteobacteria; Enterobacterales; Enterobacteriaceae; Salmonella; Salmonella enterica MNNEETFYQAMRRKGVTRRSFLKFCSLAATSLGLGAGMTPKIAWALENKPRIPVVWIHGL WP_001058312.1 hydrogenase [Salmonella enterica] Length: 372\nNumber of predicted TMHs: 2\nExp number of AAs in TMHs: 36.51738\nExp number, first 60 AAs: 19.72849\nTotal prob of N-in: 0.96170\nPOSSIBLE N-term signal sequence\ninside 1 19\nTMhelix 20 39\noutside 40 325\nTMhelix 326 348\ninside 349 372

15680 GCF_001006525.1_ASM100652v1 Salmonella enterica subsp. enterica Proteobacteria; Gammaproteobacteria; Enterobacterales; Enterobacteriaceae; Salmonella; Salmonella enterica MNNEETFYQAMRRKGVTRRSFLKFCSLAATSLGLGAGMTPKIAWALENKPRIPVVWIHGL WP_001058312.1 hydrogenase [Salmonella enterica] Length: 372\nNumber of predicted TMHs: 2\nExp number of AAs in TMHs: 36.51738\nExp number, first 60 AAs: 19.72849\nTotal prob of N-in: 0.96170\nPOSSIBLE N-term signal sequence\ninside 1 19\nTMhelix 20 39\noutside 40 325\nTMhelix 326 348\ninside 349 372

15681 GCF_000806665.1_CFSAN024768_02.0Salmonella enterica subsp. enterica Proteobacteria; Gammaproteobacteria; Enterobacterales; Enterobacteriaceae; Salmonella; Salmonella enterica MTGDNTLITSHGINRRDFMKLCAALAATMGLSSKAAAEMAESVSNPQRPPVIWIGAQECT WP_000145428.1 hydrogenase 2 small subunit [Salmonella enterica] Length: 372\nNumber of predicted TMHs: 1\nExp number of AAs in TMHs: 24.50677\nExp number, first 60 AAs: 0.2682\nTotal prob of N-in: 0.14390\noutside 1 331\nTMhelix 332 354\ninside 355 372

15682 GCF_000806985.1_CFSAN024780_01.0Salmonella enterica subsp. enterica Proteobacteria; Gammaproteobacteria; Enterobacterales; Enterobacteriaceae; Salmonella; Salmonella enterica MTGDNTLITSHGINRRDFMKLCAALAATMGLSSKAAAEMAESVSNPQRPPVIWIGAQECT WP_000145429.1 hydrogenase 2 small subunit [Salmonella enterica] Length: 372\nNumber of predicted TMHs: 1\nExp number of AAs in TMHs: 24.51309\nExp number, first 60 AAs: 0.2682\nTotal prob of N-in: 0.14402\noutside 1 331\nTMhelix 332 354\ninside 355 372

15683 GCF_001716305.1_ASM171630v1 Salmonella enterica subsp. enterica Proteobacteria; Gammaproteobacteria; Enterobacterales; Enterobacteriaceae; Salmonella; Salmonella enterica MTGDNTLITSHGINRRDFMKLCAALAATMGLSSKAAAEMAESVSNPQRPPVIWIGAQECT WP_000145429.1 hydrogenase 2 small subunit [Salmonella enterica] Length: 372\nNumber of predicted TMHs: 1\nExp number of AAs in TMHs: 24.51309\nExp number, first 60 AAs: 0.2682\nTotal prob of N-in: 0.14402\noutside 1 331\nTMhelix 332 354\ninside 355 372

15684 GCF_002201855.1_ASM220185v1 Salmonella enterica subsp. enterica Proteobacteria; Gammaproteobacteria; Enterobacterales; Enterobacteriaceae; Salmonella; Salmonella enterica MNNEETFYQAMRRKGVTRRSFLKFCSLAATSLGLGAGMTPKIAWALENKPRIPVVWIHGL WP_001058312.1 hydrogenase [Salmonella enterica] Length: 372\nNumber of predicted TMHs: 2\nExp number of AAs in TMHs: 36.51738\nExp number, first 60 AAs: 19.72849\nTotal prob of N-in: 0.96170\nPOSSIBLE N-term signal sequence\ninside 1 19\nTMhelix 20 39\noutside 40 325\nTMhelix 326 348\ninside 349 372

15685 GCF_001716715.1_ASM171671v1 Salmonella enterica subsp. enterica Proteobacteria; Gammaproteobacteria; Enterobacterales; Enterobacteriaceae; Salmonella; Salmonella enterica MTGDNTLITSHGINRRDFMKLCAALAATMGLSSKAAAEMAESVSNPQRPPVIWIGAQECT WP_000145429.1 hydrogenase 2 small subunit [Salmonella enterica] Length: 372\nNumber of predicted TMHs: 1\nExp number of AAs in TMHs: 24.51309\nExp number, first 60 AAs: 0.2682\nTotal prob of N-in: 0.14402\noutside 1 331\nTMhelix 332 354\ninside 355 372

15686 GCF_001714545.1_ASM171454v1 Salmonella enterica subsp. enterica Proteobacteria; Gammaproteobacteria; Enterobacterales; Enterobacteriaceae; Salmonella; Salmonella enterica MNNEETFYQAMRRKGVTRRSFLKFCSLAATSLGLGAGMTPKIAWALENKPRIPVVWIHGL WP_001058312.1 hydrogenase [Salmonella enterica] Length: 372\nNumber of predicted TMHs: 2\nExp number of AAs in TMHs: 36.51738\nExp number, first 60 AAs: 19.72849\nTotal prob of N-in: 0.96170\nPOSSIBLE N-term signal sequence\ninside 1 19\nTMhelix 20 39\noutside 40 325\nTMhelix 326 348\ninside 349 372

15687 GCF_001716275.1_ASM171627v1 Salmonella enterica subsp. enterica Proteobacteria; Gammaproteobacteria; Enterobacterales; Enterobacteriaceae; Salmonella; Salmonella enterica MNNEETFYQAMRRKGVTRRSFLKFCSLAATSLGLGAGMTPKIAWALENKPRIPVVWIHGL WP_001058311.1 hydrogenase [Salmonella enterica] Length: 372\nNumber of predicted TMHs: 2\nExp number of AAs in TMHs: 37.4699899999999\nExp number, first 60 AAs: 19.72699\nTotal prob of N-in: 0.96171\nPOSSIBLE N-term signal sequence\ninside 1 19\nTMhelix 20 39\noutside 40 325\nTMhelix 326 348\ninside 349 372

15688 GCF_001716135.1_ASM171613v1 Salmonella enterica subsp. enterica Proteobacteria; Gammaproteobacteria; Enterobacterales; Enterobacteriaceae; Salmonella; Salmonella enterica MTGDNTLITSHGINRRDFMKLCAALAATMGLSSKAAAEMAESVSNPQRPPVIWIGAQECT WP_000145429.1 hydrogenase 2 small subunit [Salmonella enterica] Length: 372\nNumber of predicted TMHs: 1\nExp number of AAs in TMHs: 24.51309\nExp number, first 60 AAs: 0.2682\nTotal prob of N-in: 0.14402\noutside 1 331\nTMhelix 332 354\ninside 355 372

15689 GCF_001716065.1_ASM171606v1 Salmonella enterica subsp. enterica Proteobacteria; Gammaproteobacteria; Enterobacterales; Enterobacteriaceae; Salmonella; Salmonella enterica MTGDNTLITSHGINRRDFMKLCAALAATMGLSSKAAAEMAESVSNPQRPPVIWIGAQECT WP_000145429.1 hydrogenase 2 small subunit [Salmonella enterica] Length: 372\nNumber of predicted TMHs: 1\nExp number of AAs in TMHs: 24.51309\nExp number, first 60 AAs: 0.2682\nTotal prob of N-in: 0.14402\noutside 1 331\nTMhelix 332 354\ninside 355 372

15690 GCF_001716665.1_ASM171666v1 Salmonella enterica subsp. enterica Proteobacteria; Gammaproteobacteria; Enterobacterales; Enterobacteriaceae; Salmonella; Salmonella enterica MNNEETFYQAMRRKGVTRRSFLKFCSLAATSLGLGAGMTPKIAWALENKPRIPVVWIHGL WP_001058311.1 hydrogenase [Salmonella enterica] Length: 372\nNumber of predicted TMHs: 2\nExp number of AAs in TMHs: 37.4699899999999\nExp number, first 60 AAs: 19.72699\nTotal prob of N-in: 0.96171\nPOSSIBLE N-term signal sequence\ninside 1 19\nTMhelix 20 39\noutside 40 325\nTMhelix 326 348\ninside 349 372

15691 GCF_001716405.1_ASM171640v1 Salmonella enterica subsp. enterica Proteobacteria; Gammaproteobacteria; Enterobacterales; Enterobacteriaceae; Salmonella; Salmonella enterica MNNEETFYQAMRRKGVTRRSFLKFCSLAATSLGLGAGMTPKIAWALENKPRIPVVWIHGL WP_001058311.1 hydrogenase [Salmonella enterica] Length: 372\nNumber of predicted TMHs: 2\nExp number of AAs in TMHs: 37.4699899999999\nExp number, first 60 AAs: 19.72699\nTotal prob of N-in: 0.96171\nPOSSIBLE N-term signal sequence\ninside 1 19\nTMhelix 20 39\noutside 40 325\nTMhelix 326 348\ninside 349 372

15692 GCF_001716035.1_ASM171603v1 Salmonella enterica subsp. enterica Proteobacteria; Gammaproteobacteria; Enterobacterales; Enterobacteriaceae; Salmonella; Salmonella enterica MTGDNTLITSHGINRRDFMKLCAALAATMGLSSKAAAEMAESVSNPQRPPVIWIGAQECT WP_000145429.1 hydrogenase 2 small subunit [Salmonella enterica] Length: 372\nNumber of predicted TMHs: 1\nExp

number of AAs in TMHs: 24.51309\nExp number, first 60 AAs: 0.2682\nTotal prob of N-in: 0.14402\noutside 1 331\nTMhelix 332 354\ninside 355 372

15693 GCF_001716625.1_ASM171662v1 Salmonella enterica subsp. enterica Proteobacteria; Gammaproteobacteria; Enterobacterales; Enterobacteriaceae; Salmonella; Salmonella enterica MNNEETFYQAMRRKGVTRRSFLKFCSLAATSLGLGAGMTPKIAWALENKPRIPVVWIHGL WP_001058311.1 hydrogenase [Salmonella enterica] Length: 372\nNumber of predicted TMHs: 2\nExp number of AAs in TMHs: 37.4699899999999\nExp number, first 60 AAs: 19.72699\nTotal prob of N-in: 0.96171\nPOSSIBLE N-term signal sequence\ninside 1 19\nTMhelix 20 39\noutside 40 325\nTMhelix 326 348\ninside 349 372

15694 GCF_001716305.1_ASM171630v1 Salmonella enterica subsp. enterica Proteobacteria; Gammaproteobacteria; Enterobacterales; Enterobacteriaceae; Salmonella; Salmonella enterica MNNEETFYQAMRRKGVTRRSFLKFCSLAATSLGLGAGMTPKIAWALENKPRIPVVWIHGL WP_001058311.1 hydrogenase [Salmonella enterica] Length: 372\nNumber of predicted TMHs: 2\nExp number of AAs in TMHs: 37.4699899999999\nExp number, first 60 AAs: 19.72699\nTotal prob of N-in: 0.96171\nPOSSIBLE N-term signal sequence\ninside 1 19\nTMhelix 20 39\noutside 40 325\nTMhelix 326 348\ninside 349 372

15695 GCF_002176765.1_ASM217676v1 Salmonella enterica subsp. enterica Proteobacteria; Gammaproteobacteria; Enterobacterales; Enterobacteriaceae; Salmonella; Salmonella enterica MTGDNTLITSHGINRRDFMKLCAALAATMGLSSKAAAEMAESVSNPQRPPVIWIGAQECT WP_000145429.1 hydrogenase 2 small subunit [Salmonella enterica] Length: 372\nNumber of predicted TMHs: 1\nExp number of AAs in TMHs: 24.51309\nExp number, first 60 AAs: 0.2682\nTotal prob of N-in: 0.14402\noutside 1 331\nTMhelix 332 354\ninside 355 372

15696 GCF_001716345.1_ASM171634v1 Salmonella enterica subsp. enterica Proteobacteria; Gammaproteobacteria; Enterobacterales; Enterobacteriaceae; Salmonella; Salmonella enterica MNNEETFYQAMRRKGVTRRSFLKFCSLAATSLGLGAGMTPKIAWALENKPRIPVVWIHGL WP_001058311.1 hydrogenase [Salmonella enterica] Length: 372\nNumber of predicted TMHs: 2\nExp number of AAs in TMHs: 37.4699899999999\nExp number, first 60 AAs: 19.72699\nTotal prob of N-in: 0.96171\nPOSSIBLE N-term signal sequence\ninside 1 19\nTMhelix 20 39\noutside 40 325\nTMhelix 326 348\ninside 349 372

15697 GCF_002176695.1_ASM217669v1 Salmonella enterica subsp. enterica Proteobacteria; Gammaproteobacteria; Enterobacterales; Enterobacteriaceae; Salmonella; Salmonella enterica MTGDNTLITSHGINRRDFMKLCAALAATMGLSSKAAAEMAESVSNPQRPPVIWIGAQECT WP_000145429.1 hydrogenase 2 small subunit [Salmonella enterica] Length: 372\nNumber of predicted TMHs: 1\nExp number of AAs in TMHs: 24.51309\nExp number, first 60 AAs: 0.2682\nTotal prob of N-in: 0.14402\noutside 1 331\nTMhelix 332 354\ninside 355 372

15698 GCF_002176765.1_ASM217676v1 Salmonella enterica subsp. enterica Proteobacteria; Gammaproteobacteria; Enterobacterales; Enterobacteriaceae; Salmonella; Salmonella enterica MNNEETFYQAMRRKGVTRRSFLKFCSLAATSLGLGAGMTPKIAWALENKPRIPVVWIHGL WP_001058307.1 hydrogenase [Salmonella enterica] Length: 372\nNumber of predicted TMHs: 2\nExp number of AAs in TMHs: 36.51738\nExp number, first 60 AAs: 19.72849\nTotal prob of N-in: 0.96170\nPOSSIBLE N-term signal sequence\ninside 1 19\nTMhelix 20 39\noutside 40 325\nTMhelix 326 348\ninside 349 372

15699 GCF_001714255.1_ASM171425v1 Salmonella enterica subsp. enterica Proteobacteria; Gammaproteobacteria; Enterobacterales; Enterobacteriaceae; Salmonella; Salmonella enterica MNNEETFYQAMRRKGVTRRSFLKFCSLAATSLGLGAGMTPKIAWALENKPRIPVVWIHGL WP_001058307.1 hydrogenase [Salmonella enterica] Length: 372\nNumber of predicted TMHs: 2\nExp number of AAs in TMHs: 36.51738\nExp number, first 60 AAs: 19.72849\nTotal prob of N-in: 0.96170\nPOSSIBLE N-term signal sequence\ninside 1 19\nTMhelix 20 39\noutside 40 325\nTMhelix 326 348\ninside 349 372

15700 GCF_001716345.1_ASM171634v1 Salmonella enterica subsp. enterica Proteobacteria; Gammaproteobacteria; Enterobacterales; Enterobacteriaceae; Salmonella; Salmonella enterica MTGDNTLITSHGINRRDFMKLCAALAATMGLSSKAAAEMAESVSNPQRPPVIWIGAQECT WP_000145429.1 hydrogenase 2 small subunit [Salmonella enterica] Length: 372\nNumber of predicted TMHs: 1\nExp number of AAs in TMHs: 24.51309\nExp number, first 60 AAs: 0.2682\nTotal prob of N-in: 0.14402\noutside 1 331\nTMhelix 332 354\ninside 355 372

15701 GCF_002201855.1_ASM220185v1 Salmonella enterica subsp. enterica Proteobacteria; Gammaproteobacteria; Enterobacterales; Enterobacteriaceae; Salmonella; Salmonella enterica

MTGDNTLITSHGINRRDFMKLCAALAATMGLSSKAAAEMAESVSNPQRPPVIWIGAQECT WP_000145429.1 hydrogenase 2 small subunit [Salmonella enterica] Length: 372\nNumber of predicted TMHs: 1\nExp number of AAs in TMHs: 24.51309\nExp number, first 60 AAs: 0.2682\nTotal prob of N-in: 0.14402\noutside 1 331\nTMhelix 332 354\ninside 355 372

15702 GCF_001713745.1_ASM171374v1 Salmonella enterica subsp. enterica Proteobacteria; Gammaproteobacteria; Enterobacterales; Enterobacteriaceae; Salmonella; Salmonella enterica MNNEETFYQAMRRKGVTRRSFLKFCSLAATSLGLGAGMTPKIAWALENKPRIPVVWIHGL WP_001058307.1 hydrogenase [Salmonella enterica] Length: 372\nNumber of predicted TMHs: 2\nExp number of AAs in TMHs: 36.51738\nExp number, first 60 AAs: 19.72849\nTotal prob of N-in: 0.96170\nPOSSIBLE N-term signal sequence\ninside 1 19\nTMhelix 20 39\noutside 40 325\nTMhelix 326 348\ninside 349 372

15703 GCF_001713745.1_ASM171374v1 Salmonella enterica subsp. enterica Proteobacteria; Gammaproteobacteria; Enterobacterales; Enterobacteriaceae; Salmonella; Salmonella enterica MTGDNTLITSHGINRRDFMKLCAALAATMGLSSKAAAEMAESVSNPQRPPVIWIGAQECT WP_000145429.1 hydrogenase 2 small subunit [Salmonella enterica] Length: 372\nNumber of predicted TMHs: 1\nExp number of AAs in TMHs: 24.51309\nExp number, first 60 AAs: 0.2682\nTotal prob of N-in: 0.14402\noutside 1 331\nTMhelix 332 354\ninside 355 372

15704 GCF_001647955.1_ASM164795v1 Salmonella enterica subsp. enterica Proteobacteria; Gammaproteobacteria; Enterobacterales; Enterobacteriaceae; Salmonella; Salmonella enterica MNNEETFYQAMRRKGVTRRSFLKFCSLAATSLGLGAGMTPKIAWALENKPRIPVVWIHGL WP_001058312.1 hydrogenase [Salmonella enterica] Length: 372\nNumber of predicted TMHs: 2\nExp number of AAs in TMHs: 36.51738\nExp number, first 60 AAs: 19.72849\nTotal prob of N-in: 0.96170\nPOSSIBLE N-term signal sequence\ninside 1 19\nTMhelix 20 39\noutside 40 325\nTMhelix 326 348\ninside 349 372

15705 GCF_001716805.1_ASM171680v1 Salmonella enterica subsp. enterica Proteobacteria; Gammaproteobacteria; Enterobacterales; Enterobacteriaceae; Salmonella; Salmonella enterica MTGDNTLITSHGINRRDFMKLCAALAATMGLSSKAAAEMAESVSNPQRPPVIWIGAQECT WP_000145429.1 hydrogenase 2 small subunit [Salmonella enterica] Length: 372\nNumber of predicted TMHs: 1\nExp number of AAs in TMHs: 24.51309\nExp number, first 60 AAs: 0.2682\nTotal prob of N-in: 0.14402\noutside 1 331\nTMhelix 332 354\ninside 355 372

15706 GCF_001716775.1_ASM171677v1 Salmonella enterica subsp. enterica Proteobacteria; Gammaproteobacteria; Enterobacterales; Enterobacteriaceae; Salmonella; Salmonella enterica MTGDNTLITSHGINRRDFMKLCAALAATMGLSSKAAAEMAESVSNPQRPPVIWIGAQECT WP_000145429.1 hydrogenase 2 small subunit [Salmonella enterica] Length: 372\nNumber of predicted TMHs: 1\nExp number of AAs in TMHs: 24.51309\nExp number, first 60 AAs: 0.2682\nTotal prob of N-in: 0.14402\noutside 1 331\nTMhelix 332 354\ninside 355 372

15707 GCF_001716455.1_ASM171645v1 Salmonella enterica subsp. enterica Proteobacteria; Gammaproteobacteria; Enterobacterales; Enterobacteriaceae; Salmonella; Salmonella enterica MNNEETFYQAMRRKGVTRRSFLKFCSLAATSLGLGAGMTPKIAWALENKPRIPVVWIHGL WP_001058311.1 hydrogenase [Salmonella enterica] Length: 372\nNumber of predicted TMHs: 2\nExp number of AAs in TMHs: 37.4699899999999\nExp number, first 60 AAs: 19.72699\nTotal prob of N-in: 0.96171\nPOSSIBLE N-term signal sequence\ninside 1 19\nTMhelix 20 39\noutside 40 325\nTMhelix 326 348\ninside 349 372

15708 GCF_001716715.1_ASM171671v1 Salmonella enterica subsp. enterica Proteobacteria; Gammaproteobacteria; Enterobacterales; Enterobacteriaceae; Salmonella; Salmonella enterica MNNEETFYQAMRRKGVTRRSFLKFCSLAATSLGLGAGMTPKIAWALENKPRIPVVWIHGL WP_001058311.1 hydrogenase [Salmonella enterica] Length: 372\nNumber of predicted TMHs: 2\nExp number of AAs in TMHs: 37.4699899999999\nExp number, first 60 AAs: 19.72699\nTotal prob of N-in: 0.96171\nPOSSIBLE N-term signal sequence\ninside 1 19\nTMhelix 20 39\noutside 40 325\nTMhelix 326 348\ninside 349 372

15709 GCF_002176695.1_ASM217669v1 Salmonella enterica subsp. enterica Proteobacteria; Gammaproteobacteria; Enterobacterales; Enterobacteriaceae; Salmonella; Salmonella enterica MNNEETFYQAMRRKGVTRRSFLKFCSLAATSLGLGAGMTPKIAWALENKPRIPVVWIHGL WP_001058307.1 hydrogenase [Salmonella enterica] Length: 372\nNumber of predicted TMHs: 2\nExp number of AAs in TMHs: 36.51738\nExp number, first 60 AAs: 19.72849\nTotal prob of N-in: 0.96170\nPOSSIBLE N-term signal sequence\ninside 1 19\nTMhelix 20 39\noutside 40 325\nTMhelix 326 348\ninside 349 372

15710 GCF_001714305.1_ASM171430v1 Salmonella enterica subsp. enterica Proteobacteria; Gammaproteobacteria; Enterobacterales; Enterobacteriaceae; Salmonella; Salmonella enterica
MTGDNTLITSHGINRRDFMKLCAALAATMGLSSKAAAEMAESVSNPQRPPVIWIGAQECT WP_000145429.1
hydrogenase 2 small subunit [Salmonella enterica] Length: 372\nNumber of predicted TMHs: 1\nExp number of AAs in TMHs: 24.51309\nExp number, first 60 AAs: 0.2682\nTotal prob of N-in: 0.14402\noutside 1 331\nTMhelix 332 354\ninside 355 372

15711 GCF_001714185.1_ASM171418v1 Salmonella enterica subsp. enterica Proteobacteria; Gammaproteobacteria; Enterobacterales; Enterobacteriaceae; Salmonella; Salmonella enterica
MNNEETFYQAMRRKGVTRRSFLKFCSLAATSLGLGAGMTPKIAWALENKPRIPVVWIHGL WP_001058307.1
hydrogenase [Salmonella enterica] Length: 372\nNumber of predicted TMHs: 2\nExp number of AAs in TMHs: 36.51738\nExp number, first 60 AAs: 19.72849\nTotal prob of N-in: 0.96170\nPOSSIBLE N-term signal sequence\ninside 1 19\nTMhelix 20 39\noutside 40 325\nTMhelix 326 348\ninside 349 372

15712 GCF_002094915.1_ASM209491v1 Salmonella enterica subsp. enterica Proteobacteria; Gammaproteobacteria; Enterobacterales; Enterobacteriaceae; Salmonella; Salmonella enterica
MTGDNTLITSHGINRRDFMKLCAALAATMGLSSKAAAEMAESVSNPQRPPVIWIGAQECT WP_000145431.1
hydrogenase 2 small subunit [Salmonella enterica] Length: 372\nNumber of predicted TMHs: 1\nExp number of AAs in TMHs: 24.51505\nExp number, first 60 AAs: 0.2682\nTotal prob of N-in: 0.14406\noutside 1 331\nTMhelix 332 354\ninside 355 372

15713 GCF_001714165.1_ASM171416v1 Salmonella enterica subsp. enterica Proteobacteria; Gammaproteobacteria; Enterobacterales; Enterobacteriaceae; Salmonella; Salmonella enterica
MNNEETFYQAMRRKGVTRRSFLKFCSLAATSLGLGAGMTPKIAWALENKPRIPVVWIHGL WP_001058307.1
hydrogenase [Salmonella enterica] Length: 372\nNumber of predicted TMHs: 2\nExp number of AAs in TMHs: 36.51738\nExp number, first 60 AAs: 19.72849\nTotal prob of N-in: 0.96170\nPOSSIBLE N-term signal sequence\ninside 1 19\nTMhelix 20 39\noutside 40 325\nTMhelix 326 348\ninside 349 372

15714 GCF_000806475.1_CFSAN024721_01.0Salmonella enterica subsp. enterica Proteobacteria; Gammaproteobacteria; Enterobacterales; Enterobacteriaceae; Salmonella; Salmonella enterica
MNNEETFYQAMRRKGVTRRSFLKFCSLAATSLGLGAGMTPKIAWALENKPRIPVVWIHGL WP_001058312.1
hydrogenase [Salmonella enterica] Length: 372\nNumber of predicted TMHs: 2\nExp number of AAs in TMHs: 36.51738\nExp number, first 60 AAs: 19.72849\nTotal prob of N-in: 0.96170\nPOSSIBLE N-term signal sequence\ninside 1 19\nTMhelix 20 39\noutside 40 325\nTMhelix 326 348\ninside 349 372

15715 GCF_001700595.1_ASM170059v1 Salmonella enterica subsp. enterica Proteobacteria; Gammaproteobacteria; Enterobacterales; Enterobacteriaceae; Salmonella; Salmonella enterica
MNNEETFYQAMRRKGVTRRSFLKFCSLAATSLGLGAGMTPKIAWALENKPRIPVVWIHGL WP_001058312.1
hydrogenase [Salmonella enterica] Length: 372\nNumber of predicted TMHs: 2\nExp number of AAs in TMHs: 36.51738\nExp number, first 60 AAs: 19.72849\nTotal prob of N-in: 0.96170\nPOSSIBLE N-term signal sequence\ninside 1 19\nTMhelix 20 39\noutside 40 325\nTMhelix 326 348\ninside 349 372

15716 GCF_001700345.1_ASM170034v1 Salmonella enterica subsp. enterica Proteobacteria; Gammaproteobacteria; Enterobacterales; Enterobacteriaceae; Salmonella; Salmonella enterica
MNNEETFYQAMRRKGVTRRSFLKFCSLAATSLGLGAGMTPKIAWALENKPRIPVVWIHGL WP_001058312.1
hydrogenase [Salmonella enterica] Length: 372\nNumber of predicted TMHs: 2\nExp number of AAs in TMHs: 36.51738\nExp number, first 60 AAs: 19.72849\nTotal prob of N-in: 0.96170\nPOSSIBLE N-term signal sequence\ninside 1 19\nTMhelix 20 39\noutside 40 325\nTMhelix 326 348\ninside 349 372

15717 GCF_001707605.1_ASM170760v1 Salmonella enterica subsp. enterica Proteobacteria; Gammaproteobacteria; Enterobacterales; Enterobacteriaceae; Salmonella; Salmonella enterica
MTGDNTLITSHGINRRDFMKLCAALAATMGLSSKAAAEMAESVSNPQRPPVIWIGAQECT WP_000145429.1
hydrogenase 2 small subunit [Salmonella enterica] Length: 372\nNumber of predicted TMHs: 1\nExp number of AAs in TMHs: 24.51309\nExp number, first 60 AAs: 0.2682\nTotal prob of N-in: 0.14402\noutside 1 331\nTMhelix 332 354\ninside 355 372

15718 GCF_002060085.1_ASM206008v1 Salmonella enterica subsp. enterica Proteobacteria; Gammaproteobacteria; Enterobacterales; Enterobacteriaceae; Salmonella; Salmonella enterica
MTGDNTLITSHGINRRDFMKLCAALAATMGLSSKAAAEMAESVSNPQRPPVIWIGAQECT WP_000145429.1
hydrogenase 2 small subunit [Salmonella enterica] Length: 372\nNumber of predicted TMHs: 1\nExp

number of AAs in TMHs: 24.51309\nExp number, first 60 AAs:  0.2682\nTotal prob of N-in:   0.14402\noutside   1   331\nTMhelix   332   354\ninside   355   372

15719 GCF_001714175.1_ASM171417v1 Salmonella enterica subsp. enterica Proteobacteria; Gammaproteobacteria; Enterobacterales; Enterobacteriaceae; Salmonella; Salmonella enterica MNNEETFYQAMRRKGVTRRSFLKFCSLAATSLGLGAGMTPKIAWALENKPRIPVVWIHGL WP_001058307.1 hydrogenase [Salmonella enterica] Length: 372\nNumber of predicted TMHs:  2\nExp number of AAs in TMHs: 36.51738\nExp number, first 60 AAs:  19.72849\nTotal prob of N-in:   0.96170\nPOSSIBLE N-term signal sequence\ninside   1   19\nTMhelix   20   39\noutside   40   325\nTMhelix   326   348\ninside   349   372

15720 GCF_000806785.1_CFSAN024776_01.0Salmonella enterica subsp. enterica Proteobacteria; Gammaproteobacteria; Enterobacterales; Enterobacteriaceae; Salmonella; Salmonella enterica MTGDNTLITSHGINRRDFMKLCAALAATMGLSSKAAAEMAESVSNPQRPPVIWIGAQECT WP_039509062.1 hydrogenase 2 small subunit [Salmonella enterica] Length: 372\nNumber of predicted TMHs:  1\nExp number of AAs in TMHs: 24.51305\nExp number, first 60 AAs:  0.26819\nTotal prob of N-in:   0.14402\noutside   1   331\nTMhelix   332   354\ninside   355   372

15721 GCF_001700345.1_ASM170034v1 Salmonella enterica subsp. enterica Proteobacteria; Gammaproteobacteria; Enterobacterales; Enterobacteriaceae; Salmonella; Salmonella enterica MTGDNTLITSHGINRRDFMKLCAALAATMGLSSKAAAEMAESVSNPQRPPVIWIGAQECT WP_000145429.1 hydrogenase 2 small subunit [Salmonella enterica] Length: 372\nNumber of predicted TMHs:  1\nExp number of AAs in TMHs: 24.51309\nExp number, first 60 AAs:  0.2682\nTotal prob of N-in:   0.14402\noutside   1   331\nTMhelix   332   354\ninside   355   372

15722 GCF_001700595.1_ASM170059v1 Salmonella enterica subsp. enterica Proteobacteria; Gammaproteobacteria; Enterobacterales; Enterobacteriaceae; Salmonella; Salmonella enterica MTGDNTLITSHGINRRDFMKLCAALAATMGLSSKAAAEMAESVSNPQRPPVIWIGAQECT WP_000145429.1 hydrogenase 2 small subunit [Salmonella enterica] Length: 372\nNumber of predicted TMHs:  1\nExp number of AAs in TMHs: 24.51309\nExp number, first 60 AAs:  0.2682\nTotal prob of N-in:   0.14402\noutside   1   331\nTMhelix   332   354\ninside   355   372

15723 GCF_001714225.1_ASM171422v1 Salmonella enterica subsp. enterica Proteobacteria; Gammaproteobacteria; Enterobacterales; Enterobacteriaceae; Salmonella; Salmonella enterica MNNEETFYQAMRRKGVTRRSFLKFCSLAATSLGLGAGMTPKIAWALENKPRIPVVWIHGL WP_001058307.1 hydrogenase [Salmonella enterica] Length: 372\nNumber of predicted TMHs:  2\nExp number of AAs in TMHs: 36.51738\nExp number, first 60 AAs:  19.72849\nTotal prob of N-in:   0.96170\nPOSSIBLE N-term signal sequence\ninside   1   19\nTMhelix   20   39\noutside   40   325\nTMhelix   326   348\ninside   349   372

15724 GCF_001707615.1_ASM170761v1 Salmonella enterica subsp. enterica Proteobacteria; Gammaproteobacteria; Enterobacterales; Enterobacteriaceae; Salmonella; Salmonella enterica MNNEETFYQAMRRKGVTRRSFLKFCSLAATSLGLGAGMTPKIAWALENKPRIPVVWIHGL WP_001058312.1 hydrogenase [Salmonella enterica] Length: 372\nNumber of predicted TMHs:  2\nExp number of AAs in TMHs: 36.51738\nExp number, first 60 AAs:  19.72849\nTotal prob of N-in:   0.96170\nPOSSIBLE N-term signal sequence\ninside   1   19\nTMhelix   20   39\noutside   40   325\nTMhelix   326   348\ninside   349   372

15725 GCF_001716045.1_ASM171604v1 Salmonella enterica subsp. enterica Proteobacteria; Gammaproteobacteria; Enterobacterales; Enterobacteriaceae; Salmonella; Salmonella enterica MNNEETFYQAMRRKGVTRRSFLKFCSLAATSLGLGAGMTPKIAWALENKPRIPVVWIHGL WP_001058311.1 hydrogenase [Salmonella enterica] Length: 372\nNumber of predicted TMHs:  2\nExp number of AAs in TMHs: 37.4699899999999\nExp number, first 60 AAs:  19.72699\nTotal prob of N-in:   0.96171\nPOSSIBLE N-term signal sequence\ninside   1   19\nTMhelix   20   39\noutside   40   325\nTMhelix   326   348\ninside   349   372

15726 GCF_001716215.1_ASM171621v1 Salmonella enterica subsp. enterica Proteobacteria; Gammaproteobacteria; Enterobacterales; Enterobacteriaceae; Salmonella; Salmonella enterica MTGDNTLITSHGINRRDFMKLCAALAATMGLSSKAAAEMAESVSNPQRPPVIWIGAQECT WP_000145429.1 hydrogenase 2 small subunit [Salmonella enterica] Length: 372\nNumber of predicted TMHs:  1\nExp number of AAs in TMHs: 24.51309\nExp number, first 60 AAs:  0.2682\nTotal prob of N-in:   0.14402\noutside   1   331\nTMhelix   332   354\ninside   355   372

15727 GCF_001716235.1_ASM171623v1 Salmonella enterica subsp. enterica Proteobacteria; Gammaproteobacteria; Enterobacterales; Enterobacteriaceae; Salmonella; Salmonella enterica

MTGDNTLITSHGINRRDFMKLCAALAATMGLSSKAAAEMAESVSNPQRPPVIWIGAQECT WP_000145429.1 hydrogenase 2 small subunit [Salmonella enterica] Length: 372\nNumber of predicted TMHs: 1\nExp number of AAs in TMHs: 24.51309\nExp number, first 60 AAs: 0.2682\nTotal prob of N-in: 0.14402\noutside 1 331\nTMhelix 332 354\ninside 355 372

15728 GCF_001716295.1_ASM171629v1 Salmonella enterica subsp. enterica Proteobacteria; Gammaproteobacteria; Enterobacterales; Enterobacteriaceae; Salmonella; Salmonella enterica MTGDNTLITSHGINRRDFMKLCAALAATMGLSSKAAAEMAESVSNPQRPPVIWIGAQECT WP_000145429.1 hydrogenase 2 small subunit [Salmonella enterica] Length: 372\nNumber of predicted TMHs: 1\nExp number of AAs in TMHs: 24.51309\nExp number, first 60 AAs: 0.2682\nTotal prob of N-in: 0.14402\noutside 1 331\nTMhelix 332 354\ninside 355 372

15729 GCF_001714175.1_ASM171417v1 Salmonella enterica subsp. enterica Proteobacteria; Gammaproteobacteria; Enterobacterales; Enterobacteriaceae; Salmonella; Salmonella enterica MTGDNTLITSHGINRRDFMKLCAALAATMGLSSKAAAEMAESVSNPQRPPVIWIGAQECT WP_000145429.1 hydrogenase 2 small subunit [Salmonella enterica] Length: 372\nNumber of predicted TMHs: 1\nExp number of AAs in TMHs: 24.51309\nExp number, first 60 AAs: 0.2682\nTotal prob of N-in: 0.14402\noutside 1 331\nTMhelix 332 354\ninside 355 372

15730 GCF_001716335.1_ASM171633v1 Salmonella enterica subsp. enterica Proteobacteria; Gammaproteobacteria; Enterobacterales; Enterobacteriaceae; Salmonella; Salmonella enterica MTGDNTLITSHGINRRDFMKLCAALAATMGLSSKAAAEMAESVSNPQRPPVIWIGAQECT WP_000145429.1 hydrogenase 2 small subunit [Salmonella enterica] Length: 372\nNumber of predicted TMHs: 1\nExp number of AAs in TMHs: 24.51309\nExp number, first 60 AAs: 0.2682\nTotal prob of N-in: 0.14402\noutside 1 331\nTMhelix 332 354\ninside 355 372

15731 GCF_001707615.1_ASM170761v1 Salmonella enterica subsp. enterica Proteobacteria; Gammaproteobacteria; Enterobacterales; Enterobacteriaceae; Salmonella; Salmonella enterica MTGDNTLITSHGINRRDFMKLCAALAATMGLSSKAAAEMAESVSNPQRPPVIWIGAQECT WP_000145429.1 hydrogenase 2 small subunit [Salmonella enterica] Length: 372\nNumber of predicted TMHs: 1\nExp number of AAs in TMHs: 24.51309\nExp number, first 60 AAs: 0.2682\nTotal prob of N-in: 0.14402\noutside 1 331\nTMhelix 332 354\ninside 355 372

15732 GCF_001716495.1_ASM171649v1 Salmonella enterica subsp. enterica Proteobacteria; Gammaproteobacteria; Enterobacterales; Enterobacteriaceae; Salmonella; Salmonella enterica MTGDNTLITSHGINRRDFMKLCAALAATMGLSSKAAAEMAESVSNPQRPPVIWIGAQECT WP_000145429.1 hydrogenase 2 small subunit [Salmonella enterica] Length: 372\nNumber of predicted TMHs: 1\nExp number of AAs in TMHs: 24.51309\nExp number, first 60 AAs: 0.2682\nTotal prob of N-in: 0.14402\noutside 1 331\nTMhelix 332 354\ninside 355 372

15733 GCF_001716535.1_ASM171653v1 Salmonella enterica subsp. enterica Proteobacteria; Gammaproteobacteria; Enterobacterales; Enterobacteriaceae; Salmonella; Salmonella enterica MTGDNTLITSHGINRRDFMKLCAALAATMGLSSKAAAEMAESVSNPQRPPVIWIGAQECT WP_000145429.1 hydrogenase 2 small subunit [Salmonella enterica] Length: 372\nNumber of predicted TMHs: 1\nExp number of AAs in TMHs: 24.51309\nExp number, first 60 AAs: 0.2682\nTotal prob of N-in: 0.14402\noutside 1 331\nTMhelix 332 354\ninside 355 372

15734 GCF_001586465.1_ASM158646v1 Salmonella enterica subsp. enterica Proteobacteria; Gammaproteobacteria; Enterobacterales; Enterobacteriaceae; Salmonella; Salmonella enterica MNNEETFYQAMRRKGVTRRSFLKFCSLAATSLGLGAGMTPKIAWALENKPRIPVVWIHGL WP_001058311.1 hydrogenase [Salmonella enterica] Length: 372\nNumber of predicted TMHs: 2\nExp number of AAs in TMHs: 37.4699899999999\nExp number, first 60 AAs: 19.72699\nTotal prob of N-in: 0.96171\nPOSSIBLE N-term signal sequence\ninside 1 19\nTMhelix 20 39\noutside 40 325\nTMhelix 326 348\ninside 349 372

15735 GCF_001586555.1_ASM158655v1 Salmonella enterica subsp. enterica Proteobacteria; Gammaproteobacteria; Enterobacterales; Enterobacteriaceae; Salmonella; Salmonella enterica MTGDNTLITSHGINRRDFMKLCAALAATMGLSSKAAAEMAESVSNPQRPPVIWIGAQECT WP_000145429.1 hydrogenase 2 small subunit [Salmonella enterica] Length: 372\nNumber of predicted TMHs: 1\nExp number of AAs in TMHs: 24.51309\nExp number, first 60 AAs: 0.2682\nTotal prob of N-in: 0.14402\noutside 1 331\nTMhelix 332 354\ninside 355 372

15736	GCF_001586615.1_ASM158661v1	Salmonella enterica subsp. enterica	Proteobacteria; Gammaproteobacteria; Enterobacterales; Enterobacteriaceae; Salmonella; Salmonella enterica
MTGDNTLITSHGINRRDFMKLCAALAATMGLSSKAAAEMAESVSNPQRPPVIWIGAQECT	WP_000145429.1
hydrogenase 2 small subunit [Salmonella enterica]	Length: 372\nNumber of predicted TMHs:  1\nExp number of AAs in TMHs: 24.51309\nExp number, first 60 AAs:  0.2682\nTotal prob of N-in:   0.14402\noutside   1   331\nTMhelix   332   354\ninside   355   372

15737	GCF_001586625.1_ASM158662v1	Salmonella enterica subsp. enterica	Proteobacteria; Gammaproteobacteria; Enterobacterales; Enterobacteriaceae; Salmonella; Salmonella enterica
MTGDNTLITSHGINRRDFMKLCAALAATMGLSSKAAAEMAESVSNPQRPPVIWIGAQECT	WP_000145429.1
hydrogenase 2 small subunit [Salmonella enterica]	Length: 372\nNumber of predicted TMHs:  1\nExp number of AAs in TMHs: 24.51309\nExp number, first 60 AAs:  0.2682\nTotal prob of N-in:   0.14402\noutside   1   331\nTMhelix   332   354\ninside   355   372

15738	GCF_001714225.1_ASM171422v1	Salmonella enterica subsp. enterica	Proteobacteria; Gammaproteobacteria; Enterobacterales; Enterobacteriaceae; Salmonella; Salmonella enterica
MTGDNTLITSHGINRRDFMKLCAALAATMGLSSKAAAEMAESVSNPQRPPVIWIGAQECT	WP_000145429.1
hydrogenase 2 small subunit [Salmonella enterica]	Length: 372\nNumber of predicted TMHs:  1\nExp number of AAs in TMHs: 24.51309\nExp number, first 60 AAs:  0.2682\nTotal prob of N-in:   0.14402\noutside   1   331\nTMhelix   332   354\ninside   355   372

15739	GCF_001714265.1_ASM171426v1	Salmonella enterica subsp. enterica	Proteobacteria; Gammaproteobacteria; Enterobacterales; Enterobacteriaceae; Salmonella; Salmonella enterica
MTGDNTLITSHGINRRDFMKLCAALAATMGLSSKAAAEMAESVSNPQRPPVIWIGAQECT	WP_000145429.1
hydrogenase 2 small subunit [Salmonella enterica]	Length: 372\nNumber of predicted TMHs:  1\nExp number of AAs in TMHs: 24.51309\nExp number, first 60 AAs:  0.2682\nTotal prob of N-in:   0.14402\noutside   1   331\nTMhelix   332   354\ninside   355   372

15740	GCF_001586555.1_ASM158655v1	Salmonella enterica subsp. enterica	Proteobacteria; Gammaproteobacteria; Enterobacterales; Enterobacteriaceae; Salmonella; Salmonella enterica
MNNEETFYQAMRRKGVTRRSFLKFCSLAATSLGLGAGMTPKIAWALENKPRIPVVWIHGL	WP_001058311.1
hydrogenase [Salmonella enterica]	Length: 372\nNumber of predicted TMHs:  2\nExp number of AAs in TMHs: 37.4699899999999\nExp number, first 60 AAs:  19.72699\nTotal prob of N-in:   0.96171\nPOSSIBLE N-term signal sequence\ninside   1   19\nTMhelix   20   39\noutside   40   325\nTMhelix   326   348\ninside   349   372

15741	GCF_001586615.1_ASM158661v1	Salmonella enterica subsp. enterica	Proteobacteria; Gammaproteobacteria; Enterobacterales; Enterobacteriaceae; Salmonella; Salmonella enterica
MNNEETFYQAMRRKGVTRRSFLKFCSLAATSLGLGAGMTPKIAWALENKPRIPVVWIHGL	WP_001058311.1
hydrogenase [Salmonella enterica]	Length: 372\nNumber of predicted TMHs:  2\nExp number of AAs in TMHs: 37.4699899999999\nExp number, first 60 AAs:  19.72699\nTotal prob of N-in:   0.96171\nPOSSIBLE N-term signal sequence\ninside   1   19\nTMhelix   20   39\noutside   40   325\nTMhelix   326   348\ninside   349   372

15742	GCF_001714245.1_ASM171424v1	Salmonella enterica subsp. enterica	Proteobacteria; Gammaproteobacteria; Enterobacterales; Enterobacteriaceae; Salmonella; Salmonella enterica
MTGDNTLITSHGINRRDFMKLCAALAATMGLSSKAAAEMAESVSNPQRPPVIWIGAQECT	WP_000145429.1
hydrogenase 2 small subunit [Salmonella enterica]	Length: 372\nNumber of predicted TMHs:  1\nExp number of AAs in TMHs: 24.51309\nExp number, first 60 AAs:  0.2682\nTotal prob of N-in:   0.14402\noutside   1   331\nTMhelix   332   354\ninside   355   372

15743	GCF_001586625.1_ASM158662v1	Salmonella enterica subsp. enterica	Proteobacteria; Gammaproteobacteria; Enterobacterales; Enterobacteriaceae; Salmonella; Salmonella enterica
MNNEETFYQAMRRKGVTRRSFLKFCSLAATSLGLGAGMTPKIAWALENKPRIPVVWIHGL	WP_001058311.1
hydrogenase [Salmonella enterica]	Length: 372\nNumber of predicted TMHs:  2\nExp number of AAs in TMHs: 37.4699899999999\nExp number, first 60 AAs:  19.72699\nTotal prob of N-in:   0.96171\nPOSSIBLE N-term signal sequence\ninside   1   19\nTMhelix   20   39\noutside   40   325\nTMhelix   326   348\ninside   349   372

15744	GCF_001716615.1_ASM171661v1	Salmonella enterica subsp. enterica	Proteobacteria; Gammaproteobacteria; Enterobacterales; Enterobacteriaceae; Salmonella; Salmonella enterica
MTGDNTLITSHGINRRDFMKLCAALAATMGLSSKAAAEMAESVSNPQRPPVIWIGAQECT	WP_000145429.1
hydrogenase 2 small subunit [Salmonella enterica]	Length: 372\nNumber of predicted TMHs:  1\nExp

number of AAs in TMHs: 24.51309\nExp number, first 60 AAs: 0.2682\nTotal prob of N-in: 0.14402\noutside 1 331\nTMhelix 332 354\ninside 355 372

15745 GCF_001716575.1_ASM171657v1 Salmonella enterica subsp. enterica Proteobacteria; Gammaproteobacteria; Enterobacterales; Enterobacteriaceae; Salmonella; Salmonella enterica MTGDNTLITSHGINRRDFMKLCAALAATMGLSSKAAAEMAESVSNPQRPPVIWIGAQECT WP_000145429.1 hydrogenase 2 small subunit [Salmonella enterica] Length: 372\nNumber of predicted TMHs: 1\nExp number of AAs in TMHs: 24.51309\nExp number, first 60 AAs: 0.2682\nTotal prob of N-in: 0.14402\noutside 1 331\nTMhelix 332 354\ninside 355 372

15746 GCF_001716475.1_ASM171647v1 Salmonella enterica subsp. enterica Proteobacteria; Gammaproteobacteria; Enterobacterales; Enterobacteriaceae; Salmonella; Salmonella enterica MNNEETFYQAMRRKGVTRRSFLKFCSLAATSLGLGAGMTPKIAWALENKPRIPVVWIHGL WP_001058311.1 hydrogenase [Salmonella enterica] Length: 372\nNumber of predicted TMHs: 2\nExp number of AAs in TMHs: 37.4699899999999\nExp number, first 60 AAs: 19.72699\nTotal prob of N-in: 0.96171\nPOSSIBLE N-term signal sequence\ninside 1 19\nTMhelix 20 39\noutside 40 325\nTMhelix 326 348\ninside 349 372

15747 GCF_001716655.1_ASM171665v1 Salmonella enterica subsp. enterica Proteobacteria; Gammaproteobacteria; Enterobacterales; Enterobacteriaceae; Salmonella; Salmonella enterica MTGDNTLITSHGINRRDFMKLCAALAATMGLSSKAAAEMAESVSNPQRPPVIWIGAQECT WP_000145429.1 hydrogenase 2 small subunit [Salmonella enterica] Length: 372\nNumber of predicted TMHs: 1\nExp number of AAs in TMHs: 24.51309\nExp number, first 60 AAs: 0.2682\nTotal prob of N-in: 0.14402\noutside 1 331\nTMhelix 332 354\ninside 355 372

15748 GCF_001720385.1_ASM172038v1 Salmonella enterica subsp. enterica Proteobacteria; Gammaproteobacteria; Enterobacterales; Enterobacteriaceae; Salmonella; Salmonella enterica MNNEETFYQAMRRKGVTRRSFLKFCSLAATSLGLGAGMTPKIAWALENKPRIPVVWIHGL WP_001058311.1 hydrogenase [Salmonella enterica] Length: 372\nNumber of predicted TMHs: 2\nExp number of AAs in TMHs: 37.4699899999999\nExp number, first 60 AAs: 19.72699\nTotal prob of N-in: 0.96171\nPOSSIBLE N-term signal sequence\ninside 1 19\nTMhelix 20 39\noutside 40 325\nTMhelix 326 348\ninside 349 372

15749 GCF_001006525.1_ASM100652v1 Salmonella enterica subsp. enterica Proteobacteria; Gammaproteobacteria; Enterobacterales; Enterobacteriaceae; Salmonella; Salmonella enterica MTGDNTLITSHGINRRDFMKLCAALAATMGLSSKAAAEMAESVSNPQRPPVIWIGAQECT WP_000145429.1 hydrogenase 2 small subunit [Salmonella enterica] Length: 372\nNumber of predicted TMHs: 1\nExp number of AAs in TMHs: 24.51309\nExp number, first 60 AAs: 0.2682\nTotal prob of N-in: 0.14402\noutside 1 331\nTMhelix 332 354\ninside 355 372

15750 GCF_000807005.1_CFSAN024781_01.0Salmonella enterica subsp. enterica Proteobacteria; Gammaproteobacteria; Enterobacterales; Enterobacteriaceae; Salmonella; Salmonella enterica MTGDNTLITSHGINRRDFMKLCAALAATMGLSSKAAAEMAESVSNPQRPPVIWIGAQECT WP_000145429.1 hydrogenase 2 small subunit [Salmonella enterica] Length: 372\nNumber of predicted TMHs: 1\nExp number of AAs in TMHs: 24.51309\nExp number, first 60 AAs: 0.2682\nTotal prob of N-in: 0.14402\noutside 1 331\nTMhelix 332 354\ninside 355 372

15751 GCF_000806475.1_CFSAN024721_01.0Salmonella enterica subsp. enterica Proteobacteria; Gammaproteobacteria; Enterobacterales; Enterobacteriaceae; Salmonella; Salmonella enterica MTGDNTLITSHGINRRDFMKLCAALAATMGLSSKAAAEMAESVSNPQRPPVIWIGAQECT WP_000145429.1 hydrogenase 2 small subunit [Salmonella enterica] Length: 372\nNumber of predicted TMHs: 1\nExp number of AAs in TMHs: 24.51309\nExp number, first 60 AAs: 0.2682\nTotal prob of N-in: 0.14402\noutside 1 331\nTMhelix 332 354\ninside 355 372

15752 GCF_000878845.1_10TTU468_a5 Salmonella enterica subsp. enterica Proteobacteria; Gammaproteobacteria; Enterobacterales; Enterobacteriaceae; Salmonella; Salmonella enterica MNNEETFYQAMRRKGVTRRSFLKFCSLAATSLGLGAGMTPKIAWALENKPRIPVVWIHGL WP_001058311.1 hydrogenase [Salmonella enterica] Length: 372\nNumber of predicted TMHs: 2\nExp number of AAs in TMHs: 37.4699899999999\nExp number, first 60 AAs: 19.72699\nTotal prob of N-in: 0.96171\nPOSSIBLE N-term signal sequence\ninside 1 19\nTMhelix 20 39\noutside 40 325\nTMhelix 326 348\ninside 349 372

15753 GCF_000806865.1_CFSAN024723_01.0Salmonella enterica subsp. enterica Proteobacteria; Gammaproteobacteria; Enterobacterales; Enterobacteriaceae; Salmonella; Salmonella enterica

MNNEETFYQAMRRKGVTRRSFLKFCSLAATSLGLGAGMTPKIAWALENKPRIPVVWIHGL WP_001058312.1 hydrogenase [Salmonella enterica] Length: 372\nNumber of predicted TMHs: 2\nExp number of AAs in TMHs: 36.51738\nExp number, first 60 AAs: 19.72849\nTotal prob of N-in: 0.96170\nPOSSIBLE N-term signal sequence\ninside 1 19\nTMhelix 20 39\noutside 40 325\nTMhelix 326 348\ninside 349 372

15754 GCF_000806925.1_CFSAN024763_02.0Salmonella enterica subsp. enterica Proteobacteria; Gammaproteobacteria; Enterobacterales; Enterobacteriaceae; Salmonella; Salmonella enterica MNNEETFYQAMRRKGVTRRSFLKFCSLAATSLGLGAGMTPKIAWALENKPRIPVVWIHGL WP_023243720.1 hydrogenase 1, small subunit [Salmonella enterica] Length: 372\nNumber of predicted TMHs: 2\nExp number of AAs in TMHs: 36.52147\nExp number, first 60 AAs: 19.72838\nTotal prob of N-in: 0.96170\nPOSSIBLE N-term signal sequence\ninside 1 19\nTMhelix 20 39\noutside 40 325\nTMhelix 326 348\ninside 349 372

15755 GCF_000806925.1_CFSAN024763_02.0Salmonella enterica subsp. enterica Proteobacteria; Gammaproteobacteria; Enterobacterales; Enterobacteriaceae; Salmonella; Salmonella enterica MTGDNTLITSHGINRRDFMKLCAALAATMGLSSKAAAEMAESVSNPQRPPVIWIGAQECT WP_000145429.1 hydrogenase 2 small subunit [Salmonella enterica] Length: 372\nNumber of predicted TMHs: 1\nExp number of AAs in TMHs: 24.51309\nExp number, first 60 AAs: 0.2682\nTotal prob of N-in: 0.14402\noutside 1 331\nTMhelix 332 354\ninside 355 372

15756 GCF_000806685.1_CFSAN024769_02.0Salmonella enterica subsp. enterica Proteobacteria; Gammaproteobacteria; Enterobacterales; Enterobacteriaceae; Salmonella; Salmonella enterica MNNEETLYQAMRRKGVTRRSFLKFCSLAATSLGLGAGMTPKIAWALENKPRIPVVWIHGL WP_001752445.1 hydrogenase-1 small chain [Salmonella enterica] Length: 372\nNumber of predicted TMHs: 2\nExp number of AAs in TMHs: 36.5123\nExp number, first 60 AAs: 19.72341\nTotal prob of N-in: 0.96144\nPOSSIBLE N-term signal sequence\ninside 1 19\nTMhelix 20 39\noutside 40 325\nTMhelix 326 348\ninside 349 372

15757 GCF_000806785.1_CFSAN024776_01.0Salmonella enterica subsp. enterica Proteobacteria; Gammaproteobacteria; Enterobacterales; Enterobacteriaceae; Salmonella; Salmonella enterica MNNEETFYQAMRRKGVTRRSFLKFCSLAATSLGLGAGMTPKIAWALENKPRIPVVWIHGL WP_001058312.1 hydrogenase [Salmonella enterica] Length: 372\nNumber of predicted TMHs: 2\nExp number of AAs in TMHs: 36.51738\nExp number, first 60 AAs: 19.72849\nTotal prob of N-in: 0.96170\nPOSSIBLE N-term signal sequence\ninside 1 19\nTMhelix 20 39\noutside 40 325\nTMhelix 326 348\ninside 349 372

15758 GCF_000806565.1_CFSAN024758_02.0Salmonella enterica subsp. enterica Proteobacteria; Gammaproteobacteria; Enterobacterales; Enterobacteriaceae; Salmonella; Salmonella enterica MTGDNTLITSHGINRRDFMKLCAALAATMGLSSKAAAEMAESVSNPQRPPVIWIGAQECT WP_000145429.1 hydrogenase 2 small subunit [Salmonella enterica] Length: 372\nNumber of predicted TMHs: 1\nExp number of AAs in TMHs: 24.51309\nExp number, first 60 AAs: 0.2682\nTotal prob of N-in: 0.14402\noutside 1 331\nTMhelix 332 354\ninside 355 372

15759 GCF_000806745.1_CFSAN024773_01.0Salmonella enterica subsp. enterica Proteobacteria; Gammaproteobacteria; Enterobacterales; Enterobacteriaceae; Salmonella; Salmonella enterica MNNEETFYQAMRRKGVTRRSFLKFCSLAATSLGLGAGMTPKIAWALENKPRIPVVWIHGL WP_001058312.1 hydrogenase [Salmonella enterica] Length: 372\nNumber of predicted TMHs: 2\nExp number of AAs in TMHs: 36.51738\nExp number, first 60 AAs: 19.72849\nTotal prob of N-in: 0.96170\nPOSSIBLE N-term signal sequence\ninside 1 19\nTMhelix 20 39\noutside 40 325\nTMhelix 326 348\ninside 349 372

15760 GCF_000806505.1_CFSAN024724_01.0Salmonella enterica subsp. enterica Proteobacteria; Gammaproteobacteria; Enterobacterales; Enterobacteriaceae; Salmonella; Salmonella enterica MTGDNTLITSHGINRRDFMKLCAALAATMGLSSKAAAEMAESVSNPQRPPVIWIGAQECT WP_000145429.1 hydrogenase 2 small subunit [Salmonella enterica] Length: 372\nNumber of predicted TMHs: 1\nExp number of AAs in TMHs: 24.51309\nExp number, first 60 AAs: 0.2682\nTotal prob of N-in: 0.14402\noutside 1 331\nTMhelix 332 354\ninside 355 372

15761 GCF_002060085.1_ASM206008v1 Salmonella enterica subsp. enterica Proteobacteria; Gammaproteobacteria; Enterobacterales; Enterobacteriaceae; Salmonella; Salmonella enterica MNNEETFYQAMRRKGVTRRSFLKFCSLAATSLGLGAGMTPKIAWALENKPRIPVVWIHGL WP_001058311.1 hydrogenase [Salmonella enterica] Length: 372\nNumber of predicted TMHs: 2\nExp number of AAs in TMHs: 37.4699899999999\nExp number, first 60 AAs: 19.72699\nTotal prob of N-in: 0.96171\nPOSSIBLE N-term signal sequence\ninside 1 19\nTMhelix 20 39\noutside 40 325\nTMhelix 326 348\ninside 349 372

15762 GCF_001188155.1_ASM118815v1 Salmonella enterica subsp. houtenae Proteobacteria; Gammaproteobacteria; Enterobacterales; Enterobacteriaceae; Salmonella; Salmonella enterica MNNEETFYQAMRRKGVTRRSFLKFCSLAATSLGLGAGMTPKIAWALENKPRIPVVWIHGL WP_023187532.1 hydrogenase 1, small subunit [Salmonella enterica] Length: 372\nNumber of predicted TMHs: 2\nExp number of AAs in TMHs: 36.9997\nExp number, first 60 AAs: 19.72759\nTotal prob of N-in: 0.96171\nPOSSIBLE N-term signal sequence\ninside 1 19\nTMhelix 20 39\noutside 40 325\nTMhelix 326 348\ninside 349 372

15763 GCF_001188155.1_ASM118815v1 Salmonella enterica subsp. houtenae Proteobacteria; Gammaproteobacteria; Enterobacterales; Enterobacteriaceae; Salmonella; Salmonella enterica MTGDNTLITSHGINRRDFMKLCAALAATMGLSSKAAAEMAESVSNPQRPPVIWIGAQECT WP_000145429.1 hydrogenase 2 small subunit [Salmonella enterica] Length: 372\nNumber of predicted TMHs: 1\nExp number of AAs in TMHs: 24.51309\nExp number, first 60 AAs: 0.2682\nTotal prob of N-in: 0.14402\noutside 1 331\nTMhelix 332 354\ninside 355 372

15764 GCF_000968635.1_ASM96863v1 Salmonella enterica subsp. salamae Proteobacteria; Gammaproteobacteria; Enterobacterales; Enterobacteriaceae; Salmonella; Salmonella enterica MTGDNTLITSHGINRRDFMKLCAALAATMGLSSKAAAEMAESVSNPQRPPVIWIGAQECT WP_000145432.1 hydrogenase 2 small subunit [Salmonella enterica] Length: 372\nNumber of predicted TMHs: 1\nExp number of AAs in TMHs: 24.5101\nExp number, first 60 AAs: 0.2682\nTotal prob of N-in: 0.14397\noutside 1 331\nTMhelix 332 354\ninside 355 372

15765 GCF_000968635.1_ASM96863v1 Salmonella enterica subsp. salamae Proteobacteria; Gammaproteobacteria; Enterobacterales; Enterobacteriaceae; Salmonella; Salmonella enterica MNNEETFYQAMRRKGVTRRSFLKFCSLAATSLGLGAGMTPKIAWALENKPRIPVVWIHGL WP_001058314.1 hydrogenase [Salmonella enterica] Length: 372\nNumber of predicted TMHs: 2\nExp number of AAs in TMHs: 36.3110299999999\nExp number, first 60 AAs: 19.72774\nTotal prob of N-in: 0.96169\nPOSSIBLE N-term signal sequence\ninside 1 19\nTMhelix 20 39\noutside 40 325\nTMhelix 326 348\ninside 349 372

15766 GCF_001479245.1_Salmonella_enterica_CVM_N31597-SQ_v1.0 Salmonella enterica subsp. arizonae serovar 18:z4,z23:- Proteobacteria; Gammaproteobacteria; Enterobacterales; Enterobacteriaceae; Salmonella; Salmonella enterica; Salmonella enterica subsp. arizonae MTGDNTLITSHGINRRDFMKLCAALAATMGLSSKAAAEMAESVSNPQRPPVIWIGAQECT WP_058819740.1 hydrogenase 2 small subunit [Salmonella enterica] Length: 372\nNumber of predicted TMHs: 1\nExp number of AAs in TMHs: 24.54195\nExp number, first 60 AAs: 0.269\nTotal prob of N-in: 0.14471\noutside 1 331\nTMhelix 332 354\ninside 355 372

15767 GCF_002032885.1_ASM203288v1 Salmonella enterica subsp. arizonae serovar 18:z4,z23:- Proteobacteria; Gammaproteobacteria; Enterobacterales; Enterobacteriaceae; Salmonella; Salmonella enterica; Salmonella enterica subsp. arizonae MTGDNTLITSHGINRRDFMKLCAALAATMGLSSKAAAEMAESVSNPQRPPVIWIGAQECT WP_058819740.1 hydrogenase 2 small subunit [Salmonella enterica] Length: 372\nNumber of predicted TMHs: 1\nExp number of AAs in TMHs: 24.54195\nExp number, first 60 AAs: 0.269\nTotal prob of N-in: 0.14471\noutside 1 331\nTMhelix 332 354\ninside 355 372

15768 GCF_002031985.1_ASM203198v1 Salmonella enterica subsp. arizonae serovar 18:z4,z23:- Proteobacteria; Gammaproteobacteria; Enterobacterales; Enterobacteriaceae; Salmonella; Salmonella enterica; Salmonella enterica subsp. arizonae MNNEETFYQAMRRKGVTRRSFLKFCSLAATSLGLGAGMTPKIAWALENKPRIPVVWIHGL WP_001058311.1 hydrogenase [Salmonella enterica] Length: 372\nNumber of predicted TMHs: 2\nExp number of AAs in TMHs: 37.4699899999999\nExp number, first 60 AAs: 19.72699\nTotal prob of N-in: 0.96171\nPOSSIBLE N-term signal sequence\ninside 1 19\nTMhelix 20 39\noutside 40 325\nTMhelix 326 348\ninside 349 372

15769 GCF_001479245.1_Salmonella_enterica_CVM_N31597-SQ_v1.0 Salmonella enterica subsp. arizonae serovar 18:z4,z23:- Proteobacteria; Gammaproteobacteria; Enterobacterales; Enterobacteriaceae; Salmonella; Salmonella enterica; Salmonella enterica subsp. arizonae MNNEETFYQAMRRKGVTRRSFLKFCSLAATSLGLGAGMTPKIAWALENKPRIPVVWIHGL WP_001058306.1 hydrogenase [Salmonella enterica] Length: 372\nNumber of predicted TMHs: 2\nExp number of AAs in TMHs: 36.5540099999999\nExp number, first 60 AAs: 19.72732\nTotal prob of N-in: 0.96171\nPOSSIBLE N-term signal sequence\ninside 1 19\nTMhelix 20 39\noutside 40 325\nTMhelix 326 348\ninside 349 372

15770 GCF_002032005.1_ASM203200v1 Salmonella enterica subsp. arizonae serovar 18:z4,z23:- Proteobacteria; Gammaproteobacteria; Enterobacterales; Enterobacteriaceae; Salmonella; Salmonella enterica;

Salmonella enterica subsp. arizonae MNNEETFYQAMRRKGVTRRSFLKFCSLAATSLGLGAGMTPKIAWALENKPRIPVVWIHGL WP_001058306.1 hydrogenase [Salmonella enterica] Length: 372\nNumber of predicted TMHs: 2\nExp number of AAs in TMHs: 36.5540099999999\nExp number, first 60 AAs: 19.72732\nTotal prob of N-in: 0.96171\nPOSSIBLE N-term signal sequence\ninside 1 19\nTMhelix 20 39\noutside 40 325\nTMhelix 326 348\ninside 349 372

15771 GCF_002032005.1_ASM203200v1 Salmonella enterica subsp. arizonae serovar 18:z4,z23:- Proteobacteria; Gammaproteobacteria; Enterobacterales; Enterobacteriaceae; Salmonella; Salmonella enterica; Salmonella enterica subsp. arizonae MTGDNTLITSHGINRRDFMKLCAALAATMGLSSKAAAEMAESVSNPQRPPVIWIGAQECT WP_058819740.1 hydrogenase 2 small subunit [Salmonella enterica] Length: 372\nNumber of predicted TMHs: 1\nExp number of AAs in TMHs: 24.54195\nExp number, first 60 AAs: 0.269\nTotal prob of N-in: 0.14471\noutside 1 331\nTMhelix 332 354\ninside 355 372

15772 GCF_002032885.1_ASM203288v1 Salmonella enterica subsp. arizonae serovar 18:z4,z23:- Proteobacteria; Gammaproteobacteria; Enterobacterales; Enterobacteriaceae; Salmonella; Salmonella enterica; Salmonella enterica subsp. arizonae MNNEETFYQAMRRKGVTRRSFLKFCSLAATSLGLGAGMTPKIAWALENKPRIPVVWIHGL WP_001058306.1 hydrogenase [Salmonella enterica] Length: 372\nNumber of predicted TMHs: 2\nExp number of AAs in TMHs: 36.5540099999999\nExp number, first 60 AAs: 19.72732\nTotal prob of N-in: 0.96171\nPOSSIBLE N-term signal sequence\ninside 1 19\nTMhelix 20 39\noutside 40 325\nTMhelix 326 348\ninside 349 372

15773 GCF_002031985.1_ASM203198v1 Salmonella enterica subsp. arizonae serovar 18:z4,z23:- Proteobacteria; Gammaproteobacteria; Enterobacterales; Enterobacteriaceae; Salmonella; Salmonella enterica; Salmonella enterica subsp. arizonae MTGDNTLITSHGINRRDFMKLCAALAATMGLSSKAAAEMAESVSNPQRPPVIWIGAQECT WP_000145429.1 hydrogenase 2 small subunit [Salmonella enterica] Length: 372\nNumber of predicted TMHs: 1\nExp number of AAs in TMHs: 24.51309\nExp number, first 60 AAs: 0.2682\nTotal prob of N-in: 0.14402\noutside 1 331\nTMhelix 332 354\ninside 355 372

15774 GCF_001478655.1_Salmonella_enterica_CVM_N29355-SQ_v1.0 Salmonella enterica subsp. arizonae serovar 18:z4,z23:- Proteobacteria; Gammaproteobacteria; Enterobacterales; Enterobacteriaceae; Salmonella; Salmonella enterica; Salmonella enterica subsp. arizonae MTGDNTLITSHGINRRDFMKLCAALAATMGLSSKAAAEMAESVSNPQRPPVIWIGAQECT WP_058819740.1 hydrogenase 2 small subunit [Salmonella enterica] Length: 372\nNumber of predicted TMHs: 1\nExp number of AAs in TMHs: 24.54195\nExp number, first 60 AAs: 0.269\nTotal prob of N-in: 0.14471\noutside 1 331\nTMhelix 332 354\ninside 355 372

15775 GCF_002030835.1_ASM203083v1 Salmonella enterica subsp. arizonae serovar 18:z4,z23:- Proteobacteria; Gammaproteobacteria; Enterobacterales; Enterobacteriaceae; Salmonella; Salmonella enterica; Salmonella enterica subsp. arizonae MTGDNTLITSHGINRRDFMKLCAALAATMGLSSKAAAEMAESVSNPQRPPVIWIGAQECT WP_058819740.1 hydrogenase 2 small subunit [Salmonella enterica] Length: 372\nNumber of predicted TMHs: 1\nExp number of AAs in TMHs: 24.54195\nExp number, first 60 AAs: 0.269\nTotal prob of N-in: 0.14471\noutside 1 331\nTMhelix 332 354\ninside 355 372

15776 GCF_001478385.1_Salmonella_enterica_CVM_N38891-SQ_v1.0 Salmonella enterica subsp. arizonae serovar 18:z4,z23:- Proteobacteria; Gammaproteobacteria; Enterobacterales; Enterobacteriaceae; Salmonella; Salmonella enterica; Salmonella enterica subsp. arizonae MTGDNTLITSHGINRRDFMKLCAALAATMGLSSKAAAEMAESVSNPQRPPVIWIGAQECT WP_058819740.1 hydrogenase 2 small subunit [Salmonella enterica] Length: 372\nNumber of predicted TMHs: 1\nExp number of AAs in TMHs: 24.54195\nExp number, first 60 AAs: 0.269\nTotal prob of N-in: 0.14471\noutside 1 331\nTMhelix 332 354\ninside 355 372

15777 GCF_001478655.1_Salmonella_enterica_CVM_N29355-SQ_v1.0 Salmonella enterica subsp. arizonae serovar 18:z4,z23:- Proteobacteria; Gammaproteobacteria; Enterobacterales; Enterobacteriaceae; Salmonella; Salmonella enterica; Salmonella enterica subsp. arizonae MNNEETFYQAMRRKGVTRRSFLKFCSLAATSLGLGAGMTPKIAWALENKPRIPVVWIHGL WP_001058306.1 hydrogenase [Salmonella enterica] Length: 372\nNumber of predicted TMHs: 2\nExp number of AAs in TMHs: 36.5540099999999\nExp number, first 60 AAs: 19.72732\nTotal prob of N-in: 0.96171\nPOSSIBLE N-term signal sequence\ninside 1 19\nTMhelix 20 39\noutside 40 325\nTMhelix 326 348\ninside 349 372

15778 GCF_001478385.1_Salmonella_enterica_CVM_N38891-SQ_v1.0 Salmonella enterica subsp. arizonae serovar 18:z4,z23:- Proteobacteria; Gammaproteobacteria; Enterobacterales; Enterobacteriaceae; Salmonella; Salmonella

enterica; Salmonella enterica subsp. arizonae
MNNEETFYQAMRRKGVTRRSFLKFCSLAATSLGLGAGMTPKIAWALENKPRIPVVWIHGL WP_001058306.1 hydrogenase [Salmonella enterica] Length: 372\nNumber of predicted TMHs: 2\nExp number of AAs in TMHs: 36.5540099999999\nExp number, first 60 AAs: 19.72732\nTotal prob of N-in: 0.96171\nPOSSIBLE N-term signal sequence\ninside 1 19\nTMhelix 20 39\noutside 40 325\nTMhelix 326 348\ninside 349 372

15779 GCF_002030835.1_ASM203083v1 Salmonella enterica subsp. arizonae serovar 18:z4,z23:- Proteobacteria; Gammaproteobacteria; Enterobacterales; Enterobacteriaceae; Salmonella; Salmonella enterica; Salmonella enterica subsp. arizonae MNNEETFYQAMRRKGVTRRSFLKFCSLAATSLGLGAGMTPKIAWALENKPRIPVVWIHGL WP_001058306.1 hydrogenase [Salmonella enterica] Length: 372\nNumber of predicted TMHs: 2\nExp number of AAs in TMHs: 36.5540099999999\nExp number, first 60 AAs: 19.72732\nTotal prob of N-in: 0.96171\nPOSSIBLE N-term signal sequence\ninside 1 19\nTMhelix 20 39\noutside 40 325\nTMhelix 326 348\ninside 349 372

15780 GCF_002031795.1_ASM203179v1 Salmonella enterica subsp. arizonae serovar 18:z4,z23:- Proteobacteria; Gammaproteobacteria; Enterobacterales; Enterobacteriaceae; Salmonella; Salmonella enterica; Salmonella enterica subsp. arizonae MTGDNTLITSHGINRRDFMKLCAALAATMGLSSKAAAEMAESVSNPQRPPVIWIGAQECT WP_058819740.1 hydrogenase 2 small subunit [Salmonella enterica] Length: 372\nNumber of predicted TMHs: 1\nExp number of AAs in TMHs: 24.54195\nExp number, first 60 AAs: 0.269\nTotal prob of N-in: 0.14471\noutside 1 331\nTMhelix 332 354\ninside 355 372

15781 GCF_002031795.1_ASM203179v1 Salmonella enterica subsp. arizonae serovar 18:z4,z23:- Proteobacteria; Gammaproteobacteria; Enterobacterales; Enterobacteriaceae; Salmonella; Salmonella enterica; Salmonella enterica subsp. arizonae MNNEETFYQAMRRKGVTRRSFLKFCSLAATSLGLGAGMTPKIAWALENKPRIPVVWIHGL WP_001058306.1 hydrogenase [Salmonella enterica] Length: 372\nNumber of predicted TMHs: 2\nExp number of AAs in TMHs: 36.5540099999999\nExp number, first 60 AAs: 19.72732\nTotal prob of N-in: 0.96171\nPOSSIBLE N-term signal sequence\ninside 1 19\nTMhelix 20 39\noutside 40 325\nTMhelix 326 348\ninside 349 372

15782 GCF_002106555.1_ASM210655v1 Salmonella enterica subsp. arizonae serovar 41:z4,z23:- Proteobacteria; Gammaproteobacteria; Enterobacterales; Enterobacteriaceae; Salmonella; Salmonella enterica; Salmonella enterica subsp. arizonae MNNEETFYQAMRRKGVTRRSFLKFCSLAATSLGLGAGMTPKIAWALENKPRIPVVWIHGL WP_001058306.1 hydrogenase [Salmonella enterica] Length: 372\nNumber of predicted TMHs: 2\nExp number of AAs in TMHs: 36.5540099999999\nExp number, first 60 AAs: 19.72732\nTotal prob of N-in: 0.96171\nPOSSIBLE N-term signal sequence\ninside 1 19\nTMhelix 20 39\noutside 40 325\nTMhelix 326 348\ninside 349 372

15783 GCF_002106555.1_ASM210655v1 Salmonella enterica subsp. arizonae serovar 41:z4,z23:- Proteobacteria; Gammaproteobacteria; Enterobacterales; Enterobacteriaceae; Salmonella; Salmonella enterica; Salmonella enterica subsp. arizonae MTGDNTLITSHGINRRDFMKLCAALAATMGLSSKAAAEMAESVSNPQRPPVIWIGAQECT WP_000145429.1 hydrogenase 2 small subunit [Salmonella enterica] Length: 372\nNumber of predicted TMHs: 1\nExp number of AAs in TMHs: 24.51309\nExp number, first 60 AAs: 0.2682\nTotal prob of N-in: 0.14402\noutside 1 331\nTMhelix 332 354\ninside 355 372

15784 GCF_002035775.1_ASM203577v1 Salmonella enterica subsp. arizonae serovar 48:g,z51:- Proteobacteria; Gammaproteobacteria; Enterobacterales; Enterobacteriaceae; Salmonella; Salmonella enterica; Salmonella enterica subsp. arizonae MNNEETFYQAMRRKGVTRRSFLKFCSLAATSLGLGAGMTPKIAWALENKPRIPVVWIHGL WP_001058306.1 hydrogenase [Salmonella enterica] Length: 372\nNumber of predicted TMHs: 2\nExp number of AAs in TMHs: 36.5540099999999\nExp number, first 60 AAs: 19.72732\nTotal prob of N-in: 0.96171\nPOSSIBLE N-term signal sequence\ninside 1 19\nTMhelix 20 39\noutside 40 325\nTMhelix 326 348\ninside 349 372

15785 GCF_002035775.1_ASM203577v1 Salmonella enterica subsp. arizonae serovar 48:g,z51:- Proteobacteria; Gammaproteobacteria; Enterobacterales; Enterobacteriaceae; Salmonella; Salmonella enterica; Salmonella enterica subsp. arizonae MTGDNTLITSHGINRRDFMKLCAALAATMGLSSKAAAEMAESVSNPQRPPVIWIGAQECT WP_000145429.1 hydrogenase 2 small subunit [Salmonella enterica] Length: 372\nNumber of predicted TMHs: 1\nExp number of AAs in TMHs: 24.51309\nExp number, first 60 AAs: 0.2682\nTotal prob of N-in: 0.14402\noutside 1 331\nTMhelix 332 354\ninside 355 372

15786 GCF_002035795.1_ASM203579v1 Salmonella enterica subsp. arizonae serovar 48:g,z51:- Proteobacteria; Gammaproteobacteria; Enterobacterales; Enterobacteriaceae; Salmonella; Salmonella enterica;

Salmonella enterica subsp. arizonae MTGDNTLITSHGINRRDFMKLCAALAATMGLSSKAAAEMAESVSNPQRPPVIWIGAQECT WP_000145429.1 hydrogenase 2 small subunit [Salmonella enterica] Length: 372\nNumber of predicted TMHs: 1\nExp number of AAs in TMHs: 24.51309\nExp number, first 60 AAs: 0.2682\nTotal prob of N-in: 0.14402\noutside 1 331\nTMhelix 332 354\ninside 355 372

15787 GCF_002035795.1_ASM203579v1 Salmonella enterica subsp. arizonae serovar 48:g,z51:- Proteobacteria; Gammaproteobacteria; Enterobacterales; Enterobacteriaceae; Salmonella; Salmonella enterica; Salmonella enterica subsp. arizonae MNNEETFYQAMRRKGVTRRSFLKFCSLAATSLGLGAGMTPKIAWALENKPRIPVVWIHGL WP_001058306.1 hydrogenase [Salmonella enterica] Length: 372\nNumber of predicted TMHs: 2\nExp number of AAs in TMHs: 36.5540099999999\nExp number, first 60 AAs: 19.72732\nTotal prob of N-in: 0.96171\nPOSSIBLE N-term signal sequence\ninside 1 19\nTMhelix 20 39\noutside 40 325\nTMhelix 326 348\ninside 349 372

15788 GCF_002035585.1_ASM203558v1 Salmonella enterica subsp. arizonae serovar 48:z4,z23,z32:- Proteobacteria; Gammaproteobacteria; Enterobacterales; Enterobacteriaceae; Salmonella; Salmonella enterica; Salmonella enterica subsp. arizonae MNNEETFYQAMRRKGVTRRSFLKFCSLAATSLGLGAGMTPKIAWALENKPRIPVVWIHGL WP_001058306.1 hydrogenase [Salmonella enterica] Length: 372\nNumber of predicted TMHs: 2\nExp number of AAs in TMHs: 36.5540099999999\nExp number, first 60 AAs: 19.72732\nTotal prob of N-in: 0.96171\nPOSSIBLE N-term signal sequence\ninside 1 19\nTMhelix 20 39\noutside 40 325\nTMhelix 326 348\ninside 349 372

15789 GCF_002035585.1_ASM203558v1 Salmonella enterica subsp. arizonae serovar 48:z4,z23,z32:- Proteobacteria; Gammaproteobacteria; Enterobacterales; Enterobacteriaceae; Salmonella; Salmonella enterica; Salmonella enterica subsp. arizonae MTGDNTLITSHGINRRDFMKLCAALAATMGLSSKAAAEMAESVSNPQRPPVIWIGAQECT WP_000145429.1 hydrogenase 2 small subunit [Salmonella enterica] Length: 372\nNumber of predicted TMHs: 1\nExp number of AAs in TMHs: 24.51309\nExp number, first 60 AAs: 0.2682\nTotal prob of N-in: 0.14402\noutside 1 331\nTMhelix 332 354\ninside 355 372

15790 GCF_002106495.1_ASM210649v1 Salmonella enterica subsp. arizonae serovar 50:r:z Proteobacteria; Gammaproteobacteria; Enterobacterales; Enterobacteriaceae; Salmonella; Salmonella enterica; Salmonella enterica subsp. arizonae MTGDNTLITSHGINRRDFMKLCAALAATMGLSSKAAAEMAESVSNPQRPPVIWIGAQECT WP_023248548.1 hydrogenase 2 small subunit [Salmonella enterica] Length: 372\nNumber of predicted TMHs: 1\nExp number of AAs in TMHs: 24.51281\nExp number, first 60 AAs: 0.26822\nTotal prob of N-in: 0.14402\noutside 1 331\nTMhelix 332 354\ninside 355 372

15791 GCF_002106495.1_ASM210649v1 Salmonella enterica subsp. arizonae serovar 50:r:z Proteobacteria; Gammaproteobacteria; Enterobacterales; Enterobacteriaceae; Salmonella; Salmonella enterica; Salmonella enterica subsp. arizonae MNNEETFYQAMRRKGVTRRSFLKFCSLAATSLGLGAGMTPKIAWALENKPRIPVVWIHGL WP_001058306.1 hydrogenase [Salmonella enterica] Length: 372\nNumber of predicted TMHs: 2\nExp number of AAs in TMHs: 36.5540099999999\nExp number, first 60 AAs: 19.72732\nTotal prob of N-in: 0.96171\nPOSSIBLE N-term signal sequence\ninside 1 19\nTMhelix 20 39\noutside 40 325\nTMhelix 326 348\ninside 349 372

15792 GCF_002065135.1_ASM206513v1 Salmonella enterica subsp. arizonae serovar 56:z4,z23,(z32):- Proteobacteria; Gammaproteobacteria; Enterobacterales; Enterobacteriaceae; Salmonella; Salmonella enterica; Salmonella enterica subsp. arizonae MTGDNTLITSHGINRRDFMKLCAALAATMGLSSKAAAEMAESVSNPQRPPVIWIGAQECT WP_000145429.1 hydrogenase 2 small subunit [Salmonella enterica] Length: 372\nNumber of predicted TMHs: 1\nExp number of AAs in TMHs: 24.51309\nExp number, first 60 AAs: 0.2682\nTotal prob of N-in: 0.14402\noutside 1 331\nTMhelix 332 354\ninside 355 372

15793 GCF_002065135.1_ASM206513v1 Salmonella enterica subsp. arizonae serovar 56:z4,z23,(z32):- Proteobacteria; Gammaproteobacteria; Enterobacterales; Enterobacteriaceae; Salmonella; Salmonella enterica; Salmonella enterica subsp. arizonae MNNEETFYQAMRRKGVTRRSFLKFCSLAATSLGLGAGMTPKIAWALENKPRIPVVWIHGL WP_079953656.1 [Ni/Fe] hydrogenase small subunit [Salmonella enterica] Length: 372\nNumber of predicted TMHs: 2\nExp number of AAs in TMHs: 36.5655299999999\nExp number, first 60 AAs: 19.72713\nTotal prob of N-in: 0.96172\nPOSSIBLE N-term signal sequence\ninside 1 19\nTMhelix 20 39\noutside 40 325\nTMhelix 326 348\ninside 349 372

15794 GCF_000018625.1_ASM1862v1 Salmonella enterica subsp. arizonae serovar 62:z4,z23:- Proteobacteria; Gammaproteobacteria; Enterobacterales; Enterobacteriaceae; Salmonella; Salmonella enterica; Salmonella enterica subsp. arizonae MNNEETFYQAMRRKGVTRRSFLKFCSLAATSLGLGAGMTPKIAWALENKPRIPVVWIHGL

WP_001058306.1 hydrogenase [Salmonella enterica] Length: 372\nNumber of predicted TMHs: 2\nExp number of AAs in TMHs: 36.5540099999999\nExp number, first 60 AAs: 19.72732\nTotal prob of N-in: 0.96171\nPOSSIBLE N-term signal sequence\ninside 1 19\nTMhelix 20 39\noutside 40 325\nTMhelix 326 348\ninside 349 372

15795 GCF_000018625.1_ASM1862v1 Salmonella enterica subsp. arizonae serovar 62:z4,z23:- Proteobacteria; Gammaproteobacteria; Enterobacterales; Enterobacteriaceae; Salmonella; Salmonella enterica; Salmonella enterica subsp. arizonae MTGDNTLITSHGINRRDFMKLCAALAATMGLSSKAAAEMAESVSNPQRPPVIWIGAQECT WP_000145429.1 hydrogenase 2 small subunit [Salmonella enterica] Length: 372\nNumber of predicted TMHs: 1\nExp number of AAs in TMHs: 24.51309\nExp number, first 60 AAs: 0.2682\nTotal prob of N-in: 0.14402\noutside 1 331\nTMhelix 332 354\ninside 355 372

15796 GCF_001952135.1_ASM195213v1 Salmonella enterica subsp. arizonae serovar 18:z4,z23:- str. CVM 32457 Proteobacteria; Gammaproteobacteria; Enterobacterales; Enterobacteriaceae; Salmonella; Salmonella enterica; Salmonella enterica subsp. arizonae; Salmonella enterica subsp. arizonae serovar 18 MNNEETFYQAMRRKGVTRRSFLKFCSLAATSLGLGAGMTPKIAWALENKPRIPVVWIHGL WP_001058306.1 hydrogenase [Salmonella enterica] Length: 372\nNumber of predicted TMHs: 2\nExp number of AAs in TMHs: 36.5540099999999\nExp number, first 60 AAs: 19.72732\nTotal prob of N-in: 0.96171\nPOSSIBLE N-term signal sequence\ninside 1 19\nTMhelix 20 39\noutside 40 325\nTMhelix 326 348\ninside 349 372

15797 GCF_001952135.1_ASM195213v1 Salmonella enterica subsp. arizonae serovar 18:z4,z23:- str. CVM 32457 Proteobacteria; Gammaproteobacteria; Enterobacterales; Enterobacteriaceae; Salmonella; Salmonella enterica; Salmonella enterica subsp. arizonae; Salmonella enterica subsp. arizonae serovar 18 MTGDNTLITSHGINRRDFMKLCAALAATMGLSSKAAAEMAESVSNPQRPPVIWIGAQECT WP_058819740.1 hydrogenase 2 small subunit [Salmonella enterica] Length: 372\nNumber of predicted TMHs: 1\nExp number of AAs in TMHs: 24.54195\nExp number, first 60 AAs: 0.269\nTotal prob of N-in: 0.14471\noutside 1 331\nTMhelix 332 354\ninside 355 372

15798 GCF_001951365.1_ASM195136v1 Salmonella enterica subsp. arizonae serovar 18:z4,z23:- str. CVM N18383 Proteobacteria; Gammaproteobacteria; Enterobacterales; Enterobacteriaceae; Salmonella; Salmonella enterica; Salmonella enterica subsp. arizonae; Salmonella enterica subsp. arizonae serovar 18 MNNEETFYQAMRRKGVTRRSFLKFCSLAATSLGLGAGMTPKIAWALENKPRIPVVWIHGL WP_001058306.1 hydrogenase [Salmonella enterica] Length: 372\nNumber of predicted TMHs: 2\nExp number of AAs in TMHs: 36.5540099999999\nExp number, first 60 AAs: 19.72732\nTotal prob of N-in: 0.96171\nPOSSIBLE N-term signal sequence\ninside 1 19\nTMhelix 20 39\noutside 40 325\nTMhelix 326 348\ninside 349 372

15799 GCF_001951365.1_ASM195136v1 Salmonella enterica subsp. arizonae serovar 18:z4,z23:- str. CVM N18383 Proteobacteria; Gammaproteobacteria; Enterobacterales; Enterobacteriaceae; Salmonella; Salmonella enterica; Salmonella enterica subsp. arizonae; Salmonella enterica subsp. arizonae serovar 18 MTGDNTLITSHGINRRDFMKLCAALAATMGLSSKAAAEMAESVSNPQRPPVIWIGAQECT WP_058819740.1 hydrogenase 2 small subunit [Salmonella enterica] Length: 372\nNumber of predicted TMHs: 1\nExp number of AAs in TMHs: 24.54195\nExp number, first 60 AAs: 0.269\nTotal prob of N-in: 0.14471\noutside 1 331\nTMhelix 332 354\ninside 355 372

15800 GCF_001951375.1_ASM195137v1 Salmonella enterica subsp. arizonae serovar 18:z4,z23:- str. CVM N18503 Proteobacteria; Gammaproteobacteria; Enterobacterales; Enterobacteriaceae; Salmonella; Salmonella enterica; Salmonella enterica subsp. arizonae; Salmonella enterica subsp. arizonae serovar 18 MTGDNTLITSHGINRRDFMKLCAALAATMGLSSKAAAEMAESVSNPQRPPVIWIGAQECT WP_058819740.1 hydrogenase 2 small subunit [Salmonella enterica] Length: 372\nNumber of predicted TMHs: 1\nExp number of AAs in TMHs: 24.54195\nExp number, first 60 AAs: 0.269\nTotal prob of N-in: 0.14471\noutside 1 331\nTMhelix 332 354\ninside 355 372

15801 GCF_001951375.1_ASM195137v1 Salmonella enterica subsp. arizonae serovar 18:z4,z23:- str. CVM N18503 Proteobacteria; Gammaproteobacteria; Enterobacterales; Enterobacteriaceae; Salmonella; Salmonella enterica; Salmonella enterica subsp. arizonae; Salmonella enterica subsp. arizonae serovar 18 MNNEETFYQAMRRKGVTRRSFLKFCSLAATSLGLGAGMTPKIAWALENKPRIPVVWIHGL WP_001058306.1 hydrogenase [Salmonella enterica] Length: 372\nNumber of predicted TMHs: 2\nExp number of AAs in TMHs: 36.5540099999999\nExp number, first 60 AAs: 19.72732\nTotal prob of N-in: 0.96171\nPOSSIBLE N-term signal sequence\ninside 1 19\nTMhelix 20 39\noutside 40 325\nTMhelix 326 348\ninside 349 372

15802 GCF_001951435.1_ASM195143v1 Salmonella enterica subsp. arizonae serovar 18:z4,z23:- str. CVM N18554 Proteobacteria; Gammaproteobacteria; Enterobacterales; Enterobacteriaceae; Salmonella; Salmonella enterica; Salmonella enterica subsp. arizonae; Salmonella enterica subsp. arizonae serovar 18 MTGDNTLITSHGINRRDFMKLCAALAATMGLSSKAAAEMAESVSNPQRPPVIWIGAQECT WP_058819740.1 hydrogenase 2 small subunit [Salmonella enterica] Length: 372\nNumber of predicted TMHs: 1\nExp number of AAs in TMHs: 24.54195\nExp number, first 60 AAs: 0.269\nTotal prob of N-in: 0.14471\noutside 1 331\nTMhelix 332 354\ninside 355 372

15803 GCF_001951435.1_ASM195143v1 Salmonella enterica subsp. arizonae serovar 18:z4,z23:- str. CVM N18554 Proteobacteria; Gammaproteobacteria; Enterobacterales; Enterobacteriaceae; Salmonella; Salmonella enterica; Salmonella enterica subsp. arizonae; Salmonella enterica subsp. arizonae serovar 18 MNNEETFYQAMRRKGVTRRSFLKFCSLAATSLGLGAGMTPKIAWALENKPRIPVVWIHGL WP_001058306.1 hydrogenase [Salmonella enterica] Length: 372\nNumber of predicted TMHs: 2\nExp number of AAs in TMHs: 36.5540099999999\nExp number, first 60 AAs: 19.72732\nTotal prob of N-in: 0.96171\nPOSSIBLE N-term signal sequence\ninside 1 19\nTMhelix 20 39\noutside 40 325\nTMhelix 326 348\ninside 349 372

15804 GCF_001951525.1_ASM195152v1 Salmonella enterica subsp. arizonae serovar 18:z4,z23:- str. CVM N20028 Proteobacteria; Gammaproteobacteria; Enterobacterales; Enterobacteriaceae; Salmonella; Salmonella enterica; Salmonella enterica subsp. arizonae; Salmonella enterica subsp. arizonae serovar 18 MNNEETFYQAMRRKGVTRRSFLKFCSLAATSLGLGAGMTPKIAWALENKPRIPVVWIHGL WP_001058306.1 hydrogenase [Salmonella enterica] Length: 372\nNumber of predicted TMHs: 2\nExp number of AAs in TMHs: 36.5540099999999\nExp number, first 60 AAs: 19.72732\nTotal prob of N-in: 0.96171\nPOSSIBLE N-term signal sequence\ninside 1 19\nTMhelix 20 39\noutside 40 325\nTMhelix 326 348\ninside 349 372

15805 GCF_001951525.1_ASM195152v1 Salmonella enterica subsp. arizonae serovar 18:z4,z23:- str. CVM N20028 Proteobacteria; Gammaproteobacteria; Enterobacterales; Enterobacteriaceae; Salmonella; Salmonella enterica; Salmonella enterica subsp. arizonae; Salmonella enterica subsp. arizonae serovar 18 MTGDNTLITSHGINRRDFMKLCAALAATMGLSSKAAAEMAESVSNPQRPPVIWIGAQECT WP_058819740.1 hydrogenase 2 small subunit [Salmonella enterica] Length: 372\nNumber of predicted TMHs: 1\nExp number of AAs in TMHs: 24.54195\nExp number, first 60 AAs: 0.269\nTotal prob of N-in: 0.14471\noutside 1 331\nTMhelix 332 354\ninside 355 372

15806 GCF_001951445.1_ASM195144v1 Salmonella enterica subsp. arizonae serovar 18:z4,z23:- str. CVM N23850 Proteobacteria; Gammaproteobacteria; Enterobacterales; Enterobacteriaceae; Salmonella; Salmonella enterica; Salmonella enterica subsp. arizonae; Salmonella enterica subsp. arizonae serovar 18 MNNEETFYQAMRRKGVTRRSFLKFCSLAATSLGLGAGMTPKIAWALENKPRIPVVWIHGL WP_001058312.1 hydrogenase [Salmonella enterica] Length: 372\nNumber of predicted TMHs: 2\nExp number of AAs in TMHs: 36.51738\nExp number, first 60 AAs: 19.72849\nTotal prob of N-in: 0.96170\nPOSSIBLE N-term signal sequence\ninside 1 19\nTMhelix 20 39\noutside 40 325\nTMhelix 326 348\ninside 349 372

15807 GCF_001951445.1_ASM195144v1 Salmonella enterica subsp. arizonae serovar 18:z4,z23:- str. CVM N23850 Proteobacteria; Gammaproteobacteria; Enterobacterales; Enterobacteriaceae; Salmonella; Salmonella enterica; Salmonella enterica subsp. arizonae; Salmonella enterica subsp. arizonae serovar 18 MTGDNTLITSHGINRRDFMKLCAALAATMGLSSKAAAEMAESVSNPQRPPVIWIGAQECT WP_000145429.1 hydrogenase 2 small subunit [Salmonella enterica] Length: 372\nNumber of predicted TMHs: 1\nExp number of AAs in TMHs: 24.51309\nExp number, first 60 AAs: 0.2682\nTotal prob of N-in: 0.14402\noutside 1 331\nTMhelix 332 354\ninside 355 372

15808 GCF_001951535.1_ASM195153v1 Salmonella enterica subsp. arizonae serovar 18:z4,z23:- str. CVM N25373 Proteobacteria; Gammaproteobacteria; Enterobacterales; Enterobacteriaceae; Salmonella; Salmonella enterica; Salmonella enterica subsp. arizonae; Salmonella enterica subsp. arizonae serovar 18 MTGDNTLITSHGINRRDFMKLCAALAATMGLSSKAAAEMAESVSNPQRPPVIWIGAQECT WP_058819740.1 hydrogenase 2 small subunit [Salmonella enterica] Length: 372\nNumber of predicted TMHs: 1\nExp number of AAs in TMHs: 24.54195\nExp number, first 60 AAs: 0.269\nTotal prob of N-in: 0.14471\noutside 1 331\nTMhelix 332 354\ninside 355 372

15809 GCF_001951535.1_ASM195153v1 Salmonella enterica subsp. arizonae serovar 18:z4,z23:- str. CVM N25373 Proteobacteria; Gammaproteobacteria; Enterobacterales; Enterobacteriaceae; Salmonella; Salmonella enterica; Salmonella enterica subsp. arizonae; Salmonella enterica subsp. arizonae serovar 18 MNNEETFYQAMRRKGVTRRSFLKFCSLAATSLGLGAGMTPKIAWALENKPRIPVVWIHGL WP_001058306.1 hydrogenase [Salmonella enterica] Length: 372\nNumber of predicted TMHs: 2\nExp number of AAs in

TMHs: 36.5540099999999\nExp number, first 60 AAs: 19.72732\nTotal prob of N-in: 0.96171\nPOSSIBLE N-term signal sequence\ninside 1 19\nTMhelix 20 39\noutside 40 325\nTMhelix 326 348\ninside 349 372

15810 GCF_001951545.1_ASM195154v1 Salmonella enterica subsp. arizonae serovar 18:z4,z23:- str. CVM N26624 Proteobacteria; Gammaproteobacteria; Enterobacterales; Enterobacteriaceae; Salmonella; Salmonella enterica; Salmonella enterica subsp. arizonae; Salmonella enterica subsp. arizonae serovar 18 MNNEETFYQAMRRKGVTRRSFLKFCSLAATSLGLGAGMTPKIAWALENKPRIPVVWIHGL WP_001058306.1 hydrogenase [Salmonella enterica] Length: 372\nNumber of predicted TMHs: 2\nExp number of AAs in TMHs: 36.5540099999999\nExp number, first 60 AAs: 19.72732\nTotal prob of N-in: 0.96171\nPOSSIBLE N-term signal sequence\ninside 1 19\nTMhelix 20 39\noutside 40 325\nTMhelix 326 348\ninside 349 372

15811 GCF_001951545.1_ASM195154v1 Salmonella enterica subsp. arizonae serovar 18:z4,z23:- str. CVM N26624 Proteobacteria; Gammaproteobacteria; Enterobacterales; Enterobacteriaceae; Salmonella; Salmonella enterica; Salmonella enterica subsp. arizonae; Salmonella enterica subsp. arizonae serovar 18 MTGDNTLITSHGINRRDFMKLCAALAATMGLSSKAAAEMAESVSNPQRPPVIWIGAQECT WP_058819740.1 hydrogenase 2 small subunit [Salmonella enterica] Length: 372\nNumber of predicted TMHs: 1\nExp number of AAs in TMHs: 24.54195\nExp number, first 60 AAs: 0.269\nTotal prob of N-in: 0.14471\noutside 1 331\nTMhelix 332 354\ninside 355 372

15812 GCF_001951625.1_ASM195162v1 Salmonella enterica subsp. arizonae serovar 18:z4,z23:- str. CVM N26625 Proteobacteria; Gammaproteobacteria; Enterobacterales; Enterobacteriaceae; Salmonella; Salmonella enterica; Salmonella enterica subsp. arizonae; Salmonella enterica subsp. arizonae serovar 18 MNNEETFYQAMRRKGVTRRSFLKFCSLAATSLGLGAGMTPKIAWALENKPRIPVVWIHGL WP_001058306.1 hydrogenase [Salmonella enterica] Length: 372\nNumber of predicted TMHs: 2\nExp number of AAs in TMHs: 36.5540099999999\nExp number, first 60 AAs: 19.72732\nTotal prob of N-in: 0.96171\nPOSSIBLE N-term signal sequence\ninside 1 19\nTMhelix 20 39\noutside 40 325\nTMhelix 326 348\ninside 349 372

15813 GCF_001951625.1_ASM195162v1 Salmonella enterica subsp. arizonae serovar 18:z4,z23:- str. CVM N26625 Proteobacteria; Gammaproteobacteria; Enterobacterales; Enterobacteriaceae; Salmonella; Salmonella enterica; Salmonella enterica subsp. arizonae; Salmonella enterica subsp. arizonae serovar 18 MTGDNTLITSHGINRRDFMKLCAALAATMGLSSKAAAEMAESVSNPQRPPVIWIGAQECT WP_058819740.1 hydrogenase 2 small subunit [Salmonella enterica] Length: 372\nNumber of predicted TMHs: 1\nExp number of AAs in TMHs: 24.54195\nExp number, first 60 AAs: 0.269\nTotal prob of N-in: 0.14471\noutside 1 331\nTMhelix 332 354\ninside 355 372

15814 GCF_001951455.1_ASM195145v1 Salmonella enterica subsp. arizonae serovar 18:z4,z23:- str. CVM N26626 Proteobacteria; Gammaproteobacteria; Enterobacterales; Enterobacteriaceae; Salmonella; Salmonella enterica; Salmonella enterica subsp. arizonae; Salmonella enterica subsp. arizonae serovar 18 MNNEETFYQAMRRKGVTRRSFLKFCSLAATSLGLGAGMTPKIAWALENKPRIPVVWIHGL WP_001058306.1 hydrogenase [Salmonella enterica] Length: 372\nNumber of predicted TMHs: 2\nExp number of AAs in TMHs: 36.5540099999999\nExp number, first 60 AAs: 19.72732\nTotal prob of N-in: 0.96171\nPOSSIBLE N-term signal sequence\ninside 1 19\nTMhelix 20 39\noutside 40 325\nTMhelix 326 348\ninside 349 372

15815 GCF_001951455.1_ASM195145v1 Salmonella enterica subsp. arizonae serovar 18:z4,z23:- str. CVM N26626 Proteobacteria; Gammaproteobacteria; Enterobacterales; Enterobacteriaceae; Salmonella; Salmonella enterica; Salmonella enterica subsp. arizonae; Salmonella enterica subsp. arizonae serovar 18 MTGDNTLITSHGINRRDFMKLCAALAATMGLSSKAAAEMAESVSNPQRPPVIWIGAQECT WP_058819740.1 hydrogenase 2 small subunit [Salmonella enterica] Length: 372\nNumber of predicted TMHs: 1\nExp number of AAs in TMHs: 24.54195\nExp number, first 60 AAs: 0.269\nTotal prob of N-in: 0.14471\noutside 1 331\nTMhelix 332 354\ninside 355 372

15816 GCF_001970905.1_ASM197090v1 Salmonella enterica subsp. arizonae serovar 18:z4,z23:- str. CVM N26928 Proteobacteria; Gammaproteobacteria; Enterobacterales; Enterobacteriaceae; Salmonella; Salmonella enterica; Salmonella enterica subsp. arizonae; Salmonella enterica subsp. arizonae serovar 18 MNNEETFYQAMRRKGVTRRSFLKFCSLAATSLGLGAGMTPKIAWALENKPRIPVVWIHGL WP_001058306.1 hydrogenase [Salmonella enterica] Length: 372\nNumber of predicted TMHs: 2\nExp number of AAs in TMHs: 36.5540099999999\nExp number, first 60 AAs: 19.72732\nTotal prob of N-in: 0.96171\nPOSSIBLE N-term signal sequence\ninside 1 19\nTMhelix 20 39\noutside 40 325\nTMhelix 326 348\ninside 349 372

15817 GCF_001970905.1_ASM197090v1 Salmonella enterica subsp. arizonae serovar 18:z4,z23:- str. CVM N26928 Proteobacteria; Gammaproteobacteria; Enterobacterales; Enterobacteriaceae; Salmonella; Salmonella enterica;

Salmonella enterica subsp. arizonae; Salmonella enterica subsp. arizonae serovar 18
MTGDNTLITSHGINRRDFMKLCAALAATMGLSSKAAAEMAESVSNPQRPPVIWIGAQECT WP_058819740.1
hydrogenase 2 small subunit [Salmonella enterica] Length: 372\nNumber of predicted TMHs: 1\nExp number of AAs in TMHs: 24.54195\nExp number, first 60 AAs: 0.269\nTotal prob of N-in: 0.14471\noutside 1 331\nTMhelix 332 354\ninside 355 372

15818 GCF_001951465.1_ASM195146v1 Salmonella enterica subsp. arizonae serovar 18:z4,z23:- str. CVM N27
Proteobacteria; Gammaproteobacteria; Enterobacterales; Enterobacteriaceae; Salmonella; Salmonella enterica; Salmonella enterica subsp. arizonae; Salmonella enterica subsp. arizonae serovar 18
MNNEETFYQAMRRKGVTRRSFLKFCSLAATSLGLGAGMTPKIAWALENKPRIPVVWIHGL WP_001058306.1
hydrogenase [Salmonella enterica] Length: 372\nNumber of predicted TMHs: 2\nExp number of AAs in TMHs: 36.5540099999999\nExp number, first 60 AAs: 19.72732\nTotal prob of N-in: 0.96171\nPOSSIBLE N-term signal sequence\ninside 1 19\nTMhelix 20 39\noutside 40 325\nTMhelix 326 348\ninside 349 372

15819 GCF_001951465.1_ASM195146v1 Salmonella enterica subsp. arizonae serovar 18:z4,z23:- str. CVM N27
Proteobacteria; Gammaproteobacteria; Enterobacterales; Enterobacteriaceae; Salmonella; Salmonella enterica; Salmonella enterica subsp. arizonae; Salmonella enterica subsp. arizonae serovar 18
MTGDNTLITSHGINRRDFMKLCAALAATMGLSSKAAAEMAESVSNPQRPPVIWIGAQECT WP_058819740.1
hydrogenase 2 small subunit [Salmonella enterica] Length: 372\nNumber of predicted TMHs: 1\nExp number of AAs in TMHs: 24.54195\nExp number, first 60 AAs: 0.269\nTotal prob of N-in: 0.14471\noutside 1 331\nTMhelix 332 354\ninside 355 372

15820 GCF_001952125.1_ASM195212v1 Salmonella enterica subsp. arizonae serovar 18:z4,z23:- str. CVM N29354
Proteobacteria; Gammaproteobacteria; Enterobacterales; Enterobacteriaceae; Salmonella; Salmonella enterica; Salmonella enterica subsp. arizonae; Salmonella enterica subsp. arizonae serovar 18
MTGDNTLITSHGINRRDFMKLCAALAATMGLSSKAAAEMAESVSNPQRPPVIWIGAQECT WP_058819740.1
hydrogenase 2 small subunit [Salmonella enterica] Length: 372\nNumber of predicted TMHs: 1\nExp number of AAs in TMHs: 24.54195\nExp number, first 60 AAs: 0.269\nTotal prob of N-in: 0.14471\noutside 1 331\nTMhelix 332 354\ninside 355 372

15821 GCF_001952125.1_ASM195212v1 Salmonella enterica subsp. arizonae serovar 18:z4,z23:- str. CVM N29354
Proteobacteria; Gammaproteobacteria; Enterobacterales; Enterobacteriaceae; Salmonella; Salmonella enterica; Salmonella enterica subsp. arizonae; Salmonella enterica subsp. arizonae serovar 18
MNNEETFYQAMRRKGVTRRSFLKFCSLAATSLGLGAGMTPKIAWALENKPRIPVVWIHGL WP_001058306.1
hydrogenase [Salmonella enterica] Length: 372\nNumber of predicted TMHs: 2\nExp number of AAs in TMHs: 36.5540099999999\nExp number, first 60 AAs: 19.72732\nTotal prob of N-in: 0.96171\nPOSSIBLE N-term signal sequence\ninside 1 19\nTMhelix 20 39\noutside 40 325\nTMhelix 326 348\ninside 349 372

15822 GCF_001952145.1_ASM195214v1 Salmonella enterica subsp. arizonae serovar 18:z4,z23:- str. CVM N31597
Proteobacteria; Gammaproteobacteria; Enterobacterales; Enterobacteriaceae; Salmonella; Salmonella enterica; Salmonella enterica subsp. arizonae; Salmonella enterica subsp. arizonae serovar 18
MTGDNTLITSHGINRRDFMKLCAALAATMGLSSKAAAEMAESVSNPQRPPVIWIGAQECT WP_058819740.1
hydrogenase 2 small subunit [Salmonella enterica] Length: 372\nNumber of predicted TMHs: 1\nExp number of AAs in TMHs: 24.54195\nExp number, first 60 AAs: 0.269\nTotal prob of N-in: 0.14471\noutside 1 331\nTMhelix 332 354\ninside 355 372

15823 GCF_001952145.1_ASM195214v1 Salmonella enterica subsp. arizonae serovar 18:z4,z23:- str. CVM N31597
Proteobacteria; Gammaproteobacteria; Enterobacterales; Enterobacteriaceae; Salmonella; Salmonella enterica; Salmonella enterica subsp. arizonae; Salmonella enterica subsp. arizonae serovar 18
MNNEETFYQAMRRKGVTRRSFLKFCSLAATSLGLGAGMTPKIAWALENKPRIPVVWIHGL WP_001058306.1
hydrogenase [Salmonella enterica] Length: 372\nNumber of predicted TMHs: 2\nExp number of AAs in TMHs: 36.5540099999999\nExp number, first 60 AAs: 19.72732\nTotal prob of N-in: 0.96171\nPOSSIBLE N-term signal sequence\ninside 1 19\nTMhelix 20 39\noutside 40 325\nTMhelix 326 348\ninside 349 372

15824 GCF_001951515.1_ASM195151v1 Salmonella enterica subsp. arizonae serovar 18:z4,z23:- str. CVM N4410
Proteobacteria; Gammaproteobacteria; Enterobacterales; Enterobacteriaceae; Salmonella; Salmonella enterica; Salmonella enterica subsp. arizonae; Salmonella enterica subsp. arizonae serovar 18
MNNEETFYQAMRRKGVTRRSFLKFCSLAATSLGLGAGMTPKIAWALENKPRIPVVWIHGL WP_001058306.1
hydrogenase [Salmonella enterica] Length: 372\nNumber of predicted TMHs: 2\nExp number of AAs in TMHs: 36.5540099999999\nExp number, first 60 AAs: 19.72732\nTotal prob of N-in: 0.96171\nPOSSIBLE N-term signal sequence\ninside 1 19\nTMhelix 20 39\noutside 40 325\nTMhelix 326 348\ninside 349 372

15825 GCF_001951515.1_ASM195151v1 Salmonella enterica subsp. arizonae serovar 18:z4,z23:- str. CVM N4410 Proteobacteria; Gammaproteobacteria; Enterobacterales; Enterobacteriaceae; Salmonella; Salmonella enterica; Salmonella enterica subsp. arizonae; Salmonella enterica subsp. arizonae serovar 18 MTGDNTLITSHGINRRDFMKLCAALAATMGLSSKAAAEMAESVSNPQRPPVIWIGAQECT WP_058819740.1 hydrogenase 2 small subunit [Salmonella enterica] Length: 372\nNumber of predicted TMHs: 1\nExp number of AAs in TMHs: 24.54195\nExp number, first 60 AAs: 0.269\nTotal prob of N-in: 0.14471\noutside 1 331\nTMhelix 332 354\ninside 355 372

15826 GCF_001951605.1_ASM195160v1 Salmonella enterica subsp. arizonae serovar 18:z4,z23:- str. CVM N5 Proteobacteria; Gammaproteobacteria; Enterobacterales; Enterobacteriaceae; Salmonella; Salmonella enterica; Salmonella enterica subsp. arizonae; Salmonella enterica subsp. arizonae serovar 18 MTGDNTLITSHGINRRDFMKLCAALAATMGLSSKAAAEMAESVSNPQRPPVIWIGAQECT WP_058819740.1 hydrogenase 2 small subunit [Salmonella enterica] Length: 372\nNumber of predicted TMHs: 1\nExp number of AAs in TMHs: 24.54195\nExp number, first 60 AAs: 0.269\nTotal prob of N-in: 0.14471\noutside 1 331\nTMhelix 332 354\ninside 355 372

15827 GCF_001951605.1_ASM195160v1 Salmonella enterica subsp. arizonae serovar 18:z4,z23:- str. CVM N5 Proteobacteria; Gammaproteobacteria; Enterobacterales; Enterobacteriaceae; Salmonella; Salmonella enterica; Salmonella enterica subsp. arizonae; Salmonella enterica subsp. arizonae serovar 18 MNNEETFYQAMRRKGVTRRSFLKFCSLAATSLGLGAGMTPKIAWALENKPRIPVVWIHGL WP_001058306.1 hydrogenase [Salmonella enterica] Length: 372\nNumber of predicted TMHs: 2\nExp number of AAs in TMHs: 36.5540099999999\nExp number, first 60 AAs: 19.72732\nTotal prob of N-in: 0.96171\nPOSSIBLE N-term signal sequence\ninside 1 19\nTMhelix 20 39\noutside 40 325\nTMhelix 326 348\ninside 349 372

15828 GCF_001951385.1_ASM195138v1 Salmonella enterica subsp. arizonae serovar 18:z4,z23:- str. CVM N6509 Proteobacteria; Gammaproteobacteria; Enterobacterales; Enterobacteriaceae; Salmonella; Salmonella enterica; Salmonella enterica subsp. arizonae; Salmonella enterica subsp. arizonae serovar 18 MNNEETFYQAMRRKGVTRRSFLKFCSLAATSLGLGAGMTPKIAWALENKPRIPVVWIHGL WP_001058306.1 hydrogenase [Salmonella enterica] Length: 372\nNumber of predicted TMHs: 2\nExp number of AAs in TMHs: 36.5540099999999\nExp number, first 60 AAs: 19.72732\nTotal prob of N-in: 0.96171\nPOSSIBLE N-term signal sequence\ninside 1 19\nTMhelix 20 39\noutside 40 325\nTMhelix 326 348\ninside 349 372

15829 GCF_001951385.1_ASM195138v1 Salmonella enterica subsp. arizonae serovar 18:z4,z23:- str. CVM N6509 Proteobacteria; Gammaproteobacteria; Enterobacterales; Enterobacteriaceae; Salmonella; Salmonella enterica; Salmonella enterica subsp. arizonae; Salmonella enterica subsp. arizonae serovar 18 MTGDNTLITSHGINRRDFMKLCAALAATMGLSSKAAAEMAESVSNPQRPPVIWIGAQECT WP_058819740.1 hydrogenase 2 small subunit [Salmonella enterica] Length: 372\nNumber of predicted TMHs: 1\nExp number of AAs in TMHs: 24.54195\nExp number, first 60 AAs: 0.269\nTotal prob of N-in: 0.14471\noutside 1 331\nTMhelix 332 354\ninside 355 372

15830 GCF_001951595.1_ASM195159v1 Salmonella enterica subsp. arizonae serovar 18:z4,z23:- str. CVM N7307 Proteobacteria; Gammaproteobacteria; Enterobacterales; Enterobacteriaceae; Salmonella; Salmonella enterica; Salmonella enterica subsp. arizonae; Salmonella enterica subsp. arizonae serovar 18 MNNEETFYQAMRRKGVTRRSFLKFCSLAATSLGLGAGMTPKIAWALENKPRIPVVWIHGL WP_001058312.1 hydrogenase [Salmonella enterica] Length: 372\nNumber of predicted TMHs: 2\nExp number of AAs in TMHs: 36.51738\nExp number, first 60 AAs: 19.72849\nTotal prob of N-in: 0.96170\nPOSSIBLE N-term signal sequence\ninside 1 19\nTMhelix 20 39\noutside 40 325\nTMhelix 326 348\ninside 349 372

15831 GCF_001951595.1_ASM195159v1 Salmonella enterica subsp. arizonae serovar 18:z4,z23:- str. CVM N7307 Proteobacteria; Gammaproteobacteria; Enterobacterales; Enterobacteriaceae; Salmonella; Salmonella enterica; Salmonella enterica subsp. arizonae; Salmonella enterica subsp. arizonae serovar 18 MTGDNTLITSHGINRRDFMKLCAALAATMGLSSKAAAEMAESVSNPQRPPVIWIGAQECT WP_000145429.1 hydrogenase 2 small subunit [Salmonella enterica] Length: 372\nNumber of predicted TMHs: 1\nExp number of AAs in TMHs: 24.51309\nExp number, first 60 AAs: 0.2682\nTotal prob of N-in: 0.14402\noutside 1 331\nTMhelix 332 354\ninside 355 372

15832 GCF_001951615.1_ASM195161v1 Salmonella enterica subsp. arizonae serovar 18:z4,z23:- str. CVM N9135 Proteobacteria; Gammaproteobacteria; Enterobacterales; Enterobacteriaceae; Salmonella; Salmonella enterica; Salmonella enterica subsp. arizonae; Salmonella enterica subsp. arizonae serovar 18 MNNEETFYQAMRRKGVTRRSFLKFCSLAATSLGLGAGMTPKIAWALENKPRIPVVWIHGL WP_001058306.1 hydrogenase [Salmonella enterica] Length: 372\nNumber of predicted TMHs: 2\nExp number of AAs in

TMHs: 36.5540099999999\nExp number, first 60 AAs: 19.72732\nTotal prob of N-in: 0.96171\nPOSSIBLE N-term signal sequence\ninside 1 19\nTMhelix 20 39\noutside 40 325\nTMhelix 326 348\ninside 349 372

15833 GCF_001951615.1_ASM195161v1 Salmonella enterica subsp. arizonae serovar 18:z4,z23:- str. CVM N9135 Proteobacteria; Gammaproteobacteria; Enterobacterales; Enterobacteriaceae; Salmonella; Salmonella enterica; Salmonella enterica subsp. arizonae; Salmonella enterica subsp. arizonae serovar 18 MTGDNTLITSHGINRRDFMKLCAALAATMGLSSKAAAEMAESVSNPQRPPVIWIGAQECT WP_058819740.1 hydrogenase 2 small subunit [Salmonella enterica] Length: 372\nNumber of predicted TMHs: 1\nExp number of AAs in TMHs: 24.54195\nExp number, first 60 AAs: 0.269\nTotal prob of N-in: 0.14471\noutside 1 331\nTMhelix 332 354\ninside 355 372

15834 GCF_001448925.1_ASM144892v1 Salmonella enterica subsp. arizonae serovar 62:z36:- str. 5335/86 Proteobacteria; Gammaproteobacteria; Enterobacterales; Enterobacteriaceae; Salmonella; Salmonella enterica; Salmonella enterica subsp. arizonae; Salmonella enterica subsp. arizonae serovar 62 MNNEETFYQAMRRKGVTRRSFLKFCSLAATSLGLGAGMTPKIAWALENKPRIPVVWIHGL WP_001058306.1 hydrogenase [Salmonella enterica] Length: 372\nNumber of predicted TMHs: 2\nExp number of AAs in TMHs: 36.5540099999999\nExp number, first 60 AAs: 19.72732\nTotal prob of N-in: 0.96171\nPOSSIBLE N-term signal sequence\ninside 1 19\nTMhelix 20 39\noutside 40 325\nTMhelix 326 348\ninside 349 372

15835 GCF_001448925.1_ASM144892v1 Salmonella enterica subsp. arizonae serovar 62:z36:- str. 5335/86 Proteobacteria; Gammaproteobacteria; Enterobacterales; Enterobacteriaceae; Salmonella; Salmonella enterica; Salmonella enterica subsp. arizonae; Salmonella enterica subsp. arizonae serovar 62 MTGDNTLITSHGINRRDFMKLCAALAATMGLSSKAAAEMAESVSNPQRPPVIWIGAQECT WP_000145429.1 hydrogenase 2 small subunit [Salmonella enterica] Length: 372\nNumber of predicted TMHs: 1\nExp number of AAs in TMHs: 24.51309\nExp number, first 60 AAs: 0.2682\nTotal prob of N-in: 0.14402\noutside 1 331\nTMhelix 332 354\ninside 355 372

15836 GCF_000756465.1_ASM75646v1 Salmonella enterica subsp. arizonae serovar 62:z36:- str. RKS2983 Proteobacteria; Gammaproteobacteria; Enterobacterales; Enterobacteriaceae; Salmonella; Salmonella enterica; Salmonella enterica subsp. arizonae; Salmonella enterica subsp. arizonae serovar 62 MTGDNTLITSHGINRRDFMKLCAALAATMGLSSKAAAEMAESVSNPQRPPVIWIGAQECT WP_000145429.1 hydrogenase 2 small subunit [Salmonella enterica] Length: 372\nNumber of predicted TMHs: 1\nExp number of AAs in TMHs: 24.51309\nExp number, first 60 AAs: 0.2682\nTotal prob of N-in: 0.14402\noutside 1 331\nTMhelix 332 354\ninside 355 372

15837 GCF_000756465.1_ASM75646v1 Salmonella enterica subsp. arizonae serovar 62:z36:- str. RKS2983 Proteobacteria; Gammaproteobacteria; Enterobacterales; Enterobacteriaceae; Salmonella; Salmonella enterica; Salmonella enterica subsp. arizonae; Salmonella enterica subsp. arizonae serovar 62 MNNEETFYQAMRRKGVTRRSFLKFCSLAATSLGLGAGMTPKIAWALENKPRIPVVWIHGL WP_001058306.1 hydrogenase [Salmonella enterica] Length: 372\nNumber of predicted TMHs: 2\nExp number of AAs in TMHs: 36.5540099999999\nExp number, first 60 AAs: 19.72732\nTotal prob of N-in: 0.96171\nPOSSIBLE N-term signal sequence\ninside 1 19\nTMhelix 20 39\noutside 40 325\nTMhelix 326 348\ninside 349 372

15838 GCF_001448945.1_ASM144894v1 Salmonella enterica subsp. arizonae serovar 63:g,z51:- str. So 20/20 Proteobacteria; Gammaproteobacteria; Enterobacterales; Enterobacteriaceae; Salmonella; Salmonella enterica; Salmonella enterica subsp. arizonae; Salmonella enterica subsp. arizonae serovar 63 MTGDNTLITSHGINRRDFMKLCAALAATMGLSSKAAAEMAESVSNPQRPPVIWIGAQECT WP_000145429.1 hydrogenase 2 small subunit [Salmonella enterica] Length: 372\nNumber of predicted TMHs: 1\nExp number of AAs in TMHs: 24.51309\nExp number, first 60 AAs: 0.2682\nTotal prob of N-in: 0.14402\noutside 1 331\nTMhelix 332 354\ninside 355 372

15839 GCF_001448945.1_ASM144894v1 Salmonella enterica subsp. arizonae serovar 63:g,z51:- str. So 20/20 Proteobacteria; Gammaproteobacteria; Enterobacterales; Enterobacteriaceae; Salmonella; Salmonella enterica; Salmonella enterica subsp. arizonae; Salmonella enterica subsp. arizonae serovar 63 MNNEETFYQAMRRKGVTRRSFLKFCSLAATSLGLGAGMTPKIAWALENKPRIPVVWIHGL WP_001058306.1 hydrogenase [Salmonella enterica] Length: 372\nNumber of predicted TMHs: 2\nExp number of AAs in TMHs: 36.5540099999999\nExp number, first 60 AAs: 19.72732\nTotal prob of N-in: 0.96171\nPOSSIBLE N-term signal sequence\ninside 1 19\nTMhelix 20 39\noutside 40 325\nTMhelix 326 348\ninside 349 372

15840 GCF_001832645.1_ASM183264v1 Salmonella enterica subsp. diarizonae serovar 16:z10:e,n,x,z15 Proteobacteria; Gammaproteobacteria; Enterobacterales; Enterobacteriaceae; Salmonella; Salmonella enterica;

Salmonella enterica subsp. diarizonae MTGDNTLITSHGINRRDFMKLCAALAATMGLSSKAAAEMAESVSNPQRPPVIWIGAQECT WP_023248548.1 hydrogenase 2 small subunit [Salmonella enterica] Length: 372\nNumber of predicted TMHs: 1\nExp number of AAs in TMHs: 24.51281\nExp number, first 60 AAs: 0.26822\nTotal prob of N-in: 0.14402\noutside 1 331\nTMhelix 332 354\ninside 355 372

15841 GCF_001832645.1_ASM183264v1 Salmonella enterica subsp. diarizonae serovar 16:z10:e,n,x,z15 Proteobacteria; Gammaproteobacteria; Enterobacterales; Enterobacteriaceae; Salmonella; Salmonella enterica; Salmonella enterica subsp. diarizonae MNNEETFYQAMRRKGVTRRSFLKFCSLAATSLGLGAGMTPKIAWALENKPRIPVVWIHGL WP_070801304.1 hydrogenase [Salmonella enterica] Length: 372\nNumber of predicted TMHs: 2\nExp number of AAs in TMHs: 36.3895799999999\nExp number, first 60 AAs: 19.727\nTotal prob of N-in: 0.96170\nPOSSIBLE N-term signal sequence\ninside 1 19\nTMhelix 20 39\noutside 40 325\nTMhelix 326 348\ninside 349 372

15842 GCF_001832605.1_ASM183260v1 Salmonella enterica subsp. diarizonae serovar 16:z10:e,n,x,z15 Proteobacteria; Gammaproteobacteria; Enterobacterales; Enterobacteriaceae; Salmonella; Salmonella enterica; Salmonella enterica subsp. diarizonae MTGDNTLITSHGINRRDFMKLCAALAATMGLSSKAAAEMAESVSNPQRPPVIWIGAQECT WP_023248548.1 hydrogenase 2 small subunit [Salmonella enterica] Length: 372\nNumber of predicted TMHs: 1\nExp number of AAs in TMHs: 24.51281\nExp number, first 60 AAs: 0.26822\nTotal prob of N-in: 0.14402\noutside 1 331\nTMhelix 332 354\ninside 355 372

15843 GCF_001832605.1_ASM183260v1 Salmonella enterica subsp. diarizonae serovar 16:z10:e,n,x,z15 Proteobacteria; Gammaproteobacteria; Enterobacterales; Enterobacteriaceae; Salmonella; Salmonella enterica; Salmonella enterica subsp. diarizonae MNNEETFYQAMRRKGVTRRSFLKFCSLAATSLGLGAGMTPKIAWALENKPRIPVVWIHGL WP_070801304.1 hydrogenase [Salmonella enterica] Length: 372\nNumber of predicted TMHs: 2\nExp number of AAs in TMHs: 36.3895799999999\nExp number, first 60 AAs: 19.727\nTotal prob of N-in: 0.96170\nPOSSIBLE N-term signal sequence\ninside 1 19\nTMhelix 20 39\noutside 40 325\nTMhelix 326 348\ninside 349 372

15844 GCF_001832665.1_ASM183266v1 Salmonella enterica subsp. diarizonae serovar 38:[k]:- Proteobacteria; Gammaproteobacteria; Enterobacterales; Enterobacteriaceae; Salmonella; Salmonella enterica; Salmonella enterica subsp. diarizonae MNNEETFYQAMRRKGVTRRSFLKFCSLAATSLGLGAGMTPKIAWALENKPRIPVVWIHGL WP_001058312.1 hydrogenase [Salmonella enterica] Length: 372\nNumber of predicted TMHs: 2\nExp number of AAs in TMHs: 36.51738\nExp number, first 60 AAs: 19.72849\nTotal prob of N-in: 0.96170\nPOSSIBLE N-term signal sequence\ninside 1 19\nTMhelix 20 39\noutside 40 325\nTMhelix 326 348\ninside 349 372

15845 GCF_001832245.1_ASM183224v1 Salmonella enterica subsp. diarizonae serovar 38:[k]:- Proteobacteria; Gammaproteobacteria; Enterobacterales; Enterobacteriaceae; Salmonella; Salmonella enterica; Salmonella enterica subsp. diarizonae MTGDNTLITSHGINRRDFMKLCAALAATMGLSSKAAAEMAESVSNPQRPPVIWIGAQECT WP_000145429.1 hydrogenase 2 small subunit [Salmonella enterica] Length: 372\nNumber of predicted TMHs: 1\nExp number of AAs in TMHs: 24.51309\nExp number, first 60 AAs: 0.2682\nTotal prob of N-in: 0.14402\noutside 1 331\nTMhelix 332 354\ninside 355 372

15846 GCF_001832665.1_ASM183266v1 Salmonella enterica subsp. diarizonae serovar 38:[k]:- Proteobacteria; Gammaproteobacteria; Enterobacterales; Enterobacteriaceae; Salmonella; Salmonella enterica; Salmonella enterica subsp. diarizonae MTGDNTLITSHGINRRDFMKLCAALAATMGLSSKAAAEMAESVSNPQRPPVIWIGAQECT WP_000145429.1 hydrogenase 2 small subunit [Salmonella enterica] Length: 372\nNumber of predicted TMHs: 1\nExp number of AAs in TMHs: 24.51309\nExp number, first 60 AAs: 0.2682\nTotal prob of N-in: 0.14402\noutside 1 331\nTMhelix 332 354\ninside 355 372

15847 GCF_001832285.1_ASM183228v1 Salmonella enterica subsp. diarizonae serovar 38:[k]:- Proteobacteria; Gammaproteobacteria; Enterobacterales; Enterobacteriaceae; Salmonella; Salmonella enterica; Salmonella enterica subsp. diarizonae MTGDNTLITSHGINRRDFMKLCAALAATMGLSSKAAAEMAESVSNPQRPPVIWIGAQECT WP_000145429.1 hydrogenase 2 small subunit [Salmonella enterica] Length: 372\nNumber of predicted TMHs: 1\nExp number of AAs in TMHs: 24.51309\nExp number, first 60 AAs: 0.2682\nTotal prob of N-in: 0.14402\noutside 1 331\nTMhelix 332 354\ninside 355 372

15848 GCF_001832525.1_ASM183252v1 Salmonella enterica subsp. diarizonae serovar 38:[k]:- Proteobacteria; Gammaproteobacteria; Enterobacterales; Enterobacteriaceae; Salmonella; Salmonella enterica; Salmonella enterica subsp. diarizonae MNNEETFYQAMRRKGVTRRSFLKFCSLAATSLGLGAGMTPKIAWALENKPRIPVVWIHGL WP_001058312.1 hydrogenase [Salmonella enterica] Length: 372\nNumber of predicted TMHs: 2\nExp number of AAs in TMHs: 36.51738\nExp number, first 60 AAs: 19.72849\nTotal prob of N-in: 0.96170\nPOSSIBLE N-term signal sequence\ninside 1 19\nTMhelix 20 39\noutside 40 325\nTMhelix 326 348\ninside 349 372

15849 GCF_001832305.1_ASM183230v1 Salmonella enterica subsp. diarizonae serovar 38:[k]:- Proteobacteria; Gammaproteobacteria; Enterobacterales; Enterobacteriaceae; Salmonella; Salmonella enterica; Salmonella enterica subsp. diarizonae MTGDNTLITSHGINRRDFMKLCAALAATMGLSSKAAAEMAESVSNPQRPPVIWIGAQECT WP_000145429.1 hydrogenase 2 small subunit [Salmonella enterica] Length: 372\nNumber of predicted TMHs: 1\nExp number of AAs in TMHs: 24.51309\nExp number, first 60 AAs: 0.2682\nTotal prob of N-in: 0.14402\noutside 1 331\nTMhelix 332 354\ninside 355 372

15850 GCF_001832225.1_ASM183222v1 Salmonella enterica subsp. diarizonae serovar 38:[k]:- Proteobacteria; Gammaproteobacteria; Enterobacterales; Enterobacteriaceae; Salmonella; Salmonella enterica; Salmonella enterica subsp. diarizonae MNNEETFYQAMRRKGVTRRSFLKFCSLAATSLGLGAGMTPKIAWALENKPRIPVVWIHGL WP_001058312.1 hydrogenase [Salmonella enterica] Length: 372\nNumber of predicted TMHs: 2\nExp number of AAs in TMHs: 36.51738\nExp number, first 60 AAs: 19.72849\nTotal prob of N-in: 0.96170\nPOSSIBLE N-term signal sequence\ninside 1 19\nTMhelix 20 39\noutside 40 325\nTMhelix 326 348\ninside 349 372

15851 GCF_001832305.1_ASM183230v1 Salmonella enterica subsp. diarizonae serovar 38:[k]:- Proteobacteria; Gammaproteobacteria; Enterobacterales; Enterobacteriaceae; Salmonella; Salmonella enterica; Salmonella enterica subsp. diarizonae MNNEETFYQAMRRKGVTRRSFLKFCSLAATSLGLGAGMTPKIAWALENKPRIPVVWIHGL WP_001058312.1 hydrogenase [Salmonella enterica] Length: 372\nNumber of predicted TMHs: 2\nExp number of AAs in TMHs: 36.51738\nExp number, first 60 AAs: 19.72849\nTotal prob of N-in: 0.96170\nPOSSIBLE N-term signal sequence\ninside 1 19\nTMhelix 20 39\noutside 40 325\nTMhelix 326 348\ninside 349 372

15852 GCF_001832445.1_ASM183244v1 Salmonella enterica subsp. diarizonae serovar 38:[k]:- Proteobacteria; Gammaproteobacteria; Enterobacterales; Enterobacteriaceae; Salmonella; Salmonella enterica; Salmonella enterica subsp. diarizonae MTGDNTLITSHGINRRDFMKLCAALAATMGLSSKAAAEMAESVSNPQRPPVIWIGAQECT WP_000145429.1 hydrogenase 2 small subunit [Salmonella enterica] Length: 372\nNumber of predicted TMHs: 1\nExp number of AAs in TMHs: 24.51309\nExp number, first 60 AAs: 0.2682\nTotal prob of N-in: 0.14402\noutside 1 331\nTMhelix 332 354\ninside 355 372

15853 GCF_001832525.1_ASM183252v1 Salmonella enterica subsp. diarizonae serovar 38:[k]:- Proteobacteria; Gammaproteobacteria; Enterobacterales; Enterobacteriaceae; Salmonella; Salmonella enterica; Salmonella enterica subsp. diarizonae MTGDNTLITSHGINRRDFMKLCAALAATMGLSSKAAAEMAESVSNPQRPPVIWIGAQECT WP_000145429.1 hydrogenase 2 small subunit [Salmonella enterica] Length: 372\nNumber of predicted TMHs: 1\nExp number of AAs in TMHs: 24.51309\nExp number, first 60 AAs: 0.2682\nTotal prob of N-in: 0.14402\noutside 1 331\nTMhelix 332 354\ninside 355 372

15854 GCF_001832465.1_ASM183246v1 Salmonella enterica subsp. diarizonae serovar 38:[k]:- Proteobacteria; Gammaproteobacteria; Enterobacterales; Enterobacteriaceae; Salmonella; Salmonella enterica; Salmonella enterica subsp. diarizonae MNNEETFYQAMRRKGVTRRSFLKFCSLAATSLGLGAGMTPKIAWALENKPRIPVVWIHGL WP_001058312.1 hydrogenase [Salmonella enterica] Length: 372\nNumber of predicted TMHs: 2\nExp number of AAs in TMHs: 36.51738\nExp number, first 60 AAs: 19.72849\nTotal prob of N-in: 0.96170\nPOSSIBLE N-term signal sequence\ninside 1 19\nTMhelix 20 39\noutside 40 325\nTMhelix 326 348\ninside 349 372

15855 GCF_001832465.1_ASM183246v1 Salmonella enterica subsp. diarizonae serovar 38:[k]:- Proteobacteria; Gammaproteobacteria; Enterobacterales; Enterobacteriaceae; Salmonella; Salmonella enterica; Salmonella enterica subsp. diarizonae MTGDNTLITSHGINRRDFMKLCAALAATMGLSSKAAAEMAESVSNPQRPPVIWIGAQECT WP_000145429.1 hydrogenase 2 small subunit [Salmonella enterica] Length: 372\nNumber of predicted TMHs: 1\nExp number of AAs in TMHs: 24.51309\nExp number, first 60 AAs: 0.2682\nTotal prob of N-in: 0.14402\noutside 1 331\nTMhelix 332 354\ninside 355 372

15856 GCF_001832805.1_ASM183280v1 Salmonella enterica subsp. diarizonae serovar 38:[k]:- Proteobacteria; Gammaproteobacteria; Enterobacterales; Enterobacteriaceae; Salmonella; Salmonella enterica; Salmonella enterica subsp. diarizonae MTGDNTLITSHGINRRDFMKLCAALAATMGLSSKAAAEMAESVSNPQRPPVIWIGAQECT WP_000145429.1 hydrogenase 2 small subunit [Salmonella enterica] Length: 372\nNumber of predicted TMHs: 1\nExp number of AAs in TMHs: 24.51309\nExp number, first 60 AAs: 0.2682\nTotal prob of N-in: 0.14402\noutside 1 331\nTMhelix 332 354\ninside 355 372

15857 GCF_001832845.1_ASM183284v1 Salmonella enterica subsp. diarizonae serovar 38:[k]:- Proteobacteria; Gammaproteobacteria; Enterobacterales; Enterobacteriaceae; Salmonella; Salmonella enterica; Salmonella enterica subsp. diarizonae MTGDNTLITSHGINRRDFMKLCAALAATMGLSSKAAAEMAESVSNPQRPPVIWIGAQECT WP_000145429.1 hydrogenase 2 small subunit [Salmonella enterica] Length: 372\nNumber of

predicted TMHs: 1\nExp number of AAs in TMHs: 24.51309\nExp number, first 60 AAs: 0.2682\nTotal prob of N-in: 0.14402\noutside 1 331\nTMhelix 332 354\ninside 355 372

15858 GCF_001832245.1_ASM183224v1 Salmonella enterica subsp. diarizonae serovar 38:[k]:- Proteobacteria; Gammaproteobacteria; Enterobacterales; Enterobacteriaceae; Salmonella; Salmonella enterica; Salmonella enterica subsp. diarizonae MNNEETFYQAMRRKGVTRRSFLKFCSLAATSLGLGAGMTPKIAWALENKPRIPVVWIHGL WP_001058312.1 hydrogenase [Salmonella enterica] Length: 372\nNumber of predicted TMHs: 2\nExp number of AAs in TMHs: 36.51738\nExp number, first 60 AAs: 19.72849\nTotal prob of N-in: 0.96170\nPOSSIBLE N-term signal sequence\ninside 1 19\nTMhelix 20 39\noutside 40 325\nTMhelix 326 348\ninside 349 372

15859 GCF_001832845.1_ASM183284v1 Salmonella enterica subsp. diarizonae serovar 38:[k]:- Proteobacteria; Gammaproteobacteria; Enterobacterales; Enterobacteriaceae; Salmonella; Salmonella enterica; Salmonella enterica subsp. diarizonae MNNEETFYQAMRRKGVTRRSFLKFCSLAATSLGLGAGMTPKIAWALENKPRIPVVWIHGL WP_001058312.1 hydrogenase [Salmonella enterica] Length: 372\nNumber of predicted TMHs: 2\nExp number of AAs in TMHs: 36.51738\nExp number, first 60 AAs: 19.72849\nTotal prob of N-in: 0.96170\nPOSSIBLE N-term signal sequence\ninside 1 19\nTMhelix 20 39\noutside 40 325\nTMhelix 326 348\ninside 349 372

15860 GCF_001832805.1_ASM183280v1 Salmonella enterica subsp. diarizonae serovar 38:[k]:- Proteobacteria; Gammaproteobacteria; Enterobacterales; Enterobacteriaceae; Salmonella; Salmonella enterica; Salmonella enterica subsp. diarizonae MNNEETFYQAMRRKGVTRRSFLKFCSLAATSLGLGAGMTPKIAWALENKPRIPVVWIHGL WP_001058312.1 hydrogenase [Salmonella enterica] Length: 372\nNumber of predicted TMHs: 2\nExp number of AAs in TMHs: 36.51738\nExp number, first 60 AAs: 19.72849\nTotal prob of N-in: 0.96170\nPOSSIBLE N-term signal sequence\ninside 1 19\nTMhelix 20 39\noutside 40 325\nTMhelix 326 348\ninside 349 372

15861 GCF_001832585.1_ASM183258v1 Salmonella enterica subsp. diarizonae serovar 38:[k]:- Proteobacteria; Gammaproteobacteria; Enterobacterales; Enterobacteriaceae; Salmonella; Salmonella enterica; Salmonella enterica subsp. diarizonae MTGDNTLITSHGINRRDFMKLCAALAATMGLSSKAAAEMAESVSNPQRPPVIWIGAQECT WP_000145429.1 hydrogenase 2 small subunit [Salmonella enterica] Length: 372\nNumber of predicted TMHs: 1\nExp number of AAs in TMHs: 24.51309\nExp number, first 60 AAs: 0.2682\nTotal prob of N-in: 0.14402\noutside 1 331\nTMhelix 332 354\ninside 355 372

15862 GCF_001832285.1_ASM183228v1 Salmonella enterica subsp. diarizonae serovar 38:[k]:- Proteobacteria; Gammaproteobacteria; Enterobacterales; Enterobacteriaceae; Salmonella; Salmonella enterica; Salmonella enterica subsp. diarizonae MNNEETFYQAMRRKGVTRRSFLKFCSLAATSLGLGAGMTPKIAWALENKPRIPVVWIHGL WP_001058312.1 hydrogenase [Salmonella enterica] Length: 372\nNumber of predicted TMHs: 2\nExp number of AAs in TMHs: 36.51738\nExp number, first 60 AAs: 19.72849\nTotal prob of N-in: 0.96170\nPOSSIBLE N-term signal sequence\ninside 1 19\nTMhelix 20 39\noutside 40 325\nTMhelix 326 348\ninside 349 372

15863 GCF_001832585.1_ASM183258v1 Salmonella enterica subsp. diarizonae serovar 38:[k]:- Proteobacteria; Gammaproteobacteria; Enterobacterales; Enterobacteriaceae; Salmonella; Salmonella enterica; Salmonella enterica subsp. diarizonae MNNEETFYQAMRRKGVTRRSFLKFCSLAATSLGLGAGMTPKIAWALENKPRIPVVWIHGL WP_001058312.1 hydrogenase [Salmonella enterica] Length: 372\nNumber of predicted TMHs: 2\nExp number of AAs in TMHs: 36.51738\nExp number, first 60 AAs: 19.72849\nTotal prob of N-in: 0.96170\nPOSSIBLE N-term signal sequence\ninside 1 19\nTMhelix 20 39\noutside 40 325\nTMhelix 326 348\ninside 349 372

15864 GCF_001832225.1_ASM183222v1 Salmonella enterica subsp. diarizonae serovar 38:[k]:- Proteobacteria; Gammaproteobacteria; Enterobacterales; Enterobacteriaceae; Salmonella; Salmonella enterica; Salmonella enterica subsp. diarizonae MTGDNTLITSHGINRRDFMKLCAALAATMGLSSKAAAEMAESVSNPQRPPVIWIGAQECT WP_000145429.1 hydrogenase 2 small subunit [Salmonella enterica] Length: 372\nNumber of predicted TMHs: 1\nExp number of AAs in TMHs: 24.51309\nExp number, first 60 AAs: 0.2682\nTotal prob of N-in: 0.14402\noutside 1 331\nTMhelix 332 354\ninside 355 372

15865 GCF_001832825.1_ASM183282v1 Salmonella enterica subsp. diarizonae serovar 38:[k]:- Proteobacteria; Gammaproteobacteria; Enterobacterales; Enterobacteriaceae; Salmonella; Salmonella enterica; Salmonella enterica subsp. diarizonae MTGDNTLITSHGINRRDFMKLCAALAATMGLSSKAAAEMAESVSNPQRPPVIWIGAQECT WP_000145429.1 hydrogenase 2 small subunit [Salmonella enterica] Length: 372\nNumber of predicted TMHs: 1\nExp number of AAs in TMHs: 24.51309\nExp number, first 60 AAs: 0.2682\nTotal prob of N-in: 0.14402\noutside 1 331\nTMhelix 332 354\ninside 355 372

15866 GCF_001832825.1_ASM183282v1 Salmonella enterica subsp. diarizonae serovar 38:[k]:- Proteobacteria; Gammaproteobacteria; Enterobacterales; Enterobacteriaceae; Salmonella; Salmonella enterica;

Salmonella enterica subsp. diarizonae	MNNEETFYQAMRRKGVTRRSFLKFCSLAATSLGLGAGMTPKIAWALENKPRIPVVWIHGL	WP_001058312.1	hydrogenase [Salmonella enterica]	Length: 372\nNumber of predicted TMHs: 2\nExp number of AAs in TMHs: 36.51738\nExp number, first 60 AAs: 19.72849\nTotal prob of N-in: 0.96170\nPOSSIBLE N-term signal sequence\ninside 1 19\nTMhelix 20 39\noutside 40 325\nTMhelix 326 348\ninside 349 372

15867	GCF_001832445.1_ASM183244v1	Salmonella enterica subsp. diarizonae serovar 38:[k]:-	Proteobacteria; Gammaproteobacteria; Enterobacterales; Enterobacteriaceae; Salmonella; Salmonella enterica; Salmonella enterica subsp. diarizonae	MNNEETFYQAMRRKGVTRRSFLKFCSLAATSLGLGAGMTPKIAWALENKPRIPVVWIHGL	WP_001058312.1	hydrogenase [Salmonella enterica]	Length: 372\nNumber of predicted TMHs: 2\nExp number of AAs in TMHs: 36.51738\nExp number, first 60 AAs: 19.72849\nTotal prob of N-in: 0.96170\nPOSSIBLE N-term signal sequence\ninside 1 19\nTMhelix 20 39\noutside 40 325\nTMhelix 326 348\ninside 349 372

15868	GCF_002035995.1_ASM203599v1	Salmonella enterica subsp. diarizonae serovar 38:z10:z53	Proteobacteria; Gammaproteobacteria; Enterobacterales; Enterobacteriaceae; Salmonella; Salmonella enterica; Salmonella enterica subsp. diarizonae	MTGDNTLITSHGINRRDFMKLCAALAATMGLSSKAAAEMAESVSNPQRPPVIWIGAQECT	WP_023248548.1	hydrogenase 2 small subunit [Salmonella enterica]	Length: 372\nNumber of predicted TMHs: 1\nExp number of AAs in TMHs: 24.51281\nExp number, first 60 AAs: 0.26822\nTotal prob of N-in: 0.14402\noutside 1 331\nTMhelix 332 354\ninside 355 372

15869	GCF_002035995.1_ASM203599v1	Salmonella enterica subsp. diarizonae serovar 38:z10:z53	Proteobacteria; Gammaproteobacteria; Enterobacterales; Enterobacteriaceae; Salmonella; Salmonella enterica; Salmonella enterica subsp. diarizonae	MNNEETFYQAMRRKGVTRRSFLKFCSLAATSLGLGAGMTPKIAWALENKPRIPVVWIHGL	WP_001058306.1	hydrogenase [Salmonella enterica]	Length: 372\nNumber of predicted TMHs: 2\nExp number of AAs in TMHs: 36.5540099999999\nExp number, first 60 AAs: 19.72732\nTotal prob of N-in: 0.96171\nPOSSIBLE N-term signal sequence\ninside 1 19\nTMhelix 20 39\noutside 40 325\nTMhelix 326 348\ninside 349 372

15870	GCF_002035685.1_ASM203568v1	Salmonella enterica subsp. diarizonae serovar 48:z10:e,n,x,z15	Proteobacteria; Gammaproteobacteria; Enterobacterales; Enterobacteriaceae; Salmonella; Salmonella enterica; Salmonella enterica subsp. diarizonae	MTGDNTLITSHGINRRDFMKLCAALAATMGLSSKAAAEMAESVSNPQRPPVIWIGAQECT	WP_079813156.1	hydrogenase 2 small subunit [Salmonella enterica]	Length: 372\nNumber of predicted TMHs: 1\nExp number of AAs in TMHs: 24.51604\nExp number, first 60 AAs: 0.26815\nTotal prob of N-in: 0.14420\noutside 1 331\nTMhelix 332 354\ninside 355 372

15871	GCF_002035685.1_ASM203568v1	Salmonella enterica subsp. diarizonae serovar 48:z10:e,n,x,z15	Proteobacteria; Gammaproteobacteria; Enterobacterales; Enterobacteriaceae; Salmonella; Salmonella enterica; Salmonella enterica subsp. diarizonae	MNNEETFYQAMRRKGVTRRSFLKFCSLAATSLGLGAGMTPKIAWALENKPRIPVVWIHGL	WP_079813856.1	[Ni/Fe] hydrogenase small subunit [Salmonella enterica]	Length: 372\nNumber of predicted TMHs: 2\nExp number of AAs in TMHs: 36.5946199999999\nExp number, first 60 AAs: 19.72725\nTotal prob of N-in: 0.96171\nPOSSIBLE N-term signal sequence\ninside 1 19\nTMhelix 20 39\noutside 40 325\nTMhelix 326 348\ninside 349 372

15872	GCF_001832725.1_ASM183272v1	Salmonella enterica subsp. diarizonae serovar 59:[k]:z35	Proteobacteria; Gammaproteobacteria; Enterobacterales; Enterobacteriaceae; Salmonella; Salmonella enterica; Salmonella enterica subsp. diarizonae	MNNEETFYQAMRRKGVTRRSFLKFCSLAATSLGLGAGMTPKIAWALENKPRIPVVWIHGL	WP_070803763.1	hydrogenase [Salmonella enterica]	Length: 372\nNumber of predicted TMHs: 2\nExp number of AAs in TMHs: 36.49463\nExp number, first 60 AAs: 19.72828\nTotal prob of N-in: 0.96168\nPOSSIBLE N-term signal sequence\ninside 1 19\nTMhelix 20 39\noutside 40 325\nTMhelix 326 348\ninside 349 372

15873	GCF_001832725.1_ASM183272v1	Salmonella enterica subsp. diarizonae serovar 59:[k]:z35	Proteobacteria; Gammaproteobacteria; Enterobacterales; Enterobacteriaceae; Salmonella; Salmonella enterica; Salmonella enterica subsp. diarizonae	MTGDNTLITSHGINRRDFMKLCAALAATMGLSSKAAAEMAESVSNPQRPPVIWIGAQECT	WP_023248548.1	hydrogenase 2 small subunit [Salmonella enterica]	Length: 372\nNumber of predicted TMHs: 1\nExp number of AAs in TMHs: 24.51281\nExp number, first 60 AAs: 0.26822\nTotal prob of N-in: 0.14402\noutside 1 331\nTMhelix 332 354\ninside 355 372

15874	GCF_001832505.1_ASM183250v1	Salmonella enterica subsp. diarizonae serovar 60:r:e,n,x,z15	Proteobacteria; Gammaproteobacteria; Enterobacterales; Enterobacteriaceae; Salmonella; Salmonella enterica; Salmonella enterica subsp. diarizonae	MTGDNTLITSHGINRRDFMKLCAALAATMGLSSKAAAEMAESVSNPQRPPVIWIGAQECT	WP_023248548.1	hydrogenase 2 small subunit [Salmonella enterica]	Length: 372\nNumber of

predicted TMHs: 1\nExp number of AAs in TMHs: 24.51281\nExp number, first 60 AAs: 0.26822\nTotal prob of N-in: 0.14402\noutside 1 331\nTMhelix 332 354\ninside 355 372

15875 GCF_001832505.1_ASM183250v1 Salmonella enterica subsp. diarizonae serovar 60:r:e,n,x,z15 Proteobacteria; Gammaproteobacteria; Enterobacterales; Enterobacteriaceae; Salmonella; Salmonella enterica; Salmonella enterica subsp. diarizonae MNNEETFYQAMRRKGVTRRSFLKFCSLAATSLGLGAGMTPKIAWALENKPRIPVVWIHGL WP_001058306.1 hydrogenase [Salmonella enterica] Length: 372\nNumber of predicted TMHs: 2\nExp number of AAs in TMHs: 36.5540099999999\nExp number, first 60 AAs: 19.72732\nTotal prob of N-in: 0.96171\nPOSSIBLE N-term signal sequence\ninside 1 19\nTMhelix 20 39\noutside 40 325\nTMhelix 326 348\ninside 349 372

15876 GCF_001832775.1_ASM183277v1 Salmonella enterica subsp. diarizonae serovar 60:r:e,n,x,z15 Proteobacteria; Gammaproteobacteria; Enterobacterales; Enterobacteriaceae; Salmonella; Salmonella enterica; Salmonella enterica subsp. diarizonae MNNEETFYQAMRRKGVTRRSFLKFCSLAATSLGLGAGMTPKIAWALENKPRIPVVWIHGL WP_001058306.1 hydrogenase [Salmonella enterica] Length: 372\nNumber of predicted TMHs: 2\nExp number of AAs in TMHs: 36.5540099999999\nExp number, first 60 AAs: 19.72732\nTotal prob of N-in: 0.96171\nPOSSIBLE N-term signal sequence\ninside 1 19\nTMhelix 20 39\noutside 40 325\nTMhelix 326 348\ninside 349 372

15877 GCF_002035545.1_ASM203554v1 Salmonella enterica subsp. diarizonae serovar 60:r:e,n,x,z15 Proteobacteria; Gammaproteobacteria; Enterobacterales; Enterobacteriaceae; Salmonella; Salmonella enterica; Salmonella enterica subsp. diarizonae MTGDNTLITSHGINRRDFMKLCAALAATMGLSSKAAAEMAESVSNPQRPPVIWIGAQECT WP_023248548.1 hydrogenase 2 small subunit [Salmonella enterica] Length: 372\nNumber of predicted TMHs: 1\nExp number of AAs in TMHs: 24.51281\nExp number, first 60 AAs: 0.26822\nTotal prob of N-in: 0.14402\noutside 1 331\nTMhelix 332 354\ninside 355 372

15878 GCF_001832255.1_ASM183225v1 Salmonella enterica subsp. diarizonae serovar 60:r:e,n,x,z15 Proteobacteria; Gammaproteobacteria; Enterobacterales; Enterobacteriaceae; Salmonella; Salmonella enterica; Salmonella enterica subsp. diarizonae MTGDNTLITSHGINRRDFMKLCAALAATMGLSSKAAAEMAESVSNPQRPPVIWIGAQECT WP_070796878.1 hydrogenase 2 small subunit [Salmonella enterica] Length: 372\nNumber of predicted TMHs: 1\nExp number of AAs in TMHs: 24.5133\nExp number, first 60 AAs: 0.26822\nTotal prob of N-in: 0.14404\noutside 1 331\nTMhelix 332 354\ninside 355 372

15879 GCF_001832565.1_ASM183256v1 Salmonella enterica subsp. diarizonae serovar 60:r:e,n,x,z15 Proteobacteria; Gammaproteobacteria; Enterobacterales; Enterobacteriaceae; Salmonella; Salmonella enterica; Salmonella enterica subsp. diarizonae MTGDNTLITSHGINRRDFMKLCAALAATMGLSSKAAAEMAESVSNPQRPPVIWIGAQECT WP_023248548.1 hydrogenase 2 small subunit [Salmonella enterica] Length: 372\nNumber of predicted TMHs: 1\nExp number of AAs in TMHs: 24.51281\nExp number, first 60 AAs: 0.26822\nTotal prob of N-in: 0.14402\noutside 1 331\nTMhelix 332 354\ninside 355 372

15880 GCF_001832765.1_ASM183276v1 Salmonella enterica subsp. diarizonae serovar 60:r:e,n,x,z15 Proteobacteria; Gammaproteobacteria; Enterobacterales; Enterobacteriaceae; Salmonella; Salmonella enterica; Salmonella enterica subsp. diarizonae MNNEETFYQAMRRKGVTRRSFLKFCSLAATSLGLGAGMTPKIAWALENKPRIPVVWIHGL WP_001058306.1 hydrogenase [Salmonella enterica] Length: 372\nNumber of predicted TMHs: 2\nExp number of AAs in TMHs: 36.5540099999999\nExp number, first 60 AAs: 19.72732\nTotal prob of N-in: 0.96171\nPOSSIBLE N-term signal sequence\ninside 1 19\nTMhelix 20 39\noutside 40 325\nTMhelix 326 348\ninside 349 372

15881 GCF_001832385.1_ASM183238v1 Salmonella enterica subsp. diarizonae serovar 60:r:e,n,x,z15 Proteobacteria; Gammaproteobacteria; Enterobacterales; Enterobacteriaceae; Salmonella; Salmonella enterica; Salmonella enterica subsp. diarizonae MTGDNTLITSHGINRRDFMKLCAALAATMGLSSKAAAEMAESVSNPQRPPVIWIGAQECT WP_023248548.1 hydrogenase 2 small subunit [Salmonella enterica] Length: 372\nNumber of predicted TMHs: 1\nExp number of AAs in TMHs: 24.51281\nExp number, first 60 AAs: 0.26822\nTotal prob of N-in: 0.14402\noutside 1 331\nTMhelix 332 354\ninside 355 372

15882 GCF_001832385.1_ASM183238v1 Salmonella enterica subsp. diarizonae serovar 60:r:e,n,x,z15 Proteobacteria; Gammaproteobacteria; Enterobacterales; Enterobacteriaceae; Salmonella; Salmonella enterica; Salmonella enterica subsp. diarizonae MNNEETFYQAMRRKGVTRRSFLKFCSLAATSLGLGAGMTPKIAWALENKPRIPVVWIHGL WP_001058306.1 hydrogenase [Salmonella enterica] Length: 372\nNumber of predicted TMHs: 2\nExp number of AAs in TMHs: 36.5540099999999\nExp number, first 60 AAs: 19.72732\nTotal prob of N-in:

0.96171\nPOSSIBLE N-term signal sequence\ninside 1 19\nTMhelix 20 39\noutside 40 325\nTMhelix 326 348\ninside 349 372

15883 GCF_001832335.1_ASM183233v1 Salmonella enterica subsp. diarizonae serovar 60:r:e,n,x,z15 Proteobacteria; Gammaproteobacteria; Enterobacterales; Enterobacteriaceae; Salmonella; Salmonella enterica; Salmonella enterica subsp. diarizonae MNNEETFYQAMRRKGVTRRSFLKFCSLAATSLGLGAGMTPKIAWALENKPRIPVVWIHGL WP_001058306.1 hydrogenase [Salmonella enterica] Length: 372\nNumber of predicted TMHs: 2\nExp number of AAs in TMHs: 36.5540099999999\nExp number, first 60 AAs: 19.72732\nTotal prob of N-in: 0.96171\nPOSSIBLE N-term signal sequence\ninside 1 19\nTMhelix 20 39\noutside 40 325\nTMhelix 326 348\ninside 349 372

15884 GCF_001832335.1_ASM183233v1 Salmonella enterica subsp. diarizonae serovar 60:r:e,n,x,z15 Proteobacteria; Gammaproteobacteria; Enterobacterales; Enterobacteriaceae; Salmonella; Salmonella enterica; Salmonella enterica subsp. diarizonae MTGDNTLITSHGINRRDFMKLCAALAATMGLSSKAAAEMAESVSNPQRPPVIWIGAQECT WP_023248548.1 hydrogenase 2 small subunit [Salmonella enterica] Length: 372\nNumber of predicted TMHs: 1\nExp number of AAs in TMHs: 24.51281\nExp number, first 60 AAs: 0.26822\nTotal prob of N-in: 0.14402\noutside 1 331\nTMhelix 332 354\ninside 355 372

15885 GCF_001832405.1_ASM183240v1 Salmonella enterica subsp. diarizonae serovar 60:r:e,n,x,z15 Proteobacteria; Gammaproteobacteria; Enterobacterales; Enterobacteriaceae; Salmonella; Salmonella enterica; Salmonella enterica subsp. diarizonae MTGDNTLITSHGINRRDFMKLCAALAATMGLSSKAAAEMAESVSNPQRPPVIWIGAQECT WP_023248548.1 hydrogenase 2 small subunit [Salmonella enterica] Length: 372\nNumber of predicted TMHs: 1\nExp number of AAs in TMHs: 24.51281\nExp number, first 60 AAs: 0.26822\nTotal prob of N-in: 0.14402\noutside 1 331\nTMhelix 332 354\ninside 355 372

15886 GCF_001832565.1_ASM183256v1 Salmonella enterica subsp. diarizonae serovar 60:r:e,n,x,z15 Proteobacteria; Gammaproteobacteria; Enterobacterales; Enterobacteriaceae; Salmonella; Salmonella enterica; Salmonella enterica subsp. diarizonae MNNEETFYQAMRRKGVTRRSFLKFCSLAATSLGLGAGMTPKIAWALENKPRIPVVWIHGL WP_001058306.1 hydrogenase [Salmonella enterica] Length: 372\nNumber of predicted TMHs: 2\nExp number of AAs in TMHs: 36.5540099999999\nExp number, first 60 AAs: 19.72732\nTotal prob of N-in: 0.96171\nPOSSIBLE N-term signal sequence\ninside 1 19\nTMhelix 20 39\noutside 40 325\nTMhelix 326 348\ninside 349 372

15887 GCF_001832255.1_ASM183225v1 Salmonella enterica subsp. diarizonae serovar 60:r:e,n,x,z15 Proteobacteria; Gammaproteobacteria; Enterobacterales; Enterobacteriaceae; Salmonella; Salmonella enterica; Salmonella enterica subsp. diarizonae MNNEETFYQAMRRKGVTRRSFLKFCSLAATSLGLGAGMTPKIAWALENKPRIPVVWIHGL WP_001058306.1 hydrogenase [Salmonella enterica] Length: 372\nNumber of predicted TMHs: 2\nExp number of AAs in TMHs: 36.5540099999999\nExp number, first 60 AAs: 19.72732\nTotal prob of N-in: 0.96171\nPOSSIBLE N-term signal sequence\ninside 1 19\nTMhelix 20 39\noutside 40 325\nTMhelix 326 348\ninside 349 372

15888 GCF_002035545.1_ASM203554v1 Salmonella enterica subsp. diarizonae serovar 60:r:e,n,x,z15 Proteobacteria; Gammaproteobacteria; Enterobacterales; Enterobacteriaceae; Salmonella; Salmonella enterica; Salmonella enterica subsp. diarizonae MNNEETFYQAMRRKGVTRRSFLKFCSLAATSLGLGAGMTPKIAWALENKPRIPVVWIHGL WP_001058306.1 hydrogenase [Salmonella enterica] Length: 372\nNumber of predicted TMHs: 2\nExp number of AAs in TMHs: 36.5540099999999\nExp number, first 60 AAs: 19.72732\nTotal prob of N-in: 0.96171\nPOSSIBLE N-term signal sequence\ninside 1 19\nTMhelix 20 39\noutside 40 325\nTMhelix 326 348\ninside 349 372

15889 GCF_001832425.1_ASM183242v1 Salmonella enterica subsp. diarizonae serovar 60:r:e,n,x,z15 Proteobacteria; Gammaproteobacteria; Enterobacterales; Enterobacteriaceae; Salmonella; Salmonella enterica; Salmonella enterica subsp. diarizonae MNNEETFYQAMRRKGVTRRSFLKFCSLAATSLGLGAGMTPKIAWALENKPRIPVVWIHGL WP_001058306.1 hydrogenase [Salmonella enterica] Length: 372\nNumber of predicted TMHs: 2\nExp number of AAs in TMHs: 36.5540099999999\nExp number, first 60 AAs: 19.72732\nTotal prob of N-in: 0.96171\nPOSSIBLE N-term signal sequence\ninside 1 19\nTMhelix 20 39\noutside 40 325\nTMhelix 326 348\ninside 349 372

15890 GCF_001832405.1_ASM183240v1 Salmonella enterica subsp. diarizonae serovar 60:r:e,n,x,z15 Proteobacteria; Gammaproteobacteria; Enterobacterales; Enterobacteriaceae; Salmonella; Salmonella enterica; Salmonella enterica subsp. diarizonae MNNEETFYQAMRRKGVTRRSFLKFCSLAATSLGLGAGMTPKIAWALENKPRIPVVWIHGL WP_001058306.1 hydrogenase [Salmonella enterica] Length: 372\nNumber of predicted TMHs: 2\nExp

number of AAs in TMHs: 36.5540099999999\nExp number, first 60 AAs:  19.72732\nTotal prob of N-in: 0.96171\nPOSSIBLE N-term signal sequence\ninside   1   19\nTMhelix   20   39\noutside   40   325\nTMhelix   326   348\ninside   349   372

15891 GCF_001832775.1_ASM183277v1 Salmonella enterica subsp. diarizonae serovar 60:r:e,n,x,z15 Proteobacteria; Gammaproteobacteria; Enterobacterales; Enterobacteriaceae; Salmonella; Salmonella enterica; Salmonella enterica subsp. diarizonae MTGDNTLITSHGINRRDFMKLCAALAATMGLSSKAAAEMAESVSNPQRPPVIWIGAQECT WP_023248548.1 hydrogenase 2 small subunit [Salmonella enterica] Length: 372\nNumber of predicted TMHs:  1\nExp number of AAs in TMHs: 24.51281\nExp number, first 60 AAs:  0.26822\nTotal prob of N-in: 0.14402\noutside   1   331\nTMhelix   332   354\ninside   355   372

15892 GCF_001832425.1_ASM183242v1 Salmonella enterica subsp. diarizonae serovar 60:r:e,n,x,z15 Proteobacteria; Gammaproteobacteria; Enterobacterales; Enterobacteriaceae; Salmonella; Salmonella enterica; Salmonella enterica subsp. diarizonae MTGDNTLITSHGINRRDFMKLCAALAATMGLSSKAAAEMAESVSNPQRPPVIWIGAQECT WP_023248548.1 hydrogenase 2 small subunit [Salmonella enterica] Length: 372\nNumber of predicted TMHs:  1\nExp number of AAs in TMHs: 24.51281\nExp number, first 60 AAs:  0.26822\nTotal prob of N-in: 0.14402\noutside   1   331\nTMhelix   332   354\ninside   355   372

15893 GCF_001832765.1_ASM183276v1 Salmonella enterica subsp. diarizonae serovar 60:r:e,n,x,z15 Proteobacteria; Gammaproteobacteria; Enterobacterales; Enterobacteriaceae; Salmonella; Salmonella enterica; Salmonella enterica subsp. diarizonae MTGDNTLITSHGINRRDFMKLCAALAATMGLSSKAAAEMAESVSNPQRPPVIWIGAQECT WP_023248548.1 hydrogenase 2 small subunit [Salmonella enterica] Length: 372\nNumber of predicted TMHs:  1\nExp number of AAs in TMHs: 24.51281\nExp number, first 60 AAs:  0.26822\nTotal prob of N-in: 0.14402\noutside   1   331\nTMhelix   332   354\ninside   355   372

15894 GCF_002035705.1_ASM203570v1 Salmonella enterica subsp. diarizonae serovar 60:r:z Proteobacteria; Gammaproteobacteria; Enterobacterales; Enterobacteriaceae; Salmonella; Salmonella enterica; Salmonella enterica subsp. diarizonae MNNEETFYQAMRRKGVTRRSFLKFCSLAATSLGLGAGMTPKIAWALENKPRIPVVWIHGL WP_023187532.1 hydrogenase 1, small subunit [Salmonella enterica] Length: 372\nNumber of predicted TMHs:  2\nExp number of AAs in TMHs: 36.9997\nExp number, first 60 AAs:  19.72759\nTotal prob of N-in: 0.96171\nPOSSIBLE N-term signal sequence\ninside   1   19\nTMhelix   20   39\noutside   40   325\nTMhelix   326   348\ninside   349   372

15895 GCF_002035705.1_ASM203570v1 Salmonella enterica subsp. diarizonae serovar 60:r:z Proteobacteria; Gammaproteobacteria; Enterobacterales; Enterobacteriaceae; Salmonella; Salmonella enterica; Salmonella enterica subsp. diarizonae MTGDNTLITSHGINRRDFMKLCAALAATMGLSSKAAAEMAESVSNPQRPPVIWIGAQECT WP_000145429.1 hydrogenase 2 small subunit [Salmonella enterica] Length: 372\nNumber of predicted TMHs:  1\nExp number of AAs in TMHs: 24.51309\nExp number, first 60 AAs:  0.2682\nTotal prob of N-in: 0.14402\noutside   1   331\nTMhelix   332   354\ninside   355   372

15896 GCF_002063885.1_ASM206388v1 Salmonella enterica subsp. diarizonae serovar 61:-:1,5,7 Proteobacteria; Gammaproteobacteria; Enterobacterales; Enterobacteriaceae; Salmonella; Salmonella enterica; Salmonella enterica subsp. diarizonae MTGDNTLITSHGINRRDFMKLCAALAATMGLSSKAAAEMAESVSNPQRPPVIWIGAQECT WP_023248548.1 hydrogenase 2 small subunit [Salmonella enterica] Length: 372\nNumber of predicted TMHs:  1\nExp number of AAs in TMHs: 24.51281\nExp number, first 60 AAs:  0.26822\nTotal prob of N-in: 0.14402\noutside   1   331\nTMhelix   332   354\ninside   355   372

15897 GCF_002063885.1_ASM206388v1 Salmonella enterica subsp. diarizonae serovar 61:-:1,5,7 Proteobacteria; Gammaproteobacteria; Enterobacterales; Enterobacteriaceae; Salmonella; Salmonella enterica; Salmonella enterica subsp. diarizonae MNNEETFYQAMRRKGVTRRSFLKFCSLAATSLGLGAGMTPKIAWALENKPRIPVVWIHGL WP_001058306.1 hydrogenase [Salmonella enterica] Length: 372\nNumber of predicted TMHs:  2\nExp number of AAs in TMHs: 36.5540099999999\nExp number, first 60 AAs:  19.72732\nTotal prob of N-in: 0.96171\nPOSSIBLE N-term signal sequence\ninside   1   19\nTMhelix   20   39\noutside   40   325\nTMhelix   326   348\ninside   349   372

15898 GCF_002035715.1_ASM203571v1 Salmonella enterica subsp. diarizonae serovar 61:k:1,5,7 Proteobacteria; Gammaproteobacteria; Enterobacterales; Enterobacteriaceae; Salmonella; Salmonella enterica; Salmonella enterica subsp. diarizonae MTGDNTLITSHGINRRDFMKLCAALAATMGLSSKAAAEMAESVSNPQRPPVIWIGAQECT WP_000145431.1 hydrogenase 2 small subunit [Salmonella enterica] Length: 372\nNumber of predicted TMHs:  1\nExp number of AAs in TMHs: 24.51505\nExp number, first 60 AAs:  0.2682\nTotal prob of N-in: 0.14406\noutside   1   331\nTMhelix   332   354\ninside   355   372

15899 GCF_002035715.1_ASM203571v1 Salmonella enterica subsp. diarizonae serovar 61:k:1,5,7 Proteobacteria; Gammaproteobacteria; Enterobacterales; Enterobacteriaceae; Salmonella; Salmonella enterica; Salmonella enterica subsp. diarizonae MNNEETFYQAMRRKGVTRRSFLKFCSLAATSLGLGAGMTPKIAWALENKPRIPVVWIHGL WP_001058312.1 hydrogenase [Salmonella enterica] Length: 372\nNumber of predicted TMHs: 2\nExp number of AAs in TMHs: 36.51738\nExp number, first 60 AAs: 19.72849\nTotal prob of N-in: 0.96170\nPOSSIBLE N-term signal sequence\ninside 1 19\nTMhelix 20 39\noutside 40 325\nTMhelix 326 348\ninside 349 372

15900 GCF_002107505.1_ASM210750v1 Salmonella enterica subsp. diarizonae serovar Rough:r:z Proteobacteria; Gammaproteobacteria; Enterobacterales; Enterobacteriaceae; Salmonella; Salmonella enterica; Salmonella enterica subsp. diarizonae MNNEETFYQAMRRKGVTRRSFLKFCSLAATSLGLGAGMTPKIAWALENKPRIPVVWIHGL WP_001058306.1 hydrogenase [Salmonella enterica] Length: 372\nNumber of predicted TMHs: 2\nExp number of AAs in TMHs: 36.5540099999999\nExp number, first 60 AAs: 19.72732\nTotal prob of N-in: 0.96171\nPOSSIBLE N-term signal sequence\ninside 1 19\nTMhelix 20 39\noutside 40 325\nTMhelix 326 348\ninside 349 372

15901 GCF_002107575.1_ASM210757v1 Salmonella enterica subsp. diarizonae serovar Rough:r:z Proteobacteria; Gammaproteobacteria; Enterobacterales; Enterobacteriaceae; Salmonella; Salmonella enterica; Salmonella enterica subsp. diarizonae MNNEETFYQAMRRKGVTRRSFLKFCSLAATSLGLGAGMTPKIAWALENKPRIPVVWIHGL WP_001058306.1 hydrogenase [Salmonella enterica] Length: 372\nNumber of predicted TMHs: 2\nExp number of AAs in TMHs: 36.5540099999999\nExp number, first 60 AAs: 19.72732\nTotal prob of N-in: 0.96171\nPOSSIBLE N-term signal sequence\ninside 1 19\nTMhelix 20 39\noutside 40 325\nTMhelix 326 348\ninside 349 372

15902 GCF_002107575.1_ASM210757v1 Salmonella enterica subsp. diarizonae serovar Rough:r:z Proteobacteria; Gammaproteobacteria; Enterobacterales; Enterobacteriaceae; Salmonella; Salmonella enterica; Salmonella enterica subsp. diarizonae MTGDNTLITSHGINRRDFMKLCAALAATMGLSSKAAAEMAESVSNPQRPPVIWIGAQECT WP_023248548.1 hydrogenase 2 small subunit [Salmonella enterica] Length: 372\nNumber of predicted TMHs: 1\nExp number of AAs in TMHs: 24.51281\nExp number, first 60 AAs: 0.26822\nTotal prob of N-in: 0.14402\noutside 1 331\nTMhelix 332 354\ninside 355 372

15903 GCF_002107505.1_ASM210750v1 Salmonella enterica subsp. diarizonae serovar Rough:r:z Proteobacteria; Gammaproteobacteria; Enterobacterales; Enterobacteriaceae; Salmonella; Salmonella enterica; Salmonella enterica subsp. diarizonae MTGDNTLITSHGINRRDFMKLCAALAATMGLSSKAAAEMAESVSNPQRPPVIWIGAQECT WP_023248548.1 hydrogenase 2 small subunit [Salmonella enterica] Length: 372\nNumber of predicted TMHs: 1\nExp number of AAs in TMHs: 24.51281\nExp number, first 60 AAs: 0.26822\nTotal prob of N-in: 0.14402\noutside 1 331\nTMhelix 332 354\ninside 355 372

15904 GCF_000487635.2_ASM48763v2 Salmonella enterica subsp. diarizonae serovar 60:r:e,n,x,z15 str. 01-0170 Proteobacteria; Gammaproteobacteria; Enterobacterales; Enterobacteriaceae; Salmonella; Salmonella enterica; Salmonella enterica subsp. diarizonae; Salmonella enterica subsp. diarizonae serovar 60 MTGDNTLITSHGINRRDFMKLCAALAATMGLSSKAAAEMAESVSNPQRPPVIWIGAQECT WP_023248548.1 hydrogenase 2 small subunit [Salmonella enterica] Length: 372\nNumber of predicted TMHs: 1\nExp number of AAs in TMHs: 24.51281\nExp number, first 60 AAs: 0.26822\nTotal prob of N-in: 0.14402\noutside 1 331\nTMhelix 332 354\ninside 355 372

15905 GCF_000487635.2_ASM48763v2 Salmonella enterica subsp. diarizonae serovar 60:r:e,n,x,z15 str. 01-0170 Proteobacteria; Gammaproteobacteria; Enterobacterales; Enterobacteriaceae; Salmonella; Salmonella enterica; Salmonella enterica subsp. diarizonae; Salmonella enterica subsp. diarizonae serovar 60 MNNEETFYQAMRRKGVTRRSFLKFCSLAATSLGLGAGMTPKIAWALENKPRIPVVWIHGL WP_001058306.1 hydrogenase [Salmonella enterica] Length: 372\nNumber of predicted TMHs: 2\nExp number of AAs in TMHs: 36.5540099999999\nExp number, first 60 AAs: 19.72732\nTotal prob of N-in: 0.96171\nPOSSIBLE N-term signal sequence\ninside 1 19\nTMhelix 20 39\noutside 40 325\nTMhelix 326 348\ninside 349 372

15906 GCF_002047565.1_ASM204756v1 Salmonella enterica subsp. enterica serovar 4,12:b:- Proteobacteria; Gammaproteobacteria; Enterobacterales; Enterobacteriaceae; Salmonella; Salmonella enterica; Salmonella enterica subsp. enterica MTGDNTLITSHGINRRDFMKLCAALAATMGLSSKAAAEMAESVSNPQRPPVIWIGAQECT WP_000145432.1 hydrogenase 2 small subunit [Salmonella enterica] Length: 372\nNumber of predicted TMHs: 1\nExp number of AAs in TMHs: 24.5101\nExp number, first 60 AAs: 0.2682\nTotal prob of N-in: 0.14397\noutside 1 331\nTMhelix 332 354\ninside 355 372

15907 GCF_002047565.1_ASM204756v1 Salmonella enterica subsp. enterica serovar 4,12:b:- Proteobacteria; Gammaproteobacteria; Enterobacterales; Enterobacteriaceae; Salmonella; Salmonella enterica; Salmonella enterica subsp. enterica MNNEETFYQAMRRKGVTRRSFLKFCSLAATSLGLGAGMTPKIAWALENKPRIPVVWIHGL WP_001058314.1 hydrogenase [Salmonella enterica] Length: 372\nNumber of predicted TMHs: 2\nExp number of AAs in TMHs: 36.3110299999999\nExp number, first 60 AAs: 19.72774\nTotal prob of N-in: 0.96169\nPOSSIBLE N-term signal sequence\ninside 1 19\nTMhelix 20 39\noutside 40 325\nTMhelix 326 348\ninside 349 372

15908 GCF_002032195.1_ASM203219v1 Salmonella enterica subsp. enterica serovar 4,12:d:- Proteobacteria; Gammaproteobacteria; Enterobacterales; Enterobacteriaceae; Salmonella; Salmonella enterica; Salmonella enterica subsp. enterica MNNEETFYQAMRRKGVTRRSFLKFCSLAATSLGLGAGMTPKIAWALENKPRIPVVWIHGL WP_001058311.1 hydrogenase [Salmonella enterica] Length: 372\nNumber of predicted TMHs: 2\nExp number of AAs in TMHs: 37.4699899999999\nExp number, first 60 AAs: 19.72699\nTotal prob of N-in: 0.96171\nPOSSIBLE N-term signal sequence\ninside 1 19\nTMhelix 20 39\noutside 40 325\nTMhelix 326 348\ninside 349 372

15909 GCF_002032195.1_ASM203219v1 Salmonella enterica subsp. enterica serovar 4,12:d:- Proteobacteria; Gammaproteobacteria; Enterobacterales; Enterobacteriaceae; Salmonella; Salmonella enterica; Salmonella enterica subsp. enterica MTGDNTLITSHGINRRDFMKLCAALAATMGLSSKAAAEMAESVSNPQRPPVIWIGAQECT WP_000145429.1 hydrogenase 2 small subunit [Salmonella enterica] Length: 372\nNumber of predicted TMHs: 1\nExp number of AAs in TMHs: 24.51309\nExp number, first 60 AAs: 0.2682\nTotal prob of N-in: 0.14402\noutside 1 331\nTMhelix 332 354\ninside 355 372

15910 GCF_001478855.1_Salmonella_enterica_CVM_N30650-SQ_v1.0 Salmonella enterica subsp. enterica serovar 4,12:d:- Proteobacteria; Gammaproteobacteria; Enterobacterales; Enterobacteriaceae; Salmonella; Salmonella enterica; Salmonella enterica subsp. enterica MTGDNTLITSHGINRRDFMKLCAALAATMGLSSKAAAEMAESVSNPQRPPVIWIGAQECT WP_000145429.1 hydrogenase 2 small subunit [Salmonella enterica] Length: 372\nNumber of predicted TMHs: 1\nExp number of AAs in TMHs: 24.51309\nExp number, first 60 AAs: 0.2682\nTotal prob of N-in: 0.14402\noutside 1 331\nTMhelix 332 354\ninside 355 372

15911 GCF_002032385.1_ASM203238v1 Salmonella enterica subsp. enterica serovar 4,12:d:- Proteobacteria; Gammaproteobacteria; Enterobacterales; Enterobacteriaceae; Salmonella; Salmonella enterica; Salmonella enterica subsp. enterica MNNEETFYQAMRRKGVTRRSFLKFCSLAATSLGLGAGMTPKIAWALENKPRIPVVWIHGL WP_001058311.1 hydrogenase [Salmonella enterica] Length: 372\nNumber of predicted TMHs: 2\nExp number of AAs in TMHs: 37.4699899999999\nExp number, first 60 AAs: 19.72699\nTotal prob of N-in: 0.96171\nPOSSIBLE N-term signal sequence\ninside 1 19\nTMhelix 20 39\noutside 40 325\nTMhelix 326 348\ninside 349 372

15912 GCF_002030235.1_ASM203023v1 Salmonella enterica subsp. enterica serovar 4,12:d:- Proteobacteria; Gammaproteobacteria; Enterobacterales; Enterobacteriaceae; Salmonella; Salmonella enterica; Salmonella enterica subsp. enterica MTGDNTLITSHGINRRDFMKLCAALAATMGLSSKAAAEMAESVSNPQRPPVIWIGAQECT WP_000145429.1 hydrogenase 2 small subunit [Salmonella enterica] Length: 372\nNumber of predicted TMHs: 1\nExp number of AAs in TMHs: 24.51309\nExp number, first 60 AAs: 0.2682\nTotal prob of N-in: 0.14402\noutside 1 331\nTMhelix 332 354\ninside 355 372

15913 GCF_002030235.1_ASM203023v1 Salmonella enterica subsp. enterica serovar 4,12:d:- Proteobacteria; Gammaproteobacteria; Enterobacterales; Enterobacteriaceae; Salmonella; Salmonella enterica; Salmonella enterica subsp. enterica MNNEETFYQAMRRKGVTRRSFLKFCSLAATSLGLGAGMTPKIAWALENKPRIPVVWIHGL WP_001058311.1 hydrogenase [Salmonella enterica] Length: 372\nNumber of predicted TMHs: 2\nExp number of AAs in TMHs: 37.4699899999999\nExp number, first 60 AAs: 19.72699\nTotal prob of N-in: 0.96171\nPOSSIBLE N-term signal sequence\ninside 1 19\nTMhelix 20 39\noutside 40 325\nTMhelix 326 348\ninside 349 372

15914 GCF_002032385.1_ASM203238v1 Salmonella enterica subsp. enterica serovar 4,12:d:- Proteobacteria; Gammaproteobacteria; Enterobacterales; Enterobacteriaceae; Salmonella; Salmonella enterica; Salmonella enterica subsp. enterica MTGDNTLITSHGINRRDFMKLCAALAATMGLSSKAAAEMAESVSNPQRPPVIWIGAQECT WP_000145429.1 hydrogenase 2 small subunit [Salmonella enterica] Length: 372\nNumber of predicted TMHs: 1\nExp number of AAs in TMHs: 24.51309\nExp number, first 60 AAs: 0.2682\nTotal prob of N-in: 0.14402\noutside 1 331\nTMhelix 332 354\ninside 355 372

15915 GCF_001478855.1_Salmonella_enterica_CVM_N30650-SQ_v1.0 Salmonella enterica subsp. enterica serovar 4,12:d:- Proteobacteria; Gammaproteobacteria; Enterobacterales; Enterobacteriaceae; Salmonella; Salmonella enterica; Salmonella enterica subsp. enterica MNNEETFYQAMRRKGVTRRSFLKFCSLAATSLGLGAGMTPKIAWALENKPRIPVVWIHGL WP_001058311.1 hydrogenase [Salmonella enterica] Length: 372\nNumber of predicted TMHs: 2\nExp number of AAs in TMHs: 37.4699899999999\nExp number, first 60 AAs: 19.72699\nTotal prob of N-in: 0.96171\nPOSSIBLE N-term signal sequence\ninside 1 19\nTMhelix 20 39\noutside 40 325\nTMhelix 326 348\ninside 349 372

15916 GCF_001244815.1_Salmonella_enterica_CVM_N48691_v1.0 Salmonella enterica subsp. enterica serovar 4,12:i:- Proteobacteria; Gammaproteobacteria; Enterobacterales; Enterobacteriaceae; Salmonella; Salmonella enterica; Salmonella enterica subsp. enterica MNNEETFYQAMRRKGVTRRSFLKFCSLAATSLGLGAGMTPKIAWALENKPRIPVVWIHGL WP_001058312.1 hydrogenase [Salmonella enterica] Length: 372\nNumber of predicted TMHs: 2\nExp number of AAs in TMHs: 36.51738\nExp number, first 60 AAs: 19.72849\nTotal prob of N-in: 0.96170\nPOSSIBLE N-term signal sequence\ninside 1 19\nTMhelix 20 39\noutside 40 325\nTMhelix 326 348\ninside 349 372

15917 GCF_001246615.1_Salmonella_enterica_CVM_N51270_v1.0 Salmonella enterica subsp. enterica serovar 4,12:i:- Proteobacteria; Gammaproteobacteria; Enterobacterales; Enterobacteriaceae; Salmonella; Salmonella enterica; Salmonella enterica subsp. enterica MNNEETFYQAMRRKGVTRRSFLKFCSLAATSLGLGAGMTPKIAWALENKPRIPVVWIHGL WP_001058312.1 hydrogenase [Salmonella enterica] Length: 372\nNumber of predicted TMHs: 2\nExp number of AAs in TMHs: 36.51738\nExp number, first 60 AAs: 19.72849\nTotal prob of N-in: 0.96170\nPOSSIBLE N-term signal sequence\ninside 1 19\nTMhelix 20 39\noutside 40 325\nTMhelix 326 348\ninside 349 372

15918 GCF_001293585.1_ASM129358v1 Salmonella enterica subsp. enterica serovar 4,12:i:- Proteobacteria; Gammaproteobacteria; Enterobacterales; Enterobacteriaceae; Salmonella; Salmonella enterica; Salmonella enterica subsp. enterica MNNEETFYQAMRRKGVTRRSFLKFCSLAATSLGLGAGMTPKIAWALENKPRIPVVWIHGL WP_001058312.1 hydrogenase [Salmonella enterica] Length: 372\nNumber of predicted TMHs: 2\nExp number of AAs in TMHs: 36.51738\nExp number, first 60 AAs: 19.72849\nTotal prob of N-in: 0.96170\nPOSSIBLE N-term signal sequence\ninside 1 19\nTMhelix 20 39\noutside 40 325\nTMhelix 326 348\ninside 349 372

15919 GCF_001244815.1_Salmonella_enterica_CVM_N48691_v1.0 Salmonella enterica subsp. enterica serovar 4,12:i:- Proteobacteria; Gammaproteobacteria; Enterobacterales; Enterobacteriaceae; Salmonella; Salmonella enterica; Salmonella enterica subsp. enterica MTGDNTLITSHGINRRDFMKLCAALAATMGLSSKAAAEMAESVSNPQRPPVIWIGAQECT WP_000145429.1 hydrogenase 2 small subunit [Salmonella enterica] Length: 372\nNumber of predicted TMHs: 1\nExp number of AAs in TMHs: 24.51309\nExp number, first 60 AAs: 0.2682\nTotal prob of N-in: 0.14402\noutside 1 331\nTMhelix 332 354\ninside 355 372

15920 GCF_001246615.1_Salmonella_enterica_CVM_N51270_v1.0 Salmonella enterica subsp. enterica serovar 4,12:i:- Proteobacteria; Gammaproteobacteria; Enterobacterales; Enterobacteriaceae; Salmonella; Salmonella enterica; Salmonella enterica subsp. enterica MTGDNTLITSHGINRRDFMKLCAALAATMGLSSKAAAEMAESVSNPQRPPVIWIGAQECT WP_000145429.1 hydrogenase 2 small subunit [Salmonella enterica] Length: 372\nNumber of predicted TMHs: 1\nExp number of AAs in TMHs: 24.51309\nExp number, first 60 AAs: 0.2682\nTotal prob of N-in: 0.14402\noutside 1 331\nTMhelix 332 354\ninside 355 372

15921 GCF_001293585.1_ASM129358v1 Salmonella enterica subsp. enterica serovar 4,12:i:- Proteobacteria; Gammaproteobacteria; Enterobacterales; Enterobacteriaceae; Salmonella; Salmonella enterica; Salmonella enterica subsp. enterica MTGDNTLITSHGINRRDFMKLCAALAATMGLSSKAAAEMAESVSNPQRPPVIWIGAQECT WP_000145429.1 hydrogenase 2 small subunit [Salmonella enterica] Length: 372\nNumber of predicted TMHs: 1\nExp number of AAs in TMHs: 24.51309\nExp number, first 60 AAs: 0.2682\nTotal prob of N-in: 0.14402\noutside 1 331\nTMhelix 332 354\ninside 355 372

15922 GCF_001240385.1_Salmonella_enterica_CVM_N43463_v1.0 Salmonella enterica subsp. enterica serovar 4,12:i:- Proteobacteria; Gammaproteobacteria; Enterobacterales; Enterobacteriaceae; Salmonella; Salmonella enterica; Salmonella enterica subsp. enterica MNNEETFYQAMRRKGVTRRSFLKFCSLAATSLGLGAGMTPKIAWALENKPRIPVVWIHGL WP_001058312.1 hydrogenase [Salmonella enterica] Length: 372\nNumber of predicted TMHs: 2\nExp number of AAs in TMHs: 36.51738\nExp number, first 60 AAs: 19.72849\nTotal prob of N-in: 0.96170\nPOSSIBLE N-term signal sequence\ninside 1 19\nTMhelix 20 39\noutside 40 325\nTMhelix 326 348\ninside 349 372

15923 GCF_001247365.1_Salmonella_enterica_CVM_N51306_v1.0 Salmonella enterica subsp. enterica serovar 4,12:i:- Proteobacteria; Gammaproteobacteria; Enterobacterales; Enterobacteriaceae; Salmonella; Salmonella enterica; Salmonella enterica subsp. enterica MNNEETFYQAMRRKGVTRRSFLKFCSLAATSLGLGAGMTPKIAWALENKPRIPVVWIHGL WP_001058312.1 hydrogenase [Salmonella enterica] Length: 372\nNumber of predicted TMHs: 2\nExp

number of AAs in TMHs: 36.51738\nExp number, first 60 AAs:  19.72849\nTotal prob of N-in:   0.96170\nPOSSIBLE N-term signal sequence\ninside   1   19\nTMhelix   20   39\noutside   40   325\nTMhelix   326   348\ninside   349   372

15924 GCF_001271395.1_Salmonella_enterica_CVM_N43477_v1.0 Salmonella enterica subsp. enterica serovar 4,12:i:- Proteobacteria; Gammaproteobacteria; Enterobacterales; Enterobacteriaceae; Salmonella; Salmonella enterica; Salmonella enterica subsp. enterica MNNEETFYQAMRRKGVTRRSFLKFCSLAATSLGLGAGMTPKIAWALENKPRIPVVWIHGL WP_001058312.1 hydrogenase [Salmonella enterica] Length: 372\nNumber of predicted TMHs:  2\nExp number of AAs in TMHs: 36.51738\nExp number, first 60 AAs:  19.72849\nTotal prob of N-in:   0.96170\nPOSSIBLE N-term signal sequence\ninside   1   19\nTMhelix   20   39\noutside   40   325\nTMhelix   326   348\ninside   349   372

15925 GCF_001271935.1_Salmonella_enterica_CVM_N43467_v1.0 Salmonella enterica subsp. enterica serovar 4,12:i:- Proteobacteria; Gammaproteobacteria; Enterobacterales; Enterobacteriaceae; Salmonella; Salmonella enterica; Salmonella enterica subsp. enterica MTGDNTLITSHGINRRDFMKLCAALAATMGLSSKAAAEMAESVSNPQRPPVIWIGAQECT WP_000145429.1 hydrogenase 2 small subunit [Salmonella enterica] Length: 372\nNumber of predicted TMHs:  1\nExp number of AAs in TMHs: 24.51309\nExp number, first 60 AAs:  0.2682\nTotal prob of N-in: 0.14402\noutside   1   331\nTMhelix   332   354\ninside   355   372

15926 GCF_001247365.1_Salmonella_enterica_CVM_N51306_v1.0 Salmonella enterica subsp. enterica serovar 4,12:i:- Proteobacteria; Gammaproteobacteria; Enterobacterales; Enterobacteriaceae; Salmonella; Salmonella enterica; Salmonella enterica subsp. enterica MTGDNTLITSHGINRRDFMKLCAALAATMGLSSKAAAEMAESVSNPQRPPVIWIGAQECT WP_000145429.1 hydrogenase 2 small subunit [Salmonella enterica] Length: 372\nNumber of predicted TMHs:  1\nExp number of AAs in TMHs: 24.51309\nExp number, first 60 AAs:  0.2682\nTotal prob of N-in: 0.14402\noutside   1   331\nTMhelix   332   354\ninside   355   372

15927 GCF_001271395.1_Salmonella_enterica_CVM_N43477_v1.0 Salmonella enterica subsp. enterica serovar 4,12:i:- Proteobacteria; Gammaproteobacteria; Enterobacterales; Enterobacteriaceae; Salmonella; Salmonella enterica; Salmonella enterica subsp. enterica MTGDNTLITSHGINRRDFMKLCAALAATMGLSSKAAAEMAESVSNPQRPPVIWIGAQECT WP_000145429.1 hydrogenase 2 small subunit [Salmonella enterica] Length: 372\nNumber of predicted TMHs:  1\nExp number of AAs in TMHs: 24.51309\nExp number, first 60 AAs:  0.2682\nTotal prob of N-in: 0.14402\noutside   1   331\nTMhelix   332   354\ninside   355   372

15928 GCF_001271935.1_Salmonella_enterica_CVM_N43467_v1.0 Salmonella enterica subsp. enterica serovar 4,12:i:- Proteobacteria; Gammaproteobacteria; Enterobacterales; Enterobacteriaceae; Salmonella; Salmonella enterica; Salmonella enterica subsp. enterica MNNEETFYQAMRRKGVTRRSFLKFCSLAATSLGLGAGMTPKIAWALENKPRIPVVWIHGL WP_001058312.1 hydrogenase [Salmonella enterica] Length: 372\nNumber of predicted TMHs:  2\nExp number of AAs in TMHs: 36.51738\nExp number, first 60 AAs:  19.72849\nTotal prob of N-in:   0.96170\nPOSSIBLE N-term signal sequence\ninside   1   19\nTMhelix   20   39\noutside   40   325\nTMhelix   326   348\ninside   349   372

15929 GCF_001240385.1_Salmonella_enterica_CVM_N43463_v1.0 Salmonella enterica subsp. enterica serovar 4,12:i:- Proteobacteria; Gammaproteobacteria; Enterobacterales; Enterobacteriaceae; Salmonella; Salmonella enterica; Salmonella enterica subsp. enterica MTGDNTLITSHGINRRDFMKLCAALAATMGLSSKAAAEMAESVSNPQRPPVIWIGAQECT WP_000145429.1 hydrogenase 2 small subunit [Salmonella enterica] Length: 372\nNumber of predicted TMHs:  1\nExp number of AAs in TMHs: 24.51309\nExp number, first 60 AAs:  0.2682\nTotal prob of N-in: 0.14402\noutside   1   331\nTMhelix   332   354\ninside   355   372

15930 GCF_002037405.1_ASM203740v1 Salmonella enterica subsp. enterica serovar 4,12:r:- Proteobacteria; Gammaproteobacteria; Enterobacterales; Enterobacteriaceae; Salmonella; Salmonella enterica; Salmonella enterica subsp. enterica MNNEETFYQAMRRKGVTRRSFLKFCSLAATSLGLGAGMTPKIAWALENKPRIPVVWIHGL WP_001058312.1 hydrogenase [Salmonella enterica] Length: 372\nNumber of predicted TMHs:  2\nExp number of AAs in TMHs: 36.51738\nExp number, first 60 AAs:  19.72849\nTotal prob of N-in:   0.96170\nPOSSIBLE N-term signal sequence\ninside   1   19\nTMhelix   20   39\noutside   40   325\nTMhelix   326   348\ninside   349   372

15931 GCF_002037385.1_ASM203738v1 Salmonella enterica subsp. enterica serovar 4,12:r:- Proteobacteria; Gammaproteobacteria; Enterobacterales; Enterobacteriaceae; Salmonella; Salmonella enterica; Salmonella enterica subsp. enterica MNNEETFYQAMRRKGVTRRSFLKFCSLAATSLGLGAGMTPKIAWALENKPRIPVVWIHGL WP_001058312.1 hydrogenase [Salmonella enterica] Length: 372\nNumber of predicted TMHs:  2\nExp number of AAs in TMHs: 36.51738\nExp number, first 60 AAs:  19.72849\nTotal prob of N-in:   0.96170\nPOSSIBLE N-term signal sequence\ninside   1   19\nTMhelix   20   39\noutside   40   325\nTMhelix   326   348\ninside   349   372

15932 GCF_002037345.1_ASM203734v1 Salmonella enterica subsp. enterica serovar 4,12:r:- Proteobacteria; Gammaproteobacteria; Enterobacterales; Enterobacteriaceae; Salmonella; Salmonella enterica;

Salmonella enterica subsp. enterica MNNEETFYQAMRRKGVTRRSFLKFCSLAATSLGLGAGMTPKIAWALENKPRIPVVWIHGL WP_001058312.1 hydrogenase [Salmonella enterica] Length: 372\nNumber of predicted TMHs: 2\nExp number of AAs in TMHs: 36.51738\nExp number, first 60 AAs: 19.72849\nTotal prob of N-in: 0.96170\nPOSSIBLE N-term signal sequence\ninside 1 19\nTMhelix 20 39\noutside 40 325\nTMhelix 326 348\ninside 349 372

15933 GCF_002037405.1_ASM203740v1 Salmonella enterica subsp. enterica serovar 4,12:r:- Proteobacteria; Gammaproteobacteria; Enterobacterales; Enterobacteriaceae; Salmonella; Salmonella enterica; Salmonella enterica subsp. enterica MTGDNTLITSHGINRRDFMKLCAALAATMGLSSKAAAEMAESVSNPQRPPVIWIGAQECT WP_000145429.1 hydrogenase 2 small subunit [Salmonella enterica] Length: 372\nNumber of predicted TMHs: 1\nExp number of AAs in TMHs: 24.51309\nExp number, first 60 AAs: 0.2682\nTotal prob of N-in: 0.14402\noutside 1 331\nTMhelix 332 354\ninside 355 372

15934 GCF_002037465.1_ASM203746v1 Salmonella enterica subsp. enterica serovar 4,12:r:- Proteobacteria; Gammaproteobacteria; Enterobacterales; Enterobacteriaceae; Salmonella; Salmonella enterica; Salmonella enterica subsp. enterica MNNEETFYQAMRRKGVTRRSFLKFCSLAATSLGLGAGMTPKIAWALENKPRIPVVWIHGL WP_001058312.1 hydrogenase [Salmonella enterica] Length: 372\nNumber of predicted TMHs: 2\nExp number of AAs in TMHs: 36.51738\nExp number, first 60 AAs: 19.72849\nTotal prob of N-in: 0.96170\nPOSSIBLE N-term signal sequence\ninside 1 19\nTMhelix 20 39\noutside 40 325\nTMhelix 326 348\ninside 349 372

15935 GCF_002037465.1_ASM203746v1 Salmonella enterica subsp. enterica serovar 4,12:r:- Proteobacteria; Gammaproteobacteria; Enterobacterales; Enterobacteriaceae; Salmonella; Salmonella enterica; Salmonella enterica subsp. enterica MTGDNTLITSHGINRRDFMKLCAALAATMGLSSKAAAEMAESVSNPQRPPVIWIGAQECT WP_000145429.1 hydrogenase 2 small subunit [Salmonella enterica] Length: 372\nNumber of predicted TMHs: 1\nExp number of AAs in TMHs: 24.51309\nExp number, first 60 AAs: 0.2682\nTotal prob of N-in: 0.14402\noutside 1 331\nTMhelix 332 354\ninside 355 372

15936 GCF_002037385.1_ASM203738v1 Salmonella enterica subsp. enterica serovar 4,12:r:- Proteobacteria; Gammaproteobacteria; Enterobacterales; Enterobacteriaceae; Salmonella; Salmonella enterica; Salmonella enterica subsp. enterica MTGDNTLITSHGINRRDFMKLCAALAATMGLSSKAAAEMAESVSNPQRPPVIWIGAQECT WP_000145429.1 hydrogenase 2 small subunit [Salmonella enterica] Length: 372\nNumber of predicted TMHs: 1\nExp number of AAs in TMHs: 24.51309\nExp number, first 60 AAs: 0.2682\nTotal prob of N-in: 0.14402\noutside 1 331\nTMhelix 332 354\ninside 355 372

15937 GCF_002037345.1_ASM203734v1 Salmonella enterica subsp. enterica serovar 4,12:r:- Proteobacteria; Gammaproteobacteria; Enterobacterales; Enterobacteriaceae; Salmonella; Salmonella enterica; Salmonella enterica subsp. enterica MTGDNTLITSHGINRRDFMKLCAALAATMGLSSKAAAEMAESVSNPQRPPVIWIGAQECT WP_000145429.1 hydrogenase 2 small subunit [Salmonella enterica] Length: 372\nNumber of predicted TMHs: 1\nExp number of AAs in TMHs: 24.51309\nExp number, first 60 AAs: 0.2682\nTotal prob of N-in: 0.14402\noutside 1 331\nTMhelix 332 354\ninside 355 372

15938 GCF_002047685.1_ASM204768v1 Salmonella enterica subsp. enterica serovar 4,5,12:b:- Proteobacteria; Gammaproteobacteria; Enterobacterales; Enterobacteriaceae; Salmonella; Salmonella enterica; Salmonella enterica subsp. enterica MTGDNTLITSHGINRRDFMKLCAALAATMGLSSKAAAEMAESVSNPQRPPVIWIGAQECT WP_000145429.1 hydrogenase 2 small subunit [Salmonella enterica] Length: 372\nNumber of predicted TMHs: 1\nExp number of AAs in TMHs: 24.51309\nExp number, first 60 AAs: 0.2682\nTotal prob of N-in: 0.14402\noutside 1 331\nTMhelix 332 354\ninside 355 372

15939 GCF_002048605.1_ASM204860v1 Salmonella enterica subsp. enterica serovar 4,5,12:b:- Proteobacteria; Gammaproteobacteria; Enterobacterales; Enterobacteriaceae; Salmonella; Salmonella enterica; Salmonella enterica subsp. enterica MNNEETFYQAMRRKGVTRRSFLKFCSLAATSLGLGAGMTPKIAWALENKPRIPVVWIHGL WP_001058307.1 hydrogenase [Salmonella enterica] Length: 372\nNumber of predicted TMHs: 2\nExp number of AAs in TMHs: 36.51738\nExp number, first 60 AAs: 19.72849\nTotal prob of N-in: 0.96170\nPOSSIBLE N-term signal sequence\ninside 1 19\nTMhelix 20 39\noutside 40 325\nTMhelix 326 348\ninside 349 372

15940 GCF_002048605.1_ASM204860v1 Salmonella enterica subsp. enterica serovar 4,5,12:b:- Proteobacteria; Gammaproteobacteria; Enterobacterales; Enterobacteriaceae; Salmonella; Salmonella enterica; Salmonella enterica subsp. enterica MTGDNTLITSHGINRRDFMKLCAALAATMGLSSKAAAEMAESVSNPQRPPVIWIGAQECT WP_000145429.1 hydrogenase 2 small subunit [Salmonella enterica] Length: 372\nNumber of predicted TMHs: 1\nExp number of AAs in TMHs: 24.51309\nExp number, first 60 AAs: 0.2682\nTotal prob of N-in: 0.14402\noutside 1 331\nTMhelix 332 354\ninside 355 372

15941 GCF_002047685.1_ASM204768v1 Salmonella enterica subsp. enterica serovar 4,5,12:b:- Proteobacteria; Gammaproteobacteria; Enterobacterales; Enterobacteriaceae; Salmonella; Salmonella enterica; Salmonella enterica subsp. enterica MNNEETFYQAMRRKGVTRRSFLKFCSLAATSLGLGAGMTPKIAWALENKPRIPVVWIHGL WP_001058312.1 hydrogenase [Salmonella enterica] Length: 372\nNumber of predicted TMHs: 2\nExp number of AAs in TMHs: 36.51738\nExp number, first 60 AAs: 19.72849\nTotal prob of N-in: 0.96170\nPOSSIBLE N-term signal sequence\ninside 1 19\nTMhelix 20 39\noutside 40 325\nTMhelix 326 348\ninside 349 372

15942 GCF_002034915.1_ASM203491v1 Salmonella enterica subsp. enterica serovar 4,[5],12:i:- Proteobacteria; Gammaproteobacteria; Enterobacterales; Enterobacteriaceae; Salmonella; Salmonella enterica; Salmonella enterica subsp. enterica MNNEETFYQAMRRKGVTRRSFLKFCSLAATSLGLGAGMTPKIAWALENKPRIPVVWIHGL WP_001058312.1 hydrogenase [Salmonella enterica] Length: 372\nNumber of predicted TMHs: 2\nExp number of AAs in TMHs: 36.51738\nExp number, first 60 AAs: 19.72849\nTotal prob of N-in: 0.96170\nPOSSIBLE N-term signal sequence\ninside 1 19\nTMhelix 20 39\noutside 40 325\nTMhelix 326 348\ninside 349 372

15943 GCF_002032235.1_ASM203223v1 Salmonella enterica subsp. enterica serovar 4,[5],12:i:- Proteobacteria; Gammaproteobacteria; Enterobacterales; Enterobacteriaceae; Salmonella; Salmonella enterica; Salmonella enterica subsp. enterica MTGDNTLITSHGINRRDFMKLCAALAATMGLSSKAAAEMAESVSNPQRPPVIWIGAQECT WP_000145429.1 hydrogenase 2 small subunit [Salmonella enterica] Length: 372\nNumber of predicted TMHs: 1\nExp number of AAs in TMHs: 24.51309\nExp number, first 60 AAs: 0.2682\nTotal prob of N-in: 0.14402\noutside 1 331\nTMhelix 332 354\ninside 355 372

15944 GCF_002032225.1_ASM203222v1 Salmonella enterica subsp. enterica serovar 4,[5],12:i:- Proteobacteria; Gammaproteobacteria; Enterobacterales; Enterobacteriaceae; Salmonella; Salmonella enterica; Salmonella enterica subsp. enterica MNNEETFYQAMRRKGVTRRSFLKFCSLAATSLGLGAGMTPKIAWALENKPRIPVVWIHGL WP_001058312.1 hydrogenase [Salmonella enterica] Length: 372\nNumber of predicted TMHs: 2\nExp number of AAs in TMHs: 36.51738\nExp number, first 60 AAs: 19.72849\nTotal prob of N-in: 0.96170\nPOSSIBLE N-term signal sequence\ninside 1 19\nTMhelix 20 39\noutside 40 325\nTMhelix 326 348\ninside 349 372

15945 GCF_001480365.1_Salmonella_enterica_CVM_N41895-SQ_v1.0 Salmonella enterica subsp. enterica serovar 4,[5],12:i:- Proteobacteria; Gammaproteobacteria; Enterobacterales; Enterobacteriaceae; Salmonella; Salmonella enterica; Salmonella enterica subsp. enterica MNNEETFYQAMRRKGVTRRSFLKFCSLAATSLGLGAGMTPKIAWALENKPRIPVVWIHGL WP_001058312.1 hydrogenase [Salmonella enterica] Length: 372\nNumber of predicted TMHs: 2\nExp number of AAs in TMHs: 36.51738\nExp number, first 60 AAs: 19.72849\nTotal prob of N-in: 0.96170\nPOSSIBLE N-term signal sequence\ninside 1 19\nTMhelix 20 39\noutside 40 325\nTMhelix 326 348\ninside 349 372

15946 GCF_002034905.1_ASM203490v1 Salmonella enterica subsp. enterica serovar 4,[5],12:i:- Proteobacteria; Gammaproteobacteria; Enterobacterales; Enterobacteriaceae; Salmonella; Salmonella enterica; Salmonella enterica subsp. enterica MTGDNTLITSHGINRRDFMKLCAALAATMGLSSKAAAEMAESVSNPQRPPVIWIGAQECT WP_000145429.1 hydrogenase 2 small subunit [Salmonella enterica] Length: 372\nNumber of predicted TMHs: 1\nExp number of AAs in TMHs: 24.51309\nExp number, first 60 AAs: 0.2682\nTotal prob of N-in: 0.14402\noutside 1 331\nTMhelix 332 354\ninside 355 372

15947 GCF_001480975.1_Salmonella_enterica_CVM_N40939-SQ_v1.0 Salmonella enterica subsp. enterica serovar 4,[5],12:i:- Proteobacteria; Gammaproteobacteria; Enterobacterales; Enterobacteriaceae; Salmonella; Salmonella enterica; Salmonella enterica subsp. enterica MNNEETFYQAMRRKGVTRRSFLKFCSLAATSLGLGAGMTPKIAWALENKPRIPVVWIHGL WP_001058312.1 hydrogenase [Salmonella enterica] Length: 372\nNumber of predicted TMHs: 2\nExp number of AAs in TMHs: 36.51738\nExp number, first 60 AAs: 19.72849\nTotal prob of N-in: 0.96170\nPOSSIBLE N-term signal sequence\ninside 1 19\nTMhelix 20 39\noutside 40 325\nTMhelix 326 348\ninside 349 372

15948 GCF_002034905.1_ASM203490v1 Salmonella enterica subsp. enterica serovar 4,[5],12:i:- Proteobacteria; Gammaproteobacteria; Enterobacterales; Enterobacteriaceae; Salmonella; Salmonella enterica; Salmonella enterica subsp. enterica MNNEETFYQAMRRKGVTRRSFLKFCSLAATSLGLGAGMTPKIAWALENKPRIPVVWIHGL WP_079812863.1 [Ni/Fe] hydrogenase small subunit [Salmonella enterica] Length: 372\nNumber of predicted TMHs: 2\nExp number of AAs in TMHs: 36.51688\nExp number, first 60 AAs: 19.7285\nTotal prob of N-in: 0.96169\nPOSSIBLE N-term signal sequence\ninside 1 19\nTMhelix 20 39\noutside 40 325\nTMhelix 326 348\ninside 349 372

15949 GCF_000974005.1_ABBSB1218-1.id7_v1.0 Salmonella enterica subsp. enterica serovar 4,[5],12:i:- Proteobacteria; Gammaproteobacteria; Enterobacterales; Enterobacteriaceae; Salmonella; Salmonella enterica;

Salmonella enterica subsp. enterica	MTGDNTLITSHGINRRDFMKLCAALAATMGLSSKAAAEMAESVSNPQRPPVIWIGAQECT	WP_000145429.1	hydrogenase 2 small subunit [Salmonella enterica]	Length: 372\nNumber of predicted TMHs:  1\nExp number of AAs in TMHs: 24.51309\nExp number, first 60 AAs:  0.2682\nTotal prob of N-in:   0.14402\noutside   1   331\nTMhelix   332   354\ninside   355   372

15950	GCF_000973815.1_ABBSB1205-1.id4_v1.0	Salmonella enterica subsp. enterica serovar 4,[5],12:i:-	Proteobacteria; Gammaproteobacteria; Enterobacterales; Enterobacteriaceae; Salmonella; Salmonella enterica; Salmonella enterica subsp. enterica	MTGDNTLITSHGINRRDFMKLCAALAATMGLSSKAAAEMAESVSNPQRPPVIWIGAQECT	WP_000145429.1	hydrogenase 2 small subunit [Salmonella enterica]	Length: 372\nNumber of predicted TMHs:  1\nExp number of AAs in TMHs: 24.51309\nExp number, first 60 AAs:  0.2682\nTotal prob of N-in:   0.14402\noutside   1   331\nTMhelix   332   354\ninside   355   372

15951	GCF_001480365.1_Salmonella_enterica_CVM_N41895-SQ_v1.0	Salmonella enterica subsp. enterica serovar 4,[5],12:i:-	Proteobacteria; Gammaproteobacteria; Enterobacterales; Enterobacteriaceae; Salmonella; Salmonella enterica; Salmonella enterica subsp. enterica	MTGDNTLITSHGINRRDFMKLCAALAATMGLSSKAAAEMAESVSNPQRPPVIWIGAQECT	WP_000145429.1	hydrogenase 2 small subunit [Salmonella enterica]	Length: 372\nNumber of predicted TMHs:  1\nExp number of AAs in TMHs: 24.51309\nExp number, first 60 AAs:  0.2682\nTotal prob of N-in:   0.14402\noutside   1   331\nTMhelix   332   354\ninside   355   372

15952	GCF_002032225.1_ASM203222v1	Salmonella enterica subsp. enterica serovar 4,[5],12:i:-	Proteobacteria; Gammaproteobacteria; Enterobacterales; Enterobacteriaceae; Salmonella; Salmonella enterica; Salmonella enterica subsp. enterica	MTGDNTLITSHGINRRDFMKLCAALAATMGLSSKAAAEMAESVSNPQRPPVIWIGAQECT	WP_000145429.1	hydrogenase 2 small subunit [Salmonella enterica]	Length: 372\nNumber of predicted TMHs:  1\nExp number of AAs in TMHs: 24.51309\nExp number, first 60 AAs:  0.2682\nTotal prob of N-in:   0.14402\noutside   1   331\nTMhelix   332   354\ninside   355   372

15953	GCF_002032235.1_ASM203223v1	Salmonella enterica subsp. enterica serovar 4,[5],12:i:-	Proteobacteria; Gammaproteobacteria; Enterobacterales; Enterobacteriaceae; Salmonella; Salmonella enterica; Salmonella enterica subsp. enterica	MNNEETFYQAMRRKGVTRRSFLKFCSLAATSLGLGAGMTPKIAWALENKPRIPVVWIHGL	WP_001058312.1	hydrogenase [Salmonella enterica]	Length: 372\nNumber of predicted TMHs:  2\nExp number of AAs in TMHs: 36.51738\nExp number, first 60 AAs:  19.72849\nTotal prob of N-in:   0.96170\nPOSSIBLE N-term signal sequence\ninside   1   19\nTMhelix   20   39\noutside   40   325\nTMhelix   326   348\ninside   349   372

15954	GCF_002034975.1_ASM203497v1	Salmonella enterica subsp. enterica serovar 4,[5],12:i:-	Proteobacteria; Gammaproteobacteria; Enterobacterales; Enterobacteriaceae; Salmonella; Salmonella enterica; Salmonella enterica subsp. enterica	MNNEETFYQAMRRKGVTRRSFLKFCSLAATSLGLGAGMTPKIAWALENKPRIPVVWIHGL	WP_001058312.1	hydrogenase [Salmonella enterica]	Length: 372\nNumber of predicted TMHs:  2\nExp number of AAs in TMHs: 36.51738\nExp number, first 60 AAs:  19.72849\nTotal prob of N-in:   0.96170\nPOSSIBLE N-term signal sequence\ninside   1   19\nTMhelix   20   39\noutside   40   325\nTMhelix   326   348\ninside   349   372

15955	GCF_002034975.1_ASM203497v1	Salmonella enterica subsp. enterica serovar 4,[5],12:i:-	Proteobacteria; Gammaproteobacteria; Enterobacterales; Enterobacteriaceae; Salmonella; Salmonella enterica; Salmonella enterica subsp. enterica	MTGDNTLITSHGINRRDFMKLCAALAATMGLSSKAAAEMAESVSNPQRPPVIWIGAQECT	WP_000145429.1	hydrogenase 2 small subunit [Salmonella enterica]	Length: 372\nNumber of predicted TMHs:  1\nExp number of AAs in TMHs: 24.51309\nExp number, first 60 AAs:  0.2682\nTotal prob of N-in:   0.14402\noutside   1   331\nTMhelix   332   354\ninside   355   372

15956	GCF_002034915.1_ASM203491v1	Salmonella enterica subsp. enterica serovar 4,[5],12:i:-	Proteobacteria; Gammaproteobacteria; Enterobacterales; Enterobacteriaceae; Salmonella; Salmonella enterica; Salmonella enterica subsp. enterica	MTGDNTLITSHGINRRDFMKLCAALAATMGLSSKAAAEMAESVSNPQRPPVIWIGAQECT	WP_000145429.1	hydrogenase 2 small subunit [Salmonella enterica]	Length: 372\nNumber of predicted TMHs:  1\nExp number of AAs in TMHs: 24.51309\nExp number, first 60 AAs:  0.2682\nTotal prob of N-in:   0.14402\noutside   1   331\nTMhelix   332   354\ninside   355   372

15957	GCF_002066795.1_ASM206679v1	Salmonella enterica subsp. enterica serovar 4,[5],12:i:-	Proteobacteria; Gammaproteobacteria; Enterobacterales; Enterobacteriaceae; Salmonella; Salmonella enterica; Salmonella enterica subsp. enterica	MTGDNTLITSHGINRRDFMKLCAALAATMGLSSKAAAEMAESVSNPQRPPVIWIGAQECT	WP_000145429.1	hydrogenase 2 small subunit [Salmonella enterica]	Length: 372\nNumber of predicted TMHs:  1\nExp number of AAs in TMHs: 24.51309\nExp number, first 60 AAs:  0.2682\nTotal prob of N-in:   0.14402\noutside   1   331\nTMhelix   332   354\ninside   355   372

15958 GCF_002066795.1_ASM206679v1 Salmonella enterica subsp. enterica serovar 4,[5],12:i:- Proteobacteria; Gammaproteobacteria; Enterobacterales; Enterobacteriaceae; Salmonella; Salmonella enterica; Salmonella enterica subsp. enterica MNNEETFYQAMRRKGVTRRSFLKFCSLAATSLGLGAGMTPKIAWALENKPRIPVVWIHGL WP_001058312.1 hydrogenase [Salmonella enterica] Length: 372\nNumber of predicted TMHs: 2\nExp number of AAs in TMHs: 36.51738\nExp number, first 60 AAs: 19.72849\nTotal prob of N-in: 0.96170\nPOSSIBLE N-term signal sequence\ninside 1 19\nTMhelix 20 39\noutside 40 325\nTMhelix 326 348\ninside 349 372

15959 GCF_000974005.1_ABBSB1218-1.id7_v1.0 Salmonella enterica subsp. enterica serovar 4,[5],12:i:- Proteobacteria; Gammaproteobacteria; Enterobacterales; Enterobacteriaceae; Salmonella; Salmonella enterica; Salmonella enterica subsp. enterica MNNEETFYQAMRRKGVTRRSFLKFCSLAATSLGLGAGMTPKIAWALENKPRIPVVWIHGL WP_001058312.1 hydrogenase [Salmonella enterica] Length: 372\nNumber of predicted TMHs: 2\nExp number of AAs in TMHs: 36.51738\nExp number, first 60 AAs: 19.72849\nTotal prob of N-in: 0.96170\nPOSSIBLE N-term signal sequence\ninside 1 19\nTMhelix 20 39\noutside 40 325\nTMhelix 326 348\ninside 349 372

15960 GCF_000974035.1_SALI-436-3.id6_v1.0 Salmonella enterica subsp. enterica serovar 4,[5],12:i:- Proteobacteria; Gammaproteobacteria; Enterobacterales; Enterobacteriaceae; Salmonella; Salmonella enterica; Salmonella enterica subsp. enterica MTGDNTLITSHGINRRDFMKLCAALAATMGLSSKAAAEMAESVSNPQRPPVIWIGAQECT WP_000145429.1 hydrogenase 2 small subunit [Salmonella enterica] Length: 372\nNumber of predicted TMHs: 1\nExp number of AAs in TMHs: 24.51309\nExp number, first 60 AAs: 0.2682\nTotal prob of N-in: 0.14402\noutside 1 331\nTMhelix 332 354\ninside 355 372

15961 GCF_000974035.1_SALI-436-3.id6_v1.0 Salmonella enterica subsp. enterica serovar 4,[5],12:i:- Proteobacteria; Gammaproteobacteria; Enterobacterales; Enterobacteriaceae; Salmonella; Salmonella enterica; Salmonella enterica subsp. enterica MNNEETFYQAMRRKGVTRRSFLKFCSLAATSLGLGAGMTPKIAWALENKPRIPVVWIHGL WP_001058312.1 hydrogenase [Salmonella enterica] Length: 372\nNumber of predicted TMHs: 2\nExp number of AAs in TMHs: 36.51738\nExp number, first 60 AAs: 19.72849\nTotal prob of N-in: 0.96170\nPOSSIBLE N-term signal sequence\ninside 1 19\nTMhelix 20 39\noutside 40 325\nTMhelix 326 348\ninside 349 372

15962 GCF_000973815.1_ABBSB1205-1.id4_v1.0 Salmonella enterica subsp. enterica serovar 4,[5],12:i:- Proteobacteria; Gammaproteobacteria; Enterobacterales; Enterobacteriaceae; Salmonella; Salmonella enterica; Salmonella enterica subsp. enterica MNNEETFYQAMRRKGVTRRSFLKFCSLAATSLGLGAGMTPKIAWALENKPRIPVVWIHGL WP_001058312.1 hydrogenase [Salmonella enterica] Length: 372\nNumber of predicted TMHs: 2\nExp number of AAs in TMHs: 36.51738\nExp number, first 60 AAs: 19.72849\nTotal prob of N-in: 0.96170\nPOSSIBLE N-term signal sequence\ninside 1 19\nTMhelix 20 39\noutside 40 325\nTMhelix 326 348\ninside 349 372

15963 GCF_001480975.1_Salmonella_enterica_CVM_N40939-SQ_v1.0 Salmonella enterica subsp. enterica serovar 4,[5],12:i:- Proteobacteria; Gammaproteobacteria; Enterobacterales; Enterobacteriaceae; Salmonella; Salmonella enterica; Salmonella enterica subsp. enterica MTGDNTLITSHGINRRDFMKLCAALAATMGLSSKAAAEMAESVSNPQRPPVIWIGAQECT WP_000145429.1 hydrogenase 2 small subunit [Salmonella enterica] Length: 372\nNumber of predicted TMHs: 1\nExp number of AAs in TMHs: 24.51309\nExp number, first 60 AAs: 0.2682\nTotal prob of N-in: 0.14402\noutside 1 331\nTMhelix 332 354\ninside 355 372

15964 GCF_002034965.1_ASM203496v1 Salmonella enterica subsp. enterica serovar 4,[5],12:i:- Proteobacteria; Gammaproteobacteria; Enterobacterales; Enterobacteriaceae; Salmonella; Salmonella enterica; Salmonella enterica subsp. enterica MNNEETFYQAMRRKGVTRRSFLKFCSLAATSLGLGAGMTPKIAWALENKPRIPVVWIHGL WP_001058312.1 hydrogenase [Salmonella enterica] Length: 372\nNumber of predicted TMHs: 2\nExp number of AAs in TMHs: 36.51738\nExp number, first 60 AAs: 19.72849\nTotal prob of N-in: 0.96170\nPOSSIBLE N-term signal sequence\ninside 1 19\nTMhelix 20 39\noutside 40 325\nTMhelix 326 348\ninside 349 372

15965 GCF_001479815.1_Salmonella_enterica_CVM_N38855-SQ_v1.0 Salmonella enterica subsp. enterica serovar 4,[5],12:i:- Proteobacteria; Gammaproteobacteria; Enterobacterales; Enterobacteriaceae; Salmonella; Salmonella enterica; Salmonella enterica subsp. enterica MNNEETFYQAMRRKGVTRRSFLKFCSLAATSLGLGAGMTPKIAWALENKPRIPVVWIHGL WP_001058312.1 hydrogenase [Salmonella enterica] Length: 372\nNumber of predicted TMHs: 2\nExp number of AAs in TMHs: 36.51738\nExp number, first 60 AAs: 19.72849\nTotal prob of N-in: 0.96170\nPOSSIBLE N-term signal sequence\ninside 1 19\nTMhelix 20 39\noutside 40 325\nTMhelix 326 348\ninside 349 372

15966 GCF_001479815.1_Salmonella_enterica_CVM_N38855-SQ_v1.0 Salmonella enterica subsp. enterica serovar 4,[5],12:i:- Proteobacteria; Gammaproteobacteria; Enterobacterales; Enterobacteriaceae; Salmonella; Salmonella enterica; Salmonella enterica subsp. enterica

MTGDNTLITSHGINRRDFMKLCAALAATMGLSSKAAAEMAESVSNPQRPPVIWIGAQECT WP_000145429.1 hydrogenase 2 small subunit [Salmonella enterica] Length: 372\nNumber of predicted TMHs: 1\nExp number of AAs in TMHs: 24.51309\nExp number, first 60 AAs: 0.2682\nTotal prob of N-in: 0.14402\noutside 1 331\nTMhelix 332 354\ninside 355 372

15967 GCF_002034965.1_ASM203496v1 Salmonella enterica subsp. enterica serovar 4,[5],12:i:- Proteobacteria; Gammaproteobacteria; Enterobacterales; Enterobacteriaceae; Salmonella; Salmonella enterica; Salmonella enterica subsp. enterica MTGDNTLITSHGINRRDFMKLCAALAATMGLSSKAAAEMAESVSNPQRPPVIWIGAQECT WP_000145429.1 hydrogenase 2 small subunit [Salmonella enterica] Length: 372\nNumber of predicted TMHs: 1\nExp number of AAs in TMHs: 24.51309\nExp number, first 60 AAs: 0.2682\nTotal prob of N-in: 0.14402\noutside 1 331\nTMhelix 332 354\ninside 355 372

15968 GCF_002106195.1_ASM210619v1 Salmonella enterica subsp. enterica serovar 4,[5],12:i:- Proteobacteria; Gammaproteobacteria; Enterobacterales; Enterobacteriaceae; Salmonella; Salmonella enterica; Salmonella enterica subsp. enterica MTGDNTLITSHGINRRDFMKLCAALAATMGLSSKAAAEMAESVSNPQRPPVIWIGAQECT WP_000145429.1 hydrogenase 2 small subunit [Salmonella enterica] Length: 372\nNumber of predicted TMHs: 1\nExp number of AAs in TMHs: 24.51309\nExp number, first 60 AAs: 0.2682\nTotal prob of N-in: 0.14402\noutside 1 331\nTMhelix 332 354\ninside 355 372

15969 GCF_002106195.1_ASM210619v1 Salmonella enterica subsp. enterica serovar 4,[5],12:i:- Proteobacteria; Gammaproteobacteria; Enterobacterales; Enterobacteriaceae; Salmonella; Salmonella enterica; Salmonella enterica subsp. enterica MNNEETFYQAMRRKGVTRRSFLKFCSLAATSLGLGAGMTPKIAWALENKPRIPVVWIHGL WP_001058312.1 hydrogenase [Salmonella enterica] Length: 372\nNumber of predicted TMHs: 2\nExp number of AAs in TMHs: 36.51738\nExp number, first 60 AAs: 19.72849\nTotal prob of N-in: 0.96170\nPOSSIBLE N-term signal sequence\ninside 1 19\nTMhelix 20 39\noutside 40 325\nTMhelix 326 348\ninside 349 372

15970 GCF_002063225.1_ASM206322v1 Salmonella enterica subsp. enterica serovar 4,[5],12:i:- Proteobacteria; Gammaproteobacteria; Enterobacterales; Enterobacteriaceae; Salmonella; Salmonella enterica; Salmonella enterica subsp. enterica MNNEETFYQAMRRKGVTRRSFLKFCSLAATSLGLGAGMTPKIAWALENKPRIPVVWIHGL WP_001058312.1 hydrogenase [Salmonella enterica] Length: 372\nNumber of predicted TMHs: 2\nExp number of AAs in TMHs: 36.51738\nExp number, first 60 AAs: 19.72849\nTotal prob of N-in: 0.96170\nPOSSIBLE N-term signal sequence\ninside 1 19\nTMhelix 20 39\noutside 40 325\nTMhelix 326 348\ninside 349 372

15971 GCF_001245735.1_Salmonella_enterica_CVM_N50442_v1.0 Salmonella enterica subsp. enterica serovar 4,5,12:i- Proteobacteria; Gammaproteobacteria; Enterobacterales; Enterobacteriaceae; Salmonella; Salmonella enterica; Salmonella enterica subsp. enterica MTGDNTLITSHGINRRDFMKLCAALAATMGLSSKAAAEMAESVSNPQRPPVIWIGAQECT WP_000145429.1 hydrogenase 2 small subunit [Salmonella enterica] Length: 372\nNumber of predicted TMHs: 1\nExp number of AAs in TMHs: 24.51309\nExp number, first 60 AAs: 0.2682\nTotal prob of N-in: 0.14402\noutside 1 331\nTMhelix 332 354\ninside 355 372

15972 GCF_001245735.1_Salmonella_enterica_CVM_N50442_v1.0 Salmonella enterica subsp. enterica serovar 4,5,12:i- Proteobacteria; Gammaproteobacteria; Enterobacterales; Enterobacteriaceae; Salmonella; Salmonella enterica; Salmonella enterica subsp. enterica MNNEETFYQAMRRKGVTRRSFLKFCSLAATSLGLGAGMTPKIAWALENKPRIPVVWIHGL WP_001058312.1 hydrogenase [Salmonella enterica] Length: 372\nNumber of predicted TMHs: 2\nExp number of AAs in TMHs: 36.51738\nExp number, first 60 AAs: 19.72849\nTotal prob of N-in: 0.96170\nPOSSIBLE N-term signal sequence\ninside 1 19\nTMhelix 20 39\noutside 40 325\nTMhelix 326 348\ninside 349 372

15973 GCF_001245255.1_Salmonella_enterica_CVM_N50420_v1.0 Salmonella enterica subsp. enterica serovar 4,5,12:i- Proteobacteria; Gammaproteobacteria; Enterobacterales; Enterobacteriaceae; Salmonella; Salmonella enterica; Salmonella enterica subsp. enterica MTGDNTLITSHGINRRDFMKLCAALAATMGLSSKAAAEMAESVSNPQRPPVIWIGAQECT WP_000145429.1 hydrogenase 2 small subunit [Salmonella enterica] Length: 372\nNumber of predicted TMHs: 1\nExp number of AAs in TMHs: 24.51309\nExp number, first 60 AAs: 0.2682\nTotal prob of N-in: 0.14402\noutside 1 331\nTMhelix 332 354\ninside 355 372

15974 GCF_001245255.1_Salmonella_enterica_CVM_N50420_v1.0 Salmonella enterica subsp. enterica serovar 4,5,12:i- Proteobacteria; Gammaproteobacteria; Enterobacterales; Enterobacteriaceae; Salmonella; Salmonella enterica; Salmonella enterica subsp. enterica MNNEETFYQAMRRKGVTRRSFLKFCSLAATSLGLGAGMTPKIAWALENKPRIPVVWIHGL WP_001058312.1 hydrogenase [Salmonella enterica] Length: 372\nNumber of predicted TMHs: 2\nExp number of AAs in TMHs: 36.51738\nExp number, first 60 AAs: 19.72849\nTotal prob of N-in: 0.96170\nPOSSIBLE N-term signal sequence\ninside 1 19\nTMhelix 20 39\noutside 40 325\nTMhelix 326 348\ninside 349 372

15975 GCF_001245865.1_Salmonella_enterica_CVM_N51240_v1.0 Salmonella enterica subsp. enterica serovar 4,5,12:i- Proteobacteria; Gammaproteobacteria; Enterobacterales; Enterobacteriaceae; Salmonella; Salmonella enterica; Salmonella enterica subsp. enterica MTGDNTLITSHGINRRDFMKLCAALAATMGLSSKAAAEMAESVSNPQRPPVIWIGAQECT WP_000145429.1 hydrogenase 2 small subunit [Salmonella enterica] Length: 372\nNumber of predicted TMHs: 1\nExp number of AAs in TMHs: 24.51309\nExp number, first 60 AAs: 0.2682\nTotal prob of N-in: 0.14402\noutside 1 331\nTMhelix 332 354\ninside 355 372

15976 GCF_001245765.1_Salmonella_enterica_CVM_N50445_v1.0 Salmonella enterica subsp. enterica serovar 4,5,12:i- Proteobacteria; Gammaproteobacteria; Enterobacterales; Enterobacteriaceae; Salmonella; Salmonella enterica; Salmonella enterica subsp. enterica MTGDNTLITSHGINRRDFMKLCAALAATMGLSSKAAAEMAESVSNPQRPPVIWIGAQECT WP_000145429.1 hydrogenase 2 small subunit [Salmonella enterica] Length: 372\nNumber of predicted TMHs: 1\nExp number of AAs in TMHs: 24.51309\nExp number, first 60 AAs: 0.2682\nTotal prob of N-in: 0.14402\noutside 1 331\nTMhelix 332 354\ninside 355 372

15977 GCF_001245765.1_Salmonella_enterica_CVM_N50445_v1.0 Salmonella enterica subsp. enterica serovar 4,5,12:i- Proteobacteria; Gammaproteobacteria; Enterobacterales; Enterobacteriaceae; Salmonella; Salmonella enterica; Salmonella enterica subsp. enterica MNNEETFYQAMRRKGVTRRSFLKFCSLAATSLGLGAGMTPKIAWALENKPRIPVVWIHGL WP_001058312.1 hydrogenase [Salmonella enterica] Length: 372\nNumber of predicted TMHs: 2\nExp number of AAs in TMHs: 36.51738\nExp number, first 60 AAs: 19.72849\nTotal prob of N-in: 0.96170\nPOSSIBLE N-term signal sequence\ninside 1 19\nTMhelix 20 39\noutside 40 325\nTMhelix 326 348\ninside 349 372

15978 GCF_001246945.1_Salmonella_enterica_CVM_N51287_v1.0 Salmonella enterica subsp. enterica serovar 4,5,12:i- Proteobacteria; Gammaproteobacteria; Enterobacterales; Enterobacteriaceae; Salmonella; Salmonella enterica; Salmonella enterica subsp. enterica MNNEETFYQAMRRKGVTRRSFLKFCSLAATSLGLGAGMTPKIAWALENKPRIPVVWIHGL WP_001058312.1 hydrogenase [Salmonella enterica] Length: 372\nNumber of predicted TMHs: 2\nExp number of AAs in TMHs: 36.51738\nExp number, first 60 AAs: 19.72849\nTotal prob of N-in: 0.96170\nPOSSIBLE N-term signal sequence\ninside 1 19\nTMhelix 20 39\noutside 40 325\nTMhelix 326 348\ninside 349 372

15979 GCF_001246945.1_Salmonella_enterica_CVM_N51287_v1.0 Salmonella enterica subsp. enterica serovar 4,5,12:i- Proteobacteria; Gammaproteobacteria; Enterobacterales; Enterobacteriaceae; Salmonella; Salmonella enterica; Salmonella enterica subsp. enterica MTGDNTLITSHGINRRDFMKLCAALAATMGLSSKAAAEMAESVSNPQRPPVIWIGAQECT WP_000145429.1 hydrogenase 2 small subunit [Salmonella enterica] Length: 372\nNumber of predicted TMHs: 1\nExp number of AAs in TMHs: 24.51309\nExp number, first 60 AAs: 0.2682\nTotal prob of N-in: 0.14402\noutside 1 331\nTMhelix 332 354\ninside 355 372

15980 GCF_001245865.1_Salmonella_enterica_CVM_N51240_v1.0 Salmonella enterica subsp. enterica serovar 4,5,12:i- Proteobacteria; Gammaproteobacteria; Enterobacterales; Enterobacteriaceae; Salmonella; Salmonella enterica; Salmonella enterica subsp. enterica MNNEETFYQAMRRKGVTRRSFLKFCSLAATSLGLGAGMTPKIAWALENKPRIPVVWIHGL WP_001058312.1 hydrogenase [Salmonella enterica] Length: 372\nNumber of predicted TMHs: 2\nExp number of AAs in TMHs: 36.51738\nExp number, first 60 AAs: 19.72849\nTotal prob of N-in: 0.96170\nPOSSIBLE N-term signal sequence\ninside 1 19\nTMhelix 20 39\noutside 40 325\nTMhelix 326 348\ninside 349 372

15981 GCF_002037285.1_ASM203728v1 Salmonella enterica subsp. enterica serovar 4,[5],12:r:- Proteobacteria; Gammaproteobacteria; Enterobacterales; Enterobacteriaceae; Salmonella; Salmonella enterica; Salmonella enterica subsp. enterica MTGDNTLITSHGINRRDFMKLCAALAATMGLSSKAAAEMAESVSNPQRPPVIWIGAQECT WP_000145429.1 hydrogenase 2 small subunit [Salmonella enterica] Length: 372\nNumber of predicted TMHs: 1\nExp number of AAs in TMHs: 24.51309\nExp number, first 60 AAs: 0.2682\nTotal prob of N-in: 0.14402\noutside 1 331\nTMhelix 332 354\ninside 355 372

15982 GCF_002033205.1_ASM203320v1 Salmonella enterica subsp. enterica serovar 4,[5],12:r:- Proteobacteria; Gammaproteobacteria; Enterobacterales; Enterobacteriaceae; Salmonella; Salmonella enterica; Salmonella enterica subsp. enterica MNNEETFYQAMRRKGVTRRSFLKFCSLAATSLGLGAGMTPKIAWALENKPRIPVVWIHGL WP_001058312.1 hydrogenase [Salmonella enterica] Length: 372\nNumber of predicted TMHs: 2\nExp number of AAs in TMHs: 36.51738\nExp number, first 60 AAs: 19.72849\nTotal prob of N-in: 0.96170\nPOSSIBLE N-term signal sequence\ninside 1 19\nTMhelix 20 39\noutside 40 325\nTMhelix 326 348\ninside 349 372

15983 GCF_002032095.1_ASM203209v1 Salmonella enterica subsp. enterica serovar 4,[5],12:r:- Proteobacteria; Gammaproteobacteria; Enterobacterales; Enterobacteriaceae; Salmonella; Salmonella enterica; Salmonella enterica subsp. enterica MNNEETFYQAMRRKGVTRRSFLKFCSLAATSLGLGAGMTPKIAWALENKPRIPVVWIHGL WP_001058312.1 hydrogenase [Salmonella enterica] Length: 372\nNumber of predicted TMHs: 2\nExp

number of AAs in TMHs: 36.51738\nExp number, first 60 AAs: 19.72849\nTotal prob of N-in: 0.96170\nPOSSIBLE N-term signal sequence\ninside 1 19\nTMhelix 20 39\noutside 40 325\nTMhelix 326 348\ninside 349 372

15984 GCF_002032095.1_ASM203209v1 Salmonella enterica subsp. enterica serovar 4,[5],12:r:- Proteobacteria; Gammaproteobacteria; Enterobacterales; Enterobacteriaceae; Salmonella; Salmonella enterica; Salmonella enterica subsp. enterica MTGDNTLITSHGINRRDFMKLCAALAATMGLSSKAAAEMAESVSNPQRPPVIWIGAQECT WP_000145429.1 hydrogenase 2 small subunit [Salmonella enterica] Length: 372\nNumber of predicted TMHs: 1\nExp number of AAs in TMHs: 24.51309\nExp number, first 60 AAs: 0.2682\nTotal prob of N-in: 0.14402\noutside 1 331\nTMhelix 332 354\ninside 355 372

15985 GCF_002032395.1_ASM203239v1 Salmonella enterica subsp. enterica serovar 4,[5],12:r:- Proteobacteria; Gammaproteobacteria; Enterobacterales; Enterobacteriaceae; Salmonella; Salmonella enterica; Salmonella enterica subsp. enterica MNNEETFYQAMRRKGVTRRSFLKFCSLAATSLGLGAGMTPKIAWALENKPRIPVVWIHGL WP_001058312.1 hydrogenase [Salmonella enterica] Length: 372\nNumber of predicted TMHs: 2\nExp number of AAs in TMHs: 36.51738\nExp number, first 60 AAs: 19.72849\nTotal prob of N-in: 0.96170\nPOSSIBLE N-term signal sequence\ninside 1 19\nTMhelix 20 39\noutside 40 325\nTMhelix 326 348\ninside 349 372

15986 GCF_002037285.1_ASM203728v1 Salmonella enterica subsp. enterica serovar 4,[5],12:r:- Proteobacteria; Gammaproteobacteria; Enterobacterales; Enterobacteriaceae; Salmonella; Salmonella enterica; Salmonella enterica subsp. enterica MNNEETFYQAMRRKGVTRRSFLKFCSLAATSLGLGAGMTPKIAWALENKPRIPVVWIHGL WP_001058312.1 hydrogenase [Salmonella enterica] Length: 372\nNumber of predicted TMHs: 2\nExp number of AAs in TMHs: 36.51738\nExp number, first 60 AAs: 19.72849\nTotal prob of N-in: 0.96170\nPOSSIBLE N-term signal sequence\ninside 1 19\nTMhelix 20 39\noutside 40 325\nTMhelix 326 348\ninside 349 372

15987 GCF_002033205.1_ASM203320v1 Salmonella enterica subsp. enterica serovar 4,[5],12:r:- Proteobacteria; Gammaproteobacteria; Enterobacterales; Enterobacteriaceae; Salmonella; Salmonella enterica; Salmonella enterica subsp. enterica MTGDNTLITSHGINRRDFMKLCAALAATMGLSSKAAAEMAESVSNPQRPPVIWIGAQECT WP_000145429.1 hydrogenase 2 small subunit [Salmonella enterica] Length: 372\nNumber of predicted TMHs: 1\nExp number of AAs in TMHs: 24.51309\nExp number, first 60 AAs: 0.2682\nTotal prob of N-in: 0.14402\noutside 1 331\nTMhelix 332 354\ninside 355 372

15988 GCF_002032395.1_ASM203239v1 Salmonella enterica subsp. enterica serovar 4,[5],12:r:- Proteobacteria; Gammaproteobacteria; Enterobacterales; Enterobacteriaceae; Salmonella; Salmonella enterica; Salmonella enterica subsp. enterica MTGDNTLITSHGINRRDFMKLCAALAATMGLSSKAAAEMAESVSNPQRPPVIWIGAQECT WP_000145429.1 hydrogenase 2 small subunit [Salmonella enterica] Length: 372\nNumber of predicted TMHs: 1\nExp number of AAs in TMHs: 24.51309\nExp number, first 60 AAs: 0.2682\nTotal prob of N-in: 0.14402\noutside 1 331\nTMhelix 332 354\ninside 355 372

15989 GCF_002037435.1_ASM203743v1 Salmonella enterica subsp. enterica serovar 4,[5],12:r:- Proteobacteria; Gammaproteobacteria; Enterobacterales; Enterobacteriaceae; Salmonella; Salmonella enterica; Salmonella enterica subsp. enterica MTGDNTLITSHGINRRDFMKLCAALAATMGLSSKAAAEMAESVSNPQRPPVIWIGAQECT WP_000145429.1 hydrogenase 2 small subunit [Salmonella enterica] Length: 372\nNumber of predicted TMHs: 1\nExp number of AAs in TMHs: 24.51309\nExp number, first 60 AAs: 0.2682\nTotal prob of N-in: 0.14402\noutside 1 331\nTMhelix 332 354\ninside 355 372

15990 GCF_002105885.1_ASM210588v1 Salmonella enterica subsp. enterica serovar 6,7:-:1,5 Proteobacteria; Gammaproteobacteria; Enterobacterales; Enterobacteriaceae; Salmonella; Salmonella enterica; Salmonella enterica subsp. enterica MNNEETFYQAMRRKGVTRRSFLKFCSLAATSLGLGAGMTPKIAWALENKPRIPVVWIHGL WP_001058310.1 hydrogenase [Salmonella enterica] Length: 372\nNumber of predicted TMHs: 2\nExp number of AAs in TMHs: 37.4567299999999\nExp number, first 60 AAs: 19.72694\nTotal prob of N-in: 0.96171\nPOSSIBLE N-term signal sequence\ninside 1 19\nTMhelix 20 39\noutside 40 325\nTMhelix 326 348\ninside 349 372

15991 GCF_002105885.1_ASM210588v1 Salmonella enterica subsp. enterica serovar 6,7:-:1,5 Proteobacteria; Gammaproteobacteria; Enterobacterales; Enterobacteriaceae; Salmonella; Salmonella enterica; Salmonella enterica subsp. enterica MTGDNTLITSHGINRRDFMKLCAALAATMGLSSKAAAEMAESVSNPQRPPVIWIGAQECT WP_000145429.1 hydrogenase 2 small subunit [Salmonella enterica] Length: 372\nNumber of predicted TMHs: 1\nExp number of AAs in TMHs: 24.51309\nExp number, first 60 AAs: 0.2682\nTotal prob of N-in: 0.14402\noutside 1 331\nTMhelix 332 354\ninside 355 372

15992 GCF_002031785.1_ASM203178v1 Salmonella enterica subsp. enterica serovar 6,7:k:- Proteobacteria; Gammaproteobacteria; Enterobacterales; Enterobacteriaceae; Salmonella; Salmonella enterica; Salmonella enterica subsp. enterica MTGDNTLITSHGINRRDFMKLCAALAATMGLSSKAAAEMAESVSNPQRPPVIWIGAQECT WP_000145429.1 hydrogenase 2 small subunit [Salmonella enterica] Length: 372\nNumber of predicted TMHs: 1\nExp number of AAs in TMHs: 24.51309\nExp number, first 60 AAs: 0.2682\nTotal prob of N-in: 0.14402\noutside 1 331\nTMhelix 332 354\ninside 355 372

15993 GCF_002031785.1_ASM203178v1 Salmonella enterica subsp. enterica serovar 6,7:k:- Proteobacteria; Gammaproteobacteria; Enterobacterales; Enterobacteriaceae; Salmonella; Salmonella enterica; Salmonella enterica subsp. enterica MNNEETFYQAMRRKGVTRRSFLKFCSLAATSLGLGAGMTPKIAWALENKPRIPVVWIHGL WP_001058310.1 hydrogenase [Salmonella enterica] Length: 372\nNumber of predicted TMHs: 2\nExp number of AAs in TMHs: 37.4567299999999\nExp number, first 60 AAs: 19.72694\nTotal prob of N-in: 0.96171\nPOSSIBLE N-term signal sequence\ninside 1 19\nTMhelix 20 39\noutside 40 325\nTMhelix 326 348\ninside 349 372

15994 GCF_002031605.1_ASM203160v1 Salmonella enterica subsp. enterica serovar 6,7:lw:- Proteobacteria; Gammaproteobacteria; Enterobacterales; Enterobacteriaceae; Salmonella; Salmonella enterica; Salmonella enterica subsp. enterica MTGDNTLITSHGINRRDFMKLCAALAATMGLSSKAAAEMAESVSNPQRPPVIWIGAQECT WP_000145429.1 hydrogenase 2 small subunit [Salmonella enterica] Length: 372\nNumber of predicted TMHs: 1\nExp number of AAs in TMHs: 24.51309\nExp number, first 60 AAs: 0.2682\nTotal prob of N-in: 0.14402\noutside 1 331\nTMhelix 332 354\ninside 355 372

15995 GCF_002031605.1_ASM203160v1 Salmonella enterica subsp. enterica serovar 6,7:lw:- Proteobacteria; Gammaproteobacteria; Enterobacterales; Enterobacteriaceae; Salmonella; Salmonella enterica; Salmonella enterica subsp. enterica MNNEETFYQAMRRKGVTRRSFLKFCSLATTSLGLGAGMTPKIAWALENKPRIPVVWIHGL WP_023259200.1 hydrogenase 1, small subunit [Salmonella enterica] Length: 372\nNumber of predicted TMHs: 2\nExp number of AAs in TMHs: 35.98973\nExp number, first 60 AAs: 19.19893\nTotal prob of N-in: 0.93662\nPOSSIBLE N-term signal sequence\ninside 1 19\nTMhelix 20 39\noutside 40 325\nTMhelix 326 348\ninside 349 372

15996 GCF_002064055.1_ASM206405v1 Salmonella enterica subsp. enterica serovar 6.8:eh:- Proteobacteria; Gammaproteobacteria; Enterobacterales; Enterobacteriaceae; Salmonella; Salmonella enterica; Salmonella enterica subsp. enterica MTGDNTLITSHGINRRDFMKLCAALAATMGLSSKAAAEMAESVSNPQRPPVIWIGAQECT WP_000145429.1 hydrogenase 2 small subunit [Salmonella enterica] Length: 372\nNumber of predicted TMHs: 1\nExp number of AAs in TMHs: 24.51309\nExp number, first 60 AAs: 0.2682\nTotal prob of N-in: 0.14402\noutside 1 331\nTMhelix 332 354\ninside 355 372

15997 GCF_002064055.1_ASM206405v1 Salmonella enterica subsp. enterica serovar 6.8:eh:- Proteobacteria; Gammaproteobacteria; Enterobacterales; Enterobacteriaceae; Salmonella; Salmonella enterica; Salmonella enterica subsp. enterica MNNEETFYQAMRRKGVTRRSFLKFCSLAATSLGLGAGMTPKIAWALENKPRIPVVWIHGL WP_001058312.1 hydrogenase [Salmonella enterica] Length: 372\nNumber of predicted TMHs: 2\nExp number of AAs in TMHs: 36.51738\nExp number, first 60 AAs: 19.72849\nTotal prob of N-in: 0.96170\nPOSSIBLE N-term signal sequence\ninside 1 19\nTMhelix 20 39\noutside 40 325\nTMhelix 326 348\ninside 349 372

15998 GCF_002107335.1_ASM210733v1 Salmonella enterica subsp. enterica serovar 6,8:i:- Proteobacteria; Gammaproteobacteria; Enterobacterales; Enterobacteriaceae; Salmonella; Salmonella enterica; Salmonella enterica subsp. enterica MTGDNTLITSHGINRRDFMKLCAALAATMGLSSKAAAEMAESVSNPQRPPVIWIGAQECT WP_000145429.1 hydrogenase 2 small subunit [Salmonella enterica] Length: 372\nNumber of predicted TMHs: 1\nExp number of AAs in TMHs: 24.51309\nExp number, first 60 AAs: 0.2682\nTotal prob of N-in: 0.14402\noutside 1 331\nTMhelix 332 354\ninside 355 372

15999 GCF_002107335.1_ASM210733v1 Salmonella enterica subsp. enterica serovar 6,8:i:- Proteobacteria; Gammaproteobacteria; Enterobacterales; Enterobacteriaceae; Salmonella; Salmonella enterica; Salmonella enterica subsp. enterica MNNEETFYQAMRRKGVTRRSFLKFCSLAATSLGLGAGMTPKIAWALENKPRIPVVWIHGL WP_001058311.1 hydrogenase [Salmonella enterica] Length: 372\nNumber of predicted TMHs: 2\nExp number of AAs in TMHs: 37.4699899999999\nExp number, first 60 AAs: 19.72699\nTotal prob of N-in: 0.96171\nPOSSIBLE N-term signal sequence\ninside 1 19\nTMhelix 20 39\noutside 40 325\nTMhelix 326 348\ninside 349 372

16000 GCF_001240505.1_Salmonella_enterica_CVM_N43471_v1.0 Salmonella enterica subsp. enterica serovar 8,(20):-:z6 Proteobacteria; Gammaproteobacteria; Enterobacterales; Enterobacteriaceae; Salmonella; Salmonella

enterica; Salmonella enterica subsp. enterica
MNNEETFYQAMRRKGVTRRSFLKFCSLAATSLGLGAGMTPKIAWALENKPRIPVVWIHGL WP_001058311.1 hydrogenase [Salmonella enterica] Length: 372\nNumber of predicted TMHs: 2\nExp number of AAs in TMHs: 37.4699899999999\nExp number, first 60 AAs: 19.72699\nTotal prob of N-in: 0.96171\nPOSSIBLE N-term signal sequence\ninside 1 19\nTMhelix 20 39\noutside 40 325\nTMhelix 326 348\ninside 349 372

16001 GCF_001240505.1_Salmonella_enterica_CVM_N43471_v1.0 Salmonella enterica subsp. enterica serovar 8,(20):-:z6 Proteobacteria; Gammaproteobacteria; Enterobacterales; Enterobacteriaceae; Salmonella; Salmonella enterica; Salmonella enterica subsp. enterica
MTGDNTLITSHGINRRDFMKLCAALAATMGLSSKAAAEMAESVSNPQRPPVIWIGAQECT WP_000145429.1 hydrogenase 2 small subunit [Salmonella enterica] Length: 372\nNumber of predicted TMHs: 1\nExp number of AAs in TMHs: 24.51309\nExp number, first 60 AAs: 0.2682\nTotal prob of N-in: 0.14402\noutside 1 331\nTMhelix 332 354\ninside 355 372

16002 GCF_002044925.1_ASM204492v1 Salmonella enterica subsp. enterica serovar 9,12:l,z28:- Proteobacteria; Gammaproteobacteria; Enterobacterales; Enterobacteriaceae; Salmonella; Salmonella enterica; Salmonella enterica subsp. enterica MTGDNTLITSHGINRRDFMKLCAALAATMGLSSKAAAEMAESVSNPQRPPVIWIGAQECT WP_000145429.1 hydrogenase 2 small subunit [Salmonella enterica] Length: 372\nNumber of predicted TMHs: 1\nExp number of AAs in TMHs: 24.51309\nExp number, first 60 AAs: 0.2682\nTotal prob of N-in: 0.14402\noutside 1 331\nTMhelix 332 354\ninside 355 372

16003 GCF_002044925.1_ASM204492v1 Salmonella enterica subsp. enterica serovar 9,12:l,z28:- Proteobacteria; Gammaproteobacteria; Enterobacterales; Enterobacteriaceae; Salmonella; Salmonella enterica; Salmonella enterica subsp. enterica MNNEETFYQAMRRKGVTRRSFLKFCSLAATSLGLGAGMTPKIAWALENKPRIPVVWIHGL WP_001058312.1 hydrogenase [Salmonella enterica] Length: 372\nNumber of predicted TMHs: 2\nExp number of AAs in TMHs: 36.51738\nExp number, first 60 AAs: 19.72849\nTotal prob of N-in: 0.96170\nPOSSIBLE N-term signal sequence\ninside 1 19\nTMhelix 20 39\noutside 40 325\nTMhelix 326 348\ninside 349 372

16004 GCF_002045195.1_ASM204519v1 Salmonella enterica subsp. enterica serovar 9,12:l,z28:- Proteobacteria; Gammaproteobacteria; Enterobacterales; Enterobacteriaceae; Salmonella; Salmonella enterica; Salmonella enterica subsp. enterica MNNEETFYQAMRRKGVTRRSFLKFCSLAATSLGLGAGMTPKIAWALENKPRIPVVWIHGL WP_001058312.1 hydrogenase [Salmonella enterica] Length: 372\nNumber of predicted TMHs: 2\nExp number of AAs in TMHs: 36.51738\nExp number, first 60 AAs: 19.72849\nTotal prob of N-in: 0.96170\nPOSSIBLE N-term signal sequence\ninside 1 19\nTMhelix 20 39\noutside 40 325\nTMhelix 326 348\ninside 349 372

16005 GCF_002045195.1_ASM204519v1 Salmonella enterica subsp. enterica serovar 9,12:l,z28:- Proteobacteria; Gammaproteobacteria; Enterobacterales; Enterobacteriaceae; Salmonella; Salmonella enterica; Salmonella enterica subsp. enterica MTGDNTLITSHGINRRDFMKLCAALAATMGLSSKAAAEMAESVSNPQRPPVIWIGAQECT WP_000145429.1 hydrogenase 2 small subunit [Salmonella enterica] Length: 372\nNumber of predicted TMHs: 1\nExp number of AAs in TMHs: 24.51309\nExp number, first 60 AAs: 0.2682\nTotal prob of N-in: 0.14402\noutside 1 331\nTMhelix 332 354\ninside 355 372

16006 GCF_002066715.1_ASM206671v1 Salmonella enterica subsp. enterica serovar Abaeetuba Proteobacteria; Gammaproteobacteria; Enterobacterales; Enterobacteriaceae; Salmonella; Salmonella enterica; Salmonella enterica subsp. enterica MNNEETFYQAMRRKGVTRRSFLKFCSLAATSLGLGAGMTPKIAWALENKPRIPVVWIHGL WP_001058312.1 hydrogenase [Salmonella enterica] Length: 372\nNumber of predicted TMHs: 2\nExp number of AAs in TMHs: 36.51738\nExp number, first 60 AAs: 19.72849\nTotal prob of N-in: 0.96170\nPOSSIBLE N-term signal sequence\ninside 1 19\nTMhelix 20 39\noutside 40 325\nTMhelix 326 348\ninside 349 372

16007 GCF_002066715.1_ASM206671v1 Salmonella enterica subsp. enterica serovar Abaeetuba Proteobacteria; Gammaproteobacteria; Enterobacterales; Enterobacteriaceae; Salmonella; Salmonella enterica; Salmonella enterica subsp. enterica MTGDNTLITSHGINRRDFMKLCAALAATMGLSSKAAAEMAESVSNPQRPPVIWIGAQECT WP_000145431.1 hydrogenase 2 small subunit [Salmonella enterica] Length: 372\nNumber of predicted TMHs: 1\nExp number of AAs in TMHs: 24.51505\nExp number, first 60 AAs: 0.2682\nTotal prob of N-in: 0.14406\noutside 1 331\nTMhelix 332 354\ninside 355 372

16008 GCF_001999405.1_ASM199940v1 Salmonella enterica subsp. enterica serovar Adelaide Proteobacteria; Gammaproteobacteria; Enterobacterales; Enterobacteriaceae; Salmonella; Salmonella enterica; Salmonella enterica subsp. enterica MTGDNTLITSHGINRRDFMKLCAALAATMGLSSKAAAEMAESVSNPQRPPVIWIGAQECT WP_000145429.1 hydrogenase 2 small subunit [Salmonella enterica] Length: 372\nNumber of

predicted TMHs:  1\nExp number of AAs in TMHs: 24.51309\nExp number, first 60 AAs:  0.2682\nTotal prob of N-in: 0.14402\noutside   1   331\nTMhelix   332   354\ninside   355   372

16009 GCF_001999405.1_ASM199940v1 Salmonella enterica subsp. enterica serovar Adelaide Proteobacteria; Gammaproteobacteria; Enterobacterales; Enterobacteriaceae; Salmonella; Salmonella enterica; Salmonella enterica subsp. enterica MNNEETFYQAMRRKGVTRRSFLKFCSLAATSLGLGAGMTPKIAWALENKPRIPVVWIHGL WP_001058311.1 hydrogenase [Salmonella enterica] Length: 372\nNumber of predicted TMHs:  2\nExp number of AAs in TMHs: 37.4699899999999\nExp number, first 60 AAs:  19.72699\nTotal prob of N-in: 0.96171\nPOSSIBLE N-term signal sequence\ninside   1   19\nTMhelix   20   39\noutside   40   325\nTMhelix   326   348\ninside   349   372

16010 GCF_002048945.1_ASM204894v1 Salmonella enterica subsp. enterica serovar Adelaide Proteobacteria; Gammaproteobacteria; Enterobacterales; Enterobacteriaceae; Salmonella; Salmonella enterica; Salmonella enterica subsp. enterica MNNEETFYQAMRRKGVTRRSFLKFCSLAATSLGLGAGMTPKIAWALENKPRIPVVWIHGL WP_001058311.1 hydrogenase [Salmonella enterica] Length: 372\nNumber of predicted TMHs:  2\nExp number of AAs in TMHs: 37.4699899999999\nExp number, first 60 AAs:  19.72699\nTotal prob of N-in: 0.96171\nPOSSIBLE N-term signal sequence\ninside   1   19\nTMhelix   20   39\noutside   40   325\nTMhelix   326   348\ninside   349   372

16011 GCF_002048945.1_ASM204894v1 Salmonella enterica subsp. enterica serovar Adelaide Proteobacteria; Gammaproteobacteria; Enterobacterales; Enterobacteriaceae; Salmonella; Salmonella enterica; Salmonella enterica subsp. enterica MTGDNTLITSHGINRRDFMKLCAALAATMGLSSKAAAEMAESVSNPQRPPVIWIGAQECT WP_000145429.1 hydrogenase 2 small subunit [Salmonella enterica] Length: 372\nNumber of predicted TMHs:  1\nExp number of AAs in TMHs: 24.51309\nExp number, first 60 AAs:  0.2682\nTotal prob of N-in: 0.14402\noutside   1   331\nTMhelix   332   354\ninside   355   372

16012 GCF_002049335.1_ASM204933v1 Salmonella enterica subsp. enterica serovar Adelaide Proteobacteria; Gammaproteobacteria; Enterobacterales; Enterobacteriaceae; Salmonella; Salmonella enterica; Salmonella enterica subsp. enterica MNNEETFYQAMRRKGVTRRSFLKFCSLAATSLGLGAGMTPKIAWALENKPRIPVVWIHGL WP_001058311.1 hydrogenase [Salmonella enterica] Length: 372\nNumber of predicted TMHs:  2\nExp number of AAs in TMHs: 37.4699899999999\nExp number, first 60 AAs:  19.72699\nTotal prob of N-in: 0.96171\nPOSSIBLE N-term signal sequence\ninside   1   19\nTMhelix   20   39\noutside   40   325\nTMhelix   326   348\ninside   349   372

16013 GCF_002049305.1_ASM204930v1 Salmonella enterica subsp. enterica serovar Adelaide Proteobacteria; Gammaproteobacteria; Enterobacterales; Enterobacteriaceae; Salmonella; Salmonella enterica; Salmonella enterica subsp. enterica MNNEETFYQAMRRKGVTRRSFLKFCSLAATSLGLGAGMTPKIAWALENKPRIPVVWIHGL WP_001058311.1 hydrogenase [Salmonella enterica] Length: 372\nNumber of predicted TMHs:  2\nExp number of AAs in TMHs: 37.4699899999999\nExp number, first 60 AAs:  19.72699\nTotal prob of N-in: 0.96171\nPOSSIBLE N-term signal sequence\ninside   1   19\nTMhelix   20   39\noutside   40   325\nTMhelix   326   348\ninside   349   372

16014 GCF_002049305.1_ASM204930v1 Salmonella enterica subsp. enterica serovar Adelaide Proteobacteria; Gammaproteobacteria; Enterobacterales; Enterobacteriaceae; Salmonella; Salmonella enterica; Salmonella enterica subsp. enterica MTGDNTLITSHGINRRDFMKLCAALAATMGLSSKAAAEMAESVSNPQRPPVIWIGAQECT WP_000145429.1 hydrogenase 2 small subunit [Salmonella enterica] Length: 372\nNumber of predicted TMHs:  1\nExp number of AAs in TMHs: 24.51309\nExp number, first 60 AAs:  0.2682\nTotal prob of N-in: 0.14402\noutside   1   331\nTMhelix   332   354\ninside   355   372

16015 GCF_002049335.1_ASM204933v1 Salmonella enterica subsp. enterica serovar Adelaide Proteobacteria; Gammaproteobacteria; Enterobacterales; Enterobacteriaceae; Salmonella; Salmonella enterica; Salmonella enterica subsp. enterica MTGDNTLITSHGINRRDFMKLCAALAATMGLSSKAAAEMAESVSNPQRPPVIWIGAQECT WP_000145429.1 hydrogenase 2 small subunit [Salmonella enterica] Length: 372\nNumber of predicted TMHs:  1\nExp number of AAs in TMHs: 24.51309\nExp number, first 60 AAs:  0.2682\nTotal prob of N-in: 0.14402\noutside   1   331\nTMhelix   332   354\ninside   355   372

16016 GCF_002048865.1_ASM204886v1 Salmonella enterica subsp. enterica serovar Adelaide Proteobacteria; Gammaproteobacteria; Enterobacterales; Enterobacteriaceae; Salmonella; Salmonella enterica; Salmonella enterica subsp. enterica MNNEETFYQAMRRKGVTRRSFLKFCSLAATSLGLGAGMTPKIAWALENKPRIPVVWIHGL WP_001058311.1 hydrogenase [Salmonella enterica] Length: 372\nNumber of predicted TMHs:  2\nExp number of AAs in TMHs: 37.4699899999999\nExp number, first 60 AAs:  19.72699\nTotal prob of N-in:

0.96171\nPOSSIBLE N-term signal sequence\ninside 1 19\nTMhelix 20 39\noutside 40 325\nTMhelix 326 348\ninside 349 372

16017 GCF_002048865.1_ASM204886v1 Salmonella enterica subsp. enterica serovar Adelaide Proteobacteria; Gammaproteobacteria; Enterobacterales; Enterobacteriaceae; Salmonella; Salmonella enterica; Salmonella enterica subsp. enterica MTGDNTLITSHGINRRDFMKLCAALAATMGLSSKAAAEMAESVSNPQRPPVIWIGAQECT WP_000145429.1 hydrogenase 2 small subunit [Salmonella enterica] Length: 372\nNumber of predicted TMHs: 1\nExp number of AAs in TMHs: 24.51309\nExp number, first 60 AAs: 0.2682\nTotal prob of N-in: 0.14402\noutside 1 331\nTMhelix 332 354\ninside 355 372

16018 GCF_002046745.1_ASM204674v1 Salmonella enterica subsp. enterica serovar Agbeni Proteobacteria; Gammaproteobacteria; Enterobacterales; Enterobacteriaceae; Salmonella; Salmonella enterica; Salmonella enterica subsp. enterica MTGDNTLITSHGINRRDFMKLCAALAATMGLSSKAAAEMAESVSNPQRPPVIWIGAQECT WP_000145429.1 hydrogenase 2 small subunit [Salmonella enterica] Length: 372\nNumber of predicted TMHs: 1\nExp number of AAs in TMHs: 24.51309\nExp number, first 60 AAs: 0.2682\nTotal prob of N-in: 0.14402\noutside 1 331\nTMhelix 332 354\ninside 355 372

16019 GCF_002046745.1_ASM204674v1 Salmonella enterica subsp. enterica serovar Agbeni Proteobacteria; Gammaproteobacteria; Enterobacterales; Enterobacteriaceae; Salmonella; Salmonella enterica; Salmonella enterica subsp. enterica MNNEETFYQAMRRKGVTRRSFLKFCSLAATSLGLGAGMTPKIAWALENKPRIPVVWIHGL WP_001058312.1 hydrogenase [Salmonella enterica] Length: 372\nNumber of predicted TMHs: 2\nExp number of AAs in TMHs: 36.51738\nExp number, first 60 AAs: 19.72849\nTotal prob of N-in: 0.96170\nPOSSIBLE N-term signal sequence\ninside 1 19\nTMhelix 20 39\noutside 40 325\nTMhelix 326 348\ninside 349 372

16020 GCF_002046615.1_ASM204661v1 Salmonella enterica subsp. enterica serovar Agbeni Proteobacteria; Gammaproteobacteria; Enterobacterales; Enterobacteriaceae; Salmonella; Salmonella enterica; Salmonella enterica subsp. enterica MNNEETFYQAMRRKGVTRRSFLKFCSLAATSLGLGAGMTPKIAWALENKPRIPVVWIHGL WP_001058312.1 hydrogenase [Salmonella enterica] Length: 372\nNumber of predicted TMHs: 2\nExp number of AAs in TMHs: 36.51738\nExp number, first 60 AAs: 19.72849\nTotal prob of N-in: 0.96170\nPOSSIBLE N-term signal sequence\ninside 1 19\nTMhelix 20 39\noutside 40 325\nTMhelix 326 348\ninside 349 372

16021 GCF_002046615.1_ASM204661v1 Salmonella enterica subsp. enterica serovar Agbeni Proteobacteria; Gammaproteobacteria; Enterobacterales; Enterobacteriaceae; Salmonella; Salmonella enterica; Salmonella enterica subsp. enterica MTGDNTLITSHGINRRDFMKLCAALAATMGLSSKAAAEMAESVSNPQRPPVIWIGAQECT WP_000145429.1 hydrogenase 2 small subunit [Salmonella enterica] Length: 372\nNumber of predicted TMHs: 1\nExp number of AAs in TMHs: 24.51309\nExp number, first 60 AAs: 0.2682\nTotal prob of N-in: 0.14402\noutside 1 331\nTMhelix 332 354\ninside 355 372

16022 GCF_002057065.1_ASM205706v1 Salmonella enterica subsp. enterica serovar Agona Proteobacteria; Gammaproteobacteria; Enterobacterales; Enterobacteriaceae; Salmonella; Salmonella enterica; Salmonella enterica subsp. enterica MTGDNTLITSHGINRRDFMKLCAALAATMGLSSKAAAEMAESVSNPQRPPVIWIGAQECT WP_000145429.1 hydrogenase 2 small subunit [Salmonella enterica] Length: 372\nNumber of predicted TMHs: 1\nExp number of AAs in TMHs: 24.51309\nExp number, first 60 AAs: 0.2682\nTotal prob of N-in: 0.14402\noutside 1 331\nTMhelix 332 354\ninside 355 372

16023 GCF_001240875.1_Salmonella_enterica_CVM_N43827_v1.0 Salmonella enterica subsp. enterica serovar Agona Proteobacteria; Gammaproteobacteria; Enterobacterales; Enterobacteriaceae; Salmonella; Salmonella enterica; Salmonella enterica subsp. enterica MTGDNTLITSHGINRRDFMKLCAALAATMGLSSKAAAEMAESVSNPQRPPVIWIGAQECT WP_000145429.1 hydrogenase 2 small subunit [Salmonella enterica] Length: 372\nNumber of predicted TMHs: 1\nExp number of AAs in TMHs: 24.51309\nExp number, first 60 AAs: 0.2682\nTotal prob of N-in: 0.14402\noutside 1 331\nTMhelix 332 354\ninside 355 372

16024 GCF_001240875.1_Salmonella_enterica_CVM_N43827_v1.0 Salmonella enterica subsp. enterica serovar Agona Proteobacteria; Gammaproteobacteria; Enterobacterales; Enterobacteriaceae; Salmonella; Salmonella enterica; Salmonella enterica subsp. enterica MNNEETFYQAMRRKGVTRRSFLKFCSLAATSLGLSAGMTPKIAWALENKPRIPVVWIHGL WP_001058316.1 hydrogenase [Salmonella enterica] Length: 372\nNumber of predicted TMHs: 2\nExp number of AAs in TMHs: 35.76397\nExp number, first 60 AAs: 18.97201\nTotal prob of N-in: 0.92785\nPOSSIBLE N-term signal sequence\ninside 1 19\nTMhelix 20 39\noutside 40 325\nTMhelix 326 348\ninside 349 372

16025 GCF_001479025.1_Salmonella_enterica_CVM_N30701-SQ_v1.0 Salmonella enterica subsp. enterica serovar Agona Proteobacteria; Gammaproteobacteria; Enterobacterales; Enterobacteriaceae; Salmonella; Salmonella enterica;

Salmonella enterica subsp. enterica MNNEETFYQAMRRKGVTRRSFLKFCSLAATSLGLSAGMTPKIAWALENKPRIPVVWIHGL WP_001058316.1 hydrogenase [Salmonella enterica] Length: 372\nNumber of predicted TMHs: 2\nExp number of AAs in TMHs: 35.76397\nExp number, first 60 AAs: 18.97201\nTotal prob of N-in: 0.92785\nPOSSIBLE N-term signal sequence\ninside 1 19\nTMhelix 20 39\noutside 40 325\nTMhelix 326 348\ninside 349 372

16026 GCF_002033225.1_ASM203322v1 Salmonella enterica subsp. enterica serovar Agona Proteobacteria; Gammaproteobacteria; Enterobacterales; Enterobacteriaceae; Salmonella; Salmonella enterica; Salmonella enterica subsp. enterica MTGDNTLITSHGINRRDFMKLCAALAATMGLSSKAAAEMAESVSNPQRPPVIWIGAQECT WP_000145429.1 hydrogenase 2 small subunit [Salmonella enterica] Length: 372\nNumber of predicted TMHs: 1\nExp number of AAs in TMHs: 24.51309\nExp number, first 60 AAs: 0.2682\nTotal prob of N-in: 0.14402\noutside 1 331\nTMhelix 332 354\ninside 355 372

16027 GCF_001478405.1_Salmonella_enterica_CVM_N38894-SQ_v1.0 Salmonella enterica subsp. enterica serovar Agona Proteobacteria; Gammaproteobacteria; Enterobacterales; Enterobacteriaceae; Salmonella; Salmonella enterica; Salmonella enterica subsp. enterica MNNEETFYQAMRRKGVTRRSFLKFCSLAATSLGLSAGMTPKIAWALENKPRIPVVWIHGL WP_001058316.1 hydrogenase [Salmonella enterica] Length: 372\nNumber of predicted TMHs: 2\nExp number of AAs in TMHs: 35.76397\nExp number, first 60 AAs: 18.97201\nTotal prob of N-in: 0.92785\nPOSSIBLE N-term signal sequence\ninside 1 19\nTMhelix 20 39\noutside 40 325\nTMhelix 326 348\ninside 349 372

16028 GCF_002031215.1_ASM203121v1 Salmonella enterica subsp. enterica serovar Agona Proteobacteria; Gammaproteobacteria; Enterobacterales; Enterobacteriaceae; Salmonella; Salmonella enterica; Salmonella enterica subsp. enterica MTGDNTLITSHGINRRDFMKLCAALAATMGLSSKAAAEMAESVSNPQRPPVIWIGAQECT WP_000145429.1 hydrogenase 2 small subunit [Salmonella enterica] Length: 372\nNumber of predicted TMHs: 1\nExp number of AAs in TMHs: 24.51309\nExp number, first 60 AAs: 0.2682\nTotal prob of N-in: 0.14402\noutside 1 331\nTMhelix 332 354\ninside 355 372

16029 GCF_001480835.1_Salmonella_enterica_CVM_N38874-SQ_v1.0 Salmonella enterica subsp. enterica serovar Agona Proteobacteria; Gammaproteobacteria; Enterobacterales; Enterobacteriaceae; Salmonella; Salmonella enterica; Salmonella enterica subsp. enterica MNNEETFYQAMRRKGVTRRSFLKFCSLAATSLGLSAGMTPKIAWALENKPRIPVVWIHGL WP_001058316.1 hydrogenase [Salmonella enterica] Length: 372\nNumber of predicted TMHs: 2\nExp number of AAs in TMHs: 35.76397\nExp number, first 60 AAs: 18.97201\nTotal prob of N-in: 0.92785\nPOSSIBLE N-term signal sequence\ninside 1 19\nTMhelix 20 39\noutside 40 325\nTMhelix 326 348\ninside 349 372

16030 GCF_002057015.1_ASM205701v1 Salmonella enterica subsp. enterica serovar Agona Proteobacteria; Gammaproteobacteria; Enterobacterales; Enterobacteriaceae; Salmonella; Salmonella enterica; Salmonella enterica subsp. enterica MTGDNTLITSHGINRRDFMKLCAALAATMGLSSKAAAEMAESVSNPQRPPVIWIGAQECT WP_000145429.1 hydrogenase 2 small subunit [Salmonella enterica] Length: 372\nNumber of predicted TMHs: 1\nExp number of AAs in TMHs: 24.51309\nExp number, first 60 AAs: 0.2682\nTotal prob of N-in: 0.14402\noutside 1 331\nTMhelix 332 354\ninside 355 372

16031 GCF_001480635.1_Salmonella_enterica_CVM_N42469_v1.0 Salmonella enterica subsp. enterica serovar Agona Proteobacteria; Gammaproteobacteria; Enterobacterales; Enterobacteriaceae; Salmonella; Salmonella enterica; Salmonella enterica subsp. enterica MTGDNTLITSHGINRRDFMKLCAALAATMGLSSKAAAEMAESVSNPQRPPVIWIGAQECT WP_000145429.1 hydrogenase 2 small subunit [Salmonella enterica] Length: 372\nNumber of predicted TMHs: 1\nExp number of AAs in TMHs: 24.51309\nExp number, first 60 AAs: 0.2682\nTotal prob of N-in: 0.14402\noutside 1 331\nTMhelix 332 354\ninside 355 372

16032 GCF_002106835.1_ASM210683v1 Salmonella enterica subsp. enterica serovar Agona Proteobacteria; Gammaproteobacteria; Enterobacterales; Enterobacteriaceae; Salmonella; Salmonella enterica; Salmonella enterica subsp. enterica MTGDNTLITSHGINRRDFMKLCAALAATMGLSSKAAAEMAESVSNPQRPPVIWIGAQECT WP_000145429.1 hydrogenase 2 small subunit [Salmonella enterica] Length: 372\nNumber of predicted TMHs: 1\nExp number of AAs in TMHs: 24.51309\nExp number, first 60 AAs: 0.2682\nTotal prob of N-in: 0.14402\noutside 1 331\nTMhelix 332 354\ninside 355 372

16033 GCF_002031435.1_ASM203143v1 Salmonella enterica subsp. enterica serovar Agona Proteobacteria; Gammaproteobacteria; Enterobacterales; Enterobacteriaceae; Salmonella; Salmonella enterica; Salmonella enterica subsp. enterica MTGDNTLITSHGINRRDFMKLCAALAATMGLSSKAAAEMAESVSNPQRPPVIWIGAQECT WP_000145429.1 hydrogenase 2 small subunit [Salmonella enterica] Length: 372\nNumber of predicted TMHs: 1\nExp number of AAs in TMHs: 24.51309\nExp number, first 60 AAs: 0.2682\nTotal prob of N-in: 0.14402\noutside 1 331\nTMhelix 332 354\ninside 355 372

16034	GCF_002031435.1_ASM203143v1	Salmonella enterica subsp. enterica serovar Agona	Proteobacteria; Gammaproteobacteria; Enterobacterales; Enterobacteriaceae; Salmonella; Salmonella enterica; Salmonella enterica subsp. enterica	MNNEETFYQAMRRKGVTRRSFLKFCSLAATSLGLSAGMTPKIAWALENKPRIPVVWIHGL	WP_001058316.1	hydrogenase [Salmonella enterica]	Length: 372\nNumber of predicted TMHs: 2\nExp number of AAs in TMHs: 35.76397\nExp number, first 60 AAs: 18.97201\nTotal prob of N-in: 0.92785\nPOSSIBLE N-term signal sequence\ninside 1 19\nTMhelix 20 39\noutside 40 325\nTMhelix 326 348\ninside 349 372

16035	GCF_002106835.1_ASM210683v1	Salmonella enterica subsp. enterica serovar Agona	Proteobacteria; Gammaproteobacteria; Enterobacterales; Enterobacteriaceae; Salmonella; Salmonella enterica; Salmonella enterica subsp. enterica	MNNEETFYQAMRRKGVTRRSFLKFCSLAATSLGLSAGMTPKIAWALENKPRIPVVWIHGL	WP_001058316.1	hydrogenase [Salmonella enterica]	Length: 372\nNumber of predicted TMHs: 2\nExp number of AAs in TMHs: 35.76397\nExp number, first 60 AAs: 18.97201\nTotal prob of N-in: 0.92785\nPOSSIBLE N-term signal sequence\ninside 1 19\nTMhelix 20 39\noutside 40 325\nTMhelix 326 348\ninside 349 372

16036	GCF_002031865.1_ASM203186v1	Salmonella enterica subsp. enterica serovar Agona	Proteobacteria; Gammaproteobacteria; Enterobacterales; Enterobacteriaceae; Salmonella; Salmonella enterica; Salmonella enterica subsp. enterica	MTGDNTLITSHGINRRDFMKLCAALAATMGLSSKAAAEMAESVSNPQRPPVIWIGAQECT	WP_000145429.1	hydrogenase 2 small subunit [Salmonella enterica]	Length: 372\nNumber of predicted TMHs: 1\nExp number of AAs in TMHs: 24.51309\nExp number, first 60 AAs: 0.2682\nTotal prob of N-in: 0.14402\noutside 1 331\nTMhelix 332 354\ninside 355 372

16037	GCF_002056815.1_ASM205681v1	Salmonella enterica subsp. enterica serovar Agona	Proteobacteria; Gammaproteobacteria; Enterobacterales; Enterobacteriaceae; Salmonella; Salmonella enterica; Salmonella enterica subsp. enterica	MNNEETFYQAMRRKGVTRRSFLKFCSLAATSLGLSAGMTPKIAWALENKPRIPVVWIHGL	WP_001058316.1	hydrogenase [Salmonella enterica]	Length: 372\nNumber of predicted TMHs: 2\nExp number of AAs in TMHs: 35.76397\nExp number, first 60 AAs: 18.97201\nTotal prob of N-in: 0.92785\nPOSSIBLE N-term signal sequence\ninside 1 19\nTMhelix 20 39\noutside 40 325\nTMhelix 326 348\ninside 349 372

16038	GCF_002058085.1_ASM205808v1	Salmonella enterica subsp. enterica serovar Agona	Proteobacteria; Gammaproteobacteria; Enterobacterales; Enterobacteriaceae; Salmonella; Salmonella enterica; Salmonella enterica subsp. enterica	MTGDNTLITSHGINRRDFMKLCAALAATMGLSSKAAAEMAESVSNPQRPPVIWIGAQECT	WP_000145429.1	hydrogenase 2 small subunit [Salmonella enterica]	Length: 372\nNumber of predicted TMHs: 1\nExp number of AAs in TMHs: 24.51309\nExp number, first 60 AAs: 0.2682\nTotal prob of N-in: 0.14402\noutside 1 331\nTMhelix 332 354\ninside 355 372

16039	GCF_001478405.1_Salmonella_enterica_CVM_N38894-SQ_v1.0	Salmonella enterica subsp. enterica serovar Agona	Proteobacteria; Gammaproteobacteria; Enterobacterales; Enterobacteriaceae; Salmonella; Salmonella enterica; Salmonella enterica subsp. enterica	MTGDNTLITSHGINRRDFMKLCAALAATMGLSSKAAAEMAESVSNPQRPPVIWIGAQECT	WP_000145429.1	hydrogenase 2 small subunit [Salmonella enterica]	Length: 372\nNumber of predicted TMHs: 1\nExp number of AAs in TMHs: 24.51309\nExp number, first 60 AAs: 0.2682\nTotal prob of N-in: 0.14402\noutside 1 331\nTMhelix 332 354\ninside 355 372

16040	GCF_002031425.1_ASM203142v1	Salmonella enterica subsp. enterica serovar Agona	Proteobacteria; Gammaproteobacteria; Enterobacterales; Enterobacteriaceae; Salmonella; Salmonella enterica; Salmonella enterica subsp. enterica	MNNEETFYQAMRRKGVTRRSFLKFCSLAATSLGLSAGMTPKIAWALENKPRIPVVWIHGL	WP_001058316.1	hydrogenase [Salmonella enterica]	Length: 372\nNumber of predicted TMHs: 2\nExp number of AAs in TMHs: 35.76397\nExp number, first 60 AAs: 18.97201\nTotal prob of N-in: 0.92785\nPOSSIBLE N-term signal sequence\ninside 1 19\nTMhelix 20 39\noutside 40 325\nTMhelix 326 348\ninside 349 372

16041	GCF_001244465.1_Salmonella_enterica_CVM_N48683_v1.0	Salmonella enterica subsp. enterica serovar Agona	Proteobacteria; Gammaproteobacteria; Enterobacterales; Enterobacteriaceae; Salmonella; Salmonella enterica; Salmonella enterica subsp. enterica	MTGDNTLITSHGINRRDFMKLCAALAATMGLSSKAAAEMAESVSNPQRPPVIWIGAQECT	WP_000145429.1	hydrogenase 2 small subunit [Salmonella enterica]	Length: 372\nNumber of predicted TMHs: 1\nExp number of AAs in TMHs: 24.51309\nExp number, first 60 AAs: 0.2682\nTotal prob of N-in: 0.14402\noutside 1 331\nTMhelix 332 354\ninside 355 372

16042	GCF_002058085.1_ASM205808v1	Salmonella enterica subsp. enterica serovar Agona	Proteobacteria; Gammaproteobacteria; Enterobacterales; Enterobacteriaceae; Salmonella; Salmonella enterica; Salmonella enterica subsp. enterica	MNNEETFYQAMRRKGVTRRSFLKFCSLAATSLGLSAGMTPKIAWALENKPRIPVVWIHGL	WP_001058316.1	hydrogenase [Salmonella enterica]	Length: 372\nNumber of predicted TMHs: 2\nExp

number of AAs in TMHs: 35.76397\nExp number, first 60 AAs: 18.97201\nTotal prob of N-in: 0.92785\nPOSSIBLE N-term signal sequence\ninside 1 19\nTMhelix 20 39\noutside 40 325\nTMhelix 326 348\ninside 349 372

16043 GCF_001244085.1_Salmonella_enterica_CVM_N47726_v1.0 Salmonella enterica subsp. enterica serovar Agona Proteobacteria; Gammaproteobacteria; Enterobacterales; Enterobacteriaceae; Salmonella; Salmonella enterica; Salmonella enterica subsp. enterica MNNEETFYQAMRRKGVTRRSFLKFCSLAATSLGLSAGMTPKIAWALENKPRIPVVWIHGL WP_001058316.1 hydrogenase [Salmonella enterica] Length: 372\nNumber of predicted TMHs: 2\nExp number of AAs in TMHs: 35.76397\nExp number, first 60 AAs: 18.97201\nTotal prob of N-in: 0.92785\nPOSSIBLE N-term signal sequence\ninside 1 19\nTMhelix 20 39\noutside 40 325\nTMhelix 326 348\ninside 349 372

16044 GCF_002057015.1_ASM205701v1 Salmonella enterica subsp. enterica serovar Agona Proteobacteria; Gammaproteobacteria; Enterobacterales; Enterobacteriaceae; Salmonella; Salmonella enterica; Salmonella enterica subsp. enterica MNNEETFYQAMRRKGVTRRSFLKFCSLAATSLGLSAGMTPKIAWALENKPRIPVVWIHGL WP_001058316.1 hydrogenase [Salmonella enterica] Length: 372\nNumber of predicted TMHs: 2\nExp number of AAs in TMHs: 35.76397\nExp number, first 60 AAs: 18.97201\nTotal prob of N-in: 0.92785\nPOSSIBLE N-term signal sequence\ninside 1 19\nTMhelix 20 39\noutside 40 325\nTMhelix 326 348\ninside 349 372

16045 GCF_002056815.1_ASM205681v1 Salmonella enterica subsp. enterica serovar Agona Proteobacteria; Gammaproteobacteria; Enterobacterales; Enterobacteriaceae; Salmonella; Salmonella enterica; Salmonella enterica subsp. enterica MTGDNTLITSHGINRRDFMKLCAALAATMGLSSKAAAEMAESVSNPQRPPVIWIGAQECT WP_000145429.1 hydrogenase 2 small subunit [Salmonella enterica] Length: 372\nNumber of predicted TMHs: 1\nExp number of AAs in TMHs: 24.51309\nExp number, first 60 AAs: 0.2682\nTotal prob of N-in: 0.14402\noutside 1 331\nTMhelix 332 354\ninside 355 372

16046 GCF_002106735.1_ASM210673v1 Salmonella enterica subsp. enterica serovar Agona Proteobacteria; Gammaproteobacteria; Enterobacterales; Enterobacteriaceae; Salmonella; Salmonella enterica; Salmonella enterica subsp. enterica MNNEETFYQAMRRKGVTRRSFLKFCSLAATSLGLSAGMTPKIAWALENKPRIPVVWIHGL WP_001058316.1 hydrogenase [Salmonella enterica] Length: 372\nNumber of predicted TMHs: 2\nExp number of AAs in TMHs: 35.76397\nExp number, first 60 AAs: 18.97201\nTotal prob of N-in: 0.92785\nPOSSIBLE N-term signal sequence\ninside 1 19\nTMhelix 20 39\noutside 40 325\nTMhelix 326 348\ninside 349 372

16047 GCF_002062875.1_ASM206287v1 Salmonella enterica subsp. enterica serovar Agona Proteobacteria; Gammaproteobacteria; Enterobacterales; Enterobacteriaceae; Salmonella; Salmonella enterica; Salmonella enterica subsp. enterica MNNEETFYQAMRRKGVTRRSFLKFCSLAATSLGLSAGMTPKIAWALENKPRIPVVWIHGL WP_001058316.1 hydrogenase [Salmonella enterica] Length: 372\nNumber of predicted TMHs: 2\nExp number of AAs in TMHs: 35.76397\nExp number, first 60 AAs: 18.97201\nTotal prob of N-in: 0.92785\nPOSSIBLE N-term signal sequence\ninside 1 19\nTMhelix 20 39\noutside 40 325\nTMhelix 326 348\ninside 349 372

16048 GCF_002062875.1_ASM206287v1 Salmonella enterica subsp. enterica serovar Agona Proteobacteria; Gammaproteobacteria; Enterobacterales; Enterobacteriaceae; Salmonella; Salmonella enterica; Salmonella enterica subsp. enterica MTGDNTLITSHGINRRDFMKLCAALAATMGLSSKAAAEMAESVSNPQRPPVIWIGAQECT WP_000145429.1 hydrogenase 2 small subunit [Salmonella enterica] Length: 372\nNumber of predicted TMHs: 1\nExp number of AAs in TMHs: 24.51309\nExp number, first 60 AAs: 0.2682\nTotal prob of N-in: 0.14402\noutside 1 331\nTMhelix 332 354\ninside 355 372

16049 GCF_002106735.1_ASM210673v1 Salmonella enterica subsp. enterica serovar Agona Proteobacteria; Gammaproteobacteria; Enterobacterales; Enterobacteriaceae; Salmonella; Salmonella enterica; Salmonella enterica subsp. enterica MTGDNTLITSHGINRRDFMKLCAALAATMGLSSKAAAEMAESVSNPQRPPVIWIGAQECT WP_000145429.1 hydrogenase 2 small subunit [Salmonella enterica] Length: 372\nNumber of predicted TMHs: 1\nExp number of AAs in TMHs: 24.51309\nExp number, first 60 AAs: 0.2682\nTotal prob of N-in: 0.14402\noutside 1 331\nTMhelix 332 354\ninside 355 372

16050 GCF_002031865.1_ASM203186v1 Salmonella enterica subsp. enterica serovar Agona Proteobacteria; Gammaproteobacteria; Enterobacterales; Enterobacteriaceae; Salmonella; Salmonella enterica; Salmonella enterica subsp. enterica MNNEETFYQAMRRKGVTRRSFLKFCSLAATSLGLSAGMTPKIAWALENKPRIPVVWIHGL WP_001058316.1 hydrogenase [Salmonella enterica] Length: 372\nNumber of predicted TMHs: 2\nExp number of AAs in TMHs: 35.76397\nExp number, first 60 AAs: 18.97201\nTotal prob of N-in: 0.92785\nPOSSIBLE N-term signal sequence\ninside 1 19\nTMhelix 20 39\noutside 40 325\nTMhelix 326 348\ninside 349 372

16051 GCF_002030065.1_ASM203006v1 Salmonella enterica subsp. enterica serovar Agona Proteobacteria; Gammaproteobacteria; Enterobacterales; Enterobacteriaceae; Salmonella; Salmonella enterica;

Salmonella enterica subsp. enterica MTGDNTLITSHGINRRDFMKLCAALAATMGLSSKAAAEMAESVSNPQRPPVIWIGAQECT WP_000145429.1 hydrogenase 2 small subunit [Salmonella enterica] Length: 372\nNumber of predicted TMHs: 1\nExp number of AAs in TMHs: 24.51309\nExp number, first 60 AAs: 0.2682\nTotal prob of N-in: 0.14402\noutside 1 331\nTMhelix 332 354\ninside 355 372

16052 GCF_002056705.1_ASM205670v1 Salmonella enterica subsp. enterica serovar Agona Proteobacteria; Gammaproteobacteria; Enterobacterales; Enterobacteriaceae; Salmonella; Salmonella enterica; Salmonella enterica subsp. enterica MNNEETFYQAMRRKGVTRRSFLKFCSLAATSLGLSAGMTPKIAWALENKPRIPVVWIHGL WP_001058316.1 hydrogenase [Salmonella enterica] Length: 372\nNumber of predicted TMHs: 2\nExp number of AAs in TMHs: 35.76397\nExp number, first 60 AAs: 18.97201\nTotal prob of N-in: 0.92785\nPOSSIBLE N-term signal sequence\ninside 1 19\nTMhelix 20 39\noutside 40 325\nTMhelix 326 348\ninside 349 372

16053 GCF_002056925.1_ASM205692v1 Salmonella enterica subsp. enterica serovar Agona Proteobacteria; Gammaproteobacteria; Enterobacterales; Enterobacteriaceae; Salmonella; Salmonella enterica; Salmonella enterica subsp. enterica MTGDNTLITSHGINRRDFMKLCAALAATMGLSSKAAAEMAESVSNPQRPPVIWIGAQECT WP_000145429.1 hydrogenase 2 small subunit [Salmonella enterica] Length: 372\nNumber of predicted TMHs: 1\nExp number of AAs in TMHs: 24.51309\nExp number, first 60 AAs: 0.2682\nTotal prob of N-in: 0.14402\noutside 1 331\nTMhelix 332 354\ninside 355 372

16054 GCF_001477685.1_Salmonella_enterica_CVM_N29309-SQ_v1.0 Salmonella enterica subsp. enterica serovar Agona Proteobacteria; Gammaproteobacteria; Enterobacterales; Enterobacteriaceae; Salmonella; Salmonella enterica; Salmonella enterica subsp. enterica MNNEETFYQAMRRKGVTRRSFLKFCSLAATSLGLSAGMTPKIAWALENKPRIPVVWIHGL WP_001058316.1 hydrogenase [Salmonella enterica] Length: 372\nNumber of predicted TMHs: 2\nExp number of AAs in TMHs: 35.76397\nExp number, first 60 AAs: 18.97201\nTotal prob of N-in: 0.92785\nPOSSIBLE N-term signal sequence\ninside 1 19\nTMhelix 20 39\noutside 40 325\nTMhelix 326 348\ninside 349 372

16055 GCF_001479085.1_Salmonella_enterica_CVM_N31394-SQ_v1.0 Salmonella enterica subsp. enterica serovar Agona Proteobacteria; Gammaproteobacteria; Enterobacterales; Enterobacteriaceae; Salmonella; Salmonella enterica; Salmonella enterica subsp. enterica MTGDNTLITSHGINRRDFMKLCAALAATMGLSSKAAAEMAESVSNPQRPPVIWIGAQECT WP_000145429.1 hydrogenase 2 small subunit [Salmonella enterica] Length: 372\nNumber of predicted TMHs: 1\nExp number of AAs in TMHs: 24.51309\nExp number, first 60 AAs: 0.2682\nTotal prob of N-in: 0.14402\noutside 1 331\nTMhelix 332 354\ninside 355 372

16056 GCF_002056865.1_ASM205686v1 Salmonella enterica subsp. enterica serovar Agona Proteobacteria; Gammaproteobacteria; Enterobacterales; Enterobacteriaceae; Salmonella; Salmonella enterica; Salmonella enterica subsp. enterica MTGDNTLITSHGINRRDFMKLCAALAATMGLSSKAAAEMAESVSNPQRPPVIWIGAQECT WP_000145429.1 hydrogenase 2 small subunit [Salmonella enterica] Length: 372\nNumber of predicted TMHs: 1\nExp number of AAs in TMHs: 24.51309\nExp number, first 60 AAs: 0.2682\nTotal prob of N-in: 0.14402\noutside 1 331\nTMhelix 332 354\ninside 355 372

16057 GCF_001477685.1_Salmonella_enterica_CVM_N29309-SQ_v1.0 Salmonella enterica subsp. enterica serovar Agona Proteobacteria; Gammaproteobacteria; Enterobacterales; Enterobacteriaceae; Salmonella; Salmonella enterica; Salmonella enterica subsp. enterica MTGDNTLITSHGINRRDFMKLCAALAATMGLSSKAAAEMAESVSNPQRPPVIWIGAQECT WP_000145429.1 hydrogenase 2 small subunit [Salmonella enterica] Length: 372\nNumber of predicted TMHs: 1\nExp number of AAs in TMHs: 24.51309\nExp number, first 60 AAs: 0.2682\nTotal prob of N-in: 0.14402\noutside 1 331\nTMhelix 332 354\ninside 355 372

16058 GCF_002030065.1_ASM203006v1 Salmonella enterica subsp. enterica serovar Agona Proteobacteria; Gammaproteobacteria; Enterobacterales; Enterobacteriaceae; Salmonella; Salmonella enterica; Salmonella enterica subsp. enterica MNNEETFYQAMRRKGVTRRSFLKFCSLAATSLGLSAGMTPKIAWALENKPRIPVVWIHGL WP_001058316.1 hydrogenase [Salmonella enterica] Length: 372\nNumber of predicted TMHs: 2\nExp number of AAs in TMHs: 35.76397\nExp number, first 60 AAs: 18.97201\nTotal prob of N-in: 0.92785\nPOSSIBLE N-term signal sequence\ninside 1 19\nTMhelix 20 39\noutside 40 325\nTMhelix 326 348\ninside 349 372

16059 GCF_001479085.1_Salmonella_enterica_CVM_N31394-SQ_v1.0 Salmonella enterica subsp. enterica serovar Agona Proteobacteria; Gammaproteobacteria; Enterobacterales; Enterobacteriaceae; Salmonella; Salmonella enterica; Salmonella enterica subsp. enterica MNNEETFYQAMRRKGVTRRSFLKFCSLAATSLGLSAGMTPKIAWALENKPRIPVVWIHGL WP_001058316.1 hydrogenase [Salmonella enterica] Length: 372\nNumber of predicted TMHs: 2\nExp number of AAs in TMHs: 35.76397\nExp number, first 60 AAs: 18.97201\nTotal prob of N-in: 0.92785\nPOSSIBLE N-term signal sequence\ninside 1 19\nTMhelix 20 39\noutside 40 325\nTMhelix 326 348\ninside 349 372

16060 GCF_002056705.1_ASM205670v1 Salmonella enterica subsp. enterica serovar Agona Proteobacteria; Gammaproteobacteria; Enterobacterales; Enterobacteriaceae; Salmonella; Salmonella enterica; Salmonella enterica subsp. enterica MTGDNTLITSHGINRRDFMKLCAALAATMGLSSKAAAEMAESVSNPQRPPVIWIGAQECT WP_000145429.1 hydrogenase 2 small subunit [Salmonella enterica] Length: 372\nNumber of predicted TMHs: 1\nExp number of AAs in TMHs: 24.51309\nExp number, first 60 AAs: 0.2682\nTotal prob of N-in: 0.14402\noutside 1 331\nTMhelix 332 354\ninside 355 372

16061 GCF_002056865.1_ASM205686v1 Salmonella enterica subsp. enterica serovar Agona Proteobacteria; Gammaproteobacteria; Enterobacterales; Enterobacteriaceae; Salmonella; Salmonella enterica; Salmonella enterica subsp. enterica MNNEETFYQAMRRKGVTRRSFLKFCSLAATSLGLSAGMTPKIAWALENKPRIPVVWIHGL WP_001058316.1 hydrogenase [Salmonella enterica] Length: 372\nNumber of predicted TMHs: 2\nExp number of AAs in TMHs: 35.76397\nExp number, first 60 AAs: 18.97201\nTotal prob of N-in: 0.92785\nPOSSIBLE N-term signal sequence\ninside 1 19\nTMhelix 20 39\noutside 40 325\nTMhelix 326 348\ninside 349 372

16062 GCF_002056925.1_ASM205692v1 Salmonella enterica subsp. enterica serovar Agona Proteobacteria; Gammaproteobacteria; Enterobacterales; Enterobacteriaceae; Salmonella; Salmonella enterica; Salmonella enterica subsp. enterica MNNEETFYQAMRRKGVTRRSFLKFCSLAATSLGLSAGMTPKIAWALENKPRIPVVWIHGL WP_001058316.1 hydrogenase [Salmonella enterica] Length: 372\nNumber of predicted TMHs: 2\nExp number of AAs in TMHs: 35.76397\nExp number, first 60 AAs: 18.97201\nTotal prob of N-in: 0.92785\nPOSSIBLE N-term signal sequence\ninside 1 19\nTMhelix 20 39\noutside 40 325\nTMhelix 326 348\ninside 349 372

16063 GCF_002031425.1_ASM203142v1 Salmonella enterica subsp. enterica serovar Agona Proteobacteria; Gammaproteobacteria; Enterobacterales; Enterobacteriaceae; Salmonella; Salmonella enterica; Salmonella enterica subsp. enterica MTGDNTLITSHGINRRDFMKLCAALAATMGLSSKAAAEMAESVSNPQRPPVIWIGAQECT WP_000145429.1 hydrogenase 2 small subunit [Salmonella enterica] Length: 372\nNumber of predicted TMHs: 1\nExp number of AAs in TMHs: 24.51309\nExp number, first 60 AAs: 0.2682\nTotal prob of N-in: 0.14402\noutside 1 331\nTMhelix 332 354\ninside 355 372

16064 GCF_002033225.1_ASM203322v1 Salmonella enterica subsp. enterica serovar Agona Proteobacteria; Gammaproteobacteria; Enterobacterales; Enterobacteriaceae; Salmonella; Salmonella enterica; Salmonella enterica subsp. enterica MNNEETFYQAMRRKGVTRRSFLKFCSLAATSLGLSAGMTPKIAWALENKPRIPVVWIHGL WP_001058316.1 hydrogenase [Salmonella enterica] Length: 372\nNumber of predicted TMHs: 2\nExp number of AAs in TMHs: 35.76397\nExp number, first 60 AAs: 18.97201\nTotal prob of N-in: 0.92785\nPOSSIBLE N-term signal sequence\ninside 1 19\nTMhelix 20 39\noutside 40 325\nTMhelix 326 348\ninside 349 372

16065 GCF_001480835.1_Salmonella_enterica_CVM_N38874-SQ_v1.0 Salmonella enterica subsp. enterica serovar Agona Proteobacteria; Gammaproteobacteria; Enterobacterales; Enterobacteriaceae; Salmonella; Salmonella enterica; Salmonella enterica subsp. enterica MTGDNTLITSHGINRRDFMKLCAALAATMGLSSKAAAEMAESVSNPQRPPVIWIGAQECT WP_000145429.1 hydrogenase 2 small subunit [Salmonella enterica] Length: 372\nNumber of predicted TMHs: 1\nExp number of AAs in TMHs: 24.51309\nExp number, first 60 AAs: 0.2682\nTotal prob of N-in: 0.14402\noutside 1 331\nTMhelix 332 354\ninside 355 372

16066 GCF_001479025.1_Salmonella_enterica_CVM_N30701-SQ_v1.0 Salmonella enterica subsp. enterica serovar Agona Proteobacteria; Gammaproteobacteria; Enterobacterales; Enterobacteriaceae; Salmonella; Salmonella enterica; Salmonella enterica subsp. enterica MTGDNTLITSHGINRRDFMKLCAALAATMGLSSKAAAEMAESVSNPQRPPVIWIGAQECT WP_000145429.1 hydrogenase 2 small subunit [Salmonella enterica] Length: 372\nNumber of predicted TMHs: 1\nExp number of AAs in TMHs: 24.51309\nExp number, first 60 AAs: 0.2682\nTotal prob of N-in: 0.14402\noutside 1 331\nTMhelix 332 354\ninside 355 372

16067 GCF_001244085.1_Salmonella_enterica_CVM_N47726_v1.0 Salmonella enterica subsp. enterica serovar Agona Proteobacteria; Gammaproteobacteria; Enterobacterales; Enterobacteriaceae; Salmonella; Salmonella enterica; Salmonella enterica subsp. enterica MTGDNTLITSHGINRRDFMKLCAALAATMGLSSKAAAEMAESVSNPQRPPVIWIGAQECT WP_000145429.1 hydrogenase 2 small subunit [Salmonella enterica] Length: 372\nNumber of predicted TMHs: 1\nExp number of AAs in TMHs: 24.51309\nExp number, first 60 AAs: 0.2682\nTotal prob of N-in: 0.14402\noutside 1 331\nTMhelix 332 354\ninside 355 372

16068 GCF_002031025.1_ASM203102v1 Salmonella enterica subsp. enterica serovar Alachua Proteobacteria; Gammaproteobacteria; Enterobacterales; Enterobacteriaceae; Salmonella; Salmonella enterica; Salmonella enterica subsp. enterica MNNEETFYQAMRRKGVTRRSFLKFCSLAATSLGLGAGMTPKIAWALENKPRIPVVWIHGL WP_001058312.1 hydrogenase [Salmonella enterica] Length: 372\nNumber of predicted TMHs: 2\nExp

number of AAs in TMHs: 36.51738\nExp number, first 60 AAs: 19.72849\nTotal prob of N-in: 0.96170\nPOSSIBLE N-term signal sequence\ninside 1 19\nTMhelix 20 39\noutside 40 325\nTMhelix 326 348\ninside 349 372

16069 GCF_002031025.1_ASM203102v1 Salmonella enterica subsp. enterica serovar Alachua Proteobacteria; Gammaproteobacteria; Enterobacterales; Enterobacteriaceae; Salmonella; Salmonella enterica; Salmonella enterica subsp. enterica MTGDNTLITSHGINRRDFMKLCAALAATMGLSSKAAAEMAESVSNPQRPPVIWIGAQECT WP_000145429.1 hydrogenase 2 small subunit [Salmonella enterica] Length: 372\nNumber of predicted TMHs: 1\nExp number of AAs in TMHs: 24.51309\nExp number, first 60 AAs: 0.2682\nTotal prob of N-in: 0.14402\noutside 1 331\nTMhelix 332 354\ninside 355 372

16070 GCF_002046115.1_ASM204611v1 Salmonella enterica subsp. enterica serovar Alachua Proteobacteria; Gammaproteobacteria; Enterobacterales; Enterobacteriaceae; Salmonella; Salmonella enterica; Salmonella enterica subsp. enterica MTGDNTLITSHGINRRDFMKLCAALAATMGLSSKAAAEMAESVSNPQRPPVIWIGAQECT WP_000145429.1 hydrogenase 2 small subunit [Salmonella enterica] Length: 372\nNumber of predicted TMHs: 1\nExp number of AAs in TMHs: 24.51309\nExp number, first 60 AAs: 0.2682\nTotal prob of N-in: 0.14402\noutside 1 331\nTMhelix 332 354\ninside 355 372

16071 GCF_002046115.1_ASM204611v1 Salmonella enterica subsp. enterica serovar Alachua Proteobacteria; Gammaproteobacteria; Enterobacterales; Enterobacteriaceae; Salmonella; Salmonella enterica; Salmonella enterica subsp. enterica MNNEETFYQAMRRKGVTRRSFLKFCSLAATSLGLGAGMTPKIAWALENKPRIPVVWIHGL WP_001058312.1 hydrogenase [Salmonella enterica] Length: 372\nNumber of predicted TMHs: 2\nExp number of AAs in TMHs: 36.51738\nExp number, first 60 AAs: 19.72849\nTotal prob of N-in: 0.96170\nPOSSIBLE N-term signal sequence\ninside 1 19\nTMhelix 20 39\noutside 40 325\nTMhelix 326 348\ninside 349 372

16072 GCF_002032315.1_ASM203231v1 Salmonella enterica subsp. enterica serovar Alachua Proteobacteria; Gammaproteobacteria; Enterobacterales; Enterobacteriaceae; Salmonella; Salmonella enterica; Salmonella enterica subsp. enterica MTGDNTLITSHGINRRDFMKLCAALAATMGLSSKAAAEMAESVSNPQRPPVIWIGAQECT WP_000145429.1 hydrogenase 2 small subunit [Salmonella enterica] Length: 372\nNumber of predicted TMHs: 1\nExp number of AAs in TMHs: 24.51309\nExp number, first 60 AAs: 0.2682\nTotal prob of N-in: 0.14402\noutside 1 331\nTMhelix 332 354\ninside 355 372

16073 GCF_002047955.1_ASM204795v1 Salmonella enterica subsp. enterica serovar Alachua Proteobacteria; Gammaproteobacteria; Enterobacterales; Enterobacteriaceae; Salmonella; Salmonella enterica; Salmonella enterica subsp. enterica MNNEETFYQAMRRKGVTRRSFLKFCSLAATSLGLGAGMTPKIAWALENKPRIPVVWIHGL WP_001058312.1 hydrogenase [Salmonella enterica] Length: 372\nNumber of predicted TMHs: 2\nExp number of AAs in TMHs: 36.51738\nExp number, first 60 AAs: 19.72849\nTotal prob of N-in: 0.96170\nPOSSIBLE N-term signal sequence\ninside 1 19\nTMhelix 20 39\noutside 40 325\nTMhelix 326 348\ninside 349 372

16074 GCF_002045465.1_ASM204546v1 Salmonella enterica subsp. enterica serovar Alachua Proteobacteria; Gammaproteobacteria; Enterobacterales; Enterobacteriaceae; Salmonella; Salmonella enterica; Salmonella enterica subsp. enterica MNNEETFYQAMRRKGVTRRSFLKFCSLAATSLGLGAGMTPKIAWALENKPRIPVVWIHGL WP_001058312.1 hydrogenase [Salmonella enterica] Length: 372\nNumber of predicted TMHs: 2\nExp number of AAs in TMHs: 36.51738\nExp number, first 60 AAs: 19.72849\nTotal prob of N-in: 0.96170\nPOSSIBLE N-term signal sequence\ninside 1 19\nTMhelix 20 39\noutside 40 325\nTMhelix 326 348\ninside 349 372

16075 GCF_002047955.1_ASM204795v1 Salmonella enterica subsp. enterica serovar Alachua Proteobacteria; Gammaproteobacteria; Enterobacterales; Enterobacteriaceae; Salmonella; Salmonella enterica; Salmonella enterica subsp. enterica MTGDNTLITSHGINRRDFMKLCAALAATMGLSSKAAAEMAESVSNPQRPPVIWIGAQECT WP_000145429.1 hydrogenase 2 small subunit [Salmonella enterica] Length: 372\nNumber of predicted TMHs: 1\nExp number of AAs in TMHs: 24.51309\nExp number, first 60 AAs: 0.2682\nTotal prob of N-in: 0.14402\noutside 1 331\nTMhelix 332 354\ninside 355 372

16076 GCF_002032315.1_ASM203231v1 Salmonella enterica subsp. enterica serovar Alachua Proteobacteria; Gammaproteobacteria; Enterobacterales; Enterobacteriaceae; Salmonella; Salmonella enterica; Salmonella enterica subsp. enterica MNNEETFYQAMRRKGVTRRSFLKFCSLAATSLGLGAGMTPKIAWALENKPRIPVVWIHGL WP_001058312.1 hydrogenase [Salmonella enterica] Length: 372\nNumber of predicted TMHs: 2\nExp number of AAs in TMHs: 36.51738\nExp number, first 60 AAs: 19.72849\nTotal prob of N-in: 0.96170\nPOSSIBLE N-term signal sequence\ninside 1 19\nTMhelix 20 39\noutside 40 325\nTMhelix 326 348\ninside 349 372

16077 GCF_002034675.1_ASM203467v1 Salmonella enterica subsp. enterica serovar Alachua Proteobacteria; Gammaproteobacteria; Enterobacterales; Enterobacteriaceae; Salmonella; Salmonella enterica;

Salmonella enterica subsp. enterica MNNEETFYQAMRRKGVTRRSFLKFCSLAATSLGLGAGMTPKIAWALENKPRIPVVWIHGL WP_001058312.1 hydrogenase [Salmonella enterica] Length: 372\nNumber of predicted TMHs: 2\nExp number of AAs in TMHs: 36.51738\nExp number, first 60 AAs: 19.72849\nTotal prob of N-in: 0.96170\nPOSSIBLE N-term signal sequence\ninside 1 19\nTMhelix 20 39\noutside 40 325\nTMhelix 326 348\ninside 349 372

16078 GCF_002034675.1_ASM203467v1 Salmonella enterica subsp. enterica serovar Alachua Proteobacteria; Gammaproteobacteria; Enterobacterales; Enterobacteriaceae; Salmonella; Salmonella enterica; Salmonella enterica subsp. enterica MTGDNTLITSHGINRRDFMKLCAALAATMGLSSKAAAEMAESVSNPQRPPVIWIGAQECT WP_000145429.1 hydrogenase 2 small subunit [Salmonella enterica] Length: 372\nNumber of predicted TMHs: 1\nExp number of AAs in TMHs: 24.51309\nExp number, first 60 AAs: 0.2682\nTotal prob of N-in: 0.14402\noutside 1 331\nTMhelix 332 354\ninside 355 372

16079 GCF_002046535.1_ASM204653v1 Salmonella enterica subsp. enterica serovar Alachua Proteobacteria; Gammaproteobacteria; Enterobacterales; Enterobacteriaceae; Salmonella; Salmonella enterica; Salmonella enterica subsp. enterica MNNEETFYQAMRRKGVTRRSFLKFCSLAATSLGLGAGMTPKIAWALENKPRIPVVWIHGL WP_001058312.1 hydrogenase [Salmonella enterica] Length: 372\nNumber of predicted TMHs: 2\nExp number of AAs in TMHs: 36.51738\nExp number, first 60 AAs: 19.72849\nTotal prob of N-in: 0.96170\nPOSSIBLE N-term signal sequence\ninside 1 19\nTMhelix 20 39\noutside 40 325\nTMhelix 326 348\ninside 349 372

16080 GCF_002046535.1_ASM204653v1 Salmonella enterica subsp. enterica serovar Alachua Proteobacteria; Gammaproteobacteria; Enterobacterales; Enterobacteriaceae; Salmonella; Salmonella enterica; Salmonella enterica subsp. enterica MTGDNTLITSHGINRRDFMKLCAALAATMGLSSKAAAEMAESVSNPQRPPVIWIGAQECT WP_000145429.1 hydrogenase 2 small subunit [Salmonella enterica] Length: 372\nNumber of predicted TMHs: 1\nExp number of AAs in TMHs: 24.51309\nExp number, first 60 AAs: 0.2682\nTotal prob of N-in: 0.14402\noutside 1 331\nTMhelix 332 354\ninside 355 372

16081 GCF_002062195.1_ASM206219v1 Salmonella enterica subsp. enterica serovar Albany Proteobacteria; Gammaproteobacteria; Enterobacterales; Enterobacteriaceae; Salmonella; Salmonella enterica; Salmonella enterica subsp. enterica MNNEETFYQAMRRKGVTRRSFLKFCSLAATSLGLGAGMTPKIAWALENKPRIPVVWIHGL WP_001058311.1 hydrogenase [Salmonella enterica] Length: 372\nNumber of predicted TMHs: 2\nExp number of AAs in TMHs: 37.4699899999999\nExp number, first 60 AAs: 19.72699\nTotal prob of N-in: 0.96171\nPOSSIBLE N-term signal sequence\ninside 1 19\nTMhelix 20 39\noutside 40 325\nTMhelix 326 348\ninside 349 372

16082 GCF_002048105.1_ASM204810v1 Salmonella enterica subsp. enterica serovar Albany Proteobacteria; Gammaproteobacteria; Enterobacterales; Enterobacteriaceae; Salmonella; Salmonella enterica; Salmonella enterica subsp. enterica MTGDNTLITSHGINRRDFMKLCAALAATMGLSSKAAAEMAESVSNPQRPPVIWIGAQECT WP_000145429.1 hydrogenase 2 small subunit [Salmonella enterica] Length: 372\nNumber of predicted TMHs: 1\nExp number of AAs in TMHs: 24.51309\nExp number, first 60 AAs: 0.2682\nTotal prob of N-in: 0.14402\noutside 1 331\nTMhelix 332 354\ninside 355 372

16083 GCF_001246755.1_Salmonella_enterica_CVM_N51278_v1.0 Salmonella enterica subsp. enterica serovar Albany Proteobacteria; Gammaproteobacteria; Enterobacterales; Enterobacteriaceae; Salmonella; Salmonella enterica; Salmonella enterica subsp. enterica MTGDNTLITSHGINRRDFMKLCAALAATMGLSSKAAAEMAESVSNPQRPPVIWIGAQECT WP_000145429.1 hydrogenase 2 small subunit [Salmonella enterica] Length: 372\nNumber of predicted TMHs: 1\nExp number of AAs in TMHs: 24.51309\nExp number, first 60 AAs: 0.2682\nTotal prob of N-in: 0.14402\noutside 1 331\nTMhelix 332 354\ninside 355 372

16084 GCF_002046575.1_ASM204657v1 Salmonella enterica subsp. enterica serovar Albany Proteobacteria; Gammaproteobacteria; Enterobacterales; Enterobacteriaceae; Salmonella; Salmonella enterica; Salmonella enterica subsp. enterica MTGDNTLITSHGINRRDFMKLCAALAATMGLSSKAAAEMAESVSNPQRPPVIWIGAQECT WP_000145429.1 hydrogenase 2 small subunit [Salmonella enterica] Length: 372\nNumber of predicted TMHs: 1\nExp number of AAs in TMHs: 24.51309\nExp number, first 60 AAs: 0.2682\nTotal prob of N-in: 0.14402\noutside 1 331\nTMhelix 332 354\ninside 355 372

16085 GCF_002062195.1_ASM206219v1 Salmonella enterica subsp. enterica serovar Albany Proteobacteria; Gammaproteobacteria; Enterobacterales; Enterobacteriaceae; Salmonella; Salmonella enterica; Salmonella enterica subsp. enterica MTGDNTLITSHGINRRDFMKLCAALAATMGLSSKAAAEMAESVSNPQRPPVIWIGAQECT WP_000145429.1 hydrogenase 2 small subunit [Salmonella enterica] Length: 372\nNumber of predicted TMHs: 1\nExp number of AAs in TMHs: 24.51309\nExp number, first 60 AAs: 0.2682\nTotal prob of N-in: 0.14402\noutside 1 331\nTMhelix 332 354\ninside 355 372

16086 GCF_001479845.1_Salmonella_enterica_CVM_N38860-SQ_v1.0 Salmonella enterica subsp. enterica serovar Albany Proteobacteria; Gammaproteobacteria; Enterobacterales; Enterobacteriaceae; Salmonella; Salmonella enterica; Salmonella enterica subsp. enterica MTGDNTLITSHGINRRDFMKLCAALAATMGLSSKAAAEMAESVSNPQRPPVIWIGAQECT WP_000145429.1 hydrogenase 2 small subunit [Salmonella enterica] Length: 372\nNumber of predicted TMHs: 1\nExp number of AAs in TMHs: 24.51309\nExp number, first 60 AAs: 0.2682\nTotal prob of N-in: 0.14402\noutside 1 331\nTMhelix 332 354\ninside 355 372

16087 GCF_002048105.1_ASM204810v1 Salmonella enterica subsp. enterica serovar Albany Proteobacteria; Gammaproteobacteria; Enterobacterales; Enterobacteriaceae; Salmonella; Salmonella enterica; Salmonella enterica subsp. enterica MNNEETFYQAMRRKGVTRRSFLKFCSLAATSLGLGAGMTPKIAWALENKPRIPVVWIHGL WP_001058311.1 hydrogenase [Salmonella enterica] Length: 372\nNumber of predicted TMHs: 2\nExp number of AAs in TMHs: 37.4699899999999\nExp number, first 60 AAs: 19.72699\nTotal prob of N-in: 0.96171\nPOSSIBLE N-term signal sequence\ninside 1 19\nTMhelix 20 39\noutside 40 325\nTMhelix 326 348\ninside 349 372

16088 GCF_002046915.1_ASM204691v1 Salmonella enterica subsp. enterica serovar Albany Proteobacteria; Gammaproteobacteria; Enterobacterales; Enterobacteriaceae; Salmonella; Salmonella enterica; Salmonella enterica subsp. enterica MNNEETFYQAMRRKGVTRRSFLKFCSLAATSLGLGAGMTPKIAWALENKPRIPVVWIHGL WP_001058311.1 hydrogenase [Salmonella enterica] Length: 372\nNumber of predicted TMHs: 2\nExp number of AAs in TMHs: 37.4699899999999\nExp number, first 60 AAs: 19.72699\nTotal prob of N-in: 0.96171\nPOSSIBLE N-term signal sequence\ninside 1 19\nTMhelix 20 39\noutside 40 325\nTMhelix 326 348\ninside 349 372

16089 GCF_002046915.1_ASM204691v1 Salmonella enterica subsp. enterica serovar Albany Proteobacteria; Gammaproteobacteria; Enterobacterales; Enterobacteriaceae; Salmonella; Salmonella enterica; Salmonella enterica subsp. enterica MTGDNTLITSHGINRRDFMKLCAALAATMGLSSKAAAEMAESVSNPQRPPVIWIGAQECT WP_000145429.1 hydrogenase 2 small subunit [Salmonella enterica] Length: 372\nNumber of predicted TMHs: 1\nExp number of AAs in TMHs: 24.51309\nExp number, first 60 AAs: 0.2682\nTotal prob of N-in: 0.14402\noutside 1 331\nTMhelix 332 354\ninside 355 372

16090 GCF_002046575.1_ASM204657v1 Salmonella enterica subsp. enterica serovar Albany Proteobacteria; Gammaproteobacteria; Enterobacterales; Enterobacteriaceae; Salmonella; Salmonella enterica; Salmonella enterica subsp. enterica MNNEETFYQAMRRKGVTRRSFLKFCSLAATSLGLGAGMTPKIAWALENKPRIPVVWIHGL WP_001058311.1 hydrogenase [Salmonella enterica] Length: 372\nNumber of predicted TMHs: 2\nExp number of AAs in TMHs: 37.4699899999999\nExp number, first 60 AAs: 19.72699\nTotal prob of N-in: 0.96171\nPOSSIBLE N-term signal sequence\ninside 1 19\nTMhelix 20 39\noutside 40 325\nTMhelix 326 348\ninside 349 372

16091 GCF_002071615.1_ASM207161v1 Salmonella enterica subsp. enterica serovar Albany Proteobacteria; Gammaproteobacteria; Enterobacterales; Enterobacteriaceae; Salmonella; Salmonella enterica; Salmonella enterica subsp. enterica MNNEETFYQAMRRKGVTRRSFLKFCSLAATSLGLGAGMTPKIAWALENKPRIPVVWIHGL WP_001058311.1 hydrogenase [Salmonella enterica] Length: 372\nNumber of predicted TMHs: 2\nExp number of AAs in TMHs: 37.4699899999999\nExp number, first 60 AAs: 19.72699\nTotal prob of N-in: 0.96171\nPOSSIBLE N-term signal sequence\ninside 1 19\nTMhelix 20 39\noutside 40 325\nTMhelix 326 348\ninside 349 372

16092 GCF_002071615.1_ASM207161v1 Salmonella enterica subsp. enterica serovar Albany Proteobacteria; Gammaproteobacteria; Enterobacterales; Enterobacteriaceae; Salmonella; Salmonella enterica; Salmonella enterica subsp. enterica MTGDNTLITSHGINRRDFMKLCAALAATMGLSSKAAAEMAESVSNPQRPPVIWIGAQECT WP_000145429.1 hydrogenase 2 small subunit [Salmonella enterica] Length: 372\nNumber of predicted TMHs: 1\nExp number of AAs in TMHs: 24.51309\nExp number, first 60 AAs: 0.2682\nTotal prob of N-in: 0.14402\noutside 1 331\nTMhelix 332 354\ninside 355 372

16093 GCF_001241765.1_Salmonella_enterica_CVM_N45398_v1.0 Salmonella enterica subsp. enterica serovar Albany Proteobacteria; Gammaproteobacteria; Enterobacterales; Enterobacteriaceae; Salmonella; Salmonella enterica; Salmonella enterica subsp. enterica MNNEETFYQAMRRKGVTRRSFLKFCSLAATSLGLGAGMTPKIAWALENKPRIPVVWIHGL WP_001058311.1 hydrogenase [Salmonella enterica] Length: 372\nNumber of predicted TMHs: 2\nExp number of AAs in TMHs: 37.4699899999999\nExp number, first 60 AAs: 19.72699\nTotal prob of N-in: 0.96171\nPOSSIBLE N-term signal sequence\ninside 1 19\nTMhelix 20 39\noutside 40 325\nTMhelix 326 348\ninside 349 372

16094	GCF_001243145.1_Salmonella_enterica_CVM_N46827-R_v1.0	Salmonella enterica subsp. enterica serovar Albany	Proteobacteria; Gammaproteobacteria; Enterobacterales; Enterobacteriaceae; Salmonella; Salmonella enterica; Salmonella enterica subsp. enterica	MTGDNTLITSHGINRRDFMKLCAALAATMGLSSKAAAEMAESVSNPQRPPVIWIGAQECT	WP_000145429.1	hydrogenase 2 small subunit [Salmonella enterica]	Length: 372\nNumber of predicted TMHs:  1\nExp number of AAs in TMHs: 24.51309\nExp number, first 60 AAs:  0.2682\nTotal prob of N-in: 0.14402\noutside	1	331\nTMhelix	332	354\ninside	355	372

16095	GCF_001243145.1_Salmonella_enterica_CVM_N46827-R_v1.0	Salmonella enterica subsp. enterica serovar Albany	Proteobacteria; Gammaproteobacteria; Enterobacterales; Enterobacteriaceae; Salmonella; Salmonella enterica; Salmonella enterica subsp. enterica	MNNEETFYQAMRRKGVTRRSFLKFCSLAATSLGLGAGMTPKIAWALENKPRIPVVWIHGL	WP_001058311.1	hydrogenase [Salmonella enterica]	Length: 372\nNumber of predicted TMHs:  2\nExp number of AAs in TMHs: 37.4699899999999\nExp number, first 60 AAs:  19.72699\nTotal prob of N-in: 0.96171\nPOSSIBLE N-term signal sequence\ninside	1	19\nTMhelix	20	39\noutside	40	325\nTMhelix	326	348\ninside	349	372

16096	GCF_001241765.1_Salmonella_enterica_CVM_N45398_v1.0	Salmonella enterica subsp. enterica serovar Albany	Proteobacteria; Gammaproteobacteria; Enterobacterales; Enterobacteriaceae; Salmonella; Salmonella enterica; Salmonella enterica subsp. enterica	MTGDNTLITSHGINRRDFMKLCAALAATMGLSSKAAAEMAESVSNPQRPPVIWIGAQECT	WP_000145429.1	hydrogenase 2 small subunit [Salmonella enterica]	Length: 372\nNumber of predicted TMHs:  1\nExp number of AAs in TMHs: 24.51309\nExp number, first 60 AAs:  0.2682\nTotal prob of N-in: 0.14402\noutside	1	331\nTMhelix	332	354\ninside	355	372

16097	GCF_002032745.1_ASM203274v1	Salmonella enterica subsp. enterica serovar Albany	Proteobacteria; Gammaproteobacteria; Enterobacterales; Enterobacteriaceae; Salmonella; Salmonella enterica; Salmonella enterica subsp. enterica	MTGDNTLITSHGINRRDFMKLCAALAATMGLSSKAAAEMAESVSNPQRPPVIWIGAQECT	WP_000145429.1	hydrogenase 2 small subunit [Salmonella enterica]	Length: 372\nNumber of predicted TMHs:  1\nExp number of AAs in TMHs: 24.51309\nExp number, first 60 AAs:  0.2682\nTotal prob of N-in: 0.14402\noutside	1	331\nTMhelix	332	354\ninside	355	372

16098	GCF_001244795.1_Salmonella_enterica_CVM_N48692_v1.0	Salmonella enterica subsp. enterica serovar Albany	Proteobacteria; Gammaproteobacteria; Enterobacterales; Enterobacteriaceae; Salmonella; Salmonella enterica; Salmonella enterica subsp. enterica	MTGDNTLITSHGINRRDFMKLCAALAATMGLSSKAAAEMAESVSNPQRPPVIWIGAQECT	WP_000145429.1	hydrogenase 2 small subunit [Salmonella enterica]	Length: 372\nNumber of predicted TMHs:  1\nExp number of AAs in TMHs: 24.51309\nExp number, first 60 AAs:  0.2682\nTotal prob of N-in: 0.14402\noutside	1	331\nTMhelix	332	354\ninside	355	372

16099	GCF_002047105.1_ASM204710v1	Salmonella enterica subsp. enterica serovar Albany	Proteobacteria; Gammaproteobacteria; Enterobacterales; Enterobacteriaceae; Salmonella; Salmonella enterica; Salmonella enterica subsp. enterica	MNNEETFYQAMRRKGVTRRSFLKFCSLAATSLGLGAGMTPKIAWALENKPRIPVVWIHGL	WP_001058311.1	hydrogenase [Salmonella enterica]	Length: 372\nNumber of predicted TMHs:  2\nExp number of AAs in TMHs: 37.4699899999999\nExp number, first 60 AAs:  19.72699\nTotal prob of N-in: 0.96171\nPOSSIBLE N-term signal sequence\ninside	1	19\nTMhelix	20	39\noutside	40	325\nTMhelix	326	348\ninside	349	372

16100	GCF_002047035.1_ASM204703v1	Salmonella enterica subsp. enterica serovar Albany	Proteobacteria; Gammaproteobacteria; Enterobacterales; Enterobacteriaceae; Salmonella; Salmonella enterica; Salmonella enterica subsp. enterica	MNNEETFYQAMRRKGVTRRSFLKFCSLAATSLGLGAGMTPKIAWALENKPRIPVVWIHGL	WP_001058311.1	hydrogenase [Salmonella enterica]	Length: 372\nNumber of predicted TMHs:  2\nExp number of AAs in TMHs: 37.4699899999999\nExp number, first 60 AAs:  19.72699\nTotal prob of N-in: 0.96171\nPOSSIBLE N-term signal sequence\ninside	1	19\nTMhelix	20	39\noutside	40	325\nTMhelix	326	348\ninside	349	372

16101	GCF_001244795.1_Salmonella_enterica_CVM_N48692_v1.0	Salmonella enterica subsp. enterica serovar Albany	Proteobacteria; Gammaproteobacteria; Enterobacterales; Enterobacteriaceae; Salmonella; Salmonella enterica; Salmonella enterica subsp. enterica	MNNEETFYQAMRRKGVTRRSFLKFCSLAATSLGLGAGMTPKIAWALENKPRIPVVWIHGL	WP_001058311.1	hydrogenase [Salmonella enterica]	Length: 372\nNumber of predicted TMHs:  2\nExp number of AAs in TMHs: 37.4699899999999\nExp number, first 60 AAs:  19.72699\nTotal prob of N-in: 0.96171\nPOSSIBLE N-term signal sequence\ninside	1	19\nTMhelix	20	39\noutside	40	325\nTMhelix	326	348\ninside	349	372

16102 GCF_002047105.1_ASM204710v1 Salmonella enterica subsp. enterica serovar Albany Proteobacteria; Gammaproteobacteria; Enterobacterales; Enterobacteriaceae; Salmonella; Salmonella enterica; Salmonella enterica subsp. enterica MTGDNTLITSHGINRRDFMKLCAALAATMGLSSKAAAEMAESVSNPQRPPVIWIGAQECT WP_000145429.1 hydrogenase 2 small subunit [Salmonella enterica] Length: 372\nNumber of predicted TMHs: 1\nExp number of AAs in TMHs: 24.51309\nExp number, first 60 AAs: 0.2682\nTotal prob of N-in: 0.14402\noutside 1 331\nTMhelix 332 354\ninside 355 372

16103 GCF_001246755.1_Salmonella_enterica_CVM_N51278_v1.0 Salmonella enterica subsp. enterica serovar Albany Proteobacteria; Gammaproteobacteria; Enterobacterales; Enterobacteriaceae; Salmonella; Salmonella enterica; Salmonella enterica subsp. enterica MNNEETFYQAMRRKGVTRRSFLKFCSLAATSLGLGAGMTPKIAWALENKPRIPVVWIHGL WP_001058311.1 hydrogenase [Salmonella enterica] Length: 372\nNumber of predicted TMHs: 2\nExp number of AAs in TMHs: 37.4699899999999\nExp number, first 60 AAs: 19.72699\nTotal prob of N-in: 0.96171\nPOSSIBLE N-term signal sequence\ninside 1 19\nTMhelix 20 39\noutside 40 325\nTMhelix 326 348\ninside 349 372

16104 GCF_001479845.1_Salmonella_enterica_CVM_N38860-SQ_v1.0 Salmonella enterica subsp. enterica serovar Albany Proteobacteria; Gammaproteobacteria; Enterobacterales; Enterobacteriaceae; Salmonella; Salmonella enterica; Salmonella enterica subsp. enterica MNNEETFYQAMRRKGVTRRSFLKFCSLAATSLGLGAGMTPKIAWALENKPRIPVVWIHGL WP_001058311.1 hydrogenase [Salmonella enterica] Length: 372\nNumber of predicted TMHs: 2\nExp number of AAs in TMHs: 37.4699899999999\nExp number, first 60 AAs: 19.72699\nTotal prob of N-in: 0.96171\nPOSSIBLE N-term signal sequence\ninside 1 19\nTMhelix 20 39\noutside 40 325\nTMhelix 326 348\ninside 349 372

16105 GCF_002047035.1_ASM204703v1 Salmonella enterica subsp. enterica serovar Albany Proteobacteria; Gammaproteobacteria; Enterobacterales; Enterobacteriaceae; Salmonella; Salmonella enterica; Salmonella enterica subsp. enterica MTGDNTLITSHGINRRDFMKLCAALAATMGLSSKAAAEMAESVSNPQRPPVIWIGAQECT WP_000145429.1 hydrogenase 2 small subunit [Salmonella enterica] Length: 372\nNumber of predicted TMHs: 1\nExp number of AAs in TMHs: 24.51309\nExp number, first 60 AAs: 0.2682\nTotal prob of N-in: 0.14402\noutside 1 331\nTMhelix 332 354\ninside 355 372

16106 GCF_002032745.1_ASM203274v1 Salmonella enterica subsp. enterica serovar Albany Proteobacteria; Gammaproteobacteria; Enterobacterales; Enterobacteriaceae; Salmonella; Salmonella enterica; Salmonella enterica subsp. enterica MNNEETFYQAMRRKGVTRRSFLKFCSLAATSLGLGAGMTPKIAWALENKPRIPVVWIHGL WP_001058311.1 hydrogenase [Salmonella enterica] Length: 372\nNumber of predicted TMHs: 2\nExp number of AAs in TMHs: 37.4699899999999\nExp number, first 60 AAs: 19.72699\nTotal prob of N-in: 0.96171\nPOSSIBLE N-term signal sequence\ninside 1 19\nTMhelix 20 39\noutside 40 325\nTMhelix 326 348\ninside 349 372

16107 GCF_002031475.1_ASM203147v1 Salmonella enterica subsp. enterica serovar Albany Proteobacteria; Gammaproteobacteria; Enterobacterales; Enterobacteriaceae; Salmonella; Salmonella enterica; Salmonella enterica subsp. enterica MTGDNTLITSHGINRRDFMKLCAALAATMGLSSKAAAEMAESVSNPQRPPVIWIGAQECT WP_000145429.1 hydrogenase 2 small subunit [Salmonella enterica] Length: 372\nNumber of predicted TMHs: 1\nExp number of AAs in TMHs: 24.51309\nExp number, first 60 AAs: 0.2682\nTotal prob of N-in: 0.14402\noutside 1 331\nTMhelix 332 354\ninside 355 372

16108 GCF_002031875.1_ASM203187v1 Salmonella enterica subsp. enterica serovar Albert Proteobacteria; Gammaproteobacteria; Enterobacterales; Enterobacteriaceae; Salmonella; Salmonella enterica; Salmonella enterica subsp. enterica MTGDNTLITSHGINRRDFMKLCAALAATMGLSSKAAAEMAESVSNPQRPPVIWIGAQECT WP_000145429.1 hydrogenase 2 small subunit [Salmonella enterica] Length: 372\nNumber of predicted TMHs: 1\nExp number of AAs in TMHs: 24.51309\nExp number, first 60 AAs: 0.2682\nTotal prob of N-in: 0.14402\noutside 1 331\nTMhelix 332 354\ninside 355 372

16109 GCF_002031875.1_ASM203187v1 Salmonella enterica subsp. enterica serovar Albert Proteobacteria; Gammaproteobacteria; Enterobacterales; Enterobacteriaceae; Salmonella; Salmonella enterica; Salmonella enterica subsp. enterica MNNEETFYQAMRRKGVTRRSFLKFCSLAATSLGLGAGMTPKIAWALENKPRIPVVWIHGL WP_001058308.1 hydrogenase [Salmonella enterica] Length: 372\nNumber of predicted TMHs: 2\nExp number of AAs in TMHs: 39.08204\nExp number, first 60 AAs: 19.72474\nTotal prob of N-in: 0.96174\nPOSSIBLE N-term signal sequence\ninside 1 19\nTMhelix 20 39\noutside 40 325\nTMhelix 326 348\ninside 349 372

16110 GCF_002066665.1_ASM206666v1 Salmonella enterica subsp. enterica serovar Amager Proteobacteria; Gammaproteobacteria; Enterobacterales; Enterobacteriaceae; Salmonella; Salmonella enterica;

Salmonella enterica subsp. enterica MNNEETFYQAMRRKGVTRRSFLKFCSLAATSLGLGAGMTPKIAWALENKPRIPVVWIHGL WP_001058312.1 hydrogenase [Salmonella enterica] Length: 372\nNumber of predicted TMHs: 2\nExp number of AAs in TMHs: 36.51738\nExp number, first 60 AAs: 19.72849\nTotal prob of N-in: 0.96170\nPOSSIBLE N-term signal sequence\ninside 1 19\nTMhelix 20 39\noutside 40 325\nTMhelix 326 348\ninside 349 372

16111 GCF_002066665.1_ASM206666v1 Salmonella enterica subsp. enterica serovar Amager Proteobacteria; Gammaproteobacteria; Enterobacterales; Enterobacteriaceae; Salmonella; Salmonella enterica; Salmonella enterica subsp. enterica MTGDNTLITSHGINRRDFMKLCAALAATMGLSSKAAAEMAESVSNPQRPPVIWIGAQECT WP_000145429.1 hydrogenase 2 small subunit [Salmonella enterica] Length: 372\nNumber of predicted TMHs: 1\nExp number of AAs in TMHs: 24.51309\nExp number, first 60 AAs: 0.2682\nTotal prob of N-in: 0.14402\noutside 1 331\nTMhelix 332 354\ninside 355 372

16112 GCF_002066265.1_ASM206626v1 Salmonella enterica subsp. enterica serovar Amager Proteobacteria; Gammaproteobacteria; Enterobacterales; Enterobacteriaceae; Salmonella; Salmonella enterica; Salmonella enterica subsp. enterica MNNEETFYQAMRRKGVTRRSFLKFCSLAATSLGLGAGMTPKIAWALENKPRIPVVWIHGL WP_001058312.1 hydrogenase [Salmonella enterica] Length: 372\nNumber of predicted TMHs: 2\nExp number of AAs in TMHs: 36.51738\nExp number, first 60 AAs: 19.72849\nTotal prob of N-in: 0.96170\nPOSSIBLE N-term signal sequence\ninside 1 19\nTMhelix 20 39\noutside 40 325\nTMhelix 326 348\ninside 349 372

16113 GCF_002066265.1_ASM206626v1 Salmonella enterica subsp. enterica serovar Amager Proteobacteria; Gammaproteobacteria; Enterobacterales; Enterobacteriaceae; Salmonella; Salmonella enterica; Salmonella enterica subsp. enterica MTGDNTLITSHGINRRDFMKLCAALAATMGLSSKAAAEMAESVSNPQRPPVIWIGAQECT WP_000145429.1 hydrogenase 2 small subunit [Salmonella enterica] Length: 372\nNumber of predicted TMHs: 1\nExp number of AAs in TMHs: 24.51309\nExp number, first 60 AAs: 0.2682\nTotal prob of N-in: 0.14402\noutside 1 331\nTMhelix 332 354\ninside 355 372

16114 GCF_002062115.1_ASM206211v1 Salmonella enterica subsp. enterica serovar Amsterdam Proteobacteria; Gammaproteobacteria; Enterobacterales; Enterobacteriaceae; Salmonella; Salmonella enterica; Salmonella enterica subsp. enterica MNNEETFYQAMRRKGVTRRSFLKFCSLATTSLGLGAGMTPKIAWALENKPRIPVVWIHGL WP_080163959.1 [Ni/Fe] hydrogenase small subunit [Salmonella enterica] Length: 372\nNumber of predicted TMHs: 2\nExp number of AAs in TMHs: 35.9899899999999\nExp number, first 60 AAs: 19.19891\nTotal prob of N-in: 0.93662\nPOSSIBLE N-term signal sequence\ninside 1 19\nTMhelix 20 39\noutside 40 325\nTMhelix 326 348\ninside 349 372

16115 GCF_002062115.1_ASM206211v1 Salmonella enterica subsp. enterica serovar Amsterdam Proteobacteria; Gammaproteobacteria; Enterobacterales; Enterobacteriaceae; Salmonella; Salmonella enterica; Salmonella enterica subsp. enterica MTGDNTLITSHGINRRDFMKLCAALAATMGLSSKAAAEMAESVSNPQRPPVIWIGAQECT WP_000145429.1 hydrogenase 2 small subunit [Salmonella enterica] Length: 372\nNumber of predicted TMHs: 1\nExp number of AAs in TMHs: 24.51309\nExp number, first 60 AAs: 0.2682\nTotal prob of N-in: 0.14402\noutside 1 331\nTMhelix 332 354\ninside 355 372

16116 GCF_002060035.1_ASM206003v1 Salmonella enterica subsp. enterica serovar Anatum Proteobacteria; Gammaproteobacteria; Enterobacterales; Enterobacteriaceae; Salmonella; Salmonella enterica; Salmonella enterica subsp. enterica MNNEETFYQAMRRKGVTRRSFLKFCSLAATSLGLGAGMTPKIAWALENKPRIPVVWIHGL WP_023243720.1 hydrogenase 1, small subunit [Salmonella enterica] Length: 372\nNumber of predicted TMHs: 2\nExp number of AAs in TMHs: 36.52147\nExp number, first 60 AAs: 19.72838\nTotal prob of N-in: 0.96170\nPOSSIBLE N-term signal sequence\ninside 1 19\nTMhelix 20 39\noutside 40 325\nTMhelix 326 348\ninside 349 372

16117 GCF_001478355.1_Salmonella_enterica_CVM_N38876-SQ_v1.0 Salmonella enterica subsp. enterica serovar Anatum Proteobacteria; Gammaproteobacteria; Enterobacterales; Enterobacteriaceae; Salmonella; Salmonella enterica; Salmonella enterica subsp. enterica MNNEETFYQAMRRKGVTRRSFLKFCSLAATSLGLGAGMTPKIAWALENKPRIPVVWIHGL WP_023243720.1 hydrogenase 1, small subunit [Salmonella enterica] Length: 372\nNumber of predicted TMHs: 2\nExp number of AAs in TMHs: 36.52147\nExp number, first 60 AAs: 19.72838\nTotal prob of N-in: 0.96170\nPOSSIBLE N-term signal sequence\ninside 1 19\nTMhelix 20 39\noutside 40 325\nTMhelix 326 348\ninside 349 372

16118 GCF_002060035.1_ASM206003v1 Salmonella enterica subsp. enterica serovar Anatum Proteobacteria; Gammaproteobacteria; Enterobacterales; Enterobacteriaceae; Salmonella; Salmonella enterica; Salmonella enterica subsp. enterica MTGDNTLITSHGINRRDFMKLCAALAATMGLSSKAAAEMAESVSNPQRPPVIWIGAQECT WP_000145429.1 hydrogenase 2 small subunit [Salmonella enterica] Length: 372\nNumber of

predicted TMHs: 1\nExp number of AAs in TMHs: 24.51309\nExp number, first 60 AAs: 0.2682\nTotal prob of N-in: 0.14402\noutside 1 331\nTMhelix 332 354\ninside 355 372

16119 GCF_002059305.1_ASM205930v1 Salmonella enterica subsp. enterica serovar Anatum Proteobacteria; Gammaproteobacteria; Enterobacterales; Enterobacteriaceae; Salmonella; Salmonella enterica; Salmonella enterica subsp. enterica MNNEETFYQAMRRKGVTRRSFLKFCSLAATSLGLGAGMTPKIAWALENKPRIPVVWIHGL WP_023243720.1 hydrogenase 1, small subunit [Salmonella enterica] Length: 372\nNumber of predicted TMHs: 2\nExp number of AAs in TMHs: 36.52147\nExp number, first 60 AAs: 19.72838\nTotal prob of N-in: 0.96170\nPOSSIBLE N-term signal sequence\ninside 1 19\nTMhelix 20 39\noutside 40 325\nTMhelix 326 348\ninside 349 372

16120 GCF_001247355.1_Salmonella_enterica_CVM_N51307_v1.0 Salmonella enterica subsp. enterica serovar Anatum Proteobacteria; Gammaproteobacteria; Enterobacterales; Enterobacteriaceae; Salmonella; Salmonella enterica; Salmonella enterica subsp. enterica MNNEETFYQAMRRKGVTRRSFLKFCSLAATSLGLGAGMTPKIAWALENKPRIPVVWIHGL WP_023243720.1 hydrogenase 1, small subunit [Salmonella enterica] Length: 372\nNumber of predicted TMHs: 2\nExp number of AAs in TMHs: 36.52147\nExp number, first 60 AAs: 19.72838\nTotal prob of N-in: 0.96170\nPOSSIBLE N-term signal sequence\ninside 1 19\nTMhelix 20 39\noutside 40 325\nTMhelix 326 348\ninside 349 372

16121 GCF_002032815.1_ASM203281v1 Salmonella enterica subsp. enterica serovar Anatum Proteobacteria; Gammaproteobacteria; Enterobacterales; Enterobacteriaceae; Salmonella; Salmonella enterica; Salmonella enterica subsp. enterica MNNEETFYQAMRRKGVTRRSFLKFCSLAATSLGLGAGMTPKIAWALENKPRIPVVWIHGL WP_023243720.1 hydrogenase 1, small subunit [Salmonella enterica] Length: 372\nNumber of predicted TMHs: 2\nExp number of AAs in TMHs: 36.52147\nExp number, first 60 AAs: 19.72838\nTotal prob of N-in: 0.96170\nPOSSIBLE N-term signal sequence\ninside 1 19\nTMhelix 20 39\noutside 40 325\nTMhelix 326 348\ninside 349 372

16122 GCF_001241205.1_Salmonella_enterica_CVM_N44698-R_v1.0 Salmonella enterica subsp. enterica serovar Anatum Proteobacteria; Gammaproteobacteria; Enterobacterales; Enterobacteriaceae; Salmonella; Salmonella enterica; Salmonella enterica subsp. enterica MNNEETFYQAMRRKGVTRRSFLKFCSLAATSLGLGAGMTPKIAWALENKPRIPVVWIHGL WP_023243720.1 hydrogenase 1, small subunit [Salmonella enterica] Length: 372\nNumber of predicted TMHs: 2\nExp number of AAs in TMHs: 36.52147\nExp number, first 60 AAs: 19.72838\nTotal prob of N-in: 0.96170\nPOSSIBLE N-term signal sequence\ninside 1 19\nTMhelix 20 39\noutside 40 325\nTMhelix 326 348\ninside 349 372

16123 GCF_001241225.1_Salmonella_enterica_CVM_N44699-R_v1.0 Salmonella enterica subsp. enterica serovar Anatum Proteobacteria; Gammaproteobacteria; Enterobacterales; Enterobacteriaceae; Salmonella; Salmonella enterica; Salmonella enterica subsp. enterica MTGDNTLITSHGINRRDFMKLCAALAATMGLSSKAAAEMAESVSNPQRPPVIWIGAQECT WP_000145429.1 hydrogenase 2 small subunit [Salmonella enterica] Length: 372\nNumber of predicted TMHs: 1\nExp number of AAs in TMHs: 24.51309\nExp number, first 60 AAs: 0.2682\nTotal prob of N-in: 0.14402\noutside 1 331\nTMhelix 332 354\ninside 355 372

16124 GCF_001241205.1_Salmonella_enterica_CVM_N44698-R_v1.0 Salmonella enterica subsp. enterica serovar Anatum Proteobacteria; Gammaproteobacteria; Enterobacterales; Enterobacteriaceae; Salmonella; Salmonella enterica; Salmonella enterica subsp. enterica MTGDNTLITSHGINRRDFMKLCAALAATMGLSSKAAAEMAESVSNPQRPPVIWIGAQECT WP_000145429.1 hydrogenase 2 small subunit [Salmonella enterica] Length: 372\nNumber of predicted TMHs: 1\nExp number of AAs in TMHs: 24.51309\nExp number, first 60 AAs: 0.2682\nTotal prob of N-in: 0.14402\noutside 1 331\nTMhelix 332 354\ninside 355 372

16125 GCF_001241225.1_Salmonella_enterica_CVM_N44699-R_v1.0 Salmonella enterica subsp. enterica serovar Anatum Proteobacteria; Gammaproteobacteria; Enterobacterales; Enterobacteriaceae; Salmonella; Salmonella enterica; Salmonella enterica subsp. enterica MNNEETFYQAMRRKGVTRRSFLKFCSLAATSLGLGAGMTPKIAWALENKPRIPVVWIHGL WP_023243720.1 hydrogenase 1, small subunit [Salmonella enterica] Length: 372\nNumber of predicted TMHs: 2\nExp number of AAs in TMHs: 36.52147\nExp number, first 60 AAs: 19.72838\nTotal prob of N-in: 0.96170\nPOSSIBLE N-term signal sequence\ninside 1 19\nTMhelix 20 39\noutside 40 325\nTMhelix 326 348\ninside 349 372

16126 GCF_002066255.1_ASM206625v1 Salmonella enterica subsp. enterica serovar Anatum Proteobacteria; Gammaproteobacteria; Enterobacterales; Enterobacteriaceae; Salmonella; Salmonella enterica; Salmonella enterica subsp. enterica MTGDNTLITSHGINRRDFMKLCAALAATMGLSSKAAAEMAESVSNPQRPPVIWIGAQECT WP_000145429.1 hydrogenase 2 small subunit [Salmonella enterica] Length: 372\nNumber of

predicted TMHs: 1\nExp number of AAs in TMHs: 24.51309\nExp number, first 60 AAs: 0.2682\nTotal prob of N-in: 0.14402\noutside 1 331\nTMhelix 332 354\ninside 355 372

16127 GCF_002066255.1_ASM206625v1 Salmonella enterica subsp. enterica serovar Anatum Proteobacteria; Gammaproteobacteria; Enterobacterales; Enterobacteriaceae; Salmonella; Salmonella enterica; Salmonella enterica subsp. enterica MNNEETFYQAMRRKGVTRRSFLKFCSLAATSLGLGAGMTPKIAWALENKPRIPVVWIHGL WP_023243720.1 hydrogenase 1, small subunit [Salmonella enterica] Length: 372\nNumber of predicted TMHs: 2\nExp number of AAs in TMHs: 36.52147\nExp number, first 60 AAs: 19.72838\nTotal prob of N-in: 0.96170\nPOSSIBLE N-term signal sequence\ninside 1 19\nTMhelix 20 39\noutside 40 325\nTMhelix 326 348\ninside 349 372

16128 GCF_002058525.1_ASM205852v1 Salmonella enterica subsp. enterica serovar Anatum Proteobacteria; Gammaproteobacteria; Enterobacterales; Enterobacteriaceae; Salmonella; Salmonella enterica; Salmonella enterica subsp. enterica MTGDNTLITSHGINRRDFMKLCAALAATMGLSSKAAAEMAESVSNPQRPPVIWIGAQECT WP_000145429.1 hydrogenase 2 small subunit [Salmonella enterica] Length: 372\nNumber of predicted TMHs: 1\nExp number of AAs in TMHs: 24.51309\nExp number, first 60 AAs: 0.2682\nTotal prob of N-in: 0.14402\noutside 1 331\nTMhelix 332 354\ninside 355 372

16129 GCF_001835345.1_ASM183534v1 Salmonella enterica subsp. enterica serovar Anatum Proteobacteria; Gammaproteobacteria; Enterobacterales; Enterobacteriaceae; Salmonella; Salmonella enterica; Salmonella enterica subsp. enterica MTGDNTLITSHGINRRDFMKLCAALAATMGLSSKAAAEMAESVSNPQRPPVIWIGAQECT WP_000145429.1 hydrogenase 2 small subunit [Salmonella enterica] Length: 372\nNumber of predicted TMHs: 1\nExp number of AAs in TMHs: 24.51309\nExp number, first 60 AAs: 0.2682\nTotal prob of N-in: 0.14402\noutside 1 331\nTMhelix 332 354\ninside 355 372

16130 GCF_002066235.1_ASM206623v1 Salmonella enterica subsp. enterica serovar Anatum Proteobacteria; Gammaproteobacteria; Enterobacterales; Enterobacteriaceae; Salmonella; Salmonella enterica; Salmonella enterica subsp. enterica MNNEETFYQAMRRKGVTRRSFLKFCSLAATSLGLGAGMTPKIAWALENKPRIPVVWIHGL WP_023243720.1 hydrogenase 1, small subunit [Salmonella enterica] Length: 372\nNumber of predicted TMHs: 2\nExp number of AAs in TMHs: 36.52147\nExp number, first 60 AAs: 19.72838\nTotal prob of N-in: 0.96170\nPOSSIBLE N-term signal sequence\ninside 1 19\nTMhelix 20 39\noutside 40 325\nTMhelix 326 348\ninside 349 372

16131 GCF_001835345.1_ASM183534v1 Salmonella enterica subsp. enterica serovar Anatum Proteobacteria; Gammaproteobacteria; Enterobacterales; Enterobacteriaceae; Salmonella; Salmonella enterica; Salmonella enterica subsp. enterica MNNEETFYQAMRRKGVTRRSFLKFCSLAATSLGLGAGMTPKIAWALENKPRIPVVWIHGL WP_023243720.1 hydrogenase 1, small subunit [Salmonella enterica] Length: 372\nNumber of predicted TMHs: 2\nExp number of AAs in TMHs: 36.52147\nExp number, first 60 AAs: 19.72838\nTotal prob of N-in: 0.96170\nPOSSIBLE N-term signal sequence\ninside 1 19\nTMhelix 20 39\noutside 40 325\nTMhelix 326 348\ninside 349 372

16132 GCF_002066235.1_ASM206623v1 Salmonella enterica subsp. enterica serovar Anatum Proteobacteria; Gammaproteobacteria; Enterobacterales; Enterobacteriaceae; Salmonella; Salmonella enterica; Salmonella enterica subsp. enterica MTGDNTLITSHGINRRDFMKLCAALAATMGLSSKAAAEMAESVSNPQRPPVIWIGAQECT WP_000145429.1 hydrogenase 2 small subunit [Salmonella enterica] Length: 372\nNumber of predicted TMHs: 1\nExp number of AAs in TMHs: 24.51309\nExp number, first 60 AAs: 0.2682\nTotal prob of N-in: 0.14402\noutside 1 331\nTMhelix 332 354\ninside 355 372

16133 GCF_002061315.1_ASM206131v1 Salmonella enterica subsp. enterica serovar Anatum Proteobacteria; Gammaproteobacteria; Enterobacterales; Enterobacteriaceae; Salmonella; Salmonella enterica; Salmonella enterica subsp. enterica MTGDNTLITSHGINRRDFMKLCAALAATMGLSSKAAAEMAESVSNPQRPPVIWIGAQECT WP_000145429.1 hydrogenase 2 small subunit [Salmonella enterica] Length: 372\nNumber of predicted TMHs: 1\nExp number of AAs in TMHs: 24.51309\nExp number, first 60 AAs: 0.2682\nTotal prob of N-in: 0.14402\noutside 1 331\nTMhelix 332 354\ninside 355 372

16134 GCF_002059305.1_ASM205930v1 Salmonella enterica subsp. enterica serovar Anatum Proteobacteria; Gammaproteobacteria; Enterobacterales; Enterobacteriaceae; Salmonella; Salmonella enterica; Salmonella enterica subsp. enterica MTGDNTLITSHGINRRDFMKLCAALAATMGLSSKAAAEMAESVSNPQRPPVIWIGAQECT WP_000145429.1 hydrogenase 2 small subunit [Salmonella enterica] Length: 372\nNumber of predicted TMHs: 1\nExp number of AAs in TMHs: 24.51309\nExp number, first 60 AAs: 0.2682\nTotal prob of N-in: 0.14402\noutside 1 331\nTMhelix 332 354\ninside 355 372

16135 GCF_001481105.1_Salmonella_enterica_CVM_N42473-SQ_v1.0 Salmonella enterica subsp. enterica serovar Anatum Proteobacteria; Gammaproteobacteria; Enterobacterales; Enterobacteriaceae; Salmonella; Salmonella enterica; Salmonella enterica subsp. enterica MNNEETFYQAMRRKGVTRRSFLKFCSLAATSLGLGAGMTPKIAWALENKPRIPVVWIHGL WP_023243720.1 hydrogenase 1, small subunit [Salmonella enterica] Length: 372\nNumber of predicted TMHs: 2\nExp number of AAs in TMHs: 36.52147\nExp number, first 60 AAs: 19.72838\nTotal prob of N-in: 0.96170\nPOSSIBLE N-term signal sequence\ninside 1 19\nTMhelix 20 39\noutside 40 325\nTMhelix 326 348\ninside 349 372

16136 GCF_002059595.1_ASM205959v1 Salmonella enterica subsp. enterica serovar Anatum Proteobacteria; Gammaproteobacteria; Enterobacterales; Enterobacteriaceae; Salmonella; Salmonella enterica; Salmonella enterica subsp. enterica MTGDNTLITSHGINRRDFMKLCAALAATMGLSSKAAAEMAESVSNPQRPPVIWIGAQECT WP_000145429.1 hydrogenase 2 small subunit [Salmonella enterica] Length: 372\nNumber of predicted TMHs: 1\nExp number of AAs in TMHs: 24.51309\nExp number, first 60 AAs: 0.2682\nTotal prob of N-in: 0.14402\noutside 1 331\nTMhelix 332 354\ninside 355 372

16137 GCF_002059595.1_ASM205959v1 Salmonella enterica subsp. enterica serovar Anatum Proteobacteria; Gammaproteobacteria; Enterobacterales; Enterobacteriaceae; Salmonella; Salmonella enterica; Salmonella enterica subsp. enterica MNNEETFYQAMRRKGVTRRSFLKFCSLAATSLGLGAGMTPKIAWALENKPRIPVVWIHGL WP_001058312.1 hydrogenase [Salmonella enterica] Length: 372\nNumber of predicted TMHs: 2\nExp number of AAs in TMHs: 36.51738\nExp number, first 60 AAs: 19.72849\nTotal prob of N-in: 0.96170\nPOSSIBLE N-term signal sequence\ninside 1 19\nTMhelix 20 39\noutside 40 325\nTMhelix 326 348\ninside 349 372

16138 GCF_001726095.1_ASM172609v1 Salmonella enterica subsp. enterica serovar Anatum Proteobacteria; Gammaproteobacteria; Enterobacterales; Enterobacteriaceae; Salmonella; Salmonella enterica; Salmonella enterica subsp. enterica MTGDNTLITSHGINRRDFMKLCAALAATMGLSSKAAAEMAESVSNPQRPPVIWIGAQECT WP_000145429.1 hydrogenase 2 small subunit [Salmonella enterica] Length: 372\nNumber of predicted TMHs: 1\nExp number of AAs in TMHs: 24.51309\nExp number, first 60 AAs: 0.2682\nTotal prob of N-in: 0.14402\noutside 1 331\nTMhelix 332 354\ninside 355 372

16139 GCF_002058585.1_ASM205858v1 Salmonella enterica subsp. enterica serovar Anatum Proteobacteria; Gammaproteobacteria; Enterobacterales; Enterobacteriaceae; Salmonella; Salmonella enterica; Salmonella enterica subsp. enterica MTGDNTLITSHGINRRDFMKLCAALAATMGLSSKAAAEMAESVSNPQRPPVIWIGAQECT WP_000145429.1 hydrogenase 2 small subunit [Salmonella enterica] Length: 372\nNumber of predicted TMHs: 1\nExp number of AAs in TMHs: 24.51309\nExp number, first 60 AAs: 0.2682\nTotal prob of N-in: 0.14402\noutside 1 331\nTMhelix 332 354\ninside 355 372

16140 GCF_001481105.1_Salmonella_enterica_CVM_N42473-SQ_v1.0 Salmonella enterica subsp. enterica serovar Anatum Proteobacteria; Gammaproteobacteria; Enterobacterales; Enterobacteriaceae; Salmonella; Salmonella enterica; Salmonella enterica subsp. enterica MTGDNTLITSHGINRRDFMKLCAALAATMGLSSKAAAEMAESVSNPQRPPVIWIGAQECT WP_000145429.1 hydrogenase 2 small subunit [Salmonella enterica] Length: 372\nNumber of predicted TMHs: 1\nExp number of AAs in TMHs: 24.51309\nExp number, first 60 AAs: 0.2682\nTotal prob of N-in: 0.14402\noutside 1 331\nTMhelix 332 354\ninside 355 372

16141 GCF_001478355.1_Salmonella_enterica_CVM_N38876-SQ_v1.0 Salmonella enterica subsp. enterica serovar Anatum Proteobacteria; Gammaproteobacteria; Enterobacterales; Enterobacteriaceae; Salmonella; Salmonella enterica; Salmonella enterica subsp. enterica MTGDNTLITSHGINRRDFMKLCAALAATMGLSSKAAAEMAESVSNPQRPPVIWIGAQECT WP_000145429.1 hydrogenase 2 small subunit [Salmonella enterica] Length: 372\nNumber of predicted TMHs: 1\nExp number of AAs in TMHs: 24.51309\nExp number, first 60 AAs: 0.2682\nTotal prob of N-in: 0.14402\noutside 1 331\nTMhelix 332 354\ninside 355 372

16142 GCF_001726095.1_ASM172609v1 Salmonella enterica subsp. enterica serovar Anatum Proteobacteria; Gammaproteobacteria; Enterobacterales; Enterobacteriaceae; Salmonella; Salmonella enterica; Salmonella enterica subsp. enterica MNNEETFYQAMRRKGVTRRSFLKFCSLAATSLGLGAGMTPKIAWALENKPRIPVVWIHGL WP_023243720.1 hydrogenase 1, small subunit [Salmonella enterica] Length: 372\nNumber of predicted TMHs: 2\nExp number of AAs in TMHs: 36.52147\nExp number, first 60 AAs: 19.72838\nTotal prob of N-in: 0.96170\nPOSSIBLE N-term signal sequence\ninside 1 19\nTMhelix 20 39\noutside 40 325\nTMhelix 326 348\ninside 349 372

16143 GCF_001479145.1_Salmonella_enterica_CVM_N31410-R_v1.0 Salmonella enterica subsp. enterica serovar Anatum Proteobacteria; Gammaproteobacteria; Enterobacterales; Enterobacteriaceae; Salmonella; Salmonella enterica; Salmonella enterica subsp. enterica MNNEETFYQAMRRKGVTRRSFLKFCSLAATSLGLGAGMTPKIAWALENKPRIPVVWIHGL

WP_023243720.1 hydrogenase 1, small subunit [Salmonella enterica] Length: 372\nNumber of predicted TMHs: 2\nExp number of AAs in TMHs: 36.52147\nExp number, first 60 AAs: 19.72838\nTotal prob of N-in: 0.96170\nPOSSIBLE N-term signal sequence\ninside 1 19\nTMhelix 20 39\noutside 40 325\nTMhelix 326 348\ninside 349 372

16144 GCF_002032815.1_ASM203281v1 Salmonella enterica subsp. enterica serovar Anatum Proteobacteria; Gammaproteobacteria; Enterobacterales; Enterobacteriaceae; Salmonella; Salmonella enterica; Salmonella enterica subsp. enterica MTGDNTLITSHGINRRDFMKLCAALAATMGLSSKAAAEMAESVSNPQRPPVIWIGAQECT WP_000145429.1 hydrogenase 2 small subunit [Salmonella enterica] Length: 372\nNumber of predicted TMHs: 1\nExp number of AAs in TMHs: 24.51309\nExp number, first 60 AAs: 0.2682\nTotal prob of N-in: 0.14402\noutside 1 331\nTMhelix 332 354\ninside 355 372

16145 GCF_001247355.1_Salmonella_enterica_CVM_N51307_v1.0 Salmonella enterica subsp. enterica serovar Anatum Proteobacteria; Gammaproteobacteria; Enterobacterales; Enterobacteriaceae; Salmonella; Salmonella enterica; Salmonella enterica subsp. enterica MTGDNTLITSHGINRRDFMKLCAALAATMGLSSKAAAEMAESVSNPQRPPVIWIGAQECT WP_000145429.1 hydrogenase 2 small subunit [Salmonella enterica] Length: 372\nNumber of predicted TMHs: 1\nExp number of AAs in TMHs: 24.51309\nExp number, first 60 AAs: 0.2682\nTotal prob of N-in: 0.14402\noutside 1 331\nTMhelix 332 354\ninside 355 372

16146 GCF_001479145.1_Salmonella_enterica_CVM_N31410-R_v1.0 Salmonella enterica subsp. enterica serovar Anatum Proteobacteria; Gammaproteobacteria; Enterobacterales; Enterobacteriaceae; Salmonella; Salmonella enterica; Salmonella enterica subsp. enterica MTGDNTLITSHGINRRDFMKLCAALAATMGLSSKAAAEMAESVSNPQRPPVIWIGAQECT WP_000145429.1 hydrogenase 2 small subunit [Salmonella enterica] Length: 372\nNumber of predicted TMHs: 1\nExp number of AAs in TMHs: 24.51309\nExp number, first 60 AAs: 0.2682\nTotal prob of N-in: 0.14402\noutside 1 331\nTMhelix 332 354\ninside 355 372

16147 GCF_002058525.1_ASM205852v1 Salmonella enterica subsp. enterica serovar Anatum Proteobacteria; Gammaproteobacteria; Enterobacterales; Enterobacteriaceae; Salmonella; Salmonella enterica; Salmonella enterica subsp. enterica MNNEETFYQAMRRKGVTRRSFLKFCSLAATSLGLGAGMTPKIAWALENKPRIPVVWIHGL WP_023243720.1 hydrogenase 1, small subunit [Salmonella enterica] Length: 372\nNumber of predicted TMHs: 2\nExp number of AAs in TMHs: 36.52147\nExp number, first 60 AAs: 19.72838\nTotal prob of N-in: 0.96170\nPOSSIBLE N-term signal sequence\ninside 1 19\nTMhelix 20 39\noutside 40 325\nTMhelix 326 348\ninside 349 372

16148 GCF_001447095.1_ASM144709v1 Salmonella enterica subsp. enterica serovar Anatum Proteobacteria; Gammaproteobacteria; Enterobacterales; Enterobacteriaceae; Salmonella; Salmonella enterica; Salmonella enterica subsp. enterica MNNEETFYQAMRRKGVTRRSFLKFCSLAATSLGLGAGMTPKIAWALENKPRIPVVWIHGL WP_023243720.1 hydrogenase 1, small subunit [Salmonella enterica] Length: 372\nNumber of predicted TMHs: 2\nExp number of AAs in TMHs: 36.52147\nExp number, first 60 AAs: 19.72838\nTotal prob of N-in: 0.96170\nPOSSIBLE N-term signal sequence\ninside 1 19\nTMhelix 20 39\noutside 40 325\nTMhelix 326 348\ninside 349 372

16149 GCF_002061315.1_ASM206131v1 Salmonella enterica subsp. enterica serovar Anatum Proteobacteria; Gammaproteobacteria; Enterobacterales; Enterobacteriaceae; Salmonella; Salmonella enterica; Salmonella enterica subsp. enterica MNNEETFYQAMRRKGVTRRSFLKFCSLAATSLGLGAGMTPKIAWALENKPRIPVVWIHGL WP_023243720.1 hydrogenase 1, small subunit [Salmonella enterica] Length: 372\nNumber of predicted TMHs: 2\nExp number of AAs in TMHs: 36.52147\nExp number, first 60 AAs: 19.72838\nTotal prob of N-in: 0.96170\nPOSSIBLE N-term signal sequence\ninside 1 19\nTMhelix 20 39\noutside 40 325\nTMhelix 326 348\ninside 349 372

16150 GCF_001447115.1_ASM144711v1 Salmonella enterica subsp. enterica serovar Anatum Proteobacteria; Gammaproteobacteria; Enterobacterales; Enterobacteriaceae; Salmonella; Salmonella enterica; Salmonella enterica subsp. enterica MNNEETFYQAMRRKGVTRRSFLKFCSLAATSLGLGAGMTPKIAWALENKPRIPVVWIHGL WP_023243720.1 hydrogenase 1, small subunit [Salmonella enterica] Length: 372\nNumber of predicted TMHs: 2\nExp number of AAs in TMHs: 36.52147\nExp number, first 60 AAs: 19.72838\nTotal prob of N-in: 0.96170\nPOSSIBLE N-term signal sequence\ninside 1 19\nTMhelix 20 39\noutside 40 325\nTMhelix 326 348\ninside 349 372

16151 GCF_001447095.1_ASM144709v1 Salmonella enterica subsp. enterica serovar Anatum Proteobacteria; Gammaproteobacteria; Enterobacterales; Enterobacteriaceae; Salmonella; Salmonella enterica; Salmonella enterica subsp. enterica MTGDNTLITSHGINRRDFMKLCAALAATMGLSSKAAAEMAESVSNPQRPPVIWIGAQECT

WP_000145429.1 hydrogenase 2 small subunit [Salmonella enterica] Length: 372\nNumber of predicted TMHs: 1\nExp number of AAs in TMHs: 24.51309\nExp number, first 60 AAs: 0.2682\nTotal prob of N-in: 0.14402\noutside 1 331\nTMhelix 332 354\ninside 355 372

16152 GCF_001447115.1_ASM144711v1 Salmonella enterica subsp. enterica serovar Anatum Proteobacteria; Gammaproteobacteria; Enterobacterales; Enterobacteriaceae; Salmonella; Salmonella enterica; Salmonella enterica subsp. enterica MTGDNTLITSHGINRRDFMKLCAALAATMGLSSKAAAEMAESVSNPQRPPVIWIGAQECT WP_000145429.1 hydrogenase 2 small subunit [Salmonella enterica] Length: 372\nNumber of predicted TMHs: 1\nExp number of AAs in TMHs: 24.51309\nExp number, first 60 AAs: 0.2682\nTotal prob of N-in: 0.14402\noutside 1 331\nTMhelix 332 354\ninside 355 372

16153 GCF_002060115.1_ASM206011v1 Salmonella enterica subsp. enterica serovar Anatum Proteobacteria; Gammaproteobacteria; Enterobacterales; Enterobacteriaceae; Salmonella; Salmonella enterica; Salmonella enterica subsp. enterica MNNEETFYQAMRRKGVTRRSFLKFCSLAATSLGLGAGMTPKIAWALENKPRIPVVWIHGL WP_023243720.1 hydrogenase 1, small subunit [Salmonella enterica] Length: 372\nNumber of predicted TMHs: 2\nExp number of AAs in TMHs: 36.52147\nExp number, first 60 AAs: 19.72838\nTotal prob of N-in: 0.96170\nPOSSIBLE N-term signal sequence\ninside 1 19\nTMhelix 20 39\noutside 40 325\nTMhelix 326 348\ninside 349 372

16154 GCF_002066295.1_ASM206629v1 Salmonella enterica subsp. enterica serovar Arizonae Proteobacteria; Gammaproteobacteria; Enterobacterales; Enterobacteriaceae; Salmonella; Salmonella enterica; Salmonella enterica subsp. enterica MNNEETFYQAMRRKGVTRRSFLKFCSLAATSLGLGAGMTPKIAWALENKPRIPVVWIHGL WP_001058306.1 hydrogenase [Salmonella enterica] Length: 372\nNumber of predicted TMHs: 2\nExp number of AAs in TMHs: 36.5540099999999\nExp number, first 60 AAs: 19.72732\nTotal prob of N-in: 0.96171\nPOSSIBLE N-term signal sequence\ninside 1 19\nTMhelix 20 39\noutside 40 325\nTMhelix 326 348\ninside 349 372

16155 GCF_002066295.1_ASM206629v1 Salmonella enterica subsp. enterica serovar Arizonae Proteobacteria; Gammaproteobacteria; Enterobacterales; Enterobacteriaceae; Salmonella; Salmonella enterica; Salmonella enterica subsp. enterica MTGDNTLITSHGINRRDFMKLCAALAATMGLSSKAAAEMAESVSNPQRPPVIWIGAQECT WP_000145429.1 hydrogenase 2 small subunit [Salmonella enterica] Length: 372\nNumber of predicted TMHs: 1\nExp number of AAs in TMHs: 24.51309\nExp number, first 60 AAs: 0.2682\nTotal prob of N-in: 0.14402\noutside 1 331\nTMhelix 332 354\ninside 355 372

16156 GCF_002066645.1_ASM206664v1 Salmonella enterica subsp. enterica serovar Augustenborg Proteobacteria; Gammaproteobacteria; Enterobacterales; Enterobacteriaceae; Salmonella; Salmonella enterica; Salmonella enterica subsp. enterica MNNEETFYQAMRRKGVTRRSFLKFCSLAATSLGLGAGMTPKIAWALENKPRIPVVWIHGL WP_001058311.1 hydrogenase [Salmonella enterica] Length: 372\nNumber of predicted TMHs: 2\nExp number of AAs in TMHs: 37.4699899999999\nExp number, first 60 AAs: 19.72699\nTotal prob of N-in: 0.96171\nPOSSIBLE N-term signal sequence\ninside 1 19\nTMhelix 20 39\noutside 40 325\nTMhelix 326 348\ninside 349 372

16157 GCF_002066645.1_ASM206664v1 Salmonella enterica subsp. enterica serovar Augustenborg Proteobacteria; Gammaproteobacteria; Enterobacterales; Enterobacteriaceae; Salmonella; Salmonella enterica; Salmonella enterica subsp. enterica MTGDNTLITSHGINRRDFMKLCAALAATMGLSSKAAAEMAESVSNPQRPPVIWIGAQECT WP_000145429.1 hydrogenase 2 small subunit [Salmonella enterica] Length: 372\nNumber of predicted TMHs: 1\nExp number of AAs in TMHs: 24.51309\nExp number, first 60 AAs: 0.2682\nTotal prob of N-in: 0.14402\noutside 1 331\nTMhelix 332 354\ninside 355 372

16158 GCF_002046175.1_ASM204617v1 Salmonella enterica subsp. enterica serovar Baildon Proteobacteria; Gammaproteobacteria; Enterobacterales; Enterobacteriaceae; Salmonella; Salmonella enterica; Salmonella enterica subsp. enterica MNNEETFYQAMRRKGVTRRSFLKFCSLAATSLGLGAGMTPKIAWALENKPRIPVVWIHGL WP_001058312.1 hydrogenase [Salmonella enterica] Length: 372\nNumber of predicted TMHs: 2\nExp number of AAs in TMHs: 36.51738\nExp number, first 60 AAs: 19.72849\nTotal prob of N-in: 0.96170\nPOSSIBLE N-term signal sequence\ninside 1 19\nTMhelix 20 39\noutside 40 325\nTMhelix 326 348\ninside 349 372

16159 GCF_002046175.1_ASM204617v1 Salmonella enterica subsp. enterica serovar Baildon Proteobacteria; Gammaproteobacteria; Enterobacterales; Enterobacteriaceae; Salmonella; Salmonella enterica; Salmonella enterica subsp. enterica MTGDNTLITSHGINRRDFMKLCAALAATMGLSSKAAAEMAESVSNPQRPPVIWIGAQECT WP_000145429.1 hydrogenase 2 small subunit [Salmonella enterica] Length: 372\nNumber of

predicted TMHs: 1\nExp number of AAs in TMHs: 24.51309\nExp number, first 60 AAs: 0.2682\nTotal prob of N-in: 0.14402\noutside 1 331\nTMhelix 332 354\ninside 355 372

16160 GCF_002046425.1_ASM204642v1 Salmonella enterica subsp. enterica serovar Baildon Proteobacteria; Gammaproteobacteria; Enterobacterales; Enterobacteriaceae; Salmonella; Salmonella enterica; Salmonella enterica subsp. enterica MNNEETFYQAMRRKGVTRRSFLKFCSLAATSLGLGAGMTPKIAWALENKPRIPVVWIHGL WP_001058312.1 hydrogenase [Salmonella enterica] Length: 372\nNumber of predicted TMHs: 2\nExp number of AAs in TMHs: 36.51738\nExp number, first 60 AAs: 19.72849\nTotal prob of N-in: 0.96170\nPOSSIBLE N-term signal sequence\ninside 1 19\nTMhelix 20 39\noutside 40 325\nTMhelix 326 348\ninside 349 372

16161 GCF_002045255.1_ASM204525v1 Salmonella enterica subsp. enterica serovar Baildon Proteobacteria; Gammaproteobacteria; Enterobacterales; Enterobacteriaceae; Salmonella; Salmonella enterica; Salmonella enterica subsp. enterica MTGDNTLITSHGINRRDFMKLCAALAATMGLSSKAAAEMAESVSNPQRPPVIWIGAQECT WP_000145429.1 hydrogenase 2 small subunit [Salmonella enterica] Length: 372\nNumber of predicted TMHs: 1\nExp number of AAs in TMHs: 24.51309\nExp number, first 60 AAs: 0.2682\nTotal prob of N-in: 0.14402\noutside 1 331\nTMhelix 332 354\ninside 355 372

16162 GCF_002046425.1_ASM204642v1 Salmonella enterica subsp. enterica serovar Baildon Proteobacteria; Gammaproteobacteria; Enterobacterales; Enterobacteriaceae; Salmonella; Salmonella enterica; Salmonella enterica subsp. enterica MTGDNTLITSHGINRRDFMKLCAALAATMGLSSKAAAEMAESVSNPQRPPVIWIGAQECT WP_000145429.1 hydrogenase 2 small subunit [Salmonella enterica] Length: 372\nNumber of predicted TMHs: 1\nExp number of AAs in TMHs: 24.51309\nExp number, first 60 AAs: 0.2682\nTotal prob of N-in: 0.14402\noutside 1 331\nTMhelix 332 354\ninside 355 372

16163 GCF_002045255.1_ASM204525v1 Salmonella enterica subsp. enterica serovar Baildon Proteobacteria; Gammaproteobacteria; Enterobacterales; Enterobacteriaceae; Salmonella; Salmonella enterica; Salmonella enterica subsp. enterica MNNEETFYQAMRRKGVTRRSFLKFCSLAATSLGLGAGMTPKIAWALENKPRIPVVWIHGL WP_001058312.1 hydrogenase [Salmonella enterica] Length: 372\nNumber of predicted TMHs: 2\nExp number of AAs in TMHs: 36.51738\nExp number, first 60 AAs: 19.72849\nTotal prob of N-in: 0.96170\nPOSSIBLE N-term signal sequence\ninside 1 19\nTMhelix 20 39\noutside 40 325\nTMhelix 326 348\ninside 349 372

16164 GCF_002046495.1_ASM204649v1 Salmonella enterica subsp. enterica serovar Baildon Proteobacteria; Gammaproteobacteria; Enterobacterales; Enterobacteriaceae; Salmonella; Salmonella enterica; Salmonella enterica subsp. enterica MNNEETFYQAMRRKGVTRRSFLKFCSLAATSLGLGAGMTPKIAWALENKPRIPVVWIHGL WP_001058312.1 hydrogenase [Salmonella enterica] Length: 372\nNumber of predicted TMHs: 2\nExp number of AAs in TMHs: 36.51738\nExp number, first 60 AAs: 19.72849\nTotal prob of N-in: 0.96170\nPOSSIBLE N-term signal sequence\ninside 1 19\nTMhelix 20 39\noutside 40 325\nTMhelix 326 348\ninside 349 372

16165 GCF_002045245.1_ASM204524v1 Salmonella enterica subsp. enterica serovar Baildon Proteobacteria; Gammaproteobacteria; Enterobacterales; Enterobacteriaceae; Salmonella; Salmonella enterica; Salmonella enterica subsp. enterica MTGDNTLITSHGINRRDFMKLCAALAATMGLSSKAAAEMAESVSNPQRPPVIWIGAQECT WP_000145429.1 hydrogenase 2 small subunit [Salmonella enterica] Length: 372\nNumber of predicted TMHs: 1\nExp number of AAs in TMHs: 24.51309\nExp number, first 60 AAs: 0.2682\nTotal prob of N-in: 0.14402\noutside 1 331\nTMhelix 332 354\ninside 355 372

16166 GCF_002047945.1_ASM204794v1 Salmonella enterica subsp. enterica serovar Baildon Proteobacteria; Gammaproteobacteria; Enterobacterales; Enterobacteriaceae; Salmonella; Salmonella enterica; Salmonella enterica subsp. enterica MTGDNTLITSHGINRRDFMKLCAALAATMGLSSKAAAEMAESVSNPQRPPVIWIGAQECT WP_000145429.1 hydrogenase 2 small subunit [Salmonella enterica] Length: 372\nNumber of predicted TMHs: 1\nExp number of AAs in TMHs: 24.51309\nExp number, first 60 AAs: 0.2682\nTotal prob of N-in: 0.14402\noutside 1 331\nTMhelix 332 354\ninside 355 372

16167 GCF_002047945.1_ASM204794v1 Salmonella enterica subsp. enterica serovar Baildon Proteobacteria; Gammaproteobacteria; Enterobacterales; Enterobacteriaceae; Salmonella; Salmonella enterica; Salmonella enterica subsp. enterica MNNEETFYQAMRRKGVTRRSFLKFCSLAATSLGLGAGMTPKIAWALENKPRIPVVWIHGL WP_001058312.1 hydrogenase [Salmonella enterica] Length: 372\nNumber of predicted TMHs: 2\nExp number of AAs in TMHs: 36.51738\nExp number, first 60 AAs: 19.72849\nTotal prob of N-in: 0.96170\nPOSSIBLE N-term signal sequence\ninside 1 19\nTMhelix 20 39\noutside 40 325\nTMhelix 326 348\ninside 349 372

16168 GCF_002105895.1_ASM210589v1 Salmonella enterica subsp. enterica serovar Baildon Proteobacteria; Gammaproteobacteria; Enterobacterales; Enterobacteriaceae; Salmonella; Salmonella enterica;

Salmonella enterica subsp. enterica	MNNEETFYQAMRRKGVTRRSFLKFCSLAATSLGLGAGMTPKIAWALENKPRIPVVWIHGL	WP_001058312.1	hydrogenase [Salmonella enterica]	Length: 372\nNumber of predicted TMHs:  2\nExp number of AAs in TMHs: 36.51738\nExp number, first 60 AAs:  19.72849\nTotal prob of N-in:   0.96170\nPOSSIBLE N-term signal sequence\ninside   1   19\nTMhelix   20   39\noutside   40   325\nTMhelix   326   348\ninside   349   372

16169	GCF_002105895.1_ASM210589v1	Salmonella enterica subsp. enterica serovar Baildon	Proteobacteria; Gammaproteobacteria; Enterobacterales; Enterobacteriaceae; Salmonella; Salmonella enterica; Salmonella enterica subsp. enterica	MTGDNTLITSHGINRRDFMKLCAALAATMGLSSKAAAEMAESVSNPQRPPVIWIGAQECT	WP_000145429.1	hydrogenase 2 small subunit [Salmonella enterica]	Length: 372\nNumber of predicted TMHs:  1\nExp number of AAs in TMHs: 24.51309\nExp number, first 60 AAs:  0.2682\nTotal prob of N-in:   0.14402\noutside   1   331\nTMhelix   332   354\ninside   355   372

16170	GCF_002046495.1_ASM204649v1	Salmonella enterica subsp. enterica serovar Baildon	Proteobacteria; Gammaproteobacteria; Enterobacterales; Enterobacteriaceae; Salmonella; Salmonella enterica; Salmonella enterica subsp. enterica	MTGDNTLITSHGINRRDFMKLCAALAATMGLSSKAAAEMAESVSNPQRPPVIWIGAQECT	WP_000145429.1	hydrogenase 2 small subunit [Salmonella enterica]	Length: 372\nNumber of predicted TMHs:  1\nExp number of AAs in TMHs: 24.51309\nExp number, first 60 AAs:  0.2682\nTotal prob of N-in:   0.14402\noutside   1   331\nTMhelix   332   354\ninside   355   372

16171	GCF_002045245.1_ASM204524v1	Salmonella enterica subsp. enterica serovar Baildon	Proteobacteria; Gammaproteobacteria; Enterobacterales; Enterobacteriaceae; Salmonella; Salmonella enterica; Salmonella enterica subsp. enterica	MNNEETFYQAMRRKGVTRRSFLKFCSLAATSLGLGAGMTPKIAWALENKPRIPVVWIHGL	WP_001058312.1	hydrogenase [Salmonella enterica]	Length: 372\nNumber of predicted TMHs:  2\nExp number of AAs in TMHs: 36.51738\nExp number, first 60 AAs:  19.72849\nTotal prob of N-in:   0.96170\nPOSSIBLE N-term signal sequence\ninside   1   19\nTMhelix   20   39\noutside   40   325\nTMhelix   326   348\ninside   349   372

16172	GCF_002156265.1_ASM215626v1	Salmonella enterica subsp. enterica serovar Banana	Proteobacteria; Gammaproteobacteria; Enterobacterales; Enterobacteriaceae; Salmonella; Salmonella enterica; Salmonella enterica subsp. enterica	MNNEETFYQAMRRKGVTRRSFLKFCSLAATSLGLGAGMTPKIAWALENKPRIPVVWIHGL	WP_001058312.1	hydrogenase [Salmonella enterica]	Length: 372\nNumber of predicted TMHs:  2\nExp number of AAs in TMHs: 36.51738\nExp number, first 60 AAs:  19.72849\nTotal prob of N-in:   0.96170\nPOSSIBLE N-term signal sequence\ninside   1   19\nTMhelix   20   39\noutside   40   325\nTMhelix   326   348\ninside   349   372

16173	GCF_002156265.1_ASM215626v1	Salmonella enterica subsp. enterica serovar Banana	Proteobacteria; Gammaproteobacteria; Enterobacterales; Enterobacteriaceae; Salmonella; Salmonella enterica; Salmonella enterica subsp. enterica	MTGDNTLITSHGINRRDFMKLCAALAATMGLSSKAAAEMAESVSNPQRPPVIWIGAQECT	WP_000145429.1	hydrogenase 2 small subunit [Salmonella enterica]	Length: 372\nNumber of predicted TMHs:  1\nExp number of AAs in TMHs: 24.51309\nExp number, first 60 AAs:  0.2682\nTotal prob of N-in:   0.14402\noutside   1   331\nTMhelix   332   354\ninside   355   372

16174	GCF_002066175.1_ASM206617v1	Salmonella enterica subsp. enterica serovar Bardo	Proteobacteria; Gammaproteobacteria; Enterobacterales; Enterobacteriaceae; Salmonella; Salmonella enterica; Salmonella enterica subsp. enterica	MTGDNTLITSHGINRRDFMKLCAALAATMGLSSKAAAEMAESVSNPQRPPVIWIGAQECT	WP_000145429.1	hydrogenase 2 small subunit [Salmonella enterica]	Length: 372\nNumber of predicted TMHs:  1\nExp number of AAs in TMHs: 24.51309\nExp number, first 60 AAs:  0.2682\nTotal prob of N-in:   0.14402\noutside   1   331\nTMhelix   332   354\ninside   355   372

16175	GCF_001975305.1_ASM197530v1	Salmonella enterica subsp. enterica serovar Bardo	Proteobacteria; Gammaproteobacteria; Enterobacterales; Enterobacteriaceae; Salmonella; Salmonella enterica; Salmonella enterica subsp. enterica	MTGDNTLITSHGINRRDFMKLCAALAATMGLSSKAAAEMAESVSNPQRPPVIWIGAQECT	WP_000145429.1	hydrogenase 2 small subunit [Salmonella enterica]	Length: 372\nNumber of predicted TMHs:  1\nExp number of AAs in TMHs: 24.51309\nExp number, first 60 AAs:  0.2682\nTotal prob of N-in:   0.14402\noutside   1   331\nTMhelix   332   354\ninside   355   372

16176	GCF_002066175.1_ASM206617v1	Salmonella enterica subsp. enterica serovar Bardo	Proteobacteria; Gammaproteobacteria; Enterobacterales; Enterobacteriaceae; Salmonella; Salmonella enterica; Salmonella enterica subsp. enterica	MNNEETFYQAMRRKGVTRRSFLKFCSLAATSLGLGAGMTPKIAWALENKPRIPVVWIHGL	WP_001058312.1	hydrogenase [Salmonella enterica]	Length: 372\nNumber of predicted TMHs:  2\nExp number of AAs in TMHs: 36.51738\nExp number, first 60 AAs:  19.72849\nTotal prob of N-in:   0.96170\nPOSSIBLE N-term signal sequence\ninside   1   19\nTMhelix   20   39\noutside   40   325\nTMhelix   326   348\ninside   349   372

16177 GCF_001975305.1_ASM197530v1 Salmonella enterica subsp. enterica serovar Bardo Proteobacteria; Gammaproteobacteria; Enterobacterales; Enterobacteriaceae; Salmonella; Salmonella enterica; Salmonella enterica subsp. enterica MNNEETFYQAMRRKGVTRRSFLKFCSLAATSLGLGAGMTPKIAWALENKPRIPVVWIHGL WP_001058312.1 hydrogenase [Salmonella enterica] Length: 372\nNumber of predicted TMHs: 2\nExp number of AAs in TMHs: 36.51738\nExp number, first 60 AAs: 19.72849\nTotal prob of N-in: 0.96170\nPOSSIBLE N-term signal sequence\ninside 1 19\nTMhelix 20 39\noutside 40 325\nTMhelix 326 348\ninside 349 372

16178 GCF_001832745.1_ASM183274v1 Salmonella enterica subsp. enterica serovar Bareilly Proteobacteria; Gammaproteobacteria; Enterobacterales; Enterobacteriaceae; Salmonella; Salmonella enterica; Salmonella enterica subsp. enterica MTGDNTLITSHGINRRDFMKLCAALAATMGLSSKAAAEMAESVSNPQRPPVIWIGAQECT WP_000145429.1 hydrogenase 2 small subunit [Salmonella enterica] Length: 372\nNumber of predicted TMHs: 1\nExp number of AAs in TMHs: 24.51309\nExp number, first 60 AAs: 0.2682\nTotal prob of N-in: 0.14402\noutside 1 331\nTMhelix 332 354\ninside 355 372

16179 GCF_001834225.1_ASM183422v1 Salmonella enterica subsp. enterica serovar Bareilly Proteobacteria; Gammaproteobacteria; Enterobacterales; Enterobacteriaceae; Salmonella; Salmonella enterica; Salmonella enterica subsp. enterica MTGDNTLITSHGINRRDFMKLCAALAATMGLSSKAAAEMAESVSNPQRPPVIWIGAQECT WP_000145429.1 hydrogenase 2 small subunit [Salmonella enterica] Length: 372\nNumber of predicted TMHs: 1\nExp number of AAs in TMHs: 24.51309\nExp number, first 60 AAs: 0.2682\nTotal prob of N-in: 0.14402\noutside 1 331\nTMhelix 332 354\ninside 355 372

16180 GCF_001834225.1_ASM183422v1 Salmonella enterica subsp. enterica serovar Bareilly Proteobacteria; Gammaproteobacteria; Enterobacterales; Enterobacteriaceae; Salmonella; Salmonella enterica; Salmonella enterica subsp. enterica MNNEETFYQAMRRKGVTRRSFLKFCSLAATSLGLGAGMTPKIAWALENKPRIPVVWIHGL WP_001058312.1 hydrogenase [Salmonella enterica] Length: 372\nNumber of predicted TMHs: 2\nExp number of AAs in TMHs: 36.51738\nExp number, first 60 AAs: 19.72849\nTotal prob of N-in: 0.96170\nPOSSIBLE N-term signal sequence\ninside 1 19\nTMhelix 20 39\noutside 40 325\nTMhelix 326 348\ninside 349 372

16181 GCF_001834255.1_ASM183425v1 Salmonella enterica subsp. enterica serovar Bareilly Proteobacteria; Gammaproteobacteria; Enterobacterales; Enterobacteriaceae; Salmonella; Salmonella enterica; Salmonella enterica subsp. enterica MTGDNTLITSHGINRRDFMKLCAALAATMGLSSKAAAEMAESVSNPQRPPVIWIGAQECT WP_000145429.1 hydrogenase 2 small subunit [Salmonella enterica] Length: 372\nNumber of predicted TMHs: 1\nExp number of AAs in TMHs: 24.51309\nExp number, first 60 AAs: 0.2682\nTotal prob of N-in: 0.14402\noutside 1 331\nTMhelix 332 354\ninside 355 372

16182 GCF_001835385.1_ASM183538v1 Salmonella enterica subsp. enterica serovar Bareilly Proteobacteria; Gammaproteobacteria; Enterobacterales; Enterobacteriaceae; Salmonella; Salmonella enterica; Salmonella enterica subsp. enterica MTGDNTLITSHGINRRDFMKLCAALAATMGLSSKAAAEMAESVSNPQRPPVIWIGAQECT WP_000145429.1 hydrogenase 2 small subunit [Salmonella enterica] Length: 372\nNumber of predicted TMHs: 1\nExp number of AAs in TMHs: 24.51309\nExp number, first 60 AAs: 0.2682\nTotal prob of N-in: 0.14402\noutside 1 331\nTMhelix 332 354\ninside 355 372

16183 GCF_001835385.1_ASM183538v1 Salmonella enterica subsp. enterica serovar Bareilly Proteobacteria; Gammaproteobacteria; Enterobacterales; Enterobacteriaceae; Salmonella; Salmonella enterica; Salmonella enterica subsp. enterica MNNEETFYQAMRRKGVTRRSFLKFCSLAATSLGLGAGMTPKIAWALENKPRIPVVWIHGL WP_001058312.1 hydrogenase [Salmonella enterica] Length: 372\nNumber of predicted TMHs: 2\nExp number of AAs in TMHs: 36.51738\nExp number, first 60 AAs: 19.72849\nTotal prob of N-in: 0.96170\nPOSSIBLE N-term signal sequence\ninside 1 19\nTMhelix 20 39\noutside 40 325\nTMhelix 326 348\ninside 349 372

16184 GCF_001835285.1_ASM183528v1 Salmonella enterica subsp. enterica serovar Bareilly Proteobacteria; Gammaproteobacteria; Enterobacterales; Enterobacteriaceae; Salmonella; Salmonella enterica; Salmonella enterica subsp. enterica MTGDNTLITSHGINRRDFMKLCAALAATMGLSSKAAAEMAESVSNPQRPPVIWIGAQECT WP_000145429.1 hydrogenase 2 small subunit [Salmonella enterica] Length: 372\nNumber of predicted TMHs: 1\nExp number of AAs in TMHs: 24.51309\nExp number, first 60 AAs: 0.2682\nTotal prob of N-in: 0.14402\noutside 1 331\nTMhelix 332 354\ninside 355 372

16185 GCF_001835125.1_ASM183512v1 Salmonella enterica subsp. enterica serovar Bareilly Proteobacteria; Gammaproteobacteria; Enterobacterales; Enterobacteriaceae; Salmonella; Salmonella enterica; Salmonella enterica subsp. enterica MNNEETFYQAMRRKGVTRRSFLKFCSLAATSLGLGAGMTPKIAWALENKPRIPVVWIHGL WP_001058312.1 hydrogenase [Salmonella enterica] Length: 372\nNumber of predicted TMHs: 2\nExp

number of AAs in TMHs: 36.51738\nExp number, first 60 AAs: 19.72849\nTotal prob of N-in: 0.96170\nPOSSIBLE N-term signal sequence\ninside 1 19\nTMhelix 20 39\noutside 40 325\nTMhelix 326 348\ninside 349 372

16186 GCF_001835285.1_ASM183528v1 Salmonella enterica subsp. enterica serovar Bareilly Proteobacteria; Gammaproteobacteria; Enterobacterales; Enterobacteriaceae; Salmonella; Salmonella enterica; Salmonella enterica subsp. enterica MNNEETFYQAMRRKGVTRRSFLKFCSLAATSLGLGAGMTPKIAWALENKPRIPVVWIHGL WP_001058312.1 hydrogenase [Salmonella enterica] Length: 372\nNumber of predicted TMHs: 2\nExp number of AAs in TMHs: 36.51738\nExp number, first 60 AAs: 19.72849\nTotal prob of N-in: 0.96170\nPOSSIBLE N-term signal sequence\ninside 1 19\nTMhelix 20 39\noutside 40 325\nTMhelix 326 348\ninside 349 372

16187 GCF_002063005.1_ASM206300v1 Salmonella enterica subsp. enterica serovar Bareilly Proteobacteria; Gammaproteobacteria; Enterobacterales; Enterobacteriaceae; Salmonella; Salmonella enterica; Salmonella enterica subsp. enterica MTGDNTLITSHGINRRDFMKLCAALAATMGLSSKAAAEMAESVSNPQRPPVIWIGAQECT WP_000145429.1 hydrogenase 2 small subunit [Salmonella enterica] Length: 372\nNumber of predicted TMHs: 1\nExp number of AAs in TMHs: 24.51309\nExp number, first 60 AAs: 0.2682\nTotal prob of N-in: 0.14402\noutside 1 331\nTMhelix 332 354\ninside 355 372

16188 GCF_001834405.1_ASM183440v1 Salmonella enterica subsp. enterica serovar Bareilly Proteobacteria; Gammaproteobacteria; Enterobacterales; Enterobacteriaceae; Salmonella; Salmonella enterica; Salmonella enterica subsp. enterica MTGDNTLITSHGINRRDFMKLCAALAATMGLSSKAAAEMAESVSNPQRPPVIWIGAQECT WP_000145429.1 hydrogenase 2 small subunit [Salmonella enterica] Length: 372\nNumber of predicted TMHs: 1\nExp number of AAs in TMHs: 24.51309\nExp number, first 60 AAs: 0.2682\nTotal prob of N-in: 0.14402\noutside 1 331\nTMhelix 332 354\ninside 355 372

16189 GCF_001834255.1_ASM183425v1 Salmonella enterica subsp. enterica serovar Bareilly Proteobacteria; Gammaproteobacteria; Enterobacterales; Enterobacteriaceae; Salmonella; Salmonella enterica; Salmonella enterica subsp. enterica MNNEETFYQAMRRKGVTRRSFLKFCSLAATSLGLGAGMTPKIAWALENKPRIPVVWIHGL WP_001058312.1 hydrogenase [Salmonella enterica] Length: 372\nNumber of predicted TMHs: 2\nExp number of AAs in TMHs: 36.51738\nExp number, first 60 AAs: 19.72849\nTotal prob of N-in: 0.96170\nPOSSIBLE N-term signal sequence\ninside 1 19\nTMhelix 20 39\noutside 40 325\nTMhelix 326 348\ninside 349 372

16190 GCF_001834405.1_ASM183440v1 Salmonella enterica subsp. enterica serovar Bareilly Proteobacteria; Gammaproteobacteria; Enterobacterales; Enterobacteriaceae; Salmonella; Salmonella enterica; Salmonella enterica subsp. enterica MNNEETFYQAMRRKGVTRRSFLKFCSLAATSLGLGAGMTPKIAWALENKPRIPVVWIHGL WP_001058312.1 hydrogenase [Salmonella enterica] Length: 372\nNumber of predicted TMHs: 2\nExp number of AAs in TMHs: 36.51738\nExp number, first 60 AAs: 19.72849\nTotal prob of N-in: 0.96170\nPOSSIBLE N-term signal sequence\ninside 1 19\nTMhelix 20 39\noutside 40 325\nTMhelix 326 348\ninside 349 372

16191 GCF_002066135.1_ASM206613v1 Salmonella enterica subsp. enterica serovar Bareilly Proteobacteria; Gammaproteobacteria; Enterobacterales; Enterobacteriaceae; Salmonella; Salmonella enterica; Salmonella enterica subsp. enterica MTGDNTLITSHGINRRDFMKLCAALAATMGLSSKAAAEMAESVSNPQRPPVIWIGAQECT WP_000145429.1 hydrogenase 2 small subunit [Salmonella enterica] Length: 372\nNumber of predicted TMHs: 1\nExp number of AAs in TMHs: 24.51309\nExp number, first 60 AAs: 0.2682\nTotal prob of N-in: 0.14402\noutside 1 331\nTMhelix 332 354\ninside 355 372

16192 GCF_001832745.1_ASM183274v1 Salmonella enterica subsp. enterica serovar Bareilly Proteobacteria; Gammaproteobacteria; Enterobacterales; Enterobacteriaceae; Salmonella; Salmonella enterica; Salmonella enterica subsp. enterica MNNEETFYQAMRRKGVTRRSFLKFCSLAATSLGLGAGMTPKIAWALENKPRIPVVWIHGL WP_001058312.1 hydrogenase [Salmonella enterica] Length: 372\nNumber of predicted TMHs: 2\nExp number of AAs in TMHs: 36.51738\nExp number, first 60 AAs: 19.72849\nTotal prob of N-in: 0.96170\nPOSSIBLE N-term signal sequence\ninside 1 19\nTMhelix 20 39\noutside 40 325\nTMhelix 326 348\ninside 349 372

16193 GCF_001833665.1_ASM183366v1 Salmonella enterica subsp. enterica serovar Bareilly Proteobacteria; Gammaproteobacteria; Enterobacterales; Enterobacteriaceae; Salmonella; Salmonella enterica; Salmonella enterica subsp. enterica MTGDNTLITSHGINRRDFMKLCAALAATMGLSSKAAAEMAESVSNPQRPPVIWIGAQECT WP_000145429.1 hydrogenase 2 small subunit [Salmonella enterica] Length: 372\nNumber of predicted TMHs: 1\nExp number of AAs in TMHs: 24.51309\nExp number, first 60 AAs: 0.2682\nTotal prob of N-in: 0.14402\noutside 1 331\nTMhelix 332 354\ninside 355 372

16194 GCF_002063005.1_ASM206300v1 Salmonella enterica subsp. enterica serovar Bareilly Proteobacteria; Gammaproteobacteria; Enterobacterales; Enterobacteriaceae; Salmonella; Salmonella enterica;

Salmonella enterica subsp. enterica MNNEETFYQAMRRKGVTRRSFLKFCSLAATSLGLGAGMTPKIAWALENKPRIPVVWIHGL WP_001058311.1 hydrogenase [Salmonella enterica] Length: 372\nNumber of predicted TMHs: 2\nExp number of AAs in TMHs: 37.4699899999999\nExp number, first 60 AAs: 19.72699\nTotal prob of N-in: 0.96171\nPOSSIBLE N-term signal sequence\ninside 1 19\nTMhelix 20 39\noutside 40 325\nTMhelix 326 348\ninside 349 372

16195 GCF_001833665.1_ASM183366v1 Salmonella enterica subsp. enterica serovar Bareilly Proteobacteria; Gammaproteobacteria; Enterobacterales; Enterobacteriaceae; Salmonella; Salmonella enterica; Salmonella enterica subsp. enterica MNNEETFYQAMRRKGVTRRSFLKFCSLAATSLGLGAGMTPKIAWALENKPRIPVVWIHGL WP_001058312.1 hydrogenase [Salmonella enterica] Length: 372\nNumber of predicted TMHs: 2\nExp number of AAs in TMHs: 36.51738\nExp number, first 60 AAs: 19.72849\nTotal prob of N-in: 0.96170\nPOSSIBLE N-term signal sequence\ninside 1 19\nTMhelix 20 39\noutside 40 325\nTMhelix 326 348\ninside 349 372

16196 GCF_002066135.1_ASM206613v1 Salmonella enterica subsp. enterica serovar Bareilly Proteobacteria; Gammaproteobacteria; Enterobacterales; Enterobacteriaceae; Salmonella; Salmonella enterica; Salmonella enterica subsp. enterica MNNEETFYQAMRRKGVTRRSFLKFCSLAATSLGLGAGMTPKIAWALENKPRIPVVWIHGL WP_001058311.1 hydrogenase [Salmonella enterica] Length: 372\nNumber of predicted TMHs: 2\nExp number of AAs in TMHs: 37.4699899999999\nExp number, first 60 AAs: 19.72699\nTotal prob of N-in: 0.96171\nPOSSIBLE N-term signal sequence\ninside 1 19\nTMhelix 20 39\noutside 40 325\nTMhelix 326 348\ninside 349 372

16197 GCF_001834415.1_ASM183441v1 Salmonella enterica subsp. enterica serovar Bareilly Proteobacteria; Gammaproteobacteria; Enterobacterales; Enterobacteriaceae; Salmonella; Salmonella enterica; Salmonella enterica subsp. enterica MNNEETFYQAMRRKGVTRRSFLKFCSLAATSLGLGAGMTPKIAWALENKPRIPVVWIHGL WP_001058312.1 hydrogenase [Salmonella enterica] Length: 372\nNumber of predicted TMHs: 2\nExp number of AAs in TMHs: 36.51738\nExp number, first 60 AAs: 19.72849\nTotal prob of N-in: 0.96170\nPOSSIBLE N-term signal sequence\ninside 1 19\nTMhelix 20 39\noutside 40 325\nTMhelix 326 348\ninside 349 372

16198 GCF_001835125.1_ASM183512v1 Salmonella enterica subsp. enterica serovar Bareilly Proteobacteria; Gammaproteobacteria; Enterobacterales; Enterobacteriaceae; Salmonella; Salmonella enterica; Salmonella enterica subsp. enterica MTGDNTLITSHGINRRDFMKLCAALAATMGLSSKAAAEMAESVSNPQRPPVIWIGAQECT WP_000145429.1 hydrogenase 2 small subunit [Salmonella enterica] Length: 372\nNumber of predicted TMHs: 1\nExp number of AAs in TMHs: 24.51309\nExp number, first 60 AAs: 0.2682\nTotal prob of N-in: 0.14402\noutside 1 331\nTMhelix 332 354\ninside 355 372

16199 GCF_001834415.1_ASM183441v1 Salmonella enterica subsp. enterica serovar Bareilly Proteobacteria; Gammaproteobacteria; Enterobacterales; Enterobacteriaceae; Salmonella; Salmonella enterica; Salmonella enterica subsp. enterica MTGDNTLITSHGINRRDFMKLCAALAATMGLSSKAAAEMAESVSNPQRPPVIWIGAQECT WP_000145429.1 hydrogenase 2 small subunit [Salmonella enterica] Length: 372\nNumber of predicted TMHs: 1\nExp number of AAs in TMHs: 24.51309\nExp number, first 60 AAs: 0.2682\nTotal prob of N-in: 0.14402\noutside 1 331\nTMhelix 332 354\ninside 355 372

16200 GCF_001480555.1_Salmonella_enterica_CVM_N40408-SQ_v1.0 Salmonella enterica subsp. enterica serovar Berta Proteobacteria; Gammaproteobacteria; Enterobacterales; Enterobacteriaceae; Salmonella; Salmonella enterica; Salmonella enterica subsp. enterica MTGDNTLITSHGINRRDFMKLCAALAATMGLSSKAAAEMAESVSNPQRPPVIWIGAQECT WP_000145429.1 hydrogenase 2 small subunit [Salmonella enterica] Length: 372\nNumber of predicted TMHs: 1\nExp number of AAs in TMHs: 24.51309\nExp number, first 60 AAs: 0.2682\nTotal prob of N-in: 0.14402\noutside 1 331\nTMhelix 332 354\ninside 355 372

16201 GCF_002030515.1_ASM203051v1 Salmonella enterica subsp. enterica serovar Berta Proteobacteria; Gammaproteobacteria; Enterobacterales; Enterobacteriaceae; Salmonella; Salmonella enterica; Salmonella enterica subsp. enterica MNNEETFYQAMRRKGVTRRSFLKFCSLAATSLGLGAGMTPKIAWALENKPRIPVVWIHGL WP_001058311.1 hydrogenase [Salmonella enterica] Length: 372\nNumber of predicted TMHs: 2\nExp number of AAs in TMHs: 37.4699899999999\nExp number, first 60 AAs: 19.72699\nTotal prob of N-in: 0.96171\nPOSSIBLE N-term signal sequence\ninside 1 19\nTMhelix 20 39\noutside 40 325\nTMhelix 326 348\ninside 349 372

16202 GCF_002030515.1_ASM203051v1 Salmonella enterica subsp. enterica serovar Berta Proteobacteria; Gammaproteobacteria; Enterobacterales; Enterobacteriaceae; Salmonella; Salmonella enterica; Salmonella enterica subsp. enterica MTGDNTLITSHGINRRDFMKLCAALAATMGLSSKAAAEMAESVSNPQRPPVIWIGAQECT WP_000145429.1 hydrogenase 2 small subunit [Salmonella enterica] Length: 372\nNumber of

predicted TMHs:  1\nExp number of AAs in TMHs: 24.51309\nExp number, first 60 AAs:  0.2682\nTotal prob of N-in: 0.14402\noutside   1   331\nTMhelix   332   354\ninside   355   372

16203 GCF_002061255.1_ASM206125v1 Salmonella enterica subsp. enterica serovar Berta Proteobacteria; Gammaproteobacteria; Enterobacterales; Enterobacteriaceae; Salmonella; Salmonella enterica; Salmonella enterica subsp. enterica MTGDNTLITSHGINRRDFMKLCAALAATMGLSSKAAAEMAESVSNPQRPPVIWIGAQECT WP_000145429.1 hydrogenase 2 small subunit [Salmonella enterica] Length: 372\nNumber of predicted TMHs:  1\nExp number of AAs in TMHs: 24.51309\nExp number, first 60 AAs:  0.2682\nTotal prob of N-in: 0.14402\noutside   1   331\nTMhelix   332   354\ninside   355   372

16204 GCF_001480555.1_Salmonella_enterica_CVM_N40408-SQ_v1.0 Salmonella enterica subsp. enterica serovar Berta Proteobacteria; Gammaproteobacteria; Enterobacterales; Enterobacteriaceae; Salmonella; Salmonella enterica; Salmonella enterica subsp. enterica MNNEETFYQAMRRKGVTRRSFLKFCSLAATSLGLGAGMTPKIAWALENKPRIPVVWIHGL WP_001058311.1 hydrogenase [Salmonella enterica] Length: 372\nNumber of predicted TMHs:  2\nExp number of AAs in TMHs: 37.4699899999999\nExp number, first 60 AAs:  19.72699\nTotal prob of N-in: 0.96171\nPOSSIBLE N-term signal sequence\ninside   1   19\nTMhelix   20   39\noutside   40   325\nTMhelix   326 348\ninside   349   372

16205 GCF_002058795.1_ASM205879v1 Salmonella enterica subsp. enterica serovar Berta Proteobacteria; Gammaproteobacteria; Enterobacterales; Enterobacteriaceae; Salmonella; Salmonella enterica; Salmonella enterica subsp. enterica MNNEETFYQAMRRKGVTRRSFLKFCSLAATSLGLGAGMTPKIAWALENKPRIPVVWIHGL WP_001058311.1 hydrogenase [Salmonella enterica] Length: 372\nNumber of predicted TMHs:  2\nExp number of AAs in TMHs: 37.4699899999999\nExp number, first 60 AAs:  19.72699\nTotal prob of N-in: 0.96171\nPOSSIBLE N-term signal sequence\ninside   1   19\nTMhelix   20   39\noutside   40   325\nTMhelix   326 348\ninside   349   372

16206 GCF_002061255.1_ASM206125v1 Salmonella enterica subsp. enterica serovar Berta Proteobacteria; Gammaproteobacteria; Enterobacterales; Enterobacteriaceae; Salmonella; Salmonella enterica; Salmonella enterica subsp. enterica MNNEETFYQAMRRKGVTRRSFLKFCSLAATSLGLGAGMTPKIAWALENKPRIPVVWIHGL WP_001058311.1 hydrogenase [Salmonella enterica] Length: 372\nNumber of predicted TMHs:  2\nExp number of AAs in TMHs: 37.4699899999999\nExp number, first 60 AAs:  19.72699\nTotal prob of N-in: 0.96171\nPOSSIBLE N-term signal sequence\ninside   1   19\nTMhelix   20   39\noutside   40   325\nTMhelix   326 348\ninside   349   372

16207 GCF_001242065.1_Salmonella_enterica_CVM_N45924_v1.0 Salmonella enterica subsp. enterica serovar Berta Proteobacteria; Gammaproteobacteria; Enterobacterales; Enterobacteriaceae; Salmonella; Salmonella enterica; Salmonella enterica subsp. enterica MTGDNTLITSHGINRRDFMKLCAALAATMGLSSKAAAEMAESVSNPQRPPVIWIGAQECT WP_000145429.1 hydrogenase 2 small subunit [Salmonella enterica] Length: 372\nNumber of predicted TMHs:  1\nExp number of AAs in TMHs: 24.51309\nExp number, first 60 AAs:  0.2682\nTotal prob of N-in: 0.14402\noutside   1   331\nTMhelix   332   354\ninside   355   372

16208 GCF_002031535.1_ASM203153v1 Salmonella enterica subsp. enterica serovar Berta Proteobacteria; Gammaproteobacteria; Enterobacterales; Enterobacteriaceae; Salmonella; Salmonella enterica; Salmonella enterica subsp. enterica MNNEETFYQAMRRKGVTRRSFLKFCSLAATSLGLGAGMTPKIAWALENKPRIPVVWIHGL WP_001058311.1 hydrogenase [Salmonella enterica] Length: 372\nNumber of predicted TMHs:  2\nExp number of AAs in TMHs: 37.4699899999999\nExp number, first 60 AAs:  19.72699\nTotal prob of N-in: 0.96171\nPOSSIBLE N-term signal sequence\ninside   1   19\nTMhelix   20   39\noutside   40   325\nTMhelix   326 348\ninside   349   372

16209 GCF_002030615.1_ASM203061v1 Salmonella enterica subsp. enterica serovar Berta Proteobacteria; Gammaproteobacteria; Enterobacterales; Enterobacteriaceae; Salmonella; Salmonella enterica; Salmonella enterica subsp. enterica MTGDNTLITSHGINRRDFMKLCAALAATMGLSSKAAAEMAESVSNPQRPPVIWIGAQECT WP_000145429.1 hydrogenase 2 small subunit [Salmonella enterica] Length: 372\nNumber of predicted TMHs:  1\nExp number of AAs in TMHs: 24.51309\nExp number, first 60 AAs:  0.2682\nTotal prob of N-in: 0.14402\noutside   1   331\nTMhelix   332   354\ninside   355   372

16210 GCF_002031535.1_ASM203153v1 Salmonella enterica subsp. enterica serovar Berta Proteobacteria; Gammaproteobacteria; Enterobacterales; Enterobacteriaceae; Salmonella; Salmonella enterica; Salmonella enterica subsp. enterica MTGDNTLITSHGINRRDFMKLCAALAATMGLSSKAAAEMAESVSNPQRPPVIWIGAQECT WP_000145429.1 hydrogenase 2 small subunit [Salmonella enterica] Length: 372\nNumber of

predicted TMHs: 1\nExp number of AAs in TMHs: 24.51309\nExp number, first 60 AAs: 0.2682\nTotal prob of N-in: 0.14402\noutside 1 331\nTMhelix 332 354\ninside 355 372

16211 GCF_002030615.1_ASM203061v1 Salmonella enterica subsp. enterica serovar Berta Proteobacteria; Gammaproteobacteria; Enterobacterales; Enterobacteriaceae; Salmonella; Salmonella enterica; Salmonella enterica subsp. enterica MNNEETFYQAMRRKGVTRRSFLKFCSLAATSLGLGAGMTPKIAWALENKPRIPVVWIHGL WP_001058311.1 hydrogenase [Salmonella enterica] Length: 372\nNumber of predicted TMHs: 2\nExp number of AAs in TMHs: 37.4699899999999\nExp number, first 60 AAs: 19.72699\nTotal prob of N-in: 0.96171\nPOSSIBLE N-term signal sequence\ninside 1 19\nTMhelix 20 39\noutside 40 325\nTMhelix 326 348\ninside 349 372

16212 GCF_002031935.1_ASM203193v1 Salmonella enterica subsp. enterica serovar Berta Proteobacteria; Gammaproteobacteria; Enterobacterales; Enterobacteriaceae; Salmonella; Salmonella enterica; Salmonella enterica subsp. enterica MNNEETFYQAMRRKGVTRRSFLKFCSLAATSLGLGAGMTPKIAWALENKPRIPVVWIHGL WP_001058311.1 hydrogenase [Salmonella enterica] Length: 372\nNumber of predicted TMHs: 2\nExp number of AAs in TMHs: 37.4699899999999\nExp number, first 60 AAs: 19.72699\nTotal prob of N-in: 0.96171\nPOSSIBLE N-term signal sequence\ninside 1 19\nTMhelix 20 39\noutside 40 325\nTMhelix 326 348\ninside 349 372

16213 GCF_001479635.1_Salmonella_enterica_CVM_N37918-SQ_v1.0 Salmonella enterica subsp. enterica serovar Berta Proteobacteria; Gammaproteobacteria; Enterobacterales; Enterobacteriaceae; Salmonella; Salmonella enterica; Salmonella enterica subsp. enterica MTGDNTLITSHGINRRDFMKLCAALAATMGLSSKAAAEMAESVSNPQRPPVIWIGAQECT WP_000145429.1 hydrogenase 2 small subunit [Salmonella enterica] Length: 372\nNumber of predicted TMHs: 1\nExp number of AAs in TMHs: 24.51309\nExp number, first 60 AAs: 0.2682\nTotal prob of N-in: 0.14402\noutside 1 331\nTMhelix 332 354\ninside 355 372

16214 GCF_001479635.1_Salmonella_enterica_CVM_N37918-SQ_v1.0 Salmonella enterica subsp. enterica serovar Berta Proteobacteria; Gammaproteobacteria; Enterobacterales; Enterobacteriaceae; Salmonella; Salmonella enterica; Salmonella enterica subsp. enterica MNNEETFYQAMRRKGVTRRSFLKFCSLAATSLGLGAGMTPKIAWALENKPRIPVVWIHGL WP_001058311.1 hydrogenase [Salmonella enterica] Length: 372\nNumber of predicted TMHs: 2\nExp number of AAs in TMHs: 37.4699899999999\nExp number, first 60 AAs: 19.72699\nTotal prob of N-in: 0.96171\nPOSSIBLE N-term signal sequence\ninside 1 19\nTMhelix 20 39\noutside 40 325\nTMhelix 326 348\ninside 349 372

16215 GCF_002059435.1_ASM205943v1 Salmonella enterica subsp. enterica serovar Berta Proteobacteria; Gammaproteobacteria; Enterobacterales; Enterobacteriaceae; Salmonella; Salmonella enterica; Salmonella enterica subsp. enterica MTGDNTLITSHGINRRDFMKLCAALAATMGLSSKAAAEMAESVSNPQRPPVIWIGAQECT WP_000145429.1 hydrogenase 2 small subunit [Salmonella enterica] Length: 372\nNumber of predicted TMHs: 1\nExp number of AAs in TMHs: 24.51309\nExp number, first 60 AAs: 0.2682\nTotal prob of N-in: 0.14402\noutside 1 331\nTMhelix 332 354\ninside 355 372

16216 GCF_002030645.1_ASM203064v1 Salmonella enterica subsp. enterica serovar Berta Proteobacteria; Gammaproteobacteria; Enterobacterales; Enterobacteriaceae; Salmonella; Salmonella enterica; Salmonella enterica subsp. enterica MTGDNTLITSHGINRRDFMKLCAALAATMGLSSKAAAEMAESVSNPQRPPVIWIGAQECT WP_000145429.1 hydrogenase 2 small subunit [Salmonella enterica] Length: 372\nNumber of predicted TMHs: 1\nExp number of AAs in TMHs: 24.51309\nExp number, first 60 AAs: 0.2682\nTotal prob of N-in: 0.14402\noutside 1 331\nTMhelix 332 354\ninside 355 372

16217 GCF_001242065.1_Salmonella_enterica_CVM_N45924_v1.0 Salmonella enterica subsp. enterica serovar Berta Proteobacteria; Gammaproteobacteria; Enterobacterales; Enterobacteriaceae; Salmonella; Salmonella enterica; Salmonella enterica subsp. enterica MNNEETFYQAMRRKGVTRRSFLKFCSLAATSLGLGAGMTPKIAWALENKPRIPVVWIHGL WP_001058311.1 hydrogenase [Salmonella enterica] Length: 372\nNumber of predicted TMHs: 2\nExp number of AAs in TMHs: 37.4699899999999\nExp number, first 60 AAs: 19.72699\nTotal prob of N-in: 0.96171\nPOSSIBLE N-term signal sequence\ninside 1 19\nTMhelix 20 39\noutside 40 325\nTMhelix 326 348\ninside 349 372

16218 GCF_002058795.1_ASM205879v1 Salmonella enterica subsp. enterica serovar Berta Proteobacteria; Gammaproteobacteria; Enterobacterales; Enterobacteriaceae; Salmonella; Salmonella enterica; Salmonella enterica subsp. enterica MTGDNTLITSHGINRRDFMKLCAALAATMGLSSKAAAEMAESVSNPQRPPVIWIGAQECT WP_000145429.1 hydrogenase 2 small subunit [Salmonella enterica] Length: 372\nNumber of

predicted TMHs: 1\nExp number of AAs in TMHs: 24.51309\nExp number, first 60 AAs: 0.2682\nTotal prob of N-in: 0.14402\noutside 1 331\nTMhelix 332 354\ninside 355 372

16219 GCF_002030645.1_ASM203064v1 Salmonella enterica subsp. enterica serovar Berta Proteobacteria; Gammaproteobacteria; Enterobacterales; Enterobacteriaceae; Salmonella; Salmonella enterica; Salmonella enterica subsp. enterica MNNEETFYQAMRRKGVTRRSFLKFCSLAATSLGLGAGMTPKIAWALENKPRIPVVWIHGL WP_001058311.1 hydrogenase [Salmonella enterica] Length: 372\nNumber of predicted TMHs: 2\nExp number of AAs in TMHs: 37.4699899999999\nExp number, first 60 AAs: 19.72699\nTotal prob of N-in: 0.96171\nPOSSIBLE N-term signal sequence\ninside 1 19\nTMhelix 20 39\noutside 40 325\nTMhelix 326 348\ninside 349 372

16220 GCF_000964145.1_ASM96414v1 Salmonella enterica subsp. enterica serovar Berta Proteobacteria; Gammaproteobacteria; Enterobacterales; Enterobacteriaceae; Salmonella; Salmonella enterica; Salmonella enterica subsp. enterica MTGDNTLITSHGINRRDFMKLCAALAATMGLSSKAAAEMAESVSNPQRPPVIWIGAQECT WP_000145429.1 hydrogenase 2 small subunit [Salmonella enterica] Length: 372\nNumber of predicted TMHs: 1\nExp number of AAs in TMHs: 24.51309\nExp number, first 60 AAs: 0.2682\nTotal prob of N-in: 0.14402\noutside 1 331\nTMhelix 332 354\ninside 355 372

16221 GCF_001242075.1_Salmonella_enterica_CVM_N45925_v1.0 Salmonella enterica subsp. enterica serovar Berta Proteobacteria; Gammaproteobacteria; Enterobacterales; Enterobacteriaceae; Salmonella; Salmonella enterica; Salmonella enterica subsp. enterica MNNEETFYQAMRRKGVTRRSFLKFCSLAATSLGLGAGMTPKIAWALENKPRIPVVWIHGL WP_001058311.1 hydrogenase [Salmonella enterica] Length: 372\nNumber of predicted TMHs: 2\nExp number of AAs in TMHs: 37.4699899999999\nExp number, first 60 AAs: 19.72699\nTotal prob of N-in: 0.96171\nPOSSIBLE N-term signal sequence\ninside 1 19\nTMhelix 20 39\noutside 40 325\nTMhelix 326 348\ninside 349 372

16222 GCF_000964145.1_ASM96414v1 Salmonella enterica subsp. enterica serovar Berta Proteobacteria; Gammaproteobacteria; Enterobacterales; Enterobacteriaceae; Salmonella; Salmonella enterica; Salmonella enterica subsp. enterica MNNEETFYQAMRRKGVTRRSFLKFCSLAATSLGLGAGMTPKIAWALENKPRIPVVWIHGL WP_001058311.1 hydrogenase [Salmonella enterica] Length: 372\nNumber of predicted TMHs: 2\nExp number of AAs in TMHs: 37.4699899999999\nExp number, first 60 AAs: 19.72699\nTotal prob of N-in: 0.96171\nPOSSIBLE N-term signal sequence\ninside 1 19\nTMhelix 20 39\noutside 40 325\nTMhelix 326 348\ninside 349 372

16223 GCF_001242075.1_Salmonella_enterica_CVM_N45925_v1.0 Salmonella enterica subsp. enterica serovar Berta Proteobacteria; Gammaproteobacteria; Enterobacterales; Enterobacteriaceae; Salmonella; Salmonella enterica; Salmonella enterica subsp. enterica MTGDNTLITSHGINRRDFMKLCAALAATMGLSSKAAAEMAESVSNPQRPPVIWIGAQECT WP_000145429.1 hydrogenase 2 small subunit [Salmonella enterica] Length: 372\nNumber of predicted TMHs: 1\nExp number of AAs in TMHs: 24.51309\nExp number, first 60 AAs: 0.2682\nTotal prob of N-in: 0.14402\noutside 1 331\nTMhelix 332 354\ninside 355 372

16224 GCF_002156305.1_ASM215630v1 Salmonella enterica subsp. enterica serovar Birkenhead Proteobacteria; Gammaproteobacteria; Enterobacterales; Enterobacteriaceae; Salmonella; Salmonella enterica; Salmonella enterica subsp. enterica MNNEETFYQAMRRKGVTRRSFLKFCSLAATSLGLGAGMTPKIAWALENKPRIPVVWIHGL WP_001058310.1 hydrogenase [Salmonella enterica] Length: 372\nNumber of predicted TMHs: 2\nExp number of AAs in TMHs: 37.4567299999999\nExp number, first 60 AAs: 19.72694\nTotal prob of N-in: 0.96171\nPOSSIBLE N-term signal sequence\ninside 1 19\nTMhelix 20 39\noutside 40 325\nTMhelix 326 348\ninside 349 372

16225 GCF_002156305.1_ASM215630v1 Salmonella enterica subsp. enterica serovar Birkenhead Proteobacteria; Gammaproteobacteria; Enterobacterales; Enterobacteriaceae; Salmonella; Salmonella enterica; Salmonella enterica subsp. enterica MTGDNTLITSHGINRRDFMKLCAALAATMGLSSKAAAEMAESVSNPQRPPVIWIGAQECT WP_000145429.1 hydrogenase 2 small subunit [Salmonella enterica] Length: 372\nNumber of predicted TMHs: 1\nExp number of AAs in TMHs: 24.51309\nExp number, first 60 AAs: 0.2682\nTotal prob of N-in: 0.14402\noutside 1 331\nTMhelix 332 354\ninside 355 372

16226 GCF_002049195.1_ASM204919v1 Salmonella enterica subsp. enterica serovar Blockley Proteobacteria; Gammaproteobacteria; Enterobacterales; Enterobacteriaceae; Salmonella; Salmonella enterica; Salmonella enterica subsp. enterica MTGDNTLITSHGINRRDFMKLCAALAATMGLSSKAAAEMAESVSNPQRPPVIWIGAQECT WP_000145429.1 hydrogenase 2 small subunit [Salmonella enterica] Length: 372\nNumber of

predicted TMHs:  1\nExp number of AAs in TMHs: 24.51309\nExp number, first 60 AAs:  0.2682\nTotal prob of N-in: 0.14402\noutside   1   331\nTMhelix   332   354\ninside   355   372

16227 GCF_002045325.1_ASM204532v1 Salmonella enterica subsp. enterica serovar Blockley Proteobacteria; Gammaproteobacteria; Enterobacterales; Enterobacteriaceae; Salmonella; Salmonella enterica; Salmonella enterica subsp. enterica MTGDNTLITSHGINRRDFMKLCAALAATMGLSSKAAAEMAESVSNPQRPPVIWIGAQECT WP_000145429.1 hydrogenase 2 small subunit [Salmonella enterica] Length: 372\nNumber of predicted TMHs:  1\nExp number of AAs in TMHs: 24.51309\nExp number, first 60 AAs:  0.2682\nTotal prob of N-in: 0.14402\noutside   1   331\nTMhelix   332   354\ninside   355   372

16228 GCF_002048695.1_ASM204869v1 Salmonella enterica subsp. enterica serovar Blockley Proteobacteria; Gammaproteobacteria; Enterobacterales; Enterobacteriaceae; Salmonella; Salmonella enterica; Salmonella enterica subsp. enterica MNNEETFYQAMRRKGVTRRSFLKFCSLAATSLGLGAGMTPKIAWALENKPRIPVVWIHGL WP_001058310.1 hydrogenase [Salmonella enterica] Length: 372\nNumber of predicted TMHs:  2\nExp number of AAs in TMHs: 37.4567299999999\nExp number, first 60 AAs:  19.72694\nTotal prob of N-in: 0.96171\nPOSSIBLE N-term signal sequence\ninside   1   19\nTMhelix   20   39\noutside   40   325\nTMhelix   326   348\ninside   349   372

16229 GCF_002062245.1_ASM206224v1 Salmonella enterica subsp. enterica serovar Blockley Proteobacteria; Gammaproteobacteria; Enterobacterales; Enterobacteriaceae; Salmonella; Salmonella enterica; Salmonella enterica subsp. enterica MTGDNTLITSHGINRRDFMKLCAALAATMGLSSKAAAEMAESVSNPQRPPVIWIGAQECT WP_000145429.1 hydrogenase 2 small subunit [Salmonella enterica] Length: 372\nNumber of predicted TMHs:  1\nExp number of AAs in TMHs: 24.51309\nExp number, first 60 AAs:  0.2682\nTotal prob of N-in: 0.14402\noutside   1   331\nTMhelix   332   354\ninside   355   372

16230 GCF_002062245.1_ASM206224v1 Salmonella enterica subsp. enterica serovar Blockley Proteobacteria; Gammaproteobacteria; Enterobacterales; Enterobacteriaceae; Salmonella; Salmonella enterica; Salmonella enterica subsp. enterica MNNEETFYQAMRRKGVTRRSFLKFCSLAATSLGLGAGMTPKIAWALENKPRIPVVWIHGL WP_001058310.1 hydrogenase [Salmonella enterica] Length: 372\nNumber of predicted TMHs:  2\nExp number of AAs in TMHs: 37.4567299999999\nExp number, first 60 AAs:  19.72694\nTotal prob of N-in: 0.96171\nPOSSIBLE N-term signal sequence\ninside   1   19\nTMhelix   20   39\noutside   40   325\nTMhelix   326   348\ninside   349   372

16231 GCF_002045305.1_ASM204530v1 Salmonella enterica subsp. enterica serovar Blockley Proteobacteria; Gammaproteobacteria; Enterobacterales; Enterobacteriaceae; Salmonella; Salmonella enterica; Salmonella enterica subsp. enterica MTGDNTLITSHGINRRDFMKLCAALAATMGLSSKAAAEMAESVSNPQRPPVIWIGAQECT WP_000145429.1 hydrogenase 2 small subunit [Salmonella enterica] Length: 372\nNumber of predicted TMHs:  1\nExp number of AAs in TMHs: 24.51309\nExp number, first 60 AAs:  0.2682\nTotal prob of N-in: 0.14402\noutside   1   331\nTMhelix   332   354\ninside   355   372

16232 GCF_002049195.1_ASM204919v1 Salmonella enterica subsp. enterica serovar Blockley Proteobacteria; Gammaproteobacteria; Enterobacterales; Enterobacteriaceae; Salmonella; Salmonella enterica; Salmonella enterica subsp. enterica MNNEETFYQAMRRKGVTRRSFLKFCSLAATSLGLGAGMTPKIAWALENKPRIPVVWIHGL WP_001058310.1 hydrogenase [Salmonella enterica] Length: 372\nNumber of predicted TMHs:  2\nExp number of AAs in TMHs: 37.4567299999999\nExp number, first 60 AAs:  19.72694\nTotal prob of N-in: 0.96171\nPOSSIBLE N-term signal sequence\ninside   1   19\nTMhelix   20   39\noutside   40   325\nTMhelix   326   348\ninside   349   372

16233 GCF_002045305.1_ASM204530v1 Salmonella enterica subsp. enterica serovar Blockley Proteobacteria; Gammaproteobacteria; Enterobacterales; Enterobacteriaceae; Salmonella; Salmonella enterica; Salmonella enterica subsp. enterica MNNEETFYQAMRRKGVTRRSFLKFCSLAATSLGLGAGMTPKIAWALENKPRIPVVWIHGL WP_001058310.1 hydrogenase [Salmonella enterica] Length: 372\nNumber of predicted TMHs:  2\nExp number of AAs in TMHs: 37.4567299999999\nExp number, first 60 AAs:  19.72694\nTotal prob of N-in: 0.96171\nPOSSIBLE N-term signal sequence\ninside   1   19\nTMhelix   20   39\noutside   40   325\nTMhelix   326   348\ninside   349   372

16234 GCF_002048695.1_ASM204869v1 Salmonella enterica subsp. enterica serovar Blockley Proteobacteria; Gammaproteobacteria; Enterobacterales; Enterobacteriaceae; Salmonella; Salmonella enterica; Salmonella enterica subsp. enterica MTGDNTLITSHGINRRDFMKLCAALAATMGLSSKAAAEMAESVSNPQRPPVIWIGAQECT WP_000145429.1 hydrogenase 2 small subunit [Salmonella enterica] Length: 372\nNumber of

predicted TMHs: 1\nExp number of AAs in TMHs: 24.51309\nExp number, first 60 AAs: 0.2682\nTotal prob of N-in: 0.14402\noutside 1 331\nTMhelix 332 354\ninside 355 372

16235 GCF_002045325.1_ASM204532v1 Salmonella enterica subsp. enterica serovar Blockley Proteobacteria; Gammaproteobacteria; Enterobacterales; Enterobacteriaceae; Salmonella; Salmonella enterica; Salmonella enterica subsp. enterica MNNEETFYQAMRRKGVTRRSFLKFCSLAATSLGLGAGMTPKIAWALENKPRIPVVWIHGL WP_001058310.1 hydrogenase [Salmonella enterica] Length: 372\nNumber of predicted TMHs: 2\nExp number of AAs in TMHs: 37.4567299999999\nExp number, first 60 AAs: 19.72694\nTotal prob of N-in: 0.96171\nPOSSIBLE N-term signal sequence\ninside 1 19\nTMhelix 20 39\noutside 40 325\nTMhelix 326 348\ninside 349 372

16236 GCF_002066145.1_ASM206614v1 Salmonella enterica subsp. enterica serovar Bonariensis Proteobacteria; Gammaproteobacteria; Enterobacterales; Enterobacteriaceae; Salmonella; Salmonella enterica; Salmonella enterica subsp. enterica MNNEETFYQAMRRKGVTRRSFLKFCSLAATSLGLGAGMTPKIAWALENKPRIPVVWIHGL WP_001058308.1 hydrogenase [Salmonella enterica] Length: 372\nNumber of predicted TMHs: 2\nExp number of AAs in TMHs: 39.08204\nExp number, first 60 AAs: 19.72474\nTotal prob of N-in: 0.96174\nPOSSIBLE N-term signal sequence\ninside 1 19\nTMhelix 20 39\noutside 40 325\nTMhelix 326 348\ninside 349 372

16237 GCF_002066145.1_ASM206614v1 Salmonella enterica subsp. enterica serovar Bonariensis Proteobacteria; Gammaproteobacteria; Enterobacterales; Enterobacteriaceae; Salmonella; Salmonella enterica; Salmonella enterica subsp. enterica MTGDNTLITSHGINRRDFMKLCAALAATMGLSSKAAAEMAESVSNPQRPPVIWIGAQECT WP_000145429.1 hydrogenase 2 small subunit [Salmonella enterica] Length: 372\nNumber of predicted TMHs: 1\nExp number of AAs in TMHs: 24.51309\nExp number, first 60 AAs: 0.2682\nTotal prob of N-in: 0.14402\noutside 1 331\nTMhelix 332 354\ninside 355 372

16238 GCF_002061775.1_ASM206177v1 Salmonella enterica subsp. enterica serovar Bovismorbificans Proteobacteria; Gammaproteobacteria; Enterobacterales; Enterobacteriaceae; Salmonella; Salmonella enterica; Salmonella enterica subsp. enterica MTGDNTLITSHGINRRDFMKLCAALAATMGLSSKAAAEMAESVSNPQRPPVIWIGAQECT WP_000145429.1 hydrogenase 2 small subunit [Salmonella enterica] Length: 372\nNumber of predicted TMHs: 1\nExp number of AAs in TMHs: 24.51309\nExp number, first 60 AAs: 0.2682\nTotal prob of N-in: 0.14402\noutside 1 331\nTMhelix 332 354\ninside 355 372

16239 GCF_002060595.1_ASM206059v1 Salmonella enterica subsp. enterica serovar Bovismorbificans Proteobacteria; Gammaproteobacteria; Enterobacterales; Enterobacteriaceae; Salmonella; Salmonella enterica; Salmonella enterica subsp. enterica MNNEETFYQAMRRKGVTRRSFLKFCSLAATSLGLGAGMTPKIAWALENKPRIPVVWIHGL WP_001058311.1 hydrogenase [Salmonella enterica] Length: 372\nNumber of predicted TMHs: 2\nExp number of AAs in TMHs: 37.4699899999999\nExp number, first 60 AAs: 19.72699\nTotal prob of N-in: 0.96171\nPOSSIBLE N-term signal sequence\ninside 1 19\nTMhelix 20 39\noutside 40 325\nTMhelix 326 348\ninside 349 372

16240 GCF_002061715.1_ASM206171v1 Salmonella enterica subsp. enterica serovar Bovismorbificans Proteobacteria; Gammaproteobacteria; Enterobacterales; Enterobacteriaceae; Salmonella; Salmonella enterica; Salmonella enterica subsp. enterica MNNEETFYQAMRRKGVTRRSFLKFCSLAATSLGLGAGMTPKIAWALENKPRIPVVWIHGL WP_001058312.1 hydrogenase [Salmonella enterica] Length: 372\nNumber of predicted TMHs: 2\nExp number of AAs in TMHs: 36.51738\nExp number, first 60 AAs: 19.72849\nTotal prob of N-in: 0.96170\nPOSSIBLE N-term signal sequence\ninside 1 19\nTMhelix 20 39\noutside 40 325\nTMhelix 326 348\ninside 349 372

16241 GCF_002060595.1_ASM206059v1 Salmonella enterica subsp. enterica serovar Bovismorbificans Proteobacteria; Gammaproteobacteria; Enterobacterales; Enterobacteriaceae; Salmonella; Salmonella enterica; Salmonella enterica subsp. enterica MTGDNTLITSHGINRRDFMKLCAALAATMGLSSKAAAEMAESVSNPQRPPVIWIGAQECT WP_000145429.1 hydrogenase 2 small subunit [Salmonella enterica] Length: 372\nNumber of predicted TMHs: 1\nExp number of AAs in TMHs: 24.51309\nExp number, first 60 AAs: 0.2682\nTotal prob of N-in: 0.14402\noutside 1 331\nTMhelix 332 354\ninside 355 372

16242 GCF_002061715.1_ASM206171v1 Salmonella enterica subsp. enterica serovar Bovismorbificans Proteobacteria; Gammaproteobacteria; Enterobacterales; Enterobacteriaceae; Salmonella; Salmonella enterica; Salmonella enterica subsp. enterica MTGDNTLITSHGINRRDFMKLCAALAATMGLSSKAAAEMAESVSNPQRPPVIWIGAQECT WP_000145431.1 hydrogenase 2 small subunit [Salmonella enterica] Length: 372\nNumber of predicted TMHs: 1\nExp number of AAs in TMHs: 24.51505\nExp number, first 60 AAs: 0.2682\nTotal prob of N-in: 0.14406\noutside 1 331\nTMhelix 332 354\ninside 355 372

16243 GCF_002050935.1_ASM205093v1 Salmonella enterica subsp. enterica serovar Bovismorbificans Proteobacteria; Gammaproteobacteria; Enterobacterales; Enterobacteriaceae; Salmonella; Salmonella enterica; Salmonella enterica subsp. enterica MNNEETFYQAMRRKGVTRRSFLKFCSLAATSLGLGAGMTPKIAWALENKPRIPVVWIHGL WP_001058311.1 hydrogenase [Salmonella enterica] Length: 372\nNumber of predicted TMHs: 2\nExp number of AAs in TMHs: 37.4699899999999\nExp number, first 60 AAs: 19.72699\nTotal prob of N-in: 0.96171\nPOSSIBLE N-term signal sequence\ninside 1 19\nTMhelix 20 39\noutside 40 325\nTMhelix 326 348\ninside 349 372

16244 GCF_002061735.1_ASM206173v1 Salmonella enterica subsp. enterica serovar Bovismorbificans Proteobacteria; Gammaproteobacteria; Enterobacterales; Enterobacteriaceae; Salmonella; Salmonella enterica; Salmonella enterica subsp. enterica MNNEETFYQAMRRKGVTRRSFLKFCSLAATSLGLGAGMTPKIAWALENKPRIPVVWIHGL WP_001058311.1 hydrogenase [Salmonella enterica] Length: 372\nNumber of predicted TMHs: 2\nExp number of AAs in TMHs: 37.4699899999999\nExp number, first 60 AAs: 19.72699\nTotal prob of N-in: 0.96171\nPOSSIBLE N-term signal sequence\ninside 1 19\nTMhelix 20 39\noutside 40 325\nTMhelix 326 348\ninside 349 372

16245 GCF_002061695.1_ASM206169v1 Salmonella enterica subsp. enterica serovar Bovismorbificans Proteobacteria; Gammaproteobacteria; Enterobacterales; Enterobacteriaceae; Salmonella; Salmonella enterica; Salmonella enterica subsp. enterica MNNEETFYQAMRRKGVTRRSFLKFCSLAATSLGLGAGMTPKIAWALENKPRIPVVWIHGL WP_001058312.1 hydrogenase [Salmonella enterica] Length: 372\nNumber of predicted TMHs: 2\nExp number of AAs in TMHs: 36.51738\nExp number, first 60 AAs: 19.72849\nTotal prob of N-in: 0.96170\nPOSSIBLE N-term signal sequence\ninside 1 19\nTMhelix 20 39\noutside 40 325\nTMhelix 326 348\ninside 349 372

16246 GCF_002060495.1_ASM206049v1 Salmonella enterica subsp. enterica serovar Bovismorbificans Proteobacteria; Gammaproteobacteria; Enterobacterales; Enterobacteriaceae; Salmonella; Salmonella enterica; Salmonella enterica subsp. enterica MTGDNTLITSHGINRRDFMKLCAALAATMGLSSKAAAEMAESVSNPQRPPVIWIGAQECT WP_000145429.1 hydrogenase 2 small subunit [Salmonella enterica] Length: 372\nNumber of predicted TMHs: 1\nExp number of AAs in TMHs: 24.51309\nExp number, first 60 AAs: 0.2682\nTotal prob of N-in: 0.14402\noutside 1 331\nTMhelix 332 354\ninside 355 372

16247 GCF_002060605.1_ASM206060v1 Salmonella enterica subsp. enterica serovar Bovismorbificans Proteobacteria; Gammaproteobacteria; Enterobacterales; Enterobacteriaceae; Salmonella; Salmonella enterica; Salmonella enterica subsp. enterica MNNEETFYQAMRRKGVTRRSFLKFCSLAATSLGLGAGMTPKIAWALENKPRIPVVWIHGL WP_001058311.1 hydrogenase [Salmonella enterica] Length: 372\nNumber of predicted TMHs: 2\nExp number of AAs in TMHs: 37.4699899999999\nExp number, first 60 AAs: 19.72699\nTotal prob of N-in: 0.96171\nPOSSIBLE N-term signal sequence\ninside 1 19\nTMhelix 20 39\noutside 40 325\nTMhelix 326 348\ninside 349 372

16248 GCF_002060575.1_ASM206057v1 Salmonella enterica subsp. enterica serovar Bovismorbificans Proteobacteria; Gammaproteobacteria; Enterobacterales; Enterobacteriaceae; Salmonella; Salmonella enterica; Salmonella enterica subsp. enterica MNNEETFYQAMRRKGVTRRSFLKFCSLAATSLGLGAGMTPKIAWALENKPRIPVVWIHGL WP_001058312.1 hydrogenase [Salmonella enterica] Length: 372\nNumber of predicted TMHs: 2\nExp number of AAs in TMHs: 36.51738\nExp number, first 60 AAs: 19.72849\nTotal prob of N-in: 0.96170\nPOSSIBLE N-term signal sequence\ninside 1 19\nTMhelix 20 39\noutside 40 325\nTMhelix 326 348\ninside 349 372

16249 GCF_002061735.1_ASM206173v1 Salmonella enterica subsp. enterica serovar Bovismorbificans Proteobacteria; Gammaproteobacteria; Enterobacterales; Enterobacteriaceae; Salmonella; Salmonella enterica; Salmonella enterica subsp. enterica MTGDNTLITSHGINRRDFMKLCAALAATMGLSSKAAAEMAESVSNPQRPPVIWIGAQECT WP_000145429.1 hydrogenase 2 small subunit [Salmonella enterica] Length: 372\nNumber of predicted TMHs: 1\nExp number of AAs in TMHs: 24.51309\nExp number, first 60 AAs: 0.2682\nTotal prob of N-in: 0.14402\noutside 1 331\nTMhelix 332 354\ninside 355 372

16250 GCF_002061695.1_ASM206169v1 Salmonella enterica subsp. enterica serovar Bovismorbificans Proteobacteria; Gammaproteobacteria; Enterobacterales; Enterobacteriaceae; Salmonella; Salmonella enterica; Salmonella enterica subsp. enterica MTGDNTLITSHGINRRDFMKLCAALAATMGLSSKAAAEMAESVSNPQRPPVIWIGAQECT WP_000145431.1 hydrogenase 2 small subunit [Salmonella enterica] Length: 372\nNumber of predicted TMHs: 1\nExp number of AAs in TMHs: 24.51505\nExp number, first 60 AAs: 0.2682\nTotal prob of N-in: 0.14406\noutside 1 331\nTMhelix 332 354\ninside 355 372

16251 GCF_002060605.1_ASM206060v1 Salmonella enterica subsp. enterica serovar Bovismorbificans Proteobacteria; Gammaproteobacteria; Enterobacterales; Enterobacteriaceae; Salmonella; Salmonella enterica;

Salmonella enterica subsp. enterica MTGDNTLITSHGINRRDFMKLCAALAATMGLSSKAAAEMAESVSNPQRPPVIWIGAQECT WP_000145429.1 hydrogenase 2 small subunit [Salmonella enterica] Length: 372\nNumber of predicted TMHs: 1\nExp number of AAs in TMHs: 24.51309\nExp number, first 60 AAs: 0.2682\nTotal prob of N-in: 0.14402\noutside 1 331\nTMhelix 332 354\ninside 355 372

16252 GCF_002060575.1_ASM206057v1 Salmonella enterica subsp. enterica serovar Bovismorbificans Proteobacteria; Gammaproteobacteria; Enterobacterales; Enterobacteriaceae; Salmonella; Salmonella enterica; Salmonella enterica subsp. enterica MTGDNTLITSHGINRRDFMKLCAALAATMGLSSKAAAEMAESVSNPQRPPVIWIGAQECT WP_000145431.1 hydrogenase 2 small subunit [Salmonella enterica] Length: 372\nNumber of predicted TMHs: 1\nExp number of AAs in TMHs: 24.51505\nExp number, first 60 AAs: 0.2682\nTotal prob of N-in: 0.14406\noutside 1 331\nTMhelix 332 354\ninside 355 372

16253 GCF_002060495.1_ASM206049v1 Salmonella enterica subsp. enterica serovar Bovismorbificans Proteobacteria; Gammaproteobacteria; Enterobacterales; Enterobacteriaceae; Salmonella; Salmonella enterica; Salmonella enterica subsp. enterica MNNEETFYQAMRRKGVTRRSFLKFCSLAATSLGLGAGMTPKIAWALENKPRIPVVWIHGL WP_001058311.1 hydrogenase [Salmonella enterica] Length: 372\nNumber of predicted TMHs: 2\nExp number of AAs in TMHs: 37.4699899999999\nExp number, first 60 AAs: 19.72699\nTotal prob of N-in: 0.96171\nPOSSIBLE N-term signal sequence\ninside 1 19\nTMhelix 20 39\noutside 40 325\nTMhelix 326 348\ninside 349 372

16254 GCF_002060515.1_ASM206051v1 Salmonella enterica subsp. enterica serovar Bovismorbificans Proteobacteria; Gammaproteobacteria; Enterobacterales; Enterobacteriaceae; Salmonella; Salmonella enterica; Salmonella enterica subsp. enterica MNNEETFYQAMRRKGVTRRSFLKFCSLAATSLGLGAGMTPKIAWALENKPRIPVVWIHGL WP_001058312.1 hydrogenase [Salmonella enterica] Length: 372\nNumber of predicted TMHs: 2\nExp number of AAs in TMHs: 36.51738\nExp number, first 60 AAs: 19.72849\nTotal prob of N-in: 0.96170\nPOSSIBLE N-term signal sequence\ninside 1 19\nTMhelix 20 39\noutside 40 325\nTMhelix 326 348\ninside 349 372

16255 GCF_002060515.1_ASM206051v1 Salmonella enterica subsp. enterica serovar Bovismorbificans Proteobacteria; Gammaproteobacteria; Enterobacterales; Enterobacteriaceae; Salmonella; Salmonella enterica; Salmonella enterica subsp. enterica MTGDNTLITSHGINRRDFMKLCAALAATMGLSSKAAAEMAESVSNPQRPPVIWIGAQECT WP_000145431.1 hydrogenase 2 small subunit [Salmonella enterica] Length: 372\nNumber of predicted TMHs: 1\nExp number of AAs in TMHs: 24.51505\nExp number, first 60 AAs: 0.2682\nTotal prob of N-in: 0.14406\noutside 1 331\nTMhelix 332 354\ninside 355 372

16256 GCF_002062655.1_ASM206265v1 Salmonella enterica subsp. enterica serovar Bovismorbificans Proteobacteria; Gammaproteobacteria; Enterobacterales; Enterobacteriaceae; Salmonella; Salmonella enterica; Salmonella enterica subsp. enterica MTGDNTLITSHGINRRDFMKLCAALAATMGLSSKAAAEMAESVSNPQRPPVIWIGAQECT WP_000145429.1 hydrogenase 2 small subunit [Salmonella enterica] Length: 372\nNumber of predicted TMHs: 1\nExp number of AAs in TMHs: 24.51309\nExp number, first 60 AAs: 0.2682\nTotal prob of N-in: 0.14402\noutside 1 331\nTMhelix 332 354\ninside 355 372

16257 GCF_002049045.1_ASM204904v1 Salmonella enterica subsp. enterica serovar Bovismorbificans Proteobacteria; Gammaproteobacteria; Enterobacterales; Enterobacteriaceae; Salmonella; Salmonella enterica; Salmonella enterica subsp. enterica MTGDNTLITSHGINRRDFMKLCAALAATMGLSSKAAAEMAESVSNPQRPPVIWIGAQECT WP_000145429.1 hydrogenase 2 small subunit [Salmonella enterica] Length: 372\nNumber of predicted TMHs: 1\nExp number of AAs in TMHs: 24.51309\nExp number, first 60 AAs: 0.2682\nTotal prob of N-in: 0.14402\noutside 1 331\nTMhelix 332 354\ninside 355 372

16258 GCF_002049045.1_ASM204904v1 Salmonella enterica subsp. enterica serovar Bovismorbificans Proteobacteria; Gammaproteobacteria; Enterobacterales; Enterobacteriaceae; Salmonella; Salmonella enterica; Salmonella enterica subsp. enterica MNNEETFYQAMRRKGVTRRSFLKFCSLAATSLGLGAGMTPKIAWALENKPRIPVVWIHGL WP_001058311.1 hydrogenase [Salmonella enterica] Length: 372\nNumber of predicted TMHs: 2\nExp number of AAs in TMHs: 37.4699899999999\nExp number, first 60 AAs: 19.72699\nTotal prob of N-in: 0.96171\nPOSSIBLE N-term signal sequence\ninside 1 19\nTMhelix 20 39\noutside 40 325\nTMhelix 326 348\ninside 349 372

16259 GCF_002062655.1_ASM206265v1 Salmonella enterica subsp. enterica serovar Bovismorbificans Proteobacteria; Gammaproteobacteria; Enterobacterales; Enterobacteriaceae; Salmonella; Salmonella enterica; Salmonella enterica subsp. enterica MNNEETFYQAMRRKGVTRRSFLKFCSLAATSLGLGAGMTPKIAWALENKPRIPVVWIHGL WP_001058311.1 hydrogenase [Salmonella enterica] Length: 372\nNumber of predicted TMHs: 2\nExp number of AAs in TMHs: 37.4699899999999\nExp number, first 60 AAs: 19.72699\nTotal prob of N-in:

0.96171\nPOSSIBLE N-term signal sequence\ninside 1 19\nTMhelix 20 39\noutside 40 325\nTMhelix 326 348\ninside 349 372

16260 GCF_002050735.1_ASM205073v1 Salmonella enterica subsp. enterica serovar Bovismorbificans Proteobacteria; Gammaproteobacteria; Enterobacterales; Enterobacteriaceae; Salmonella; Salmonella enterica; Salmonella enterica subsp. enterica MTGDNTLITSHGINRRDFMKLCAALAATMGLSSKAAAEMAESVSNPQRPPVIWIGAQECT WP_000145429.1 hydrogenase 2 small subunit [Salmonella enterica] Length: 372\nNumber of predicted TMHs: 1\nExp number of AAs in TMHs: 24.51309\nExp number, first 60 AAs: 0.2682\nTotal prob of N-in: 0.14402\noutside 1 331\nTMhelix 332 354\ninside 355 372

16261 GCF_002050735.1_ASM205073v1 Salmonella enterica subsp. enterica serovar Bovismorbificans Proteobacteria; Gammaproteobacteria; Enterobacterales; Enterobacteriaceae; Salmonella; Salmonella enterica; Salmonella enterica subsp. enterica MNNEETFYQAMRRKGVTRRSFLKFCSLAATSLGLGAGMTPKIAWALENKPRIPVVWIHGL WP_001058311.1 hydrogenase [Salmonella enterica] Length: 372\nNumber of predicted TMHs: 2\nExp number of AAs in TMHs: 37.4699899999999\nExp number, first 60 AAs: 19.72699\nTotal prob of N-in: 0.96171\nPOSSIBLE N-term signal sequence\ninside 1 19\nTMhelix 20 39\noutside 40 325\nTMhelix 326 348\ninside 349 372

16262 GCF_002060635.1_ASM206063v1 Salmonella enterica subsp. enterica serovar Bovismorbificans Proteobacteria; Gammaproteobacteria; Enterobacterales; Enterobacteriaceae; Salmonella; Salmonella enterica; Salmonella enterica subsp. enterica MTGDNTLITSHGINRRDFMKLCAALAATMGLSSKAAAEMAESVSNPQRPPVIWIGAQECT WP_000145429.1 hydrogenase 2 small subunit [Salmonella enterica] Length: 372\nNumber of predicted TMHs: 1\nExp number of AAs in TMHs: 24.51309\nExp number, first 60 AAs: 0.2682\nTotal prob of N-in: 0.14402\noutside 1 331\nTMhelix 332 354\ninside 355 372

16263 GCF_002061775.1_ASM206177v1 Salmonella enterica subsp. enterica serovar Bovismorbificans Proteobacteria; Gammaproteobacteria; Enterobacterales; Enterobacteriaceae; Salmonella; Salmonella enterica; Salmonella enterica subsp. enterica MNNEETFYQAMRRKGVTRRSFLKFCSLAATSLGLGAGMTPKIAWALENKPRIPVVWIHGL WP_001058311.1 hydrogenase [Salmonella enterica] Length: 372\nNumber of predicted TMHs: 2\nExp number of AAs in TMHs: 37.4699899999999\nExp number, first 60 AAs: 19.72699\nTotal prob of N-in: 0.96171\nPOSSIBLE N-term signal sequence\ninside 1 19\nTMhelix 20 39\noutside 40 325\nTMhelix 326 348\ninside 349 372

16264 GCF_002060635.1_ASM206063v1 Salmonella enterica subsp. enterica serovar Bovismorbificans Proteobacteria; Gammaproteobacteria; Enterobacterales; Enterobacteriaceae; Salmonella; Salmonella enterica; Salmonella enterica subsp. enterica MNNEETFYQAMRRKGVTRRSFLKFCSLAATSLGLGAGMTPKIAWALENKPRIPVVWIHGL WP_001058311.1 hydrogenase [Salmonella enterica] Length: 372\nNumber of predicted TMHs: 2\nExp number of AAs in TMHs: 37.4699899999999\nExp number, first 60 AAs: 19.72699\nTotal prob of N-in: 0.96171\nPOSSIBLE N-term signal sequence\ninside 1 19\nTMhelix 20 39\noutside 40 325\nTMhelix 326 348\ninside 349 372

16265 GCF_002050885.1_ASM205088v1 Salmonella enterica subsp. enterica serovar Bovismorbificans Proteobacteria; Gammaproteobacteria; Enterobacterales; Enterobacteriaceae; Salmonella; Salmonella enterica; Salmonella enterica subsp. enterica MTGDNTLITSHGINRRDFMKLCAALAATMGLSSKAAAEMAESVSNPQRPPVIWIGAQECT WP_000145429.1 hydrogenase 2 small subunit [Salmonella enterica] Length: 372\nNumber of predicted TMHs: 1\nExp number of AAs in TMHs: 24.51309\nExp number, first 60 AAs: 0.2682\nTotal prob of N-in: 0.14402\noutside 1 331\nTMhelix 332 354\ninside 355 372

16266 GCF_002050935.1_ASM205093v1 Salmonella enterica subsp. enterica serovar Bovismorbificans Proteobacteria; Gammaproteobacteria; Enterobacterales; Enterobacteriaceae; Salmonella; Salmonella enterica; Salmonella enterica subsp. enterica MTGDNTLITSHGINRRDFMKLCAALAATMGLSSKAAAEMAESVSNPQRPPVIWIGAQECT WP_000145429.1 hydrogenase 2 small subunit [Salmonella enterica] Length: 372\nNumber of predicted TMHs: 1\nExp number of AAs in TMHs: 24.51309\nExp number, first 60 AAs: 0.2682\nTotal prob of N-in: 0.14402\noutside 1 331\nTMhelix 332 354\ninside 355 372

16267 GCF_002050885.1_ASM205088v1 Salmonella enterica subsp. enterica serovar Bovismorbificans Proteobacteria; Gammaproteobacteria; Enterobacterales; Enterobacteriaceae; Salmonella; Salmonella enterica; Salmonella enterica subsp. enterica MNNEETFYQAMRRKGVTRRSFLKFCSLAATSLGLGAGMTPKIAWALENKPRIPVVWIHGL WP_001058311.1 hydrogenase [Salmonella enterica] Length: 372\nNumber of predicted TMHs: 2\nExp number of AAs in TMHs: 37.4699899999999\nExp number, first 60 AAs: 19.72699\nTotal prob of N-in:

0.96171\nPOSSIBLE N-term signal sequence\ninside 1 19\nTMhelix 20 39\noutside 40 325\nTMhelix 326 348\ninside 349 372

16268 GCF_002062565.1_ASM206256v1 Salmonella enterica subsp. enterica serovar Braenddrup Proteobacteria; Gammaproteobacteria; Enterobacterales; Enterobacteriaceae; Salmonella; Salmonella enterica; Salmonella enterica subsp. enterica MTGDNTLITSHGINRRDFMKLCAALAATMGLSSKAAAEMAESVSNPQRPPVIWIGAQECT WP_000145429.1 hydrogenase 2 small subunit [Salmonella enterica] Length: 372\nNumber of predicted TMHs: 1\nExp number of AAs in TMHs: 24.51309\nExp number, first 60 AAs: 0.2682\nTotal prob of N-in: 0.14402\noutside 1 331\nTMhelix 332 354\ninside 355 372

16269 GCF_002062565.1_ASM206256v1 Salmonella enterica subsp. enterica serovar Braenddrup Proteobacteria; Gammaproteobacteria; Enterobacterales; Enterobacteriaceae; Salmonella; Salmonella enterica; Salmonella enterica subsp. enterica MNNEETFYQAMRRKGVTRRSFLKFCSLAATSLGLGAGMTPKIAWALENKPRIPVVWIHGL WP_001058312.1 hydrogenase [Salmonella enterica] Length: 372\nNumber of predicted TMHs: 2\nExp number of AAs in TMHs: 36.51738\nExp number, first 60 AAs: 19.72849\nTotal prob of N-in: 0.96170\nPOSSIBLE N-term signal sequence\ninside 1 19\nTMhelix 20 39\noutside 40 325\nTMhelix 326 348\ninside 349 372

16270 GCF_001833895.1_ASM183389v1 Salmonella enterica subsp. enterica serovar Braenderup Proteobacteria; Gammaproteobacteria; Enterobacterales; Enterobacteriaceae; Salmonella; Salmonella enterica; Salmonella enterica subsp. enterica MNNEETFYQAMRRKGVTRRSFLKFCSLAATSLGLGAGMTPKIAWALENKPRIPVVWIHGL WP_001058312.1 hydrogenase [Salmonella enterica] Length: 372\nNumber of predicted TMHs: 2\nExp number of AAs in TMHs: 36.51738\nExp number, first 60 AAs: 19.72849\nTotal prob of N-in: 0.96170\nPOSSIBLE N-term signal sequence\ninside 1 19\nTMhelix 20 39\noutside 40 325\nTMhelix 326 348\ninside 349 372

16271 GCF_002036245.1_ASM203624v1 Salmonella enterica subsp. enterica serovar Braenderup Proteobacteria; Gammaproteobacteria; Enterobacterales; Enterobacteriaceae; Salmonella; Salmonella enterica; Salmonella enterica subsp. enterica MTGDNTLITSHGINRRDFMKLCAALAATMGLSSKAAAEMAESVSNPQRPPVIWIGAQECT WP_000145429.1 hydrogenase 2 small subunit [Salmonella enterica] Length: 372\nNumber of predicted TMHs: 1\nExp number of AAs in TMHs: 24.51309\nExp number, first 60 AAs: 0.2682\nTotal prob of N-in: 0.14402\noutside 1 331\nTMhelix 332 354\ninside 355 372

16272 GCF_001245445.1_Salmonella_enterica_CVM_N50427_v1.0 Salmonella enterica subsp. enterica serovar Braenderup Proteobacteria; Gammaproteobacteria; Enterobacterales; Enterobacteriaceae; Salmonella; Salmonella enterica; Salmonella enterica subsp. enterica MTGDNTLITSHGINRRDFMKLCAALAATMGLSSKAAAEMAESVSNPQRPPVIWIGAQECT WP_000145429.1 hydrogenase 2 small subunit [Salmonella enterica] Length: 372\nNumber of predicted TMHs: 1\nExp number of AAs in TMHs: 24.51309\nExp number, first 60 AAs: 0.2682\nTotal prob of N-in: 0.14402\noutside 1 331\nTMhelix 332 354\ninside 355 372

16273 GCF_002059815.1_ASM205981v1 Salmonella enterica subsp. enterica serovar Braenderup Proteobacteria; Gammaproteobacteria; Enterobacterales; Enterobacteriaceae; Salmonella; Salmonella enterica; Salmonella enterica subsp. enterica MNNEETFYQAMRRKGVTRRSFLKFCSLAATSLGLGAGMTPKIAWALENKPRIPVVWIHGL WP_001058312.1 hydrogenase [Salmonella enterica] Length: 372\nNumber of predicted TMHs: 2\nExp number of AAs in TMHs: 36.51738\nExp number, first 60 AAs: 19.72849\nTotal prob of N-in: 0.96170\nPOSSIBLE N-term signal sequence\ninside 1 19\nTMhelix 20 39\noutside 40 325\nTMhelix 326 348\ninside 349 372

16274 GCF_001245025.1_Salmonella_enterica_CVM_N48701_v1.0 Salmonella enterica subsp. enterica serovar Braenderup Proteobacteria; Gammaproteobacteria; Enterobacterales; Enterobacteriaceae; Salmonella; Salmonella enterica; Salmonella enterica subsp. enterica MNNEETFYQAMRRKGVTRRSFLKFCSLAATSLGLGAGMTPKIAWALENKPRIPVVWIHGL WP_001058312.1 hydrogenase [Salmonella enterica] Length: 372\nNumber of predicted TMHs: 2\nExp number of AAs in TMHs: 36.51738\nExp number, first 60 AAs: 19.72849\nTotal prob of N-in: 0.96170\nPOSSIBLE N-term signal sequence\ninside 1 19\nTMhelix 20 39\noutside 40 325\nTMhelix 326 348\ninside 349 372

16275 GCF_001245445.1_Salmonella_enterica_CVM_N50427_v1.0 Salmonella enterica subsp. enterica serovar Braenderup Proteobacteria; Gammaproteobacteria; Enterobacterales; Enterobacteriaceae; Salmonella; Salmonella enterica; Salmonella enterica subsp. enterica MNNEETFYQAMRRKGVTRRSFLKFCSLAATSLGLGAGMTPKIAWALENKPRIPVVWIHGL WP_001058312.1 hydrogenase [Salmonella enterica] Length: 372\nNumber of predicted TMHs: 2\nExp number of AAs in TMHs: 36.51738\nExp number, first 60 AAs: 19.72849\nTotal prob of N-in: 0.96170\nPOSSIBLE N-term signal sequence\ninside 1 19\nTMhelix 20 39\noutside 40 325\nTMhelix 326 348\ninside 349 372

16276 GCF_002059815.1_ASM205981v1 Salmonella enterica subsp. enterica serovar Braenderup Proteobacteria; Gammaproteobacteria; Enterobacterales; Enterobacteriaceae; Salmonella; Salmonella enterica; Salmonella enterica subsp. enterica MTGDNTLITSHGINRRDFMKLCAALAATMGLSSKAAAEMAESVSNPQRPPVIWIGAQECT WP_000145429.1 hydrogenase 2 small subunit [Salmonella enterica] Length: 372\nNumber of predicted TMHs: 1\nExp number of AAs in TMHs: 24.51309\nExp number, first 60 AAs: 0.2682\nTotal prob of N-in: 0.14402\noutside 1 331\nTMhelix 332 354\ninside 355 372

16277 GCF_001480345.1_Salmonella_enterica_CVM_N41747-SQ_v1.0 Salmonella enterica subsp. enterica serovar Braenderup Proteobacteria; Gammaproteobacteria; Enterobacterales; Enterobacteriaceae; Salmonella; Salmonella enterica; Salmonella enterica subsp. enterica MTGDNTLITSHGINRRDFMKLCAALAATMGLSSKAAAEMAESVSNPQRPPVIWIGAQECT WP_000145429.1 hydrogenase 2 small subunit [Salmonella enterica] Length: 372\nNumber of predicted TMHs: 1\nExp number of AAs in TMHs: 24.51309\nExp number, first 60 AAs: 0.2682\nTotal prob of N-in: 0.14402\noutside 1 331\nTMhelix 332 354\ninside 355 372

16278 GCF_002059465.1_ASM205946v1 Salmonella enterica subsp. enterica serovar Braenderup Proteobacteria; Gammaproteobacteria; Enterobacterales; Enterobacteriaceae; Salmonella; Salmonella enterica; Salmonella enterica subsp. enterica MTGDNTLITSHGINRRDFMKLCAALAATMGLSSKAAAEMAESVSNPQRPPVIWIGAQECT WP_000145429.1 hydrogenase 2 small subunit [Salmonella enterica] Length: 372\nNumber of predicted TMHs: 1\nExp number of AAs in TMHs: 24.51309\nExp number, first 60 AAs: 0.2682\nTotal prob of N-in: 0.14402\noutside 1 331\nTMhelix 332 354\ninside 355 372

16279 GCF_002036245.1_ASM203624v1 Salmonella enterica subsp. enterica serovar Braenderup Proteobacteria; Gammaproteobacteria; Enterobacterales; Enterobacteriaceae; Salmonella; Salmonella enterica; Salmonella enterica subsp. enterica MNNEETFYQAMRRKGVTRRSFLKFCSLAATSLGLGAGMTPKIAWALENKPRIPVVWIHGL WP_001058312.1 hydrogenase [Salmonella enterica] Length: 372\nNumber of predicted TMHs: 2\nExp number of AAs in TMHs: 36.51738\nExp number, first 60 AAs: 19.72849\nTotal prob of N-in: 0.96170\nPOSSIBLE N-term signal sequence\ninside 1 19\nTMhelix 20 39\noutside 40 325\nTMhelix 326 348\ninside 349 372

16280 GCF_001480425.1_Salmonella_enterica_CVM_N42235-SQ_v1.0 Salmonella enterica subsp. enterica serovar Braenderup Proteobacteria; Gammaproteobacteria; Enterobacterales; Enterobacteriaceae; Salmonella; Salmonella enterica; Salmonella enterica subsp. enterica MTGDNTLITSHGINRRDFMKLCAALAATMGLSSKAAAEMAESVSNPQRPPVIWIGAQECT WP_000145429.1 hydrogenase 2 small subunit [Salmonella enterica] Length: 372\nNumber of predicted TMHs: 1\nExp number of AAs in TMHs: 24.51309\nExp number, first 60 AAs: 0.2682\nTotal prob of N-in: 0.14402\noutside 1 331\nTMhelix 332 354\ninside 355 372

16281 GCF_001480345.1_Salmonella_enterica_CVM_N41747-SQ_v1.0 Salmonella enterica subsp. enterica serovar Braenderup Proteobacteria; Gammaproteobacteria; Enterobacterales; Enterobacteriaceae; Salmonella; Salmonella enterica; Salmonella enterica subsp. enterica MNNEETFYQAMRRKGVTRRSFLKFCSLAATSLGLGAGMTPKIAWALENKPRIPVVWIHGL WP_001058312.1 hydrogenase [Salmonella enterica] Length: 372\nNumber of predicted TMHs: 2\nExp number of AAs in TMHs: 36.51738\nExp number, first 60 AAs: 19.72849\nTotal prob of N-in: 0.96170\nPOSSIBLE N-term signal sequence\ninside 1 19\nTMhelix 20 39\noutside 40 325\nTMhelix 326 348\ninside 349 372

16282 GCF_001833895.1_ASM183389v1 Salmonella enterica subsp. enterica serovar Braenderup Proteobacteria; Gammaproteobacteria; Enterobacterales; Enterobacteriaceae; Salmonella; Salmonella enterica; Salmonella enterica subsp. enterica MTGDNTLITSHGINRRDFMKLCAALAATMGLSSKAAAEMAESVSNPQRPPVIWIGAQECT WP_000145429.1 hydrogenase 2 small subunit [Salmonella enterica] Length: 372\nNumber of predicted TMHs: 1\nExp number of AAs in TMHs: 24.51309\nExp number, first 60 AAs: 0.2682\nTotal prob of N-in: 0.14402\noutside 1 331\nTMhelix 332 354\ninside 355 372

16283 GCF_001480425.1_Salmonella_enterica_CVM_N42235-SQ_v1.0 Salmonella enterica subsp. enterica serovar Braenderup Proteobacteria; Gammaproteobacteria; Enterobacterales; Enterobacteriaceae; Salmonella; Salmonella enterica; Salmonella enterica subsp. enterica MNNEETFYQAMRRKGVTRRSFLKFCSLAATSLGLGAGMTPKIAWALENKPRIPVVWIHGL WP_001058312.1 hydrogenase [Salmonella enterica] Length: 372\nNumber of predicted TMHs: 2\nExp number of AAs in TMHs: 36.51738\nExp number, first 60 AAs: 19.72849\nTotal prob of N-in: 0.96170\nPOSSIBLE N-term signal sequence\ninside 1 19\nTMhelix 20 39\noutside 40 325\nTMhelix 326 348\ninside 349 372

16284 GCF_002059465.1_ASM205946v1 Salmonella enterica subsp. enterica serovar Braenderup Proteobacteria; Gammaproteobacteria; Enterobacterales; Enterobacteriaceae; Salmonella; Salmonella enterica; Salmonella enterica subsp. enterica MNNEETFYQAMRRKGVTRRSFLKFCSLAATSLGLGAGMTPKIAWALENKPRIPVVWIHGL WP_001058312.1 hydrogenase [Salmonella enterica] Length: 372\nNumber of predicted TMHs: 2\nExp number of AAs in TMHs: 36.51738\nExp number, first 60 AAs: 19.72849\nTotal prob of N-in: 0.96170\nPOSSIBLE N-term signal sequence\ninside 1 19\nTMhelix 20 39\noutside 40 325\nTMhelix 326 348\ninside 349 372

16285 GCF_001833885.1_ASM183388v1 Salmonella enterica subsp. enterica serovar Braenderup Proteobacteria; Gammaproteobacteria; Enterobacterales; Enterobacteriaceae; Salmonella; Salmonella enterica; Salmonella enterica subsp. enterica MNNEETFYQAMRRKGVTRRSFLKFCSLAATSLGLGAGMTPKIAWALENKPRIPVVWIHGL WP_001058312.1 hydrogenase [Salmonella enterica] Length: 372\nNumber of predicted TMHs: 2\nExp number of AAs in TMHs: 36.51738\nExp number, first 60 AAs: 19.72849\nTotal prob of N-in: 0.96170\nPOSSIBLE N-term signal sequence\ninside 1 19\nTMhelix 20 39\noutside 40 325\nTMhelix 326 348\ninside 349 372

16286 GCF_001833885.1_ASM183388v1 Salmonella enterica subsp. enterica serovar Braenderup Proteobacteria; Gammaproteobacteria; Enterobacterales; Enterobacteriaceae; Salmonella; Salmonella enterica; Salmonella enterica subsp. enterica MTGDNTLITSHGINRRDFMKLCAALAATMGLSSKAAAEMAESVSNPQRPPVIWIGAQECT WP_000145429.1 hydrogenase 2 small subunit [Salmonella enterica] Length: 372\nNumber of predicted TMHs: 1\nExp number of AAs in TMHs: 24.51309\nExp number, first 60 AAs: 0.2682\nTotal prob of N-in: 0.14402\noutside 1 331\nTMhelix 332 354\ninside 355 372

16287 GCF_001243855.1_Salmonella_enterica_CVM_N47716_v1.0 Salmonella enterica subsp. enterica serovar Brandenburg Proteobacteria; Gammaproteobacteria; Enterobacterales; Enterobacteriaceae; Salmonella; Salmonella enterica; Salmonella enterica subsp. enterica MTGDNTLITSHGINRRDFMKLCAALAATMGLSSKAAAEMAESVSNPQRPPVIWIGAQECT WP_000145429.1 hydrogenase 2 small subunit [Salmonella enterica] Length: 372\nNumber of predicted TMHs: 1\nExp number of AAs in TMHs: 24.51309\nExp number, first 60 AAs: 0.2682\nTotal prob of N-in: 0.14402\noutside 1 331\nTMhelix 332 354\ninside 355 372

16288 GCF_001243855.1_Salmonella_enterica_CVM_N47716_v1.0 Salmonella enterica subsp. enterica serovar Brandenburg Proteobacteria; Gammaproteobacteria; Enterobacterales; Enterobacteriaceae; Salmonella; Salmonella enterica; Salmonella enterica subsp. enterica MNNEETFYQAMRRKGVTRRSFLKFCSLAATSLGLGAGMTPKIAWALENKPRIPVVWIHGL WP_001058312.1 hydrogenase [Salmonella enterica] Length: 372\nNumber of predicted TMHs: 2\nExp number of AAs in TMHs: 36.51738\nExp number, first 60 AAs: 19.72849\nTotal prob of N-in: 0.96170\nPOSSIBLE N-term signal sequence\ninside 1 19\nTMhelix 20 39\noutside 40 325\nTMhelix 326 348\ninside 349 372

16289 GCF_001480545.1_Salmonella_enterica_CVM_N40367-SQ_v1.0 Salmonella enterica subsp. enterica serovar Brandenburg Proteobacteria; Gammaproteobacteria; Enterobacterales; Enterobacteriaceae; Salmonella; Salmonella enterica; Salmonella enterica subsp. enterica MTGDNTLITSHGINRRDFMKLCAALAATMGLSSKAAAEMAESVSNPQRPPVIWIGAQECT WP_000145429.1 hydrogenase 2 small subunit [Salmonella enterica] Length: 372\nNumber of predicted TMHs: 1\nExp number of AAs in TMHs: 24.51309\nExp number, first 60 AAs: 0.2682\nTotal prob of N-in: 0.14402\noutside 1 331\nTMhelix 332 354\ninside 355 372

16290 GCF_002030445.1_ASM203044v1 Salmonella enterica subsp. enterica serovar Brandenburg Proteobacteria; Gammaproteobacteria; Enterobacterales; Enterobacteriaceae; Salmonella; Salmonella enterica; Salmonella enterica subsp. enterica MNNEETFYQAMRRKGVTRRSFLKFCSLAATSLGLGAGMTPKIAWALENKPRIPVVWIHGL WP_001058312.1 hydrogenase [Salmonella enterica] Length: 372\nNumber of predicted TMHs: 2\nExp number of AAs in TMHs: 36.51738\nExp number, first 60 AAs: 19.72849\nTotal prob of N-in: 0.96170\nPOSSIBLE N-term signal sequence\ninside 1 19\nTMhelix 20 39\noutside 40 325\nTMhelix 326 348\ninside 349 372

16291 GCF_001480545.1_Salmonella_enterica_CVM_N40367-SQ_v1.0 Salmonella enterica subsp. enterica serovar Brandenburg Proteobacteria; Gammaproteobacteria; Enterobacterales; Enterobacteriaceae; Salmonella; Salmonella enterica; Salmonella enterica subsp. enterica MNNEETFYQAMRRKGVTRRSFLKFCSLAATSLGLGAGMTPKIAWALENKPRIPVVWIHGL WP_001058312.1 hydrogenase [Salmonella enterica] Length: 372\nNumber of predicted TMHs: 2\nExp number of AAs in TMHs: 36.51738\nExp number, first 60 AAs: 19.72849\nTotal prob of N-in: 0.96170\nPOSSIBLE N-term signal sequence\ninside 1 19\nTMhelix 20 39\noutside 40 325\nTMhelix 326 348\ninside 349 372

16292 GCF_002035925.1_ASM203592v1 Salmonella enterica subsp. enterica serovar Brandenburg Proteobacteria; Gammaproteobacteria; Enterobacterales; Enterobacteriaceae; Salmonella; Salmonella enterica; Salmonella enterica subsp. enterica MNNEETFYQAMRRKGVTRRSFLKFCSLAATSLGLGAGMTPKIAWALENKPRIPVVWIHGL WP_001058312.1 hydrogenase [Salmonella enterica] Length: 372\nNumber of predicted TMHs: 2\nExp number of AAs in TMHs: 36.51738\nExp number, first 60 AAs: 19.72849\nTotal prob of N-in: 0.96170\nPOSSIBLE N-term signal sequence\ninside 1 19\nTMhelix 20 39\noutside 40 325\nTMhelix 326 348\ninside 349 372

16293 GCF_002066025.1_ASM206602v1 Salmonella enterica subsp. enterica serovar Brandenburg Proteobacteria; Gammaproteobacteria; Enterobacterales; Enterobacteriaceae; Salmonella; Salmonella enterica; Salmonella enterica subsp. enterica MNNEETFYQAMRRKGVTRRSFLKFCSLAATSLGLGAGMTPKIAWALENKPRIPVVWIHGL WP_001058312.1 hydrogenase [Salmonella enterica] Length: 372\nNumber of predicted TMHs: 2\nExp number of AAs in TMHs: 36.51738\nExp number, first 60 AAs: 19.72849\nTotal prob of N-in: 0.96170\nPOSSIBLE N-term signal sequence\ninside 1 19\nTMhelix 20 39\noutside 40 325\nTMhelix 326 348\ninside 349 372

16294 GCF_002030445.1_ASM203044v1 Salmonella enterica subsp. enterica serovar Brandenburg Proteobacteria; Gammaproteobacteria; Enterobacterales; Enterobacteriaceae; Salmonella; Salmonella enterica; Salmonella enterica subsp. enterica MTGDNTLITSHGINRRDFMKLCAALAATMGLSSKAAAEMAESVSNPQRPPVIWIGAQECT WP_000145429.1 hydrogenase 2 small subunit [Salmonella enterica] Length: 372\nNumber of predicted TMHs: 1\nExp number of AAs in TMHs: 24.51309\nExp number, first 60 AAs: 0.2682\nTotal prob of N-in: 0.14402\noutside 1 331\nTMhelix 332 354\ninside 355 372

16295 GCF_002035925.1_ASM203592v1 Salmonella enterica subsp. enterica serovar Brandenburg Proteobacteria; Gammaproteobacteria; Enterobacterales; Enterobacteriaceae; Salmonella; Salmonella enterica; Salmonella enterica subsp. enterica MTGDNTLITSHGINRRDFMKLCAALAATMGLSSKAAAEMAESVSNPQRPPVIWIGAQECT WP_000145429.1 hydrogenase 2 small subunit [Salmonella enterica] Length: 372\nNumber of predicted TMHs: 1\nExp number of AAs in TMHs: 24.51309\nExp number, first 60 AAs: 0.2682\nTotal prob of N-in: 0.14402\noutside 1 331\nTMhelix 332 354\ninside 355 372

16296 GCF_002066025.1_ASM206602v1 Salmonella enterica subsp. enterica serovar Brandenburg Proteobacteria; Gammaproteobacteria; Enterobacterales; Enterobacteriaceae; Salmonella; Salmonella enterica; Salmonella enterica subsp. enterica MTGDNTLITSHGINRRDFMKLCAALAATMGLSSKAAAEMAESVSNPQRPPVIWIGAQECT WP_000145429.1 hydrogenase 2 small subunit [Salmonella enterica] Length: 372\nNumber of predicted TMHs: 1\nExp number of AAs in TMHs: 24.51309\nExp number, first 60 AAs: 0.2682\nTotal prob of N-in: 0.14402\noutside 1 331\nTMhelix 332 354\ninside 355 372

16297 GCF_001242575.1_Salmonella_enterica_CVM_N45949_v1.0 Salmonella enterica subsp. enterica serovar Brandenburg Proteobacteria; Gammaproteobacteria; Enterobacterales; Enterobacteriaceae; Salmonella; Salmonella enterica; Salmonella enterica subsp. enterica MTGDNTLITSHGINRRDFMKLCAALAATMGLSSKAAAEMAESVSNPQRPPVIWIGAQECT WP_000145429.1 hydrogenase 2 small subunit [Salmonella enterica] Length: 372\nNumber of predicted TMHs: 1\nExp number of AAs in TMHs: 24.51309\nExp number, first 60 AAs: 0.2682\nTotal prob of N-in: 0.14402\noutside 1 331\nTMhelix 332 354\ninside 355 372

16298 GCF_001242575.1_Salmonella_enterica_CVM_N45949_v1.0 Salmonella enterica subsp. enterica serovar Brandenburg Proteobacteria; Gammaproteobacteria; Enterobacterales; Enterobacteriaceae; Salmonella; Salmonella enterica; Salmonella enterica subsp. enterica MNNEETFYQAMRRKGVTRRSFLKFCSLAATSLGLGAGMTPKIAWALENKPRIPVVWIHGL WP_001058312.1 hydrogenase [Salmonella enterica] Length: 372\nNumber of predicted TMHs: 2\nExp number of AAs in TMHs: 36.51738\nExp number, first 60 AAs: 19.72849\nTotal prob of N-in: 0.96170\nPOSSIBLE N-term signal sequence\ninside 1 19\nTMhelix 20 39\noutside 40 325\nTMhelix 326 348\ninside 349 372

16299 GCF_001246205.1_Salmonella_enterica_CVM_N51253_v1.0 Salmonella enterica subsp. enterica serovar Brandenburg Proteobacteria; Gammaproteobacteria; Enterobacterales; Enterobacteriaceae; Salmonella; Salmonella enterica; Salmonella enterica subsp. enterica MNNEETFYQAMRRKGVTRRSFLKFCSLAATSLGLGAGMTPKIAWALENKPRIPVVWIHGL WP_001058312.1 hydrogenase [Salmonella enterica] Length: 372\nNumber of predicted TMHs: 2\nExp number of AAs in TMHs: 36.51738\nExp number, first 60 AAs: 19.72849\nTotal prob of N-in: 0.96170\nPOSSIBLE N-term signal sequence\ninside 1 19\nTMhelix 20 39\noutside 40 325\nTMhelix 326 348\ninside 349 372

16300 GCF_001246205.1_Salmonella_enterica_CVM_N51253_v1.0 Salmonella enterica subsp. enterica serovar Brandenburg Proteobacteria; Gammaproteobacteria; Enterobacterales; Enterobacteriaceae; Salmonella; Salmonella

enterica; Salmonella enterica subsp. enterica
MTGDNTLITSHGINRRDFMKLCAALAATMGLSSKAAAEMAESVSNPQRPPVIWIGAQECT WP_000145429.1 hydrogenase 2 small subunit [Salmonella enterica] Length: 372\nNumber of predicted TMHs: 1\nExp number of AAs in TMHs: 24.51309\nExp number, first 60 AAs: 0.2682\nTotal prob of N-in: 0.14402\noutside 1 331\nTMhelix 332 354\ninside 355 372

16301 GCF_002063195.1_ASM206319v1 Salmonella enterica subsp. enterica serovar Brandenburg Proteobacteria; Gammaproteobacteria; Enterobacterales; Enterobacteriaceae; Salmonella; Salmonella enterica; Salmonella enterica subsp. enterica MTGDNTLITSHGINRRDFMKLCAALAATMGLSSKAAAEMAESVSNPQRPPVIWIGAQECT WP_000145429.1 hydrogenase 2 small subunit [Salmonella enterica] Length: 372\nNumber of predicted TMHs: 1\nExp number of AAs in TMHs: 24.51309\nExp number, first 60 AAs: 0.2682\nTotal prob of N-in: 0.14402\noutside 1 331\nTMhelix 332 354\ninside 355 372

16302 GCF_002047245.1_ASM204724v1 Salmonella enterica subsp. enterica serovar Bredeney Proteobacteria; Gammaproteobacteria; Enterobacterales; Enterobacteriaceae; Salmonella; Salmonella enterica; Salmonella enterica subsp. enterica MTGDNTLITSHGINRRDFMKLCAALAATMGLSSKAAAEMAESVSNPQRPPVIWIGAQECT WP_000145429.1 hydrogenase 2 small subunit [Salmonella enterica] Length: 372\nNumber of predicted TMHs: 1\nExp number of AAs in TMHs: 24.51309\nExp number, first 60 AAs: 0.2682\nTotal prob of N-in: 0.14402\noutside 1 331\nTMhelix 332 354\ninside 355 372

16303 GCF_001478645.1_Salmonella_enterica_CVM_N29351-R_v1.0 Salmonella enterica subsp. enterica serovar Bredeney Proteobacteria; Gammaproteobacteria; Enterobacterales; Enterobacteriaceae; Salmonella; Salmonella enterica; Salmonella enterica subsp. enterica MTGDNTLITSHGINRRDFMKLCAALAATMGLSSKAAAEMAESVSNPQRPPVIWIGAQECT WP_000145429.1 hydrogenase 2 small subunit [Salmonella enterica] Length: 372\nNumber of predicted TMHs: 1\nExp number of AAs in TMHs: 24.51309\nExp number, first 60 AAs: 0.2682\nTotal prob of N-in: 0.14402\noutside 1 331\nTMhelix 332 354\ninside 355 372

16304 GCF_002047245.1_ASM204724v1 Salmonella enterica subsp. enterica serovar Bredeney Proteobacteria; Gammaproteobacteria; Enterobacterales; Enterobacteriaceae; Salmonella; Salmonella enterica; Salmonella enterica subsp. enterica MNNEETFYQAMRRKGVTRRSFLKFCSLAATSLGLGAGMTPKIAWALENKPRIPVVWIHGL WP_001058312.1 hydrogenase [Salmonella enterica] Length: 372\nNumber of predicted TMHs: 2\nExp number of AAs in TMHs: 36.51738\nExp number, first 60 AAs: 19.72849\nTotal prob of N-in: 0.96170\nPOSSIBLE N-term signal sequence\ninside 1 19\nTMhelix 20 39\noutside 40 325\nTMhelix 326 348\ninside 349 372

16305 GCF_002047265.1_ASM204726v1 Salmonella enterica subsp. enterica serovar Bredeney Proteobacteria; Gammaproteobacteria; Enterobacterales; Enterobacteriaceae; Salmonella; Salmonella enterica; Salmonella enterica subsp. enterica MNNEETFYQAMRRKGVTRRSFLKFCSLAATSLGLGAGMTPKIAWALENKPRIPVVWIHGL WP_001058312.1 hydrogenase [Salmonella enterica] Length: 372\nNumber of predicted TMHs: 2\nExp number of AAs in TMHs: 36.51738\nExp number, first 60 AAs: 19.72849\nTotal prob of N-in: 0.96170\nPOSSIBLE N-term signal sequence\ninside 1 19\nTMhelix 20 39\noutside 40 325\nTMhelix 326 348\ninside 349 372

16306 GCF_001478645.1_Salmonella_enterica_CVM_N29351-R_v1.0 Salmonella enterica subsp. enterica serovar Bredeney Proteobacteria; Gammaproteobacteria; Enterobacterales; Enterobacteriaceae; Salmonella; Salmonella enterica; Salmonella enterica subsp. enterica MNNEETFYQAMRRKGVTRRSFLKFCSLAATSLGLGAGMTPKIAWALENKPRIPVVWIHGL WP_001058312.1 hydrogenase [Salmonella enterica] Length: 372\nNumber of predicted TMHs: 2\nExp number of AAs in TMHs: 36.51738\nExp number, first 60 AAs: 19.72849\nTotal prob of N-in: 0.96170\nPOSSIBLE N-term signal sequence\ninside 1 19\nTMhelix 20 39\noutside 40 325\nTMhelix 326 348\ninside 349 372

16307 GCF_001271405.1_Salmonella_enterica_CVM_N43475_v1.0 Salmonella enterica subsp. enterica serovar Bredeney Proteobacteria; Gammaproteobacteria; Enterobacterales; Enterobacteriaceae; Salmonella; Salmonella enterica; Salmonella enterica subsp. enterica MNNEETFYQAMRRKGVTRRSFLKFCSLAATSLGLGAGMTPKIAWALENKPRIPVVWIHGL WP_001058312.1 hydrogenase [Salmonella enterica] Length: 372\nNumber of predicted TMHs: 2\nExp number of AAs in TMHs: 36.51738\nExp number, first 60 AAs: 19.72849\nTotal prob of N-in: 0.96170\nPOSSIBLE N-term signal sequence\ninside 1 19\nTMhelix 20 39\noutside 40 325\nTMhelix 326 348\ninside 349 372

16308 GCF_002048365.1_ASM204836v1 Salmonella enterica subsp. enterica serovar Bredeney Proteobacteria; Gammaproteobacteria; Enterobacterales; Enterobacteriaceae; Salmonella; Salmonella enterica; Salmonella enterica subsp. enterica MNNEETFYQAMRRKGVTRRSFLKFCSLAATSLGLGAGMTPKIAWALENKPRIPVVWIHGL WP_001058312.1 hydrogenase [Salmonella enterica] Length: 372\nNumber of predicted TMHs: 2\nExp number of AAs in TMHs: 36.51738\nExp number, first 60 AAs: 19.72849\nTotal prob of N-in: 0.96170\nPOSSIBLE N-term signal sequence\ninside 1 19\nTMhelix 20 39\noutside 40 325\nTMhelix 326 348\ninside 349 372

16309 GCF_002048505.1_ASM204850v1 Salmonella enterica subsp. enterica serovar Bredeney Proteobacteria; Gammaproteobacteria; Enterobacterales; Enterobacteriaceae; Salmonella; Salmonella enterica; Salmonella enterica subsp. enterica MNNEETFYQAMRRKGVTRRSFLKFCSLAATSLGLGAGMTPKIAWALENKPRIPVVWIHGL WP_001058312.1 hydrogenase [Salmonella enterica] Length: 372\nNumber of predicted TMHs: 2\nExp number of AAs in TMHs: 36.51738\nExp number, first 60 AAs: 19.72849\nTotal prob of N-in: 0.96170\nPOSSIBLE N-term signal sequence\ninside 1 19\nTMhelix 20 39\noutside 40 325\nTMhelix 326 348\ninside 349 372

16310 GCF_002048505.1_ASM204850v1 Salmonella enterica subsp. enterica serovar Bredeney Proteobacteria; Gammaproteobacteria; Enterobacterales; Enterobacteriaceae; Salmonella; Salmonella enterica; Salmonella enterica subsp. enterica MTGDNTLITSHGINRRDFMKLCAALAATMGLSSKAAAEMAESVSNPQRPPVIWIGAQECT WP_000145429.1 hydrogenase 2 small subunit [Salmonella enterica] Length: 372\nNumber of predicted TMHs: 1\nExp number of AAs in TMHs: 24.51309\nExp number, first 60 AAs: 0.2682\nTotal prob of N-in: 0.14402\noutside 1 331\nTMhelix 332 354\ninside 355 372

16311 GCF_002048365.1_ASM204836v1 Salmonella enterica subsp. enterica serovar Bredeney Proteobacteria; Gammaproteobacteria; Enterobacterales; Enterobacteriaceae; Salmonella; Salmonella enterica; Salmonella enterica subsp. enterica MTGDNTLITSHGINRRDFMKLCAALAATMGLSSKAAAEMAESVSNPQRPPVIWIGAQECT WP_000145429.1 hydrogenase 2 small subunit [Salmonella enterica] Length: 372\nNumber of predicted TMHs: 1\nExp number of AAs in TMHs: 24.51309\nExp number, first 60 AAs: 0.2682\nTotal prob of N-in: 0.14402\noutside 1 331\nTMhelix 332 354\ninside 355 372

16312 GCF_002048415.1_ASM204841v1 Salmonella enterica subsp. enterica serovar Bredeney Proteobacteria; Gammaproteobacteria; Enterobacterales; Enterobacteriaceae; Salmonella; Salmonella enterica; Salmonella enterica subsp. enterica MTGDNTLITSHGINRRDFMKLCAALAATMGLSSKAAAEMAESVSNPQRPPVIWIGAQECT WP_000145429.1 hydrogenase 2 small subunit [Salmonella enterica] Length: 372\nNumber of predicted TMHs: 1\nExp number of AAs in TMHs: 24.51309\nExp number, first 60 AAs: 0.2682\nTotal prob of N-in: 0.14402\noutside 1 331\nTMhelix 332 354\ninside 355 372

16313 GCF_002063755.1_ASM206375v1 Salmonella enterica subsp. enterica serovar Bredeney Proteobacteria; Gammaproteobacteria; Enterobacterales; Enterobacteriaceae; Salmonella; Salmonella enterica; Salmonella enterica subsp. enterica MNNEETFYQAMRRKGVTRRSFLKFCSLAATSLGLGAGMTPKIAWALENKPRIPVVWIHGL WP_080153733.1 [Ni/Fe] hydrogenase small subunit [Salmonella enterica] Length: 372\nNumber of predicted TMHs: 2\nExp number of AAs in TMHs: 36.51949\nExp number, first 60 AAs: 19.72829\nTotal prob of N-in: 0.96170\nPOSSIBLE N-term signal sequence\ninside 1 19\nTMhelix 20 39\noutside 40 325\nTMhelix 326 348\ninside 349 372

16314 GCF_002047285.1_ASM204728v1 Salmonella enterica subsp. enterica serovar Bredeney Proteobacteria; Gammaproteobacteria; Enterobacterales; Enterobacteriaceae; Salmonella; Salmonella enterica; Salmonella enterica subsp. enterica MTGDNTLITSHGINRRDFMKLCAALAATMGLSSKAAAEMAESVSNPQRPPVIWIGAQECT WP_000145429.1 hydrogenase 2 small subunit [Salmonella enterica] Length: 372\nNumber of predicted TMHs: 1\nExp number of AAs in TMHs: 24.51309\nExp number, first 60 AAs: 0.2682\nTotal prob of N-in: 0.14402\noutside 1 331\nTMhelix 332 354\ninside 355 372

16315 GCF_002047265.1_ASM204726v1 Salmonella enterica subsp. enterica serovar Bredeney Proteobacteria; Gammaproteobacteria; Enterobacterales; Enterobacteriaceae; Salmonella; Salmonella enterica; Salmonella enterica subsp. enterica MTGDNTLITSHGINRRDFMKLCAALAATMGLSSKAAAEMAESVSNPQRPPVIWIGAQECT WP_000145429.1 hydrogenase 2 small subunit [Salmonella enterica] Length: 372\nNumber of predicted TMHs: 1\nExp number of AAs in TMHs: 24.51309\nExp number, first 60 AAs: 0.2682\nTotal prob of N-in: 0.14402\noutside 1 331\nTMhelix 332 354\ninside 355 372

16316 GCF_002048415.1_ASM204841v1 Salmonella enterica subsp. enterica serovar Bredeney Proteobacteria; Gammaproteobacteria; Enterobacterales; Enterobacteriaceae; Salmonella; Salmonella enterica; Salmonella enterica subsp. enterica MNNEETFYQAMRRKGVTRRSFLKFCSLAATSLGLGAGMTPKIAWALENKPRIPVVWIHGL WP_001058312.1 hydrogenase [Salmonella enterica] Length: 372\nNumber of predicted TMHs: 2\nExp number of AAs in TMHs: 36.51738\nExp number, first 60 AAs: 19.72849\nTotal prob of N-in: 0.96170\nPOSSIBLE N-term signal sequence\ninside 1 19\nTMhelix 20 39\noutside 40 325\nTMhelix 326 348\ninside 349 372

16317 GCF_002063755.1_ASM206375v1 Salmonella enterica subsp. enterica serovar Bredeney Proteobacteria; Gammaproteobacteria; Enterobacterales; Enterobacteriaceae; Salmonella; Salmonella enterica; Salmonella enterica subsp. enterica MTGDNTLITSHGINRRDFMKLCAALAATMGLSSKAAAEMAESVSNPQRPPVIWIGAQECT WP_000145429.1 hydrogenase 2 small subunit [Salmonella enterica] Length: 372\nNumber of

predicted TMHs:  1\nExp number of AAs in TMHs: 24.51309\nExp number, first 60 AAs:  0.2682\nTotal prob of N-in:  0.14402\noutside   1   331\nTMhelix   332   354\ninside   355   372

16318 GCF_002047285.1_ASM204728v1 Salmonella enterica subsp. enterica serovar Bredeney Proteobacteria; Gammaproteobacteria; Enterobacterales; Enterobacteriaceae; Salmonella; Salmonella enterica; Salmonella enterica subsp. enterica MNNEETFYQAMRRKGVTRRSFLKFCSLAATSLGLGAGMTPKIAWALENKPRIPVVWIHGL WP_001058312.1 hydrogenase [Salmonella enterica] Length: 372\nNumber of predicted TMHs:  2\nExp number of AAs in TMHs: 36.51738\nExp number, first 60 AAs:  19.72849\nTotal prob of N-in:   0.96170\nPOSSIBLE N-term signal sequence\ninside   1   19\nTMhelix   20   39\noutside   40   325\nTMhelix   326   348\ninside   349   372

16319 GCF_001271405.1_Salmonella_enterica_CVM_N43475_v1.0 Salmonella enterica subsp. enterica serovar Bredeney Proteobacteria; Gammaproteobacteria; Enterobacterales; Enterobacteriaceae; Salmonella; Salmonella enterica; Salmonella enterica subsp. enterica MTGDNTLITSHGINRRDFMKLCAALAATMGLSSKAAAEMAESVSNPQRPPVIWIGAQECT WP_000145429.1 hydrogenase 2 small subunit [Salmonella enterica] Length: 372\nNumber of predicted TMHs:  1\nExp number of AAs in TMHs: 24.51309\nExp number, first 60 AAs:  0.2682\nTotal prob of N-in:  0.14402\noutside   1   331\nTMhelix   332   354\ninside   355   372

16320 GCF_002071655.1_ASM207165v1 Salmonella enterica subsp. enterica serovar Bredeny Proteobacteria; Gammaproteobacteria; Enterobacterales; Enterobacteriaceae; Salmonella; Salmonella enterica; Salmonella enterica subsp. enterica MTGDNTLITSHGINRRDFMKLCAALAATMGLSSKAAAEMAESVSNPQRPPVIWIGAQECT WP_000145429.1 hydrogenase 2 small subunit [Salmonella enterica] Length: 372\nNumber of predicted TMHs:  1\nExp number of AAs in TMHs: 24.51309\nExp number, first 60 AAs:  0.2682\nTotal prob of N-in:  0.14402\noutside   1   331\nTMhelix   332   354\ninside   355   372

16321 GCF_002062945.1_ASM206294v1 Salmonella enterica subsp. enterica serovar Bredeny Proteobacteria; Gammaproteobacteria; Enterobacterales; Enterobacteriaceae; Salmonella; Salmonella enterica; Salmonella enterica subsp. enterica MTGDNTLITSHGINRRDFMKLCAALAATMGLSSKAAAEMAESVSNPQRPPVIWIGAQECT WP_000145429.1 hydrogenase 2 small subunit [Salmonella enterica] Length: 372\nNumber of predicted TMHs:  1\nExp number of AAs in TMHs: 24.51309\nExp number, first 60 AAs:  0.2682\nTotal prob of N-in:  0.14402\noutside   1   331\nTMhelix   332   354\ninside   355   372

16322 GCF_002071655.1_ASM207165v1 Salmonella enterica subsp. enterica serovar Bredeny Proteobacteria; Gammaproteobacteria; Enterobacterales; Enterobacteriaceae; Salmonella; Salmonella enterica; Salmonella enterica subsp. enterica MNNEETFYQAMRRKGVTRRSFLKFCSLAATSLGLGAGMTPKIAWALENKPRIPVVWIHGL WP_001058312.1 hydrogenase [Salmonella enterica] Length: 372\nNumber of predicted TMHs:  2\nExp number of AAs in TMHs: 36.51738\nExp number, first 60 AAs:  19.72849\nTotal prob of N-in:   0.96170\nPOSSIBLE N-term signal sequence\ninside   1   19\nTMhelix   20   39\noutside   40   325\nTMhelix   326   348\ninside   349   372

16323 GCF_002062945.1_ASM206294v1 Salmonella enterica subsp. enterica serovar Bredeny Proteobacteria; Gammaproteobacteria; Enterobacterales; Enterobacteriaceae; Salmonella; Salmonella enterica; Salmonella enterica subsp. enterica MNNEETFYQAMRRKGVTRRSFLKFCSLAATSLGLGAGMTPKIAWALENKPRIPVVWIHGL WP_001058312.1 hydrogenase [Salmonella enterica] Length: 372\nNumber of predicted TMHs:  2\nExp number of AAs in TMHs: 36.51738\nExp number, first 60 AAs:  19.72849\nTotal prob of N-in:   0.96170\nPOSSIBLE N-term signal sequence\ninside   1   19\nTMhelix   20   39\noutside   40   325\nTMhelix   326   348\ninside   349   372

16324 GCF_002034805.1_ASM203480v1 Salmonella enterica subsp. enterica serovar Brisbane Proteobacteria; Gammaproteobacteria; Enterobacterales; Enterobacteriaceae; Salmonella; Salmonella enterica; Salmonella enterica subsp. enterica MNNEETFYQAMRRKGVTRRSFLKFCSLAATSLGLGAGMTPKIAWALENKPRIPVVWIHGL WP_001058312.1 hydrogenase [Salmonella enterica] Length: 372\nNumber of predicted TMHs:  2\nExp number of AAs in TMHs: 36.51738\nExp number, first 60 AAs:  19.72849\nTotal prob of N-in:   0.96170\nPOSSIBLE N-term signal sequence\ninside   1   19\nTMhelix   20   39\noutside   40   325\nTMhelix   326   348\ninside   349   372

16325 GCF_002034805.1_ASM203480v1 Salmonella enterica subsp. enterica serovar Brisbane Proteobacteria; Gammaproteobacteria; Enterobacterales; Enterobacteriaceae; Salmonella; Salmonella enterica; Salmonella enterica subsp. enterica MTGDNTLITSHGINRRDFMKLCAALAATMGLSSKAAAEMAESVSNPQRPPVIWIGAQECT WP_000145431.1 hydrogenase 2 small subunit [Salmonella enterica] Length: 372\nNumber of predicted TMHs:  1\nExp number of AAs in TMHs: 24.51505\nExp number, first 60 AAs:  0.2682\nTotal prob of N-in:  0.14406\noutside   1   331\nTMhelix   332   354\ninside   355   372

16326 GCF_002156235.1_ASM215623v1 Salmonella enterica subsp. enterica serovar Bristol Proteobacteria; Gammaproteobacteria; Enterobacterales; Enterobacteriaceae; Salmonella; Salmonella enterica;

Salmonella enterica subsp. enterica MNNEETFYQAMRRKGVTRRSFLKFCSLAATSLGLGAGMTPKIAWALENKPRIPVVWIHGL
WP_001058312.1 hydrogenase [Salmonella enterica] Length: 372\nNumber of predicted TMHs: 2\nExp number of AAs in TMHs: 36.51738\nExp number, first 60 AAs: 19.72849\nTotal prob of N-in: 0.96170\nPOSSIBLE N-term signal sequence\ninside 1 19\nTMhelix 20 39\noutside 40 325\nTMhelix 326 348\ninside 349 372

16327 GCF_002156235.1_ASM215623v1 Salmonella enterica subsp. enterica serovar Bristol
Proteobacteria; Gammaproteobacteria; Enterobacterales; Enterobacteriaceae; Salmonella; Salmonella enterica; Salmonella enterica subsp. enterica MTGDNTLITSHGINRRDFMKLCAALAATMGLSSKAAAEMAESVSNPQRPPVIWIGAQECT
WP_000145429.1 hydrogenase 2 small subunit [Salmonella enterica] Length: 372\nNumber of predicted TMHs: 1\nExp number of AAs in TMHs: 24.51309\nExp number, first 60 AAs: 0.2682\nTotal prob of N-in: 0.14402\noutside 1 331\nTMhelix 332 354\ninside 355 372

16328 GCF_002106295.1_ASM210629v1 Salmonella enterica subsp. enterica serovar Cerro
Proteobacteria; Gammaproteobacteria; Enterobacterales; Enterobacteriaceae; Salmonella; Salmonella enterica; Salmonella enterica subsp. enterica MNNEETFYQAMRRKGVTRRSFLKFCSLAATSLGLGAGMTPKIAWALENKPRIPVVWIHGL
WP_042847606.1 hydrogenase [Salmonella enterica] Length: 372\nNumber of predicted TMHs: 2\nExp number of AAs in TMHs: 37.4207199999999\nExp number, first 60 AAs: 19.72732\nTotal prob of N-in: 0.96168\nPOSSIBLE N-term signal sequence\ninside 1 19\nTMhelix 20 39\noutside 40 325\nTMhelix 326 348\ninside 349 372

16329 GCF_002107175.1_ASM210717v1 Salmonella enterica subsp. enterica serovar Cerro
Proteobacteria; Gammaproteobacteria; Enterobacterales; Enterobacteriaceae; Salmonella; Salmonella enterica; Salmonella enterica subsp. enterica MNNEETFYQAMRRKGVTRRSFLKFCSLAATSLGLGAGMTPKIAWALENKPRIPVVWIHGL
WP_042847606.1 hydrogenase [Salmonella enterica] Length: 372\nNumber of predicted TMHs: 2\nExp number of AAs in TMHs: 37.4207199999999\nExp number, first 60 AAs: 19.72732\nTotal prob of N-in: 0.96168\nPOSSIBLE N-term signal sequence\ninside 1 19\nTMhelix 20 39\noutside 40 325\nTMhelix 326 348\ninside 349 372

16330 GCF_002105795.1_ASM210579v1 Salmonella enterica subsp. enterica serovar Cerro
Proteobacteria; Gammaproteobacteria; Enterobacterales; Enterobacteriaceae; Salmonella; Salmonella enterica; Salmonella enterica subsp. enterica MTGDNTLITSHGINRRDFMKLCAALAATMGLSSKAAAEMAESVSNPQRPPVIWIGAQECT
WP_000145429.1 hydrogenase 2 small subunit [Salmonella enterica] Length: 372\nNumber of predicted TMHs: 1\nExp number of AAs in TMHs: 24.51309\nExp number, first 60 AAs: 0.2682\nTotal prob of N-in: 0.14402\noutside 1 331\nTMhelix 332 354\ninside 355 372

16331 GCF_002107455.1_ASM210745v1 Salmonella enterica subsp. enterica serovar Cerro
Proteobacteria; Gammaproteobacteria; Enterobacterales; Enterobacteriaceae; Salmonella; Salmonella enterica; Salmonella enterica subsp. enterica MNNEETFYQAMRRKGVTRRSFLKFCSLAATSLGLGAGMTPKIAWALENKPRIPVVWIHGL
WP_042847606.1 hydrogenase [Salmonella enterica] Length: 372\nNumber of predicted TMHs: 2\nExp number of AAs in TMHs: 37.4207199999999\nExp number, first 60 AAs: 19.72732\nTotal prob of N-in: 0.96168\nPOSSIBLE N-term signal sequence\ninside 1 19\nTMhelix 20 39\noutside 40 325\nTMhelix 326 348\ninside 349 372

16332 GCF_002107175.1_ASM210717v1 Salmonella enterica subsp. enterica serovar Cerro
Proteobacteria; Gammaproteobacteria; Enterobacterales; Enterobacteriaceae; Salmonella; Salmonella enterica; Salmonella enterica subsp. enterica MTGDNTLITSHGINRRDFMKLCAALAATMGLSSKAAAEMAESVSNPQRPPVIWIGAQECT
WP_000145429.1 hydrogenase 2 small subunit [Salmonella enterica] Length: 372\nNumber of predicted TMHs: 1\nExp number of AAs in TMHs: 24.51309\nExp number, first 60 AAs: 0.2682\nTotal prob of N-in: 0.14402\noutside 1 331\nTMhelix 332 354\ninside 355 372

16333 GCF_002106055.1_ASM210605v1 Salmonella enterica subsp. enterica serovar Cerro
Proteobacteria; Gammaproteobacteria; Enterobacterales; Enterobacteriaceae; Salmonella; Salmonella enterica; Salmonella enterica subsp. enterica MTGDNTLITSHGINRRDFMKLCAALAATMGLSSKAAAEMAESVSNPQRPPVIWIGAQECT
WP_000145429.1 hydrogenase 2 small subunit [Salmonella enterica] Length: 372\nNumber of predicted TMHs: 1\nExp number of AAs in TMHs: 24.51309\nExp number, first 60 AAs: 0.2682\nTotal prob of N-in: 0.14402\noutside 1 331\nTMhelix 332 354\ninside 355 372

16334 GCF_002105825.1_ASM210582v1 Salmonella enterica subsp. enterica serovar Cerro
Proteobacteria; Gammaproteobacteria; Enterobacterales; Enterobacteriaceae; Salmonella; Salmonella enterica; Salmonella enterica subsp. enterica MTGDNTLITSHGINRRDFMKLCAALAATMGLSSKAAAEMAESVSNPQRPPVIWIGAQECT
WP_000145429.1 hydrogenase 2 small subunit [Salmonella enterica] Length: 372\nNumber of

predicted TMHs: 1\nExp number of AAs in TMHs: 24.51309\nExp number, first 60 AAs: 0.2682\nTotal prob of N-in: 0.14402\noutside 1 331\nTMhelix 332 354\ninside 355 372

16335 GCF_002105795.1_ASM210579v1 Salmonella enterica subsp. enterica serovar Cerro Proteobacteria; Gammaproteobacteria; Enterobacterales; Enterobacteriaceae; Salmonella; Salmonella enterica; Salmonella enterica subsp. enterica MNNEETFYQAMRRKGVTRRSFLKFCSLAATSLGLGAGMTPKIAWALENKPRIPVVWIHGL WP_042847606.1 hydrogenase [Salmonella enterica] Length: 372\nNumber of predicted TMHs: 2\nExp number of AAs in TMHs: 37.4207199999999\nExp number, first 60 AAs: 19.72732\nTotal prob of N-in: 0.96168\nPOSSIBLE N-term signal sequence\ninside 1 19\nTMhelix 20 39\noutside 40 325\nTMhelix 326 348\ninside 349 372

16336 GCF_002105825.1_ASM210582v1 Salmonella enterica subsp. enterica serovar Cerro Proteobacteria; Gammaproteobacteria; Enterobacterales; Enterobacteriaceae; Salmonella; Salmonella enterica; Salmonella enterica subsp. enterica MNNEETFYQAMRRKGVTRRSFLKFCSLAATSLGLGAGMTPKIAWALENKPRIPVVWIHGL WP_001058312.1 hydrogenase [Salmonella enterica] Length: 372\nNumber of predicted TMHs: 2\nExp number of AAs in TMHs: 36.51738\nExp number, first 60 AAs: 19.72849\nTotal prob of N-in: 0.96170\nPOSSIBLE N-term signal sequence\ninside 1 19\nTMhelix 20 39\noutside 40 325\nTMhelix 326 348\ninside 349 372

16337 GCF_002106055.1_ASM210605v1 Salmonella enterica subsp. enterica serovar Cerro Proteobacteria; Gammaproteobacteria; Enterobacterales; Enterobacteriaceae; Salmonella; Salmonella enterica; Salmonella enterica subsp. enterica MNNEETFYQAMRRKGVTRRSFLKFCSLAATSLGLGAGMTPKIAWALENKPRIPVVWIHGL WP_042847606.1 hydrogenase [Salmonella enterica] Length: 372\nNumber of predicted TMHs: 2\nExp number of AAs in TMHs: 37.4207199999999\nExp number, first 60 AAs: 19.72732\nTotal prob of N-in: 0.96168\nPOSSIBLE N-term signal sequence\ninside 1 19\nTMhelix 20 39\noutside 40 325\nTMhelix 326 348\ninside 349 372

16338 GCF_002106295.1_ASM210629v1 Salmonella enterica subsp. enterica serovar Cerro Proteobacteria; Gammaproteobacteria; Enterobacterales; Enterobacteriaceae; Salmonella; Salmonella enterica; Salmonella enterica subsp. enterica MTGDNTLITSHGINRRDFMKLCAALAATMGLSSKAAAEMAESVSNPQRPPVIWIGAQECT WP_000145429.1 hydrogenase 2 small subunit [Salmonella enterica] Length: 372\nNumber of predicted TMHs: 1\nExp number of AAs in TMHs: 24.51309\nExp number, first 60 AAs: 0.2682\nTotal prob of N-in: 0.14402\noutside 1 331\nTMhelix 332 354\ninside 355 372

16339 GCF_002063315.1_ASM206331v1 Salmonella enterica subsp. enterica serovar Cerro Proteobacteria; Gammaproteobacteria; Enterobacterales; Enterobacteriaceae; Salmonella; Salmonella enterica; Salmonella enterica subsp. enterica MNNEETFYQAMRRKGVTRRSFLKFCSLAATSLGLGAGMTPKIAWALENKPRIPVVWIHGL WP_023233644.1 hydrogenase 1, small subunit [Salmonella enterica] Length: 372\nNumber of predicted TMHs: 2\nExp number of AAs in TMHs: 37.4208299999999\nExp number, first 60 AAs: 19.72732\nTotal prob of N-in: 0.96168\nPOSSIBLE N-term signal sequence\ninside 1 19\nTMhelix 20 39\noutside 40 325\nTMhelix 326 348\ninside 349 372

16340 GCF_002106095.1_ASM210609v1 Salmonella enterica subsp. enterica serovar Cerro Proteobacteria; Gammaproteobacteria; Enterobacterales; Enterobacteriaceae; Salmonella; Salmonella enterica; Salmonella enterica subsp. enterica MTGDNTLITSHGINRRDFMKLCAALAATMGLSSKAAAEMAESVSNPQRPPVIWIGAQECT WP_000145429.1 hydrogenase 2 small subunit [Salmonella enterica] Length: 372\nNumber of predicted TMHs: 1\nExp number of AAs in TMHs: 24.51309\nExp number, first 60 AAs: 0.2682\nTotal prob of N-in: 0.14402\noutside 1 331\nTMhelix 332 354\ninside 355 372

16341 GCF_002063315.1_ASM206331v1 Salmonella enterica subsp. enterica serovar Cerro Proteobacteria; Gammaproteobacteria; Enterobacterales; Enterobacteriaceae; Salmonella; Salmonella enterica; Salmonella enterica subsp. enterica MTGDNTLITSHGINRRDFMKLCAALAATMGLSSKAAAEMAESVSNPQRPPVIWIGAQECT WP_000145429.1 hydrogenase 2 small subunit [Salmonella enterica] Length: 372\nNumber of predicted TMHs: 1\nExp number of AAs in TMHs: 24.51309\nExp number, first 60 AAs: 0.2682\nTotal prob of N-in: 0.14402\noutside 1 331\nTMhelix 332 354\ninside 355 372

16342 GCF_002106095.1_ASM210609v1 Salmonella enterica subsp. enterica serovar Cerro Proteobacteria; Gammaproteobacteria; Enterobacterales; Enterobacteriaceae; Salmonella; Salmonella enterica; Salmonella enterica subsp. enterica MNNEETFYQAMRRKGVTRRSFLKFCSLAATSLGLGAGMTPKIAWALENKPRIPVVWIHGL WP_042847606.1 hydrogenase [Salmonella enterica] Length: 372\nNumber of predicted TMHs: 2\nExp number of AAs in TMHs: 37.4207199999999\nExp number, first 60 AAs: 19.72732\nTotal prob of N-in:

0.96168\nPOSSIBLE N-term signal sequence\ninside 1 19\nTMhelix 20 39\noutside 40 325\nTMhelix 326 348\ninside 349 372

16343 GCF_002107455.1_ASM210745v1 Salmonella enterica subsp. enterica serovar Cerro Proteobacteria; Gammaproteobacteria; Enterobacterales; Enterobacteriaceae; Salmonella; Salmonella enterica; Salmonella enterica subsp. enterica MTGDNTLITSHGINRRDFMKLCAALAATMGLSSKAAAEMAESVSNPQRPPVIWIGAQECT WP_000145429.1 hydrogenase 2 small subunit [Salmonella enterica] Length: 372\nNumber of predicted TMHs: 1\nExp number of AAs in TMHs: 24.51309\nExp number, first 60 AAs: 0.2682\nTotal prob of N-in: 0.14402\noutside 1 331\nTMhelix 332 354\ninside 355 372

16344 GCF_001043095.1_ASM104309v1 Salmonella enterica subsp. enterica serovar Cerro Proteobacteria; Gammaproteobacteria; Enterobacterales; Enterobacteriaceae; Salmonella; Salmonella enterica; Salmonella enterica subsp. enterica MTGDNTLITSHGINRRDFMKLCAALAATMGLSSKAAAEMAESVSNPQRPPVIWIGAQECT WP_000145429.1 hydrogenase 2 small subunit [Salmonella enterica] Length: 372\nNumber of predicted TMHs: 1\nExp number of AAs in TMHs: 24.51309\nExp number, first 60 AAs: 0.2682\nTotal prob of N-in: 0.14402\noutside 1 331\nTMhelix 332 354\ninside 355 372

16345 GCF_001941405.1_ASM194140v1 Salmonella enterica subsp. enterica serovar Cerro Proteobacteria; Gammaproteobacteria; Enterobacterales; Enterobacteriaceae; Salmonella; Salmonella enterica; Salmonella enterica subsp. enterica MTGDNTLITSHGINRRDFMKLCAALAATMGLSSKAAAEMAESVSNPQRPPVIWIGAQECT WP_000145429.1 hydrogenase 2 small subunit [Salmonella enterica] Length: 372\nNumber of predicted TMHs: 1\nExp number of AAs in TMHs: 24.51309\nExp number, first 60 AAs: 0.2682\nTotal prob of N-in: 0.14402\noutside 1 331\nTMhelix 332 354\ninside 355 372

16346 GCF_002047115.1_ASM204711v1 Salmonella enterica subsp. enterica serovar Cerro Proteobacteria; Gammaproteobacteria; Enterobacterales; Enterobacteriaceae; Salmonella; Salmonella enterica; Salmonella enterica subsp. enterica MTGDNTLITSHGINRRDFMKLCAALAATMGLSSKAAAEMAESVSNPQRPPVIWIGAQECT WP_000145429.1 hydrogenase 2 small subunit [Salmonella enterica] Length: 372\nNumber of predicted TMHs: 1\nExp number of AAs in TMHs: 24.51309\nExp number, first 60 AAs: 0.2682\nTotal prob of N-in: 0.14402\noutside 1 331\nTMhelix 332 354\ninside 355 372

16347 GCF_002047115.1_ASM204711v1 Salmonella enterica subsp. enterica serovar Cerro Proteobacteria; Gammaproteobacteria; Enterobacterales; Enterobacteriaceae; Salmonella; Salmonella enterica; Salmonella enterica subsp. enterica MNNEETFYQAMRRKGVTRRSFLKFCSLAATSLGLGAGMTPKIAWALENKPRIPVVWIHGL WP_023233644.1 hydrogenase 1, small subunit [Salmonella enterica] Length: 372\nNumber of predicted TMHs: 2\nExp number of AAs in TMHs: 37.4208299999999\nExp number, first 60 AAs: 19.72732\nTotal prob of N-in: 0.96168\nPOSSIBLE N-term signal sequence\ninside 1 19\nTMhelix 20 39\noutside 40 325\nTMhelix 326 348\ninside 349 372

16348 GCF_002034285.1_ASM203428v1 Salmonella enterica subsp. enterica serovar Cerro Proteobacteria; Gammaproteobacteria; Enterobacterales; Enterobacteriaceae; Salmonella; Salmonella enterica; Salmonella enterica subsp. enterica MTGDNTLITSHGINRRDFMKLCAALAATMGLSSKAAAEMAESVSNPQRPPVIWIGAQECT WP_000145429.1 hydrogenase 2 small subunit [Salmonella enterica] Length: 372\nNumber of predicted TMHs: 1\nExp number of AAs in TMHs: 24.51309\nExp number, first 60 AAs: 0.2682\nTotal prob of N-in: 0.14402\noutside 1 331\nTMhelix 332 354\ninside 355 372

16349 GCF_001941405.1_ASM194140v1 Salmonella enterica subsp. enterica serovar Cerro Proteobacteria; Gammaproteobacteria; Enterobacterales; Enterobacteriaceae; Salmonella; Salmonella enterica; Salmonella enterica subsp. enterica MNNEETFYQAMRRKGVTRRSFLKFCSLAATSLGLGAGMTPKIAWALENKPRIPVVWIHGL WP_023233644.1 hydrogenase 1, small subunit [Salmonella enterica] Length: 372\nNumber of predicted TMHs: 2\nExp number of AAs in TMHs: 37.4208299999999\nExp number, first 60 AAs: 19.72732\nTotal prob of N-in: 0.96168\nPOSSIBLE N-term signal sequence\ninside 1 19\nTMhelix 20 39\noutside 40 325\nTMhelix 326 348\ninside 349 372

16350 GCF_001043095.1_ASM104309v1 Salmonella enterica subsp. enterica serovar Cerro Proteobacteria; Gammaproteobacteria; Enterobacterales; Enterobacteriaceae; Salmonella; Salmonella enterica; Salmonella enterica subsp. enterica MNNEETFYQAMRRKGVTRRSFLKFCSLAATSLGLGAGMTPKIAWALENKPRIPVVWIHGL WP_023233644.1 hydrogenase 1, small subunit [Salmonella enterica] Length: 372\nNumber of predicted TMHs: 2\nExp number of AAs in TMHs: 37.4208299999999\nExp number, first 60 AAs: 19.72732\nTotal prob of N-in: 0.96168\nPOSSIBLE N-term signal sequence\ninside 1 19\nTMhelix 20 39\noutside 40 325\nTMhelix 326 348\ninside 349 372

16351 GCF_002047325.1_ASM204732v1 Salmonella enterica subsp. enterica serovar Cerro Proteobacteria; Gammaproteobacteria; Enterobacterales; Enterobacteriaceae; Salmonella; Salmonella enterica; Salmonella enterica subsp. enterica MNNEETFYQAMRRKGVTRRSFLKFCSLAATSLGLGAGMTPKIAWALENKPRIPVVWIHGL WP_023233644.1 hydrogenase 1, small subunit [Salmonella enterica] Length: 372\nNumber of predicted TMHs: 2\nExp number of AAs in TMHs: 37.4208299999999\nExp number, first 60 AAs: 19.72732\nTotal prob of N-in: 0.96168\nPOSSIBLE N-term signal sequence\ninside 1 19\nTMhelix 20 39\noutside 40 325\nTMhelix 326 348\ninside 349 372

16352 GCF_002106035.1_ASM210603v1 Salmonella enterica subsp. enterica serovar Cerro Proteobacteria; Gammaproteobacteria; Enterobacterales; Enterobacteriaceae; Salmonella; Salmonella enterica; Salmonella enterica subsp. enterica MNNEETFYQAMRRKGVTRRSFLKFCSLAATSLGLGAGMTPKIAWALENKPRIPVVWIHGL WP_042847606.1 hydrogenase [Salmonella enterica] Length: 372\nNumber of predicted TMHs: 2\nExp number of AAs in TMHs: 37.4207199999999\nExp number, first 60 AAs: 19.72732\nTotal prob of N-in: 0.96168\nPOSSIBLE N-term signal sequence\ninside 1 19\nTMhelix 20 39\noutside 40 325\nTMhelix 326 348\ninside 349 372

16353 GCF_002106765.1_ASM210676v1 Salmonella enterica subsp. enterica serovar Cerro Proteobacteria; Gammaproteobacteria; Enterobacterales; Enterobacteriaceae; Salmonella; Salmonella enterica; Salmonella enterica subsp. enterica MNNEETFYQAMRRKGVTRRSFLKFCSLAATSLGLGAGMTPKIAWALENKPRIPVVWIHGL WP_042847606.1 hydrogenase [Salmonella enterica] Length: 372\nNumber of predicted TMHs: 2\nExp number of AAs in TMHs: 37.4207199999999\nExp number, first 60 AAs: 19.72732\nTotal prob of N-in: 0.96168\nPOSSIBLE N-term signal sequence\ninside 1 19\nTMhelix 20 39\noutside 40 325\nTMhelix 326 348\ninside 349 372

16354 GCF_002106325.1_ASM210632v1 Salmonella enterica subsp. enterica serovar Cerro Proteobacteria; Gammaproteobacteria; Enterobacterales; Enterobacteriaceae; Salmonella; Salmonella enterica; Salmonella enterica subsp. enterica MTGDNTLITSHGINRRDFMKLCAALAATMGLSSKAAAEMAESVSNPQRPPVIWIGAQECT WP_000145429.1 hydrogenase 2 small subunit [Salmonella enterica] Length: 372\nNumber of predicted TMHs: 1\nExp number of AAs in TMHs: 24.51309\nExp number, first 60 AAs: 0.2682\nTotal prob of N-in: 0.14402\noutside 1 331\nTMhelix 332 354\ninside 355 372

16355 GCF_002105875.1_ASM210587v1 Salmonella enterica subsp. enterica serovar Cerro Proteobacteria; Gammaproteobacteria; Enterobacterales; Enterobacteriaceae; Salmonella; Salmonella enterica; Salmonella enterica subsp. enterica MNNEETFYQAMRRKGVTRRSFLKFCSLAATSLGLGAGMTPKIAWALENKPRIPVVWIHGL WP_042847606.1 hydrogenase [Salmonella enterica] Length: 372\nNumber of predicted TMHs: 2\nExp number of AAs in TMHs: 37.4207199999999\nExp number, first 60 AAs: 19.72732\nTotal prob of N-in: 0.96168\nPOSSIBLE N-term signal sequence\ninside 1 19\nTMhelix 20 39\noutside 40 325\nTMhelix 326 348\ninside 349 372

16356 GCF_002106035.1_ASM210603v1 Salmonella enterica subsp. enterica serovar Cerro Proteobacteria; Gammaproteobacteria; Enterobacterales; Enterobacteriaceae; Salmonella; Salmonella enterica; Salmonella enterica subsp. enterica MTGDNTLITSHGINRRDFMKLCAALAATMGLSSKAAAEMAESVSNPQRPPVIWIGAQECT WP_000145429.1 hydrogenase 2 small subunit [Salmonella enterica] Length: 372\nNumber of predicted TMHs: 1\nExp number of AAs in TMHs: 24.51309\nExp number, first 60 AAs: 0.2682\nTotal prob of N-in: 0.14402\noutside 1 331\nTMhelix 332 354\ninside 355 372

16357 GCF_002066035.1_ASM206603v1 Salmonella enterica subsp. enterica serovar Cerro Proteobacteria; Gammaproteobacteria; Enterobacterales; Enterobacteriaceae; Salmonella; Salmonella enterica; Salmonella enterica subsp. enterica MNNEETFYQAMRRKGVTRRSFLKFCSLAATSLGLGAGMTPKIAWALENKPRIPVVWIHGL WP_023233644.1 hydrogenase 1, small subunit [Salmonella enterica] Length: 372\nNumber of predicted TMHs: 2\nExp number of AAs in TMHs: 37.4208299999999\nExp number, first 60 AAs: 19.72732\nTotal prob of N-in: 0.96168\nPOSSIBLE N-term signal sequence\ninside 1 19\nTMhelix 20 39\noutside 40 325\nTMhelix 326 348\ninside 349 372

16358 GCF_002066035.1_ASM206603v1 Salmonella enterica subsp. enterica serovar Cerro Proteobacteria; Gammaproteobacteria; Enterobacterales; Enterobacteriaceae; Salmonella; Salmonella enterica; Salmonella enterica subsp. enterica MTGDNTLITSHGINRRDFMKLCAALAATMGLSSKAAAEMAESVSNPQRPPVIWIGAQECT WP_000145429.1 hydrogenase 2 small subunit [Salmonella enterica] Length: 372\nNumber of predicted TMHs: 1\nExp number of AAs in TMHs: 24.51309\nExp number, first 60 AAs: 0.2682\nTotal prob of N-in: 0.14402\noutside 1 331\nTMhelix 332 354\ninside 355 372

16359 GCF_002106325.1_ASM210632v1 Salmonella enterica subsp. enterica serovar Cerro Proteobacteria; Gammaproteobacteria; Enterobacterales; Enterobacteriaceae; Salmonella; Salmonella enterica; Salmonella enterica subsp. enterica MNNEETFYQAMRRKGVTRRSFLKFCSLAATSLGLGAGMTPKIAWALENKPRIPVVWIHGL WP_042847606.1 hydrogenase [Salmonella enterica] Length: 372\nNumber of predicted TMHs: 2\nExp number of AAs in TMHs: 37.4207199999999\nExp number, first 60 AAs: 19.72732\nTotal prob of N-in: 0.96168\nPOSSIBLE N-term signal sequence\ninside 1 19\nTMhelix 20 39\noutside 40 325\nTMhelix 326 348\ninside 349 372

16360 GCF_002105875.1_ASM210587v1 Salmonella enterica subsp. enterica serovar Cerro Proteobacteria; Gammaproteobacteria; Enterobacterales; Enterobacteriaceae; Salmonella; Salmonella enterica; Salmonella enterica subsp. enterica MTGDNTLITSHGINRRDFMKLCAALAATMGLSSKAAAEMAESVSNPQRPPVIWIGAQECT WP_000145429.1 hydrogenase 2 small subunit [Salmonella enterica] Length: 372\nNumber of predicted TMHs: 1\nExp number of AAs in TMHs: 24.51309\nExp number, first 60 AAs: 0.2682\nTotal prob of N-in: 0.14402\noutside 1 331\nTMhelix 332 354\ninside 355 372

16361 GCF_002106765.1_ASM210676v1 Salmonella enterica subsp. enterica serovar Cerro Proteobacteria; Gammaproteobacteria; Enterobacterales; Enterobacteriaceae; Salmonella; Salmonella enterica; Salmonella enterica subsp. enterica MTGDNTLITSHGINRRDFMKLCAALAATMGLSSKAAAEMAESVSNPQRPPVIWIGAQECT WP_000145429.1 hydrogenase 2 small subunit [Salmonella enterica] Length: 372\nNumber of predicted TMHs: 1\nExp number of AAs in TMHs: 24.51309\nExp number, first 60 AAs: 0.2682\nTotal prob of N-in: 0.14402\noutside 1 331\nTMhelix 332 354\ninside 355 372

16362 GCF_002066015.1_ASM206601v1 Salmonella enterica subsp. enterica serovar Cerro Proteobacteria; Gammaproteobacteria; Enterobacterales; Enterobacteriaceae; Salmonella; Salmonella enterica; Salmonella enterica subsp. enterica MTGDNTLITSHGINRRDFMKLCAALAATMGLSSKAAAEMAESVSNPQRPPVIWIGAQECT WP_000145429.1 hydrogenase 2 small subunit [Salmonella enterica] Length: 372\nNumber of predicted TMHs: 1\nExp number of AAs in TMHs: 24.51309\nExp number, first 60 AAs: 0.2682\nTotal prob of N-in: 0.14402\noutside 1 331\nTMhelix 332 354\ninside 355 372

16363 GCF_002066015.1_ASM206601v1 Salmonella enterica subsp. enterica serovar Cerro Proteobacteria; Gammaproteobacteria; Enterobacterales; Enterobacteriaceae; Salmonella; Salmonella enterica; Salmonella enterica subsp. enterica MNNEETFYQAMRRKGVTRRSFLKFCSLAATSLGLGAGMTPKIAWALENKPRIPVVWIHGL WP_023233644.1 hydrogenase 1, small subunit [Salmonella enterica] Length: 372\nNumber of predicted TMHs: 2\nExp number of AAs in TMHs: 37.4208299999999\nExp number, first 60 AAs: 19.72732\nTotal prob of N-in: 0.96168\nPOSSIBLE N-term signal sequence\ninside 1 19\nTMhelix 20 39\noutside 40 325\nTMhelix 326 348\ninside 349 372

16364 GCF_002047325.1_ASM204732v1 Salmonella enterica subsp. enterica serovar Cerro Proteobacteria; Gammaproteobacteria; Enterobacterales; Enterobacteriaceae; Salmonella; Salmonella enterica; Salmonella enterica subsp. enterica MTGDNTLITSHGINRRDFMKLCAALAATMGLSSKAAAEMAESVSNPQRPPVIWIGAQECT WP_000145429.1 hydrogenase 2 small subunit [Salmonella enterica] Length: 372\nNumber of predicted TMHs: 1\nExp number of AAs in TMHs: 24.51309\nExp number, first 60 AAs: 0.2682\nTotal prob of N-in: 0.14402\noutside 1 331\nTMhelix 332 354\ninside 355 372

16365 GCF_002047405.1_ASM204740v1 Salmonella enterica subsp. enterica serovar Cerro Proteobacteria; Gammaproteobacteria; Enterobacterales; Enterobacteriaceae; Salmonella; Salmonella enterica; Salmonella enterica subsp. enterica MNNEETFYQAMRRKGVTRRSFLKFCSLAATSLGLGAGMTPKIAWALENKPRIPVVWIHGL WP_023233644.1 hydrogenase 1, small subunit [Salmonella enterica] Length: 372\nNumber of predicted TMHs: 2\nExp number of AAs in TMHs: 37.4208299999999\nExp number, first 60 AAs: 19.72732\nTotal prob of N-in: 0.96168\nPOSSIBLE N-term signal sequence\ninside 1 19\nTMhelix 20 39\noutside 40 325\nTMhelix 326 348\ninside 349 372

16366 GCF_002047405.1_ASM204740v1 Salmonella enterica subsp. enterica serovar Cerro Proteobacteria; Gammaproteobacteria; Enterobacterales; Enterobacteriaceae; Salmonella; Salmonella enterica; Salmonella enterica subsp. enterica MTGDNTLITSHGINRRDFMKLCAALAATMGLSSKAAAEMAESVSNPQRPPVIWIGAQECT WP_000145429.1 hydrogenase 2 small subunit [Salmonella enterica] Length: 372\nNumber of predicted TMHs: 1\nExp number of AAs in TMHs: 24.51309\nExp number, first 60 AAs: 0.2682\nTotal prob of N-in: 0.14402\noutside 1 331\nTMhelix 332 354\ninside 355 372

16367 GCF_002065995.1_ASM206599v1 Salmonella enterica subsp. enterica serovar Chandans Proteobacteria; Gammaproteobacteria; Enterobacterales; Enterobacteriaceae; Salmonella; Salmonella enterica;

Salmonella enterica subsp. enterica MNNEETFYQAMRRKGVTRRSFLKFCSLAATSLGLGAGMTPKIAWALENKPRIPVVWIHGL WP_001058312.1 hydrogenase [Salmonella enterica] Length: 372\nNumber of predicted TMHs: 2\nExp number of AAs in TMHs: 36.51738\nExp number, first 60 AAs: 19.72849\nTotal prob of N-in: 0.96170\nPOSSIBLE N-term signal sequence\ninside 1 19\nTMhelix 20 39\noutside 40 325\nTMhelix 326 348\ninside 349 372

16368 GCF_002065995.1_ASM206599v1 Salmonella enterica subsp. enterica serovar Chandans Proteobacteria; Gammaproteobacteria; Enterobacterales; Enterobacteriaceae; Salmonella; Salmonella enterica; Salmonella enterica subsp. enterica MTGDNTLITSHGINRRDFMKLCAALAATMGLSSKAAAEMAESVSNPQRPPVIWIGAQECT WP_000145429.1 hydrogenase 2 small subunit [Salmonella enterica] Length: 372\nNumber of predicted TMHs: 1\nExp number of AAs in TMHs: 24.51309\nExp number, first 60 AAs: 0.2682\nTotal prob of N-in: 0.14402\noutside 1 331\nTMhelix 332 354\ninside 355 372

16369 GCF_002047045.1_ASM204704v1 Salmonella enterica subsp. enterica serovar Chester Proteobacteria; Gammaproteobacteria; Enterobacterales; Enterobacteriaceae; Salmonella; Salmonella enterica; Salmonella enterica subsp. enterica MTGDNTLITSHGINRRDFMKLCAALAATMGLSSKAAAEMAESVSNPQRPPVIWIGAQECT WP_079965169.1 hydrogenase 2 small subunit [Salmonella enterica] Length: 372\nNumber of predicted TMHs: 1\nExp number of AAs in TMHs: 24.51333\nExp number, first 60 AAs: 0.2682\nTotal prob of N-in: 0.14403\noutside 1 331\nTMhelix 332 354\ninside 355 372

16370 GCF_002047045.1_ASM204704v1 Salmonella enterica subsp. enterica serovar Chester Proteobacteria; Gammaproteobacteria; Enterobacterales; Enterobacteriaceae; Salmonella; Salmonella enterica; Salmonella enterica subsp. enterica MNNEETFYQAMRRKGVTRRSFLKFCSLAATSLGLGAGMTPKIAWALENKPRIPVVWIHGL WP_001058312.1 hydrogenase [Salmonella enterica] Length: 372\nNumber of predicted TMHs: 2\nExp number of AAs in TMHs: 36.51738\nExp number, first 60 AAs: 19.72849\nTotal prob of N-in: 0.96170\nPOSSIBLE N-term signal sequence\ninside 1 19\nTMhelix 20 39\noutside 40 325\nTMhelix 326 348\ninside 349 372

16371 GCF_002046865.1_ASM204686v1 Salmonella enterica subsp. enterica serovar Chester Proteobacteria; Gammaproteobacteria; Enterobacterales; Enterobacteriaceae; Salmonella; Salmonella enterica; Salmonella enterica subsp. enterica MNNEETFYQAMRRKGVTRRSFLKFCSLAATSLGLGAGMTPKIAWALENKPRIPVVWIHGL WP_001058312.1 hydrogenase [Salmonella enterica] Length: 372\nNumber of predicted TMHs: 2\nExp number of AAs in TMHs: 36.51738\nExp number, first 60 AAs: 19.72849\nTotal prob of N-in: 0.96170\nPOSSIBLE N-term signal sequence\ninside 1 19\nTMhelix 20 39\noutside 40 325\nTMhelix 326 348\ninside 349 372

16372 GCF_002046725.1_ASM204672v1 Salmonella enterica subsp. enterica serovar Chester Proteobacteria; Gammaproteobacteria; Enterobacterales; Enterobacteriaceae; Salmonella; Salmonella enterica; Salmonella enterica subsp. enterica MTGDNTLITSHGINRRDFMKLCAALAATMGLSSKAAAEMAESVSNPQRPPVIWIGAQECT WP_000145429.1 hydrogenase 2 small subunit [Salmonella enterica] Length: 372\nNumber of predicted TMHs: 1\nExp number of AAs in TMHs: 24.51309\nExp number, first 60 AAs: 0.2682\nTotal prob of N-in: 0.14402\noutside 1 331\nTMhelix 332 354\ninside 355 372

16373 GCF_002046725.1_ASM204672v1 Salmonella enterica subsp. enterica serovar Chester Proteobacteria; Gammaproteobacteria; Enterobacterales; Enterobacteriaceae; Salmonella; Salmonella enterica; Salmonella enterica subsp. enterica MNNEETFYQAMRRKGVTRRSFLKFCSLAATSLGLGAGMTPKIAWALENKPRIPVVWIHGL WP_001058312.1 hydrogenase [Salmonella enterica] Length: 372\nNumber of predicted TMHs: 2\nExp number of AAs in TMHs: 36.51738\nExp number, first 60 AAs: 19.72849\nTotal prob of N-in: 0.96170\nPOSSIBLE N-term signal sequence\ninside 1 19\nTMhelix 20 39\noutside 40 325\nTMhelix 326 348\ninside 349 372

16374 GCF_002046965.1_ASM204696v1 Salmonella enterica subsp. enterica serovar Chester Proteobacteria; Gammaproteobacteria; Enterobacterales; Enterobacteriaceae; Salmonella; Salmonella enterica; Salmonella enterica subsp. enterica MNNEETFYQAMRRKGVTRRSFLKFCSLAATSLGLGAGMTPKIAWALENKPRIPVVWIHGL WP_001058312.1 hydrogenase [Salmonella enterica] Length: 372\nNumber of predicted TMHs: 2\nExp number of AAs in TMHs: 36.51738\nExp number, first 60 AAs: 19.72849\nTotal prob of N-in: 0.96170\nPOSSIBLE N-term signal sequence\ninside 1 19\nTMhelix 20 39\noutside 40 325\nTMhelix 326 348\ninside 349 372

16375 GCF_002048125.1_ASM204812v1 Salmonella enterica subsp. enterica serovar Chester Proteobacteria; Gammaproteobacteria; Enterobacterales; Enterobacteriaceae; Salmonella; Salmonella enterica; Salmonella enterica subsp. enterica MNNEETFYQAMRRKGVTRRSFLKFCSLAATSLGLGAGMTPKIAWALENKPRIPVVWIHGL WP_001058312.1 hydrogenase [Salmonella enterica] Length: 372\nNumber of predicted TMHs: 2\nExp number of AAs in TMHs: 36.51738\nExp number, first 60 AAs: 19.72849\nTotal prob of N-in: 0.96170\nPOSSIBLE N-term signal sequence\ninside 1 19\nTMhelix 20 39\noutside 40 325\nTMhelix 326 348\ninside 349 372

16376 GCF_002048125.1_ASM204812v1 Salmonella enterica subsp. enterica serovar Chester Proteobacteria; Gammaproteobacteria; Enterobacterales; Enterobacteriaceae; Salmonella; Salmonella enterica; Salmonella enterica subsp. enterica MTGDNTLITSHGINRRDFMKLCAALAATMGLSSKAAAEMAESVSNPQRPPVIWIGAQECT WP_000145429.1 hydrogenase 2 small subunit [Salmonella enterica] Length: 372\nNumber of predicted TMHs: 1\nExp number of AAs in TMHs: 24.51309\nExp number, first 60 AAs: 0.2682\nTotal prob of N-in: 0.14402\noutside 1 331\nTMhelix 332 354\ninside 355 372

16377 GCF_002062475.1_ASM206247v1 Salmonella enterica subsp. enterica serovar Chester Proteobacteria; Gammaproteobacteria; Enterobacterales; Enterobacteriaceae; Salmonella; Salmonella enterica; Salmonella enterica subsp. enterica MTGDNTLITSHGINRRDFMKLCAALAATMGLSSKAAAEMAESVSNPQRPPVIWIGAQECT WP_000145429.1 hydrogenase 2 small subunit [Salmonella enterica] Length: 372\nNumber of predicted TMHs: 1\nExp number of AAs in TMHs: 24.51309\nExp number, first 60 AAs: 0.2682\nTotal prob of N-in: 0.14402\noutside 1 331\nTMhelix 332 354\ninside 355 372

16378 GCF_002046965.1_ASM204696v1 Salmonella enterica subsp. enterica serovar Chester Proteobacteria; Gammaproteobacteria; Enterobacterales; Enterobacteriaceae; Salmonella; Salmonella enterica; Salmonella enterica subsp. enterica MTGDNTLITSHGINRRDFMKLCAALAATMGLSSKAAAEMAESVSNPQRPPVIWIGAQECT WP_000145429.1 hydrogenase 2 small subunit [Salmonella enterica] Length: 372\nNumber of predicted TMHs: 1\nExp number of AAs in TMHs: 24.51309\nExp number, first 60 AAs: 0.2682\nTotal prob of N-in: 0.14402\noutside 1 331\nTMhelix 332 354\ninside 355 372

16379 GCF_002062475.1_ASM206247v1 Salmonella enterica subsp. enterica serovar Chester Proteobacteria; Gammaproteobacteria; Enterobacterales; Enterobacteriaceae; Salmonella; Salmonella enterica; Salmonella enterica subsp. enterica MNNEETFYQAMRRKGVTRRSFLKFCSLAATSLGLGAGMTPKIAWALENKPRIPVVWIHGL WP_001058312.1 hydrogenase [Salmonella enterica] Length: 372\nNumber of predicted TMHs: 2\nExp number of AAs in TMHs: 36.51738\nExp number, first 60 AAs: 19.72849\nTotal prob of N-in: 0.96170\nPOSSIBLE N-term signal sequence\ninside 1 19\nTMhelix 20 39\noutside 40 325\nTMhelix 326 348\ninside 349 372

16380 GCF_002048115.1_ASM204811v1 Salmonella enterica subsp. enterica serovar Chester Proteobacteria; Gammaproteobacteria; Enterobacterales; Enterobacteriaceae; Salmonella; Salmonella enterica; Salmonella enterica subsp. enterica MNNEETFYQAMRRKGVTRRSFLKFCSLAATSLGLGAGMTPKIAWALENKPRIPVVWIHGL WP_001058312.1 hydrogenase [Salmonella enterica] Length: 372\nNumber of predicted TMHs: 2\nExp number of AAs in TMHs: 36.51738\nExp number, first 60 AAs: 19.72849\nTotal prob of N-in: 0.96170\nPOSSIBLE N-term signal sequence\ninside 1 19\nTMhelix 20 39\noutside 40 325\nTMhelix 326 348\ninside 349 372

16381 GCF_002048115.1_ASM204811v1 Salmonella enterica subsp. enterica serovar Chester Proteobacteria; Gammaproteobacteria; Enterobacterales; Enterobacteriaceae; Salmonella; Salmonella enterica; Salmonella enterica subsp. enterica MTGDNTLITSHGINRRDFMKLCAALAATMGLSSKAAAEMAESVSNPQRPPVIWIGAQECT WP_000145429.1 hydrogenase 2 small subunit [Salmonella enterica] Length: 372\nNumber of predicted TMHs: 1\nExp number of AAs in TMHs: 24.51309\nExp number, first 60 AAs: 0.2682\nTotal prob of N-in: 0.14402\noutside 1 331\nTMhelix 332 354\ninside 355 372

16382 GCF_002046865.1_ASM204686v1 Salmonella enterica subsp. enterica serovar Chester Proteobacteria; Gammaproteobacteria; Enterobacterales; Enterobacteriaceae; Salmonella; Salmonella enterica; Salmonella enterica subsp. enterica MTGDNTLITSHGINRRDFMKLCAALAATMGLSSKAAAEMAESVSNPQRPPVIWIGAQECT WP_000145429.1 hydrogenase 2 small subunit [Salmonella enterica] Length: 372\nNumber of predicted TMHs: 1\nExp number of AAs in TMHs: 24.51309\nExp number, first 60 AAs: 0.2682\nTotal prob of N-in: 0.14402\noutside 1 331\nTMhelix 332 354\ninside 355 372

16383 GCF_002044825.1_ASM204482v1 Salmonella enterica subsp. enterica serovar Choleraesuis Proteobacteria; Gammaproteobacteria; Enterobacterales; Enterobacteriaceae; Salmonella; Salmonella enterica; Salmonella enterica subsp. enterica MTGDNTLITSHGINRRDFMKLCAALAATMGLSSKAAAEMAESVSNPQRPPVIWIGAQECT WP_000145429.1 hydrogenase 2 small subunit [Salmonella enterica] Length: 372\nNumber of predicted TMHs: 1\nExp number of AAs in TMHs: 24.51309\nExp number, first 60 AAs: 0.2682\nTotal prob of N-in: 0.14402\noutside 1 331\nTMhelix 332 354\ninside 355 372

16384 GCF_002044825.1_ASM204482v1 Salmonella enterica subsp. enterica serovar Choleraesuis Proteobacteria; Gammaproteobacteria; Enterobacterales; Enterobacteriaceae; Salmonella; Salmonella enterica; Salmonella enterica subsp. enterica MNNEETFYQAMRRKGVTRRSFLKFCSLAATSLGLGAGMTPKIAWALENKPRIPVVWIHGL WP_023234992.1 hydrogenase 1, small subunit [Salmonella enterica] Length: 372\nNumber of predicted TMHs: 2\nExp number of AAs in TMHs: 37.41309\nExp number, first 60 AAs: 19.72865\nTotal prob of N-in:

0.96169\nPOSSIBLE N-term signal sequence\ninside 1 19\nTMhelix 20 39\noutside 40 325\nTMhelix 326 348\ninside 349 372

16385 GCF_000742815.1_ASM74281v1 Salmonella enterica subsp. enterica serovar Choleraesuis Proteobacteria; Gammaproteobacteria; Enterobacterales; Enterobacteriaceae; Salmonella; Salmonella enterica; Salmonella enterica subsp. enterica MNNEETFYQAMRRKGVTRRSFLKFCSLAATSLGLGAGMTPKIAWALENKPRIPVVWIHGL WP_023234992.1 hydrogenase 1, small subunit [Salmonella enterica] Length: 372\nNumber of predicted TMHs: 2\nExp number of AAs in TMHs: 37.41309\nExp number, first 60 AAs: 19.72865\nTotal prob of N-in: 0.96169\nPOSSIBLE N-term signal sequence\ninside 1 19\nTMhelix 20 39\noutside 40 325\nTMhelix 326 348\ninside 349 372

16386 GCF_000742815.1_ASM74281v1 Salmonella enterica subsp. enterica serovar Choleraesuis Proteobacteria; Gammaproteobacteria; Enterobacterales; Enterobacteriaceae; Salmonella; Salmonella enterica; Salmonella enterica subsp. enterica MTGDNTLITSHGINRRDFMKLCAALAATMGLSSKAAAEMAESVSNPQRPPVIWIGAQECT WP_000145429.1 hydrogenase 2 small subunit [Salmonella enterica] Length: 372\nNumber of predicted TMHs: 1\nExp number of AAs in TMHs: 24.51309\nExp number, first 60 AAs: 0.2682\nTotal prob of N-in: 0.14402\noutside 1 331\nTMhelix 332 354\ninside 355 372

16387 GCF_002066615.1_ASM206661v1 Salmonella enterica subsp. enterica serovar Colindale Proteobacteria; Gammaproteobacteria; Enterobacterales; Enterobacteriaceae; Salmonella; Salmonella enterica; Salmonella enterica subsp. enterica MNNEETFYQAMRRKGVTRRSFLKFCSLAATSLGLGAGMTPKIAWALENKPRIPVVWIHGL WP_001058311.1 hydrogenase [Salmonella enterica] Length: 372\nNumber of predicted TMHs: 2\nExp number of AAs in TMHs: 37.4699899999999\nExp number, first 60 AAs: 19.72699\nTotal prob of N-in: 0.96171\nPOSSIBLE N-term signal sequence\ninside 1 19\nTMhelix 20 39\noutside 40 325\nTMhelix 326 348\ninside 349 372

16388 GCF_002066615.1_ASM206661v1 Salmonella enterica subsp. enterica serovar Colindale Proteobacteria; Gammaproteobacteria; Enterobacterales; Enterobacteriaceae; Salmonella; Salmonella enterica; Salmonella enterica subsp. enterica MTGDNTLITSHGINRRDFMKLCAALAATMGLSSKAAAEMAESVSNPQRPPVIWIGAQECT WP_000145429.1 hydrogenase 2 small subunit [Salmonella enterica] Length: 372\nNumber of predicted TMHs: 1\nExp number of AAs in TMHs: 24.51309\nExp number, first 60 AAs: 0.2682\nTotal prob of N-in: 0.14402\noutside 1 331\nTMhelix 332 354\ninside 355 372

16389 GCF_002061015.1_ASM206101v1 Salmonella enterica subsp. enterica serovar Colindale Proteobacteria; Gammaproteobacteria; Enterobacterales; Enterobacteriaceae; Salmonella; Salmonella enterica; Salmonella enterica subsp. enterica MTGDNTLITSHGINRRDFMKLCAALAATMGLSSKAAAEMAESVSNPQRPPVIWIGAQECT WP_000145429.1 hydrogenase 2 small subunit [Salmonella enterica] Length: 372\nNumber of predicted TMHs: 1\nExp number of AAs in TMHs: 24.51309\nExp number, first 60 AAs: 0.2682\nTotal prob of N-in: 0.14402\noutside 1 331\nTMhelix 332 354\ninside 355 372

16390 GCF_002061015.1_ASM206101v1 Salmonella enterica subsp. enterica serovar Colindale Proteobacteria; Gammaproteobacteria; Enterobacterales; Enterobacteriaceae; Salmonella; Salmonella enterica; Salmonella enterica subsp. enterica MNNEETFYQAMRRKGVTRRSFLKFCSLAATSLGLGAGMTPKIAWALENKPRIPVVWIHGL WP_001058311.1 hydrogenase [Salmonella enterica] Length: 372\nNumber of predicted TMHs: 2\nExp number of AAs in TMHs: 37.4699899999999\nExp number, first 60 AAs: 19.72699\nTotal prob of N-in: 0.96171\nPOSSIBLE N-term signal sequence\ninside 1 19\nTMhelix 20 39\noutside 40 325\nTMhelix 326 348\ninside 349 372

16391 GCF_002065935.1_ASM206593v1 Salmonella enterica subsp. enterica serovar Concord Proteobacteria; Gammaproteobacteria; Enterobacterales; Enterobacteriaceae; Salmonella; Salmonella enterica; Salmonella enterica subsp. enterica MTGDNTLITSHGINRRDFMKLCAALAATMGLSSKAAAEMAESVSNPQRPPVIWIGAQECT WP_000145429.1 hydrogenase 2 small subunit [Salmonella enterica] Length: 372\nNumber of predicted TMHs: 1\nExp number of AAs in TMHs: 24.51309\nExp number, first 60 AAs: 0.2682\nTotal prob of N-in: 0.14402\noutside 1 331\nTMhelix 332 354\ninside 355 372

16392 GCF_002065935.1_ASM206593v1 Salmonella enterica subsp. enterica serovar Concord Proteobacteria; Gammaproteobacteria; Enterobacterales; Enterobacteriaceae; Salmonella; Salmonella enterica; Salmonella enterica subsp. enterica MNNEETFYQAMRRKGVTRRSFLKFCSLAATSLGLGAGMTPKIAWALENKPRIPVVWIHGL WP_001058312.1 hydrogenase [Salmonella enterica] Length: 372\nNumber of predicted TMHs: 2\nExp number of AAs in TMHs: 36.51738\nExp number, first 60 AAs: 19.72849\nTotal prob of N-in: 0.96170\nPOSSIBLE N-term signal sequence\ninside 1 19\nTMhelix 20 39\noutside 40 325\nTMhelix 326 348\ninside 349 372

16393	GCF_002065885.1_ASM206588v1	Salmonella enterica subsp. enterica serovar Concord	Proteobacteria; Gammaproteobacteria; Enterobacterales; Enterobacteriaceae; Salmonella; Salmonella enterica; Salmonella enterica subsp. enterica	MTGDNTLITSHGINRRDFMKLCAALAATMGLSSKAAAEMAESVSNPQRPPVIWIGAQECT	WP_000145429.1	hydrogenase 2 small subunit [Salmonella enterica]	Length: 372\nNumber of predicted TMHs: 1\nExp number of AAs in TMHs: 24.51309\nExp number, first 60 AAs: 0.2682\nTotal prob of N-in: 0.14402\noutside 1 331\nTMhelix 332 354\ninside 355 372

16394	GCF_002065885.1_ASM206588v1	Salmonella enterica subsp. enterica serovar Concord	Proteobacteria; Gammaproteobacteria; Enterobacterales; Enterobacteriaceae; Salmonella; Salmonella enterica; Salmonella enterica subsp. enterica	MNNEETFYQAMRRKGVTRRSFLKFCSLAATSLGLGAGMTPKIAWALENKPRIPVVWIHGL	WP_001058312.1	hydrogenase [Salmonella enterica]	Length: 372\nNumber of predicted TMHs: 2\nExp number of AAs in TMHs: 36.51738\nExp number, first 60 AAs: 19.72849\nTotal prob of N-in: 0.96170\nPOSSIBLE N-term signal sequence\ninside 1 19\nTMhelix 20 39\noutside 40 325\nTMhelix 326 348\ninside 349 372

16395	GCF_002065915.1_ASM206591v1	Salmonella enterica subsp. enterica serovar Concord	Proteobacteria; Gammaproteobacteria; Enterobacterales; Enterobacteriaceae; Salmonella; Salmonella enterica; Salmonella enterica subsp. enterica	MNNEETFYQAMRRKGVTRRSFLKFCSLAATSLGLGAGMTPKIAWALENKPRIPVVWIHGL	WP_001058312.1	hydrogenase [Salmonella enterica]	Length: 372\nNumber of predicted TMHs: 2\nExp number of AAs in TMHs: 36.51738\nExp number, first 60 AAs: 19.72849\nTotal prob of N-in: 0.96170\nPOSSIBLE N-term signal sequence\ninside 1 19\nTMhelix 20 39\noutside 40 325\nTMhelix 326 348\ninside 349 372

16396	GCF_002065915.1_ASM206591v1	Salmonella enterica subsp. enterica serovar Concord	Proteobacteria; Gammaproteobacteria; Enterobacterales; Enterobacteriaceae; Salmonella; Salmonella enterica; Salmonella enterica subsp. enterica	MTGDNTLITSHGINRRDFMKLCAALAATMGLSSKAAAEMAESVSNPQRPPVIWIGAQECT	WP_000145429.1	hydrogenase 2 small subunit [Salmonella enterica]	Length: 372\nNumber of predicted TMHs: 1\nExp number of AAs in TMHs: 24.51309\nExp number, first 60 AAs: 0.2682\nTotal prob of N-in: 0.14402\noutside 1 331\nTMhelix 332 354\ninside 355 372

16397	GCF_002062415.1_ASM206241v1	Salmonella enterica subsp. enterica serovar Concord	Proteobacteria; Gammaproteobacteria; Enterobacterales; Enterobacteriaceae; Salmonella; Salmonella enterica; Salmonella enterica subsp. enterica	MNNEETFYQAMRRKGVTRRSFLKFCSLAATSLGLGAGMTPKIAWALENKPRIPVVWIHGL	WP_001058310.1	hydrogenase [Salmonella enterica]	Length: 372\nNumber of predicted TMHs: 2\nExp number of AAs in TMHs: 37.4567299999999\nExp number, first 60 AAs: 19.72694\nTotal prob of N-in: 0.96171\nPOSSIBLE N-term signal sequence\ninside 1 19\nTMhelix 20 39\noutside 40 325\nTMhelix 326 348\ninside 349 372

16398	GCF_002062415.1_ASM206241v1	Salmonella enterica subsp. enterica serovar Concord	Proteobacteria; Gammaproteobacteria; Enterobacterales; Enterobacteriaceae; Salmonella; Salmonella enterica; Salmonella enterica subsp. enterica	MTGDNTLITSHGINRRDFMKLCAALAATMGLSSKAAAEMAESVSNPQRPPVIWIGAQECT	WP_000145429.1	hydrogenase 2 small subunit [Salmonella enterica]	Length: 372\nNumber of predicted TMHs: 1\nExp number of AAs in TMHs: 24.51309\nExp number, first 60 AAs: 0.2682\nTotal prob of N-in: 0.14402\noutside 1 331\nTMhelix 332 354\ninside 355 372

16399	GCF_002061625.1_ASM206162v1	Salmonella enterica subsp. enterica serovar Copenhagen	Proteobacteria; Gammaproteobacteria; Enterobacterales; Enterobacteriaceae; Salmonella; Salmonella enterica; Salmonella enterica subsp. enterica	MTGDNTLITSHGINRRDFMKLCAALAATMGLSSKAAAEMAESVSNPQRPPVIWIGAQECT	WP_000145429.1	hydrogenase 2 small subunit [Salmonella enterica]	Length: 372\nNumber of predicted TMHs: 1\nExp number of AAs in TMHs: 24.51309\nExp number, first 60 AAs: 0.2682\nTotal prob of N-in: 0.14402\noutside 1 331\nTMhelix 332 354\ninside 355 372

16400	GCF_002105815.1_ASM210581v1	Salmonella enterica subsp. enterica serovar Copenhagen	Proteobacteria; Gammaproteobacteria; Enterobacterales; Enterobacteriaceae; Salmonella; Salmonella enterica; Salmonella enterica subsp. enterica	MNNEETFYQAMRRKGVTRRSFLKFCSLAATSLGLGAGMTPKIAWALENKPRIPVVWIHGL	WP_001058312.1	hydrogenase [Salmonella enterica]	Length: 372\nNumber of predicted TMHs: 2\nExp number of AAs in TMHs: 36.51738\nExp number, first 60 AAs: 19.72849\nTotal prob of N-in: 0.96170\nPOSSIBLE N-term signal sequence\ninside 1 19\nTMhelix 20 39\noutside 40 325\nTMhelix 326 348\ninside 349 372

16401	GCF_002060305.1_ASM206030v1	Salmonella enterica subsp. enterica serovar Copenhagen	Proteobacteria; Gammaproteobacteria; Enterobacterales; Enterobacteriaceae; Salmonella; Salmonella enterica; Salmonella enterica subsp. enterica	MNNEETFYQAMRRKGVTRRSFLKFCSLAATSLGLGAGMTPKIAWALENKPRIPVVWIHGL	WP_001058312.1	hydrogenase [Salmonella enterica]	Length: 372\nNumber of predicted TMHs: 2\nExp

number of AAs in TMHs: 36.51738\nExp number, first 60 AAs: 19.72849\nTotal prob of N-in: 0.96170\nPOSSIBLE N-term signal sequence\ninside 1 19\nTMhelix 20 39\noutside 40 325\nTMhelix 326 348\ninside 349 372

16402 GCF_002036765.1_ASM203676v1 Salmonella enterica subsp. enterica serovar Copenhagen Proteobacteria; Gammaproteobacteria; Enterobacterales; Enterobacteriaceae; Salmonella; Salmonella enterica; Salmonella enterica subsp. enterica MNNEETFYQAMRRKGVTRRSFLKFCSLAATSLGLGAGMTPKIAWALENKPRIPVVWIHGL WP_001058312.1 hydrogenase [Salmonella enterica] Length: 372\nNumber of predicted TMHs: 2\nExp number of AAs in TMHs: 36.51738\nExp number, first 60 AAs: 19.72849\nTotal prob of N-in: 0.96170\nPOSSIBLE N-term signal sequence\ninside 1 19\nTMhelix 20 39\noutside 40 325\nTMhelix 326 348\ninside 349 372

16403 GCF_002061625.1_ASM206162v1 Salmonella enterica subsp. enterica serovar Copenhagen Proteobacteria; Gammaproteobacteria; Enterobacterales; Enterobacteriaceae; Salmonella; Salmonella enterica; Salmonella enterica subsp. enterica MNNEETFYQAMRRKGVTRRSFLKFCSLAATSLGLGAGMTPKIAWALENKPRIPVVWIHGL WP_001058312.1 hydrogenase [Salmonella enterica] Length: 372\nNumber of predicted TMHs: 2\nExp number of AAs in TMHs: 36.51738\nExp number, first 60 AAs: 19.72849\nTotal prob of N-in: 0.96170\nPOSSIBLE N-term signal sequence\ninside 1 19\nTMhelix 20 39\noutside 40 325\nTMhelix 326 348\ninside 349 372

16404 GCF_002105815.1_ASM210581v1 Salmonella enterica subsp. enterica serovar Copenhagen Proteobacteria; Gammaproteobacteria; Enterobacterales; Enterobacteriaceae; Salmonella; Salmonella enterica; Salmonella enterica subsp. enterica MTGDNTLITSHGINRRDFMKLCAALAATMGLSSKAAAEMAESVSNPQRPPVIWIGAQECT WP_000145429.1 hydrogenase 2 small subunit [Salmonella enterica] Length: 372\nNumber of predicted TMHs: 1\nExp number of AAs in TMHs: 24.51309\nExp number, first 60 AAs: 0.2682\nTotal prob of N-in: 0.14402\noutside 1 331\nTMhelix 332 354\ninside 355 372

16405 GCF_002060305.1_ASM206030v1 Salmonella enterica subsp. enterica serovar Copenhagen Proteobacteria; Gammaproteobacteria; Enterobacterales; Enterobacteriaceae; Salmonella; Salmonella enterica; Salmonella enterica subsp. enterica MTGDNTLITSHGINRRDFMKLCAALAATMGLSSKAAAEMAESVSNPQRPPVIWIGAQECT WP_000145429.1 hydrogenase 2 small subunit [Salmonella enterica] Length: 372\nNumber of predicted TMHs: 1\nExp number of AAs in TMHs: 24.51309\nExp number, first 60 AAs: 0.2682\nTotal prob of N-in: 0.14402\noutside 1 331\nTMhelix 332 354\ninside 355 372

16406 GCF_002036765.1_ASM203676v1 Salmonella enterica subsp. enterica serovar Copenhagen Proteobacteria; Gammaproteobacteria; Enterobacterales; Enterobacteriaceae; Salmonella; Salmonella enterica; Salmonella enterica subsp. enterica MTGDNTLITSHGINRRDFMKLCAALAATMGLSSKAAAEMAESVSNPQRPPVIWIGAQECT WP_000145429.1 hydrogenase 2 small subunit [Salmonella enterica] Length: 372\nNumber of predicted TMHs: 1\nExp number of AAs in TMHs: 24.51309\nExp number, first 60 AAs: 0.2682\nTotal prob of N-in: 0.14402\noutside 1 331\nTMhelix 332 354\ninside 355 372

16407 GCF_002066565.1_ASM206656v1 Salmonella enterica subsp. enterica serovar Corvallis Proteobacteria; Gammaproteobacteria; Enterobacterales; Enterobacteriaceae; Salmonella; Salmonella enterica; Salmonella enterica subsp. enterica MTGDNTLITSHGINRRDFMKLCAALAATMGLSSKAAAEMAESVSNPQRPPVIWIGAQECT WP_000145429.1 hydrogenase 2 small subunit [Salmonella enterica] Length: 372\nNumber of predicted TMHs: 1\nExp number of AAs in TMHs: 24.51309\nExp number, first 60 AAs: 0.2682\nTotal prob of N-in: 0.14402\noutside 1 331\nTMhelix 332 354\ninside 355 372

16408 GCF_002066565.1_ASM206656v1 Salmonella enterica subsp. enterica serovar Corvallis Proteobacteria; Gammaproteobacteria; Enterobacterales; Enterobacteriaceae; Salmonella; Salmonella enterica; Salmonella enterica subsp. enterica MNNEETFYQAMRRKGVTRRSFLKFCSLAATSLGLGAGMTPKIAWALENKPRIPVVWIHGL WP_001058311.1 hydrogenase [Salmonella enterica] Length: 372\nNumber of predicted TMHs: 2\nExp number of AAs in TMHs: 37.4699899999999\nExp number, first 60 AAs: 19.72699\nTotal prob of N-in: 0.96171\nPOSSIBLE N-term signal sequence\ninside 1 19\nTMhelix 20 39\noutside 40 325\nTMhelix 326 348\ninside 349 372

16409 GCF_002045955.1_ASM204595v1 Salmonella enterica subsp. enterica serovar Corvallis Proteobacteria; Gammaproteobacteria; Enterobacterales; Enterobacteriaceae; Salmonella; Salmonella enterica; Salmonella enterica subsp. enterica MTGDNTLITSHGINRRDFMKLCAALAATMGLSSKAAAEMAESVSNPQRPPVIWIGAQECT WP_000145429.1 hydrogenase 2 small subunit [Salmonella enterica] Length: 372\nNumber of predicted TMHs: 1\nExp number of AAs in TMHs: 24.51309\nExp number, first 60 AAs: 0.2682\nTotal prob of N-in: 0.14402\noutside 1 331\nTMhelix 332 354\ninside 355 372

16410 GCF_002045955.1_ASM204595v1 Salmonella enterica subsp. enterica serovar Corvallis Proteobacteria; Gammaproteobacteria; Enterobacterales; Enterobacteriaceae; Salmonella; Salmonella enterica; Salmonella enterica subsp. enterica MNNEETFYQAMRRKGVTRRSFLKFCSLAATSLGLGAGMTPKIAWALENKPRIPVVWIHGL WP_001058311.1 hydrogenase [Salmonella enterica] Length: 372\nNumber of predicted TMHs: 2\nExp number of AAs in TMHs: 37.4699899999999\nExp number, first 60 AAs: 19.72699\nTotal prob of N-in: 0.96171\nPOSSIBLE N-term signal sequence\ninside 1 19\nTMhelix 20 39\noutside 40 325\nTMhelix 326 348\ninside 349 372

16411 GCF_002045985.1_ASM204598v1 Salmonella enterica subsp. enterica serovar Corvallis Proteobacteria; Gammaproteobacteria; Enterobacterales; Enterobacteriaceae; Salmonella; Salmonella enterica; Salmonella enterica subsp. enterica MTGDNTLITSHGINRRDFMKLCAALAATMGLSSKAAAEMAESVSNPQRPPVIWIGAQECT WP_000145429.1 hydrogenase 2 small subunit [Salmonella enterica] Length: 372\nNumber of predicted TMHs: 1\nExp number of AAs in TMHs: 24.51309\nExp number, first 60 AAs: 0.2682\nTotal prob of N-in: 0.14402\noutside 1 331\nTMhelix 332 354\ninside 355 372

16412 GCF_002062535.1_ASM206253v1 Salmonella enterica subsp. enterica serovar Corvallis Proteobacteria; Gammaproteobacteria; Enterobacterales; Enterobacteriaceae; Salmonella; Salmonella enterica; Salmonella enterica subsp. enterica MNNEETFYQAMRRKGVTRRSFLKFCSLAATSLGLGAGMTPKIAWALENKPRIPVVWIHGL WP_001058311.1 hydrogenase [Salmonella enterica] Length: 372\nNumber of predicted TMHs: 2\nExp number of AAs in TMHs: 37.4699899999999\nExp number, first 60 AAs: 19.72699\nTotal prob of N-in: 0.96171\nPOSSIBLE N-term signal sequence\ninside 1 19\nTMhelix 20 39\noutside 40 325\nTMhelix 326 348\ninside 349 372

16413 GCF_002063295.1_ASM206329v1 Salmonella enterica subsp. enterica serovar Corvallis Proteobacteria; Gammaproteobacteria; Enterobacterales; Enterobacteriaceae; Salmonella; Salmonella enterica; Salmonella enterica subsp. enterica MNNEETFYQAMRRKGVTRRSFLKFCSLAATSLGLGAGMTPKIAWALENKPRIPVVWIHGL WP_001058311.1 hydrogenase [Salmonella enterica] Length: 372\nNumber of predicted TMHs: 2\nExp number of AAs in TMHs: 37.4699899999999\nExp number, first 60 AAs: 19.72699\nTotal prob of N-in: 0.96171\nPOSSIBLE N-term signal sequence\ninside 1 19\nTMhelix 20 39\noutside 40 325\nTMhelix 326 348\ninside 349 372

16414 GCF_002062535.1_ASM206253v1 Salmonella enterica subsp. enterica serovar Corvallis Proteobacteria; Gammaproteobacteria; Enterobacterales; Enterobacteriaceae; Salmonella; Salmonella enterica; Salmonella enterica subsp. enterica MTGDNTLITSHGINRRDFMKLCAALAATMGLSSKAAAEMAESVSNPQRPPVIWIGAQECT WP_000145429.1 hydrogenase 2 small subunit [Salmonella enterica] Length: 372\nNumber of predicted TMHs: 1\nExp number of AAs in TMHs: 24.51309\nExp number, first 60 AAs: 0.2682\nTotal prob of N-in: 0.14402\noutside 1 331\nTMhelix 332 354\ninside 355 372

16415 GCF_002063295.1_ASM206329v1 Salmonella enterica subsp. enterica serovar Corvallis Proteobacteria; Gammaproteobacteria; Enterobacterales; Enterobacteriaceae; Salmonella; Salmonella enterica; Salmonella enterica subsp. enterica MTGDNTLITSHGINRRDFMKLCAALAATMGLSSKAAAEMAESVSNPQRPPVIWIGAQECT WP_000145429.1 hydrogenase 2 small subunit [Salmonella enterica] Length: 372\nNumber of predicted TMHs: 1\nExp number of AAs in TMHs: 24.51309\nExp number, first 60 AAs: 0.2682\nTotal prob of N-in: 0.14402\noutside 1 331\nTMhelix 332 354\ninside 355 372

16416 GCF_002045985.1_ASM204598v1 Salmonella enterica subsp. enterica serovar Corvallis Proteobacteria; Gammaproteobacteria; Enterobacterales; Enterobacteriaceae; Salmonella; Salmonella enterica; Salmonella enterica subsp. enterica MNNEETFYQAMRRKGVTRRSFLKFCSLAATSLGLGAGMTPKIAWALENKPRIPVVWIHGL WP_001058311.1 hydrogenase [Salmonella enterica] Length: 372\nNumber of predicted TMHs: 2\nExp number of AAs in TMHs: 37.4699899999999\nExp number, first 60 AAs: 19.72699\nTotal prob of N-in: 0.96171\nPOSSIBLE N-term signal sequence\ninside 1 19\nTMhelix 20 39\noutside 40 325\nTMhelix 326 348\ninside 349 372

16417 GCF_002057115.1_ASM205711v1 Salmonella enterica subsp. enterica serovar Cubana Proteobacteria; Gammaproteobacteria; Enterobacterales; Enterobacteriaceae; Salmonella; Salmonella enterica; Salmonella enterica subsp. enterica MTGDNTLITSHGINRRDFMKLCAALAATMGLSSKAAAEMAESVSNPQRPPVIWIGAQECT WP_000145429.1 hydrogenase 2 small subunit [Salmonella enterica] Length: 372\nNumber of predicted TMHs: 1\nExp number of AAs in TMHs: 24.51309\nExp number, first 60 AAs: 0.2682\nTotal prob of N-in: 0.14402\noutside 1 331\nTMhelix 332 354\ninside 355 372

16418	GCF_002046285.1_ASM204628v1	Salmonella enterica subsp. enterica serovar Cubana	Proteobacteria; Gammaproteobacteria; Enterobacterales; Enterobacteriaceae; Salmonella; Salmonella enterica; Salmonella enterica subsp. enterica	MNNEETFYQAMRRKGVTRRSFLKFCSLAATSLGLGAGMTPKIAWALENKPRIPVVWIHGL	WP_001058311.1	hydrogenase [Salmonella enterica]	Length: 372\nNumber of predicted TMHs:  2\nExp number of AAs in TMHs: 37.4699899999999\nExp number, first 60 AAs:  19.72699\nTotal prob of N-in:  0.96171\nPOSSIBLE N-term signal sequence\ninside   1   19\nTMhelix   20   39\noutside   40   325\nTMhelix   326   348\ninside   349   372

16419	GCF_002045355.1_ASM204535v1	Salmonella enterica subsp. enterica serovar Cubana	Proteobacteria; Gammaproteobacteria; Enterobacterales; Enterobacteriaceae; Salmonella; Salmonella enterica; Salmonella enterica subsp. enterica	MTGDNTLITSHGINRRDFMKLCAALAATMGLSSKAAAEMAESVSNPQRPPVIWIGAQECT	WP_000145429.1	hydrogenase 2 small subunit [Salmonella enterica]	Length: 372\nNumber of predicted TMHs:  1\nExp number of AAs in TMHs: 24.51309\nExp number, first 60 AAs:  0.2682\nTotal prob of N-in:  0.14402\noutside   1   331\nTMhelix   332   354\ninside   355   372

16420	GCF_002034725.1_ASM203472v1	Salmonella enterica subsp. enterica serovar Cubana	Proteobacteria; Gammaproteobacteria; Enterobacterales; Enterobacteriaceae; Salmonella; Salmonella enterica; Salmonella enterica subsp. enterica	MTGDNTLITSHGINRRDFMKLCAALAATMGLSSKAAAEMAESVSNPQRPPVIWIGAQECT	WP_000145429.1	hydrogenase 2 small subunit [Salmonella enterica]	Length: 372\nNumber of predicted TMHs:  1\nExp number of AAs in TMHs: 24.51309\nExp number, first 60 AAs:  0.2682\nTotal prob of N-in:  0.14402\noutside   1   331\nTMhelix   332   354\ninside   355   372

16421	GCF_002034725.1_ASM203472v1	Salmonella enterica subsp. enterica serovar Cubana	Proteobacteria; Gammaproteobacteria; Enterobacterales; Enterobacteriaceae; Salmonella; Salmonella enterica; Salmonella enterica subsp. enterica	MNNEETFYQAMRRKGVTRRSFLKFCSLAATSLGLGAGMTPKIAWALENKPRIPVVWIHGL	WP_001058311.1	hydrogenase [Salmonella enterica]	Length: 372\nNumber of predicted TMHs:  2\nExp number of AAs in TMHs: 37.4699899999999\nExp number, first 60 AAs:  19.72699\nTotal prob of N-in:  0.96171\nPOSSIBLE N-term signal sequence\ninside   1   19\nTMhelix   20   39\noutside   40   325\nTMhelix   326   348\ninside   349   372

16422	GCF_002057115.1_ASM205711v1	Salmonella enterica subsp. enterica serovar Cubana	Proteobacteria; Gammaproteobacteria; Enterobacterales; Enterobacteriaceae; Salmonella; Salmonella enterica; Salmonella enterica subsp. enterica	MNNEETFYQAMRRKGVTRRSFLKFCSLAATSLGLGAGMTPKIAWALENKPRIPVVWIHGL	WP_001058311.1	hydrogenase [Salmonella enterica]	Length: 372\nNumber of predicted TMHs:  2\nExp number of AAs in TMHs: 37.4699899999999\nExp number, first 60 AAs:  19.72699\nTotal prob of N-in:  0.96171\nPOSSIBLE N-term signal sequence\ninside   1   19\nTMhelix   20   39\noutside   40   325\nTMhelix   326   348\ninside   349   372

16423	GCF_002046215.1_ASM204621v1	Salmonella enterica subsp. enterica serovar Cubana	Proteobacteria; Gammaproteobacteria; Enterobacterales; Enterobacteriaceae; Salmonella; Salmonella enterica; Salmonella enterica subsp. enterica	MNNEETFYQAMRRKGVTRRSFLKFCSLAATSLGLGAGMTPKIAWALENKPRIPVVWIHGL	WP_001058312.1	hydrogenase [Salmonella enterica]	Length: 372\nNumber of predicted TMHs:  2\nExp number of AAs in TMHs: 36.51738\nExp number, first 60 AAs:  19.72849\nTotal prob of N-in:   0.96170\nPOSSIBLE N-term signal sequence\ninside   1   19\nTMhelix   20   39\noutside   40   325\nTMhelix   326   348\ninside   349   372

16424	GCF_002046285.1_ASM204628v1	Salmonella enterica subsp. enterica serovar Cubana	Proteobacteria; Gammaproteobacteria; Enterobacterales; Enterobacteriaceae; Salmonella; Salmonella enterica; Salmonella enterica subsp. enterica	MTGDNTLITSHGINRRDFMKLCAALAATMGLSSKAAAEMAESVSNPQRPPVIWIGAQECT	WP_000145429.1	hydrogenase 2 small subunit [Salmonella enterica]	Length: 372\nNumber of predicted TMHs:  1\nExp number of AAs in TMHs: 24.51309\nExp number, first 60 AAs:  0.2682\nTotal prob of N-in:  0.14402\noutside   1   331\nTMhelix   332   354\ninside   355   372

16425	GCF_002046215.1_ASM204621v1	Salmonella enterica subsp. enterica serovar Cubana	Proteobacteria; Gammaproteobacteria; Enterobacterales; Enterobacteriaceae; Salmonella; Salmonella enterica; Salmonella enterica subsp. enterica	MTGDNTLITSHGINRRDFMKLCAALAATMGLSSKAAAEMAESVSNPQRPPVIWIGAQECT	WP_000145429.1	hydrogenase 2 small subunit [Salmonella enterica]	Length: 372\nNumber of predicted TMHs:  1\nExp number of AAs in TMHs: 24.51309\nExp number, first 60 AAs:  0.2682\nTotal prob of N-in:  0.14402\noutside   1   331\nTMhelix   332   354\ninside   355   372

16426	GCF_002045355.1_ASM204535v1	Salmonella enterica subsp. enterica serovar Cubana	Proteobacteria; Gammaproteobacteria; Enterobacterales; Enterobacteriaceae; Salmonella; Salmonella enterica;

Salmonella enterica subsp. enterica	MNNEETFYQAMRRKGVTRRSFLKFCSLAATSLGLGAGMTPKIAWALENKPRIPVVWIHGL	WP_001058311.1	hydrogenase [Salmonella enterica]	Length: 372\nNumber of predicted TMHs: 2\nExp number of AAs in TMHs: 37.4699899999999\nExp number, first 60 AAs: 19.72699\nTotal prob of N-in: 0.96171\nPOSSIBLE N-term signal sequence\ninside 1 19\nTMhelix 20 39\noutside 40 325\nTMhelix 326 348\ninside 349 372

16427	GCF_002048065.1_ASM204806v1	Salmonella enterica subsp. enterica serovar Cubana	Proteobacteria; Gammaproteobacteria; Enterobacterales; Enterobacteriaceae; Salmonella; Salmonella enterica; Salmonella enterica subsp. enterica	MTGDNTLITSHGINRRDFMKLCAALAATMGLSSKAAAEMAESVSNPQRPPVIWIGAQECT	WP_000145429.1	hydrogenase 2 small subunit [Salmonella enterica]	Length: 372\nNumber of predicted TMHs: 1\nExp number of AAs in TMHs: 24.51309\nExp number, first 60 AAs: 0.2682\nTotal prob of N-in: 0.14402\noutside 1 331\nTMhelix 332 354\ninside 355 372

16428	GCF_002045265.1_ASM204526v1	Salmonella enterica subsp. enterica serovar Cubana	Proteobacteria; Gammaproteobacteria; Enterobacterales; Enterobacteriaceae; Salmonella; Salmonella enterica; Salmonella enterica subsp. enterica	MTGDNTLITSHGINRRDFMKLCAALAATMGLSSKAAAEMAESVSNPQRPPVIWIGAQECT	WP_000145429.1	hydrogenase 2 small subunit [Salmonella enterica]	Length: 372\nNumber of predicted TMHs: 1\nExp number of AAs in TMHs: 24.51309\nExp number, first 60 AAs: 0.2682\nTotal prob of N-in: 0.14402\noutside 1 331\nTMhelix 332 354\ninside 355 372

16429	GCF_002048065.1_ASM204806v1	Salmonella enterica subsp. enterica serovar Cubana	Proteobacteria; Gammaproteobacteria; Enterobacterales; Enterobacteriaceae; Salmonella; Salmonella enterica; Salmonella enterica subsp. enterica	MNNEETFYQAMRRKGVTRRSFLKFCSLAATSLGLGAGMTPKIAWALENKPRIPVVWIHGL	WP_001058312.1	hydrogenase [Salmonella enterica]	Length: 372\nNumber of predicted TMHs: 2\nExp number of AAs in TMHs: 36.51738\nExp number, first 60 AAs: 19.72849\nTotal prob of N-in: 0.96170\nPOSSIBLE N-term signal sequence\ninside 1 19\nTMhelix 20 39\noutside 40 325\nTMhelix 326 348\ninside 349 372

16430	GCF_002045265.1_ASM204526v1	Salmonella enterica subsp. enterica serovar Cubana	Proteobacteria; Gammaproteobacteria; Enterobacterales; Enterobacteriaceae; Salmonella; Salmonella enterica; Salmonella enterica subsp. enterica	MNNEETFYQAMRRKGVTRRSFLKFCSLAATSLGLGAGMTPKIAWALENKPRIPVVWIHGL	WP_001058311.1	hydrogenase [Salmonella enterica]	Length: 372\nNumber of predicted TMHs: 2\nExp number of AAs in TMHs: 37.4699899999999\nExp number, first 60 AAs: 19.72699\nTotal prob of N-in: 0.96171\nPOSSIBLE N-term signal sequence\ninside 1 19\nTMhelix 20 39\noutside 40 325\nTMhelix 326 348\ninside 349 372

16431	GCF_002156345.1_ASM215634v1	Salmonella enterica subsp. enterica serovar Daytona	Proteobacteria; Gammaproteobacteria; Enterobacterales; Enterobacteriaceae; Salmonella; Salmonella enterica; Salmonella enterica subsp. enterica	MTGDNTLITSHGINRRDFMKLCAALAATMGLSSKAAAEMAESVSNPQRPPVIWIGAQECT	WP_000145429.1	hydrogenase 2 small subunit [Salmonella enterica]	Length: 372\nNumber of predicted TMHs: 1\nExp number of AAs in TMHs: 24.51309\nExp number, first 60 AAs: 0.2682\nTotal prob of N-in: 0.14402\noutside 1 331\nTMhelix 332 354\ninside 355 372

16432	GCF_002065855.1_ASM206585v1	Salmonella enterica subsp. enterica serovar Derby	Proteobacteria; Gammaproteobacteria; Enterobacterales; Enterobacteriaceae; Salmonella; Salmonella enterica; Salmonella enterica subsp. enterica	MNNEETFYQAMRRKGVTRRSFLKFCSLAATSLGLGAGMTPKIAWALENKPRIPVVWIHGL	WP_001058312.1	hydrogenase [Salmonella enterica]	Length: 372\nNumber of predicted TMHs: 2\nExp number of AAs in TMHs: 36.51738\nExp number, first 60 AAs: 19.72849\nTotal prob of N-in: 0.96170\nPOSSIBLE N-term signal sequence\ninside 1 19\nTMhelix 20 39\noutside 40 325\nTMhelix 326 348\ninside 349 372

16433	GCF_002065855.1_ASM206585v1	Salmonella enterica subsp. enterica serovar Derby	Proteobacteria; Gammaproteobacteria; Enterobacterales; Enterobacteriaceae; Salmonella; Salmonella enterica; Salmonella enterica subsp. enterica	MTGDNTLITSHGINRRDFMKLCAALAATMGLSSKAAAEMAESVSNPQRPPVIWIGAQECT	WP_000145429.1	hydrogenase 2 small subunit [Salmonella enterica]	Length: 372\nNumber of predicted TMHs: 1\nExp number of AAs in TMHs: 24.51309\nExp number, first 60 AAs: 0.2682\nTotal prob of N-in: 0.14402\noutside 1 331\nTMhelix 332 354\ninside 355 372

16434	GCF_002063355.1_ASM206335v1	Salmonella enterica subsp. enterica serovar Derby	Proteobacteria; Gammaproteobacteria; Enterobacterales; Enterobacteriaceae; Salmonella; Salmonella enterica; Salmonella enterica subsp. enterica	MTGDNTLITSHGINRRDFMKLCAALAATMGLSSKAAAEMAESVSNPQRPPVIWIGAQECT	WP_000145429.1	hydrogenase 2 small subunit [Salmonella enterica]	Length: 372\nNumber of

predicted TMHs: 1\nExp number of AAs in TMHs: 24.51309\nExp number, first 60 AAs: 0.2682\nTotal prob of N-in: 0.14402\noutside 1 331\nTMhelix 332 354\ninside 355 372

16435 GCF_002030215.1_ASM203021v1 Salmonella enterica subsp. enterica serovar Derby Proteobacteria; Gammaproteobacteria; Enterobacterales; Enterobacteriaceae; Salmonella; Salmonella enterica; Salmonella enterica subsp. enterica MTGDNTLITSHGINRRDFMKLCAALAATMGLSSKAAAEMAESVSNPQRPPVIWIGAQECT WP_000145429.1 hydrogenase 2 small subunit [Salmonella enterica] Length: 372\nNumber of predicted TMHs: 1\nExp number of AAs in TMHs: 24.51309\nExp number, first 60 AAs: 0.2682\nTotal prob of N-in: 0.14402\noutside 1 331\nTMhelix 332 354\ninside 355 372

16436 GCF_002063355.1_ASM206335v1 Salmonella enterica subsp. enterica serovar Derby Proteobacteria; Gammaproteobacteria; Enterobacterales; Enterobacteriaceae; Salmonella; Salmonella enterica; Salmonella enterica subsp. enterica MNNEETFYQAMRRKGVTRRSFLKFCSLAATSLGLGAGMTPKIAWALENKPRIPVVWIHGL WP_001058312.1 hydrogenase [Salmonella enterica] Length: 372\nNumber of predicted TMHs: 2\nExp number of AAs in TMHs: 36.51738\nExp number, first 60 AAs: 19.72849\nTotal prob of N-in: 0.96170\nPOSSIBLE N-term signal sequence\ninside 1 19\nTMhelix 20 39\noutside 40 325\nTMhelix 326 348\ninside 349 372

16437 GCF_001478085.1_Salmonella_enterica_CVM_N30705-R_v1.0 Salmonella enterica subsp. enterica serovar Derby Proteobacteria; Gammaproteobacteria; Enterobacterales; Enterobacteriaceae; Salmonella; Salmonella enterica; Salmonella enterica subsp. enterica MNNEETFYQAMRRKGVTRRSFLKFCSLAATSLGLGAGMTPKIAWALENKPRIPVVWIHGL WP_001058312.1 hydrogenase [Salmonella enterica] Length: 372\nNumber of predicted TMHs: 2\nExp number of AAs in TMHs: 36.51738\nExp number, first 60 AAs: 19.72849\nTotal prob of N-in: 0.96170\nPOSSIBLE N-term signal sequence\ninside 1 19\nTMhelix 20 39\noutside 40 325\nTMhelix 326 348\ninside 349 372

16438 GCF_002061135.1_ASM206113v1 Salmonella enterica subsp. enterica serovar Derby Proteobacteria; Gammaproteobacteria; Enterobacterales; Enterobacteriaceae; Salmonella; Salmonella enterica; Salmonella enterica subsp. enterica MNNEETFYQAMRRKGVTRRSFLKFCSLAATSLGLGAGMTPKIAWALENKPRIPVVWIHGL WP_001058312.1 hydrogenase [Salmonella enterica] Length: 372\nNumber of predicted TMHs: 2\nExp number of AAs in TMHs: 36.51738\nExp number, first 60 AAs: 19.72849\nTotal prob of N-in: 0.96170\nPOSSIBLE N-term signal sequence\ninside 1 19\nTMhelix 20 39\noutside 40 325\nTMhelix 326 348\ninside 349 372

16439 GCF_002031835.1_ASM203183v1 Salmonella enterica subsp. enterica serovar Derby Proteobacteria; Gammaproteobacteria; Enterobacterales; Enterobacteriaceae; Salmonella; Salmonella enterica; Salmonella enterica subsp. enterica MTGDNTLITSHGINRRDFMKLCAALAATMGLSSKAAAEMAESVSNPQRPPVIWIGAQECT WP_000145429.1 hydrogenase 2 small subunit [Salmonella enterica] Length: 372\nNumber of predicted TMHs: 1\nExp number of AAs in TMHs: 24.51309\nExp number, first 60 AAs: 0.2682\nTotal prob of N-in: 0.14402\noutside 1 331\nTMhelix 332 354\ninside 355 372

16440 GCF_002061135.1_ASM206113v1 Salmonella enterica subsp. enterica serovar Derby Proteobacteria; Gammaproteobacteria; Enterobacterales; Enterobacteriaceae; Salmonella; Salmonella enterica; Salmonella enterica subsp. enterica MTGDNTLITSHGINRRDFMKLCAALAATMGLSSKAAAEMAESVSNPQRPPVIWIGAQECT WP_000145429.1 hydrogenase 2 small subunit [Salmonella enterica] Length: 372\nNumber of predicted TMHs: 1\nExp number of AAs in TMHs: 24.51309\nExp number, first 60 AAs: 0.2682\nTotal prob of N-in: 0.14402\noutside 1 331\nTMhelix 332 354\ninside 355 372

16441 GCF_002030965.1_ASM203096v1 Salmonella enterica subsp. enterica serovar Derby Proteobacteria; Gammaproteobacteria; Enterobacterales; Enterobacteriaceae; Salmonella; Salmonella enterica; Salmonella enterica subsp. enterica MTGDNTLITSHGINRRDFMKLCAALAATMGLSSKAAAEMAESVSNPQRPPVIWIGAQECT WP_000145429.1 hydrogenase 2 small subunit [Salmonella enterica] Length: 372\nNumber of predicted TMHs: 1\nExp number of AAs in TMHs: 24.51309\nExp number, first 60 AAs: 0.2682\nTotal prob of N-in: 0.14402\noutside 1 331\nTMhelix 332 354\ninside 355 372

16442 GCF_001262435.1_Salmonella_enterica_CVM_N46856_v1.0 Salmonella enterica subsp. enterica serovar Derby Proteobacteria; Gammaproteobacteria; Enterobacterales; Enterobacteriaceae; Salmonella; Salmonella enterica; Salmonella enterica subsp. enterica MTGDNTLITSHGINRRDFMKLCAALAATMGLSSKAAAEMAESVSNPQRPPVIWIGAQECT WP_000145429.1 hydrogenase 2 small subunit [Salmonella enterica] Length: 372\nNumber of predicted TMHs: 1\nExp number of AAs in TMHs: 24.51309\nExp number, first 60 AAs: 0.2682\nTotal prob of N-in: 0.14402\noutside 1 331\nTMhelix 332 354\ninside 355 372

16443 GCF_001262435.1_Salmonella_enterica_CVM_N46856_v1.0 Salmonella enterica subsp. enterica serovar Derby Proteobacteria; Gammaproteobacteria; Enterobacterales; Enterobacteriaceae; Salmonella; Salmonella enterica;

Salmonella enterica subsp. enterica	MNNEETFYQAMRRKGVTRRSFLKFCSLAATSLGLGAGMTPKIAWALENKPRIPVVWIHGL	WP_001058312.1	hydrogenase [Salmonella enterica]	Length: 372\nNumber of predicted TMHs:  2\nExp number of AAs in TMHs: 36.51738\nExp number, first 60 AAs:  19.72849\nTotal prob of N-in:   0.96170\nPOSSIBLE N-term signal sequence\ninside   1   19\nTMhelix   20   39\noutside   40   325\nTMhelix   326   348\ninside   349   372

16444	GCF_002032085.1_ASM203208v1	Salmonella enterica subsp. enterica serovar Derby	Proteobacteria; Gammaproteobacteria; Enterobacterales; Enterobacteriaceae; Salmonella; Salmonella enterica; Salmonella enterica subsp. enterica	MTGDNTLITSHGINRRDFMKLCAALAATMGLSSKAAAEMAESVSNPQRPPVIWIGAQECT	WP_000145429.1	hydrogenase 2 small subunit [Salmonella enterica]	Length: 372\nNumber of predicted TMHs:  1\nExp number of AAs in TMHs: 24.51309\nExp number, first 60 AAs:  0.2682\nTotal prob of N-in:   0.14402\noutside   1   331\nTMhelix   332   354\ninside   355   372

16445	GCF_002032085.1_ASM203208v1	Salmonella enterica subsp. enterica serovar Derby	Proteobacteria; Gammaproteobacteria; Enterobacterales; Enterobacteriaceae; Salmonella; Salmonella enterica; Salmonella enterica subsp. enterica	MNNEETFYQAMRRKGVTRRSFLKFCSLAATSLGLGAGMTPKIAWALENKPRIPVVWIHGL	WP_001058312.1	hydrogenase [Salmonella enterica]	Length: 372\nNumber of predicted TMHs:  2\nExp number of AAs in TMHs: 36.51738\nExp number, first 60 AAs:  19.72849\nTotal prob of N-in:   0.96170\nPOSSIBLE N-term signal sequence\ninside   1   19\nTMhelix   20   39\noutside   40   325\nTMhelix   326   348\ninside   349   372

16446	GCF_001613955.1_ASM161395v1	Salmonella enterica subsp. enterica serovar Derby	Proteobacteria; Gammaproteobacteria; Enterobacterales; Enterobacteriaceae; Salmonella; Salmonella enterica; Salmonella enterica subsp. enterica	MTGDNTLITSHGINRRDFMKLCAALAATMGLSSKAAAEMAESVSNPQRPPVIWIGAQECT	WP_000145429.1	hydrogenase 2 small subunit [Salmonella enterica]	Length: 372\nNumber of predicted TMHs:  1\nExp number of AAs in TMHs: 24.51309\nExp number, first 60 AAs:  0.2682\nTotal prob of N-in:   0.14402\noutside   1   331\nTMhelix   332   354\ninside   355   372

16447	GCF_001242475.1_Salmonella_enterica_CVM_N45941_v1.0	Salmonella enterica subsp. enterica serovar Derby	Proteobacteria; Gammaproteobacteria; Enterobacterales; Enterobacteriaceae; Salmonella; Salmonella enterica; Salmonella enterica subsp. enterica	MTGDNTLITSHGINRRDFMKLCAALAATMGLSSKAAAEMAESVSNPQRPPVIWIGAQECT	WP_000145429.1	hydrogenase 2 small subunit [Salmonella enterica]	Length: 372\nNumber of predicted TMHs:  1\nExp number of AAs in TMHs: 24.51309\nExp number, first 60 AAs:  0.2682\nTotal prob of N-in:   0.14402\noutside   1   331\nTMhelix   332   354\ninside   355   372

16448	GCF_001478085.1_Salmonella_enterica_CVM_N30705-R_v1.0	Salmonella enterica subsp. enterica serovar Derby	Proteobacteria; Gammaproteobacteria; Enterobacterales; Enterobacteriaceae; Salmonella; Salmonella enterica; Salmonella enterica subsp. enterica	MTGDNTLITSHGINRRDFMKLCAALAATMGLSSKAAAEMAESVSNPQRPPVIWIGAQECT	WP_000145429.1	hydrogenase 2 small subunit [Salmonella enterica]	Length: 372\nNumber of predicted TMHs:  1\nExp number of AAs in TMHs: 24.51309\nExp number, first 60 AAs:  0.2682\nTotal prob of N-in:   0.14402\noutside   1   331\nTMhelix   332   354\ninside   355   372

16449	GCF_001242475.1_Salmonella_enterica_CVM_N45941_v1.0	Salmonella enterica subsp. enterica serovar Derby	Proteobacteria; Gammaproteobacteria; Enterobacterales; Enterobacteriaceae; Salmonella; Salmonella enterica; Salmonella enterica subsp. enterica	MNNEETFYQAMRRKGVTRRSFLKFCSLAATSLGLGAGMTPKIAWALENKPRIPVVWIHGL	WP_001058312.1	hydrogenase [Salmonella enterica]	Length: 372\nNumber of predicted TMHs:  2\nExp number of AAs in TMHs: 36.51738\nExp number, first 60 AAs:  19.72849\nTotal prob of N-in:   0.96170\nPOSSIBLE N-term signal sequence\ninside   1   19\nTMhelix   20   39\noutside   40   325\nTMhelix   326   348\ninside   349   372

16450	GCF_001244515.1_Salmonella_enterica_CVM_N48685_v1.0	Salmonella enterica subsp. enterica serovar Derby	Proteobacteria; Gammaproteobacteria; Enterobacterales; Enterobacteriaceae; Salmonella; Salmonella enterica; Salmonella enterica subsp. enterica	MTGDNTLITSHGINRRDFMKLCAALAATMGLSSKAAAEMAESVSNPQRPPVIWIGAQECT	WP_000145429.1	hydrogenase 2 small subunit [Salmonella enterica]	Length: 372\nNumber of predicted TMHs:  1\nExp number of AAs in TMHs: 24.51309\nExp number, first 60 AAs:  0.2682\nTotal prob of N-in:   0.14402\noutside   1   331\nTMhelix   332   354\ninside   355   372

16451	GCF_001244515.1_Salmonella_enterica_CVM_N48685_v1.0	Salmonella enterica subsp. enterica serovar Derby	Proteobacteria; Gammaproteobacteria; Enterobacterales; Enterobacteriaceae; Salmonella; Salmonella enterica; Salmonella enterica subsp. enterica	MNNEETFYQAMRRKGVTRRSFLKFCSLAATSLGLGAGMTPKIAWALENKPRIPVVWIHGL	WP_001058312.1	hydrogenase [Salmonella enterica]	Length: 372\nNumber of predicted TMHs:  2\nExp number of AAs in TMHs: 36.51738\nExp number, first 60 AAs:  19.72849\nTotal prob of N-in:   0.96170\nPOSSIBLE N-term signal sequence\ninside   1   19\nTMhelix   20   39\noutside   40   325\nTMhelix   326   348\ninside   349   372

16452 GCF_001246825.1_Salmonella_enterica_CVM_N51280_v1.0 Salmonella enterica subsp. enterica serovar Derby Proteobacteria; Gammaproteobacteria; Enterobacterales; Enterobacteriaceae; Salmonella; Salmonella enterica; Salmonella enterica subsp. enterica MTGDNTLITSHGINRRDFMKLCAALAATMGLSSKAAAEMAESVSNPQRPPVIWIGAQECT WP_000145429.1 hydrogenase 2 small subunit [Salmonella enterica] Length: 372\nNumber of predicted TMHs: 1\nExp number of AAs in TMHs: 24.51309\nExp number, first 60 AAs: 0.2682\nTotal prob of N-in: 0.14402\noutside 1 331\nTMhelix 332 354\ninside 355 372

16453 GCF_001040825.1_ASM104082v1 Salmonella enterica subsp. enterica serovar Derby Proteobacteria; Gammaproteobacteria; Enterobacterales; Enterobacteriaceae; Salmonella; Salmonella enterica; Salmonella enterica subsp. enterica MTGDNTLITSHGINRRDFMKLCAALAATMGLSSKAAAEMAESVSNPQRPPVIWIGAQECT WP_000145429.1 hydrogenase 2 small subunit [Salmonella enterica] Length: 372\nNumber of predicted TMHs: 1\nExp number of AAs in TMHs: 24.51309\nExp number, first 60 AAs: 0.2682\nTotal prob of N-in: 0.14402\noutside 1 331\nTMhelix 332 354\ninside 355 372

16454 GCF_001244725.1_Salmonella_enterica_CVM_N48686_v1.0 Salmonella enterica subsp. enterica serovar Derby Proteobacteria; Gammaproteobacteria; Enterobacterales; Enterobacteriaceae; Salmonella; Salmonella enterica; Salmonella enterica subsp. enterica MTGDNTLITSHGINRRDFMKLCAALAATMGLSSKAAAEMAESVSNPQRPPVIWIGAQECT WP_000145429.1 hydrogenase 2 small subunit [Salmonella enterica] Length: 372\nNumber of predicted TMHs: 1\nExp number of AAs in TMHs: 24.51309\nExp number, first 60 AAs: 0.2682\nTotal prob of N-in: 0.14402\noutside 1 331\nTMhelix 332 354\ninside 355 372

16455 GCF_001040825.1_ASM104082v1 Salmonella enterica subsp. enterica serovar Derby Proteobacteria; Gammaproteobacteria; Enterobacterales; Enterobacteriaceae; Salmonella; Salmonella enterica; Salmonella enterica subsp. enterica MNNEETFYQAMRRKGVTRRSFLKFCSLAATSLGLGAGMTPKIAWALENKPRIPVVWIHGL WP_001058312.1 hydrogenase [Salmonella enterica] Length: 372\nNumber of predicted TMHs: 2\nExp number of AAs in TMHs: 36.51738\nExp number, first 60 AAs: 19.72849\nTotal prob of N-in: 0.96170\nPOSSIBLE N-term signal sequence\ninside 1 19\nTMhelix 20 39\noutside 40 325\nTMhelix 326 348\ninside 349 372

16456 GCF_001244725.1_Salmonella_enterica_CVM_N48686_v1.0 Salmonella enterica subsp. enterica serovar Derby Proteobacteria; Gammaproteobacteria; Enterobacterales; Enterobacteriaceae; Salmonella; Salmonella enterica; Salmonella enterica subsp. enterica MNNEETFYQAMRRKGVTRRSFLKFCSLAATSLGLGAGMTPKIAWALENKPRIPVVWIHGL WP_001058312.1 hydrogenase [Salmonella enterica] Length: 372\nNumber of predicted TMHs: 2\nExp number of AAs in TMHs: 36.51738\nExp number, first 60 AAs: 19.72849\nTotal prob of N-in: 0.96170\nPOSSIBLE N-term signal sequence\ninside 1 19\nTMhelix 20 39\noutside 40 325\nTMhelix 326 348\ninside 349 372

16457 GCF_001246825.1_Salmonella_enterica_CVM_N51280_v1.0 Salmonella enterica subsp. enterica serovar Derby Proteobacteria; Gammaproteobacteria; Enterobacterales; Enterobacteriaceae; Salmonella; Salmonella enterica; Salmonella enterica subsp. enterica MNNEETFYQAMRRKGVTRRSFLKFCSLAATSLGLGAGMTPKIAWALENKPRIPVVWIHGL WP_001058312.1 hydrogenase [Salmonella enterica] Length: 372\nNumber of predicted TMHs: 2\nExp number of AAs in TMHs: 36.51738\nExp number, first 60 AAs: 19.72849\nTotal prob of N-in: 0.96170\nPOSSIBLE N-term signal sequence\ninside 1 19\nTMhelix 20 39\noutside 40 325\nTMhelix 326 348\ninside 349 372

16458 GCF_002066555.1_ASM206655v1 Salmonella enterica subsp. enterica serovar Derby Proteobacteria; Gammaproteobacteria; Enterobacterales; Enterobacteriaceae; Salmonella; Salmonella enterica; Salmonella enterica subsp. enterica MTGDNTLITSHGINRRDFMKLCAALAATMGLSSKAAAEMAESVSNPQRPPVIWIGAQECT WP_000145429.1 hydrogenase 2 small subunit [Salmonella enterica] Length: 372\nNumber of predicted TMHs: 1\nExp number of AAs in TMHs: 24.51309\nExp number, first 60 AAs: 0.2682\nTotal prob of N-in: 0.14402\noutside 1 331\nTMhelix 332 354\ninside 355 372

16459 GCF_002065865.1_ASM206586v1 Salmonella enterica subsp. enterica serovar Derby Proteobacteria; Gammaproteobacteria; Enterobacterales; Enterobacteriaceae; Salmonella; Salmonella enterica; Salmonella enterica subsp. enterica MTGDNTLITSHGINRRDFMKLCAALAATMGLSSKAAAEMAESVSNPQRPPVIWIGAQECT WP_000145429.1 hydrogenase 2 small subunit [Salmonella enterica] Length: 372\nNumber of predicted TMHs: 1\nExp number of AAs in TMHs: 24.51309\nExp number, first 60 AAs: 0.2682\nTotal prob of N-in: 0.14402\noutside 1 331\nTMhelix 332 354\ninside 355 372

16460 GCF_002066555.1_ASM206655v1 Salmonella enterica subsp. enterica serovar Derby Proteobacteria; Gammaproteobacteria; Enterobacterales; Enterobacteriaceae; Salmonella; Salmonella enterica; Salmonella enterica subsp. enterica MNNEETFYQAMRRKGVTRRSFLKFCSLAATSLGLGAGMTPKIAWALENKPRIPVVWIHGL WP_001058312.1 hydrogenase [Salmonella enterica] Length: 372\nNumber of predicted TMHs: 2\nExp

number of AAs in TMHs: 36.51738\nExp number, first 60 AAs: 19.72849\nTotal prob of N-in: 0.96170\nPOSSIBLE N-term signal sequence\ninside 1 19\nTMhelix 20 39\noutside 40 325\nTMhelix 326 348\ninside 349 372

16461 GCF_002065865.1_ASM206586v1 Salmonella enterica subsp. enterica serovar Derby Proteobacteria; Gammaproteobacteria; Enterobacterales; Enterobacteriaceae; Salmonella; Salmonella enterica; Salmonella enterica subsp. enterica MNNEETFYQAMRRKGVTRRSFLKFCSLAATSLGLGAGMTPKIAWALENKPRIPVVWIHGL WP_001058312.1 hydrogenase [Salmonella enterica] Length: 372\nNumber of predicted TMHs: 2\nExp number of AAs in TMHs: 36.51738\nExp number, first 60 AAs: 19.72849\nTotal prob of N-in: 0.96170\nPOSSIBLE N-term signal sequence\ninside 1 19\nTMhelix 20 39\noutside 40 325\nTMhelix 326 348\ninside 349 372

16462 GCF_002031835.1_ASM203183v1 Salmonella enterica subsp. enterica serovar Derby Proteobacteria; Gammaproteobacteria; Enterobacterales; Enterobacteriaceae; Salmonella; Salmonella enterica; Salmonella enterica subsp. enterica MNNEETFYQAMRRKGVTRRSFLKFCSLAATSLGLGAGMTPKIAWALENKPRIPVVWIHGL WP_001058312.1 hydrogenase [Salmonella enterica] Length: 372\nNumber of predicted TMHs: 2\nExp number of AAs in TMHs: 36.51738\nExp number, first 60 AAs: 19.72849\nTotal prob of N-in: 0.96170\nPOSSIBLE N-term signal sequence\ninside 1 19\nTMhelix 20 39\noutside 40 325\nTMhelix 326 348\ninside 349 372

16463 GCF_002030965.1_ASM203096v1 Salmonella enterica subsp. enterica serovar Derby Proteobacteria; Gammaproteobacteria; Enterobacterales; Enterobacteriaceae; Salmonella; Salmonella enterica; Salmonella enterica subsp. enterica MNNEETFYQAMRRKGVTRRSFLKFCSLAATSLGLGAGMTPKIAWALENKPRIPVVWIHGL WP_001058312.1 hydrogenase [Salmonella enterica] Length: 372\nNumber of predicted TMHs: 2\nExp number of AAs in TMHs: 36.51738\nExp number, first 60 AAs: 19.72849\nTotal prob of N-in: 0.96170\nPOSSIBLE N-term signal sequence\ninside 1 19\nTMhelix 20 39\noutside 40 325\nTMhelix 326 348\ninside 349 372

16464 GCF_001613955.1_ASM161395v1 Salmonella enterica subsp. enterica serovar Derby Proteobacteria; Gammaproteobacteria; Enterobacterales; Enterobacteriaceae; Salmonella; Salmonella enterica; Salmonella enterica subsp. enterica MNNEETFYQAMRRKGVTRRSFLKFCSLAATSLGLGAGMTPKIAWALENKPRIPVVWIHGL WP_001058312.1 hydrogenase [Salmonella enterica] Length: 372\nNumber of predicted TMHs: 2\nExp number of AAs in TMHs: 36.51738\nExp number, first 60 AAs: 19.72849\nTotal prob of N-in: 0.96170\nPOSSIBLE N-term signal sequence\ninside 1 19\nTMhelix 20 39\noutside 40 325\nTMhelix 326 348\ninside 349 372

16465 GCF_002032345.1_ASM203234v1 Salmonella enterica subsp. enterica serovar Derby Proteobacteria; Gammaproteobacteria; Enterobacterales; Enterobacteriaceae; Salmonella; Salmonella enterica; Salmonella enterica subsp. enterica MTGDNTLITSHGINRRDFMKLCAALAATMGLSSKAAAEMAESVSNPQRPPVIWIGAQECT WP_000145429.1 hydrogenase 2 small subunit [Salmonella enterica] Length: 372\nNumber of predicted TMHs: 1\nExp number of AAs in TMHs: 24.51309\nExp number, first 60 AAs: 0.2682\nTotal prob of N-in: 0.14402\noutside 1 331\nTMhelix 332 354\ninside 355 372

16466 GCF_002032345.1_ASM203234v1 Salmonella enterica subsp. enterica serovar Derby Proteobacteria; Gammaproteobacteria; Enterobacterales; Enterobacteriaceae; Salmonella; Salmonella enterica; Salmonella enterica subsp. enterica MNNEETFYQAMRRKGVTRRSFLKFCSLAATSLGLGAGMTPKIAWALENKPRIPVVWIHGL WP_001058312.1 hydrogenase [Salmonella enterica] Length: 372\nNumber of predicted TMHs: 2\nExp number of AAs in TMHs: 36.51738\nExp number, first 60 AAs: 19.72849\nTotal prob of N-in: 0.96170\nPOSSIBLE N-term signal sequence\ninside 1 19\nTMhelix 20 39\noutside 40 325\nTMhelix 326 348\ninside 349 372

16467 GCF_001478025.1_Salmonella_enterica_CVM_N30679-SQ_v1.0 Salmonella enterica subsp. enterica serovar Derby Proteobacteria; Gammaproteobacteria; Enterobacterales; Enterobacteriaceae; Salmonella; Salmonella enterica; Salmonella enterica subsp. enterica MNNEETFYQAMRRKGVTRRSFLKFCSLAATSLGLGAGMTPKIAWALENKPRIPVVWIHGL WP_001058312.1 hydrogenase [Salmonella enterica] Length: 372\nNumber of predicted TMHs: 2\nExp number of AAs in TMHs: 36.51738\nExp number, first 60 AAs: 19.72849\nTotal prob of N-in: 0.96170\nPOSSIBLE N-term signal sequence\ninside 1 19\nTMhelix 20 39\noutside 40 325\nTMhelix 326 348\ninside 349 372

16468 GCF_002030215.1_ASM203021v1 Salmonella enterica subsp. enterica serovar Derby Proteobacteria; Gammaproteobacteria; Enterobacterales; Enterobacteriaceae; Salmonella; Salmonella enterica; Salmonella enterica subsp. enterica MNNEETFYQAMRRKGVTRRSFLKFCSLAATSLGLGAGMTPKIAWALENKPRIPVVWIHGL WP_001058312.1 hydrogenase [Salmonella enterica] Length: 372\nNumber of predicted TMHs: 2\nExp number of AAs in TMHs: 36.51738\nExp number, first 60 AAs: 19.72849\nTotal prob of N-in: 0.96170\nPOSSIBLE N-term signal sequence\ninside 1 19\nTMhelix 20 39\noutside 40 325\nTMhelix 326 348\ninside 349 372

16469 GCF_001478465.1_Salmonella_enterica_CVM_N40386-R_v1.0 Salmonella enterica subsp. enterica serovar Derby Proteobacteria; Gammaproteobacteria; Enterobacterales; Enterobacteriaceae; Salmonella; Salmonella enterica;

Salmonella enterica subsp. enterica	MNNEETFYQAMRRKGVTRRSFLKFCSLAATSLGLGAGMTPKIAWALENKPRIPVVWIHGL
	WP_001058312.1	hydrogenase [Salmonella enterica]	Length: 372\nNumber of predicted TMHs:  2\nExp number of AAs in TMHs: 36.51738\nExp number, first 60 AAs:  19.72849\nTotal prob of N-in:   0.96170\nPOSSIBLE N-term signal sequence\ninside   1   19\nTMhelix   20   39\noutside   40   325\nTMhelix   326   348\ninside   349   372

16470	GCF_001475515.1_ASM147551v1	Salmonella enterica subsp. enterica serovar Derby	Proteobacteria; Gammaproteobacteria; Enterobacterales; Enterobacteriaceae; Salmonella; Salmonella enterica; Salmonella enterica subsp. enterica	MTGDNTLITSHGINRRDFMKLCAALAATMGLSSKAAAEMAESVSNPQRPPVIWIGAQECT	WP_000145429.1	hydrogenase 2 small subunit [Salmonella enterica]	Length: 372\nNumber of predicted TMHs:  1\nExp number of AAs in TMHs: 24.51309\nExp number, first 60 AAs:  0.2682\nTotal prob of N-in:   0.14402\noutside   1   331\nTMhelix   332   354\ninside   355   372

16471	GCF_001475515.1_ASM147551v1	Salmonella enterica subsp. enterica serovar Derby	Proteobacteria; Gammaproteobacteria; Enterobacterales; Enterobacteriaceae; Salmonella; Salmonella enterica; Salmonella enterica subsp. enterica	MNNEETFYQAMRRKGVTRRSFLKFCSLAATSLGLGAGMTPKIAWALENKPRIPVVWIHGL	WP_001058312.1	hydrogenase [Salmonella enterica]	Length: 372\nNumber of predicted TMHs:  2\nExp number of AAs in TMHs: 36.51738\nExp number, first 60 AAs:  19.72849\nTotal prob of N-in:   0.96170\nPOSSIBLE N-term signal sequence\ninside   1   19\nTMhelix   20   39\noutside   40   325\nTMhelix   326   348\ninside   349   372

16472	GCF_001478465.1_Salmonella_enterica_CVM_N40386-R_v1.0	Salmonella enterica subsp. enterica serovar Derby	Proteobacteria; Gammaproteobacteria; Enterobacterales; Enterobacteriaceae; Salmonella; Salmonella enterica; Salmonella enterica subsp. enterica	MTGDNTLITSHGINRRDFMKLCAALAATMGLSSKAAAEMAESVSNPQRPPVIWIGAQECT	WP_000145429.1	hydrogenase 2 small subunit [Salmonella enterica]	Length: 372\nNumber of predicted TMHs:  1\nExp number of AAs in TMHs: 24.51309\nExp number, first 60 AAs:  0.2682\nTotal prob of N-in:   0.14402\noutside   1   331\nTMhelix   332   354\ninside   355   372

16473	GCF_001481215.1_Salmonella_enterica_CVM_N42507_v1.0	Salmonella enterica subsp. enterica serovar Derby	Proteobacteria; Gammaproteobacteria; Enterobacterales; Enterobacteriaceae; Salmonella; Salmonella enterica; Salmonella enterica subsp. enterica	MNNEETFYQAMRRKGVTRRSFLKFCSLAATSLGLGAGMTPKIAWALENKPRIPVVWIHGL	WP_001058312.1	hydrogenase [Salmonella enterica]	Length: 372\nNumber of predicted TMHs:  2\nExp number of AAs in TMHs: 36.51738\nExp number, first 60 AAs:  19.72849\nTotal prob of N-in:   0.96170\nPOSSIBLE N-term signal sequence\ninside   1   19\nTMhelix   20   39\noutside   40   325\nTMhelix   326   348\ninside   349   372

16474	GCF_001478025.1_Salmonella_enterica_CVM_N30679-SQ_v1.0	Salmonella enterica subsp. enterica serovar Derby	Proteobacteria; Gammaproteobacteria; Enterobacterales; Enterobacteriaceae; Salmonella; Salmonella enterica; Salmonella enterica subsp. enterica	MTGDNTLITSHGINRRDFMKLCAALAATMGLSSKAAAEMAESVSNPQRPPVIWIGAQECT	WP_000145429.1	hydrogenase 2 small subunit [Salmonella enterica]	Length: 372\nNumber of predicted TMHs:  1\nExp number of AAs in TMHs: 24.51309\nExp number, first 60 AAs:  0.2682\nTotal prob of N-in:   0.14402\noutside   1   331\nTMhelix   332   354\ninside   355   372

16475	GCF_001613945.1_ASM161394v1	Salmonella enterica subsp. enterica serovar Derby	Proteobacteria; Gammaproteobacteria; Enterobacterales; Enterobacteriaceae; Salmonella; Salmonella enterica; Salmonella enterica subsp. enterica	MNNEETFYQAMRRKGVTRRSFLKFCSLAATSLGLGAGMTPKIAWALENKPRIPVVWIHGL	WP_001058312.1	hydrogenase [Salmonella enterica]	Length: 372\nNumber of predicted TMHs:  2\nExp number of AAs in TMHs: 36.51738\nExp number, first 60 AAs:  19.72849\nTotal prob of N-in:   0.96170\nPOSSIBLE N-term signal sequence\ninside   1   19\nTMhelix   20   39\noutside   40   325\nTMhelix   326   348\ninside   349   372

16476	GCF_001613945.1_ASM161394v1	Salmonella enterica subsp. enterica serovar Derby	Proteobacteria; Gammaproteobacteria; Enterobacterales; Enterobacteriaceae; Salmonella; Salmonella enterica; Salmonella enterica subsp. enterica	MTGDNTLITSHGINRRDFMKLCAALAATMGLSSKAAAEMAESVSNPQRPPVIWIGAQECT	WP_000145429.1	hydrogenase 2 small subunit [Salmonella enterica]	Length: 372\nNumber of predicted TMHs:  1\nExp number of AAs in TMHs: 24.51309\nExp number, first 60 AAs:  0.2682\nTotal prob of N-in:   0.14402\noutside   1   331\nTMhelix   332   354\ninside   355   372

16477	GCF_001481215.1_Salmonella_enterica_CVM_N42507_v1.0	Salmonella enterica subsp. enterica serovar Derby	Proteobacteria; Gammaproteobacteria; Enterobacterales; Enterobacteriaceae; Salmonella; Salmonella enterica; Salmonella enterica subsp. enterica	MTGDNTLITSHGINRRDFMKLCAALAATMGLSSKAAAEMAESVSNPQRPPVIWIGAQECT	WP_000145429.1	hydrogenase 2 small subunit [Salmonella enterica]	Length: 372\nNumber of predicted TMHs:  1\nExp number of AAs in TMHs: 24.51309\nExp number, first 60 AAs:  0.2682\nTotal prob of N-in:   0.14402\noutside   1   331\nTMhelix   332   354\ninside   355   372

16478 GCF_001480725.1_Salmonella_enterica_CVM_N42512_v1.0 Salmonella enterica subsp. enterica serovar Dublin Proteobacteria; Gammaproteobacteria; Enterobacterales; Enterobacteriaceae; Salmonella; Salmonella enterica; Salmonella enterica subsp. enterica MTGDNTLITSHGINRRDFMKLCAALAATMGLSSKAAAEMAESVSNPQRPPVIWIGAQECT WP_000145428.1 hydrogenase 2 small subunit [Salmonella enterica] Length: 372\nNumber of predicted TMHs: 1\nExp number of AAs in TMHs: 24.50677\nExp number, first 60 AAs: 0.2682\nTotal prob of N-in: 0.14390\noutside 1 331\nTMhelix 332 354\ninside 355 372

16479 GCF_002057025.1_ASM205702v1 Salmonella enterica subsp. enterica serovar Dublin Proteobacteria; Gammaproteobacteria; Enterobacterales; Enterobacteriaceae; Salmonella; Salmonella enterica; Salmonella enterica subsp. enterica MNNEETLYQAMRRKGVTRRSFLKFCSLAATSLGLGAGMTPKIAWALENKPRIPVVWIHGL WP_001752445.1 hydrogenase-1 small chain [Salmonella enterica] Length: 372\nNumber of predicted TMHs: 2\nExp number of AAs in TMHs: 36.5123\nExp number, first 60 AAs: 19.72341\nTotal prob of N-in: 0.96144\nPOSSIBLE N-term signal sequence\ninside 1 19\nTMhelix 20 39\noutside 40 325\nTMhelix 326 348\ninside 349 372

16480 GCF_001480245.1_Salmonella_enterica_CVM_N40387-SQ_v1.0 Salmonella enterica subsp. enterica serovar Dublin Proteobacteria; Gammaproteobacteria; Enterobacterales; Enterobacteriaceae; Salmonella; Salmonella enterica; Salmonella enterica subsp. enterica MNNEETLYQAMRRKGVTRRSFLKFCSLAATSLGLGAGMTPKIAWALENKPRIPVVWIHGL WP_001752445.1 hydrogenase-1 small chain [Salmonella enterica] Length: 372\nNumber of predicted TMHs: 2\nExp number of AAs in TMHs: 36.5123\nExp number, first 60 AAs: 19.72341\nTotal prob of N-in: 0.96144\nPOSSIBLE N-term signal sequence\ninside 1 19\nTMhelix 20 39\noutside 40 325\nTMhelix 326 348\ninside 349 372

16481 GCF_001480725.1_Salmonella_enterica_CVM_N42512_v1.0 Salmonella enterica subsp. enterica serovar Dublin Proteobacteria; Gammaproteobacteria; Enterobacterales; Enterobacteriaceae; Salmonella; Salmonella enterica; Salmonella enterica subsp. enterica MNNEETLYQAMRRKGVTRRSFLKFCSLAATSLGLGAGMTPKIAWALENKPRIPVVWIHGL WP_001752445.1 hydrogenase-1 small chain [Salmonella enterica] Length: 372\nNumber of predicted TMHs: 2\nExp number of AAs in TMHs: 36.5123\nExp number, first 60 AAs: 19.72341\nTotal prob of N-in: 0.96144\nPOSSIBLE N-term signal sequence\ninside 1 19\nTMhelix 20 39\noutside 40 325\nTMhelix 326 348\ninside 349 372

16482 GCF_001954995.1_ASM195499v1 Salmonella enterica subsp. enterica serovar Dublin Proteobacteria; Gammaproteobacteria; Enterobacterales; Enterobacteriaceae; Salmonella; Salmonella enterica; Salmonella enterica subsp. enterica MTGDNTLITSHGINRRDFMKLCAALAATMGLSSKAAAEMAESVSNPQRPPVIWIGAQECT WP_000145428.1 hydrogenase 2 small subunit [Salmonella enterica] Length: 372\nNumber of predicted TMHs: 1\nExp number of AAs in TMHs: 24.50677\nExp number, first 60 AAs: 0.2682\nTotal prob of N-in: 0.14390\noutside 1 331\nTMhelix 332 354\ninside 355 372

16483 GCF_001954985.1_ASM195498v1 Salmonella enterica subsp. enterica serovar Dublin Proteobacteria; Gammaproteobacteria; Enterobacterales; Enterobacteriaceae; Salmonella; Salmonella enterica; Salmonella enterica subsp. enterica MNNEETLYQAMRRKGVTRRSFLKFCSLAATSLGLGAGMTPKIAWALENKPRIPVVWIHGL WP_001752445.1 hydrogenase-1 small chain [Salmonella enterica] Length: 372\nNumber of predicted TMHs: 2\nExp number of AAs in TMHs: 36.5123\nExp number, first 60 AAs: 19.72341\nTotal prob of N-in: 0.96144\nPOSSIBLE N-term signal sequence\ninside 1 19\nTMhelix 20 39\noutside 40 325\nTMhelix 326 348\ninside 349 372

16484 GCF_001480245.1_Salmonella_enterica_CVM_N40387-SQ_v1.0 Salmonella enterica subsp. enterica serovar Dublin Proteobacteria; Gammaproteobacteria; Enterobacterales; Enterobacteriaceae; Salmonella; Salmonella enterica; Salmonella enterica subsp. enterica MTGDNTLITSHGINRRDFMKLCAALAATMGLSSKAAAEMAESVSNPQRPPVIWIGAQECT WP_000145428.1 hydrogenase 2 small subunit [Salmonella enterica] Length: 372\nNumber of predicted TMHs: 1\nExp number of AAs in TMHs: 24.50677\nExp number, first 60 AAs: 0.2682\nTotal prob of N-in: 0.14390\noutside 1 331\nTMhelix 332 354\ninside 355 372

16485 GCF_001954735.1_ASM195473v1 Salmonella enterica subsp. enterica serovar Dublin Proteobacteria; Gammaproteobacteria; Enterobacterales; Enterobacteriaceae; Salmonella; Salmonella enterica; Salmonella enterica subsp. enterica MNNEETLYQAMRRKGVTRRSFLKFCSLAATSLGLGAGMTPKIAWALENKPRIPVVWIHGL WP_001752445.1 hydrogenase-1 small chain [Salmonella enterica] Length: 372\nNumber of predicted TMHs: 2\nExp number of AAs in TMHs: 36.5123\nExp number, first 60 AAs: 19.72341\nTotal prob of N-in: 0.96144\nPOSSIBLE N-term signal sequence\ninside 1 19\nTMhelix 20 39\noutside 40 325\nTMhelix 326 348\ninside 349 372

16486 GCF_001954735.1_ASM195473v1 Salmonella enterica subsp. enterica serovar Dublin Proteobacteria; Gammaproteobacteria; Enterobacterales; Enterobacteriaceae; Salmonella; Salmonella enterica; Salmonella enterica subsp. enterica MTGDNTLITSHGINRRDFMKLCAALAATMGLSSKAAAEMAESVSNPQRPPVIWIGAQECT WP_000145428.1 hydrogenase 2 small subunit [Salmonella enterica] Length: 372\nNumber of

predicted TMHs: 1\nExp number of AAs in TMHs: 24.50677\nExp number, first 60 AAs: 0.2682\nTotal prob of N-in: 0.14390\noutside 1 331\nTMhelix 332 354\ninside 355 372

16487 GCF_001954705.1_ASM195470v1 Salmonella enterica subsp. enterica serovar Dublin Proteobacteria; Gammaproteobacteria; Enterobacterales; Enterobacteriaceae; Salmonella; Salmonella enterica; Salmonella enterica subsp. enterica MNNEETLYQAMRRKGVTRRSFLKFCSLAATSLGLGAGMTPKIAWALENKPRIPVVWIHGL WP_001752445.1 hydrogenase-1 small chain [Salmonella enterica] Length: 372\nNumber of predicted TMHs: 2\nExp number of AAs in TMHs: 36.5123\nExp number, first 60 AAs: 19.72341\nTotal prob of N-in: 0.96144\nPOSSIBLE N-term signal sequence\ninside 1 19\nTMhelix 20 39\noutside 40 325\nTMhelix 326 348\ninside 349 372

16488 GCF_001245745.1_Salmonella_enterica_CVM_N50443_v1.0 Salmonella enterica subsp. enterica serovar Dublin Proteobacteria; Gammaproteobacteria; Enterobacterales; Enterobacteriaceae; Salmonella; Salmonella enterica; Salmonella enterica subsp. enterica MTGDNTLITSHGINRRDFMKLCAALAATMGLSSKAAAEMAESVSNPQRPPVIWIGAQECT WP_000145428.1 hydrogenase 2 small subunit [Salmonella enterica] Length: 372\nNumber of predicted TMHs: 1\nExp number of AAs in TMHs: 24.50677\nExp number, first 60 AAs: 0.2682\nTotal prob of N-in: 0.14390\noutside 1 331\nTMhelix 332 354\ninside 355 372

16489 GCF_002030155.1_ASM203015v1 Salmonella enterica subsp. enterica serovar Dublin Proteobacteria; Gammaproteobacteria; Enterobacterales; Enterobacteriaceae; Salmonella; Salmonella enterica; Salmonella enterica subsp. enterica MTGDNTLITSHGINRRDFMKLCAALAATMGLSSKAAAEMAESVSNPQRPPVIWIGAQECT WP_079823063.1 hydrogenase 2 small subunit [Salmonella enterica] Length: 372\nNumber of predicted TMHs: 1\nExp number of AAs in TMHs: 24.51705\nExp number, first 60 AAs: 0.2682\nTotal prob of N-in: 0.14411\noutside 1 331\nTMhelix 332 354\ninside 355 372

16490 GCF_002030155.1_ASM203015v1 Salmonella enterica subsp. enterica serovar Dublin Proteobacteria; Gammaproteobacteria; Enterobacterales; Enterobacteriaceae; Salmonella; Salmonella enterica; Salmonella enterica subsp. enterica MNNEETLYQAMRRKGVTRRSFLKFCSLAATSLGLGAGMTPKIAWALENKPRIPVVWIHGL WP_001752445.1 hydrogenase-1 small chain [Salmonella enterica] Length: 372\nNumber of predicted TMHs: 2\nExp number of AAs in TMHs: 36.5123\nExp number, first 60 AAs: 19.72341\nTotal prob of N-in: 0.96144\nPOSSIBLE N-term signal sequence\ninside 1 19\nTMhelix 20 39\noutside 40 325\nTMhelix 326 348\ninside 349 372

16491 GCF_001954905.1_ASM195490v1 Salmonella enterica subsp. enterica serovar Dublin Proteobacteria; Gammaproteobacteria; Enterobacterales; Enterobacteriaceae; Salmonella; Salmonella enterica; Salmonella enterica subsp. enterica MNNEETLYQAMRRKGVTRRSFLKFCSLAATSLGLGAGMTPKIAWALENKPRIPVVWIHGL WP_001752445.1 hydrogenase-1 small chain [Salmonella enterica] Length: 372\nNumber of predicted TMHs: 2\nExp number of AAs in TMHs: 36.5123\nExp number, first 60 AAs: 19.72341\nTotal prob of N-in: 0.96144\nPOSSIBLE N-term signal sequence\ninside 1 19\nTMhelix 20 39\noutside 40 325\nTMhelix 326 348\ninside 349 372

16492 GCF_001480305.1_Salmonella_enterica_CVM_N41733-SQ_v1.0 Salmonella enterica subsp. enterica serovar Dublin Proteobacteria; Gammaproteobacteria; Enterobacterales; Enterobacteriaceae; Salmonella; Salmonella enterica; Salmonella enterica subsp. enterica MNNEETLYQAMRRKGVTRRSFLKFCSLAATSLGLGAGMTPKIAWALENKPRIPVVWIHGL WP_001752445.1 hydrogenase-1 small chain [Salmonella enterica] Length: 372\nNumber of predicted TMHs: 2\nExp number of AAs in TMHs: 36.5123\nExp number, first 60 AAs: 19.72341\nTotal prob of N-in: 0.96144\nPOSSIBLE N-term signal sequence\ninside 1 19\nTMhelix 20 39\noutside 40 325\nTMhelix 326 348\ninside 349 372

16493 GCF_001242765.1_Salmonella_enterica_CVM_N45955_v1.0 Salmonella enterica subsp. enterica serovar Dublin Proteobacteria; Gammaproteobacteria; Enterobacterales; Enterobacteriaceae; Salmonella; Salmonella enterica; Salmonella enterica subsp. enterica MNNEETLYQAMRRKGVTRRSFLKFCSLAATSLGLGAGMTPKIAWALENKPRIPVVWIHGL WP_001752445.1 hydrogenase-1 small chain [Salmonella enterica] Length: 372\nNumber of predicted TMHs: 2\nExp number of AAs in TMHs: 36.5123\nExp number, first 60 AAs: 19.72341\nTotal prob of N-in: 0.96144\nPOSSIBLE N-term signal sequence\ninside 1 19\nTMhelix 20 39\noutside 40 325\nTMhelix 326 348\ninside 349 372

16494 GCF_001954935.1_ASM195493v1 Salmonella enterica subsp. enterica serovar Dublin Proteobacteria; Gammaproteobacteria; Enterobacterales; Enterobacteriaceae; Salmonella; Salmonella enterica; Salmonella enterica subsp. enterica MNNEETLYQAMRRKGVTRRSFLKFCSLAATSLGLGAGMTPKIAWALENKPRIPVVWIHGL WP_001752445.1 hydrogenase-1 small chain [Salmonella enterica] Length: 372\nNumber of predicted TMHs: 2\nExp number of AAs in TMHs: 36.5123\nExp number, first 60 AAs: 19.72341\nTotal prob of N-in: 0.96144\nPOSSIBLE N-term signal sequence\ninside 1 19\nTMhelix 20 39\noutside 40 325\nTMhelix 326 348\ninside 349 372

16495 GCF_000336035.1_ASM33603v1 Salmonella enterica subsp. enterica serovar Dublin Proteobacteria; Gammaproteobacteria; Enterobacterales; Enterobacteriaceae; Salmonella; Salmonella enterica;

Salmonella enterica subsp. enterica	MNNEETFYQAMRRKGVTRRSFLKFCSLAATSLGLGAGMTPKIAWALENKPRIPVVWIHGL
	WP_001058312.1	hydrogenase [Salmonella enterica]	Length: 372\nNumber of predicted TMHs: 2\nExp number of AAs in TMHs: 36.51738\nExp number, first 60 AAs: 19.72849\nTotal prob of N-in: 0.96170\nPOSSIBLE N-term signal sequence\ninside 1 19\nTMhelix 20 39\noutside 40 325\nTMhelix 326 348\ninside 349 372

16496	GCF_002090485.1_ASM209048v1	Salmonella enterica subsp. enterica serovar Dublin
	Proteobacteria; Gammaproteobacteria; Enterobacterales; Enterobacteriaceae; Salmonella; Salmonella enterica; Salmonella enterica subsp. enterica	MNNEETLYQAMRRKGVTRRSFLKFCSLAATSLGLGAGMTPKIAWALENKPRIPVVWIHGL
	WP_001752445.1	hydrogenase-1 small chain [Salmonella enterica]	Length: 372\nNumber of predicted TMHs: 2\nExp number of AAs in TMHs: 36.5123\nExp number, first 60 AAs: 19.72341\nTotal prob of N-in: 0.96144\nPOSSIBLE N-term signal sequence\ninside 1 19\nTMhelix 20 39\noutside 40 325\nTMhelix 326 348\ninside 349 372

16497	GCF_002090565.1_ASM209056v1	Salmonella enterica subsp. enterica serovar Dublin
	Proteobacteria; Gammaproteobacteria; Enterobacterales; Enterobacteriaceae; Salmonella; Salmonella enterica; Salmonella enterica subsp. enterica	MNNEETLYQAMRRKGVTRRSFLKFCSLAATSLGLGAGMTPKIAWALENKPRIPVVWIHGL
	WP_001752445.1	hydrogenase-1 small chain [Salmonella enterica]	Length: 372\nNumber of predicted TMHs: 2\nExp number of AAs in TMHs: 36.5123\nExp number, first 60 AAs: 19.72341\nTotal prob of N-in: 0.96144\nPOSSIBLE N-term signal sequence\ninside 1 19\nTMhelix 20 39\noutside 40 325\nTMhelix 326 348\ninside 349 372

16498	GCF_002090755.1_ASM209075v1	Salmonella enterica subsp. enterica serovar Dublin
	Proteobacteria; Gammaproteobacteria; Enterobacterales; Enterobacteriaceae; Salmonella; Salmonella enterica; Salmonella enterica subsp. enterica	MTGDNTLITSHGINRRDFMKLCAALAATMGLSSKAAAEMAESVSNPQRPPVIWIGAQECT
	WP_000145428.1	hydrogenase 2 small subunit [Salmonella enterica]	Length: 372\nNumber of predicted TMHs: 1\nExp number of AAs in TMHs: 24.50677\nExp number, first 60 AAs: 0.2682\nTotal prob of N-in: 0.14390\noutside 1 331\nTMhelix 332 354\ninside 355 372

16499	GCF_002090775.1_ASM209077v1	Salmonella enterica subsp. enterica serovar Dublin
	Proteobacteria; Gammaproteobacteria; Enterobacterales; Enterobacteriaceae; Salmonella; Salmonella enterica; Salmonella enterica subsp. enterica	MTGDNTLITSHGINRRDFMKLCAALAATMGLSSKAAAEMAESVSNPQRPPVIWIGAQECT
	WP_000145428.1	hydrogenase 2 small subunit [Salmonella enterica]	Length: 372\nNumber of predicted TMHs: 1\nExp number of AAs in TMHs: 24.50677\nExp number, first 60 AAs: 0.2682\nTotal prob of N-in: 0.14390\noutside 1 331\nTMhelix 332 354\ninside 355 372

16500	GCF_001954745.1_ASM195474v1	Salmonella enterica subsp. enterica serovar Dublin
	Proteobacteria; Gammaproteobacteria; Enterobacterales; Enterobacteriaceae; Salmonella; Salmonella enterica; Salmonella enterica subsp. enterica	MTGDNTLITSHGINRRDFMKLCAALAATMGLSSKAAAEMAESVSNPQRPPVIWIGAQECT
	WP_000145428.1	hydrogenase 2 small subunit [Salmonella enterica]	Length: 372\nNumber of predicted TMHs: 1\nExp number of AAs in TMHs: 24.50677\nExp number, first 60 AAs: 0.2682\nTotal prob of N-in: 0.14390\noutside 1 331\nTMhelix 332 354\ninside 355 372

16501	GCF_002061665.1_ASM206166v1	Salmonella enterica subsp. enterica serovar Dublin
	Proteobacteria; Gammaproteobacteria; Enterobacterales; Enterobacteriaceae; Salmonella; Salmonella enterica; Salmonella enterica subsp. enterica	MNNEETFYQAMRRKGVTRRSFLKFCSLAATSLGLGAGMTPKIAWALENKPRIPVVWIHGL
	WP_001058312.1	hydrogenase [Salmonella enterica]	Length: 372\nNumber of predicted TMHs: 2\nExp number of AAs in TMHs: 36.51738\nExp number, first 60 AAs: 19.72849\nTotal prob of N-in: 0.96170\nPOSSIBLE N-term signal sequence\ninside 1 19\nTMhelix 20 39\noutside 40 325\nTMhelix 326 348\ninside 349 372

16502	GCF_002061665.1_ASM206166v1	Salmonella enterica subsp. enterica serovar Dublin
	Proteobacteria; Gammaproteobacteria; Enterobacterales; Enterobacteriaceae; Salmonella; Salmonella enterica; Salmonella enterica subsp. enterica	MTGDNTLITSHGINRRDFMKLCAALAATMGLSSKAAAEMAESVSNPQRPPVIWIGAQECT
	WP_000145429.1	hydrogenase 2 small subunit [Salmonella enterica]	Length: 372\nNumber of predicted TMHs: 1\nExp number of AAs in TMHs: 24.51309\nExp number, first 60 AAs: 0.2682\nTotal prob of N-in: 0.14402\noutside 1 331\nTMhelix 332 354\ninside 355 372

16503	GCF_001954705.1_ASM195470v1	Salmonella enterica subsp. enterica serovar Dublin
	Proteobacteria; Gammaproteobacteria; Enterobacterales; Enterobacteriaceae; Salmonella; Salmonella enterica; Salmonella enterica subsp. enterica	MTGDNTLITSHGINRRDFMKLCAALAATMGLSSKAAAEMAESVSNPQRPPVIWIGAQECT
	WP_000145428.1	hydrogenase 2 small subunit [Salmonella enterica]	Length: 372\nNumber of predicted TMHs: 1\nExp number of AAs in TMHs: 24.50677\nExp number, first 60 AAs: 0.2682\nTotal prob of N-in: 0.14390\noutside 1 331\nTMhelix 332 354\ninside 355 372

16504 GCF_001242765.1_Salmonella_enterica_CVM_N45955_v1.0 Salmonella enterica subsp. enterica serovar Dublin Proteobacteria; Gammaproteobacteria; Enterobacterales; Enterobacteriaceae; Salmonella; Salmonella enterica; Salmonella enterica subsp. enterica MTGDNTLITSHGINRRDFMKLCAALAATMGLSSKAAAEMAESVSNPQRPPVIWIGAQECT WP_000145428.1 hydrogenase 2 small subunit [Salmonella enterica] Length: 372\nNumber of predicted TMHs: 1\nExp number of AAs in TMHs: 24.50677\nExp number, first 60 AAs: 0.2682\nTotal prob of N-in: 0.14390\noutside 1 331\nTMhelix 332 354\ninside 355 372

16505 GCF_001481195.1_Salmonella_enterica_CVM_N42502_v1.0 Salmonella enterica subsp. enterica serovar Dublin Proteobacteria; Gammaproteobacteria; Enterobacterales; Enterobacteriaceae; Salmonella; Salmonella enterica; Salmonella enterica subsp. enterica MTGDNTLITSHGINRRDFMKLCAALAATMGLSSKAAAEMAESVSNPQRPPVIWIGAQECT WP_000145428.1 hydrogenase 2 small subunit [Salmonella enterica] Length: 372\nNumber of predicted TMHs: 1\nExp number of AAs in TMHs: 24.50677\nExp number, first 60 AAs: 0.2682\nTotal prob of N-in: 0.14390\noutside 1 331\nTMhelix 332 354\ninside 355 372

16506 GCF_001954745.1_ASM195474v1 Salmonella enterica subsp. enterica serovar Dublin Proteobacteria; Gammaproteobacteria; Enterobacterales; Enterobacteriaceae; Salmonella; Salmonella enterica; Salmonella enterica subsp. enterica MNNEETLYQAMRRKGVTRRSFLKFCSLAATSLGLGAGMTPKIAWALENKPRIPVVWIHGL WP_001752445.1 hydrogenase-1 small chain [Salmonella enterica] Length: 372\nNumber of predicted TMHs: 2\nExp number of AAs in TMHs: 36.5123\nExp number, first 60 AAs: 19.72341\nTotal prob of N-in: 0.96144\nPOSSIBLE N-term signal sequence\ninside 1 19\nTMhelix 20 39\noutside 40 325\nTMhelix 326 348\ninside 349 372

16507 GCF_001480305.1_Salmonella_enterica_CVM_N41733-SQ_v1.0 Salmonella enterica subsp. enterica serovar Dublin Proteobacteria; Gammaproteobacteria; Enterobacterales; Enterobacteriaceae; Salmonella; Salmonella enterica; Salmonella enterica subsp. enterica MTGDNTLITSHGINRRDFMKLCAALAATMGLSSKAAAEMAESVSNPQRPPVIWIGAQECT WP_000145428.1 hydrogenase 2 small subunit [Salmonella enterica] Length: 372\nNumber of predicted TMHs: 1\nExp number of AAs in TMHs: 24.50677\nExp number, first 60 AAs: 0.2682\nTotal prob of N-in: 0.14390\noutside 1 331\nTMhelix 332 354\ninside 355 372

16508 GCF_001245745.1_Salmonella_enterica_CVM_N50443_v1.0 Salmonella enterica subsp. enterica serovar Dublin Proteobacteria; Gammaproteobacteria; Enterobacterales; Enterobacteriaceae; Salmonella; Salmonella enterica; Salmonella enterica subsp. enterica MNNEETLYQAMRRKGVTRRSFLKFCSLAATSLGLGAGMTPKIAWALENKPRIPVVWIHGL WP_001752445.1 hydrogenase-1 small chain [Salmonella enterica] Length: 372\nNumber of predicted TMHs: 2\nExp number of AAs in TMHs: 36.5123\nExp number, first 60 AAs: 19.72341\nTotal prob of N-in: 0.96144\nPOSSIBLE N-term signal sequence\ninside 1 19\nTMhelix 20 39\noutside 40 325\nTMhelix 326 348\ninside 349 372

16509 GCF_002090735.1_ASM209073v1 Salmonella enterica subsp. enterica serovar Dublin Proteobacteria; Gammaproteobacteria; Enterobacterales; Enterobacteriaceae; Salmonella; Salmonella enterica; Salmonella enterica subsp. enterica MNNEETLYQAMRRKGVTRRSFLKFCSLAATSLGLGAGMTPKIAWALENKPRIPVVWIHGL WP_001752445.1 hydrogenase-1 small chain [Salmonella enterica] Length: 372\nNumber of predicted TMHs: 2\nExp number of AAs in TMHs: 36.5123\nExp number, first 60 AAs: 19.72341\nTotal prob of N-in: 0.96144\nPOSSIBLE N-term signal sequence\ninside 1 19\nTMhelix 20 39\noutside 40 325\nTMhelix 326 348\ninside 349 372

16510 GCF_002061495.1_ASM206149v1 Salmonella enterica subsp. enterica serovar Dublin Proteobacteria; Gammaproteobacteria; Enterobacterales; Enterobacteriaceae; Salmonella; Salmonella enterica; Salmonella enterica subsp. enterica MTGDNTLITSHGINRRDFMKLCAALAATMGLSSKAAAEMAESVSNPQRPPVIWIGAQECT WP_000145428.1 hydrogenase 2 small subunit [Salmonella enterica] Length: 372\nNumber of predicted TMHs: 1\nExp number of AAs in TMHs: 24.50677\nExp number, first 60 AAs: 0.2682\nTotal prob of N-in: 0.14390\noutside 1 331\nTMhelix 332 354\ninside 355 372

16511 GCF_002090735.1_ASM209073v1 Salmonella enterica subsp. enterica serovar Dublin Proteobacteria; Gammaproteobacteria; Enterobacterales; Enterobacteriaceae; Salmonella; Salmonella enterica; Salmonella enterica subsp. enterica MTGDNTLITSHGINRRDFMKLCAALAATMGLSSKAAAEMAESVSNPQRPPVIWIGAQECT WP_000145428.1 hydrogenase 2 small subunit [Salmonella enterica] Length: 372\nNumber of predicted TMHs: 1\nExp number of AAs in TMHs: 24.50677\nExp number, first 60 AAs: 0.2682\nTotal prob of N-in: 0.14390\noutside 1 331\nTMhelix 332 354\ninside 355 372

16512 GCF_002066575.1_ASM206657v1 Salmonella enterica subsp. enterica serovar Dublin Proteobacteria; Gammaproteobacteria; Enterobacterales; Enterobacteriaceae; Salmonella; Salmonella enterica; Salmonella enterica subsp. enterica MNNEETLYQAMRRKGVTRRSFLKFCSLAATSLGLGAGMTPKIAWALENKPRIPVVWIHGL WP_001752445.1 hydrogenase-1 small chain [Salmonella enterica] Length: 372\nNumber of predicted TMHs:

2\nExp number of AAs in TMHs: 36.5123\nExp number, first 60 AAs: 19.72341\nTotal prob of N-in: 0.96144\nPOSSIBLE N-term signal sequence\ninside 1 19\nTMhelix 20 39\noutside 40 325\nTMhelix 326 348\ninside 349 372

16513 GCF_002090385.1_ASM209038v1 Salmonella enterica subsp. enterica serovar Dublin Proteobacteria; Gammaproteobacteria; Enterobacterales; Enterobacteriaceae; Salmonella; Salmonella enterica; Salmonella enterica subsp. enterica MNNEETLYQAMRRKGVTRRSFLKFCSLAATSLGLGAGMTPKIAWALENKPRIPVVWIHGL WP_001752445.1 hydrogenase-1 small chain [Salmonella enterica] Length: 372\nNumber of predicted TMHs: 2\nExp number of AAs in TMHs: 36.5123\nExp number, first 60 AAs: 19.72341\nTotal prob of N-in: 0.96144\nPOSSIBLE N-term signal sequence\ninside 1 19\nTMhelix 20 39\noutside 40 325\nTMhelix 326 348\ninside 349 372

16514 GCF_002030115.1_ASM203011v1 Salmonella enterica subsp. enterica serovar Dublin Proteobacteria; Gammaproteobacteria; Enterobacterales; Enterobacteriaceae; Salmonella; Salmonella enterica; Salmonella enterica subsp. enterica MTGDNTLITSHGINRRDFMKLCAALAATMGLSSKAAAEMAESVSNPQRPPVIWIGAQECT WP_000145429.1 hydrogenase 2 small subunit [Salmonella enterica] Length: 372\nNumber of predicted TMHs: 1\nExp number of AAs in TMHs: 24.51309\nExp number, first 60 AAs: 0.2682\nTotal prob of N-in: 0.14402\noutside 1 331\nTMhelix 332 354\ninside 355 372

16515 GCF_002030115.1_ASM203011v1 Salmonella enterica subsp. enterica serovar Dublin Proteobacteria; Gammaproteobacteria; Enterobacterales; Enterobacteriaceae; Salmonella; Salmonella enterica; Salmonella enterica subsp. enterica MNNEETFYQAMRRKGVTRRSFLKFCSLAATSLGLGAGMTPKIAWALENKPRIPVVWIHGL WP_001058312.1 hydrogenase [Salmonella enterica] Length: 372\nNumber of predicted TMHs: 2\nExp number of AAs in TMHs: 36.51738\nExp number, first 60 AAs: 19.72849\nTotal prob of N-in: 0.96170\nPOSSIBLE N-term signal sequence\ninside 1 19\nTMhelix 20 39\noutside 40 325\nTMhelix 326 348\ninside 349 372

16516 GCF_002090475.1_ASM209047v1 Salmonella enterica subsp. enterica serovar Dublin Proteobacteria; Gammaproteobacteria; Enterobacterales; Enterobacteriaceae; Salmonella; Salmonella enterica; Salmonella enterica subsp. enterica MTGDNTLITSHGINRRDFMKLCAALAATMGLSSKAAAEMAESVSNPQRPPVIWIGAQECT WP_000145428.1 hydrogenase 2 small subunit [Salmonella enterica] Length: 372\nNumber of predicted TMHs: 1\nExp number of AAs in TMHs: 24.50677\nExp number, first 60 AAs: 0.2682\nTotal prob of N-in: 0.14390\noutside 1 331\nTMhelix 332 354\ninside 355 372

16517 GCF_001954925.1_ASM195492v1 Salmonella enterica subsp. enterica serovar Dublin Proteobacteria; Gammaproteobacteria; Enterobacterales; Enterobacteriaceae; Salmonella; Salmonella enterica; Salmonella enterica subsp. enterica MTGDNTLITSHGINRRDFMKLCAALAATMGLSSKAAAEMAESVSNPQRPPVIWIGAQECT WP_000145428.1 hydrogenase 2 small subunit [Salmonella enterica] Length: 372\nNumber of predicted TMHs: 1\nExp number of AAs in TMHs: 24.50677\nExp number, first 60 AAs: 0.2682\nTotal prob of N-in: 0.14390\noutside 1 331\nTMhelix 332 354\ninside 355 372

16518 GCF_001954975.1_ASM195497v1 Salmonella enterica subsp. enterica serovar Dublin Proteobacteria; Gammaproteobacteria; Enterobacterales; Enterobacteriaceae; Salmonella; Salmonella enterica; Salmonella enterica subsp. enterica MTGDNTLITSHGINRRDFMKLCAALAATMGLSSKAAAEMAESVSNPQRPPVIWIGAQECT WP_000145428.1 hydrogenase 2 small subunit [Salmonella enterica] Length: 372\nNumber of predicted TMHs: 1\nExp number of AAs in TMHs: 24.50677\nExp number, first 60 AAs: 0.2682\nTotal prob of N-in: 0.14390\noutside 1 331\nTMhelix 332 354\ninside 355 372

16519 GCF_002090385.1_ASM209038v1 Salmonella enterica subsp. enterica serovar Dublin Proteobacteria; Gammaproteobacteria; Enterobacterales; Enterobacteriaceae; Salmonella; Salmonella enterica; Salmonella enterica subsp. enterica MTGDNTLITSHGINRRDFMKLCAALAATMGLSSKAAAEMAESVSNPQRPPVIWIGAQECT WP_000145428.1 hydrogenase 2 small subunit [Salmonella enterica] Length: 372\nNumber of predicted TMHs: 1\nExp number of AAs in TMHs: 24.50677\nExp number, first 60 AAs: 0.2682\nTotal prob of N-in: 0.14390\noutside 1 331\nTMhelix 332 354\ninside 355 372

16520 GCF_002066575.1_ASM206657v1 Salmonella enterica subsp. enterica serovar Dublin Proteobacteria; Gammaproteobacteria; Enterobacterales; Enterobacteriaceae; Salmonella; Salmonella enterica; Salmonella enterica subsp. enterica MTGDNTLITSHGINRRDFMKLCAALAATMGLSSKAAAEMAESVSNPQRPPVIWIGAQECT WP_000145428.1 hydrogenase 2 small subunit [Salmonella enterica] Length: 372\nNumber of predicted TMHs: 1\nExp number of AAs in TMHs: 24.50677\nExp number, first 60 AAs: 0.2682\nTotal prob of N-in: 0.14390\noutside 1 331\nTMhelix 332 354\ninside 355 372

16521 GCF_002061495.1_ASM206149v1 Salmonella enterica subsp. enterica serovar Dublin Proteobacteria; Gammaproteobacteria; Enterobacterales; Enterobacteriaceae; Salmonella; Salmonella enterica;

Salmonella enterica subsp. enterica	MNNEETLYQAMRRKGVTRRSFLKFCSLAATSLGLGAGMTPKIAWALENKPRIPVVWIHGL	WP_001752445.1	hydrogenase-1 small chain [Salmonella enterica]	Length: 372\nNumber of predicted TMHs: 2\nExp number of AAs in TMHs: 36.5123\nExp number, first 60 AAs: 19.72341\nTotal prob of N-in: 0.96144\nPOSSIBLE N-term signal sequence\ninside 1 19\nTMhelix 20 39\noutside 40 325\nTMhelix 326 348\ninside 349 372

16522	GCF_001954985.1_ASM195498v1	Salmonella enterica subsp. enterica serovar Dublin	Proteobacteria; Gammaproteobacteria; Enterobacterales; Enterobacteriaceae; Salmonella; Salmonella enterica; Salmonella enterica subsp. enterica	MTGDNTLITSHGINRRDFMKLCAALAATMGLSSKAAAEMAESVSNPQRPPVIWIGAQECT	WP_000145428.1	hydrogenase 2 small subunit [Salmonella enterica]	Length: 372\nNumber of predicted TMHs: 1\nExp number of AAs in TMHs: 24.50677\nExp number, first 60 AAs: 0.2682\nTotal prob of N-in: 0.14390\noutside 1 331\nTMhelix 332 354\ninside 355 372

16523	GCF_002057025.1_ASM205702v1	Salmonella enterica subsp. enterica serovar Dublin	Proteobacteria; Gammaproteobacteria; Enterobacterales; Enterobacteriaceae; Salmonella; Salmonella enterica; Salmonella enterica subsp. enterica	MTGDNTLITSHGINRRDFMKLCAALAATMGLSSKAAAEMAESVSNPQRPPVIWIGAQECT	WP_000145428.1	hydrogenase 2 small subunit [Salmonella enterica]	Length: 372\nNumber of predicted TMHs: 1\nExp number of AAs in TMHs: 24.50677\nExp number, first 60 AAs: 0.2682\nTotal prob of N-in: 0.14390\noutside 1 331\nTMhelix 332 354\ninside 355 372

16524	GCF_002090455.1_ASM209045v1	Salmonella enterica subsp. enterica serovar Dublin	Proteobacteria; Gammaproteobacteria; Enterobacterales; Enterobacteriaceae; Salmonella; Salmonella enterica; Salmonella enterica subsp. enterica	MTGDNTLITSHGINRRDFMKLCAALAATMGLSSKAAAEMAESVSNPQRPPVIWIGAQECT	WP_000145428.1	hydrogenase 2 small subunit [Salmonella enterica]	Length: 372\nNumber of predicted TMHs: 1\nExp number of AAs in TMHs: 24.50677\nExp number, first 60 AAs: 0.2682\nTotal prob of N-in: 0.14390\noutside 1 331\nTMhelix 332 354\ninside 355 372

16525	GCF_002090475.1_ASM209047v1	Salmonella enterica subsp. enterica serovar Dublin	Proteobacteria; Gammaproteobacteria; Enterobacterales; Enterobacteriaceae; Salmonella; Salmonella enterica; Salmonella enterica subsp. enterica	MNNEETLYQAMRRKGVTRRSFLKFCSLAATSLGLGAGMTPKIAWALENKPRIPVVWIHGL	WP_001752445.1	hydrogenase-1 small chain [Salmonella enterica]	Length: 372\nNumber of predicted TMHs: 2\nExp number of AAs in TMHs: 36.5123\nExp number, first 60 AAs: 19.72341\nTotal prob of N-in: 0.96144\nPOSSIBLE N-term signal sequence\ninside 1 19\nTMhelix 20 39\noutside 40 325\nTMhelix 326 348\ninside 349 372

16526	GCF_001954995.1_ASM195499v1	Salmonella enterica subsp. enterica serovar Dublin	Proteobacteria; Gammaproteobacteria; Enterobacterales; Enterobacteriaceae; Salmonella; Salmonella enterica; Salmonella enterica subsp. enterica	MNNEETLYQAMRRKGVTRRSFLKFCSLAATSLGLGAGMTPKIAWALENKPRIPVVWIHGL	WP_001752445.1	hydrogenase-1 small chain [Salmonella enterica]	Length: 372\nNumber of predicted TMHs: 2\nExp number of AAs in TMHs: 36.5123\nExp number, first 60 AAs: 19.72341\nTotal prob of N-in: 0.96144\nPOSSIBLE N-term signal sequence\ninside 1 19\nTMhelix 20 39\noutside 40 325\nTMhelix 326 348\ninside 349 372

16527	GCF_001954975.1_ASM195497v1	Salmonella enterica subsp. enterica serovar Dublin	Proteobacteria; Gammaproteobacteria; Enterobacterales; Enterobacteriaceae; Salmonella; Salmonella enterica; Salmonella enterica subsp. enterica	MNNEETLYQAMRRKGVTRRSFLKFCSLAATSLGLGAGMTPKIAWALENKPRIPVVWIHGL	WP_001752445.1	hydrogenase-1 small chain [Salmonella enterica]	Length: 372\nNumber of predicted TMHs: 2\nExp number of AAs in TMHs: 36.5123\nExp number, first 60 AAs: 19.72341\nTotal prob of N-in: 0.96144\nPOSSIBLE N-term signal sequence\ninside 1 19\nTMhelix 20 39\noutside 40 325\nTMhelix 326 348\ninside 349 372

16528	GCF_001954925.1_ASM195492v1	Salmonella enterica subsp. enterica serovar Dublin	Proteobacteria; Gammaproteobacteria; Enterobacterales; Enterobacteriaceae; Salmonella; Salmonella enterica; Salmonella enterica subsp. enterica	MNNEETLYQAMRRKGVTRRSFLKFCSLAATSLGLGAGMTPKIAWALENKPRIPVVWIHGL	WP_001752445.1	hydrogenase-1 small chain [Salmonella enterica]	Length: 372\nNumber of predicted TMHs: 2\nExp number of AAs in TMHs: 36.5123\nExp number, first 60 AAs: 19.72341\nTotal prob of N-in: 0.96144\nPOSSIBLE N-term signal sequence\ninside 1 19\nTMhelix 20 39\noutside 40 325\nTMhelix 326 348\ninside 349 372

16529	GCF_002090455.1_ASM209045v1	Salmonella enterica subsp. enterica serovar Dublin	Proteobacteria; Gammaproteobacteria; Enterobacterales; Enterobacteriaceae; Salmonella; Salmonella enterica; Salmonella enterica subsp. enterica	MNNEETLYQAMRRKGVTRRSFLKFCSLAATSLGLGAGMTPKIAWALENKPRIPVVWIHGL	WP_001752445.1	hydrogenase-1 small chain [Salmonella enterica]	Length: 372\nNumber of predicted TMHs: 2\nExp number of AAs in TMHs: 36.5123\nExp number, first 60 AAs: 19.72341\nTotal prob of N-in: 0.96144\nPOSSIBLE N-term signal sequence\ninside 1 19\nTMhelix 20 39\noutside 40 325\nTMhelix 326 348\ninside 349 372

16530 GCF_000336035.1_ASM33603v1 Salmonella enterica subsp. enterica serovar Dublin Proteobacteria; Gammaproteobacteria; Enterobacterales; Enterobacteriaceae; Salmonella; Salmonella enterica; Salmonella enterica subsp. enterica MTGDNTLITSHGINRRDFMKLCAALAATMGLSSKAAAEMAESVSNPQRPPVIWIGAQECT WP_000145428.1 hydrogenase 2 small subunit [Salmonella enterica] Length: 372\nNumber of predicted TMHs: 1\nExp number of AAs in TMHs: 24.50677\nExp number, first 60 AAs: 0.2682\nTotal prob of N-in: 0.14390\noutside 1 331\nTMhelix 332 354\ninside 355 372

16531 GCF_002090565.1_ASM209056v1 Salmonella enterica subsp. enterica serovar Dublin Proteobacteria; Gammaproteobacteria; Enterobacterales; Enterobacteriaceae; Salmonella; Salmonella enterica; Salmonella enterica subsp. enterica MTGDNTLITSHGINRRDFMKLCAALAATMGLSSKAAAEMAESVSNPQRPPVIWIGAQECT WP_000145428.1 hydrogenase 2 small subunit [Salmonella enterica] Length: 372\nNumber of predicted TMHs: 1\nExp number of AAs in TMHs: 24.50677\nExp number, first 60 AAs: 0.2682\nTotal prob of N-in: 0.14390\noutside 1 331\nTMhelix 332 354\ninside 355 372

16532 GCF_002063385.1_ASM206338v1 Salmonella enterica subsp. enterica serovar Dublin Proteobacteria; Gammaproteobacteria; Enterobacterales; Enterobacteriaceae; Salmonella; Salmonella enterica; Salmonella enterica subsp. enterica MNNEETFYQAMRRKGVTRRSFLKFCSLAATSLGLGAGMTPKIAWALENKPRIPVVWIHGL WP_001058312.1 hydrogenase [Salmonella enterica] Length: 372\nNumber of predicted TMHs: 2\nExp number of AAs in TMHs: 36.51738\nExp number, first 60 AAs: 19.72849\nTotal prob of N-in: 0.96170\nPOSSIBLE N-term signal sequence\ninside 1 19\nTMhelix 20 39\noutside 40 325\nTMhelix 326 348\ninside 349 372

16533 GCF_002065815.1_ASM206581v1 Salmonella enterica subsp. enterica serovar Dublin Proteobacteria; Gammaproteobacteria; Enterobacterales; Enterobacteriaceae; Salmonella; Salmonella enterica; Salmonella enterica subsp. enterica MTGDNTLITSHGINRRDFMKLCAALAATMGLSSKAAAEMAESVSNPQRPPVIWIGAQECT WP_000145428.1 hydrogenase 2 small subunit [Salmonella enterica] Length: 372\nNumber of predicted TMHs: 1\nExp number of AAs in TMHs: 24.50677\nExp number, first 60 AAs: 0.2682\nTotal prob of N-in: 0.14390\noutside 1 331\nTMhelix 332 354\ninside 355 372

16534 GCF_002090755.1_ASM209075v1 Salmonella enterica subsp. enterica serovar Dublin Proteobacteria; Gammaproteobacteria; Enterobacterales; Enterobacteriaceae; Salmonella; Salmonella enterica; Salmonella enterica subsp. enterica MNNEETLYQAMRRKGVTRRSFLKFCSLAATSLGLGAGMTPKIAWALENKPRIPVVWIHGL WP_001752445.1 hydrogenase-1 small chain [Salmonella enterica] Length: 372\nNumber of predicted TMHs: 2\nExp number of AAs in TMHs: 36.5123\nExp number, first 60 AAs: 19.72341\nTotal prob of N-in: 0.96144\nPOSSIBLE N-term signal sequence\ninside 1 19\nTMhelix 20 39\noutside 40 325\nTMhelix 326 348\ninside 349 372

16535 GCF_002090395.1_ASM209039v1 Salmonella enterica subsp. enterica serovar Dublin Proteobacteria; Gammaproteobacteria; Enterobacterales; Enterobacteriaceae; Salmonella; Salmonella enterica; Salmonella enterica subsp. enterica MNNEETLYQAMRRKGVTRRSFLKFCSLAATSLGLGAGMTPKIAWALENKPRIPVVWIHGL WP_001752445.1 hydrogenase-1 small chain [Salmonella enterica] Length: 372\nNumber of predicted TMHs: 2\nExp number of AAs in TMHs: 36.5123\nExp number, first 60 AAs: 19.72341\nTotal prob of N-in: 0.96144\nPOSSIBLE N-term signal sequence\ninside 1 19\nTMhelix 20 39\noutside 40 325\nTMhelix 326 348\ninside 349 372

16536 GCF_002090775.1_ASM209077v1 Salmonella enterica subsp. enterica serovar Dublin Proteobacteria; Gammaproteobacteria; Enterobacterales; Enterobacteriaceae; Salmonella; Salmonella enterica; Salmonella enterica subsp. enterica MNNEETLYQAMRRKGVTRRSFLKFCSLAATSLGLGAGMTPKIAWALENKPRIPVVWIHGL WP_001752445.1 hydrogenase-1 small chain [Salmonella enterica] Length: 372\nNumber of predicted TMHs: 2\nExp number of AAs in TMHs: 36.5123\nExp number, first 60 AAs: 19.72341\nTotal prob of N-in: 0.96144\nPOSSIBLE N-term signal sequence\ninside 1 19\nTMhelix 20 39\noutside 40 325\nTMhelix 326 348\ninside 349 372

16537 GCF_002065815.1_ASM206581v1 Salmonella enterica subsp. enterica serovar Dublin Proteobacteria; Gammaproteobacteria; Enterobacterales; Enterobacteriaceae; Salmonella; Salmonella enterica; Salmonella enterica subsp. enterica MNNEETFYQAMRRKGVTRRSFLKFCSLAATSLGLGAGMTPKIAWALENKPRIPVVWIHGL WP_001058303.1 hydrogenase [Salmonella enterica] Length: 372\nNumber of predicted TMHs: 2\nExp number of AAs in TMHs: 36.518\nExp number, first 60 AAs: 19.72854\nTotal prob of N-in: 0.96169\nPOSSIBLE N-term signal sequence\ninside 1 19\nTMhelix 20 39\noutside 40 325\nTMhelix 326 348\ninside 349 372

16538 GCF_002090395.1_ASM209039v1 Salmonella enterica subsp. enterica serovar Dublin Proteobacteria; Gammaproteobacteria; Enterobacterales; Enterobacteriaceae; Salmonella; Salmonella enterica; Salmonella enterica subsp. enterica MTGDNTLITSHGINRRDFMKLCAALAATMGLSSKAAAEMAESVSNPQRPPVIWIGAQECT WP_000145428.1 hydrogenase 2 small subunit [Salmonella enterica] Length: 372\nNumber of

predicted TMHs: 1\nExp number of AAs in TMHs: 24.50677\nExp number, first 60 AAs: 0.2682\nTotal prob of N-in: 0.14390\noutside 1 331\nTMhelix 332 354\ninside 355 372

16539 GCF_002090465.1_ASM209046v1 Salmonella enterica subsp. enterica serovar Dublin Proteobacteria; Gammaproteobacteria; Enterobacterales; Enterobacteriaceae; Salmonella; Salmonella enterica; Salmonella enterica subsp. enterica MTGDNTLITSHGINRRDFMKLCAALAATMGLSSKAAAEMAESVSNPQRPPVIWIGAQECT WP_000145428.1 hydrogenase 2 small subunit [Salmonella enterica] Length: 372\nNumber of predicted TMHs: 1\nExp number of AAs in TMHs: 24.50677\nExp number, first 60 AAs: 0.2682\nTotal prob of N-in: 0.14390\noutside 1 331\nTMhelix 332 354\ninside 355 372

16540 GCF_002063385.1_ASM206338v1 Salmonella enterica subsp. enterica serovar Dublin Proteobacteria; Gammaproteobacteria; Enterobacterales; Enterobacteriaceae; Salmonella; Salmonella enterica; Salmonella enterica subsp. enterica MTGDNTLITSHGINRRDFMKLCAALAATMGLSSKAAAEMAESVSNPQRPPVIWIGAQECT WP_000145428.1 hydrogenase 2 small subunit [Salmonella enterica] Length: 372\nNumber of predicted TMHs: 1\nExp number of AAs in TMHs: 24.50677\nExp number, first 60 AAs: 0.2682\nTotal prob of N-in: 0.14390\noutside 1 331\nTMhelix 332 354\ninside 355 372

16541 GCF_002090375.1_ASM209037v1 Salmonella enterica subsp. enterica serovar Dublin Proteobacteria; Gammaproteobacteria; Enterobacterales; Enterobacteriaceae; Salmonella; Salmonella enterica; Salmonella enterica subsp. enterica MNNEETLYQAMRRKGVTRRSFLKFCSLAATSLGLGAGMTPKIAWALENKPRIPVVWIHGL WP_001752445.1 hydrogenase-1 small chain [Salmonella enterica] Length: 372\nNumber of predicted TMHs: 2\nExp number of AAs in TMHs: 36.5123\nExp number, first 60 AAs: 19.72341\nTotal prob of N-in: 0.96144\nPOSSIBLE N-term signal sequence\ninside 1 19\nTMhelix 20 39\noutside 40 325\nTMhelix 326 348\ninside 349 372

16542 GCF_002090465.1_ASM209046v1 Salmonella enterica subsp. enterica serovar Dublin Proteobacteria; Gammaproteobacteria; Enterobacterales; Enterobacteriaceae; Salmonella; Salmonella enterica; Salmonella enterica subsp. enterica MNNEETLYQAMRRKGVTRRSFLKFCSLAATSLGLGAGMTPKIAWALENKPRIPVVWIHGL WP_001752445.1 hydrogenase-1 small chain [Salmonella enterica] Length: 372\nNumber of predicted TMHs: 2\nExp number of AAs in TMHs: 36.5123\nExp number, first 60 AAs: 19.72341\nTotal prob of N-in: 0.96144\nPOSSIBLE N-term signal sequence\ninside 1 19\nTMhelix 20 39\noutside 40 325\nTMhelix 326 348\ninside 349 372

16543 GCF_002090375.1_ASM209037v1 Salmonella enterica subsp. enterica serovar Dublin Proteobacteria; Gammaproteobacteria; Enterobacterales; Enterobacteriaceae; Salmonella; Salmonella enterica; Salmonella enterica subsp. enterica MTGDNTLITSHGINRRDFMKLCAALAATMGLSSKAAAEMAESVSNPQRPPVIWIGAQECT WP_000145428.1 hydrogenase 2 small subunit [Salmonella enterica] Length: 372\nNumber of predicted TMHs: 1\nExp number of AAs in TMHs: 24.50677\nExp number, first 60 AAs: 0.2682\nTotal prob of N-in: 0.14390\noutside 1 331\nTMhelix 332 354\ninside 355 372

16544 GCF_002090485.1_ASM209048v1 Salmonella enterica subsp. enterica serovar Dublin Proteobacteria; Gammaproteobacteria; Enterobacterales; Enterobacteriaceae; Salmonella; Salmonella enterica; Salmonella enterica subsp. enterica MTGDNTLITSHGINRRDFMKLCAALAATMGLSSKAAAEMAESVSNPQRPPVIWIGAQECT WP_000145428.1 hydrogenase 2 small subunit [Salmonella enterica] Length: 372\nNumber of predicted TMHs: 1\nExp number of AAs in TMHs: 24.50677\nExp number, first 60 AAs: 0.2682\nTotal prob of N-in: 0.14390\noutside 1 331\nTMhelix 332 354\ninside 355 372

16545 GCF_001954935.1_ASM195493v1 Salmonella enterica subsp. enterica serovar Dublin Proteobacteria; Gammaproteobacteria; Enterobacterales; Enterobacteriaceae; Salmonella; Salmonella enterica; Salmonella enterica subsp. enterica MTGDNTLITSHGINRRDFMKLCAALAATMGLSSKAAAEMAESVSNPQRPPVIWIGAQECT WP_000145428.1 hydrogenase 2 small subunit [Salmonella enterica] Length: 372\nNumber of predicted TMHs: 1\nExp number of AAs in TMHs: 24.50677\nExp number, first 60 AAs: 0.2682\nTotal prob of N-in: 0.14390\noutside 1 331\nTMhelix 332 354\ninside 355 372

16546 GCF_001481195.1_Salmonella_enterica_CVM_N42502_v1.0 Salmonella enterica subsp. enterica serovar Dublin Proteobacteria; Gammaproteobacteria; Enterobacterales; Enterobacteriaceae; Salmonella; Salmonella enterica; Salmonella enterica subsp. enterica MNNEETLYQAMRRKGVTRRSFLKFCSLAATSLGLGAGMTPKIAWALENKPRIPVVWIHGL WP_001752445.1 hydrogenase-1 small chain [Salmonella enterica] Length: 372\nNumber of predicted TMHs: 2\nExp number of AAs in TMHs: 36.5123\nExp number, first 60 AAs: 19.72341\nTotal prob of N-in: 0.96144\nPOSSIBLE N-term signal sequence\ninside 1 19\nTMhelix 20 39\noutside 40 325\nTMhelix 326 348\ninside 349 372

16547 GCF_001954905.1_ASM195490v1 Salmonella enterica subsp. enterica serovar Dublin Proteobacteria; Gammaproteobacteria; Enterobacterales; Enterobacteriaceae; Salmonella; Salmonella enterica;

Salmonella enterica subsp. enterica MTGDNTLITSHGINRRDFMKLCAALAATMGLSSKAAAEMAESVSNPQRPPVIWIGAQECT WP_000145428.1 hydrogenase 2 small subunit [Salmonella enterica] Length: 372\nNumber of predicted TMHs: 1\nExp number of AAs in TMHs: 24.50677\nExp number, first 60 AAs: 0.2682\nTotal prob of N-in: 0.14390\noutside 1 331\nTMhelix 332 354\ninside 355 372

16548 GCF_001245835.1_Salmonella_enterica_CVM_N50446_v1.0 Salmonella enterica subsp. enterica serovar Dublin Proteobacteria; Gammaproteobacteria; Enterobacterales; Enterobacteriaceae; Salmonella; Salmonella enterica; Salmonella enterica subsp. enterica MTGDNTLITSHGINRRDFMKLCAALAATMGLSSKAAAEMAESVSNPQRPPVIWIGAQECT WP_000145428.1 hydrogenase 2 small subunit [Salmonella enterica] Length: 372\nNumber of predicted TMHs: 1\nExp number of AAs in TMHs: 24.50677\nExp number, first 60 AAs: 0.2682\nTotal prob of N-in: 0.14390\noutside 1 331\nTMhelix 332 354\ninside 355 372

16549 GCF_001247195.1_Salmonella_enterica_CVM_N51298_v1.0 Salmonella enterica subsp. enterica serovar Dublin Proteobacteria; Gammaproteobacteria; Enterobacterales; Enterobacteriaceae; Salmonella; Salmonella enterica; Salmonella enterica subsp. enterica MTGDNTLITSHGINRRDFMKLCAALAATMGLSSKAAAEMAESVSNPQRPPVIWIGAQECT WP_000145428.1 hydrogenase 2 small subunit [Salmonella enterica] Length: 372\nNumber of predicted TMHs: 1\nExp number of AAs in TMHs: 24.50677\nExp number, first 60 AAs: 0.2682\nTotal prob of N-in: 0.14390\noutside 1 331\nTMhelix 332 354\ninside 355 372

16550 GCF_001245835.1_Salmonella_enterica_CVM_N50446_v1.0 Salmonella enterica subsp. enterica serovar Dublin Proteobacteria; Gammaproteobacteria; Enterobacterales; Enterobacteriaceae; Salmonella; Salmonella enterica; Salmonella enterica subsp. enterica MNNEETLYQAMRRKGVTRRSFLKFCSLAATSLGLGAGMTPKIAWALENKPRIPVVWIHGL WP_001752445.1 hydrogenase-1 small chain [Salmonella enterica] Length: 372\nNumber of predicted TMHs: 2\nExp number of AAs in TMHs: 36.5123\nExp number, first 60 AAs: 19.72341\nTotal prob of N-in: 0.96144\nPOSSIBLE N-term signal sequence\ninside 1 19\nTMhelix 20 39\noutside 40 325\nTMhelix 326 348\ninside 349 372

16551 GCF_001246165.1_Salmonella_enterica_CVM_N51251_v1.0 Salmonella enterica subsp. enterica serovar Dublin Proteobacteria; Gammaproteobacteria; Enterobacterales; Enterobacteriaceae; Salmonella; Salmonella enterica; Salmonella enterica subsp. enterica MNNEETLYQAMRRKGVTRRSFLKFCSLAATSLGLGAGMTPKIAWALENKPRIPVVWIHGL WP_001752445.1 hydrogenase-1 small chain [Salmonella enterica] Length: 372\nNumber of predicted TMHs: 2\nExp number of AAs in TMHs: 36.5123\nExp number, first 60 AAs: 19.72341\nTotal prob of N-in: 0.96144\nPOSSIBLE N-term signal sequence\ninside 1 19\nTMhelix 20 39\noutside 40 325\nTMhelix 326 348\ninside 349 372

16552 GCF_001247195.1_Salmonella_enterica_CVM_N51298_v1.0 Salmonella enterica subsp. enterica serovar Dublin Proteobacteria; Gammaproteobacteria; Enterobacterales; Enterobacteriaceae; Salmonella; Salmonella enterica; Salmonella enterica subsp. enterica MNNEETLYQAMRRKGVTRRSFLKFCSLAATSLGLGAGMTPKIAWALENKPRIPVVWIHGL WP_001752445.1 hydrogenase-1 small chain [Salmonella enterica] Length: 372\nNumber of predicted TMHs: 2\nExp number of AAs in TMHs: 36.5123\nExp number, first 60 AAs: 19.72341\nTotal prob of N-in: 0.96144\nPOSSIBLE N-term signal sequence\ninside 1 19\nTMhelix 20 39\noutside 40 325\nTMhelix 326 348\ninside 349 372

16553 GCF_001246165.1_Salmonella_enterica_CVM_N51251_v1.0 Salmonella enterica subsp. enterica serovar Dublin Proteobacteria; Gammaproteobacteria; Enterobacterales; Enterobacteriaceae; Salmonella; Salmonella enterica; Salmonella enterica subsp. enterica MTGDNTLITSHGINRRDFMKLCAALAATMGLSSKAAAEMAESVSNPQRPPVIWIGAQECT WP_000145428.1 hydrogenase 2 small subunit [Salmonella enterica] Length: 372\nNumber of predicted TMHs: 1\nExp number of AAs in TMHs: 24.50677\nExp number, first 60 AAs: 0.2682\nTotal prob of N-in: 0.14390\noutside 1 331\nTMhelix 332 354\ninside 355 372

16554 GCF_002156225.1_ASM215622v1 Salmonella enterica subsp. enterica serovar Ealing Proteobacteria; Gammaproteobacteria; Enterobacterales; Enterobacteriaceae; Salmonella; Salmonella enterica; Salmonella enterica subsp. enterica MTGDNTLITSHGINRRDFMKLCAALAATMGLSSKAAAEMAESVSNPQRPPVIWIGAQECT WP_000145429.1 hydrogenase 2 small subunit [Salmonella enterica] Length: 372\nNumber of predicted TMHs: 1\nExp number of AAs in TMHs: 24.51309\nExp number, first 60 AAs: 0.2682\nTotal prob of N-in: 0.14402\noutside 1 331\nTMhelix 332 354\ninside 355 372

16555 GCF_002156225.1_ASM215622v1 Salmonella enterica subsp. enterica serovar Ealing Proteobacteria; Gammaproteobacteria; Enterobacterales; Enterobacteriaceae; Salmonella; Salmonella enterica; Salmonella enterica subsp. enterica MNNEETFYQAMRRKGVTRRSFLKFCSLAATSLGLGAGMTPKIAWALENKPRIPVVWIHGL WP_001058311.1 hydrogenase [Salmonella enterica] Length: 372\nNumber of predicted TMHs: 2\nExp number of AAs in TMHs: 37.4699899999999\nExp number, first 60 AAs: 19.72699\nTotal prob of N-in: 0.96171\nPOSSIBLE N-term signal sequence\ninside 1 19\nTMhelix 20 39\noutside 40 325\nTMhelix 326 348\ninside 349 372

16556	GCF_002048035.1_ASM204803v1	Salmonella enterica subsp. enterica serovar Ealing	Proteobacteria; Gammaproteobacteria; Enterobacterales; Enterobacteriaceae; Salmonella; Salmonella enterica; Salmonella enterica subsp. enterica	MTGDNTLITSHGINRRDFMKLCAALAATMGLSSKAAAEMAESVSNPQRPPVIWIGAQECT	WP_000145429.1	hydrogenase 2 small subunit [Salmonella enterica]	Length: 372\nNumber of predicted TMHs: 1\nExp number of AAs in TMHs: 24.51309\nExp number, first 60 AAs: 0.2682\nTotal prob of N-in: 0.14402\noutside 1 331\nTMhelix 332 354\ninside 355 372

16557	GCF_002046005.1_ASM204600v1	Salmonella enterica subsp. enterica serovar Ealing	Proteobacteria; Gammaproteobacteria; Enterobacterales; Enterobacteriaceae; Salmonella; Salmonella enterica; Salmonella enterica subsp. enterica	MNNEETFYQAMRRKGVTRRSFLKFCSLAATSLGLGAGMTPKIAWALENKPRIPVVWIHGL	WP_001058311.1	hydrogenase [Salmonella enterica]	Length: 372\nNumber of predicted TMHs: 2\nExp number of AAs in TMHs: 37.4699899999999\nExp number, first 60 AAs: 19.72699\nTotal prob of N-in: 0.96171\nPOSSIBLE N-term signal sequence\ninside 1 19\nTMhelix 20 39\noutside 40 325\nTMhelix 326 348\ninside 349 372

16558	GCF_002048035.1_ASM204803v1	Salmonella enterica subsp. enterica serovar Ealing	Proteobacteria; Gammaproteobacteria; Enterobacterales; Enterobacteriaceae; Salmonella; Salmonella enterica; Salmonella enterica subsp. enterica	MNNEETFYQAMRRKGVTRRSFLKFCSLAATSLGLGAGMTPKIAWALENKPRIPVVWIHGL	WP_001058311.1	hydrogenase [Salmonella enterica]	Length: 372\nNumber of predicted TMHs: 2\nExp number of AAs in TMHs: 37.4699899999999\nExp number, first 60 AAs: 19.72699\nTotal prob of N-in: 0.96171\nPOSSIBLE N-term signal sequence\ninside 1 19\nTMhelix 20 39\noutside 40 325\nTMhelix 326 348\ninside 349 372

16559	GCF_002046005.1_ASM204600v1	Salmonella enterica subsp. enterica serovar Ealing	Proteobacteria; Gammaproteobacteria; Enterobacterales; Enterobacteriaceae; Salmonella; Salmonella enterica; Salmonella enterica subsp. enterica	MTGDNTLITSHGINRRDFMKLCAALAATMGLSSKAAAEMAESVSNPQRPPVIWIGAQECT	WP_000145429.1	hydrogenase 2 small subunit [Salmonella enterica]	Length: 372\nNumber of predicted TMHs: 1\nExp number of AAs in TMHs: 24.51309\nExp number, first 60 AAs: 0.2682\nTotal prob of N-in: 0.14402\noutside 1 331\nTMhelix 332 354\ninside 355 372

16560	GCF_002046055.1_ASM204605v1	Salmonella enterica subsp. enterica serovar Eastbourne	Proteobacteria; Gammaproteobacteria; Enterobacterales; Enterobacteriaceae; Salmonella; Salmonella enterica; Salmonella enterica subsp. enterica	MNNEETFYQAMRRKGVTRRSFLKFCSLAATSLGLGAGMTPKIAWALENKPRIPVVWIHGL	WP_001058312.1	hydrogenase [Salmonella enterica]	Length: 372\nNumber of predicted TMHs: 2\nExp number of AAs in TMHs: 36.51738\nExp number, first 60 AAs: 19.72849\nTotal prob of N-in: 0.96170\nPOSSIBLE N-term signal sequence\ninside 1 19\nTMhelix 20 39\noutside 40 325\nTMhelix 326 348\ninside 349 372

16561	GCF_002045425.1_ASM204542v1	Salmonella enterica subsp. enterica serovar Eastbourne	Proteobacteria; Gammaproteobacteria; Enterobacterales; Enterobacteriaceae; Salmonella; Salmonella enterica; Salmonella enterica subsp. enterica	MNNEETFYQAMRRKGVTRRSFLKFCSLAATSLGLGAGMTPKIAWALENKPRIPVVWIHGL	WP_001058312.1	hydrogenase [Salmonella enterica]	Length: 372\nNumber of predicted TMHs: 2\nExp number of AAs in TMHs: 36.51738\nExp number, first 60 AAs: 19.72849\nTotal prob of N-in: 0.96170\nPOSSIBLE N-term signal sequence\ninside 1 19\nTMhelix 20 39\noutside 40 325\nTMhelix 326 348\ninside 349 372

16562	GCF_002045425.1_ASM204542v1	Salmonella enterica subsp. enterica serovar Eastbourne	Proteobacteria; Gammaproteobacteria; Enterobacterales; Enterobacteriaceae; Salmonella; Salmonella enterica; Salmonella enterica subsp. enterica	MTGDNTLITSHGINRRDFMKLCAALAATMGLSSKAAAEMAESVSNPQRPPVIWIGAQECT	WP_000145429.1	hydrogenase 2 small subunit [Salmonella enterica]	Length: 372\nNumber of predicted TMHs: 1\nExp number of AAs in TMHs: 24.51309\nExp number, first 60 AAs: 0.2682\nTotal prob of N-in: 0.14402\noutside 1 331\nTMhelix 332 354\ninside 355 372

16563	GCF_002046055.1_ASM204605v1	Salmonella enterica subsp. enterica serovar Eastbourne	Proteobacteria; Gammaproteobacteria; Enterobacterales; Enterobacteriaceae; Salmonella; Salmonella enterica; Salmonella enterica subsp. enterica	MTGDNTLITSHGINRRDFMKLCAALAATMGLSSKAAAEMAESVSNPQRPPVIWIGAQECT	WP_000145431.1	hydrogenase 2 small subunit [Salmonella enterica]	Length: 372\nNumber of predicted TMHs: 1\nExp number of AAs in TMHs: 24.51505\nExp number, first 60 AAs: 0.2682\nTotal prob of N-in: 0.14406\noutside 1 331\nTMhelix 332 354\ninside 355 372

16564	GCF_002047975.1_ASM204797v1	Salmonella enterica subsp. enterica serovar Eastbourne	Proteobacteria; Gammaproteobacteria; Enterobacterales; Enterobacteriaceae; Salmonella; Salmonella enterica; Salmonella enterica subsp. enterica	MTGDNTLITSHGINRRDFMKLCAALAATMGLSSKAAAEMAESVSNPQRPPVIWIGAQECT

WP_000145429.1 hydrogenase 2 small subunit [Salmonella enterica] Length: 372\nNumber of predicted TMHs: 1\nExp number of AAs in TMHs: 24.51309\nExp number, first 60 AAs: 0.2682\nTotal prob of N-in: 0.14402\noutside 1 331\nTMhelix 332 354\ninside 355 372

16565 GCF_002046145.1_ASM204614v1 Salmonella enterica subsp. enterica serovar Eastbourne Proteobacteria; Gammaproteobacteria; Enterobacterales; Enterobacteriaceae; Salmonella; Salmonella enterica; Salmonella enterica subsp. enterica MTGDNTLITSHGINRRDFMKLCAALAATMGLSSKAAAEMAESVSNPQRPPVIWIGAQECT WP_000145429.1 hydrogenase 2 small subunit [Salmonella enterica] Length: 372\nNumber of predicted TMHs: 1\nExp number of AAs in TMHs: 24.51309\nExp number, first 60 AAs: 0.2682\nTotal prob of N-in: 0.14402\noutside 1 331\nTMhelix 332 354\ninside 355 372

16566 GCF_002046145.1_ASM204614v1 Salmonella enterica subsp. enterica serovar Eastbourne Proteobacteria; Gammaproteobacteria; Enterobacterales; Enterobacteriaceae; Salmonella; Salmonella enterica; Salmonella enterica subsp. enterica MNNEETFYQAMRRKGVTRRSFLKFCSLAATSLGLGAGMTPKIAWALENKPRIPVVWIHGL WP_001058312.1 hydrogenase [Salmonella enterica] Length: 372\nNumber of predicted TMHs: 2\nExp number of AAs in TMHs: 36.51738\nExp number, first 60 AAs: 19.72849\nTotal prob of N-in: 0.96170\nPOSSIBLE N-term signal sequence\ninside 1 19\nTMhelix 20 39\noutside 40 325\nTMhelix 326 348\ninside 349 372

16567 GCF_002047975.1_ASM204797v1 Salmonella enterica subsp. enterica serovar Eastbourne Proteobacteria; Gammaproteobacteria; Enterobacterales; Enterobacteriaceae; Salmonella; Salmonella enterica; Salmonella enterica subsp. enterica MNNEETFYQAMRRKGVTRRSFLKFCSLAATSLGLGAGMTPKIAWALENKPRIPVVWIHGL WP_001058312.1 hydrogenase [Salmonella enterica] Length: 372\nNumber of predicted TMHs: 2\nExp number of AAs in TMHs: 36.51738\nExp number, first 60 AAs: 19.72849\nTotal prob of N-in: 0.96170\nPOSSIBLE N-term signal sequence\ninside 1 19\nTMhelix 20 39\noutside 40 325\nTMhelix 326 348\ninside 349 372

16568 GCF_002048025.1_ASM204802v1 Salmonella enterica subsp. enterica serovar Eastbourne Proteobacteria; Gammaproteobacteria; Enterobacterales; Enterobacteriaceae; Salmonella; Salmonella enterica; Salmonella enterica subsp. enterica MTGDNTLITSHGINRRDFMKLCAALAATMGLSSKAAAEMAESVSNPQRPPVIWIGAQECT WP_000145429.1 hydrogenase 2 small subunit [Salmonella enterica] Length: 372\nNumber of predicted TMHs: 1\nExp number of AAs in TMHs: 24.51309\nExp number, first 60 AAs: 0.2682\nTotal prob of N-in: 0.14402\noutside 1 331\nTMhelix 332 354\ninside 355 372

16569 GCF_002048025.1_ASM204802v1 Salmonella enterica subsp. enterica serovar Eastbourne Proteobacteria; Gammaproteobacteria; Enterobacterales; Enterobacteriaceae; Salmonella; Salmonella enterica; Salmonella enterica subsp. enterica MNNEETFYQAMRRKGVTRRSFLKFCSLAATSLGLGAGMTPKIAWALENKPRIPVVWIHGL WP_001058312.1 hydrogenase [Salmonella enterica] Length: 372\nNumber of predicted TMHs: 2\nExp number of AAs in TMHs: 36.51738\nExp number, first 60 AAs: 19.72849\nTotal prob of N-in: 0.96170\nPOSSIBLE N-term signal sequence\ninside 1 19\nTMhelix 20 39\noutside 40 325\nTMhelix 326 348\ninside 349 372

16570 GCF_002045435.1_ASM204543v1 Salmonella enterica subsp. enterica serovar Edinburgh Proteobacteria; Gammaproteobacteria; Enterobacterales; Enterobacteriaceae; Salmonella; Salmonella enterica; Salmonella enterica subsp. enterica MNNEETFYQAMRRKGVTRRSFLKFCSLAATSLGLGAGMTPKIAWALENKPRIPVVWIHGL WP_001058312.1 hydrogenase [Salmonella enterica] Length: 372\nNumber of predicted TMHs: 2\nExp number of AAs in TMHs: 36.51738\nExp number, first 60 AAs: 19.72849\nTotal prob of N-in: 0.96170\nPOSSIBLE N-term signal sequence\ninside 1 19\nTMhelix 20 39\noutside 40 325\nTMhelix 326 348\ninside 349 372

16571 GCF_002047965.1_ASM204796v1 Salmonella enterica subsp. enterica serovar Edinburgh Proteobacteria; Gammaproteobacteria; Enterobacterales; Enterobacteriaceae; Salmonella; Salmonella enterica; Salmonella enterica subsp. enterica MTGDNTLITSHGINRRDFMKLCAALAATMGLSSKAAAEMAESVSNPQRPPVIWIGAQECT WP_000145429.1 hydrogenase 2 small subunit [Salmonella enterica] Length: 372\nNumber of predicted TMHs: 1\nExp number of AAs in TMHs: 24.51309\nExp number, first 60 AAs: 0.2682\nTotal prob of N-in: 0.14402\noutside 1 331\nTMhelix 332 354\ninside 355 372

16572 GCF_002047965.1_ASM204796v1 Salmonella enterica subsp. enterica serovar Edinburgh Proteobacteria; Gammaproteobacteria; Enterobacterales; Enterobacteriaceae; Salmonella; Salmonella enterica; Salmonella enterica subsp. enterica MNNEETFYQAMRRKGVTRRSFLKFCSLAATSLGLGAGMTPKIAWALENKPRIPVVWIHGL WP_001058312.1 hydrogenase [Salmonella enterica] Length: 372\nNumber of predicted TMHs: 2\nExp number of AAs in TMHs: 36.51738\nExp number, first 60 AAs: 19.72849\nTotal prob of N-in: 0.96170\nPOSSIBLE N-term signal sequence\ninside 1 19\nTMhelix 20 39\noutside 40 325\nTMhelix 326 348\ninside 349 372

16573 GCF_002046605.1_ASM204660v1 Salmonella enterica subsp. enterica serovar Edinburgh Proteobacteria; Gammaproteobacteria; Enterobacterales; Enterobacteriaceae; Salmonella; Salmonella enterica; Salmonella enterica subsp. enterica MTGDNTLITSHGINRRDFMKLCAALAATMGLSSKAAAEMAESVSNPQRPPVIWIGAQECT WP_000145429.1 hydrogenase 2 small subunit [Salmonella enterica] Length: 372\nNumber of predicted TMHs: 1\nExp number of AAs in TMHs: 24.51309\nExp number, first 60 AAs: 0.2682\nTotal prob of N-in: 0.14402\noutside 1 331\nTMhelix 332 354\ninside 355 372

16574 GCF_002046605.1_ASM204660v1 Salmonella enterica subsp. enterica serovar Edinburgh Proteobacteria; Gammaproteobacteria; Enterobacterales; Enterobacteriaceae; Salmonella; Salmonella enterica; Salmonella enterica subsp. enterica MNNEETFYQAMRRKGVTRRSFLKFCSLAATSLGLGAGMTPKIAWALENKPRIPVVWIHGL WP_001058312.1 hydrogenase [Salmonella enterica] Length: 372\nNumber of predicted TMHs: 2\nExp number of AAs in TMHs: 36.51738\nExp number, first 60 AAs: 19.72849\nTotal prob of N-in: 0.96170\nPOSSIBLE N-term signal sequence\ninside 1 19\nTMhelix 20 39\noutside 40 325\nTMhelix 326 348\ninside 349 372

16575 GCF_002046365.1_ASM204636v1 Salmonella enterica subsp. enterica serovar Edinburgh Proteobacteria; Gammaproteobacteria; Enterobacterales; Enterobacteriaceae; Salmonella; Salmonella enterica; Salmonella enterica subsp. enterica MNNEETFYQAMRRKGVTRRSFLKFCSLAATSLGLGAGMTPKIAWALENKPRIPVVWIHGL WP_001058312.1 hydrogenase [Salmonella enterica] Length: 372\nNumber of predicted TMHs: 2\nExp number of AAs in TMHs: 36.51738\nExp number, first 60 AAs: 19.72849\nTotal prob of N-in: 0.96170\nPOSSIBLE N-term signal sequence\ninside 1 19\nTMhelix 20 39\noutside 40 325\nTMhelix 326 348\ninside 349 372

16576 GCF_002045435.1_ASM204543v1 Salmonella enterica subsp. enterica serovar Edinburgh Proteobacteria; Gammaproteobacteria; Enterobacterales; Enterobacteriaceae; Salmonella; Salmonella enterica; Salmonella enterica subsp. enterica MTGDNTLITSHGINRRDFMKLCAALAATMGLSSKAAAEMAESVSNPQRPPVIWIGAQECT WP_000145429.1 hydrogenase 2 small subunit [Salmonella enterica] Length: 372\nNumber of predicted TMHs: 1\nExp number of AAs in TMHs: 24.51309\nExp number, first 60 AAs: 0.2682\nTotal prob of N-in: 0.14402\noutside 1 331\nTMhelix 332 354\ninside 355 372

16577 GCF_002065785.1_ASM206578v1 Salmonella enterica subsp. enterica serovar Elizabethville Proteobacteria; Gammaproteobacteria; Enterobacterales; Enterobacteriaceae; Salmonella; Salmonella enterica; Salmonella enterica subsp. enterica MTGDNTLITSHGINRRDFMKLCAALAATMGLSSKAAAEMAESVSNPQRPPVIWIGAQECT WP_000145429.1 hydrogenase 2 small subunit [Salmonella enterica] Length: 372\nNumber of predicted TMHs: 1\nExp number of AAs in TMHs: 24.51309\nExp number, first 60 AAs: 0.2682\nTotal prob of N-in: 0.14402\noutside 1 331\nTMhelix 332 354\ninside 355 372

16578 GCF_002063115.1_ASM206311v1 Salmonella enterica subsp. enterica serovar Elizabethville Proteobacteria; Gammaproteobacteria; Enterobacterales; Enterobacteriaceae; Salmonella; Salmonella enterica; Salmonella enterica subsp. enterica MTGDNTLITSHGINRRDFMKLCAALAATMGLSSKAAAEMAESVSNPQRPPVIWIGAQECT WP_000145429.1 hydrogenase 2 small subunit [Salmonella enterica] Length: 372\nNumber of predicted TMHs: 1\nExp number of AAs in TMHs: 24.51309\nExp number, first 60 AAs: 0.2682\nTotal prob of N-in: 0.14402\noutside 1 331\nTMhelix 332 354\ninside 355 372

16579 GCF_002065785.1_ASM206578v1 Salmonella enterica subsp. enterica serovar Elizabethville Proteobacteria; Gammaproteobacteria; Enterobacterales; Enterobacteriaceae; Salmonella; Salmonella enterica; Salmonella enterica subsp. enterica MNNEETFYQAMRRKGVTRRSFLKFCSLAATSLGLGAGMTPKIAWALENKPRIPVVWIHGL WP_001058311.1 hydrogenase [Salmonella enterica] Length: 372\nNumber of predicted TMHs: 2\nExp number of AAs in TMHs: 37.4699899999999\nExp number, first 60 AAs: 19.72699\nTotal prob of N-in: 0.96171\nPOSSIBLE N-term signal sequence\ninside 1 19\nTMhelix 20 39\noutside 40 325\nTMhelix 326 348\ninside 349 372

16580 GCF_002077015.1_ASM207701v1 Salmonella enterica subsp. enterica serovar Enteritidis Proteobacteria; Gammaproteobacteria; Enterobacterales; Enterobacteriaceae; Salmonella; Salmonella enterica; Salmonella enterica subsp. enterica MNNEETFYQAMRRKGVTRRSFLKFCSLAATSLGLGAGMTPKIAWALENKPRIPVVWIHGL WP_001058307.1 hydrogenase [Salmonella enterica] Length: 372\nNumber of predicted TMHs: 2\nExp number of AAs in TMHs: 36.51738\nExp number, first 60 AAs: 19.72849\nTotal prob of N-in: 0.96170\nPOSSIBLE N-term signal sequence\ninside 1 19\nTMhelix 20 39\noutside 40 325\nTMhelix 326 348\ninside 349 372

16581 GCF_002076975.1_ASM207697v1 Salmonella enterica subsp. enterica serovar Enteritidis Proteobacteria; Gammaproteobacteria; Enterobacterales; Enterobacteriaceae; Salmonella; Salmonella enterica; Salmonella enterica subsp. enterica MNNEETFYQAMRRKGVTRRSFLKFCSLAATSLGLGAGMTPKIAWALENKPRIPVVWIHGL WP_001058307.1 hydrogenase [Salmonella enterica] Length: 372\nNumber of predicted TMHs: 2\nExp

number of AAs in TMHs: 36.51738\nExp number, first 60 AAs: 19.72849\nTotal prob of N-in: 0.96170\nPOSSIBLE N-term signal sequence\ninside 1 19\nTMhelix 20 39\noutside 40 325\nTMhelix 326 348\ninside 349 372

16582 GCF_002066435.1_ASM206643v1 Salmonella enterica subsp. enterica serovar Enteritidis Proteobacteria; Gammaproteobacteria; Enterobacterales; Enterobacteriaceae; Salmonella; Salmonella enterica; Salmonella enterica subsp. enterica MNNEETFYQAMRRKGVTRRSFLKFCSLAATSLGLGAGMTPKIAWALENKPRIPVVWIHGL WP_001058307.1 hydrogenase [Salmonella enterica] Length: 372\nNumber of predicted TMHs: 2\nExp number of AAs in TMHs: 36.51738\nExp number, first 60 AAs: 19.72849\nTotal prob of N-in: 0.96170\nPOSSIBLE N-term signal sequence\ninside 1 19\nTMhelix 20 39\noutside 40 325\nTMhelix 326 348\ninside 349 372

16583 GCF_002066435.1_ASM206643v1 Salmonella enterica subsp. enterica serovar Enteritidis Proteobacteria; Gammaproteobacteria; Enterobacterales; Enterobacteriaceae; Salmonella; Salmonella enterica; Salmonella enterica subsp. enterica MTGDNTLITSHGINRRDFMKLCAALAATMGLSSKAAAEMAESVSNPQRPPVIWIGAQECT WP_000145429.1 hydrogenase 2 small subunit [Salmonella enterica] Length: 372\nNumber of predicted TMHs: 1\nExp number of AAs in TMHs: 24.51309\nExp number, first 60 AAs: 0.2682\nTotal prob of N-in: 0.14402\noutside 1 331\nTMhelix 332 354\ninside 355 372

16584 GCF_002076975.1_ASM207697v1 Salmonella enterica subsp. enterica serovar Enteritidis Proteobacteria; Gammaproteobacteria; Enterobacterales; Enterobacteriaceae; Salmonella; Salmonella enterica; Salmonella enterica subsp. enterica MTGDNTLITSHGINRRDFMKLCAALAATMGLSSKAAAEMAESVSNPQRPPVIWIGAQECT WP_000145429.1 hydrogenase 2 small subunit [Salmonella enterica] Length: 372\nNumber of predicted TMHs: 1\nExp number of AAs in TMHs: 24.51309\nExp number, first 60 AAs: 0.2682\nTotal prob of N-in: 0.14402\noutside 1 331\nTMhelix 332 354\ninside 355 372

16585 GCF_002063265.1_ASM206326v1 Salmonella enterica subsp. enterica serovar Enteritidis Proteobacteria; Gammaproteobacteria; Enterobacterales; Enterobacteriaceae; Salmonella; Salmonella enterica; Salmonella enterica subsp. enterica MTGDNTLITSHGINRRDFMKLCAALAATMGLSSKAAAEMAESVSNPQRPPVIWIGAQECT WP_000145429.1 hydrogenase 2 small subunit [Salmonella enterica] Length: 372\nNumber of predicted TMHs: 1\nExp number of AAs in TMHs: 24.51309\nExp number, first 60 AAs: 0.2682\nTotal prob of N-in: 0.14402\noutside 1 331\nTMhelix 332 354\ninside 355 372

16586 GCF_002065735.1_ASM206573v1 Salmonella enterica subsp. enterica serovar Enteritidis Proteobacteria; Gammaproteobacteria; Enterobacterales; Enterobacteriaceae; Salmonella; Salmonella enterica; Salmonella enterica subsp. enterica MNNEETFYQAMRRKGVTRRSFLKFCSLAATSLGLGAGMTPKIAWALENKPRIPVVWIHGL WP_001058307.1 hydrogenase [Salmonella enterica] Length: 372\nNumber of predicted TMHs: 2\nExp number of AAs in TMHs: 36.51738\nExp number, first 60 AAs: 19.72849\nTotal prob of N-in: 0.96170\nPOSSIBLE N-term signal sequence\ninside 1 19\nTMhelix 20 39\noutside 40 325\nTMhelix 326 348\ninside 349 372

16587 GCF_002107355.1_ASM210735v1 Salmonella enterica subsp. enterica serovar Enteritidis Proteobacteria; Gammaproteobacteria; Enterobacterales; Enterobacteriaceae; Salmonella; Salmonella enterica; Salmonella enterica subsp. enterica MNNEETFYQAMRRKGVTRRSFLKFCSLAATSLGLGAGMTPKIAWALENKPRIPVVWIHGL WP_001058307.1 hydrogenase [Salmonella enterica] Length: 372\nNumber of predicted TMHs: 2\nExp number of AAs in TMHs: 36.51738\nExp number, first 60 AAs: 19.72849\nTotal prob of N-in: 0.96170\nPOSSIBLE N-term signal sequence\ninside 1 19\nTMhelix 20 39\noutside 40 325\nTMhelix 326 348\ninside 349 372

16588 GCF_002063265.1_ASM206326v1 Salmonella enterica subsp. enterica serovar Enteritidis Proteobacteria; Gammaproteobacteria; Enterobacterales; Enterobacteriaceae; Salmonella; Salmonella enterica; Salmonella enterica subsp. enterica MNNEETFYQAMRRKGVTRRSFLKFCSLAATSLGLGAGMTPKIAWALENKPRIPVVWIHGL WP_001058307.1 hydrogenase [Salmonella enterica] Length: 372\nNumber of predicted TMHs: 2\nExp number of AAs in TMHs: 36.51738\nExp number, first 60 AAs: 19.72849\nTotal prob of N-in: 0.96170\nPOSSIBLE N-term signal sequence\ninside 1 19\nTMhelix 20 39\noutside 40 325\nTMhelix 326 348\ninside 349 372

16589 GCF_002107355.1_ASM210735v1 Salmonella enterica subsp. enterica serovar Enteritidis Proteobacteria; Gammaproteobacteria; Enterobacterales; Enterobacteriaceae; Salmonella; Salmonella enterica; Salmonella enterica subsp. enterica MTGDNTLITSHGINRRDFMKLCAALAATMGLSSKAAAEMAESVSNPQRPPVIWIGAQECT WP_000145429.1 hydrogenase 2 small subunit [Salmonella enterica] Length: 372\nNumber of predicted TMHs: 1\nExp number of AAs in TMHs: 24.51309\nExp number, first 60 AAs: 0.2682\nTotal prob of N-in: 0.14402\noutside 1 331\nTMhelix 332 354\ninside 355 372

16590 GCF_002065735.1_ASM206573v1 Salmonella enterica subsp. enterica serovar Enteritidis Proteobacteria; Gammaproteobacteria; Enterobacterales; Enterobacteriaceae; Salmonella; Salmonella enterica;

Salmonella enterica subsp. enterica	MTGDNTLITSHGINRRDFMKLCAALAATMGLSSKAAAEMAESVSNPQRPPVIWIGAQECT	WP_000145429.1	hydrogenase 2 small subunit [Salmonella enterica]	Length: 372\nNumber of predicted TMHs:  1\nExp number of AAs in TMHs: 24.51309\nExp number, first 60 AAs:  0.2682\nTotal prob of N-in:   0.14402\noutside   1   331\nTMhelix   332   354\ninside   355   372

16591	GCF_002107235.1_ASM210723v1	Salmonella enterica subsp. enterica serovar Enteritidis	Proteobacteria; Gammaproteobacteria; Enterobacterales; Enterobacteriaceae; Salmonella; Salmonella enterica; Salmonella enterica subsp. enterica	MTGDNTLITSHGINRRDFMKLCAALAATMGLSSKAAAEMAESVSNPQRPPVIWIGAQECT	WP_000145429.1	hydrogenase 2 small subunit [Salmonella enterica]	Length: 372\nNumber of predicted TMHs:  1\nExp number of AAs in TMHs: 24.51309\nExp number, first 60 AAs:  0.2682\nTotal prob of N-in:   0.14402\noutside   1   331\nTMhelix   332   354\ninside   355   372

16592	GCF_002077015.1_ASM207701v1	Salmonella enterica subsp. enterica serovar Enteritidis	Proteobacteria; Gammaproteobacteria; Enterobacterales; Enterobacteriaceae; Salmonella; Salmonella enterica; Salmonella enterica subsp. enterica	MTGDNTLITSHGINRRDFMKLCAALAATMGLSSKAAAEMAESVSNPQRPPVIWIGAQECT	WP_000145429.1	hydrogenase 2 small subunit [Salmonella enterica]	Length: 372\nNumber of predicted TMHs:  1\nExp number of AAs in TMHs: 24.51309\nExp number, first 60 AAs:  0.2682\nTotal prob of N-in:   0.14402\noutside   1   331\nTMhelix   332   354\ninside   355   372

16593	GCF_001271895.1_Salmonella_enterica_CVM_N43451_v1.0	Salmonella enterica subsp. enterica serovar Enteritidis	Proteobacteria; Gammaproteobacteria; Enterobacterales; Enterobacteriaceae; Salmonella; Salmonella enterica; Salmonella enterica subsp. enterica	MTGDNTLITSHGINRRDFMKLCAALAATMGLSSKAAAEMAESVSNPQRPPVIWIGAQECT	WP_000145429.1	hydrogenase 2 small subunit [Salmonella enterica]	Length: 372\nNumber of predicted TMHs:  1\nExp number of AAs in TMHs: 24.51309\nExp number, first 60 AAs:  0.2682\nTotal prob of N-in:   0.14402\noutside   1   331\nTMhelix   332   354\ninside   355   372

16594	GCF_001271735.1_Salmonella_enterica_CVM_N46839_v1.0	Salmonella enterica subsp. enterica serovar Enteritidis	Proteobacteria; Gammaproteobacteria; Enterobacterales; Enterobacteriaceae; Salmonella; Salmonella enterica; Salmonella enterica subsp. enterica	MTGDNTLITSHGINRRDFMKLCAALAATMGLSSKAAAEMAESVSNPQRPPVIWIGAQECT	WP_000145429.1	hydrogenase 2 small subunit [Salmonella enterica]	Length: 372\nNumber of predicted TMHs:  1\nExp number of AAs in TMHs: 24.51309\nExp number, first 60 AAs:  0.2682\nTotal prob of N-in:   0.14402\noutside   1   331\nTMhelix   332   354\ninside   355   372

16595	GCF_001271685.1_Salmonella_enterica_CVM_N46822_v1.0	Salmonella enterica subsp. enterica serovar Enteritidis	Proteobacteria; Gammaproteobacteria; Enterobacterales; Enterobacteriaceae; Salmonella; Salmonella enterica; Salmonella enterica subsp. enterica	MTGDNTLITSHGINRRDFMKLCAALAATMGLSSKAAAEMAESVSNPQRPPVIWIGAQECT	WP_000145429.1	hydrogenase 2 small subunit [Salmonella enterica]	Length: 372\nNumber of predicted TMHs:  1\nExp number of AAs in TMHs: 24.51309\nExp number, first 60 AAs:  0.2682\nTotal prob of N-in:   0.14402\noutside   1   331\nTMhelix   332   354\ninside   355   372

16596	GCF_001271615.1_Salmonella_enterica_CVM_N45409_v1.0	Salmonella enterica subsp. enterica serovar Enteritidis	Proteobacteria; Gammaproteobacteria; Enterobacterales; Enterobacteriaceae; Salmonella; Salmonella enterica; Salmonella enterica subsp. enterica	MNNEETFYQAMRRKGVTRRSFLKFCSLAATSLGLGAGMTPKIAWALENKPRIPVVWIHGL	WP_001058307.1	hydrogenase [Salmonella enterica]	Length: 372\nNumber of predicted TMHs:  2\nExp number of AAs in TMHs: 36.51738\nExp number, first 60 AAs:  19.72849\nTotal prob of N-in:   0.96170\nPOSSIBLE N-term signal sequence\ninside   1   19\nTMhelix   20   39\noutside   40   325\nTMhelix   326   348\ninside   349   372

16597	GCF_001271615.1_Salmonella_enterica_CVM_N45409_v1.0	Salmonella enterica subsp. enterica serovar Enteritidis	Proteobacteria; Gammaproteobacteria; Enterobacterales; Enterobacteriaceae; Salmonella; Salmonella enterica; Salmonella enterica subsp. enterica	MTGDNTLITSHGINRRDFMKLCAALAATMGLSSKAAAEMAESVSNPQRPPVIWIGAQECT	WP_000145429.1	hydrogenase 2 small subunit [Salmonella enterica]	Length: 372\nNumber of predicted TMHs:  1\nExp number of AAs in TMHs: 24.51309\nExp number, first 60 AAs:  0.2682\nTotal prob of N-in:   0.14402\noutside   1   331\nTMhelix   332   354\ninside   355   372

16598	GCF_001271575.1_Salmonella_enterica_CVM_N45407_v1.0	Salmonella enterica subsp. enterica serovar Enteritidis	Proteobacteria; Gammaproteobacteria; Enterobacterales; Enterobacteriaceae; Salmonella; Salmonella

enterica; Salmonella enterica subsp. enterica
MNNEETFYQAMRRKGVTRRSFLKFCSLAATSLGLGAGMTPKIAWALENKPRIPVVWIHGL WP_001058307.1 hydrogenase [Salmonella enterica] Length: 372\nNumber of predicted TMHs: 2\nExp number of AAs in TMHs: 36.51738\nExp number, first 60 AAs: 19.72849\nTotal prob of N-in: 0.96170\nPOSSIBLE N-term signal sequence\ninside 1 19\nTMhelix 20 39\noutside 40 325\nTMhelix 326 348\ninside 349 372

16599 GCF_001271575.1_Salmonella_enterica_CVM_N45407_v1.0 Salmonella enterica subsp. enterica serovar Enteritidis Proteobacteria; Gammaproteobacteria; Enterobacterales; Enterobacteriaceae; Salmonella; Salmonella enterica; Salmonella enterica subsp. enterica
MTGDNTLITSHGINRRDFMKLCAALAATMGLSSKAAAEMAESVSNPQRPPVIWIGAQECT WP_000145429.1 hydrogenase 2 small subunit [Salmonella enterica] Length: 372\nNumber of predicted TMHs: 1\nExp number of AAs in TMHs: 24.51309\nExp number, first 60 AAs: 0.2682\nTotal prob of N-in: 0.14402\noutside 1 331\nTMhelix 332 354\ninside 355 372

16600 GCF_001271415.1_Salmonella_enterica_CVM_N43468_v1.0 Salmonella enterica subsp. enterica serovar Enteritidis Proteobacteria; Gammaproteobacteria; Enterobacterales; Enterobacteriaceae; Salmonella; Salmonella enterica; Salmonella enterica subsp. enterica
MNNEETFYQAMRRKGVTRRSFLKFCSLAATSLGLGAGMTPKIAWALENKPRIPVVWIHGL WP_001058307.1 hydrogenase [Salmonella enterica] Length: 372\nNumber of predicted TMHs: 2\nExp number of AAs in TMHs: 36.51738\nExp number, first 60 AAs: 19.72849\nTotal prob of N-in: 0.96170\nPOSSIBLE N-term signal sequence\ninside 1 19\nTMhelix 20 39\noutside 40 325\nTMhelix 326 348\ninside 349 372

16601 GCF_001271415.1_Salmonella_enterica_CVM_N43468_v1.0 Salmonella enterica subsp. enterica serovar Enteritidis Proteobacteria; Gammaproteobacteria; Enterobacterales; Enterobacteriaceae; Salmonella; Salmonella enterica; Salmonella enterica subsp. enterica
MTGDNTLITSHGINRRDFMKLCAALAATMGLSSKAAAEMAESVSNPQRPPVIWIGAQECT WP_000145429.1 hydrogenase 2 small subunit [Salmonella enterica] Length: 372\nNumber of predicted TMHs: 1\nExp number of AAs in TMHs: 24.51309\nExp number, first 60 AAs: 0.2682\nTotal prob of N-in: 0.14402\noutside 1 331\nTMhelix 332 354\ninside 355 372

16602 GCF_001245355.1_Salmonella_enterica_CVM_N50424_v1.0 Salmonella enterica subsp. enterica serovar Enteritidis Proteobacteria; Gammaproteobacteria; Enterobacterales; Enterobacteriaceae; Salmonella; Salmonella enterica; Salmonella enterica subsp. enterica
MNNEETFYQAMRRKGVTRRSFLKFCSLAATSLGLGAGMTPKIAWALENKPRIPVVWIHGL WP_001058307.1 hydrogenase [Salmonella enterica] Length: 372\nNumber of predicted TMHs: 2\nExp number of AAs in TMHs: 36.51738\nExp number, first 60 AAs: 19.72849\nTotal prob of N-in: 0.96170\nPOSSIBLE N-term signal sequence\ninside 1 19\nTMhelix 20 39\noutside 40 325\nTMhelix 326 348\ninside 349 372

16603 GCF_001245355.1_Salmonella_enterica_CVM_N50424_v1.0 Salmonella enterica subsp. enterica serovar Enteritidis Proteobacteria; Gammaproteobacteria; Enterobacterales; Enterobacteriaceae; Salmonella; Salmonella enterica; Salmonella enterica subsp. enterica
MTGDNTLITSHGINRRDFMKLCAALAATMGLSSKAAAEMAESVSNPQRPPVIWIGAQECT WP_000145429.1 hydrogenase 2 small subunit [Salmonella enterica] Length: 372\nNumber of predicted TMHs: 1\nExp number of AAs in TMHs: 24.51309\nExp number, first 60 AAs: 0.2682\nTotal prob of N-in: 0.14402\noutside 1 331\nTMhelix 332 354\ninside 355 372

16604 GCF_001240255.1_Salmonella_enterica_CVM_N43457_v1.0 Salmonella enterica subsp. enterica serovar Enteritidis Proteobacteria; Gammaproteobacteria; Enterobacterales; Enterobacteriaceae; Salmonella; Salmonella enterica; Salmonella enterica subsp. enterica
MTGDNTLITSHGINRRDFMKLCAALAATMGLSSKAAAEMAESVSNPQRPPVIWIGAQECT WP_000145429.1 hydrogenase 2 small subunit [Salmonella enterica] Length: 372\nNumber of predicted TMHs: 1\nExp number of AAs in TMHs: 24.51309\nExp number, first 60 AAs: 0.2682\nTotal prob of N-in: 0.14402\noutside 1 331\nTMhelix 332 354\ninside 355 372

16605 GCF_001102865.1_8616_4_55 Salmonella enterica subsp. enterica serovar Enteritidis Proteobacteria; Gammaproteobacteria; Enterobacterales; Enterobacteriaceae; Salmonella; Salmonella enterica; Salmonella enterica subsp. enterica MTGDNTLITSHGINRRDFMKLCAALAATMGLSSKAAAEMAESVSNPQRPPVIWIGAQECT WP_000145429.1 hydrogenase 2 small subunit [Salmonella enterica] Length: 372\nNumber of predicted TMHs: 1\nExp number of AAs in TMHs: 24.51309\nExp number, first 60 AAs: 0.2682\nTotal prob of N-in: 0.14402\noutside 1 331\nTMhelix 332 354\ninside 355 372

16606 GCF_001240255.1_Salmonella_enterica_CVM_N43457_v1.0 Salmonella enterica subsp. enterica serovar Enteritidis Proteobacteria; Gammaproteobacteria; Enterobacterales; Enterobacteriaceae; Salmonella; Salmonella enterica; Salmonella enterica subsp. enterica MNNEETFYQAMRRKGVTRRSFLKFCSLAATSLGLGAGMTPKIAWALENKPRIPVVWIHGL WP_001058307.1 hydrogenase [Salmonella enterica] Length: 372\nNumber of predicted TMHs: 2\nExp number of AAs in TMHs: 36.51738\nExp number, first 60 AAs: 19.72849\nTotal prob of N-in: 0.96170\nPOSSIBLE N-term signal sequence\ninside 1 19\nTMhelix 20 39\noutside 40 325\nTMhelix 326 348\ninside 349 372

16607 GCF_001185245.1_ASM118524v1 Salmonella enterica subsp. enterica serovar Enteritidis Proteobacteria; Gammaproteobacteria; Enterobacterales; Enterobacteriaceae; Salmonella; Salmonella enterica; Salmonella enterica subsp. enterica MNNEETFYQAMRRKGVTRRSFLKFCSLAATSLGLGAGMTPKIAWALENKPRIPVVWIHGL WP_001058307.1 hydrogenase [Salmonella enterica] Length: 372\nNumber of predicted TMHs: 2\nExp number of AAs in TMHs: 36.51738\nExp number, first 60 AAs: 19.72849\nTotal prob of N-in: 0.96170\nPOSSIBLE N-term signal sequence\ninside 1 19\nTMhelix 20 39\noutside 40 325\nTMhelix 326 348\ninside 349 372

16608 GCF_001185245.1_ASM118524v1 Salmonella enterica subsp. enterica serovar Enteritidis Proteobacteria; Gammaproteobacteria; Enterobacterales; Enterobacteriaceae; Salmonella; Salmonella enterica; Salmonella enterica subsp. enterica MTGDNTLITSHGINRRDFMKLCAALAATMGLSSKAAAEMAESVSNPQRPPVIWIGAQECT WP_000145429.1 hydrogenase 2 small subunit [Salmonella enterica] Length: 372\nNumber of predicted TMHs: 1\nExp number of AAs in TMHs: 24.51309\nExp number, first 60 AAs: 0.2682\nTotal prob of N-in: 0.14402\noutside 1 331\nTMhelix 332 354\ninside 355 372

16609 GCF_000750495.1_ASM75049v1 Salmonella enterica subsp. enterica serovar Enteritidis Proteobacteria; Gammaproteobacteria; Enterobacterales; Enterobacteriaceae; Salmonella; Salmonella enterica; Salmonella enterica subsp. enterica MNNEETFYQAMRRKGVTRRSFLKFCSLAATSLGLGAGMTPKIAWALENKPRIPVVWIHGL WP_001058307.1 hydrogenase [Salmonella enterica] Length: 372\nNumber of predicted TMHs: 2\nExp number of AAs in TMHs: 36.51738\nExp number, first 60 AAs: 19.72849\nTotal prob of N-in: 0.96170\nPOSSIBLE N-term signal sequence\ninside 1 19\nTMhelix 20 39\noutside 40 325\nTMhelix 326 348\ninside 349 372

16610 GCF_000750455.1_ASM75045v1 Salmonella enterica subsp. enterica serovar Enteritidis Proteobacteria; Gammaproteobacteria; Enterobacterales; Enterobacteriaceae; Salmonella; Salmonella enterica; Salmonella enterica subsp. enterica MNNEETFYQAMRRKGVTRRSFLKFCSLAATSLGLGAGMTPKIAWALENKPRIPVVWIHGL WP_001058307.1 hydrogenase [Salmonella enterica] Length: 372\nNumber of predicted TMHs: 2\nExp number of AAs in TMHs: 36.51738\nExp number, first 60 AAs: 19.72849\nTotal prob of N-in: 0.96170\nPOSSIBLE N-term signal sequence\ninside 1 19\nTMhelix 20 39\noutside 40 325\nTMhelix 326 348\ninside 349 372

16611 GCF_000750395.2_ASM75039v2 Salmonella enterica subsp. enterica serovar Enteritidis Proteobacteria; Gammaproteobacteria; Enterobacterales; Enterobacteriaceae; Salmonella; Salmonella enterica; Salmonella enterica subsp. enterica MNNEETFYQAMRRKGVTRRSFLKFCSLAATSLGLGAGMTPKIAWALENKPRIPVVWIHGL WP_001058307.1 hydrogenase [Salmonella enterica] Length: 372\nNumber of predicted TMHs: 2\nExp number of AAs in TMHs: 36.51738\nExp number, first 60 AAs: 19.72849\nTotal prob of N-in: 0.96170\nPOSSIBLE N-term signal sequence\ninside 1 19\nTMhelix 20 39\noutside 40 325\nTMhelix 326 348\ninside 349 372

16612 GCF_000750335.1_ASM75033v1 Salmonella enterica subsp. enterica serovar Enteritidis Proteobacteria; Gammaproteobacteria; Enterobacterales; Enterobacteriaceae; Salmonella; Salmonella enterica; Salmonella enterica subsp. enterica MNNEETFYQAMRRKGVTRRSFLKFCSLAATSLGLGAGMTPKIAWALENKPRIPVVWIHGL WP_001058307.1 hydrogenase [Salmonella enterica] Length: 372\nNumber of predicted TMHs: 2\nExp number of AAs in TMHs: 36.51738\nExp number, first 60 AAs: 19.72849\nTotal prob of N-in: 0.96170\nPOSSIBLE N-term signal sequence\ninside 1 19\nTMhelix 20 39\noutside 40 325\nTMhelix 326 348\ninside 349 372

16613 GCF_000750215.1_ASM75021v1 Salmonella enterica subsp. enterica serovar Enteritidis Proteobacteria; Gammaproteobacteria; Enterobacterales; Enterobacteriaceae; Salmonella; Salmonella enterica; Salmonella enterica subsp. enterica MNNEETFYQAMRRKGVTRRSFLKFCSLAATSLGLGAGMTPKIAWALENKPRIPVVWIHGL WP_001058307.1 hydrogenase [Salmonella enterica] Length: 372\nNumber of predicted TMHs: 2\nExp number of AAs in TMHs: 36.51738\nExp number, first 60 AAs: 19.72849\nTotal prob of N-in: 0.96170\nPOSSIBLE N-term signal sequence\ninside 1 19\nTMhelix 20 39\noutside 40 325\nTMhelix 326 348\ninside 349 372

16614 GCF_000750455.1_ASM75045v1 Salmonella enterica subsp. enterica serovar Enteritidis Proteobacteria; Gammaproteobacteria; Enterobacterales; Enterobacteriaceae; Salmonella; Salmonella enterica; Salmonella enterica subsp. enterica MTGDNTLITSHGINRRDFMKLCAALAATMGLSSKAAAEMAESVSNPQRPPVIWIGAQECT WP_000145429.1 hydrogenase 2 small subunit [Salmonella enterica] Length: 372\nNumber of

predicted TMHs:  1\nExp number of AAs in TMHs: 24.51309\nExp number, first 60 AAs:  0.2682\nTotal prob of N-in: 0.14402\noutside   1   331\nTMhelix   332   354\ninside   355   372

16615 GCF_000750295.1_ASM75029v1 Salmonella enterica subsp. enterica serovar Enteritidis Proteobacteria; Gammaproteobacteria; Enterobacterales; Enterobacteriaceae; Salmonella; Salmonella enterica; Salmonella enterica subsp. enterica MNNEETFYQAMRRKGVTRRSFLKFCSLAATSLGLGAGMTPKIAWALENKPRIPVVWIHGL WP_001058307.1 hydrogenase [Salmonella enterica] Length: 372\nNumber of predicted TMHs:  2\nExp number of AAs in TMHs: 36.51738\nExp number, first 60 AAs:  19.72849\nTotal prob of N-in:   0.96170\nPOSSIBLE N-term signal sequence\ninside   1   19\nTMhelix   20   39\noutside   40   325\nTMhelix   326   348\ninside   349   372

16616 GCF_000750335.1_ASM75033v1 Salmonella enterica subsp. enterica serovar Enteritidis Proteobacteria; Gammaproteobacteria; Enterobacterales; Enterobacteriaceae; Salmonella; Salmonella enterica; Salmonella enterica subsp. enterica MTGDNTLITSHGINRRDFMKLCAALAATMGLSSKAAAEMAESVSNPQRPPVIWIGAQECT WP_000145429.1 hydrogenase 2 small subunit [Salmonella enterica] Length: 372\nNumber of predicted TMHs:  1\nExp number of AAs in TMHs: 24.51309\nExp number, first 60 AAs:  0.2682\nTotal prob of N-in: 0.14402\noutside   1   331\nTMhelix   332   354\ninside   355   372

16617 GCF_000750215.1_ASM75021v1 Salmonella enterica subsp. enterica serovar Enteritidis Proteobacteria; Gammaproteobacteria; Enterobacterales; Enterobacteriaceae; Salmonella; Salmonella enterica; Salmonella enterica subsp. enterica MTGDNTLITSHGINRRDFMKLCAALAATMGLSSKAAAEMAESVSNPQRPPVIWIGAQECT WP_000145429.1 hydrogenase 2 small subunit [Salmonella enterica] Length: 372\nNumber of predicted TMHs:  1\nExp number of AAs in TMHs: 24.51309\nExp number, first 60 AAs:  0.2682\nTotal prob of N-in: 0.14402\noutside   1   331\nTMhelix   332   354\ninside   355   372

16618 GCF_001245855.1_Salmonella_enterica_CVM_N50448_v1.0 Salmonella enterica subsp. enterica serovar Enteritidis Proteobacteria; Gammaproteobacteria; Enterobacterales; Enterobacteriaceae; Salmonella; Salmonella enterica; Salmonella enterica subsp. enterica MTGDNTLITSHGINRRDFMKLCAALAATMGLSSKAAAEMAESVSNPQRPPVIWIGAQECT WP_000145429.1 hydrogenase 2 small subunit [Salmonella enterica] Length: 372\nNumber of predicted TMHs:  1\nExp number of AAs in TMHs: 24.51309\nExp number, first 60 AAs:  0.2682\nTotal prob of N-in:   0.14402\noutside   1   331\nTMhelix   332   354\ninside   355   372

16619 GCF_001245855.1_Salmonella_enterica_CVM_N50448_v1.0 Salmonella enterica subsp. enterica serovar Enteritidis Proteobacteria; Gammaproteobacteria; Enterobacterales; Enterobacteriaceae; Salmonella; Salmonella enterica; Salmonella enterica subsp. enterica MNNEETFYQAMRRKGVTRRSFLKFCSLAATSLGLGAGMTPKIAWALENKPRIPVVWIHGL WP_001058307.1 hydrogenase [Salmonella enterica] Length: 372\nNumber of predicted TMHs:  2\nExp number of AAs in TMHs: 36.51738\nExp number, first 60 AAs:  19.72849\nTotal prob of N-in:   0.96170\nPOSSIBLE N-term signal sequence\ninside   1   19\nTMhelix   20   39\noutside   40   325\nTMhelix   326   348\ninside   349   372

16620 GCF_000750395.2_ASM75039v2 Salmonella enterica subsp. enterica serovar Enteritidis Proteobacteria; Gammaproteobacteria; Enterobacterales; Enterobacteriaceae; Salmonella; Salmonella enterica; Salmonella enterica subsp. enterica MTGDNTLITSHGINRRDFMKLCAALAATMGLSSKAAAEMAESVSNPQRPPVIWIGAQECT WP_000145429.1 hydrogenase 2 small subunit [Salmonella enterica] Length: 372\nNumber of predicted TMHs:  1\nExp number of AAs in TMHs: 24.51309\nExp number, first 60 AAs:  0.2682\nTotal prob of N-in: 0.14402\noutside   1   331\nTMhelix   332   354\ninside   355   372

16621 GCF_001280185.1_ASM128018v1 Salmonella enterica subsp. enterica serovar Enteritidis Proteobacteria; Gammaproteobacteria; Enterobacterales; Enterobacteriaceae; Salmonella; Salmonella enterica; Salmonella enterica subsp. enterica MNNEETFYQAMRRKGVTRRSFLKFCSLAATSLGLGAGMTPKIAWALENKPRIPVVWIHGL WP_001058307.1 hydrogenase [Salmonella enterica] Length: 372\nNumber of predicted TMHs:  2\nExp number of AAs in TMHs: 36.51738\nExp number, first 60 AAs:  19.72849\nTotal prob of N-in:   0.96170\nPOSSIBLE N-term signal sequence\ninside   1   19\nTMhelix   20   39\noutside   40   325\nTMhelix   326   348\ninside   349   372

16622 GCF_000750495.1_ASM75049v1 Salmonella enterica subsp. enterica serovar Enteritidis Proteobacteria; Gammaproteobacteria; Enterobacterales; Enterobacteriaceae; Salmonella; Salmonella enterica; Salmonella enterica subsp. enterica MTGDNTLITSHGINRRDFMKLCAALAATMGLSSKAAAEMAESVSNPQRPPVIWIGAQECT WP_000145429.1 hydrogenase 2 small subunit [Salmonella enterica] Length: 372\nNumber of predicted TMHs:  1\nExp number of AAs in TMHs: 24.51309\nExp number, first 60 AAs:  0.2682\nTotal prob of N-in: 0.14402\noutside   1   331\nTMhelix   332   354\ninside   355   372

16623 GCF_001272455.1_Salmonella_enterica_CVM_N51309_v1.0 Salmonella enterica subsp. enterica serovar Enteritidis Proteobacteria; Gammaproteobacteria; Enterobacterales; Enterobacteriaceae; Salmonella; Salmonella enterica; Salmonella enterica subsp. enterica
MNNEETFYQAMRRKGVTRRSFLKFCSLAATSLGLGAGMTPKIAWALENKPRIPVVWIHGL WP_001058307.1
hydrogenase [Salmonella enterica] Length: 372\nNumber of predicted TMHs: 2\nExp number of AAs in TMHs: 36.51738\nExp number, first 60 AAs: 19.72849\nTotal prob of N-in: 0.96170\nPOSSIBLE N-term signal sequence\ninside 1 19\nTMhelix 20 39\noutside 40 325\nTMhelix 326 348\ninside 349 372

16624 GCF_001272455.1_Salmonella_enterica_CVM_N51309_v1.0 Salmonella enterica subsp. enterica serovar Enteritidis Proteobacteria; Gammaproteobacteria; Enterobacterales; Enterobacteriaceae; Salmonella; Salmonella enterica; Salmonella enterica subsp. enterica
MTGDNTLITSHGINRRDFMKLCAALAATMGLSSKAAAEMAESVSNPQRPPVIWIGAQECT WP_000145429.1
hydrogenase 2 small subunit [Salmonella enterica] Length: 372\nNumber of predicted TMHs: 1\nExp number of AAs in TMHs: 24.51309\nExp number, first 60 AAs: 0.2682\nTotal prob of N-in: 0.14402\noutside 1 331\nTMhelix 332 354\ninside 355 372

16625 GCF_001272125.1_Salmonella_enterica_CVM_N48681_v1.0 Salmonella enterica subsp. enterica serovar Enteritidis Proteobacteria; Gammaproteobacteria; Enterobacterales; Enterobacteriaceae; Salmonella; Salmonella enterica; Salmonella enterica subsp. enterica
MNNEETFYQAMRRKGVTRRSFLKFCSLAATSLGLGAGMTPKIAWALENKPRIPVVWIHGL WP_001058307.1
hydrogenase [Salmonella enterica] Length: 372\nNumber of predicted TMHs: 2\nExp number of AAs in TMHs: 36.51738\nExp number, first 60 AAs: 19.72849\nTotal prob of N-in: 0.96170\nPOSSIBLE N-term signal sequence\ninside 1 19\nTMhelix 20 39\noutside 40 325\nTMhelix 326 348\ninside 349 372

16626 GCF_001272285.1_Salmonella_enterica_CVM_N50447_v1.0 Salmonella enterica subsp. enterica serovar Enteritidis Proteobacteria; Gammaproteobacteria; Enterobacterales; Enterobacteriaceae; Salmonella; Salmonella enterica; Salmonella enterica subsp. enterica
MNNEETFYQAMRRKGVTRRSFLKFCSLAATSLGLGAGMTPKIAWALENKPRIPVVWIHGL WP_001058307.1
hydrogenase [Salmonella enterica] Length: 372\nNumber of predicted TMHs: 2\nExp number of AAs in TMHs: 36.51738\nExp number, first 60 AAs: 19.72849\nTotal prob of N-in: 0.96170\nPOSSIBLE N-term signal sequence\ninside 1 19\nTMhelix 20 39\noutside 40 325\nTMhelix 326 348\ninside 349 372

16627 GCF_001272285.1_Salmonella_enterica_CVM_N50447_v1.0 Salmonella enterica subsp. enterica serovar Enteritidis Proteobacteria; Gammaproteobacteria; Enterobacterales; Enterobacteriaceae; Salmonella; Salmonella enterica; Salmonella enterica subsp. enterica
MTGDNTLITSHGINRRDFMKLCAALAATMGLSSKAAAEMAESVSNPQRPPVIWIGAQECT WP_000145429.1
hydrogenase 2 small subunit [Salmonella enterica] Length: 372\nNumber of predicted TMHs: 1\nExp number of AAs in TMHs: 24.51309\nExp number, first 60 AAs: 0.2682\nTotal prob of N-in: 0.14402\noutside 1 331\nTMhelix 332 354\ninside 355 372

16628 GCF_001272255.1_Salmonella_enterica_CVM_N50431_v1.0 Salmonella enterica subsp. enterica serovar Enteritidis Proteobacteria; Gammaproteobacteria; Enterobacterales; Enterobacteriaceae; Salmonella; Salmonella enterica; Salmonella enterica subsp. enterica
MNNEETFYQAMRRKGVTRRSFLKFCSLAATSLGLGAGMTPKIAWALENKPRIPVVWIHGL WP_001058307.1
hydrogenase [Salmonella enterica] Length: 372\nNumber of predicted TMHs: 2\nExp number of AAs in TMHs: 36.51738\nExp number, first 60 AAs: 19.72849\nTotal prob of N-in: 0.96170\nPOSSIBLE N-term signal sequence\ninside 1 19\nTMhelix 20 39\noutside 40 325\nTMhelix 326 348\ninside 349 372

16629 GCF_001272255.1_Salmonella_enterica_CVM_N50431_v1.0 Salmonella enterica subsp. enterica serovar Enteritidis Proteobacteria; Gammaproteobacteria; Enterobacterales; Enterobacteriaceae; Salmonella; Salmonella enterica; Salmonella enterica subsp. enterica
MTGDNTLITSHGINRRDFMKLCAALAATMGLSSKAAAEMAESVSNPQRPPVIWIGAQECT WP_000145429.1
hydrogenase 2 small subunit [Salmonella enterica] Length: 372\nNumber of predicted TMHs: 1\nExp number of AAs in TMHs: 24.51309\nExp number, first 60 AAs: 0.2682\nTotal prob of N-in: 0.14402\noutside 1 331\nTMhelix 332 354\ninside 355 372

16630 GCF_001272015.1_Salmonella_enterica_CVM_N43833_v1.0 Salmonella enterica subsp. enterica serovar Enteritidis Proteobacteria; Gammaproteobacteria; Enterobacterales; Enterobacteriaceae; Salmonella; Salmonella enterica; Salmonella enterica subsp. enterica
MNNEETFYQAMRRKGVTRRSFLKFCSLAATSLGLGAGMTPKIAWALENKPRIPVVWIHGL WP_001058307.1
hydrogenase [Salmonella enterica] Length: 372\nNumber of predicted TMHs: 2\nExp number of AAs in

TMHs: 36.51738\nExp number, first 60 AAs: 19.72849\nTotal prob of N-in: 0.96170\nPOSSIBLE N-term signal sequence\ninside 1 19\nTMhelix 20 39\noutside 40 325\nTMhelix 326 348\ninside 349 372

16631 GCF_001272125.1_Salmonella_enterica_CVM_N48681_v1.0 Salmonella enterica subsp. enterica serovar Enteritidis Proteobacteria; Gammaproteobacteria; Enterobacterales; Enterobacteriaceae; Salmonella; Salmonella enterica; Salmonella enterica subsp. enterica MTGDNTLITSHGINRRDFMKLCAALAATMGLSSKAAAEMAESVSNPQRPPVIWIGAQECT WP_000145429.1 hydrogenase 2 small subunit [Salmonella enterica] Length: 372\nNumber of predicted TMHs: 1\nExp number of AAs in TMHs: 24.51309\nExp number, first 60 AAs: 0.2682\nTotal prob of N-in: 0.14402\noutside 1 331\nTMhelix 332 354\ninside 355 372

16632 GCF_001272015.1_Salmonella_enterica_CVM_N43833_v1.0 Salmonella enterica subsp. enterica serovar Enteritidis Proteobacteria; Gammaproteobacteria; Enterobacterales; Enterobacteriaceae; Salmonella; Salmonella enterica; Salmonella enterica subsp. enterica MTGDNTLITSHGINRRDFMKLCAALAATMGLSSKAAAEMAESVSNPQRPPVIWIGAQECT WP_000145429.1 hydrogenase 2 small subunit [Salmonella enterica] Length: 372\nNumber of predicted TMHs: 1\nExp number of AAs in TMHs: 24.51309\nExp number, first 60 AAs: 0.2682\nTotal prob of N-in: 0.14402\noutside 1 331\nTMhelix 332 354\ninside 355 372

16633 GCF_001271895.1_Salmonella_enterica_CVM_N43451_v1.0 Salmonella enterica subsp. enterica serovar Enteritidis Proteobacteria; Gammaproteobacteria; Enterobacterales; Enterobacteriaceae; Salmonella; Salmonella enterica; Salmonella enterica subsp. enterica MNNEETFYQAMRRKGVTRRSFLKFCSLAATSLGLGAGMTPKIAWALENKPRIPVVWIHGL WP_001058307.1 hydrogenase [Salmonella enterica] Length: 372\nNumber of predicted TMHs: 2\nExp number of AAs in TMHs: 36.51738\nExp number, first 60 AAs: 19.72849\nTotal prob of N-in: 0.96170\nPOSSIBLE N-term signal sequence\ninside 1 19\nTMhelix 20 39\noutside 40 325\nTMhelix 326 348\ninside 349 372

16634 GCF_001271735.1_Salmonella_enterica_CVM_N46839_v1.0 Salmonella enterica subsp. enterica serovar Enteritidis Proteobacteria; Gammaproteobacteria; Enterobacterales; Enterobacteriaceae; Salmonella; Salmonella enterica; Salmonella enterica subsp. enterica MNNEETFYQAMRRKGVTRRSFLKFCSLAATSLGLGAGMTPKIAWALENKPRIPVVWIHGL WP_001058307.1 hydrogenase [Salmonella enterica] Length: 372\nNumber of predicted TMHs: 2\nExp number of AAs in TMHs: 36.51738\nExp number, first 60 AAs: 19.72849\nTotal prob of N-in: 0.96170\nPOSSIBLE N-term signal sequence\ninside 1 19\nTMhelix 20 39\noutside 40 325\nTMhelix 326 348\ninside 349 372

16635 GCF_001271685.1_Salmonella_enterica_CVM_N46822_v1.0 Salmonella enterica subsp. enterica serovar Enteritidis Proteobacteria; Gammaproteobacteria; Enterobacterales; Enterobacteriaceae; Salmonella; Salmonella enterica; Salmonella enterica subsp. enterica MNNEETFYQAMRRKGVTRRSFLKFCSLAATSLGLGAGMTPKIAWALENKPRIPVVWIHGL WP_001058307.1 hydrogenase [Salmonella enterica] Length: 372\nNumber of predicted TMHs: 2\nExp number of AAs in TMHs: 36.51738\nExp number, first 60 AAs: 19.72849\nTotal prob of N-in: 0.96170\nPOSSIBLE N-term signal sequence\ninside 1 19\nTMhelix 20 39\noutside 40 325\nTMhelix 326 348\ninside 349 372

16636 GCF_001713955.1_ASM171395v1 Salmonella enterica subsp. enterica serovar Enteritidis Proteobacteria; Gammaproteobacteria; Enterobacterales; Enterobacteriaceae; Salmonella; Salmonella enterica; Salmonella enterica subsp. enterica MNNEETFYQAMRRKGVTRRSFLKFCSLAATSLGLGAGMTPKIAWALENKPRIPVVWIHGL WP_001058307.1 hydrogenase [Salmonella enterica] Length: 372\nNumber of predicted TMHs: 2\nExp number of AAs in TMHs: 36.51738\nExp number, first 60 AAs: 19.72849\nTotal prob of N-in: 0.96170\nPOSSIBLE N-term signal sequence\ninside 1 19\nTMhelix 20 39\noutside 40 325\nTMhelix 326 348\ninside 349 372

16637 GCF_002041955.1_ASM204195v1 Salmonella enterica subsp. enterica serovar Enteritidis Proteobacteria; Gammaproteobacteria; Enterobacterales; Enterobacteriaceae; Salmonella; Salmonella enterica; Salmonella enterica subsp. enterica MNNEETFYQAMRRKGVTRRSFLKFCSLAATSLGLGAGMTPKIAWALENKPRIPVVWIHGL WP_001058307.1 hydrogenase [Salmonella enterica] Length: 372\nNumber of predicted TMHs: 2\nExp number of AAs in TMHs: 36.51738\nExp number, first 60 AAs: 19.72849\nTotal prob of N-in: 0.96170\nPOSSIBLE N-term signal sequence\ninside 1 19\nTMhelix 20 39\noutside 40 325\nTMhelix 326 348\ninside 349 372

16638 GCF_002042675.1_ASM204267v1 Salmonella enterica subsp. enterica serovar Enteritidis Proteobacteria; Gammaproteobacteria; Enterobacterales; Enterobacteriaceae; Salmonella; Salmonella enterica; Salmonella enterica subsp. enterica MTGDNTLITSHGINRRDFMKLCAALAATMGLSSKAAAEMAESVSNPQRPPVIWIGAQECT WP_000145429.1 hydrogenase 2 small subunit [Salmonella enterica] Length: 372\nNumber of

predicted TMHs: 1\nExp number of AAs in TMHs: 24.51309\nExp number, first 60 AAs: 0.2682\nTotal prob of N-in: 0.14402\noutside 1 331\nTMhelix 332 354\ninside 355 372

16639 GCF_001973105.1_ASM197310v1 Salmonella enterica subsp. enterica serovar Enteritidis Proteobacteria; Gammaproteobacteria; Enterobacterales; Enterobacteriaceae; Salmonella; Salmonella enterica; Salmonella enterica subsp. enterica MNNEETFYQAMRRKGVTRRSFLKFCSLAATSLGLGAGMTPKIAWALENKPRIPVVWIHGL WP_001058307.1 hydrogenase [Salmonella enterica] Length: 372\nNumber of predicted TMHs: 2\nExp number of AAs in TMHs: 36.51738\nExp number, first 60 AAs: 19.72849\nTotal prob of N-in: 0.96170\nPOSSIBLE N-term signal sequence\ninside 1 19\nTMhelix 20 39\noutside 40 325\nTMhelix 326 348\ninside 349 372

16640 GCF_002041775.1_ASM204177v1 Salmonella enterica subsp. enterica serovar Enteritidis Proteobacteria; Gammaproteobacteria; Enterobacterales; Enterobacteriaceae; Salmonella; Salmonella enterica; Salmonella enterica subsp. enterica MNNEETFYQAMRRKGVTRRSFLKFCSLAATSLGLGAGMTPKIAWALENKPRIPVVWIHGL WP_001058307.1 hydrogenase [Salmonella enterica] Length: 372\nNumber of predicted TMHs: 2\nExp number of AAs in TMHs: 36.51738\nExp number, first 60 AAs: 19.72849\nTotal prob of N-in: 0.96170\nPOSSIBLE N-term signal sequence\ninside 1 19\nTMhelix 20 39\noutside 40 325\nTMhelix 326 348\ninside 349 372

16641 GCF_002041775.1_ASM204177v1 Salmonella enterica subsp. enterica serovar Enteritidis Proteobacteria; Gammaproteobacteria; Enterobacterales; Enterobacteriaceae; Salmonella; Salmonella enterica; Salmonella enterica subsp. enterica MTGDNTLITSHGINRRDFMKLCAALAATMGLSSKAAAEMAESVSNPQRPPVIWIGAQECT WP_000145429.1 hydrogenase 2 small subunit [Salmonella enterica] Length: 372\nNumber of predicted TMHs: 1\nExp number of AAs in TMHs: 24.51309\nExp number, first 60 AAs: 0.2682\nTotal prob of N-in: 0.14402\noutside 1 331\nTMhelix 332 354\ninside 355 372

16642 GCF_002041625.1_ASM204162v1 Salmonella enterica subsp. enterica serovar Enteritidis Proteobacteria; Gammaproteobacteria; Enterobacterales; Enterobacteriaceae; Salmonella; Salmonella enterica; Salmonella enterica subsp. enterica MNNEETFYQAMRRKGVTRRSFLKFCSLAATSLGLGAGMTPKIAWALENKPRIPVVWIHGL WP_001058307.1 hydrogenase [Salmonella enterica] Length: 372\nNumber of predicted TMHs: 2\nExp number of AAs in TMHs: 36.51738\nExp number, first 60 AAs: 19.72849\nTotal prob of N-in: 0.96170\nPOSSIBLE N-term signal sequence\ninside 1 19\nTMhelix 20 39\noutside 40 325\nTMhelix 326 348\ninside 349 372

16643 GCF_002041755.1_ASM204175v1 Salmonella enterica subsp. enterica serovar Enteritidis Proteobacteria; Gammaproteobacteria; Enterobacterales; Enterobacteriaceae; Salmonella; Salmonella enterica; Salmonella enterica subsp. enterica MNNEETFYQAMRRKGVTRRSFLKFCSLAATSLGLGAGMTPKIAWALENKPRIPVVWIHGL WP_001058307.1 hydrogenase [Salmonella enterica] Length: 372\nNumber of predicted TMHs: 2\nExp number of AAs in TMHs: 36.51738\nExp number, first 60 AAs: 19.72849\nTotal prob of N-in: 0.96170\nPOSSIBLE N-term signal sequence\ninside 1 19\nTMhelix 20 39\noutside 40 325\nTMhelix 326 348\ninside 349 372

16644 GCF_002041755.1_ASM204175v1 Salmonella enterica subsp. enterica serovar Enteritidis Proteobacteria; Gammaproteobacteria; Enterobacterales; Enterobacteriaceae; Salmonella; Salmonella enterica; Salmonella enterica subsp. enterica MTGDNTLITSHGINRRDFMKLCAALAATMGLSSKAAAEMAESVSNPQRPPVIWIGAQECT WP_000145429.1 hydrogenase 2 small subunit [Salmonella enterica] Length: 372\nNumber of predicted TMHs: 1\nExp number of AAs in TMHs: 24.51309\nExp number, first 60 AAs: 0.2682\nTotal prob of N-in: 0.14402\noutside 1 331\nTMhelix 332 354\ninside 355 372

16645 GCF_002041605.1_ASM204160v1 Salmonella enterica subsp. enterica serovar Enteritidis Proteobacteria; Gammaproteobacteria; Enterobacterales; Enterobacteriaceae; Salmonella; Salmonella enterica; Salmonella enterica subsp. enterica MNNEETFYQAMRRKGVTRRSFLKFCSLAATSLGLGAGMTPKIAWALENKPRIPVVWIHGL WP_001058307.1 hydrogenase [Salmonella enterica] Length: 372\nNumber of predicted TMHs: 2\nExp number of AAs in TMHs: 36.51738\nExp number, first 60 AAs: 19.72849\nTotal prob of N-in: 0.96170\nPOSSIBLE N-term signal sequence\ninside 1 19\nTMhelix 20 39\noutside 40 325\nTMhelix 326 348\ninside 349 372

16646 GCF_002041365.1_ASM204136v1 Salmonella enterica subsp. enterica serovar Enteritidis Proteobacteria; Gammaproteobacteria; Enterobacterales; Enterobacteriaceae; Salmonella; Salmonella enterica; Salmonella enterica subsp. enterica MNNEETFYQAMRRKGVTRRSFLKFCSLAATSLGLGAGMTPKIAWALENKPRIPVVWIHGL WP_001058307.1 hydrogenase [Salmonella enterica] Length: 372\nNumber of predicted TMHs: 2\nExp number of AAs in TMHs: 36.51738\nExp number, first 60 AAs: 19.72849\nTotal prob of N-in: 0.96170\nPOSSIBLE N-term signal sequence\ninside 1 19\nTMhelix 20 39\noutside 40 325\nTMhelix 326 348\ninside 349 372

16647 GCF_002041385.1_ASM204138v1 Salmonella enterica subsp. enterica serovar Enteritidis Proteobacteria; Gammaproteobacteria; Enterobacterales; Enterobacteriaceae; Salmonella; Salmonella enterica;

Salmonella enterica subsp. enterica MTGDNTLITSHGINRRDFMKLCAALAATMGLSSKAAAEMAESVSNPQRPPVIWIGAQECT WP_000145429.1 hydrogenase 2 small subunit [Salmonella enterica] Length: 372\nNumber of predicted TMHs: 1\nExp number of AAs in TMHs: 24.51309\nExp number, first 60 AAs: 0.2682\nTotal prob of N-in: 0.14402\noutside 1 331\nTMhelix 332 354\ninside 355 372

16648 GCF_000612325.1_ASM61232v1 Salmonella enterica subsp. enterica serovar Enteritidis Proteobacteria; Gammaproteobacteria; Enterobacterales; Enterobacteriaceae; Salmonella; Salmonella enterica; Salmonella enterica subsp. enterica MTGDNTLITSHGINRRDFMKLCAALAATMGLSSKAAAEMAESVSNPQRPPVIWIGAQECT WP_000145429.1 hydrogenase 2 small subunit [Salmonella enterica] Length: 372\nNumber of predicted TMHs: 1\nExp number of AAs in TMHs: 24.51309\nExp number, first 60 AAs: 0.2682\nTotal prob of N-in: 0.14402\noutside 1 331\nTMhelix 332 354\ninside 355 372

16649 GCF_002041365.1_ASM204136v1 Salmonella enterica subsp. enterica serovar Enteritidis Proteobacteria; Gammaproteobacteria; Enterobacterales; Enterobacteriaceae; Salmonella; Salmonella enterica; Salmonella enterica subsp. enterica MTGDNTLITSHGINRRDFMKLCAALAATMGLSSKAAAEMAESVSNPQRPPVIWIGAQECT WP_000145429.1 hydrogenase 2 small subunit [Salmonella enterica] Length: 372\nNumber of predicted TMHs: 1\nExp number of AAs in TMHs: 24.51309\nExp number, first 60 AAs: 0.2682\nTotal prob of N-in: 0.14402\noutside 1 331\nTMhelix 332 354\ninside 355 372

16650 GCF_002041245.1_ASM204124v1 Salmonella enterica subsp. enterica serovar Enteritidis Proteobacteria; Gammaproteobacteria; Enterobacterales; Enterobacteriaceae; Salmonella; Salmonella enterica; Salmonella enterica subsp. enterica MNNEETFYQAMRRKGVTRRSFLKFCSLAATSLGLGAGMTPKIAWALENKPRIPVVWIHGL WP_001058307.1 hydrogenase [Salmonella enterica] Length: 372\nNumber of predicted TMHs: 2\nExp number of AAs in TMHs: 36.51738\nExp number, first 60 AAs: 19.72849\nTotal prob of N-in: 0.96170\nPOSSIBLE N-term signal sequence\ninside 1 19\nTMhelix 20 39\noutside 40 325\nTMhelix 326 348\ninside 349 372

16651 GCF_002041205.1_ASM204120v1 Salmonella enterica subsp. enterica serovar Enteritidis Proteobacteria; Gammaproteobacteria; Enterobacterales; Enterobacteriaceae; Salmonella; Salmonella enterica; Salmonella enterica subsp. enterica MNNEETFYQAMRRKGVTRRSFLKFCSLAATSLGLGAGMTPKIAWALENKPRIPVVWIHGL WP_001058307.1 hydrogenase [Salmonella enterica] Length: 372\nNumber of predicted TMHs: 2\nExp number of AAs in TMHs: 36.51738\nExp number, first 60 AAs: 19.72849\nTotal prob of N-in: 0.96170\nPOSSIBLE N-term signal sequence\ninside 1 19\nTMhelix 20 39\noutside 40 325\nTMhelix 326 348\ninside 349 372

16652 GCF_002040975.1_ASM204097v1 Salmonella enterica subsp. enterica serovar Enteritidis Proteobacteria; Gammaproteobacteria; Enterobacterales; Enterobacteriaceae; Salmonella; Salmonella enterica; Salmonella enterica subsp. enterica MNNEETFYQAMRRKGVTRRSFLKFCSLAATSLGLGAGMTPKIAWALENKPRIPVVWIHGL WP_001058307.1 hydrogenase [Salmonella enterica] Length: 372\nNumber of predicted TMHs: 2\nExp number of AAs in TMHs: 36.51738\nExp number, first 60 AAs: 19.72849\nTotal prob of N-in: 0.96170\nPOSSIBLE N-term signal sequence\ninside 1 19\nTMhelix 20 39\noutside 40 325\nTMhelix 326 348\ninside 349 372

16653 GCF_002041095.1_ASM204109v1 Salmonella enterica subsp. enterica serovar Enteritidis Proteobacteria; Gammaproteobacteria; Enterobacterales; Enterobacteriaceae; Salmonella; Salmonella enterica; Salmonella enterica subsp. enterica MTGDNTLITSHGINRRDFMKLCAALAATMGLSSKAAAEMAESVSNPQRPPVIWIGAQECT WP_000145429.1 hydrogenase 2 small subunit [Salmonella enterica] Length: 372\nNumber of predicted TMHs: 1\nExp number of AAs in TMHs: 24.51309\nExp number, first 60 AAs: 0.2682\nTotal prob of N-in: 0.14402\noutside 1 331\nTMhelix 332 354\ninside 355 372

16654 GCF_001238365.1_Salmonella_enterica_CVM_N43449_v1.0 Salmonella enterica subsp. enterica serovar Enteritidis Proteobacteria; Gammaproteobacteria; Enterobacterales; Enterobacteriaceae; Salmonella; Salmonella enterica; Salmonella enterica subsp. enterica MTGDNTLITSHGINRRDFMKLCAALAATMGLSSKAAAEMAESVSNPQRPPVIWIGAQECT WP_000145429.1 hydrogenase 2 small subunit [Salmonella enterica] Length: 372\nNumber of predicted TMHs: 1\nExp number of AAs in TMHs: 24.51309\nExp number, first 60 AAs: 0.2682\nTotal prob of N-in: 0.14402\noutside 1 331\nTMhelix 332 354\ninside 355 372

16655 GCF_002041065.1_ASM204106v1 Salmonella enterica subsp. enterica serovar Enteritidis Proteobacteria; Gammaproteobacteria; Enterobacterales; Enterobacteriaceae; Salmonella; Salmonella enterica; Salmonella enterica subsp. enterica MTGDNTLITSHGINRRDFMKLCAALAATMGLSSKAAAEMAESVSNPQRPPVIWIGAQECT WP_000145429.1 hydrogenase 2 small subunit [Salmonella enterica] Length: 372\nNumber of predicted TMHs: 1\nExp number of AAs in TMHs: 24.51309\nExp number, first 60 AAs: 0.2682\nTotal prob of N-in: 0.14402\noutside 1 331\nTMhelix 332 354\ninside 355 372

16656 GCF_001973515.1_ASM197351v1 Salmonella enterica subsp. enterica serovar Enteritidis Proteobacteria; Gammaproteobacteria; Enterobacterales; Enterobacteriaceae; Salmonella; Salmonella enterica; Salmonella enterica subsp. enterica MNNEETFYQAMRRKGVTRRSFLKFCSLAATSLGLGAGMTPKIAWALENKPRIPVVWIHGL WP_001058307.1 hydrogenase [Salmonella enterica] Length: 372\nNumber of predicted TMHs: 2\nExp number of AAs in TMHs: 36.51738\nExp number, first 60 AAs: 19.72849\nTotal prob of N-in: 0.96170\nPOSSIBLE N-term signal sequence\ninside 1 19\nTMhelix 20 39\noutside 40 325\nTMhelix 326 348\ninside 349 372

16657 GCF_002041025.1_ASM204102v1 Salmonella enterica subsp. enterica serovar Enteritidis Proteobacteria; Gammaproteobacteria; Enterobacterales; Enterobacteriaceae; Salmonella; Salmonella enterica; Salmonella enterica subsp. enterica MTGDNTLITSHGINRRDFMKLCAALAATMGLSSKAAAEMAESVSNPQRPPVIWIGAQECT WP_000145429.1 hydrogenase 2 small subunit [Salmonella enterica] Length: 372\nNumber of predicted TMHs: 1\nExp number of AAs in TMHs: 24.51309\nExp number, first 60 AAs: 0.2682\nTotal prob of N-in: 0.14402\noutside 1 331\nTMhelix 332 354\ninside 355 372

16658 GCF_001238365.1_Salmonella_enterica_CVM_N43449_v1.0 Salmonella enterica subsp. enterica serovar Enteritidis Proteobacteria; Gammaproteobacteria; Enterobacterales; Enterobacteriaceae; Salmonella; Salmonella enterica; Salmonella enterica subsp. enterica MNNEETFYQAMRRKGVTRRSFLKFCSLAATSLGLGAGMTPKIAWALENKPRIPVVWIHGL WP_001058307.1 hydrogenase [Salmonella enterica] Length: 372\nNumber of predicted TMHs: 2\nExp number of AAs in TMHs: 36.51738\nExp number, first 60 AAs: 19.72849\nTotal prob of N-in: 0.96170\nPOSSIBLE N-term signal sequence\ninside 1 19\nTMhelix 20 39\noutside 40 325\nTMhelix 326 348\ninside 349 372

16659 GCF_001972545.1_ASM197254v1 Salmonella enterica subsp. enterica serovar Enteritidis Proteobacteria; Gammaproteobacteria; Enterobacterales; Enterobacteriaceae; Salmonella; Salmonella enterica; Salmonella enterica subsp. enterica MNNEETFYQAMRRKGVTRRSFLKFCSLAATSLGLGAGMTPKIAWALENKPRIPVVWIHGL WP_001058307.1 hydrogenase [Salmonella enterica] Length: 372\nNumber of predicted TMHs: 2\nExp number of AAs in TMHs: 36.51738\nExp number, first 60 AAs: 19.72849\nTotal prob of N-in: 0.96170\nPOSSIBLE N-term signal sequence\ninside 1 19\nTMhelix 20 39\noutside 40 325\nTMhelix 326 348\ninside 349 372

16660 GCF_002040975.1_ASM204097v1 Salmonella enterica subsp. enterica serovar Enteritidis Proteobacteria; Gammaproteobacteria; Enterobacterales; Enterobacteriaceae; Salmonella; Salmonella enterica; Salmonella enterica subsp. enterica MTGDNTLITSHGINRRDFMKLCAALAATMGLSSKAAAEMAESVSNPQRPPVIWIGAQECT WP_000145429.1 hydrogenase 2 small subunit [Salmonella enterica] Length: 372\nNumber of predicted TMHs: 1\nExp number of AAs in TMHs: 24.51309\nExp number, first 60 AAs: 0.2682\nTotal prob of N-in: 0.14402\noutside 1 331\nTMhelix 332 354\ninside 355 372

16661 GCF_002038705.1_ASM203870v1 Salmonella enterica subsp. enterica serovar Enteritidis Proteobacteria; Gammaproteobacteria; Enterobacterales; Enterobacteriaceae; Salmonella; Salmonella enterica; Salmonella enterica subsp. enterica MTGDNTLITSHGINRRDFMKLCAALAATMGLSSKAAAEMAESVSNPQRPPVIWIGAQECT WP_000145429.1 hydrogenase 2 small subunit [Salmonella enterica] Length: 372\nNumber of predicted TMHs: 1\nExp number of AAs in TMHs: 24.51309\nExp number, first 60 AAs: 0.2682\nTotal prob of N-in: 0.14402\noutside 1 331\nTMhelix 332 354\ninside 355 372

16662 GCF_002042485.1_ASM204248v1 Salmonella enterica subsp. enterica serovar Enteritidis Proteobacteria; Gammaproteobacteria; Enterobacterales; Enterobacteriaceae; Salmonella; Salmonella enterica; Salmonella enterica subsp. enterica MNNEETFYQAMRRKGVTRRSFLKFCSLAATSLGLGAGMTPKIAWALENKPRIPVVWIHGL WP_001058307.1 hydrogenase [Salmonella enterica] Length: 372\nNumber of predicted TMHs: 2\nExp number of AAs in TMHs: 36.51738\nExp number, first 60 AAs: 19.72849\nTotal prob of N-in: 0.96170\nPOSSIBLE N-term signal sequence\ninside 1 19\nTMhelix 20 39\noutside 40 325\nTMhelix 326 348\ninside 349 372

16663 GCF_001973755.1_ASM197375v1 Salmonella enterica subsp. enterica serovar Enteritidis Proteobacteria; Gammaproteobacteria; Enterobacterales; Enterobacteriaceae; Salmonella; Salmonella enterica; Salmonella enterica subsp. enterica MNNEETFYQAMRRKGVTRRSFLKFCSLAATSLGLGAGMTPKIAWALENKPRIPVVWIHGL WP_001058307.1 hydrogenase [Salmonella enterica] Length: 372\nNumber of predicted TMHs: 2\nExp number of AAs in TMHs: 36.51738\nExp number, first 60 AAs: 19.72849\nTotal prob of N-in: 0.96170\nPOSSIBLE N-term signal sequence\ninside 1 19\nTMhelix 20 39\noutside 40 325\nTMhelix 326 348\ninside 349 372

16664 GCF_001973275.1_ASM197327v1 Salmonella enterica subsp. enterica serovar Enteritidis Proteobacteria; Gammaproteobacteria; Enterobacterales; Enterobacteriaceae; Salmonella; Salmonella enterica; Salmonella enterica subsp. enterica MNNEETFYQAMRRKGVTRRSFLKFCSLAATSLGLGAGMTPKIAWALENKPRIPVVWIHGL WP_001058307.1 hydrogenase [Salmonella enterica] Length: 372\nNumber of predicted TMHs: 2\nExp

number of AAs in TMHs: 36.51738\nExp number, first 60 AAs: 19.72849\nTotal prob of N-in: 0.96170\nPOSSIBLE N-term signal sequence\ninside 1 19\nTMhelix 20 39\noutside 40 325\nTMhelix 326 348\ninside 349 372

16665 GCF_001973325.1_ASM197332v1 Salmonella enterica subsp. enterica serovar Enteritidis Proteobacteria; Gammaproteobacteria; Enterobacterales; Enterobacteriaceae; Salmonella; Salmonella enterica; Salmonella enterica subsp. enterica MNNEETFYQAMRRKGVTRRSFLKFCSLAATSLGLGAGMTPKIAWALENKPRIPVVWIHGL WP_001058307.1 hydrogenase [Salmonella enterica] Length: 372\nNumber of predicted TMHs: 2\nExp number of AAs in TMHs: 36.51738\nExp number, first 60 AAs: 19.72849\nTotal prob of N-in: 0.96170\nPOSSIBLE N-term signal sequence\ninside 1 19\nTMhelix 20 39\noutside 40 325\nTMhelix 326 348\ninside 349 372

16666 GCF_001973065.1_ASM197306v1 Salmonella enterica subsp. enterica serovar Enteritidis Proteobacteria; Gammaproteobacteria; Enterobacterales; Enterobacteriaceae; Salmonella; Salmonella enterica; Salmonella enterica subsp. enterica MNNEETFYQAMRRKGVTRRSFLKFCSLAATSLGLGAGMTPKIAWALENKPRIPVVWIHGL WP_001058307.1 hydrogenase [Salmonella enterica] Length: 372\nNumber of predicted TMHs: 2\nExp number of AAs in TMHs: 36.51738\nExp number, first 60 AAs: 19.72849\nTotal prob of N-in: 0.96170\nPOSSIBLE N-term signal sequence\ninside 1 19\nTMhelix 20 39\noutside 40 325\nTMhelix 326 348\ninside 349 372

16667 GCF_001973355.1_ASM197335v1 Salmonella enterica subsp. enterica serovar Enteritidis Proteobacteria; Gammaproteobacteria; Enterobacterales; Enterobacteriaceae; Salmonella; Salmonella enterica; Salmonella enterica subsp. enterica MTGDNTLITSHGINRRDFMKLCAALAATMGLSSKAAAEMAESVSNPQRPPVIWIGAQECT WP_000145429.1 hydrogenase 2 small subunit [Salmonella enterica] Length: 372\nNumber of predicted TMHs: 1\nExp number of AAs in TMHs: 24.51309\nExp number, first 60 AAs: 0.2682\nTotal prob of N-in: 0.14402\noutside 1 331\nTMhelix 332 354\ninside 355 372

16668 GCF_002042525.1_ASM204252v1 Salmonella enterica subsp. enterica serovar Enteritidis Proteobacteria; Gammaproteobacteria; Enterobacterales; Enterobacteriaceae; Salmonella; Salmonella enterica; Salmonella enterica subsp. enterica MNNEETFYQAMRRKGVTRRSFLKFCSLAATSLGLGAGMTPKIAWALENKPRIPVVWIHGL WP_001058307.1 hydrogenase [Salmonella enterica] Length: 372\nNumber of predicted TMHs: 2\nExp number of AAs in TMHs: 36.51738\nExp number, first 60 AAs: 19.72849\nTotal prob of N-in: 0.96170\nPOSSIBLE N-term signal sequence\ninside 1 19\nTMhelix 20 39\noutside 40 325\nTMhelix 326 348\ninside 349 372

16669 GCF_000612325.1_ASM61232v1 Salmonella enterica subsp. enterica serovar Enteritidis Proteobacteria; Gammaproteobacteria; Enterobacterales; Enterobacteriaceae; Salmonella; Salmonella enterica; Salmonella enterica subsp. enterica MNNEETFYQAMRRKGVTRRSFLKFCSLAATSLGLGAGMTPKIAWALENKPRIPVVWIHGL WP_001058307.1 hydrogenase [Salmonella enterica] Length: 372\nNumber of predicted TMHs: 2\nExp number of AAs in TMHs: 36.51738\nExp number, first 60 AAs: 19.72849\nTotal prob of N-in: 0.96170\nPOSSIBLE N-term signal sequence\ninside 1 19\nTMhelix 20 39\noutside 40 325\nTMhelix 326 348\ninside 349 372

16670 GCF_002042565.1_ASM204256v1 Salmonella enterica subsp. enterica serovar Enteritidis Proteobacteria; Gammaproteobacteria; Enterobacterales; Enterobacteriaceae; Salmonella; Salmonella enterica; Salmonella enterica subsp. enterica MTGDNTLITSHGINRRDFMKLCAALAATMGLSSKAAAEMAESVSNPQRPPVIWIGAQECT WP_000145429.1 hydrogenase 2 small subunit [Salmonella enterica] Length: 372\nNumber of predicted TMHs: 1\nExp number of AAs in TMHs: 24.51309\nExp number, first 60 AAs: 0.2682\nTotal prob of N-in: 0.14402\noutside 1 331\nTMhelix 332 354\ninside 355 372

16671 GCF_001973805.1_ASM197380v1 Salmonella enterica subsp. enterica serovar Enteritidis Proteobacteria; Gammaproteobacteria; Enterobacterales; Enterobacteriaceae; Salmonella; Salmonella enterica; Salmonella enterica subsp. enterica MNNEETFYQAMRRKGVTRRSFLKFCSLAATSLGLGAGMTPKIAWALENKPRIPVVWIHGL WP_001058307.1 hydrogenase [Salmonella enterica] Length: 372\nNumber of predicted TMHs: 2\nExp number of AAs in TMHs: 36.51738\nExp number, first 60 AAs: 19.72849\nTotal prob of N-in: 0.96170\nPOSSIBLE N-term signal sequence\ninside 1 19\nTMhelix 20 39\noutside 40 325\nTMhelix 326 348\ninside 349 372

16672 GCF_001271675.1_Salmonella_enterica_CVM_N46817_v1.0 Salmonella enterica subsp. enterica serovar Enteritidis Proteobacteria; Gammaproteobacteria; Enterobacterales; Enterobacteriaceae; Salmonella; Salmonella enterica; Salmonella enterica subsp. enterica MTGDNTLITSHGINRRDFMKLCAALAATMGLSSKAAAEMAESVSNPQRPPVIWIGAQECT WP_000145429.1 hydrogenase 2 small subunit [Salmonella enterica] Length: 372\nNumber of predicted TMHs: 1\nExp number of AAs in TMHs: 24.51309\nExp number, first 60 AAs: 0.2682\nTotal prob of N-in: 0.14402\noutside 1 331\nTMhelix 332 354\ninside 355 372

16673 GCF_002042525.1_ASM204252v1 Salmonella enterica subsp. enterica serovar Enteritidis Proteobacteria; Gammaproteobacteria; Enterobacterales; Enterobacteriaceae; Salmonella; Salmonella enterica; Salmonella enterica subsp. enterica MTGDNTLITSHGINRRDFMKLCAALAATMGLSSKAAAEMAESVSNPQRPPVIWIGAQECT WP_000145429.1 hydrogenase 2 small subunit [Salmonella enterica] Length: 372\nNumber of predicted TMHs: 1\nExp number of AAs in TMHs: 24.51309\nExp number, first 60 AAs: 0.2682\nTotal prob of N-in: 0.14402\noutside 1 331\nTMhelix 332 354\ninside 355 372

16674 GCF_002042485.1_ASM204248v1 Salmonella enterica subsp. enterica serovar Enteritidis Proteobacteria; Gammaproteobacteria; Enterobacterales; Enterobacteriaceae; Salmonella; Salmonella enterica; Salmonella enterica subsp. enterica MTGDNTLITSHGINRRDFMKLCAALAATMGLSSKAAAEMAESVSNPQRPPVIWIGAQECT WP_000145429.1 hydrogenase 2 small subunit [Salmonella enterica] Length: 372\nNumber of predicted TMHs: 1\nExp number of AAs in TMHs: 24.51309\nExp number, first 60 AAs: 0.2682\nTotal prob of N-in: 0.14402\noutside 1 331\nTMhelix 332 354\ninside 355 372

16675 GCF_001973725.1_ASM197372v1 Salmonella enterica subsp. enterica serovar Enteritidis Proteobacteria; Gammaproteobacteria; Enterobacterales; Enterobacteriaceae; Salmonella; Salmonella enterica; Salmonella enterica subsp. enterica MNNEETFYQAMRRKGVTRRSFLKFCSLAATSLGLGAGMTPKIAWALENKPRIPVVWIHGL WP_001058307.1 hydrogenase [Salmonella enterica] Length: 372\nNumber of predicted TMHs: 2\nExp number of AAs in TMHs: 36.51738\nExp number, first 60 AAs: 19.72849\nTotal prob of N-in: 0.96170\nPOSSIBLE N-term signal sequence\ninside 1 19\nTMhelix 20 39\noutside 40 325\nTMhelix 326 348\ninside 349 372

16676 GCF_001973805.1_ASM197380v1 Salmonella enterica subsp. enterica serovar Enteritidis Proteobacteria; Gammaproteobacteria; Enterobacterales; Enterobacteriaceae; Salmonella; Salmonella enterica; Salmonella enterica subsp. enterica MTGDNTLITSHGINRRDFMKLCAALAATMGLSSKAAAEMAESVSNPQRPPVIWIGAQECT WP_000145429.1 hydrogenase 2 small subunit [Salmonella enterica] Length: 372\nNumber of predicted TMHs: 1\nExp number of AAs in TMHs: 24.51309\nExp number, first 60 AAs: 0.2682\nTotal prob of N-in: 0.14402\noutside 1 331\nTMhelix 332 354\ninside 355 372

16677 GCF_001271975.1_Salmonella_enterica_CVM_N43470_v1.0 Salmonella enterica subsp. enterica serovar Enteritidis Proteobacteria; Gammaproteobacteria; Enterobacterales; Enterobacteriaceae; Salmonella; Salmonella enterica; Salmonella enterica subsp. enterica MTGDNTLITSHGINRRDFMKLCAALAATMGLSSKAAAEMAESVSNPQRPPVIWIGAQECT WP_000145429.1 hydrogenase 2 small subunit [Salmonella enterica] Length: 372\nNumber of predicted TMHs: 1\nExp number of AAs in TMHs: 24.51309\nExp number, first 60 AAs: 0.2682\nTotal prob of N-in: 0.14402\noutside 1 331\nTMhelix 332 354\ninside 355 372

16678 GCF_001973755.1_ASM197375v1 Salmonella enterica subsp. enterica serovar Enteritidis Proteobacteria; Gammaproteobacteria; Enterobacterales; Enterobacteriaceae; Salmonella; Salmonella enterica; Salmonella enterica subsp. enterica MTGDNTLITSHGINRRDFMKLCAALAATMGLSSKAAAEMAESVSNPQRPPVIWIGAQECT WP_000145429.1 hydrogenase 2 small subunit [Salmonella enterica] Length: 372\nNumber of predicted TMHs: 1\nExp number of AAs in TMHs: 24.51309\nExp number, first 60 AAs: 0.2682\nTotal prob of N-in: 0.14402\noutside 1 331\nTMhelix 332 354\ninside 355 372

16679 GCF_002036335.1_ASM203633v1 Salmonella enterica subsp. enterica serovar Enteritidis Proteobacteria; Gammaproteobacteria; Enterobacterales; Enterobacteriaceae; Salmonella; Salmonella enterica; Salmonella enterica subsp. enterica MNNEETFYQAMRRKGVTRRSFLKFCSLAATSLGLGAGMTPKIAWALENKPRIPVVWIHGL WP_001058307.1 hydrogenase [Salmonella enterica] Length: 372\nNumber of predicted TMHs: 2\nExp number of AAs in TMHs: 36.51738\nExp number, first 60 AAs: 19.72849\nTotal prob of N-in: 0.96170\nPOSSIBLE N-term signal sequence\ninside 1 19\nTMhelix 20 39\noutside 40 325\nTMhelix 326 348\ninside 349 372

16680 GCF_001271975.1_Salmonella_enterica_CVM_N43470_v1.0 Salmonella enterica subsp. enterica serovar Enteritidis Proteobacteria; Gammaproteobacteria; Enterobacterales; Enterobacteriaceae; Salmonella; Salmonella enterica; Salmonella enterica subsp. enterica MNNEETFYQAMRRKGVTRRSFLKFCSLAATSLGLGAGMTPKIAWALENKPRIPVVWIHGL WP_001058307.1 hydrogenase [Salmonella enterica] Length: 372\nNumber of predicted TMHs: 2\nExp number of AAs in TMHs: 36.51738\nExp number, first 60 AAs: 19.72849\nTotal prob of N-in: 0.96170\nPOSSIBLE N-term signal sequence\ninside 1 19\nTMhelix 20 39\noutside 40 325\nTMhelix 326 348\ninside 349 372

16681 GCF_002042305.1_ASM204230v1 Salmonella enterica subsp. enterica serovar Enteritidis Proteobacteria; Gammaproteobacteria; Enterobacterales; Enterobacteriaceae; Salmonella; Salmonella enterica; Salmonella enterica subsp. enterica MNNEETFYQAMRRKGVTRRSFLKFCSLAATSLGLGAGMTPKIAWALENKPRIPVVWIHGL

WP_001058307.1 hydrogenase [Salmonella enterica] Length: 372\nNumber of predicted TMHs: 2\nExp number of AAs in TMHs: 36.51738\nExp number, first 60 AAs: 19.72849\nTotal prob of N-in: 0.96170\nPOSSIBLE N-term signal sequence\ninside 1 19\nTMhelix 20 39\noutside 40 325\nTMhelix 326 348\ninside 349 372

16682 GCF_002042225.1_ASM204222v1 Salmonella enterica subsp. enterica serovar Enteritidis Proteobacteria; Gammaproteobacteria; Enterobacterales; Enterobacteriaceae; Salmonella; Salmonella enterica; Salmonella enterica subsp. enterica MTGDNTLITSHGINRRDFMKLCAALAATMGLSSKAAAEMAESVSNPQRPPVIWIGAQECT WP_000145429.1 hydrogenase 2 small subunit [Salmonella enterica] Length: 372\nNumber of predicted TMHs: 1\nExp number of AAs in TMHs: 24.51309\nExp number, first 60 AAs: 0.2682\nTotal prob of N-in: 0.14402\noutside 1 331\nTMhelix 332 354\ninside 355 372

16683 GCF_001973355.1_ASM197335v1 Salmonella enterica subsp. enterica serovar Enteritidis Proteobacteria; Gammaproteobacteria; Enterobacterales; Enterobacteriaceae; Salmonella; Salmonella enterica; Salmonella enterica subsp. enterica MNNEETFYQAMRRKGVTRRSFLKFCSLAATSLGLGAGMTPKIAWALENKPRIPVVWIHGL WP_001058307.1 hydrogenase [Salmonella enterica] Length: 372\nNumber of predicted TMHs: 2\nExp number of AAs in TMHs: 36.51738\nExp number, first 60 AAs: 19.72849\nTotal prob of N-in: 0.96170\nPOSSIBLE N-term signal sequence\ninside 1 19\nTMhelix 20 39\noutside 40 325\nTMhelix 326 348\ninside 349 372

16684 GCF_001272215.1_Salmonella_enterica_CVM_N50426_v1.0 Salmonella enterica subsp. enterica serovar Enteritidis Proteobacteria; Gammaproteobacteria; Enterobacterales; Enterobacteriaceae; Salmonella; Salmonella enterica; Salmonella enterica subsp. enterica MNNEETFYQAMRRKGVTRRSFLKFCSLAATSLGLGAGMTPKIAWALENKPRIPVVWIHGL WP_001058307.1 hydrogenase [Salmonella enterica] Length: 372\nNumber of predicted TMHs: 2\nExp number of AAs in TMHs: 36.51738\nExp number, first 60 AAs: 19.72849\nTotal prob of N-in: 0.96170\nPOSSIBLE N-term signal sequence\ninside 1 19\nTMhelix 20 39\noutside 40 325\nTMhelix 326 348\ninside 349 372

16685 GCF_002042145.1_ASM204214v1 Salmonella enterica subsp. enterica serovar Enteritidis Proteobacteria; Gammaproteobacteria; Enterobacterales; Enterobacteriaceae; Salmonella; Salmonella enterica; Salmonella enterica subsp. enterica MTGDNTLITSHGINRRDFMKLCAALAATMGLSSKAAAEMAESVSNPQRPPVIWIGAQECT WP_000145429.1 hydrogenase 2 small subunit [Salmonella enterica] Length: 372\nNumber of predicted TMHs: 1\nExp number of AAs in TMHs: 24.51309\nExp number, first 60 AAs: 0.2682\nTotal prob of N-in: 0.14402\noutside 1 331\nTMhelix 332 354\ninside 355 372

16686 GCF_002042105.1_ASM204210v1 Salmonella enterica subsp. enterica serovar Enteritidis Proteobacteria; Gammaproteobacteria; Enterobacterales; Enterobacteriaceae; Salmonella; Salmonella enterica; Salmonella enterica subsp. enterica MTGDNTLITSHGINRRDFMKLCAALAATMGLSSKAAAEMAESVSNPQRPPVIWIGAQECT WP_000145429.1 hydrogenase 2 small subunit [Salmonella enterica] Length: 372\nNumber of predicted TMHs: 1\nExp number of AAs in TMHs: 24.51309\nExp number, first 60 AAs: 0.2682\nTotal prob of N-in: 0.14402\noutside 1 331\nTMhelix 332 354\ninside 355 372

16687 GCF_002042015.1_ASM204201v1 Salmonella enterica subsp. enterica serovar Enteritidis Proteobacteria; Gammaproteobacteria; Enterobacterales; Enterobacteriaceae; Salmonella; Salmonella enterica; Salmonella enterica subsp. enterica MNNEETFYQAMRRKGVTRRSFLKFCSLAATSLGLGAGMTPKIAWALENKPRIPVVWIHGL WP_001058307.1 hydrogenase [Salmonella enterica] Length: 372\nNumber of predicted TMHs: 2\nExp number of AAs in TMHs: 36.51738\nExp number, first 60 AAs: 19.72849\nTotal prob of N-in: 0.96170\nPOSSIBLE N-term signal sequence\ninside 1 19\nTMhelix 20 39\noutside 40 325\nTMhelix 326 348\ninside 349 372

16688 GCF_002039245.1_ASM203924v1 Salmonella enterica subsp. enterica serovar Enteritidis Proteobacteria; Gammaproteobacteria; Enterobacterales; Enterobacteriaceae; Salmonella; Salmonella enterica; Salmonella enterica subsp. enterica MNNEETFYQAMRRKGVTRRSFLKFCSLAATSLGLGAGMTPKIAWALENKPRIPVVWIHGL WP_001058307.1 hydrogenase [Salmonella enterica] Length: 372\nNumber of predicted TMHs: 2\nExp number of AAs in TMHs: 36.51738\nExp number, first 60 AAs: 19.72849\nTotal prob of N-in: 0.96170\nPOSSIBLE N-term signal sequence\ninside 1 19\nTMhelix 20 39\noutside 40 325\nTMhelix 326 348\ninside 349 372

16689 GCF_002039325.1_ASM203932v1 Salmonella enterica subsp. enterica serovar Enteritidis Proteobacteria; Gammaproteobacteria; Enterobacterales; Enterobacteriaceae; Salmonella; Salmonella enterica; Salmonella enterica subsp. enterica MNNEETFYQAMRRKGVTRRSFLKFCSLAATSLGLGAGMTPKIAWALENKPRIPVVWIHGL WP_001058307.1 hydrogenase [Salmonella enterica] Length: 372\nNumber of predicted TMHs: 2\nExp number of AAs in TMHs: 36.51738\nExp number, first 60 AAs: 19.72849\nTotal prob of N-in: 0.96170\nPOSSIBLE N-term signal sequence\ninside 1 19\nTMhelix 20 39\noutside 40 325\nTMhelix 326 348\ninside 349 372

16690 GCF_001243345.1_Salmonella_enterica_CVM_N46838_v1.0 Salmonella enterica subsp. enterica serovar Enteritidis Proteobacteria; Gammaproteobacteria; Enterobacterales; Enterobacteriaceae; Salmonella; Salmonella enterica; Salmonella enterica subsp. enterica MTGDNTLITSHGINRRDFMKLCAALAATMGLSSKAAAEMAESVSNPQRPPVIWIGAQECT WP_000145429.1 hydrogenase 2 small subunit [Salmonella enterica] Length: 372\nNumber of predicted TMHs: 1\nExp number of AAs in TMHs: 24.51309\nExp number, first 60 AAs: 0.2682\nTotal prob of N-in: 0.14402\noutside 1 331\nTMhelix 332 354\ninside 355 372

16691 GCF_000973935.1_ABBSB1004-1.id11_v1.0 Salmonella enterica subsp. enterica serovar Enteritidis Proteobacteria; Gammaproteobacteria; Enterobacterales; Enterobacteriaceae; Salmonella; Salmonella enterica; Salmonella enterica subsp. enterica MTGDNTLITSHGINRRDFMKLCAALAATMGLSSKAAAEMAESVSNPQRPPVIWIGAQECT WP_000145429.1 hydrogenase 2 small subunit [Salmonella enterica] Length: 372\nNumber of predicted TMHs: 1\nExp number of AAs in TMHs: 24.51309\nExp number, first 60 AAs: 0.2682\nTotal prob of N-in: 0.14402\noutside 1 331\nTMhelix 332 354\ninside 355 372

16692 GCF_001713955.1_ASM171395v1 Salmonella enterica subsp. enterica serovar Enteritidis Proteobacteria; Gammaproteobacteria; Enterobacterales; Enterobacteriaceae; Salmonella; Salmonella enterica; Salmonella enterica subsp. enterica MTGDNTLITSHGINRRDFMKLCAALAATMGLSSKAAAEMAESVSNPQRPPVIWIGAQECT WP_000145429.1 hydrogenase 2 small subunit [Salmonella enterica] Length: 372\nNumber of predicted TMHs: 1\nExp number of AAs in TMHs: 24.51309\nExp number, first 60 AAs: 0.2682\nTotal prob of N-in: 0.14402\noutside 1 331\nTMhelix 332 354\ninside 355 372

16693 GCF_002039285.1_ASM203928v1 Salmonella enterica subsp. enterica serovar Enteritidis Proteobacteria; Gammaproteobacteria; Enterobacterales; Enterobacteriaceae; Salmonella; Salmonella enterica; Salmonella enterica subsp. enterica MNNEETFYQAMRRKGVTRRSFLKFCSLAATSLGLGAGMTPKIAWALENKPRIPVVWIHGL WP_001058307.1 hydrogenase [Salmonella enterica] Length: 372\nNumber of predicted TMHs: 2\nExp number of AAs in TMHs: 36.51738\nExp number, first 60 AAs: 19.72849\nTotal prob of N-in: 0.96170\nPOSSIBLE N-term signal sequence\ninside 1 19\nTMhelix 20 39\noutside 40 325\nTMhelix 326 348\ninside 349 372

16694 GCF_002039445.1_ASM203944v1 Salmonella enterica subsp. enterica serovar Enteritidis Proteobacteria; Gammaproteobacteria; Enterobacterales; Enterobacteriaceae; Salmonella; Salmonella enterica; Salmonella enterica subsp. enterica MTGDNTLITSHGINRRDFMKLCAALAATMGLSSKAAAEMAESVSNPQRPPVIWIGAQECT WP_000145429.1 hydrogenase 2 small subunit [Salmonella enterica] Length: 372\nNumber of predicted TMHs: 1\nExp number of AAs in TMHs: 24.51309\nExp number, first 60 AAs: 0.2682\nTotal prob of N-in: 0.14402\noutside 1 331\nTMhelix 332 354\ninside 355 372

16695 GCF_002038905.1_ASM203890v1 Salmonella enterica subsp. enterica serovar Enteritidis Proteobacteria; Gammaproteobacteria; Enterobacterales; Enterobacteriaceae; Salmonella; Salmonella enterica; Salmonella enterica subsp. enterica MNNEETFYQAMRRKGVTRRSFLKFCSLAATSLGLGAGMTPKIAWALENKPRIPVVWIHGL WP_001058307.1 hydrogenase [Salmonella enterica] Length: 372\nNumber of predicted TMHs: 2\nExp number of AAs in TMHs: 36.51738\nExp number, first 60 AAs: 19.72849\nTotal prob of N-in: 0.96170\nPOSSIBLE N-term signal sequence\ninside 1 19\nTMhelix 20 39\noutside 40 325\nTMhelix 326 348\ninside 349 372

16696 GCF_002038975.1_ASM203897v1 Salmonella enterica subsp. enterica serovar Enteritidis Proteobacteria; Gammaproteobacteria; Enterobacterales; Enterobacteriaceae; Salmonella; Salmonella enterica; Salmonella enterica subsp. enterica MNNEETFYQAMRRKGVTRRSFLKFCSLAATSLGLGAGMTPKIAWALENKPRIPVVWIHGL WP_001058307.1 hydrogenase [Salmonella enterica] Length: 372\nNumber of predicted TMHs: 2\nExp number of AAs in TMHs: 36.51738\nExp number, first 60 AAs: 19.72849\nTotal prob of N-in: 0.96170\nPOSSIBLE N-term signal sequence\ninside 1 19\nTMhelix 20 39\noutside 40 325\nTMhelix 326 348\ninside 349 372

16697 GCF_002039025.1_ASM203902v1 Salmonella enterica subsp. enterica serovar Enteritidis Proteobacteria; Gammaproteobacteria; Enterobacterales; Enterobacteriaceae; Salmonella; Salmonella enterica; Salmonella enterica subsp. enterica MNNEETFYQAMRRKGVTRRSFLKFCSLAATSLGLGAGMTPKIAWALENKPRIPVVWIHGL WP_001058307.1 hydrogenase [Salmonella enterica] Length: 372\nNumber of predicted TMHs: 2\nExp number of AAs in TMHs: 36.51738\nExp number, first 60 AAs: 19.72849\nTotal prob of N-in: 0.96170\nPOSSIBLE N-term signal sequence\ninside 1 19\nTMhelix 20 39\noutside 40 325\nTMhelix 326 348\ninside 349 372

16698 GCF_002039025.1_ASM203902v1 Salmonella enterica subsp. enterica serovar Enteritidis Proteobacteria; Gammaproteobacteria; Enterobacterales; Enterobacteriaceae; Salmonella; Salmonella enterica; Salmonella enterica subsp. enterica MTGDNTLITSHGINRRDFMKLCAALAATMGLSSKAAAEMAESVSNPQRPPVIWIGAQECT WP_000145429.1 hydrogenase 2 small subunit [Salmonella enterica] Length: 372\nNumber of

predicted TMHs: 1\nExp number of AAs in TMHs: 24.51309\nExp number, first 60 AAs: 0.2682\nTotal prob of N-in: 0.14402\noutside 1 331\nTMhelix 332 354\ninside 355 372

16699 GCF_002038975.1_ASM203897v1 Salmonella enterica subsp. enterica serovar Enteritidis Proteobacteria; Gammaproteobacteria; Enterobacterales; Enterobacteriaceae; Salmonella; Salmonella enterica; Salmonella enterica subsp. enterica MTGDNTLITSHGINRRDFMKLCAALAATMGLSSKAAAEMAESVSNPQRPPVIWIGAQECT WP_000145429.1 hydrogenase 2 small subunit [Salmonella enterica] Length: 372\nNumber of predicted TMHs: 1\nExp number of AAs in TMHs: 24.51309\nExp number, first 60 AAs: 0.2682\nTotal prob of N-in: 0.14402\noutside 1 331\nTMhelix 332 354\ninside 355 372

16700 GCF_002038825.1_ASM203882v1 Salmonella enterica subsp. enterica serovar Enteritidis Proteobacteria; Gammaproteobacteria; Enterobacterales; Enterobacteriaceae; Salmonella; Salmonella enterica; Salmonella enterica subsp. enterica MNNEETFYQAMRRKGVTRRSFLKFCSLAATSLGLGAGMTPKIAWALENKPRIPVVWIHGL WP_001058307.1 hydrogenase [Salmonella enterica] Length: 372\nNumber of predicted TMHs: 2\nExp number of AAs in TMHs: 36.51738\nExp number, first 60 AAs: 19.72849\nTotal prob of N-in: 0.96170\nPOSSIBLE N-term signal sequence\ninside 1 19\nTMhelix 20 39\noutside 40 325\nTMhelix 326 348\ninside 349 372

16701 GCF_002061465.1_ASM206146v1 Salmonella enterica subsp. enterica serovar Enteritidis Proteobacteria; Gammaproteobacteria; Enterobacterales; Enterobacteriaceae; Salmonella; Salmonella enterica; Salmonella enterica subsp. enterica MTGDNTLITSHGINRRDFMKLCAALAATMGLSSKAAAEMAESVSNPQRPPVIWIGAQECT WP_000145429.1 hydrogenase 2 small subunit [Salmonella enterica] Length: 372\nNumber of predicted TMHs: 1\nExp number of AAs in TMHs: 24.51309\nExp number, first 60 AAs: 0.2682\nTotal prob of N-in: 0.14402\noutside 1 331\nTMhelix 332 354\ninside 355 372

16702 GCF_002038665.1_ASM203866v1 Salmonella enterica subsp. enterica serovar Enteritidis Proteobacteria; Gammaproteobacteria; Enterobacterales; Enterobacteriaceae; Salmonella; Salmonella enterica; Salmonella enterica subsp. enterica MNNEETFYQAMRRKGVTRRSFLKFCSLAATSLGLGAGMTPKIAWALENKPRIPVVWIHGL WP_001058307.1 hydrogenase [Salmonella enterica] Length: 372\nNumber of predicted TMHs: 2\nExp number of AAs in TMHs: 36.51738\nExp number, first 60 AAs: 19.72849\nTotal prob of N-in: 0.96170\nPOSSIBLE N-term signal sequence\ninside 1 19\nTMhelix 20 39\noutside 40 325\nTMhelix 326 348\ninside 349 372

16703 GCF_002042595.1_ASM204259v1 Salmonella enterica subsp. enterica serovar Enteritidis Proteobacteria; Gammaproteobacteria; Enterobacterales; Enterobacteriaceae; Salmonella; Salmonella enterica; Salmonella enterica subsp. enterica MTGDNTLITSHGINRRDFMKLCAALAATMGLSSKAAAEMAESVSNPQRPPVIWIGAQECT WP_000145429.1 hydrogenase 2 small subunit [Salmonella enterica] Length: 372\nNumber of predicted TMHs: 1\nExp number of AAs in TMHs: 24.51309\nExp number, first 60 AAs: 0.2682\nTotal prob of N-in: 0.14402\noutside 1 331\nTMhelix 332 354\ninside 355 372

16704 GCF_002038665.1_ASM203866v1 Salmonella enterica subsp. enterica serovar Enteritidis Proteobacteria; Gammaproteobacteria; Enterobacterales; Enterobacteriaceae; Salmonella; Salmonella enterica; Salmonella enterica subsp. enterica MTGDNTLITSHGINRRDFMKLCAALAATMGLSSKAAAEMAESVSNPQRPPVIWIGAQECT WP_000145429.1 hydrogenase 2 small subunit [Salmonella enterica] Length: 372\nNumber of predicted TMHs: 1\nExp number of AAs in TMHs: 24.51309\nExp number, first 60 AAs: 0.2682\nTotal prob of N-in: 0.14402\noutside 1 331\nTMhelix 332 354\ninside 355 372

16705 GCF_001713995.1_ASM171399v1 Salmonella enterica subsp. enterica serovar Enteritidis Proteobacteria; Gammaproteobacteria; Enterobacterales; Enterobacteriaceae; Salmonella; Salmonella enterica; Salmonella enterica subsp. enterica MTGDNTLITSHGINRRDFMKLCAALAATMGLSSKAAAEMAESVSNPQRPPVIWIGAQECT WP_000145429.1 hydrogenase 2 small subunit [Salmonella enterica] Length: 372\nNumber of predicted TMHs: 1\nExp number of AAs in TMHs: 24.51309\nExp number, first 60 AAs: 0.2682\nTotal prob of N-in: 0.14402\noutside 1 331\nTMhelix 332 354\ninside 355 372

16706 GCF_002038625.1_ASM203862v1 Salmonella enterica subsp. enterica serovar Enteritidis Proteobacteria; Gammaproteobacteria; Enterobacterales; Enterobacteriaceae; Salmonella; Salmonella enterica; Salmonella enterica subsp. enterica MTGDNTLITSHGINRRDFMKLCAALAATMGLSSKAAAEMAESVSNPQRPPVIWIGAQECT WP_000145429.1 hydrogenase 2 small subunit [Salmonella enterica] Length: 372\nNumber of predicted TMHs: 1\nExp number of AAs in TMHs: 24.51309\nExp number, first 60 AAs: 0.2682\nTotal prob of N-in: 0.14402\noutside 1 331\nTMhelix 332 354\ninside 355 372

16707 GCF_001714025.1_ASM171402v1 Salmonella enterica subsp. enterica serovar Enteritidis Proteobacteria; Gammaproteobacteria; Enterobacterales; Enterobacteriaceae; Salmonella; Salmonella enterica;

Salmonella enterica subsp. enterica MTGDNTLITSHGINRRDFMKLCAALAATMGLSSKAAAEMAESVSNPQRPPVIWIGAQECT WP_000145429.1 hydrogenase 2 small subunit [Salmonella enterica] Length: 372\nNumber of predicted TMHs: 1\nExp number of AAs in TMHs: 24.51309\nExp number, first 60 AAs: 0.2682\nTotal prob of N-in: 0.14402\noutside 1 331\nTMhelix 332 354\ninside 355 372

16708 GCF_002038585.1_ASM203858v1 Salmonella enterica subsp. enterica serovar Enteritidis Proteobacteria; Gammaproteobacteria; Enterobacterales; Enterobacteriaceae; Salmonella; Salmonella enterica; Salmonella enterica subsp. enterica MTGDNTLITSHGINRRDFMKLCAALAATMGLSSKAAAEMAESVSNPQRPPVIWIGAQECT WP_000145429.1 hydrogenase 2 small subunit [Salmonella enterica] Length: 372\nNumber of predicted TMHs: 1\nExp number of AAs in TMHs: 24.51309\nExp number, first 60 AAs: 0.2682\nTotal prob of N-in: 0.14402\noutside 1 331\nTMhelix 332 354\ninside 355 372

16709 GCF_002038455.1_ASM203845v1 Salmonella enterica subsp. enterica serovar Enteritidis Proteobacteria; Gammaproteobacteria; Enterobacterales; Enterobacteriaceae; Salmonella; Salmonella enterica; Salmonella enterica subsp. enterica MNNEETFYQAMRRKGVTRRSFLKFCSLAATSLGLGAGMTPKIAWALENKPRIPVVWIHGL WP_001058307.1 hydrogenase [Salmonella enterica] Length: 372\nNumber of predicted TMHs: 2\nExp number of AAs in TMHs: 36.51738\nExp number, first 60 AAs: 19.72849\nTotal prob of N-in: 0.96170\nPOSSIBLE N-term signal sequence\ninside 1 19\nTMhelix 20 39\noutside 40 325\nTMhelix 326 348\ninside 349 372

16710 GCF_002038505.1_ASM203850v1 Salmonella enterica subsp. enterica serovar Enteritidis Proteobacteria; Gammaproteobacteria; Enterobacterales; Enterobacteriaceae; Salmonella; Salmonella enterica; Salmonella enterica subsp. enterica MTGDNTLITSHGINRRDFMKLCAALAATMGLSSKAAAEMAESVSNPQRPPVIWIGAQECT WP_000145429.1 hydrogenase 2 small subunit [Salmonella enterica] Length: 372\nNumber of predicted TMHs: 1\nExp number of AAs in TMHs: 24.51309\nExp number, first 60 AAs: 0.2682\nTotal prob of N-in: 0.14402\noutside 1 331\nTMhelix 332 354\ninside 355 372

16711 GCF_002038185.1_ASM203818v1 Salmonella enterica subsp. enterica serovar Enteritidis Proteobacteria; Gammaproteobacteria; Enterobacterales; Enterobacteriaceae; Salmonella; Salmonella enterica; Salmonella enterica subsp. enterica MTGDNTLITSHGINRRDFMKLCAALAATMGLSSKAAAEMAESVSNPQRPPVIWIGAQECT WP_000145429.1 hydrogenase 2 small subunit [Salmonella enterica] Length: 372\nNumber of predicted TMHs: 1\nExp number of AAs in TMHs: 24.51309\nExp number, first 60 AAs: 0.2682\nTotal prob of N-in: 0.14402\noutside 1 331\nTMhelix 332 354\ninside 355 372

16712 GCF_001714075.1_ASM171407v1 Salmonella enterica subsp. enterica serovar Enteritidis Proteobacteria; Gammaproteobacteria; Enterobacterales; Enterobacteriaceae; Salmonella; Salmonella enterica; Salmonella enterica subsp. enterica MTGDNTLITSHGINRRDFMKLCAALAATMGLSSKAAAEMAESVSNPQRPPVIWIGAQECT WP_000145429.1 hydrogenase 2 small subunit [Salmonella enterica] Length: 372\nNumber of predicted TMHs: 1\nExp number of AAs in TMHs: 24.51309\nExp number, first 60 AAs: 0.2682\nTotal prob of N-in: 0.14402\noutside 1 331\nTMhelix 332 354\ninside 355 372

16713 GCF_002038135.1_ASM203813v1 Salmonella enterica subsp. enterica serovar Enteritidis Proteobacteria; Gammaproteobacteria; Enterobacterales; Enterobacteriaceae; Salmonella; Salmonella enterica; Salmonella enterica subsp. enterica MTGDNTLITSHGINRRDFMKLCAALAATMGLSSKAAAEMAESVSNPQRPPVIWIGAQECT WP_000145429.1 hydrogenase 2 small subunit [Salmonella enterica] Length: 372\nNumber of predicted TMHs: 1\nExp number of AAs in TMHs: 24.51309\nExp number, first 60 AAs: 0.2682\nTotal prob of N-in: 0.14402\noutside 1 331\nTMhelix 332 354\ninside 355 372

16714 GCF_002038105.1_ASM203810v1 Salmonella enterica subsp. enterica serovar Enteritidis Proteobacteria; Gammaproteobacteria; Enterobacterales; Enterobacteriaceae; Salmonella; Salmonella enterica; Salmonella enterica subsp. enterica MTGDNTLITSHGINRRDFMKLCAALAATMGLSSKAAAEMAESVSNPQRPPVIWIGAQECT WP_000145429.1 hydrogenase 2 small subunit [Salmonella enterica] Length: 372\nNumber of predicted TMHs: 1\nExp number of AAs in TMHs: 24.51309\nExp number, first 60 AAs: 0.2682\nTotal prob of N-in: 0.14402\noutside 1 331\nTMhelix 332 354\ninside 355 372

16715 GCF_002038055.1_ASM203805v1 Salmonella enterica subsp. enterica serovar Enteritidis Proteobacteria; Gammaproteobacteria; Enterobacterales; Enterobacteriaceae; Salmonella; Salmonella enterica; Salmonella enterica subsp. enterica MTGDNTLITSHGINRRDFMKLCAALAATMGLSSKAAAEMAESVSNPQRPPVIWIGAQECT WP_000145429.1 hydrogenase 2 small subunit [Salmonella enterica] Length: 372\nNumber of predicted TMHs: 1\nExp number of AAs in TMHs: 24.51309\nExp number, first 60 AAs: 0.2682\nTotal prob of N-in: 0.14402\noutside 1 331\nTMhelix 332 354\ninside 355 372

16716 GCF_002037945.1_ASM203794v1 Salmonella enterica subsp. enterica serovar Enteritidis Proteobacteria; Gammaproteobacteria; Enterobacterales; Enterobacteriaceae; Salmonella; Salmonella enterica; Salmonella enterica subsp. enterica MNNEETFYQAMRRKGVTRRSFLKFCSLAATSLGLGAGMTPKIAWALENKPRIPVVWIHGL WP_001058307.1 hydrogenase [Salmonella enterica] Length: 372\nNumber of predicted TMHs: 2\nExp number of AAs in TMHs: 36.51738\nExp number, first 60 AAs: 19.72849\nTotal prob of N-in: 0.96170\nPOSSIBLE N-term signal sequence\ninside 1 19\nTMhelix 20 39\noutside 40 325\nTMhelix 326 348\ninside 349 372

16717 GCF_002037895.1_ASM203789v1 Salmonella enterica subsp. enterica serovar Enteritidis Proteobacteria; Gammaproteobacteria; Enterobacterales; Enterobacteriaceae; Salmonella; Salmonella enterica; Salmonella enterica subsp. enterica MNNEETFYQAMRRKGVTRRSFLKFCSLAATSLGLGAGMTPKIAWALENKPRIPVVWIHGL WP_001058307.1 hydrogenase [Salmonella enterica] Length: 372\nNumber of predicted TMHs: 2\nExp number of AAs in TMHs: 36.51738\nExp number, first 60 AAs: 19.72849\nTotal prob of N-in: 0.96170\nPOSSIBLE N-term signal sequence\ninside 1 19\nTMhelix 20 39\noutside 40 325\nTMhelix 326 348\ninside 349 372

16718 GCF_002038025.1_ASM203802v1 Salmonella enterica subsp. enterica serovar Enteritidis Proteobacteria; Gammaproteobacteria; Enterobacterales; Enterobacteriaceae; Salmonella; Salmonella enterica; Salmonella enterica subsp. enterica MNNEETFYQAMRRKGVTRRSFLKFCSLAATSLGLGAGMTPKIAWALENKPRIPVVWIHGL WP_001058307.1 hydrogenase [Salmonella enterica] Length: 372\nNumber of predicted TMHs: 2\nExp number of AAs in TMHs: 36.51738\nExp number, first 60 AAs: 19.72849\nTotal prob of N-in: 0.96170\nPOSSIBLE N-term signal sequence\ninside 1 19\nTMhelix 20 39\noutside 40 325\nTMhelix 326 348\ninside 349 372

16719 GCF_000750375.1_ASM75037v1 Salmonella enterica subsp. enterica serovar Enteritidis Proteobacteria; Gammaproteobacteria; Enterobacterales; Enterobacteriaceae; Salmonella; Salmonella enterica; Salmonella enterica subsp. enterica MTGDNTLITSHGINRRDFMKLCAALAATMGLSSKAAAEMAESVSNPQRPPVIWIGAQECT WP_000145429.1 hydrogenase 2 small subunit [Salmonella enterica] Length: 372\nNumber of predicted TMHs: 1\nExp number of AAs in TMHs: 24.51309\nExp number, first 60 AAs: 0.2682\nTotal prob of N-in: 0.14402\noutside 1 331\nTMhelix 332 354\ninside 355 372

16720 GCF_000750415.2_ASM75041v2 Salmonella enterica subsp. enterica serovar Enteritidis Proteobacteria; Gammaproteobacteria; Enterobacterales; Enterobacteriaceae; Salmonella; Salmonella enterica; Salmonella enterica subsp. enterica MTGDNTLITSHGINRRDFMKLCAALAATMGLSSKAAAEMAESVSNPQRPPVIWIGAQECT WP_000145429.1 hydrogenase 2 small subunit [Salmonella enterica] Length: 372\nNumber of predicted TMHs: 1\nExp number of AAs in TMHs: 24.51309\nExp number, first 60 AAs: 0.2682\nTotal prob of N-in: 0.14402\noutside 1 331\nTMhelix 332 354\ninside 355 372

16721 GCF_001973195.1_ASM197319v1 Salmonella enterica subsp. enterica serovar Enteritidis Proteobacteria; Gammaproteobacteria; Enterobacterales; Enterobacteriaceae; Salmonella; Salmonella enterica; Salmonella enterica subsp. enterica MTGDNTLITSHGINRRDFMKLCAALAATMGLSSKAAAEMAESVSNPQRPPVIWIGAQECT WP_000145429.1 hydrogenase 2 small subunit [Salmonella enterica] Length: 372\nNumber of predicted TMHs: 1\nExp number of AAs in TMHs: 24.51309\nExp number, first 60 AAs: 0.2682\nTotal prob of N-in: 0.14402\noutside 1 331\nTMhelix 332 354\ninside 355 372

16722 GCF_000750435.1_ASM75043v1 Salmonella enterica subsp. enterica serovar Enteritidis Proteobacteria; Gammaproteobacteria; Enterobacterales; Enterobacteriaceae; Salmonella; Salmonella enterica; Salmonella enterica subsp. enterica MTGDNTLITSHGINRRDFMKLCAALAATMGLSSKAAAEMAESVSNPQRPPVIWIGAQECT WP_000145429.1 hydrogenase 2 small subunit [Salmonella enterica] Length: 372\nNumber of predicted TMHs: 1\nExp number of AAs in TMHs: 24.51309\nExp number, first 60 AAs: 0.2682\nTotal prob of N-in: 0.14402\noutside 1 331\nTMhelix 332 354\ninside 355 372

16723 GCF_001997445.1_ASM199744v1 Salmonella enterica subsp. enterica serovar Enteritidis Proteobacteria; Gammaproteobacteria; Enterobacterales; Enterobacteriaceae; Salmonella; Salmonella enterica; Salmonella enterica subsp. enterica MNNEETFYQAMRRKGVTRRSFLKFCSLAATSLGLGAGMTPKIAWALENKPRIPVVWIHGL WP_001058307.1 hydrogenase [Salmonella enterica] Length: 372\nNumber of predicted TMHs: 2\nExp number of AAs in TMHs: 36.51738\nExp number, first 60 AAs: 19.72849\nTotal prob of N-in: 0.96170\nPOSSIBLE N-term signal sequence\ninside 1 19\nTMhelix 20 39\noutside 40 325\nTMhelix 326 348\ninside 349 372

16724 GCF_001972945.1_ASM197294v1 Salmonella enterica subsp. enterica serovar Enteritidis Proteobacteria; Gammaproteobacteria; Enterobacterales; Enterobacteriaceae; Salmonella; Salmonella enterica; Salmonella enterica subsp. enterica MNNEETFYQAMRRKGVTRRSFLKFCSLAATSLGLGAGMTPKIAWALGNKPRIPVVWIHGL WP_076926151.1 [Ni/Fe] hydrogenase small subunit [Salmonella enterica] Length: 372\nNumber of predicted TMHs: 2\nExp number of AAs in TMHs: 36.7858799999999\nExp number, first 60 AAs: 19.99153\nTotal prob of

N-in: 0.97261\nPOSSIBLE N-term signal sequence\ninside 1 19\nTMhelix 20 39\noutside 40 325\nTMhelix 326 348\ninside 349 372

16725 GCF_001972985.1_ASM197298v1 Salmonella enterica subsp. enterica serovar Enteritidis Proteobacteria; Gammaproteobacteria; Enterobacterales; Enterobacteriaceae; Salmonella; Salmonella enterica; Salmonella enterica subsp. enterica MNNEETFYQAMRRKGVTRRSFLKFCSLAATSLGLGAGMTPKIAWALENKPRIPVVWIHGL WP_001058307.1 hydrogenase [Salmonella enterica] Length: 372\nNumber of predicted TMHs: 2\nExp number of AAs in TMHs: 36.51738\nExp number, first 60 AAs: 19.72849\nTotal prob of N-in: 0.96170\nPOSSIBLE N-term signal sequence\ninside 1 19\nTMhelix 20 39\noutside 40 325\nTMhelix 326 348\ninside 349 372

16726 GCF_002062265.1_ASM206226v1 Salmonella enterica subsp. enterica serovar Enteritidis Proteobacteria; Gammaproteobacteria; Enterobacterales; Enterobacteriaceae; Salmonella; Salmonella enterica; Salmonella enterica subsp. enterica MNNEETFYQAMRRKGVTRRSFLKFCSLAATSLGLGAGMTPKIAWALENKPRIPVVWIHGL WP_001058307.1 hydrogenase [Salmonella enterica] Length: 372\nNumber of predicted TMHs: 2\nExp number of AAs in TMHs: 36.51738\nExp number, first 60 AAs: 19.72849\nTotal prob of N-in: 0.96170\nPOSSIBLE N-term signal sequence\ninside 1 19\nTMhelix 20 39\noutside 40 325\nTMhelix 326 348\ninside 349 372

16727 GCF_001972985.1_ASM197298v1 Salmonella enterica subsp. enterica serovar Enteritidis Proteobacteria; Gammaproteobacteria; Enterobacterales; Enterobacteriaceae; Salmonella; Salmonella enterica; Salmonella enterica subsp. enterica MTGDNTLITSHGINRRDFMKLCAALAATMGLSSKAAAEMAESVSNPQRPPVIWIGAQECT WP_000145429.1 hydrogenase 2 small subunit [Salmonella enterica] Length: 372\nNumber of predicted TMHs: 1\nExp number of AAs in TMHs: 24.51309\nExp number, first 60 AAs: 0.2682\nTotal prob of N-in: 0.14402\noutside 1 331\nTMhelix 332 354\ninside 355 372

16728 GCF_001972945.1_ASM197294v1 Salmonella enterica subsp. enterica serovar Enteritidis Proteobacteria; Gammaproteobacteria; Enterobacterales; Enterobacteriaceae; Salmonella; Salmonella enterica; Salmonella enterica subsp. enterica MTGDNTLITSHGINRRDFMKLCAALAATMGLSSKAAAEMAESVSNPQRPPVIWIGAQECT WP_000145429.1 hydrogenase 2 small subunit [Salmonella enterica] Length: 372\nNumber of predicted TMHs: 1\nExp number of AAs in TMHs: 24.51309\nExp number, first 60 AAs: 0.2682\nTotal prob of N-in: 0.14402\noutside 1 331\nTMhelix 332 354\ninside 355 372

16729 GCF_001997445.1_ASM199744v1 Salmonella enterica subsp. enterica serovar Enteritidis Proteobacteria; Gammaproteobacteria; Enterobacterales; Enterobacteriaceae; Salmonella; Salmonella enterica; Salmonella enterica subsp. enterica MTGDNTLITSHGINRRDFMKLCAALAATMGLSSKAAAEMAESVSNPQRPPVIWIGAQECT WP_000145429.1 hydrogenase 2 small subunit [Salmonella enterica] Length: 372\nNumber of predicted TMHs: 1\nExp number of AAs in TMHs: 24.51309\nExp number, first 60 AAs: 0.2682\nTotal prob of N-in: 0.14402\noutside 1 331\nTMhelix 332 354\ninside 355 372

16730 GCF_001972785.1_ASM197278v1 Salmonella enterica subsp. enterica serovar Enteritidis Proteobacteria; Gammaproteobacteria; Enterobacterales; Enterobacteriaceae; Salmonella; Salmonella enterica; Salmonella enterica subsp. enterica MNNEETFYQAMRRKGVTRRSFLKFCSLAATSLGLGAGMTPKIAWALENKPRIPVVWIHGL WP_001058307.1 hydrogenase [Salmonella enterica] Length: 372\nNumber of predicted TMHs: 2\nExp number of AAs in TMHs: 36.51738\nExp number, first 60 AAs: 19.72849\nTotal prob of N-in: 0.96170\nPOSSIBLE N-term signal sequence\ninside 1 19\nTMhelix 20 39\noutside 40 325\nTMhelix 326 348\ninside 349 372

16731 GCF_001972545.1_ASM197254v1 Salmonella enterica subsp. enterica serovar Enteritidis Proteobacteria; Gammaproteobacteria; Enterobacterales; Enterobacteriaceae; Salmonella; Salmonella enterica; Salmonella enterica subsp. enterica MTGDNTLITSHGINRRDFMKLCAALAATMGLSSKAAAEMAESVSNPQRPPVIWIGAQECT WP_000145429.1 hydrogenase 2 small subunit [Salmonella enterica] Length: 372\nNumber of predicted TMHs: 1\nExp number of AAs in TMHs: 24.51309\nExp number, first 60 AAs: 0.2682\nTotal prob of N-in: 0.14402\noutside 1 331\nTMhelix 332 354\ninside 355 372

16732 GCF_001972705.1_ASM197270v1 Salmonella enterica subsp. enterica serovar Enteritidis Proteobacteria; Gammaproteobacteria; Enterobacterales; Enterobacteriaceae; Salmonella; Salmonella enterica; Salmonella enterica subsp. enterica MNNEETFYQAMRRKGVTRRSFLKFCSLAATSLGLGAGMTPKIAWALENKPRIPVVWIHGL WP_001058307.1 hydrogenase [Salmonella enterica] Length: 372\nNumber of predicted TMHs: 2\nExp number of AAs in TMHs: 36.51738\nExp number, first 60 AAs: 19.72849\nTotal prob of N-in: 0.96170\nPOSSIBLE N-term signal sequence\ninside 1 19\nTMhelix 20 39\noutside 40 325\nTMhelix 326 348\ninside 349 372

16733 GCF_001972565.1_ASM197256v1 Salmonella enterica subsp. enterica serovar Enteritidis Proteobacteria; Gammaproteobacteria; Enterobacterales; Enterobacteriaceae; Salmonella; Salmonella enterica;

Salmonella enterica subsp. enterica	MTGDNTLITSHGINRRDFMKLCAALAATMGLSSKAAAEMAESVSNPQRPPVIWIGAQECT	WP_000145429.1	hydrogenase 2 small subunit [Salmonella enterica]	Length: 372\nNumber of predicted TMHs:  1\nExp number of AAs in TMHs: 24.51309\nExp number, first 60 AAs:  0.2682\nTotal prob of N-in:  0.14402\noutside   1   331\nTMhelix   332   354\ninside   355   372

16734	GCF_002062265.1_ASM206226v1	Salmonella enterica subsp. enterica serovar Enteritidis	Proteobacteria; Gammaproteobacteria; Enterobacterales; Enterobacteriaceae; Salmonella; Salmonella enterica; Salmonella enterica subsp. enterica	MTGDNTLITSHGINRRDFMKLCAALAATMGLSSKAAAEMAESVSNPQRPPVIWIGAQECT	WP_000145429.1	hydrogenase 2 small subunit [Salmonella enterica]	Length: 372\nNumber of predicted TMHs:  1\nExp number of AAs in TMHs: 24.51309\nExp number, first 60 AAs:  0.2682\nTotal prob of N-in:  0.14402\noutside   1   331\nTMhelix   332   354\ninside   355   372

16735	GCF_001484025.1_ASM148402v1	Salmonella enterica subsp. enterica serovar Enteritidis	Proteobacteria; Gammaproteobacteria; Enterobacterales; Enterobacteriaceae; Salmonella; Salmonella enterica; Salmonella enterica subsp. enterica	MNNEETFYQAMRRKGVTRRSFLKFCSLAATSLGLGAGMTPKIAWALENKPRIPVVWIHGL	WP_001058307.1	hydrogenase [Salmonella enterica]	Length: 372\nNumber of predicted TMHs:  2\nExp number of AAs in TMHs: 36.51738\nExp number, first 60 AAs:  19.72849\nTotal prob of N-in:   0.96170\nPOSSIBLE N-term signal sequence\ninside   1   19\nTMhelix   20   39\noutside   40   325\nTMhelix   326   348\ninside   349   372

16736	GCF_002040785.1_ASM204078v1	Salmonella enterica subsp. enterica serovar Enteritidis	Proteobacteria; Gammaproteobacteria; Enterobacterales; Enterobacteriaceae; Salmonella; Salmonella enterica; Salmonella enterica subsp. enterica	MNNEETFYQAMRRKGVTRRSFLKFCSLAATSLGLGAGMTPKIAWALENKPRIPVVWIHGL	WP_001058307.1	hydrogenase [Salmonella enterica]	Length: 372\nNumber of predicted TMHs:  2\nExp number of AAs in TMHs: 36.51738\nExp number, first 60 AAs:  19.72849\nTotal prob of N-in:   0.96170\nPOSSIBLE N-term signal sequence\ninside   1   19\nTMhelix   20   39\noutside   40   325\nTMhelix   326   348\ninside   349   372

16737	GCF_002040825.1_ASM204082v1	Salmonella enterica subsp. enterica serovar Enteritidis	Proteobacteria; Gammaproteobacteria; Enterobacterales; Enterobacteriaceae; Salmonella; Salmonella enterica; Salmonella enterica subsp. enterica	MNNEETFYQAMRRKGVTRRSFLKFCSLAATSLGLGAGMTPKIAWALENKPRIPVVWIHGL	WP_001058307.1	hydrogenase [Salmonella enterica]	Length: 372\nNumber of predicted TMHs:  2\nExp number of AAs in TMHs: 36.51738\nExp number, first 60 AAs:  19.72849\nTotal prob of N-in:   0.96170\nPOSSIBLE N-term signal sequence\ninside   1   19\nTMhelix   20   39\noutside   40   325\nTMhelix   326   348\ninside   349   372

16738	GCF_002040525.1_ASM204052v1	Salmonella enterica subsp. enterica serovar Enteritidis	Proteobacteria; Gammaproteobacteria; Enterobacterales; Enterobacteriaceae; Salmonella; Salmonella enterica; Salmonella enterica subsp. enterica	MNNEETFYQAMRRKGVTRRSFLKFCSLAATSLGLGAGMTPKIAWALENKPRIPVVWIHGL	WP_001058307.1	hydrogenase [Salmonella enterica]	Length: 372\nNumber of predicted TMHs:  2\nExp number of AAs in TMHs: 36.51738\nExp number, first 60 AAs:  19.72849\nTotal prob of N-in:   0.96170\nPOSSIBLE N-term signal sequence\ninside   1   19\nTMhelix   20   39\noutside   40   325\nTMhelix   326   348\ninside   349   372

16739	GCF_002040485.1_ASM204048v1	Salmonella enterica subsp. enterica serovar Enteritidis	Proteobacteria; Gammaproteobacteria; Enterobacterales; Enterobacteriaceae; Salmonella; Salmonella enterica; Salmonella enterica subsp. enterica	MNNEETFYQAMRRKGVTRRSFLKFCSLAATSLGLGAGMTPKIAWALENKPRIPVVWIHGL	WP_001058307.1	hydrogenase [Salmonella enterica]	Length: 372\nNumber of predicted TMHs:  2\nExp number of AAs in TMHs: 36.51738\nExp number, first 60 AAs:  19.72849\nTotal prob of N-in:   0.96170\nPOSSIBLE N-term signal sequence\ninside   1   19\nTMhelix   20   39\noutside   40   325\nTMhelix   326   348\ninside   349   372

16740	GCF_002040615.1_ASM204061v1	Salmonella enterica subsp. enterica serovar Enteritidis	Proteobacteria; Gammaproteobacteria; Enterobacterales; Enterobacteriaceae; Salmonella; Salmonella enterica; Salmonella enterica subsp. enterica	MTGDNTLITSHGINRRDFMKLCAALAATMGLSSKAAAEMAESVSNPQRPPVIWIGAQECT	WP_000145429.1	hydrogenase 2 small subunit [Salmonella enterica]	Length: 372\nNumber of predicted TMHs:  1\nExp number of AAs in TMHs: 24.51309\nExp number, first 60 AAs:  0.2682\nTotal prob of N-in:  0.14402\noutside   1   331\nTMhelix   332   354\ninside   355   372

16741	GCF_002040595.1_ASM204059v1	Salmonella enterica subsp. enterica serovar Enteritidis	Proteobacteria; Gammaproteobacteria; Enterobacterales; Enterobacteriaceae; Salmonella; Salmonella enterica; Salmonella enterica subsp. enterica	MTGDNTLITSHGINRRDFMKLCAALAATMGLSSKAAAEMAESVSNPQRPPVIWIGAQECT	WP_000145429.1	hydrogenase 2 small subunit [Salmonella enterica]	Length: 372\nNumber of predicted TMHs:  1\nExp number of AAs in TMHs: 24.51309\nExp number, first 60 AAs:  0.2682\nTotal prob of N-in:  0.14402\noutside   1   331\nTMhelix   332   354\ninside   355   372

16742 GCF_002040565.1_ASM204056v1 Salmonella enterica subsp. enterica serovar Enteritidis Proteobacteria; Gammaproteobacteria; Enterobacterales; Enterobacteriaceae; Salmonella; Salmonella enterica; Salmonella enterica subsp. enterica MTGDNTLITSHGINRRDFMKLCAALAATMGLSSKAAAEMAESVSNPQRPPVIWIGAQECT WP_000145429.1 hydrogenase 2 small subunit [Salmonella enterica] Length: 372\nNumber of predicted TMHs: 1\nExp number of AAs in TMHs: 24.51309\nExp number, first 60 AAs: 0.2682\nTotal prob of N-in: 0.14402\noutside 1 331\nTMhelix 332 354\ninside 355 372

16743 GCF_002040525.1_ASM204052v1 Salmonella enterica subsp. enterica serovar Enteritidis Proteobacteria; Gammaproteobacteria; Enterobacterales; Enterobacteriaceae; Salmonella; Salmonella enterica; Salmonella enterica subsp. enterica MTGDNTLITSHGINRRDFMKLCAALAATMGLSSKAAAEMAESVSNPQRPPVIWIGAQECT WP_000145429.1 hydrogenase 2 small subunit [Salmonella enterica] Length: 372\nNumber of predicted TMHs: 1\nExp number of AAs in TMHs: 24.51309\nExp number, first 60 AAs: 0.2682\nTotal prob of N-in: 0.14402\noutside 1 331\nTMhelix 332 354\ninside 355 372

16744 GCF_002040485.1_ASM204048v1 Salmonella enterica subsp. enterica serovar Enteritidis Proteobacteria; Gammaproteobacteria; Enterobacterales; Enterobacteriaceae; Salmonella; Salmonella enterica; Salmonella enterica subsp. enterica MTGDNTLITSHGINRRDFMKLCAALAATMGLSSKAAAEMAESVSNPQRPPVIWIGAQECT WP_000145429.1 hydrogenase 2 small subunit [Salmonella enterica] Length: 372\nNumber of predicted TMHs: 1\nExp number of AAs in TMHs: 24.51309\nExp number, first 60 AAs: 0.2682\nTotal prob of N-in: 0.14402\noutside 1 331\nTMhelix 332 354\ninside 355 372

16745 GCF_002040255.1_ASM204025v1 Salmonella enterica subsp. enterica serovar Enteritidis Proteobacteria; Gammaproteobacteria; Enterobacterales; Enterobacteriaceae; Salmonella; Salmonella enterica; Salmonella enterica subsp. enterica MTGDNTLITSHGINRRDFMKLCAALAATMGLSSKAAAEMAESVSNPQRPPVIWIGAQECT WP_000145429.1 hydrogenase 2 small subunit [Salmonella enterica] Length: 372\nNumber of predicted TMHs: 1\nExp number of AAs in TMHs: 24.51309\nExp number, first 60 AAs: 0.2682\nTotal prob of N-in: 0.14402\noutside 1 331\nTMhelix 332 354\ninside 355 372

16746 GCF_002040095.1_ASM204009v1 Salmonella enterica subsp. enterica serovar Enteritidis Proteobacteria; Gammaproteobacteria; Enterobacterales; Enterobacteriaceae; Salmonella; Salmonella enterica; Salmonella enterica subsp. enterica MNNEETFYQAMRRKGVTRRSFLKFCSLAATSLGLGAGMTPKIAWALENKPRIPVVWIHGL WP_001058307.1 hydrogenase [Salmonella enterica] Length: 372\nNumber of predicted TMHs: 2\nExp number of AAs in TMHs: 36.51738\nExp number, first 60 AAs: 19.72849\nTotal prob of N-in: 0.96170\nPOSSIBLE N-term signal sequence\ninside 1 19\nTMhelix 20 39\noutside 40 325\nTMhelix 326 348\ninside 349 372

16747 GCF_002040065.1_ASM204006v1 Salmonella enterica subsp. enterica serovar Enteritidis Proteobacteria; Gammaproteobacteria; Enterobacterales; Enterobacteriaceae; Salmonella; Salmonella enterica; Salmonella enterica subsp. enterica MNNEETFYQAMRRKGVTRRSFLKFCSLAATSLGLGAGMTPKIAWALENKPRIPVVWIHGL WP_001058307.1 hydrogenase [Salmonella enterica] Length: 372\nNumber of predicted TMHs: 2\nExp number of AAs in TMHs: 36.51738\nExp number, first 60 AAs: 19.72849\nTotal prob of N-in: 0.96170\nPOSSIBLE N-term signal sequence\ninside 1 19\nTMhelix 20 39\noutside 40 325\nTMhelix 326 348\ninside 349 372

16748 GCF_002040015.1_ASM204001v1 Salmonella enterica subsp. enterica serovar Enteritidis Proteobacteria; Gammaproteobacteria; Enterobacterales; Enterobacteriaceae; Salmonella; Salmonella enterica; Salmonella enterica subsp. enterica MNNEETFYQAMRRKGVTRRSFLKFCSLAATSLGLGAGMTPKIAWALENKPRIPVVWIHGL WP_001058307.1 hydrogenase [Salmonella enterica] Length: 372\nNumber of predicted TMHs: 2\nExp number of AAs in TMHs: 36.51738\nExp number, first 60 AAs: 19.72849\nTotal prob of N-in: 0.96170\nPOSSIBLE N-term signal sequence\ninside 1 19\nTMhelix 20 39\noutside 40 325\nTMhelix 326 348\ninside 349 372

16749 GCF_002039985.1_ASM203998v1 Salmonella enterica subsp. enterica serovar Enteritidis Proteobacteria; Gammaproteobacteria; Enterobacterales; Enterobacteriaceae; Salmonella; Salmonella enterica; Salmonella enterica subsp. enterica MNNEETFYQAMRRKGVTRRSFLKFCSLAATSLGLGAGMTPKIAWALENKPRIPVVWIHGL WP_001058307.1 hydrogenase [Salmonella enterica] Length: 372\nNumber of predicted TMHs: 2\nExp number of AAs in TMHs: 36.51738\nExp number, first 60 AAs: 19.72849\nTotal prob of N-in: 0.96170\nPOSSIBLE N-term signal sequence\ninside 1 19\nTMhelix 20 39\noutside 40 325\nTMhelix 326 348\ninside 349 372

16750 GCF_002040135.1_ASM204013v1 Salmonella enterica subsp. enterica serovar Enteritidis Proteobacteria; Gammaproteobacteria; Enterobacterales; Enterobacteriaceae; Salmonella; Salmonella enterica; Salmonella enterica subsp. enterica MNNEETFYQAMRRKGVTRRSFLKFCSLAATSLGLGAGMTPKIAWALENKPRIPVVWIHGL WP_001058307.1 hydrogenase [Salmonella enterica] Length: 372\nNumber of predicted TMHs: 2\nExp

number of AAs in TMHs: 36.51738\nExp number, first 60 AAs: 19.72849\nTotal prob of N-in: 0.96170\nPOSSIBLE N-term signal sequence\ninside 1 19\nTMhelix 20 39\noutside 40 325\nTMhelix 326 348\ninside 349 372

16751 GCF_002039815.1_ASM203981v1 Salmonella enterica subsp. enterica serovar Enteritidis Proteobacteria; Gammaproteobacteria; Enterobacterales; Enterobacteriaceae; Salmonella; Salmonella enterica; Salmonella enterica subsp. enterica MNNEETFYQAMRRKGVTRRSFLKFCSLAATSLGLGAGMTPKIAWALENKPRIPVVWIHGL WP_001058307.1 hydrogenase [Salmonella enterica] Length: 372\nNumber of predicted TMHs: 2\nExp number of AAs in TMHs: 36.51738\nExp number, first 60 AAs: 19.72849\nTotal prob of N-in: 0.96170\nPOSSIBLE N-term signal sequence\ninside 1 19\nTMhelix 20 39\noutside 40 325\nTMhelix 326 348\ninside 349 372

16752 GCF_002039855.1_ASM203985v1 Salmonella enterica subsp. enterica serovar Enteritidis Proteobacteria; Gammaproteobacteria; Enterobacterales; Enterobacteriaceae; Salmonella; Salmonella enterica; Salmonella enterica subsp. enterica MTGDNTLITSHGINRRDFMKLCAALAATMGLSSKAAAEMAESVSNPQRPPVIWIGAQECT WP_000145429.1 hydrogenase 2 small subunit [Salmonella enterica] Length: 372\nNumber of predicted TMHs: 1\nExp number of AAs in TMHs: 24.51309\nExp number, first 60 AAs: 0.2682\nTotal prob of N-in: 0.14402\noutside 1 331\nTMhelix 332 354\ninside 355 372

16753 GCF_002039815.1_ASM203981v1 Salmonella enterica subsp. enterica serovar Enteritidis Proteobacteria; Gammaproteobacteria; Enterobacterales; Enterobacteriaceae; Salmonella; Salmonella enterica; Salmonella enterica subsp. enterica MTGDNTLITSHGINRRDFMKLCAALAATMGLSSKAAAEMAESVSNPQRPPVIWIGAQECT WP_000145429.1 hydrogenase 2 small subunit [Salmonella enterica] Length: 372\nNumber of predicted TMHs: 1\nExp number of AAs in TMHs: 24.51309\nExp number, first 60 AAs: 0.2682\nTotal prob of N-in: 0.14402\noutside 1 331\nTMhelix 332 354\ninside 355 372

16754 GCF_001713915.1_ASM171391v1 Salmonella enterica subsp. enterica serovar Enteritidis Proteobacteria; Gammaproteobacteria; Enterobacterales; Enterobacteriaceae; Salmonella; Salmonella enterica; Salmonella enterica subsp. enterica MTGDNTLITSHGINRRDFMKLCAALAATMGLSSKAAAEMAESVSNPQRPPVIWIGAQECT WP_000145429.1 hydrogenase 2 small subunit [Salmonella enterica] Length: 372\nNumber of predicted TMHs: 1\nExp number of AAs in TMHs: 24.51309\nExp number, first 60 AAs: 0.2682\nTotal prob of N-in: 0.14402\noutside 1 331\nTMhelix 332 354\ninside 355 372

16755 GCF_002039715.1_ASM203971v1 Salmonella enterica subsp. enterica serovar Enteritidis Proteobacteria; Gammaproteobacteria; Enterobacterales; Enterobacteriaceae; Salmonella; Salmonella enterica; Salmonella enterica subsp. enterica MNNEETFYQAMRRKGVTRRSFLKFCSLAATSLGLGAGMTPKIAWALENKPRIPVVWIHGL WP_001058307.1 hydrogenase [Salmonella enterica] Length: 372\nNumber of predicted TMHs: 2\nExp number of AAs in TMHs: 36.51738\nExp number, first 60 AAs: 19.72849\nTotal prob of N-in: 0.96170\nPOSSIBLE N-term signal sequence\ninside 1 19\nTMhelix 20 39\noutside 40 325\nTMhelix 326 348\ninside 349 372

16756 GCF_002039685.1_ASM203968v1 Salmonella enterica subsp. enterica serovar Enteritidis Proteobacteria; Gammaproteobacteria; Enterobacterales; Enterobacteriaceae; Salmonella; Salmonella enterica; Salmonella enterica subsp. enterica MNNEETFYQAMRRKGVTRRSFLKFCSLAATSLGLGAGMTPKIAWALENKPRIPVVWIHGL WP_001058307.1 hydrogenase [Salmonella enterica] Length: 372\nNumber of predicted TMHs: 2\nExp number of AAs in TMHs: 36.51738\nExp number, first 60 AAs: 19.72849\nTotal prob of N-in: 0.96170\nPOSSIBLE N-term signal sequence\ninside 1 19\nTMhelix 20 39\noutside 40 325\nTMhelix 326 348\ninside 349 372

16757 GCF_002039785.1_ASM203978v1 Salmonella enterica subsp. enterica serovar Enteritidis Proteobacteria; Gammaproteobacteria; Enterobacterales; Enterobacteriaceae; Salmonella; Salmonella enterica; Salmonella enterica subsp. enterica MTGDNTLITSHGINRRDFMKLCAALAATMGLSSKAAAEMAESVSNPQRPPVIWIGAQECT WP_000145429.1 hydrogenase 2 small subunit [Salmonella enterica] Length: 372\nNumber of predicted TMHs: 1\nExp number of AAs in TMHs: 24.51309\nExp number, first 60 AAs: 0.2682\nTotal prob of N-in: 0.14402\noutside 1 331\nTMhelix 332 354\ninside 355 372

16758 GCF_001713915.1_ASM171391v1 Salmonella enterica subsp. enterica serovar Enteritidis Proteobacteria; Gammaproteobacteria; Enterobacterales; Enterobacteriaceae; Salmonella; Salmonella enterica; Salmonella enterica subsp. enterica MNNEETFYQAMRRKGVTRRSFLKFCSLAATSLGLGAGMTPKIAWALENKPRIPVVWIHGL WP_001058307.1 hydrogenase [Salmonella enterica] Length: 372\nNumber of predicted TMHs: 2\nExp number of AAs in TMHs: 36.51738\nExp number, first 60 AAs: 19.72849\nTotal prob of N-in: 0.96170\nPOSSIBLE N-term signal sequence\ninside 1 19\nTMhelix 20 39\noutside 40 325\nTMhelix 326 348\ninside 349 372

16759 GCF_001713755.1_ASM171375v1 Salmonella enterica subsp. enterica serovar Enteritidis Proteobacteria; Gammaproteobacteria; Enterobacterales; Enterobacteriaceae; Salmonella; Salmonella enterica;

Salmonella enterica subsp. enterica	MNNEETFYQAMRRKGVTRRSFLKFCSLAATSLGLGAGMTPKIAWALENKPRIPVVWIHGL	WP_001058307.1	hydrogenase [Salmonella enterica]	Length: 372\nNumber of predicted TMHs:  2\nExp number of AAs in TMHs: 36.51738\nExp number, first 60 AAs:  19.72849\nTotal prob of N-in:   0.96170\nPOSSIBLE N-term signal sequence\ninside   1   19\nTMhelix   20   39\noutside   40   325\nTMhelix   326   348\ninside   349   372

16760	GCF_002039755.1_ASM203975v1	Salmonella enterica subsp. enterica serovar Enteritidis	Proteobacteria; Gammaproteobacteria; Enterobacterales; Enterobacteriaceae; Salmonella; Salmonella enterica; Salmonella enterica subsp. enterica	MTGDNTLITSHGINRRDFMKLCAALAATMGLSSKAAAEMAESVSNPQRPPVIWIGAQECT	WP_000145429.1	hydrogenase 2 small subunit [Salmonella enterica]	Length: 372\nNumber of predicted TMHs:  1\nExp number of AAs in TMHs: 24.51309\nExp number, first 60 AAs:  0.2682\nTotal prob of N-in:  0.14402\noutside   1   331\nTMhelix   332   354\ninside   355   372

16761	GCF_002039395.1_ASM203939v1	Salmonella enterica subsp. enterica serovar Enteritidis	Proteobacteria; Gammaproteobacteria; Enterobacterales; Enterobacteriaceae; Salmonella; Salmonella enterica; Salmonella enterica subsp. enterica	MTGDNTLITSHGINRRDFMKLCAALAATMGLSSKAAAEMAESVSNPQRPPVIWIGAQECT	WP_000145429.1	hydrogenase 2 small subunit [Salmonella enterica]	Length: 372\nNumber of predicted TMHs:  1\nExp number of AAs in TMHs: 24.51309\nExp number, first 60 AAs:  0.2682\nTotal prob of N-in:  0.14402\noutside   1   331\nTMhelix   332   354\ninside   355   372

16762	GCF_002039365.1_ASM203936v1	Salmonella enterica subsp. enterica serovar Enteritidis	Proteobacteria; Gammaproteobacteria; Enterobacterales; Enterobacteriaceae; Salmonella; Salmonella enterica; Salmonella enterica subsp. enterica	MTGDNTLITSHGINRRDFMKLCAALAATMGLSSKAAAEMAESVSNPQRPPVIWIGAQECT	WP_000145429.1	hydrogenase 2 small subunit [Salmonella enterica]	Length: 372\nNumber of predicted TMHs:  1\nExp number of AAs in TMHs: 24.51309\nExp number, first 60 AAs:  0.2682\nTotal prob of N-in:  0.14402\noutside   1   331\nTMhelix   332   354\ninside   355   372

16763	GCF_002039205.1_ASM203920v1	Salmonella enterica subsp. enterica serovar Enteritidis	Proteobacteria; Gammaproteobacteria; Enterobacterales; Enterobacteriaceae; Salmonella; Salmonella enterica; Salmonella enterica subsp. enterica	MNNEETFYQAMRRKGVTRRSFLKFCSLAATSLGLGAGMTPKIAWALENKPRIPVVWIHGL	WP_001058307.1	hydrogenase [Salmonella enterica]	Length: 372\nNumber of predicted TMHs:  2\nExp number of AAs in TMHs: 36.51738\nExp number, first 60 AAs:  19.72849\nTotal prob of N-in:   0.96170\nPOSSIBLE N-term signal sequence\ninside   1   19\nTMhelix   20   39\noutside   40   325\nTMhelix   326   348\ninside   349   372

16764	GCF_002038155.1_ASM203815v1	Salmonella enterica subsp. enterica serovar Enteritidis	Proteobacteria; Gammaproteobacteria; Enterobacterales; Enterobacteriaceae; Salmonella; Salmonella enterica; Salmonella enterica subsp. enterica	MNNEETFYQAMRRKGVTRRSFLKFCSLAATSLGLGAGMTPKIAWALENKPRIPVVWIHGL	WP_001058307.1	hydrogenase [Salmonella enterica]	Length: 372\nNumber of predicted TMHs:  2\nExp number of AAs in TMHs: 36.51738\nExp number, first 60 AAs:  19.72849\nTotal prob of N-in:   0.96170\nPOSSIBLE N-term signal sequence\ninside   1   19\nTMhelix   20   39\noutside   40   325\nTMhelix   326   348\ninside   349   372

16765	GCF_002038115.1_ASM203811v1	Salmonella enterica subsp. enterica serovar Enteritidis	Proteobacteria; Gammaproteobacteria; Enterobacterales; Enterobacteriaceae; Salmonella; Salmonella enterica; Salmonella enterica subsp. enterica	MNNEETFYQAMRRKGVTRRSFLKFCSLAATSLGLGAGMTPKIAWALENKPRIPVVWIHGL	WP_001058307.1	hydrogenase [Salmonella enterica]	Length: 372\nNumber of predicted TMHs:  2\nExp number of AAs in TMHs: 36.51738\nExp number, first 60 AAs:  19.72849\nTotal prob of N-in:   0.96170\nPOSSIBLE N-term signal sequence\ninside   1   19\nTMhelix   20   39\noutside   40   325\nTMhelix   326   348\ninside   349   372

16766	GCF_002038035.1_ASM203803v1	Salmonella enterica subsp. enterica serovar Enteritidis	Proteobacteria; Gammaproteobacteria; Enterobacterales; Enterobacteriaceae; Salmonella; Salmonella enterica; Salmonella enterica subsp. enterica	MTGDNTLITSHGINRRDFMKLCAALAATMGLSSKAAAEMAESVSNPQRPPVIWIGAQECT	WP_000145429.1	hydrogenase 2 small subunit [Salmonella enterica]	Length: 372\nNumber of predicted TMHs:  1\nExp number of AAs in TMHs: 24.51309\nExp number, first 60 AAs:  0.2682\nTotal prob of N-in:  0.14402\noutside   1   331\nTMhelix   332   354\ninside   355   372

16767	GCF_002037865.1_ASM203786v1	Salmonella enterica subsp. enterica serovar Enteritidis	Proteobacteria; Gammaproteobacteria; Enterobacterales; Enterobacteriaceae; Salmonella; Salmonella enterica; Salmonella enterica subsp. enterica	MNNEETFYQAMRRKGVTRRSFLKFCSLAATSLGLGAGMTPKIAWALENKPRIPVVWIHGL	WP_001058307.1	hydrogenase [Salmonella enterica]	Length: 372\nNumber of predicted TMHs:  2\nExp number of AAs in TMHs: 36.51738\nExp number, first 60 AAs:  19.72849\nTotal prob of N-in:   0.96170\nPOSSIBLE N-term signal sequence\ninside   1   19\nTMhelix   20   39\noutside   40   325\nTMhelix   326   348\ninside   349   372

16768	GCF_002037995.1_ASM203799v1	Salmonella enterica subsp. enterica serovar Enteritidis	Proteobacteria; Gammaproteobacteria; Enterobacterales; Enterobacteriaceae; Salmonella; Salmonella enterica; Salmonella enterica subsp. enterica	MTGDNTLITSHGINRRDFMKLCAALAATMGLSSKAAAEMAESVSNPQRPPVIWIGAQECT	WP_000145429.1	hydrogenase 2 small subunit [Salmonella enterica]	Length: 372\nNumber of predicted TMHs: 1\nExp number of AAs in TMHs: 24.51309\nExp number, first 60 AAs: 0.2682\nTotal prob of N-in: 0.14402\noutside 1 331\nTMhelix 332 354\ninside 355 372

16769	GCF_001713905.1_ASM171390v1	Salmonella enterica subsp. enterica serovar Enteritidis	Proteobacteria; Gammaproteobacteria; Enterobacterales; Enterobacteriaceae; Salmonella; Salmonella enterica; Salmonella enterica subsp. enterica	MNNEETFYQAMRRKGVTRRSFLKFCSLAATSLGLGAGMTPKIAWALENKPRIPVVWIHGL	WP_001058307.1	hydrogenase [Salmonella enterica]	Length: 372\nNumber of predicted TMHs: 2\nExp number of AAs in TMHs: 36.51738\nExp number, first 60 AAs: 19.72849\nTotal prob of N-in: 0.96170\nPOSSIBLE N-term signal sequence\ninside 1 19\nTMhelix 20 39\noutside 40 325\nTMhelix 326 348\ninside 349 372

16770	GCF_002030505.1_ASM203050v1	Salmonella enterica subsp. enterica serovar Enteritidis	Proteobacteria; Gammaproteobacteria; Enterobacterales; Enterobacteriaceae; Salmonella; Salmonella enterica; Salmonella enterica subsp. enterica	MNNEETFYQAMRRKGVTRRSFLKFCSLAATSLGLGAGMTPKIAWALENKPRIPVVWIHGL	WP_001058307.1	hydrogenase [Salmonella enterica]	Length: 372\nNumber of predicted TMHs: 2\nExp number of AAs in TMHs: 36.51738\nExp number, first 60 AAs: 19.72849\nTotal prob of N-in: 0.96170\nPOSSIBLE N-term signal sequence\ninside 1 19\nTMhelix 20 39\noutside 40 325\nTMhelix 326 348\ninside 349 372

16771	GCF_001713855.1_ASM171385v1	Salmonella enterica subsp. enterica serovar Enteritidis	Proteobacteria; Gammaproteobacteria; Enterobacterales; Enterobacteriaceae; Salmonella; Salmonella enterica; Salmonella enterica subsp. enterica	MNNEETFYQAMRRKGVTRRSFLKFCSLAATSLGLGAGMTPKIAWALENKPRIPVVWIHGL	WP_001058307.1	hydrogenase [Salmonella enterica]	Length: 372\nNumber of predicted TMHs: 2\nExp number of AAs in TMHs: 36.51738\nExp number, first 60 AAs: 19.72849\nTotal prob of N-in: 0.96170\nPOSSIBLE N-term signal sequence\ninside 1 19\nTMhelix 20 39\noutside 40 325\nTMhelix 326 348\ninside 349 372

16772	GCF_001713835.1_ASM171383v1	Salmonella enterica subsp. enterica serovar Enteritidis	Proteobacteria; Gammaproteobacteria; Enterobacterales; Enterobacteriaceae; Salmonella; Salmonella enterica; Salmonella enterica subsp. enterica	MNNEETFYQAMRRKGVTRRSFLKFCSLAATSLGLGAGMTPKIAWALENKPRIPVVWIHGL	WP_001058307.1	hydrogenase [Salmonella enterica]	Length: 372\nNumber of predicted TMHs: 2\nExp number of AAs in TMHs: 36.51738\nExp number, first 60 AAs: 19.72849\nTotal prob of N-in: 0.96170\nPOSSIBLE N-term signal sequence\ninside 1 19\nTMhelix 20 39\noutside 40 325\nTMhelix 326 348\ninside 349 372

16773	GCF_002037965.1_ASM203796v1	Salmonella enterica subsp. enterica serovar Enteritidis	Proteobacteria; Gammaproteobacteria; Enterobacterales; Enterobacteriaceae; Salmonella; Salmonella enterica; Salmonella enterica subsp. enterica	MTGDNTLITSHGINRRDFMKLCAALAATMGLSSKAAAEMAESVSNPQRPPVIWIGAQECT	WP_000145429.1	hydrogenase 2 small subunit [Salmonella enterica]	Length: 372\nNumber of predicted TMHs: 1\nExp number of AAs in TMHs: 24.51309\nExp number, first 60 AAs: 0.2682\nTotal prob of N-in: 0.14402\noutside 1 331\nTMhelix 332 354\ninside 355 372

16774	GCF_001714065.1_ASM171406v1	Salmonella enterica subsp. enterica serovar Enteritidis	Proteobacteria; Gammaproteobacteria; Enterobacterales; Enterobacteriaceae; Salmonella; Salmonella enterica; Salmonella enterica subsp. enterica	MNNEETFYQAMRRKGVTRRSFLKFCSLAATSLGLGAGMTPKIAWALENKPRIPVVWIHGL	WP_001058307.1	hydrogenase [Salmonella enterica]	Length: 372\nNumber of predicted TMHs: 2\nExp number of AAs in TMHs: 36.51738\nExp number, first 60 AAs: 19.72849\nTotal prob of N-in: 0.96170\nPOSSIBLE N-term signal sequence\ninside 1 19\nTMhelix 20 39\noutside 40 325\nTMhelix 326 348\ninside 349 372

16775	GCF_001714015.1_ASM171401v1	Salmonella enterica subsp. enterica serovar Enteritidis	Proteobacteria; Gammaproteobacteria; Enterobacterales; Enterobacteriaceae; Salmonella; Salmonella enterica; Salmonella enterica subsp. enterica	MNNEETFYQAMRRKGVTRRSFLKFCSLAATSLGLGAGMTPKIAWALENKPRIPVVWIHGL	WP_001058307.1	hydrogenase [Salmonella enterica]	Length: 372\nNumber of predicted TMHs: 2\nExp number of AAs in TMHs: 36.51738\nExp number, first 60 AAs: 19.72849\nTotal prob of N-in: 0.96170\nPOSSIBLE N-term signal sequence\ninside 1 19\nTMhelix 20 39\noutside 40 325\nTMhelix 326 348\ninside 349 372

16776	GCF_001713905.1_ASM171390v1	Salmonella enterica subsp. enterica serovar Enteritidis	Proteobacteria; Gammaproteobacteria; Enterobacterales; Enterobacteriaceae; Salmonella; Salmonella enterica; Salmonella enterica subsp. enterica	MTGDNTLITSHGINRRDFMKLCAALAATMGLSSKAAAEMAESVSNPQRPPVIWIGAQECT	WP_000145429.1	hydrogenase 2 small subunit [Salmonella enterica]	Length: 372\nNumber of

predicted TMHs: 1\nExp number of AAs in TMHs: 24.51309\nExp number, first 60 AAs: 0.2682\nTotal prob of N-in: 0.14402\noutside 1 331\nTMhelix 332 354\ninside 355 372

16777 GCF_001713775.1_ASM171377v1 Salmonella enterica subsp. enterica serovar Enteritidis Proteobacteria; Gammaproteobacteria; Enterobacterales; Enterobacteriaceae; Salmonella; Salmonella enterica; Salmonella enterica subsp. enterica MNNEETFYQAMRRKGVTRRSFLKFCSLAATSLGLGAGMTPKIAWALENKPRIPVVWIHGL WP_001058307.1 hydrogenase [Salmonella enterica] Length: 372\nNumber of predicted TMHs: 2\nExp number of AAs in TMHs: 36.51738\nExp number, first 60 AAs: 19.72849\nTotal prob of N-in: 0.96170\nPOSSIBLE N-term signal sequence\ninside 1 19\nTMhelix 20 39\noutside 40 325\nTMhelix 326 348\ninside 349 372

16778 GCF_001713855.1_ASM171385v1 Salmonella enterica subsp. enterica serovar Enteritidis Proteobacteria; Gammaproteobacteria; Enterobacterales; Enterobacteriaceae; Salmonella; Salmonella enterica; Salmonella enterica subsp. enterica MTGDNTLITSHGINRRDFMKLCAALAATMGLSSKAAAEMAESVSNPQRPPVIWIGAQECT WP_000145429.1 hydrogenase 2 small subunit [Salmonella enterica] Length: 372\nNumber of predicted TMHs: 1\nExp number of AAs in TMHs: 24.51309\nExp number, first 60 AAs: 0.2682\nTotal prob of N-in: 0.14402\noutside 1 331\nTMhelix 332 354\ninside 355 372

16779 GCF_001713835.1_ASM171383v1 Salmonella enterica subsp. enterica serovar Enteritidis Proteobacteria; Gammaproteobacteria; Enterobacterales; Enterobacteriaceae; Salmonella; Salmonella enterica; Salmonella enterica subsp. enterica MTGDNTLITSHGINRRDFMKLCAALAATMGLSSKAAAEMAESVSNPQRPPVIWIGAQECT WP_000145429.1 hydrogenase 2 small subunit [Salmonella enterica] Length: 372\nNumber of predicted TMHs: 1\nExp number of AAs in TMHs: 24.51309\nExp number, first 60 AAs: 0.2682\nTotal prob of N-in: 0.14402\noutside 1 331\nTMhelix 332 354\ninside 355 372

16780 GCF_002037905.1_ASM203790v1 Salmonella enterica subsp. enterica serovar Enteritidis Proteobacteria; Gammaproteobacteria; Enterobacterales; Enterobacteriaceae; Salmonella; Salmonella enterica; Salmonella enterica subsp. enterica MTGDNTLITSHGINRRDFMKLCAALAATMGLSSKAAAEMAESVSNPQRPPVIWIGAQECT WP_000145429.1 hydrogenase 2 small subunit [Salmonella enterica] Length: 372\nNumber of predicted TMHs: 1\nExp number of AAs in TMHs: 24.51309\nExp number, first 60 AAs: 0.2682\nTotal prob of N-in: 0.14402\noutside 1 331\nTMhelix 332 354\ninside 355 372

16781 GCF_001713825.1_ASM171382v1 Salmonella enterica subsp. enterica serovar Enteritidis Proteobacteria; Gammaproteobacteria; Enterobacterales; Enterobacteriaceae; Salmonella; Salmonella enterica; Salmonella enterica subsp. enterica MTGDNTLITSHGINRRDFMKLCAALAATMGLSSKAAAEMAESVSNPQRPPVIWIGAQECT WP_000145429.1 hydrogenase 2 small subunit [Salmonella enterica] Length: 372\nNumber of predicted TMHs: 1\nExp number of AAs in TMHs: 24.51309\nExp number, first 60 AAs: 0.2682\nTotal prob of N-in: 0.14402\noutside 1 331\nTMhelix 332 354\ninside 355 372

16782 GCF_001478145.1_Salmonella_enterica_CVM_N31397-SQ_v1.0 Salmonella enterica subsp. enterica serovar Enteritidis Proteobacteria; Gammaproteobacteria; Enterobacterales; Enterobacteriaceae; Salmonella; Salmonella enterica; Salmonella enterica subsp. enterica MTGDNTLITSHGINRRDFMKLCAALAATMGLSSKAAAEMAESVSNPQRPPVIWIGAQECT WP_000145429.1 hydrogenase 2 small subunit [Salmonella enterica] Length: 372\nNumber of predicted TMHs: 1\nExp number of AAs in TMHs: 24.51309\nExp number, first 60 AAs: 0.2682\nTotal prob of N-in: 0.14402\noutside 1 331\nTMhelix 332 354\ninside 355 372

16783 GCF_002040545.1_ASM204054v1 Salmonella enterica subsp. enterica serovar Enteritidis Proteobacteria; Gammaproteobacteria; Enterobacterales; Enterobacteriaceae; Salmonella; Salmonella enterica; Salmonella enterica subsp. enterica MTGDNTLITSHGINRRDFMKLCAALAATMGLSSKAAAEMAESVSNPQRPPVIWIGAQECT WP_000145429.1 hydrogenase 2 small subunit [Salmonella enterica] Length: 372\nNumber of predicted TMHs: 1\nExp number of AAs in TMHs: 24.51309\nExp number, first 60 AAs: 0.2682\nTotal prob of N-in: 0.14402\noutside 1 331\nTMhelix 332 354\ninside 355 372

16784 GCF_001713775.1_ASM171377v1 Salmonella enterica subsp. enterica serovar Enteritidis Proteobacteria; Gammaproteobacteria; Enterobacterales; Enterobacteriaceae; Salmonella; Salmonella enterica; Salmonella enterica subsp. enterica MTGDNTLITSHGINRRDFMKLCAALAATMGLSSKAAAEMAESVSNPQRPPVIWIGAQECT WP_000145429.1 hydrogenase 2 small subunit [Salmonella enterica] Length: 372\nNumber of predicted TMHs: 1\nExp number of AAs in TMHs: 24.51309\nExp number, first 60 AAs: 0.2682\nTotal prob of N-in: 0.14402\noutside 1 331\nTMhelix 332 354\ninside 355 372

16785 GCF_002041175.1_ASM204117v1 Salmonella enterica subsp. enterica serovar Enteritidis Proteobacteria; Gammaproteobacteria; Enterobacterales; Enterobacteriaceae; Salmonella; Salmonella enterica; Salmonella enterica subsp. enterica MNNEETFYQAMRRKGVTRRSFLKFCSLAATSLGLGAGMTPKIAWALENKPRIPVVWIHGL WP_001058307.1 hydrogenase [Salmonella enterica] Length: 372\nNumber of predicted TMHs: 2\nExp number of AAs in TMHs: 36.51738\nExp number, first 60 AAs: 19.72849\nTotal prob of N-in: 0.96170\nPOSSIBLE N-term signal sequence\ninside 1 19\nTMhelix 20 39\noutside 40 325\nTMhelix 326 348\ninside 349 372

16786 GCF_002030505.1_ASM203050v1 Salmonella enterica subsp. enterica serovar Enteritidis Proteobacteria; Gammaproteobacteria; Enterobacterales; Enterobacteriaceae; Salmonella; Salmonella enterica; Salmonella enterica subsp. enterica MTGDNTLITSHGINRRDFMKLCAALAATMGLSSKAAAEMAESVSNPQRPPVIWIGAQECT WP_000145429.1 hydrogenase 2 small subunit [Salmonella enterica] Length: 372\nNumber of predicted TMHs: 1\nExp number of AAs in TMHs: 24.51309\nExp number, first 60 AAs: 0.2682\nTotal prob of N-in: 0.14402\noutside 1 331\nTMhelix 332 354\ninside 355 372

16787 GCF_002040245.1_ASM204024v1 Salmonella enterica subsp. enterica serovar Enteritidis Proteobacteria; Gammaproteobacteria; Enterobacterales; Enterobacteriaceae; Salmonella; Salmonella enterica; Salmonella enterica subsp. enterica MNNEETFYQAMRRKGVTRRSFLKFCSLAATSLGLGAGMTPKIAWALENKPRIPVVWIHGL WP_001058307.1 hydrogenase [Salmonella enterica] Length: 372\nNumber of predicted TMHs: 2\nExp number of AAs in TMHs: 36.51738\nExp number, first 60 AAs: 19.72849\nTotal prob of N-in: 0.96170\nPOSSIBLE N-term signal sequence\ninside 1 19\nTMhelix 20 39\noutside 40 325\nTMhelix 326 348\ninside 349 372

16788 GCF_002040365.1_ASM204036v1 Salmonella enterica subsp. enterica serovar Enteritidis Proteobacteria; Gammaproteobacteria; Enterobacterales; Enterobacteriaceae; Salmonella; Salmonella enterica; Salmonella enterica subsp. enterica MTGDNTLITSHGINRRDFMKLCAALAATMGLSSKAAAEMAESVSNPQRPPVIWIGAQECT WP_000145429.1 hydrogenase 2 small subunit [Salmonella enterica] Length: 372\nNumber of predicted TMHs: 1\nExp number of AAs in TMHs: 24.51309\nExp number, first 60 AAs: 0.2682\nTotal prob of N-in: 0.14402\noutside 1 331\nTMhelix 332 354\ninside 355 372

16789 GCF_002037865.1_ASM203786v1 Salmonella enterica subsp. enterica serovar Enteritidis Proteobacteria; Gammaproteobacteria; Enterobacterales; Enterobacteriaceae; Salmonella; Salmonella enterica; Salmonella enterica subsp. enterica MTGDNTLITSHGINRRDFMKLCAALAATMGLSSKAAAEMAESVSNPQRPPVIWIGAQECT WP_000145429.1 hydrogenase 2 small subunit [Salmonella enterica] Length: 372\nNumber of predicted TMHs: 1\nExp number of AAs in TMHs: 24.51309\nExp number, first 60 AAs: 0.2682\nTotal prob of N-in: 0.14402\noutside 1 331\nTMhelix 332 354\ninside 355 372

16790 GCF_002041305.1_ASM204130v1 Salmonella enterica subsp. enterica serovar Enteritidis Proteobacteria; Gammaproteobacteria; Enterobacterales; Enterobacteriaceae; Salmonella; Salmonella enterica; Salmonella enterica subsp. enterica MNNEETFYQAMRRKGVTRRSFLKFCSLAATSLGLGAGMTPKIAWALENKPRIPVVWIHGL WP_001058307.1 hydrogenase [Salmonella enterica] Length: 372\nNumber of predicted TMHs: 2\nExp number of AAs in TMHs: 36.51738\nExp number, first 60 AAs: 19.72849\nTotal prob of N-in: 0.96170\nPOSSIBLE N-term signal sequence\ninside 1 19\nTMhelix 20 39\noutside 40 325\nTMhelix 326 348\ninside 349 372

16791 GCF_001478145.1_Salmonella_enterica_CVM_N31397-SQ_v1.0 Salmonella enterica subsp. enterica serovar Enteritidis Proteobacteria; Gammaproteobacteria; Enterobacterales; Enterobacteriaceae; Salmonella; Salmonella enterica; Salmonella enterica subsp. enterica MNNEETFYQAMRRKGVTRRSFLKFCSLAATSLGLGAGMTPKIAWALENKPRIPVVWIHGL WP_001058307.1 hydrogenase [Salmonella enterica] Length: 372\nNumber of predicted TMHs: 2\nExp number of AAs in TMHs: 36.51738\nExp number, first 60 AAs: 19.72849\nTotal prob of N-in: 0.96170\nPOSSIBLE N-term signal sequence\ninside 1 19\nTMhelix 20 39\noutside 40 325\nTMhelix 326 348\ninside 349 372

16792 GCF_002039045.1_ASM203904v1 Salmonella enterica subsp. enterica serovar Enteritidis Proteobacteria; Gammaproteobacteria; Enterobacterales; Enterobacteriaceae; Salmonella; Salmonella enterica; Salmonella enterica subsp. enterica MTGDNTLITSHGINRRDFMKLCAALAATMGLSSKAAAEMAESVSNPQRPPVIWIGAQECT WP_000145429.1 hydrogenase 2 small subunit [Salmonella enterica] Length: 372\nNumber of predicted TMHs: 1\nExp number of AAs in TMHs: 24.51309\nExp number, first 60 AAs: 0.2682\nTotal prob of N-in: 0.14402\noutside 1 331\nTMhelix 332 354\ninside 355 372

16793 GCF_002032355.1_ASM203235v1 Salmonella enterica subsp. enterica serovar Enteritidis Proteobacteria; Gammaproteobacteria; Enterobacterales; Enterobacteriaceae; Salmonella; Salmonella enterica; Salmonella enterica subsp. enterica MTGDNTLITSHGINRRDFMKLCAALAATMGLSSKAAAEMAESVSNPQRPPVIWIGAQECT WP_000145429.1 hydrogenase 2 small subunit [Salmonella enterica] Length: 372\nNumber of

predicted TMHs: 1\nExp number of AAs in TMHs: 24.51309\nExp number, first 60 AAs: 0.2682\nTotal prob of N-in: 0.14402\noutside 1 331\nTMhelix 332 354\ninside 355 372

16794 GCF_002039705.1_ASM203970v1 Salmonella enterica subsp. enterica serovar Enteritidis Proteobacteria; Gammaproteobacteria; Enterobacterales; Enterobacteriaceae; Salmonella; Salmonella enterica; Salmonella enterica subsp. enterica MNNEETFYQAMRRKGVTRRSFLKFCSLAATSLGLGAGMTPKIAWALENKPRIPVVWIHGL WP_001058307.1 hydrogenase [Salmonella enterica] Length: 372\nNumber of predicted TMHs: 2\nExp number of AAs in TMHs: 36.51738\nExp number, first 60 AAs: 19.72849\nTotal prob of N-in: 0.96170\nPOSSIBLE N-term signal sequence\ninside 1 19\nTMhelix 20 39\noutside 40 325\nTMhelix 326 348\ninside 349 372

16795 GCF_002041525.1_ASM204152v1 Salmonella enterica subsp. enterica serovar Enteritidis Proteobacteria; Gammaproteobacteria; Enterobacterales; Enterobacteriaceae; Salmonella; Salmonella enterica; Salmonella enterica subsp. enterica MTGDNTLITSHGINRRDFMKLCAALAATMGLSSKAAAEMAESVSNPQRPPVIWIGAQECT WP_000145429.1 hydrogenase 2 small subunit [Salmonella enterica] Length: 372\nNumber of predicted TMHs: 1\nExp number of AAs in TMHs: 24.51309\nExp number, first 60 AAs: 0.2682\nTotal prob of N-in: 0.14402\noutside 1 331\nTMhelix 332 354\ninside 355 372

16796 GCF_002041535.1_ASM204153v1 Salmonella enterica subsp. enterica serovar Enteritidis Proteobacteria; Gammaproteobacteria; Enterobacterales; Enterobacteriaceae; Salmonella; Salmonella enterica; Salmonella enterica subsp. enterica MTGDNTLITSHGINRRDFMKLCAALAATMGLSSKAAAEMAESVSNPQRPPVIWIGAQECT WP_000145429.1 hydrogenase 2 small subunit [Salmonella enterica] Length: 372\nNumber of predicted TMHs: 1\nExp number of AAs in TMHs: 24.51309\nExp number, first 60 AAs: 0.2682\nTotal prob of N-in: 0.14402\noutside 1 331\nTMhelix 332 354\ninside 355 372

16797 GCF_002040395.1_ASM204039v1 Salmonella enterica subsp. enterica serovar Enteritidis Proteobacteria; Gammaproteobacteria; Enterobacterales; Enterobacteriaceae; Salmonella; Salmonella enterica; Salmonella enterica subsp. enterica MTGDNTLITSHGINRRDFMKLCAALAATMGLSSKAAAEMAESVSNPQRPPVIWIGAQECT WP_000145429.1 hydrogenase 2 small subunit [Salmonella enterica] Length: 372\nNumber of predicted TMHs: 1\nExp number of AAs in TMHs: 24.51309\nExp number, first 60 AAs: 0.2682\nTotal prob of N-in: 0.14402\noutside 1 331\nTMhelix 332 354\ninside 355 372

16798 GCF_002032355.1_ASM203235v1 Salmonella enterica subsp. enterica serovar Enteritidis Proteobacteria; Gammaproteobacteria; Enterobacterales; Enterobacteriaceae; Salmonella; Salmonella enterica; Salmonella enterica subsp. enterica MNNEETFYQAMRRKGVTRRSFLKFCSLAATSLGLGAGMTPKIAWALENKPRIPVVWIHGL WP_001058307.1 hydrogenase [Salmonella enterica] Length: 372\nNumber of predicted TMHs: 2\nExp number of AAs in TMHs: 36.51738\nExp number, first 60 AAs: 19.72849\nTotal prob of N-in: 0.96170\nPOSSIBLE N-term signal sequence\ninside 1 19\nTMhelix 20 39\noutside 40 325\nTMhelix 326 348\ninside 349 372

16799 GCF_002040315.1_ASM204031v1 Salmonella enterica subsp. enterica serovar Enteritidis Proteobacteria; Gammaproteobacteria; Enterobacterales; Enterobacteriaceae; Salmonella; Salmonella enterica; Salmonella enterica subsp. enterica MTGDNTLITSHGINRRDFMKLCAALAATMGLSSKAAAEMAESVSNPQRPPVIWIGAQECT WP_000145429.1 hydrogenase 2 small subunit [Salmonella enterica] Length: 372\nNumber of predicted TMHs: 1\nExp number of AAs in TMHs: 24.51309\nExp number, first 60 AAs: 0.2682\nTotal prob of N-in: 0.14402\noutside 1 331\nTMhelix 332 354\ninside 355 372

16800 GCF_002042315.1_ASM204231v1 Salmonella enterica subsp. enterica serovar Enteritidis Proteobacteria; Gammaproteobacteria; Enterobacterales; Enterobacteriaceae; Salmonella; Salmonella enterica; Salmonella enterica subsp. enterica MNNEETFYQAMRRKGVTRRSFLKFCSLAATSLGLGAGMTPKIAWALENKPRIPVVWIHGL WP_001058307.1 hydrogenase [Salmonella enterica] Length: 372\nNumber of predicted TMHs: 2\nExp number of AAs in TMHs: 36.51738\nExp number, first 60 AAs: 19.72849\nTotal prob of N-in: 0.96170\nPOSSIBLE N-term signal sequence\ninside 1 19\nTMhelix 20 39\noutside 40 325\nTMhelix 326 348\ninside 349 372

16801 GCF_002040285.1_ASM204028v1 Salmonella enterica subsp. enterica serovar Enteritidis Proteobacteria; Gammaproteobacteria; Enterobacterales; Enterobacteriaceae; Salmonella; Salmonella enterica; Salmonella enterica subsp. enterica MTGDNTLITSHGINRRDFMKLCAALAATMGLSSKAAAEMAESVSNPQRPPVIWIGAQECT WP_000145429.1 hydrogenase 2 small subunit [Salmonella enterica] Length: 372\nNumber of predicted TMHs: 1\nExp number of AAs in TMHs: 24.51309\nExp number, first 60 AAs: 0.2682\nTotal prob of N-in: 0.14402\noutside 1 331\nTMhelix 332 354\ninside 355 372

16802 GCF_002040205.1_ASM204020v1 Salmonella enterica subsp. enterica serovar Enteritidis Proteobacteria; Gammaproteobacteria; Enterobacterales; Enterobacteriaceae; Salmonella; Salmonella enterica;

Salmonella enterica subsp. enterica	MTGDNTLITSHGINRRDFMKLCAALAATMGLSSKAAAEMAESVSNPQRPPVIWIGAQECT	WP_000145429.1	hydrogenase 2 small subunit [Salmonella enterica]	Length: 372\nNumber of predicted TMHs:  1\nExp number of AAs in TMHs: 24.51309\nExp number, first 60 AAs:  0.2682\nTotal prob of N-in:  0.14402\noutside   1   331\nTMhelix   332   354\ninside   355   372

16803	GCF_002036395.1_ASM203639v1	Salmonella enterica subsp. enterica serovar Enteritidis	Proteobacteria; Gammaproteobacteria; Enterobacterales; Enterobacteriaceae; Salmonella; Salmonella enterica; Salmonella enterica subsp. enterica	MTGDNTLITSHGINRRDFMKLCAALAATMGLSSKAAAEMAESVSNPQRPPVIWIGAQECT	WP_079830008.1	hydrogenase 2 small subunit, partial [Salmonella enterica]	Length: 357\nNumber of predicted TMHs:  1\nExp number of AAs in TMHs: 24.14871\nExp number, first 60 AAs:  0.27617\nTotal prob of N-in:  0.22797\noutside   1   332\nTMhelix   333   355\ninside   356   357

16804	GCF_002040155.1_ASM204015v1	Salmonella enterica subsp. enterica serovar Enteritidis	Proteobacteria; Gammaproteobacteria; Enterobacterales; Enterobacteriaceae; Salmonella; Salmonella enterica; Salmonella enterica subsp. enterica	MTGDNTLITSHGINRRDFMKLCAALAATMGLSSKAAAEMAESVSNPQRPPVIWIGAQECT	WP_000145429.1	hydrogenase 2 small subunit [Salmonella enterica]	Length: 372\nNumber of predicted TMHs:  1\nExp number of AAs in TMHs: 24.51309\nExp number, first 60 AAs:  0.2682\nTotal prob of N-in:  0.14402\noutside   1   331\nTMhelix   332   354\ninside   355   372

16805	GCF_002040145.1_ASM204014v1	Salmonella enterica subsp. enterica serovar Enteritidis	Proteobacteria; Gammaproteobacteria; Enterobacterales; Enterobacteriaceae; Salmonella; Salmonella enterica; Salmonella enterica subsp. enterica	MTGDNTLITSHGINRRDFMKLCAALAATMGLSSKAAAEMAESVSNPQRPPVIWIGAQECT	WP_000145429.1	hydrogenase 2 small subunit [Salmonella enterica]	Length: 372\nNumber of predicted TMHs:  1\nExp number of AAs in TMHs: 24.51309\nExp number, first 60 AAs:  0.2682\nTotal prob of N-in:  0.14402\noutside   1   331\nTMhelix   332   354\ninside   355   372

16806	GCF_002039965.1_ASM203996v1	Salmonella enterica subsp. enterica serovar Enteritidis	Proteobacteria; Gammaproteobacteria; Enterobacterales; Enterobacteriaceae; Salmonella; Salmonella enterica; Salmonella enterica subsp. enterica	MNNEETFYQAMRRKGVTRRSFLKFCSLAATSLGLGAGMTPKIAWALENKPRIPVVWIHGL	WP_001058307.1	hydrogenase [Salmonella enterica]	Length: 372\nNumber of predicted TMHs:  2\nExp number of AAs in TMHs: 36.51738\nExp number, first 60 AAs:  19.72849\nTotal prob of N-in:   0.96170\nPOSSIBLE N-term signal sequence\ninside   1   19\nTMhelix   20   39\noutside   40   325\nTMhelix   326   348\ninside   349   372

16807	GCF_002040045.1_ASM204004v1	Salmonella enterica subsp. enterica serovar Enteritidis	Proteobacteria; Gammaproteobacteria; Enterobacterales; Enterobacteriaceae; Salmonella; Salmonella enterica; Salmonella enterica subsp. enterica	MNNEETFYQAMRRKGVTRRSFLKFCSLAATSLGLGAGMTPKIAWALENKPRIPVVWIHGL	WP_001058307.1	hydrogenase [Salmonella enterica]	Length: 372\nNumber of predicted TMHs:  2\nExp number of AAs in TMHs: 36.51738\nExp number, first 60 AAs:  19.72849\nTotal prob of N-in:   0.96170\nPOSSIBLE N-term signal sequence\ninside   1   19\nTMhelix   20   39\noutside   40   325\nTMhelix   326   348\ninside   349   372

16808	GCF_002040005.1_ASM204000v1	Salmonella enterica subsp. enterica serovar Enteritidis	Proteobacteria; Gammaproteobacteria; Enterobacterales; Enterobacteriaceae; Salmonella; Salmonella enterica; Salmonella enterica subsp. enterica	MNNEETFYQAMRRKGVTRRSFLKFCSLAATSLGLGAGMTPKIAWALENKPRIPVVWIHGL	WP_001058307.1	hydrogenase [Salmonella enterica]	Length: 372\nNumber of predicted TMHs:  2\nExp number of AAs in TMHs: 36.51738\nExp number, first 60 AAs:  19.72849\nTotal prob of N-in:   0.96170\nPOSSIBLE N-term signal sequence\ninside   1   19\nTMhelix   20   39\noutside   40   325\nTMhelix   326   348\ninside   349   372

16809	GCF_001478845.1_Salmonella_enterica_CVM_N29385-SQ_v1.0	Salmonella enterica subsp. enterica serovar Enteritidis	Proteobacteria; Gammaproteobacteria; Enterobacterales; Enterobacteriaceae; Salmonella; Salmonella enterica; Salmonella enterica subsp. enterica	MTGDNTLITSHGINRRDFMKLCAALAATMGLSSKAAAEMAESVSNPQRPPVIWIGAQECT	WP_000145429.1	hydrogenase 2 small subunit [Salmonella enterica]	Length: 372\nNumber of predicted TMHs:  1\nExp number of AAs in TMHs: 24.51309\nExp number, first 60 AAs:  0.2682\nTotal prob of N-in:   0.14402\noutside   1   331\nTMhelix   332   354\ninside   355   372

16810	GCF_002036435.1_ASM203643v1	Salmonella enterica subsp. enterica serovar Enteritidis	Proteobacteria; Gammaproteobacteria; Enterobacterales; Enterobacteriaceae; Salmonella; Salmonella enterica; Salmonella enterica subsp. enterica	MTGDNTLITSHGINRRDFMKLCAALAATMGLSSKAAAEMAESVSNPQRPPVIWIGAQECT	WP_000145429.1	hydrogenase 2 small subunit [Salmonella enterica]	Length: 372\nNumber of predicted TMHs:  1\nExp number of AAs in TMHs: 24.51309\nExp number, first 60 AAs:  0.2682\nTotal prob of N-in:  0.14402\noutside   1   331\nTMhelix   332   354\ninside   355   372

16811 GCF_001478845.1_Salmonella_enterica_CVM_N29385-SQ_v1.0 Salmonella enterica subsp. enterica serovar Enteritidis Proteobacteria; Gammaproteobacteria; Enterobacterales; Enterobacteriaceae; Salmonella; Salmonella enterica; Salmonella enterica subsp. enterica MNNEETFYQAMRRKGVTRRSFLKFCSLAATSLGLGAGMTPKIAWALENKPRIPVVWIHGL WP_001058307.1 hydrogenase [Salmonella enterica] Length: 372\nNumber of predicted TMHs: 2\nExp number of AAs in TMHs: 36.51738\nExp number, first 60 AAs: 19.72849\nTotal prob of N-in: 0.96170\nPOSSIBLE N-term signal sequence\ninside 1 19\nTMhelix 20 39\noutside 40 325\nTMhelix 326 348\ninside 349 372

16812 GCF_001973665.1_ASM197366v1 Salmonella enterica subsp. enterica serovar Enteritidis Proteobacteria; Gammaproteobacteria; Enterobacterales; Enterobacteriaceae; Salmonella; Salmonella enterica; Salmonella enterica subsp. enterica MTGDNTLITSHGINRRDFMKLCAALAATMGLSSKAAAEMAESVSNPQRPPVIWIGAQECT WP_000145429.1 hydrogenase 2 small subunit [Salmonella enterica] Length: 372\nNumber of predicted TMHs: 1\nExp number of AAs in TMHs: 24.51309\nExp number, first 60 AAs: 0.2682\nTotal prob of N-in: 0.14402\noutside 1 331\nTMhelix 332 354\ninside 355 372

16813 GCF_002041575.1_ASM204157v1 Salmonella enterica subsp. enterica serovar Enteritidis Proteobacteria; Gammaproteobacteria; Enterobacterales; Enterobacteriaceae; Salmonella; Salmonella enterica; Salmonella enterica subsp. enterica MTGDNTLITSHGINRRDFMKLCAALAATMGLSSKAAAEMAESVSNPQRPPVIWIGAQECT WP_000145429.1 hydrogenase 2 small subunit [Salmonella enterica] Length: 372\nNumber of predicted TMHs: 1\nExp number of AAs in TMHs: 24.51309\nExp number, first 60 AAs: 0.2682\nTotal prob of N-in: 0.14402\noutside 1 331\nTMhelix 332 354\ninside 355 372

16814 GCF_001647045.1_ASM164704v1 Salmonella enterica subsp. enterica serovar Enteritidis Proteobacteria; Gammaproteobacteria; Enterobacterales; Enterobacteriaceae; Salmonella; Salmonella enterica; Salmonella enterica subsp. enterica MNNEETFYQAMRRKGVTRRSFLKFCSLAATSLGLGAGMTPKIAWALENKPRIPVVWIHGL WP_001058307.1 hydrogenase [Salmonella enterica] Length: 372\nNumber of predicted TMHs: 2\nExp number of AAs in TMHs: 36.51738\nExp number, first 60 AAs: 19.72849\nTotal prob of N-in: 0.96170\nPOSSIBLE N-term signal sequence\ninside 1 19\nTMhelix 20 39\noutside 40 325\nTMhelix 326 348\ninside 349 372

16815 GCF_001655705.1_ASM165570v1 Salmonella enterica subsp. enterica serovar Enteritidis Proteobacteria; Gammaproteobacteria; Enterobacterales; Enterobacteriaceae; Salmonella; Salmonella enterica; Salmonella enterica subsp. enterica MNNEETFYQAMRRKGVTRRSFLKFCSLAATSLGLGAGMTPKIAWALENKPRIPVVWIHGL WP_001058307.1 hydrogenase [Salmonella enterica] Length: 372\nNumber of predicted TMHs: 2\nExp number of AAs in TMHs: 36.51738\nExp number, first 60 AAs: 19.72849\nTotal prob of N-in: 0.96170\nPOSSIBLE N-term signal sequence\ninside 1 19\nTMhelix 20 39\noutside 40 325\nTMhelix 326 348\ninside 349 372

16816 GCF_002040545.1_ASM204054v1 Salmonella enterica subsp. enterica serovar Enteritidis Proteobacteria; Gammaproteobacteria; Enterobacterales; Enterobacteriaceae; Salmonella; Salmonella enterica; Salmonella enterica subsp. enterica MNNEETFYQAMRRKGVTRRSFLKFCSLAATSLGLGAGMTPKIAWALENKPRIPVVWIHGL WP_001058307.1 hydrogenase [Salmonella enterica] Length: 372\nNumber of predicted TMHs: 2\nExp number of AAs in TMHs: 36.51738\nExp number, first 60 AAs: 19.72849\nTotal prob of N-in: 0.96170\nPOSSIBLE N-term signal sequence\ninside 1 19\nTMhelix 20 39\noutside 40 325\nTMhelix 326 348\ninside 349 372

16817 GCF_001655705.1_ASM165570v1 Salmonella enterica subsp. enterica serovar Enteritidis Proteobacteria; Gammaproteobacteria; Enterobacterales; Enterobacteriaceae; Salmonella; Salmonella enterica; Salmonella enterica subsp. enterica MTGDNTLITSHGINRRDFMKLCAALAATMGLSSKAAAEMAESVSNPQRPPVIWIGAQECT WP_000145429.1 hydrogenase 2 small subunit [Salmonella enterica] Length: 372\nNumber of predicted TMHs: 1\nExp number of AAs in TMHs: 24.51309\nExp number, first 60 AAs: 0.2682\nTotal prob of N-in: 0.14402\noutside 1 331\nTMhelix 332 354\ninside 355 372

16818 GCF_001643395.1_ASM164339v1 Salmonella enterica subsp. enterica serovar Enteritidis Proteobacteria; Gammaproteobacteria; Enterobacterales; Enterobacteriaceae; Salmonella; Salmonella enterica; Salmonella enterica subsp. enterica MNNEETFYQAMRRKGVTRRSFLKFCSLAATSLGLGAGMTPKIAWALENKPRIPVVWIHGL WP_001058307.1 hydrogenase [Salmonella enterica] Length: 372\nNumber of predicted TMHs: 2\nExp number of AAs in TMHs: 36.51738\nExp number, first 60 AAs: 19.72849\nTotal prob of N-in: 0.96170\nPOSSIBLE N-term signal sequence\ninside 1 19\nTMhelix 20 39\noutside 40 325\nTMhelix 326 348\ninside 349 372

16819 GCF_002040985.1_ASM204098v1 Salmonella enterica subsp. enterica serovar Enteritidis Proteobacteria; Gammaproteobacteria; Enterobacterales; Enterobacteriaceae; Salmonella; Salmonella enterica; Salmonella enterica subsp. enterica MTGDNTLITSHGINRRDFMKLCAALAATMGLSSKAAAEMAESVSNPQRPPVIWIGAQECT WP_000145429.1 hydrogenase 2 small subunit [Salmonella enterica] Length: 372\nNumber of

predicted TMHs: 1\nExp number of AAs in TMHs: 24.51309\nExp number, first 60 AAs: 0.2682\nTotal prob of N-in: 0.14402\noutside 1 331\nTMhelix 332 354\ninside 355 372

16820 GCF_001643395.1_ASM164339v1 Salmonella enterica subsp. enterica serovar Enteritidis Proteobacteria; Gammaproteobacteria; Enterobacterales; Enterobacteriaceae; Salmonella; Salmonella enterica; Salmonella enterica subsp. enterica MTGDNTLITSHGINRRDFMKLCAALAATMGLSSKAAAEMAESVSNPQRPPVIWIGAQECT WP_000145429.1 hydrogenase 2 small subunit [Salmonella enterica] Length: 372\nNumber of predicted TMHs: 1\nExp number of AAs in TMHs: 24.51309\nExp number, first 60 AAs: 0.2682\nTotal prob of N-in: 0.14402\noutside 1 331\nTMhelix 332 354\ninside 355 372

16821 GCF_002039805.1_ASM203980v1 Salmonella enterica subsp. enterica serovar Enteritidis Proteobacteria; Gammaproteobacteria; Enterobacterales; Enterobacteriaceae; Salmonella; Salmonella enterica; Salmonella enterica subsp. enterica MTGDNTLITSHGINRRDFMKLCAALAATMGLSSKAAAEMAESVSNPQRPPVIWIGAQECT WP_000145429.1 hydrogenase 2 small subunit [Salmonella enterica] Length: 372\nNumber of predicted TMHs: 1\nExp number of AAs in TMHs: 24.51309\nExp number, first 60 AAs: 0.2682\nTotal prob of N-in: 0.14402\noutside 1 331\nTMhelix 332 354\ninside 355 372

16822 GCF_002041615.1_ASM204161v1 Salmonella enterica subsp. enterica serovar Enteritidis Proteobacteria; Gammaproteobacteria; Enterobacterales; Enterobacteriaceae; Salmonella; Salmonella enterica; Salmonella enterica subsp. enterica MTGDNTLITSHGINRRDFMKLCAALAATMGLSSKAAAEMAESVSNPQRPPVIWIGAQECT WP_000145429.1 hydrogenase 2 small subunit [Salmonella enterica] Length: 372\nNumber of predicted TMHs: 1\nExp number of AAs in TMHs: 24.51309\nExp number, first 60 AAs: 0.2682\nTotal prob of N-in: 0.14402\noutside 1 331\nTMhelix 332 354\ninside 355 372

16823 GCF_002039865.1_ASM203986v1 Salmonella enterica subsp. enterica serovar Enteritidis Proteobacteria; Gammaproteobacteria; Enterobacterales; Enterobacteriaceae; Salmonella; Salmonella enterica; Salmonella enterica subsp. enterica MTGDNTLITSHGINRRDFMKLCAALAATMGLSSKAAAEMAESVSNPQRPPVIWIGAQECT WP_000145429.1 hydrogenase 2 small subunit [Salmonella enterica] Length: 372\nNumber of predicted TMHs: 1\nExp number of AAs in TMHs: 24.51309\nExp number, first 60 AAs: 0.2682\nTotal prob of N-in: 0.14402\noutside 1 331\nTMhelix 332 354\ninside 355 372

16824 GCF_002041525.1_ASM204152v1 Salmonella enterica subsp. enterica serovar Enteritidis Proteobacteria; Gammaproteobacteria; Enterobacterales; Enterobacteriaceae; Salmonella; Salmonella enterica; Salmonella enterica subsp. enterica MNNEETFYQAMRRKGVTRRSFLKFCSLAATSLGLGAGMTPKIAWALENKPRIPVVWIHGL WP_001058307.1 hydrogenase [Salmonella enterica] Length: 372\nNumber of predicted TMHs: 2\nExp number of AAs in TMHs: 36.51738\nExp number, first 60 AAs: 19.72849\nTotal prob of N-in: 0.96170\nPOSSIBLE N-term signal sequence\ninside 1 19\nTMhelix 20 39\noutside 40 325\nTMhelix 326 348\ninside 349 372

16825 GCF_002039845.1_ASM203984v1 Salmonella enterica subsp. enterica serovar Enteritidis Proteobacteria; Gammaproteobacteria; Enterobacterales; Enterobacteriaceae; Salmonella; Salmonella enterica; Salmonella enterica subsp. enterica MTGDNTLITSHGINRRDFMKLCAALAATMGLSSKAAAEMAESVSNPQRPPVIWIGAQECT WP_000145429.1 hydrogenase 2 small subunit [Salmonella enterica] Length: 372\nNumber of predicted TMHs: 1\nExp number of AAs in TMHs: 24.51309\nExp number, first 60 AAs: 0.2682\nTotal prob of N-in: 0.14402\noutside 1 331\nTMhelix 332 354\ninside 355 372

16826 GCF_002040605.1_ASM204060v1 Salmonella enterica subsp. enterica serovar Enteritidis Proteobacteria; Gammaproteobacteria; Enterobacterales; Enterobacteriaceae; Salmonella; Salmonella enterica; Salmonella enterica subsp. enterica MNNEETFYQAMRRKGVTRRSFLKFCSLAATSLGLGAGMTPKIAWALENKPRIPVVWIHGL WP_001058307.1 hydrogenase [Salmonella enterica] Length: 372\nNumber of predicted TMHs: 2\nExp number of AAs in TMHs: 36.51738\nExp number, first 60 AAs: 19.72849\nTotal prob of N-in: 0.96170\nPOSSIBLE N-term signal sequence\ninside 1 19\nTMhelix 20 39\noutside 40 325\nTMhelix 326 348\ninside 349 372

16827 GCF_002039865.1_ASM203986v1 Salmonella enterica subsp. enterica serovar Enteritidis Proteobacteria; Gammaproteobacteria; Enterobacterales; Enterobacteriaceae; Salmonella; Salmonella enterica; Salmonella enterica subsp. enterica MNNEETFYQAMRRKGVTRRSFLKFCSLAATSLGLGAGMTPKIAWALENKPRIPVVWIHGL WP_001058307.1 hydrogenase [Salmonella enterica] Length: 372\nNumber of predicted TMHs: 2\nExp number of AAs in TMHs: 36.51738\nExp number, first 60 AAs: 19.72849\nTotal prob of N-in: 0.96170\nPOSSIBLE N-term signal sequence\ninside 1 19\nTMhelix 20 39\noutside 40 325\nTMhelix 326 348\ninside 349 372

16828 GCF_002041535.1_ASM204153v1 Salmonella enterica subsp. enterica serovar Enteritidis Proteobacteria; Gammaproteobacteria; Enterobacterales; Enterobacteriaceae; Salmonella; Salmonella enterica;

Salmonella enterica subsp. enterica MNNEETFYQAMRRKGVTRRSFLKFCSLAATSLGLGAGMTPKIAWALENKPRIPVVWIHGL WP_001058307.1 hydrogenase [Salmonella enterica] Length: 372\nNumber of predicted TMHs: 2\nExp number of AAs in TMHs: 36.51738\nExp number, first 60 AAs: 19.72849\nTotal prob of N-in: 0.96170\nPOSSIBLE N-term signal sequence\ninside 1 19\nTMhelix 20 39\noutside 40 325\nTMhelix 326 348\ninside 349 372

16829 GCF_002041615.1_ASM204161v1 Salmonella enterica subsp. enterica serovar Enteritidis Proteobacteria; Gammaproteobacteria; Enterobacterales; Enterobacteriaceae; Salmonella; Salmonella enterica; Salmonella enterica subsp. enterica MNNEETFYQAMRRKGVTRRSFLKFCSLAATSLGLGAGMTPKIAWALENKPRIPVVWIHGL WP_001058307.1 hydrogenase [Salmonella enterica] Length: 372\nNumber of predicted TMHs: 2\nExp number of AAs in TMHs: 36.51738\nExp number, first 60 AAs: 19.72849\nTotal prob of N-in: 0.96170\nPOSSIBLE N-term signal sequence\ninside 1 19\nTMhelix 20 39\noutside 40 325\nTMhelix 326 348\ninside 349 372

16830 GCF_002039805.1_ASM203980v1 Salmonella enterica subsp. enterica serovar Enteritidis Proteobacteria; Gammaproteobacteria; Enterobacterales; Enterobacteriaceae; Salmonella; Salmonella enterica; Salmonella enterica subsp. enterica MNNEETFYQAMRRKGVTRRSFLKFCSLAATSLGLGAGMTPKIAWALENKPRIPVVWIHGL WP_001058307.1 hydrogenase [Salmonella enterica] Length: 372\nNumber of predicted TMHs: 2\nExp number of AAs in TMHs: 36.51738\nExp number, first 60 AAs: 19.72849\nTotal prob of N-in: 0.96170\nPOSSIBLE N-term signal sequence\ninside 1 19\nTMhelix 20 39\noutside 40 325\nTMhelix 326 348\ninside 349 372

16831 GCF_002041885.1_ASM204188v1 Salmonella enterica subsp. enterica serovar Enteritidis Proteobacteria; Gammaproteobacteria; Enterobacterales; Enterobacteriaceae; Salmonella; Salmonella enterica; Salmonella enterica subsp. enterica MTGDNTLITSHGINRRDFMKLCAALAATMGLSSKAAAEMAESVSNPQRPPVIWIGAQECT WP_000145429.1 hydrogenase 2 small subunit [Salmonella enterica] Length: 372\nNumber of predicted TMHs: 1\nExp number of AAs in TMHs: 24.51309\nExp number, first 60 AAs: 0.2682\nTotal prob of N-in: 0.14402\noutside 1 331\nTMhelix 332 354\ninside 355 372

16832 GCF_002041885.1_ASM204188v1 Salmonella enterica subsp. enterica serovar Enteritidis Proteobacteria; Gammaproteobacteria; Enterobacterales; Enterobacteriaceae; Salmonella; Salmonella enterica; Salmonella enterica subsp. enterica MNNEETFYQAMRRKGVTRRSFLKFCSLAATSLGLGAGMTPKIAWALENKPRIPVVWIHGL WP_001058307.1 hydrogenase [Salmonella enterica] Length: 372\nNumber of predicted TMHs: 2\nExp number of AAs in TMHs: 36.51738\nExp number, first 60 AAs: 19.72849\nTotal prob of N-in: 0.96170\nPOSSIBLE N-term signal sequence\ninside 1 19\nTMhelix 20 39\noutside 40 325\nTMhelix 326 348\ninside 349 372

16833 GCF_001973705.1_ASM197370v1 Salmonella enterica subsp. enterica serovar Enteritidis Proteobacteria; Gammaproteobacteria; Enterobacterales; Enterobacteriaceae; Salmonella; Salmonella enterica; Salmonella enterica subsp. enterica MTGDNTLITSHGINRRDFMKLCAALAATMGLSSKAAAEMAESVSNPQRPPVIWIGAQECT WP_000145429.1 hydrogenase 2 small subunit [Salmonella enterica] Length: 372\nNumber of predicted TMHs: 1\nExp number of AAs in TMHs: 24.51309\nExp number, first 60 AAs: 0.2682\nTotal prob of N-in: 0.14402\noutside 1 331\nTMhelix 332 354\ninside 355 372

16834 GCF_001973745.1_ASM197374v1 Salmonella enterica subsp. enterica serovar Enteritidis Proteobacteria; Gammaproteobacteria; Enterobacterales; Enterobacteriaceae; Salmonella; Salmonella enterica; Salmonella enterica subsp. enterica MNNEETFYQAMRRKGVTRRSFLKFCSLAATSLGLGAGMTPKIAWALENKPRIPVVWIHGL WP_001058307.1 hydrogenase [Salmonella enterica] Length: 372\nNumber of predicted TMHs: 2\nExp number of AAs in TMHs: 36.51738\nExp number, first 60 AAs: 19.72849\nTotal prob of N-in: 0.96170\nPOSSIBLE N-term signal sequence\ninside 1 19\nTMhelix 20 39\noutside 40 325\nTMhelix 326 348\ninside 349 372

16835 GCF_002039915.1_ASM203991v1 Salmonella enterica subsp. enterica serovar Enteritidis Proteobacteria; Gammaproteobacteria; Enterobacterales; Enterobacteriaceae; Salmonella; Salmonella enterica; Salmonella enterica subsp. enterica MNNEETFYQAMRRKGVTRRSFLKFCSLAATSLGLGAGMTPKIAWALENKPRIPVVWIHGL WP_001058307.1 hydrogenase [Salmonella enterica] Length: 372\nNumber of predicted TMHs: 2\nExp number of AAs in TMHs: 36.51738\nExp number, first 60 AAs: 19.72849\nTotal prob of N-in: 0.96170\nPOSSIBLE N-term signal sequence\ninside 1 19\nTMhelix 20 39\noutside 40 325\nTMhelix 326 348\ninside 349 372

16836 GCF_002041695.1_ASM204169v1 Salmonella enterica subsp. enterica serovar Enteritidis Proteobacteria; Gammaproteobacteria; Enterobacterales; Enterobacteriaceae; Salmonella; Salmonella enterica; Salmonella enterica subsp. enterica MNNEETFYQAMRRKGVTRRSFLKFCSLAATSLGLGAGMTPKIAWALENKPRIPVVWIHGL WP_001058307.1 hydrogenase [Salmonella enterica] Length: 372\nNumber of predicted TMHs: 2\nExp number of AAs in TMHs: 36.51738\nExp number, first 60 AAs: 19.72849\nTotal prob of N-in: 0.96170\nPOSSIBLE N-term signal sequence\ninside 1 19\nTMhelix 20 39\noutside 40 325\nTMhelix 326 348\ninside 349 372

16837 GCF_002034445.1_ASM203444v1 Salmonella enterica subsp. enterica serovar Enteritidis Proteobacteria; Gammaproteobacteria; Enterobacterales; Enterobacteriaceae; Salmonella; Salmonella enterica; Salmonella enterica subsp. enterica MTGDNTLITSHGINRRDFMKLCAALAATMGLSSKAAAEMAESVSNPQRPPVIWIGAQECT WP_000145429.1 hydrogenase 2 small subunit [Salmonella enterica] Length: 372\nNumber of predicted TMHs: 1\nExp number of AAs in TMHs: 24.51309\nExp number, first 60 AAs: 0.2682\nTotal prob of N-in: 0.14402\noutside 1 331\nTMhelix 332 354\ninside 355 372

16838 GCF_002041925.1_ASM204192v1 Salmonella enterica subsp. enterica serovar Enteritidis Proteobacteria; Gammaproteobacteria; Enterobacterales; Enterobacteriaceae; Salmonella; Salmonella enterica; Salmonella enterica subsp. enterica MTGDNTLITSHGINRRDFMKLCAALAATMGLSSKAAAEMAESVSNPQRPPVIWIGAQECT WP_000145429.1 hydrogenase 2 small subunit [Salmonella enterica] Length: 372\nNumber of predicted TMHs: 1\nExp number of AAs in TMHs: 24.51309\nExp number, first 60 AAs: 0.2682\nTotal prob of N-in: 0.14402\noutside 1 331\nTMhelix 332 354\ninside 355 372

16839 GCF_002041925.1_ASM204192v1 Salmonella enterica subsp. enterica serovar Enteritidis Proteobacteria; Gammaproteobacteria; Enterobacterales; Enterobacteriaceae; Salmonella; Salmonella enterica; Salmonella enterica subsp. enterica MNNEETFYQAMRRKGVTRRSFLKFCSLAATSLGLGAGMTPKIAWALENKPRIPVVWIHGL WP_001058307.1 hydrogenase [Salmonella enterica] Length: 372\nNumber of predicted TMHs: 2\nExp number of AAs in TMHs: 36.51738\nExp number, first 60 AAs: 19.72849\nTotal prob of N-in: 0.96170\nPOSSIBLE N-term signal sequence\ninside 1 19\nTMhelix 20 39\noutside 40 325\nTMhelix 326 348\ninside 349 372

16840 GCF_002042175.1_ASM204217v1 Salmonella enterica subsp. enterica serovar Enteritidis Proteobacteria; Gammaproteobacteria; Enterobacterales; Enterobacteriaceae; Salmonella; Salmonella enterica; Salmonella enterica subsp. enterica MTGDNTLITSHGINRRDFMKLCAALAATMGLSSKAAAEMAESVSNPQRPPVIWIGAQECT WP_000145429.1 hydrogenase 2 small subunit [Salmonella enterica] Length: 372\nNumber of predicted TMHs: 1\nExp number of AAs in TMHs: 24.51309\nExp number, first 60 AAs: 0.2682\nTotal prob of N-in: 0.14402\noutside 1 331\nTMhelix 332 354\ninside 355 372

16841 GCF_001973745.1_ASM197374v1 Salmonella enterica subsp. enterica serovar Enteritidis Proteobacteria; Gammaproteobacteria; Enterobacterales; Enterobacteriaceae; Salmonella; Salmonella enterica; Salmonella enterica subsp. enterica MTGDNTLITSHGINRRDFMKLCAALAATMGLSSKAAAEMAESVSNPQRPPVIWIGAQECT WP_000145429.1 hydrogenase 2 small subunit [Salmonella enterica] Length: 372\nNumber of predicted TMHs: 1\nExp number of AAs in TMHs: 24.51309\nExp number, first 60 AAs: 0.2682\nTotal prob of N-in: 0.14402\noutside 1 331\nTMhelix 332 354\ninside 355 372

16842 GCF_002041945.1_ASM204194v1 Salmonella enterica subsp. enterica serovar Enteritidis Proteobacteria; Gammaproteobacteria; Enterobacterales; Enterobacteriaceae; Salmonella; Salmonella enterica; Salmonella enterica subsp. enterica MNNEETFYQAMRRKGVTRRSFLKFCSLAATSLGLGAGMTPKIAWALENKPRIPVVWIHGL WP_001058307.1 hydrogenase [Salmonella enterica] Length: 372\nNumber of predicted TMHs: 2\nExp number of AAs in TMHs: 36.51738\nExp number, first 60 AAs: 19.72849\nTotal prob of N-in: 0.96170\nPOSSIBLE N-term signal sequence\ninside 1 19\nTMhelix 20 39\noutside 40 325\nTMhelix 326 348\ninside 349 372

16843 GCF_001973785.1_ASM197378v1 Salmonella enterica subsp. enterica serovar Enteritidis Proteobacteria; Gammaproteobacteria; Enterobacterales; Enterobacteriaceae; Salmonella; Salmonella enterica; Salmonella enterica subsp. enterica MTGDNTLITSHGINRRDFMKLCAALAATMGLSSKAAAEMAESVSNPQRPPVIWIGAQECT WP_000145429.1 hydrogenase 2 small subunit [Salmonella enterica] Length: 372\nNumber of predicted TMHs: 1\nExp number of AAs in TMHs: 24.51309\nExp number, first 60 AAs: 0.2682\nTotal prob of N-in: 0.14402\noutside 1 331\nTMhelix 332 354\ninside 355 372

16844 GCF_002042005.1_ASM204200v1 Salmonella enterica subsp. enterica serovar Enteritidis Proteobacteria; Gammaproteobacteria; Enterobacterales; Enterobacteriaceae; Salmonella; Salmonella enterica; Salmonella enterica subsp. enterica MNNEETFYQAMRRKGVTRRSFLKFCSLAATSLGLGAGMTPKIAWALENKPRIPVVWIHGL WP_001058307.1 hydrogenase [Salmonella enterica] Length: 372\nNumber of predicted TMHs: 2\nExp number of AAs in TMHs: 36.51738\nExp number, first 60 AAs: 19.72849\nTotal prob of N-in: 0.96170\nPOSSIBLE N-term signal sequence\ninside 1 19\nTMhelix 20 39\noutside 40 325\nTMhelix 326 348\ninside 349 372

16845 GCF_002042115.1_ASM204211v1 Salmonella enterica subsp. enterica serovar Enteritidis Proteobacteria; Gammaproteobacteria; Enterobacterales; Enterobacteriaceae; Salmonella; Salmonella enterica; Salmonella enterica subsp. enterica MTGDNTLITSHGINRRDFMKLCAALAATMGLSSKAAAEMAESVSNPQRPPVIWIGAQECT WP_000145429.1 hydrogenase 2 small subunit [Salmonella enterica] Length: 372\nNumber of

predicted TMHs:  1\nExp number of AAs in TMHs: 24.51309\nExp number, first 60 AAs:  0.2682\nTotal prob of N-in:  0.14402\noutside   1   331\nTMhelix   332   354\ninside   355   372

16846 GCF_002040705.1_ASM204070v1 Salmonella enterica subsp. enterica serovar Enteritidis Proteobacteria; Gammaproteobacteria; Enterobacterales; Enterobacteriaceae; Salmonella; Salmonella enterica; Salmonella enterica subsp. enterica MNNEETFYQAMRRKGVTRRSFLKFCSLAATSLGLGAGMTPKIAWALENKPRIPVVWIHGL WP_001058307.1 hydrogenase [Salmonella enterica] Length: 372\nNumber of predicted TMHs:  2\nExp number of AAs in TMHs: 36.51738\nExp number, first 60 AAs:  19.72849\nTotal prob of N-in:   0.96170\nPOSSIBLE N-term signal sequence\ninside   1   19\nTMhelix   20   39\noutside   40   325\nTMhelix   326   348\ninside   349   372

16847 GCF_002042115.1_ASM204211v1 Salmonella enterica subsp. enterica serovar Enteritidis Proteobacteria; Gammaproteobacteria; Enterobacterales; Enterobacteriaceae; Salmonella; Salmonella enterica; Salmonella enterica subsp. enterica MNNEETFYQAMRRKGVTRRSFLKFCSLAATSLGLGAGMTPKIAWALENKPRIPVVWIHGL WP_001058307.1 hydrogenase [Salmonella enterica] Length: 372\nNumber of predicted TMHs:  2\nExp number of AAs in TMHs: 36.51738\nExp number, first 60 AAs:  19.72849\nTotal prob of N-in:   0.96170\nPOSSIBLE N-term signal sequence\ninside   1   19\nTMhelix   20   39\noutside   40   325\nTMhelix   326   348\ninside   349   372

16848 GCF_002042025.1_ASM204202v1 Salmonella enterica subsp. enterica serovar Enteritidis Proteobacteria; Gammaproteobacteria; Enterobacterales; Enterobacteriaceae; Salmonella; Salmonella enterica; Salmonella enterica subsp. enterica MNNEETFYQAMRRKGVTRRSFLKFCSLAATSLGLGAGMTPKIAWALENKPRIPVVWIHGL WP_001058307.1 hydrogenase [Salmonella enterica] Length: 372\nNumber of predicted TMHs:  2\nExp number of AAs in TMHs: 36.51738\nExp number, first 60 AAs:  19.72849\nTotal prob of N-in:   0.96170\nPOSSIBLE N-term signal sequence\ninside   1   19\nTMhelix   20   39\noutside   40   325\nTMhelix   326   348\ninside   349   372

16849 GCF_002040765.1_ASM204076v1 Salmonella enterica subsp. enterica serovar Enteritidis Proteobacteria; Gammaproteobacteria; Enterobacterales; Enterobacteriaceae; Salmonella; Salmonella enterica; Salmonella enterica subsp. enterica MNNEETFYQAMRRKGVTRRSFLKFCSLAATSLGLGAGMTPKIAWALENKPRIPVVWIHGL WP_001058307.1 hydrogenase [Salmonella enterica] Length: 372\nNumber of predicted TMHs:  2\nExp number of AAs in TMHs: 36.51738\nExp number, first 60 AAs:  19.72849\nTotal prob of N-in:   0.96170\nPOSSIBLE N-term signal sequence\ninside   1   19\nTMhelix   20   39\noutside   40   325\nTMhelix   326   348\ninside   349   372

16850 GCF_002042075.1_ASM204207v1 Salmonella enterica subsp. enterica serovar Enteritidis Proteobacteria; Gammaproteobacteria; Enterobacterales; Enterobacteriaceae; Salmonella; Salmonella enterica; Salmonella enterica subsp. enterica MNNEETFYQAMRRKGVTRRSFLKFCSLAATSLGLGAGMTPKIAWALENKPRIPVVWIHGL WP_001058307.1 hydrogenase [Salmonella enterica] Length: 372\nNumber of predicted TMHs:  2\nExp number of AAs in TMHs: 36.51738\nExp number, first 60 AAs:  19.72849\nTotal prob of N-in:   0.96170\nPOSSIBLE N-term signal sequence\ninside   1   19\nTMhelix   20   39\noutside   40   325\nTMhelix   326   348\ninside   349   372

16851 GCF_001597875.1_ASM159787v1 Salmonella enterica subsp. enterica serovar Enteritidis Proteobacteria; Gammaproteobacteria; Enterobacterales; Enterobacteriaceae; Salmonella; Salmonella enterica; Salmonella enterica subsp. enterica MNNEETFYQAMRRKGVTRRSFLKFCSLAATSLGLGAGMTPKIAWALENKPRIPVVWIHGL WP_001058307.1 hydrogenase [Salmonella enterica] Length: 372\nNumber of predicted TMHs:  2\nExp number of AAs in TMHs: 36.51738\nExp number, first 60 AAs:  19.72849\nTotal prob of N-in:   0.96170\nPOSSIBLE N-term signal sequence\ninside   1   19\nTMhelix   20   39\noutside   40   325\nTMhelix   326   348\ninside   349   372

16852 GCF_002040805.1_ASM204080v1 Salmonella enterica subsp. enterica serovar Enteritidis Proteobacteria; Gammaproteobacteria; Enterobacterales; Enterobacteriaceae; Salmonella; Salmonella enterica; Salmonella enterica subsp. enterica MNNEETFYQAMRRKGVTRRSFLKFCSLAATSLGLGAGMTPKIAWALENKPRIPVVWIHGL WP_001058307.1 hydrogenase [Salmonella enterica] Length: 372\nNumber of predicted TMHs:  2\nExp number of AAs in TMHs: 36.51738\nExp number, first 60 AAs:  19.72849\nTotal prob of N-in:   0.96170\nPOSSIBLE N-term signal sequence\ninside   1   19\nTMhelix   20   39\noutside   40   325\nTMhelix   326   348\ninside   349   372

16853 GCF_001597875.1_ASM159787v1 Salmonella enterica subsp. enterica serovar Enteritidis Proteobacteria; Gammaproteobacteria; Enterobacterales; Enterobacteriaceae; Salmonella; Salmonella enterica; Salmonella enterica subsp. enterica MTGDNTLITSHGINRRDFMKLCAALAATMGLSSKAAAEMAESVSNPQRPPVIWIGAQECT WP_000145429.1 hydrogenase 2 small subunit [Salmonella enterica] Length: 372\nNumber of predicted TMHs:  1\nExp number of AAs in TMHs: 24.51309\nExp number, first 60 AAs:  0.2682\nTotal prob of N-in:  0.14402\noutside   1   331\nTMhelix   332   354\ninside   355   372

16854 GCF_002040765.1_ASM204076v1 Salmonella enterica subsp. enterica serovar Enteritidis Proteobacteria; Gammaproteobacteria; Enterobacterales; Enterobacteriaceae; Salmonella; Salmonella enterica;

Salmonella enterica subsp. enterica MTGDNTLITSHGINRRDFMKLCAALAATMGLSSKAAAEMAESVSNPQRPPVIWIGAQECT WP_000145429.1 hydrogenase 2 small subunit [Salmonella enterica] Length: 372\nNumber of predicted TMHs: 1\nExp number of AAs in TMHs: 24.51309\nExp number, first 60 AAs: 0.2682\nTotal prob of N-in: 0.14402\noutside 1 331\nTMhelix 332 354\ninside 355 372

16855 GCF_002040805.1_ASM204080v1 Salmonella enterica subsp. enterica serovar Enteritidis Proteobacteria; Gammaproteobacteria; Enterobacterales; Enterobacteriaceae; Salmonella; Salmonella enterica; Salmonella enterica subsp. enterica MTGDNTLITSHGINRRDFMKLCAALAATMGLSSKAAAEMAESVSNPQRPPVIWIGAQECT WP_000145429.1 hydrogenase 2 small subunit [Salmonella enterica] Length: 372\nNumber of predicted TMHs: 1\nExp number of AAs in TMHs: 24.51309\nExp number, first 60 AAs: 0.2682\nTotal prob of N-in: 0.14402\noutside 1 331\nTMhelix 332 354\ninside 355 372

16856 GCF_002034445.1_ASM203444v1 Salmonella enterica subsp. enterica serovar Enteritidis Proteobacteria; Gammaproteobacteria; Enterobacterales; Enterobacteriaceae; Salmonella; Salmonella enterica; Salmonella enterica subsp. enterica MNNEETFYQAMRRKGVTRRSFLKFCSLAATSLGLGAGMTPKIAWALENKPRIPVVWIHGL WP_001058307.1 hydrogenase [Salmonella enterica] Length: 372\nNumber of predicted TMHs: 2\nExp number of AAs in TMHs: 36.51738\nExp number, first 60 AAs: 19.72849\nTotal prob of N-in: 0.96170\nPOSSIBLE N-term signal sequence\ninside 1 19\nTMhelix 20 39\noutside 40 325\nTMhelix 326 348\ninside 349 372

16857 GCF_002042845.1_ASM204284v1 Salmonella enterica subsp. enterica serovar Enteritidis Proteobacteria; Gammaproteobacteria; Enterobacterales; Enterobacteriaceae; Salmonella; Salmonella enterica; Salmonella enterica subsp. enterica MTGDNTLITSHGINRRDFMKLCAALAATMGLSSKAAAEMAESVSNPQRPPVIWIGAQECT WP_000145429.1 hydrogenase 2 small subunit [Salmonella enterica] Length: 372\nNumber of predicted TMHs: 1\nExp number of AAs in TMHs: 24.51309\nExp number, first 60 AAs: 0.2682\nTotal prob of N-in: 0.14402\noutside 1 331\nTMhelix 332 354\ninside 355 372

16858 GCF_002042535.1_ASM204253v1 Salmonella enterica subsp. enterica serovar Enteritidis Proteobacteria; Gammaproteobacteria; Enterobacterales; Enterobacteriaceae; Salmonella; Salmonella enterica; Salmonella enterica subsp. enterica MNNEETFYQAMRRKGVTRRSFLKFCSLAATSLGLGAGMTPKIAWALENKPRIPVVWIHGL WP_001058307.1 hydrogenase [Salmonella enterica] Length: 372\nNumber of predicted TMHs: 2\nExp number of AAs in TMHs: 36.51738\nExp number, first 60 AAs: 19.72849\nTotal prob of N-in: 0.96170\nPOSSIBLE N-term signal sequence\ninside 1 19\nTMhelix 20 39\noutside 40 325\nTMhelix 326 348\ninside 349 372

16859 GCF_002040495.1_ASM204049v1 Salmonella enterica subsp. enterica serovar Enteritidis Proteobacteria; Gammaproteobacteria; Enterobacterales; Enterobacteriaceae; Salmonella; Salmonella enterica; Salmonella enterica subsp. enterica MTGDNTLITSHGINRRDFMKLCAALAATMGLSSKAAAEMAESVSNPQRPPVIWIGAQECT WP_000145429.1 hydrogenase 2 small subunit [Salmonella enterica] Length: 372\nNumber of predicted TMHs: 1\nExp number of AAs in TMHs: 24.51309\nExp number, first 60 AAs: 0.2682\nTotal prob of N-in: 0.14402\noutside 1 331\nTMhelix 332 354\ninside 355 372

16860 GCF_002042685.1_ASM204268v1 Salmonella enterica subsp. enterica serovar Enteritidis Proteobacteria; Gammaproteobacteria; Enterobacterales; Enterobacteriaceae; Salmonella; Salmonella enterica; Salmonella enterica subsp. enterica MNNEETFYQAMRRKGVTRRSFLKFCSLAATSLGLGAGMTPKIAWALENKPRIPVVWIHGL WP_001058307.1 hydrogenase [Salmonella enterica] Length: 372\nNumber of predicted TMHs: 2\nExp number of AAs in TMHs: 36.51738\nExp number, first 60 AAs: 19.72849\nTotal prob of N-in: 0.96170\nPOSSIBLE N-term signal sequence\ninside 1 19\nTMhelix 20 39\noutside 40 325\nTMhelix 326 348\ninside 349 372

16861 GCF_002042665.1_ASM204266v1 Salmonella enterica subsp. enterica serovar Enteritidis Proteobacteria; Gammaproteobacteria; Enterobacterales; Enterobacteriaceae; Salmonella; Salmonella enterica; Salmonella enterica subsp. enterica MNNEETFYQAMRRKGVTRRSFLKFCSLAATSLGLGAGMTPKIAWALENKPRIPVVWIHGL WP_001058307.1 hydrogenase [Salmonella enterica] Length: 372\nNumber of predicted TMHs: 2\nExp number of AAs in TMHs: 36.51738\nExp number, first 60 AAs: 19.72849\nTotal prob of N-in: 0.96170\nPOSSIBLE N-term signal sequence\ninside 1 19\nTMhelix 20 39\noutside 40 325\nTMhelix 326 348\ninside 349 372

16862 GCF_002042625.1_ASM204262v1 Salmonella enterica subsp. enterica serovar Enteritidis Proteobacteria; Gammaproteobacteria; Enterobacterales; Enterobacteriaceae; Salmonella; Salmonella enterica; Salmonella enterica subsp. enterica MNNEETFYQAMRRKGVTRRSFLKFCSLAATSLGLGAGMTPKIAWALENKPRIPVVWIHGL WP_001058307.1 hydrogenase [Salmonella enterica] Length: 372\nNumber of predicted TMHs: 2\nExp number of AAs in TMHs: 36.51738\nExp number, first 60 AAs: 19.72849\nTotal prob of N-in: 0.96170\nPOSSIBLE N-term signal sequence\ninside 1 19\nTMhelix 20 39\noutside 40 325\nTMhelix 326 348\ninside 349 372

16863 GCF_002042665.1_ASM204266v1 Salmonella enterica subsp. enterica serovar Enteritidis Proteobacteria; Gammaproteobacteria; Enterobacterales; Enterobacteriaceae; Salmonella; Salmonella enterica; Salmonella enterica subsp. enterica MTGDNTLITSHGINRRDFMKLCAALAATMGLSSKAAAEMAESVSNPQRPPVIWIGAQECT WP_000145429.1 hydrogenase 2 small subunit [Salmonella enterica] Length: 372\nNumber of predicted TMHs: 1\nExp number of AAs in TMHs: 24.51309\nExp number, first 60 AAs: 0.2682\nTotal prob of N-in: 0.14402\noutside 1 331\nTMhelix 332 354\ninside 355 372

16864 GCF_002042685.1_ASM204268v1 Salmonella enterica subsp. enterica serovar Enteritidis Proteobacteria; Gammaproteobacteria; Enterobacterales; Enterobacteriaceae; Salmonella; Salmonella enterica; Salmonella enterica subsp. enterica MTGDNTLITSHGINRRDFMKLCAALAATMGLSSKAAAEMAESVSNPQRPPVIWIGAQECT WP_000145429.1 hydrogenase 2 small subunit [Salmonella enterica] Length: 372\nNumber of predicted TMHs: 1\nExp number of AAs in TMHs: 24.51309\nExp number, first 60 AAs: 0.2682\nTotal prob of N-in: 0.14402\noutside 1 331\nTMhelix 332 354\ninside 355 372

16865 GCF_002042625.1_ASM204262v1 Salmonella enterica subsp. enterica serovar Enteritidis Proteobacteria; Gammaproteobacteria; Enterobacterales; Enterobacteriaceae; Salmonella; Salmonella enterica; Salmonella enterica subsp. enterica MTGDNTLITSHGINRRDFMKLCAALAATMGLSSKAAAEMAESVSNPQRPPVIWIGAQECT WP_000145429.1 hydrogenase 2 small subunit [Salmonella enterica] Length: 372\nNumber of predicted TMHs: 1\nExp number of AAs in TMHs: 24.51309\nExp number, first 60 AAs: 0.2682\nTotal prob of N-in: 0.14402\noutside 1 331\nTMhelix 332 354\ninside 355 372

16866 GCF_002040285.1_ASM204028v1 Salmonella enterica subsp. enterica serovar Enteritidis Proteobacteria; Gammaproteobacteria; Enterobacterales; Enterobacteriaceae; Salmonella; Salmonella enterica; Salmonella enterica subsp. enterica MNNEETFYQAMRRKGVTRRSFLKFCSLAATSLGLGAGMTPKIAWALENKPRIPVVWIHGL WP_001058307.1 hydrogenase [Salmonella enterica] Length: 372\nNumber of predicted TMHs: 2\nExp number of AAs in TMHs: 36.51738\nExp number, first 60 AAs: 19.72849\nTotal prob of N-in: 0.96170\nPOSSIBLE N-term signal sequence\ninside 1 19\nTMhelix 20 39\noutside 40 325\nTMhelix 326 348\ninside 349 372

16867 GCF_002040315.1_ASM204031v1 Salmonella enterica subsp. enterica serovar Enteritidis Proteobacteria; Gammaproteobacteria; Enterobacterales; Enterobacteriaceae; Salmonella; Salmonella enterica; Salmonella enterica subsp. enterica MNNEETFYQAMRRKGVTRRSFLKFCSLAATSLGLGAGMTPKIAWALENKPRIPVVWIHGL WP_001058307.1 hydrogenase [Salmonella enterica] Length: 372\nNumber of predicted TMHs: 2\nExp number of AAs in TMHs: 36.51738\nExp number, first 60 AAs: 19.72849\nTotal prob of N-in: 0.96170\nPOSSIBLE N-term signal sequence\ninside 1 19\nTMhelix 20 39\noutside 40 325\nTMhelix 326 348\ninside 349 372

16868 GCF_002040445.1_ASM204044v1 Salmonella enterica subsp. enterica serovar Enteritidis Proteobacteria; Gammaproteobacteria; Enterobacterales; Enterobacteriaceae; Salmonella; Salmonella enterica; Salmonella enterica subsp. enterica MNNEETFYQAMRRKGVTRRSFLKFCSLAATSLGLGAGMTPKIAWALENKPRIPVVWIHGL WP_001058307.1 hydrogenase [Salmonella enterica] Length: 372\nNumber of predicted TMHs: 2\nExp number of AAs in TMHs: 36.51738\nExp number, first 60 AAs: 19.72849\nTotal prob of N-in: 0.96170\nPOSSIBLE N-term signal sequence\ninside 1 19\nTMhelix 20 39\noutside 40 325\nTMhelix 326 348\ninside 349 372

16869 GCF_002030735.1_ASM203073v1 Salmonella enterica subsp. enterica serovar Enteritidis Proteobacteria; Gammaproteobacteria; Enterobacterales; Enterobacteriaceae; Salmonella; Salmonella enterica; Salmonella enterica subsp. enterica MTGDNTLITSHGINRRDFMKLCAALAATMGLSSKAAAEMAESVSNPQRPPVIWIGAQECT WP_000145429.1 hydrogenase 2 small subunit [Salmonella enterica] Length: 372\nNumber of predicted TMHs: 1\nExp number of AAs in TMHs: 24.51309\nExp number, first 60 AAs: 0.2682\nTotal prob of N-in: 0.14402\noutside 1 331\nTMhelix 332 354\ninside 355 372

16870 GCF_002042405.1_ASM204240v1 Salmonella enterica subsp. enterica serovar Enteritidis Proteobacteria; Gammaproteobacteria; Enterobacterales; Enterobacteriaceae; Salmonella; Salmonella enterica; Salmonella enterica subsp. enterica MNNEETFYQAMRRKGVTRRSFLKFCSLAATSLGLGAGMTPKIAWALENKPRIPVVWIHGL WP_001058307.1 hydrogenase [Salmonella enterica] Length: 372\nNumber of predicted TMHs: 2\nExp number of AAs in TMHs: 36.51738\nExp number, first 60 AAs: 19.72849\nTotal prob of N-in: 0.96170\nPOSSIBLE N-term signal sequence\ninside 1 19\nTMhelix 20 39\noutside 40 325\nTMhelix 326 348\ninside 349 372

16871 GCF_002042405.1_ASM204240v1 Salmonella enterica subsp. enterica serovar Enteritidis Proteobacteria; Gammaproteobacteria; Enterobacterales; Enterobacteriaceae; Salmonella; Salmonella enterica; Salmonella enterica subsp. enterica MTGDNTLITSHGINRRDFMKLCAALAATMGLSSKAAAEMAESVSNPQRPPVIWIGAQECT WP_000145429.1 hydrogenase 2 small subunit [Salmonella enterica] Length: 372\nNumber of

predicted TMHs: 1\nExp number of AAs in TMHs: 24.51309\nExp number, first 60 AAs: 0.2682\nTotal prob of N-in: 0.14402\noutside 1 331\nTMhelix 332 354\ninside 355 372

16872 GCF_002030735.1_ASM203073v1 Salmonella enterica subsp. enterica serovar Enteritidis Proteobacteria; Gammaproteobacteria; Enterobacterales; Enterobacteriaceae; Salmonella; Salmonella enterica; Salmonella enterica subsp. enterica MNNEETFYQAMRRKGVTRRSFLKFCSLAATSLGLGAGMTPKIAWALENKPRIPVVWIHGL WP_001058307.1 hydrogenase [Salmonella enterica] Length: 372\nNumber of predicted TMHs: 2\nExp number of AAs in TMHs: 36.51738\nExp number, first 60 AAs: 19.72849\nTotal prob of N-in: 0.96170\nPOSSIBLE N-term signal sequence\ninside 1 19\nTMhelix 20 39\noutside 40 325\nTMhelix 326 348\ninside 349 372

16873 GCF_002042855.1_ASM204285v1 Salmonella enterica subsp. enterica serovar Enteritidis Proteobacteria; Gammaproteobacteria; Enterobacterales; Enterobacteriaceae; Salmonella; Salmonella enterica; Salmonella enterica subsp. enterica MTGDNTLITSHGINRRDFMKLCAALAATMGLSSKAAAEMAESVSNPQRPPVIWIGAQECT WP_000145429.1 hydrogenase 2 small subunit [Salmonella enterica] Length: 372\nNumber of predicted TMHs: 1\nExp number of AAs in TMHs: 24.51309\nExp number, first 60 AAs: 0.2682\nTotal prob of N-in: 0.14402\noutside 1 331\nTMhelix 332 354\ninside 355 372

16874 GCF_002042235.1_ASM204223v1 Salmonella enterica subsp. enterica serovar Enteritidis Proteobacteria; Gammaproteobacteria; Enterobacterales; Enterobacteriaceae; Salmonella; Salmonella enterica; Salmonella enterica subsp. enterica MNNEETFYQAMRRKGVTRRSFLKFCSLAATSLGLGAGMTPKIAWALENKPRIPVVWIHGL WP_001058307.1 hydrogenase [Salmonella enterica] Length: 372\nNumber of predicted TMHs: 2\nExp number of AAs in TMHs: 36.51738\nExp number, first 60 AAs: 19.72849\nTotal prob of N-in: 0.96170\nPOSSIBLE N-term signal sequence\ninside 1 19\nTMhelix 20 39\noutside 40 325\nTMhelix 326 348\ninside 349 372

16875 GCF_002040495.1_ASM204049v1 Salmonella enterica subsp. enterica serovar Enteritidis Proteobacteria; Gammaproteobacteria; Enterobacterales; Enterobacteriaceae; Salmonella; Salmonella enterica; Salmonella enterica subsp. enterica MNNEETFYQAMRRKGVTRRSFLKFCSLAATSLGLGAGMTPKIAWALENKPRIPVVWIHGL WP_001058307.1 hydrogenase [Salmonella enterica] Length: 372\nNumber of predicted TMHs: 2\nExp number of AAs in TMHs: 36.51738\nExp number, first 60 AAs: 19.72849\nTotal prob of N-in: 0.96170\nPOSSIBLE N-term signal sequence\ninside 1 19\nTMhelix 20 39\noutside 40 325\nTMhelix 326 348\ninside 349 372

16876 GCF_002042185.1_ASM204218v1 Salmonella enterica subsp. enterica serovar Enteritidis Proteobacteria; Gammaproteobacteria; Enterobacterales; Enterobacteriaceae; Salmonella; Salmonella enterica; Salmonella enterica subsp. enterica MNNEETFYQAMRRKGVTRRSFLKFCSLAATSLGLGAGMTPKIAWALENKPRIPVVWIHGL WP_001058307.1 hydrogenase [Salmonella enterica] Length: 372\nNumber of predicted TMHs: 2\nExp number of AAs in TMHs: 36.51738\nExp number, first 60 AAs: 19.72849\nTotal prob of N-in: 0.96170\nPOSSIBLE N-term signal sequence\ninside 1 19\nTMhelix 20 39\noutside 40 325\nTMhelix 326 348\ninside 349 372

16877 GCF_002040445.1_ASM204044v1 Salmonella enterica subsp. enterica serovar Enteritidis Proteobacteria; Gammaproteobacteria; Enterobacterales; Enterobacteriaceae; Salmonella; Salmonella enterica; Salmonella enterica subsp. enterica MTGDNTLITSHGINRRDFMKLCAALAATMGLSSKAAAEMAESVSNPQRPPVIWIGAQECT WP_000145429.1 hydrogenase 2 small subunit [Salmonella enterica] Length: 372\nNumber of predicted TMHs: 1\nExp number of AAs in TMHs: 24.51309\nExp number, first 60 AAs: 0.2682\nTotal prob of N-in: 0.14402\noutside 1 331\nTMhelix 332 354\ninside 355 372

16878 GCF_002042375.1_ASM204237v1 Salmonella enterica subsp. enterica serovar Enteritidis Proteobacteria; Gammaproteobacteria; Enterobacterales; Enterobacteriaceae; Salmonella; Salmonella enterica; Salmonella enterica subsp. enterica MNNEETFYQAMRRKGVTRRSFLKFCSLAATSLGLGAGMTPKIAWALENKPRIPVVWIHGL WP_001058307.1 hydrogenase [Salmonella enterica] Length: 372\nNumber of predicted TMHs: 2\nExp number of AAs in TMHs: 36.51738\nExp number, first 60 AAs: 19.72849\nTotal prob of N-in: 0.96170\nPOSSIBLE N-term signal sequence\ninside 1 19\nTMhelix 20 39\noutside 40 325\nTMhelix 326 348\ninside 349 372

16879 GCF_002041425.1_ASM204142v1 Salmonella enterica subsp. enterica serovar Enteritidis Proteobacteria; Gammaproteobacteria; Enterobacterales; Enterobacteriaceae; Salmonella; Salmonella enterica; Salmonella enterica subsp. enterica MNNEETFYQAMRRKGVTRRSFLKFCSLAATSLGLGAGMTPKIAWALENKPRIPVVWIHGL WP_079786335.1 [Ni/Fe] hydrogenase small subunit [Salmonella enterica] Length: 372\nNumber of predicted TMHs: 2\nExp number of AAs in TMHs: 36.51716\nExp number, first 60 AAs: 19.7285\nTotal prob of N-in: 0.96169\nPOSSIBLE N-term signal sequence\ninside 1 19\nTMhelix 20 39\noutside 40 325\nTMhelix 326 348\ninside 349 372

16880 GCF_002041375.1_ASM204137v1 Salmonella enterica subsp. enterica serovar Enteritidis Proteobacteria; Gammaproteobacteria; Enterobacterales; Enterobacteriaceae; Salmonella; Salmonella enterica; Salmonella enterica subsp. enterica MNNEETFYQAMRRKGVTRRSFLKFCSLAATSLGLGAGMTPKIAWALENKPRIPVVWIHGL WP_001058307.1 hydrogenase [Salmonella enterica] Length: 372\nNumber of predicted TMHs: 2\nExp number of AAs in TMHs: 36.51738\nExp number, first 60 AAs: 19.72849\nTotal prob of N-in: 0.96170\nPOSSIBLE N-term signal sequence\ninside 1 19\nTMhelix 20 39\noutside 40 325\nTMhelix 326 348\ninside 349 372

16881 GCF_002041345.1_ASM204134v1 Salmonella enterica subsp. enterica serovar Enteritidis Proteobacteria; Gammaproteobacteria; Enterobacterales; Enterobacteriaceae; Salmonella; Salmonella enterica; Salmonella enterica subsp. enterica MNNEETFYQAMRRKGVTRRSFLKFCSLAATSLGLGAGMTPKIAWALENKPRIPVVWIHGL WP_001058307.1 hydrogenase [Salmonella enterica] Length: 372\nNumber of predicted TMHs: 2\nExp number of AAs in TMHs: 36.51738\nExp number, first 60 AAs: 19.72849\nTotal prob of N-in: 0.96170\nPOSSIBLE N-term signal sequence\ninside 1 19\nTMhelix 20 39\noutside 40 325\nTMhelix 326 348\ninside 349 372

16882 GCF_002041375.1_ASM204137v1 Salmonella enterica subsp. enterica serovar Enteritidis Proteobacteria; Gammaproteobacteria; Enterobacterales; Enterobacteriaceae; Salmonella; Salmonella enterica; Salmonella enterica subsp. enterica MTGDNTLITSHGINRRDFMKLCAALAATMGLSSKAAAEMAESVSNPQRPPVIWIGAQECT WP_000145429.1 hydrogenase 2 small subunit [Salmonella enterica] Length: 372\nNumber of predicted TMHs: 1\nExp number of AAs in TMHs: 24.51309\nExp number, first 60 AAs: 0.2682\nTotal prob of N-in: 0.14402\noutside 1 331\nTMhelix 332 354\ninside 355 372

16883 GCF_002041345.1_ASM204134v1 Salmonella enterica subsp. enterica serovar Enteritidis Proteobacteria; Gammaproteobacteria; Enterobacterales; Enterobacteriaceae; Salmonella; Salmonella enterica; Salmonella enterica subsp. enterica MTGDNTLITSHGINRRDFMKLCAALAATMGLSSKAAAEMAESVSNPQRPPVIWIGAQECT WP_000145429.1 hydrogenase 2 small subunit [Salmonella enterica] Length: 372\nNumber of predicted TMHs: 1\nExp number of AAs in TMHs: 24.51309\nExp number, first 60 AAs: 0.2682\nTotal prob of N-in: 0.14402\noutside 1 331\nTMhelix 332 354\ninside 355 372

16884 GCF_002041425.1_ASM204142v1 Salmonella enterica subsp. enterica serovar Enteritidis Proteobacteria; Gammaproteobacteria; Enterobacterales; Enterobacteriaceae; Salmonella; Salmonella enterica; Salmonella enterica subsp. enterica MTGDNTLITSHGINRRDFMKLCAALAATMGLSSKAAAEMAESVSNPQRPPVIWIGAQECT WP_000145429.1 hydrogenase 2 small subunit [Salmonella enterica] Length: 372\nNumber of predicted TMHs: 1\nExp number of AAs in TMHs: 24.51309\nExp number, first 60 AAs: 0.2682\nTotal prob of N-in: 0.14402\noutside 1 331\nTMhelix 332 354\ninside 355 372

16885 GCF_002041305.1_ASM204130v1 Salmonella enterica subsp. enterica serovar Enteritidis Proteobacteria; Gammaproteobacteria; Enterobacterales; Enterobacteriaceae; Salmonella; Salmonella enterica; Salmonella enterica subsp. enterica MTGDNTLITSHGINRRDFMKLCAALAATMGLSSKAAAEMAESVSNPQRPPVIWIGAQECT WP_000145429.1 hydrogenase 2 small subunit [Salmonella enterica] Length: 372\nNumber of predicted TMHs: 1\nExp number of AAs in TMHs: 24.51309\nExp number, first 60 AAs: 0.2682\nTotal prob of N-in: 0.14402\noutside 1 331\nTMhelix 332 354\ninside 355 372

16886 GCF_002041085.1_ASM204108v1 Salmonella enterica subsp. enterica serovar Enteritidis Proteobacteria; Gammaproteobacteria; Enterobacterales; Enterobacteriaceae; Salmonella; Salmonella enterica; Salmonella enterica subsp. enterica MTGDNTLITSHGINRRDFMKLCAALAATMGLSSKAAAEMAESVSNPQRPPVIWIGAQECT WP_000145429.1 hydrogenase 2 small subunit [Salmonella enterica] Length: 372\nNumber of predicted TMHs: 1\nExp number of AAs in TMHs: 24.51309\nExp number, first 60 AAs: 0.2682\nTotal prob of N-in: 0.14402\noutside 1 331\nTMhelix 332 354\ninside 355 372

16887 GCF_002042845.1_ASM204284v1 Salmonella enterica subsp. enterica serovar Enteritidis Proteobacteria; Gammaproteobacteria; Enterobacterales; Enterobacteriaceae; Salmonella; Salmonella enterica; Salmonella enterica subsp. enterica MNNEETFYQAMRRKGVTRRSFLKFCSLAATSLGLGAGMTPKIAWALENKPRIPVVWIHGL WP_001058307.1 hydrogenase [Salmonella enterica] Length: 372\nNumber of predicted TMHs: 2\nExp number of AAs in TMHs: 36.51738\nExp number, first 60 AAs: 19.72849\nTotal prob of N-in: 0.96170\nPOSSIBLE N-term signal sequence\ninside 1 19\nTMhelix 20 39\noutside 40 325\nTMhelix 326 348\ninside 349 372

16888 GCF_002042855.1_ASM204285v1 Salmonella enterica subsp. enterica serovar Enteritidis Proteobacteria; Gammaproteobacteria; Enterobacterales; Enterobacteriaceae; Salmonella; Salmonella enterica; Salmonella enterica subsp. enterica MNNEETFYQAMRRKGVTRRSFLKFCSLAATSLGLGAGMTPKIAWALENKPRIPVVWIHGL WP_001058307.1 hydrogenase [Salmonella enterica] Length: 372\nNumber of predicted TMHs: 2\nExp

number of AAs in TMHs: 36.51738\nExp number, first 60 AAs:  19.72849\nTotal prob of N-in:   0.96170\nPOSSIBLE N-term signal sequence\ninside   1   19\nTMhelix   20   39\noutside   40   325\nTMhelix   326   348\ninside   349   372

16889 GCF_002041035.1_ASM204103v1 Salmonella enterica subsp. enterica serovar Enteritidis Proteobacteria; Gammaproteobacteria; Enterobacterales; Enterobacteriaceae; Salmonella; Salmonella enterica; Salmonella enterica subsp. enterica MTGDNTLITSHGINRRDFMKLCAALAATMGLSSKAAAEMAESVSNPQRPPVIWIGAQECT WP_000145429.1 hydrogenase 2 small subunit [Salmonella enterica] Length: 372\nNumber of predicted TMHs:  1\nExp number of AAs in TMHs: 24.51309\nExp number, first 60 AAs:  0.2682\nTotal prob of N-in: 0.14402\noutside   1   331\nTMhelix   332   354\ninside   355   372

16890 GCF_002040915.1_ASM204091v1 Salmonella enterica subsp. enterica serovar Enteritidis Proteobacteria; Gammaproteobacteria; Enterobacterales; Enterobacteriaceae; Salmonella; Salmonella enterica; Salmonella enterica subsp. enterica MNNEETFYQAMRRKGVTRRSFLKFCSLAATSLGLGAGMTPKIAWALENKPRIPVVWIHGL WP_001058307.1 hydrogenase [Salmonella enterica] Length: 372\nNumber of predicted TMHs:  2\nExp number of AAs in TMHs: 36.51738\nExp number, first 60 AAs:  19.72849\nTotal prob of N-in:   0.96170\nPOSSIBLE N-term signal sequence\ninside   1   19\nTMhelix   20   39\noutside   40   325\nTMhelix   326   348\ninside   349   372

16891 GCF_002040885.1_ASM204088v1 Salmonella enterica subsp. enterica serovar Enteritidis Proteobacteria; Gammaproteobacteria; Enterobacterales; Enterobacteriaceae; Salmonella; Salmonella enterica; Salmonella enterica subsp. enterica MNNEETFYQAMRRKGVTRRSFLKFCSLAATSLGLGAGMTPKIAWALENKPRIPVVWIHGL WP_001058307.1 hydrogenase [Salmonella enterica] Length: 372\nNumber of predicted TMHs:  2\nExp number of AAs in TMHs: 36.51738\nExp number, first 60 AAs:  19.72849\nTotal prob of N-in:   0.96170\nPOSSIBLE N-term signal sequence\ninside   1   19\nTMhelix   20   39\noutside   40   325\nTMhelix   326   348\ninside   349   372

16892 GCF_002040845.1_ASM204084v1 Salmonella enterica subsp. enterica serovar Enteritidis Proteobacteria; Gammaproteobacteria; Enterobacterales; Enterobacteriaceae; Salmonella; Salmonella enterica; Salmonella enterica subsp. enterica MNNEETFYQAMRRKGVTRRSFLKFCSLAATSLGLGAGMTPKIAWALENKPRIPVVWIHGL WP_001058307.1 hydrogenase [Salmonella enterica] Length: 372\nNumber of predicted TMHs:  2\nExp number of AAs in TMHs: 36.51738\nExp number, first 60 AAs:  19.72849\nTotal prob of N-in:   0.96170\nPOSSIBLE N-term signal sequence\ninside   1   19\nTMhelix   20   39\noutside   40   325\nTMhelix   326   348\ninside   349   372

16893 GCF_002040985.1_ASM204098v1 Salmonella enterica subsp. enterica serovar Enteritidis Proteobacteria; Gammaproteobacteria; Enterobacterales; Enterobacteriaceae; Salmonella; Salmonella enterica; Salmonella enterica subsp. enterica MNNEETFYQAMRRKGVTRRSFLKFCSLAATSLGLGAGMTPKIAWALENKPRIPVVWIHGL WP_001058307.1 hydrogenase [Salmonella enterica] Length: 372\nNumber of predicted TMHs:  2\nExp number of AAs in TMHs: 36.51738\nExp number, first 60 AAs:  19.72849\nTotal prob of N-in:   0.96170\nPOSSIBLE N-term signal sequence\ninside   1   19\nTMhelix   20   39\noutside   40   325\nTMhelix   326   348\ninside   349   372

16894 GCF_002041265.1_ASM204126v1 Salmonella enterica subsp. enterica serovar Enteritidis Proteobacteria; Gammaproteobacteria; Enterobacterales; Enterobacteriaceae; Salmonella; Salmonella enterica; Salmonella enterica subsp. enterica MTGDNTLITSHGINRRDFMKLCAALAATMGLSSKAAAEMAESVSNPQRPPVIWIGAQECT WP_000145429.1 hydrogenase 2 small subunit [Salmonella enterica] Length: 372\nNumber of predicted TMHs:  1\nExp number of AAs in TMHs: 24.51309\nExp number, first 60 AAs:  0.2682\nTotal prob of N-in: 0.14402\noutside   1   331\nTMhelix   332   354\ninside   355   372

16895 GCF_001972555.1_ASM197255v1 Salmonella enterica subsp. enterica serovar Enteritidis Proteobacteria; Gammaproteobacteria; Enterobacterales; Enterobacteriaceae; Salmonella; Salmonella enterica; Salmonella enterica subsp. enterica MTGDNTLITSHGINRRDFMKLCAALAATMGLSSKAAAEMAESVSNPQRPPVIWIGAQECT WP_000145429.1 hydrogenase 2 small subunit [Salmonella enterica] Length: 372\nNumber of predicted TMHs:  1\nExp number of AAs in TMHs: 24.51309\nExp number, first 60 AAs:  0.2682\nTotal prob of N-in: 0.14402\noutside   1   331\nTMhelix   332   354\ninside   355   372

16896 GCF_001972635.1_ASM197263v1 Salmonella enterica subsp. enterica serovar Enteritidis Proteobacteria; Gammaproteobacteria; Enterobacterales; Enterobacteriaceae; Salmonella; Salmonella enterica; Salmonella enterica subsp. enterica MNNEETFYQAMRRKGVTRRSFLKFCSLAATSLGLGAGMTPKIAWALENKPRIPVVWIHGL WP_001058307.1 hydrogenase [Salmonella enterica] Length: 372\nNumber of predicted TMHs:  2\nExp number of AAs in TMHs: 36.51738\nExp number, first 60 AAs:  19.72849\nTotal prob of N-in:   0.96170\nPOSSIBLE N-term signal sequence\ninside   1   19\nTMhelix   20   39\noutside   40   325\nTMhelix   326   348\ninside   349   372

16897 GCF_001972715.1_ASM197271v1 Salmonella enterica subsp. enterica serovar Enteritidis Proteobacteria; Gammaproteobacteria; Enterobacterales; Enterobacteriaceae; Salmonella; Salmonella enterica;

Salmonella enterica subsp. enterica MNNEETFYQAMRRKGVTRRSFLKFCSLAATSLGLGAGMTPKIAWALENKPRIPVVWIHGL WP_001058307.1 hydrogenase [Salmonella enterica] Length: 372\nNumber of predicted TMHs: 2\nExp number of AAs in TMHs: 36.51738\nExp number, first 60 AAs: 19.72849\nTotal prob of N-in: 0.96170\nPOSSIBLE N-term signal sequence\ninside 1 19\nTMhelix 20 39\noutside 40 325\nTMhelix 326 348\ninside 349 372

16898 GCF_001972735.1_ASM197273v1 Salmonella enterica subsp. enterica serovar Enteritidis Proteobacteria; Gammaproteobacteria; Enterobacterales; Enterobacteriaceae; Salmonella; Salmonella enterica; Salmonella enterica subsp. enterica MNNEETFYQAMRRKGVTRRSFLKFCSLAATSLGLGAGMTPKIAWALENKPRIPVVWIHGL WP_001058307.1 hydrogenase [Salmonella enterica] Length: 372\nNumber of predicted TMHs: 2\nExp number of AAs in TMHs: 36.51738\nExp number, first 60 AAs: 19.72849\nTotal prob of N-in: 0.96170\nPOSSIBLE N-term signal sequence\ninside 1 19\nTMhelix 20 39\noutside 40 325\nTMhelix 326 348\ninside 349 372

16899 GCF_001972835.1_ASM197283v1 Salmonella enterica subsp. enterica serovar Enteritidis Proteobacteria; Gammaproteobacteria; Enterobacterales; Enterobacteriaceae; Salmonella; Salmonella enterica; Salmonella enterica subsp. enterica MTGDNTLITSHGINRRDFMKLCAALAATMGLSSKAAAEMAESVSNPQRPPVIWIGAQECT WP_000145429.1 hydrogenase 2 small subunit [Salmonella enterica] Length: 372\nNumber of predicted TMHs: 1\nExp number of AAs in TMHs: 24.51309\nExp number, first 60 AAs: 0.2682\nTotal prob of N-in: 0.14402\noutside 1 331\nTMhelix 332 354\ninside 355 372

16900 GCF_001972795.1_ASM197279v1 Salmonella enterica subsp. enterica serovar Enteritidis Proteobacteria; Gammaproteobacteria; Enterobacterales; Enterobacteriaceae; Salmonella; Salmonella enterica; Salmonella enterica subsp. enterica MTGDNTLITSHGINRRDFMKLCAALAATMGLSSKAAAEMAESVSNPQRPPVIWIGAQECT WP_000145429.1 hydrogenase 2 small subunit [Salmonella enterica] Length: 372\nNumber of predicted TMHs: 1\nExp number of AAs in TMHs: 24.51309\nExp number, first 60 AAs: 0.2682\nTotal prob of N-in: 0.14402\noutside 1 331\nTMhelix 332 354\ninside 355 372

16901 GCF_001972875.1_ASM197287v1 Salmonella enterica subsp. enterica serovar Enteritidis Proteobacteria; Gammaproteobacteria; Enterobacterales; Enterobacteriaceae; Salmonella; Salmonella enterica; Salmonella enterica subsp. enterica MTGDNTLITSHGINRRDFMKLCAALAATMGLSSKAAAEMAESVSNPQRPPVIWIGAQECT WP_000145429.1 hydrogenase 2 small subunit [Salmonella enterica] Length: 372\nNumber of predicted TMHs: 1\nExp number of AAs in TMHs: 24.51309\nExp number, first 60 AAs: 0.2682\nTotal prob of N-in: 0.14402\noutside 1 331\nTMhelix 332 354\ninside 355 372

16902 GCF_001972915.1_ASM197291v1 Salmonella enterica subsp. enterica serovar Enteritidis Proteobacteria; Gammaproteobacteria; Enterobacterales; Enterobacteriaceae; Salmonella; Salmonella enterica; Salmonella enterica subsp. enterica MTGDNTLITSHGINRRDFMKLCAALAATMGLSSKAAAEMAESVSNPQRPPVIWIGAQECT WP_000145429.1 hydrogenase 2 small subunit [Salmonella enterica] Length: 372\nNumber of predicted TMHs: 1\nExp number of AAs in TMHs: 24.51309\nExp number, first 60 AAs: 0.2682\nTotal prob of N-in: 0.14402\noutside 1 331\nTMhelix 332 354\ninside 355 372

16903 GCF_001972835.1_ASM197283v1 Salmonella enterica subsp. enterica serovar Enteritidis Proteobacteria; Gammaproteobacteria; Enterobacterales; Enterobacteriaceae; Salmonella; Salmonella enterica; Salmonella enterica subsp. enterica MNNEETFYQAMRRKGVTRRSFLKFCSLAATSLGLGAGMTPKIAWALENKPRIPVVWIHGL WP_001058307.1 hydrogenase [Salmonella enterica] Length: 372\nNumber of predicted TMHs: 2\nExp number of AAs in TMHs: 36.51738\nExp number, first 60 AAs: 19.72849\nTotal prob of N-in: 0.96170\nPOSSIBLE N-term signal sequence\ninside 1 19\nTMhelix 20 39\noutside 40 325\nTMhelix 326 348\ninside 349 372

16904 GCF_001972795.1_ASM197279v1 Salmonella enterica subsp. enterica serovar Enteritidis Proteobacteria; Gammaproteobacteria; Enterobacterales; Enterobacteriaceae; Salmonella; Salmonella enterica; Salmonella enterica subsp. enterica MNNEETFYQAMRRKGVTRRSFLKFCSLAATSLGLGAGMTPKIAWALENKPRIPVVWIHGL WP_001058307.1 hydrogenase [Salmonella enterica] Length: 372\nNumber of predicted TMHs: 2\nExp number of AAs in TMHs: 36.51738\nExp number, first 60 AAs: 19.72849\nTotal prob of N-in: 0.96170\nPOSSIBLE N-term signal sequence\ninside 1 19\nTMhelix 20 39\noutside 40 325\nTMhelix 326 348\ninside 349 372

16905 GCF_001972955.1_ASM197295v1 Salmonella enterica subsp. enterica serovar Enteritidis Proteobacteria; Gammaproteobacteria; Enterobacterales; Enterobacteriaceae; Salmonella; Salmonella enterica; Salmonella enterica subsp. enterica MTGDNTLITSHGINRRDFMKLCAALAATMGLSSKAAAEMAESVSNPQRPPVIWIGAQECT WP_000145429.1 hydrogenase 2 small subunit [Salmonella enterica] Length: 372\nNumber of predicted TMHs: 1\nExp number of AAs in TMHs: 24.51309\nExp number, first 60 AAs: 0.2682\nTotal prob of N-in: 0.14402\noutside 1 331\nTMhelix 332 354\ninside 355 372

16906 GCF_001972955.1_ASM197295v1 Salmonella enterica subsp. enterica serovar Enteritidis Proteobacteria; Gammaproteobacteria; Enterobacterales; Enterobacteriaceae; Salmonella; Salmonella enterica; Salmonella enterica subsp. enterica MNNEETFYQAMRRKGVTRRSFLKFCSLAATSLGLGAGMTPKIAWALENKPRIPVVWIHGL WP_001058307.1 hydrogenase [Salmonella enterica] Length: 372\nNumber of predicted TMHs: 2\nExp number of AAs in TMHs: 36.51738\nExp number, first 60 AAs: 19.72849\nTotal prob of N-in: 0.96170\nPOSSIBLE N-term signal sequence\ninside 1 19\nTMhelix 20 39\noutside 40 325\nTMhelix 326 348\ninside 349 372

16907 GCF_001972875.1_ASM197287v1 Salmonella enterica subsp. enterica serovar Enteritidis Proteobacteria; Gammaproteobacteria; Enterobacterales; Enterobacteriaceae; Salmonella; Salmonella enterica; Salmonella enterica subsp. enterica MNNEETFYQAMRRKGVTRRSFLKFCSLAATSLGLGAGMTPKIAWALENKPRIPVVWIHGL WP_001058307.1 hydrogenase [Salmonella enterica] Length: 372\nNumber of predicted TMHs: 2\nExp number of AAs in TMHs: 36.51738\nExp number, first 60 AAs: 19.72849\nTotal prob of N-in: 0.96170\nPOSSIBLE N-term signal sequence\ninside 1 19\nTMhelix 20 39\noutside 40 325\nTMhelix 326 348\ninside 349 372

16908 GCF_001972995.1_ASM197299v1 Salmonella enterica subsp. enterica serovar Enteritidis Proteobacteria; Gammaproteobacteria; Enterobacterales; Enterobacteriaceae; Salmonella; Salmonella enterica; Salmonella enterica subsp. enterica MTGDNTLITSHGINRRDFMKLCAALAATMGLSSKAAAEMAESVSNPQRPPVIWIGAQECT WP_000145429.1 hydrogenase 2 small subunit [Salmonella enterica] Length: 372\nNumber of predicted TMHs: 1\nExp number of AAs in TMHs: 24.51309\nExp number, first 60 AAs: 0.2682\nTotal prob of N-in: 0.14402\noutside 1 331\nTMhelix 332 354\ninside 355 372

16909 GCF_001973035.1_ASM197303v1 Salmonella enterica subsp. enterica serovar Enteritidis Proteobacteria; Gammaproteobacteria; Enterobacterales; Enterobacteriaceae; Salmonella; Salmonella enterica; Salmonella enterica subsp. enterica MTGDNTLITSHGINRRDFMKLCAALAATMGLSSKAAAEMAESVSNPQRPPVIWIGAQECT WP_000145429.1 hydrogenase 2 small subunit [Salmonella enterica] Length: 372\nNumber of predicted TMHs: 1\nExp number of AAs in TMHs: 24.51309\nExp number, first 60 AAs: 0.2682\nTotal prob of N-in: 0.14402\noutside 1 331\nTMhelix 332 354\ninside 355 372

16910 GCF_001973165.1_ASM197316v1 Salmonella enterica subsp. enterica serovar Enteritidis Proteobacteria; Gammaproteobacteria; Enterobacterales; Enterobacteriaceae; Salmonella; Salmonella enterica; Salmonella enterica subsp. enterica MNNEETFYQAMRRKGVTRRSFLKFCSLAATSLGLGAGMTPKIAWALENKPRIPVVWIHGL WP_001058307.1 hydrogenase [Salmonella enterica] Length: 372\nNumber of predicted TMHs: 2\nExp number of AAs in TMHs: 36.51738\nExp number, first 60 AAs: 19.72849\nTotal prob of N-in: 0.96170\nPOSSIBLE N-term signal sequence\ninside 1 19\nTMhelix 20 39\noutside 40 325\nTMhelix 326 348\ninside 349 372

16911 GCF_001973185.1_ASM197318v1 Salmonella enterica subsp. enterica serovar Enteritidis Proteobacteria; Gammaproteobacteria; Enterobacterales; Enterobacteriaceae; Salmonella; Salmonella enterica; Salmonella enterica subsp. enterica MNNEETFYQAMRRKGVTRRSFLKFCSLAATSLGLGAGMTPKIAWALENKPRIPVVWIHGL WP_001058307.1 hydrogenase [Salmonella enterica] Length: 372\nNumber of predicted TMHs: 2\nExp number of AAs in TMHs: 36.51738\nExp number, first 60 AAs: 19.72849\nTotal prob of N-in: 0.96170\nPOSSIBLE N-term signal sequence\ninside 1 19\nTMhelix 20 39\noutside 40 325\nTMhelix 326 348\ninside 349 372

16912 GCF_001973085.1_ASM197308v1 Salmonella enterica subsp. enterica serovar Enteritidis Proteobacteria; Gammaproteobacteria; Enterobacterales; Enterobacteriaceae; Salmonella; Salmonella enterica; Salmonella enterica subsp. enterica MNNEETFYQAMRRKGVTRRSFLKFCSLAATSLGLGAGMTPKIAWALENKPRIPVVWIHGL WP_001058307.1 hydrogenase [Salmonella enterica] Length: 372\nNumber of predicted TMHs: 2\nExp number of AAs in TMHs: 36.51738\nExp number, first 60 AAs: 19.72849\nTotal prob of N-in: 0.96170\nPOSSIBLE N-term signal sequence\ninside 1 19\nTMhelix 20 39\noutside 40 325\nTMhelix 326 348\ninside 349 372

16913 GCF_001973115.1_ASM197311v1 Salmonella enterica subsp. enterica serovar Enteritidis Proteobacteria; Gammaproteobacteria; Enterobacterales; Enterobacteriaceae; Salmonella; Salmonella enterica; Salmonella enterica subsp. enterica MNNEETFYQAMRRKGVTRRSFLKFCSLAATSLGLGAGMTPKIAWALENKPRIPVVWIHGL WP_001058307.1 hydrogenase [Salmonella enterica] Length: 372\nNumber of predicted TMHs: 2\nExp number of AAs in TMHs: 36.51738\nExp number, first 60 AAs: 19.72849\nTotal prob of N-in: 0.96170\nPOSSIBLE N-term signal sequence\ninside 1 19\nTMhelix 20 39\noutside 40 325\nTMhelix 326 348\ninside 349 372

16914 GCF_001973225.1_ASM197322v1 Salmonella enterica subsp. enterica serovar Enteritidis Proteobacteria; Gammaproteobacteria; Enterobacterales; Enterobacteriaceae; Salmonella; Salmonella enterica; Salmonella enterica subsp. enterica MTGDNTLITSHGINRRDFMKLCAALAATMGLSSKAAAEMAESVSNPQRPPVIWIGAQECT WP_000145429.1 hydrogenase 2 small subunit [Salmonella enterica] Length: 372\nNumber of

predicted TMHs:  1\nExp number of AAs in TMHs: 24.51309\nExp number, first 60 AAs:  0.2682\nTotal prob of N-in: 0.14402\noutside   1   331\nTMhelix   332   354\ninside   355   372

16915 GCF_001973265.1_ASM197326v1 Salmonella enterica subsp. enterica serovar Enteritidis Proteobacteria; Gammaproteobacteria; Enterobacterales; Enterobacteriaceae; Salmonella; Salmonella enterica; Salmonella enterica subsp. enterica MTGDNTLITSHGINRRDFMKLCAALAATMGLSSKAAAEMAESVSNPQRPPVIWIGAQECT WP_000145429.1 hydrogenase 2 small subunit [Salmonella enterica] Length: 372\nNumber of predicted TMHs:  1\nExp number of AAs in TMHs: 24.51309\nExp number, first 60 AAs:  0.2682\nTotal prob of N-in: 0.14402\noutside   1   331\nTMhelix   332   354\ninside   355   372

16916 GCF_001973345.1_ASM197334v1 Salmonella enterica subsp. enterica serovar Enteritidis Proteobacteria; Gammaproteobacteria; Enterobacterales; Enterobacteriaceae; Salmonella; Salmonella enterica; Salmonella enterica subsp. enterica MTGDNTLITSHGINRRDFMKLCAALAATMGLSSKAAAEMAESVSNPQRPPVIWIGAQECT WP_000145429.1 hydrogenase 2 small subunit [Salmonella enterica] Length: 372\nNumber of predicted TMHs:  1\nExp number of AAs in TMHs: 24.51309\nExp number, first 60 AAs:  0.2682\nTotal prob of N-in: 0.14402\noutside   1   331\nTMhelix   332   354\ninside   355   372

16917 GCF_001973545.1_ASM197354v1 Salmonella enterica subsp. enterica serovar Enteritidis Proteobacteria; Gammaproteobacteria; Enterobacterales; Enterobacteriaceae; Salmonella; Salmonella enterica; Salmonella enterica subsp. enterica MTGDNTLITSHGINRRDFMKLCAALAATMGLSSKAAAEMAESVSNPQRPPVIWIGAQECT WP_000145429.1 hydrogenase 2 small subunit [Salmonella enterica] Length: 372\nNumber of predicted TMHs:  1\nExp number of AAs in TMHs: 24.51309\nExp number, first 60 AAs:  0.2682\nTotal prob of N-in: 0.14402\noutside   1   331\nTMhelix   332   354\ninside   355   372

16918 GCF_001973505.1_ASM197350v1 Salmonella enterica subsp. enterica serovar Enteritidis Proteobacteria; Gammaproteobacteria; Enterobacterales; Enterobacteriaceae; Salmonella; Salmonella enterica; Salmonella enterica subsp. enterica MNNEETFYQAMRRKGVTRRSFLKFCSLAATSLGLGAGMTPKIAWALENKPRIPVVWIHGL WP_001058307.1 hydrogenase [Salmonella enterica] Length: 372\nNumber of predicted TMHs:  2\nExp number of AAs in TMHs: 36.51738\nExp number, first 60 AAs:  19.72849\nTotal prob of N-in:   0.96170\nPOSSIBLE N-term signal sequence\ninside   1   19\nTMhelix   20   39\noutside   40   325\nTMhelix   326   348\ninside   349   372

16919 GCF_001973465.1_ASM197346v1 Salmonella enterica subsp. enterica serovar Enteritidis Proteobacteria; Gammaproteobacteria; Enterobacterales; Enterobacteriaceae; Salmonella; Salmonella enterica; Salmonella enterica subsp. enterica MNNEETFYQAMRRKGVTRRSFLKFCSLAATSLGLGAGMTPKIAWALENKPRIPVVWIHGL WP_001058307.1 hydrogenase [Salmonella enterica] Length: 372\nNumber of predicted TMHs:  2\nExp number of AAs in TMHs: 36.51738\nExp number, first 60 AAs:  19.72849\nTotal prob of N-in:   0.96170\nPOSSIBLE N-term signal sequence\ninside   1   19\nTMhelix   20   39\noutside   40   325\nTMhelix   326   348\ninside   349   372

16920 GCF_001973345.1_ASM197334v1 Salmonella enterica subsp. enterica serovar Enteritidis Proteobacteria; Gammaproteobacteria; Enterobacterales; Enterobacteriaceae; Salmonella; Salmonella enterica; Salmonella enterica subsp. enterica MNNEETFYQAMRRKGVTRRSFLKFCSLAATSLGLGAGMTPKIAWALENKPRIPVVWIHGL WP_001058307.1 hydrogenase [Salmonella enterica] Length: 372\nNumber of predicted TMHs:  2\nExp number of AAs in TMHs: 36.51738\nExp number, first 60 AAs:  19.72849\nTotal prob of N-in:   0.96170\nPOSSIBLE N-term signal sequence\ninside   1   19\nTMhelix   20   39\noutside   40   325\nTMhelix   326   348\ninside   349   372

16921 GCF_001973585.1_ASM197358v1 Salmonella enterica subsp. enterica serovar Enteritidis Proteobacteria; Gammaproteobacteria; Enterobacterales; Enterobacteriaceae; Salmonella; Salmonella enterica; Salmonella enterica subsp. enterica MTGDNTLITSHGINRRDFMKLCAALAATMGLSSKAAAEMAESVSNPQRPPVIWIGAQECT WP_000145429.1 hydrogenase 2 small subunit [Salmonella enterica] Length: 372\nNumber of predicted TMHs:  1\nExp number of AAs in TMHs: 24.51309\nExp number, first 60 AAs:  0.2682\nTotal prob of N-in: 0.14402\noutside   1   331\nTMhelix   332   354\ninside   355   372

16922 GCF_001973585.1_ASM197358v1 Salmonella enterica subsp. enterica serovar Enteritidis Proteobacteria; Gammaproteobacteria; Enterobacterales; Enterobacteriaceae; Salmonella; Salmonella enterica; Salmonella enterica subsp. enterica MNNEETFYQAMRRKGVTRRSFLKFCSLAATSLGLGAGMTPKIAWALENKPRIPVVWIHGL WP_001058307.1 hydrogenase [Salmonella enterica] Length: 372\nNumber of predicted TMHs:  2\nExp number of AAs in TMHs: 36.51738\nExp number, first 60 AAs:  19.72849\nTotal prob of N-in:   0.96170\nPOSSIBLE N-term signal sequence\ninside   1   19\nTMhelix   20   39\noutside   40   325\nTMhelix   326   348\ninside   349   372

16923 GCF_002039665.1_ASM203966v1 Salmonella enterica subsp. enterica serovar Enteritidis Proteobacteria; Gammaproteobacteria; Enterobacterales; Enterobacteriaceae; Salmonella; Salmonella enterica;

Salmonella enterica subsp. enterica MTGDNTLITSHGINRRDFMKLCAALAATMGLSSKAAAEMAESVSNPQRPPVIWIGAQECT WP_000145429.1 hydrogenase 2 small subunit [Salmonella enterica] Length: 372\nNumber of predicted TMHs: 1\nExp number of AAs in TMHs: 24.51309\nExp number, first 60 AAs: 0.2682\nTotal prob of N-in: 0.14402\noutside 1 331\nTMhelix 332 354\ninside 355 372

16924 GCF_002039525.1_ASM203952v1 Salmonella enterica subsp. enterica serovar Enteritidis Proteobacteria; Gammaproteobacteria; Enterobacterales; Enterobacteriaceae; Salmonella; Salmonella enterica; Salmonella enterica subsp. enterica MNNEETFYQAMRRKGVTRRSFLKFCSLAATSLGLGAGMTPKIAWALENKPRIPVVWIHGL WP_001058307.1 hydrogenase [Salmonella enterica] Length: 372\nNumber of predicted TMHs: 2\nExp number of AAs in TMHs: 36.51738\nExp number, first 60 AAs: 19.72849\nTotal prob of N-in: 0.96170\nPOSSIBLE N-term signal sequence\ninside 1 19\nTMhelix 20 39\noutside 40 325\nTMhelix 326 348\ninside 349 372

16925 GCF_002039505.1_ASM203950v1 Salmonella enterica subsp. enterica serovar Enteritidis Proteobacteria; Gammaproteobacteria; Enterobacterales; Enterobacteriaceae; Salmonella; Salmonella enterica; Salmonella enterica subsp. enterica MNNEETFYQAMRRKGVTRRSFLKFCSLAATSLGLGAGMTPKIAWALENKPRIPVVWIHGL WP_001058307.1 hydrogenase [Salmonella enterica] Length: 372\nNumber of predicted TMHs: 2\nExp number of AAs in TMHs: 36.51738\nExp number, first 60 AAs: 19.72849\nTotal prob of N-in: 0.96170\nPOSSIBLE N-term signal sequence\ninside 1 19\nTMhelix 20 39\noutside 40 325\nTMhelix 326 348\ninside 349 372

16926 GCF_002039625.1_ASM203962v1 Salmonella enterica subsp. enterica serovar Enteritidis Proteobacteria; Gammaproteobacteria; Enterobacterales; Enterobacteriaceae; Salmonella; Salmonella enterica; Salmonella enterica subsp. enterica MTGDNTLITSHGINRRDFMKLCAALAATMGLSSKAAAEMAESVSNPQRPPVIWIGAQECT WP_000145429.1 hydrogenase 2 small subunit [Salmonella enterica] Length: 372\nNumber of predicted TMHs: 1\nExp number of AAs in TMHs: 24.51309\nExp number, first 60 AAs: 0.2682\nTotal prob of N-in: 0.14402\noutside 1 331\nTMhelix 332 354\ninside 355 372

16927 GCF_002039585.1_ASM203958v1 Salmonella enterica subsp. enterica serovar Enteritidis Proteobacteria; Gammaproteobacteria; Enterobacterales; Enterobacteriaceae; Salmonella; Salmonella enterica; Salmonella enterica subsp. enterica MTGDNTLITSHGINRRDFMKLCAALAATMGLSSKAAAEMAESVSNPQRPPVIWIGAQECT WP_000145429.1 hydrogenase 2 small subunit [Salmonella enterica] Length: 372\nNumber of predicted TMHs: 1\nExp number of AAs in TMHs: 24.51309\nExp number, first 60 AAs: 0.2682\nTotal prob of N-in: 0.14402\noutside 1 331\nTMhelix 332 354\ninside 355 372

16928 GCF_002039455.1_ASM203945v1 Salmonella enterica subsp. enterica serovar Enteritidis Proteobacteria; Gammaproteobacteria; Enterobacterales; Enterobacteriaceae; Salmonella; Salmonella enterica; Salmonella enterica subsp. enterica MNNEETFYQAMRRKGVTRRSFLKFCSLAATSLGLGAGMTPKIAWALENKPRIPVVWIHGL WP_001058307.1 hydrogenase [Salmonella enterica] Length: 372\nNumber of predicted TMHs: 2\nExp number of AAs in TMHs: 36.51738\nExp number, first 60 AAs: 19.72849\nTotal prob of N-in: 0.96170\nPOSSIBLE N-term signal sequence\ninside 1 19\nTMhelix 20 39\noutside 40 325\nTMhelix 326 348\ninside 349 372

16929 GCF_002039545.1_ASM203954v1 Salmonella enterica subsp. enterica serovar Enteritidis Proteobacteria; Gammaproteobacteria; Enterobacterales; Enterobacteriaceae; Salmonella; Salmonella enterica; Salmonella enterica subsp. enterica MTGDNTLITSHGINRRDFMKLCAALAATMGLSSKAAAEMAESVSNPQRPPVIWIGAQECT WP_000145429.1 hydrogenase 2 small subunit [Salmonella enterica] Length: 372\nNumber of predicted TMHs: 1\nExp number of AAs in TMHs: 24.51309\nExp number, first 60 AAs: 0.2682\nTotal prob of N-in: 0.14402\noutside 1 331\nTMhelix 332 354\ninside 355 372

16930 GCF_002039335.1_ASM203933v1 Salmonella enterica subsp. enterica serovar Enteritidis Proteobacteria; Gammaproteobacteria; Enterobacterales; Enterobacteriaceae; Salmonella; Salmonella enterica; Salmonella enterica subsp. enterica MNNEETFYQAMRRKGVTRRSFLKFCSLAATSLGLGAGMTPKIAWALENKPRIPVVWIHGL WP_001058307.1 hydrogenase [Salmonella enterica] Length: 372\nNumber of predicted TMHs: 2\nExp number of AAs in TMHs: 36.51738\nExp number, first 60 AAs: 19.72849\nTotal prob of N-in: 0.96170\nPOSSIBLE N-term signal sequence\ninside 1 19\nTMhelix 20 39\noutside 40 325\nTMhelix 326 348\ninside 349 372

16931 GCF_002039385.1_ASM203938v1 Salmonella enterica subsp. enterica serovar Enteritidis Proteobacteria; Gammaproteobacteria; Enterobacterales; Enterobacteriaceae; Salmonella; Salmonella enterica; Salmonella enterica subsp. enterica MTGDNTLITSHGINRRDFMKLCAALAATMGLSSKAAAEMAESVSNPQRPPVIWIGAQECT WP_000145429.1 hydrogenase 2 small subunit [Salmonella enterica] Length: 372\nNumber of predicted TMHs: 1\nExp number of AAs in TMHs: 24.51309\nExp number, first 60 AAs: 0.2682\nTotal prob of N-in: 0.14402\noutside 1 331\nTMhelix 332 354\ninside 355 372

16932 GCF_002039335.1_ASM203933v1 Salmonella enterica subsp. enterica serovar Enteritidis Proteobacteria; Gammaproteobacteria; Enterobacterales; Enterobacteriaceae; Salmonella; Salmonella enterica; Salmonella enterica subsp. enterica MTGDNTLITSHGINRRDFMKLCAALAATMGLSSKAAAEMAESVSNPQRPPVIWIGAQECT WP_000145429.1 hydrogenase 2 small subunit [Salmonella enterica] Length: 372\nNumber of predicted TMHs: 1\nExp number of AAs in TMHs: 24.51309\nExp number, first 60 AAs: 0.2682\nTotal prob of N-in: 0.14402\noutside 1 331\nTMhelix 332 354\ninside 355 372

16933 GCF_002039185.1_ASM203918v1 Salmonella enterica subsp. enterica serovar Enteritidis Proteobacteria; Gammaproteobacteria; Enterobacterales; Enterobacteriaceae; Salmonella; Salmonella enterica; Salmonella enterica subsp. enterica MNNEETFYQAMRRKGVTRRSFLKFCSLAATSLGLGAGMTPKIAWALENKPRIPVVWIHGL WP_001058307.1 hydrogenase [Salmonella enterica] Length: 372\nNumber of predicted TMHs: 2\nExp number of AAs in TMHs: 36.51738\nExp number, first 60 AAs: 19.72849\nTotal prob of N-in: 0.96170\nPOSSIBLE N-term signal sequence\ninside 1 19\nTMhelix 20 39\noutside 40 325\nTMhelix 326 348\ninside 349 372

16934 GCF_002039295.1_ASM203929v1 Salmonella enterica subsp. enterica serovar Enteritidis Proteobacteria; Gammaproteobacteria; Enterobacterales; Enterobacteriaceae; Salmonella; Salmonella enterica; Salmonella enterica subsp. enterica MNNEETFYQAMRRKGVTRRSFLKFCSLAATSLGLGAGMTPKIAWALENKPRIPVVWIHGL WP_001058307.1 hydrogenase [Salmonella enterica] Length: 372\nNumber of predicted TMHs: 2\nExp number of AAs in TMHs: 36.51738\nExp number, first 60 AAs: 19.72849\nTotal prob of N-in: 0.96170\nPOSSIBLE N-term signal sequence\ninside 1 19\nTMhelix 20 39\noutside 40 325\nTMhelix 326 348\ninside 349 372

16935 GCF_002039295.1_ASM203929v1 Salmonella enterica subsp. enterica serovar Enteritidis Proteobacteria; Gammaproteobacteria; Enterobacterales; Enterobacteriaceae; Salmonella; Salmonella enterica; Salmonella enterica subsp. enterica MTGDNTLITSHGINRRDFMKLCAALAATMGLSSKAAAEMAESVSNPQRPPVIWIGAQECT WP_000145429.1 hydrogenase 2 small subunit [Salmonella enterica] Length: 372\nNumber of predicted TMHs: 1\nExp number of AAs in TMHs: 24.51309\nExp number, first 60 AAs: 0.2682\nTotal prob of N-in: 0.14402\noutside 1 331\nTMhelix 332 354\ninside 355 372

16936 GCF_002039145.1_ASM203914v1 Salmonella enterica subsp. enterica serovar Enteritidis Proteobacteria; Gammaproteobacteria; Enterobacterales; Enterobacteriaceae; Salmonella; Salmonella enterica; Salmonella enterica subsp. enterica MNNEETFYQAMRRKGVTRRSFLKFCSLAATSLGLGAGMTPKIAWALENKPRIPVVWIHGL WP_001058307.1 hydrogenase [Salmonella enterica] Length: 372\nNumber of predicted TMHs: 2\nExp number of AAs in TMHs: 36.51738\nExp number, first 60 AAs: 19.72849\nTotal prob of N-in: 0.96170\nPOSSIBLE N-term signal sequence\ninside 1 19\nTMhelix 20 39\noutside 40 325\nTMhelix 326 348\ninside 349 372

16937 GCF_002038965.1_ASM203896v1 Salmonella enterica subsp. enterica serovar Enteritidis Proteobacteria; Gammaproteobacteria; Enterobacterales; Enterobacteriaceae; Salmonella; Salmonella enterica; Salmonella enterica subsp. enterica MNNEETFYQAMRRKGVTRRSFLKFCSLAATSLGLGAGMTPKIAWALENKPRIPVVWIHGL WP_001058307.1 hydrogenase [Salmonella enterica] Length: 372\nNumber of predicted TMHs: 2\nExp number of AAs in TMHs: 36.51738\nExp number, first 60 AAs: 19.72849\nTotal prob of N-in: 0.96170\nPOSSIBLE N-term signal sequence\ninside 1 19\nTMhelix 20 39\noutside 40 325\nTMhelix 326 348\ninside 349 372

16938 GCF_002039005.1_ASM203900v1 Salmonella enterica subsp. enterica serovar Enteritidis Proteobacteria; Gammaproteobacteria; Enterobacterales; Enterobacteriaceae; Salmonella; Salmonella enterica; Salmonella enterica subsp. enterica MNNEETFYQAMRRKGVTRRSFLKFCSLAATSLGLGAGMTPKIAWALENKPRIPVVWIHGL WP_001058307.1 hydrogenase [Salmonella enterica] Length: 372\nNumber of predicted TMHs: 2\nExp number of AAs in TMHs: 36.51738\nExp number, first 60 AAs: 19.72849\nTotal prob of N-in: 0.96170\nPOSSIBLE N-term signal sequence\ninside 1 19\nTMhelix 20 39\noutside 40 325\nTMhelix 326 348\ninside 349 372

16939 GCF_002039005.1_ASM203900v1 Salmonella enterica subsp. enterica serovar Enteritidis Proteobacteria; Gammaproteobacteria; Enterobacterales; Enterobacteriaceae; Salmonella; Salmonella enterica; Salmonella enterica subsp. enterica MTGDNTLITSHGINRRDFMKLCAALAATMGLSSKAAAEMAESVSNPQRPPVIWIGAQECT WP_000145429.1 hydrogenase 2 small subunit [Salmonella enterica] Length: 372\nNumber of predicted TMHs: 1\nExp number of AAs in TMHs: 24.51309\nExp number, first 60 AAs: 0.2682\nTotal prob of N-in: 0.14402\noutside 1 331\nTMhelix 332 354\ninside 355 372

16940 GCF_002038915.1_ASM203891v1 Salmonella enterica subsp. enterica serovar Enteritidis Proteobacteria; Gammaproteobacteria; Enterobacterales; Enterobacteriaceae; Salmonella; Salmonella enterica; Salmonella enterica subsp. enterica MNNEETFYQAMRRKGVTRRSFLKFCSLAATSLGLGAGMTPKIAWALENKPRIPVVWIHGL WP_001058307.1 hydrogenase [Salmonella enterica] Length: 372\nNumber of predicted TMHs: 2\nExp

number of AAs in TMHs: 36.51738\nExp number, first 60 AAs: 19.72849\nTotal prob of N-in: 0.96170\nPOSSIBLE N-term signal sequence\ninside 1 19\nTMhelix 20 39\noutside 40 325\nTMhelix 326 348\ninside 349 372

16941 GCF_002038885.1_ASM203888v1 Salmonella enterica subsp. enterica serovar Enteritidis Proteobacteria; Gammaproteobacteria; Enterobacterales; Enterobacteriaceae; Salmonella; Salmonella enterica; Salmonella enterica subsp. enterica MNNEETFYQAMRRKGVTRRSFLKFCSLAATSLGLGAGMTPKIAWALENKPRIPVVWIHGL WP_001058307.1 hydrogenase [Salmonella enterica] Length: 372\nNumber of predicted TMHs: 2\nExp number of AAs in TMHs: 36.51738\nExp number, first 60 AAs: 19.72849\nTotal prob of N-in: 0.96170\nPOSSIBLE N-term signal sequence\ninside 1 19\nTMhelix 20 39\noutside 40 325\nTMhelix 326 348\ninside 349 372

16942 GCF_002038865.1_ASM203886v1 Salmonella enterica subsp. enterica serovar Enteritidis Proteobacteria; Gammaproteobacteria; Enterobacterales; Enterobacteriaceae; Salmonella; Salmonella enterica; Salmonella enterica subsp. enterica MNNEETFYQAMRRKGVTRRSFLKFCSLAATSLGLGAGMTPKIAWALENKPRIPVVWIHGL WP_001058307.1 hydrogenase [Salmonella enterica] Length: 372\nNumber of predicted TMHs: 2\nExp number of AAs in TMHs: 36.51738\nExp number, first 60 AAs: 19.72849\nTotal prob of N-in: 0.96170\nPOSSIBLE N-term signal sequence\ninside 1 19\nTMhelix 20 39\noutside 40 325\nTMhelix 326 348\ninside 349 372

16943 GCF_002038805.1_ASM203880v1 Salmonella enterica subsp. enterica serovar Enteritidis Proteobacteria; Gammaproteobacteria; Enterobacterales; Enterobacteriaceae; Salmonella; Salmonella enterica; Salmonella enterica subsp. enterica MNNEETFYQAMRRKGVTRRSFLKFCSLAATSLGLGAGMTPKIAWALENKPRIPVVWIHGL WP_001058307.1 hydrogenase [Salmonella enterica] Length: 372\nNumber of predicted TMHs: 2\nExp number of AAs in TMHs: 36.51738\nExp number, first 60 AAs: 19.72849\nTotal prob of N-in: 0.96170\nPOSSIBLE N-term signal sequence\ninside 1 19\nTMhelix 20 39\noutside 40 325\nTMhelix 326 348\ninside 349 372

16944 GCF_002038765.1_ASM203876v1 Salmonella enterica subsp. enterica serovar Enteritidis Proteobacteria; Gammaproteobacteria; Enterobacterales; Enterobacteriaceae; Salmonella; Salmonella enterica; Salmonella enterica subsp. enterica MNNEETFYQAMRRKGVTRRSFLKFCSLAATSLGLGAGMTPKIAWALENKPRIPVVWIHGL WP_001058307.1 hydrogenase [Salmonella enterica] Length: 372\nNumber of predicted TMHs: 2\nExp number of AAs in TMHs: 36.51738\nExp number, first 60 AAs: 19.72849\nTotal prob of N-in: 0.96170\nPOSSIBLE N-term signal sequence\ninside 1 19\nTMhelix 20 39\noutside 40 325\nTMhelix 326 348\ninside 349 372

16945 GCF_002038715.1_ASM203871v1 Salmonella enterica subsp. enterica serovar Enteritidis Proteobacteria; Gammaproteobacteria; Enterobacterales; Enterobacteriaceae; Salmonella; Salmonella enterica; Salmonella enterica subsp. enterica MNNEETFYQAMRRKGVTRRSFLKFCSLAATSLGLGAGMTPKIAWALENKPRIPVVWIHGL WP_001058307.1 hydrogenase [Salmonella enterica] Length: 372\nNumber of predicted TMHs: 2\nExp number of AAs in TMHs: 36.51738\nExp number, first 60 AAs: 19.72849\nTotal prob of N-in: 0.96170\nPOSSIBLE N-term signal sequence\ninside 1 19\nTMhelix 20 39\noutside 40 325\nTMhelix 326 348\ninside 349 372

16946 GCF_002038805.1_ASM203880v1 Salmonella enterica subsp. enterica serovar Enteritidis Proteobacteria; Gammaproteobacteria; Enterobacterales; Enterobacteriaceae; Salmonella; Salmonella enterica; Salmonella enterica subsp. enterica MTGDNTLITSHGINRRDFMKLCAALAATMGLSSKAAAEMAESVSNPQRPPVIWIGAQECT WP_000145429.1 hydrogenase 2 small subunit [Salmonella enterica] Length: 372\nNumber of predicted TMHs: 1\nExp number of AAs in TMHs: 24.51309\nExp number, first 60 AAs: 0.2682\nTotal prob of N-in: 0.14402\noutside 1 331\nTMhelix 332 354\ninside 355 372

16947 GCF_002038765.1_ASM203876v1 Salmonella enterica subsp. enterica serovar Enteritidis Proteobacteria; Gammaproteobacteria; Enterobacterales; Enterobacteriaceae; Salmonella; Salmonella enterica; Salmonella enterica subsp. enterica MTGDNTLITSHGINRRDFMKLCAALAATMGLSSKAAAEMAESVSNPQRPPVIWIGAQECT WP_000145429.1 hydrogenase 2 small subunit [Salmonella enterica] Length: 372\nNumber of predicted TMHs: 1\nExp number of AAs in TMHs: 24.51309\nExp number, first 60 AAs: 0.2682\nTotal prob of N-in: 0.14402\noutside 1 331\nTMhelix 332 354\ninside 355 372

16948 GCF_002038715.1_ASM203871v1 Salmonella enterica subsp. enterica serovar Enteritidis Proteobacteria; Gammaproteobacteria; Enterobacterales; Enterobacteriaceae; Salmonella; Salmonella enterica; Salmonella enterica subsp. enterica MTGDNTLITSHGINRRDFMKLCAALAATMGLSSKAAAEMAESVSNPQRPPVIWIGAQECT WP_000145429.1 hydrogenase 2 small subunit [Salmonella enterica] Length: 372\nNumber of predicted TMHs: 1\nExp number of AAs in TMHs: 24.51309\nExp number, first 60 AAs: 0.2682\nTotal prob of N-in: 0.14402\noutside 1 331\nTMhelix 332 354\ninside 355 372

16949 GCF_002038525.1_ASM203852v1 Salmonella enterica subsp. enterica serovar Enteritidis Proteobacteria; Gammaproteobacteria; Enterobacterales; Enterobacteriaceae; Salmonella; Salmonella enterica;

Salmonella enterica subsp. enterica	MNNEETFYQAMRRKGVTRRSFLKFCSLAATSLGLGAGMTPKIAWALENKPRIPVVWIHGL
WP_001058307.1	hydrogenase [Salmonella enterica]	Length: 372\nNumber of predicted TMHs:  2\nExp number of AAs in TMHs: 36.51738\nExp number, first 60 AAs:  19.72849\nTotal prob of N-in:   0.96170\nPOSSIBLE N-term signal sequence\ninside   1   19\nTMhelix   20   39\noutside   40   325\nTMhelix   326   348\ninside   349   372

16950	GCF_002038485.1_ASM203848v1	Salmonella enterica subsp. enterica serovar Enteritidis
Proteobacteria; Gammaproteobacteria; Enterobacterales; Enterobacteriaceae; Salmonella; Salmonella enterica; Salmonella enterica subsp. enterica	MNNEETFYQAMRRKGVTRRSFLKFCSLAATSLGLGAGMTPKIAWALENKPRIPVVWIHGL
WP_001058307.1	hydrogenase [Salmonella enterica]	Length: 372\nNumber of predicted TMHs:  2\nExp number of AAs in TMHs: 36.51738\nExp number, first 60 AAs:  19.72849\nTotal prob of N-in:   0.96170\nPOSSIBLE N-term signal sequence\ninside   1   19\nTMhelix   20   39\noutside   40   325\nTMhelix   326   348\ninside   349   372

16951	GCF_002038555.1_ASM203855v1	Salmonella enterica subsp. enterica serovar Enteritidis
Proteobacteria; Gammaproteobacteria; Enterobacterales; Enterobacteriaceae; Salmonella; Salmonella enterica; Salmonella enterica subsp. enterica	MTGDNTLITSHGINRRDFMKLCAALAATMGLSSKAAAEMAESVSNPQRPPVIWIGAQECT
WP_000145429.1	hydrogenase 2 small subunit [Salmonella enterica]	Length: 372\nNumber of predicted TMHs:  1\nExp number of AAs in TMHs: 24.51309\nExp number, first 60 AAs:  0.2682\nTotal prob of N-in:   0.14402\noutside   1   331\nTMhelix   332   354\ninside   355   372

16952	GCF_002038525.1_ASM203852v1	Salmonella enterica subsp. enterica serovar Enteritidis
Proteobacteria; Gammaproteobacteria; Enterobacterales; Enterobacteriaceae; Salmonella; Salmonella enterica; Salmonella enterica subsp. enterica	MTGDNTLITSHGINRRDFMKLCAALAATMGLSSKAAAEMAESVSNPQRPPVIWIGAQECT
WP_000145429.1	hydrogenase 2 small subunit [Salmonella enterica]	Length: 372\nNumber of predicted TMHs:  1\nExp number of AAs in TMHs: 24.51309\nExp number, first 60 AAs:  0.2682\nTotal prob of N-in:   0.14402\noutside   1   331\nTMhelix   332   354\ninside   355   372

16953	GCF_002038485.1_ASM203848v1	Salmonella enterica subsp. enterica serovar Enteritidis
Proteobacteria; Gammaproteobacteria; Enterobacterales; Enterobacteriaceae; Salmonella; Salmonella enterica; Salmonella enterica subsp. enterica	MTGDNTLITSHGINRRDFMKLCAALAATMGLSSKAAAEMAESVSNPQRPPVIWIGAQECT
WP_000145429.1	hydrogenase 2 small subunit [Salmonella enterica]	Length: 372\nNumber of predicted TMHs:  1\nExp number of AAs in TMHs: 24.51309\nExp number, first 60 AAs:  0.2682\nTotal prob of N-in:   0.14402\noutside   1   331\nTMhelix   332   354\ninside   355   372

16954	GCF_002038445.1_ASM203844v1	Salmonella enterica subsp. enterica serovar Enteritidis
Proteobacteria; Gammaproteobacteria; Enterobacterales; Enterobacteriaceae; Salmonella; Salmonella enterica; Salmonella enterica subsp. enterica	MTGDNTLITSHGINRRDFMKLCAALAATMGLSSKAAAEMAESVSNPQRPPVIWIGAQECT
WP_000145429.1	hydrogenase 2 small subunit [Salmonella enterica]	Length: 372\nNumber of predicted TMHs:  1\nExp number of AAs in TMHs: 24.51309\nExp number, first 60 AAs:  0.2682\nTotal prob of N-in:   0.14402\noutside   1   331\nTMhelix   332   354\ninside   355   372

16955	GCF_002038325.1_ASM203832v1	Salmonella enterica subsp. enterica serovar Enteritidis
Proteobacteria; Gammaproteobacteria; Enterobacterales; Enterobacteriaceae; Salmonella; Salmonella enterica; Salmonella enterica subsp. enterica	MNNEETFYQAMRRKGVTRRSFLKFCSLAATSLGLGAGMTPKIAWALENKPRIPVVWIHGL
WP_001058307.1	hydrogenase [Salmonella enterica]	Length: 372\nNumber of predicted TMHs:  2\nExp number of AAs in TMHs: 36.51738\nExp number, first 60 AAs:  19.72849\nTotal prob of N-in:   0.96170\nPOSSIBLE N-term signal sequence\ninside   1   19\nTMhelix   20   39\noutside   40   325\nTMhelix   326   348\ninside   349   372

16956	GCF_002038285.1_ASM203828v1	Salmonella enterica subsp. enterica serovar Enteritidis
Proteobacteria; Gammaproteobacteria; Enterobacterales; Enterobacteriaceae; Salmonella; Salmonella enterica; Salmonella enterica subsp. enterica	MNNEETFYQAMRRKGVTRRSFLKFCSLAATSLGLGAGMTPKIAWALENKPRIPVVWIHGL
WP_001058307.1	hydrogenase [Salmonella enterica]	Length: 372\nNumber of predicted TMHs:  2\nExp number of AAs in TMHs: 36.51738\nExp number, first 60 AAs:  19.72849\nTotal prob of N-in:   0.96170\nPOSSIBLE N-term signal sequence\ninside   1   19\nTMhelix   20   39\noutside   40   325\nTMhelix   326   348\ninside   349   372

16957	GCF_002038325.1_ASM203832v1	Salmonella enterica subsp. enterica serovar Enteritidis
Proteobacteria; Gammaproteobacteria; Enterobacterales; Enterobacteriaceae; Salmonella; Salmonella enterica; Salmonella enterica subsp. enterica	MTGDNTLITSHGINRRDFMKLCAALAATMGLSSKAAAEMAESVSNPQRPPVIWIGAQECT
WP_000145429.1	hydrogenase 2 small subunit [Salmonella enterica]	Length: 372\nNumber of predicted TMHs:  1\nExp number of AAs in TMHs: 24.51309\nExp number, first 60 AAs:  0.2682\nTotal prob of N-in:   0.14402\noutside   1   331\nTMhelix   332   354\ninside   355   372

16958	GCF_002038285.1_ASM203828v1	Salmonella enterica subsp. enterica serovar Enteritidis	Proteobacteria; Gammaproteobacteria; Enterobacterales; Enterobacteriaceae; Salmonella; Salmonella enterica; Salmonella enterica subsp. enterica	MTGDNTLITSHGINRRDFMKLCAALAATMGLSSKAAAEMAESVSNPQRPPVIWIGAQECT	WP_000145429.1	hydrogenase 2 small subunit [Salmonella enterica]	Length: 372\nNumber of predicted TMHs:  1\nExp number of AAs in TMHs: 24.51309\nExp number, first 60 AAs:  0.2682\nTotal prob of N-in:  0.14402\noutside   1   331\nTMhelix   332   354\ninside   355   372

16959	GCF_002038215.1_ASM203821v1	Salmonella enterica subsp. enterica serovar Enteritidis	Proteobacteria; Gammaproteobacteria; Enterobacterales; Enterobacteriaceae; Salmonella; Salmonella enterica; Salmonella enterica subsp. enterica	MTGDNTLITSHGINRRDFMKLCAALAATMGLSSKAAAEMAESVSNPQRPPVIWIGAQECT	WP_000145429.1	hydrogenase 2 small subunit [Salmonella enterica]	Length: 372\nNumber of predicted TMHs:  1\nExp number of AAs in TMHs: 24.51309\nExp number, first 60 AAs:  0.2682\nTotal prob of N-in:  0.14402\noutside   1   331\nTMhelix   332   354\ninside   355   372

16960	GCF_002038205.1_ASM203820v1	Salmonella enterica subsp. enterica serovar Enteritidis	Proteobacteria; Gammaproteobacteria; Enterobacterales; Enterobacteriaceae; Salmonella; Salmonella enterica; Salmonella enterica subsp. enterica	MTGDNTLITSHGINRRDFMKLCAALAATMGLSSKAAAEMAESVSNPQRPPVIWIGAQECT	WP_000145429.1	hydrogenase 2 small subunit [Salmonella enterica]	Length: 372\nNumber of predicted TMHs:  1\nExp number of AAs in TMHs: 24.51309\nExp number, first 60 AAs:  0.2682\nTotal prob of N-in:  0.14402\noutside   1   331\nTMhelix   332   354\ninside   355   372

16961	GCF_002038035.1_ASM203803v1	Salmonella enterica subsp. enterica serovar Enteritidis	Proteobacteria; Gammaproteobacteria; Enterobacterales; Enterobacteriaceae; Salmonella; Salmonella enterica; Salmonella enterica subsp. enterica	MNNEETFYQAMRRKGVTRRSFLKFCSLAATSLGLGAGMTPKIAWALENKPRIPVVWIHGL	WP_001058307.1	hydrogenase [Salmonella enterica]	Length: 372\nNumber of predicted TMHs:  2\nExp number of AAs in TMHs: 36.51738\nExp number, first 60 AAs:  19.72849\nTotal prob of N-in:   0.96170\nPOSSIBLE N-term signal sequence\ninside   1   19\nTMhelix   20   39\noutside   40   325\nTMhelix   326   348\ninside   349   372

16962	GCF_002107235.1_ASM210723v1	Salmonella enterica subsp. enterica serovar Enteritidis	Proteobacteria; Gammaproteobacteria; Enterobacterales; Enterobacteriaceae; Salmonella; Salmonella enterica; Salmonella enterica subsp. enterica	MNNEETFYQAMRRKGVTRRSFLKFCSLAATSLGLGAGMTPKIAWALENKPRIPVVWIHGL	WP_001058307.1	hydrogenase [Salmonella enterica]	Length: 372\nNumber of predicted TMHs:  2\nExp number of AAs in TMHs: 36.51738\nExp number, first 60 AAs:  19.72849\nTotal prob of N-in:   0.96170\nPOSSIBLE N-term signal sequence\ninside   1   19\nTMhelix   20   39\noutside   40   325\nTMhelix   326   348\ninside   349   372

16963	GCF_001479035.1_Salmonella_enterica_CVM_N31380-SQ_v1.0	Salmonella enterica subsp. enterica serovar Enteritidis	Proteobacteria; Gammaproteobacteria; Enterobacterales; Enterobacteriaceae; Salmonella; Salmonella enterica; Salmonella enterica subsp. enterica	MTGDNTLITSHGINRRDFMKLCAALAATMGLSSKAAAEMAESVSNPQRPPVIWIGAQECT	WP_000145429.1	hydrogenase 2 small subunit [Salmonella enterica]	Length: 372\nNumber of predicted TMHs:  1\nExp number of AAs in TMHs: 24.51309\nExp number, first 60 AAs:  0.2682\nTotal prob of N-in:   0.14402\noutside   1   331\nTMhelix   332   354\ninside   355   372

16964	GCF_000750475.1_ASM75047v1	Salmonella enterica subsp. enterica serovar Enteritidis	Proteobacteria; Gammaproteobacteria; Enterobacterales; Enterobacteriaceae; Salmonella; Salmonella enterica; Salmonella enterica subsp. enterica	MTGDNTLITSHGINRRDFMKLCAALAATMGLSSKAAAEMAESVSNPQRPPVIWIGAQECT	WP_000145429.1	hydrogenase 2 small subunit [Salmonella enterica]	Length: 372\nNumber of predicted TMHs:  1\nExp number of AAs in TMHs: 24.51309\nExp number, first 60 AAs:  0.2682\nTotal prob of N-in:  0.14402\noutside   1   331\nTMhelix   332   354\ninside   355   372

16965	GCF_002038785.1_ASM203878v1	Salmonella enterica subsp. enterica serovar Enteritidis	Proteobacteria; Gammaproteobacteria; Enterobacterales; Enterobacteriaceae; Salmonella; Salmonella enterica; Salmonella enterica subsp. enterica	MNNEETFYQAMRRKGVTRRSFLKFCSLAATSLGLGAGMTPKIAWALENKPRIPVVWIHGL	WP_001058307.1	hydrogenase [Salmonella enterica]	Length: 372\nNumber of predicted TMHs:  2\nExp number of AAs in TMHs: 36.51738\nExp number, first 60 AAs:  19.72849\nTotal prob of N-in:   0.96170\nPOSSIBLE N-term signal sequence\ninside   1   19\nTMhelix   20   39\noutside   40   325\nTMhelix   326   348\ninside   349   372

16966	GCF_002038545.1_ASM203854v1	Salmonella enterica subsp. enterica serovar Enteritidis	Proteobacteria; Gammaproteobacteria; Enterobacterales; Enterobacteriaceae; Salmonella; Salmonella enterica; Salmonella enterica subsp. enterica	MTGDNTLITSHGINRRDFMKLCAALAATMGLSSKAAAEMAESVSNPQRPPVIWIGAQECT	WP_000145429.1	hydrogenase 2 small subunit [Salmonella enterica]	Length: 372\nNumber of

predicted TMHs: 1\nExp number of AAs in TMHs: 24.51309\nExp number, first 60 AAs: 0.2682\nTotal prob of N-in: 0.14402\noutside 1 331\nTMhelix 332 354\ninside 355 372

16967 GCF_002038455.1_ASM203845v1 Salmonella enterica subsp. enterica serovar Enteritidis Proteobacteria; Gammaproteobacteria; Enterobacterales; Enterobacteriaceae; Salmonella; Salmonella enterica; Salmonella enterica subsp. enterica MTGDNTLITSHGINRRDFMKLCAALAATMGLSSKAAAEMAESVSNPQRPPVIWIGAQECT WP_000145429.1 hydrogenase 2 small subunit [Salmonella enterica] Length: 372\nNumber of predicted TMHs: 1\nExp number of AAs in TMHs: 24.51309\nExp number, first 60 AAs: 0.2682\nTotal prob of N-in: 0.14402\noutside 1 331\nTMhelix 332 354\ninside 355 372

16968 GCF_002038425.1_ASM203842v1 Salmonella enterica subsp. enterica serovar Enteritidis Proteobacteria; Gammaproteobacteria; Enterobacterales; Enterobacteriaceae; Salmonella; Salmonella enterica; Salmonella enterica subsp. enterica MNNEETFYQAMRRKGVTRRSFLKFCSLAATSLGLGAGMTPKIAWALENKPRIPVVWIHGL WP_001058307.1 hydrogenase [Salmonella enterica] Length: 372\nNumber of predicted TMHs: 2\nExp number of AAs in TMHs: 36.51738\nExp number, first 60 AAs: 19.72849\nTotal prob of N-in: 0.96170\nPOSSIBLE N-term signal sequence\ninside 1 19\nTMhelix 20 39\noutside 40 325\nTMhelix 326 348\ninside 349 372

16969 GCF_002038385.1_ASM203838v1 Salmonella enterica subsp. enterica serovar Enteritidis Proteobacteria; Gammaproteobacteria; Enterobacterales; Enterobacteriaceae; Salmonella; Salmonella enterica; Salmonella enterica subsp. enterica MNNEETFYQAMRRKGVTRRSFLKFCSLAATSLGLGAGMTPKIAWALENKPRIPVVWIHGL WP_001058307.1 hydrogenase [Salmonella enterica] Length: 372\nNumber of predicted TMHs: 2\nExp number of AAs in TMHs: 36.51738\nExp number, first 60 AAs: 19.72849\nTotal prob of N-in: 0.96170\nPOSSIBLE N-term signal sequence\ninside 1 19\nTMhelix 20 39\noutside 40 325\nTMhelix 326 348\ninside 349 372

16970 GCF_002038425.1_ASM203842v1 Salmonella enterica subsp. enterica serovar Enteritidis Proteobacteria; Gammaproteobacteria; Enterobacterales; Enterobacteriaceae; Salmonella; Salmonella enterica; Salmonella enterica subsp. enterica MTGDNTLITSHGINRRDFMKLCAALAATMGLSSKAAAEMAESVSNPQRPPVIWIGAQECT WP_000145429.1 hydrogenase 2 small subunit [Salmonella enterica] Length: 372\nNumber of predicted TMHs: 1\nExp number of AAs in TMHs: 24.51309\nExp number, first 60 AAs: 0.2682\nTotal prob of N-in: 0.14402\noutside 1 331\nTMhelix 332 354\ninside 355 372

16971 GCF_002038385.1_ASM203838v1 Salmonella enterica subsp. enterica serovar Enteritidis Proteobacteria; Gammaproteobacteria; Enterobacterales; Enterobacteriaceae; Salmonella; Salmonella enterica; Salmonella enterica subsp. enterica MTGDNTLITSHGINRRDFMKLCAALAATMGLSSKAAAEMAESVSNPQRPPVIWIGAQECT WP_000145429.1 hydrogenase 2 small subunit [Salmonella enterica] Length: 372\nNumber of predicted TMHs: 1\nExp number of AAs in TMHs: 24.51309\nExp number, first 60 AAs: 0.2682\nTotal prob of N-in: 0.14402\noutside 1 331\nTMhelix 332 354\ninside 355 372

16972 GCF_002038305.1_ASM203830v1 Salmonella enterica subsp. enterica serovar Enteritidis Proteobacteria; Gammaproteobacteria; Enterobacterales; Enterobacteriaceae; Salmonella; Salmonella enterica; Salmonella enterica subsp. enterica MTGDNTLITSHGINRRDFMKLCAALAATMGLSSKAAAEMAESVSNPQRPPVIWIGAQECT WP_000145429.1 hydrogenase 2 small subunit [Salmonella enterica] Length: 372\nNumber of predicted TMHs: 1\nExp number of AAs in TMHs: 24.51309\nExp number, first 60 AAs: 0.2682\nTotal prob of N-in: 0.14402\noutside 1 331\nTMhelix 332 354\ninside 355 372

16973 GCF_002038105.1_ASM203810v1 Salmonella enterica subsp. enterica serovar Enteritidis Proteobacteria; Gammaproteobacteria; Enterobacterales; Enterobacteriaceae; Salmonella; Salmonella enterica; Salmonella enterica subsp. enterica MNNEETFYQAMRRKGVTRRSFLKFCSLAATSLGLGAGMTPKIAWALENKPRIPVVWIHGL WP_001058307.1 hydrogenase [Salmonella enterica] Length: 372\nNumber of predicted TMHs: 2\nExp number of AAs in TMHs: 36.51738\nExp number, first 60 AAs: 19.72849\nTotal prob of N-in: 0.96170\nPOSSIBLE N-term signal sequence\ninside 1 19\nTMhelix 20 39\noutside 40 325\nTMhelix 326 348\ninside 349 372

16974 GCF_002038255.1_ASM203825v1 Salmonella enterica subsp. enterica serovar Enteritidis Proteobacteria; Gammaproteobacteria; Enterobacterales; Enterobacteriaceae; Salmonella; Salmonella enterica; Salmonella enterica subsp. enterica MNNEETFYQAMRRKGVTRRSFLKFCSLAATSLGLGAGMTPKIAWALENKPRIPVVWIHGL WP_001058307.1 hydrogenase [Salmonella enterica] Length: 372\nNumber of predicted TMHs: 2\nExp number of AAs in TMHs: 36.51738\nExp number, first 60 AAs: 19.72849\nTotal prob of N-in: 0.96170\nPOSSIBLE N-term signal sequence\ninside 1 19\nTMhelix 20 39\noutside 40 325\nTMhelix 326 348\ninside 349 372

16975 GCF_002042795.1_ASM204279v1 Salmonella enterica subsp. enterica serovar Enteritidis Proteobacteria; Gammaproteobacteria; Enterobacterales; Enterobacteriaceae; Salmonella; Salmonella enterica;

Salmonella enterica subsp. enterica MTGDNTLITSHGINRRDFMKLCAALAATMGLSSKAAAEMAESVSNPQRPPVIWIGAQECT WP_000145429.1 hydrogenase 2 small subunit [Salmonella enterica] Length: 372\nNumber of predicted TMHs: 1\nExp number of AAs in TMHs: 24.51309\nExp number, first 60 AAs: 0.2682\nTotal prob of N-in: 0.14402\noutside 1 331\nTMhelix 332 354\ninside 355 372

16976 GCF_002038255.1_ASM203825v1 Salmonella enterica subsp. enterica serovar Enteritidis Proteobacteria; Gammaproteobacteria; Enterobacterales; Enterobacteriaceae; Salmonella; Salmonella enterica; Salmonella enterica subsp. enterica MTGDNTLITSHGINRRDFMKLCAALAATMGLSSKAAAEMAESVSNPQRPPVIWIGAQECT WP_000145429.1 hydrogenase 2 small subunit [Salmonella enterica] Length: 372\nNumber of predicted TMHs: 1\nExp number of AAs in TMHs: 24.51309\nExp number, first 60 AAs: 0.2682\nTotal prob of N-in: 0.14402\noutside 1 331\nTMhelix 332 354\ninside 355 372

16977 GCF_002038055.1_ASM203805v1 Salmonella enterica subsp. enterica serovar Enteritidis Proteobacteria; Gammaproteobacteria; Enterobacterales; Enterobacteriaceae; Salmonella; Salmonella enterica; Salmonella enterica subsp. enterica MNNEETFYQAMRRKGVTRRSFLKFCSLAATSLGLGAGMTPKIAWALENKPRIPVVWIHGL WP_001058307.1 hydrogenase [Salmonella enterica] Length: 372\nNumber of predicted TMHs: 2\nExp number of AAs in TMHs: 36.51738\nExp number, first 60 AAs: 19.72849\nTotal prob of N-in: 0.96170\nPOSSIBLE N-term signal sequence\ninside 1 19\nTMhelix 20 39\noutside 40 325\nTMhelix 326 348\ninside 349 372

16978 GCF_002038225.1_ASM203822v1 Salmonella enterica subsp. enterica serovar Enteritidis Proteobacteria; Gammaproteobacteria; Enterobacterales; Enterobacteriaceae; Salmonella; Salmonella enterica; Salmonella enterica subsp. enterica MTGDNTLITSHGINRRDFMKLCAALAATMGLSSKAAAEMAESVSNPQRPPVIWIGAQECT WP_000145429.1 hydrogenase 2 small subunit [Salmonella enterica] Length: 372\nNumber of predicted TMHs: 1\nExp number of AAs in TMHs: 24.51309\nExp number, first 60 AAs: 0.2682\nTotal prob of N-in: 0.14402\noutside 1 331\nTMhelix 332 354\ninside 355 372

16979 GCF_000750255.1_ASM75025v1 Salmonella enterica subsp. enterica serovar Enteritidis Proteobacteria; Gammaproteobacteria; Enterobacterales; Enterobacteriaceae; Salmonella; Salmonella enterica; Salmonella enterica subsp. enterica MTGDNTLITSHGINRRDFMKLCAALAATMGLSSKAAAEMAESVSNPQRPPVIWIGAQECT WP_000145429.1 hydrogenase 2 small subunit [Salmonella enterica] Length: 372\nNumber of predicted TMHs: 1\nExp number of AAs in TMHs: 24.51309\nExp number, first 60 AAs: 0.2682\nTotal prob of N-in: 0.14402\noutside 1 331\nTMhelix 332 354\ninside 355 372

16980 GCF_000750255.1_ASM75025v1 Salmonella enterica subsp. enterica serovar Enteritidis Proteobacteria; Gammaproteobacteria; Enterobacterales; Enterobacteriaceae; Salmonella; Salmonella enterica; Salmonella enterica subsp. enterica MNNEETFYQAMRRKGVTRRSFLKFCSLAATSLGLGAGMTPKIAWALENKPRIPVVWIHGL WP_001058307.1 hydrogenase [Salmonella enterica] Length: 372\nNumber of predicted TMHs: 2\nExp number of AAs in TMHs: 36.51738\nExp number, first 60 AAs: 19.72849\nTotal prob of N-in: 0.96170\nPOSSIBLE N-term signal sequence\ninside 1 19\nTMhelix 20 39\noutside 40 325\nTMhelix 326 348\ninside 349 372

16981 GCF_000750475.1_ASM75047v1 Salmonella enterica subsp. enterica serovar Enteritidis Proteobacteria; Gammaproteobacteria; Enterobacterales; Enterobacteriaceae; Salmonella; Salmonella enterica; Salmonella enterica subsp. enterica MNNEETFYQAMRRKGVTRRSFLKFCSLAATSLGLGAGMTPKIAWALENKPRIPVVWIHGL WP_001058307.1 hydrogenase [Salmonella enterica] Length: 372\nNumber of predicted TMHs: 2\nExp number of AAs in TMHs: 36.51738\nExp number, first 60 AAs: 19.72849\nTotal prob of N-in: 0.96170\nPOSSIBLE N-term signal sequence\ninside 1 19\nTMhelix 20 39\noutside 40 325\nTMhelix 326 348\ninside 349 372

16982 GCF_000754375.1_ASM75437v1 Salmonella enterica subsp. enterica serovar Enteritidis Proteobacteria; Gammaproteobacteria; Enterobacterales; Enterobacteriaceae; Salmonella; Salmonella enterica; Salmonella enterica subsp. enterica MNNEETFYQAMRRKGVTRRSFLKFCSLAATSLGLGAGMTPKIAWALENKPRIPVVWIHGL WP_001058307.1 hydrogenase [Salmonella enterica] Length: 372\nNumber of predicted TMHs: 2\nExp number of AAs in TMHs: 36.51738\nExp number, first 60 AAs: 19.72849\nTotal prob of N-in: 0.96170\nPOSSIBLE N-term signal sequence\ninside 1 19\nTMhelix 20 39\noutside 40 325\nTMhelix 326 348\ninside 349 372

16983 GCF_002038825.1_ASM203882v1 Salmonella enterica subsp. enterica serovar Enteritidis Proteobacteria; Gammaproteobacteria; Enterobacterales; Enterobacteriaceae; Salmonella; Salmonella enterica; Salmonella enterica subsp. enterica MTGDNTLITSHGINRRDFMKLCAALAATMGLSSKAAAEMAESVSNPQRPPVIWIGAQECT WP_000145429.1 hydrogenase 2 small subunit [Salmonella enterica] Length: 372\nNumber of predicted TMHs: 1\nExp number of AAs in TMHs: 24.51309\nExp number, first 60 AAs: 0.2682\nTotal prob of N-in: 0.14402\noutside 1 331\nTMhelix 332 354\ninside 355 372

16984 GCF_002037885.1_ASM203788v1 Salmonella enterica subsp. enterica serovar Enteritidis Proteobacteria; Gammaproteobacteria; Enterobacterales; Enterobacteriaceae; Salmonella; Salmonella enterica; Salmonella enterica subsp. enterica MTGDNTLITSHGINRRDFMKLCAALAATMGLSSKAAAEMAESVSNPQRPPVIWIGAQECT WP_000145429.1 hydrogenase 2 small subunit [Salmonella enterica] Length: 372\nNumber of predicted TMHs: 1\nExp number of AAs in TMHs: 24.51309\nExp number, first 60 AAs: 0.2682\nTotal prob of N-in: 0.14402\noutside 1 331\nTMhelix 332 354\ninside 355 372

16985 GCF_002037835.1_ASM203783v1 Salmonella enterica subsp. enterica serovar Enteritidis Proteobacteria; Gammaproteobacteria; Enterobacterales; Enterobacteriaceae; Salmonella; Salmonella enterica; Salmonella enterica subsp. enterica MTGDNTLITSHGINRRDFMKLCAALAATMGLSSKAAAEMAESVSNPQRPPVIWIGAQECT WP_000145429.1 hydrogenase 2 small subunit [Salmonella enterica] Length: 372\nNumber of predicted TMHs: 1\nExp number of AAs in TMHs: 24.51309\nExp number, first 60 AAs: 0.2682\nTotal prob of N-in: 0.14402\noutside 1 331\nTMhelix 332 354\ninside 355 372

16986 GCF_002035985.1_ASM203598v1 Salmonella enterica subsp. enterica serovar Enteritidis Proteobacteria; Gammaproteobacteria; Enterobacterales; Enterobacteriaceae; Salmonella; Salmonella enterica; Salmonella enterica subsp. enterica MNNEETFYQAMRRKGVTRRSFLKFCSLAATSLGLGAGMTPKIAWALENKPRIPVVWIHGL WP_001058312.1 hydrogenase [Salmonella enterica] Length: 372\nNumber of predicted TMHs: 2\nExp number of AAs in TMHs: 36.51738\nExp number, first 60 AAs: 19.72849\nTotal prob of N-in: 0.96170\nPOSSIBLE N-term signal sequence\ninside 1 19\nTMhelix 20 39\noutside 40 325\nTMhelix 326 348\ninside 349 372

16987 GCF_002035985.1_ASM203598v1 Salmonella enterica subsp. enterica serovar Enteritidis Proteobacteria; Gammaproteobacteria; Enterobacterales; Enterobacteriaceae; Salmonella; Salmonella enterica; Salmonella enterica subsp. enterica MTGDNTLITSHGINRRDFMKLCAALAATMGLSSKAAAEMAESVSNPQRPPVIWIGAQECT WP_000145429.1 hydrogenase 2 small subunit [Salmonella enterica] Length: 372\nNumber of predicted TMHs: 1\nExp number of AAs in TMHs: 24.51309\nExp number, first 60 AAs: 0.2682\nTotal prob of N-in: 0.14402\noutside 1 331\nTMhelix 332 354\ninside 355 372

16988 GCF_000974045.1_ABB07-SB3071.id12_v1.0 Salmonella enterica subsp. enterica serovar Enteritidis Proteobacteria; Gammaproteobacteria; Enterobacterales; Enterobacteriaceae; Salmonella; Salmonella enterica; Salmonella enterica subsp. enterica MTGDNTLITSHGINRRDFMKLCAALAATMGLSSKAAAEMAESVSNPQRPPVIWIGAQECT WP_000145429.1 hydrogenase 2 small subunit [Salmonella enterica] Length: 372\nNumber of predicted TMHs: 1\nExp number of AAs in TMHs: 24.51309\nExp number, first 60 AAs: 0.2682\nTotal prob of N-in: 0.14402\noutside 1 331\nTMhelix 332 354\ninside 355 372

16989 GCF_000974045.1_ABB07-SB3071.id12_v1.0 Salmonella enterica subsp. enterica serovar Enteritidis Proteobacteria; Gammaproteobacteria; Enterobacterales; Enterobacteriaceae; Salmonella; Salmonella enterica; Salmonella enterica subsp. enterica MNNEETFYQAMRRKGVTRRSFLKFCSLAATSLGLGAGMTPKIAWALENKPRIPVVWIHGL WP_001058307.1 hydrogenase [Salmonella enterica] Length: 372\nNumber of predicted TMHs: 2\nExp number of AAs in TMHs: 36.51738\nExp number, first 60 AAs: 19.72849\nTotal prob of N-in: 0.96170\nPOSSIBLE N-term signal sequence\ninside 1 19\nTMhelix 20 39\noutside 40 325\nTMhelix 326 348\ninside 349 372

16990 GCF_000973935.1_ABBSB1004-1.id11_v1.0 Salmonella enterica subsp. enterica serovar Enteritidis Proteobacteria; Gammaproteobacteria; Enterobacterales; Enterobacteriaceae; Salmonella; Salmonella enterica; Salmonella enterica subsp. enterica MNNEETFYQAMRRKGVTRRSFLKFCSLAATSLGLGAGMTPKIAWALENKPRIPVVWIHGL WP_001058307.1 hydrogenase [Salmonella enterica] Length: 372\nNumber of predicted TMHs: 2\nExp number of AAs in TMHs: 36.51738\nExp number, first 60 AAs: 19.72849\nTotal prob of N-in: 0.96170\nPOSSIBLE N-term signal sequence\ninside 1 19\nTMhelix 20 39\noutside 40 325\nTMhelix 326 348\ninside 349 372

16991 GCF_002042755.1_ASM204275v1 Salmonella enterica subsp. enterica serovar Enteritidis Proteobacteria; Gammaproteobacteria; Enterobacterales; Enterobacteriaceae; Salmonella; Salmonella enterica; Salmonella enterica subsp. enterica MTGDNTLITSHGINRRDFMKLCAALAATMGLSSKAAAEMAESVSNPQRPPVIWIGAQECT WP_000145429.1 hydrogenase 2 small subunit [Salmonella enterica] Length: 372\nNumber of predicted TMHs: 1\nExp number of AAs in TMHs: 24.51309\nExp number, first 60 AAs: 0.2682\nTotal prob of N-in: 0.14402\noutside 1 331\nTMhelix 332 354\ninside 355 372

16992 GCF_002042725.1_ASM204272v1 Salmonella enterica subsp. enterica serovar Enteritidis Proteobacteria; Gammaproteobacteria; Enterobacterales; Enterobacteriaceae; Salmonella; Salmonella enterica; Salmonella enterica subsp. enterica MTGDNTLITSHGINRRDFMKLCAALAATMGLSSKAAAEMAESVSNPQRPPVIWIGAQECT WP_000145429.1 hydrogenase 2 small subunit [Salmonella enterica] Length: 372\nNumber of

predicted TMHs: 1\nExp number of AAs in TMHs: 24.51309\nExp number, first 60 AAs: 0.2682\nTotal prob of N-in: 0.14402\noutside 1 331\nTMhelix 332 354\ninside 355 372

16993 GCF_002042595.1_ASM204259v1 Salmonella enterica subsp. enterica serovar Enteritidis Proteobacteria; Gammaproteobacteria; Enterobacterales; Enterobacteriaceae; Salmonella; Salmonella enterica; Salmonella enterica subsp. enterica MNNEETFYQAMRRKGVTRRSFLKFCSLAATSLGLGAGMTPKIAWALENKPRIPVVWIHGL WP_001058307.1 hydrogenase [Salmonella enterica] Length: 372\nNumber of predicted TMHs: 2\nExp number of AAs in TMHs: 36.51738\nExp number, first 60 AAs: 19.72849\nTotal prob of N-in: 0.96170\nPOSSIBLE N-term signal sequence\ninside 1 19\nTMhelix 20 39\noutside 40 325\nTMhelix 326 348\ninside 349 372

16994 GCF_002042565.1_ASM204256v1 Salmonella enterica subsp. enterica serovar Enteritidis Proteobacteria; Gammaproteobacteria; Enterobacterales; Enterobacteriaceae; Salmonella; Salmonella enterica; Salmonella enterica subsp. enterica MNNEETFYQAMRRKGVTRRSFLKFCSLAATSLGLGAGMTPKIAWALENKPRIPVVWIHGL WP_001058307.1 hydrogenase [Salmonella enterica] Length: 372\nNumber of predicted TMHs: 2\nExp number of AAs in TMHs: 36.51738\nExp number, first 60 AAs: 19.72849\nTotal prob of N-in: 0.96170\nPOSSIBLE N-term signal sequence\ninside 1 19\nTMhelix 20 39\noutside 40 325\nTMhelix 326 348\ninside 349 372

16995 GCF_002042675.1_ASM204267v1 Salmonella enterica subsp. enterica serovar Enteritidis Proteobacteria; Gammaproteobacteria; Enterobacterales; Enterobacteriaceae; Salmonella; Salmonella enterica; Salmonella enterica subsp. enterica MNNEETFYQAMRRKGVTRRSFLKFCSLAATSLGLGAGMTPKIAWALENKPRIPVVWIHGL WP_001058307.1 hydrogenase [Salmonella enterica] Length: 372\nNumber of predicted TMHs: 2\nExp number of AAs in TMHs: 36.51738\nExp number, first 60 AAs: 19.72849\nTotal prob of N-in: 0.96170\nPOSSIBLE N-term signal sequence\ninside 1 19\nTMhelix 20 39\noutside 40 325\nTMhelix 326 348\ninside 349 372

16996 GCF_002042635.1_ASM204263v1 Salmonella enterica subsp. enterica serovar Enteritidis Proteobacteria; Gammaproteobacteria; Enterobacterales; Enterobacteriaceae; Salmonella; Salmonella enterica; Salmonella enterica subsp. enterica MNNEETFYQAMRRKGVTRRSFLKFCSLAATSLGLGAGMTPKIAWALENKPRIPVVWIHGL WP_001058307.1 hydrogenase [Salmonella enterica] Length: 372\nNumber of predicted TMHs: 2\nExp number of AAs in TMHs: 36.51738\nExp number, first 60 AAs: 19.72849\nTotal prob of N-in: 0.96170\nPOSSIBLE N-term signal sequence\ninside 1 19\nTMhelix 20 39\noutside 40 325\nTMhelix 326 348\ninside 349 372

16997 GCF_002042325.1_ASM204232v1 Salmonella enterica subsp. enterica serovar Enteritidis Proteobacteria; Gammaproteobacteria; Enterobacterales; Enterobacteriaceae; Salmonella; Salmonella enterica; Salmonella enterica subsp. enterica MNNEETFYQAMRRKGVTRRSFLKFCSLAATSLGLGAGMTPKIAWALENKPRIPVVWIHGL WP_001058307.1 hydrogenase [Salmonella enterica] Length: 372\nNumber of predicted TMHs: 2\nExp number of AAs in TMHs: 36.51738\nExp number, first 60 AAs: 19.72849\nTotal prob of N-in: 0.96170\nPOSSIBLE N-term signal sequence\ninside 1 19\nTMhelix 20 39\noutside 40 325\nTMhelix 326 348\ninside 349 372

16998 GCF_002042245.1_ASM204224v1 Salmonella enterica subsp. enterica serovar Enteritidis Proteobacteria; Gammaproteobacteria; Enterobacterales; Enterobacteriaceae; Salmonella; Salmonella enterica; Salmonella enterica subsp. enterica MNNEETFYQAMRRKGVTRRSFLKFCSLAATSLGLGAGMTPKIAWALENKPRIPVVWIHGL WP_001058307.1 hydrogenase [Salmonella enterica] Length: 372\nNumber of predicted TMHs: 2\nExp number of AAs in TMHs: 36.51738\nExp number, first 60 AAs: 19.72849\nTotal prob of N-in: 0.96170\nPOSSIBLE N-term signal sequence\ninside 1 19\nTMhelix 20 39\noutside 40 325\nTMhelix 326 348\ninside 349 372

16999 GCF_002042325.1_ASM204232v1 Salmonella enterica subsp. enterica serovar Enteritidis Proteobacteria; Gammaproteobacteria; Enterobacterales; Enterobacteriaceae; Salmonella; Salmonella enterica; Salmonella enterica subsp. enterica MTGDNTLITSHGINRRDFMKLCAALAATMGLSSKAAAEMAESVSNPQRPPVIWIGAQECT WP_000145429.1 hydrogenase 2 small subunit [Salmonella enterica] Length: 372\nNumber of predicted TMHs: 1\nExp number of AAs in TMHs: 24.51309\nExp number, first 60 AAs: 0.2682\nTotal prob of N-in: 0.14402\noutside 1 331\nTMhelix 332 354\ninside 355 372

17000 GCF_002042245.1_ASM204224v1 Salmonella enterica subsp. enterica serovar Enteritidis Proteobacteria; Gammaproteobacteria; Enterobacterales; Enterobacteriaceae; Salmonella; Salmonella enterica; Salmonella enterica subsp. enterica MTGDNTLITSHGINRRDFMKLCAALAATMGLSSKAAAEMAESVSNPQRPPVIWIGAQECT WP_000145429.1 hydrogenase 2 small subunit [Salmonella enterica] Length: 372\nNumber of predicted TMHs: 1\nExp number of AAs in TMHs: 24.51309\nExp number, first 60 AAs: 0.2682\nTotal prob of N-in: 0.14402\noutside 1 331\nTMhelix 332 354\ninside 355 372

17001 GCF_002042305.1_ASM204230v1 Salmonella enterica subsp. enterica serovar Enteritidis Proteobacteria; Gammaproteobacteria; Enterobacterales; Enterobacteriaceae; Salmonella; Salmonella enterica;

Salmonella enterica subsp. enterica MTGDNTLITSHGINRRDFMKLCAALAATMGLSSKAAAEMAESVSNPQRPPVIWIGAQECT WP_000145429.1 hydrogenase 2 small subunit [Salmonella enterica] Length: 372\nNumber of predicted TMHs: 1\nExp number of AAs in TMHs: 24.51309\nExp number, first 60 AAs: 0.2682\nTotal prob of N-in: 0.14402\noutside 1 331\nTMhelix 332 354\ninside 355 372

17002 GCF_002042145.1_ASM204214v1 Salmonella enterica subsp. enterica serovar Enteritidis Proteobacteria; Gammaproteobacteria; Enterobacterales; Enterobacteriaceae; Salmonella; Salmonella enterica; Salmonella enterica subsp. enterica MNNEETFYQAMRRKGVTRRSFLKFCSLAATSLGLGAGMTPKIAWALENKPRIPVVWIHGL WP_001058307.1 hydrogenase [Salmonella enterica] Length: 372\nNumber of predicted TMHs: 2\nExp number of AAs in TMHs: 36.51738\nExp number, first 60 AAs: 19.72849\nTotal prob of N-in: 0.96170\nPOSSIBLE N-term signal sequence\ninside 1 19\nTMhelix 20 39\noutside 40 325\nTMhelix 326 348\ninside 349 372

17003 GCF_002042105.1_ASM204210v1 Salmonella enterica subsp. enterica serovar Enteritidis Proteobacteria; Gammaproteobacteria; Enterobacterales; Enterobacteriaceae; Salmonella; Salmonella enterica; Salmonella enterica subsp. enterica MNNEETFYQAMRRKGVTRRSFLKFCSLAATSLGLGAGMTPKIAWALENKPRIPVVWIHGL WP_001058307.1 hydrogenase [Salmonella enterica] Length: 372\nNumber of predicted TMHs: 2\nExp number of AAs in TMHs: 36.51738\nExp number, first 60 AAs: 19.72849\nTotal prob of N-in: 0.96170\nPOSSIBLE N-term signal sequence\ninside 1 19\nTMhelix 20 39\noutside 40 325\nTMhelix 326 348\ninside 349 372

17004 GCF_002041955.1_ASM204195v1 Salmonella enterica subsp. enterica serovar Enteritidis Proteobacteria; Gammaproteobacteria; Enterobacterales; Enterobacteriaceae; Salmonella; Salmonella enterica; Salmonella enterica subsp. enterica MTGDNTLITSHGINRRDFMKLCAALAATMGLSSKAAAEMAESVSNPQRPPVIWIGAQECT WP_000145429.1 hydrogenase 2 small subunit [Salmonella enterica] Length: 372\nNumber of predicted TMHs: 1\nExp number of AAs in TMHs: 24.51309\nExp number, first 60 AAs: 0.2682\nTotal prob of N-in: 0.14402\noutside 1 331\nTMhelix 332 354\ninside 355 372

17005 GCF_002041935.1_ASM204193v1 Salmonella enterica subsp. enterica serovar Enteritidis Proteobacteria; Gammaproteobacteria; Enterobacterales; Enterobacteriaceae; Salmonella; Salmonella enterica; Salmonella enterica subsp. enterica MTGDNTLITSHGINRRDFMKLCAALAATMGLSSKAAAEMAESVSNPQRPPVIWIGAQECT WP_000145429.1 hydrogenase 2 small subunit [Salmonella enterica] Length: 372\nNumber of predicted TMHs: 1\nExp number of AAs in TMHs: 24.51309\nExp number, first 60 AAs: 0.2682\nTotal prob of N-in: 0.14402\noutside 1 331\nTMhelix 332 354\ninside 355 372

17006 GCF_002041905.1_ASM204190v1 Salmonella enterica subsp. enterica serovar Enteritidis Proteobacteria; Gammaproteobacteria; Enterobacterales; Enterobacteriaceae; Salmonella; Salmonella enterica; Salmonella enterica subsp. enterica MTGDNTLITSHGINRRDFMKLCAALAATMGLSSKAAAEMAESVSNPQRPPVIWIGAQECT WP_000145429.1 hydrogenase 2 small subunit [Salmonella enterica] Length: 372\nNumber of predicted TMHs: 1\nExp number of AAs in TMHs: 24.51309\nExp number, first 60 AAs: 0.2682\nTotal prob of N-in: 0.14402\noutside 1 331\nTMhelix 332 354\ninside 355 372

17007 GCF_002041845.1_ASM204184v1 Salmonella enterica subsp. enterica serovar Enteritidis Proteobacteria; Gammaproteobacteria; Enterobacterales; Enterobacteriaceae; Salmonella; Salmonella enterica; Salmonella enterica subsp. enterica MTGDNTLITSHGINRRDFMKLCAALAATMGLSSKAAAEMAESVSNPQRPPVIWIGAQECT WP_000145429.1 hydrogenase 2 small subunit [Salmonella enterica] Length: 372\nNumber of predicted TMHs: 1\nExp number of AAs in TMHs: 24.51309\nExp number, first 60 AAs: 0.2682\nTotal prob of N-in: 0.14402\noutside 1 331\nTMhelix 332 354\ninside 355 372

17008 GCF_002041825.1_ASM204182v1 Salmonella enterica subsp. enterica serovar Enteritidis Proteobacteria; Gammaproteobacteria; Enterobacterales; Enterobacteriaceae; Salmonella; Salmonella enterica; Salmonella enterica subsp. enterica MTGDNTLITSHGINRRDFMKLCAALAATMGLSSKAAAEMAESVSNPQRPPVIWIGAQECT WP_000145429.1 hydrogenase 2 small subunit [Salmonella enterica] Length: 372\nNumber of predicted TMHs: 1\nExp number of AAs in TMHs: 24.51309\nExp number, first 60 AAs: 0.2682\nTotal prob of N-in: 0.14402\noutside 1 331\nTMhelix 332 354\ninside 355 372

17009 GCF_002041685.1_ASM204168v1 Salmonella enterica subsp. enterica serovar Enteritidis Proteobacteria; Gammaproteobacteria; Enterobacterales; Enterobacteriaceae; Salmonella; Salmonella enterica; Salmonella enterica subsp. enterica MNNEETFYQAMRRKGVTRRSFLKFCSLAATSLGLGAGMTPKIAWALENKPRIPVVWIHGL WP_001058307.1 hydrogenase [Salmonella enterica] Length: 372\nNumber of predicted TMHs: 2\nExp number of AAs in TMHs: 36.51738\nExp number, first 60 AAs: 19.72849\nTotal prob of N-in: 0.96170\nPOSSIBLE N-term signal sequence\ninside 1 19\nTMhelix 20 39\noutside 40 325\nTMhelix 326 348\ninside 349 372

17010 GCF_002041705.1_ASM204170v1 Salmonella enterica subsp. enterica serovar Enteritidis Proteobacteria; Gammaproteobacteria; Enterobacterales; Enterobacteriaceae; Salmonella; Salmonella enterica; Salmonella enterica subsp. enterica MNNEETFYQAMRRKGVTRRSFLKFCSLAATSLGLGAGMTPKIAWALENKPRIPVVWIHGL WP_001058307.1 hydrogenase [Salmonella enterica] Length: 372\nNumber of predicted TMHs: 2\nExp number of AAs in TMHs: 36.51738\nExp number, first 60 AAs: 19.72849\nTotal prob of N-in: 0.96170\nPOSSIBLE N-term signal sequence\ninside 1 19\nTMhelix 20 39\noutside 40 325\nTMhelix 326 348\ninside 349 372

17011 GCF_002042635.1_ASM204263v1 Salmonella enterica subsp. enterica serovar Enteritidis Proteobacteria; Gammaproteobacteria; Enterobacterales; Enterobacteriaceae; Salmonella; Salmonella enterica; Salmonella enterica subsp. enterica MTGDNTLITSHGINRRDFMKLCAALAATMGLSSKAAAEMAESVSNPQRPPVIWIGAQECT WP_000145429.1 hydrogenase 2 small subunit [Salmonella enterica] Length: 372\nNumber of predicted TMHs: 1\nExp number of AAs in TMHs: 24.51309\nExp number, first 60 AAs: 0.2682\nTotal prob of N-in: 0.14402\noutside 1 331\nTMhelix 332 354\ninside 355 372

17012 GCF_002041465.1_ASM204146v1 Salmonella enterica subsp. enterica serovar Enteritidis Proteobacteria; Gammaproteobacteria; Enterobacterales; Enterobacteriaceae; Salmonella; Salmonella enterica; Salmonella enterica subsp. enterica MNNEETFYQAMRRKGVTRRSFLKFCSLAATSLGLGAGMTPKIAWALENKPRIPVVWIHGL WP_001058307.1 hydrogenase [Salmonella enterica] Length: 372\nNumber of predicted TMHs: 2\nExp number of AAs in TMHs: 36.51738\nExp number, first 60 AAs: 19.72849\nTotal prob of N-in: 0.96170\nPOSSIBLE N-term signal sequence\ninside 1 19\nTMhelix 20 39\noutside 40 325\nTMhelix 326 348\ninside 349 372

17013 GCF_002041445.1_ASM204144v1 Salmonella enterica subsp. enterica serovar Enteritidis Proteobacteria; Gammaproteobacteria; Enterobacterales; Enterobacteriaceae; Salmonella; Salmonella enterica; Salmonella enterica subsp. enterica MNNEETFYQAMRRKGVTRRSFLKFCSLAATSLGLGAGMTPKIAWALENKPRIPVVWIHGL WP_001058307.1 hydrogenase [Salmonella enterica] Length: 372\nNumber of predicted TMHs: 2\nExp number of AAs in TMHs: 36.51738\nExp number, first 60 AAs: 19.72849\nTotal prob of N-in: 0.96170\nPOSSIBLE N-term signal sequence\ninside 1 19\nTMhelix 20 39\noutside 40 325\nTMhelix 326 348\ninside 349 372

17014 GCF_002041465.1_ASM204146v1 Salmonella enterica subsp. enterica serovar Enteritidis Proteobacteria; Gammaproteobacteria; Enterobacterales; Enterobacteriaceae; Salmonella; Salmonella enterica; Salmonella enterica subsp. enterica MTGDNTLITSHGINRRDFMKLCAALAATMGLSSKAAAEMAESVSNPQRPPVIWIGAQECT WP_000145429.1 hydrogenase 2 small subunit [Salmonella enterica] Length: 372\nNumber of predicted TMHs: 1\nExp number of AAs in TMHs: 24.51309\nExp number, first 60 AAs: 0.2682\nTotal prob of N-in: 0.14402\noutside 1 331\nTMhelix 332 354\ninside 355 372

17015 GCF_002041445.1_ASM204144v1 Salmonella enterica subsp. enterica serovar Enteritidis Proteobacteria; Gammaproteobacteria; Enterobacterales; Enterobacteriaceae; Salmonella; Salmonella enterica; Salmonella enterica subsp. enterica MTGDNTLITSHGINRRDFMKLCAALAATMGLSSKAAAEMAESVSNPQRPPVIWIGAQECT WP_000145429.1 hydrogenase 2 small subunit [Salmonella enterica] Length: 372\nNumber of predicted TMHs: 1\nExp number of AAs in TMHs: 24.51309\nExp number, first 60 AAs: 0.2682\nTotal prob of N-in: 0.14402\noutside 1 331\nTMhelix 332 354\ninside 355 372

17016 GCF_002041275.1_ASM204127v1 Salmonella enterica subsp. enterica serovar Enteritidis Proteobacteria; Gammaproteobacteria; Enterobacterales; Enterobacteriaceae; Salmonella; Salmonella enterica; Salmonella enterica subsp. enterica MNNEETFYQAMRRKGVTRRSFLKFCSLAATSLGLGAGMTPKIAWALENKPRIPVVWIHGL WP_001058307.1 hydrogenase [Salmonella enterica] Length: 372\nNumber of predicted TMHs: 2\nExp number of AAs in TMHs: 36.51738\nExp number, first 60 AAs: 19.72849\nTotal prob of N-in: 0.96170\nPOSSIBLE N-term signal sequence\ninside 1 19\nTMhelix 20 39\noutside 40 325\nTMhelix 326 348\ninside 349 372

17017 GCF_002041385.1_ASM204138v1 Salmonella enterica subsp. enterica serovar Enteritidis Proteobacteria; Gammaproteobacteria; Enterobacterales; Enterobacteriaceae; Salmonella; Salmonella enterica; Salmonella enterica subsp. enterica MNNEETFYQAMRRKGVTRRSFLKFCSLAATSLGLGAGMTPKIAWALENKPRIPVVWIHGL WP_001058307.1 hydrogenase [Salmonella enterica] Length: 372\nNumber of predicted TMHs: 2\nExp number of AAs in TMHs: 36.51738\nExp number, first 60 AAs: 19.72849\nTotal prob of N-in: 0.96170\nPOSSIBLE N-term signal sequence\ninside 1 19\nTMhelix 20 39\noutside 40 325\nTMhelix 326 348\ninside 349 372

17018 GCF_001714075.1_ASM171407v1 Salmonella enterica subsp. enterica serovar Enteritidis Proteobacteria; Gammaproteobacteria; Enterobacterales; Enterobacteriaceae; Salmonella; Salmonella enterica; Salmonella enterica subsp. enterica MNNEETFYQAMRRKGVTRRSFLKFCSLAATSLGLGAGMTPKIAWALENKPRIPVVWIHGL WP_001058307.1 hydrogenase [Salmonella enterica] Length: 372\nNumber of predicted TMHs: 2\nExp

number of AAs in TMHs: 36.51738\nExp number, first 60 AAs: 19.72849\nTotal prob of N-in: 0.96170\nPOSSIBLE N-term signal sequence\ninside 1 19\nTMhelix 20 39\noutside 40 325\nTMhelix 326 348\ninside 349 372

17019 GCF_001714025.1_ASM171402v1 Salmonella enterica subsp. enterica serovar Enteritidis Proteobacteria; Gammaproteobacteria; Enterobacterales; Enterobacteriaceae; Salmonella; Salmonella enterica; Salmonella enterica subsp. enterica MNNEETFYQAMRRKGVTRRSFLKFCSLAATSLGLGAGMTPKIAWALENKPRIPVVWIHGL WP_001058307.1 hydrogenase [Salmonella enterica] Length: 372\nNumber of predicted TMHs: 2\nExp number of AAs in TMHs: 36.51738\nExp number, first 60 AAs: 19.72849\nTotal prob of N-in: 0.96170\nPOSSIBLE N-term signal sequence\ninside 1 19\nTMhelix 20 39\noutside 40 325\nTMhelix 326 348\ninside 349 372

17020 GCF_002041065.1_ASM204106v1 Salmonella enterica subsp. enterica serovar Enteritidis Proteobacteria; Gammaproteobacteria; Enterobacterales; Enterobacteriaceae; Salmonella; Salmonella enterica; Salmonella enterica subsp. enterica MNNEETFYQAMRRKGVTRRSFLKFCSLAATSLGLGAGMTPKIAWALENKPRIPVVWIHGL WP_001058307.1 hydrogenase [Salmonella enterica] Length: 372\nNumber of predicted TMHs: 2\nExp number of AAs in TMHs: 36.51738\nExp number, first 60 AAs: 19.72849\nTotal prob of N-in: 0.96170\nPOSSIBLE N-term signal sequence\ninside 1 19\nTMhelix 20 39\noutside 40 325\nTMhelix 326 348\ninside 349 372

17021 GCF_001713755.1_ASM171375v1 Salmonella enterica subsp. enterica serovar Enteritidis Proteobacteria; Gammaproteobacteria; Enterobacterales; Enterobacteriaceae; Salmonella; Salmonella enterica; Salmonella enterica subsp. enterica MTGDNTLITSHGINRRDFMKLCAALAATMGLSSKAAAEMAESVSNPQRPPVIWIGAQECT WP_000145429.1 hydrogenase 2 small subunit [Salmonella enterica] Length: 372\nNumber of predicted TMHs: 1\nExp number of AAs in TMHs: 24.51309\nExp number, first 60 AAs: 0.2682\nTotal prob of N-in: 0.14402\noutside 1 331\nTMhelix 332 354\ninside 355 372

17022 GCF_002040925.1_ASM204092v1 Salmonella enterica subsp. enterica serovar Enteritidis Proteobacteria; Gammaproteobacteria; Enterobacterales; Enterobacteriaceae; Salmonella; Salmonella enterica; Salmonella enterica subsp. enterica MNNEETFYQAMRRKGVTRRSFLKFCSLAATSLGLGAGMTPKIAWALENKPRIPVVWIHGL WP_001058307.1 hydrogenase [Salmonella enterica] Length: 372\nNumber of predicted TMHs: 2\nExp number of AAs in TMHs: 36.51738\nExp number, first 60 AAs: 19.72849\nTotal prob of N-in: 0.96170\nPOSSIBLE N-term signal sequence\ninside 1 19\nTMhelix 20 39\noutside 40 325\nTMhelix 326 348\ninside 349 372

17023 GCF_002041145.1_ASM204114v1 Salmonella enterica subsp. enterica serovar Enteritidis Proteobacteria; Gammaproteobacteria; Enterobacterales; Enterobacteriaceae; Salmonella; Salmonella enterica; Salmonella enterica subsp. enterica MNNEETFYQAMRRKGVTRRSFLKFCSLAATSLGLGAGMTPKIAWALENKPRIPVVWIHGL WP_001058307.1 hydrogenase [Salmonella enterica] Length: 372\nNumber of predicted TMHs: 2\nExp number of AAs in TMHs: 36.51738\nExp number, first 60 AAs: 19.72849\nTotal prob of N-in: 0.96170\nPOSSIBLE N-term signal sequence\ninside 1 19\nTMhelix 20 39\noutside 40 325\nTMhelix 326 348\ninside 349 372

17024 GCF_001691655.1_ASM169165v1 Salmonella enterica subsp. enterica serovar Enteritidis Proteobacteria; Gammaproteobacteria; Enterobacterales; Enterobacteriaceae; Salmonella; Salmonella enterica; Salmonella enterica subsp. enterica MNNEETFYQAMRRKGVTRRSFLKFCSLAATSLGLGAGMTPKIAWALENKPRIPVVWIHGL WP_001058307.1 hydrogenase [Salmonella enterica] Length: 372\nNumber of predicted TMHs: 2\nExp number of AAs in TMHs: 36.51738\nExp number, first 60 AAs: 19.72849\nTotal prob of N-in: 0.96170\nPOSSIBLE N-term signal sequence\ninside 1 19\nTMhelix 20 39\noutside 40 325\nTMhelix 326 348\ninside 349 372

17025 GCF_001691675.1_ASM169167v1 Salmonella enterica subsp. enterica serovar Enteritidis Proteobacteria; Gammaproteobacteria; Enterobacterales; Enterobacteriaceae; Salmonella; Salmonella enterica; Salmonella enterica subsp. enterica MNNEETFYQAMRRKGVTRRSFLKFCSLAATSLGLGAGMTPKIAWALENKPRIPVVWIHGL WP_001058307.1 hydrogenase [Salmonella enterica] Length: 372\nNumber of predicted TMHs: 2\nExp number of AAs in TMHs: 36.51738\nExp number, first 60 AAs: 19.72849\nTotal prob of N-in: 0.96170\nPOSSIBLE N-term signal sequence\ninside 1 19\nTMhelix 20 39\noutside 40 325\nTMhelix 326 348\ninside 349 372

17026 GCF_001691655.1_ASM169165v1 Salmonella enterica subsp. enterica serovar Enteritidis Proteobacteria; Gammaproteobacteria; Enterobacterales; Enterobacteriaceae; Salmonella; Salmonella enterica; Salmonella enterica subsp. enterica MTGDNTLITSHGINRRDFMKLCAALAATMGLSSKAAAEMAESVSNPQRPPVIWIGAQECT WP_000145429.1 hydrogenase 2 small subunit [Salmonella enterica] Length: 372\nNumber of predicted TMHs: 1\nExp number of AAs in TMHs: 24.51309\nExp number, first 60 AAs: 0.2682\nTotal prob of N-in: 0.14402\noutside 1 331\nTMhelix 332 354\ninside 355 372

17027 GCF_001691675.1_ASM169167v1 Salmonella enterica subsp. enterica serovar Enteritidis Proteobacteria; Gammaproteobacteria; Enterobacterales; Enterobacteriaceae; Salmonella; Salmonella enterica;

Salmonella enterica subsp. enterica	MTGDNTLITSHGINRRDFMKLCAALAATMGLSSKAAAEMAESVSNPQRPPVIWIGAQECT	WP_000145429.1	hydrogenase 2 small subunit [Salmonella enterica]	Length: 372\nNumber of predicted TMHs:  1\nExp number of AAs in TMHs: 24.51309\nExp number, first 60 AAs:  0.2682\nTotal prob of N-in:  0.14402\noutside   1   331\nTMhelix   332   354\ninside   355   372

17028	GCF_002041145.1_ASM204114v1	Salmonella enterica subsp. enterica serovar Enteritidis	Proteobacteria; Gammaproteobacteria; Enterobacterales; Enterobacteriaceae; Salmonella; Salmonella enterica; Salmonella enterica subsp. enterica	MTGDNTLITSHGINRRDFMKLCAALAATMGLSSKAAAEMAESVSNPQRPPVIWIGAQECT	WP_000145429.1	hydrogenase 2 small subunit [Salmonella enterica]	Length: 372\nNumber of predicted TMHs:  1\nExp number of AAs in TMHs: 24.51309\nExp number, first 60 AAs:  0.2682\nTotal prob of N-in:  0.14402\noutside   1   331\nTMhelix   332   354\ninside   355   372

17029	GCF_002040905.1_ASM204090v1	Salmonella enterica subsp. enterica serovar Enteritidis	Proteobacteria; Gammaproteobacteria; Enterobacterales; Enterobacteriaceae; Salmonella; Salmonella enterica; Salmonella enterica subsp. enterica	MNNEETFYQAMRRKGVTRRSFLKFCSLAATSLGLGAGMTPKIAWALENKPRIPVVWIHGL	WP_001058307.1	hydrogenase [Salmonella enterica]	Length: 372\nNumber of predicted TMHs:  2\nExp number of AAs in TMHs: 36.51738\nExp number, first 60 AAs:  19.72849\nTotal prob of N-in:   0.96170\nPOSSIBLE N-term signal sequence\ninside   1   19\nTMhelix   20   39\noutside   40   325\nTMhelix   326   348\ninside   349   372

17030	GCF_002041025.1_ASM204102v1	Salmonella enterica subsp. enterica serovar Enteritidis	Proteobacteria; Gammaproteobacteria; Enterobacterales; Enterobacteriaceae; Salmonella; Salmonella enterica; Salmonella enterica subsp. enterica	MNNEETFYQAMRRKGVTRRSFLKFCSLAATSLGLGAGMTPKIAWALENKPRIPVVWIHGL	WP_001058307.1	hydrogenase [Salmonella enterica]	Length: 372\nNumber of predicted TMHs:  2\nExp number of AAs in TMHs: 36.51738\nExp number, first 60 AAs:  19.72849\nTotal prob of N-in:   0.96170\nPOSSIBLE N-term signal sequence\ninside   1   19\nTMhelix   20   39\noutside   40   325\nTMhelix   326   348\ninside   349   372

17031	GCF_002040925.1_ASM204092v1	Salmonella enterica subsp. enterica serovar Enteritidis	Proteobacteria; Gammaproteobacteria; Enterobacterales; Enterobacteriaceae; Salmonella; Salmonella enterica; Salmonella enterica subsp. enterica	MTGDNTLITSHGINRRDFMKLCAALAATMGLSSKAAAEMAESVSNPQRPPVIWIGAQECT	WP_000145429.1	hydrogenase 2 small subunit [Salmonella enterica]	Length: 372\nNumber of predicted TMHs:  1\nExp number of AAs in TMHs: 24.51309\nExp number, first 60 AAs:  0.2682\nTotal prob of N-in:  0.14402\noutside   1   331\nTMhelix   332   354\ninside   355   372

17032	GCF_002040905.1_ASM204090v1	Salmonella enterica subsp. enterica serovar Enteritidis	Proteobacteria; Gammaproteobacteria; Enterobacterales; Enterobacteriaceae; Salmonella; Salmonella enterica; Salmonella enterica subsp. enterica	MTGDNTLITSHGINRRDFMKLCAALAATMGLSSKAAAEMAESVSNPQRPPVIWIGAQECT	WP_000145429.1	hydrogenase 2 small subunit [Salmonella enterica]	Length: 372\nNumber of predicted TMHs:  1\nExp number of AAs in TMHs: 24.51309\nExp number, first 60 AAs:  0.2682\nTotal prob of N-in:  0.14402\noutside   1   331\nTMhelix   332   354\ninside   355   372

17033	GCF_002040785.1_ASM204078v1	Salmonella enterica subsp. enterica serovar Enteritidis	Proteobacteria; Gammaproteobacteria; Enterobacterales; Enterobacteriaceae; Salmonella; Salmonella enterica; Salmonella enterica subsp. enterica	MTGDNTLITSHGINRRDFMKLCAALAATMGLSSKAAAEMAESVSNPQRPPVIWIGAQECT	WP_000145429.1	hydrogenase 2 small subunit [Salmonella enterica]	Length: 372\nNumber of predicted TMHs:  1\nExp number of AAs in TMHs: 24.51309\nExp number, first 60 AAs:  0.2682\nTotal prob of N-in:  0.14402\noutside   1   331\nTMhelix   332   354\ninside   355   372

17034	GCF_002040825.1_ASM204082v1	Salmonella enterica subsp. enterica serovar Enteritidis	Proteobacteria; Gammaproteobacteria; Enterobacterales; Enterobacteriaceae; Salmonella; Salmonella enterica; Salmonella enterica subsp. enterica	MTGDNTLITSHGINRRDFMKLCAALAATMGLSSKAAAEMAESVSNPQRPPVIWIGAQECT	WP_000145429.1	hydrogenase 2 small subunit [Salmonella enterica]	Length: 372\nNumber of predicted TMHs:  1\nExp number of AAs in TMHs: 24.51309\nExp number, first 60 AAs:  0.2682\nTotal prob of N-in:  0.14402\noutside   1   331\nTMhelix   332   354\ninside   355   372

17035	GCF_002040615.1_ASM204061v1	Salmonella enterica subsp. enterica serovar Enteritidis	Proteobacteria; Gammaproteobacteria; Enterobacterales; Enterobacteriaceae; Salmonella; Salmonella enterica; Salmonella enterica subsp. enterica	MNNEETFYQAMRRKGVTRRSFLKFCSLAATSLGLGAGMTPKIAWALENKPRIPVVWIHGL	WP_001058307.1	hydrogenase [Salmonella enterica]	Length: 372\nNumber of predicted TMHs:  2\nExp number of AAs in TMHs: 36.51738\nExp number, first 60 AAs:  19.72849\nTotal prob of N-in:   0.96170\nPOSSIBLE N-term signal sequence\ninside   1   19\nTMhelix   20   39\noutside   40   325\nTMhelix   326   348\ninside   349   372

17036 GCF_002040325.1_ASM204032v1 Salmonella enterica subsp. enterica serovar Enteritidis Proteobacteria; Gammaproteobacteria; Enterobacterales; Enterobacteriaceae; Salmonella; Salmonella enterica; Salmonella enterica subsp. enterica MNNEETFYQAMRRKGVTRRSFLKFCSLAATSLGLGAGMTPKIAWALENKPRIPVVWIHGL WP_001058307.1 hydrogenase [Salmonella enterica] Length: 372\nNumber of predicted TMHs: 2\nExp number of AAs in TMHs: 36.51738\nExp number, first 60 AAs: 19.72849\nTotal prob of N-in: 0.96170\nPOSSIBLE N-term signal sequence\ninside 1 19\nTMhelix 20 39\noutside 40 325\nTMhelix 326 348\ninside 349 372

17037 GCF_002040465.1_ASM204046v1 Salmonella enterica subsp. enterica serovar Enteritidis Proteobacteria; Gammaproteobacteria; Enterobacterales; Enterobacteriaceae; Salmonella; Salmonella enterica; Salmonella enterica subsp. enterica MNNEETFYQAMRRKGVTRRSFLKFCSLAATSLGLGAGMTPKIAWALENKPRIPVVWIHGL WP_001058307.1 hydrogenase [Salmonella enterica] Length: 372\nNumber of predicted TMHs: 2\nExp number of AAs in TMHs: 36.51738\nExp number, first 60 AAs: 19.72849\nTotal prob of N-in: 0.96170\nPOSSIBLE N-term signal sequence\ninside 1 19\nTMhelix 20 39\noutside 40 325\nTMhelix 326 348\ninside 349 372

17038 GCF_002040465.1_ASM204046v1 Salmonella enterica subsp. enterica serovar Enteritidis Proteobacteria; Gammaproteobacteria; Enterobacterales; Enterobacteriaceae; Salmonella; Salmonella enterica; Salmonella enterica subsp. enterica MTGDNTLITSHGINRRDFMKLCAALAATMGLSSKAAAEMAESVSNPQRPPVIWIGAQECT WP_000145429.1 hydrogenase 2 small subunit [Salmonella enterica] Length: 372\nNumber of predicted TMHs: 1\nExp number of AAs in TMHs: 24.51309\nExp number, first 60 AAs: 0.2682\nTotal prob of N-in: 0.14402\noutside 1 331\nTMhelix 332 354\ninside 355 372

17039 GCF_002040305.1_ASM204030v1 Salmonella enterica subsp. enterica serovar Enteritidis Proteobacteria; Gammaproteobacteria; Enterobacterales; Enterobacteriaceae; Salmonella; Salmonella enterica; Salmonella enterica subsp. enterica MNNEETFYQAMRRKGVTRRSFLKFCSLAATSLGLGAGMTPKIAWALENKPRIPVVWIHGL WP_001058307.1 hydrogenase [Salmonella enterica] Length: 372\nNumber of predicted TMHs: 2\nExp number of AAs in TMHs: 36.51738\nExp number, first 60 AAs: 19.72849\nTotal prob of N-in: 0.96170\nPOSSIBLE N-term signal sequence\ninside 1 19\nTMhelix 20 39\noutside 40 325\nTMhelix 326 348\ninside 349 372

17040 GCF_002040405.1_ASM204040v1 Salmonella enterica subsp. enterica serovar Enteritidis Proteobacteria; Gammaproteobacteria; Enterobacterales; Enterobacteriaceae; Salmonella; Salmonella enterica; Salmonella enterica subsp. enterica MNNEETFYQAMRRKGVTRRSFLKFCSLAATSLGLGAGMTPKIAWALENKPRIPVVWIHGL WP_001058307.1 hydrogenase [Salmonella enterica] Length: 372\nNumber of predicted TMHs: 2\nExp number of AAs in TMHs: 36.51738\nExp number, first 60 AAs: 19.72849\nTotal prob of N-in: 0.96170\nPOSSIBLE N-term signal sequence\ninside 1 19\nTMhelix 20 39\noutside 40 325\nTMhelix 326 348\ninside 349 372

17041 GCF_002040385.1_ASM204038v1 Salmonella enterica subsp. enterica serovar Enteritidis Proteobacteria; Gammaproteobacteria; Enterobacterales; Enterobacteriaceae; Salmonella; Salmonella enterica; Salmonella enterica subsp. enterica MNNEETFYQAMRRKGVTRRSFLKFCSLAATSLGLGAGMTPKIAWALENKPRIPVVWIHGL WP_001058307.1 hydrogenase [Salmonella enterica] Length: 372\nNumber of predicted TMHs: 2\nExp number of AAs in TMHs: 36.51738\nExp number, first 60 AAs: 19.72849\nTotal prob of N-in: 0.96170\nPOSSIBLE N-term signal sequence\ninside 1 19\nTMhelix 20 39\noutside 40 325\nTMhelix 326 348\ninside 349 372

17042 GCF_002040135.1_ASM204013v1 Salmonella enterica subsp. enterica serovar Enteritidis Proteobacteria; Gammaproteobacteria; Enterobacterales; Enterobacteriaceae; Salmonella; Salmonella enterica; Salmonella enterica subsp. enterica MTGDNTLITSHGINRRDFMKLCAALAATMGLSSKAAAEMAESVSNPQRPPVIWIGAQECT WP_000145429.1 hydrogenase 2 small subunit [Salmonella enterica] Length: 372\nNumber of predicted TMHs: 1\nExp number of AAs in TMHs: 24.51309\nExp number, first 60 AAs: 0.2682\nTotal prob of N-in: 0.14402\noutside 1 331\nTMhelix 332 354\ninside 355 372

17043 GCF_002039905.1_ASM203990v1 Salmonella enterica subsp. enterica serovar Enteritidis Proteobacteria; Gammaproteobacteria; Enterobacterales; Enterobacteriaceae; Salmonella; Salmonella enterica; Salmonella enterica subsp. enterica MNNEETFYQAMRRKGVTRRSFLKFCSLAATSLGLGAGMTPKIAWALENKPRIPVVWIHGL WP_001058307.1 hydrogenase [Salmonella enterica] Length: 372\nNumber of predicted TMHs: 2\nExp number of AAs in TMHs: 36.51738\nExp number, first 60 AAs: 19.72849\nTotal prob of N-in: 0.96170\nPOSSIBLE N-term signal sequence\ninside 1 19\nTMhelix 20 39\noutside 40 325\nTMhelix 326 348\ninside 349 372

17044 GCF_002039925.1_ASM203992v1 Salmonella enterica subsp. enterica serovar Enteritidis Proteobacteria; Gammaproteobacteria; Enterobacterales; Enterobacteriaceae; Salmonella; Salmonella enterica; Salmonella enterica subsp. enterica MNNEETFYQAMRRKGVTRRSFLKFCSLAATSLGLGAGMTPKIAWALENKPRIPVVWIHGL WP_001058307.1 hydrogenase [Salmonella enterica] Length: 372\nNumber of predicted TMHs: 2\nExp

number of AAs in TMHs: 36.51738\nExp number, first 60 AAs: 19.72849\nTotal prob of N-in: 0.96170\nPOSSIBLE N-term signal sequence\ninside 1 19\nTMhelix 20 39\noutside 40 325\nTMhelix 326 348\ninside 349 372

17045 GCF_001479035.1_Salmonella_enterica_CVM_N31380-SQ_v1.0 Salmonella enterica subsp. enterica serovar Enteritidis Proteobacteria; Gammaproteobacteria; Enterobacterales; Enterobacteriaceae; Salmonella; Salmonella enterica; Salmonella enterica subsp. enterica MNNEETFYQAMRRKGVTRRSFLKFCSLAATSLGLGAGMTPKIAWALENKPRIPVVWIHGL WP_001058307.1 hydrogenase [Salmonella enterica] Length: 372\nNumber of predicted TMHs: 2\nExp number of AAs in TMHs: 36.51738\nExp number, first 60 AAs: 19.72849\nTotal prob of N-in: 0.96170\nPOSSIBLE N-term signal sequence\ninside 1 19\nTMhelix 20 39\noutside 40 325\nTMhelix 326 348\ninside 349 372

17046 GCF_002039245.1_ASM203924v1 Salmonella enterica subsp. enterica serovar Enteritidis Proteobacteria; Gammaproteobacteria; Enterobacterales; Enterobacteriaceae; Salmonella; Salmonella enterica; Salmonella enterica subsp. enterica MTGDNTLITSHGINRRDFMKLCAALAATMGLSSKAAAEMAESVSNPQRPPVIWIGAQECT WP_000145429.1 hydrogenase 2 small subunit [Salmonella enterica] Length: 372\nNumber of predicted TMHs: 1\nExp number of AAs in TMHs: 24.51309\nExp number, first 60 AAs: 0.2682\nTotal prob of N-in: 0.14402\noutside 1 331\nTMhelix 332 354\ninside 355 372

17047 GCF_002039595.1_ASM203959v1 Salmonella enterica subsp. enterica serovar Enteritidis Proteobacteria; Gammaproteobacteria; Enterobacterales; Enterobacteriaceae; Salmonella; Salmonella enterica; Salmonella enterica subsp. enterica MNNEETFYQAMRRKGVTRRSFLKFCSLAATSLGLGAGMTPKIAWALENKPRIPVVWIHGL WP_001058307.1 hydrogenase [Salmonella enterica] Length: 372\nNumber of predicted TMHs: 2\nExp number of AAs in TMHs: 36.51738\nExp number, first 60 AAs: 19.72849\nTotal prob of N-in: 0.96170\nPOSSIBLE N-term signal sequence\ninside 1 19\nTMhelix 20 39\noutside 40 325\nTMhelix 326 348\ninside 349 372

17048 GCF_002039645.1_ASM203964v1 Salmonella enterica subsp. enterica serovar Enteritidis Proteobacteria; Gammaproteobacteria; Enterobacterales; Enterobacteriaceae; Salmonella; Salmonella enterica; Salmonella enterica subsp. enterica MNNEETFYQAMRRKGVTRRSFLKFCSLAATSLGLGAGMTPKIAWALENKPRIPVVWIHGL WP_001058307.1 hydrogenase [Salmonella enterica] Length: 372\nNumber of predicted TMHs: 2\nExp number of AAs in TMHs: 36.51738\nExp number, first 60 AAs: 19.72849\nTotal prob of N-in: 0.96170\nPOSSIBLE N-term signal sequence\ninside 1 19\nTMhelix 20 39\noutside 40 325\nTMhelix 326 348\ninside 349 372

17049 GCF_002039445.1_ASM203944v1 Salmonella enterica subsp. enterica serovar Enteritidis Proteobacteria; Gammaproteobacteria; Enterobacterales; Enterobacteriaceae; Salmonella; Salmonella enterica; Salmonella enterica subsp. enterica MNNEETFYQAMRRKGVTRRSFLKFCSLAATSLGLGAGMTPKIAWALENKPRIPVVWIHGL WP_001058307.1 hydrogenase [Salmonella enterica] Length: 372\nNumber of predicted TMHs: 2\nExp number of AAs in TMHs: 36.51738\nExp number, first 60 AAs: 19.72849\nTotal prob of N-in: 0.96170\nPOSSIBLE N-term signal sequence\ninside 1 19\nTMhelix 20 39\noutside 40 325\nTMhelix 326 348\ninside 349 372

17050 GCF_002039485.1_ASM203948v1 Salmonella enterica subsp. enterica serovar Enteritidis Proteobacteria; Gammaproteobacteria; Enterobacterales; Enterobacteriaceae; Salmonella; Salmonella enterica; Salmonella enterica subsp. enterica MNNEETFYQAMRRKGVTRRSFLKFCSLAATSLGLGAGMTPKIAWALENKPRIPVVWIHGL WP_001058307.1 hydrogenase [Salmonella enterica] Length: 372\nNumber of predicted TMHs: 2\nExp number of AAs in TMHs: 36.51738\nExp number, first 60 AAs: 19.72849\nTotal prob of N-in: 0.96170\nPOSSIBLE N-term signal sequence\ninside 1 19\nTMhelix 20 39\noutside 40 325\nTMhelix 326 348\ninside 349 372

17051 GCF_002039685.1_ASM203968v1 Salmonella enterica subsp. enterica serovar Enteritidis Proteobacteria; Gammaproteobacteria; Enterobacterales; Enterobacteriaceae; Salmonella; Salmonella enterica; Salmonella enterica subsp. enterica MTGDNTLITSHGINRRDFMKLCAALAATMGLSSKAAAEMAESVSNPQRPPVIWIGAQECT WP_000145429.1 hydrogenase 2 small subunit [Salmonella enterica] Length: 372\nNumber of predicted TMHs: 1\nExp number of AAs in TMHs: 24.51309\nExp number, first 60 AAs: 0.2682\nTotal prob of N-in: 0.14402\noutside 1 331\nTMhelix 332 354\ninside 355 372

17052 GCF_002040065.1_ASM204006v1 Salmonella enterica subsp. enterica serovar Enteritidis Proteobacteria; Gammaproteobacteria; Enterobacterales; Enterobacteriaceae; Salmonella; Salmonella enterica; Salmonella enterica subsp. enterica MTGDNTLITSHGINRRDFMKLCAALAATMGLSSKAAAEMAESVSNPQRPPVIWIGAQECT WP_000145429.1 hydrogenase 2 small subunit [Salmonella enterica] Length: 372\nNumber of predicted TMHs: 1\nExp number of AAs in TMHs: 24.51309\nExp number, first 60 AAs: 0.2682\nTotal prob of N-in: 0.14402\noutside 1 331\nTMhelix 332 354\ninside 355 372

17053 GCF_002040095.1_ASM204009v1 Salmonella enterica subsp. enterica serovar Enteritidis Proteobacteria; Gammaproteobacteria; Enterobacterales; Enterobacteriaceae; Salmonella; Salmonella enterica; Salmonella enterica subsp. enterica MTGDNTLITSHGINRRDFMKLCAALAATMGLSSKAAAEMAESVSNPQRPPVIWIGAQECT WP_000145429.1 hydrogenase 2 small subunit [Salmonella enterica] Length: 372\nNumber of predicted TMHs: 1\nExp number of AAs in TMHs: 24.51309\nExp number, first 60 AAs: 0.2682\nTotal prob of N-in: 0.14402\noutside 1 331\nTMhelix 332 354\ninside 355 372

17054 GCF_002038625.1_ASM203862v1 Salmonella enterica subsp. enterica serovar Enteritidis Proteobacteria; Gammaproteobacteria; Enterobacterales; Enterobacteriaceae; Salmonella; Salmonella enterica; Salmonella enterica subsp. enterica MNNEETFYQAMRRKGVTRRSFLKFCSLAATSLGLGAGMTPKIAWALENKPRIPVVWIHGL WP_001058307.1 hydrogenase [Salmonella enterica] Length: 372\nNumber of predicted TMHs: 2\nExp number of AAs in TMHs: 36.51738\nExp number, first 60 AAs: 19.72849\nTotal prob of N-in: 0.96170\nPOSSIBLE N-term signal sequence\ninside 1 19\nTMhelix 20 39\noutside 40 325\nTMhelix 326 348\ninside 349 372

17055 GCF_001185215.1_ASM118521v1 Salmonella enterica subsp. enterica serovar Enteritidis Proteobacteria; Gammaproteobacteria; Enterobacterales; Enterobacteriaceae; Salmonella; Salmonella enterica; Salmonella enterica subsp. enterica MTGDNTLITSHGINRRDFMKLCAALAATMGLSSKAAAEMAESVSNPQRPPVIWIGAQECT WP_000145429.1 hydrogenase 2 small subunit [Salmonella enterica] Length: 372\nNumber of predicted TMHs: 1\nExp number of AAs in TMHs: 24.51309\nExp number, first 60 AAs: 0.2682\nTotal prob of N-in: 0.14402\noutside 1 331\nTMhelix 332 354\ninside 355 372

17056 GCF_002032765.1_ASM203276v1 Salmonella enterica subsp. enterica serovar Enteritidis Proteobacteria; Gammaproteobacteria; Enterobacterales; Enterobacteriaceae; Salmonella; Salmonella enterica; Salmonella enterica subsp. enterica MTGDNTLITSHGINRRDFMKLCAALAATMGLSSKAAAEMAESVSNPQRPPVIWIGAQECT WP_000145429.1 hydrogenase 2 small subunit [Salmonella enterica] Length: 372\nNumber of predicted TMHs: 1\nExp number of AAs in TMHs: 24.51309\nExp number, first 60 AAs: 0.2682\nTotal prob of N-in: 0.14402\noutside 1 331\nTMhelix 332 354\ninside 355 372

17057 GCF_002038585.1_ASM203858v1 Salmonella enterica subsp. enterica serovar Enteritidis Proteobacteria; Gammaproteobacteria; Enterobacterales; Enterobacteriaceae; Salmonella; Salmonella enterica; Salmonella enterica subsp. enterica MNNEETFYQAMRRKGVTRRSFLKFCSLAATSLGLGAGMTPKIAWALENKPRIPVVWIHGL WP_001058307.1 hydrogenase [Salmonella enterica] Length: 372\nNumber of predicted TMHs: 2\nExp number of AAs in TMHs: 36.51738\nExp number, first 60 AAs: 19.72849\nTotal prob of N-in: 0.96170\nPOSSIBLE N-term signal sequence\ninside 1 19\nTMhelix 20 39\noutside 40 325\nTMhelix 326 348\ninside 349 372

17058 GCF_001973725.1_ASM197372v1 Salmonella enterica subsp. enterica serovar Enteritidis Proteobacteria; Gammaproteobacteria; Enterobacterales; Enterobacteriaceae; Salmonella; Salmonella enterica; Salmonella enterica subsp. enterica MTGDNTLITSHGINRRDFMKLCAALAATMGLSSKAAAEMAESVSNPQRPPVIWIGAQECT WP_000145429.1 hydrogenase 2 small subunit [Salmonella enterica] Length: 372\nNumber of predicted TMHs: 1\nExp number of AAs in TMHs: 24.51309\nExp number, first 60 AAs: 0.2682\nTotal prob of N-in: 0.14402\noutside 1 331\nTMhelix 332 354\ninside 355 372

17059 GCF_001973475.1_ASM197347v1 Salmonella enterica subsp. enterica serovar Enteritidis Proteobacteria; Gammaproteobacteria; Enterobacterales; Enterobacteriaceae; Salmonella; Salmonella enterica; Salmonella enterica subsp. enterica MNNEETFYQAMRRKGVTRRSFLKFCSLAATSLGLGAGMTPKIAWALENKPRIPVVWIHGL WP_001058307.1 hydrogenase [Salmonella enterica] Length: 372\nNumber of predicted TMHs: 2\nExp number of AAs in TMHs: 36.51738\nExp number, first 60 AAs: 19.72849\nTotal prob of N-in: 0.96170\nPOSSIBLE N-term signal sequence\ninside 1 19\nTMhelix 20 39\noutside 40 325\nTMhelix 326 348\ninside 349 372

17060 GCF_001973675.1_ASM197367v1 Salmonella enterica subsp. enterica serovar Enteritidis Proteobacteria; Gammaproteobacteria; Enterobacterales; Enterobacteriaceae; Salmonella; Salmonella enterica; Salmonella enterica subsp. enterica MNNEETFYQAMRRKGVTRRSFLKFCSLAATSLGLGAGMTPKIAWALENKPRIPVVWIHGL WP_001058307.1 hydrogenase [Salmonella enterica] Length: 372\nNumber of predicted TMHs: 2\nExp number of AAs in TMHs: 36.51738\nExp number, first 60 AAs: 19.72849\nTotal prob of N-in: 0.96170\nPOSSIBLE N-term signal sequence\ninside 1 19\nTMhelix 20 39\noutside 40 325\nTMhelix 326 348\ninside 349 372

17061 GCF_001973645.1_ASM197364v1 Salmonella enterica subsp. enterica serovar Enteritidis Proteobacteria; Gammaproteobacteria; Enterobacterales; Enterobacteriaceae; Salmonella; Salmonella enterica; Salmonella enterica subsp. enterica MNNEETFYQAMRRKGVTRRSFLKFCSLAATSLGLGAGMTPKIAWALENKPRIPVVWIHGL WP_001058307.1 hydrogenase [Salmonella enterica] Length: 372\nNumber of predicted TMHs: 2\nExp

number of AAs in TMHs: 36.51738\nExp number, first 60 AAs: 19.72849\nTotal prob of N-in: 0.96170\nPOSSIBLE N-term signal sequence\ninside 1 19\nTMhelix 20 39\noutside 40 325\nTMhelix 326 348\ninside 349 372

17062 GCF_001973565.1_ASM197356v1 Salmonella enterica subsp. enterica serovar Enteritidis Proteobacteria; Gammaproteobacteria; Enterobacterales; Enterobacteriaceae; Salmonella; Salmonella enterica; Salmonella enterica subsp. enterica MNNEETFYQAMRRKGVTRRSFLKFCSLAATSLGLGAGMTPKIAWALENKPRIPVVWIHGL WP_001058307.1 hydrogenase [Salmonella enterica] Length: 372\nNumber of predicted TMHs: 2\nExp number of AAs in TMHs: 36.51738\nExp number, first 60 AAs: 19.72849\nTotal prob of N-in: 0.96170\nPOSSIBLE N-term signal sequence\ninside 1 19\nTMhelix 20 39\noutside 40 325\nTMhelix 326 348\ninside 349 372

17063 GCF_001973675.1_ASM197367v1 Salmonella enterica subsp. enterica serovar Enteritidis Proteobacteria; Gammaproteobacteria; Enterobacterales; Enterobacteriaceae; Salmonella; Salmonella enterica; Salmonella enterica subsp. enterica MTGDNTLITSHGINRRDFMKLCAALAATMGLSSKAAAEMAESVSNPQRPPVIWIGAQECT WP_000145429.1 hydrogenase 2 small subunit [Salmonella enterica] Length: 372\nNumber of predicted TMHs: 1\nExp number of AAs in TMHs: 24.51309\nExp number, first 60 AAs: 0.2682\nTotal prob of N-in: 0.14402\noutside 1 331\nTMhelix 332 354\ninside 355 372

17064 GCF_001973645.1_ASM197364v1 Salmonella enterica subsp. enterica serovar Enteritidis Proteobacteria; Gammaproteobacteria; Enterobacterales; Enterobacteriaceae; Salmonella; Salmonella enterica; Salmonella enterica subsp. enterica MTGDNTLITSHGINRRDFMKLCAALAATMGLSSKAAAEMAESVSNPQRPPVIWIGAQECT WP_000145429.1 hydrogenase 2 small subunit [Salmonella enterica] Length: 372\nNumber of predicted TMHs: 1\nExp number of AAs in TMHs: 24.51309\nExp number, first 60 AAs: 0.2682\nTotal prob of N-in: 0.14402\noutside 1 331\nTMhelix 332 354\ninside 355 372

17065 GCF_001973325.1_ASM197332v1 Salmonella enterica subsp. enterica serovar Enteritidis Proteobacteria; Gammaproteobacteria; Enterobacterales; Enterobacteriaceae; Salmonella; Salmonella enterica; Salmonella enterica subsp. enterica MTGDNTLITSHGINRRDFMKLCAALAATMGLSSKAAAEMAESVSNPQRPPVIWIGAQECT WP_000145429.1 hydrogenase 2 small subunit [Salmonella enterica] Length: 372\nNumber of predicted TMHs: 1\nExp number of AAs in TMHs: 24.51309\nExp number, first 60 AAs: 0.2682\nTotal prob of N-in: 0.14402\noutside 1 331\nTMhelix 332 354\ninside 355 372

17066 GCF_001973275.1_ASM197327v1 Salmonella enterica subsp. enterica serovar Enteritidis Proteobacteria; Gammaproteobacteria; Enterobacterales; Enterobacteriaceae; Salmonella; Salmonella enterica; Salmonella enterica subsp. enterica MTGDNTLITSHGINRRDFMKLCAALAATMGLSSKAAAEMAESVSNPQRPPVIWIGAQECT WP_000145429.1 hydrogenase 2 small subunit [Salmonella enterica] Length: 372\nNumber of predicted TMHs: 1\nExp number of AAs in TMHs: 24.51309\nExp number, first 60 AAs: 0.2682\nTotal prob of N-in: 0.14402\noutside 1 331\nTMhelix 332 354\ninside 355 372

17067 GCF_001973145.1_ASM197314v1 Salmonella enterica subsp. enterica serovar Enteritidis Proteobacteria; Gammaproteobacteria; Enterobacterales; Enterobacteriaceae; Salmonella; Salmonella enterica; Salmonella enterica subsp. enterica MNNEETFYQAMRRKGVTRRSFLKFCSLAATSLGLGAGMTPKIAWALENKPRIPVVWIHGL WP_001058307.1 hydrogenase [Salmonella enterica] Length: 372\nNumber of predicted TMHs: 2\nExp number of AAs in TMHs: 36.51738\nExp number, first 60 AAs: 19.72849\nTotal prob of N-in: 0.96170\nPOSSIBLE N-term signal sequence\ninside 1 19\nTMhelix 20 39\noutside 40 325\nTMhelix 326 348\ninside 349 372

17068 GCF_001973245.1_ASM197324v1 Salmonella enterica subsp. enterica serovar Enteritidis Proteobacteria; Gammaproteobacteria; Enterobacterales; Enterobacteriaceae; Salmonella; Salmonella enterica; Salmonella enterica subsp. enterica MNNEETFYQAMRRKGVTRRSFLKFCSLAATSLGLGAGMTPKIAWALENKPRIPVVWIHGL WP_001058307.1 hydrogenase [Salmonella enterica] Length: 372\nNumber of predicted TMHs: 2\nExp number of AAs in TMHs: 36.51738\nExp number, first 60 AAs: 19.72849\nTotal prob of N-in: 0.96170\nPOSSIBLE N-term signal sequence\ninside 1 19\nTMhelix 20 39\noutside 40 325\nTMhelix 326 348\ninside 349 372

17069 GCF_001973195.1_ASM197319v1 Salmonella enterica subsp. enterica serovar Enteritidis Proteobacteria; Gammaproteobacteria; Enterobacterales; Enterobacteriaceae; Salmonella; Salmonella enterica; Salmonella enterica subsp. enterica MNNEETFYQAMRRKGVTRRSFLKFCSLAATSLGLGAGMTPKIAWALENKPRIPVVWIHGL WP_001058307.1 hydrogenase [Salmonella enterica] Length: 372\nNumber of predicted TMHs: 2\nExp number of AAs in TMHs: 36.51738\nExp number, first 60 AAs: 19.72849\nTotal prob of N-in: 0.96170\nPOSSIBLE N-term signal sequence\ninside 1 19\nTMhelix 20 39\noutside 40 325\nTMhelix 326 348\ninside 349 372

17070 GCF_001973245.1_ASM197324v1 Salmonella enterica subsp. enterica serovar Enteritidis Proteobacteria; Gammaproteobacteria; Enterobacterales; Enterobacteriaceae; Salmonella; Salmonella enterica;

Salmonella enterica subsp. enterica MTGDNTLITSHGINRRDFMKLCAALAATMGLSSKAAAEMAESVSNPQRPPVIWIGAQECT WP_000145429.1 hydrogenase 2 small subunit [Salmonella enterica] Length: 372\nNumber of predicted TMHs: 1\nExp number of AAs in TMHs: 24.51309\nExp number, first 60 AAs: 0.2682\nTotal prob of N-in: 0.14402\noutside 1 331\nTMhelix 332 354\ninside 355 372

17071 GCF_001972725.1_ASM197272v1 Salmonella enterica subsp. enterica serovar Enteritidis Proteobacteria; Gammaproteobacteria; Enterobacterales; Enterobacteriaceae; Salmonella; Salmonella enterica; Salmonella enterica subsp. enterica MNNEETFYQAMRRKGVTRRSFLKFCSLAATSLGLGAGMTPKIAWALENKPRIPVVWIHGL WP_001058307.1 hydrogenase [Salmonella enterica] Length: 372\nNumber of predicted TMHs: 2\nExp number of AAs in TMHs: 36.51738\nExp number, first 60 AAs: 19.72849\nTotal prob of N-in: 0.96170\nPOSSIBLE N-term signal sequence\ninside 1 19\nTMhelix 20 39\noutside 40 325\nTMhelix 326 348\ninside 349 372

17072 GCF_001972645.1_ASM197264v1 Salmonella enterica subsp. enterica serovar Enteritidis Proteobacteria; Gammaproteobacteria; Enterobacterales; Enterobacteriaceae; Salmonella; Salmonella enterica; Salmonella enterica subsp. enterica MNNEETFYQAMRRKGVTRRSFLKFCSLAATSLGLGAGMTPKIAWALENKPRIPVVWIHGL WP_001058307.1 hydrogenase [Salmonella enterica] Length: 372\nNumber of predicted TMHs: 2\nExp number of AAs in TMHs: 36.51738\nExp number, first 60 AAs: 19.72849\nTotal prob of N-in: 0.96170\nPOSSIBLE N-term signal sequence\ninside 1 19\nTMhelix 20 39\noutside 40 325\nTMhelix 326 348\ninside 349 372

17073 GCF_001972905.1_ASM197290v1 Salmonella enterica subsp. enterica serovar Enteritidis Proteobacteria; Gammaproteobacteria; Enterobacterales; Enterobacteriaceae; Salmonella; Salmonella enterica; Salmonella enterica subsp. enterica MNNEETFYQAMRRKGVTRRSFLKFCSLAATSLGLGAGMTPKIAWALENKPRIPVVWIHGL WP_001058307.1 hydrogenase [Salmonella enterica] Length: 372\nNumber of predicted TMHs: 2\nExp number of AAs in TMHs: 36.51738\nExp number, first 60 AAs: 19.72849\nTotal prob of N-in: 0.96170\nPOSSIBLE N-term signal sequence\ninside 1 19\nTMhelix 20 39\noutside 40 325\nTMhelix 326 348\ninside 349 372

17074 GCF_001972905.1_ASM197290v1 Salmonella enterica subsp. enterica serovar Enteritidis Proteobacteria; Gammaproteobacteria; Enterobacterales; Enterobacteriaceae; Salmonella; Salmonella enterica; Salmonella enterica subsp. enterica MTGDNTLITSHGINRRDFMKLCAALAATMGLSSKAAAEMAESVSNPQRPPVIWIGAQECT WP_000145429.1 hydrogenase 2 small subunit [Salmonella enterica] Length: 372\nNumber of predicted TMHs: 1\nExp number of AAs in TMHs: 24.51309\nExp number, first 60 AAs: 0.2682\nTotal prob of N-in: 0.14402\noutside 1 331\nTMhelix 332 354\ninside 355 372

17075 GCF_001972625.1_ASM197262v1 Salmonella enterica subsp. enterica serovar Enteritidis Proteobacteria; Gammaproteobacteria; Enterobacterales; Enterobacteriaceae; Salmonella; Salmonella enterica; Salmonella enterica subsp. enterica MNNEETFYQAMRRKGVTRRSFLKFCSLAATSLGLGAGMTPKIAWALENKPRIPVVWIHGL WP_001058307.1 hydrogenase [Salmonella enterica] Length: 372\nNumber of predicted TMHs: 2\nExp number of AAs in TMHs: 36.51738\nExp number, first 60 AAs: 19.72849\nTotal prob of N-in: 0.96170\nPOSSIBLE N-term signal sequence\ninside 1 19\nTMhelix 20 39\noutside 40 325\nTMhelix 326 348\ninside 349 372

17076 GCF_001972565.1_ASM197256v1 Salmonella enterica subsp. enterica serovar Enteritidis Proteobacteria; Gammaproteobacteria; Enterobacterales; Enterobacteriaceae; Salmonella; Salmonella enterica; Salmonella enterica subsp. enterica MNNEETFYQAMRRKGVTRRSFLKFCSLAATSLGLGAGMTPKIAWALENKPRIPVVWIHGL WP_001058307.1 hydrogenase [Salmonella enterica] Length: 372\nNumber of predicted TMHs: 2\nExp number of AAs in TMHs: 36.51738\nExp number, first 60 AAs: 19.72849\nTotal prob of N-in: 0.96170\nPOSSIBLE N-term signal sequence\ninside 1 19\nTMhelix 20 39\noutside 40 325\nTMhelix 326 348\ninside 349 372

17077 GCF_002038905.1_ASM203890v1 Salmonella enterica subsp. enterica serovar Enteritidis Proteobacteria; Gammaproteobacteria; Enterobacterales; Enterobacteriaceae; Salmonella; Salmonella enterica; Salmonella enterica subsp. enterica MTGDNTLITSHGINRRDFMKLCAALAATMGLSSKAAAEMAESVSNPQRPPVIWIGAQECT WP_000145429.1 hydrogenase 2 small subunit [Salmonella enterica] Length: 372\nNumber of predicted TMHs: 1\nExp number of AAs in TMHs: 24.51309\nExp number, first 60 AAs: 0.2682\nTotal prob of N-in: 0.14402\noutside 1 331\nTMhelix 332 354\ninside 355 372

17078 GCF_001240265.1_Salmonella_enterica_CVM_N43458_v1.0 Salmonella enterica subsp. enterica serovar Enteritidis Proteobacteria; Gammaproteobacteria; Enterobacterales; Enterobacteriaceae; Salmonella; Salmonella enterica; Salmonella enterica subsp. enterica MNNEETFYQAMRRKGVTRRSFLKFCSLAATSLGLGAGMTPKIAWALENKPRIPVVWIHGL WP_001058307.1 hydrogenase [Salmonella enterica] Length: 372\nNumber of predicted TMHs: 2\nExp number of AAs in TMHs: 36.51738\nExp number, first 60 AAs: 19.72849\nTotal prob of N-in: 0.96170\nPOSSIBLE N-term signal sequence\ninside 1 19\nTMhelix 20 39\noutside 40 325\nTMhelix 326 348\ninside 349 372

17079 GCF_001243345.1_Salmonella_enterica_CVM_N46838_v1.0 Salmonella enterica subsp. enterica serovar Enteritidis Proteobacteria; Gammaproteobacteria; Enterobacterales; Enterobacteriaceae; Salmonella; Salmonella enterica; Salmonella enterica subsp. enterica MNNEETFYQAMRRKGVTRRSFLKFCSLAATSLGLGAGMTPKIAWALENKPRIPVVWIHGL WP_001058307.1 hydrogenase [Salmonella enterica] Length: 372\nNumber of predicted TMHs: 2\nExp number of AAs in TMHs: 36.51738\nExp number, first 60 AAs: 19.72849\nTotal prob of N-in: 0.96170\nPOSSIBLE N-term signal sequence\ninside 1 19\nTMhelix 20 39\noutside 40 325\nTMhelix 326 348\ninside 349 372

17080 GCF_002038945.1_ASM203894v1 Salmonella enterica subsp. enterica serovar Enteritidis Proteobacteria; Gammaproteobacteria; Enterobacterales; Enterobacteriaceae; Salmonella; Salmonella enterica; Salmonella enterica subsp. enterica MNNEETFYQAMRRKGVTRRSFLKFCSLAATSLGLGAGMTPKIAWALENKPRIPVVWIHGL WP_001058307.1 hydrogenase [Salmonella enterica] Length: 372\nNumber of predicted TMHs: 2\nExp number of AAs in TMHs: 36.51738\nExp number, first 60 AAs: 19.72849\nTotal prob of N-in: 0.96170\nPOSSIBLE N-term signal sequence\ninside 1 19\nTMhelix 20 39\noutside 40 325\nTMhelix 326 348\ninside 349 372

17081 GCF_002039065.1_ASM203906v1 Salmonella enterica subsp. enterica serovar Enteritidis Proteobacteria; Gammaproteobacteria; Enterobacterales; Enterobacteriaceae; Salmonella; Salmonella enterica; Salmonella enterica subsp. enterica MTGDNTLITSHGINRRDFMKLCAALAATMGLSSKAAAEMAESVSNPQRPPVIWIGAQECT WP_000145429.1 hydrogenase 2 small subunit [Salmonella enterica] Length: 372\nNumber of predicted TMHs: 1\nExp number of AAs in TMHs: 24.51309\nExp number, first 60 AAs: 0.2682\nTotal prob of N-in: 0.14402\noutside 1 331\nTMhelix 332 354\ninside 355 372

17082 GCF_001271425.1_Salmonella_enterica_CVM_N43464_v1.0 Salmonella enterica subsp. enterica serovar Enteritidis Proteobacteria; Gammaproteobacteria; Enterobacterales; Enterobacteriaceae; Salmonella; Salmonella enterica; Salmonella enterica subsp. enterica MNNEETFYQAMRRKGVTRRSFLKFCSLAATSLGLGAGMTPKIAWALENKPRIPVVWIHGL WP_001058307.1 hydrogenase [Salmonella enterica] Length: 372\nNumber of predicted TMHs: 2\nExp number of AAs in TMHs: 36.51738\nExp number, first 60 AAs: 19.72849\nTotal prob of N-in: 0.96170\nPOSSIBLE N-term signal sequence\ninside 1 19\nTMhelix 20 39\noutside 40 325\nTMhelix 326 348\ninside 349 372

17083 GCF_001271485.1_Salmonella_enterica_CVM_N43834_v1.0 Salmonella enterica subsp. enterica serovar Enteritidis Proteobacteria; Gammaproteobacteria; Enterobacterales; Enterobacteriaceae; Salmonella; Salmonella enterica; Salmonella enterica subsp. enterica MTGDNTLITSHGINRRDFMKLCAALAATMGLSSKAAAEMAESVSNPQRPPVIWIGAQECT WP_000145429.1 hydrogenase 2 small subunit [Salmonella enterica] Length: 372\nNumber of predicted TMHs: 1\nExp number of AAs in TMHs: 24.51309\nExp number, first 60 AAs: 0.2682\nTotal prob of N-in: 0.14402\noutside 1 331\nTMhelix 332 354\ninside 355 372

17084 GCF_001271495.1_Salmonella_enterica_CVM_N44704_v1.0 Salmonella enterica subsp. enterica serovar Enteritidis Proteobacteria; Gammaproteobacteria; Enterobacterales; Enterobacteriaceae; Salmonella; Salmonella enterica; Salmonella enterica subsp. enterica MTGDNTLITSHGINRRDFMKLCAALAATMGLSSKAAAEMAESVSNPQRPPVIWIGAQECT WP_000145429.1 hydrogenase 2 small subunit [Salmonella enterica] Length: 372\nNumber of predicted TMHs: 1\nExp number of AAs in TMHs: 24.51309\nExp number, first 60 AAs: 0.2682\nTotal prob of N-in: 0.14402\noutside 1 331\nTMhelix 332 354\ninside 355 372

17085 GCF_001271485.1_Salmonella_enterica_CVM_N43834_v1.0 Salmonella enterica subsp. enterica serovar Enteritidis Proteobacteria; Gammaproteobacteria; Enterobacterales; Enterobacteriaceae; Salmonella; Salmonella enterica; Salmonella enterica subsp. enterica MNNEETFYQAMRRKGVTRRSFLKFCSLAATSLGLGAGMTPKIAWALENKPRIPVVWIHGL WP_001058307.1 hydrogenase [Salmonella enterica] Length: 372\nNumber of predicted TMHs: 2\nExp number of AAs in TMHs: 36.51738\nExp number, first 60 AAs: 19.72849\nTotal prob of N-in: 0.96170\nPOSSIBLE N-term signal sequence\ninside 1 19\nTMhelix 20 39\noutside 40 325\nTMhelix 326 348\ninside 349 372

17086 GCF_001271495.1_Salmonella_enterica_CVM_N44704_v1.0 Salmonella enterica subsp. enterica serovar Enteritidis Proteobacteria; Gammaproteobacteria; Enterobacterales; Enterobacteriaceae; Salmonella; Salmonella enterica; Salmonella enterica subsp. enterica MNNEETFYQAMRRKGVTRRSFLKFCSLAATSLGLGAGMTPKIAWALENKPRIPVVWIHGL WP_001058307.1 hydrogenase [Salmonella enterica] Length: 372\nNumber of predicted TMHs: 2\nExp number of AAs in TMHs: 36.51738\nExp number, first 60 AAs: 19.72849\nTotal prob of N-in: 0.96170\nPOSSIBLE N-term signal sequence\ninside 1 19\nTMhelix 20 39\noutside 40 325\nTMhelix 326 348\ninside 349 372

17087 GCF_002039085.1_ASM203908v1 Salmonella enterica subsp. enterica serovar Enteritidis Proteobacteria; Gammaproteobacteria; Enterobacterales; Enterobacteriaceae; Salmonella; Salmonella enterica; Salmonella enterica subsp. enterica MTGDNTLITSHGINRRDFMKLCAALAATMGLSSKAAAEMAESVSNPQRPPVIWIGAQECT WP_000145429.1 hydrogenase 2 small subunit [Salmonella enterica] Length: 372\nNumber of predicted TMHs: 1\nExp number of AAs in TMHs: 24.51309\nExp number, first 60 AAs: 0.2682\nTotal prob of N-in: 0.14402\noutside 1 331\nTMhelix 332 354\ninside 355 372

17088 GCF_001271675.1_Salmonella_enterica_CVM_N46817_v1.0 Salmonella enterica subsp. enterica serovar Enteritidis Proteobacteria; Gammaproteobacteria; Enterobacterales; Enterobacteriaceae; Salmonella; Salmonella enterica; Salmonella enterica subsp. enterica MNNEETFYQAMRRKGVTRRSFLKFCSLAATSLGLGAGMTPKIAWALENKPRIPVVWIHGL WP_001058307.1 hydrogenase [Salmonella enterica] Length: 372\nNumber of predicted TMHs: 2\nExp number of AAs in TMHs: 36.51738\nExp number, first 60 AAs: 19.72849\nTotal prob of N-in: 0.96170\nPOSSIBLE N-term signal sequence\ninside 1 19\nTMhelix 20 39\noutside 40 325\nTMhelix 326 348\ninside 349 372

17089 GCF_001271905.1_Salmonella_enterica_CVM_N47727_v1.0 Salmonella enterica subsp. enterica serovar Enteritidis Proteobacteria; Gammaproteobacteria; Enterobacterales; Enterobacteriaceae; Salmonella; Salmonella enterica; Salmonella enterica subsp. enterica MTGDNTLITSHGINRRDFMKLCAALAATMGLSSKAAAEMAESVSNPQRPPVIWIGAQECT WP_000145429.1 hydrogenase 2 small subunit [Salmonella enterica] Length: 372\nNumber of predicted TMHs: 1\nExp number of AAs in TMHs: 24.51309\nExp number, first 60 AAs: 0.2682\nTotal prob of N-in: 0.14402\noutside 1 331\nTMhelix 332 354\ninside 355 372

17090 GCF_001271955.1_Salmonella_enterica_CVM_N43476_v1.0 Salmonella enterica subsp. enterica serovar Enteritidis Proteobacteria; Gammaproteobacteria; Enterobacterales; Enterobacteriaceae; Salmonella; Salmonella enterica; Salmonella enterica subsp. enterica MTGDNTLITSHGINRRDFMKLCAALAATMGLSSKAAAEMAESVSNPQRPPVIWIGAQECT WP_000145429.1 hydrogenase 2 small subunit [Salmonella enterica] Length: 372\nNumber of predicted TMHs: 1\nExp number of AAs in TMHs: 24.51309\nExp number, first 60 AAs: 0.2682\nTotal prob of N-in: 0.14402\noutside 1 331\nTMhelix 332 354\ninside 355 372

17091 GCF_001272175.1_Salmonella_enterica_CVM_N48702_v1.0 Salmonella enterica subsp. enterica serovar Enteritidis Proteobacteria; Gammaproteobacteria; Enterobacterales; Enterobacteriaceae; Salmonella; Salmonella enterica; Salmonella enterica subsp. enterica MTGDNTLITSHGINRRDFMKLCAALAATMGLSSKAAAEMAESVSNPQRPPVIWIGAQECT WP_000145429.1 hydrogenase 2 small subunit [Salmonella enterica] Length: 372\nNumber of predicted TMHs: 1\nExp number of AAs in TMHs: 24.51309\nExp number, first 60 AAs: 0.2682\nTotal prob of N-in: 0.14402\noutside 1 331\nTMhelix 332 354\ninside 355 372

17092 GCF_001272215.1_Salmonella_enterica_CVM_N50426_v1.0 Salmonella enterica subsp. enterica serovar Enteritidis Proteobacteria; Gammaproteobacteria; Enterobacterales; Enterobacteriaceae; Salmonella; Salmonella enterica; Salmonella enterica subsp. enterica MTGDNTLITSHGINRRDFMKLCAALAATMGLSSKAAAEMAESVSNPQRPPVIWIGAQECT WP_000145429.1 hydrogenase 2 small subunit [Salmonella enterica] Length: 372\nNumber of predicted TMHs: 1\nExp number of AAs in TMHs: 24.51309\nExp number, first 60 AAs: 0.2682\nTotal prob of N-in: 0.14402\noutside 1 331\nTMhelix 332 354\ninside 355 372

17093 GCF_001272175.1_Salmonella_enterica_CVM_N48702_v1.0 Salmonella enterica subsp. enterica serovar Enteritidis Proteobacteria; Gammaproteobacteria; Enterobacterales; Enterobacteriaceae; Salmonella; Salmonella enterica; Salmonella enterica subsp. enterica MNNEETFYQAMRRKGVTRRSFLKFCSLAATSLGLGAGMTPKIAWALENKPRIPVVWIHGL WP_001058307.1 hydrogenase [Salmonella enterica] Length: 372\nNumber of predicted TMHs: 2\nExp number of AAs in TMHs: 36.51738\nExp number, first 60 AAs: 19.72849\nTotal prob of N-in: 0.96170\nPOSSIBLE N-term signal sequence\ninside 1 19\nTMhelix 20 39\noutside 40 325\nTMhelix 326 348\ninside 349 372

17094 GCF_001272485.1_Salmonella_enterica_CVM_N51310_v1.0 Salmonella enterica subsp. enterica serovar Enteritidis Proteobacteria; Gammaproteobacteria; Enterobacterales; Enterobacteriaceae; Salmonella; Salmonella enterica; Salmonella enterica subsp. enterica MNNEETFYQAMRRKGVTRRSFLKFCSLAATSLGLGAGMTPKIAWALENKPRIPVVWIHGL WP_001058307.1 hydrogenase [Salmonella enterica] Length: 372\nNumber of predicted TMHs: 2\nExp number of AAs in

TMHs: 36.51738\nExp number, first 60 AAs: 19.72849\nTotal prob of N-in: 0.96170\nPOSSIBLE N-term signal sequence\ninside 1 19\nTMhelix 20 39\noutside 40 325\nTMhelix 326 348\ninside 349 372

17095 GCF_002039125.1_ASM203912v1 Salmonella enterica subsp. enterica serovar Enteritidis Proteobacteria; Gammaproteobacteria; Enterobacterales; Enterobacteriaceae; Salmonella; Salmonella enterica; Salmonella enterica subsp. enterica MTGDNTLITSHGINRRDFMKLCAALAATMGLSSKAAAEMAESVSNPQRPPVIWIGAQECT WP_000145429.1 hydrogenase 2 small subunit [Salmonella enterica] Length: 372\nNumber of predicted TMHs: 1\nExp number of AAs in TMHs: 24.51309\nExp number, first 60 AAs: 0.2682\nTotal prob of N-in: 0.14402\noutside 1 331\nTMhelix 332 354\ninside 355 372

17096 GCF_002039165.1_ASM203916v1 Salmonella enterica subsp. enterica serovar Enteritidis Proteobacteria; Gammaproteobacteria; Enterobacterales; Enterobacteriaceae; Salmonella; Salmonella enterica; Salmonella enterica subsp. enterica MTGDNTLITSHGINRRDFMKLCAALAATMGLSSKAAAEMAESVSNPQRPPVIWIGAQECT WP_000145429.1 hydrogenase 2 small subunit [Salmonella enterica] Length: 372\nNumber of predicted TMHs: 1\nExp number of AAs in TMHs: 24.51309\nExp number, first 60 AAs: 0.2682\nTotal prob of N-in: 0.14402\noutside 1 331\nTMhelix 332 354\ninside 355 372

17097 GCF_002039205.1_ASM203920v1 Salmonella enterica subsp. enterica serovar Enteritidis Proteobacteria; Gammaproteobacteria; Enterobacterales; Enterobacteriaceae; Salmonella; Salmonella enterica; Salmonella enterica subsp. enterica MTGDNTLITSHGINRRDFMKLCAALAATMGLSSKAAAEMAESVSNPQRPPVIWIGAQECT WP_000145429.1 hydrogenase 2 small subunit [Salmonella enterica] Length: 372\nNumber of predicted TMHs: 1\nExp number of AAs in TMHs: 24.51309\nExp number, first 60 AAs: 0.2682\nTotal prob of N-in: 0.14402\noutside 1 331\nTMhelix 332 354\ninside 355 372

17098 GCF_002041275.1_ASM204127v1 Salmonella enterica subsp. enterica serovar Enteritidis Proteobacteria; Gammaproteobacteria; Enterobacterales; Enterobacteriaceae; Salmonella; Salmonella enterica; Salmonella enterica subsp. enterica MTGDNTLITSHGINRRDFMKLCAALAATMGLSSKAAAEMAESVSNPQRPPVIWIGAQECT WP_000145429.1 hydrogenase 2 small subunit [Salmonella enterica] Length: 372\nNumber of predicted TMHs: 1\nExp number of AAs in TMHs: 24.51309\nExp number, first 60 AAs: 0.2682\nTotal prob of N-in: 0.14402\noutside 1 331\nTMhelix 332 354\ninside 355 372

17099 GCF_002041245.1_ASM204124v1 Salmonella enterica subsp. enterica serovar Enteritidis Proteobacteria; Gammaproteobacteria; Enterobacterales; Enterobacteriaceae; Salmonella; Salmonella enterica; Salmonella enterica subsp. enterica MTGDNTLITSHGINRRDFMKLCAALAATMGLSSKAAAEMAESVSNPQRPPVIWIGAQECT WP_000145429.1 hydrogenase 2 small subunit [Salmonella enterica] Length: 372\nNumber of predicted TMHs: 1\nExp number of AAs in TMHs: 24.51309\nExp number, first 60 AAs: 0.2682\nTotal prob of N-in: 0.14402\noutside 1 331\nTMhelix 332 354\ninside 355 372

17100 GCF_002041205.1_ASM204120v1 Salmonella enterica subsp. enterica serovar Enteritidis Proteobacteria; Gammaproteobacteria; Enterobacterales; Enterobacteriaceae; Salmonella; Salmonella enterica; Salmonella enterica subsp. enterica MTGDNTLITSHGINRRDFMKLCAALAATMGLSSKAAAEMAESVSNPQRPPVIWIGAQECT WP_000145429.1 hydrogenase 2 small subunit [Salmonella enterica] Length: 372\nNumber of predicted TMHs: 1\nExp number of AAs in TMHs: 24.51309\nExp number, first 60 AAs: 0.2682\nTotal prob of N-in: 0.14402\noutside 1 331\nTMhelix 332 354\ninside 355 372

17101 GCF_002038185.1_ASM203818v1 Salmonella enterica subsp. enterica serovar Enteritidis Proteobacteria; Gammaproteobacteria; Enterobacterales; Enterobacteriaceae; Salmonella; Salmonella enterica; Salmonella enterica subsp. enterica MNNEETFYQAMRRKGVTRRSFLKFCSLAATSLGLGAGMTPKIAWALENKPRIPVVWIHGL WP_001058307.1 hydrogenase [Salmonella enterica] Length: 372\nNumber of predicted TMHs: 2\nExp number of AAs in TMHs: 36.51738\nExp number, first 60 AAs: 19.72849\nTotal prob of N-in: 0.96170\nPOSSIBLE N-term signal sequence\ninside 1 19\nTMhelix 20 39\noutside 40 325\nTMhelix 326 348\ninside 349 372

17102 GCF_002038135.1_ASM203813v1 Salmonella enterica subsp. enterica serovar Enteritidis Proteobacteria; Gammaproteobacteria; Enterobacterales; Enterobacteriaceae; Salmonella; Salmonella enterica; Salmonella enterica subsp. enterica MNNEETFYQAMRRKGVTRRSFLKFCSLAATSLGLGAGMTPKIAWALENKPRIPVVWIHGL WP_001058307.1 hydrogenase [Salmonella enterica] Length: 372\nNumber of predicted TMHs: 2\nExp number of AAs in TMHs: 36.51738\nExp number, first 60 AAs: 19.72849\nTotal prob of N-in: 0.96170\nPOSSIBLE N-term signal sequence\ninside 1 19\nTMhelix 20 39\noutside 40 325\nTMhelix 326 348\ninside 349 372

17103 GCF_002038305.1_ASM203830v1 Salmonella enterica subsp. enterica serovar Enteritidis Proteobacteria; Gammaproteobacteria; Enterobacterales; Enterobacteriaceae; Salmonella; Salmonella enterica;

Salmonella enterica subsp. enterica	MNNEETFYQAMRRKGVTRRSFLKFCSLAATSLGLGAGMTPKIAWALENKPRIPVVWIHGL
WP_001058307.1	hydrogenase [Salmonella enterica]	Length: 372\nNumber of predicted TMHs: 2\nExp number of AAs in TMHs: 36.51738\nExp number, first 60 AAs: 19.72849\nTotal prob of N-in: 0.96170\nPOSSIBLE N-term signal sequence\ninside 1 19\nTMhelix 20 39\noutside 40 325\nTMhelix 326 348\ninside 349 372

17104	GCF_002038345.1_ASM203834v1	Salmonella enterica subsp. enterica serovar Enteritidis
Proteobacteria; Gammaproteobacteria; Enterobacterales; Enterobacteriaceae; Salmonella; Salmonella enterica; Salmonella enterica subsp. enterica	MNNEETFYQAMRRKGVTRRSFLKFCSLAATSLGLGAGMTPKIAWALENKPRIPVVWIHGL
WP_001058307.1	hydrogenase [Salmonella enterica]	Length: 372\nNumber of predicted TMHs: 2\nExp number of AAs in TMHs: 36.51738\nExp number, first 60 AAs: 19.72849\nTotal prob of N-in: 0.96170\nPOSSIBLE N-term signal sequence\ninside 1 19\nTMhelix 20 39\noutside 40 325\nTMhelix 326 348\ninside 349 372

17105	GCF_002038225.1_ASM203822v1	Salmonella enterica subsp. enterica serovar Enteritidis
Proteobacteria; Gammaproteobacteria; Enterobacterales; Enterobacteriaceae; Salmonella; Salmonella enterica; Salmonella enterica subsp. enterica	MNNEETFYQAMRRKGVTRRSFLKFCSLAATSLGLGAGMTPKIAWALENKPRIPVVWIHGL
WP_001058307.1	hydrogenase [Salmonella enterica]	Length: 372\nNumber of predicted TMHs: 2\nExp number of AAs in TMHs: 36.51738\nExp number, first 60 AAs: 19.72849\nTotal prob of N-in: 0.96170\nPOSSIBLE N-term signal sequence\ninside 1 19\nTMhelix 20 39\noutside 40 325\nTMhelix 326 348\ninside 349 372

17106	GCF_002037985.1_ASM203798v1	Salmonella enterica subsp. enterica serovar Enteritidis
Proteobacteria; Gammaproteobacteria; Enterobacterales; Enterobacteriaceae; Salmonella; Salmonella enterica; Salmonella enterica subsp. enterica	MNNEETFYQAMRRKGVTRRSFLKFCSLAATSLGLGAGMTPKIAWALENKPRIPVVWIHGL
WP_001058307.1	hydrogenase [Salmonella enterica]	Length: 372\nNumber of predicted TMHs: 2\nExp number of AAs in TMHs: 36.51738\nExp number, first 60 AAs: 19.72849\nTotal prob of N-in: 0.96170\nPOSSIBLE N-term signal sequence\ninside 1 19\nTMhelix 20 39\noutside 40 325\nTMhelix 326 348\ninside 349 372

17107	GCF_002038025.1_ASM203802v1	Salmonella enterica subsp. enterica serovar Enteritidis
Proteobacteria; Gammaproteobacteria; Enterobacterales; Enterobacteriaceae; Salmonella; Salmonella enterica; Salmonella enterica subsp. enterica	MTGDNTLITSHGINRRDFMKLCAALAATMGLSSKAAAEMAESVSNPQRPPVIWIGAQECT
WP_000145429.1	hydrogenase 2 small subunit [Salmonella enterica]	Length: 372\nNumber of predicted TMHs: 1\nExp number of AAs in TMHs: 24.51309\nExp number, first 60 AAs: 0.2682\nTotal prob of N-in: 0.14402\noutside 1 331\nTMhelix 332 354\ninside 355 372

17108	GCF_000750375.1_ASM75037v1	Salmonella enterica subsp. enterica serovar Enteritidis
Proteobacteria; Gammaproteobacteria; Enterobacterales; Enterobacteriaceae; Salmonella; Salmonella enterica; Salmonella enterica subsp. enterica	MNNEETFYQAMRRKGVTRRSFLKFCSLAATSLGLGAGMTPKIAWALENKPRIPVVWIHGL
WP_001058307.1	hydrogenase [Salmonella enterica]	Length: 372\nNumber of predicted TMHs: 2\nExp number of AAs in TMHs: 36.51738\nExp number, first 60 AAs: 19.72849\nTotal prob of N-in: 0.96170\nPOSSIBLE N-term signal sequence\ninside 1 19\nTMhelix 20 39\noutside 40 325\nTMhelix 326 348\ninside 349 372

17109	GCF_002037895.1_ASM203789v1	Salmonella enterica subsp. enterica serovar Enteritidis
Proteobacteria; Gammaproteobacteria; Enterobacterales; Enterobacteriaceae; Salmonella; Salmonella enterica; Salmonella enterica subsp. enterica	MTGDNTLITSHGINRRDFMKLCAALAATMGLSSKAAAEMAESVSNPQRPPVIWIGAQECT
WP_000145429.1	hydrogenase 2 small subunit [Salmonella enterica]	Length: 372\nNumber of predicted TMHs: 1\nExp number of AAs in TMHs: 24.51309\nExp number, first 60 AAs: 0.2682\nTotal prob of N-in: 0.14402\noutside 1 331\nTMhelix 332 354\ninside 355 372

17110	GCF_002037945.1_ASM203794v1	Salmonella enterica subsp. enterica serovar Enteritidis
Proteobacteria; Gammaproteobacteria; Enterobacterales; Enterobacteriaceae; Salmonella; Salmonella enterica; Salmonella enterica subsp. enterica	MTGDNTLITSHGINRRDFMKLCAALAATMGLSSKAAAEMAESVSNPQRPPVIWIGAQECT
WP_000145429.1	hydrogenase 2 small subunit [Salmonella enterica]	Length: 372\nNumber of predicted TMHs: 1\nExp number of AAs in TMHs: 24.51309\nExp number, first 60 AAs: 0.2682\nTotal prob of N-in: 0.14402\noutside 1 331\nTMhelix 332 354\ninside 355 372

17111	GCF_002037835.1_ASM203783v1	Salmonella enterica subsp. enterica serovar Enteritidis
Proteobacteria; Gammaproteobacteria; Enterobacterales; Enterobacteriaceae; Salmonella; Salmonella enterica; Salmonella enterica subsp. enterica	MNNEETFYQAMRRKGVTRRSFLKFCSLAATSLGLGAGMTPKIAWALENKPRIPVVWIHGL
WP_001058307.1	hydrogenase [Salmonella enterica]	Length: 372\nNumber of predicted TMHs: 2\nExp number of AAs in TMHs: 36.51738\nExp number, first 60 AAs: 19.72849\nTotal prob of N-in: 0.96170\nPOSSIBLE N-term signal sequence\ninside 1 19\nTMhelix 20 39\noutside 40 325\nTMhelix 326 348\ninside 349 372

17112 GCF_002037885.1_ASM203788v1 Salmonella enterica subsp. enterica serovar Enteritidis Proteobacteria; Gammaproteobacteria; Enterobacterales; Enterobacteriaceae; Salmonella; Salmonella enterica; Salmonella enterica subsp. enterica MNNEETFYQAMRRKGVTRRSFLKFCSLAATSLGLGAGMTPKIAWALENKPRIPVVWIHGL WP_001058307.1 hydrogenase [Salmonella enterica] Length: 372\nNumber of predicted TMHs: 2\nExp number of AAs in TMHs: 36.51738\nExp number, first 60 AAs: 19.72849\nTotal prob of N-in: 0.96170\nPOSSIBLE N-term signal sequence\ninside 1 19\nTMhelix 20 39\noutside 40 325\nTMhelix 326 348\ninside 349 372

17113 GCF_000750435.1_ASM75043v1 Salmonella enterica subsp. enterica serovar Enteritidis Proteobacteria; Gammaproteobacteria; Enterobacterales; Enterobacteriaceae; Salmonella; Salmonella enterica; Salmonella enterica subsp. enterica MNNEETFYQAMRRKGVTRRSFLKFCSLAATSLGLGAGMTPKIAWALENKPRIPVVWIHGL WP_001058307.1 hydrogenase [Salmonella enterica] Length: 372\nNumber of predicted TMHs: 2\nExp number of AAs in TMHs: 36.51738\nExp number, first 60 AAs: 19.72849\nTotal prob of N-in: 0.96170\nPOSSIBLE N-term signal sequence\ninside 1 19\nTMhelix 20 39\noutside 40 325\nTMhelix 326 348\ninside 349 372

17114 GCF_002037985.1_ASM203798v1 Salmonella enterica subsp. enterica serovar Enteritidis Proteobacteria; Gammaproteobacteria; Enterobacterales; Enterobacteriaceae; Salmonella; Salmonella enterica; Salmonella enterica subsp. enterica MTGDNTLITSHGINRRDFMKLCAALAATMGLSSKAAAEMAESVSNPQRPPVIWIGAQECT WP_000145429.1 hydrogenase 2 small subunit [Salmonella enterica] Length: 372\nNumber of predicted TMHs: 1\nExp number of AAs in TMHs: 24.51309\nExp number, first 60 AAs: 0.2682\nTotal prob of N-in: 0.14402\noutside 1 331\nTMhelix 332 354\ninside 355 372

17115 GCF_000750415.2_ASM75041v2 Salmonella enterica subsp. enterica serovar Enteritidis Proteobacteria; Gammaproteobacteria; Enterobacterales; Enterobacteriaceae; Salmonella; Salmonella enterica; Salmonella enterica subsp. enterica MNNEETFYQAMRRKGVTRRSFLKFCSLAATSLGLGAGMTPKIAWALENKPRIPVVWIHGL WP_001058307.1 hydrogenase [Salmonella enterica] Length: 372\nNumber of predicted TMHs: 2\nExp number of AAs in TMHs: 36.51738\nExp number, first 60 AAs: 19.72849\nTotal prob of N-in: 0.96170\nPOSSIBLE N-term signal sequence\ninside 1 19\nTMhelix 20 39\noutside 40 325\nTMhelix 326 348\ninside 349 372

17116 GCF_002038705.1_ASM203870v1 Salmonella enterica subsp. enterica serovar Enteritidis Proteobacteria; Gammaproteobacteria; Enterobacterales; Enterobacteriaceae; Salmonella; Salmonella enterica; Salmonella enterica subsp. enterica MNNEETFYQAMRRKGVTRRSFLKFCSLAATSLGLGAGMTPKIAWALENKPRIPVVWIHGL WP_001058307.1 hydrogenase [Salmonella enterica] Length: 372\nNumber of predicted TMHs: 2\nExp number of AAs in TMHs: 36.51738\nExp number, first 60 AAs: 19.72849\nTotal prob of N-in: 0.96170\nPOSSIBLE N-term signal sequence\ninside 1 19\nTMhelix 20 39\noutside 40 325\nTMhelix 326 348\ninside 349 372

17117 GCF_001713995.1_ASM171399v1 Salmonella enterica subsp. enterica serovar Enteritidis Proteobacteria; Gammaproteobacteria; Enterobacterales; Enterobacteriaceae; Salmonella; Salmonella enterica; Salmonella enterica subsp. enterica MNNEETFYQAMRRKGVTRRSFLKFCSLAATSLGLGAGMTPKIAWALENKPRIPVVWIHGL WP_001058307.1 hydrogenase [Salmonella enterica] Length: 372\nNumber of predicted TMHs: 2\nExp number of AAs in TMHs: 36.51738\nExp number, first 60 AAs: 19.72849\nTotal prob of N-in: 0.96170\nPOSSIBLE N-term signal sequence\ninside 1 19\nTMhelix 20 39\noutside 40 325\nTMhelix 326 348\ninside 349 372

17118 GCF_001484025.1_ASM148402v1 Salmonella enterica subsp. enterica serovar Enteritidis Proteobacteria; Gammaproteobacteria; Enterobacterales; Enterobacteriaceae; Salmonella; Salmonella enterica; Salmonella enterica subsp. enterica MTGDNTLITSHGINRRDFMKLCAALAATMGLSSKAAAEMAESVSNPQRPPVIWIGAQECT WP_000145429.1 hydrogenase 2 small subunit [Salmonella enterica] Length: 372\nNumber of predicted TMHs: 1\nExp number of AAs in TMHs: 24.51309\nExp number, first 60 AAs: 0.2682\nTotal prob of N-in: 0.14402\noutside 1 331\nTMhelix 332 354\ninside 355 372

17119 GCF_002038545.1_ASM203854v1 Salmonella enterica subsp. enterica serovar Enteritidis Proteobacteria; Gammaproteobacteria; Enterobacterales; Enterobacteriaceae; Salmonella; Salmonella enterica; Salmonella enterica subsp. enterica MNNEETFYQAMRRKGVTRRSFLKFCSLAATSLGLGAGMTPKIAWALENKPRIPVVWIHGL WP_001058307.1 hydrogenase [Salmonella enterica] Length: 372\nNumber of predicted TMHs: 2\nExp number of AAs in TMHs: 36.51738\nExp number, first 60 AAs: 19.72849\nTotal prob of N-in: 0.96170\nPOSSIBLE N-term signal sequence\ninside 1 19\nTMhelix 20 39\noutside 40 325\nTMhelix 326 348\ninside 349 372

17120 GCF_001479885.1_Salmonella_enterica_CVM_N38869-SQ_v1.0 Salmonella enterica subsp. enterica serovar Enteritidis Proteobacteria; Gammaproteobacteria; Enterobacterales; Enterobacteriaceae; Salmonella; Salmonella enterica; Salmonella enterica subsp. enterica MNNEETFYQAMRRKGVTRRSFLKFCSLAATSLGLGAGMTPKIAWALENKPRIPVVWIHGL WP_001058308.1 hydrogenase [Salmonella enterica] Length: 372\nNumber of predicted TMHs: 2\nExp number of AAs in

TMHs: 39.08204\nExp number, first 60 AAs: 19.72474\nTotal prob of N-in: 0.96174\nPOSSIBLE N-term signal sequence\ninside 1 19\nTMhelix 20 39\noutside 40 325\nTMhelix 326 348\ninside 349 372

17121 GCF_001479885.1_Salmonella_enterica_CVM_N38869-SQ_v1.0 Salmonella enterica subsp. enterica serovar Enteritidis Proteobacteria; Gammaproteobacteria; Enterobacterales; Enterobacteriaceae; Salmonella; Salmonella enterica; Salmonella enterica subsp. enterica MTGDNTLITSHGINRRDFMKLCAALAATMGLSSKAAAEMAESVSNPQRPPVIWIGAQECT WP_000145429.1 hydrogenase 2 small subunit [Salmonella enterica] Length: 372\nNumber of predicted TMHs: 1\nExp number of AAs in TMHs: 24.51309\nExp number, first 60 AAs: 0.2682\nTotal prob of N-in: 0.14402\noutside 1 331\nTMhelix 332 354\ninside 355 372

17122 GCF_002038505.1_ASM203850v1 Salmonella enterica subsp. enterica serovar Enteritidis Proteobacteria; Gammaproteobacteria; Enterobacterales; Enterobacteriaceae; Salmonella; Salmonella enterica; Salmonella enterica subsp. enterica MNNEETFYQAMRRKGVTRRSFLKFCSLAATSLGLGAGMTPKIAWALENKPRIPVVWIHGL WP_001058307.1 hydrogenase [Salmonella enterica] Length: 372\nNumber of predicted TMHs: 2\nExp number of AAs in TMHs: 36.51738\nExp number, first 60 AAs: 19.72849\nTotal prob of N-in: 0.96170\nPOSSIBLE N-term signal sequence\ninside 1 19\nTMhelix 20 39\noutside 40 325\nTMhelix 326 348\ninside 349 372

17123 GCF_001185215.1_ASM118521v1 Salmonella enterica subsp. enterica serovar Enteritidis Proteobacteria; Gammaproteobacteria; Enterobacterales; Enterobacteriaceae; Salmonella; Salmonella enterica; Salmonella enterica subsp. enterica MNNEETFYQAMRRKGVTRRSFLKFCSLAATSLGLGAGMTPKIAWALENKPRIPVVWIHGL WP_001058307.1 hydrogenase [Salmonella enterica] Length: 372\nNumber of predicted TMHs: 2\nExp number of AAs in TMHs: 36.51738\nExp number, first 60 AAs: 19.72849\nTotal prob of N-in: 0.96170\nPOSSIBLE N-term signal sequence\ninside 1 19\nTMhelix 20 39\noutside 40 325\nTMhelix 326 348\ninside 349 372

17124 GCF_002036335.1_ASM203633v1 Salmonella enterica subsp. enterica serovar Enteritidis Proteobacteria; Gammaproteobacteria; Enterobacterales; Enterobacteriaceae; Salmonella; Salmonella enterica; Salmonella enterica subsp. enterica MTGDNTLITSHGINRRDFMKLCAALAATMGLSSKAAAEMAESVSNPQRPPVIWIGAQECT WP_000145429.1 hydrogenase 2 small subunit [Salmonella enterica] Length: 372\nNumber of predicted TMHs: 1\nExp number of AAs in TMHs: 24.51309\nExp number, first 60 AAs: 0.2682\nTotal prob of N-in: 0.14402\noutside 1 331\nTMhelix 332 354\ninside 355 372

17125 GCF_002038745.1_ASM203874v1 Salmonella enterica subsp. enterica serovar Enteritidis Proteobacteria; Gammaproteobacteria; Enterobacterales; Enterobacteriaceae; Salmonella; Salmonella enterica; Salmonella enterica subsp. enterica MTGDNTLITSHGINRRDFMKLCAALAATMGLSSKAAAEMAESVSNPQRPPVIWIGAQECT WP_000145429.1 hydrogenase 2 small subunit [Salmonella enterica] Length: 372\nNumber of predicted TMHs: 1\nExp number of AAs in TMHs: 24.51309\nExp number, first 60 AAs: 0.2682\nTotal prob of N-in: 0.14402\noutside 1 331\nTMhelix 332 354\ninside 355 372

17126 GCF_002038785.1_ASM203878v1 Salmonella enterica subsp. enterica serovar Enteritidis Proteobacteria; Gammaproteobacteria; Enterobacterales; Enterobacteriaceae; Salmonella; Salmonella enterica; Salmonella enterica subsp. enterica MTGDNTLITSHGINRRDFMKLCAALAATMGLSSKAAAEMAESVSNPQRPPVIWIGAQECT WP_000145429.1 hydrogenase 2 small subunit [Salmonella enterica] Length: 372\nNumber of predicted TMHs: 1\nExp number of AAs in TMHs: 24.51309\nExp number, first 60 AAs: 0.2682\nTotal prob of N-in: 0.14402\noutside 1 331\nTMhelix 332 354\ninside 355 372

17127 GCF_002038745.1_ASM203874v1 Salmonella enterica subsp. enterica serovar Enteritidis Proteobacteria; Gammaproteobacteria; Enterobacterales; Enterobacteriaceae; Salmonella; Salmonella enterica; Salmonella enterica subsp. enterica MNNEETFYQAMRRKGVTRRSFLKFCSLAATSLGLGAGMTPKIAWALENKPRIPVVWIHGL WP_001058307.1 hydrogenase [Salmonella enterica] Length: 372\nNumber of predicted TMHs: 2\nExp number of AAs in TMHs: 36.51738\nExp number, first 60 AAs: 19.72849\nTotal prob of N-in: 0.96170\nPOSSIBLE N-term signal sequence\ninside 1 19\nTMhelix 20 39\noutside 40 325\nTMhelix 326 348\ninside 349 372

17128 GCF_002038945.1_ASM203894v1 Salmonella enterica subsp. enterica serovar Enteritidis Proteobacteria; Gammaproteobacteria; Enterobacterales; Enterobacteriaceae; Salmonella; Salmonella enterica; Salmonella enterica subsp. enterica MTGDNTLITSHGINRRDFMKLCAALAATMGLSSKAAAEMAESVSNPQRPPVIWIGAQECT WP_000145429.1 hydrogenase 2 small subunit [Salmonella enterica] Length: 372\nNumber of predicted TMHs: 1\nExp number of AAs in TMHs: 24.51309\nExp number, first 60 AAs: 0.2682\nTotal prob of N-in: 0.14402\noutside 1 331\nTMhelix 332 354\ninside 355 372

17129 GCF_002039065.1_ASM203906v1 Salmonella enterica subsp. enterica serovar Enteritidis Proteobacteria; Gammaproteobacteria; Enterobacterales; Enterobacteriaceae; Salmonella; Salmonella enterica; Salmonella enterica subsp. enterica MNNEETFYQAMRRKGVTRRSFLKFCSLAATSLGLGAGMTPKIAWALENKPRIPVVWIHGL WP_001058307.1 hydrogenase [Salmonella enterica] Length: 372\nNumber of predicted TMHs: 2\nExp number of AAs in TMHs: 36.51738\nExp number, first 60 AAs: 19.72849\nTotal prob of N-in: 0.96170\nPOSSIBLE N-term signal sequence\ninside 1 19\nTMhelix 20 39\noutside 40 325\nTMhelix 326 348\ninside 349 372

17130 GCF_002039125.1_ASM203912v1 Salmonella enterica subsp. enterica serovar Enteritidis Proteobacteria; Gammaproteobacteria; Enterobacterales; Enterobacteriaceae; Salmonella; Salmonella enterica; Salmonella enterica subsp. enterica MNNEETFYQAMRRKGVTRRSFLKFCSLAATSLGLGAGMTPKIAWALENKPRIPVVWIHGL WP_001058307.1 hydrogenase [Salmonella enterica] Length: 372\nNumber of predicted TMHs: 2\nExp number of AAs in TMHs: 36.51738\nExp number, first 60 AAs: 19.72849\nTotal prob of N-in: 0.96170\nPOSSIBLE N-term signal sequence\ninside 1 19\nTMhelix 20 39\noutside 40 325\nTMhelix 326 348\ninside 349 372

17131 GCF_002039085.1_ASM203908v1 Salmonella enterica subsp. enterica serovar Enteritidis Proteobacteria; Gammaproteobacteria; Enterobacterales; Enterobacteriaceae; Salmonella; Salmonella enterica; Salmonella enterica subsp. enterica MNNEETFYQAMRRKGVTRRSFLKFCSLAATSLGLGAGMTPKIAWALENKPRIPVVWIHGL WP_001058307.1 hydrogenase [Salmonella enterica] Length: 372\nNumber of predicted TMHs: 2\nExp number of AAs in TMHs: 36.51738\nExp number, first 60 AAs: 19.72849\nTotal prob of N-in: 0.96170\nPOSSIBLE N-term signal sequence\ninside 1 19\nTMhelix 20 39\noutside 40 325\nTMhelix 326 348\ninside 349 372

17132 GCF_002038345.1_ASM203834v1 Salmonella enterica subsp. enterica serovar Enteritidis Proteobacteria; Gammaproteobacteria; Enterobacterales; Enterobacteriaceae; Salmonella; Salmonella enterica; Salmonella enterica subsp. enterica MTGDNTLITSHGINRRDFMKLCAALAATMGLSSKAAAEMAESVSNPQRPPVIWIGAQECT WP_000145429.1 hydrogenase 2 small subunit [Salmonella enterica] Length: 372\nNumber of predicted TMHs: 1\nExp number of AAs in TMHs: 24.51309\nExp number, first 60 AAs: 0.2682\nTotal prob of N-in: 0.14402\noutside 1 331\nTMhelix 332 354\ninside 355 372

17133 GCF_002039165.1_ASM203916v1 Salmonella enterica subsp. enterica serovar Enteritidis Proteobacteria; Gammaproteobacteria; Enterobacterales; Enterobacteriaceae; Salmonella; Salmonella enterica; Salmonella enterica subsp. enterica MNNEETFYQAMRRKGVTRRSFLKFCSLAATSLGLGAGMTPKIAWALENKPRIPVVWIHGL WP_001058307.1 hydrogenase [Salmonella enterica] Length: 372\nNumber of predicted TMHs: 2\nExp number of AAs in TMHs: 36.51738\nExp number, first 60 AAs: 19.72849\nTotal prob of N-in: 0.96170\nPOSSIBLE N-term signal sequence\ninside 1 19\nTMhelix 20 39\noutside 40 325\nTMhelix 326 348\ninside 349 372

17134 GCF_002039285.1_ASM203928v1 Salmonella enterica subsp. enterica serovar Enteritidis Proteobacteria; Gammaproteobacteria; Enterobacterales; Enterobacteriaceae; Salmonella; Salmonella enterica; Salmonella enterica subsp. enterica MTGDNTLITSHGINRRDFMKLCAALAATMGLSSKAAAEMAESVSNPQRPPVIWIGAQECT WP_000145429.1 hydrogenase 2 small subunit [Salmonella enterica] Length: 372\nNumber of predicted TMHs: 1\nExp number of AAs in TMHs: 24.51309\nExp number, first 60 AAs: 0.2682\nTotal prob of N-in: 0.14402\noutside 1 331\nTMhelix 332 354\ninside 355 372

17135 GCF_001713875.1_ASM171387v1 Salmonella enterica subsp. enterica serovar Enteritidis Proteobacteria; Gammaproteobacteria; Enterobacterales; Enterobacteriaceae; Salmonella; Salmonella enterica; Salmonella enterica subsp. enterica MNNEETFYQAMRRKGVTRRSFLKFCSLAATSLGLGAGMTPKIAWALENKPRIPVVWIHGL WP_001058307.1 hydrogenase [Salmonella enterica] Length: 372\nNumber of predicted TMHs: 2\nExp number of AAs in TMHs: 36.51738\nExp number, first 60 AAs: 19.72849\nTotal prob of N-in: 0.96170\nPOSSIBLE N-term signal sequence\ninside 1 19\nTMhelix 20 39\noutside 40 325\nTMhelix 326 348\ninside 349 372

17136 GCF_001713785.1_ASM171378v1 Salmonella enterica subsp. enterica serovar Enteritidis Proteobacteria; Gammaproteobacteria; Enterobacterales; Enterobacteriaceae; Salmonella; Salmonella enterica; Salmonella enterica subsp. enterica MNNEETFYQAMRRKGVTRRSFLKFCSLAATSLGLGAGMTPKIAWALENKPRIPVVWIHGL WP_001058307.1 hydrogenase [Salmonella enterica] Length: 372\nNumber of predicted TMHs: 2\nExp number of AAs in TMHs: 36.51738\nExp number, first 60 AAs: 19.72849\nTotal prob of N-in: 0.96170\nPOSSIBLE N-term signal sequence\ninside 1 19\nTMhelix 20 39\noutside 40 325\nTMhelix 326 348\ninside 349 372

17137 GCF_002039325.1_ASM203932v1 Salmonella enterica subsp. enterica serovar Enteritidis Proteobacteria; Gammaproteobacteria; Enterobacterales; Enterobacteriaceae; Salmonella; Salmonella enterica; Salmonella enterica subsp. enterica MTGDNTLITSHGINRRDFMKLCAALAATMGLSSKAAAEMAESVSNPQRPPVIWIGAQECT WP_000145429.1 hydrogenase 2 small subunit [Salmonella enterica] Length: 372\nNumber of

predicted TMHs: 1\nExp number of AAs in TMHs: 24.51309\nExp number, first 60 AAs: 0.2682\nTotal prob of N-in: 0.14402\noutside 1 331\nTMhelix 332 354\ninside 355 372

17138 GCF_001713875.1_ASM171387v1 Salmonella enterica subsp. enterica serovar Enteritidis Proteobacteria; Gammaproteobacteria; Enterobacterales; Enterobacteriaceae; Salmonella; Salmonella enterica; Salmonella enterica subsp. enterica MTGDNTLITSHGINRRDFMKLCAALAATMGLSSKAAAEMAESVSNPQRPPVIWIGAQECT WP_000145429.1 hydrogenase 2 small subunit [Salmonella enterica] Length: 372\nNumber of predicted TMHs: 1\nExp number of AAs in TMHs: 24.51309\nExp number, first 60 AAs: 0.2682\nTotal prob of N-in: 0.14402\noutside 1 331\nTMhelix 332 354\ninside 355 372

17139 GCF_001713785.1_ASM171378v1 Salmonella enterica subsp. enterica serovar Enteritidis Proteobacteria; Gammaproteobacteria; Enterobacterales; Enterobacteriaceae; Salmonella; Salmonella enterica; Salmonella enterica subsp. enterica MTGDNTLITSHGINRRDFMKLCAALAATMGLSSKAAAEMAESVSNPQRPPVIWIGAQECT WP_000145429.1 hydrogenase 2 small subunit [Salmonella enterica] Length: 372\nNumber of predicted TMHs: 1\nExp number of AAs in TMHs: 24.51309\nExp number, first 60 AAs: 0.2682\nTotal prob of N-in: 0.14402\noutside 1 331\nTMhelix 332 354\ninside 355 372

17140 GCF_002039715.1_ASM203971v1 Salmonella enterica subsp. enterica serovar Enteritidis Proteobacteria; Gammaproteobacteria; Enterobacterales; Enterobacteriaceae; Salmonella; Salmonella enterica; Salmonella enterica subsp. enterica MTGDNTLITSHGINRRDFMKLCAALAATMGLSSKAAAEMAESVSNPQRPPVIWIGAQECT WP_000145429.1 hydrogenase 2 small subunit [Salmonella enterica] Length: 372\nNumber of predicted TMHs: 1\nExp number of AAs in TMHs: 24.51309\nExp number, first 60 AAs: 0.2682\nTotal prob of N-in: 0.14402\noutside 1 331\nTMhelix 332 354\ninside 355 372

17141 GCF_001240265.1_Salmonella_enterica_CVM_N43458_v1.0 Salmonella enterica subsp. enterica serovar Enteritidis Proteobacteria; Gammaproteobacteria; Enterobacterales; Enterobacteriaceae; Salmonella; Salmonella enterica; Salmonella enterica subsp. enterica MTGDNTLITSHGINRRDFMKLCAALAATMGLSSKAAAEMAESVSNPQRPPVIWIGAQECT WP_000145429.1 hydrogenase 2 small subunit [Salmonella enterica] Length: 372\nNumber of predicted TMHs: 1\nExp number of AAs in TMHs: 24.51309\nExp number, first 60 AAs: 0.2682\nTotal prob of N-in: 0.14402\noutside 1 331\nTMhelix 332 354\ninside 355 372

17142 GCF_002039365.1_ASM203936v1 Salmonella enterica subsp. enterica serovar Enteritidis Proteobacteria; Gammaproteobacteria; Enterobacterales; Enterobacteriaceae; Salmonella; Salmonella enterica; Salmonella enterica subsp. enterica MNNEETFYQAMRRKGVTRRSFLKFCSLAATSLGLGAGMTPKIAWALENKPRIPVVWIHGL WP_001058307.1 hydrogenase [Salmonella enterica] Length: 372\nNumber of predicted TMHs: 2\nExp number of AAs in TMHs: 36.51738\nExp number, first 60 AAs: 19.72849\nTotal prob of N-in: 0.96170\nPOSSIBLE N-term signal sequence\ninside 1 19\nTMhelix 20 39\noutside 40 325\nTMhelix 326 348\ninside 349 372

17143 GCF_002039485.1_ASM203948v1 Salmonella enterica subsp. enterica serovar Enteritidis Proteobacteria; Gammaproteobacteria; Enterobacterales; Enterobacteriaceae; Salmonella; Salmonella enterica; Salmonella enterica subsp. enterica MTGDNTLITSHGINRRDFMKLCAALAATMGLSSKAAAEMAESVSNPQRPPVIWIGAQECT WP_000145429.1 hydrogenase 2 small subunit [Salmonella enterica] Length: 372\nNumber of predicted TMHs: 1\nExp number of AAs in TMHs: 24.51309\nExp number, first 60 AAs: 0.2682\nTotal prob of N-in: 0.14402\noutside 1 331\nTMhelix 332 354\ninside 355 372

17144 GCF_002039395.1_ASM203939v1 Salmonella enterica subsp. enterica serovar Enteritidis Proteobacteria; Gammaproteobacteria; Enterobacterales; Enterobacteriaceae; Salmonella; Salmonella enterica; Salmonella enterica subsp. enterica MNNEETFYQAMRRKGVTRRSFLKFCSLAATSLGLGAGMTPKIAWALENKPRIPVVWIHGL WP_001058307.1 hydrogenase [Salmonella enterica] Length: 372\nNumber of predicted TMHs: 2\nExp number of AAs in TMHs: 36.51738\nExp number, first 60 AAs: 19.72849\nTotal prob of N-in: 0.96170\nPOSSIBLE N-term signal sequence\ninside 1 19\nTMhelix 20 39\noutside 40 325\nTMhelix 326 348\ninside 349 372

17145 GCF_002039565.1_ASM203956v1 Salmonella enterica subsp. enterica serovar Enteritidis Proteobacteria; Gammaproteobacteria; Enterobacterales; Enterobacteriaceae; Salmonella; Salmonella enterica; Salmonella enterica subsp. enterica MTGDNTLITSHGINRRDFMKLCAALAATMGLSSKAAAEMAESVSNPQRPPVIWIGAQECT WP_000145429.1 hydrogenase 2 small subunit [Salmonella enterica] Length: 372\nNumber of predicted TMHs: 1\nExp number of AAs in TMHs: 24.51309\nExp number, first 60 AAs: 0.2682\nTotal prob of N-in: 0.14402\noutside 1 331\nTMhelix 332 354\ninside 355 372

17146 GCF_001973595.1_ASM197359v1 Salmonella enterica subsp. enterica serovar Enteritidis Proteobacteria; Gammaproteobacteria; Enterobacterales; Enterobacteriaceae; Salmonella; Salmonella enterica; Salmonella enterica subsp. enterica MTGDNTLITSHGINRRDFMKLCAALAATMGLSSKAAAEMAESVSNPQRPPVIWIGAQECT WP_076935543.1 hydrogenase 2 small subunit [Salmonella enterica] Length: 372\nNumber of predicted TMHs: 1\nExp number of AAs in TMHs: 24.16349\nExp number, first 60 AAs: 0.26819\nTotal prob of N-in: 0.12948\noutside 1 331\nTMhelix 332 354\ninside 355 372

17147 GCF_002039595.1_ASM203959v1 Salmonella enterica subsp. enterica serovar Enteritidis Proteobacteria; Gammaproteobacteria; Enterobacterales; Enterobacteriaceae; Salmonella; Salmonella enterica; Salmonella enterica subsp. enterica MTGDNTLITSHGINRRDFMKLCAALAATMGLSSKAAAEMAESVSNPQRPPVIWIGAQECT WP_000145429.1 hydrogenase 2 small subunit [Salmonella enterica] Length: 372\nNumber of predicted TMHs: 1\nExp number of AAs in TMHs: 24.51309\nExp number, first 60 AAs: 0.2682\nTotal prob of N-in: 0.14402\noutside 1 331\nTMhelix 332 354\ninside 355 372

17148 GCF_001973595.1_ASM197359v1 Salmonella enterica subsp. enterica serovar Enteritidis Proteobacteria; Gammaproteobacteria; Enterobacterales; Enterobacteriaceae; Salmonella; Salmonella enterica; Salmonella enterica subsp. enterica MNNEETFYQAMRRKGVTRRSFLKFCSLAATSLGLGAGMTPKIAWALENKPRIPVVWIHGL WP_001058307.1 hydrogenase [Salmonella enterica] Length: 372\nNumber of predicted TMHs: 2\nExp number of AAs in TMHs: 36.51738\nExp number, first 60 AAs: 19.72849\nTotal prob of N-in: 0.96170\nPOSSIBLE N-term signal sequence\ninside 1 19\nTMhelix 20 39\noutside 40 325\nTMhelix 326 348\ninside 349 372

17149 GCF_002039645.1_ASM203964v1 Salmonella enterica subsp. enterica serovar Enteritidis Proteobacteria; Gammaproteobacteria; Enterobacterales; Enterobacteriaceae; Salmonella; Salmonella enterica; Salmonella enterica subsp. enterica MTGDNTLITSHGINRRDFMKLCAALAATMGLSSKAAAEMAESVSNPQRPPVIWIGAQECT WP_000145429.1 hydrogenase 2 small subunit [Salmonella enterica] Length: 372\nNumber of predicted TMHs: 1\nExp number of AAs in TMHs: 24.51309\nExp number, first 60 AAs: 0.2682\nTotal prob of N-in: 0.14402\noutside 1 331\nTMhelix 332 354\ninside 355 372

17150 GCF_002039565.1_ASM203956v1 Salmonella enterica subsp. enterica serovar Enteritidis Proteobacteria; Gammaproteobacteria; Enterobacterales; Enterobacteriaceae; Salmonella; Salmonella enterica; Salmonella enterica subsp. enterica MNNEETFYQAMRRKGVTRRSFLKFCSLAATSLGLGAGMTPKIAWALENKPRIPVVWIHGL WP_001058307.1 hydrogenase [Salmonella enterica] Length: 372\nNumber of predicted TMHs: 2\nExp number of AAs in TMHs: 36.51738\nExp number, first 60 AAs: 19.72849\nTotal prob of N-in: 0.96170\nPOSSIBLE N-term signal sequence\ninside 1 19\nTMhelix 20 39\noutside 40 325\nTMhelix 326 348\ninside 349 372

17151 GCF_001973565.1_ASM197356v1 Salmonella enterica subsp. enterica serovar Enteritidis Proteobacteria; Gammaproteobacteria; Enterobacterales; Enterobacteriaceae; Salmonella; Salmonella enterica; Salmonella enterica subsp. enterica MTGDNTLITSHGINRRDFMKLCAALAATMGLSSKAAAEMAESVSNPQRPPVIWIGAQECT WP_000145429.1 hydrogenase 2 small subunit [Salmonella enterica] Length: 372\nNumber of predicted TMHs: 1\nExp number of AAs in TMHs: 24.51309\nExp number, first 60 AAs: 0.2682\nTotal prob of N-in: 0.14402\noutside 1 331\nTMhelix 332 354\ninside 355 372

17152 GCF_002039855.1_ASM203985v1 Salmonella enterica subsp. enterica serovar Enteritidis Proteobacteria; Gammaproteobacteria; Enterobacterales; Enterobacteriaceae; Salmonella; Salmonella enterica; Salmonella enterica subsp. enterica MNNEETFYQAMRRKGVTRRSFLKFCSLAATSLGLGAGMTPKIAWALENKPRIPVVWIHGL WP_001058307.1 hydrogenase [Salmonella enterica] Length: 372\nNumber of predicted TMHs: 2\nExp number of AAs in TMHs: 36.51738\nExp number, first 60 AAs: 19.72849\nTotal prob of N-in: 0.96170\nPOSSIBLE N-term signal sequence\ninside 1 19\nTMhelix 20 39\noutside 40 325\nTMhelix 326 348\ninside 349 372

17153 GCF_001973515.1_ASM197351v1 Salmonella enterica subsp. enterica serovar Enteritidis Proteobacteria; Gammaproteobacteria; Enterobacterales; Enterobacteriaceae; Salmonella; Salmonella enterica; Salmonella enterica subsp. enterica MTGDNTLITSHGINRRDFMKLCAALAATMGLSSKAAAEMAESVSNPQRPPVIWIGAQECT WP_000145429.1 hydrogenase 2 small subunit [Salmonella enterica] Length: 372\nNumber of predicted TMHs: 1\nExp number of AAs in TMHs: 24.51309\nExp number, first 60 AAs: 0.2682\nTotal prob of N-in: 0.14402\noutside 1 331\nTMhelix 332 354\ninside 355 372

17154 GCF_001973435.1_ASM197343v1 Salmonella enterica subsp. enterica serovar Enteritidis Proteobacteria; Gammaproteobacteria; Enterobacterales; Enterobacteriaceae; Salmonella; Salmonella enterica; Salmonella enterica subsp. enterica MNNEETFYQAMRRKGVTRRSFLKFCSLAATSLGLGAGMTPKIAWALENKPRIPVVWIHGL WP_001058307.1 hydrogenase [Salmonella enterica] Length: 372\nNumber of predicted TMHs: 2\nExp

number of AAs in TMHs: 36.51738\nExp number, first 60 AAs:  19.72849\nTotal prob of N-in:   0.96170\nPOSSIBLE N-term signal sequence\ninside   1   19\nTMhelix   20   39\noutside   40   325\nTMhelix   326   348\ninside   349   372

17155	GCF_001973405.1_ASM197340v1	Salmonella enterica subsp. enterica serovar Enteritidis	Proteobacteria; Gammaproteobacteria; Enterobacterales; Enterobacteriaceae; Salmonella; Salmonella enterica; Salmonella enterica subsp. enterica	MNNEETFYQAMRRKGVTRRSFLKFCSLAATSLGLGAGMTPKIAWALENKPRIPVVWIHGL	WP_001058307.1	hydrogenase [Salmonella enterica]	Length: 372\nNumber of predicted TMHs:  2\nExp number of AAs in TMHs: 36.51738\nExp number, first 60 AAs:  19.72849\nTotal prob of N-in:   0.96170\nPOSSIBLE N-term signal sequence\ninside   1   19\nTMhelix   20   39\noutside   40   325\nTMhelix   326   348\ninside   349   372

17156	GCF_002039755.1_ASM203975v1	Salmonella enterica subsp. enterica serovar Enteritidis	Proteobacteria; Gammaproteobacteria; Enterobacterales; Enterobacteriaceae; Salmonella; Salmonella enterica; Salmonella enterica subsp. enterica	MNNEETFYQAMRRKGVTRRSFLKFCSLAATSLGLGAGMTPKIAWALENKPRIPVVWIHGL	WP_001058307.1	hydrogenase [Salmonella enterica]	Length: 372\nNumber of predicted TMHs:  2\nExp number of AAs in TMHs: 36.51738\nExp number, first 60 AAs:  19.72849\nTotal prob of N-in:   0.96170\nPOSSIBLE N-term signal sequence\ninside   1   19\nTMhelix   20   39\noutside   40   325\nTMhelix   326   348\ninside   349   372

17157	GCF_001973475.1_ASM197347v1	Salmonella enterica subsp. enterica serovar Enteritidis	Proteobacteria; Gammaproteobacteria; Enterobacterales; Enterobacteriaceae; Salmonella; Salmonella enterica; Salmonella enterica subsp. enterica	MTGDNTLITSHGINRRDFMKLCAALAATMGLSSKAAAEMAESVSNPQRPPVIWIGAQECT	WP_000145429.1	hydrogenase 2 small subunit [Salmonella enterica]	Length: 372\nNumber of predicted TMHs:  1\nExp number of AAs in TMHs: 24.51309\nExp number, first 60 AAs:  0.2682\nTotal prob of N-in:   0.14402\noutside   1   331\nTMhelix   332   354\ninside   355   372

17158	GCF_002039785.1_ASM203978v1	Salmonella enterica subsp. enterica serovar Enteritidis	Proteobacteria; Gammaproteobacteria; Enterobacterales; Enterobacteriaceae; Salmonella; Salmonella enterica; Salmonella enterica subsp. enterica	MNNEETFYQAMRRKGVTRRSFLKFCSLAATSLGLGAGMTPKIAWALENKPRIPVVWIHGL	WP_001058307.1	hydrogenase [Salmonella enterica]	Length: 372\nNumber of predicted TMHs:  2\nExp number of AAs in TMHs: 36.51738\nExp number, first 60 AAs:  19.72849\nTotal prob of N-in:   0.96170\nPOSSIBLE N-term signal sequence\ninside   1   19\nTMhelix   20   39\noutside   40   325\nTMhelix   326   348\ninside   349   372

17159	GCF_001973435.1_ASM197343v1	Salmonella enterica subsp. enterica serovar Enteritidis	Proteobacteria; Gammaproteobacteria; Enterobacterales; Enterobacteriaceae; Salmonella; Salmonella enterica; Salmonella enterica subsp. enterica	MTGDNTLITSHGINRRDFMKLCAALAATMGLSSKAAAEMAESVSNPQRPPVIWIGAQECT	WP_000145429.1	hydrogenase 2 small subunit [Salmonella enterica]	Length: 372\nNumber of predicted TMHs:  1\nExp number of AAs in TMHs: 24.51309\nExp number, first 60 AAs:  0.2682\nTotal prob of N-in:   0.14402\noutside   1   331\nTMhelix   332   354\ninside   355   372

17160	GCF_002061465.1_ASM206146v1	Salmonella enterica subsp. enterica serovar Enteritidis	Proteobacteria; Gammaproteobacteria; Enterobacterales; Enterobacteriaceae; Salmonella; Salmonella enterica; Salmonella enterica subsp. enterica	MNNEETFYQAMRRKGVTRRSFLKFCSLAATSLGLGAGMTPKIAWALENKPRIPVVWIHGL	WP_001058307.1	hydrogenase [Salmonella enterica]	Length: 372\nNumber of predicted TMHs:  2\nExp number of AAs in TMHs: 36.51738\nExp number, first 60 AAs:  19.72849\nTotal prob of N-in:   0.96170\nPOSSIBLE N-term signal sequence\ninside   1   19\nTMhelix   20   39\noutside   40   325\nTMhelix   326   348\ninside   349   372

17161	GCF_001973405.1_ASM197340v1	Salmonella enterica subsp. enterica serovar Enteritidis	Proteobacteria; Gammaproteobacteria; Enterobacterales; Enterobacteriaceae; Salmonella; Salmonella enterica; Salmonella enterica subsp. enterica	MTGDNTLITSHGINRRDFMKLCAALAATMGLSSKAAAEMAESVSNPQRPPVIWIGAQECT	WP_000145429.1	hydrogenase 2 small subunit [Salmonella enterica]	Length: 372\nNumber of predicted TMHs:  1\nExp number of AAs in TMHs: 24.51309\nExp number, first 60 AAs:  0.2682\nTotal prob of N-in:   0.14402\noutside   1   331\nTMhelix   332   354\ninside   355   372

17162	GCF_001973025.1_ASM197302v1	Salmonella enterica subsp. enterica serovar Enteritidis	Proteobacteria; Gammaproteobacteria; Enterobacterales; Enterobacteriaceae; Salmonella; Salmonella enterica; Salmonella enterica subsp. enterica	MNNEETFYQAMRRKGVTRRSFLKFCSLAATSLGLGAGMTPKIAWALENKPRIPVVWIHGL	WP_001058307.1	hydrogenase [Salmonella enterica]	Length: 372\nNumber of predicted TMHs:  2\nExp number of AAs in TMHs: 36.51738\nExp number, first 60 AAs:  19.72849\nTotal prob of N-in:   0.96170\nPOSSIBLE N-term signal sequence\ninside   1   19\nTMhelix   20   39\noutside   40   325\nTMhelix   326   348\ninside   349   372

17163	GCF_002039905.1_ASM203990v1	Salmonella enterica subsp. enterica serovar Enteritidis	Proteobacteria; Gammaproteobacteria; Enterobacterales; Enterobacteriaceae; Salmonella; Salmonella enterica;

Salmonella enterica subsp. enterica	MTGDNTLITSHGINRRDFMKLCAALAATMGLSSKAAAEMAESVSNPQRPPVIWIGAQECT	WP_000145429.1	hydrogenase 2 small subunit [Salmonella enterica]	Length: 372\nNumber of predicted TMHs: 1\nExp number of AAs in TMHs: 24.51309\nExp number, first 60 AAs: 0.2682\nTotal prob of N-in: 0.14402\noutside 1 331\nTMhelix 332 354\ninside 355 372

17164	GCF_001973105.1_ASM197310v1	Salmonella enterica subsp. enterica serovar Enteritidis	Proteobacteria; Gammaproteobacteria; Enterobacterales; Enterobacteriaceae; Salmonella; Salmonella enterica; Salmonella enterica subsp. enterica	MTGDNTLITSHGINRRDFMKLCAALAATMGLSSKAAAEMAESVSNPQRPPVIWIGAQECT	WP_000145429.1	hydrogenase 2 small subunit [Salmonella enterica]	Length: 372\nNumber of predicted TMHs: 1\nExp number of AAs in TMHs: 24.51309\nExp number, first 60 AAs: 0.2682\nTotal prob of N-in: 0.14402\noutside 1 331\nTMhelix 332 354\ninside 355 372

17165	GCF_001973065.1_ASM197306v1	Salmonella enterica subsp. enterica serovar Enteritidis	Proteobacteria; Gammaproteobacteria; Enterobacterales; Enterobacteriaceae; Salmonella; Salmonella enterica; Salmonella enterica subsp. enterica	MTGDNTLITSHGINRRDFMKLCAALAATMGLSSKAAAEMAESVSNPQRPPVIWIGAQECT	WP_000145429.1	hydrogenase 2 small subunit [Salmonella enterica]	Length: 372\nNumber of predicted TMHs: 1\nExp number of AAs in TMHs: 24.51309\nExp number, first 60 AAs: 0.2682\nTotal prob of N-in: 0.14402\noutside 1 331\nTMhelix 332 354\ninside 355 372

17166	GCF_002039925.1_ASM203992v1	Salmonella enterica subsp. enterica serovar Enteritidis	Proteobacteria; Gammaproteobacteria; Enterobacterales; Enterobacteriaceae; Salmonella; Salmonella enterica; Salmonella enterica subsp. enterica	MTGDNTLITSHGINRRDFMKLCAALAATMGLSSKAAAEMAESVSNPQRPPVIWIGAQECT	WP_000145429.1	hydrogenase 2 small subunit [Salmonella enterica]	Length: 372\nNumber of predicted TMHs: 1\nExp number of AAs in TMHs: 24.51309\nExp number, first 60 AAs: 0.2682\nTotal prob of N-in: 0.14402\noutside 1 331\nTMhelix 332 354\ninside 355 372

17167	GCF_001973145.1_ASM197314v1	Salmonella enterica subsp. enterica serovar Enteritidis	Proteobacteria; Gammaproteobacteria; Enterobacterales; Enterobacteriaceae; Salmonella; Salmonella enterica; Salmonella enterica subsp. enterica	MTGDNTLITSHGINRRDFMKLCAALAATMGLSSKAAAEMAESVSNPQRPPVIWIGAQECT	WP_000145429.1	hydrogenase 2 small subunit [Salmonella enterica]	Length: 372\nNumber of predicted TMHs: 1\nExp number of AAs in TMHs: 24.51309\nExp number, first 60 AAs: 0.2682\nTotal prob of N-in: 0.14402\noutside 1 331\nTMhelix 332 354\ninside 355 372

17168	GCF_002039985.1_ASM203998v1	Salmonella enterica subsp. enterica serovar Enteritidis	Proteobacteria; Gammaproteobacteria; Enterobacterales; Enterobacteriaceae; Salmonella; Salmonella enterica; Salmonella enterica subsp. enterica	MTGDNTLITSHGINRRDFMKLCAALAATMGLSSKAAAEMAESVSNPQRPPVIWIGAQECT	WP_000145429.1	hydrogenase 2 small subunit [Salmonella enterica]	Length: 372\nNumber of predicted TMHs: 1\nExp number of AAs in TMHs: 24.51309\nExp number, first 60 AAs: 0.2682\nTotal prob of N-in: 0.14402\noutside 1 331\nTMhelix 332 354\ninside 355 372

17169	GCF_001271425.1_Salmonella_enterica_CVM_N43464_v1.0	Salmonella enterica subsp. enterica serovar Enteritidis	Proteobacteria; Gammaproteobacteria; Enterobacterales; Enterobacteriaceae; Salmonella; Salmonella enterica; Salmonella enterica subsp. enterica	MTGDNTLITSHGINRRDFMKLCAALAATMGLSSKAAAEMAESVSNPQRPPVIWIGAQECT	WP_000145429.1	hydrogenase 2 small subunit [Salmonella enterica]	Length: 372\nNumber of predicted TMHs: 1\nExp number of AAs in TMHs: 24.51309\nExp number, first 60 AAs: 0.2682\nTotal prob of N-in: 0.14402\noutside 1 331\nTMhelix 332 354\ninside 355 372

17170	GCF_001973025.1_ASM197302v1	Salmonella enterica subsp. enterica serovar Enteritidis	Proteobacteria; Gammaproteobacteria; Enterobacterales; Enterobacteriaceae; Salmonella; Salmonella enterica; Salmonella enterica subsp. enterica	MTGDNTLITSHGINRRDFMKLCAALAATMGLSSKAAAEMAESVSNPQRPPVIWIGAQECT	WP_000145429.1	hydrogenase 2 small subunit [Salmonella enterica]	Length: 372\nNumber of predicted TMHs: 1\nExp number of AAs in TMHs: 24.51309\nExp number, first 60 AAs: 0.2682\nTotal prob of N-in: 0.14402\noutside 1 331\nTMhelix 332 354\ninside 355 372

17171	GCF_001972865.1_ASM197286v1	Salmonella enterica subsp. enterica serovar Enteritidis	Proteobacteria; Gammaproteobacteria; Enterobacterales; Enterobacteriaceae; Salmonella; Salmonella enterica; Salmonella enterica subsp. enterica	MNNEETFYQAMRRKGVTRRSFLKFCSLAATSLGLGAGMTPKIAWALENKPRIPVVWIHGL	WP_001058307.1	hydrogenase [Salmonella enterica]	Length: 372\nNumber of predicted TMHs: 2\nExp number of AAs in TMHs: 36.51738\nExp number, first 60 AAs: 19.72849\nTotal prob of N-in: 0.96170\nPOSSIBLE N-term signal sequence\ninside 1 19\nTMhelix 20 39\noutside 40 325\nTMhelix 326 348\ninside 349 372

17172 GCF_001972815.1_ASM197281v1 Salmonella enterica subsp. enterica serovar Enteritidis Proteobacteria; Gammaproteobacteria; Enterobacterales; Enterobacteriaceae; Salmonella; Salmonella enterica; Salmonella enterica subsp. enterica MNNEETFYQAMRRKGVTRRSFLKFCSLAATSLGLGAGMTPKIAWALENKPRIPVVWIHGL WP_001058307.1 hydrogenase [Salmonella enterica] Length: 372\nNumber of predicted TMHs: 2\nExp number of AAs in TMHs: 36.51738\nExp number, first 60 AAs: 19.72849\nTotal prob of N-in: 0.96170\nPOSSIBLE N-term signal sequence\ninside 1 19\nTMhelix 20 39\noutside 40 325\nTMhelix 326 348\ninside 349 372

17173 GCF_001972815.1_ASM197281v1 Salmonella enterica subsp. enterica serovar Enteritidis Proteobacteria; Gammaproteobacteria; Enterobacterales; Enterobacteriaceae; Salmonella; Salmonella enterica; Salmonella enterica subsp. enterica MTGDNTLITSHGINRRDFMKLCAALAATMGLSSKAAAEMAESVSNPQRPPVIWIGAQECT WP_000145429.1 hydrogenase 2 small subunit [Salmonella enterica] Length: 372\nNumber of predicted TMHs: 1\nExp number of AAs in TMHs: 24.51309\nExp number, first 60 AAs: 0.2682\nTotal prob of N-in: 0.14402\noutside 1 331\nTMhelix 332 354\ninside 355 372

17174 GCF_001271905.1_Salmonella_enterica_CVM_N47727_v1.0 Salmonella enterica subsp. enterica serovar Enteritidis Proteobacteria; Gammaproteobacteria; Enterobacterales; Enterobacteriaceae; Salmonella; Salmonella enterica; Salmonella enterica subsp. enterica MNNEETFYQAMRRKGVTRRSFLKFCSLAATSLGLGAGMTPKIAWALENKPRIPVVWIHGL WP_001058307.1 hydrogenase [Salmonella enterica] Length: 372\nNumber of predicted TMHs: 2\nExp number of AAs in TMHs: 36.51738\nExp number, first 60 AAs: 19.72849\nTotal prob of N-in: 0.96170\nPOSSIBLE N-term signal sequence\ninside 1 19\nTMhelix 20 39\noutside 40 325\nTMhelix 326 348\ninside 349 372

17175 GCF_001271955.1_Salmonella_enterica_CVM_N43476_v1.0 Salmonella enterica subsp. enterica serovar Enteritidis Proteobacteria; Gammaproteobacteria; Enterobacterales; Enterobacteriaceae; Salmonella; Salmonella enterica; Salmonella enterica subsp. enterica MNNEETFYQAMRRKGVTRRSFLKFCSLAATSLGLGAGMTPKIAWALENKPRIPVVWIHGL WP_001058307.1 hydrogenase [Salmonella enterica] Length: 372\nNumber of predicted TMHs: 2\nExp number of AAs in TMHs: 36.51738\nExp number, first 60 AAs: 19.72849\nTotal prob of N-in: 0.96170\nPOSSIBLE N-term signal sequence\ninside 1 19\nTMhelix 20 39\noutside 40 325\nTMhelix 326 348\ninside 349 372

17176 GCF_001972785.1_ASM197278v1 Salmonella enterica subsp. enterica serovar Enteritidis Proteobacteria; Gammaproteobacteria; Enterobacterales; Enterobacteriaceae; Salmonella; Salmonella enterica; Salmonella enterica subsp. enterica MTGDNTLITSHGINRRDFMKLCAALAATMGLSSKAAAEMAESVSNPQRPPVIWIGAQECT WP_000145429.1 hydrogenase 2 small subunit [Salmonella enterica] Length: 372\nNumber of predicted TMHs: 1\nExp number of AAs in TMHs: 24.51309\nExp number, first 60 AAs: 0.2682\nTotal prob of N-in: 0.14402\noutside 1 331\nTMhelix 332 354\ninside 355 372

17177 GCF_001972705.1_ASM197270v1 Salmonella enterica subsp. enterica serovar Enteritidis Proteobacteria; Gammaproteobacteria; Enterobacterales; Enterobacteriaceae; Salmonella; Salmonella enterica; Salmonella enterica subsp. enterica MTGDNTLITSHGINRRDFMKLCAALAATMGLSSKAAAEMAESVSNPQRPPVIWIGAQECT WP_000145429.1 hydrogenase 2 small subunit [Salmonella enterica] Length: 372\nNumber of predicted TMHs: 1\nExp number of AAs in TMHs: 24.51309\nExp number, first 60 AAs: 0.2682\nTotal prob of N-in: 0.14402\noutside 1 331\nTMhelix 332 354\ninside 355 372

17178 GCF_001272485.1_Salmonella_enterica_CVM_N51310_v1.0 Salmonella enterica subsp. enterica serovar Enteritidis Proteobacteria; Gammaproteobacteria; Enterobacterales; Enterobacteriaceae; Salmonella; Salmonella enterica; Salmonella enterica subsp. enterica MTGDNTLITSHGINRRDFMKLCAALAATMGLSSKAAAEMAESVSNPQRPPVIWIGAQECT WP_000145429.1 hydrogenase 2 small subunit [Salmonella enterica] Length: 372\nNumber of predicted TMHs: 1\nExp number of AAs in TMHs: 24.51309\nExp number, first 60 AAs: 0.2682\nTotal prob of N-in: 0.14402\noutside 1 331\nTMhelix 332 354\ninside 355 372

17179 GCF_001972725.1_ASM197272v1 Salmonella enterica subsp. enterica serovar Enteritidis Proteobacteria; Gammaproteobacteria; Enterobacterales; Enterobacteriaceae; Salmonella; Salmonella enterica; Salmonella enterica subsp. enterica MTGDNTLITSHGINRRDFMKLCAALAATMGLSSKAAAEMAESVSNPQRPPVIWIGAQECT WP_000145429.1 hydrogenase 2 small subunit [Salmonella enterica] Length: 372\nNumber of predicted TMHs: 1\nExp number of AAs in TMHs: 24.51309\nExp number, first 60 AAs: 0.2682\nTotal prob of N-in: 0.14402\noutside 1 331\nTMhelix 332 354\ninside 355 372

17180 GCF_001972645.1_ASM197264v1 Salmonella enterica subsp. enterica serovar Enteritidis Proteobacteria; Gammaproteobacteria; Enterobacterales; Enterobacteriaceae; Salmonella; Salmonella enterica;

Salmonella enterica subsp. enterica	MTGDNTLITSHGINRRDFMKLCAALAATMGLSSKAAAEMAESVSNPQRPPVIWIGAQECT
	WP_000145429.1	hydrogenase 2 small subunit [Salmonella enterica]	Length: 372\nNumber of predicted TMHs:  1\nExp number of AAs in TMHs: 24.51309\nExp number, first 60 AAs:  0.2682\nTotal prob of N-in: 0.14402\noutside	1	331\nTMhelix	332	354\ninside	355	372

17181	GCF_002040015.1_ASM204001v1	Salmonella enterica subsp. enterica serovar Enteritidis
	Proteobacteria; Gammaproteobacteria; Enterobacterales; Enterobacteriaceae; Salmonella; Salmonella enterica; Salmonella enterica subsp. enterica	MTGDNTLITSHGINRRDFMKLCAALAATMGLSSKAAAEMAESVSNPQRPPVIWIGAQECT
	WP_000145429.1	hydrogenase 2 small subunit [Salmonella enterica]	Length: 372\nNumber of predicted TMHs:  1\nExp number of AAs in TMHs: 24.51309\nExp number, first 60 AAs:  0.2682\nTotal prob of N-in: 0.14402\noutside	1	331\nTMhelix	332	354\ninside	355	372

17182	GCF_002040215.1_ASM204021v1	Salmonella enterica subsp. enterica serovar Enteritidis
	Proteobacteria; Gammaproteobacteria; Enterobacterales; Enterobacteriaceae; Salmonella; Salmonella enterica; Salmonella enterica subsp. enterica	MTGDNTLITSHGINRRDFMKLCAALAATMGLSSKAAAEMAESVSNPQRPPVIWIGAQECT
	WP_000145429.1	hydrogenase 2 small subunit [Salmonella enterica]	Length: 372\nNumber of predicted TMHs:  1\nExp number of AAs in TMHs: 24.51309\nExp number, first 60 AAs:  0.2682\nTotal prob of N-in: 0.14402\noutside	1	331\nTMhelix	332	354\ninside	355	372

17183	GCF_001305235.1_ASM130523v1	Salmonella enterica subsp. enterica serovar Enteritidis
	Proteobacteria; Gammaproteobacteria; Enterobacterales; Enterobacteriaceae; Salmonella; Salmonella enterica; Salmonella enterica subsp. enterica	MTGDNTLITSHGINRRDFMKLCAALAATMGLSSKAAAEMAESVSNPQRPPVIWIGAQECT
	WP_000145429.1	hydrogenase 2 small subunit [Salmonella enterica]	Length: 372\nNumber of predicted TMHs:  1\nExp number of AAs in TMHs: 24.51309\nExp number, first 60 AAs:  0.2682\nTotal prob of N-in: 0.14402\noutside	1	331\nTMhelix	332	354\ninside	355	372

17184	GCF_001305235.1_ASM130523v1	Salmonella enterica subsp. enterica serovar Enteritidis
	Proteobacteria; Gammaproteobacteria; Enterobacterales; Enterobacteriaceae; Salmonella; Salmonella enterica; Salmonella enterica subsp. enterica	MNNEETFYQAMRRKGVTRRSFLKFCSLAATSLGLGAGMTPKIAWALENKPRIPVVWIHGL
	WP_001058307.1	hydrogenase [Salmonella enterica]	Length: 372\nNumber of predicted TMHs:  2\nExp number of AAs in TMHs: 36.51738\nExp number, first 60 AAs:  19.72849\nTotal prob of N-in:	0.96170\nPOSSIBLE N-term signal sequence\ninside	1	19\nTMhelix	20	39\noutside	40	325\nTMhelix	326	348\ninside	349	372

17185	GCF_001972625.1_ASM197262v1	Salmonella enterica subsp. enterica serovar Enteritidis
	Proteobacteria; Gammaproteobacteria; Enterobacterales; Enterobacteriaceae; Salmonella; Salmonella enterica; Salmonella enterica subsp. enterica	MTGDNTLITSHGINRRDFMKLCAALAATMGLSSKAAAEMAESVSNPQRPPVIWIGAQECT
	WP_000145429.1	hydrogenase 2 small subunit [Salmonella enterica]	Length: 372\nNumber of predicted TMHs:  1\nExp number of AAs in TMHs: 24.51309\nExp number, first 60 AAs:  0.2682\nTotal prob of N-in: 0.14402\noutside	1	331\nTMhelix	332	354\ninside	355	372

17186	GCF_001972865.1_ASM197286v1	Salmonella enterica subsp. enterica serovar Enteritidis
	Proteobacteria; Gammaproteobacteria; Enterobacterales; Enterobacteriaceae; Salmonella; Salmonella enterica; Salmonella enterica subsp. enterica	MTGDNTLITSHGINRRDFMKLCAALAATMGLSSKAAAEMAESVSNPQRPPVIWIGAQECT
	WP_000145429.1	hydrogenase 2 small subunit [Salmonella enterica]	Length: 372\nNumber of predicted TMHs:  1\nExp number of AAs in TMHs: 24.51309\nExp number, first 60 AAs:  0.2682\nTotal prob of N-in: 0.14402\noutside	1	331\nTMhelix	332	354\ninside	355	372

17187	GCF_002040305.1_ASM204030v1	Salmonella enterica subsp. enterica serovar Enteritidis
	Proteobacteria; Gammaproteobacteria; Enterobacterales; Enterobacteriaceae; Salmonella; Salmonella enterica; Salmonella enterica subsp. enterica	MTGDNTLITSHGINRRDFMKLCAALAATMGLSSKAAAEMAESVSNPQRPPVIWIGAQECT
	WP_000145429.1	hydrogenase 2 small subunit [Salmonella enterica]	Length: 372\nNumber of predicted TMHs:  1\nExp number of AAs in TMHs: 24.51309\nExp number, first 60 AAs:  0.2682\nTotal prob of N-in: 0.14402\noutside	1	331\nTMhelix	332	354\ninside	355	372

17188	GCF_002040325.1_ASM204032v1	Salmonella enterica subsp. enterica serovar Enteritidis
	Proteobacteria; Gammaproteobacteria; Enterobacterales; Enterobacteriaceae; Salmonella; Salmonella enterica; Salmonella enterica subsp. enterica	MTGDNTLITSHGINRRDFMKLCAALAATMGLSSKAAAEMAESVSNPQRPPVIWIGAQECT
	WP_000145429.1	hydrogenase 2 small subunit [Salmonella enterica]	Length: 372\nNumber of predicted TMHs:  1\nExp number of AAs in TMHs: 24.51309\nExp number, first 60 AAs:  0.2682\nTotal prob of N-in: 0.14402\noutside	1	331\nTMhelix	332	354\ninside	355	372

17189 GCF_002040215.1_ASM204021v1 Salmonella enterica subsp. enterica serovar Enteritidis Proteobacteria; Gammaproteobacteria; Enterobacterales; Enterobacteriaceae; Salmonella; Salmonella enterica; Salmonella enterica subsp. enterica MNNEETFYQAMRRKGVTRRSFLKFCSLAATSLGLGAGMTPKIAWALENKPRIPVVWIHGL WP_001058307.1 hydrogenase [Salmonella enterica] Length: 372\nNumber of predicted TMHs: 2\nExp number of AAs in TMHs: 36.51738\nExp number, first 60 AAs: 19.72849\nTotal prob of N-in: 0.96170\nPOSSIBLE N-term signal sequence\ninside 1 19\nTMhelix 20 39\noutside 40 325\nTMhelix 326 348\ninside 349 372

17190 GCF_002040255.1_ASM204025v1 Salmonella enterica subsp. enterica serovar Enteritidis Proteobacteria; Gammaproteobacteria; Enterobacterales; Enterobacteriaceae; Salmonella; Salmonella enterica; Salmonella enterica subsp. enterica MNNEETFYQAMRRKGVTRRSFLKFCSLAATSLGLGAGMTPKIAWALENKPRIPVVWIHGL WP_001058307.1 hydrogenase [Salmonella enterica] Length: 372\nNumber of predicted TMHs: 2\nExp number of AAs in TMHs: 36.51738\nExp number, first 60 AAs: 19.72849\nTotal prob of N-in: 0.96170\nPOSSIBLE N-term signal sequence\ninside 1 19\nTMhelix 20 39\noutside 40 325\nTMhelix 326 348\ninside 349 372

17191 GCF_002040385.1_ASM204038v1 Salmonella enterica subsp. enterica serovar Enteritidis Proteobacteria; Gammaproteobacteria; Enterobacterales; Enterobacteriaceae; Salmonella; Salmonella enterica; Salmonella enterica subsp. enterica MTGDNTLITSHGINRRDFMKLCAALAATMGLSSKAAAEMAESVSNPQRPPVIWIGAQECT WP_000145429.1 hydrogenase 2 small subunit [Salmonella enterica] Length: 372\nNumber of predicted TMHs: 1\nExp number of AAs in TMHs: 24.51309\nExp number, first 60 AAs: 0.2682\nTotal prob of N-in: 0.14402\noutside 1 331\nTMhelix 332 354\ninside 355 372

17192 GCF_002040405.1_ASM204040v1 Salmonella enterica subsp. enterica serovar Enteritidis Proteobacteria; Gammaproteobacteria; Enterobacterales; Enterobacteriaceae; Salmonella; Salmonella enterica; Salmonella enterica subsp. enterica MTGDNTLITSHGINRRDFMKLCAALAATMGLSSKAAAEMAESVSNPQRPPVIWIGAQECT WP_000145429.1 hydrogenase 2 small subunit [Salmonella enterica] Length: 372\nNumber of predicted TMHs: 1\nExp number of AAs in TMHs: 24.51309\nExp number, first 60 AAs: 0.2682\nTotal prob of N-in: 0.14402\noutside 1 331\nTMhelix 332 354\ninside 355 372

17193 GCF_002040565.1_ASM204056v1 Salmonella enterica subsp. enterica serovar Enteritidis Proteobacteria; Gammaproteobacteria; Enterobacterales; Enterobacteriaceae; Salmonella; Salmonella enterica; Salmonella enterica subsp. enterica MNNEETFYQAMRRKGVTRRSFLKFCSLAATSLGLGAGMTPKIAWALENKPRIPVVWIHGL WP_001058307.1 hydrogenase [Salmonella enterica] Length: 372\nNumber of predicted TMHs: 2\nExp number of AAs in TMHs: 36.51738\nExp number, first 60 AAs: 19.72849\nTotal prob of N-in: 0.96170\nPOSSIBLE N-term signal sequence\ninside 1 19\nTMhelix 20 39\noutside 40 325\nTMhelix 326 348\ninside 349 372

17194 GCF_000754375.1_ASM75437v1 Salmonella enterica subsp. enterica serovar Enteritidis Proteobacteria; Gammaproteobacteria; Enterobacterales; Enterobacteriaceae; Salmonella; Salmonella enterica; Salmonella enterica subsp. enterica MTGDNTLITSHGINRRDFMKLCAALAATMGLSSKAAAEMAESVSNPQRPPVIWIGAQECT WP_000145429.1 hydrogenase 2 small subunit [Salmonella enterica] Length: 372\nNumber of predicted TMHs: 1\nExp number of AAs in TMHs: 24.51309\nExp number, first 60 AAs: 0.2682\nTotal prob of N-in: 0.14402\noutside 1 331\nTMhelix 332 354\ninside 355 372

17195 GCF_002042795.1_ASM204279v1 Salmonella enterica subsp. enterica serovar Enteritidis Proteobacteria; Gammaproteobacteria; Enterobacterales; Enterobacteriaceae; Salmonella; Salmonella enterica; Salmonella enterica subsp. enterica MNNEETFYQAMRRKGVTRRSFLKFCSLAATSLGLGAGMTPKIAWALENKPRIPVVWIHGL WP_001058307.1 hydrogenase [Salmonella enterica] Length: 372\nNumber of predicted TMHs: 2\nExp number of AAs in TMHs: 36.51738\nExp number, first 60 AAs: 19.72849\nTotal prob of N-in: 0.96170\nPOSSIBLE N-term signal sequence\ninside 1 19\nTMhelix 20 39\noutside 40 325\nTMhelix 326 348\ninside 349 372

17196 GCF_002042755.1_ASM204275v1 Salmonella enterica subsp. enterica serovar Enteritidis Proteobacteria; Gammaproteobacteria; Enterobacterales; Enterobacteriaceae; Salmonella; Salmonella enterica; Salmonella enterica subsp. enterica MNNEETFYQAMRRKGVTRRSFLKFCSLAATSLGLGAGMTPKIAWALENKPRIPVVWIHGL WP_001058307.1 hydrogenase [Salmonella enterica] Length: 372\nNumber of predicted TMHs: 2\nExp number of AAs in TMHs: 36.51738\nExp number, first 60 AAs: 19.72849\nTotal prob of N-in: 0.96170\nPOSSIBLE N-term signal sequence\ninside 1 19\nTMhelix 20 39\noutside 40 325\nTMhelix 326 348\ninside 349 372

17197 GCF_002042725.1_ASM204272v1 Salmonella enterica subsp. enterica serovar Enteritidis Proteobacteria; Gammaproteobacteria; Enterobacterales; Enterobacteriaceae; Salmonella; Salmonella enterica; Salmonella enterica subsp. enterica MNNEETFYQAMRRKGVTRRSFLKFCSLAATSLGLGAGMTPKIAWALENKPRIPVVWIHGL WP_001058307.1 hydrogenase [Salmonella enterica] Length: 372\nNumber of predicted TMHs: 2\nExp

number of AAs in TMHs: 36.51738\nExp number, first 60 AAs: 19.72849\nTotal prob of N-in: 0.96170\nPOSSIBLE N-term signal sequence\ninside 1 19\nTMhelix 20 39\noutside 40 325\nTMhelix 326 348\ninside 349 372

17198 GCF_002042395.1_ASM204239v1 Salmonella enterica subsp. enterica serovar Enteritidis Proteobacteria; Gammaproteobacteria; Enterobacterales; Enterobacteriaceae; Salmonella; Salmonella enterica; Salmonella enterica subsp. enterica MNNEETFYQAMRRKGVTRRSFLKFCSLAATSLGLGAGMTPKIAWALENKPRIPVVWIHGL WP_001058307.1 hydrogenase [Salmonella enterica] Length: 372\nNumber of predicted TMHs: 2\nExp number of AAs in TMHs: 36.51738\nExp number, first 60 AAs: 19.72849\nTotal prob of N-in: 0.96170\nPOSSIBLE N-term signal sequence\ninside 1 19\nTMhelix 20 39\noutside 40 325\nTMhelix 326 348\ninside 349 372

17199 GCF_002042445.1_ASM204244v1 Salmonella enterica subsp. enterica serovar Enteritidis Proteobacteria; Gammaproteobacteria; Enterobacterales; Enterobacteriaceae; Salmonella; Salmonella enterica; Salmonella enterica subsp. enterica MNNEETFYQAMRRKGVTRRSFLKFCSLAATSLGLGAGMTPKIAWALENKPRIPVVWIHGL WP_001058307.1 hydrogenase [Salmonella enterica] Length: 372\nNumber of predicted TMHs: 2\nExp number of AAs in TMHs: 36.51738\nExp number, first 60 AAs: 19.72849\nTotal prob of N-in: 0.96170\nPOSSIBLE N-term signal sequence\ninside 1 19\nTMhelix 20 39\noutside 40 325\nTMhelix 326 348\ninside 349 372

17200 GCF_002042365.1_ASM204236v1 Salmonella enterica subsp. enterica serovar Enteritidis Proteobacteria; Gammaproteobacteria; Enterobacterales; Enterobacteriaceae; Salmonella; Salmonella enterica; Salmonella enterica subsp. enterica MNNEETFYQAMRRKGVTRRSFLKFCSLAATSLGLGAGMTPKIAWALENKPRIPVVWIHGL WP_001058307.1 hydrogenase [Salmonella enterica] Length: 372\nNumber of predicted TMHs: 2\nExp number of AAs in TMHs: 36.51738\nExp number, first 60 AAs: 19.72849\nTotal prob of N-in: 0.96170\nPOSSIBLE N-term signal sequence\ninside 1 19\nTMhelix 20 39\noutside 40 325\nTMhelix 326 348\ninside 349 372

17201 GCF_002042445.1_ASM204244v1 Salmonella enterica subsp. enterica serovar Enteritidis Proteobacteria; Gammaproteobacteria; Enterobacterales; Enterobacteriaceae; Salmonella; Salmonella enterica; Salmonella enterica subsp. enterica MTGDNTLITSHGINRRDFMKLCAALAATMGLSSKAAAEMAESVSNPQRPPVIWIGAQECT WP_000145429.1 hydrogenase 2 small subunit [Salmonella enterica] Length: 372\nNumber of predicted TMHs: 1\nExp number of AAs in TMHs: 24.51309\nExp number, first 60 AAs: 0.2682\nTotal prob of N-in: 0.14402\noutside 1 331\nTMhelix 332 354\ninside 355 372

17202 GCF_002042395.1_ASM204239v1 Salmonella enterica subsp. enterica serovar Enteritidis Proteobacteria; Gammaproteobacteria; Enterobacterales; Enterobacteriaceae; Salmonella; Salmonella enterica; Salmonella enterica subsp. enterica MTGDNTLITSHGINRRDFMKLCAALAATMGLSSKAAAEMAESVSNPQRPPVIWIGAQECT WP_000145429.1 hydrogenase 2 small subunit [Salmonella enterica] Length: 372\nNumber of predicted TMHs: 1\nExp number of AAs in TMHs: 24.51309\nExp number, first 60 AAs: 0.2682\nTotal prob of N-in: 0.14402\noutside 1 331\nTMhelix 332 354\ninside 355 372

17203 GCF_002042365.1_ASM204236v1 Salmonella enterica subsp. enterica serovar Enteritidis Proteobacteria; Gammaproteobacteria; Enterobacterales; Enterobacteriaceae; Salmonella; Salmonella enterica; Salmonella enterica subsp. enterica MTGDNTLITSHGINRRDFMKLCAALAATMGLSSKAAAEMAESVSNPQRPPVIWIGAQECT WP_000145429.1 hydrogenase 2 small subunit [Salmonella enterica] Length: 372\nNumber of predicted TMHs: 1\nExp number of AAs in TMHs: 24.51309\nExp number, first 60 AAs: 0.2682\nTotal prob of N-in: 0.14402\noutside 1 331\nTMhelix 332 354\ninside 355 372

17204 GCF_002042225.1_ASM204222v1 Salmonella enterica subsp. enterica serovar Enteritidis Proteobacteria; Gammaproteobacteria; Enterobacterales; Enterobacteriaceae; Salmonella; Salmonella enterica; Salmonella enterica subsp. enterica MNNEETFYQAMRRKGVTRRSFLKFCSLAATSLGLGAGMTPKIAWALENKPRIPVVWIHGL WP_001058307.1 hydrogenase [Salmonella enterica] Length: 372\nNumber of predicted TMHs: 2\nExp number of AAs in TMHs: 36.51738\nExp number, first 60 AAs: 19.72849\nTotal prob of N-in: 0.96170\nPOSSIBLE N-term signal sequence\ninside 1 19\nTMhelix 20 39\noutside 40 325\nTMhelix 326 348\ninside 349 372

17205 GCF_002041935.1_ASM204193v1 Salmonella enterica subsp. enterica serovar Enteritidis Proteobacteria; Gammaproteobacteria; Enterobacterales; Enterobacteriaceae; Salmonella; Salmonella enterica; Salmonella enterica subsp. enterica MNNEETFYQAMRRKGVTRRSFLKFCSLAATSLGLGAGMTPKIAWALENKPRIPVVWIHGL WP_001058307.1 hydrogenase [Salmonella enterica] Length: 372\nNumber of predicted TMHs: 2\nExp number of AAs in TMHs: 36.51738\nExp number, first 60 AAs: 19.72849\nTotal prob of N-in: 0.96170\nPOSSIBLE N-term signal sequence\ninside 1 19\nTMhelix 20 39\noutside 40 325\nTMhelix 326 348\ninside 349 372

17206 GCF_002040675.1_ASM204067v1 Salmonella enterica subsp. enterica serovar Enteritidis Proteobacteria; Gammaproteobacteria; Enterobacterales; Enterobacteriaceae; Salmonella; Salmonella enterica;

Salmonella enterica subsp. enterica	MTGDNTLITSHGINRRDFMKLCAALAATMGLSSKAAAEMAESVSNPQRPPVIWIGAQECT	WP_000145429.1	hydrogenase 2 small subunit [Salmonella enterica]	Length: 372\nNumber of predicted TMHs:  1\nExp number of AAs in TMHs: 24.51309\nExp number, first 60 AAs:  0.2682\nTotal prob of N-in:  0.14402\noutside   1   331\nTMhelix   332   354\ninside   355   372

17207	GCF_002041905.1_ASM204190v1	Salmonella enterica subsp. enterica serovar Enteritidis	Proteobacteria; Gammaproteobacteria; Enterobacterales; Enterobacteriaceae; Salmonella; Salmonella enterica; Salmonella enterica subsp. enterica	MNNEETFYQAMRRKGVTRRSFLKFCSLAATSLGLGAGMTPKIAWALENKPRIPVVWIHGL	WP_001058307.1	hydrogenase [Salmonella enterica]	Length: 372\nNumber of predicted TMHs:  2\nExp number of AAs in TMHs: 36.51738\nExp number, first 60 AAs:  19.72849\nTotal prob of N-in:   0.96170\nPOSSIBLE N-term signal sequence\ninside   1   19\nTMhelix   20   39\noutside   40   325\nTMhelix   326   348\ninside   349   372

17208	GCF_002042065.1_ASM204206v1	Salmonella enterica subsp. enterica serovar Enteritidis	Proteobacteria; Gammaproteobacteria; Enterobacterales; Enterobacteriaceae; Salmonella; Salmonella enterica; Salmonella enterica subsp. enterica	MNNEETFYQAMRRKGVTRRSFLKFCSLAATSLGLGAGMTPKIAWALENKPRIPVVWIHGL	WP_001058307.1	hydrogenase [Salmonella enterica]	Length: 372\nNumber of predicted TMHs:  2\nExp number of AAs in TMHs: 36.51738\nExp number, first 60 AAs:  19.72849\nTotal prob of N-in:   0.96170\nPOSSIBLE N-term signal sequence\ninside   1   19\nTMhelix   20   39\noutside   40   325\nTMhelix   326   348\ninside   349   372

17209	GCF_002042065.1_ASM204206v1	Salmonella enterica subsp. enterica serovar Enteritidis	Proteobacteria; Gammaproteobacteria; Enterobacterales; Enterobacteriaceae; Salmonella; Salmonella enterica; Salmonella enterica subsp. enterica	MTGDNTLITSHGINRRDFMKLCAALAATMGLSSKAAAEMAESVSNPQRPPVIWIGAQECT	WP_000145429.1	hydrogenase 2 small subunit [Salmonella enterica]	Length: 372\nNumber of predicted TMHs:  1\nExp number of AAs in TMHs: 24.51309\nExp number, first 60 AAs:  0.2682\nTotal prob of N-in:  0.14402\noutside   1   331\nTMhelix   332   354\ninside   355   372

17210	GCF_002041845.1_ASM204184v1	Salmonella enterica subsp. enterica serovar Enteritidis	Proteobacteria; Gammaproteobacteria; Enterobacterales; Enterobacteriaceae; Salmonella; Salmonella enterica; Salmonella enterica subsp. enterica	MNNEETFYQAMRRKGVTRRSFLKFCSLAATSLGLGAGMTPKIAWALENKPRIPVVWIHGL	WP_001058307.1	hydrogenase [Salmonella enterica]	Length: 372\nNumber of predicted TMHs:  2\nExp number of AAs in TMHs: 36.51738\nExp number, first 60 AAs:  19.72849\nTotal prob of N-in:   0.96170\nPOSSIBLE N-term signal sequence\ninside   1   19\nTMhelix   20   39\noutside   40   325\nTMhelix   326   348\ninside   349   372

17211	GCF_002041825.1_ASM204182v1	Salmonella enterica subsp. enterica serovar Enteritidis	Proteobacteria; Gammaproteobacteria; Enterobacterales; Enterobacteriaceae; Salmonella; Salmonella enterica; Salmonella enterica subsp. enterica	MNNEETFYQAMRRKGVTRRSFLKFCSLAATSLGLGAGMTPKIAWALENKPRIPVVWIHGL	WP_001058307.1	hydrogenase [Salmonella enterica]	Length: 372\nNumber of predicted TMHs:  2\nExp number of AAs in TMHs: 36.51738\nExp number, first 60 AAs:  19.72849\nTotal prob of N-in:   0.96170\nPOSSIBLE N-term signal sequence\ninside   1   19\nTMhelix   20   39\noutside   40   325\nTMhelix   326   348\ninside   349   372

17212	GCF_002041685.1_ASM204168v1	Salmonella enterica subsp. enterica serovar Enteritidis	Proteobacteria; Gammaproteobacteria; Enterobacterales; Enterobacteriaceae; Salmonella; Salmonella enterica; Salmonella enterica subsp. enterica	MTGDNTLITSHGINRRDFMKLCAALAATMGLSSKAAAEMAESVSNPQRPPVIWIGAQECT	WP_000145429.1	hydrogenase 2 small subunit [Salmonella enterica]	Length: 372\nNumber of predicted TMHs:  1\nExp number of AAs in TMHs: 24.51309\nExp number, first 60 AAs:  0.2682\nTotal prob of N-in:  0.14402\noutside   1   331\nTMhelix   332   354\ninside   355   372

17213	GCF_002041625.1_ASM204162v1	Salmonella enterica subsp. enterica serovar Enteritidis	Proteobacteria; Gammaproteobacteria; Enterobacterales; Enterobacteriaceae; Salmonella; Salmonella enterica; Salmonella enterica subsp. enterica	MTGDNTLITSHGINRRDFMKLCAALAATMGLSSKAAAEMAESVSNPQRPPVIWIGAQECT	WP_000145429.1	hydrogenase 2 small subunit [Salmonella enterica]	Length: 372\nNumber of predicted TMHs:  1\nExp number of AAs in TMHs: 24.51309\nExp number, first 60 AAs:  0.2682\nTotal prob of N-in:  0.14402\noutside   1   331\nTMhelix   332   354\ninside   355   372

17214	GCF_002041705.1_ASM204170v1	Salmonella enterica subsp. enterica serovar Enteritidis	Proteobacteria; Gammaproteobacteria; Enterobacterales; Enterobacteriaceae; Salmonella; Salmonella enterica; Salmonella enterica subsp. enterica	MTGDNTLITSHGINRRDFMKLCAALAATMGLSSKAAAEMAESVSNPQRPPVIWIGAQECT	WP_000145429.1	hydrogenase 2 small subunit [Salmonella enterica]	Length: 372\nNumber of predicted TMHs:  1\nExp number of AAs in TMHs: 24.51309\nExp number, first 60 AAs:  0.2682\nTotal prob of N-in:  0.14402\noutside   1   331\nTMhelix   332   354\ninside   355   372

17215 GCF_002040675.1_ASM204067v1 Salmonella enterica subsp. enterica serovar Enteritidis Proteobacteria; Gammaproteobacteria; Enterobacterales; Enterobacteriaceae; Salmonella; Salmonella enterica; Salmonella enterica subsp. enterica MNNEETFYQAMRRKGVTRRSFLKFCSLAATSLGLGAGMTPKIAWALENKPRIPVVWIHGL WP_001058307.1 hydrogenase [Salmonella enterica] Length: 372\nNumber of predicted TMHs: 2\nExp number of AAs in TMHs: 36.51738\nExp number, first 60 AAs: 19.72849\nTotal prob of N-in: 0.96170\nPOSSIBLE N-term signal sequence\ninside 1 19\nTMhelix 20 39\noutside 40 325\nTMhelix 326 348\ninside 349 372

17216 GCF_002041545.1_ASM204154v1 Salmonella enterica subsp. enterica serovar Enteritidis Proteobacteria; Gammaproteobacteria; Enterobacterales; Enterobacteriaceae; Salmonella; Salmonella enterica; Salmonella enterica subsp. enterica MNNEETFYQAMRRKGVTRRSFLKFCSLAATSLGLGAGMTPKIAWALENKPRIPVVWIHGL WP_001058307.1 hydrogenase [Salmonella enterica] Length: 372\nNumber of predicted TMHs: 2\nExp number of AAs in TMHs: 36.51738\nExp number, first 60 AAs: 19.72849\nTotal prob of N-in: 0.96170\nPOSSIBLE N-term signal sequence\ninside 1 19\nTMhelix 20 39\noutside 40 325\nTMhelix 326 348\ninside 349 372

17217 GCF_002041605.1_ASM204160v1 Salmonella enterica subsp. enterica serovar Enteritidis Proteobacteria; Gammaproteobacteria; Enterobacterales; Enterobacteriaceae; Salmonella; Salmonella enterica; Salmonella enterica subsp. enterica MTGDNTLITSHGINRRDFMKLCAALAATMGLSSKAAAEMAESVSNPQRPPVIWIGAQECT WP_000145429.1 hydrogenase 2 small subunit [Salmonella enterica] Length: 372\nNumber of predicted TMHs: 1\nExp number of AAs in TMHs: 24.51309\nExp number, first 60 AAs: 0.2682\nTotal prob of N-in: 0.14402\noutside 1 331\nTMhelix 332 354\ninside 355 372

17218 GCF_002041545.1_ASM204154v1 Salmonella enterica subsp. enterica serovar Enteritidis Proteobacteria; Gammaproteobacteria; Enterobacterales; Enterobacteriaceae; Salmonella; Salmonella enterica; Salmonella enterica subsp. enterica MTGDNTLITSHGINRRDFMKLCAALAATMGLSSKAAAEMAESVSNPQRPPVIWIGAQECT WP_000145429.1 hydrogenase 2 small subunit [Salmonella enterica] Length: 372\nNumber of predicted TMHs: 1\nExp number of AAs in TMHs: 24.51309\nExp number, first 60 AAs: 0.2682\nTotal prob of N-in: 0.14402\noutside 1 331\nTMhelix 332 354\ninside 355 372

17219 GCF_002041325.1_ASM204132v1 Salmonella enterica subsp. enterica serovar Enteritidis Proteobacteria; Gammaproteobacteria; Enterobacterales; Enterobacteriaceae; Salmonella; Salmonella enterica; Salmonella enterica subsp. enterica MNNEETFYQAMRRKGVTRRSFLKFCSLAATSLGLGAGMTPKIAWALENKPRIPVVWIHGL WP_001058307.1 hydrogenase [Salmonella enterica] Length: 372\nNumber of predicted TMHs: 2\nExp number of AAs in TMHs: 36.51738\nExp number, first 60 AAs: 19.72849\nTotal prob of N-in: 0.96170\nPOSSIBLE N-term signal sequence\ninside 1 19\nTMhelix 20 39\noutside 40 325\nTMhelix 326 348\ninside 349 372

17220 GCF_002041165.1_ASM204116v1 Salmonella enterica subsp. enterica serovar Enteritidis Proteobacteria; Gammaproteobacteria; Enterobacterales; Enterobacteriaceae; Salmonella; Salmonella enterica; Salmonella enterica subsp. enterica MNNEETFYQAMRRKGVTRRSFLKFCSLAATSLGLGAGMTPKIAWALENKPRIPVVWIHGL WP_001058307.1 hydrogenase [Salmonella enterica] Length: 372\nNumber of predicted TMHs: 2\nExp number of AAs in TMHs: 36.51738\nExp number, first 60 AAs: 19.72849\nTotal prob of N-in: 0.96170\nPOSSIBLE N-term signal sequence\ninside 1 19\nTMhelix 20 39\noutside 40 325\nTMhelix 326 348\ninside 349 372

17221 GCF_002040595.1_ASM204059v1 Salmonella enterica subsp. enterica serovar Enteritidis Proteobacteria; Gammaproteobacteria; Enterobacterales; Enterobacteriaceae; Salmonella; Salmonella enterica; Salmonella enterica subsp. enterica MNNEETFYQAMRRKGVTRRSFLKFCSLAATSLGLGAGMTPKIAWALENKPRIPVVWIHGL WP_001058307.1 hydrogenase [Salmonella enterica] Length: 372\nNumber of predicted TMHs: 2\nExp number of AAs in TMHs: 36.51738\nExp number, first 60 AAs: 19.72849\nTotal prob of N-in: 0.96170\nPOSSIBLE N-term signal sequence\ninside 1 19\nTMhelix 20 39\noutside 40 325\nTMhelix 326 348\ninside 349 372

17222 GCF_002040735.1_ASM204073v1 Salmonella enterica subsp. enterica serovar Enteritidis Proteobacteria; Gammaproteobacteria; Enterobacterales; Enterobacteriaceae; Salmonella; Salmonella enterica; Salmonella enterica subsp. enterica MTGDNTLITSHGINRRDFMKLCAALAATMGLSSKAAAEMAESVSNPQRPPVIWIGAQECT WP_000145429.1 hydrogenase 2 small subunit [Salmonella enterica] Length: 372\nNumber of predicted TMHs: 1\nExp number of AAs in TMHs: 24.51309\nExp number, first 60 AAs: 0.2682\nTotal prob of N-in: 0.14402\noutside 1 331\nTMhelix 332 354\ninside 355 372

17223 GCF_002040735.1_ASM204073v1 Salmonella enterica subsp. enterica serovar Enteritidis Proteobacteria; Gammaproteobacteria; Enterobacterales; Enterobacteriaceae; Salmonella; Salmonella enterica; Salmonella enterica subsp. enterica MNNEETFYQAMRRKGVTRRSFLKFCSLAATSLGLGAGMTPKIAWALENKPRIPVVWIHGL WP_001058307.1 hydrogenase [Salmonella enterica] Length: 372\nNumber of predicted TMHs: 2\nExp

number of AAs in TMHs: 36.51738\nExp number, first 60 AAs: 19.72849\nTotal prob of N-in: 0.96170\nPOSSIBLE N-term signal sequence\ninside 1 19\nTMhelix 20 39\noutside 40 325\nTMhelix 326 348\ninside 349 372

17224 GCF_002042015.1_ASM204201v1 Salmonella enterica subsp. enterica serovar Enteritidis Proteobacteria; Gammaproteobacteria; Enterobacterales; Enterobacteriaceae; Salmonella; Salmonella enterica; Salmonella enterica subsp. enterica MTGDNTLITSHGINRRDFMKLCAALAATMGLSSKAAAEMAESVSNPQRPPVIWIGAQECT WP_000145429.1 hydrogenase 2 small subunit [Salmonella enterica] Length: 372\nNumber of predicted TMHs: 1\nExp number of AAs in TMHs: 24.51309\nExp number, first 60 AAs: 0.2682\nTotal prob of N-in: 0.14402\noutside 1 331\nTMhelix 332 354\ninside 355 372

17225 GCF_002041095.1_ASM204109v1 Salmonella enterica subsp. enterica serovar Enteritidis Proteobacteria; Gammaproteobacteria; Enterobacterales; Enterobacteriaceae; Salmonella; Salmonella enterica; Salmonella enterica subsp. enterica MNNEETFYQAMRRKGVTRRSFLKFCSLAATSLGLGAGMTPKIAWALENKPRIPVVWIHGL WP_001058307.1 hydrogenase [Salmonella enterica] Length: 372\nNumber of predicted TMHs: 2\nExp number of AAs in TMHs: 36.51738\nExp number, first 60 AAs: 19.72849\nTotal prob of N-in: 0.96170\nPOSSIBLE N-term signal sequence\ninside 1 19\nTMhelix 20 39\noutside 40 325\nTMhelix 326 348\ninside 349 372

17226 GCF_002041165.1_ASM204116v1 Salmonella enterica subsp. enterica serovar Enteritidis Proteobacteria; Gammaproteobacteria; Enterobacterales; Enterobacteriaceae; Salmonella; Salmonella enterica; Salmonella enterica subsp. enterica MTGDNTLITSHGINRRDFMKLCAALAATMGLSSKAAAEMAESVSNPQRPPVIWIGAQECT WP_000145429.1 hydrogenase 2 small subunit [Salmonella enterica] Length: 372\nNumber of predicted TMHs: 1\nExp number of AAs in TMHs: 24.51309\nExp number, first 60 AAs: 0.2682\nTotal prob of N-in: 0.14402\noutside 1 331\nTMhelix 332 354\ninside 355 372

17227 GCF_002041325.1_ASM204132v1 Salmonella enterica subsp. enterica serovar Enteritidis Proteobacteria; Gammaproteobacteria; Enterobacterales; Enterobacteriaceae; Salmonella; Salmonella enterica; Salmonella enterica subsp. enterica MTGDNTLITSHGINRRDFMKLCAALAATMGLSSKAAAEMAESVSNPQRPPVIWIGAQECT WP_000145429.1 hydrogenase 2 small subunit [Salmonella enterica] Length: 372\nNumber of predicted TMHs: 1\nExp number of AAs in TMHs: 24.51309\nExp number, first 60 AAs: 0.2682\nTotal prob of N-in: 0.14402\noutside 1 331\nTMhelix 332 354\ninside 355 372

17228 GCF_002039845.1_ASM203984v1 Salmonella enterica subsp. enterica serovar Enteritidis Proteobacteria; Gammaproteobacteria; Enterobacterales; Enterobacteriaceae; Salmonella; Salmonella enterica; Salmonella enterica subsp. enterica MNNEETFYQAMRRKGVTRRSFLKFCSLAATSLGLGAGMTPKIAWALENKPRIPVVWIHGL WP_001058307.1 hydrogenase [Salmonella enterica] Length: 372\nNumber of predicted TMHs: 2\nExp number of AAs in TMHs: 36.51738\nExp number, first 60 AAs: 19.72849\nTotal prob of N-in: 0.96170\nPOSSIBLE N-term signal sequence\ninside 1 19\nTMhelix 20 39\noutside 40 325\nTMhelix 326 348\ninside 349 372

17229 GCF_002039745.1_ASM203974v1 Salmonella enterica subsp. enterica serovar Enteritidis Proteobacteria; Gammaproteobacteria; Enterobacterales; Enterobacteriaceae; Salmonella; Salmonella enterica; Salmonella enterica subsp. enterica MNNEETFYQAMRRKGVTRRSFLKFCSLAATSLGLGAGMTPKIAWALENKPRIPVVWIHGL WP_001058307.1 hydrogenase [Salmonella enterica] Length: 372\nNumber of predicted TMHs: 2\nExp number of AAs in TMHs: 36.51738\nExp number, first 60 AAs: 19.72849\nTotal prob of N-in: 0.96170\nPOSSIBLE N-term signal sequence\ninside 1 19\nTMhelix 20 39\noutside 40 325\nTMhelix 326 348\ninside 349 372

17230 GCF_002039915.1_ASM203991v1 Salmonella enterica subsp. enterica serovar Enteritidis Proteobacteria; Gammaproteobacteria; Enterobacterales; Enterobacteriaceae; Salmonella; Salmonella enterica; Salmonella enterica subsp. enterica MTGDNTLITSHGINRRDFMKLCAALAATMGLSSKAAAEMAESVSNPQRPPVIWIGAQECT WP_000145429.1 hydrogenase 2 small subunit [Salmonella enterica] Length: 372\nNumber of predicted TMHs: 1\nExp number of AAs in TMHs: 24.51309\nExp number, first 60 AAs: 0.2682\nTotal prob of N-in: 0.14402\noutside 1 331\nTMhelix 332 354\ninside 355 372

17231 GCF_002039965.1_ASM203996v1 Salmonella enterica subsp. enterica serovar Enteritidis Proteobacteria; Gammaproteobacteria; Enterobacterales; Enterobacteriaceae; Salmonella; Salmonella enterica; Salmonella enterica subsp. enterica MTGDNTLITSHGINRRDFMKLCAALAATMGLSSKAAAEMAESVSNPQRPPVIWIGAQECT WP_000145429.1 hydrogenase 2 small subunit [Salmonella enterica] Length: 372\nNumber of predicted TMHs: 1\nExp number of AAs in TMHs: 24.51309\nExp number, first 60 AAs: 0.2682\nTotal prob of N-in: 0.14402\noutside 1 331\nTMhelix 332 354\ninside 355 372

17232 GCF_002040005.1_ASM204000v1 Salmonella enterica subsp. enterica serovar Enteritidis Proteobacteria; Gammaproteobacteria; Enterobacterales; Enterobacteriaceae; Salmonella; Salmonella enterica;

Salmonella enterica subsp. enterica MTGDNTLITSHGINRRDFMKLCAALAATMGLSSKAAAEMAESVSNPQRPPVIWIGAQECT WP_000145429.1 hydrogenase 2 small subunit [Salmonella enterica] Length: 372\nNumber of predicted TMHs: 1\nExp number of AAs in TMHs: 24.51309\nExp number, first 60 AAs: 0.2682\nTotal prob of N-in: 0.14402\noutside 1 331\nTMhelix 332 354\ninside 355 372

17233 GCF_002040045.1_ASM204004v1 Salmonella enterica subsp. enterica serovar Enteritidis Proteobacteria; Gammaproteobacteria; Enterobacterales; Enterobacteriaceae; Salmonella; Salmonella enterica; Salmonella enterica subsp. enterica MTGDNTLITSHGINRRDFMKLCAALAATMGLSSKAAAEMAESVSNPQRPPVIWIGAQECT WP_000145429.1 hydrogenase 2 small subunit [Salmonella enterica] Length: 372\nNumber of predicted TMHs: 1\nExp number of AAs in TMHs: 24.51309\nExp number, first 60 AAs: 0.2682\nTotal prob of N-in: 0.14402\noutside 1 331\nTMhelix 332 354\ninside 355 372

17234 GCF_002040085.1_ASM204008v1 Salmonella enterica subsp. enterica serovar Enteritidis Proteobacteria; Gammaproteobacteria; Enterobacterales; Enterobacteriaceae; Salmonella; Salmonella enterica; Salmonella enterica subsp. enterica MTGDNTLITSHGINRRDFMKLCAALAATMGLSSKAAAEMAESVSNPQRPPVIWIGAQECT WP_000145429.1 hydrogenase 2 small subunit [Salmonella enterica] Length: 372\nNumber of predicted TMHs: 1\nExp number of AAs in TMHs: 24.51309\nExp number, first 60 AAs: 0.2682\nTotal prob of N-in: 0.14402\noutside 1 331\nTMhelix 332 354\ninside 355 372

17235 GCF_002040155.1_ASM204015v1 Salmonella enterica subsp. enterica serovar Enteritidis Proteobacteria; Gammaproteobacteria; Enterobacterales; Enterobacteriaceae; Salmonella; Salmonella enterica; Salmonella enterica subsp. enterica MNNEETFYQAMRRKGVTRRSFLKFCSLAATSLGLGAGMTPKIAWALENKPRIPVVWIHGL WP_001058307.1 hydrogenase [Salmonella enterica] Length: 372\nNumber of predicted TMHs: 2\nExp number of AAs in TMHs: 36.51738\nExp number, first 60 AAs: 19.72849\nTotal prob of N-in: 0.96170\nPOSSIBLE N-term signal sequence\ninside 1 19\nTMhelix 20 39\noutside 40 325\nTMhelix 326 348\ninside 349 372

17236 GCF_002040145.1_ASM204014v1 Salmonella enterica subsp. enterica serovar Enteritidis Proteobacteria; Gammaproteobacteria; Enterobacterales; Enterobacteriaceae; Salmonella; Salmonella enterica; Salmonella enterica subsp. enterica MNNEETFYQAMRRKGVTRRSFLKFCSLAATSLGLGAGMTPKIAWALENKPRIPVVWIHGL WP_001058307.1 hydrogenase [Salmonella enterica] Length: 372\nNumber of predicted TMHs: 2\nExp number of AAs in TMHs: 36.51738\nExp number, first 60 AAs: 19.72849\nTotal prob of N-in: 0.96170\nPOSSIBLE N-term signal sequence\ninside 1 19\nTMhelix 20 39\noutside 40 325\nTMhelix 326 348\ninside 349 372

17237 GCF_002040205.1_ASM204020v1 Salmonella enterica subsp. enterica serovar Enteritidis Proteobacteria; Gammaproteobacteria; Enterobacterales; Enterobacteriaceae; Salmonella; Salmonella enterica; Salmonella enterica subsp. enterica MNNEETFYQAMRRKGVTRRSFLKFCSLAATSLGLGAGMTPKIAWALENKPRIPVVWIHGL WP_001058307.1 hydrogenase [Salmonella enterica] Length: 372\nNumber of predicted TMHs: 2\nExp number of AAs in TMHs: 36.51738\nExp number, first 60 AAs: 19.72849\nTotal prob of N-in: 0.96170\nPOSSIBLE N-term signal sequence\ninside 1 19\nTMhelix 20 39\noutside 40 325\nTMhelix 326 348\ninside 349 372

17238 GCF_002040085.1_ASM204008v1 Salmonella enterica subsp. enterica serovar Enteritidis Proteobacteria; Gammaproteobacteria; Enterobacterales; Enterobacteriaceae; Salmonella; Salmonella enterica; Salmonella enterica subsp. enterica MNNEETFYQAMRRKGVTRRSFLKFCSLAATSLGLGAGMTPKIAWALENKPRIPVVWIHGL WP_001058307.1 hydrogenase [Salmonella enterica] Length: 372\nNumber of predicted TMHs: 2\nExp number of AAs in TMHs: 36.51738\nExp number, first 60 AAs: 19.72849\nTotal prob of N-in: 0.96170\nPOSSIBLE N-term signal sequence\ninside 1 19\nTMhelix 20 39\noutside 40 325\nTMhelix 326 348\ninside 349 372

17239 GCF_002040125.1_ASM204012v1 Salmonella enterica subsp. enterica serovar Enteritidis Proteobacteria; Gammaproteobacteria; Enterobacterales; Enterobacteriaceae; Salmonella; Salmonella enterica; Salmonella enterica subsp. enterica MNNEETFYQAMRRKGVTRRSFLKFCSLAATSLGLGAGMTPKIAWALENKPRIPVVWIHGL WP_001058307.1 hydrogenase [Salmonella enterica] Length: 372\nNumber of predicted TMHs: 2\nExp number of AAs in TMHs: 36.51738\nExp number, first 60 AAs: 19.72849\nTotal prob of N-in: 0.96170\nPOSSIBLE N-term signal sequence\ninside 1 19\nTMhelix 20 39\noutside 40 325\nTMhelix 326 348\ninside 349 372

17240 GCF_002040245.1_ASM204024v1 Salmonella enterica subsp. enterica serovar Enteritidis Proteobacteria; Gammaproteobacteria; Enterobacterales; Enterobacteriaceae; Salmonella; Salmonella enterica; Salmonella enterica subsp. enterica MTGDNTLITSHGINRRDFMKLCAALAATMGLSSKAAAEMAESVSNPQRPPVIWIGAQECT WP_000145429.1 hydrogenase 2 small subunit [Salmonella enterica] Length: 372\nNumber of predicted TMHs: 1\nExp number of AAs in TMHs: 24.51309\nExp number, first 60 AAs: 0.2682\nTotal prob of N-in: 0.14402\noutside 1 331\nTMhelix 332 354\ninside 355 372

17241 GCF_002038365.1_ASM203836v1 Salmonella enterica subsp. enterica serovar Enteritidis Proteobacteria; Gammaproteobacteria; Enterobacterales; Enterobacteriaceae; Salmonella; Salmonella enterica; Salmonella enterica subsp. enterica MTGDNTLITSHGINRRDFMKLCAALAATMGLSSKAAAEMAESVSNPQRPPVIWIGAQECT WP_000145429.1 hydrogenase 2 small subunit [Salmonella enterica] Length: 372\nNumber of predicted TMHs: 1\nExp number of AAs in TMHs: 24.51309\nExp number, first 60 AAs: 0.2682\nTotal prob of N-in: 0.14402\noutside 1 331\nTMhelix 332 354\ninside 355 372

17242 GCF_002040395.1_ASM204039v1 Salmonella enterica subsp. enterica serovar Enteritidis Proteobacteria; Gammaproteobacteria; Enterobacterales; Enterobacteriaceae; Salmonella; Salmonella enterica; Salmonella enterica subsp. enterica MNNEETFYQAMRRKGVTRRSFLKFCSLAATSLGLGAGMTPKIAWALENKPRIPVVWIHGL WP_001058307.1 hydrogenase [Salmonella enterica] Length: 372\nNumber of predicted TMHs: 2\nExp number of AAs in TMHs: 36.51738\nExp number, first 60 AAs: 19.72849\nTotal prob of N-in: 0.96170\nPOSSIBLE N-term signal sequence\ninside 1 19\nTMhelix 20 39\noutside 40 325\nTMhelix 326 348\ninside 349 372

17243 GCF_002040365.1_ASM204036v1 Salmonella enterica subsp. enterica serovar Enteritidis Proteobacteria; Gammaproteobacteria; Enterobacterales; Enterobacteriaceae; Salmonella; Salmonella enterica; Salmonella enterica subsp. enterica MNNEETFYQAMRRKGVTRRSFLKFCSLAATSLGLGAGMTPKIAWALENKPRIPVVWIHGL WP_001058307.1 hydrogenase [Salmonella enterica] Length: 372\nNumber of predicted TMHs: 2\nExp number of AAs in TMHs: 36.51738\nExp number, first 60 AAs: 19.72849\nTotal prob of N-in: 0.96170\nPOSSIBLE N-term signal sequence\ninside 1 19\nTMhelix 20 39\noutside 40 325\nTMhelix 326 348\ninside 349 372

17244 GCF_002040585.1_ASM204058v1 Salmonella enterica subsp. enterica serovar Enteritidis Proteobacteria; Gammaproteobacteria; Enterobacterales; Enterobacteriaceae; Salmonella; Salmonella enterica; Salmonella enterica subsp. enterica MTGDNTLITSHGINRRDFMKLCAALAATMGLSSKAAAEMAESVSNPQRPPVIWIGAQECT WP_000145429.1 hydrogenase 2 small subunit [Salmonella enterica] Length: 372\nNumber of predicted TMHs: 1\nExp number of AAs in TMHs: 24.51309\nExp number, first 60 AAs: 0.2682\nTotal prob of N-in: 0.14402\noutside 1 331\nTMhelix 332 354\ninside 355 372

17245 GCF_002040605.1_ASM204060v1 Salmonella enterica subsp. enterica serovar Enteritidis Proteobacteria; Gammaproteobacteria; Enterobacterales; Enterobacteriaceae; Salmonella; Salmonella enterica; Salmonella enterica subsp. enterica MTGDNTLITSHGINRRDFMKLCAALAATMGLSSKAAAEMAESVSNPQRPPVIWIGAQECT WP_000145429.1 hydrogenase 2 small subunit [Salmonella enterica] Length: 372\nNumber of predicted TMHs: 1\nExp number of AAs in TMHs: 24.51309\nExp number, first 60 AAs: 0.2682\nTotal prob of N-in: 0.14402\noutside 1 331\nTMhelix 332 354\ninside 355 372

17246 GCF_002040665.1_ASM204066v1 Salmonella enterica subsp. enterica serovar Enteritidis Proteobacteria; Gammaproteobacteria; Enterobacterales; Enterobacteriaceae; Salmonella; Salmonella enterica; Salmonella enterica subsp. enterica MTGDNTLITSHGINRRDFMKLCAALAATMGLSSKAAAEMAESVSNPQRPPVIWIGAQECT WP_000145429.1 hydrogenase 2 small subunit [Salmonella enterica] Length: 372\nNumber of predicted TMHs: 1\nExp number of AAs in TMHs: 24.51309\nExp number, first 60 AAs: 0.2682\nTotal prob of N-in: 0.14402\noutside 1 331\nTMhelix 332 354\ninside 355 372

17247 GCF_002040705.1_ASM204070v1 Salmonella enterica subsp. enterica serovar Enteritidis Proteobacteria; Gammaproteobacteria; Enterobacterales; Enterobacteriaceae; Salmonella; Salmonella enterica; Salmonella enterica subsp. enterica MTGDNTLITSHGINRRDFMKLCAALAATMGLSSKAAAEMAESVSNPQRPPVIWIGAQECT WP_000145429.1 hydrogenase 2 small subunit [Salmonella enterica] Length: 372\nNumber of predicted TMHs: 1\nExp number of AAs in TMHs: 24.51309\nExp number, first 60 AAs: 0.2682\nTotal prob of N-in: 0.14402\noutside 1 331\nTMhelix 332 354\ninside 355 372

17248 GCF_002040585.1_ASM204058v1 Salmonella enterica subsp. enterica serovar Enteritidis Proteobacteria; Gammaproteobacteria; Enterobacterales; Enterobacteriaceae; Salmonella; Salmonella enterica; Salmonella enterica subsp. enterica MNNEETFYQAMRRKGVTRRSFLKFCSLAATSLGLGAGMTPKIAWALENKPRIPVVWIHGL WP_001058307.1 hydrogenase [Salmonella enterica] Length: 372\nNumber of predicted TMHs: 2\nExp number of AAs in TMHs: 36.51738\nExp number, first 60 AAs: 19.72849\nTotal prob of N-in: 0.96170\nPOSSIBLE N-term signal sequence\ninside 1 19\nTMhelix 20 39\noutside 40 325\nTMhelix 326 348\ninside 349 372

17249 GCF_002040665.1_ASM204066v1 Salmonella enterica subsp. enterica serovar Enteritidis Proteobacteria; Gammaproteobacteria; Enterobacterales; Enterobacteriaceae; Salmonella; Salmonella enterica; Salmonella enterica subsp. enterica MNNEETFYQAMRRKGVTRRSFLKFCSLAATSLGLGAGMTPKIAWALENKPRIPVVWIHGL WP_001058307.1 hydrogenase [Salmonella enterica] Length: 372\nNumber of predicted TMHs: 2\nExp

number of AAs in TMHs: 36.51738\nExp number, first 60 AAs: 19.72849\nTotal prob of N-in: 0.96170\nPOSSIBLE N-term signal sequence\ninside 1 19\nTMhelix 20 39\noutside 40 325\nTMhelix 326 348\ninside 349 372

17250 GCF_002040835.1_ASM204083v1 Salmonella enterica subsp. enterica serovar Enteritidis Proteobacteria; Gammaproteobacteria; Enterobacterales; Enterobacteriaceae; Salmonella; Salmonella enterica; Salmonella enterica subsp. enterica MTGDNTLITSHGINRRDFMKLCAALAATMGLSSKAAAEMAESVSNPQRPPVIWIGAQECT WP_000145429.1 hydrogenase 2 small subunit [Salmonella enterica] Length: 372\nNumber of predicted TMHs: 1\nExp number of AAs in TMHs: 24.51309\nExp number, first 60 AAs: 0.2682\nTotal prob of N-in: 0.14402\noutside 1 331\nTMhelix 332 354\ninside 355 372

17251 GCF_002040845.1_ASM204084v1 Salmonella enterica subsp. enterica serovar Enteritidis Proteobacteria; Gammaproteobacteria; Enterobacterales; Enterobacteriaceae; Salmonella; Salmonella enterica; Salmonella enterica subsp. enterica MTGDNTLITSHGINRRDFMKLCAALAATMGLSSKAAAEMAESVSNPQRPPVIWIGAQECT WP_000145429.1 hydrogenase 2 small subunit [Salmonella enterica] Length: 372\nNumber of predicted TMHs: 1\nExp number of AAs in TMHs: 24.51309\nExp number, first 60 AAs: 0.2682\nTotal prob of N-in: 0.14402\noutside 1 331\nTMhelix 332 354\ninside 355 372

17252 GCF_002040885.1_ASM204088v1 Salmonella enterica subsp. enterica serovar Enteritidis Proteobacteria; Gammaproteobacteria; Enterobacterales; Enterobacteriaceae; Salmonella; Salmonella enterica; Salmonella enterica subsp. enterica MTGDNTLITSHGINRRDFMKLCAALAATMGLSSKAAAEMAESVSNPQRPPVIWIGAQECT WP_000145429.1 hydrogenase 2 small subunit [Salmonella enterica] Length: 372\nNumber of predicted TMHs: 1\nExp number of AAs in TMHs: 24.51309\nExp number, first 60 AAs: 0.2682\nTotal prob of N-in: 0.14402\noutside 1 331\nTMhelix 332 354\ninside 355 372

17253 GCF_002040915.1_ASM204091v1 Salmonella enterica subsp. enterica serovar Enteritidis Proteobacteria; Gammaproteobacteria; Enterobacterales; Enterobacteriaceae; Salmonella; Salmonella enterica; Salmonella enterica subsp. enterica MTGDNTLITSHGINRRDFMKLCAALAATMGLSSKAAAEMAESVSNPQRPPVIWIGAQECT WP_000145429.1 hydrogenase 2 small subunit [Salmonella enterica] Length: 372\nNumber of predicted TMHs: 1\nExp number of AAs in TMHs: 24.51309\nExp number, first 60 AAs: 0.2682\nTotal prob of N-in: 0.14402\noutside 1 331\nTMhelix 332 354\ninside 355 372

17254 GCF_002040965.1_ASM204096v1 Salmonella enterica subsp. enterica serovar Enteritidis Proteobacteria; Gammaproteobacteria; Enterobacterales; Enterobacteriaceae; Salmonella; Salmonella enterica; Salmonella enterica subsp. enterica MTGDNTLITSHGINRRDFMKLCAALAATMGLSSKAAAEMAESVSNPQRPPVIWIGAQECT WP_000145429.1 hydrogenase 2 small subunit [Salmonella enterica] Length: 372\nNumber of predicted TMHs: 1\nExp number of AAs in TMHs: 24.51309\nExp number, first 60 AAs: 0.2682\nTotal prob of N-in: 0.14402\noutside 1 331\nTMhelix 332 354\ninside 355 372

17255 GCF_002040835.1_ASM204083v1 Salmonella enterica subsp. enterica serovar Enteritidis Proteobacteria; Gammaproteobacteria; Enterobacterales; Enterobacteriaceae; Salmonella; Salmonella enterica; Salmonella enterica subsp. enterica MNNEETFYQAMRRKGVTRRSFLKFCSLAATSLGLGAGMTPKIAWALENKPRIPVVWIHGL WP_001058307.1 hydrogenase [Salmonella enterica] Length: 372\nNumber of predicted TMHs: 2\nExp number of AAs in TMHs: 36.51738\nExp number, first 60 AAs: 19.72849\nTotal prob of N-in: 0.96170\nPOSSIBLE N-term signal sequence\ninside 1 19\nTMhelix 20 39\noutside 40 325\nTMhelix 326 348\ninside 349 372

17256 GCF_002041035.1_ASM204103v1 Salmonella enterica subsp. enterica serovar Enteritidis Proteobacteria; Gammaproteobacteria; Enterobacterales; Enterobacteriaceae; Salmonella; Salmonella enterica; Salmonella enterica subsp. enterica MNNEETFYQAMRRKGVTRRSFLKFCSLAATSLGLGAGMTPKIAWALENKPRIPVVWIHGL WP_001058307.1 hydrogenase [Salmonella enterica] Length: 372\nNumber of predicted TMHs: 2\nExp number of AAs in TMHs: 36.51738\nExp number, first 60 AAs: 19.72849\nTotal prob of N-in: 0.96170\nPOSSIBLE N-term signal sequence\ninside 1 19\nTMhelix 20 39\noutside 40 325\nTMhelix 326 348\ninside 349 372

17257 GCF_002041085.1_ASM204108v1 Salmonella enterica subsp. enterica serovar Enteritidis Proteobacteria; Gammaproteobacteria; Enterobacterales; Enterobacteriaceae; Salmonella; Salmonella enterica; Salmonella enterica subsp. enterica MNNEETFYQAMRRKGVTRRSFLKFCSLAATSLGLGAGMTPKIAWALENKPRIPVVWIHGL WP_001058307.1 hydrogenase [Salmonella enterica] Length: 372\nNumber of predicted TMHs: 2\nExp number of AAs in TMHs: 36.51738\nExp number, first 60 AAs: 19.72849\nTotal prob of N-in: 0.96170\nPOSSIBLE N-term signal sequence\ninside 1 19\nTMhelix 20 39\noutside 40 325\nTMhelix 326 348\ninside 349 372

17258 GCF_002040965.1_ASM204096v1 Salmonella enterica subsp. enterica serovar Enteritidis Proteobacteria; Gammaproteobacteria; Enterobacterales; Enterobacteriaceae; Salmonella; Salmonella enterica;

Salmonella enterica subsp. enterica	MNNEETFYQAMRRKGVTRRSFLKFCSLAATSLGLGAGMTPKIAWALENKPRIPVVWIHGL	WP_001058307.1	hydrogenase [Salmonella enterica]	Length: 372\nNumber of predicted TMHs:  2\nExp number of AAs in TMHs: 36.51738\nExp number, first 60 AAs:  19.72849\nTotal prob of N-in:   0.96170\nPOSSIBLE N-term signal sequence\ninside   1   19\nTMhelix   20   39\noutside   40   325\nTMhelix   326   348\ninside   349   372

17259	GCF_002041125.1_ASM204112v1	Salmonella enterica subsp. enterica serovar Enteritidis	Proteobacteria; Gammaproteobacteria; Enterobacterales; Enterobacteriaceae; Salmonella; Salmonella enterica; Salmonella enterica subsp. enterica	MTGDNTLITSHGINRRDFMKLCAALAATMGLSSKAAAEMAESVSNPQRPPVIWIGAQECT	WP_000145429.1	hydrogenase 2 small subunit [Salmonella enterica]	Length: 372\nNumber of predicted TMHs:  1\nExp number of AAs in TMHs: 24.51309\nExp number, first 60 AAs:  0.2682\nTotal prob of N-in:  0.14402\noutside   1   331\nTMhelix   332   354\ninside   355   372

17260	GCF_002041175.1_ASM204117v1	Salmonella enterica subsp. enterica serovar Enteritidis	Proteobacteria; Gammaproteobacteria; Enterobacterales; Enterobacteriaceae; Salmonella; Salmonella enterica; Salmonella enterica subsp. enterica	MTGDNTLITSHGINRRDFMKLCAALAATMGLSSKAAAEMAESVSNPQRPPVIWIGAQECT	WP_000145429.1	hydrogenase 2 small subunit [Salmonella enterica]	Length: 372\nNumber of predicted TMHs:  1\nExp number of AAs in TMHs: 24.51309\nExp number, first 60 AAs:  0.2682\nTotal prob of N-in:  0.14402\noutside   1   331\nTMhelix   332   354\ninside   355   372

17261	GCF_002041215.1_ASM204121v1	Salmonella enterica subsp. enterica serovar Enteritidis	Proteobacteria; Gammaproteobacteria; Enterobacterales; Enterobacteriaceae; Salmonella; Salmonella enterica; Salmonella enterica subsp. enterica	MTGDNTLITSHGINRRDFMKLCAALAATMGLSSKAAAEMAESVSNPQRPPVIWIGAQECT	WP_000145429.1	hydrogenase 2 small subunit [Salmonella enterica]	Length: 372\nNumber of predicted TMHs:  1\nExp number of AAs in TMHs: 24.51309\nExp number, first 60 AAs:  0.2682\nTotal prob of N-in:  0.14402\noutside   1   331\nTMhelix   332   354\ninside   355   372

17262	GCF_002041125.1_ASM204112v1	Salmonella enterica subsp. enterica serovar Enteritidis	Proteobacteria; Gammaproteobacteria; Enterobacterales; Enterobacteriaceae; Salmonella; Salmonella enterica; Salmonella enterica subsp. enterica	MNNEETFYQAMRRKGVTRRSFLKFCSLAATSLGLGAGMTPKIAWALENKPRIPVVWIHGL	WP_001058307.1	hydrogenase [Salmonella enterica]	Length: 372\nNumber of predicted TMHs:  2\nExp number of AAs in TMHs: 36.51738\nExp number, first 60 AAs:  19.72849\nTotal prob of N-in:   0.96170\nPOSSIBLE N-term signal sequence\ninside   1   19\nTMhelix   20   39\noutside   40   325\nTMhelix   326   348\ninside   349   372

17263	GCF_002041215.1_ASM204121v1	Salmonella enterica subsp. enterica serovar Enteritidis	Proteobacteria; Gammaproteobacteria; Enterobacterales; Enterobacteriaceae; Salmonella; Salmonella enterica; Salmonella enterica subsp. enterica	MNNEETFYQAMRRKGVTRRSFLKFCSLAATSLGLGAGMTPKIAWALENKPRIPVVWIHGL	WP_001058307.1	hydrogenase [Salmonella enterica]	Length: 372\nNumber of predicted TMHs:  2\nExp number of AAs in TMHs: 36.51738\nExp number, first 60 AAs:  19.72849\nTotal prob of N-in:   0.96170\nPOSSIBLE N-term signal sequence\ninside   1   19\nTMhelix   20   39\noutside   40   325\nTMhelix   326   348\ninside   349   372

17264	GCF_002041455.1_ASM204145v1	Salmonella enterica subsp. enterica serovar Enteritidis	Proteobacteria; Gammaproteobacteria; Enterobacterales; Enterobacteriaceae; Salmonella; Salmonella enterica; Salmonella enterica subsp. enterica	MTGDNTLITSHGINRRDFMKLCAALAATMGLSSKAAAEMAESVSNPQRPPVIWIGAQECT	WP_000145429.1	hydrogenase 2 small subunit [Salmonella enterica]	Length: 372\nNumber of predicted TMHs:  1\nExp number of AAs in TMHs: 24.51309\nExp number, first 60 AAs:  0.2682\nTotal prob of N-in:  0.14402\noutside   1   331\nTMhelix   332   354\ninside   355   372

17265	GCF_002041455.1_ASM204145v1	Salmonella enterica subsp. enterica serovar Enteritidis	Proteobacteria; Gammaproteobacteria; Enterobacterales; Enterobacteriaceae; Salmonella; Salmonella enterica; Salmonella enterica subsp. enterica	MNNEETFYQAMRRKGVTRRSFLKFCSLAATSLGLGAGMTPKIAWALENKPRIPVVWIHGL	WP_001058307.1	hydrogenase [Salmonella enterica]	Length: 372\nNumber of predicted TMHs:  2\nExp number of AAs in TMHs: 36.51738\nExp number, first 60 AAs:  19.72849\nTotal prob of N-in:   0.96170\nPOSSIBLE N-term signal sequence\ninside   1   19\nTMhelix   20   39\noutside   40   325\nTMhelix   326   348\ninside   349   372

17266	GCF_002041265.1_ASM204126v1	Salmonella enterica subsp. enterica serovar Enteritidis	Proteobacteria; Gammaproteobacteria; Enterobacterales; Enterobacteriaceae; Salmonella; Salmonella enterica; Salmonella enterica subsp. enterica	MNNEETFYQAMRRKGVTRRSFLKFCSLAATSLGLGAGMTPKIAWALENKPRIPVVWIHGL	WP_001058307.1	hydrogenase [Salmonella enterica]	Length: 372\nNumber of predicted TMHs:  2\nExp number of AAs in TMHs: 36.51738\nExp number, first 60 AAs:  19.72849\nTotal prob of N-in:   0.96170\nPOSSIBLE N-term signal sequence\ninside   1   19\nTMhelix   20   39\noutside   40   325\nTMhelix   326   348\ninside   349   372

17267 GCF_002041505.1_ASM204150v1 Salmonella enterica subsp. enterica serovar Enteritidis Proteobacteria; Gammaproteobacteria; Enterobacterales; Enterobacteriaceae; Salmonella; Salmonella enterica; Salmonella enterica subsp. enterica MTGDNTLITSHGINRRDFMKLCAALAATMGLSSKAAAEMAESVSNPQRPPVIWIGAQECT WP_000145429.1 hydrogenase 2 small subunit [Salmonella enterica] Length: 372\nNumber of predicted TMHs: 1\nExp number of AAs in TMHs: 24.51309\nExp number, first 60 AAs: 0.2682\nTotal prob of N-in: 0.14402\noutside 1 331\nTMhelix 332 354\ninside 355 372

17268 GCF_002041505.1_ASM204150v1 Salmonella enterica subsp. enterica serovar Enteritidis Proteobacteria; Gammaproteobacteria; Enterobacterales; Enterobacteriaceae; Salmonella; Salmonella enterica; Salmonella enterica subsp. enterica MNNEETFYQAMRRKGVTRRSFLKFCSLAATSLGLGAGMTPKIAWALENKPRIPVVWIHGL WP_001058307.1 hydrogenase [Salmonella enterica] Length: 372\nNumber of predicted TMHs: 2\nExp number of AAs in TMHs: 36.51738\nExp number, first 60 AAs: 19.72849\nTotal prob of N-in: 0.96170\nPOSSIBLE N-term signal sequence\ninside 1 19\nTMhelix 20 39\noutside 40 325\nTMhelix 326 348\ninside 349 372

17269 GCF_002042165.1_ASM204216v1 Salmonella enterica subsp. enterica serovar Enteritidis Proteobacteria; Gammaproteobacteria; Enterobacterales; Enterobacteriaceae; Salmonella; Salmonella enterica; Salmonella enterica subsp. enterica MTGDNTLITSHGINRRDFMKLCAALAATMGLSSKAAAEMAESVSNPQRPPVIWIGAQECT WP_000145429.1 hydrogenase 2 small subunit [Salmonella enterica] Length: 372\nNumber of predicted TMHs: 1\nExp number of AAs in TMHs: 24.51309\nExp number, first 60 AAs: 0.2682\nTotal prob of N-in: 0.14402\noutside 1 331\nTMhelix 332 354\ninside 355 372

17270 GCF_002041575.1_ASM204157v1 Salmonella enterica subsp. enterica serovar Enteritidis Proteobacteria; Gammaproteobacteria; Enterobacterales; Enterobacteriaceae; Salmonella; Salmonella enterica; Salmonella enterica subsp. enterica MNNEETFYQAMRRKGVTRRSFLKFCSLAATSLGLGAGMTPKIAWALENKPRIPVVWIHGL WP_001058307.1 hydrogenase [Salmonella enterica] Length: 372\nNumber of predicted TMHs: 2\nExp number of AAs in TMHs: 36.51738\nExp number, first 60 AAs: 19.72849\nTotal prob of N-in: 0.96170\nPOSSIBLE N-term signal sequence\ninside 1 19\nTMhelix 20 39\noutside 40 325\nTMhelix 326 348\ninside 349 372

17271 GCF_002041665.1_ASM204166v1 Salmonella enterica subsp. enterica serovar Enteritidis Proteobacteria; Gammaproteobacteria; Enterobacterales; Enterobacteriaceae; Salmonella; Salmonella enterica; Salmonella enterica subsp. enterica MTGDNTLITSHGINRRDFMKLCAALAATMGLSSKAAAEMAESVSNPQRPPVIWIGAQECT WP_000145429.1 hydrogenase 2 small subunit [Salmonella enterica] Length: 372\nNumber of predicted TMHs: 1\nExp number of AAs in TMHs: 24.51309\nExp number, first 60 AAs: 0.2682\nTotal prob of N-in: 0.14402\noutside 1 331\nTMhelix 332 354\ninside 355 372

17272 GCF_002041695.1_ASM204169v1 Salmonella enterica subsp. enterica serovar Enteritidis Proteobacteria; Gammaproteobacteria; Enterobacterales; Enterobacteriaceae; Salmonella; Salmonella enterica; Salmonella enterica subsp. enterica MTGDNTLITSHGINRRDFMKLCAALAATMGLSSKAAAEMAESVSNPQRPPVIWIGAQECT WP_000145429.1 hydrogenase 2 small subunit [Salmonella enterica] Length: 372\nNumber of predicted TMHs: 1\nExp number of AAs in TMHs: 24.51309\nExp number, first 60 AAs: 0.2682\nTotal prob of N-in: 0.14402\noutside 1 331\nTMhelix 332 354\ninside 355 372

17273 GCF_002041745.1_ASM204174v1 Salmonella enterica subsp. enterica serovar Enteritidis Proteobacteria; Gammaproteobacteria; Enterobacterales; Enterobacteriaceae; Salmonella; Salmonella enterica; Salmonella enterica subsp. enterica MTGDNTLITSHGINRRDFMKLCAALAATMGLSSKAAAEMAESVSNPQRPPVIWIGAQECT WP_000145429.1 hydrogenase 2 small subunit [Salmonella enterica] Length: 372\nNumber of predicted TMHs: 1\nExp number of AAs in TMHs: 24.51309\nExp number, first 60 AAs: 0.2682\nTotal prob of N-in: 0.14402\noutside 1 331\nTMhelix 332 354\ninside 355 372

17274 GCF_002041765.1_ASM204176v1 Salmonella enterica subsp. enterica serovar Enteritidis Proteobacteria; Gammaproteobacteria; Enterobacterales; Enterobacteriaceae; Salmonella; Salmonella enterica; Salmonella enterica subsp. enterica MTGDNTLITSHGINRRDFMKLCAALAATMGLSSKAAAEMAESVSNPQRPPVIWIGAQECT WP_000145429.1 hydrogenase 2 small subunit [Salmonella enterica] Length: 372\nNumber of predicted TMHs: 1\nExp number of AAs in TMHs: 24.51309\nExp number, first 60 AAs: 0.2682\nTotal prob of N-in: 0.14402\noutside 1 331\nTMhelix 332 354\ninside 355 372

17275 GCF_002041835.1_ASM204183v1 Salmonella enterica subsp. enterica serovar Enteritidis Proteobacteria; Gammaproteobacteria; Enterobacterales; Enterobacteriaceae; Salmonella; Salmonella enterica; Salmonella enterica subsp. enterica MTGDNTLITSHGINRRDFMKLCAALAATMGLSSKAAAEMAESVSNPQRPPVIWIGAQECT WP_000145429.1 hydrogenase 2 small subunit [Salmonella enterica] Length: 372\nNumber of

predicted TMHs:  1\nExp number of AAs in TMHs: 24.51309\nExp number, first 60 AAs:  0.2682\nTotal prob of N-in: 0.14402\noutside   1   331\nTMhelix   332   354\ninside   355   372

17276 GCF_002041665.1_ASM204166v1 Salmonella enterica subsp. enterica serovar Enteritidis Proteobacteria; Gammaproteobacteria; Enterobacterales; Enterobacteriaceae; Salmonella; Salmonella enterica; Salmonella enterica subsp. enterica MNNEETFYQAMRRKGVTRRSFLKFCSLAATSLGLGAGMTPKIAWALENKPRIPVVWIHGL WP_001058307.1 hydrogenase [Salmonella enterica] Length: 372\nNumber of predicted TMHs:  2\nExp number of AAs in TMHs: 36.51738\nExp number, first 60 AAs:  19.72849\nTotal prob of N-in:   0.96170\nPOSSIBLE N-term signal sequence\ninside   1   19\nTMhelix   20   39\noutside   40   325\nTMhelix   326   348\ninside   349   372

17277 GCF_002041745.1_ASM204174v1 Salmonella enterica subsp. enterica serovar Enteritidis Proteobacteria; Gammaproteobacteria; Enterobacterales; Enterobacteriaceae; Salmonella; Salmonella enterica; Salmonella enterica subsp. enterica MNNEETFYQAMRRKGVTRRSFLKFCSLAATSLGLGAGMTPKIAWALENKPRIPVVWIHGL WP_001058307.1 hydrogenase [Salmonella enterica] Length: 372\nNumber of predicted TMHs:  2\nExp number of AAs in TMHs: 36.51738\nExp number, first 60 AAs:  19.72849\nTotal prob of N-in:   0.96170\nPOSSIBLE N-term signal sequence\ninside   1   19\nTMhelix   20   39\noutside   40   325\nTMhelix   326   348\ninside   349   372

17278 GCF_002041765.1_ASM204176v1 Salmonella enterica subsp. enterica serovar Enteritidis Proteobacteria; Gammaproteobacteria; Enterobacterales; Enterobacteriaceae; Salmonella; Salmonella enterica; Salmonella enterica subsp. enterica MNNEETFYQAMRRKGVTRRSFLKFCSLAATSLGLGAGMTPKIAWALENKPRIPVVWIHGL WP_001058307.1 hydrogenase [Salmonella enterica] Length: 372\nNumber of predicted TMHs:  2\nExp number of AAs in TMHs: 36.51738\nExp number, first 60 AAs:  19.72849\nTotal prob of N-in:   0.96170\nPOSSIBLE N-term signal sequence\ninside   1   19\nTMhelix   20   39\noutside   40   325\nTMhelix   326   348\ninside   349   372

17279 GCF_002041835.1_ASM204183v1 Salmonella enterica subsp. enterica serovar Enteritidis Proteobacteria; Gammaproteobacteria; Enterobacterales; Enterobacteriaceae; Salmonella; Salmonella enterica; Salmonella enterica subsp. enterica MNNEETFYQAMRRKGVTRRSFLKFCSLAATSLGLGAGMTPKIAWALENKPRIPVVWIHGL WP_001058307.1 hydrogenase [Salmonella enterica] Length: 372\nNumber of predicted TMHs:  2\nExp number of AAs in TMHs: 36.51738\nExp number, first 60 AAs:  19.72849\nTotal prob of N-in:   0.96170\nPOSSIBLE N-term signal sequence\ninside   1   19\nTMhelix   20   39\noutside   40   325\nTMhelix   326   348\ninside   349   372

17280 GCF_002041945.1_ASM204194v1 Salmonella enterica subsp. enterica serovar Enteritidis Proteobacteria; Gammaproteobacteria; Enterobacterales; Enterobacteriaceae; Salmonella; Salmonella enterica; Salmonella enterica subsp. enterica MTGDNTLITSHGINRRDFMKLCAALAATMGLSSKAAAEMAESVSNPQRPPVIWIGAQECT WP_000145429.1 hydrogenase 2 small subunit [Salmonella enterica] Length: 372\nNumber of predicted TMHs:  1\nExp number of AAs in TMHs: 24.51309\nExp number, first 60 AAs:  0.2682\nTotal prob of N-in: 0.14402\noutside   1   331\nTMhelix   332   354\ninside   355   372

17281 GCF_002042005.1_ASM204200v1 Salmonella enterica subsp. enterica serovar Enteritidis Proteobacteria; Gammaproteobacteria; Enterobacterales; Enterobacteriaceae; Salmonella; Salmonella enterica; Salmonella enterica subsp. enterica MTGDNTLITSHGINRRDFMKLCAALAATMGLSSKAAAEMAESVSNPQRPPVIWIGAQECT WP_000145429.1 hydrogenase 2 small subunit [Salmonella enterica] Length: 372\nNumber of predicted TMHs:  1\nExp number of AAs in TMHs: 24.51309\nExp number, first 60 AAs:  0.2682\nTotal prob of N-in: 0.14402\noutside   1   331\nTMhelix   332   354\ninside   355   372

17282 GCF_002042025.1_ASM204202v1 Salmonella enterica subsp. enterica serovar Enteritidis Proteobacteria; Gammaproteobacteria; Enterobacterales; Enterobacteriaceae; Salmonella; Salmonella enterica; Salmonella enterica subsp. enterica MTGDNTLITSHGINRRDFMKLCAALAATMGLSSKAAAEMAESVSNPQRPPVIWIGAQECT WP_000145429.1 hydrogenase 2 small subunit [Salmonella enterica] Length: 372\nNumber of predicted TMHs:  1\nExp number of AAs in TMHs: 24.51309\nExp number, first 60 AAs:  0.2682\nTotal prob of N-in: 0.14402\noutside   1   331\nTMhelix   332   354\ninside   355   372

17283 GCF_002042075.1_ASM204207v1 Salmonella enterica subsp. enterica serovar Enteritidis Proteobacteria; Gammaproteobacteria; Enterobacterales; Enterobacteriaceae; Salmonella; Salmonella enterica; Salmonella enterica subsp. enterica MTGDNTLITSHGINRRDFMKLCAALAATMGLSSKAAAEMAESVSNPQRPPVIWIGAQECT WP_000145429.1 hydrogenase 2 small subunit [Salmonella enterica] Length: 372\nNumber of predicted TMHs:  1\nExp number of AAs in TMHs: 24.51309\nExp number, first 60 AAs:  0.2682\nTotal prob of N-in: 0.14402\noutside   1   331\nTMhelix   332   354\ninside   355   372

17284 GCF_002042185.1_ASM204218v1 Salmonella enterica subsp. enterica serovar Enteritidis Proteobacteria; Gammaproteobacteria; Enterobacterales; Enterobacteriaceae; Salmonella; Salmonella enterica;

Salmonella enterica subsp. enterica MTGDNTLITSHGINRRDFMKLCAALAATMGLSSKAAAEMAESVSNPQRPPVIWIGAQECT WP_000145429.1 hydrogenase 2 small subunit [Salmonella enterica] Length: 372\nNumber of predicted TMHs: 1\nExp number of AAs in TMHs: 24.51309\nExp number, first 60 AAs: 0.2682\nTotal prob of N-in: 0.14402\noutside 1 331\nTMhelix 332 354\ninside 355 372

17285 GCF_002042235.1_ASM204223v1 Salmonella enterica subsp. enterica serovar Enteritidis Proteobacteria; Gammaproteobacteria; Enterobacterales; Enterobacteriaceae; Salmonella; Salmonella enterica; Salmonella enterica subsp. enterica MTGDNTLITSHGINRRDFMKLCAALAATMGLSSKAAAEMAESVSNPQRPPVIWIGAQECT WP_000145429.1 hydrogenase 2 small subunit [Salmonella enterica] Length: 372\nNumber of predicted TMHs: 1\nExp number of AAs in TMHs: 24.51309\nExp number, first 60 AAs: 0.2682\nTotal prob of N-in: 0.14402\noutside 1 331\nTMhelix 332 354\ninside 355 372

17286 GCF_002042255.1_ASM204225v1 Salmonella enterica subsp. enterica serovar Enteritidis Proteobacteria; Gammaproteobacteria; Enterobacterales; Enterobacteriaceae; Salmonella; Salmonella enterica; Salmonella enterica subsp. enterica MTGDNTLITSHGINRRDFMKLCAALAATMGLSSKAAAEMAESVSNPQRPPVIWIGAQECT WP_000145429.1 hydrogenase 2 small subunit [Salmonella enterica] Length: 372\nNumber of predicted TMHs: 1\nExp number of AAs in TMHs: 24.51309\nExp number, first 60 AAs: 0.2682\nTotal prob of N-in: 0.14402\noutside 1 331\nTMhelix 332 354\ninside 355 372

17287 GCF_002042165.1_ASM204216v1 Salmonella enterica subsp. enterica serovar Enteritidis Proteobacteria; Gammaproteobacteria; Enterobacterales; Enterobacteriaceae; Salmonella; Salmonella enterica; Salmonella enterica subsp. enterica MNNEETFYQAMRRKGVTRRSFLKFCSLAATSLGLGAGMTPKIAWALENKPRIPVVWIHGL WP_001058307.1 hydrogenase [Salmonella enterica] Length: 372\nNumber of predicted TMHs: 2\nExp number of AAs in TMHs: 36.51738\nExp number, first 60 AAs: 19.72849\nTotal prob of N-in: 0.96170\nPOSSIBLE N-term signal sequence\ninside 1 19\nTMhelix 20 39\noutside 40 325\nTMhelix 326 348\ninside 349 372

17288 GCF_002042175.1_ASM204217v1 Salmonella enterica subsp. enterica serovar Enteritidis Proteobacteria; Gammaproteobacteria; Enterobacterales; Enterobacteriaceae; Salmonella; Salmonella enterica; Salmonella enterica subsp. enterica MNNEETFYQAMRRKGVTRRSFLKFCSLAATSLGLGAGMTPKIAWALENKPRIPVVWIHGL WP_001058307.1 hydrogenase [Salmonella enterica] Length: 372\nNumber of predicted TMHs: 2\nExp number of AAs in TMHs: 36.51738\nExp number, first 60 AAs: 19.72849\nTotal prob of N-in: 0.96170\nPOSSIBLE N-term signal sequence\ninside 1 19\nTMhelix 20 39\noutside 40 325\nTMhelix 326 348\ninside 349 372

17289 GCF_002042375.1_ASM204237v1 Salmonella enterica subsp. enterica serovar Enteritidis Proteobacteria; Gammaproteobacteria; Enterobacterales; Enterobacteriaceae; Salmonella; Salmonella enterica; Salmonella enterica subsp. enterica MTGDNTLITSHGINRRDFMKLCAALAATMGLSSKAAAEMAESVSNPQRPPVIWIGAQECT WP_000145429.1 hydrogenase 2 small subunit [Salmonella enterica] Length: 372\nNumber of predicted TMHs: 1\nExp number of AAs in TMHs: 24.51309\nExp number, first 60 AAs: 0.2682\nTotal prob of N-in: 0.14402\noutside 1 331\nTMhelix 332 354\ninside 355 372

17290 GCF_002042315.1_ASM204231v1 Salmonella enterica subsp. enterica serovar Enteritidis Proteobacteria; Gammaproteobacteria; Enterobacterales; Enterobacteriaceae; Salmonella; Salmonella enterica; Salmonella enterica subsp. enterica MTGDNTLITSHGINRRDFMKLCAALAATMGLSSKAAAEMAESVSNPQRPPVIWIGAQECT WP_000145429.1 hydrogenase 2 small subunit [Salmonella enterica] Length: 372\nNumber of predicted TMHs: 1\nExp number of AAs in TMHs: 24.51309\nExp number, first 60 AAs: 0.2682\nTotal prob of N-in: 0.14402\noutside 1 331\nTMhelix 332 354\ninside 355 372

17291 GCF_002042255.1_ASM204225v1 Salmonella enterica subsp. enterica serovar Enteritidis Proteobacteria; Gammaproteobacteria; Enterobacterales; Enterobacteriaceae; Salmonella; Salmonella enterica; Salmonella enterica subsp. enterica MNNEETFYQAMRRKGVTRRSFLKFCSLAATSLGLGAGMTPKIAWALENKPRIPVVWIHGL WP_001058307.1 hydrogenase [Salmonella enterica] Length: 372\nNumber of predicted TMHs: 2\nExp number of AAs in TMHs: 36.51738\nExp number, first 60 AAs: 19.72849\nTotal prob of N-in: 0.96170\nPOSSIBLE N-term signal sequence\ninside 1 19\nTMhelix 20 39\noutside 40 325\nTMhelix 326 348\ninside 349 372

17292 GCF_002042455.1_ASM204245v1 Salmonella enterica subsp. enterica serovar Enteritidis Proteobacteria; Gammaproteobacteria; Enterobacterales; Enterobacteriaceae; Salmonella; Salmonella enterica; Salmonella enterica subsp. enterica MTGDNTLITSHGINRRDFMKLCAALAATMGLSSKAAAEMAESVSNPQRPPVIWIGAQECT WP_000145429.1 hydrogenase 2 small subunit [Salmonella enterica] Length: 372\nNumber of predicted TMHs: 1\nExp number of AAs in TMHs: 24.51309\nExp number, first 60 AAs: 0.2682\nTotal prob of N-in: 0.14402\noutside 1 331\nTMhelix 332 354\ninside 355 372

17293	GCF_002042495.1_ASM204249v1	Salmonella enterica subsp. enterica serovar Enteritidis	Proteobacteria; Gammaproteobacteria; Enterobacterales; Enterobacteriaceae; Salmonella; Salmonella enterica; Salmonella enterica subsp. enterica	MTGDNTLITSHGINRRDFMKLCAALAATMGLSSKAAAEMAESVSNPQRPPVIWIGAQECT	WP_000145429.1	hydrogenase 2 small subunit [Salmonella enterica]	Length: 372\nNumber of predicted TMHs:  1\nExp number of AAs in TMHs: 24.51309\nExp number, first 60 AAs:  0.2682\nTotal prob of N-in:  0.14402\noutside   1   331\nTMhelix   332   354\ninside   355   372

17294	GCF_002042535.1_ASM204253v1	Salmonella enterica subsp. enterica serovar Enteritidis	Proteobacteria; Gammaproteobacteria; Enterobacterales; Enterobacteriaceae; Salmonella; Salmonella enterica; Salmonella enterica subsp. enterica	MTGDNTLITSHGINRRDFMKLCAALAATMGLSSKAAAEMAESVSNPQRPPVIWIGAQECT	WP_000145429.1	hydrogenase 2 small subunit [Salmonella enterica]	Length: 372\nNumber of predicted TMHs:  1\nExp number of AAs in TMHs: 24.51309\nExp number, first 60 AAs:  0.2682\nTotal prob of N-in:  0.14402\noutside   1   331\nTMhelix   332   354\ninside   355   372

17295	GCF_002042455.1_ASM204245v1	Salmonella enterica subsp. enterica serovar Enteritidis	Proteobacteria; Gammaproteobacteria; Enterobacterales; Enterobacteriaceae; Salmonella; Salmonella enterica; Salmonella enterica subsp. enterica	MNNEETFYQAMRRKGVTRRSFLKFCSLAATSLGLGAGMTPKIAWALENKPRIPVVWIHGL	WP_001058307.1	hydrogenase [Salmonella enterica]	Length: 372\nNumber of predicted TMHs:  2\nExp number of AAs in TMHs: 36.51738\nExp number, first 60 AAs:  19.72849\nTotal prob of N-in:   0.96170\nPOSSIBLE N-term signal sequence\ninside   1   19\nTMhelix   20   39\noutside   40   325\nTMhelix   326   348\ninside   349   372

17296	GCF_002042495.1_ASM204249v1	Salmonella enterica subsp. enterica serovar Enteritidis	Proteobacteria; Gammaproteobacteria; Enterobacterales; Enterobacteriaceae; Salmonella; Salmonella enterica; Salmonella enterica subsp. enterica	MNNEETFYQAMRRKGVTRRSFLKFCSLAATSLGLGAGMTPKIAWALENKPRIPVVWIHGL	WP_001058307.1	hydrogenase [Salmonella enterica]	Length: 372\nNumber of predicted TMHs:  2\nExp number of AAs in TMHs: 36.51738\nExp number, first 60 AAs:  19.72849\nTotal prob of N-in:   0.96170\nPOSSIBLE N-term signal sequence\ninside   1   19\nTMhelix   20   39\noutside   40   325\nTMhelix   326   348\ninside   349   372

17297	GCF_002042585.1_ASM204258v1	Salmonella enterica subsp. enterica serovar Enteritidis	Proteobacteria; Gammaproteobacteria; Enterobacterales; Enterobacteriaceae; Salmonella; Salmonella enterica; Salmonella enterica subsp. enterica	MTGDNTLITSHGINRRDFMKLCAALAATMGLSSKAAAEMAESVSNPQRPPVIWIGAQECT	WP_000145429.1	hydrogenase 2 small subunit [Salmonella enterica]	Length: 372\nNumber of predicted TMHs:  1\nExp number of AAs in TMHs: 24.51309\nExp number, first 60 AAs:  0.2682\nTotal prob of N-in:  0.14402\noutside   1   331\nTMhelix   332   354\ninside   355   372

17298	GCF_002042745.1_ASM204274v1	Salmonella enterica subsp. enterica serovar Enteritidis	Proteobacteria; Gammaproteobacteria; Enterobacterales; Enterobacteriaceae; Salmonella; Salmonella enterica; Salmonella enterica subsp. enterica	MTGDNTLITSHGINRRDFMKLCAALAATMGLSSKAAAEMAESVSNPQRPPVIWIGAQECT	WP_000145429.1	hydrogenase 2 small subunit [Salmonella enterica]	Length: 372\nNumber of predicted TMHs:  1\nExp number of AAs in TMHs: 24.51309\nExp number, first 60 AAs:  0.2682\nTotal prob of N-in:  0.14402\noutside   1   331\nTMhelix   332   354\ninside   355   372

17299	GCF_002042785.1_ASM204278v1	Salmonella enterica subsp. enterica serovar Enteritidis	Proteobacteria; Gammaproteobacteria; Enterobacterales; Enterobacteriaceae; Salmonella; Salmonella enterica; Salmonella enterica subsp. enterica	MTGDNTLITSHGINRRDFMKLCAALAATMGLSSKAAAEMAESVSNPQRPPVIWIGAQECT	WP_000145429.1	hydrogenase 2 small subunit [Salmonella enterica]	Length: 372\nNumber of predicted TMHs:  1\nExp number of AAs in TMHs: 24.51309\nExp number, first 60 AAs:  0.2682\nTotal prob of N-in:  0.14402\noutside   1   331\nTMhelix   332   354\ninside   355   372

17300	GCF_002042825.1_ASM204282v1	Salmonella enterica subsp. enterica serovar Enteritidis	Proteobacteria; Gammaproteobacteria; Enterobacterales; Enterobacteriaceae; Salmonella; Salmonella enterica; Salmonella enterica subsp. enterica	MTGDNTLITSHGINRRDFMKLCAALAATMGLSSKAAAEMAESVSNPQRPPVIWIGAQECT	WP_000145429.1	hydrogenase 2 small subunit [Salmonella enterica]	Length: 372\nNumber of predicted TMHs:  1\nExp number of AAs in TMHs: 24.51309\nExp number, first 60 AAs:  0.2682\nTotal prob of N-in:  0.14402\noutside   1   331\nTMhelix   332   354\ninside   355   372

17301	GCF_002042745.1_ASM204274v1	Salmonella enterica subsp. enterica serovar Enteritidis	Proteobacteria; Gammaproteobacteria; Enterobacterales; Enterobacteriaceae; Salmonella; Salmonella enterica; Salmonella enterica subsp. enterica	MNNEETFYQAMRRKGVTRRSFLKFCSLAATSLGLGAGMTPKIAWALENKPRIPVVWIHGL	WP_001058307.1	hydrogenase [Salmonella enterica]	Length: 372\nNumber of predicted TMHs:  2\nExp

number of AAs in TMHs: 36.51738\nExp number, first 60 AAs:  19.72849\nTotal prob of N-in:   0.96170\nPOSSIBLE N-term signal sequence\ninside   1   19\nTMhelix   20   39\noutside   40   325\nTMhelix   326   348\ninside   349   372

17302 GCF_002042785.1_ASM204278v1 Salmonella enterica subsp. enterica serovar Enteritidis Proteobacteria; Gammaproteobacteria; Enterobacterales; Enterobacteriaceae; Salmonella; Salmonella enterica; Salmonella enterica subsp. enterica MNNEETFYQAMRRKGVTRRSFLKFCSLAATSLGLGAGMTPKIAWALENKPRIPVVWIHGL WP_001058307.1 hydrogenase [Salmonella enterica] Length: 372\nNumber of predicted TMHs:  2\nExp number of AAs in TMHs: 36.51738\nExp number, first 60 AAs:  19.72849\nTotal prob of N-in:   0.96170\nPOSSIBLE N-term signal sequence\ninside   1   19\nTMhelix   20   39\noutside   40   325\nTMhelix   326   348\ninside   349   372

17303 GCF_002042825.1_ASM204282v1 Salmonella enterica subsp. enterica serovar Enteritidis Proteobacteria; Gammaproteobacteria; Enterobacterales; Enterobacteriaceae; Salmonella; Salmonella enterica; Salmonella enterica subsp. enterica MNNEETFYQAMRRKGVTRRSFLKFCSLAATSLGLGAGMTPKIAWALENKPRIPVVWIHGL WP_001058307.1 hydrogenase [Salmonella enterica] Length: 372\nNumber of predicted TMHs:  2\nExp number of AAs in TMHs: 36.51738\nExp number, first 60 AAs:  19.72849\nTotal prob of N-in:   0.96170\nPOSSIBLE N-term signal sequence\ninside   1   19\nTMhelix   20   39\noutside   40   325\nTMhelix   326   348\ninside   349   372

17304 GCF_002042585.1_ASM204258v1 Salmonella enterica subsp. enterica serovar Enteritidis Proteobacteria; Gammaproteobacteria; Enterobacterales; Enterobacteriaceae; Salmonella; Salmonella enterica; Salmonella enterica subsp. enterica MNNEETFYQAMRRKGVTRRSFLKFCSLAATSLGLGAGMTPKIAWALENKPRIPVVWIHGL WP_001058307.1 hydrogenase [Salmonella enterica] Length: 372\nNumber of predicted TMHs:  2\nExp number of AAs in TMHs: 36.51738\nExp number, first 60 AAs:  19.72849\nTotal prob of N-in:   0.96170\nPOSSIBLE N-term signal sequence\ninside   1   19\nTMhelix   20   39\noutside   40   325\nTMhelix   326   348\ninside   349   372

17305 GCF_002038215.1_ASM203821v1 Salmonella enterica subsp. enterica serovar Enteritidis Proteobacteria; Gammaproteobacteria; Enterobacterales; Enterobacteriaceae; Salmonella; Salmonella enterica; Salmonella enterica subsp. enterica MNNEETFYQAMRRKGVTRRSFLKFCSLAATSLGLGAGMTPKIAWALENKPRIPVVWIHGL WP_001058307.1 hydrogenase [Salmonella enterica] Length: 372\nNumber of predicted TMHs:  2\nExp number of AAs in TMHs: 36.51738\nExp number, first 60 AAs:  19.72849\nTotal prob of N-in:   0.96170\nPOSSIBLE N-term signal sequence\ninside   1   19\nTMhelix   20   39\noutside   40   325\nTMhelix   326   348\ninside   349   372

17306 GCF_002038205.1_ASM203820v1 Salmonella enterica subsp. enterica serovar Enteritidis Proteobacteria; Gammaproteobacteria; Enterobacterales; Enterobacteriaceae; Salmonella; Salmonella enterica; Salmonella enterica subsp. enterica MNNEETFYQAMRRKGVTRRSFLKFCSLAATSLGLGAGMTPKIAWALENKPRIPVVWIHGL WP_001058307.1 hydrogenase [Salmonella enterica] Length: 372\nNumber of predicted TMHs:  2\nExp number of AAs in TMHs: 36.51738\nExp number, first 60 AAs:  19.72849\nTotal prob of N-in:   0.96170\nPOSSIBLE N-term signal sequence\ninside   1   19\nTMhelix   20   39\noutside   40   325\nTMhelix   326   348\ninside   349   372

17307 GCF_002038155.1_ASM203815v1 Salmonella enterica subsp. enterica serovar Enteritidis Proteobacteria; Gammaproteobacteria; Enterobacterales; Enterobacteriaceae; Salmonella; Salmonella enterica; Salmonella enterica subsp. enterica MTGDNTLITSHGINRRDFMKLCAALAATMGLSSKAAAEMAESVSNPQRPPVIWIGAQECT WP_000145429.1 hydrogenase 2 small subunit [Salmonella enterica] Length: 372\nNumber of predicted TMHs:  1\nExp number of AAs in TMHs: 24.51309\nExp number, first 60 AAs:  0.2682\nTotal prob of N-in: 0.14402\noutside   1   331\nTMhelix   332   354\ninside   355   372

17308 GCF_002038115.1_ASM203811v1 Salmonella enterica subsp. enterica serovar Enteritidis Proteobacteria; Gammaproteobacteria; Enterobacterales; Enterobacteriaceae; Salmonella; Salmonella enterica; Salmonella enterica subsp. enterica MTGDNTLITSHGINRRDFMKLCAALAATMGLSSKAAAEMAESVSNPQRPPVIWIGAQECT WP_000145429.1 hydrogenase 2 small subunit [Salmonella enterica] Length: 372\nNumber of predicted TMHs:  1\nExp number of AAs in TMHs: 24.51309\nExp number, first 60 AAs:  0.2682\nTotal prob of N-in: 0.14402\noutside   1   331\nTMhelix   332   354\ninside   355   372

17309 GCF_002037995.1_ASM203799v1 Salmonella enterica subsp. enterica serovar Enteritidis Proteobacteria; Gammaproteobacteria; Enterobacterales; Enterobacteriaceae; Salmonella; Salmonella enterica; Salmonella enterica subsp. enterica MNNEETFYQAMRRKGVTRRSFLKFCSLAATSLGLGAGMTPKIAWALENKPRIPVVWIHGL WP_001058307.1 hydrogenase [Salmonella enterica] Length: 372\nNumber of predicted TMHs:  2\nExp number of AAs in TMHs: 36.51738\nExp number, first 60 AAs:  19.72849\nTotal prob of N-in:   0.96170\nPOSSIBLE N-term signal sequence\ninside   1   19\nTMhelix   20   39\noutside   40   325\nTMhelix   326   348\ninside   349   372

17310 GCF_002038065.1_ASM203806v1 Salmonella enterica subsp. enterica serovar Enteritidis Proteobacteria; Gammaproteobacteria; Enterobacterales; Enterobacteriaceae; Salmonella; Salmonella enterica;

Salmonella enterica subsp. enterica MNNEETFYQAMRRKGVTRRSFLKFCSLAATSLGLGAGMTPKIAWALENKPRIPVVWIHGL WP_001058307.1 hydrogenase [Salmonella enterica] Length: 372\nNumber of predicted TMHs: 2\nExp number of AAs in TMHs: 36.51738\nExp number, first 60 AAs: 19.72849\nTotal prob of N-in: 0.96170\nPOSSIBLE N-term signal sequence\ninside 1 19\nTMhelix 20 39\noutside 40 325\nTMhelix 326 348\ninside 349 372

17311 GCF_002038065.1_ASM203806v1 Salmonella enterica subsp. enterica serovar Enteritidis Proteobacteria; Gammaproteobacteria; Enterobacterales; Enterobacteriaceae; Salmonella; Salmonella enterica; Salmonella enterica subsp. enterica MTGDNTLITSHGINRRDFMKLCAALAATMGLSSKAAAEMAESVSNPQRPPVIWIGAQECT WP_000145429.1 hydrogenase 2 small subunit [Salmonella enterica] Length: 372\nNumber of predicted TMHs: 1\nExp number of AAs in TMHs: 24.51309\nExp number, first 60 AAs: 0.2682\nTotal prob of N-in: 0.14402\noutside 1 331\nTMhelix 332 354\ninside 355 372

17312 GCF_002037965.1_ASM203796v1 Salmonella enterica subsp. enterica serovar Enteritidis Proteobacteria; Gammaproteobacteria; Enterobacterales; Enterobacteriaceae; Salmonella; Salmonella enterica; Salmonella enterica subsp. enterica MNNEETFYQAMRRKGVTRRSFLKFCSLAATSLGLGAGMTPKIAWALENKPRIPVVWIHGL WP_001058307.1 hydrogenase [Salmonella enterica] Length: 372\nNumber of predicted TMHs: 2\nExp number of AAs in TMHs: 36.51738\nExp number, first 60 AAs: 19.72849\nTotal prob of N-in: 0.96170\nPOSSIBLE N-term signal sequence\ninside 1 19\nTMhelix 20 39\noutside 40 325\nTMhelix 326 348\ninside 349 372

17313 GCF_002037905.1_ASM203790v1 Salmonella enterica subsp. enterica serovar Enteritidis Proteobacteria; Gammaproteobacteria; Enterobacterales; Enterobacteriaceae; Salmonella; Salmonella enterica; Salmonella enterica subsp. enterica MNNEETFYQAMRRKGVTRRSFLKFCSLAATSLGLGAGMTPKIAWALENKPRIPVVWIHGL WP_001058307.1 hydrogenase [Salmonella enterica] Length: 372\nNumber of predicted TMHs: 2\nExp number of AAs in TMHs: 36.51738\nExp number, first 60 AAs: 19.72849\nTotal prob of N-in: 0.96170\nPOSSIBLE N-term signal sequence\ninside 1 19\nTMhelix 20 39\noutside 40 325\nTMhelix 326 348\ninside 349 372

17314 GCF_002037825.1_ASM203782v1 Salmonella enterica subsp. enterica serovar Enteritidis Proteobacteria; Gammaproteobacteria; Enterobacterales; Enterobacteriaceae; Salmonella; Salmonella enterica; Salmonella enterica subsp. enterica MNNEETFYQAMRRKGVTRRSFLKFCSLAATSLGLGAGMTPKIAWALENKPRIPVVWIHGL WP_001058307.1 hydrogenase [Salmonella enterica] Length: 372\nNumber of predicted TMHs: 2\nExp number of AAs in TMHs: 36.51738\nExp number, first 60 AAs: 19.72849\nTotal prob of N-in: 0.96170\nPOSSIBLE N-term signal sequence\ninside 1 19\nTMhelix 20 39\noutside 40 325\nTMhelix 326 348\ninside 349 372

17315 GCF_002037825.1_ASM203782v1 Salmonella enterica subsp. enterica serovar Enteritidis Proteobacteria; Gammaproteobacteria; Enterobacterales; Enterobacteriaceae; Salmonella; Salmonella enterica; Salmonella enterica subsp. enterica MTGDNTLITSHGINRRDFMKLCAALAATMGLSSKAAAEMAESVSNPQRPPVIWIGAQECT WP_000145429.1 hydrogenase 2 small subunit [Salmonella enterica] Length: 372\nNumber of predicted TMHs: 1\nExp number of AAs in TMHs: 24.51309\nExp number, first 60 AAs: 0.2682\nTotal prob of N-in: 0.14402\noutside 1 331\nTMhelix 332 354\ninside 355 372

17316 GCF_002066535.1_ASM206653v1 Salmonella enterica subsp. enterica serovar Enteritidis Proteobacteria; Gammaproteobacteria; Enterobacterales; Enterobacteriaceae; Salmonella; Salmonella enterica; Salmonella enterica subsp. enterica MNNEETFYQAMRRKGVTRRSFLKFCSLAATSLGLGAGMTPKIAWALENKPRIPVVWIHGL WP_001058307.1 hydrogenase [Salmonella enterica] Length: 372\nNumber of predicted TMHs: 2\nExp number of AAs in TMHs: 36.51738\nExp number, first 60 AAs: 19.72849\nTotal prob of N-in: 0.96170\nPOSSIBLE N-term signal sequence\ninside 1 19\nTMhelix 20 39\noutside 40 325\nTMhelix 326 348\ninside 349 372

17317 GCF_002036435.1_ASM203643v1 Salmonella enterica subsp. enterica serovar Enteritidis Proteobacteria; Gammaproteobacteria; Enterobacterales; Enterobacteriaceae; Salmonella; Salmonella enterica; Salmonella enterica subsp. enterica MNNEETFYQAMRRKGVTRRSFLKFCSLAATSLGLGAGMTPKIAWALENKPRIPVVWIHGL WP_001058307.1 hydrogenase [Salmonella enterica] Length: 372\nNumber of predicted TMHs: 2\nExp number of AAs in TMHs: 36.51738\nExp number, first 60 AAs: 19.72849\nTotal prob of N-in: 0.96170\nPOSSIBLE N-term signal sequence\ninside 1 19\nTMhelix 20 39\noutside 40 325\nTMhelix 326 348\ninside 349 372

17318 GCF_002036395.1_ASM203639v1 Salmonella enterica subsp. enterica serovar Enteritidis Proteobacteria; Gammaproteobacteria; Enterobacterales; Enterobacteriaceae; Salmonella; Salmonella enterica; Salmonella enterica subsp. enterica MNNEETFYQAMRRKGVTRRSFLKFCSLAATSLGLGAGMTPKIAWALENKPRIPVVWIHGL WP_001058307.1 hydrogenase [Salmonella enterica] Length: 372\nNumber of predicted TMHs: 2\nExp number of AAs in TMHs: 36.51738\nExp number, first 60 AAs: 19.72849\nTotal prob of N-in: 0.96170\nPOSSIBLE N-term signal sequence\ninside 1 19\nTMhelix 20 39\noutside 40 325\nTMhelix 326 348\ninside 349 372

17319 GCF_002034885.1_ASM203488v1 Salmonella enterica subsp. enterica serovar Enteritidis Proteobacteria; Gammaproteobacteria; Enterobacterales; Enterobacteriaceae; Salmonella; Salmonella enterica; Salmonella enterica subsp. enterica MNNEETFYQAMRRKGVTRRSFLKFCSLAATSLGLGAGMTPKIAWALENKPRIPVVWIHGL WP_001058307.1 hydrogenase [Salmonella enterica] Length: 372\nNumber of predicted TMHs: 2\nExp number of AAs in TMHs: 36.51738\nExp number, first 60 AAs: 19.72849\nTotal prob of N-in: 0.96170\nPOSSIBLE N-term signal sequence\ninside 1 19\nTMhelix 20 39\noutside 40 325\nTMhelix 326 348\ninside 349 372

17320 GCF_002034885.1_ASM203488v1 Salmonella enterica subsp. enterica serovar Enteritidis Proteobacteria; Gammaproteobacteria; Enterobacterales; Enterobacteriaceae; Salmonella; Salmonella enterica; Salmonella enterica subsp. enterica MTGDNTLITSHGINRRDFMKLCAALAATMGLSSKAAAEMAESVSNPQRPPVIWIGAQECT WP_000145429.1 hydrogenase 2 small subunit [Salmonella enterica] Length: 372\nNumber of predicted TMHs: 1\nExp number of AAs in TMHs: 24.51309\nExp number, first 60 AAs: 0.2682\nTotal prob of N-in: 0.14402\noutside 1 331\nTMhelix 332 354\ninside 355 372

17321 GCF_002034005.1_ASM203400v1 Salmonella enterica subsp. enterica serovar Enteritidis Proteobacteria; Gammaproteobacteria; Enterobacterales; Enterobacteriaceae; Salmonella; Salmonella enterica; Salmonella enterica subsp. enterica MNNEETFYQAMRRKGVTRRSFLKFCSLAATSLGLGAGMTPKIAWALENKPRIPVVWIHGL WP_001728654.1 hydrogenase 1, small subunit [Salmonella enterica] Length: 372\nNumber of predicted TMHs: 2\nExp number of AAs in TMHs: 37.4248699999999\nExp number, first 60 AAs: 19.72699\nTotal prob of N-in: 0.96171\nPOSSIBLE N-term signal sequence\ninside 1 19\nTMhelix 20 39\noutside 40 325\nTMhelix 326 348\ninside 349 372

17322 GCF_002034005.1_ASM203400v1 Salmonella enterica subsp. enterica serovar Enteritidis Proteobacteria; Gammaproteobacteria; Enterobacterales; Enterobacteriaceae; Salmonella; Salmonella enterica; Salmonella enterica subsp. enterica MTGDNTLITSHGINRRDFMKLCAALAATMGLSSKAAAEMAESVSNPQRPPVIWIGAQECT WP_000145429.1 hydrogenase 2 small subunit [Salmonella enterica] Length: 372\nNumber of predicted TMHs: 1\nExp number of AAs in TMHs: 24.51309\nExp number, first 60 AAs: 0.2682\nTotal prob of N-in: 0.14402\noutside 1 331\nTMhelix 332 354\ninside 355 372

17323 GCF_002033105.1_ASM203310v1 Salmonella enterica subsp. enterica serovar Enteritidis Proteobacteria; Gammaproteobacteria; Enterobacterales; Enterobacteriaceae; Salmonella; Salmonella enterica; Salmonella enterica subsp. enterica MNNEETFYQAMRRKGVTRRSFLKFCSLAATSLGLGAGMTPKIAWALENKPRIPVVWIHGL WP_001058307.1 hydrogenase [Salmonella enterica] Length: 372\nNumber of predicted TMHs: 2\nExp number of AAs in TMHs: 36.51738\nExp number, first 60 AAs: 19.72849\nTotal prob of N-in: 0.96170\nPOSSIBLE N-term signal sequence\ninside 1 19\nTMhelix 20 39\noutside 40 325\nTMhelix 326 348\ninside 349 372

17324 GCF_002033105.1_ASM203310v1 Salmonella enterica subsp. enterica serovar Enteritidis Proteobacteria; Gammaproteobacteria; Enterobacterales; Enterobacteriaceae; Salmonella; Salmonella enterica; Salmonella enterica subsp. enterica MTGDNTLITSHGINRRDFMKLCAALAATMGLSSKAAAEMAESVSNPQRPPVIWIGAQECT WP_000145429.1 hydrogenase 2 small subunit [Salmonella enterica] Length: 372\nNumber of predicted TMHs: 1\nExp number of AAs in TMHs: 24.51309\nExp number, first 60 AAs: 0.2682\nTotal prob of N-in: 0.14402\noutside 1 331\nTMhelix 332 354\ninside 355 372

17325 GCF_002062685.1_ASM206268v1 Salmonella enterica subsp. enterica serovar Enteritidis Proteobacteria; Gammaproteobacteria; Enterobacterales; Enterobacteriaceae; Salmonella; Salmonella enterica; Salmonella enterica subsp. enterica MTGDNTLITSHGINRRDFMKLCAALAATMGLSSKAAAEMAESVSNPQRPPVIWIGAQECT WP_000145429.1 hydrogenase 2 small subunit [Salmonella enterica] Length: 372\nNumber of predicted TMHs: 1\nExp number of AAs in TMHs: 24.51309\nExp number, first 60 AAs: 0.2682\nTotal prob of N-in: 0.14402\noutside 1 331\nTMhelix 332 354\ninside 355 372

17326 GCF_002062685.1_ASM206268v1 Salmonella enterica subsp. enterica serovar Enteritidis Proteobacteria; Gammaproteobacteria; Enterobacterales; Enterobacteriaceae; Salmonella; Salmonella enterica; Salmonella enterica subsp. enterica MNNEETFYQAMRRKGVTRRSFLKFCSLAATSLGLGAGMTPKIAWALENKPRIPVVWIHGL WP_001058307.1 hydrogenase [Salmonella enterica] Length: 372\nNumber of predicted TMHs: 2\nExp number of AAs in TMHs: 36.51738\nExp number, first 60 AAs: 19.72849\nTotal prob of N-in: 0.96170\nPOSSIBLE N-term signal sequence\ninside 1 19\nTMhelix 20 39\noutside 40 325\nTMhelix 326 348\ninside 349 372

17327 GCF_001973115.1_ASM197311v1 Salmonella enterica subsp. enterica serovar Enteritidis Proteobacteria; Gammaproteobacteria; Enterobacterales; Enterobacteriaceae; Salmonella; Salmonella enterica; Salmonella enterica subsp. enterica MTGDNTLITSHGINRRDFMKLCAALAATMGLSSKAAAEMAESVSNPQRPPVIWIGAQECT WP_000145429.1 hydrogenase 2 small subunit [Salmonella enterica] Length: 372\nNumber of

predicted TMHs: 1\nExp number of AAs in TMHs: 24.51309\nExp number, first 60 AAs: 0.2682\nTotal prob of N-in: 0.14402\noutside 1 331\nTMhelix 332 354\ninside 355 372

17328 GCF_002038365.1_ASM203836v1 Salmonella enterica subsp. enterica serovar Enteritidis Proteobacteria; Gammaproteobacteria; Enterobacterales; Enterobacteriaceae; Salmonella; Salmonella enterica; Salmonella enterica subsp. enterica MNNEETFYQAMRRKGVTRRSFLKFCSLAATSLGLGAGMTPKIAWALENKPRIPVVWIHGL WP_001058307.1 hydrogenase [Salmonella enterica] Length: 372\nNumber of predicted TMHs: 2\nExp number of AAs in TMHs: 36.51738\nExp number, first 60 AAs: 19.72849\nTotal prob of N-in: 0.96170\nPOSSIBLE N-term signal sequence\ninside 1 19\nTMhelix 20 39\noutside 40 325\nTMhelix 326 348\ninside 349 372

17329 GCF_002038395.1_ASM203839v1 Salmonella enterica subsp. enterica serovar Enteritidis Proteobacteria; Gammaproteobacteria; Enterobacterales; Enterobacteriaceae; Salmonella; Salmonella enterica; Salmonella enterica subsp. enterica MNNEETFYQAMRRKGVTRRSFLKFCSLAATSLGLGAGMTPKIAWALENKPRIPVVWIHGL WP_001058307.1 hydrogenase [Salmonella enterica] Length: 372\nNumber of predicted TMHs: 2\nExp number of AAs in TMHs: 36.51738\nExp number, first 60 AAs: 19.72849\nTotal prob of N-in: 0.96170\nPOSSIBLE N-term signal sequence\ninside 1 19\nTMhelix 20 39\noutside 40 325\nTMhelix 326 348\ninside 349 372

17330 GCF_002039105.1_ASM203910v1 Salmonella enterica subsp. enterica serovar Enteritidis Proteobacteria; Gammaproteobacteria; Enterobacterales; Enterobacteriaceae; Salmonella; Salmonella enterica; Salmonella enterica subsp. enterica MTGDNTLITSHGINRRDFMKLCAALAATMGLSSKAAAEMAESVSNPQRPPVIWIGAQECT WP_000145429.1 hydrogenase 2 small subunit [Salmonella enterica] Length: 372\nNumber of predicted TMHs: 1\nExp number of AAs in TMHs: 24.51309\nExp number, first 60 AAs: 0.2682\nTotal prob of N-in: 0.14402\noutside 1 331\nTMhelix 332 354\ninside 355 372

17331 GCF_002038445.1_ASM203844v1 Salmonella enterica subsp. enterica serovar Enteritidis Proteobacteria; Gammaproteobacteria; Enterobacterales; Enterobacteriaceae; Salmonella; Salmonella enterica; Salmonella enterica subsp. enterica MNNEETFYQAMRRKGVTRRSFLKFCSLAATSLGLGAGMTPKIAWALENKPRIPVVWIHGL WP_001058307.1 hydrogenase [Salmonella enterica] Length: 372\nNumber of predicted TMHs: 2\nExp number of AAs in TMHs: 36.51738\nExp number, first 60 AAs: 19.72849\nTotal prob of N-in: 0.96170\nPOSSIBLE N-term signal sequence\ninside 1 19\nTMhelix 20 39\noutside 40 325\nTMhelix 326 348\ninside 349 372

17332 GCF_002038645.1_ASM203864v1 Salmonella enterica subsp. enterica serovar Enteritidis Proteobacteria; Gammaproteobacteria; Enterobacterales; Enterobacteriaceae; Salmonella; Salmonella enterica; Salmonella enterica subsp. enterica MTGDNTLITSHGINRRDFMKLCAALAATMGLSSKAAAEMAESVSNPQRPPVIWIGAQECT WP_000145429.1 hydrogenase 2 small subunit [Salmonella enterica] Length: 372\nNumber of predicted TMHs: 1\nExp number of AAs in TMHs: 24.51309\nExp number, first 60 AAs: 0.2682\nTotal prob of N-in: 0.14402\noutside 1 331\nTMhelix 332 354\ninside 355 372

17333 GCF_002038605.1_ASM203860v1 Salmonella enterica subsp. enterica serovar Enteritidis Proteobacteria; Gammaproteobacteria; Enterobacterales; Enterobacteriaceae; Salmonella; Salmonella enterica; Salmonella enterica subsp. enterica MTGDNTLITSHGINRRDFMKLCAALAATMGLSSKAAAEMAESVSNPQRPPVIWIGAQECT WP_000145429.1 hydrogenase 2 small subunit [Salmonella enterica] Length: 372\nNumber of predicted TMHs: 1\nExp number of AAs in TMHs: 24.51309\nExp number, first 60 AAs: 0.2682\nTotal prob of N-in: 0.14402\noutside 1 331\nTMhelix 332 354\ninside 355 372

17334 GCF_002038685.1_ASM203868v1 Salmonella enterica subsp. enterica serovar Enteritidis Proteobacteria; Gammaproteobacteria; Enterobacterales; Enterobacteriaceae; Salmonella; Salmonella enterica; Salmonella enterica subsp. enterica MTGDNTLITSHGINRRDFMKLCAALAATMGLSSKAAAEMAESVSNPQRPPVIWIGAQECT WP_000145429.1 hydrogenase 2 small subunit [Salmonella enterica] Length: 372\nNumber of predicted TMHs: 1\nExp number of AAs in TMHs: 24.51309\nExp number, first 60 AAs: 0.2682\nTotal prob of N-in: 0.14402\noutside 1 331\nTMhelix 332 354\ninside 355 372

17335 GCF_002038645.1_ASM203864v1 Salmonella enterica subsp. enterica serovar Enteritidis Proteobacteria; Gammaproteobacteria; Enterobacterales; Enterobacteriaceae; Salmonella; Salmonella enterica; Salmonella enterica subsp. enterica MNNEETFYQAMRRKGVTRRSFLKFCSLAATSLGLGAGMTPKIAWALENKPRIPVVWIHGL WP_001058307.1 hydrogenase [Salmonella enterica] Length: 372\nNumber of predicted TMHs: 2\nExp number of AAs in TMHs: 36.51738\nExp number, first 60 AAs: 19.72849\nTotal prob of N-in: 0.96170\nPOSSIBLE N-term signal sequence\ninside 1 19\nTMhelix 20 39\noutside 40 325\nTMhelix 326 348\ninside 349 372

17336 GCF_002038605.1_ASM203860v1 Salmonella enterica subsp. enterica serovar Enteritidis Proteobacteria; Gammaproteobacteria; Enterobacterales; Enterobacteriaceae; Salmonella; Salmonella enterica;

Salmonella enterica subsp. enterica	MNNEETFYQAMRRKGVTRRSFLKFCSLAATSLGLGAGMTPKIAWALENKPRIPVVWIHGL	WP_001058307.1	hydrogenase [Salmonella enterica]	Length: 372\nNumber of predicted TMHs:  2\nExp number of AAs in TMHs: 36.51738\nExp number, first 60 AAs:  19.72849\nTotal prob of N-in:   0.96170\nPOSSIBLE N-term signal sequence\ninside   1   19\nTMhelix   20   39\noutside   40   325\nTMhelix   326   348\ninside   349   372

17337	GCF_002038685.1_ASM203868v1	Salmonella enterica subsp. enterica serovar Enteritidis	Proteobacteria; Gammaproteobacteria; Enterobacterales; Enterobacteriaceae; Salmonella; Salmonella enterica; Salmonella enterica subsp. enterica	MNNEETFYQAMRRKGVTRRSFLKFCSLAATSLGLGAGMTPKIAWALENKPRIPVVWIHGL	WP_001058307.1	hydrogenase [Salmonella enterica]	Length: 372\nNumber of predicted TMHs:  2\nExp number of AAs in TMHs: 36.51738\nExp number, first 60 AAs:  19.72849\nTotal prob of N-in:   0.96170\nPOSSIBLE N-term signal sequence\ninside   1   19\nTMhelix   20   39\noutside   40   325\nTMhelix   326   348\ninside   349   372

17338	GCF_002038555.1_ASM203855v1	Salmonella enterica subsp. enterica serovar Enteritidis	Proteobacteria; Gammaproteobacteria; Enterobacterales; Enterobacteriaceae; Salmonella; Salmonella enterica; Salmonella enterica subsp. enterica	MNNEETFYQAMRRKGVTRRSFLKFCSLAATSLGLGAGMTPKIAWALENKPRIPVVWIHGL	WP_001058307.1	hydrogenase [Salmonella enterica]	Length: 372\nNumber of predicted TMHs:  2\nExp number of AAs in TMHs: 36.51738\nExp number, first 60 AAs:  19.72849\nTotal prob of N-in:   0.96170\nPOSSIBLE N-term signal sequence\ninside   1   19\nTMhelix   20   39\noutside   40   325\nTMhelix   326   348\ninside   349   372

17339	GCF_002038835.1_ASM203883v1	Salmonella enterica subsp. enterica serovar Enteritidis	Proteobacteria; Gammaproteobacteria; Enterobacterales; Enterobacteriaceae; Salmonella; Salmonella enterica; Salmonella enterica subsp. enterica	MTGDNTLITSHGINRRDFMKLCAALAATMGLSSKAAAEMAESVSNPQRPPVIWIGAQECT	WP_000145429.1	hydrogenase 2 small subunit [Salmonella enterica]	Length: 372\nNumber of predicted TMHs:  1\nExp number of AAs in TMHs: 24.51309\nExp number, first 60 AAs:  0.2682\nTotal prob of N-in:   0.14402\noutside   1   331\nTMhelix   332   354\ninside   355   372

17340	GCF_002038865.1_ASM203886v1	Salmonella enterica subsp. enterica serovar Enteritidis	Proteobacteria; Gammaproteobacteria; Enterobacterales; Enterobacteriaceae; Salmonella; Salmonella enterica; Salmonella enterica subsp. enterica	MTGDNTLITSHGINRRDFMKLCAALAATMGLSSKAAAEMAESVSNPQRPPVIWIGAQECT	WP_000145429.1	hydrogenase 2 small subunit [Salmonella enterica]	Length: 372\nNumber of predicted TMHs:  1\nExp number of AAs in TMHs: 24.51309\nExp number, first 60 AAs:  0.2682\nTotal prob of N-in:   0.14402\noutside   1   331\nTMhelix   332   354\ninside   355   372

17341	GCF_002038885.1_ASM203888v1	Salmonella enterica subsp. enterica serovar Enteritidis	Proteobacteria; Gammaproteobacteria; Enterobacterales; Enterobacteriaceae; Salmonella; Salmonella enterica; Salmonella enterica subsp. enterica	MTGDNTLITSHGINRRDFMKLCAALAATMGLSSKAAAEMAESVSNPQRPPVIWIGAQECT	WP_000145429.1	hydrogenase 2 small subunit [Salmonella enterica]	Length: 372\nNumber of predicted TMHs:  1\nExp number of AAs in TMHs: 24.51309\nExp number, first 60 AAs:  0.2682\nTotal prob of N-in:   0.14402\noutside   1   331\nTMhelix   332   354\ninside   355   372

17342	GCF_002038915.1_ASM203891v1	Salmonella enterica subsp. enterica serovar Enteritidis	Proteobacteria; Gammaproteobacteria; Enterobacterales; Enterobacteriaceae; Salmonella; Salmonella enterica; Salmonella enterica subsp. enterica	MTGDNTLITSHGINRRDFMKLCAALAATMGLSSKAAAEMAESVSNPQRPPVIWIGAQECT	WP_000145429.1	hydrogenase 2 small subunit [Salmonella enterica]	Length: 372\nNumber of predicted TMHs:  1\nExp number of AAs in TMHs: 24.51309\nExp number, first 60 AAs:  0.2682\nTotal prob of N-in:   0.14402\noutside   1   331\nTMhelix   332   354\ninside   355   372

17343	GCF_002038835.1_ASM203883v1	Salmonella enterica subsp. enterica serovar Enteritidis	Proteobacteria; Gammaproteobacteria; Enterobacterales; Enterobacteriaceae; Salmonella; Salmonella enterica; Salmonella enterica subsp. enterica	MNNEETFYQAMRRKGVTRRSFLKFCSLAATSLGLGAGMTPKIAWALENKPRIPVVWIHGL	WP_001058307.1	hydrogenase [Salmonella enterica]	Length: 372\nNumber of predicted TMHs:  2\nExp number of AAs in TMHs: 36.51738\nExp number, first 60 AAs:  19.72849\nTotal prob of N-in:   0.96170\nPOSSIBLE N-term signal sequence\ninside   1   19\nTMhelix   20   39\noutside   40   325\nTMhelix   326   348\ninside   349   372

17344	GCF_002038965.1_ASM203896v1	Salmonella enterica subsp. enterica serovar Enteritidis	Proteobacteria; Gammaproteobacteria; Enterobacterales; Enterobacteriaceae; Salmonella; Salmonella enterica; Salmonella enterica subsp. enterica	MTGDNTLITSHGINRRDFMKLCAALAATMGLSSKAAAEMAESVSNPQRPPVIWIGAQECT	WP_000145429.1	hydrogenase 2 small subunit [Salmonella enterica]	Length: 372\nNumber of predicted TMHs:  1\nExp number of AAs in TMHs: 24.51309\nExp number, first 60 AAs:  0.2682\nTotal prob of N-in:   0.14402\noutside   1   331\nTMhelix   332   354\ninside   355   372

17345 GCF_002039145.1_ASM203914v1 Salmonella enterica subsp. enterica serovar Enteritidis Proteobacteria; Gammaproteobacteria; Enterobacterales; Enterobacteriaceae; Salmonella; Salmonella enterica; Salmonella enterica subsp. enterica MTGDNTLITSHGINRRDFMKLCAALAATMGLSSKAAAEMAESVSNPQRPPVIWIGAQECT WP_000145429.1 hydrogenase 2 small subunit [Salmonella enterica] Length: 372\nNumber of predicted TMHs: 1\nExp number of AAs in TMHs: 24.51309\nExp number, first 60 AAs: 0.2682\nTotal prob of N-in: 0.14402\noutside 1 331\nTMhelix 332 354\ninside 355 372

17346 GCF_002039185.1_ASM203918v1 Salmonella enterica subsp. enterica serovar Enteritidis Proteobacteria; Gammaproteobacteria; Enterobacterales; Enterobacteriaceae; Salmonella; Salmonella enterica; Salmonella enterica subsp. enterica MTGDNTLITSHGINRRDFMKLCAALAATMGLSSKAAAEMAESVSNPQRPPVIWIGAQECT WP_000145429.1 hydrogenase 2 small subunit [Salmonella enterica] Length: 372\nNumber of predicted TMHs: 1\nExp number of AAs in TMHs: 24.51309\nExp number, first 60 AAs: 0.2682\nTotal prob of N-in: 0.14402\noutside 1 331\nTMhelix 332 354\ninside 355 372

17347 GCF_002039105.1_ASM203910v1 Salmonella enterica subsp. enterica serovar Enteritidis Proteobacteria; Gammaproteobacteria; Enterobacterales; Enterobacteriaceae; Salmonella; Salmonella enterica; Salmonella enterica subsp. enterica MNNEETFYQAMRRKGVTRRSFLKFCSLAATSLGLGAGMTPKIAWALENKPRIPVVWIHGL WP_001058307.1 hydrogenase [Salmonella enterica] Length: 372\nNumber of predicted TMHs: 2\nExp number of AAs in TMHs: 36.51738\nExp number, first 60 AAs: 19.72849\nTotal prob of N-in: 0.96170\nPOSSIBLE N-term signal sequence\ninside 1 19\nTMhelix 20 39\noutside 40 325\nTMhelix 326 348\ninside 349 372

17348 GCF_002039215.1_ASM203921v1 Salmonella enterica subsp. enterica serovar Enteritidis Proteobacteria; Gammaproteobacteria; Enterobacterales; Enterobacteriaceae; Salmonella; Salmonella enterica; Salmonella enterica subsp. enterica MTGDNTLITSHGINRRDFMKLCAALAATMGLSSKAAAEMAESVSNPQRPPVIWIGAQECT WP_000145429.1 hydrogenase 2 small subunit [Salmonella enterica] Length: 372\nNumber of predicted TMHs: 1\nExp number of AAs in TMHs: 24.51309\nExp number, first 60 AAs: 0.2682\nTotal prob of N-in: 0.14402\noutside 1 331\nTMhelix 332 354\ninside 355 372

17349 GCF_002039045.1_ASM203904v1 Salmonella enterica subsp. enterica serovar Enteritidis Proteobacteria; Gammaproteobacteria; Enterobacterales; Enterobacteriaceae; Salmonella; Salmonella enterica; Salmonella enterica subsp. enterica MNNEETFYQAMRRKGVTRRSFLKFCSLAATSLGLGAGMTPKIAWALENKPRIPVVWIHGL WP_001058307.1 hydrogenase [Salmonella enterica] Length: 372\nNumber of predicted TMHs: 2\nExp number of AAs in TMHs: 36.51738\nExp number, first 60 AAs: 19.72849\nTotal prob of N-in: 0.96170\nPOSSIBLE N-term signal sequence\ninside 1 19\nTMhelix 20 39\noutside 40 325\nTMhelix 326 348\ninside 349 372

17350 GCF_002039265.1_ASM203926v1 Salmonella enterica subsp. enterica serovar Enteritidis Proteobacteria; Gammaproteobacteria; Enterobacterales; Enterobacteriaceae; Salmonella; Salmonella enterica; Salmonella enterica subsp. enterica MTGDNTLITSHGINRRDFMKLCAALAATMGLSSKAAAEMAESVSNPQRPPVIWIGAQECT WP_000145429.1 hydrogenase 2 small subunit [Salmonella enterica] Length: 372\nNumber of predicted TMHs: 1\nExp number of AAs in TMHs: 24.51309\nExp number, first 60 AAs: 0.2682\nTotal prob of N-in: 0.14402\noutside 1 331\nTMhelix 332 354\ninside 355 372

17351 GCF_002039265.1_ASM203926v1 Salmonella enterica subsp. enterica serovar Enteritidis Proteobacteria; Gammaproteobacteria; Enterobacterales; Enterobacteriaceae; Salmonella; Salmonella enterica; Salmonella enterica subsp. enterica MNNEETFYQAMRRKGVTRRSFLKFCSLAATSLGLGAGMTPKIAWALENKPRIPVVWIHGL WP_001058307.1 hydrogenase [Salmonella enterica] Length: 372\nNumber of predicted TMHs: 2\nExp number of AAs in TMHs: 36.51738\nExp number, first 60 AAs: 19.72849\nTotal prob of N-in: 0.96170\nPOSSIBLE N-term signal sequence\ninside 1 19\nTMhelix 20 39\noutside 40 325\nTMhelix 326 348\ninside 349 372

17352 GCF_002039385.1_ASM203938v1 Salmonella enterica subsp. enterica serovar Enteritidis Proteobacteria; Gammaproteobacteria; Enterobacterales; Enterobacteriaceae; Salmonella; Salmonella enterica; Salmonella enterica subsp. enterica MNNEETFYQAMRRKGVTRRSFLKFCSLAATSLGLGAGMTPKIAWALENKPRIPVVWIHGL WP_001058307.1 hydrogenase [Salmonella enterica] Length: 372\nNumber of predicted TMHs: 2\nExp number of AAs in TMHs: 36.51738\nExp number, first 60 AAs: 19.72849\nTotal prob of N-in: 0.96170\nPOSSIBLE N-term signal sequence\ninside 1 19\nTMhelix 20 39\noutside 40 325\nTMhelix 326 348\ninside 349 372

17353 GCF_002039215.1_ASM203921v1 Salmonella enterica subsp. enterica serovar Enteritidis Proteobacteria; Gammaproteobacteria; Enterobacterales; Enterobacteriaceae; Salmonella; Salmonella enterica; Salmonella enterica subsp. enterica MNNEETFYQAMRRKGVTRRSFLKFCSLAATSLGLGAGMTPKIAWALENKPRIPVVWIHGL WP_001058307.1 hydrogenase [Salmonella enterica] Length: 372\nNumber of predicted TMHs: 2\nExp

number of AAs in TMHs: 36.51738\nExp number, first 60 AAs: 19.72849\nTotal prob of N-in: 0.96170\nPOSSIBLE N-term signal sequence\ninside 1 19\nTMhelix 20 39\noutside 40 325\nTMhelix 326 348\ninside 349 372

17354 GCF_002039425.1_ASM203942v1 Salmonella enterica subsp. enterica serovar Enteritidis Proteobacteria; Gammaproteobacteria; Enterobacterales; Enterobacteriaceae; Salmonella; Salmonella enterica; Salmonella enterica subsp. enterica MTGDNTLITSHGINRRDFMKLCAALAATMGLSSKAAAEMAESVSNPQRPPVIWIGAQECT WP_000145429.1 hydrogenase 2 small subunit [Salmonella enterica] Length: 372\nNumber of predicted TMHs: 1\nExp number of AAs in TMHs: 24.51309\nExp number, first 60 AAs: 0.2682\nTotal prob of N-in: 0.14402\noutside 1 331\nTMhelix 332 354\ninside 355 372

17355 GCF_002039455.1_ASM203945v1 Salmonella enterica subsp. enterica serovar Enteritidis Proteobacteria; Gammaproteobacteria; Enterobacterales; Enterobacteriaceae; Salmonella; Salmonella enterica; Salmonella enterica subsp. enterica MTGDNTLITSHGINRRDFMKLCAALAATMGLSSKAAAEMAESVSNPQRPPVIWIGAQECT WP_000145429.1 hydrogenase 2 small subunit [Salmonella enterica] Length: 372\nNumber of predicted TMHs: 1\nExp number of AAs in TMHs: 24.51309\nExp number, first 60 AAs: 0.2682\nTotal prob of N-in: 0.14402\noutside 1 331\nTMhelix 332 354\ninside 355 372

17356 GCF_002039505.1_ASM203950v1 Salmonella enterica subsp. enterica serovar Enteritidis Proteobacteria; Gammaproteobacteria; Enterobacterales; Enterobacteriaceae; Salmonella; Salmonella enterica; Salmonella enterica subsp. enterica MTGDNTLITSHGINRRDFMKLCAALAATMGLSSKAAAEMAESVSNPQRPPVIWIGAQECT WP_000145429.1 hydrogenase 2 small subunit [Salmonella enterica] Length: 372\nNumber of predicted TMHs: 1\nExp number of AAs in TMHs: 24.51309\nExp number, first 60 AAs: 0.2682\nTotal prob of N-in: 0.14402\noutside 1 331\nTMhelix 332 354\ninside 355 372

17357 GCF_002039425.1_ASM203942v1 Salmonella enterica subsp. enterica serovar Enteritidis Proteobacteria; Gammaproteobacteria; Enterobacterales; Enterobacteriaceae; Salmonella; Salmonella enterica; Salmonella enterica subsp. enterica MNNEETFYQAMRRKGVTRRSFLKFCSLAATSLGLGAGMTPKIAWALENKPRIPVVWIHGL WP_001058307.1 hydrogenase [Salmonella enterica] Length: 372\nNumber of predicted TMHs: 2\nExp number of AAs in TMHs: 36.51738\nExp number, first 60 AAs: 19.72849\nTotal prob of N-in: 0.96170\nPOSSIBLE N-term signal sequence\ninside 1 19\nTMhelix 20 39\noutside 40 325\nTMhelix 326 348\ninside 349 372

17358 GCF_002039525.1_ASM203952v1 Salmonella enterica subsp. enterica serovar Enteritidis Proteobacteria; Gammaproteobacteria; Enterobacterales; Enterobacteriaceae; Salmonella; Salmonella enterica; Salmonella enterica subsp. enterica MTGDNTLITSHGINRRDFMKLCAALAATMGLSSKAAAEMAESVSNPQRPPVIWIGAQECT WP_000145429.1 hydrogenase 2 small subunit [Salmonella enterica] Length: 372\nNumber of predicted TMHs: 1\nExp number of AAs in TMHs: 24.51309\nExp number, first 60 AAs: 0.2682\nTotal prob of N-in: 0.14402\noutside 1 331\nTMhelix 332 354\ninside 355 372

17359 GCF_002039705.1_ASM203970v1 Salmonella enterica subsp. enterica serovar Enteritidis Proteobacteria; Gammaproteobacteria; Enterobacterales; Enterobacteriaceae; Salmonella; Salmonella enterica; Salmonella enterica subsp. enterica MTGDNTLITSHGINRRDFMKLCAALAATMGLSSKAAAEMAESVSNPQRPPVIWIGAQECT WP_000145429.1 hydrogenase 2 small subunit [Salmonella enterica] Length: 372\nNumber of predicted TMHs: 1\nExp number of AAs in TMHs: 24.51309\nExp number, first 60 AAs: 0.2682\nTotal prob of N-in: 0.14402\noutside 1 331\nTMhelix 332 354\ninside 355 372

17360 GCF_002039545.1_ASM203954v1 Salmonella enterica subsp. enterica serovar Enteritidis Proteobacteria; Gammaproteobacteria; Enterobacterales; Enterobacteriaceae; Salmonella; Salmonella enterica; Salmonella enterica subsp. enterica MNNEETFYQAMRRKGVTRRSFLKFCSLAATSLGLGAGMTPKIAWALENKPRIPVVWIHGL WP_001058307.1 hydrogenase [Salmonella enterica] Length: 372\nNumber of predicted TMHs: 2\nExp number of AAs in TMHs: 36.51738\nExp number, first 60 AAs: 19.72849\nTotal prob of N-in: 0.96170\nPOSSIBLE N-term signal sequence\ninside 1 19\nTMhelix 20 39\noutside 40 325\nTMhelix 326 348\ninside 349 372

17361 GCF_002039585.1_ASM203958v1 Salmonella enterica subsp. enterica serovar Enteritidis Proteobacteria; Gammaproteobacteria; Enterobacterales; Enterobacteriaceae; Salmonella; Salmonella enterica; Salmonella enterica subsp. enterica MNNEETFYQAMRRKGVTRRSFLKFCSLAATSLGLGAGMTPKIAWALENKPRIPVVWIHGL WP_001058307.1 hydrogenase [Salmonella enterica] Length: 372\nNumber of predicted TMHs: 2\nExp number of AAs in TMHs: 36.51738\nExp number, first 60 AAs: 19.72849\nTotal prob of N-in: 0.96170\nPOSSIBLE N-term signal sequence\ninside 1 19\nTMhelix 20 39\noutside 40 325\nTMhelix 326 348\ninside 349 372

17362 GCF_002039745.1_ASM203974v1 Salmonella enterica subsp. enterica serovar Enteritidis Proteobacteria; Gammaproteobacteria; Enterobacterales; Enterobacteriaceae; Salmonella; Salmonella enterica;

Salmonella enterica subsp. enterica	MTGDNTLITSHGINRRDFMKLCAALAATMGLSSKAAAEMAESVSNPQRPPVIWIGAQECT	WP_000145429.1	hydrogenase 2 small subunit [Salmonella enterica]	Length: 372\nNumber of predicted TMHs: 1\nExp number of AAs in TMHs: 24.51309\nExp number, first 60 AAs: 0.2682\nTotal prob of N-in: 0.14402\noutside 1 331\nTMhelix 332 354\ninside 355 372

17363	GCF_002039625.1_ASM203962v1	Salmonella enterica subsp. enterica serovar Enteritidis	Proteobacteria; Gammaproteobacteria; Enterobacterales; Enterobacteriaceae; Salmonella; Salmonella enterica; Salmonella enterica subsp. enterica	MNNEETFYQAMRRKGVTRRSFLKFCSLAATSLGLGAGMTPKIAWALENKPRIPVVWIHGL	WP_001058307.1	hydrogenase [Salmonella enterica]	Length: 372\nNumber of predicted TMHs: 2\nExp number of AAs in TMHs: 36.51738\nExp number, first 60 AAs: 19.72849\nTotal prob of N-in: 0.96170\nPOSSIBLE N-term signal sequence\ninside 1 19\nTMhelix 20 39\noutside 40 325\nTMhelix 326 348\ninside 349 372

17364	GCF_002039665.1_ASM203966v1	Salmonella enterica subsp. enterica serovar Enteritidis	Proteobacteria; Gammaproteobacteria; Enterobacterales; Enterobacteriaceae; Salmonella; Salmonella enterica; Salmonella enterica subsp. enterica	MNNEETFYQAMRRKGVTRRSFLKFCSLAATSLGLGAGMTPKIAWALENKPRIPVVWIHGL	WP_001058307.1	hydrogenase [Salmonella enterica]	Length: 372\nNumber of predicted TMHs: 2\nExp number of AAs in TMHs: 36.51738\nExp number, first 60 AAs: 19.72849\nTotal prob of N-in: 0.96170\nPOSSIBLE N-term signal sequence\ninside 1 19\nTMhelix 20 39\noutside 40 325\nTMhelix 326 348\ninside 349 372

17365	GCF_002040125.1_ASM204012v1	Salmonella enterica subsp. enterica serovar Enteritidis	Proteobacteria; Gammaproteobacteria; Enterobacterales; Enterobacteriaceae; Salmonella; Salmonella enterica; Salmonella enterica subsp. enterica	MTGDNTLITSHGINRRDFMKLCAALAATMGLSSKAAAEMAESVSNPQRPPVIWIGAQECT	WP_000145429.1	hydrogenase 2 small subunit [Salmonella enterica]	Length: 372\nNumber of predicted TMHs: 1\nExp number of AAs in TMHs: 24.51309\nExp number, first 60 AAs: 0.2682\nTotal prob of N-in: 0.14402\noutside 1 331\nTMhelix 332 354\ninside 355 372

17366	GCF_001691665.1_ASM169166v1	Salmonella enterica subsp. enterica serovar Enteritidis	Proteobacteria; Gammaproteobacteria; Enterobacterales; Enterobacteriaceae; Salmonella; Salmonella enterica; Salmonella enterica subsp. enterica	MNNEETFYQAMRRKGVTRRSFLKFCSLAATSLGLGAGMTPKIAWALENKPRIPVVWIHGL	WP_001058307.1	hydrogenase [Salmonella enterica]	Length: 372\nNumber of predicted TMHs: 2\nExp number of AAs in TMHs: 36.51738\nExp number, first 60 AAs: 19.72849\nTotal prob of N-in: 0.96170\nPOSSIBLE N-term signal sequence\ninside 1 19\nTMhelix 20 39\noutside 40 325\nTMhelix 326 348\ninside 349 372

17367	GCF_001691665.1_ASM169166v1	Salmonella enterica subsp. enterica serovar Enteritidis	Proteobacteria; Gammaproteobacteria; Enterobacterales; Enterobacteriaceae; Salmonella; Salmonella enterica; Salmonella enterica subsp. enterica	MTGDNTLITSHGINRRDFMKLCAALAATMGLSSKAAAEMAESVSNPQRPPVIWIGAQECT	WP_000145429.1	hydrogenase 2 small subunit [Salmonella enterica]	Length: 372\nNumber of predicted TMHs: 1\nExp number of AAs in TMHs: 24.51309\nExp number, first 60 AAs: 0.2682\nTotal prob of N-in: 0.14402\noutside 1 331\nTMhelix 332 354\ninside 355 372

17368	GCF_001647045.1_ASM164704v1	Salmonella enterica subsp. enterica serovar Enteritidis	Proteobacteria; Gammaproteobacteria; Enterobacterales; Enterobacteriaceae; Salmonella; Salmonella enterica; Salmonella enterica subsp. enterica	MTGDNTLITSHGINRRDFMKLCAALAATMGLSSKAAAEMAESVSNPQRPPVIWIGAQECT	WP_000145429.1	hydrogenase 2 small subunit [Salmonella enterica]	Length: 372\nNumber of predicted TMHs: 1\nExp number of AAs in TMHs: 24.51309\nExp number, first 60 AAs: 0.2682\nTotal prob of N-in: 0.14402\noutside 1 331\nTMhelix 332 354\ninside 355 372

17369	GCF_001973785.1_ASM197378v1	Salmonella enterica subsp. enterica serovar Enteritidis	Proteobacteria; Gammaproteobacteria; Enterobacterales; Enterobacteriaceae; Salmonella; Salmonella enterica; Salmonella enterica subsp. enterica	MNNEETFYQAMRRKGVTRRSFLKFCSLAATSLGLGAGMTPKIAWALENKPRIPVVWIHGL	WP_001058307.1	hydrogenase [Salmonella enterica]	Length: 372\nNumber of predicted TMHs: 2\nExp number of AAs in TMHs: 36.51738\nExp number, first 60 AAs: 19.72849\nTotal prob of N-in: 0.96170\nPOSSIBLE N-term signal sequence\ninside 1 19\nTMhelix 20 39\noutside 40 325\nTMhelix 326 348\ninside 349 372

17370	GCF_001973665.1_ASM197366v1	Salmonella enterica subsp. enterica serovar Enteritidis	Proteobacteria; Gammaproteobacteria; Enterobacterales; Enterobacteriaceae; Salmonella; Salmonella enterica; Salmonella enterica subsp. enterica	MNNEETFYQAMRRKGVTRRSFLKFCSLAATSLGLGAGMTPKIAWALENKPRIPVVWIHGL	WP_001058307.1	hydrogenase [Salmonella enterica]	Length: 372\nNumber of predicted TMHs: 2\nExp number of AAs in TMHs: 36.51738\nExp number, first 60 AAs: 19.72849\nTotal prob of N-in: 0.96170\nPOSSIBLE N-term signal sequence\ninside 1 19\nTMhelix 20 39\noutside 40 325\nTMhelix 326 348\ninside 349 372

17371	GCF_001973705.1_ASM197370v1	Salmonella enterica subsp. enterica serovar Enteritidis	Proteobacteria; Gammaproteobacteria; Enterobacterales; Enterobacteriaceae; Salmonella; Salmonella enterica; Salmonella enterica subsp. enterica	MNNEETFYQAMRRKGVTRRSFLKFCSLAATSLGLGAGMTPKIAWALENKPRIPVVWIHGL	WP_001058307.1	hydrogenase [Salmonella enterica]	Length: 372\nNumber of predicted TMHs:  2\nExp number of AAs in TMHs: 36.51738\nExp number, first 60 AAs:  19.72849\nTotal prob of N-in:   0.96170\nPOSSIBLE N-term signal sequence\ninside   1   19\nTMhelix   20   39\noutside   40   325\nTMhelix   326   348\ninside   349   372

17372	GCF_001973545.1_ASM197354v1	Salmonella enterica subsp. enterica serovar Enteritidis	Proteobacteria; Gammaproteobacteria; Enterobacterales; Enterobacteriaceae; Salmonella; Salmonella enterica; Salmonella enterica subsp. enterica	MNNEETFYQAMRRKGVTRRSFLKFCSLAATSLGLGAGMTPKIAWALENKPRIPVVWIHGL	WP_001058307.1	hydrogenase [Salmonella enterica]	Length: 372\nNumber of predicted TMHs:  2\nExp number of AAs in TMHs: 36.51738\nExp number, first 60 AAs:  19.72849\nTotal prob of N-in:   0.96170\nPOSSIBLE N-term signal sequence\ninside   1   19\nTMhelix   20   39\noutside   40   325\nTMhelix   326   348\ninside   349   372

17373	GCF_001973625.1_ASM197362v1	Salmonella enterica subsp. enterica serovar Enteritidis	Proteobacteria; Gammaproteobacteria; Enterobacterales; Enterobacteriaceae; Salmonella; Salmonella enterica; Salmonella enterica subsp. enterica	MNNEETFYQAMRRKGVTRRSFLKFCSLAATSLGLGAGMTPKIAWALENKPRIPVVWIHGL	WP_001058307.1	hydrogenase [Salmonella enterica]	Length: 372\nNumber of predicted TMHs:  2\nExp number of AAs in TMHs: 36.51738\nExp number, first 60 AAs:  19.72849\nTotal prob of N-in:   0.96170\nPOSSIBLE N-term signal sequence\ninside   1   19\nTMhelix   20   39\noutside   40   325\nTMhelix   326   348\ninside   349   372

17374	GCF_001973625.1_ASM197362v1	Salmonella enterica subsp. enterica serovar Enteritidis	Proteobacteria; Gammaproteobacteria; Enterobacterales; Enterobacteriaceae; Salmonella; Salmonella enterica; Salmonella enterica subsp. enterica	MTGDNTLITSHGINRRDFMKLCAALAATMGLSSKAAAEMAESVSNPQRPPVIWIGAQECT	WP_000145429.1	hydrogenase 2 small subunit [Salmonella enterica]	Length: 372\nNumber of predicted TMHs:  1\nExp number of AAs in TMHs: 24.51309\nExp number, first 60 AAs:  0.2682\nTotal prob of N-in: 0.14402\noutside   1   331\nTMhelix   332   354\ninside   355   372

17375	GCF_001973425.1_ASM197342v1	Salmonella enterica subsp. enterica serovar Enteritidis	Proteobacteria; Gammaproteobacteria; Enterobacterales; Enterobacteriaceae; Salmonella; Salmonella enterica; Salmonella enterica subsp. enterica	MNNEETFYQAMRRKGVTRRSFLKFCSLAATSLGLGAGMTPKIAWALENKPRIPVVWIHGL	WP_001058307.1	hydrogenase [Salmonella enterica]	Length: 372\nNumber of predicted TMHs:  2\nExp number of AAs in TMHs: 36.51738\nExp number, first 60 AAs:  19.72849\nTotal prob of N-in:   0.96170\nPOSSIBLE N-term signal sequence\ninside   1   19\nTMhelix   20   39\noutside   40   325\nTMhelix   326   348\ninside   349   372

17376	GCF_001973385.1_ASM197338v1	Salmonella enterica subsp. enterica serovar Enteritidis	Proteobacteria; Gammaproteobacteria; Enterobacterales; Enterobacteriaceae; Salmonella; Salmonella enterica; Salmonella enterica subsp. enterica	MNNEETFYQAMRRKGVTRRSFLKFCSLAATSLGLGAGMTPKIAWALENKPRIPVVWIHGL	WP_001058307.1	hydrogenase [Salmonella enterica]	Length: 372\nNumber of predicted TMHs:  2\nExp number of AAs in TMHs: 36.51738\nExp number, first 60 AAs:  19.72849\nTotal prob of N-in:   0.96170\nPOSSIBLE N-term signal sequence\ninside   1   19\nTMhelix   20   39\noutside   40   325\nTMhelix   326   348\ninside   349   372

17377	GCF_001973305.1_ASM197330v1	Salmonella enterica subsp. enterica serovar Enteritidis	Proteobacteria; Gammaproteobacteria; Enterobacterales; Enterobacteriaceae; Salmonella; Salmonella enterica; Salmonella enterica subsp. enterica	MNNEETFYQAMRRKGVTRRSFLKFCSLAATSLGLGAGMTPKIAWALENKPRIPVVWIHGL	WP_001058307.1	hydrogenase [Salmonella enterica]	Length: 372\nNumber of predicted TMHs:  2\nExp number of AAs in TMHs: 36.51738\nExp number, first 60 AAs:  19.72849\nTotal prob of N-in:   0.96170\nPOSSIBLE N-term signal sequence\ninside   1   19\nTMhelix   20   39\noutside   40   325\nTMhelix   326   348\ninside   349   372

17378	GCF_001973505.1_ASM197350v1	Salmonella enterica subsp. enterica serovar Enteritidis	Proteobacteria; Gammaproteobacteria; Enterobacterales; Enterobacteriaceae; Salmonella; Salmonella enterica; Salmonella enterica subsp. enterica	MTGDNTLITSHGINRRDFMKLCAALAATMGLSSKAAAEMAESVSNPQRPPVIWIGAQECT	WP_000145429.1	hydrogenase 2 small subunit [Salmonella enterica]	Length: 372\nNumber of predicted TMHs:  1\nExp number of AAs in TMHs: 24.51309\nExp number, first 60 AAs:  0.2682\nTotal prob of N-in: 0.14402\noutside   1   331\nTMhelix   332   354\ninside   355   372

17379	GCF_001973425.1_ASM197342v1	Salmonella enterica subsp. enterica serovar Enteritidis	Proteobacteria; Gammaproteobacteria; Enterobacterales; Enterobacteriaceae; Salmonella; Salmonella enterica; Salmonella enterica subsp. enterica	MTGDNTLITSHGINRRDFMKLCAALAATMGLSSKAAAEMAESVSNPQRPPVIWIGAQECT	WP_000145429.1	hydrogenase 2 small subunit [Salmonella enterica]	Length: 372\nNumber of

predicted TMHs:  1\nExp number of AAs in TMHs: 24.51309\nExp number, first 60 AAs:  0.2682\nTotal prob of N-in: 0.14402\noutside   1   331\nTMhelix   332   354\ninside   355   372

17380 GCF_001973465.1_ASM197346v1 Salmonella enterica subsp. enterica serovar Enteritidis Proteobacteria; Gammaproteobacteria; Enterobacterales; Enterobacteriaceae; Salmonella; Salmonella enterica; Salmonella enterica subsp. enterica MTGDNTLITSHGINRRDFMKLCAALAATMGLSSKAAAEMAESVSNPQRPPVIWIGAQECT WP_000145429.1 hydrogenase 2 small subunit [Salmonella enterica] Length: 372\nNumber of predicted TMHs:  1\nExp number of AAs in TMHs: 24.51309\nExp number, first 60 AAs:  0.2682\nTotal prob of N-in: 0.14402\noutside   1   331\nTMhelix   332   354\ninside   355   372

17381 GCF_001973265.1_ASM197326v1 Salmonella enterica subsp. enterica serovar Enteritidis Proteobacteria; Gammaproteobacteria; Enterobacterales; Enterobacteriaceae; Salmonella; Salmonella enterica; Salmonella enterica subsp. enterica MNNEETFYQAMRRKGVTRRSFLKFCSLAATSLGLGAGMTPKIAWALENKPRIPVVWIHGL WP_001058307.1 hydrogenase [Salmonella enterica] Length: 372\nNumber of predicted TMHs:  2\nExp number of AAs in TMHs: 36.51738\nExp number, first 60 AAs:  19.72849\nTotal prob of N-in:   0.96170\nPOSSIBLE N-term signal sequence\ninside   1   19\nTMhelix   20   39\noutside   40   325\nTMhelix   326   348\ninside   349   372

17382 GCF_001973225.1_ASM197322v1 Salmonella enterica subsp. enterica serovar Enteritidis Proteobacteria; Gammaproteobacteria; Enterobacterales; Enterobacteriaceae; Salmonella; Salmonella enterica; Salmonella enterica subsp. enterica MNNEETFYQAMRRKGVTRRSFLKFCSLAATSLGLGAGMTPKIAWALENKPRIPVVWIHGL WP_001058307.1 hydrogenase [Salmonella enterica] Length: 372\nNumber of predicted TMHs:  2\nExp number of AAs in TMHs: 36.51738\nExp number, first 60 AAs:  19.72849\nTotal prob of N-in:   0.96170\nPOSSIBLE N-term signal sequence\ninside   1   19\nTMhelix   20   39\noutside   40   325\nTMhelix   326   348\ninside   349   372

17383 GCF_001973385.1_ASM197338v1 Salmonella enterica subsp. enterica serovar Enteritidis Proteobacteria; Gammaproteobacteria; Enterobacterales; Enterobacteriaceae; Salmonella; Salmonella enterica; Salmonella enterica subsp. enterica MTGDNTLITSHGINRRDFMKLCAALAATMGLSSKAAAEMAESVSNPQRPPVIWIGAQECT WP_000145429.1 hydrogenase 2 small subunit [Salmonella enterica] Length: 372\nNumber of predicted TMHs:  1\nExp number of AAs in TMHs: 24.51309\nExp number, first 60 AAs:  0.2682\nTotal prob of N-in: 0.14402\noutside   1   331\nTMhelix   332   354\ninside   355   372

17384 GCF_001973305.1_ASM197330v1 Salmonella enterica subsp. enterica serovar Enteritidis Proteobacteria; Gammaproteobacteria; Enterobacterales; Enterobacteriaceae; Salmonella; Salmonella enterica; Salmonella enterica subsp. enterica MTGDNTLITSHGINRRDFMKLCAALAATMGLSSKAAAEMAESVSNPQRPPVIWIGAQECT WP_000145429.1 hydrogenase 2 small subunit [Salmonella enterica] Length: 372\nNumber of predicted TMHs:  1\nExp number of AAs in TMHs: 24.51309\nExp number, first 60 AAs:  0.2682\nTotal prob of N-in: 0.14402\noutside   1   331\nTMhelix   332   354\ninside   355   372

17385 GCF_001973185.1_ASM197318v1 Salmonella enterica subsp. enterica serovar Enteritidis Proteobacteria; Gammaproteobacteria; Enterobacterales; Enterobacteriaceae; Salmonella; Salmonella enterica; Salmonella enterica subsp. enterica MTGDNTLITSHGINRRDFMKLCAALAATMGLSSKAAAEMAESVSNPQRPPVIWIGAQECT WP_000145429.1 hydrogenase 2 small subunit [Salmonella enterica] Length: 372\nNumber of predicted TMHs:  1\nExp number of AAs in TMHs: 24.51309\nExp number, first 60 AAs:  0.2682\nTotal prob of N-in: 0.14402\noutside   1   331\nTMhelix   332   354\ninside   355   372

17386 GCF_002060155.1_ASM206015v1 Salmonella enterica subsp. enterica serovar Enteritidis Proteobacteria; Gammaproteobacteria; Enterobacterales; Enterobacteriaceae; Salmonella; Salmonella enterica; Salmonella enterica subsp. enterica MNNEETFYQAMRRKGVTRRSFLKFCSLAATSLGLGAGMTPKIAWALENKPRIPVVWIHGL WP_001058307.1 hydrogenase [Salmonella enterica] Length: 372\nNumber of predicted TMHs:  2\nExp number of AAs in TMHs: 36.51738\nExp number, first 60 AAs:  19.72849\nTotal prob of N-in:   0.96170\nPOSSIBLE N-term signal sequence\ninside   1   19\nTMhelix   20   39\noutside   40   325\nTMhelix   326   348\ninside   349   372

17387 GCF_002060155.1_ASM206015v1 Salmonella enterica subsp. enterica serovar Enteritidis Proteobacteria; Gammaproteobacteria; Enterobacterales; Enterobacteriaceae; Salmonella; Salmonella enterica; Salmonella enterica subsp. enterica MTGDNTLITSHGINRRDFMKLCAALAATMGLSSKAAAEMAESVSNPQRPPVIWIGAQECT WP_000145429.1 hydrogenase 2 small subunit [Salmonella enterica] Length: 372\nNumber of predicted TMHs:  1\nExp number of AAs in TMHs: 24.51309\nExp number, first 60 AAs:  0.2682\nTotal prob of N-in: 0.14402\noutside   1   331\nTMhelix   332   354\ninside   355   372

17388 GCF_001973165.1_ASM197316v1 Salmonella enterica subsp. enterica serovar Enteritidis Proteobacteria; Gammaproteobacteria; Enterobacterales; Enterobacteriaceae; Salmonella; Salmonella enterica;

Salmonella enterica subsp. enterica MTGDNTLITSHGINRRDFMKLCAALAATMGLSSKAAAEMAESVSNPQRPPVIWIGAQECT WP_000145429.1 hydrogenase 2 small subunit [Salmonella enterica] Length: 372\nNumber of predicted TMHs: 1\nExp number of AAs in TMHs: 24.51309\nExp number, first 60 AAs: 0.2682\nTotal prob of N-in: 0.14402\noutside 1 331\nTMhelix 332 354\ninside 355 372

17389 GCF_001972915.1_ASM197291v1 Salmonella enterica subsp. enterica serovar Enteritidis Proteobacteria; Gammaproteobacteria; Enterobacterales; Enterobacteriaceae; Salmonella; Salmonella enterica; Salmonella enterica subsp. enterica MNNEETFYQAMRRKGVTRRSFLKFCSLAATSLGLGAGMTPKIAWALENKPRIPVVWIHGL WP_001058307.1 hydrogenase [Salmonella enterica] Length: 372\nNumber of predicted TMHs: 2\nExp number of AAs in TMHs: 36.51738\nExp number, first 60 AAs: 19.72849\nTotal prob of N-in: 0.96170\nPOSSIBLE N-term signal sequence\ninside 1 19\nTMhelix 20 39\noutside 40 325\nTMhelix 326 348\ninside 349 372

17390 GCF_001973035.1_ASM197303v1 Salmonella enterica subsp. enterica serovar Enteritidis Proteobacteria; Gammaproteobacteria; Enterobacterales; Enterobacteriaceae; Salmonella; Salmonella enterica; Salmonella enterica subsp. enterica MNNEETFYQAMRRKGVTRRSFLKFCSLAATSLGLGAGMTPKIAWALENKPRIPVVWIHGL WP_001058307.1 hydrogenase [Salmonella enterica] Length: 372\nNumber of predicted TMHs: 2\nExp number of AAs in TMHs: 36.51738\nExp number, first 60 AAs: 19.72849\nTotal prob of N-in: 0.96170\nPOSSIBLE N-term signal sequence\ninside 1 19\nTMhelix 20 39\noutside 40 325\nTMhelix 326 348\ninside 349 372

17391 GCF_001972995.1_ASM197299v1 Salmonella enterica subsp. enterica serovar Enteritidis Proteobacteria; Gammaproteobacteria; Enterobacterales; Enterobacteriaceae; Salmonella; Salmonella enterica; Salmonella enterica subsp. enterica MNNEETFYQAMRRKGVTRRSFLKFCSLAATSLGLGAGMTPKIAWALENKPRIPVVWIHGL WP_001058307.1 hydrogenase [Salmonella enterica] Length: 372\nNumber of predicted TMHs: 2\nExp number of AAs in TMHs: 36.51738\nExp number, first 60 AAs: 19.72849\nTotal prob of N-in: 0.96170\nPOSSIBLE N-term signal sequence\ninside 1 19\nTMhelix 20 39\noutside 40 325\nTMhelix 326 348\ninside 349 372

17392 GCF_002038395.1_ASM203839v1 Salmonella enterica subsp. enterica serovar Enteritidis Proteobacteria; Gammaproteobacteria; Enterobacterales; Enterobacteriaceae; Salmonella; Salmonella enterica; Salmonella enterica subsp. enterica MTGDNTLITSHGINRRDFMKLCAALAATMGLSSKAAAEMAESVSNPQRPPVIWIGAQECT WP_000145429.1 hydrogenase 2 small subunit [Salmonella enterica] Length: 372\nNumber of predicted TMHs: 1\nExp number of AAs in TMHs: 24.51309\nExp number, first 60 AAs: 0.2682\nTotal prob of N-in: 0.14402\noutside 1 331\nTMhelix 332 354\ninside 355 372

17393 GCF_001973085.1_ASM197308v1 Salmonella enterica subsp. enterica serovar Enteritidis Proteobacteria; Gammaproteobacteria; Enterobacterales; Enterobacteriaceae; Salmonella; Salmonella enterica; Salmonella enterica subsp. enterica MTGDNTLITSHGINRRDFMKLCAALAATMGLSSKAAAEMAESVSNPQRPPVIWIGAQECT WP_000145429.1 hydrogenase 2 small subunit [Salmonella enterica] Length: 372\nNumber of predicted TMHs: 1\nExp number of AAs in TMHs: 24.51309\nExp number, first 60 AAs: 0.2682\nTotal prob of N-in: 0.14402\noutside 1 331\nTMhelix 332 354\ninside 355 372

17394 GCF_001972675.1_ASM197267v1 Salmonella enterica subsp. enterica serovar Enteritidis Proteobacteria; Gammaproteobacteria; Enterobacterales; Enterobacteriaceae; Salmonella; Salmonella enterica; Salmonella enterica subsp. enterica MNNEETFYQAMRRKGVTRRSFLKFCSLAATSLGLGAGMTPKIAWALENKPRIPVVWIHGL WP_001058307.1 hydrogenase [Salmonella enterica] Length: 372\nNumber of predicted TMHs: 2\nExp number of AAs in TMHs: 36.51738\nExp number, first 60 AAs: 19.72849\nTotal prob of N-in: 0.96170\nPOSSIBLE N-term signal sequence\ninside 1 19\nTMhelix 20 39\noutside 40 325\nTMhelix 326 348\ninside 349 372

17395 GCF_001972575.1_ASM197257v1 Salmonella enterica subsp. enterica serovar Enteritidis Proteobacteria; Gammaproteobacteria; Enterobacterales; Enterobacteriaceae; Salmonella; Salmonella enterica; Salmonella enterica subsp. enterica MNNEETFYQAMRRKGVTRRSFLKFCSLAATSLGLGAGMTPKIAWALENKPRIPVVWIHGL WP_001058307.1 hydrogenase [Salmonella enterica] Length: 372\nNumber of predicted TMHs: 2\nExp number of AAs in TMHs: 36.51738\nExp number, first 60 AAs: 19.72849\nTotal prob of N-in: 0.96170\nPOSSIBLE N-term signal sequence\ninside 1 19\nTMhelix 20 39\noutside 40 325\nTMhelix 326 348\ninside 349 372

17396 GCF_001972555.1_ASM197255v1 Salmonella enterica subsp. enterica serovar Enteritidis Proteobacteria; Gammaproteobacteria; Enterobacterales; Enterobacteriaceae; Salmonella; Salmonella enterica; Salmonella enterica subsp. enterica MNNEETFYQAMRRKGVTRRSFLKFCSLAATSLGLGAGMTPKIAWALENKPRIPVVWIHGL WP_001058307.1 hydrogenase [Salmonella enterica] Length: 372\nNumber of predicted TMHs: 2\nExp number of AAs in TMHs: 36.51738\nExp number, first 60 AAs: 19.72849\nTotal prob of N-in: 0.96170\nPOSSIBLE N-term signal sequence\ninside 1 19\nTMhelix 20 39\noutside 40 325\nTMhelix 326 348\ninside 349 372

17397 GCF_001972735.1_ASM197273v1 Salmonella enterica subsp. enterica serovar Enteritidis Proteobacteria; Gammaproteobacteria; Enterobacterales; Enterobacteriaceae; Salmonella; Salmonella enterica; Salmonella enterica subsp. enterica MTGDNTLITSHGINRRDFMKLCAALAATMGLSSKAAAEMAESVSNPQRPPVIWIGAQECT WP_000145429.1 hydrogenase 2 small subunit [Salmonella enterica] Length: 372\nNumber of predicted TMHs: 1\nExp number of AAs in TMHs: 24.51309\nExp number, first 60 AAs: 0.2682\nTotal prob of N-in: 0.14402\noutside 1 331\nTMhelix 332 354\ninside 355 372

17398 GCF_001972715.1_ASM197271v1 Salmonella enterica subsp. enterica serovar Enteritidis Proteobacteria; Gammaproteobacteria; Enterobacterales; Enterobacteriaceae; Salmonella; Salmonella enterica; Salmonella enterica subsp. enterica MTGDNTLITSHGINRRDFMKLCAALAATMGLSSKAAAEMAESVSNPQRPPVIWIGAQECT WP_000145429.1 hydrogenase 2 small subunit [Salmonella enterica] Length: 372\nNumber of predicted TMHs: 1\nExp number of AAs in TMHs: 24.51309\nExp number, first 60 AAs: 0.2682\nTotal prob of N-in: 0.14402\noutside 1 331\nTMhelix 332 354\ninside 355 372

17399 GCF_001972635.1_ASM197263v1 Salmonella enterica subsp. enterica serovar Enteritidis Proteobacteria; Gammaproteobacteria; Enterobacterales; Enterobacteriaceae; Salmonella; Salmonella enterica; Salmonella enterica subsp. enterica MTGDNTLITSHGINRRDFMKLCAALAATMGLSSKAAAEMAESVSNPQRPPVIWIGAQECT WP_000145429.1 hydrogenase 2 small subunit [Salmonella enterica] Length: 372\nNumber of predicted TMHs: 1\nExp number of AAs in TMHs: 24.51309\nExp number, first 60 AAs: 0.2682\nTotal prob of N-in: 0.14402\noutside 1 331\nTMhelix 332 354\ninside 355 372

17400 GCF_001972675.1_ASM197267v1 Salmonella enterica subsp. enterica serovar Enteritidis Proteobacteria; Gammaproteobacteria; Enterobacterales; Enterobacteriaceae; Salmonella; Salmonella enterica; Salmonella enterica subsp. enterica MTGDNTLITSHGINRRDFMKLCAALAATMGLSSKAAAEMAESVSNPQRPPVIWIGAQECT WP_000145429.1 hydrogenase 2 small subunit [Salmonella enterica] Length: 372\nNumber of predicted TMHs: 1\nExp number of AAs in TMHs: 24.51309\nExp number, first 60 AAs: 0.2682\nTotal prob of N-in: 0.14402\noutside 1 331\nTMhelix 332 354\ninside 355 372

17401 GCF_001972575.1_ASM197257v1 Salmonella enterica subsp. enterica serovar Enteritidis Proteobacteria; Gammaproteobacteria; Enterobacterales; Enterobacteriaceae; Salmonella; Salmonella enterica; Salmonella enterica subsp. enterica MTGDNTLITSHGINRRDFMKLCAALAATMGLSSKAAAEMAESVSNPQRPPVIWIGAQECT WP_000145429.1 hydrogenase 2 small subunit [Salmonella enterica] Length: 372\nNumber of predicted TMHs: 1\nExp number of AAs in TMHs: 24.51309\nExp number, first 60 AAs: 0.2682\nTotal prob of N-in: 0.14402\noutside 1 331\nTMhelix 332 354\ninside 355 372

17402 GCF_002077315.1_ASM207731v1 Salmonella enterica subsp. enterica serovar Enteritidis Proteobacteria; Gammaproteobacteria; Enterobacterales; Enterobacteriaceae; Salmonella; Salmonella enterica; Salmonella enterica subsp. enterica MNNEETFYQAMRRKGVTRRSFLKFCSLAATSLGLGAGMTPKIAWALENKPRIPVVWIHGL WP_001058307.1 hydrogenase [Salmonella enterica] Length: 372\nNumber of predicted TMHs: 2\nExp number of AAs in TMHs: 36.51738\nExp number, first 60 AAs: 19.72849\nTotal prob of N-in: 0.96170\nPOSSIBLE N-term signal sequence\ninside 1 19\nTMhelix 20 39\noutside 40 325\nTMhelix 326 348\ninside 349 372

17403 GCF_002076985.1_ASM207698v1 Salmonella enterica subsp. enterica serovar Enteritidis Proteobacteria; Gammaproteobacteria; Enterobacterales; Enterobacteriaceae; Salmonella; Salmonella enterica; Salmonella enterica subsp. enterica MNNEETFYQAMRRKGVTRRSFLKFCSLAATSLGLGAGMTPKIAWALENKPRIPVVWIHGL WP_001058307.1 hydrogenase [Salmonella enterica] Length: 372\nNumber of predicted TMHs: 2\nExp number of AAs in TMHs: 36.51738\nExp number, first 60 AAs: 19.72849\nTotal prob of N-in: 0.96170\nPOSSIBLE N-term signal sequence\ninside 1 19\nTMhelix 20 39\noutside 40 325\nTMhelix 326 348\ninside 349 372

17404 GCF_002077315.1_ASM207731v1 Salmonella enterica subsp. enterica serovar Enteritidis Proteobacteria; Gammaproteobacteria; Enterobacterales; Enterobacteriaceae; Salmonella; Salmonella enterica; Salmonella enterica subsp. enterica MTGDNTLITSHGINRRDFMKLCAALAATMGLSSKAAAEMAESVSNPQRPPVIWIGAQECT WP_000145429.1 hydrogenase 2 small subunit [Salmonella enterica] Length: 372\nNumber of predicted TMHs: 1\nExp number of AAs in TMHs: 24.51309\nExp number, first 60 AAs: 0.2682\nTotal prob of N-in: 0.14402\noutside 1 331\nTMhelix 332 354\ninside 355 372

17405 GCF_002076985.1_ASM207698v1 Salmonella enterica subsp. enterica serovar Enteritidis Proteobacteria; Gammaproteobacteria; Enterobacterales; Enterobacteriaceae; Salmonella; Salmonella enterica; Salmonella enterica subsp. enterica MTGDNTLITSHGINRRDFMKLCAALAATMGLSSKAAAEMAESVSNPQRPPVIWIGAQECT WP_000145429.1 hydrogenase 2 small subunit [Salmonella enterica] Length: 372\nNumber of

predicted TMHs: 1\nExp number of AAs in TMHs: 24.51309\nExp number, first 60 AAs: 0.2682\nTotal prob of N-in: 0.14402\noutside 1 331\nTMhelix 332 354\ninside 355 372

17406 GCF_002065685.1_ASM206568v1 Salmonella enterica subsp. enterica serovar Enteritidis Proteobacteria; Gammaproteobacteria; Enterobacterales; Enterobacteriaceae; Salmonella; Salmonella enterica; Salmonella enterica subsp. enterica MTGDNTLITSHGINRRDFMKLCAALAATMGLSSKAAAEMAESVSNPQRPPVIWIGAQECT WP_000145429.1 hydrogenase 2 small subunit [Salmonella enterica] Length: 372\nNumber of predicted TMHs: 1\nExp number of AAs in TMHs: 24.51309\nExp number, first 60 AAs: 0.2682\nTotal prob of N-in: 0.14402\noutside 1 331\nTMhelix 332 354\ninside 355 372

17407 GCF_002065725.1_ASM206572v1 Salmonella enterica subsp. enterica serovar Enteritidis Proteobacteria; Gammaproteobacteria; Enterobacterales; Enterobacteriaceae; Salmonella; Salmonella enterica; Salmonella enterica subsp. enterica MTGDNTLITSHGINRRDFMKLCAALAATMGLSSKAAAEMAESVSNPQRPPVIWIGAQECT WP_000145429.1 hydrogenase 2 small subunit [Salmonella enterica] Length: 372\nNumber of predicted TMHs: 1\nExp number of AAs in TMHs: 24.51309\nExp number, first 60 AAs: 0.2682\nTotal prob of N-in: 0.14402\noutside 1 331\nTMhelix 332 354\ninside 355 372

17408 GCF_002065775.1_ASM206577v1 Salmonella enterica subsp. enterica serovar Enteritidis Proteobacteria; Gammaproteobacteria; Enterobacterales; Enterobacteriaceae; Salmonella; Salmonella enterica; Salmonella enterica subsp. enterica MTGDNTLITSHGINRRDFMKLCAALAATMGLSSKAAAEMAESVSNPQRPPVIWIGAQECT WP_000145429.1 hydrogenase 2 small subunit [Salmonella enterica] Length: 372\nNumber of predicted TMHs: 1\nExp number of AAs in TMHs: 24.51309\nExp number, first 60 AAs: 0.2682\nTotal prob of N-in: 0.14402\noutside 1 331\nTMhelix 332 354\ninside 355 372

17409 GCF_002065805.1_ASM206580v1 Salmonella enterica subsp. enterica serovar Enteritidis Proteobacteria; Gammaproteobacteria; Enterobacterales; Enterobacteriaceae; Salmonella; Salmonella enterica; Salmonella enterica subsp. enterica MTGDNTLITSHGINRRDFMKLCAALAATMGLSSKAAAEMAESVSNPQRPPVIWIGAQECT WP_000145429.1 hydrogenase 2 small subunit [Salmonella enterica] Length: 372\nNumber of predicted TMHs: 1\nExp number of AAs in TMHs: 24.51309\nExp number, first 60 AAs: 0.2682\nTotal prob of N-in: 0.14402\noutside 1 331\nTMhelix 332 354\ninside 355 372

17410 GCF_002065685.1_ASM206568v1 Salmonella enterica subsp. enterica serovar Enteritidis Proteobacteria; Gammaproteobacteria; Enterobacterales; Enterobacteriaceae; Salmonella; Salmonella enterica; Salmonella enterica subsp. enterica MNNEETFYQAMRRKGVTRRSFLKFCSLAATSLGLGAGMTPKIAWALENKPRIPVVWIHGL WP_001058307.1 hydrogenase [Salmonella enterica] Length: 372\nNumber of predicted TMHs: 2\nExp number of AAs in TMHs: 36.51738\nExp number, first 60 AAs: 19.72849\nTotal prob of N-in: 0.96170\nPOSSIBLE N-term signal sequence\ninside 1 19\nTMhelix 20 39\noutside 40 325\nTMhelix 326 348\ninside 349 372

17411 GCF_002065725.1_ASM206572v1 Salmonella enterica subsp. enterica serovar Enteritidis Proteobacteria; Gammaproteobacteria; Enterobacterales; Enterobacteriaceae; Salmonella; Salmonella enterica; Salmonella enterica subsp. enterica MNNEETFYQAMRRKGVTRRSFLKFCSLAATSLGLGAGMTPKIAWALENKPRIPVVWIHGL WP_001058307.1 hydrogenase [Salmonella enterica] Length: 372\nNumber of predicted TMHs: 2\nExp number of AAs in TMHs: 36.51738\nExp number, first 60 AAs: 19.72849\nTotal prob of N-in: 0.96170\nPOSSIBLE N-term signal sequence\ninside 1 19\nTMhelix 20 39\noutside 40 325\nTMhelix 326 348\ninside 349 372

17412 GCF_002065775.1_ASM206577v1 Salmonella enterica subsp. enterica serovar Enteritidis Proteobacteria; Gammaproteobacteria; Enterobacterales; Enterobacteriaceae; Salmonella; Salmonella enterica; Salmonella enterica subsp. enterica MNNEETFYQAMRRKGVTRRSFLKFCSLAATSLGLGAGMTPKIAWALENKPRIPVVWIHGL WP_001058307.1 hydrogenase [Salmonella enterica] Length: 372\nNumber of predicted TMHs: 2\nExp number of AAs in TMHs: 36.51738\nExp number, first 60 AAs: 19.72849\nTotal prob of N-in: 0.96170\nPOSSIBLE N-term signal sequence\ninside 1 19\nTMhelix 20 39\noutside 40 325\nTMhelix 326 348\ninside 349 372

17413 GCF_002065805.1_ASM206580v1 Salmonella enterica subsp. enterica serovar Enteritidis Proteobacteria; Gammaproteobacteria; Enterobacterales; Enterobacteriaceae; Salmonella; Salmonella enterica; Salmonella enterica subsp. enterica MNNEETFYQAMRRKGVTRRSFLKFCSLAATSLGLGAGMTPKIAWALENKPRIPVVWIHGL WP_001058307.1 hydrogenase [Salmonella enterica] Length: 372\nNumber of predicted TMHs: 2\nExp number of AAs in TMHs: 36.51738\nExp number, first 60 AAs: 19.72849\nTotal prob of N-in: 0.96170\nPOSSIBLE N-term signal sequence\ninside 1 19\nTMhelix 20 39\noutside 40 325\nTMhelix 326 348\ninside 349 372

17414 GCF_002066415.1_ASM206641v1 Salmonella enterica subsp. enterica serovar Enteritidis Proteobacteria; Gammaproteobacteria; Enterobacterales; Enterobacteriaceae; Salmonella; Salmonella enterica;

Salmonella enterica subsp. enterica MTGDNTLITSHGINRRDFMKLCAALAATMGLSSKAAAEMAESVSNPQRPPVIWIGAQECT WP_000145429.1 hydrogenase 2 small subunit [Salmonella enterica] Length: 372\nNumber of predicted TMHs: 1\nExp number of AAs in TMHs: 24.51309\nExp number, first 60 AAs: 0.2682\nTotal prob of N-in: 0.14402\noutside 1 331\nTMhelix 332 354\ninside 355 372

17415 GCF_002066495.1_ASM206649v1 Salmonella enterica subsp. enterica serovar Enteritidis Proteobacteria; Gammaproteobacteria; Enterobacterales; Enterobacteriaceae; Salmonella; Salmonella enterica; Salmonella enterica subsp. enterica MTGDNTLITSHGINRRDFMKLCAALAATMGLSSKAAAEMAESVSNPQRPPVIWIGAQECT WP_000145429.1 hydrogenase 2 small subunit [Salmonella enterica] Length: 372\nNumber of predicted TMHs: 1\nExp number of AAs in TMHs: 24.51309\nExp number, first 60 AAs: 0.2682\nTotal prob of N-in: 0.14402\noutside 1 331\nTMhelix 332 354\ninside 355 372

17416 GCF_002066415.1_ASM206641v1 Salmonella enterica subsp. enterica serovar Enteritidis Proteobacteria; Gammaproteobacteria; Enterobacterales; Enterobacteriaceae; Salmonella; Salmonella enterica; Salmonella enterica subsp. enterica MNNEETFYQAMRRKGVTRRSFLKFCSLAATSLGLGAGMTPKIAWALENKPRIPVVWIHGL WP_001058307.1 hydrogenase [Salmonella enterica] Length: 372\nNumber of predicted TMHs: 2\nExp number of AAs in TMHs: 36.51738\nExp number, first 60 AAs: 19.72849\nTotal prob of N-in: 0.96170\nPOSSIBLE N-term signal sequence\ninside 1 19\nTMhelix 20 39\noutside 40 325\nTMhelix 326 348\ninside 349 372

17417 GCF_002066535.1_ASM206653v1 Salmonella enterica subsp. enterica serovar Enteritidis Proteobacteria; Gammaproteobacteria; Enterobacterales; Enterobacteriaceae; Salmonella; Salmonella enterica; Salmonella enterica subsp. enterica MTGDNTLITSHGINRRDFMKLCAALAATMGLSSKAAAEMAESVSNPQRPPVIWIGAQECT WP_000145429.1 hydrogenase 2 small subunit [Salmonella enterica] Length: 372\nNumber of predicted TMHs: 1\nExp number of AAs in TMHs: 24.51309\nExp number, first 60 AAs: 0.2682\nTotal prob of N-in: 0.14402\noutside 1 331\nTMhelix 332 354\ninside 355 372

17418 GCF_002066495.1_ASM206649v1 Salmonella enterica subsp. enterica serovar Enteritidis Proteobacteria; Gammaproteobacteria; Enterobacterales; Enterobacteriaceae; Salmonella; Salmonella enterica; Salmonella enterica subsp. enterica MNNEETFYQAMRRKGVTRRSFLKFCSLAATSLGLGAGMTPKIAWALENKPRIPVVWIHGL WP_001058307.1 hydrogenase [Salmonella enterica] Length: 372\nNumber of predicted TMHs: 2\nExp number of AAs in TMHs: 36.51738\nExp number, first 60 AAs: 19.72849\nTotal prob of N-in: 0.96170\nPOSSIBLE N-term signal sequence\ninside 1 19\nTMhelix 20 39\noutside 40 325\nTMhelix 326 348\ninside 349 372

17419 GCF_001714065.1_ASM171406v1 Salmonella enterica subsp. enterica serovar Enteritidis Proteobacteria; Gammaproteobacteria; Enterobacterales; Enterobacteriaceae; Salmonella; Salmonella enterica; Salmonella enterica subsp. enterica MTGDNTLITSHGINRRDFMKLCAALAATMGLSSKAAAEMAESVSNPQRPPVIWIGAQECT WP_000145429.1 hydrogenase 2 small subunit [Salmonella enterica] Length: 372\nNumber of predicted TMHs: 1\nExp number of AAs in TMHs: 24.51309\nExp number, first 60 AAs: 0.2682\nTotal prob of N-in: 0.14402\noutside 1 331\nTMhelix 332 354\ninside 355 372

17420 GCF_001714015.1_ASM171401v1 Salmonella enterica subsp. enterica serovar Enteritidis Proteobacteria; Gammaproteobacteria; Enterobacterales; Enterobacteriaceae; Salmonella; Salmonella enterica; Salmonella enterica subsp. enterica MTGDNTLITSHGINRRDFMKLCAALAATMGLSSKAAAEMAESVSNPQRPPVIWIGAQECT WP_000145429.1 hydrogenase 2 small subunit [Salmonella enterica] Length: 372\nNumber of predicted TMHs: 1\nExp number of AAs in TMHs: 24.51309\nExp number, first 60 AAs: 0.2682\nTotal prob of N-in: 0.14402\noutside 1 331\nTMhelix 332 354\ninside 355 372

17421 GCF_001713825.1_ASM171382v1 Salmonella enterica subsp. enterica serovar Enteritidis Proteobacteria; Gammaproteobacteria; Enterobacterales; Enterobacteriaceae; Salmonella; Salmonella enterica; Salmonella enterica subsp. enterica MNNEETFYQAMRRKGVTRRSFLKFCSLAATSLGLGAGMTPKIAWALENKPRIPVVWIHGL WP_001058307.1 hydrogenase [Salmonella enterica] Length: 372\nNumber of predicted TMHs: 2\nExp number of AAs in TMHs: 36.51738\nExp number, first 60 AAs: 19.72849\nTotal prob of N-in: 0.96170\nPOSSIBLE N-term signal sequence\ninside 1 19\nTMhelix 20 39\noutside 40 325\nTMhelix 326 348\ninside 349 372

17422 GCF_001713985.1_ASM171398v1 Salmonella enterica subsp. enterica serovar Enteritidis Proteobacteria; Gammaproteobacteria; Enterobacterales; Enterobacteriaceae; Salmonella; Salmonella enterica; Salmonella enterica subsp. enterica MNNEETFYQAMRRKGVTRRSFLKFCSLAATSLGLGAGMTPKIAWALENKPRIPVVWIHGL WP_001058307.1 hydrogenase [Salmonella enterica] Length: 372\nNumber of predicted TMHs: 2\nExp number of AAs in TMHs: 36.51738\nExp number, first 60 AAs: 19.72849\nTotal prob of N-in: 0.96170\nPOSSIBLE N-term signal sequence\ninside 1 19\nTMhelix 20 39\noutside 40 325\nTMhelix 326 348\ninside 349 372

17423 GCF_001713945.1_ASM171394v1 Salmonella enterica subsp. enterica serovar Enteritidis Proteobacteria; Gammaproteobacteria; Enterobacterales; Enterobacteriaceae; Salmonella; Salmonella enterica; Salmonella enterica subsp. enterica MNNEETFYQAMRRKGVTRRSFLKFCSLAATSLGLGAGMTPKIAWALENKPRIPVVWIHGL WP_001058307.1 hydrogenase [Salmonella enterica] Length: 372\nNumber of predicted TMHs: 2\nExp number of AAs in TMHs: 36.51738\nExp number, first 60 AAs: 19.72849\nTotal prob of N-in: 0.96170\nPOSSIBLE N-term signal sequence\ninside 1 19\nTMhelix 20 39\noutside 40 325\nTMhelix 326 348\ninside 349 372

17424 GCF_001713985.1_ASM171398v1 Salmonella enterica subsp. enterica serovar Enteritidis Proteobacteria; Gammaproteobacteria; Enterobacterales; Enterobacteriaceae; Salmonella; Salmonella enterica; Salmonella enterica subsp. enterica MTGDNTLITSHGINRRDFMKLCAALAATMGLSSKAAAEMAESVSNPQRPPVIWIGAQECT WP_000145429.1 hydrogenase 2 small subunit [Salmonella enterica] Length: 372\nNumber of predicted TMHs: 1\nExp number of AAs in TMHs: 24.51309\nExp number, first 60 AAs: 0.2682\nTotal prob of N-in: 0.14402\noutside 1 331\nTMhelix 332 354\ninside 355 372

17425 GCF_001713945.1_ASM171394v1 Salmonella enterica subsp. enterica serovar Enteritidis Proteobacteria; Gammaproteobacteria; Enterobacterales; Enterobacteriaceae; Salmonella; Salmonella enterica; Salmonella enterica subsp. enterica MTGDNTLITSHGINRRDFMKLCAALAATMGLSSKAAAEMAESVSNPQRPPVIWIGAQECT WP_000145429.1 hydrogenase 2 small subunit [Salmonella enterica] Length: 372\nNumber of predicted TMHs: 1\nExp number of AAs in TMHs: 24.51309\nExp number, first 60 AAs: 0.2682\nTotal prob of N-in: 0.14402\noutside 1 331\nTMhelix 332 354\ninside 355 372

17426 GCF_002057075.1_ASM205707v1 Salmonella enterica subsp. enterica serovar Enteritidis Proteobacteria; Gammaproteobacteria; Enterobacterales; Enterobacteriaceae; Salmonella; Salmonella enterica; Salmonella enterica subsp. enterica MTGDNTLITSHGINRRDFMKLCAALAATMGLSSKAAAEMAESVSNPQRPPVIWIGAQECT WP_000145428.1 hydrogenase 2 small subunit [Salmonella enterica] Length: 372\nNumber of predicted TMHs: 1\nExp number of AAs in TMHs: 24.50677\nExp number, first 60 AAs: 0.2682\nTotal prob of N-in: 0.14390\noutside 1 331\nTMhelix 332 354\ninside 355 372

17427 GCF_002156255.1_ASM215625v1 Salmonella enterica subsp. enterica serovar Gallinarum Proteobacteria; Gammaproteobacteria; Enterobacterales; Enterobacteriaceae; Salmonella; Salmonella enterica; Salmonella enterica subsp. enterica MNNEETFYQAMRRKGVTRRSFLKFCSLAATSLGLGAGMTPKIAWALENKPRIPVVWIHGL WP_001058304.1 hydrogenase [Salmonella enterica] Length: 372\nNumber of predicted TMHs: 2\nExp number of AAs in TMHs: 36.51703\nExp number, first 60 AAs: 19.7285\nTotal prob of N-in: 0.96169\nPOSSIBLE N-term signal sequence\ninside 1 19\nTMhelix 20 39\noutside 40 325\nTMhelix 326 348\ninside 349 372

17428 GCF_001484835.1_ASM148483v1 Salmonella enterica subsp. enterica serovar Gallinarum Proteobacteria; Gammaproteobacteria; Enterobacterales; Enterobacteriaceae; Salmonella; Salmonella enterica; Salmonella enterica subsp. enterica MTGDNTLITSHGINRRDFMKLCAALAATMGLSSKAAAEMAESVSNPQRPPVIWIGAQECT WP_000145429.1 hydrogenase 2 small subunit [Salmonella enterica] Length: 372\nNumber of predicted TMHs: 1\nExp number of AAs in TMHs: 24.51309\nExp number, first 60 AAs: 0.2682\nTotal prob of N-in: 0.14402\noutside 1 331\nTMhelix 332 354\ninside 355 372

17429 GCF_001448635.1_ASM144863v1 Salmonella enterica subsp. enterica serovar Gallinarum Proteobacteria; Gammaproteobacteria; Enterobacterales; Enterobacteriaceae; Salmonella; Salmonella enterica; Salmonella enterica subsp. enterica MNNEETFYQAMRRKGVTRRSFLKFCSLAATSLGLGAGMTPKIAWALENKPRIPVVWIHGL WP_001058304.1 hydrogenase [Salmonella enterica] Length: 372\nNumber of predicted TMHs: 2\nExp number of AAs in TMHs: 36.51703\nExp number, first 60 AAs: 19.7285\nTotal prob of N-in: 0.96169\nPOSSIBLE N-term signal sequence\ninside 1 19\nTMhelix 20 39\noutside 40 325\nTMhelix 326 348\ninside 349 372

17430 GCF_001484835.1_ASM148483v1 Salmonella enterica subsp. enterica serovar Gallinarum Proteobacteria; Gammaproteobacteria; Enterobacterales; Enterobacteriaceae; Salmonella; Salmonella enterica; Salmonella enterica subsp. enterica MNNEETFYQAMRRKGVTRRSFLKFCSLAATSLGLGAGMTPKIAWALENKPRIPVVWIHGL WP_001058304.1 hydrogenase [Salmonella enterica] Length: 372\nNumber of predicted TMHs: 2\nExp number of AAs in TMHs: 36.51703\nExp number, first 60 AAs: 19.7285\nTotal prob of N-in: 0.96169\nPOSSIBLE N-term signal sequence\ninside 1 19\nTMhelix 20 39\noutside 40 325\nTMhelix 326 348\ninside 349 372

17431 GCF_001448635.1_ASM144863v1 Salmonella enterica subsp. enterica serovar Gallinarum Proteobacteria; Gammaproteobacteria; Enterobacterales; Enterobacteriaceae; Salmonella; Salmonella enterica; Salmonella enterica subsp. enterica MTGDNTLITSHGINRRDFMKLCAALAATMGLSSKAAAEMAESVSNPQRPPVIWIGAQECT WP_000145429.1 hydrogenase 2 small subunit [Salmonella enterica] Length: 372\nNumber of

predicted TMHs: 1\nExp number of AAs in TMHs: 24.51309\nExp number, first 60 AAs: 0.2682\nTotal prob of N-in: 0.14402\noutside 1 331\nTMhelix 332 354\ninside 355 372

17432 GCF_002156255.1_ASM215625v1 Salmonella enterica subsp. enterica serovar Gallinarum Proteobacteria; Gammaproteobacteria; Enterobacterales; Enterobacteriaceae; Salmonella; Salmonella enterica; Salmonella enterica subsp. enterica MTGDNTLITSHGINRRDFMKLCAALAATMGLSSKAAAEMAESVSNPQRPPVIWIGAQECT WP_000145429.1 hydrogenase 2 small subunit [Salmonella enterica] Length: 372\nNumber of predicted TMHs: 1\nExp number of AAs in TMHs: 24.51309\nExp number, first 60 AAs: 0.2682\nTotal prob of N-in: 0.14402\noutside 1 331\nTMhelix 332 354\ninside 355 372

17433 GCF_002049445.1_ASM204944v1 Salmonella enterica subsp. enterica serovar Gaminara Proteobacteria; Gammaproteobacteria; Enterobacterales; Enterobacteriaceae; Salmonella; Salmonella enterica; Salmonella enterica subsp. enterica MNNEETFYQAMRRKGVTRRSFLKFCSLAATSLGLGAGMTPKIAWALENKPRIPVVWIHGL WP_001662439.1 hydrogenase [Salmonella enterica] Length: 372\nNumber of predicted TMHs: 2\nExp number of AAs in TMHs: 36.5172\nExp number, first 60 AAs: 19.72849\nTotal prob of N-in: 0.96169\nPOSSIBLE N-term signal sequence\ninside 1 19\nTMhelix 20 39\noutside 40 325\nTMhelix 326 348\ninside 349 372

17434 GCF_002050675.1_ASM205067v1 Salmonella enterica subsp. enterica serovar Gaminara Proteobacteria; Gammaproteobacteria; Enterobacterales; Enterobacteriaceae; Salmonella; Salmonella enterica; Salmonella enterica subsp. enterica MTGDNTLITSHGINRRDFMKLCAALAATMGLSSKAAAEMAESVSNPQRPPVIWIGAQECT WP_000145429.1 hydrogenase 2 small subunit [Salmonella enterica] Length: 372\nNumber of predicted TMHs: 1\nExp number of AAs in TMHs: 24.51309\nExp number, first 60 AAs: 0.2682\nTotal prob of N-in: 0.14402\noutside 1 331\nTMhelix 332 354\ninside 355 372

17435 GCF_002049145.1_ASM204914v1 Salmonella enterica subsp. enterica serovar Gaminara Proteobacteria; Gammaproteobacteria; Enterobacterales; Enterobacteriaceae; Salmonella; Salmonella enterica; Salmonella enterica subsp. enterica MTGDNTLITSHGINRRDFMKLCAALAATMGLSSKAAAEMAESVSNPQRPPVIWIGAQECT WP_000145429.1 hydrogenase 2 small subunit [Salmonella enterica] Length: 372\nNumber of predicted TMHs: 1\nExp number of AAs in TMHs: 24.51309\nExp number, first 60 AAs: 0.2682\nTotal prob of N-in: 0.14402\noutside 1 331\nTMhelix 332 354\ninside 355 372

17436 GCF_002049145.1_ASM204914v1 Salmonella enterica subsp. enterica serovar Gaminara Proteobacteria; Gammaproteobacteria; Enterobacterales; Enterobacteriaceae; Salmonella; Salmonella enterica; Salmonella enterica subsp. enterica MNNEETFYQAMRRKGVTRRSFLKFCSLAATSLGLGAGMTPKIAWALENKPRIPVVWIHGL WP_001662439.1 hydrogenase [Salmonella enterica] Length: 372\nNumber of predicted TMHs: 2\nExp number of AAs in TMHs: 36.5172\nExp number, first 60 AAs: 19.72849\nTotal prob of N-in: 0.96169\nPOSSIBLE N-term signal sequence\ninside 1 19\nTMhelix 20 39\noutside 40 325\nTMhelix 326 348\ninside 349 372

17437 GCF_002106395.1_ASM210639v1 Salmonella enterica subsp. enterica serovar Gaminara Proteobacteria; Gammaproteobacteria; Enterobacterales; Enterobacteriaceae; Salmonella; Salmonella enterica; Salmonella enterica subsp. enterica MTGDNTLITSHGINRRDFMKLCAALAATMGLSSKAAAEMAESVSNPQRPPVIWIGAQECT WP_000145429.1 hydrogenase 2 small subunit [Salmonella enterica] Length: 372\nNumber of predicted TMHs: 1\nExp number of AAs in TMHs: 24.51309\nExp number, first 60 AAs: 0.2682\nTotal prob of N-in: 0.14402\noutside 1 331\nTMhelix 332 354\ninside 355 372

17438 GCF_002049005.1_ASM204900v1 Salmonella enterica subsp. enterica serovar Gaminara Proteobacteria; Gammaproteobacteria; Enterobacterales; Enterobacteriaceae; Salmonella; Salmonella enterica; Salmonella enterica subsp. enterica MNNEETFYQAMRRKGVTRRSFLKFCSLAATSLGLGAGMTPKIAWALENKPRIPVVWIHGL WP_001662439.1 hydrogenase [Salmonella enterica] Length: 372\nNumber of predicted TMHs: 2\nExp number of AAs in TMHs: 36.5172\nExp number, first 60 AAs: 19.72849\nTotal prob of N-in: 0.96169\nPOSSIBLE N-term signal sequence\ninside 1 19\nTMhelix 20 39\noutside 40 325\nTMhelix 326 348\ninside 349 372

17439 GCF_002049005.1_ASM204900v1 Salmonella enterica subsp. enterica serovar Gaminara Proteobacteria; Gammaproteobacteria; Enterobacterales; Enterobacteriaceae; Salmonella; Salmonella enterica; Salmonella enterica subsp. enterica MTGDNTLITSHGINRRDFMKLCAALAATMGLSSKAAAEMAESVSNPQRPPVIWIGAQECT WP_000145429.1 hydrogenase 2 small subunit [Salmonella enterica] Length: 372\nNumber of predicted TMHs: 1\nExp number of AAs in TMHs: 24.51309\nExp number, first 60 AAs: 0.2682\nTotal prob of N-in: 0.14402\noutside 1 331\nTMhelix 332 354\ninside 355 372

17440 GCF_002106395.1_ASM210639v1 Salmonella enterica subsp. enterica serovar Gaminara Proteobacteria; Gammaproteobacteria; Enterobacterales; Enterobacteriaceae; Salmonella; Salmonella enterica;

Salmonella enterica subsp. enterica MNNEETFYQAMRRKGVTRRSFLKFCSLAATSLGLGAGMTPKIAWALENKPRIPVVWIHGL WP_001662439.1 hydrogenase [Salmonella enterica] Length: 372\nNumber of predicted TMHs: 2\nExp number of AAs in TMHs: 36.5172\nExp number, first 60 AAs: 19.72849\nTotal prob of N-in: 0.96169\nPOSSIBLE N-term signal sequence\ninside 1 19\nTMhelix 20 39\noutside 40 325\nTMhelix 326 348\ninside 349 372

17441 GCF_002050675.1_ASM205067v1 Salmonella enterica subsp. enterica serovar Gaminara Proteobacteria; Gammaproteobacteria; Enterobacterales; Enterobacteriaceae; Salmonella; Salmonella enterica; Salmonella enterica subsp. enterica MNNEETFYQAMRRKGVTRRSFLKFCSLAATSLGLGAGMTPKIAWALENKPRIPVVWIHGL WP_001662439.1 hydrogenase [Salmonella enterica] Length: 372\nNumber of predicted TMHs: 2\nExp number of AAs in TMHs: 36.5172\nExp number, first 60 AAs: 19.72849\nTotal prob of N-in: 0.96169\nPOSSIBLE N-term signal sequence\ninside 1 19\nTMhelix 20 39\noutside 40 325\nTMhelix 326 348\ninside 349 372

17442 GCF_002049445.1_ASM204944v1 Salmonella enterica subsp. enterica serovar Gaminara Proteobacteria; Gammaproteobacteria; Enterobacterales; Enterobacteriaceae; Salmonella; Salmonella enterica; Salmonella enterica subsp. enterica MTGDNTLITSHGINRRDFMKLCAALAATMGLSSKAAAEMAESVSNPQRPPVIWIGAQECT WP_000145429.1 hydrogenase 2 small subunit [Salmonella enterica] Length: 372\nNumber of predicted TMHs: 1\nExp number of AAs in TMHs: 24.51309\nExp number, first 60 AAs: 0.2682\nTotal prob of N-in: 0.14402\noutside 1 331\nTMhelix 332 354\ninside 355 372

17443 GCF_002050505.1_ASM205050v1 Salmonella enterica subsp. enterica serovar Gaminara Proteobacteria; Gammaproteobacteria; Enterobacterales; Enterobacteriaceae; Salmonella; Salmonella enterica; Salmonella enterica subsp. enterica MNNEETFYQAMRRKGVTRRSFLKFCSLAATSLGLGAGMTPKIAWALENKPRIPVVWIHGL WP_001662439.1 hydrogenase [Salmonella enterica] Length: 372\nNumber of predicted TMHs: 2\nExp number of AAs in TMHs: 36.5172\nExp number, first 60 AAs: 19.72849\nTotal prob of N-in: 0.96169\nPOSSIBLE N-term signal sequence\ninside 1 19\nTMhelix 20 39\noutside 40 325\nTMhelix 326 348\ninside 349 372

17444 GCF_002050505.1_ASM205050v1 Salmonella enterica subsp. enterica serovar Gaminara Proteobacteria; Gammaproteobacteria; Enterobacterales; Enterobacteriaceae; Salmonella; Salmonella enterica; Salmonella enterica subsp. enterica MTGDNTLITSHGINRRDFMKLCAALAATMGLSSKAAAEMAESVSNPQRPPVIWIGAQECT WP_080102385.1 hydrogenase 2 small subunit [Salmonella enterica] Length: 372\nNumber of predicted TMHs: 1\nExp number of AAs in TMHs: 24.48766\nExp number, first 60 AAs: 0.26839\nTotal prob of N-in: 0.14782\noutside 1 331\nTMhelix 332 354\ninside 355 372

17445 GCF_002034385.1_ASM203438v1 Salmonella enterica subsp. enterica serovar Gera Proteobacteria; Gammaproteobacteria; Enterobacterales; Enterobacteriaceae; Salmonella; Salmonella enterica; Salmonella enterica subsp. enterica MNNEETFYQAMRRKGVTRRSFLKFCSLAATSLGLGAGMTPKIAWALENKPRIPVVWIHGL WP_001058311.1 hydrogenase [Salmonella enterica] Length: 372\nNumber of predicted TMHs: 2\nExp number of AAs in TMHs: 37.4699899999999\nExp number, first 60 AAs: 19.72699\nTotal prob of N-in: 0.96171\nPOSSIBLE N-term signal sequence\ninside 1 19\nTMhelix 20 39\noutside 40 325\nTMhelix 326 348\ninside 349 372

17446 GCF_002034385.1_ASM203438v1 Salmonella enterica subsp. enterica serovar Gera Proteobacteria; Gammaproteobacteria; Enterobacterales; Enterobacteriaceae; Salmonella; Salmonella enterica; Salmonella enterica subsp. enterica MTGDNTLITSHGINRRDFMKLCAALAATMGLSSKAAAEMAESVSNPQRPPVIWIGAQECT WP_000145429.1 hydrogenase 2 small subunit [Salmonella enterica] Length: 372\nNumber of predicted TMHs: 1\nExp number of AAs in TMHs: 24.51309\nExp number, first 60 AAs: 0.2682\nTotal prob of N-in: 0.14402\noutside 1 331\nTMhelix 332 354\ninside 355 372

17447 GCF_000732045.1_CFSAN012622_01.0Salmonella enterica subsp. enterica serovar Give Proteobacteria; Gammaproteobacteria; Enterobacterales; Enterobacteriaceae; Salmonella; Salmonella enterica; Salmonella enterica subsp. enterica MNNEETFYQAMRRKGVTRRSFLKFCSLAATSLGLGAGMTPKIAWALENKPRIPVVWIHGL WP_001058312.1 hydrogenase [Salmonella enterica] Length: 372\nNumber of predicted TMHs: 2\nExp number of AAs in TMHs: 36.51738\nExp number, first 60 AAs: 19.72849\nTotal prob of N-in: 0.96170\nPOSSIBLE N-term signal sequence\ninside 1 19\nTMhelix 20 39\noutside 40 325\nTMhelix 326 348\ninside 349 372

17448 GCF_002106825.1_ASM210682v1 Salmonella enterica subsp. enterica serovar Give Proteobacteria; Gammaproteobacteria; Enterobacterales; Enterobacteriaceae; Salmonella; Salmonella enterica; Salmonella enterica subsp. enterica MTGDNTLITSHGINRRDFMKLCAALAATMGLSSKAAAEMAESVSNPQRPPVIWIGAQECT WP_000145429.1 hydrogenase 2 small subunit [Salmonella enterica] Length: 372\nNumber of predicted TMHs: 1\nExp number of AAs in TMHs: 24.51309\nExp number, first 60 AAs: 0.2682\nTotal prob of N-in: 0.14402\noutside 1 331\nTMhelix 332 354\ninside 355 372

17449 GCF_000732045.1_CFSAN012622_01.0Salmonella enterica subsp. enterica serovar Give Proteobacteria; Gammaproteobacteria; Enterobacterales; Enterobacteriaceae; Salmonella; Salmonella enterica; Salmonella enterica subsp. enterica MTGDNTLITSHGINRRDFMKLCAALAATMGLSSKAAAEMAESVSNPQRPPVIWIGAQECT WP_000145429.1 hydrogenase 2 small subunit [Salmonella enterica] Length: 372\nNumber of predicted TMHs: 1\nExp number of AAs in TMHs: 24.51309\nExp number, first 60 AAs: 0.2682\nTotal prob of N-in: 0.14402\noutside 1 331\nTMhelix 332 354\ninside 355 372

17450 GCF_002062555.1_ASM206255v1 Salmonella enterica subsp. enterica serovar Give Proteobacteria; Gammaproteobacteria; Enterobacterales; Enterobacteriaceae; Salmonella; Salmonella enterica; Salmonella enterica subsp. enterica MNNEETFYQAMRRKGVTRRSFLKFCSLAATSLGLGAGMTPKIAWALENKPRIPVVWIHGL WP_080191388.1 [Ni/Fe] hydrogenase small subunit [Salmonella enterica] Length: 372\nNumber of predicted TMHs: 2\nExp number of AAs in TMHs: 36.51738\nExp number, first 60 AAs: 19.72849\nTotal prob of N-in: 0.96170\nPOSSIBLE N-term signal sequence\ninside 1 19\nTMhelix 20 39\noutside 40 325\nTMhelix 326 348\ninside 349 372

17451 GCF_002062555.1_ASM206255v1 Salmonella enterica subsp. enterica serovar Give Proteobacteria; Gammaproteobacteria; Enterobacterales; Enterobacteriaceae; Salmonella; Salmonella enterica; Salmonella enterica subsp. enterica MTGDNTLITSHGINRRDFMKLCAALAATMGLSSKAAAEMAESVSNPQRPPVIWIGAQECT WP_000145429.1 hydrogenase 2 small subunit [Salmonella enterica] Length: 372\nNumber of predicted TMHs: 1\nExp number of AAs in TMHs: 24.51309\nExp number, first 60 AAs: 0.2682\nTotal prob of N-in: 0.14402\noutside 1 331\nTMhelix 332 354\ninside 355 372

17452 GCF_002105985.1_ASM210598v1 Salmonella enterica subsp. enterica serovar Give Proteobacteria; Gammaproteobacteria; Enterobacterales; Enterobacteriaceae; Salmonella; Salmonella enterica; Salmonella enterica subsp. enterica MTGDNTLITSHGINRRDFMKLCAALAATMGLSSKAAAEMAESVSNPQRPPVIWIGAQECT WP_000145429.1 hydrogenase 2 small subunit [Salmonella enterica] Length: 372\nNumber of predicted TMHs: 1\nExp number of AAs in TMHs: 24.51309\nExp number, first 60 AAs: 0.2682\nTotal prob of N-in: 0.14402\noutside 1 331\nTMhelix 332 354\ninside 355 372

17453 GCF_002105965.1_ASM210596v1 Salmonella enterica subsp. enterica serovar Give Proteobacteria; Gammaproteobacteria; Enterobacterales; Enterobacteriaceae; Salmonella; Salmonella enterica; Salmonella enterica subsp. enterica MTGDNTLITSHGINRRDFMKLCAALAATMGLSSKAAAEMAESVSNPQRPPVIWIGAQECT WP_000145429.1 hydrogenase 2 small subunit [Salmonella enterica] Length: 372\nNumber of predicted TMHs: 1\nExp number of AAs in TMHs: 24.51309\nExp number, first 60 AAs: 0.2682\nTotal prob of N-in: 0.14402\noutside 1 331\nTMhelix 332 354\ninside 355 372

17454 GCF_002106825.1_ASM210682v1 Salmonella enterica subsp. enterica serovar Give Proteobacteria; Gammaproteobacteria; Enterobacterales; Enterobacteriaceae; Salmonella; Salmonella enterica; Salmonella enterica subsp. enterica MNNEETFYQAMRRKGVTRRSFLKFCSLAATSLGLGAGMTPKIAWALENKPRIPVVWIHGL WP_001058312.1 hydrogenase [Salmonella enterica] Length: 372\nNumber of predicted TMHs: 2\nExp number of AAs in TMHs: 36.51738\nExp number, first 60 AAs: 19.72849\nTotal prob of N-in: 0.96170\nPOSSIBLE N-term signal sequence\ninside 1 19\nTMhelix 20 39\noutside 40 325\nTMhelix 326 348\ninside 349 372

17455 GCF_002105985.1_ASM210598v1 Salmonella enterica subsp. enterica serovar Give Proteobacteria; Gammaproteobacteria; Enterobacterales; Enterobacteriaceae; Salmonella; Salmonella enterica; Salmonella enterica subsp. enterica MNNEETFYQAMRRKGVTRRSFLKFCSLAATSLGLGAGMTPKIAWALENKPRIPVVWIHGL WP_001058312.1 hydrogenase [Salmonella enterica] Length: 372\nNumber of predicted TMHs: 2\nExp number of AAs in TMHs: 36.51738\nExp number, first 60 AAs: 19.72849\nTotal prob of N-in: 0.96170\nPOSSIBLE N-term signal sequence\ninside 1 19\nTMhelix 20 39\noutside 40 325\nTMhelix 326 348\ninside 349 372

17456 GCF_002105965.1_ASM210596v1 Salmonella enterica subsp. enterica serovar Give Proteobacteria; Gammaproteobacteria; Enterobacterales; Enterobacteriaceae; Salmonella; Salmonella enterica; Salmonella enterica subsp. enterica MNNEETFYQAMRRKGVTRRSFLKFCSLAATSLGLGAGMTPKIAWALENKPRIPVVWIHGL WP_001058312.1 hydrogenase [Salmonella enterica] Length: 372\nNumber of predicted TMHs: 2\nExp number of AAs in TMHs: 36.51738\nExp number, first 60 AAs: 19.72849\nTotal prob of N-in: 0.96170\nPOSSIBLE N-term signal sequence\ninside 1 19\nTMhelix 20 39\noutside 40 325\nTMhelix 326 348\ninside 349 372

17457 GCF_002106715.1_ASM210671v1 Salmonella enterica subsp. enterica serovar Give Proteobacteria; Gammaproteobacteria; Enterobacterales; Enterobacteriaceae; Salmonella; Salmonella enterica; Salmonella enterica subsp. enterica MTGDNTLITSHGINRRDFMKLCAALAATMGLSSKAAAEMAESVSNPQRPPVIWIGAQECT WP_000145429.1 hydrogenase 2 small subunit [Salmonella enterica] Length: 372\nNumber of predicted TMHs: 1\nExp

number of AAs in TMHs: 24.51309\nExp number, first 60 AAs: 0.2682\nTotal prob of N-in: 0.14402\noutside 1 331\nTMhelix 332 354\ninside 355 372

17458 GCF_002107545.1_ASM210754v1 Salmonella enterica subsp. enterica serovar Give Proteobacteria; Gammaproteobacteria; Enterobacterales; Enterobacteriaceae; Salmonella; Salmonella enterica; Salmonella enterica subsp. enterica MNNEETFYQAMRRKGVTRRSFLKFCSLAATSLGLGAGMTPKIAWALENKPRIPVVWIHGL WP_001058312.1 hydrogenase [Salmonella enterica] Length: 372\nNumber of predicted TMHs: 2\nExp number of AAs in TMHs: 36.51738\nExp number, first 60 AAs: 19.72849\nTotal prob of N-in: 0.96170\nPOSSIBLE N-term signal sequence\ninside 1 19\nTMhelix 20 39\noutside 40 325\nTMhelix 326 348\ninside 349 372

17459 GCF_002107545.1_ASM210754v1 Salmonella enterica subsp. enterica serovar Give Proteobacteria; Gammaproteobacteria; Enterobacterales; Enterobacteriaceae; Salmonella; Salmonella enterica; Salmonella enterica subsp. enterica MTGDNTLITSHGINRRDFMKLCAALAATMGLSSKAAAEMAESVSNPQRPPVIWIGAQECT WP_000145429.1 hydrogenase 2 small subunit [Salmonella enterica] Length: 372\nNumber of predicted TMHs: 1\nExp number of AAs in TMHs: 24.51309\nExp number, first 60 AAs: 0.2682\nTotal prob of N-in: 0.14402\noutside 1 331\nTMhelix 332 354\ninside 355 372

17460 GCF_002063135.1_ASM206313v1 Salmonella enterica subsp. enterica serovar Give Proteobacteria; Gammaproteobacteria; Enterobacterales; Enterobacteriaceae; Salmonella; Salmonella enterica; Salmonella enterica subsp. enterica MNNEETFYQAMRRKGVTRRSFLKFCSLAATSLGLGAGMTPKIAWALENKPRIPVVWIHGL WP_001058312.1 hydrogenase [Salmonella enterica] Length: 372\nNumber of predicted TMHs: 2\nExp number of AAs in TMHs: 36.51738\nExp number, first 60 AAs: 19.72849\nTotal prob of N-in: 0.96170\nPOSSIBLE N-term signal sequence\ninside 1 19\nTMhelix 20 39\noutside 40 325\nTMhelix 326 348\ninside 349 372

17461 GCF_002107415.1_ASM210741v1 Salmonella enterica subsp. enterica serovar Give Proteobacteria; Gammaproteobacteria; Enterobacterales; Enterobacteriaceae; Salmonella; Salmonella enterica; Salmonella enterica subsp. enterica MNNEETFYQAMRRKGVTRRSFLKFCSLAATSLGLGAGMTPKIAWALENKPRIPVVWIHGL WP_001058312.1 hydrogenase [Salmonella enterica] Length: 372\nNumber of predicted TMHs: 2\nExp number of AAs in TMHs: 36.51738\nExp number, first 60 AAs: 19.72849\nTotal prob of N-in: 0.96170\nPOSSIBLE N-term signal sequence\ninside 1 19\nTMhelix 20 39\noutside 40 325\nTMhelix 326 348\ninside 349 372

17462 GCF_002106715.1_ASM210671v1 Salmonella enterica subsp. enterica serovar Give Proteobacteria; Gammaproteobacteria; Enterobacterales; Enterobacteriaceae; Salmonella; Salmonella enterica; Salmonella enterica subsp. enterica MNNEETFYQAMRRKGVTRRSFLKFCSLAATSLGLGAGMTPKIAWALENKPRIPVVWIHGL WP_001058312.1 hydrogenase [Salmonella enterica] Length: 372\nNumber of predicted TMHs: 2\nExp number of AAs in TMHs: 36.51738\nExp number, first 60 AAs: 19.72849\nTotal prob of N-in: 0.96170\nPOSSIBLE N-term signal sequence\ninside 1 19\nTMhelix 20 39\noutside 40 325\nTMhelix 326 348\ninside 349 372

17463 GCF_002065645.1_ASM206564v1 Salmonella enterica subsp. enterica serovar Give Proteobacteria; Gammaproteobacteria; Enterobacterales; Enterobacteriaceae; Salmonella; Salmonella enterica; Salmonella enterica subsp. enterica MTGDNTLITSHGINRRDFMKLCAALAATMGLSSKAAAEMAESVSNPQRPPVIWIGAQECT WP_000145429.1 hydrogenase 2 small subunit [Salmonella enterica] Length: 372\nNumber of predicted TMHs: 1\nExp number of AAs in TMHs: 24.51309\nExp number, first 60 AAs: 0.2682\nTotal prob of N-in: 0.14402\noutside 1 331\nTMhelix 332 354\ninside 355 372

17464 GCF_002063135.1_ASM206313v1 Salmonella enterica subsp. enterica serovar Give Proteobacteria; Gammaproteobacteria; Enterobacterales; Enterobacteriaceae; Salmonella; Salmonella enterica; Salmonella enterica subsp. enterica MTGDNTLITSHGINRRDFMKLCAALAATMGLSSKAAAEMAESVSNPQRPPVIWIGAQECT WP_000145429.1 hydrogenase 2 small subunit [Salmonella enterica] Length: 372\nNumber of predicted TMHs: 1\nExp number of AAs in TMHs: 24.51309\nExp number, first 60 AAs: 0.2682\nTotal prob of N-in: 0.14402\noutside 1 331\nTMhelix 332 354\ninside 355 372

17465 GCF_002107415.1_ASM210741v1 Salmonella enterica subsp. enterica serovar Give Proteobacteria; Gammaproteobacteria; Enterobacterales; Enterobacteriaceae; Salmonella; Salmonella enterica; Salmonella enterica subsp. enterica MTGDNTLITSHGINRRDFMKLCAALAATMGLSSKAAAEMAESVSNPQRPPVIWIGAQECT WP_000145429.1 hydrogenase 2 small subunit [Salmonella enterica] Length: 372\nNumber of predicted TMHs: 1\nExp number of AAs in TMHs: 24.51309\nExp number, first 60 AAs: 0.2682\nTotal prob of N-in: 0.14402\noutside 1 331\nTMhelix 332 354\ninside 355 372

17466 GCF_002065645.1_ASM206564v1 Salmonella enterica subsp. enterica serovar Give Proteobacteria; Gammaproteobacteria; Enterobacterales; Enterobacteriaceae; Salmonella; Salmonella enterica; Salmonella enterica subsp.

enterica MNNEETFYQAMRRKGVTRRSFLKFCSLAATSLGLGAGMTPKIAWALENKPRIPVVWIHGL WP_080191388.1 [Ni/Fe] hydrogenase small subunit [Salmonella enterica] Length: 372\nNumber of predicted TMHs: 2\nExp number of AAs in TMHs: 36.51738\nExp number, first 60 AAs: 19.72849\nTotal prob of N-in: 0.96170\nPOSSIBLE N-term signal sequence\ninside 1 19\nTMhelix 20 39\noutside 40 325\nTMhelix 326 348\ninside 349 372

17467 GCF_002046655.1_ASM204665v1 Salmonella enterica subsp. enterica serovar Grumpensis Proteobacteria; Gammaproteobacteria; Enterobacterales; Enterobacteriaceae; Salmonella; Salmonella enterica; Salmonella enterica subsp. enterica MTGDNTLITSHGINRRDFMKLCAALAATMGLSSKAAAEMAESVSNPQRPPVIWIGAQECT WP_000145429.1 hydrogenase 2 small subunit [Salmonella enterica] Length: 372\nNumber of predicted TMHs: 1\nExp number of AAs in TMHs: 24.51309\nExp number, first 60 AAs: 0.2682\nTotal prob of N-in: 0.14402\noutside 1 331\nTMhelix 332 354\ninside 355 372

17468 GCF_002048265.1_ASM204826v1 Salmonella enterica subsp. enterica serovar Grumpensis Proteobacteria; Gammaproteobacteria; Enterobacterales; Enterobacteriaceae; Salmonella; Salmonella enterica; Salmonella enterica subsp. enterica MTGDNTLITSHGINRRDFMKLCAALAATMGLSSKAAAEMAESVSNPQRPPVIWIGAQECT WP_000145429.1 hydrogenase 2 small subunit [Salmonella enterica] Length: 372\nNumber of predicted TMHs: 1\nExp number of AAs in TMHs: 24.51309\nExp number, first 60 AAs: 0.2682\nTotal prob of N-in: 0.14402\noutside 1 331\nTMhelix 332 354\ninside 355 372

17469 GCF_002046655.1_ASM204665v1 Salmonella enterica subsp. enterica serovar Grumpensis Proteobacteria; Gammaproteobacteria; Enterobacterales; Enterobacteriaceae; Salmonella; Salmonella enterica; Salmonella enterica subsp. enterica MNNEETFYQAMRRKGVTRRSFLKFCSLAATSLGLGAGMTPKIAWALENKPRIPVVWIHGL WP_001662439.1 hydrogenase [Salmonella enterica] Length: 372\nNumber of predicted TMHs: 2\nExp number of AAs in TMHs: 36.5172\nExp number, first 60 AAs: 19.72849\nTotal prob of N-in: 0.96169\nPOSSIBLE N-term signal sequence\ninside 1 19\nTMhelix 20 39\noutside 40 325\nTMhelix 326 348\ninside 349 372

17470 GCF_002046695.1_ASM204669v1 Salmonella enterica subsp. enterica serovar Grumpensis Proteobacteria; Gammaproteobacteria; Enterobacterales; Enterobacteriaceae; Salmonella; Salmonella enterica; Salmonella enterica subsp. enterica MTGDNTLITSHGINRRDFMKLCAALAATMGLSSKAAAEMAESVSNPQRPPVIWIGAQECT WP_000145429.1 hydrogenase 2 small subunit [Salmonella enterica] Length: 372\nNumber of predicted TMHs: 1\nExp number of AAs in TMHs: 24.51309\nExp number, first 60 AAs: 0.2682\nTotal prob of N-in: 0.14402\noutside 1 331\nTMhelix 332 354\ninside 355 372

17471 GCF_002046945.1_ASM204694v1 Salmonella enterica subsp. enterica serovar Grumpensis Proteobacteria; Gammaproteobacteria; Enterobacterales; Enterobacteriaceae; Salmonella; Salmonella enterica; Salmonella enterica subsp. enterica MTGDNTLITSHGINRRDFMKLCAALAATMGLSSKAAAEMAESVSNPQRPPVIWIGAQECT WP_000145429.1 hydrogenase 2 small subunit [Salmonella enterica] Length: 372\nNumber of predicted TMHs: 1\nExp number of AAs in TMHs: 24.51309\nExp number, first 60 AAs: 0.2682\nTotal prob of N-in: 0.14402\noutside 1 331\nTMhelix 332 354\ninside 355 372

17472 GCF_002046695.1_ASM204669v1 Salmonella enterica subsp. enterica serovar Grumpensis Proteobacteria; Gammaproteobacteria; Enterobacterales; Enterobacteriaceae; Salmonella; Salmonella enterica; Salmonella enterica subsp. enterica MNNEETFYQAMRRKGVTRRSFLKFCSLAATSLGLGAGMTPKIAWALENKPRIPVVWIHGL WP_001662439.1 hydrogenase [Salmonella enterica] Length: 372\nNumber of predicted TMHs: 2\nExp number of AAs in TMHs: 36.5172\nExp number, first 60 AAs: 19.72849\nTotal prob of N-in: 0.96169\nPOSSIBLE N-term signal sequence\ninside 1 19\nTMhelix 20 39\noutside 40 325\nTMhelix 326 348\ninside 349 372

17473 GCF_001246475.1_Salmonella_enterica_CVM_N51263_v1.0 Salmonella enterica subsp. enterica serovar Hadar Proteobacteria; Gammaproteobacteria; Enterobacterales; Enterobacteriaceae; Salmonella; Salmonella enterica; Salmonella enterica subsp. enterica MTGDNTLITSHGINRRDFMKLCAALAATMGLSSKAAAEMAESVSNPQRPPVIWIGAQECT WP_000145429.1 hydrogenase 2 small subunit [Salmonella enterica] Length: 372\nNumber of predicted TMHs: 1\nExp number of AAs in TMHs: 24.51309\nExp number, first 60 AAs: 0.2682\nTotal prob of N-in: 0.14402\noutside 1 331\nTMhelix 332 354\ninside 355 372

17474 GCF_002030685.1_ASM203068v1 Salmonella enterica subsp. enterica serovar Hadar Proteobacteria; Gammaproteobacteria; Enterobacterales; Enterobacteriaceae; Salmonella; Salmonella enterica; Salmonella enterica subsp. enterica MNNEETFYQAMRRKGVTRRSFLKFCSLAATSLGLGAGMTPKIAWALENKPRIPVVWIHGL WP_079831006.1 [Ni/Fe] hydrogenase small subunit [Salmonella enterica] Length: 372\nNumber of predicted TMHs: 2\nExp number of AAs in TMHs: 39.08204\nExp number, first 60 AAs: 19.72474\nTotal prob of N-in: 0.96174\nPOSSIBLE N-term signal sequence\ninside 1 19\nTMhelix 20 39\noutside 40 325\nTMhelix 326 348\ninside 349 372

17475 GCF_001243705.1_Salmonella_enterica_CVM_N46859_v1.0 Salmonella enterica subsp. enterica serovar Hadar Proteobacteria; Gammaproteobacteria; Enterobacterales; Enterobacteriaceae; Salmonella; Salmonella enterica; Salmonella enterica subsp. enterica MTGDNTLITSHGINRRDFMKLCAALAATMGLSSKAAAEMAESVSNPQRPPVIWIGAQECT WP_000145429.1 hydrogenase 2 small subunit [Salmonella enterica] Length: 372\nNumber of predicted TMHs: 1\nExp number of AAs in TMHs: 24.51309\nExp number, first 60 AAs: 0.2682\nTotal prob of N-in: 0.14402\noutside 1 331\nTMhelix 332 354\ninside 355 372

17476 GCF_002032945.1_ASM203294v1 Salmonella enterica subsp. enterica serovar Hadar Proteobacteria; Gammaproteobacteria; Enterobacterales; Enterobacteriaceae; Salmonella; Salmonella enterica; Salmonella enterica subsp. enterica MTGDNTLITSHGINRRDFMKLCAALAATMGLSSKAAAEMAESVSNPQRPPVIWIGAQECT WP_000145429.1 hydrogenase 2 small subunit [Salmonella enterica] Length: 372\nNumber of predicted TMHs: 1\nExp number of AAs in TMHs: 24.51309\nExp number, first 60 AAs: 0.2682\nTotal prob of N-in: 0.14402\noutside 1 331\nTMhelix 332 354\ninside 355 372

17477 GCF_001245215.1_Salmonella_enterica_CVM_N48712_v1.0 Salmonella enterica subsp. enterica serovar Hadar Proteobacteria; Gammaproteobacteria; Enterobacterales; Enterobacteriaceae; Salmonella; Salmonella enterica; Salmonella enterica subsp. enterica MNNEETFYQAMRRKGVTRRSFLKFCSLAATSLGLGAGMTPKIAWALENKPRIPVVWIHGL WP_001058308.1 hydrogenase [Salmonella enterica] Length: 372\nNumber of predicted TMHs: 2\nExp number of AAs in TMHs: 39.08204\nExp number, first 60 AAs: 19.72474\nTotal prob of N-in: 0.96174\nPOSSIBLE N-term signal sequence\ninside 1 19\nTMhelix 20 39\noutside 40 325\nTMhelix 326 348\ninside 349 372

17478 GCF_002030685.1_ASM203068v1 Salmonella enterica subsp. enterica serovar Hadar Proteobacteria; Gammaproteobacteria; Enterobacterales; Enterobacteriaceae; Salmonella; Salmonella enterica; Salmonella enterica subsp. enterica MTGDNTLITSHGINRRDFMKLCAALAATMGLSSKAAAEMAESVSNPQRPPVIWIGAQECT WP_000145429.1 hydrogenase 2 small subunit [Salmonella enterica] Length: 372\nNumber of predicted TMHs: 1\nExp number of AAs in TMHs: 24.51309\nExp number, first 60 AAs: 0.2682\nTotal prob of N-in: 0.14402\noutside 1 331\nTMhelix 332 354\ninside 355 372

17479 GCF_001479565.1_Salmonella_enterica_CVM_N32788-R_v1.0 Salmonella enterica subsp. enterica serovar Hadar Proteobacteria; Gammaproteobacteria; Enterobacterales; Enterobacteriaceae; Salmonella; Salmonella enterica; Salmonella enterica subsp. enterica MTGDNTLITSHGINRRDFMKLCAALAATMGLSSKAAAEMAESVSNPQRPPVIWIGAQECT WP_000145429.1 hydrogenase 2 small subunit [Salmonella enterica] Length: 372\nNumber of predicted TMHs: 1\nExp number of AAs in TMHs: 24.51309\nExp number, first 60 AAs: 0.2682\nTotal prob of N-in: 0.14402\noutside 1 331\nTMhelix 332 354\ninside 355 372

17480 GCF_001246485.1_Salmonella_enterica_CVM_N51265_v1.0 Salmonella enterica subsp. enterica serovar Hadar Proteobacteria; Gammaproteobacteria; Enterobacterales; Enterobacteriaceae; Salmonella; Salmonella enterica; Salmonella enterica subsp. enterica MTGDNTLITSHGINRRDFMKLCAALAATMGLSSKAAAEMAESVSNPQRPPVIWIGAQECT WP_000145429.1 hydrogenase 2 small subunit [Salmonella enterica] Length: 372\nNumber of predicted TMHs: 1\nExp number of AAs in TMHs: 24.51309\nExp number, first 60 AAs: 0.2682\nTotal prob of N-in: 0.14402\noutside 1 331\nTMhelix 332 354\ninside 355 372

17481 GCF_001479565.1_Salmonella_enterica_CVM_N32788-R_v1.0 Salmonella enterica subsp. enterica serovar Hadar Proteobacteria; Gammaproteobacteria; Enterobacterales; Enterobacteriaceae; Salmonella; Salmonella enterica; Salmonella enterica subsp. enterica MNNEETFYQAMRRKGVTRRSFLKFCSLAATSLGLGAGMTPKIAWALENKPRIPVVWIHGL WP_001058308.1 hydrogenase [Salmonella enterica] Length: 372\nNumber of predicted TMHs: 2\nExp number of AAs in TMHs: 39.08204\nExp number, first 60 AAs: 19.72474\nTotal prob of N-in: 0.96174\nPOSSIBLE N-term signal sequence\ninside 1 19\nTMhelix 20 39\noutside 40 325\nTMhelix 326 348\ninside 349 372

17482 GCF_002065715.1_ASM206571v1 Salmonella enterica subsp. enterica serovar Hadar Proteobacteria; Gammaproteobacteria; Enterobacterales; Enterobacteriaceae; Salmonella; Salmonella enterica; Salmonella enterica subsp. enterica MLRFRTIFLKKIFTLKNSILARRGAVIVIVSAVFTSIMFFAHSWASDKEVAMTLSVLNSN WP_080151465.1 hypothetical protein [Salmonella enterica] Length: 204\nNumber of predicted TMHs: 2\nExp number of AAs in TMHs: 44.99538\nExp number, first 60 AAs: 22.34317\nTotal prob of N-in: 0.89744\nPOSSIBLE N-term signal sequence\ninside 1 19\nTMhelix 20 42\noutside 43 171\nTMhelix 172 194\ninside 195 204

17483 GCF_002065715.1_ASM206571v1 Salmonella enterica subsp. enterica serovar Hadar Proteobacteria; Gammaproteobacteria; Enterobacterales; Enterobacteriaceae; Salmonella; Salmonella enterica; Salmonella enterica subsp. enterica MTGDNTLITSHGINRRDFMKLCAALAATMGLSSKAAAEMAESVSNPQRPPVIWIGAQECT WP_000145429.1 hydrogenase 2 small subunit [Salmonella enterica] Length: 372\nNumber of

predicted TMHs: 1\nExp number of AAs in TMHs: 24.51309\nExp number, first 60 AAs: 0.2682\nTotal prob of N-in: 0.14402\noutside 1 331\nTMhelix 332 354\ninside 355 372

17484 GCF_002065635.1_ASM206563v1 Salmonella enterica subsp. enterica serovar Hadar Proteobacteria; Gammaproteobacteria; Enterobacterales; Enterobacteriaceae; Salmonella; Salmonella enterica; Salmonella enterica subsp. enterica MNNEETFYQAMRRKGVTRRSFLKFCSLAATSLGLGAGMTPKIAWALENKPRIPVVWIHGL WP_001058308.1 hydrogenase [Salmonella enterica] Length: 372\nNumber of predicted TMHs: 2\nExp number of AAs in TMHs: 39.08204\nExp number, first 60 AAs: 19.72474\nTotal prob of N-in: 0.96174\nPOSSIBLE N-term signal sequence\ninside 1 19\nTMhelix 20 39\noutside 40 325\nTMhelix 326 348\ninside 349 372

17485 GCF_002065715.1_ASM206571v1 Salmonella enterica subsp. enterica serovar Hadar Proteobacteria; Gammaproteobacteria; Enterobacterales; Enterobacteriaceae; Salmonella; Salmonella enterica; Salmonella enterica subsp. enterica MNNEETFYQAMRRKGVTRRSFLKFCSLAATSLGLGAGMTPKIAWALENKPRIPVVWIHGL WP_001058311.1 hydrogenase [Salmonella enterica] Length: 372\nNumber of predicted TMHs: 2\nExp number of AAs in TMHs: 37.4699899999999\nExp number, first 60 AAs: 19.72699\nTotal prob of N-in: 0.96171\nPOSSIBLE N-term signal sequence\ninside 1 19\nTMhelix 20 39\noutside 40 325\nTMhelix 326 348\ninside 349 372

17486 GCF_002065635.1_ASM206563v1 Salmonella enterica subsp. enterica serovar Hadar Proteobacteria; Gammaproteobacteria; Enterobacterales; Enterobacteriaceae; Salmonella; Salmonella enterica; Salmonella enterica subsp. enterica MTGDNTLITSHGINRRDFMKLCAALAATMGLSSKAAAEMAESVSNPQRPPVIWIGAQECT WP_000145429.1 hydrogenase 2 small subunit [Salmonella enterica] Length: 372\nNumber of predicted TMHs: 1\nExp number of AAs in TMHs: 24.51309\nExp number, first 60 AAs: 0.2682\nTotal prob of N-in: 0.14402\noutside 1 331\nTMhelix 332 354\ninside 355 372

17487 GCF_001243355.1_Salmonella_enterica_CVM_N46842_v1.0 Salmonella enterica subsp. enterica serovar Hadar Proteobacteria; Gammaproteobacteria; Enterobacterales; Enterobacteriaceae; Salmonella; Salmonella enterica; Salmonella enterica subsp. enterica MTGDNTLITSHGINRRDFMKLCAALAATMGLSSKAAAEMAESVSNPQRPPVIWIGAQECT WP_000145429.1 hydrogenase 2 small subunit [Salmonella enterica] Length: 372\nNumber of predicted TMHs: 1\nExp number of AAs in TMHs: 24.51309\nExp number, first 60 AAs: 0.2682\nTotal prob of N-in: 0.14402\noutside 1 331\nTMhelix 332 354\ninside 355 372

17488 GCF_001245555.1_Salmonella_enterica_CVM_N50436_v1.0 Salmonella enterica subsp. enterica serovar Hadar Proteobacteria; Gammaproteobacteria; Enterobacterales; Enterobacteriaceae; Salmonella; Salmonella enterica; Salmonella enterica subsp. enterica MTGDNTLITSHGINRRDFMKLCAALAATMGLSSKAAAEMAESVSNPQRPPVIWIGAQECT WP_000145429.1 hydrogenase 2 small subunit [Salmonella enterica] Length: 372\nNumber of predicted TMHs: 1\nExp number of AAs in TMHs: 24.51309\nExp number, first 60 AAs: 0.2682\nTotal prob of N-in: 0.14402\noutside 1 331\nTMhelix 332 354\ninside 355 372

17489 GCF_001246765.1_Salmonella_enterica_CVM_N51279_v1.0 Salmonella enterica subsp. enterica serovar Hadar Proteobacteria; Gammaproteobacteria; Enterobacterales; Enterobacteriaceae; Salmonella; Salmonella enterica; Salmonella enterica subsp. enterica MNNEETFYQAMRRKGVTRRSFLKFCSLAATSLGLGAGMTPKIAWALENKPRIPVVWIHGL WP_001058308.1 hydrogenase [Salmonella enterica] Length: 372\nNumber of predicted TMHs: 2\nExp number of AAs in TMHs: 39.08204\nExp number, first 60 AAs: 19.72474\nTotal prob of N-in: 0.96174\nPOSSIBLE N-term signal sequence\ninside 1 19\nTMhelix 20 39\noutside 40 325\nTMhelix 326 348\ninside 349 372

17490 GCF_001246355.1_Salmonella_enterica_CVM_N51260_v1.0 Salmonella enterica subsp. enterica serovar Hadar Proteobacteria; Gammaproteobacteria; Enterobacterales; Enterobacteriaceae; Salmonella; Salmonella enterica; Salmonella enterica subsp. enterica MNNEETFYQAMRRKGVTRRSFLKFCSLAATSLGLGAGMTPKIAWALENKPRIPVVWIHGL WP_001058308.1 hydrogenase [Salmonella enterica] Length: 372\nNumber of predicted TMHs: 2\nExp number of AAs in TMHs: 39.08204\nExp number, first 60 AAs: 19.72474\nTotal prob of N-in: 0.96174\nPOSSIBLE N-term signal sequence\ninside 1 19\nTMhelix 20 39\noutside 40 325\nTMhelix 326 348\ninside 349 372

17491 GCF_001245555.1_Salmonella_enterica_CVM_N50436_v1.0 Salmonella enterica subsp. enterica serovar Hadar Proteobacteria; Gammaproteobacteria; Enterobacterales; Enterobacteriaceae; Salmonella; Salmonella enterica; Salmonella enterica subsp. enterica MNNEETFYQAMRRKGVTRRSFLKFCSLAATSLGLGAGMTPKIAWALENKPRIPVVWIHGL WP_001058308.1 hydrogenase [Salmonella enterica] Length: 372\nNumber of predicted TMHs: 2\nExp number of AAs in TMHs: 39.08204\nExp number, first 60 AAs: 19.72474\nTotal prob of N-in: 0.96174\nPOSSIBLE N-term signal sequence\ninside 1 19\nTMhelix 20 39\noutside 40 325\nTMhelix 326 348\ninside 349 372

17492 GCF_001241055.1_Salmonella_enterica_CVM_N43836_v1.0 Salmonella enterica subsp. enterica serovar Hadar Proteobacteria; Gammaproteobacteria; Enterobacterales; Enterobacteriaceae; Salmonella; Salmonella enterica; Salmonella enterica subsp. enterica MTGDNTLITSHGINRRDFMKLCAALAATMGLSSKAAAEMAESVSNPQRPPVIWIGAQECT WP_000145429.1 hydrogenase 2 small subunit [Salmonella enterica] Length: 372\nNumber of predicted TMHs: 1\nExp number of AAs in TMHs: 24.51309\nExp number, first 60 AAs: 0.2682\nTotal prob of N-in: 0.14402\noutside 1 331\nTMhelix 332 354\ninside 355 372

17493 GCF_001246765.1_Salmonella_enterica_CVM_N51279_v1.0 Salmonella enterica subsp. enterica serovar Hadar Proteobacteria; Gammaproteobacteria; Enterobacterales; Enterobacteriaceae; Salmonella; Salmonella enterica; Salmonella enterica subsp. enterica MTGDNTLITSHGINRRDFMKLCAALAATMGLSSKAAAEMAESVSNPQRPPVIWIGAQECT WP_000145429.1 hydrogenase 2 small subunit [Salmonella enterica] Length: 372\nNumber of predicted TMHs: 1\nExp number of AAs in TMHs: 24.51309\nExp number, first 60 AAs: 0.2682\nTotal prob of N-in: 0.14402\noutside 1 331\nTMhelix 332 354\ninside 355 372

17494 GCF_001243355.1_Salmonella_enterica_CVM_N46842_v1.0 Salmonella enterica subsp. enterica serovar Hadar Proteobacteria; Gammaproteobacteria; Enterobacterales; Enterobacteriaceae; Salmonella; Salmonella enterica; Salmonella enterica subsp. enterica MNNEETFYQAMRRKGVTRRSFLKFCSLAATSLGLGAGMTPKIAWALENKPRIPVVWIHGL WP_001058308.1 hydrogenase [Salmonella enterica] Length: 372\nNumber of predicted TMHs: 2\nExp number of AAs in TMHs: 39.08204\nExp number, first 60 AAs: 19.72474\nTotal prob of N-in: 0.96174\nPOSSIBLE N-term signal sequence\ninside 1 19\nTMhelix 20 39\noutside 40 325\nTMhelix 326 348\ninside 349 372

17495 GCF_001241055.1_Salmonella_enterica_CVM_N43836_v1.0 Salmonella enterica subsp. enterica serovar Hadar Proteobacteria; Gammaproteobacteria; Enterobacterales; Enterobacteriaceae; Salmonella; Salmonella enterica; Salmonella enterica subsp. enterica MNNEETFYQAMRRKGVTRRSFLKFCSLAATSLGLGAGMTPKIAWALENKPRIPVVWIHGL WP_001058308.1 hydrogenase [Salmonella enterica] Length: 372\nNumber of predicted TMHs: 2\nExp number of AAs in TMHs: 39.08204\nExp number, first 60 AAs: 19.72474\nTotal prob of N-in: 0.96174\nPOSSIBLE N-term signal sequence\ninside 1 19\nTMhelix 20 39\noutside 40 325\nTMhelix 326 348\ninside 349 372

17496 GCF_001246435.1_Salmonella_enterica_CVM_N51262_v1.0 Salmonella enterica subsp. enterica serovar Hadar Proteobacteria; Gammaproteobacteria; Enterobacterales; Enterobacteriaceae; Salmonella; Salmonella enterica; Salmonella enterica subsp. enterica MTGDNTLITSHGINRRDFMKLCAALAATMGLSSKAAAEMAESVSNPQRPPVIWIGAQECT WP_000145429.1 hydrogenase 2 small subunit [Salmonella enterica] Length: 372\nNumber of predicted TMHs: 1\nExp number of AAs in TMHs: 24.51309\nExp number, first 60 AAs: 0.2682\nTotal prob of N-in: 0.14402\noutside 1 331\nTMhelix 332 354\ninside 355 372

17497 GCF_001242445.1_Salmonella_enterica_CVM_N45940_v1.0 Salmonella enterica subsp. enterica serovar Hadar Proteobacteria; Gammaproteobacteria; Enterobacterales; Enterobacteriaceae; Salmonella; Salmonella enterica; Salmonella enterica subsp. enterica MTGDNTLITSHGINRRDFMKLCAALAATMGLSSKAAAEMAESVSNPQRPPVIWIGAQECT WP_000145429.1 hydrogenase 2 small subunit [Salmonella enterica] Length: 372\nNumber of predicted TMHs: 1\nExp number of AAs in TMHs: 24.51309\nExp number, first 60 AAs: 0.2682\nTotal prob of N-in: 0.14402\noutside 1 331\nTMhelix 332 354\ninside 355 372

17498 GCF_001246495.1_Salmonella_enterica_CVM_N51264_v1.0 Salmonella enterica subsp. enterica serovar Hadar Proteobacteria; Gammaproteobacteria; Enterobacterales; Enterobacteriaceae; Salmonella; Salmonella enterica; Salmonella enterica subsp. enterica MTGDNTLITSHGINRRDFMKLCAALAATMGLSSKAAAEMAESVSNPQRPPVIWIGAQECT WP_000145429.1 hydrogenase 2 small subunit [Salmonella enterica] Length: 372\nNumber of predicted TMHs: 1\nExp number of AAs in TMHs: 24.51309\nExp number, first 60 AAs: 0.2682\nTotal prob of N-in: 0.14402\noutside 1 331\nTMhelix 332 354\ninside 355 372

17499 GCF_001246435.1_Salmonella_enterica_CVM_N51262_v1.0 Salmonella enterica subsp. enterica serovar Hadar Proteobacteria; Gammaproteobacteria; Enterobacterales; Enterobacteriaceae; Salmonella; Salmonella enterica; Salmonella enterica subsp. enterica MNNEETFYQAMRRKGVTRRSFLKFCSLAATSLGLGAGMTPKIAWALENKPRIPVVWIHGL WP_001058308.1 hydrogenase [Salmonella enterica] Length: 372\nNumber of predicted TMHs: 2\nExp number of AAs in TMHs: 39.08204\nExp number, first 60 AAs: 19.72474\nTotal prob of N-in: 0.96174\nPOSSIBLE N-term signal sequence\ninside 1 19\nTMhelix 20 39\noutside 40 325\nTMhelix 326 348\ninside 349 372

17500 GCF_001242005.1_Salmonella_enterica_CVM_N45411_v1.0 Salmonella enterica subsp. enterica serovar Hadar Proteobacteria; Gammaproteobacteria; Enterobacterales; Enterobacteriaceae; Salmonella; Salmonella enterica; Salmonella enterica subsp. enterica MNNEETFYQAMRRKGVTRRSFLKFCSLAATSLGLGAGMTPKIAWALENKPRIPVVWIHGL WP_001058308.1 hydrogenase [Salmonella enterica] Length: 372\nNumber of predicted TMHs: 2\nExp

number of AAs in TMHs: 39.08204\nExp number, first 60 AAs: 19.72474\nTotal prob of N-in: 0.96174\nPOSSIBLE N-term signal sequence\ninside 1 19\nTMhelix 20 39\noutside 40 325\nTMhelix 326 348\ninside 349 372

17501 GCF_001246495.1_Salmonella_enterica_CVM_N51264_v1.0 Salmonella enterica subsp. enterica serovar Hadar Proteobacteria; Gammaproteobacteria; Enterobacterales; Enterobacteriaceae; Salmonella; Salmonella enterica; Salmonella enterica subsp. enterica MNNEETFYQAMRRKGVTRRSFLKFCSLAATSLGLGAGMTPKIAWALENKPRIPVVWIHGL WP_001058308.1 hydrogenase [Salmonella enterica] Length: 372\nNumber of predicted TMHs: 2\nExp number of AAs in TMHs: 39.08204\nExp number, first 60 AAs: 19.72474\nTotal prob of N-in: 0.96174\nPOSSIBLE N-term signal sequence\ninside 1 19\nTMhelix 20 39\noutside 40 325\nTMhelix 326 348\ninside 349 372

17502 GCF_001242445.1_Salmonella_enterica_CVM_N45940_v1.0 Salmonella enterica subsp. enterica serovar Hadar Proteobacteria; Gammaproteobacteria; Enterobacterales; Enterobacteriaceae; Salmonella; Salmonella enterica; Salmonella enterica subsp. enterica MNNEETFYQAMRRKGVTRRSFLKFCSLAATSLGLGAGMTPKIAWALENKPRIPVVWIHGL WP_001058308.1 hydrogenase [Salmonella enterica] Length: 372\nNumber of predicted TMHs: 2\nExp number of AAs in TMHs: 39.08204\nExp number, first 60 AAs: 19.72474\nTotal prob of N-in: 0.96174\nPOSSIBLE N-term signal sequence\ninside 1 19\nTMhelix 20 39\noutside 40 325\nTMhelix 326 348\ninside 349 372

17503 GCF_001246355.1_Salmonella_enterica_CVM_N51260_v1.0 Salmonella enterica subsp. enterica serovar Hadar Proteobacteria; Gammaproteobacteria; Enterobacterales; Enterobacteriaceae; Salmonella; Salmonella enterica; Salmonella enterica subsp. enterica MTGDNTLITSHGINRRDFMKLCAALAATMGLSSKAAAEMAESVSNPQRPPVIWIGAQECT WP_000145429.1 hydrogenase 2 small subunit [Salmonella enterica] Length: 372\nNumber of predicted TMHs: 1\nExp number of AAs in TMHs: 24.51309\nExp number, first 60 AAs: 0.2682\nTotal prob of N-in: 0.14402\noutside 1 331\nTMhelix 332 354\ninside 355 372

17504 GCF_001242005.1_Salmonella_enterica_CVM_N45411_v1.0 Salmonella enterica subsp. enterica serovar Hadar Proteobacteria; Gammaproteobacteria; Enterobacterales; Enterobacteriaceae; Salmonella; Salmonella enterica; Salmonella enterica subsp. enterica MTGDNTLITSHGINRRDFMKLCAALAATMGLSSKAAAEMAESVSNPQRPPVIWIGAQECT WP_000145429.1 hydrogenase 2 small subunit [Salmonella enterica] Length: 372\nNumber of predicted TMHs: 1\nExp number of AAs in TMHs: 24.51309\nExp number, first 60 AAs: 0.2682\nTotal prob of N-in: 0.14402\noutside 1 331\nTMhelix 332 354\ninside 355 372

17505 GCF_000973745.1_ABBSB1020-2.id16_v1.0 Salmonella enterica subsp. enterica serovar Hadar Proteobacteria; Gammaproteobacteria; Enterobacterales; Enterobacteriaceae; Salmonella; Salmonella enterica; Salmonella enterica subsp. enterica MNNEETFYQAMRRKGVTRRSFLKFCSLAATSLGLGAGMTPKIAWALENKPRIPVVWIHGL WP_001058308.1 hydrogenase [Salmonella enterica] Length: 372\nNumber of predicted TMHs: 2\nExp number of AAs in TMHs: 39.08204\nExp number, first 60 AAs: 19.72474\nTotal prob of N-in: 0.96174\nPOSSIBLE N-term signal sequence\ninside 1 19\nTMhelix 20 39\noutside 40 325\nTMhelix 326 348\ninside 349 372

17506 GCF_002033235.1_ASM203323v1 Salmonella enterica subsp. enterica serovar Hadar Proteobacteria; Gammaproteobacteria; Enterobacterales; Enterobacteriaceae; Salmonella; Salmonella enterica; Salmonella enterica subsp. enterica MTGDNTLITSHGINRRDFMKLCAALAATMGLSSKAAAEMAESVSNPQRPPVIWIGAQECT WP_000145429.1 hydrogenase 2 small subunit [Salmonella enterica] Length: 372\nNumber of predicted TMHs: 1\nExp number of AAs in TMHs: 24.51309\nExp number, first 60 AAs: 0.2682\nTotal prob of N-in: 0.14402\noutside 1 331\nTMhelix 332 354\ninside 355 372

17507 GCF_001243605.1_Salmonella_enterica_CVM_N46854_v1.0 Salmonella enterica subsp. enterica serovar Hadar Proteobacteria; Gammaproteobacteria; Enterobacterales; Enterobacteriaceae; Salmonella; Salmonella enterica; Salmonella enterica subsp. enterica MNNEETFYQAMRRKGVTRRSFLKFCSLAATSLGLGAGMTPKIAWALENKPRIPVVWIHGL WP_001058308.1 hydrogenase [Salmonella enterica] Length: 372\nNumber of predicted TMHs: 2\nExp number of AAs in TMHs: 39.08204\nExp number, first 60 AAs: 19.72474\nTotal prob of N-in: 0.96174\nPOSSIBLE N-term signal sequence\ninside 1 19\nTMhelix 20 39\noutside 40 325\nTMhelix 326 348\ninside 349 372

17508 GCF_001246485.1_Salmonella_enterica_CVM_N51265_v1.0 Salmonella enterica subsp. enterica serovar Hadar Proteobacteria; Gammaproteobacteria; Enterobacterales; Enterobacteriaceae; Salmonella; Salmonella enterica; Salmonella enterica subsp. enterica MNNEETFYQAMRRKGVTRRSFLKFCSLAATSLGLGAGMTPKIAWALENKPRIPVVWIHGL WP_001058308.1 hydrogenase [Salmonella enterica] Length: 372\nNumber of predicted TMHs: 2\nExp number of AAs in TMHs: 39.08204\nExp number, first 60 AAs: 19.72474\nTotal prob of N-in: 0.96174\nPOSSIBLE N-term signal sequence\ninside 1 19\nTMhelix 20 39\noutside 40 325\nTMhelix 326 348\ninside 349 372

17509 GCF_001246475.1_Salmonella_enterica_CVM_N51263_v1.0 Salmonella enterica subsp. enterica serovar Hadar Proteobacteria; Gammaproteobacteria; Enterobacterales; Enterobacteriaceae; Salmonella; Salmonella enterica;

Salmonella enterica subsp. enterica MNNEETFYQAMRRKGVTRRSFLKFCSLAATSLGLGAGMTPKIAWALENKPRIPVVWIHGL WP_001058308.1 hydrogenase [Salmonella enterica] Length: 372\nNumber of predicted TMHs: 2\nExp number of AAs in TMHs: 39.08204\nExp number, first 60 AAs: 19.72474\nTotal prob of N-in: 0.96174\nPOSSIBLE N-term signal sequence\ninside 1 19\nTMhelix 20 39\noutside 40 325\nTMhelix 326 348\ninside 349 372

17510 GCF_000973775.1_ABB1048-1.id15_v1.0 Salmonella enterica subsp. enterica serovar Hadar Proteobacteria; Gammaproteobacteria; Enterobacterales; Enterobacteriaceae; Salmonella; Salmonella enterica; Salmonella enterica subsp. enterica MNNEETFYQAMRRKGVTRRSFLKFCSLAATSLGLGAGMTPKIAWALENKPRIPVVWIHGL WP_001058308.1 hydrogenase [Salmonella enterica] Length: 372\nNumber of predicted TMHs: 2\nExp number of AAs in TMHs: 39.08204\nExp number, first 60 AAs: 19.72474\nTotal prob of N-in: 0.96174\nPOSSIBLE N-term signal sequence\ninside 1 19\nTMhelix 20 39\noutside 40 325\nTMhelix 326 348\ninside 349 372

17511 GCF_001477835.1_Salmonella_enterica_CVM_N29339-R_v1.0 Salmonella enterica subsp. enterica serovar Hadar Proteobacteria; Gammaproteobacteria; Enterobacterales; Enterobacteriaceae; Salmonella; Salmonella enterica; Salmonella enterica subsp. enterica MTGDNTLITSHGINRRDFMKLCAALAATMGLSSKAAAEMAESVSNPQRPPVIWIGAQECT WP_000145429.1 hydrogenase 2 small subunit [Salmonella enterica] Length: 372\nNumber of predicted TMHs: 1\nExp number of AAs in TMHs: 24.51309\nExp number, first 60 AAs: 0.2682\nTotal prob of N-in: 0.14402\noutside 1 331\nTMhelix 332 354\ninside 355 372

17512 GCF_000973775.1_ABB1048-1.id15_v1.0 Salmonella enterica subsp. enterica serovar Hadar Proteobacteria; Gammaproteobacteria; Enterobacterales; Enterobacteriaceae; Salmonella; Salmonella enterica; Salmonella enterica subsp. enterica MTGDNTLITSHGINRRDFMKLCAALAATMGLSSKAAAEMAESVSNPQRPPVIWIGAQECT WP_000145429.1 hydrogenase 2 small subunit [Salmonella enterica] Length: 372\nNumber of predicted TMHs: 1\nExp number of AAs in TMHs: 24.51309\nExp number, first 60 AAs: 0.2682\nTotal prob of N-in: 0.14402\noutside 1 331\nTMhelix 332 354\ninside 355 372

17513 GCF_002031825.1_ASM203182v1 Salmonella enterica subsp. enterica serovar Hadar Proteobacteria; Gammaproteobacteria; Enterobacterales; Enterobacteriaceae; Salmonella; Salmonella enterica; Salmonella enterica subsp. enterica MNNEETFYQAMRRKGVTRRSFLKFCSLAATSLGLGAGMTPKIAWALENKPRIPVVWIHGL WP_001058308.1 hydrogenase [Salmonella enterica] Length: 372\nNumber of predicted TMHs: 2\nExp number of AAs in TMHs: 39.08204\nExp number, first 60 AAs: 19.72474\nTotal prob of N-in: 0.96174\nPOSSIBLE N-term signal sequence\ninside 1 19\nTMhelix 20 39\noutside 40 325\nTMhelix 326 348\ninside 349 372

17514 GCF_002033235.1_ASM203323v1 Salmonella enterica subsp. enterica serovar Hadar Proteobacteria; Gammaproteobacteria; Enterobacterales; Enterobacteriaceae; Salmonella; Salmonella enterica; Salmonella enterica subsp. enterica MNNEETFYQAMRRKGVTRRSFLKFCSLAATSLGLGAGMTPKIAWALENKPRIPVVWIHGL WP_001058308.1 hydrogenase [Salmonella enterica] Length: 372\nNumber of predicted TMHs: 2\nExp number of AAs in TMHs: 39.08204\nExp number, first 60 AAs: 19.72474\nTotal prob of N-in: 0.96174\nPOSSIBLE N-term signal sequence\ninside 1 19\nTMhelix 20 39\noutside 40 325\nTMhelix 326 348\ninside 349 372

17515 GCF_002062815.1_ASM206281v1 Salmonella enterica subsp. enterica serovar Hadar Proteobacteria; Gammaproteobacteria; Enterobacterales; Enterobacteriaceae; Salmonella; Salmonella enterica; Salmonella enterica subsp. enterica MTGDNTLITSHGINRRDFMKLCAALAATMGLSSKAAAEMAESVSNPQRPPVIWIGAQECT WP_000145429.1 hydrogenase 2 small subunit [Salmonella enterica] Length: 372\nNumber of predicted TMHs: 1\nExp number of AAs in TMHs: 24.51309\nExp number, first 60 AAs: 0.2682\nTotal prob of N-in: 0.14402\noutside 1 331\nTMhelix 332 354\ninside 355 372

17516 GCF_002036665.1_ASM203666v1 Salmonella enterica subsp. enterica serovar Hadar Proteobacteria; Gammaproteobacteria; Enterobacterales; Enterobacteriaceae; Salmonella; Salmonella enterica; Salmonella enterica subsp. enterica MNNEETFYQAMRRKGVTRRSFLKFCSLAATSLGLGAGMTPKIAWALENKPRIPVVWIHGL WP_001058308.1 hydrogenase [Salmonella enterica] Length: 372\nNumber of predicted TMHs: 2\nExp number of AAs in TMHs: 39.08204\nExp number, first 60 AAs: 19.72474\nTotal prob of N-in: 0.96174\nPOSSIBLE N-term signal sequence\ninside 1 19\nTMhelix 20 39\noutside 40 325\nTMhelix 326 348\ninside 349 372

17517 GCF_002062815.1_ASM206281v1 Salmonella enterica subsp. enterica serovar Hadar Proteobacteria; Gammaproteobacteria; Enterobacterales; Enterobacteriaceae; Salmonella; Salmonella enterica; Salmonella enterica subsp. enterica MNNEETFYQAMRRKGVTRRSFLKFCSLAATSLGLGAGMTPKIAWALENKPRIPVVWIHGL WP_001058308.1 hydrogenase [Salmonella enterica] Length: 372\nNumber of predicted TMHs: 2\nExp number of AAs in TMHs: 39.08204\nExp number, first 60 AAs: 19.72474\nTotal prob of N-in: 0.96174\nPOSSIBLE N-term signal sequence\ninside 1 19\nTMhelix 20 39\noutside 40 325\nTMhelix 326 348\ninside 349 372

17518 GCF_002030665.1_ASM203066v1 Salmonella enterica subsp. enterica serovar Hadar Proteobacteria; Gammaproteobacteria; Enterobacterales; Enterobacteriaceae; Salmonella; Salmonella enterica; Salmonella enterica subsp. enterica MNNEETFYQAMRRKGVTRRSFLKFCSLAATSLGLGAGMTPKIAWALENKPRIPVVWIHGL WP_001058308.1 hydrogenase [Salmonella enterica] Length: 372\nNumber of predicted TMHs: 2\nExp number of AAs in TMHs: 39.08204\nExp number, first 60 AAs: 19.72474\nTotal prob of N-in: 0.96174\nPOSSIBLE N-term signal sequence\ninside 1 19\nTMhelix 20 39\noutside 40 325\nTMhelix 326 348\ninside 349 372

17519 GCF_002030665.1_ASM203066v1 Salmonella enterica subsp. enterica serovar Hadar Proteobacteria; Gammaproteobacteria; Enterobacterales; Enterobacteriaceae; Salmonella; Salmonella enterica; Salmonella enterica subsp. enterica MTGDNTLITSHGINRRDFMKLCAALAATMGLSSKAAAEMAESVSNPQRPPVIWIGAQECT WP_000145429.1 hydrogenase 2 small subunit [Salmonella enterica] Length: 372\nNumber of predicted TMHs: 1\nExp number of AAs in TMHs: 24.51309\nExp number, first 60 AAs: 0.2682\nTotal prob of N-in: 0.14402\noutside 1 331\nTMhelix 332 354\ninside 355 372

17520 GCF_002031055.1_ASM203105v1 Salmonella enterica subsp. enterica serovar Hadar Proteobacteria; Gammaproteobacteria; Enterobacterales; Enterobacteriaceae; Salmonella; Salmonella enterica; Salmonella enterica subsp. enterica MTGDNTLITSHGINRRDFMKLCAALAATMGLSSKAAAEMAESVSNPQRPPVIWIGAQECT WP_000145429.1 hydrogenase 2 small subunit [Salmonella enterica] Length: 372\nNumber of predicted TMHs: 1\nExp number of AAs in TMHs: 24.51309\nExp number, first 60 AAs: 0.2682\nTotal prob of N-in: 0.14402\noutside 1 331\nTMhelix 332 354\ninside 355 372

17521 GCF_002058685.1_ASM205868v1 Salmonella enterica subsp. enterica serovar Hadar Proteobacteria; Gammaproteobacteria; Enterobacterales; Enterobacteriaceae; Salmonella; Salmonella enterica; Salmonella enterica subsp. enterica MNNEETFYQAMRRKGVTRRSFLKFCSLAATSLGLGAGMTPKIAWALENKPRIPVVWIHGL WP_001058308.1 hydrogenase [Salmonella enterica] Length: 372\nNumber of predicted TMHs: 2\nExp number of AAs in TMHs: 39.08204\nExp number, first 60 AAs: 19.72474\nTotal prob of N-in: 0.96174\nPOSSIBLE N-term signal sequence\ninside 1 19\nTMhelix 20 39\noutside 40 325\nTMhelix 326 348\ninside 349 372

17522 GCF_000974135.1_ABBSB1121-1.id17_v1.0 Salmonella enterica subsp. enterica serovar Hadar Proteobacteria; Gammaproteobacteria; Enterobacterales; Enterobacteriaceae; Salmonella; Salmonella enterica; Salmonella enterica subsp. enterica MTGDNTLITSHGINRRDFMKLCAALAATMGLSSKAAAEMAESVSNPQRPPVIWIGAQECT WP_000145429.1 hydrogenase 2 small subunit [Salmonella enterica] Length: 372\nNumber of predicted TMHs: 1\nExp number of AAs in TMHs: 24.51309\nExp number, first 60 AAs: 0.2682\nTotal prob of N-in: 0.14402\noutside 1 331\nTMhelix 332 354\ninside 355 372

17523 GCF_000974135.1_ABBSB1121-1.id17_v1.0 Salmonella enterica subsp. enterica serovar Hadar Proteobacteria; Gammaproteobacteria; Enterobacterales; Enterobacteriaceae; Salmonella; Salmonella enterica; Salmonella enterica subsp. enterica MNNEETFYQAMRRKGVTRRSFLKFCSLAATSLGLGAGMTPKIAWALENKPRIPVVWIHGL WP_001058308.1 hydrogenase [Salmonella enterica] Length: 372\nNumber of predicted TMHs: 2\nExp number of AAs in TMHs: 39.08204\nExp number, first 60 AAs: 19.72474\nTotal prob of N-in: 0.96174\nPOSSIBLE N-term signal sequence\ninside 1 19\nTMhelix 20 39\noutside 40 325\nTMhelix 326 348\ninside 349 372

17524 GCF_001479865.1_Salmonella_enterica_CVM_N38861-SQ_v1.0 Salmonella enterica subsp. enterica serovar Hadar Proteobacteria; Gammaproteobacteria; Enterobacterales; Enterobacteriaceae; Salmonella; Salmonella enterica; Salmonella enterica subsp. enterica MNNEETFYQAMRRKGVTRRSFLKFCSLAATSLGLGAGMTPKIAWALENKPRIPVVWIHGL WP_001058308.1 hydrogenase [Salmonella enterica] Length: 372\nNumber of predicted TMHs: 2\nExp number of AAs in TMHs: 39.08204\nExp number, first 60 AAs: 19.72474\nTotal prob of N-in: 0.96174\nPOSSIBLE N-term signal sequence\ninside 1 19\nTMhelix 20 39\noutside 40 325\nTMhelix 326 348\ninside 349 372

17525 GCF_001479865.1_Salmonella_enterica_CVM_N38861-SQ_v1.0 Salmonella enterica subsp. enterica serovar Hadar Proteobacteria; Gammaproteobacteria; Enterobacterales; Enterobacteriaceae; Salmonella; Salmonella enterica; Salmonella enterica subsp. enterica MTGDNTLITSHGINRRDFMKLCAALAATMGLSSKAAAEMAESVSNPQRPPVIWIGAQECT WP_000145429.1 hydrogenase 2 small subunit [Salmonella enterica] Length: 372\nNumber of predicted TMHs: 1\nExp number of AAs in TMHs: 24.51309\nExp number, first 60 AAs: 0.2682\nTotal prob of N-in: 0.14402\noutside 1 331\nTMhelix 332 354\ninside 355 372

17526 GCF_001479605.1_Salmonella_enterica_CVM_N37910-SQ_v1.0 Salmonella enterica subsp. enterica serovar Hadar Proteobacteria; Gammaproteobacteria; Enterobacterales; Enterobacteriaceae; Salmonella; Salmonella enterica; Salmonella enterica subsp. enterica MNNEETFYQAMRRKGVTRRSFLKFCSLAATSLGLGAGMTPKIAWALENKPRIPVVWIHGL WP_001058308.1 hydrogenase [Salmonella enterica] Length: 372\nNumber of predicted TMHs: 2\nExp

number of AAs in TMHs: 39.08204\nExp number, first 60 AAs: 19.72474\nTotal prob of N-in: 0.96174\nPOSSIBLE N-term signal sequence\ninside 1 19\nTMhelix 20 39\noutside 40 325\nTMhelix 326 348\ninside 349 372

17527 GCF_001479605.1_Salmonella_enterica_CVM_N37910-SQ_v1.0 Salmonella enterica subsp. enterica serovar Hadar Proteobacteria; Gammaproteobacteria; Enterobacterales; Enterobacteriaceae; Salmonella; Salmonella enterica; Salmonella enterica subsp. enterica MTGDNTLITSHGINRRDFMKLCAALAATMGLSSKAAAEMAESVSNPQRPPVIWIGAQECT WP_000145429.1 hydrogenase 2 small subunit [Salmonella enterica] Length: 372\nNumber of predicted TMHs: 1\nExp number of AAs in TMHs: 24.51309\nExp number, first 60 AAs: 0.2682\nTotal prob of N-in: 0.14402\noutside 1 331\nTMhelix 332 354\ninside 355 372

17528 GCF_002035955.1_ASM203595v1 Salmonella enterica subsp. enterica serovar Hadar Proteobacteria; Gammaproteobacteria; Enterobacterales; Enterobacteriaceae; Salmonella; Salmonella enterica; Salmonella enterica subsp. enterica MTGDNTLITSHGINRRDFMKLCAALAATMGLSSKAAAEMAESVSNPQRPPVIWIGAQECT WP_000145429.1 hydrogenase 2 small subunit [Salmonella enterica] Length: 372\nNumber of predicted TMHs: 1\nExp number of AAs in TMHs: 24.51309\nExp number, first 60 AAs: 0.2682\nTotal prob of N-in: 0.14402\noutside 1 331\nTMhelix 332 354\ninside 355 372

17529 GCF_002035955.1_ASM203595v1 Salmonella enterica subsp. enterica serovar Hadar Proteobacteria; Gammaproteobacteria; Enterobacterales; Enterobacteriaceae; Salmonella; Salmonella enterica; Salmonella enterica subsp. enterica MNNEETFYQAMRRKGVTRRSFLKFCSLAATSLGLGAGMTPKIAWALENKPRIPVVWIHGL WP_001058308.1 hydrogenase [Salmonella enterica] Length: 372\nNumber of predicted TMHs: 2\nExp number of AAs in TMHs: 39.08204\nExp number, first 60 AAs: 19.72474\nTotal prob of N-in: 0.96174\nPOSSIBLE N-term signal sequence\ninside 1 19\nTMhelix 20 39\noutside 40 325\nTMhelix 326 348\ninside 349 372

17530 GCF_001480605.1_Salmonella_enterica_CVM_N42468_v1.0 Salmonella enterica subsp. enterica serovar Hadar Proteobacteria; Gammaproteobacteria; Enterobacterales; Enterobacteriaceae; Salmonella; Salmonella enterica; Salmonella enterica subsp. enterica MTGDNTLITSHGINRRDFMKLCAALAATMGLSSKAAAEMAESVSNPQRPPVIWIGAQECT WP_000145429.1 hydrogenase 2 small subunit [Salmonella enterica] Length: 372\nNumber of predicted TMHs: 1\nExp number of AAs in TMHs: 24.51309\nExp number, first 60 AAs: 0.2682\nTotal prob of N-in: 0.14402\noutside 1 331\nTMhelix 332 354\ninside 355 372

17531 GCF_002036665.1_ASM203666v1 Salmonella enterica subsp. enterica serovar Hadar Proteobacteria; Gammaproteobacteria; Enterobacterales; Enterobacteriaceae; Salmonella; Salmonella enterica; Salmonella enterica subsp. enterica MTGDNTLITSHGINRRDFMKLCAALAATMGLSSKAAAEMAESVSNPQRPPVIWIGAQECT WP_000145429.1 hydrogenase 2 small subunit [Salmonella enterica] Length: 372\nNumber of predicted TMHs: 1\nExp number of AAs in TMHs: 24.51309\nExp number, first 60 AAs: 0.2682\nTotal prob of N-in: 0.14402\noutside 1 331\nTMhelix 332 354\ninside 355 372

17532 GCF_001480605.1_Salmonella_enterica_CVM_N42468_v1.0 Salmonella enterica subsp. enterica serovar Hadar Proteobacteria; Gammaproteobacteria; Enterobacterales; Enterobacteriaceae; Salmonella; Salmonella enterica; Salmonella enterica subsp. enterica MNNEETFYQAMRRKGVTRRSFLKFCSLAATSLGLGAGMTPKIAWALENKPRIPVVWIHGL WP_001058308.1 hydrogenase [Salmonella enterica] Length: 372\nNumber of predicted TMHs: 2\nExp number of AAs in TMHs: 39.08204\nExp number, first 60 AAs: 19.72474\nTotal prob of N-in: 0.96174\nPOSSIBLE N-term signal sequence\ninside 1 19\nTMhelix 20 39\noutside 40 325\nTMhelix 326 348\ninside 349 372

17533 GCF_002031055.1_ASM203105v1 Salmonella enterica subsp. enterica serovar Hadar Proteobacteria; Gammaproteobacteria; Enterobacterales; Enterobacteriaceae; Salmonella; Salmonella enterica; Salmonella enterica subsp. enterica MNNEETFYQAMRRKGVTRRSFLKFCSLAATSLGLGAGMTPKIAWALENKPRIPVVWIHGL WP_001058308.1 hydrogenase [Salmonella enterica] Length: 372\nNumber of predicted TMHs: 2\nExp number of AAs in TMHs: 39.08204\nExp number, first 60 AAs: 19.72474\nTotal prob of N-in: 0.96174\nPOSSIBLE N-term signal sequence\ninside 1 19\nTMhelix 20 39\noutside 40 325\nTMhelix 326 348\ninside 349 372

17534 GCF_001478535.1_Salmonella_enterica_CVM_N41909-SQ_v1.0 Salmonella enterica subsp. enterica serovar Hadar Proteobacteria; Gammaproteobacteria; Enterobacterales; Enterobacteriaceae; Salmonella; Salmonella enterica; Salmonella enterica subsp. enterica MNNEETFYQAMRRKGVTRRSFLKFCSLAATSLGLGAGMTPKIAWALENKPRIPVVWIHGL WP_001058308.1 hydrogenase [Salmonella enterica] Length: 372\nNumber of predicted TMHs: 2\nExp number of AAs in TMHs: 39.08204\nExp number, first 60 AAs: 19.72474\nTotal prob of N-in: 0.96174\nPOSSIBLE N-term signal sequence\ninside 1 19\nTMhelix 20 39\noutside 40 325\nTMhelix 326 348\ninside 349 372

17535 GCF_001478535.1_Salmonella_enterica_CVM_N41909-SQ_v1.0 Salmonella enterica subsp. enterica serovar Hadar Proteobacteria; Gammaproteobacteria; Enterobacterales; Enterobacteriaceae; Salmonella; Salmonella enterica;

Salmonella enterica subsp. enterica MTGDNTLITSHGINRRDFMKLCAALAATMGLSSKAAAEMAESVSNPQRPPVIWIGAQECT WP_000145429.1 hydrogenase 2 small subunit [Salmonella enterica] Length: 372\nNumber of predicted TMHs: 1\nExp number of AAs in TMHs: 24.51309\nExp number, first 60 AAs: 0.2682\nTotal prob of N-in: 0.14402\noutside 1 331\nTMhelix 332 354\ninside 355 372

17536 GCF_001477965.1_Salmonella_enterica_CVM_N30645-SQ_v1.0 Salmonella enterica subsp. enterica serovar Hadar Proteobacteria; Gammaproteobacteria; Enterobacterales; Enterobacteriaceae; Salmonella; Salmonella enterica; Salmonella enterica subsp. enterica MNNEETFYQAMRRKGVTRRSFLKFCSLAATSLGLGAGMTPKIAWALENKPRIPVVWIHGL WP_001058308.1 hydrogenase [Salmonella enterica] Length: 372\nNumber of predicted TMHs: 2\nExp number of AAs in TMHs: 39.08204\nExp number, first 60 AAs: 19.72474\nTotal prob of N-in: 0.96174\nPOSSIBLE N-term signal sequence\ninside 1 19\nTMhelix 20 39\noutside 40 325\nTMhelix 326 348\ninside 349 372

17537 GCF_001477965.1_Salmonella_enterica_CVM_N30645-SQ_v1.0 Salmonella enterica subsp. enterica serovar Hadar Proteobacteria; Gammaproteobacteria; Enterobacterales; Enterobacteriaceae; Salmonella; Salmonella enterica; Salmonella enterica subsp. enterica MTGDNTLITSHGINRRDFMKLCAALAATMGLSSKAAAEMAESVSNPQRPPVIWIGAQECT WP_000145429.1 hydrogenase 2 small subunit [Salmonella enterica] Length: 372\nNumber of predicted TMHs: 1\nExp number of AAs in TMHs: 24.51309\nExp number, first 60 AAs: 0.2682\nTotal prob of N-in: 0.14402\noutside 1 331\nTMhelix 332 354\ninside 355 372

17538 GCF_001477755.1_Salmonella_enterica_CVM_N29317-SQ_v1.0 Salmonella enterica subsp. enterica serovar Hadar Proteobacteria; Gammaproteobacteria; Enterobacterales; Enterobacteriaceae; Salmonella; Salmonella enterica; Salmonella enterica subsp. enterica MNNEETFYQAMRRKGVTRRSFLKFCSLAATSLGLGAGMTPKIAWALENKPRIPVVWIHGL WP_001058308.1 hydrogenase [Salmonella enterica] Length: 372\nNumber of predicted TMHs: 2\nExp number of AAs in TMHs: 39.08204\nExp number, first 60 AAs: 19.72474\nTotal prob of N-in: 0.96174\nPOSSIBLE N-term signal sequence\ninside 1 19\nTMhelix 20 39\noutside 40 325\nTMhelix 326 348\ninside 349 372

17539 GCF_001477755.1_Salmonella_enterica_CVM_N29317-SQ_v1.0 Salmonella enterica subsp. enterica serovar Hadar Proteobacteria; Gammaproteobacteria; Enterobacterales; Enterobacteriaceae; Salmonella; Salmonella enterica; Salmonella enterica subsp. enterica MTGDNTLITSHGINRRDFMKLCAALAATMGLSSKAAAEMAESVSNPQRPPVIWIGAQECT WP_000145429.1 hydrogenase 2 small subunit [Salmonella enterica] Length: 372\nNumber of predicted TMHs: 1\nExp number of AAs in TMHs: 24.51309\nExp number, first 60 AAs: 0.2682\nTotal prob of N-in: 0.14402\noutside 1 331\nTMhelix 332 354\ninside 355 372

17540 GCF_002061195.1_ASM206119v1 Salmonella enterica subsp. enterica serovar Hadar Proteobacteria; Gammaproteobacteria; Enterobacterales; Enterobacteriaceae; Salmonella; Salmonella enterica; Salmonella enterica subsp. enterica MNNEETFYQAMRRKGVTRRSFLKFCSLAATSLGLGAGMTPKIAWALENKPRIPVVWIHGL WP_001058308.1 hydrogenase [Salmonella enterica] Length: 372\nNumber of predicted TMHs: 2\nExp number of AAs in TMHs: 39.08204\nExp number, first 60 AAs: 19.72474\nTotal prob of N-in: 0.96174\nPOSSIBLE N-term signal sequence\ninside 1 19\nTMhelix 20 39\noutside 40 325\nTMhelix 326 348\ninside 349 372

17541 GCF_002061195.1_ASM206119v1 Salmonella enterica subsp. enterica serovar Hadar Proteobacteria; Gammaproteobacteria; Enterobacterales; Enterobacteriaceae; Salmonella; Salmonella enterica; Salmonella enterica subsp. enterica MTGDNTLITSHGINRRDFMKLCAALAATMGLSSKAAAEMAESVSNPQRPPVIWIGAQECT WP_000145429.1 hydrogenase 2 small subunit [Salmonella enterica] Length: 372\nNumber of predicted TMHs: 1\nExp number of AAs in TMHs: 24.51309\nExp number, first 60 AAs: 0.2682\nTotal prob of N-in: 0.14402\noutside 1 331\nTMhelix 332 354\ninside 355 372

17542 GCF_002058685.1_ASM205868v1 Salmonella enterica subsp. enterica serovar Hadar Proteobacteria; Gammaproteobacteria; Enterobacterales; Enterobacteriaceae; Salmonella; Salmonella enterica; Salmonella enterica subsp. enterica MTGDNTLITSHGINRRDFMKLCAALAATMGLSSKAAAEMAESVSNPQRPPVIWIGAQECT WP_000145429.1 hydrogenase 2 small subunit [Salmonella enterica] Length: 372\nNumber of predicted TMHs: 1\nExp number of AAs in TMHs: 24.51309\nExp number, first 60 AAs: 0.2682\nTotal prob of N-in: 0.14402\noutside 1 331\nTMhelix 332 354\ninside 355 372

17543 GCF_000973745.1_ABBSB1020-2.id16_v1.0 Salmonella enterica subsp. enterica serovar Hadar Proteobacteria; Gammaproteobacteria; Enterobacterales; Enterobacteriaceae; Salmonella; Salmonella enterica; Salmonella enterica subsp. enterica MTGDNTLITSHGINRRDFMKLCAALAATMGLSSKAAAEMAESVSNPQRPPVIWIGAQECT WP_000145429.1 hydrogenase 2 small subunit [Salmonella enterica] Length: 372\nNumber of predicted TMHs: 1\nExp number of AAs in TMHs: 24.51309\nExp number, first 60 AAs: 0.2682\nTotal prob of N-in: 0.14402\noutside 1 331\nTMhelix 332 354\ninside 355 372

17544 GCF_001477835.1_Salmonella_enterica_CVM_N29339-R_v1.0 Salmonella enterica subsp. enterica serovar Hadar Proteobacteria; Gammaproteobacteria; Enterobacterales; Enterobacteriaceae; Salmonella; Salmonella enterica; Salmonella enterica subsp. enterica MNNEETFYQAMRRKGVTRRSFLKFCSLAATSLGLGAGMTPKIAWALENKPRIPVVWIHGL WP_001058308.1 hydrogenase [Salmonella enterica] Length: 372\nNumber of predicted TMHs: 2\nExp number of AAs in TMHs: 39.08204\nExp number, first 60 AAs: 19.72474\nTotal prob of N-in: 0.96174\nPOSSIBLE N-term signal sequence\ninside 1 19\nTMhelix 20 39\noutside 40 325\nTMhelix 326 348\ninside 349 372

17545 GCF_001243705.1_Salmonella_enterica_CVM_N46859_v1.0 Salmonella enterica subsp. enterica serovar Hadar Proteobacteria; Gammaproteobacteria; Enterobacterales; Enterobacteriaceae; Salmonella; Salmonella enterica; Salmonella enterica subsp. enterica MNNEETFYQAMRRKGVTRRSFLKFCSLAATSLGLGAGMTPKIAWALENKPRIPVVWIHGL WP_001058308.1 hydrogenase [Salmonella enterica] Length: 372\nNumber of predicted TMHs: 2\nExp number of AAs in TMHs: 39.08204\nExp number, first 60 AAs: 19.72474\nTotal prob of N-in: 0.96174\nPOSSIBLE N-term signal sequence\ninside 1 19\nTMhelix 20 39\noutside 40 325\nTMhelix 326 348\ninside 349 372

17546 GCF_002031825.1_ASM203182v1 Salmonella enterica subsp. enterica serovar Hadar Proteobacteria; Gammaproteobacteria; Enterobacterales; Enterobacteriaceae; Salmonella; Salmonella enterica; Salmonella enterica subsp. enterica MTGDNTLITSHGINRRDFMKLCAALAATMGLSSKAAAEMAESVSNPQRPPVIWIGAQECT WP_000145429.1 hydrogenase 2 small subunit [Salmonella enterica] Length: 372\nNumber of predicted TMHs: 1\nExp number of AAs in TMHs: 24.51309\nExp number, first 60 AAs: 0.2682\nTotal prob of N-in: 0.14402\noutside 1 331\nTMhelix 332 354\ninside 355 372

17547 GCF_001241115.1_Salmonella_enterica_CVM_N44694_v1.0 Salmonella enterica subsp. enterica serovar Hadar Proteobacteria; Gammaproteobacteria; Enterobacterales; Enterobacteriaceae; Salmonella; Salmonella enterica; Salmonella enterica subsp. enterica MTGDNTLITSHGINRRDFMKLCAALAATMGLSSKAAAEMAESVSNPQRPPVIWIGAQECT WP_000145429.1 hydrogenase 2 small subunit [Salmonella enterica] Length: 372\nNumber of predicted TMHs: 1\nExp number of AAs in TMHs: 24.51309\nExp number, first 60 AAs: 0.2682\nTotal prob of N-in: 0.14402\noutside 1 331\nTMhelix 332 354\ninside 355 372

17548 GCF_001243605.1_Salmonella_enterica_CVM_N46854_v1.0 Salmonella enterica subsp. enterica serovar Hadar Proteobacteria; Gammaproteobacteria; Enterobacterales; Enterobacteriaceae; Salmonella; Salmonella enterica; Salmonella enterica subsp. enterica MTGDNTLITSHGINRRDFMKLCAALAATMGLSSKAAAEMAESVSNPQRPPVIWIGAQECT WP_000145429.1 hydrogenase 2 small subunit [Salmonella enterica] Length: 372\nNumber of predicted TMHs: 1\nExp number of AAs in TMHs: 24.51309\nExp number, first 60 AAs: 0.2682\nTotal prob of N-in: 0.14402\noutside 1 331\nTMhelix 332 354\ninside 355 372

17549 GCF_001241115.1_Salmonella_enterica_CVM_N44694_v1.0 Salmonella enterica subsp. enterica serovar Hadar Proteobacteria; Gammaproteobacteria; Enterobacterales; Enterobacteriaceae; Salmonella; Salmonella enterica; Salmonella enterica subsp. enterica MNNEETFYQAMRRKGVTRRSFLKFCSLAATSLGLGAGMTPKIAWALENKPRIPVVWIHGL WP_001058308.1 hydrogenase [Salmonella enterica] Length: 372\nNumber of predicted TMHs: 2\nExp number of AAs in TMHs: 39.08204\nExp number, first 60 AAs: 19.72474\nTotal prob of N-in: 0.96174\nPOSSIBLE N-term signal sequence\ninside 1 19\nTMhelix 20 39\noutside 40 325\nTMhelix 326 348\ninside 349 372

17550 GCF_002032945.1_ASM203294v1 Salmonella enterica subsp. enterica serovar Hadar Proteobacteria; Gammaproteobacteria; Enterobacterales; Enterobacteriaceae; Salmonella; Salmonella enterica; Salmonella enterica subsp. enterica MNNEETFYQAMRRKGVTRRSFLKFCSLAATSLGLGAGMTPKIAWALENKPRIPVVWIHGL WP_001058308.1 hydrogenase [Salmonella enterica] Length: 372\nNumber of predicted TMHs: 2\nExp number of AAs in TMHs: 39.08204\nExp number, first 60 AAs: 19.72474\nTotal prob of N-in: 0.96174\nPOSSIBLE N-term signal sequence\ninside 1 19\nTMhelix 20 39\noutside 40 325\nTMhelix 326 348\ninside 349 372

17551 GCF_001245215.1_Salmonella_enterica_CVM_N48712_v1.0 Salmonella enterica subsp. enterica serovar Hadar Proteobacteria; Gammaproteobacteria; Enterobacterales; Enterobacteriaceae; Salmonella; Salmonella enterica; Salmonella enterica subsp. enterica MTGDNTLITSHGINRRDFMKLCAALAATMGLSSKAAAEMAESVSNPQRPPVIWIGAQECT WP_000145429.1 hydrogenase 2 small subunit [Salmonella enterica] Length: 372\nNumber of predicted TMHs: 1\nExp number of AAs in TMHs: 24.51309\nExp number, first 60 AAs: 0.2682\nTotal prob of N-in: 0.14402\noutside 1 331\nTMhelix 332 354\ninside 355 372

17552 GCF_002065675.1_ASM206567v1 Salmonella enterica subsp. enterica serovar Haifa Proteobacteria; Gammaproteobacteria; Enterobacterales; Enterobacteriaceae; Salmonella; Salmonella enterica; Salmonella enterica subsp. enterica MTGDNTLITSHGINRRDFMKLCAALAATMGLSSKAAAEMAESVSNPQRPPVIWIGAQECT WP_000145429.1 hydrogenase 2 small subunit [Salmonella enterica] Length: 372\nNumber of

predicted TMHs: 1\nExp number of AAs in TMHs: 24.51309\nExp number, first 60 AAs: 0.2682\nTotal prob of N-in: 0.14402\noutside 1 331\nTMhelix 332 354\ninside 355 372

17553 GCF_002065675.1_ASM206567v1 Salmonella enterica subsp. enterica serovar Haifa Proteobacteria; Gammaproteobacteria; Enterobacterales; Enterobacteriaceae; Salmonella; Salmonella enterica; Salmonella enterica subsp. enterica MNNEETFYQAMRRKGVTRRSFLKFCSLAATSLGLGAGMTPKIAWALENKPRIPVVWIHGL WP_001058312.1 hydrogenase [Salmonella enterica] Length: 372\nNumber of predicted TMHs: 2\nExp number of AAs in TMHs: 36.51738\nExp number, first 60 AAs: 19.72849\nTotal prob of N-in: 0.96170\nPOSSIBLE N-term signal sequence\ninside 1 19\nTMhelix 20 39\noutside 40 325\nTMhelix 326 348\ninside 349 372

17554 GCF_002065015.1_ASM206501v1 Salmonella enterica subsp. enterica serovar Haifa Proteobacteria; Gammaproteobacteria; Enterobacterales; Enterobacteriaceae; Salmonella; Salmonella enterica; Salmonella enterica subsp. enterica MTGDNTLITSHGINRRDFMKLCAALAATMGLSSKAAAEMAESVSNPQRPPVIWIGAQECT WP_000145429.1 hydrogenase 2 small subunit [Salmonella enterica] Length: 372\nNumber of predicted TMHs: 1\nExp number of AAs in TMHs: 24.51309\nExp number, first 60 AAs: 0.2682\nTotal prob of N-in: 0.14402\noutside 1 331\nTMhelix 332 354\ninside 355 372

17555 GCF_002065015.1_ASM206501v1 Salmonella enterica subsp. enterica serovar Haifa Proteobacteria; Gammaproteobacteria; Enterobacterales; Enterobacteriaceae; Salmonella; Salmonella enterica; Salmonella enterica subsp. enterica MNNEETFYQAMRRKGVTRRSFLKFCSLAATSLGLGAGMTPKIAWALENKPRIPVVWIHGL WP_001058312.1 hydrogenase [Salmonella enterica] Length: 372\nNumber of predicted TMHs: 2\nExp number of AAs in TMHs: 36.51738\nExp number, first 60 AAs: 19.72849\nTotal prob of N-in: 0.96170\nPOSSIBLE N-term signal sequence\ninside 1 19\nTMhelix 20 39\noutside 40 325\nTMhelix 326 348\ninside 349 372

17556 GCF_002061155.1_ASM206115v1 Salmonella enterica subsp. enterica serovar Haifa Proteobacteria; Gammaproteobacteria; Enterobacterales; Enterobacteriaceae; Salmonella; Salmonella enterica; Salmonella enterica subsp. enterica MTGDNTLITSHGINRRDFMKLCAALAATMGLSSKAAAEMAESVSNPQRPPVIWIGAQECT WP_000145429.1 hydrogenase 2 small subunit [Salmonella enterica] Length: 372\nNumber of predicted TMHs: 1\nExp number of AAs in TMHs: 24.51309\nExp number, first 60 AAs: 0.2682\nTotal prob of N-in: 0.14402\noutside 1 331\nTMhelix 332 354\ninside 355 372

17557 GCF_002061155.1_ASM206115v1 Salmonella enterica subsp. enterica serovar Haifa Proteobacteria; Gammaproteobacteria; Enterobacterales; Enterobacteriaceae; Salmonella; Salmonella enterica; Salmonella enterica subsp. enterica MNNEETFYQAMRRKGVTRRSFLKFCSLAATSLGLGAGMTPKIAWALENKPRIPVVWIHGL WP_001058312.1 hydrogenase [Salmonella enterica] Length: 372\nNumber of predicted TMHs: 2\nExp number of AAs in TMHs: 36.51738\nExp number, first 60 AAs: 19.72849\nTotal prob of N-in: 0.96170\nPOSSIBLE N-term signal sequence\ninside 1 19\nTMhelix 20 39\noutside 40 325\nTMhelix 326 348\ninside 349 372

17558 GCF_002058515.1_ASM205851v1 Salmonella enterica subsp. enterica serovar Hartford Proteobacteria; Gammaproteobacteria; Enterobacterales; Enterobacteriaceae; Salmonella; Salmonella enterica; Salmonella enterica subsp. enterica MTGDNTLITSHGINRRDFMKLCAALAATMGLSSKAAAEMAESVSNPQRPPVIWIGAQECT WP_000145429.1 hydrogenase 2 small subunit [Salmonella enterica] Length: 372\nNumber of predicted TMHs: 1\nExp number of AAs in TMHs: 24.51309\nExp number, first 60 AAs: 0.2682\nTotal prob of N-in: 0.14402\noutside 1 331\nTMhelix 332 354\ninside 355 372

17559 GCF_002058735.1_ASM205873v1 Salmonella enterica subsp. enterica serovar Hartford Proteobacteria; Gammaproteobacteria; Enterobacterales; Enterobacteriaceae; Salmonella; Salmonella enterica; Salmonella enterica subsp. enterica MNNEETFYQAMRRKGVTRRSFLKFCSLAATSLGLGAGMTPKIAWALENKPRIPVVWIHGL WP_001058308.1 hydrogenase [Salmonella enterica] Length: 372\nNumber of predicted TMHs: 2\nExp number of AAs in TMHs: 39.08204\nExp number, first 60 AAs: 19.72474\nTotal prob of N-in: 0.96174\nPOSSIBLE N-term signal sequence\ninside 1 19\nTMhelix 20 39\noutside 40 325\nTMhelix 326 348\ninside 349 372

17560 GCF_002106515.1_ASM210651v1 Salmonella enterica subsp. enterica serovar Hartford Proteobacteria; Gammaproteobacteria; Enterobacterales; Enterobacteriaceae; Salmonella; Salmonella enterica; Salmonella enterica subsp. enterica MTGDNTLITSHGINRRDFMKLCAALAATMGLSSKAAAEMAESVSNPQRPPVIWIGAQECT WP_000145429.1 hydrogenase 2 small subunit [Salmonella enterica] Length: 372\nNumber of predicted TMHs: 1\nExp number of AAs in TMHs: 24.51309\nExp number, first 60 AAs: 0.2682\nTotal prob of N-in: 0.14402\noutside 1 331\nTMhelix 332 354\ninside 355 372

17561 GCF_002106515.1_ASM210651v1 Salmonella enterica subsp. enterica serovar Hartford Proteobacteria; Gammaproteobacteria; Enterobacterales; Enterobacteriaceae; Salmonella; Salmonella enterica;

Salmonella enterica subsp. enterica MNNEETFYQAMRRKGVTRRSFLKFCSLAATSLGLGAGMTPKIAWALENKPRIPVVWIHGL WP_001058312.1 hydrogenase [Salmonella enterica] Length: 372\nNumber of predicted TMHs: 2\nExp number of AAs in TMHs: 36.51738\nExp number, first 60 AAs: 19.72849\nTotal prob of N-in: 0.96170\nPOSSIBLE N-term signal sequence\ninside 1 19\nTMhelix 20 39\noutside 40 325\nTMhelix 326 348\ninside 349 372

17562 GCF_002058515.1_ASM205851v1 Salmonella enterica subsp. enterica serovar Hartford Proteobacteria; Gammaproteobacteria; Enterobacterales; Enterobacteriaceae; Salmonella; Salmonella enterica; Salmonella enterica subsp. enterica MNNEETFYQAMRRKGVTRRSFLKFCSLAATSLGLGAGMTPKIAWALENKPRIPVVWIHGL WP_001058312.1 hydrogenase [Salmonella enterica] Length: 372\nNumber of predicted TMHs: 2\nExp number of AAs in TMHs: 36.51738\nExp number, first 60 AAs: 19.72849\nTotal prob of N-in: 0.96170\nPOSSIBLE N-term signal sequence\ninside 1 19\nTMhelix 20 39\noutside 40 325\nTMhelix 326 348\ninside 349 372

17563 GCF_002058735.1_ASM205873v1 Salmonella enterica subsp. enterica serovar Hartford Proteobacteria; Gammaproteobacteria; Enterobacterales; Enterobacteriaceae; Salmonella; Salmonella enterica; Salmonella enterica subsp. enterica MTGDNTLITSHGINRRDFMKLCAALAATMGLSSKAAAEMAESVSNPQRPPVIWIGAQECT WP_000145429.1 hydrogenase 2 small subunit [Salmonella enterica] Length: 372\nNumber of predicted TMHs: 1\nExp number of AAs in TMHs: 24.51309\nExp number, first 60 AAs: 0.2682\nTotal prob of N-in: 0.14402\noutside 1 331\nTMhelix 332 354\ninside 355 372

17564 GCF_002064975.1_ASM206497v1 Salmonella enterica subsp. enterica serovar Havana Proteobacteria; Gammaproteobacteria; Enterobacterales; Enterobacteriaceae; Salmonella; Salmonella enterica; Salmonella enterica subsp. enterica MNNEETFYQAMRRKGVTRRSFLKFCSLAATSLGLGAGMTPKIAWALENKPRIPVVWIHGL WP_001058312.1 hydrogenase [Salmonella enterica] Length: 372\nNumber of predicted TMHs: 2\nExp number of AAs in TMHs: 36.51738\nExp number, first 60 AAs: 19.72849\nTotal prob of N-in: 0.96170\nPOSSIBLE N-term signal sequence\ninside 1 19\nTMhelix 20 39\noutside 40 325\nTMhelix 326 348\ninside 349 372

17565 GCF_002063345.1_ASM206334v1 Salmonella enterica subsp. enterica serovar Havana Proteobacteria; Gammaproteobacteria; Enterobacterales; Enterobacteriaceae; Salmonella; Salmonella enterica; Salmonella enterica subsp. enterica MTGDNTLITSHGINRRDFMKLCAALAATMGLSSKAAAEMAESVSNPQRPPVIWIGAQECT WP_000145429.1 hydrogenase 2 small subunit [Salmonella enterica] Length: 372\nNumber of predicted TMHs: 1\nExp number of AAs in TMHs: 24.51309\nExp number, first 60 AAs: 0.2682\nTotal prob of N-in: 0.14402\noutside 1 331\nTMhelix 332 354\ninside 355 372

17566 GCF_002063345.1_ASM206334v1 Salmonella enterica subsp. enterica serovar Havana Proteobacteria; Gammaproteobacteria; Enterobacterales; Enterobacteriaceae; Salmonella; Salmonella enterica; Salmonella enterica subsp. enterica MNNEETFYQAMRRKGVTRRSFLKFCSLAATSLGLGAGMTPKIAWALENKPRIPVVWIHGL WP_001058312.1 hydrogenase [Salmonella enterica] Length: 372\nNumber of predicted TMHs: 2\nExp number of AAs in TMHs: 36.51738\nExp number, first 60 AAs: 19.72849\nTotal prob of N-in: 0.96170\nPOSSIBLE N-term signal sequence\ninside 1 19\nTMhelix 20 39\noutside 40 325\nTMhelix 326 348\ninside 349 372

17567 GCF_002064955.1_ASM206495v1 Salmonella enterica subsp. enterica serovar Havana Proteobacteria; Gammaproteobacteria; Enterobacterales; Enterobacteriaceae; Salmonella; Salmonella enterica; Salmonella enterica subsp. enterica MTGDNTLITSHGINRRDFMKLCAALAATMGLSSKAAAEMAESVSNPQRPPVIWIGAQECT WP_000145429.1 hydrogenase 2 small subunit [Salmonella enterica] Length: 372\nNumber of predicted TMHs: 1\nExp number of AAs in TMHs: 24.51309\nExp number, first 60 AAs: 0.2682\nTotal prob of N-in: 0.14402\noutside 1 331\nTMhelix 332 354\ninside 355 372

17568 GCF_002064975.1_ASM206497v1 Salmonella enterica subsp. enterica serovar Havana Proteobacteria; Gammaproteobacteria; Enterobacterales; Enterobacteriaceae; Salmonella; Salmonella enterica; Salmonella enterica subsp. enterica MTGDNTLITSHGINRRDFMKLCAALAATMGLSSKAAAEMAESVSNPQRPPVIWIGAQECT WP_000145429.1 hydrogenase 2 small subunit [Salmonella enterica] Length: 372\nNumber of predicted TMHs: 1\nExp number of AAs in TMHs: 24.51309\nExp number, first 60 AAs: 0.2682\nTotal prob of N-in: 0.14402\noutside 1 331\nTMhelix 332 354\ninside 355 372

17569 GCF_002064955.1_ASM206495v1 Salmonella enterica subsp. enterica serovar Havana Proteobacteria; Gammaproteobacteria; Enterobacterales; Enterobacteriaceae; Salmonella; Salmonella enterica; Salmonella enterica subsp. enterica MNNEETFYQAMRRKGVTRRSFLKFCSLAATSLGLGAGMTPKIAWALENKPRIPVVWIHGL WP_001058311.1 hydrogenase [Salmonella enterica] Length: 372\nNumber of predicted TMHs: 2\nExp number of AAs in TMHs: 37.4699899999999\nExp number, first 60 AAs: 19.72699\nTotal prob of N-in: 0.96171\nPOSSIBLE N-term signal sequence\ninside 1 19\nTMhelix 20 39\noutside 40 325\nTMhelix 326 348\ninside 349 372

17570 GCF_002048515.1_ASM204851v1 Salmonella enterica subsp. enterica serovar Havana Proteobacteria; Gammaproteobacteria; Enterobacterales; Enterobacteriaceae; Salmonella; Salmonella enterica; Salmonella enterica subsp. enterica MNNEETFYQAMRRKGVTRRSFLKFCSLAATSLGLGAGMTPKIAWALENKPRIPVVWIHGL WP_001058311.1 hydrogenase [Salmonella enterica] Length: 372\nNumber of predicted TMHs: 2\nExp number of AAs in TMHs: 37.4699899999999\nExp number, first 60 AAs: 19.72699\nTotal prob of N-in: 0.96171\nPOSSIBLE N-term signal sequence\ninside 1 19\nTMhelix 20 39\noutside 40 325\nTMhelix 326 348\ninside 349 372

17571 GCF_002048515.1_ASM204851v1 Salmonella enterica subsp. enterica serovar Havana Proteobacteria; Gammaproteobacteria; Enterobacterales; Enterobacteriaceae; Salmonella; Salmonella enterica; Salmonella enterica subsp. enterica MTGDNTLITSHGINRRDFMKLCAALAATMGLSSKAAAEMAESVSNPQRPPVIWIGAQECT WP_000145429.1 hydrogenase 2 small subunit [Salmonella enterica] Length: 372\nNumber of predicted TMHs: 1\nExp number of AAs in TMHs: 24.51309\nExp number, first 60 AAs: 0.2682\nTotal prob of N-in: 0.14402\noutside 1 331\nTMhelix 332 354\ninside 355 372

17572 GCF_002047095.1_ASM204709v1 Salmonella enterica subsp. enterica serovar Havana Proteobacteria; Gammaproteobacteria; Enterobacterales; Enterobacteriaceae; Salmonella; Salmonella enterica; Salmonella enterica subsp. enterica MNNEETFYQAMRRKGVTRRSFLKFCSLAATSLGLGAGMTPKIAWALENKPRIPVVWIHGL WP_001058312.1 hydrogenase [Salmonella enterica] Length: 372\nNumber of predicted TMHs: 2\nExp number of AAs in TMHs: 36.51738\nExp number, first 60 AAs: 19.72849\nTotal prob of N-in: 0.96170\nPOSSIBLE N-term signal sequence\ninside 1 19\nTMhelix 20 39\noutside 40 325\nTMhelix 326 348\ninside 349 372

17573 GCF_002047345.1_ASM204734v1 Salmonella enterica subsp. enterica serovar Havana Proteobacteria; Gammaproteobacteria; Enterobacterales; Enterobacteriaceae; Salmonella; Salmonella enterica; Salmonella enterica subsp. enterica MNNEETFYQAMRRKGVTRRSFLKFCSLAATSLGLGAGMTPKIAWALENKPRIPVVWIHGL WP_001058312.1 hydrogenase [Salmonella enterica] Length: 372\nNumber of predicted TMHs: 2\nExp number of AAs in TMHs: 36.51738\nExp number, first 60 AAs: 19.72849\nTotal prob of N-in: 0.96170\nPOSSIBLE N-term signal sequence\ninside 1 19\nTMhelix 20 39\noutside 40 325\nTMhelix 326 348\ninside 349 372

17574 GCF_002064965.1_ASM206496v1 Salmonella enterica subsp. enterica serovar Havana Proteobacteria; Gammaproteobacteria; Enterobacterales; Enterobacteriaceae; Salmonella; Salmonella enterica; Salmonella enterica subsp. enterica MNNEETFYQAMRRKGVTRRSFLKFCSLAATSLGLGAGMTPKIAWALENKPRIPVVWIHGL WP_001058311.1 hydrogenase [Salmonella enterica] Length: 372\nNumber of predicted TMHs: 2\nExp number of AAs in TMHs: 37.4699899999999\nExp number, first 60 AAs: 19.72699\nTotal prob of N-in: 0.96171\nPOSSIBLE N-term signal sequence\ninside 1 19\nTMhelix 20 39\noutside 40 325\nTMhelix 326 348\ninside 349 372

17575 GCF_002047095.1_ASM204709v1 Salmonella enterica subsp. enterica serovar Havana Proteobacteria; Gammaproteobacteria; Enterobacterales; Enterobacteriaceae; Salmonella; Salmonella enterica; Salmonella enterica subsp. enterica MTGDNTLITSHGINRRDFMKLCAALAATMGLSSKAAAEMAESVSNPQRPPVIWIGAQECT WP_000145429.1 hydrogenase 2 small subunit [Salmonella enterica] Length: 372\nNumber of predicted TMHs: 1\nExp number of AAs in TMHs: 24.51309\nExp number, first 60 AAs: 0.2682\nTotal prob of N-in: 0.14402\noutside 1 331\nTMhelix 332 354\ninside 355 372

17576 GCF_002047345.1_ASM204734v1 Salmonella enterica subsp. enterica serovar Havana Proteobacteria; Gammaproteobacteria; Enterobacterales; Enterobacteriaceae; Salmonella; Salmonella enterica; Salmonella enterica subsp. enterica MTGDNTLITSHGINRRDFMKLCAALAATMGLSSKAAAEMAESVSNPQRPPVIWIGAQECT WP_000145429.1 hydrogenase 2 small subunit [Salmonella enterica] Length: 372\nNumber of predicted TMHs: 1\nExp number of AAs in TMHs: 24.51309\nExp number, first 60 AAs: 0.2682\nTotal prob of N-in: 0.14402\noutside 1 331\nTMhelix 332 354\ninside 355 372

17577 GCF_002048335.1_ASM204833v1 Salmonella enterica subsp. enterica serovar Havana Proteobacteria; Gammaproteobacteria; Enterobacterales; Enterobacteriaceae; Salmonella; Salmonella enterica; Salmonella enterica subsp. enterica MNNEETFYQAMRRKGVTRRSFLKFCSLAATSLGLGAGMTPKIAWALENKPRIPVVWIHGL WP_001058311.1 hydrogenase [Salmonella enterica] Length: 372\nNumber of predicted TMHs: 2\nExp number of AAs in TMHs: 37.4699899999999\nExp number, first 60 AAs: 19.72699\nTotal prob of N-in: 0.96171\nPOSSIBLE N-term signal sequence\ninside 1 19\nTMhelix 20 39\noutside 40 325\nTMhelix 326 348\ninside 349 372

17578 GCF_002064965.1_ASM206496v1 Salmonella enterica subsp. enterica serovar Havana Proteobacteria; Gammaproteobacteria; Enterobacterales; Enterobacteriaceae; Salmonella; Salmonella enterica;

Salmonella enterica subsp. enterica	MTGDNTLITSHGINRRDFMKLCAALAATMGLSSKAAAEMAESVSNPQRPPVIWIGAQECT	WP_000145429.1	hydrogenase 2 small subunit [Salmonella enterica]	Length: 372\nNumber of predicted TMHs: 1\nExp number of AAs in TMHs: 24.51309\nExp number, first 60 AAs: 0.2682\nTotal prob of N-in: 0.14402\noutside 1 331\nTMhelix 332 354\ninside 355 372

17579	GCF_002048335.1_ASM204833v1	Salmonella enterica subsp. enterica serovar Havana	Proteobacteria; Gammaproteobacteria; Enterobacterales; Enterobacteriaceae; Salmonella; Salmonella enterica; Salmonella enterica subsp. enterica	MTGDNTLITSHGINRRDFMKLCAALAATMGLSSKAAAEMAESVSNPQRPPVIWIGAQECT	WP_000145429.1	hydrogenase 2 small subunit [Salmonella enterica]	Length: 372\nNumber of predicted TMHs: 1\nExp number of AAs in TMHs: 24.51309\nExp number, first 60 AAs: 0.2682\nTotal prob of N-in: 0.14402\noutside 1 331\nTMhelix 332 354\ninside 355 372

17580	GCF_002048245.1_ASM204824v1	Salmonella enterica subsp. enterica serovar Havana	Proteobacteria; Gammaproteobacteria; Enterobacterales; Enterobacteriaceae; Salmonella; Salmonella enterica; Salmonella enterica subsp. enterica	MNNEETFYQAMRRKGVTRRSFLKFCSLAATSLGLGAGMTPKIAWALENKPRIPVVWIHGL	WP_001058311.1	hydrogenase [Salmonella enterica]	Length: 372\nNumber of predicted TMHs: 2\nExp number of AAs in TMHs: 37.4699899999999\nExp number, first 60 AAs: 19.72699\nTotal prob of N-in: 0.96171\nPOSSIBLE N-term signal sequence\ninside 1 19\nTMhelix 20 39\noutside 40 325\nTMhelix 326 348\ninside 349 372

17581	GCF_002062035.1_ASM206203v1	Salmonella enterica subsp. enterica serovar Havana	Proteobacteria; Gammaproteobacteria; Enterobacterales; Enterobacteriaceae; Salmonella; Salmonella enterica; Salmonella enterica subsp. enterica	MNNEETFYQAMRRKGVTRRSFLKFCSLAATSLGLGAGMTPKIAWALENKPRIPVVWIHGL	WP_001058312.1	hydrogenase [Salmonella enterica]	Length: 372\nNumber of predicted TMHs: 2\nExp number of AAs in TMHs: 36.51738\nExp number, first 60 AAs: 19.72849\nTotal prob of N-in: 0.96170\nPOSSIBLE N-term signal sequence\ninside 1 19\nTMhelix 20 39\noutside 40 325\nTMhelix 326 348\ninside 349 372

17582	GCF_002062035.1_ASM206203v1	Salmonella enterica subsp. enterica serovar Havana	Proteobacteria; Gammaproteobacteria; Enterobacterales; Enterobacteriaceae; Salmonella; Salmonella enterica; Salmonella enterica subsp. enterica	MTGDNTLITSHGINRRDFMKLCAALAATMGLSSKAAAEMAESVSNPQRPPVIWIGAQECT	WP_000145429.1	hydrogenase 2 small subunit [Salmonella enterica]	Length: 372\nNumber of predicted TMHs: 1\nExp number of AAs in TMHs: 24.51309\nExp number, first 60 AAs: 0.2682\nTotal prob of N-in: 0.14402\noutside 1 331\nTMhelix 332 354\ninside 355 372

17583	GCF_002048245.1_ASM204824v1	Salmonella enterica subsp. enterica serovar Havana	Proteobacteria; Gammaproteobacteria; Enterobacterales; Enterobacteriaceae; Salmonella; Salmonella enterica; Salmonella enterica subsp. enterica	MTGDNTLITSHGINRRDFMKLCAALAATMGLSSKAAAEMAESVSNPQRPPVIWIGAQECT	WP_000145429.1	hydrogenase 2 small subunit [Salmonella enterica]	Length: 372\nNumber of predicted TMHs: 1\nExp number of AAs in TMHs: 24.51309\nExp number, first 60 AAs: 0.2682\nTotal prob of N-in: 0.14402\noutside 1 331\nTMhelix 332 354\ninside 355 372

17584	GCF_001242675.1_Salmonella_enterica_CVM_N45954_v1.0	Salmonella enterica subsp. enterica serovar Heidelberg	Proteobacteria; Gammaproteobacteria; Enterobacterales; Enterobacteriaceae; Salmonella; Salmonella enterica; Salmonella enterica subsp. enterica	MNNEETFYQAMRRKGVTRRSFLKFCSLAATSLGLGAGMTPKIAWALENKPRIPVVWIHGL	WP_001058312.1	hydrogenase [Salmonella enterica]	Length: 372\nNumber of predicted TMHs: 2\nExp number of AAs in TMHs: 36.51738\nExp number, first 60 AAs: 19.72849\nTotal prob of N-in: 0.96170\nPOSSIBLE N-term signal sequence\ninside 1 19\nTMhelix 20 39\noutside 40 325\nTMhelix 326 348\ninside 349 372

17585	GCF_001244225.1_Salmonella_enterica_CVM_N48578_v1.0	Salmonella enterica subsp. enterica serovar Heidelberg	Proteobacteria; Gammaproteobacteria; Enterobacterales; Enterobacteriaceae; Salmonella; Salmonella enterica; Salmonella enterica subsp. enterica	MTGDNTLITSHGINRRDFMKLCAALAATMGLSSKAAAEMAESVSNPQRPPVIWIGAQECT	WP_000145429.1	hydrogenase 2 small subunit [Salmonella enterica]	Length: 372\nNumber of predicted TMHs: 1\nExp number of AAs in TMHs: 24.51309\nExp number, first 60 AAs: 0.2682\nTotal prob of N-in: 0.14402\noutside 1 331\nTMhelix 332 354\ninside 355 372

17586	GCF_001242965.1_Salmonella_enterica_CVM_N46818_v1.0	Salmonella enterica subsp. enterica serovar Heidelberg	Proteobacteria; Gammaproteobacteria; Enterobacterales; Enterobacteriaceae; Salmonella; Salmonella enterica; Salmonella enterica subsp. enterica	MNNEETFYQAMRRKGVTRRSFLKFCSLAATSLGLGAGMTPKIAWALENKPRIPVVWIHGL	WP_001058312.1

hydrogenase [Salmonella enterica] Length: 372\nNumber of predicted TMHs: 2\nExp number of AAs in TMHs: 36.51738\nExp number, first 60 AAs: 19.72849\nTotal prob of N-in: 0.96170\nPOSSIBLE N-term signal sequence\ninside 1 19\nTMhelix 20 39\noutside 40 325\nTMhelix 326 348\ninside 349 372

17587 GCF_001241365.1_Salmonella_enterica_CVM_N44707_v1.0 Salmonella enterica subsp. enterica serovar Heidelberg Proteobacteria; Gammaproteobacteria; Enterobacterales; Enterobacteriaceae; Salmonella; Salmonella enterica; Salmonella enterica subsp. enterica MTGDNTLITSHGINRRDFMKLCAALAATMGLSSKAAAEMAESVSNPQRPPVIWIGAQECT WP_000145429.1 hydrogenase 2 small subunit [Salmonella enterica] Length: 372\nNumber of predicted TMHs: 1\nExp number of AAs in TMHs: 24.51309\nExp number, first 60 AAs: 0.2682\nTotal prob of N-in: 0.14402\noutside 1 331\nTMhelix 332 354\ninside 355 372

17588 GCF_001478505.1_Salmonella_enterica_CVM_N41746-SQ_v1.0 Salmonella enterica subsp. enterica serovar Heidelberg Proteobacteria; Gammaproteobacteria; Enterobacterales; Enterobacteriaceae; Salmonella; Salmonella enterica; Salmonella enterica subsp. enterica MTGDNTLITSHGINRRDFMKLCAALAATMGLSSKAAAEMAESVSNPQRPPVIWIGAQECT WP_000145429.1 hydrogenase 2 small subunit [Salmonella enterica] Length: 372\nNumber of predicted TMHs: 1\nExp number of AAs in TMHs: 24.51309\nExp number, first 60 AAs: 0.2682\nTotal prob of N-in: 0.14402\noutside 1 331\nTMhelix 332 354\ninside 355 372

17589 GCF_001241725.1_Salmonella_enterica_CVM_N45397_v1.0 Salmonella enterica subsp. enterica serovar Heidelberg Proteobacteria; Gammaproteobacteria; Enterobacterales; Enterobacteriaceae; Salmonella; Salmonella enterica; Salmonella enterica subsp. enterica MNNEETFYQAMRRKGVTRRSFLKFCSLAATSLGLGAGMTPKIAWALENKPRIPVVWIHGL WP_001058312.1 hydrogenase [Salmonella enterica] Length: 372\nNumber of predicted TMHs: 2\nExp number of AAs in TMHs: 36.51738\nExp number, first 60 AAs: 19.72849\nTotal prob of N-in: 0.96170\nPOSSIBLE N-term signal sequence\ninside 1 19\nTMhelix 20 39\noutside 40 325\nTMhelix 326 348\ninside 349 372

17590 GCF_001692655.1_ASM169265v1 Salmonella enterica subsp. enterica serovar Heidelberg Proteobacteria; Gammaproteobacteria; Enterobacterales; Enterobacteriaceae; Salmonella; Salmonella enterica; Salmonella enterica subsp. enterica MNNEETFYQAMRRKGVTRRSFLKFCSLAATSLGLGAGMTPKIAWALENKPRIPVVWIHGL WP_001058312.1 hydrogenase [Salmonella enterica] Length: 372\nNumber of predicted TMHs: 2\nExp number of AAs in TMHs: 36.51738\nExp number, first 60 AAs: 19.72849\nTotal prob of N-in: 0.96170\nPOSSIBLE N-term signal sequence\ninside 1 19\nTMhelix 20 39\noutside 40 325\nTMhelix 326 348\ninside 349 372

17591 GCF_001478975.1_Salmonella_enterica_CVM_N30683-SQ_v1.0 Salmonella enterica subsp. enterica serovar Heidelberg Proteobacteria; Gammaproteobacteria; Enterobacterales; Enterobacteriaceae; Salmonella; Salmonella enterica; Salmonella enterica subsp. enterica MNNEETFYQAMRRKGVTRRSFLKFCSLAATSLGLGAGMTPKIAWALENKPRIPVVWIHGL WP_001058312.1 hydrogenase [Salmonella enterica] Length: 372\nNumber of predicted TMHs: 2\nExp number of AAs in TMHs: 36.51738\nExp number, first 60 AAs: 19.72849\nTotal prob of N-in: 0.96170\nPOSSIBLE N-term signal sequence\ninside 1 19\nTMhelix 20 39\noutside 40 325\nTMhelix 326 348\ninside 349 372

17592 GCF_002032425.1_ASM203242v1 Salmonella enterica subsp. enterica serovar Heidelberg Proteobacteria; Gammaproteobacteria; Enterobacterales; Enterobacteriaceae; Salmonella; Salmonella enterica; Salmonella enterica subsp. enterica MNNEETFYQAMRRKGVTRRSFLKFCSLAATSLGLGAGMTPKIAWALENKPRIPVVWIHGL WP_001058312.1 hydrogenase [Salmonella enterica] Length: 372\nNumber of predicted TMHs: 2\nExp number of AAs in TMHs: 36.51738\nExp number, first 60 AAs: 19.72849\nTotal prob of N-in: 0.96170\nPOSSIBLE N-term signal sequence\ninside 1 19\nTMhelix 20 39\noutside 40 325\nTMhelix 326 348\ninside 349 372

17593 GCF_001241305.1_Salmonella_enterica_CVM_N44703_v1.0 Salmonella enterica subsp. enterica serovar Heidelberg Proteobacteria; Gammaproteobacteria; Enterobacterales; Enterobacteriaceae; Salmonella; Salmonella enterica; Salmonella enterica subsp. enterica MNNEETFYQAMRRKGVTRRSFLKFCSLAATSLGLGAGMTPKIAWALENKPRIPVVWIHGL WP_001058312.1 hydrogenase [Salmonella enterica] Length: 372\nNumber of predicted TMHs: 2\nExp number of AAs in TMHs: 36.51738\nExp number, first 60 AAs: 19.72849\nTotal prob of N-in: 0.96170\nPOSSIBLE N-term signal sequence\ninside 1 19\nTMhelix 20 39\noutside 40 325\nTMhelix 326 348\ninside 349 372

17594 GCF_001478505.1_Salmonella_enterica_CVM_N41746-SQ_v1.0 Salmonella enterica subsp. enterica serovar Heidelberg Proteobacteria; Gammaproteobacteria; Enterobacterales; Enterobacteriaceae; Salmonella; Salmonella enterica; Salmonella enterica subsp. enterica

MNNEETFYQAMRRKGVTRRSFLKFCSLAATSLGLGAGMTPKIAWALENKPRIPVVWIHGL WP_001058312.1 hydrogenase [Salmonella enterica] Length: 372\nNumber of predicted TMHs: 2\nExp number of AAs in TMHs: 36.51738\nExp number, first 60 AAs: 19.72849\nTotal prob of N-in: 0.96170\nPOSSIBLE N-term signal sequence\ninside 1 19\nTMhelix 20 39\noutside 40 325\nTMhelix 326 348\ninside 349 372

17595 GCF_001952325.1_ASM195232v1 Salmonella enterica subsp. enterica serovar Heidelberg Proteobacteria; Gammaproteobacteria; Enterobacterales; Enterobacteriaceae; Salmonella; Salmonella enterica; Salmonella enterica subsp. enterica MTGDNTLITSHGINRRDFMKLCAALAATMGLSSKAAAEMAESVSNPQRPPVIWIGAQECT WP_000145429.1 hydrogenase 2 small subunit [Salmonella enterica] Length: 372\nNumber of predicted TMHs: 1\nExp number of AAs in TMHs: 24.51309\nExp number, first 60 AAs: 0.2682\nTotal prob of N-in: 0.14402\noutside 1 331\nTMhelix 332 354\ninside 355 372

17596 GCF_001479485.1_Salmonella_enterica_CVM_N32772-SQ_v1.0 Salmonella enterica subsp. enterica serovar Heidelberg Proteobacteria; Gammaproteobacteria; Enterobacterales; Enterobacteriaceae; Salmonella; Salmonella enterica; Salmonella enterica subsp. enterica MNNEETFYQAMRRKGVTRRSFLKFCSLAATSLGLGAGMTPKIAWALENKPRIPVVWIHGL WP_001058312.1 hydrogenase [Salmonella enterica] Length: 372\nNumber of predicted TMHs: 2\nExp number of AAs in TMHs: 36.51738\nExp number, first 60 AAs: 19.72849\nTotal prob of N-in: 0.96170\nPOSSIBLE N-term signal sequence\ninside 1 19\nTMhelix 20 39\noutside 40 325\nTMhelix 326 348\ninside 349 372

17597 GCF_001480665.1_Salmonella_enterica_CVM_N42482_v1.0 Salmonella enterica subsp. enterica serovar Heidelberg Proteobacteria; Gammaproteobacteria; Enterobacterales; Enterobacteriaceae; Salmonella; Salmonella enterica; Salmonella enterica subsp. enterica MTGDNTLITSHGINRRDFMKLCAALAATMGLSSKAAAEMAESVSNPQRPPVIWIGAQECT WP_000145429.1 hydrogenase 2 small subunit [Salmonella enterica] Length: 372\nNumber of predicted TMHs: 1\nExp number of AAs in TMHs: 24.51309\nExp number, first 60 AAs: 0.2682\nTotal prob of N-in: 0.14402\noutside 1 331\nTMhelix 332 354\ninside 355 372

17598 GCF_001480665.1_Salmonella_enterica_CVM_N42482_v1.0 Salmonella enterica subsp. enterica serovar Heidelberg Proteobacteria; Gammaproteobacteria; Enterobacterales; Enterobacteriaceae; Salmonella; Salmonella enterica; Salmonella enterica subsp. enterica MNNEETFYQAMRRKGVTRRSFLKFCSLAATSLGLGAGMTPKIAWALENKPRIPVVWIHGL WP_001058312.1 hydrogenase [Salmonella enterica] Length: 372\nNumber of predicted TMHs: 2\nExp number of AAs in TMHs: 36.51738\nExp number, first 60 AAs: 19.72849\nTotal prob of N-in: 0.96170\nPOSSIBLE N-term signal sequence\ninside 1 19\nTMhelix 20 39\noutside 40 325\nTMhelix 326 348\ninside 349 372

17599 GCF_001247075.1_Salmonella_enterica_CVM_N51292_v1.0 Salmonella enterica subsp. enterica serovar Heidelberg Proteobacteria; Gammaproteobacteria; Enterobacterales; Enterobacteriaceae; Salmonella; Salmonella enterica; Salmonella enterica subsp. enterica MNNEETFYQAMRRKGVTRRSFLKFCSLAATSLGLGAGMTPKIAWALENKPRIPVVWIHGL WP_001058312.1 hydrogenase [Salmonella enterica] Length: 372\nNumber of predicted TMHs: 2\nExp number of AAs in TMHs: 36.51738\nExp number, first 60 AAs: 19.72849\nTotal prob of N-in: 0.96170\nPOSSIBLE N-term signal sequence\ninside 1 19\nTMhelix 20 39\noutside 40 325\nTMhelix 326 348\ninside 349 372

17600 GCF_002037355.1_ASM203735v1 Salmonella enterica subsp. enterica serovar Heidelberg Proteobacteria; Gammaproteobacteria; Enterobacterales; Enterobacteriaceae; Salmonella; Salmonella enterica; Salmonella enterica subsp. enterica MTGDNTLITSHGINRRDFMKLCAALAATMGLSSKAAAEMAESVSNPQRPPVIWIGAQECT WP_000145429.1 hydrogenase 2 small subunit [Salmonella enterica] Length: 372\nNumber of predicted TMHs: 1\nExp number of AAs in TMHs: 24.51309\nExp number, first 60 AAs: 0.2682\nTotal prob of N-in: 0.14402\noutside 1 331\nTMhelix 332 354\ninside 355 372

17601 GCF_001241455.1_Salmonella_enterica_CVM_N44710_v1.0 Salmonella enterica subsp. enterica serovar Heidelberg Proteobacteria; Gammaproteobacteria; Enterobacterales; Enterobacteriaceae; Salmonella; Salmonella enterica; Salmonella enterica subsp. enterica MTGDNTLITSHGINRRDFMKLCAALAATMGLSSKAAAEMAESVSNPQRPPVIWIGAQECT WP_000145429.1 hydrogenase 2 small subunit [Salmonella enterica] Length: 372\nNumber of predicted TMHs: 1\nExp number of AAs in TMHs: 24.51309\nExp number, first 60 AAs: 0.2682\nTotal prob of N-in: 0.14402\noutside 1 331\nTMhelix 332 354\ninside 355 372

17602 GCF_001481095.1_Salmonella_enterica_CVM_N42472_v1.0 Salmonella enterica subsp. enterica serovar Heidelberg Proteobacteria; Gammaproteobacteria; Enterobacterales; Enterobacteriaceae; Salmonella; Salmonella

enterica; Salmonella enterica subsp. enterica MTGDNTLITSHGINRRDFMKLCAALAATMGLSSKAAAEMAESVSNPQRPPVIWIGAQECT WP_000145429.1 hydrogenase 2 small subunit [Salmonella enterica] Length: 372\nNumber of predicted TMHs: 1\nExp number of AAs in TMHs: 24.51309\nExp number, first 60 AAs: 0.2682\nTotal prob of N-in: 0.14402\noutside 1 331\nTMhelix 332 354\ninside 355 372

17603 GCF_001481095.1_Salmonella_enterica_CVM_N42472_v1.0 Salmonella enterica subsp. enterica serovar Heidelberg Proteobacteria; Gammaproteobacteria; Enterobacterales; Enterobacteriaceae; Salmonella; Salmonella enterica; Salmonella enterica subsp. enterica MNNEETFYQAMRRKGVTRRSFLKFCSLAATSLGLGAGMTPKIAWALENKPRIPVVWIHGL WP_001058312.1 hydrogenase [Salmonella enterica] Length: 372\nNumber of predicted TMHs: 2\nExp number of AAs in TMHs: 36.51738\nExp number, first 60 AAs: 19.72849\nTotal prob of N-in: 0.96170\nPOSSIBLE N-term signal sequence\ninside 1 19\nTMhelix 20 39\noutside 40 325\nTMhelix 326 348\ninside 349 372

17604 GCF_001247585.1_Salmonella_enterica_CVM_N51984_v1.0 Salmonella enterica subsp. enterica serovar Heidelberg Proteobacteria; Gammaproteobacteria; Enterobacterales; Enterobacteriaceae; Salmonella; Salmonella enterica; Salmonella enterica subsp. enterica MTGDNTLITSHGINRRDFMKLCAALAATMGLSSKAAAEMAESVSNPQRPPVIWIGAQECT WP_000145429.1 hydrogenase 2 small subunit [Salmonella enterica] Length: 372\nNumber of predicted TMHs: 1\nExp number of AAs in TMHs: 24.51309\nExp number, first 60 AAs: 0.2682\nTotal prob of N-in: 0.14402\noutside 1 331\nTMhelix 332 354\ninside 355 372

17605 GCF_001242865.1_Salmonella_enterica_CVM_N46812_v1.0 Salmonella enterica subsp. enterica serovar Heidelberg Proteobacteria; Gammaproteobacteria; Enterobacterales; Enterobacteriaceae; Salmonella; Salmonella enterica; Salmonella enterica subsp. enterica MTGDNTLITSHGINRRDFMKLCAALAATMGLSSKAAAEMAESVSNPQRPPVIWIGAQECT WP_000145429.1 hydrogenase 2 small subunit [Salmonella enterica] Length: 372\nNumber of predicted TMHs: 1\nExp number of AAs in TMHs: 24.51309\nExp number, first 60 AAs: 0.2682\nTotal prob of N-in: 0.14402\noutside 1 331\nTMhelix 332 354\ninside 355 372

17606 GCF_001271555.1_Salmonella_enterica_CVM_N45393_v1.0 Salmonella enterica subsp. enterica serovar Heidelberg Proteobacteria; Gammaproteobacteria; Enterobacterales; Enterobacteriaceae; Salmonella; Salmonella enterica; Salmonella enterica subsp. enterica MNNEETFYQAMRRKGVTRRSFLKFCSLAATSLGLGAGMTPKIAWALENKPRIPVVWIHGL WP_001058312.1 hydrogenase [Salmonella enterica] Length: 372\nNumber of predicted TMHs: 2\nExp number of AAs in TMHs: 36.51738\nExp number, first 60 AAs: 19.72849\nTotal prob of N-in: 0.96170\nPOSSIBLE N-term signal sequence\ninside 1 19\nTMhelix 20 39\noutside 40 325\nTMhelix 326 348\ninside 349 372

17607 GCF_000974165.1_SALB-159-4.id19_v1.0 Salmonella enterica subsp. enterica serovar Heidelberg Proteobacteria; Gammaproteobacteria; Enterobacterales; Enterobacteriaceae; Salmonella; Salmonella enterica; Salmonella enterica subsp. enterica MTGDNTLITSHGINRRDFMKLCAALAATMGLSSKAAAEMAESVSNPQRPPVIWIGAQECT WP_000145429.1 hydrogenase 2 small subunit [Salmonella enterica] Length: 372\nNumber of predicted TMHs: 1\nExp number of AAs in TMHs: 24.51309\nExp number, first 60 AAs: 0.2682\nTotal prob of N-in: 0.14402\noutside 1 331\nTMhelix 332 354\ninside 355 372

17608 GCF_001479585.1_Salmonella_enterica_CVM_N37907-SQ_v1.0 Salmonella enterica subsp. enterica serovar Heidelberg Proteobacteria; Gammaproteobacteria; Enterobacterales; Enterobacteriaceae; Salmonella; Salmonella enterica; Salmonella enterica subsp. enterica MNNEETFYQAMRRKGVTRRSFLKFCSLAATSLGLGAGMTPKIAWALENKPRIPVVWIHGL WP_001058312.1 hydrogenase [Salmonella enterica] Length: 372\nNumber of predicted TMHs: 2\nExp number of AAs in TMHs: 36.51738\nExp number, first 60 AAs: 19.72849\nTotal prob of N-in: 0.96170\nPOSSIBLE N-term signal sequence\ninside 1 19\nTMhelix 20 39\noutside 40 325\nTMhelix 326 348\ninside 349 372

17609 GCF_001272055.1_Salmonella_enterica_CVM_N44702_v1.0 Salmonella enterica subsp. enterica serovar Heidelberg Proteobacteria; Gammaproteobacteria; Enterobacterales; Enterobacteriaceae; Salmonella; Salmonella enterica; Salmonella enterica subsp. enterica MNNEETFYQAMRRKGVTRRSFLKFCSLAATSLGLGAGMTPKIAWALENKPRIPVVWIHGL WP_001058312.1 hydrogenase [Salmonella enterica] Length: 372\nNumber of predicted TMHs: 2\nExp number of AAs in TMHs: 36.51738\nExp number, first 60 AAs: 19.72849\nTotal prob of N-in: 0.96170\nPOSSIBLE N-term signal sequence\ninside 1 19\nTMhelix 20 39\noutside 40 325\nTMhelix 326 348\ninside 349 372

17610 GCF_001952575.1_ASM195257v1 Salmonella enterica subsp. enterica serovar Heidelberg Proteobacteria; Gammaproteobacteria; Enterobacterales; Enterobacteriaceae; Salmonella; Salmonella enterica; Salmonella enterica subsp. enterica MNNEETFYQAMRRKGVTRRSFLKFCSLAATSLGLGAGMTPKIAWALENKPRIPVVWIHGL WP_001058312.1 hydrogenase [Salmonella enterica] Length: 372\nNumber of predicted TMHs: 2\nExp number of AAs in TMHs: 36.51738\nExp number, first 60 AAs: 19.72849\nTotal prob of N-in: 0.96170\nPOSSIBLE N-term signal sequence\ninside 1 19\nTMhelix 20 39\noutside 40 325\nTMhelix 326 348\ninside 349 372

17611 GCF_001240155.1_Salmonella_enterica_CVM_N43452_v1.0 Salmonella enterica subsp. enterica serovar Heidelberg Proteobacteria; Gammaproteobacteria; Enterobacterales; Enterobacteriaceae; Salmonella; Salmonella enterica; Salmonella enterica subsp. enterica MTGDNTLITSHGINRRDFMKLCAALAATMGLSSKAAAEMAESVSNPQRPPVIWIGAQECT WP_000145429.1 hydrogenase 2 small subunit [Salmonella enterica] Length: 372\nNumber of predicted TMHs: 1\nExp number of AAs in TMHs: 24.51309\nExp number, first 60 AAs: 0.2682\nTotal prob of N-in: 0.14402\noutside 1 331\nTMhelix 332 354\ninside 355 372

17612 GCF_001478945.1_Salmonella_enterica_CVM_N30671-SQ_v1.0 Salmonella enterica subsp. enterica serovar Heidelberg Proteobacteria; Gammaproteobacteria; Enterobacterales; Enterobacteriaceae; Salmonella; Salmonella enterica; Salmonella enterica subsp. enterica MTGDNTLITSHGINRRDFMKLCAALAATMGLSSKAAAEMAESVSNPQRPPVIWIGAQECT WP_000145429.1 hydrogenase 2 small subunit [Salmonella enterica] Length: 372\nNumber of predicted TMHs: 1\nExp number of AAs in TMHs: 24.51309\nExp number, first 60 AAs: 0.2682\nTotal prob of N-in: 0.14402\noutside 1 331\nTMhelix 332 354\ninside 355 372

17613 GCF_001241305.1_Salmonella_enterica_CVM_N44703_v1.0 Salmonella enterica subsp. enterica serovar Heidelberg Proteobacteria; Gammaproteobacteria; Enterobacterales; Enterobacteriaceae; Salmonella; Salmonella enterica; Salmonella enterica subsp. enterica MTGDNTLITSHGINRRDFMKLCAALAATMGLSSKAAAEMAESVSNPQRPPVIWIGAQECT WP_000145429.1 hydrogenase 2 small subunit [Salmonella enterica] Length: 372\nNumber of predicted TMHs: 1\nExp number of AAs in TMHs: 24.51309\nExp number, first 60 AAs: 0.2682\nTotal prob of N-in: 0.14402\noutside 1 331\nTMhelix 332 354\ninside 355 372

17614 GCF_001242675.1_Salmonella_enterica_CVM_N45954_v1.0 Salmonella enterica subsp. enterica serovar Heidelberg Proteobacteria; Gammaproteobacteria; Enterobacterales; Enterobacteriaceae; Salmonella; Salmonella enterica; Salmonella enterica subsp. enterica MTGDNTLITSHGINRRDFMKLCAALAATMGLSSKAAAEMAESVSNPQRPPVIWIGAQECT WP_000145429.1 hydrogenase 2 small subunit [Salmonella enterica] Length: 372\nNumber of predicted TMHs: 1\nExp number of AAs in TMHs: 24.51309\nExp number, first 60 AAs: 0.2682\nTotal prob of N-in: 0.14402\noutside 1 331\nTMhelix 332 354\ninside 355 372

17615 GCF_001952555.1_ASM195255v1 Salmonella enterica subsp. enterica serovar Heidelberg Proteobacteria; Gammaproteobacteria; Enterobacterales; Enterobacteriaceae; Salmonella; Salmonella enterica; Salmonella enterica subsp. enterica MNNEETFYQAMRRKGVTRRSFLKFCSLAATSLGLGAGMTPKIAWALENKPRIPVVWIHGL WP_001058312.1 hydrogenase [Salmonella enterica] Length: 372\nNumber of predicted TMHs: 2\nExp number of AAs in TMHs: 36.51738\nExp number, first 60 AAs: 19.72849\nTotal prob of N-in: 0.96170\nPOSSIBLE N-term signal sequence\ninside 1 19\nTMhelix 20 39\noutside 40 325\nTMhelix 326 348\ninside 349 372

17616 GCF_002060235.1_ASM206023v1 Salmonella enterica subsp. enterica serovar Heidelberg Proteobacteria; Gammaproteobacteria; Enterobacterales; Enterobacteriaceae; Salmonella; Salmonella enterica; Salmonella enterica subsp. enterica MTGDNTLITSHGINRRDFMKLCAALAATMGLSSKAAAEMAESVSNPQRPPVIWIGAQECT WP_000145429.1 hydrogenase 2 small subunit [Salmonella enterica] Length: 372\nNumber of predicted TMHs: 1\nExp number of AAs in TMHs: 24.51309\nExp number, first 60 AAs: 0.2682\nTotal prob of N-in: 0.14402\noutside 1 331\nTMhelix 332 354\ninside 355 372

17617 GCF_002061535.1_ASM206153v1 Salmonella enterica subsp. enterica serovar Heidelberg Proteobacteria; Gammaproteobacteria; Enterobacterales; Enterobacteriaceae; Salmonella; Salmonella enterica; Salmonella enterica subsp. enterica MTGDNTLITSHGINRRDFMKLCAALAATMGLSSKAAAEMAESVSNPQRPPVIWIGAQECT WP_000145429.1 hydrogenase 2 small subunit [Salmonella enterica] Length: 372\nNumber of predicted TMHs: 1\nExp number of AAs in TMHs: 24.51309\nExp number, first 60 AAs: 0.2682\nTotal prob of N-in: 0.14402\noutside 1 331\nTMhelix 332 354\ninside 355 372

17618 GCF_001952325.1_ASM195232v1 Salmonella enterica subsp. enterica serovar Heidelberg Proteobacteria; Gammaproteobacteria; Enterobacterales; Enterobacteriaceae; Salmonella; Salmonella enterica; Salmonella enterica subsp. enterica MNNEETFYQAMRRKGVTRRSFLKFCSLAATSLGLGAGMTPKIAWALENKPRIPVVWIHGL WP_001058312.1 hydrogenase [Salmonella enterica] Length: 372\nNumber of predicted TMHs: 2\nExp number of AAs in TMHs: 36.51738\nExp number, first 60 AAs: 19.72849\nTotal prob of N-in: 0.96170\nPOSSIBLE N-term signal sequence\ninside 1 19\nTMhelix 20 39\noutside 40 325\nTMhelix 326 348\ninside 349 372

17619 GCF_002060215.1_ASM206021v1 Salmonella enterica subsp. enterica serovar Heidelberg Proteobacteria; Gammaproteobacteria; Enterobacterales; Enterobacteriaceae; Salmonella; Salmonella enterica; Salmonella enterica subsp. enterica MNNEETFYQAMRRKGVTRRSFLKFCSLAATSLGLGAGMTPKIAWALENKPRIPVVWIHGL WP_001058312.1 hydrogenase [Salmonella enterica] Length: 372\nNumber of predicted TMHs: 2\nExp number of AAs in TMHs: 36.51738\nExp number, first 60 AAs: 19.72849\nTotal prob of N-in: 0.96170\nPOSSIBLE N-term signal sequence\ninside 1 19\nTMhelix 20 39\noutside 40 325\nTMhelix 326 348\ninside 349 372

17620 GCF_002037295.1_ASM203729v1 Salmonella enterica subsp. enterica serovar Heidelberg Proteobacteria; Gammaproteobacteria; Enterobacterales; Enterobacteriaceae; Salmonella; Salmonella enterica; Salmonella enterica subsp. enterica MNNEETFYQAMRRKGVTRRSFLKFCSLAATSLGLGAGMTPKIAWALENKPRIPVVWIHGL WP_001058312.1 hydrogenase [Salmonella enterica] Length: 372\nNumber of predicted TMHs: 2\nExp number of AAs in TMHs: 36.51738\nExp number, first 60 AAs: 19.72849\nTotal prob of N-in: 0.96170\nPOSSIBLE N-term signal sequence\ninside 1 19\nTMhelix 20 39\noutside 40 325\nTMhelix 326 348\ninside 349 372

17621 GCF_002037505.1_ASM203750v1 Salmonella enterica subsp. enterica serovar Heidelberg Proteobacteria; Gammaproteobacteria; Enterobacterales; Enterobacteriaceae; Salmonella; Salmonella enterica; Salmonella enterica subsp. enterica MNNEETFYQAMRRKGVTRRSFLKFCSLAATSLGLGAGMTPKIAWALENKPRIPVVWIHGL WP_001058312.1 hydrogenase [Salmonella enterica] Length: 372\nNumber of predicted TMHs: 2\nExp number of AAs in TMHs: 36.51738\nExp number, first 60 AAs: 19.72849\nTotal prob of N-in: 0.96170\nPOSSIBLE N-term signal sequence\ninside 1 19\nTMhelix 20 39\noutside 40 325\nTMhelix 326 348\ninside 349 372

17622 GCF_002037325.1_ASM203732v1 Salmonella enterica subsp. enterica serovar Heidelberg Proteobacteria; Gammaproteobacteria; Enterobacterales; Enterobacteriaceae; Salmonella; Salmonella enterica; Salmonella enterica subsp. enterica MTGDNTLITSHGINRRDFMKLCAALAATMGLSSKAAAEMAESVSNPQRPPVIWIGAQECT WP_000145429.1 hydrogenase 2 small subunit [Salmonella enterica] Length: 372\nNumber of predicted TMHs: 1\nExp number of AAs in TMHs: 24.51309\nExp number, first 60 AAs: 0.2682\nTotal prob of N-in: 0.14402\noutside 1 331\nTMhelix 332 354\ninside 355 372

17623 GCF_001692595.1_ASM169259v1 Salmonella enterica subsp. enterica serovar Heidelberg Proteobacteria; Gammaproteobacteria; Enterobacterales; Enterobacteriaceae; Salmonella; Salmonella enterica; Salmonella enterica subsp. enterica MNNEETFYQAMRRKGVTRRSFLKFCSLAATSLGLGAGMTPKIAWALENKPRIPVVWIHGL WP_001058312.1 hydrogenase [Salmonella enterica] Length: 372\nNumber of predicted TMHs: 2\nExp number of AAs in TMHs: 36.51738\nExp number, first 60 AAs: 19.72849\nTotal prob of N-in: 0.96170\nPOSSIBLE N-term signal sequence\ninside 1 19\nTMhelix 20 39\noutside 40 325\nTMhelix 326 348\ninside 349 372

17624 GCF_002062895.1_ASM206289v1 Salmonella enterica subsp. enterica serovar Heidelberg Proteobacteria; Gammaproteobacteria; Enterobacterales; Enterobacteriaceae; Salmonella; Salmonella enterica; Salmonella enterica subsp. enterica MNNEETFYQAMRRKGVTRRSFLKFCSLAATSLGLGAGMTPKIAWALENKPRIPVVWIHGL WP_001058312.1 hydrogenase [Salmonella enterica] Length: 372\nNumber of predicted TMHs: 2\nExp number of AAs in TMHs: 36.51738\nExp number, first 60 AAs: 19.72849\nTotal prob of N-in: 0.96170\nPOSSIBLE N-term signal sequence\ninside 1 19\nTMhelix 20 39\noutside 40 325\nTMhelix 326 348\ninside 349 372

17625 GCF_002062895.1_ASM206289v1 Salmonella enterica subsp. enterica serovar Heidelberg Proteobacteria; Gammaproteobacteria; Enterobacterales; Enterobacteriaceae; Salmonella; Salmonella enterica; Salmonella enterica subsp. enterica MTGDNTLITSHGINRRDFMKLCAALAATMGLSSKAAAEMAESVSNPQRPPVIWIGAQECT WP_000145429.1 hydrogenase 2 small subunit [Salmonella enterica] Length: 372\nNumber of predicted TMHs: 1\nExp number of AAs in TMHs: 24.51309\nExp number, first 60 AAs: 0.2682\nTotal prob of N-in: 0.14402\noutside 1 331\nTMhelix 332 354\ninside 355 372

17626 GCF_001690135.1_ASM169013v1 Salmonella enterica subsp. enterica serovar Heidelberg Proteobacteria; Gammaproteobacteria; Enterobacterales; Enterobacteriaceae; Salmonella; Salmonella enterica; Salmonella enterica subsp. enterica MTGDNTLITSHGINRRDFMKLCAALAATMGLSSKAAAEMAESVSNPQRPPVIWIGAQECT WP_000145429.1 hydrogenase 2 small subunit [Salmonella enterica] Length: 372\nNumber of

predicted TMHs:  1\nExp number of AAs in TMHs: 24.51309\nExp number, first 60 AAs:  0.2682\nTotal prob of N-in: 0.14402\noutside   1   331\nTMhelix   332   354\ninside   355   372

17627 GCF_002061515.1_ASM206151v1 Salmonella enterica subsp. enterica serovar Heidelberg Proteobacteria; Gammaproteobacteria; Enterobacterales; Enterobacteriaceae; Salmonella; Salmonella enterica; Salmonella enterica subsp. enterica MTGDNTLITSHGINRRDFMKLCAALAATMGLSSKAAAEMAESVSNPQRPPVIWIGAQECT WP_000145429.1 hydrogenase 2 small subunit [Salmonella enterica] Length: 372\nNumber of predicted TMHs:  1\nExp number of AAs in TMHs: 24.51309\nExp number, first 60 AAs:  0.2682\nTotal prob of N-in: 0.14402\noutside   1   331\nTMhelix   332   354\ninside   355   372

17628 GCF_002061515.1_ASM206151v1 Salmonella enterica subsp. enterica serovar Heidelberg Proteobacteria; Gammaproteobacteria; Enterobacterales; Enterobacteriaceae; Salmonella; Salmonella enterica; Salmonella enterica subsp. enterica MNNEETFYQAMRRKGVTRRSFLKFCSLAATSLGLGAGMTPKIAWALENKPRIPVVWIHGL WP_001058312.1 hydrogenase [Salmonella enterica] Length: 372\nNumber of predicted TMHs:  2\nExp number of AAs in TMHs: 36.51738\nExp number, first 60 AAs:  19.72849\nTotal prob of N-in:   0.96170\nPOSSIBLE N-term signal sequence\ninside   1   19\nTMhelix   20   39\noutside   40   325\nTMhelix   326   348\ninside   349   372

17629 GCF_001690135.1_ASM169013v1 Salmonella enterica subsp. enterica serovar Heidelberg Proteobacteria; Gammaproteobacteria; Enterobacterales; Enterobacteriaceae; Salmonella; Salmonella enterica; Salmonella enterica subsp. enterica MNNEETFYQAMRRKGVTRRSFLKFCSLAATSLGLGAGMTPKIAWALENKPRIPVVWIHGL WP_001058312.1 hydrogenase [Salmonella enterica] Length: 372\nNumber of predicted TMHs:  2\nExp number of AAs in TMHs: 36.51738\nExp number, first 60 AAs:  19.72849\nTotal prob of N-in:   0.96170\nPOSSIBLE N-term signal sequence\ninside   1   19\nTMhelix   20   39\noutside   40   325\nTMhelix   326   348\ninside   349   372

17630 GCF_001952305.1_ASM195230v1 Salmonella enterica subsp. enterica serovar Heidelberg Proteobacteria; Gammaproteobacteria; Enterobacterales; Enterobacteriaceae; Salmonella; Salmonella enterica; Salmonella enterica subsp. enterica MNNEETFYQAMRRKGVTRRSFLKFCSLAATSLGLGAGMTPKIAWALENKPRIPVVWIHGL WP_001058312.1 hydrogenase [Salmonella enterica] Length: 372\nNumber of predicted TMHs:  2\nExp number of AAs in TMHs: 36.51738\nExp number, first 60 AAs:  19.72849\nTotal prob of N-in:   0.96170\nPOSSIBLE N-term signal sequence\ninside   1   19\nTMhelix   20   39\noutside   40   325\nTMhelix   326   348\ninside   349   372

17631 GCF_001952475.1_ASM195247v1 Salmonella enterica subsp. enterica serovar Heidelberg Proteobacteria; Gammaproteobacteria; Enterobacterales; Enterobacteriaceae; Salmonella; Salmonella enterica; Salmonella enterica subsp. enterica MTGDNTLITSHGINRRDFMKLCAALAATMGLSSKAAAEMAESVSNPQRPPVIWIGAQECT WP_000145429.1 hydrogenase 2 small subunit [Salmonella enterica] Length: 372\nNumber of predicted TMHs:  1\nExp number of AAs in TMHs: 24.51309\nExp number, first 60 AAs:  0.2682\nTotal prob of N-in: 0.14402\noutside   1   331\nTMhelix   332   354\ninside   355   372

17632 GCF_001690005.1_ASM169000v1 Salmonella enterica subsp. enterica serovar Heidelberg Proteobacteria; Gammaproteobacteria; Enterobacterales; Enterobacteriaceae; Salmonella; Salmonella enterica; Salmonella enterica subsp. enterica MTGDNTLITSHGINRRDFMKLCAALAATMGLSSKAAAEMAESVSNPQRPPVIWIGAQECT WP_000145429.1 hydrogenase 2 small subunit [Salmonella enterica] Length: 372\nNumber of predicted TMHs:  1\nExp number of AAs in TMHs: 24.51309\nExp number, first 60 AAs:  0.2682\nTotal prob of N-in: 0.14402\noutside   1   331\nTMhelix   332   354\ninside   355   372

17633 GCF_001952405.1_ASM195240v1 Salmonella enterica subsp. enterica serovar Heidelberg Proteobacteria; Gammaproteobacteria; Enterobacterales; Enterobacteriaceae; Salmonella; Salmonella enterica; Salmonella enterica subsp. enterica MTGDNTLITSHGINRRDFMKLCAALAATMGLSSKAAAEMAESVSNPQRPPVIWIGAQECT WP_000145429.1 hydrogenase 2 small subunit [Salmonella enterica] Length: 372\nNumber of predicted TMHs:  1\nExp number of AAs in TMHs: 24.51309\nExp number, first 60 AAs:  0.2682\nTotal prob of N-in: 0.14402\noutside   1   331\nTMhelix   332   354\ninside   355   372

17634 GCF_001952305.1_ASM195230v1 Salmonella enterica subsp. enterica serovar Heidelberg Proteobacteria; Gammaproteobacteria; Enterobacterales; Enterobacteriaceae; Salmonella; Salmonella enterica; Salmonella enterica subsp. enterica MTGDNTLITSHGINRRDFMKLCAALAATMGLSSKAAAEMAESVSNPQRPPVIWIGAQECT WP_000145429.1 hydrogenase 2 small subunit [Salmonella enterica] Length: 372\nNumber of predicted TMHs:  1\nExp number of AAs in TMHs: 24.51309\nExp number, first 60 AAs:  0.2682\nTotal prob of N-in: 0.14402\noutside   1   331\nTMhelix   332   354\ninside   355   372

17635 GCF_002036185.1_ASM203618v1 Salmonella enterica subsp. enterica serovar Heidelberg Proteobacteria; Gammaproteobacteria; Enterobacterales; Enterobacteriaceae; Salmonella; Salmonella enterica;

Salmonella enterica subsp. enterica MTGDNTLITSHGINRRDFMKLCAALAATMGLSSKAAAEMAESVSNPQRPPVIWIGAQECT WP_000145429.1 hydrogenase 2 small subunit [Salmonella enterica] Length: 372\nNumber of predicted TMHs: 1\nExp number of AAs in TMHs: 24.51309\nExp number, first 60 AAs: 0.2682\nTotal prob of N-in: 0.14402\noutside 1 331\nTMhelix 332 354\ninside 355 372

17636 GCF_002037295.1_ASM203729v1 Salmonella enterica subsp. enterica serovar Heidelberg Proteobacteria; Gammaproteobacteria; Enterobacterales; Enterobacteriaceae; Salmonella; Salmonella enterica; Salmonella enterica subsp. enterica MTGDNTLITSHGINRRDFMKLCAALAATMGLSSKAAAEMAESVSNPQRPPVIWIGAQECT WP_000145429.1 hydrogenase 2 small subunit [Salmonella enterica] Length: 372\nNumber of predicted TMHs: 1\nExp number of AAs in TMHs: 24.51309\nExp number, first 60 AAs: 0.2682\nTotal prob of N-in: 0.14402\noutside 1 331\nTMhelix 332 354\ninside 355 372

17637 GCF_002064935.1_ASM206493v1 Salmonella enterica subsp. enterica serovar Heidelberg Proteobacteria; Gammaproteobacteria; Enterobacterales; Enterobacteriaceae; Salmonella; Salmonella enterica; Salmonella enterica subsp. enterica MTGDNTLITSHGINRRDFMKLCAALAATMGLSSKAAAEMAESVSNPQRPPVIWIGAQECT WP_000145429.1 hydrogenase 2 small subunit [Salmonella enterica] Length: 372\nNumber of predicted TMHs: 1\nExp number of AAs in TMHs: 24.51309\nExp number, first 60 AAs: 0.2682\nTotal prob of N-in: 0.14402\noutside 1 331\nTMhelix 332 354\ninside 355 372

17638 GCF_002060275.1_ASM206027v1 Salmonella enterica subsp. enterica serovar Heidelberg Proteobacteria; Gammaproteobacteria; Enterobacterales; Enterobacteriaceae; Salmonella; Salmonella enterica; Salmonella enterica subsp. enterica MNNEETFYQAMRRKGVTRRSFLKFCSLAATSLGLGAGMTPKIAWALENKPRIPVVWIHGL WP_001058312.1 hydrogenase [Salmonella enterica] Length: 372\nNumber of predicted TMHs: 2\nExp number of AAs in TMHs: 36.51738\nExp number, first 60 AAs: 19.72849\nTotal prob of N-in: 0.96170\nPOSSIBLE N-term signal sequence\ninside 1 19\nTMhelix 20 39\noutside 40 325\nTMhelix 326 348\ninside 349 372

17639 GCF_001692555.1_ASM169255v1 Salmonella enterica subsp. enterica serovar Heidelberg Proteobacteria; Gammaproteobacteria; Enterobacterales; Enterobacteriaceae; Salmonella; Salmonella enterica; Salmonella enterica subsp. enterica MNNEETFYQAMRRKGVTRRSFLKFCSLAATSLGLGAGMTPKIAWALENKPRIPVVWIHGL WP_001058312.1 hydrogenase [Salmonella enterica] Length: 372\nNumber of predicted TMHs: 2\nExp number of AAs in TMHs: 36.51738\nExp number, first 60 AAs: 19.72849\nTotal prob of N-in: 0.96170\nPOSSIBLE N-term signal sequence\ninside 1 19\nTMhelix 20 39\noutside 40 325\nTMhelix 326 348\ninside 349 372

17640 GCF_002064935.1_ASM206493v1 Salmonella enterica subsp. enterica serovar Heidelberg Proteobacteria; Gammaproteobacteria; Enterobacterales; Enterobacteriaceae; Salmonella; Salmonella enterica; Salmonella enterica subsp. enterica MNNEETFYQAMRRKGVTRRSFLKFCSLAATSLGLGAGMTPKIAWALENKPRIPVVWIHGL WP_001058312.1 hydrogenase [Salmonella enterica] Length: 372\nNumber of predicted TMHs: 2\nExp number of AAs in TMHs: 36.51738\nExp number, first 60 AAs: 19.72849\nTotal prob of N-in: 0.96170\nPOSSIBLE N-term signal sequence\ninside 1 19\nTMhelix 20 39\noutside 40 325\nTMhelix 326 348\ninside 349 372

17641 GCF_002060275.1_ASM206027v1 Salmonella enterica subsp. enterica serovar Heidelberg Proteobacteria; Gammaproteobacteria; Enterobacterales; Enterobacteriaceae; Salmonella; Salmonella enterica; Salmonella enterica subsp. enterica MTGDNTLITSHGINRRDFMKLCAALAATMGLSSKAAAEMAESVSNPQRPPVIWIGAQECT WP_000145429.1 hydrogenase 2 small subunit [Salmonella enterica] Length: 372\nNumber of predicted TMHs: 1\nExp number of AAs in TMHs: 24.51309\nExp number, first 60 AAs: 0.2682\nTotal prob of N-in: 0.14402\noutside 1 331\nTMhelix 332 354\ninside 355 372

17642 GCF_001723795.1_ASM172379v1 Salmonella enterica subsp. enterica serovar Heidelberg Proteobacteria; Gammaproteobacteria; Enterobacterales; Enterobacteriaceae; Salmonella; Salmonella enterica; Salmonella enterica subsp. enterica MTGDNTLITSHGINRRDFMKLCAALAATMGLSSKAAAEMAESVSNPQRPPVIWIGAQECT WP_039509062.1 hydrogenase 2 small subunit [Salmonella enterica] Length: 372\nNumber of predicted TMHs: 1\nExp number of AAs in TMHs: 24.51305\nExp number, first 60 AAs: 0.26819\nTotal prob of N-in: 0.14402\noutside 1 331\nTMhelix 332 354\ninside 355 372

17643 GCF_002036185.1_ASM203618v1 Salmonella enterica subsp. enterica serovar Heidelberg Proteobacteria; Gammaproteobacteria; Enterobacterales; Enterobacteriaceae; Salmonella; Salmonella enterica; Salmonella enterica subsp. enterica MNNEETFYQAMRRKGVTRRSFLKFCSLAATSLGLGAGMTPKIAWALENKPRIPVVWIHGL WP_001058312.1 hydrogenase [Salmonella enterica] Length: 372\nNumber of predicted TMHs: 2\nExp number of AAs in TMHs: 36.51738\nExp number, first 60 AAs: 19.72849\nTotal prob of N-in: 0.96170\nPOSSIBLE N-term signal sequence\ninside 1 19\nTMhelix 20 39\noutside 40 325\nTMhelix 326 348\ninside 349 372

17644 GCF_002060215.1_ASM206021v1 Salmonella enterica subsp. enterica serovar Heidelberg Proteobacteria; Gammaproteobacteria; Enterobacterales; Enterobacteriaceae; Salmonella; Salmonella enterica; Salmonella enterica subsp. enterica MTGDNTLITSHGINRRDFMKLCAALAATMGLSSKAAAEMAESVSNPQRPPVIWIGAQECT WP_000145429.1 hydrogenase 2 small subunit [Salmonella enterica] Length: 372\nNumber of predicted TMHs: 1\nExp number of AAs in TMHs: 24.51309\nExp number, first 60 AAs: 0.2682\nTotal prob of N-in: 0.14402\noutside 1 331\nTMhelix 332 354\ninside 355 372

17645 GCF_001723795.1_ASM172379v1 Salmonella enterica subsp. enterica serovar Heidelberg Proteobacteria; Gammaproteobacteria; Enterobacterales; Enterobacteriaceae; Salmonella; Salmonella enterica; Salmonella enterica subsp. enterica MNNEETFYQAMRRKGVTRRSFLKFCSLAATSLGLGAGMTPKIAWALENKPRIPVVWIHGL WP_001058312.1 hydrogenase [Salmonella enterica] Length: 372\nNumber of predicted TMHs: 2\nExp number of AAs in TMHs: 36.51738\nExp number, first 60 AAs: 19.72849\nTotal prob of N-in: 0.96170\nPOSSIBLE N-term signal sequence\ninside 1 19\nTMhelix 20 39\noutside 40 325\nTMhelix 326 348\ninside 349 372

17646 GCF_000974175.1_SALB-46.id18_v1.0 Salmonella enterica subsp. enterica serovar Heidelberg Proteobacteria; Gammaproteobacteria; Enterobacterales; Enterobacteriaceae; Salmonella; Salmonella enterica; Salmonella enterica subsp. enterica MNNEETFYQAMRRKGVTRRSFLKFCSLAATSLGLGAGMTPKIAWALENKPRIPVVWIHGL WP_001058312.1 hydrogenase [Salmonella enterica] Length: 372\nNumber of predicted TMHs: 2\nExp number of AAs in TMHs: 36.51738\nExp number, first 60 AAs: 19.72849\nTotal prob of N-in: 0.96170\nPOSSIBLE N-term signal sequence\ninside 1 19\nTMhelix 20 39\noutside 40 325\nTMhelix 326 348\ninside 349 372

17647 GCF_000974175.1_SALB-46.id18_v1.0 Salmonella enterica subsp. enterica serovar Heidelberg Proteobacteria; Gammaproteobacteria; Enterobacterales; Enterobacteriaceae; Salmonella; Salmonella enterica; Salmonella enterica subsp. enterica MTGDNTLITSHGINRRDFMKLCAALAATMGLSSKAAAEMAESVSNPQRPPVIWIGAQECT WP_000145429.1 hydrogenase 2 small subunit [Salmonella enterica] Length: 372\nNumber of predicted TMHs: 1\nExp number of AAs in TMHs: 24.51309\nExp number, first 60 AAs: 0.2682\nTotal prob of N-in: 0.14402\noutside 1 331\nTMhelix 332 354\ninside 355 372

17648 GCF_000973785.1_ABB07-SB3031.id20_v1.0 Salmonella enterica subsp. enterica serovar Heidelberg Proteobacteria; Gammaproteobacteria; Enterobacterales; Enterobacteriaceae; Salmonella; Salmonella enterica; Salmonella enterica subsp. enterica MTGDNTLITSHGINRRDFMKLCAALAATMGLSSKAAAEMAESVSNPQRPPVIWIGAQECT WP_000145429.1 hydrogenase 2 small subunit [Salmonella enterica] Length: 372\nNumber of predicted TMHs: 1\nExp number of AAs in TMHs: 24.51309\nExp number, first 60 AAs: 0.2682\nTotal prob of N-in: 0.14402\noutside 1 331\nTMhelix 332 354\ninside 355 372

17649 GCF_001690115.1_ASM169011v1 Salmonella enterica subsp. enterica serovar Heidelberg Proteobacteria; Gammaproteobacteria; Enterobacterales; Enterobacteriaceae; Salmonella; Salmonella enterica; Salmonella enterica subsp. enterica MNNEETFYQAMRRKGVTRRSFLKFCSLAATSLGLGAGMTPKIAWALENKPRIPVVWIHGL WP_001058312.1 hydrogenase [Salmonella enterica] Length: 372\nNumber of predicted TMHs: 2\nExp number of AAs in TMHs: 36.51738\nExp number, first 60 AAs: 19.72849\nTotal prob of N-in: 0.96170\nPOSSIBLE N-term signal sequence\ninside 1 19\nTMhelix 20 39\noutside 40 325\nTMhelix 326 348\ninside 349 372

17650 GCF_001690075.1_ASM169007v1 Salmonella enterica subsp. enterica serovar Heidelberg Proteobacteria; Gammaproteobacteria; Enterobacterales; Enterobacteriaceae; Salmonella; Salmonella enterica; Salmonella enterica subsp. enterica MNNEETFYQAMRRKGVTRRSFLKFCSLAATSLGLGAGMTPKIAWALENKPRIPVVWIHGL WP_001058312.1 hydrogenase [Salmonella enterica] Length: 372\nNumber of predicted TMHs: 2\nExp number of AAs in TMHs: 36.51738\nExp number, first 60 AAs: 19.72849\nTotal prob of N-in: 0.96170\nPOSSIBLE N-term signal sequence\ninside 1 19\nTMhelix 20 39\noutside 40 325\nTMhelix 326 348\ninside 349 372

17651 GCF_001690115.1_ASM169011v1 Salmonella enterica subsp. enterica serovar Heidelberg Proteobacteria; Gammaproteobacteria; Enterobacterales; Enterobacteriaceae; Salmonella; Salmonella enterica; Salmonella enterica subsp. enterica MTGDNTLITSHGINRRDFMKLCAALAATMGLSSKAAAEMAESVSNPQRPPVIWIGAQECT WP_000145429.1 hydrogenase 2 small subunit [Salmonella enterica] Length: 372\nNumber of predicted TMHs: 1\nExp number of AAs in TMHs: 24.51309\nExp number, first 60 AAs: 0.2682\nTotal prob of N-in: 0.14402\noutside 1 331\nTMhelix 332 354\ninside 355 372

17652 GCF_001690075.1_ASM169007v1 Salmonella enterica subsp. enterica serovar Heidelberg Proteobacteria; Gammaproteobacteria; Enterobacterales; Enterobacteriaceae; Salmonella; Salmonella enterica; Salmonella enterica subsp. enterica MTGDNTLITSHGINRRDFMKLCAALAATMGLSSKAAAEMAESVSNPQRPPVIWIGAQECT WP_000145429.1 hydrogenase 2 small subunit [Salmonella enterica] Length: 372\nNumber of

predicted TMHs: 1\nExp number of AAs in TMHs: 24.51309\nExp number, first 60 AAs: 0.2682\nTotal prob of N-in: 0.14402\noutside 1 331\nTMhelix 332 354\ninside 355 372

17653 GCF_001272375.1_Salmonella_enterica_CVM_N51271_v1.0 Salmonella enterica subsp. enterica serovar Heidelberg Proteobacteria; Gammaproteobacteria; Enterobacterales; Enterobacteriaceae; Salmonella; Salmonella enterica; Salmonella enterica subsp. enterica MNNEETFYQAMRRKGVTRRSFLKFCSLAATSLGLGAGMTPKIAWALENKPRIPVVWIHGL WP_001058312.1 hydrogenase [Salmonella enterica] Length: 372\nNumber of predicted TMHs: 2\nExp number of AAs in TMHs: 36.51738\nExp number, first 60 AAs: 19.72849\nTotal prob of N-in: 0.96170\nPOSSIBLE N-term signal sequence\ninside 1 19\nTMhelix 20 39\noutside 40 325\nTMhelix 326 348\ninside 349 372

17654 GCF_001690035.1_ASM169003v1 Salmonella enterica subsp. enterica serovar Heidelberg Proteobacteria; Gammaproteobacteria; Enterobacterales; Enterobacteriaceae; Salmonella; Salmonella enterica; Salmonella enterica subsp. enterica MNNEETFYQAMRRKGVTRRSFLKFCSLAATSLGLGAGMTPKIAWALENKPRIPVVWIHGL WP_001058312.1 hydrogenase [Salmonella enterica] Length: 372\nNumber of predicted TMHs: 2\nExp number of AAs in TMHs: 36.51738\nExp number, first 60 AAs: 19.72849\nTotal prob of N-in: 0.96170\nPOSSIBLE N-term signal sequence\ninside 1 19\nTMhelix 20 39\noutside 40 325\nTMhelix 326 348\ninside 349 372

17655 GCF_001242885.1_Salmonella_enterica_CVM_N46811_v1.0 Salmonella enterica subsp. enterica serovar Heidelberg Proteobacteria; Gammaproteobacteria; Enterobacterales; Enterobacteriaceae; Salmonella; Salmonella enterica; Salmonella enterica subsp. enterica MTGDNTLITSHGINRRDFMKLCAALAATMGLSSKAAAEMAESVSNPQRPPVIWIGAQECT WP_000145429.1 hydrogenase 2 small subunit [Salmonella enterica] Length: 372\nNumber of predicted TMHs: 1\nExp number of AAs in TMHs: 24.51309\nExp number, first 60 AAs: 0.2682\nTotal prob of N-in: 0.14402\noutside 1 331\nTMhelix 332 354\ninside 355 372

17656 GCF_001242965.1_Salmonella_enterica_CVM_N46818_v1.0 Salmonella enterica subsp. enterica serovar Heidelberg Proteobacteria; Gammaproteobacteria; Enterobacterales; Enterobacteriaceae; Salmonella; Salmonella enterica; Salmonella enterica subsp. enterica MTGDNTLITSHGINRRDFMKLCAALAATMGLSSKAAAEMAESVSNPQRPPVIWIGAQECT WP_000145429.1 hydrogenase 2 small subunit [Salmonella enterica] Length: 372\nNumber of predicted TMHs: 1\nExp number of AAs in TMHs: 24.51309\nExp number, first 60 AAs: 0.2682\nTotal prob of N-in: 0.14402\noutside 1 331\nTMhelix 332 354\ninside 355 372

17657 GCF_002062495.1_ASM206249v1 Salmonella enterica subsp. enterica serovar Heidelberg Proteobacteria; Gammaproteobacteria; Enterobacterales; Enterobacteriaceae; Salmonella; Salmonella enterica; Salmonella enterica subsp. enterica MTGDNTLITSHGINRRDFMKLCAALAATMGLSSKAAAEMAESVSNPQRPPVIWIGAQECT WP_000145429.1 hydrogenase 2 small subunit [Salmonella enterica] Length: 372\nNumber of predicted TMHs: 1\nExp number of AAs in TMHs: 24.51309\nExp number, first 60 AAs: 0.2682\nTotal prob of N-in: 0.14402\noutside 1 331\nTMhelix 332 354\ninside 355 372

17658 GCF_002060235.1_ASM206023v1 Salmonella enterica subsp. enterica serovar Heidelberg Proteobacteria; Gammaproteobacteria; Enterobacterales; Enterobacteriaceae; Salmonella; Salmonella enterica; Salmonella enterica subsp. enterica MNNEETFYQAMRRKGVTRRSFLKFCSLAATSLGLGAGMTPKIAWALENKPRIPVVWIHGL WP_001058312.1 hydrogenase [Salmonella enterica] Length: 372\nNumber of predicted TMHs: 2\nExp number of AAs in TMHs: 36.51738\nExp number, first 60 AAs: 19.72849\nTotal prob of N-in: 0.96170\nPOSSIBLE N-term signal sequence\ninside 1 19\nTMhelix 20 39\noutside 40 325\nTMhelix 326 348\ninside 349 372

17659 GCF_001479585.1_Salmonella_enterica_CVM_N37907-SQ_v1.0 Salmonella enterica subsp. enterica serovar Heidelberg Proteobacteria; Gammaproteobacteria; Enterobacterales; Enterobacteriaceae; Salmonella; Salmonella enterica; Salmonella enterica subsp. enterica MTGDNTLITSHGINRRDFMKLCAALAATMGLSSKAAAEMAESVSNPQRPPVIWIGAQECT WP_000145429.1 hydrogenase 2 small subunit [Salmonella enterica] Length: 372\nNumber of predicted TMHs: 1\nExp number of AAs in TMHs: 24.51309\nExp number, first 60 AAs: 0.2682\nTotal prob of N-in: 0.14402\noutside 1 331\nTMhelix 332 354\ninside 355 372

17660 GCF_001246855.1_Salmonella_enterica_CVM_N51282_v1.0 Salmonella enterica subsp. enterica serovar Heidelberg Proteobacteria; Gammaproteobacteria; Enterobacterales; Enterobacteriaceae; Salmonella; Salmonella enterica; Salmonella enterica subsp. enterica MNNEETFYQAMRRKGVTRRSFLKFCSLAATSLGLGAGMTPKIAWALENKPRIPVVWIHGL WP_001058312.1 hydrogenase [Salmonella enterica] Length: 372\nNumber of predicted TMHs: 2\nExp number of AAs in

TMHs: 36.51738\nExp number, first 60 AAs: 19.72849\nTotal prob of N-in: 0.96170\nPOSSIBLE N-term signal sequence\ninside 1 19\nTMhelix 20 39\noutside 40 325\nTMhelix 326 348\ninside 349 372

17661 GCF_001245845.1_Salmonella_enterica_CVM_N50449_v1.0 Salmonella enterica subsp. enterica serovar Heidelberg Proteobacteria; Gammaproteobacteria; Enterobacterales; Enterobacteriaceae; Salmonella; Salmonella enterica; Salmonella enterica subsp. enterica MNNEETFYQAMRRKGVTRRSFLKFCSLAATSLGLGAGMTPKIAWALENKPRIPVVWIHGL WP_001058312.1 hydrogenase [Salmonella enterica] Length: 372\nNumber of predicted TMHs: 2\nExp number of AAs in TMHs: 36.51738\nExp number, first 60 AAs: 19.72849\nTotal prob of N-in: 0.96170\nPOSSIBLE N-term signal sequence\ninside 1 19\nTMhelix 20 39\noutside 40 325\nTMhelix 326 348\ninside 349 372

17662 GCF_002037355.1_ASM203735v1 Salmonella enterica subsp. enterica serovar Heidelberg Proteobacteria; Gammaproteobacteria; Enterobacterales; Enterobacteriaceae; Salmonella; Salmonella enterica; Salmonella enterica subsp. enterica MNNEETFYQAMRRKGVTRRSFLKFCSLAATSLGLGAGMTPKIAWALENKPRIPVVWIHGL WP_001058312.1 hydrogenase [Salmonella enterica] Length: 372\nNumber of predicted TMHs: 2\nExp number of AAs in TMHs: 36.51738\nExp number, first 60 AAs: 19.72849\nTotal prob of N-in: 0.96170\nPOSSIBLE N-term signal sequence\ninside 1 19\nTMhelix 20 39\noutside 40 325\nTMhelix 326 348\ninside 349 372

17663 GCF_001272055.1_Salmonella_enterica_CVM_N44702_v1.0 Salmonella enterica subsp. enterica serovar Heidelberg Proteobacteria; Gammaproteobacteria; Enterobacterales; Enterobacteriaceae; Salmonella; Salmonella enterica; Salmonella enterica subsp. enterica MTGDNTLITSHGINRRDFMKLCAALAATMGLSSKAAAEMAESVSNPQRPPVIWIGAQECT WP_000145429.1 hydrogenase 2 small subunit [Salmonella enterica] Length: 372\nNumber of predicted TMHs: 1\nExp number of AAs in TMHs: 24.51309\nExp number, first 60 AAs: 0.2682\nTotal prob of N-in: 0.14402\noutside 1 331\nTMhelix 332 354\ninside 355 372

17664 GCF_001479435.1_Salmonella_enterica_CVM_N32754-SQ_v1.0 Salmonella enterica subsp. enterica serovar Heidelberg Proteobacteria; Gammaproteobacteria; Enterobacterales; Enterobacteriaceae; Salmonella; Salmonella enterica; Salmonella enterica subsp. enterica MTGDNTLITSHGINRRDFMKLCAALAATMGLSSKAAAEMAESVSNPQRPPVIWIGAQECT WP_000145429.1 hydrogenase 2 small subunit [Salmonella enterica] Length: 372\nNumber of predicted TMHs: 1\nExp number of AAs in TMHs: 24.51309\nExp number, first 60 AAs: 0.2682\nTotal prob of N-in: 0.14402\noutside 1 331\nTMhelix 332 354\ninside 355 372

17665 GCF_001241275.1_Salmonella_enterica_CVM_N44701-R_v1.0 Salmonella enterica subsp. enterica serovar Heidelberg Proteobacteria; Gammaproteobacteria; Enterobacterales; Enterobacteriaceae; Salmonella; Salmonella enterica; Salmonella enterica subsp. enterica MTGDNTLITSHGINRRDFMKLCAALAATMGLSSKAAAEMAESVSNPQRPPVIWIGAQECT WP_000145429.1 hydrogenase 2 small subunit [Salmonella enterica] Length: 372\nNumber of predicted TMHs: 1\nExp number of AAs in TMHs: 24.51309\nExp number, first 60 AAs: 0.2682\nTotal prob of N-in: 0.14402\noutside 1 331\nTMhelix 332 354\ninside 355 372

17666 GCF_001952465.1_Salmonella_enterica_CVM_N44695_v1.0 Salmonella enterica subsp. enterica serovar Heidelberg Proteobacteria; Gammaproteobacteria; Enterobacterales; Enterobacteriaceae; Salmonella; Salmonella enterica; Salmonella enterica subsp. enterica MTGDNTLITSHGINRRDFMKLCAALAATMGLSSKAAAEMAESVSNPQRPPVIWIGAQECT WP_000145429.1 hydrogenase 2 small subunit [Salmonella enterica] Length: 372\nNumber of predicted TMHs: 1\nExp number of AAs in TMHs: 24.51309\nExp number, first 60 AAs: 0.2682\nTotal prob of N-in: 0.14402\noutside 1 331\nTMhelix 332 354\ninside 355 372

17667 GCF_001241365.1_Salmonella_enterica_CVM_N44707_v1.0 Salmonella enterica subsp. enterica serovar Heidelberg Proteobacteria; Gammaproteobacteria; Enterobacterales; Enterobacteriaceae; Salmonella; Salmonella enterica; Salmonella enterica subsp. enterica MNNEETFYQAMRRKGVTRRSFLKFCSLAATSLGLGAGMTPKIAWALENKPRIPVVWIHGL WP_001058312.1 hydrogenase [Salmonella enterica] Length: 372\nNumber of predicted TMHs: 2\nExp number of AAs in TMHs: 36.51738\nExp number, first 60 AAs: 19.72849\nTotal prob of N-in: 0.96170\nPOSSIBLE N-term signal sequence\ninside 1 19\nTMhelix 20 39\noutside 40 325\nTMhelix 326 348\ninside 349 372

17668 GCF_001247585.1_Salmonella_enterica_CVM_N51984_v1.0 Salmonella enterica subsp. enterica serovar Heidelberg Proteobacteria; Gammaproteobacteria; Enterobacterales; Enterobacteriaceae; Salmonella; Salmonella enterica; Salmonella enterica subsp. enterica

MNNEETFYQAMRRKGVTRRSFLKFCSLAATSLGLGAGMTPKIAWALENKPRIPVVWIHGL WP_001058312.1 hydrogenase [Salmonella enterica] Length: 372\nNumber of predicted TMHs: 2\nExp number of AAs in TMHs: 36.51738\nExp number, first 60 AAs: 19.72849\nTotal prob of N-in: 0.96170\nPOSSIBLE N-term signal sequence\ninside 1 19\nTMhelix 20 39\noutside 40 325\nTMhelix 326 348\ninside 349 372

17669 GCF_002062495.1_ASM206249v1 Salmonella enterica subsp. enterica serovar Heidelberg Proteobacteria; Gammaproteobacteria; Enterobacterales; Enterobacteriaceae; Salmonella; Salmonella enterica; Salmonella enterica subsp. enterica MNNEETFYQAMRRKGVTRRSFLKFCSLAATSLGLGAGMTPKIAWALENKPRIPVVWIHGL WP_001058312.1 hydrogenase [Salmonella enterica] Length: 372\nNumber of predicted TMHs: 2\nExp number of AAs in TMHs: 36.51738\nExp number, first 60 AAs: 19.72849\nTotal prob of N-in: 0.96170\nPOSSIBLE N-term signal sequence\ninside 1 19\nTMhelix 20 39\noutside 40 325\nTMhelix 326 348\ninside 349 372

17670 GCF_001245845.1_Salmonella_enterica_CVM_N50449_v1.0 Salmonella enterica subsp. enterica serovar Heidelberg Proteobacteria; Gammaproteobacteria; Enterobacterales; Enterobacteriaceae; Salmonella; Salmonella enterica; Salmonella enterica subsp. enterica MTGDNTLITSHGINRRDFMKLCAALAATMGLSSKAAAEMAESVSNPQRPPVIWIGAQECT WP_000145429.1 hydrogenase 2 small subunit [Salmonella enterica] Length: 372\nNumber of predicted TMHs: 1\nExp number of AAs in TMHs: 24.51309\nExp number, first 60 AAs: 0.2682\nTotal prob of N-in: 0.14402\noutside 1 331\nTMhelix 332 354\ninside 355 372

17671 GCF_001477675.1_Salmonella_enterica_CVM_N29307-SQ_v1.0 Salmonella enterica subsp. enterica serovar Heidelberg Proteobacteria; Gammaproteobacteria; Enterobacterales; Enterobacteriaceae; Salmonella; Salmonella enterica; Salmonella enterica subsp. enterica MTGDNTLITSHGINRRDFMKLCAALAATMGLSSKAAAEMAESVSNPQRPPVIWIGAQECT WP_000145429.1 hydrogenase 2 small subunit [Salmonella enterica] Length: 372\nNumber of predicted TMHs: 1\nExp number of AAs in TMHs: 24.51309\nExp number, first 60 AAs: 0.2682\nTotal prob of N-in: 0.14402\noutside 1 331\nTMhelix 332 354\ninside 355 372

17672 GCF_001480445.1_Salmonella_enterica_CVM_N42239-SQ_v1.0 Salmonella enterica subsp. enterica serovar Heidelberg Proteobacteria; Gammaproteobacteria; Enterobacterales; Enterobacteriaceae; Salmonella; Salmonella enterica; Salmonella enterica subsp. enterica MNNEETFYQAMRRKGVTRRSFLKFCSLAATSLGLGAGMTPKIAWALENKPRIPVVWIHGL WP_001058312.1 hydrogenase [Salmonella enterica] Length: 372\nNumber of predicted TMHs: 2\nExp number of AAs in TMHs: 36.51738\nExp number, first 60 AAs: 19.72849\nTotal prob of N-in: 0.96170\nPOSSIBLE N-term signal sequence\ninside 1 19\nTMhelix 20 39\noutside 40 325\nTMhelix 326 348\ninside 349 372

17673 GCF_001479485.1_Salmonella_enterica_CVM_N32772-SQ_v1.0 Salmonella enterica subsp. enterica serovar Heidelberg Proteobacteria; Gammaproteobacteria; Enterobacterales; Enterobacteriaceae; Salmonella; Salmonella enterica; Salmonella enterica subsp. enterica MTGDNTLITSHGINRRDFMKLCAALAATMGLSSKAAAEMAESVSNPQRPPVIWIGAQECT WP_000145429.1 hydrogenase 2 small subunit [Salmonella enterica] Length: 372\nNumber of predicted TMHs: 1\nExp number of AAs in TMHs: 24.51309\nExp number, first 60 AAs: 0.2682\nTotal prob of N-in: 0.14402\noutside 1 331\nTMhelix 332 354\ninside 355 372

17674 GCF_001692615.1_ASM169261v1 Salmonella enterica subsp. enterica serovar Heidelberg Proteobacteria; Gammaproteobacteria; Enterobacterales; Enterobacteriaceae; Salmonella; Salmonella enterica; Salmonella enterica subsp. enterica MTGDNTLITSHGINRRDFMKLCAALAATMGLSSKAAAEMAESVSNPQRPPVIWIGAQECT WP_000145429.1 hydrogenase 2 small subunit [Salmonella enterica] Length: 372\nNumber of predicted TMHs: 1\nExp number of AAs in TMHs: 24.51309\nExp number, first 60 AAs: 0.2682\nTotal prob of N-in: 0.14402\noutside 1 331\nTMhelix 332 354\ninside 355 372

17675 GCF_001692495.1_ASM169249v1 Salmonella enterica subsp. enterica serovar Heidelberg Proteobacteria; Gammaproteobacteria; Enterobacterales; Enterobacteriaceae; Salmonella; Salmonella enterica; Salmonella enterica subsp. enterica MTGDNTLITSHGINRRDFMKLCAALAATMGLSSKAAAEMAESVSNPQRPPVIWIGAQECT WP_000145429.1 hydrogenase 2 small subunit [Salmonella enterica] Length: 372\nNumber of predicted TMHs: 1\nExp number of AAs in TMHs: 24.51309\nExp number, first 60 AAs: 0.2682\nTotal prob of N-in: 0.14402\noutside 1 331\nTMhelix 332 354\ninside 355 372

17676 GCF_001271555.1_Salmonella_enterica_CVM_N45393_v1.0 Salmonella enterica subsp. enterica serovar Heidelberg Proteobacteria; Gammaproteobacteria; Enterobacterales; Enterobacteriaceae; Salmonella; Salmonella enterica; Salmonella enterica subsp. enterica

MTGDNTLITSHGINRRDFMKLCAALAATMGLSSKAAAEMAESVSNPQRPPVIWIGAQECT WP_000145429.1 hydrogenase 2 small subunit [Salmonella enterica] Length: 372\nNumber of predicted TMHs: 1\nExp number of AAs in TMHs: 24.51309\nExp number, first 60 AAs: 0.2682\nTotal prob of N-in: 0.14402\noutside 1 331\nTMhelix 332 354\ninside 355 372

17677 GCF_001247075.1_Salmonella_enterica_CVM_N51292_v1.0 Salmonella enterica subsp. enterica serovar Heidelberg Proteobacteria; Gammaproteobacteria; Enterobacterales; Enterobacteriaceae; Salmonella; Salmonella enterica; Salmonella enterica subsp. enterica MTGDNTLITSHGINRRDFMKLCAALAATMGLSSKAAAEMAESVSNPQRPPVIWIGAQECT WP_000145429.1 hydrogenase 2 small subunit [Salmonella enterica] Length: 372\nNumber of predicted TMHs: 1\nExp number of AAs in TMHs: 24.51309\nExp number, first 60 AAs: 0.2682\nTotal prob of N-in: 0.14402\noutside 1 331\nTMhelix 332 354\ninside 355 372

17678 GCF_001243875.1_Salmonella_enterica_CVM_N47717_v1.0 Salmonella enterica subsp. enterica serovar Heidelberg Proteobacteria; Gammaproteobacteria; Enterobacterales; Enterobacteriaceae; Salmonella; Salmonella enterica; Salmonella enterica subsp. enterica MTGDNTLITSHGINRRDFMKLCAALAATMGLSSKAAAEMAESVSNPQRPPVIWIGAQECT WP_000145429.1 hydrogenase 2 small subunit [Salmonella enterica] Length: 372\nNumber of predicted TMHs: 1\nExp number of AAs in TMHs: 24.51309\nExp number, first 60 AAs: 0.2682\nTotal prob of N-in: 0.14402\noutside 1 331\nTMhelix 332 354\ninside 355 372

17679 GCF_001245655.1_Salmonella_enterica_CVM_N50440_v1.0 Salmonella enterica subsp. enterica serovar Heidelberg Proteobacteria; Gammaproteobacteria; Enterobacterales; Enterobacteriaceae; Salmonella; Salmonella enterica; Salmonella enterica subsp. enterica MTGDNTLITSHGINRRDFMKLCAALAATMGLSSKAAAEMAESVSNPQRPPVIWIGAQECT WP_000145429.1 hydrogenase 2 small subunit [Salmonella enterica] Length: 372\nNumber of predicted TMHs: 1\nExp number of AAs in TMHs: 24.51309\nExp number, first 60 AAs: 0.2682\nTotal prob of N-in: 0.14402\noutside 1 331\nTMhelix 332 354\ninside 355 372

17680 GCF_001245035.1_Salmonella_enterica_CVM_N48704_v1.0 Salmonella enterica subsp. enterica serovar Heidelberg Proteobacteria; Gammaproteobacteria; Enterobacterales; Enterobacteriaceae; Salmonella; Salmonella enterica; Salmonella enterica subsp. enterica MNNEETFYQAMRRKGVTRRSFLKFCSLAATSLGLGAGMTPKIAWALENKPRIPVVWIHGL WP_001058312.1 hydrogenase [Salmonella enterica] Length: 372\nNumber of predicted TMHs: 2\nExp number of AAs in TMHs: 36.51738\nExp number, first 60 AAs: 19.72849\nTotal prob of N-in: 0.96170\nPOSSIBLE N-term signal sequence\ninside 1 19\nTMhelix 20 39\noutside 40 325\nTMhelix 326 348\ninside 349 372

17681 GCF_001241355.1_Salmonella_enterica_CVM_N44706-R_v1.0 Salmonella enterica subsp. enterica serovar Heidelberg Proteobacteria; Gammaproteobacteria; Enterobacterales; Enterobacteriaceae; Salmonella; Salmonella enterica; Salmonella enterica subsp. enterica MNNEETFYQAMRRKGVTRRSFLKFCSLAATSLGLGAGMTPKIAWALENKPRIPVVWIHGL WP_001058312.1 hydrogenase [Salmonella enterica] Length: 372\nNumber of predicted TMHs: 2\nExp number of AAs in TMHs: 36.51738\nExp number, first 60 AAs: 19.72849\nTotal prob of N-in: 0.96170\nPOSSIBLE N-term signal sequence\ninside 1 19\nTMhelix 20 39\noutside 40 325\nTMhelix 326 348\ninside 349 372

17682 GCF_001241355.1_Salmonella_enterica_CVM_N44706-R_v1.0 Salmonella enterica subsp. enterica serovar Heidelberg Proteobacteria; Gammaproteobacteria; Enterobacterales; Enterobacteriaceae; Salmonella; Salmonella enterica; Salmonella enterica subsp. enterica MTGDNTLITSHGINRRDFMKLCAALAATMGLSSKAAAEMAESVSNPQRPPVIWIGAQECT WP_000145429.1 hydrogenase 2 small subunit [Salmonella enterica] Length: 372\nNumber of predicted TMHs: 1\nExp number of AAs in TMHs: 24.51309\nExp number, first 60 AAs: 0.2682\nTotal prob of N-in: 0.14402\noutside 1 331\nTMhelix 332 354\ninside 355 372

17683 GCF_001241105.1_Salmonella_enterica_CVM_N44695-R_v1.0 Salmonella enterica subsp. enterica serovar Heidelberg Proteobacteria; Gammaproteobacteria; Enterobacterales; Enterobacteriaceae; Salmonella; Salmonella enterica; Salmonella enterica subsp. enterica MTGDNTLITSHGINRRDFMKLCAALAATMGLSSKAAAEMAESVSNPQRPPVIWIGAQECT WP_000145429.1 hydrogenase 2 small subunit [Salmonella enterica] Length: 372\nNumber of predicted TMHs: 1\nExp number of AAs in TMHs: 24.51309\nExp number, first 60 AAs: 0.2682\nTotal prob of N-in: 0.14402\noutside 1 331\nTMhelix 332 354\ninside 355 372

17684 GCF_000974205.1_SALB-47-2.id25_v1.0 Salmonella enterica subsp. enterica serovar Heidelberg Proteobacteria; Gammaproteobacteria; Enterobacterales; Enterobacteriaceae; Salmonella; Salmonella enterica; Salmonella enterica subsp. enterica MNNEETFYQAMRRKGVTRRSFLKFCSLAATSLGLGAGMTPKIAWALENKPRIPVVWIHGL WP_001058312.1 hydrogenase [Salmonella enterica] Length: 372\nNumber of predicted TMHs: 2\nExp number of AAs in TMHs: 36.51738\nExp number, first 60 AAs: 19.72849\nTotal prob of N-in: 0.96170\nPOSSIBLE N-term signal sequence\ninside 1 19\nTMhelix 20 39\noutside 40 325\nTMhelix 326 348\ninside 349 372

17685 GCF_000974205.1_SALB-47-2.id25_v1.0 Salmonella enterica subsp. enterica serovar Heidelberg Proteobacteria; Gammaproteobacteria; Enterobacterales; Enterobacteriaceae; Salmonella; Salmonella enterica; Salmonella enterica subsp. enterica MTGDNTLITSHGINRRDFMKLCAALAATMGLSSKAAAEMAESVSNPQRPPVIWIGAQECT WP_000145429.1 hydrogenase 2 small subunit [Salmonella enterica] Length: 372\nNumber of predicted TMHs: 1\nExp number of AAs in TMHs: 24.51309\nExp number, first 60 AAs: 0.2682\nTotal prob of N-in: 0.14402\noutside 1 331\nTMhelix 332 354\ninside 355 372

17686 GCF_001242195.1_Salmonella_enterica_CVM_N45930_v1.0 Salmonella enterica subsp. enterica serovar Heidelberg Proteobacteria; Gammaproteobacteria; Enterobacterales; Enterobacteriaceae; Salmonella; Salmonella enterica; Salmonella enterica subsp. enterica MTGDNTLITSHGINRRDFMKLCAALAATMGLSSKAAAEMAESVSNPQRPPVIWIGAQECT WP_000145429.1 hydrogenase 2 small subunit [Salmonella enterica] Length: 372\nNumber of predicted TMHs: 1\nExp number of AAs in TMHs: 24.51309\nExp number, first 60 AAs: 0.2682\nTotal prob of N-in: 0.14402\noutside 1 331\nTMhelix 332 354\ninside 355 372

17687 GCF_001246915.1_Salmonella_enterica_CVM_N51285_v1.0 Salmonella enterica subsp. enterica serovar Heidelberg Proteobacteria; Gammaproteobacteria; Enterobacterales; Enterobacteriaceae; Salmonella; Salmonella enterica; Salmonella enterica subsp. enterica MTGDNTLITSHGINRRDFMKLCAALAATMGLSSKAAAEMAESVSNPQRPPVIWIGAQECT WP_000145429.1 hydrogenase 2 small subunit [Salmonella enterica] Length: 372\nNumber of predicted TMHs: 1\nExp number of AAs in TMHs: 24.51309\nExp number, first 60 AAs: 0.2682\nTotal prob of N-in: 0.14402\noutside 1 331\nTMhelix 332 354\ninside 355 372

17688 GCF_001241625.1_Salmonella_enterica_CVM_N45392_v1.0 Salmonella enterica subsp. enterica serovar Heidelberg Proteobacteria; Gammaproteobacteria; Enterobacterales; Enterobacteriaceae; Salmonella; Salmonella enterica; Salmonella enterica subsp. enterica MTGDNTLITSHGINRRDFMKLCAALAATMGLSSKAAAEMAESVSNPQRPPVIWIGAQECT WP_000145429.1 hydrogenase 2 small subunit [Salmonella enterica] Length: 372\nNumber of predicted TMHs: 1\nExp number of AAs in TMHs: 24.51309\nExp number, first 60 AAs: 0.2682\nTotal prob of N-in: 0.14402\noutside 1 331\nTMhelix 332 354\ninside 355 372

17689 GCF_001243875.1_Salmonella_enterica_CVM_N47717_v1.0 Salmonella enterica subsp. enterica serovar Heidelberg Proteobacteria; Gammaproteobacteria; Enterobacterales; Enterobacteriaceae; Salmonella; Salmonella enterica; Salmonella enterica subsp. enterica MNNEETFYQAMRRKGVTRRSFLKFCSLAATSLGLGAGMTPKIAWALENKPRIPVVWIHGL WP_001058312.1 hydrogenase [Salmonella enterica] Length: 372\nNumber of predicted TMHs: 2\nExp number of AAs in TMHs: 36.51738\nExp number, first 60 AAs: 19.72849\nTotal prob of N-in: 0.96170\nPOSSIBLE N-term signal sequence\ninside 1 19\nTMhelix 20 39\noutside 40 325\nTMhelix 326 348\ninside 349 372

17690 GCF_001240635.1_Salmonella_enterica_CVM_N43480_v1.0 Salmonella enterica subsp. enterica serovar Heidelberg Proteobacteria; Gammaproteobacteria; Enterobacterales; Enterobacteriaceae; Salmonella; Salmonella enterica; Salmonella enterica subsp. enterica MNNEETFYQAMRRKGVTRRSFLKFCSLAATSLGLGAGMTPKIAWALENKPRIPVVWIHGL WP_001058312.1 hydrogenase [Salmonella enterica] Length: 372\nNumber of predicted TMHs: 2\nExp number of AAs in TMHs: 36.51738\nExp number, first 60 AAs: 19.72849\nTotal prob of N-in: 0.96170\nPOSSIBLE N-term signal sequence\ninside 1 19\nTMhelix 20 39\noutside 40 325\nTMhelix 326 348\ninside 349 372

17691 GCF_001243065.1_Salmonella_enterica_CVM_N46821_v1.0 Salmonella enterica subsp. enterica serovar Heidelberg Proteobacteria; Gammaproteobacteria; Enterobacterales; Enterobacteriaceae; Salmonella; Salmonella enterica; Salmonella enterica subsp. enterica MNNEETFYQAMRRKGVTRRSFLKFCSLAATSLGLGAGMTPKIAWALENKPRIPVVWIHGL WP_001058312.1 hydrogenase [Salmonella enterica] Length: 372\nNumber of predicted TMHs: 2\nExp number of AAs in TMHs: 36.51738\nExp number, first 60 AAs: 19.72849\nTotal prob of N-in: 0.96170\nPOSSIBLE N-term signal sequence\ninside 1 19\nTMhelix 20 39\noutside 40 325\nTMhelix 326 348\ninside 349 372

17692 GCF_001240675.1_Salmonella_enterica_CVM_N43481_v1.0 Salmonella enterica subsp. enterica serovar Heidelberg Proteobacteria; Gammaproteobacteria; Enterobacterales; Enterobacteriaceae; Salmonella; Salmonella enterica; Salmonella enterica subsp. enterica
MTGDNTLITSHGINRRDFMKLCAALAATMGLSSKAAAEMAESVSNPQRPPVIWIGAQECT WP_000145429.1
hydrogenase 2 small subunit [Salmonella enterica] Length: 372\nNumber of predicted TMHs: 1\nExp number of AAs in TMHs: 24.51309\nExp number, first 60 AAs: 0.2682\nTotal prob of N-in: 0.14402\noutside 1 331\nTMhelix 332 354\ninside 355 372

17693 GCF_001240635.1_Salmonella_enterica_CVM_N43480_v1.0 Salmonella enterica subsp. enterica serovar Heidelberg Proteobacteria; Gammaproteobacteria; Enterobacterales; Enterobacteriaceae; Salmonella; Salmonella enterica; Salmonella enterica subsp. enterica
MTGDNTLITSHGINRRDFMKLCAALAATMGLSSKAAAEMAESVSNPQRPPVIWIGAQECT WP_000145429.1
hydrogenase 2 small subunit [Salmonella enterica] Length: 372\nNumber of predicted TMHs: 1\nExp number of AAs in TMHs: 24.51309\nExp number, first 60 AAs: 0.2682\nTotal prob of N-in: 0.14402\noutside 1 331\nTMhelix 332 354\ninside 355 372

17694 GCF_001240545.1_Salmonella_enterica_CVM_N43473_v1.0 Salmonella enterica subsp. enterica serovar Heidelberg Proteobacteria; Gammaproteobacteria; Enterobacterales; Enterobacteriaceae; Salmonella; Salmonella enterica; Salmonella enterica subsp. enterica
MTGDNTLITSHGINRRDFMKLCAALAATMGLSSKAAAEMAESVSNPQRPPVIWIGAQECT WP_000145429.1
hydrogenase 2 small subunit [Salmonella enterica] Length: 372\nNumber of predicted TMHs: 1\nExp number of AAs in TMHs: 24.51309\nExp number, first 60 AAs: 0.2682\nTotal prob of N-in: 0.14402\noutside 1 331\nTMhelix 332 354\ninside 355 372

17695 GCF_001240145.1_Salmonella_enterica_CVM_N43453_v1.0 Salmonella enterica subsp. enterica serovar Heidelberg Proteobacteria; Gammaproteobacteria; Enterobacterales; Enterobacteriaceae; Salmonella; Salmonella enterica; Salmonella enterica subsp. enterica
MNNEETFYQAMRRKGVTRRSFLKFCSLAATSLGLGAGMTPKIAWALENKPRIPVVWIHGL WP_001058312.1
hydrogenase [Salmonella enterica] Length: 372\nNumber of predicted TMHs: 2\nExp number of AAs in TMHs: 36.51738\nExp number, first 60 AAs: 19.72849\nTotal prob of N-in: 0.96170\nPOSSIBLE N-term signal sequence\ninside 1 19\nTMhelix 20 39\noutside 40 325\nTMhelix 326 348\ninside 349 372

17696 GCF_001241625.1_Salmonella_enterica_CVM_N45392_v1.0 Salmonella enterica subsp. enterica serovar Heidelberg Proteobacteria; Gammaproteobacteria; Enterobacterales; Enterobacteriaceae; Salmonella; Salmonella enterica; Salmonella enterica subsp. enterica
MNNEETFYQAMRRKGVTRRSFLKFCSLAATSLGLGAGMTPKIAWALENKPRIPVVWIHGL WP_001058312.1
hydrogenase [Salmonella enterica] Length: 372\nNumber of predicted TMHs: 2\nExp number of AAs in TMHs: 36.51738\nExp number, first 60 AAs: 19.72849\nTotal prob of N-in: 0.96170\nPOSSIBLE N-term signal sequence\ninside 1 19\nTMhelix 20 39\noutside 40 325\nTMhelix 326 348\ninside 349 372

17697 GCF_001246915.1_Salmonella_enterica_CVM_N51285_v1.0 Salmonella enterica subsp. enterica serovar Heidelberg Proteobacteria; Gammaproteobacteria; Enterobacterales; Enterobacteriaceae; Salmonella; Salmonella enterica; Salmonella enterica subsp. enterica
MNNEETFYQAMRRKGVTRRSFLKFCSLAATSLGLGAGMTPKIAWALENKPRIPVVWIHGL WP_001058312.1
hydrogenase [Salmonella enterica] Length: 372\nNumber of predicted TMHs: 2\nExp number of AAs in TMHs: 36.51738\nExp number, first 60 AAs: 19.72849\nTotal prob of N-in: 0.96170\nPOSSIBLE N-term signal sequence\ninside 1 19\nTMhelix 20 39\noutside 40 325\nTMhelix 326 348\ninside 349 372

17698 GCF_001245005.1_Salmonella_enterica_CVM_N48699_v1.0 Salmonella enterica subsp. enterica serovar Heidelberg Proteobacteria; Gammaproteobacteria; Enterobacterales; Enterobacteriaceae; Salmonella; Salmonella enterica; Salmonella enterica subsp. enterica
MNNEETFYQAMRRKGVTRRSFLKFCSLAATSLGLGAGMTPKIAWALENKPRIPVVWIHGL WP_001058312.1
hydrogenase [Salmonella enterica] Length: 372\nNumber of predicted TMHs: 2\nExp number of AAs in TMHs: 36.51738\nExp number, first 60 AAs: 19.72849\nTotal prob of N-in: 0.96170\nPOSSIBLE N-term signal sequence\ninside 1 19\nTMhelix 20 39\noutside 40 325\nTMhelix 326 348\ninside 349 372

17699 GCF_001245005.1_Salmonella_enterica_CVM_N48699_v1.0 Salmonella enterica subsp. enterica serovar Heidelberg Proteobacteria; Gammaproteobacteria; Enterobacterales; Enterobacteriaceae; Salmonella; Salmonella enterica; Salmonella enterica subsp. enterica
MTGDNTLITSHGINRRDFMKLCAALAATMGLSSKAAAEMAESVSNPQRPPVIWIGAQECT WP_000145429.1
hydrogenase 2 small subunit [Salmonella enterica] Length: 372\nNumber of predicted TMHs: 1\nExp

number of AAs in TMHs: 24.51309\nExp number, first 60 AAs: 0.2682\nTotal prob of N-in: 0.14402\noutside 1 331\nTMhelix 332 354\ninside 355 372

17700 GCF_001242755.1_Salmonella_enterica_CVM_N45958_v1.0 Salmonella enterica subsp. enterica serovar Heidelberg Proteobacteria; Gammaproteobacteria; Enterobacterales; Enterobacteriaceae; Salmonella; Salmonella enterica; Salmonella enterica subsp. enterica MTGDNTLITSHGINRRDFMKLCAALAATMGLSSKAAAEMAESVSNPQRPPVIWIGAQECT WP_000145429.1 hydrogenase 2 small subunit [Salmonella enterica] Length: 372\nNumber of predicted TMHs: 1\nExp number of AAs in TMHs: 24.51309\nExp number, first 60 AAs: 0.2682\nTotal prob of N-in: 0.14402\noutside 1 331\nTMhelix 332 354\ninside 355 372

17701 GCF_001245435.1_Salmonella_enterica_CVM_N50428_v1.0 Salmonella enterica subsp. enterica serovar Heidelberg Proteobacteria; Gammaproteobacteria; Enterobacterales; Enterobacteriaceae; Salmonella; Salmonella enterica; Salmonella enterica subsp. enterica MTGDNTLITSHGINRRDFMKLCAALAATMGLSSKAAAEMAESVSNPQRPPVIWIGAQECT WP_000145429.1 hydrogenase 2 small subunit [Salmonella enterica] Length: 372\nNumber of predicted TMHs: 1\nExp number of AAs in TMHs: 24.51309\nExp number, first 60 AAs: 0.2682\nTotal prob of N-in: 0.14402\noutside 1 331\nTMhelix 332 354\ninside 355 372

17702 GCF_001245035.1_Salmonella_enterica_CVM_N48704_v1.0 Salmonella enterica subsp. enterica serovar Heidelberg Proteobacteria; Gammaproteobacteria; Enterobacterales; Enterobacteriaceae; Salmonella; Salmonella enterica; Salmonella enterica subsp. enterica MTGDNTLITSHGINRRDFMKLCAALAATMGLSSKAAAEMAESVSNPQRPPVIWIGAQECT WP_000145429.1 hydrogenase 2 small subunit [Salmonella enterica] Length: 372\nNumber of predicted TMHs: 1\nExp number of AAs in TMHs: 24.51309\nExp number, first 60 AAs: 0.2682\nTotal prob of N-in: 0.14402\noutside 1 331\nTMhelix 332 354\ninside 355 372

17703 GCF_001240885.1_Salmonella_enterica_CVM_N43828_v1.0 Salmonella enterica subsp. enterica serovar Heidelberg Proteobacteria; Gammaproteobacteria; Enterobacterales; Enterobacteriaceae; Salmonella; Salmonella enterica; Salmonella enterica subsp. enterica MTGDNTLITSHGINRRDFMKLCAALAATMGLSSKAAAEMAESVSNPQRPPVIWIGAQECT WP_000145429.1 hydrogenase 2 small subunit [Salmonella enterica] Length: 372\nNumber of predicted TMHs: 1\nExp number of AAs in TMHs: 24.51309\nExp number, first 60 AAs: 0.2682\nTotal prob of N-in: 0.14402\noutside 1 331\nTMhelix 332 354\ninside 355 372

17704 GCF_001241845.1_Salmonella_enterica_CVM_N45401_v1.0 Salmonella enterica subsp. enterica serovar Heidelberg Proteobacteria; Gammaproteobacteria; Enterobacterales; Enterobacteriaceae; Salmonella; Salmonella enterica; Salmonella enterica subsp. enterica MNNEETFYQAMRRKGVTRRSFLKFCSLAATSLGLGAGMTPKIAWALENKPRIPVVWIHGL WP_001058312.1 hydrogenase [Salmonella enterica] Length: 372\nNumber of predicted TMHs: 2\nExp number of AAs in TMHs: 36.51738\nExp number, first 60 AAs: 19.72849\nTotal prob of N-in: 0.96170\nPOSSIBLE N-term signal sequence\ninside 1 19\nTMhelix 20 39\noutside 40 325\nTMhelix 326 348\ninside 349 372

17705 GCF_001272325.1_Salmonella_enterica_CVM_N51243_v1.0 Salmonella enterica subsp. enterica serovar Heidelberg Proteobacteria; Gammaproteobacteria; Enterobacterales; Enterobacteriaceae; Salmonella; Salmonella enterica; Salmonella enterica subsp. enterica MNNEETFYQAMRRKGVTRRSFLKFCSLAATSLGLGAGMTPKIAWALENKPRIPVVWIHGL WP_001058312.1 hydrogenase [Salmonella enterica] Length: 372\nNumber of predicted TMHs: 2\nExp number of AAs in TMHs: 36.51738\nExp number, first 60 AAs: 19.72849\nTotal prob of N-in: 0.96170\nPOSSIBLE N-term signal sequence\ninside 1 19\nTMhelix 20 39\noutside 40 325\nTMhelix 326 348\ninside 349 372

17706 GCF_001245435.1_Salmonella_enterica_CVM_N50428_v1.0 Salmonella enterica subsp. enterica serovar Heidelberg Proteobacteria; Gammaproteobacteria; Enterobacterales; Enterobacteriaceae; Salmonella; Salmonella enterica; Salmonella enterica subsp. enterica MNNEETFYQAMRRKGVTRRSFLKFCSLAATSLGLGAGMTPKIAWALENKPRIPVVWIHGL WP_001058312.1 hydrogenase [Salmonella enterica] Length: 372\nNumber of predicted TMHs: 2\nExp number of AAs in TMHs: 36.51738\nExp number, first 60 AAs: 19.72849\nTotal prob of N-in: 0.96170\nPOSSIBLE N-term signal sequence\ninside 1 19\nTMhelix 20 39\noutside 40 325\nTMhelix 326 348\ninside 349 372

17707 GCF_001272325.1_Salmonella_enterica_CVM_N51243_v1.0 Salmonella enterica subsp. enterica serovar Heidelberg Proteobacteria; Gammaproteobacteria; Enterobacterales; Enterobacteriaceae; Salmonella; Salmonella

enterica; Salmonella enterica subsp. enterica
MTGDNTLITSHGINRRDFMKLCAALAATMGLSSKAAAEMAESVSNPQRPPVIWIGAQECT WP_000145429.1
hydrogenase 2 small subunit [Salmonella enterica] Length: 372\nNumber of predicted TMHs: 1\nExp number of AAs in TMHs: 24.51309\nExp number, first 60 AAs: 0.2682\nTotal prob of N-in: 0.14402\noutside 1 331\nTMhelix 332 354\ninside 355 372

17708 GCF_001241845.1_Salmonella_enterica_CVM_N45401_v1.0 Salmonella enterica subsp. enterica serovar Heidelberg Proteobacteria; Gammaproteobacteria; Enterobacterales; Enterobacteriaceae; Salmonella; Salmonella enterica; Salmonella enterica subsp. enterica
MTGDNTLITSHGINRRDFMKLCAALAATMGLSSKAAAEMAESVSNPQRPPVIWIGAQECT WP_000145429.1
hydrogenase 2 small subunit [Salmonella enterica] Length: 372\nNumber of predicted TMHs: 1\nExp number of AAs in TMHs: 24.51309\nExp number, first 60 AAs: 0.2682\nTotal prob of N-in: 0.14402\noutside 1 331\nTMhelix 332 354\ninside 355 372

17709 GCF_001247145.1_Salmonella_enterica_CVM_N51297_v1.0 Salmonella enterica subsp. enterica serovar Heidelberg Proteobacteria; Gammaproteobacteria; Enterobacterales; Enterobacteriaceae; Salmonella; Salmonella enterica; Salmonella enterica subsp. enterica
MNNEETFYQAMRRKGVTRRSFLKFCSLAATSLGLGAGMTPKIAWALENKPRIPVVWIHGL WP_001058312.1
hydrogenase [Salmonella enterica] Length: 372\nNumber of predicted TMHs: 2\nExp number of AAs in TMHs: 36.51738\nExp number, first 60 AAs: 19.72849\nTotal prob of N-in: 0.96170\nPOSSIBLE N-term signal sequence\ninside 1 19\nTMhelix 20 39\noutside 40 325\nTMhelix 326 348\ninside 349 372

17710 GCF_001245965.1_Salmonella_enterica_CVM_N51244_v1.0 Salmonella enterica subsp. enterica serovar Heidelberg Proteobacteria; Gammaproteobacteria; Enterobacterales; Enterobacteriaceae; Salmonella; Salmonella enterica; Salmonella enterica subsp. enterica
MTGDNTLITSHGINRRDFMKLCAALAATMGLSSKAAAEMAESVSNPQRPPVIWIGAQECT WP_000145429.1
hydrogenase 2 small subunit [Salmonella enterica] Length: 372\nNumber of predicted TMHs: 1\nExp number of AAs in TMHs: 24.51309\nExp number, first 60 AAs: 0.2682\nTotal prob of N-in: 0.14402\noutside 1 331\nTMhelix 332 354\ninside 355 372

17711 GCF_001246035.1_Salmonella_enterica_CVM_N51246_v1.0 Salmonella enterica subsp. enterica serovar Heidelberg Proteobacteria; Gammaproteobacteria; Enterobacterales; Enterobacteriaceae; Salmonella; Salmonella enterica; Salmonella enterica subsp. enterica
MTGDNTLITSHGINRRDFMKLCAALAATMGLSSKAAAEMAESVSNPQRPPVIWIGAQECT WP_000145429.1
hydrogenase 2 small subunit [Salmonella enterica] Length: 372\nNumber of predicted TMHs: 1\nExp number of AAs in TMHs: 24.51309\nExp number, first 60 AAs: 0.2682\nTotal prob of N-in: 0.14402\noutside 1 331\nTMhelix 332 354\ninside 355 372

17712 GCF_001240145.1_Salmonella_enterica_CVM_N43453_v1.0 Salmonella enterica subsp. enterica serovar Heidelberg Proteobacteria; Gammaproteobacteria; Enterobacterales; Enterobacteriaceae; Salmonella; Salmonella enterica; Salmonella enterica subsp. enterica
MTGDNTLITSHGINRRDFMKLCAALAATMGLSSKAAAEMAESVSNPQRPPVIWIGAQECT WP_000145429.1
hydrogenase 2 small subunit [Salmonella enterica] Length: 372\nNumber of predicted TMHs: 1\nExp number of AAs in TMHs: 24.51309\nExp number, first 60 AAs: 0.2682\nTotal prob of N-in: 0.14402\noutside 1 331\nTMhelix 332 354\ninside 355 372

17713 GCF_001242195.1_Salmonella_enterica_CVM_N45930_v1.0 Salmonella enterica subsp. enterica serovar Heidelberg Proteobacteria; Gammaproteobacteria; Enterobacterales; Enterobacteriaceae; Salmonella; Salmonella enterica; Salmonella enterica subsp. enterica
MNNEETFYQAMRRKGVTRRSFLKFCSLAATSLGLGAGMTPKIAWALENKPRIPVVWIHGL WP_001058312.1
hydrogenase [Salmonella enterica] Length: 372\nNumber of predicted TMHs: 2\nExp number of AAs in TMHs: 36.51738\nExp number, first 60 AAs: 19.72849\nTotal prob of N-in: 0.96170\nPOSSIBLE N-term signal sequence\ninside 1 19\nTMhelix 20 39\noutside 40 325\nTMhelix 326 348\ninside 349 372

17714 GCF_001245965.1_Salmonella_enterica_CVM_N51244_v1.0 Salmonella enterica subsp. enterica serovar Heidelberg Proteobacteria; Gammaproteobacteria; Enterobacterales; Enterobacteriaceae; Salmonella; Salmonella enterica; Salmonella enterica subsp. enterica
MNNEETFYQAMRRKGVTRRSFLKFCSLAATSLGLGAGMTPKIAWALENKPRIPVVWIHGL WP_001058312.1
hydrogenase [Salmonella enterica] Length: 372\nNumber of predicted TMHs: 2\nExp number of AAs in TMHs: 36.51738\nExp number, first 60 AAs: 19.72849\nTotal prob of N-in: 0.96170\nPOSSIBLE N-term signal sequence\ninside 1 19\nTMhelix 20 39\noutside 40 325\nTMhelix 326 348\ninside 349 372

17715 GCF_001243065.1_Salmonella_enterica_CVM_N46821_v1.0 Salmonella enterica subsp. enterica serovar Heidelberg Proteobacteria; Gammaproteobacteria; Enterobacterales; Enterobacteriaceae; Salmonella; Salmonella enterica; Salmonella enterica subsp. enterica
MTGDNTLITSHGINRRDFMKLCAALAATMGLSSKAAAEMAESVSNPQRPPVIWIGAQECT WP_000145429.1
hydrogenase 2 small subunit [Salmonella enterica] Length: 372\nNumber of predicted TMHs: 1\nExp number of AAs in TMHs: 24.51309\nExp number, first 60 AAs: 0.2682\nTotal prob of N-in: 0.14402\noutside 1 331\nTMhelix 332 354\ninside 355 372

17716 GCF_001245655.1_Salmonella_enterica_CVM_N50440_v1.0 Salmonella enterica subsp. enterica serovar Heidelberg Proteobacteria; Gammaproteobacteria; Enterobacterales; Enterobacteriaceae; Salmonella; Salmonella enterica; Salmonella enterica subsp. enterica
MNNEETFYQAMRRKGVTRRSFLKFCSLAATSLGLGAGMTPKIAWALENKPRIPVVWIHGL WP_001058312.1
hydrogenase [Salmonella enterica] Length: 372\nNumber of predicted TMHs: 2\nExp number of AAs in TMHs: 36.51738\nExp number, first 60 AAs: 19.72849\nTotal prob of N-in: 0.96170\nPOSSIBLE N-term signal sequence\ninside 1 19\nTMhelix 20 39\noutside 40 325\nTMhelix 326 348\ninside 349 372

17717 GCF_001242755.1_Salmonella_enterica_CVM_N45958_v1.0 Salmonella enterica subsp. enterica serovar Heidelberg Proteobacteria; Gammaproteobacteria; Enterobacterales; Enterobacteriaceae; Salmonella; Salmonella enterica; Salmonella enterica subsp. enterica
MNNEETFYQAMRRKGVTRRSFLKFCSLAATSLGLGAGMTPKIAWALENKPRIPVVWIHGL WP_001058312.1
hydrogenase [Salmonella enterica] Length: 372\nNumber of predicted TMHs: 2\nExp number of AAs in TMHs: 36.51738\nExp number, first 60 AAs: 19.72849\nTotal prob of N-in: 0.96170\nPOSSIBLE N-term signal sequence\ninside 1 19\nTMhelix 20 39\noutside 40 325\nTMhelix 326 348\ninside 349 372

17718 GCF_001246035.1_Salmonella_enterica_CVM_N51246_v1.0 Salmonella enterica subsp. enterica serovar Heidelberg Proteobacteria; Gammaproteobacteria; Enterobacterales; Enterobacteriaceae; Salmonella; Salmonella enterica; Salmonella enterica subsp. enterica
MNNEETFYQAMRRKGVTRRSFLKFCSLAATSLGLGAGMTPKIAWALENKPRIPVVWIHGL WP_001058312.1
hydrogenase [Salmonella enterica] Length: 372\nNumber of predicted TMHs: 2\nExp number of AAs in TMHs: 36.51738\nExp number, first 60 AAs: 19.72849\nTotal prob of N-in: 0.96170\nPOSSIBLE N-term signal sequence\ninside 1 19\nTMhelix 20 39\noutside 40 325\nTMhelix 326 348\ninside 349 372

17719 GCF_001241105.1_Salmonella_enterica_CVM_N44695-R_v1.0 Salmonella enterica subsp. enterica serovar Heidelberg Proteobacteria; Gammaproteobacteria; Enterobacterales; Enterobacteriaceae; Salmonella; Salmonella enterica; Salmonella enterica subsp. enterica
MNNEETFYQAMRRKGVTRRSFLKFCSLAATSLGLGAGMTPKIAWALENKPRIPVVWIHGL WP_001058312.1
hydrogenase [Salmonella enterica] Length: 372\nNumber of predicted TMHs: 2\nExp number of AAs in TMHs: 36.51738\nExp number, first 60 AAs: 19.72849\nTotal prob of N-in: 0.96170\nPOSSIBLE N-term signal sequence\ninside 1 19\nTMhelix 20 39\noutside 40 325\nTMhelix 326 348\ninside 349 372

17720 GCF_001240675.1_Salmonella_enterica_CVM_N43481_v1.0 Salmonella enterica subsp. enterica serovar Heidelberg Proteobacteria; Gammaproteobacteria; Enterobacterales; Enterobacteriaceae; Salmonella; Salmonella enterica; Salmonella enterica subsp. enterica
MNNEETFYQAMRRKGVTRRSFLKFCSLAATSLGLGAGMTPKIAWALENKPRIPVVWIHGL WP_001058312.1
hydrogenase [Salmonella enterica] Length: 372\nNumber of predicted TMHs: 2\nExp number of AAs in TMHs: 36.51738\nExp number, first 60 AAs: 19.72849\nTotal prob of N-in: 0.96170\nPOSSIBLE N-term signal sequence\ninside 1 19\nTMhelix 20 39\noutside 40 325\nTMhelix 326 348\ninside 349 372

17721 GCF_001246965.1_Salmonella_enterica_CVM_N51286_v1.0 Salmonella enterica subsp. enterica serovar Heidelberg Proteobacteria; Gammaproteobacteria; Enterobacterales; Enterobacteriaceae; Salmonella; Salmonella enterica; Salmonella enterica subsp. enterica
MTGDNTLITSHGINRRDFMKLCAALAATMGLSSKAAAEMAESVSNPQRPPVIWIGAQECT WP_000145429.1
hydrogenase 2 small subunit [Salmonella enterica] Length: 372\nNumber of predicted TMHs: 1\nExp number of AAs in TMHs: 24.51309\nExp number, first 60 AAs: 0.2682\nTotal prob of N-in: 0.14402\noutside 1 331\nTMhelix 332 354\ninside 355 372

17722 GCF_001240885.1_Salmonella_enterica_CVM_N43828_v1.0 Salmonella enterica subsp. enterica serovar Heidelberg Proteobacteria; Gammaproteobacteria; Enterobacterales; Enterobacteriaceae; Salmonella; Salmonella enterica; Salmonella enterica subsp. enterica
MNNEETFYQAMRRKGVTRRSFLKFCSLAATSLGLGAGMTPKIAWALENKPRIPVVWIHGL WP_001058312.1
hydrogenase [Salmonella enterica] Length: 372\nNumber of predicted TMHs: 2\nExp number of AAs in

TMHs: 36.51738\nExp number, first 60 AAs: 19.72849\nTotal prob of N-in: 0.96170\nPOSSIBLE N-term signal sequence\ninside 1 19\nTMhelix 20 39\noutside 40 325\nTMhelix 326 348\ninside 349 372

17723 GCF_001240545.1_Salmonella_enterica_CVM_N43473_v1.0 Salmonella enterica subsp. enterica serovar Heidelberg Proteobacteria; Gammaproteobacteria; Enterobacterales; Enterobacteriaceae; Salmonella; Salmonella enterica; Salmonella enterica subsp. enterica MNNEETFYQAMRRKGVTRRSFLKFCSLAATSLGLGAGMTPKIAWALENKPRIPVVWIHGL WP_001058312.1 hydrogenase [Salmonella enterica] Length: 372\nNumber of predicted TMHs: 2\nExp number of AAs in TMHs: 36.51738\nExp number, first 60 AAs: 19.72849\nTotal prob of N-in: 0.96170\nPOSSIBLE N-term signal sequence\ninside 1 19\nTMhelix 20 39\noutside 40 325\nTMhelix 326 348\ninside 349 372

17724 GCF_001247145.1_Salmonella_enterica_CVM_N51297_v1.0 Salmonella enterica subsp. enterica serovar Heidelberg Proteobacteria; Gammaproteobacteria; Enterobacterales; Enterobacteriaceae; Salmonella; Salmonella enterica; Salmonella enterica subsp. enterica MTGDNTLITSHGINRRDFMKLCAALAATMGLSSKAAAEMAESVSNPQRPPVIWIGAQECT WP_000145429.1 hydrogenase 2 small subunit [Salmonella enterica] Length: 372\nNumber of predicted TMHs: 1\nExp number of AAs in TMHs: 24.51309\nExp number, first 60 AAs: 0.2682\nTotal prob of N-in: 0.14402\noutside 1 331\nTMhelix 332 354\ninside 355 372

17725 GCF_001246965.1_Salmonella_enterica_CVM_N51286_v1.0 Salmonella enterica subsp. enterica serovar Heidelberg Proteobacteria; Gammaproteobacteria; Enterobacterales; Enterobacteriaceae; Salmonella; Salmonella enterica; Salmonella enterica subsp. enterica MNNEETFYQAMRRKGVTRRSFLKFCSLAATSLGLGAGMTPKIAWALENKPRIPVVWIHGL WP_001058312.1 hydrogenase [Salmonella enterica] Length: 372\nNumber of predicted TMHs: 2\nExp number of AAs in TMHs: 36.51738\nExp number, first 60 AAs: 19.72849\nTotal prob of N-in: 0.96170\nPOSSIBLE N-term signal sequence\ninside 1 19\nTMhelix 20 39\noutside 40 325\nTMhelix 326 348\ninside 349 372

17726 GCF_001441205.1_ASM144120v1 Salmonella enterica subsp. enterica serovar Heidelberg Proteobacteria; Gammaproteobacteria; Enterobacterales; Enterobacteriaceae; Salmonella; Salmonella enterica; Salmonella enterica subsp. enterica MNNEETFYQAMRRKGVTRRSFLKFCSLAATSLGLGAGMTPKIAWALENKPRIPVVWIHGL WP_001058312.1 hydrogenase [Salmonella enterica] Length: 372\nNumber of predicted TMHs: 2\nExp number of AAs in TMHs: 36.51738\nExp number, first 60 AAs: 19.72849\nTotal prob of N-in: 0.96170\nPOSSIBLE N-term signal sequence\ninside 1 19\nTMhelix 20 39\noutside 40 325\nTMhelix 326 348\ninside 349 372

17727 GCF_001441245.1_ASM144124v1 Salmonella enterica subsp. enterica serovar Heidelberg Proteobacteria; Gammaproteobacteria; Enterobacterales; Enterobacteriaceae; Salmonella; Salmonella enterica; Salmonella enterica subsp. enterica MTGDNTLITSHGINRRDFMKLCAALAATMGLSSKAAAEMAESVSNPQRPPVIWIGAQECT WP_000145429.1 hydrogenase 2 small subunit [Salmonella enterica] Length: 372\nNumber of predicted TMHs: 1\nExp number of AAs in TMHs: 24.51309\nExp number, first 60 AAs: 0.2682\nTotal prob of N-in: 0.14402\noutside 1 331\nTMhelix 332 354\ninside 355 372

17728 GCF_001479665.1_Salmonella_enterica_CVM_N37938-SQ_v1.0 Salmonella enterica subsp. enterica serovar Heidelberg Proteobacteria; Gammaproteobacteria; Enterobacterales; Enterobacteriaceae; Salmonella; Salmonella enterica; Salmonella enterica subsp. enterica MNNEETFYQAMRRKGVTRRSFLKFCSLAATSLGLGAGMTPKIAWALENKPRIPVVWIHGL WP_001058312.1 hydrogenase [Salmonella enterica] Length: 372\nNumber of predicted TMHs: 2\nExp number of AAs in TMHs: 36.51738\nExp number, first 60 AAs: 19.72849\nTotal prob of N-in: 0.96170\nPOSSIBLE N-term signal sequence\ninside 1 19\nTMhelix 20 39\noutside 40 325\nTMhelix 326 348\ninside 349 372

17729 GCF_001692515.1_ASM169251v1 Salmonella enterica subsp. enterica serovar Heidelberg Proteobacteria; Gammaproteobacteria; Enterobacterales; Enterobacteriaceae; Salmonella; Salmonella enterica; Salmonella enterica subsp. enterica MTGDNTLITSHGINRRDFMKLCAALAATMGLSSKAAAEMAESVSNPQRPPVIWIGAQECT WP_000145429.1 hydrogenase 2 small subunit [Salmonella enterica] Length: 372\nNumber of predicted TMHs: 1\nExp number of AAs in TMHs: 24.51309\nExp number, first 60 AAs: 0.2682\nTotal prob of N-in: 0.14402\noutside 1 331\nTMhelix 332 354\ninside 355 372

17730 GCF_001692555.1_ASM169255v1 Salmonella enterica subsp. enterica serovar Heidelberg Proteobacteria; Gammaproteobacteria; Enterobacterales; Enterobacteriaceae; Salmonella; Salmonella enterica; Salmonella enterica subsp. enterica MTGDNTLITSHGINRRDFMKLCAALAATMGLSSKAAAEMAESVSNPQRPPVIWIGAQECT WP_000145429.1 hydrogenase 2 small subunit [Salmonella enterica] Length: 372\nNumber of

predicted TMHs: 1\nExp number of AAs in TMHs: 24.51309\nExp number, first 60 AAs: 0.2682\nTotal prob of N-in: 0.14402\noutside 1 331\nTMhelix 332 354\ninside 355 372

17731 GCF_002037425.1_ASM203742v1 Salmonella enterica subsp. enterica serovar Heidelberg Proteobacteria; Gammaproteobacteria; Enterobacterales; Enterobacteriaceae; Salmonella; Salmonella enterica; Salmonella enterica subsp. enterica MTGDNTLITSHGINRRDFMKLCAALAATMGLSSKAAAEMAESVSNPQRPPVIWIGAQECT WP_000145429.1 hydrogenase 2 small subunit [Salmonella enterica] Length: 372\nNumber of predicted TMHs: 1\nExp number of AAs in TMHs: 24.51309\nExp number, first 60 AAs: 0.2682\nTotal prob of N-in: 0.14402\noutside 1 331\nTMhelix 332 354\ninside 355 372

17732 GCF_001692595.1_ASM169259v1 Salmonella enterica subsp. enterica serovar Heidelberg Proteobacteria; Gammaproteobacteria; Enterobacterales; Enterobacteriaceae; Salmonella; Salmonella enterica; Salmonella enterica subsp. enterica MTGDNTLITSHGINRRDFMKLCAALAATMGLSSKAAAEMAESVSNPQRPPVIWIGAQECT WP_000145429.1 hydrogenase 2 small subunit [Salmonella enterica] Length: 372\nNumber of predicted TMHs: 1\nExp number of AAs in TMHs: 24.51309\nExp number, first 60 AAs: 0.2682\nTotal prob of N-in: 0.14402\noutside 1 331\nTMhelix 332 354\ninside 355 372

17733 GCF_001690005.1_ASM169000v1 Salmonella enterica subsp. enterica serovar Heidelberg Proteobacteria; Gammaproteobacteria; Enterobacterales; Enterobacteriaceae; Salmonella; Salmonella enterica; Salmonella enterica subsp. enterica MNNEETFYQAMRRKGVTRRSFLKFCSLAATSLGLGAGMTPKIAWALENKPRIPVVWIHGL WP_001058312.1 hydrogenase [Salmonella enterica] Length: 372\nNumber of predicted TMHs: 2\nExp number of AAs in TMHs: 36.51738\nExp number, first 60 AAs: 19.72849\nTotal prob of N-in: 0.96170\nPOSSIBLE N-term signal sequence\ninside 1 19\nTMhelix 20 39\noutside 40 325\nTMhelix 326 348\ninside 349 372

17734 GCF_002037325.1_ASM203732v1 Salmonella enterica subsp. enterica serovar Heidelberg Proteobacteria; Gammaproteobacteria; Enterobacterales; Enterobacteriaceae; Salmonella; Salmonella enterica; Salmonella enterica subsp. enterica MNNEETFYQAMRRKGVTRRSFLKFCSLAATSLGLGAGMTPKIAWALENKPRIPVVWIHGL WP_001058312.1 hydrogenase [Salmonella enterica] Length: 372\nNumber of predicted TMHs: 2\nExp number of AAs in TMHs: 36.51738\nExp number, first 60 AAs: 19.72849\nTotal prob of N-in: 0.96170\nPOSSIBLE N-term signal sequence\ninside 1 19\nTMhelix 20 39\noutside 40 325\nTMhelix 326 348\ninside 349 372

17735 GCF_002037505.1_ASM203750v1 Salmonella enterica subsp. enterica serovar Heidelberg Proteobacteria; Gammaproteobacteria; Enterobacterales; Enterobacteriaceae; Salmonella; Salmonella enterica; Salmonella enterica subsp. enterica MTGDNTLITSHGINRRDFMKLCAALAATMGLSSKAAAEMAESVSNPQRPPVIWIGAQECT WP_000145429.1 hydrogenase 2 small subunit [Salmonella enterica] Length: 372\nNumber of predicted TMHs: 1\nExp number of AAs in TMHs: 24.51309\nExp number, first 60 AAs: 0.2682\nTotal prob of N-in: 0.14402\noutside 1 331\nTMhelix 332 354\ninside 355 372

17736 GCF_002037425.1_ASM203742v1 Salmonella enterica subsp. enterica serovar Heidelberg Proteobacteria; Gammaproteobacteria; Enterobacterales; Enterobacteriaceae; Salmonella; Salmonella enterica; Salmonella enterica subsp. enterica MNNEETFYQAMRRKGVTRRSFLKFCSLAATSLGLGAGMTPKIAWALENKPRIPVVWIHGL WP_001058312.1 hydrogenase [Salmonella enterica] Length: 372\nNumber of predicted TMHs: 2\nExp number of AAs in TMHs: 36.51738\nExp number, first 60 AAs: 19.72849\nTotal prob of N-in: 0.96170\nPOSSIBLE N-term signal sequence\ninside 1 19\nTMhelix 20 39\noutside 40 325\nTMhelix 326 348\ninside 349 372

17737 GCF_001692635.1_ASM169263v1 Salmonella enterica subsp. enterica serovar Heidelberg Proteobacteria; Gammaproteobacteria; Enterobacterales; Enterobacteriaceae; Salmonella; Salmonella enterica; Salmonella enterica subsp. enterica MNNEETFYQAMRRKGVTRRSFLKFCSLAATSLGLGAGMTPKIAWALENKPRIPVVWIHGL WP_001058312.1 hydrogenase [Salmonella enterica] Length: 372\nNumber of predicted TMHs: 2\nExp number of AAs in TMHs: 36.51738\nExp number, first 60 AAs: 19.72849\nTotal prob of N-in: 0.96170\nPOSSIBLE N-term signal sequence\ninside 1 19\nTMhelix 20 39\noutside 40 325\nTMhelix 326 348\ninside 349 372

17738 GCF_001478185.1_Salmonella_enterica_CVM_N32052-SQ_v1.0 Salmonella enterica subsp. enterica serovar Heidelberg Proteobacteria; Gammaproteobacteria; Enterobacterales; Enterobacteriaceae; Salmonella; Salmonella enterica; Salmonella enterica subsp. enterica MNNEETFYQAMRRKGVTRRSFLKFCSLAATSLGLGAGMTPKIAWALENKPRIPVVWIHGL WP_001058312.1 hydrogenase [Salmonella enterica] Length: 372\nNumber of predicted TMHs: 2\nExp number of AAs in TMHs: 36.51738\nExp number, first 60 AAs: 19.72849\nTotal prob of N-in: 0.96170\nPOSSIBLE N-term signal sequence\ninside 1 19\nTMhelix 20 39\noutside 40 325\nTMhelix 326 348\ninside 349 372

17739 GCF_001692515.1_ASM169251v1 Salmonella enterica subsp. enterica serovar Heidelberg Proteobacteria; Gammaproteobacteria; Enterobacterales; Enterobacteriaceae; Salmonella; Salmonella enterica; Salmonella enterica subsp. enterica MNNEETFYQAMRRKGVTRRSFLKFCSLAATSLGLGAGMTPKIAWALENKPRIPVVWIHGL WP_001058312.1 hydrogenase [Salmonella enterica] Length: 372\nNumber of predicted TMHs: 2\nExp number of AAs in TMHs: 36.51738\nExp number, first 60 AAs: 19.72849\nTotal prob of N-in: 0.96170\nPOSSIBLE N-term signal sequence\ninside 1 19\nTMhelix 20 39\noutside 40 325\nTMhelix 326 348\ninside 349 372

17740 GCF_001478185.1_Salmonella_enterica_CVM_N32052-SQ_v1.0 Salmonella enterica subsp. enterica serovar Heidelberg Proteobacteria; Gammaproteobacteria; Enterobacterales; Enterobacteriaceae; Salmonella; Salmonella enterica; Salmonella enterica subsp. enterica MTGDNTLITSHGINRRDFMKLCAALAATMGLSSKAAAEMAESVSNPQRPPVIWIGAQECT WP_000145429.1 hydrogenase 2 small subunit [Salmonella enterica] Length: 372\nNumber of predicted TMHs: 1\nExp number of AAs in TMHs: 24.51309\nExp number, first 60 AAs: 0.2682\nTotal prob of N-in: 0.14402\noutside 1 331\nTMhelix 332 354\ninside 355 372

17741 GCF_001441225.1_ASM144122v1 Salmonella enterica subsp. enterica serovar Heidelberg Proteobacteria; Gammaproteobacteria; Enterobacterales; Enterobacteriaceae; Salmonella; Salmonella enterica; Salmonella enterica subsp. enterica MNNEETFYQAMRRKGVTRRSFLKFCSLAATSLGLGAGMTPKIAWALENKPRIPVVWIHGL WP_001058312.1 hydrogenase [Salmonella enterica] Length: 372\nNumber of predicted TMHs: 2\nExp number of AAs in TMHs: 36.51738\nExp number, first 60 AAs: 19.72849\nTotal prob of N-in: 0.96170\nPOSSIBLE N-term signal sequence\ninside 1 19\nTMhelix 20 39\noutside 40 325\nTMhelix 326 348\ninside 349 372

17742 GCF_001952405.1_ASM195240v1 Salmonella enterica subsp. enterica serovar Heidelberg Proteobacteria; Gammaproteobacteria; Enterobacterales; Enterobacteriaceae; Salmonella; Salmonella enterica; Salmonella enterica subsp. enterica MNNEETFYQAMRRKGVTRRSFLKFCSLAATSLGLGAGMTPKIAWALENKPRIPVVWIHGL WP_001058312.1 hydrogenase [Salmonella enterica] Length: 372\nNumber of predicted TMHs: 2\nExp number of AAs in TMHs: 36.51738\nExp number, first 60 AAs: 19.72849\nTotal prob of N-in: 0.96170\nPOSSIBLE N-term signal sequence\ninside 1 19\nTMhelix 20 39\noutside 40 325\nTMhelix 326 348\ninside 349 372

17743 GCF_001952565.1_ASM195256v1 Salmonella enterica subsp. enterica serovar Heidelberg Proteobacteria; Gammaproteobacteria; Enterobacterales; Enterobacteriaceae; Salmonella; Salmonella enterica; Salmonella enterica subsp. enterica MNNEETFYQAMRRKGVTRRSFLKFCSLAATSLGLGAGMTPKIAWALENKPRIPVVWIHGL WP_001058312.1 hydrogenase [Salmonella enterica] Length: 372\nNumber of predicted TMHs: 2\nExp number of AAs in TMHs: 36.51738\nExp number, first 60 AAs: 19.72849\nTotal prob of N-in: 0.96170\nPOSSIBLE N-term signal sequence\ninside 1 19\nTMhelix 20 39\noutside 40 325\nTMhelix 326 348\ninside 349 372

17744 GCF_001952475.1_ASM195247v1 Salmonella enterica subsp. enterica serovar Heidelberg Proteobacteria; Gammaproteobacteria; Enterobacterales; Enterobacteriaceae; Salmonella; Salmonella enterica; Salmonella enterica subsp. enterica MNNEETFYQAMRRKGVTRRSFLKFCSLAATSLGLGAGMTPKIAWALENKPRIPVVWIHGL WP_001058312.1 hydrogenase [Salmonella enterica] Length: 372\nNumber of predicted TMHs: 2\nExp number of AAs in TMHs: 36.51738\nExp number, first 60 AAs: 19.72849\nTotal prob of N-in: 0.96170\nPOSSIBLE N-term signal sequence\ninside 1 19\nTMhelix 20 39\noutside 40 325\nTMhelix 326 348\ninside 349 372

17745 GCF_001952565.1_ASM195256v1 Salmonella enterica subsp. enterica serovar Heidelberg Proteobacteria; Gammaproteobacteria; Enterobacterales; Enterobacteriaceae; Salmonella; Salmonella enterica; Salmonella enterica subsp. enterica MTGDNTLITSHGINRRDFMKLCAALAATMGLSSKAAAEMAESVSNPQRPPVIWIGAQECT WP_000145429.1 hydrogenase 2 small subunit [Salmonella enterica] Length: 372\nNumber of predicted TMHs: 1\nExp number of AAs in TMHs: 24.51309\nExp number, first 60 AAs: 0.2682\nTotal prob of N-in: 0.14402\noutside 1 331\nTMhelix 332 354\ninside 355 372

17746 GCF_001692635.1_ASM169263v1 Salmonella enterica subsp. enterica serovar Heidelberg Proteobacteria; Gammaproteobacteria; Enterobacterales; Enterobacteriaceae; Salmonella; Salmonella enterica; Salmonella enterica subsp. enterica MTGDNTLITSHGINRRDFMKLCAALAATMGLSSKAAAEMAESVSNPQRPPVIWIGAQECT WP_000145429.1 hydrogenase 2 small subunit [Salmonella enterica] Length: 372\nNumber of predicted TMHs: 1\nExp number of AAs in TMHs: 24.51309\nExp number, first 60 AAs: 0.2682\nTotal prob of N-in: 0.14402\noutside 1 331\nTMhelix 332 354\ninside 355 372

17747 GCF_001478705.1_Salmonella_enterica_CVM_N29362-R_v1.0 Salmonella enterica subsp. enterica serovar Heidelberg Proteobacteria; Gammaproteobacteria; Enterobacterales; Enterobacteriaceae; Salmonella; Salmonella enterica; Salmonella enterica subsp. enterica MTGDNTLITSHGINRRDFMKLCAALAATMGLSSKAAAEMAESVSNPQRPPVIWIGAQECT WP_000145429.1

hydrogenase 2 small subunit [Salmonella enterica] Length: 372\nNumber of predicted TMHs: 1\nExp number of AAs in TMHs: 24.51309\nExp number, first 60 AAs: 0.2682\nTotal prob of N-in: 0.14402\noutside 1 331\nTMhelix 332 354\ninside 355 372

17748 GCF_001441205.1_ASM144120v1 Salmonella enterica subsp. enterica serovar Heidelberg Proteobacteria; Gammaproteobacteria; Enterobacterales; Enterobacteriaceae; Salmonella; Salmonella enterica; Salmonella enterica subsp. enterica MTGDNTLITSHGINRRDFMKLCAALAATMGLSSKAAAEMAESVSNPQRPPVIWIGAQECT WP_000145429.1 hydrogenase 2 small subunit [Salmonella enterica] Length: 372\nNumber of predicted TMHs: 1\nExp number of AAs in TMHs: 24.51309\nExp number, first 60 AAs: 0.2682\nTotal prob of N-in: 0.14402\noutside 1 331\nTMhelix 332 354\ninside 355 372

17749 GCF_001441225.1_ASM144122v1 Salmonella enterica subsp. enterica serovar Heidelberg Proteobacteria; Gammaproteobacteria; Enterobacterales; Enterobacteriaceae; Salmonella; Salmonella enterica; Salmonella enterica subsp. enterica MTGDNTLITSHGINRRDFMKLCAALAATMGLSSKAAAEMAESVSNPQRPPVIWIGAQECT WP_000145429.1 hydrogenase 2 small subunit [Salmonella enterica] Length: 372\nNumber of predicted TMHs: 1\nExp number of AAs in TMHs: 24.51309\nExp number, first 60 AAs: 0.2682\nTotal prob of N-in: 0.14402\noutside 1 331\nTMhelix 332 354\ninside 355 372

17750 GCF_001478965.1_Salmonella_enterica_CVM_N30677-SQ_v1.0 Salmonella enterica subsp. enterica serovar Heidelberg Proteobacteria; Gammaproteobacteria; Enterobacterales; Enterobacteriaceae; Salmonella; Salmonella enterica; Salmonella enterica subsp. enterica MNNEETFYQAMRRKGVTRRSFLKFCSLAATSLGLGAGMTPKIAWALENKPRIPVVWIHGL WP_001058312.1 hydrogenase [Salmonella enterica] Length: 372\nNumber of predicted TMHs: 2\nExp number of AAs in TMHs: 36.51738\nExp number, first 60 AAs: 19.72849\nTotal prob of N-in: 0.96170\nPOSSIBLE N-term signal sequence\ninside 1 19\nTMhelix 20 39\noutside 40 325\nTMhelix 326 348\ninside 349 372

17751 GCF_002036305.1_ASM203630v1 Salmonella enterica subsp. enterica serovar Heidelberg Proteobacteria; Gammaproteobacteria; Enterobacterales; Enterobacteriaceae; Salmonella; Salmonella enterica; Salmonella enterica subsp. enterica MTGDNTLITSHGINRRDFMKLCAALAATMGLSSKAAAEMAESVSNPQRPPVIWIGAQECT WP_000145429.1 hydrogenase 2 small subunit [Salmonella enterica] Length: 372\nNumber of predicted TMHs: 1\nExp number of AAs in TMHs: 24.51309\nExp number, first 60 AAs: 0.2682\nTotal prob of N-in: 0.14402\noutside 1 331\nTMhelix 332 354\ninside 355 372

17752 GCF_002036305.1_ASM203630v1 Salmonella enterica subsp. enterica serovar Heidelberg Proteobacteria; Gammaproteobacteria; Enterobacterales; Enterobacteriaceae; Salmonella; Salmonella enterica; Salmonella enterica subsp. enterica MNNEETFYQAMRRKGVTRRSFLKFCSLAATSLGLGAGMTPKIAWALENKPRIPVVWIHGL WP_001058312.1 hydrogenase [Salmonella enterica] Length: 372\nNumber of predicted TMHs: 2\nExp number of AAs in TMHs: 36.51738\nExp number, first 60 AAs: 19.72849\nTotal prob of N-in: 0.96170\nPOSSIBLE N-term signal sequence\ninside 1 19\nTMhelix 20 39\noutside 40 325\nTMhelix 326 348\ninside 349 372

17753 GCF_001690055.1_ASM169005v1 Salmonella enterica subsp. enterica serovar Heidelberg Proteobacteria; Gammaproteobacteria; Enterobacterales; Enterobacteriaceae; Salmonella; Salmonella enterica; Salmonella enterica subsp. enterica MTGDNTLITSHGINRRDFMKLCAALAATMGLSSKAAAEMAESVSNPQRPPVIWIGAQECT WP_000145429.1 hydrogenase 2 small subunit [Salmonella enterica] Length: 372\nNumber of predicted TMHs: 1\nExp number of AAs in TMHs: 24.51309\nExp number, first 60 AAs: 0.2682\nTotal prob of N-in: 0.14402\noutside 1 331\nTMhelix 332 354\ninside 355 372

17754 GCF_001478965.1_Salmonella_enterica_CVM_N30677-SQ_v1.0 Salmonella enterica subsp. enterica serovar Heidelberg Proteobacteria; Gammaproteobacteria; Enterobacterales; Enterobacteriaceae; Salmonella; Salmonella enterica; Salmonella enterica subsp. enterica MTGDNTLITSHGINRRDFMKLCAALAATMGLSSKAAAEMAESVSNPQRPPVIWIGAQECT WP_000145429.1 hydrogenase 2 small subunit [Salmonella enterica] Length: 372\nNumber of predicted TMHs: 1\nExp number of AAs in TMHs: 24.51309\nExp number, first 60 AAs: 0.2682\nTotal prob of N-in: 0.14402\noutside 1 331\nTMhelix 332 354\ninside 355 372

17755 GCF_001690095.1_ASM169009v1 Salmonella enterica subsp. enterica serovar Heidelberg Proteobacteria; Gammaproteobacteria; Enterobacterales; Enterobacteriaceae; Salmonella; Salmonella enterica; Salmonella enterica subsp. enterica MTGDNTLITSHGINRRDFMKLCAALAATMGLSSKAAAEMAESVSNPQRPPVIWIGAQECT WP_000145429.1 hydrogenase 2 small subunit [Salmonella enterica] Length: 372\nNumber of predicted TMHs: 1\nExp number of AAs in TMHs: 24.51309\nExp number, first 60 AAs: 0.2682\nTotal prob of N-in: 0.14402\noutside 1 331\nTMhelix 332 354\ninside 355 372

17756 GCF_001690095.1_ASM169009v1 Salmonella enterica subsp. enterica serovar Heidelberg Proteobacteria; Gammaproteobacteria; Enterobacterales; Enterobacteriaceae; Salmonella; Salmonella enterica; Salmonella enterica subsp. enterica MNNEETFYQAMRRKGVTRRSFLKFCSLAATSLGLGAGMTPKIAWALENKPRIPVVWIHGL WP_001058312.1 hydrogenase [Salmonella enterica] Length: 372\nNumber of predicted TMHs: 2\nExp number of AAs in TMHs: 36.51738\nExp number, first 60 AAs: 19.72849\nTotal prob of N-in: 0.96170\nPOSSIBLE N-term signal sequence\ninside 1 19\nTMhelix 20 39\noutside 40 325\nTMhelix 326 348\ninside 349 372

17757 GCF_001690055.1_ASM169005v1 Salmonella enterica subsp. enterica serovar Heidelberg Proteobacteria; Gammaproteobacteria; Enterobacterales; Enterobacteriaceae; Salmonella; Salmonella enterica; Salmonella enterica subsp. enterica MNNEETFYQAMRRKGVTRRSFLKFCSLAATSLGLGAGMTPKIAWALENKPRIPVVWIHGL WP_001058312.1 hydrogenase [Salmonella enterica] Length: 372\nNumber of predicted TMHs: 2\nExp number of AAs in TMHs: 36.51738\nExp number, first 60 AAs: 19.72849\nTotal prob of N-in: 0.96170\nPOSSIBLE N-term signal sequence\ninside 1 19\nTMhelix 20 39\noutside 40 325\nTMhelix 326 348\ninside 349 372

17758 GCF_001478705.1_Salmonella_enterica_CVM_N29362-R_v1.0 Salmonella enterica subsp. enterica serovar Heidelberg Proteobacteria; Gammaproteobacteria; Enterobacterales; Enterobacteriaceae; Salmonella; Salmonella enterica; Salmonella enterica subsp. enterica MNNEETFYQAMRRKGVTRRSFLKFCSLAATSLGLGAGMTPKIAWALENKPRIPVVWIHGL WP_001058312.1 hydrogenase [Salmonella enterica] Length: 372\nNumber of predicted TMHs: 2\nExp number of AAs in TMHs: 36.51738\nExp number, first 60 AAs: 19.72849\nTotal prob of N-in: 0.96170\nPOSSIBLE N-term signal sequence\ninside 1 19\nTMhelix 20 39\noutside 40 325\nTMhelix 326 348\ninside 349 372

17759 GCF_001441245.1_ASM144124v1 Salmonella enterica subsp. enterica serovar Heidelberg Proteobacteria; Gammaproteobacteria; Enterobacterales; Enterobacteriaceae; Salmonella; Salmonella enterica; Salmonella enterica subsp. enterica MNNEETFYQAMRRKGVTRRSFLKFCSLAATSLGLGAGMTPKIAWALENKPRIPVVWIHGL WP_001058312.1 hydrogenase [Salmonella enterica] Length: 372\nNumber of predicted TMHs: 2\nExp number of AAs in TMHs: 36.51738\nExp number, first 60 AAs: 19.72849\nTotal prob of N-in: 0.96170\nPOSSIBLE N-term signal sequence\ninside 1 19\nTMhelix 20 39\noutside 40 325\nTMhelix 326 348\ninside 349 372

17760 GCF_002060365.1_ASM206036v1 Salmonella enterica subsp. enterica serovar Heidelberg Proteobacteria; Gammaproteobacteria; Enterobacterales; Enterobacteriaceae; Salmonella; Salmonella enterica; Salmonella enterica subsp. enterica MTGDNTLITSHGINRRDFMKLCAALAATMGLSSKAAAEMAESVSNPQRPPVIWIGAQECT WP_000145429.1 hydrogenase 2 small subunit [Salmonella enterica] Length: 372\nNumber of predicted TMHs: 1\nExp number of AAs in TMHs: 24.51309\nExp number, first 60 AAs: 0.2682\nTotal prob of N-in: 0.14402\noutside 1 331\nTMhelix 332 354\ninside 355 372

17761 GCF_001479665.1_Salmonella_enterica_CVM_N37938-SQ_v1.0 Salmonella enterica subsp. enterica serovar Heidelberg Proteobacteria; Gammaproteobacteria; Enterobacterales; Enterobacteriaceae; Salmonella; Salmonella enterica; Salmonella enterica subsp. enterica MTGDNTLITSHGINRRDFMKLCAALAATMGLSSKAAAEMAESVSNPQRPPVIWIGAQECT WP_000145429.1 hydrogenase 2 small subunit [Salmonella enterica] Length: 372\nNumber of predicted TMHs: 1\nExp number of AAs in TMHs: 24.51309\nExp number, first 60 AAs: 0.2682\nTotal prob of N-in: 0.14402\noutside 1 331\nTMhelix 332 354\ninside 355 372

17762 GCF_001479265.1_Salmonella_enterica_CVM_N31844-SQ_v1.0 Salmonella enterica subsp. enterica serovar Heidelberg Proteobacteria; Gammaproteobacteria; Enterobacterales; Enterobacteriaceae; Salmonella; Salmonella enterica; Salmonella enterica subsp. enterica MNNEETFYQAMRRKGVTRRSFLKFCSLAATSLGLGAGMTPKIAWALENKPRIPVVWIHGL WP_001058312.1 hydrogenase [Salmonella enterica] Length: 372\nNumber of predicted TMHs: 2\nExp number of AAs in TMHs: 36.51738\nExp number, first 60 AAs: 19.72849\nTotal prob of N-in: 0.96170\nPOSSIBLE N-term signal sequence\ninside 1 19\nTMhelix 20 39\noutside 40 325\nTMhelix 326 348\ninside 349 372

17763 GCF_001479265.1_Salmonella_enterica_CVM_N31844-SQ_v1.0 Salmonella enterica subsp. enterica serovar Heidelberg Proteobacteria; Gammaproteobacteria; Enterobacterales; Enterobacteriaceae; Salmonella; Salmonella enterica; Salmonella enterica subsp. enterica MTGDNTLITSHGINRRDFMKLCAALAATMGLSSKAAAEMAESVSNPQRPPVIWIGAQECT WP_000145429.1 hydrogenase 2 small subunit [Salmonella enterica] Length: 372\nNumber of predicted TMHs: 1\nExp number of AAs in TMHs: 24.51309\nExp number, first 60 AAs: 0.2682\nTotal prob of N-in: 0.14402\noutside 1 331\nTMhelix 332 354\ninside 355 372

17764 GCF_001692575.1_ASM169257v1 Salmonella enterica subsp. enterica serovar Heidelberg Proteobacteria; Gammaproteobacteria; Enterobacterales; Enterobacteriaceae; Salmonella; Salmonella enterica; Salmonella enterica subsp. enterica MTGDNTLITSHGINRRDFMKLCAALAATMGLSSKAAAEMAESVSNPQRPPVIWIGAQECT WP_000145429.1 hydrogenase 2 small subunit [Salmonella enterica] Length: 372\nNumber of predicted TMHs: 1\nExp number of AAs in TMHs: 24.51309\nExp number, first 60 AAs: 0.2682\nTotal prob of N-in: 0.14402\noutside 1 331\nTMhelix 332 354\ninside 355 372

17765 GCF_001692535.1_ASM169253v1 Salmonella enterica subsp. enterica serovar Heidelberg Proteobacteria; Gammaproteobacteria; Enterobacterales; Enterobacteriaceae; Salmonella; Salmonella enterica; Salmonella enterica subsp. enterica MTGDNTLITSHGINRRDFMKLCAALAATMGLSSKAAAEMAESVSNPQRPPVIWIGAQECT WP_000145429.1 hydrogenase 2 small subunit [Salmonella enterica] Length: 372\nNumber of predicted TMHs: 1\nExp number of AAs in TMHs: 24.51309\nExp number, first 60 AAs: 0.2682\nTotal prob of N-in: 0.14402\noutside 1 331\nTMhelix 332 354\ninside 355 372

17766 GCF_001692655.1_ASM169265v1 Salmonella enterica subsp. enterica serovar Heidelberg Proteobacteria; Gammaproteobacteria; Enterobacterales; Enterobacteriaceae; Salmonella; Salmonella enterica; Salmonella enterica subsp. enterica MTGDNTLITSHGINRRDFMKLCAALAATMGLSSKAAAEMAESVSNPQRPPVIWIGAQECT WP_000145429.1 hydrogenase 2 small subunit [Salmonella enterica] Length: 372\nNumber of predicted TMHs: 1\nExp number of AAs in TMHs: 24.51309\nExp number, first 60 AAs: 0.2682\nTotal prob of N-in: 0.14402\noutside 1 331\nTMhelix 332 354\ninside 355 372

17767 GCF_001692495.1_ASM169249v1 Salmonella enterica subsp. enterica serovar Heidelberg Proteobacteria; Gammaproteobacteria; Enterobacterales; Enterobacteriaceae; Salmonella; Salmonella enterica; Salmonella enterica subsp. enterica MNNEETFYQAMRRKGVTRRSFLKFCSLAATSLGLGAGMTPKIAWALENKPRIPVVWIHGL WP_001058312.1 hydrogenase [Salmonella enterica] Length: 372\nNumber of predicted TMHs: 2\nExp number of AAs in TMHs: 36.51738\nExp number, first 60 AAs: 19.72849\nTotal prob of N-in: 0.96170\nPOSSIBLE N-term signal sequence\ninside 1 19\nTMhelix 20 39\noutside 40 325\nTMhelix 326 348\ninside 349 372

17768 GCF_001692575.1_ASM169257v1 Salmonella enterica subsp. enterica serovar Heidelberg Proteobacteria; Gammaproteobacteria; Enterobacterales; Enterobacteriaceae; Salmonella; Salmonella enterica; Salmonella enterica subsp. enterica MNNEETFYQAMRRKGVTRRSFLKFCSLAATSLGLGAGMTPKIAWALENKPRIPVVWIHGL WP_001058312.1 hydrogenase [Salmonella enterica] Length: 372\nNumber of predicted TMHs: 2\nExp number of AAs in TMHs: 36.51738\nExp number, first 60 AAs: 19.72849\nTotal prob of N-in: 0.96170\nPOSSIBLE N-term signal sequence\ninside 1 19\nTMhelix 20 39\noutside 40 325\nTMhelix 326 348\ninside 349 372

17769 GCF_001244225.1_Salmonella_enterica_CVM_N48578_v1.0 Salmonella enterica subsp. enterica serovar Heidelberg Proteobacteria; Gammaproteobacteria; Enterobacterales; Enterobacteriaceae; Salmonella; Salmonella enterica; Salmonella enterica subsp. enterica MNNEETFYQAMRRKGVTRRSFLKFCSLAATSLGLGAGMTPKIAWALENKPRIPVVWIHGL WP_001058312.1 hydrogenase [Salmonella enterica] Length: 372\nNumber of predicted TMHs: 2\nExp number of AAs in TMHs: 36.51738\nExp number, first 60 AAs: 19.72849\nTotal prob of N-in: 0.96170\nPOSSIBLE N-term signal sequence\ninside 1 19\nTMhelix 20 39\noutside 40 325\nTMhelix 326 348\ninside 349 372

17770 GCF_001242885.1_Salmonella_enterica_CVM_N46811_v1.0 Salmonella enterica subsp. enterica serovar Heidelberg Proteobacteria; Gammaproteobacteria; Enterobacterales; Enterobacteriaceae; Salmonella; Salmonella enterica; Salmonella enterica subsp. enterica MNNEETFYQAMRRKGVTRRSFLKFCSLAATSLGLGAGMTPKIAWALENKPRIPVVWIHGL WP_001058312.1 hydrogenase [Salmonella enterica] Length: 372\nNumber of predicted TMHs: 2\nExp number of AAs in TMHs: 36.51738\nExp number, first 60 AAs: 19.72849\nTotal prob of N-in: 0.96170\nPOSSIBLE N-term signal sequence\ninside 1 19\nTMhelix 20 39\noutside 40 325\nTMhelix 326 348\ninside 349 372

17771 GCF_001272375.1_Salmonella_enterica_CVM_N51271_v1.0 Salmonella enterica subsp. enterica serovar Heidelberg Proteobacteria; Gammaproteobacteria; Enterobacterales; Enterobacteriaceae; Salmonella; Salmonella enterica; Salmonella enterica subsp. enterica MTGDNTLITSHGINRRDFMKLCAALAATMGLSSKAAAEMAESVSNPQRPPVIWIGAQECT WP_000145429.1 hydrogenase 2 small subunit [Salmonella enterica] Length: 372\nNumber of predicted TMHs: 1\nExp number of AAs in TMHs: 24.51309\nExp number, first 60 AAs: 0.2682\nTotal prob of N-in: 0.14402\noutside 1 331\nTMhelix 332 354\ninside 355 372

17772 GCF_001246805.1_Salmonella_enterica_CVM_N51281_v1.0 Salmonella enterica subsp. enterica serovar Heidelberg Proteobacteria; Gammaproteobacteria; Enterobacterales; Enterobacteriaceae; Salmonella; Salmonella

enterica; Salmonella enterica subsp. enterica
MTGDNTLITSHGINRRDFMKLCAALAATMGLSSKAAAEMAESVSNPQRPPVIWIGAQECT WP_000145429.1 hydrogenase 2 small subunit [Salmonella enterica] Length: 372\nNumber of predicted TMHs: 1\nExp number of AAs in TMHs: 24.51309\nExp number, first 60 AAs: 0.2682\nTotal prob of N-in: 0.14402\noutside 1 331\nTMhelix 332 354\ninside 355 372

17773 GCF_001952555.1_ASM195255v1 Salmonella enterica subsp. enterica serovar Heidelberg Proteobacteria; Gammaproteobacteria; Enterobacterales; Enterobacteriaceae; Salmonella; Salmonella enterica; Salmonella enterica subsp. enterica MTGDNTLITSHGINRRDFMKLCAALAATMGLSSKAAAEMAESVSNPQRPPVIWIGAQECT WP_000145429.1 hydrogenase 2 small subunit [Salmonella enterica] Length: 372\nNumber of predicted TMHs: 1\nExp number of AAs in TMHs: 24.51309\nExp number, first 60 AAs: 0.2682\nTotal prob of N-in: 0.14402\noutside 1 331\nTMhelix 332 354\ninside 355 372

17774 GCF_001952575.1_ASM195257v1 Salmonella enterica subsp. enterica serovar Heidelberg Proteobacteria; Gammaproteobacteria; Enterobacterales; Enterobacteriaceae; Salmonella; Salmonella enterica; Salmonella enterica subsp. enterica MTGDNTLITSHGINRRDFMKLCAALAATMGLSSKAAAEMAESVSNPQRPPVIWIGAQECT WP_000145429.1 hydrogenase 2 small subunit [Salmonella enterica] Length: 372\nNumber of predicted TMHs: 1\nExp number of AAs in TMHs: 24.51309\nExp number, first 60 AAs: 0.2682\nTotal prob of N-in: 0.14402\noutside 1 331\nTMhelix 332 354\ninside 355 372

17775 GCF_001952465.1_Salmonella_enterica_CVM_N44695_v1.0 Salmonella enterica subsp. enterica serovar Heidelberg Proteobacteria; Gammaproteobacteria; Enterobacterales; Enterobacteriaceae; Salmonella; Salmonella enterica; Salmonella enterica subsp. enterica
MNNEETFYQAMRRKGVTRRSFLKFCSLAATSLGLGAGMTPKIAWALENKPRIPVVWIHGL WP_001058312.1 hydrogenase [Salmonella enterica] Length: 372\nNumber of predicted TMHs: 2\nExp number of AAs in TMHs: 36.51738\nExp number, first 60 AAs: 19.72849\nTotal prob of N-in: 0.96170\nPOSSIBLE N-term signal sequence\ninside 1 19\nTMhelix 20 39\noutside 40 325\nTMhelix 326 348\ninside 349 372

17776 GCF_000974165.1_SALB-159-4.id19_v1.0 Salmonella enterica subsp. enterica serovar Heidelberg Proteobacteria; Gammaproteobacteria; Enterobacterales; Enterobacteriaceae; Salmonella; Salmonella enterica; Salmonella enterica subsp. enterica MNNEETFYQAMRRKGVTRRSFLKFCSLAATSLGLGAGMTPKIAWALENKPRIPVVWIHGL WP_001058312.1 hydrogenase [Salmonella enterica] Length: 372\nNumber of predicted TMHs: 2\nExp number of AAs in TMHs: 36.51738\nExp number, first 60 AAs: 19.72849\nTotal prob of N-in: 0.96170\nPOSSIBLE N-term signal sequence\ninside 1 19\nTMhelix 20 39\noutside 40 325\nTMhelix 326 348\ninside 349 372

17777 GCF_001479435.1_Salmonella_enterica_CVM_N32754-SQ_v1.0 Salmonella enterica subsp. enterica serovar Heidelberg Proteobacteria; Gammaproteobacteria; Enterobacterales; Enterobacteriaceae; Salmonella; Salmonella enterica; Salmonella enterica subsp. enterica
MNNEETFYQAMRRKGVTRRSFLKFCSLAATSLGLGAGMTPKIAWALENKPRIPVVWIHGL WP_001058312.1 hydrogenase [Salmonella enterica] Length: 372\nNumber of predicted TMHs: 2\nExp number of AAs in TMHs: 36.51738\nExp number, first 60 AAs: 19.72849\nTotal prob of N-in: 0.96170\nPOSSIBLE N-term signal sequence\ninside 1 19\nTMhelix 20 39\noutside 40 325\nTMhelix 326 348\ninside 349 372

17778 GCF_001480445.1_Salmonella_enterica_CVM_N42239-SQ_v1.0 Salmonella enterica subsp. enterica serovar Heidelberg Proteobacteria; Gammaproteobacteria; Enterobacterales; Enterobacteriaceae; Salmonella; Salmonella enterica; Salmonella enterica subsp. enterica
MTGDNTLITSHGINRRDFMKLCAALAATMGLSSKAAAEMAESVSNPQRPPVIWIGAQECT WP_000145429.1 hydrogenase 2 small subunit [Salmonella enterica] Length: 372\nNumber of predicted TMHs: 1\nExp number of AAs in TMHs: 24.51309\nExp number, first 60 AAs: 0.2682\nTotal prob of N-in: 0.14402\noutside 1 331\nTMhelix 332 354\ninside 355 372

17779 GCF_001246095.1_Salmonella_enterica_CVM_N51248_v1.0 Salmonella enterica subsp. enterica serovar Heidelberg Proteobacteria; Gammaproteobacteria; Enterobacterales; Enterobacteriaceae; Salmonella; Salmonella enterica; Salmonella enterica subsp. enterica
MNNEETFYQAMRRKGVTRRSFLKFCSLAATSLGLGAGMTPKIAWALENKPRIPVVWIHGL WP_001058312.1 hydrogenase [Salmonella enterica] Length: 372\nNumber of predicted TMHs: 2\nExp number of AAs in TMHs: 36.51738\nExp number, first 60 AAs: 19.72849\nTotal prob of N-in: 0.96170\nPOSSIBLE N-term signal sequence\ninside 1 19\nTMhelix 20 39\noutside 40 325\nTMhelix 326 348\ninside 349 372

17780 GCF_001246095.1_Salmonella_enterica_CVM_N51248_v1.0 Salmonella enterica subsp. enterica serovar Heidelberg Proteobacteria; Gammaproteobacteria; Enterobacterales; Enterobacteriaceae; Salmonella; Salmonella

enterica; Salmonella enterica subsp. enterica
MTGDNTLITSHGINRRDFMKLCAALAATMGLSSKAAAEMAESVSNPQRPPVIWIGAQECT WP_000145429.1 hydrogenase 2 small subunit [Salmonella enterica] Length: 372\nNumber of predicted TMHs: 1\nExp number of AAs in TMHs: 24.51309\nExp number, first 60 AAs: 0.2682\nTotal prob of N-in: 0.14402\noutside 1 331\nTMhelix 332 354\ninside 355 372

17781 GCF_001689935.1_ASM168993v1 Salmonella enterica subsp. enterica serovar Heidelberg Proteobacteria; Gammaproteobacteria; Enterobacterales; Enterobacteriaceae; Salmonella; Salmonella enterica; Salmonella enterica subsp. enterica MTGDNTLITSHGINRRDFMKLCAALAATMGLSSKAAAEMAESVSNPQRPPVIWIGAQECT WP_000145429.1 hydrogenase 2 small subunit [Salmonella enterica] Length: 372\nNumber of predicted TMHs: 1\nExp number of AAs in TMHs: 24.51309\nExp number, first 60 AAs: 0.2682\nTotal prob of N-in: 0.14402\noutside 1 331\nTMhelix 332 354\ninside 355 372

17782 GCF_001246855.1_Salmonella_enterica_CVM_N51282_v1.0 Salmonella enterica subsp. enterica serovar Heidelberg Proteobacteria; Gammaproteobacteria; Enterobacterales; Enterobacteriaceae; Salmonella; Salmonella enterica; Salmonella enterica subsp. enterica
MTGDNTLITSHGINRRDFMKLCAALAATMGLSSKAAAEMAESVSNPQRPPVIWIGAQECT WP_000145429.1 hydrogenase 2 small subunit [Salmonella enterica] Length: 372\nNumber of predicted TMHs: 1\nExp number of AAs in TMHs: 24.51309\nExp number, first 60 AAs: 0.2682\nTotal prob of N-in: 0.14402\noutside 1 331\nTMhelix 332 354\ninside 355 372

17783 GCF_001690035.1_ASM169003v1 Salmonella enterica subsp. enterica serovar Heidelberg Proteobacteria; Gammaproteobacteria; Enterobacterales; Enterobacteriaceae; Salmonella; Salmonella enterica; Salmonella enterica subsp. enterica MTGDNTLITSHGINRRDFMKLCAALAATMGLSSKAAAEMAESVSNPQRPPVIWIGAQECT WP_000145429.1 hydrogenase 2 small subunit [Salmonella enterica] Length: 372\nNumber of predicted TMHs: 1\nExp number of AAs in TMHs: 24.51309\nExp number, first 60 AAs: 0.2682\nTotal prob of N-in: 0.14402\noutside 1 331\nTMhelix 332 354\ninside 355 372

17784 GCF_001689935.1_ASM168993v1 Salmonella enterica subsp. enterica serovar Heidelberg Proteobacteria; Gammaproteobacteria; Enterobacterales; Enterobacteriaceae; Salmonella; Salmonella enterica; Salmonella enterica subsp. enterica MNNEETFYQAMRRKGVTRRSFLKFCSLAATSLGLGAGMTPKIAWALENKPRIPVVWIHGL WP_001058312.1 hydrogenase [Salmonella enterica] Length: 372\nNumber of predicted TMHs: 2\nExp number of AAs in TMHs: 36.51738\nExp number, first 60 AAs: 19.72849\nTotal prob of N-in: 0.96170\nPOSSIBLE N-term signal sequence\ninside 1 19\nTMhelix 20 39\noutside 40 325\nTMhelix 326 348\ninside 349 372

17785 GCF_002032425.1_ASM203242v1 Salmonella enterica subsp. enterica serovar Heidelberg Proteobacteria; Gammaproteobacteria; Enterobacterales; Enterobacteriaceae; Salmonella; Salmonella enterica; Salmonella enterica subsp. enterica MTGDNTLITSHGINRRDFMKLCAALAATMGLSSKAAAEMAESVSNPQRPPVIWIGAQECT WP_000145429.1 hydrogenase 2 small subunit [Salmonella enterica] Length: 372\nNumber of predicted TMHs: 1\nExp number of AAs in TMHs: 24.51309\nExp number, first 60 AAs: 0.2682\nTotal prob of N-in: 0.14402\noutside 1 331\nTMhelix 332 354\ninside 355 372

17786 GCF_001692615.1_ASM169261v1 Salmonella enterica subsp. enterica serovar Heidelberg Proteobacteria; Gammaproteobacteria; Enterobacterales; Enterobacteriaceae; Salmonella; Salmonella enterica; Salmonella enterica subsp. enterica MNNEETFYQAMRRKGVTRRSFLKFCSLAATSLGLGAGMTPKIAWALENKPRIPVVWIHGL WP_001058312.1 hydrogenase [Salmonella enterica] Length: 372\nNumber of predicted TMHs: 2\nExp number of AAs in TMHs: 36.51738\nExp number, first 60 AAs: 19.72849\nTotal prob of N-in: 0.96170\nPOSSIBLE N-term signal sequence\ninside 1 19\nTMhelix 20 39\noutside 40 325\nTMhelix 326 348\ninside 349 372

17787 GCF_001692535.1_ASM169253v1 Salmonella enterica subsp. enterica serovar Heidelberg Proteobacteria; Gammaproteobacteria; Enterobacterales; Enterobacteriaceae; Salmonella; Salmonella enterica; Salmonella enterica subsp. enterica MNNEETFYQAMRRKGVTRRSFLKFCSLAATSLGLGAGMTPKIAWALENKPRIPVVWIHGL WP_001058312.1 hydrogenase [Salmonella enterica] Length: 372\nNumber of predicted TMHs: 2\nExp number of AAs in TMHs: 36.51738\nExp number, first 60 AAs: 19.72849\nTotal prob of N-in: 0.96170\nPOSSIBLE N-term signal sequence\ninside 1 19\nTMhelix 20 39\noutside 40 325\nTMhelix 326 348\ninside 349 372

17788 GCF_001243585.1_Salmonella_enterica_CVM_N46853_v1.0 Salmonella enterica subsp. enterica serovar Heidelberg Proteobacteria; Gammaproteobacteria; Enterobacterales; Enterobacteriaceae; Salmonella; Salmonella enterica; Salmonella enterica subsp. enterica
MNNEETFYQAMRRKGVTRRSFLKFCSLAATSLGLGAGMTPKIAWALENKPRIPVVWIHGL WP_001058312.1 hydrogenase [Salmonella enterica] Length: 372\nNumber of predicted TMHs: 2\nExp number of AAs in

TMHs: 36.51738\nExp number, first 60 AAs: 19.72849\nTotal prob of N-in: 0.96170\nPOSSIBLE N-term signal sequence\ninside 1 19\nTMhelix 20 39\noutside 40 325\nTMhelix 326 348\ninside 349 372

17789 GCF_001243585.1_Salmonella_enterica_CVM_N46853_v1.0 Salmonella enterica subsp. enterica serovar Heidelberg Proteobacteria; Gammaproteobacteria; Enterobacterales; Enterobacteriaceae; Salmonella; Salmonella enterica; Salmonella enterica subsp. enterica
MTGDNTLITSHGINRRDFMKLCAALAATMGLSSKAAAEMAESVSNPQRPPVIWIGAQECT WP_000145429.1
hydrogenase 2 small subunit [Salmonella enterica] Length: 372\nNumber of predicted TMHs: 1\nExp number of AAs in TMHs: 24.51309\nExp number, first 60 AAs: 0.2682\nTotal prob of N-in: 0.14402\noutside 1 331\nTMhelix 332 354\ninside 355 372

17790 GCF_001242865.1_Salmonella_enterica_CVM_N46812_v1.0 Salmonella enterica subsp. enterica serovar Heidelberg Proteobacteria; Gammaproteobacteria; Enterobacterales; Enterobacteriaceae; Salmonella; Salmonella enterica; Salmonella enterica subsp. enterica
MNNEETFYQAMRRKGVTRRSFLKFCSLAATSLGLGAGMTPKIAWALENKPRIPVVWIHGL WP_001058312.1
hydrogenase [Salmonella enterica] Length: 372\nNumber of predicted TMHs: 2\nExp number of AAs in TMHs: 36.51738\nExp number, first 60 AAs: 19.72849\nTotal prob of N-in: 0.96170\nPOSSIBLE N-term signal sequence\ninside 1 19\nTMhelix 20 39\noutside 40 325\nTMhelix 326 348\ninside 349 372

17791 GCF_001478975.1_Salmonella_enterica_CVM_N30683-SQ_v1.0 Salmonella enterica subsp. enterica serovar Heidelberg Proteobacteria; Gammaproteobacteria; Enterobacterales; Enterobacteriaceae; Salmonella; Salmonella enterica; Salmonella enterica subsp. enterica
MTGDNTLITSHGINRRDFMKLCAALAATMGLSSKAAAEMAESVSNPQRPPVIWIGAQECT WP_000145429.1
hydrogenase 2 small subunit [Salmonella enterica] Length: 372\nNumber of predicted TMHs: 1\nExp number of AAs in TMHs: 24.51309\nExp number, first 60 AAs: 0.2682\nTotal prob of N-in: 0.14402\noutside 1 331\nTMhelix 332 354\ninside 355 372

17792 GCF_001241725.1_Salmonella_enterica_CVM_N45397_v1.0 Salmonella enterica subsp. enterica serovar Heidelberg Proteobacteria; Gammaproteobacteria; Enterobacterales; Enterobacteriaceae; Salmonella; Salmonella enterica; Salmonella enterica subsp. enterica
MTGDNTLITSHGINRRDFMKLCAALAATMGLSSKAAAEMAESVSNPQRPPVIWIGAQECT WP_000145429.1
hydrogenase 2 small subunit [Salmonella enterica] Length: 372\nNumber of predicted TMHs: 1\nExp number of AAs in TMHs: 24.51309\nExp number, first 60 AAs: 0.2682\nTotal prob of N-in: 0.14402\noutside 1 331\nTMhelix 332 354\ninside 355 372

17793 GCF_001241455.1_Salmonella_enterica_CVM_N44710_v1.0 Salmonella enterica subsp. enterica serovar Heidelberg Proteobacteria; Gammaproteobacteria; Enterobacterales; Enterobacteriaceae; Salmonella; Salmonella enterica; Salmonella enterica subsp. enterica
MNNEETFYQAMRRKGVTRRSFLKFCSLAATSLGLGAGMTPKIAWALENKPRIPVVWIHGL WP_001058312.1
hydrogenase [Salmonella enterica] Length: 372\nNumber of predicted TMHs: 2\nExp number of AAs in TMHs: 36.51738\nExp number, first 60 AAs: 19.72849\nTotal prob of N-in: 0.96170\nPOSSIBLE N-term signal sequence\ninside 1 19\nTMhelix 20 39\noutside 40 325\nTMhelix 326 348\ninside 349 372

17794 GCF_001478945.1_Salmonella_enterica_CVM_N30671-SQ_v1.0 Salmonella enterica subsp. enterica serovar Heidelberg Proteobacteria; Gammaproteobacteria; Enterobacterales; Enterobacteriaceae; Salmonella; Salmonella enterica; Salmonella enterica subsp. enterica
MNNEETFYQAMRRKGVTRRSFLKFCSLAATSLGLGAGMTPKIAWALENKPRIPVVWIHGL WP_001058312.1
hydrogenase [Salmonella enterica] Length: 372\nNumber of predicted TMHs: 2\nExp number of AAs in TMHs: 36.51738\nExp number, first 60 AAs: 19.72849\nTotal prob of N-in: 0.96170\nPOSSIBLE N-term signal sequence\ninside 1 19\nTMhelix 20 39\noutside 40 325\nTMhelix 326 348\ninside 349 372

17795 GCF_001241155.1_Salmonella_enterica_CVM_N44696-R_v1.0 Salmonella enterica subsp. enterica serovar Heidelberg Proteobacteria; Gammaproteobacteria; Enterobacterales; Enterobacteriaceae; Salmonella; Salmonella enterica; Salmonella enterica subsp. enterica
MNNEETFYQAMRRKGVTRRSFLKFCSLAATSLGLGAGMTPKIAWALENKPRIPVVWIHGL WP_001058312.1
hydrogenase [Salmonella enterica] Length: 372\nNumber of predicted TMHs: 2\nExp number of AAs in TMHs: 36.51738\nExp number, first 60 AAs: 19.72849\nTotal prob of N-in: 0.96170\nPOSSIBLE N-term signal sequence\ninside 1 19\nTMhelix 20 39\noutside 40 325\nTMhelix 326 348\ninside 349 372

17796 GCF_001477675.1_Salmonella_enterica_CVM_N29307-SQ_v1.0 Salmonella enterica subsp. enterica serovar Heidelberg Proteobacteria; Gammaproteobacteria; Enterobacterales; Enterobacteriaceae; Salmonella; Salmonella

enterica; Salmonella enterica subsp. enterica
MNNEETFYQAMRRKGVTRRSFLKFCSLAATSLGLGAGMTPKIAWALENKPRIPVVWIHGL WP_001058312.1 hydrogenase [Salmonella enterica] Length: 372\nNumber of predicted TMHs: 2\nExp number of AAs in TMHs: 36.51738\nExp number, first 60 AAs: 19.72849\nTotal prob of N-in: 0.96170\nPOSSIBLE N-term signal sequence\ninside 1 19\nTMhelix 20 39\noutside 40 325\nTMhelix 326 348\ninside 349 372

17797 GCF_001241275.1_Salmonella_enterica_CVM_N44701-R_v1.0 Salmonella enterica subsp. enterica serovar Heidelberg Proteobacteria; Gammaproteobacteria; Enterobacterales; Enterobacteriaceae; Salmonella; Salmonella enterica; Salmonella enterica subsp. enterica
MNNEETFYQAMRRKGVTRRSFLKFCSLAATSLGLGAGMTPKIAWALENKPRIPVVWIHGL WP_001058312.1 hydrogenase [Salmonella enterica] Length: 372\nNumber of predicted TMHs: 2\nExp number of AAs in TMHs: 36.51738\nExp number, first 60 AAs: 19.72849\nTotal prob of N-in: 0.96170\nPOSSIBLE N-term signal sequence\ninside 1 19\nTMhelix 20 39\noutside 40 325\nTMhelix 326 348\ninside 349 372

17798 GCF_001241155.1_Salmonella_enterica_CVM_N44696-R_v1.0 Salmonella enterica subsp. enterica serovar Heidelberg Proteobacteria; Gammaproteobacteria; Enterobacterales; Enterobacteriaceae; Salmonella; Salmonella enterica; Salmonella enterica subsp. enterica
MTGDNTLITSHGINRRDFMKLCAALAATMGLSSKAAAEMAESVSNPQRPPVIWIGAQECT WP_000145429.1 hydrogenase 2 small subunit [Salmonella enterica] Length: 372\nNumber of predicted TMHs: 1\nExp number of AAs in TMHs: 24.51309\nExp number, first 60 AAs: 0.2682\nTotal prob of N-in: 0.14402\noutside 1 331\nTMhelix 332 354\ninside 355 372

17799 GCF_001240155.1_Salmonella_enterica_CVM_N43452_v1.0 Salmonella enterica subsp. enterica serovar Heidelberg Proteobacteria; Gammaproteobacteria; Enterobacterales; Enterobacteriaceae; Salmonella; Salmonella enterica; Salmonella enterica subsp. enterica
MNNEETFYQAMRRKGVTRRSFLKFCSLAATSLGLGAGMTPKIAWALENKPRIPVVWIHGL WP_001058312.1 hydrogenase [Salmonella enterica] Length: 372\nNumber of predicted TMHs: 2\nExp number of AAs in TMHs: 36.51738\nExp number, first 60 AAs: 19.72849\nTotal prob of N-in: 0.96170\nPOSSIBLE N-term signal sequence\ninside 1 19\nTMhelix 20 39\noutside 40 325\nTMhelix 326 348\ninside 349 372

17800 GCF_001246805.1_Salmonella_enterica_CVM_N51281_v1.0 Salmonella enterica subsp. enterica serovar Heidelberg Proteobacteria; Gammaproteobacteria; Enterobacterales; Enterobacteriaceae; Salmonella; Salmonella enterica; Salmonella enterica subsp. enterica
MNNEETFYQAMRRKGVTRRSFLKFCSLAATSLGLGAGMTPKIAWALENKPRIPVVWIHGL WP_001058312.1 hydrogenase [Salmonella enterica] Length: 372\nNumber of predicted TMHs: 2\nExp number of AAs in TMHs: 36.51738\nExp number, first 60 AAs: 19.72849\nTotal prob of N-in: 0.96170\nPOSSIBLE N-term signal sequence\ninside 1 19\nTMhelix 20 39\noutside 40 325\nTMhelix 326 348\ninside 349 372

17801 GCF_002063205.1_ASM206320v1 Salmonella enterica subsp. enterica serovar Hiduddify Proteobacteria; Gammaproteobacteria; Enterobacterales; Enterobacteriaceae; Salmonella; Salmonella enterica; Salmonella enterica subsp. enterica MNNEETFYQAMRRKGVTRRSFLKFCSLAATSLGLGAGMTPKIAWALENKPRIPVVWIHGL WP_001058311.1 hydrogenase [Salmonella enterica] Length: 372\nNumber of predicted TMHs: 2\nExp number of AAs in TMHs: 37.4699899999999\nExp number, first 60 AAs: 19.72699\nTotal prob of N-in: 0.96171\nPOSSIBLE N-term signal sequence\ninside 1 19\nTMhelix 20 39\noutside 40 325\nTMhelix 326 348\ninside 349 372

17802 GCF_002063205.1_ASM206320v1 Salmonella enterica subsp. enterica serovar Hiduddify Proteobacteria; Gammaproteobacteria; Enterobacterales; Enterobacteriaceae; Salmonella; Salmonella enterica; Salmonella enterica subsp. enterica MTGDNTLITSHGINRRDFMKLCAALAATMGLSSKAAAEMAESVSNPQRPPVIWIGAQECT WP_000145429.1 hydrogenase 2 small subunit [Salmonella enterica] Length: 372\nNumber of predicted TMHs: 1\nExp number of AAs in TMHs: 24.51309\nExp number, first 60 AAs: 0.2682\nTotal prob of N-in: 0.14402\noutside 1 331\nTMhelix 332 354\ninside 355 372

17803 GCF_002031165.1_ASM203116v1 Salmonella enterica subsp. enterica serovar Hindmarsh Proteobacteria; Gammaproteobacteria; Enterobacterales; Enterobacteriaceae; Salmonella; Salmonella enterica; Salmonella enterica subsp. enterica MTGDNTLITSHGINRRDFMKLCAALAATMGLSSKAAAEMAESVSNPQRPPVIWIGAQECT WP_000145429.1 hydrogenase 2 small subunit [Salmonella enterica] Length: 372\nNumber of predicted TMHs: 1\nExp number of AAs in TMHs: 24.51309\nExp number, first 60 AAs: 0.2682\nTotal prob of N-in: 0.14402\noutside 1 331\nTMhelix 332 354\ninside 355 372

17804 GCF_001478005.1_Salmonella_enterica_CVM_N30673-SQ_v1.0 Salmonella enterica subsp. enterica serovar Hindmarsh Proteobacteria; Gammaproteobacteria; Enterobacterales; Enterobacteriaceae; Salmonella; Salmonella enterica; Salmonella enterica subsp. enterica MTGDNTLITSHGINRRDFMKLCAALAATMGLSSKAAAEMAESVSNPQRPPVIWIGAQECT WP_000145429.1 hydrogenase 2 small subunit [Salmonella enterica] Length: 372\nNumber of predicted TMHs: 1\nExp number of AAs in TMHs: 24.51309\nExp number, first 60 AAs: 0.2682\nTotal prob of N-in: 0.14402\noutside 1 331\nTMhelix 332 354\ninside 355 372

17805 GCF_002031165.1_ASM203116v1 Salmonella enterica subsp. enterica serovar Hindmarsh Proteobacteria; Gammaproteobacteria; Enterobacterales; Enterobacteriaceae; Salmonella; Salmonella enterica; Salmonella enterica subsp. enterica MNNEETFYQAMRRKGVTRRSFLKFCSLAATSLGLGAGMTPKIAWALENKPRIPVVWIHGL WP_001058311.1 hydrogenase [Salmonella enterica] Length: 372\nNumber of predicted TMHs: 2\nExp number of AAs in TMHs: 37.4699899999999\nExp number, first 60 AAs: 19.72699\nTotal prob of N-in: 0.96171\nPOSSIBLE N-term signal sequence\ninside 1 19\nTMhelix 20 39\noutside 40 325\nTMhelix 326 348\ninside 349 372

17806 GCF_001478005.1_Salmonella_enterica_CVM_N30673-SQ_v1.0 Salmonella enterica subsp. enterica serovar Hindmarsh Proteobacteria; Gammaproteobacteria; Enterobacterales; Enterobacteriaceae; Salmonella; Salmonella enterica; Salmonella enterica subsp. enterica MNNEETFYQAMRRKGVTRRSFLKFCSLAATSLGLGAGMTPKIAWALENKPRIPVVWIHGL WP_001058311.1 hydrogenase [Salmonella enterica] Length: 372\nNumber of predicted TMHs: 2\nExp number of AAs in TMHs: 37.4699899999999\nExp number, first 60 AAs: 19.72699\nTotal prob of N-in: 0.96171\nPOSSIBLE N-term signal sequence\ninside 1 19\nTMhelix 20 39\noutside 40 325\nTMhelix 326 348\ninside 349 372

17807 GCF_002106505.1_ASM210650v1 Salmonella enterica subsp. enterica serovar Holcomb Proteobacteria; Gammaproteobacteria; Enterobacterales; Enterobacteriaceae; Salmonella; Salmonella enterica; Salmonella enterica subsp. enterica MTGDNTLITSHGINRRDFMKLCAALAATMGLSSKAAAEMAESVSNPQRPPVIWIGAQECT WP_000145429.1 hydrogenase 2 small subunit [Salmonella enterica] Length: 372\nNumber of predicted TMHs: 1\nExp number of AAs in TMHs: 24.51309\nExp number, first 60 AAs: 0.2682\nTotal prob of N-in: 0.14402\noutside 1 331\nTMhelix 332 354\ninside 355 372

17808 GCF_002106505.1_ASM210650v1 Salmonella enterica subsp. enterica serovar Holcomb Proteobacteria; Gammaproteobacteria; Enterobacterales; Enterobacteriaceae; Salmonella; Salmonella enterica; Salmonella enterica subsp. enterica MNNEETFYQAMRRKGVTRRSFLKFCSLAATSLGLGAGMTPKIAWALENKPRIPVVWIHGL WP_023201174.1 hydrogenase 1, small subunit [Salmonella enterica] Length: 372\nNumber of predicted TMHs: 2\nExp number of AAs in TMHs: 38.09509\nExp number, first 60 AAs: 19.72914\nTotal prob of N-in: 0.96171\nPOSSIBLE N-term signal sequence\ninside 1 19\nTMhelix 20 39\noutside 40 325\nTMhelix 326 348\ninside 349 372

17809 GCF_002106355.1_ASM210635v1 Salmonella enterica subsp. enterica serovar Holcomb Proteobacteria; Gammaproteobacteria; Enterobacterales; Enterobacteriaceae; Salmonella; Salmonella enterica; Salmonella enterica subsp. enterica MNNEETFYQAMRRKGVTRRSFLKFCSLAATSLGLGAGMTPKIAWALENKPRIPVVWIHGL WP_023201174.1 hydrogenase 1, small subunit [Salmonella enterica] Length: 372\nNumber of predicted TMHs: 2\nExp number of AAs in TMHs: 38.09509\nExp number, first 60 AAs: 19.72914\nTotal prob of N-in: 0.96171\nPOSSIBLE N-term signal sequence\ninside 1 19\nTMhelix 20 39\noutside 40 325\nTMhelix 326 348\ninside 349 372

17810 GCF_002106355.1_ASM210635v1 Salmonella enterica subsp. enterica serovar Holcomb Proteobacteria; Gammaproteobacteria; Enterobacterales; Enterobacteriaceae; Salmonella; Salmonella enterica; Salmonella enterica subsp. enterica MTGDNTLITSHGINRRDFMKLCAALAATMGLSSKAAAEMAESVSNPQRPPVIWIGAQECT WP_000145429.1 hydrogenase 2 small subunit [Salmonella enterica] Length: 372\nNumber of predicted TMHs: 1\nExp number of AAs in TMHs: 24.51309\nExp number, first 60 AAs: 0.2682\nTotal prob of N-in: 0.14402\noutside 1 331\nTMhelix 332 354\ninside 355 372

17811 GCF_002047625.1_ASM204762v1 Salmonella enterica subsp. enterica serovar Hvittingfoss Proteobacteria; Gammaproteobacteria; Enterobacterales; Enterobacteriaceae; Salmonella; Salmonella enterica; Salmonella enterica subsp. enterica MTGDNTLITSHGINRRDFMKLCAALAATMGLSSKAAAEMAESVSNPQRPPVIWIGAQECT WP_000145429.1 hydrogenase 2 small subunit [Salmonella enterica] Length: 372\nNumber of predicted TMHs: 1\nExp number of AAs in TMHs: 24.51309\nExp number, first 60 AAs: 0.2682\nTotal prob of N-in: 0.14402\noutside 1 331\nTMhelix 332 354\ninside 355 372

17812 GCF_002047875.1_ASM204787v1 Salmonella enterica subsp. enterica serovar Hvittingfoss Proteobacteria; Gammaproteobacteria; Enterobacterales; Enterobacteriaceae; Salmonella; Salmonella enterica; Salmonella enterica subsp. enterica MTGDNTLITSHGINRRDFMKLCAALAATMGLSSKAAAEMAESVSNPQRPPVIWIGAQECT WP_000145429.1 hydrogenase 2 small subunit [Salmonella enterica] Length: 372\nNumber of predicted TMHs: 1\nExp number of AAs in TMHs: 24.51309\nExp number, first 60 AAs: 0.2682\nTotal prob of N-in: 0.14402\noutside 1 331\nTMhelix 332 354\ninside 355 372

17813 GCF_002047725.1_ASM204772v1 Salmonella enterica subsp. enterica serovar Hvittingfoss Proteobacteria; Gammaproteobacteria; Enterobacterales; Enterobacteriaceae; Salmonella; Salmonella enterica; Salmonella enterica subsp. enterica MNNEETFYQAMRRKGVTRRSFLKFCSLAATSLGLGAGMTPKIAWALENKPRIPVVWIHGL WP_001058312.1 hydrogenase [Salmonella enterica] Length: 372\nNumber of predicted TMHs: 2\nExp number of AAs in TMHs: 36.51738\nExp number, first 60 AAs: 19.72849\nTotal prob of N-in: 0.96170\nPOSSIBLE N-term signal sequence\ninside 1 19\nTMhelix 20 39\noutside 40 325\nTMhelix 326 348\ninside 349 372

17814 GCF_002047725.1_ASM204772v1 Salmonella enterica subsp. enterica serovar Hvittingfoss Proteobacteria; Gammaproteobacteria; Enterobacterales; Enterobacteriaceae; Salmonella; Salmonella enterica; Salmonella enterica subsp. enterica MTGDNTLITSHGINRRDFMKLCAALAATMGLSSKAAAEMAESVSNPQRPPVIWIGAQECT WP_000145429.1 hydrogenase 2 small subunit [Salmonella enterica] Length: 372\nNumber of predicted TMHs: 1\nExp number of AAs in TMHs: 24.51309\nExp number, first 60 AAs: 0.2682\nTotal prob of N-in: 0.14402\noutside 1 331\nTMhelix 332 354\ninside 355 372

17815 GCF_002047625.1_ASM204762v1 Salmonella enterica subsp. enterica serovar Hvittingfoss Proteobacteria; Gammaproteobacteria; Enterobacterales; Enterobacteriaceae; Salmonella; Salmonella enterica; Salmonella enterica subsp. enterica MNNEETFYQAMRRKGVTRRSFLKFCSLAATSLGLGAGMTPKIAWALENKPRIPVVWIHGL WP_001058307.1 hydrogenase [Salmonella enterica] Length: 372\nNumber of predicted TMHs: 2\nExp number of AAs in TMHs: 36.51738\nExp number, first 60 AAs: 19.72849\nTotal prob of N-in: 0.96170\nPOSSIBLE N-term signal sequence\ninside 1 19\nTMhelix 20 39\noutside 40 325\nTMhelix 326 348\ninside 349 372

17816 GCF_002047875.1_ASM204787v1 Salmonella enterica subsp. enterica serovar Hvittingfoss Proteobacteria; Gammaproteobacteria; Enterobacterales; Enterobacteriaceae; Salmonella; Salmonella enterica; Salmonella enterica subsp. enterica MNNEETFYQAMRRKGVTRRSFLKFCSLAATSLGLGAGMTPKIAWALENKPRIPVVWIHGL WP_001058307.1 hydrogenase [Salmonella enterica] Length: 372\nNumber of predicted TMHs: 2\nExp number of AAs in TMHs: 36.51738\nExp number, first 60 AAs: 19.72849\nTotal prob of N-in: 0.96170\nPOSSIBLE N-term signal sequence\ninside 1 19\nTMhelix 20 39\noutside 40 325\nTMhelix 326 348\ninside 349 372

17817 GCF_002047475.1_ASM204747v1 Salmonella enterica subsp. enterica serovar Hvittingfoss Proteobacteria; Gammaproteobacteria; Enterobacterales; Enterobacteriaceae; Salmonella; Salmonella enterica; Salmonella enterica subsp. enterica MTGDNTLITSHGINRRDFMKLCAALAATMGLSSKAAAEMAESVSNPQRPPVIWIGAQECT WP_000145429.1 hydrogenase 2 small subunit [Salmonella enterica] Length: 372\nNumber of predicted TMHs: 1\nExp number of AAs in TMHs: 24.51309\nExp number, first 60 AAs: 0.2682\nTotal prob of N-in: 0.14402\noutside 1 331\nTMhelix 332 354\ninside 355 372

17818 GCF_002047475.1_ASM204747v1 Salmonella enterica subsp. enterica serovar Hvittingfoss Proteobacteria; Gammaproteobacteria; Enterobacterales; Enterobacteriaceae; Salmonella; Salmonella enterica; Salmonella enterica subsp. enterica MNNEETFYQAMRRKGVTRRSFLKFCSLAATSLGLGAGMTPKIAWALENKPRIPVVWIHGL WP_001058312.1 hydrogenase [Salmonella enterica] Length: 372\nNumber of predicted TMHs: 2\nExp number of AAs in TMHs: 36.51738\nExp number, first 60 AAs: 19.72849\nTotal prob of N-in: 0.96170\nPOSSIBLE N-term signal sequence\ninside 1 19\nTMhelix 20 39\noutside 40 325\nTMhelix 326 348\ninside 349 372

17819 GCF_002045025.1_ASM204502v1 Salmonella enterica subsp. enterica serovar Ibadan Proteobacteria; Gammaproteobacteria; Enterobacterales; Enterobacteriaceae; Salmonella; Salmonella enterica; Salmonella enterica subsp. enterica MNNEETFYQAMRRKGVTRRSFLKFCSLAATSLGLGAGMTPKIAWALENKPRIPVVWIHGL WP_001058312.1 hydrogenase [Salmonella enterica] Length: 372\nNumber of predicted TMHs: 2\nExp number of AAs in TMHs: 36.51738\nExp number, first 60 AAs: 19.72849\nTotal prob of N-in: 0.96170\nPOSSIBLE N-term signal sequence\ninside 1 19\nTMhelix 20 39\noutside 40 325\nTMhelix 326 348\ninside 349 372

17820 GCF_002045725.1_ASM204572v1 Salmonella enterica subsp. enterica serovar Ibadan Proteobacteria; Gammaproteobacteria; Enterobacterales; Enterobacteriaceae; Salmonella; Salmonella enterica; Salmonella enterica subsp. enterica MTGDNTLITSHGINRRDFMKLCAALAATMGLSSKAAAEMAESVSNPQRPPVIWIGAQECT WP_000145431.1 hydrogenase 2 small subunit [Salmonella enterica] Length: 372\nNumber of

predicted TMHs:  1\nExp number of AAs in TMHs: 24.51505\nExp number, first 60 AAs:  0.2682\nTotal prob of N-in:  0.14406\noutside   1   331\nTMhelix   332   354\ninside   355   372

17821	GCF_002045025.1_ASM204502v1	Salmonella enterica subsp. enterica serovar Ibadan	Proteobacteria; Gammaproteobacteria; Enterobacterales; Enterobacteriaceae; Salmonella; Salmonella enterica; Salmonella enterica subsp. enterica	MTGDNTLITSHGINRRDFMKLCAALAATMGLSSKAAAEMAESVSNPQRPPVIWIGAQECT	WP_000145431.1	hydrogenase 2 small subunit [Salmonella enterica]	Length: 372\nNumber of predicted TMHs:  1\nExp number of AAs in TMHs: 24.51505\nExp number, first 60 AAs:  0.2682\nTotal prob of N-in:  0.14406\noutside   1   331\nTMhelix   332   354\ninside   355   372

17822	GCF_002045725.1_ASM204572v1	Salmonella enterica subsp. enterica serovar Ibadan	Proteobacteria; Gammaproteobacteria; Enterobacterales; Enterobacteriaceae; Salmonella; Salmonella enterica; Salmonella enterica subsp. enterica	MNNEETFYQAMRRKGVTRRSFLKFCSLAATSLGLGAGMTPKIAWALENKPRIPVVWIHGL	WP_001058312.1	hydrogenase [Salmonella enterica]	Length: 372\nNumber of predicted TMHs:  2\nExp number of AAs in TMHs: 36.51738\nExp number, first 60 AAs:  19.72849\nTotal prob of N-in:   0.96170\nPOSSIBLE N-term signal sequence\ninside   1   19\nTMhelix   20   39\noutside   40   325\nTMhelix   326   348\ninside   349   372

17823	GCF_002046685.1_ASM204668v1	Salmonella enterica subsp. enterica serovar Indiana	Proteobacteria; Gammaproteobacteria; Enterobacterales; Enterobacteriaceae; Salmonella; Salmonella enterica; Salmonella enterica subsp. enterica	MNNEETFYQAMRRKGVTRRSFLKFCSLAATSLGLGAGMTPQIAWALENKPRIPVVWIHGL	WP_023227615.1	hydrogenase 1, small subunit [Salmonella enterica]	Length: 372\nNumber of predicted TMHs:  2\nExp number of AAs in TMHs: 39.8591699999999\nExp number, first 60 AAs:  22.13093\nTotal prob of N-in:   0.98639\nPOSSIBLE N-term signal sequence\ninside   1   20\nTMhelix   21   43\noutside   44   325\nTMhelix   326   348\ninside   349   372

17824	GCF_002048205.1_ASM204820v1	Salmonella enterica subsp. enterica serovar Indiana	Proteobacteria; Gammaproteobacteria; Enterobacterales; Enterobacteriaceae; Salmonella; Salmonella enterica; Salmonella enterica subsp. enterica	MLKFRTIFLKKIFTLKNSILARRGAVIVIVSAVFTSIMFFAHSWASDKEVAMTLSVLNSN	WP_023226644.1	hypothetical protein [Salmonella enterica]	Length: 204\nNumber of predicted TMHs:  2\nExp number of AAs in TMHs: 44.99526\nExp number, first 60 AAs:  22.33786\nTotal prob of N-in:   0.87083\nPOSSIBLE N-term signal sequence\ninside   1   19\nTMhelix   20   42\noutside   43   171\nTMhelix   172   194\ninside   195   204

17825	GCF_002048205.1_ASM204820v1	Salmonella enterica subsp. enterica serovar Indiana	Proteobacteria; Gammaproteobacteria; Enterobacterales; Enterobacteriaceae; Salmonella; Salmonella enterica; Salmonella enterica subsp. enterica	MNNEETFYQAMRRKGVTRRSFLKFCSLAATSLGLGAGMTPQIAWALENKPRIPVVWIHGL	WP_023227615.1	hydrogenase 1, small subunit [Salmonella enterica]	Length: 372\nNumber of predicted TMHs:  2\nExp number of AAs in TMHs: 39.8591699999999\nExp number, first 60 AAs:  22.13093\nTotal prob of N-in:   0.98639\nPOSSIBLE N-term signal sequence\ninside   1   20\nTMhelix   21   43\noutside   44   325\nTMhelix   326   348\ninside   349   372

17826	GCF_002048175.1_ASM204817v1	Salmonella enterica subsp. enterica serovar Indiana	Proteobacteria; Gammaproteobacteria; Enterobacterales; Enterobacteriaceae; Salmonella; Salmonella enterica; Salmonella enterica subsp. enterica	MNNEETFYQAMRRKGVTRRSFLKFCSLAATSLGLGAGMTPQIAWALENKPRIPVVWIHGL	WP_023227615.1	hydrogenase 1, small subunit [Salmonella enterica]	Length: 372\nNumber of predicted TMHs:  2\nExp number of AAs in TMHs: 39.8591699999999\nExp number, first 60 AAs:  22.13093\nTotal prob of N-in:   0.98639\nPOSSIBLE N-term signal sequence\ninside   1   20\nTMhelix   21   43\noutside   44   325\nTMhelix   326   348\ninside   349   372

17827	GCF_002047005.1_ASM204700v1	Salmonella enterica subsp. enterica serovar Indiana	Proteobacteria; Gammaproteobacteria; Enterobacterales; Enterobacteriaceae; Salmonella; Salmonella enterica; Salmonella enterica subsp. enterica	MTGDNTLITSHGINRRDFMKLCAALAATMGLSSKAAAEMAESVSNPQRPPVIWIGAQECT	WP_000145429.1	hydrogenase 2 small subunit [Salmonella enterica]	Length: 372\nNumber of predicted TMHs:  1\nExp number of AAs in TMHs: 24.51309\nExp number, first 60 AAs:  0.2682\nTotal prob of N-in:  0.14402\noutside   1   331\nTMhelix   332   354\ninside   355   372

17828	GCF_002046685.1_ASM204668v1	Salmonella enterica subsp. enterica serovar Indiana	Proteobacteria; Gammaproteobacteria; Enterobacterales; Enterobacteriaceae; Salmonella; Salmonella enterica; Salmonella enterica subsp. enterica	MTGDNTLITSHGINRRDFMKLCAALAATMGLSSKAAAEMAESVSNPQRPPVIWIGAQECT	WP_000145429.1	hydrogenase 2 small subunit [Salmonella enterica]	Length: 372\nNumber of predicted TMHs:  1\nExp number of AAs in TMHs: 24.51309\nExp number, first 60 AAs:  0.2682\nTotal prob of N-in:  0.14402\noutside   1   331\nTMhelix   332   354\ninside   355   372

17829 GCF_002046685.1_ASM204668v1 Salmonella enterica subsp. enterica serovar Indiana Proteobacteria; Gammaproteobacteria; Enterobacterales; Enterobacteriaceae; Salmonella; Salmonella enterica; Salmonella enterica subsp. enterica MLKFRTIFLKKIFTLKNSILARRGAVIVIVSAVFTSIMFFAHSWASDKEVAMTLSVLNSN WP_023226644.1 hypothetical protein [Salmonella enterica] Length: 204\nNumber of predicted TMHs: 2\nExp number of AAs in TMHs: 44.99526\nExp number, first 60 AAs: 22.33786\nTotal prob of N-in: 0.87083\nPOSSIBLE N-term signal sequence\ninside 1 19\nTMhelix 20 42\noutside 43 171\nTMhelix 172 194\ninside 195 204

17830 GCF_002046785.1_ASM204678v1 Salmonella enterica subsp. enterica serovar Indiana Proteobacteria; Gammaproteobacteria; Enterobacterales; Enterobacteriaceae; Salmonella; Salmonella enterica; Salmonella enterica subsp. enterica MLKFRTIFLKKIFTLKNSILARRGAVIVIVSAVFTSIMFFAHSWASDKEVAMTLSVLNSN WP_023226644.1 hypothetical protein [Salmonella enterica] Length: 204\nNumber of predicted TMHs: 2\nExp number of AAs in TMHs: 44.99526\nExp number, first 60 AAs: 22.33786\nTotal prob of N-in: 0.87083\nPOSSIBLE N-term signal sequence\ninside 1 19\nTMhelix 20 42\noutside 43 171\nTMhelix 172 194\ninside 195 204

17831 GCF_002046785.1_ASM204678v1 Salmonella enterica subsp. enterica serovar Indiana Proteobacteria; Gammaproteobacteria; Enterobacterales; Enterobacteriaceae; Salmonella; Salmonella enterica; Salmonella enterica subsp. enterica MNNEETFYQAMRRKGVTRRSFLKFCSLAATSLGLGAGMTPQIAWALENKPRIPVVWIHGL WP_023227615.1 hydrogenase 1, small subunit [Salmonella enterica] Length: 372\nNumber of predicted TMHs: 2\nExp number of AAs in TMHs: 39.8591699999999\nExp number, first 60 AAs: 22.13093\nTotal prob of N-in: 0.98639\nPOSSIBLE N-term signal sequence\ninside 1 20\nTMhelix 21 43\noutside 44 325\nTMhelix 326 348\ninside 349 372

17832 GCF_002047005.1_ASM204700v1 Salmonella enterica subsp. enterica serovar Indiana Proteobacteria; Gammaproteobacteria; Enterobacterales; Enterobacteriaceae; Salmonella; Salmonella enterica; Salmonella enterica subsp. enterica MNNEETFYQAMRRKGVTRRSFLKFCSLAATSLGLGAGMTPQIAWALENKPRIPVVWIHGL WP_023227615.1 hydrogenase 1, small subunit [Salmonella enterica] Length: 372\nNumber of predicted TMHs: 2\nExp number of AAs in TMHs: 39.8591699999999\nExp number, first 60 AAs: 22.13093\nTotal prob of N-in: 0.98639\nPOSSIBLE N-term signal sequence\ninside 1 20\nTMhelix 21 43\noutside 44 325\nTMhelix 326 348\ninside 349 372

17833 GCF_002048175.1_ASM204817v1 Salmonella enterica subsp. enterica serovar Indiana Proteobacteria; Gammaproteobacteria; Enterobacterales; Enterobacteriaceae; Salmonella; Salmonella enterica; Salmonella enterica subsp. enterica MTGDNTLITSHGINRRDFMKLCAALAATMGLSSKAAAEMAESVSNPQRPPVIWIGAQECT WP_000145429.1 hydrogenase 2 small subunit [Salmonella enterica] Length: 372\nNumber of predicted TMHs: 1\nExp number of AAs in TMHs: 24.51309\nExp number, first 60 AAs: 0.2682\nTotal prob of N-in: 0.14402\noutside 1 331\nTMhelix 332 354\ninside 355 372

17834 GCF_002048175.1_ASM204817v1 Salmonella enterica subsp. enterica serovar Indiana Proteobacteria; Gammaproteobacteria; Enterobacterales; Enterobacteriaceae; Salmonella; Salmonella enterica; Salmonella enterica subsp. enterica MLKFRTIFLKKIFTLKNSILARRGAVIVIVSAVFTSIMFFAHSWASDKEVAMTLSVLNSN WP_023226644.1 hypothetical protein [Salmonella enterica] Length: 204\nNumber of predicted TMHs: 2\nExp number of AAs in TMHs: 44.99526\nExp number, first 60 AAs: 22.33786\nTotal prob of N-in: 0.87083\nPOSSIBLE N-term signal sequence\ninside 1 19\nTMhelix 20 42\noutside 43 171\nTMhelix 172 194\ninside 195 204

17835 GCF_002047005.1_ASM204700v1 Salmonella enterica subsp. enterica serovar Indiana Proteobacteria; Gammaproteobacteria; Enterobacterales; Enterobacteriaceae; Salmonella; Salmonella enterica; Salmonella enterica subsp. enterica MLKFRTIFLKKIFTLKNSILARRGAVIVIVSAVFTSIMFFAHSWASDKEVAMTLSVLNSN WP_023226644.1 hypothetical protein [Salmonella enterica] Length: 204\nNumber of predicted TMHs: 2\nExp number of AAs in TMHs: 44.99526\nExp number, first 60 AAs: 22.33786\nTotal prob of N-in: 0.87083\nPOSSIBLE N-term signal sequence\ninside 1 19\nTMhelix 20 42\noutside 43 171\nTMhelix 172 194\ninside 195 204

17836 GCF_002048205.1_ASM204820v1 Salmonella enterica subsp. enterica serovar Indiana Proteobacteria; Gammaproteobacteria; Enterobacterales; Enterobacteriaceae; Salmonella; Salmonella enterica; Salmonella enterica subsp. enterica MTGDNTLITSHGINRRDFMKLCAALAATMGLSSKAAAEMAESVSNPQRPPVIWIGAQECT WP_000145429.1 hydrogenase 2 small subunit [Salmonella enterica] Length: 372\nNumber of predicted TMHs: 1\nExp number of AAs in TMHs: 24.51309\nExp number, first 60 AAs: 0.2682\nTotal prob of N-in: 0.14402\noutside 1 331\nTMhelix 332 354\ninside 355 372

17837 GCF_002046785.1_ASM204678v1 Salmonella enterica subsp. enterica serovar Indiana Proteobacteria; Gammaproteobacteria; Enterobacterales; Enterobacteriaceae; Salmonella; Salmonella enterica; Salmonella enterica subsp. enterica MTGDNTLITSHGINRRDFMKLCAALAATMGLSSKAAAEMAESVSNPQRPPVIWIGAQECT

WP_000145429.1 hydrogenase 2 small subunit [Salmonella enterica] Length: 372\nNumber of predicted TMHs: 1\nExp number of AAs in TMHs: 24.51309\nExp number, first 60 AAs: 0.2682\nTotal prob of N-in: 0.14402\noutside 1 331\nTMhelix 332 354\ninside 355 372

17838 GCF_002062235.1_ASM206223v1 Salmonella enterica subsp. enterica serovar Indiana Proteobacteria; Gammaproteobacteria; Enterobacterales; Enterobacteriaceae; Salmonella; Salmonella enterica; Salmonella enterica subsp. enterica MTGDNTLITSHGINRRDFMKLCAALAATMGLSSKAAAEMAESVSNPQRPPVIWIGAQECT WP_000145429.1 hydrogenase 2 small subunit [Salmonella enterica] Length: 372\nNumber of predicted TMHs: 1\nExp number of AAs in TMHs: 24.51309\nExp number, first 60 AAs: 0.2682\nTotal prob of N-in: 0.14402\noutside 1 331\nTMhelix 332 354\ninside 355 372

17839 GCF_002062235.1_ASM206223v1 Salmonella enterica subsp. enterica serovar Indiana Proteobacteria; Gammaproteobacteria; Enterobacterales; Enterobacteriaceae; Salmonella; Salmonella enterica; Salmonella enterica subsp. enterica MNNEETFYQAMRRKGVTRRSFLKFCSLAATSLGLGAGMTPQIAWALENKPRIPVVWIHGL WP_023227615.1 hydrogenase 1, small subunit [Salmonella enterica] Length: 372\nNumber of predicted TMHs: 2\nExp number of AAs in TMHs: 39.8591699999999\nExp number, first 60 AAs: 22.13093\nTotal prob of N-in: 0.98639\nPOSSIBLE N-term signal sequence\ninside 1 20\nTMhelix 21 43\noutside 44 325\nTMhelix 326 348\ninside 349 372

17840 GCF_002062235.1_ASM206223v1 Salmonella enterica subsp. enterica serovar Indiana Proteobacteria; Gammaproteobacteria; Enterobacterales; Enterobacteriaceae; Salmonella; Salmonella enterica; Salmonella enterica subsp. enterica MLKFRTIFLKKIFTLKNSILARRGAVIVIVSAVFTSIMFFAHSWASDKEVAMTLSVLNSN WP_023226644.1 hypothetical protein [Salmonella enterica] Length: 204\nNumber of predicted TMHs: 2\nExp number of AAs in TMHs: 44.99526\nExp number, first 60 AAs: 22.33786\nTotal prob of N-in: 0.87083\nPOSSIBLE N-term signal sequence\ninside 1 19\nTMhelix 20 42\noutside 43 171\nTMhelix 172 194\ninside 195 204

17841 GCF_001276865.1_ASM127686v1 Salmonella enterica subsp. enterica serovar Indiana Proteobacteria; Gammaproteobacteria; Enterobacterales; Enterobacteriaceae; Salmonella; Salmonella enterica; Salmonella enterica subsp. enterica MLKFRTIFLKKIFTLKNSILARRGAVIVIVSAVFTSIMFFAHSWASDKEVAMTLSVLNSN WP_023226644.1 hypothetical protein [Salmonella enterica] Length: 204\nNumber of predicted TMHs: 2\nExp number of AAs in TMHs: 44.99526\nExp number, first 60 AAs: 22.33786\nTotal prob of N-in: 0.87083\nPOSSIBLE N-term signal sequence\ninside 1 19\nTMhelix 20 42\noutside 43 171\nTMhelix 172 194\ninside 195 204

17842 GCF_001276865.1_ASM127686v1 Salmonella enterica subsp. enterica serovar Indiana Proteobacteria; Gammaproteobacteria; Enterobacterales; Enterobacteriaceae; Salmonella; Salmonella enterica; Salmonella enterica subsp. enterica MNNEETFYQAMRRKGVTRRSFLKFCSLAATSLGLGAGMTPQIAWALENKPRIPVVWIHGL WP_023227615.1 hydrogenase 1, small subunit [Salmonella enterica] Length: 372\nNumber of predicted TMHs: 2\nExp number of AAs in TMHs: 39.8591699999999\nExp number, first 60 AAs: 22.13093\nTotal prob of N-in: 0.98639\nPOSSIBLE N-term signal sequence\ninside 1 20\nTMhelix 21 43\noutside 44 325\nTMhelix 326 348\ninside 349 372

17843 GCF_001276865.1_ASM127686v1 Salmonella enterica subsp. enterica serovar Indiana Proteobacteria; Gammaproteobacteria; Enterobacterales; Enterobacteriaceae; Salmonella; Salmonella enterica; Salmonella enterica subsp. enterica MTGDNTLITSHGINRRDFMKLCAALAATMGLSSKAAAEMAESVSNPQRPPVIWIGAQECT WP_000145429.1 hydrogenase 2 small subunit [Salmonella enterica] Length: 372\nNumber of predicted TMHs: 1\nExp number of AAs in TMHs: 24.51309\nExp number, first 60 AAs: 0.2682\nTotal prob of N-in: 0.14402\noutside 1 331\nTMhelix 332 354\ninside 355 372

17844 GCF_001242165.1_Salmonella_enterica_CVM_N45928_v1.0 Salmonella enterica subsp. enterica serovar Infantis Proteobacteria; Gammaproteobacteria; Enterobacterales; Enterobacteriaceae; Salmonella; Salmonella enterica; Salmonella enterica subsp. enterica MTGDNTLITSHGINRRDFMKLCAALAATMGLSSKAAAEMAESVSNPQRPPVIWIGAQECT WP_000145429.1 hydrogenase 2 small subunit [Salmonella enterica] Length: 372\nNumber of predicted TMHs: 1\nExp number of AAs in TMHs: 24.51309\nExp number, first 60 AAs: 0.2682\nTotal prob of N-in: 0.14402\noutside 1 331\nTMhelix 332 354\ninside 355 372

17845 GCF_001241325.1_Salmonella_enterica_CVM_N44705_v1.0 Salmonella enterica subsp. enterica serovar Infantis Proteobacteria; Gammaproteobacteria; Enterobacterales; Enterobacteriaceae; Salmonella; Salmonella enterica; Salmonella enterica subsp. enterica MTGDNTLITSHGINRRDFMKLCAALAATMGLSSKAAAEMAESVSNPQRPPVIWIGAQECT WP_000145429.1 hydrogenase 2 small subunit [Salmonella enterica] Length: 372\nNumber of predicted TMHs: 1\nExp number of AAs in TMHs: 24.51309\nExp number, first 60 AAs: 0.2682\nTotal prob of N-in: 0.14402\noutside 1 331\nTMhelix 332 354\ninside 355 372

17846 GCF_001246465.1_Salmonella_enterica_CVM_N51266_v1.0 Salmonella enterica subsp. enterica serovar Infantis Proteobacteria; Gammaproteobacteria; Enterobacterales; Enterobacteriaceae; Salmonella; Salmonella enterica; Salmonella enterica subsp. enterica MNNEETFYQAMRRKGVTRRSFLKFCSLAATSLGLGAGMTPKIAWALENKPRIPVVWIHGL WP_001058311.1 hydrogenase [Salmonella enterica] Length: 372\nNumber of predicted TMHs: 2\nExp number of AAs in TMHs: 37.4699899999999\nExp number, first 60 AAs: 19.72699\nTotal prob of N-in: 0.96171\nPOSSIBLE N-term signal sequence\ninside 1 19\nTMhelix 20 39\noutside 40 325\nTMhelix 326 348\ninside 349 372

17847 GCF_001246295.1_Salmonella_enterica_CVM_N51258_v1.0 Salmonella enterica subsp. enterica serovar Infantis Proteobacteria; Gammaproteobacteria; Enterobacterales; Enterobacteriaceae; Salmonella; Salmonella enterica; Salmonella enterica subsp. enterica MNNEETFYQAMRRKGVTRRSFLKFCSLAATSLGLGAGMTPKIAWALENKPRIPVVWIHGL WP_001058311.1 hydrogenase [Salmonella enterica] Length: 372\nNumber of predicted TMHs: 2\nExp number of AAs in TMHs: 37.4699899999999\nExp number, first 60 AAs: 19.72699\nTotal prob of N-in: 0.96171\nPOSSIBLE N-term signal sequence\ninside 1 19\nTMhelix 20 39\noutside 40 325\nTMhelix 326 348\ninside 349 372

17848 GCF_000953495.1_SINFA Salmonella enterica subsp. enterica serovar Infantis Proteobacteria; Gammaproteobacteria; Enterobacterales; Enterobacteriaceae; Salmonella; Salmonella enterica; Salmonella enterica subsp. enterica MNNEETFYQAMRRKGVTRRSFLKFCSLAATSLGLGAGMTPKIAWALENKPRIPVVWIHGL WP_001058311.1 hydrogenase [Salmonella enterica] Length: 372\nNumber of predicted TMHs: 2\nExp number of AAs in TMHs: 37.4699899999999\nExp number, first 60 AAs: 19.72699\nTotal prob of N-in: 0.96171\nPOSSIBLE N-term signal sequence\ninside 1 19\nTMhelix 20 39\noutside 40 325\nTMhelix 326 348\ninside 349 372

17849 GCF_001244865.1_Salmonella_enterica_CVM_N48694_v1.0 Salmonella enterica subsp. enterica serovar Infantis Proteobacteria; Gammaproteobacteria; Enterobacterales; Enterobacteriaceae; Salmonella; Salmonella enterica; Salmonella enterica subsp. enterica MTGDNTLITSHGINRRDFMKLCAALAATMGLSSKAAAEMAESVSNPQRPPVIWIGAQECT WP_000145429.1 hydrogenase 2 small subunit [Salmonella enterica] Length: 372\nNumber of predicted TMHs: 1\nExp number of AAs in TMHs: 24.51309\nExp number, first 60 AAs: 0.2682\nTotal prob of N-in: 0.14402\noutside 1 331\nTMhelix 332 354\ninside 355 372

17850 GCF_001242875.1_Salmonella_enterica_CVM_N46813_v1.0 Salmonella enterica subsp. enterica serovar Infantis Proteobacteria; Gammaproteobacteria; Enterobacterales; Enterobacteriaceae; Salmonella; Salmonella enterica; Salmonella enterica subsp. enterica MTGDNTLITSHGINRRDFMKLCAALAATMGLSSKAAAEMAESVSNPQRPPVIWIGAQECT WP_000145429.1 hydrogenase 2 small subunit [Salmonella enterica] Length: 372\nNumber of predicted TMHs: 1\nExp number of AAs in TMHs: 24.51309\nExp number, first 60 AAs: 0.2682\nTotal prob of N-in: 0.14402\noutside 1 331\nTMhelix 332 354\ninside 355 372

17851 GCF_001242165.1_Salmonella_enterica_CVM_N45928_v1.0 Salmonella enterica subsp. enterica serovar Infantis Proteobacteria; Gammaproteobacteria; Enterobacterales; Enterobacteriaceae; Salmonella; Salmonella enterica; Salmonella enterica subsp. enterica MNNEETFYQAMRRKGVTRRSFLKFCSLAATSLGLGAGMTPKIAWALENKPRIPVVWIHGL WP_001058311.1 hydrogenase [Salmonella enterica] Length: 372\nNumber of predicted TMHs: 2\nExp number of AAs in TMHs: 37.4699899999999\nExp number, first 60 AAs: 19.72699\nTotal prob of N-in: 0.96171\nPOSSIBLE N-term signal sequence\ninside 1 19\nTMhelix 20 39\noutside 40 325\nTMhelix 326 348\ninside 349 372

17852 GCF_001244905.1_Salmonella_enterica_CVM_N48695_v1.0 Salmonella enterica subsp. enterica serovar Infantis Proteobacteria; Gammaproteobacteria; Enterobacterales; Enterobacteriaceae; Salmonella; Salmonella enterica; Salmonella enterica subsp. enterica MTGDNTLITSHGINRRDFMKLCAALAATMGLSSKAAAEMAESVSNPQRPPVIWIGAQECT WP_000145429.1 hydrogenase 2 small subunit [Salmonella enterica] Length: 372\nNumber of predicted TMHs: 1\nExp number of AAs in TMHs: 24.51309\nExp number, first 60 AAs: 0.2682\nTotal prob of N-in: 0.14402\noutside 1 331\nTMhelix 332 354\ninside 355 372

17853 GCF_001244905.1_Salmonella_enterica_CVM_N48695_v1.0 Salmonella enterica subsp. enterica serovar Infantis Proteobacteria; Gammaproteobacteria; Enterobacterales; Enterobacteriaceae; Salmonella; Salmonella enterica; Salmonella enterica subsp. enterica MNNEETFYQAMRRKGVTRRSFLKFCSLAATSLGLGAGMTPKIAWALENKPRIPVVWIHGL WP_001058311.1 hydrogenase [Salmonella enterica] Length: 372\nNumber of predicted TMHs: 2\nExp number of AAs in TMHs: 37.4699899999999\nExp number, first 60 AAs: 19.72699\nTotal prob of N-in: 0.96171\nPOSSIBLE N-term signal sequence\ninside 1 19\nTMhelix 20 39\noutside 40 325\nTMhelix 326 348\ninside 349 372

17854 GCF_001244865.1_Salmonella_enterica_CVM_N48694_v1.0 Salmonella enterica subsp. enterica serovar Infantis Proteobacteria; Gammaproteobacteria; Enterobacterales; Enterobacteriaceae; Salmonella; Salmonella enterica; Salmonella enterica subsp. enterica MNNEETFYQAMRRKGVTRRSFLKFCSLAATSLGLGAGMTPKIAWALENKPRIPVVWIHGL WP_001058311.1 hydrogenase [Salmonella enterica] Length: 372\nNumber of predicted TMHs: 2\nExp number of AAs in TMHs: 37.4699899999999\nExp number, first 60 AAs: 19.72699\nTotal prob of N-in: 0.96171\nPOSSIBLE N-term signal sequence\ninside 1 19\nTMhelix 20 39\noutside 40 325\nTMhelix 326 348\ninside 349 372

17855 GCF_001241325.1_Salmonella_enterica_CVM_N44705_v1.0 Salmonella enterica subsp. enterica serovar Infantis Proteobacteria; Gammaproteobacteria; Enterobacterales; Enterobacteriaceae; Salmonella; Salmonella enterica; Salmonella enterica subsp. enterica MNNEETFYQAMRRKGVTRRSFLKFCSLAATSLGLGAGMTPKIAWALENKPRIPVVWIHGL WP_001058311.1 hydrogenase [Salmonella enterica] Length: 372\nNumber of predicted TMHs: 2\nExp number of AAs in TMHs: 37.4699899999999\nExp number, first 60 AAs: 19.72699\nTotal prob of N-in: 0.96171\nPOSSIBLE N-term signal sequence\ninside 1 19\nTMhelix 20 39\noutside 40 325\nTMhelix 326 348\ninside 349 372

17856 GCF_001242875.1_Salmonella_enterica_CVM_N46813_v1.0 Salmonella enterica subsp. enterica serovar Infantis Proteobacteria; Gammaproteobacteria; Enterobacterales; Enterobacteriaceae; Salmonella; Salmonella enterica; Salmonella enterica subsp. enterica MNNEETFYQAMRRKGVTRRSFLKFCSLAATSLGLGAGMTPKIAWALENKPRIPVVWIHGL WP_001058311.1 hydrogenase [Salmonella enterica] Length: 372\nNumber of predicted TMHs: 2\nExp number of AAs in TMHs: 37.4699899999999\nExp number, first 60 AAs: 19.72699\nTotal prob of N-in: 0.96171\nPOSSIBLE N-term signal sequence\ninside 1 19\nTMhelix 20 39\noutside 40 325\nTMhelix 326 348\ninside 349 372

17857 GCF_001242545.1_Salmonella_enterica_CVM_N45943_v1.0 Salmonella enterica subsp. enterica serovar Infantis Proteobacteria; Gammaproteobacteria; Enterobacterales; Enterobacteriaceae; Salmonella; Salmonella enterica; Salmonella enterica subsp. enterica MNNEETFYQAMRRKGVTRRSFLKFCSLAATSLGLGAGMTPKIAWALENKPRIPVVWIHGL WP_001058311.1 hydrogenase [Salmonella enterica] Length: 372\nNumber of predicted TMHs: 2\nExp number of AAs in TMHs: 37.4699899999999\nExp number, first 60 AAs: 19.72699\nTotal prob of N-in: 0.96171\nPOSSIBLE N-term signal sequence\ninside 1 19\nTMhelix 20 39\noutside 40 325\nTMhelix 326 348\ninside 349 372

17858 GCF_001244215.1_Salmonella_enterica_CVM_N48577_v1.0 Salmonella enterica subsp. enterica serovar Infantis Proteobacteria; Gammaproteobacteria; Enterobacterales; Enterobacteriaceae; Salmonella; Salmonella enterica; Salmonella enterica subsp. enterica MTGDNTLITSHGINRRDFMKLCAALAATMGLSSKAAAEMAESVSNPQRPPVIWIGAQECT WP_000145429.1 hydrogenase 2 small subunit [Salmonella enterica] Length: 372\nNumber of predicted TMHs: 1\nExp number of AAs in TMHs: 24.51309\nExp number, first 60 AAs: 0.2682\nTotal prob of N-in: 0.14402\noutside 1 331\nTMhelix 332 354\ninside 355 372

17859 GCF_001246295.1_Salmonella_enterica_CVM_N51258_v1.0 Salmonella enterica subsp. enterica serovar Infantis Proteobacteria; Gammaproteobacteria; Enterobacterales; Enterobacteriaceae; Salmonella; Salmonella enterica; Salmonella enterica subsp. enterica MTGDNTLITSHGINRRDFMKLCAALAATMGLSSKAAAEMAESVSNPQRPPVIWIGAQECT WP_000145429.1 hydrogenase 2 small subunit [Salmonella enterica] Length: 372\nNumber of predicted TMHs: 1\nExp number of AAs in TMHs: 24.51309\nExp number, first 60 AAs: 0.2682\nTotal prob of N-in: 0.14402\noutside 1 331\nTMhelix 332 354\ninside 355 372

17860 GCF_000812595.1_ASM81259v1 Salmonella enterica subsp. enterica serovar Infantis Proteobacteria; Gammaproteobacteria; Enterobacterales; Enterobacteriaceae; Salmonella; Salmonella enterica; Salmonella enterica subsp. enterica MNNEETFYQAMRRKGVTRRSFLKFCSLAATSLGLGAGMTPKIAWALENKPRIPVVWIHGL WP_001058311.1 hydrogenase [Salmonella enterica] Length: 372\nNumber of predicted TMHs: 2\nExp number of AAs in TMHs: 37.4699899999999\nExp number, first 60 AAs: 19.72699\nTotal prob of N-in: 0.96171\nPOSSIBLE N-term signal sequence\ninside 1 19\nTMhelix 20 39\noutside 40 325\nTMhelix 326 348\ninside 349 372

17861 GCF_001244215.1_Salmonella_enterica_CVM_N48577_v1.0 Salmonella enterica subsp. enterica serovar Infantis Proteobacteria; Gammaproteobacteria; Enterobacterales; Enterobacteriaceae; Salmonella; Salmonella enterica; Salmonella enterica subsp. enterica MNNEETFYQAMRRKGVTRRSFLKFCSLAATSLGLGAGMTPKIAWALENKPRIPVVWIHGL WP_001058311.1 hydrogenase [Salmonella enterica] Length: 372\nNumber of predicted TMHs: 2\nExp number of AAs in TMHs: 37.4699899999999\nExp number, first 60 AAs: 19.72699\nTotal prob of N-in: 0.96171\nPOSSIBLE N-term signal sequence\ninside 1 19\nTMhelix 20 39\noutside 40 325\nTMhelix 326 348\ninside 349 372

17862 GCF_000953495.1_SINFA Salmonella enterica subsp. enterica serovar Infantis Proteobacteria; Gammaproteobacteria; Enterobacterales; Enterobacteriaceae; Salmonella; Salmonella enterica; Salmonella enterica subsp. enterica MTGDNTLITSHGINRRDFMKLCAALAATMGLSSKAAAEMAESVSNPQRPPVIWIGAQECT WP_000145429.1 hydrogenase 2 small subunit [Salmonella enterica] Length: 372\nNumber of predicted TMHs: 1\nExp number of AAs in TMHs: 24.51309\nExp number, first 60 AAs: 0.2682\nTotal prob of N-in: 0.14402\noutside 1 331\nTMhelix 332 354\ninside 355 372

17863 GCF_001244805.1_Salmonella_enterica_CVM_N48693_v1.0 Salmonella enterica subsp. enterica serovar Infantis Proteobacteria; Gammaproteobacteria; Enterobacterales; Enterobacteriaceae; Salmonella; Salmonella enterica; Salmonella enterica subsp. enterica MTGDNTLITSHGINRRDFMKLCAALAATMGLSSKAAAEMAESVSNPQRPPVIWIGAQECT WP_000145429.1 hydrogenase 2 small subunit [Salmonella enterica] Length: 372\nNumber of predicted TMHs: 1\nExp number of AAs in TMHs: 24.51309\nExp number, first 60 AAs: 0.2682\nTotal prob of N-in: 0.14402\noutside 1 331\nTMhelix 332 354\ninside 355 372

17864 GCF_001242545.1_Salmonella_enterica_CVM_N45943_v1.0 Salmonella enterica subsp. enterica serovar Infantis Proteobacteria; Gammaproteobacteria; Enterobacterales; Enterobacteriaceae; Salmonella; Salmonella enterica; Salmonella enterica subsp. enterica MTGDNTLITSHGINRRDFMKLCAALAATMGLSSKAAAEMAESVSNPQRPPVIWIGAQECT WP_000145429.1 hydrogenase 2 small subunit [Salmonella enterica] Length: 372\nNumber of predicted TMHs: 1\nExp number of AAs in TMHs: 24.51309\nExp number, first 60 AAs: 0.2682\nTotal prob of N-in: 0.14402\noutside 1 331\nTMhelix 332 354\ninside 355 372

17865 GCF_001246465.1_Salmonella_enterica_CVM_N51266_v1.0 Salmonella enterica subsp. enterica serovar Infantis Proteobacteria; Gammaproteobacteria; Enterobacterales; Enterobacteriaceae; Salmonella; Salmonella enterica; Salmonella enterica subsp. enterica MTGDNTLITSHGINRRDFMKLCAALAATMGLSSKAAAEMAESVSNPQRPPVIWIGAQECT WP_000145429.1 hydrogenase 2 small subunit [Salmonella enterica] Length: 372\nNumber of predicted TMHs: 1\nExp number of AAs in TMHs: 24.51309\nExp number, first 60 AAs: 0.2682\nTotal prob of N-in: 0.14402\noutside 1 331\nTMhelix 332 354\ninside 355 372

17866 GCF_001244805.1_Salmonella_enterica_CVM_N48693_v1.0 Salmonella enterica subsp. enterica serovar Infantis Proteobacteria; Gammaproteobacteria; Enterobacterales; Enterobacteriaceae; Salmonella; Salmonella enterica; Salmonella enterica subsp. enterica MNNEETFYQAMRRKGVTRRSFLKFCSLAATSLGLGAGMTPKIAWALENKPRIPVVWIHGL WP_001058311.1 hydrogenase [Salmonella enterica] Length: 372\nNumber of predicted TMHs: 2\nExp number of AAs in TMHs: 37.4699899999999\nExp number, first 60 AAs: 19.72699\nTotal prob of N-in: 0.96171\nPOSSIBLE N-term signal sequence\ninside 1 19\nTMhelix 20 39\noutside 40 325\nTMhelix 326 348\ninside 349 372

17867 GCF_002200415.1_ASM220041v1 Salmonella enterica subsp. enterica serovar Infantis Proteobacteria; Gammaproteobacteria; Enterobacterales; Enterobacteriaceae; Salmonella; Salmonella enterica; Salmonella enterica subsp. enterica MTGDNTLITSHGINRRDFMKLCAALAATMGLSSKAAAEMAESVSNPQRPPVIWIGAQECT WP_000145429.1 hydrogenase 2 small subunit [Salmonella enterica] Length: 372\nNumber of predicted TMHs: 1\nExp number of AAs in TMHs: 24.51309\nExp number, first 60 AAs: 0.2682\nTotal prob of N-in: 0.14402\noutside 1 331\nTMhelix 332 354\ninside 355 372

17868 GCF_002200395.1_ASM220039v1 Salmonella enterica subsp. enterica serovar Infantis Proteobacteria; Gammaproteobacteria; Enterobacterales; Enterobacteriaceae; Salmonella; Salmonella enterica; Salmonella enterica subsp. enterica MTGDNTLITSHGINRRDFMKLCAALAATMGLSSKAAAEMAESVSNPQRPPVIWIGAQECT WP_000145429.1 hydrogenase 2 small subunit [Salmonella enterica] Length: 372\nNumber of predicted TMHs: 1\nExp number of AAs in TMHs: 24.51309\nExp number, first 60 AAs: 0.2682\nTotal prob of N-in: 0.14402\noutside 1 331\nTMhelix 332 354\ninside 355 372

17869 GCF_002200335.1_ASM220033v1 Salmonella enterica subsp. enterica serovar Infantis Proteobacteria; Gammaproteobacteria; Enterobacterales; Enterobacteriaceae; Salmonella; Salmonella enterica; Salmonella enterica subsp. enterica MTGDNTLITSHGINRRDFMKLCAALAATMGLSSKAAAEMAESVSNPQRPPVIWIGAQECT WP_000145429.1 hydrogenase 2 small subunit [Salmonella enterica] Length: 372\nNumber of predicted TMHs: 1\nExp number of AAs in TMHs: 24.51309\nExp number, first 60 AAs: 0.2682\nTotal prob of N-in: 0.14402\noutside 1 331\nTMhelix 332 354\ninside 355 372

17870 GCF_002200275.1_ASM220027v1 Salmonella enterica subsp. enterica serovar Infantis Proteobacteria; Gammaproteobacteria; Enterobacterales; Enterobacteriaceae; Salmonella; Salmonella enterica; Salmonella enterica subsp. enterica MTGDNTLITSHGINRRDFMKLCAALAATMGLSSKAAAEMAESVSNPQRPPVIWIGAQECT WP_000145429.1 hydrogenase 2 small subunit [Salmonella enterica] Length: 372\nNumber of

predicted TMHs:  1\nExp number of AAs in TMHs: 24.51309\nExp number, first 60 AAs:  0.2682\nTotal prob of N-in: 0.14402\noutside   1   331\nTMhelix   332   354\ninside   355   372

17871 GCF_002064905.1_ASM206490v1 Salmonella enterica subsp. enterica serovar Infantis Proteobacteria; Gammaproteobacteria; Enterobacterales; Enterobacteriaceae; Salmonella; Salmonella enterica; Salmonella enterica subsp. enterica MTGDNTLITSHGINRRDFMKLCAALAATMGLSSKAAAEMAESVSNPQRPPVIWIGAQECT WP_000145429.1 hydrogenase 2 small subunit [Salmonella enterica] Length: 372\nNumber of predicted TMHs:  1\nExp number of AAs in TMHs: 24.51309\nExp number, first 60 AAs:  0.2682\nTotal prob of N-in: 0.14402\noutside   1   331\nTMhelix   332   354\ninside   355   372

17872 GCF_002064875.1_ASM206487v1 Salmonella enterica subsp. enterica serovar Infantis Proteobacteria; Gammaproteobacteria; Enterobacterales; Enterobacteriaceae; Salmonella; Salmonella enterica; Salmonella enterica subsp. enterica MNNEETFYQAMRRKGVTRRSFLKFCSLAATSLGLGAGMTPKIAWALENKPRIPVVWIHGL WP_001058311.1 hydrogenase [Salmonella enterica] Length: 372\nNumber of predicted TMHs:  2\nExp number of AAs in TMHs: 37.4699899999999\nExp number, first 60 AAs:  19.72699\nTotal prob of N-in: 0.96171\nPOSSIBLE N-term signal sequence\ninside   1   19\nTMhelix   20   39\noutside   40   325\nTMhelix   326 348\ninside   349   372

17873 GCF_002110515.1_ASM211051v1 Salmonella enterica subsp. enterica serovar Infantis Proteobacteria; Gammaproteobacteria; Enterobacterales; Enterobacteriaceae; Salmonella; Salmonella enterica; Salmonella enterica subsp. enterica MTGDNTLITSHGINRRDFMKLCAALAATMGLSSKAAAEMAESVSNPQRPPVIWIGAQECT WP_000145429.1 hydrogenase 2 small subunit [Salmonella enterica] Length: 372\nNumber of predicted TMHs:  1\nExp number of AAs in TMHs: 24.51309\nExp number, first 60 AAs:  0.2682\nTotal prob of N-in: 0.14402\noutside   1   331\nTMhelix   332   354\ninside   355   372

17874 GCF_002200735.1_ASM220073v1 Salmonella enterica subsp. enterica serovar Infantis Proteobacteria; Gammaproteobacteria; Enterobacterales; Enterobacteriaceae; Salmonella; Salmonella enterica; Salmonella enterica subsp. enterica MNNEETFYQAMRRKGVTRRSFLKFCSLAATSLGLGAGMTPKIAWALENKPRIPVVWIHGL WP_001058311.1 hydrogenase [Salmonella enterica] Length: 372\nNumber of predicted TMHs:  2\nExp number of AAs in TMHs: 37.4699899999999\nExp number, first 60 AAs:  19.72699\nTotal prob of N-in: 0.96171\nPOSSIBLE N-term signal sequence\ninside   1   19\nTMhelix   20   39\noutside   40   325\nTMhelix   326 348\ninside   349   372

17875 GCF_002110515.1_ASM211051v1 Salmonella enterica subsp. enterica serovar Infantis Proteobacteria; Gammaproteobacteria; Enterobacterales; Enterobacteriaceae; Salmonella; Salmonella enterica; Salmonella enterica subsp. enterica MNNEETFYQAMRRKGVTRRSFLKFCSLAATSLGLGAGMTPKIAWALENKPRIPVVWIHGL WP_001058311.1 hydrogenase [Salmonella enterica] Length: 372\nNumber of predicted TMHs:  2\nExp number of AAs in TMHs: 37.4699899999999\nExp number, first 60 AAs:  19.72699\nTotal prob of N-in: 0.96171\nPOSSIBLE N-term signal sequence\ninside   1   19\nTMhelix   20   39\noutside   40   325\nTMhelix   326 348\ninside   349   372

17876 GCF_002200635.1_ASM220063v1 Salmonella enterica subsp. enterica serovar Infantis Proteobacteria; Gammaproteobacteria; Enterobacterales; Enterobacteriaceae; Salmonella; Salmonella enterica; Salmonella enterica subsp. enterica MTGDNTLITSHGINRRDFMKLCAALAATMGLSSKAAAEMAESVSNPQRPPVIWIGAQECT WP_000145429.1 hydrogenase 2 small subunit [Salmonella enterica] Length: 372\nNumber of predicted TMHs:  1\nExp number of AAs in TMHs: 24.51309\nExp number, first 60 AAs:  0.2682\nTotal prob of N-in: 0.14402\noutside   1   331\nTMhelix   332   354\ninside   355   372

17877 GCF_002064875.1_ASM206487v1 Salmonella enterica subsp. enterica serovar Infantis Proteobacteria; Gammaproteobacteria; Enterobacterales; Enterobacteriaceae; Salmonella; Salmonella enterica; Salmonella enterica subsp. enterica MTGDNTLITSHGINRRDFMKLCAALAATMGLSSKAAAEMAESVSNPQRPPVIWIGAQECT WP_000145429.1 hydrogenase 2 small subunit [Salmonella enterica] Length: 372\nNumber of predicted TMHs:  1\nExp number of AAs in TMHs: 24.51309\nExp number, first 60 AAs:  0.2682\nTotal prob of N-in: 0.14402\noutside   1   331\nTMhelix   332   354\ninside   355   372

17878 GCF_002200415.1_ASM220041v1 Salmonella enterica subsp. enterica serovar Infantis Proteobacteria; Gammaproteobacteria; Enterobacterales; Enterobacteriaceae; Salmonella; Salmonella enterica; Salmonella enterica subsp. enterica MNNEETFYQAMRRKGVTRRSFLKFCSLAATSLGLGAGMTPKIAWALENKPRIPVVWIHGL WP_001058311.1 hydrogenase [Salmonella enterica] Length: 372\nNumber of predicted TMHs:  2\nExp number of AAs in TMHs: 37.4699899999999\nExp number, first 60 AAs:  19.72699\nTotal prob of N-in:

0.96171\nPOSSIBLE N-term signal sequence\ninside 1 19\nTMhelix 20 39\noutside 40 325\nTMhelix 326 348\ninside 349 372

17879 GCF_002200475.1_ASM220047v1 Salmonella enterica subsp. enterica serovar Infantis Proteobacteria; Gammaproteobacteria; Enterobacterales; Enterobacteriaceae; Salmonella; Salmonella enterica; Salmonella enterica subsp. enterica MTGDNTLITSHGINRRDFMKLCAALAATMGLSSKAAAEMAESVSNPQRPPVIWIGAQECT WP_000145429.1 hydrogenase 2 small subunit [Salmonella enterica] Length: 372\nNumber of predicted TMHs: 1\nExp number of AAs in TMHs: 24.51309\nExp number, first 60 AAs: 0.2682\nTotal prob of N-in: 0.14402\noutside 1 331\nTMhelix 332 354\ninside 355 372

17880 GCF_002200715.1_ASM220071v1 Salmonella enterica subsp. enterica serovar Infantis Proteobacteria; Gammaproteobacteria; Enterobacterales; Enterobacteriaceae; Salmonella; Salmonella enterica; Salmonella enterica subsp. enterica MNNEETFYQAMRRKGVTRRSFLKFCSLAATSLGLGAGMTPKIAWALENKPRIPVVWIHGL WP_001058311.1 hydrogenase [Salmonella enterica] Length: 372\nNumber of predicted TMHs: 2\nExp number of AAs in TMHs: 37.4699899999999\nExp number, first 60 AAs: 19.72699\nTotal prob of N-in: 0.96171\nPOSSIBLE N-term signal sequence\ninside 1 19\nTMhelix 20 39\noutside 40 325\nTMhelix 326 348\ninside 349 372

17881 GCF_002200575.1_ASM220057v1 Salmonella enterica subsp. enterica serovar Infantis Proteobacteria; Gammaproteobacteria; Enterobacterales; Enterobacteriaceae; Salmonella; Salmonella enterica; Salmonella enterica subsp. enterica MTGDNTLITSHGINRRDFMKLCAALAATMGLSSKAAAEMAESVSNPQRPPVIWIGAQECT WP_000145429.1 hydrogenase 2 small subunit [Salmonella enterica] Length: 372\nNumber of predicted TMHs: 1\nExp number of AAs in TMHs: 24.51309\nExp number, first 60 AAs: 0.2682\nTotal prob of N-in: 0.14402\noutside 1 331\nTMhelix 332 354\ninside 355 372

17882 GCF_002200555.1_ASM220055v1 Salmonella enterica subsp. enterica serovar Infantis Proteobacteria; Gammaproteobacteria; Enterobacterales; Enterobacteriaceae; Salmonella; Salmonella enterica; Salmonella enterica subsp. enterica MTGDNTLITSHGINRRDFMKLCAALAATMGLSSKAAAEMAESVSNPQRPPVIWIGAQECT WP_000145429.1 hydrogenase 2 small subunit [Salmonella enterica] Length: 372\nNumber of predicted TMHs: 1\nExp number of AAs in TMHs: 24.51309\nExp number, first 60 AAs: 0.2682\nTotal prob of N-in: 0.14402\noutside 1 331\nTMhelix 332 354\ninside 355 372

17883 GCF_002064905.1_ASM206490v1 Salmonella enterica subsp. enterica serovar Infantis Proteobacteria; Gammaproteobacteria; Enterobacterales; Enterobacteriaceae; Salmonella; Salmonella enterica; Salmonella enterica subsp. enterica MNNEETFYQAMRRKGVTRRSFLKFCSLAATSLGLGAGMTPKIAWALENKPRIPVVWIHGL WP_001058312.1 hydrogenase [Salmonella enterica] Length: 372\nNumber of predicted TMHs: 2\nExp number of AAs in TMHs: 36.51738\nExp number, first 60 AAs: 19.72849\nTotal prob of N-in: 0.96170\nPOSSIBLE N-term signal sequence\ninside 1 19\nTMhelix 20 39\noutside 40 325\nTMhelix 326 348\ninside 349 372

17884 GCF_002200665.1_ASM220066v1 Salmonella enterica subsp. enterica serovar Infantis Proteobacteria; Gammaproteobacteria; Enterobacterales; Enterobacteriaceae; Salmonella; Salmonella enterica; Salmonella enterica subsp. enterica MTGDNTLITSHGINRRDFMKLCAALAATMGLSSKAAAEMAESVSNPQRPPVIWIGAQECT WP_000145429.1 hydrogenase 2 small subunit [Salmonella enterica] Length: 372\nNumber of predicted TMHs: 1\nExp number of AAs in TMHs: 24.51309\nExp number, first 60 AAs: 0.2682\nTotal prob of N-in: 0.14402\noutside 1 331\nTMhelix 332 354\ninside 355 372

17885 GCF_002200555.1_ASM220055v1 Salmonella enterica subsp. enterica serovar Infantis Proteobacteria; Gammaproteobacteria; Enterobacterales; Enterobacteriaceae; Salmonella; Salmonella enterica; Salmonella enterica subsp. enterica MNNEETFYQAMRRKGVTRRSFLKFCSLAATSLGLGAGMTPKIAWALENKPRIPVVWIHGL WP_001058311.1 hydrogenase [Salmonella enterica] Length: 372\nNumber of predicted TMHs: 2\nExp number of AAs in TMHs: 37.4699899999999\nExp number, first 60 AAs: 19.72699\nTotal prob of N-in: 0.96171\nPOSSIBLE N-term signal sequence\ninside 1 19\nTMhelix 20 39\noutside 40 325\nTMhelix 326 348\ninside 349 372

17886 GCF_002065585.1_ASM206558v1 Salmonella enterica subsp. enterica serovar Infantis Proteobacteria; Gammaproteobacteria; Enterobacterales; Enterobacteriaceae; Salmonella; Salmonella enterica; Salmonella enterica subsp. enterica MNNEETFYQAMRRKGVTRRSFLKFCSLAATSLGLGAGMTPKIAWALENKPRIPVVWIHGL WP_001058311.1 hydrogenase [Salmonella enterica] Length: 372\nNumber of predicted TMHs: 2\nExp number of AAs in TMHs: 37.4699899999999\nExp number, first 60 AAs: 19.72699\nTotal prob of N-in: 0.96171\nPOSSIBLE N-term signal sequence\ninside 1 19\nTMhelix 20 39\noutside 40 325\nTMhelix 326 348\ninside 349 372

17887	GCF_002200575.1_ASM220057v1	Salmonella enterica subsp. enterica serovar Infantis	Proteobacteria; Gammaproteobacteria; Enterobacterales; Enterobacteriaceae; Salmonella; Salmonella enterica; Salmonella enterica subsp. enterica	MNNEETFYQAMRRKGVTRRSFLKFCSLAATSLGLGAGMTPKIAWALENKPRIPVVWIHGL	WP_001058311.1	hydrogenase [Salmonella enterica]	Length: 372\nNumber of predicted TMHs: 2\nExp number of AAs in TMHs: 37.4699899999999\nExp number, first 60 AAs: 19.72699\nTotal prob of N-in: 0.96171\nPOSSIBLE N-term signal sequence\ninside 1 19\nTMhelix 20 39\noutside 40 325\nTMhelix 326 348\ninside 349 372

17888	GCF_002065585.1_ASM206558v1	Salmonella enterica subsp. enterica serovar Infantis	Proteobacteria; Gammaproteobacteria; Enterobacterales; Enterobacteriaceae; Salmonella; Salmonella enterica; Salmonella enterica subsp. enterica	MTGDNTLITSHGINRRDFMKLCAALAATMGLSSKAAAEMAESVSNPQRPPVIWIGAQECT	WP_000145429.1	hydrogenase 2 small subunit [Salmonella enterica]	Length: 372\nNumber of predicted TMHs: 1\nExp number of AAs in TMHs: 24.51309\nExp number, first 60 AAs: 0.2682\nTotal prob of N-in: 0.14402\noutside 1 331\nTMhelix 332 354\ninside 355 372

17889	GCF_001478785.1_Salmonella_enterica_CVM_N29379-SQ_v1.0	Salmonella enterica subsp. enterica serovar Infantis	Proteobacteria; Gammaproteobacteria; Enterobacterales; Enterobacteriaceae; Salmonella; Salmonella enterica; Salmonella enterica subsp. enterica	MTGDNTLITSHGINRRDFMKLCAALAATMGLSSKAAAEMAESVSNPQRPPVIWIGAQECT	WP_000145429.1	hydrogenase 2 small subunit [Salmonella enterica]	Length: 372\nNumber of predicted TMHs: 1\nExp number of AAs in TMHs: 24.51309\nExp number, first 60 AAs: 0.2682\nTotal prob of N-in: 0.14402\noutside 1 331\nTMhelix 332 354\ninside 355 372

17890	GCF_001906535.1_ASM190653v1	Salmonella enterica subsp. enterica serovar Infantis	Proteobacteria; Gammaproteobacteria; Enterobacterales; Enterobacteriaceae; Salmonella; Salmonella enterica; Salmonella enterica subsp. enterica	MTGDNTLITSHGINRRDFMKLCAALAATMGLSSKAAAEMAESVSNPQRPPVIWIGAQECT	WP_000145429.1	hydrogenase 2 small subunit [Salmonella enterica]	Length: 372\nNumber of predicted TMHs: 1\nExp number of AAs in TMHs: 24.51309\nExp number, first 60 AAs: 0.2682\nTotal prob of N-in: 0.14402\noutside 1 331\nTMhelix 332 354\ninside 355 372

17891	GCF_001906515.1_ASM190651v1	Salmonella enterica subsp. enterica serovar Infantis	Proteobacteria; Gammaproteobacteria; Enterobacterales; Enterobacteriaceae; Salmonella; Salmonella enterica; Salmonella enterica subsp. enterica	MNNEETFYQAMRRKGVTRRSFLKFCSLAATSLGLGAGMTPKIAWALENKPRIPVVWIHGL	WP_001058311.1	hydrogenase [Salmonella enterica]	Length: 372\nNumber of predicted TMHs: 2\nExp number of AAs in TMHs: 37.4699899999999\nExp number, first 60 AAs: 19.72699\nTotal prob of N-in: 0.96171\nPOSSIBLE N-term signal sequence\ninside 1 19\nTMhelix 20 39\noutside 40 325\nTMhelix 326 348\ninside 349 372

17892	GCF_001906535.1_ASM190653v1	Salmonella enterica subsp. enterica serovar Infantis	Proteobacteria; Gammaproteobacteria; Enterobacterales; Enterobacteriaceae; Salmonella; Salmonella enterica; Salmonella enterica subsp. enterica	MNNEETFYQAMRRKGVTRRSFLKFCSLAATSLGLGAGMTPKIAWALENKPRIPVVWIHGL	WP_001058311.1	hydrogenase [Salmonella enterica]	Length: 372\nNumber of predicted TMHs: 2\nExp number of AAs in TMHs: 37.4699899999999\nExp number, first 60 AAs: 19.72699\nTotal prob of N-in: 0.96171\nPOSSIBLE N-term signal sequence\ninside 1 19\nTMhelix 20 39\noutside 40 325\nTMhelix 326 348\ninside 349 372

17893	GCF_002059955.1_ASM205995v1	Salmonella enterica subsp. enterica serovar Infantis	Proteobacteria; Gammaproteobacteria; Enterobacterales; Enterobacteriaceae; Salmonella; Salmonella enterica; Salmonella enterica subsp. enterica	MNNEETFYQAMRRKGVTRRSFLKFCSLAATSLGLGAGMTPKIAWALENKPRIPVVWIHGL	WP_001058311.1	hydrogenase [Salmonella enterica]	Length: 372\nNumber of predicted TMHs: 2\nExp number of AAs in TMHs: 37.4699899999999\nExp number, first 60 AAs: 19.72699\nTotal prob of N-in: 0.96171\nPOSSIBLE N-term signal sequence\ninside 1 19\nTMhelix 20 39\noutside 40 325\nTMhelix 326 348\ninside 349 372

17894	GCF_002059955.1_ASM205995v1	Salmonella enterica subsp. enterica serovar Infantis	Proteobacteria; Gammaproteobacteria; Enterobacterales; Enterobacteriaceae; Salmonella; Salmonella enterica; Salmonella enterica subsp. enterica	MTGDNTLITSHGINRRDFMKLCAALAATMGLSSKAAAEMAESVSNPQRPPVIWIGAQECT	WP_000145429.1	hydrogenase 2 small subunit [Salmonella enterica]	Length: 372\nNumber of predicted TMHs: 1\nExp number of AAs in TMHs: 24.51309\nExp number, first 60 AAs: 0.2682\nTotal prob of N-in: 0.14402\noutside 1 331\nTMhelix 332 354\ninside 355 372

17895 GCF_001187365.1_ASM118736v1 Salmonella enterica subsp. enterica serovar Infantis Proteobacteria; Gammaproteobacteria; Enterobacterales; Enterobacteriaceae; Salmonella; Salmonella enterica; Salmonella enterica subsp. enterica MTGDNTLITSHGINRRDFMKLCAALAATMGLSSKAAAEMAESVSNPQRPPVIWIGAQECT WP_000145429.1 hydrogenase 2 small subunit [Salmonella enterica] Length: 372\nNumber of predicted TMHs: 1\nExp number of AAs in TMHs: 24.51309\nExp number, first 60 AAs: 0.2682\nTotal prob of N-in: 0.14402\noutside 1 331\nTMhelix 332 354\ninside 355 372

17896 GCF_001766515.1_ASM176651v1 Salmonella enterica subsp. enterica serovar Infantis Proteobacteria; Gammaproteobacteria; Enterobacterales; Enterobacteriaceae; Salmonella; Salmonella enterica; Salmonella enterica subsp. enterica MNNEETFYQAMRRKGVTRRSFLKFCSLAATSLGLGAGMTPKIAWALENKPRIPVVWIHGL WP_001058311.1 hydrogenase [Salmonella enterica] Length: 372\nNumber of predicted TMHs: 2\nExp number of AAs in TMHs: 37.4699899999999\nExp number, first 60 AAs: 19.72699\nTotal prob of N-in: 0.96171\nPOSSIBLE N-term signal sequence\ninside 1 19\nTMhelix 20 39\noutside 40 325\nTMhelix 326 348\ninside 349 372

17897 GCF_001766495.1_ASM176649v1 Salmonella enterica subsp. enterica serovar Infantis Proteobacteria; Gammaproteobacteria; Enterobacterales; Enterobacteriaceae; Salmonella; Salmonella enterica; Salmonella enterica subsp. enterica MNNEETFYQAMRRKGVTRRSFLKFCSLAATSLGLGAGMTPKIAWALENKPRIPVVWIHGL WP_001058311.1 hydrogenase [Salmonella enterica] Length: 372\nNumber of predicted TMHs: 2\nExp number of AAs in TMHs: 37.4699899999999\nExp number, first 60 AAs: 19.72699\nTotal prob of N-in: 0.96171\nPOSSIBLE N-term signal sequence\ninside 1 19\nTMhelix 20 39\noutside 40 325\nTMhelix 326 348\ninside 349 372

17898 GCF_001242985.1_Salmonella_enterica_CVM_N46819_v1.0 Salmonella enterica subsp. enterica serovar Infantis Proteobacteria; Gammaproteobacteria; Enterobacterales; Enterobacteriaceae; Salmonella; Salmonella enterica; Salmonella enterica subsp. enterica MTGDNTLITSHGINRRDFMKLCAALAATMGLSSKAAAEMAESVSNPQRPPVIWIGAQECT WP_000145429.1 hydrogenase 2 small subunit [Salmonella enterica] Length: 372\nNumber of predicted TMHs: 1\nExp number of AAs in TMHs: 24.51309\nExp number, first 60 AAs: 0.2682\nTotal prob of N-in: 0.14402\noutside 1 331\nTMhelix 332 354\ninside 355 372

17899 GCF_000812575.1_ASM81257v1 Salmonella enterica subsp. enterica serovar Infantis Proteobacteria; Gammaproteobacteria; Enterobacterales; Enterobacteriaceae; Salmonella; Salmonella enterica; Salmonella enterica subsp. enterica MTGDNTLITSHGINRRDFMKLCAALAATMGLSSKAAAEMAESVSNPQRPPVIWIGAQECT WP_000145429.1 hydrogenase 2 small subunit [Salmonella enterica] Length: 372\nNumber of predicted TMHs: 1\nExp number of AAs in TMHs: 24.51309\nExp number, first 60 AAs: 0.2682\nTotal prob of N-in: 0.14402\noutside 1 331\nTMhelix 332 354\ninside 355 372

17900 GCF_001242985.1_Salmonella_enterica_CVM_N46819_v1.0 Salmonella enterica subsp. enterica serovar Infantis Proteobacteria; Gammaproteobacteria; Enterobacterales; Enterobacteriaceae; Salmonella; Salmonella enterica; Salmonella enterica subsp. enterica MNNEETFYQAMRRKGVTRRSFLKFCSLAATSLGLGAGMTPKIAWALENKPRIPVVWIHGL WP_052901769.1 hydrogenase [Salmonella enterica] Length: 372\nNumber of predicted TMHs: 2\nExp number of AAs in TMHs: 37.42632\nExp number, first 60 AAs: 19.72508\nTotal prob of N-in: 0.96168\nPOSSIBLE N-term signal sequence\ninside 1 19\nTMhelix 20 39\noutside 40 325\nTMhelix 326 348\ninside 349 372

17901 GCF_002032265.1_ASM203226v1 Salmonella enterica subsp. enterica serovar Infantis Proteobacteria; Gammaproteobacteria; Enterobacterales; Enterobacteriaceae; Salmonella; Salmonella enterica; Salmonella enterica subsp. enterica MNNEETFYQAMRRKGVTRRSFLKFCSLAATSLGLGAGMTPKIAWALENKPRIPVVWIHGL WP_052901769.1 hydrogenase [Salmonella enterica] Length: 372\nNumber of predicted TMHs: 2\nExp number of AAs in TMHs: 37.42632\nExp number, first 60 AAs: 19.72508\nTotal prob of N-in: 0.96168\nPOSSIBLE N-term signal sequence\ninside 1 19\nTMhelix 20 39\noutside 40 325\nTMhelix 326 348\ninside 349 372

17902 GCF_001243785.1_Salmonella_enterica_CVM_N47714_v1.0 Salmonella enterica subsp. enterica serovar Infantis Proteobacteria; Gammaproteobacteria; Enterobacterales; Enterobacteriaceae; Salmonella; Salmonella enterica; Salmonella enterica subsp. enterica MTGDNTLITSHGINRRDFMKLCAALAATMGLSSKAAAEMAESVSNPQRPPVIWIGAQECT WP_000145429.1 hydrogenase 2 small subunit [Salmonella enterica] Length: 372\nNumber of predicted TMHs: 1\nExp number of AAs in TMHs: 24.51309\nExp number, first 60 AAs: 0.2682\nTotal prob of N-in: 0.14402\noutside 1 331\nTMhelix 332 354\ninside 355 372

17903 GCF_001479785.1_Salmonella_enterica_CVM_N38848-SQ_v1.0 Salmonella enterica subsp. enterica serovar Infantis Proteobacteria; Gammaproteobacteria; Enterobacterales; Enterobacteriaceae; Salmonella; Salmonella enterica; Salmonella enterica subsp. enterica MNNEETFYQAMRRKGVTRRSFLKFCSLAATSLGLGAGMTPKIAWALENKPRIPVVWIHGL

WP_001058311.1 hydrogenase [Salmonella enterica] Length: 372\nNumber of predicted TMHs: 2\nExp number of AAs in TMHs: 37.4699899999999\nExp number, first 60 AAs: 19.72699\nTotal prob of N-in: 0.96171\nPOSSIBLE N-term signal sequence\ninside 1 19\nTMhelix 20 39\noutside 40 325\nTMhelix 326 348\ninside 349 372

17904 GCF_001906515.1_ASM190651v1 Salmonella enterica subsp. enterica serovar Infantis Proteobacteria; Gammaproteobacteria; Enterobacterales; Enterobacteriaceae; Salmonella; Salmonella enterica; Salmonella enterica subsp. enterica MTGDNTLITSHGINRRDFMKLCAALAATMGLSSKAAAEMAESVSNPQRPPVIWIGAQECT WP_000145429.1 hydrogenase 2 small subunit [Salmonella enterica] Length: 372\nNumber of predicted TMHs: 1\nExp number of AAs in TMHs: 24.51309\nExp number, first 60 AAs: 0.2682\nTotal prob of N-in: 0.14402\noutside 1 331\nTMhelix 332 354\ninside 355 372

17905 GCF_001931575.1_ASM193157v1 Salmonella enterica subsp. enterica serovar Infantis Proteobacteria; Gammaproteobacteria; Enterobacterales; Enterobacteriaceae; Salmonella; Salmonella enterica; Salmonella enterica subsp. enterica MTGDNTLITSHGINRRDFMKLCAALAATMGLSSKAAAEMAESVSNPQRPPVIWIGAQECT WP_000145429.1 hydrogenase 2 small subunit [Salmonella enterica] Length: 372\nNumber of predicted TMHs: 1\nExp number of AAs in TMHs: 24.51309\nExp number, first 60 AAs: 0.2682\nTotal prob of N-in: 0.14402\noutside 1 331\nTMhelix 332 354\ninside 355 372

17906 GCF_001518495.1_ASM151849v1 Salmonella enterica subsp. enterica serovar Infantis Proteobacteria; Gammaproteobacteria; Enterobacterales; Enterobacteriaceae; Salmonella; Salmonella enterica; Salmonella enterica subsp. enterica MNNEETFYQAMRRKGVTRRSFLKFCSLAATSLGLGAGMTPKIAWALENKPRIPVVWIHGL WP_001058311.1 hydrogenase [Salmonella enterica] Length: 372\nNumber of predicted TMHs: 2\nExp number of AAs in TMHs: 37.4699899999999\nExp number, first 60 AAs: 19.72699\nTotal prob of N-in: 0.96171\nPOSSIBLE N-term signal sequence\ninside 1 19\nTMhelix 20 39\noutside 40 325\nTMhelix 326 348\ninside 349 372

17907 GCF_001242185.1_Salmonella_enterica_CVM_N45929_v1.0 Salmonella enterica subsp. enterica serovar Infantis Proteobacteria; Gammaproteobacteria; Enterobacterales; Enterobacteriaceae; Salmonella; Salmonella enterica; Salmonella enterica subsp. enterica MTGDNTLITSHGINRRDFMKLCAALAATMGLSSKAAAEMAESVSNPQRPPVIWIGAQECT WP_000145429.1 hydrogenase 2 small subunit [Salmonella enterica] Length: 372\nNumber of predicted TMHs: 1\nExp number of AAs in TMHs: 24.51309\nExp number, first 60 AAs: 0.2682\nTotal prob of N-in: 0.14402\noutside 1 331\nTMhelix 332 354\ninside 355 372

17908 GCF_002032265.1_ASM203226v1 Salmonella enterica subsp. enterica serovar Infantis Proteobacteria; Gammaproteobacteria; Enterobacterales; Enterobacteriaceae; Salmonella; Salmonella enterica; Salmonella enterica subsp. enterica MTGDNTLITSHGINRRDFMKLCAALAATMGLSSKAAAEMAESVSNPQRPPVIWIGAQECT WP_000145429.1 hydrogenase 2 small subunit [Salmonella enterica] Length: 372\nNumber of predicted TMHs: 1\nExp number of AAs in TMHs: 24.51309\nExp number, first 60 AAs: 0.2682\nTotal prob of N-in: 0.14402\noutside 1 331\nTMhelix 332 354\ninside 355 372

17909 GCF_001931575.1_ASM193157v1 Salmonella enterica subsp. enterica serovar Infantis Proteobacteria; Gammaproteobacteria; Enterobacterales; Enterobacteriaceae; Salmonella; Salmonella enterica; Salmonella enterica subsp. enterica MNNEETFYQAMRRKGVTRRSFLKFCSLAATSLGLGAGMTPKIAWALENKPRIPVVWIHGL WP_001058311.1 hydrogenase [Salmonella enterica] Length: 372\nNumber of predicted TMHs: 2\nExp number of AAs in TMHs: 37.4699899999999\nExp number, first 60 AAs: 19.72699\nTotal prob of N-in: 0.96171\nPOSSIBLE N-term signal sequence\ninside 1 19\nTMhelix 20 39\noutside 40 325\nTMhelix 326 348\ninside 349 372

17910 GCF_002030855.1_ASM203085v1 Salmonella enterica subsp. enterica serovar Infantis Proteobacteria; Gammaproteobacteria; Enterobacterales; Enterobacteriaceae; Salmonella; Salmonella enterica; Salmonella enterica subsp. enterica MTGDNTLITSHGINRRDFMKLCAALAATMGLSSKAAAEMAESVSNPQRPPVIWIGAQECT WP_000145429.1 hydrogenase 2 small subunit [Salmonella enterica] Length: 372\nNumber of predicted TMHs: 1\nExp number of AAs in TMHs: 24.51309\nExp number, first 60 AAs: 0.2682\nTotal prob of N-in: 0.14402\noutside 1 331\nTMhelix 332 354\ninside 355 372

17911 GCF_001931595.1_ASM193159v1 Salmonella enterica subsp. enterica serovar Infantis Proteobacteria; Gammaproteobacteria; Enterobacterales; Enterobacteriaceae; Salmonella; Salmonella enterica; Salmonella enterica subsp. enterica MTGDNTLITSHGINRRDFMKLCAALAATMGLSSKAAAEMAESVSNPQRPPVIWIGAQECT WP_000145429.1 hydrogenase 2 small subunit [Salmonella enterica] Length: 372\nNumber of

predicted TMHs: 1\nExp number of AAs in TMHs: 24.51309\nExp number, first 60 AAs: 0.2682\nTotal prob of N-in: 0.14402\noutside 1 331\nTMhelix 332 354\ninside 355 372

17912 GCF_002059515.1_ASM205951v1 Salmonella enterica subsp. enterica serovar Infantis Proteobacteria; Gammaproteobacteria; Enterobacterales; Enterobacteriaceae; Salmonella; Salmonella enterica; Salmonella enterica subsp. enterica MNNEETFYQAMRRKGVTRRSFLKFCSLAATSLGLGAGMTPKIAWALENKPRIPVVWIHGL WP_001058311.1 hydrogenase [Salmonella enterica] Length: 372\nNumber of predicted TMHs: 2\nExp number of AAs in TMHs: 37.4699899999999\nExp number, first 60 AAs: 19.72699\nTotal prob of N-in: 0.96171\nPOSSIBLE N-term signal sequence\ninside 1 19\nTMhelix 20 39\noutside 40 325\nTMhelix 326 348\ninside 349 372

17913 GCF_002059515.1_ASM205951v1 Salmonella enterica subsp. enterica serovar Infantis Proteobacteria; Gammaproteobacteria; Enterobacterales; Enterobacteriaceae; Salmonella; Salmonella enterica; Salmonella enterica subsp. enterica MTGDNTLITSHGINRRDFMKLCAALAATMGLSSKAAAEMAESVSNPQRPPVIWIGAQECT WP_000145429.1 hydrogenase 2 small subunit [Salmonella enterica] Length: 372\nNumber of predicted TMHs: 1\nExp number of AAs in TMHs: 24.51309\nExp number, first 60 AAs: 0.2682\nTotal prob of N-in: 0.14402\noutside 1 331\nTMhelix 332 354\ninside 355 372

17914 GCF_001518495.1_ASM151849v1 Salmonella enterica subsp. enterica serovar Infantis Proteobacteria; Gammaproteobacteria; Enterobacterales; Enterobacteriaceae; Salmonella; Salmonella enterica; Salmonella enterica subsp. enterica MTGDNTLITSHGINRRDFMKLCAALAATMGLSSKAAAEMAESVSNPQRPPVIWIGAQECT WP_000145429.1 hydrogenase 2 small subunit [Salmonella enterica] Length: 372\nNumber of predicted TMHs: 1\nExp number of AAs in TMHs: 24.51309\nExp number, first 60 AAs: 0.2682\nTotal prob of N-in: 0.14402\noutside 1 331\nTMhelix 332 354\ninside 355 372

17915 GCF_002032145.1_ASM203214v1 Salmonella enterica subsp. enterica serovar Infantis Proteobacteria; Gammaproteobacteria; Enterobacterales; Enterobacteriaceae; Salmonella; Salmonella enterica; Salmonella enterica subsp. enterica MTGDNTLITSHGINRRDFMKLCAALAATMGLSSKAAAEMAESVSNPQRPPVIWIGAQECT WP_000145429.1 hydrogenase 2 small subunit [Salmonella enterica] Length: 372\nNumber of predicted TMHs: 1\nExp number of AAs in TMHs: 24.51309\nExp number, first 60 AAs: 0.2682\nTotal prob of N-in: 0.14402\noutside 1 331\nTMhelix 332 354\ninside 355 372

17916 GCF_002032145.1_ASM203214v1 Salmonella enterica subsp. enterica serovar Infantis Proteobacteria; Gammaproteobacteria; Enterobacterales; Enterobacteriaceae; Salmonella; Salmonella enterica; Salmonella enterica subsp. enterica MNNEETFYQAMRRKGVTRRSFLKFCSLAATSLGLGAGMTPKIAWALENKPRIPVVWIHGL WP_001058311.1 hydrogenase [Salmonella enterica] Length: 372\nNumber of predicted TMHs: 2\nExp number of AAs in TMHs: 37.4699899999999\nExp number, first 60 AAs: 19.72699\nTotal prob of N-in: 0.96171\nPOSSIBLE N-term signal sequence\ninside 1 19\nTMhelix 20 39\noutside 40 325\nTMhelix 326 348\ninside 349 372

17917 GCF_001766495.1_ASM176649v1 Salmonella enterica subsp. enterica serovar Infantis Proteobacteria; Gammaproteobacteria; Enterobacterales; Enterobacteriaceae; Salmonella; Salmonella enterica; Salmonella enterica subsp. enterica MTGDNTLITSHGINRRDFMKLCAALAATMGLSSKAAAEMAESVSNPQRPPVIWIGAQECT WP_000145429.1 hydrogenase 2 small subunit [Salmonella enterica] Length: 372\nNumber of predicted TMHs: 1\nExp number of AAs in TMHs: 24.51309\nExp number, first 60 AAs: 0.2682\nTotal prob of N-in: 0.14402\noutside 1 331\nTMhelix 332 354\ninside 355 372

17918 GCF_001766515.1_ASM176651v1 Salmonella enterica subsp. enterica serovar Infantis Proteobacteria; Gammaproteobacteria; Enterobacterales; Enterobacteriaceae; Salmonella; Salmonella enterica; Salmonella enterica subsp. enterica MTGDNTLITSHGINRRDFMKLCAALAATMGLSSKAAAEMAESVSNPQRPPVIWIGAQECT WP_000145429.1 hydrogenase 2 small subunit [Salmonella enterica] Length: 372\nNumber of predicted TMHs: 1\nExp number of AAs in TMHs: 24.51309\nExp number, first 60 AAs: 0.2682\nTotal prob of N-in: 0.14402\noutside 1 331\nTMhelix 332 354\ninside 355 372

17919 GCF_001931595.1_ASM193159v1 Salmonella enterica subsp. enterica serovar Infantis Proteobacteria; Gammaproteobacteria; Enterobacterales; Enterobacteriaceae; Salmonella; Salmonella enterica; Salmonella enterica subsp. enterica MNNEETFYQAMRRKGVTRRSFLKFCSLAATSLGLGAGMTPKIAWALENKPRIPVVWIHGL WP_001058311.1 hydrogenase [Salmonella enterica] Length: 372\nNumber of predicted TMHs: 2\nExp number of AAs in TMHs: 37.4699899999999\nExp number, first 60 AAs: 19.72699\nTotal prob of N-in: 0.96171\nPOSSIBLE N-term signal sequence\ninside 1 19\nTMhelix 20 39\noutside 40 325\nTMhelix 326 348\ninside 349 372

17920 GCF_002066895.1_ASM206689v1 Salmonella enterica subsp. enterica serovar Infantis Proteobacteria; Gammaproteobacteria; Enterobacterales; Enterobacteriaceae; Salmonella; Salmonella enterica; Salmonella enterica subsp. enterica MTGDNTLITSHGINRRDFMKLCAALAATMGLSSKAAAEMAESVSNPQRPPVIWIGAQECT WP_000145429.1 hydrogenase 2 small subunit [Salmonella enterica] Length: 372\nNumber of predicted TMHs: 1\nExp number of AAs in TMHs: 24.51309\nExp number, first 60 AAs: 0.2682\nTotal prob of N-in: 0.14402\noutside 1 331\nTMhelix 332 354\ninside 355 372

17921 GCF_002066895.1_ASM206689v1 Salmonella enterica subsp. enterica serovar Infantis Proteobacteria; Gammaproteobacteria; Enterobacterales; Enterobacteriaceae; Salmonella; Salmonella enterica; Salmonella enterica subsp. enterica MNNEETFYQAMRRKGVTRRSFLKFCSLAATSLGLGAGMTPKIAWALENKPRIPVVWIHGL WP_001058311.1 hydrogenase [Salmonella enterica] Length: 372\nNumber of predicted TMHs: 2\nExp number of AAs in TMHs: 37.4699899999999\nExp number, first 60 AAs: 19.72699\nTotal prob of N-in: 0.96171\nPOSSIBLE N-term signal sequence\ninside 1 19\nTMhelix 20 39\noutside 40 325\nTMhelix 326 348\ninside 349 372

17922 GCF_001691885.1_ASM169188v1 Salmonella enterica subsp. enterica serovar Infantis Proteobacteria; Gammaproteobacteria; Enterobacterales; Enterobacteriaceae; Salmonella; Salmonella enterica; Salmonella enterica subsp. enterica MTGDNTLITSHGINRRDFMKLCAALAATMGLSSKAAAEMAESVSNPQRPPVIWIGAQECT WP_000145429.1 hydrogenase 2 small subunit [Salmonella enterica] Length: 372\nNumber of predicted TMHs: 1\nExp number of AAs in TMHs: 24.51309\nExp number, first 60 AAs: 0.2682\nTotal prob of N-in: 0.14402\noutside 1 331\nTMhelix 332 354\ninside 355 372

17923 GCF_001691915.1_ASM169191v1 Salmonella enterica subsp. enterica serovar Infantis Proteobacteria; Gammaproteobacteria; Enterobacterales; Enterobacteriaceae; Salmonella; Salmonella enterica; Salmonella enterica subsp. enterica MTGDNTLITSHGINRRDFMKLCAALAATMGLSSKAAAEMAESVSNPQRPPVIWIGAQECT WP_000145429.1 hydrogenase 2 small subunit [Salmonella enterica] Length: 372\nNumber of predicted TMHs: 1\nExp number of AAs in TMHs: 24.51309\nExp number, first 60 AAs: 0.2682\nTotal prob of N-in: 0.14402\noutside 1 331\nTMhelix 332 354\ninside 355 372

17924 GCF_001691915.1_ASM169191v1 Salmonella enterica subsp. enterica serovar Infantis Proteobacteria; Gammaproteobacteria; Enterobacterales; Enterobacteriaceae; Salmonella; Salmonella enterica; Salmonella enterica subsp. enterica MNNEETFYQAMRRKGVTRRSFLKFCSLAATSLGLGAGMTPKIAWALENKPRIPVVWIHGL WP_001058311.1 hydrogenase [Salmonella enterica] Length: 372\nNumber of predicted TMHs: 2\nExp number of AAs in TMHs: 37.4699899999999\nExp number, first 60 AAs: 19.72699\nTotal prob of N-in: 0.96171\nPOSSIBLE N-term signal sequence\ninside 1 19\nTMhelix 20 39\noutside 40 325\nTMhelix 326 348\ninside 349 372

17925 GCF_002206125.1_ASM220612v1 Salmonella enterica subsp. enterica serovar Infantis Proteobacteria; Gammaproteobacteria; Enterobacterales; Enterobacteriaceae; Salmonella; Salmonella enterica; Salmonella enterica subsp. enterica MNNEETFYQAMRRKGVTRRSFLKFCSLAATSLGLGAGMTPKIAWALENKPRIPVVWIHGL WP_001058311.1 hydrogenase [Salmonella enterica] Length: 372\nNumber of predicted TMHs: 2\nExp number of AAs in TMHs: 37.4699899999999\nExp number, first 60 AAs: 19.72699\nTotal prob of N-in: 0.96171\nPOSSIBLE N-term signal sequence\ninside 1 19\nTMhelix 20 39\noutside 40 325\nTMhelix 326 348\ninside 349 372

17926 GCF_001766505.1_ASM176650v1 Salmonella enterica subsp. enterica serovar Infantis Proteobacteria; Gammaproteobacteria; Enterobacterales; Enterobacteriaceae; Salmonella; Salmonella enterica; Salmonella enterica subsp. enterica MTGDNTLITSHGINRRDFMKLCAALAATMGLSSKAAAEMAESVSNPQRPPVIWIGAQECT WP_000145429.1 hydrogenase 2 small subunit [Salmonella enterica] Length: 372\nNumber of predicted TMHs: 1\nExp number of AAs in TMHs: 24.51309\nExp number, first 60 AAs: 0.2682\nTotal prob of N-in: 0.14402\noutside 1 331\nTMhelix 332 354\ninside 355 372

17927 GCF_002206125.1_ASM220612v1 Salmonella enterica subsp. enterica serovar Infantis Proteobacteria; Gammaproteobacteria; Enterobacterales; Enterobacteriaceae; Salmonella; Salmonella enterica; Salmonella enterica subsp. enterica MTGDNTLITSHGINRRDFMKLCAALAATMGLSSKAAAEMAESVSNPQRPPVIWIGAQECT WP_000145429.1 hydrogenase 2 small subunit [Salmonella enterica] Length: 372\nNumber of predicted TMHs: 1\nExp number of AAs in TMHs: 24.51309\nExp number, first 60 AAs: 0.2682\nTotal prob of N-in: 0.14402\noutside 1 331\nTMhelix 332 354\ninside 355 372

17928 GCF_002200795.1_ASM220079v1 Salmonella enterica subsp. enterica serovar Infantis Proteobacteria; Gammaproteobacteria; Enterobacterales; Enterobacteriaceae; Salmonella; Salmonella enterica;

Salmonella enterica subsp. enterica	MNNEETFYQAMRRKGVTRRSFLKFCSLAATSLGLGAGMTPKIAWALENKPRIPVVWIHGL	WP_001058311.1	hydrogenase [Salmonella enterica]	Length: 372\nNumber of predicted TMHs:  2\nExp number of AAs in TMHs: 37.4699899999999\nExp number, first 60 AAs:  19.72699\nTotal prob of N-in: 0.96171\nPOSSIBLE N-term signal sequence\ninside   1   19\nTMhelix   20   39\noutside   40   325\nTMhelix   326   348\ninside   349   372

17929	GCF_001766505.1_ASM176650v1	Salmonella enterica subsp. enterica serovar Infantis	Proteobacteria; Gammaproteobacteria; Enterobacterales; Enterobacteriaceae; Salmonella; Salmonella enterica; Salmonella enterica subsp. enterica	MNNEETFYQAMRRKGVTRRSFLKFCSLAATSLGLGAGMTPKIAWALENKPRIPVVWIHGL	WP_001058311.1	hydrogenase [Salmonella enterica]	Length: 372\nNumber of predicted TMHs:  2\nExp number of AAs in TMHs: 37.4699899999999\nExp number, first 60 AAs:  19.72699\nTotal prob of N-in: 0.96171\nPOSSIBLE N-term signal sequence\ninside   1   19\nTMhelix   20   39\noutside   40   325\nTMhelix   326   348\ninside   349   372

17930	GCF_001906505.1_ASM190650v1	Salmonella enterica subsp. enterica serovar Infantis	Proteobacteria; Gammaproteobacteria; Enterobacterales; Enterobacteriaceae; Salmonella; Salmonella enterica; Salmonella enterica subsp. enterica	MTGDNTLITSHGINRRDFMKLCAALAATMGLSSKAAAEMAESVSNPQRPPVIWIGAQECT	WP_000145429.1	hydrogenase 2 small subunit [Salmonella enterica]	Length: 372\nNumber of predicted TMHs:  1\nExp number of AAs in TMHs: 24.51309\nExp number, first 60 AAs:  0.2682\nTotal prob of N-in: 0.14402\noutside   1   331\nTMhelix   332   354\ninside   355   372

17931	GCF_001906505.1_ASM190650v1	Salmonella enterica subsp. enterica serovar Infantis	Proteobacteria; Gammaproteobacteria; Enterobacterales; Enterobacteriaceae; Salmonella; Salmonella enterica; Salmonella enterica subsp. enterica	MNNEETFYQAMRRKGVTRRSFLKFCSLAATSLGLGAGMTPKIAWALENKPRIPVVWIHGL	WP_001058311.1	hydrogenase [Salmonella enterica]	Length: 372\nNumber of predicted TMHs:  2\nExp number of AAs in TMHs: 37.4699899999999\nExp number, first 60 AAs:  19.72699\nTotal prob of N-in: 0.96171\nPOSSIBLE N-term signal sequence\ninside   1   19\nTMhelix   20   39\noutside   40   325\nTMhelix   326   348\ninside   349   372

17932	GCF_001931555.1_ASM193155v1	Salmonella enterica subsp. enterica serovar Infantis	Proteobacteria; Gammaproteobacteria; Enterobacterales; Enterobacteriaceae; Salmonella; Salmonella enterica; Salmonella enterica subsp. enterica	MNNEETFYQAMRRKGVTRRSFLKFCSLAATSLGLGAGMTPKIAWALENKPRIPVVWIHGL	WP_001058311.1	hydrogenase [Salmonella enterica]	Length: 372\nNumber of predicted TMHs:  2\nExp number of AAs in TMHs: 37.4699899999999\nExp number, first 60 AAs:  19.72699\nTotal prob of N-in: 0.96171\nPOSSIBLE N-term signal sequence\ninside   1   19\nTMhelix   20   39\noutside   40   325\nTMhelix   326   348\ninside   349   372

17933	GCF_002200655.1_ASM220065v1	Salmonella enterica subsp. enterica serovar Infantis	Proteobacteria; Gammaproteobacteria; Enterobacterales; Enterobacteriaceae; Salmonella; Salmonella enterica; Salmonella enterica subsp. enterica	MNNEETFYQAMRRKGVTRRSFLKFCSLAATSLGLGAGMTPKIAWALENKPRIPVVWIHGL	WP_001058311.1	hydrogenase [Salmonella enterica]	Length: 372\nNumber of predicted TMHs:  2\nExp number of AAs in TMHs: 37.4699899999999\nExp number, first 60 AAs:  19.72699\nTotal prob of N-in: 0.96171\nPOSSIBLE N-term signal sequence\ninside   1   19\nTMhelix   20   39\noutside   40   325\nTMhelix   326   348\ninside   349   372

17934	GCF_002200745.1_ASM220074v1	Salmonella enterica subsp. enterica serovar Infantis	Proteobacteria; Gammaproteobacteria; Enterobacterales; Enterobacteriaceae; Salmonella; Salmonella enterica; Salmonella enterica subsp. enterica	MNNEETFYQAMRRKGVTRRSFLKFCSLAATSLGLGAGMTPKIAWALENKPRIPVVWIHGL	WP_001058311.1	hydrogenase [Salmonella enterica]	Length: 372\nNumber of predicted TMHs:  2\nExp number of AAs in TMHs: 37.4699899999999\nExp number, first 60 AAs:  19.72699\nTotal prob of N-in: 0.96171\nPOSSIBLE N-term signal sequence\ninside   1   19\nTMhelix   20   39\noutside   40   325\nTMhelix   326   348\ninside   349   372

17935	GCF_002200675.1_ASM220067v1	Salmonella enterica subsp. enterica serovar Infantis	Proteobacteria; Gammaproteobacteria; Enterobacterales; Enterobacteriaceae; Salmonella; Salmonella enterica; Salmonella enterica subsp. enterica	MNNEETFYQAMRRKGVTRRSFLKFCSLAATSLGLGAGMTPKIAWALENKPRIPVVWIHGL	WP_001058311.1	hydrogenase [Salmonella enterica]	Length: 372\nNumber of predicted TMHs:  2\nExp number of AAs in TMHs: 37.4699899999999\nExp number, first 60 AAs:  19.72699\nTotal prob of N-in: 0.96171\nPOSSIBLE N-term signal sequence\ninside   1   19\nTMhelix   20   39\noutside   40   325\nTMhelix   326   348\ninside   349   372

17936 GCF_002200795.1_ASM220079v1 Salmonella enterica subsp. enterica serovar Infantis Proteobacteria; Gammaproteobacteria; Enterobacterales; Enterobacteriaceae; Salmonella; Salmonella enterica; Salmonella enterica subsp. enterica MTGDNTLITSHGINRRDFMKLCAALAATMGLSSKAAAEMAESVSNPQRPPVIWIGAQECT WP_000145429.1 hydrogenase 2 small subunit [Salmonella enterica] Length: 372\nNumber of predicted TMHs: 1\nExp number of AAs in TMHs: 24.51309\nExp number, first 60 AAs: 0.2682\nTotal prob of N-in: 0.14402\noutside 1 331\nTMhelix 332 354\ninside 355 372

17937 GCF_002200745.1_ASM220074v1 Salmonella enterica subsp. enterica serovar Infantis Proteobacteria; Gammaproteobacteria; Enterobacterales; Enterobacteriaceae; Salmonella; Salmonella enterica; Salmonella enterica subsp. enterica MTGDNTLITSHGINRRDFMKLCAALAATMGLSSKAAAEMAESVSNPQRPPVIWIGAQECT WP_000145429.1 hydrogenase 2 small subunit [Salmonella enterica] Length: 372\nNumber of predicted TMHs: 1\nExp number of AAs in TMHs: 24.51309\nExp number, first 60 AAs: 0.2682\nTotal prob of N-in: 0.14402\noutside 1 331\nTMhelix 332 354\ninside 355 372

17938 GCF_002200655.1_ASM220065v1 Salmonella enterica subsp. enterica serovar Infantis Proteobacteria; Gammaproteobacteria; Enterobacterales; Enterobacteriaceae; Salmonella; Salmonella enterica; Salmonella enterica subsp. enterica MTGDNTLITSHGINRRDFMKLCAALAATMGLSSKAAAEMAESVSNPQRPPVIWIGAQECT WP_000145429.1 hydrogenase 2 small subunit [Salmonella enterica] Length: 372\nNumber of predicted TMHs: 1\nExp number of AAs in TMHs: 24.51309\nExp number, first 60 AAs: 0.2682\nTotal prob of N-in: 0.14402\noutside 1 331\nTMhelix 332 354\ninside 355 372

17939 GCF_002200675.1_ASM220067v1 Salmonella enterica subsp. enterica serovar Infantis Proteobacteria; Gammaproteobacteria; Enterobacterales; Enterobacteriaceae; Salmonella; Salmonella enterica; Salmonella enterica subsp. enterica MTGDNTLITSHGINRRDFMKLCAALAATMGLSSKAAAEMAESVSNPQRPPVIWIGAQECT WP_000145429.1 hydrogenase 2 small subunit [Salmonella enterica] Length: 372\nNumber of predicted TMHs: 1\nExp number of AAs in TMHs: 24.51309\nExp number, first 60 AAs: 0.2682\nTotal prob of N-in: 0.14402\noutside 1 331\nTMhelix 332 354\ninside 355 372

17940 GCF_002200435.1_ASM220043v1 Salmonella enterica subsp. enterica serovar Infantis Proteobacteria; Gammaproteobacteria; Enterobacterales; Enterobacteriaceae; Salmonella; Salmonella enterica; Salmonella enterica subsp. enterica MNNEETFYQAMRRKGVTRRSFLKFCSLAATSLGLGAGMTPKIAWALENKPRIPVVWIHGL WP_001058311.1 hydrogenase [Salmonella enterica] Length: 372\nNumber of predicted TMHs: 2\nExp number of AAs in TMHs: 37.4699899999999\nExp number, first 60 AAs: 19.72699\nTotal prob of N-in: 0.96171\nPOSSIBLE N-term signal sequence\ninside 1 19\nTMhelix 20 39\noutside 40 325\nTMhelix 326 348\ninside 349 372

17941 GCF_002200605.1_ASM220060v1 Salmonella enterica subsp. enterica serovar Infantis Proteobacteria; Gammaproteobacteria; Enterobacterales; Enterobacteriaceae; Salmonella; Salmonella enterica; Salmonella enterica subsp. enterica MNNEETFYQAMRRKGVTRRSFLKFCSLAATSLGLGAGMTPKIAWALENKPRIPVVWIHGL WP_001058311.1 hydrogenase [Salmonella enterica] Length: 372\nNumber of predicted TMHs: 2\nExp number of AAs in TMHs: 37.4699899999999\nExp number, first 60 AAs: 19.72699\nTotal prob of N-in: 0.96171\nPOSSIBLE N-term signal sequence\ninside 1 19\nTMhelix 20 39\noutside 40 325\nTMhelix 326 348\ninside 349 372

17942 GCF_002200565.1_ASM220056v1 Salmonella enterica subsp. enterica serovar Infantis Proteobacteria; Gammaproteobacteria; Enterobacterales; Enterobacteriaceae; Salmonella; Salmonella enterica; Salmonella enterica subsp. enterica MNNEETFYQAMRRKGVTRRSFLKFCSLAATSLGLGAGMTPKIAWALENKPRIPVVWIHGL WP_001058311.1 hydrogenase [Salmonella enterica] Length: 372\nNumber of predicted TMHs: 2\nExp number of AAs in TMHs: 37.4699899999999\nExp number, first 60 AAs: 19.72699\nTotal prob of N-in: 0.96171\nPOSSIBLE N-term signal sequence\ninside 1 19\nTMhelix 20 39\noutside 40 325\nTMhelix 326 348\ninside 349 372

17943 GCF_002200485.1_ASM220048v1 Salmonella enterica subsp. enterica serovar Infantis Proteobacteria; Gammaproteobacteria; Enterobacterales; Enterobacteriaceae; Salmonella; Salmonella enterica; Salmonella enterica subsp. enterica MNNEETFYQAMRRKGVTRRSFLKFCSLAATSLGLGAGMTPKIAWALENKPRIPVVWIHGL WP_001058311.1 hydrogenase [Salmonella enterica] Length: 372\nNumber of predicted TMHs: 2\nExp number of AAs in TMHs: 37.4699899999999\nExp number, first 60 AAs: 19.72699\nTotal prob of N-in: 0.96171\nPOSSIBLE N-term signal sequence\ninside 1 19\nTMhelix 20 39\noutside 40 325\nTMhelix 326 348\ninside 349 372

17944 GCF_002200605.1_ASM220060v1 Salmonella enterica subsp. enterica serovar Infantis Proteobacteria; Gammaproteobacteria; Enterobacterales; Enterobacteriaceae; Salmonella; Salmonella enterica; Salmonella enterica subsp. enterica MTGDNTLITSHGINRRDFMKLCAALAATMGLSSKAAAEMAESVSNPQRPPVIWIGAQECT WP_000145429.1 hydrogenase 2 small subunit [Salmonella enterica] Length: 372\nNumber of predicted TMHs: 1\nExp number of AAs in TMHs: 24.51309\nExp number, first 60 AAs: 0.2682\nTotal prob of N-in: 0.14402\noutside 1 331\nTMhelix 332 354\ninside 355 372

17945 GCF_002200565.1_ASM220056v1 Salmonella enterica subsp. enterica serovar Infantis Proteobacteria; Gammaproteobacteria; Enterobacterales; Enterobacteriaceae; Salmonella; Salmonella enterica; Salmonella enterica subsp. enterica MTGDNTLITSHGINRRDFMKLCAALAATMGLSSKAAAEMAESVSNPQRPPVIWIGAQECT WP_000145429.1 hydrogenase 2 small subunit [Salmonella enterica] Length: 372\nNumber of predicted TMHs: 1\nExp number of AAs in TMHs: 24.51309\nExp number, first 60 AAs: 0.2682\nTotal prob of N-in: 0.14402\noutside 1 331\nTMhelix 332 354\ninside 355 372

17946 GCF_002200485.1_ASM220048v1 Salmonella enterica subsp. enterica serovar Infantis Proteobacteria; Gammaproteobacteria; Enterobacterales; Enterobacteriaceae; Salmonella; Salmonella enterica; Salmonella enterica subsp. enterica MTGDNTLITSHGINRRDFMKLCAALAATMGLSSKAAAEMAESVSNPQRPPVIWIGAQECT WP_000145429.1 hydrogenase 2 small subunit [Salmonella enterica] Length: 372\nNumber of predicted TMHs: 1\nExp number of AAs in TMHs: 24.51309\nExp number, first 60 AAs: 0.2682\nTotal prob of N-in: 0.14402\noutside 1 331\nTMhelix 332 354\ninside 355 372

17947 GCF_002200405.1_ASM220040v1 Salmonella enterica subsp. enterica serovar Infantis Proteobacteria; Gammaproteobacteria; Enterobacterales; Enterobacteriaceae; Salmonella; Salmonella enterica; Salmonella enterica subsp. enterica MNNEETFYQAMRRKGVTRRSFLKFCSLAATSLGLGAGMTPKIAWALENKPRIPVVWIHGL WP_001058311.1 hydrogenase [Salmonella enterica] Length: 372\nNumber of predicted TMHs: 2\nExp number of AAs in TMHs: 37.4699899999999\nExp number, first 60 AAs: 19.72699\nTotal prob of N-in: 0.96171\nPOSSIBLE N-term signal sequence\ninside 1 19\nTMhelix 20 39\noutside 40 325\nTMhelix 326 348\ninside 349 372

17948 GCF_002200525.1_ASM220052v1 Salmonella enterica subsp. enterica serovar Infantis Proteobacteria; Gammaproteobacteria; Enterobacterales; Enterobacteriaceae; Salmonella; Salmonella enterica; Salmonella enterica subsp. enterica MNNEETFYQAMRRKGVTRRSFLKFCSLAATSLGLGAGMTPKIAWALENKPRIPVVWIHGL WP_001058311.1 hydrogenase [Salmonella enterica] Length: 372\nNumber of predicted TMHs: 2\nExp number of AAs in TMHs: 37.4699899999999\nExp number, first 60 AAs: 19.72699\nTotal prob of N-in: 0.96171\nPOSSIBLE N-term signal sequence\ninside 1 19\nTMhelix 20 39\noutside 40 325\nTMhelix 326 348\ninside 349 372

17949 GCF_002200525.1_ASM220052v1 Salmonella enterica subsp. enterica serovar Infantis Proteobacteria; Gammaproteobacteria; Enterobacterales; Enterobacteriaceae; Salmonella; Salmonella enterica; Salmonella enterica subsp. enterica MTGDNTLITSHGINRRDFMKLCAALAATMGLSSKAAAEMAESVSNPQRPPVIWIGAQECT WP_000145429.1 hydrogenase 2 small subunit [Salmonella enterica] Length: 372\nNumber of predicted TMHs: 1\nExp number of AAs in TMHs: 24.51309\nExp number, first 60 AAs: 0.2682\nTotal prob of N-in: 0.14402\noutside 1 331\nTMhelix 332 354\ninside 355 372

17950 GCF_002200435.1_ASM220043v1 Salmonella enterica subsp. enterica serovar Infantis Proteobacteria; Gammaproteobacteria; Enterobacterales; Enterobacteriaceae; Salmonella; Salmonella enterica; Salmonella enterica subsp. enterica MTGDNTLITSHGINRRDFMKLCAALAATMGLSSKAAAEMAESVSNPQRPPVIWIGAQECT WP_000145429.1 hydrogenase 2 small subunit [Salmonella enterica] Length: 372\nNumber of predicted TMHs: 1\nExp number of AAs in TMHs: 24.51309\nExp number, first 60 AAs: 0.2682\nTotal prob of N-in: 0.14402\noutside 1 331\nTMhelix 332 354\ninside 355 372

17951 GCF_002200365.1_ASM220036v1 Salmonella enterica subsp. enterica serovar Infantis Proteobacteria; Gammaproteobacteria; Enterobacterales; Enterobacteriaceae; Salmonella; Salmonella enterica; Salmonella enterica subsp. enterica MNNEETFYQAMRRKGVTRRSFLKFCSLAATSLGLGAGMTPKIAWALENKPRIPVVWIHGL WP_001058311.1 hydrogenase [Salmonella enterica] Length: 372\nNumber of predicted TMHs: 2\nExp number of AAs in TMHs: 37.4699899999999\nExp number, first 60 AAs: 19.72699\nTotal prob of N-in: 0.96171\nPOSSIBLE N-term signal sequence\ninside 1 19\nTMhelix 20 39\noutside 40 325\nTMhelix 326 348\ninside 349 372

17952 GCF_002200405.1_ASM220040v1 Salmonella enterica subsp. enterica serovar Infantis Proteobacteria; Gammaproteobacteria; Enterobacterales; Enterobacteriaceae; Salmonella; Salmonella enterica;

Salmonella enterica subsp. enterica MTGDNTLITSHGINRRDFMKLCAALAATMGLSSKAAAEMAESVSNPQRPPVIWIGAQECT WP_000145429.1 hydrogenase 2 small subunit [Salmonella enterica] Length: 372\nNumber of predicted TMHs: 1\nExp number of AAs in TMHs: 24.51309\nExp number, first 60 AAs: 0.2682\nTotal prob of N-in: 0.14402\noutside 1 331\nTMhelix 332 354\ninside 355 372

17953 GCF_002200325.1_ASM220032v1 Salmonella enterica subsp. enterica serovar Infantis Proteobacteria; Gammaproteobacteria; Enterobacterales; Enterobacteriaceae; Salmonella; Salmonella enterica; Salmonella enterica subsp. enterica MNNEETFYQAMRRKGVTRRSFLKFCSLAATSLGLGAGMTPKIAWALENKPRIPVVWIHGL WP_001058311.1 hydrogenase [Salmonella enterica] Length: 372\nNumber of predicted TMHs: 2\nExp number of AAs in TMHs: 37.4699899999999\nExp number, first 60 AAs: 19.72699\nTotal prob of N-in: 0.96171\nPOSSIBLE N-term signal sequence\ninside 1 19\nTMhelix 20 39\noutside 40 325\nTMhelix 326 348\ninside 349 372

17954 GCF_002200365.1_ASM220036v1 Salmonella enterica subsp. enterica serovar Infantis Proteobacteria; Gammaproteobacteria; Enterobacterales; Enterobacteriaceae; Salmonella; Salmonella enterica; Salmonella enterica subsp. enterica MTGDNTLITSHGINRRDFMKLCAALAATMGLSSKAAAEMAESVSNPQRPPVIWIGAQECT WP_000145429.1 hydrogenase 2 small subunit [Salmonella enterica] Length: 372\nNumber of predicted TMHs: 1\nExp number of AAs in TMHs: 24.51309\nExp number, first 60 AAs: 0.2682\nTotal prob of N-in: 0.14402\noutside 1 331\nTMhelix 332 354\ninside 355 372

17955 GCF_002200325.1_ASM220032v1 Salmonella enterica subsp. enterica serovar Infantis Proteobacteria; Gammaproteobacteria; Enterobacterales; Enterobacteriaceae; Salmonella; Salmonella enterica; Salmonella enterica subsp. enterica MTGDNTLITSHGINRRDFMKLCAALAATMGLSSKAAAEMAESVSNPQRPPVIWIGAQECT WP_000145429.1 hydrogenase 2 small subunit [Salmonella enterica] Length: 372\nNumber of predicted TMHs: 1\nExp number of AAs in TMHs: 24.51309\nExp number, first 60 AAs: 0.2682\nTotal prob of N-in: 0.14402\noutside 1 331\nTMhelix 332 354\ninside 355 372

17956 GCF_002200315.1_ASM220031v1 Salmonella enterica subsp. enterica serovar Infantis Proteobacteria; Gammaproteobacteria; Enterobacterales; Enterobacteriaceae; Salmonella; Salmonella enterica; Salmonella enterica subsp. enterica MNNEETFYQAMRRKGVTRRSFLKFCSLAATSLGLGAGMTPKIAWALENKPRIPVVWIHGL WP_001058311.1 hydrogenase [Salmonella enterica] Length: 372\nNumber of predicted TMHs: 2\nExp number of AAs in TMHs: 37.4699899999999\nExp number, first 60 AAs: 19.72699\nTotal prob of N-in: 0.96171\nPOSSIBLE N-term signal sequence\ninside 1 19\nTMhelix 20 39\noutside 40 325\nTMhelix 326 348\ninside 349 372

17957 GCF_002200285.1_ASM220028v1 Salmonella enterica subsp. enterica serovar Infantis Proteobacteria; Gammaproteobacteria; Enterobacterales; Enterobacteriaceae; Salmonella; Salmonella enterica; Salmonella enterica subsp. enterica MNNEETFYQAMRRKGVTRRSFLKFCSLAATSLGLGAGMTPKIAWALENKPRIPVVWIHGL WP_001058311.1 hydrogenase [Salmonella enterica] Length: 372\nNumber of predicted TMHs: 2\nExp number of AAs in TMHs: 37.4699899999999\nExp number, first 60 AAs: 19.72699\nTotal prob of N-in: 0.96171\nPOSSIBLE N-term signal sequence\ninside 1 19\nTMhelix 20 39\noutside 40 325\nTMhelix 326 348\ninside 349 372

17958 GCF_002198745.1_ASM219874v1 Salmonella enterica subsp. enterica serovar Infantis Proteobacteria; Gammaproteobacteria; Enterobacterales; Enterobacteriaceae; Salmonella; Salmonella enterica; Salmonella enterica subsp. enterica MNNEETFYQAMRRKGVTRRSFLKFCSLAATSLGLGAGMTPKIAWALENKPRIPVVWIHGL WP_001058311.1 hydrogenase [Salmonella enterica] Length: 372\nNumber of predicted TMHs: 2\nExp number of AAs in TMHs: 37.4699899999999\nExp number, first 60 AAs: 19.72699\nTotal prob of N-in: 0.96171\nPOSSIBLE N-term signal sequence\ninside 1 19\nTMhelix 20 39\noutside 40 325\nTMhelix 326 348\ninside 349 372

17959 GCF_001997485.1_ASM199748v1 Salmonella enterica subsp. enterica serovar Infantis Proteobacteria; Gammaproteobacteria; Enterobacterales; Enterobacteriaceae; Salmonella; Salmonella enterica; Salmonella enterica subsp. enterica MTGDNTLITSHGINRRDFMKLCAALAATMGLSSKAAAEMAESVSNPQRPPVIWIGAQECT WP_000145429.1 hydrogenase 2 small subunit [Salmonella enterica] Length: 372\nNumber of predicted TMHs: 1\nExp number of AAs in TMHs: 24.51309\nExp number, first 60 AAs: 0.2682\nTotal prob of N-in: 0.14402\noutside 1 331\nTMhelix 332 354\ninside 355 372

17960 GCF_002200315.1_ASM220031v1 Salmonella enterica subsp. enterica serovar Infantis Proteobacteria; Gammaproteobacteria; Enterobacterales; Enterobacteriaceae; Salmonella; Salmonella enterica; Salmonella enterica subsp. enterica MTGDNTLITSHGINRRDFMKLCAALAATMGLSSKAAAEMAESVSNPQRPPVIWIGAQECT

WP_000145429.1	hydrogenase 2 small subunit [Salmonella enterica]	Length: 372\nNumber of predicted TMHs: 1\nExp number of AAs in TMHs: 24.51309\nExp number, first 60 AAs: 0.2682\nTotal prob of N-in: 0.14402\noutside 1 331\nTMhelix 332 354\ninside 355 372

17961	GCF_002200285.1_ASM220028v1	Salmonella enterica subsp. enterica serovar Infantis	Proteobacteria; Gammaproteobacteria; Enterobacterales; Enterobacteriaceae; Salmonella; Salmonella enterica; Salmonella enterica subsp. enterica	MTGDNTLITSHGINRRDFMKLCAALAATMGLSSKAAAEMAESVSNPQRPPVIWIGAQECT	WP_000145429.1	hydrogenase 2 small subunit [Salmonella enterica]	Length: 372\nNumber of predicted TMHs: 1\nExp number of AAs in TMHs: 24.51309\nExp number, first 60 AAs: 0.2682\nTotal prob of N-in: 0.14402\noutside 1 331\nTMhelix 332 354\ninside 355 372

17962	GCF_002198745.1_ASM219874v1	Salmonella enterica subsp. enterica serovar Infantis	Proteobacteria; Gammaproteobacteria; Enterobacterales; Enterobacteriaceae; Salmonella; Salmonella enterica; Salmonella enterica subsp. enterica	MTGDNTLITSHGINRRDFMKLCAALAATMGLSSKAAAEMAESVSNPQRPPVIWIGAQECT	WP_000145429.1	hydrogenase 2 small subunit [Salmonella enterica]	Length: 372\nNumber of predicted TMHs: 1\nExp number of AAs in TMHs: 24.51309\nExp number, first 60 AAs: 0.2682\nTotal prob of N-in: 0.14402\noutside 1 331\nTMhelix 332 354\ninside 355 372

17963	GCF_002030305.1_ASM203030v1	Salmonella enterica subsp. enterica serovar Infantis	Proteobacteria; Gammaproteobacteria; Enterobacterales; Enterobacteriaceae; Salmonella; Salmonella enterica; Salmonella enterica subsp. enterica	MTGDNTLITSHGINRRDFMKLCAALAATMGLSSKAAAEMAESVSNPQRPPVIWIGAQECT	WP_000145429.1	hydrogenase 2 small subunit [Salmonella enterica]	Length: 372\nNumber of predicted TMHs: 1\nExp number of AAs in TMHs: 24.51309\nExp number, first 60 AAs: 0.2682\nTotal prob of N-in: 0.14402\noutside 1 331\nTMhelix 332 354\ninside 355 372

17964	GCF_002030305.1_ASM203030v1	Salmonella enterica subsp. enterica serovar Infantis	Proteobacteria; Gammaproteobacteria; Enterobacterales; Enterobacteriaceae; Salmonella; Salmonella enterica; Salmonella enterica subsp. enterica	MNNEETFYQAMRRKGVTRRSFLKFCSLAATSLGLGAGMTPKIAWALENKPRIPVVWIHGL	WP_001058311.1	hydrogenase [Salmonella enterica]	Length: 372\nNumber of predicted TMHs: 2\nExp number of AAs in TMHs: 37.4699899999999\nExp number, first 60 AAs: 19.72699\nTotal prob of N-in: 0.96171\nPOSSIBLE N-term signal sequence\ninside 1 19\nTMhelix 20 39\noutside 40 325\nTMhelix 326 348\ninside 349 372

17965	GCF_002030975.1_ASM203097v1	Salmonella enterica subsp. enterica serovar Infantis	Proteobacteria; Gammaproteobacteria; Enterobacterales; Enterobacteriaceae; Salmonella; Salmonella enterica; Salmonella enterica subsp. enterica	MTGDNTLITSHGINRRDFMKLCAALAATMGLSSKAAAEMAESVSNPQRPPVIWIGAQECT	WP_000145429.1	hydrogenase 2 small subunit [Salmonella enterica]	Length: 372\nNumber of predicted TMHs: 1\nExp number of AAs in TMHs: 24.51309\nExp number, first 60 AAs: 0.2682\nTotal prob of N-in: 0.14402\noutside 1 331\nTMhelix 332 354\ninside 355 372

17966	GCF_002030975.1_ASM203097v1	Salmonella enterica subsp. enterica serovar Infantis	Proteobacteria; Gammaproteobacteria; Enterobacterales; Enterobacteriaceae; Salmonella; Salmonella enterica; Salmonella enterica subsp. enterica	MNNEETFYQAMRRKGVTRRSFLKFCSLAATSLGLGAGMTPKIAWALENKPRIPVVWIHGL	WP_001058311.1	hydrogenase [Salmonella enterica]	Length: 372\nNumber of predicted TMHs: 2\nExp number of AAs in TMHs: 37.4699899999999\nExp number, first 60 AAs: 19.72699\nTotal prob of N-in: 0.96171\nPOSSIBLE N-term signal sequence\ninside 1 19\nTMhelix 20 39\noutside 40 325\nTMhelix 326 348\ninside 349 372

17967	GCF_002119135.1_ASM211913v1	Salmonella enterica subsp. enterica serovar Infantis	Proteobacteria; Gammaproteobacteria; Enterobacterales; Enterobacteriaceae; Salmonella; Salmonella enterica; Salmonella enterica subsp. enterica	MNNEETFYQAMRRKGVTRRSFLKFCSLAATSLGLGAGMTPKIAWALENKPRIPVVWIHGL	WP_001058311.1	hydrogenase [Salmonella enterica]	Length: 372\nNumber of predicted TMHs: 2\nExp number of AAs in TMHs: 37.4699899999999\nExp number, first 60 AAs: 19.72699\nTotal prob of N-in: 0.96171\nPOSSIBLE N-term signal sequence\ninside 1 19\nTMhelix 20 39\noutside 40 325\nTMhelix 326 348\ninside 349 372

17968	GCF_002119135.1_ASM211913v1	Salmonella enterica subsp. enterica serovar Infantis	Proteobacteria; Gammaproteobacteria; Enterobacterales; Enterobacteriaceae; Salmonella; Salmonella enterica; Salmonella enterica subsp. enterica	MTGDNTLITSHGINRRDFMKLCAALAATMGLSSKAAAEMAESVSNPQRPPVIWIGAQECT	WP_000145429.1	hydrogenase 2 small subunit [Salmonella enterica]	Length: 372\nNumber of

predicted TMHs: 1\nExp number of AAs in TMHs: 24.51309\nExp number, first 60 AAs: 0.2682\nTotal prob of N-in: 0.14402\noutside 1 331\nTMhelix 332 354\ninside 355 372

17969 GCF_002107535.1_ASM210753v1 Salmonella enterica subsp. enterica serovar Infantis Proteobacteria; Gammaproteobacteria; Enterobacterales; Enterobacteriaceae; Salmonella; Salmonella enterica; Salmonella enterica subsp. enterica MNNEETFYQAMRRKGVTRRSFLKFCSLAATSLGLGAGMTPKIAWALENKPRIPVVWIHGL WP_001058311.1 hydrogenase [Salmonella enterica] Length: 372\nNumber of predicted TMHs: 2\nExp number of AAs in TMHs: 37.4699899999999\nExp number, first 60 AAs: 19.72699\nTotal prob of N-in: 0.96171\nPOSSIBLE N-term signal sequence\ninside 1 19\nTMhelix 20 39\noutside 40 325\nTMhelix 326 348\ninside 349 372

17970 GCF_002107535.1_ASM210753v1 Salmonella enterica subsp. enterica serovar Infantis Proteobacteria; Gammaproteobacteria; Enterobacterales; Enterobacteriaceae; Salmonella; Salmonella enterica; Salmonella enterica subsp. enterica MTGDNTLITSHGINRRDFMKLCAALAATMGLSSKAAAEMAESVSNPQRPPVIWIGAQECT WP_000145429.1 hydrogenase 2 small subunit [Salmonella enterica] Length: 372\nNumber of predicted TMHs: 1\nExp number of AAs in TMHs: 24.51309\nExp number, first 60 AAs: 0.2682\nTotal prob of N-in: 0.14402\noutside 1 331\nTMhelix 332 354\ninside 355 372

17971 GCF_002107195.1_ASM210719v1 Salmonella enterica subsp. enterica serovar Infantis Proteobacteria; Gammaproteobacteria; Enterobacterales; Enterobacteriaceae; Salmonella; Salmonella enterica; Salmonella enterica subsp. enterica MNNEETFYQAMRRKGVTRRSFLKFCSLAATSLGLGAGMTPKIAWALENKPRIPVVWIHGL WP_001058311.1 hydrogenase [Salmonella enterica] Length: 372\nNumber of predicted TMHs: 2\nExp number of AAs in TMHs: 37.4699899999999\nExp number, first 60 AAs: 19.72699\nTotal prob of N-in: 0.96171\nPOSSIBLE N-term signal sequence\ninside 1 19\nTMhelix 20 39\noutside 40 325\nTMhelix 326 348\ninside 349 372

17972 GCF_002107195.1_ASM210719v1 Salmonella enterica subsp. enterica serovar Infantis Proteobacteria; Gammaproteobacteria; Enterobacterales; Enterobacteriaceae; Salmonella; Salmonella enterica; Salmonella enterica subsp. enterica MTGDNTLITSHGINRRDFMKLCAALAATMGLSSKAAAEMAESVSNPQRPPVIWIGAQECT WP_000145429.1 hydrogenase 2 small subunit [Salmonella enterica] Length: 372\nNumber of predicted TMHs: 1\nExp number of AAs in TMHs: 24.51309\nExp number, first 60 AAs: 0.2682\nTotal prob of N-in: 0.14402\noutside 1 331\nTMhelix 332 354\ninside 355 372

17973 GCF_002059795.1_ASM205979v1 Salmonella enterica subsp. enterica serovar Infantis Proteobacteria; Gammaproteobacteria; Enterobacterales; Enterobacteriaceae; Salmonella; Salmonella enterica; Salmonella enterica subsp. enterica MNNEETFYQAMRRKGVTRRSFLKFCSLAATSLGLGAGMTPKIAWALENKPRIPVVWIHGL WP_001058311.1 hydrogenase [Salmonella enterica] Length: 372\nNumber of predicted TMHs: 2\nExp number of AAs in TMHs: 37.4699899999999\nExp number, first 60 AAs: 19.72699\nTotal prob of N-in: 0.96171\nPOSSIBLE N-term signal sequence\ninside 1 19\nTMhelix 20 39\noutside 40 325\nTMhelix 326 348\ninside 349 372

17974 GCF_002062835.1_ASM206283v1 Salmonella enterica subsp. enterica serovar Infantis Proteobacteria; Gammaproteobacteria; Enterobacterales; Enterobacteriaceae; Salmonella; Salmonella enterica; Salmonella enterica subsp. enterica MTGDNTLITSHGINRRDFMKLCAALAATMGLSSKAAAEMAESVSNPQRPPVIWIGAQECT WP_000145429.1 hydrogenase 2 small subunit [Salmonella enterica] Length: 372\nNumber of predicted TMHs: 1\nExp number of AAs in TMHs: 24.51309\nExp number, first 60 AAs: 0.2682\nTotal prob of N-in: 0.14402\noutside 1 331\nTMhelix 332 354\ninside 355 372

17975 GCF_002062835.1_ASM206283v1 Salmonella enterica subsp. enterica serovar Infantis Proteobacteria; Gammaproteobacteria; Enterobacterales; Enterobacteriaceae; Salmonella; Salmonella enterica; Salmonella enterica subsp. enterica MNNEETFYQAMRRKGVTRRSFLKFCSLAATSLGLGAGMTPKIAWALENKPRIPVVWIHGL WP_001058311.1 hydrogenase [Salmonella enterica] Length: 372\nNumber of predicted TMHs: 2\nExp number of AAs in TMHs: 37.4699899999999\nExp number, first 60 AAs: 19.72699\nTotal prob of N-in: 0.96171\nPOSSIBLE N-term signal sequence\ninside 1 19\nTMhelix 20 39\noutside 40 325\nTMhelix 326 348\ninside 349 372

17976 GCF_002064835.1_ASM206483v1 Salmonella enterica subsp. enterica serovar Infantis Proteobacteria; Gammaproteobacteria; Enterobacterales; Enterobacteriaceae; Salmonella; Salmonella enterica; Salmonella enterica subsp. enterica MTGDNTLITSHGINRRDFMKLCAALAATMGLSSKAAAEMAESVSNPQRPPVIWIGAQECT WP_000145429.1 hydrogenase 2 small subunit [Salmonella enterica] Length: 372\nNumber of

predicted TMHs:  1\nExp number of AAs in TMHs: 24.51309\nExp number, first 60 AAs:  0.2682\nTotal prob of N-in: 0.14402\noutside   1   331\nTMhelix   332   354\ninside   355   372

17977 GCF_002064895.1_ASM206489v1 Salmonella enterica subsp. enterica serovar Infantis Proteobacteria; Gammaproteobacteria; Enterobacterales; Enterobacteriaceae; Salmonella; Salmonella enterica; Salmonella enterica subsp. enterica MTGDNTLITSHGINRRDFMKLCAALAATMGLSSKAAAEMAESVSNPQRPPVIWIGAQECT WP_000145429.1 hydrogenase 2 small subunit [Salmonella enterica] Length: 372\nNumber of predicted TMHs:  1\nExp number of AAs in TMHs: 24.51309\nExp number, first 60 AAs:  0.2682\nTotal prob of N-in: 0.14402\noutside   1   331\nTMhelix   332   354\ninside   355   372

17978 GCF_002064835.1_ASM206483v1 Salmonella enterica subsp. enterica serovar Infantis Proteobacteria; Gammaproteobacteria; Enterobacterales; Enterobacteriaceae; Salmonella; Salmonella enterica; Salmonella enterica subsp. enterica MNNEETFYQAMRRKGVTRRSFLKFCSLAATSLGLGAGMTPKIAWALENKPRIPVVWIHGL WP_001058311.1 hydrogenase [Salmonella enterica] Length: 372\nNumber of predicted TMHs:  2\nExp number of AAs in TMHs: 37.4699899999999\nExp number, first 60 AAs:  19.72699\nTotal prob of N-in: 0.96171\nPOSSIBLE N-term signal sequence\ninside   1   19\nTMhelix   20   39\noutside   40   325\nTMhelix   326   348\ninside   349   372

17979 GCF_002064895.1_ASM206489v1 Salmonella enterica subsp. enterica serovar Infantis Proteobacteria; Gammaproteobacteria; Enterobacterales; Enterobacteriaceae; Salmonella; Salmonella enterica; Salmonella enterica subsp. enterica MNNEETFYQAMRRKGVTRRSFLKFCSLAATSLGLGAGMTPKIAWALENKPRIPVVWIHGL WP_001058311.1 hydrogenase [Salmonella enterica] Length: 372\nNumber of predicted TMHs:  2\nExp number of AAs in TMHs: 37.4699899999999\nExp number, first 60 AAs:  19.72699\nTotal prob of N-in: 0.96171\nPOSSIBLE N-term signal sequence\ninside   1   19\nTMhelix   20   39\noutside   40   325\nTMhelix   326   348\ninside   349   372

17980 GCF_000812595.1_ASM81259v1 Salmonella enterica subsp. enterica serovar Infantis Proteobacteria; Gammaproteobacteria; Enterobacterales; Enterobacteriaceae; Salmonella; Salmonella enterica; Salmonella enterica subsp. enterica MTGDNTLITSHGINRRDFMKLCAALAATMGLSSKAAAEMAESVSNPQRPPVIWIGAQECT WP_000145429.1 hydrogenase 2 small subunit [Salmonella enterica] Length: 372\nNumber of predicted TMHs:  1\nExp number of AAs in TMHs: 24.51309\nExp number, first 60 AAs:  0.2682\nTotal prob of N-in: 0.14402\noutside   1   331\nTMhelix   332   354\ninside   355   372

17981 GCF_001187365.1_ASM118736v1 Salmonella enterica subsp. enterica serovar Infantis Proteobacteria; Gammaproteobacteria; Enterobacterales; Enterobacteriaceae; Salmonella; Salmonella enterica; Salmonella enterica subsp. enterica MNNEETFYQAMRRKGVTRRSFLKFCSLAATSLGLSAGMTPKIAWALENKPRIPVVWIHGL WP_001058316.1 hydrogenase [Salmonella enterica] Length: 372\nNumber of predicted TMHs:  2\nExp number of AAs in TMHs: 35.76397\nExp number, first 60 AAs:  18.97201\nTotal prob of N-in:   0.92785\nPOSSIBLE N-term signal sequence\ninside   1   19\nTMhelix   20   39\noutside   40   325\nTMhelix   326   348\ninside   349   372

17982 GCF_001243785.1_Salmonella_enterica_CVM_N47714_v1.0 Salmonella enterica subsp. enterica serovar Infantis Proteobacteria; Gammaproteobacteria; Enterobacterales; Enterobacteriaceae; Salmonella; Salmonella enterica; Salmonella enterica subsp. enterica MNNEETFYQAMRRKGVTRRSFLKFCSLAATSLGLGAGMTPKIAWALENKPRIPVVWIHGL WP_001058311.1 hydrogenase [Salmonella enterica] Length: 372\nNumber of predicted TMHs:  2\nExp number of AAs in TMHs: 37.4699899999999\nExp number, first 60 AAs:  19.72699\nTotal prob of N-in: 0.96171\nPOSSIBLE N-term signal sequence\ninside   1   19\nTMhelix   20   39\noutside   40   325\nTMhelix   326   348\ninside   349   372

17983 GCF_002030855.1_ASM203085v1 Salmonella enterica subsp. enterica serovar Infantis Proteobacteria; Gammaproteobacteria; Enterobacterales; Enterobacteriaceae; Salmonella; Salmonella enterica; Salmonella enterica subsp. enterica MNNEETFYQAMRRKGVTRRSFLKFCSLAATSLGLGAGMTPKIAWALENKPRIPVVWIHGL WP_001058311.1 hydrogenase [Salmonella enterica] Length: 372\nNumber of predicted TMHs:  2\nExp number of AAs in TMHs: 37.4699899999999\nExp number, first 60 AAs:  19.72699\nTotal prob of N-in: 0.96171\nPOSSIBLE N-term signal sequence\ninside   1   19\nTMhelix   20   39\noutside   40   325\nTMhelix   326   348\ninside   349   372

17984 GCF_001479785.1_Salmonella_enterica_CVM_N38848-SQ_v1.0 Salmonella enterica subsp. enterica serovar Infantis Proteobacteria; Gammaproteobacteria; Enterobacterales; Enterobacteriaceae; Salmonella; Salmonella enterica; Salmonella enterica subsp. enterica MTGDNTLITSHGINRRDFMKLCAALAATMGLSSKAAAEMAESVSNPQRPPVIWIGAQECT WP_000145429.1 hydrogenase 2 small subunit [Salmonella enterica] Length: 372\nNumber of

predicted TMHs:  1\nExp number of AAs in TMHs: 24.51309\nExp number, first 60 AAs:  0.2682\nTotal prob of N-in: 0.14402\noutside   1   331\nTMhelix   332   354\ninside   355   372

17985 GCF_001242185.1_Salmonella_enterica_CVM_N45929_v1.0 Salmonella enterica subsp. enterica serovar Infantis Proteobacteria; Gammaproteobacteria; Enterobacterales; Enterobacteriaceae; Salmonella; Salmonella enterica; Salmonella enterica subsp. enterica MNNEETFYQAMRRKGVTRRSFLKFCSLAATSLGLGAGMTPKIAWALENKPRIPVVWIHGL WP_001058311.1 hydrogenase [Salmonella enterica] Length: 372\nNumber of predicted TMHs:  2\nExp number of AAs in TMHs: 37.4699899999999\nExp number, first 60 AAs:  19.72699\nTotal prob of N-in: 0.96171\nPOSSIBLE N-term signal sequence\ninside   1   19\nTMhelix   20   39\noutside   40   325\nTMhelix   326   348\ninside   349   372

17986 GCF_000812575.1_ASM81257v1 Salmonella enterica subsp. enterica serovar Infantis Proteobacteria; Gammaproteobacteria; Enterobacterales; Enterobacteriaceae; Salmonella; Salmonella enterica; Salmonella enterica subsp. enterica MNNEETFYQAMRRKGVTRRSFLKFCSLAATSLGLGAGMTPKIAWALENKPRIPVVWIHGL WP_001058311.1 hydrogenase [Salmonella enterica] Length: 372\nNumber of predicted TMHs:  2\nExp number of AAs in TMHs: 37.4699899999999\nExp number, first 60 AAs:  19.72699\nTotal prob of N-in: 0.96171\nPOSSIBLE N-term signal sequence\ninside   1   19\nTMhelix   20   39\noutside   40   325\nTMhelix   326   348\ninside   349   372

17987 GCF_001478785.1_Salmonella_enterica_CVM_N29379-SQ_v1.0 Salmonella enterica subsp. enterica serovar Infantis Proteobacteria; Gammaproteobacteria; Enterobacterales; Enterobacteriaceae; Salmonella; Salmonella enterica; Salmonella enterica subsp. enterica MNNEETFYQAMRRKGVTRRSFLKFCSLAATSLGLGAGMTPKIAWALENKPRIPVVWIHGL WP_052901769.1 hydrogenase [Salmonella enterica] Length: 372\nNumber of predicted TMHs:  2\nExp number of AAs in TMHs: 37.42632\nExp number, first 60 AAs:  19.72508\nTotal prob of N-in:   0.96168\nPOSSIBLE N-term signal sequence\ninside   1   19\nTMhelix   20   39\noutside   40   325\nTMhelix   326   348\ninside   349   372

17988 GCF_002200635.1_ASM220063v1 Salmonella enterica subsp. enterica serovar Infantis Proteobacteria; Gammaproteobacteria; Enterobacterales; Enterobacteriaceae; Salmonella; Salmonella enterica; Salmonella enterica subsp. enterica MNNEETFYQAMRRKGVTRRSFLKFCSLAATSLGLGAGMTPKIAWALENKPRIPVVWIHGL WP_001058311.1 hydrogenase [Salmonella enterica] Length: 372\nNumber of predicted TMHs:  2\nExp number of AAs in TMHs: 37.4699899999999\nExp number, first 60 AAs:  19.72699\nTotal prob of N-in: 0.96171\nPOSSIBLE N-term signal sequence\ninside   1   19\nTMhelix   20   39\noutside   40   325\nTMhelix   326   348\ninside   349   372

17989 GCF_002200515.1_ASM220051v1 Salmonella enterica subsp. enterica serovar Infantis Proteobacteria; Gammaproteobacteria; Enterobacterales; Enterobacteriaceae; Salmonella; Salmonella enterica; Salmonella enterica subsp. enterica MNNEETFYQAMRRKGVTRRSFLKFCSLAATSLGLGAGMTPKIAWALENKPRIPVVWIHGL WP_001058311.1 hydrogenase [Salmonella enterica] Length: 372\nNumber of predicted TMHs:  2\nExp number of AAs in TMHs: 37.4699899999999\nExp number, first 60 AAs:  19.72699\nTotal prob of N-in: 0.96171\nPOSSIBLE N-term signal sequence\ninside   1   19\nTMhelix   20   39\noutside   40   325\nTMhelix   326   348\ninside   349   372

17990 GCF_002110505.1_ASM211050v1 Salmonella enterica subsp. enterica serovar Infantis Proteobacteria; Gammaproteobacteria; Enterobacterales; Enterobacteriaceae; Salmonella; Salmonella enterica; Salmonella enterica subsp. enterica MNNEETFYQAMRRKGVTRRSFLKFCSLAATSLGLGAGMTPKIAWALENKPRIPVVWIHGL WP_001058311.1 hydrogenase [Salmonella enterica] Length: 372\nNumber of predicted TMHs:  2\nExp number of AAs in TMHs: 37.4699899999999\nExp number, first 60 AAs:  19.72699\nTotal prob of N-in: 0.96171\nPOSSIBLE N-term signal sequence\ninside   1   19\nTMhelix   20   39\noutside   40   325\nTMhelix   326   348\ninside   349   372

17991 GCF_002200805.1_ASM220080v1 Salmonella enterica subsp. enterica serovar Infantis Proteobacteria; Gammaproteobacteria; Enterobacterales; Enterobacteriaceae; Salmonella; Salmonella enterica; Salmonella enterica subsp. enterica MTGDNTLITSHGINRRDFMKLCAALAATMGLSSKAAAEMAESVSNPQRPPVIWIGAQECT WP_000145429.1 hydrogenase 2 small subunit [Salmonella enterica] Length: 372\nNumber of predicted TMHs:  1\nExp number of AAs in TMHs: 24.51309\nExp number, first 60 AAs:  0.2682\nTotal prob of N-in: 0.14402\noutside   1   331\nTMhelix   332   354\ninside   355   372

17992 GCF_002200755.1_ASM220075v1 Salmonella enterica subsp. enterica serovar Infantis Proteobacteria; Gammaproteobacteria; Enterobacterales; Enterobacteriaceae; Salmonella; Salmonella enterica; Salmonella enterica subsp. enterica MTGDNTLITSHGINRRDFMKLCAALAATMGLSSKAAAEMAESVSNPQRPPVIWIGAQECT WP_000145429.1 hydrogenase 2 small subunit [Salmonella enterica] Length: 372\nNumber of

predicted TMHs: 1\nExp number of AAs in TMHs: 24.51309\nExp number, first 60 AAs: 0.2682\nTotal prob of N-in: 0.14402\noutside 1 331\nTMhelix 332 354\ninside 355 372

17993 GCF_002200735.1_ASM220073v1 Salmonella enterica subsp. enterica serovar Infantis Proteobacteria; Gammaproteobacteria; Enterobacterales; Enterobacteriaceae; Salmonella; Salmonella enterica; Salmonella enterica subsp. enterica MTGDNTLITSHGINRRDFMKLCAALAATMGLSSKAAAEMAESVSNPQRPPVIWIGAQECT WP_000145429.1 hydrogenase 2 small subunit [Salmonella enterica] Length: 372\nNumber of predicted TMHs: 1\nExp number of AAs in TMHs: 24.51309\nExp number, first 60 AAs: 0.2682\nTotal prob of N-in: 0.14402\noutside 1 331\nTMhelix 332 354\ninside 355 372

17994 GCF_002200715.1_ASM220071v1 Salmonella enterica subsp. enterica serovar Infantis Proteobacteria; Gammaproteobacteria; Enterobacterales; Enterobacteriaceae; Salmonella; Salmonella enterica; Salmonella enterica subsp. enterica MTGDNTLITSHGINRRDFMKLCAALAATMGLSSKAAAEMAESVSNPQRPPVIWIGAQECT WP_000145429.1 hydrogenase 2 small subunit [Salmonella enterica] Length: 372\nNumber of predicted TMHs: 1\nExp number of AAs in TMHs: 24.51309\nExp number, first 60 AAs: 0.2682\nTotal prob of N-in: 0.14402\noutside 1 331\nTMhelix 332 354\ninside 355 372

17995 GCF_002200665.1_ASM220066v1 Salmonella enterica subsp. enterica serovar Infantis Proteobacteria; Gammaproteobacteria; Enterobacterales; Enterobacteriaceae; Salmonella; Salmonella enterica; Salmonella enterica subsp. enterica MNNEETFYQAMRRKGVTRRSFLKFCSLAATSLGLGAGMTPKIAWALENKPRIPVVWIHGL WP_001058311.1 hydrogenase [Salmonella enterica] Length: 372\nNumber of predicted TMHs: 2\nExp number of AAs in TMHs: 37.4699899999999\nExp number, first 60 AAs: 19.72699\nTotal prob of N-in: 0.96171\nPOSSIBLE N-term signal sequence\ninside 1 19\nTMhelix 20 39\noutside 40 325\nTMhelix 326 348\ninside 349 372

17996 GCF_002200475.1_ASM220047v1 Salmonella enterica subsp. enterica serovar Infantis Proteobacteria; Gammaproteobacteria; Enterobacterales; Enterobacteriaceae; Salmonella; Salmonella enterica; Salmonella enterica subsp. enterica MNNEETFYQAMRRKGVTRRSFLKFCSLAATSLGLGAGMTPKIAWALENKPRIPVVWIHGL WP_001058311.1 hydrogenase [Salmonella enterica] Length: 372\nNumber of predicted TMHs: 2\nExp number of AAs in TMHs: 37.4699899999999\nExp number, first 60 AAs: 19.72699\nTotal prob of N-in: 0.96171\nPOSSIBLE N-term signal sequence\ninside 1 19\nTMhelix 20 39\noutside 40 325\nTMhelix 326 348\ninside 349 372

17997 GCF_002200515.1_ASM220051v1 Salmonella enterica subsp. enterica serovar Infantis Proteobacteria; Gammaproteobacteria; Enterobacterales; Enterobacteriaceae; Salmonella; Salmonella enterica; Salmonella enterica subsp. enterica MTGDNTLITSHGINRRDFMKLCAALAATMGLSSKAAAEMAESVSNPQRPPVIWIGAQECT WP_000145429.1 hydrogenase 2 small subunit [Salmonella enterica] Length: 372\nNumber of predicted TMHs: 1\nExp number of AAs in TMHs: 24.51309\nExp number, first 60 AAs: 0.2682\nTotal prob of N-in: 0.14402\noutside 1 331\nTMhelix 332 354\ninside 355 372

17998 GCF_002110505.1_ASM211050v1 Salmonella enterica subsp. enterica serovar Infantis Proteobacteria; Gammaproteobacteria; Enterobacterales; Enterobacteriaceae; Salmonella; Salmonella enterica; Salmonella enterica subsp. enterica MTGDNTLITSHGINRRDFMKLCAALAATMGLSSKAAAEMAESVSNPQRPPVIWIGAQECT WP_000145429.1 hydrogenase 2 small subunit [Salmonella enterica] Length: 372\nNumber of predicted TMHs: 1\nExp number of AAs in TMHs: 24.51309\nExp number, first 60 AAs: 0.2682\nTotal prob of N-in: 0.14402\noutside 1 331\nTMhelix 332 354\ninside 355 372

17999 GCF_002200335.1_ASM220033v1 Salmonella enterica subsp. enterica serovar Infantis Proteobacteria; Gammaproteobacteria; Enterobacterales; Enterobacteriaceae; Salmonella; Salmonella enterica; Salmonella enterica subsp. enterica MNNEETFYQAMRRKGVTRRSFLKFCSLAATSLGLGAGMTPKIAWALENKPRIPVVWIHGL WP_001058311.1 hydrogenase [Salmonella enterica] Length: 372\nNumber of predicted TMHs: 2\nExp number of AAs in TMHs: 37.4699899999999\nExp number, first 60 AAs: 19.72699\nTotal prob of N-in: 0.96171\nPOSSIBLE N-term signal sequence\ninside 1 19\nTMhelix 20 39\noutside 40 325\nTMhelix 326 348\ninside 349 372

18000 GCF_002200395.1_ASM220039v1 Salmonella enterica subsp. enterica serovar Infantis Proteobacteria; Gammaproteobacteria; Enterobacterales; Enterobacteriaceae; Salmonella; Salmonella enterica; Salmonella enterica subsp. enterica MNNEETFYQAMRRKGVTRRSFLKFCSLAATSLGLGAGMTPKIAWALENKPRIPVVWIHGL WP_001058311.1 hydrogenase [Salmonella enterica] Length: 372\nNumber of predicted TMHs: 2\nExp number of AAs in TMHs: 37.4699899999999\nExp number, first 60 AAs: 19.72699\nTotal prob of N-in:

0.96171\nPOSSIBLE N-term signal sequence\ninside 1 19\nTMhelix 20 39\noutside 40 325\nTMhelix 326 348\ninside 349 372

18001 GCF_002200275.1_ASM220027v1 Salmonella enterica subsp. enterica serovar Infantis Proteobacteria; Gammaproteobacteria; Enterobacterales; Enterobacteriaceae; Salmonella; Salmonella enterica; Salmonella enterica subsp. enterica MNNEETFYQAMRRKGVTRRSFLKFCSLAATSLGLGAGMTPKIAWALENKPRIPVVWIHGL WP_001058311.1 hydrogenase [Salmonella enterica] Length: 372\nNumber of predicted TMHs: 2\nExp number of AAs in TMHs: 37.4699899999999\nExp number, first 60 AAs: 19.72699\nTotal prob of N-in: 0.96171\nPOSSIBLE N-term signal sequence\ninside 1 19\nTMhelix 20 39\noutside 40 325\nTMhelix 326 348\ninside 349 372

18002 GCF_002200805.1_ASM220080v1 Salmonella enterica subsp. enterica serovar Infantis Proteobacteria; Gammaproteobacteria; Enterobacterales; Enterobacteriaceae; Salmonella; Salmonella enterica; Salmonella enterica subsp. enterica MNNEETFYQAMRRKGVTRRSFLKFCSLAATSLGLGAGMTPKIAWALENKPRIPVVWIHGL WP_001058311.1 hydrogenase [Salmonella enterica] Length: 372\nNumber of predicted TMHs: 2\nExp number of AAs in TMHs: 37.4699899999999\nExp number, first 60 AAs: 19.72699\nTotal prob of N-in: 0.96171\nPOSSIBLE N-term signal sequence\ninside 1 19\nTMhelix 20 39\noutside 40 325\nTMhelix 326 348\ninside 349 372

18003 GCF_002200755.1_ASM220075v1 Salmonella enterica subsp. enterica serovar Infantis Proteobacteria; Gammaproteobacteria; Enterobacterales; Enterobacteriaceae; Salmonella; Salmonella enterica; Salmonella enterica subsp. enterica MNNEETFYQAMRRKGVTRRSFLKFCSLAATSLGLGAGMTPKIAWALENKPRIPVVWIHGL WP_001058311.1 hydrogenase [Salmonella enterica] Length: 372\nNumber of predicted TMHs: 2\nExp number of AAs in TMHs: 37.4699899999999\nExp number, first 60 AAs: 19.72699\nTotal prob of N-in: 0.96171\nPOSSIBLE N-term signal sequence\ninside 1 19\nTMhelix 20 39\noutside 40 325\nTMhelix 326 348\ninside 349 372

18004 GCF_001478585.1_Salmonella_enterica_CVM_N42237-SQ_v1.0 Salmonella enterica subsp. enterica serovar Infantis Proteobacteria; Gammaproteobacteria; Enterobacterales; Enterobacteriaceae; Salmonella; Salmonella enterica; Salmonella enterica subsp. enterica MTGDNTLITSHGINRRDFMKLCAALAATMGLSSKAAAEMAESVSNPQRPPVIWIGAQECT WP_000145429.1 hydrogenase 2 small subunit [Salmonella enterica] Length: 372\nNumber of predicted TMHs: 1\nExp number of AAs in TMHs: 24.51309\nExp number, first 60 AAs: 0.2682\nTotal prob of N-in: 0.14402\noutside 1 331\nTMhelix 332 354\ninside 355 372

18005 GCF_002059795.1_ASM205979v1 Salmonella enterica subsp. enterica serovar Infantis Proteobacteria; Gammaproteobacteria; Enterobacterales; Enterobacteriaceae; Salmonella; Salmonella enterica; Salmonella enterica subsp. enterica MTGDNTLITSHGINRRDFMKLCAALAATMGLSSKAAAEMAESVSNPQRPPVIWIGAQECT WP_000145429.1 hydrogenase 2 small subunit [Salmonella enterica] Length: 372\nNumber of predicted TMHs: 1\nExp number of AAs in TMHs: 24.51309\nExp number, first 60 AAs: 0.2682\nTotal prob of N-in: 0.14402\noutside 1 331\nTMhelix 332 354\ninside 355 372

18006 GCF_001478585.1_Salmonella_enterica_CVM_N42237-SQ_v1.0 Salmonella enterica subsp. enterica serovar Infantis Proteobacteria; Gammaproteobacteria; Enterobacterales; Enterobacteriaceae; Salmonella; Salmonella enterica; Salmonella enterica subsp. enterica MNNEETFYQAMRRKGVTRRSFLKFCSLAATSLGLGAGMTPKIAWALENKPRIPVVWIHGL WP_001058311.1 hydrogenase [Salmonella enterica] Length: 372\nNumber of predicted TMHs: 2\nExp number of AAs in TMHs: 37.4699899999999\nExp number, first 60 AAs: 19.72699\nTotal prob of N-in: 0.96171\nPOSSIBLE N-term signal sequence\ninside 1 19\nTMhelix 20 39\noutside 40 325\nTMhelix 326 348\ninside 349 372

18007 GCF_001691885.1_ASM169188v1 Salmonella enterica subsp. enterica serovar Infantis Proteobacteria; Gammaproteobacteria; Enterobacterales; Enterobacteriaceae; Salmonella; Salmonella enterica; Salmonella enterica subsp. enterica MNNEETFYQAMRRKGVTRRSFLKFCSLAATSLGLGAGMTPKIAWALENKPRIPVVWIHGL WP_001058311.1 hydrogenase [Salmonella enterica] Length: 372\nNumber of predicted TMHs: 2\nExp number of AAs in TMHs: 37.4699899999999\nExp number, first 60 AAs: 19.72699\nTotal prob of N-in: 0.96171\nPOSSIBLE N-term signal sequence\ninside 1 19\nTMhelix 20 39\noutside 40 325\nTMhelix 326 348\ninside 349 372

18008 GCF_001906525.1_ASM190652v1 Salmonella enterica subsp. enterica serovar Infantis Proteobacteria; Gammaproteobacteria; Enterobacterales; Enterobacteriaceae; Salmonella; Salmonella enterica; Salmonella enterica subsp. enterica MTGDNTLITSHGINRRDFMKLCAALAATMGLSSKAAAEMAESVSNPQRPPVIWIGAQECT WP_000145429.1 hydrogenase 2 small subunit [Salmonella enterica] Length: 372\nNumber of

predicted TMHs: 1\nExp number of AAs in TMHs: 24.51309\nExp number, first 60 AAs: 0.2682\nTotal prob of N-in: 0.14402\noutside 1 331\nTMhelix 332 354\ninside 355 372

18009 GCF_001931555.1_ASM193155v1 Salmonella enterica subsp. enterica serovar Infantis Proteobacteria; Gammaproteobacteria; Enterobacterales; Enterobacteriaceae; Salmonella; Salmonella enterica; Salmonella enterica subsp. enterica MTGDNTLITSHGINRRDFMKLCAALAATMGLSSKAAAEMAESVSNPQRPPVIWIGAQECT WP_000145429.1 hydrogenase 2 small subunit [Salmonella enterica] Length: 372\nNumber of predicted TMHs: 1\nExp number of AAs in TMHs: 24.51309\nExp number, first 60 AAs: 0.2682\nTotal prob of N-in: 0.14402\noutside 1 331\nTMhelix 332 354\ninside 355 372

18010 GCF_001931615.1_ASM193161v1 Salmonella enterica subsp. enterica serovar Infantis Proteobacteria; Gammaproteobacteria; Enterobacterales; Enterobacteriaceae; Salmonella; Salmonella enterica; Salmonella enterica subsp. enterica MTGDNTLITSHGINRRDFMKLCAALAATMGLSSKAAAEMAESVSNPQRPPVIWIGAQECT WP_000145429.1 hydrogenase 2 small subunit [Salmonella enterica] Length: 372\nNumber of predicted TMHs: 1\nExp number of AAs in TMHs: 24.51309\nExp number, first 60 AAs: 0.2682\nTotal prob of N-in: 0.14402\noutside 1 331\nTMhelix 332 354\ninside 355 372

18011 GCF_001906525.1_ASM190652v1 Salmonella enterica subsp. enterica serovar Infantis Proteobacteria; Gammaproteobacteria; Enterobacterales; Enterobacteriaceae; Salmonella; Salmonella enterica; Salmonella enterica subsp. enterica MNNEETFYQAMRRKGVTRRSFLKFCSLAATSLGLGAGMTPKIAWALENKPRIPVVWIHGL WP_001058311.1 hydrogenase [Salmonella enterica] Length: 372\nNumber of predicted TMHs: 2\nExp number of AAs in TMHs: 37.4699899999999\nExp number, first 60 AAs: 19.72699\nTotal prob of N-in: 0.96171\nPOSSIBLE N-term signal sequence\ninside 1 19\nTMhelix 20 39\noutside 40 325\nTMhelix 326 348\ninside 349 372

18012 GCF_001931615.1_ASM193161v1 Salmonella enterica subsp. enterica serovar Infantis Proteobacteria; Gammaproteobacteria; Enterobacterales; Enterobacteriaceae; Salmonella; Salmonella enterica; Salmonella enterica subsp. enterica MNNEETFYQAMRRKGVTRRSFLKFCSLAATSLGLGAGMTPKIAWALENKPRIPVVWIHGL WP_001058311.1 hydrogenase [Salmonella enterica] Length: 372\nNumber of predicted TMHs: 2\nExp number of AAs in TMHs: 37.4699899999999\nExp number, first 60 AAs: 19.72699\nTotal prob of N-in: 0.96171\nPOSSIBLE N-term signal sequence\ninside 1 19\nTMhelix 20 39\noutside 40 325\nTMhelix 326 348\ninside 349 372

18013 GCF_001997485.1_ASM199748v1 Salmonella enterica subsp. enterica serovar Infantis Proteobacteria; Gammaproteobacteria; Enterobacterales; Enterobacteriaceae; Salmonella; Salmonella enterica; Salmonella enterica subsp. enterica MNNEETFYQAMRRKGVTRRSFLKFCSLAATSLGLGAGMTPKIAWALENKPRIPVVWIHGL WP_001058311.1 hydrogenase [Salmonella enterica] Length: 372\nNumber of predicted TMHs: 2\nExp number of AAs in TMHs: 37.4699899999999\nExp number, first 60 AAs: 19.72699\nTotal prob of N-in: 0.96171\nPOSSIBLE N-term signal sequence\ninside 1 19\nTMhelix 20 39\noutside 40 325\nTMhelix 326 348\ninside 349 372

18014 GCF_001834735.1_ASM183473v1 Salmonella enterica subsp. enterica serovar Inverness Proteobacteria; Gammaproteobacteria; Enterobacterales; Enterobacteriaceae; Salmonella; Salmonella enterica; Salmonella enterica subsp. enterica MTGDNTLITSHGINRRDFMKLCAALAATMGLSSKAAAEMAESVSNPQRPPVIWIGAQECT WP_000145429.1 hydrogenase 2 small subunit [Salmonella enterica] Length: 372\nNumber of predicted TMHs: 1\nExp number of AAs in TMHs: 24.51309\nExp number, first 60 AAs: 0.2682\nTotal prob of N-in: 0.14402\noutside 1 331\nTMhelix 332 354\ninside 355 372

18015 GCF_001833785.1_ASM183378v1 Salmonella enterica subsp. enterica serovar Inverness Proteobacteria; Gammaproteobacteria; Enterobacterales; Enterobacteriaceae; Salmonella; Salmonella enterica; Salmonella enterica subsp. enterica MNNEETFYQAMRRKGVTRRSFLKFCSLAATSLGLGAGMTPKIAWALENKPRIPVVWIHGL WP_001058312.1 hydrogenase [Salmonella enterica] Length: 372\nNumber of predicted TMHs: 2\nExp number of AAs in TMHs: 36.51738\nExp number, first 60 AAs: 19.72849\nTotal prob of N-in: 0.96170\nPOSSIBLE N-term signal sequence\ninside 1 19\nTMhelix 20 39\noutside 40 325\nTMhelix 326 348\ninside 349 372

18016 GCF_002047715.1_ASM204771v1 Salmonella enterica subsp. enterica serovar Inverness Proteobacteria; Gammaproteobacteria; Enterobacterales; Enterobacteriaceae; Salmonella; Salmonella enterica; Salmonella enterica subsp. enterica MTGDNTLITSHGINRRDFMKLCAALAATMGLSSKAAAEMAESVSNPQRPPVIWIGAQECT WP_000145429.1 hydrogenase 2 small subunit [Salmonella enterica] Length: 372\nNumber of predicted TMHs: 1\nExp number of AAs in TMHs: 24.51309\nExp number, first 60 AAs: 0.2682\nTotal prob of N-in: 0.14402\noutside 1 331\nTMhelix 332 354\ninside 355 372

18017	GCF_001835225.1_ASM183522v1	Salmonella enterica subsp. enterica serovar Inverness	Proteobacteria; Gammaproteobacteria; Enterobacterales; Enterobacteriaceae; Salmonella; Salmonella enterica; Salmonella enterica subsp. enterica	MNNEETFYQAMRRKGVTRRSFLKFCSLAATSLGLGAGMTPKIAWALENKPRIPVVWIHGL	WP_001058311.1	hydrogenase [Salmonella enterica]	Length: 372\nNumber of predicted TMHs:  2\nExp number of AAs in TMHs: 37.4699899999999\nExp number, first 60 AAs:  19.72699\nTotal prob of N-in:  0.96171\nPOSSIBLE N-term signal sequence\ninside   1   19\nTMhelix   20   39\noutside   40   325\nTMhelix   326   348\ninside   349   372

18018	GCF_001834735.1_ASM183473v1	Salmonella enterica subsp. enterica serovar Inverness	Proteobacteria; Gammaproteobacteria; Enterobacterales; Enterobacteriaceae; Salmonella; Salmonella enterica; Salmonella enterica subsp. enterica	MNNEETFYQAMRRKGVTRRSFLKFCSLAATSLGLGAGMTPKIAWALENKPRIPVVWIHGL	WP_001058312.1	hydrogenase [Salmonella enterica]	Length: 372\nNumber of predicted TMHs:  2\nExp number of AAs in TMHs: 36.51738\nExp number, first 60 AAs:  19.72849\nTotal prob of N-in:   0.96170\nPOSSIBLE N-term signal sequence\ninside   1   19\nTMhelix   20   39\noutside   40   325\nTMhelix   326   348\ninside   349   372

18019	GCF_002047715.1_ASM204771v1	Salmonella enterica subsp. enterica serovar Inverness	Proteobacteria; Gammaproteobacteria; Enterobacterales; Enterobacteriaceae; Salmonella; Salmonella enterica; Salmonella enterica subsp. enterica	MNNEETFYQAMRRKGVTRRSFLKFCSLAATSLGLGAGMTPKIAWALENKPRIPVVWIHGL	WP_001058312.1	hydrogenase [Salmonella enterica]	Length: 372\nNumber of predicted TMHs:  2\nExp number of AAs in TMHs: 36.51738\nExp number, first 60 AAs:  19.72849\nTotal prob of N-in:   0.96170\nPOSSIBLE N-term signal sequence\ninside   1   19\nTMhelix   20   39\noutside   40   325\nTMhelix   326   348\ninside   349   372

18020	GCF_001835205.1_ASM183520v1	Salmonella enterica subsp. enterica serovar Inverness	Proteobacteria; Gammaproteobacteria; Enterobacterales; Enterobacteriaceae; Salmonella; Salmonella enterica; Salmonella enterica subsp. enterica	MNNEETFYQAMRRKGVTRRSFLKFCSLAATSLGLGAGMTPKIAWALENKPRIPVVWIHGL	WP_001058312.1	hydrogenase [Salmonella enterica]	Length: 372\nNumber of predicted TMHs:  2\nExp number of AAs in TMHs: 36.51738\nExp number, first 60 AAs:  19.72849\nTotal prob of N-in:   0.96170\nPOSSIBLE N-term signal sequence\ninside   1   19\nTMhelix   20   39\noutside   40   325\nTMhelix   326   348\ninside   349   372

18021	GCF_001833745.1_ASM183374v1	Salmonella enterica subsp. enterica serovar Inverness	Proteobacteria; Gammaproteobacteria; Enterobacterales; Enterobacteriaceae; Salmonella; Salmonella enterica; Salmonella enterica subsp. enterica	MNNEETFYQAMRRKGVTRRSFLKFCSLAATSLGLGAGMTPKIAWALENKPRIPVVWIHGL	WP_001058312.1	hydrogenase [Salmonella enterica]	Length: 372\nNumber of predicted TMHs:  2\nExp number of AAs in TMHs: 36.51738\nExp number, first 60 AAs:  19.72849\nTotal prob of N-in:   0.96170\nPOSSIBLE N-term signal sequence\ninside   1   19\nTMhelix   20   39\noutside   40   325\nTMhelix   326   348\ninside   349   372

18022	GCF_001834205.1_ASM183420v1	Salmonella enterica subsp. enterica serovar Inverness	Proteobacteria; Gammaproteobacteria; Enterobacterales; Enterobacteriaceae; Salmonella; Salmonella enterica; Salmonella enterica subsp. enterica	MTGDNTLITSHGINRRDFMKLCAALAATMGLSSKAAAEMAESVSNPQRPPVIWIGAQECT	WP_000145429.1	hydrogenase 2 small subunit [Salmonella enterica]	Length: 372\nNumber of predicted TMHs:  1\nExp number of AAs in TMHs: 24.51309\nExp number, first 60 AAs:  0.2682\nTotal prob of N-in:  0.14402\noutside   1   331\nTMhelix   332   354\ninside   355   372

18023	GCF_001834205.1_ASM183420v1	Salmonella enterica subsp. enterica serovar Inverness	Proteobacteria; Gammaproteobacteria; Enterobacterales; Enterobacteriaceae; Salmonella; Salmonella enterica; Salmonella enterica subsp. enterica	MNNEETFYQAMRRKGVTRRSFLKFCSLAATSLGLGAGMTPKIAWALENKPRIPVVWIHGL	WP_001058312.1	hydrogenase [Salmonella enterica]	Length: 372\nNumber of predicted TMHs:  2\nExp number of AAs in TMHs: 36.51738\nExp number, first 60 AAs:  19.72849\nTotal prob of N-in:   0.96170\nPOSSIBLE N-term signal sequence\ninside   1   19\nTMhelix   20   39\noutside   40   325\nTMhelix   326   348\ninside   349   372

18024	GCF_002049115.1_ASM204911v1	Salmonella enterica subsp. enterica serovar Inverness	Proteobacteria; Gammaproteobacteria; Enterobacterales; Enterobacteriaceae; Salmonella; Salmonella enterica; Salmonella enterica subsp. enterica	MNNEETFYQAMRRKGVTRRSFLKFCSLAATSLGLGAGMTPKIAWALENKPRIPVVWIHGL	WP_001058312.1	hydrogenase [Salmonella enterica]	Length: 372\nNumber of predicted TMHs:  2\nExp number of AAs in TMHs: 36.51738\nExp number, first 60 AAs:  19.72849\nTotal prob of N-in:   0.96170\nPOSSIBLE N-term signal sequence\ninside   1   19\nTMhelix   20   39\noutside   40   325\nTMhelix   326   348\ninside   349   372

18025	GCF_001833745.1_ASM183374v1	Salmonella enterica subsp. enterica serovar Inverness	Proteobacteria; Gammaproteobacteria; Enterobacterales; Enterobacteriaceae; Salmonella; Salmonella enterica; Salmonella enterica subsp. enterica	MTGDNTLITSHGINRRDFMKLCAALAATMGLSSKAAAEMAESVSNPQRPPVIWIGAQECT	WP_000145429.1	hydrogenase 2 small subunit [Salmonella enterica]	Length: 372\nNumber of

predicted TMHs: 1\nExp number of AAs in TMHs: 24.51309\nExp number, first 60 AAs: 0.2682\nTotal prob of N-in: 0.14402\noutside 1 331\nTMhelix 332 354\ninside 355 372

18026 GCF_002049115.1_ASM204911v1 Salmonella enterica subsp. enterica serovar Inverness Proteobacteria; Gammaproteobacteria; Enterobacterales; Enterobacteriaceae; Salmonella; Salmonella enterica; Salmonella enterica subsp. enterica MTGDNTLITSHGINRRDFMKLCAALAATMGLSSKAAAEMAESVSNPQRPPVIWIGAQECT WP_000145429.1 hydrogenase 2 small subunit [Salmonella enterica] Length: 372\nNumber of predicted TMHs: 1\nExp number of AAs in TMHs: 24.51309\nExp number, first 60 AAs: 0.2682\nTotal prob of N-in: 0.14402\noutside 1 331\nTMhelix 332 354\ninside 355 372

18027 GCF_002047815.1_ASM204781v1 Salmonella enterica subsp. enterica serovar Inverness Proteobacteria; Gammaproteobacteria; Enterobacterales; Enterobacteriaceae; Salmonella; Salmonella enterica; Salmonella enterica subsp. enterica MTGDNTLITSHGINRRDFMKLCAALAATMGLSSKAAAEMAESVSNPQRPPVIWIGAQECT WP_000145429.1 hydrogenase 2 small subunit [Salmonella enterica] Length: 372\nNumber of predicted TMHs: 1\nExp number of AAs in TMHs: 24.51309\nExp number, first 60 AAs: 0.2682\nTotal prob of N-in: 0.14402\noutside 1 331\nTMhelix 332 354\ninside 355 372

18028 GCF_001835175.1_ASM183517v1 Salmonella enterica subsp. enterica serovar Inverness Proteobacteria; Gammaproteobacteria; Enterobacterales; Enterobacteriaceae; Salmonella; Salmonella enterica; Salmonella enterica subsp. enterica MTGDNTLITSHGINRRDFMKLCAALAATMGLSSKAAAEMAESVSNPQRPPVIWIGAQECT WP_000145429.1 hydrogenase 2 small subunit [Salmonella enterica] Length: 372\nNumber of predicted TMHs: 1\nExp number of AAs in TMHs: 24.51309\nExp number, first 60 AAs: 0.2682\nTotal prob of N-in: 0.14402\noutside 1 331\nTMhelix 332 354\ninside 355 372

18029 GCF_001835225.1_ASM183522v1 Salmonella enterica subsp. enterica serovar Inverness Proteobacteria; Gammaproteobacteria; Enterobacterales; Enterobacteriaceae; Salmonella; Salmonella enterica; Salmonella enterica subsp. enterica MTGDNTLITSHGINRRDFMKLCAALAATMGLSSKAAAEMAESVSNPQRPPVIWIGAQECT WP_000145429.1 hydrogenase 2 small subunit [Salmonella enterica] Length: 372\nNumber of predicted TMHs: 1\nExp number of AAs in TMHs: 24.51309\nExp number, first 60 AAs: 0.2682\nTotal prob of N-in: 0.14402\noutside 1 331\nTMhelix 332 354\ninside 355 372

18030 GCF_001833785.1_ASM183378v1 Salmonella enterica subsp. enterica serovar Inverness Proteobacteria; Gammaproteobacteria; Enterobacterales; Enterobacteriaceae; Salmonella; Salmonella enterica; Salmonella enterica subsp. enterica MTGDNTLITSHGINRRDFMKLCAALAATMGLSSKAAAEMAESVSNPQRPPVIWIGAQECT WP_000145429.1 hydrogenase 2 small subunit [Salmonella enterica] Length: 372\nNumber of predicted TMHs: 1\nExp number of AAs in TMHs: 24.51309\nExp number, first 60 AAs: 0.2682\nTotal prob of N-in: 0.14402\noutside 1 331\nTMhelix 332 354\ninside 355 372

18031 GCF_001835175.1_ASM183517v1 Salmonella enterica subsp. enterica serovar Inverness Proteobacteria; Gammaproteobacteria; Enterobacterales; Enterobacteriaceae; Salmonella; Salmonella enterica; Salmonella enterica subsp. enterica MNNEETFYQAMRRKGVTRRSFLKFCSLAATSLGLGAGMTPKIAWALENKPRIPVVWIHGL WP_001058312.1 hydrogenase [Salmonella enterica] Length: 372\nNumber of predicted TMHs: 2\nExp number of AAs in TMHs: 36.51738\nExp number, first 60 AAs: 19.72849\nTotal prob of N-in: 0.96170\nPOSSIBLE N-term signal sequence\ninside 1 19\nTMhelix 20 39\noutside 40 325\nTMhelix 326 348\ninside 349 372

18032 GCF_002047815.1_ASM204781v1 Salmonella enterica subsp. enterica serovar Inverness Proteobacteria; Gammaproteobacteria; Enterobacterales; Enterobacteriaceae; Salmonella; Salmonella enterica; Salmonella enterica subsp. enterica MNNEETFYQAMRRKGVTRRSFLKFCSLAATSLGLGAGMTPKIAWALENKPRIPVVWIHGL WP_001058312.1 hydrogenase [Salmonella enterica] Length: 372\nNumber of predicted TMHs: 2\nExp number of AAs in TMHs: 36.51738\nExp number, first 60 AAs: 19.72849\nTotal prob of N-in: 0.96170\nPOSSIBLE N-term signal sequence\ninside 1 19\nTMhelix 20 39\noutside 40 325\nTMhelix 326 348\ninside 349 372

18033 GCF_001835205.1_ASM183520v1 Salmonella enterica subsp. enterica serovar Inverness Proteobacteria; Gammaproteobacteria; Enterobacterales; Enterobacteriaceae; Salmonella; Salmonella enterica; Salmonella enterica subsp. enterica MTGDNTLITSHGINRRDFMKLCAALAATMGLSSKAAAEMAESVSNPQRPPVIWIGAQECT WP_000145429.1 hydrogenase 2 small subunit [Salmonella enterica] Length: 372\nNumber of predicted TMHs: 1\nExp number of AAs in TMHs: 24.51309\nExp number, first 60 AAs: 0.2682\nTotal prob of N-in: 0.14402\noutside 1 331\nTMhelix 332 354\ninside 355 372

18034 GCF_002047515.1_ASM204751v1 Salmonella enterica subsp. enterica serovar Inverness Proteobacteria; Gammaproteobacteria; Enterobacterales; Enterobacteriaceae; Salmonella; Salmonella enterica;

Salmonella enterica subsp. enterica MNNEETFYQAMRRKGVTRRSFLKFCSLAATSLGLGAGMTPKIAWALENKPRIPVVWIHGL WP_001058312.1 hydrogenase [Salmonella enterica] Length: 372\nNumber of predicted TMHs: 2\nExp number of AAs in TMHs: 36.51738\nExp number, first 60 AAs: 19.72849\nTotal prob of N-in: 0.96170\nPOSSIBLE N-term signal sequence\ninside 1 19\nTMhelix 20 39\noutside 40 325\nTMhelix 326 348\ninside 349 372

18035 GCF_002034735.1_ASM203473v1 Salmonella enterica subsp. enterica serovar Inverness Proteobacteria; Gammaproteobacteria; Enterobacterales; Enterobacteriaceae; Salmonella; Salmonella enterica; Salmonella enterica subsp. enterica MNNEETFYQAMRRKGVTRRSFLKFCSLAATSLGLGAGMTPKIAWALENKPRIPVVWIHGL WP_001058312.1 hydrogenase [Salmonella enterica] Length: 372\nNumber of predicted TMHs: 2\nExp number of AAs in TMHs: 36.51738\nExp number, first 60 AAs: 19.72849\nTotal prob of N-in: 0.96170\nPOSSIBLE N-term signal sequence\ninside 1 19\nTMhelix 20 39\noutside 40 325\nTMhelix 326 348\ninside 349 372

18036 GCF_002034735.1_ASM203473v1 Salmonella enterica subsp. enterica serovar Inverness Proteobacteria; Gammaproteobacteria; Enterobacterales; Enterobacteriaceae; Salmonella; Salmonella enterica; Salmonella enterica subsp. enterica MTGDNTLITSHGINRRDFMKLCAALAATMGLSSKAAAEMAESVSNPQRPPVIWIGAQECT WP_000145429.1 hydrogenase 2 small subunit [Salmonella enterica] Length: 372\nNumber of predicted TMHs: 1\nExp number of AAs in TMHs: 24.51309\nExp number, first 60 AAs: 0.2682\nTotal prob of N-in: 0.14402\noutside 1 331\nTMhelix 332 354\ninside 355 372

18037 GCF_002047515.1_ASM204751v1 Salmonella enterica subsp. enterica serovar Inverness Proteobacteria; Gammaproteobacteria; Enterobacterales; Enterobacteriaceae; Salmonella; Salmonella enterica; Salmonella enterica subsp. enterica MTGDNTLITSHGINRRDFMKLCAALAATMGLSSKAAAEMAESVSNPQRPPVIWIGAQECT WP_000145429.1 hydrogenase 2 small subunit [Salmonella enterica] Length: 372\nNumber of predicted TMHs: 1\nExp number of AAs in TMHs: 24.51309\nExp number, first 60 AAs: 0.2682\nTotal prob of N-in: 0.14402\noutside 1 331\nTMhelix 332 354\ninside 355 372

18038 GCF_002064825.1_ASM206482v1 Salmonella enterica subsp. enterica serovar Isangi Proteobacteria; Gammaproteobacteria; Enterobacterales; Enterobacteriaceae; Salmonella; Salmonella enterica; Salmonella enterica subsp. enterica MNNEETFYQAMRRKGVTRRSFLKFCSLAATSLGLGAGMTPKIAWALENKPRIPVVWIHGL WP_001058310.1 hydrogenase [Salmonella enterica] Length: 372\nNumber of predicted TMHs: 2\nExp number of AAs in TMHs: 37.4567299999999\nExp number, first 60 AAs: 19.72694\nTotal prob of N-in: 0.96171\nPOSSIBLE N-term signal sequence\ninside 1 19\nTMhelix 20 39\noutside 40 325\nTMhelix 326 348\ninside 349 372

18039 GCF_002064825.1_ASM206482v1 Salmonella enterica subsp. enterica serovar Isangi Proteobacteria; Gammaproteobacteria; Enterobacterales; Enterobacteriaceae; Salmonella; Salmonella enterica; Salmonella enterica subsp. enterica MTGDNTLITSHGINRRDFMKLCAALAATMGLSSKAAAEMAESVSNPQRPPVIWIGAQECT WP_000145429.1 hydrogenase 2 small subunit [Salmonella enterica] Length: 372\nNumber of predicted TMHs: 1\nExp number of AAs in TMHs: 24.51309\nExp number, first 60 AAs: 0.2682\nTotal prob of N-in: 0.14402\noutside 1 331\nTMhelix 332 354\ninside 355 372

18040 GCF_002064795.1_ASM206479v1 Salmonella enterica subsp. enterica serovar Istanbul Proteobacteria; Gammaproteobacteria; Enterobacterales; Enterobacteriaceae; Salmonella; Salmonella enterica; Salmonella enterica subsp. enterica MTGDNTLITSHGINRRDFMKLCAALAATMGLSSKAAAEMAESVSNPQRPPVIWIGAQECT WP_000145429.1 hydrogenase 2 small subunit [Salmonella enterica] Length: 372\nNumber of predicted TMHs: 1\nExp number of AAs in TMHs: 24.51309\nExp number, first 60 AAs: 0.2682\nTotal prob of N-in: 0.14402\noutside 1 331\nTMhelix 332 354\ninside 355 372

18041 GCF_002064795.1_ASM206479v1 Salmonella enterica subsp. enterica serovar Istanbul Proteobacteria; Gammaproteobacteria; Enterobacterales; Enterobacteriaceae; Salmonella; Salmonella enterica; Salmonella enterica subsp. enterica MNNEETFYQAMRRKGVTRRSFLKFCSLAATSLGLGAGMTPKIAWALENKPRIPVVWIHGL WP_001058308.1 hydrogenase [Salmonella enterica] Length: 372\nNumber of predicted TMHs: 2\nExp number of AAs in TMHs: 39.08204\nExp number, first 60 AAs: 19.72474\nTotal prob of N-in: 0.96174\nPOSSIBLE N-term signal sequence\ninside 1 19\nTMhelix 20 39\noutside 40 325\nTMhelix 326 348\ninside 349 372

18042 GCF_002064815.1_ASM206481v1 Salmonella enterica subsp. enterica serovar Itami Proteobacteria; Gammaproteobacteria; Enterobacterales; Enterobacteriaceae; Salmonella; Salmonella enterica; Salmonella enterica subsp. enterica MTGDNTLITSHGINRRDFMKLCAALAATMGLSSKAAAEMAESVSNPQRPPVIWIGAQECT WP_000145429.1 hydrogenase 2 small subunit [Salmonella enterica] Length: 372\nNumber of predicted TMHs: 1\nExp number of AAs in TMHs: 24.51309\nExp number, first 60 AAs: 0.2682\nTotal prob of N-in: 0.14402\noutside 1 331\nTMhelix 332 354\ninside 355 372

18043 GCF_002064815.1_ASM206481v1 Salmonella enterica subsp. enterica serovar Itami Proteobacteria; Gammaproteobacteria; Enterobacterales; Enterobacteriaceae; Salmonella; Salmonella enterica; Salmonella enterica subsp. enterica MNNEETFYQAMRRKGVTRRSFLKFCSLAATSLGLGAGMTPKIAWALENKPRIPVVWIHGL WP_001058312.1 hydrogenase [Salmonella enterica] Length: 372\nNumber of predicted TMHs: 2\nExp number of AAs in TMHs: 36.51738\nExp number, first 60 AAs: 19.72849\nTotal prob of N-in: 0.96170\nPOSSIBLE N-term signal sequence\ninside 1 19\nTMhelix 20 39\noutside 40 325\nTMhelix 326 348\ninside 349 372

18044 GCF_002061385.1_ASM206138v1 Salmonella enterica subsp. enterica serovar Javiana Proteobacteria; Gammaproteobacteria; Enterobacterales; Enterobacteriaceae; Salmonella; Salmonella enterica; Salmonella enterica subsp. enterica MTGDNTLITSHGINRRDFMKLCAALAATMGLSSKAAAEMAESVSNPQRPPVIWIGAQECT WP_000145429.1 hydrogenase 2 small subunit [Salmonella enterica] Length: 372\nNumber of predicted TMHs: 1\nExp number of AAs in TMHs: 24.51309\nExp number, first 60 AAs: 0.2682\nTotal prob of N-in: 0.14402\noutside 1 331\nTMhelix 332 354\ninside 355 372

18045 GCF_002061385.1_ASM206138v1 Salmonella enterica subsp. enterica serovar Javiana Proteobacteria; Gammaproteobacteria; Enterobacterales; Enterobacteriaceae; Salmonella; Salmonella enterica; Salmonella enterica subsp. enterica MNNEETFYQAMRRKGVTRRSFLKFCSLAATSLGLGAGMTPKIAWALENKPRIPVVWIHGL WP_001058312.1 hydrogenase [Salmonella enterica] Length: 372\nNumber of predicted TMHs: 2\nExp number of AAs in TMHs: 36.51738\nExp number, first 60 AAs: 19.72849\nTotal prob of N-in: 0.96170\nPOSSIBLE N-term signal sequence\ninside 1 19\nTMhelix 20 39\noutside 40 325\nTMhelix 326 348\ninside 349 372

18046 GCF_002063095.1_ASM206309v1 Salmonella enterica subsp. enterica serovar Javiana Proteobacteria; Gammaproteobacteria; Enterobacterales; Enterobacteriaceae; Salmonella; Salmonella enterica; Salmonella enterica subsp. enterica MTGDNTLITSHGINRRDFMKLCAALAATMGLSSKAAAEMAESVSNPQRPPVIWIGAQECT WP_000145429.1 hydrogenase 2 small subunit [Salmonella enterica] Length: 372\nNumber of predicted TMHs: 1\nExp number of AAs in TMHs: 24.51309\nExp number, first 60 AAs: 0.2682\nTotal prob of N-in: 0.14402\noutside 1 331\nTMhelix 332 354\ninside 355 372

18047 GCF_002059825.1_ASM205982v1 Salmonella enterica subsp. enterica serovar Javiana Proteobacteria; Gammaproteobacteria; Enterobacterales; Enterobacteriaceae; Salmonella; Salmonella enterica; Salmonella enterica subsp. enterica MTGDNTLITSHGINRRDFMKLCAALAATMGLSSKAAAEMAESVSNPQRPPVIWIGAQECT WP_000145429.1 hydrogenase 2 small subunit [Salmonella enterica] Length: 372\nNumber of predicted TMHs: 1\nExp number of AAs in TMHs: 24.51309\nExp number, first 60 AAs: 0.2682\nTotal prob of N-in: 0.14402\noutside 1 331\nTMhelix 332 354\ninside 355 372

18048 GCF_002059295.1_ASM205929v1 Salmonella enterica subsp. enterica serovar Javiana Proteobacteria; Gammaproteobacteria; Enterobacterales; Enterobacteriaceae; Salmonella; Salmonella enterica; Salmonella enterica subsp. enterica MTGDNTLITSHGINRRDFMKLCAALAATMGLSSKAAAEMAESVSNPQRPPVIWIGAQECT WP_000145429.1 hydrogenase 2 small subunit [Salmonella enterica] Length: 372\nNumber of predicted TMHs: 1\nExp number of AAs in TMHs: 24.51309\nExp number, first 60 AAs: 0.2682\nTotal prob of N-in: 0.14402\noutside 1 331\nTMhelix 332 354\ninside 355 372

18049 GCF_002064765.1_ASM206476v1 Salmonella enterica subsp. enterica serovar Javiana Proteobacteria; Gammaproteobacteria; Enterobacterales; Enterobacteriaceae; Salmonella; Salmonella enterica; Salmonella enterica subsp. enterica MNNEETFYQAMRRKGVTRRSFLKFCSLAATSLGLGAGMTPKIAWALENKPRIPVVWIHGL WP_001058312.1 hydrogenase [Salmonella enterica] Length: 372\nNumber of predicted TMHs: 2\nExp number of AAs in TMHs: 36.51738\nExp number, first 60 AAs: 19.72849\nTotal prob of N-in: 0.96170\nPOSSIBLE N-term signal sequence\ninside 1 19\nTMhelix 20 39\noutside 40 325\nTMhelix 326 348\ninside 349 372

18050 GCF_002064765.1_ASM206476v1 Salmonella enterica subsp. enterica serovar Javiana Proteobacteria; Gammaproteobacteria; Enterobacterales; Enterobacteriaceae; Salmonella; Salmonella enterica; Salmonella enterica subsp. enterica MTGDNTLITSHGINRRDFMKLCAALAATMGLSSKAAAEMAESVSNPQRPPVIWIGAQECT WP_000145431.1 hydrogenase 2 small subunit [Salmonella enterica] Length: 372\nNumber of predicted TMHs: 1\nExp number of AAs in TMHs: 24.51505\nExp number, first 60 AAs: 0.2682\nTotal prob of N-in: 0.14406\noutside 1 331\nTMhelix 332 354\ninside 355 372

18051 GCF_002034745.1_ASM203474v1 Salmonella enterica subsp. enterica serovar Javiana Proteobacteria; Gammaproteobacteria; Enterobacterales; Enterobacteriaceae; Salmonella; Salmonella enterica; Salmonella enterica subsp. enterica MTGDNTLITSHGINRRDFMKLCAALAATMGLSSKAAAEMAESVSNPQRPPVIWIGAQECT WP_000145429.1 hydrogenase 2 small subunit [Salmonella enterica] Length: 372\nNumber of

predicted TMHs: 1\nExp number of AAs in TMHs: 24.51309\nExp number, first 60 AAs: 0.2682\nTotal prob of N-in: 0.14402\noutside 1 331\nTMhelix 332 354\ninside 355 372

18052 GCF_002034745.1_ASM203474v1 Salmonella enterica subsp. enterica serovar Javiana Proteobacteria; Gammaproteobacteria; Enterobacterales; Enterobacteriaceae; Salmonella; Salmonella enterica; Salmonella enterica subsp. enterica MNNEETFYQAMRRKGVTRRSFLKFCSLAATSLGLGAGMTPKIAWALENKPRIPVVWIHGL WP_001058312.1 hydrogenase [Salmonella enterica] Length: 372\nNumber of predicted TMHs: 2\nExp number of AAs in TMHs: 36.51738\nExp number, first 60 AAs: 19.72849\nTotal prob of N-in: 0.96170\nPOSSIBLE N-term signal sequence\ninside 1 19\nTMhelix 20 39\noutside 40 325\nTMhelix 326 348\ninside 349 372

18053 GCF_002059295.1_ASM205929v1 Salmonella enterica subsp. enterica serovar Javiana Proteobacteria; Gammaproteobacteria; Enterobacterales; Enterobacteriaceae; Salmonella; Salmonella enterica; Salmonella enterica subsp. enterica MNNEETFYQAMRRKGVTRRSFLKFCSLAATSLGLGAGMTPKIAWALENKPRIPVVWIHGL WP_001058312.1 hydrogenase [Salmonella enterica] Length: 372\nNumber of predicted TMHs: 2\nExp number of AAs in TMHs: 36.51738\nExp number, first 60 AAs: 19.72849\nTotal prob of N-in: 0.96170\nPOSSIBLE N-term signal sequence\ninside 1 19\nTMhelix 20 39\noutside 40 325\nTMhelix 326 348\ninside 349 372

18054 GCF_002059385.1_ASM205938v1 Salmonella enterica subsp. enterica serovar Javiana Proteobacteria; Gammaproteobacteria; Enterobacterales; Enterobacteriaceae; Salmonella; Salmonella enterica; Salmonella enterica subsp. enterica MTGDNTLITSHGINRRDFMKLCAALAATMGLSSKAAAEMAESVSNPQRPPVIWIGAQECT WP_000145429.1 hydrogenase 2 small subunit [Salmonella enterica] Length: 372\nNumber of predicted TMHs: 1\nExp number of AAs in TMHs: 24.51309\nExp number, first 60 AAs: 0.2682\nTotal prob of N-in: 0.14402\noutside 1 331\nTMhelix 332 354\ninside 355 372

18055 GCF_002059665.1_ASM205966v1 Salmonella enterica subsp. enterica serovar Javiana Proteobacteria; Gammaproteobacteria; Enterobacterales; Enterobacteriaceae; Salmonella; Salmonella enterica; Salmonella enterica subsp. enterica MTGDNTLITSHGINRRDFMKLCAALAATMGLSSKAAAEMAESVSNPQRPPVIWIGAQECT WP_000145429.1 hydrogenase 2 small subunit [Salmonella enterica] Length: 372\nNumber of predicted TMHs: 1\nExp number of AAs in TMHs: 24.51309\nExp number, first 60 AAs: 0.2682\nTotal prob of N-in: 0.14402\noutside 1 331\nTMhelix 332 354\ninside 355 372

18056 GCF_001938885.1_ASM193888v1 Salmonella enterica subsp. enterica serovar Javiana Proteobacteria; Gammaproteobacteria; Enterobacterales; Enterobacteriaceae; Salmonella; Salmonella enterica; Salmonella enterica subsp. enterica MNNEETFYQAMRRKGVTRRSFLKFCSLAATSLGLGAGMTPKIAWALENKPRIPVVWIHGL WP_001058312.1 hydrogenase [Salmonella enterica] Length: 372\nNumber of predicted TMHs: 2\nExp number of AAs in TMHs: 36.51738\nExp number, first 60 AAs: 19.72849\nTotal prob of N-in: 0.96170\nPOSSIBLE N-term signal sequence\ninside 1 19\nTMhelix 20 39\noutside 40 325\nTMhelix 326 348\ninside 349 372

18057 GCF_001938885.1_ASM193888v1 Salmonella enterica subsp. enterica serovar Javiana Proteobacteria; Gammaproteobacteria; Enterobacterales; Enterobacteriaceae; Salmonella; Salmonella enterica; Salmonella enterica subsp. enterica MTGDNTLITSHGINRRDFMKLCAALAATMGLSSKAAAEMAESVSNPQRPPVIWIGAQECT WP_000145429.1 hydrogenase 2 small subunit [Salmonella enterica] Length: 372\nNumber of predicted TMHs: 1\nExp number of AAs in TMHs: 24.51309\nExp number, first 60 AAs: 0.2682\nTotal prob of N-in: 0.14402\noutside 1 331\nTMhelix 332 354\ninside 355 372

18058 GCF_002063095.1_ASM206309v1 Salmonella enterica subsp. enterica serovar Javiana Proteobacteria; Gammaproteobacteria; Enterobacterales; Enterobacteriaceae; Salmonella; Salmonella enterica; Salmonella enterica subsp. enterica MNNEETFYQAMRRKGVTRRSFLKFCSLAATSLGLGAGMTPKIAWALENKPRIPVVWIHGL WP_001058312.1 hydrogenase [Salmonella enterica] Length: 372\nNumber of predicted TMHs: 2\nExp number of AAs in TMHs: 36.51738\nExp number, first 60 AAs: 19.72849\nTotal prob of N-in: 0.96170\nPOSSIBLE N-term signal sequence\ninside 1 19\nTMhelix 20 39\noutside 40 325\nTMhelix 326 348\ninside 349 372

18059 GCF_002061335.1_ASM206133v1 Salmonella enterica subsp. enterica serovar Javiana Proteobacteria; Gammaproteobacteria; Enterobacterales; Enterobacteriaceae; Salmonella; Salmonella enterica; Salmonella enterica subsp. enterica MNNEETFYQAMRRKGVTRRSFLKFCSLAATSLGLGAGMTPKIAWALENKPRIPVVWIHGL WP_001058312.1 hydrogenase [Salmonella enterica] Length: 372\nNumber of predicted TMHs: 2\nExp number of AAs in TMHs: 36.51738\nExp number, first 60 AAs: 19.72849\nTotal prob of N-in: 0.96170\nPOSSIBLE N-term signal sequence\ninside 1 19\nTMhelix 20 39\noutside 40 325\nTMhelix 326 348\ninside 349 372

18060 GCF_002060175.1_ASM206017v1 Salmonella enterica subsp. enterica serovar Javiana Proteobacteria; Gammaproteobacteria; Enterobacterales; Enterobacteriaceae; Salmonella; Salmonella enterica;

Salmonella enterica subsp. enterica MTGDNTLITSHGINRRDFMKLCAALAATMGLSSKAAAEMAESVSNPQRPPVIWIGAQECT WP_000145429.1 hydrogenase 2 small subunit [Salmonella enterica] Length: 372\nNumber of predicted TMHs: 1\nExp number of AAs in TMHs: 24.51309\nExp number, first 60 AAs: 0.2682\nTotal prob of N-in: 0.14402\noutside 1 331\nTMhelix 332 354\ninside 355 372

18061 GCF_001938795.1_ASM193879v1 Salmonella enterica subsp. enterica serovar Javiana Proteobacteria; Gammaproteobacteria; Enterobacterales; Enterobacteriaceae; Salmonella; Salmonella enterica; Salmonella enterica subsp. enterica MTGDNTLITSHGINRRDFMKLCAALAATMGLSSKAAAEMAESVSNPQRPPVIWIGAQECT WP_000145429.1 hydrogenase 2 small subunit [Salmonella enterica] Length: 372\nNumber of predicted TMHs: 1\nExp number of AAs in TMHs: 24.51309\nExp number, first 60 AAs: 0.2682\nTotal prob of N-in: 0.14402\noutside 1 331\nTMhelix 332 354\ninside 355 372

18062 GCF_001478595.1_Salmonella_enterica_CVM_N42337-SQ_v1.0 Salmonella enterica subsp. enterica serovar Javiana Proteobacteria; Gammaproteobacteria; Enterobacterales; Enterobacteriaceae; Salmonella; Salmonella enterica; Salmonella enterica subsp. enterica MTGDNTLITSHGINRRDFMKLCAALAATMGLSSKAAAEMAESVSNPQRPPVIWIGAQECT WP_000145429.1 hydrogenase 2 small subunit [Salmonella enterica] Length: 372\nNumber of predicted TMHs: 1\nExp number of AAs in TMHs: 24.51309\nExp number, first 60 AAs: 0.2682\nTotal prob of N-in: 0.14402\noutside 1 331\nTMhelix 332 354\ninside 355 372

18063 GCF_001938925.1_ASM193892v1 Salmonella enterica subsp. enterica serovar Javiana Proteobacteria; Gammaproteobacteria; Enterobacterales; Enterobacteriaceae; Salmonella; Salmonella enterica; Salmonella enterica subsp. enterica MNNEETFYQAMRRKGVTRRSFLKFCSLAATSLGLGAGMTPKIAWALENKPRIPVVWIHGL WP_001058312.1 hydrogenase [Salmonella enterica] Length: 372\nNumber of predicted TMHs: 2\nExp number of AAs in TMHs: 36.51738\nExp number, first 60 AAs: 19.72849\nTotal prob of N-in: 0.96170\nPOSSIBLE N-term signal sequence\ninside 1 19\nTMhelix 20 39\noutside 40 325\nTMhelix 326 348\ninside 349 372

18064 GCF_001938865.1_ASM193886v1 Salmonella enterica subsp. enterica serovar Javiana Proteobacteria; Gammaproteobacteria; Enterobacterales; Enterobacteriaceae; Salmonella; Salmonella enterica; Salmonella enterica subsp. enterica MTGDNTLITSHGINRRDFMKLCAALAATMGLSSKAAAEMAESVSNPQRPPVIWIGAQECT WP_000145429.1 hydrogenase 2 small subunit [Salmonella enterica] Length: 372\nNumber of predicted TMHs: 1\nExp number of AAs in TMHs: 24.51309\nExp number, first 60 AAs: 0.2682\nTotal prob of N-in: 0.14402\noutside 1 331\nTMhelix 332 354\ninside 355 372

18065 GCF_001938865.1_ASM193886v1 Salmonella enterica subsp. enterica serovar Javiana Proteobacteria; Gammaproteobacteria; Enterobacterales; Enterobacteriaceae; Salmonella; Salmonella enterica; Salmonella enterica subsp. enterica MNNEETFYQAMRRKGVTRRSFLKFCSLAATSLGLGAGMTPKIAWALENKPRIPVVWIHGL WP_001058312.1 hydrogenase [Salmonella enterica] Length: 372\nNumber of predicted TMHs: 2\nExp number of AAs in TMHs: 36.51738\nExp number, first 60 AAs: 19.72849\nTotal prob of N-in: 0.96170\nPOSSIBLE N-term signal sequence\ninside 1 19\nTMhelix 20 39\noutside 40 325\nTMhelix 326 348\ninside 349 372

18066 GCF_001938925.1_ASM193892v1 Salmonella enterica subsp. enterica serovar Javiana Proteobacteria; Gammaproteobacteria; Enterobacterales; Enterobacteriaceae; Salmonella; Salmonella enterica; Salmonella enterica subsp. enterica MTGDNTLITSHGINRRDFMKLCAALAATMGLSSKAAAEMAESVSNPQRPPVIWIGAQECT WP_000145429.1 hydrogenase 2 small subunit [Salmonella enterica] Length: 372\nNumber of predicted TMHs: 1\nExp number of AAs in TMHs: 24.51309\nExp number, first 60 AAs: 0.2682\nTotal prob of N-in: 0.14402\noutside 1 331\nTMhelix 332 354\ninside 355 372

18067 GCF_001938855.1_ASM193885v1 Salmonella enterica subsp. enterica serovar Javiana Proteobacteria; Gammaproteobacteria; Enterobacterales; Enterobacteriaceae; Salmonella; Salmonella enterica; Salmonella enterica subsp. enterica MTGDNTLITSHGINRRDFMKLCAALAATMGLSSKAAAEMAESVSNPQRPPVIWIGAQECT WP_000145429.1 hydrogenase 2 small subunit [Salmonella enterica] Length: 372\nNumber of predicted TMHs: 1\nExp number of AAs in TMHs: 24.51309\nExp number, first 60 AAs: 0.2682\nTotal prob of N-in: 0.14402\noutside 1 331\nTMhelix 332 354\ninside 355 372

18068 GCF_002059825.1_ASM205982v1 Salmonella enterica subsp. enterica serovar Javiana Proteobacteria; Gammaproteobacteria; Enterobacterales; Enterobacteriaceae; Salmonella; Salmonella enterica; Salmonella enterica subsp. enterica MNNEETFYQAMRRKGVTRRSFLKFCSLAATSLGLGAGMTPKIAWALENKPRIPVVWIHGL WP_001058312.1 hydrogenase [Salmonella enterica] Length: 372\nNumber of predicted TMHs: 2\nExp number of AAs in TMHs: 36.51738\nExp number, first 60 AAs: 19.72849\nTotal prob of N-in: 0.96170\nPOSSIBLE N-term signal sequence\ninside 1 19\nTMhelix 20 39\noutside 40 325\nTMhelix 326 348\ninside 349 372

18069 GCF_002061445.1_ASM206144v1 Salmonella enterica subsp. enterica serovar Javiana Proteobacteria; Gammaproteobacteria; Enterobacterales; Enterobacteriaceae; Salmonella; Salmonella enterica; Salmonella enterica subsp. enterica MTGDNTLITSHGINRRDFMKLCAALAATMGLSSKAAAEMAESVSNPQRPPVIWIGAQECT WP_000145429.1 hydrogenase 2 small subunit [Salmonella enterica] Length: 372\nNumber of predicted TMHs: 1\nExp number of AAs in TMHs: 24.51309\nExp number, first 60 AAs: 0.2682\nTotal prob of N-in: 0.14402\noutside 1 331\nTMhelix 332 354\ninside 355 372

18070 GCF_002061445.1_ASM206144v1 Salmonella enterica subsp. enterica serovar Javiana Proteobacteria; Gammaproteobacteria; Enterobacterales; Enterobacteriaceae; Salmonella; Salmonella enterica; Salmonella enterica subsp. enterica MNNEETFYQAMRRKGVTRRSFLKFCSLAATSLGLGAGMTPKIAWALENKPRIPVVWIHGL WP_001058312.1 hydrogenase [Salmonella enterica] Length: 372\nNumber of predicted TMHs: 2\nExp number of AAs in TMHs: 36.51738\nExp number, first 60 AAs: 19.72849\nTotal prob of N-in: 0.96170\nPOSSIBLE N-term signal sequence\ninside 1 19\nTMhelix 20 39\noutside 40 325\nTMhelix 326 348\ninside 349 372

18071 GCF_002059385.1_ASM205938v1 Salmonella enterica subsp. enterica serovar Javiana Proteobacteria; Gammaproteobacteria; Enterobacterales; Enterobacteriaceae; Salmonella; Salmonella enterica; Salmonella enterica subsp. enterica MNNEETFYQAMRRKGVTRRSFLKFCSLAATSLGLGAGMTPKIAWALENKPRIPVVWIHGL WP_001058312.1 hydrogenase [Salmonella enterica] Length: 372\nNumber of predicted TMHs: 2\nExp number of AAs in TMHs: 36.51738\nExp number, first 60 AAs: 19.72849\nTotal prob of N-in: 0.96170\nPOSSIBLE N-term signal sequence\ninside 1 19\nTMhelix 20 39\noutside 40 325\nTMhelix 326 348\ninside 349 372

18072 GCF_001938855.1_ASM193885v1 Salmonella enterica subsp. enterica serovar Javiana Proteobacteria; Gammaproteobacteria; Enterobacterales; Enterobacteriaceae; Salmonella; Salmonella enterica; Salmonella enterica subsp. enterica MNNEETFYQAMRRKGVTRRSFLKFCSLAATSLGLGAGMTPKIAWALENKPRIPVVWIHGL WP_001058312.1 hydrogenase [Salmonella enterica] Length: 372\nNumber of predicted TMHs: 2\nExp number of AAs in TMHs: 36.51738\nExp number, first 60 AAs: 19.72849\nTotal prob of N-in: 0.96170\nPOSSIBLE N-term signal sequence\ninside 1 19\nTMhelix 20 39\noutside 40 325\nTMhelix 326 348\ninside 349 372

18073 GCF_002059665.1_ASM205966v1 Salmonella enterica subsp. enterica serovar Javiana Proteobacteria; Gammaproteobacteria; Enterobacterales; Enterobacteriaceae; Salmonella; Salmonella enterica; Salmonella enterica subsp. enterica MNNEETFYQAMRRKGVTRRSFLKFCSLAATSLGLGAGMTPKIAWALENKPRIPVVWIHGL WP_001058312.1 hydrogenase [Salmonella enterica] Length: 372\nNumber of predicted TMHs: 2\nExp number of AAs in TMHs: 36.51738\nExp number, first 60 AAs: 19.72849\nTotal prob of N-in: 0.96170\nPOSSIBLE N-term signal sequence\ninside 1 19\nTMhelix 20 39\noutside 40 325\nTMhelix 326 348\ninside 349 372

18074 GCF_002064715.1_ASM206471v1 Salmonella enterica subsp. enterica serovar Javiana Proteobacteria; Gammaproteobacteria; Enterobacterales; Enterobacteriaceae; Salmonella; Salmonella enterica; Salmonella enterica subsp. enterica MTGDNTLITSHGINRRDFMKLCAALAATMGLSSKAAAEMAESVSNPQRPPVIWIGAQECT WP_000145429.1 hydrogenase 2 small subunit [Salmonella enterica] Length: 372\nNumber of predicted TMHs: 1\nExp number of AAs in TMHs: 24.51309\nExp number, first 60 AAs: 0.2682\nTotal prob of N-in: 0.14402\noutside 1 331\nTMhelix 332 354\ninside 355 372

18075 GCF_002064715.1_ASM206471v1 Salmonella enterica subsp. enterica serovar Javiana Proteobacteria; Gammaproteobacteria; Enterobacterales; Enterobacteriaceae; Salmonella; Salmonella enterica; Salmonella enterica subsp. enterica MNNEETFYQAMRRKGVTRRSFLKFCSLAATSLGLGAGMTPKIAWALENKPRIPVVWIHGL WP_001058312.1 hydrogenase [Salmonella enterica] Length: 372\nNumber of predicted TMHs: 2\nExp number of AAs in TMHs: 36.51738\nExp number, first 60 AAs: 19.72849\nTotal prob of N-in: 0.96170\nPOSSIBLE N-term signal sequence\ninside 1 19\nTMhelix 20 39\noutside 40 325\nTMhelix 326 348\ninside 349 372

18076 GCF_001478595.1_Salmonella_enterica_CVM_N42337-SQ_v1.0 Salmonella enterica subsp. enterica serovar Javiana Proteobacteria; Gammaproteobacteria; Enterobacterales; Enterobacteriaceae; Salmonella; Salmonella enterica; Salmonella enterica subsp. enterica MNNEETFYQAMRRKGVTRRSFLKFCSLAATSLGLGAGMTPKIAWALENKPRIPVVWIHGL WP_001058312.1 hydrogenase [Salmonella enterica] Length: 372\nNumber of predicted TMHs: 2\nExp number of AAs in TMHs: 36.51738\nExp number, first 60 AAs: 19.72849\nTotal prob of N-in: 0.96170\nPOSSIBLE N-term signal sequence\ninside 1 19\nTMhelix 20 39\noutside 40 325\nTMhelix 326 348\ninside 349 372

18077 GCF_002060175.1_ASM206017v1 Salmonella enterica subsp. enterica serovar Javiana Proteobacteria; Gammaproteobacteria; Enterobacterales; Enterobacteriaceae; Salmonella; Salmonella enterica; Salmonella enterica subsp. enterica MNNEETFYQAMRRKGVTRRSFLKFCSLAATSLGLGAGMTPKIAWALENKPRIPVVWIHGL WP_001058312.1 hydrogenase [Salmonella enterica] Length: 372\nNumber of predicted TMHs: 2\nExp

number of AAs in TMHs: 36.51738\nExp number, first 60 AAs: 19.72849\nTotal prob of N-in: 0.96170\nPOSSIBLE N-term signal sequence\ninside 1 19\nTMhelix 20 39\noutside 40 325\nTMhelix 326 348\ninside 349 372

18078 GCF_002059715.1_ASM205971v1 Salmonella enterica subsp. enterica serovar Javiana Proteobacteria; Gammaproteobacteria; Enterobacterales; Enterobacteriaceae; Salmonella; Salmonella enterica; Salmonella enterica subsp. enterica MTGDNTLITSHGINRRDFMKLCAALAATMGLSSKAAAEMAESVSNPQRPPVIWIGAQECT WP_000145429.1 hydrogenase 2 small subunit [Salmonella enterica] Length: 372\nNumber of predicted TMHs: 1\nExp number of AAs in TMHs: 24.51309\nExp number, first 60 AAs: 0.2682\nTotal prob of N-in: 0.14402\noutside 1 331\nTMhelix 332 354\ninside 355 372

18079 GCF_002059715.1_ASM205971v1 Salmonella enterica subsp. enterica serovar Javiana Proteobacteria; Gammaproteobacteria; Enterobacterales; Enterobacteriaceae; Salmonella; Salmonella enterica; Salmonella enterica subsp. enterica MNNEETFYQAMRRKGVTRRSFLKFCSLAATSLGLGAGMTPKIAWALENKPRIPVVWIHGL WP_001058312.1 hydrogenase [Salmonella enterica] Length: 372\nNumber of predicted TMHs: 2\nExp number of AAs in TMHs: 36.51738\nExp number, first 60 AAs: 19.72849\nTotal prob of N-in: 0.96170\nPOSSIBLE N-term signal sequence\ninside 1 19\nTMhelix 20 39\noutside 40 325\nTMhelix 326 348\ninside 349 372

18080 GCF_001271635.1_Salmonella_enterica_CVM_N45933_v1.0 Salmonella enterica subsp. enterica serovar Johannesburg Proteobacteria; Gammaproteobacteria; Enterobacterales; Enterobacteriaceae; Salmonella; Salmonella enterica; Salmonella enterica subsp. enterica MTGDNTLITSHGINRRDFMKLCAALAATMGLSSKAAAEMAESVSNPQRPPVIWIGAQECT WP_000145429.1 hydrogenase 2 small subunit [Salmonella enterica] Length: 372\nNumber of predicted TMHs: 1\nExp number of AAs in TMHs: 24.51309\nExp number, first 60 AAs: 0.2682\nTotal prob of N-in: 0.14402\noutside 1 331\nTMhelix 332 354\ninside 355 372

18081 GCF_001242975.1_Salmonella_enterica_CVM_N46815_v1.0 Salmonella enterica subsp. enterica serovar Johannesburg Proteobacteria; Gammaproteobacteria; Enterobacterales; Enterobacteriaceae; Salmonella; Salmonella enterica; Salmonella enterica subsp. enterica MTGDNTLITSHGINRRDFMKLCAALAATMGLSSKAAAEMAESVSNPQRPPVIWIGAQECT WP_000145429.1 hydrogenase 2 small subunit [Salmonella enterica] Length: 372\nNumber of predicted TMHs: 1\nExp number of AAs in TMHs: 24.51309\nExp number, first 60 AAs: 0.2682\nTotal prob of N-in: 0.14402\noutside 1 331\nTMhelix 332 354\ninside 355 372

18082 GCF_001247345.1_Salmonella_enterica_CVM_N51305_v1.0 Salmonella enterica subsp. enterica serovar Johannesburg Proteobacteria; Gammaproteobacteria; Enterobacterales; Enterobacteriaceae; Salmonella; Salmonella enterica; Salmonella enterica subsp. enterica MNNEETFYQAMRRKGVTRRSFLKFCSLAATSLGLGAGMTPKIAWALENKPRIPVVWIHGL WP_023198237.1 hydrogenase 1, small subunit [Salmonella enterica] Length: 372\nNumber of predicted TMHs: 2\nExp number of AAs in TMHs: 36.39423\nExp number, first 60 AAs: 19.72877\nTotal prob of N-in: 0.96162\nPOSSIBLE N-term signal sequence\ninside 1 19\nTMhelix 20 39\noutside 40 325\nTMhelix 326 348\ninside 349 372

18083 GCF_001247345.1_Salmonella_enterica_CVM_N51305_v1.0 Salmonella enterica subsp. enterica serovar Johannesburg Proteobacteria; Gammaproteobacteria; Enterobacterales; Enterobacteriaceae; Salmonella; Salmonella enterica; Salmonella enterica subsp. enterica MTGDNTLITSHGINRRDFMKLCAALAATMGLSSKAAAEMAESVSNPQRPPVIWIGAQECT WP_000145429.1 hydrogenase 2 small subunit [Salmonella enterica] Length: 372\nNumber of predicted TMHs: 1\nExp number of AAs in TMHs: 24.51309\nExp number, first 60 AAs: 0.2682\nTotal prob of N-in: 0.14402\noutside 1 331\nTMhelix 332 354\ninside 355 372

18084 GCF_001242975.1_Salmonella_enterica_CVM_N46815_v1.0 Salmonella enterica subsp. enterica serovar Johannesburg Proteobacteria; Gammaproteobacteria; Enterobacterales; Enterobacteriaceae; Salmonella; Salmonella enterica; Salmonella enterica subsp. enterica MNNEETFYQAMRRKGVTRRSFLKFCSLAATSLGLGAGMTPKIAWALENKPRIPVVWIHGL WP_023198237.1 hydrogenase 1, small subunit [Salmonella enterica] Length: 372\nNumber of predicted TMHs: 2\nExp number of AAs in TMHs: 36.39423\nExp number, first 60 AAs: 19.72877\nTotal prob of N-in: 0.96162\nPOSSIBLE N-term signal sequence\ninside 1 19\nTMhelix 20 39\noutside 40 325\nTMhelix 326 348\ninside 349 372

18085 GCF_002048395.1_ASM204839v1 Salmonella enterica subsp. enterica serovar Johannesburg Proteobacteria; Gammaproteobacteria; Enterobacterales; Enterobacteriaceae; Salmonella; Salmonella enterica; Salmonella enterica subsp. enterica MNNEETFYQAMRRKGVTRRSFLKFCSLAATSLGLGAGMTPKIAWALENKPRIPVVWIHGL WP_023198237.1 hydrogenase 1, small subunit [Salmonella enterica] Length: 372\nNumber of

predicted TMHs: 2\nExp number of AAs in TMHs: 36.39423\nExp number, first 60 AAs: 19.72877\nTotal prob of N-in: 0.96162\nPOSSIBLE N-term signal sequence\ninside 1 19\nTMhelix 20 39\noutside 40 325\nTMhelix 326 348\ninside 349 372

18086 GCF_002034785.1_ASM203478v1 Salmonella enterica subsp. enterica serovar Johannesburg Proteobacteria; Gammaproteobacteria; Enterobacterales; Enterobacteriaceae; Salmonella; Salmonella enterica; Salmonella enterica subsp. enterica MTGDNTLITSHGINRRDFMKLCAALAATMGLSSKAAAEMAESVSNPQRPPVIWIGAQECT WP_000145429.1 hydrogenase 2 small subunit [Salmonella enterica] Length: 372\nNumber of predicted TMHs: 1\nExp number of AAs in TMHs: 24.51309\nExp number, first 60 AAs: 0.2682\nTotal prob of N-in: 0.14402\noutside 1 331\nTMhelix 332 354\ninside 355 372

18087 GCF_001271635.1_Salmonella_enterica_CVM_N45933_v1.0 Salmonella enterica subsp. enterica serovar Johannesburg Proteobacteria; Gammaproteobacteria; Enterobacterales; Enterobacteriaceae; Salmonella; Salmonella enterica; Salmonella enterica subsp. enterica MNNEETFYQAMRRKGVTRRSFLKFCSLAATSLGLGAGMTPKIAWALENKPRIPVVWIHGL WP_023198237.1 hydrogenase 1, small subunit [Salmonella enterica] Length: 372\nNumber of predicted TMHs: 2\nExp number of AAs in TMHs: 36.39423\nExp number, first 60 AAs: 19.72877\nTotal prob of N-in: 0.96162\nPOSSIBLE N-term signal sequence\ninside 1 19\nTMhelix 20 39\noutside 40 325\nTMhelix 326 348\ninside 349 372

18088 GCF_002048385.1_ASM204838v1 Salmonella enterica subsp. enterica serovar Johannesburg Proteobacteria; Gammaproteobacteria; Enterobacterales; Enterobacteriaceae; Salmonella; Salmonella enterica; Salmonella enterica subsp. enterica MTGDNTLITSHGINRRDFMKLCAALAATMGLSSKAAAEMAESVSNPQRPPVIWIGAQECT WP_000145429.1 hydrogenase 2 small subunit [Salmonella enterica] Length: 372\nNumber of predicted TMHs: 1\nExp number of AAs in TMHs: 24.51309\nExp number, first 60 AAs: 0.2682\nTotal prob of N-in: 0.14402\noutside 1 331\nTMhelix 332 354\ninside 355 372

18089 GCF_002048385.1_ASM204838v1 Salmonella enterica subsp. enterica serovar Johannesburg Proteobacteria; Gammaproteobacteria; Enterobacterales; Enterobacteriaceae; Salmonella; Salmonella enterica; Salmonella enterica subsp. enterica MNNEETFYQAMRRKGVTRRSFLKFCSLAATSLGLGAGMTPKIAWALENKPRIPVVWIHGL WP_023198237.1 hydrogenase 1, small subunit [Salmonella enterica] Length: 372\nNumber of predicted TMHs: 2\nExp number of AAs in TMHs: 36.39423\nExp number, first 60 AAs: 19.72877\nTotal prob of N-in: 0.96162\nPOSSIBLE N-term signal sequence\ninside 1 19\nTMhelix 20 39\noutside 40 325\nTMhelix 326 348\ninside 349 372

18090 GCF_002048395.1_ASM204839v1 Salmonella enterica subsp. enterica serovar Johannesburg Proteobacteria; Gammaproteobacteria; Enterobacterales; Enterobacteriaceae; Salmonella; Salmonella enterica; Salmonella enterica subsp. enterica MTGDNTLITSHGINRRDFMKLCAALAATMGLSSKAAAEMAESVSNPQRPPVIWIGAQECT WP_000145429.1 hydrogenase 2 small subunit [Salmonella enterica] Length: 372\nNumber of predicted TMHs: 1\nExp number of AAs in TMHs: 24.51309\nExp number, first 60 AAs: 0.2682\nTotal prob of N-in: 0.14402\noutside 1 331\nTMhelix 332 354\ninside 355 372

18091 GCF_002034785.1_ASM203478v1 Salmonella enterica subsp. enterica serovar Johannesburg Proteobacteria; Gammaproteobacteria; Enterobacterales; Enterobacteriaceae; Salmonella; Salmonella enterica; Salmonella enterica subsp. enterica MNNEETFYQAMRRKGVTRRSFLKFCSLAATSLGLGAGMTPKIAWALENKPRIPVVWIHGL WP_023198237.1 hydrogenase 1, small subunit [Salmonella enterica] Length: 372\nNumber of predicted TMHs: 2\nExp number of AAs in TMHs: 36.39423\nExp number, first 60 AAs: 19.72877\nTotal prob of N-in: 0.96162\nPOSSIBLE N-term signal sequence\ninside 1 19\nTMhelix 20 39\noutside 40 325\nTMhelix 326 348\ninside 349 372

18092 GCF_002047165.1_ASM204716v1 Salmonella enterica subsp. enterica serovar Johannesburg Proteobacteria; Gammaproteobacteria; Enterobacterales; Enterobacteriaceae; Salmonella; Salmonella enterica; Salmonella enterica subsp. enterica MNNEETFYQAMRRKGVTRRSFLKFCSLAATSLGLGAGMTPKIAWALENKPRIPVVWIHGL WP_023198237.1 hydrogenase 1, small subunit [Salmonella enterica] Length: 372\nNumber of predicted TMHs: 2\nExp number of AAs in TMHs: 36.39423\nExp number, first 60 AAs: 19.72877\nTotal prob of N-in: 0.96162\nPOSSIBLE N-term signal sequence\ninside 1 19\nTMhelix 20 39\noutside 40 325\nTMhelix 326 348\ninside 349 372

18093 GCF_002047165.1_ASM204716v1 Salmonella enterica subsp. enterica serovar Johannesburg Proteobacteria; Gammaproteobacteria; Enterobacterales; Enterobacteriaceae; Salmonella; Salmonella enterica; Salmonella enterica subsp. enterica MTGDNTLITSHGINRRDFMKLCAALAATMGLSSKAAAEMAESVSNPQRPPVIWIGAQECT WP_000145429.1 hydrogenase 2 small subunit [Salmonella enterica] Length: 372\nNumber of

predicted TMHs: 1\nExp number of AAs in TMHs: 24.51309\nExp number, first 60 AAs: 0.2682\nTotal prob of N-in: 0.14402\noutside 1 331\nTMhelix 332 354\ninside 355 372

18094 GCF_002064755.1_ASM206475v1 Salmonella enterica subsp. enterica serovar Jukestown Proteobacteria; Gammaproteobacteria; Enterobacterales; Enterobacteriaceae; Salmonella; Salmonella enterica; Salmonella enterica subsp. enterica MNNEETFYQAMRRKGVTRRSFLKFCSLAATSLGLGAGMTPKIAWALENKPRIPVVWIHGL WP_001058311.1 hydrogenase [Salmonella enterica] Length: 372\nNumber of predicted TMHs: 2\nExp number of AAs in TMHs: 37.4699899999999\nExp number, first 60 AAs: 19.72699\nTotal prob of N-in: 0.96171\nPOSSIBLE N-term signal sequence\ninside 1 19\nTMhelix 20 39\noutside 40 325\nTMhelix 326 348\ninside 349 372

18095 GCF_002064755.1_ASM206475v1 Salmonella enterica subsp. enterica serovar Jukestown Proteobacteria; Gammaproteobacteria; Enterobacterales; Enterobacteriaceae; Salmonella; Salmonella enterica; Salmonella enterica subsp. enterica MTGDNTLITSHGINRRDFMKLCAALAATMGLSSKAAAEMAESVSNPQRPPVIWIGAQECT WP_000145429.1 hydrogenase 2 small subunit [Salmonella enterica] Length: 372\nNumber of predicted TMHs: 1\nExp number of AAs in TMHs: 24.51309\nExp number, first 60 AAs: 0.2682\nTotal prob of N-in: 0.14402\noutside 1 331\nTMhelix 332 354\ninside 355 372

18096 GCF_002065595.1_ASM206559v1 Salmonella enterica subsp. enterica serovar Kambole Proteobacteria; Gammaproteobacteria; Enterobacterales; Enterobacteriaceae; Salmonella; Salmonella enterica; Salmonella enterica subsp. enterica MNNEETFYQAMRRKGVTRRSFLKFCSLAATSLGLGAGMTPKIAWALENKPRIPVVWIHGL WP_001058312.1 hydrogenase [Salmonella enterica] Length: 372\nNumber of predicted TMHs: 2\nExp number of AAs in TMHs: 36.51738\nExp number, first 60 AAs: 19.72849\nTotal prob of N-in: 0.96170\nPOSSIBLE N-term signal sequence\ninside 1 19\nTMhelix 20 39\noutside 40 325\nTMhelix 326 348\ninside 349 372

18097 GCF_002065595.1_ASM206559v1 Salmonella enterica subsp. enterica serovar Kambole Proteobacteria; Gammaproteobacteria; Enterobacterales; Enterobacteriaceae; Salmonella; Salmonella enterica; Salmonella enterica subsp. enterica MTGDNTLITSHGINRRDFMKLCAALAATMGLSSKAAAEMAESVSNPQRPPVIWIGAQECT WP_000145429.1 hydrogenase 2 small subunit [Salmonella enterica] Length: 372\nNumber of predicted TMHs: 1\nExp number of AAs in TMHs: 24.51309\nExp number, first 60 AAs: 0.2682\nTotal prob of N-in: 0.14402\noutside 1 331\nTMhelix 332 354\ninside 355 372

18098 GCF_002064695.1_ASM206469v1 Salmonella enterica subsp. enterica serovar Kedougou Proteobacteria; Gammaproteobacteria; Enterobacterales; Enterobacteriaceae; Salmonella; Salmonella enterica; Salmonella enterica subsp. enterica MNNEETFYQAMRRKGVTRRSFLKFCSLAATSLGLGAGMTPKIAWALENKPRIPVVWIHGL WP_001058312.1 hydrogenase [Salmonella enterica] Length: 372\nNumber of predicted TMHs: 2\nExp number of AAs in TMHs: 36.51738\nExp number, first 60 AAs: 19.72849\nTotal prob of N-in: 0.96170\nPOSSIBLE N-term signal sequence\ninside 1 19\nTMhelix 20 39\noutside 40 325\nTMhelix 326 348\ninside 349 372

18099 GCF_002064635.1_ASM206463v1 Salmonella enterica subsp. enterica serovar Kedougou Proteobacteria; Gammaproteobacteria; Enterobacterales; Enterobacteriaceae; Salmonella; Salmonella enterica; Salmonella enterica subsp. enterica MTGDNTLITSHGINRRDFMKLCAALAATMGLSSKAAAEMAESVSNPQRPPVIWIGAQECT WP_000145429.1 hydrogenase 2 small subunit [Salmonella enterica] Length: 372\nNumber of predicted TMHs: 1\nExp number of AAs in TMHs: 24.51309\nExp number, first 60 AAs: 0.2682\nTotal prob of N-in: 0.14402\noutside 1 331\nTMhelix 332 354\ninside 355 372

18100 GCF_002065575.1_ASM206557v1 Salmonella enterica subsp. enterica serovar Kedougou Proteobacteria; Gammaproteobacteria; Enterobacterales; Enterobacteriaceae; Salmonella; Salmonella enterica; Salmonella enterica subsp. enterica MTGDNTLITSHGINRRDFMKLCAALAATMGLSSKAAAEMAESVSNPQRPPVIWIGAQECT WP_000145429.1 hydrogenase 2 small subunit [Salmonella enterica] Length: 372\nNumber of predicted TMHs: 1\nExp number of AAs in TMHs: 24.51309\nExp number, first 60 AAs: 0.2682\nTotal prob of N-in: 0.14402\noutside 1 331\nTMhelix 332 354\ninside 355 372

18101 GCF_002065575.1_ASM206557v1 Salmonella enterica subsp. enterica serovar Kedougou Proteobacteria; Gammaproteobacteria; Enterobacterales; Enterobacteriaceae; Salmonella; Salmonella enterica; Salmonella enterica subsp. enterica MNNEETFYQAMRRKGVTRRSFLKFCSLAATSLGLGAGMTPKIAWALENKPRIPVVWIHGL WP_001058312.1 hydrogenase [Salmonella enterica] Length: 372\nNumber of predicted TMHs: 2\nExp number of AAs in TMHs: 36.51738\nExp number, first 60 AAs: 19.72849\nTotal prob of N-in: 0.96170\nPOSSIBLE N-term signal sequence\ninside 1 19\nTMhelix 20 39\noutside 40 325\nTMhelix 326 348\ninside 349 372

18102 GCF_002064725.1_ASM206472v1 Salmonella enterica subsp. enterica serovar Kedougou Proteobacteria; Gammaproteobacteria; Enterobacterales; Enterobacteriaceae; Salmonella; Salmonella enterica; Salmonella enterica subsp. enterica MNNEETFYQAMRRKGVTRRSFLKFCSLAATSLGLGAGMTPKIAWALENKPRIPVVWIHGL WP_001058312.1 hydrogenase [Salmonella enterica] Length: 372\nNumber of predicted TMHs: 2\nExp number of AAs in TMHs: 36.51738\nExp number, first 60 AAs: 19.72849\nTotal prob of N-in: 0.96170\nPOSSIBLE N-term signal sequence\ninside 1 19\nTMhelix 20 39\noutside 40 325\nTMhelix 326 348\ninside 349 372

18103 GCF_002064725.1_ASM206472v1 Salmonella enterica subsp. enterica serovar Kedougou Proteobacteria; Gammaproteobacteria; Enterobacterales; Enterobacteriaceae; Salmonella; Salmonella enterica; Salmonella enterica subsp. enterica MTGDNTLITSHGINRRDFMKLCAALAATMGLSSKAAAEMAESVSNPQRPPVIWIGAQECT WP_000145429.1 hydrogenase 2 small subunit [Salmonella enterica] Length: 372\nNumber of predicted TMHs: 1\nExp number of AAs in TMHs: 24.51309\nExp number, first 60 AAs: 0.2682\nTotal prob of N-in: 0.14402\noutside 1 331\nTMhelix 332 354\ninside 355 372

18104 GCF_002064635.1_ASM206463v1 Salmonella enterica subsp. enterica serovar Kedougou Proteobacteria; Gammaproteobacteria; Enterobacterales; Enterobacteriaceae; Salmonella; Salmonella enterica; Salmonella enterica subsp. enterica MNNEETFYQAMRRKGVTRRSFLKFCSLAATSLGLGAGMTPKIAWALENKPRIPVVWIHGL WP_001058312.1 hydrogenase [Salmonella enterica] Length: 372\nNumber of predicted TMHs: 2\nExp number of AAs in TMHs: 36.51738\nExp number, first 60 AAs: 19.72849\nTotal prob of N-in: 0.96170\nPOSSIBLE N-term signal sequence\ninside 1 19\nTMhelix 20 39\noutside 40 325\nTMhelix 326 348\ninside 349 372

18105 GCF_002064695.1_ASM206469v1 Salmonella enterica subsp. enterica serovar Kedougou Proteobacteria; Gammaproteobacteria; Enterobacterales; Enterobacteriaceae; Salmonella; Salmonella enterica; Salmonella enterica subsp. enterica MTGDNTLITSHGINRRDFMKLCAALAATMGLSSKAAAEMAESVSNPQRPPVIWIGAQECT WP_000145429.1 hydrogenase 2 small subunit [Salmonella enterica] Length: 372\nNumber of predicted TMHs: 1\nExp number of AAs in TMHs: 24.51309\nExp number, first 60 AAs: 0.2682\nTotal prob of N-in: 0.14402\noutside 1 331\nTMhelix 332 354\ninside 355 372

18106 GCF_002062135.1_ASM206213v1 Salmonella enterica subsp. enterica serovar Kedougou Proteobacteria; Gammaproteobacteria; Enterobacterales; Enterobacteriaceae; Salmonella; Salmonella enterica; Salmonella enterica subsp. enterica TRRSFLKFCSLAATSLGLGAGMTPKIAWALENKPRIPVVWIHGLECTCCTESFIRSSHPL WP_080236668.1 [Ni/Fe] hydrogenase small subunit, partial [Salmonella enterica] Length: 356\nNumber of predicted TMHs: 1\nExp number of AAs in TMHs: 18.07289\nExp number, first 60 AAs: 1.24286\nTotal prob of N-in: 0.08098\noutside 1 309\nTMhelix 310 332\ninside 333 356

18107 GCF_002062135.1_ASM206213v1 Salmonella enterica subsp. enterica serovar Kedougou Proteobacteria; Gammaproteobacteria; Enterobacterales; Enterobacteriaceae; Salmonella; Salmonella enterica; Salmonella enterica subsp. enterica MTGDNTLITSHGINRRDFMKLCAALAATMGLSSKAAAEMAESVSNPQRPPVIWIGAQECT WP_000145429.1 hydrogenase 2 small subunit [Salmonella enterica] Length: 372\nNumber of predicted TMHs: 1\nExp number of AAs in TMHs: 24.51309\nExp number, first 60 AAs: 0.2682\nTotal prob of N-in: 0.14402\noutside 1 331\nTMhelix 332 354\ninside 355 372

18108 GCF_002064645.1_ASM206464v1 Salmonella enterica subsp. enterica serovar Kedougou Proteobacteria; Gammaproteobacteria; Enterobacterales; Enterobacteriaceae; Salmonella; Salmonella enterica; Salmonella enterica subsp. enterica MTGDNTLITSHGINRRDFMKLCAALAATMGLSSKAAAEMAESVSNPQRPPVIWIGAQECT WP_000145429.1 hydrogenase 2 small subunit [Salmonella enterica] Length: 372\nNumber of predicted TMHs: 1\nExp number of AAs in TMHs: 24.51309\nExp number, first 60 AAs: 0.2682\nTotal prob of N-in: 0.14402\noutside 1 331\nTMhelix 332 354\ninside 355 372

18109 GCF_002064645.1_ASM206464v1 Salmonella enterica subsp. enterica serovar Kedougou Proteobacteria; Gammaproteobacteria; Enterobacterales; Enterobacteriaceae; Salmonella; Salmonella enterica; Salmonella enterica subsp. enterica MNNEETFYQAMRRKGVTRRSFLKFCSLAATSLGLGAGMTPKIAWALENKPRIPVVWIHGL WP_001058312.1 hydrogenase [Salmonella enterica] Length: 372\nNumber of predicted TMHs: 2\nExp number of AAs in TMHs: 36.51738\nExp number, first 60 AAs: 19.72849\nTotal prob of N-in: 0.96170\nPOSSIBLE N-term signal sequence\ninside 1 19\nTMhelix 20 39\noutside 40 325\nTMhelix 326 348\ninside 349 372

18110 GCF_002065555.1_ASM206555v1 Salmonella enterica subsp. enterica serovar Kedougou Proteobacteria; Gammaproteobacteria; Enterobacterales; Enterobacteriaceae; Salmonella; Salmonella enterica; Salmonella enterica subsp. enterica MTGDNTLITSHGINRRDFMKLCAALAATMGLSSKAAAEMAESVSNPQRPPVIWIGAQECT WP_000145429.1 hydrogenase 2 small subunit [Salmonella enterica] Length: 372\nNumber of

predicted TMHs: 1\nExp number of AAs in TMHs: 24.51309\nExp number, first 60 AAs: 0.2682\nTotal prob of N-in: 0.14402\noutside 1 331\nTMhelix 332 354\ninside 355 372

18111 GCF_002065555.1_ASM206555v1 Salmonella enterica subsp. enterica serovar Kedougou Proteobacteria; Gammaproteobacteria; Enterobacterales; Enterobacteriaceae; Salmonella; Salmonella enterica; Salmonella enterica subsp. enterica MNNEETFYQAMRRKGVTRRSFLKFCSLAATSLGLGAGMTPKIAWALENKPRIPVVWIHGL WP_001058312.1 hydrogenase [Salmonella enterica] Length: 372\nNumber of predicted TMHs: 2\nExp number of AAs in TMHs: 36.51738\nExp number, first 60 AAs: 19.72849\nTotal prob of N-in: 0.96170\nPOSSIBLE N-term signal sequence\ninside 1 19\nTMhelix 20 39\noutside 40 325\nTMhelix 326 348\ninside 349 372

18112 GCF_002065505.1_ASM206550v1 Salmonella enterica subsp. enterica serovar Kentucky Proteobacteria; Gammaproteobacteria; Enterobacterales; Enterobacteriaceae; Salmonella; Salmonella enterica; Salmonella enterica subsp. enterica MNNEETFYQAMRRKGVTRRSFLKFCSLAATSLGLGAGMTPKIAWALENKPRIPVVWIHGL WP_001058311.1 hydrogenase [Salmonella enterica] Length: 372\nNumber of predicted TMHs: 2\nExp number of AAs in TMHs: 37.4699899999999\nExp number, first 60 AAs: 19.72699\nTotal prob of N-in: 0.96171\nPOSSIBLE N-term signal sequence\ninside 1 19\nTMhelix 20 39\noutside 40 325\nTMhelix 326 348\ninside 349 372

18113 GCF_002065505.1_ASM206550v1 Salmonella enterica subsp. enterica serovar Kentucky Proteobacteria; Gammaproteobacteria; Enterobacterales; Enterobacteriaceae; Salmonella; Salmonella enterica; Salmonella enterica subsp. enterica MTGDNTLITSHGINRRDFMKLCAALAATMGLSSKAAAEMAESVSNPQRPPVIWIGAQECT WP_000145429.1 hydrogenase 2 small subunit [Salmonella enterica] Length: 372\nNumber of predicted TMHs: 1\nExp number of AAs in TMHs: 24.51309\nExp number, first 60 AAs: 0.2682\nTotal prob of N-in: 0.14402\noutside 1 331\nTMhelix 332 354\ninside 355 372

18114 GCF_002064625.1_ASM206462v1 Salmonella enterica subsp. enterica serovar Kentucky Proteobacteria; Gammaproteobacteria; Enterobacterales; Enterobacteriaceae; Salmonella; Salmonella enterica; Salmonella enterica subsp. enterica MTGDNTLITSHGINRRDFMKLCAALAATMGLSSKAAAEMAESVSNPQRPPVIWIGAQECT WP_000145429.1 hydrogenase 2 small subunit [Salmonella enterica] Length: 372\nNumber of predicted TMHs: 1\nExp number of AAs in TMHs: 24.51309\nExp number, first 60 AAs: 0.2682\nTotal prob of N-in: 0.14402\noutside 1 331\nTMhelix 332 354\ninside 355 372

18115 GCF_002064625.1_ASM206462v1 Salmonella enterica subsp. enterica serovar Kentucky Proteobacteria; Gammaproteobacteria; Enterobacterales; Enterobacteriaceae; Salmonella; Salmonella enterica; Salmonella enterica subsp. enterica MNNEETFYQAMRRKGVTRRSFLKFCSLAATSLGLGAGMTPKIAWALENKPRIPVVWIHGL WP_001058307.1 hydrogenase [Salmonella enterica] Length: 372\nNumber of predicted TMHs: 2\nExp number of AAs in TMHs: 36.51738\nExp number, first 60 AAs: 19.72849\nTotal prob of N-in: 0.96170\nPOSSIBLE N-term signal sequence\ninside 1 19\nTMhelix 20 39\noutside 40 325\nTMhelix 326 348\ninside 349 372

18116 GCF_002064595.1_ASM206459v1 Salmonella enterica subsp. enterica serovar Kentucky Proteobacteria; Gammaproteobacteria; Enterobacterales; Enterobacteriaceae; Salmonella; Salmonella enterica; Salmonella enterica subsp. enterica MNNEETFYQAMRRKGVTRRSFLKFCSLAATSLGLGAGMTPKIAWALENKPRIPVVWIHGL WP_001058307.1 hydrogenase [Salmonella enterica] Length: 372\nNumber of predicted TMHs: 2\nExp number of AAs in TMHs: 36.51738\nExp number, first 60 AAs: 19.72849\nTotal prob of N-in: 0.96170\nPOSSIBLE N-term signal sequence\ninside 1 19\nTMhelix 20 39\noutside 40 325\nTMhelix 326 348\ninside 349 372

18117 GCF_002064595.1_ASM206459v1 Salmonella enterica subsp. enterica serovar Kentucky Proteobacteria; Gammaproteobacteria; Enterobacterales; Enterobacteriaceae; Salmonella; Salmonella enterica; Salmonella enterica subsp. enterica MTGDNTLITSHGINRRDFMKLCAALAATMGLSSKAAAEMAESVSNPQRPPVIWIGAQECT WP_000145429.1 hydrogenase 2 small subunit [Salmonella enterica] Length: 372\nNumber of predicted TMHs: 1\nExp number of AAs in TMHs: 24.51309\nExp number, first 60 AAs: 0.2682\nTotal prob of N-in: 0.14402\noutside 1 331\nTMhelix 332 354\ninside 355 372

18118 GCF_002062995.1_ASM206299v1 Salmonella enterica subsp. enterica serovar Kentucky Proteobacteria; Gammaproteobacteria; Enterobacterales; Enterobacteriaceae; Salmonella; Salmonella enterica; Salmonella enterica subsp. enterica MNNEETFYQAMRRKGVTRRSFLKFCSLAATSLGLGAGMTPKIAWALENKPRIPVVWIHGL WP_001058307.1 hydrogenase [Salmonella enterica] Length: 372\nNumber of predicted TMHs: 2\nExp number of AAs in TMHs: 36.51738\nExp number, first 60 AAs: 19.72849\nTotal prob of N-in: 0.96170\nPOSSIBLE N-term signal sequence\ninside 1 19\nTMhelix 20 39\noutside 40 325\nTMhelix 326 348\ninside 349 372

18119 GCF_002062995.1_ASM206299v1 Salmonella enterica subsp. enterica serovar Kentucky Proteobacteria; Gammaproteobacteria; Enterobacterales; Enterobacteriaceae; Salmonella; Salmonella enterica; Salmonella enterica subsp. enterica MTGDNTLITSHGINRRDFMKLCAALAATMGLSSKAAAEMAESVSNPQRPPVIWIGAQECT WP_000145429.1 hydrogenase 2 small subunit [Salmonella enterica] Length: 372\nNumber of predicted TMHs: 1\nExp number of AAs in TMHs: 24.51309\nExp number, first 60 AAs: 0.2682\nTotal prob of N-in: 0.14402\noutside 1 331\nTMhelix 332 354\ninside 355 372

18120 GCF_001241395.1_Salmonella_enterica_CVM_N44708_v1.0 Salmonella enterica subsp. enterica serovar Kentucky Proteobacteria; Gammaproteobacteria; Enterobacterales; Enterobacteriaceae; Salmonella; Salmonella enterica; Salmonella enterica subsp. enterica MNNEETFYQAMRRKGVTRRSFLKFCSLAATSLGLGAGMTPKIAWALENKPRIPVVWIHGL WP_001058311.1 hydrogenase [Salmonella enterica] Length: 372\nNumber of predicted TMHs: 2\nExp number of AAs in TMHs: 37.4699899999999\nExp number, first 60 AAs: 19.72699\nTotal prob of N-in: 0.96171\nPOSSIBLE N-term signal sequence\ninside 1 19\nTMhelix 20 39\noutside 40 325\nTMhelix 326 348\ninside 349 372

18121 GCF_001245135.1_Salmonella_enterica_CVM_N48710_v1.0 Salmonella enterica subsp. enterica serovar Kentucky Proteobacteria; Gammaproteobacteria; Enterobacterales; Enterobacteriaceae; Salmonella; Salmonella enterica; Salmonella enterica subsp. enterica MTGDNTLITSHGINRRDFMKLCAALAATMGLSSKAAAEMAESVSNPQRPPVIWIGAQECT WP_000145429.1 hydrogenase 2 small subunit [Salmonella enterica] Length: 372\nNumber of predicted TMHs: 1\nExp number of AAs in TMHs: 24.51309\nExp number, first 60 AAs: 0.2682\nTotal prob of N-in: 0.14402\noutside 1 331\nTMhelix 332 354\ninside 355 372

18122 GCF_001244155.1_Salmonella_enterica_CVM_N47729_v1.0 Salmonella enterica subsp. enterica serovar Kentucky Proteobacteria; Gammaproteobacteria; Enterobacterales; Enterobacteriaceae; Salmonella; Salmonella enterica; Salmonella enterica subsp. enterica MNNEETFYQAMRRKGVTRRSFLKFCSLAATSLGLGAGMTPKIAWALENKPRIPVVWIHGL WP_001058311.1 hydrogenase [Salmonella enterica] Length: 372\nNumber of predicted TMHs: 2\nExp number of AAs in TMHs: 37.4699899999999\nExp number, first 60 AAs: 19.72699\nTotal prob of N-in: 0.96171\nPOSSIBLE N-term signal sequence\ninside 1 19\nTMhelix 20 39\noutside 40 325\nTMhelix 326 348\ninside 349 372

18123 GCF_001244155.1_Salmonella_enterica_CVM_N47729_v1.0 Salmonella enterica subsp. enterica serovar Kentucky Proteobacteria; Gammaproteobacteria; Enterobacterales; Enterobacteriaceae; Salmonella; Salmonella enterica; Salmonella enterica subsp. enterica MTGDNTLITSHGINRRDFMKLCAALAATMGLSSKAAAEMAESVSNPQRPPVIWIGAQECT WP_000145429.1 hydrogenase 2 small subunit [Salmonella enterica] Length: 372\nNumber of predicted TMHs: 1\nExp number of AAs in TMHs: 24.51309\nExp number, first 60 AAs: 0.2682\nTotal prob of N-in: 0.14402\noutside 1 331\nTMhelix 332 354\ninside 355 372

18124 GCF_001243955.1_Salmonella_enterica_CVM_N47720_v1.0 Salmonella enterica subsp. enterica serovar Kentucky Proteobacteria; Gammaproteobacteria; Enterobacterales; Enterobacteriaceae; Salmonella; Salmonella enterica; Salmonella enterica subsp. enterica MNNEETFYQAMRRKGVTRRSFLKFCSLAATSLGLGAGMTPKIAWALENKPRIPVVWIHGL WP_001058311.1 hydrogenase [Salmonella enterica] Length: 372\nNumber of predicted TMHs: 2\nExp number of AAs in TMHs: 37.4699899999999\nExp number, first 60 AAs: 19.72699\nTotal prob of N-in: 0.96171\nPOSSIBLE N-term signal sequence\ninside 1 19\nTMhelix 20 39\noutside 40 325\nTMhelix 326 348\ninside 349 372

18125 GCF_001243915.1_Salmonella_enterica_CVM_N47718_v1.0 Salmonella enterica subsp. enterica serovar Kentucky Proteobacteria; Gammaproteobacteria; Enterobacterales; Enterobacteriaceae; Salmonella; Salmonella enterica; Salmonella enterica subsp. enterica MNNEETFYQAMRRKGVTRRSFLKFCSLAATSLGLGAGMTPKIAWALENKPRIPVVWIHGL WP_001058311.1 hydrogenase [Salmonella enterica] Length: 372\nNumber of predicted TMHs: 2\nExp number of AAs in TMHs: 37.4699899999999\nExp number, first 60 AAs: 19.72699\nTotal prob of N-in: 0.96171\nPOSSIBLE N-term signal sequence\ninside 1 19\nTMhelix 20 39\noutside 40 325\nTMhelix 326 348\ninside 349 372

18126 GCF_000973755.1_ABBSB1008-2.id24_v1.0 Salmonella enterica subsp. enterica serovar Kentucky Proteobacteria; Gammaproteobacteria; Enterobacterales; Enterobacteriaceae; Salmonella; Salmonella enterica; Salmonella enterica subsp. enterica MTGDNTLITSHGINRRDFMKLCAALAATMGLSSKAAAEMAESVSNPQRPPVIWIGAQECT WP_000145429.1 hydrogenase 2 small subunit [Salmonella enterica] Length: 372\nNumber of predicted TMHs: 1\nExp number of AAs in TMHs: 24.51309\nExp number, first 60 AAs: 0.2682\nTotal prob of N-in: 0.14402\noutside 1 331\nTMhelix 332 354\ninside 355 372

18127 GCF_000973835.1_SALC-205-3.id22_v1.0 Salmonella enterica subsp. enterica serovar Kentucky Proteobacteria; Gammaproteobacteria; Enterobacterales; Enterobacteriaceae; Salmonella; Salmonella enterica; Salmonella enterica subsp. enterica MTGDNTLITSHGINRRDFMKLCAALAATMGLSSKAAAEMAESVSNPQRPPVIWIGAQECT WP_000145429.1 hydrogenase 2 small subunit [Salmonella enterica] Length: 372\nNumber of predicted TMHs: 1\nExp number of AAs in TMHs: 24.51309\nExp number, first 60 AAs: 0.2682\nTotal prob of N-in: 0.14402\noutside 1 331\nTMhelix 332 354\ninside 355 372

18128 GCF_000973755.1_ABBSB1008-2.id24_v1.0 Salmonella enterica subsp. enterica serovar Kentucky Proteobacteria; Gammaproteobacteria; Enterobacterales; Enterobacteriaceae; Salmonella; Salmonella enterica; Salmonella enterica subsp. enterica MNNEETFYQAMRRKGVTRRSFLKFCSLAATSLGLGAGMTPKIAWALENKPRIPVVWIHGL WP_001058311.1 hydrogenase [Salmonella enterica] Length: 372\nNumber of predicted TMHs: 2\nExp number of AAs in TMHs: 37.4699899999999\nExp number, first 60 AAs: 19.72699\nTotal prob of N-in: 0.96171\nPOSSIBLE N-term signal sequence\ninside 1 19\nTMhelix 20 39\noutside 40 325\nTMhelix 326 348\ninside 349 372

18129 GCF_001243955.1_Salmonella_enterica_CVM_N47720_v1.0 Salmonella enterica subsp. enterica serovar Kentucky Proteobacteria; Gammaproteobacteria; Enterobacterales; Enterobacteriaceae; Salmonella; Salmonella enterica; Salmonella enterica subsp. enterica MTGDNTLITSHGINRRDFMKLCAALAATMGLSSKAAAEMAESVSNPQRPPVIWIGAQECT WP_000145429.1 hydrogenase 2 small subunit [Salmonella enterica] Length: 372\nNumber of predicted TMHs: 1\nExp number of AAs in TMHs: 24.51309\nExp number, first 60 AAs: 0.2682\nTotal prob of N-in: 0.14402\noutside 1 331\nTMhelix 332 354\ninside 355 372

18130 GCF_001243915.1_Salmonella_enterica_CVM_N47718_v1.0 Salmonella enterica subsp. enterica serovar Kentucky Proteobacteria; Gammaproteobacteria; Enterobacterales; Enterobacteriaceae; Salmonella; Salmonella enterica; Salmonella enterica subsp. enterica MTGDNTLITSHGINRRDFMKLCAALAATMGLSSKAAAEMAESVSNPQRPPVIWIGAQECT WP_000145429.1 hydrogenase 2 small subunit [Salmonella enterica] Length: 372\nNumber of predicted TMHs: 1\nExp number of AAs in TMHs: 24.51309\nExp number, first 60 AAs: 0.2682\nTotal prob of N-in: 0.14402\noutside 1 331\nTMhelix 332 354\ninside 355 372

18131 GCF_001242405.1_Salmonella_enterica_CVM_N45939_v1.0 Salmonella enterica subsp. enterica serovar Kentucky Proteobacteria; Gammaproteobacteria; Enterobacterales; Enterobacteriaceae; Salmonella; Salmonella enterica; Salmonella enterica subsp. enterica MNNEETFYQAMRRKGVTRRSFLKFCSLAATSLGLGAGMTPKIAWALENKPRIPVVWIHGL WP_001058311.1 hydrogenase [Salmonella enterica] Length: 372\nNumber of predicted TMHs: 2\nExp number of AAs in TMHs: 37.4699899999999\nExp number, first 60 AAs: 19.72699\nTotal prob of N-in: 0.96171\nPOSSIBLE N-term signal sequence\ninside 1 19\nTMhelix 20 39\noutside 40 325\nTMhelix 326 348\ninside 349 372

18132 GCF_001242495.1_Salmonella_enterica_CVM_N45944_v1.0 Salmonella enterica subsp. enterica serovar Kentucky Proteobacteria; Gammaproteobacteria; Enterobacterales; Enterobacteriaceae; Salmonella; Salmonella enterica; Salmonella enterica subsp. enterica MTGDNTLITSHGINRRDFMKLCAALAATMGLSSKAAAEMAESVSNPQRPPVIWIGAQECT WP_000145429.1 hydrogenase 2 small subunit [Salmonella enterica] Length: 372\nNumber of predicted TMHs: 1\nExp number of AAs in TMHs: 24.51309\nExp number, first 60 AAs: 0.2682\nTotal prob of N-in: 0.14402\noutside 1 331\nTMhelix 332 354\ninside 355 372

18133 GCF_001242405.1_Salmonella_enterica_CVM_N45939_v1.0 Salmonella enterica subsp. enterica serovar Kentucky Proteobacteria; Gammaproteobacteria; Enterobacterales; Enterobacteriaceae; Salmonella; Salmonella enterica; Salmonella enterica subsp. enterica MTGDNTLITSHGINRRDFMKLCAALAATMGLSSKAAAEMAESVSNPQRPPVIWIGAQECT WP_000145429.1 hydrogenase 2 small subunit [Salmonella enterica] Length: 372\nNumber of predicted TMHs: 1\nExp number of AAs in TMHs: 24.51309\nExp number, first 60 AAs: 0.2682\nTotal prob of N-in: 0.14402\noutside 1 331\nTMhelix 332 354\ninside 355 372

18134 GCF_001242375.1_Salmonella_enterica_CVM_N45937_v1.0 Salmonella enterica subsp. enterica serovar Kentucky Proteobacteria; Gammaproteobacteria; Enterobacterales; Enterobacteriaceae; Salmonella; Salmonella enterica; Salmonella enterica subsp. enterica MNNEETFYQAMRRKGVTRRSFLKFCSLAATSLGLGAGMTPKIAWALENKPRIPVVWIHGL WP_001058311.1 hydrogenase [Salmonella enterica] Length: 372\nNumber of predicted TMHs: 2\nExp number of AAs in TMHs: 37.4699899999999\nExp number, first 60 AAs: 19.72699\nTotal prob of N-in: 0.96171\nPOSSIBLE N-term signal sequence\ninside 1 19\nTMhelix 20 39\noutside 40 325\nTMhelix 326 348\ninside 349 372

18135 GCF_001242375.1_Salmonella_enterica_CVM_N45937_v1.0 Salmonella enterica subsp. enterica serovar Kentucky Proteobacteria; Gammaproteobacteria; Enterobacterales; Enterobacteriaceae; Salmonella; Salmonella enterica;

Salmonella enterica subsp. enterica	MTGDNTLITSHGINRRDFMKLCAALAATMGLSSKAAAEMAESVSNPQRPPVIWIGAQECT	WP_000145429.1	hydrogenase 2 small subunit [Salmonella enterica]	Length: 372\nNumber of predicted TMHs: 1\nExp number of AAs in TMHs: 24.51309\nExp number, first 60 AAs: 0.2682\nTotal prob of N-in: 0.14402\noutside 1 331\nTMhelix 332 354\ninside 355 372

18136	GCF_001245205.1_Salmonella_enterica_CVM_N50419_v1.0	Salmonella enterica subsp. enterica serovar Kentucky Proteobacteria; Gammaproteobacteria; Enterobacterales; Enterobacteriaceae; Salmonella; Salmonella enterica; Salmonella enterica subsp. enterica	MTGDNTLITSHGINRRDFMKLCAALAATMGLSSKAAAEMAESVSNPQRPPVIWIGAQECT	WP_000145429.1	hydrogenase 2 small subunit [Salmonella enterica]	Length: 372\nNumber of predicted TMHs: 1\nExp number of AAs in TMHs: 24.51309\nExp number, first 60 AAs: 0.2682\nTotal prob of N-in: 0.14402\noutside 1 331\nTMhelix 332 354\ninside 355 372

18137	GCF_001272205.1_Salmonella_enterica_CVM_N48705_v1.0	Salmonella enterica subsp. enterica serovar Kentucky Proteobacteria; Gammaproteobacteria; Enterobacterales; Enterobacteriaceae; Salmonella; Salmonella enterica; Salmonella enterica subsp. enterica	MNNEETFYQAMRRKGVTRRSFLKFCSLAATSLGLGAGMTPKIAWALENKPRIPVVWIHGL	WP_001058311.1	hydrogenase [Salmonella enterica]	Length: 372\nNumber of predicted TMHs: 2\nExp number of AAs in TMHs: 37.4699899999999\nExp number, first 60 AAs: 19.72699\nTotal prob of N-in: 0.96171\nPOSSIBLE N-term signal sequence\ninside 1 19\nTMhelix 20 39\noutside 40 325\nTMhelix 326 348\ninside 349 372

18138	GCF_001272205.1_Salmonella_enterica_CVM_N48705_v1.0	Salmonella enterica subsp. enterica serovar Kentucky Proteobacteria; Gammaproteobacteria; Enterobacterales; Enterobacteriaceae; Salmonella; Salmonella enterica; Salmonella enterica subsp. enterica	MTGDNTLITSHGINRRDFMKLCAALAATMGLSSKAAAEMAESVSNPQRPPVIWIGAQECT	WP_000145429.1	hydrogenase 2 small subunit [Salmonella enterica]	Length: 372\nNumber of predicted TMHs: 1\nExp number of AAs in TMHs: 24.51309\nExp number, first 60 AAs: 0.2682\nTotal prob of N-in: 0.14402\noutside 1 331\nTMhelix 332 354\ninside 355 372

18139	GCF_001271475.1_Salmonella_enterica_CVM_N44693_v1.0	Salmonella enterica subsp. enterica serovar Kentucky Proteobacteria; Gammaproteobacteria; Enterobacterales; Enterobacteriaceae; Salmonella; Salmonella enterica; Salmonella enterica subsp. enterica	MNNEETFYQAMRRKGVTRRSFLKFCSLAATSLGLGAGMTPKIAWALENKPRIPVVWIHGL	WP_001058311.1	hydrogenase [Salmonella enterica]	Length: 372\nNumber of predicted TMHs: 2\nExp number of AAs in TMHs: 37.4699899999999\nExp number, first 60 AAs: 19.72699\nTotal prob of N-in: 0.96171\nPOSSIBLE N-term signal sequence\ninside 1 19\nTMhelix 20 39\noutside 40 325\nTMhelix 326 348\ninside 349 372

18140	GCF_001271475.1_Salmonella_enterica_CVM_N44693_v1.0	Salmonella enterica subsp. enterica serovar Kentucky Proteobacteria; Gammaproteobacteria; Enterobacterales; Enterobacteriaceae; Salmonella; Salmonella enterica; Salmonella enterica subsp. enterica	MTGDNTLITSHGINRRDFMKLCAALAATMGLSSKAAAEMAESVSNPQRPPVIWIGAQECT	WP_000145429.1	hydrogenase 2 small subunit [Salmonella enterica]	Length: 372\nNumber of predicted TMHs: 1\nExp number of AAs in TMHs: 24.51309\nExp number, first 60 AAs: 0.2682\nTotal prob of N-in: 0.14402\noutside 1 331\nTMhelix 332 354\ninside 355 372

18141	GCF_001247545.1_Salmonella_enterica_CVM_N51982_v1.0	Salmonella enterica subsp. enterica serovar Kentucky Proteobacteria; Gammaproteobacteria; Enterobacterales; Enterobacteriaceae; Salmonella; Salmonella enterica; Salmonella enterica subsp. enterica	MNNEETFYQAMRRKGVTRRSFLKFCSLAATSLGLGAGMTPKIAWALENKPRIPVVWIHGL	WP_001058311.1	hydrogenase [Salmonella enterica]	Length: 372\nNumber of predicted TMHs: 2\nExp number of AAs in TMHs: 37.4699899999999\nExp number, first 60 AAs: 19.72699\nTotal prob of N-in: 0.96171\nPOSSIBLE N-term signal sequence\ninside 1 19\nTMhelix 20 39\noutside 40 325\nTMhelix 326 348\ninside 349 372

18142	GCF_001247545.1_Salmonella_enterica_CVM_N51982_v1.0	Salmonella enterica subsp. enterica serovar Kentucky Proteobacteria; Gammaproteobacteria; Enterobacterales; Enterobacteriaceae; Salmonella; Salmonella enterica; Salmonella enterica subsp. enterica	MTGDNTLITSHGINRRDFMKLCAALAATMGLSSKAAAEMAESVSNPQRPPVIWIGAQECT	WP_000145429.1	hydrogenase 2 small subunit [Salmonella enterica]	Length: 372\nNumber of predicted TMHs: 1\nExp number of AAs in TMHs: 24.51309\nExp number, first 60 AAs: 0.2682\nTotal prob of N-in: 0.14402\noutside 1 331\nTMhelix 332 354\ninside 355 372

18143	GCF_001247085.1_Salmonella_enterica_CVM_N51294_v1.0	Salmonella enterica subsp. enterica serovar Kentucky Proteobacteria; Gammaproteobacteria; Enterobacterales; Enterobacteriaceae; Salmonella; Salmonella enterica; Salmonella enterica subsp. enterica	MNNEETFYQAMRRKGVTRRSFLKFCSLAATSLGLGAGMTPKIAWALENKPRIPVVWIHGL	WP_001058311.1	hydrogenase [Salmonella enterica]	Length: 372\nNumber of predicted TMHs: 2\nExp

number of AAs in TMHs: 37.4699899999999\nExp number, first 60 AAs: 19.72699\nTotal prob of N-in: 0.96171\nPOSSIBLE N-term signal sequence\ninside 1 19\nTMhelix 20 39\noutside 40 325\nTMhelix 326 348\ninside 349 372

18144 GCF_001247085.1_Salmonella_enterica_CVM_N51294_v1.0 Salmonella enterica subsp. enterica serovar Kentucky Proteobacteria; Gammaproteobacteria; Enterobacterales; Enterobacteriaceae; Salmonella; Salmonella enterica; Salmonella enterica subsp. enterica MTGDNTLITSHGINRRDFMKLCAALAATMGLSSKAAAEMAESVSNPQRPPVIWIGAQECT WP_000145429.1 hydrogenase 2 small subunit [Salmonella enterica] Length: 372\nNumber of predicted TMHs: 1\nExp number of AAs in TMHs: 24.51309\nExp number, first 60 AAs: 0.2682\nTotal prob of N-in: 0.14402\noutside 1 331\nTMhelix 332 354\ninside 355 372

18145 GCF_001247055.1_Salmonella_enterica_CVM_N51290_v1.0 Salmonella enterica subsp. enterica serovar Kentucky Proteobacteria; Gammaproteobacteria; Enterobacterales; Enterobacteriaceae; Salmonella; Salmonella enterica; Salmonella enterica subsp. enterica MNNEETFYQAMRRKGVTRRSFLKFCSLAATSLGLGAGMTPKIAWALENKPRIPVVWIHGL WP_001058307.1 hydrogenase [Salmonella enterica] Length: 372\nNumber of predicted TMHs: 2\nExp number of AAs in TMHs: 36.51738\nExp number, first 60 AAs: 19.72849\nTotal prob of N-in: 0.96170\nPOSSIBLE N-term signal sequence\ninside 1 19\nTMhelix 20 39\noutside 40 325\nTMhelix 326 348\ninside 349 372

18146 GCF_001247055.1_Salmonella_enterica_CVM_N51290_v1.0 Salmonella enterica subsp. enterica serovar Kentucky Proteobacteria; Gammaproteobacteria; Enterobacterales; Enterobacteriaceae; Salmonella; Salmonella enterica; Salmonella enterica subsp. enterica MTGDNTLITSHGINRRDFMKLCAALAATMGLSSKAAAEMAESVSNPQRPPVIWIGAQECT WP_000145429.1 hydrogenase 2 small subunit [Salmonella enterica] Length: 372\nNumber of predicted TMHs: 1\nExp number of AAs in TMHs: 24.51309\nExp number, first 60 AAs: 0.2682\nTotal prob of N-in: 0.14402\noutside 1 331\nTMhelix 332 354\ninside 355 372

18147 GCF_001246665.1_Salmonella_enterica_CVM_N51273_v1.0 Salmonella enterica subsp. enterica serovar Kentucky Proteobacteria; Gammaproteobacteria; Enterobacterales; Enterobacteriaceae; Salmonella; Salmonella enterica; Salmonella enterica subsp. enterica MNNEETFYQAMRRKGVTRRSFLKFCSLAATSLGLGAGMTPKIAWALENKPRIPVVWIHGL WP_001058311.1 hydrogenase [Salmonella enterica] Length: 372\nNumber of predicted TMHs: 2\nExp number of AAs in TMHs: 37.4699899999999\nExp number, first 60 AAs: 19.72699\nTotal prob of N-in: 0.96171\nPOSSIBLE N-term signal sequence\ninside 1 19\nTMhelix 20 39\noutside 40 325\nTMhelix 326 348\ninside 349 372

18148 GCF_001246665.1_Salmonella_enterica_CVM_N51273_v1.0 Salmonella enterica subsp. enterica serovar Kentucky Proteobacteria; Gammaproteobacteria; Enterobacterales; Enterobacteriaceae; Salmonella; Salmonella enterica; Salmonella enterica subsp. enterica MTGDNTLITSHGINRRDFMKLCAALAATMGLSSKAAAEMAESVSNPQRPPVIWIGAQECT WP_000145429.1 hydrogenase 2 small subunit [Salmonella enterica] Length: 372\nNumber of predicted TMHs: 1\nExp number of AAs in TMHs: 24.51309\nExp number, first 60 AAs: 0.2682\nTotal prob of N-in: 0.14402\noutside 1 331\nTMhelix 332 354\ninside 355 372

18149 GCF_001246255.1_Salmonella_enterica_CVM_N51256_v1.0 Salmonella enterica subsp. enterica serovar Kentucky Proteobacteria; Gammaproteobacteria; Enterobacterales; Enterobacteriaceae; Salmonella; Salmonella enterica; Salmonella enterica subsp. enterica MNNEETFYQAMRRKGVTRRSFLKFCSLAATSLGLGAGMTPKIAWALENKPRIPVVWIHGL WP_001058311.1 hydrogenase [Salmonella enterica] Length: 372\nNumber of predicted TMHs: 2\nExp number of AAs in TMHs: 37.4699899999999\nExp number, first 60 AAs: 19.72699\nTotal prob of N-in: 0.96171\nPOSSIBLE N-term signal sequence\ninside 1 19\nTMhelix 20 39\noutside 40 325\nTMhelix 326 348\ninside 349 372

18150 GCF_001246255.1_Salmonella_enterica_CVM_N51256_v1.0 Salmonella enterica subsp. enterica serovar Kentucky Proteobacteria; Gammaproteobacteria; Enterobacterales; Enterobacteriaceae; Salmonella; Salmonella enterica; Salmonella enterica subsp. enterica MTGDNTLITSHGINRRDFMKLCAALAATMGLSSKAAAEMAESVSNPQRPPVIWIGAQECT WP_000145429.1 hydrogenase 2 small subunit [Salmonella enterica] Length: 372\nNumber of predicted TMHs: 1\nExp number of AAs in TMHs: 24.51309\nExp number, first 60 AAs: 0.2682\nTotal prob of N-in: 0.14402\noutside 1 331\nTMhelix 332 354\ninside 355 372

18151 GCF_001246115.1_Salmonella_enterica_CVM_N51249_v1.0 Salmonella enterica subsp. enterica serovar Kentucky Proteobacteria; Gammaproteobacteria; Enterobacterales; Enterobacteriaceae; Salmonella; Salmonella enterica; Salmonella enterica subsp. enterica MNNEETFYQAMRRKGVTRRSFLKFCSLAATSLGLGAGMTPKIAWALENKPRIPVVWIHGL WP_001058311.1 hydrogenase [Salmonella enterica] Length: 372\nNumber of predicted TMHs: 2\nExp number of AAs in TMHs: 37.4699899999999\nExp number, first 60 AAs: 19.72699\nTotal prob of N-in:

0.96171\nPOSSIBLE N-term signal sequence\ninside 1 19\nTMhelix 20 39\noutside 40 325\nTMhelix 326 348\ninside 349 372

18152 GCF_001246115.1_Salmonella_enterica_CVM_N51249_v1.0 Salmonella enterica subsp. enterica serovar Kentucky Proteobacteria; Gammaproteobacteria; Enterobacterales; Enterobacteriaceae; Salmonella; Salmonella enterica; Salmonella enterica subsp. enterica MTGDNTLITSHGINRRDFMKLCAALAATMGLSSKAAAEMAESVSNPQRPPVIWIGAQECT WP_000145429.1 hydrogenase 2 small subunit [Salmonella enterica] Length: 372\nNumber of predicted TMHs: 1\nExp number of AAs in TMHs: 24.51309\nExp number, first 60 AAs: 0.2682\nTotal prob of N-in: 0.14402\noutside 1 331\nTMhelix 332 354\ninside 355 372

18153 GCF_001245465.1_Salmonella_enterica_CVM_N50429_v1.0 Salmonella enterica subsp. enterica serovar Kentucky Proteobacteria; Gammaproteobacteria; Enterobacterales; Enterobacteriaceae; Salmonella; Salmonella enterica; Salmonella enterica subsp. enterica MNNEETFYQAMRRKGVTRRSFLKFCSLAATSLGLGAGMTPKIAWALENKPRIPVVWIHGL WP_001058311.1 hydrogenase [Salmonella enterica] Length: 372\nNumber of predicted TMHs: 2\nExp number of AAs in TMHs: 37.4699899999999\nExp number, first 60 AAs: 19.72699\nTotal prob of N-in: 0.96171\nPOSSIBLE N-term signal sequence\ninside 1 19\nTMhelix 20 39\noutside 40 325\nTMhelix 326 348\ninside 349 372

18154 GCF_001245465.1_Salmonella_enterica_CVM_N50429_v1.0 Salmonella enterica subsp. enterica serovar Kentucky Proteobacteria; Gammaproteobacteria; Enterobacterales; Enterobacteriaceae; Salmonella; Salmonella enterica; Salmonella enterica subsp. enterica MTGDNTLITSHGINRRDFMKLCAALAATMGLSSKAAAEMAESVSNPQRPPVIWIGAQECT WP_000145429.1 hydrogenase 2 small subunit [Salmonella enterica] Length: 372\nNumber of predicted TMHs: 1\nExp number of AAs in TMHs: 24.51309\nExp number, first 60 AAs: 0.2682\nTotal prob of N-in: 0.14402\noutside 1 331\nTMhelix 332 354\ninside 355 372

18155 GCF_001242495.1_Salmonella_enterica_CVM_N45944_v1.0 Salmonella enterica subsp. enterica serovar Kentucky Proteobacteria; Gammaproteobacteria; Enterobacterales; Enterobacteriaceae; Salmonella; Salmonella enterica; Salmonella enterica subsp. enterica MNNEETFYQAMRRKGVTRRSFLKFCSLAATSLGLGAGMTPKIAWALENKPRIPVVWIHGL WP_001058311.1 hydrogenase [Salmonella enterica] Length: 372\nNumber of predicted TMHs: 2\nExp number of AAs in TMHs: 37.4699899999999\nExp number, first 60 AAs: 19.72699\nTotal prob of N-in: 0.96171\nPOSSIBLE N-term signal sequence\ninside 1 19\nTMhelix 20 39\noutside 40 325\nTMhelix 326 348\ninside 349 372

18156 GCF_001241395.1_Salmonella_enterica_CVM_N44708_v1.0 Salmonella enterica subsp. enterica serovar Kentucky Proteobacteria; Gammaproteobacteria; Enterobacterales; Enterobacteriaceae; Salmonella; Salmonella enterica; Salmonella enterica subsp. enterica MTGDNTLITSHGINRRDFMKLCAALAATMGLSSKAAAEMAESVSNPQRPPVIWIGAQECT WP_000145429.1 hydrogenase 2 small subunit [Salmonella enterica] Length: 372\nNumber of predicted TMHs: 1\nExp number of AAs in TMHs: 24.51309\nExp number, first 60 AAs: 0.2682\nTotal prob of N-in: 0.14402\noutside 1 331\nTMhelix 332 354\ninside 355 372

18157 GCF_001241025.1_Salmonella_enterica_CVM_N43835_v1.0 Salmonella enterica subsp. enterica serovar Kentucky Proteobacteria; Gammaproteobacteria; Enterobacterales; Enterobacteriaceae; Salmonella; Salmonella enterica; Salmonella enterica subsp. enterica MNNEETFYQAMRRKGVTRRSFLKFCSLAATSLGLGAGMTPKIAWALENKPRIPVVWIHGL WP_001058311.1 hydrogenase [Salmonella enterica] Length: 372\nNumber of predicted TMHs: 2\nExp number of AAs in TMHs: 37.4699899999999\nExp number, first 60 AAs: 19.72699\nTotal prob of N-in: 0.96171\nPOSSIBLE N-term signal sequence\ninside 1 19\nTMhelix 20 39\noutside 40 325\nTMhelix 326 348\ninside 349 372

18158 GCF_001241025.1_Salmonella_enterica_CVM_N43835_v1.0 Salmonella enterica subsp. enterica serovar Kentucky Proteobacteria; Gammaproteobacteria; Enterobacterales; Enterobacteriaceae; Salmonella; Salmonella enterica; Salmonella enterica subsp. enterica MTGDNTLITSHGINRRDFMKLCAALAATMGLSSKAAAEMAESVSNPQRPPVIWIGAQECT WP_000145429.1 hydrogenase 2 small subunit [Salmonella enterica] Length: 372\nNumber of predicted TMHs: 1\nExp number of AAs in TMHs: 24.51309\nExp number, first 60 AAs: 0.2682\nTotal prob of N-in: 0.14402\noutside 1 331\nTMhelix 332 354\ninside 355 372

18159 GCF_001240565.1_Salmonella_enterica_CVM_N43478_v1.0 Salmonella enterica subsp. enterica serovar Kentucky Proteobacteria; Gammaproteobacteria; Enterobacterales; Enterobacteriaceae; Salmonella; Salmonella enterica; Salmonella enterica subsp. enterica MNNEETFYQAMRRKGVTRRSFLKFCSLAATSLGLGAGMTPKIAWALENKPRIPVVWIHGL WP_001058311.1 hydrogenase [Salmonella enterica] Length: 372\nNumber of predicted TMHs: 2\nExp number of AAs in TMHs: 37.4699899999999\nExp number, first 60 AAs: 19.72699\nTotal prob of N-in:

0.96171\nPOSSIBLE N-term signal sequence\ninside 1 19\nTMhelix 20 39\noutside 40 325\nTMhelix 326 348\ninside 349 372

18160 GCF_001245135.1_Salmonella_enterica_CVM_N48710_v1.0 Salmonella enterica subsp. enterica serovar Kentucky Proteobacteria; Gammaproteobacteria; Enterobacterales; Enterobacteriaceae; Salmonella; Salmonella enterica; Salmonella enterica subsp. enterica MNNEETFYQAMRRKGVTRRSFLKFCSLAATSLGLGAGMTPKIAWALENKPRIPVVWIHGL WP_001058311.1 hydrogenase [Salmonella enterica] Length: 372\nNumber of predicted TMHs: 2\nExp number of AAs in TMHs: 37.4699899999999\nExp number, first 60 AAs: 19.72699\nTotal prob of N-in: 0.96171\nPOSSIBLE N-term signal sequence\ninside 1 19\nTMhelix 20 39\noutside 40 325\nTMhelix 326 348\ninside 349 372

18161 GCF_001240565.1_Salmonella_enterica_CVM_N43478_v1.0 Salmonella enterica subsp. enterica serovar Kentucky Proteobacteria; Gammaproteobacteria; Enterobacterales; Enterobacteriaceae; Salmonella; Salmonella enterica; Salmonella enterica subsp. enterica MTGDNTLITSHGINRRDFMKLCAALAATMGLSSKAAAEMAESVSNPQRPPVIWIGAQECT WP_000145429.1 hydrogenase 2 small subunit [Salmonella enterica] Length: 372\nNumber of predicted TMHs: 1\nExp number of AAs in TMHs: 24.51309\nExp number, first 60 AAs: 0.2682\nTotal prob of N-in: 0.14402\noutside 1 331\nTMhelix 332 354\ninside 355 372

18162 GCF_001240425.1_Salmonella_enterica_CVM_N43465_v1.0 Salmonella enterica subsp. enterica serovar Kentucky Proteobacteria; Gammaproteobacteria; Enterobacterales; Enterobacteriaceae; Salmonella; Salmonella enterica; Salmonella enterica subsp. enterica MNNEETFYQAMRRKGVTRRSFLKFCSLAATSLGLGAGMTPKIAWALENKPRIPVVWIHGL WP_001058311.1 hydrogenase [Salmonella enterica] Length: 372\nNumber of predicted TMHs: 2\nExp number of AAs in TMHs: 37.4699899999999\nExp number, first 60 AAs: 19.72699\nTotal prob of N-in: 0.96171\nPOSSIBLE N-term signal sequence\ninside 1 19\nTMhelix 20 39\noutside 40 325\nTMhelix 326 348\ninside 349 372

18163 GCF_001240425.1_Salmonella_enterica_CVM_N43465_v1.0 Salmonella enterica subsp. enterica serovar Kentucky Proteobacteria; Gammaproteobacteria; Enterobacterales; Enterobacteriaceae; Salmonella; Salmonella enterica; Salmonella enterica subsp. enterica MTGDNTLITSHGINRRDFMKLCAALAATMGLSSKAAAEMAESVSNPQRPPVIWIGAQECT WP_000145429.1 hydrogenase 2 small subunit [Salmonella enterica] Length: 372\nNumber of predicted TMHs: 1\nExp number of AAs in TMHs: 24.51309\nExp number, first 60 AAs: 0.2682\nTotal prob of N-in: 0.14402\noutside 1 331\nTMhelix 332 354\ninside 355 372

18164 GCF_001238375.1_Salmonella_enterica_CVM_N43447-R_v1.0 Salmonella enterica subsp. enterica serovar Kentucky Proteobacteria; Gammaproteobacteria; Enterobacterales; Enterobacteriaceae; Salmonella; Salmonella enterica; Salmonella enterica subsp. enterica MNNEETFYQAMRRKGVTRRSFLKFCSLAATSLGLGAGMTPKIAWALENKPRIPVVWIHGL WP_001058311.1 hydrogenase [Salmonella enterica] Length: 372\nNumber of predicted TMHs: 2\nExp number of AAs in TMHs: 37.4699899999999\nExp number, first 60 AAs: 19.72699\nTotal prob of N-in: 0.96171\nPOSSIBLE N-term signal sequence\ninside 1 19\nTMhelix 20 39\noutside 40 325\nTMhelix 326 348\ninside 349 372

18165 GCF_001238285.1_Salmonella_enterica_CVM_N43450_v1.0 Salmonella enterica subsp. enterica serovar Kentucky Proteobacteria; Gammaproteobacteria; Enterobacterales; Enterobacteriaceae; Salmonella; Salmonella enterica; Salmonella enterica subsp. enterica MNNEETFYQAMRRKGVTRRSFLKFCSLAATSLGLGAGMTPKIAWALENKPRIPVVWIHGL WP_001058311.1 hydrogenase [Salmonella enterica] Length: 372\nNumber of predicted TMHs: 2\nExp number of AAs in TMHs: 37.4699899999999\nExp number, first 60 AAs: 19.72699\nTotal prob of N-in: 0.96171\nPOSSIBLE N-term signal sequence\ninside 1 19\nTMhelix 20 39\noutside 40 325\nTMhelix 326 348\ninside 349 372

18166 GCF_001238285.1_Salmonella_enterica_CVM_N43450_v1.0 Salmonella enterica subsp. enterica serovar Kentucky Proteobacteria; Gammaproteobacteria; Enterobacterales; Enterobacteriaceae; Salmonella; Salmonella enterica; Salmonella enterica subsp. enterica MTGDNTLITSHGINRRDFMKLCAALAATMGLSSKAAAEMAESVSNPQRPPVIWIGAQECT WP_000145429.1 hydrogenase 2 small subunit [Salmonella enterica] Length: 372\nNumber of predicted TMHs: 1\nExp number of AAs in TMHs: 24.51309\nExp number, first 60 AAs: 0.2682\nTotal prob of N-in: 0.14402\noutside 1 331\nTMhelix 332 354\ninside 355 372

18167 GCF_001238375.1_Salmonella_enterica_CVM_N43447-R_v1.0 Salmonella enterica subsp. enterica serovar Kentucky Proteobacteria; Gammaproteobacteria; Enterobacterales; Enterobacteriaceae; Salmonella; Salmonella enterica; Salmonella enterica subsp. enterica MTGDNTLITSHGINRRDFMKLCAALAATMGLSSKAAAEMAESVSNPQRPPVIWIGAQECT WP_000145429.1 hydrogenase 2 small subunit [Salmonella enterica] Length: 372\nNumber of

predicted TMHs: 1\nExp number of AAs in TMHs: 24.51309\nExp number, first 60 AAs: 0.2682\nTotal prob of N-in: 0.14402\noutside 1 331\nTMhelix 332 354\ninside 355 372

18168 GCF_001244185.1_Salmonella_enterica_CVM_N47730_v1.0 Salmonella enterica subsp. enterica serovar Kentucky Proteobacteria; Gammaproteobacteria; Enterobacterales; Enterobacteriaceae; Salmonella; Salmonella enterica; Salmonella enterica subsp. enterica MNNEETFYQAMRRKGVTRRSFLKFCSLAATSLGLGAGMTPKIAWALENKPRIPVVWIHGL WP_001058311.1 hydrogenase [Salmonella enterica] Length: 372\nNumber of predicted TMHs: 2\nExp number of AAs in TMHs: 37.4699899999999\nExp number, first 60 AAs: 19.72699\nTotal prob of N-in: 0.96171\nPOSSIBLE N-term signal sequence\ninside 1 19\nTMhelix 20 39\noutside 40 325\nTMhelix 326 348\ninside 349 372

18169 GCF_001245295.1_Salmonella_enterica_CVM_N50421_v1.0 Salmonella enterica subsp. enterica serovar Kentucky Proteobacteria; Gammaproteobacteria; Enterobacterales; Enterobacteriaceae; Salmonella; Salmonella enterica; Salmonella enterica subsp. enterica MNNEETFYQAMRRKGVTRRSFLKFCSLAATSLGLGAGMTPKIAWALENKPRIPVVWIHGL WP_001058311.1 hydrogenase [Salmonella enterica] Length: 372\nNumber of predicted TMHs: 2\nExp number of AAs in TMHs: 37.4699899999999\nExp number, first 60 AAs: 19.72699\nTotal prob of N-in: 0.96171\nPOSSIBLE N-term signal sequence\ninside 1 19\nTMhelix 20 39\noutside 40 325\nTMhelix 326 348\ninside 349 372

18170 GCF_002050605.1_ASM205060v1 Salmonella enterica subsp. enterica serovar Kentucky Proteobacteria; Gammaproteobacteria; Enterobacterales; Enterobacteriaceae; Salmonella; Salmonella enterica; Salmonella enterica subsp. enterica MNNEETFYQAMRRKGVTRRSFLKFCSLAATSLGLGAGMTPKIAWALENKPRIPVVWIHGL WP_001058311.1 hydrogenase [Salmonella enterica] Length: 372\nNumber of predicted TMHs: 2\nExp number of AAs in TMHs: 37.4699899999999\nExp number, first 60 AAs: 19.72699\nTotal prob of N-in: 0.96171\nPOSSIBLE N-term signal sequence\ninside 1 19\nTMhelix 20 39\noutside 40 325\nTMhelix 326 348\ninside 349 372

18171 GCF_001245955.1_Salmonella_enterica_CVM_N51241_v1.0 Salmonella enterica subsp. enterica serovar Kentucky Proteobacteria; Gammaproteobacteria; Enterobacterales; Enterobacteriaceae; Salmonella; Salmonella enterica; Salmonella enterica subsp. enterica MTGDNTLITSHGINRRDFMKLCAALAATMGLSSKAAAEMAESVSNPQRPPVIWIGAQECT WP_000145429.1 hydrogenase 2 small subunit [Salmonella enterica] Length: 372\nNumber of predicted TMHs: 1\nExp number of AAs in TMHs: 24.51309\nExp number, first 60 AAs: 0.2682\nTotal prob of N-in: 0.14402\noutside 1 331\nTMhelix 332 354\ninside 355 372

18172 GCF_001242995.1_Salmonella_enterica_CVM_N46820_v1.0 Salmonella enterica subsp. enterica serovar Kentucky Proteobacteria; Gammaproteobacteria; Enterobacterales; Enterobacteriaceae; Salmonella; Salmonella enterica; Salmonella enterica subsp. enterica MTGDNTLITSHGINRRDFMKLCAALAATMGLSSKAAAEMAESVSNPQRPPVIWIGAQECT WP_000145429.1 hydrogenase 2 small subunit [Salmonella enterica] Length: 372\nNumber of predicted TMHs: 1\nExp number of AAs in TMHs: 24.51309\nExp number, first 60 AAs: 0.2682\nTotal prob of N-in: 0.14402\noutside 1 331\nTMhelix 332 354\ninside 355 372

18173 GCF_001478565.1_Salmonella_enterica_CVM_N41924-SQ_v1.0 Salmonella enterica subsp. enterica serovar Kentucky Proteobacteria; Gammaproteobacteria; Enterobacterales; Enterobacteriaceae; Salmonella; Salmonella enterica; Salmonella enterica subsp. enterica MTGDNTLITSHGINRRDFMKLCAALAATMGLSSKAAAEMAESVSNPQRPPVIWIGAQECT WP_000145429.1 hydrogenase 2 small subunit [Salmonella enterica] Length: 372\nNumber of predicted TMHs: 1\nExp number of AAs in TMHs: 24.51309\nExp number, first 60 AAs: 0.2682\nTotal prob of N-in: 0.14402\noutside 1 331\nTMhelix 332 354\ninside 355 372

18174 GCF_001478715.1_Salmonella_enterica_CVM_N29363-SQ_v1.0 Salmonella enterica subsp. enterica serovar Kentucky Proteobacteria; Gammaproteobacteria; Enterobacterales; Enterobacteriaceae; Salmonella; Salmonella enterica; Salmonella enterica subsp. enterica MTGDNTLITSHGINRRDFMKLCAALAATMGLSSKAAAEMAESVSNPQRPPVIWIGAQECT WP_000145429.1 hydrogenase 2 small subunit [Salmonella enterica] Length: 372\nNumber of predicted TMHs: 1\nExp number of AAs in TMHs: 24.51309\nExp number, first 60 AAs: 0.2682\nTotal prob of N-in: 0.14402\noutside 1 331\nTMhelix 332 354\ninside 355 372

18175 GCF_001272405.1_Salmonella_enterica_CVM_N51277_v1.0 Salmonella enterica subsp. enterica serovar Kentucky Proteobacteria; Gammaproteobacteria; Enterobacterales; Enterobacteriaceae; Salmonella; Salmonella enterica; Salmonella enterica subsp. enterica MNNEETFYQAMRRKGVTRRSFLKFCSLAATSLGLGAGMTPKIAWALENKPRIPVVWIHGL WP_001058311.1 hydrogenase [Salmonella enterica] Length: 372\nNumber of predicted TMHs: 2\nExp number of AAs in TMHs: 37.4699899999999\nExp number, first 60 AAs: 19.72699\nTotal prob of N-in:

0.96171\nPOSSIBLE N-term signal sequence\ninside 1 19\nTMhelix 20 39\noutside 40 325\nTMhelix 326 348\ninside 349 372

18176 GCF_002036235.1_ASM203623v1 Salmonella enterica subsp. enterica serovar Kentucky Proteobacteria; Gammaproteobacteria; Enterobacterales; Enterobacteriaceae; Salmonella; Salmonella enterica; Salmonella enterica subsp. enterica MNNEETFYQAMRRKGVTRRSFLKFCSLAATSLGLGAGMTPKIAWALENKPRIPVVWIHGL WP_001058311.1 hydrogenase [Salmonella enterica] Length: 372\nNumber of predicted TMHs: 2\nExp number of AAs in TMHs: 37.4699899999999\nExp number, first 60 AAs: 19.72699\nTotal prob of N-in: 0.96171\nPOSSIBLE N-term signal sequence\ninside 1 19\nTMhelix 20 39\noutside 40 325\nTMhelix 326 348\ninside 349 372

18177 GCF_001479615.1_Salmonella_enterica_CVM_N37917-SQ_v1.0 Salmonella enterica subsp. enterica serovar Kentucky Proteobacteria; Gammaproteobacteria; Enterobacterales; Enterobacteriaceae; Salmonella; Salmonella enterica; Salmonella enterica subsp. enterica MTGDNTLITSHGINRRDFMKLCAALAATMGLSSKAAAEMAESVSNPQRPPVIWIGAQECT WP_000145429.1 hydrogenase 2 small subunit [Salmonella enterica] Length: 372\nNumber of predicted TMHs: 1\nExp number of AAs in TMHs: 24.51309\nExp number, first 60 AAs: 0.2682\nTotal prob of N-in: 0.14402\noutside 1 331\nTMhelix 332 354\ninside 355 372

18178 GCF_001247535.1_Salmonella_enterica_CVM_N51981_v1.0 Salmonella enterica subsp. enterica serovar Kentucky Proteobacteria; Gammaproteobacteria; Enterobacterales; Enterobacteriaceae; Salmonella; Salmonella enterica; Salmonella enterica subsp. enterica MNNEETFYQAMRRKGVTRRSFLKFCSLAATSLGLGAGMTPKIAWALENKPRIPVVWIHGL WP_052908125.1 hydrogenase [Salmonella enterica] Length: 372\nNumber of predicted TMHs: 2\nExp number of AAs in TMHs: 37.4698099999999\nExp number, first 60 AAs: 19.72699\nTotal prob of N-in: 0.96171\nPOSSIBLE N-term signal sequence\ninside 1 19\nTMhelix 20 39\noutside 40 325\nTMhelix 326 348\ninside 349 372

18179 GCF_001247535.1_Salmonella_enterica_CVM_N51981_v1.0 Salmonella enterica subsp. enterica serovar Kentucky Proteobacteria; Gammaproteobacteria; Enterobacterales; Enterobacteriaceae; Salmonella; Salmonella enterica; Salmonella enterica subsp. enterica MTGDNTLITSHGINRRDFMKLCAALAATMGLSSKAAAEMAESVSNPQRPPVIWIGAQECT WP_000145429.1 hydrogenase 2 small subunit [Salmonella enterica] Length: 372\nNumber of predicted TMHs: 1\nExp number of AAs in TMHs: 24.51309\nExp number, first 60 AAs: 0.2682\nTotal prob of N-in: 0.14402\noutside 1 331\nTMhelix 332 354\ninside 355 372

18180 GCF_001247465.1_Salmonella_enterica_CVM_N51313_v1.0 Salmonella enterica subsp. enterica serovar Kentucky Proteobacteria; Gammaproteobacteria; Enterobacterales; Enterobacteriaceae; Salmonella; Salmonella enterica; Salmonella enterica subsp. enterica MTGDNTLITSHGINRRDFMKLCAALAATMGLSSKAAAEMAESVSNPQRPPVIWIGAQECT WP_000145429.1 hydrogenase 2 small subunit [Salmonella enterica] Length: 372\nNumber of predicted TMHs: 1\nExp number of AAs in TMHs: 24.51309\nExp number, first 60 AAs: 0.2682\nTotal prob of N-in: 0.14402\noutside 1 331\nTMhelix 332 354\ninside 355 372

18181 GCF_001479745.1_Salmonella_enterica_CVM_N38238-SQ_v1.0 Salmonella enterica subsp. enterica serovar Kentucky Proteobacteria; Gammaproteobacteria; Enterobacterales; Enterobacteriaceae; Salmonella; Salmonella enterica; Salmonella enterica subsp. enterica MNNEETFYQAMRRKGVTRRSFLKFCSLAATSLGLGAGMTPKIAWALENKPRIPVVWIHGL WP_001058311.1 hydrogenase [Salmonella enterica] Length: 372\nNumber of predicted TMHs: 2\nExp number of AAs in TMHs: 37.4699899999999\nExp number, first 60 AAs: 19.72699\nTotal prob of N-in: 0.96171\nPOSSIBLE N-term signal sequence\ninside 1 19\nTMhelix 20 39\noutside 40 325\nTMhelix 326 348\ninside 349 372

18182 GCF_001480405.1_Salmonella_enterica_CVM_N42233-SQ_v1.0 Salmonella enterica subsp. enterica serovar Kentucky Proteobacteria; Gammaproteobacteria; Enterobacterales; Enterobacteriaceae; Salmonella; Salmonella enterica; Salmonella enterica subsp. enterica MNNEETFYQAMRRKGVTRRSFLKFCSLAATSLGLGAGMTPKIAWALENKPRIPVVWIHGL WP_001058311.1 hydrogenase [Salmonella enterica] Length: 372\nNumber of predicted TMHs: 2\nExp number of AAs in TMHs: 37.4699899999999\nExp number, first 60 AAs: 19.72699\nTotal prob of N-in: 0.96171\nPOSSIBLE N-term signal sequence\ninside 1 19\nTMhelix 20 39\noutside 40 325\nTMhelix 326 348\ninside 349 372

18183 GCF_001480715.1_Salmonella_enterica_CVM_N42508_v1.0 Salmonella enterica subsp. enterica serovar Kentucky Proteobacteria; Gammaproteobacteria; Enterobacterales; Enterobacteriaceae; Salmonella; Salmonella enterica; Salmonella enterica subsp. enterica MTGDNTLITSHGINRRDFMKLCAALAATMGLSSKAAAEMAESVSNPQRPPVIWIGAQECT WP_000145429.1 hydrogenase 2 small subunit [Salmonella enterica] Length: 372\nNumber of

predicted TMHs:  1\nExp number of AAs in TMHs: 24.51309\nExp number, first 60 AAs:  0.2682\nTotal prob of N-in: 0.14402\noutside   1   331\nTMhelix   332   354\ninside   355   372

18184 GCF_001480715.1_Salmonella_enterica_CVM_N42508_v1.0 Salmonella enterica subsp. enterica serovar Kentucky Proteobacteria; Gammaproteobacteria; Enterobacterales; Enterobacteriaceae; Salmonella; Salmonella enterica; Salmonella enterica subsp. enterica MNNEETFYQAMRRKGVTRRSFLKFCSLAATSLGLGAGMTPKIAWALENKPRIPVVWIHGL WP_001058311.1 hydrogenase [Salmonella enterica] Length: 372\nNumber of predicted TMHs:  2\nExp number of AAs in TMHs: 37.4699899999999\nExp number, first 60 AAs:  19.72699\nTotal prob of N-in: 0.96171\nPOSSIBLE N-term signal sequence\ninside   1   19\nTMhelix   20   39\noutside   40   325\nTMhelix   326   348\ninside   349   372

18185 GCF_001480595.1_Salmonella_enterica_CVM_N42453-SQ_v1.0 Salmonella enterica subsp. enterica serovar Kentucky Proteobacteria; Gammaproteobacteria; Enterobacterales; Enterobacteriaceae; Salmonella; Salmonella enterica; Salmonella enterica subsp. enterica MNNEETFYQAMRRKGVTRRSFLKFCSLAATSLGLGAGMTPKIAWALENKPRIPVVWIHGL WP_001058307.1 hydrogenase [Salmonella enterica] Length: 372\nNumber of predicted TMHs:  2\nExp number of AAs in TMHs: 36.51738\nExp number, first 60 AAs:  19.72849\nTotal prob of N-in:   0.96170\nPOSSIBLE N-term signal sequence\ninside   1   19\nTMhelix   20   39\noutside   40   325\nTMhelix   326   348\ninside   349   372

18186 GCF_001481015.1_Salmonella_enterica_CVM_N41913-SQ_v1.0 Salmonella enterica subsp. enterica serovar Kentucky Proteobacteria; Gammaproteobacteria; Enterobacterales; Enterobacteriaceae; Salmonella; Salmonella enterica; Salmonella enterica subsp. enterica MTGDNTLITSHGINRRDFMKLCAALAATMGLSSKAAAEMAESVSNPQRPPVIWIGAQECT WP_000145429.1 hydrogenase 2 small subunit [Salmonella enterica] Length: 372\nNumber of predicted TMHs:  1\nExp number of AAs in TMHs: 24.51309\nExp number, first 60 AAs:  0.2682\nTotal prob of N-in: 0.14402\noutside   1   331\nTMhelix   332   354\ninside   355   372

18187 GCF_001480925.1_Salmonella_enterica_CVM_N40395-SQ_v1.0 Salmonella enterica subsp. enterica serovar Kentucky Proteobacteria; Gammaproteobacteria; Enterobacterales; Enterobacteriaceae; Salmonella; Salmonella enterica; Salmonella enterica subsp. enterica MTGDNTLITSHGINRRDFMKLCAALAATMGLSSKAAAEMAESVSNPQRPPVIWIGAQECT WP_000145429.1 hydrogenase 2 small subunit [Salmonella enterica] Length: 372\nNumber of predicted TMHs:  1\nExp number of AAs in TMHs: 24.51309\nExp number, first 60 AAs:  0.2682\nTotal prob of N-in: 0.14402\noutside   1   331\nTMhelix   332   354\ninside   355   372

18188 GCF_001480925.1_Salmonella_enterica_CVM_N40395-SQ_v1.0 Salmonella enterica subsp. enterica serovar Kentucky Proteobacteria; Gammaproteobacteria; Enterobacterales; Enterobacteriaceae; Salmonella; Salmonella enterica; Salmonella enterica subsp. enterica MNNEETFYQAMRRKGVTRRSFLKFCSLAATSLGLGAGMTPKIAWALENKPRIPVVWIHGL WP_001058311.1 hydrogenase [Salmonella enterica] Length: 372\nNumber of predicted TMHs:  2\nExp number of AAs in TMHs: 37.4699899999999\nExp number, first 60 AAs:  19.72699\nTotal prob of N-in: 0.96171\nPOSSIBLE N-term signal sequence\ninside   1   19\nTMhelix   20   39\noutside   40   325\nTMhelix   326   348\ninside   349   372

18189 GCF_001480755.1_Salmonella_enterica_CVM_N42521-SQ_v1.0 Salmonella enterica subsp. enterica serovar Kentucky Proteobacteria; Gammaproteobacteria; Enterobacterales; Enterobacteriaceae; Salmonella; Salmonella enterica; Salmonella enterica subsp. enterica MNNEETFYQAMRRKGVTRRSFLKFCSLAATSLGLGAGMTPKIAWALENKPRIPVVWIHGL WP_001058311.1 hydrogenase [Salmonella enterica] Length: 372\nNumber of predicted TMHs:  2\nExp number of AAs in TMHs: 37.4699899999999\nExp number, first 60 AAs:  19.72699\nTotal prob of N-in: 0.96171\nPOSSIBLE N-term signal sequence\ninside   1   19\nTMhelix   20   39\noutside   40   325\nTMhelix   326   348\ninside   349   372

18190 GCF_002031485.1_ASM203148v1 Salmonella enterica subsp. enterica serovar Kentucky Proteobacteria; Gammaproteobacteria; Enterobacterales; Enterobacteriaceae; Salmonella; Salmonella enterica; Salmonella enterica subsp. enterica MNNEETFYQAMRRKGVTRRSFLKFCSLAATSLGLGAGMTPKIAWALENKPRIPVVWIHGL WP_001058311.1 hydrogenase [Salmonella enterica] Length: 372\nNumber of predicted TMHs:  2\nExp number of AAs in TMHs: 37.4699899999999\nExp number, first 60 AAs:  19.72699\nTotal prob of N-in: 0.96171\nPOSSIBLE N-term signal sequence\ninside   1   19\nTMhelix   20   39\noutside   40   325\nTMhelix   326   348\ninside   349   372

18191 GCF_002031485.1_ASM203148v1 Salmonella enterica subsp. enterica serovar Kentucky Proteobacteria; Gammaproteobacteria; Enterobacterales; Enterobacteriaceae; Salmonella; Salmonella enterica; Salmonella enterica subsp. enterica MTGDNTLITSHGINRRDFMKLCAALAATMGLSSKAAAEMAESVSNPQRPPVIWIGAQECT WP_000145429.1 hydrogenase 2 small subunit [Salmonella enterica] Length: 372\nNumber of

predicted TMHs: 1\nExp number of AAs in TMHs: 24.51309\nExp number, first 60 AAs: 0.2682\nTotal prob of N-in: 0.14402\noutside 1 331\nTMhelix 332 354\ninside 355 372

18192 GCF_002031045.1_ASM203104v1 Salmonella enterica subsp. enterica serovar Kentucky Proteobacteria; Gammaproteobacteria; Enterobacterales; Enterobacteriaceae; Salmonella; Salmonella enterica; Salmonella enterica subsp. enterica MNNEETFYQAMRRKGVTRRSFLKFCSLAATSLGLGAGMTPKIAWALENKPRIPVVWIHGL WP_001058311.1 hydrogenase [Salmonella enterica] Length: 372\nNumber of predicted TMHs: 2\nExp number of AAs in TMHs: 37.4699899999999\nExp number, first 60 AAs: 19.72699\nTotal prob of N-in: 0.96171\nPOSSIBLE N-term signal sequence\ninside 1 19\nTMhelix 20 39\noutside 40 325\nTMhelix 326 348\ninside 349 372

18193 GCF_002031045.1_ASM203104v1 Salmonella enterica subsp. enterica serovar Kentucky Proteobacteria; Gammaproteobacteria; Enterobacterales; Enterobacteriaceae; Salmonella; Salmonella enterica; Salmonella enterica subsp. enterica MTGDNTLITSHGINRRDFMKLCAALAATMGLSSKAAAEMAESVSNPQRPPVIWIGAQECT WP_000145429.1 hydrogenase 2 small subunit [Salmonella enterica] Length: 372\nNumber of predicted TMHs: 1\nExp number of AAs in TMHs: 24.51309\nExp number, first 60 AAs: 0.2682\nTotal prob of N-in: 0.14402\noutside 1 331\nTMhelix 332 354\ninside 355 372

18194 GCF_001238265.1_Salmonella_enterica_CVM_N43448_v1.0 Salmonella enterica subsp. enterica serovar Kentucky Proteobacteria; Gammaproteobacteria; Enterobacterales; Enterobacteriaceae; Salmonella; Salmonella enterica; Salmonella enterica subsp. enterica MNNEETFYQAMRRKGVTRRSFLKFCSLAATSLGLGAGMTPKIAWALENKPRIPVVWIHGL WP_001058311.1 hydrogenase [Salmonella enterica] Length: 372\nNumber of predicted TMHs: 2\nExp number of AAs in TMHs: 37.4699899999999\nExp number, first 60 AAs: 19.72699\nTotal prob of N-in: 0.96171\nPOSSIBLE N-term signal sequence\ninside 1 19\nTMhelix 20 39\noutside 40 325\nTMhelix 326 348\ninside 349 372

18195 GCF_001890855.1_ASM189085v1 Salmonella enterica subsp. enterica serovar Kentucky Proteobacteria; Gammaproteobacteria; Enterobacterales; Enterobacteriaceae; Salmonella; Salmonella enterica; Salmonella enterica subsp. enterica MNNEETFYQAMRRKGVTRRSFLKFCSLAATSLGLGAGMTPKIAWALENKPRIPVVWIHGL WP_001058307.1 hydrogenase [Salmonella enterica] Length: 372\nNumber of predicted TMHs: 2\nExp number of AAs in TMHs: 36.51738\nExp number, first 60 AAs: 19.72849\nTotal prob of N-in: 0.96170\nPOSSIBLE N-term signal sequence\ninside 1 19\nTMhelix 20 39\noutside 40 325\nTMhelix 326 348\ninside 349 372

18196 GCF_001240465.1_Salmonella_enterica_CVM_N43466_v1.0 Salmonella enterica subsp. enterica serovar Kentucky Proteobacteria; Gammaproteobacteria; Enterobacterales; Enterobacteriaceae; Salmonella; Salmonella enterica; Salmonella enterica subsp. enterica MNNEETFYQAMRRKGVTRRSFLKFCSLAATSLGLGAGMTPKIAWALENKPRIPVVWIHGL WP_001058311.1 hydrogenase [Salmonella enterica] Length: 372\nNumber of predicted TMHs: 2\nExp number of AAs in TMHs: 37.4699899999999\nExp number, first 60 AAs: 19.72699\nTotal prob of N-in: 0.96171\nPOSSIBLE N-term signal sequence\ninside 1 19\nTMhelix 20 39\noutside 40 325\nTMhelix 326 348\ninside 349 372

18197 GCF_001242015.1_Salmonella_enterica_CVM_N45412_v1.0 Salmonella enterica subsp. enterica serovar Kentucky Proteobacteria; Gammaproteobacteria; Enterobacterales; Enterobacteriaceae; Salmonella; Salmonella enterica; Salmonella enterica subsp. enterica MTGDNTLITSHGINRRDFMKLCAALAATMGLSSKAAAEMAESVSNPQRPPVIWIGAQECT WP_000145429.1 hydrogenase 2 small subunit [Salmonella enterica] Length: 372\nNumber of predicted TMHs: 1\nExp number of AAs in TMHs: 24.51309\nExp number, first 60 AAs: 0.2682\nTotal prob of N-in: 0.14402\noutside 1 331\nTMhelix 332 354\ninside 355 372

18198 GCF_002030345.1_ASM203034v1 Salmonella enterica subsp. enterica serovar Kentucky Proteobacteria; Gammaproteobacteria; Enterobacterales; Enterobacteriaceae; Salmonella; Salmonella enterica; Salmonella enterica subsp. enterica MTGDNTLITSHGINRRDFMKLCAALAATMGLSSKAAAEMAESVSNPQRPPVIWIGAQECT WP_000145429.1 hydrogenase 2 small subunit [Salmonella enterica] Length: 372\nNumber of predicted TMHs: 1\nExp number of AAs in TMHs: 24.51309\nExp number, first 60 AAs: 0.2682\nTotal prob of N-in: 0.14402\noutside 1 331\nTMhelix 332 354\ninside 355 372

18199 GCF_002030185.1_ASM203018v1 Salmonella enterica subsp. enterica serovar Kentucky Proteobacteria; Gammaproteobacteria; Enterobacterales; Enterobacteriaceae; Salmonella; Salmonella enterica; Salmonella enterica subsp. enterica MTGDNTLITSHGINRRDFMKLCAALAATMGLSSKAAAEMAESVSNPQRPPVIWIGAQECT WP_000145429.1 hydrogenase 2 small subunit [Salmonella enterica] Length: 372\nNumber of predicted TMHs: 1\nExp number of AAs in TMHs: 24.51309\nExp number, first 60 AAs: 0.2682\nTotal prob of N-in: 0.14402\noutside 1 331\nTMhelix 332 354\ninside 355 372

18200	GCF_001243485.1_Salmonella_enterica_CVM_N46849-R_v1.0	Salmonella enterica subsp. enterica serovar Kentucky Proteobacteria; Gammaproteobacteria; Enterobacterales; Enterobacteriaceae; Salmonella; Salmonella enterica; Salmonella enterica subsp. enterica	MTGDNTLITSHGINRRDFMKLCAALAATMGLSSKAAAEMAESVSNPQRPPVIWIGAQECT	WP_000145429.1	hydrogenase 2 small subunit [Salmonella enterica]	Length: 372\nNumber of predicted TMHs: 1\nExp number of AAs in TMHs: 24.51309\nExp number, first 60 AAs: 0.2682\nTotal prob of N-in: 0.14402\noutside 1 331\nTMhelix 332 354\ninside 355 372

18201	GCF_001244185.1_Salmonella_enterica_CVM_N47730_v1.0	Salmonella enterica subsp. enterica serovar Kentucky Proteobacteria; Gammaproteobacteria; Enterobacterales; Enterobacteriaceae; Salmonella; Salmonella enterica; Salmonella enterica subsp. enterica	MTGDNTLITSHGINRRDFMKLCAALAATMGLSSKAAAEMAESVSNPQRPPVIWIGAQECT	WP_000145429.1	hydrogenase 2 small subunit [Salmonella enterica]	Length: 372\nNumber of predicted TMHs: 1\nExp number of AAs in TMHs: 24.51309\nExp number, first 60 AAs: 0.2682\nTotal prob of N-in: 0.14402\noutside 1 331\nTMhelix 332 354\ninside 355 372

18202	GCF_001244005.1_Salmonella_enterica_CVM_N47723_v1.0	Salmonella enterica subsp. enterica serovar Kentucky Proteobacteria; Gammaproteobacteria; Enterobacterales; Enterobacteriaceae; Salmonella; Salmonella enterica; Salmonella enterica subsp. enterica	MNNEETFYQAMRRKGVTRRSFLKFCSLAATSLGLGAGMTPKIAWALENKPRIPVVWIHGL	WP_001058311.1	hydrogenase [Salmonella enterica]	Length: 372\nNumber of predicted TMHs: 2\nExp number of AAs in TMHs: 37.4699899999999\nExp number, first 60 AAs: 19.72699\nTotal prob of N-in: 0.96171\nPOSSIBLE N-term signal sequence\ninside 1 19\nTMhelix 20 39\noutside 40 325\nTMhelix 326 348\ninside 349 372

18203	GCF_002032275.1_ASM203227v1	Salmonella enterica subsp. enterica serovar Kentucky	Proteobacteria; Gammaproteobacteria; Enterobacterales; Enterobacteriaceae; Salmonella; Salmonella enterica; Salmonella enterica subsp. enterica	MNNEETFYQAMRRKGVTRRSFLKFCSLAATSLGLGAGMTPKIAWALENKPRIPVVWIHGL	WP_001058307.1	hydrogenase [Salmonella enterica]	Length: 372\nNumber of predicted TMHs: 2\nExp number of AAs in TMHs: 36.51738\nExp number, first 60 AAs: 19.72849\nTotal prob of N-in: 0.96170\nPOSSIBLE N-term signal sequence\ninside 1 19\nTMhelix 20 39\noutside 40 325\nTMhelix 326 348\ninside 349 372

18204	GCF_002030365.1_ASM203036v1	Salmonella enterica subsp. enterica serovar Kentucky	Proteobacteria; Gammaproteobacteria; Enterobacterales; Enterobacteriaceae; Salmonella; Salmonella enterica; Salmonella enterica subsp. enterica	MNNEETFYQAMRRKGVTRRSFLKFCSLAATSLGLGAGMTPKIAWALENKPRIPVVWIHGL	WP_001058311.1	hydrogenase [Salmonella enterica]	Length: 372\nNumber of predicted TMHs: 2\nExp number of AAs in TMHs: 37.4699899999999\nExp number, first 60 AAs: 19.72699\nTotal prob of N-in: 0.96171\nPOSSIBLE N-term signal sequence\ninside 1 19\nTMhelix 20 39\noutside 40 325\nTMhelix 326 348\ninside 349 372

18205	GCF_002030365.1_ASM203036v1	Salmonella enterica subsp. enterica serovar Kentucky	Proteobacteria; Gammaproteobacteria; Enterobacterales; Enterobacteriaceae; Salmonella; Salmonella enterica; Salmonella enterica subsp. enterica	MTGDNTLITSHGINRRDFMKLCAALAATMGLSSKAAAEMAESVSNPQRPPVIWIGAQECT	WP_000145429.1	hydrogenase 2 small subunit [Salmonella enterica]	Length: 372\nNumber of predicted TMHs: 1\nExp number of AAs in TMHs: 24.51309\nExp number, first 60 AAs: 0.2682\nTotal prob of N-in: 0.14402\noutside 1 331\nTMhelix 332 354\ninside 355 372

18206	GCF_002064615.1_ASM206461v1	Salmonella enterica subsp. enterica serovar Kentucky	Proteobacteria; Gammaproteobacteria; Enterobacterales; Enterobacteriaceae; Salmonella; Salmonella enterica; Salmonella enterica subsp. enterica	MNNEETFYQAMRRKGVTRRSFLKFCSLAATSLGLGAGMTPKIAWALENKPRIPVVWIHGL	WP_001058307.1	hydrogenase [Salmonella enterica]	Length: 372\nNumber of predicted TMHs: 2\nExp number of AAs in TMHs: 36.51738\nExp number, first 60 AAs: 19.72849\nTotal prob of N-in: 0.96170\nPOSSIBLE N-term signal sequence\ninside 1 19\nTMhelix 20 39\noutside 40 325\nTMhelix 326 348\ninside 349 372

18207	GCF_002064615.1_ASM206461v1	Salmonella enterica subsp. enterica serovar Kentucky	Proteobacteria; Gammaproteobacteria; Enterobacterales; Enterobacteriaceae; Salmonella; Salmonella enterica; Salmonella enterica subsp. enterica	MTGDNTLITSHGINRRDFMKLCAALAATMGLSSKAAAEMAESVSNPQRPPVIWIGAQECT	WP_000145429.1	hydrogenase 2 small subunit [Salmonella enterica]	Length: 372\nNumber of predicted TMHs: 1\nExp number of AAs in TMHs: 24.51309\nExp number, first 60 AAs: 0.2682\nTotal prob of N-in: 0.14402\noutside 1 331\nTMhelix 332 354\ninside 355 372

18208	GCF_002036325.1_ASM203632v1	Salmonella enterica subsp. enterica serovar Kentucky	Proteobacteria; Gammaproteobacteria; Enterobacterales; Enterobacteriaceae; Salmonella; Salmonella enterica; Salmonella enterica subsp. enterica	MNNEETFYQAMRRKGVTRRSFLKFCSLAATSLGLGAGMTPKIAWALENKPRIPVVWIHGL

WP_001058311.1 hydrogenase [Salmonella enterica] Length: 372\nNumber of predicted TMHs: 2\nExp number of AAs in TMHs: 37.4699899999999\nExp number, first 60 AAs: 19.72699\nTotal prob of N-in: 0.96171\nPOSSIBLE N-term signal sequence\ninside 1 19\nTMhelix 20 39\noutside 40 325\nTMhelix 326 348\ninside 349 372

18209 GCF_002050645.1_ASM205064v1 Salmonella enterica subsp. enterica serovar Kentucky Proteobacteria; Gammaproteobacteria; Enterobacterales; Enterobacteriaceae; Salmonella; Salmonella enterica; Salmonella enterica subsp. enterica MTGDNTLITSHGINRRDFMKLCAALAATMGLSSKAAAEMAESVSNPQRPPVIWIGAQECT WP_000145429.1 hydrogenase 2 small subunit [Salmonella enterica] Length: 372\nNumber of predicted TMHs: 1\nExp number of AAs in TMHs: 24.51309\nExp number, first 60 AAs: 0.2682\nTotal prob of N-in: 0.14402\noutside 1 331\nTMhelix 332 354\ninside 355 372

18210 GCF_002065535.1_ASM206553v1 Salmonella enterica subsp. enterica serovar Kentucky Proteobacteria; Gammaproteobacteria; Enterobacterales; Enterobacteriaceae; Salmonella; Salmonella enterica; Salmonella enterica subsp. enterica MNNEETFYQAMRRKGVTRRSFLKFCSLAATSLGLGAGMTPKIAWALENKPRIPVVWIHGL WP_001058307.1 hydrogenase [Salmonella enterica] Length: 372\nNumber of predicted TMHs: 2\nExp number of AAs in TMHs: 36.51738\nExp number, first 60 AAs: 19.72849\nTotal prob of N-in: 0.96170\nPOSSIBLE N-term signal sequence\ninside 1 19\nTMhelix 20 39\noutside 40 325\nTMhelix 326 348\ninside 349 372

18211 GCF_002065495.1_ASM206549v1 Salmonella enterica subsp. enterica serovar Kentucky Proteobacteria; Gammaproteobacteria; Enterobacterales; Enterobacteriaceae; Salmonella; Salmonella enterica; Salmonella enterica subsp. enterica MNNEETFYQAMRRKGVTRRSFLKFCSLAATSLGLGAGMTPKIAWALENKPRIPVVWIHGL WP_001058307.1 hydrogenase [Salmonella enterica] Length: 372\nNumber of predicted TMHs: 2\nExp number of AAs in TMHs: 36.51738\nExp number, first 60 AAs: 19.72849\nTotal prob of N-in: 0.96170\nPOSSIBLE N-term signal sequence\ninside 1 19\nTMhelix 20 39\noutside 40 325\nTMhelix 326 348\ninside 349 372

18212 GCF_002065535.1_ASM206553v1 Salmonella enterica subsp. enterica serovar Kentucky Proteobacteria; Gammaproteobacteria; Enterobacterales; Enterobacteriaceae; Salmonella; Salmonella enterica; Salmonella enterica subsp. enterica MTGDNTLITSHGINRRDFMKLCAALAATMGLSSKAAAEMAESVSNPQRPPVIWIGAQECT WP_000145429.1 hydrogenase 2 small subunit [Salmonella enterica] Length: 372\nNumber of predicted TMHs: 1\nExp number of AAs in TMHs: 24.51309\nExp number, first 60 AAs: 0.2682\nTotal prob of N-in: 0.14402\noutside 1 331\nTMhelix 332 354\ninside 355 372

18213 GCF_002032275.1_ASM203227v1 Salmonella enterica subsp. enterica serovar Kentucky Proteobacteria; Gammaproteobacteria; Enterobacterales; Enterobacteriaceae; Salmonella; Salmonella enterica; Salmonella enterica subsp. enterica MTGDNTLITSHGINRRDFMKLCAALAATMGLSSKAAAEMAESVSNPQRPPVIWIGAQECT WP_000145429.1 hydrogenase 2 small subunit [Salmonella enterica] Length: 372\nNumber of predicted TMHs: 1\nExp number of AAs in TMHs: 24.51309\nExp number, first 60 AAs: 0.2682\nTotal prob of N-in: 0.14402\noutside 1 331\nTMhelix 332 354\ninside 355 372

18214 GCF_002065495.1_ASM206549v1 Salmonella enterica subsp. enterica serovar Kentucky Proteobacteria; Gammaproteobacteria; Enterobacterales; Enterobacteriaceae; Salmonella; Salmonella enterica; Salmonella enterica subsp. enterica MTGDNTLITSHGINRRDFMKLCAALAATMGLSSKAAAEMAESVSNPQRPPVIWIGAQECT WP_000145429.1 hydrogenase 2 small subunit [Salmonella enterica] Length: 372\nNumber of predicted TMHs: 1\nExp number of AAs in TMHs: 24.51309\nExp number, first 60 AAs: 0.2682\nTotal prob of N-in: 0.14402\noutside 1 331\nTMhelix 332 354\ninside 355 372

18215 GCF_002031615.1_ASM203161v1 Salmonella enterica subsp. enterica serovar Kentucky Proteobacteria; Gammaproteobacteria; Enterobacterales; Enterobacteriaceae; Salmonella; Salmonella enterica; Salmonella enterica subsp. enterica MNNEETFYQAMRRKGVTRRSFLKFCSLAATSLGLGAGMTPKIAWALENKPRIPVVWIHGL WP_001058311.1 hydrogenase [Salmonella enterica] Length: 372\nNumber of predicted TMHs: 2\nExp number of AAs in TMHs: 37.4699899999999\nExp number, first 60 AAs: 19.72699\nTotal prob of N-in: 0.96171\nPOSSIBLE N-term signal sequence\ninside 1 19\nTMhelix 20 39\noutside 40 325\nTMhelix 326 348\ninside 349 372

18216 GCF_002050645.1_ASM205064v1 Salmonella enterica subsp. enterica serovar Kentucky Proteobacteria; Gammaproteobacteria; Enterobacterales; Enterobacteriaceae; Salmonella; Salmonella enterica; Salmonella enterica subsp. enterica MNNEETFYQAMRRKGVTRRSFLKFCSLAATSLGLGAGMTPKIAWALENKPRIPVVWIHGL WP_001058311.1 hydrogenase [Salmonella enterica] Length: 372\nNumber of predicted TMHs: 2\nExp number of AAs in TMHs: 37.4699899999999\nExp number, first 60 AAs: 19.72699\nTotal prob of N-in:

0.96171\nPOSSIBLE N-term signal sequence\ninside 1 19\nTMhelix 20 39\noutside 40 325\nTMhelix 326 348\ninside 349 372

18217 GCF_001890825.1_ASM189082v1 Salmonella enterica subsp. enterica serovar Kentucky Proteobacteria; Gammaproteobacteria; Enterobacterales; Enterobacteriaceae; Salmonella; Salmonella enterica; Salmonella enterica subsp. enterica MTGDNTLITSHGINRRDFMKLCAALAATMGLSSKAAAEMAESVSNPQRPPVIWIGAQECT WP_000145429.1 hydrogenase 2 small subunit [Salmonella enterica] Length: 372\nNumber of predicted TMHs: 1\nExp number of AAs in TMHs: 24.51309\nExp number, first 60 AAs: 0.2682\nTotal prob of N-in: 0.14402\noutside 1 331\nTMhelix 332 354\ninside 355 372

18218 GCF_002031345.1_ASM203134v1 Salmonella enterica subsp. enterica serovar Kentucky Proteobacteria; Gammaproteobacteria; Enterobacterales; Enterobacteriaceae; Salmonella; Salmonella enterica; Salmonella enterica subsp. enterica MTGDNTLITSHGINRRDFMKLCAALAATMGLSSKAAAEMAESVSNPQRPPVIWIGAQECT WP_000145429.1 hydrogenase 2 small subunit [Salmonella enterica] Length: 372\nNumber of predicted TMHs: 1\nExp number of AAs in TMHs: 24.51309\nExp number, first 60 AAs: 0.2682\nTotal prob of N-in: 0.14402\noutside 1 331\nTMhelix 332 354\ninside 355 372

18219 GCF_001890825.1_ASM189082v1 Salmonella enterica subsp. enterica serovar Kentucky Proteobacteria; Gammaproteobacteria; Enterobacterales; Enterobacteriaceae; Salmonella; Salmonella enterica; Salmonella enterica subsp. enterica MNNEETFYQAMRRKGVTRRSFLKFCSLAATSLGLGAGMTPKIAWALENKPRIPVVWIHGL WP_001058307.1 hydrogenase [Salmonella enterica] Length: 372\nNumber of predicted TMHs: 2\nExp number of AAs in TMHs: 36.51738\nExp number, first 60 AAs: 19.72849\nTotal prob of N-in: 0.96170\nPOSSIBLE N-term signal sequence\ninside 1 19\nTMhelix 20 39\noutside 40 325\nTMhelix 326 348\ninside 349 372

18220 GCF_001890845.1_ASM189084v1 Salmonella enterica subsp. enterica serovar Kentucky Proteobacteria; Gammaproteobacteria; Enterobacterales; Enterobacteriaceae; Salmonella; Salmonella enterica; Salmonella enterica subsp. enterica MTGDNTLITSHGINRRDFMKLCAALAATMGLSSKAAAEMAESVSNPQRPPVIWIGAQECT WP_000145429.1 hydrogenase 2 small subunit [Salmonella enterica] Length: 372\nNumber of predicted TMHs: 1\nExp number of AAs in TMHs: 24.51309\nExp number, first 60 AAs: 0.2682\nTotal prob of N-in: 0.14402\noutside 1 331\nTMhelix 332 354\ninside 355 372

18221 GCF_002050525.1_ASM205052v1 Salmonella enterica subsp. enterica serovar Kentucky Proteobacteria; Gammaproteobacteria; Enterobacterales; Enterobacteriaceae; Salmonella; Salmonella enterica; Salmonella enterica subsp. enterica MNNEETFYQAMRRKGVTRRSFLKFCSLAATSLGLGAGMTPKIAWALENKPRIPVVWIHGL WP_001058311.1 hydrogenase [Salmonella enterica] Length: 372\nNumber of predicted TMHs: 2\nExp number of AAs in TMHs: 37.4699899999999\nExp number, first 60 AAs: 19.72699\nTotal prob of N-in: 0.96171\nPOSSIBLE N-term signal sequence\ninside 1 19\nTMhelix 20 39\noutside 40 325\nTMhelix 326 348\ninside 349 372

18222 GCF_001890845.1_ASM189084v1 Salmonella enterica subsp. enterica serovar Kentucky Proteobacteria; Gammaproteobacteria; Enterobacterales; Enterobacteriaceae; Salmonella; Salmonella enterica; Salmonella enterica subsp. enterica MNNEETFYQAMRRKGVTRRSFLKFCSLAATSLGLGAGMTPKIAWALENKPRIPVVWIHGL WP_001058307.1 hydrogenase [Salmonella enterica] Length: 372\nNumber of predicted TMHs: 2\nExp number of AAs in TMHs: 36.51738\nExp number, first 60 AAs: 19.72849\nTotal prob of N-in: 0.96170\nPOSSIBLE N-term signal sequence\ninside 1 19\nTMhelix 20 39\noutside 40 325\nTMhelix 326 348\ninside 349 372

18223 GCF_001243485.1_Salmonella_enterica_CVM_N46849-R_v1.0 Salmonella enterica subsp. enterica serovar Kentucky Proteobacteria; Gammaproteobacteria; Enterobacterales; Enterobacteriaceae; Salmonella; Salmonella enterica; Salmonella enterica subsp. enterica MNNEETFYQAMRRKGVTRRSFLKFCSLAATSLGLGAGMTPKIAWALENKPRIPVVWIHGL WP_001058311.1 hydrogenase [Salmonella enterica] Length: 372\nNumber of predicted TMHs: 2\nExp number of AAs in TMHs: 37.4699899999999\nExp number, first 60 AAs: 19.72699\nTotal prob of N-in: 0.96171\nPOSSIBLE N-term signal sequence\ninside 1 19\nTMhelix 20 39\noutside 40 325\nTMhelix 326 348\ninside 349 372

18224 GCF_001243655.1_Salmonella_enterica_CVM_N46857_v1.0 Salmonella enterica subsp. enterica serovar Kentucky Proteobacteria; Gammaproteobacteria; Enterobacterales; Enterobacteriaceae; Salmonella; Salmonella enterica; Salmonella enterica subsp. enterica MNNEETFYQAMRRKGVTRRSFLKFCSLAATSLGLGAGMTPKIAWALENKPRIPVVWIHGL WP_001058311.1 hydrogenase [Salmonella enterica] Length: 372\nNumber of predicted TMHs: 2\nExp number of AAs in TMHs: 37.4699899999999\nExp number, first 60 AAs: 19.72699\nTotal prob of N-in: 0.96171\nPOSSIBLE N-term signal sequence\ninside 1 19\nTMhelix 20 39\noutside 40 325\nTMhelix 326 348\ninside 349 372

18225 GCF_001479615.1_Salmonella_enterica_CVM_N37917-SQ_v1.0 Salmonella enterica subsp. enterica serovar Kentucky Proteobacteria; Gammaproteobacteria; Enterobacterales; Enterobacteriaceae; Salmonella; Salmonella enterica; Salmonella enterica subsp. enterica MNNEETFYQAMRRKGVTRRSFLKFCSLAATSLGLGAGMTPKIAWALENKPRIPVVWIHGL WP_001058311.1 hydrogenase [Salmonella enterica] Length: 372\nNumber of predicted TMHs: 2\nExp number of AAs in TMHs: 37.4699899999999\nExp number, first 60 AAs: 19.72699\nTotal prob of N-in: 0.96171\nPOSSIBLE N-term signal sequence\ninside 1 19\nTMhelix 20 39\noutside 40 325\nTMhelix 326 348\ninside 349 372

18226 GCF_001243995.1_Salmonella_enterica_CVM_N47721_v1.0 Salmonella enterica subsp. enterica serovar Kentucky Proteobacteria; Gammaproteobacteria; Enterobacterales; Enterobacteriaceae; Salmonella; Salmonella enterica; Salmonella enterica subsp. enterica MNNEETFYQAMRRKGVTRRSFLKFCSLAATSLGLGAGMTPKIAWALENKPRIPVVWIHGL WP_001058311.1 hydrogenase [Salmonella enterica] Length: 372\nNumber of predicted TMHs: 2\nExp number of AAs in TMHs: 37.4699899999999\nExp number, first 60 AAs: 19.72699\nTotal prob of N-in: 0.96171\nPOSSIBLE N-term signal sequence\ninside 1 19\nTMhelix 20 39\noutside 40 325\nTMhelix 326 348\ninside 349 372

18227 GCF_001244715.1_Salmonella_enterica_CVM_N48688_v1.0 Salmonella enterica subsp. enterica serovar Kentucky Proteobacteria; Gammaproteobacteria; Enterobacterales; Enterobacteriaceae; Salmonella; Salmonella enterica; Salmonella enterica subsp. enterica MTGDNTLITSHGINRRDFMKLCAALAATMGLSSKAAAEMAESVSNPQRPPVIWIGAQECT WP_000145429.1 hydrogenase 2 small subunit [Salmonella enterica] Length: 372\nNumber of predicted TMHs: 1\nExp number of AAs in TMHs: 24.51309\nExp number, first 60 AAs: 0.2682\nTotal prob of N-in: 0.14402\noutside 1 331\nTMhelix 332 354\ninside 355 372

18228 GCF_001244715.1_Salmonella_enterica_CVM_N48688_v1.0 Salmonella enterica subsp. enterica serovar Kentucky Proteobacteria; Gammaproteobacteria; Enterobacterales; Enterobacteriaceae; Salmonella; Salmonella enterica; Salmonella enterica subsp. enterica MNNEETFYQAMRRKGVTRRSFLKFCSLAATSLGLGAGMTPKIAWALENKPRIPVVWIHGL WP_001058311.1 hydrogenase [Salmonella enterica] Length: 372\nNumber of predicted TMHs: 2\nExp number of AAs in TMHs: 37.4699899999999\nExp number, first 60 AAs: 19.72699\nTotal prob of N-in: 0.96171\nPOSSIBLE N-term signal sequence\ninside 1 19\nTMhelix 20 39\noutside 40 325\nTMhelix 326 348\ninside 349 372

18229 GCF_001245295.1_Salmonella_enterica_CVM_N50421_v1.0 Salmonella enterica subsp. enterica serovar Kentucky Proteobacteria; Gammaproteobacteria; Enterobacterales; Enterobacteriaceae; Salmonella; Salmonella enterica; Salmonella enterica subsp. enterica MTGDNTLITSHGINRRDFMKLCAALAATMGLSSKAAAEMAESVSNPQRPPVIWIGAQECT WP_000145429.1 hydrogenase 2 small subunit [Salmonella enterica] Length: 372\nNumber of predicted TMHs: 1\nExp number of AAs in TMHs: 24.51309\nExp number, first 60 AAs: 0.2682\nTotal prob of N-in: 0.14402\noutside 1 331\nTMhelix 332 354\ninside 355 372

18230 GCF_001245195.1_Salmonella_enterica_CVM_N48711_v1.0 Salmonella enterica subsp. enterica serovar Kentucky Proteobacteria; Gammaproteobacteria; Enterobacterales; Enterobacteriaceae; Salmonella; Salmonella enterica; Salmonella enterica subsp. enterica MNNEETFYQAMRRKGVTRRSFLKFCSLAATSLGLGAGMTPKIAWALENKPRIPVVWIHGL WP_001058311.1 hydrogenase [Salmonella enterica] Length: 372\nNumber of predicted TMHs: 2\nExp number of AAs in TMHs: 37.4699899999999\nExp number, first 60 AAs: 19.72699\nTotal prob of N-in: 0.96171\nPOSSIBLE N-term signal sequence\ninside 1 19\nTMhelix 20 39\noutside 40 325\nTMhelix 326 348\ninside 349 372

18231 GCF_001480045.1_Salmonella_enterica_CVM_N38909-SQ_v1.0 Salmonella enterica subsp. enterica serovar Kentucky Proteobacteria; Gammaproteobacteria; Enterobacterales; Enterobacteriaceae; Salmonella; Salmonella enterica; Salmonella enterica subsp. enterica MTGDNTLITSHGINRRDFMKLCAALAATMGLSSKAAAEMAESVSNPQRPPVIWIGAQECT WP_000145429.1 hydrogenase 2 small subunit [Salmonella enterica] Length: 372\nNumber of predicted TMHs: 1\nExp number of AAs in TMHs: 24.51309\nExp number, first 60 AAs: 0.2682\nTotal prob of N-in: 0.14402\noutside 1 331\nTMhelix 332 354\ninside 355 372

18232 GCF_001480595.1_Salmonella_enterica_CVM_N42453-SQ_v1.0 Salmonella enterica subsp. enterica serovar Kentucky Proteobacteria; Gammaproteobacteria; Enterobacterales; Enterobacteriaceae; Salmonella; Salmonella enterica; Salmonella enterica subsp. enterica MTGDNTLITSHGINRRDFMKLCAALAATMGLSSKAAAEMAESVSNPQRPPVIWIGAQECT WP_000145429.1 hydrogenase 2 small subunit [Salmonella enterica] Length: 372\nNumber of predicted TMHs: 1\nExp number of AAs in TMHs: 24.51309\nExp number, first 60 AAs: 0.2682\nTotal prob of N-in: 0.14402\noutside 1 331\nTMhelix 332 354\ninside 355 372

18233 GCF_001245565.1_Salmonella_enterica_CVM_N50435_v1.0 Salmonella enterica subsp. enterica serovar Kentucky Proteobacteria; Gammaproteobacteria; Enterobacterales; Enterobacteriaceae; Salmonella; Salmonella enterica; Salmonella enterica subsp. enterica MNNEETFYQAMRRKGVTRRSFLKFCSLAATSLGLGAGMTPKIAWALENKPRIPVVWIHGL WP_001058311.1 hydrogenase [Salmonella enterica] Length: 372\nNumber of predicted TMHs: 2\nExp number of AAs in TMHs: 37.4699899999999\nExp number, first 60 AAs: 19.72699\nTotal prob of N-in: 0.96171\nPOSSIBLE N-term signal sequence\ninside 1 19\nTMhelix 20 39\noutside 40 325\nTMhelix 326 348\ninside 349 372

18234 GCF_001246195.1_Salmonella_enterica_CVM_N51252_v1.0 Salmonella enterica subsp. enterica serovar Kentucky Proteobacteria; Gammaproteobacteria; Enterobacterales; Enterobacteriaceae; Salmonella; Salmonella enterica; Salmonella enterica subsp. enterica MTGDNTLITSHGINRRDFMKLCAALAATMGLSSKAAAEMAESVSNPQRPPVIWIGAQECT WP_000145429.1 hydrogenase 2 small subunit [Salmonella enterica] Length: 372\nNumber of predicted TMHs: 1\nExp number of AAs in TMHs: 24.51309\nExp number, first 60 AAs: 0.2682\nTotal prob of N-in: 0.14402\noutside 1 331\nTMhelix 332 354\ninside 355 372

18235 GCF_001246195.1_Salmonella_enterica_CVM_N51252_v1.0 Salmonella enterica subsp. enterica serovar Kentucky Proteobacteria; Gammaproteobacteria; Enterobacterales; Enterobacteriaceae; Salmonella; Salmonella enterica; Salmonella enterica subsp. enterica MNNEETFYQAMRRKGVTRRSFLKFCSLAATSLGLGAGMTPKIAWALENKPRIPVVWIHGL WP_001058311.1 hydrogenase [Salmonella enterica] Length: 372\nNumber of predicted TMHs: 2\nExp number of AAs in TMHs: 37.4699899999999\nExp number, first 60 AAs: 19.72699\nTotal prob of N-in: 0.96171\nPOSSIBLE N-term signal sequence\ninside 1 19\nTMhelix 20 39\noutside 40 325\nTMhelix 326 348\ninside 349 372

18236 GCF_001245955.1_Salmonella_enterica_CVM_N51241_v1.0 Salmonella enterica subsp. enterica serovar Kentucky Proteobacteria; Gammaproteobacteria; Enterobacterales; Enterobacteriaceae; Salmonella; Salmonella enterica; Salmonella enterica subsp. enterica MNNEETFYQAMRRKGVTRRSFLKFCSLAATSLGLGAGMTPKIAWALENKPRIPVVWIHGL WP_001058311.1 hydrogenase [Salmonella enterica] Length: 372\nNumber of predicted TMHs: 2\nExp number of AAs in TMHs: 37.4699899999999\nExp number, first 60 AAs: 19.72699\nTotal prob of N-in: 0.96171\nPOSSIBLE N-term signal sequence\ninside 1 19\nTMhelix 20 39\noutside 40 325\nTMhelix 326 348\ninside 349 372

18237 GCF_001481015.1_Salmonella_enterica_CVM_N41913-SQ_v1.0 Salmonella enterica subsp. enterica serovar Kentucky Proteobacteria; Gammaproteobacteria; Enterobacterales; Enterobacteriaceae; Salmonella; Salmonella enterica; Salmonella enterica subsp. enterica MNNEETFYQAMRRKGVTRRSFLKFCSLAATSLGLGAGMTPKIAWALENKPRIPVVWIHGL WP_001058307.1 hydrogenase [Salmonella enterica] Length: 372\nNumber of predicted TMHs: 2\nExp number of AAs in TMHs: 36.51738\nExp number, first 60 AAs: 19.72849\nTotal prob of N-in: 0.96170\nPOSSIBLE N-term signal sequence\ninside 1 19\nTMhelix 20 39\noutside 40 325\nTMhelix 326 348\ninside 349 372

18238 GCF_001243655.1_Salmonella_enterica_CVM_N46857_v1.0 Salmonella enterica subsp. enterica serovar Kentucky Proteobacteria; Gammaproteobacteria; Enterobacterales; Enterobacteriaceae; Salmonella; Salmonella enterica; Salmonella enterica subsp. enterica MTGDNTLITSHGINRRDFMKLCAALAATMGLSSKAAAEMAESVSNPQRPPVIWIGAQECT WP_000145429.1 hydrogenase 2 small subunit [Salmonella enterica] Length: 372\nNumber of predicted TMHs: 1\nExp number of AAs in TMHs: 24.51309\nExp number, first 60 AAs: 0.2682\nTotal prob of N-in: 0.14402\noutside 1 331\nTMhelix 332 354\ninside 355 372

18239 GCF_001247465.1_Salmonella_enterica_CVM_N51313_v1.0 Salmonella enterica subsp. enterica serovar Kentucky Proteobacteria; Gammaproteobacteria; Enterobacterales; Enterobacteriaceae; Salmonella; Salmonella enterica; Salmonella enterica subsp. enterica MNNEETFYQAMRRKGVTRRSFLKFCSLAATSLGLGAGMTPKIAWALENKPRIPVVWIHGL WP_001058311.1 hydrogenase [Salmonella enterica] Length: 372\nNumber of predicted TMHs: 2\nExp number of AAs in TMHs: 37.4699899999999\nExp number, first 60 AAs: 19.72699\nTotal prob of N-in: 0.96171\nPOSSIBLE N-term signal sequence\ninside 1 19\nTMhelix 20 39\noutside 40 325\nTMhelix 326 348\ninside 349 372

18240 GCF_001272135.1_Salmonella_enterica_CVM_N48687_v1.0 Salmonella enterica subsp. enterica serovar Kentucky Proteobacteria; Gammaproteobacteria; Enterobacterales; Enterobacteriaceae; Salmonella; Salmonella enterica; Salmonella enterica subsp. enterica MTGDNTLITSHGINRRDFMKLCAALAATMGLSSKAAAEMAESVSNPQRPPVIWIGAQECT WP_000145429.1 hydrogenase 2 small subunit [Salmonella enterica] Length: 372\nNumber of predicted TMHs: 1\nExp number of AAs in TMHs: 24.51309\nExp number, first 60 AAs: 0.2682\nTotal prob of N-in: 0.14402\noutside 1 331\nTMhelix 332 354\ninside 355 372

18241 GCF_001271875.1_Salmonella_enterica_CVM_N47722_v1.0 Salmonella enterica subsp. enterica serovar Kentucky Proteobacteria; Gammaproteobacteria; Enterobacterales; Enterobacteriaceae; Salmonella; Salmonella enterica; Salmonella enterica subsp. enterica MNNEETFYQAMRRKGVTRRSFLKFCSLAATSLGLGAGMTPKIAWALENKPRIPVVWIHGL WP_001058311.1 hydrogenase [Salmonella enterica] Length: 372\nNumber of predicted TMHs: 2\nExp number of AAs in TMHs: 37.4699899999999\nExp number, first 60 AAs: 19.72699\nTotal prob of N-in: 0.96171\nPOSSIBLE N-term signal sequence\ninside 1 19\nTMhelix 20 39\noutside 40 325\nTMhelix 326 348\ninside 349 372

18242 GCF_001240465.1_Salmonella_enterica_CVM_N43466_v1.0 Salmonella enterica subsp. enterica serovar Kentucky Proteobacteria; Gammaproteobacteria; Enterobacterales; Enterobacteriaceae; Salmonella; Salmonella enterica; Salmonella enterica subsp. enterica MTGDNTLITSHGINRRDFMKLCAALAATMGLSSKAAAEMAESVSNPQRPPVIWIGAQECT WP_000145429.1 hydrogenase 2 small subunit [Salmonella enterica] Length: 372\nNumber of predicted TMHs: 1\nExp number of AAs in TMHs: 24.51309\nExp number, first 60 AAs: 0.2682\nTotal prob of N-in: 0.14402\noutside 1 331\nTMhelix 332 354\ninside 355 372

18243 GCF_000973905.1_ABB07-SB3057-2.id23_v1.0 Salmonella enterica subsp. enterica serovar Kentucky Proteobacteria; Gammaproteobacteria; Enterobacterales; Enterobacteriaceae; Salmonella; Salmonella enterica; Salmonella enterica subsp. enterica MTGDNTLITSHGINRRDFMKLCAALAATMGLSSKAAAEMAESVSNPQRPPVIWIGAQECT WP_000145429.1 hydrogenase 2 small subunit [Salmonella enterica] Length: 372\nNumber of predicted TMHs: 1\nExp number of AAs in TMHs: 24.51309\nExp number, first 60 AAs: 0.2682\nTotal prob of N-in: 0.14402\noutside 1 331\nTMhelix 332 354\ninside 355 372

18244 GCF_002036235.1_ASM203623v1 Salmonella enterica subsp. enterica serovar Kentucky Proteobacteria; Gammaproteobacteria; Enterobacterales; Enterobacteriaceae; Salmonella; Salmonella enterica; Salmonella enterica subsp. enterica MTGDNTLITSHGINRRDFMKLCAALAATMGLSSKAAAEMAESVSNPQRPPVIWIGAQECT WP_000145429.1 hydrogenase 2 small subunit [Salmonella enterica] Length: 372\nNumber of predicted TMHs: 1\nExp number of AAs in TMHs: 24.51309\nExp number, first 60 AAs: 0.2682\nTotal prob of N-in: 0.14402\noutside 1 331\nTMhelix 332 354\ninside 355 372

18245 GCF_001242285.1_Salmonella_enterica_CVM_N45934_v1.0 Salmonella enterica subsp. enterica serovar Kentucky Proteobacteria; Gammaproteobacteria; Enterobacterales; Enterobacteriaceae; Salmonella; Salmonella enterica; Salmonella enterica subsp. enterica MTGDNTLITSHGINRRDFMKLCAALAATMGLSSKAAAEMAESVSNPQRPPVIWIGAQECT WP_000145429.1 hydrogenase 2 small subunit [Salmonella enterica] Length: 372\nNumber of predicted TMHs: 1\nExp number of AAs in TMHs: 24.51309\nExp number, first 60 AAs: 0.2682\nTotal prob of N-in: 0.14402\noutside 1 331\nTMhelix 332 354\ninside 355 372

18246 GCF_002030345.1_ASM203034v1 Salmonella enterica subsp. enterica serovar Kentucky Proteobacteria; Gammaproteobacteria; Enterobacterales; Enterobacteriaceae; Salmonella; Salmonella enterica; Salmonella enterica subsp. enterica MNNEETFYQAMRRKGVTRRSFLKFCSLAATSLGLGAGMTPKIAWALENKPRIPVVWIHGL WP_001058311.1 hydrogenase [Salmonella enterica] Length: 372\nNumber of predicted TMHs: 2\nExp number of AAs in TMHs: 37.4699899999999\nExp number, first 60 AAs: 19.72699\nTotal prob of N-in: 0.96171\nPOSSIBLE N-term signal sequence\ninside 1 19\nTMhelix 20 39\noutside 40 325\nTMhelix 326 348\ninside 349 372

18247 GCF_002033015.1_ASM203301v1 Salmonella enterica subsp. enterica serovar Kentucky Proteobacteria; Gammaproteobacteria; Enterobacterales; Enterobacteriaceae; Salmonella; Salmonella enterica; Salmonella enterica subsp. enterica MTGDNTLITSHGINRRDFMKLCAALAATMGLSSKAAAEMAESVSNPQRPPVIWIGAQECT WP_000145429.1 hydrogenase 2 small subunit [Salmonella enterica] Length: 372\nNumber of predicted TMHs: 1\nExp number of AAs in TMHs: 24.51309\nExp number, first 60 AAs: 0.2682\nTotal prob of N-in: 0.14402\noutside 1 331\nTMhelix 332 354\ninside 355 372

18248 GCF_002030185.1_ASM203018v1 Salmonella enterica subsp. enterica serovar Kentucky Proteobacteria; Gammaproteobacteria; Enterobacterales; Enterobacteriaceae; Salmonella; Salmonella enterica; Salmonella enterica subsp. enterica MNNEETFYQAMRRKGVTRRSFLKFCSLAATSLGLGAGMTPKIAWALENKPRIPVVWIHGL WP_001058311.1 hydrogenase [Salmonella enterica] Length: 372\nNumber of predicted TMHs: 2\nExp number of AAs in TMHs: 37.4699899999999\nExp number, first 60 AAs: 19.72699\nTotal prob of N-in: 0.96171\nPOSSIBLE N-term signal sequence\ninside 1 19\nTMhelix 20 39\noutside 40 325\nTMhelix 326 348\ninside 349 372

18249 GCF_002031745.1_ASM203174v1 Salmonella enterica subsp. enterica serovar Kentucky Proteobacteria; Gammaproteobacteria; Enterobacterales; Enterobacteriaceae; Salmonella; Salmonella enterica;

Salmonella enterica subsp. enterica MTGDNTLITSHGINRRDFMKLCAALAATMGLSSKAAAEMAESVSNPQRPPVIWIGAQECT WP_000145429.1 hydrogenase 2 small subunit [Salmonella enterica] Length: 372\nNumber of predicted TMHs: 1\nExp number of AAs in TMHs: 24.51309\nExp number, first 60 AAs: 0.2682\nTotal prob of N-in: 0.14402\noutside 1 331\nTMhelix 332 354\ninside 355 372

18250 GCF_002031145.1_ASM203114v1 Salmonella enterica subsp. enterica serovar Kentucky Proteobacteria; Gammaproteobacteria; Enterobacterales; Enterobacteriaceae; Salmonella; Salmonella enterica; Salmonella enterica subsp. enterica MNNEETFYQAMRRKGVTRRSFLKFCSLAATSLGLGAGMTPKIAWALENKPRIPVVWIHGL WP_001058311.1 hydrogenase [Salmonella enterica] Length: 372\nNumber of predicted TMHs: 2\nExp number of AAs in TMHs: 37.4699899999999\nExp number, first 60 AAs: 19.72699\nTotal prob of N-in: 0.96171\nPOSSIBLE N-term signal sequence\ninside 1 19\nTMhelix 20 39\noutside 40 325\nTMhelix 326 348\ninside 349 372

18251 GCF_002031305.1_ASM203130v1 Salmonella enterica subsp. enterica serovar Kentucky Proteobacteria; Gammaproteobacteria; Enterobacterales; Enterobacteriaceae; Salmonella; Salmonella enterica; Salmonella enterica subsp. enterica MNNEETFYQAMRRKGVTRRSFLKFCSLAATSLGLGAGMTPKIAWALENKPRIPVVWIHGL WP_001058311.1 hydrogenase [Salmonella enterica] Length: 372\nNumber of predicted TMHs: 2\nExp number of AAs in TMHs: 37.4699899999999\nExp number, first 60 AAs: 19.72699\nTotal prob of N-in: 0.96171\nPOSSIBLE N-term signal sequence\ninside 1 19\nTMhelix 20 39\noutside 40 325\nTMhelix 326 348\ninside 349 372

18252 GCF_002031305.1_ASM203130v1 Salmonella enterica subsp. enterica serovar Kentucky Proteobacteria; Gammaproteobacteria; Enterobacterales; Enterobacteriaceae; Salmonella; Salmonella enterica; Salmonella enterica subsp. enterica MTGDNTLITSHGINRRDFMKLCAALAATMGLSSKAAAEMAESVSNPQRPPVIWIGAQECT WP_000145429.1 hydrogenase 2 small subunit [Salmonella enterica] Length: 372\nNumber of predicted TMHs: 1\nExp number of AAs in TMHs: 24.51309\nExp number, first 60 AAs: 0.2682\nTotal prob of N-in: 0.14402\noutside 1 331\nTMhelix 332 354\ninside 355 372

18253 GCF_001542055.1_ASM154205v1 Salmonella enterica subsp. enterica serovar Kentucky Proteobacteria; Gammaproteobacteria; Enterobacterales; Enterobacteriaceae; Salmonella; Salmonella enterica; Salmonella enterica subsp. enterica MTGDNTLITSHGINRRDFMKLCAALAATMGLSSKAAAEMAESVSNPQRPPVIWIGAQECT WP_000145429.1 hydrogenase 2 small subunit [Salmonella enterica] Length: 372\nNumber of predicted TMHs: 1\nExp number of AAs in TMHs: 24.51309\nExp number, first 60 AAs: 0.2682\nTotal prob of N-in: 0.14402\noutside 1 331\nTMhelix 332 354\ninside 355 372

18254 GCF_001478565.1_Salmonella_enterica_CVM_N41924-SQ_v1.0 Salmonella enterica subsp. enterica serovar Kentucky Proteobacteria; Gammaproteobacteria; Enterobacterales; Enterobacteriaceae; Salmonella; Salmonella enterica; Salmonella enterica subsp. enterica MNNEETFYQAMRRKGVTRRSFLKFCSLAATSLGLGAGMTPKIAWALENKPRIPVVWIHGL WP_001058311.1 hydrogenase [Salmonella enterica] Length: 372\nNumber of predicted TMHs: 2\nExp number of AAs in TMHs: 37.4699899999999\nExp number, first 60 AAs: 19.72699\nTotal prob of N-in: 0.96171\nPOSSIBLE N-term signal sequence\ninside 1 19\nTMhelix 20 39\noutside 40 325\nTMhelix 326 348\ninside 349 372

18255 GCF_002030765.1_ASM203076v1 Salmonella enterica subsp. enterica serovar Kentucky Proteobacteria; Gammaproteobacteria; Enterobacterales; Enterobacteriaceae; Salmonella; Salmonella enterica; Salmonella enterica subsp. enterica MTGDNTLITSHGINRRDFMKLCAALAATMGLSSKAAAEMAESVSNPQRPPVIWIGAQECT WP_000145429.1 hydrogenase 2 small subunit [Salmonella enterica] Length: 372\nNumber of predicted TMHs: 1\nExp number of AAs in TMHs: 24.51309\nExp number, first 60 AAs: 0.2682\nTotal prob of N-in: 0.14402\noutside 1 331\nTMhelix 332 354\ninside 355 372

18256 GCF_002030465.1_ASM203046v1 Salmonella enterica subsp. enterica serovar Kentucky Proteobacteria; Gammaproteobacteria; Enterobacterales; Enterobacteriaceae; Salmonella; Salmonella enterica; Salmonella enterica subsp. enterica MNNEETFYQAMRRKGVTRRSFLKFCSLAATSLGLGAGMTPKIAWALENKPRIPVVWIHGL WP_001058311.1 hydrogenase [Salmonella enterica] Length: 372\nNumber of predicted TMHs: 2\nExp number of AAs in TMHs: 37.4699899999999\nExp number, first 60 AAs: 19.72699\nTotal prob of N-in: 0.96171\nPOSSIBLE N-term signal sequence\ninside 1 19\nTMhelix 20 39\noutside 40 325\nTMhelix 326 348\ninside 349 372

18257 GCF_002030465.1_ASM203046v1 Salmonella enterica subsp. enterica serovar Kentucky Proteobacteria; Gammaproteobacteria; Enterobacterales; Enterobacteriaceae; Salmonella; Salmonella enterica; Salmonella enterica subsp. enterica MTGDNTLITSHGINRRDFMKLCAALAATMGLSSKAAAEMAESVSNPQRPPVIWIGAQECT

WP_000145429.1 hydrogenase 2 small subunit [Salmonella enterica] Length: 372\nNumber of predicted TMHs: 1\nExp number of AAs in TMHs: 24.51309\nExp number, first 60 AAs: 0.2682\nTotal prob of N-in: 0.14402\noutside 1 331\nTMhelix 332 354\ninside 355 372

18258 GCF_001478825.1_Salmonella_enterica_CVM_N29383-SQ_v1.0 Salmonella enterica subsp. enterica serovar Kentucky Proteobacteria; Gammaproteobacteria; Enterobacterales; Enterobacteriaceae; Salmonella; Salmonella enterica; Salmonella enterica subsp. enterica MNNEETFYQAMRRKGVTRRSFLKFCSLAATSLGLGAGMTPKIAWALENKPRIPVVWIHGL
WP_001058311.1 hydrogenase [Salmonella enterica] Length: 372\nNumber of predicted TMHs: 2\nExp number of AAs in TMHs: 37.4699899999999\nExp number, first 60 AAs: 19.72699\nTotal prob of N-in: 0.96171\nPOSSIBLE N-term signal sequence\ninside 1 19\nTMhelix 20 39\noutside 40 325\nTMhelix 326 348\ninside 349 372

18259 GCF_001240135.1_Salmonella_enterica_CVM_N43455_v1.0 Salmonella enterica subsp. enterica serovar Kentucky Proteobacteria; Gammaproteobacteria; Enterobacterales; Enterobacteriaceae; Salmonella; Salmonella enterica; Salmonella enterica subsp. enterica MTGDNTLITSHGINRRDFMKLCAALAATMGLSSKAAAEMAESVSNPQRPPVIWIGAQECT
WP_000145429.1 hydrogenase 2 small subunit [Salmonella enterica] Length: 372\nNumber of predicted TMHs: 1\nExp number of AAs in TMHs: 24.51309\nExp number, first 60 AAs: 0.2682\nTotal prob of N-in: 0.14402\noutside 1 331\nTMhelix 332 354\ninside 355 372

18260 GCF_001480755.1_Salmonella_enterica_CVM_N42521-SQ_v1.0 Salmonella enterica subsp. enterica serovar Kentucky Proteobacteria; Gammaproteobacteria; Enterobacterales; Enterobacteriaceae; Salmonella; Salmonella enterica; Salmonella enterica subsp. enterica MTGDNTLITSHGINRRDFMKLCAALAATMGLSSKAAAEMAESVSNPQRPPVIWIGAQECT
WP_000145429.1 hydrogenase 2 small subunit [Salmonella enterica] Length: 372\nNumber of predicted TMHs: 1\nExp number of AAs in TMHs: 24.51309\nExp number, first 60 AAs: 0.2682\nTotal prob of N-in: 0.14402\noutside 1 331\nTMhelix 332 354\ninside 355 372

18261 GCF_001478825.1_Salmonella_enterica_CVM_N29383-SQ_v1.0 Salmonella enterica subsp. enterica serovar Kentucky Proteobacteria; Gammaproteobacteria; Enterobacterales; Enterobacteriaceae; Salmonella; Salmonella enterica; Salmonella enterica subsp. enterica MTGDNTLITSHGINRRDFMKLCAALAATMGLSSKAAAEMAESVSNPQRPPVIWIGAQECT
WP_000145429.1 hydrogenase 2 small subunit [Salmonella enterica] Length: 372\nNumber of predicted TMHs: 1\nExp number of AAs in TMHs: 24.51309\nExp number, first 60 AAs: 0.2682\nTotal prob of N-in: 0.14402\noutside 1 331\nTMhelix 332 354\ninside 355 372

18262 GCF_001478715.1_Salmonella_enterica_CVM_N29363-SQ_v1.0 Salmonella enterica subsp. enterica serovar Kentucky Proteobacteria; Gammaproteobacteria; Enterobacterales; Enterobacteriaceae; Salmonella; Salmonella enterica; Salmonella enterica subsp. enterica MNNEETFYQAMRRKGVTRRSFLKFCSLAATSLGLGAGMTPKIAWALENKPRIPVVWIHGL
WP_001058311.1 hydrogenase [Salmonella enterica] Length: 372\nNumber of predicted TMHs: 2\nExp number of AAs in TMHs: 37.4699899999999\nExp number, first 60 AAs: 19.72699\nTotal prob of N-in: 0.96171\nPOSSIBLE N-term signal sequence\ninside 1 19\nTMhelix 20 39\noutside 40 325\nTMhelix 326 348\ninside 349 372

18263 GCF_002033015.1_ASM203301v1 Salmonella enterica subsp. enterica serovar Kentucky Proteobacteria; Gammaproteobacteria; Enterobacterales; Enterobacteriaceae; Salmonella; Salmonella enterica; Salmonella enterica subsp. enterica MNNEETFYQAMRRKGVTRRSFLKFCSLAATSLGLGAGMTPKIAWALENKPRIPVVWIHGL
WP_001058311.1 hydrogenase [Salmonella enterica] Length: 372\nNumber of predicted TMHs: 2\nExp number of AAs in TMHs: 37.4699899999999\nExp number, first 60 AAs: 19.72699\nTotal prob of N-in: 0.96171\nPOSSIBLE N-term signal sequence\ninside 1 19\nTMhelix 20 39\noutside 40 325\nTMhelix 326 348\ninside 349 372

18264 GCF_002031745.1_ASM203174v1 Salmonella enterica subsp. enterica serovar Kentucky Proteobacteria; Gammaproteobacteria; Enterobacterales; Enterobacteriaceae; Salmonella; Salmonella enterica; Salmonella enterica subsp. enterica MNNEETFYQAMRRKGVTRRSFLKFCSLAATSLGLGAGMTPKIAWALENKPRIPVVWIHGL
WP_001058311.1 hydrogenase [Salmonella enterica] Length: 372\nNumber of predicted TMHs: 2\nExp number of AAs in TMHs: 37.4699899999999\nExp number, first 60 AAs: 19.72699\nTotal prob of N-in: 0.96171\nPOSSIBLE N-term signal sequence\ninside 1 19\nTMhelix 20 39\noutside 40 325\nTMhelix 326 348\ninside 349 372

18265 GCF_002031145.1_ASM203114v1 Salmonella enterica subsp. enterica serovar Kentucky Proteobacteria; Gammaproteobacteria; Enterobacterales; Enterobacteriaceae; Salmonella; Salmonella enterica; Salmonella enterica subsp. enterica MTGDNTLITSHGINRRDFMKLCAALAATMGLSSKAAAEMAESVSNPQRPPVIWIGAQECT
WP_000145429.1 hydrogenase 2 small subunit [Salmonella enterica] Length: 372\nNumber of

predicted TMHs: 1\nExp number of AAs in TMHs: 24.51309\nExp number, first 60 AAs: 0.2682\nTotal prob of N-in: 0.14402\noutside 1 331\nTMhelix 332 354\ninside 355 372

18266 GCF_002030765.1_ASM203076v1 Salmonella enterica subsp. enterica serovar Kentucky Proteobacteria; Gammaproteobacteria; Enterobacterales; Enterobacteriaceae; Salmonella; Salmonella enterica; Salmonella enterica subsp. enterica MNNEETFYQAMRRKGVTRRSFLKFCSLAATSLGLGAGMTPKIAWALENKPRIPVVWIHGL WP_001058311.1 hydrogenase [Salmonella enterica] Length: 372\nNumber of predicted TMHs: 2\nExp number of AAs in TMHs: 37.4699899999999\nExp number, first 60 AAs: 19.72699\nTotal prob of N-in: 0.96171\nPOSSIBLE N-term signal sequence\ninside 1 19\nTMhelix 20 39\noutside 40 325\nTMhelix 326 348\ninside 349 372

18267 GCF_002050605.1_ASM205060v1 Salmonella enterica subsp. enterica serovar Kentucky Proteobacteria; Gammaproteobacteria; Enterobacterales; Enterobacteriaceae; Salmonella; Salmonella enterica; Salmonella enterica subsp. enterica MTGDNTLITSHGINRRDFMKLCAALAATMGLSSKAAAEMAESVSNPQRPPVIWIGAQECT WP_000145429.1 hydrogenase 2 small subunit [Salmonella enterica] Length: 372\nNumber of predicted TMHs: 1\nExp number of AAs in TMHs: 24.51309\nExp number, first 60 AAs: 0.2682\nTotal prob of N-in: 0.14402\noutside 1 331\nTMhelix 332 354\ninside 355 372

18268 GCF_001272405.1_Salmonella_enterica_CVM_N51277_v1.0 Salmonella enterica subsp. enterica serovar Kentucky Proteobacteria; Gammaproteobacteria; Enterobacterales; Enterobacteriaceae; Salmonella; Salmonella enterica; Salmonella enterica subsp. enterica MTGDNTLITSHGINRRDFMKLCAALAATMGLSSKAAAEMAESVSNPQRPPVIWIGAQECT WP_000145429.1 hydrogenase 2 small subunit [Salmonella enterica] Length: 372\nNumber of predicted TMHs: 1\nExp number of AAs in TMHs: 24.51309\nExp number, first 60 AAs: 0.2682\nTotal prob of N-in: 0.14402\noutside 1 331\nTMhelix 332 354\ninside 355 372

18269 GCF_001271875.1_Salmonella_enterica_CVM_N47722_v1.0 Salmonella enterica subsp. enterica serovar Kentucky Proteobacteria; Gammaproteobacteria; Enterobacterales; Enterobacteriaceae; Salmonella; Salmonella enterica; Salmonella enterica subsp. enterica MTGDNTLITSHGINRRDFMKLCAALAATMGLSSKAAAEMAESVSNPQRPPVIWIGAQECT WP_000145429.1 hydrogenase 2 small subunit [Salmonella enterica] Length: 372\nNumber of predicted TMHs: 1\nExp number of AAs in TMHs: 24.51309\nExp number, first 60 AAs: 0.2682\nTotal prob of N-in: 0.14402\noutside 1 331\nTMhelix 332 354\ninside 355 372

18270 GCF_002049015.1_ASM204901v1 Salmonella enterica subsp. enterica serovar Kentucky Proteobacteria; Gammaproteobacteria; Enterobacterales; Enterobacteriaceae; Salmonella; Salmonella enterica; Salmonella enterica subsp. enterica MNNEETFYQAMRRKGVTRRSFLKFCSLAATSLGLGAGMTPKIAWALENKPRIPVVWIHGL WP_001058307.1 hydrogenase [Salmonella enterica] Length: 372\nNumber of predicted TMHs: 2\nExp number of AAs in TMHs: 36.51738\nExp number, first 60 AAs: 19.72849\nTotal prob of N-in: 0.96170\nPOSSIBLE N-term signal sequence\ninside 1 19\nTMhelix 20 39\noutside 40 325\nTMhelix 326 348\ninside 349 372

18271 GCF_002049015.1_ASM204901v1 Salmonella enterica subsp. enterica serovar Kentucky Proteobacteria; Gammaproteobacteria; Enterobacterales; Enterobacteriaceae; Salmonella; Salmonella enterica; Salmonella enterica subsp. enterica MTGDNTLITSHGINRRDFMKLCAALAATMGLSSKAAAEMAESVSNPQRPPVIWIGAQECT WP_000145429.1 hydrogenase 2 small subunit [Salmonella enterica] Length: 372\nNumber of predicted TMHs: 1\nExp number of AAs in TMHs: 24.51309\nExp number, first 60 AAs: 0.2682\nTotal prob of N-in: 0.14402\noutside 1 331\nTMhelix 332 354\ninside 355 372

18272 GCF_000974085.1_ABB1087-1.id21_v1.0 Salmonella enterica subsp. enterica serovar Kentucky Proteobacteria; Gammaproteobacteria; Enterobacterales; Enterobacteriaceae; Salmonella; Salmonella enterica; Salmonella enterica subsp. enterica MTGDNTLITSHGINRRDFMKLCAALAATMGLSSKAAAEMAESVSNPQRPPVIWIGAQECT WP_000145429.1 hydrogenase 2 small subunit [Salmonella enterica] Length: 372\nNumber of predicted TMHs: 1\nExp number of AAs in TMHs: 24.51309\nExp number, first 60 AAs: 0.2682\nTotal prob of N-in: 0.14402\noutside 1 331\nTMhelix 332 354\ninside 355 372

18273 GCF_000973905.1_ABB07-SB3057-2.id23_v1.0 Salmonella enterica subsp. enterica serovar Kentucky Proteobacteria; Gammaproteobacteria; Enterobacterales; Enterobacteriaceae; Salmonella; Salmonella enterica; Salmonella enterica subsp. enterica MNNEETFYQAMRRKGVTRRSFLKFCSLAATSLGLGAGMTPKIAWALENKPRIPVVWIHGL WP_001058311.1 hydrogenase [Salmonella enterica] Length: 372\nNumber of predicted TMHs: 2\nExp number of AAs in TMHs: 37.4699899999999\nExp number, first 60 AAs: 19.72699\nTotal prob of N-in: 0.96171\nPOSSIBLE N-term signal sequence\ninside 1 19\nTMhelix 20 39\noutside 40 325\nTMhelix 326 348\ninside 349 372

18274 GCF_001238265.1_Salmonella_enterica_CVM_N43448_v1.0 Salmonella enterica subsp. enterica serovar Kentucky Proteobacteria; Gammaproteobacteria; Enterobacterales; Enterobacteriaceae; Salmonella; Salmonella enterica; Salmonella enterica subsp. enterica MTGDNTLITSHGINRRDFMKLCAALAATMGLSSKAAAEMAESVSNPQRPPVIWIGAQECT WP_000145429.1 hydrogenase 2 small subunit [Salmonella enterica] Length: 372\nNumber of predicted TMHs: 1\nExp number of AAs in TMHs: 24.51309\nExp number, first 60 AAs: 0.2682\nTotal prob of N-in: 0.14402\noutside 1 331\nTMhelix 332 354\ninside 355 372

18275 GCF_001240135.1_Salmonella_enterica_CVM_N43455_v1.0 Salmonella enterica subsp. enterica serovar Kentucky Proteobacteria; Gammaproteobacteria; Enterobacterales; Enterobacteriaceae; Salmonella; Salmonella enterica; Salmonella enterica subsp. enterica MNNEETFYQAMRRKGVTRRSFLKFCSLAATSLGLGAGMTPKIAWALENKPRIPVVWIHGL WP_001058311.1 hydrogenase [Salmonella enterica] Length: 372\nNumber of predicted TMHs: 2\nExp number of AAs in TMHs: 37.4699899999999\nExp number, first 60 AAs: 19.72699\nTotal prob of N-in: 0.96171\nPOSSIBLE N-term signal sequence\ninside 1 19\nTMhelix 20 39\noutside 40 325\nTMhelix 326 348\ninside 349 372

18276 GCF_001240725.1_Salmonella_enterica_CVM_N43820_v1.0 Salmonella enterica subsp. enterica serovar Kentucky Proteobacteria; Gammaproteobacteria; Enterobacterales; Enterobacteriaceae; Salmonella; Salmonella enterica; Salmonella enterica subsp. enterica MTGDNTLITSHGINRRDFMKLCAALAATMGLSSKAAAEMAESVSNPQRPPVIWIGAQECT WP_000145429.1 hydrogenase 2 small subunit [Salmonella enterica] Length: 372\nNumber of predicted TMHs: 1\nExp number of AAs in TMHs: 24.51309\nExp number, first 60 AAs: 0.2682\nTotal prob of N-in: 0.14402\noutside 1 331\nTMhelix 332 354\ninside 355 372

18277 GCF_001240725.1_Salmonella_enterica_CVM_N43820_v1.0 Salmonella enterica subsp. enterica serovar Kentucky Proteobacteria; Gammaproteobacteria; Enterobacterales; Enterobacteriaceae; Salmonella; Salmonella enterica; Salmonella enterica subsp. enterica MNNEETFYQAMRRKGVTRRSFLKFCSLAATSLGLGAGMTPKIAWALENKPRIPVVWIHGL WP_001058311.1 hydrogenase [Salmonella enterica] Length: 372\nNumber of predicted TMHs: 2\nExp number of AAs in TMHs: 37.4699899999999\nExp number, first 60 AAs: 19.72699\nTotal prob of N-in: 0.96171\nPOSSIBLE N-term signal sequence\ninside 1 19\nTMhelix 20 39\noutside 40 325\nTMhelix 326 348\ninside 349 372

18278 GCF_001890835.1_ASM189083v1 Salmonella enterica subsp. enterica serovar Kentucky Proteobacteria; Gammaproteobacteria; Enterobacterales; Enterobacteriaceae; Salmonella; Salmonella enterica; Salmonella enterica subsp. enterica MNNEETFYQAMRRKGVTRRSFLKFCSLAATSLGLGAGMTPKIAWALENKPRIPVVWIHGL WP_001058307.1 hydrogenase [Salmonella enterica] Length: 372\nNumber of predicted TMHs: 2\nExp number of AAs in TMHs: 36.51738\nExp number, first 60 AAs: 19.72849\nTotal prob of N-in: 0.96170\nPOSSIBLE N-term signal sequence\ninside 1 19\nTMhelix 20 39\noutside 40 325\nTMhelix 326 348\ninside 349 372

18279 GCF_001890855.1_ASM189085v1 Salmonella enterica subsp. enterica serovar Kentucky Proteobacteria; Gammaproteobacteria; Enterobacterales; Enterobacteriaceae; Salmonella; Salmonella enterica; Salmonella enterica subsp. enterica MTGDNTLITSHGINRRDFMKLCAALAATMGLSSKAAAEMAESVSNPQRPPVIWIGAQECT WP_000145429.1 hydrogenase 2 small subunit [Salmonella enterica] Length: 372\nNumber of predicted TMHs: 1\nExp number of AAs in TMHs: 24.51309\nExp number, first 60 AAs: 0.2682\nTotal prob of N-in: 0.14402\noutside 1 331\nTMhelix 332 354\ninside 355 372

18280 GCF_001242015.1_Salmonella_enterica_CVM_N45412_v1.0 Salmonella enterica subsp. enterica serovar Kentucky Proteobacteria; Gammaproteobacteria; Enterobacterales; Enterobacteriaceae; Salmonella; Salmonella enterica; Salmonella enterica subsp. enterica MNNEETFYQAMRRKGVTRRSFLKFCSLAATSLGLGAGMTPKIAWALENKPRIPVVWIHGL WP_001058311.1 hydrogenase [Salmonella enterica] Length: 372\nNumber of predicted TMHs: 2\nExp number of AAs in TMHs: 37.4699899999999\nExp number, first 60 AAs: 19.72699\nTotal prob of N-in: 0.96171\nPOSSIBLE N-term signal sequence\ninside 1 19\nTMhelix 20 39\noutside 40 325\nTMhelix 326 348\ninside 349 372

18281 GCF_001242285.1_Salmonella_enterica_CVM_N45934_v1.0 Salmonella enterica subsp. enterica serovar Kentucky Proteobacteria; Gammaproteobacteria; Enterobacterales; Enterobacteriaceae; Salmonella; Salmonella enterica; Salmonella enterica subsp. enterica MNNEETFYQAMRRKGVTRRSFLKFCSLAATSLGLGAGMTPKIAWALENKPRIPVVWIHGL WP_001058311.1 hydrogenase [Salmonella enterica] Length: 372\nNumber of predicted TMHs: 2\nExp number of AAs in TMHs: 37.4699899999999\nExp number, first 60 AAs: 19.72699\nTotal prob of N-in: 0.96171\nPOSSIBLE N-term signal sequence\ninside 1 19\nTMhelix 20 39\noutside 40 325\nTMhelix 326 348\ninside 349 372

18282 GCF_001890835.1_ASM189083v1 Salmonella enterica subsp. enterica serovar Kentucky Proteobacteria; Gammaproteobacteria; Enterobacterales; Enterobacteriaceae; Salmonella; Salmonella enterica; Salmonella enterica subsp. enterica MTGDNTLITSHGINRRDFMKLCAALAATMGLSSKAAAEMAESVSNPQRPPVIWIGAQECT WP_000145429.1 hydrogenase 2 small subunit [Salmonella enterica] Length: 372\nNumber of predicted TMHs: 1\nExp number of AAs in TMHs: 24.51309\nExp number, first 60 AAs: 0.2682\nTotal prob of N-in: 0.14402\noutside 1 331\nTMhelix 332 354\ninside 355 372

18283 GCF_001242995.1_Salmonella_enterica_CVM_N46820_v1.0 Salmonella enterica subsp. enterica serovar Kentucky Proteobacteria; Gammaproteobacteria; Enterobacterales; Enterobacteriaceae; Salmonella; Salmonella enterica; Salmonella enterica subsp. enterica MNNEETFYQAMRRKGVTRRSFLKFCSLAATSLGLGAGMTPKIAWALENKPRIPVVWIHGL WP_001058311.1 hydrogenase [Salmonella enterica] Length: 372\nNumber of predicted TMHs: 2\nExp number of AAs in TMHs: 37.4699899999999\nExp number, first 60 AAs: 19.72699\nTotal prob of N-in: 0.96171\nPOSSIBLE N-term signal sequence\ninside 1 19\nTMhelix 20 39\noutside 40 325\nTMhelix 326 348\ninside 349 372

18284 GCF_001272135.1_Salmonella_enterica_CVM_N48687_v1.0 Salmonella enterica subsp. enterica serovar Kentucky Proteobacteria; Gammaproteobacteria; Enterobacterales; Enterobacteriaceae; Salmonella; Salmonella enterica; Salmonella enterica subsp. enterica MNNEETFYQAMRRKGVTRRSFLKFCSLAATSLGLGAGMTPKIAWALENKPRIPVVWIHGL WP_001058311.1 hydrogenase [Salmonella enterica] Length: 372\nNumber of predicted TMHs: 2\nExp number of AAs in TMHs: 37.4699899999999\nExp number, first 60 AAs: 19.72699\nTotal prob of N-in: 0.96171\nPOSSIBLE N-term signal sequence\ninside 1 19\nTMhelix 20 39\noutside 40 325\nTMhelix 326 348\ninside 349 372

18285 GCF_001243995.1_Salmonella_enterica_CVM_N47721_v1.0 Salmonella enterica subsp. enterica serovar Kentucky Proteobacteria; Gammaproteobacteria; Enterobacterales; Enterobacteriaceae; Salmonella; Salmonella enterica; Salmonella enterica subsp. enterica MTGDNTLITSHGINRRDFMKLCAALAATMGLSSKAAAEMAESVSNPQRPPVIWIGAQECT WP_000145429.1 hydrogenase 2 small subunit [Salmonella enterica] Length: 372\nNumber of predicted TMHs: 1\nExp number of AAs in TMHs: 24.51309\nExp number, first 60 AAs: 0.2682\nTotal prob of N-in: 0.14402\noutside 1 331\nTMhelix 332 354\ninside 355 372

18286 GCF_001244005.1_Salmonella_enterica_CVM_N47723_v1.0 Salmonella enterica subsp. enterica serovar Kentucky Proteobacteria; Gammaproteobacteria; Enterobacterales; Enterobacteriaceae; Salmonella; Salmonella enterica; Salmonella enterica subsp. enterica MTGDNTLITSHGINRRDFMKLCAALAATMGLSSKAAAEMAESVSNPQRPPVIWIGAQECT WP_000145429.1 hydrogenase 2 small subunit [Salmonella enterica] Length: 372\nNumber of predicted TMHs: 1\nExp number of AAs in TMHs: 24.51309\nExp number, first 60 AAs: 0.2682\nTotal prob of N-in: 0.14402\noutside 1 331\nTMhelix 332 354\ninside 355 372

18287 GCF_001245105.1_Salmonella_enterica_CVM_N48707_v1.0 Salmonella enterica subsp. enterica serovar Kentucky Proteobacteria; Gammaproteobacteria; Enterobacterales; Enterobacteriaceae; Salmonella; Salmonella enterica; Salmonella enterica subsp. enterica MTGDNTLITSHGINRRDFMKLCAALAATMGLSSKAAAEMAESVSNPQRPPVIWIGAQECT WP_000145429.1 hydrogenase 2 small subunit [Salmonella enterica] Length: 372\nNumber of predicted TMHs: 1\nExp number of AAs in TMHs: 24.51309\nExp number, first 60 AAs: 0.2682\nTotal prob of N-in: 0.14402\noutside 1 331\nTMhelix 332 354\ninside 355 372

18288 GCF_001245105.1_Salmonella_enterica_CVM_N48707_v1.0 Salmonella enterica subsp. enterica serovar Kentucky Proteobacteria; Gammaproteobacteria; Enterobacterales; Enterobacteriaceae; Salmonella; Salmonella enterica; Salmonella enterica subsp. enterica MNNEETFYQAMRRKGVTRRSFLKFCSLAATSLGLGAGMTPKIAWALENKPRIPVVWIHGL WP_001058311.1 hydrogenase [Salmonella enterica] Length: 372\nNumber of predicted TMHs: 2\nExp number of AAs in TMHs: 37.4699899999999\nExp number, first 60 AAs: 19.72699\nTotal prob of N-in: 0.96171\nPOSSIBLE N-term signal sequence\ninside 1 19\nTMhelix 20 39\noutside 40 325\nTMhelix 326 348\ninside 349 372

18289 GCF_001245195.1_Salmonella_enterica_CVM_N48711_v1.0 Salmonella enterica subsp. enterica serovar Kentucky Proteobacteria; Gammaproteobacteria; Enterobacterales; Enterobacteriaceae; Salmonella; Salmonella enterica; Salmonella enterica subsp. enterica MTGDNTLITSHGINRRDFMKLCAALAATMGLSSKAAAEMAESVSNPQRPPVIWIGAQECT WP_000145429.1 hydrogenase 2 small subunit [Salmonella enterica] Length: 372\nNumber of predicted TMHs: 1\nExp number of AAs in TMHs: 24.51309\nExp number, first 60 AAs: 0.2682\nTotal prob of N-in: 0.14402\noutside 1 331\nTMhelix 332 354\ninside 355 372

18290 GCF_001245565.1_Salmonella_enterica_CVM_N50435_v1.0 Salmonella enterica subsp. enterica serovar Kentucky Proteobacteria; Gammaproteobacteria; Enterobacterales; Enterobacteriaceae; Salmonella; Salmonella enterica;

Salmonella enterica subsp. enterica MTGDNTLITSHGINRRDFMKLCAALAATMGLSSKAAAEMAESVSNPQRPPVIWIGAQECT WP_000145429.1 hydrogenase 2 small subunit [Salmonella enterica] Length: 372\nNumber of predicted TMHs: 1\nExp number of AAs in TMHs: 24.51309\nExp number, first 60 AAs: 0.2682\nTotal prob of N-in: 0.14402\noutside 1 331\nTMhelix 332 354\ninside 355 372

18291 GCF_001245665.1_Salmonella_enterica_CVM_N50437_v1.0 Salmonella enterica subsp. enterica serovar Kentucky Proteobacteria; Gammaproteobacteria; Enterobacterales; Enterobacteriaceae; Salmonella; Salmonella enterica; Salmonella enterica subsp. enterica MTGDNTLITSHGINRRDFMKLCAALAATMGLSSKAAAEMAESVSNPQRPPVIWIGAQECT WP_000145429.1 hydrogenase 2 small subunit [Salmonella enterica] Length: 372\nNumber of predicted TMHs: 1\nExp number of AAs in TMHs: 24.51309\nExp number, first 60 AAs: 0.2682\nTotal prob of N-in: 0.14402\noutside 1 331\nTMhelix 332 354\ninside 355 372

18292 GCF_001245665.1_Salmonella_enterica_CVM_N50437_v1.0 Salmonella enterica subsp. enterica serovar Kentucky Proteobacteria; Gammaproteobacteria; Enterobacterales; Enterobacteriaceae; Salmonella; Salmonella enterica; Salmonella enterica subsp. enterica MNNEETFYQAMRRKGVTRRSFLKFCSLAATSLGLGAGMTPKIAWALENKPRIPVVWIHGL WP_001058311.1 hydrogenase [Salmonella enterica] Length: 372\nNumber of predicted TMHs: 2\nExp number of AAs in TMHs: 37.4699899999999\nExp number, first 60 AAs: 19.72699\nTotal prob of N-in: 0.96171\nPOSSIBLE N-term signal sequence\ninside 1 19\nTMhelix 20 39\noutside 40 325\nTMhelix 326 348\ninside 349 372

18293 GCF_001542055.1_ASM154205v1 Salmonella enterica subsp. enterica serovar Kentucky Proteobacteria; Gammaproteobacteria; Enterobacterales; Enterobacteriaceae; Salmonella; Salmonella enterica; Salmonella enterica subsp. enterica MNNEETFYQAMRRKGVTRRSFLKFCSLAATSLGLGAGMTPKIAWALENKPRIPVVWIHGL WP_001058311.1 hydrogenase [Salmonella enterica] Length: 372\nNumber of predicted TMHs: 2\nExp number of AAs in TMHs: 37.4699899999999\nExp number, first 60 AAs: 19.72699\nTotal prob of N-in: 0.96171\nPOSSIBLE N-term signal sequence\ninside 1 19\nTMhelix 20 39\noutside 40 325\nTMhelix 326 348\ninside 349 372

18294 GCF_001542175.1_ASM154217v1 Salmonella enterica subsp. enterica serovar Kentucky Proteobacteria; Gammaproteobacteria; Enterobacterales; Enterobacteriaceae; Salmonella; Salmonella enterica; Salmonella enterica subsp. enterica MNNEETFYQAMRRKGVTRRSFLKFCSLAATSLGLGAGMTPKIAWALENKPRIPVVWIHGL WP_001058311.1 hydrogenase [Salmonella enterica] Length: 372\nNumber of predicted TMHs: 2\nExp number of AAs in TMHs: 37.4699899999999\nExp number, first 60 AAs: 19.72699\nTotal prob of N-in: 0.96171\nPOSSIBLE N-term signal sequence\ninside 1 19\nTMhelix 20 39\noutside 40 325\nTMhelix 326 348\ninside 349 372

18295 GCF_001542095.1_ASM154209v1 Salmonella enterica subsp. enterica serovar Kentucky Proteobacteria; Gammaproteobacteria; Enterobacterales; Enterobacteriaceae; Salmonella; Salmonella enterica; Salmonella enterica subsp. enterica MNNEETFYQAMRRKGVTRRSFLKFCSLAATSLGLGAGMTPKIAWALENKPRIPVVWIHGL WP_001058311.1 hydrogenase [Salmonella enterica] Length: 372\nNumber of predicted TMHs: 2\nExp number of AAs in TMHs: 37.4699899999999\nExp number, first 60 AAs: 19.72699\nTotal prob of N-in: 0.96171\nPOSSIBLE N-term signal sequence\ninside 1 19\nTMhelix 20 39\noutside 40 325\nTMhelix 326 348\ninside 349 372

18296 GCF_001542135.1_ASM154213v1 Salmonella enterica subsp. enterica serovar Kentucky Proteobacteria; Gammaproteobacteria; Enterobacterales; Enterobacteriaceae; Salmonella; Salmonella enterica; Salmonella enterica subsp. enterica MNNEETFYQAMRRKGVTRRSFLKFCSLAATSLGLGAGMTPKIAWALENKPRIPVVWIHGL WP_001058311.1 hydrogenase [Salmonella enterica] Length: 372\nNumber of predicted TMHs: 2\nExp number of AAs in TMHs: 37.4699899999999\nExp number, first 60 AAs: 19.72699\nTotal prob of N-in: 0.96171\nPOSSIBLE N-term signal sequence\ninside 1 19\nTMhelix 20 39\noutside 40 325\nTMhelix 326 348\ninside 349 372

18297 GCF_001542175.1_ASM154217v1 Salmonella enterica subsp. enterica serovar Kentucky Proteobacteria; Gammaproteobacteria; Enterobacterales; Enterobacteriaceae; Salmonella; Salmonella enterica; Salmonella enterica subsp. enterica MTGDNTLITSHGINRRDFMKLCAALAATMGLSSKAAAEMAESVSNPQRPPVIWIGAQECT WP_000145429.1 hydrogenase 2 small subunit [Salmonella enterica] Length: 372\nNumber of predicted TMHs: 1\nExp number of AAs in TMHs: 24.51309\nExp number, first 60 AAs: 0.2682\nTotal prob of N-in: 0.14402\noutside 1 331\nTMhelix 332 354\ninside 355 372

18298 GCF_001542135.1_ASM154213v1 Salmonella enterica subsp. enterica serovar Kentucky Proteobacteria; Gammaproteobacteria; Enterobacterales; Enterobacteriaceae; Salmonella; Salmonella enterica;

Salmonella enterica subsp. enterica	MTGDNTLITSHGINRRDFMKLCAALAATMGLSSKAAAEMAESVSNPQRPPVIWIGAQECT	WP_000145429.1	hydrogenase 2 small subunit [Salmonella enterica]	Length: 372\nNumber of predicted TMHs:  1\nExp number of AAs in TMHs: 24.51309\nExp number, first 60 AAs:  0.2682\nTotal prob of N-in: 0.14402\noutside   1   331\nTMhelix   332   354\ninside   355   372

18299	GCF_001480405.1_Salmonella_enterica_CVM_N42233-SQ_v1.0	Salmonella enterica subsp. enterica serovar Kentucky	Proteobacteria; Gammaproteobacteria; Enterobacterales; Enterobacteriaceae; Salmonella; Salmonella enterica; Salmonella enterica subsp. enterica	MTGDNTLITSHGINRRDFMKLCAALAATMGLSSKAAAEMAESVSNPQRPPVIWIGAQECT	WP_000145429.1	hydrogenase 2 small subunit [Salmonella enterica]	Length: 372\nNumber of predicted TMHs:  1\nExp number of AAs in TMHs: 24.51309\nExp number, first 60 AAs:  0.2682\nTotal prob of N-in: 0.14402\noutside   1   331\nTMhelix   332   354\ninside   355   372

18300	GCF_001542095.1_ASM154209v1	Salmonella enterica subsp. enterica serovar Kentucky	Proteobacteria; Gammaproteobacteria; Enterobacterales; Enterobacteriaceae; Salmonella; Salmonella enterica; Salmonella enterica subsp. enterica	MTGDNTLITSHGINRRDFMKLCAALAATMGLSSKAAAEMAESVSNPQRPPVIWIGAQECT	WP_000145429.1	hydrogenase 2 small subunit [Salmonella enterica]	Length: 372\nNumber of predicted TMHs:  1\nExp number of AAs in TMHs: 24.51309\nExp number, first 60 AAs:  0.2682\nTotal prob of N-in: 0.14402\noutside   1   331\nTMhelix   332   354\ninside   355   372

18301	GCF_001480045.1_Salmonella_enterica_CVM_N38909-SQ_v1.0	Salmonella enterica subsp. enterica serovar Kentucky	Proteobacteria; Gammaproteobacteria; Enterobacterales; Enterobacteriaceae; Salmonella; Salmonella enterica; Salmonella enterica subsp. enterica	MNNEETFYQAMRRKGVTRRSFLKFCSLAATSLGLGAGMTPKIAWALENKPRIPVVWIHGL	WP_001058311.1	hydrogenase [Salmonella enterica]	Length: 372\nNumber of predicted TMHs:  2\nExp number of AAs in TMHs: 37.4699899999999\nExp number, first 60 AAs:  19.72699\nTotal prob of N-in: 0.96171\nPOSSIBLE N-term signal sequence\ninside   1   19\nTMhelix   20   39\noutside   40   325\nTMhelix   326   348\ninside   349   372

18302	GCF_001479745.1_Salmonella_enterica_CVM_N38238-SQ_v1.0	Salmonella enterica subsp. enterica serovar Kentucky	Proteobacteria; Gammaproteobacteria; Enterobacterales; Enterobacteriaceae; Salmonella; Salmonella enterica; Salmonella enterica subsp. enterica	MTGDNTLITSHGINRRDFMKLCAALAATMGLSSKAAAEMAESVSNPQRPPVIWIGAQECT	WP_000145429.1	hydrogenase 2 small subunit [Salmonella enterica]	Length: 372\nNumber of predicted TMHs:  1\nExp number of AAs in TMHs: 24.51309\nExp number, first 60 AAs:  0.2682\nTotal prob of N-in: 0.14402\noutside   1   331\nTMhelix   332   354\ninside   355   372

18303	GCF_002050525.1_ASM205052v1	Salmonella enterica subsp. enterica serovar Kentucky	Proteobacteria; Gammaproteobacteria; Enterobacterales; Enterobacteriaceae; Salmonella; Salmonella enterica; Salmonella enterica subsp. enterica	MTGDNTLITSHGINRRDFMKLCAALAATMGLSSKAAAEMAESVSNPQRPPVIWIGAQECT	WP_000145429.1	hydrogenase 2 small subunit [Salmonella enterica]	Length: 372\nNumber of predicted TMHs:  1\nExp number of AAs in TMHs: 24.51309\nExp number, first 60 AAs:  0.2682\nTotal prob of N-in: 0.14402\noutside   1   331\nTMhelix   332   354\ninside   355   372

18304	GCF_002036325.1_ASM203632v1	Salmonella enterica subsp. enterica serovar Kentucky	Proteobacteria; Gammaproteobacteria; Enterobacterales; Enterobacteriaceae; Salmonella; Salmonella enterica; Salmonella enterica subsp. enterica	MTGDNTLITSHGINRRDFMKLCAALAATMGLSSKAAAEMAESVSNPQRPPVIWIGAQECT	WP_000145429.1	hydrogenase 2 small subunit [Salmonella enterica]	Length: 372\nNumber of predicted TMHs:  1\nExp number of AAs in TMHs: 24.51309\nExp number, first 60 AAs:  0.2682\nTotal prob of N-in: 0.14402\noutside   1   331\nTMhelix   332   354\ninside   355   372

18305	GCF_001479905.1_Salmonella_enterica_CVM_N38870-SQ_v1.0	Salmonella enterica subsp. enterica serovar Kentucky	Proteobacteria; Gammaproteobacteria; Enterobacterales; Enterobacteriaceae; Salmonella; Salmonella enterica; Salmonella enterica subsp. enterica	MNNEETFYQAMRRKGVTRRSFLKFCSLAATSLGLGAGMTPKIAWALENKPRIPVVWIHGL	WP_023243720.1	hydrogenase 1, small subunit [Salmonella enterica]	Length: 372\nNumber of predicted TMHs:  2\nExp number of AAs in TMHs: 36.52147\nExp number, first 60 AAs:  19.72838\nTotal prob of N-in: 0.96170\nPOSSIBLE N-term signal sequence\ninside   1   19\nTMhelix   20   39\noutside   40   325\nTMhelix   326   348\ninside   349   372

18306	GCF_001479725.1_Salmonella_enterica_CVM_N38236-SQ_v1.0	Salmonella enterica subsp. enterica serovar Kentucky	Proteobacteria; Gammaproteobacteria; Enterobacterales; Enterobacteriaceae; Salmonella; Salmonella enterica; Salmonella enterica subsp. enterica	MNNEETFYQAMRRKGVTRRSFLKFCSLAATSLGLGAGMTPKIAWALENKPRIPVVWIHGL	WP_001058311.1	hydrogenase [Salmonella enterica]	Length: 372\nNumber of predicted TMHs:  2\nExp number of AAs in TMHs: 37.4699899999999\nExp number, first 60 AAs:  19.72699\nTotal prob of N-in:

0.96171\nPOSSIBLE N-term signal sequence\ninside 1 19\nTMhelix 20 39\noutside 40 325\nTMhelix 326 348\ninside 349 372

18307 GCF_001478475.1_Salmonella_enterica_CVM_N40925-SQ_v1.0 Salmonella enterica subsp. enterica serovar Kentucky Proteobacteria; Gammaproteobacteria; Enterobacterales; Enterobacteriaceae; Salmonella; Salmonella enterica; Salmonella enterica subsp. enterica MNNEETFYQAMRRKGVTRRSFLKFCSLAATSLGLGAGMTPKIAWALENKPRIPVVWIHGL WP_001058311.1 hydrogenase [Salmonella enterica] Length: 372\nNumber of predicted TMHs: 2\nExp number of AAs in TMHs: 37.4699899999999\nExp number, first 60 AAs: 19.72699\nTotal prob of N-in: 0.96171\nPOSSIBLE N-term signal sequence\ninside 1 19\nTMhelix 20 39\noutside 40 325\nTMhelix 326 348\ninside 349 372

18308 GCF_001480865.1_Salmonella_enterica_CVM_N39863-SQ_v1.0 Salmonella enterica subsp. enterica serovar Kentucky Proteobacteria; Gammaproteobacteria; Enterobacterales; Enterobacteriaceae; Salmonella; Salmonella enterica; Salmonella enterica subsp. enterica MNNEETFYQAMRRKGVTRRSFLKFCSLAATSLGLGAGMTPKIAWALENKPRIPVVWIHGL WP_001058311.1 hydrogenase [Salmonella enterica] Length: 372\nNumber of predicted TMHs: 2\nExp number of AAs in TMHs: 37.4699899999999\nExp number, first 60 AAs: 19.72699\nTotal prob of N-in: 0.96171\nPOSSIBLE N-term signal sequence\ninside 1 19\nTMhelix 20 39\noutside 40 325\nTMhelix 326 348\ninside 349 372

18309 GCF_001542155.1_ASM154215v1 Salmonella enterica subsp. enterica serovar Kentucky Proteobacteria; Gammaproteobacteria; Enterobacterales; Enterobacteriaceae; Salmonella; Salmonella enterica; Salmonella enterica subsp. enterica MNNEETFYQAMRRKGVTRRSFLKFCSLAATSLGLGAGMTPKIAWALENKPRIPVVWIHGL WP_001058311.1 hydrogenase [Salmonella enterica] Length: 372\nNumber of predicted TMHs: 2\nExp number of AAs in TMHs: 37.4699899999999\nExp number, first 60 AAs: 19.72699\nTotal prob of N-in: 0.96171\nPOSSIBLE N-term signal sequence\ninside 1 19\nTMhelix 20 39\noutside 40 325\nTMhelix 326 348\ninside 349 372

18310 GCF_001478475.1_Salmonella_enterica_CVM_N40925-SQ_v1.0 Salmonella enterica subsp. enterica serovar Kentucky Proteobacteria; Gammaproteobacteria; Enterobacterales; Enterobacteriaceae; Salmonella; Salmonella enterica; Salmonella enterica subsp. enterica MTGDNTLITSHGINRRDFMKLCAALAATMGLSSKAAAEMAESVSNPQRPPVIWIGAQECT WP_000145429.1 hydrogenase 2 small subunit [Salmonella enterica] Length: 372\nNumber of predicted TMHs: 1\nExp number of AAs in TMHs: 24.51309\nExp number, first 60 AAs: 0.2682\nTotal prob of N-in: 0.14402\noutside 1 331\nTMhelix 332 354\ninside 355 372

18311 GCF_001480865.1_Salmonella_enterica_CVM_N39863-SQ_v1.0 Salmonella enterica subsp. enterica serovar Kentucky Proteobacteria; Gammaproteobacteria; Enterobacterales; Enterobacteriaceae; Salmonella; Salmonella enterica; Salmonella enterica subsp. enterica MTGDNTLITSHGINRRDFMKLCAALAATMGLSSKAAAEMAESVSNPQRPPVIWIGAQECT WP_000145429.1 hydrogenase 2 small subunit [Salmonella enterica] Length: 372\nNumber of predicted TMHs: 1\nExp number of AAs in TMHs: 24.51309\nExp number, first 60 AAs: 0.2682\nTotal prob of N-in: 0.14402\noutside 1 331\nTMhelix 332 354\ninside 355 372

18312 GCF_001478105.1_Salmonella_enterica_CVM_N31384-SQ_v1.0 Salmonella enterica subsp. enterica serovar Kentucky Proteobacteria; Gammaproteobacteria; Enterobacterales; Enterobacteriaceae; Salmonella; Salmonella enterica; Salmonella enterica subsp. enterica MNNEETFYQAMRRKGVTRRSFLKFCSLAATSLGLGAGMTPKIAWALENKPRIPVVWIHGL WP_001058311.1 hydrogenase [Salmonella enterica] Length: 372\nNumber of predicted TMHs: 2\nExp number of AAs in TMHs: 37.4699899999999\nExp number, first 60 AAs: 19.72699\nTotal prob of N-in: 0.96171\nPOSSIBLE N-term signal sequence\ninside 1 19\nTMhelix 20 39\noutside 40 325\nTMhelix 326 348\ninside 349 372

18313 GCF_001478105.1_Salmonella_enterica_CVM_N31384-SQ_v1.0 Salmonella enterica subsp. enterica serovar Kentucky Proteobacteria; Gammaproteobacteria; Enterobacterales; Enterobacteriaceae; Salmonella; Salmonella enterica; Salmonella enterica subsp. enterica MTGDNTLITSHGINRRDFMKLCAALAATMGLSSKAAAEMAESVSNPQRPPVIWIGAQECT WP_000145429.1 hydrogenase 2 small subunit [Salmonella enterica] Length: 372\nNumber of predicted TMHs: 1\nExp number of AAs in TMHs: 24.51309\nExp number, first 60 AAs: 0.2682\nTotal prob of N-in: 0.14402\noutside 1 331\nTMhelix 332 354\ninside 355 372

18314 GCF_002031175.1_ASM203117v1 Salmonella enterica subsp. enterica serovar Kentucky Proteobacteria; Gammaproteobacteria; Enterobacterales; Enterobacteriaceae; Salmonella; Salmonella enterica; Salmonella enterica subsp. enterica MTGDNTLITSHGINRRDFMKLCAALAATMGLSSKAAAEMAESVSNPQRPPVIWIGAQECT WP_000145429.1 hydrogenase 2 small subunit [Salmonella enterica] Length: 372\nNumber of

predicted TMHs:  1\nExp number of AAs in TMHs: 24.51309\nExp number, first 60 AAs:  0.2682\nTotal prob of N-in: 0.14402\noutside   1   331\nTMhelix   332   354\ninside   355   372

18315 GCF_001542125.1_ASM154212v1 Salmonella enterica subsp. enterica serovar Kentucky Proteobacteria; Gammaproteobacteria; Enterobacterales; Enterobacteriaceae; Salmonella; Salmonella enterica; Salmonella enterica subsp. enterica MNNEETFYQAMRRKGVTRRSFLKFCSLAATSLGLGAGMTPKIAWALENKPRIPVVWIHGL WP_001058311.1 hydrogenase [Salmonella enterica] Length: 372\nNumber of predicted TMHs:  2\nExp number of AAs in TMHs: 37.4699899999999\nExp number, first 60 AAs:  19.72699\nTotal prob of N-in: 0.96171\nPOSSIBLE N-term signal sequence\ninside   1   19\nTMhelix   20   39\noutside   40   325\nTMhelix   326   348\ninside   349   372

18316 GCF_001479685.1_Salmonella_enterica_CVM_N38232-SQ_v1.0 Salmonella enterica subsp. enterica serovar Kentucky Proteobacteria; Gammaproteobacteria; Enterobacterales; Enterobacteriaceae; Salmonella; Salmonella enterica; Salmonella enterica subsp. enterica MNNEETFYQAMRRKGVTRRSFLKFCSLAATSLGLGAGMTPKIAWALENKPRIPVVWIHGL WP_001058311.1 hydrogenase [Salmonella enterica] Length: 372\nNumber of predicted TMHs:  2\nExp number of AAs in TMHs: 37.4699899999999\nExp number, first 60 AAs:  19.72699\nTotal prob of N-in: 0.96171\nPOSSIBLE N-term signal sequence\ninside   1   19\nTMhelix   20   39\noutside   40   325\nTMhelix   326   348\ninside   349   372

18317 GCF_001542215.1_ASM154221v1 Salmonella enterica subsp. enterica serovar Kentucky Proteobacteria; Gammaproteobacteria; Enterobacterales; Enterobacteriaceae; Salmonella; Salmonella enterica; Salmonella enterica subsp. enterica MTGDNTLITSHGINRRDFMKLCAALAATMGLSSKAAAEMAESVSNPQRPPVIWIGAQECT WP_000145429.1 hydrogenase 2 small subunit [Salmonella enterica] Length: 372\nNumber of predicted TMHs:  1\nExp number of AAs in TMHs: 24.51309\nExp number, first 60 AAs:  0.2682\nTotal prob of N-in: 0.14402\noutside   1   331\nTMhelix   332   354\ninside   355   372

18318 GCF_001542155.1_ASM154215v1 Salmonella enterica subsp. enterica serovar Kentucky Proteobacteria; Gammaproteobacteria; Enterobacterales; Enterobacteriaceae; Salmonella; Salmonella enterica; Salmonella enterica subsp. enterica MTGDNTLITSHGINRRDFMKLCAALAATMGLSSKAAAEMAESVSNPQRPPVIWIGAQECT WP_000145429.1 hydrogenase 2 small subunit [Salmonella enterica] Length: 372\nNumber of predicted TMHs:  1\nExp number of AAs in TMHs: 24.51309\nExp number, first 60 AAs:  0.2682\nTotal prob of N-in: 0.14402\noutside   1   331\nTMhelix   332   354\ninside   355   372

18319 GCF_001479425.1_Salmonella_enterica_CVM_N32064-SQ_v1.0 Salmonella enterica subsp. enterica serovar Kentucky Proteobacteria; Gammaproteobacteria; Enterobacterales; Enterobacteriaceae; Salmonella; Salmonella enterica; Salmonella enterica subsp. enterica MTGDNTLITSHGINRRDFMKLCAALAATMGLSSKAAAEMAESVSNPQRPPVIWIGAQECT WP_000145429.1 hydrogenase 2 small subunit [Salmonella enterica] Length: 372\nNumber of predicted TMHs:  1\nExp number of AAs in TMHs: 24.51309\nExp number, first 60 AAs:  0.2682\nTotal prob of N-in: 0.14402\noutside   1   331\nTMhelix   332   354\ninside   355   372

18320 GCF_001479425.1_Salmonella_enterica_CVM_N32064-SQ_v1.0 Salmonella enterica subsp. enterica serovar Kentucky Proteobacteria; Gammaproteobacteria; Enterobacterales; Enterobacteriaceae; Salmonella; Salmonella enterica; Salmonella enterica subsp. enterica MNNEETFYQAMRRKGVTRRSFLKFCSLAATSLGLGAGMTPKIAWALENKPRIPVVWIHGL WP_001058311.1 hydrogenase [Salmonella enterica] Length: 372\nNumber of predicted TMHs:  2\nExp number of AAs in TMHs: 37.4699899999999\nExp number, first 60 AAs:  19.72699\nTotal prob of N-in: 0.96171\nPOSSIBLE N-term signal sequence\ninside   1   19\nTMhelix   20   39\noutside   40   325\nTMhelix   326   348\ninside   349   372

18321 GCF_001479685.1_Salmonella_enterica_CVM_N38232-SQ_v1.0 Salmonella enterica subsp. enterica serovar Kentucky Proteobacteria; Gammaproteobacteria; Enterobacterales; Enterobacteriaceae; Salmonella; Salmonella enterica; Salmonella enterica subsp. enterica MTGDNTLITSHGINRRDFMKLCAALAATMGLSSKAAAEMAESVSNPQRPPVIWIGAQECT WP_000145429.1 hydrogenase 2 small subunit [Salmonella enterica] Length: 372\nNumber of predicted TMHs:  1\nExp number of AAs in TMHs: 24.51309\nExp number, first 60 AAs:  0.2682\nTotal prob of N-in: 0.14402\noutside   1   331\nTMhelix   332   354\ninside   355   372

18322 GCF_001479755.1_Salmonella_enterica_CVM_N38240-SQ_v1.0 Salmonella enterica subsp. enterica serovar Kentucky Proteobacteria; Gammaproteobacteria; Enterobacterales; Enterobacteriaceae; Salmonella; Salmonella enterica; Salmonella enterica subsp. enterica MTGDNTLITSHGINRRDFMKLCAALAATMGLSSKAAAEMAESVSNPQRPPVIWIGAQECT WP_000145429.1 hydrogenase 2 small subunit [Salmonella enterica] Length: 372\nNumber of predicted TMHs:  1\nExp number of AAs in TMHs: 24.51309\nExp number, first 60 AAs:  0.2682\nTotal prob of N-in: 0.14402\noutside   1   331\nTMhelix   332   354\ninside   355   372

18323 GCF_001479725.1_Salmonella_enterica_CVM_N38236-SQ_v1.0 Salmonella enterica subsp. enterica serovar Kentucky Proteobacteria; Gammaproteobacteria; Enterobacterales; Enterobacteriaceae; Salmonella; Salmonella enterica; Salmonella enterica subsp. enterica MTGDNTLITSHGINRRDFMKLCAALAATMGLSSKAAAEMAESVSNPQRPPVIWIGAQECT WP_000145429.1 hydrogenase 2 small subunit [Salmonella enterica] Length: 372\nNumber of predicted TMHs: 1\nExp number of AAs in TMHs: 24.51309\nExp number, first 60 AAs: 0.2682\nTotal prob of N-in: 0.14402\noutside 1 331\nTMhelix 332 354\ninside 355 372

18324 GCF_001479755.1_Salmonella_enterica_CVM_N38240-SQ_v1.0 Salmonella enterica subsp. enterica serovar Kentucky Proteobacteria; Gammaproteobacteria; Enterobacterales; Enterobacteriaceae; Salmonella; Salmonella enterica; Salmonella enterica subsp. enterica MNNEETFYQAMRRKGVTRRSFLKFCSLAATSLGLGAGMTPKIAWALENKPRIPVVWIHGL WP_001058311.1 hydrogenase [Salmonella enterica] Length: 372\nNumber of predicted TMHs: 2\nExp number of AAs in TMHs: 37.4699899999999\nExp number, first 60 AAs: 19.72699\nTotal prob of N-in: 0.96171\nPOSSIBLE N-term signal sequence\ninside 1 19\nTMhelix 20 39\noutside 40 325\nTMhelix 326 348\ninside 349 372

18325 GCF_001542065.1_ASM154206v1 Salmonella enterica subsp. enterica serovar Kentucky Proteobacteria; Gammaproteobacteria; Enterobacterales; Enterobacteriaceae; Salmonella; Salmonella enterica; Salmonella enterica subsp. enterica MNNEETFYQAMRRKGVTRRSFLKFCSLAATSLGLGAGMTPKIAWALENKPRIPVVWIHGL WP_001058311.1 hydrogenase [Salmonella enterica] Length: 372\nNumber of predicted TMHs: 2\nExp number of AAs in TMHs: 37.4699899999999\nExp number, first 60 AAs: 19.72699\nTotal prob of N-in: 0.96171\nPOSSIBLE N-term signal sequence\ninside 1 19\nTMhelix 20 39\noutside 40 325\nTMhelix 326 348\ninside 349 372

18326 GCF_001479905.1_Salmonella_enterica_CVM_N38870-SQ_v1.0 Salmonella enterica subsp. enterica serovar Kentucky Proteobacteria; Gammaproteobacteria; Enterobacterales; Enterobacteriaceae; Salmonella; Salmonella enterica; Salmonella enterica subsp. enterica MTGDNTLITSHGINRRDFMKLCAALAATMGLSSKAAAEMAESVSNPQRPPVIWIGAQECT WP_000145429.1 hydrogenase 2 small subunit [Salmonella enterica] Length: 372\nNumber of predicted TMHs: 1\nExp number of AAs in TMHs: 24.51309\nExp number, first 60 AAs: 0.2682\nTotal prob of N-in: 0.14402\noutside 1 331\nTMhelix 332 354\ninside 355 372

18327 GCF_002030455.1_ASM203045v1 Salmonella enterica subsp. enterica serovar Kentucky Proteobacteria; Gammaproteobacteria; Enterobacterales; Enterobacteriaceae; Salmonella; Salmonella enterica; Salmonella enterica subsp. enterica MTGDNTLITSHGINRRDFMKLCAALAATMGLSSKAAAEMAESVSNPQRPPVIWIGAQECT WP_000145429.1 hydrogenase 2 small subunit [Salmonella enterica] Length: 372\nNumber of predicted TMHs: 1\nExp number of AAs in TMHs: 24.51309\nExp number, first 60 AAs: 0.2682\nTotal prob of N-in: 0.14402\noutside 1 331\nTMhelix 332 354\ninside 355 372

18328 GCF_002030455.1_ASM203045v1 Salmonella enterica subsp. enterica serovar Kentucky Proteobacteria; Gammaproteobacteria; Enterobacterales; Enterobacteriaceae; Salmonella; Salmonella enterica; Salmonella enterica subsp. enterica MNNEETFYQAMRRKGVTRRSFLKFCSLAATSLGLGAGMTPKIAWALENKPRIPVVWIHGL WP_001058311.1 hydrogenase [Salmonella enterica] Length: 372\nNumber of predicted TMHs: 2\nExp number of AAs in TMHs: 37.4699899999999\nExp number, first 60 AAs: 19.72699\nTotal prob of N-in: 0.96171\nPOSSIBLE N-term signal sequence\ninside 1 19\nTMhelix 20 39\noutside 40 325\nTMhelix 326 348\ninside 349 372

18329 GCF_001480695.1_Salmonella_enterica_CVM_N42500_v1.0 Salmonella enterica subsp. enterica serovar Kentucky Proteobacteria; Gammaproteobacteria; Enterobacterales; Enterobacteriaceae; Salmonella; Salmonella enterica; Salmonella enterica subsp. enterica MTGDNTLITSHGINRRDFMKLCAALAATMGLSSKAAAEMAESVSNPQRPPVIWIGAQECT WP_000145429.1 hydrogenase 2 small subunit [Salmonella enterica] Length: 372\nNumber of predicted TMHs: 1\nExp number of AAs in TMHs: 24.51309\nExp number, first 60 AAs: 0.2682\nTotal prob of N-in: 0.14402\noutside 1 331\nTMhelix 332 354\ninside 355 372

18330 GCF_001480805.1_Salmonella_enterica_CVM_N38868-SQ_v1.0 Salmonella enterica subsp. enterica serovar Kentucky Proteobacteria; Gammaproteobacteria; Enterobacterales; Enterobacteriaceae; Salmonella; Salmonella enterica; Salmonella enterica subsp. enterica MTGDNTLITSHGINRRDFMKLCAALAATMGLSSKAAAEMAESVSNPQRPPVIWIGAQECT WP_000145429.1 hydrogenase 2 small subunit [Salmonella enterica] Length: 372\nNumber of predicted TMHs: 1\nExp number of AAs in TMHs: 24.51309\nExp number, first 60 AAs: 0.2682\nTotal prob of N-in: 0.14402\noutside 1 331\nTMhelix 332 354\ninside 355 372

18331 GCF_001457355.1_ASM145735v1 Salmonella enterica subsp. enterica serovar Kentucky Proteobacteria; Gammaproteobacteria; Enterobacterales; Enterobacteriaceae; Salmonella; Salmonella enterica;

Salmonella enterica subsp. enterica	MTGDNTLITSHGINRRDFMKLCAALAATMGLSSKAAAEMAESVSNPQRPPVIWIGAQECT	WP_000145429.1	hydrogenase 2 small subunit [Salmonella enterica]	Length: 372\nNumber of predicted TMHs:  1\nExp number of AAs in TMHs: 24.51309\nExp number, first 60 AAs:  0.2682\nTotal prob of N-in:  0.14402\noutside   1   331\nTMhelix   332   354\ninside   355   372

18332	GCF_001457355.1_ASM145735v1	Salmonella enterica subsp. enterica serovar Kentucky	Proteobacteria; Gammaproteobacteria; Enterobacterales; Enterobacteriaceae; Salmonella; Salmonella enterica; Salmonella enterica subsp. enterica	MNNEETFYQAMRRKGVTRRSFLKFCSLAATSLGLGAGMTPKIAWALENKPRIPVVWIHGL	WP_001058311.1	hydrogenase [Salmonella enterica]	Length: 372\nNumber of predicted TMHs:  2\nExp number of AAs in TMHs: 37.4699899999999\nExp number, first 60 AAs:  19.72699\nTotal prob of N-in:  0.96171\nPOSSIBLE N-term signal sequence\ninside   1   19\nTMhelix   20   39\noutside   40   325\nTMhelix   326   348\ninside   349   372

18333	GCF_001480805.1_Salmonella_enterica_CVM_N38868-SQ_v1.0	Salmonella enterica subsp. enterica serovar Kentucky	Proteobacteria; Gammaproteobacteria; Enterobacterales; Enterobacteriaceae; Salmonella; Salmonella enterica; Salmonella enterica subsp. enterica	MNNEETFYQAMRRKGVTRRSFLKFCSLAATSLGLGAGMTPKIAWALENKPRIPVVWIHGL	WP_001058312.1	hydrogenase [Salmonella enterica]	Length: 372\nNumber of predicted TMHs:  2\nExp number of AAs in TMHs: 36.51738\nExp number, first 60 AAs:  19.72849\nTotal prob of N-in:   0.96170\nPOSSIBLE N-term signal sequence\ninside   1   19\nTMhelix   20   39\noutside   40   325\nTMhelix   326   348\ninside   349   372

18334	GCF_001480695.1_Salmonella_enterica_CVM_N42500_v1.0	Salmonella enterica subsp. enterica serovar Kentucky	Proteobacteria; Gammaproteobacteria; Enterobacterales; Enterobacteriaceae; Salmonella; Salmonella enterica; Salmonella enterica subsp. enterica	MNNEETFYQAMRRKGVTRRSFLKFCSLAATSLGLGAGMTPKIAWALENKPRIPVVWIHGL	WP_001058311.1	hydrogenase [Salmonella enterica]	Length: 372\nNumber of predicted TMHs:  2\nExp number of AAs in TMHs: 37.4699899999999\nExp number, first 60 AAs:  19.72699\nTotal prob of N-in:  0.96171\nPOSSIBLE N-term signal sequence\ninside   1   19\nTMhelix   20   39\noutside   40   325\nTMhelix   326   348\ninside   349   372

18335	GCF_001542215.1_ASM154221v1	Salmonella enterica subsp. enterica serovar Kentucky	Proteobacteria; Gammaproteobacteria; Enterobacterales; Enterobacteriaceae; Salmonella; Salmonella enterica; Salmonella enterica subsp. enterica	MNNEETFYQAMRRKGVTRRSFLKFCSLAATSLGLGAGMTPKIAWALENKPRIPVVWIHGL	WP_001058311.1	hydrogenase [Salmonella enterica]	Length: 372\nNumber of predicted TMHs:  2\nExp number of AAs in TMHs: 37.4699899999999\nExp number, first 60 AAs:  19.72699\nTotal prob of N-in:  0.96171\nPOSSIBLE N-term signal sequence\ninside   1   19\nTMhelix   20   39\noutside   40   325\nTMhelix   326   348\ninside   349   372

18336	GCF_001542125.1_ASM154212v1	Salmonella enterica subsp. enterica serovar Kentucky	Proteobacteria; Gammaproteobacteria; Enterobacterales; Enterobacteriaceae; Salmonella; Salmonella enterica; Salmonella enterica subsp. enterica	MTGDNTLITSHGINRRDFMKLCAALAATMGLSSKAAAEMAESVSNPQRPPVIWIGAQECT	WP_000145429.1	hydrogenase 2 small subunit [Salmonella enterica]	Length: 372\nNumber of predicted TMHs:  1\nExp number of AAs in TMHs: 24.51309\nExp number, first 60 AAs:  0.2682\nTotal prob of N-in:  0.14402\noutside   1   331\nTMhelix   332   354\ninside   355   372

18337	GCF_001542065.1_ASM154206v1	Salmonella enterica subsp. enterica serovar Kentucky	Proteobacteria; Gammaproteobacteria; Enterobacterales; Enterobacteriaceae; Salmonella; Salmonella enterica; Salmonella enterica subsp. enterica	MTGDNTLITSHGINRRDFMKLCAALAATMGLSSKAAAEMAESVSNPQRPPVIWIGAQECT	WP_000145429.1	hydrogenase 2 small subunit [Salmonella enterica]	Length: 372\nNumber of predicted TMHs:  1\nExp number of AAs in TMHs: 24.51309\nExp number, first 60 AAs:  0.2682\nTotal prob of N-in:  0.14402\noutside   1   331\nTMhelix   332   354\ninside   355   372

18338	GCF_002031345.1_ASM203134v1	Salmonella enterica subsp. enterica serovar Kentucky	Proteobacteria; Gammaproteobacteria; Enterobacterales; Enterobacteriaceae; Salmonella; Salmonella enterica; Salmonella enterica subsp. enterica	MNNEETFYQAMRRKGVTRRSFLKFCSLAATSLGLGAGMTPKIAWALENKPRIPVVWIHGL	WP_001058312.1	hydrogenase [Salmonella enterica]	Length: 372\nNumber of predicted TMHs:  2\nExp number of AAs in TMHs: 36.51738\nExp number, first 60 AAs:  19.72849\nTotal prob of N-in:   0.96170\nPOSSIBLE N-term signal sequence\ninside   1   19\nTMhelix   20   39\noutside   40   325\nTMhelix   326   348\ninside   349   372

18339	GCF_002031615.1_ASM203161v1	Salmonella enterica subsp. enterica serovar Kentucky	Proteobacteria; Gammaproteobacteria; Enterobacterales; Enterobacteriaceae; Salmonella; Salmonella enterica; Salmonella enterica subsp. enterica	MTGDNTLITSHGINRRDFMKLCAALAATMGLSSKAAAEMAESVSNPQRPPVIWIGAQECT	WP_000145429.1	hydrogenase 2 small subunit [Salmonella enterica]	Length: 372\nNumber of

predicted TMHs:  1\nExp number of AAs in TMHs: 24.51309\nExp number, first 60 AAs:  0.2682\nTotal prob of N-in: 0.14402\noutside   1   331\nTMhelix   332   354\ninside   355   372

18340 GCF_002031175.1_ASM203117v1 Salmonella enterica subsp. enterica serovar Kentucky Proteobacteria; Gammaproteobacteria; Enterobacterales; Enterobacteriaceae; Salmonella; Salmonella enterica; Salmonella enterica subsp. enterica MNNEETFYQAMRRKGVTRRSFLKFCSLAATSLGLGAGMTPKIAWALENKPRIPVVWIHGL WP_001058311.1 hydrogenase [Salmonella enterica] Length: 372\nNumber of predicted TMHs:  2\nExp number of AAs in TMHs: 37.4699899999999\nExp number, first 60 AAs:  19.72699\nTotal prob of N-in: 0.96171\nPOSSIBLE N-term signal sequence\ninside   1   19\nTMhelix   20   39\noutside   40   325\nTMhelix   326   348\ninside   349   372

18341 GCF_002048595.1_ASM204859v1 Salmonella enterica subsp. enterica serovar Kiambu Proteobacteria; Gammaproteobacteria; Enterobacterales; Enterobacteriaceae; Salmonella; Salmonella enterica; Salmonella enterica subsp. enterica MNNEETFYQAMRRKGVTRRSFLKFCSLAATSLGLGAGMTPKIAWALENKPRIPVVWIHGL WP_064000476.1 hydrogenase [Salmonella enterica] Length: 372\nNumber of predicted TMHs:  2\nExp number of AAs in TMHs: 37.4700099999999\nExp number, first 60 AAs:  19.72698\nTotal prob of N-in: 0.96171\nPOSSIBLE N-term signal sequence\ninside   1   19\nTMhelix   20   39\noutside   40   325\nTMhelix   326   348\ninside   349   372

18342 GCF_002048595.1_ASM204859v1 Salmonella enterica subsp. enterica serovar Kiambu Proteobacteria; Gammaproteobacteria; Enterobacterales; Enterobacteriaceae; Salmonella; Salmonella enterica; Salmonella enterica subsp. enterica MTGDNTLITSHGINRRDFMKLCAALAATMGLSSKAAAEMAESVSNPQRPPVIWIGAQECT WP_000145429.1 hydrogenase 2 small subunit [Salmonella enterica] Length: 372\nNumber of predicted TMHs:  1\nExp number of AAs in TMHs: 24.51309\nExp number, first 60 AAs:  0.2682\nTotal prob of N-in: 0.14402\noutside   1   331\nTMhelix   332   354\ninside   355   372

18343 GCF_002048645.1_ASM204864v1 Salmonella enterica subsp. enterica serovar Kiambu Proteobacteria; Gammaproteobacteria; Enterobacterales; Enterobacteriaceae; Salmonella; Salmonella enterica; Salmonella enterica subsp. enterica MTGDNTLITSHGINRRDFMKLCAALAATMGLSSKAAAEMAESVSNPQRPPVIWIGAQECT WP_000145429.1 hydrogenase 2 small subunit [Salmonella enterica] Length: 372\nNumber of predicted TMHs:  1\nExp number of AAs in TMHs: 24.51309\nExp number, first 60 AAs:  0.2682\nTotal prob of N-in: 0.14402\noutside   1   331\nTMhelix   332   354\ninside   355   372

18344 GCF_002047545.1_ASM204754v1 Salmonella enterica subsp. enterica serovar Kiambu Proteobacteria; Gammaproteobacteria; Enterobacterales; Enterobacteriaceae; Salmonella; Salmonella enterica; Salmonella enterica subsp. enterica MTGDNTLITSHGINRRDFMKLCAALAATMGLSSKAAAEMAESVSNPQRPPVIWIGAQECT WP_000145429.1 hydrogenase 2 small subunit [Salmonella enterica] Length: 372\nNumber of predicted TMHs:  1\nExp number of AAs in TMHs: 24.51309\nExp number, first 60 AAs:  0.2682\nTotal prob of N-in: 0.14402\noutside   1   331\nTMhelix   332   354\ninside   355   372

18345 GCF_002048595.1_ASM204859v1 Salmonella enterica subsp. enterica serovar Kiambu Proteobacteria; Gammaproteobacteria; Enterobacterales; Enterobacteriaceae; Salmonella; Salmonella enterica; Salmonella enterica subsp. enterica MLKFRTIFLKKIFTLKNSILARRGAVIVIVSAVFTSIMFFAHSWASDKEVAMTLSVLNSN WP_023226644.1 hypothetical protein [Salmonella enterica] Length: 204\nNumber of predicted TMHs: 2\nExp number of AAs in TMHs: 44.99526\nExp number, first 60 AAs:  22.33786\nTotal prob of N-in:   0.87083\nPOSSIBLE N-term signal sequence\ninside   1   19\nTMhelix   20   42\noutside   43   171\nTMhelix   172   194\ninside   195   204

18346 GCF_002047805.1_ASM204780v1 Salmonella enterica subsp. enterica serovar Kiambu Proteobacteria; Gammaproteobacteria; Enterobacterales; Enterobacteriaceae; Salmonella; Salmonella enterica; Salmonella enterica subsp. enterica MLKFRTIFLKKIFTLKNSILARRGAVIVIVSAVFTSIMFFAHSWASDKEVAMTLSVLNSN WP_023226644.1 hypothetical protein [Salmonella enterica] Length: 204\nNumber of predicted TMHs: 2\nExp number of AAs in TMHs: 44.99526\nExp number, first 60 AAs:  22.33786\nTotal prob of N-in:   0.87083\nPOSSIBLE N-term signal sequence\ninside   1   19\nTMhelix   20   42\noutside   43   171\nTMhelix   172   194\ninside   195   204

18347 GCF_002047805.1_ASM204780v1 Salmonella enterica subsp. enterica serovar Kiambu Proteobacteria; Gammaproteobacteria; Enterobacterales; Enterobacteriaceae; Salmonella; Salmonella enterica; Salmonella enterica subsp. enterica MNNEETFYQAMRRKGVTRRSFLKFCSLAATSLGLGAGMTPKIAWALENKPRIPVVWIHGL WP_064000476.1 hydrogenase [Salmonella enterica] Length: 372\nNumber of predicted TMHs:  2\nExp number of AAs in TMHs: 37.4700099999999\nExp number, first 60 AAs:  19.72698\nTotal prob of N-in: 0.96171\nPOSSIBLE N-term signal sequence\ninside   1   19\nTMhelix   20   39\noutside   40   325\nTMhelix   326   348\ninside   349   372

18348 GCF_002047555.1_ASM204755v1 Salmonella enterica subsp. enterica serovar Kiambu Proteobacteria; Gammaproteobacteria; Enterobacterales; Enterobacteriaceae; Salmonella; Salmonella enterica; Salmonella enterica subsp. enterica MNNEETFYQAMRRKGVTRRSFLKFCSLAATSLGLGAGMTPKIAWALENKPRIPVVWIHGL WP_064000476.1 hydrogenase [Salmonella enterica] Length: 372\nNumber of predicted TMHs: 2\nExp number of AAs in TMHs: 37.4700099999999\nExp number, first 60 AAs: 19.72698\nTotal prob of N-in: 0.96171\nPOSSIBLE N-term signal sequence\ninside 1 19\nTMhelix 20 39\noutside 40 325\nTMhelix 326 348\ninside 349 372

18349 GCF_002047555.1_ASM204755v1 Salmonella enterica subsp. enterica serovar Kiambu Proteobacteria; Gammaproteobacteria; Enterobacterales; Enterobacteriaceae; Salmonella; Salmonella enterica; Salmonella enterica subsp. enterica MTGDNTLITSHGINRRDFMKLCAALAATMGLSSKAAAEMAESVSNPQRPPVIWIGAQECT WP_000145429.1 hydrogenase 2 small subunit [Salmonella enterica] Length: 372\nNumber of predicted TMHs: 1\nExp number of AAs in TMHs: 24.51309\nExp number, first 60 AAs: 0.2682\nTotal prob of N-in: 0.14402\noutside 1 331\nTMhelix 332 354\ninside 355 372

18350 GCF_002047555.1_ASM204755v1 Salmonella enterica subsp. enterica serovar Kiambu Proteobacteria; Gammaproteobacteria; Enterobacterales; Enterobacteriaceae; Salmonella; Salmonella enterica; Salmonella enterica subsp. enterica MLKFRTIFLKKIFTLKNSILARRGAVIVIVSAVFTSIMFFAHSWASDKEVAMTLSVLNSN WP_023226644.1 hypothetical protein [Salmonella enterica] Length: 204\nNumber of predicted TMHs: 2\nExp number of AAs in TMHs: 44.99526\nExp number, first 60 AAs: 22.33786\nTotal prob of N-in: 0.87083\nPOSSIBLE N-term signal sequence\ninside 1 19\nTMhelix 20 42\noutside 43 171\nTMhelix 172 194\ninside 195 204

18351 GCF_002047805.1_ASM204780v1 Salmonella enterica subsp. enterica serovar Kiambu Proteobacteria; Gammaproteobacteria; Enterobacterales; Enterobacteriaceae; Salmonella; Salmonella enterica; Salmonella enterica subsp. enterica MTGDNTLITSHGINRRDFMKLCAALAATMGLSSKAAAEMAESVSNPQRPPVIWIGAQECT WP_000145429.1 hydrogenase 2 small subunit [Salmonella enterica] Length: 372\nNumber of predicted TMHs: 1\nExp number of AAs in TMHs: 24.51309\nExp number, first 60 AAs: 0.2682\nTotal prob of N-in: 0.14402\noutside 1 331\nTMhelix 332 354\ninside 355 372

18352 GCF_002047545.1_ASM204754v1 Salmonella enterica subsp. enterica serovar Kiambu Proteobacteria; Gammaproteobacteria; Enterobacterales; Enterobacteriaceae; Salmonella; Salmonella enterica; Salmonella enterica subsp. enterica MNNEETFYQAMRRKGVTRRSFLKFCSLAATSLGLGAGMTPKIAWALENKPRIPVVWIHGL WP_064000476.1 hydrogenase [Salmonella enterica] Length: 372\nNumber of predicted TMHs: 2\nExp number of AAs in TMHs: 37.4700099999999\nExp number, first 60 AAs: 19.72698\nTotal prob of N-in: 0.96171\nPOSSIBLE N-term signal sequence\ninside 1 19\nTMhelix 20 39\noutside 40 325\nTMhelix 326 348\ninside 349 372

18353 GCF_002047545.1_ASM204754v1 Salmonella enterica subsp. enterica serovar Kiambu Proteobacteria; Gammaproteobacteria; Enterobacterales; Enterobacteriaceae; Salmonella; Salmonella enterica; Salmonella enterica subsp. enterica MLKFRTIFLKKIFTLKNSILARRGAVIVIVSAVFTSIMFFAHSWASDKEVAMTLSVLNSN WP_023226644.1 hypothetical protein [Salmonella enterica] Length: 204\nNumber of predicted TMHs: 2\nExp number of AAs in TMHs: 44.99526\nExp number, first 60 AAs: 22.33786\nTotal prob of N-in: 0.87083\nPOSSIBLE N-term signal sequence\ninside 1 19\nTMhelix 20 42\noutside 43 171\nTMhelix 172 194\ninside 195 204

18354 GCF_002048645.1_ASM204864v1 Salmonella enterica subsp. enterica serovar Kiambu Proteobacteria; Gammaproteobacteria; Enterobacterales; Enterobacteriaceae; Salmonella; Salmonella enterica; Salmonella enterica subsp. enterica MLKFRTIFLKKIFTLKNSILARRGAVIVIVSAVFTSIMFFAHSWASDKEVAMTLSVLNSN WP_023226644.1 hypothetical protein [Salmonella enterica] Length: 204\nNumber of predicted TMHs: 2\nExp number of AAs in TMHs: 44.99526\nExp number, first 60 AAs: 22.33786\nTotal prob of N-in: 0.87083\nPOSSIBLE N-term signal sequence\ninside 1 19\nTMhelix 20 42\noutside 43 171\nTMhelix 172 194\ninside 195 204

18355 GCF_002048645.1_ASM204864v1 Salmonella enterica subsp. enterica serovar Kiambu Proteobacteria; Gammaproteobacteria; Enterobacterales; Enterobacteriaceae; Salmonella; Salmonella enterica; Salmonella enterica subsp. enterica MNNEETFYQAMRRKGVTRRSFLKFCSLAATSLGLGAGMTPKIAWALENKPRIPVVWIHGL WP_064000476.1 hydrogenase [Salmonella enterica] Length: 372\nNumber of predicted TMHs: 2\nExp number of AAs in TMHs: 37.4700099999999\nExp number, first 60 AAs: 19.72698\nTotal prob of N-in: 0.96171\nPOSSIBLE N-term signal sequence\ninside 1 19\nTMhelix 20 39\noutside 40 325\nTMhelix 326 348\ninside 349 372

18356 GCF_002045095.1_ASM204509v1 Salmonella enterica subsp. enterica serovar Kintambo Proteobacteria; Gammaproteobacteria; Enterobacterales; Enterobacteriaceae; Salmonella; Salmonella enterica;

Salmonella enterica subsp. enterica MNNEETFYQAMRRKGVTRRSFLKFCSLAATSLGLGAGMTPKIAWALENKPRIPVVWIHGL WP_001058312.1 hydrogenase [Salmonella enterica] Length: 372\nNumber of predicted TMHs: 2\nExp number of AAs in TMHs: 36.51738\nExp number, first 60 AAs: 19.72849\nTotal prob of N-in: 0.96170\nPOSSIBLE N-term signal sequence\ninside 1 19\nTMhelix 20 39\noutside 40 325\nTMhelix 326 348\ninside 349 372

18357 GCF_002044735.1_ASM204473v1 Salmonella enterica subsp. enterica serovar Kintambo Proteobacteria; Gammaproteobacteria; Enterobacterales; Enterobacteriaceae; Salmonella; Salmonella enterica; Salmonella enterica subsp. enterica MNNEETFYQAMRRKGVTRRSFLKFCSLAATSLGLGAGMTPKIAWALENKPRIPVVWIHGL WP_001058311.1 hydrogenase [Salmonella enterica] Length: 372\nNumber of predicted TMHs: 2\nExp number of AAs in TMHs: 37.4699899999999\nExp number, first 60 AAs: 19.72699\nTotal prob of N-in: 0.96171\nPOSSIBLE N-term signal sequence\ninside 1 19\nTMhelix 20 39\noutside 40 325\nTMhelix 326 348\ninside 349 372

18358 GCF_002044735.1_ASM204473v1 Salmonella enterica subsp. enterica serovar Kintambo Proteobacteria; Gammaproteobacteria; Enterobacterales; Enterobacteriaceae; Salmonella; Salmonella enterica; Salmonella enterica subsp. enterica MTGDNTLITSHGINRRDFMKLCAALAATMGLSSKAAAEMAESVSNPQRPPVIWIGAQECT WP_000145429.1 hydrogenase 2 small subunit [Salmonella enterica] Length: 372\nNumber of predicted TMHs: 1\nExp number of AAs in TMHs: 24.51309\nExp number, first 60 AAs: 0.2682\nTotal prob of N-in: 0.14402\noutside 1 331\nTMhelix 332 354\ninside 355 372

18359 GCF_002045095.1_ASM204509v1 Salmonella enterica subsp. enterica serovar Kintambo Proteobacteria; Gammaproteobacteria; Enterobacterales; Enterobacteriaceae; Salmonella; Salmonella enterica; Salmonella enterica subsp. enterica MTGDNTLITSHGINRRDFMKLCAALAATMGLSSKAAAEMAESVSNPQRPPVIWIGAQECT WP_000145429.1 hydrogenase 2 small subunit [Salmonella enterica] Length: 372\nNumber of predicted TMHs: 1\nExp number of AAs in TMHs: 24.51309\nExp number, first 60 AAs: 0.2682\nTotal prob of N-in: 0.14402\noutside 1 331\nTMhelix 332 354\ninside 355 372

18360 GCF_002045875.1_ASM204587v1 Salmonella enterica subsp. enterica serovar Kintambo Proteobacteria; Gammaproteobacteria; Enterobacterales; Enterobacteriaceae; Salmonella; Salmonella enterica; Salmonella enterica subsp. enterica MNNEETFYQAMRRKGVTRRSFLKFCSLAATSLGLGAGMTPKIAWALENKPRIPVVWIHGL WP_001058311.1 hydrogenase [Salmonella enterica] Length: 372\nNumber of predicted TMHs: 2\nExp number of AAs in TMHs: 37.4699899999999\nExp number, first 60 AAs: 19.72699\nTotal prob of N-in: 0.96171\nPOSSIBLE N-term signal sequence\ninside 1 19\nTMhelix 20 39\noutside 40 325\nTMhelix 326 348\ninside 349 372

18361 GCF_002045875.1_ASM204587v1 Salmonella enterica subsp. enterica serovar Kintambo Proteobacteria; Gammaproteobacteria; Enterobacterales; Enterobacteriaceae; Salmonella; Salmonella enterica; Salmonella enterica subsp. enterica MTGDNTLITSHGINRRDFMKLCAALAATMGLSSKAAAEMAESVSNPQRPPVIWIGAQECT WP_000145429.1 hydrogenase 2 small subunit [Salmonella enterica] Length: 372\nNumber of predicted TMHs: 1\nExp number of AAs in TMHs: 24.51309\nExp number, first 60 AAs: 0.2682\nTotal prob of N-in: 0.14402\noutside 1 331\nTMhelix 332 354\ninside 355 372

18362 GCF_002045995.1_ASM204599v1 Salmonella enterica subsp. enterica serovar Kintambo Proteobacteria; Gammaproteobacteria; Enterobacterales; Enterobacteriaceae; Salmonella; Salmonella enterica; Salmonella enterica subsp. enterica MNNEETFYQAMRRKGVTRRSFLKFCSLAATSLGLGAGMTPKIAWALENKPRIPVVWIHGL WP_001058312.1 hydrogenase [Salmonella enterica] Length: 372\nNumber of predicted TMHs: 2\nExp number of AAs in TMHs: 36.51738\nExp number, first 60 AAs: 19.72849\nTotal prob of N-in: 0.96170\nPOSSIBLE N-term signal sequence\ninside 1 19\nTMhelix 20 39\noutside 40 325\nTMhelix 326 348\ninside 349 372

18363 GCF_002045995.1_ASM204599v1 Salmonella enterica subsp. enterica serovar Kintambo Proteobacteria; Gammaproteobacteria; Enterobacterales; Enterobacteriaceae; Salmonella; Salmonella enterica; Salmonella enterica subsp. enterica MTGDNTLITSHGINRRDFMKLCAALAATMGLSSKAAAEMAESVSNPQRPPVIWIGAQECT WP_000145429.1 hydrogenase 2 small subunit [Salmonella enterica] Length: 372\nNumber of predicted TMHs: 1\nExp number of AAs in TMHs: 24.51309\nExp number, first 60 AAs: 0.2682\nTotal prob of N-in: 0.14402\noutside 1 331\nTMhelix 332 354\ninside 355 372

18364 GCF_002062825.1_ASM206282v1 Salmonella enterica subsp. enterica serovar Kohbu Proteobacteria; Gammaproteobacteria; Enterobacterales; Enterobacteriaceae; Salmonella; Salmonella enterica; Salmonella enterica subsp. enterica MTGDNTLITSHGINRRDFMKLCAALAATMGLSSKAAAEMAESVSNPQRPPVIWIGAQECT WP_000145429.1 hydrogenase 2 small subunit [Salmonella enterica] Length: 372\nNumber of

predicted TMHs: 1\nExp number of AAs in TMHs: 24.51309\nExp number, first 60 AAs: 0.2682\nTotal prob of N-in: 0.14402\noutside 1 331\nTMhelix 332 354\ninside 355 372

18365 GCF_002062825.1_ASM206282v1 Salmonella enterica subsp. enterica serovar Kohbu Proteobacteria; Gammaproteobacteria; Enterobacterales; Enterobacteriaceae; Salmonella; Salmonella enterica; Salmonella enterica subsp. enterica MNNEETFYQAMRRKGVTRRSFLKFCSLAATSLGLGAGMTPKIAWALENKPRIPVVWIHGL WP_001058308.1 hydrogenase [Salmonella enterica] Length: 372\nNumber of predicted TMHs: 2\nExp number of AAs in TMHs: 39.08204\nExp number, first 60 AAs: 19.72474\nTotal prob of N-in: 0.96174\nPOSSIBLE N-term signal sequence\ninside 1 19\nTMhelix 20 39\noutside 40 325\nTMhelix 326 348\ninside 349 372

18366 GCF_002044865.1_ASM204486v1 Salmonella enterica subsp. enterica serovar Kottbus Proteobacteria; Gammaproteobacteria; Enterobacterales; Enterobacteriaceae; Salmonella; Salmonella enterica; Salmonella enterica subsp. enterica MTGDNTLITSHGINRRDFMKLCAALAATMGLSSKAAAEMAESVSNPQRPPVIWIGAQECT WP_000145429.1 hydrogenase 2 small subunit [Salmonella enterica] Length: 372\nNumber of predicted TMHs: 1\nExp number of AAs in TMHs: 24.51309\nExp number, first 60 AAs: 0.2682\nTotal prob of N-in: 0.14402\noutside 1 331\nTMhelix 332 354\ninside 355 372

18367 GCF_002045115.1_ASM204511v1 Salmonella enterica subsp. enterica serovar Kottbus Proteobacteria; Gammaproteobacteria; Enterobacterales; Enterobacteriaceae; Salmonella; Salmonella enterica; Salmonella enterica subsp. enterica MNNEETFYQAMRRKGVTRRSFLKFCSLAATSLGLGAGMTPKIAWALENKPRIPVVWIHGL WP_001058308.1 hydrogenase [Salmonella enterica] Length: 372\nNumber of predicted TMHs: 2\nExp number of AAs in TMHs: 39.08204\nExp number, first 60 AAs: 19.72474\nTotal prob of N-in: 0.96174\nPOSSIBLE N-term signal sequence\ninside 1 19\nTMhelix 20 39\noutside 40 325\nTMhelix 326 348\ninside 349 372

18368 GCF_002045115.1_ASM204511v1 Salmonella enterica subsp. enterica serovar Kottbus Proteobacteria; Gammaproteobacteria; Enterobacterales; Enterobacteriaceae; Salmonella; Salmonella enterica; Salmonella enterica subsp. enterica MTGDNTLITSHGINRRDFMKLCAALAATMGLSSKAAAEMAESVSNPQRPPVIWIGAQECT WP_000145429.1 hydrogenase 2 small subunit [Salmonella enterica] Length: 372\nNumber of predicted TMHs: 1\nExp number of AAs in TMHs: 24.51309\nExp number, first 60 AAs: 0.2682\nTotal prob of N-in: 0.14402\noutside 1 331\nTMhelix 332 354\ninside 355 372

18369 GCF_002044865.1_ASM204486v1 Salmonella enterica subsp. enterica serovar Kottbus Proteobacteria; Gammaproteobacteria; Enterobacterales; Enterobacteriaceae; Salmonella; Salmonella enterica; Salmonella enterica subsp. enterica MNNEETFYQAMRRKGVTRRSFLKFCSLAATSLGLGAGMTPKIAWALENKPRIPVVWIHGL WP_001058312.1 hydrogenase [Salmonella enterica] Length: 372\nNumber of predicted TMHs: 2\nExp number of AAs in TMHs: 36.51738\nExp number, first 60 AAs: 19.72849\nTotal prob of N-in: 0.96170\nPOSSIBLE N-term signal sequence\ninside 1 19\nTMhelix 20 39\noutside 40 325\nTMhelix 326 348\ninside 349 372

18370 GCF_002045035.1_ASM204503v1 Salmonella enterica subsp. enterica serovar Kottbus Proteobacteria; Gammaproteobacteria; Enterobacterales; Enterobacteriaceae; Salmonella; Salmonella enterica; Salmonella enterica subsp. enterica MTGDNTLITSHGINRRDFMKLCAALAATMGLSSKAAAEMAESVSNPQRPPVIWIGAQECT WP_000145429.1 hydrogenase 2 small subunit [Salmonella enterica] Length: 372\nNumber of predicted TMHs: 1\nExp number of AAs in TMHs: 24.51309\nExp number, first 60 AAs: 0.2682\nTotal prob of N-in: 0.14402\noutside 1 331\nTMhelix 332 354\ninside 355 372

18371 GCF_002045035.1_ASM204503v1 Salmonella enterica subsp. enterica serovar Kottbus Proteobacteria; Gammaproteobacteria; Enterobacterales; Enterobacteriaceae; Salmonella; Salmonella enterica; Salmonella enterica subsp. enterica MNNEETFYQAMRRKGVTRRSFLKFCSLAATSLGLGAGMTPKIAWALENKPRIPVVWIHGL WP_001058312.1 hydrogenase [Salmonella enterica] Length: 372\nNumber of predicted TMHs: 2\nExp number of AAs in TMHs: 36.51738\nExp number, first 60 AAs: 19.72849\nTotal prob of N-in: 0.96170\nPOSSIBLE N-term signal sequence\ninside 1 19\nTMhelix 20 39\noutside 40 325\nTMhelix 326 348\ninside 349 372

18372 GCF_002107495.1_ASM210749v1 Salmonella enterica subsp. enterica serovar Litchfield Proteobacteria; Gammaproteobacteria; Enterobacterales; Enterobacteriaceae; Salmonella; Salmonella enterica; Salmonella enterica subsp. enterica MNNEETFYQAMRRKGVTRRSFLKFCSLAATSLGLGAGMTPKIAWALENKPRIPVVWIHGL WP_001058312.1 hydrogenase [Salmonella enterica] Length: 372\nNumber of predicted TMHs: 2\nExp number of AAs in TMHs: 36.51738\nExp number, first 60 AAs: 19.72849\nTotal prob of N-in: 0.96170\nPOSSIBLE N-term signal sequence\ninside 1 19\nTMhelix 20 39\noutside 40 325\nTMhelix 326 348\ninside 349 372

18373 GCF_002059605.1_ASM205960v1 Salmonella enterica subsp. enterica serovar Litchfield Proteobacteria; Gammaproteobacteria; Enterobacterales; Enterobacteriaceae; Salmonella; Salmonella enterica;

Salmonella enterica subsp. enterica MTGDNTLITSHGINRRDFMKLCAALAATMGLSSKAAAEMAESVSNPQRPPVIWIGAQECT WP_000145429.1 hydrogenase 2 small subunit [Salmonella enterica] Length: 372\nNumber of predicted TMHs: 1\nExp number of AAs in TMHs: 24.51309\nExp number, first 60 AAs: 0.2682\nTotal prob of N-in: 0.14402\noutside 1 331\nTMhelix 332 354\ninside 355 372

18374 GCF_002064575.1_ASM206457v1 Salmonella enterica subsp. enterica serovar Litchfield Proteobacteria; Gammaproteobacteria; Enterobacterales; Enterobacteriaceae; Salmonella; Salmonella enterica; Salmonella enterica subsp. enterica MTGDNTLITSHGINRRDFMKLCAALAATMGLSSKAAAEMAESVSNPQRPPVIWIGAQECT WP_000145429.1 hydrogenase 2 small subunit [Salmonella enterica] Length: 372\nNumber of predicted TMHs: 1\nExp number of AAs in TMHs: 24.51309\nExp number, first 60 AAs: 0.2682\nTotal prob of N-in: 0.14402\noutside 1 331\nTMhelix 332 354\ninside 355 372

18375 GCF_002064525.1_ASM206452v1 Salmonella enterica subsp. enterica serovar Litchfield Proteobacteria; Gammaproteobacteria; Enterobacterales; Enterobacteriaceae; Salmonella; Salmonella enterica; Salmonella enterica subsp. enterica MTGDNTLITSHGINRRDFMKLCAALAATMGLSSKAAAEMAESVSNPQRPPVIWIGAQECT WP_000145429.1 hydrogenase 2 small subunit [Salmonella enterica] Length: 372\nNumber of predicted TMHs: 1\nExp number of AAs in TMHs: 24.51309\nExp number, first 60 AAs: 0.2682\nTotal prob of N-in: 0.14402\noutside 1 331\nTMhelix 332 354\ninside 355 372

18376 GCF_002064575.1_ASM206457v1 Salmonella enterica subsp. enterica serovar Litchfield Proteobacteria; Gammaproteobacteria; Enterobacterales; Enterobacteriaceae; Salmonella; Salmonella enterica; Salmonella enterica subsp. enterica MNNEETFYQAMRRKGVTRRSFLKFCSLAATSLGLGAGMTPKIAWALENKPRIPVVWIHGL WP_001058312.1 hydrogenase [Salmonella enterica] Length: 372\nNumber of predicted TMHs: 2\nExp number of AAs in TMHs: 36.51738\nExp number, first 60 AAs: 19.72849\nTotal prob of N-in: 0.96170\nPOSSIBLE N-term signal sequence\ninside 1 19\nTMhelix 20 39\noutside 40 325\nTMhelix 326 348\ninside 349 372

18377 GCF_002064525.1_ASM206452v1 Salmonella enterica subsp. enterica serovar Litchfield Proteobacteria; Gammaproteobacteria; Enterobacterales; Enterobacteriaceae; Salmonella; Salmonella enterica; Salmonella enterica subsp. enterica MNNEETFYQAMRRKGVTRRSFLKFCSLAATSLGLGAGMTPKIAWALENKPRIPVVWIHGL WP_001058312.1 hydrogenase [Salmonella enterica] Length: 372\nNumber of predicted TMHs: 2\nExp number of AAs in TMHs: 36.51738\nExp number, first 60 AAs: 19.72849\nTotal prob of N-in: 0.96170\nPOSSIBLE N-term signal sequence\ninside 1 19\nTMhelix 20 39\noutside 40 325\nTMhelix 326 348\ninside 349 372

18378 GCF_002107495.1_ASM210749v1 Salmonella enterica subsp. enterica serovar Litchfield Proteobacteria; Gammaproteobacteria; Enterobacterales; Enterobacteriaceae; Salmonella; Salmonella enterica; Salmonella enterica subsp. enterica MTGDNTLITSHGINRRDFMKLCAALAATMGLSSKAAAEMAESVSNPQRPPVIWIGAQECT WP_000145429.1 hydrogenase 2 small subunit [Salmonella enterica] Length: 372\nNumber of predicted TMHs: 1\nExp number of AAs in TMHs: 24.51309\nExp number, first 60 AAs: 0.2682\nTotal prob of N-in: 0.14402\noutside 1 331\nTMhelix 332 354\ninside 355 372

18379 GCF_001243305.1_Salmonella_enterica_CVM_N46836_v1.0 Salmonella enterica subsp. enterica serovar Litchfield Proteobacteria; Gammaproteobacteria; Enterobacterales; Enterobacteriaceae; Salmonella; Salmonella enterica; Salmonella enterica subsp. enterica MTGDNTLITSHGINRRDFMKLCAALAATMGLSSKAAAEMAESVSNPQRPPVIWIGAQECT WP_000145429.1 hydrogenase 2 small subunit [Salmonella enterica] Length: 372\nNumber of predicted TMHs: 1\nExp number of AAs in TMHs: 24.51309\nExp number, first 60 AAs: 0.2682\nTotal prob of N-in: 0.14402\noutside 1 331\nTMhelix 332 354\ninside 355 372

18380 GCF_001243305.1_Salmonella_enterica_CVM_N46836_v1.0 Salmonella enterica subsp. enterica serovar Litchfield Proteobacteria; Gammaproteobacteria; Enterobacterales; Enterobacteriaceae; Salmonella; Salmonella enterica; Salmonella enterica subsp. enterica MNNEETFYQAMRRKGVTRRSFLKFCSLAATSLGLGAGMTPKIAWALENKPRIPVVWIHGL WP_001058312.1 hydrogenase [Salmonella enterica] Length: 372\nNumber of predicted TMHs: 2\nExp number of AAs in TMHs: 36.51738\nExp number, first 60 AAs: 19.72849\nTotal prob of N-in: 0.96170\nPOSSIBLE N-term signal sequence\ninside 1 19\nTMhelix 20 39\noutside 40 325\nTMhelix 326 348\ninside 349 372

18381 GCF_001243175.1_Salmonella_enterica_CVM_N46829_v1.0 Salmonella enterica subsp. enterica serovar Litchfield Proteobacteria; Gammaproteobacteria; Enterobacterales; Enterobacteriaceae; Salmonella; Salmonella enterica; Salmonella enterica subsp. enterica MTGDNTLITSHGINRRDFMKLCAALAATMGLSSKAAAEMAESVSNPQRPPVIWIGAQECT WP_000145429.1 hydrogenase 2 small subunit [Salmonella enterica] Length: 372\nNumber of predicted TMHs: 1\nExp number of AAs in TMHs: 24.51309\nExp number, first 60 AAs: 0.2682\nTotal prob of N-in: 0.14402\noutside 1 331\nTMhelix 332 354\ninside 355 372

18382	GCF_002063035.1_ASM206303v1	Salmonella enterica subsp. enterica serovar Litchfield	Proteobacteria; Gammaproteobacteria; Enterobacterales; Enterobacteriaceae; Salmonella; Salmonella enterica; Salmonella enterica subsp. enterica	MNNEETFYQAMRRKGVTRRSFLKFCSLAATSLGLGAGMTPKIAWALENKPRIPVVWIHGL	WP_001058312.1	hydrogenase [Salmonella enterica]	Length: 372\nNumber of predicted TMHs:  2\nExp number of AAs in TMHs: 36.51738\nExp number, first 60 AAs:  19.72849\nTotal prob of N-in:   0.96170\nPOSSIBLE N-term signal sequence\ninside   1   19\nTMhelix   20   39\noutside   40   325\nTMhelix   326   348\ninside   349   372

18383	GCF_002063035.1_ASM206303v1	Salmonella enterica subsp. enterica serovar Litchfield	Proteobacteria; Gammaproteobacteria; Enterobacterales; Enterobacteriaceae; Salmonella; Salmonella enterica; Salmonella enterica subsp. enterica	MTGDNTLITSHGINRRDFMKLCAALAATMGLSSKAAAEMAESVSNPQRPPVIWIGAQECT	WP_000145429.1	hydrogenase 2 small subunit [Salmonella enterica]	Length: 372\nNumber of predicted TMHs:  1\nExp number of AAs in TMHs: 24.51309\nExp number, first 60 AAs:  0.2682\nTotal prob of N-in:   0.14402\noutside   1   331\nTMhelix   332   354\ninside   355   372

18384	GCF_001241815.1_Salmonella_enterica_CVM_N45400_v1.0	Salmonella enterica subsp. enterica serovar Litchfield	Proteobacteria; Gammaproteobacteria; Enterobacterales; Enterobacteriaceae; Salmonella; Salmonella enterica; Salmonella enterica subsp. enterica	MNNEETFYQAMRRKGVTRRSFLKFCSLAATSLGLGAGMTPKIAWALENKPRIPVVWIHGL	WP_001058312.1	hydrogenase [Salmonella enterica]	Length: 372\nNumber of predicted TMHs:  2\nExp number of AAs in TMHs: 36.51738\nExp number, first 60 AAs:  19.72849\nTotal prob of N-in:   0.96170\nPOSSIBLE N-term signal sequence\ninside   1   19\nTMhelix   20   39\noutside   40   325\nTMhelix   326   348\ninside   349   372

18385	GCF_001243325.1_Salmonella_enterica_CVM_N46837_v1.0	Salmonella enterica subsp. enterica serovar Litchfield	Proteobacteria; Gammaproteobacteria; Enterobacterales; Enterobacteriaceae; Salmonella; Salmonella enterica; Salmonella enterica subsp. enterica	MTGDNTLITSHGINRRDFMKLCAALAATMGLSSKAAAEMAESVSNPQRPPVIWIGAQECT	WP_000145429.1	hydrogenase 2 small subunit [Salmonella enterica]	Length: 372\nNumber of predicted TMHs:  1\nExp number of AAs in TMHs: 24.51309\nExp number, first 60 AAs:  0.2682\nTotal prob of N-in:   0.14402\noutside   1   331\nTMhelix   332   354\ninside   355   372

18386	GCF_001241815.1_Salmonella_enterica_CVM_N45400_v1.0	Salmonella enterica subsp. enterica serovar Litchfield	Proteobacteria; Gammaproteobacteria; Enterobacterales; Enterobacteriaceae; Salmonella; Salmonella enterica; Salmonella enterica subsp. enterica	MTGDNTLITSHGINRRDFMKLCAALAATMGLSSKAAAEMAESVSNPQRPPVIWIGAQECT	WP_000145429.1	hydrogenase 2 small subunit [Salmonella enterica]	Length: 372\nNumber of predicted TMHs:  1\nExp number of AAs in TMHs: 24.51309\nExp number, first 60 AAs:  0.2682\nTotal prob of N-in:   0.14402\noutside   1   331\nTMhelix   332   354\ninside   355   372

18387	GCF_001243325.1_Salmonella_enterica_CVM_N46837_v1.0	Salmonella enterica subsp. enterica serovar Litchfield	Proteobacteria; Gammaproteobacteria; Enterobacterales; Enterobacteriaceae; Salmonella; Salmonella enterica; Salmonella enterica subsp. enterica	MNNEETFYQAMRRKGVTRRSFLKFCSLAATSLGLGAGMTPKIAWALENKPRIPVVWIHGL	WP_001058312.1	hydrogenase [Salmonella enterica]	Length: 372\nNumber of predicted TMHs:  2\nExp number of AAs in TMHs: 36.51738\nExp number, first 60 AAs:  19.72849\nTotal prob of N-in:   0.96170\nPOSSIBLE N-term signal sequence\ninside   1   19\nTMhelix   20   39\noutside   40   325\nTMhelix   326   348\ninside   349   372

18388	GCF_001243175.1_Salmonella_enterica_CVM_N46829_v1.0	Salmonella enterica subsp. enterica serovar Litchfield	Proteobacteria; Gammaproteobacteria; Enterobacterales; Enterobacteriaceae; Salmonella; Salmonella enterica; Salmonella enterica subsp. enterica	MNNEETFYQAMRRKGVTRRSFLKFCSLAATSLGLGAGMTPKIAWALENKPRIPVVWIHGL	WP_001058312.1	hydrogenase [Salmonella enterica]	Length: 372\nNumber of predicted TMHs:  2\nExp number of AAs in TMHs: 36.51738\nExp number, first 60 AAs:  19.72849\nTotal prob of N-in:   0.96170\nPOSSIBLE N-term signal sequence\ninside   1   19\nTMhelix   20   39\noutside   40   325\nTMhelix   326   348\ninside   349   372

18389	GCF_002058745.1_ASM205874v1	Salmonella enterica subsp. enterica serovar Litchfield	Proteobacteria; Gammaproteobacteria; Enterobacterales; Enterobacteriaceae; Salmonella; Salmonella enterica; Salmonella enterica subsp. enterica	MTGDNTLITSHGINRRDFMKLCAALAATMGLSSKAAAEMAESVSNPQRPPVIWIGAQECT	WP_000145429.1	hydrogenase 2 small subunit [Salmonella enterica]	Length: 372\nNumber of predicted TMHs:  1\nExp number of AAs in TMHs: 24.51309\nExp number, first 60 AAs:  0.2682\nTotal prob of N-in:   0.14402\noutside   1   331\nTMhelix   332   354\ninside   355   372

18390	GCF_002030935.1_ASM203093v1	Salmonella enterica subsp. enterica serovar Litchfield	Proteobacteria; Gammaproteobacteria; Enterobacterales; Enterobacteriaceae; Salmonella; Salmonella enterica; Salmonella enterica subsp. enterica	MTGDNTLITSHGINRRDFMKLCAALAATMGLSSKAAAEMAESVSNPQRPPVIWIGAQECT	WP_000145429.1	hydrogenase 2 small subunit [Salmonella enterica]	Length: 372\nNumber of

predicted TMHs:  1\nExp number of AAs in TMHs: 24.51309\nExp number, first 60 AAs:  0.2682\nTotal prob of N-in:  0.14402\noutside   1   331\nTMhelix   332   354\ninside   355   372

18391 GCF_002030935.1_ASM203093v1 Salmonella enterica subsp. enterica serovar Litchfield Proteobacteria; Gammaproteobacteria; Enterobacterales; Enterobacteriaceae; Salmonella; Salmonella enterica; Salmonella enterica subsp. enterica MNNEETFYQAMRRKGVTRRSFLKFCSLAATSLGLGAGMTPKIAWALENKPRIPVVWIHGL WP_001058312.1 hydrogenase [Salmonella enterica] Length: 372\nNumber of predicted TMHs:  2\nExp number of AAs in TMHs: 36.51738\nExp number, first 60 AAs:  19.72849\nTotal prob of N-in:   0.96170\nPOSSIBLE N-term signal sequence\ninside   1   19\nTMhelix   20   39\noutside   40   325\nTMhelix   326   348\ninside   349   372

18392 GCF_001477885.1_Salmonella_enterica_CVM_N29367_v1.0 Salmonella enterica subsp. enterica serovar Litchfield Proteobacteria; Gammaproteobacteria; Enterobacterales; Enterobacteriaceae; Salmonella; Salmonella enterica; Salmonella enterica subsp. enterica MTGDNTLITSHGINRRDFMKLCAALAATMGLSSKAAAEMAESVSNPQRPPVIWIGAQECT WP_000145429.1 hydrogenase 2 small subunit [Salmonella enterica] Length: 372\nNumber of predicted TMHs:  1\nExp number of AAs in TMHs: 24.51309\nExp number, first 60 AAs:  0.2682\nTotal prob of N-in:  0.14402\noutside   1   331\nTMhelix   332   354\ninside   355   372

18393 GCF_001479305.1_Salmonella_enterica_CVM_N32042-SQ_v1.0 Salmonella enterica subsp. enterica serovar Litchfield Proteobacteria; Gammaproteobacteria; Enterobacterales; Enterobacteriaceae; Salmonella; Salmonella enterica; Salmonella enterica subsp. enterica MNNEETFYQAMRRKGVTRRSFLKFCSLAATSLGLGAGMTPKIAWALENKPRIPVVWIHGL WP_001058312.1 hydrogenase [Salmonella enterica] Length: 372\nNumber of predicted TMHs:  2\nExp number of AAs in TMHs: 36.51738\nExp number, first 60 AAs:  19.72849\nTotal prob of N-in:   0.96170\nPOSSIBLE N-term signal sequence\ninside   1   19\nTMhelix   20   39\noutside   40   325\nTMhelix   326   348\ninside   349   372

18394 GCF_002058745.1_ASM205874v1 Salmonella enterica subsp. enterica serovar Litchfield Proteobacteria; Gammaproteobacteria; Enterobacterales; Enterobacteriaceae; Salmonella; Salmonella enterica; Salmonella enterica subsp. enterica MNNEETFYQAMRRKGVTRRSFLKFCSLAATSLGLGAGMTPKIAWALENKPRIPVVWIHGL WP_001058312.1 hydrogenase [Salmonella enterica] Length: 372\nNumber of predicted TMHs:  2\nExp number of AAs in TMHs: 36.51738\nExp number, first 60 AAs:  19.72849\nTotal prob of N-in:   0.96170\nPOSSIBLE N-term signal sequence\ninside   1   19\nTMhelix   20   39\noutside   40   325\nTMhelix   326   348\ninside   349   372

18395 GCF_001479305.1_Salmonella_enterica_CVM_N32042-SQ_v1.0 Salmonella enterica subsp. enterica serovar Litchfield Proteobacteria; Gammaproteobacteria; Enterobacterales; Enterobacteriaceae; Salmonella; Salmonella enterica; Salmonella enterica subsp. enterica MTGDNTLITSHGINRRDFMKLCAALAATMGLSSKAAAEMAESVSNPQRPPVIWIGAQECT WP_000145429.1 hydrogenase 2 small subunit [Salmonella enterica] Length: 372\nNumber of predicted TMHs:  1\nExp number of AAs in TMHs: 24.51309\nExp number, first 60 AAs:  0.2682\nTotal prob of N-in:  0.14402\noutside   1   331\nTMhelix   332   354\ninside   355   372

18396 GCF_002059605.1_ASM205960v1 Salmonella enterica subsp. enterica serovar Litchfield Proteobacteria; Gammaproteobacteria; Enterobacterales; Enterobacteriaceae; Salmonella; Salmonella enterica; Salmonella enterica subsp. enterica MNNEETFYQAMRRKGVTRRSFLKFCSLAATSLGLGAGMTPKIAWALENKPRIPVVWIHGL WP_001058312.1 hydrogenase [Salmonella enterica] Length: 372\nNumber of predicted TMHs:  2\nExp number of AAs in TMHs: 36.51738\nExp number, first 60 AAs:  19.72849\nTotal prob of N-in:   0.96170\nPOSSIBLE N-term signal sequence\ninside   1   19\nTMhelix   20   39\noutside   40   325\nTMhelix   326   348\ninside   349   372

18397 GCF_001477885.1_Salmonella_enterica_CVM_N29367_v1.0 Salmonella enterica subsp. enterica serovar Litchfield Proteobacteria; Gammaproteobacteria; Enterobacterales; Enterobacteriaceae; Salmonella; Salmonella enterica; Salmonella enterica subsp. enterica MNNEETFYQAMRRKGVTRRSFLKFCSLAATSLGLGAGMTPKIAWALENKPRIPVVWIHGL WP_001058312.1 hydrogenase [Salmonella enterica] Length: 372\nNumber of predicted TMHs:  2\nExp number of AAs in TMHs: 36.51738\nExp number, first 60 AAs:  19.72849\nTotal prob of N-in:   0.96170\nPOSSIBLE N-term signal sequence\ninside   1   19\nTMhelix   20   39\noutside   40   325\nTMhelix   326   348\ninside   349   372

18398 GCF_002061095.1_ASM206109v1 Salmonella enterica subsp. enterica serovar Liverpool Proteobacteria; Gammaproteobacteria; Enterobacterales; Enterobacteriaceae; Salmonella; Salmonella enterica; Salmonella enterica subsp. enterica MNNEETFYQAMRRKGVTRRSFLKFCSLAATSLGLGAGMTPKIAWALENKPRIPVVWIHGL WP_001058312.1 hydrogenase [Salmonella enterica] Length: 372\nNumber of predicted TMHs:  2\nExp number of AAs in TMHs: 36.51738\nExp number, first 60 AAs:  19.72849\nTotal prob of N-in:   0.96170\nPOSSIBLE N-term signal sequence\ninside   1   19\nTMhelix   20   39\noutside   40   325\nTMhelix   326   348\ninside   349   372

18399 GCF_002061095.1_ASM206109v1 Salmonella enterica subsp. enterica serovar Liverpool Proteobacteria; Gammaproteobacteria; Enterobacterales; Enterobacteriaceae; Salmonella; Salmonella enterica;

Salmonella enterica subsp. enterica MTGDNTLITSHGINRRDFMKLCAALAATMGLSSKAAAEMAESVSNPQRPPVIWIGAQECT WP_000145429.1 hydrogenase 2 small subunit [Salmonella enterica] Length: 372\nNumber of predicted TMHs: 1\nExp number of AAs in TMHs: 24.51309\nExp number, first 60 AAs: 0.2682\nTotal prob of N-in: 0.14402\noutside 1 331\nTMhelix 332 354\ninside 355 372

18400 GCF_001242905.1_Salmonella_enterica_CVM_N46814_v1.0 Salmonella enterica subsp. enterica serovar Liverpool Proteobacteria; Gammaproteobacteria; Enterobacterales; Enterobacteriaceae; Salmonella; Salmonella enterica; Salmonella enterica subsp. enterica MTGDNTLITSHGINRRDFMKLCAALAATMGLSSKAAAEMAESVSNPQRPPVIWIGAQECT WP_000145429.1 hydrogenase 2 small subunit [Salmonella enterica] Length: 372\nNumber of predicted TMHs: 1\nExp number of AAs in TMHs: 24.51309\nExp number, first 60 AAs: 0.2682\nTotal prob of N-in: 0.14402\noutside 1 331\nTMhelix 332 354\ninside 355 372

18401 GCF_001245755.1_Salmonella_enterica_CVM_N50444_v1.0 Salmonella enterica subsp. enterica serovar Liverpool Proteobacteria; Gammaproteobacteria; Enterobacterales; Enterobacteriaceae; Salmonella; Salmonella enterica; Salmonella enterica subsp. enterica MTGDNTLITSHGINRRDFMKLCAALAATMGLSSKAAAEMAESVSNPQRPPVIWIGAQECT WP_000145429.1 hydrogenase 2 small subunit [Salmonella enterica] Length: 372\nNumber of predicted TMHs: 1\nExp number of AAs in TMHs: 24.51309\nExp number, first 60 AAs: 0.2682\nTotal prob of N-in: 0.14402\noutside 1 331\nTMhelix 332 354\ninside 355 372

18402 GCF_001242905.1_Salmonella_enterica_CVM_N46814_v1.0 Salmonella enterica subsp. enterica serovar Liverpool Proteobacteria; Gammaproteobacteria; Enterobacterales; Enterobacteriaceae; Salmonella; Salmonella enterica; Salmonella enterica subsp. enterica MNNEETFYQAMRRKGVTRRSFLKFCSLAATSLGLGAGMTPKIAWALENKPRIPVVWIHGL WP_001058312.1 hydrogenase [Salmonella enterica] Length: 372\nNumber of predicted TMHs: 2\nExp number of AAs in TMHs: 36.51738\nExp number, first 60 AAs: 19.72849\nTotal prob of N-in: 0.96170\nPOSSIBLE N-term signal sequence\ninside 1 19\nTMhelix 20 39\noutside 40 325\nTMhelix 326 348\ninside 349 372

18403 GCF_001245755.1_Salmonella_enterica_CVM_N50444_v1.0 Salmonella enterica subsp. enterica serovar Liverpool Proteobacteria; Gammaproteobacteria; Enterobacterales; Enterobacteriaceae; Salmonella; Salmonella enterica; Salmonella enterica subsp. enterica MNNEETFYQAMRRKGVTRRSFLKFCSLAATSLGLGAGMTPKIAWALENKPRIPVVWIHGL WP_001058312.1 hydrogenase [Salmonella enterica] Length: 372\nNumber of predicted TMHs: 2\nExp number of AAs in TMHs: 36.51738\nExp number, first 60 AAs: 19.72849\nTotal prob of N-in: 0.96170\nPOSSIBLE N-term signal sequence\ninside 1 19\nTMhelix 20 39\noutside 40 325\nTMhelix 326 348\ninside 349 372

18404 GCF_000956125.1_ASM95612v1 Salmonella enterica subsp. enterica serovar Livingstone Proteobacteria; Gammaproteobacteria; Enterobacterales; Enterobacteriaceae; Salmonella; Salmonella enterica; Salmonella enterica subsp. enterica MTGDNTLITSHGINRRDFMKLCAALAATMGLSSKAAAEMAESVSNPQRPPVIWIGAQECT WP_000145429.1 hydrogenase 2 small subunit [Salmonella enterica] Length: 372\nNumber of predicted TMHs: 1\nExp number of AAs in TMHs: 24.51309\nExp number, first 60 AAs: 0.2682\nTotal prob of N-in: 0.14402\noutside 1 331\nTMhelix 332 354\ninside 355 372

18405 GCF_000956125.1_ASM95612v1 Salmonella enterica subsp. enterica serovar Livingstone Proteobacteria; Gammaproteobacteria; Enterobacterales; Enterobacteriaceae; Salmonella; Salmonella enterica; Salmonella enterica subsp. enterica MNNEETFYQAMRRKGVTRRSFLKFCSLAATSLGLGAGMTPKIAWALENKPRIPVVWIHGL WP_001058311.1 hydrogenase [Salmonella enterica] Length: 372\nNumber of predicted TMHs: 2\nExp number of AAs in TMHs: 37.4699899999999\nExp number, first 60 AAs: 19.72699\nTotal prob of N-in: 0.96171\nPOSSIBLE N-term signal sequence\ninside 1 19\nTMhelix 20 39\noutside 40 325\nTMhelix 326 348\ninside 349 372

18406 GCF_001241775.1_Salmonella_enterica_CVM_N45399_v1.0 Salmonella enterica subsp. enterica serovar Livingstone Proteobacteria; Gammaproteobacteria; Enterobacterales; Enterobacteriaceae; Salmonella; Salmonella enterica; Salmonella enterica subsp. enterica MTGDNTLITSHGINRRDFMKLCAALAATMGLSSKAAAEMAESVSNPQRPPVIWIGAQECT WP_000145429.1 hydrogenase 2 small subunit [Salmonella enterica] Length: 372\nNumber of predicted TMHs: 1\nExp number of AAs in TMHs: 24.51309\nExp number, first 60 AAs: 0.2682\nTotal prob of N-in: 0.14402\noutside 1 331\nTMhelix 332 354\ninside 355 372

18407 GCF_002062375.1_ASM206237v1 Salmonella enterica subsp. enterica serovar Livingstone Proteobacteria; Gammaproteobacteria; Enterobacterales; Enterobacteriaceae; Salmonella; Salmonella enterica; Salmonella enterica subsp. enterica MNNEETFYQAMRRKGVTRRSFLKFCSLATTSLGLGAGMTPKIAWALENKPRIPVVWIHGL WP_023259200.1 hydrogenase 1, small subunit [Salmonella enterica] Length: 372\nNumber of predicted TMHs: 2\nExp number of AAs in TMHs: 35.98973\nExp number, first 60 AAs: 19.19893\nTotal prob of N-in:

0.93662\nPOSSIBLE N-term signal sequence\ninside 1 19\nTMhelix 20 39\noutside 40 325\nTMhelix 326 348\ninside 349 372

18408 GCF_002062375.1_ASM206237v1 Salmonella enterica subsp. enterica serovar Livingstone Proteobacteria; Gammaproteobacteria; Enterobacterales; Enterobacteriaceae; Salmonella; Salmonella enterica; Salmonella enterica subsp. enterica MTGDNTLITSHGINRRDFMKLCAALAATMGLSSKAAAEMAESVSNPQRPPVIWIGAQECT WP_000145429.1 hydrogenase 2 small subunit [Salmonella enterica] Length: 372\nNumber of predicted TMHs: 1\nExp number of AAs in TMHs: 24.51309\nExp number, first 60 AAs: 0.2682\nTotal prob of N-in: 0.14402\noutside 1 331\nTMhelix 332 354\ninside 355 372

18409 GCF_001241775.1_Salmonella_enterica_CVM_N45399_v1.0 Salmonella enterica subsp. enterica serovar Livingstone Proteobacteria; Gammaproteobacteria; Enterobacterales; Enterobacteriaceae; Salmonella; Salmonella enterica; Salmonella enterica subsp. enterica MNNEETFYQAMRRKGVTRRSFLKFCSLAATSLGLGAGMTPKIAWALENKPRIPVVWIHGL WP_001058311.1 hydrogenase [Salmonella enterica] Length: 372\nNumber of predicted TMHs: 2\nExp number of AAs in TMHs: 37.4699899999999\nExp number, first 60 AAs: 19.72699\nTotal prob of N-in: 0.96171\nPOSSIBLE N-term signal sequence\ninside 1 19\nTMhelix 20 39\noutside 40 325\nTMhelix 326 348\ninside 349 372

18410 GCF_002045405.1_ASM204540v1 Salmonella enterica subsp. enterica serovar Lomalinda Proteobacteria; Gammaproteobacteria; Enterobacterales; Enterobacteriaceae; Salmonella; Salmonella enterica; Salmonella enterica subsp. enterica MTGDNTLITSHGINRRDFMKLCAALAATMGLSSKAAAEMAESVSNPQRPPVIWIGAQECT WP_000145429.1 hydrogenase 2 small subunit [Salmonella enterica] Length: 372\nNumber of predicted TMHs: 1\nExp number of AAs in TMHs: 24.51309\nExp number, first 60 AAs: 0.2682\nTotal prob of N-in: 0.14402\noutside 1 331\nTMhelix 332 354\ninside 355 372

18411 GCF_002046245.1_ASM204624v1 Salmonella enterica subsp. enterica serovar Lomalinda Proteobacteria; Gammaproteobacteria; Enterobacterales; Enterobacteriaceae; Salmonella; Salmonella enterica; Salmonella enterica subsp. enterica MTGDNTLITSHGINRRDFMKLCAALAATMGLSSKAAAEMAESVSNPQRPPVIWIGAQECT WP_000145429.1 hydrogenase 2 small subunit [Salmonella enterica] Length: 372\nNumber of predicted TMHs: 1\nExp number of AAs in TMHs: 24.51309\nExp number, first 60 AAs: 0.2682\nTotal prob of N-in: 0.14402\noutside 1 331\nTMhelix 332 354\ninside 355 372

18412 GCF_002045405.1_ASM204540v1 Salmonella enterica subsp. enterica serovar Lomalinda Proteobacteria; Gammaproteobacteria; Enterobacterales; Enterobacteriaceae; Salmonella; Salmonella enterica; Salmonella enterica subsp. enterica MNNEETFYQAMRRKGVTRRSFLKFCSLAATSLGLGAGMTPKIAWALENKPRIPVVWIHGL WP_001058312.1 hydrogenase [Salmonella enterica] Length: 372\nNumber of predicted TMHs: 2\nExp number of AAs in TMHs: 36.51738\nExp number, first 60 AAs: 19.72849\nTotal prob of N-in: 0.96170\nPOSSIBLE N-term signal sequence\ninside 1 19\nTMhelix 20 39\noutside 40 325\nTMhelix 326 348\ninside 349 372

18413 GCF_002045385.1_ASM204538v1 Salmonella enterica subsp. enterica serovar Lomalinda Proteobacteria; Gammaproteobacteria; Enterobacterales; Enterobacteriaceae; Salmonella; Salmonella enterica; Salmonella enterica subsp. enterica MTGDNTLITSHGINRRDFMKLCAALAATMGLSSKAAAEMAESVSNPQRPPVIWIGAQECT WP_000145429.1 hydrogenase 2 small subunit [Salmonella enterica] Length: 372\nNumber of predicted TMHs: 1\nExp number of AAs in TMHs: 24.51309\nExp number, first 60 AAs: 0.2682\nTotal prob of N-in: 0.14402\noutside 1 331\nTMhelix 332 354\ninside 355 372

18414 GCF_002046525.1_ASM204652v1 Salmonella enterica subsp. enterica serovar Lomalinda Proteobacteria; Gammaproteobacteria; Enterobacterales; Enterobacteriaceae; Salmonella; Salmonella enterica; Salmonella enterica subsp. enterica MTGDNTLITSHGINRRDFMKLCAALAATMGLSSKAAAEMAESVSNPQRPPVIWIGAQECT WP_000145429.1 hydrogenase 2 small subunit [Salmonella enterica] Length: 372\nNumber of predicted TMHs: 1\nExp number of AAs in TMHs: 24.51309\nExp number, first 60 AAs: 0.2682\nTotal prob of N-in: 0.14402\noutside 1 331\nTMhelix 332 354\ninside 355 372

18415 GCF_002046245.1_ASM204624v1 Salmonella enterica subsp. enterica serovar Lomalinda Proteobacteria; Gammaproteobacteria; Enterobacterales; Enterobacteriaceae; Salmonella; Salmonella enterica; Salmonella enterica subsp. enterica MNNEETFYQAMRRKGVTRRSFLKFCSLAATSLGLGAGMTPKIAWALENKPRIPVVWIHGL WP_001058312.1 hydrogenase [Salmonella enterica] Length: 372\nNumber of predicted TMHs: 2\nExp number of AAs in TMHs: 36.51738\nExp number, first 60 AAs: 19.72849\nTotal prob of N-in: 0.96170\nPOSSIBLE N-term signal sequence\ninside 1 19\nTMhelix 20 39\noutside 40 325\nTMhelix 326 348\ninside 349 372

18416 GCF_002046275.1_ASM204627v1 Salmonella enterica subsp. enterica serovar Lomalinda Proteobacteria; Gammaproteobacteria; Enterobacterales; Enterobacteriaceae; Salmonella; Salmonella enterica; Salmonella enterica subsp. enterica MTGDNTLITSHGINRRDFMKLCAALAATMGLSSKAAAEMAESVSNPQRPPVIWIGAQECT WP_000145429.1 hydrogenase 2 small subunit [Salmonella enterica] Length: 372\nNumber of predicted TMHs: 1\nExp number of AAs in TMHs: 24.51309\nExp number, first 60 AAs: 0.2682\nTotal prob of N-in: 0.14402\noutside 1 331\nTMhelix 332 354\ninside 355 372

18417 GCF_002045385.1_ASM204538v1 Salmonella enterica subsp. enterica serovar Lomalinda Proteobacteria; Gammaproteobacteria; Enterobacterales; Enterobacteriaceae; Salmonella; Salmonella enterica; Salmonella enterica subsp. enterica MNNEETFYQAMRRKGVTRRSFLKFCSLAATSLGLGAGMTPKIAWALENKPRIPVVWIHGL WP_001058312.1 hydrogenase [Salmonella enterica] Length: 372\nNumber of predicted TMHs: 2\nExp number of AAs in TMHs: 36.51738\nExp number, first 60 AAs: 19.72849\nTotal prob of N-in: 0.96170\nPOSSIBLE N-term signal sequence\ninside 1 19\nTMhelix 20 39\noutside 40 325\nTMhelix 326 348\ninside 349 372

18418 GCF_002046525.1_ASM204652v1 Salmonella enterica subsp. enterica serovar Lomalinda Proteobacteria; Gammaproteobacteria; Enterobacterales; Enterobacteriaceae; Salmonella; Salmonella enterica; Salmonella enterica subsp. enterica MNNEETFYQAMRRKGVTRRSFLKFCSLAATSLGLGAGMTPKIAWALENKPRIPVVWIHGL WP_001058312.1 hydrogenase [Salmonella enterica] Length: 372\nNumber of predicted TMHs: 2\nExp number of AAs in TMHs: 36.51738\nExp number, first 60 AAs: 19.72849\nTotal prob of N-in: 0.96170\nPOSSIBLE N-term signal sequence\ninside 1 19\nTMhelix 20 39\noutside 40 325\nTMhelix 326 348\ninside 349 372

18419 GCF_002048075.1_ASM204807v1 Salmonella enterica subsp. enterica serovar Lomalinda Proteobacteria; Gammaproteobacteria; Enterobacterales; Enterobacteriaceae; Salmonella; Salmonella enterica; Salmonella enterica subsp. enterica MNNEETFYQAMRRKGVTRRSFLKFCSLAATSLGLGAGMTPKIAWALENKPRIPVVWIHGL WP_001058312.1 hydrogenase [Salmonella enterica] Length: 372\nNumber of predicted TMHs: 2\nExp number of AAs in TMHs: 36.51738\nExp number, first 60 AAs: 19.72849\nTotal prob of N-in: 0.96170\nPOSSIBLE N-term signal sequence\ninside 1 19\nTMhelix 20 39\noutside 40 325\nTMhelix 326 348\ninside 349 372

18420 GCF_002048075.1_ASM204807v1 Salmonella enterica subsp. enterica serovar Lomalinda Proteobacteria; Gammaproteobacteria; Enterobacterales; Enterobacteriaceae; Salmonella; Salmonella enterica; Salmonella enterica subsp. enterica MTGDNTLITSHGINRRDFMKLCAALAATMGLSSKAAAEMAESVSNPQRPPVIWIGAQECT WP_000145429.1 hydrogenase 2 small subunit [Salmonella enterica] Length: 372\nNumber of predicted TMHs: 1\nExp number of AAs in TMHs: 24.51309\nExp number, first 60 AAs: 0.2682\nTotal prob of N-in: 0.14402\noutside 1 331\nTMhelix 332 354\ninside 355 372

18421 GCF_002047355.1_ASM204735v1 Salmonella enterica subsp. enterica serovar London Proteobacteria; Gammaproteobacteria; Enterobacterales; Enterobacteriaceae; Salmonella; Salmonella enterica; Salmonella enterica subsp. enterica MTGDNTLITSHGINRRDFMKLCAALAATMGLSSKAAAEMAESVSNPQRPPVIWIGAQECT WP_000145429.1 hydrogenase 2 small subunit [Salmonella enterica] Length: 372\nNumber of predicted TMHs: 1\nExp number of AAs in TMHs: 24.51309\nExp number, first 60 AAs: 0.2682\nTotal prob of N-in: 0.14402\noutside 1 331\nTMhelix 332 354\ninside 355 372

18422 GCF_002047085.1_ASM204708v1 Salmonella enterica subsp. enterica serovar London Proteobacteria; Gammaproteobacteria; Enterobacterales; Enterobacteriaceae; Salmonella; Salmonella enterica; Salmonella enterica subsp. enterica MNNEETFYQAMRRKGVTRRSFLKFCSLAATSLGLGAGMTPKIAWALENKPRIPVVWIHGL WP_001058311.1 hydrogenase [Salmonella enterica] Length: 372\nNumber of predicted TMHs: 2\nExp number of AAs in TMHs: 37.4699899999999\nExp number, first 60 AAs: 19.72699\nTotal prob of N-in: 0.96171\nPOSSIBLE N-term signal sequence\ninside 1 19\nTMhelix 20 39\noutside 40 325\nTMhelix 326 348\ninside 349 372

18423 GCF_002047355.1_ASM204735v1 Salmonella enterica subsp. enterica serovar London Proteobacteria; Gammaproteobacteria; Enterobacterales; Enterobacteriaceae; Salmonella; Salmonella enterica; Salmonella enterica subsp. enterica MNNEETFYQAMRRKGVTRRSFLKFCSLAATSLGLGAGMTPKIAWALENKPRIPVVWIHGL WP_001058311.1 hydrogenase [Salmonella enterica] Length: 372\nNumber of predicted TMHs: 2\nExp number of AAs in TMHs: 37.4699899999999\nExp number, first 60 AAs: 19.72699\nTotal prob of N-in: 0.96171\nPOSSIBLE N-term signal sequence\ninside 1 19\nTMhelix 20 39\noutside 40 325\nTMhelix 326 348\ninside 349 372

18424 GCF_001479935.1_Salmonella_enterica_CVM_N38882-SQ_v1.0 Salmonella enterica subsp. enterica serovar London Proteobacteria; Gammaproteobacteria; Enterobacterales; Enterobacteriaceae; Salmonella; Salmonella enterica; Salmonella enterica subsp. enterica MNNEETFYQAMRRKGVTRRSFLKFCSLAATSLGLGAGMTPKIAWALENKPRIPVVWIHGL

WP_001058311.1 hydrogenase [Salmonella enterica] Length: 372\nNumber of predicted TMHs: 2\nExp number of AAs in TMHs: 37.4699899999999\nExp number, first 60 AAs: 19.72699\nTotal prob of N-in: 0.96171\nPOSSIBLE N-term signal sequence\ninside 1 19\nTMhelix 20 39\noutside 40 325\nTMhelix 326 348\ninside 349 372

18425 GCF_002064515.1_ASM206451v1 Salmonella enterica subsp. enterica serovar London Proteobacteria; Gammaproteobacteria; Enterobacterales; Enterobacteriaceae; Salmonella; Salmonella enterica; Salmonella enterica subsp. enterica MNNEETFYQAMRRKGVTRRSFLKFCSLAATSLGLGAGMTPKIAWALENKPRIPVVWIHGL WP_001058311.1 hydrogenase [Salmonella enterica] Length: 372\nNumber of predicted TMHs: 2\nExp number of AAs in TMHs: 37.4699899999999\nExp number, first 60 AAs: 19.72699\nTotal prob of N-in: 0.96171\nPOSSIBLE N-term signal sequence\ninside 1 19\nTMhelix 20 39\noutside 40 325\nTMhelix 326 348\ninside 349 372

18426 GCF_002064515.1_ASM206451v1 Salmonella enterica subsp. enterica serovar London Proteobacteria; Gammaproteobacteria; Enterobacterales; Enterobacteriaceae; Salmonella; Salmonella enterica; Salmonella enterica subsp. enterica MTGDNTLITSHGINRRDFMKLCAALAATMGLSSKAAAEMAESVSNPQRPPVIWIGAQECT WP_000145429.1 hydrogenase 2 small subunit [Salmonella enterica] Length: 372\nNumber of predicted TMHs: 1\nExp number of AAs in TMHs: 24.51309\nExp number, first 60 AAs: 0.2682\nTotal prob of N-in: 0.14402\noutside 1 331\nTMhelix 332 354\ninside 355 372

18427 GCF_002048305.1_ASM204830v1 Salmonella enterica subsp. enterica serovar London Proteobacteria; Gammaproteobacteria; Enterobacterales; Enterobacteriaceae; Salmonella; Salmonella enterica; Salmonella enterica subsp. enterica MTGDNTLITSHGINRRDFMKLCAALAATMGLSSKAAAEMAESVSNPQRPPVIWIGAQECT WP_000145429.1 hydrogenase 2 small subunit [Salmonella enterica] Length: 372\nNumber of predicted TMHs: 1\nExp number of AAs in TMHs: 24.51309\nExp number, first 60 AAs: 0.2682\nTotal prob of N-in: 0.14402\noutside 1 331\nTMhelix 332 354\ninside 355 372

18428 GCF_002048305.1_ASM204830v1 Salmonella enterica subsp. enterica serovar London Proteobacteria; Gammaproteobacteria; Enterobacterales; Enterobacteriaceae; Salmonella; Salmonella enterica; Salmonella enterica subsp. enterica MNNEETFYQAMRRKGVTRRSFLKFCSLAATSLGLGAGMTPKIAWALENKPRIPVVWIHGL WP_001058311.1 hydrogenase [Salmonella enterica] Length: 372\nNumber of predicted TMHs: 2\nExp number of AAs in TMHs: 37.4699899999999\nExp number, first 60 AAs: 19.72699\nTotal prob of N-in: 0.96171\nPOSSIBLE N-term signal sequence\ninside 1 19\nTMhelix 20 39\noutside 40 325\nTMhelix 326 348\ninside 349 372

18429 GCF_002063165.1_ASM206316v1 Salmonella enterica subsp. enterica serovar London Proteobacteria; Gammaproteobacteria; Enterobacterales; Enterobacteriaceae; Salmonella; Salmonella enterica; Salmonella enterica subsp. enterica MTGDNTLITSHGINRRDFMKLCAALAATMGLSSKAAAEMAESVSNPQRPPVIWIGAQECT WP_000145429.1 hydrogenase 2 small subunit [Salmonella enterica] Length: 372\nNumber of predicted TMHs: 1\nExp number of AAs in TMHs: 24.51309\nExp number, first 60 AAs: 0.2682\nTotal prob of N-in: 0.14402\noutside 1 331\nTMhelix 332 354\ninside 355 372

18430 GCF_002063165.1_ASM206316v1 Salmonella enterica subsp. enterica serovar London Proteobacteria; Gammaproteobacteria; Enterobacterales; Enterobacteriaceae; Salmonella; Salmonella enterica; Salmonella enterica subsp. enterica MNNEETFYQAMRRKGVTRRSFLKFCSLAATSLGLGAGMTPKIAWALENKPRIPVVWIHGL WP_001058311.1 hydrogenase [Salmonella enterica] Length: 372\nNumber of predicted TMHs: 2\nExp number of AAs in TMHs: 37.4699899999999\nExp number, first 60 AAs: 19.72699\nTotal prob of N-in: 0.96171\nPOSSIBLE N-term signal sequence\ninside 1 19\nTMhelix 20 39\noutside 40 325\nTMhelix 326 348\ninside 349 372

18431 GCF_001479935.1_Salmonella_enterica_CVM_N38882-SQ_v1.0 Salmonella enterica subsp. enterica serovar London Proteobacteria; Gammaproteobacteria; Enterobacterales; Enterobacteriaceae; Salmonella; Salmonella enterica; Salmonella enterica subsp. enterica MTGDNTLITSHGINRRDFMKLCAALAATMGLSSKAAAEMAESVSNPQRPPVIWIGAQECT WP_000145429.1 hydrogenase 2 small subunit [Salmonella enterica] Length: 372\nNumber of predicted TMHs: 1\nExp number of AAs in TMHs: 24.51309\nExp number, first 60 AAs: 0.2682\nTotal prob of N-in: 0.14402\noutside 1 331\nTMhelix 332 354\ninside 355 372

18432 GCF_002047085.1_ASM204708v1 Salmonella enterica subsp. enterica serovar London Proteobacteria; Gammaproteobacteria; Enterobacterales; Enterobacteriaceae; Salmonella; Salmonella enterica; Salmonella enterica subsp. enterica MTGDNTLITSHGINRRDFMKLCAALAATMGLSSKAAAEMAESVSNPQRPPVIWIGAQECT WP_000145429.1 hydrogenase 2 small subunit [Salmonella enterica] Length: 372\nNumber of

predicted TMHs:  1\nExp number of AAs in TMHs: 24.51309\nExp number, first 60 AAs:  0.2682\nTotal prob of N-in: 0.14402\noutside   1   331\nTMhelix   332   354\ninside   355   372

18433 GCF_002049245.1_ASM204924v1 Salmonella enterica subsp. enterica serovar Manhattan Proteobacteria; Gammaproteobacteria; Enterobacterales; Enterobacteriaceae; Salmonella; Salmonella enterica; Salmonella enterica subsp. enterica MTGDNTLITSHGINRRDFMKLCAALAATMGLSSKAAAEMAESVSNPQRPPVIWIGAQECT WP_000145429.1 hydrogenase 2 small subunit [Salmonella enterica] Length: 372\nNumber of predicted TMHs:  1\nExp number of AAs in TMHs: 24.51309\nExp number, first 60 AAs:  0.2682\nTotal prob of N-in: 0.14402\noutside   1   331\nTMhelix   332   354\ninside   355   372

18434 GCF_002049245.1_ASM204924v1 Salmonella enterica subsp. enterica serovar Manhattan Proteobacteria; Gammaproteobacteria; Enterobacterales; Enterobacteriaceae; Salmonella; Salmonella enterica; Salmonella enterica subsp. enterica MNNEETFYQAMRRKGVTRRSFLKFCSLAATSLGLGAGMTPKIAWALENKPRIPVVWIHGL WP_001058311.1 hydrogenase [Salmonella enterica] Length: 372\nNumber of predicted TMHs:  2\nExp number of AAs in TMHs: 37.4699899999999\nExp number, first 60 AAs:  19.72699\nTotal prob of N-in: 0.96171\nPOSSIBLE N-term signal sequence\ninside   1   19\nTMhelix   20   39\noutside   40   325\nTMhelix   326   348\ninside   349   372

18435 GCF_002049165.1_ASM204916v1 Salmonella enterica subsp. enterica serovar Manhattan Proteobacteria; Gammaproteobacteria; Enterobacterales; Enterobacteriaceae; Salmonella; Salmonella enterica; Salmonella enterica subsp. enterica MTGDNTLITSHGINRRDFMKLCAALAATMGLSSKAAAEMAESVSNPQRPPVIWIGAQECT WP_000145429.1 hydrogenase 2 small subunit [Salmonella enterica] Length: 372\nNumber of predicted TMHs:  1\nExp number of AAs in TMHs: 24.51309\nExp number, first 60 AAs:  0.2682\nTotal prob of N-in: 0.14402\noutside   1   331\nTMhelix   332   354\ninside   355   372

18436 GCF_002049165.1_ASM204916v1 Salmonella enterica subsp. enterica serovar Manhattan Proteobacteria; Gammaproteobacteria; Enterobacterales; Enterobacteriaceae; Salmonella; Salmonella enterica; Salmonella enterica subsp. enterica MNNEETFYQAMRRKGVTRRSFLKFCSLAATSLGLGAGMTPKIAWALENKPRIPVVWIHGL WP_001058311.1 hydrogenase [Salmonella enterica] Length: 372\nNumber of predicted TMHs:  2\nExp number of AAs in TMHs: 37.4699899999999\nExp number, first 60 AAs:  19.72699\nTotal prob of N-in: 0.96171\nPOSSIBLE N-term signal sequence\ninside   1   19\nTMhelix   20   39\noutside   40   325\nTMhelix   326   348\ninside   349   372

18437 GCF_002064535.1_ASM206453v1 Salmonella enterica subsp. enterica serovar Manhattan Proteobacteria; Gammaproteobacteria; Enterobacterales; Enterobacteriaceae; Salmonella; Salmonella enterica; Salmonella enterica subsp. enterica MTGDNTLITSHGINRRDFMKLCAALAATMGLSSKAAAEMAESVSNPQRPPVIWIGAQECT WP_000145429.1 hydrogenase 2 small subunit [Salmonella enterica] Length: 372\nNumber of predicted TMHs:  1\nExp number of AAs in TMHs: 24.51309\nExp number, first 60 AAs:  0.2682\nTotal prob of N-in: 0.14402\noutside   1   331\nTMhelix   332   354\ninside   355   372

18438 GCF_002064535.1_ASM206453v1 Salmonella enterica subsp. enterica serovar Manhattan Proteobacteria; Gammaproteobacteria; Enterobacterales; Enterobacteriaceae; Salmonella; Salmonella enterica; Salmonella enterica subsp. enterica MNNEETFYQAMRRKGVTRRSFLKFCSLAATSLGLGAGMTPKIAWALENKPRIPVVWIHGL WP_001058311.1 hydrogenase [Salmonella enterica] Length: 372\nNumber of predicted TMHs:  2\nExp number of AAs in TMHs: 37.4699899999999\nExp number, first 60 AAs:  19.72699\nTotal prob of N-in: 0.96171\nPOSSIBLE N-term signal sequence\ninside   1   19\nTMhelix   20   39\noutside   40   325\nTMhelix   326   348\ninside   349   372

18439 GCF_002062715.1_ASM206271v1 Salmonella enterica subsp. enterica serovar Manhatten Proteobacteria; Gammaproteobacteria; Enterobacterales; Enterobacteriaceae; Salmonella; Salmonella enterica; Salmonella enterica subsp. enterica MTGDNTLITSHGINRRDFMKLCAALAATMGLSSKAAAEMAESVSNPQRPPVIWIGAQECT WP_000145429.1 hydrogenase 2 small subunit [Salmonella enterica] Length: 372\nNumber of predicted TMHs:  1\nExp number of AAs in TMHs: 24.51309\nExp number, first 60 AAs:  0.2682\nTotal prob of N-in: 0.14402\noutside   1   331\nTMhelix   332   354\ninside   355   372

18440 GCF_002062715.1_ASM206271v1 Salmonella enterica subsp. enterica serovar Manhatten Proteobacteria; Gammaproteobacteria; Enterobacterales; Enterobacteriaceae; Salmonella; Salmonella enterica; Salmonella enterica subsp. enterica MNNEETFYQAMRRKGVTRRSFLKFCSLAATSLGLGAGMTPKIAWALENKPRIPVVWIHGL WP_001058311.1 hydrogenase [Salmonella enterica] Length: 372\nNumber of predicted TMHs:  2\nExp number of AAs in TMHs: 37.4699899999999\nExp number, first 60 AAs:  19.72699\nTotal prob of N-in:

0.96171\nPOSSIBLE N-term signal sequence\ninside 1 19\nTMhelix 20 39\noutside 40 325\nTMhelix 326 348\ninside 349 372

18441 GCF_002062345.1_ASM206234v1 Salmonella enterica subsp. enterica serovar Mbadaka Proteobacteria; Gammaproteobacteria; Enterobacterales; Enterobacteriaceae; Salmonella; Salmonella enterica; Salmonella enterica subsp. enterica MNNEETFYQAMRRKGVTRRSFLKFCSLAATSLGLGAGMTPKIAWALENKPRIPVVWIHGL WP_001058311.1 hydrogenase [Salmonella enterica] Length: 372\nNumber of predicted TMHs: 2\nExp number of AAs in TMHs: 37.4699899999999\nExp number, first 60 AAs: 19.72699\nTotal prob of N-in: 0.96171\nPOSSIBLE N-term signal sequence\ninside 1 19\nTMhelix 20 39\noutside 40 325\nTMhelix 326 348\ninside 349 372

18442 GCF_002062345.1_ASM206234v1 Salmonella enterica subsp. enterica serovar Mbadaka Proteobacteria; Gammaproteobacteria; Enterobacterales; Enterobacteriaceae; Salmonella; Salmonella enterica; Salmonella enterica subsp. enterica MTGDNTLITSHGINRRDFMKLCAALAATMGLSSKAAAEMAESVSNPQRPPVIWIGAQECT WP_000145429.1 hydrogenase 2 small subunit [Salmonella enterica] Length: 372\nNumber of predicted TMHs: 1\nExp number of AAs in TMHs: 24.51309\nExp number, first 60 AAs: 0.2682\nTotal prob of N-in: 0.14402\noutside 1 331\nTMhelix 332 354\ninside 355 372

18443 GCF_002065465.1_ASM206546v1 Salmonella enterica subsp. enterica serovar Mbadaka Proteobacteria; Gammaproteobacteria; Enterobacterales; Enterobacteriaceae; Salmonella; Salmonella enterica; Salmonella enterica subsp. enterica MTGDNTLITSHGINRRDFMKLCAALAATMGLSSKAAAEMAESVSNPQRPPVIWIGAQECT WP_000145429.1 hydrogenase 2 small subunit [Salmonella enterica] Length: 372\nNumber of predicted TMHs: 1\nExp number of AAs in TMHs: 24.51309\nExp number, first 60 AAs: 0.2682\nTotal prob of N-in: 0.14402\noutside 1 331\nTMhelix 332 354\ninside 355 372

18444 GCF_002065465.1_ASM206546v1 Salmonella enterica subsp. enterica serovar Mbadaka Proteobacteria; Gammaproteobacteria; Enterobacterales; Enterobacteriaceae; Salmonella; Salmonella enterica; Salmonella enterica subsp. enterica MNNEETFYQAMRRKGVTRRSFLKFCSLAATSLGLGAGMTPKIAWALENKPRIPVVWIHGL WP_001058311.1 hydrogenase [Salmonella enterica] Length: 372\nNumber of predicted TMHs: 2\nExp number of AAs in TMHs: 37.4699899999999\nExp number, first 60 AAs: 19.72699\nTotal prob of N-in: 0.96171\nPOSSIBLE N-term signal sequence\ninside 1 19\nTMhelix 20 39\noutside 40 325\nTMhelix 326 348\ninside 349 372

18445 GCF_002036225.1_ASM203622v1 Salmonella enterica subsp. enterica serovar Mbandaka Proteobacteria; Gammaproteobacteria; Enterobacterales; Enterobacteriaceae; Salmonella; Salmonella enterica; Salmonella enterica subsp. enterica MTGDNTLITSHGINRRDFMKLCAALAATMGLSSKAAAEMAESVSNPQRPPVIWIGAQECT WP_000145429.1 hydrogenase 2 small subunit [Salmonella enterica] Length: 372\nNumber of predicted TMHs: 1\nExp number of AAs in TMHs: 24.51309\nExp number, first 60 AAs: 0.2682\nTotal prob of N-in: 0.14402\noutside 1 331\nTMhelix 332 354\ninside 355 372

18446 GCF_002064495.1_ASM206449v1 Salmonella enterica subsp. enterica serovar Mbandaka Proteobacteria; Gammaproteobacteria; Enterobacterales; Enterobacteriaceae; Salmonella; Salmonella enterica; Salmonella enterica subsp. enterica MNNEETFYQAMRRKGVTRRSFLKFCSLAATSLGLGAGMTPKIAWALENKPRIPVVWIHGL WP_001058311.1 hydrogenase [Salmonella enterica] Length: 372\nNumber of predicted TMHs: 2\nExp number of AAs in TMHs: 37.4699899999999\nExp number, first 60 AAs: 19.72699\nTotal prob of N-in: 0.96171\nPOSSIBLE N-term signal sequence\ninside 1 19\nTMhelix 20 39\noutside 40 325\nTMhelix 326 348\ninside 349 372

18447 GCF_002064495.1_ASM206449v1 Salmonella enterica subsp. enterica serovar Mbandaka Proteobacteria; Gammaproteobacteria; Enterobacterales; Enterobacteriaceae; Salmonella; Salmonella enterica; Salmonella enterica subsp. enterica MTGDNTLITSHGINRRDFMKLCAALAATMGLSSKAAAEMAESVSNPQRPPVIWIGAQECT WP_000145429.1 hydrogenase 2 small subunit [Salmonella enterica] Length: 372\nNumber of predicted TMHs: 1\nExp number of AAs in TMHs: 24.51309\nExp number, first 60 AAs: 0.2682\nTotal prob of N-in: 0.14402\noutside 1 331\nTMhelix 332 354\ninside 355 372

18448 GCF_002064445.1_ASM206444v1 Salmonella enterica subsp. enterica serovar Mbandaka Proteobacteria; Gammaproteobacteria; Enterobacterales; Enterobacteriaceae; Salmonella; Salmonella enterica; Salmonella enterica subsp. enterica MTGDNTLITSHGINRRDFMKLCAALAATMGLSSKAAAEMAESVSNPQRPPVIWIGAQECT WP_000145429.1 hydrogenase 2 small subunit [Salmonella enterica] Length: 372\nNumber of predicted TMHs: 1\nExp number of AAs in TMHs: 24.51309\nExp number, first 60 AAs: 0.2682\nTotal prob of N-in: 0.14402\noutside 1 331\nTMhelix 332 354\ninside 355 372

18449 GCF_002036225.1_ASM203622v1 Salmonella enterica subsp. enterica serovar Mbandaka Proteobacteria; Gammaproteobacteria; Enterobacterales; Enterobacteriaceae; Salmonella; Salmonella enterica; Salmonella enterica subsp. enterica MNNEETFYQAMRRKGVTRRSFLKFCSLAATSLGLGAGMTPKIAWALENKPRIPVVWIHGL WP_001058311.1 hydrogenase [Salmonella enterica] Length: 372\nNumber of predicted TMHs: 2\nExp number of AAs in TMHs: 37.4699899999999\nExp number, first 60 AAs: 19.72699\nTotal prob of N-in: 0.96171\nPOSSIBLE N-term signal sequence\ninside 1 19\nTMhelix 20 39\noutside 40 325\nTMhelix 326 348\ninside 349 372

18450 GCF_002030575.1_ASM203057v1 Salmonella enterica subsp. enterica serovar Mbandaka Proteobacteria; Gammaproteobacteria; Enterobacterales; Enterobacteriaceae; Salmonella; Salmonella enterica; Salmonella enterica subsp. enterica MTGDNTLITSHGINRRDFMKLCAALAATMGLSSKAAAEMAESVSNPQRPPVIWIGAQECT WP_000145429.1 hydrogenase 2 small subunit [Salmonella enterica] Length: 372\nNumber of predicted TMHs: 1\nExp number of AAs in TMHs: 24.51309\nExp number, first 60 AAs: 0.2682\nTotal prob of N-in: 0.14402\noutside 1 331\nTMhelix 332 354\ninside 355 372

18451 GCF_001833595.1_ASM183359v1 Salmonella enterica subsp. enterica serovar Mbandaka Proteobacteria; Gammaproteobacteria; Enterobacterales; Enterobacteriaceae; Salmonella; Salmonella enterica; Salmonella enterica subsp. enterica MTGDNTLITSHGINRRDFMKLCAALAATMGLSSKAAAEMAESVSNPQRPPVIWIGAQECT WP_023248548.1 hydrogenase 2 small subunit [Salmonella enterica] Length: 372\nNumber of predicted TMHs: 1\nExp number of AAs in TMHs: 24.51281\nExp number, first 60 AAs: 0.26822\nTotal prob of N-in: 0.14402\noutside 1 331\nTMhelix 332 354\ninside 355 372

18452 GCF_002064445.1_ASM206444v1 Salmonella enterica subsp. enterica serovar Mbandaka Proteobacteria; Gammaproteobacteria; Enterobacterales; Enterobacteriaceae; Salmonella; Salmonella enterica; Salmonella enterica subsp. enterica MNNEETFYQAMRRKGVTRRSFLKFCSLAATSLGLGAGMTPKIAWALENKPRIPVVWIHGL WP_001058311.1 hydrogenase [Salmonella enterica] Length: 372\nNumber of predicted TMHs: 2\nExp number of AAs in TMHs: 37.4699899999999\nExp number, first 60 AAs: 19.72699\nTotal prob of N-in: 0.96171\nPOSSIBLE N-term signal sequence\ninside 1 19\nTMhelix 20 39\noutside 40 325\nTMhelix 326 348\ninside 349 372

18453 GCF_001833595.1_ASM183359v1 Salmonella enterica subsp. enterica serovar Mbandaka Proteobacteria; Gammaproteobacteria; Enterobacterales; Enterobacteriaceae; Salmonella; Salmonella enterica; Salmonella enterica subsp. enterica MNNEETFYQAMRRKGVTRRSFLKFCSLAATSLGLGAGMTPKIAWALENKPRIPVVWIHGL WP_070801304.1 hydrogenase [Salmonella enterica] Length: 372\nNumber of predicted TMHs: 2\nExp number of AAs in TMHs: 36.3895799999999\nExp number, first 60 AAs: 19.727\nTotal prob of N-in: 0.96170\nPOSSIBLE N-term signal sequence\ninside 1 19\nTMhelix 20 39\noutside 40 325\nTMhelix 326 348\ninside 349 372

18454 GCF_001835085.1_ASM183508v1 Salmonella enterica subsp. enterica serovar Mbandaka Proteobacteria; Gammaproteobacteria; Enterobacterales; Enterobacteriaceae; Salmonella; Salmonella enterica; Salmonella enterica subsp. enterica MTGDNTLITSHGINRRDFMKLCAALAATMGLSSKAAAEMAESVSNPQRPPVIWIGAQECT WP_023248548.1 hydrogenase 2 small subunit [Salmonella enterica] Length: 372\nNumber of predicted TMHs: 1\nExp number of AAs in TMHs: 24.51281\nExp number, first 60 AAs: 0.26822\nTotal prob of N-in: 0.14402\noutside 1 331\nTMhelix 332 354\ninside 355 372

18455 GCF_001478205.1_Salmonella_enterica_CVM_N32055-SQ_v1.0 Salmonella enterica subsp. enterica serovar Mbandaka Proteobacteria; Gammaproteobacteria; Enterobacterales; Enterobacteriaceae; Salmonella; Salmonella enterica; Salmonella enterica subsp. enterica MTGDNTLITSHGINRRDFMKLCAALAATMGLSSKAAAEMAESVSNPQRPPVIWIGAQECT WP_000145429.1 hydrogenase 2 small subunit [Salmonella enterica] Length: 372\nNumber of predicted TMHs: 1\nExp number of AAs in TMHs: 24.51309\nExp number, first 60 AAs: 0.2682\nTotal prob of N-in: 0.14402\noutside 1 331\nTMhelix 332 354\ninside 355 372

18456 GCF_001835085.1_ASM183508v1 Salmonella enterica subsp. enterica serovar Mbandaka Proteobacteria; Gammaproteobacteria; Enterobacterales; Enterobacteriaceae; Salmonella; Salmonella enterica; Salmonella enterica subsp. enterica MNNEETFYQAMRRKGVTRRSFLKFCSLAATSLGLGAGMTPKIAWALENKPRIPVVWIHGL WP_070801304.1 hydrogenase [Salmonella enterica] Length: 372\nNumber of predicted TMHs: 2\nExp number of AAs in TMHs: 36.3895799999999\nExp number, first 60 AAs: 19.727\nTotal prob of N-in: 0.96170\nPOSSIBLE N-term signal sequence\ninside 1 19\nTMhelix 20 39\noutside 40 325\nTMhelix 326 348\ninside 349 372

18457 GCF_001247475.1_Salmonella_enterica_CVM_N51315_v1.0 Salmonella enterica subsp. enterica serovar Mbandaka Proteobacteria; Gammaproteobacteria; Enterobacterales; Enterobacteriaceae; Salmonella; Salmonella

enterica; Salmonella enterica subsp. enterica
MNNEETFYQAMRRKGVTRRSFLKFCSLAATSLGLGAGMTPKIAWALENKPRIPVVWIHGL WP_001058311.1 hydrogenase [Salmonella enterica] Length: 372\nNumber of predicted TMHs: 2\nExp number of AAs in TMHs: 37.4699899999999\nExp number, first 60 AAs: 19.72699\nTotal prob of N-in: 0.96171\nPOSSIBLE N-term signal sequence\ninside 1 19\nTMhelix 20 39\noutside 40 325\nTMhelix 326 348\ninside 349 372

18458 GCF_001247475.1_Salmonella_enterica_CVM_N51315_v1.0 Salmonella enterica subsp. enterica serovar Mbandaka Proteobacteria; Gammaproteobacteria; Enterobacterales; Enterobacteriaceae; Salmonella; Salmonella enterica; Salmonella enterica subsp. enterica
MTGDNTLITSHGINRRDFMKLCAALAATMGLSSKAAAEMAESVSNPQRPPVIWIGAQECT WP_000145429.1 hydrogenase 2 small subunit [Salmonella enterica] Length: 372\nNumber of predicted TMHs: 1\nExp number of AAs in TMHs: 24.51309\nExp number, first 60 AAs: 0.2682\nTotal prob of N-in: 0.14402\noutside 1 331\nTMhelix 332 354\ninside 355 372

18459 GCF_001242775.1_Salmonella_enterica_CVM_N45957_v1.0 Salmonella enterica subsp. enterica serovar Mbandaka Proteobacteria; Gammaproteobacteria; Enterobacterales; Enterobacteriaceae; Salmonella; Salmonella enterica; Salmonella enterica subsp. enterica
MNNEETFYQAMRRKGVTRRSFLKFCSLAATSLGLGAGMTPKIAWALENKPRIPVVWIHGL WP_001058311.1 hydrogenase [Salmonella enterica] Length: 372\nNumber of predicted TMHs: 2\nExp number of AAs in TMHs: 37.4699899999999\nExp number, first 60 AAs: 19.72699\nTotal prob of N-in: 0.96171\nPOSSIBLE N-term signal sequence\ninside 1 19\nTMhelix 20 39\noutside 40 325\nTMhelix 326 348\ninside 349 372

18460 GCF_001247255.1_Salmonella_enterica_CVM_N51302_v1.0 Salmonella enterica subsp. enterica serovar Mbandaka Proteobacteria; Gammaproteobacteria; Enterobacterales; Enterobacteriaceae; Salmonella; Salmonella enterica; Salmonella enterica subsp. enterica
MTGDNTLITSHGINRRDFMKLCAALAATMGLSSKAAAEMAESVSNPQRPPVIWIGAQECT WP_000145429.1 hydrogenase 2 small subunit [Salmonella enterica] Length: 372\nNumber of predicted TMHs: 1\nExp number of AAs in TMHs: 24.51309\nExp number, first 60 AAs: 0.2682\nTotal prob of N-in: 0.14402\noutside 1 331\nTMhelix 332 354\ninside 355 372

18461 GCF_001247255.1_Salmonella_enterica_CVM_N51302_v1.0 Salmonella enterica subsp. enterica serovar Mbandaka Proteobacteria; Gammaproteobacteria; Enterobacterales; Enterobacteriaceae; Salmonella; Salmonella enterica; Salmonella enterica subsp. enterica
MNNEETFYQAMRRKGVTRRSFLKFCSLAATSLGLGAGMTPKIAWALENKPRIPVVWIHGL WP_001058311.1 hydrogenase [Salmonella enterica] Length: 372\nNumber of predicted TMHs: 2\nExp number of AAs in TMHs: 37.4699899999999\nExp number, first 60 AAs: 19.72699\nTotal prob of N-in: 0.96171\nPOSSIBLE N-term signal sequence\ninside 1 19\nTMhelix 20 39\noutside 40 325\nTMhelix 326 348\ninside 349 372

18462 GCF_001272395.1_Salmonella_enterica_CVM_N51293_v1.0 Salmonella enterica subsp. enterica serovar Mbandaka Proteobacteria; Gammaproteobacteria; Enterobacterales; Enterobacteriaceae; Salmonella; Salmonella enterica; Salmonella enterica subsp. enterica
MNNEETFYQAMRRKGVTRRSFLKFCSLAATSLGLGAGMTPKIAWALENKPRIPVVWIHGL WP_001058311.1 hydrogenase [Salmonella enterica] Length: 372\nNumber of predicted TMHs: 2\nExp number of AAs in TMHs: 37.4699899999999\nExp number, first 60 AAs: 19.72699\nTotal prob of N-in: 0.96171\nPOSSIBLE N-term signal sequence\ninside 1 19\nTMhelix 20 39\noutside 40 325\nTMhelix 326 348\ninside 349 372

18463 GCF_001272395.1_Salmonella_enterica_CVM_N51293_v1.0 Salmonella enterica subsp. enterica serovar Mbandaka Proteobacteria; Gammaproteobacteria; Enterobacterales; Enterobacteriaceae; Salmonella; Salmonella enterica; Salmonella enterica subsp. enterica
MTGDNTLITSHGINRRDFMKLCAALAATMGLSSKAAAEMAESVSNPQRPPVIWIGAQECT WP_000145429.1 hydrogenase 2 small subunit [Salmonella enterica] Length: 372\nNumber of predicted TMHs: 1\nExp number of AAs in TMHs: 24.51309\nExp number, first 60 AAs: 0.2682\nTotal prob of N-in: 0.14402\noutside 1 331\nTMhelix 332 354\ninside 355 372

18464 GCF_001242775.1_Salmonella_enterica_CVM_N45957_v1.0 Salmonella enterica subsp. enterica serovar Mbandaka Proteobacteria; Gammaproteobacteria; Enterobacterales; Enterobacteriaceae; Salmonella; Salmonella enterica; Salmonella enterica subsp. enterica
MTGDNTLITSHGINRRDFMKLCAALAATMGLSSKAAAEMAESVSNPQRPPVIWIGAQECT WP_000145429.1 hydrogenase 2 small subunit [Salmonella enterica] Length: 372\nNumber of predicted TMHs: 1\nExp number of AAs in TMHs: 24.51309\nExp number, first 60 AAs: 0.2682\nTotal prob of N-in: 0.14402\noutside 1 331\nTMhelix 332 354\ninside 355 372

18465 GCF_002064435.1_ASM206443v1 Salmonella enterica subsp. enterica serovar Mbandaka Proteobacteria; Gammaproteobacteria; Enterobacterales; Enterobacteriaceae; Salmonella; Salmonella enterica; Salmonella enterica subsp. enterica MTGDNTLITSHGINRRDFMKLCAALAATMGLSSKAAAEMAESVSNPQRPPVIWIGAQECT WP_000145429.1 hydrogenase 2 small subunit [Salmonella enterica] Length: 372\nNumber of predicted TMHs: 1\nExp number of AAs in TMHs: 24.51309\nExp number, first 60 AAs: 0.2682\nTotal prob of N-in: 0.14402\noutside 1 331\nTMhelix 332 354\ninside 355 372

18466 GCF_002066805.1_ASM206680v1 Salmonella enterica subsp. enterica serovar Mbandaka Proteobacteria; Gammaproteobacteria; Enterobacterales; Enterobacteriaceae; Salmonella; Salmonella enterica; Salmonella enterica subsp. enterica MTGDNTLITSHGINRRDFMKLCAALAATMGLSSKAAAEMAESVSNPQRPPVIWIGAQECT WP_000145429.1 hydrogenase 2 small subunit [Salmonella enterica] Length: 372\nNumber of predicted TMHs: 1\nExp number of AAs in TMHs: 24.51309\nExp number, first 60 AAs: 0.2682\nTotal prob of N-in: 0.14402\noutside 1 331\nTMhelix 332 354\ninside 355 372

18467 GCF_002066805.1_ASM206680v1 Salmonella enterica subsp. enterica serovar Mbandaka Proteobacteria; Gammaproteobacteria; Enterobacterales; Enterobacteriaceae; Salmonella; Salmonella enterica; Salmonella enterica subsp. enterica MNNEETFYQAMRRKGVTRRSFLKFCSLAATSLGLGAGMTPKIAWALENKPRIPVVWIHGL WP_001058311.1 hydrogenase [Salmonella enterica] Length: 372\nNumber of predicted TMHs: 2\nExp number of AAs in TMHs: 37.4699899999999\nExp number, first 60 AAs: 19.72699\nTotal prob of N-in: 0.96171\nPOSSIBLE N-term signal sequence\ninside 1 19\nTMhelix 20 39\noutside 40 325\nTMhelix 326 348\ninside 349 372

18468 GCF_002064455.1_ASM206445v1 Salmonella enterica subsp. enterica serovar Mbandaka Proteobacteria; Gammaproteobacteria; Enterobacterales; Enterobacteriaceae; Salmonella; Salmonella enterica; Salmonella enterica subsp. enterica MNNEETFYQAMRRKGVTRRSFLKFCSLAATSLGLGAGMTPKIAWALENKPRIPVVWIHGL WP_001058311.1 hydrogenase [Salmonella enterica] Length: 372\nNumber of predicted TMHs: 2\nExp number of AAs in TMHs: 37.4699899999999\nExp number, first 60 AAs: 19.72699\nTotal prob of N-in: 0.96171\nPOSSIBLE N-term signal sequence\ninside 1 19\nTMhelix 20 39\noutside 40 325\nTMhelix 326 348\ninside 349 372

18469 GCF_002064455.1_ASM206445v1 Salmonella enterica subsp. enterica serovar Mbandaka Proteobacteria; Gammaproteobacteria; Enterobacterales; Enterobacteriaceae; Salmonella; Salmonella enterica; Salmonella enterica subsp. enterica MTGDNTLITSHGINRRDFMKLCAALAATMGLSSKAAAEMAESVSNPQRPPVIWIGAQECT WP_000145429.1 hydrogenase 2 small subunit [Salmonella enterica] Length: 372\nNumber of predicted TMHs: 1\nExp number of AAs in TMHs: 24.51309\nExp number, first 60 AAs: 0.2682\nTotal prob of N-in: 0.14402\noutside 1 331\nTMhelix 332 354\ninside 355 372

18470 GCF_002064435.1_ASM206443v1 Salmonella enterica subsp. enterica serovar Mbandaka Proteobacteria; Gammaproteobacteria; Enterobacterales; Enterobacteriaceae; Salmonella; Salmonella enterica; Salmonella enterica subsp. enterica MNNEETFYQAMRRKGVTRRSFLKFCSLAATSLGLGAGMTPKIAWALENKPRIPVVWIHGL WP_001058311.1 hydrogenase [Salmonella enterica] Length: 372\nNumber of predicted TMHs: 2\nExp number of AAs in TMHs: 37.4699899999999\nExp number, first 60 AAs: 19.72699\nTotal prob of N-in: 0.96171\nPOSSIBLE N-term signal sequence\ninside 1 19\nTMhelix 20 39\noutside 40 325\nTMhelix 326 348\ninside 349 372

18471 GCF_002058425.1_ASM205842v1 Salmonella enterica subsp. enterica serovar Mbandaka Proteobacteria; Gammaproteobacteria; Enterobacterales; Enterobacteriaceae; Salmonella; Salmonella enterica; Salmonella enterica subsp. enterica MTGDNTLITSHGINRRDFMKLCAALAATMGLSSKAAAEMAESVSNPQRPPVIWIGAQECT WP_000145429.1 hydrogenase 2 small subunit [Salmonella enterica] Length: 372\nNumber of predicted TMHs: 1\nExp number of AAs in TMHs: 24.51309\nExp number, first 60 AAs: 0.2682\nTotal prob of N-in: 0.14402\noutside 1 331\nTMhelix 332 354\ninside 355 372

18472 GCF_001479985.1_Salmonella_enterica_CVM_N38886-SQ_v1.0 Salmonella enterica subsp. enterica serovar Mbandaka Proteobacteria; Gammaproteobacteria; Enterobacterales; Enterobacteriaceae; Salmonella; Salmonella enterica; Salmonella enterica subsp. enterica MNNEETFYQAMRRKGVTRRSFLKFCSLAATSLGLGAGMTPKIAWALENKPRIPVVWIHGL WP_001058311.1 hydrogenase [Salmonella enterica] Length: 372\nNumber of predicted TMHs: 2\nExp number of AAs in TMHs: 37.4699899999999\nExp number, first 60 AAs: 19.72699\nTotal prob of N-in: 0.96171\nPOSSIBLE N-term signal sequence\ninside 1 19\nTMhelix 20 39\noutside 40 325\nTMhelix 326 348\ninside 349 372

18473 GCF_001480505.1_Salmonella_enterica_CVM_N42451-SQ_v1.0 Salmonella enterica subsp. enterica serovar Mbandaka Proteobacteria; Gammaproteobacteria; Enterobacterales; Enterobacteriaceae; Salmonella; Salmonella enterica; Salmonella enterica subsp. enterica MTGDNTLITSHGINRRDFMKLCAALAATMGLSSKAAAEMAESVSNPQRPPVIWIGAQECT WP_000145429.1 hydrogenase 2 small subunit [Salmonella enterica] Length: 372\nNumber of predicted TMHs: 1\nExp number of AAs in TMHs: 24.51309\nExp number, first 60 AAs: 0.2682\nTotal prob of N-in: 0.14402\noutside 1 331\nTMhelix 332 354\ninside 355 372

18474 GCF_001479985.1_Salmonella_enterica_CVM_N38886-SQ_v1.0 Salmonella enterica subsp. enterica serovar Mbandaka Proteobacteria; Gammaproteobacteria; Enterobacterales; Enterobacteriaceae; Salmonella; Salmonella enterica; Salmonella enterica subsp. enterica MTGDNTLITSHGINRRDFMKLCAALAATMGLSSKAAAEMAESVSNPQRPPVIWIGAQECT WP_000145429.1 hydrogenase 2 small subunit [Salmonella enterica] Length: 372\nNumber of predicted TMHs: 1\nExp number of AAs in TMHs: 24.51309\nExp number, first 60 AAs: 0.2682\nTotal prob of N-in: 0.14402\noutside 1 331\nTMhelix 332 354\ninside 355 372

18475 GCF_002030575.1_ASM203057v1 Salmonella enterica subsp. enterica serovar Mbandaka Proteobacteria; Gammaproteobacteria; Enterobacterales; Enterobacteriaceae; Salmonella; Salmonella enterica; Salmonella enterica subsp. enterica MNNEETFYQAMRRKGVTRRSFLKFCSLAATSLGLGAGMTPKIAWALENKPRIPVVWIHGL WP_001058311.1 hydrogenase [Salmonella enterica] Length: 372\nNumber of predicted TMHs: 2\nExp number of AAs in TMHs: 37.4699899999999\nExp number, first 60 AAs: 19.72699\nTotal prob of N-in: 0.96171\nPOSSIBLE N-term signal sequence\ninside 1 19\nTMhelix 20 39\noutside 40 325\nTMhelix 326 348\ninside 349 372

18476 GCF_001480505.1_Salmonella_enterica_CVM_N42451-SQ_v1.0 Salmonella enterica subsp. enterica serovar Mbandaka Proteobacteria; Gammaproteobacteria; Enterobacterales; Enterobacteriaceae; Salmonella; Salmonella enterica; Salmonella enterica subsp. enterica MNNEETFYQAMRRKGVTRRSFLKFCSLAATSLGLGAGMTPKIAWALENKPRIPVVWIHGL WP_001058311.1 hydrogenase [Salmonella enterica] Length: 372\nNumber of predicted TMHs: 2\nExp number of AAs in TMHs: 37.4699899999999\nExp number, first 60 AAs: 19.72699\nTotal prob of N-in: 0.96171\nPOSSIBLE N-term signal sequence\ninside 1 19\nTMhelix 20 39\noutside 40 325\nTMhelix 326 348\ninside 349 372

18477 GCF_001481155.1_Salmonella_enterica_CVM_N42487-R_v1.0 Salmonella enterica subsp. enterica serovar Mbandaka Proteobacteria; Gammaproteobacteria; Enterobacterales; Enterobacteriaceae; Salmonella; Salmonella enterica; Salmonella enterica subsp. enterica MTGDNTLITSHGINRRDFMKLCAALAATMGLSSKAAAEMAESVSNPQRPPVIWIGAQECT WP_000145429.1 hydrogenase 2 small subunit [Salmonella enterica] Length: 372\nNumber of predicted TMHs: 1\nExp number of AAs in TMHs: 24.51309\nExp number, first 60 AAs: 0.2682\nTotal prob of N-in: 0.14402\noutside 1 331\nTMhelix 332 354\ninside 355 372

18478 GCF_001481155.1_Salmonella_enterica_CVM_N42487-R_v1.0 Salmonella enterica subsp. enterica serovar Mbandaka Proteobacteria; Gammaproteobacteria; Enterobacterales; Enterobacteriaceae; Salmonella; Salmonella enterica; Salmonella enterica subsp. enterica MNNEETFYQAMRRKGVTRRSFLKFCSLAATSLGLGAGMTPKIAWALENKPRIPVVWIHGL WP_001058311.1 hydrogenase [Salmonella enterica] Length: 372\nNumber of predicted TMHs: 2\nExp number of AAs in TMHs: 37.4699899999999\nExp number, first 60 AAs: 19.72699\nTotal prob of N-in: 0.96171\nPOSSIBLE N-term signal sequence\ninside 1 19\nTMhelix 20 39\noutside 40 325\nTMhelix 326 348\ninside 349 372

18479 GCF_001478205.1_Salmonella_enterica_CVM_N32055-SQ_v1.0 Salmonella enterica subsp. enterica serovar Mbandaka Proteobacteria; Gammaproteobacteria; Enterobacterales; Enterobacteriaceae; Salmonella; Salmonella enterica; Salmonella enterica subsp. enterica MNNEETFYQAMRRKGVTRRSFLKFCSLAATSLGLGAGMTPKIAWALENKPRIPVVWIHGL WP_001058311.1 hydrogenase [Salmonella enterica] Length: 372\nNumber of predicted TMHs: 2\nExp number of AAs in TMHs: 37.4699899999999\nExp number, first 60 AAs: 19.72699\nTotal prob of N-in: 0.96171\nPOSSIBLE N-term signal sequence\ninside 1 19\nTMhelix 20 39\noutside 40 325\nTMhelix 326 348\ninside 349 372

18480 GCF_002046085.1_ASM204608v1 Salmonella enterica subsp. enterica serovar Meleagridis Proteobacteria; Gammaproteobacteria; Enterobacterales; Enterobacteriaceae; Salmonella; Salmonella enterica; Salmonella enterica subsp. enterica MTGDNTLITSHGINRRDFMKLCAALAATMGLSSKAAAEMAESVSNPQRPPVIWIGAQECT WP_000145429.1 hydrogenase 2 small subunit [Salmonella enterica] Length: 372\nNumber of

predicted TMHs:  1\nExp number of AAs in TMHs: 24.51309\nExp number, first 60 AAs:  0.2682\nTotal prob of N-in: 0.14402\noutside   1   331\nTMhelix   332   354\ninside   355   372

18481 GCF_002044745.1_ASM204474v1 Salmonella enterica subsp. enterica serovar Meleagridis Proteobacteria; Gammaproteobacteria; Enterobacterales; Enterobacteriaceae; Salmonella; Salmonella enterica; Salmonella enterica subsp. enterica MNNEETFYQAMRRKGVTRRSFLKFCSLAATSLGLGAGMTPKIAWALENKPRIPVVWIHGL WP_001058311.1 hydrogenase [Salmonella enterica] Length: 372\nNumber of predicted TMHs:  2\nExp number of AAs in TMHs: 37.4699899999999\nExp number, first 60 AAs:  19.72699\nTotal prob of N-in: 0.96171\nPOSSIBLE N-term signal sequence\ninside   1   19\nTMhelix   20   39\noutside   40   325\nTMhelix   326   348\ninside   349   372

18482 GCF_002046085.1_ASM204608v1 Salmonella enterica subsp. enterica serovar Meleagridis Proteobacteria; Gammaproteobacteria; Enterobacterales; Enterobacteriaceae; Salmonella; Salmonella enterica; Salmonella enterica subsp. enterica MNNEETFYQAMRRKGVTRRSFLKFCSLAATSLGLGAGMTPKIAWALENKPRIPVVWIHGL WP_001058311.1 hydrogenase [Salmonella enterica] Length: 372\nNumber of predicted TMHs:  2\nExp number of AAs in TMHs: 37.4699899999999\nExp number, first 60 AAs:  19.72699\nTotal prob of N-in: 0.96171\nPOSSIBLE N-term signal sequence\ninside   1   19\nTMhelix   20   39\noutside   40   325\nTMhelix   326   348\ninside   349   372

18483 GCF_002044885.1_ASM204488v1 Salmonella enterica subsp. enterica serovar Meleagridis Proteobacteria; Gammaproteobacteria; Enterobacterales; Enterobacteriaceae; Salmonella; Salmonella enterica; Salmonella enterica subsp. enterica MTGDNTLITSHGINRRDFMKLCAALAATMGLSSKAAAEMAESVSNPQRPPVIWIGAQECT WP_000145429.1 hydrogenase 2 small subunit [Salmonella enterica] Length: 372\nNumber of predicted TMHs:  1\nExp number of AAs in TMHs: 24.51309\nExp number, first 60 AAs:  0.2682\nTotal prob of N-in: 0.14402\noutside   1   331\nTMhelix   332   354\ninside   355   372

18484 GCF_002045015.1_ASM204501v1 Salmonella enterica subsp. enterica serovar Meleagridis Proteobacteria; Gammaproteobacteria; Enterobacterales; Enterobacteriaceae; Salmonella; Salmonella enterica; Salmonella enterica subsp. enterica MNNEETFYQAMRRKGVTRRSFLKFCSLAATSLGLGAGMTPKIAWALENKPRIPVVWIHGL WP_079941068.1 [Ni/Fe] hydrogenase small subunit [Salmonella enterica] Length: 372\nNumber of predicted TMHs:  2\nExp number of AAs in TMHs: 37.37321\nExp number, first 60 AAs:  19.7296\nTotal prob of N-in: 0.96168\nPOSSIBLE N-term signal sequence\ninside   1   19\nTMhelix   20   39\noutside   40   325\nTMhelix   326   348\ninside   349   372

18485 GCF_002045015.1_ASM204501v1 Salmonella enterica subsp. enterica serovar Meleagridis Proteobacteria; Gammaproteobacteria; Enterobacterales; Enterobacteriaceae; Salmonella; Salmonella enterica; Salmonella enterica subsp. enterica MTGDNTLITSHGINRRDFMKLCAALAATMGLSSKAAAEMAESVSNPQRPPVIWIGAQECT WP_000145429.1 hydrogenase 2 small subunit [Salmonella enterica] Length: 372\nNumber of predicted TMHs:  1\nExp number of AAs in TMHs: 24.51309\nExp number, first 60 AAs:  0.2682\nTotal prob of N-in: 0.14402\noutside   1   331\nTMhelix   332   354\ninside   355   372

18486 GCF_002044885.1_ASM204488v1 Salmonella enterica subsp. enterica serovar Meleagridis Proteobacteria; Gammaproteobacteria; Enterobacterales; Enterobacteriaceae; Salmonella; Salmonella enterica; Salmonella enterica subsp. enterica MNNEETFYQAMRRKGVTRRSFLKFCSLAATSLGLGAGMTPKIAWALENKPRIPVVWIHGL WP_001058311.1 hydrogenase [Salmonella enterica] Length: 372\nNumber of predicted TMHs:  2\nExp number of AAs in TMHs: 37.4699899999999\nExp number, first 60 AAs:  19.72699\nTotal prob of N-in: 0.96171\nPOSSIBLE N-term signal sequence\ninside   1   19\nTMhelix   20   39\noutside   40   325\nTMhelix   326   348\ninside   349   372

18487 GCF_002044805.1_ASM204480v1 Salmonella enterica subsp. enterica serovar Meleagridis Proteobacteria; Gammaproteobacteria; Enterobacterales; Enterobacteriaceae; Salmonella; Salmonella enterica; Salmonella enterica subsp. enterica MNNEETFYQAMRRKGVTRRSFLKFCSLAATSLGLGAGMTPKIAWALENKPRIPVVWIHGL WP_001058311.1 hydrogenase [Salmonella enterica] Length: 372\nNumber of predicted TMHs:  2\nExp number of AAs in TMHs: 37.4699899999999\nExp number, first 60 AAs:  19.72699\nTotal prob of N-in: 0.96171\nPOSSIBLE N-term signal sequence\ninside   1   19\nTMhelix   20   39\noutside   40   325\nTMhelix   326   348\ninside   349   372

18488 GCF_002044805.1_ASM204480v1 Salmonella enterica subsp. enterica serovar Meleagridis Proteobacteria; Gammaproteobacteria; Enterobacterales; Enterobacteriaceae; Salmonella; Salmonella enterica; Salmonella enterica subsp. enterica MTGDNTLITSHGINRRDFMKLCAALAATMGLSSKAAAEMAESVSNPQRPPVIWIGAQECT WP_000145429.1 hydrogenase 2 small subunit [Salmonella enterica] Length: 372\nNumber of

predicted TMHs: 1\nExp number of AAs in TMHs: 24.51309\nExp number, first 60 AAs: 0.2682\nTotal prob of N-in: 0.14402\noutside 1 331\nTMhelix 332 354\ninside 355 372

18489 GCF_002063075.1_ASM206307v1 Salmonella enterica subsp. enterica serovar Meleagridis Proteobacteria; Gammaproteobacteria; Enterobacterales; Enterobacteriaceae; Salmonella; Salmonella enterica; Salmonella enterica subsp. enterica MNNEETFYQAMRRKGVTRRSFLKFCSLAATSLGLGAGMTPKIAWALENKPRIPVVWIHGL WP_001058311.1 hydrogenase [Salmonella enterica] Length: 372\nNumber of predicted TMHs: 2\nExp number of AAs in TMHs: 37.4699899999999\nExp number, first 60 AAs: 19.72699\nTotal prob of N-in: 0.96171\nPOSSIBLE N-term signal sequence\ninside 1 19\nTMhelix 20 39\noutside 40 325\nTMhelix 326 348\ninside 349 372

18490 GCF_002063075.1_ASM206307v1 Salmonella enterica subsp. enterica serovar Meleagridis Proteobacteria; Gammaproteobacteria; Enterobacterales; Enterobacteriaceae; Salmonella; Salmonella enterica; Salmonella enterica subsp. enterica MTGDNTLITSHGINRRDFMKLCAALAATMGLSSKAAAEMAESVSNPQRPPVIWIGAQECT WP_000145429.1 hydrogenase 2 small subunit [Salmonella enterica] Length: 372\nNumber of predicted TMHs: 1\nExp number of AAs in TMHs: 24.51309\nExp number, first 60 AAs: 0.2682\nTotal prob of N-in: 0.14402\noutside 1 331\nTMhelix 332 354\ninside 355 372

18491 GCF_002064405.1_ASM206440v1 Salmonella enterica subsp. enterica serovar Mgulani Proteobacteria; Gammaproteobacteria; Enterobacterales; Enterobacteriaceae; Salmonella; Salmonella enterica; Salmonella enterica subsp. enterica MNNEETFYQAMRRKGVTRRSFLKFCSLAATSLGLGAGMTPKIAWALENKPRIPVVWIHGL WP_001058311.1 hydrogenase [Salmonella enterica] Length: 372\nNumber of predicted TMHs: 2\nExp number of AAs in TMHs: 37.4699899999999\nExp number, first 60 AAs: 19.72699\nTotal prob of N-in: 0.96171\nPOSSIBLE N-term signal sequence\ninside 1 19\nTMhelix 20 39\noutside 40 325\nTMhelix 326 348\ninside 349 372

18492 GCF_002064405.1_ASM206440v1 Salmonella enterica subsp. enterica serovar Mgulani Proteobacteria; Gammaproteobacteria; Enterobacterales; Enterobacteriaceae; Salmonella; Salmonella enterica; Salmonella enterica subsp. enterica MTGDNTLITSHGINRRDFMKLCAALAATMGLSSKAAAEMAESVSNPQRPPVIWIGAQECT WP_000145429.1 hydrogenase 2 small subunit [Salmonella enterica] Length: 372\nNumber of predicted TMHs: 1\nExp number of AAs in TMHs: 24.51309\nExp number, first 60 AAs: 0.2682\nTotal prob of N-in: 0.14402\noutside 1 331\nTMhelix 332 354\ninside 355 372

18493 GCF_002049065.1_ASM204906v1 Salmonella enterica subsp. enterica serovar Miami Proteobacteria; Gammaproteobacteria; Enterobacterales; Enterobacteriaceae; Salmonella; Salmonella enterica; Salmonella enterica subsp. enterica MTGDNTLITSHGINRRDFMKLCAALAATMGLSSKAAAEMAESVSNPQRPPVIWIGAQECT WP_000145429.1 hydrogenase 2 small subunit [Salmonella enterica] Length: 372\nNumber of predicted TMHs: 1\nExp number of AAs in TMHs: 24.51309\nExp number, first 60 AAs: 0.2682\nTotal prob of N-in: 0.14402\noutside 1 331\nTMhelix 332 354\ninside 355 372

18494 GCF_002050945.1_ASM205094v1 Salmonella enterica subsp. enterica serovar Miami Proteobacteria; Gammaproteobacteria; Enterobacterales; Enterobacteriaceae; Salmonella; Salmonella enterica; Salmonella enterica subsp. enterica MNNEETFYQAMRRKGVTRRSFLKFCSLAATSLGLGAGMTPKIAWALENKPRIPVVWIHGL WP_001662439.1 hydrogenase [Salmonella enterica] Length: 372\nNumber of predicted TMHs: 2\nExp number of AAs in TMHs: 36.5172\nExp number, first 60 AAs: 19.72849\nTotal prob of N-in: 0.96169\nPOSSIBLE N-term signal sequence\ninside 1 19\nTMhelix 20 39\noutside 40 325\nTMhelix 326 348\ninside 349 372

18495 GCF_002049065.1_ASM204906v1 Salmonella enterica subsp. enterica serovar Miami Proteobacteria; Gammaproteobacteria; Enterobacterales; Enterobacteriaceae; Salmonella; Salmonella enterica; Salmonella enterica subsp. enterica MNNEETFYQAMRRKGVTRRSFLKFCSLAATSLGLGAGMTPKIAWALENKPRIPVVWIHGL WP_001662439.1 hydrogenase [Salmonella enterica] Length: 372\nNumber of predicted TMHs: 2\nExp number of AAs in TMHs: 36.5172\nExp number, first 60 AAs: 19.72849\nTotal prob of N-in: 0.96169\nPOSSIBLE N-term signal sequence\ninside 1 19\nTMhelix 20 39\noutside 40 325\nTMhelix 326 348\ninside 349 372

18496 GCF_002033985.1_ASM203398v1 Salmonella enterica subsp. enterica serovar Miami Proteobacteria; Gammaproteobacteria; Enterobacterales; Enterobacteriaceae; Salmonella; Salmonella enterica; Salmonella enterica subsp. enterica MNNEETFYQAMRRKGVTRRSFLKFCSLAATSLGLGAGMTPKIAWALENKPRIPVVWIHGL WP_001058312.1 hydrogenase [Salmonella enterica] Length: 372\nNumber of predicted TMHs: 2\nExp number of AAs in TMHs: 36.51738\nExp number, first 60 AAs: 19.72849\nTotal prob of N-in: 0.96170\nPOSSIBLE N-term signal sequence\ninside 1 19\nTMhelix 20 39\noutside 40 325\nTMhelix 326 348\ninside 349 372

18497 GCF_002050915.1_ASM205091v1 Salmonella enterica subsp. enterica serovar Miami Proteobacteria; Gammaproteobacteria; Enterobacterales; Enterobacteriaceae; Salmonella; Salmonella enterica; Salmonella enterica subsp. enterica MNNEETFYQAMRRKGVTRRSFLKFCSLAATSLGLGAGMTPKIAWALENKPRIPVVWIHGL WP_001662439.1 hydrogenase [Salmonella enterica] Length: 372\nNumber of predicted TMHs: 2\nExp number of AAs in TMHs: 36.5172\nExp number, first 60 AAs: 19.72849\nTotal prob of N-in: 0.96169\nPOSSIBLE N-term signal sequence\ninside 1 19\nTMhelix 20 39\noutside 40 325\nTMhelix 326 348\ninside 349 372

18498 GCF_002033985.1_ASM203398v1 Salmonella enterica subsp. enterica serovar Miami Proteobacteria; Gammaproteobacteria; Enterobacterales; Enterobacteriaceae; Salmonella; Salmonella enterica; Salmonella enterica subsp. enterica MTGDNTLITSHGINRRDFMKLCAALAATMGLSSKAAAEMAESVSNPQRPPVIWIGAQECT WP_000145429.1 hydrogenase 2 small subunit [Salmonella enterica] Length: 372\nNumber of predicted TMHs: 1\nExp number of AAs in TMHs: 24.51309\nExp number, first 60 AAs: 0.2682\nTotal prob of N-in: 0.14402\noutside 1 331\nTMhelix 332 354\ninside 355 372

18499 GCF_002050855.1_ASM205085v1 Salmonella enterica subsp. enterica serovar Miami Proteobacteria; Gammaproteobacteria; Enterobacterales; Enterobacteriaceae; Salmonella; Salmonella enterica; Salmonella enterica subsp. enterica MTGDNTLITSHGINRRDFMKLCAALAATMGLSSKAAAEMAESVSNPQRPPVIWIGAQECT WP_000145429.1 hydrogenase 2 small subunit [Salmonella enterica] Length: 372\nNumber of predicted TMHs: 1\nExp number of AAs in TMHs: 24.51309\nExp number, first 60 AAs: 0.2682\nTotal prob of N-in: 0.14402\noutside 1 331\nTMhelix 332 354\ninside 355 372

18500 GCF_002050915.1_ASM205091v1 Salmonella enterica subsp. enterica serovar Miami Proteobacteria; Gammaproteobacteria; Enterobacterales; Enterobacteriaceae; Salmonella; Salmonella enterica; Salmonella enterica subsp. enterica MTGDNTLITSHGINRRDFMKLCAALAATMGLSSKAAAEMAESVSNPQRPPVIWIGAQECT WP_000145429.1 hydrogenase 2 small subunit [Salmonella enterica] Length: 372\nNumber of predicted TMHs: 1\nExp number of AAs in TMHs: 24.51309\nExp number, first 60 AAs: 0.2682\nTotal prob of N-in: 0.14402\noutside 1 331\nTMhelix 332 354\ninside 355 372

18501 GCF_002050945.1_ASM205094v1 Salmonella enterica subsp. enterica serovar Miami Proteobacteria; Gammaproteobacteria; Enterobacterales; Enterobacteriaceae; Salmonella; Salmonella enterica; Salmonella enterica subsp. enterica MTGDNTLITSHGINRRDFMKLCAALAATMGLSSKAAAEMAESVSNPQRPPVIWIGAQECT WP_000145429.1 hydrogenase 2 small subunit [Salmonella enterica] Length: 372\nNumber of predicted TMHs: 1\nExp number of AAs in TMHs: 24.51309\nExp number, first 60 AAs: 0.2682\nTotal prob of N-in: 0.14402\noutside 1 331\nTMhelix 332 354\ninside 355 372

18502 GCF_002050975.1_ASM205097v1 Salmonella enterica subsp. enterica serovar Miami Proteobacteria; Gammaproteobacteria; Enterobacterales; Enterobacteriaceae; Salmonella; Salmonella enterica; Salmonella enterica subsp. enterica MTGDNTLITSHGINRRDFMKLCAALAATMGLSSKAAAEMAESVSNPQRPPVIWIGAQECT WP_000145429.1 hydrogenase 2 small subunit [Salmonella enterica] Length: 372\nNumber of predicted TMHs: 1\nExp number of AAs in TMHs: 24.51309\nExp number, first 60 AAs: 0.2682\nTotal prob of N-in: 0.14402\noutside 1 331\nTMhelix 332 354\ninside 355 372

18503 GCF_002050975.1_ASM205097v1 Salmonella enterica subsp. enterica serovar Miami Proteobacteria; Gammaproteobacteria; Enterobacterales; Enterobacteriaceae; Salmonella; Salmonella enterica; Salmonella enterica subsp. enterica MNNEETFYQAMRRKGVTRRSFLKFCSLAATSLGLGAGMTPKIAWALENKPRIPVVWIHGL WP_001662439.1 hydrogenase [Salmonella enterica] Length: 372\nNumber of predicted TMHs: 2\nExp number of AAs in TMHs: 36.5172\nExp number, first 60 AAs: 19.72849\nTotal prob of N-in: 0.96169\nPOSSIBLE N-term signal sequence\ninside 1 19\nTMhelix 20 39\noutside 40 325\nTMhelix 326 348\ninside 349 372

18504 GCF_002050855.1_ASM205085v1 Salmonella enterica subsp. enterica serovar Miami Proteobacteria; Gammaproteobacteria; Enterobacterales; Enterobacteriaceae; Salmonella; Salmonella enterica; Salmonella enterica subsp. enterica MNNEETFYQAMRRKGVTRRSFLKFCSLAATSLGLGAGMTPKIAWALENKPRIPVVWIHGL WP_001662439.1 hydrogenase [Salmonella enterica] Length: 372\nNumber of predicted TMHs: 2\nExp number of AAs in TMHs: 36.5172\nExp number, first 60 AAs: 19.72849\nTotal prob of N-in: 0.96169\nPOSSIBLE N-term signal sequence\ninside 1 19\nTMhelix 20 39\noutside 40 325\nTMhelix 326 348\ninside 349 372

18505 GCF_002034405.1_ASM203440v1 Salmonella enterica subsp. enterica serovar Michigan Proteobacteria; Gammaproteobacteria; Enterobacterales; Enterobacteriaceae; Salmonella; Salmonella enterica; Salmonella enterica subsp. enterica MTGDNTLITSHGINRRDFMKLCAALAATMGLSSKAAAEMAESVSNPQRPPVIWIGAQECT WP_000145429.1 hydrogenase 2 small subunit [Salmonella enterica] Length: 372\nNumber of

predicted TMHs:  1\nExp number of AAs in TMHs: 24.51309\nExp number, first 60 AAs:  0.2682\nTotal prob of N-in: 0.14402\noutside   1   331\nTMhelix   332   354\ninside   355   372

18506 GCF_002034405.1_ASM203440v1 Salmonella enterica subsp. enterica serovar Michigan Proteobacteria; Gammaproteobacteria; Enterobacterales; Enterobacteriaceae; Salmonella; Salmonella enterica; Salmonella enterica subsp. enterica MNNEETFYQAMRRKGVTRRSFLKFCSLAATSLGLGAGMTPKIAWALENKPRIPVVWIHGL WP_079840579.1 [Ni/Fe] hydrogenase small subunit [Salmonella enterica] Length: 372\nNumber of predicted TMHs:  2\nExp number of AAs in TMHs: 36.50242\nExp number, first 60 AAs:  19.72845\nTotal prob of N-in: 0.96169\nPOSSIBLE N-term signal sequence\ninside   1   19\nTMhelix   20   39\noutside   40   325\nTMhelix   326   348\ninside   349   372

18507 GCF_002047885.1_ASM204788v1 Salmonella enterica subsp. enterica serovar Minnesota Proteobacteria; Gammaproteobacteria; Enterobacterales; Enterobacteriaceae; Salmonella; Salmonella enterica; Salmonella enterica subsp. enterica MTGDNTLITSHGINRRDFMKLCAALAATMGLSSKAAAEMAESVSNPQRPPVIWIGAQECT WP_000145429.1 hydrogenase 2 small subunit [Salmonella enterica] Length: 372\nNumber of predicted TMHs:  1\nExp number of AAs in TMHs: 24.51309\nExp number, first 60 AAs:  0.2682\nTotal prob of N-in: 0.14402\noutside   1   331\nTMhelix   332   354\ninside   355   372

18508 GCF_001680585.1_ASM168058v1 Salmonella enterica subsp. enterica serovar Minnesota Proteobacteria; Gammaproteobacteria; Enterobacterales; Enterobacteriaceae; Salmonella; Salmonella enterica; Salmonella enterica subsp. enterica MTGDNTLITSHGINRRDFMKLCAALAATMGLSSKAAAEMAESVSNPQRPPVIWIGAQECT WP_000145429.1 hydrogenase 2 small subunit [Salmonella enterica] Length: 372\nNumber of predicted TMHs:  1\nExp number of AAs in TMHs: 24.51309\nExp number, first 60 AAs:  0.2682\nTotal prob of N-in: 0.14402\noutside   1   331\nTMhelix   332   354\ninside   355   372

18509 GCF_002047795.1_ASM204779v1 Salmonella enterica subsp. enterica serovar Minnesota Proteobacteria; Gammaproteobacteria; Enterobacterales; Enterobacteriaceae; Salmonella; Salmonella enterica; Salmonella enterica subsp. enterica MTGDNTLITSHGINRRDFMKLCAALAATMGLSSKAAAEMAESVSNPQRPPVIWIGAQECT WP_000145429.1 hydrogenase 2 small subunit [Salmonella enterica] Length: 372\nNumber of predicted TMHs:  1\nExp number of AAs in TMHs: 24.51309\nExp number, first 60 AAs:  0.2682\nTotal prob of N-in: 0.14402\noutside   1   331\nTMhelix   332   354\ninside   355   372

18510 GCF_002048475.1_ASM204847v1 Salmonella enterica subsp. enterica serovar Minnesota Proteobacteria; Gammaproteobacteria; Enterobacterales; Enterobacteriaceae; Salmonella; Salmonella enterica; Salmonella enterica subsp. enterica MTGDNTLITSHGINRRDFMKLCAALAATMGLSSKAAAEMAESVSNPQRPPVIWIGAQECT WP_000145429.1 hydrogenase 2 small subunit [Salmonella enterica] Length: 372\nNumber of predicted TMHs:  1\nExp number of AAs in TMHs: 24.51309\nExp number, first 60 AAs:  0.2682\nTotal prob of N-in: 0.14402\noutside   1   331\nTMhelix   332   354\ninside   355   372

18511 GCF_002196535.1_ASM219653v1 Salmonella enterica subsp. enterica serovar Minnesota Proteobacteria; Gammaproteobacteria; Enterobacterales; Enterobacteriaceae; Salmonella; Salmonella enterica; Salmonella enterica subsp. enterica MTGDNTLITSHGINRRDFMKLCAALAATMGLSSKAAAEMAESVSNPQRPPVIWIGAQECT WP_000145429.1 hydrogenase 2 small subunit [Salmonella enterica] Length: 372\nNumber of predicted TMHs:  1\nExp number of AAs in TMHs: 24.51309\nExp number, first 60 AAs:  0.2682\nTotal prob of N-in: 0.14402\noutside   1   331\nTMhelix   332   354\ninside   355   372

18512 GCF_002064375.1_ASM206437v1 Salmonella enterica subsp. enterica serovar Minnesota Proteobacteria; Gammaproteobacteria; Enterobacterales; Enterobacteriaceae; Salmonella; Salmonella enterica; Salmonella enterica subsp. enterica MTGDNTLITSHGINRRDFMKLCAALAATMGLSSKAAAEMAESVSNPQRPPVIWIGAQECT WP_000145429.1 hydrogenase 2 small subunit [Salmonella enterica] Length: 372\nNumber of predicted TMHs:  1\nExp number of AAs in TMHs: 24.51309\nExp number, first 60 AAs:  0.2682\nTotal prob of N-in: 0.14402\noutside   1   331\nTMhelix   332   354\ninside   355   372

18513 GCF_002048585.1_ASM204858v1 Salmonella enterica subsp. enterica serovar Minnesota Proteobacteria; Gammaproteobacteria; Enterobacterales; Enterobacteriaceae; Salmonella; Salmonella enterica; Salmonella enterica subsp. enterica MTGDNTLITSHGINRRDFMKLCAALAATMGLSSKAAAEMAESVSNPQRPPVIWIGAQECT WP_000145429.1 hydrogenase 2 small subunit [Salmonella enterica] Length: 372\nNumber of predicted TMHs:  1\nExp number of AAs in TMHs: 24.51309\nExp number, first 60 AAs:  0.2682\nTotal prob of N-in: 0.14402\noutside   1   331\nTMhelix   332   354\ninside   355   372

18514	GCF_002048655.1_ASM204865v1	Salmonella enterica subsp. enterica serovar Minnesota	Proteobacteria; Gammaproteobacteria; Enterobacterales; Enterobacteriaceae; Salmonella; Salmonella enterica; Salmonella enterica subsp. enterica	MTGDNTLITSHGINRRDFMKLCAALAATMGLSSKAAAEMAESVSNPQRPPVIWIGAQECT	WP_000145429.1	hydrogenase 2 small subunit [Salmonella enterica]	Length: 372\nNumber of predicted TMHs:  1\nExp number of AAs in TMHs: 24.51309\nExp number, first 60 AAs:  0.2682\nTotal prob of N-in:  0.14402\noutside  1  331\nTMhelix  332  354\ninside  355  372

18515	GCF_000964185.1_ASM96418v1	Salmonella enterica subsp. enterica serovar Mishmarhaemek	Proteobacteria; Gammaproteobacteria; Enterobacterales; Enterobacteriaceae; Salmonella; Salmonella enterica; Salmonella enterica subsp. enterica	MTGDNTLITSHGINRRDFMKLCAALAATMGLSSKAAAEMAESVSNPQRPPVIWIGAQECT	WP_045902238.1	hydrogenase 2 small subunit [Salmonella enterica]	Length: 372\nNumber of predicted TMHs:  1\nExp number of AAs in TMHs: 24.51562\nExp number, first 60 AAs:  0.26822\nTotal prob of N-in:  0.14409\noutside  1  331\nTMhelix  332  354\ninside  355  372

18516	GCF_000964185.1_ASM96418v1	Salmonella enterica subsp. enterica serovar Mishmarhaemek	Proteobacteria; Gammaproteobacteria; Enterobacterales; Enterobacteriaceae; Salmonella; Salmonella enterica; Salmonella enterica subsp. enterica	MNNEETFYQAMRRKGVTRRSFLKFCSLAATSLGLGAGMTPKIAWALENKPRIPVVWIHGL	WP_001058312.1	hydrogenase [Salmonella enterica]	Length: 372\nNumber of predicted TMHs:  2\nExp number of AAs in TMHs: 36.51738\nExp number, first 60 AAs:  19.72849\nTotal prob of N-in:  0.96170\nPOSSIBLE N-term signal sequence\ninside  1  19\nTMhelix  20  39\noutside  40  325\nTMhelix  326  348\ninside  349  372

18517	GCF_002127805.1_ASM212780v1	Salmonella enterica subsp. enterica serovar Mississippi	Proteobacteria; Gammaproteobacteria; Enterobacterales; Enterobacteriaceae; Salmonella; Salmonella enterica; Salmonella enterica subsp. enterica	MTGDNTLITSHGINRRDFMKLCAALAATMGLSSKAAAEMAESVSNPQRPPVIWIGAQECT	WP_000145429.1	hydrogenase 2 small subunit [Salmonella enterica]	Length: 372\nNumber of predicted TMHs:  1\nExp number of AAs in TMHs: 24.51309\nExp number, first 60 AAs:  0.2682\nTotal prob of N-in:  0.14402\noutside  1  331\nTMhelix  332  354\ninside  355  372

18518	GCF_002127765.1_ASM212776v1	Salmonella enterica subsp. enterica serovar Mississippi	Proteobacteria; Gammaproteobacteria; Enterobacterales; Enterobacteriaceae; Salmonella; Salmonella enterica; Salmonella enterica subsp. enterica	MTGDNTLITSHGINRRDFMKLCAALAATMGLSSKAAAEMAESVSNPQRPPVIWIGAQECT	WP_000145429.1	hydrogenase 2 small subunit [Salmonella enterica]	Length: 372\nNumber of predicted TMHs:  1\nExp number of AAs in TMHs: 24.51309\nExp number, first 60 AAs:  0.2682\nTotal prob of N-in:  0.14402\noutside  1  331\nTMhelix  332  354\ninside  355  372

18519	GCF_002127845.1_ASM212784v1	Salmonella enterica subsp. enterica serovar Mississippi	Proteobacteria; Gammaproteobacteria; Enterobacterales; Enterobacteriaceae; Salmonella; Salmonella enterica; Salmonella enterica subsp. enterica	MTGDNTLITSHGINRRDFMKLCAALAATMGLSSKAAAEMAESVSNPQRPPVIWIGAQECT	WP_000145429.1	hydrogenase 2 small subunit [Salmonella enterica]	Length: 372\nNumber of predicted TMHs:  1\nExp number of AAs in TMHs: 24.51309\nExp number, first 60 AAs:  0.2682\nTotal prob of N-in:  0.14402\noutside  1  331\nTMhelix  332  354\ninside  355  372

18520	GCF_002127865.1_ASM212786v1	Salmonella enterica subsp. enterica serovar Mississippi	Proteobacteria; Gammaproteobacteria; Enterobacterales; Enterobacteriaceae; Salmonella; Salmonella enterica; Salmonella enterica subsp. enterica	MTGDNTLITSHGINRRDFMKLCAALAATMGLSSKAAAEMAESVSNPQRPPVIWIGAQECT	WP_000145429.1	hydrogenase 2 small subunit [Salmonella enterica]	Length: 372\nNumber of predicted TMHs:  1\nExp number of AAs in TMHs: 24.51309\nExp number, first 60 AAs:  0.2682\nTotal prob of N-in:  0.14402\noutside  1  331\nTMhelix  332  354\ninside  355  372

18521	GCF_002059325.1_ASM205932v1	Salmonella enterica subsp. enterica serovar Mississippi	Proteobacteria; Gammaproteobacteria; Enterobacterales; Enterobacteriaceae; Salmonella; Salmonella enterica; Salmonella enterica subsp. enterica	MNNEETFYQAMRRKGVTRRSFLKFCSLAATSLGLGAGMTPKIAWALENKPRIPVVWIHGL	WP_001058312.1	hydrogenase [Salmonella enterica]	Length: 372\nNumber of predicted TMHs:  2\nExp number of AAs in TMHs: 36.51738\nExp number, first 60 AAs:  19.72849\nTotal prob of N-in:  0.96170\nPOSSIBLE N-term signal sequence\ninside  1  19\nTMhelix  20  39\noutside  40  325\nTMhelix  326  348\ninside  349  372

18522	GCF_002059325.1_ASM205932v1	Salmonella enterica subsp. enterica serovar Mississippi	Proteobacteria; Gammaproteobacteria; Enterobacterales; Enterobacteriaceae; Salmonella; Salmonella enterica; Salmonella enterica subsp. enterica	MTGDNTLITSHGINRRDFMKLCAALAATMGLSSKAAAEMAESVSNPQRPPVIWIGAQECT	WP_000145431.1	hydrogenase 2 small subunit [Salmonella enterica]	Length: 372\nNumber of

predicted TMHs: 1\nExp number of AAs in TMHs: 24.51505\nExp number, first 60 AAs: 0.2682\nTotal prob of N-in: 0.14406\noutside 1 331\nTMhelix 332 354\ninside 355 372

18523 GCF_002127855.1_ASM212785v1 Salmonella enterica subsp. enterica serovar Mississippi Proteobacteria; Gammaproteobacteria; Enterobacterales; Enterobacteriaceae; Salmonella; Salmonella enterica; Salmonella enterica subsp. enterica MTGDNTLITSHGINRRDFMKLCAALAATMGLSSKAAAEMAESVSNPQRPPVIWIGAQECT WP_000145429.1 hydrogenase 2 small subunit [Salmonella enterica] Length: 372\nNumber of predicted TMHs: 1\nExp number of AAs in TMHs: 24.51309\nExp number, first 60 AAs: 0.2682\nTotal prob of N-in: 0.14402\noutside 1 331\nTMhelix 332 354\ninside 355 372

18524 GCF_002127925.1_ASM212792v1 Salmonella enterica subsp. enterica serovar Mississippi Proteobacteria; Gammaproteobacteria; Enterobacterales; Enterobacteriaceae; Salmonella; Salmonella enterica; Salmonella enterica subsp. enterica MNNEETFYQAMRRKGVTRRSFLKFCSLAATSLGLGAGMTPKIAWALENKPRIPVVWIHGL WP_001058312.1 hydrogenase [Salmonella enterica] Length: 372\nNumber of predicted TMHs: 2\nExp number of AAs in TMHs: 36.51738\nExp number, first 60 AAs: 19.72849\nTotal prob of N-in: 0.96170\nPOSSIBLE N-term signal sequence\ninside 1 19\nTMhelix 20 39\noutside 40 325\nTMhelix 326 348\ninside 349 372

18525 GCF_002127775.1_ASM212777v1 Salmonella enterica subsp. enterica serovar Mississippi Proteobacteria; Gammaproteobacteria; Enterobacterales; Enterobacteriaceae; Salmonella; Salmonella enterica; Salmonella enterica subsp. enterica MTGDNTLITSHGINRRDFMKLCAALAATMGLSSKAAAEMAESVSNPQRPPVIWIGAQECT WP_000145429.1 hydrogenase 2 small subunit [Salmonella enterica] Length: 372\nNumber of predicted TMHs: 1\nExp number of AAs in TMHs: 24.51309\nExp number, first 60 AAs: 0.2682\nTotal prob of N-in: 0.14402\noutside 1 331\nTMhelix 332 354\ninside 355 372

18526 GCF_002127935.1_ASM212793v1 Salmonella enterica subsp. enterica serovar Mississippi Proteobacteria; Gammaproteobacteria; Enterobacterales; Enterobacteriaceae; Salmonella; Salmonella enterica; Salmonella enterica subsp. enterica MNNEETFYQAMRRKGVTRRSFLKFCSLAATSLGLGAGMTPKIAWALENKPRIPVVWIHGL WP_001058312.1 hydrogenase [Salmonella enterica] Length: 372\nNumber of predicted TMHs: 2\nExp number of AAs in TMHs: 36.51738\nExp number, first 60 AAs: 19.72849\nTotal prob of N-in: 0.96170\nPOSSIBLE N-term signal sequence\ninside 1 19\nTMhelix 20 39\noutside 40 325\nTMhelix 326 348\ninside 349 372

18527 GCF_002127875.1_ASM212787v1 Salmonella enterica subsp. enterica serovar Mississippi Proteobacteria; Gammaproteobacteria; Enterobacterales; Enterobacteriaceae; Salmonella; Salmonella enterica; Salmonella enterica subsp. enterica MTGDNTLITSHGINRRDFMKLCAALAATMGLSSKAAAEMAESVSNPQRPPVIWIGAQECT WP_000145429.1 hydrogenase 2 small subunit [Salmonella enterica] Length: 372\nNumber of predicted TMHs: 1\nExp number of AAs in TMHs: 24.51309\nExp number, first 60 AAs: 0.2682\nTotal prob of N-in: 0.14402\noutside 1 331\nTMhelix 332 354\ninside 355 372

18528 GCF_002127925.1_ASM212792v1 Salmonella enterica subsp. enterica serovar Mississippi Proteobacteria; Gammaproteobacteria; Enterobacterales; Enterobacteriaceae; Salmonella; Salmonella enterica; Salmonella enterica subsp. enterica MTGDNTLITSHGINRRDFMKLCAALAATMGLSSKAAAEMAESVSNPQRPPVIWIGAQECT WP_000145429.1 hydrogenase 2 small subunit [Salmonella enterica] Length: 372\nNumber of predicted TMHs: 1\nExp number of AAs in TMHs: 24.51309\nExp number, first 60 AAs: 0.2682\nTotal prob of N-in: 0.14402\noutside 1 331\nTMhelix 332 354\ninside 355 372

18529 GCF_002127855.1_ASM212785v1 Salmonella enterica subsp. enterica serovar Mississippi Proteobacteria; Gammaproteobacteria; Enterobacterales; Enterobacteriaceae; Salmonella; Salmonella enterica; Salmonella enterica subsp. enterica MNNEETFYQAMRRKGVTRRSFLKFCSLAATSLGLGAGMTPKIAWALENKPRIPVVWIHGL WP_001058312.1 hydrogenase [Salmonella enterica] Length: 372\nNumber of predicted TMHs: 2\nExp number of AAs in TMHs: 36.51738\nExp number, first 60 AAs: 19.72849\nTotal prob of N-in: 0.96170\nPOSSIBLE N-term signal sequence\ninside 1 19\nTMhelix 20 39\noutside 40 325\nTMhelix 326 348\ninside 349 372

18530 GCF_002127775.1_ASM212777v1 Salmonella enterica subsp. enterica serovar Mississippi Proteobacteria; Gammaproteobacteria; Enterobacterales; Enterobacteriaceae; Salmonella; Salmonella enterica; Salmonella enterica subsp. enterica MNNEETFYQAMRRKGVTRRSFLKFCSLAATSLGLGAGMTPKIAWALENKPRIPVVWIHGL WP_001058312.1 hydrogenase [Salmonella enterica] Length: 372\nNumber of predicted TMHs: 2\nExp number of AAs in TMHs: 36.51738\nExp number, first 60 AAs: 19.72849\nTotal prob of N-in: 0.96170\nPOSSIBLE N-term signal sequence\ninside 1 19\nTMhelix 20 39\noutside 40 325\nTMhelix 326 348\ninside 349 372

18531 GCF_002127935.1_ASM212793v1 Salmonella enterica subsp. enterica serovar Mississippi Proteobacteria; Gammaproteobacteria; Enterobacterales; Enterobacteriaceae; Salmonella; Salmonella enterica;

Salmonella enterica subsp. enterica MTGDNTLITSHGINRRDFMKLCAALAATMGLSSKAAAEMAESVSNPQRPPVIWIGAQECT WP_000145429.1 hydrogenase 2 small subunit [Salmonella enterica] Length: 372\nNumber of predicted TMHs: 1\nExp number of AAs in TMHs: 24.51309\nExp number, first 60 AAs: 0.2682\nTotal prob of N-in: 0.14402\noutside 1 331\nTMhelix 332 354\ninside 355 372

18532 GCF_002127875.1_ASM212787v1 Salmonella enterica subsp. enterica serovar Mississippi Proteobacteria; Gammaproteobacteria; Enterobacterales; Enterobacteriaceae; Salmonella; Salmonella enterica; Salmonella enterica subsp. enterica MNNEETFYQAMRRKGVTRRSFLKFCSLAATSLGLGAGMTPKIAWALENKPRIPVVWIHGL WP_001058312.1 hydrogenase [Salmonella enterica] Length: 372\nNumber of predicted TMHs: 2\nExp number of AAs in TMHs: 36.51738\nExp number, first 60 AAs: 19.72849\nTotal prob of N-in: 0.96170\nPOSSIBLE N-term signal sequence\ninside 1 19\nTMhelix 20 39\noutside 40 325\nTMhelix 326 348\ninside 349 372

18533 GCF_002127765.1_ASM212776v1 Salmonella enterica subsp. enterica serovar Mississippi Proteobacteria; Gammaproteobacteria; Enterobacterales; Enterobacteriaceae; Salmonella; Salmonella enterica; Salmonella enterica subsp. enterica MNNEETFYQAMRRKGVTRRSFLKFCSLAATSLGLGAGMTPKIAWALENKPRIPVVWIHGL WP_001058312.1 hydrogenase [Salmonella enterica] Length: 372\nNumber of predicted TMHs: 2\nExp number of AAs in TMHs: 36.51738\nExp number, first 60 AAs: 19.72849\nTotal prob of N-in: 0.96170\nPOSSIBLE N-term signal sequence\ninside 1 19\nTMhelix 20 39\noutside 40 325\nTMhelix 326 348\ninside 349 372

18534 GCF_002127845.1_ASM212784v1 Salmonella enterica subsp. enterica serovar Mississippi Proteobacteria; Gammaproteobacteria; Enterobacterales; Enterobacteriaceae; Salmonella; Salmonella enterica; Salmonella enterica subsp. enterica MNNEETFYQAMRRKGVTRRSFLKFCSLAATSLGLGAGMTPKIAWALENKPRIPVVWIHGL WP_001058312.1 hydrogenase [Salmonella enterica] Length: 372\nNumber of predicted TMHs: 2\nExp number of AAs in TMHs: 36.51738\nExp number, first 60 AAs: 19.72849\nTotal prob of N-in: 0.96170\nPOSSIBLE N-term signal sequence\ninside 1 19\nTMhelix 20 39\noutside 40 325\nTMhelix 326 348\ninside 349 372

18535 GCF_002127865.1_ASM212786v1 Salmonella enterica subsp. enterica serovar Mississippi Proteobacteria; Gammaproteobacteria; Enterobacterales; Enterobacteriaceae; Salmonella; Salmonella enterica; Salmonella enterica subsp. enterica MNNEETFYQAMRRKGVTRRSFLKFCSLAATSLGLGAGMTPKIAWALENKPRIPVVWIHGL WP_001058312.1 hydrogenase [Salmonella enterica] Length: 372\nNumber of predicted TMHs: 2\nExp number of AAs in TMHs: 36.51738\nExp number, first 60 AAs: 19.72849\nTotal prob of N-in: 0.96170\nPOSSIBLE N-term signal sequence\ninside 1 19\nTMhelix 20 39\noutside 40 325\nTMhelix 326 348\ninside 349 372

18536 GCF_002127805.1_ASM212780v1 Salmonella enterica subsp. enterica serovar Mississippi Proteobacteria; Gammaproteobacteria; Enterobacterales; Enterobacteriaceae; Salmonella; Salmonella enterica; Salmonella enterica subsp. enterica MNNEETFYQAMRRKGVTRRSFLKFCSLAATSLGLGAGMTPKIAWALENKPRIPVVWIHGL WP_001058312.1 hydrogenase [Salmonella enterica] Length: 372\nNumber of predicted TMHs: 2\nExp number of AAs in TMHs: 36.51738\nExp number, first 60 AAs: 19.72849\nTotal prob of N-in: 0.96170\nPOSSIBLE N-term signal sequence\ninside 1 19\nTMhelix 20 39\noutside 40 325\nTMhelix 326 348\ninside 349 372

18537 GCF_002045845.1_ASM204584v1 Salmonella enterica subsp. enterica serovar Monschaui Proteobacteria; Gammaproteobacteria; Enterobacterales; Enterobacteriaceae; Salmonella; Salmonella enterica; Salmonella enterica subsp. enterica MNNEETFYQAMRRKGVTRRSFLKFCSLAATSLGLGAGMTPKIAWALENKPRIPVVWIHGL WP_001058311.1 hydrogenase [Salmonella enterica] Length: 372\nNumber of predicted TMHs: 2\nExp number of AAs in TMHs: 37.4699899999999\nExp number, first 60 AAs: 19.72699\nTotal prob of N-in: 0.96171\nPOSSIBLE N-term signal sequence\ninside 1 19\nTMhelix 20 39\noutside 40 325\nTMhelix 326 348\ninside 349 372

18538 GCF_002045005.1_ASM204500v1 Salmonella enterica subsp. enterica serovar Monschaui Proteobacteria; Gammaproteobacteria; Enterobacterales; Enterobacteriaceae; Salmonella; Salmonella enterica; Salmonella enterica subsp. enterica MNNEETFYQAMRRKGVTRRSFLKFCSLAATSLGLSAGMTPKIAWALENKPRIPVVWIHGL WP_001058316.1 hydrogenase [Salmonella enterica] Length: 372\nNumber of predicted TMHs: 2\nExp number of AAs in TMHs: 35.76397\nExp number, first 60 AAs: 18.97201\nTotal prob of N-in: 0.92785\nPOSSIBLE N-term signal sequence\ninside 1 19\nTMhelix 20 39\noutside 40 325\nTMhelix 326 348\ninside 349 372

18539 GCF_002045805.1_ASM204580v1 Salmonella enterica subsp. enterica serovar Monschaui Proteobacteria; Gammaproteobacteria; Enterobacterales; Enterobacteriaceae; Salmonella; Salmonella enterica; Salmonella enterica subsp. enterica MNNEETFYQAMRRKGVTRRSFLKFCSLAATSLGLSAGMTPKIAWALENKPRIPVVWIHGL WP_001058316.1 hydrogenase [Salmonella enterica] Length: 372\nNumber of predicted TMHs: 2\nExp number of AAs in TMHs: 35.76397\nExp number, first 60 AAs: 18.97201\nTotal prob of N-in: 0.92785\nPOSSIBLE N-term signal sequence\ninside 1 19\nTMhelix 20 39\noutside 40 325\nTMhelix 326 348\ninside 349 372

18540 GCF_002045705.1_ASM204570v1 Salmonella enterica subsp. enterica serovar Monschaui Proteobacteria; Gammaproteobacteria; Enterobacterales; Enterobacteriaceae; Salmonella; Salmonella enterica; Salmonella enterica subsp. enterica MTGDNTLITSHGINRRDFMKLCAALAATMGLSSKAAAEMAESVSNPQRPPVIWIGAQECT WP_000145429.1 hydrogenase 2 small subunit [Salmonella enterica] Length: 372\nNumber of predicted TMHs: 1\nExp number of AAs in TMHs: 24.51309\nExp number, first 60 AAs: 0.2682\nTotal prob of N-in: 0.14402\noutside 1 331\nTMhelix 332 354\ninside 355 372

18541 GCF_002045705.1_ASM204570v1 Salmonella enterica subsp. enterica serovar Monschaui Proteobacteria; Gammaproteobacteria; Enterobacterales; Enterobacteriaceae; Salmonella; Salmonella enterica; Salmonella enterica subsp. enterica MNNEETFYQAMRRKGVTRRSFLKFCSLAATSLGLSAGMTPKIAWALENKPRIPVVWIHGL WP_001058316.1 hydrogenase [Salmonella enterica] Length: 372\nNumber of predicted TMHs: 2\nExp number of AAs in TMHs: 35.76397\nExp number, first 60 AAs: 18.97201\nTotal prob of N-in: 0.92785\nPOSSIBLE N-term signal sequence\ninside 1 19\nTMhelix 20 39\noutside 40 325\nTMhelix 326 348\ninside 349 372

18542 GCF_002065435.1_ASM206543v1 Salmonella enterica subsp. enterica serovar Monschaui Proteobacteria; Gammaproteobacteria; Enterobacterales; Enterobacteriaceae; Salmonella; Salmonella enterica; Salmonella enterica subsp. enterica MNNEETFYQAMRRKGVTRRSFLKFCSLAATSLGLGAGMTPKIAWALENKPRIPVVWIHGL WP_001058311.1 hydrogenase [Salmonella enterica] Length: 372\nNumber of predicted TMHs: 2\nExp number of AAs in TMHs: 37.4699899999999\nExp number, first 60 AAs: 19.72699\nTotal prob of N-in: 0.96171\nPOSSIBLE N-term signal sequence\ninside 1 19\nTMhelix 20 39\noutside 40 325\nTMhelix 326 348\ninside 349 372

18543 GCF_002045005.1_ASM204500v1 Salmonella enterica subsp. enterica serovar Monschaui Proteobacteria; Gammaproteobacteria; Enterobacterales; Enterobacteriaceae; Salmonella; Salmonella enterica; Salmonella enterica subsp. enterica MTGDNTLITSHGINRRDFMKLCAALAATMGLSSKAAAEMAESVSNPQRPPVIWIGAQECT WP_000145429.1 hydrogenase 2 small subunit [Salmonella enterica] Length: 372\nNumber of predicted TMHs: 1\nExp number of AAs in TMHs: 24.51309\nExp number, first 60 AAs: 0.2682\nTotal prob of N-in: 0.14402\noutside 1 331\nTMhelix 332 354\ninside 355 372

18544 GCF_002065435.1_ASM206543v1 Salmonella enterica subsp. enterica serovar Monschaui Proteobacteria; Gammaproteobacteria; Enterobacterales; Enterobacteriaceae; Salmonella; Salmonella enterica; Salmonella enterica subsp. enterica MTGDNTLITSHGINRRDFMKLCAALAATMGLSSKAAAEMAESVSNPQRPPVIWIGAQECT WP_000145429.1 hydrogenase 2 small subunit [Salmonella enterica] Length: 372\nNumber of predicted TMHs: 1\nExp number of AAs in TMHs: 24.51309\nExp number, first 60 AAs: 0.2682\nTotal prob of N-in: 0.14402\noutside 1 331\nTMhelix 332 354\ninside 355 372

18545 GCF_002045805.1_ASM204580v1 Salmonella enterica subsp. enterica serovar Monschaui Proteobacteria; Gammaproteobacteria; Enterobacterales; Enterobacteriaceae; Salmonella; Salmonella enterica; Salmonella enterica subsp. enterica MTGDNTLITSHGINRRDFMKLCAALAATMGLSSKAAAEMAESVSNPQRPPVIWIGAQECT WP_000145429.1 hydrogenase 2 small subunit [Salmonella enterica] Length: 372\nNumber of predicted TMHs: 1\nExp number of AAs in TMHs: 24.51309\nExp number, first 60 AAs: 0.2682\nTotal prob of N-in: 0.14402\noutside 1 331\nTMhelix 332 354\ninside 355 372

18546 GCF_002045845.1_ASM204584v1 Salmonella enterica subsp. enterica serovar Monschaui Proteobacteria; Gammaproteobacteria; Enterobacterales; Enterobacteriaceae; Salmonella; Salmonella enterica; Salmonella enterica subsp. enterica MTGDNTLITSHGINRRDFMKLCAALAATMGLSSKAAAEMAESVSNPQRPPVIWIGAQECT WP_000145429.1 hydrogenase 2 small subunit [Salmonella enterica] Length: 372\nNumber of predicted TMHs: 1\nExp number of AAs in TMHs: 24.51309\nExp number, first 60 AAs: 0.2682\nTotal prob of N-in: 0.14402\noutside 1 331\nTMhelix 332 354\ninside 355 372

18547 GCF_001272295.1_Salmonella_enterica_CVM_N50432_v1.0 Salmonella enterica subsp. enterica serovar Montevideo Proteobacteria; Gammaproteobacteria; Enterobacterales; Enterobacteriaceae; Salmonella; Salmonella enterica; Salmonella enterica subsp. enterica MNNEETFYQAMRRKGVTRRSFLKFCSLAATSLGLGAGMTPKIAWALENKPRIPVVWIHGL WP_001058312.1 hydrogenase [Salmonella enterica] Length: 372\nNumber of predicted TMHs: 2\nExp number of AAs in TMHs: 36.51738\nExp number, first 60 AAs: 19.72849\nTotal prob of N-in: 0.96170\nPOSSIBLE N-term signal sequence\ninside 1 19\nTMhelix 20 39\noutside 40 325\nTMhelix 326 348\ninside 349 372

18548 GCF_001246955.1_Salmonella_enterica_CVM_N51288_v1.0 Salmonella enterica subsp. enterica serovar Montevideo Proteobacteria; Gammaproteobacteria; Enterobacterales; Enterobacteriaceae; Salmonella; Salmonella enterica; Salmonella enterica subsp. enterica

MNNEETFYQAMRRKGVTRRSFLKFCSLAATSLGLGAGMTPKIAWALENKPRIPVVWIHGL WP_001058312.1 hydrogenase [Salmonella enterica] Length: 372\nNumber of predicted TMHs: 2\nExp number of AAs in TMHs: 36.51738\nExp number, first 60 AAs: 19.72849\nTotal prob of N-in: 0.96170\nPOSSIBLE N-term signal sequence\ninside 1 19\nTMhelix 20 39\noutside 40 325\nTMhelix 326 348\ninside 349 372

18549 GCF_001241855.1_Salmonella_enterica_CVM_N45402_v1.0 Salmonella enterica subsp. enterica serovar Montevideo Proteobacteria; Gammaproteobacteria; Enterobacterales; Enterobacteriaceae; Salmonella; Salmonella enterica; Salmonella enterica subsp. enterica MTGDNTLITSHGINRRDFMKLCAALAATMGLSSKAAAEMAESVSNPQRPPVIWIGAQECT WP_000145429.1 hydrogenase 2 small subunit [Salmonella enterica] Length: 372\nNumber of predicted TMHs: 1\nExp number of AAs in TMHs: 24.51309\nExp number, first 60 AAs: 0.2682\nTotal prob of N-in: 0.14402\noutside 1 331\nTMhelix 332 354\ninside 355 372

18550 GCF_001241855.1_Salmonella_enterica_CVM_N45402_v1.0 Salmonella enterica subsp. enterica serovar Montevideo Proteobacteria; Gammaproteobacteria; Enterobacterales; Enterobacteriaceae; Salmonella; Salmonella enterica; Salmonella enterica subsp. enterica MNNEETFYQAMRRKGVTRRSFLKFCSLAATSLGLGAGMTPKIAWALENKPRIPVVWIHGL WP_044782462.1 hydrogenase [Salmonella enterica] Length: 372\nNumber of predicted TMHs: 2\nExp number of AAs in TMHs: 36.51702\nExp number, first 60 AAs: 19.72849\nTotal prob of N-in: 0.96170\nPOSSIBLE N-term signal sequence\ninside 1 19\nTMhelix 20 39\noutside 40 325\nTMhelix 326 348\ninside 349 372

18551 GCF_002059705.1_ASM205970v1 Salmonella enterica subsp. enterica serovar Montevideo Proteobacteria; Gammaproteobacteria; Enterobacterales; Enterobacteriaceae; Salmonella; Salmonella enterica; Salmonella enterica subsp. enterica MNNEETFYQAMRRKGVTRRSFLKFCSLAATSLGLGAGMTPKIAWALENKPRIPVVWIHGL WP_001058312.1 hydrogenase [Salmonella enterica] Length: 372\nNumber of predicted TMHs: 2\nExp number of AAs in TMHs: 36.51738\nExp number, first 60 AAs: 19.72849\nTotal prob of N-in: 0.96170\nPOSSIBLE N-term signal sequence\ninside 1 19\nTMhelix 20 39\noutside 40 325\nTMhelix 326 348\ninside 349 372

18552 GCF_002037075.1_ASM203707v1 Salmonella enterica subsp. enterica serovar Montevideo Proteobacteria; Gammaproteobacteria; Enterobacterales; Enterobacteriaceae; Salmonella; Salmonella enterica; Salmonella enterica subsp. enterica MTGDNTLITSHGINRRDFMKLCAALAATMGLSSKAAAEMAESVSNPQRPPVIWIGAQECT WP_000145429.1 hydrogenase 2 small subunit [Salmonella enterica] Length: 372\nNumber of predicted TMHs: 1\nExp number of AAs in TMHs: 24.51309\nExp number, first 60 AAs: 0.2682\nTotal prob of N-in: 0.14402\noutside 1 331\nTMhelix 332 354\ninside 355 372

18553 GCF_002037075.1_ASM203707v1 Salmonella enterica subsp. enterica serovar Montevideo Proteobacteria; Gammaproteobacteria; Enterobacterales; Enterobacteriaceae; Salmonella; Salmonella enterica; Salmonella enterica subsp. enterica MNNEETFYQAMRRKGVTRRSFLKFCSLAATSLGLGAGMTPKIAWALENKPRIPVVWIHGL WP_001058312.1 hydrogenase [Salmonella enterica] Length: 372\nNumber of predicted TMHs: 2\nExp number of AAs in TMHs: 36.51738\nExp number, first 60 AAs: 19.72849\nTotal prob of N-in: 0.96170\nPOSSIBLE N-term signal sequence\ninside 1 19\nTMhelix 20 39\noutside 40 325\nTMhelix 326 348\ninside 349 372

18554 GCF_001692045.1_ASM169204v1 Salmonella enterica subsp. enterica serovar Montevideo Proteobacteria; Gammaproteobacteria; Enterobacterales; Enterobacteriaceae; Salmonella; Salmonella enterica; Salmonella enterica subsp. enterica MTGDNTLITSHGINRRDFMKLCAALAATMGLSSKAAAEMAESVSNPQRPPVIWIGAQECT WP_000145429.1 hydrogenase 2 small subunit [Salmonella enterica] Length: 372\nNumber of predicted TMHs: 1\nExp number of AAs in TMHs: 24.51309\nExp number, first 60 AAs: 0.2682\nTotal prob of N-in: 0.14402\noutside 1 331\nTMhelix 332 354\ninside 355 372

18555 GCF_002066305.1_ASM206630v1 Salmonella enterica subsp. enterica serovar Montevideo Proteobacteria; Gammaproteobacteria; Enterobacterales; Enterobacteriaceae; Salmonella; Salmonella enterica; Salmonella enterica subsp. enterica MNNEETFYQAMRRKGVTRRSFLKFCSLAATSLGLGAGMTPKIAWALENKPRIPVVWIHGL WP_001058312.1 hydrogenase [Salmonella enterica] Length: 372\nNumber of predicted TMHs: 2\nExp number of AAs in TMHs: 36.51738\nExp number, first 60 AAs: 19.72849\nTotal prob of N-in: 0.96170\nPOSSIBLE N-term signal sequence\ninside 1 19\nTMhelix 20 39\noutside 40 325\nTMhelix 326 348\ninside 349 372

18556 GCF_002066305.1_ASM206630v1 Salmonella enterica subsp. enterica serovar Montevideo Proteobacteria; Gammaproteobacteria; Enterobacterales; Enterobacteriaceae; Salmonella; Salmonella enterica; Salmonella enterica subsp. enterica MTGDNTLITSHGINRRDFMKLCAALAATMGLSSKAAAEMAESVSNPQRPPVIWIGAQECT WP_000145429.1 hydrogenase 2 small subunit [Salmonella enterica] Length: 372\nNumber of

predicted TMHs: 1\nExp number of AAs in TMHs: 24.51309\nExp number, first 60 AAs: 0.2682\nTotal prob of N-in: 0.14402\noutside 1 331\nTMhelix 332 354\ninside 355 372

18557 GCF_001692045.1_ASM169204v1 Salmonella enterica subsp. enterica serovar Montevideo Proteobacteria; Gammaproteobacteria; Enterobacterales; Enterobacteriaceae; Salmonella; Salmonella enterica; Salmonella enterica subsp. enterica MNNEETFYQAMRRKGVTRRSFLKFCSLAATSLGLGAGMTPKIAWALENKPRIPVVWIHGL WP_001058312.1 hydrogenase [Salmonella enterica] Length: 372\nNumber of predicted TMHs: 2\nExp number of AAs in TMHs: 36.51738\nExp number, first 60 AAs: 19.72849\nTotal prob of N-in: 0.96170\nPOSSIBLE N-term signal sequence\ninside 1 19\nTMhelix 20 39\noutside 40 325\nTMhelix 326 348\ninside 349 372

18558 GCF_002059705.1_ASM205970v1 Salmonella enterica subsp. enterica serovar Montevideo Proteobacteria; Gammaproteobacteria; Enterobacterales; Enterobacteriaceae; Salmonella; Salmonella enterica; Salmonella enterica subsp. enterica MTGDNTLITSHGINRRDFMKLCAALAATMGLSSKAAAEMAESVSNPQRPPVIWIGAQECT WP_000145429.1 hydrogenase 2 small subunit [Salmonella enterica] Length: 372\nNumber of predicted TMHs: 1\nExp number of AAs in TMHs: 24.51309\nExp number, first 60 AAs: 0.2682\nTotal prob of N-in: 0.14402\noutside 1 331\nTMhelix 332 354\ninside 355 372

18559 GCF_001478915.1_Salmonella_enterica_CVM_N30669-SQ_v1.0 Salmonella enterica subsp. enterica serovar Montevideo Proteobacteria; Gammaproteobacteria; Enterobacterales; Enterobacteriaceae; Salmonella; Salmonella enterica; Salmonella enterica subsp. enterica MNNEETFYQAMRRKGVTRRSFLKFCSLAATSLGLGAGMTPKIAWALENKPRIPVVWIHGL WP_001058312.1 hydrogenase [Salmonella enterica] Length: 372\nNumber of predicted TMHs: 2\nExp number of AAs in TMHs: 36.51738\nExp number, first 60 AAs: 19.72849\nTotal prob of N-in: 0.96170\nPOSSIBLE N-term signal sequence\ninside 1 19\nTMhelix 20 39\noutside 40 325\nTMhelix 326 348\ninside 349 372

18560 GCF_001480465.1_Salmonella_enterica_CVM_N42336-SQ_v1.0 Salmonella enterica subsp. enterica serovar Montevideo Proteobacteria; Gammaproteobacteria; Enterobacterales; Enterobacteriaceae; Salmonella; Salmonella enterica; Salmonella enterica subsp. enterica MTGDNTLITSHGINRRDFMKLCAALAATMGLSSKAAAEMAESVSNPQRPPVIWIGAQECT WP_000145429.1 hydrogenase 2 small subunit [Salmonella enterica] Length: 372\nNumber of predicted TMHs: 1\nExp number of AAs in TMHs: 24.51309\nExp number, first 60 AAs: 0.2682\nTotal prob of N-in: 0.14402\noutside 1 331\nTMhelix 332 354\ninside 355 372

18561 GCF_002061225.1_ASM206122v1 Salmonella enterica subsp. enterica serovar Montevideo Proteobacteria; Gammaproteobacteria; Enterobacterales; Enterobacteriaceae; Salmonella; Salmonella enterica; Salmonella enterica subsp. enterica MNNEETFYQAMRRKGVTRRSFLKFCSLAATSLGLGAGMTPKIAWALENKPRIPVVWIHGL WP_001058312.1 hydrogenase [Salmonella enterica] Length: 372\nNumber of predicted TMHs: 2\nExp number of AAs in TMHs: 36.51738\nExp number, first 60 AAs: 19.72849\nTotal prob of N-in: 0.96170\nPOSSIBLE N-term signal sequence\ninside 1 19\nTMhelix 20 39\noutside 40 325\nTMhelix 326 348\ninside 349 372

18562 GCF_002061225.1_ASM206122v1 Salmonella enterica subsp. enterica serovar Montevideo Proteobacteria; Gammaproteobacteria; Enterobacterales; Enterobacteriaceae; Salmonella; Salmonella enterica; Salmonella enterica subsp. enterica MTGDNTLITSHGINRRDFMKLCAALAATMGLSSKAAAEMAESVSNPQRPPVIWIGAQECT WP_000145429.1 hydrogenase 2 small subunit [Salmonella enterica] Length: 372\nNumber of predicted TMHs: 1\nExp number of AAs in TMHs: 24.51309\nExp number, first 60 AAs: 0.2682\nTotal prob of N-in: 0.14402\noutside 1 331\nTMhelix 332 354\ninside 355 372

18563 GCF_002034305.1_ASM203430v1 Salmonella enterica subsp. enterica serovar Montevideo Proteobacteria; Gammaproteobacteria; Enterobacterales; Enterobacteriaceae; Salmonella; Salmonella enterica; Salmonella enterica subsp. enterica MTGDNTLITSHGINRRDFMKLCAALAATMGLSSKAAAEMAESVSNPQRPPVIWIGAQECT WP_000145429.1 hydrogenase 2 small subunit [Salmonella enterica] Length: 372\nNumber of predicted TMHs: 1\nExp number of AAs in TMHs: 24.51309\nExp number, first 60 AAs: 0.2682\nTotal prob of N-in: 0.14402\noutside 1 331\nTMhelix 332 354\ninside 355 372

18564 GCF_001691975.1_ASM169197v1 Salmonella enterica subsp. enterica serovar Montevideo Proteobacteria; Gammaproteobacteria; Enterobacterales; Enterobacteriaceae; Salmonella; Salmonella enterica; Salmonella enterica subsp. enterica MNNEETFYQAMRRKGVTRRSFLKFCSLAATSLGLGAGMTPKIAWALENKPRIPVVWIHGL WP_001058312.1 hydrogenase [Salmonella enterica] Length: 372\nNumber of predicted TMHs: 2\nExp number of AAs in TMHs: 36.51738\nExp number, first 60 AAs: 19.72849\nTotal prob of N-in: 0.96170\nPOSSIBLE N-term signal sequence\ninside 1 19\nTMhelix 20 39\noutside 40 325\nTMhelix 326 348\ninside 349 372

18565 GCF_001480465.1_Salmonella_enterica_CVM_N42336-SQ_v1.0 Salmonella enterica subsp. enterica serovar Montevideo Proteobacteria; Gammaproteobacteria; Enterobacterales; Enterobacteriaceae; Salmonella; Salmonella enterica; Salmonella enterica subsp. enterica MNNEETFYQAMRRKGVTRRSFLKFCSLAATSLGLGAGMTPKIAWALENKPRIPVVWIHGL WP_001058312.1 hydrogenase [Salmonella enterica] Length: 372\nNumber of predicted TMHs: 2\nExp number of AAs in TMHs: 36.51738\nExp number, first 60 AAs: 19.72849\nTotal prob of N-in: 0.96170\nPOSSIBLE N-term signal sequence\ninside 1 19\nTMhelix 20 39\noutside 40 325\nTMhelix 326 348\ninside 349 372

18566 GCF_002059315.1_ASM205931v1 Salmonella enterica subsp. enterica serovar Montevideo Proteobacteria; Gammaproteobacteria; Enterobacterales; Enterobacteriaceae; Salmonella; Salmonella enterica; Salmonella enterica subsp. enterica MNNEETFYQAMRRKGVTRRSFLKFCSLAATSLGLGAGMTPKIAWALENKPRIPVVWIHGL WP_001058312.1 hydrogenase [Salmonella enterica] Length: 372\nNumber of predicted TMHs: 2\nExp number of AAs in TMHs: 36.51738\nExp number, first 60 AAs: 19.72849\nTotal prob of N-in: 0.96170\nPOSSIBLE N-term signal sequence\ninside 1 19\nTMhelix 20 39\noutside 40 325\nTMhelix 326 348\ninside 349 372

18567 GCF_002059315.1_ASM205931v1 Salmonella enterica subsp. enterica serovar Montevideo Proteobacteria; Gammaproteobacteria; Enterobacterales; Enterobacteriaceae; Salmonella; Salmonella enterica; Salmonella enterica subsp. enterica MTGDNTLITSHGINRRDFMKLCAALAATMGLSSKAAAEMAESVSNPQRPPVIWIGAQECT WP_000145429.1 hydrogenase 2 small subunit [Salmonella enterica] Length: 372\nNumber of predicted TMHs: 1\nExp number of AAs in TMHs: 24.51309\nExp number, first 60 AAs: 0.2682\nTotal prob of N-in: 0.14402\noutside 1 331\nTMhelix 332 354\ninside 355 372

18568 GCF_001691975.1_ASM169197v1 Salmonella enterica subsp. enterica serovar Montevideo Proteobacteria; Gammaproteobacteria; Enterobacterales; Enterobacteriaceae; Salmonella; Salmonella enterica; Salmonella enterica subsp. enterica MTGDNTLITSHGINRRDFMKLCAALAATMGLSSKAAAEMAESVSNPQRPPVIWIGAQECT WP_000145429.1 hydrogenase 2 small subunit [Salmonella enterica] Length: 372\nNumber of predicted TMHs: 1\nExp number of AAs in TMHs: 24.51309\nExp number, first 60 AAs: 0.2682\nTotal prob of N-in: 0.14402\noutside 1 331\nTMhelix 332 354\ninside 355 372

18569 GCF_002034305.1_ASM203430v1 Salmonella enterica subsp. enterica serovar Montevideo Proteobacteria; Gammaproteobacteria; Enterobacterales; Enterobacteriaceae; Salmonella; Salmonella enterica; Salmonella enterica subsp. enterica MNNEETFYQAMRRKGVTRRSFLKFCSLAATSLGLGAGMTPKIAWALENKPRIPVVWIHGL WP_001058312.1 hydrogenase [Salmonella enterica] Length: 372\nNumber of predicted TMHs: 2\nExp number of AAs in TMHs: 36.51738\nExp number, first 60 AAs: 19.72849\nTotal prob of N-in: 0.96170\nPOSSIBLE N-term signal sequence\ninside 1 19\nTMhelix 20 39\noutside 40 325\nTMhelix 326 348\ninside 349 372

18570 GCF_001478915.1_Salmonella_enterica_CVM_N30669-SQ_v1.0 Salmonella enterica subsp. enterica serovar Montevideo Proteobacteria; Gammaproteobacteria; Enterobacterales; Enterobacteriaceae; Salmonella; Salmonella enterica; Salmonella enterica subsp. enterica MTGDNTLITSHGINRRDFMKLCAALAATMGLSSKAAAEMAESVSNPQRPPVIWIGAQECT WP_000145429.1 hydrogenase 2 small subunit [Salmonella enterica] Length: 372\nNumber of predicted TMHs: 1\nExp number of AAs in TMHs: 24.51309\nExp number, first 60 AAs: 0.2682\nTotal prob of N-in: 0.14402\noutside 1 331\nTMhelix 332 354\ninside 355 372

18571 GCF_001272295.1_Salmonella_enterica_CVM_N50432_v1.0 Salmonella enterica subsp. enterica serovar Montevideo Proteobacteria; Gammaproteobacteria; Enterobacterales; Enterobacteriaceae; Salmonella; Salmonella enterica; Salmonella enterica subsp. enterica MTGDNTLITSHGINRRDFMKLCAALAATMGLSSKAAAEMAESVSNPQRPPVIWIGAQECT WP_000145429.1 hydrogenase 2 small subunit [Salmonella enterica] Length: 372\nNumber of predicted TMHs: 1\nExp number of AAs in TMHs: 24.51309\nExp number, first 60 AAs: 0.2682\nTotal prob of N-in: 0.14402\noutside 1 331\nTMhelix 332 354\ninside 355 372

18572 GCF_001246955.1_Salmonella_enterica_CVM_N51288_v1.0 Salmonella enterica subsp. enterica serovar Montevideo Proteobacteria; Gammaproteobacteria; Enterobacterales; Enterobacteriaceae; Salmonella; Salmonella enterica; Salmonella enterica subsp. enterica MTGDNTLITSHGINRRDFMKLCAALAATMGLSSKAAAEMAESVSNPQRPPVIWIGAQECT WP_000145429.1 hydrogenase 2 small subunit [Salmonella enterica] Length: 372\nNumber of predicted TMHs: 1\nExp number of AAs in TMHs: 24.51309\nExp number, first 60 AAs: 0.2682\nTotal prob of N-in: 0.14402\noutside 1 331\nTMhelix 332 354\ninside 355 372

18573 GCF_001245015.1_Salmonella_enterica_CVM_N48700_v1.0 Salmonella enterica subsp. enterica serovar Montevideo Proteobacteria; Gammaproteobacteria; Enterobacterales; Enterobacteriaceae; Salmonella; Salmonella enterica; Salmonella enterica subsp. enterica MTGDNTLITSHGINRRDFMKLCAALAATMGLSSKAAAEMAESVSNPQRPPVIWIGAQECT WP_000145429.1 hydrogenase 2 small subunit [Salmonella enterica] Length: 372\nNumber of predicted TMHs: 1\nExp number of AAs in TMHs: 24.51309\nExp number, first 60 AAs: 0.2682\nTotal prob of N-in: 0.14402\noutside 1 331\nTMhelix 332 354\ninside 355 372

18574 GCF_001245015.1_Salmonella_enterica_CVM_N48700_v1.0 Salmonella enterica subsp. enterica serovar Montevideo Proteobacteria; Gammaproteobacteria; Enterobacterales; Enterobacteriaceae; Salmonella; Salmonella enterica; Salmonella enterica subsp. enterica MNNEETFYQAMRRKGVTRRSFLKFCSLAATSLGLGAGMTPKIAWALENKPRIPVVWIHGL WP_001058312.1 hydrogenase [Salmonella enterica] Length: 372\nNumber of predicted TMHs: 2\nExp number of AAs in TMHs: 36.51738\nExp number, first 60 AAs: 19.72849\nTotal prob of N-in: 0.96170\nPOSSIBLE N-term signal sequence\ninside 1 19\nTMhelix 20 39\noutside 40 325\nTMhelix 326 348\ninside 349 372

18575 GCF_001245115.1_Salmonella_enterica_CVM_N48706_v1.0 Salmonella enterica subsp. enterica serovar Montevideo Proteobacteria; Gammaproteobacteria; Enterobacterales; Enterobacteriaceae; Salmonella; Salmonella enterica; Salmonella enterica subsp. enterica MNNEETFYQAMRRKGVTRRSFLKFCSLAATSLGLGAGMTPKIAWALENKPRIPVVWIHGL WP_001058312.1 hydrogenase [Salmonella enterica] Length: 372\nNumber of predicted TMHs: 2\nExp number of AAs in TMHs: 36.51738\nExp number, first 60 AAs: 19.72849\nTotal prob of N-in: 0.96170\nPOSSIBLE N-term signal sequence\ninside 1 19\nTMhelix 20 39\noutside 40 325\nTMhelix 326 348\ninside 349 372

18576 GCF_001276695.1_ASM127669v1 Salmonella enterica subsp. enterica serovar Montevideo Proteobacteria; Gammaproteobacteria; Enterobacterales; Enterobacteriaceae; Salmonella; Salmonella enterica; Salmonella enterica subsp. enterica MTGDNTLITSHGINRRDFMKLCAALAATMGLSSKAAAEMAESVSNPQRPPVIWIGAQECT WP_000145429.1 hydrogenase 2 small subunit [Salmonella enterica] Length: 372\nNumber of predicted TMHs: 1\nExp number of AAs in TMHs: 24.51309\nExp number, first 60 AAs: 0.2682\nTotal prob of N-in: 0.14402\noutside 1 331\nTMhelix 332 354\ninside 355 372

18577 GCF_001276905.1_ASM127690v1 Salmonella enterica subsp. enterica serovar Montevideo Proteobacteria; Gammaproteobacteria; Enterobacterales; Enterobacteriaceae; Salmonella; Salmonella enterica; Salmonella enterica subsp. enterica MNNEETFYQAMRRKGVTRRSFLKFCSLAATSLGLGAGMTPKIAWALENKPRIPVVWIHGL WP_001058312.1 hydrogenase [Salmonella enterica] Length: 372\nNumber of predicted TMHs: 2\nExp number of AAs in TMHs: 36.51738\nExp number, first 60 AAs: 19.72849\nTotal prob of N-in: 0.96170\nPOSSIBLE N-term signal sequence\ninside 1 19\nTMhelix 20 39\noutside 40 325\nTMhelix 326 348\ninside 349 372

18578 GCF_001276905.1_ASM127690v1 Salmonella enterica subsp. enterica serovar Montevideo Proteobacteria; Gammaproteobacteria; Enterobacterales; Enterobacteriaceae; Salmonella; Salmonella enterica; Salmonella enterica subsp. enterica MTGDNTLITSHGINRRDFMKLCAALAATMGLSSKAAAEMAESVSNPQRPPVIWIGAQECT WP_000145429.1 hydrogenase 2 small subunit [Salmonella enterica] Length: 372\nNumber of predicted TMHs: 1\nExp number of AAs in TMHs: 24.51309\nExp number, first 60 AAs: 0.2682\nTotal prob of N-in: 0.14402\noutside 1 331\nTMhelix 332 354\ninside 355 372

18579 GCF_001245115.1_Salmonella_enterica_CVM_N48706_v1.0 Salmonella enterica subsp. enterica serovar Montevideo Proteobacteria; Gammaproteobacteria; Enterobacterales; Enterobacteriaceae; Salmonella; Salmonella enterica; Salmonella enterica subsp. enterica MTGDNTLITSHGINRRDFMKLCAALAATMGLSSKAAAEMAESVSNPQRPPVIWIGAQECT WP_000145429.1 hydrogenase 2 small subunit [Salmonella enterica] Length: 372\nNumber of predicted TMHs: 1\nExp number of AAs in TMHs: 24.51309\nExp number, first 60 AAs: 0.2682\nTotal prob of N-in: 0.14402\noutside 1 331\nTMhelix 332 354\ninside 355 372

18580 GCF_001276695.1_ASM127669v1 Salmonella enterica subsp. enterica serovar Montevideo Proteobacteria; Gammaproteobacteria; Enterobacterales; Enterobacteriaceae; Salmonella; Salmonella enterica; Salmonella enterica subsp. enterica MNNEETFYQAMRRKGVTRRSFLKFCSLAATSLGLGAGMTPKIAWALENKPRIPVVWIHGL WP_001058312.1 hydrogenase [Salmonella enterica] Length: 372\nNumber of predicted TMHs: 2\nExp number of AAs in TMHs: 36.51738\nExp number, first 60 AAs: 19.72849\nTotal prob of N-in: 0.96170\nPOSSIBLE N-term signal sequence\ninside 1 19\nTMhelix 20 39\noutside 40 325\nTMhelix 326 348\ninside 349 372

18581 GCF_002062665.1_ASM206266v1 Salmonella enterica subsp. enterica serovar Montevideo Proteobacteria; Gammaproteobacteria; Enterobacterales; Enterobacteriaceae; Salmonella; Salmonella enterica; Salmonella enterica subsp. enterica MTGDNTLITSHGINRRDFMKLCAALAATMGLSSKAAAEMAESVSNPQRPPVIWIGAQECT WP_000145429.1 hydrogenase 2 small subunit [Salmonella enterica] Length: 372\nNumber of predicted TMHs: 1\nExp number of AAs in TMHs: 24.51309\nExp number, first 60 AAs: 0.2682\nTotal prob of N-in: 0.14402\noutside 1 331\nTMhelix 332 354\ninside 355 372

18582 GCF_002062665.1_ASM206266v1 Salmonella enterica subsp. enterica serovar Montevideo Proteobacteria; Gammaproteobacteria; Enterobacterales; Enterobacteriaceae; Salmonella; Salmonella enterica; Salmonella enterica subsp. enterica MNNEETFYQAMRRKGVTRRSFLKFCSLAATSLGLGAGMTPKIAWALENKPRIPVVWIHGL WP_001058312.1 hydrogenase [Salmonella enterica] Length: 372\nNumber of predicted TMHs: 2\nExp number of AAs in TMHs: 36.51738\nExp number, first 60 AAs: 19.72849\nTotal prob of N-in: 0.96170\nPOSSIBLE N-term signal sequence\ninside 1 19\nTMhelix 20 39\noutside 40 325\nTMhelix 326 348\ninside 349 372

18583 GCF_002065395.1_ASM206539v1 Salmonella enterica subsp. enterica serovar Montevideo Proteobacteria; Gammaproteobacteria; Enterobacterales; Enterobacteriaceae; Salmonella; Salmonella enterica; Salmonella enterica subsp. enterica MNNEETFYQAMRRKGVTRRSFLKFCSLAATSLGLGAGMTPKIAWALENKPRIPVVWIHGL WP_001058312.1 hydrogenase [Salmonella enterica] Length: 372\nNumber of predicted TMHs: 2\nExp number of AAs in TMHs: 36.51738\nExp number, first 60 AAs: 19.72849\nTotal prob of N-in: 0.96170\nPOSSIBLE N-term signal sequence\ninside 1 19\nTMhelix 20 39\noutside 40 325\nTMhelix 326 348\ninside 349 372

18584 GCF_002065395.1_ASM206539v1 Salmonella enterica subsp. enterica serovar Montevideo Proteobacteria; Gammaproteobacteria; Enterobacterales; Enterobacteriaceae; Salmonella; Salmonella enterica; Salmonella enterica subsp. enterica MTGDNTLITSHGINRRDFMKLCAALAATMGLSSKAAAEMAESVSNPQRPPVIWIGAQECT WP_000145429.1 hydrogenase 2 small subunit [Salmonella enterica] Length: 372\nNumber of predicted TMHs: 1\nExp number of AAs in TMHs: 24.51309\nExp number, first 60 AAs: 0.2682\nTotal prob of N-in: 0.14402\noutside 1 331\nTMhelix 332 354\ninside 355 372

18585 GCF_002064355.1_ASM206435v1 Salmonella enterica subsp. enterica serovar Muenchen Proteobacteria; Gammaproteobacteria; Enterobacterales; Enterobacteriaceae; Salmonella; Salmonella enterica; Salmonella enterica subsp. enterica MTGDNTLITSHGINRRDFMKLCAALAATMGLSSKAAAEMAESVSNPQRPPVIWIGAQECT WP_000145429.1 hydrogenase 2 small subunit [Salmonella enterica] Length: 372\nNumber of predicted TMHs: 1\nExp number of AAs in TMHs: 24.51309\nExp number, first 60 AAs: 0.2682\nTotal prob of N-in: 0.14402\noutside 1 331\nTMhelix 332 354\ninside 355 372

18586 GCF_002064355.1_ASM206435v1 Salmonella enterica subsp. enterica serovar Muenchen Proteobacteria; Gammaproteobacteria; Enterobacterales; Enterobacteriaceae; Salmonella; Salmonella enterica; Salmonella enterica subsp. enterica MNNEETFYQAMRRKGVTRRSFLKFCSLAATSLGLGAGMTPKIAWALENKPRIPVVWIHGL WP_001058311.1 hydrogenase [Salmonella enterica] Length: 372\nNumber of predicted TMHs: 2\nExp number of AAs in TMHs: 37.4699899999999\nExp number, first 60 AAs: 19.72699\nTotal prob of N-in: 0.96171\nPOSSIBLE N-term signal sequence\ninside 1 19\nTMhelix 20 39\noutside 40 325\nTMhelix 326 348\ninside 349 372

18587 GCF_001240705.1_Salmonella_enterica_CVM_N43482_v1.0 Salmonella enterica subsp. enterica serovar Muenchen Proteobacteria; Gammaproteobacteria; Enterobacterales; Enterobacteriaceae; Salmonella; Salmonella enterica; Salmonella enterica subsp. enterica MTGDNTLITSHGINRRDFMKLCAALAATMGLSSKAAAEMAESVSNPQRPPVIWIGAQECT WP_000145429.1 hydrogenase 2 small subunit [Salmonella enterica] Length: 372\nNumber of predicted TMHs: 1\nExp number of AAs in TMHs: 24.51309\nExp number, first 60 AAs: 0.2682\nTotal prob of N-in: 0.14402\noutside 1 331\nTMhelix 332 354\ninside 355 372

18588 GCF_001244895.1_Salmonella_enterica_CVM_N48696_v1.0 Salmonella enterica subsp. enterica serovar Muenchen Proteobacteria; Gammaproteobacteria; Enterobacterales; Enterobacteriaceae; Salmonella; Salmonella enterica; Salmonella enterica subsp. enterica MNNEETFYQAMRRKGVTRRSFLKFCSLAATSLGLGAGMTPKIAWALENKPRIPVVWIHGL WP_001058312.1 hydrogenase [Salmonella enterica] Length: 372\nNumber of predicted TMHs: 2\nExp number of AAs in TMHs: 36.51738\nExp number, first 60 AAs: 19.72849\nTotal prob of N-in: 0.96170\nPOSSIBLE N-term signal sequence\ninside 1 19\nTMhelix 20 39\noutside 40 325\nTMhelix 326 348\ninside 349 372

18589 GCF_001242505.1_Salmonella_enterica_CVM_N45942_v1.0 Salmonella enterica subsp. enterica serovar Muenchen Proteobacteria; Gammaproteobacteria; Enterobacterales; Enterobacteriaceae; Salmonella; Salmonella

enterica; Salmonella enterica subsp. enterica
MTGDNTLITSHGINRRDFMKLCAALAATMGLSSKAAAEMAESVSNPQRPPVIWIGAQECT WP_000145429.1
hydrogenase 2 small subunit [Salmonella enterica] Length: 372\nNumber of predicted TMHs: 1\nExp number of AAs in TMHs: 24.51309\nExp number, first 60 AAs: 0.2682\nTotal prob of N-in: 0.14402\noutside 1 331\nTMhelix 332 354\ninside 355 372

18590 GCF_002059935.1_ASM205993v1 Salmonella enterica subsp. enterica serovar Muenchen
Proteobacteria; Gammaproteobacteria; Enterobacterales; Enterobacteriaceae; Salmonella; Salmonella enterica; Salmonella enterica subsp. enterica MNNEETFYQAMRRKGVTRRSFLKFCSLAATSLGLGAGMTPKIAWALENKPRIPVVWIHGL
WP_001058311.1 hydrogenase [Salmonella enterica] Length: 372\nNumber of predicted TMHs: 2\nExp number of AAs in TMHs: 37.4699899999999\nExp number, first 60 AAs: 19.72699\nTotal prob of N-in: 0.96171\nPOSSIBLE N-term signal sequence\ninside 1 19\nTMhelix 20 39\noutside 40 325\nTMhelix 326 348\ninside 349 372

18591 GCF_001834155.1_ASM183415v1 Salmonella enterica subsp. enterica serovar Muenchen
Proteobacteria; Gammaproteobacteria; Enterobacterales; Enterobacteriaceae; Salmonella; Salmonella enterica; Salmonella enterica subsp. enterica MNNEETFYQAMRRKGVTRRSFLKFCSLAATSLGLGAGMTPKIAWALENKPRIPVVWIHGL
WP_001058312.1 hydrogenase [Salmonella enterica] Length: 372\nNumber of predicted TMHs: 2\nExp number of AAs in TMHs: 36.51738\nExp number, first 60 AAs: 19.72849\nTotal prob of N-in: 0.96170\nPOSSIBLE N-term signal sequence\ninside 1 19\nTMhelix 20 39\noutside 40 325\nTMhelix 326 348\ninside 349 372

18592 GCF_001247375.1_Salmonella_enterica_CVM_N51308_v1.0 Salmonella enterica subsp. enterica serovar Muenchen Proteobacteria; Gammaproteobacteria; Enterobacterales; Enterobacteriaceae; Salmonella; Salmonella enterica; Salmonella enterica subsp. enterica
MNNEETFYQAMRRKGVTRRSFLKFCSLAATSLGLGAGMTPKIAWALENKPRIPVVWIHGL WP_001058312.1
hydrogenase [Salmonella enterica] Length: 372\nNumber of predicted TMHs: 2\nExp number of AAs in TMHs: 36.51738\nExp number, first 60 AAs: 19.72849\nTotal prob of N-in: 0.96170\nPOSSIBLE N-term signal sequence\ninside 1 19\nTMhelix 20 39\noutside 40 325\nTMhelix 326 348\ninside 349 372

18593 GCF_001834045.1_ASM183404v1 Salmonella enterica subsp. enterica serovar Muenchen
Proteobacteria; Gammaproteobacteria; Enterobacterales; Enterobacteriaceae; Salmonella; Salmonella enterica; Salmonella enterica subsp. enterica MNNEETFYQAMRRKGVTRRSFLKFCSLAATSLGLGAGMTPKIAWALENKPRIPVVWIHGL
WP_070809206.1 hydrogenase [Salmonella enterica] Length: 372\nNumber of predicted TMHs: 2\nExp number of AAs in TMHs: 37.52613\nExp number, first 60 AAs: 19.72559\nTotal prob of N-in: 0.96174\nPOSSIBLE N-term signal sequence\ninside 1 19\nTMhelix 20 39\noutside 40 325\nTMhelix 326 348\ninside 349 372

18594 GCF_001834575.1_ASM183457v1 Salmonella enterica subsp. enterica serovar Muenchen
Proteobacteria; Gammaproteobacteria; Enterobacterales; Enterobacteriaceae; Salmonella; Salmonella enterica; Salmonella enterica subsp. enterica MNNEETFYQAMRRKGVTRRSFLKFCSLAATSLGLGAGMTPKIAWALENKPRIPVVWIHGL
WP_001058311.1 hydrogenase [Salmonella enterica] Length: 372\nNumber of predicted TMHs: 2\nExp number of AAs in TMHs: 37.4699899999999\nExp number, first 60 AAs: 19.72699\nTotal prob of N-in: 0.96171\nPOSSIBLE N-term signal sequence\ninside 1 19\nTMhelix 20 39\noutside 40 325\nTMhelix 326 348\ninside 349 372

18595 GCF_001242125.1_Salmonella_enterica_CVM_N45927_v1.0 Salmonella enterica subsp. enterica serovar Muenchen Proteobacteria; Gammaproteobacteria; Enterobacterales; Enterobacteriaceae; Salmonella; Salmonella enterica; Salmonella enterica subsp. enterica
MTGDNTLITSHGINRRDFMKLCAALAATMGLSSKAAAEMAESVSNPQRPPVIWIGAQECT WP_000145429.1
hydrogenase 2 small subunit [Salmonella enterica] Length: 372\nNumber of predicted TMHs: 1\nExp number of AAs in TMHs: 24.51309\nExp number, first 60 AAs: 0.2682\nTotal prob of N-in: 0.14402\noutside 1 331\nTMhelix 332 354\ninside 355 372

18596 GCF_001242505.1_Salmonella_enterica_CVM_N45942_v1.0 Salmonella enterica subsp. enterica serovar Muenchen Proteobacteria; Gammaproteobacteria; Enterobacterales; Enterobacteriaceae; Salmonella; Salmonella enterica; Salmonella enterica subsp. enterica
MNNEETFYQAMRRKGVTRRSFLKFCSLAATSLGLGAGMTPKIAWALENKPRIPVVWIHGL WP_001058312.1
hydrogenase [Salmonella enterica] Length: 372\nNumber of predicted TMHs: 2\nExp number of AAs in TMHs: 36.51738\nExp number, first 60 AAs: 19.72849\nTotal prob of N-in: 0.96170\nPOSSIBLE N-term signal sequence\ninside 1 19\nTMhelix 20 39\noutside 40 325\nTMhelix 326 348\ninside 349 372

18597 GCF_001834155.1_ASM183415v1 Salmonella enterica subsp. enterica serovar Muenchen Proteobacteria; Gammaproteobacteria; Enterobacterales; Enterobacteriaceae; Salmonella; Salmonella enterica; Salmonella enterica subsp. enterica MTGDNTLITSHGINRRDFMKLCAALAATMGLSSKAAAEMAESVSNPQRPPVIWIGAQECT WP_000145429.1 hydrogenase 2 small subunit [Salmonella enterica] Length: 372\nNumber of predicted TMHs: 1\nExp number of AAs in TMHs: 24.51309\nExp number, first 60 AAs: 0.2682\nTotal prob of N-in: 0.14402\noutside 1 331\nTMhelix 332 354\ninside 355 372

18598 GCF_001246885.1_Salmonella_enterica_CVM_N51284_v1.0 Salmonella enterica subsp. enterica serovar Muenchen Proteobacteria; Gammaproteobacteria; Enterobacterales; Enterobacteriaceae; Salmonella; Salmonella enterica; Salmonella enterica subsp. enterica MNNEETFYQAMRRKGVTRRSFLKFCSLAATSLGLGAGMTPKIAWALENKPRIPVVWIHGL WP_001058312.1 hydrogenase [Salmonella enterica] Length: 372\nNumber of predicted TMHs: 2\nExp number of AAs in TMHs: 36.51738\nExp number, first 60 AAs: 19.72849\nTotal prob of N-in: 0.96170\nPOSSIBLE N-term signal sequence\ninside 1 19\nTMhelix 20 39\noutside 40 325\nTMhelix 326 348\ninside 349 372

18599 GCF_001240705.1_Salmonella_enterica_CVM_N43482_v1.0 Salmonella enterica subsp. enterica serovar Muenchen Proteobacteria; Gammaproteobacteria; Enterobacterales; Enterobacteriaceae; Salmonella; Salmonella enterica; Salmonella enterica subsp. enterica MNNEETFYQAMRRKGVTRRSFLKFCSLAATSLGLGAGMTPKIAWALENKPRIPVVWIHGL WP_001058312.1 hydrogenase [Salmonella enterica] Length: 372\nNumber of predicted TMHs: 2\nExp number of AAs in TMHs: 36.51738\nExp number, first 60 AAs: 19.72849\nTotal prob of N-in: 0.96170\nPOSSIBLE N-term signal sequence\ninside 1 19\nTMhelix 20 39\noutside 40 325\nTMhelix 326 348\ninside 349 372

18600 GCF_001834725.1_ASM183472v1 Salmonella enterica subsp. enterica serovar Muenchen Proteobacteria; Gammaproteobacteria; Enterobacterales; Enterobacteriaceae; Salmonella; Salmonella enterica; Salmonella enterica subsp. enterica MTGDNTLITSHGINRRDFMKLCAALAATMGLSSKAAAEMAESVSNPQRPPVIWIGAQECT WP_000145429.1 hydrogenase 2 small subunit [Salmonella enterica] Length: 372\nNumber of predicted TMHs: 1\nExp number of AAs in TMHs: 24.51309\nExp number, first 60 AAs: 0.2682\nTotal prob of N-in: 0.14402\noutside 1 331\nTMhelix 332 354\ninside 355 372

18601 GCF_001240125.1_Salmonella_enterica_CVM_N43454_v1.0 Salmonella enterica subsp. enterica serovar Muenchen Proteobacteria; Gammaproteobacteria; Enterobacterales; Enterobacteriaceae; Salmonella; Salmonella enterica; Salmonella enterica subsp. enterica MNNEETFYQAMRRKGVTRRSFLKFCSLAATSLGLGAGMTPKIAWALENKPRIPVVWIHGL WP_001058312.1 hydrogenase [Salmonella enterica] Length: 372\nNumber of predicted TMHs: 2\nExp number of AAs in TMHs: 36.51738\nExp number, first 60 AAs: 19.72849\nTotal prob of N-in: 0.96170\nPOSSIBLE N-term signal sequence\ninside 1 19\nTMhelix 20 39\noutside 40 325\nTMhelix 326 348\ninside 349 372

18602 GCF_001241675.1_Salmonella_enterica_CVM_N45395_v1.0 Salmonella enterica subsp. enterica serovar Muenchen Proteobacteria; Gammaproteobacteria; Enterobacterales; Enterobacteriaceae; Salmonella; Salmonella enterica; Salmonella enterica subsp. enterica MNNEETFYQAMRRKGVTRRSFLKFCSLAATSLGLGAGMTPKIAWALENKPRIPVVWIHGL WP_001058312.1 hydrogenase [Salmonella enterica] Length: 372\nNumber of predicted TMHs: 2\nExp number of AAs in TMHs: 36.51738\nExp number, first 60 AAs: 19.72849\nTotal prob of N-in: 0.96170\nPOSSIBLE N-term signal sequence\ninside 1 19\nTMhelix 20 39\noutside 40 325\nTMhelix 326 348\ninside 349 372

18603 GCF_001243235.1_Salmonella_enterica_CVM_N46833_v1.0 Salmonella enterica subsp. enterica serovar Muenchen Proteobacteria; Gammaproteobacteria; Enterobacterales; Enterobacteriaceae; Salmonella; Salmonella enterica; Salmonella enterica subsp. enterica MNNEETFYQAMRRKGVTRRSFLKFCSLAATSLGLGAGMTPKIAWALENKPRIPVVWIHGL WP_001058312.1 hydrogenase [Salmonella enterica] Length: 372\nNumber of predicted TMHs: 2\nExp number of AAs in TMHs: 36.51738\nExp number, first 60 AAs: 19.72849\nTotal prob of N-in: 0.96170\nPOSSIBLE N-term signal sequence\ninside 1 19\nTMhelix 20 39\noutside 40 325\nTMhelix 326 348\ninside 349 372

18604 GCF_001243615.1_Salmonella_enterica_CVM_N46855_v1.0 Salmonella enterica subsp. enterica serovar Muenchen Proteobacteria; Gammaproteobacteria; Enterobacterales; Enterobacteriaceae; Salmonella; Salmonella enterica; Salmonella enterica subsp. enterica MTGDNTLITSHGINRRDFMKLCAALAATMGLSSKAAAEMAESVSNPQRPPVIWIGAQECT WP_000145429.1 hydrogenase 2 small subunit [Salmonella enterica] Length: 372\nNumber of predicted TMHs: 1\nExp number of AAs in TMHs: 24.51309\nExp number, first 60 AAs: 0.2682\nTotal prob of N-in: 0.14402\noutside 1 331\nTMhelix 332 354\ninside 355 372

18605 GCF_001243615.1_Salmonella_enterica_CVM_N46855_v1.0 Salmonella enterica subsp. enterica serovar Muenchen Proteobacteria; Gammaproteobacteria; Enterobacterales; Enterobacteriaceae; Salmonella; Salmonella enterica; Salmonella enterica subsp. enterica
MNNEETFYQAMRRKGVTRRSFLKFCSLAATSLGLGAGMTPKIAWALENKPRIPVVWIHGL WP_001058312.1
hydrogenase [Salmonella enterica] Length: 372\nNumber of predicted TMHs: 2\nExp number of AAs in TMHs: 36.51738\nExp number, first 60 AAs: 19.72849\nTotal prob of N-in: 0.96170\nPOSSIBLE N-term signal sequence\ninside 1 19\nTMhelix 20 39\noutside 40 325\nTMhelix 326 348\ninside 349 372

18606 GCF_001241675.1_Salmonella_enterica_CVM_N45395_v1.0 Salmonella enterica subsp. enterica serovar Muenchen Proteobacteria; Gammaproteobacteria; Enterobacterales; Enterobacteriaceae; Salmonella; Salmonella enterica; Salmonella enterica subsp. enterica
MTGDNTLITSHGINRRDFMKLCAALAATMGLSSKAAAEMAESVSNPQRPPVIWIGAQECT WP_000145429.1
hydrogenase 2 small subunit [Salmonella enterica] Length: 372\nNumber of predicted TMHs: 1\nExp number of AAs in TMHs: 24.51309\nExp number, first 60 AAs: 0.2682\nTotal prob of N-in: 0.14402\noutside 1 331\nTMhelix 332 354\ninside 355 372

18607 GCF_001245945.1_Salmonella_enterica_CVM_N51242_v1.0 Salmonella enterica subsp. enterica serovar Muenchen Proteobacteria; Gammaproteobacteria; Enterobacterales; Enterobacteriaceae; Salmonella; Salmonella enterica; Salmonella enterica subsp. enterica
MNNEETFYQAMRRKGVTRRSFLKFCSLAATSLGLGAGMTPKIAWALENKPRIPVVWIHGL WP_001058312.1
hydrogenase [Salmonella enterica] Length: 372\nNumber of predicted TMHs: 2\nExp number of AAs in TMHs: 36.51738\nExp number, first 60 AAs: 19.72849\nTotal prob of N-in: 0.96170\nPOSSIBLE N-term signal sequence\ninside 1 19\nTMhelix 20 39\noutside 40 325\nTMhelix 326 348\ninside 349 372

18608 GCF_001245945.1_Salmonella_enterica_CVM_N51242_v1.0 Salmonella enterica subsp. enterica serovar Muenchen Proteobacteria; Gammaproteobacteria; Enterobacterales; Enterobacteriaceae; Salmonella; Salmonella enterica; Salmonella enterica subsp. enterica
MTGDNTLITSHGINRRDFMKLCAALAATMGLSSKAAAEMAESVSNPQRPPVIWIGAQECT WP_000145429.1
hydrogenase 2 small subunit [Salmonella enterica] Length: 372\nNumber of predicted TMHs: 1\nExp number of AAs in TMHs: 24.51309\nExp number, first 60 AAs: 0.2682\nTotal prob of N-in: 0.14402\noutside 1 331\nTMhelix 332 354\ninside 355 372

18609 GCF_001245635.1_Salmonella_enterica_CVM_N50441_v1.0 Salmonella enterica subsp. enterica serovar Muenchen Proteobacteria; Gammaproteobacteria; Enterobacterales; Enterobacteriaceae; Salmonella; Salmonella enterica; Salmonella enterica subsp. enterica
MNNEETFYQAMRRKGVTRRSFLKFCSLAATSLGLGAGMTPKIAWALENKPRIPVVWIHGL WP_001058312.1
hydrogenase [Salmonella enterica] Length: 372\nNumber of predicted TMHs: 2\nExp number of AAs in TMHs: 36.51738\nExp number, first 60 AAs: 19.72849\nTotal prob of N-in: 0.96170\nPOSSIBLE N-term signal sequence\ninside 1 19\nTMhelix 20 39\noutside 40 325\nTMhelix 326 348\ninside 349 372

18610 GCF_001245635.1_Salmonella_enterica_CVM_N50441_v1.0 Salmonella enterica subsp. enterica serovar Muenchen Proteobacteria; Gammaproteobacteria; Enterobacterales; Enterobacteriaceae; Salmonella; Salmonella enterica; Salmonella enterica subsp. enterica
MTGDNTLITSHGINRRDFMKLCAALAATMGLSSKAAAEMAESVSNPQRPPVIWIGAQECT WP_000145429.1
hydrogenase 2 small subunit [Salmonella enterica] Length: 372\nNumber of predicted TMHs: 1\nExp number of AAs in TMHs: 24.51309\nExp number, first 60 AAs: 0.2682\nTotal prob of N-in: 0.14402\noutside 1 331\nTMhelix 332 354\ninside 355 372

18611 GCF_001240125.1_Salmonella_enterica_CVM_N43454_v1.0 Salmonella enterica subsp. enterica serovar Muenchen Proteobacteria; Gammaproteobacteria; Enterobacterales; Enterobacteriaceae; Salmonella; Salmonella enterica; Salmonella enterica subsp. enterica
MTGDNTLITSHGINRRDFMKLCAALAATMGLSSKAAAEMAESVSNPQRPPVIWIGAQECT WP_000145429.1
hydrogenase 2 small subunit [Salmonella enterica] Length: 372\nNumber of predicted TMHs: 1\nExp number of AAs in TMHs: 24.51309\nExp number, first 60 AAs: 0.2682\nTotal prob of N-in: 0.14402\noutside 1 331\nTMhelix 332 354\ninside 355 372

18612 GCF_001272185.1_Salmonella_enterica_CVM_N48703_v1.0 Salmonella enterica subsp. enterica serovar Muenchen Proteobacteria; Gammaproteobacteria; Enterobacterales; Enterobacteriaceae; Salmonella; Salmonella enterica; Salmonella enterica subsp. enterica
MTGDNTLITSHGINRRDFMKLCAALAATMGLSSKAAAEMAESVSNPQRPPVIWIGAQECT WP_000145429.1
hydrogenase 2 small subunit [Salmonella enterica] Length: 372\nNumber of predicted TMHs: 1\nExp

number of AAs in TMHs: 24.51309\nExp number, first 60 AAs: 0.2682\nTotal prob of N-in: 0.14402\noutside 1 331\nTMhelix 332 354\ninside 355 372

18613 GCF_001247595.1_Salmonella_enterica_CVM_N51983_v1.0 Salmonella enterica subsp. enterica serovar Muenchen Proteobacteria; Gammaproteobacteria; Enterobacterales; Enterobacteriaceae; Salmonella; Salmonella enterica; Salmonella enterica subsp. enterica MNNEETFYQAMRRKGVTRRSFLKFCSLAATSLGLGAGMTPKIAWALENKPRIPVVWIHGL WP_001058312.1 hydrogenase [Salmonella enterica] Length: 372\nNumber of predicted TMHs: 2\nExp number of AAs in TMHs: 36.51738\nExp number, first 60 AAs: 19.72849\nTotal prob of N-in: 0.96170\nPOSSIBLE N-term signal sequence\ninside 1 19\nTMhelix 20 39\noutside 40 325\nTMhelix 326 348\ninside 349 372

18614 GCF_001247595.1_Salmonella_enterica_CVM_N51983_v1.0 Salmonella enterica subsp. enterica serovar Muenchen Proteobacteria; Gammaproteobacteria; Enterobacterales; Enterobacteriaceae; Salmonella; Salmonella enterica; Salmonella enterica subsp. enterica MTGDNTLITSHGINRRDFMKLCAALAATMGLSSKAAAEMAESVSNPQRPPVIWIGAQECT WP_000145429.1 hydrogenase 2 small subunit [Salmonella enterica] Length: 372\nNumber of predicted TMHs: 1\nExp number of AAs in TMHs: 24.51309\nExp number, first 60 AAs: 0.2682\nTotal prob of N-in: 0.14402\noutside 1 331\nTMhelix 332 354\ninside 355 372

18615 GCF_001272185.1_Salmonella_enterica_CVM_N48703_v1.0 Salmonella enterica subsp. enterica serovar Muenchen Proteobacteria; Gammaproteobacteria; Enterobacterales; Enterobacteriaceae; Salmonella; Salmonella enterica; Salmonella enterica subsp. enterica MNNEETFYQAMRRKGVTRRSFLKFCSLAATSLGLGAGMTPKIAWALENKPRIPVVWIHGL WP_001058312.1 hydrogenase [Salmonella enterica] Length: 372\nNumber of predicted TMHs: 2\nExp number of AAs in TMHs: 36.51738\nExp number, first 60 AAs: 19.72849\nTotal prob of N-in: 0.96170\nPOSSIBLE N-term signal sequence\ninside 1 19\nTMhelix 20 39\noutside 40 325\nTMhelix 326 348\ninside 349 372

18616 GCF_001243235.1_Salmonella_enterica_CVM_N46833_v1.0 Salmonella enterica subsp. enterica serovar Muenchen Proteobacteria; Gammaproteobacteria; Enterobacterales; Enterobacteriaceae; Salmonella; Salmonella enterica; Salmonella enterica subsp. enterica MTGDNTLITSHGINRRDFMKLCAALAATMGLSSKAAAEMAESVSNPQRPPVIWIGAQECT WP_000145429.1 hydrogenase 2 small subunit [Salmonella enterica] Length: 372\nNumber of predicted TMHs: 1\nExp number of AAs in TMHs: 24.51309\nExp number, first 60 AAs: 0.2682\nTotal prob of N-in: 0.14402\noutside 1 331\nTMhelix 332 354\ninside 355 372

18617 GCF_001692085.1_ASM169208v1 Salmonella enterica subsp. enterica serovar Muenchen Proteobacteria; Gammaproteobacteria; Enterobacterales; Enterobacteriaceae; Salmonella; Salmonella enterica; Salmonella enterica subsp. enterica MTGDNTLITSHGINRRDFMKLCAALAATMGLSSKAAAEMAESVSNPQRPPVIWIGAQECT WP_000145429.1 hydrogenase 2 small subunit [Salmonella enterica] Length: 372\nNumber of predicted TMHs: 1\nExp number of AAs in TMHs: 24.51309\nExp number, first 60 AAs: 0.2682\nTotal prob of N-in: 0.14402\noutside 1 331\nTMhelix 332 354\ninside 355 372

18618 GCF_002031245.1_ASM203124v1 Salmonella enterica subsp. enterica serovar Muenchen Proteobacteria; Gammaproteobacteria; Enterobacterales; Enterobacteriaceae; Salmonella; Salmonella enterica; Salmonella enterica subsp. enterica MNNEETFYQAMRRKGVTRRSFLKFCSLAATSLGLGAGMTPKIAWALENKPRIPVVWIHGL WP_001058312.1 hydrogenase [Salmonella enterica] Length: 372\nNumber of predicted TMHs: 2\nExp number of AAs in TMHs: 36.51738\nExp number, first 60 AAs: 19.72849\nTotal prob of N-in: 0.96170\nPOSSIBLE N-term signal sequence\ninside 1 19\nTMhelix 20 39\noutside 40 325\nTMhelix 326 348\ninside 349 372

18619 GCF_002059525.1_ASM205952v1 Salmonella enterica subsp. enterica serovar Muenchen Proteobacteria; Gammaproteobacteria; Enterobacterales; Enterobacteriaceae; Salmonella; Salmonella enterica; Salmonella enterica subsp. enterica MTGDNTLITSHGINRRDFMKLCAALAATMGLSSKAAAEMAESVSNPQRPPVIWIGAQECT WP_000145429.1 hydrogenase 2 small subunit [Salmonella enterica] Length: 372\nNumber of predicted TMHs: 1\nExp number of AAs in TMHs: 24.51309\nExp number, first 60 AAs: 0.2682\nTotal prob of N-in: 0.14402\noutside 1 331\nTMhelix 332 354\ninside 355 372

18620 GCF_002059525.1_ASM205952v1 Salmonella enterica subsp. enterica serovar Muenchen Proteobacteria; Gammaproteobacteria; Enterobacterales; Enterobacteriaceae; Salmonella; Salmonella enterica; Salmonella enterica subsp. enterica MNNEETFYQAMRRKGVTRRSFLKFCSLAATSLGLGAGMTPKIAWALENKPRIPVVWIHGL WP_001058312.1 hydrogenase [Salmonella enterica] Length: 372\nNumber of predicted TMHs: 2\nExp

number of AAs in TMHs: 36.51738\nExp number, first 60 AAs:  19.72849\nTotal prob of N-in:   0.96170\nPOSSIBLE N-term signal sequence\ninside   1   19\nTMhelix   20   39\noutside   40   325\nTMhelix   326   348\ninside   349   372

18621 GCF_002059695.1_ASM205969v1 Salmonella enterica subsp. enterica serovar Muenchen Proteobacteria; Gammaproteobacteria; Enterobacterales; Enterobacteriaceae; Salmonella; Salmonella enterica; Salmonella enterica subsp. enterica MNNEETFYQAMRRKGVTRRSFLKFCSLAATSLGLGAGMTPKIAWALENKPRIPVVWIHGL WP_001058312.1 hydrogenase [Salmonella enterica] Length: 372\nNumber of predicted TMHs:  2\nExp number of AAs in TMHs: 36.51738\nExp number, first 60 AAs:  19.72849\nTotal prob of N-in:   0.96170\nPOSSIBLE N-term signal sequence\ninside   1   19\nTMhelix   20   39\noutside   40   325\nTMhelix   326   348\ninside   349   372

18622 GCF_002065415.1_ASM206541v1 Salmonella enterica subsp. enterica serovar Muenchen Proteobacteria; Gammaproteobacteria; Enterobacterales; Enterobacteriaceae; Salmonella; Salmonella enterica; Salmonella enterica subsp. enterica MTGDNTLITSHGINRRDFMKLCAALAATMGLSSKAAAEMAESVSNPQRPPVIWIGAQECT WP_000145429.1 hydrogenase 2 small subunit [Salmonella enterica] Length: 372\nNumber of predicted TMHs:  1\nExp number of AAs in TMHs: 24.51309\nExp number, first 60 AAs:  0.2682\nTotal prob of N-in: 0.14402\noutside   1   331\nTMhelix   332   354\ninside   355   372

18623 GCF_002064365.1_ASM206436v1 Salmonella enterica subsp. enterica serovar Muenchen Proteobacteria; Gammaproteobacteria; Enterobacterales; Enterobacteriaceae; Salmonella; Salmonella enterica; Salmonella enterica subsp. enterica MNNEETFYQAMRRKGVTRRSFLKFCSLAATSLGLGAGMTPKIAWALENKPRIPVVWIHGL WP_001058311.1 hydrogenase [Salmonella enterica] Length: 372\nNumber of predicted TMHs:  2\nExp number of AAs in TMHs: 37.4699899999999\nExp number, first 60 AAs:  19.72699\nTotal prob of N-in: 0.96171\nPOSSIBLE N-term signal sequence\ninside   1   19\nTMhelix   20   39\noutside   40   325\nTMhelix   326 348\ninside   349   372

18624 GCF_001833235.1_ASM183323v1 Salmonella enterica subsp. enterica serovar Muenchen Proteobacteria; Gammaproteobacteria; Enterobacterales; Enterobacteriaceae; Salmonella; Salmonella enterica; Salmonella enterica subsp. enterica MNNEETFYQAMRRKGVTRRSFLKFCSLAATSLGLGAGMTPKIAWALENKPRIPVVWIHGL WP_001058311.1 hydrogenase [Salmonella enterica] Length: 372\nNumber of predicted TMHs:  2\nExp number of AAs in TMHs: 37.4699899999999\nExp number, first 60 AAs:  19.72699\nTotal prob of N-in: 0.96171\nPOSSIBLE N-term signal sequence\ninside   1   19\nTMhelix   20   39\noutside   40   325\nTMhelix   326 348\ninside   349   372

18625 GCF_002064365.1_ASM206436v1 Salmonella enterica subsp. enterica serovar Muenchen Proteobacteria; Gammaproteobacteria; Enterobacterales; Enterobacteriaceae; Salmonella; Salmonella enterica; Salmonella enterica subsp. enterica MTGDNTLITSHGINRRDFMKLCAALAATMGLSSKAAAEMAESVSNPQRPPVIWIGAQECT WP_000145429.1 hydrogenase 2 small subunit [Salmonella enterica] Length: 372\nNumber of predicted TMHs:  1\nExp number of AAs in TMHs: 24.51309\nExp number, first 60 AAs:  0.2682\nTotal prob of N-in: 0.14402\noutside   1   331\nTMhelix   332   354\ninside   355   372

18626 GCF_002065415.1_ASM206541v1 Salmonella enterica subsp. enterica serovar Muenchen Proteobacteria; Gammaproteobacteria; Enterobacterales; Enterobacteriaceae; Salmonella; Salmonella enterica; Salmonella enterica subsp. enterica MNNEETFYQAMRRKGVTRRSFLKFCSLAATSLGLGAGMTPKIAWALENKPRIPVVWIHGL WP_001058311.1 hydrogenase [Salmonella enterica] Length: 372\nNumber of predicted TMHs:  2\nExp number of AAs in TMHs: 37.4699899999999\nExp number, first 60 AAs:  19.72699\nTotal prob of N-in: 0.96171\nPOSSIBLE N-term signal sequence\ninside   1   19\nTMhelix   20   39\noutside   40   325\nTMhelix   326 348\ninside   349   372

18627 GCF_001834365.1_ASM183436v1 Salmonella enterica subsp. enterica serovar Muenchen Proteobacteria; Gammaproteobacteria; Enterobacterales; Enterobacteriaceae; Salmonella; Salmonella enterica; Salmonella enterica subsp. enterica MTGDNTLITSHGINRRDFMKLCAALAATMGLSSKAAAEMAESVSNPQRPPVIWIGAQECT WP_000145429.1 hydrogenase 2 small subunit [Salmonella enterica] Length: 372\nNumber of predicted TMHs:  1\nExp number of AAs in TMHs: 24.51309\nExp number, first 60 AAs:  0.2682\nTotal prob of N-in: 0.14402\noutside   1   331\nTMhelix   332   354\ninside   355   372

18628 GCF_001834365.1_ASM183436v1 Salmonella enterica subsp. enterica serovar Muenchen Proteobacteria; Gammaproteobacteria; Enterobacterales; Enterobacteriaceae; Salmonella; Salmonella enterica; Salmonella enterica subsp. enterica MNNEETFYQAMRRKGVTRRSFLKFCSLAATSLGLGAGMTPKIAWALENKPRIPVVWIHGL WP_001058311.1 hydrogenase [Salmonella enterica] Length: 372\nNumber of predicted TMHs:  2\nExp number of AAs in TMHs: 37.4699899999999\nExp number, first 60 AAs:  19.72699\nTotal prob of N-in:

0.96171\nPOSSIBLE N-term signal sequence\ninside 1 19\nTMhelix 20 39\noutside 40 325\nTMhelix 326 348\ninside 349 372

18629 GCF_002059695.1_ASM205969v1 Salmonella enterica subsp. enterica serovar Muenchen Proteobacteria; Gammaproteobacteria; Enterobacterales; Enterobacteriaceae; Salmonella; Salmonella enterica; Salmonella enterica subsp. enterica MTGDNTLITSHGINRRDFMKLCAALAATMGLSSKAAAEMAESVSNPQRPPVIWIGAQECT WP_000145429.1 hydrogenase 2 small subunit [Salmonella enterica] Length: 372\nNumber of predicted TMHs: 1\nExp number of AAs in TMHs: 24.51309\nExp number, first 60 AAs: 0.2682\nTotal prob of N-in: 0.14402\noutside 1 331\nTMhelix 332 354\ninside 355 372

18630 GCF_001692005.1_ASM169200v1 Salmonella enterica subsp. enterica serovar Muenchen Proteobacteria; Gammaproteobacteria; Enterobacterales; Enterobacteriaceae; Salmonella; Salmonella enterica; Salmonella enterica subsp. enterica MNNEETFYQAMRRKGVTRRSFLKFCSLAATSLGLGAGMTPKIAWALENKPRIPVVWIHGL WP_001058311.1 hydrogenase [Salmonella enterica] Length: 372\nNumber of predicted TMHs: 2\nExp number of AAs in TMHs: 37.4699899999999\nExp number, first 60 AAs: 19.72699\nTotal prob of N-in: 0.96171\nPOSSIBLE N-term signal sequence\ninside 1 19\nTMhelix 20 39\noutside 40 325\nTMhelix 326 348\ninside 349 372

18631 GCF_001834085.1_ASM183408v1 Salmonella enterica subsp. enterica serovar Muenchen Proteobacteria; Gammaproteobacteria; Enterobacterales; Enterobacteriaceae; Salmonella; Salmonella enterica; Salmonella enterica subsp. enterica MTGDNTLITSHGINRRDFMKLCAALAATMGLSSKAAAEMAESVSNPQRPPVIWIGAQECT WP_000145429.1 hydrogenase 2 small subunit [Salmonella enterica] Length: 372\nNumber of predicted TMHs: 1\nExp number of AAs in TMHs: 24.51309\nExp number, first 60 AAs: 0.2682\nTotal prob of N-in: 0.14402\noutside 1 331\nTMhelix 332 354\ninside 355 372

18632 GCF_001244895.1_Salmonella_enterica_CVM_N48696_v1.0 Salmonella enterica subsp. enterica serovar Muenchen Proteobacteria; Gammaproteobacteria; Enterobacterales; Enterobacteriaceae; Salmonella; Salmonella enterica; Salmonella enterica subsp. enterica MTGDNTLITSHGINRRDFMKLCAALAATMGLSSKAAAEMAESVSNPQRPPVIWIGAQECT WP_000145429.1 hydrogenase 2 small subunit [Salmonella enterica] Length: 372\nNumber of predicted TMHs: 1\nExp number of AAs in TMHs: 24.51309\nExp number, first 60 AAs: 0.2682\nTotal prob of N-in: 0.14402\noutside 1 331\nTMhelix 332 354\ninside 355 372

18633 GCF_001481325.1_Salmonella_enterica_CVM_N42480_v1.0 Salmonella enterica subsp. enterica serovar Muenchen Proteobacteria; Gammaproteobacteria; Enterobacterales; Enterobacteriaceae; Salmonella; Salmonella enterica; Salmonella enterica subsp. enterica MNNEETFYQAMRRKGVTRRSFLKFCSLAATSLGLGAGMTPKIAWALENKPRIPVVWIHGL WP_001058312.1 hydrogenase [Salmonella enterica] Length: 372\nNumber of predicted TMHs: 2\nExp number of AAs in TMHs: 36.51738\nExp number, first 60 AAs: 19.72849\nTotal prob of N-in: 0.96170\nPOSSIBLE N-term signal sequence\ninside 1 19\nTMhelix 20 39\noutside 40 325\nTMhelix 326 348\ninside 349 372

18634 GCF_001692005.1_ASM169200v1 Salmonella enterica subsp. enterica serovar Muenchen Proteobacteria; Gammaproteobacteria; Enterobacterales; Enterobacteriaceae; Salmonella; Salmonella enterica; Salmonella enterica subsp. enterica MTGDNTLITSHGINRRDFMKLCAALAATMGLSSKAAAEMAESVSNPQRPPVIWIGAQECT WP_000145429.1 hydrogenase 2 small subunit [Salmonella enterica] Length: 372\nNumber of predicted TMHs: 1\nExp number of AAs in TMHs: 24.51309\nExp number, first 60 AAs: 0.2682\nTotal prob of N-in: 0.14402\noutside 1 331\nTMhelix 332 354\ninside 355 372

18635 GCF_001834045.1_ASM183404v1 Salmonella enterica subsp. enterica serovar Muenchen Proteobacteria; Gammaproteobacteria; Enterobacterales; Enterobacteriaceae; Salmonella; Salmonella enterica; Salmonella enterica subsp. enterica MTGDNTLITSHGINRRDFMKLCAALAATMGLSSKAAAEMAESVSNPQRPPVIWIGAQECT WP_000145429.1 hydrogenase 2 small subunit [Salmonella enterica] Length: 372\nNumber of predicted TMHs: 1\nExp number of AAs in TMHs: 24.51309\nExp number, first 60 AAs: 0.2682\nTotal prob of N-in: 0.14402\noutside 1 331\nTMhelix 332 354\ninside 355 372

18636 GCF_001834325.1_ASM183432v1 Salmonella enterica subsp. enterica serovar Muenchen Proteobacteria; Gammaproteobacteria; Enterobacterales; Enterobacteriaceae; Salmonella; Salmonella enterica; Salmonella enterica subsp. enterica MNNEETFYQAMRRKGVTRRSFLKFCSLAATSLGLGAGMTPKIAWALENKPRIPVVWIHGL WP_001058311.1 hydrogenase [Salmonella enterica] Length: 372\nNumber of predicted TMHs: 2\nExp number of AAs in TMHs: 37.4699899999999\nExp number, first 60 AAs: 19.72699\nTotal prob of N-in:

0.96171\nPOSSIBLE N-term signal sequence\ninside 1 19\nTMhelix 20 39\noutside 40 325\nTMhelix 326 348\ninside 349 372

18637 GCF_001481325.1_Salmonella_enterica_CVM_N42480_v1.0 Salmonella enterica subsp. enterica serovar Muenchen Proteobacteria; Gammaproteobacteria; Enterobacterales; Enterobacteriaceae; Salmonella; Salmonella enterica; Salmonella enterica subsp. enterica MTGDNTLITSHGINRRDFMKLCAALAATMGLSSKAAAEMAESVSNPQRPPVIWIGAQECT WP_000145429.1 hydrogenase 2 small subunit [Salmonella enterica] Length: 372\nNumber of predicted TMHs: 1\nExp number of AAs in TMHs: 24.51309\nExp number, first 60 AAs: 0.2682\nTotal prob of N-in: 0.14402\noutside 1 331\nTMhelix 332 354\ninside 355 372

18638 GCF_001240835.1_Salmonella_enterica_CVM_N43826_v1.0 Salmonella enterica subsp. enterica serovar Muenchen Proteobacteria; Gammaproteobacteria; Enterobacterales; Enterobacteriaceae; Salmonella; Salmonella enterica; Salmonella enterica subsp. enterica MNNEETFYQAMRRKGVTRRSFLKFCSLAATSLGLGAGMTPKIAWALENKPRIPVVWIHGL WP_001058312.1 hydrogenase [Salmonella enterica] Length: 372\nNumber of predicted TMHs: 2\nExp number of AAs in TMHs: 36.51738\nExp number, first 60 AAs: 19.72849\nTotal prob of N-in: 0.96170\nPOSSIBLE N-term signal sequence\ninside 1 19\nTMhelix 20 39\noutside 40 325\nTMhelix 326 348\ninside 349 372

18639 GCF_002059935.1_ASM205993v1 Salmonella enterica subsp. enterica serovar Muenchen Proteobacteria; Gammaproteobacteria; Enterobacterales; Enterobacteriaceae; Salmonella; Salmonella enterica; Salmonella enterica subsp. enterica MTGDNTLITSHGINRRDFMKLCAALAATMGLSSKAAAEMAESVSNPQRPPVIWIGAQECT WP_000145429.1 hydrogenase 2 small subunit [Salmonella enterica] Length: 372\nNumber of predicted TMHs: 1\nExp number of AAs in TMHs: 24.51309\nExp number, first 60 AAs: 0.2682\nTotal prob of N-in: 0.14402\noutside 1 331\nTMhelix 332 354\ninside 355 372

18640 GCF_001479965.1_Salmonella_enterica_CVM_N38885-SQ_v1.0 Salmonella enterica subsp. enterica serovar Muenchen Proteobacteria; Gammaproteobacteria; Enterobacterales; Enterobacteriaceae; Salmonella; Salmonella enterica; Salmonella enterica subsp. enterica MTGDNTLITSHGINRRDFMKLCAALAATMGLSSKAAAEMAESVSNPQRPPVIWIGAQECT WP_000145429.1 hydrogenase 2 small subunit [Salmonella enterica] Length: 372\nNumber of predicted TMHs: 1\nExp number of AAs in TMHs: 24.51309\nExp number, first 60 AAs: 0.2682\nTotal prob of N-in: 0.14402\noutside 1 331\nTMhelix 332 354\ninside 355 372

18641 GCF_001834725.1_ASM183472v1 Salmonella enterica subsp. enterica serovar Muenchen Proteobacteria; Gammaproteobacteria; Enterobacterales; Enterobacteriaceae; Salmonella; Salmonella enterica; Salmonella enterica subsp. enterica MNNEETFYQAMRRKGVTRRSFLKFCSLAATSLGLGAGMTPKIAWALENKPRIPVVWIHGL WP_001058311.1 hydrogenase [Salmonella enterica] Length: 372\nNumber of predicted TMHs: 2\nExp number of AAs in TMHs: 37.4699899999999\nExp number, first 60 AAs: 19.72699\nTotal prob of N-in: 0.96171\nPOSSIBLE N-term signal sequence\ninside 1 19\nTMhelix 20 39\noutside 40 325\nTMhelix 326 348\ninside 349 372

18642 GCF_001246885.1_Salmonella_enterica_CVM_N51284_v1.0 Salmonella enterica subsp. enterica serovar Muenchen Proteobacteria; Gammaproteobacteria; Enterobacterales; Enterobacteriaceae; Salmonella; Salmonella enterica; Salmonella enterica subsp. enterica MTGDNTLITSHGINRRDFMKLCAALAATMGLSSKAAAEMAESVSNPQRPPVIWIGAQECT WP_000145429.1 hydrogenase 2 small subunit [Salmonella enterica] Length: 372\nNumber of predicted TMHs: 1\nExp number of AAs in TMHs: 24.51309\nExp number, first 60 AAs: 0.2682\nTotal prob of N-in: 0.14402\noutside 1 331\nTMhelix 332 354\ninside 355 372

18643 GCF_001240345.1_Salmonella_enterica_CVM_N43460_v1.0 Salmonella enterica subsp. enterica serovar Muenchen Proteobacteria; Gammaproteobacteria; Enterobacterales; Enterobacteriaceae; Salmonella; Salmonella enterica; Salmonella enterica subsp. enterica MTGDNTLITSHGINRRDFMKLCAALAATMGLSSKAAAEMAESVSNPQRPPVIWIGAQECT WP_000145429.1 hydrogenase 2 small subunit [Salmonella enterica] Length: 372\nNumber of predicted TMHs: 1\nExp number of AAs in TMHs: 24.51309\nExp number, first 60 AAs: 0.2682\nTotal prob of N-in: 0.14402\noutside 1 331\nTMhelix 332 354\ninside 355 372

18644 GCF_001240345.1_Salmonella_enterica_CVM_N43460_v1.0 Salmonella enterica subsp. enterica serovar Muenchen Proteobacteria; Gammaproteobacteria; Enterobacterales; Enterobacteriaceae; Salmonella; Salmonella enterica; Salmonella enterica subsp. enterica

MNNEETFYQAMRRKGVTRRSFLKFCSLAATSLGLGAGMTPKIAWALENKPRIPVVWIHGL WP_001058312.1 hydrogenase [Salmonella enterica] Length: 372\nNumber of predicted TMHs: 2\nExp number of AAs in TMHs: 36.51738\nExp number, first 60 AAs: 19.72849\nTotal prob of N-in: 0.96170\nPOSSIBLE N-term signal sequence\ninside 1 19\nTMhelix 20 39\noutside 40 325\nTMhelix 326 348\ninside 349 372

18645 GCF_001247375.1_Salmonella_enterica_CVM_N51308_v1.0 Salmonella enterica subsp. enterica serovar Muenchen Proteobacteria; Gammaproteobacteria; Enterobacterales; Enterobacteriaceae; Salmonella; Salmonella enterica; Salmonella enterica subsp. enterica MTGDNTLITSHGINRRDFMKLCAALAATMGLSSKAAAEMAESVSNPQRPPVIWIGAQECT WP_000145429.1 hydrogenase 2 small subunit [Salmonella enterica] Length: 372\nNumber of predicted TMHs: 1\nExp number of AAs in TMHs: 24.51309\nExp number, first 60 AAs: 0.2682\nTotal prob of N-in: 0.14402\noutside 1 331\nTMhelix 332 354\ninside 355 372

18646 GCF_001834245.1_ASM183424v1 Salmonella enterica subsp. enterica serovar Muenchen Proteobacteria; Gammaproteobacteria; Enterobacterales; Enterobacteriaceae; Salmonella; Salmonella enterica; Salmonella enterica subsp. enterica MNNEETFYQAMRRKGVTRRSFLKFCSLAATSLGLGAGMTPKIAWALENKPRIPVVWIHGL WP_001058311.1 hydrogenase [Salmonella enterica] Length: 372\nNumber of predicted TMHs: 2\nExp number of AAs in TMHs: 37.4699899999999\nExp number, first 60 AAs: 19.72699\nTotal prob of N-in: 0.96171\nPOSSIBLE N-term signal sequence\ninside 1 19\nTMhelix 20 39\noutside 40 325\nTMhelix 326 348\ninside 349 372

18647 GCF_001240835.1_Salmonella_enterica_CVM_N43826_v1.0 Salmonella enterica subsp. enterica serovar Muenchen Proteobacteria; Gammaproteobacteria; Enterobacterales; Enterobacteriaceae; Salmonella; Salmonella enterica; Salmonella enterica subsp. enterica MTGDNTLITSHGINRRDFMKLCAALAATMGLSSKAAAEMAESVSNPQRPPVIWIGAQECT WP_000145429.1 hydrogenase 2 small subunit [Salmonella enterica] Length: 372\nNumber of predicted TMHs: 1\nExp number of AAs in TMHs: 24.51309\nExp number, first 60 AAs: 0.2682\nTotal prob of N-in: 0.14402\noutside 1 331\nTMhelix 332 354\ninside 355 372

18648 GCF_001479965.1_Salmonella_enterica_CVM_N38885-SQ_v1.0 Salmonella enterica subsp. enterica serovar Muenchen Proteobacteria; Gammaproteobacteria; Enterobacterales; Enterobacteriaceae; Salmonella; Salmonella enterica; Salmonella enterica subsp. enterica MNNEETFYQAMRRKGVTRRSFLKFCSLAATSLGLGAGMTPKIAWALENKPRIPVVWIHGL WP_001058312.1 hydrogenase [Salmonella enterica] Length: 372\nNumber of predicted TMHs: 2\nExp number of AAs in TMHs: 36.51738\nExp number, first 60 AAs: 19.72849\nTotal prob of N-in: 0.96170\nPOSSIBLE N-term signal sequence\ninside 1 19\nTMhelix 20 39\noutside 40 325\nTMhelix 326 348\ninside 349 372

18649 GCF_001834575.1_ASM183457v1 Salmonella enterica subsp. enterica serovar Muenchen Proteobacteria; Gammaproteobacteria; Enterobacterales; Enterobacteriaceae; Salmonella; Salmonella enterica; Salmonella enterica subsp. enterica MTGDNTLITSHGINRRDFMKLCAALAATMGLSSKAAAEMAESVSNPQRPPVIWIGAQECT WP_000145429.1 hydrogenase 2 small subunit [Salmonella enterica] Length: 372\nNumber of predicted TMHs: 1\nExp number of AAs in TMHs: 24.51309\nExp number, first 60 AAs: 0.2682\nTotal prob of N-in: 0.14402\noutside 1 331\nTMhelix 332 354\ninside 355 372

18650 GCF_001242125.1_Salmonella_enterica_CVM_N45927_v1.0 Salmonella enterica subsp. enterica serovar Muenchen Proteobacteria; Gammaproteobacteria; Enterobacterales; Enterobacteriaceae; Salmonella; Salmonella enterica; Salmonella enterica subsp. enterica MNNEETFYQAMRRKGVTRRSFLKFCSLAATSLGLGAGMTPKIAWALENKPRIPVVWIHGL WP_001058312.1 hydrogenase [Salmonella enterica] Length: 372\nNumber of predicted TMHs: 2\nExp number of AAs in TMHs: 36.51738\nExp number, first 60 AAs: 19.72849\nTotal prob of N-in: 0.96170\nPOSSIBLE N-term signal sequence\ninside 1 19\nTMhelix 20 39\noutside 40 325\nTMhelix 326 348\ninside 349 372

18651 GCF_001834325.1_ASM183432v1 Salmonella enterica subsp. enterica serovar Muenchen Proteobacteria; Gammaproteobacteria; Enterobacterales; Enterobacteriaceae; Salmonella; Salmonella enterica; Salmonella enterica subsp. enterica MTGDNTLITSHGINRRDFMKLCAALAATMGLSSKAAAEMAESVSNPQRPPVIWIGAQECT WP_000145429.1 hydrogenase 2 small subunit [Salmonella enterica] Length: 372\nNumber of predicted TMHs: 1\nExp number of AAs in TMHs: 24.51309\nExp number, first 60 AAs: 0.2682\nTotal prob of N-in: 0.14402\noutside 1 331\nTMhelix 332 354\ninside 355 372

18652 GCF_001834085.1_ASM183408v1 Salmonella enterica subsp. enterica serovar Muenchen Proteobacteria; Gammaproteobacteria; Enterobacterales; Enterobacteriaceae; Salmonella; Salmonella enterica;

Salmonella enterica subsp. enterica	MNNEETFYQAMRRKGVTRRSFLKFCSLAATSLGLGAGMTPKIAWALENKPRIPVVWIHGL	WP_070809206.1	hydrogenase [Salmonella enterica]	Length: 372\nNumber of predicted TMHs:  2\nExp number of AAs in TMHs: 37.52613\nExp number, first 60 AAs:  19.72559\nTotal prob of N-in:   0.96174\nPOSSIBLE N-term signal sequence\ninside   1   19\nTMhelix   20   39\noutside   40   325\nTMhelix   326   348\ninside   349   372

18653	GCF_001834245.1_ASM183424v1	Salmonella enterica subsp. enterica serovar Muenchen	Proteobacteria; Gammaproteobacteria; Enterobacterales; Enterobacteriaceae; Salmonella; Salmonella enterica; Salmonella enterica subsp. enterica	MTGDNTLITSHGINRRDFMKLCAALAATMGLSSKAAAEMAESVSNPQRPPVIWIGAQECT	WP_000145429.1	hydrogenase 2 small subunit [Salmonella enterica]	Length: 372\nNumber of predicted TMHs:  1\nExp number of AAs in TMHs: 24.51309\nExp number, first 60 AAs:  0.2682\nTotal prob of N-in:   0.14402\noutside   1   331\nTMhelix   332   354\ninside   355   372

18654	GCF_001692085.1_ASM169208v1	Salmonella enterica subsp. enterica serovar Muenchen	Proteobacteria; Gammaproteobacteria; Enterobacterales; Enterobacteriaceae; Salmonella; Salmonella enterica; Salmonella enterica subsp. enterica	MNNEETFYQAMRRKGVTRRSFLKFCSLAATSLGLGAGMTPKIAWALENKPRIPVVWIHGL	WP_001058311.1	hydrogenase [Salmonella enterica]	Length: 372\nNumber of predicted TMHs:  2\nExp number of AAs in TMHs: 37.4699899999999\nExp number, first 60 AAs:  19.72699\nTotal prob of N-in:   0.96171\nPOSSIBLE N-term signal sequence\ninside   1   19\nTMhelix   20   39\noutside   40   325\nTMhelix   326   348\ninside   349   372

18655	GCF_001834095.1_ASM183409v1	Salmonella enterica subsp. enterica serovar Muenchen	Proteobacteria; Gammaproteobacteria; Enterobacterales; Enterobacteriaceae; Salmonella; Salmonella enterica; Salmonella enterica subsp. enterica	MTGDNTLITSHGINRRDFMKLCAALAATMGLSSKAAAEMAESVSNPQRPPVIWIGAQECT	WP_000145429.1	hydrogenase 2 small subunit [Salmonella enterica]	Length: 372\nNumber of predicted TMHs:  1\nExp number of AAs in TMHs: 24.51309\nExp number, first 60 AAs:  0.2682\nTotal prob of N-in:   0.14402\noutside   1   331\nTMhelix   332   354\ninside   355   372

18656	GCF_001832485.1_ASM183248v1	Salmonella enterica subsp. enterica serovar Muenchen	Proteobacteria; Gammaproteobacteria; Enterobacterales; Enterobacteriaceae; Salmonella; Salmonella enterica; Salmonella enterica subsp. enterica	MNNEETFYQAMRRKGVTRRSFLKFCSLAATSLGLGAGMTPKIAWALENKPRIPVVWIHGL	WP_001058311.1	hydrogenase [Salmonella enterica]	Length: 372\nNumber of predicted TMHs:  2\nExp number of AAs in TMHs: 37.4699899999999\nExp number, first 60 AAs:  19.72699\nTotal prob of N-in:   0.96171\nPOSSIBLE N-term signal sequence\ninside   1   19\nTMhelix   20   39\noutside   40   325\nTMhelix   326   348\ninside   349   372

18657	GCF_001834055.1_ASM183405v1	Salmonella enterica subsp. enterica serovar Muenchen	Proteobacteria; Gammaproteobacteria; Enterobacterales; Enterobacteriaceae; Salmonella; Salmonella enterica; Salmonella enterica subsp. enterica	MNNEETFYQAMRRKGVTRRSFLKFCSLAATSLGLGAGMTPKIAWALENKPRIPVVWIHGL	WP_070809206.1	hydrogenase [Salmonella enterica]	Length: 372\nNumber of predicted TMHs:  2\nExp number of AAs in TMHs: 37.52613\nExp number, first 60 AAs:  19.72559\nTotal prob of N-in:   0.96174\nPOSSIBLE N-term signal sequence\ninside   1   19\nTMhelix   20   39\noutside   40   325\nTMhelix   326   348\ninside   349   372

18658	GCF_002059945.1_ASM205994v1	Salmonella enterica subsp. enterica serovar Muenchen	Proteobacteria; Gammaproteobacteria; Enterobacterales; Enterobacteriaceae; Salmonella; Salmonella enterica; Salmonella enterica subsp. enterica	MNNEETFYQAMRRKGVTRRSFLKFCSLAATSLGLGAGMTPKIAWALENKPRIPVVWIHGL	WP_001058311.1	hydrogenase [Salmonella enterica]	Length: 372\nNumber of predicted TMHs:  2\nExp number of AAs in TMHs: 37.4699899999999\nExp number, first 60 AAs:  19.72699\nTotal prob of N-in:   0.96171\nPOSSIBLE N-term signal sequence\ninside   1   19\nTMhelix   20   39\noutside   40   325\nTMhelix   326   348\ninside   349   372

18659	GCF_001834055.1_ASM183405v1	Salmonella enterica subsp. enterica serovar Muenchen	Proteobacteria; Gammaproteobacteria; Enterobacterales; Enterobacteriaceae; Salmonella; Salmonella enterica; Salmonella enterica subsp. enterica	MTGDNTLITSHGINRRDFMKLCAALAATMGLSSKAAAEMAESVSNPQRPPVIWIGAQECT	WP_000145429.1	hydrogenase 2 small subunit [Salmonella enterica]	Length: 372\nNumber of predicted TMHs:  1\nExp number of AAs in TMHs: 24.51309\nExp number, first 60 AAs:  0.2682\nTotal prob of N-in:   0.14402\noutside   1   331\nTMhelix   332   354\ninside   355   372

18660	GCF_002031245.1_ASM203124v1	Salmonella enterica subsp. enterica serovar Muenchen	Proteobacteria; Gammaproteobacteria; Enterobacterales; Enterobacteriaceae; Salmonella; Salmonella enterica; Salmonella enterica subsp. enterica	MTGDNTLITSHGINRRDFMKLCAALAATMGLSSKAAAEMAESVSNPQRPPVIWIGAQECT	WP_000145429.1	hydrogenase 2 small subunit [Salmonella enterica]	Length: 372\nNumber of

predicted TMHs:  1\nExp number of AAs in TMHs: 24.51309\nExp number, first 60 AAs:  0.2682\nTotal prob of N-in: 0.14402\noutside   1   331\nTMhelix   332   354\ninside   355   372

18661 GCF_002032075.1_ASM203207v1 Salmonella enterica subsp. enterica serovar Muenchen Proteobacteria; Gammaproteobacteria; Enterobacterales; Enterobacteriaceae; Salmonella; Salmonella enterica; Salmonella enterica subsp. enterica MTGDNTLITSHGINRRDFMKLCAALAATMGLSSKAAAEMAESVSNPQRPPVIWIGAQECT WP_000145429.1 hydrogenase 2 small subunit [Salmonella enterica] Length: 372\nNumber of predicted TMHs:  1\nExp number of AAs in TMHs: 24.51309\nExp number, first 60 AAs:  0.2682\nTotal prob of N-in: 0.14402\noutside   1   331\nTMhelix   332   354\ninside   355   372

18662 GCF_002060015.1_ASM206001v1 Salmonella enterica subsp. enterica serovar Muenchen Proteobacteria; Gammaproteobacteria; Enterobacterales; Enterobacteriaceae; Salmonella; Salmonella enterica; Salmonella enterica subsp. enterica MNNEETFYQAMRRKGVTRRSFLKFCSLAATSLGLGAGMTPKIAWALENKPRIPVVWIHGL WP_001058311.1 hydrogenase [Salmonella enterica] Length: 372\nNumber of predicted TMHs:  2\nExp number of AAs in TMHs: 37.4699899999999\nExp number, first 60 AAs:  19.72699\nTotal prob of N-in: 0.96171\nPOSSIBLE N-term signal sequence\ninside   1   19\nTMhelix   20   39\noutside   40   325\nTMhelix   326   348\ninside   349   372

18663 GCF_002060015.1_ASM206001v1 Salmonella enterica subsp. enterica serovar Muenchen Proteobacteria; Gammaproteobacteria; Enterobacterales; Enterobacteriaceae; Salmonella; Salmonella enterica; Salmonella enterica subsp. enterica MTGDNTLITSHGINRRDFMKLCAALAATMGLSSKAAAEMAESVSNPQRPPVIWIGAQECT WP_000145429.1 hydrogenase 2 small subunit [Salmonella enterica] Length: 372\nNumber of predicted TMHs:  1\nExp number of AAs in TMHs: 24.51309\nExp number, first 60 AAs:  0.2682\nTotal prob of N-in: 0.14402\noutside   1   331\nTMhelix   332   354\ninside   355   372

18664 GCF_001835095.1_ASM183509v1 Salmonella enterica subsp. enterica serovar Muenchen Proteobacteria; Gammaproteobacteria; Enterobacterales; Enterobacteriaceae; Salmonella; Salmonella enterica; Salmonella enterica subsp. enterica MNNEETFYQAMRRKGVTRRSFLKFCSLAATSLGLGAGMTPKIAWALENKPRIPVVWIHGL WP_001058311.1 hydrogenase [Salmonella enterica] Length: 372\nNumber of predicted TMHs:  2\nExp number of AAs in TMHs: 37.4699899999999\nExp number, first 60 AAs:  19.72699\nTotal prob of N-in: 0.96171\nPOSSIBLE N-term signal sequence\ninside   1   19\nTMhelix   20   39\noutside   40   325\nTMhelix   326   348\ninside   349   372

18665 GCF_002059945.1_ASM205994v1 Salmonella enterica subsp. enterica serovar Muenchen Proteobacteria; Gammaproteobacteria; Enterobacterales; Enterobacteriaceae; Salmonella; Salmonella enterica; Salmonella enterica subsp. enterica MTGDNTLITSHGINRRDFMKLCAALAATMGLSSKAAAEMAESVSNPQRPPVIWIGAQECT WP_000145429.1 hydrogenase 2 small subunit [Salmonella enterica] Length: 372\nNumber of predicted TMHs:  1\nExp number of AAs in TMHs: 24.51309\nExp number, first 60 AAs:  0.2682\nTotal prob of N-in: 0.14402\noutside   1   331\nTMhelix   332   354\ninside   355   372

18666 GCF_001835095.1_ASM183509v1 Salmonella enterica subsp. enterica serovar Muenchen Proteobacteria; Gammaproteobacteria; Enterobacterales; Enterobacteriaceae; Salmonella; Salmonella enterica; Salmonella enterica subsp. enterica MTGDNTLITSHGINRRDFMKLCAALAATMGLSSKAAAEMAESVSNPQRPPVIWIGAQECT WP_000145429.1 hydrogenase 2 small subunit [Salmonella enterica] Length: 372\nNumber of predicted TMHs:  1\nExp number of AAs in TMHs: 24.51309\nExp number, first 60 AAs:  0.2682\nTotal prob of N-in: 0.14402\noutside   1   331\nTMhelix   332   354\ninside   355   372

18667 GCF_001833235.1_ASM183323v1 Salmonella enterica subsp. enterica serovar Muenchen Proteobacteria; Gammaproteobacteria; Enterobacterales; Enterobacteriaceae; Salmonella; Salmonella enterica; Salmonella enterica subsp. enterica MTGDNTLITSHGINRRDFMKLCAALAATMGLSSKAAAEMAESVSNPQRPPVIWIGAQECT WP_000145429.1 hydrogenase 2 small subunit [Salmonella enterica] Length: 372\nNumber of predicted TMHs:  1\nExp number of AAs in TMHs: 24.51309\nExp number, first 60 AAs:  0.2682\nTotal prob of N-in: 0.14402\noutside   1   331\nTMhelix   332   354\ninside   355   372

18668 GCF_001832485.1_ASM183248v1 Salmonella enterica subsp. enterica serovar Muenchen Proteobacteria; Gammaproteobacteria; Enterobacterales; Enterobacteriaceae; Salmonella; Salmonella enterica; Salmonella enterica subsp. enterica MTGDNTLITSHGINRRDFMKLCAALAATMGLSSKAAAEMAESVSNPQRPPVIWIGAQECT WP_000145429.1 hydrogenase 2 small subunit [Salmonella enterica] Length: 372\nNumber of predicted TMHs:  1\nExp number of AAs in TMHs: 24.51309\nExp number, first 60 AAs:  0.2682\nTotal prob of N-in: 0.14402\noutside   1   331\nTMhelix   332   354\ninside   355   372

18669 GCF_001834175.1_ASM183417v1 Salmonella enterica subsp. enterica serovar Muenchen Proteobacteria; Gammaproteobacteria; Enterobacterales; Enterobacteriaceae; Salmonella; Salmonella enterica; Salmonella enterica subsp. enterica MNNEETFYQAMRRKGVTRRSFLKFCSLAATSLGLGAGMTPKIAWALENKPRIPVVWIHGL WP_001058312.1 hydrogenase [Salmonella enterica] Length: 372\nNumber of predicted TMHs: 2\nExp number of AAs in TMHs: 36.51738\nExp number, first 60 AAs: 19.72849\nTotal prob of N-in: 0.96170\nPOSSIBLE N-term signal sequence\ninside 1 19\nTMhelix 20 39\noutside 40 325\nTMhelix 326 348\ninside 349 372

18670 GCF_001834175.1_ASM183417v1 Salmonella enterica subsp. enterica serovar Muenchen Proteobacteria; Gammaproteobacteria; Enterobacterales; Enterobacteriaceae; Salmonella; Salmonella enterica; Salmonella enterica subsp. enterica MTGDNTLITSHGINRRDFMKLCAALAATMGLSSKAAAEMAESVSNPQRPPVIWIGAQECT WP_000145429.1 hydrogenase 2 small subunit [Salmonella enterica] Length: 372\nNumber of predicted TMHs: 1\nExp number of AAs in TMHs: 24.51309\nExp number, first 60 AAs: 0.2682\nTotal prob of N-in: 0.14402\noutside 1 331\nTMhelix 332 354\ninside 355 372

18671 GCF_002034335.1_ASM203433v1 Salmonella enterica subsp. enterica serovar Muenchen Proteobacteria; Gammaproteobacteria; Enterobacterales; Enterobacteriaceae; Salmonella; Salmonella enterica; Salmonella enterica subsp. enterica MTGDNTLITSHGINRRDFMKLCAALAATMGLSSKAAAEMAESVSNPQRPPVIWIGAQECT WP_000145429.1 hydrogenase 2 small subunit [Salmonella enterica] Length: 372\nNumber of predicted TMHs: 1\nExp number of AAs in TMHs: 24.51309\nExp number, first 60 AAs: 0.2682\nTotal prob of N-in: 0.14402\noutside 1 331\nTMhelix 332 354\ninside 355 372

18672 GCF_001481135.1_Salmonella_enterica_CVM_N42479_v1.0 Salmonella enterica subsp. enterica serovar Muenchen Proteobacteria; Gammaproteobacteria; Enterobacterales; Enterobacteriaceae; Salmonella; Salmonella enterica; Salmonella enterica subsp. enterica MNNEETFYQAMRRKGVTRRSFLKFCSLAATSLGLGAGMTPKIAWALENKPRIPVVWIHGL WP_001058312.1 hydrogenase [Salmonella enterica] Length: 372\nNumber of predicted TMHs: 2\nExp number of AAs in TMHs: 36.51738\nExp number, first 60 AAs: 19.72849\nTotal prob of N-in: 0.96170\nPOSSIBLE N-term signal sequence\ninside 1 19\nTMhelix 20 39\noutside 40 325\nTMhelix 326 348\ninside 349 372

18673 GCF_002034335.1_ASM203433v1 Salmonella enterica subsp. enterica serovar Muenchen Proteobacteria; Gammaproteobacteria; Enterobacterales; Enterobacteriaceae; Salmonella; Salmonella enterica; Salmonella enterica subsp. enterica MNNEETFYQAMRRKGVTRRSFLKFCSLAATSLGLGAGMTPKIAWALENKPRIPVVWIHGL WP_001058312.1 hydrogenase [Salmonella enterica] Length: 372\nNumber of predicted TMHs: 2\nExp number of AAs in TMHs: 36.51738\nExp number, first 60 AAs: 19.72849\nTotal prob of N-in: 0.96170\nPOSSIBLE N-term signal sequence\ninside 1 19\nTMhelix 20 39\noutside 40 325\nTMhelix 326 348\ninside 349 372

18674 GCF_001834145.1_ASM183414v1 Salmonella enterica subsp. enterica serovar Muenchen Proteobacteria; Gammaproteobacteria; Enterobacterales; Enterobacteriaceae; Salmonella; Salmonella enterica; Salmonella enterica subsp. enterica MNNEETFYQAMRRKGVTRRSFLKFCSLAATSLGLGAGMTPKIAWALENKPRIPVVWIHGL WP_001058311.1 hydrogenase [Salmonella enterica] Length: 372\nNumber of predicted TMHs: 2\nExp number of AAs in TMHs: 37.4699899999999\nExp number, first 60 AAs: 19.72699\nTotal prob of N-in: 0.96171\nPOSSIBLE N-term signal sequence\ninside 1 19\nTMhelix 20 39\noutside 40 325\nTMhelix 326 348\ninside 349 372

18675 GCF_001834095.1_ASM183409v1 Salmonella enterica subsp. enterica serovar Muenchen Proteobacteria; Gammaproteobacteria; Enterobacterales; Enterobacteriaceae; Salmonella; Salmonella enterica; Salmonella enterica subsp. enterica MNNEETFYQAMRRKGVTRRSFLKFCSLAATSLGLGAGMTPKIAWALENKPRIPVVWIHGL WP_001058311.1 hydrogenase [Salmonella enterica] Length: 372\nNumber of predicted TMHs: 2\nExp number of AAs in TMHs: 37.4699899999999\nExp number, first 60 AAs: 19.72699\nTotal prob of N-in: 0.96171\nPOSSIBLE N-term signal sequence\ninside 1 19\nTMhelix 20 39\noutside 40 325\nTMhelix 326 348\ninside 349 372

18676 GCF_001834145.1_ASM183414v1 Salmonella enterica subsp. enterica serovar Muenchen Proteobacteria; Gammaproteobacteria; Enterobacterales; Enterobacteriaceae; Salmonella; Salmonella enterica; Salmonella enterica subsp. enterica MTGDNTLITSHGINRRDFMKLCAALAATMGLSSKAAAEMAESVSNPQRPPVIWIGAQECT WP_000145429.1 hydrogenase 2 small subunit [Salmonella enterica] Length: 372\nNumber of predicted TMHs: 1\nExp number of AAs in TMHs: 24.51309\nExp number, first 60 AAs: 0.2682\nTotal prob of N-in: 0.14402\noutside 1 331\nTMhelix 332 354\ninside 355 372

18677 GCF_001481135.1_Salmonella_enterica_CVM_N42479_v1.0 Salmonella enterica subsp. enterica serovar Muenchen Proteobacteria; Gammaproteobacteria; Enterobacterales; Enterobacteriaceae; Salmonella; Salmonella

enterica; Salmonella enterica subsp. enterica
MTGDNTLITSHGINRRDFMKLCAALAATMGLSSKAAAEMAESVSNPQRPPVIWIGAQECT WP_000145429.1 hydrogenase 2 small subunit [Salmonella enterica] Length: 372\nNumber of predicted TMHs: 1\nExp number of AAs in TMHs: 24.51309\nExp number, first 60 AAs: 0.2682\nTotal prob of N-in: 0.14402\noutside 1 331\nTMhelix 332 354\ninside 355 372

18678 GCF_002050685.1_ASM205068v1 Salmonella enterica subsp. enterica serovar Muenster Proteobacteria; Gammaproteobacteria; Enterobacterales; Enterobacteriaceae; Salmonella; Salmonella enterica; Salmonella enterica subsp. enterica MNNEETFYQAMRRKGVTRRSFLKFCSLAATSLGLGAGMTPKIAWALENKPRIPVVWIHGL WP_001058312.1 hydrogenase [Salmonella enterica] Length: 372\nNumber of predicted TMHs: 2\nExp number of AAs in TMHs: 36.51738\nExp number, first 60 AAs: 19.72849\nTotal prob of N-in: 0.96170\nPOSSIBLE N-term signal sequence\ninside 1 19\nTMhelix 20 39\noutside 40 325\nTMhelix 326 348\ninside 349 372

18679 GCF_002050685.1_ASM205068v1 Salmonella enterica subsp. enterica serovar Muenster Proteobacteria; Gammaproteobacteria; Enterobacterales; Enterobacteriaceae; Salmonella; Salmonella enterica; Salmonella enterica subsp. enterica MTGDNTLITSHGINRRDFMKLCAALAATMGLSSKAAAEMAESVSNPQRPPVIWIGAQECT WP_000145431.1 hydrogenase 2 small subunit [Salmonella enterica] Length: 372\nNumber of predicted TMHs: 1\nExp number of AAs in TMHs: 24.51505\nExp number, first 60 AAs: 0.2682\nTotal prob of N-in: 0.14406\noutside 1 331\nTMhelix 332 354\ninside 355 372

18680 GCF_002050875.1_ASM205087v1 Salmonella enterica subsp. enterica serovar Muenster Proteobacteria; Gammaproteobacteria; Enterobacterales; Enterobacteriaceae; Salmonella; Salmonella enterica; Salmonella enterica subsp. enterica MNNEETFYQAMRRKGVTRRSFLKFCSLAATSLGLGAGMTPKIAWALENKPRIPVVWIHGL WP_001058312.1 hydrogenase [Salmonella enterica] Length: 372\nNumber of predicted TMHs: 2\nExp number of AAs in TMHs: 36.51738\nExp number, first 60 AAs: 19.72849\nTotal prob of N-in: 0.96170\nPOSSIBLE N-term signal sequence\ninside 1 19\nTMhelix 20 39\noutside 40 325\nTMhelix 326 348\ninside 349 372

18681 GCF_002050495.1_ASM205049v1 Salmonella enterica subsp. enterica serovar Muenster Proteobacteria; Gammaproteobacteria; Enterobacterales; Enterobacteriaceae; Salmonella; Salmonella enterica; Salmonella enterica subsp. enterica MNNEETFYQAMRRKGVTRRSFLKFCSLAATSLGLGAGMTPKIAWALENKPRIPVVWIHGL WP_001058312.1 hydrogenase [Salmonella enterica] Length: 372\nNumber of predicted TMHs: 2\nExp number of AAs in TMHs: 36.51738\nExp number, first 60 AAs: 19.72849\nTotal prob of N-in: 0.96170\nPOSSIBLE N-term signal sequence\ninside 1 19\nTMhelix 20 39\noutside 40 325\nTMhelix 326 348\ninside 349 372

18682 GCF_002050495.1_ASM205049v1 Salmonella enterica subsp. enterica serovar Muenster Proteobacteria; Gammaproteobacteria; Enterobacterales; Enterobacteriaceae; Salmonella; Salmonella enterica; Salmonella enterica subsp. enterica MTGDNTLITSHGINRRDFMKLCAALAATMGLSSKAAAEMAESVSNPQRPPVIWIGAQECT WP_000145431.1 hydrogenase 2 small subunit [Salmonella enterica] Length: 372\nNumber of predicted TMHs: 1\nExp number of AAs in TMHs: 24.51505\nExp number, first 60 AAs: 0.2682\nTotal prob of N-in: 0.14406\noutside 1 331\nTMhelix 332 354\ninside 355 372

18683 GCF_002050875.1_ASM205087v1 Salmonella enterica subsp. enterica serovar Muenster Proteobacteria; Gammaproteobacteria; Enterobacterales; Enterobacteriaceae; Salmonella; Salmonella enterica; Salmonella enterica subsp. enterica MTGDNTLITSHGINRRDFMKLCAALAATMGLSSKAAAEMAESVSNPQRPPVIWIGAQECT WP_000145431.1 hydrogenase 2 small subunit [Salmonella enterica] Length: 372\nNumber of predicted TMHs: 1\nExp number of AAs in TMHs: 24.51505\nExp number, first 60 AAs: 0.2682\nTotal prob of N-in: 0.14406\noutside 1 331\nTMhelix 332 354\ninside 355 372

18684 GCF_001246125.1_Salmonella_enterica_CVM_N51250_v1.0 Salmonella enterica subsp. enterica serovar MuensterProteobacteria; Gammaproteobacteria; Enterobacterales; Enterobacteriaceae; Salmonella; Salmonella enterica; Salmonella enterica subsp. enterica MNNEETFYQAMRRKGVTRRSFLKFCSLAATSLGLGAGMTPKIAWALENKPRIPVVWIHGL WP_001058312.1 hydrogenase [Salmonella enterica] Length: 372\nNumber of predicted TMHs: 2\nExp number of AAs in TMHs: 36.51738\nExp number, first 60 AAs: 19.72849\nTotal prob of N-in: 0.96170\nPOSSIBLE N-term signal sequence\ninside 1 19\nTMhelix 20 39\noutside 40 325\nTMhelix 326 348\ninside 349 372

18685 GCF_001246125.1_Salmonella_enterica_CVM_N51250_v1.0 Salmonella enterica subsp. enterica serovar MuensterProteobacteria; Gammaproteobacteria; Enterobacterales; Enterobacteriaceae; Salmonella; Salmonella enterica; Salmonella enterica subsp. enterica MTGDNTLITSHGINRRDFMKLCAALAATMGLSSKAAAEMAESVSNPQRPPVIWIGAQECT WP_000145431.1 hydrogenase 2 small subunit [Salmonella enterica] Length: 372\nNumber of predicted TMHs: 1\nExp number of AAs in TMHs: 24.51505\nExp number, first 60 AAs: 0.2682\nTotal prob of N-in: 0.14406\noutside 1 331\nTMhelix 332 354\ninside 355 372

18686 GCF_002050585.1_ASM205058v1 Salmonella enterica subsp. enterica serovar Muenster Proteobacteria; Gammaproteobacteria; Enterobacterales; Enterobacteriaceae; Salmonella; Salmonella enterica; Salmonella enterica subsp. enterica MNNEETFYQAMRRKGVTRRSFLKFCSLAATSLGLGAGMTPKIAWALENKPRIPVVWIHGL WP_001058312.1 hydrogenase [Salmonella enterica] Length: 372\nNumber of predicted TMHs: 2\nExp number of AAs in TMHs: 36.51738\nExp number, first 60 AAs: 19.72849\nTotal prob of N-in: 0.96170\nPOSSIBLE N-term signal sequence\ninside 1 19\nTMhelix 20 39\noutside 40 325\nTMhelix 326 348\ninside 349 372

18687 GCF_002065425.1_ASM206542v1 Salmonella enterica subsp. enterica serovar Muenster Proteobacteria; Gammaproteobacteria; Enterobacterales; Enterobacteriaceae; Salmonella; Salmonella enterica; Salmonella enterica subsp. enterica MTGDNTLITSHGINRRDFMKLCAALAATMGLSSKAAAEMAESVSNPQRPPVIWIGAQECT WP_000145431.1 hydrogenase 2 small subunit [Salmonella enterica] Length: 372\nNumber of predicted TMHs: 1\nExp number of AAs in TMHs: 24.51505\nExp number, first 60 AAs: 0.2682\nTotal prob of N-in: 0.14406\noutside 1 331\nTMhelix 332 354\ninside 355 372

18688 GCF_002065425.1_ASM206542v1 Salmonella enterica subsp. enterica serovar Muenster Proteobacteria; Gammaproteobacteria; Enterobacterales; Enterobacteriaceae; Salmonella; Salmonella enterica; Salmonella enterica subsp. enterica MNNEETFYQAMRRKGVTRRSFLKFCSLAATSLGLGAGMTPKIAWALENKPRIPVVWIHGL WP_001662439.1 hydrogenase [Salmonella enterica] Length: 372\nNumber of predicted TMHs: 2\nExp number of AAs in TMHs: 36.5172\nExp number, first 60 AAs: 19.72849\nTotal prob of N-in: 0.96169\nPOSSIBLE N-term signal sequence\ninside 1 19\nTMhelix 20 39\noutside 40 325\nTMhelix 326 348\ninside 349 372

18689 GCF_002062055.1_ASM206205v1 Salmonella enterica subsp. enterica serovar Muenster Proteobacteria; Gammaproteobacteria; Enterobacterales; Enterobacteriaceae; Salmonella; Salmonella enterica; Salmonella enterica subsp. enterica MNNEETFYQAMRRKGVTRRSFLKFCSLAATSLGLGAGMTPKIAWALENKPRIPVVWIHGL WP_001058311.1 hydrogenase [Salmonella enterica] Length: 372\nNumber of predicted TMHs: 2\nExp number of AAs in TMHs: 37.4699899999999\nExp number, first 60 AAs: 19.72699\nTotal prob of N-in: 0.96171\nPOSSIBLE N-term signal sequence\ninside 1 19\nTMhelix 20 39\noutside 40 325\nTMhelix 326 348\ninside 349 372

18690 GCF_001477695.1_Salmonella_enterica_CVM_N29310-SQ_v1.0 Salmonella enterica subsp. enterica serovar MuensterProteobacteria; Gammaproteobacteria; Enterobacterales; Enterobacteriaceae; Salmonella; Salmonella enterica; Salmonella enterica subsp. enterica MTGDNTLITSHGINRRDFMKLCAALAATMGLSSKAAAEMAESVSNPQRPPVIWIGAQECT WP_000145431.1 hydrogenase 2 small subunit [Salmonella enterica] Length: 372\nNumber of predicted TMHs: 1\nExp number of AAs in TMHs: 24.51505\nExp number, first 60 AAs: 0.2682\nTotal prob of N-in: 0.14406\noutside 1 331\nTMhelix 332 354\ninside 355 372

18691 GCF_002062055.1_ASM206205v1 Salmonella enterica subsp. enterica serovar Muenster Proteobacteria; Gammaproteobacteria; Enterobacterales; Enterobacteriaceae; Salmonella; Salmonella enterica; Salmonella enterica subsp. enterica MTGDNTLITSHGINRRDFMKLCAALAATMGLSSKAAAEMAESVSNPQRPPVIWIGAQECT WP_000145429.1 hydrogenase 2 small subunit [Salmonella enterica] Length: 372\nNumber of predicted TMHs: 1\nExp number of AAs in TMHs: 24.51309\nExp number, first 60 AAs: 0.2682\nTotal prob of N-in: 0.14402\noutside 1 331\nTMhelix 332 354\ninside 355 372

18692 GCF_002050585.1_ASM205058v1 Salmonella enterica subsp. enterica serovar Muenster Proteobacteria; Gammaproteobacteria; Enterobacterales; Enterobacteriaceae; Salmonella; Salmonella enterica; Salmonella enterica subsp. enterica MTGDNTLITSHGINRRDFMKLCAALAATMGLSSKAAAEMAESVSNPQRPPVIWIGAQECT WP_000145431.1 hydrogenase 2 small subunit [Salmonella enterica] Length: 372\nNumber of predicted TMHs: 1\nExp number of AAs in TMHs: 24.51505\nExp number, first 60 AAs: 0.2682\nTotal prob of N-in: 0.14406\noutside 1 331\nTMhelix 332 354\ninside 355 372

18693 GCF_002034315.1_ASM203431v1 Salmonella enterica subsp. enterica serovar Muenster Proteobacteria; Gammaproteobacteria; Enterobacterales; Enterobacteriaceae; Salmonella; Salmonella enterica; Salmonella enterica subsp. enterica MNNEETFYQAMRRKGVTRRSFLKFCSLAATSLGLGAGMTPKIAWALENKPRIPVVWIHGL WP_001058312.1 hydrogenase [Salmonella enterica] Length: 372\nNumber of predicted TMHs: 2\nExp number of AAs in TMHs: 36.51738\nExp number, first 60 AAs: 19.72849\nTotal prob of N-in: 0.96170\nPOSSIBLE N-term signal sequence\ninside 1 19\nTMhelix 20 39\noutside 40 325\nTMhelix 326 348\ninside 349 372

18694 GCF_001477695.1_Salmonella_enterica_CVM_N29310-SQ_v1.0 Salmonella enterica subsp. enterica serovar MuensterProteobacteria; Gammaproteobacteria; Enterobacterales; Enterobacteriaceae; Salmonella; Salmonella enterica; Salmonella enterica subsp. enterica MNNEETFYQAMRRKGVTRRSFLKFCSLAATSLGLGAGMTPKIAWALENKPRIPVVWIHGL WP_001058312.1 hydrogenase [Salmonella enterica] Length: 372\nNumber of predicted TMHs: 2\nExp

number of AAs in TMHs: 36.51738\nExp number, first 60 AAs:  19.72849\nTotal prob of N-in:   0.96170\nPOSSIBLE N-term signal sequence\ninside   1   19\nTMhelix   20   39\noutside   40   325\nTMhelix   326   348\ninside   349   372

18695 GCF_002034315.1_ASM203431v1 Salmonella enterica subsp. enterica serovar Muenster Proteobacteria; Gammaproteobacteria; Enterobacterales; Enterobacteriaceae; Salmonella; Salmonella enterica; Salmonella enterica subsp. enterica MTGDNTLITSHGINRRDFMKLCAALAATMGLSSKAAAEMAESVSNPQRPPVIWIGAQECT WP_000145431.1 hydrogenase 2 small subunit [Salmonella enterica] Length: 372\nNumber of predicted TMHs:  1\nExp number of AAs in TMHs: 24.51505\nExp number, first 60 AAs:  0.2682\nTotal prob of N-in: 0.14406\noutside   1   331\nTMhelix   332   354\ninside   355   372

18696 GCF_002062165.1_ASM206216v1 Salmonella enterica subsp. enterica serovar Muenster Proteobacteria; Gammaproteobacteria; Enterobacterales; Enterobacteriaceae; Salmonella; Salmonella enterica; Salmonella enterica subsp. enterica MTGDNTLITSHGINRRDFMKLCAALAATMGLSSKAAAEMAESVSNPQRPPVIWIGAQECT WP_000145431.1 hydrogenase 2 small subunit [Salmonella enterica] Length: 372\nNumber of predicted TMHs:  1\nExp number of AAs in TMHs: 24.51505\nExp number, first 60 AAs:  0.2682\nTotal prob of N-in: 0.14406\noutside   1   331\nTMhelix   332   354\ninside   355   372

18697 GCF_002062205.1_ASM206220v1 Salmonella enterica subsp. enterica serovar Muenster Proteobacteria; Gammaproteobacteria; Enterobacterales; Enterobacteriaceae; Salmonella; Salmonella enterica; Salmonella enterica subsp. enterica MTGDNTLITSHGINRRDFMKLCAALAATMGLSSKAAAEMAESVSNPQRPPVIWIGAQECT WP_000145431.1 hydrogenase 2 small subunit [Salmonella enterica] Length: 372\nNumber of predicted TMHs:  1\nExp number of AAs in TMHs: 24.51505\nExp number, first 60 AAs:  0.2682\nTotal prob of N-in: 0.14406\noutside   1   331\nTMhelix   332   354\ninside   355   372

18698 GCF_002064315.1_ASM206431v1 Salmonella enterica subsp. enterica serovar Muenster Proteobacteria; Gammaproteobacteria; Enterobacterales; Enterobacteriaceae; Salmonella; Salmonella enterica; Salmonella enterica subsp. enterica MTGDNTLITSHGINRRDFMKLCAALAATMGLSSKAAAEMAESVSNPQRPPVIWIGAQECT WP_000145431.1 hydrogenase 2 small subunit [Salmonella enterica] Length: 372\nNumber of predicted TMHs:  1\nExp number of AAs in TMHs: 24.51505\nExp number, first 60 AAs:  0.2682\nTotal prob of N-in: 0.14406\noutside   1   331\nTMhelix   332   354\ninside   355   372

18699 GCF_002062165.1_ASM206216v1 Salmonella enterica subsp. enterica serovar Muenster Proteobacteria; Gammaproteobacteria; Enterobacterales; Enterobacteriaceae; Salmonella; Salmonella enterica; Salmonella enterica subsp. enterica MNNEETFYQAMRRKGVTRRSFLKFCSLAATSLGLGAGMTPKIAWALENKPRIPVVWIHGL WP_001662439.1 hydrogenase [Salmonella enterica] Length: 372\nNumber of predicted TMHs:  2\nExp number of AAs in TMHs: 36.5172\nExp number, first 60 AAs:  19.72849\nTotal prob of N-in:   0.96169\nPOSSIBLE N-term signal sequence\ninside   1   19\nTMhelix   20   39\noutside   40   325\nTMhelix   326   348\ninside   349   372

18700 GCF_002062205.1_ASM206220v1 Salmonella enterica subsp. enterica serovar Muenster Proteobacteria; Gammaproteobacteria; Enterobacterales; Enterobacteriaceae; Salmonella; Salmonella enterica; Salmonella enterica subsp. enterica MNNEETFYQAMRRKGVTRRSFLKFCSLAATSLGLGAGMTPKIAWALENKPRIPVVWIHGL WP_001662439.1 hydrogenase [Salmonella enterica] Length: 372\nNumber of predicted TMHs:  2\nExp number of AAs in TMHs: 36.5172\nExp number, first 60 AAs:  19.72849\nTotal prob of N-in:   0.96169\nPOSSIBLE N-term signal sequence\ninside   1   19\nTMhelix   20   39\noutside   40   325\nTMhelix   326   348\ninside   349   372

18701 GCF_002064315.1_ASM206431v1 Salmonella enterica subsp. enterica serovar Muenster Proteobacteria; Gammaproteobacteria; Enterobacterales; Enterobacteriaceae; Salmonella; Salmonella enterica; Salmonella enterica subsp. enterica MNNEETFYQAMRRKGVTRRSFLKFCSLAATSLGLGAGMTPKIAWALENKPRIPVVWIHGL WP_001058312.1 hydrogenase [Salmonella enterica] Length: 372\nNumber of predicted TMHs:  2\nExp number of AAs in TMHs: 36.51738\nExp number, first 60 AAs:  19.72849\nTotal prob of N-in:   0.96170\nPOSSIBLE N-term signal sequence\ninside   1   19\nTMhelix   20   39\noutside   40   325\nTMhelix   326   348\ninside   349   372

18702 GCF_001047615.1_ASM104761v1 Salmonella enterica subsp. enterica serovar Napoli Proteobacteria; Gammaproteobacteria; Enterobacterales; Enterobacteriaceae; Salmonella; Salmonella enterica; Salmonella enterica subsp. enterica MTGDNTLITSHGINRRDFMKLCAALAATMGLSSKAAAEMAESVSNPQRPPVIWIGAQECT WP_000145429.1 hydrogenase 2 small subunit [Salmonella enterica] Length: 372\nNumber of predicted TMHs:  1\nExp number of AAs in TMHs: 24.51309\nExp number, first 60 AAs:  0.2682\nTotal prob of N-in: 0.14402\noutside   1   331\nTMhelix   332   354\ninside   355   372

18703 GCF_001047615.1_ASM104761v1 Salmonella enterica subsp. enterica serovar Napoli Proteobacteria; Gammaproteobacteria; Enterobacterales; Enterobacteriaceae; Salmonella; Salmonella enterica;

Salmonella enterica subsp. enterica MNNEETFYQAMRRKGVTRRSFLKFCSLAATSLGLGAGMTPKIAWALENKPRIPVVWIHGL WP_001058312.1 hydrogenase [Salmonella enterica] Length: 372\nNumber of predicted TMHs: 2\nExp number of AAs in TMHs: 36.51738\nExp number, first 60 AAs: 19.72849\nTotal prob of N-in: 0.96170\nPOSSIBLE N-term signal sequence\ninside 1 19\nTMhelix 20 39\noutside 40 325\nTMhelix 326 348\ninside 349 372

18704 GCF_001835415.1_ASM183541v1 Salmonella enterica subsp. enterica serovar Newport Proteobacteria; Gammaproteobacteria; Enterobacterales; Enterobacteriaceae; Salmonella; Salmonella enterica; Salmonella enterica subsp. enterica MTGDNTLITSHGINRRDFMKLCAALAATMGLSSKAAAEMAESVSNPQRPPVIWIGAQECT WP_000145429.1 hydrogenase 2 small subunit [Salmonella enterica] Length: 372\nNumber of predicted TMHs: 1\nExp number of AAs in TMHs: 24.51309\nExp number, first 60 AAs: 0.2682\nTotal prob of N-in: 0.14402\noutside 1 331\nTMhelix 332 354\ninside 355 372

18705 GCF_002060195.1_ASM206019v1 Salmonella enterica subsp. enterica serovar Newport Proteobacteria; Gammaproteobacteria; Enterobacterales; Enterobacteriaceae; Salmonella; Salmonella enterica; Salmonella enterica subsp. enterica MNNEETFYQAMRRKGVTRRSFLKFCSLAATSLGLGAGMTPKIAWALENKPRIPVVWIHGL WP_001058311.1 hydrogenase [Salmonella enterica] Length: 372\nNumber of predicted TMHs: 2\nExp number of AAs in TMHs: 37.4699899999999\nExp number, first 60 AAs: 19.72699\nTotal prob of N-in: 0.96171\nPOSSIBLE N-term signal sequence\ninside 1 19\nTMhelix 20 39\noutside 40 325\nTMhelix 326 348\ninside 349 372

18706 GCF_001954765.1_ASM195476v1 Salmonella enterica subsp. enterica serovar Newport Proteobacteria; Gammaproteobacteria; Enterobacterales; Enterobacteriaceae; Salmonella; Salmonella enterica; Salmonella enterica subsp. enterica MTGDNTLITSHGINRRDFMKLCAALAATMGLSSKAAAEMAESVSNPQRPPVIWIGAQECT WP_000145429.1 hydrogenase 2 small subunit [Salmonella enterica] Length: 372\nNumber of predicted TMHs: 1\nExp number of AAs in TMHs: 24.51309\nExp number, first 60 AAs: 0.2682\nTotal prob of N-in: 0.14402\noutside 1 331\nTMhelix 332 354\ninside 355 372

18707 GCF_001954845.1_ASM195484v1 Salmonella enterica subsp. enterica serovar Newport Proteobacteria; Gammaproteobacteria; Enterobacterales; Enterobacteriaceae; Salmonella; Salmonella enterica; Salmonella enterica subsp. enterica MTGDNTLITSHGINRRDFMKLCAALAATMGLSSKAAAEMAESVSNPQRPPVIWIGAQECT WP_000145429.1 hydrogenase 2 small subunit [Salmonella enterica] Length: 372\nNumber of predicted TMHs: 1\nExp number of AAs in TMHs: 24.51309\nExp number, first 60 AAs: 0.2682\nTotal prob of N-in: 0.14402\noutside 1 331\nTMhelix 332 354\ninside 355 372

18708 GCF_001954825.1_ASM195482v1 Salmonella enterica subsp. enterica serovar Newport Proteobacteria; Gammaproteobacteria; Enterobacterales; Enterobacteriaceae; Salmonella; Salmonella enterica; Salmonella enterica subsp. enterica MTGDNTLITSHGINRRDFMKLCAALAATMGLSSKAAAEMAESVSNPQRPPVIWIGAQECT WP_000145429.1 hydrogenase 2 small subunit [Salmonella enterica] Length: 372\nNumber of predicted TMHs: 1\nExp number of AAs in TMHs: 24.51309\nExp number, first 60 AAs: 0.2682\nTotal prob of N-in: 0.14402\noutside 1 331\nTMhelix 332 354\ninside 355 372

18709 GCF_002056845.1_ASM205684v1 Salmonella enterica subsp. enterica serovar Newport Proteobacteria; Gammaproteobacteria; Enterobacterales; Enterobacteriaceae; Salmonella; Salmonella enterica; Salmonella enterica subsp. enterica MTGDNTLITSHGINRRDFMKLCAALAATMGLSSKAAAEMAESVSNPQRPPVIWIGAQECT WP_000145429.1 hydrogenase 2 small subunit [Salmonella enterica] Length: 372\nNumber of predicted TMHs: 1\nExp number of AAs in TMHs: 24.51309\nExp number, first 60 AAs: 0.2682\nTotal prob of N-in: 0.14402\noutside 1 331\nTMhelix 332 354\ninside 355 372

18710 GCF_002056915.1_ASM205691v1 Salmonella enterica subsp. enterica serovar Newport Proteobacteria; Gammaproteobacteria; Enterobacterales; Enterobacteriaceae; Salmonella; Salmonella enterica; Salmonella enterica subsp. enterica MTGDNTLITSHGINRRDFMKLCAALAATMGLSSKAAAEMAESVSNPQRPPVIWIGAQECT WP_000145429.1 hydrogenase 2 small subunit [Salmonella enterica] Length: 372\nNumber of predicted TMHs: 1\nExp number of AAs in TMHs: 24.51309\nExp number, first 60 AAs: 0.2682\nTotal prob of N-in: 0.14402\noutside 1 331\nTMhelix 332 354\ninside 355 372

18711 GCF_002057125.1_ASM205712v1 Salmonella enterica subsp. enterica serovar Newport Proteobacteria; Gammaproteobacteria; Enterobacterales; Enterobacteriaceae; Salmonella; Salmonella enterica; Salmonella enterica subsp. enterica MNNEETFYQAMRRKGVTRRSFLKFCSLAATSLGLGAGMTPKIAWALENKPRIPVVWIHGL WP_001058312.1 hydrogenase [Salmonella enterica] Length: 372\nNumber of predicted TMHs: 2\nExp number of AAs in TMHs: 36.51738\nExp number, first 60 AAs: 19.72849\nTotal prob of N-in: 0.96170\nPOSSIBLE N-term signal sequence\ninside 1 19\nTMhelix 20 39\noutside 40 325\nTMhelix 326 348\ninside 349 372

18712 GCF_002032465.1_ASM203246v1 Salmonella enterica subsp. enterica serovar Newport Proteobacteria; Gammaproteobacteria; Enterobacterales; Enterobacteriaceae; Salmonella; Salmonella enterica; Salmonella enterica subsp. enterica MNNEETFYQAMRRKGVTRRSFLKFCSLAATSLGLGAGMTPKIAWALENKPRIPVVWIHGL WP_001058312.1 hydrogenase [Salmonella enterica] Length: 372\nNumber of predicted TMHs: 2\nExp number of AAs in TMHs: 36.51738\nExp number, first 60 AAs: 19.72849\nTotal prob of N-in: 0.96170\nPOSSIBLE N-term signal sequence\ninside 1 19\nTMhelix 20 39\noutside 40 325\nTMhelix 326 348\ninside 349 372

18713 GCF_002032465.1_ASM203246v1 Salmonella enterica subsp. enterica serovar Newport Proteobacteria; Gammaproteobacteria; Enterobacterales; Enterobacteriaceae; Salmonella; Salmonella enterica; Salmonella enterica subsp. enterica MTGDNTLITSHGINRRDFMKLCAALAATMGLSSKAAAEMAESVSNPQRPPVIWIGAQECT WP_000145429.1 hydrogenase 2 small subunit [Salmonella enterica] Length: 372\nNumber of predicted TMHs: 1\nExp number of AAs in TMHs: 24.51309\nExp number, first 60 AAs: 0.2682\nTotal prob of N-in: 0.14402\noutside 1 331\nTMhelix 332 354\ninside 355 372

18714 GCF_001243515.1_Salmonella_enterica_CVM_N46850-R_v1.0 Salmonella enterica subsp. enterica serovar Newport Proteobacteria; Gammaproteobacteria; Enterobacterales; Enterobacteriaceae; Salmonella; Salmonella enterica; Salmonella enterica subsp. enterica MNNEETFYQAMRRKGVTRRSFLKFCSLAATSLGLGAGMTPKIAWALENKPRIPVVWIHGL WP_001058312.1 hydrogenase [Salmonella enterica] Length: 372\nNumber of predicted TMHs: 2\nExp number of AAs in TMHs: 36.51738\nExp number, first 60 AAs: 19.72849\nTotal prob of N-in: 0.96170\nPOSSIBLE N-term signal sequence\ninside 1 19\nTMhelix 20 39\noutside 40 325\nTMhelix 326 348\ninside 349 372

18715 GCF_002057865.1_ASM205786v1 Salmonella enterica subsp. enterica serovar Newport Proteobacteria; Gammaproteobacteria; Enterobacterales; Enterobacteriaceae; Salmonella; Salmonella enterica; Salmonella enterica subsp. enterica MTGDNTLITSHGINRRDFMKLCAALAATMGLSSKAAAEMAESVSNPQRPPVIWIGAQECT WP_000145429.1 hydrogenase 2 small subunit [Salmonella enterica] Length: 372\nNumber of predicted TMHs: 1\nExp number of AAs in TMHs: 24.51309\nExp number, first 60 AAs: 0.2682\nTotal prob of N-in: 0.14402\noutside 1 331\nTMhelix 332 354\ninside 355 372

18716 GCF_002058285.1_ASM205828v1 Salmonella enterica subsp. enterica serovar Newport Proteobacteria; Gammaproteobacteria; Enterobacterales; Enterobacteriaceae; Salmonella; Salmonella enterica; Salmonella enterica subsp. enterica MTGDNTLITSHGINRRDFMKLCAALAATMGLSSKAAAEMAESVSNPQRPPVIWIGAQECT WP_000145429.1 hydrogenase 2 small subunit [Salmonella enterica] Length: 372\nNumber of predicted TMHs: 1\nExp number of AAs in TMHs: 24.51309\nExp number, first 60 AAs: 0.2682\nTotal prob of N-in: 0.14402\noutside 1 331\nTMhelix 332 354\ninside 355 372

18717 GCF_002058285.1_ASM205828v1 Salmonella enterica subsp. enterica serovar Newport Proteobacteria; Gammaproteobacteria; Enterobacterales; Enterobacteriaceae; Salmonella; Salmonella enterica; Salmonella enterica subsp. enterica MNNEETFYQAMRRKGVTRRSFLKFCSLAATSLGLGAGMTPKIAWALENKPRIPVVWIHGL WP_001058312.1 hydrogenase [Salmonella enterica] Length: 372\nNumber of predicted TMHs: 2\nExp number of AAs in TMHs: 36.51738\nExp number, first 60 AAs: 19.72849\nTotal prob of N-in: 0.96170\nPOSSIBLE N-term signal sequence\ninside 1 19\nTMhelix 20 39\noutside 40 325\nTMhelix 326 348\ninside 349 372

18718 GCF_002058605.1_ASM205860v1 Salmonella enterica subsp. enterica serovar Newport Proteobacteria; Gammaproteobacteria; Enterobacterales; Enterobacteriaceae; Salmonella; Salmonella enterica; Salmonella enterica subsp. enterica MTGDNTLITSHGINRRDFMKLCAALAATMGLSSKAAAEMAESVSNPQRPPVIWIGAQECT WP_000145429.1 hydrogenase 2 small subunit [Salmonella enterica] Length: 372\nNumber of predicted TMHs: 1\nExp number of AAs in TMHs: 24.51309\nExp number, first 60 AAs: 0.2682\nTotal prob of N-in: 0.14402\noutside 1 331\nTMhelix 332 354\ninside 355 372

18719 GCF_002058605.1_ASM205860v1 Salmonella enterica subsp. enterica serovar Newport Proteobacteria; Gammaproteobacteria; Enterobacterales; Enterobacteriaceae; Salmonella; Salmonella enterica; Salmonella enterica subsp. enterica MNNEETFYQAMRRKGVTRRSFLKFCSLAATSLGLGAGMTPKIAWALENKPRIPVVWIHGL WP_001058312.1 hydrogenase [Salmonella enterica] Length: 372\nNumber of predicted TMHs: 2\nExp number of AAs in TMHs: 36.51738\nExp number, first 60 AAs: 19.72849\nTotal prob of N-in: 0.96170\nPOSSIBLE N-term signal sequence\ninside 1 19\nTMhelix 20 39\noutside 40 325\nTMhelix 326 348\ninside 349 372

18720 GCF_001278315.1_ASM127831v1 Salmonella enterica subsp. enterica serovar Newport Proteobacteria; Gammaproteobacteria; Enterobacterales; Enterobacteriaceae; Salmonella; Salmonella enterica; Salmonella enterica subsp. enterica MTGDNTLITSHGINRRDFMKLCAALAATMGLSSKAAAEMAESVSNPQRPPVIWIGAQECT WP_000145429.1 hydrogenase 2 small subunit [Salmonella enterica] Length: 372\nNumber of

predicted TMHs:  1\nExp number of AAs in TMHs: 24.51309\nExp number, first 60 AAs:  0.2682\nTotal prob of N-in:  0.14402\noutside   1   331\nTMhelix   332   354\ninside   355   372

18721 GCF_001278315.1_ASM127831v1 Salmonella enterica subsp. enterica serovar Newport Proteobacteria; Gammaproteobacteria; Enterobacterales; Enterobacteriaceae; Salmonella; Salmonella enterica; Salmonella enterica subsp. enterica MNNEETFYQAMRRKGVTRRSFLKFCSLAATSLGLGAGMTPKIAWALENKPRIPVVWIHGL WP_001058312.1 hydrogenase [Salmonella enterica] Length: 372\nNumber of predicted TMHs:  2\nExp number of AAs in TMHs: 36.51738\nExp number, first 60 AAs:  19.72849\nTotal prob of N-in:   0.96170\nPOSSIBLE N-term signal sequence\ninside   1   19\nTMhelix   20   39\noutside   40   325\nTMhelix   326   348\ninside   349   372

18722 GCF_002090545.1_ASM209054v1 Salmonella enterica subsp. enterica serovar Newport Proteobacteria; Gammaproteobacteria; Enterobacterales; Enterobacteriaceae; Salmonella; Salmonella enterica; Salmonella enterica subsp. enterica MNNEETFYQAMRRKGVTRRSFLKFCSLAATSLGLGAGMTPKIAWALENKPRIPVVWIHGL WP_001058312.1 hydrogenase [Salmonella enterica] Length: 372\nNumber of predicted TMHs:  2\nExp number of AAs in TMHs: 36.51738\nExp number, first 60 AAs:  19.72849\nTotal prob of N-in:   0.96170\nPOSSIBLE N-term signal sequence\ninside   1   19\nTMhelix   20   39\noutside   40   325\nTMhelix   326   348\ninside   349   372

18723 GCF_002090835.1_ASM209083v1 Salmonella enterica subsp. enterica serovar Newport Proteobacteria; Gammaproteobacteria; Enterobacterales; Enterobacteriaceae; Salmonella; Salmonella enterica; Salmonella enterica subsp. enterica MTGDNTLITSHGINRRDFMKLCAALAATMGLSSKAAAEMAESVSNPQRPPVIWIGAQECT WP_000145429.1 hydrogenase 2 small subunit [Salmonella enterica] Length: 372\nNumber of predicted TMHs:  1\nExp number of AAs in TMHs: 24.51309\nExp number, first 60 AAs:  0.2682\nTotal prob of N-in:  0.14402\noutside   1   331\nTMhelix   332   354\ninside   355   372

18724 GCF_002064275.1_ASM206427v1 Salmonella enterica subsp. enterica serovar Newport Proteobacteria; Gammaproteobacteria; Enterobacterales; Enterobacteriaceae; Salmonella; Salmonella enterica; Salmonella enterica subsp. enterica MTGDNTLITSHGINRRDFMKLCAALAATMGLSSKAAAEMAESVSNPQRPPVIWIGAQECT WP_000145429.1 hydrogenase 2 small subunit [Salmonella enterica] Length: 372\nNumber of predicted TMHs:  1\nExp number of AAs in TMHs: 24.51309\nExp number, first 60 AAs:  0.2682\nTotal prob of N-in:  0.14402\noutside   1   331\nTMhelix   332   354\ninside   355   372

18725 GCF_002107465.1_ASM210746v1 Salmonella enterica subsp. enterica serovar Newport Proteobacteria; Gammaproteobacteria; Enterobacterales; Enterobacteriaceae; Salmonella; Salmonella enterica; Salmonella enterica subsp. enterica MTGDNTLITSHGINRRDFMKLCAALAATMGLSSKAAAEMAESVSNPQRPPVIWIGAQECT WP_000145429.1 hydrogenase 2 small subunit [Salmonella enterica] Length: 372\nNumber of predicted TMHs:  1\nExp number of AAs in TMHs: 24.51309\nExp number, first 60 AAs:  0.2682\nTotal prob of N-in:  0.14402\noutside   1   331\nTMhelix   332   354\ninside   355   372

18726 GCF_002064305.1_ASM206430v1 Salmonella enterica subsp. enterica serovar Newport Proteobacteria; Gammaproteobacteria; Enterobacterales; Enterobacteriaceae; Salmonella; Salmonella enterica; Salmonella enterica subsp. enterica MTGDNTLITSHGINRRDFMKLCAALAATMGLSSKAAAEMAESVSNPQRPPVIWIGAQECT WP_000145429.1 hydrogenase 2 small subunit [Salmonella enterica] Length: 372\nNumber of predicted TMHs:  1\nExp number of AAs in TMHs: 24.51309\nExp number, first 60 AAs:  0.2682\nTotal prob of N-in:  0.14402\noutside   1   331\nTMhelix   332   354\ninside   355   372

18727 GCF_002064275.1_ASM206427v1 Salmonella enterica subsp. enterica serovar Newport Proteobacteria; Gammaproteobacteria; Enterobacterales; Enterobacteriaceae; Salmonella; Salmonella enterica; Salmonella enterica subsp. enterica MNNEETFYQAMRRKGVTRRSFLKFCSLAATSLGLGAGMTPKIAWALENKPRIPVVWIHGL WP_001058312.1 hydrogenase [Salmonella enterica] Length: 372\nNumber of predicted TMHs:  2\nExp number of AAs in TMHs: 36.51738\nExp number, first 60 AAs:  19.72849\nTotal prob of N-in:   0.96170\nPOSSIBLE N-term signal sequence\ninside   1   19\nTMhelix   20   39\noutside   40   325\nTMhelix   326   348\ninside   349   372

18728 GCF_002065345.1_ASM206534v1 Salmonella enterica subsp. enterica serovar Newport Proteobacteria; Gammaproteobacteria; Enterobacterales; Enterobacteriaceae; Salmonella; Salmonella enterica; Salmonella enterica subsp. enterica MNNEETFYQAMRRKGVTRRSFLKFCSLAATSLGLGAGMTPKIAWALENKPRIPVVWIHGL WP_001058312.1 hydrogenase [Salmonella enterica] Length: 372\nNumber of predicted TMHs:  2\nExp number of AAs in TMHs: 36.51738\nExp number, first 60 AAs:  19.72849\nTotal prob of N-in:   0.96170\nPOSSIBLE N-term signal sequence\ninside   1   19\nTMhelix   20   39\noutside   40   325\nTMhelix   326   348\ninside   349   372

18729 GCF_002065345.1_ASM206534v1 Salmonella enterica subsp. enterica serovar Newport Proteobacteria; Gammaproteobacteria; Enterobacterales; Enterobacteriaceae; Salmonella; Salmonella enterica;

Salmonella enterica subsp. enterica	MTGDNTLITSHGINRRDFMKLCAALAATMGLSSKAAAEMAESVSNPQRPPVIWIGAQECT	WP_000145429.1	hydrogenase 2 small subunit [Salmonella enterica]	Length: 372\nNumber of predicted TMHs:  1\nExp number of AAs in TMHs: 24.51309\nExp number, first 60 AAs:  0.2682\nTotal prob of N-in:  0.14402\noutside   1   331\nTMhelix   332   354\ninside   355   372

18730	GCF_002106475.1_ASM210647v1	Salmonella enterica subsp. enterica serovar Newport	Proteobacteria; Gammaproteobacteria; Enterobacterales; Enterobacteriaceae; Salmonella; Salmonella enterica; Salmonella enterica subsp. enterica	MNNEETFYQAMRRKGVTRRSFLKFCSLAATSLGLGAGMTPKIAWALENKPRIPVVWIHGL	WP_001058312.1	hydrogenase [Salmonella enterica]	Length: 372\nNumber of predicted TMHs:  2\nExp number of AAs in TMHs: 36.51738\nExp number, first 60 AAs:  19.72849\nTotal prob of N-in:   0.96170\nPOSSIBLE N-term signal sequence\ninside   1   19\nTMhelix   20   39\noutside   40   325\nTMhelix   326   348\ninside   349   372

18731	GCF_002106475.1_ASM210647v1	Salmonella enterica subsp. enterica serovar Newport	Proteobacteria; Gammaproteobacteria; Enterobacterales; Enterobacteriaceae; Salmonella; Salmonella enterica; Salmonella enterica subsp. enterica	MTGDNTLITSHGINRRDFMKLCAALAATMGLSSKAAAEMAESVSNPQRPPVIWIGAQECT	WP_000145429.1	hydrogenase 2 small subunit [Salmonella enterica]	Length: 372\nNumber of predicted TMHs:  1\nExp number of AAs in TMHs: 24.51309\nExp number, first 60 AAs:  0.2682\nTotal prob of N-in:  0.14402\noutside   1   331\nTMhelix   332   354\ninside   355   372

18732	GCF_002106275.1_ASM210627v1	Salmonella enterica subsp. enterica serovar Newport	Proteobacteria; Gammaproteobacteria; Enterobacterales; Enterobacteriaceae; Salmonella; Salmonella enterica; Salmonella enterica subsp. enterica	MNNEETFYQAMRRKGVTRRSFLKFCSLAATSLGLGAGMTPKIAWALENKPRIPVVWIHGL	WP_001058312.1	hydrogenase [Salmonella enterica]	Length: 372\nNumber of predicted TMHs:  2\nExp number of AAs in TMHs: 36.51738\nExp number, first 60 AAs:  19.72849\nTotal prob of N-in:   0.96170\nPOSSIBLE N-term signal sequence\ninside   1   19\nTMhelix   20   39\noutside   40   325\nTMhelix   326   348\ninside   349   372

18733	GCF_002107585.1_ASM210758v1	Salmonella enterica subsp. enterica serovar Newport	Proteobacteria; Gammaproteobacteria; Enterobacterales; Enterobacteriaceae; Salmonella; Salmonella enterica; Salmonella enterica subsp. enterica	MTGDNTLITSHGINRRDFMKLCAALAATMGLSSKAAAEMAESVSNPQRPPVIWIGAQECT	WP_000145429.1	hydrogenase 2 small subunit [Salmonella enterica]	Length: 372\nNumber of predicted TMHs:  1\nExp number of AAs in TMHs: 24.51309\nExp number, first 60 AAs:  0.2682\nTotal prob of N-in:  0.14402\noutside   1   331\nTMhelix   332   354\ninside   355   372

18734	GCF_002090705.1_ASM209070v1	Salmonella enterica subsp. enterica serovar Newport	Proteobacteria; Gammaproteobacteria; Enterobacterales; Enterobacteriaceae; Salmonella; Salmonella enterica; Salmonella enterica subsp. enterica	MNNEETFYQAMRRKGVTRRSFLKFCSLAATSLGLGAGMTPKIAWALENKPRIPVVWIHGL	WP_001058312.1	hydrogenase [Salmonella enterica]	Length: 372\nNumber of predicted TMHs:  2\nExp number of AAs in TMHs: 36.51738\nExp number, first 60 AAs:  19.72849\nTotal prob of N-in:   0.96170\nPOSSIBLE N-term signal sequence\ninside   1   19\nTMhelix   20   39\noutside   40   325\nTMhelix   326   348\ninside   349   372

18735	GCF_002048325.1_ASM204832v1	Salmonella enterica subsp. enterica serovar Newport	Proteobacteria; Gammaproteobacteria; Enterobacterales; Enterobacteriaceae; Salmonella; Salmonella enterica; Salmonella enterica subsp. enterica	MNNEETFYQAMRRKGVTRRSFLKFCSLAATSLGLGAGMTPKIAWALENKPRIPVVWIHGL	WP_001058312.1	hydrogenase [Salmonella enterica]	Length: 372\nNumber of predicted TMHs:  2\nExp number of AAs in TMHs: 36.51738\nExp number, first 60 AAs:  19.72849\nTotal prob of N-in:   0.96170\nPOSSIBLE N-term signal sequence\ninside   1   19\nTMhelix   20   39\noutside   40   325\nTMhelix   326   348\ninside   349   372

18736	GCF_002033925.1_ASM203392v1	Salmonella enterica subsp. enterica serovar Newport	Proteobacteria; Gammaproteobacteria; Enterobacterales; Enterobacteriaceae; Salmonella; Salmonella enterica; Salmonella enterica subsp. enterica	MNNEETFYQAMRRKGVTRRSFLKFCSLAATSLGLGAGMTPKIAWALENKPRIPVVWIHGL	WP_001058312.1	hydrogenase [Salmonella enterica]	Length: 372\nNumber of predicted TMHs:  2\nExp number of AAs in TMHs: 36.51738\nExp number, first 60 AAs:  19.72849\nTotal prob of N-in:   0.96170\nPOSSIBLE N-term signal sequence\ninside   1   19\nTMhelix   20   39\noutside   40   325\nTMhelix   326   348\ninside   349   372

18737	GCF_001955135.1_ASM195513v1	Salmonella enterica subsp. enterica serovar Newport	Proteobacteria; Gammaproteobacteria; Enterobacterales; Enterobacteriaceae; Salmonella; Salmonella enterica; Salmonella enterica subsp. enterica	MTGDNTLITSHGINRRDFMKLCAALAATMGLSSKAAAEMAESVSNPQRPPVIWIGAQECT	WP_000145429.1	hydrogenase 2 small subunit [Salmonella enterica]	Length: 372\nNumber of predicted TMHs:  1\nExp number of AAs in TMHs: 24.51309\nExp number, first 60 AAs:  0.2682\nTotal prob of N-in:  0.14402\noutside   1   331\nTMhelix   332   354\ninside   355   372

18738	GCF_001954845.1_ASM195484v1	Salmonella enterica subsp. enterica serovar Newport	Proteobacteria; Gammaproteobacteria; Enterobacterales; Enterobacteriaceae; Salmonella; Salmonella enterica; Salmonella enterica subsp. enterica	MNNEETFYQAMRRKGVTRRSFLKFCSLAATSLGLGAGMTPKIAWALENKPRIPVVWIHGL	WP_001058312.1	hydrogenase [Salmonella enterica]	Length: 372\nNumber of predicted TMHs: 2\nExp number of AAs in TMHs: 36.51738\nExp number, first 60 AAs: 19.72849\nTotal prob of N-in: 0.96170\nPOSSIBLE N-term signal sequence\ninside 1 19\nTMhelix 20 39\noutside 40 325\nTMhelix 326 348\ninside 349 372

18739	GCF_001955035.1_ASM195503v1	Salmonella enterica subsp. enterica serovar Newport	Proteobacteria; Gammaproteobacteria; Enterobacterales; Enterobacteriaceae; Salmonella; Salmonella enterica; Salmonella enterica subsp. enterica	MTGDNTLITSHGINRRDFMKLCAALAATMGLSSKAAAEMAESVSNPQRPPVIWIGAQECT	WP_000145429.1	hydrogenase 2 small subunit [Salmonella enterica]	Length: 372\nNumber of predicted TMHs: 1\nExp number of AAs in TMHs: 24.51309\nExp number, first 60 AAs: 0.2682\nTotal prob of N-in: 0.14402\noutside 1 331\nTMhelix 332 354\ninside 355 372

18740	GCF_001955165.1_ASM195516v1	Salmonella enterica subsp. enterica serovar Newport	Proteobacteria; Gammaproteobacteria; Enterobacterales; Enterobacteriaceae; Salmonella; Salmonella enterica; Salmonella enterica subsp. enterica	MTGDNTLITSHGINRRDFMKLCAALAATMGLSSKAAAEMAESVSNPQRPPVIWIGAQECT	WP_000145429.1	hydrogenase 2 small subunit [Salmonella enterica]	Length: 372\nNumber of predicted TMHs: 1\nExp number of AAs in TMHs: 24.51309\nExp number, first 60 AAs: 0.2682\nTotal prob of N-in: 0.14402\noutside 1 331\nTMhelix 332 354\ninside 355 372

18741	GCF_001955165.1_ASM195516v1	Salmonella enterica subsp. enterica serovar Newport	Proteobacteria; Gammaproteobacteria; Enterobacterales; Enterobacteriaceae; Salmonella; Salmonella enterica; Salmonella enterica subsp. enterica	MNNEETFYQAMRRKGVTRRSFLKFCSLAATSLGLGAGMTPKIAWALENKPRIPVVWIHGL	WP_001058312.1	hydrogenase [Salmonella enterica]	Length: 372\nNumber of predicted TMHs: 2\nExp number of AAs in TMHs: 36.51738\nExp number, first 60 AAs: 19.72849\nTotal prob of N-in: 0.96170\nPOSSIBLE N-term signal sequence\ninside 1 19\nTMhelix 20 39\noutside 40 325\nTMhelix 326 348\ninside 349 372

18742	GCF_001243515.1_Salmonella_enterica_CVM_N46850-R_v1.0	Salmonella enterica subsp. enterica serovar Newport	Proteobacteria; Gammaproteobacteria; Enterobacterales; Enterobacteriaceae; Salmonella; Salmonella enterica; Salmonella enterica subsp. enterica	MTGDNTLITSHGINRRDFMKLCAALAATMGLSSKAAAEMAESVSNPQRPPVIWIGAQECT	WP_000145429.1	hydrogenase 2 small subunit [Salmonella enterica]	Length: 372\nNumber of predicted TMHs: 1\nExp number of AAs in TMHs: 24.51309\nExp number, first 60 AAs: 0.2682\nTotal prob of N-in: 0.14402\noutside 1 331\nTMhelix 332 354\ninside 355 372

18743	GCF_002036025.1_ASM203602v1	Salmonella enterica subsp. enterica serovar Newport	Proteobacteria; Gammaproteobacteria; Enterobacterales; Enterobacteriaceae; Salmonella; Salmonella enterica; Salmonella enterica subsp. enterica	MTGDNTLITSHGINRRDFMKLCAALAATMGLSSKAAAEMAESVSNPQRPPVIWIGAQECT	WP_000145429.1	hydrogenase 2 small subunit [Salmonella enterica]	Length: 372\nNumber of predicted TMHs: 1\nExp number of AAs in TMHs: 24.51309\nExp number, first 60 AAs: 0.2682\nTotal prob of N-in: 0.14402\noutside 1 331\nTMhelix 332 354\ninside 355 372

18744	GCF_001890165.1_ASM189016v1	Salmonella enterica subsp. enterica serovar Newport	Proteobacteria; Gammaproteobacteria; Enterobacterales; Enterobacteriaceae; Salmonella; Salmonella enterica; Salmonella enterica subsp. enterica	MTGDNTLITSHGINRRDFMKLCAALAATMGLSSKAAAEMAESVSNPQRPPVIWIGAQECT	WP_000145429.1	hydrogenase 2 small subunit [Salmonella enterica]	Length: 372\nNumber of predicted TMHs: 1\nExp number of AAs in TMHs: 24.51309\nExp number, first 60 AAs: 0.2682\nTotal prob of N-in: 0.14402\noutside 1 331\nTMhelix 332 354\ninside 355 372

18745	GCF_001479995.1_Salmonella_enterica_CVM_N38893-SQ_v1.0	Salmonella enterica subsp. enterica serovar Newport	Proteobacteria; Gammaproteobacteria; Enterobacterales; Enterobacteriaceae; Salmonella; Salmonella enterica; Salmonella enterica subsp. enterica	MNNEETFYQAMRRKGVTRRSFLKFCSLAATSLGLGAGMTPKIAWALENKPRIPVVWIHGL	WP_001058312.1	hydrogenase [Salmonella enterica]	Length: 372\nNumber of predicted TMHs: 2\nExp number of AAs in TMHs: 36.51738\nExp number, first 60 AAs: 19.72849\nTotal prob of N-in: 0.96170\nPOSSIBLE N-term signal sequence\ninside 1 19\nTMhelix 20 39\noutside 40 325\nTMhelix 326 348\ninside 349 372

18746	GCF_001890165.1_ASM189016v1	Salmonella enterica subsp. enterica serovar Newport	Proteobacteria; Gammaproteobacteria; Enterobacterales; Enterobacteriaceae; Salmonella; Salmonella enterica; Salmonella enterica subsp. enterica	MNNEETFYQAMRRKGVTRRSFLKFCSLAATSLGLGAGMTPKIAWALENKPRIPVVWIHGL	WP_001058312.1	hydrogenase [Salmonella enterica]	Length: 372\nNumber of predicted TMHs: 2\nExp

number of AAs in TMHs: 36.51738\nExp number, first 60 AAs: 19.72849\nTotal prob of N-in: 0.96170\nPOSSIBLE N-term signal sequence\ninside 1 19\nTMhelix 20 39\noutside 40 325\nTMhelix 326 348\ninside 349 372

18747 GCF_001955075.1_ASM195507v1 Salmonella enterica subsp. enterica serovar Newport Proteobacteria; Gammaproteobacteria; Enterobacterales; Enterobacteriaceae; Salmonella; Salmonella enterica; Salmonella enterica subsp. enterica MTGDNTLITSHGINRRDFMKLCAALAATMGLSSKAAAEMAESVSNPQRPPVIWIGAQECT WP_000145429.1 hydrogenase 2 small subunit [Salmonella enterica] Length: 372\nNumber of predicted TMHs: 1\nExp number of AAs in TMHs: 24.51309\nExp number, first 60 AAs: 0.2682\nTotal prob of N-in: 0.14402\noutside 1 331\nTMhelix 332 354\ninside 355 372

18748 GCF_001272275.1_Salmonella_enterica_CVM_N50439_v1.0 Salmonella enterica subsp. enterica serovar Newport Proteobacteria; Gammaproteobacteria; Enterobacterales; Enterobacteriaceae; Salmonella; Salmonella enterica; Salmonella enterica subsp. enterica MTGDNTLITSHGINRRDFMKLCAALAATMGLSSKAAAEMAESVSNPQRPPVIWIGAQECT WP_000145429.1 hydrogenase 2 small subunit [Salmonella enterica] Length: 372\nNumber of predicted TMHs: 1\nExp number of AAs in TMHs: 24.51309\nExp number, first 60 AAs: 0.2682\nTotal prob of N-in: 0.14402\noutside 1 331\nTMhelix 332 354\ninside 355 372

18749 GCF_002090915.1_ASM209091v1 Salmonella enterica subsp. enterica serovar Newport Proteobacteria; Gammaproteobacteria; Enterobacterales; Enterobacteriaceae; Salmonella; Salmonella enterica; Salmonella enterica subsp. enterica MTGDNTLITSHGINRRDFMKLCAALAATMGLSSKAAAEMAESVSNPQRPPVIWIGAQECT WP_000145429.1 hydrogenase 2 small subunit [Salmonella enterica] Length: 372\nNumber of predicted TMHs: 1\nExp number of AAs in TMHs: 24.51309\nExp number, first 60 AAs: 0.2682\nTotal prob of N-in: 0.14402\noutside 1 331\nTMhelix 332 354\ninside 355 372

18750 GCF_002107185.1_ASM210718v1 Salmonella enterica subsp. enterica serovar Newport Proteobacteria; Gammaproteobacteria; Enterobacterales; Enterobacteriaceae; Salmonella; Salmonella enterica; Salmonella enterica subsp. enterica MNNEETFYQAMRRKGVTRRSFLKFCSLAATSLGLGAGMTPKIAWALENKPRIPVVWIHGL WP_001058312.1 hydrogenase [Salmonella enterica] Length: 372\nNumber of predicted TMHs: 2\nExp number of AAs in TMHs: 36.51738\nExp number, first 60 AAs: 19.72849\nTotal prob of N-in: 0.96170\nPOSSIBLE N-term signal sequence\ninside 1 19\nTMhelix 20 39\noutside 40 325\nTMhelix 326 348\ninside 349 372

18751 GCF_002090705.1_ASM209070v1 Salmonella enterica subsp. enterica serovar Newport Proteobacteria; Gammaproteobacteria; Enterobacterales; Enterobacteriaceae; Salmonella; Salmonella enterica; Salmonella enterica subsp. enterica MTGDNTLITSHGINRRDFMKLCAALAATMGLSSKAAAEMAESVSNPQRPPVIWIGAQECT WP_000145429.1 hydrogenase 2 small subunit [Salmonella enterica] Length: 372\nNumber of predicted TMHs: 1\nExp number of AAs in TMHs: 24.51309\nExp number, first 60 AAs: 0.2682\nTotal prob of N-in: 0.14402\noutside 1 331\nTMhelix 332 354\ninside 355 372

18752 GCF_002106275.1_ASM210627v1 Salmonella enterica subsp. enterica serovar Newport Proteobacteria; Gammaproteobacteria; Enterobacterales; Enterobacteriaceae; Salmonella; Salmonella enterica; Salmonella enterica subsp. enterica MTGDNTLITSHGINRRDFMKLCAALAATMGLSSKAAAEMAESVSNPQRPPVIWIGAQECT WP_000145429.1 hydrogenase 2 small subunit [Salmonella enterica] Length: 372\nNumber of predicted TMHs: 1\nExp number of AAs in TMHs: 24.51309\nExp number, first 60 AAs: 0.2682\nTotal prob of N-in: 0.14402\noutside 1 331\nTMhelix 332 354\ninside 355 372

18753 GCF_002091155.1_ASM209115v1 Salmonella enterica subsp. enterica serovar Newport Proteobacteria; Gammaproteobacteria; Enterobacterales; Enterobacteriaceae; Salmonella; Salmonella enterica; Salmonella enterica subsp. enterica MNNEETFYQAMRRKGVTRRSFLKFCSLAATSLGLGAGMTPKIAWALENKPRIPVVWIHGL WP_001058312.1 hydrogenase [Salmonella enterica] Length: 372\nNumber of predicted TMHs: 2\nExp number of AAs in TMHs: 36.51738\nExp number, first 60 AAs: 19.72849\nTotal prob of N-in: 0.96170\nPOSSIBLE N-term signal sequence\ninside 1 19\nTMhelix 20 39\noutside 40 325\nTMhelix 326 348\ninside 349 372

18754 GCF_002090665.1_ASM209066v1 Salmonella enterica subsp. enterica serovar Newport Proteobacteria; Gammaproteobacteria; Enterobacterales; Enterobacteriaceae; Salmonella; Salmonella enterica; Salmonella enterica subsp. enterica MTGDNTLITSHGINRRDFMKLCAALAATMGLSSKAAAEMAESVSNPQRPPVIWIGAQECT WP_000145429.1 hydrogenase 2 small subunit [Salmonella enterica] Length: 372\nNumber of predicted TMHs: 1\nExp number of AAs in TMHs: 24.51309\nExp number, first 60 AAs: 0.2682\nTotal prob of N-in: 0.14402\noutside 1 331\nTMhelix 332 354\ninside 355 372

18755 GCF_002107465.1_ASM210746v1 Salmonella enterica subsp. enterica serovar Newport Proteobacteria; Gammaproteobacteria; Enterobacterales; Enterobacteriaceae; Salmonella; Salmonella enterica;

Salmonella enterica subsp. enterica MNNEETFYQAMRRKGVTRRSFLKFCSLAATSLGLGAGMTPKIAWALENKPRIPVVWIHGL WP_001058312.1 hydrogenase [Salmonella enterica] Length: 372\nNumber of predicted TMHs: 2\nExp number of AAs in TMHs: 36.51738\nExp number, first 60 AAs: 19.72849\nTotal prob of N-in: 0.96170\nPOSSIBLE N-term signal sequence\ninside 1 19\nTMhelix 20 39\noutside 40 325\nTMhelix 326 348\ninside 349 372

18756 GCF_002106655.1_ASM210665v1 Salmonella enterica subsp. enterica serovar Newport Proteobacteria; Gammaproteobacteria; Enterobacterales; Enterobacteriaceae; Salmonella; Salmonella enterica; Salmonella enterica subsp. enterica MTGDNTLITSHGINRRDFMKLCAALAATMGLSSKAAAEMAESVSNPQRPPVIWIGAQECT WP_000145429.1 hydrogenase 2 small subunit [Salmonella enterica] Length: 372\nNumber of predicted TMHs: 1\nExp number of AAs in TMHs: 24.51309\nExp number, first 60 AAs: 0.2682\nTotal prob of N-in: 0.14402\noutside 1 331\nTMhelix 332 354\ninside 355 372

18757 GCF_002091155.1_ASM209115v1 Salmonella enterica subsp. enterica serovar Newport Proteobacteria; Gammaproteobacteria; Enterobacterales; Enterobacteriaceae; Salmonella; Salmonella enterica; Salmonella enterica subsp. enterica MTGDNTLITSHGINRRDFMKLCAALAATMGLSSKAAAEMAESVSNPQRPPVIWIGAQECT WP_000145429.1 hydrogenase 2 small subunit [Salmonella enterica] Length: 372\nNumber of predicted TMHs: 1\nExp number of AAs in TMHs: 24.51309\nExp number, first 60 AAs: 0.2682\nTotal prob of N-in: 0.14402\noutside 1 331\nTMhelix 332 354\ninside 355 372

18758 GCF_002107585.1_ASM210758v1 Salmonella enterica subsp. enterica serovar Newport Proteobacteria; Gammaproteobacteria; Enterobacterales; Enterobacteriaceae; Salmonella; Salmonella enterica; Salmonella enterica subsp. enterica MNNEETFYQAMRRKGVTRRSFLKFCSLAATSLGLGAGMTPKIAWALENKPRIPVVWIHGL WP_001058312.1 hydrogenase [Salmonella enterica] Length: 372\nNumber of predicted TMHs: 2\nExp number of AAs in TMHs: 36.51738\nExp number, first 60 AAs: 19.72849\nTotal prob of N-in: 0.96170\nPOSSIBLE N-term signal sequence\ninside 1 19\nTMhelix 20 39\noutside 40 325\nTMhelix 326 348\ninside 349 372

18759 GCF_002106655.1_ASM210665v1 Salmonella enterica subsp. enterica serovar Newport Proteobacteria; Gammaproteobacteria; Enterobacterales; Enterobacteriaceae; Salmonella; Salmonella enterica; Salmonella enterica subsp. enterica MNNEETFYQAMRRKGVTRRSFLKFCSLAATSLGLGAGMTPKIAWALENKPRIPVVWIHGL WP_001058312.1 hydrogenase [Salmonella enterica] Length: 372\nNumber of predicted TMHs: 2\nExp number of AAs in TMHs: 36.51738\nExp number, first 60 AAs: 19.72849\nTotal prob of N-in: 0.96170\nPOSSIBLE N-term signal sequence\ninside 1 19\nTMhelix 20 39\noutside 40 325\nTMhelix 326 348\ninside 349 372

18760 GCF_002107155.1_ASM210715v1 Salmonella enterica subsp. enterica serovar Newport Proteobacteria; Gammaproteobacteria; Enterobacterales; Enterobacteriaceae; Salmonella; Salmonella enterica; Salmonella enterica subsp. enterica MTGDNTLITSHGINRRDFMKLCAALAATMGLSSKAAAEMAESVSNPQRPPVIWIGAQECT WP_000145429.1 hydrogenase 2 small subunit [Salmonella enterica] Length: 372\nNumber of predicted TMHs: 1\nExp number of AAs in TMHs: 24.51309\nExp number, first 60 AAs: 0.2682\nTotal prob of N-in: 0.14402\noutside 1 331\nTMhelix 332 354\ninside 355 372

18761 GCF_002107185.1_ASM210718v1 Salmonella enterica subsp. enterica serovar Newport Proteobacteria; Gammaproteobacteria; Enterobacterales; Enterobacteriaceae; Salmonella; Salmonella enterica; Salmonella enterica subsp. enterica MTGDNTLITSHGINRRDFMKLCAALAATMGLSSKAAAEMAESVSNPQRPPVIWIGAQECT WP_000145429.1 hydrogenase 2 small subunit [Salmonella enterica] Length: 372\nNumber of predicted TMHs: 1\nExp number of AAs in TMHs: 24.51309\nExp number, first 60 AAs: 0.2682\nTotal prob of N-in: 0.14402\noutside 1 331\nTMhelix 332 354\ninside 355 372

18762 GCF_002107265.1_ASM210726v1 Salmonella enterica subsp. enterica serovar Newport Proteobacteria; Gammaproteobacteria; Enterobacterales; Enterobacteriaceae; Salmonella; Salmonella enterica; Salmonella enterica subsp. enterica MTGDNTLITSHGINRRDFMKLCAALAATMGLSSKAAAEMAESVSNPQRPPVIWIGAQECT WP_000145429.1 hydrogenase 2 small subunit [Salmonella enterica] Length: 372\nNumber of predicted TMHs: 1\nExp number of AAs in TMHs: 24.51309\nExp number, first 60 AAs: 0.2682\nTotal prob of N-in: 0.14402\noutside 1 331\nTMhelix 332 354\ninside 355 372

18763 GCF_002090545.1_ASM209054v1 Salmonella enterica subsp. enterica serovar Newport Proteobacteria; Gammaproteobacteria; Enterobacterales; Enterobacteriaceae; Salmonella; Salmonella enterica; Salmonella enterica subsp. enterica MTGDNTLITSHGINRRDFMKLCAALAATMGLSSKAAAEMAESVSNPQRPPVIWIGAQECT WP_000145429.1 hydrogenase 2 small subunit [Salmonella enterica] Length: 372\nNumber of predicted TMHs: 1\nExp number of AAs in TMHs: 24.51309\nExp number, first 60 AAs: 0.2682\nTotal prob of N-in: 0.14402\noutside 1 331\nTMhelix 332 354\ninside 355 372

18764 GCF_002090625.1_ASM209062v1 Salmonella enterica subsp. enterica serovar Newport Proteobacteria; Gammaproteobacteria; Enterobacterales; Enterobacteriaceae; Salmonella; Salmonella enterica; Salmonella enterica subsp. enterica MTGDNTLITSHGINRRDFMKLCAALAATMGLSSKAAAEMAESVSNPQRPPVIWIGAQECT WP_000145429.1 hydrogenase 2 small subunit [Salmonella enterica] Length: 372\nNumber of predicted TMHs: 1\nExp number of AAs in TMHs: 24.51309\nExp number, first 60 AAs: 0.2682\nTotal prob of N-in: 0.14402\noutside 1 331\nTMhelix 332 354\ninside 355 372

18765 GCF_002107265.1_ASM210726v1 Salmonella enterica subsp. enterica serovar Newport Proteobacteria; Gammaproteobacteria; Enterobacterales; Enterobacteriaceae; Salmonella; Salmonella enterica; Salmonella enterica subsp. enterica MNNEETFYQAMRRKGVTRRSFLKFCSLAATSLGLGAGMTPKIAWALENKPRIPVVWIHGL WP_001058312.1 hydrogenase [Salmonella enterica] Length: 372\nNumber of predicted TMHs: 2\nExp number of AAs in TMHs: 36.51738\nExp number, first 60 AAs: 19.72849\nTotal prob of N-in: 0.96170\nPOSSIBLE N-term signal sequence\ninside 1 19\nTMhelix 20 39\noutside 40 325\nTMhelix 326 348\ninside 349 372

18766 GCF_002107155.1_ASM210715v1 Salmonella enterica subsp. enterica serovar Newport Proteobacteria; Gammaproteobacteria; Enterobacterales; Enterobacteriaceae; Salmonella; Salmonella enterica; Salmonella enterica subsp. enterica MNNEETFYQAMRRKGVTRRSFLKFCSLAATSLGLGAGMTPKIAWALENKPRIPVVWIHGL WP_001058312.1 hydrogenase [Salmonella enterica] Length: 372\nNumber of predicted TMHs: 2\nExp number of AAs in TMHs: 36.51738\nExp number, first 60 AAs: 19.72849\nTotal prob of N-in: 0.96170\nPOSSIBLE N-term signal sequence\ninside 1 19\nTMhelix 20 39\noutside 40 325\nTMhelix 326 348\ninside 349 372

18767 GCF_002090625.1_ASM209062v1 Salmonella enterica subsp. enterica serovar Newport Proteobacteria; Gammaproteobacteria; Enterobacterales; Enterobacteriaceae; Salmonella; Salmonella enterica; Salmonella enterica subsp. enterica MNNEETFYQAMRRKGVTRRSFLKFCSLAATSLGLGAGMTPKIAWALENKPRIPVVWIHGL WP_001058312.1 hydrogenase [Salmonella enterica] Length: 372\nNumber of predicted TMHs: 2\nExp number of AAs in TMHs: 36.51738\nExp number, first 60 AAs: 19.72849\nTotal prob of N-in: 0.96170\nPOSSIBLE N-term signal sequence\ninside 1 19\nTMhelix 20 39\noutside 40 325\nTMhelix 326 348\ninside 349 372

18768 GCF_002090915.1_ASM209091v1 Salmonella enterica subsp. enterica serovar Newport Proteobacteria; Gammaproteobacteria; Enterobacterales; Enterobacteriaceae; Salmonella; Salmonella enterica; Salmonella enterica subsp. enterica MNNEETFYQAMRRKGVTRRSFLKFCSLAATSLGLGAGMTPKIAWALENKPRIPVVWIHGL WP_001058312.1 hydrogenase [Salmonella enterica] Length: 372\nNumber of predicted TMHs: 2\nExp number of AAs in TMHs: 36.51738\nExp number, first 60 AAs: 19.72849\nTotal prob of N-in: 0.96170\nPOSSIBLE N-term signal sequence\ninside 1 19\nTMhelix 20 39\noutside 40 325\nTMhelix 326 348\ninside 349 372

18769 GCF_002090835.1_ASM209083v1 Salmonella enterica subsp. enterica serovar Newport Proteobacteria; Gammaproteobacteria; Enterobacterales; Enterobacteriaceae; Salmonella; Salmonella enterica; Salmonella enterica subsp. enterica MNNEETFYQAMRRKGVTRRSFLKFCSLAATSLGLGAGMTPKIAWALENKPRIPVVWIHGL WP_001058312.1 hydrogenase [Salmonella enterica] Length: 372\nNumber of predicted TMHs: 2\nExp number of AAs in TMHs: 36.51738\nExp number, first 60 AAs: 19.72849\nTotal prob of N-in: 0.96170\nPOSSIBLE N-term signal sequence\ninside 1 19\nTMhelix 20 39\noutside 40 325\nTMhelix 326 348\ninside 349 372

18770 GCF_002090665.1_ASM209066v1 Salmonella enterica subsp. enterica serovar Newport Proteobacteria; Gammaproteobacteria; Enterobacterales; Enterobacteriaceae; Salmonella; Salmonella enterica; Salmonella enterica subsp. enterica MNNEETFYQAMRRKGVTRRSFLKFCSLAATSLGLGAGMTPKIAWALENKPRIPVVWIHGL WP_001058312.1 hydrogenase [Salmonella enterica] Length: 372\nNumber of predicted TMHs: 2\nExp number of AAs in TMHs: 36.51738\nExp number, first 60 AAs: 19.72849\nTotal prob of N-in: 0.96170\nPOSSIBLE N-term signal sequence\ninside 1 19\nTMhelix 20 39\noutside 40 325\nTMhelix 326 348\ninside 349 372

18771 GCF_002056835.1_ASM205683v1 Salmonella enterica subsp. enterica serovar Newport Proteobacteria; Gammaproteobacteria; Enterobacterales; Enterobacteriaceae; Salmonella; Salmonella enterica; Salmonella enterica subsp. enterica MTGDNTLITSHGINRRDFMKLCAALAATMGLSSKAAAEMAESVSNPQRPPVIWIGAQECT WP_000145429.1 hydrogenase 2 small subunit [Salmonella enterica] Length: 372\nNumber of predicted TMHs: 1\nExp number of AAs in TMHs: 24.51309\nExp number, first 60 AAs: 0.2682\nTotal prob of N-in: 0.14402\noutside 1 331\nTMhelix 332 354\ninside 355 372

18772 GCF_002056835.1_ASM205683v1 Salmonella enterica subsp. enterica serovar Newport Proteobacteria; Gammaproteobacteria; Enterobacterales; Enterobacteriaceae; Salmonella; Salmonella enterica; Salmonella enterica subsp. enterica MNNEETFYQAMRRKGVTRRSFLKFCSLAATSLGLGAGMTPKIAWALENKPRIPVVWIHGL WP_001058312.1 hydrogenase [Salmonella enterica] Length: 372\nNumber of predicted TMHs: 2\nExp

number of AAs in TMHs: 36.51738\nExp number, first 60 AAs: 19.72849\nTotal prob of N-in: 0.96170\nPOSSIBLE N-term signal sequence\ninside 1 19\nTMhelix 20 39\noutside 40 325\nTMhelix 326 348\ninside 349 372

18773 GCF_002057935.1_ASM205793v1 Salmonella enterica subsp. enterica serovar Newport Proteobacteria; Gammaproteobacteria; Enterobacterales; Enterobacteriaceae; Salmonella; Salmonella enterica; Salmonella enterica subsp. enterica MTGDNTLITSHGINRRDFMKLCAALAATMGLSSKAAAEMAESVSNPQRPPVIWIGAQECT WP_000145429.1 hydrogenase 2 small subunit [Salmonella enterica] Length: 372\nNumber of predicted TMHs: 1\nExp number of AAs in TMHs: 24.51309\nExp number, first 60 AAs: 0.2682\nTotal prob of N-in: 0.14402\noutside 1 331\nTMhelix 332 354\ninside 355 372

18774 GCF_002058635.1_ASM205863v1 Salmonella enterica subsp. enterica serovar Newport Proteobacteria; Gammaproteobacteria; Enterobacterales; Enterobacteriaceae; Salmonella; Salmonella enterica; Salmonella enterica subsp. enterica MTGDNTLITSHGINRRDFMKLCAALAATMGLSSKAAAEMAESVSNPQRPPVIWIGAQECT WP_000145429.1 hydrogenase 2 small subunit [Salmonella enterica] Length: 372\nNumber of predicted TMHs: 1\nExp number of AAs in TMHs: 24.51309\nExp number, first 60 AAs: 0.2682\nTotal prob of N-in: 0.14402\noutside 1 331\nTMhelix 332 354\ninside 355 372

18775 GCF_002059255.1_ASM205925v1 Salmonella enterica subsp. enterica serovar Newport Proteobacteria; Gammaproteobacteria; Enterobacterales; Enterobacteriaceae; Salmonella; Salmonella enterica; Salmonella enterica subsp. enterica MNNEETFYQAMRRKGVTRRSFLKFCSLAATSLGLGAGMTPKIAWALENKPRIPVVWIHGL WP_001058312.1 hydrogenase [Salmonella enterica] Length: 372\nNumber of predicted TMHs: 2\nExp number of AAs in TMHs: 36.51738\nExp number, first 60 AAs: 19.72849\nTotal prob of N-in: 0.96170\nPOSSIBLE N-term signal sequence\ninside 1 19\nTMhelix 20 39\noutside 40 325\nTMhelix 326 348\ninside 349 372

18776 GCF_001834665.1_ASM183466v1 Salmonella enterica subsp. enterica serovar Newport Proteobacteria; Gammaproteobacteria; Enterobacterales; Enterobacteriaceae; Salmonella; Salmonella enterica; Salmonella enterica subsp. enterica MNNEETFYQAMRRKGVTRRSFLKFCSLAATSLGLGAGMTPKIAWALENKPRIPVVWIHGL WP_001058312.1 hydrogenase [Salmonella enterica] Length: 372\nNumber of predicted TMHs: 2\nExp number of AAs in TMHs: 36.51738\nExp number, first 60 AAs: 19.72849\nTotal prob of N-in: 0.96170\nPOSSIBLE N-term signal sequence\ninside 1 19\nTMhelix 20 39\noutside 40 325\nTMhelix 326 348\ninside 349 372

18777 GCF_001834705.1_ASM183470v1 Salmonella enterica subsp. enterica serovar Newport Proteobacteria; Gammaproteobacteria; Enterobacterales; Enterobacteriaceae; Salmonella; Salmonella enterica; Salmonella enterica subsp. enterica MNNEETFYQAMRRKGVTRRSFLKFCSLAATSLGLGAGMTPKIAWALENKPRIPVVWIHGL WP_001058312.1 hydrogenase [Salmonella enterica] Length: 372\nNumber of predicted TMHs: 2\nExp number of AAs in TMHs: 36.51738\nExp number, first 60 AAs: 19.72849\nTotal prob of N-in: 0.96170\nPOSSIBLE N-term signal sequence\ninside 1 19\nTMhelix 20 39\noutside 40 325\nTMhelix 326 348\ninside 349 372

18778 GCF_001835255.1_ASM183525v1 Salmonella enterica subsp. enterica serovar Newport Proteobacteria; Gammaproteobacteria; Enterobacterales; Enterobacteriaceae; Salmonella; Salmonella enterica; Salmonella enterica subsp. enterica MTGDNTLITSHGINRRDFMKLCAALAATMGLSSKAAAEMAESVSNPQRPPVIWIGAQECT WP_000145429.1 hydrogenase 2 small subunit [Salmonella enterica] Length: 372\nNumber of predicted TMHs: 1\nExp number of AAs in TMHs: 24.51309\nExp number, first 60 AAs: 0.2682\nTotal prob of N-in: 0.14402\noutside 1 331\nTMhelix 332 354\ninside 355 372

18779 GCF_001922165.1_ASM192216v1 Salmonella enterica subsp. enterica serovar Newport Proteobacteria; Gammaproteobacteria; Enterobacterales; Enterobacteriaceae; Salmonella; Salmonella enterica; Salmonella enterica subsp. enterica MTGDNTLITSHGINRRDFMKLCAALAATMGLSSKAAAEMAESVSNPQRPPVIWIGAQECT WP_000145429.1 hydrogenase 2 small subunit [Salmonella enterica] Length: 372\nNumber of predicted TMHs: 1\nExp number of AAs in TMHs: 24.51309\nExp number, first 60 AAs: 0.2682\nTotal prob of N-in: 0.14402\noutside 1 331\nTMhelix 332 354\ninside 355 372

18780 GCF_001922165.1_ASM192216v1 Salmonella enterica subsp. enterica serovar Newport Proteobacteria; Gammaproteobacteria; Enterobacterales; Enterobacteriaceae; Salmonella; Salmonella enterica; Salmonella enterica subsp. enterica MNNEETFYQAMRRKGVTRRSFLKFCSLAATSLGLGAGMTPKIAWALENKPRIPVVWIHGL WP_001058312.1 hydrogenase [Salmonella enterica] Length: 372\nNumber of predicted TMHs: 2\nExp number of AAs in TMHs: 36.51738\nExp number, first 60 AAs: 19.72849\nTotal prob of N-in: 0.96170\nPOSSIBLE N-term signal sequence\ninside 1 19\nTMhelix 20 39\noutside 40 325\nTMhelix 326 348\ninside 349 372

18781 GCF_001954755.1_ASM195475v1 Salmonella enterica subsp. enterica serovar Newport Proteobacteria; Gammaproteobacteria; Enterobacterales; Enterobacteriaceae; Salmonella; Salmonella enterica;

Salmonella enterica subsp. enterica MNNEETFYQAMRRKGVTRRSFLKFCSLAATSLGLGAGMTPKIAWALENKPRIPVVWIHGL WP_001058312.1 hydrogenase [Salmonella enterica] Length: 372\nNumber of predicted TMHs: 2\nExp number of AAs in TMHs: 36.51738\nExp number, first 60 AAs: 19.72849\nTotal prob of N-in: 0.96170\nPOSSIBLE N-term signal sequence\ninside 1 19\nTMhelix 20 39\noutside 40 325\nTMhelix 326 348\ninside 349 372

18782 GCF_001954815.1_ASM195481v1 Salmonella enterica subsp. enterica serovar Newport Proteobacteria; Gammaproteobacteria; Enterobacterales; Enterobacteriaceae; Salmonella; Salmonella enterica; Salmonella enterica subsp. enterica MNNEETFYQAMRRKGVTRRSFLKFCSLAATSLGLGAGMTPKIAWALENKPRIPVVWIHGL WP_001058312.1 hydrogenase [Salmonella enterica] Length: 372\nNumber of predicted TMHs: 2\nExp number of AAs in TMHs: 36.51738\nExp number, first 60 AAs: 19.72849\nTotal prob of N-in: 0.96170\nPOSSIBLE N-term signal sequence\ninside 1 19\nTMhelix 20 39\noutside 40 325\nTMhelix 326 348\ninside 349 372

18783 GCF_001954835.1_ASM195483v1 Salmonella enterica subsp. enterica serovar Newport Proteobacteria; Gammaproteobacteria; Enterobacterales; Enterobacteriaceae; Salmonella; Salmonella enterica; Salmonella enterica subsp. enterica MNNEETFYQAMRRKGVTRRSFLKFCSLAATSLGLGAGMTPKIAWALENKPRIPVVWIHGL WP_001058312.1 hydrogenase [Salmonella enterica] Length: 372\nNumber of predicted TMHs: 2\nExp number of AAs in TMHs: 36.51738\nExp number, first 60 AAs: 19.72849\nTotal prob of N-in: 0.96170\nPOSSIBLE N-term signal sequence\ninside 1 19\nTMhelix 20 39\noutside 40 325\nTMhelix 326 348\ninside 349 372

18784 GCF_002031995.1_ASM203199v1 Salmonella enterica subsp. enterica serovar Newport Proteobacteria; Gammaproteobacteria; Enterobacterales; Enterobacteriaceae; Salmonella; Salmonella enterica; Salmonella enterica subsp. enterica MTGDNTLITSHGINRRDFMKLCAALAATMGLSSKAAAEMAESVSNPQRPPVIWIGAQECT WP_000145429.1 hydrogenase 2 small subunit [Salmonella enterica] Length: 372\nNumber of predicted TMHs: 1\nExp number of AAs in TMHs: 24.51309\nExp number, first 60 AAs: 0.2682\nTotal prob of N-in: 0.14402\noutside 1 331\nTMhelix 332 354\ninside 355 372

18785 GCF_001480655.1_Salmonella_enterica_CVM_N42477_v1.0 Salmonella enterica subsp. enterica serovar Newport Proteobacteria; Gammaproteobacteria; Enterobacterales; Enterobacteriaceae; Salmonella; Salmonella enterica; Salmonella enterica subsp. enterica MNNEETFYQAMRRKGVTRRSFLKFCSLAATSLGLGAGMTPKIAWALENKPRIPVVWIHGL WP_001058312.1 hydrogenase [Salmonella enterica] Length: 372\nNumber of predicted TMHs: 2\nExp number of AAs in TMHs: 36.51738\nExp number, first 60 AAs: 19.72849\nTotal prob of N-in: 0.96170\nPOSSIBLE N-term signal sequence\ninside 1 19\nTMhelix 20 39\noutside 40 325\nTMhelix 326 348\ninside 349 372

18786 GCF_002061655.1_ASM206165v1 Salmonella enterica subsp. enterica serovar Newport Proteobacteria; Gammaproteobacteria; Enterobacterales; Enterobacteriaceae; Salmonella; Salmonella enterica; Salmonella enterica subsp. enterica MNNEETFYQAMRRKGVTRRSFLKFCSLAATSLGLGAGMTPKIAWALENKPRIPVVWIHGL WP_001058312.1 hydrogenase [Salmonella enterica] Length: 372\nNumber of predicted TMHs: 2\nExp number of AAs in TMHs: 36.51738\nExp number, first 60 AAs: 19.72849\nTotal prob of N-in: 0.96170\nPOSSIBLE N-term signal sequence\ninside 1 19\nTMhelix 20 39\noutside 40 325\nTMhelix 326 348\ninside 349 372

18787 GCF_002047445.1_ASM204744v1 Salmonella enterica subsp. enterica serovar Newport Proteobacteria; Gammaproteobacteria; Enterobacterales; Enterobacteriaceae; Salmonella; Salmonella enterica; Salmonella enterica subsp. enterica MTGDNTLITSHGINRRDFMKLCAALAATMGLSSKAAAEMAESVSNPQRPPVIWIGAQECT WP_000145429.1 hydrogenase 2 small subunit [Salmonella enterica] Length: 372\nNumber of predicted TMHs: 1\nExp number of AAs in TMHs: 24.51309\nExp number, first 60 AAs: 0.2682\nTotal prob of N-in: 0.14402\noutside 1 331\nTMhelix 332 354\ninside 355 372

18788 GCF_002047445.1_ASM204744v1 Salmonella enterica subsp. enterica serovar Newport Proteobacteria; Gammaproteobacteria; Enterobacterales; Enterobacteriaceae; Salmonella; Salmonella enterica; Salmonella enterica subsp. enterica MNNEETFYQAMRRKGVTRRSFLKFCSLAATSLGLGAGMTPKIAWALENKPRIPVVWIHGL WP_001058312.1 hydrogenase [Salmonella enterica] Length: 372\nNumber of predicted TMHs: 2\nExp number of AAs in TMHs: 36.51738\nExp number, first 60 AAs: 19.72849\nTotal prob of N-in: 0.96170\nPOSSIBLE N-term signal sequence\ninside 1 19\nTMhelix 20 39\noutside 40 325\nTMhelix 326 348\ninside 349 372

18789 GCF_001480655.1_Salmonella_enterica_CVM_N42477_v1.0 Salmonella enterica subsp. enterica serovar Newport Proteobacteria; Gammaproteobacteria; Enterobacterales; Enterobacteriaceae; Salmonella; Salmonella enterica; Salmonella enterica subsp. enterica MTGDNTLITSHGINRRDFMKLCAALAATMGLSSKAAAEMAESVSNPQRPPVIWIGAQECT WP_000145429.1 hydrogenase 2 small subunit [Salmonella enterica] Length: 372\nNumber of predicted TMHs: 1\nExp number of AAs in TMHs: 24.51309\nExp number, first 60 AAs: 0.2682\nTotal prob of N-in: 0.14402\noutside 1 331\nTMhelix 332 354\ninside 355 372

18790 GCF_001955135.1_ASM195513v1 Salmonella enterica subsp. enterica serovar Newport Proteobacteria; Gammaproteobacteria; Enterobacterales; Enterobacteriaceae; Salmonella; Salmonella enterica; Salmonella enterica subsp. enterica MNNEETFYQAMRRKGVTRRSFLKFCSLAATSLGLGAGMTPKIAWALENKPRIPVVWIHGL WP_001058312.1 hydrogenase [Salmonella enterica] Length: 372\nNumber of predicted TMHs: 2\nExp number of AAs in TMHs: 36.51738\nExp number, first 60 AAs: 19.72849\nTotal prob of N-in: 0.96170\nPOSSIBLE N-term signal sequence\ninside 1 19\nTMhelix 20 39\noutside 40 325\nTMhelix 326 348\ninside 349 372

18791 GCF_002058335.1_ASM205833v1 Salmonella enterica subsp. enterica serovar Newport Proteobacteria; Gammaproteobacteria; Enterobacterales; Enterobacteriaceae; Salmonella; Salmonella enterica; Salmonella enterica subsp. enterica MNNEETFYQAMRRKGVTRRSFLKFCSLAATSLGLGAGMTPKIAWALENKPRIPVVWIHGL WP_001058312.1 hydrogenase [Salmonella enterica] Length: 372\nNumber of predicted TMHs: 2\nExp number of AAs in TMHs: 36.51738\nExp number, first 60 AAs: 19.72849\nTotal prob of N-in: 0.96170\nPOSSIBLE N-term signal sequence\ninside 1 19\nTMhelix 20 39\noutside 40 325\nTMhelix 326 348\ninside 349 372

18792 GCF_001272275.1_Salmonella_enterica_CVM_N50439_v1.0 Salmonella enterica subsp. enterica serovar Newport Proteobacteria; Gammaproteobacteria; Enterobacterales; Enterobacteriaceae; Salmonella; Salmonella enterica; Salmonella enterica subsp. enterica MNNEETFYQAMRRKGVTRRSFLKFCSLAATSLGLGAGMTPKIAWALENKPRIPVVWIHGL WP_001058312.1 hydrogenase [Salmonella enterica] Length: 372\nNumber of predicted TMHs: 2\nExp number of AAs in TMHs: 36.51738\nExp number, first 60 AAs: 19.72849\nTotal prob of N-in: 0.96170\nPOSSIBLE N-term signal sequence\ninside 1 19\nTMhelix 20 39\noutside 40 325\nTMhelix 326 348\ninside 349 372

18793 GCF_001955035.1_ASM195503v1 Salmonella enterica subsp. enterica serovar Newport Proteobacteria; Gammaproteobacteria; Enterobacterales; Enterobacteriaceae; Salmonella; Salmonella enterica; Salmonella enterica subsp. enterica MNNEETFYQAMRRKGVTRRSFLKFCSLAATSLGLGAGMTPKIAWALENKPRIPVVWIHGL WP_001058312.1 hydrogenase [Salmonella enterica] Length: 372\nNumber of predicted TMHs: 2\nExp number of AAs in TMHs: 36.51738\nExp number, first 60 AAs: 19.72849\nTotal prob of N-in: 0.96170\nPOSSIBLE N-term signal sequence\ninside 1 19\nTMhelix 20 39\noutside 40 325\nTMhelix 326 348\ninside 349 372

18794 GCF_001479995.1_Salmonella_enterica_CVM_N38893-SQ_v1.0 Salmonella enterica subsp. enterica serovar Newport Proteobacteria; Gammaproteobacteria; Enterobacterales; Enterobacteriaceae; Salmonella; Salmonella enterica; Salmonella enterica subsp. enterica MTGDNTLITSHGINRRDFMKLCAALAATMGLSSKAAAEMAESVSNPQRPPVIWIGAQECT WP_000145429.1 hydrogenase 2 small subunit [Salmonella enterica] Length: 372\nNumber of predicted TMHs: 1\nExp number of AAs in TMHs: 24.51309\nExp number, first 60 AAs: 0.2682\nTotal prob of N-in: 0.14402\noutside 1 331\nTMhelix 332 354\ninside 355 372

18795 GCF_002059855.1_ASM205985v1 Salmonella enterica subsp. enterica serovar Newport Proteobacteria; Gammaproteobacteria; Enterobacterales; Enterobacteriaceae; Salmonella; Salmonella enterica; Salmonella enterica subsp. enterica MTGDNTLITSHGINRRDFMKLCAALAATMGLSSKAAAEMAESVSNPQRPPVIWIGAQECT WP_000145429.1 hydrogenase 2 small subunit [Salmonella enterica] Length: 372\nNumber of predicted TMHs: 1\nExp number of AAs in TMHs: 24.51309\nExp number, first 60 AAs: 0.2682\nTotal prob of N-in: 0.14402\noutside 1 331\nTMhelix 332 354\ninside 355 372

18796 GCF_002059855.1_ASM205985v1 Salmonella enterica subsp. enterica serovar Newport Proteobacteria; Gammaproteobacteria; Enterobacterales; Enterobacteriaceae; Salmonella; Salmonella enterica; Salmonella enterica subsp. enterica MNNEETFYQAMRRKGVTRRSFLKFCSLAATSLGLGAGMTPKIAWALENKPRIPVVWIHGL WP_001058312.1 hydrogenase [Salmonella enterica] Length: 372\nNumber of predicted TMHs: 2\nExp number of AAs in TMHs: 36.51738\nExp number, first 60 AAs: 19.72849\nTotal prob of N-in: 0.96170\nPOSSIBLE N-term signal sequence\ninside 1 19\nTMhelix 20 39\noutside 40 325\nTMhelix 326 348\ninside 349 372

18797 GCF_001955075.1_ASM195507v1 Salmonella enterica subsp. enterica serovar Newport Proteobacteria; Gammaproteobacteria; Enterobacterales; Enterobacteriaceae; Salmonella; Salmonella enterica; Salmonella enterica subsp. enterica MNNEETFYQAMRRKGVTRRSFLKFCSLAATSLGLGAGMTPKIAWALENKPRIPVVWIHGL WP_001058312.1 hydrogenase [Salmonella enterica] Length: 372\nNumber of predicted TMHs: 2\nExp number of AAs in TMHs: 36.51738\nExp number, first 60 AAs: 19.72849\nTotal prob of N-in: 0.96170\nPOSSIBLE N-term signal sequence\ninside 1 19\nTMhelix 20 39\noutside 40 325\nTMhelix 326 348\ninside 349 372

18798 GCF_001954765.1_ASM195476v1 Salmonella enterica subsp. enterica serovar Newport Proteobacteria; Gammaproteobacteria; Enterobacterales; Enterobacteriaceae; Salmonella; Salmonella enterica; Salmonella enterica subsp. enterica MNNEETFYQAMRRKGVTRRSFLKFCSLAATSLGLGAGMTPKIAWALENKPRIPVVWIHGL WP_001058312.1 hydrogenase [Salmonella enterica] Length: 372\nNumber of predicted TMHs: 2\nExp

number of AAs in TMHs: 36.51738\nExp number, first 60 AAs: 19.72849\nTotal prob of N-in: 0.96170\nPOSSIBLE N-term signal sequence\ninside 1 19\nTMhelix 20 39\noutside 40 325\nTMhelix 326 348\ninside 349 372

18799 GCF_002060195.1_ASM206019v1 Salmonella enterica subsp. enterica serovar Newport Proteobacteria; Gammaproteobacteria; Enterobacterales; Enterobacteriaceae; Salmonella; Salmonella enterica; Salmonella enterica subsp. enterica MTGDNTLITSHGINRRDFMKLCAALAATMGLSSKAAAEMAESVSNPQRPPVIWIGAQECT WP_000145429.1 hydrogenase 2 small subunit [Salmonella enterica] Length: 372\nNumber of predicted TMHs: 1\nExp number of AAs in TMHs: 24.51309\nExp number, first 60 AAs: 0.2682\nTotal prob of N-in: 0.14402\noutside 1 331\nTMhelix 332 354\ninside 355 372

18800 GCF_002056745.1_ASM205674v1 Salmonella enterica subsp. enterica serovar Newport Proteobacteria; Gammaproteobacteria; Enterobacterales; Enterobacteriaceae; Salmonella; Salmonella enterica; Salmonella enterica subsp. enterica MTGDNTLITSHGINRRDFMKLCAALAATMGLSSKAAAEMAESVSNPQRPPVIWIGAQECT WP_000145429.1 hydrogenase 2 small subunit [Salmonella enterica] Length: 372\nNumber of predicted TMHs: 1\nExp number of AAs in TMHs: 24.51309\nExp number, first 60 AAs: 0.2682\nTotal prob of N-in: 0.14402\noutside 1 331\nTMhelix 332 354\ninside 355 372

18801 GCF_002056845.1_ASM205684v1 Salmonella enterica subsp. enterica serovar Newport Proteobacteria; Gammaproteobacteria; Enterobacterales; Enterobacteriaceae; Salmonella; Salmonella enterica; Salmonella enterica subsp. enterica MNNEETFYQAMRRKGVTRRSFLKFCSLAATSLGLGAGMTPKIAWALENKPRIPVVWIHGL WP_001058312.1 hydrogenase [Salmonella enterica] Length: 372\nNumber of predicted TMHs: 2\nExp number of AAs in TMHs: 36.51738\nExp number, first 60 AAs: 19.72849\nTotal prob of N-in: 0.96170\nPOSSIBLE N-term signal sequence\ninside 1 19\nTMhelix 20 39\noutside 40 325\nTMhelix 326 348\ninside 349 372

18802 GCF_001954825.1_ASM195482v1 Salmonella enterica subsp. enterica serovar Newport Proteobacteria; Gammaproteobacteria; Enterobacterales; Enterobacteriaceae; Salmonella; Salmonella enterica; Salmonella enterica subsp. enterica MNNEETFYQAMRRKGVTRRSFLKFCSLAATSLGLGAGMTPKIAWALENKPRIPVVWIHGL WP_001058312.1 hydrogenase [Salmonella enterica] Length: 372\nNumber of predicted TMHs: 2\nExp number of AAs in TMHs: 36.51738\nExp number, first 60 AAs: 19.72849\nTotal prob of N-in: 0.96170\nPOSSIBLE N-term signal sequence\ninside 1 19\nTMhelix 20 39\noutside 40 325\nTMhelix 326 348\ninside 349 372

18803 GCF_002033925.1_ASM203392v1 Salmonella enterica subsp. enterica serovar Newport Proteobacteria; Gammaproteobacteria; Enterobacterales; Enterobacteriaceae; Salmonella; Salmonella enterica; Salmonella enterica subsp. enterica MTGDNTLITSHGINRRDFMKLCAALAATMGLSSKAAAEMAESVSNPQRPPVIWIGAQECT WP_000145429.1 hydrogenase 2 small subunit [Salmonella enterica] Length: 372\nNumber of predicted TMHs: 1\nExp number of AAs in TMHs: 24.51309\nExp number, first 60 AAs: 0.2682\nTotal prob of N-in: 0.14402\noutside 1 331\nTMhelix 332 354\ninside 355 372

18804 GCF_002057085.1_ASM205708v1 Salmonella enterica subsp. enterica serovar Newport Proteobacteria; Gammaproteobacteria; Enterobacterales; Enterobacteriaceae; Salmonella; Salmonella enterica; Salmonella enterica subsp. enterica MTGDNTLITSHGINRRDFMKLCAALAATMGLSSKAAAEMAESVSNPQRPPVIWIGAQECT WP_000145429.1 hydrogenase 2 small subunit [Salmonella enterica] Length: 372\nNumber of predicted TMHs: 1\nExp number of AAs in TMHs: 24.51309\nExp number, first 60 AAs: 0.2682\nTotal prob of N-in: 0.14402\noutside 1 331\nTMhelix 332 354\ninside 355 372

18805 GCF_001835415.1_ASM183541v1 Salmonella enterica subsp. enterica serovar Newport Proteobacteria; Gammaproteobacteria; Enterobacterales; Enterobacteriaceae; Salmonella; Salmonella enterica; Salmonella enterica subsp. enterica MNNEETFYQAMRRKGVTRRSFLKFCSLAATSLGLGAGMTPKIAWALENKPRIPVVWIHGL WP_001058312.1 hydrogenase [Salmonella enterica] Length: 372\nNumber of predicted TMHs: 2\nExp number of AAs in TMHs: 36.51738\nExp number, first 60 AAs: 19.72849\nTotal prob of N-in: 0.96170\nPOSSIBLE N-term signal sequence\ninside 1 19\nTMhelix 20 39\noutside 40 325\nTMhelix 326 348\ninside 349 372

18806 GCF_002057085.1_ASM205708v1 Salmonella enterica subsp. enterica serovar Newport Proteobacteria; Gammaproteobacteria; Enterobacterales; Enterobacteriaceae; Salmonella; Salmonella enterica; Salmonella enterica subsp. enterica MNNEETFYQAMRRKGVTRRSFLKFCSLAATSLGLGAGMTPKIAWALENKPRIPVVWIHGL WP_001058312.1 hydrogenase [Salmonella enterica] Length: 372\nNumber of predicted TMHs: 2\nExp number of AAs in TMHs: 36.51738\nExp number, first 60 AAs: 19.72849\nTotal prob of N-in: 0.96170\nPOSSIBLE N-term signal sequence\ninside 1 19\nTMhelix 20 39\noutside 40 325\nTMhelix 326 348\ninside 349 372

18807 GCF_002056915.1_ASM205691v1 Salmonella enterica subsp. enterica serovar Newport Proteobacteria; Gammaproteobacteria; Enterobacterales; Enterobacteriaceae; Salmonella; Salmonella enterica;

Salmonella enterica subsp. enterica MNNEETFYQAMRRKGVTRRSFLKFCSLAATSLGLGAGMTPKIAWALENKPRIPVVWIHGL
WP_001058312.1 hydrogenase [Salmonella enterica] Length: 372\nNumber of predicted TMHs: 2\nExp number of AAs in TMHs: 36.51738\nExp number, first 60 AAs: 19.72849\nTotal prob of N-in: 0.96170\nPOSSIBLE N-term signal sequence\ninside 1 19\nTMhelix 20 39\noutside 40 325\nTMhelix 326 348\ninside 349 372

18808 GCF_002058035.1_ASM205803v1 Salmonella enterica subsp. enterica serovar Newport
Proteobacteria; Gammaproteobacteria; Enterobacterales; Enterobacteriaceae; Salmonella; Salmonella enterica; Salmonella enterica subsp. enterica MTGDNTLITSHGINRRDFMKLCAALAATMGLSSKAAAEMAESVSNPQRPPVIWIGAQECT
WP_000145429.1 hydrogenase 2 small subunit [Salmonella enterica] Length: 372\nNumber of predicted TMHs: 1\nExp number of AAs in TMHs: 24.51309\nExp number, first 60 AAs: 0.2682\nTotal prob of N-in: 0.14402\noutside 1 331\nTMhelix 332 354\ninside 355 372

18809 GCF_002048325.1_ASM204832v1 Salmonella enterica subsp. enterica serovar Newport
Proteobacteria; Gammaproteobacteria; Enterobacterales; Enterobacteriaceae; Salmonella; Salmonella enterica; Salmonella enterica subsp. enterica MTGDNTLITSHGINRRDFMKLCAALAATMGLSSKAAAEMAESVSNPQRPPVIWIGAQECT
WP_000145429.1 hydrogenase 2 small subunit [Salmonella enterica] Length: 372\nNumber of predicted TMHs: 1\nExp number of AAs in TMHs: 24.51309\nExp number, first 60 AAs: 0.2682\nTotal prob of N-in: 0.14402\noutside 1 331\nTMhelix 332 354\ninside 355 372

18810 GCF_002058105.1_ASM205810v1 Salmonella enterica subsp. enterica serovar Newport
Proteobacteria; Gammaproteobacteria; Enterobacterales; Enterobacteriaceae; Salmonella; Salmonella enterica; Salmonella enterica subsp. enterica MTGDNTLITSHGINRRDFMKLCAALAATMGLSSKAAAEMAESVSNPQRPPVIWIGAQECT
WP_000145429.1 hydrogenase 2 small subunit [Salmonella enterica] Length: 372\nNumber of predicted TMHs: 1\nExp number of AAs in TMHs: 24.51309\nExp number, first 60 AAs: 0.2682\nTotal prob of N-in: 0.14402\noutside 1 331\nTMhelix 332 354\ninside 355 372

18811 GCF_002058105.1_ASM205810v1 Salmonella enterica subsp. enterica serovar Newport
Proteobacteria; Gammaproteobacteria; Enterobacterales; Enterobacteriaceae; Salmonella; Salmonella enterica; Salmonella enterica subsp. enterica MNNEETFYQAMRRKGVTRRSFLKFCSLAATSLGLGAGMTPKIAWALENKPRIPVVWIHGL
WP_001058312.1 hydrogenase [Salmonella enterica] Length: 372\nNumber of predicted TMHs: 2\nExp number of AAs in TMHs: 36.51738\nExp number, first 60 AAs: 19.72849\nTotal prob of N-in: 0.96170\nPOSSIBLE N-term signal sequence\ninside 1 19\nTMhelix 20 39\noutside 40 325\nTMhelix 326 348\ninside 349 372

18812 GCF_002036025.1_ASM203602v1 Salmonella enterica subsp. enterica serovar Newport
Proteobacteria; Gammaproteobacteria; Enterobacterales; Enterobacteriaceae; Salmonella; Salmonella enterica; Salmonella enterica subsp. enterica MNNEETFYQAMRRKGVTRRSFLKFCSLAATSLGLGAGMTPKIAWALENKPRIPVVWIHGL
WP_001058312.1 hydrogenase [Salmonella enterica] Length: 372\nNumber of predicted TMHs: 2\nExp number of AAs in TMHs: 36.51738\nExp number, first 60 AAs: 19.72849\nTotal prob of N-in: 0.96170\nPOSSIBLE N-term signal sequence\ninside 1 19\nTMhelix 20 39\noutside 40 325\nTMhelix 326 348\ninside 349 372

18813 GCF_002057865.1_ASM205786v1 Salmonella enterica subsp. enterica serovar Newport
Proteobacteria; Gammaproteobacteria; Enterobacterales; Enterobacteriaceae; Salmonella; Salmonella enterica; Salmonella enterica subsp. enterica MNNEETFYQAMRRKGVTRRSFLKFCSLAATSLGLGAGMTPKIAWALENKPRIPVVWIHGL
WP_001058312.1 hydrogenase [Salmonella enterica] Length: 372\nNumber of predicted TMHs: 2\nExp number of AAs in TMHs: 36.51738\nExp number, first 60 AAs: 19.72849\nTotal prob of N-in: 0.96170\nPOSSIBLE N-term signal sequence\ninside 1 19\nTMhelix 20 39\noutside 40 325\nTMhelix 326 348\ninside 349 372

18814 GCF_002058035.1_ASM205803v1 Salmonella enterica subsp. enterica serovar Newport
Proteobacteria; Gammaproteobacteria; Enterobacterales; Enterobacteriaceae; Salmonella; Salmonella enterica; Salmonella enterica subsp. enterica MNNEETFYQAMRRKGVTRRSFLKFCSLAATSLGLGAGMTPKIAWALENKPRIPVVWIHGL
WP_001058312.1 hydrogenase [Salmonella enterica] Length: 372\nNumber of predicted TMHs: 2\nExp number of AAs in TMHs: 36.51738\nExp number, first 60 AAs: 19.72849\nTotal prob of N-in: 0.96170\nPOSSIBLE N-term signal sequence\ninside 1 19\nTMhelix 20 39\noutside 40 325\nTMhelix 326 348\ninside 349 372

18815 GCF_002058335.1_ASM205833v1 Salmonella enterica subsp. enterica serovar Newport
Proteobacteria; Gammaproteobacteria; Enterobacterales; Enterobacteriaceae; Salmonella; Salmonella enterica; Salmonella enterica subsp. enterica MTGDNTLITSHGINRRDFMKLCAALAATMGLSSKAAAEMAESVSNPQRPPVIWIGAQECT
WP_000145429.1 hydrogenase 2 small subunit [Salmonella enterica] Length: 372\nNumber of predicted TMHs: 1\nExp number of AAs in TMHs: 24.51309\nExp number, first 60 AAs: 0.2682\nTotal prob of N-in: 0.14402\noutside 1 331\nTMhelix 332 354\ninside 355 372

18816	GCF_002056745.1_ASM205674v1	Salmonella enterica subsp. enterica serovar Newport	Proteobacteria; Gammaproteobacteria; Enterobacterales; Enterobacteriaceae; Salmonella; Salmonella enterica; Salmonella enterica subsp. enterica	MNNEETFYQAMRRKGVTRRSFLKFCSLAATSLGLGAGMTPKIAWALENKPRIPVVWIHGL	WP_001058312.1	hydrogenase [Salmonella enterica]	Length: 372\nNumber of predicted TMHs:  2\nExp number of AAs in TMHs: 36.51738\nExp number, first 60 AAs:  19.72849\nTotal prob of N-in:   0.96170\nPOSSIBLE N-term signal sequence\ninside   1   19\nTMhelix   20   39\noutside   40   325\nTMhelix   326   348\ninside   349   372

18817	GCF_001834665.1_ASM183466v1	Salmonella enterica subsp. enterica serovar Newport	Proteobacteria; Gammaproteobacteria; Enterobacterales; Enterobacteriaceae; Salmonella; Salmonella enterica; Salmonella enterica subsp. enterica	MTGDNTLITSHGINRRDFMKLCAALAATMGLSSKAAAEMAESVSNPQRPPVIWIGAQECT	WP_000145429.1	hydrogenase 2 small subunit [Salmonella enterica]	Length: 372\nNumber of predicted TMHs:  1\nExp number of AAs in TMHs: 24.51309\nExp number, first 60 AAs:  0.2682\nTotal prob of N-in:   0.14402\noutside   1   331\nTMhelix   332   354\ninside   355   372

18818	GCF_002106575.1_ASM210657v1	Salmonella enterica subsp. enterica serovar Newport	Proteobacteria; Gammaproteobacteria; Enterobacterales; Enterobacteriaceae; Salmonella; Salmonella enterica; Salmonella enterica subsp. enterica	MTGDNTLITSHGINRRDFMKLCAALAATMGLSSKAAAEMAESVSNPQRPPVIWIGAQECT	WP_000145429.1	hydrogenase 2 small subunit [Salmonella enterica]	Length: 372\nNumber of predicted TMHs:  1\nExp number of AAs in TMHs: 24.51309\nExp number, first 60 AAs:  0.2682\nTotal prob of N-in:   0.14402\noutside   1   331\nTMhelix   332   354\ninside   355   372

18819	GCF_002058635.1_ASM205863v1	Salmonella enterica subsp. enterica serovar Newport	Proteobacteria; Gammaproteobacteria; Enterobacterales; Enterobacteriaceae; Salmonella; Salmonella enterica; Salmonella enterica subsp. enterica	MNNEETFYQAMRRKGVTRRSFLKFCSLAATSLGLGAGMTPKIAWALENKPRIPVVWIHGL	WP_001058312.1	hydrogenase [Salmonella enterica]	Length: 372\nNumber of predicted TMHs:  2\nExp number of AAs in TMHs: 36.51738\nExp number, first 60 AAs:  19.72849\nTotal prob of N-in:   0.96170\nPOSSIBLE N-term signal sequence\ninside   1   19\nTMhelix   20   39\noutside   40   325\nTMhelix   326   348\ninside   349   372

18820	GCF_002106595.1_ASM210659v1	Salmonella enterica subsp. enterica serovar Newport	Proteobacteria; Gammaproteobacteria; Enterobacterales; Enterobacteriaceae; Salmonella; Salmonella enterica; Salmonella enterica subsp. enterica	MTGDNTLITSHGINRRDFMKLCAALAATMGLSSKAAAEMAESVSNPQRPPVIWIGAQECT	WP_000145429.1	hydrogenase 2 small subunit [Salmonella enterica]	Length: 372\nNumber of predicted TMHs:  1\nExp number of AAs in TMHs: 24.51309\nExp number, first 60 AAs:  0.2682\nTotal prob of N-in:   0.14402\noutside   1   331\nTMhelix   332   354\ninside   355   372

18821	GCF_002059255.1_ASM205925v1	Salmonella enterica subsp. enterica serovar Newport	Proteobacteria; Gammaproteobacteria; Enterobacterales; Enterobacteriaceae; Salmonella; Salmonella enterica; Salmonella enterica subsp. enterica	MTGDNTLITSHGINRRDFMKLCAALAATMGLSSKAAAEMAESVSNPQRPPVIWIGAQECT	WP_000145429.1	hydrogenase 2 small subunit [Salmonella enterica]	Length: 372\nNumber of predicted TMHs:  1\nExp number of AAs in TMHs: 24.51309\nExp number, first 60 AAs:  0.2682\nTotal prob of N-in:   0.14402\noutside   1   331\nTMhelix   332   354\ninside   355   372

18822	GCF_002106215.1_ASM210621v1	Salmonella enterica subsp. enterica serovar Newport	Proteobacteria; Gammaproteobacteria; Enterobacterales; Enterobacteriaceae; Salmonella; Salmonella enterica; Salmonella enterica subsp. enterica	MNNEETFYQAMRRKGVTRRSFLKFCSLAATSLGLGAGMTPKIAWALENKPRIPVVWIHGL	WP_001058312.1	hydrogenase [Salmonella enterica]	Length: 372\nNumber of predicted TMHs:  2\nExp number of AAs in TMHs: 36.51738\nExp number, first 60 AAs:  19.72849\nTotal prob of N-in:   0.96170\nPOSSIBLE N-term signal sequence\ninside   1   19\nTMhelix   20   39\noutside   40   325\nTMhelix   326   348\ninside   349   372

18823	GCF_002106135.1_ASM210613v1	Salmonella enterica subsp. enterica serovar Newport	Proteobacteria; Gammaproteobacteria; Enterobacterales; Enterobacteriaceae; Salmonella; Salmonella enterica; Salmonella enterica subsp. enterica	MNNEETFYQAMRRKGVTRRSFLKFCSLAATSLGLGAGMTPKIAWALENKPRIPVVWIHGL	WP_001058312.1	hydrogenase [Salmonella enterica]	Length: 372\nNumber of predicted TMHs:  2\nExp number of AAs in TMHs: 36.51738\nExp number, first 60 AAs:  19.72849\nTotal prob of N-in:   0.96170\nPOSSIBLE N-term signal sequence\ninside   1   19\nTMhelix   20   39\noutside   40   325\nTMhelix   326   348\ninside   349   372

18824	GCF_002106215.1_ASM210621v1	Salmonella enterica subsp. enterica serovar Newport	Proteobacteria; Gammaproteobacteria; Enterobacterales; Enterobacteriaceae; Salmonella; Salmonella enterica; Salmonella enterica subsp. enterica	MTGDNTLITSHGINRRDFMKLCAALAATMGLSSKAAAEMAESVSNPQRPPVIWIGAQECT	WP_000145429.1	hydrogenase 2 small subunit [Salmonella enterica]	Length: 372\nNumber of

predicted TMHs: 1\nExp number of AAs in TMHs: 24.51309\nExp number, first 60 AAs: 0.2682\nTotal prob of N-in: 0.14402\noutside 1 331\nTMhelix 332 354\ninside 355 372

18825 GCF_002106135.1_ASM210613v1 Salmonella enterica subsp. enterica serovar Newport Proteobacteria; Gammaproteobacteria; Enterobacterales; Enterobacteriaceae; Salmonella; Salmonella enterica; Salmonella enterica subsp. enterica MTGDNTLITSHGINRRDFMKLCAALAATMGLSSKAAAEMAESVSNPQRPPVIWIGAQECT WP_000145429.1 hydrogenase 2 small subunit [Salmonella enterica] Length: 372\nNumber of predicted TMHs: 1\nExp number of AAs in TMHs: 24.51309\nExp number, first 60 AAs: 0.2682\nTotal prob of N-in: 0.14402\noutside 1 331\nTMhelix 332 354\ninside 355 372

18826 GCF_002106745.1_ASM210674v1 Salmonella enterica subsp. enterica serovar Newport Proteobacteria; Gammaproteobacteria; Enterobacterales; Enterobacteriaceae; Salmonella; Salmonella enterica; Salmonella enterica subsp. enterica MTGDNTLITSHGINRRDFMKLCAALAATMGLSSKAAAEMAESVSNPQRPPVIWIGAQECT WP_000145429.1 hydrogenase 2 small subunit [Salmonella enterica] Length: 372\nNumber of predicted TMHs: 1\nExp number of AAs in TMHs: 24.51309\nExp number, first 60 AAs: 0.2682\nTotal prob of N-in: 0.14402\noutside 1 331\nTMhelix 332 354\ninside 355 372

18827 GCF_002106595.1_ASM210659v1 Salmonella enterica subsp. enterica serovar Newport Proteobacteria; Gammaproteobacteria; Enterobacterales; Enterobacteriaceae; Salmonella; Salmonella enterica; Salmonella enterica subsp. enterica MNNEETFYQAMRRKGVTRRSFLKFCSLAATSLGLGAGMTPKIAWALENKPRIPVVWIHGL WP_001058312.1 hydrogenase [Salmonella enterica] Length: 372\nNumber of predicted TMHs: 2\nExp number of AAs in TMHs: 36.51738\nExp number, first 60 AAs: 19.72849\nTotal prob of N-in: 0.96170\nPOSSIBLE N-term signal sequence\ninside 1 19\nTMhelix 20 39\noutside 40 325\nTMhelix 326 348\ninside 349 372

18828 GCF_002106745.1_ASM210674v1 Salmonella enterica subsp. enterica serovar Newport Proteobacteria; Gammaproteobacteria; Enterobacterales; Enterobacteriaceae; Salmonella; Salmonella enterica; Salmonella enterica subsp. enterica MNNEETFYQAMRRKGVTRRSFLKFCSLAATSLGLGAGMTPKIAWALENKPRIPVVWIHGL WP_001058312.1 hydrogenase [Salmonella enterica] Length: 372\nNumber of predicted TMHs: 2\nExp number of AAs in TMHs: 36.51738\nExp number, first 60 AAs: 19.72849\nTotal prob of N-in: 0.96170\nPOSSIBLE N-term signal sequence\ninside 1 19\nTMhelix 20 39\noutside 40 325\nTMhelix 326 348\ninside 349 372

18829 GCF_002106575.1_ASM210657v1 Salmonella enterica subsp. enterica serovar Newport Proteobacteria; Gammaproteobacteria; Enterobacterales; Enterobacteriaceae; Salmonella; Salmonella enterica; Salmonella enterica subsp. enterica MNNEETFYQAMRRKGVTRRSFLKFCSLAATSLGLGAGMTPKIAWALENKPRIPVVWIHGL WP_001058312.1 hydrogenase [Salmonella enterica] Length: 372\nNumber of predicted TMHs: 2\nExp number of AAs in TMHs: 36.51738\nExp number, first 60 AAs: 19.72849\nTotal prob of N-in: 0.96170\nPOSSIBLE N-term signal sequence\ninside 1 19\nTMhelix 20 39\noutside 40 325\nTMhelix 326 348\ninside 349 372

18830 GCF_002106815.1_ASM210681v1 Salmonella enterica subsp. enterica serovar Newport Proteobacteria; Gammaproteobacteria; Enterobacterales; Enterobacteriaceae; Salmonella; Salmonella enterica; Salmonella enterica subsp. enterica MTGDNTLITSHGINRRDFMKLCAALAATMGLSSKAAAEMAESVSNPQRPPVIWIGAQECT WP_000145429.1 hydrogenase 2 small subunit [Salmonella enterica] Length: 372\nNumber of predicted TMHs: 1\nExp number of AAs in TMHs: 24.51309\nExp number, first 60 AAs: 0.2682\nTotal prob of N-in: 0.14402\noutside 1 331\nTMhelix 332 354\ninside 355 372

18831 GCF_002106815.1_ASM210681v1 Salmonella enterica subsp. enterica serovar Newport Proteobacteria; Gammaproteobacteria; Enterobacterales; Enterobacteriaceae; Salmonella; Salmonella enterica; Salmonella enterica subsp. enterica MNNEETFYQAMRRKGVTRRSFLKFCSLAATSLGLGAGMTPKIAWALENKPRIPVVWIHGL WP_001058312.1 hydrogenase [Salmonella enterica] Length: 372\nNumber of predicted TMHs: 2\nExp number of AAs in TMHs: 36.51738\nExp number, first 60 AAs: 19.72849\nTotal prob of N-in: 0.96170\nPOSSIBLE N-term signal sequence\ninside 1 19\nTMhelix 20 39\noutside 40 325\nTMhelix 326 348\ninside 349 372

18832 GCF_002107255.1_ASM210725v1 Salmonella enterica subsp. enterica serovar Newport Proteobacteria; Gammaproteobacteria; Enterobacterales; Enterobacteriaceae; Salmonella; Salmonella enterica; Salmonella enterica subsp. enterica MTGDNTLITSHGINRRDFMKLCAALAATMGLSSKAAAEMAESVSNPQRPPVIWIGAQECT WP_000145429.1 hydrogenase 2 small subunit [Salmonella enterica] Length: 372\nNumber of predicted TMHs: 1\nExp number of AAs in TMHs: 24.51309\nExp number, first 60 AAs: 0.2682\nTotal prob of N-in: 0.14402\noutside 1 331\nTMhelix 332 354\ninside 355 372

18833 GCF_002107295.1_ASM210729v1 Salmonella enterica subsp. enterica serovar Newport Proteobacteria; Gammaproteobacteria; Enterobacterales; Enterobacteriaceae; Salmonella; Salmonella enterica;

Salmonella enterica subsp. enterica	MTGDNTLITSHGINRRDFMKLCAALAATMGLSSKAAAEMAESVSNPQRPPVIWIGAQECT	WP_000145429.1	hydrogenase 2 small subunit [Salmonella enterica]	Length: 372\nNumber of predicted TMHs:  1\nExp number of AAs in TMHs: 24.51309\nExp number, first 60 AAs:  0.2682\nTotal prob of N-in:  0.14402\noutside   1   331\nTMhelix   332   354\ninside   355   372

18834	GCF_002107295.1_ASM210729v1	Salmonella enterica subsp. enterica serovar Newport	Proteobacteria; Gammaproteobacteria; Enterobacterales; Enterobacteriaceae; Salmonella; Salmonella enterica; Salmonella enterica subsp. enterica	MNNEETFYQAMRRKGVTRRSFLKFCSLAATSLGLGAGMTPKIAWALENKPRIPVVWIHGL	WP_001058312.1	hydrogenase [Salmonella enterica]	Length: 372\nNumber of predicted TMHs:  2\nExp number of AAs in TMHs: 36.51738\nExp number, first 60 AAs:  19.72849\nTotal prob of N-in:   0.96170\nPOSSIBLE N-term signal sequence\ninside   1   19\nTMhelix   20   39\noutside   40   325\nTMhelix   326   348\ninside   349   372

18835	GCF_002107375.1_ASM210737v1	Salmonella enterica subsp. enterica serovar Newport	Proteobacteria; Gammaproteobacteria; Enterobacterales; Enterobacteriaceae; Salmonella; Salmonella enterica; Salmonella enterica subsp. enterica	MTGDNTLITSHGINRRDFMKLCAALAATMGLSSKAAAEMAESVSNPQRPPVIWIGAQECT	WP_000145429.1	hydrogenase 2 small subunit [Salmonella enterica]	Length: 372\nNumber of predicted TMHs:  1\nExp number of AAs in TMHs: 24.51309\nExp number, first 60 AAs:  0.2682\nTotal prob of N-in:  0.14402\noutside   1   331\nTMhelix   332   354\ninside   355   372

18836	GCF_002061655.1_ASM206165v1	Salmonella enterica subsp. enterica serovar Newport	Proteobacteria; Gammaproteobacteria; Enterobacterales; Enterobacteriaceae; Salmonella; Salmonella enterica; Salmonella enterica subsp. enterica	MTGDNTLITSHGINRRDFMKLCAALAATMGLSSKAAAEMAESVSNPQRPPVIWIGAQECT	WP_000145429.1	hydrogenase 2 small subunit [Salmonella enterica]	Length: 372\nNumber of predicted TMHs:  1\nExp number of AAs in TMHs: 24.51309\nExp number, first 60 AAs:  0.2682\nTotal prob of N-in:  0.14402\noutside   1   331\nTMhelix   332   354\ninside   355   372

18837	GCF_002107255.1_ASM210725v1	Salmonella enterica subsp. enterica serovar Newport	Proteobacteria; Gammaproteobacteria; Enterobacterales; Enterobacteriaceae; Salmonella; Salmonella enterica; Salmonella enterica subsp. enterica	MNNEETFYQAMRRKGVTRRSFLKFCSLAATSLGLGAGMTPKIAWALENKPRIPVVWIHGL	WP_001058312.1	hydrogenase [Salmonella enterica]	Length: 372\nNumber of predicted TMHs:  2\nExp number of AAs in TMHs: 36.51738\nExp number, first 60 AAs:  19.72849\nTotal prob of N-in:   0.96170\nPOSSIBLE N-term signal sequence\ninside   1   19\nTMhelix   20   39\noutside   40   325\nTMhelix   326   348\ninside   349   372

18838	GCF_002107375.1_ASM210737v1	Salmonella enterica subsp. enterica serovar Newport	Proteobacteria; Gammaproteobacteria; Enterobacterales; Enterobacteriaceae; Salmonella; Salmonella enterica; Salmonella enterica subsp. enterica	MNNEETFYQAMRRKGVTRRSFLKFCSLAATSLGLGAGMTPKIAWALENKPRIPVVWIHGL	WP_001058312.1	hydrogenase [Salmonella enterica]	Length: 372\nNumber of predicted TMHs:  2\nExp number of AAs in TMHs: 36.51738\nExp number, first 60 AAs:  19.72849\nTotal prob of N-in:   0.96170\nPOSSIBLE N-term signal sequence\ninside   1   19\nTMhelix   20   39\noutside   40   325\nTMhelix   326   348\ninside   349   372

18839	GCF_002091175.1_ASM209117v1	Salmonella enterica subsp. enterica serovar Newport	Proteobacteria; Gammaproteobacteria; Enterobacterales; Enterobacteriaceae; Salmonella; Salmonella enterica; Salmonella enterica subsp. enterica	MNNEETFYQAMRRKGVTRRSFLKFCSLAATSLGLGAGMTPKIAWALENKPRIPVVWIHGL	WP_001058312.1	hydrogenase [Salmonella enterica]	Length: 372\nNumber of predicted TMHs:  2\nExp number of AAs in TMHs: 36.51738\nExp number, first 60 AAs:  19.72849\nTotal prob of N-in:   0.96170\nPOSSIBLE N-term signal sequence\ninside   1   19\nTMhelix   20   39\noutside   40   325\nTMhelix   326   348\ninside   349   372

18840	GCF_002031995.1_ASM203199v1	Salmonella enterica subsp. enterica serovar Newport	Proteobacteria; Gammaproteobacteria; Enterobacterales; Enterobacteriaceae; Salmonella; Salmonella enterica; Salmonella enterica subsp. enterica	MNNEETFYQAMRRKGVTRRSFLKFCSLAATSLGLGAGMTPKIAWALENKPRIPVVWIHGL	WP_001058312.1	hydrogenase [Salmonella enterica]	Length: 372\nNumber of predicted TMHs:  2\nExp number of AAs in TMHs: 36.51738\nExp number, first 60 AAs:  19.72849\nTotal prob of N-in:   0.96170\nPOSSIBLE N-term signal sequence\ninside   1   19\nTMhelix   20   39\noutside   40   325\nTMhelix   326   348\ninside   349   372

18841	GCF_002091175.1_ASM209117v1	Salmonella enterica subsp. enterica serovar Newport	Proteobacteria; Gammaproteobacteria; Enterobacterales; Enterobacteriaceae; Salmonella; Salmonella enterica; Salmonella enterica subsp. enterica	MTGDNTLITSHGINRRDFMKLCAALAATMGLSSKAAAEMAESVSNPQRPPVIWIGAQECT	WP_000145429.1	hydrogenase 2 small subunit [Salmonella enterica]	Length: 372\nNumber of predicted TMHs:  1\nExp number of AAs in TMHs: 24.51309\nExp number, first 60 AAs:  0.2682\nTotal prob of N-in:  0.14402\noutside   1   331\nTMhelix   332   354\ninside   355   372

18842 GCF_002090695.1_ASM209069v1 Salmonella enterica subsp. enterica serovar Newport Proteobacteria; Gammaproteobacteria; Enterobacterales; Enterobacteriaceae; Salmonella; Salmonella enterica; Salmonella enterica subsp. enterica MNNEETFYQAMRRKGVTRRSFLKFCSLAATSLGLGAGMTPKIAWALENKPRIPVVWIHGL WP_001058312.1 hydrogenase [Salmonella enterica] Length: 372\nNumber of predicted TMHs: 2\nExp number of AAs in TMHs: 36.51738\nExp number, first 60 AAs: 19.72849\nTotal prob of N-in: 0.96170\nPOSSIBLE N-term signal sequence\ninside 1 19\nTMhelix 20 39\noutside 40 325\nTMhelix 326 348\ninside 349 372

18843 GCF_002090655.1_ASM209065v1 Salmonella enterica subsp. enterica serovar Newport Proteobacteria; Gammaproteobacteria; Enterobacterales; Enterobacteriaceae; Salmonella; Salmonella enterica; Salmonella enterica subsp. enterica MNNEETFYQAMRRKGVTRRSFLKFCSLAATSLGLGAGMTPKIAWALENKPRIPVVWIHGL WP_001058312.1 hydrogenase [Salmonella enterica] Length: 372\nNumber of predicted TMHs: 2\nExp number of AAs in TMHs: 36.51738\nExp number, first 60 AAs: 19.72849\nTotal prob of N-in: 0.96170\nPOSSIBLE N-term signal sequence\ninside 1 19\nTMhelix 20 39\noutside 40 325\nTMhelix 326 348\ninside 349 372

18844 GCF_002090865.1_ASM209086v1 Salmonella enterica subsp. enterica serovar Newport Proteobacteria; Gammaproteobacteria; Enterobacterales; Enterobacteriaceae; Salmonella; Salmonella enterica; Salmonella enterica subsp. enterica MNNEETFYQAMRRKGVTRRSFLKFCSLAATSLGLGAGMTPKIAWALENKPRIPVVWIHGL WP_001058312.1 hydrogenase [Salmonella enterica] Length: 372\nNumber of predicted TMHs: 2\nExp number of AAs in TMHs: 36.51738\nExp number, first 60 AAs: 19.72849\nTotal prob of N-in: 0.96170\nPOSSIBLE N-term signal sequence\ninside 1 19\nTMhelix 20 39\noutside 40 325\nTMhelix 326 348\ninside 349 372

18845 GCF_002090865.1_ASM209086v1 Salmonella enterica subsp. enterica serovar Newport Proteobacteria; Gammaproteobacteria; Enterobacterales; Enterobacteriaceae; Salmonella; Salmonella enterica; Salmonella enterica subsp. enterica MTGDNTLITSHGINRRDFMKLCAALAATMGLSSKAAAEMAESVSNPQRPPVIWIGAQECT WP_000145429.1 hydrogenase 2 small subunit [Salmonella enterica] Length: 372\nNumber of predicted TMHs: 1\nExp number of AAs in TMHs: 24.51309\nExp number, first 60 AAs: 0.2682\nTotal prob of N-in: 0.14402\noutside 1 331\nTMhelix 332 354\ninside 355 372

18846 GCF_002090615.1_ASM209061v1 Salmonella enterica subsp. enterica serovar Newport Proteobacteria; Gammaproteobacteria; Enterobacterales; Enterobacteriaceae; Salmonella; Salmonella enterica; Salmonella enterica subsp. enterica MNNEETFYQAMRRKGVTRRSFLKFCSLAATSLGLGAGMTPKIAWALENKPRIPVVWIHGL WP_001058312.1 hydrogenase [Salmonella enterica] Length: 372\nNumber of predicted TMHs: 2\nExp number of AAs in TMHs: 36.51738\nExp number, first 60 AAs: 19.72849\nTotal prob of N-in: 0.96170\nPOSSIBLE N-term signal sequence\ninside 1 19\nTMhelix 20 39\noutside 40 325\nTMhelix 326 348\ninside 349 372

18847 GCF_002090875.1_ASM209087v1 Salmonella enterica subsp. enterica serovar Newport Proteobacteria; Gammaproteobacteria; Enterobacterales; Enterobacteriaceae; Salmonella; Salmonella enterica; Salmonella enterica subsp. enterica MNNEETFYQAMRRKGVTRRSFLKFCSLAATSLGLGAGMTPKIAWALENKPRIPVVWIHGL WP_001058312.1 hydrogenase [Salmonella enterica] Length: 372\nNumber of predicted TMHs: 2\nExp number of AAs in TMHs: 36.51738\nExp number, first 60 AAs: 19.72849\nTotal prob of N-in: 0.96170\nPOSSIBLE N-term signal sequence\ninside 1 19\nTMhelix 20 39\noutside 40 325\nTMhelix 326 348\ninside 349 372

18848 GCF_002090845.1_ASM209084v1 Salmonella enterica subsp. enterica serovar Newport Proteobacteria; Gammaproteobacteria; Enterobacterales; Enterobacteriaceae; Salmonella; Salmonella enterica; Salmonella enterica subsp. enterica MNNEETFYQAMRRKGVTRRSFLKFCSLAATSLGLGAGMTPKIAWALENKPRIPVVWIHGL WP_001058312.1 hydrogenase [Salmonella enterica] Length: 372\nNumber of predicted TMHs: 2\nExp number of AAs in TMHs: 36.51738\nExp number, first 60 AAs: 19.72849\nTotal prob of N-in: 0.96170\nPOSSIBLE N-term signal sequence\ninside 1 19\nTMhelix 20 39\noutside 40 325\nTMhelix 326 348\ninside 349 372

18849 GCF_002090875.1_ASM209087v1 Salmonella enterica subsp. enterica serovar Newport Proteobacteria; Gammaproteobacteria; Enterobacterales; Enterobacteriaceae; Salmonella; Salmonella enterica; Salmonella enterica subsp. enterica MTGDNTLITSHGINRRDFMKLCAALAATMGLSSKAAAEMAESVSNPQRPPVIWIGAQECT WP_000145429.1 hydrogenase 2 small subunit [Salmonella enterica] Length: 372\nNumber of predicted TMHs: 1\nExp number of AAs in TMHs: 24.51309\nExp number, first 60 AAs: 0.2682\nTotal prob of N-in: 0.14402\noutside 1 331\nTMhelix 332 354\ninside 355 372

18850 GCF_002090845.1_ASM209084v1 Salmonella enterica subsp. enterica serovar Newport Proteobacteria; Gammaproteobacteria; Enterobacterales; Enterobacteriaceae; Salmonella; Salmonella enterica; Salmonella enterica subsp. enterica MTGDNTLITSHGINRRDFMKLCAALAATMGLSSKAAAEMAESVSNPQRPPVIWIGAQECT WP_000145429.1 hydrogenase 2 small subunit [Salmonella enterica] Length: 372\nNumber of

predicted TMHs: 1\nExp number of AAs in TMHs: 24.51309\nExp number, first 60 AAs: 0.2682\nTotal prob of N-in: 0.14402\noutside 1 331\nTMhelix 332 354\ninside 355 372

18851 GCF_002090555.1_ASM209055v1 Salmonella enterica subsp. enterica serovar Newport Proteobacteria; Gammaproteobacteria; Enterobacterales; Enterobacteriaceae; Salmonella; Salmonella enterica; Salmonella enterica subsp. enterica MNNEETFYQAMRRKGVTRRSFLKFCSLAATSLGLGAGMTPKIAWALENKPRIPVVWIHGL WP_001058312.1 hydrogenase [Salmonella enterica] Length: 372\nNumber of predicted TMHs: 2\nExp number of AAs in TMHs: 36.51738\nExp number, first 60 AAs: 19.72849\nTotal prob of N-in: 0.96170\nPOSSIBLE N-term signal sequence\ninside 1 19\nTMhelix 20 39\noutside 40 325\nTMhelix 326 348\ninside 349 372

18852 GCF_002090815.1_ASM209081v1 Salmonella enterica subsp. enterica serovar Newport Proteobacteria; Gammaproteobacteria; Enterobacterales; Enterobacteriaceae; Salmonella; Salmonella enterica; Salmonella enterica subsp. enterica MNNEETFYQAMRRKGVTRRSFLKFCSLAATSLGLGAGMTPKIAWALENKPRIPVVWIHGL WP_001058312.1 hydrogenase [Salmonella enterica] Length: 372\nNumber of predicted TMHs: 2\nExp number of AAs in TMHs: 36.51738\nExp number, first 60 AAs: 19.72849\nTotal prob of N-in: 0.96170\nPOSSIBLE N-term signal sequence\ninside 1 19\nTMhelix 20 39\noutside 40 325\nTMhelix 326 348\ninside 349 372

18853 GCF_002090815.1_ASM209081v1 Salmonella enterica subsp. enterica serovar Newport Proteobacteria; Gammaproteobacteria; Enterobacterales; Enterobacteriaceae; Salmonella; Salmonella enterica; Salmonella enterica subsp. enterica MTGDNTLITSHGINRRDFMKLCAALAATMGLSSKAAAEMAESVSNPQRPPVIWIGAQECT WP_000145429.1 hydrogenase 2 small subunit [Salmonella enterica] Length: 372\nNumber of predicted TMHs: 1\nExp number of AAs in TMHs: 24.51309\nExp number, first 60 AAs: 0.2682\nTotal prob of N-in: 0.14402\noutside 1 331\nTMhelix 332 354\ninside 355 372

18854 GCF_002090535.1_ASM209053v1 Salmonella enterica subsp. enterica serovar Newport Proteobacteria; Gammaproteobacteria; Enterobacterales; Enterobacteriaceae; Salmonella; Salmonella enterica; Salmonella enterica subsp. enterica MNNEETFYQAMRRKGVTRRSFLKFCSLAATSLGLGAGMTPKIAWALENKPRIPVVWIHGL WP_001058312.1 hydrogenase [Salmonella enterica] Length: 372\nNumber of predicted TMHs: 2\nExp number of AAs in TMHs: 36.51738\nExp number, first 60 AAs: 19.72849\nTotal prob of N-in: 0.96170\nPOSSIBLE N-term signal sequence\ninside 1 19\nTMhelix 20 39\noutside 40 325\nTMhelix 326 348\ninside 349 372

18855 GCF_002090765.1_ASM209076v1 Salmonella enterica subsp. enterica serovar Newport Proteobacteria; Gammaproteobacteria; Enterobacterales; Enterobacteriaceae; Salmonella; Salmonella enterica; Salmonella enterica subsp. enterica MNNEETFYQAMRRKGVTRRSFLKFCSLAATSLGLGAGMTPKIAWALENKPRIPVVWIHGL WP_001058312.1 hydrogenase [Salmonella enterica] Length: 372\nNumber of predicted TMHs: 2\nExp number of AAs in TMHs: 36.51738\nExp number, first 60 AAs: 19.72849\nTotal prob of N-in: 0.96170\nPOSSIBLE N-term signal sequence\ninside 1 19\nTMhelix 20 39\noutside 40 325\nTMhelix 326 348\ninside 349 372

18856 GCF_002090765.1_ASM209076v1 Salmonella enterica subsp. enterica serovar Newport Proteobacteria; Gammaproteobacteria; Enterobacterales; Enterobacteriaceae; Salmonella; Salmonella enterica; Salmonella enterica subsp. enterica MTGDNTLITSHGINRRDFMKLCAALAATMGLSSKAAAEMAESVSNPQRPPVIWIGAQECT WP_000145429.1 hydrogenase 2 small subunit [Salmonella enterica] Length: 372\nNumber of predicted TMHs: 1\nExp number of AAs in TMHs: 24.51309\nExp number, first 60 AAs: 0.2682\nTotal prob of N-in: 0.14402\noutside 1 331\nTMhelix 332 354\ninside 355 372

18857 GCF_002090655.1_ASM209065v1 Salmonella enterica subsp. enterica serovar Newport Proteobacteria; Gammaproteobacteria; Enterobacterales; Enterobacteriaceae; Salmonella; Salmonella enterica; Salmonella enterica subsp. enterica MTGDNTLITSHGINRRDFMKLCAALAATMGLSSKAAAEMAESVSNPQRPPVIWIGAQECT WP_000145429.1 hydrogenase 2 small subunit [Salmonella enterica] Length: 372\nNumber of predicted TMHs: 1\nExp number of AAs in TMHs: 24.51309\nExp number, first 60 AAs: 0.2682\nTotal prob of N-in: 0.14402\noutside 1 331\nTMhelix 332 354\ninside 355 372

18858 GCF_002090695.1_ASM209069v1 Salmonella enterica subsp. enterica serovar Newport Proteobacteria; Gammaproteobacteria; Enterobacterales; Enterobacteriaceae; Salmonella; Salmonella enterica; Salmonella enterica subsp. enterica MTGDNTLITSHGINRRDFMKLCAALAATMGLSSKAAAEMAESVSNPQRPPVIWIGAQECT WP_000145429.1 hydrogenase 2 small subunit [Salmonella enterica] Length: 372\nNumber of predicted TMHs: 1\nExp number of AAs in TMHs: 24.51309\nExp number, first 60 AAs: 0.2682\nTotal prob of N-in: 0.14402\noutside 1 331\nTMhelix 332 354\ninside 355 372

18859 GCF_001955145.1_ASM195514v1 Salmonella enterica subsp. enterica serovar Newport Proteobacteria; Gammaproteobacteria; Enterobacterales; Enterobacteriaceae; Salmonella; Salmonella enterica;

Salmonella enterica subsp. enterica MNNEETFYQAMRRKGVTRRSFLKFCSLAATSLGLGAGMTPKIAWALENKPRIPVVWIHGL WP_001058312.1 hydrogenase [Salmonella enterica] Length: 372\nNumber of predicted TMHs: 2\nExp number of AAs in TMHs: 36.51738\nExp number, first 60 AAs: 19.72849\nTotal prob of N-in: 0.96170\nPOSSIBLE N-term signal sequence\ninside 1 19\nTMhelix 20 39\noutside 40 325\nTMhelix 326 348\ninside 349 372

18860 GCF_001955105.1_ASM195510v1 Salmonella enterica subsp. enterica serovar Newport Proteobacteria; Gammaproteobacteria; Enterobacterales; Enterobacteriaceae; Salmonella; Salmonella enterica; Salmonella enterica subsp. enterica MNNEETFYQAMRRKGVTRRSFLKFCSLAATSLGLGAGMTPKIAWALENKPRIPVVWIHGL WP_001058312.1 hydrogenase [Salmonella enterica] Length: 372\nNumber of predicted TMHs: 2\nExp number of AAs in TMHs: 36.51738\nExp number, first 60 AAs: 19.72849\nTotal prob of N-in: 0.96170\nPOSSIBLE N-term signal sequence\ninside 1 19\nTMhelix 20 39\noutside 40 325\nTMhelix 326 348\ninside 349 372

18861 GCF_001955145.1_ASM195514v1 Salmonella enterica subsp. enterica serovar Newport Proteobacteria; Gammaproteobacteria; Enterobacterales; Enterobacteriaceae; Salmonella; Salmonella enterica; Salmonella enterica subsp. enterica MTGDNTLITSHGINRRDFMKLCAALAATMGLSSKAAAEMAESVSNPQRPPVIWIGAQECT WP_000145429.1 hydrogenase 2 small subunit [Salmonella enterica] Length: 372\nNumber of predicted TMHs: 1\nExp number of AAs in TMHs: 24.51309\nExp number, first 60 AAs: 0.2682\nTotal prob of N-in: 0.14402\noutside 1 331\nTMhelix 332 354\ninside 355 372

18862 GCF_002090615.1_ASM209061v1 Salmonella enterica subsp. enterica serovar Newport Proteobacteria; Gammaproteobacteria; Enterobacterales; Enterobacteriaceae; Salmonella; Salmonella enterica; Salmonella enterica subsp. enterica MTGDNTLITSHGINRRDFMKLCAALAATMGLSSKAAAEMAESVSNPQRPPVIWIGAQECT WP_000145429.1 hydrogenase 2 small subunit [Salmonella enterica] Length: 372\nNumber of predicted TMHs: 1\nExp number of AAs in TMHs: 24.51309\nExp number, first 60 AAs: 0.2682\nTotal prob of N-in: 0.14402\noutside 1 331\nTMhelix 332 354\ninside 355 372

18863 GCF_002090555.1_ASM209055v1 Salmonella enterica subsp. enterica serovar Newport Proteobacteria; Gammaproteobacteria; Enterobacterales; Enterobacteriaceae; Salmonella; Salmonella enterica; Salmonella enterica subsp. enterica MTGDNTLITSHGINRRDFMKLCAALAATMGLSSKAAAEMAESVSNPQRPPVIWIGAQECT WP_000145429.1 hydrogenase 2 small subunit [Salmonella enterica] Length: 372\nNumber of predicted TMHs: 1\nExp number of AAs in TMHs: 24.51309\nExp number, first 60 AAs: 0.2682\nTotal prob of N-in: 0.14402\noutside 1 331\nTMhelix 332 354\ninside 355 372

18864 GCF_001955105.1_ASM195510v1 Salmonella enterica subsp. enterica serovar Newport Proteobacteria; Gammaproteobacteria; Enterobacterales; Enterobacteriaceae; Salmonella; Salmonella enterica; Salmonella enterica subsp. enterica MTGDNTLITSHGINRRDFMKLCAALAATMGLSSKAAAEMAESVSNPQRPPVIWIGAQECT WP_000145429.1 hydrogenase 2 small subunit [Salmonella enterica] Length: 372\nNumber of predicted TMHs: 1\nExp number of AAs in TMHs: 24.51309\nExp number, first 60 AAs: 0.2682\nTotal prob of N-in: 0.14402\noutside 1 331\nTMhelix 332 354\ninside 355 372

18865 GCF_001955065.1_ASM195506v1 Salmonella enterica subsp. enterica serovar Newport Proteobacteria; Gammaproteobacteria; Enterobacterales; Enterobacteriaceae; Salmonella; Salmonella enterica; Salmonella enterica subsp. enterica MNNEETFYQAMRRKGVTRRSFLKFCSLAATSLGLGAGMTPKIAWALENKPRIPVVWIHGL WP_001058312.1 hydrogenase [Salmonella enterica] Length: 372\nNumber of predicted TMHs: 2\nExp number of AAs in TMHs: 36.51738\nExp number, first 60 AAs: 19.72849\nTotal prob of N-in: 0.96170\nPOSSIBLE N-term signal sequence\ninside 1 19\nTMhelix 20 39\noutside 40 325\nTMhelix 326 348\ninside 349 372

18866 GCF_002090535.1_ASM209053v1 Salmonella enterica subsp. enterica serovar Newport Proteobacteria; Gammaproteobacteria; Enterobacterales; Enterobacteriaceae; Salmonella; Salmonella enterica; Salmonella enterica subsp. enterica MTGDNTLITSHGINRRDFMKLCAALAATMGLSSKAAAEMAESVSNPQRPPVIWIGAQECT WP_000145429.1 hydrogenase 2 small subunit [Salmonella enterica] Length: 372\nNumber of predicted TMHs: 1\nExp number of AAs in TMHs: 24.51309\nExp number, first 60 AAs: 0.2682\nTotal prob of N-in: 0.14402\noutside 1 331\nTMhelix 332 354\ninside 355 372

18867 GCF_001955065.1_ASM195506v1 Salmonella enterica subsp. enterica serovar Newport Proteobacteria; Gammaproteobacteria; Enterobacterales; Enterobacteriaceae; Salmonella; Salmonella enterica; Salmonella enterica subsp. enterica MTGDNTLITSHGINRRDFMKLCAALAATMGLSSKAAAEMAESVSNPQRPPVIWIGAQECT WP_000145429.1 hydrogenase 2 small subunit [Salmonella enterica] Length: 372\nNumber of predicted TMHs: 1\nExp number of AAs in TMHs: 24.51309\nExp number, first 60 AAs: 0.2682\nTotal prob of N-in: 0.14402\noutside 1 331\nTMhelix 332 354\ninside 355 372

18868 GCF_001954835.1_ASM195483v1 Salmonella enterica subsp. enterica serovar Newport Proteobacteria; Gammaproteobacteria; Enterobacterales; Enterobacteriaceae; Salmonella; Salmonella enterica; Salmonella enterica subsp. enterica MTGDNTLITSHGINRRDFMKLCAALAATMGLSSKAAAEMAESVSNPQRPPVIWIGAQECT WP_000145429.1 hydrogenase 2 small subunit [Salmonella enterica] Length: 372\nNumber of predicted TMHs: 1\nExp number of AAs in TMHs: 24.51309\nExp number, first 60 AAs: 0.2682\nTotal prob of N-in: 0.14402\noutside 1 331\nTMhelix 332 354\ninside 355 372

18869 GCF_001954815.1_ASM195481v1 Salmonella enterica subsp. enterica serovar Newport Proteobacteria; Gammaproteobacteria; Enterobacterales; Enterobacteriaceae; Salmonella; Salmonella enterica; Salmonella enterica subsp. enterica MTGDNTLITSHGINRRDFMKLCAALAATMGLSSKAAAEMAESVSNPQRPPVIWIGAQECT WP_000145429.1 hydrogenase 2 small subunit [Salmonella enterica] Length: 372\nNumber of predicted TMHs: 1\nExp number of AAs in TMHs: 24.51309\nExp number, first 60 AAs: 0.2682\nTotal prob of N-in: 0.14402\noutside 1 331\nTMhelix 332 354\ninside 355 372

18870 GCF_001954755.1_ASM195475v1 Salmonella enterica subsp. enterica serovar Newport Proteobacteria; Gammaproteobacteria; Enterobacterales; Enterobacteriaceae; Salmonella; Salmonella enterica; Salmonella enterica subsp. enterica MTGDNTLITSHGINRRDFMKLCAALAATMGLSSKAAAEMAESVSNPQRPPVIWIGAQECT WP_000145429.1 hydrogenase 2 small subunit [Salmonella enterica] Length: 372\nNumber of predicted TMHs: 1\nExp number of AAs in TMHs: 24.51309\nExp number, first 60 AAs: 0.2682\nTotal prob of N-in: 0.14402\noutside 1 331\nTMhelix 332 354\ninside 355 372

18871 GCF_002064235.1_ASM206423v1 Salmonella enterica subsp. enterica serovar Newport Proteobacteria; Gammaproteobacteria; Enterobacterales; Enterobacteriaceae; Salmonella; Salmonella enterica; Salmonella enterica subsp. enterica MTGDNTLITSHGINRRDFMKLCAALAATMGLSSKAAAEMAESVSNPQRPPVIWIGAQECT WP_000145429.1 hydrogenase 2 small subunit [Salmonella enterica] Length: 372\nNumber of predicted TMHs: 1\nExp number of AAs in TMHs: 24.51309\nExp number, first 60 AAs: 0.2682\nTotal prob of N-in: 0.14402\noutside 1 331\nTMhelix 332 354\ninside 355 372

18872 GCF_002064235.1_ASM206423v1 Salmonella enterica subsp. enterica serovar Newport Proteobacteria; Gammaproteobacteria; Enterobacterales; Enterobacteriaceae; Salmonella; Salmonella enterica; Salmonella enterica subsp. enterica MNNEETFYQAMRRKGVTRRSFLKFCSLAATSLGLGAGMTPKIAWALENKPRIPVVWIHGL WP_001058312.1 hydrogenase [Salmonella enterica] Length: 372\nNumber of predicted TMHs: 2\nExp number of AAs in TMHs: 36.51738\nExp number, first 60 AAs: 19.72849\nTotal prob of N-in: 0.96170\nPOSSIBLE N-term signal sequence\ninside 1 19\nTMhelix 20 39\noutside 40 325\nTMhelix 326 348\ninside 349 372

18873 GCF_002064285.1_ASM206428v1 Salmonella enterica subsp. enterica serovar Newport Proteobacteria; Gammaproteobacteria; Enterobacterales; Enterobacteriaceae; Salmonella; Salmonella enterica; Salmonella enterica subsp. enterica MNNEETFYQAMRRKGVTRRSFLKFCSLAATSLGLGAGMTPKIAWALENKPRIPVVWIHGL WP_001058312.1 hydrogenase [Salmonella enterica] Length: 372\nNumber of predicted TMHs: 2\nExp number of AAs in TMHs: 36.51738\nExp number, first 60 AAs: 19.72849\nTotal prob of N-in: 0.96170\nPOSSIBLE N-term signal sequence\ninside 1 19\nTMhelix 20 39\noutside 40 325\nTMhelix 326 348\ninside 349 372

18874 GCF_001890185.1_ASM189018v1 Salmonella enterica subsp. enterica serovar Newport Proteobacteria; Gammaproteobacteria; Enterobacterales; Enterobacteriaceae; Salmonella; Salmonella enterica; Salmonella enterica subsp. enterica MNNEETFYQAMRRKGVTRRSFLKFCSLAATSLGLGAGMTPKIAWALENKPRIPVVWIHGL WP_001058312.1 hydrogenase [Salmonella enterica] Length: 372\nNumber of predicted TMHs: 2\nExp number of AAs in TMHs: 36.51738\nExp number, first 60 AAs: 19.72849\nTotal prob of N-in: 0.96170\nPOSSIBLE N-term signal sequence\ninside 1 19\nTMhelix 20 39\noutside 40 325\nTMhelix 326 348\ninside 349 372

18875 GCF_001890185.1_ASM189018v1 Salmonella enterica subsp. enterica serovar Newport Proteobacteria; Gammaproteobacteria; Enterobacterales; Enterobacteriaceae; Salmonella; Salmonella enterica; Salmonella enterica subsp. enterica MTGDNTLITSHGINRRDFMKLCAALAATMGLSSKAAAEMAESVSNPQRPPVIWIGAQECT WP_000145429.1 hydrogenase 2 small subunit [Salmonella enterica] Length: 372\nNumber of predicted TMHs: 1\nExp number of AAs in TMHs: 24.51309\nExp number, first 60 AAs: 0.2682\nTotal prob of N-in: 0.14402\noutside 1 331\nTMhelix 332 354\ninside 355 372

18876 GCF_001890445.1_ASM189044v1 Salmonella enterica subsp. enterica serovar Newport Proteobacteria; Gammaproteobacteria; Enterobacterales; Enterobacteriaceae; Salmonella; Salmonella enterica; Salmonella enterica subsp. enterica MNNEETFYQAMRRKGVTRRSFLKFCSLAATSLGLGAGMTPKIAWALENKPRIPVVWIHGL WP_001058312.1 hydrogenase [Salmonella enterica] Length: 372\nNumber of predicted TMHs: 2\nExp

number of AAs in TMHs: 36.51738\nExp number, first 60 AAs: 19.72849\nTotal prob of N-in: 0.96170\nPOSSIBLE N-term signal sequence\ninside 1 19\nTMhelix 20 39\noutside 40 325\nTMhelix 326 348\ninside 349 372

18877 GCF_001890445.1_ASM189044v1 Salmonella enterica subsp. enterica serovar Newport Proteobacteria; Gammaproteobacteria; Enterobacterales; Enterobacteriaceae; Salmonella; Salmonella enterica; Salmonella enterica subsp. enterica MTGDNTLITSHGINRRDFMKLCAALAATMGLSSKAAAEMAESVSNPQRPPVIWIGAQECT WP_000145429.1 hydrogenase 2 small subunit [Salmonella enterica] Length: 372\nNumber of predicted TMHs: 1\nExp number of AAs in TMHs: 24.51309\nExp number, first 60 AAs: 0.2682\nTotal prob of N-in: 0.14402\noutside 1 331\nTMhelix 332 354\ninside 355 372

18878 GCF_002065335.1_ASM206533v1 Salmonella enterica subsp. enterica serovar Newport Proteobacteria; Gammaproteobacteria; Enterobacterales; Enterobacteriaceae; Salmonella; Salmonella enterica; Salmonella enterica subsp. enterica MTGDNTLITSHGINRRDFMKLCAALAATMGLSSKAAAEMAESVSNPQRPPVIWIGAQECT WP_000145429.1 hydrogenase 2 small subunit [Salmonella enterica] Length: 372\nNumber of predicted TMHs: 1\nExp number of AAs in TMHs: 24.51309\nExp number, first 60 AAs: 0.2682\nTotal prob of N-in: 0.14402\noutside 1 331\nTMhelix 332 354\ninside 355 372

18879 GCF_002065335.1_ASM206533v1 Salmonella enterica subsp. enterica serovar Newport Proteobacteria; Gammaproteobacteria; Enterobacterales; Enterobacteriaceae; Salmonella; Salmonella enterica; Salmonella enterica subsp. enterica MNNEETFYQAMRRKGVTRRSFLKFCSLAATSLGLGAGMTPKIAWALENKPRIPVVWIHGL WP_001058312.1 hydrogenase [Salmonella enterica] Length: 372\nNumber of predicted TMHs: 2\nExp number of AAs in TMHs: 36.51738\nExp number, first 60 AAs: 19.72849\nTotal prob of N-in: 0.96170\nPOSSIBLE N-term signal sequence\ninside 1 19\nTMhelix 20 39\noutside 40 325\nTMhelix 326 348\ninside 349 372

18880 GCF_002065355.1_ASM206535v1 Salmonella enterica subsp. enterica serovar Newport Proteobacteria; Gammaproteobacteria; Enterobacterales; Enterobacteriaceae; Salmonella; Salmonella enterica; Salmonella enterica subsp. enterica MTGDNTLITSHGINRRDFMKLCAALAATMGLSSKAAAEMAESVSNPQRPPVIWIGAQECT WP_000145429.1 hydrogenase 2 small subunit [Salmonella enterica] Length: 372\nNumber of predicted TMHs: 1\nExp number of AAs in TMHs: 24.51309\nExp number, first 60 AAs: 0.2682\nTotal prob of N-in: 0.14402\noutside 1 331\nTMhelix 332 354\ninside 355 372

18881 GCF_002065355.1_ASM206535v1 Salmonella enterica subsp. enterica serovar Newport Proteobacteria; Gammaproteobacteria; Enterobacterales; Enterobacteriaceae; Salmonella; Salmonella enterica; Salmonella enterica subsp. enterica MNNEETFYQAMRRKGVTRRSFLKFCSLAATSLGLGAGMTPKIAWALENKPRIPVVWIHGL WP_001058312.1 hydrogenase [Salmonella enterica] Length: 372\nNumber of predicted TMHs: 2\nExp number of AAs in TMHs: 36.51738\nExp number, first 60 AAs: 19.72849\nTotal prob of N-in: 0.96170\nPOSSIBLE N-term signal sequence\ninside 1 19\nTMhelix 20 39\noutside 40 325\nTMhelix 326 348\ninside 349 372

18882 GCF_001835445.1_ASM183544v1 Salmonella enterica subsp. enterica serovar Newport Proteobacteria; Gammaproteobacteria; Enterobacterales; Enterobacteriaceae; Salmonella; Salmonella enterica; Salmonella enterica subsp. enterica MNNEETFYQAMRRKGVTRRSFLKFCSLAATSLGLGAGMTPKIAWALENKPRIPVVWIHGL WP_001058312.1 hydrogenase [Salmonella enterica] Length: 372\nNumber of predicted TMHs: 2\nExp number of AAs in TMHs: 36.51738\nExp number, first 60 AAs: 19.72849\nTotal prob of N-in: 0.96170\nPOSSIBLE N-term signal sequence\ninside 1 19\nTMhelix 20 39\noutside 40 325\nTMhelix 326 348\ninside 349 372

18883 GCF_001835445.1_ASM183544v1 Salmonella enterica subsp. enterica serovar Newport Proteobacteria; Gammaproteobacteria; Enterobacterales; Enterobacteriaceae; Salmonella; Salmonella enterica; Salmonella enterica subsp. enterica MTGDNTLITSHGINRRDFMKLCAALAATMGLSSKAAAEMAESVSNPQRPPVIWIGAQECT WP_000145429.1 hydrogenase 2 small subunit [Salmonella enterica] Length: 372\nNumber of predicted TMHs: 1\nExp number of AAs in TMHs: 24.51309\nExp number, first 60 AAs: 0.2682\nTotal prob of N-in: 0.14402\noutside 1 331\nTMhelix 332 354\ninside 355 372

18884 GCF_001835255.1_ASM183525v1 Salmonella enterica subsp. enterica serovar Newport Proteobacteria; Gammaproteobacteria; Enterobacterales; Enterobacteriaceae; Salmonella; Salmonella enterica; Salmonella enterica subsp. enterica MNNEETFYQAMRRKGVTRRSFLKFCSLAATSLGLGAGMTPKIAWALENKPRIPVVWIHGL WP_001058312.1 hydrogenase [Salmonella enterica] Length: 372\nNumber of predicted TMHs: 2\nExp number of AAs in TMHs: 36.51738\nExp number, first 60 AAs: 19.72849\nTotal prob of N-in: 0.96170\nPOSSIBLE N-term signal sequence\ninside 1 19\nTMhelix 20 39\noutside 40 325\nTMhelix 326 348\ninside 349 372

18885 GCF_001834705.1_ASM183470v1 Salmonella enterica subsp. enterica serovar Newport Proteobacteria; Gammaproteobacteria; Enterobacterales; Enterobacteriaceae; Salmonella; Salmonella enterica;

Salmonella enterica subsp. enterica MTGDNTLITSHGINRRDFMKLCAALAATMGLSSKAAAEMAESVSNPQRPPVIWIGAQECT WP_000145429.1 hydrogenase 2 small subunit [Salmonella enterica] Length: 372\nNumber of predicted TMHs: 1\nExp number of AAs in TMHs: 24.51309\nExp number, first 60 AAs: 0.2682\nTotal prob of N-in: 0.14402\noutside 1 331\nTMhelix 332 354\ninside 355 372

18886 GCF_001834645.1_ASM183464v1 Salmonella enterica subsp. enterica serovar Newport Proteobacteria; Gammaproteobacteria; Enterobacterales; Enterobacteriaceae; Salmonella; Salmonella enterica; Salmonella enterica subsp. enterica MNNEETFYQAMRRKGVTRRSFLKFCSLAATSLGLGAGMTPKIAWALENKPRIPVVWIHGL WP_001058312.1 hydrogenase [Salmonella enterica] Length: 372\nNumber of predicted TMHs: 2\nExp number of AAs in TMHs: 36.51738\nExp number, first 60 AAs: 19.72849\nTotal prob of N-in: 0.96170\nPOSSIBLE N-term signal sequence\ninside 1 19\nTMhelix 20 39\noutside 40 325\nTMhelix 326 348\ninside 349 372

18887 GCF_001834645.1_ASM183464v1 Salmonella enterica subsp. enterica serovar Newport Proteobacteria; Gammaproteobacteria; Enterobacterales; Enterobacteriaceae; Salmonella; Salmonella enterica; Salmonella enterica subsp. enterica MTGDNTLITSHGINRRDFMKLCAALAATMGLSSKAAAEMAESVSNPQRPPVIWIGAQECT WP_000145429.1 hydrogenase 2 small subunit [Salmonella enterica] Length: 372\nNumber of predicted TMHs: 1\nExp number of AAs in TMHs: 24.51309\nExp number, first 60 AAs: 0.2682\nTotal prob of N-in: 0.14402\noutside 1 331\nTMhelix 332 354\ninside 355 372

18888 GCF_002056755.1_ASM205675v1 Salmonella enterica subsp. enterica serovar Newport Proteobacteria; Gammaproteobacteria; Enterobacterales; Enterobacteriaceae; Salmonella; Salmonella enterica; Salmonella enterica subsp. enterica MNNEETFYQAMRRKGVTRRSFLKFCSLAATSLGLGAGMTPKIAWALENKPRIPVVWIHGL WP_001058312.1 hydrogenase [Salmonella enterica] Length: 372\nNumber of predicted TMHs: 2\nExp number of AAs in TMHs: 36.51738\nExp number, first 60 AAs: 19.72849\nTotal prob of N-in: 0.96170\nPOSSIBLE N-term signal sequence\ninside 1 19\nTMhelix 20 39\noutside 40 325\nTMhelix 326 348\ninside 349 372

18889 GCF_002056755.1_ASM205675v1 Salmonella enterica subsp. enterica serovar Newport Proteobacteria; Gammaproteobacteria; Enterobacterales; Enterobacteriaceae; Salmonella; Salmonella enterica; Salmonella enterica subsp. enterica MTGDNTLITSHGINRRDFMKLCAALAATMGLSSKAAAEMAESVSNPQRPPVIWIGAQECT WP_000145429.1 hydrogenase 2 small subunit [Salmonella enterica] Length: 372\nNumber of predicted TMHs: 1\nExp number of AAs in TMHs: 24.51309\nExp number, first 60 AAs: 0.2682\nTotal prob of N-in: 0.14402\noutside 1 331\nTMhelix 332 354\ninside 355 372

18890 GCF_002057935.1_ASM205793v1 Salmonella enterica subsp. enterica serovar Newport Proteobacteria; Gammaproteobacteria; Enterobacterales; Enterobacteriaceae; Salmonella; Salmonella enterica; Salmonella enterica subsp. enterica MNNEETFYQAMRRKGVTRRSFLKFCSLAATSLGLGAGMTPKIAWALENKPRIPVVWIHGL WP_001058312.1 hydrogenase [Salmonella enterica] Length: 372\nNumber of predicted TMHs: 2\nExp number of AAs in TMHs: 36.51738\nExp number, first 60 AAs: 19.72849\nTotal prob of N-in: 0.96170\nPOSSIBLE N-term signal sequence\ninside 1 19\nTMhelix 20 39\noutside 40 325\nTMhelix 326 348\ninside 349 372

18891 GCF_001975605.1_ASM197560v1 Salmonella enterica subsp. enterica serovar Nitra Proteobacteria; Gammaproteobacteria; Enterobacterales; Enterobacteriaceae; Salmonella; Salmonella enterica; Salmonella enterica subsp. enterica MTGDNTLITSHGINRRDFMKLCAALAATMGLSSKAAAEMAESVSNPQRPPVIWIGAQECT WP_000145429.1 hydrogenase 2 small subunit [Salmonella enterica] Length: 372\nNumber of predicted TMHs: 1\nExp number of AAs in TMHs: 24.51309\nExp number, first 60 AAs: 0.2682\nTotal prob of N-in: 0.14402\noutside 1 331\nTMhelix 332 354\ninside 355 372

18892 GCF_001975605.1_ASM197560v1 Salmonella enterica subsp. enterica serovar Nitra Proteobacteria; Gammaproteobacteria; Enterobacterales; Enterobacteriaceae; Salmonella; Salmonella enterica; Salmonella enterica subsp. enterica MNNEETFYQAMRRKGVTRRSFLKFCSLAATSLGLGAGMTPKIAWALENKPRIPVVWIHGL WP_001058312.1 hydrogenase [Salmonella enterica] Length: 372\nNumber of predicted TMHs: 2\nExp number of AAs in TMHs: 36.51738\nExp number, first 60 AAs: 19.72849\nTotal prob of N-in: 0.96170\nPOSSIBLE N-term signal sequence\ninside 1 19\nTMhelix 20 39\noutside 40 325\nTMhelix 326 348\ninside 349 372

18893 GCF_001642895.1_CFSAN006803_01.0 Salmonella enterica subsp. enterica serovar Nottingham Proteobacteria; Gammaproteobacteria; Enterobacterales; Enterobacteriaceae; Salmonella; Salmonella enterica; Salmonella enterica subsp. enterica MTGDNTLITSHGINRRDFMKLCAALAATMGLSSKAAAEMAESVSNPQRPPVIWIGAQECT WP_000145429.1 hydrogenase 2 small subunit [Salmonella enterica] Length: 372\nNumber of predicted TMHs: 1\nExp number of AAs in TMHs: 24.51309\nExp number, first 60 AAs: 0.2682\nTotal prob of N-in: 0.14402\noutside 1 331\nTMhelix 332 354\ninside 355 372

18894 GCF_001642895.1_CFSAN006803_01.0Salmonella enterica subsp. enterica serovar Nottingham Proteobacteria; Gammaproteobacteria; Enterobacterales; Enterobacteriaceae; Salmonella; Salmonella enterica; Salmonella enterica subsp. enterica MNNEETFYQAMRRKGVTRRSFLKFCSLAATSLGLGAGMTPKIAWALENKPRIPVVWIHGL WP_001058312.1 hydrogenase [Salmonella enterica] Length: 372\nNumber of predicted TMHs: 2\nExp number of AAs in TMHs: 36.51738\nExp number, first 60 AAs: 19.72849\nTotal prob of N-in: 0.96170\nPOSSIBLE N-term signal sequence\ninside 1 19\nTMhelix 20 39\noutside 40 325\nTMhelix 326 348\ninside 349 372

18895 GCF_002048935.1_ASM204893v1 Salmonella enterica subsp. enterica serovar Ohio Proteobacteria; Gammaproteobacteria; Enterobacterales; Enterobacteriaceae; Salmonella; Salmonella enterica; Salmonella enterica subsp. enterica MTGDNTLITSHGINRRDFMKLCAALAATMGLSSKAAAEMAESVSNPQRPPVIWIGAQECT WP_000145429.1 hydrogenase 2 small subunit [Salmonella enterica] Length: 372\nNumber of predicted TMHs: 1\nExp number of AAs in TMHs: 24.51309\nExp number, first 60 AAs: 0.2682\nTotal prob of N-in: 0.14402\noutside 1 331\nTMhelix 332 354\ninside 355 372

18896 GCF_002048935.1_ASM204893v1 Salmonella enterica subsp. enterica serovar Ohio Proteobacteria; Gammaproteobacteria; Enterobacterales; Enterobacteriaceae; Salmonella; Salmonella enterica; Salmonella enterica subsp. enterica MNNEETFYQAMRRKGVTRRSFLKFCSLATTSLGLGAGMTPKIAWALENKPRIPVVWIHGL WP_023259200.1 hydrogenase 1, small subunit [Salmonella enterica] Length: 372\nNumber of predicted TMHs: 2\nExp number of AAs in TMHs: 35.98973\nExp number, first 60 AAs: 19.19893\nTotal prob of N-in: 0.93662\nPOSSIBLE N-term signal sequence\ninside 1 19\nTMhelix 20 39\noutside 40 325\nTMhelix 326 348\ninside 349 372

18897 GCF_002048905.1_ASM204890v1 Salmonella enterica subsp. enterica serovar Ohio Proteobacteria; Gammaproteobacteria; Enterobacterales; Enterobacteriaceae; Salmonella; Salmonella enterica; Salmonella enterica subsp. enterica MTGDNTLITSHGINRRDFMKLCAALAATMGLSSKAAAEMAESVSNPQRPPVIWIGAQECT WP_000145429.1 hydrogenase 2 small subunit [Salmonella enterica] Length: 372\nNumber of predicted TMHs: 1\nExp number of AAs in TMHs: 24.51309\nExp number, first 60 AAs: 0.2682\nTotal prob of N-in: 0.14402\noutside 1 331\nTMhelix 332 354\ninside 355 372

18898 GCF_002031335.1_ASM203133v1 Salmonella enterica subsp. enterica serovar Ohio Proteobacteria; Gammaproteobacteria; Enterobacterales; Enterobacteriaceae; Salmonella; Salmonella enterica; Salmonella enterica subsp. enterica MNNEETFYQAMRRKGVTRRSFLKFCSLATTSLGLGAGMTPKIAWALENKPRIPVVWIHGL WP_023259200.1 hydrogenase 1, small subunit [Salmonella enterica] Length: 372\nNumber of predicted TMHs: 2\nExp number of AAs in TMHs: 35.98973\nExp number, first 60 AAs: 19.19893\nTotal prob of N-in: 0.93662\nPOSSIBLE N-term signal sequence\ninside 1 19\nTMhelix 20 39\noutside 40 325\nTMhelix 326 348\ninside 349 372

18899 GCF_002031335.1_ASM203133v1 Salmonella enterica subsp. enterica serovar Ohio Proteobacteria; Gammaproteobacteria; Enterobacterales; Enterobacteriaceae; Salmonella; Salmonella enterica; Salmonella enterica subsp. enterica MTGDNTLITSHGINRRDFMKLCAALAATMGLSSKAAAEMAESVSNPQRPPVIWIGAQECT WP_000145429.1 hydrogenase 2 small subunit [Salmonella enterica] Length: 372\nNumber of predicted TMHs: 1\nExp number of AAs in TMHs: 24.51309\nExp number, first 60 AAs: 0.2682\nTotal prob of N-in: 0.14402\noutside 1 331\nTMhelix 332 354\ninside 355 372

18900 GCF_002048745.1_ASM204874v1 Salmonella enterica subsp. enterica serovar Ohio Proteobacteria; Gammaproteobacteria; Enterobacterales; Enterobacteriaceae; Salmonella; Salmonella enterica; Salmonella enterica subsp. enterica MTGDNTLITSHGINRRDFMKLCAALAATMGLSSKAAAEMAESVSNPQRPPVIWIGAQECT WP_000145429.1 hydrogenase 2 small subunit [Salmonella enterica] Length: 372\nNumber of predicted TMHs: 1\nExp number of AAs in TMHs: 24.51309\nExp number, first 60 AAs: 0.2682\nTotal prob of N-in: 0.14402\noutside 1 331\nTMhelix 332 354\ninside 355 372

18901 GCF_002048805.1_ASM204880v1 Salmonella enterica subsp. enterica serovar Ohio Proteobacteria; Gammaproteobacteria; Enterobacterales; Enterobacteriaceae; Salmonella; Salmonella enterica; Salmonella enterica subsp. enterica MTGDNTLITSHGINRRDFMKLCAALAATMGLSSKAAAEMAESVSNPQRPPVIWIGAQECT WP_000145429.1 hydrogenase 2 small subunit [Salmonella enterica] Length: 372\nNumber of predicted TMHs: 1\nExp number of AAs in TMHs: 24.51309\nExp number, first 60 AAs: 0.2682\nTotal prob of N-in: 0.14402\noutside 1 331\nTMhelix 332 354\ninside 355 372

18902 GCF_002048905.1_ASM204890v1 Salmonella enterica subsp. enterica serovar Ohio Proteobacteria; Gammaproteobacteria; Enterobacterales; Enterobacteriaceae; Salmonella; Salmonella enterica; Salmonella enterica subsp. enterica MNNEETFYQAMRRKGVTRRSFLKFCSLATTSLGLGAGMTPKIAWALENKPRIPVVWIHGL WP_023259200.1 hydrogenase 1, small subunit [Salmonella enterica] Length: 372\nNumber of predicted TMHs: 2\nExp

number of AAs in TMHs: 35.98973\nExp number, first 60 AAs: 19.19893\nTotal prob of N-in: 0.93662\nPOSSIBLE N-term signal sequence\ninside 1 19\nTMhelix 20 39\noutside 40 325\nTMhelix 326 348\ninside 349 372

18903 GCF_002048805.1_ASM204880v1 Salmonella enterica subsp. enterica serovar Ohio Proteobacteria; Gammaproteobacteria; Enterobacterales; Enterobacteriaceae; Salmonella; Salmonella enterica; Salmonella enterica subsp. enterica MNNEETFYQAMRRKGVTRRSFLKFCSLATTSLGLGAGMTPKIAWALENKPRIPVVWIHGL WP_023259200.1 hydrogenase 1, small subunit [Salmonella enterica] Length: 372\nNumber of predicted TMHs: 2\nExp number of AAs in TMHs: 35.98973\nExp number, first 60 AAs: 19.19893\nTotal prob of N-in: 0.93662\nPOSSIBLE N-term signal sequence\ninside 1 19\nTMhelix 20 39\noutside 40 325\nTMhelix 326 348\ninside 349 372

18904 GCF_001241475.1_Salmonella_enterica_CVM_N44711_v1.0 Salmonella enterica subsp. enterica serovar Ohio Proteobacteria; Gammaproteobacteria; Enterobacterales; Enterobacteriaceae; Salmonella; Salmonella enterica; Salmonella enterica subsp. enterica MTGDNTLITSHGINRRDFMKLCAALAATMGLSSKAAAEMAESVSNPQRPPVIWIGAQECT WP_000145429.1 hydrogenase 2 small subunit [Salmonella enterica] Length: 372\nNumber of predicted TMHs: 1\nExp number of AAs in TMHs: 24.51309\nExp number, first 60 AAs: 0.2682\nTotal prob of N-in: 0.14402\noutside 1 331\nTMhelix 332 354\ninside 355 372

18905 GCF_001241475.1_Salmonella_enterica_CVM_N44711_v1.0 Salmonella enterica subsp. enterica serovar Ohio Proteobacteria; Gammaproteobacteria; Enterobacterales; Enterobacteriaceae; Salmonella; Salmonella enterica; Salmonella enterica subsp. enterica MNNEETFYQAMRRKGVTRRSFLKFCSLATTSLGLGAGMTPKIAWALENKPRIPVVWIHGL WP_023259200.1 hydrogenase 1, small subunit [Salmonella enterica] Length: 372\nNumber of predicted TMHs: 2\nExp number of AAs in TMHs: 35.98973\nExp number, first 60 AAs: 19.19893\nTotal prob of N-in: 0.93662\nPOSSIBLE N-term signal sequence\ninside 1 19\nTMhelix 20 39\noutside 40 325\nTMhelix 326 348\ninside 349 372

18906 GCF_002031325.1_ASM203132v1 Salmonella enterica subsp. enterica serovar Ohio Proteobacteria; Gammaproteobacteria; Enterobacterales; Enterobacteriaceae; Salmonella; Salmonella enterica; Salmonella enterica subsp. enterica MNNEETFYQAMRRKGVTRRSFLKFCSLATTSLGLGAGMTPKIAWALENKPRIPVVWIHGL WP_023259200.1 hydrogenase 1, small subunit [Salmonella enterica] Length: 372\nNumber of predicted TMHs: 2\nExp number of AAs in TMHs: 35.98973\nExp number, first 60 AAs: 19.19893\nTotal prob of N-in: 0.93662\nPOSSIBLE N-term signal sequence\ninside 1 19\nTMhelix 20 39\noutside 40 325\nTMhelix 326 348\ninside 349 372

18907 GCF_001478795.1_Salmonella_enterica_CVM_N29382-R_v1.0 Salmonella enterica subsp. enterica serovar Ohio Proteobacteria; Gammaproteobacteria; Enterobacterales; Enterobacteriaceae; Salmonella; Salmonella enterica; Salmonella enterica subsp. enterica MNNEETFYQAMRRKGVTRRSFLKFCSLATTSLGLGAGMTPKIAWALENKPRIPVVWIHGL WP_023259200.1 hydrogenase 1, small subunit [Salmonella enterica] Length: 372\nNumber of predicted TMHs: 2\nExp number of AAs in TMHs: 35.98973\nExp number, first 60 AAs: 19.19893\nTotal prob of N-in: 0.93662\nPOSSIBLE N-term signal sequence\ninside 1 19\nTMhelix 20 39\noutside 40 325\nTMhelix 326 348\ninside 349 372

18908 GCF_001478795.1_Salmonella_enterica_CVM_N29382-R_v1.0 Salmonella enterica subsp. enterica serovar Ohio Proteobacteria; Gammaproteobacteria; Enterobacterales; Enterobacteriaceae; Salmonella; Salmonella enterica; Salmonella enterica subsp. enterica MTGDNTLITSHGINRRDFMKLCAALAATMGLSSKAAAEMAESVSNPQRPPVIWIGAQECT WP_000145429.1 hydrogenase 2 small subunit [Salmonella enterica] Length: 372\nNumber of predicted TMHs: 1\nExp number of AAs in TMHs: 24.51309\nExp number, first 60 AAs: 0.2682\nTotal prob of N-in: 0.14402\noutside 1 331\nTMhelix 332 354\ninside 355 372

18909 GCF_002048745.1_ASM204874v1 Salmonella enterica subsp. enterica serovar Ohio Proteobacteria; Gammaproteobacteria; Enterobacterales; Enterobacteriaceae; Salmonella; Salmonella enterica; Salmonella enterica subsp. enterica MNNEETFYQAMRRKGVTRRSFLKFCSLATTSLGLGAGMTPKIAWALENKPRIPVVWIHGL WP_023259200.1 hydrogenase 1, small subunit [Salmonella enterica] Length: 372\nNumber of predicted TMHs: 2\nExp number of AAs in TMHs: 35.98973\nExp number, first 60 AAs: 19.19893\nTotal prob of N-in: 0.93662\nPOSSIBLE N-term signal sequence\ninside 1 19\nTMhelix 20 39\noutside 40 325\nTMhelix 326 348\ninside 349 372

18910 GCF_002061365.1_ASM206136v1 Salmonella enterica subsp. enterica serovar Onireke Proteobacteria; Gammaproteobacteria; Enterobacterales; Enterobacteriaceae; Salmonella; Salmonella enterica; Salmonella enterica subsp. enterica MTGDNTLITSHGINRRDFMKLCAALAATMGLSSKAAAEMAESVSNPQRPPVIWIGAQECT WP_000145429.1 hydrogenase 2 small subunit [Salmonella enterica] Length: 372\nNumber of predicted TMHs: 1\nExp number of AAs in TMHs: 24.51309\nExp number, first 60 AAs: 0.2682\nTotal prob of N-in: 0.14402\noutside 1 331\nTMhelix 332 354\ninside 355 372

18911 GCF_002064225.1_ASM206422v1 Salmonella enterica subsp. enterica serovar Onireke Proteobacteria; Gammaproteobacteria; Enterobacterales; Enterobacteriaceae; Salmonella; Salmonella enterica; Salmonella enterica subsp. enterica MNNEETFYQAMRRKGVTRRSFLKFCSLAATSLGLGAGMTPKIAWALENKPRIPVVWIHGL WP_001058311.1 hydrogenase [Salmonella enterica] Length: 372\nNumber of predicted TMHs: 2\nExp number of AAs in TMHs: 37.4699899999999\nExp number, first 60 AAs: 19.72699\nTotal prob of N-in: 0.96171\nPOSSIBLE N-term signal sequence\ninside 1 19\nTMhelix 20 39\noutside 40 325\nTMhelix 326 348\ninside 349 372

18912 GCF_002065285.1_ASM206528v1 Salmonella enterica subsp. enterica serovar Oranienburg Proteobacteria; Gammaproteobacteria; Enterobacterales; Enterobacteriaceae; Salmonella; Salmonella enterica; Salmonella enterica subsp. enterica MTGDNTLITSHGINRRDFMKLCAALAATMGLSSKAAAEMAESVSNPQRPPVIWIGAQECT WP_000145429.1 hydrogenase 2 small subunit [Salmonella enterica] Length: 372\nNumber of predicted TMHs: 1\nExp number of AAs in TMHs: 24.51309\nExp number, first 60 AAs: 0.2682\nTotal prob of N-in: 0.14402\noutside 1 331\nTMhelix 332 354\ninside 355 372

18913 GCF_002065285.1_ASM206528v1 Salmonella enterica subsp. enterica serovar Oranienburg Proteobacteria; Gammaproteobacteria; Enterobacterales; Enterobacteriaceae; Salmonella; Salmonella enterica; Salmonella enterica subsp. enterica MNNEETFYQAMRRKGVTRRSFLKFCSLAATSLGLGAGMTPKIAWALENKPRIPVVWIHGL WP_001058311.1 hydrogenase [Salmonella enterica] Length: 372\nNumber of predicted TMHs: 2\nExp number of AAs in TMHs: 37.4699899999999\nExp number, first 60 AAs: 19.72699\nTotal prob of N-in: 0.96171\nPOSSIBLE N-term signal sequence\ninside 1 19\nTMhelix 20 39\noutside 40 325\nTMhelix 326 348\ninside 349 372

18914 GCF_002156335.1_ASM215633v1 Salmonella enterica subsp. enterica serovar Oranienburg Proteobacteria; Gammaproteobacteria; Enterobacterales; Enterobacteriaceae; Salmonella; Salmonella enterica; Salmonella enterica subsp. enterica MNNEETFYQAMRRKGVTRRSFLKFCSLAATSLGLGAGMTPKIAWALENKPRIPVVWIHGL WP_001058312.1 hydrogenase [Salmonella enterica] Length: 372\nNumber of predicted TMHs: 2\nExp number of AAs in TMHs: 36.51738\nExp number, first 60 AAs: 19.72849\nTotal prob of N-in: 0.96170\nPOSSIBLE N-term signal sequence\ninside 1 19\nTMhelix 20 39\noutside 40 325\nTMhelix 326 348\ninside 349 372

18915 GCF_001879975.1_ASM187997v1 Salmonella enterica subsp. enterica serovar Oranienburg Proteobacteria; Gammaproteobacteria; Enterobacterales; Enterobacteriaceae; Salmonella; Salmonella enterica; Salmonella enterica subsp. enterica MTGDNTLITSHGINRRDFMKLCAALAATMGLSSKAAAEMAESVSNPQRPPVIWIGAQECT WP_000145431.1 hydrogenase 2 small subunit [Salmonella enterica] Length: 372\nNumber of predicted TMHs: 1\nExp number of AAs in TMHs: 24.51505\nExp number, first 60 AAs: 0.2682\nTotal prob of N-in: 0.14406\noutside 1 331\nTMhelix 332 354\ninside 355 372

18916 GCF_002062305.1_ASM206230v1 Salmonella enterica subsp. enterica serovar Oranienburg Proteobacteria; Gammaproteobacteria; Enterobacterales; Enterobacteriaceae; Salmonella; Salmonella enterica; Salmonella enterica subsp. enterica MNNEETFYQAMRRKGVTRRSFLKFCSLAATSLGLGAGMTPKIAWALENKPRIPVVWIHGL WP_001058312.1 hydrogenase [Salmonella enterica] Length: 372\nNumber of predicted TMHs: 2\nExp number of AAs in TMHs: 36.51738\nExp number, first 60 AAs: 19.72849\nTotal prob of N-in: 0.96170\nPOSSIBLE N-term signal sequence\ninside 1 19\nTMhelix 20 39\noutside 40 325\nTMhelix 326 348\ninside 349 372

18917 GCF_002059755.1_ASM205975v1 Salmonella enterica subsp. enterica serovar Oranienburg Proteobacteria; Gammaproteobacteria; Enterobacterales; Enterobacteriaceae; Salmonella; Salmonella enterica; Salmonella enterica subsp. enterica MTGDNTLITSHGINRRDFMKLCAALAATMGLSSKAAAEMAESVSNPQRPPVIWIGAQECT WP_000145431.1 hydrogenase 2 small subunit [Salmonella enterica] Length: 372\nNumber of predicted TMHs: 1\nExp number of AAs in TMHs: 24.51505\nExp number, first 60 AAs: 0.2682\nTotal prob of N-in: 0.14406\noutside 1 331\nTMhelix 332 354\ninside 355 372

18918 GCF_001879795.1_ASM187979v1 Salmonella enterica subsp. enterica serovar Oranienburg Proteobacteria; Gammaproteobacteria; Enterobacterales; Enterobacteriaceae; Salmonella; Salmonella enterica; Salmonella enterica subsp. enterica MNNEETFYQAMRRKGVTRRSFLKFCSLAATSLGLGAGMTPKIAWALENKPRIPVVWIHGL WP_001058312.1 hydrogenase [Salmonella enterica] Length: 372\nNumber of predicted TMHs: 2\nExp number of AAs in TMHs: 36.51738\nExp number, first 60 AAs: 19.72849\nTotal prob of N-in: 0.96170\nPOSSIBLE N-term signal sequence\ninside 1 19\nTMhelix 20 39\noutside 40 325\nTMhelix 326 348\ninside 349 372

18919 GCF_001878895.1_ASM187889v1 Salmonella enterica subsp. enterica serovar Oranienburg Proteobacteria; Gammaproteobacteria; Enterobacterales; Enterobacteriaceae; Salmonella; Salmonella enterica; Salmonella enterica subsp. enterica MNNEETFYQAMRRKGVTRRSFLKFCSLAATSLGLGAGMTPKIAWALENKPRIPVVWIHGL

WP_001058312.1	hydrogenase [Salmonella enterica]	Length: 372\nNumber of predicted TMHs:  2\nExp number of AAs in TMHs: 36.51738\nExp number, first 60 AAs:  19.72849\nTotal prob of N-in:   0.96170\nPOSSIBLE N-term signal sequence\ninside   1   19\nTMhelix   20   39\noutside   40   325\nTMhelix   326   348\ninside   349   372

18920	GCF_001879975.1_ASM187997v1	Salmonella enterica subsp. enterica serovar Oranienburg	Proteobacteria; Gammaproteobacteria; Enterobacterales; Enterobacteriaceae; Salmonella; Salmonella enterica; Salmonella enterica subsp. enterica	MNNEETFYQAMRRKGVTRRSFLKFCSLAATSLGLGAGMTPKIAWALENKPRIPVVWIHGL	WP_001058312.1	hydrogenase [Salmonella enterica]	Length: 372\nNumber of predicted TMHs:  2\nExp number of AAs in TMHs: 36.51738\nExp number, first 60 AAs:  19.72849\nTotal prob of N-in:   0.96170\nPOSSIBLE N-term signal sequence\ninside   1   19\nTMhelix   20   39\noutside   40   325\nTMhelix   326   348\ninside   349   372

18921	GCF_002062305.1_ASM206230v1	Salmonella enterica subsp. enterica serovar Oranienburg	Proteobacteria; Gammaproteobacteria; Enterobacterales; Enterobacteriaceae; Salmonella; Salmonella enterica; Salmonella enterica subsp. enterica	MTGDNTLITSHDINRRDFMKLCAALAATMGLSSKAAAEMAESVSNPQRPPVIWIGAQECT	WP_080191561.1	hydrogenase 2 small subunit [Salmonella enterica]	Length: 372\nNumber of predicted TMHs:  1\nExp number of AAs in TMHs: 24.53892\nExp number, first 60 AAs:  0.29527\nTotal prob of N-in:   0.14517\noutside   1   331\nTMhelix   332   354\ninside   355   372

18922	GCF_001878945.1_ASM187894v1	Salmonella enterica subsp. enterica serovar Oranienburg	Proteobacteria; Gammaproteobacteria; Enterobacterales; Enterobacteriaceae; Salmonella; Salmonella enterica; Salmonella enterica subsp. enterica	MNNEETFYQAMRRKGVTRRSFLKFCSLAATSLGLGAGMTPKIAWALENKPRIPVVWIHGL	WP_023187532.1	hydrogenase 1, small subunit [Salmonella enterica]	Length: 372\nNumber of predicted TMHs:  2\nExp number of AAs in TMHs: 36.9997\nExp number, first 60 AAs:  19.72759\nTotal prob of N-in:   0.96171\nPOSSIBLE N-term signal sequence\ninside   1   19\nTMhelix   20   39\noutside   40   325\nTMhelix   326   348\ninside   349   372

18923	GCF_001879935.1_ASM187993v1	Salmonella enterica subsp. enterica serovar Oranienburg	Proteobacteria; Gammaproteobacteria; Enterobacterales; Enterobacteriaceae; Salmonella; Salmonella enterica; Salmonella enterica subsp. enterica	MNNEETFYQAMRRKGVTRRSFLKFCSLAATSLGLGAGMTPKIAWALENKPRIPVVWIHGL	WP_001058312.1	hydrogenase [Salmonella enterica]	Length: 372\nNumber of predicted TMHs:  2\nExp number of AAs in TMHs: 36.51738\nExp number, first 60 AAs:  19.72849\nTotal prob of N-in:   0.96170\nPOSSIBLE N-term signal sequence\ninside   1   19\nTMhelix   20   39\noutside   40   325\nTMhelix   326   348\ninside   349   372

18924	GCF_002059755.1_ASM205975v1	Salmonella enterica subsp. enterica serovar Oranienburg	Proteobacteria; Gammaproteobacteria; Enterobacterales; Enterobacteriaceae; Salmonella; Salmonella enterica; Salmonella enterica subsp. enterica	MNNEETFYQAMRRKGVTRRSFLKFCSLAATSLGLGAGMTPKIAWALENKPRIPVVWIHGL	WP_001058312.1	hydrogenase [Salmonella enterica]	Length: 372\nNumber of predicted TMHs:  2\nExp number of AAs in TMHs: 36.51738\nExp number, first 60 AAs:  19.72849\nTotal prob of N-in:   0.96170\nPOSSIBLE N-term signal sequence\ninside   1   19\nTMhelix   20   39\noutside   40   325\nTMhelix   326   348\ninside   349   372

18925	GCF_001880105.1_ASM188010v1	Salmonella enterica subsp. enterica serovar Oranienburg	Proteobacteria; Gammaproteobacteria; Enterobacterales; Enterobacteriaceae; Salmonella; Salmonella enterica; Salmonella enterica subsp. enterica	MNNEETFYQAMRRKGVTRRSFLKFCSLAATSLGLGAGMTPKIAWALENKPRIPVVWIHGL	WP_001058312.1	hydrogenase [Salmonella enterica]	Length: 372\nNumber of predicted TMHs:  2\nExp number of AAs in TMHs: 36.51738\nExp number, first 60 AAs:  19.72849\nTotal prob of N-in:   0.96170\nPOSSIBLE N-term signal sequence\ninside   1   19\nTMhelix   20   39\noutside   40   325\nTMhelix   326   348\ninside   349   372

18926	GCF_001879935.1_ASM187993v1	Salmonella enterica subsp. enterica serovar Oranienburg	Proteobacteria; Gammaproteobacteria; Enterobacterales; Enterobacteriaceae; Salmonella; Salmonella enterica; Salmonella enterica subsp. enterica	MTGDNTLITSHGINRRDFMKLCAALAATMGLSSKAAAEMAESVSNPQRPPVIWIGAQECT	WP_000145431.1	hydrogenase 2 small subunit [Salmonella enterica]	Length: 372\nNumber of predicted TMHs:  1\nExp number of AAs in TMHs: 24.51505\nExp number, first 60 AAs:  0.2682\nTotal prob of N-in:   0.14406\noutside   1   331\nTMhelix   332   354\ninside   355   372

18927	GCF_001879855.1_ASM187985v1	Salmonella enterica subsp. enterica serovar Oranienburg	Proteobacteria; Gammaproteobacteria; Enterobacterales; Enterobacteriaceae; Salmonella; Salmonella enterica; Salmonella enterica subsp. enterica	MTGDNTLITSHGINRRDFMKLCAALAATMGLSSKAAAEMAESVSNPQRPPVIWIGAQECT	WP_000145431.1	hydrogenase 2 small subunit [Salmonella enterica]	Length: 372\nNumber of predicted TMHs:  1\nExp number of AAs in TMHs: 24.51505\nExp number, first 60 AAs:  0.2682\nTotal prob of N-in:   0.14406\noutside   1   331\nTMhelix   332   354\ninside   355   372

18928 GCF_001879905.1_ASM187990v1 Salmonella enterica subsp. enterica serovar Oranienburg Proteobacteria; Gammaproteobacteria; Enterobacterales; Enterobacteriaceae; Salmonella; Salmonella enterica; Salmonella enterica subsp. enterica MTGDNTLITSHGINRRDFMKLCAALAATMGLSSKAAAEMAESVSNPQRPPVIWIGAQECT WP_000145431.1 hydrogenase 2 small subunit [Salmonella enterica] Length: 372\nNumber of predicted TMHs: 1\nExp number of AAs in TMHs: 24.51505\nExp number, first 60 AAs: 0.2682\nTotal prob of N-in: 0.14406\noutside 1 331\nTMhelix 332 354\ninside 355 372

18929 GCF_001879795.1_ASM187979v1 Salmonella enterica subsp. enterica serovar Oranienburg Proteobacteria; Gammaproteobacteria; Enterobacterales; Enterobacteriaceae; Salmonella; Salmonella enterica; Salmonella enterica subsp. enterica MTGDNTLITSHGINRRDFMKLCAALAATMGLSSKAAAEMAESVSNPQRPPVIWIGAQECT WP_000145431.1 hydrogenase 2 small subunit [Salmonella enterica] Length: 372\nNumber of predicted TMHs: 1\nExp number of AAs in TMHs: 24.51505\nExp number, first 60 AAs: 0.2682\nTotal prob of N-in: 0.14406\noutside 1 331\nTMhelix 332 354\ninside 355 372

18930 GCF_001879855.1_ASM187985v1 Salmonella enterica subsp. enterica serovar Oranienburg Proteobacteria; Gammaproteobacteria; Enterobacterales; Enterobacteriaceae; Salmonella; Salmonella enterica; Salmonella enterica subsp. enterica MNNEETFYQAMRRKGVTRRSFLKFCSLAATSLGLGAGMTPKIAWALENKPRIPVVWIHGL WP_001058312.1 hydrogenase [Salmonella enterica] Length: 372\nNumber of predicted TMHs: 2\nExp number of AAs in TMHs: 36.51738\nExp number, first 60 AAs: 19.72849\nTotal prob of N-in: 0.96170\nPOSSIBLE N-term signal sequence\ninside 1 19\nTMhelix 20 39\noutside 40 325\nTMhelix 326 348\ninside 349 372

18931 GCF_001878895.1_ASM187889v1 Salmonella enterica subsp. enterica serovar Oranienburg Proteobacteria; Gammaproteobacteria; Enterobacterales; Enterobacteriaceae; Salmonella; Salmonella enterica; Salmonella enterica subsp. enterica MTGDNTLITSHGINRRDFMKLCAALAATMGLSSKAAAEMAESVSNPQRPPVIWIGAQECT WP_000145429.1 hydrogenase 2 small subunit [Salmonella enterica] Length: 372\nNumber of predicted TMHs: 1\nExp number of AAs in TMHs: 24.51309\nExp number, first 60 AAs: 0.2682\nTotal prob of N-in: 0.14402\noutside 1 331\nTMhelix 332 354\ninside 355 372

18932 GCF_001879785.1_ASM187978v1 Salmonella enterica subsp. enterica serovar Oranienburg Proteobacteria; Gammaproteobacteria; Enterobacterales; Enterobacteriaceae; Salmonella; Salmonella enterica; Salmonella enterica subsp. enterica MTGDNTLITSHGINRRDFMKLCAALAATMGLSSKAAAEMAESVSNPQRPPVIWIGAQECT WP_000145431.1 hydrogenase 2 small subunit [Salmonella enterica] Length: 372\nNumber of predicted TMHs: 1\nExp number of AAs in TMHs: 24.51505\nExp number, first 60 AAs: 0.2682\nTotal prob of N-in: 0.14406\noutside 1 331\nTMhelix 332 354\ninside 355 372

18933 GCF_002060075.1_ASM206007v1 Salmonella enterica subsp. enterica serovar Oranienburg Proteobacteria; Gammaproteobacteria; Enterobacterales; Enterobacteriaceae; Salmonella; Salmonella enterica; Salmonella enterica subsp. enterica MTGDNTLITSHGINRRDFMKLCAALAATMGLSSKAAAEMAESVSNPQRPPVIWIGAQECT WP_000145431.1 hydrogenase 2 small subunit [Salmonella enterica] Length: 372\nNumber of predicted TMHs: 1\nExp number of AAs in TMHs: 24.51505\nExp number, first 60 AAs: 0.2682\nTotal prob of N-in: 0.14406\noutside 1 331\nTMhelix 332 354\ninside 355 372

18934 GCF_002032155.1_ASM203215v1 Salmonella enterica subsp. enterica serovar Oranienburg Proteobacteria; Gammaproteobacteria; Enterobacterales; Enterobacteriaceae; Salmonella; Salmonella enterica; Salmonella enterica subsp. enterica MTGDNTLITSHGINRRDFMKLCAALAATMGLSSKAAAEMAESVSNPQRPPVIWIGAQECT WP_000145431.1 hydrogenase 2 small subunit [Salmonella enterica] Length: 372\nNumber of predicted TMHs: 1\nExp number of AAs in TMHs: 24.51505\nExp number, first 60 AAs: 0.2682\nTotal prob of N-in: 0.14406\noutside 1 331\nTMhelix 332 354\ninside 355 372

18935 GCF_001879925.1_ASM187992v1 Salmonella enterica subsp. enterica serovar Oranienburg Proteobacteria; Gammaproteobacteria; Enterobacterales; Enterobacteriaceae; Salmonella; Salmonella enterica; Salmonella enterica subsp. enterica MTGDNTLITSHGINRRDFMKLCAALAATMGLSSKAAAEMAESVSNPQRPPVIWIGAQECT WP_000145431.1 hydrogenase 2 small subunit [Salmonella enterica] Length: 372\nNumber of predicted TMHs: 1\nExp number of AAs in TMHs: 24.51505\nExp number, first 60 AAs: 0.2682\nTotal prob of N-in: 0.14406\noutside 1 331\nTMhelix 332 354\ninside 355 372

18936 GCF_001880025.1_ASM188002v1 Salmonella enterica subsp. enterica serovar Oranienburg Proteobacteria; Gammaproteobacteria; Enterobacterales; Enterobacteriaceae; Salmonella; Salmonella enterica; Salmonella enterica subsp. enterica MNNEETFYQAMRRKGVTRRSFLKFCSLAATSLGLGAGMTPKIAWALENKPRIPVVWIHGL WP_001058312.1 hydrogenase [Salmonella enterica] Length: 372\nNumber of predicted TMHs: 2\nExp

number of AAs in TMHs: 36.51738\nExp number, first 60 AAs: 19.72849\nTotal prob of N-in: 0.96170\nPOSSIBLE N-term signal sequence\ninside 1 19\nTMhelix 20 39\noutside 40 325\nTMhelix 326 348\ninside 349 372

18937 GCF_001879955.1_ASM187995v1 Salmonella enterica subsp. enterica serovar Oranienburg Proteobacteria; Gammaproteobacteria; Enterobacterales; Enterobacteriaceae; Salmonella; Salmonella enterica; Salmonella enterica subsp. enterica MTGDNTLITSHGINRRDFMKLCAALAATMGLSSKAAAEMAESVSNPQRPPVIWIGAQECT WP_000145431.1 hydrogenase 2 small subunit [Salmonella enterica] Length: 372\nNumber of predicted TMHs: 1\nExp number of AAs in TMHs: 24.51505\nExp number, first 60 AAs: 0.2682\nTotal prob of N-in: 0.14406\noutside 1 331\nTMhelix 332 354\ninside 355 372

18938 GCF_001878935.1_ASM187893v1 Salmonella enterica subsp. enterica serovar Oranienburg Proteobacteria; Gammaproteobacteria; Enterobacterales; Enterobacteriaceae; Salmonella; Salmonella enterica; Salmonella enterica subsp. enterica MNNEETFYQAMRRKGVTRRSFLKFCSLAATSLGLGAGMMPKIAWALENKPRIPVVWIHGL WP_057393961.1 hydrogenase [Salmonella enterica] Length: 372\nNumber of predicted TMHs: 2\nExp number of AAs in TMHs: 37.03037\nExp number, first 60 AAs: 20.25216\nTotal prob of N-in: 0.98577\nPOSSIBLE N-term signal sequence\ninside 1 19\nTMhelix 20 39\noutside 40 325\nTMhelix 326 348\ninside 349 372

18939 GCF_001880005.1_ASM188000v1 Salmonella enterica subsp. enterica serovar Oranienburg Proteobacteria; Gammaproteobacteria; Enterobacterales; Enterobacteriaceae; Salmonella; Salmonella enterica; Salmonella enterica subsp. enterica MTGDNTLITSHGINRRDFMKLCAALAATMGLSSKAAAEMAESVSNPQRPPVIWIGAQECT WP_000145431.1 hydrogenase 2 small subunit [Salmonella enterica] Length: 372\nNumber of predicted TMHs: 1\nExp number of AAs in TMHs: 24.51505\nExp number, first 60 AAs: 0.2682\nTotal prob of N-in: 0.14406\noutside 1 331\nTMhelix 332 354\ninside 355 372

18940 GCF_001880025.1_ASM188002v1 Salmonella enterica subsp. enterica serovar Oranienburg Proteobacteria; Gammaproteobacteria; Enterobacterales; Enterobacteriaceae; Salmonella; Salmonella enterica; Salmonella enterica subsp. enterica MTGDNTLITSHGINRRDFMKLCAALAATMGLSSKAAAEMAESVSNPQRPPVIWIGAQECT WP_000145431.1 hydrogenase 2 small subunit [Salmonella enterica] Length: 372\nNumber of predicted TMHs: 1\nExp number of AAs in TMHs: 24.51505\nExp number, first 60 AAs: 0.2682\nTotal prob of N-in: 0.14406\noutside 1 331\nTMhelix 332 354\ninside 355 372

18941 GCF_001879955.1_ASM187995v1 Salmonella enterica subsp. enterica serovar Oranienburg Proteobacteria; Gammaproteobacteria; Enterobacterales; Enterobacteriaceae; Salmonella; Salmonella enterica; Salmonella enterica subsp. enterica MNNEETFYQAMRRKGVTRRSFLKFCSLAATSLGLGAGMTPKIAWALENKPRIPVVWIHGL WP_001058312.1 hydrogenase [Salmonella enterica] Length: 372\nNumber of predicted TMHs: 2\nExp number of AAs in TMHs: 36.51738\nExp number, first 60 AAs: 19.72849\nTotal prob of N-in: 0.96170\nPOSSIBLE N-term signal sequence\ninside 1 19\nTMhelix 20 39\noutside 40 325\nTMhelix 326 348\ninside 349 372

18942 GCF_001878885.1_ASM187888v1 Salmonella enterica subsp. enterica serovar Oranienburg Proteobacteria; Gammaproteobacteria; Enterobacterales; Enterobacteriaceae; Salmonella; Salmonella enterica; Salmonella enterica subsp. enterica MTGDNTLITSHGINRRDFMKLCAALAATMGLSSKAAAEMAESVSNPQRPPVIWIGAQECT WP_000145431.1 hydrogenase 2 small subunit [Salmonella enterica] Length: 372\nNumber of predicted TMHs: 1\nExp number of AAs in TMHs: 24.51505\nExp number, first 60 AAs: 0.2682\nTotal prob of N-in: 0.14406\noutside 1 331\nTMhelix 332 354\ninside 355 372

18943 GCF_001880005.1_ASM188000v1 Salmonella enterica subsp. enterica serovar Oranienburg Proteobacteria; Gammaproteobacteria; Enterobacterales; Enterobacteriaceae; Salmonella; Salmonella enterica; Salmonella enterica subsp. enterica MNNEETFYQAMRRKGVTRRSFLKFCSLAATSLGLGAGMTPKIAWALENKPRIPVVWIHGL WP_001058312.1 hydrogenase [Salmonella enterica] Length: 372\nNumber of predicted TMHs: 2\nExp number of AAs in TMHs: 36.51738\nExp number, first 60 AAs: 19.72849\nTotal prob of N-in: 0.96170\nPOSSIBLE N-term signal sequence\ninside 1 19\nTMhelix 20 39\noutside 40 325\nTMhelix 326 348\ninside 349 372

18944 GCF_001878965.1_ASM187896v1 Salmonella enterica subsp. enterica serovar Oranienburg Proteobacteria; Gammaproteobacteria; Enterobacterales; Enterobacteriaceae; Salmonella; Salmonella enterica; Salmonella enterica subsp. enterica MNNEETFYQAMRRKGVTRRSFLKFCSLAATSLGLGAGMTPKIAWALENKPRIPVVWIHGL WP_001058312.1 hydrogenase [Salmonella enterica] Length: 372\nNumber of predicted TMHs: 2\nExp number of AAs in TMHs: 36.51738\nExp number, first 60 AAs: 19.72849\nTotal prob of N-in: 0.96170\nPOSSIBLE N-term signal sequence\ninside 1 19\nTMhelix 20 39\noutside 40 325\nTMhelix 326 348\ninside 349 372

18945 GCF_001879865.1_ASM187986v1 Salmonella enterica subsp. enterica serovar Oranienburg Proteobacteria; Gammaproteobacteria; Enterobacterales; Enterobacteriaceae; Salmonella; Salmonella enterica;

Salmonella enterica subsp. enterica MNNEETFYQAMRRKGVTRRSFLKFCSLAATSLGLGAGMTPKIAWALENKPRIPVVWIHGL WP_001058312.1 hydrogenase [Salmonella enterica] Length: 372\nNumber of predicted TMHs: 2\nExp number of AAs in TMHs: 36.51738\nExp number, first 60 AAs: 19.72849\nTotal prob of N-in: 0.96170\nPOSSIBLE N-term signal sequence\ninside 1 19\nTMhelix 20 39\noutside 40 325\nTMhelix 326 348\ninside 349 372

18946 GCF_002060075.1_ASM206007v1 Salmonella enterica subsp. enterica serovar Oranienburg Proteobacteria; Gammaproteobacteria; Enterobacterales; Enterobacteriaceae; Salmonella; Salmonella enterica; Salmonella enterica subsp. enterica MNNEETFYQAMRRKGVTRRSFLKFCSLAATSLGLGAGMTPKIAWALENKPRIPVVWIHGL WP_001058312.1 hydrogenase [Salmonella enterica] Length: 372\nNumber of predicted TMHs: 2\nExp number of AAs in TMHs: 36.51738\nExp number, first 60 AAs: 19.72849\nTotal prob of N-in: 0.96170\nPOSSIBLE N-term signal sequence\ninside 1 19\nTMhelix 20 39\noutside 40 325\nTMhelix 326 348\ninside 349 372

18947 GCF_001879845.1_ASM187984v1 Salmonella enterica subsp. enterica serovar Oranienburg Proteobacteria; Gammaproteobacteria; Enterobacterales; Enterobacteriaceae; Salmonella; Salmonella enterica; Salmonella enterica subsp. enterica MTGDNTLITSHGINRRDFMKLCAALAATMGLSSKAAAEMAESVSNPQRPPVIWIGAQECT WP_000145431.1 hydrogenase 2 small subunit [Salmonella enterica] Length: 372\nNumber of predicted TMHs: 1\nExp number of AAs in TMHs: 24.51505\nExp number, first 60 AAs: 0.2682\nTotal prob of N-in: 0.14406\noutside 1 331\nTMhelix 332 354\ninside 355 372

18948 GCF_002156335.1_ASM215633v1 Salmonella enterica subsp. enterica serovar Oranienburg Proteobacteria; Gammaproteobacteria; Enterobacterales; Enterobacteriaceae; Salmonella; Salmonella enterica; Salmonella enterica subsp. enterica MTGDNTLITSHGINRRDFMKLCAALAATMGLSSKAAAEMAESVSNPQRPPVIWIGAQECT WP_000145429.1 hydrogenase 2 small subunit [Salmonella enterica] Length: 372\nNumber of predicted TMHs: 1\nExp number of AAs in TMHs: 24.51309\nExp number, first 60 AAs: 0.2682\nTotal prob of N-in: 0.14402\noutside 1 331\nTMhelix 332 354\ninside 355 372

18949 GCF_001878985.1_ASM187898v1 Salmonella enterica subsp. enterica serovar Oranienburg Proteobacteria; Gammaproteobacteria; Enterobacterales; Enterobacteriaceae; Salmonella; Salmonella enterica; Salmonella enterica subsp. enterica MTGDNTLITSHGINRRDFMKLCAALAATMGLSSKAAAEMAESVSNPQRPPVIWIGAQECT WP_000145429.1 hydrogenase 2 small subunit [Salmonella enterica] Length: 372\nNumber of predicted TMHs: 1\nExp number of AAs in TMHs: 24.51309\nExp number, first 60 AAs: 0.2682\nTotal prob of N-in: 0.14402\noutside 1 331\nTMhelix 332 354\ninside 355 372

18950 GCF_001880015.1_ASM188001v1 Salmonella enterica subsp. enterica serovar Oranienburg Proteobacteria; Gammaproteobacteria; Enterobacterales; Enterobacteriaceae; Salmonella; Salmonella enterica; Salmonella enterica subsp. enterica MNNEETFYQAMRRKGVTRRSFLKFCSLAATSLGLGAGMTPKIAWALENKPRIPVVWIHGL WP_001058312.1 hydrogenase [Salmonella enterica] Length: 372\nNumber of predicted TMHs: 2\nExp number of AAs in TMHs: 36.51738\nExp number, first 60 AAs: 19.72849\nTotal prob of N-in: 0.96170\nPOSSIBLE N-term signal sequence\ninside 1 19\nTMhelix 20 39\noutside 40 325\nTMhelix 326 348\ninside 349 372

18951 GCF_001879905.1_ASM187990v1 Salmonella enterica subsp. enterica serovar Oranienburg Proteobacteria; Gammaproteobacteria; Enterobacterales; Enterobacteriaceae; Salmonella; Salmonella enterica; Salmonella enterica subsp. enterica MNNEETFYQAMRRKGVTRRSFLKFCSLAATSLGLGAGMTPKIAWALENKPRIPVVWIHGL WP_001058312.1 hydrogenase [Salmonella enterica] Length: 372\nNumber of predicted TMHs: 2\nExp number of AAs in TMHs: 36.51738\nExp number, first 60 AAs: 19.72849\nTotal prob of N-in: 0.96170\nPOSSIBLE N-term signal sequence\ninside 1 19\nTMhelix 20 39\noutside 40 325\nTMhelix 326 348\ninside 349 372

18952 GCF_001880035.1_ASM188003v1 Salmonella enterica subsp. enterica serovar Oranienburg Proteobacteria; Gammaproteobacteria; Enterobacterales; Enterobacteriaceae; Salmonella; Salmonella enterica; Salmonella enterica subsp. enterica MTGDNTLITSHGINRRDFMKLCAALAATMGLSSKAAAEMAESVSNPQRPPVIWIGAQECT WP_000145431.1 hydrogenase 2 small subunit [Salmonella enterica] Length: 372\nNumber of predicted TMHs: 1\nExp number of AAs in TMHs: 24.51505\nExp number, first 60 AAs: 0.2682\nTotal prob of N-in: 0.14406\noutside 1 331\nTMhelix 332 354\ninside 355 372

18953 GCF_001880035.1_ASM188003v1 Salmonella enterica subsp. enterica serovar Oranienburg Proteobacteria; Gammaproteobacteria; Enterobacterales; Enterobacteriaceae; Salmonella; Salmonella enterica; Salmonella enterica subsp. enterica MNNEETFYQAMRRKGVTRRSFLKFCSLAATSLGLGAGMTPKIAWALENKPRIPVVWIHGL WP_001058312.1 hydrogenase [Salmonella enterica] Length: 372\nNumber of predicted TMHs: 2\nExp number of AAs in TMHs: 36.51738\nExp number, first 60 AAs: 19.72849\nTotal prob of N-in: 0.96170\nPOSSIBLE N-term signal sequence\ninside 1 19\nTMhelix 20 39\noutside 40 325\nTMhelix 326 348\ninside 349 372

18954 GCF_001880105.1_ASM188010v1 Salmonella enterica subsp. enterica serovar Oranienburg Proteobacteria; Gammaproteobacteria; Enterobacterales; Enterobacteriaceae; Salmonella; Salmonella enterica; Salmonella enterica subsp. enterica MTGDNTLITSHGINRRDFMKLCAALAATMGLSSKAAAEMAESVSNPQRPPVIWIGAQECT WP_000145431.1 hydrogenase 2 small subunit [Salmonella enterica] Length: 372\nNumber of predicted TMHs: 1\nExp number of AAs in TMHs: 24.51505\nExp number, first 60 AAs: 0.2682\nTotal prob of N-in: 0.14406\noutside 1 331\nTMhelix 332 354\ninside 355 372

18955 GCF_001878985.1_ASM187898v1 Salmonella enterica subsp. enterica serovar Oranienburg Proteobacteria; Gammaproteobacteria; Enterobacterales; Enterobacteriaceae; Salmonella; Salmonella enterica; Salmonella enterica subsp. enterica MNNEETFYQAMRRKGVTRRSFLKFCSLAATSLGLGAGMTPKIAWALENKPRIPVVWIHGL WP_001058311.1 hydrogenase [Salmonella enterica] Length: 372\nNumber of predicted TMHs: 2\nExp number of AAs in TMHs: 37.4699899999999\nExp number, first 60 AAs: 19.72699\nTotal prob of N-in: 0.96171\nPOSSIBLE N-term signal sequence\ninside 1 19\nTMhelix 20 39\noutside 40 325\nTMhelix 326 348\ninside 349 372

18956 GCF_002061275.1_ASM206127v1 Salmonella enterica subsp. enterica serovar Oranienburg Proteobacteria; Gammaproteobacteria; Enterobacterales; Enterobacteriaceae; Salmonella; Salmonella enterica; Salmonella enterica subsp. enterica MNNEETFYQAMRRKGVTRRSFLKFCSLAATSLGLGAGMTPKIAWALENKPRIPVVWIHGL WP_001058312.1 hydrogenase [Salmonella enterica] Length: 372\nNumber of predicted TMHs: 2\nExp number of AAs in TMHs: 36.51738\nExp number, first 60 AAs: 19.72849\nTotal prob of N-in: 0.96170\nPOSSIBLE N-term signal sequence\ninside 1 19\nTMhelix 20 39\noutside 40 325\nTMhelix 326 348\ninside 349 372

18957 GCF_001878875.1_ASM187887v1 Salmonella enterica subsp. enterica serovar Oranienburg Proteobacteria; Gammaproteobacteria; Enterobacterales; Enterobacteriaceae; Salmonella; Salmonella enterica; Salmonella enterica subsp. enterica MTGDNTLITSHGINRRDFMKLCAALAATMGLSSKAAAEMAESVSNPQRPPVIWIGAQECT WP_000145429.1 hydrogenase 2 small subunit [Salmonella enterica] Length: 372\nNumber of predicted TMHs: 1\nExp number of AAs in TMHs: 24.51309\nExp number, first 60 AAs: 0.2682\nTotal prob of N-in: 0.14402\noutside 1 331\nTMhelix 332 354\ninside 355 372

18958 GCF_001878945.1_ASM187894v1 Salmonella enterica subsp. enterica serovar Oranienburg Proteobacteria; Gammaproteobacteria; Enterobacterales; Enterobacteriaceae; Salmonella; Salmonella enterica; Salmonella enterica subsp. enterica MTGDNTLITSHGINRRDFMKLCAALAATMGLSSKAAAEMAESVSNPQRPPVIWIGAQECT WP_000145429.1 hydrogenase 2 small subunit [Salmonella enterica] Length: 372\nNumber of predicted TMHs: 1\nExp number of AAs in TMHs: 24.51309\nExp number, first 60 AAs: 0.2682\nTotal prob of N-in: 0.14402\noutside 1 331\nTMhelix 332 354\ninside 355 372

18959 GCF_001880015.1_ASM188001v1 Salmonella enterica subsp. enterica serovar Oranienburg Proteobacteria; Gammaproteobacteria; Enterobacterales; Enterobacteriaceae; Salmonella; Salmonella enterica; Salmonella enterica subsp. enterica MTGDNTLITSHGINRRDFMKLCAALAATMGLSSKAAAEMAESVSNPQRPPVIWIGAQECT WP_000145431.1 hydrogenase 2 small subunit [Salmonella enterica] Length: 372\nNumber of predicted TMHs: 1\nExp number of AAs in TMHs: 24.51505\nExp number, first 60 AAs: 0.2682\nTotal prob of N-in: 0.14406\noutside 1 331\nTMhelix 332 354\ninside 355 372

18960 GCF_002061275.1_ASM206127v1 Salmonella enterica subsp. enterica serovar Oranienburg Proteobacteria; Gammaproteobacteria; Enterobacterales; Enterobacteriaceae; Salmonella; Salmonella enterica; Salmonella enterica subsp. enterica MTGDNTLITSHGINRRDFMKLCAALAATMGLSSKAAAEMAESVSNPQRPPVIWIGAQECT WP_000145431.1 hydrogenase 2 small subunit [Salmonella enterica] Length: 372\nNumber of predicted TMHs: 1\nExp number of AAs in TMHs: 24.51505\nExp number, first 60 AAs: 0.2682\nTotal prob of N-in: 0.14406\noutside 1 331\nTMhelix 332 354\ninside 355 372

18961 GCF_001878875.1_ASM187887v1 Salmonella enterica subsp. enterica serovar Oranienburg Proteobacteria; Gammaproteobacteria; Enterobacterales; Enterobacteriaceae; Salmonella; Salmonella enterica; Salmonella enterica subsp. enterica MNNEETFYQAMRRKGVTRRSFLKFCSLAATSLGLGAGMTPKIAWALENKPRIPVVWIHGL WP_001058310.1 hydrogenase [Salmonella enterica] Length: 372\nNumber of predicted TMHs: 2\nExp number of AAs in TMHs: 37.4567299999999\nExp number, first 60 AAs: 19.72694\nTotal prob of N-in: 0.96171\nPOSSIBLE N-term signal sequence\ninside 1 19\nTMhelix 20 39\noutside 40 325\nTMhelix 326 348\ninside 349 372

18962 GCF_001880085.1_ASM188008v1 Salmonella enterica subsp. enterica serovar Oranienburg Proteobacteria; Gammaproteobacteria; Enterobacterales; Enterobacteriaceae; Salmonella; Salmonella enterica; Salmonella enterica subsp. enterica MNNEETFYQAMRRKGVTRRSFLKFCSLAATSLGLGAGMTPKIAWALENKPRIPVVWIHGL

WP_001058312.1	hydrogenase [Salmonella enterica]	Length: 372\nNumber of predicted TMHs:  2\nExp number of AAs in TMHs: 36.51738\nExp number, first 60 AAs:  19.72849\nTotal prob of N-in:   0.96170\nPOSSIBLE N-term signal sequence\ninside   1   19\nTMhelix   20   39\noutside   40   325\nTMhelix   326   348\ninside   349   372

18963	GCF_002064215.1_ASM206421v1	Salmonella enterica subsp. enterica serovar Oranienburg	Proteobacteria; Gammaproteobacteria; Enterobacterales; Enterobacteriaceae; Salmonella; Salmonella enterica; Salmonella enterica subsp. enterica	MTGDNTLITSHDINRRDFMKLCAALAATMGLSSKAAAEMAESVSNPQRPPVIWIGAQECT	WP_080191561.1	hydrogenase 2 small subunit [Salmonella enterica]	Length: 372\nNumber of predicted TMHs:  1\nExp number of AAs in TMHs: 24.53892\nExp number, first 60 AAs:  0.29527\nTotal prob of N-in:   0.14517\noutside   1   331\nTMhelix   332   354\ninside   355   372

18964	GCF_002032155.1_ASM203215v1	Salmonella enterica subsp. enterica serovar Oranienburg	Proteobacteria; Gammaproteobacteria; Enterobacterales; Enterobacteriaceae; Salmonella; Salmonella enterica; Salmonella enterica subsp. enterica	MNNEETFYQAMRRKGVTRRSFLKFCSLAATSLGLGAGMTPKIAWALENKPRIPVVWIHGL	WP_001058312.1	hydrogenase [Salmonella enterica]	Length: 372\nNumber of predicted TMHs:  2\nExp number of AAs in TMHs: 36.51738\nExp number, first 60 AAs:  19.72849\nTotal prob of N-in:   0.96170\nPOSSIBLE N-term signal sequence\ninside   1   19\nTMhelix   20   39\noutside   40   325\nTMhelix   326   348\ninside   349   372

18965	GCF_001878935.1_ASM187893v1	Salmonella enterica subsp. enterica serovar Oranienburg	Proteobacteria; Gammaproteobacteria; Enterobacterales; Enterobacteriaceae; Salmonella; Salmonella enterica; Salmonella enterica subsp. enterica	MTGDNTLITSHGINRRDFMKLCAALAATMGLSSKAAAEMAESVSNPQRPPVIWIGAQECT	WP_000145429.1	hydrogenase 2 small subunit [Salmonella enterica]	Length: 372\nNumber of predicted TMHs:  1\nExp number of AAs in TMHs: 24.51309\nExp number, first 60 AAs:  0.2682\nTotal prob of N-in:   0.14402\noutside   1   331\nTMhelix   332   354\ninside   355   372

18966	GCF_001878965.1_ASM187896v1	Salmonella enterica subsp. enterica serovar Oranienburg	Proteobacteria; Gammaproteobacteria; Enterobacterales; Enterobacteriaceae; Salmonella; Salmonella enterica; Salmonella enterica subsp. enterica	MTGDNTLITSHGINRRDFMKLCAALAATMGLSSKAAAEMAESVSNPQRPPVIWIGAQECT	WP_000145429.1	hydrogenase 2 small subunit [Salmonella enterica]	Length: 372\nNumber of predicted TMHs:  1\nExp number of AAs in TMHs: 24.51309\nExp number, first 60 AAs:  0.2682\nTotal prob of N-in:   0.14402\noutside   1   331\nTMhelix   332   354\ninside   355   372

18967	GCF_002064215.1_ASM206421v1	Salmonella enterica subsp. enterica serovar Oranienburg	Proteobacteria; Gammaproteobacteria; Enterobacterales; Enterobacteriaceae; Salmonella; Salmonella enterica; Salmonella enterica subsp. enterica	MNNEETFYQAMRRKGVTRRSFLKFCSLAATSLGLGAGMTPKIAWALENKPRIPVVWIHGL	WP_001058312.1	hydrogenase [Salmonella enterica]	Length: 372\nNumber of predicted TMHs:  2\nExp number of AAs in TMHs: 36.51738\nExp number, first 60 AAs:  19.72849\nTotal prob of N-in:   0.96170\nPOSSIBLE N-term signal sequence\ninside   1   19\nTMhelix   20   39\noutside   40   325\nTMhelix   326   348\ninside   349   372

18968	GCF_001879845.1_ASM187984v1	Salmonella enterica subsp. enterica serovar Oranienburg	Proteobacteria; Gammaproteobacteria; Enterobacterales; Enterobacteriaceae; Salmonella; Salmonella enterica; Salmonella enterica subsp. enterica	MNNEETFYQAMRRKGVTRRSFLKFCSLAATSLGLGAGMTPKIAWALENKPRIPVVWIHGL	WP_001058312.1	hydrogenase [Salmonella enterica]	Length: 372\nNumber of predicted TMHs:  2\nExp number of AAs in TMHs: 36.51738\nExp number, first 60 AAs:  19.72849\nTotal prob of N-in:   0.96170\nPOSSIBLE N-term signal sequence\ninside   1   19\nTMhelix   20   39\noutside   40   325\nTMhelix   326   348\ninside   349   372

18969	GCF_001879785.1_ASM187978v1	Salmonella enterica subsp. enterica serovar Oranienburg	Proteobacteria; Gammaproteobacteria; Enterobacterales; Enterobacteriaceae; Salmonella; Salmonella enterica; Salmonella enterica subsp. enterica	MNNEETFYQAMRRKGVTRRSFLKFCSLAATSLGLGAGMTPKIAWALENKPRIPVVWIHGL	WP_001058312.1	hydrogenase [Salmonella enterica]	Length: 372\nNumber of predicted TMHs:  2\nExp number of AAs in TMHs: 36.51738\nExp number, first 60 AAs:  19.72849\nTotal prob of N-in:   0.96170\nPOSSIBLE N-term signal sequence\ninside   1   19\nTMhelix   20   39\noutside   40   325\nTMhelix   326   348\ninside   349   372

18970	GCF_001879925.1_ASM187992v1	Salmonella enterica subsp. enterica serovar Oranienburg	Proteobacteria; Gammaproteobacteria; Enterobacterales; Enterobacteriaceae; Salmonella; Salmonella enterica; Salmonella enterica subsp. enterica	MNNEETFYQAMRRKGVTRRSFLKFCSLAATSLGLGAGMTPKIAWALENKPRIPVVWIHGL	WP_001058312.1	hydrogenase [Salmonella enterica]	Length: 372\nNumber of predicted TMHs:  2\nExp number of AAs in TMHs: 36.51738\nExp number, first 60 AAs:  19.72849\nTotal prob of N-in:   0.96170\nPOSSIBLE N-term signal sequence\ninside   1   19\nTMhelix   20   39\noutside   40   325\nTMhelix   326   348\ninside   349   372

18971 GCF_001880085.1_ASM188008v1 Salmonella enterica subsp. enterica serovar Oranienburg Proteobacteria; Gammaproteobacteria; Enterobacterales; Enterobacteriaceae; Salmonella; Salmonella enterica; Salmonella enterica subsp. enterica MTGDNTLITSHGINRRDFMKLCAALAATMGLSSKAAAEMAESVSNPQRPPVIWIGAQECT WP_000145431.1 hydrogenase 2 small subunit [Salmonella enterica] Length: 372\nNumber of predicted TMHs: 1\nExp number of AAs in TMHs: 24.51505\nExp number, first 60 AAs: 0.2682\nTotal prob of N-in: 0.14406\noutside 1 331\nTMhelix 332 354\ninside 355 372

18972 GCF_001879765.1_ASM187976v1 Salmonella enterica subsp. enterica serovar Oranienburg Proteobacteria; Gammaproteobacteria; Enterobacterales; Enterobacteriaceae; Salmonella; Salmonella enterica; Salmonella enterica subsp. enterica MNNEETFYQAMRRKGVTRRSFLKFCSLAATSLGLGAGMTPKIAWALENKPRIPVVWIHGL WP_001058312.1 hydrogenase [Salmonella enterica] Length: 372\nNumber of predicted TMHs: 2\nExp number of AAs in TMHs: 36.51738\nExp number, first 60 AAs: 19.72849\nTotal prob of N-in: 0.96170\nPOSSIBLE N-term signal sequence\ninside 1 19\nTMhelix 20 39\noutside 40 325\nTMhelix 326 348\ninside 349 372

18973 GCF_001879765.1_ASM187976v1 Salmonella enterica subsp. enterica serovar Oranienburg Proteobacteria; Gammaproteobacteria; Enterobacterales; Enterobacteriaceae; Salmonella; Salmonella enterica; Salmonella enterica subsp. enterica MTGDNTLITSHGINRRDFMKLCAALAATMGLSSKAAAEMAESVSNPQRPPVIWIGAQECT WP_000145431.1 hydrogenase 2 small subunit [Salmonella enterica] Length: 372\nNumber of predicted TMHs: 1\nExp number of AAs in TMHs: 24.51505\nExp number, first 60 AAs: 0.2682\nTotal prob of N-in: 0.14406\noutside 1 331\nTMhelix 332 354\ninside 355 372

18974 GCF_001879865.1_ASM187986v1 Salmonella enterica subsp. enterica serovar Oranienburg Proteobacteria; Gammaproteobacteria; Enterobacterales; Enterobacteriaceae; Salmonella; Salmonella enterica; Salmonella enterica subsp. enterica MTGDNTLITSHGINRRDFMKLCAALAATMGLSSKAAAEMAESVSNPQRPPVIWIGAQECT WP_000145429.1 hydrogenase 2 small subunit [Salmonella enterica] Length: 372\nNumber of predicted TMHs: 1\nExp number of AAs in TMHs: 24.51309\nExp number, first 60 AAs: 0.2682\nTotal prob of N-in: 0.14402\noutside 1 331\nTMhelix 332 354\ninside 355 372

18975 GCF_002064195.1_ASM206419v1 Salmonella enterica subsp. enterica serovar Oritamerin Proteobacteria; Gammaproteobacteria; Enterobacterales; Enterobacteriaceae; Salmonella; Salmonella enterica; Salmonella enterica subsp. enterica MTGDNTLITSHGINRRDFMKLCAALAATMGLSSKAAAEMAESVSNPQRPPVIWIGAQECT WP_000145429.1 hydrogenase 2 small subunit [Salmonella enterica] Length: 372\nNumber of predicted TMHs: 1\nExp number of AAs in TMHs: 24.51309\nExp number, first 60 AAs: 0.2682\nTotal prob of N-in: 0.14402\noutside 1 331\nTMhelix 332 354\ninside 355 372

18976 GCF_002064195.1_ASM206419v1 Salmonella enterica subsp. enterica serovar Oritamerin Proteobacteria; Gammaproteobacteria; Enterobacterales; Enterobacteriaceae; Salmonella; Salmonella enterica; Salmonella enterica subsp. enterica MNNEETFYQAMRRKGVTRRSFLKFCSLAATSLGLGAGMTPKIAWALENKPRIPVVWIHGL WP_001058312.1 hydrogenase [Salmonella enterica] Length: 372\nNumber of predicted TMHs: 2\nExp number of AAs in TMHs: 36.51738\nExp number, first 60 AAs: 19.72849\nTotal prob of N-in: 0.96170\nPOSSIBLE N-term signal sequence\ninside 1 19\nTMhelix 20 39\noutside 40 325\nTMhelix 326 348\ninside 349 372

18977 GCF_002048195.1_ASM204819v1 Salmonella enterica subsp. enterica serovar Oslo Proteobacteria; Gammaproteobacteria; Enterobacterales; Enterobacteriaceae; Salmonella; Salmonella enterica; Salmonella enterica subsp. enterica MTGDNTLITSHGINRRDFMKLCAALAATMGLSSKAAAEMAESVSNPQRPPVIWIGAQECT WP_000145429.1 hydrogenase 2 small subunit [Salmonella enterica] Length: 372\nNumber of predicted TMHs: 1\nExp number of AAs in TMHs: 24.51309\nExp number, first 60 AAs: 0.2682\nTotal prob of N-in: 0.14402\noutside 1 331\nTMhelix 332 354\ninside 355 372

18978 GCF_002046975.1_ASM204697v1 Salmonella enterica subsp. enterica serovar Oslo Proteobacteria; Gammaproteobacteria; Enterobacterales; Enterobacteriaceae; Salmonella; Salmonella enterica; Salmonella enterica subsp. enterica MTGDNTLITSHGINRRDFMKLCAALAATMGLSSKAAAEMAESVSNPQRPPVIWIGAQECT WP_000145429.1 hydrogenase 2 small subunit [Salmonella enterica] Length: 372\nNumber of predicted TMHs: 1\nExp number of AAs in TMHs: 24.51309\nExp number, first 60 AAs: 0.2682\nTotal prob of N-in: 0.14402\noutside 1 331\nTMhelix 332 354\ninside 355 372

18979 GCF_002047025.1_ASM204702v1 Salmonella enterica subsp. enterica serovar Oslo Proteobacteria; Gammaproteobacteria; Enterobacterales; Enterobacteriaceae; Salmonella; Salmonella enterica; Salmonella enterica subsp. enterica MNNEETFYQAMRRKGVTRRSFLKFCSLAATSLGLGAGMTPKIAWALENKPRIPVVWIHGL WP_001058312.1 hydrogenase [Salmonella enterica] Length: 372\nNumber of predicted TMHs: 2\nExp number of AAs in

TMHs: 36.51738\nExp number, first 60 AAs: 19.72849\nTotal prob of N-in: 0.96170\nPOSSIBLE N-term signal sequence\ninside 1 19\nTMhelix 20 39\noutside 40 325\nTMhelix 326 348\ninside 349 372

18980 GCF_002047025.1_ASM204702v1 Salmonella enterica subsp. enterica serovar Oslo Proteobacteria; Gammaproteobacteria; Enterobacterales; Enterobacteriaceae; Salmonella; Salmonella enterica; Salmonella enterica subsp. enterica MTGDNTLITSHGINRRDFMKLCAALAATMGLSSKAAAEMAESVSNPQRPPVIWIGAQECT WP_000145429.1 hydrogenase 2 small subunit [Salmonella enterica] Length: 372\nNumber of predicted TMHs: 1\nExp number of AAs in TMHs: 24.51309\nExp number, first 60 AAs: 0.2682\nTotal prob of N-in: 0.14402\noutside 1 331\nTMhelix 332 354\ninside 355 372

18981 GCF_002048195.1_ASM204819v1 Salmonella enterica subsp. enterica serovar Oslo Proteobacteria; Gammaproteobacteria; Enterobacterales; Enterobacteriaceae; Salmonella; Salmonella enterica; Salmonella enterica subsp. enterica MNNEETFYQAMRRKGVTRRSFLKFCSLAATSLGLGAGMTPKIAWALENKPRIPVVWIHGL WP_001058312.1 hydrogenase [Salmonella enterica] Length: 372\nNumber of predicted TMHs: 2\nExp number of AAs in TMHs: 36.51738\nExp number, first 60 AAs: 19.72849\nTotal prob of N-in: 0.96170\nPOSSIBLE N-term signal sequence\ninside 1 19\nTMhelix 20 39\noutside 40 325\nTMhelix 326 348\ninside 349 372

18982 GCF_002046815.1_ASM204681v1 Salmonella enterica subsp. enterica serovar Oslo Proteobacteria; Gammaproteobacteria; Enterobacterales; Enterobacteriaceae; Salmonella; Salmonella enterica; Salmonella enterica subsp. enterica MNNEETFYQAMRRKGVTRRSFLKFCSLAATSLGLGAGMTPKIAWALENKPRIPVVWIHGL WP_001058312.1 hydrogenase [Salmonella enterica] Length: 372\nNumber of predicted TMHs: 2\nExp number of AAs in TMHs: 36.51738\nExp number, first 60 AAs: 19.72849\nTotal prob of N-in: 0.96170\nPOSSIBLE N-term signal sequence\ninside 1 19\nTMhelix 20 39\noutside 40 325\nTMhelix 326 348\ninside 349 372

18983 GCF_002046815.1_ASM204681v1 Salmonella enterica subsp. enterica serovar Oslo Proteobacteria; Gammaproteobacteria; Enterobacterales; Enterobacteriaceae; Salmonella; Salmonella enterica; Salmonella enterica subsp. enterica MTGDNTLITSHGINRRDFMKLCAALAATMGLSSKAAAEMAESVSNPQRPPVIWIGAQECT WP_000145429.1 hydrogenase 2 small subunit [Salmonella enterica] Length: 372\nNumber of predicted TMHs: 1\nExp number of AAs in TMHs: 24.51309\nExp number, first 60 AAs: 0.2682\nTotal prob of N-in: 0.14402\noutside 1 331\nTMhelix 332 354\ninside 355 372

18984 GCF_001077835.1_ASM107783v1 Salmonella enterica subsp. enterica serovar Ouakam Proteobacteria; Gammaproteobacteria; Enterobacterales; Enterobacteriaceae; Salmonella; Salmonella enterica; Salmonella enterica subsp. enterica MTGDNTLITSHGINRRDFMKLCAALAATMGLSSKAAAEMAESVSNPQRPPVIWIGAQECT WP_000145429.1 hydrogenase 2 small subunit [Salmonella enterica] Length: 372\nNumber of predicted TMHs: 1\nExp number of AAs in TMHs: 24.51309\nExp number, first 60 AAs: 0.2682\nTotal prob of N-in: 0.14402\noutside 1 331\nTMhelix 332 354\ninside 355 372

18985 GCF_001077835.1_ASM107783v1 Salmonella enterica subsp. enterica serovar Ouakam Proteobacteria; Gammaproteobacteria; Enterobacterales; Enterobacteriaceae; Salmonella; Salmonella enterica; Salmonella enterica subsp. enterica MNNEETFYQAMRRKGVTRRSFLKFCSLAATSLGLGAGMTPKIAWALENKPRIPVVWIHGL WP_001058311.1 hydrogenase [Salmonella enterica] Length: 372\nNumber of predicted TMHs: 2\nExp number of AAs in TMHs: 37.4699899999999\nExp number, first 60 AAs: 19.72699\nTotal prob of N-in: 0.96171\nPOSSIBLE N-term signal sequence\ninside 1 19\nTMhelix 20 39\noutside 40 325\nTMhelix 326 348\ninside 349 372

18986 GCF_002062625.1_ASM206262v1 Salmonella enterica subsp. enterica serovar Panama Proteobacteria; Gammaproteobacteria; Enterobacterales; Enterobacteriaceae; Salmonella; Salmonella enterica; Salmonella enterica subsp. enterica MTGDNTLITSHGINRRDFMKLCAALAATMGLSSKAAAEMAESVSNPQRPPVIWIGAQECT WP_080195303.1 hydrogenase 2 small subunit [Salmonella enterica] Length: 372\nNumber of predicted TMHs: 1\nExp number of AAs in TMHs: 24.51336\nExp number, first 60 AAs: 0.26822\nTotal prob of N-in: 0.14403\noutside 1 331\nTMhelix 332 354\ninside 355 372

18987 GCF_002062625.1_ASM206262v1 Salmonella enterica subsp. enterica serovar Panama Proteobacteria; Gammaproteobacteria; Enterobacterales; Enterobacteriaceae; Salmonella; Salmonella enterica; Salmonella enterica subsp. enterica MNNEETFYQAMRRKGVTRRSFLKFCSLAATSLGLGAGMTPKIAWALENKPRIPVVWIHGL WP_001058312.1 hydrogenase [Salmonella enterica] Length: 372\nNumber of predicted TMHs: 2\nExp number of AAs in TMHs: 36.51738\nExp number, first 60 AAs: 19.72849\nTotal prob of N-in: 0.96170\nPOSSIBLE N-term signal sequence\ninside 1 19\nTMhelix 20 39\noutside 40 325\nTMhelix 326 348\ninside 349 372

18988	GCF_002064145.1_ASM206414v1	Salmonella enterica subsp. enterica serovar Panama	Proteobacteria; Gammaproteobacteria; Enterobacterales; Enterobacteriaceae; Salmonella; Salmonella enterica; Salmonella enterica subsp. enterica	MNNEETFYQAMRRKGVTRRSFLKFCSLAATSLGLGAGMTPKIAWALENKPRIPVVWIHGL	WP_001058312.1	hydrogenase [Salmonella enterica]	Length: 372\nNumber of predicted TMHs: 2\nExp number of AAs in TMHs: 36.51738\nExp number, first 60 AAs: 19.72849\nTotal prob of N-in: 0.96170\nPOSSIBLE N-term signal sequence\ninside 1 19\nTMhelix 20 39\noutside 40 325\nTMhelix 326 348\ninside 349 372

18989	GCF_002064145.1_ASM206414v1	Salmonella enterica subsp. enterica serovar Panama	Proteobacteria; Gammaproteobacteria; Enterobacterales; Enterobacteriaceae; Salmonella; Salmonella enterica; Salmonella enterica subsp. enterica	MTGDNTLITSHGINRRDFMKLCAALAATMGLSSKAAAEMAESVSNPQRPPVIWIGAQECT	WP_080195303.1	hydrogenase 2 small subunit [Salmonella enterica]	Length: 372\nNumber of predicted TMHs: 1\nExp number of AAs in TMHs: 24.51336\nExp number, first 60 AAs: 0.26822\nTotal prob of N-in: 0.14403\noutside 1 331\nTMhelix 332 354\ninside 355 372

18990	GCF_000984075.1_PA132	Salmonella enterica subsp. enterica serovar Paratyphi A	Proteobacteria; Gammaproteobacteria; Enterobacterales; Enterobacteriaceae; Salmonella; Salmonella enterica; Salmonella enterica subsp. enterica	MTGDNTLITSHGINRRDFMKLCAALAATMGLSSKAAAEMAESVSNPQRPPVIWIGAQECT	WP_000145427.1	hydrogenase 2 small subunit [Salmonella enterica]	Length: 372\nNumber of predicted TMHs: 1\nExp number of AAs in TMHs: 24.48576\nExp number, first 60 AAs: 0.2692\nTotal prob of N-in: 0.14251\noutside 1 331\nTMhelix 332 354\ninside 355 372

18991	GCF_000984195.1_PA102	Salmonella enterica subsp. enterica serovar Paratyphi A	Proteobacteria; Gammaproteobacteria; Enterobacterales; Enterobacteriaceae; Salmonella; Salmonella enterica; Salmonella enterica subsp. enterica	MTGDNTLITSHGINRRDFMKLCAALAATMGLSSKAAAEMAESVSNPQRPPVIWIGAQECT	WP_000145427.1	hydrogenase 2 small subunit [Salmonella enterica]	Length: 372\nNumber of predicted TMHs: 1\nExp number of AAs in TMHs: 24.48576\nExp number, first 60 AAs: 0.2692\nTotal prob of N-in: 0.14251\noutside 1 331\nTMhelix 332 354\ninside 355 372

18992	GCF_000984275.1_PA017	Salmonella enterica subsp. enterica serovar Paratyphi A	Proteobacteria; Gammaproteobacteria; Enterobacterales; Enterobacteriaceae; Salmonella; Salmonella enterica; Salmonella enterica subsp. enterica	MTGDNTLITSHGINRRDFMKLCAALAATMGLSSKAAAEMAESVSNPQRPPVIWIGAQECT	WP_000145427.1	hydrogenase 2 small subunit [Salmonella enterica]	Length: 372\nNumber of predicted TMHs: 1\nExp number of AAs in TMHs: 24.48576\nExp number, first 60 AAs: 0.2692\nTotal prob of N-in: 0.14251\noutside 1 331\nTMhelix 332 354\ninside 355 372

18993	GCF_000984235.1_PA068	Salmonella enterica subsp. enterica serovar Paratyphi A	Proteobacteria; Gammaproteobacteria; Enterobacterales; Enterobacteriaceae; Salmonella; Salmonella enterica; Salmonella enterica subsp. enterica	MTGDNTLITSHGINRRDFMKLCAALAATMGLSSKAAAEMAESVSNPQRPPVIWIGAQECT	WP_000145427.1	hydrogenase 2 small subunit [Salmonella enterica]	Length: 372\nNumber of predicted TMHs: 1\nExp number of AAs in TMHs: 24.48576\nExp number, first 60 AAs: 0.2692\nTotal prob of N-in: 0.14251\noutside 1 331\nTMhelix 332 354\ninside 355 372

18994	GCF_000984155.1_PA030	Salmonella enterica subsp. enterica serovar Paratyphi A	Proteobacteria; Gammaproteobacteria; Enterobacterales; Enterobacteriaceae; Salmonella; Salmonella enterica; Salmonella enterica subsp. enterica	MTGDNTLITSHGINRRDFMKLCAALAATMGLSSKAAAEMAESVSNPQRPPVIWIGAQECT	WP_046595276.1	hydrogenase 2 small subunit [Salmonella enterica]	Length: 372\nNumber of predicted TMHs: 1\nExp number of AAs in TMHs: 24.40967\nExp number, first 60 AAs: 0.26399\nTotal prob of N-in: 0.14296\noutside 1 331\nTMhelix 332 354\ninside 355 372

18995	GCF_000984895.1_PA070	Salmonella enterica subsp. enterica serovar Paratyphi A	Proteobacteria; Gammaproteobacteria; Enterobacterales; Enterobacteriaceae; Salmonella; Salmonella enterica; Salmonella enterica subsp. enterica	MTGDNTLITSHGINRRDFMKLCAALAATMGLSSKAAAEMAESVSNPQRPPVIWIGAQECT	WP_000145427.1	hydrogenase 2 small subunit [Salmonella enterica]	Length: 372\nNumber of predicted TMHs: 1\nExp number of AAs in TMHs: 24.48576\nExp number, first 60 AAs: 0.2692\nTotal prob of N-in: 0.14251\noutside 1 331\nTMhelix 332 354\ninside 355 372

18996	GCF_000984935.1_PA015	Salmonella enterica subsp. enterica serovar Paratyphi A	Proteobacteria; Gammaproteobacteria; Enterobacterales; Enterobacteriaceae; Salmonella; Salmonella enterica; Salmonella enterica subsp. enterica	MTGDNTLITSHGINRRDFMKLCAALAATMGLSSKAAAEMAESVSNPQRPPVIWIGAQECT	WP_000145427.1	hydrogenase 2 small subunit [Salmonella enterica]	Length: 372\nNumber of predicted TMHs: 1\nExp

number of AAs in TMHs: 24.48576\nExp number, first 60 AAs:  0.2692\nTotal prob of N-in:   0.14251\noutside   1   331\nTMhelix   332   354\ninside   355   372

18997 GCF_000984975.1_PA086 Salmonella enterica subsp. enterica serovar Paratyphi A Proteobacteria; Gammaproteobacteria; Enterobacterales; Enterobacteriaceae; Salmonella; Salmonella enterica; Salmonella enterica subsp. enterica MTGDNTLITSHGINRRDFMKLCAALAATMGLSSKAAAEMAESVSNPQRPPVIWIGAQECT WP_000145427.1 hydrogenase 2 small subunit [Salmonella enterica] Length: 372\nNumber of predicted TMHs:  1\nExp number of AAs in TMHs: 24.48576\nExp number, first 60 AAs:  0.2692\nTotal prob of N-in:   0.14251\noutside   1   331\nTMhelix   332   354\ninside   355   372

18998 GCF_000985015.1_PA053 Salmonella enterica subsp. enterica serovar Paratyphi A Proteobacteria; Gammaproteobacteria; Enterobacterales; Enterobacteriaceae; Salmonella; Salmonella enterica; Salmonella enterica subsp. enterica MTGDNTLITSHGINRRDFMKLCAALAATMGLSSKAAAEMAESVSNPQRPPVIWIGAQECT WP_000145427.1 hydrogenase 2 small subunit [Salmonella enterica] Length: 372\nNumber of predicted TMHs:  1\nExp number of AAs in TMHs: 24.48576\nExp number, first 60 AAs:  0.2692\nTotal prob of N-in:   0.14251\noutside   1   331\nTMhelix   332   354\ninside   355   372

18999 GCF_000985775.1_PA144 Salmonella enterica subsp. enterica serovar Paratyphi A Proteobacteria; Gammaproteobacteria; Enterobacterales; Enterobacteriaceae; Salmonella; Salmonella enterica; Salmonella enterica subsp. enterica MTGDNTLITSHGINRRDFMKLCAALAATMGLSSKAAAEMAESVSNPQRPPVIWIGAQECT WP_000145427.1 hydrogenase 2 small subunit [Salmonella enterica] Length: 372\nNumber of predicted TMHs:  1\nExp number of AAs in TMHs: 24.48576\nExp number, first 60 AAs:  0.2692\nTotal prob of N-in:   0.14251\noutside   1   331\nTMhelix   332   354\ninside   355   372

19000 GCF_000985655.1_PA073 Salmonella enterica subsp. enterica serovar Paratyphi A Proteobacteria; Gammaproteobacteria; Enterobacterales; Enterobacteriaceae; Salmonella; Salmonella enterica; Salmonella enterica subsp. enterica MTGDNTLITSHGINRRDFMKLCAALAATMGLSSKAAAEMAESVSNPQRPPVIWIGAQECT WP_000145427.1 hydrogenase 2 small subunit [Salmonella enterica] Length: 372\nNumber of predicted TMHs:  1\nExp number of AAs in TMHs: 24.48576\nExp number, first 60 AAs:  0.2692\nTotal prob of N-in:   0.14251\noutside   1   331\nTMhelix   332   354\ninside   355   372

19001 GCF_000985195.1_PA029 Salmonella enterica subsp. enterica serovar Paratyphi A Proteobacteria; Gammaproteobacteria; Enterobacterales; Enterobacteriaceae; Salmonella; Salmonella enterica; Salmonella enterica subsp. enterica MTGDNTLITSHGINRRDFMKLCAALAATMGLSSKAAAEMAESVSNPQRPPVIWIGAQECT WP_000145427.1 hydrogenase 2 small subunit [Salmonella enterica] Length: 372\nNumber of predicted TMHs:  1\nExp number of AAs in TMHs: 24.48576\nExp number, first 60 AAs:  0.2692\nTotal prob of N-in:   0.14251\noutside   1   331\nTMhelix   332   354\ninside   355   372

19002 GCF_000983705.1_PA142 Salmonella enterica subsp. enterica serovar Paratyphi A Proteobacteria; Gammaproteobacteria; Enterobacterales; Enterobacteriaceae; Salmonella; Salmonella enterica; Salmonella enterica subsp. enterica MTGDNTLITSHGINRRDFMKLCAALAATMGLSSKAAAEMAESVSNPQRPPVIWIGAQECT WP_000145427.1 hydrogenase 2 small subunit [Salmonella enterica] Length: 372\nNumber of predicted TMHs:  1\nExp number of AAs in TMHs: 24.48576\nExp number, first 60 AAs:  0.2692\nTotal prob of N-in:   0.14251\noutside   1   331\nTMhelix   332   354\ninside   355   372

19003 GCF_000985835.1_PA055 Salmonella enterica subsp. enterica serovar Paratyphi A Proteobacteria; Gammaproteobacteria; Enterobacterales; Enterobacteriaceae; Salmonella; Salmonella enterica; Salmonella enterica subsp. enterica MTGDNTLITSHGINRRDFMKLCAALAATMGLSSKAAAEMAESVSNPQRPPVIWIGAQECT WP_000145427.1 hydrogenase 2 small subunit [Salmonella enterica] Length: 372\nNumber of predicted TMHs:  1\nExp number of AAs in TMHs: 24.48576\nExp number, first 60 AAs:  0.2692\nTotal prob of N-in:   0.14251\noutside   1   331\nTMhelix   332   354\ninside   355   372

19004 GCF_000983155.1_PA094 Salmonella enterica subsp. enterica serovar Paratyphi A Proteobacteria; Gammaproteobacteria; Enterobacterales; Enterobacteriaceae; Salmonella; Salmonella enterica; Salmonella enterica subsp. enterica MTGDNTLITSHGINRRDFMKLCAALAATMGLSSKAAAEMAESVSNPQRPPVIWIGAQECT WP_000145427.1 hydrogenase 2 small subunit [Salmonella enterica] Length: 372\nNumber of predicted TMHs:  1\nExp number of AAs in TMHs: 24.48576\nExp number, first 60 AAs:  0.2692\nTotal prob of N-in:   0.14251\noutside   1   331\nTMhelix   332   354\ninside   355   372

19005 GCF_000985155.1_PA111 Salmonella enterica subsp. enterica serovar Paratyphi A Proteobacteria; Gammaproteobacteria; Enterobacterales; Enterobacteriaceae; Salmonella; Salmonella enterica; Salmonella enterica subsp.

enterica MTGDNTLITSHGINRRDFMKLCAALAATMGLSSKAAAEMAESVSNPQRPPVIWIGAQECT WP_000145427.1 hydrogenase 2 small subunit [Salmonella enterica] Length: 372\nNumber of predicted TMHs: 1\nExp number of AAs in TMHs: 24.48576\nExp number, first 60 AAs: 0.2692\nTotal prob of N-in: 0.14251\noutside 1 331\nTMhelix 332 354\ninside 355 372

19006 GCF_000983195.1_PA079 Salmonella enterica subsp. enterica serovar Paratyphi A Proteobacteria; Gammaproteobacteria; Enterobacterales; Enterobacteriaceae; Salmonella; Salmonella enterica; Salmonella enterica subsp. enterica MTGDNTLITSHGINRRDFMKLCAALAATMGLSSKAAAEMAESVSNPQRPPVIWIGAQECT WP_000145427.1 hydrogenase 2 small subunit [Salmonella enterica] Length: 372\nNumber of predicted TMHs: 1\nExp number of AAs in TMHs: 24.48576\nExp number, first 60 AAs: 0.2692\nTotal prob of N-in: 0.14251\noutside 1 331\nTMhelix 332 354\ninside 355 372

19007 GCF_000984115.1_PA067 Salmonella enterica subsp. enterica serovar Paratyphi A Proteobacteria; Gammaproteobacteria; Enterobacterales; Enterobacteriaceae; Salmonella; Salmonella enterica; Salmonella enterica subsp. enterica MTGDNTLITSHGINRRDFMKLCAALAATMGLSSKAAAEMAESVSNPQRPPVIWIGAQECT WP_000145427.1 hydrogenase 2 small subunit [Salmonella enterica] Length: 372\nNumber of predicted TMHs: 1\nExp number of AAs in TMHs: 24.48576\nExp number, first 60 AAs: 0.2692\nTotal prob of N-in: 0.14251\noutside 1 331\nTMhelix 332 354\ninside 355 372

19008 GCF_000985055.1_PA074 Salmonella enterica subsp. enterica serovar Paratyphi A Proteobacteria; Gammaproteobacteria; Enterobacterales; Enterobacteriaceae; Salmonella; Salmonella enterica; Salmonella enterica subsp. enterica MTGDNTLITSHGINRRDFMKLCAALAATMGLSSKAAAEMAESVSNPQRPPVIWIGAQECT WP_000145427.1 hydrogenase 2 small subunit [Salmonella enterica] Length: 372\nNumber of predicted TMHs: 1\nExp number of AAs in TMHs: 24.48576\nExp number, first 60 AAs: 0.2692\nTotal prob of N-in: 0.14251\noutside 1 331\nTMhelix 332 354\ninside 355 372

19009 GCF_000983435.1_PA146 Salmonella enterica subsp. enterica serovar Paratyphi A Proteobacteria; Gammaproteobacteria; Enterobacterales; Enterobacteriaceae; Salmonella; Salmonella enterica; Salmonella enterica subsp. enterica MTGDNTLITSHGINRRDFMKLCAALAATMGLSSKAAAEMAESVSNPQRPPVIWIGAQECT WP_000145427.1 hydrogenase 2 small subunit [Salmonella enterica] Length: 372\nNumber of predicted TMHs: 1\nExp number of AAs in TMHs: 24.48576\nExp number, first 60 AAs: 0.2692\nTotal prob of N-in: 0.14251\noutside 1 331\nTMhelix 332 354\ninside 355 372

19010 GCF_000983685.1_PA116 Salmonella enterica subsp. enterica serovar Paratyphi A Proteobacteria; Gammaproteobacteria; Enterobacterales; Enterobacteriaceae; Salmonella; Salmonella enterica; Salmonella enterica subsp. enterica MTGDNTLITSHGINRRDFMKLCAALAATMGLSSKAAAEMAESVSNPQRPPVIWIGAQECT WP_000145427.1 hydrogenase 2 small subunit [Salmonella enterica] Length: 372\nNumber of predicted TMHs: 1\nExp number of AAs in TMHs: 24.48576\nExp number, first 60 AAs: 0.2692\nTotal prob of N-in: 0.14251\noutside 1 331\nTMhelix 332 354\ninside 355 372

19011 GCF_000983565.1_PA063 Salmonella enterica subsp. enterica serovar Paratyphi A Proteobacteria; Gammaproteobacteria; Enterobacterales; Enterobacteriaceae; Salmonella; Salmonella enterica; Salmonella enterica subsp. enterica MTGDNTLITSHGINRRDFMKLCAALAATMGLSSKAAAEMAESVSNPQRPPVIWIGAQECT WP_000145427.1 hydrogenase 2 small subunit [Salmonella enterica] Length: 372\nNumber of predicted TMHs: 1\nExp number of AAs in TMHs: 24.48576\nExp number, first 60 AAs: 0.2692\nTotal prob of N-in: 0.14251\noutside 1 331\nTMhelix 332 354\ninside 355 372

19012 GCF_000983775.1_PA139 Salmonella enterica subsp. enterica serovar Paratyphi A Proteobacteria; Gammaproteobacteria; Enterobacterales; Enterobacteriaceae; Salmonella; Salmonella enterica; Salmonella enterica subsp. enterica MTGDNTLITSHGINRRDFMKLCAALAATMGLSSKAAAEMAESVSNPQRPPVIWIGAQECT WP_000145427.1 hydrogenase 2 small subunit [Salmonella enterica] Length: 372\nNumber of predicted TMHs: 1\nExp number of AAs in TMHs: 24.48576\nExp number, first 60 AAs: 0.2692\nTotal prob of N-in: 0.14251\noutside 1 331\nTMhelix 332 354\ninside 355 372

19013 GCF_000983625.1_PA039 Salmonella enterica subsp. enterica serovar Paratyphi A Proteobacteria; Gammaproteobacteria; Enterobacterales; Enterobacteriaceae; Salmonella; Salmonella enterica; Salmonella enterica subsp. enterica MTGDNTLITSHGINRRDFMKLCAALAATMGLSSKAAAEMAESVSNPQRPPVIWIGAQECT WP_000145427.1 hydrogenase 2 small subunit [Salmonella enterica] Length: 372\nNumber of predicted TMHs: 1\nExp number of AAs in TMHs: 24.48576\nExp number, first 60 AAs: 0.2692\nTotal prob of N-in: 0.14251\noutside 1 331\nTMhelix 332 354\ninside 355 372

19014 GCF_000983135.1_PA005 Salmonella enterica subsp. enterica serovar Paratyphi A Proteobacteria; Gammaproteobacteria; Enterobacterales; Enterobacteriaceae; Salmonella; Salmonella enterica; Salmonella enterica subsp. enterica MTGDNTLITSHGINRRDFMKLCAALAATMGLSSKAAAEMAESVSNPQRPPVIWIGAQECT WP_000145427.1 hydrogenase 2 small subunit [Salmonella enterica] Length: 372\nNumber of predicted TMHs: 1\nExp number of AAs in TMHs: 24.48576\nExp number, first 60 AAs: 0.2692\nTotal prob of N-in: 0.14251\noutside 1 331\nTMhelix 332 354\ninside 355 372

19015 GCF_000984855.1_PA020 Salmonella enterica subsp. enterica serovar Paratyphi A Proteobacteria; Gammaproteobacteria; Enterobacterales; Enterobacteriaceae; Salmonella; Salmonella enterica; Salmonella enterica subsp. enterica MTGDNTLITSHGINRRDFMKLCAALAATMGLSSKAAAEMAESVSNPQRPPVIWIGAQECT WP_000145427.1 hydrogenase 2 small subunit [Salmonella enterica] Length: 372\nNumber of predicted TMHs: 1\nExp number of AAs in TMHs: 24.48576\nExp number, first 60 AAs: 0.2692\nTotal prob of N-in: 0.14251\noutside 1 331\nTMhelix 332 354\ninside 355 372

19016 GCF_000985095.1_PA114 Salmonella enterica subsp. enterica serovar Paratyphi A Proteobacteria; Gammaproteobacteria; Enterobacterales; Enterobacteriaceae; Salmonella; Salmonella enterica; Salmonella enterica subsp. enterica MTGDNTLITSHGINRRDFMKLCAALAATMGLSSKAAAEMAESVSNPQRPPVIWIGAQECT WP_000145427.1 hydrogenase 2 small subunit [Salmonella enterica] Length: 372\nNumber of predicted TMHs: 1\nExp number of AAs in TMHs: 24.48576\nExp number, first 60 AAs: 0.2692\nTotal prob of N-in: 0.14251\noutside 1 331\nTMhelix 332 354\ninside 355 372

19017 GCF_000983275.1_PA010 Salmonella enterica subsp. enterica serovar Paratyphi A Proteobacteria; Gammaproteobacteria; Enterobacterales; Enterobacteriaceae; Salmonella; Salmonella enterica; Salmonella enterica subsp. enterica MTGDNTLITSHGINRRDFMKLCAALAATMGLSSKAAAEMAESVSNPQRPPVIWIGAQECT WP_000145427.1 hydrogenase 2 small subunit [Salmonella enterica] Length: 372\nNumber of predicted TMHs: 1\nExp number of AAs in TMHs: 24.48576\nExp number, first 60 AAs: 0.2692\nTotal prob of N-in: 0.14251\noutside 1 331\nTMhelix 332 354\ninside 355 372

19018 GCF_000985355.1_PA016 Salmonella enterica subsp. enterica serovar Paratyphi A Proteobacteria; Gammaproteobacteria; Enterobacterales; Enterobacteriaceae; Salmonella; Salmonella enterica; Salmonella enterica subsp. enterica MTGDNTLITSHGINRRDFMKLCAALAATMGLSSKAAAEMAESVSNPQRPPVIWIGAQECT WP_000145427.1 hydrogenase 2 small subunit [Salmonella enterica] Length: 372\nNumber of predicted TMHs: 1\nExp number of AAs in TMHs: 24.48576\nExp number, first 60 AAs: 0.2692\nTotal prob of N-in: 0.14251\noutside 1 331\nTMhelix 332 354\ninside 355 372

19019 GCF_000985315.1_PA088 Salmonella enterica subsp. enterica serovar Paratyphi A Proteobacteria; Gammaproteobacteria; Enterobacterales; Enterobacteriaceae; Salmonella; Salmonella enterica; Salmonella enterica subsp. enterica MTGDNTLITSHGINRRDFMKLCAALAATMGLSSKAAAEMAESVSNPQRPPVIWIGAQECT WP_000145427.1 hydrogenase 2 small subunit [Salmonella enterica] Length: 372\nNumber of predicted TMHs: 1\nExp number of AAs in TMHs: 24.48576\nExp number, first 60 AAs: 0.2692\nTotal prob of N-in: 0.14251\noutside 1 331\nTMhelix 332 354\ninside 355 372

19020 GCF_000983315.1_PA100 Salmonella enterica subsp. enterica serovar Paratyphi A Proteobacteria; Gammaproteobacteria; Enterobacterales; Enterobacteriaceae; Salmonella; Salmonella enterica; Salmonella enterica subsp. enterica MTGDNTLITSHGINRRDFMKLCAALAATMGLSSKAAAEMAESVSNPQRPPVIWIGAQECT WP_000145427.1 hydrogenase 2 small subunit [Salmonella enterica] Length: 372\nNumber of predicted TMHs: 1\nExp number of AAs in TMHs: 24.48576\nExp number, first 60 AAs: 0.2692\nTotal prob of N-in: 0.14251\noutside 1 331\nTMhelix 332 354\ninside 355 372

19021 GCF_000984775.1_PA089 Salmonella enterica subsp. enterica serovar Paratyphi A Proteobacteria; Gammaproteobacteria; Enterobacterales; Enterobacteriaceae; Salmonella; Salmonella enterica; Salmonella enterica subsp. enterica MTGDNTLITSHGINRRDFMKLCAALAATMGLSSKAAAEMAESVSNPQRPPVIWIGAQECT WP_000145429.1 hydrogenase 2 small subunit [Salmonella enterica] Length: 372\nNumber of predicted TMHs: 1\nExp number of AAs in TMHs: 24.51309\nExp number, first 60 AAs: 0.2682\nTotal prob of N-in: 0.14402\noutside 1 331\nTMhelix 332 354\ninside 355 372

19022 GCF_000983235.1_PA035 Salmonella enterica subsp. enterica serovar Paratyphi A Proteobacteria; Gammaproteobacteria; Enterobacterales; Enterobacteriaceae; Salmonella; Salmonella enterica; Salmonella enterica subsp. enterica MTGDNTLITSHGINRRDFMKLCAALAATMGLSSKAAAEMAESVSNPQRPPVIWIGAQECT WP_000145427.1 hydrogenase 2 small subunit [Salmonella enterica] Length: 372\nNumber of predicted TMHs: 1\nExp

number of AAs in TMHs: 24.48576\nExp number, first 60 AAs:  0.2692\nTotal prob of N-in:   0.14251\noutside   1   331\nTMhelix   332   354\ninside   355   372

19023 GCF_000985235.1_PA090 Salmonella enterica subsp. enterica serovar Paratyphi A Proteobacteria; Gammaproteobacteria; Enterobacterales; Enterobacteriaceae; Salmonella; Salmonella enterica; Salmonella enterica subsp. enterica MTGDNTLITSHGINRRDFMKLCAALAATMGLSSKAAAEMAESVSNPQRPPVIWIGAQECT WP_000145427.1 hydrogenase 2 small subunit [Salmonella enterica] Length: 372\nNumber of predicted TMHs:  1\nExp number of AAs in TMHs: 24.48576\nExp number, first 60 AAs:  0.2692\nTotal prob of N-in:   0.14251\noutside   1   331\nTMhelix   332   354\ninside   355   372

19024 GCF_000985275.1_PA148 Salmonella enterica subsp. enterica serovar Paratyphi A Proteobacteria; Gammaproteobacteria; Enterobacterales; Enterobacteriaceae; Salmonella; Salmonella enterica; Salmonella enterica subsp. enterica MTGDNTLITSHGINRRDFMKLCAALAATMGLSSKAAAEMAESVSNPQRPPVIWIGAQECT WP_000145427.1 hydrogenase 2 small subunit [Salmonella enterica] Length: 372\nNumber of predicted TMHs:  1\nExp number of AAs in TMHs: 24.48576\nExp number, first 60 AAs:  0.2692\nTotal prob of N-in:   0.14251\noutside   1   331\nTMhelix   332   354\ninside   355   372

19025 GCF_000984695.1_PA131 Salmonella enterica subsp. enterica serovar Paratyphi A Proteobacteria; Gammaproteobacteria; Enterobacterales; Enterobacteriaceae; Salmonella; Salmonella enterica; Salmonella enterica subsp. enterica MTGDNTLITSHGINRRDFMKLCAALAATMGLSSKAAAEMAESVSNPQRPPVIWIGAQECT WP_000145427.1 hydrogenase 2 small subunit [Salmonella enterica] Length: 372\nNumber of predicted TMHs:  1\nExp number of AAs in TMHs: 24.48576\nExp number, first 60 AAs:  0.2692\nTotal prob of N-in:   0.14251\noutside   1   331\nTMhelix   332   354\ninside   355   372

19026 GCF_000984735.1_PA140 Salmonella enterica subsp. enterica serovar Paratyphi A Proteobacteria; Gammaproteobacteria; Enterobacterales; Enterobacteriaceae; Salmonella; Salmonella enterica; Salmonella enterica subsp. enterica MTGDNTLITSHGINRRDFMKLCAALAATMGLSSKAAAEMAESVSNPQRPPVIWIGAQECT WP_000145427.1 hydrogenase 2 small subunit [Salmonella enterica] Length: 372\nNumber of predicted TMHs:  1\nExp number of AAs in TMHs: 24.48576\nExp number, first 60 AAs:  0.2692\nTotal prob of N-in:   0.14251\noutside   1   331\nTMhelix   332   354\ninside   355   372

19027 GCF_000983535.1_PA057 Salmonella enterica subsp. enterica serovar Paratyphi A Proteobacteria; Gammaproteobacteria; Enterobacterales; Enterobacteriaceae; Salmonella; Salmonella enterica; Salmonella enterica subsp. enterica MTGDNTLITSHGINRRDFMKLCAALAATMGLSSKAAAEMAESVSNPQRPPVIWIGAQECT WP_000145427.1 hydrogenase 2 small subunit [Salmonella enterica] Length: 372\nNumber of predicted TMHs:  1\nExp number of AAs in TMHs: 24.48576\nExp number, first 60 AAs:  0.2692\nTotal prob of N-in:   0.14251\noutside   1   331\nTMhelix   332   354\ninside   355   372

19028 GCF_000985435.1_PA072 Salmonella enterica subsp. enterica serovar Paratyphi A Proteobacteria; Gammaproteobacteria; Enterobacterales; Enterobacteriaceae; Salmonella; Salmonella enterica; Salmonella enterica subsp. enterica MTGDNTLITSHGINRRDFMKLCAALAATMGLSSKAAAEMAESVSNPQRPPVIWIGAQECT WP_000145427.1 hydrogenase 2 small subunit [Salmonella enterica] Length: 372\nNumber of predicted TMHs:  1\nExp number of AAs in TMHs: 24.48576\nExp number, first 60 AAs:  0.2692\nTotal prob of N-in:   0.14251\noutside   1   331\nTMhelix   332   354\ninside   355   372

19029 GCF_000983355.1_PA027 Salmonella enterica subsp. enterica serovar Paratyphi A Proteobacteria; Gammaproteobacteria; Enterobacterales; Enterobacteriaceae; Salmonella; Salmonella enterica; Salmonella enterica subsp. enterica MTGDNTLITSHGINRRDFMKLCAALAATMGLSSKAAAEMAESVSNPQRPPVIWIGAQECT WP_000145427.1 hydrogenase 2 small subunit [Salmonella enterica] Length: 372\nNumber of predicted TMHs:  1\nExp number of AAs in TMHs: 24.48576\nExp number, first 60 AAs:  0.2692\nTotal prob of N-in:   0.14251\noutside   1   331\nTMhelix   332   354\ninside   355   372

19030 GCF_000983415.1_PA064 Salmonella enterica subsp. enterica serovar Paratyphi A Proteobacteria; Gammaproteobacteria; Enterobacterales; Enterobacteriaceae; Salmonella; Salmonella enterica; Salmonella enterica subsp. enterica MTGDNTLITSHGINRRDFMKLCAALAATMGLSSKAAAEMAESVSNPQRPPVIWIGAQECT WP_000145427.1 hydrogenase 2 small subunit [Salmonella enterica] Length: 372\nNumber of predicted TMHs:  1\nExp number of AAs in TMHs: 24.48576\nExp number, first 60 AAs:  0.2692\nTotal prob of N-in:   0.14251\noutside   1   331\nTMhelix   332   354\ninside   355   372

19031 GCF_000983645.1_PA122 Salmonella enterica subsp. enterica serovar Paratyphi A Proteobacteria; Gammaproteobacteria; Enterobacterales; Enterobacteriaceae; Salmonella; Salmonella enterica; Salmonella enterica subsp.

enterica MTGDNTLITSHGINRRDFMKLCAALAATMGLSSKAAAEMAESVSNPQRPPVIWIGAQECT WP_000145427.1 hydrogenase 2 small subunit [Salmonella enterica] Length: 372\nNumber of predicted TMHs: 1\nExp number of AAs in TMHs: 24.48576\nExp number, first 60 AAs: 0.2692\nTotal prob of N-in: 0.14251\noutside 1 331\nTMhelix 332 354\ninside 355 372

19032 GCF_000984815.1_PA130 Salmonella enterica subsp. enterica serovar Paratyphi A Proteobacteria; Gammaproteobacteria; Enterobacterales; Enterobacteriaceae; Salmonella; Salmonella enterica; Salmonella enterica subsp. enterica MTGDNTLITSHGINRRDFMKLCAALAATMGLSSKAAAEMAESVSNPQRPPVIWIGAQECT WP_000145427.1 hydrogenase 2 small subunit [Salmonella enterica] Length: 372\nNumber of predicted TMHs: 1\nExp number of AAs in TMHs: 24.48576\nExp number, first 60 AAs: 0.2692\nTotal prob of N-in: 0.14251\noutside 1 331\nTMhelix 332 354\ninside 355 372

19033 GCF_000983735.1_PA126 Salmonella enterica subsp. enterica serovar Paratyphi A Proteobacteria; Gammaproteobacteria; Enterobacterales; Enterobacteriaceae; Salmonella; Salmonella enterica; Salmonella enterica subsp. enterica MTGDNTLITSHGINRRDFMKLCAALAATMGLSSKAAAEMAESVSNPQRPPVIWIGAQECT WP_000145427.1 hydrogenase 2 small subunit [Salmonella enterica] Length: 372\nNumber of predicted TMHs: 1\nExp number of AAs in TMHs: 24.48576\nExp number, first 60 AAs: 0.2692\nTotal prob of N-in: 0.14251\noutside 1 331\nTMhelix 332 354\ninside 355 372

19034 GCF_000983875.1_PA105 Salmonella enterica subsp. enterica serovar Paratyphi A Proteobacteria; Gammaproteobacteria; Enterobacterales; Enterobacteriaceae; Salmonella; Salmonella enterica; Salmonella enterica subsp. enterica MTGDNTLITSHGINRRDFMKLCAALAATMGLSSKAAAEMAESVSNPQRPPVIWIGAQECT WP_000145427.1 hydrogenase 2 small subunit [Salmonella enterica] Length: 372\nNumber of predicted TMHs: 1\nExp number of AAs in TMHs: 24.48576\nExp number, first 60 AAs: 0.2692\nTotal prob of N-in: 0.14251\noutside 1 331\nTMhelix 332 354\ninside 355 372

19035 GCF_000983505.1_PA147 Salmonella enterica subsp. enterica serovar Paratyphi A Proteobacteria; Gammaproteobacteria; Enterobacterales; Enterobacteriaceae; Salmonella; Salmonella enterica; Salmonella enterica subsp. enterica MTGDNTLITSHGINRRDFMKLCAALAATMGLSSKAAAEMAESVSNPQRPPVIWIGAQECT WP_000145427.1 hydrogenase 2 small subunit [Salmonella enterica] Length: 372\nNumber of predicted TMHs: 1\nExp number of AAs in TMHs: 24.48576\nExp number, first 60 AAs: 0.2692\nTotal prob of N-in: 0.14251\noutside 1 331\nTMhelix 332 354\ninside 355 372

19036 GCF_000983935.1_PA031 Salmonella enterica subsp. enterica serovar Paratyphi A Proteobacteria; Gammaproteobacteria; Enterobacterales; Enterobacteriaceae; Salmonella; Salmonella enterica; Salmonella enterica subsp. enterica MTGDNTLITSHGINRRDFMKLCAALAATMGLSSKAAAEMAESVSNPQRPPVIWIGAQECT WP_000145427.1 hydrogenase 2 small subunit [Salmonella enterica] Length: 372\nNumber of predicted TMHs: 1\nExp number of AAs in TMHs: 24.48576\nExp number, first 60 AAs: 0.2692\nTotal prob of N-in: 0.14251\noutside 1 331\nTMhelix 332 354\ninside 355 372

19037 GCF_000983755.1_PA037 Salmonella enterica subsp. enterica serovar Paratyphi A Proteobacteria; Gammaproteobacteria; Enterobacterales; Enterobacteriaceae; Salmonella; Salmonella enterica; Salmonella enterica subsp. enterica MTGDNTLITSHGINRRDFMKLCAALAATMGLSSKAAAEMAESVSNPQRPPVIWIGAQECT WP_000145427.1 hydrogenase 2 small subunit [Salmonella enterica] Length: 372\nNumber of predicted TMHs: 1\nExp number of AAs in TMHs: 24.48576\nExp number, first 60 AAs: 0.2692\nTotal prob of N-in: 0.14251\noutside 1 331\nTMhelix 332 354\ninside 355 372

19038 GCF_000818075.1_ASM81807v1 Salmonella enterica subsp. enterica serovar Paratyphi A Proteobacteria; Gammaproteobacteria; Enterobacterales; Enterobacteriaceae; Salmonella; Salmonella enterica; Salmonella enterica subsp. enterica MTGDNTLITSHGINRRDFMKLCAALAATMGLSSKAAAEMAESVSNPQRPPVIWIGAQECT WP_000145427.1 hydrogenase 2 small subunit [Salmonella enterica] Length: 372\nNumber of predicted TMHs: 1\nExp number of AAs in TMHs: 24.48576\nExp number, first 60 AAs: 0.2692\nTotal prob of N-in: 0.14251\noutside 1 331\nTMhelix 332 354\ninside 355 372

19039 GCF_000983485.1_PA023 Salmonella enterica subsp. enterica serovar Paratyphi A Proteobacteria; Gammaproteobacteria; Enterobacterales; Enterobacteriaceae; Salmonella; Salmonella enterica; Salmonella enterica subsp. enterica MTGDNTLITSHGINRRDFMKLCAALAATMGLSSKAAAEMAESVSNPQRPPVIWIGAQECT WP_000145427.1 hydrogenase 2 small subunit [Salmonella enterica] Length: 372\nNumber of predicted TMHs: 1\nExp number of AAs in TMHs: 24.48576\nExp number, first 60 AAs: 0.2692\nTotal prob of N-in: 0.14251\noutside 1 331\nTMhelix 332 354\ninside 355 372

19040 GCF_000983795.1_PA128 Salmonella enterica subsp. enterica serovar Paratyphi A Proteobacteria; Gammaproteobacteria; Enterobacterales; Enterobacteriaceae; Salmonella; Salmonella enterica; Salmonella enterica subsp. enterica MTGDNTLITSHGINRRDFMKLCAALAATMGLSSKAAAEMAESVSNPQRPPVIWIGAQECT WP_000145427.1 hydrogenase 2 small subunit [Salmonella enterica] Length: 372\nNumber of predicted TMHs: 1\nExp number of AAs in TMHs: 24.48576\nExp number, first 60 AAs: 0.2692\nTotal prob of N-in: 0.14251\noutside 1 331\nTMhelix 332 354\ninside 355 372

19041 GCF_000983855.1_PA051 Salmonella enterica subsp. enterica serovar Paratyphi A Proteobacteria; Gammaproteobacteria; Enterobacterales; Enterobacteriaceae; Salmonella; Salmonella enterica; Salmonella enterica subsp. enterica MTGDNTLITSHGINRRDFMKLCAALAATMGLSSKAAAEMAESVSNPQRPPVIWIGAQECT WP_000145427.1 hydrogenase 2 small subunit [Salmonella enterica] Length: 372\nNumber of predicted TMHs: 1\nExp number of AAs in TMHs: 24.48576\nExp number, first 60 AAs: 0.2692\nTotal prob of N-in: 0.14251\noutside 1 331\nTMhelix 332 354\ninside 355 372

19042 GCF_000983975.1_PA059 Salmonella enterica subsp. enterica serovar Paratyphi A Proteobacteria; Gammaproteobacteria; Enterobacterales; Enterobacteriaceae; Salmonella; Salmonella enterica; Salmonella enterica subsp. enterica MTGDNTLITSHGINRRDFMKLCAALAATMGLSSKAAAEMAESVSNPQRPPVIWIGAQECT WP_000145427.1 hydrogenase 2 small subunit [Salmonella enterica] Length: 372\nNumber of predicted TMHs: 1\nExp number of AAs in TMHs: 24.48576\nExp number, first 60 AAs: 0.2692\nTotal prob of N-in: 0.14251\noutside 1 331\nTMhelix 332 354\ninside 355 372

19043 GCF_000984025.1_PA112 Salmonella enterica subsp. enterica serovar Paratyphi A Proteobacteria; Gammaproteobacteria; Enterobacterales; Enterobacteriaceae; Salmonella; Salmonella enterica; Salmonella enterica subsp. enterica MTGDNTLITSHGINRRDFMKLCAALAATMGLSSKAAAEMAESVSNPQRPPVIWIGAQECT WP_000145427.1 hydrogenase 2 small subunit [Salmonella enterica] Length: 372\nNumber of predicted TMHs: 1\nExp number of AAs in TMHs: 24.48576\nExp number, first 60 AAs: 0.2692\nTotal prob of N-in: 0.14251\noutside 1 331\nTMhelix 332 354\ninside 355 372

19044 GCF_000984375.1_PA127 Salmonella enterica subsp. enterica serovar Paratyphi A Proteobacteria; Gammaproteobacteria; Enterobacterales; Enterobacteriaceae; Salmonella; Salmonella enterica; Salmonella enterica subsp. enterica MTGDNTLITSHGINRRDFMKLCAALAATMGLSSKAAAEMAESVSNPQRPPVIWIGAQECT WP_000145427.1 hydrogenase 2 small subunit [Salmonella enterica] Length: 372\nNumber of predicted TMHs: 1\nExp number of AAs in TMHs: 24.48576\nExp number, first 60 AAs: 0.2692\nTotal prob of N-in: 0.14251\noutside 1 331\nTMhelix 332 354\ninside 355 372

19045 GCF_000984415.1_PA011 Salmonella enterica subsp. enterica serovar Paratyphi A Proteobacteria; Gammaproteobacteria; Enterobacterales; Enterobacteriaceae; Salmonella; Salmonella enterica; Salmonella enterica subsp. enterica MTGDNTLITSHGINRRDFMKLCAALAATMGLSSKAAAEMAESVSNPQRPPVIWIGAQECT WP_000145427.1 hydrogenase 2 small subunit [Salmonella enterica] Length: 372\nNumber of predicted TMHs: 1\nExp number of AAs in TMHs: 24.48576\nExp number, first 60 AAs: 0.2692\nTotal prob of N-in: 0.14251\noutside 1 331\nTMhelix 332 354\ninside 355 372

19046 GCF_000984455.1_PA136 Salmonella enterica subsp. enterica serovar Paratyphi A Proteobacteria; Gammaproteobacteria; Enterobacterales; Enterobacteriaceae; Salmonella; Salmonella enterica; Salmonella enterica subsp. enterica MTGDNTLITSHGINRRDFMKLCAALAATMGLSSKAAAEMAESVSNPQRPPVIWIGAQECT WP_000145427.1 hydrogenase 2 small subunit [Salmonella enterica] Length: 372\nNumber of predicted TMHs: 1\nExp number of AAs in TMHs: 24.48576\nExp number, first 60 AAs: 0.2692\nTotal prob of N-in: 0.14251\noutside 1 331\nTMhelix 332 354\ninside 355 372

19047 GCF_000984535.1_PA044 Salmonella enterica subsp. enterica serovar Paratyphi A Proteobacteria; Gammaproteobacteria; Enterobacterales; Enterobacteriaceae; Salmonella; Salmonella enterica; Salmonella enterica subsp. enterica MTGDNTLITSHGINRRDFMKLCAALAATMGLSSKAAAEMAESVSNPQRPPVIWIGAQECT WP_000145427.1 hydrogenase 2 small subunit [Salmonella enterica] Length: 372\nNumber of predicted TMHs: 1\nExp number of AAs in TMHs: 24.48576\nExp number, first 60 AAs: 0.2692\nTotal prob of N-in: 0.14251\noutside 1 331\nTMhelix 332 354\ninside 355 372

19048 GCF_000984495.1_PA137 Salmonella enterica subsp. enterica serovar Paratyphi A Proteobacteria; Gammaproteobacteria; Enterobacterales; Enterobacteriaceae; Salmonella; Salmonella enterica; Salmonella enterica subsp. enterica MTGDNTLITSHGINRRDFMKLCAALAATMGLSSKAAAEMAESVSNPQRPPVIWIGAQECT WP_000145427.1 hydrogenase 2 small subunit [Salmonella enterica] Length: 372\nNumber of predicted TMHs: 1\nExp

number of AAs in TMHs: 24.48576\nExp number, first 60 AAs: 0.2692\nTotal prob of N-in: 0.14251\noutside 1 331\nTMhelix 332 354\ninside 355 372

19049 GCF_000984575.1_PA052 Salmonella enterica subsp. enterica serovar Paratyphi A Proteobacteria; Gammaproteobacteria; Enterobacterales; Enterobacteriaceae; Salmonella; Salmonella enterica; Salmonella enterica subsp. enterica MTGDNTLITSHGINRRDFMKLCAALAATMGLSSKAAAEMAESVSNPQRPPVIWIGAQECT WP_000145427.1 hydrogenase 2 small subunit [Salmonella enterica] Length: 372\nNumber of predicted TMHs: 1\nExp number of AAs in TMHs: 24.48576\nExp number, first 60 AAs: 0.2692\nTotal prob of N-in: 0.14251\noutside 1 331\nTMhelix 332 354\ninside 355 372

19050 GCF_000984615.1_PA103A Salmonella enterica subsp. enterica serovar Paratyphi A Proteobacteria; Gammaproteobacteria; Enterobacterales; Enterobacteriaceae; Salmonella; Salmonella enterica; Salmonella enterica subsp. enterica MTGDNTLITSHGINRRDFMKLCAALAATMGLSSKAAAEMAESVSNPQRPPVIWIGAQECT WP_000145427.1 hydrogenase 2 small subunit [Salmonella enterica] Length: 372\nNumber of predicted TMHs: 1\nExp number of AAs in TMHs: 24.48576\nExp number, first 60 AAs: 0.2692\nTotal prob of N-in: 0.14251\noutside 1 331\nTMhelix 332 354\ninside 355 372

19051 GCF_000985475.1_PA125 Salmonella enterica subsp. enterica serovar Paratyphi A Proteobacteria; Gammaproteobacteria; Enterobacterales; Enterobacteriaceae; Salmonella; Salmonella enterica; Salmonella enterica subsp. enterica MTGDNTLITSHGINRRDFMKLCAALAATMGLSSKAAAEMAESVSNPQRPPVIWIGAQECT WP_000145427.1 hydrogenase 2 small subunit [Salmonella enterica] Length: 372\nNumber of predicted TMHs: 1\nExp number of AAs in TMHs: 24.48576\nExp number, first 60 AAs: 0.2692\nTotal prob of N-in: 0.14251\noutside 1 331\nTMhelix 332 354\ninside 355 372

19052 GCF_000985515.1_PA097 Salmonella enterica subsp. enterica serovar Paratyphi A Proteobacteria; Gammaproteobacteria; Enterobacterales; Enterobacteriaceae; Salmonella; Salmonella enterica; Salmonella enterica subsp. enterica MTGDNTLITSHGINRRDFMKLCAALAATMGLSSKAAAEMAESVSNPQRPPVIWIGAQECT WP_000145427.1 hydrogenase 2 small subunit [Salmonella enterica] Length: 372\nNumber of predicted TMHs: 1\nExp number of AAs in TMHs: 24.48576\nExp number, first 60 AAs: 0.2692\nTotal prob of N-in: 0.14251\noutside 1 331\nTMhelix 332 354\ninside 355 372

19053 GCF_000985555.1_PA075 Salmonella enterica subsp. enterica serovar Paratyphi A Proteobacteria; Gammaproteobacteria; Enterobacterales; Enterobacteriaceae; Salmonella; Salmonella enterica; Salmonella enterica subsp. enterica MTGDNTLITSHGINRRDFMKLCAALAATMGLSSKAAAEMAESVSNPQRPPVIWIGAQECT WP_000145427.1 hydrogenase 2 small subunit [Salmonella enterica] Length: 372\nNumber of predicted TMHs: 1\nExp number of AAs in TMHs: 24.48576\nExp number, first 60 AAs: 0.2692\nTotal prob of N-in: 0.14251\noutside 1 331\nTMhelix 332 354\ninside 355 372

19054 GCF_000985695.1_PA082 Salmonella enterica subsp. enterica serovar Paratyphi A Proteobacteria; Gammaproteobacteria; Enterobacterales; Enterobacteriaceae; Salmonella; Salmonella enterica; Salmonella enterica subsp. enterica MTGDNTLITSHGINRRDFMKLCAALAATMGLSSKAAAEMAESVSNPQRPPVIWIGAQECT WP_000145427.1 hydrogenase 2 small subunit [Salmonella enterica] Length: 372\nNumber of predicted TMHs: 1\nExp number of AAs in TMHs: 24.48576\nExp number, first 60 AAs: 0.2692\nTotal prob of N-in: 0.14251\noutside 1 331\nTMhelix 332 354\ninside 355 372

19055 GCF_000985595.1_PA040 Salmonella enterica subsp. enterica serovar Paratyphi A Proteobacteria; Gammaproteobacteria; Enterobacterales; Enterobacteriaceae; Salmonella; Salmonella enterica; Salmonella enterica subsp. enterica MTGDNTLITSHGINRRDFMKLCAALAATMGLSSKAAAEMAESVSNPQRPPVIWIGAQECT WP_000145427.1 hydrogenase 2 small subunit [Salmonella enterica] Length: 372\nNumber of predicted TMHs: 1\nExp number of AAs in TMHs: 24.48576\nExp number, first 60 AAs: 0.2692\nTotal prob of N-in: 0.14251\noutside 1 331\nTMhelix 332 354\ninside 355 372

19056 GCF_000985735.1_PA104 Salmonella enterica subsp. enterica serovar Paratyphi A Proteobacteria; Gammaproteobacteria; Enterobacterales; Enterobacteriaceae; Salmonella; Salmonella enterica; Salmonella enterica subsp. enterica MTGDNTLITSHGINRRDFMKLCAALAATMGLSSKAAAEMAESVSNPQRPPVIWIGAQECT WP_000145427.1 hydrogenase 2 small subunit [Salmonella enterica] Length: 372\nNumber of predicted TMHs: 1\nExp number of AAs in TMHs: 24.48576\nExp number, first 60 AAs: 0.2692\nTotal prob of N-in: 0.14251\noutside 1 331\nTMhelix 332 354\ninside 355 372

19057 GCF_000983595.1_PA083 Salmonella enterica subsp. enterica serovar Paratyphi A Proteobacteria; Gammaproteobacteria; Enterobacterales; Enterobacteriaceae; Salmonella; Salmonella enterica; Salmonella enterica subsp.

enterica	MTGDNTLITSHGINRRDFMKLCAALAATMGLSSKAAAEMAESVSNPQRPPVIWIGAQECT	WP_000145427.1	hydrogenase 2 small subunit [Salmonella enterica]	Length: 372\nNumber of predicted TMHs: 1\nExp number of AAs in TMHs: 24.48576\nExp number, first 60 AAs: 0.2692\nTotal prob of N-in: 0.14251\noutside 1 331\nTMhelix 332 354\ninside 355 372

19058	GCF_000985575.1_PA007	Salmonella enterica subsp. enterica serovar Paratyphi A	Proteobacteria; Gammaproteobacteria; Enterobacterales; Enterobacteriaceae; Salmonella; Salmonella enterica; Salmonella enterica subsp. enterica	MTGDNTLITSHGINRRDFMKLCAALAATMGLSSKAAAEMAESVSNPQRPPVIWIGAQECT	WP_000145427.1	hydrogenase 2 small subunit [Salmonella enterica]	Length: 372\nNumber of predicted TMHs: 1\nExp number of AAs in TMHs: 24.48576\nExp number, first 60 AAs: 0.2692\nTotal prob of N-in: 0.14251\noutside 1 331\nTMhelix 332 354\ninside 355 372

19059	GCF_000985615.1_PA098	Salmonella enterica subsp. enterica serovar Paratyphi A	Proteobacteria; Gammaproteobacteria; Enterobacterales; Enterobacteriaceae; Salmonella; Salmonella enterica; Salmonella enterica subsp. enterica	MTGDNTLITSHGINRRDFMKLCAALAATMGLSSKAAAEMAESVSNPQRPPVIWIGAQECT	WP_000145427.1	hydrogenase 2 small subunit [Salmonella enterica]	Length: 372\nNumber of predicted TMHs: 1\nExp number of AAs in TMHs: 24.48576\nExp number, first 60 AAs: 0.2692\nTotal prob of N-in: 0.14251\noutside 1 331\nTMhelix 332 354\ninside 355 372

19060	GCF_000985535.1_PA046	Salmonella enterica subsp. enterica serovar Paratyphi A	Proteobacteria; Gammaproteobacteria; Enterobacterales; Enterobacteriaceae; Salmonella; Salmonella enterica; Salmonella enterica subsp. enterica	MTGDNTLITSHGINRRDFMKLCAALAATMGLSSKAAAEMAESVSNPQRPPVIWIGAQECT	WP_000145427.1	hydrogenase 2 small subunit [Salmonella enterica]	Length: 372\nNumber of predicted TMHs: 1\nExp number of AAs in TMHs: 24.48576\nExp number, first 60 AAs: 0.2692\nTotal prob of N-in: 0.14251\noutside 1 331\nTMhelix 332 354\ninside 355 372

19061	GCF_000985855.1_PA124	Salmonella enterica subsp. enterica serovar Paratyphi A	Proteobacteria; Gammaproteobacteria; Enterobacterales; Enterobacteriaceae; Salmonella; Salmonella enterica; Salmonella enterica subsp. enterica	MTGDNTLITSHGINRRDFMKLCAALAATMGLSSKAAAEMAESVSNPQRPPVIWIGAQECT	WP_000145427.1	hydrogenase 2 small subunit [Salmonella enterica]	Length: 372\nNumber of predicted TMHs: 1\nExp number of AAs in TMHs: 24.48576\nExp number, first 60 AAs: 0.2692\nTotal prob of N-in: 0.14251\noutside 1 331\nTMhelix 332 354\ninside 355 372

19062	GCF_000985495.1_PA006	Salmonella enterica subsp. enterica serovar Paratyphi A	Proteobacteria; Gammaproteobacteria; Enterobacterales; Enterobacteriaceae; Salmonella; Salmonella enterica; Salmonella enterica subsp. enterica	MTGDNTLITSHGINRRDFMKLCAALAATMGLSSKAAAEMAESVSNPQRPPVIWIGAQECT	WP_000145427.1	hydrogenase 2 small subunit [Salmonella enterica]	Length: 372\nNumber of predicted TMHs: 1\nExp number of AAs in TMHs: 24.48576\nExp number, first 60 AAs: 0.2692\nTotal prob of N-in: 0.14251\noutside 1 331\nTMhelix 332 354\ninside 355 372

19063	GCF_000985455.1_PA050	Salmonella enterica subsp. enterica serovar Paratyphi A	Proteobacteria; Gammaproteobacteria; Enterobacterales; Enterobacteriaceae; Salmonella; Salmonella enterica; Salmonella enterica subsp. enterica	MTGDNTLITSHGINRRDFMKLCAALAATMGLSSKAAAEMAESVSNPQRPPVIWIGAQECT	WP_000145427.1	hydrogenase 2 small subunit [Salmonella enterica]	Length: 372\nNumber of predicted TMHs: 1\nExp number of AAs in TMHs: 24.48576\nExp number, first 60 AAs: 0.2692\nTotal prob of N-in: 0.14251\noutside 1 331\nTMhelix 332 354\ninside 355 372

19064	GCF_000985415.1_PA021	Salmonella enterica subsp. enterica serovar Paratyphi A	Proteobacteria; Gammaproteobacteria; Enterobacterales; Enterobacteriaceae; Salmonella; Salmonella enterica; Salmonella enterica subsp. enterica	MTGDNTLITSHGINRRDFMKLCAALAATMGLSSKAAAEMAESVSNPQRPPVIWIGAQECT	WP_000145427.1	hydrogenase 2 small subunit [Salmonella enterica]	Length: 372\nNumber of predicted TMHs: 1\nExp number of AAs in TMHs: 24.48576\nExp number, first 60 AAs: 0.2692\nTotal prob of N-in: 0.14251\noutside 1 331\nTMhelix 332 354\ninside 355 372

19065	GCF_000985395.1_PA058	Salmonella enterica subsp. enterica serovar Paratyphi A	Proteobacteria; Gammaproteobacteria; Enterobacterales; Enterobacteriaceae; Salmonella; Salmonella enterica; Salmonella enterica subsp. enterica	MTGDNTLITSHGINRRDFMKLCAALAATMGLSSKAAAEMAESVSNPQRPPVIWIGAQECT	WP_000145427.1	hydrogenase 2 small subunit [Salmonella enterica]	Length: 372\nNumber of predicted TMHs: 1\nExp number of AAs in TMHs: 24.48576\nExp number, first 60 AAs: 0.2692\nTotal prob of N-in: 0.14251\noutside 1 331\nTMhelix 332 354\ninside 355 372

19066 GCF_000985375.1_PA121 Salmonella enterica subsp. enterica serovar Paratyphi A Proteobacteria; Gammaproteobacteria; Enterobacterales; Enterobacteriaceae; Salmonella; Salmonella enterica; Salmonella enterica subsp. enterica MTGDNTLITSHGINRRDFMKLCAALAATMGLSSKAAAEMAESVSNPQRPPVIWIGAQECT WP_000145427.1 hydrogenase 2 small subunit [Salmonella enterica] Length: 372\nNumber of predicted TMHs: 1\nExp number of AAs in TMHs: 24.48576\nExp number, first 60 AAs: 0.2692\nTotal prob of N-in: 0.14251\noutside 1 331\nTMhelix 332 354\ninside 355 372

19067 GCF_000985335.1_PA018 Salmonella enterica subsp. enterica serovar Paratyphi A Proteobacteria; Gammaproteobacteria; Enterobacterales; Enterobacteriaceae; Salmonella; Salmonella enterica; Salmonella enterica subsp. enterica MTGDNTLITSHGINRRDFMKLCAALAATMGLSSKAAAEMAESVSNPQRPPVIWIGAQECT WP_000145427.1 hydrogenase 2 small subunit [Salmonella enterica] Length: 372\nNumber of predicted TMHs: 1\nExp number of AAs in TMHs: 24.48576\nExp number, first 60 AAs: 0.2692\nTotal prob of N-in: 0.14251\noutside 1 331\nTMhelix 332 354\ninside 355 372

19068 GCF_000985215.1_PA042 Salmonella enterica subsp. enterica serovar Paratyphi A Proteobacteria; Gammaproteobacteria; Enterobacterales; Enterobacteriaceae; Salmonella; Salmonella enterica; Salmonella enterica subsp. enterica MTGDNTLITSHGINRRDFMKLCAALAATMGLSSKAAAEMAESVSNPQRPPVIWIGAQECT WP_000145427.1 hydrogenase 2 small subunit [Salmonella enterica] Length: 372\nNumber of predicted TMHs: 1\nExp number of AAs in TMHs: 24.48576\nExp number, first 60 AAs: 0.2692\nTotal prob of N-in: 0.14251\noutside 1 331\nTMhelix 332 354\ninside 355 372

19069 GCF_000985255.1_PA138 Salmonella enterica subsp. enterica serovar Paratyphi A Proteobacteria; Gammaproteobacteria; Enterobacterales; Enterobacteriaceae; Salmonella; Salmonella enterica; Salmonella enterica subsp. enterica MTGDNTLITSHGINRRDFMKLCAALAATMGLSSKAAAEMAESVSNPQRPPVIWIGAQECT WP_000145427.1 hydrogenase 2 small subunit [Salmonella enterica] Length: 372\nNumber of predicted TMHs: 1\nExp number of AAs in TMHs: 24.48576\nExp number, first 60 AAs: 0.2692\nTotal prob of N-in: 0.14251\noutside 1 331\nTMhelix 332 354\ninside 355 372

19070 GCF_000985295.1_PA034 Salmonella enterica subsp. enterica serovar Paratyphi A Proteobacteria; Gammaproteobacteria; Enterobacterales; Enterobacteriaceae; Salmonella; Salmonella enterica; Salmonella enterica subsp. enterica MTGDNTLITSHGINRRDFMKLCAALAATMGLSSKAAAEMAESVSNPQRPPVIWIGAQECT WP_000145427.1 hydrogenase 2 small subunit [Salmonella enterica] Length: 372\nNumber of predicted TMHs: 1\nExp number of AAs in TMHs: 24.48576\nExp number, first 60 AAs: 0.2692\nTotal prob of N-in: 0.14251\noutside 1 331\nTMhelix 332 354\ninside 355 372

19071 GCF_000985175.1_PA085 Salmonella enterica subsp. enterica serovar Paratyphi A Proteobacteria; Gammaproteobacteria; Enterobacterales; Enterobacteriaceae; Salmonella; Salmonella enterica; Salmonella enterica subsp. enterica MTGDNTLITSHGINRRDFMKLCAALAATMGLSSKAAAEMAESVSNPQRPPVIWIGAQECT WP_000145427.1 hydrogenase 2 small subunit [Salmonella enterica] Length: 372\nNumber of predicted TMHs: 1\nExp number of AAs in TMHs: 24.48576\nExp number, first 60 AAs: 0.2692\nTotal prob of N-in: 0.14251\noutside 1 331\nTMhelix 332 354\ninside 355 372

19072 GCF_000985135.1_PA041 Salmonella enterica subsp. enterica serovar Paratyphi A Proteobacteria; Gammaproteobacteria; Enterobacterales; Enterobacteriaceae; Salmonella; Salmonella enterica; Salmonella enterica subsp. enterica MTGDNTLITSHGINRRDFMKLCAALAATMGLSSKAAAEMAESVSNPQRPPVIWIGAQECT WP_000145427.1 hydrogenase 2 small subunit [Salmonella enterica] Length: 372\nNumber of predicted TMHs: 1\nExp number of AAs in TMHs: 24.48576\nExp number, first 60 AAs: 0.2692\nTotal prob of N-in: 0.14251\noutside 1 331\nTMhelix 332 354\ninside 355 372

19073 GCF_000985115.1_PA084 Salmonella enterica subsp. enterica serovar Paratyphi A Proteobacteria; Gammaproteobacteria; Enterobacterales; Enterobacteriaceae; Salmonella; Salmonella enterica; Salmonella enterica subsp. enterica MTGDNTLITSHGINRRDFMKLCAALAATMGLSSKAAAEMAESVSNPQRPPVIWIGAQECT WP_000145427.1 hydrogenase 2 small subunit [Salmonella enterica] Length: 372\nNumber of predicted TMHs: 1\nExp number of AAs in TMHs: 24.48576\nExp number, first 60 AAs: 0.2692\nTotal prob of N-in: 0.14251\noutside 1 331\nTMhelix 332 354\ninside 355 372

19074 GCF_000985075.1_PA022 Salmonella enterica subsp. enterica serovar Paratyphi A Proteobacteria; Gammaproteobacteria; Enterobacterales; Enterobacteriaceae; Salmonella; Salmonella enterica; Salmonella enterica subsp. enterica MTGDNTLITSHGINRRDFMKLCAALAATMGLSSKAAAEMAESVSNPQRPPVIWIGAQECT WP_000145427.1 hydrogenase 2 small subunit [Salmonella enterica] Length: 372\nNumber of predicted TMHs: 1\nExp

number of AAs in TMHs: 24.48576\nExp number, first 60 AAs: 0.2692\nTotal prob of N-in: 0.14251\noutside 1 331\nTMhelix 332 354\ninside 355 372

19075 GCF_000983295.1_PA038 Salmonella enterica subsp. enterica serovar Paratyphi A Proteobacteria; Gammaproteobacteria; Enterobacterales; Enterobacteriaceae; Salmonella; Salmonella enterica; Salmonella enterica subsp. enterica MTGDNTLITSHGINRRDFMKLCAALAATMGLSSKAAAEMAESVSNPQRPPVIWIGAQECT WP_000145427.1 hydrogenase 2 small subunit [Salmonella enterica] Length: 372\nNumber of predicted TMHs: 1\nExp number of AAs in TMHs: 24.48576\nExp number, first 60 AAs: 0.2692\nTotal prob of N-in: 0.14251\noutside 1 331\nTMhelix 332 354\ninside 355 372

19076 GCF_000983255.1_PA024 Salmonella enterica subsp. enterica serovar Paratyphi A Proteobacteria; Gammaproteobacteria; Enterobacterales; Enterobacteriaceae; Salmonella; Salmonella enterica; Salmonella enterica subsp. enterica MTGDNTLITSHGINRRDFMKLCAALAATMGLSSKAAAEMAESVSNPQRPPVIWIGAQECT WP_000145427.1 hydrogenase 2 small subunit [Salmonella enterica] Length: 372\nNumber of predicted TMHs: 1\nExp number of AAs in TMHs: 24.48576\nExp number, first 60 AAs: 0.2692\nTotal prob of N-in: 0.14251\noutside 1 331\nTMhelix 332 354\ninside 355 372

19077 GCF_000983175.1_PA115 Salmonella enterica subsp. enterica serovar Paratyphi A Proteobacteria; Gammaproteobacteria; Enterobacterales; Enterobacteriaceae; Salmonella; Salmonella enterica; Salmonella enterica subsp. enterica MTGDNTLITSHGINRRDFMKLCAALAATMGLSSKAAAEMAESVSNPQRPPVIWIGAQECT WP_000145427.1 hydrogenase 2 small subunit [Salmonella enterica] Length: 372\nNumber of predicted TMHs: 1\nExp number of AAs in TMHs: 24.48576\nExp number, first 60 AAs: 0.2692\nTotal prob of N-in: 0.14251\noutside 1 331\nTMhelix 332 354\ninside 355 372

19078 GCF_000985675.1_PA123 Salmonella enterica subsp. enterica serovar Paratyphi A Proteobacteria; Gammaproteobacteria; Enterobacterales; Enterobacteriaceae; Salmonella; Salmonella enterica; Salmonella enterica subsp. enterica MTGDNTLITSHGINRRDFMKLCAALAATMGLSSKAAAEMAESVSNPQRPPVIWIGAQECT WP_000145427.1 hydrogenase 2 small subunit [Salmonella enterica] Length: 372\nNumber of predicted TMHs: 1\nExp number of AAs in TMHs: 24.48576\nExp number, first 60 AAs: 0.2692\nTotal prob of N-in: 0.14251\noutside 1 331\nTMhelix 332 354\ninside 355 372

19079 GCF_000983375.1_PA060 Salmonella enterica subsp. enterica serovar Paratyphi A Proteobacteria; Gammaproteobacteria; Enterobacterales; Enterobacteriaceae; Salmonella; Salmonella enterica; Salmonella enterica subsp. enterica MTGDNTLITSHGINRRDFMKLCAALAATMGLSSKAAAEMAESVSNPQRPPVIWIGAQECT WP_000145427.1 hydrogenase 2 small subunit [Salmonella enterica] Length: 372\nNumber of predicted TMHs: 1\nExp number of AAs in TMHs: 24.48576\nExp number, first 60 AAs: 0.2692\nTotal prob of N-in: 0.14251\noutside 1 331\nTMhelix 332 354\ninside 355 372

19080 GCF_000983335.1_PA049 Salmonella enterica subsp. enterica serovar Paratyphi A Proteobacteria; Gammaproteobacteria; Enterobacterales; Enterobacteriaceae; Salmonella; Salmonella enterica; Salmonella enterica subsp. enterica MTGDNTLITSHGINRRDFMKLCAALAATMGLSSKAAAEMAESVSNPQRPPVIWIGAQECT WP_000145427.1 hydrogenase 2 small subunit [Salmonella enterica] Length: 372\nNumber of predicted TMHs: 1\nExp number of AAs in TMHs: 24.48576\nExp number, first 60 AAs: 0.2692\nTotal prob of N-in: 0.14251\noutside 1 331\nTMhelix 332 354\ninside 355 372

19081 GCF_000985755.1_PA120 Salmonella enterica subsp. enterica serovar Paratyphi A Proteobacteria; Gammaproteobacteria; Enterobacterales; Enterobacteriaceae; Salmonella; Salmonella enterica; Salmonella enterica subsp. enterica MTGDNTLITSHGINRRDFMKLCAALAATMGLSSKAAAEMAESVSNPQRPPVIWIGAQECT WP_000145427.1 hydrogenase 2 small subunit [Salmonella enterica] Length: 372\nNumber of predicted TMHs: 1\nExp number of AAs in TMHs: 24.48576\nExp number, first 60 AAs: 0.2692\nTotal prob of N-in: 0.14251\noutside 1 331\nTMhelix 332 354\ninside 355 372

19082 GCF_000983215.1_04_9176 Salmonella enterica subsp. enterica serovar Paratyphi A Proteobacteria; Gammaproteobacteria; Enterobacterales; Enterobacteriaceae; Salmonella; Salmonella enterica; Salmonella enterica subsp. enterica MTGDNTLITSHGINRRDFMKLCAALAATMGLSSKAAAEMAESVSNPQRPPVIWIGAQECT WP_000145427.1 hydrogenase 2 small subunit [Salmonella enterica] Length: 372\nNumber of predicted TMHs: 1\nExp number of AAs in TMHs: 24.48576\nExp number, first 60 AAs: 0.2692\nTotal prob of N-in: 0.14251\noutside 1 331\nTMhelix 332 354\ninside 355 372

19083 GCF_000985035.1_PA026 Salmonella enterica subsp. enterica serovar Paratyphi A Proteobacteria; Gammaproteobacteria; Enterobacterales; Enterobacteriaceae; Salmonella; Salmonella enterica; Salmonella enterica subsp.

enterica MTGDNTLITSHGINRRDFMKLCAALAATMGLSSKAAAEMAESVSNPQRPPVIWIGAQECT WP_000145427.1 hydrogenase 2 small subunit [Salmonella enterica] Length: 372\nNumber of predicted TMHs: 1\nExp number of AAs in TMHs: 24.48576\nExp number, first 60 AAs: 0.2692\nTotal prob of N-in: 0.14251\noutside 1 331\nTMhelix 332 354\ninside 355 372

19084 GCF_000984995.1_PA047 Salmonella enterica subsp. enterica serovar Paratyphi A Proteobacteria; Gammaproteobacteria; Enterobacterales; Enterobacteriaceae; Salmonella; Salmonella enterica; Salmonella enterica subsp. enterica MTGDNTLITSHGINRRDFMKLCAALAATMGLSSKAAAEMAESVSNPQRPPVIWIGAQECT WP_000145427.1 hydrogenase 2 small subunit [Salmonella enterica] Length: 372\nNumber of predicted TMHs: 1\nExp number of AAs in TMHs: 24.48576\nExp number, first 60 AAs: 0.2692\nTotal prob of N-in: 0.14251\noutside 1 331\nTMhelix 332 354\ninside 355 372

19085 GCF_000984955.1_PA019 Salmonella enterica subsp. enterica serovar Paratyphi A Proteobacteria; Gammaproteobacteria; Enterobacterales; Enterobacteriaceae; Salmonella; Salmonella enterica; Salmonella enterica subsp. enterica MTGDNTLITSHGINRRDFMKLCAALAATMGLSSKAAAEMAESVSNPQRPPVIWIGAQECT WP_000145427.1 hydrogenase 2 small subunit [Salmonella enterica] Length: 372\nNumber of predicted TMHs: 1\nExp number of AAs in TMHs: 24.48576\nExp number, first 60 AAs: 0.2692\nTotal prob of N-in: 0.14251\noutside 1 331\nTMhelix 332 354\ninside 355 372

19086 GCF_000984915.1_PA054 Salmonella enterica subsp. enterica serovar Paratyphi A Proteobacteria; Gammaproteobacteria; Enterobacterales; Enterobacteriaceae; Salmonella; Salmonella enterica; Salmonella enterica subsp. enterica MTGDNTLITSHGINRRDFMKLCAALAATMGLSSKAAAEMAESVSNPQRPPVIWIGAQECT WP_000145427.1 hydrogenase 2 small subunit [Salmonella enterica] Length: 372\nNumber of predicted TMHs: 1\nExp number of AAs in TMHs: 24.48576\nExp number, first 60 AAs: 0.2692\nTotal prob of N-in: 0.14251\noutside 1 331\nTMhelix 332 354\ninside 355 372

19087 GCF_000984875.1_PA107 Salmonella enterica subsp. enterica serovar Paratyphi A Proteobacteria; Gammaproteobacteria; Enterobacterales; Enterobacteriaceae; Salmonella; Salmonella enterica; Salmonella enterica subsp. enterica MTGDNTLITSHGINRRDFMKLCAALAATMGLSSKAAAEMAESVSNPQRPPVIWIGAQECT WP_000145427.1 hydrogenase 2 small subunit [Salmonella enterica] Length: 372\nNumber of predicted TMHs: 1\nExp number of AAs in TMHs: 24.48576\nExp number, first 60 AAs: 0.2692\nTotal prob of N-in: 0.14251\noutside 1 331\nTMhelix 332 354\ninside 355 372

19088 GCF_000984835.1_PA076 Salmonella enterica subsp. enterica serovar Paratyphi A Proteobacteria; Gammaproteobacteria; Enterobacterales; Enterobacteriaceae; Salmonella; Salmonella enterica; Salmonella enterica subsp. enterica MTGDNTLITSHGINRRDFMKLCAALAATMGLSSKAAAEMAESVSNPQRPPVIWIGAQECT WP_000145427.1 hydrogenase 2 small subunit [Salmonella enterica] Length: 372\nNumber of predicted TMHs: 1\nExp number of AAs in TMHs: 24.48576\nExp number, first 60 AAs: 0.2692\nTotal prob of N-in: 0.14251\noutside 1 331\nTMhelix 332 354\ninside 355 372

19089 GCF_000984795.1_PA087 Salmonella enterica subsp. enterica serovar Paratyphi A Proteobacteria; Gammaproteobacteria; Enterobacterales; Enterobacteriaceae; Salmonella; Salmonella enterica; Salmonella enterica subsp. enterica MTGDNTLITSHGINRRDFMKLCAALAATMGLSSKAAAEMAESVSNPQRPPVIWIGAQECT WP_000145427.1 hydrogenase 2 small subunit [Salmonella enterica] Length: 372\nNumber of predicted TMHs: 1\nExp number of AAs in TMHs: 24.48576\nExp number, first 60 AAs: 0.2692\nTotal prob of N-in: 0.14251\noutside 1 331\nTMhelix 332 354\ninside 355 372

19090 GCF_000984755.1_PA008 Salmonella enterica subsp. enterica serovar Paratyphi A Proteobacteria; Gammaproteobacteria; Enterobacterales; Enterobacteriaceae; Salmonella; Salmonella enterica; Salmonella enterica subsp. enterica MTGDNTLITSHGINRRDFMKLCAALAATMGLSSKAAAEMAESVSNPQRPPVIWIGAQECT WP_000145427.1 hydrogenase 2 small subunit [Salmonella enterica] Length: 372\nNumber of predicted TMHs: 1\nExp number of AAs in TMHs: 24.48576\nExp number, first 60 AAs: 0.2692\nTotal prob of N-in: 0.14251\noutside 1 331\nTMhelix 332 354\ninside 355 372

19091 GCF_000984715.1_PA118 Salmonella enterica subsp. enterica serovar Paratyphi A Proteobacteria; Gammaproteobacteria; Enterobacterales; Enterobacteriaceae; Salmonella; Salmonella enterica; Salmonella enterica subsp. enterica MTGDNTLITSHGINRRDFMKLCAALAATMGLSSKAAAEMAESVSNPQRPPVIWIGAQECT WP_000145427.1 hydrogenase 2 small subunit [Salmonella enterica] Length: 372\nNumber of predicted TMHs: 1\nExp number of AAs in TMHs: 24.48576\nExp number, first 60 AAs: 0.2692\nTotal prob of N-in: 0.14251\noutside 1 331\nTMhelix 332 354\ninside 355 372

19092 GCF_000984675.1_PA110 Salmonella enterica subsp. enterica serovar Paratyphi A Proteobacteria; Gammaproteobacteria; Enterobacterales; Enterobacteriaceae; Salmonella; Salmonella enterica; Salmonella enterica subsp. enterica MTGDNTLITSHGINRRDFMKLCAALAATMGLSSKAAAEMAESVSNPQRPPVIWIGAQECT WP_000145427.1 hydrogenase 2 small subunit [Salmonella enterica] Length: 372\nNumber of predicted TMHs: 1\nExp number of AAs in TMHs: 24.48576\nExp number, first 60 AAs: 0.2692\nTotal prob of N-in: 0.14251\noutside 1 331\nTMhelix 332 354\ninside 355 372

19093 GCF_000984655.1_PA133 Salmonella enterica subsp. enterica serovar Paratyphi A Proteobacteria; Gammaproteobacteria; Enterobacterales; Enterobacteriaceae; Salmonella; Salmonella enterica; Salmonella enterica subsp. enterica MTGDNTLITSHGINRRDFMKLCAALAATMGLSSKAAAEMAESVSNPQRPPVIWIGAQECT WP_000145427.1 hydrogenase 2 small subunit [Salmonella enterica] Length: 372\nNumber of predicted TMHs: 1\nExp number of AAs in TMHs: 24.48576\nExp number, first 60 AAs: 0.2692\nTotal prob of N-in: 0.14251\noutside 1 331\nTMhelix 332 354\ninside 355 372

19094 GCF_000984635.1_PA141 Salmonella enterica subsp. enterica serovar Paratyphi A Proteobacteria; Gammaproteobacteria; Enterobacterales; Enterobacteriaceae; Salmonella; Salmonella enterica; Salmonella enterica subsp. enterica MTGDNTLITSHGINRRDFMKLCAALAATMGLSSKAAAEMAESVSNPQRPPVIWIGAQECT WP_000145427.1 hydrogenase 2 small subunit [Salmonella enterica] Length: 372\nNumber of predicted TMHs: 1\nExp number of AAs in TMHs: 24.48576\nExp number, first 60 AAs: 0.2692\nTotal prob of N-in: 0.14251\noutside 1 331\nTMhelix 332 354\ninside 355 372

19095 GCF_000984595.1_PA113 Salmonella enterica subsp. enterica serovar Paratyphi A Proteobacteria; Gammaproteobacteria; Enterobacterales; Enterobacteriaceae; Salmonella; Salmonella enterica; Salmonella enterica subsp. enterica MTGDNTLITSHGINRRDFMKLCAALAATMGLSSKAAAEMAESVSNPQRPPVIWIGAQECT WP_000145427.1 hydrogenase 2 small subunit [Salmonella enterica] Length: 372\nNumber of predicted TMHs: 1\nExp number of AAs in TMHs: 24.48576\nExp number, first 60 AAs: 0.2692\nTotal prob of N-in: 0.14251\noutside 1 331\nTMhelix 332 354\ninside 355 372

19096 GCF_000984555.1_PA145 Salmonella enterica subsp. enterica serovar Paratyphi A Proteobacteria; Gammaproteobacteria; Enterobacterales; Enterobacteriaceae; Salmonella; Salmonella enterica; Salmonella enterica subsp. enterica MTGDNTLITSHGINRRDFMKLCAALAATMGLSSKAAAEMAESVSNPQRPPVIWIGAQECT WP_000145427.1 hydrogenase 2 small subunit [Salmonella enterica] Length: 372\nNumber of predicted TMHs: 1\nExp number of AAs in TMHs: 24.48576\nExp number, first 60 AAs: 0.2692\nTotal prob of N-in: 0.14251\noutside 1 331\nTMhelix 332 354\ninside 355 372

19097 GCF_000984515.1_PA129 Salmonella enterica subsp. enterica serovar Paratyphi A Proteobacteria; Gammaproteobacteria; Enterobacterales; Enterobacteriaceae; Salmonella; Salmonella enterica; Salmonella enterica subsp. enterica MTGDNTLITSHGINRRDFMKLCAALAATMGLSSKAAAEMAESVSNPQRPPVIWIGAQECT WP_000145427.1 hydrogenase 2 small subunit [Salmonella enterica] Length: 372\nNumber of predicted TMHs: 1\nExp number of AAs in TMHs: 24.48576\nExp number, first 60 AAs: 0.2692\nTotal prob of N-in: 0.14251\noutside 1 331\nTMhelix 332 354\ninside 355 372

19098 GCF_001045395.1_ASM104539v1 Salmonella enterica subsp. enterica serovar Paratyphi A Proteobacteria; Gammaproteobacteria; Enterobacterales; Enterobacteriaceae; Salmonella; Salmonella enterica; Salmonella enterica subsp. enterica MTGDNTLITSHGINRRDFMKLCAALAATMGLSSKAAAEMAESVSNPQRPPVIWIGAQECT WP_000145427.1 hydrogenase 2 small subunit [Salmonella enterica] Length: 372\nNumber of predicted TMHs: 1\nExp number of AAs in TMHs: 24.48576\nExp number, first 60 AAs: 0.2692\nTotal prob of N-in: 0.14251\noutside 1 331\nTMhelix 332 354\ninside 355 372

19099 GCF_000984475.1_PA013 Salmonella enterica subsp. enterica serovar Paratyphi A Proteobacteria; Gammaproteobacteria; Enterobacterales; Enterobacteriaceae; Salmonella; Salmonella enterica; Salmonella enterica subsp. enterica MTGDNTLITSHGINRRDFMKLCAALAATMGLSSKAAAEMAESVSNPQRPPVIWIGAQECT WP_000145427.1 hydrogenase 2 small subunit [Salmonella enterica] Length: 372\nNumber of predicted TMHs: 1\nExp number of AAs in TMHs: 24.48576\nExp number, first 60 AAs: 0.2692\nTotal prob of N-in: 0.14251\noutside 1 331\nTMhelix 332 354\ninside 355 372

19100 GCF_000985795.1_PA117 Salmonella enterica subsp. enterica serovar Paratyphi A Proteobacteria; Gammaproteobacteria; Enterobacterales; Enterobacteriaceae; Salmonella; Salmonella enterica; Salmonella enterica subsp. enterica MTGDNTLITSHGINRRDFMKLCAALAATMGLSSKAAAEMAESVSNPQRPPVIWIGAQECT WP_000145427.1 hydrogenase 2 small subunit [Salmonella enterica] Length: 372\nNumber of predicted TMHs: 1\nExp

number of AAs in TMHs: 24.48576\nExp number, first 60 AAs:  0.2692\nTotal prob of N-in:   0.14251\noutside   1   331\nTMhelix   332   354\ninside   355   372

19101 GCF_000984435.1_PA134 Salmonella enterica subsp. enterica serovar Paratyphi A Proteobacteria; Gammaproteobacteria; Enterobacterales; Enterobacteriaceae; Salmonella; Salmonella enterica; Salmonella enterica subsp. enterica MTGDNTLITSHGINRRDFMKLCAALAATMGLSSKAAAEMAESVSNPQRPPVIWIGAQECT WP_000145427.1 hydrogenase 2 small subunit [Salmonella enterica] Length: 372\nNumber of predicted TMHs:  1\nExp number of AAs in TMHs: 24.48576\nExp number, first 60 AAs:  0.2692\nTotal prob of N-in:   0.14251\noutside   1   331\nTMhelix   332   354\ninside   355   372

19102 GCF_000984395.1_PA033 Salmonella enterica subsp. enterica serovar Paratyphi A Proteobacteria; Gammaproteobacteria; Enterobacterales; Enterobacteriaceae; Salmonella; Salmonella enterica; Salmonella enterica subsp. enterica MTGDNTLITSHGINRRDFMKLCAALAATMGLSSKAAAEMAESVSNPQRPPVIWIGAQECT WP_000145427.1 hydrogenase 2 small subunit [Salmonella enterica] Length: 372\nNumber of predicted TMHs:  1\nExp number of AAs in TMHs: 24.48576\nExp number, first 60 AAs:  0.2692\nTotal prob of N-in:   0.14251\noutside   1   331\nTMhelix   332   354\ninside   355   372

19103 GCF_000983835.1_PA014 Salmonella enterica subsp. enterica serovar Paratyphi A Proteobacteria; Gammaproteobacteria; Enterobacterales; Enterobacteriaceae; Salmonella; Salmonella enterica; Salmonella enterica subsp. enterica MTGDNTLITSHGINRRDFMKLCAALAATMGLSSKAAAEMAESVSNPQRPPVIWIGAQECT WP_000145427.1 hydrogenase 2 small subunit [Salmonella enterica] Length: 372\nNumber of predicted TMHs:  1\nExp number of AAs in TMHs: 24.48576\nExp number, first 60 AAs:  0.2692\nTotal prob of N-in:   0.14251\noutside   1   331\nTMhelix   332   354\ninside   355   372

19104 GCF_000984355.1_PA096 Salmonella enterica subsp. enterica serovar Paratyphi A Proteobacteria; Gammaproteobacteria; Enterobacterales; Enterobacteriaceae; Salmonella; Salmonella enterica; Salmonella enterica subsp. enterica MTGDNTLITSHGINRRDFMKLCAALAATMGLSSKAAAEMAESVSNPQRPPVIWIGAQECT WP_000145427.1 hydrogenase 2 small subunit [Salmonella enterica] Length: 372\nNumber of predicted TMHs:  1\nExp number of AAs in TMHs: 24.48576\nExp number, first 60 AAs:  0.2692\nTotal prob of N-in:   0.14251\noutside   1   331\nTMhelix   332   354\ninside   355   372

19105 GCF_000985715.1_PA032 Salmonella enterica subsp. enterica serovar Paratyphi A Proteobacteria; Gammaproteobacteria; Enterobacterales; Enterobacteriaceae; Salmonella; Salmonella enterica; Salmonella enterica subsp. enterica MTGDNTLITSHGINRRDFMKLCAALAATMGLSSKAAAEMAESVSNPQRPPVIWIGAQECT WP_000145427.1 hydrogenase 2 small subunit [Salmonella enterica] Length: 372\nNumber of predicted TMHs:  1\nExp number of AAs in TMHs: 24.48576\nExp number, first 60 AAs:  0.2692\nTotal prob of N-in:   0.14251\noutside   1   331\nTMhelix   332   354\ninside   355   372

19106 GCF_000984295.1_PA135 Salmonella enterica subsp. enterica serovar Paratyphi A Proteobacteria; Gammaproteobacteria; Enterobacterales; Enterobacteriaceae; Salmonella; Salmonella enterica; Salmonella enterica subsp. enterica MTGDNTLITSHGINRRDFMKLCAALAATMGLSSKAAAEMAESVSNPQRPPVIWIGAQECT WP_000145427.1 hydrogenase 2 small subunit [Salmonella enterica] Length: 372\nNumber of predicted TMHs:  1\nExp number of AAs in TMHs: 24.48576\nExp number, first 60 AAs:  0.2692\nTotal prob of N-in:   0.14251\noutside   1   331\nTMhelix   332   354\ninside   355   372

19107 GCF_000984335.1_PA056 Salmonella enterica subsp. enterica serovar Paratyphi A Proteobacteria; Gammaproteobacteria; Enterobacterales; Enterobacteriaceae; Salmonella; Salmonella enterica; Salmonella enterica subsp. enterica MTGDNTLITSHGINRRDFMKLCAALAATMGLSSKAAAEMAESVSNPQRPPVIWIGAQECT WP_000145427.1 hydrogenase 2 small subunit [Salmonella enterica] Length: 372\nNumber of predicted TMHs:  1\nExp number of AAs in TMHs: 24.48576\nExp number, first 60 AAs:  0.2692\nTotal prob of N-in:   0.14251\noutside   1   331\nTMhelix   332   354\ninside   355   372

19108 GCF_000984255.1_PA036 Salmonella enterica subsp. enterica serovar Paratyphi A Proteobacteria; Gammaproteobacteria; Enterobacterales; Enterobacteriaceae; Salmonella; Salmonella enterica; Salmonella enterica subsp. enterica MTGDNTLITSHGINRRDFMKLCAALAATMGLSSKAAAEMAESVSNPQRPPVIWIGAQECT WP_000145427.1 hydrogenase 2 small subunit [Salmonella enterica] Length: 372\nNumber of predicted TMHs:  1\nExp number of AAs in TMHs: 24.48576\nExp number, first 60 AAs:  0.2692\nTotal prob of N-in:   0.14251\noutside   1   331\nTMhelix   332   354\ninside   355   372

19109 GCF_000983895.1_PA062 Salmonella enterica subsp. enterica serovar Paratyphi A Proteobacteria; Gammaproteobacteria; Enterobacterales; Enterobacteriaceae; Salmonella; Salmonella enterica; Salmonella enterica subsp.

enterica MTGDNTLITSHGINRRDFMKLCAALAATMGLSSKAAAEMAESVSNPQRPPVIWIGAQECT WP_000145427.1 hydrogenase 2 small subunit [Salmonella enterica] Length: 372\nNumber of predicted TMHs: 1\nExp number of AAs in TMHs: 24.48576\nExp number, first 60 AAs: 0.2692\nTotal prob of N-in: 0.14251\noutside 1 331\nTMhelix 332 354\ninside 355 372

19110 GCF_000984215.1_PA101 Salmonella enterica subsp. enterica serovar Paratyphi A Proteobacteria; Gammaproteobacteria; Enterobacterales; Enterobacteriaceae; Salmonella; Salmonella enterica; Salmonella enterica subsp. enterica MTGDNTLITSHGINRRDFMKLCAALAATMGLSSKAAAEMAESVSNPQRPPVIWIGAQECT WP_000145427.1 hydrogenase 2 small subunit [Salmonella enterica] Length: 372\nNumber of predicted TMHs: 1\nExp number of AAs in TMHs: 24.48576\nExp number, first 60 AAs: 0.2692\nTotal prob of N-in: 0.14251\noutside 1 331\nTMhelix 332 354\ninside 355 372

19111 GCF_000984175.1_PA069 Salmonella enterica subsp. enterica serovar Paratyphi A Proteobacteria; Gammaproteobacteria; Enterobacterales; Enterobacteriaceae; Salmonella; Salmonella enterica; Salmonella enterica subsp. enterica MTGDNTLITSHGINRRDFMKLCAALAATMGLSSKAAAEMAESVSNPQRPPVIWIGAQECT WP_000145427.1 hydrogenase 2 small subunit [Salmonella enterica] Length: 372\nNumber of predicted TMHs: 1\nExp number of AAs in TMHs: 24.48576\nExp number, first 60 AAs: 0.2692\nTotal prob of N-in: 0.14251\noutside 1 331\nTMhelix 332 354\ninside 355 372

19112 GCF_000984135.1_PA028 Salmonella enterica subsp. enterica serovar Paratyphi A Proteobacteria; Gammaproteobacteria; Enterobacterales; Enterobacteriaceae; Salmonella; Salmonella enterica; Salmonella enterica subsp. enterica MTGDNTLITSHGINRRDFMKLCAALAATMGLSSKAAAEMAESVSNPQRPPVIWIGAQECT WP_000145427.1 hydrogenase 2 small subunit [Salmonella enterica] Length: 372\nNumber of predicted TMHs: 1\nExp number of AAs in TMHs: 24.48576\nExp number, first 60 AAs: 0.2692\nTotal prob of N-in: 0.14251\noutside 1 331\nTMhelix 332 354\ninside 355 372

19113 GCF_000984095.1_PA009 Salmonella enterica subsp. enterica serovar Paratyphi A Proteobacteria; Gammaproteobacteria; Enterobacterales; Enterobacteriaceae; Salmonella; Salmonella enterica; Salmonella enterica subsp. enterica MTGDNTLITSHGINRRDFMKLCAALAATMGLSSKAAAEMAESVSNPQRPPVIWIGAQECT WP_000145427.1 hydrogenase 2 small subunit [Salmonella enterica] Length: 372\nNumber of predicted TMHs: 1\nExp number of AAs in TMHs: 24.48576\nExp number, first 60 AAs: 0.2692\nTotal prob of N-in: 0.14251\noutside 1 331\nTMhelix 332 354\ninside 355 372

19114 GCF_000818115.1_ASM81811v1 Salmonella enterica subsp. enterica serovar Paratyphi A Proteobacteria; Gammaproteobacteria; Enterobacterales; Enterobacteriaceae; Salmonella; Salmonella enterica; Salmonella enterica subsp. enterica MTGDNTLITSHGINRRDFMKLCAALAATMGLSSKAAAEMAESVSNPQRPPVIWIGAQECT WP_000145427.1 hydrogenase 2 small subunit [Salmonella enterica] Length: 372\nNumber of predicted TMHs: 1\nExp number of AAs in TMHs: 24.48576\nExp number, first 60 AAs: 0.2692\nTotal prob of N-in: 0.14251\noutside 1 331\nTMhelix 332 354\ninside 355 372

19115 GCF_000984045.1_PA012 Salmonella enterica subsp. enterica serovar Paratyphi A Proteobacteria; Gammaproteobacteria; Enterobacterales; Enterobacteriaceae; Salmonella; Salmonella enterica; Salmonella enterica subsp. enterica MTGDNTLITSHGINRRDFMKLCAALAATMGLSSKAAAEMAESVSNPQRPPVIWIGAQECT WP_000145427.1 hydrogenase 2 small subunit [Salmonella enterica] Length: 372\nNumber of predicted TMHs: 1\nExp number of AAs in TMHs: 24.48576\nExp number, first 60 AAs: 0.2692\nTotal prob of N-in: 0.14251\noutside 1 331\nTMhelix 332 354\ninside 355 372

19116 GCF_000985815.1_PA061 Salmonella enterica subsp. enterica serovar Paratyphi A Proteobacteria; Gammaproteobacteria; Enterobacterales; Enterobacteriaceae; Salmonella; Salmonella enterica; Salmonella enterica subsp. enterica MTGDNTLITSHGINRRDFMKLCAALAATMGLSSKAAAEMAESVSNPQRPPVIWIGAQECT WP_000145427.1 hydrogenase 2 small subunit [Salmonella enterica] Length: 372\nNumber of predicted TMHs: 1\nExp number of AAs in TMHs: 24.48576\nExp number, first 60 AAs: 0.2692\nTotal prob of N-in: 0.14251\noutside 1 331\nTMhelix 332 354\ninside 355 372

19117 GCF_000983955.1_PA045 Salmonella enterica subsp. enterica serovar Paratyphi A Proteobacteria; Gammaproteobacteria; Enterobacterales; Enterobacteriaceae; Salmonella; Salmonella enterica; Salmonella enterica subsp. enterica MTGDNTLITSHGINRRDFMKLCAALAATMGLSSKAAAEMAESVSNPQRPPVIWIGAQECT WP_000145427.1 hydrogenase 2 small subunit [Salmonella enterica] Length: 372\nNumber of predicted TMHs: 1\nExp number of AAs in TMHs: 24.48576\nExp number, first 60 AAs: 0.2692\nTotal prob of N-in: 0.14251\noutside 1 331\nTMhelix 332 354\ninside 355 372

19118 GCF_000983915.1_PA048 Salmonella enterica subsp. enterica serovar Paratyphi A Proteobacteria; Gammaproteobacteria; Enterobacterales; Enterobacteriaceae; Salmonella; Salmonella enterica; Salmonella enterica subsp. enterica MTGDNTLITSHGINRRDFMKLCAALAATMGLSSKAAAEMAESVSNPQRPPVIWIGAQECT WP_000145427.1 hydrogenase 2 small subunit [Salmonella enterica] Length: 372\nNumber of predicted TMHs: 1\nExp number of AAs in TMHs: 24.48576\nExp number, first 60 AAs: 0.2692\nTotal prob of N-in: 0.14251\noutside 1 331\nTMhelix 332 354\ninside 355 372

19119 GCF_000983395.1_PA093 Salmonella enterica subsp. enterica serovar Paratyphi A Proteobacteria; Gammaproteobacteria; Enterobacterales; Enterobacteriaceae; Salmonella; Salmonella enterica; Salmonella enterica subsp. enterica MTGDNTLITSHGINRRDFMKLCAALAATMGLSSKAAAEMAESVSNPQRPPVIWIGAQECT WP_000145427.1 hydrogenase 2 small subunit [Salmonella enterica] Length: 372\nNumber of predicted TMHs: 1\nExp number of AAs in TMHs: 24.48576\nExp number, first 60 AAs: 0.2692\nTotal prob of N-in: 0.14251\noutside 1 331\nTMhelix 332 354\ninside 355 372

19120 GCF_000983995.1_PA095 Salmonella enterica subsp. enterica serovar Paratyphi A Proteobacteria; Gammaproteobacteria; Enterobacterales; Enterobacteriaceae; Salmonella; Salmonella enterica; Salmonella enterica subsp. enterica MTGDNTLITSHGINRRDFMKLCAALAATMGLSSKAAAEMAESVSNPQRPPVIWIGAQECT WP_000145427.1 hydrogenase 2 small subunit [Salmonella enterica] Length: 372\nNumber of predicted TMHs: 1\nExp number of AAs in TMHs: 24.48576\nExp number, first 60 AAs: 0.2692\nTotal prob of N-in: 0.14251\noutside 1 331\nTMhelix 332 354\ninside 355 372

19121 GCF_002058475.1_ASM205847v1 Salmonella enterica subsp. enterica serovar Paratyphi B Proteobacteria; Gammaproteobacteria; Enterobacterales; Enterobacteriaceae; Salmonella; Salmonella enterica; Salmonella enterica subsp. enterica MNNEETFYQAMRRKGVTRRSFLKFCSLAATSLGLGAGMTPKIAWALENKPRIPVVWIHGL WP_001058312.1 hydrogenase [Salmonella enterica] Length: 372\nNumber of predicted TMHs: 2\nExp number of AAs in TMHs: 36.51738\nExp number, first 60 AAs: 19.72849\nTotal prob of N-in: 0.96170\nPOSSIBLE N-term signal sequence\ninside 1 19\nTMhelix 20 39\noutside 40 325\nTMhelix 326 348\ninside 349 372

19122 GCF_002034145.1_ASM203414v1 Salmonella enterica subsp. enterica serovar Paratyphi B Proteobacteria; Gammaproteobacteria; Enterobacterales; Enterobacteriaceae; Salmonella; Salmonella enterica; Salmonella enterica subsp. enterica MTGDNTLITSHGINRRDFMKLCAALAATMGLSSKAAAEMAESVSNPQRPPVIWIGAQECT WP_000145429.1 hydrogenase 2 small subunit [Salmonella enterica] Length: 372\nNumber of predicted TMHs: 1\nExp number of AAs in TMHs: 24.51309\nExp number, first 60 AAs: 0.2682\nTotal prob of N-in: 0.14402\noutside 1 331\nTMhelix 332 354\ninside 355 372

19123 GCF_002058475.1_ASM205847v1 Salmonella enterica subsp. enterica serovar Paratyphi B Proteobacteria; Gammaproteobacteria; Enterobacterales; Enterobacteriaceae; Salmonella; Salmonella enterica; Salmonella enterica subsp. enterica MTGDNTLITSHGINRRDFMKLCAALAATMGLSSKAAAEMAESVSNPQRPPVIWIGAQECT WP_000145429.1 hydrogenase 2 small subunit [Salmonella enterica] Length: 372\nNumber of predicted TMHs: 1\nExp number of AAs in TMHs: 24.51309\nExp number, first 60 AAs: 0.2682\nTotal prob of N-in: 0.14402\noutside 1 331\nTMhelix 332 354\ninside 355 372

19124 GCF_001614125.1_ASM161412v1 Salmonella enterica subsp. enterica serovar Paratyphi B Proteobacteria; Gammaproteobacteria; Enterobacterales; Enterobacteriaceae; Salmonella; Salmonella enterica; Salmonella enterica subsp. enterica MTGDNTLITSHGINRRDFMKLCAALAATMGLSSKAAAEMAESVSNPQRPPVIWIGAQECT WP_000145429.1 hydrogenase 2 small subunit [Salmonella enterica] Length: 372\nNumber of predicted TMHs: 1\nExp number of AAs in TMHs: 24.51309\nExp number, first 60 AAs: 0.2682\nTotal prob of N-in: 0.14402\noutside 1 331\nTMhelix 332 354\ninside 355 372

19125 GCF_001614015.1_ASM161401v1 Salmonella enterica subsp. enterica serovar Paratyphi B Proteobacteria; Gammaproteobacteria; Enterobacterales; Enterobacteriaceae; Salmonella; Salmonella enterica; Salmonella enterica subsp. enterica MTGDNTLITSHGINRRDFMKLCAALAATMGLSSKAAAEMAESVSNPQRPPVIWIGAQECT WP_000145429.1 hydrogenase 2 small subunit [Salmonella enterica] Length: 372\nNumber of predicted TMHs: 1\nExp number of AAs in TMHs: 24.51309\nExp number, first 60 AAs: 0.2682\nTotal prob of N-in: 0.14402\noutside 1 331\nTMhelix 332 354\ninside 355 372

19126 GCF_001614125.1_ASM161412v1 Salmonella enterica subsp. enterica serovar Paratyphi B Proteobacteria; Gammaproteobacteria; Enterobacterales; Enterobacteriaceae; Salmonella; Salmonella enterica; Salmonella enterica subsp. enterica MNNEETFYQAMRRKGVTRRSFLKFCSLAATSLGLGAGMTPKIAWALENKPRIPVVWIHGL WP_001058311.1 hydrogenase [Salmonella enterica] Length: 372\nNumber of predicted TMHs: 2\nExp number of AAs in TMHs: 37.4699899999999\nExp number, first 60 AAs: 19.72699\nTotal prob of N-in:

0.96171\nPOSSIBLE N-term signal sequence\ninside 1 19\nTMhelix 20 39\noutside 40 325\nTMhelix 326 348\ninside 349 372

19127 GCF_001614015.1_ASM161401v1 Salmonella enterica subsp. enterica serovar Paratyphi B Proteobacteria; Gammaproteobacteria; Enterobacterales; Enterobacteriaceae; Salmonella; Salmonella enterica; Salmonella enterica subsp. enterica MNNEETFYQAMRRKGVTRRSFLKFCSLAATSLGLGAGMTPKIAWALENKPRIPVVWIHGL WP_001058311.1 hydrogenase [Salmonella enterica] Length: 372\nNumber of predicted TMHs: 2\nExp number of AAs in TMHs: 37.4699899999999\nExp number, first 60 AAs: 19.72699\nTotal prob of N-in: 0.96171\nPOSSIBLE N-term signal sequence\ninside 1 19\nTMhelix 20 39\noutside 40 325\nTMhelix 326 348\ninside 349 372

19128 GCF_002034145.1_ASM203414v1 Salmonella enterica subsp. enterica serovar Paratyphi B Proteobacteria; Gammaproteobacteria; Enterobacterales; Enterobacteriaceae; Salmonella; Salmonella enterica; Salmonella enterica subsp. enterica MNNEETFYQAMRRKGVTRRSFLKFCSLAATSLGLGAGMTPKIAWALENKPRIPVVWIHGL WP_001058312.1 hydrogenase [Salmonella enterica] Length: 372\nNumber of predicted TMHs: 2\nExp number of AAs in TMHs: 36.51738\nExp number, first 60 AAs: 19.72849\nTotal prob of N-in: 0.96170\nPOSSIBLE N-term signal sequence\ninside 1 19\nTMhelix 20 39\noutside 40 325\nTMhelix 326 348\ninside 349 372

19129 GCF_002065255.1_ASM206525v1 Salmonella enterica subsp. enterica serovar Paratyphi B Proteobacteria; Gammaproteobacteria; Enterobacterales; Enterobacteriaceae; Salmonella; Salmonella enterica; Salmonella enterica subsp. enterica MNNEETFYQAMRRKGVTRRSFLKFCSLAATSLGLGAGMTPKIAWALENKPRIPVVWIHGL WP_001058312.1 hydrogenase [Salmonella enterica] Length: 372\nNumber of predicted TMHs: 2\nExp number of AAs in TMHs: 36.51738\nExp number, first 60 AAs: 19.72849\nTotal prob of N-in: 0.96170\nPOSSIBLE N-term signal sequence\ninside 1 19\nTMhelix 20 39\noutside 40 325\nTMhelix 326 348\ninside 349 372

19130 GCF_002033905.1_ASM203390v1 Salmonella enterica subsp. enterica serovar Paratyphi B Proteobacteria; Gammaproteobacteria; Enterobacterales; Enterobacteriaceae; Salmonella; Salmonella enterica; Salmonella enterica subsp. enterica MTGDNTLITSHGINRRDFMKLCAALAATMGLSSKAAAEMAESVSNPQRPPVIWIGAQECT WP_000145429.1 hydrogenase 2 small subunit [Salmonella enterica] Length: 372\nNumber of predicted TMHs: 1\nExp number of AAs in TMHs: 24.51309\nExp number, first 60 AAs: 0.2682\nTotal prob of N-in: 0.14402\noutside 1 331\nTMhelix 332 354\ninside 355 372

19131 GCF_002065255.1_ASM206525v1 Salmonella enterica subsp. enterica serovar Paratyphi B Proteobacteria; Gammaproteobacteria; Enterobacterales; Enterobacteriaceae; Salmonella; Salmonella enterica; Salmonella enterica subsp. enterica MTGDNTLITSHGINRRDFMKLCAALAATMGLSSKAAAEMAESVSNPQRPPVIWIGAQECT WP_000145429.1 hydrogenase 2 small subunit [Salmonella enterica] Length: 372\nNumber of predicted TMHs: 1\nExp number of AAs in TMHs: 24.51309\nExp number, first 60 AAs: 0.2682\nTotal prob of N-in: 0.14402\noutside 1 331\nTMhelix 332 354\ninside 355 372

19132 GCF_002033905.1_ASM203390v1 Salmonella enterica subsp. enterica serovar Paratyphi B Proteobacteria; Gammaproteobacteria; Enterobacterales; Enterobacteriaceae; Salmonella; Salmonella enterica; Salmonella enterica subsp. enterica MNNEETFYQAMRRKGVTRRSFLKFCSLAATSLGLGAGMTPKIAWALENKPRIPVVWIHGL WP_001058311.1 hydrogenase [Salmonella enterica] Length: 372\nNumber of predicted TMHs: 2\nExp number of AAs in TMHs: 37.4699899999999\nExp number, first 60 AAs: 19.72699\nTotal prob of N-in: 0.96171\nPOSSIBLE N-term signal sequence\ninside 1 19\nTMhelix 20 39\noutside 40 325\nTMhelix 326 348\ninside 349 372

19133 GCF_001613965.1_ASM161396v1 Salmonella enterica subsp. enterica serovar Paratyphi B Proteobacteria; Gammaproteobacteria; Enterobacterales; Enterobacteriaceae; Salmonella; Salmonella enterica; Salmonella enterica subsp. enterica MTGDNTLITSHGINRRDFMKLCAALAATMGLSSKAAAEMAESVSNPQRPPVIWIGAQECT WP_000145429.1 hydrogenase 2 small subunit [Salmonella enterica] Length: 372\nNumber of predicted TMHs: 1\nExp number of AAs in TMHs: 24.51309\nExp number, first 60 AAs: 0.2682\nTotal prob of N-in: 0.14402\noutside 1 331\nTMhelix 332 354\ninside 355 372

19134 GCF_001466355.1_ASM146635v1 Salmonella enterica subsp. enterica serovar Paratyphi B Proteobacteria; Gammaproteobacteria; Enterobacterales; Enterobacteriaceae; Salmonella; Salmonella enterica; Salmonella enterica subsp. enterica MNNEETFYQAMRRKGVTRRSFLKFCSLAATSLGLGAGMTPKIAWALENKPRIPVVWIHGL WP_001058311.1 hydrogenase [Salmonella enterica] Length: 372\nNumber of predicted TMHs: 2\nExp number of AAs in TMHs: 37.4699899999999\nExp number, first 60 AAs: 19.72699\nTotal prob of N-in: 0.96171\nPOSSIBLE N-term signal sequence\ninside 1 19\nTMhelix 20 39\noutside 40 325\nTMhelix 326 348\ninside 349 372

19135 GCF_001466365.1_ASM146636v1 Salmonella enterica subsp. enterica serovar Paratyphi B Proteobacteria; Gammaproteobacteria; Enterobacterales; Enterobacteriaceae; Salmonella; Salmonella enterica; Salmonella enterica subsp. enterica MTGDNTLITSHGINRRDFMKLCAALAATMGLSSKAAAEMAESVSNPQRPPVIWIGAQECT WP_000145429.1 hydrogenase 2 small subunit [Salmonella enterica] Length: 372\nNumber of predicted TMHs: 1\nExp number of AAs in TMHs: 24.51309\nExp number, first 60 AAs: 0.2682\nTotal prob of N-in: 0.14402\noutside 1 331\nTMhelix 332 354\ninside 355 372

19136 GCF_001466365.1_ASM146636v1 Salmonella enterica subsp. enterica serovar Paratyphi B Proteobacteria; Gammaproteobacteria; Enterobacterales; Enterobacteriaceae; Salmonella; Salmonella enterica; Salmonella enterica subsp. enterica MNNEETFYQAMRRKGVTRRSFLKFCSLAATSLGLGAGMTPKIAWALENKPRIPVVWIHGL WP_001058311.1 hydrogenase [Salmonella enterica] Length: 372\nNumber of predicted TMHs: 2\nExp number of AAs in TMHs: 37.4699899999999\nExp number, first 60 AAs: 19.72699\nTotal prob of N-in: 0.96171\nPOSSIBLE N-term signal sequence\ninside 1 19\nTMhelix 20 39\noutside 40 325\nTMhelix 326 348\ninside 349 372

19137 GCF_001466355.1_ASM146635v1 Salmonella enterica subsp. enterica serovar Paratyphi B Proteobacteria; Gammaproteobacteria; Enterobacterales; Enterobacteriaceae; Salmonella; Salmonella enterica; Salmonella enterica subsp. enterica MTGDNTLITSHGINRRDFMKLCAALAATMGLSSKAAAEMAESVSNPQRPPVIWIGAQECT WP_000145429.1 hydrogenase 2 small subunit [Salmonella enterica] Length: 372\nNumber of predicted TMHs: 1\nExp number of AAs in TMHs: 24.51309\nExp number, first 60 AAs: 0.2682\nTotal prob of N-in: 0.14402\noutside 1 331\nTMhelix 332 354\ninside 355 372

19138 GCF_001614145.1_ASM161414v1 Salmonella enterica subsp. enterica serovar Paratyphi B Proteobacteria; Gammaproteobacteria; Enterobacterales; Enterobacteriaceae; Salmonella; Salmonella enterica; Salmonella enterica subsp. enterica MNNEETFYQAMRRKGVTRRSFLKFCSLAATSLGLGAGMTPKIAWALENKPRIPVVWIHGL WP_001058311.1 hydrogenase [Salmonella enterica] Length: 372\nNumber of predicted TMHs: 2\nExp number of AAs in TMHs: 37.4699899999999\nExp number, first 60 AAs: 19.72699\nTotal prob of N-in: 0.96171\nPOSSIBLE N-term signal sequence\ninside 1 19\nTMhelix 20 39\noutside 40 325\nTMhelix 326 348\ninside 349 372

19139 GCF_002034115.1_ASM203411v1 Salmonella enterica subsp. enterica serovar Paratyphi B Proteobacteria; Gammaproteobacteria; Enterobacterales; Enterobacteriaceae; Salmonella; Salmonella enterica; Salmonella enterica subsp. enterica MTGDNTLITSHGINRRDFMKLCAALAATMGLSSKAAAEMAESVSNPQRPPVIWIGAQECT WP_000145429.1 hydrogenase 2 small subunit [Salmonella enterica] Length: 372\nNumber of predicted TMHs: 1\nExp number of AAs in TMHs: 24.51309\nExp number, first 60 AAs: 0.2682\nTotal prob of N-in: 0.14402\noutside 1 331\nTMhelix 332 354\ninside 355 372

19140 GCF_001614145.1_ASM161414v1 Salmonella enterica subsp. enterica serovar Paratyphi B Proteobacteria; Gammaproteobacteria; Enterobacterales; Enterobacteriaceae; Salmonella; Salmonella enterica; Salmonella enterica subsp. enterica MTGDNTLITSHGINRRDFMKLCAALAATMGLSSKAAAEMAESVSNPQRPPVIWIGAQECT WP_000145429.1 hydrogenase 2 small subunit [Salmonella enterica] Length: 372\nNumber of predicted TMHs: 1\nExp number of AAs in TMHs: 24.51309\nExp number, first 60 AAs: 0.2682\nTotal prob of N-in: 0.14402\noutside 1 331\nTMhelix 332 354\ninside 355 372

19141 GCF_002034115.1_ASM203411v1 Salmonella enterica subsp. enterica serovar Paratyphi B Proteobacteria; Gammaproteobacteria; Enterobacterales; Enterobacteriaceae; Salmonella; Salmonella enterica; Salmonella enterica subsp. enterica MNNEETFYQAMRRKGVTRRSFLKFCSLAATSLGLGAGMTPKIAWALENKPRIPVVWIHGL WP_001058307.1 hydrogenase [Salmonella enterica] Length: 372\nNumber of predicted TMHs: 2\nExp number of AAs in TMHs: 36.51738\nExp number, first 60 AAs: 19.72849\nTotal prob of N-in: 0.96170\nPOSSIBLE N-term signal sequence\ninside 1 19\nTMhelix 20 39\noutside 40 325\nTMhelix 326 348\ninside 349 372

19142 GCF_001613965.1_ASM161396v1 Salmonella enterica subsp. enterica serovar Paratyphi B Proteobacteria; Gammaproteobacteria; Enterobacterales; Enterobacteriaceae; Salmonella; Salmonella enterica; Salmonella enterica subsp. enterica MNNEETFYQAMRRKGVTRRSFLKFCSLAATSLGLGAGMTPKIAWALENKPRIPVVWIHGL WP_001058311.1 hydrogenase [Salmonella enterica] Length: 372\nNumber of predicted TMHs: 2\nExp number of AAs in TMHs: 37.4699899999999\nExp number, first 60 AAs: 19.72699\nTotal prob of N-in: 0.96171\nPOSSIBLE N-term signal sequence\ninside 1 19\nTMhelix 20 39\noutside 40 325\nTMhelix 326 348\ninside 349 372

19143 GCF_002034065.1_ASM203406v1 Salmonella enterica subsp. enterica serovar Paratyphi B Proteobacteria; Gammaproteobacteria; Enterobacterales; Enterobacteriaceae; Salmonella; Salmonella enterica;

Salmonella enterica subsp. enterica MTGDNTLITSHGINRRDFMKLCAALAATMGLSSKAAAEMAESVSNPQRPPVIWIGAQECT WP_000145429.1 hydrogenase 2 small subunit [Salmonella enterica] Length: 372\nNumber of predicted TMHs: 1\nExp number of AAs in TMHs: 24.51309\nExp number, first 60 AAs: 0.2682\nTotal prob of N-in: 0.14402\noutside 1 331\nTMhelix 332 354\ninside 355 372

19144 GCF_002048725.1_ASM204872v1 Salmonella enterica subsp. enterica serovar Pomona Proteobacteria; Gammaproteobacteria; Enterobacterales; Enterobacteriaceae; Salmonella; Salmonella enterica; Salmonella enterica subsp. enterica MTGDNTLITSHGINRRDFMKLCAALAATMGLSSKAAAEMAESVSNPQRPPVIWIGAQECT WP_000145431.1 hydrogenase 2 small subunit [Salmonella enterica] Length: 372\nNumber of predicted TMHs: 1\nExp number of AAs in TMHs: 24.51505\nExp number, first 60 AAs: 0.2682\nTotal prob of N-in: 0.14406\noutside 1 331\nTMhelix 332 354\ninside 355 372

19145 GCF_002065215.1_ASM206521v1 Salmonella enterica subsp. enterica serovar Pomona Proteobacteria; Gammaproteobacteria; Enterobacterales; Enterobacteriaceae; Salmonella; Salmonella enterica; Salmonella enterica subsp. enterica MTGDNTLITSHGINRRDFMKLCAALAATMGLSSKAAAEMAESVSNPQRPPVIWIGAQECT WP_000145431.1 hydrogenase 2 small subunit [Salmonella enterica] Length: 372\nNumber of predicted TMHs: 1\nExp number of AAs in TMHs: 24.51505\nExp number, first 60 AAs: 0.2682\nTotal prob of N-in: 0.14406\noutside 1 331\nTMhelix 332 354\ninside 355 372

19146 GCF_002049235.1_ASM204923v1 Salmonella enterica subsp. enterica serovar Pomona Proteobacteria; Gammaproteobacteria; Enterobacterales; Enterobacteriaceae; Salmonella; Salmonella enterica; Salmonella enterica subsp. enterica MTGDNTLITSHGINRRDFMKLCAALAATMGLSSKAAAEMAESVSNPQRPPVIWIGAQECT WP_000145431.1 hydrogenase 2 small subunit [Salmonella enterica] Length: 372\nNumber of predicted TMHs: 1\nExp number of AAs in TMHs: 24.51505\nExp number, first 60 AAs: 0.2682\nTotal prob of N-in: 0.14406\noutside 1 331\nTMhelix 332 354\ninside 355 372

19147 GCF_002049225.1_ASM204922v1 Salmonella enterica subsp. enterica serovar Pomona Proteobacteria; Gammaproteobacteria; Enterobacterales; Enterobacteriaceae; Salmonella; Salmonella enterica; Salmonella enterica subsp. enterica MTGDNTLITSHGINRRDFMKLCAALAATMGLSSKAAAEMAESVSNPQRPPVIWIGAQECT WP_000145431.1 hydrogenase 2 small subunit [Salmonella enterica] Length: 372\nNumber of predicted TMHs: 1\nExp number of AAs in TMHs: 24.51505\nExp number, first 60 AAs: 0.2682\nTotal prob of N-in: 0.14406\noutside 1 331\nTMhelix 332 354\ninside 355 372

19148 GCF_002049325.1_ASM204932v1 Salmonella enterica subsp. enterica serovar Pomona Proteobacteria; Gammaproteobacteria; Enterobacterales; Enterobacteriaceae; Salmonella; Salmonella enterica; Salmonella enterica subsp. enterica MTGDNTLITSHGINRRDFMKLCAALAATMGLSSKAAAEMAESVSNPQRPPVIWIGAQECT WP_000145431.1 hydrogenase 2 small subunit [Salmonella enterica] Length: 372\nNumber of predicted TMHs: 1\nExp number of AAs in TMHs: 24.51505\nExp number, first 60 AAs: 0.2682\nTotal prob of N-in: 0.14406\noutside 1 331\nTMhelix 332 354\ninside 355 372

19149 GCF_002062335.1_ASM206233v1 Salmonella enterica subsp. enterica serovar Poona Proteobacteria; Gammaproteobacteria; Enterobacterales; Enterobacteriaceae; Salmonella; Salmonella enterica; Salmonella enterica subsp. enterica MNNEETFYQAMRRKGVTRRSFLKFCSLAATSLGLGAGMTPKIAWALENKPRIPVVWIHGL WP_001058312.1 hydrogenase [Salmonella enterica] Length: 372\nNumber of predicted TMHs: 2\nExp number of AAs in TMHs: 36.51738\nExp number, first 60 AAs: 19.72849\nTotal prob of N-in: 0.96170\nPOSSIBLE N-term signal sequence\ninside 1 19\nTMhelix 20 39\noutside 40 325\nTMhelix 326 348\ninside 349 372

19150 GCF_002061265.1_ASM206126v1 Salmonella enterica subsp. enterica serovar Poona Proteobacteria; Gammaproteobacteria; Enterobacterales; Enterobacteriaceae; Salmonella; Salmonella enterica; Salmonella enterica subsp. enterica MNNEETFYQAMRRKGVTRRSFLKFCSLAATSLGLGAGMTPKIAWALENKPRIPVVWIHGL WP_023213673.1 hydrogenase 1, small subunit [Salmonella enterica] Length: 372\nNumber of predicted TMHs: 2\nExp number of AAs in TMHs: 36.51753\nExp number, first 60 AAs: 19.72849\nTotal prob of N-in: 0.96170\nPOSSIBLE N-term signal sequence\ninside 1 19\nTMhelix 20 39\noutside 40 325\nTMhelix 326 348\ninside 349 372

19151 GCF_002065205.1_ASM206520v1 Salmonella enterica subsp. enterica serovar Poona Proteobacteria; Gammaproteobacteria; Enterobacterales; Enterobacteriaceae; Salmonella; Salmonella enterica; Salmonella enterica subsp. enterica MTGDNTLITSHGINRRDFMKLCAALAATMGLSSKAAAEMAESVSNPQRPPVIWIGAQECT WP_000145429.1 hydrogenase 2 small subunit [Salmonella enterica] Length: 372\nNumber of predicted TMHs: 1\nExp number of AAs in TMHs: 24.51309\nExp number, first 60 AAs: 0.2682\nTotal prob of N-in: 0.14402\noutside 1 331\nTMhelix 332 354\ninside 355 372

19152 GCF_002065205.1_ASM206520v1 Salmonella enterica subsp. enterica serovar Poona Proteobacteria; Gammaproteobacteria; Enterobacterales; Enterobacteriaceae; Salmonella; Salmonella enterica; Salmonella enterica subsp. enterica MNNEETFYQAMRRKGVTRRSFLKFCSLAATSLGLGAGMTPKIAWALENKPRIPVVWIHGL WP_001058312.1 hydrogenase [Salmonella enterica] Length: 372\nNumber of predicted TMHs: 2\nExp number of AAs in TMHs: 36.51738\nExp number, first 60 AAs: 19.72849\nTotal prob of N-in: 0.96170\nPOSSIBLE N-term signal sequence\ninside 1 19\nTMhelix 20 39\noutside 40 325\nTMhelix 326 348\ninside 349 372

19153 GCF_002062335.1_ASM206233v1 Salmonella enterica subsp. enterica serovar Poona Proteobacteria; Gammaproteobacteria; Enterobacterales; Enterobacteriaceae; Salmonella; Salmonella enterica; Salmonella enterica subsp. enterica MTGDNTLITSHGINRRDFMKLCAALAATMGLSSKAAAEMAESVSNPQRPPVIWIGAQECT WP_000145429.1 hydrogenase 2 small subunit [Salmonella enterica] Length: 372\nNumber of predicted TMHs: 1\nExp number of AAs in TMHs: 24.51309\nExp number, first 60 AAs: 0.2682\nTotal prob of N-in: 0.14402\noutside 1 331\nTMhelix 332 354\ninside 355 372

19154 GCF_002061145.1_ASM206114v1 Salmonella enterica subsp. enterica serovar Poona Proteobacteria; Gammaproteobacteria; Enterobacterales; Enterobacteriaceae; Salmonella; Salmonella enterica; Salmonella enterica subsp. enterica MTGDNTLITSHGINRRDFMKLCAALAATMGLSSKAAAEMAESVSNPQRPPVIWIGAQECT WP_000145429.1 hydrogenase 2 small subunit [Salmonella enterica] Length: 372\nNumber of predicted TMHs: 1\nExp number of AAs in TMHs: 24.51309\nExp number, first 60 AAs: 0.2682\nTotal prob of N-in: 0.14402\noutside 1 331\nTMhelix 332 354\ninside 355 372

19155 GCF_002061265.1_ASM206126v1 Salmonella enterica subsp. enterica serovar Poona Proteobacteria; Gammaproteobacteria; Enterobacterales; Enterobacteriaceae; Salmonella; Salmonella enterica; Salmonella enterica subsp. enterica MTGDNTLITSHGINRRDFMKLCAALAATMGLSSKAAAEMAESVSNPQRPPVIWIGAQECT WP_000145429.1 hydrogenase 2 small subunit [Salmonella enterica] Length: 372\nNumber of predicted TMHs: 1\nExp number of AAs in TMHs: 24.51309\nExp number, first 60 AAs: 0.2682\nTotal prob of N-in: 0.14402\noutside 1 331\nTMhelix 332 354\ninside 355 372

19156 GCF_002061145.1_ASM206114v1 Salmonella enterica subsp. enterica serovar Poona Proteobacteria; Gammaproteobacteria; Enterobacterales; Enterobacteriaceae; Salmonella; Salmonella enterica; Salmonella enterica subsp. enterica MNNEETFYQAMRRKGVTRRSFLKFCSLAATSLGLGAGMTPKIAWALENKPRIPVVWIHGL WP_023213673.1 hydrogenase 1, small subunit [Salmonella enterica] Length: 372\nNumber of predicted TMHs: 2\nExp number of AAs in TMHs: 36.51753\nExp number, first 60 AAs: 19.72849\nTotal prob of N-in: 0.96170\nPOSSIBLE N-term signal sequence\ninside 1 19\nTMhelix 20 39\noutside 40 325\nTMhelix 326 348\ninside 349 372

19157 GCF_002062095.1_ASM206209v1 Salmonella enterica subsp. enterica serovar Potsdam Proteobacteria; Gammaproteobacteria; Enterobacterales; Enterobacteriaceae; Salmonella; Salmonella enterica; Salmonella enterica subsp. enterica MNNEETFYQAMRRKGVTRRSFLKFCSLAATSLGLGAGMTPKIAWALENKPRIPVVWIHGL WP_001058311.1 hydrogenase [Salmonella enterica] Length: 372\nNumber of predicted TMHs: 2\nExp number of AAs in TMHs: 37.4699899999999\nExp number, first 60 AAs: 19.72699\nTotal prob of N-in: 0.96171\nPOSSIBLE N-term signal sequence\ninside 1 19\nTMhelix 20 39\noutside 40 325\nTMhelix 326 348\ninside 349 372

19158 GCF_002062095.1_ASM206209v1 Salmonella enterica subsp. enterica serovar Potsdam Proteobacteria; Gammaproteobacteria; Enterobacterales; Enterobacteriaceae; Salmonella; Salmonella enterica; Salmonella enterica subsp. enterica MTGDNTLITSHGINRRDFMKLCAALAATMGLSSKAAAEMAESVSNPQRPPVIWIGAQECT WP_000145429.1 hydrogenase 2 small subunit [Salmonella enterica] Length: 372\nNumber of predicted TMHs: 1\nExp number of AAs in TMHs: 24.51309\nExp number, first 60 AAs: 0.2682\nTotal prob of N-in: 0.14402\noutside 1 331\nTMhelix 332 354\ninside 355 372

19159 GCF_002065195.1_ASM206519v1 Salmonella enterica subsp. enterica serovar Potsdam Proteobacteria; Gammaproteobacteria; Enterobacterales; Enterobacteriaceae; Salmonella; Salmonella enterica; Salmonella enterica subsp. enterica MNNEETFYQAMRRKGVTRRSFLKFCSLAATSLGLGAGMTPKIAWALENKPRIPVVWIHGL WP_001058311.1 hydrogenase [Salmonella enterica] Length: 372\nNumber of predicted TMHs: 2\nExp number of AAs in TMHs: 37.4699899999999\nExp number, first 60 AAs: 19.72699\nTotal prob of N-in: 0.96171\nPOSSIBLE N-term signal sequence\ninside 1 19\nTMhelix 20 39\noutside 40 325\nTMhelix 326 348\ninside 349 372

19160 GCF_002065195.1_ASM206519v1 Salmonella enterica subsp. enterica serovar Potsdam Proteobacteria; Gammaproteobacteria; Enterobacterales; Enterobacteriaceae; Salmonella; Salmonella enterica;

Salmonella enterica subsp. enterica	MTGDNTLITSHGINRRDFMKLCAALAATMGLSSKAAAEMAESVSNPQRPPVIWIGAQECT	WP_000145429.1	hydrogenase 2 small subunit [Salmonella enterica]	Length: 372\nNumber of predicted TMHs:  1\nExp number of AAs in TMHs: 24.51309\nExp number, first 60 AAs:  0.2682\nTotal prob of N-in:   0.14402\noutside   1   331\nTMhelix   332   354\ninside   355   372

19161	GCF_000953595.1_S44987_scaffold1	Salmonella enterica subsp. enterica serovar Pullorum	Proteobacteria; Gammaproteobacteria; Enterobacterales; Enterobacteriaceae; Salmonella; Salmonella enterica; Salmonella enterica subsp. enterica	MNNEETFYQAMRRKGVTRRSFLKFCSLAATSLGLGAGMTPKIAWALENKPRIPVVWIHGL	WP_001058309.1	hydrogenase [Salmonella enterica]	Length: 372\nNumber of predicted TMHs:  2\nExp number of AAs in TMHs: 36.51725\nExp number, first 60 AAs:  19.7285\nTotal prob of N-in:   0.96170\nPOSSIBLE N-term signal sequence\ninside   1   19\nTMhelix   20   39\noutside   40   325\nTMhelix   326   348\ninside   349   372

19162	GCF_001448575.1_ASM144857v1	Salmonella enterica subsp. enterica serovar Pullorum	Proteobacteria; Gammaproteobacteria; Enterobacterales; Enterobacteriaceae; Salmonella; Salmonella enterica; Salmonella enterica subsp. enterica	MTGDNTLITSHGINRRDFMKLCAALAATMGLSSKAAAEMAESVSNPQRPPVIWIGAQECT	WP_000145430.1	hydrogenase 2 small subunit [Salmonella enterica]	Length: 372\nNumber of predicted TMHs:  1\nExp number of AAs in TMHs: 24.51306\nExp number, first 60 AAs:  0.2682\nTotal prob of N-in:   0.14402\noutside   1   331\nTMhelix   332   354\ninside   355   372

19163	GCF_001448575.1_ASM144857v1	Salmonella enterica subsp. enterica serovar Pullorum	Proteobacteria; Gammaproteobacteria; Enterobacterales; Enterobacteriaceae; Salmonella; Salmonella enterica; Salmonella enterica subsp. enterica	MNNEETFYQAMRRKGVTRRSFLKFCSLAATSLGLGAGMTPKIAWALENKPRIPVVWIHGL	WP_001058309.1	hydrogenase [Salmonella enterica]	Length: 372\nNumber of predicted TMHs:  2\nExp number of AAs in TMHs: 36.51725\nExp number, first 60 AAs:  19.7285\nTotal prob of N-in:   0.96170\nPOSSIBLE N-term signal sequence\ninside   1   19\nTMhelix   20   39\noutside   40   325\nTMhelix   326   348\ninside   349   372

19164	GCF_900067255.1_NeoSp11	Salmonella enterica subsp. enterica serovar Pullorum	Proteobacteria; Gammaproteobacteria; Enterobacterales; Enterobacteriaceae; Salmonella; Salmonella enterica; Salmonella enterica subsp. enterica	MNNEETFYQAMRRKGVTRRSFLKFCSLAATSLGLGAGMTPKIAWALENKPRIPVVWIHGL	WP_001058309.1	hydrogenase [Salmonella enterica]	Length: 372\nNumber of predicted TMHs:  2\nExp number of AAs in TMHs: 36.51725\nExp number, first 60 AAs:  19.7285\nTotal prob of N-in:   0.96170\nPOSSIBLE N-term signal sequence\ninside   1   19\nTMhelix   20   39\noutside   40   325\nTMhelix   326   348\ninside   349   372

19165	GCF_900067255.1_NeoSp11	Salmonella enterica subsp. enterica serovar Pullorum	Proteobacteria; Gammaproteobacteria; Enterobacterales; Enterobacteriaceae; Salmonella; Salmonella enterica; Salmonella enterica subsp. enterica	MTGDNTLITSHGINRRDFMKLCAALAATMGLSSKAAAEMAESVSNPQRPPVIWIGAQECT	WP_000145430.1	hydrogenase 2 small subunit [Salmonella enterica]	Length: 372\nNumber of predicted TMHs:  1\nExp number of AAs in TMHs: 24.51306\nExp number, first 60 AAs:  0.2682\nTotal prob of N-in:   0.14402\noutside   1   331\nTMhelix   332   354\ninside   355   372

19166	GCF_000953595.1_S44987_scaffold1	Salmonella enterica subsp. enterica serovar Pullorum	Proteobacteria; Gammaproteobacteria; Enterobacterales; Enterobacteriaceae; Salmonella; Salmonella enterica; Salmonella enterica subsp. enterica	MTGDNTLITSHGINRRDFMKLCAALAATMGLSSKAAAEMAESVSNPQRPPVIWIGAQECT	WP_000145430.1	hydrogenase 2 small subunit [Salmonella enterica]	Length: 372\nNumber of predicted TMHs:  1\nExp number of AAs in TMHs: 24.51306\nExp number, first 60 AAs:  0.2682\nTotal prob of N-in:   0.14402\noutside   1   331\nTMhelix   332   354\ninside   355   372

19167	GCF_001244455.1_Salmonella_enterica_CVM_N48684_v1.0	Salmonella enterica subsp. enterica serovar Reading	Proteobacteria; Gammaproteobacteria; Enterobacterales; Enterobacteriaceae; Salmonella; Salmonella enterica; Salmonella enterica subsp. enterica	MTGDNTLITSHGINRRDFMKLCAALAATMGLSSKAAAEMAESVSNPQRPPVIWIGAQECT	WP_000145429.1	hydrogenase 2 small subunit [Salmonella enterica]	Length: 372\nNumber of predicted TMHs:  1\nExp number of AAs in TMHs: 24.51309\nExp number, first 60 AAs:  0.2682\nTotal prob of N-in:   0.14402\noutside   1   331\nTMhelix   332   354\ninside   355   372

19168	GCF_002030715.1_ASM203071v1	Salmonella enterica subsp. enterica serovar Reading	Proteobacteria; Gammaproteobacteria; Enterobacterales; Enterobacteriaceae; Salmonella; Salmonella enterica; Salmonella enterica subsp. enterica	MNNEETFYQAMRRKGVTRRSFLKFCSLAATSLGLGAGMTPKIAWALENKPRIPVVWIHGL	WP_001058312.1	hydrogenase [Salmonella enterica]	Length: 372\nNumber of predicted TMHs:  2\nExp number of AAs in TMHs: 36.51738\nExp number, first 60 AAs:  19.72849\nTotal prob of N-in:   0.96170\nPOSSIBLE N-term signal sequence\ninside   1   19\nTMhelix   20   39\noutside   40   325\nTMhelix   326   348\ninside   349   372

19169	GCF_001480185.1_Salmonella_enterica_CVM_N40375-R_v1.0	Salmonella enterica subsp. enterica serovar Reading	Proteobacteria; Gammaproteobacteria; Enterobacterales; Enterobacteriaceae; Salmonella; Salmonella enterica; Salmonella enterica subsp. enterica	MNNEETFYQAMRRKGVTRRSFLKFCSLAATSLGLGAGMTPKIAWALENKPRIPVVWIHGL	WP_001058312.1	hydrogenase [Salmonella enterica]	Length: 372\nNumber of predicted TMHs: 2\nExp number of AAs in TMHs: 36.51738\nExp number, first 60 AAs: 19.72849\nTotal prob of N-in: 0.96170\nPOSSIBLE N-term signal sequence\ninside 1 19\nTMhelix 20 39\noutside 40 325\nTMhelix 326 348\ninside 349 372

19170	GCF_002048855.1_ASM204885v1	Salmonella enterica subsp. enterica serovar Reading	Proteobacteria; Gammaproteobacteria; Enterobacterales; Enterobacteriaceae; Salmonella; Salmonella enterica; Salmonella enterica subsp. enterica	MTGDNTLITSHGINRRDFMKLCAALAATMGLSSKAAAEMAESVSNPQRPPVIWIGAQECT	WP_000145429.1	hydrogenase 2 small subunit [Salmonella enterica]	Length: 372\nNumber of predicted TMHs: 1\nExp number of AAs in TMHs: 24.51309\nExp number, first 60 AAs: 0.2682\nTotal prob of N-in: 0.14402\noutside 1 331\nTMhelix 332 354\ninside 355 372

19171	GCF_002066775.1_ASM206677v1	Salmonella enterica subsp. enterica serovar Reading	Proteobacteria; Gammaproteobacteria; Enterobacterales; Enterobacteriaceae; Salmonella; Salmonella enterica; Salmonella enterica subsp. enterica	MNNEETFYQAMRRKGVTRRSFLKFCSLAATSLGLGAGMTPKIAWALENKPRIPVVWIHGL	WP_001058312.1	hydrogenase [Salmonella enterica]	Length: 372\nNumber of predicted TMHs: 2\nExp number of AAs in TMHs: 36.51738\nExp number, first 60 AAs: 19.72849\nTotal prob of N-in: 0.96170\nPOSSIBLE N-term signal sequence\ninside 1 19\nTMhelix 20 39\noutside 40 325\nTMhelix 326 348\ninside 349 372

19172	GCF_002066775.1_ASM206677v1	Salmonella enterica subsp. enterica serovar Reading	Proteobacteria; Gammaproteobacteria; Enterobacterales; Enterobacteriaceae; Salmonella; Salmonella enterica; Salmonella enterica subsp. enterica	MTGDNTLITSHGINRRDFMKLCAALAATMGLSSKAAAEMAESVSNPQRPPVIWIGAQECT	WP_000145429.1	hydrogenase 2 small subunit [Salmonella enterica]	Length: 372\nNumber of predicted TMHs: 1\nExp number of AAs in TMHs: 24.51309\nExp number, first 60 AAs: 0.2682\nTotal prob of N-in: 0.14402\noutside 1 331\nTMhelix 332 354\ninside 355 372

19173	GCF_002048825.1_ASM204882v1	Salmonella enterica subsp. enterica serovar Reading	Proteobacteria; Gammaproteobacteria; Enterobacterales; Enterobacteriaceae; Salmonella; Salmonella enterica; Salmonella enterica subsp. enterica	MTGDNTLITSHGINRRDFMKLCAALAATMGLSSKAAAEMAESVSNPQRPPVIWIGAQECT	WP_000145429.1	hydrogenase 2 small subunit [Salmonella enterica]	Length: 372\nNumber of predicted TMHs: 1\nExp number of AAs in TMHs: 24.51309\nExp number, first 60 AAs: 0.2682\nTotal prob of N-in: 0.14402\noutside 1 331\nTMhelix 332 354\ninside 355 372

19174	GCF_001244455.1_Salmonella_enterica_CVM_N48684_v1.0	Salmonella enterica subsp. enterica serovar Reading	Proteobacteria; Gammaproteobacteria; Enterobacterales; Enterobacteriaceae; Salmonella; Salmonella enterica; Salmonella enterica subsp. enterica	MNNEETFYQAMRRKGVTRRSFLKFCSLAATSLGLGAGMTPKIAWALENKPRIPVVWIHGL	WP_001058312.1	hydrogenase [Salmonella enterica]	Length: 372\nNumber of predicted TMHs: 2\nExp number of AAs in TMHs: 36.51738\nExp number, first 60 AAs: 19.72849\nTotal prob of N-in: 0.96170\nPOSSIBLE N-term signal sequence\ninside 1 19\nTMhelix 20 39\noutside 40 325\nTMhelix 326 348\ninside 349 372

19175	GCF_001477955.1_Salmonella_enterica_CVM_N29378-SQ_v1.0	Salmonella enterica subsp. enterica serovar Reading	Proteobacteria; Gammaproteobacteria; Enterobacterales; Enterobacteriaceae; Salmonella; Salmonella enterica; Salmonella enterica subsp. enterica	MNNEETFYQAMRRKGVTRRSFLKFCSLAATSLGLGAGMTPKIAWALENKPRIPVVWIHGL	WP_001058312.1	hydrogenase [Salmonella enterica]	Length: 372\nNumber of predicted TMHs: 2\nExp number of AAs in TMHs: 36.51738\nExp number, first 60 AAs: 19.72849\nTotal prob of N-in: 0.96170\nPOSSIBLE N-term signal sequence\ninside 1 19\nTMhelix 20 39\noutside 40 325\nTMhelix 326 348\ninside 349 372

19176	GCF_002030715.1_ASM203071v1	Salmonella enterica subsp. enterica serovar Reading	Proteobacteria; Gammaproteobacteria; Enterobacterales; Enterobacteriaceae; Salmonella; Salmonella enterica; Salmonella enterica subsp. enterica	MTGDNTLITSHGINRRDFMKLCAALAATMGLSSKAAAEMAESVSNPQRPPVIWIGAQECT	WP_000145429.1	hydrogenase 2 small subunit [Salmonella enterica]	Length: 372\nNumber of predicted TMHs: 1\nExp number of AAs in TMHs: 24.51309\nExp number, first 60 AAs: 0.2682\nTotal prob of N-in: 0.14402\noutside 1 331\nTMhelix 332 354\ninside 355 372

19177	GCF_001246075.1_Salmonella_enterica_CVM_N51247_v1.0	Salmonella enterica subsp. enterica serovar Reading	Proteobacteria; Gammaproteobacteria; Enterobacterales; Enterobacteriaceae; Salmonella; Salmonella enterica; Salmonella enterica subsp. enterica	MTGDNTLITSHGINRRDFMKLCAALAATMGLSSKAAAEMAESVSNPQRPPVIWIGAQECT	WP_000145429.1	hydrogenase 2 small subunit [Salmonella enterica]	Length: 372\nNumber of

predicted TMHs:  1\nExp number of AAs in TMHs: 24.51309\nExp number, first 60 AAs:  0.2682\nTotal prob of N-in: 0.14402\noutside   1   331\nTMhelix   332   354\ninside   355   372

19178 GCF_001480185.1_Salmonella_enterica_CVM_N40375-R_v1.0 Salmonella enterica subsp. enterica serovar Reading Proteobacteria; Gammaproteobacteria; Enterobacterales; Enterobacteriaceae; Salmonella; Salmonella enterica; Salmonella enterica subsp. enterica MTGDNTLITSHGINRRDFMKLCAALAATMGLSSKAAAEMAESVSNPQRPPVIWIGAQECT WP_000145429.1 hydrogenase 2 small subunit [Salmonella enterica] Length: 372\nNumber of predicted TMHs:  1\nExp number of AAs in TMHs: 24.51309\nExp number, first 60 AAs:  0.2682\nTotal prob of N-in: 0.14402\noutside   1   331\nTMhelix   332   354\ninside   355   372

19179 GCF_002064115.1_ASM206411v1 Salmonella enterica subsp. enterica serovar Reading Proteobacteria; Gammaproteobacteria; Enterobacterales; Enterobacteriaceae; Salmonella; Salmonella enterica; Salmonella enterica subsp. enterica MTGDNTLITSHGINRRDFMKLCAALAATMGLSSKAAAEMAESVSNPQRPPVIWIGAQECT WP_000145429.1 hydrogenase 2 small subunit [Salmonella enterica] Length: 372\nNumber of predicted TMHs:  1\nExp number of AAs in TMHs: 24.51309\nExp number, first 60 AAs:  0.2682\nTotal prob of N-in: 0.14402\noutside   1   331\nTMhelix   332   354\ninside   355   372

19180 GCF_002064115.1_ASM206411v1 Salmonella enterica subsp. enterica serovar Reading Proteobacteria; Gammaproteobacteria; Enterobacterales; Enterobacteriaceae; Salmonella; Salmonella enterica; Salmonella enterica subsp. enterica MNNEETFYQAMRRKGVTRRSFLKFCSLAATSLGLGAGMTPKIAWALENKPRIPVVWIHGL WP_001058312.1 hydrogenase [Salmonella enterica] Length: 372\nNumber of predicted TMHs:  2\nExp number of AAs in TMHs: 36.51738\nExp number, first 60 AAs:  19.72849\nTotal prob of N-in:   0.96170\nPOSSIBLE N-term signal sequence\ninside   1   19\nTMhelix   20   39\noutside   40   325\nTMhelix   326   348\ninside   349   372

19181 GCF_001243985.1_Salmonella_enterica_CVM_N46858_v1.0 Salmonella enterica subsp. enterica serovar Reading Proteobacteria; Gammaproteobacteria; Enterobacterales; Enterobacteriaceae; Salmonella; Salmonella enterica; Salmonella enterica subsp. enterica MTGDNTLITSHGINRRDFMKLCAALAATMGLSSKAAAEMAESVSNPQRPPVIWIGAQECT WP_000145429.1 hydrogenase 2 small subunit [Salmonella enterica] Length: 372\nNumber of predicted TMHs:  1\nExp number of AAs in TMHs: 24.51309\nExp number, first 60 AAs:  0.2682\nTotal prob of N-in: 0.14402\noutside   1   331\nTMhelix   332   354\ninside   355   372

19182 GCF_001241715.1_Salmonella_enterica_CVM_N45396_v1.0 Salmonella enterica subsp. enterica serovar Reading Proteobacteria; Gammaproteobacteria; Enterobacterales; Enterobacteriaceae; Salmonella; Salmonella enterica; Salmonella enterica subsp. enterica MNNEETFYQAMRRKGVTRRSFLKFCSLAATSLGLGAGMTPKIAWALENKPRIPVVWIHGL WP_001058312.1 hydrogenase [Salmonella enterica] Length: 372\nNumber of predicted TMHs:  2\nExp number of AAs in TMHs: 36.51738\nExp number, first 60 AAs:  19.72849\nTotal prob of N-in:   0.96170\nPOSSIBLE N-term signal sequence\ninside   1   19\nTMhelix   20   39\noutside   40   325\nTMhelix   326   348\ninside   349   372

19183 GCF_001243985.1_Salmonella_enterica_CVM_N46858_v1.0 Salmonella enterica subsp. enterica serovar Reading Proteobacteria; Gammaproteobacteria; Enterobacterales; Enterobacteriaceae; Salmonella; Salmonella enterica; Salmonella enterica subsp. enterica MNNEETFYQAMRRKGVTRRSFLKFCSLAATSLGLGAGMTPKIAWALENKPRIPVVWIHGL WP_001058312.1 hydrogenase [Salmonella enterica] Length: 372\nNumber of predicted TMHs:  2\nExp number of AAs in TMHs: 36.51738\nExp number, first 60 AAs:  19.72849\nTotal prob of N-in:   0.96170\nPOSSIBLE N-term signal sequence\ninside   1   19\nTMhelix   20   39\noutside   40   325\nTMhelix   326   348\ninside   349   372

19184 GCF_001241715.1_Salmonella_enterica_CVM_N45396_v1.0 Salmonella enterica subsp. enterica serovar Reading Proteobacteria; Gammaproteobacteria; Enterobacterales; Enterobacteriaceae; Salmonella; Salmonella enterica; Salmonella enterica subsp. enterica MTGDNTLITSHGINRRDFMKLCAALAATMGLSSKAAAEMAESVSNPQRPPVIWIGAQECT WP_000145429.1 hydrogenase 2 small subunit [Salmonella enterica] Length: 372\nNumber of predicted TMHs:  1\nExp number of AAs in TMHs: 24.51309\nExp number, first 60 AAs:  0.2682\nTotal prob of N-in: 0.14402\noutside   1   331\nTMhelix   332   354\ninside   355   372

19185 GCF_001481335.1_Salmonella_enterica_CVM_N42528-SQ_v1.0 Salmonella enterica subsp. enterica serovar Reading Proteobacteria; Gammaproteobacteria; Enterobacterales; Enterobacteriaceae; Salmonella; Salmonella enterica; Salmonella enterica subsp. enterica MTGDNTLITSHGINRRDFMKLCAALAATMGLSSKAAAEMAESVSNPQRPPVIWIGAQECT WP_000145429.1 hydrogenase 2 small subunit [Salmonella enterica] Length: 372\nNumber of predicted TMHs:  1\nExp number of AAs in TMHs: 24.51309\nExp number, first 60 AAs:  0.2682\nTotal prob of N-in: 0.14402\noutside   1   331\nTMhelix   332   354\ninside   355   372

19186 GCF_001481255.1_Salmonella_enterica_CVM_N43241_v1.0 Salmonella enterica subsp. enterica serovar Reading Proteobacteria; Gammaproteobacteria; Enterobacterales; Enterobacteriaceae; Salmonella; Salmonella enterica;

Salmonella enterica subsp. enterica	MNNEETFYQAMRRKGVTRRSFLKFCSLAATSLGLGAGMTPKIAWALENKPRIPVVWIHGL	WP_001058312.1	hydrogenase [Salmonella enterica]	Length: 372\nNumber of predicted TMHs: 2\nExp number of AAs in TMHs: 36.51738\nExp number, first 60 AAs: 19.72849\nTotal prob of N-in: 0.96170\nPOSSIBLE N-term signal sequence\ninside 1 19\nTMhelix 20 39\noutside 40 325\nTMhelix 326 348\ninside 349 372

19187	GCF_001481335.1_Salmonella_enterica_CVM_N42528-SQ_v1.0	Salmonella enterica subsp. enterica serovar Reading	Proteobacteria; Gammaproteobacteria; Enterobacterales; Enterobacteriaceae; Salmonella; Salmonella enterica; Salmonella enterica subsp. enterica	MNNEETFYQAMRRKGVTRRSFLKFCSLAATSLGLGAGMTPKIAWALENKPRIPVVWIHGL	WP_001058312.1	hydrogenase [Salmonella enterica]	Length: 372\nNumber of predicted TMHs: 2\nExp number of AAs in TMHs: 36.51738\nExp number, first 60 AAs: 19.72849\nTotal prob of N-in: 0.96170\nPOSSIBLE N-term signal sequence\ninside 1 19\nTMhelix 20 39\noutside 40 325\nTMhelix 326 348\ninside 349 372

19188	GCF_002048825.1_ASM204882v1	Salmonella enterica subsp. enterica serovar Reading	Proteobacteria; Gammaproteobacteria; Enterobacterales; Enterobacteriaceae; Salmonella; Salmonella enterica; Salmonella enterica subsp. enterica	MNNEETFYQAMRRKGVTRRSFLKFCSLAATSLGLGAGMTPKIAWALENKPRIPVVWIHGL	WP_001058312.1	hydrogenase [Salmonella enterica]	Length: 372\nNumber of predicted TMHs: 2\nExp number of AAs in TMHs: 36.51738\nExp number, first 60 AAs: 19.72849\nTotal prob of N-in: 0.96170\nPOSSIBLE N-term signal sequence\ninside 1 19\nTMhelix 20 39\noutside 40 325\nTMhelix 326 348\ninside 349 372

19189	GCF_002031205.1_ASM203120v1	Salmonella enterica subsp. enterica serovar Reading	Proteobacteria; Gammaproteobacteria; Enterobacterales; Enterobacteriaceae; Salmonella; Salmonella enterica; Salmonella enterica subsp. enterica	MNNEETFYQAMRRKGVTRRSFLKFCSLAATSLGLGAGMTPKIAWALENKPRIPVVWIHGL	WP_001058312.1	hydrogenase [Salmonella enterica]	Length: 372\nNumber of predicted TMHs: 2\nExp number of AAs in TMHs: 36.51738\nExp number, first 60 AAs: 19.72849\nTotal prob of N-in: 0.96170\nPOSSIBLE N-term signal sequence\ninside 1 19\nTMhelix 20 39\noutside 40 325\nTMhelix 326 348\ninside 349 372

19190	GCF_001481255.1_Salmonella_enterica_CVM_N43241_v1.0	Salmonella enterica subsp. enterica serovar Reading	Proteobacteria; Gammaproteobacteria; Enterobacterales; Enterobacteriaceae; Salmonella; Salmonella enterica; Salmonella enterica subsp. enterica	MTGDNTLITSHGINRRDFMKLCAALAATMGLSSKAAAEMAESVSNPQRPPVIWIGAQECT	WP_000145429.1	hydrogenase 2 small subunit [Salmonella enterica]	Length: 372\nNumber of predicted TMHs: 1\nExp number of AAs in TMHs: 24.51309\nExp number, first 60 AAs: 0.2682\nTotal prob of N-in: 0.14402\noutside 1 331\nTMhelix 332 354\ninside 355 372

19191	GCF_002031205.1_ASM203120v1	Salmonella enterica subsp. enterica serovar Reading	Proteobacteria; Gammaproteobacteria; Enterobacterales; Enterobacteriaceae; Salmonella; Salmonella enterica; Salmonella enterica subsp. enterica	MTGDNTLITSHGINRRDFMKLCAALAATMGLSSKAAAEMAESVSNPQRPPVIWIGAQECT	WP_000145429.1	hydrogenase 2 small subunit [Salmonella enterica]	Length: 372\nNumber of predicted TMHs: 1\nExp number of AAs in TMHs: 24.51309\nExp number, first 60 AAs: 0.2682\nTotal prob of N-in: 0.14402\noutside 1 331\nTMhelix 332 354\ninside 355 372

19192	GCF_001246075.1_Salmonella_enterica_CVM_N51247_v1.0	Salmonella enterica subsp. enterica serovar Reading	Proteobacteria; Gammaproteobacteria; Enterobacterales; Enterobacteriaceae; Salmonella; Salmonella enterica; Salmonella enterica subsp. enterica	MNNEETFYQAMRRKGVTRRSFLKFCSLAATSLGLGAGMTPKIAWALENKPRIPVVWIHGL	WP_001058312.1	hydrogenase [Salmonella enterica]	Length: 372\nNumber of predicted TMHs: 2\nExp number of AAs in TMHs: 36.51738\nExp number, first 60 AAs: 19.72849\nTotal prob of N-in: 0.96170\nPOSSIBLE N-term signal sequence\ninside 1 19\nTMhelix 20 39\noutside 40 325\nTMhelix 326 348\ninside 349 372

19193	GCF_001477955.1_Salmonella_enterica_CVM_N29378-SQ_v1.0	Salmonella enterica subsp. enterica serovar Reading	Proteobacteria; Gammaproteobacteria; Enterobacterales; Enterobacteriaceae; Salmonella; Salmonella enterica; Salmonella enterica subsp. enterica	MTGDNTLITSHGINRRDFMKLCAALAATMGLSSKAAAEMAESVSNPQRPPVIWIGAQECT	WP_000145429.1	hydrogenase 2 small subunit [Salmonella enterica]	Length: 372\nNumber of predicted TMHs: 1\nExp number of AAs in TMHs: 24.51309\nExp number, first 60 AAs: 0.2682\nTotal prob of N-in: 0.14402\noutside 1 331\nTMhelix 332 354\ninside 355 372

19194	GCF_001246365.1_Salmonella_enterica_CVM_N51261_v1.0	Salmonella enterica subsp. enterica serovar Reading	Proteobacteria; Gammaproteobacteria; Enterobacterales; Enterobacteriaceae; Salmonella; Salmonella enterica; Salmonella enterica subsp. enterica	MNNEETFYQAMRRKGVTRRSFLKFCSLAATSLGLGAGMTPKIAWALENKPRIPVVWIHGL	WP_001058312.1	hydrogenase [Salmonella enterica]	Length: 372\nNumber of predicted TMHs: 2\nExp number of AAs in TMHs: 36.51738\nExp number, first 60 AAs: 19.72849\nTotal prob of N-in: 0.96170\nPOSSIBLE N-term signal sequence\ninside 1 19\nTMhelix 20 39\noutside 40 325\nTMhelix 326 348\ninside 349 372

19195 GCF_001246365.1_Salmonella_enterica_CVM_N51261_v1.0 Salmonella enterica subsp. enterica serovar Reading Proteobacteria; Gammaproteobacteria; Enterobacterales; Enterobacteriaceae; Salmonella; Salmonella enterica; Salmonella enterica subsp. enterica MTGDNTLITSHGINRRDFMKLCAALAATMGLSSKAAAEMAESVSNPQRPPVIWIGAQECT WP_000145429.1 hydrogenase 2 small subunit [Salmonella enterica] Length: 372\nNumber of predicted TMHs: 1\nExp number of AAs in TMHs: 24.51309\nExp number, first 60 AAs: 0.2682\nTotal prob of N-in: 0.14402\noutside 1 331\nTMhelix 332 354\ninside 355 372

19196 GCF_002048855.1_ASM204885v1 Salmonella enterica subsp. enterica serovar Reading Proteobacteria; Gammaproteobacteria; Enterobacterales; Enterobacteriaceae; Salmonella; Salmonella enterica; Salmonella enterica subsp. enterica MNNEETFYQAMRRKGVTRRSFLKFCSLAATSLGLGAGMTPKIAWALENKPRIPVVWIHGL WP_001058312.1 hydrogenase [Salmonella enterica] Length: 372\nNumber of predicted TMHs: 2\nExp number of AAs in TMHs: 36.51738\nExp number, first 60 AAs: 19.72849\nTotal prob of N-in: 0.96170\nPOSSIBLE N-term signal sequence\ninside 1 19\nTMhelix 20 39\noutside 40 325\nTMhelix 326 348\ninside 349 372

19197 GCF_002045825.1_ASM204582v1 Salmonella enterica subsp. enterica serovar Rissen Proteobacteria; Gammaproteobacteria; Enterobacterales; Enterobacteriaceae; Salmonella; Salmonella enterica; Salmonella enterica subsp. enterica MTGDNTLITSHGINRRDFMKLCAALAATMGLSSKAAAEMAESVSNPQRPPVIWIGAQECT WP_000145429.1 hydrogenase 2 small subunit [Salmonella enterica] Length: 372\nNumber of predicted TMHs: 1\nExp number of AAs in TMHs: 24.51309\nExp number, first 60 AAs: 0.2682\nTotal prob of N-in: 0.14402\noutside 1 331\nTMhelix 332 354\ninside 355 372

19198 GCF_002065155.1_ASM206515v1 Salmonella enterica subsp. enterica serovar Rissen Proteobacteria; Gammaproteobacteria; Enterobacterales; Enterobacteriaceae; Salmonella; Salmonella enterica; Salmonella enterica subsp. enterica MNNEETFYQAMRRKGVTRRSFLKFCSLAATSLGLSAGMTPKIAWALENKPRIPVVWIHGL WP_001058316.1 hydrogenase [Salmonella enterica] Length: 372\nNumber of predicted TMHs: 2\nExp number of AAs in TMHs: 35.76397\nExp number, first 60 AAs: 18.97201\nTotal prob of N-in: 0.92785\nPOSSIBLE N-term signal sequence\ninside 1 19\nTMhelix 20 39\noutside 40 325\nTMhelix 326 348\ninside 349 372

19199 GCF_002062455.1_ASM206245v1 Salmonella enterica subsp. enterica serovar Rissen Proteobacteria; Gammaproteobacteria; Enterobacterales; Enterobacteriaceae; Salmonella; Salmonella enterica; Salmonella enterica subsp. enterica MTGDNTLITSHGINRRDFMKLCAALAATMGLSSKAAAEMAESVSNPQRPPVIWIGAQECT WP_000145429.1 hydrogenase 2 small subunit [Salmonella enterica] Length: 372\nNumber of predicted TMHs: 1\nExp number of AAs in TMHs: 24.51309\nExp number, first 60 AAs: 0.2682\nTotal prob of N-in: 0.14402\noutside 1 331\nTMhelix 332 354\ninside 355 372

19200 GCF_002064035.1_ASM206403v1 Salmonella enterica subsp. enterica serovar Rissen Proteobacteria; Gammaproteobacteria; Enterobacterales; Enterobacteriaceae; Salmonella; Salmonella enterica; Salmonella enterica subsp. enterica MTGDNTLITSHGINRRDFMKLCAALAATMGLSSKAAAEMAESVSNPQRPPVIWIGAQECT WP_000145429.1 hydrogenase 2 small subunit [Salmonella enterica] Length: 372\nNumber of predicted TMHs: 1\nExp number of AAs in TMHs: 24.51309\nExp number, first 60 AAs: 0.2682\nTotal prob of N-in: 0.14402\noutside 1 331\nTMhelix 332 354\ninside 355 372

19201 GCF_002064085.1_ASM206408v1 Salmonella enterica subsp. enterica serovar Rissen Proteobacteria; Gammaproteobacteria; Enterobacterales; Enterobacteriaceae; Salmonella; Salmonella enterica; Salmonella enterica subsp. enterica MTGDNTLITSHGINRRDFMKLCAALAATMGLSSKAAAEMAESVSNPQRPPVIWIGAQECT WP_000145429.1 hydrogenase 2 small subunit [Salmonella enterica] Length: 372\nNumber of predicted TMHs: 1\nExp number of AAs in TMHs: 24.51309\nExp number, first 60 AAs: 0.2682\nTotal prob of N-in: 0.14402\noutside 1 331\nTMhelix 332 354\ninside 355 372

19202 GCF_002064085.1_ASM206408v1 Salmonella enterica subsp. enterica serovar Rissen Proteobacteria; Gammaproteobacteria; Enterobacterales; Enterobacteriaceae; Salmonella; Salmonella enterica; Salmonella enterica subsp. enterica MNNEETFYQAMRRKGVTRRSFLKFCSLAATSLGLSAGMTPKIAWALENKPRIPVVWIHGL WP_080202861.1 [Ni/Fe] hydrogenase small subunit, partial [Salmonella enterica] Length: 353\nNumber of predicted TMHs: 2\nExp number of AAs in TMHs: 36.97941\nExp number, first 60 AAs: 18.9528\nTotal prob of N-in: 0.92802\nPOSSIBLE N-term signal sequence\ninside 1 19\nTMhelix 20 39\noutside 40 325\nTMhelix 326 348\ninside 349 353

19203 GCF_002065155.1_ASM206515v1 Salmonella enterica subsp. enterica serovar Rissen Proteobacteria; Gammaproteobacteria; Enterobacterales; Enterobacteriaceae; Salmonella; Salmonella enterica; Salmonella enterica subsp. enterica MTGDNTLITSHGINRRDFMKLCAALAATMGLSSKAAAEMAESVSNPQRPPVIWIGAQECT WP_000145429.1 hydrogenase 2 small subunit [Salmonella enterica] Length: 372\nNumber of

predicted TMHs: 1\nExp number of AAs in TMHs: 24.51309\nExp number, first 60 AAs: 0.2682\nTotal prob of N-in: 0.14402\noutside 1 331\nTMhelix 332 354\ninside 355 372

19204 GCF_002062455.1_ASM206245v1 Salmonella enterica subsp. enterica serovar Rissen Proteobacteria; Gammaproteobacteria; Enterobacterales; Enterobacteriaceae; Salmonella; Salmonella enterica; Salmonella enterica subsp. enterica MNNEETFYQAMRRKGVTRRSFLKFCSLAATSLGLSAGMTPKIAWALENKPRIPVVWIHGL WP_001058316.1 hydrogenase [Salmonella enterica] Length: 372\nNumber of predicted TMHs: 2\nExp number of AAs in TMHs: 35.76397\nExp number, first 60 AAs: 18.97201\nTotal prob of N-in: 0.92785\nPOSSIBLE N-term signal sequence\ninside 1 19\nTMhelix 20 39\noutside 40 325\nTMhelix 326 348\ninside 349 372

19205 GCF_002045825.1_ASM204582v1 Salmonella enterica subsp. enterica serovar Rissen Proteobacteria; Gammaproteobacteria; Enterobacterales; Enterobacteriaceae; Salmonella; Salmonella enterica; Salmonella enterica subsp. enterica MNNEETFYQAMRRKGVTRRSFLKFCSLAATSLGLSAGMTPKIAWALENKPRIPVVWIHGL WP_001058316.1 hydrogenase [Salmonella enterica] Length: 372\nNumber of predicted TMHs: 2\nExp number of AAs in TMHs: 35.76397\nExp number, first 60 AAs: 18.97201\nTotal prob of N-in: 0.92785\nPOSSIBLE N-term signal sequence\ninside 1 19\nTMhelix 20 39\noutside 40 325\nTMhelix 326 348\ninside 349 372

19206 GCF_002044955.1_ASM204495v1 Salmonella enterica subsp. enterica serovar Rissen Proteobacteria; Gammaproteobacteria; Enterobacterales; Enterobacteriaceae; Salmonella; Salmonella enterica; Salmonella enterica subsp. enterica MTGDNTLITSHGINRRDFMKLCAALAATMGLSSKAAAEMAESVSNPQRPPVIWIGAQECT WP_000145429.1 hydrogenase 2 small subunit [Salmonella enterica] Length: 372\nNumber of predicted TMHs: 1\nExp number of AAs in TMHs: 24.51309\nExp number, first 60 AAs: 0.2682\nTotal prob of N-in: 0.14402\noutside 1 331\nTMhelix 332 354\ninside 355 372

19207 GCF_002044955.1_ASM204495v1 Salmonella enterica subsp. enterica serovar Rissen Proteobacteria; Gammaproteobacteria; Enterobacterales; Enterobacteriaceae; Salmonella; Salmonella enterica; Salmonella enterica subsp. enterica MNNEETFYQAMRRKGVTRRSFLKFCSLAATSLGLSAGMTPKIAWALENKPRIPVVWIHGL WP_001058316.1 hydrogenase [Salmonella enterica] Length: 372\nNumber of predicted TMHs: 2\nExp number of AAs in TMHs: 35.76397\nExp number, first 60 AAs: 18.97201\nTotal prob of N-in: 0.92785\nPOSSIBLE N-term signal sequence\ninside 1 19\nTMhelix 20 39\noutside 40 325\nTMhelix 326 348\ninside 349 372

19208 GCF_001835155.1_ASM183515v1 Salmonella enterica subsp. enterica serovar Rubislaw Proteobacteria; Gammaproteobacteria; Enterobacterales; Enterobacteriaceae; Salmonella; Salmonella enterica; Salmonella enterica subsp. enterica MTGDNTLITSHGINRRDFMKLCAALAATMGLSSKAAAEMAESVSNPQRPPVIWIGAQECT WP_000145429.1 hydrogenase 2 small subunit [Salmonella enterica] Length: 372\nNumber of predicted TMHs: 1\nExp number of AAs in TMHs: 24.51309\nExp number, first 60 AAs: 0.2682\nTotal prob of N-in: 0.14402\noutside 1 331\nTMhelix 332 354\ninside 355 372

19209 GCF_001833255.1_ASM183325v1 Salmonella enterica subsp. enterica serovar Rubislaw Proteobacteria; Gammaproteobacteria; Enterobacterales; Enterobacteriaceae; Salmonella; Salmonella enterica; Salmonella enterica subsp. enterica MTGDNTLITSHGINRRDFMKLCAALAATMGLSSKAAAEMAESVSNPQRPPVIWIGAQECT WP_000145429.1 hydrogenase 2 small subunit [Salmonella enterica] Length: 372\nNumber of predicted TMHs: 1\nExp number of AAs in TMHs: 24.51309\nExp number, first 60 AAs: 0.2682\nTotal prob of N-in: 0.14402\noutside 1 331\nTMhelix 332 354\ninside 355 372

19210 GCF_001834655.1_ASM183465v1 Salmonella enterica subsp. enterica serovar Rubislaw Proteobacteria; Gammaproteobacteria; Enterobacterales; Enterobacteriaceae; Salmonella; Salmonella enterica; Salmonella enterica subsp. enterica MNNEETFYQAMRRKGVTRRSFLKFCSLAATSLGLGAGMTPKIAWALENKPRIPVVWIHGL WP_001058312.1 hydrogenase [Salmonella enterica] Length: 372\nNumber of predicted TMHs: 2\nExp number of AAs in TMHs: 36.51738\nExp number, first 60 AAs: 19.72849\nTotal prob of N-in: 0.96170\nPOSSIBLE N-term signal sequence\ninside 1 19\nTMhelix 20 39\noutside 40 325\nTMhelix 326 348\ninside 349 372

19211 GCF_001833925.1_ASM183392v1 Salmonella enterica subsp. enterica serovar Rubislaw Proteobacteria; Gammaproteobacteria; Enterobacterales; Enterobacteriaceae; Salmonella; Salmonella enterica; Salmonella enterica subsp. enterica MNNEETFYQAMRRKGVTRRSFLKFCSLAATSLGLGAGMTPKIAWALENKPRIPVVWIHGL WP_001058311.1 hydrogenase [Salmonella enterica] Length: 372\nNumber of predicted TMHs: 2\nExp number of AAs in TMHs: 37.4699899999999\nExp number, first 60 AAs: 19.72699\nTotal prob of N-in: 0.96171\nPOSSIBLE N-term signal sequence\ninside 1 19\nTMhelix 20 39\noutside 40 325\nTMhelix 326 348\ninside 349 372

19212 GCF_001833255.1_ASM183325v1 Salmonella enterica subsp. enterica serovar Rubislaw Proteobacteria; Gammaproteobacteria; Enterobacterales; Enterobacteriaceae; Salmonella; Salmonella enterica; Salmonella enterica subsp. enterica MNNEETFYQAMRRKGVTRRSFLKFCSLAATSLGLGAGMTPKIAWALENKPRIPVVWIHGL WP_001058312.1 hydrogenase [Salmonella enterica] Length: 372\nNumber of predicted TMHs: 2\nExp number of AAs in TMHs: 36.51738\nExp number, first 60 AAs: 19.72849\nTotal prob of N-in: 0.96170\nPOSSIBLE N-term signal sequence\ninside 1 19\nTMhelix 20 39\noutside 40 325\nTMhelix 326 348\ninside 349 372

19213 GCF_001833825.1_ASM183382v1 Salmonella enterica subsp. enterica serovar Rubislaw Proteobacteria; Gammaproteobacteria; Enterobacterales; Enterobacteriaceae; Salmonella; Salmonella enterica; Salmonella enterica subsp. enterica MNNEETFYQAMRRKGVTRRSFLKFCSLAATSLGLGAGMTPKIAWALENKPRIPVVWIHGL WP_001058312.1 hydrogenase [Salmonella enterica] Length: 372\nNumber of predicted TMHs: 2\nExp number of AAs in TMHs: 36.51738\nExp number, first 60 AAs: 19.72849\nTotal prob of N-in: 0.96170\nPOSSIBLE N-term signal sequence\ninside 1 19\nTMhelix 20 39\noutside 40 325\nTMhelix 326 348\ninside 349 372

19214 GCF_002107635.1_ASM210763v1 Salmonella enterica subsp. enterica serovar Rubislaw Proteobacteria; Gammaproteobacteria; Enterobacterales; Enterobacteriaceae; Salmonella; Salmonella enterica; Salmonella enterica subsp. enterica MTGDNTLITSHGINRRDFMKLCAALAATMGLSSKAAAEMAESVSNPQRPPVIWIGAQECT WP_000145429.1 hydrogenase 2 small subunit [Salmonella enterica] Length: 372\nNumber of predicted TMHs: 1\nExp number of AAs in TMHs: 24.51309\nExp number, first 60 AAs: 0.2682\nTotal prob of N-in: 0.14402\noutside 1 331\nTMhelix 332 354\ninside 355 372

19215 GCF_001835135.1_ASM183513v1 Salmonella enterica subsp. enterica serovar Rubislaw Proteobacteria; Gammaproteobacteria; Enterobacterales; Enterobacteriaceae; Salmonella; Salmonella enterica; Salmonella enterica subsp. enterica MNNEETFYQAMRRKGVTRRSFLKFCSLAATSLGLGAGMTPKIAWALENKPRIPVVWIHGL WP_001058312.1 hydrogenase [Salmonella enterica] Length: 372\nNumber of predicted TMHs: 2\nExp number of AAs in TMHs: 36.51738\nExp number, first 60 AAs: 19.72849\nTotal prob of N-in: 0.96170\nPOSSIBLE N-term signal sequence\ninside 1 19\nTMhelix 20 39\noutside 40 325\nTMhelix 326 348\ninside 349 372

19216 GCF_002107635.1_ASM210763v1 Salmonella enterica subsp. enterica serovar Rubislaw Proteobacteria; Gammaproteobacteria; Enterobacterales; Enterobacteriaceae; Salmonella; Salmonella enterica; Salmonella enterica subsp. enterica MNNEETFYQAMRRKGVTRRSFLKFCSLAATSLGLGAGMTPKIAWALENKPRIPVVWIHGL WP_001058312.1 hydrogenase [Salmonella enterica] Length: 372\nNumber of predicted TMHs: 2\nExp number of AAs in TMHs: 36.51738\nExp number, first 60 AAs: 19.72849\nTotal prob of N-in: 0.96170\nPOSSIBLE N-term signal sequence\ninside 1 19\nTMhelix 20 39\noutside 40 325\nTMhelix 326 348\ninside 349 372

19217 GCF_002034845.1_ASM203484v1 Salmonella enterica subsp. enterica serovar Rubislaw Proteobacteria; Gammaproteobacteria; Enterobacterales; Enterobacteriaceae; Salmonella; Salmonella enterica; Salmonella enterica subsp. enterica MNNEETFYQAMRRKGVTRRSFLKFCSLAATSLGLGAGMTPKIAWALENKPRIPVVWIHGL WP_001058312.1 hydrogenase [Salmonella enterica] Length: 372\nNumber of predicted TMHs: 2\nExp number of AAs in TMHs: 36.51738\nExp number, first 60 AAs: 19.72849\nTotal prob of N-in: 0.96170\nPOSSIBLE N-term signal sequence\ninside 1 19\nTMhelix 20 39\noutside 40 325\nTMhelix 326 348\ninside 349 372

19218 GCF_001835405.1_ASM183540v1 Salmonella enterica subsp. enterica serovar Rubislaw Proteobacteria; Gammaproteobacteria; Enterobacterales; Enterobacteriaceae; Salmonella; Salmonella enterica; Salmonella enterica subsp. enterica MTGDNTLITSHGINRRDFMKLCAALAATMGLSSKAAAEMAESVSNPQRPPVIWIGAQECT WP_000145429.1 hydrogenase 2 small subunit [Salmonella enterica] Length: 372\nNumber of predicted TMHs: 1\nExp number of AAs in TMHs: 24.51309\nExp number, first 60 AAs: 0.2682\nTotal prob of N-in: 0.14402\noutside 1 331\nTMhelix 332 354\ninside 355 372

19219 GCF_001833735.1_ASM183373v1 Salmonella enterica subsp. enterica serovar Rubislaw Proteobacteria; Gammaproteobacteria; Enterobacterales; Enterobacteriaceae; Salmonella; Salmonella enterica; Salmonella enterica subsp. enterica MTGDNTLITSHGINRRDFMKLCAALAATMGLSSKAAAEMAESVSNPQRPPVIWIGAQECT WP_000145429.1 hydrogenase 2 small subunit [Salmonella enterica] Length: 372\nNumber of predicted TMHs: 1\nExp number of AAs in TMHs: 24.51309\nExp number, first 60 AAs: 0.2682\nTotal prob of N-in: 0.14402\noutside 1 331\nTMhelix 332 354\ninside 355 372

19220 GCF_002034845.1_ASM203484v1 Salmonella enterica subsp. enterica serovar Rubislaw Proteobacteria; Gammaproteobacteria; Enterobacterales; Enterobacteriaceae; Salmonella; Salmonella enterica; Salmonella enterica subsp. enterica MTGDNTLITSHGINRRDFMKLCAALAATMGLSSKAAAEMAESVSNPQRPPVIWIGAQECT WP_000145429.1 hydrogenase 2 small subunit [Salmonella enterica] Length: 372\nNumber of

predicted TMHs: 1\nExp number of AAs in TMHs: 24.51309\nExp number, first 60 AAs: 0.2682\nTotal prob of N-in: 0.14402\noutside 1 331\nTMhelix 332 354\ninside 355 372

19221 GCF_001835305.1_ASM183530v1 Salmonella enterica subsp. enterica serovar Rubislaw Proteobacteria; Gammaproteobacteria; Enterobacterales; Enterobacteriaceae; Salmonella; Salmonella enterica; Salmonella enterica subsp. enterica MTGDNTLITSHGINRRDFMKLCAALAATMGLSSKAAAEMAESVSNPQRPPVIWIGAQECT WP_000145429.1 hydrogenase 2 small subunit [Salmonella enterica] Length: 372\nNumber of predicted TMHs: 1\nExp number of AAs in TMHs: 24.51309\nExp number, first 60 AAs: 0.2682\nTotal prob of N-in: 0.14402\noutside 1 331\nTMhelix 332 354\ninside 355 372

19222 GCF_001835405.1_ASM183540v1 Salmonella enterica subsp. enterica serovar Rubislaw Proteobacteria; Gammaproteobacteria; Enterobacterales; Enterobacteriaceae; Salmonella; Salmonella enterica; Salmonella enterica subsp. enterica MNNEETFYQAMRRKGVTRRSFLKFCSLAATSLGLGAGMTPKIAWALENKPRIPVVWIHGL WP_001058312.1 hydrogenase [Salmonella enterica] Length: 372\nNumber of predicted TMHs: 2\nExp number of AAs in TMHs: 36.51738\nExp number, first 60 AAs: 19.72849\nTotal prob of N-in: 0.96170\nPOSSIBLE N-term signal sequence\ninside 1 19\nTMhelix 20 39\noutside 40 325\nTMhelix 326 348\ninside 349 372

19223 GCF_001835305.1_ASM183530v1 Salmonella enterica subsp. enterica serovar Rubislaw Proteobacteria; Gammaproteobacteria; Enterobacterales; Enterobacteriaceae; Salmonella; Salmonella enterica; Salmonella enterica subsp. enterica MNNEETFYQAMRRKGVTRRSFLKFCSLAATSLGLGAGMTPKIAWALENKPRIPVVWIHGL WP_001058312.1 hydrogenase [Salmonella enterica] Length: 372\nNumber of predicted TMHs: 2\nExp number of AAs in TMHs: 36.51738\nExp number, first 60 AAs: 19.72849\nTotal prob of N-in: 0.96170\nPOSSIBLE N-term signal sequence\ninside 1 19\nTMhelix 20 39\noutside 40 325\nTMhelix 326 348\ninside 349 372

19224 GCF_001835135.1_ASM183513v1 Salmonella enterica subsp. enterica serovar Rubislaw Proteobacteria; Gammaproteobacteria; Enterobacterales; Enterobacteriaceae; Salmonella; Salmonella enterica; Salmonella enterica subsp. enterica MTGDNTLITSHGINRRDFMKLCAALAATMGLSSKAAAEMAESVSNPQRPPVIWIGAQECT WP_000145429.1 hydrogenase 2 small subunit [Salmonella enterica] Length: 372\nNumber of predicted TMHs: 1\nExp number of AAs in TMHs: 24.51309\nExp number, first 60 AAs: 0.2682\nTotal prob of N-in: 0.14402\noutside 1 331\nTMhelix 332 354\ninside 355 372

19225 GCF_001833735.1_ASM183373v1 Salmonella enterica subsp. enterica serovar Rubislaw Proteobacteria; Gammaproteobacteria; Enterobacterales; Enterobacteriaceae; Salmonella; Salmonella enterica; Salmonella enterica subsp. enterica MNNEETFYQAMRRKGVTRRSFLKFCSLAATSLGLGAGMTPKIAWALENKPRIPVVWIHGL WP_001058312.1 hydrogenase [Salmonella enterica] Length: 372\nNumber of predicted TMHs: 2\nExp number of AAs in TMHs: 36.51738\nExp number, first 60 AAs: 19.72849\nTotal prob of N-in: 0.96170\nPOSSIBLE N-term signal sequence\ninside 1 19\nTMhelix 20 39\noutside 40 325\nTMhelix 326 348\ninside 349 372

19226 GCF_001833925.1_ASM183392v1 Salmonella enterica subsp. enterica serovar Rubislaw Proteobacteria; Gammaproteobacteria; Enterobacterales; Enterobacteriaceae; Salmonella; Salmonella enterica; Salmonella enterica subsp. enterica MTGDNTLITSHGINRRDFMKLCAALAATMGLSSKAAAEMAESVSNPQRPPVIWIGAQECT WP_000145429.1 hydrogenase 2 small subunit [Salmonella enterica] Length: 372\nNumber of predicted TMHs: 1\nExp number of AAs in TMHs: 24.51309\nExp number, first 60 AAs: 0.2682\nTotal prob of N-in: 0.14402\noutside 1 331\nTMhelix 332 354\ninside 355 372

19227 GCF_001835365.1_ASM183536v1 Salmonella enterica subsp. enterica serovar Rubislaw Proteobacteria; Gammaproteobacteria; Enterobacterales; Enterobacteriaceae; Salmonella; Salmonella enterica; Salmonella enterica subsp. enterica MTGDNTLITSHGINRRDFMKLCAALAATMGLSSKAAAEMAESVSNPQRPPVIWIGAQECT WP_000145429.1 hydrogenase 2 small subunit [Salmonella enterica] Length: 372\nNumber of predicted TMHs: 1\nExp number of AAs in TMHs: 24.51309\nExp number, first 60 AAs: 0.2682\nTotal prob of N-in: 0.14402\noutside 1 331\nTMhelix 332 354\ninside 355 372

19228 GCF_001834285.1_ASM183428v1 Salmonella enterica subsp. enterica serovar Rubislaw Proteobacteria; Gammaproteobacteria; Enterobacterales; Enterobacteriaceae; Salmonella; Salmonella enterica; Salmonella enterica subsp. enterica MTGDNTLITSHGINRRDFMKLCAALAATMGLSSKAAAEMAESVSNPQRPPVIWIGAQECT WP_000145429.1 hydrogenase 2 small subunit [Salmonella enterica] Length: 372\nNumber of predicted TMHs: 1\nExp number of AAs in TMHs: 24.51309\nExp number, first 60 AAs: 0.2682\nTotal prob of N-in: 0.14402\noutside 1 331\nTMhelix 332 354\ninside 355 372

19229 GCF_001835155.1_ASM183515v1 Salmonella enterica subsp. enterica serovar Rubislaw Proteobacteria; Gammaproteobacteria; Enterobacterales; Enterobacteriaceae; Salmonella; Salmonella enterica;

Salmonella enterica subsp. enterica MNNEETFYQAMRRKGVTRRSFLKFCSLAATSLGLGAGMTPKIAWALENKPRIPVVWIHGL WP_001058312.1 hydrogenase [Salmonella enterica] Length: 372\nNumber of predicted TMHs: 2\nExp number of AAs in TMHs: 36.51738\nExp number, first 60 AAs: 19.72849\nTotal prob of N-in: 0.96170\nPOSSIBLE N-term signal sequence\ninside 1 19\nTMhelix 20 39\noutside 40 325\nTMhelix 326 348\ninside 349 372

19230 GCF_001832545.1_ASM183254v1 Salmonella enterica subsp. enterica serovar Rubislaw Proteobacteria; Gammaproteobacteria; Enterobacterales; Enterobacteriaceae; Salmonella; Salmonella enterica; Salmonella enterica subsp. enterica MNNEETFYQAMRRKGVTRRSFLKFCSLAATSLGLGAGMTPKIAWALENKPRIPVVWIHGL WP_001058312.1 hydrogenase [Salmonella enterica] Length: 372\nNumber of predicted TMHs: 2\nExp number of AAs in TMHs: 36.51738\nExp number, first 60 AAs: 19.72849\nTotal prob of N-in: 0.96170\nPOSSIBLE N-term signal sequence\ninside 1 19\nTMhelix 20 39\noutside 40 325\nTMhelix 326 348\ninside 349 372

19231 GCF_001832545.1_ASM183254v1 Salmonella enterica subsp. enterica serovar Rubislaw Proteobacteria; Gammaproteobacteria; Enterobacterales; Enterobacteriaceae; Salmonella; Salmonella enterica; Salmonella enterica subsp. enterica MTGDNTLITSHGINRRDFMKLCAALAATMGLSSKAAAEMAESVSNPQRPPVIWIGAQECT WP_000145429.1 hydrogenase 2 small subunit [Salmonella enterica] Length: 372\nNumber of predicted TMHs: 1\nExp number of AAs in TMHs: 24.51309\nExp number, first 60 AAs: 0.2682\nTotal prob of N-in: 0.14402\noutside 1 331\nTMhelix 332 354\ninside 355 372

19232 GCF_001834625.1_ASM183462v1 Salmonella enterica subsp. enterica serovar Rubislaw Proteobacteria; Gammaproteobacteria; Enterobacterales; Enterobacteriaceae; Salmonella; Salmonella enterica; Salmonella enterica subsp. enterica MTGDNTLITSHGINRRDFMKLCAALAATMGLSSKAAAEMAESVSNPQRPPVIWIGAQECT WP_000145429.1 hydrogenase 2 small subunit [Salmonella enterica] Length: 372\nNumber of predicted TMHs: 1\nExp number of AAs in TMHs: 24.51309\nExp number, first 60 AAs: 0.2682\nTotal prob of N-in: 0.14402\noutside 1 331\nTMhelix 332 354\ninside 355 372

19233 GCF_001833725.1_ASM183372v1 Salmonella enterica subsp. enterica serovar Rubislaw Proteobacteria; Gammaproteobacteria; Enterobacterales; Enterobacteriaceae; Salmonella; Salmonella enterica; Salmonella enterica subsp. enterica MNNEETFYQAMRRKGVTRRSFLKFCSLAATSLGLGAGMTPKIAWALENKPRIPVVWIHGL WP_001058312.1 hydrogenase [Salmonella enterica] Length: 372\nNumber of predicted TMHs: 2\nExp number of AAs in TMHs: 36.51738\nExp number, first 60 AAs: 19.72849\nTotal prob of N-in: 0.96170\nPOSSIBLE N-term signal sequence\ninside 1 19\nTMhelix 20 39\noutside 40 325\nTMhelix 326 348\ninside 349 372

19234 GCF_001834285.1_ASM183428v1 Salmonella enterica subsp. enterica serovar Rubislaw Proteobacteria; Gammaproteobacteria; Enterobacterales; Enterobacteriaceae; Salmonella; Salmonella enterica; Salmonella enterica subsp. enterica MNNEETFYQAMRRKGVTRRSFLKFCSLAATSLGLGAGMTPKIAWALENKPRIPVVWIHGL WP_001058312.1 hydrogenase [Salmonella enterica] Length: 372\nNumber of predicted TMHs: 2\nExp number of AAs in TMHs: 36.51738\nExp number, first 60 AAs: 19.72849\nTotal prob of N-in: 0.96170\nPOSSIBLE N-term signal sequence\ninside 1 19\nTMhelix 20 39\noutside 40 325\nTMhelix 326 348\ninside 349 372

19235 GCF_001834625.1_ASM183462v1 Salmonella enterica subsp. enterica serovar Rubislaw Proteobacteria; Gammaproteobacteria; Enterobacterales; Enterobacteriaceae; Salmonella; Salmonella enterica; Salmonella enterica subsp. enterica MNNEETFYQAMRRKGVTRRSFLKFCSLAATSLGLGAGMTPKIAWALENKPRIPVVWIHGL WP_001058312.1 hydrogenase [Salmonella enterica] Length: 372\nNumber of predicted TMHs: 2\nExp number of AAs in TMHs: 36.51738\nExp number, first 60 AAs: 19.72849\nTotal prob of N-in: 0.96170\nPOSSIBLE N-term signal sequence\ninside 1 19\nTMhelix 20 39\noutside 40 325\nTMhelix 326 348\ninside 349 372

19236 GCF_001835365.1_ASM183536v1 Salmonella enterica subsp. enterica serovar Rubislaw Proteobacteria; Gammaproteobacteria; Enterobacterales; Enterobacteriaceae; Salmonella; Salmonella enterica; Salmonella enterica subsp. enterica MNNEETFYQAMRRKGVTRRSFLKFCSLAATSLGLGAGMTPKIAWALENKPRIPVVWIHGL WP_001058312.1 hydrogenase [Salmonella enterica] Length: 372\nNumber of predicted TMHs: 2\nExp number of AAs in TMHs: 36.51738\nExp number, first 60 AAs: 19.72849\nTotal prob of N-in: 0.96170\nPOSSIBLE N-term signal sequence\ninside 1 19\nTMhelix 20 39\noutside 40 325\nTMhelix 326 348\ninside 349 372

19237 GCF_001833705.1_ASM183370v1 Salmonella enterica subsp. enterica serovar Rubislaw Proteobacteria; Gammaproteobacteria; Enterobacterales; Enterobacteriaceae; Salmonella; Salmonella enterica; Salmonella enterica subsp. enterica MNNEETFYQAMRRKGVTRRSFLKFCSLAATSLGLGAGMTPKIAWALENKPRIPVVWIHGL WP_001058312.1 hydrogenase [Salmonella enterica] Length: 372\nNumber of predicted TMHs: 2\nExp number of AAs in TMHs: 36.51738\nExp number, first 60 AAs: 19.72849\nTotal prob of N-in: 0.96170\nPOSSIBLE N-term signal sequence\ninside 1 19\nTMhelix 20 39\noutside 40 325\nTMhelix 326 348\ninside 349 372

19238 GCF_001833705.1_ASM183370v1 Salmonella enterica subsp. enterica serovar Rubislaw Proteobacteria; Gammaproteobacteria; Enterobacterales; Enterobacteriaceae; Salmonella; Salmonella enterica; Salmonella enterica subsp. enterica MTGDNTLITSHGINRRDFMKLCAALAATMGLSSKAAAEMAESVSNPQRPPVIWIGAQECT WP_000145429.1 hydrogenase 2 small subunit [Salmonella enterica] Length: 372\nNumber of predicted TMHs: 1\nExp number of AAs in TMHs: 24.51309\nExp number, first 60 AAs: 0.2682\nTotal prob of N-in: 0.14402\noutside 1 331\nTMhelix 332 354\ninside 355 372

19239 GCF_001833825.1_ASM183382v1 Salmonella enterica subsp. enterica serovar Rubislaw Proteobacteria; Gammaproteobacteria; Enterobacterales; Enterobacteriaceae; Salmonella; Salmonella enterica; Salmonella enterica subsp. enterica MTGDNTLITSHGINRRDFMKLCAALAATMGLSSKAAAEMAESVSNPQRPPVIWIGAQECT WP_000145429.1 hydrogenase 2 small subunit [Salmonella enterica] Length: 372\nNumber of predicted TMHs: 1\nExp number of AAs in TMHs: 24.51309\nExp number, first 60 AAs: 0.2682\nTotal prob of N-in: 0.14402\noutside 1 331\nTMhelix 332 354\ninside 355 372

19240 GCF_001833725.1_ASM183372v1 Salmonella enterica subsp. enterica serovar Rubislaw Proteobacteria; Gammaproteobacteria; Enterobacterales; Enterobacteriaceae; Salmonella; Salmonella enterica; Salmonella enterica subsp. enterica MTGDNTLITSHGINRRDFMKLCAALAATMGLSSKAAAEMAESVSNPQRPPVIWIGAQECT WP_000145429.1 hydrogenase 2 small subunit [Salmonella enterica] Length: 372\nNumber of predicted TMHs: 1\nExp number of AAs in TMHs: 24.51309\nExp number, first 60 AAs: 0.2682\nTotal prob of N-in: 0.14402\noutside 1 331\nTMhelix 332 354\ninside 355 372

19241 GCF_001834655.1_ASM183465v1 Salmonella enterica subsp. enterica serovar Rubislaw Proteobacteria; Gammaproteobacteria; Enterobacterales; Enterobacteriaceae; Salmonella; Salmonella enterica; Salmonella enterica subsp. enterica MTGDNTLITSHGINRRDFMKLCAALAATMGLSSKAAAEMAESVSNPQRPPVIWIGAQECT WP_000145429.1 hydrogenase 2 small subunit [Salmonella enterica] Length: 372\nNumber of predicted TMHs: 1\nExp number of AAs in TMHs: 24.51309\nExp number, first 60 AAs: 0.2682\nTotal prob of N-in: 0.14402\noutside 1 331\nTMhelix 332 354\ninside 355 372

19242 GCF_001244935.1_Salmonella_enterica_CVM_N48698_v1.0 Salmonella enterica subsp. enterica serovar Saintpaul Proteobacteria; Gammaproteobacteria; Enterobacterales; Enterobacteriaceae; Salmonella; Salmonella enterica; Salmonella enterica subsp. enterica MNNEETFYQAMRRKGVTRRSFLKFCSLAATSLGLGAGMTPKIAWALENKPRIPVVWIHGL WP_001058312.1 hydrogenase [Salmonella enterica] Length: 372\nNumber of predicted TMHs: 2\nExp number of AAs in TMHs: 36.51738\nExp number, first 60 AAs: 19.72849\nTotal prob of N-in: 0.96170\nPOSSIBLE N-term signal sequence\ninside 1 19\nTMhelix 20 39\noutside 40 325\nTMhelix 326 348\ninside 349 372

19243 GCF_001245345.1_Salmonella_enterica_CVM_N50422_v1.0 Salmonella enterica subsp. enterica serovar Saintpaul Proteobacteria; Gammaproteobacteria; Enterobacterales; Enterobacteriaceae; Salmonella; Salmonella enterica; Salmonella enterica subsp. enterica MNNEETFYQAMRRKGVTRRSFLKFCSLAATSLGLGAGMTPKIAWALENKPRIPVVWIHGL WP_001058312.1 hydrogenase [Salmonella enterica] Length: 372\nNumber of predicted TMHs: 2\nExp number of AAs in TMHs: 36.51738\nExp number, first 60 AAs: 19.72849\nTotal prob of N-in: 0.96170\nPOSSIBLE N-term signal sequence\ninside 1 19\nTMhelix 20 39\noutside 40 325\nTMhelix 326 348\ninside 349 372

19244 GCF_001834125.1_ASM183412v1 Salmonella enterica subsp. enterica serovar Saintpaul Proteobacteria; Gammaproteobacteria; Enterobacterales; Enterobacteriaceae; Salmonella; Salmonella enterica; Salmonella enterica subsp. enterica MTGDNTLITSHGINRRDFMKLCAALAATMGLSSKAAAEMAESVSNPQRPPVIWIGAQECT WP_000145429.1 hydrogenase 2 small subunit [Salmonella enterica] Length: 372\nNumber of predicted TMHs: 1\nExp number of AAs in TMHs: 24.51309\nExp number, first 60 AAs: 0.2682\nTotal prob of N-in: 0.14402\noutside 1 331\nTMhelix 332 354\ninside 355 372

19245 GCF_001833345.1_ASM183334v1 Salmonella enterica subsp. enterica serovar Saintpaul Proteobacteria; Gammaproteobacteria; Enterobacterales; Enterobacteriaceae; Salmonella; Salmonella enterica; Salmonella enterica subsp. enterica MTGDNTLITSHGINRRDFMKLCAALAATMGLSSKAAAEMAESVSNPQRPPVIWIGAQECT WP_000145429.1 hydrogenase 2 small subunit [Salmonella enterica] Length: 372\nNumber of predicted TMHs: 1\nExp number of AAs in TMHs: 24.51309\nExp number, first 60 AAs: 0.2682\nTotal prob of N-in: 0.14402\noutside 1 331\nTMhelix 332 354\ninside 355 372

19246 GCF_001953675.1_ASM195367v1 Salmonella enterica subsp. enterica serovar Saintpaul Proteobacteria; Gammaproteobacteria; Enterobacterales; Enterobacteriaceae; Salmonella; Salmonella enterica; Salmonella enterica subsp. enterica MNNEETFYQAMRRKGVTRRSFLKFCSLAATSLGLGAGMTPKIAWALENKPRIPVVWIHGL WP_001058312.1 hydrogenase [Salmonella enterica] Length: 372\nNumber of predicted TMHs: 2\nExp

number of AAs in TMHs: 36.51738\nExp number, first 60 AAs: 19.72849\nTotal prob of N-in: 0.96170\nPOSSIBLE N-term signal sequence\ninside 1 19\nTMhelix 20 39\noutside 40 325\nTMhelix 326 348\ninside 349 372

19247 GCF_002031105.1_ASM203110v1 Salmonella enterica subsp. enterica serovar Saintpaul Proteobacteria; Gammaproteobacteria; Enterobacterales; Enterobacteriaceae; Salmonella; Salmonella enterica; Salmonella enterica subsp. enterica MTGDNTLITSHGINRRDFMKLCAALAATMGLSSKAAAEMAESVSNPQRPPVIWIGAQECT WP_000145429.1 hydrogenase 2 small subunit [Salmonella enterica] Length: 372\nNumber of predicted TMHs: 1\nExp number of AAs in TMHs: 24.51309\nExp number, first 60 AAs: 0.2682\nTotal prob of N-in: 0.14402\noutside 1 331\nTMhelix 332 354\ninside 355 372

19248 GCF_002031105.1_ASM203110v1 Salmonella enterica subsp. enterica serovar Saintpaul Proteobacteria; Gammaproteobacteria; Enterobacterales; Enterobacteriaceae; Salmonella; Salmonella enterica; Salmonella enterica subsp. enterica MNNEETFYQAMRRKGVTRRSFLKFCSLAATSLGLGAGMTPKIAWALENKPRIPVVWIHGL WP_001058312.1 hydrogenase [Salmonella enterica] Length: 372\nNumber of predicted TMHs: 2\nExp number of AAs in TMHs: 36.51738\nExp number, first 60 AAs: 19.72849\nTotal prob of N-in: 0.96170\nPOSSIBLE N-term signal sequence\ninside 1 19\nTMhelix 20 39\noutside 40 325\nTMhelix 326 348\ninside 349 372

19249 GCF_001241665.1_Salmonella_enterica_CVM_N45394_v1.0 Salmonella enterica subsp. enterica serovar Saintpaul Proteobacteria; Gammaproteobacteria; Enterobacterales; Enterobacteriaceae; Salmonella; Salmonella enterica; Salmonella enterica subsp. enterica MTGDNTLITSHGINRRDFMKLCAALAATMGLSSKAAAEMAESVSNPQRPPVIWIGAQECT WP_000145429.1 hydrogenase 2 small subunit [Salmonella enterica] Length: 372\nNumber of predicted TMHs: 1\nExp number of AAs in TMHs: 24.51309\nExp number, first 60 AAs: 0.2682\nTotal prob of N-in: 0.14402\noutside 1 331\nTMhelix 332 354\ninside 355 372

19250 GCF_002031465.1_ASM203146v1 Salmonella enterica subsp. enterica serovar Saintpaul Proteobacteria; Gammaproteobacteria; Enterobacterales; Enterobacteriaceae; Salmonella; Salmonella enterica; Salmonella enterica subsp. enterica MNNEETFYQAMRRKGVTRRSFLKFCSLAATSLGLGAGMTPKIAWALENKPRIPVVWIHGL WP_001058312.1 hydrogenase [Salmonella enterica] Length: 372\nNumber of predicted TMHs: 2\nExp number of AAs in TMHs: 36.51738\nExp number, first 60 AAs: 19.72849\nTotal prob of N-in: 0.96170\nPOSSIBLE N-term signal sequence\ninside 1 19\nTMhelix 20 39\noutside 40 325\nTMhelix 326 348\ninside 349 372

19251 GCF_001243085.1_Salmonella_enterica_CVM_N46823_v1.0 Salmonella enterica subsp. enterica serovar Saintpaul Proteobacteria; Gammaproteobacteria; Enterobacterales; Enterobacteriaceae; Salmonella; Salmonella enterica; Salmonella enterica subsp. enterica MTGDNTLITSHGINRRDFMKLCAALAATMGLSSKAAAEMAESVSNPQRPPVIWIGAQECT WP_000145429.1 hydrogenase 2 small subunit [Salmonella enterica] Length: 372\nNumber of predicted TMHs: 1\nExp number of AAs in TMHs: 24.51309\nExp number, first 60 AAs: 0.2682\nTotal prob of N-in: 0.14402\noutside 1 331\nTMhelix 332 354\ninside 355 372

19252 GCF_001243445.1_Salmonella_enterica_CVM_N46846-R_v1.0 Salmonella enterica subsp. enterica serovar Saintpaul Proteobacteria; Gammaproteobacteria; Enterobacterales; Enterobacteriaceae; Salmonella; Salmonella enterica; Salmonella enterica subsp. enterica MTGDNTLITSHGINRRDFMKLCAALAATMGLSSKAAAEMAESVSNPQRPPVIWIGAQECT WP_000145429.1 hydrogenase 2 small subunit [Salmonella enterica] Length: 372\nNumber of predicted TMHs: 1\nExp number of AAs in TMHs: 24.51309\nExp number, first 60 AAs: 0.2682\nTotal prob of N-in: 0.14402\noutside 1 331\nTMhelix 332 354\ninside 355 372

19253 GCF_001243775.1_Salmonella_enterica_CVM_N47713_v1.0 Salmonella enterica subsp. enterica serovar Saintpaul Proteobacteria; Gammaproteobacteria; Enterobacterales; Enterobacteriaceae; Salmonella; Salmonella enterica; Salmonella enterica subsp. enterica MNNEETFYQAMRRKGVTRRSFLKFCSLAATSLGLGAGMTPKIAWALENKPRIPVVWIHGL WP_001058312.1 hydrogenase [Salmonella enterica] Length: 372\nNumber of predicted TMHs: 2\nExp number of AAs in TMHs: 36.51738\nExp number, first 60 AAs: 19.72849\nTotal prob of N-in: 0.96170\nPOSSIBLE N-term signal sequence\ninside 1 19\nTMhelix 20 39\noutside 40 325\nTMhelix 326 348\ninside 349 372

19254 GCF_001243905.1_Salmonella_enterica_CVM_N47719_v1.0 Salmonella enterica subsp. enterica serovar Saintpaul Proteobacteria; Gammaproteobacteria; Enterobacterales; Enterobacteriaceae; Salmonella; Salmonella enterica; Salmonella enterica subsp. enterica MTGDNTLITSHGINRRDFMKLCAALAATMGLSSKAAAEMAESVSNPQRPPVIWIGAQECT WP_000145429.1 hydrogenase 2 small subunit [Salmonella enterica] Length: 372\nNumber of predicted TMHs: 1\nExp number of AAs in TMHs: 24.51309\nExp number, first 60 AAs: 0.2682\nTotal prob of N-in: 0.14402\noutside 1 331\nTMhelix 332 354\ninside 355 372

19255 GCF_001243905.1_Salmonella_enterica_CVM_N47719_v1.0 Salmonella enterica subsp. enterica serovar Saintpaul Proteobacteria; Gammaproteobacteria; Enterobacterales; Enterobacteriaceae; Salmonella; Salmonella enterica;

Salmonella enterica subsp. enterica MNNEETFYQAMRRKGVTRRSFLKFCSLAATSLGLGAGMTPKIAWALENKPRIPVVWIHGL WP_001058312.1 hydrogenase [Salmonella enterica] Length: 372\nNumber of predicted TMHs: 2\nExp number of AAs in TMHs: 36.51738\nExp number, first 60 AAs: 19.72849\nTotal prob of N-in: 0.96170\nPOSSIBLE N-term signal sequence\ninside 1 19\nTMhelix 20 39\noutside 40 325\nTMhelix 326 348\ninside 349 372

19256 GCF_002032725.1_ASM203272v1 Salmonella enterica subsp. enterica serovar Saintpaul Proteobacteria; Gammaproteobacteria; Enterobacterales; Enterobacteriaceae; Salmonella; Salmonella enterica; Salmonella enterica subsp. enterica MNNEETFYQAMRRKGVTRRSFLKFCSLAATSLGLGAGMTPKIAWALENKPRIPVVWIHGL WP_001058312.1 hydrogenase [Salmonella enterica] Length: 372\nNumber of predicted TMHs: 2\nExp number of AAs in TMHs: 36.51738\nExp number, first 60 AAs: 19.72849\nTotal prob of N-in: 0.96170\nPOSSIBLE N-term signal sequence\ninside 1 19\nTMhelix 20 39\noutside 40 325\nTMhelix 326 348\ninside 349 372

19257 GCF_001246025.1_Salmonella_enterica_CVM_N51245_v1.0 Salmonella enterica subsp. enterica serovar Saintpaul Proteobacteria; Gammaproteobacteria; Enterobacterales; Enterobacteriaceae; Salmonella; Salmonella enterica; Salmonella enterica subsp. enterica MTGDNTLITSHGINRRDFMKLCAALAATMGLSSKAAAEMAESVSNPQRPPVIWIGAQECT WP_000145429.1 hydrogenase 2 small subunit [Salmonella enterica] Length: 372\nNumber of predicted TMHs: 1\nExp number of AAs in TMHs: 24.51309\nExp number, first 60 AAs: 0.2682\nTotal prob of N-in: 0.14402\noutside 1 331\nTMhelix 332 354\ninside 355 372

19258 GCF_001478155.1_Salmonella_enterica_CVM_N31414-SQ_v1.0 Salmonella enterica subsp. enterica serovar Saintpaul Proteobacteria; Gammaproteobacteria; Enterobacterales; Enterobacteriaceae; Salmonella; Salmonella enterica; Salmonella enterica subsp. enterica MNNEETFYQAMRRKGVTRRSFLKFCSLAATSLGLGAGMTPKIAWALENKPRIPVVWIHGL WP_001058312.1 hydrogenase [Salmonella enterica] Length: 372\nNumber of predicted TMHs: 2\nExp number of AAs in TMHs: 36.51738\nExp number, first 60 AAs: 19.72849\nTotal prob of N-in: 0.96170\nPOSSIBLE N-term signal sequence\ninside 1 19\nTMhelix 20 39\noutside 40 325\nTMhelix 326 348\ninside 349 372

19259 GCF_001247105.1_Salmonella_enterica_CVM_N51291_v1.0 Salmonella enterica subsp. enterica serovar Saintpaul Proteobacteria; Gammaproteobacteria; Enterobacterales; Enterobacteriaceae; Salmonella; Salmonella enterica; Salmonella enterica subsp. enterica MTGDNTLITSHGINRRDFMKLCAALAATMGLSSKAAAEMAESVSNPQRPPVIWIGAQECT WP_000145429.1 hydrogenase 2 small subunit [Salmonella enterica] Length: 372\nNumber of predicted TMHs: 1\nExp number of AAs in TMHs: 24.51309\nExp number, first 60 AAs: 0.2682\nTotal prob of N-in: 0.14402\noutside 1 331\nTMhelix 332 354\ninside 355 372

19260 GCF_001834125.1_ASM183412v1 Salmonella enterica subsp. enterica serovar Saintpaul Proteobacteria; Gammaproteobacteria; Enterobacterales; Enterobacteriaceae; Salmonella; Salmonella enterica; Salmonella enterica subsp. enterica MNNEETFYQAMRRKGVTRRSFLKFCSLAATSLGLGAGMTPKIAWALENKPRIPVVWIHGL WP_001058311.1 hydrogenase [Salmonella enterica] Length: 372\nNumber of predicted TMHs: 2\nExp number of AAs in TMHs: 37.4699899999999\nExp number, first 60 AAs: 19.72699\nTotal prob of N-in: 0.96171\nPOSSIBLE N-term signal sequence\ninside 1 19\nTMhelix 20 39\noutside 40 325\nTMhelix 326 348\ninside 349 372

19261 GCF_001479185.1_Salmonella_enterica_CVM_N31412-SQ_v1.0 Salmonella enterica subsp. enterica serovar Saintpaul Proteobacteria; Gammaproteobacteria; Enterobacterales; Enterobacteriaceae; Salmonella; Salmonella enterica; Salmonella enterica subsp. enterica MNNEETFYQAMRRKGVTRRSFLKFCSLAATSLGLGAGMTPKIAWALENKPRIPVVWIHGL WP_001058312.1 hydrogenase [Salmonella enterica] Length: 372\nNumber of predicted TMHs: 2\nExp number of AAs in TMHs: 36.51738\nExp number, first 60 AAs: 19.72849\nTotal prob of N-in: 0.96170\nPOSSIBLE N-term signal sequence\ninside 1 19\nTMhelix 20 39\noutside 40 325\nTMhelix 326 348\ninside 349 372

19262 GCF_001479185.1_Salmonella_enterica_CVM_N31412-SQ_v1.0 Salmonella enterica subsp. enterica serovar Saintpaul Proteobacteria; Gammaproteobacteria; Enterobacterales; Enterobacteriaceae; Salmonella; Salmonella enterica; Salmonella enterica subsp. enterica MTGDNTLITSHGINRRDFMKLCAALAATMGLSSKAAAEMAESVSNPQRPPVIWIGAQECT WP_000145429.1 hydrogenase 2 small subunit [Salmonella enterica] Length: 372\nNumber of predicted TMHs: 1\nExp number of AAs in TMHs: 24.51309\nExp number, first 60 AAs: 0.2682\nTotal prob of N-in: 0.14402\noutside 1 331\nTMhelix 332 354\ninside 355 372

19263 GCF_001833805.1_ASM183380v1 Salmonella enterica subsp. enterica serovar Saintpaul Proteobacteria; Gammaproteobacteria; Enterobacterales; Enterobacteriaceae; Salmonella; Salmonella enterica; Salmonella enterica subsp. enterica MTGDNTLITSHGINRRDFMKLCAALAATMGLSSKAAAEMAESVSNPQRPPVIWIGAQECT WP_000145429.1 hydrogenase 2 small subunit [Salmonella enterica] Length: 372\nNumber of predicted TMHs: 1\nExp number of AAs in TMHs: 24.51309\nExp number, first 60 AAs: 0.2682\nTotal prob of N-in: 0.14402\noutside 1 331\nTMhelix 332 354\ninside 355 372

19264 GCF_001478765.1_Salmonella_enterica_CVM_N29377-SQ_v1.0 Salmonella enterica subsp. enterica serovar Saintpaul Proteobacteria; Gammaproteobacteria; Enterobacterales; Enterobacteriaceae; Salmonella; Salmonella enterica; Salmonella enterica subsp. enterica MTGDNTLITSHGINRRDFMKLCAALAATMGLSSKAAAEMAESVSNPQRPPVIWIGAQECT WP_000145429.1 hydrogenase 2 small subunit [Salmonella enterica] Length: 372\nNumber of predicted TMHs: 1\nExp number of AAs in TMHs: 24.51309\nExp number, first 60 AAs: 0.2682\nTotal prob of N-in: 0.14402\noutside 1 331\nTMhelix 332 354\ninside 355 372

19265 GCF_001240225.1_Salmonella_enterica_CVM_N43456_v1.0 Salmonella enterica subsp. enterica serovar Saintpaul Proteobacteria; Gammaproteobacteria; Enterobacterales; Enterobacteriaceae; Salmonella; Salmonella enterica; Salmonella enterica subsp. enterica MNNEETFYQAMRRKGVTRRSFLKFCSLAATSLGLGAGMTPKIAWALENKPRIPVVWIHGL WP_001058312.1 hydrogenase [Salmonella enterica] Length: 372\nNumber of predicted TMHs: 2\nExp number of AAs in TMHs: 36.51738\nExp number, first 60 AAs: 19.72849\nTotal prob of N-in: 0.96170\nPOSSIBLE N-term signal sequence\ninside 1 19\nTMhelix 20 39\noutside 40 325\nTMhelix 326 348\ninside 349 372

19266 GCF_001242115.1_Salmonella_enterica_CVM_N45926_v1.0 Salmonella enterica subsp. enterica serovar Saintpaul Proteobacteria; Gammaproteobacteria; Enterobacterales; Enterobacteriaceae; Salmonella; Salmonella enterica; Salmonella enterica subsp. enterica MNNEETFYQAMRRKGVTRRSFLKFCSLAATSLGLGAGMTPKIAWALENKPRIPVVWIHGL WP_001058312.1 hydrogenase [Salmonella enterica] Length: 372\nNumber of predicted TMHs: 2\nExp number of AAs in TMHs: 36.51738\nExp number, first 60 AAs: 19.72849\nTotal prob of N-in: 0.96170\nPOSSIBLE N-term signal sequence\ninside 1 19\nTMhelix 20 39\noutside 40 325\nTMhelix 326 348\ninside 349 372

19267 GCF_001242115.1_Salmonella_enterica_CVM_N45926_v1.0 Salmonella enterica subsp. enterica serovar Saintpaul Proteobacteria; Gammaproteobacteria; Enterobacterales; Enterobacteriaceae; Salmonella; Salmonella enterica; Salmonella enterica subsp. enterica MTGDNTLITSHGINRRDFMKLCAALAATMGLSSKAAAEMAESVSNPQRPPVIWIGAQECT WP_000145429.1 hydrogenase 2 small subunit [Salmonella enterica] Length: 372\nNumber of predicted TMHs: 1\nExp number of AAs in TMHs: 24.51309\nExp number, first 60 AAs: 0.2682\nTotal prob of N-in: 0.14402\noutside 1 331\nTMhelix 332 354\ninside 355 372

19268 GCF_001245335.1_Salmonella_enterica_CVM_N50423_v1.0 Salmonella enterica subsp. enterica serovar Saintpaul Proteobacteria; Gammaproteobacteria; Enterobacterales; Enterobacteriaceae; Salmonella; Salmonella enterica; Salmonella enterica subsp. enterica MTGDNTLITSHGINRRDFMKLCAALAATMGLSSKAAAEMAESVSNPQRPPVIWIGAQECT WP_000145429.1 hydrogenase 2 small subunit [Salmonella enterica] Length: 372\nNumber of predicted TMHs: 1\nExp number of AAs in TMHs: 24.51309\nExp number, first 60 AAs: 0.2682\nTotal prob of N-in: 0.14402\noutside 1 331\nTMhelix 332 354\ninside 355 372

19269 GCF_001240335.1_Salmonella_enterica_CVM_N43461_v1.0 Salmonella enterica subsp. enterica serovar Saintpaul Proteobacteria; Gammaproteobacteria; Enterobacterales; Enterobacteriaceae; Salmonella; Salmonella enterica; Salmonella enterica subsp. enterica MTGDNTLITSHGINRRDFMKLCAALAATMGLSSKAAAEMAESVSNPQRPPVIWIGAQECT WP_000145429.1 hydrogenase 2 small subunit [Salmonella enterica] Length: 372\nNumber of predicted TMHs: 1\nExp number of AAs in TMHs: 24.51309\nExp number, first 60 AAs: 0.2682\nTotal prob of N-in: 0.14402\noutside 1 331\nTMhelix 332 354\ninside 355 372

19270 GCF_001243455.1_Salmonella_enterica_CVM_N46847-R_v1.0 Salmonella enterica subsp. enterica serovar Saintpaul Proteobacteria; Gammaproteobacteria; Enterobacterales; Enterobacteriaceae; Salmonella; Salmonella enterica; Salmonella enterica subsp. enterica MNNEETFYQAMRRKGVTRRSFLKFCSLAATSLGLGAGMTPKIAWALENKPRIPVVWIHGL WP_001058312.1 hydrogenase [Salmonella enterica] Length: 372\nNumber of predicted TMHs: 2\nExp number of AAs in TMHs: 36.51738\nExp number, first 60 AAs: 19.72849\nTotal prob of N-in: 0.96170\nPOSSIBLE N-term signal sequence\ninside 1 19\nTMhelix 20 39\noutside 40 325\nTMhelix 326 348\ninside 349 372

19271 GCF_001243455.1_Salmonella_enterica_CVM_N46847-R_v1.0 Salmonella enterica subsp. enterica serovar Saintpaul Proteobacteria; Gammaproteobacteria; Enterobacterales; Enterobacteriaceae; Salmonella; Salmonella enterica; Salmonella enterica subsp. enterica MTGDNTLITSHGINRRDFMKLCAALAATMGLSSKAAAEMAESVSNPQRPPVIWIGAQECT WP_000145429.1 hydrogenase 2 small subunit [Salmonella enterica] Length: 372\nNumber of predicted TMHs: 1\nExp number of AAs in TMHs: 24.51309\nExp number, first 60 AAs: 0.2682\nTotal prob of N-in: 0.14402\noutside 1 331\nTMhelix 332 354\ninside 355 372

19272 GCF_001240225.1_Salmonella_enterica_CVM_N43456_v1.0 Salmonella enterica subsp. enterica serovar Saintpaul Proteobacteria; Gammaproteobacteria; Enterobacterales; Enterobacteriaceae; Salmonella; Salmonella enterica; Salmonella enterica subsp. enterica MTGDNTLITSHGINRRDFMKLCAALAATMGLSSKAAAEMAESVSNPQRPPVIWIGAQECT WP_000145429.1 hydrogenase 2 small subunit [Salmonella enterica] Length: 372\nNumber of

predicted TMHs: 1\nExp number of AAs in TMHs: 24.51309\nExp number, first 60 AAs: 0.2682\nTotal prob of N-in: 0.14402\noutside 1 331\nTMhelix 332 354\ninside 355 372

19273 GCF_001244055.1_Salmonella_enterica_CVM_N47724_v1.0 Salmonella enterica subsp. enterica serovar Saintpaul Proteobacteria; Gammaproteobacteria; Enterobacterales; Enterobacteriaceae; Salmonella; Salmonella enterica; Salmonella enterica subsp. enterica MTGDNTLITSHGINRRDFMKLCAALAATMGLSSKAAAEMAESVSNPQRPPVIWIGAQECT WP_000145429.1 hydrogenase 2 small subunit [Salmonella enterica] Length: 372\nNumber of predicted TMHs: 1\nExp number of AAs in TMHs: 24.51309\nExp number, first 60 AAs: 0.2682\nTotal prob of N-in: 0.14402\noutside 1 331\nTMhelix 332 354\ninside 355 372

19274 GCF_001244095.1_Salmonella_enterica_CVM_N47725_v1.0 Salmonella enterica subsp. enterica serovar Saintpaul Proteobacteria; Gammaproteobacteria; Enterobacterales; Enterobacteriaceae; Salmonella; Salmonella enterica; Salmonella enterica subsp. enterica MTGDNTLITSHGINRRDFMKLCAALAATMGLSSKAAAEMAESVSNPQRPPVIWIGAQECT WP_000145429.1 hydrogenase 2 small subunit [Salmonella enterica] Length: 372\nNumber of predicted TMHs: 1\nExp number of AAs in TMHs: 24.51309\nExp number, first 60 AAs: 0.2682\nTotal prob of N-in: 0.14402\noutside 1 331\nTMhelix 332 354\ninside 355 372

19275 GCF_001244095.1_Salmonella_enterica_CVM_N47725_v1.0 Salmonella enterica subsp. enterica serovar Saintpaul Proteobacteria; Gammaproteobacteria; Enterobacterales; Enterobacteriaceae; Salmonella; Salmonella enterica; Salmonella enterica subsp. enterica MNNEETFYQAMRRKGVTRRSFLKFCSLAATSLGLGAGMTPKIAWALENKPRIPVVWIHGL WP_001058312.1 hydrogenase [Salmonella enterica] Length: 372\nNumber of predicted TMHs: 2\nExp number of AAs in TMHs: 36.51738\nExp number, first 60 AAs: 19.72849\nTotal prob of N-in: 0.96170\nPOSSIBLE N-term signal sequence\ninside 1 19\nTMhelix 20 39\noutside 40 325\nTMhelix 326 348\ninside 349 372

19276 GCF_001244055.1_Salmonella_enterica_CVM_N47724_v1.0 Salmonella enterica subsp. enterica serovar Saintpaul Proteobacteria; Gammaproteobacteria; Enterobacterales; Enterobacteriaceae; Salmonella; Salmonella enterica; Salmonella enterica subsp. enterica MNNEETFYQAMRRKGVTRRSFLKFCSLAATSLGLGAGMTPKIAWALENKPRIPVVWIHGL WP_001058312.1 hydrogenase [Salmonella enterica] Length: 372\nNumber of predicted TMHs: 2\nExp number of AAs in TMHs: 36.51738\nExp number, first 60 AAs: 19.72849\nTotal prob of N-in: 0.96170\nPOSSIBLE N-term signal sequence\ninside 1 19\nTMhelix 20 39\noutside 40 325\nTMhelix 326 348\ninside 349 372

19277 GCF_001244925.1_Salmonella_enterica_CVM_N48697_v1.0 Salmonella enterica subsp. enterica serovar Saintpaul Proteobacteria; Gammaproteobacteria; Enterobacterales; Enterobacteriaceae; Salmonella; Salmonella enterica; Salmonella enterica subsp. enterica MNNEETFYQAMRRKGVTRRSFLKFCSLAATSLGLGAGMTPKIAWALENKPRIPVVWIHGL WP_001058312.1 hydrogenase [Salmonella enterica] Length: 372\nNumber of predicted TMHs: 2\nExp number of AAs in TMHs: 36.51738\nExp number, first 60 AAs: 19.72849\nTotal prob of N-in: 0.96170\nPOSSIBLE N-term signal sequence\ninside 1 19\nTMhelix 20 39\noutside 40 325\nTMhelix 326 348\ninside 349 372

19278 GCF_001244325.1_Salmonella_enterica_CVM_N48678_v1.0 Salmonella enterica subsp. enterica serovar Saintpaul Proteobacteria; Gammaproteobacteria; Enterobacterales; Enterobacteriaceae; Salmonella; Salmonella enterica; Salmonella enterica subsp. enterica MTGDNTLITSHGINRRDFMKLCAALAATMGLSSKAAAEMAESVSNPQRPPVIWIGAQECT WP_000145429.1 hydrogenase 2 small subunit [Salmonella enterica] Length: 372\nNumber of predicted TMHs: 1\nExp number of AAs in TMHs: 24.51309\nExp number, first 60 AAs: 0.2682\nTotal prob of N-in: 0.14402\noutside 1 331\nTMhelix 332 354\ninside 355 372

19279 GCF_001245335.1_Salmonella_enterica_CVM_N50423_v1.0 Salmonella enterica subsp. enterica serovar Saintpaul Proteobacteria; Gammaproteobacteria; Enterobacterales; Enterobacteriaceae; Salmonella; Salmonella enterica; Salmonella enterica subsp. enterica MNNEETFYQAMRRKGVTRRSFLKFCSLAATSLGLGAGMTPKIAWALENKPRIPVVWIHGL WP_001058312.1 hydrogenase [Salmonella enterica] Length: 372\nNumber of predicted TMHs: 2\nExp number of AAs in TMHs: 36.51738\nExp number, first 60 AAs: 19.72849\nTotal prob of N-in: 0.96170\nPOSSIBLE N-term signal sequence\ninside 1 19\nTMhelix 20 39\noutside 40 325\nTMhelix 326 348\ninside 349 372

19280 GCF_001271755.1_Salmonella_enterica_CVM_N46831_v1.0 Salmonella enterica subsp. enterica serovar Saintpaul Proteobacteria; Gammaproteobacteria; Enterobacterales; Enterobacteriaceae; Salmonella; Salmonella enterica; Salmonella enterica subsp. enterica MTGDNTLITSHGINRRDFMKLCAALAATMGLSSKAAAEMAESVSNPQRPPVIWIGAQECT WP_000145429.1 hydrogenase 2 small subunit [Salmonella enterica] Length: 372\nNumber of predicted TMHs: 1\nExp number of AAs in TMHs: 24.51309\nExp number, first 60 AAs: 0.2682\nTotal prob of N-in: 0.14402\noutside 1 331\nTMhelix 332 354\ninside 355 372

19281 GCF_001271755.1_Salmonella_enterica_CVM_N46831_v1.0 Salmonella enterica subsp. enterica serovar Saintpaul Proteobacteria; Gammaproteobacteria; Enterobacterales; Enterobacteriaceae; Salmonella; Salmonella enterica;

Salmonella enterica subsp. enterica MNNEETFYQAMRRKGVTRRSFLKFCSLAATSLGLGAGMTPKIAWALENKPRIPVVWIHGL WP_001058312.1 hydrogenase [Salmonella enterica] Length: 372\nNumber of predicted TMHs: 2\nExp number of AAs in TMHs: 36.51738\nExp number, first 60 AAs: 19.72849\nTotal prob of N-in: 0.96170\nPOSSIBLE N-term signal sequence\ninside 1 19\nTMhelix 20 39\noutside 40 325\nTMhelix 326 348\ninside 349 372

19282 GCF_001243195.1_Salmonella_enterica_CVM_N46830_v1.0 Salmonella enterica subsp. enterica serovar Saintpaul Proteobacteria; Gammaproteobacteria; Enterobacterales; Enterobacteriaceae; Salmonella; Salmonella enterica; Salmonella enterica subsp. enterica MTGDNTLITSHGINRRDFMKLCAALAATMGLSSKAAAEMAESVSNPQRPPVIWIGAQECT WP_000145429.1 hydrogenase 2 small subunit [Salmonella enterica] Length: 372\nNumber of predicted TMHs: 1\nExp number of AAs in TMHs: 24.51309\nExp number, first 60 AAs: 0.2682\nTotal prob of N-in: 0.14402\noutside 1 331\nTMhelix 332 354\ninside 355 372

19283 GCF_001243205.1_Salmonella_enterica_CVM_N46832_v1.0 Salmonella enterica subsp. enterica serovar Saintpaul Proteobacteria; Gammaproteobacteria; Enterobacterales; Enterobacteriaceae; Salmonella; Salmonella enterica; Salmonella enterica subsp. enterica MTGDNTLITSHGINRRDFMKLCAALAATMGLSSKAAAEMAESVSNPQRPPVIWIGAQECT WP_000145429.1 hydrogenase 2 small subunit [Salmonella enterica] Length: 372\nNumber of predicted TMHs: 1\nExp number of AAs in TMHs: 24.51309\nExp number, first 60 AAs: 0.2682\nTotal prob of N-in: 0.14402\noutside 1 331\nTMhelix 332 354\ninside 355 372

19284 GCF_001244325.1_Salmonella_enterica_CVM_N48678_v1.0 Salmonella enterica subsp. enterica serovar Saintpaul Proteobacteria; Gammaproteobacteria; Enterobacterales; Enterobacteriaceae; Salmonella; Salmonella enterica; Salmonella enterica subsp. enterica MNNEETFYQAMRRKGVTRRSFLKFCSLAATSLGLGAGMTPKIAWALENKPRIPVVWIHGL WP_001058312.1 hydrogenase [Salmonella enterica] Length: 372\nNumber of predicted TMHs: 2\nExp number of AAs in TMHs: 36.51738\nExp number, first 60 AAs: 19.72849\nTotal prob of N-in: 0.96170\nPOSSIBLE N-term signal sequence\ninside 1 19\nTMhelix 20 39\noutside 40 325\nTMhelix 326 348\ninside 349 372

19285 GCF_001240335.1_Salmonella_enterica_CVM_N43461_v1.0 Salmonella enterica subsp. enterica serovar Saintpaul Proteobacteria; Gammaproteobacteria; Enterobacterales; Enterobacteriaceae; Salmonella; Salmonella enterica; Salmonella enterica subsp. enterica MNNEETFYQAMRRKGVTRRSFLKFCSLAATSLGLGAGMTPKIAWALENKPRIPVVWIHGL WP_001058312.1 hydrogenase [Salmonella enterica] Length: 372\nNumber of predicted TMHs: 2\nExp number of AAs in TMHs: 36.51738\nExp number, first 60 AAs: 19.72849\nTotal prob of N-in: 0.96170\nPOSSIBLE N-term signal sequence\ninside 1 19\nTMhelix 20 39\noutside 40 325\nTMhelix 326 348\ninside 349 372

19286 GCF_001243205.1_Salmonella_enterica_CVM_N46832_v1.0 Salmonella enterica subsp. enterica serovar Saintpaul Proteobacteria; Gammaproteobacteria; Enterobacterales; Enterobacteriaceae; Salmonella; Salmonella enterica; Salmonella enterica subsp. enterica MNNEETFYQAMRRKGVTRRSFLKFCSLAATSLGLGAGMTPKIAWALENKPRIPVVWIHGL WP_001058312.1 hydrogenase [Salmonella enterica] Length: 372\nNumber of predicted TMHs: 2\nExp number of AAs in TMHs: 36.51738\nExp number, first 60 AAs: 19.72849\nTotal prob of N-in: 0.96170\nPOSSIBLE N-term signal sequence\ninside 1 19\nTMhelix 20 39\noutside 40 325\nTMhelix 326 348\ninside 349 372

19287 GCF_001243195.1_Salmonella_enterica_CVM_N46830_v1.0 Salmonella enterica subsp. enterica serovar Saintpaul Proteobacteria; Gammaproteobacteria; Enterobacterales; Enterobacteriaceae; Salmonella; Salmonella enterica; Salmonella enterica subsp. enterica MNNEETFYQAMRRKGVTRRSFLKFCSLAATSLGLGAGMTPKIAWALENKPRIPVVWIHGL WP_001058312.1 hydrogenase [Salmonella enterica] Length: 372\nNumber of predicted TMHs: 2\nExp number of AAs in TMHs: 36.51738\nExp number, first 60 AAs: 19.72849\nTotal prob of N-in: 0.96170\nPOSSIBLE N-term signal sequence\ninside 1 19\nTMhelix 20 39\noutside 40 325\nTMhelix 326 348\ninside 349 372

19288 GCF_001244925.1_Salmonella_enterica_CVM_N48697_v1.0 Salmonella enterica subsp. enterica serovar Saintpaul Proteobacteria; Gammaproteobacteria; Enterobacterales; Enterobacteriaceae; Salmonella; Salmonella enterica; Salmonella enterica subsp. enterica MTGDNTLITSHGINRRDFMKLCAALAATMGLSSKAAAEMAESVSNPQRPPVIWIGAQECT WP_000145429.1 hydrogenase 2 small subunit [Salmonella enterica] Length: 372\nNumber of predicted TMHs: 1\nExp number of AAs in TMHs: 24.51309\nExp number, first 60 AAs: 0.2682\nTotal prob of N-in: 0.14402\noutside 1 331\nTMhelix 332 354\ninside 355 372

19289 GCF_002063995.1_ASM206399v1 Salmonella enterica subsp. enterica serovar Saintpaul Proteobacteria; Gammaproteobacteria; Enterobacterales; Enterobacteriaceae; Salmonella; Salmonella enterica; Salmonella enterica subsp. enterica MTGDNTLITSHGINRRDFMKLCAALAATMGLSSKAAAEMAESVSNPQRPPVIWIGAQECT WP_000145429.1 hydrogenase 2 small subunit [Salmonella enterica] Length: 372\nNumber of predicted TMHs: 1\nExp number of AAs in TMHs: 24.51309\nExp number, first 60 AAs: 0.2682\nTotal prob of N-in: 0.14402\noutside 1 331\nTMhelix 332 354\ninside 355 372

19290 GCF_002063995.1_ASM206399v1 Salmonella enterica subsp. enterica serovar Saintpaul Proteobacteria; Gammaproteobacteria; Enterobacterales; Enterobacteriaceae; Salmonella; Salmonella enterica; Salmonella enterica subsp. enterica MNNEETFYQAMRRKGVTRRSFLKFCSLAATSLGLGAGMTPKIAWALENKPRIPVVWIHGL WP_001058312.1 hydrogenase [Salmonella enterica] Length: 372\nNumber of predicted TMHs: 2\nExp number of AAs in TMHs: 36.51738\nExp number, first 60 AAs: 19.72849\nTotal prob of N-in: 0.96170\nPOSSIBLE N-term signal sequence\ninside 1 19\nTMhelix 20 39\noutside 40 325\nTMhelix 326 348\ninside 349 372

19291 GCF_002064045.1_ASM206404v1 Salmonella enterica subsp. enterica serovar Saintpaul Proteobacteria; Gammaproteobacteria; Enterobacterales; Enterobacteriaceae; Salmonella; Salmonella enterica; Salmonella enterica subsp. enterica MTGDNTLITSHGINRRDFMKLCAALAATMGLSSKAAAEMAESVSNPQRPPVIWIGAQECT WP_000145429.1 hydrogenase 2 small subunit [Salmonella enterica] Length: 372\nNumber of predicted TMHs: 1\nExp number of AAs in TMHs: 24.51309\nExp number, first 60 AAs: 0.2682\nTotal prob of N-in: 0.14402\noutside 1 331\nTMhelix 332 354\ninside 355 372

19292 GCF_002064045.1_ASM206404v1 Salmonella enterica subsp. enterica serovar Saintpaul Proteobacteria; Gammaproteobacteria; Enterobacterales; Enterobacteriaceae; Salmonella; Salmonella enterica; Salmonella enterica subsp. enterica MNNEETFYQAMRRKGVTRRSFLKFCSLAATSLGLGAGMTPKIAWALENKPRIPVVWIHGL WP_001058312.1 hydrogenase [Salmonella enterica] Length: 372\nNumber of predicted TMHs: 2\nExp number of AAs in TMHs: 36.51738\nExp number, first 60 AAs: 19.72849\nTotal prob of N-in: 0.96170\nPOSSIBLE N-term signal sequence\ninside 1 19\nTMhelix 20 39\noutside 40 325\nTMhelix 326 348\ninside 349 372

19293 GCF_001834005.1_ASM183400v1 Salmonella enterica subsp. enterica serovar Saintpaul Proteobacteria; Gammaproteobacteria; Enterobacterales; Enterobacteriaceae; Salmonella; Salmonella enterica; Salmonella enterica subsp. enterica MTGDNTLITSHGINRRDFMKLCAALAATMGLSSKAAAEMAESVSNPQRPPVIWIGAQECT WP_000145429.1 hydrogenase 2 small subunit [Salmonella enterica] Length: 372\nNumber of predicted TMHs: 1\nExp number of AAs in TMHs: 24.51309\nExp number, first 60 AAs: 0.2682\nTotal prob of N-in: 0.14402\noutside 1 331\nTMhelix 332 354\ninside 355 372

19294 GCF_002031275.1_ASM203127v1 Salmonella enterica subsp. enterica serovar Saintpaul Proteobacteria; Gammaproteobacteria; Enterobacterales; Enterobacteriaceae; Salmonella; Salmonella enterica; Salmonella enterica subsp. enterica MTGDNTLITSHGINRRDFMKLCAALAATMGLSSKAAAEMAESVSNPQRPPVIWIGAQECT WP_000145429.1 hydrogenase 2 small subunit [Salmonella enterica] Length: 372\nNumber of predicted TMHs: 1\nExp number of AAs in TMHs: 24.51309\nExp number, first 60 AAs: 0.2682\nTotal prob of N-in: 0.14402\noutside 1 331\nTMhelix 332 354\ninside 355 372

19295 GCF_001832855.1_ASM183285v1 Salmonella enterica subsp. enterica serovar Saintpaul Proteobacteria; Gammaproteobacteria; Enterobacterales; Enterobacteriaceae; Salmonella; Salmonella enterica; Salmonella enterica subsp. enterica MNNEETFYQAMRRKGVTRRSFLKFCSLAATSLGLGAGMTPKIAWALENKPRIPVVWIHGL WP_001058311.1 hydrogenase [Salmonella enterica] Length: 372\nNumber of predicted TMHs: 2\nExp number of AAs in TMHs: 37.4699899999999\nExp number, first 60 AAs: 19.72699\nTotal prob of N-in: 0.96171\nPOSSIBLE N-term signal sequence\ninside 1 19\nTMhelix 20 39\noutside 40 325\nTMhelix 326 348\ninside 349 372

19296 GCF_002030545.1_ASM203054v1 Salmonella enterica subsp. enterica serovar Saintpaul Proteobacteria; Gammaproteobacteria; Enterobacterales; Enterobacteriaceae; Salmonella; Salmonella enterica; Salmonella enterica subsp. enterica MNNEETFYQAMRRKGVTRRSFLKFCSLAATSLGLGAGMTPKIAWALENKPRIPVVWIHGL WP_001058312.1 hydrogenase [Salmonella enterica] Length: 372\nNumber of predicted TMHs: 2\nExp number of AAs in TMHs: 36.51738\nExp number, first 60 AAs: 19.72849\nTotal prob of N-in: 0.96170\nPOSSIBLE N-term signal sequence\ninside 1 19\nTMhelix 20 39\noutside 40 325\nTMhelix 326 348\ninside 349 372

19297 GCF_001833315.1_ASM183331v1 Salmonella enterica subsp. enterica serovar Saintpaul Proteobacteria; Gammaproteobacteria; Enterobacterales; Enterobacteriaceae; Salmonella; Salmonella enterica; Salmonella enterica subsp. enterica MTGDNTLITSHGINRRDFMKLCAALAATMGLSSKAAAEMAESVSNPQRPPVIWIGAQECT WP_000145429.1 hydrogenase 2 small subunit [Salmonella enterica] Length: 372\nNumber of predicted TMHs: 1\nExp number of AAs in TMHs: 24.51309\nExp number, first 60 AAs: 0.2682\nTotal prob of N-in: 0.14402\noutside 1 331\nTMhelix 332 354\ninside 355 372

19298 GCF_001834605.1_ASM183460v1 Salmonella enterica subsp. enterica serovar Saintpaul Proteobacteria; Gammaproteobacteria; Enterobacterales; Enterobacteriaceae; Salmonella; Salmonella enterica; Salmonella enterica subsp. enterica MTGDNTLITSHGINRRDFMKLCAALAATMGLSSKAAAEMAESVSNPQRPPVIWIGAQECT WP_000145429.1 hydrogenase 2 small subunit [Salmonella enterica] Length: 372\nNumber of

predicted TMHs:  1\nExp number of AAs in TMHs: 24.51309\nExp number, first 60 AAs:  0.2682\nTotal prob of N-in: 0.14402\noutside   1   331\nTMhelix   332   354\ninside   355   372

19299 GCF_001834565.1_ASM183456v1 Salmonella enterica subsp. enterica serovar Saintpaul Proteobacteria; Gammaproteobacteria; Enterobacterales; Enterobacteriaceae; Salmonella; Salmonella enterica; Salmonella enterica subsp. enterica MTGDNTLITSHGINRRDFMKLCAALAATMGLSSKAAAEMAESVSNPQRPPVIWIGAQECT WP_000145429.1 hydrogenase 2 small subunit [Salmonella enterica] Length: 372\nNumber of predicted TMHs:  1\nExp number of AAs in TMHs: 24.51309\nExp number, first 60 AAs:  0.2682\nTotal prob of N-in: 0.14402\noutside   1   331\nTMhelix   332   354\ninside   355   372

19300 GCF_001834525.1_ASM183452v1 Salmonella enterica subsp. enterica serovar Saintpaul Proteobacteria; Gammaproteobacteria; Enterobacterales; Enterobacteriaceae; Salmonella; Salmonella enterica; Salmonella enterica subsp. enterica MTGDNTLITSHGINRRDFMKLCAALAATMGLSSKAAAEMAESVSNPQRPPVIWIGAQECT WP_000145429.1 hydrogenase 2 small subunit [Salmonella enterica] Length: 372\nNumber of predicted TMHs:  1\nExp number of AAs in TMHs: 24.51309\nExp number, first 60 AAs:  0.2682\nTotal prob of N-in: 0.14402\noutside   1   331\nTMhelix   332   354\ninside   355   372

19301 GCF_001834485.1_ASM183448v1 Salmonella enterica subsp. enterica serovar Saintpaul Proteobacteria; Gammaproteobacteria; Enterobacterales; Enterobacteriaceae; Salmonella; Salmonella enterica; Salmonella enterica subsp. enterica MTGDNTLITSHGINRRDFMKLCAALAATMGLSSKAAAEMAESVSNPQRPPVIWIGAQECT WP_000145429.1 hydrogenase 2 small subunit [Salmonella enterica] Length: 372\nNumber of predicted TMHs:  1\nExp number of AAs in TMHs: 24.51309\nExp number, first 60 AAs:  0.2682\nTotal prob of N-in: 0.14402\noutside   1   331\nTMhelix   332   354\ninside   355   372

19302 GCF_001833415.1_ASM183341v1 Salmonella enterica subsp. enterica serovar Saintpaul Proteobacteria; Gammaproteobacteria; Enterobacterales; Enterobacteriaceae; Salmonella; Salmonella enterica; Salmonella enterica subsp. enterica MTGDNTLITSHGINRRDFMKLCAALAATMGLSSKAAAEMAESVSNPQRPPVIWIGAQECT WP_000145429.1 hydrogenase 2 small subunit [Salmonella enterica] Length: 372\nNumber of predicted TMHs:  1\nExp number of AAs in TMHs: 24.51309\nExp number, first 60 AAs:  0.2682\nTotal prob of N-in: 0.14402\noutside   1   331\nTMhelix   332   354\ninside   355   372

19303 GCF_001834305.1_ASM183430v1 Salmonella enterica subsp. enterica serovar Saintpaul Proteobacteria; Gammaproteobacteria; Enterobacterales; Enterobacteriaceae; Salmonella; Salmonella enterica; Salmonella enterica subsp. enterica MTGDNTLITSHGINRRDFMKLCAALAATMGLSSKAAAEMAESVSNPQRPPVIWIGAQECT WP_000145429.1 hydrogenase 2 small subunit [Salmonella enterica] Length: 372\nNumber of predicted TMHs:  1\nExp number of AAs in TMHs: 24.51309\nExp number, first 60 AAs:  0.2682\nTotal prob of N-in: 0.14402\noutside   1   331\nTMhelix   332   354\ninside   355   372

19304 GCF_001834445.1_ASM183444v1 Salmonella enterica subsp. enterica serovar Saintpaul Proteobacteria; Gammaproteobacteria; Enterobacterales; Enterobacteriaceae; Salmonella; Salmonella enterica; Salmonella enterica subsp. enterica MTGDNTLITSHGINRRDFMKLCAALAATMGLSSKAAAEMAESVSNPQRPPVIWIGAQECT WP_000145429.1 hydrogenase 2 small subunit [Salmonella enterica] Length: 372\nNumber of predicted TMHs:  1\nExp number of AAs in TMHs: 24.51309\nExp number, first 60 AAs:  0.2682\nTotal prob of N-in: 0.14402\noutside   1   331\nTMhelix   332   354\ninside   355   372

19305 GCF_001834015.1_ASM183401v1 Salmonella enterica subsp. enterica serovar Saintpaul Proteobacteria; Gammaproteobacteria; Enterobacterales; Enterobacteriaceae; Salmonella; Salmonella enterica; Salmonella enterica subsp. enterica MNNEETFYQAMRRKGVTRRSFLKFCSLAATSLGLGAGMTPKIAWALENKPRIPVVWIHGL WP_001058311.1 hydrogenase [Salmonella enterica] Length: 372\nNumber of predicted TMHs:  2\nExp number of AAs in TMHs: 37.4699899999999\nExp number, first 60 AAs:  19.72699\nTotal prob of N-in: 0.96171\nPOSSIBLE N-term signal sequence\ninside   1   19\nTMhelix   20   39\noutside   40   325\nTMhelix   326   348\ninside   349   372

19306 GCF_001834015.1_ASM183401v1 Salmonella enterica subsp. enterica serovar Saintpaul Proteobacteria; Gammaproteobacteria; Enterobacterales; Enterobacteriaceae; Salmonella; Salmonella enterica; Salmonella enterica subsp. enterica MTGDNTLITSHGINRRDFMKLCAALAATMGLSSKAAAEMAESVSNPQRPPVIWIGAQECT WP_000145429.1 hydrogenase 2 small subunit [Salmonella enterica] Length: 372\nNumber of predicted TMHs:  1\nExp number of AAs in TMHs: 24.51309\nExp number, first 60 AAs:  0.2682\nTotal prob of N-in: 0.14402\noutside   1   331\nTMhelix   332   354\ninside   355   372

19307 GCF_001833855.1_ASM183385v1 Salmonella enterica subsp. enterica serovar Saintpaul Proteobacteria; Gammaproteobacteria; Enterobacterales; Enterobacteriaceae; Salmonella; Salmonella enterica; Salmonella enterica subsp. enterica MNNEETFYQAMRRKGVTRRSFLKFCSLAATSLGLGAGMTPKIAWALENKPRIPVVWIHGL WP_001058311.1 hydrogenase [Salmonella enterica] Length: 372\nNumber of predicted TMHs: 2\nExp number of AAs in TMHs: 37.4699899999999\nExp number, first 60 AAs: 19.72699\nTotal prob of N-in: 0.96171\nPOSSIBLE N-term signal sequence\ninside 1 19\nTMhelix 20 39\noutside 40 325\nTMhelix 326 348\ninside 349 372

19308 GCF_001833815.1_ASM183381v1 Salmonella enterica subsp. enterica serovar Saintpaul Proteobacteria; Gammaproteobacteria; Enterobacterales; Enterobacteriaceae; Salmonella; Salmonella enterica; Salmonella enterica subsp. enterica MNNEETFYQAMRRKGVTRRSFLKFCSLAATSLGLGAGMTPKIAWALENKPRIPVVWIHGL WP_001058311.1 hydrogenase [Salmonella enterica] Length: 372\nNumber of predicted TMHs: 2\nExp number of AAs in TMHs: 37.4699899999999\nExp number, first 60 AAs: 19.72699\nTotal prob of N-in: 0.96171\nPOSSIBLE N-term signal sequence\ninside 1 19\nTMhelix 20 39\noutside 40 325\nTMhelix 326 348\ninside 349 372

19309 GCF_001833655.1_ASM183365v1 Salmonella enterica subsp. enterica serovar Saintpaul Proteobacteria; Gammaproteobacteria; Enterobacterales; Enterobacteriaceae; Salmonella; Salmonella enterica; Salmonella enterica subsp. enterica MNNEETFYQAMRRKGVTRRSFLKFCSLAATSLGLGAGMTPKIAWALENKPRIPVVWIHGL WP_001058311.1 hydrogenase [Salmonella enterica] Length: 372\nNumber of predicted TMHs: 2\nExp number of AAs in TMHs: 37.4699899999999\nExp number, first 60 AAs: 19.72699\nTotal prob of N-in: 0.96171\nPOSSIBLE N-term signal sequence\ninside 1 19\nTMhelix 20 39\noutside 40 325\nTMhelix 326 348\ninside 349 372

19310 GCF_002063255.1_ASM206325v1 Salmonella enterica subsp. enterica serovar Saintpaul Proteobacteria; Gammaproteobacteria; Enterobacterales; Enterobacteriaceae; Salmonella; Salmonella enterica; Salmonella enterica subsp. enterica MTGDNTLITSHGINRRDFMKLCAALAATMGLSSKAAAEMAESVSNPQRPPVIWIGAQECT WP_000145429.1 hydrogenase 2 small subunit [Salmonella enterica] Length: 372\nNumber of predicted TMHs: 1\nExp number of AAs in TMHs: 24.51309\nExp number, first 60 AAs: 0.2682\nTotal prob of N-in: 0.14402\noutside 1 331\nTMhelix 332 354\ninside 355 372

19311 GCF_001833465.1_ASM183346v1 Salmonella enterica subsp. enterica serovar Saintpaul Proteobacteria; Gammaproteobacteria; Enterobacterales; Enterobacteriaceae; Salmonella; Salmonella enterica; Salmonella enterica subsp. enterica MTGDNTLITSHGINRRDFMKLCAALAATMGLSSKAAAEMAESVSNPQRPPVIWIGAQECT WP_000145429.1 hydrogenase 2 small subunit [Salmonella enterica] Length: 372\nNumber of predicted TMHs: 1\nExp number of AAs in TMHs: 24.51309\nExp number, first 60 AAs: 0.2682\nTotal prob of N-in: 0.14402\noutside 1 331\nTMhelix 332 354\ninside 355 372

19312 GCF_001833415.1_ASM183341v1 Salmonella enterica subsp. enterica serovar Saintpaul Proteobacteria; Gammaproteobacteria; Enterobacterales; Enterobacteriaceae; Salmonella; Salmonella enterica; Salmonella enterica subsp. enterica MNNEETFYQAMRRKGVTRRSFLKFCSLAATSLGLGAGMTPKIAWALENKPRIPVVWIHGL WP_001058311.1 hydrogenase [Salmonella enterica] Length: 372\nNumber of predicted TMHs: 2\nExp number of AAs in TMHs: 37.4699899999999\nExp number, first 60 AAs: 19.72699\nTotal prob of N-in: 0.96171\nPOSSIBLE N-term signal sequence\ninside 1 19\nTMhelix 20 39\noutside 40 325\nTMhelix 326 348\ninside 349 372

19313 GCF_002063255.1_ASM206325v1 Salmonella enterica subsp. enterica serovar Saintpaul Proteobacteria; Gammaproteobacteria; Enterobacterales; Enterobacteriaceae; Salmonella; Salmonella enterica; Salmonella enterica subsp. enterica MNNEETFYQAMRRKGVTRRSFLKFCSLAATSLGLGAGMTPKIAWALENKPRIPVVWIHGL WP_001058312.1 hydrogenase [Salmonella enterica] Length: 372\nNumber of predicted TMHs: 2\nExp number of AAs in TMHs: 36.51738\nExp number, first 60 AAs: 19.72849\nTotal prob of N-in: 0.96170\nPOSSIBLE N-term signal sequence\ninside 1 19\nTMhelix 20 39\noutside 40 325\nTMhelix 326 348\ninside 349 372

19314 GCF_001952995.1_ASM195299v1 Salmonella enterica subsp. enterica serovar Saintpaul Proteobacteria; Gammaproteobacteria; Enterobacterales; Enterobacteriaceae; Salmonella; Salmonella enterica; Salmonella enterica subsp. enterica MTGDNTLITSHGINRRDFMKLCAALAATMGLSSKAAAEMAESVSNPQRPPVIWIGAQECT WP_000145429.1 hydrogenase 2 small subunit [Salmonella enterica] Length: 372\nNumber of predicted TMHs: 1\nExp number of AAs in TMHs: 24.51309\nExp number, first 60 AAs: 0.2682\nTotal prob of N-in: 0.14402\noutside 1 331\nTMhelix 332 354\ninside 355 372

19315 GCF_002063415.1_ASM206341v1 Salmonella enterica subsp. enterica serovar Saintpaul Proteobacteria; Gammaproteobacteria; Enterobacterales; Enterobacteriaceae; Salmonella; Salmonella enterica; Salmonella enterica subsp. enterica MNNEETFYQAMRRKGVTRRSFLKFCSLAATSLGLGAGMTPKIAWALENKPRIPVVWIHGL WP_001058312.1 hydrogenase [Salmonella enterica] Length: 372\nNumber of predicted TMHs: 2\nExp number of AAs in TMHs: 36.51738\nExp number, first 60 AAs: 19.72849\nTotal prob of N-in: 0.96170\nPOSSIBLE N-term signal sequence\ninside 1 19\nTMhelix 20 39\noutside 40 325\nTMhelix 326 348\ninside 349 372

19316 GCF_002063415.1_ASM206341v1 Salmonella enterica subsp. enterica serovar Saintpaul Proteobacteria; Gammaproteobacteria; Enterobacterales; Enterobacteriaceae; Salmonella; Salmonella enterica; Salmonella enterica subsp. enterica MTGDNTLITSHGINRRDFMKLCAALAATMGLSSKAAAEMAESVSNPQRPPVIWIGAQECT WP_000145429.1 hydrogenase 2 small subunit [Salmonella enterica] Length: 372\nNumber of predicted TMHs: 1\nExp number of AAs in TMHs: 24.51309\nExp number, first 60 AAs: 0.2682\nTotal prob of N-in: 0.14402\noutside 1 331\nTMhelix 332 354\ninside 355 372

19317 GCF_002030545.1_ASM203054v1 Salmonella enterica subsp. enterica serovar Saintpaul Proteobacteria; Gammaproteobacteria; Enterobacterales; Enterobacteriaceae; Salmonella; Salmonella enterica; Salmonella enterica subsp. enterica MTGDNTLITSHGINRRDFMKLCAALAATMGLSSKAAAEMAESVSNPQRPPVIWIGAQECT WP_000145429.1 hydrogenase 2 small subunit [Salmonella enterica] Length: 372\nNumber of predicted TMHs: 1\nExp number of AAs in TMHs: 24.51309\nExp number, first 60 AAs: 0.2682\nTotal prob of N-in: 0.14402\noutside 1 331\nTMhelix 332 354\ninside 355 372

19318 GCF_001832595.1_ASM183259v1 Salmonella enterica subsp. enterica serovar Saintpaul Proteobacteria; Gammaproteobacteria; Enterobacterales; Enterobacteriaceae; Salmonella; Salmonella enterica; Salmonella enterica subsp. enterica MTGDNTLITSHGINRRDFMKLCAALAATMGLSSKAAAEMAESVSNPQRPPVIWIGAQECT WP_000145429.1 hydrogenase 2 small subunit [Salmonella enterica] Length: 372\nNumber of predicted TMHs: 1\nExp number of AAs in TMHs: 24.51309\nExp number, first 60 AAs: 0.2682\nTotal prob of N-in: 0.14402\noutside 1 331\nTMhelix 332 354\ninside 355 372

19319 GCF_001834465.1_ASM183446v1 Salmonella enterica subsp. enterica serovar Saintpaul Proteobacteria; Gammaproteobacteria; Enterobacterales; Enterobacteriaceae; Salmonella; Salmonella enterica; Salmonella enterica subsp. enterica MNNEETFYQAMRRKGVTRRSFLKFCSLAATSLGLGAGMTPKIAWALENKPRIPVVWIHGL WP_001058311.1 hydrogenase [Salmonella enterica] Length: 372\nNumber of predicted TMHs: 2\nExp number of AAs in TMHs: 37.4699899999999\nExp number, first 60 AAs: 19.72699\nTotal prob of N-in: 0.96171\nPOSSIBLE N-term signal sequence\ninside 1 19\nTMhelix 20 39\noutside 40 325\nTMhelix 326 348\ninside 349 372

19320 GCF_001246025.1_Salmonella_enterica_CVM_N51245_v1.0 Salmonella enterica subsp. enterica serovar Saintpaul Proteobacteria; Gammaproteobacteria; Enterobacterales; Enterobacteriaceae; Salmonella; Salmonella enterica; Salmonella enterica subsp. enterica MNNEETFYQAMRRKGVTRRSFLKFCSLAATSLGLGAGMTPKIAWALENKPRIPVVWIHGL WP_001058312.1 hydrogenase [Salmonella enterica] Length: 372\nNumber of predicted TMHs: 2\nExp number of AAs in TMHs: 36.51738\nExp number, first 60 AAs: 19.72849\nTotal prob of N-in: 0.96170\nPOSSIBLE N-term signal sequence\ninside 1 19\nTMhelix 20 39\noutside 40 325\nTMhelix 326 348\ninside 349 372

19321 GCF_001834385.1_ASM183438v1 Salmonella enterica subsp. enterica serovar Saintpaul Proteobacteria; Gammaproteobacteria; Enterobacterales; Enterobacteriaceae; Salmonella; Salmonella enterica; Salmonella enterica subsp. enterica MNNEETFYQAMRRKGVTRRSFLKFCSLAATSLGLGAGMTPKIAWALENKPRIPVVWIHGL WP_001058311.1 hydrogenase [Salmonella enterica] Length: 372\nNumber of predicted TMHs: 2\nExp number of AAs in TMHs: 37.4699899999999\nExp number, first 60 AAs: 19.72699\nTotal prob of N-in: 0.96171\nPOSSIBLE N-term signal sequence\ninside 1 19\nTMhelix 20 39\noutside 40 325\nTMhelix 326 348\ninside 349 372

19322 GCF_001835315.1_ASM183531v1 Salmonella enterica subsp. enterica serovar Saintpaul Proteobacteria; Gammaproteobacteria; Enterobacterales; Enterobacteriaceae; Salmonella; Salmonella enterica; Salmonella enterica subsp. enterica MTGDNTLITSHGINRRDFMKLCAALAATMGLSSKAAAEMAESVSNPQRPPVIWIGAQECT WP_000145429.1 hydrogenase 2 small subunit [Salmonella enterica] Length: 372\nNumber of predicted TMHs: 1\nExp number of AAs in TMHs: 24.51309\nExp number, first 60 AAs: 0.2682\nTotal prob of N-in: 0.14402\noutside 1 331\nTMhelix 332 354\ninside 355 372

19323 GCF_001478765.1_Salmonella_enterica_CVM_N29377-SQ_v1.0 Salmonella enterica subsp. enterica serovar Saintpaul Proteobacteria; Gammaproteobacteria; Enterobacterales; Enterobacteriaceae; Salmonella; Salmonella enterica; Salmonella enterica subsp. enterica MNNEETFYQAMRRKGVTRRSFLKFCSLAATSLGLGAGMTPKIAWALENKPRIPVVWIHGL

WP_001058312.1 hydrogenase [Salmonella enterica] Length: 372\nNumber of predicted TMHs: 2\nExp number of AAs in TMHs: 36.51738\nExp number, first 60 AAs: 19.72849\nTotal prob of N-in: 0.96170\nPOSSIBLE N-term signal sequence\ninside 1 19\nTMhelix 20 39\noutside 40 325\nTMhelix 326 348\ninside 349 372

19324 GCF_001479275.1_Salmonella_enterica_CVM_N31846-SQ_v1.0 Salmonella enterica subsp. enterica serovar Saintpaul Proteobacteria; Gammaproteobacteria; Enterobacterales; Enterobacteriaceae; Salmonella; Salmonella enterica; Salmonella enterica subsp. enterica MNNEETFYQAMRRKGVTRRSFLKFCSLAATSLGLGAGMTPKIAWALENKPRIPVVWIHGL WP_001058312.1 hydrogenase [Salmonella enterica] Length: 372\nNumber of predicted TMHs: 2\nExp number of AAs in TMHs: 36.51738\nExp number, first 60 AAs: 19.72849\nTotal prob of N-in: 0.96170\nPOSSIBLE N-term signal sequence\ninside 1 19\nTMhelix 20 39\noutside 40 325\nTMhelix 326 348\ninside 349 372

19325 GCF_001479205.1_Salmonella_enterica_CVM_N31415-SQ_v1.0 Salmonella enterica subsp. enterica serovar Saintpaul Proteobacteria; Gammaproteobacteria; Enterobacterales; Enterobacteriaceae; Salmonella; Salmonella enterica; Salmonella enterica subsp. enterica MNNEETFYQAMRRKGVTRRSFLKFCSLAATSLGLGAGMTPKIAWALENKPRIPVVWIHGL WP_001058312.1 hydrogenase [Salmonella enterica] Length: 372\nNumber of predicted TMHs: 2\nExp number of AAs in TMHs: 36.51738\nExp number, first 60 AAs: 19.72849\nTotal prob of N-in: 0.96170\nPOSSIBLE N-term signal sequence\ninside 1 19\nTMhelix 20 39\noutside 40 325\nTMhelix 326 348\ninside 349 372

19326 GCF_001479275.1_Salmonella_enterica_CVM_N31846-SQ_v1.0 Salmonella enterica subsp. enterica serovar Saintpaul Proteobacteria; Gammaproteobacteria; Enterobacterales; Enterobacteriaceae; Salmonella; Salmonella enterica; Salmonella enterica subsp. enterica MTGDNTLITSHGINRRDFMKLCAALAATMGLSSKAAAEMAESVSNPQRPPVIWIGAQECT WP_000145429.1 hydrogenase 2 small subunit [Salmonella enterica] Length: 372\nNumber of predicted TMHs: 1\nExp number of AAs in TMHs: 24.51309\nExp number, first 60 AAs: 0.2682\nTotal prob of N-in: 0.14402\noutside 1 331\nTMhelix 332 354\ninside 355 372

19327 GCF_002031635.1_ASM203163v1 Salmonella enterica subsp. enterica serovar Saintpaul Proteobacteria; Gammaproteobacteria; Enterobacterales; Enterobacteriaceae; Salmonella; Salmonella enterica; Salmonella enterica subsp. enterica MTGDNTLITSHGINRRDFMKLCAALAATMGLSSKAAAEMAESVSNPQRPPVIWIGAQECT WP_000145429.1 hydrogenase 2 small subunit [Salmonella enterica] Length: 372\nNumber of predicted TMHs: 1\nExp number of AAs in TMHs: 24.51309\nExp number, first 60 AAs: 0.2682\nTotal prob of N-in: 0.14402\noutside 1 331\nTMhelix 332 354\ninside 355 372

19328 GCF_001479205.1_Salmonella_enterica_CVM_N31415-SQ_v1.0 Salmonella enterica subsp. enterica serovar Saintpaul Proteobacteria; Gammaproteobacteria; Enterobacterales; Enterobacteriaceae; Salmonella; Salmonella enterica; Salmonella enterica subsp. enterica MTGDNTLITSHGINRRDFMKLCAALAATMGLSSKAAAEMAESVSNPQRPPVIWIGAQECT WP_000145429.1 hydrogenase 2 small subunit [Salmonella enterica] Length: 372\nNumber of predicted TMHs: 1\nExp number of AAs in TMHs: 24.51309\nExp number, first 60 AAs: 0.2682\nTotal prob of N-in: 0.14402\noutside 1 331\nTMhelix 332 354\ninside 355 372

19329 GCF_001953675.1_ASM195367v1 Salmonella enterica subsp. enterica serovar Saintpaul Proteobacteria; Gammaproteobacteria; Enterobacterales; Enterobacteriaceae; Salmonella; Salmonella enterica; Salmonella enterica subsp. enterica MTGDNTLITSHGINRRDFMKLCAALAATMGLSSKAAAEMAESVSNPQRPPVIWIGAQECT WP_000145429.1 hydrogenase 2 small subunit [Salmonella enterica] Length: 372\nNumber of predicted TMHs: 1\nExp number of AAs in TMHs: 24.51309\nExp number, first 60 AAs: 0.2682\nTotal prob of N-in: 0.14402\noutside 1 331\nTMhelix 332 354\ninside 355 372

19330 GCF_001243775.1_Salmonella_enterica_CVM_N47713_v1.0 Salmonella enterica subsp. enterica serovar Saintpaul Proteobacteria; Gammaproteobacteria; Enterobacterales; Enterobacteriaceae; Salmonella; Salmonella enterica; Salmonella enterica subsp. enterica MTGDNTLITSHGINRRDFMKLCAALAATMGLSSKAAAEMAESVSNPQRPPVIWIGAQECT WP_000145429.1 hydrogenase 2 small subunit [Salmonella enterica] Length: 372\nNumber of predicted TMHs: 1\nExp number of AAs in TMHs: 24.51309\nExp number, first 60 AAs: 0.2682\nTotal prob of N-in: 0.14402\noutside 1 331\nTMhelix 332 354\ninside 355 372

19331 GCF_001244935.1_Salmonella_enterica_CVM_N48698_v1.0 Salmonella enterica subsp. enterica serovar Saintpaul Proteobacteria; Gammaproteobacteria; Enterobacterales; Enterobacteriaceae; Salmonella; Salmonella enterica; Salmonella enterica subsp. enterica MTGDNTLITSHGINRRDFMKLCAALAATMGLSSKAAAEMAESVSNPQRPPVIWIGAQECT WP_000145429.1 hydrogenase 2 small subunit [Salmonella enterica] Length: 372\nNumber of predicted TMHs: 1\nExp number of AAs in TMHs: 24.51309\nExp number, first 60 AAs: 0.2682\nTotal prob of N-in: 0.14402\noutside 1 331\nTMhelix 332 354\ninside 355 372

19332 GCF_001247105.1_Salmonella_enterica_CVM_N51291_v1.0 Salmonella enterica subsp. enterica serovar Saintpaul Proteobacteria; Gammaproteobacteria; Enterobacterales; Enterobacteriaceae; Salmonella; Salmonella enterica; Salmonella enterica subsp. enterica MNNEETFYQAMRRKGVTRRSFLKFCSLAATSLGLGAGMTPKIAWALENKPRIPVVWIHGL WP_001058312.1 hydrogenase [Salmonella enterica] Length: 372\nNumber of predicted TMHs: 2\nExp number of AAs in TMHs: 36.51738\nExp number, first 60 AAs: 19.72849\nTotal prob of N-in: 0.96170\nPOSSIBLE N-term signal sequence\ninside 1 19\nTMhelix 20 39\noutside 40 325\nTMhelix 326 348\ninside 349 372

19333 GCF_001245345.1_Salmonella_enterica_CVM_N50422_v1.0 Salmonella enterica subsp. enterica serovar Saintpaul Proteobacteria; Gammaproteobacteria; Enterobacterales; Enterobacteriaceae; Salmonella; Salmonella enterica; Salmonella enterica subsp. enterica MTGDNTLITSHGINRRDFMKLCAALAATMGLSSKAAAEMAESVSNPQRPPVIWIGAQECT WP_000145429.1 hydrogenase 2 small subunit [Salmonella enterica] Length: 372\nNumber of predicted TMHs: 1\nExp number of AAs in TMHs: 24.51309\nExp number, first 60 AAs: 0.2682\nTotal prob of N-in: 0.14402\noutside 1 331\nTMhelix 332 354\ninside 355 372

19334 GCF_001834385.1_ASM183438v1 Salmonella enterica subsp. enterica serovar Saintpaul Proteobacteria; Gammaproteobacteria; Enterobacterales; Enterobacteriaceae; Salmonella; Salmonella enterica; Salmonella enterica subsp. enterica MTGDNTLITSHGINRRDFMKLCAALAATMGLSSKAAAEMAESVSNPQRPPVIWIGAQECT WP_000145429.1 hydrogenase 2 small subunit [Salmonella enterica] Length: 372\nNumber of predicted TMHs: 1\nExp number of AAs in TMHs: 24.51309\nExp number, first 60 AAs: 0.2682\nTotal prob of N-in: 0.14402\noutside 1 331\nTMhelix 332 354\ninside 355 372

19335 GCF_001478245.1_Salmonella_enterica_CVM_N32763-SQ_v1.0 Salmonella enterica subsp. enterica serovar Saintpaul Proteobacteria; Gammaproteobacteria; Enterobacterales; Enterobacteriaceae; Salmonella; Salmonella enterica; Salmonella enterica subsp. enterica MTGDNTLITSHGINRRDFMKLCAALAATMGLSSKAAAEMAESVSNPQRPPVIWIGAQECT WP_000145429.1 hydrogenase 2 small subunit [Salmonella enterica] Length: 372\nNumber of predicted TMHs: 1\nExp number of AAs in TMHs: 24.51309\nExp number, first 60 AAs: 0.2682\nTotal prob of N-in: 0.14402\noutside 1 331\nTMhelix 332 354\ninside 355 372

19336 GCF_001833225.1_ASM183322v1 Salmonella enterica subsp. enterica serovar Saintpaul Proteobacteria; Gammaproteobacteria; Enterobacterales; Enterobacteriaceae; Salmonella; Salmonella enterica; Salmonella enterica subsp. enterica MNNEETFYQAMRRKGVTRRSFLKFCSLAATSLGLGAGMTPKIAWALENKPRIPVVWIHGL WP_001058311.1 hydrogenase [Salmonella enterica] Length: 372\nNumber of predicted TMHs: 2\nExp number of AAs in TMHs: 37.4699899999999\nExp number, first 60 AAs: 19.72699\nTotal prob of N-in: 0.96171\nPOSSIBLE N-term signal sequence\ninside 1 19\nTMhelix 20 39\noutside 40 325\nTMhelix 326 348\ninside 349 372

19337 GCF_001833965.1_ASM183396v1 Salmonella enterica subsp. enterica serovar Saintpaul Proteobacteria; Gammaproteobacteria; Enterobacterales; Enterobacteriaceae; Salmonella; Salmonella enterica; Salmonella enterica subsp. enterica MNNEETFYQAMRRKGVTRRSFLKFCSLAATSLGLGAGMTPKIAWALENKPRIPVVWIHGL WP_001058311.1 hydrogenase [Salmonella enterica] Length: 372\nNumber of predicted TMHs: 2\nExp number of AAs in TMHs: 37.4699899999999\nExp number, first 60 AAs: 19.72699\nTotal prob of N-in: 0.96171\nPOSSIBLE N-term signal sequence\ninside 1 19\nTMhelix 20 39\noutside 40 325\nTMhelix 326 348\ninside 349 372

19338 GCF_001478155.1_Salmonella_enterica_CVM_N31414-SQ_v1.0 Salmonella enterica subsp. enterica serovar Saintpaul Proteobacteria; Gammaproteobacteria; Enterobacterales; Enterobacteriaceae; Salmonella; Salmonella enterica; Salmonella enterica subsp. enterica MTGDNTLITSHGINRRDFMKLCAALAATMGLSSKAAAEMAESVSNPQRPPVIWIGAQECT WP_000145429.1 hydrogenase 2 small subunit [Salmonella enterica] Length: 372\nNumber of predicted TMHs: 1\nExp number of AAs in TMHs: 24.51309\nExp number, first 60 AAs: 0.2682\nTotal prob of N-in: 0.14402\noutside 1 331\nTMhelix 332 354\ninside 355 372

19339 GCF_001834465.1_ASM183446v1 Salmonella enterica subsp. enterica serovar Saintpaul Proteobacteria; Gammaproteobacteria; Enterobacterales; Enterobacteriaceae; Salmonella; Salmonella enterica; Salmonella enterica subsp. enterica MTGDNTLITSHGINRRDFMKLCAALAATMGLSSKAAAEMAESVSNPQRPPVIWIGAQECT WP_000145429.1 hydrogenase 2 small subunit [Salmonella enterica] Length: 372\nNumber of predicted TMHs: 1\nExp number of AAs in TMHs: 24.51309\nExp number, first 60 AAs: 0.2682\nTotal prob of N-in: 0.14402\noutside 1 331\nTMhelix 332 354\ninside 355 372

19340 GCF_001477945.1_Salmonella_enterica_CVM_N30642-R_v1.0 Salmonella enterica subsp. enterica serovar Saintpaul Proteobacteria; Gammaproteobacteria; Enterobacterales; Enterobacteriaceae; Salmonella; Salmonella enterica; Salmonella enterica subsp. enterica MTGDNTLITSHGINRRDFMKLCAALAATMGLSSKAAAEMAESVSNPQRPPVIWIGAQECT

WP_000145429.1 hydrogenase 2 small subunit [Salmonella enterica] Length: 372\nNumber of predicted TMHs: 1\nExp number of AAs in TMHs: 24.51309\nExp number, first 60 AAs: 0.2682\nTotal prob of N-in: 0.14402\noutside 1 331\nTMhelix 332 354\ninside 355 372

19341 GCF_001243085.1_Salmonella_enterica_CVM_N46823_v1.0 Salmonella enterica subsp. enterica serovar Saintpaul Proteobacteria; Gammaproteobacteria; Enterobacterales; Enterobacteriaceae; Salmonella; Salmonella enterica; Salmonella enterica subsp. enterica MNNEETFYQAMRRKGVTRRSFLKFCSLAATSLGLGAGMTPKIAWALENKPRIPVVWIHGL WP_001058312.1 hydrogenase [Salmonella enterica] Length: 372\nNumber of predicted TMHs: 2\nExp number of AAs in TMHs: 36.51738\nExp number, first 60 AAs: 19.72849\nTotal prob of N-in: 0.96170\nPOSSIBLE N-term signal sequence\ninside 1 19\nTMhelix 20 39\noutside 40 325\nTMhelix 326 348\ninside 349 372

19342 GCF_001478245.1_Salmonella_enterica_CVM_N32763-SQ_v1.0 Salmonella enterica subsp. enterica serovar Saintpaul Proteobacteria; Gammaproteobacteria; Enterobacterales; Enterobacteriaceae; Salmonella; Salmonella enterica; Salmonella enterica subsp. enterica MNNEETFYQAMRRKGVTRRSFLKFCSLAATSLGLGAGMTPKIAWALENKPRIPVVWIHGL WP_001058312.1 hydrogenase [Salmonella enterica] Length: 372\nNumber of predicted TMHs: 2\nExp number of AAs in TMHs: 36.51738\nExp number, first 60 AAs: 19.72849\nTotal prob of N-in: 0.96170\nPOSSIBLE N-term signal sequence\ninside 1 19\nTMhelix 20 39\noutside 40 325\nTMhelix 326 348\ninside 349 372

19343 GCF_001243445.1_Salmonella_enterica_CVM_N46846-R_v1.0 Salmonella enterica subsp. enterica serovar Saintpaul Proteobacteria; Gammaproteobacteria; Enterobacterales; Enterobacteriaceae; Salmonella; Salmonella enterica; Salmonella enterica subsp. enterica MNNEETFYQAMRRKGVTRRSFLKFCSLAATSLGLGAGMTPKIAWALENKPRIPVVWIHGL WP_001058312.1 hydrogenase [Salmonella enterica] Length: 372\nNumber of predicted TMHs: 2\nExp number of AAs in TMHs: 36.51738\nExp number, first 60 AAs: 19.72849\nTotal prob of N-in: 0.96170\nPOSSIBLE N-term signal sequence\ninside 1 19\nTMhelix 20 39\noutside 40 325\nTMhelix 326 348\ninside 349 372

19344 GCF_001477945.1_Salmonella_enterica_CVM_N30642-R_v1.0 Salmonella enterica subsp. enterica serovar Saintpaul Proteobacteria; Gammaproteobacteria; Enterobacterales; Enterobacteriaceae; Salmonella; Salmonella enterica; Salmonella enterica subsp. enterica MNNEETFYQAMRRKGVTRRSFLKFCSLAATSLGLGAGMTPKIAWALENKPRIPVVWIHGL WP_001058312.1 hydrogenase [Salmonella enterica] Length: 372\nNumber of predicted TMHs: 2\nExp number of AAs in TMHs: 36.51738\nExp number, first 60 AAs: 19.72849\nTotal prob of N-in: 0.96170\nPOSSIBLE N-term signal sequence\ninside 1 19\nTMhelix 20 39\noutside 40 325\nTMhelix 326 348\ninside 349 372

19345 GCF_001835315.1_ASM183531v1 Salmonella enterica subsp. enterica serovar Saintpaul Proteobacteria; Gammaproteobacteria; Enterobacterales; Enterobacteriaceae; Salmonella; Salmonella enterica; Salmonella enterica subsp. enterica MNNEETFYQAMRRKGVTRRSFLKFCSLAATSLGLGAGMTPKIAWALENKPRIPVVWIHGL WP_001058311.1 hydrogenase [Salmonella enterica] Length: 372\nNumber of predicted TMHs: 2\nExp number of AAs in TMHs: 37.4699899999999\nExp number, first 60 AAs: 19.72699\nTotal prob of N-in: 0.96171\nPOSSIBLE N-term signal sequence\ninside 1 19\nTMhelix 20 39\noutside 40 325\nTMhelix 326 348\ninside 349 372

19346 GCF_001833805.1_ASM183380v1 Salmonella enterica subsp. enterica serovar Saintpaul Proteobacteria; Gammaproteobacteria; Enterobacterales; Enterobacteriaceae; Salmonella; Salmonella enterica; Salmonella enterica subsp. enterica MNNEETFYQAMRRKGVTRRSFLKFCSLAATSLGLGAGMTPKIAWALENKPRIPVVWIHGL WP_001058311.1 hydrogenase [Salmonella enterica] Length: 372\nNumber of predicted TMHs: 2\nExp number of AAs in TMHs: 37.4699899999999\nExp number, first 60 AAs: 19.72699\nTotal prob of N-in: 0.96171\nPOSSIBLE N-term signal sequence\ninside 1 19\nTMhelix 20 39\noutside 40 325\nTMhelix 326 348\ninside 349 372

19347 GCF_002032725.1_ASM203272v1 Salmonella enterica subsp. enterica serovar Saintpaul Proteobacteria; Gammaproteobacteria; Enterobacterales; Enterobacteriaceae; Salmonella; Salmonella enterica; Salmonella enterica subsp. enterica MTGDNTLITSHGINRRDFMKLCAALAATMGLSSKAAAEMAESVSNPQRPPVIWIGAQECT WP_000145429.1 hydrogenase 2 small subunit [Salmonella enterica] Length: 372\nNumber of predicted TMHs: 1\nExp number of AAs in TMHs: 24.51309\nExp number, first 60 AAs: 0.2682\nTotal prob of N-in: 0.14402\noutside 1 331\nTMhelix 332 354\ninside 355 372

19348 GCF_001834535.1_ASM183453v1 Salmonella enterica subsp. enterica serovar Saintpaul Proteobacteria; Gammaproteobacteria; Enterobacterales; Enterobacteriaceae; Salmonella; Salmonella enterica; Salmonella enterica subsp. enterica MTGDNTLITSHGINRRDFMKLCAALAATMGLSSKAAAEMAESVSNPQRPPVIWIGAQECT WP_000145429.1 hydrogenase 2 small subunit [Salmonella enterica] Length: 372\nNumber of predicted TMHs: 1\nExp number of AAs in TMHs: 24.51309\nExp number, first 60 AAs: 0.2682\nTotal prob of N-in: 0.14402\noutside 1 331\nTMhelix 332 354\ninside 355 372

19349 GCF_001478685.1_Salmonella_enterica_CVM_N29357-SQ_v1.0 Salmonella enterica subsp. enterica serovar Saintpaul Proteobacteria; Gammaproteobacteria; Enterobacterales; Enterobacteriaceae; Salmonella; Salmonella enterica; Salmonella enterica subsp. enterica MNNEETFYQAMRRKGVTRRSFLKFCSLAATSLGLGAGMTPKIAWALENKPRIPVVWIHGL WP_001058312.1 hydrogenase [Salmonella enterica] Length: 372\nNumber of predicted TMHs: 2\nExp number of AAs in TMHs: 36.51738\nExp number, first 60 AAs: 19.72849\nTotal prob of N-in: 0.96170\nPOSSIBLE N-term signal sequence\ninside 1 19\nTMhelix 20 39\noutside 40 325\nTMhelix 326 348\ninside 349 372

19350 GCF_001481045.1_Salmonella_enterica_CVM_N42458_v1.0 Salmonella enterica subsp. enterica serovar Saintpaul Proteobacteria; Gammaproteobacteria; Enterobacterales; Enterobacteriaceae; Salmonella; Salmonella enterica; Salmonella enterica subsp. enterica MNNEETFYQAMRRKGVTRRSFLKFCSLAATSLGLGAGMTPKIAWALENKPRIPVVWIHGL WP_001058312.1 hydrogenase [Salmonella enterica] Length: 372\nNumber of predicted TMHs: 2\nExp number of AAs in TMHs: 36.51738\nExp number, first 60 AAs: 19.72849\nTotal prob of N-in: 0.96170\nPOSSIBLE N-term signal sequence\ninside 1 19\nTMhelix 20 39\noutside 40 325\nTMhelix 326 348\ninside 349 372

19351 GCF_001833225.1_ASM183322v1 Salmonella enterica subsp. enterica serovar Saintpaul Proteobacteria; Gammaproteobacteria; Enterobacterales; Enterobacteriaceae; Salmonella; Salmonella enterica; Salmonella enterica subsp. enterica MTGDNTLITSHGINRRDFMKLCAALAATMGLSSKAAAEMAESVSNPQRPPVIWIGAQECT WP_000145429.1 hydrogenase 2 small subunit [Salmonella enterica] Length: 372\nNumber of predicted TMHs: 1\nExp number of AAs in TMHs: 24.51309\nExp number, first 60 AAs: 0.2682\nTotal prob of N-in: 0.14402\noutside 1 331\nTMhelix 332 354\ninside 355 372

19352 GCF_001481045.1_Salmonella_enterica_CVM_N42458_v1.0 Salmonella enterica subsp. enterica serovar Saintpaul Proteobacteria; Gammaproteobacteria; Enterobacterales; Enterobacteriaceae; Salmonella; Salmonella enterica; Salmonella enterica subsp. enterica MTGDNTLITSHGINRRDFMKLCAALAATMGLSSKAAAEMAESVSNPQRPPVIWIGAQECT WP_000145429.1 hydrogenase 2 small subunit [Salmonella enterica] Length: 372\nNumber of predicted TMHs: 1\nExp number of AAs in TMHs: 24.51309\nExp number, first 60 AAs: 0.2682\nTotal prob of N-in: 0.14402\noutside 1 331\nTMhelix 332 354\ninside 355 372

19353 GCF_001833345.1_ASM183334v1 Salmonella enterica subsp. enterica serovar Saintpaul Proteobacteria; Gammaproteobacteria; Enterobacterales; Enterobacteriaceae; Salmonella; Salmonella enterica; Salmonella enterica subsp. enterica MNNEETFYQAMRRKGVTRRSFLKFCSLAATSLGLGAGMTPKIAWALENKPRIPVVWIHGL WP_001058311.1 hydrogenase [Salmonella enterica] Length: 372\nNumber of predicted TMHs: 2\nExp number of AAs in TMHs: 37.4699899999999\nExp number, first 60 AAs: 19.72699\nTotal prob of N-in: 0.96171\nPOSSIBLE N-term signal sequence\ninside 1 19\nTMhelix 20 39\noutside 40 325\nTMhelix 326 348\ninside 349 372

19354 GCF_001834535.1_ASM183453v1 Salmonella enterica subsp. enterica serovar Saintpaul Proteobacteria; Gammaproteobacteria; Enterobacterales; Enterobacteriaceae; Salmonella; Salmonella enterica; Salmonella enterica subsp. enterica MNNEETFYQAMRRKGVTRRSFLKFCSLAATSLGLGAGMTPKIAWALENKPRIPVVWIHGL WP_001058311.1 hydrogenase [Salmonella enterica] Length: 372\nNumber of predicted TMHs: 2\nExp number of AAs in TMHs: 37.4699899999999\nExp number, first 60 AAs: 19.72699\nTotal prob of N-in: 0.96171\nPOSSIBLE N-term signal sequence\ninside 1 19\nTMhelix 20 39\noutside 40 325\nTMhelix 326 348\ninside 349 372

19355 GCF_001834495.1_ASM183449v1 Salmonella enterica subsp. enterica serovar Saintpaul Proteobacteria; Gammaproteobacteria; Enterobacterales; Enterobacteriaceae; Salmonella; Salmonella enterica; Salmonella enterica subsp. enterica MTGDNTLITSHGINRRDFMKLCAALAATMGLSSKAAAEMAESVSNPQRPPVIWIGAQECT WP_000145429.1 hydrogenase 2 small subunit [Salmonella enterica] Length: 372\nNumber of predicted TMHs: 1\nExp number of AAs in TMHs: 24.51309\nExp number, first 60 AAs: 0.2682\nTotal prob of N-in: 0.14402\noutside 1 331\nTMhelix 332 354\ninside 355 372

19356 GCF_001242655.1_Salmonella_enterica_CVM_N45953_v1.0 Salmonella enterica subsp. enterica serovar Saintpaul Proteobacteria; Gammaproteobacteria; Enterobacterales; Enterobacteriaceae; Salmonella; Salmonella enterica; Salmonella enterica subsp. enterica MNNEETFYQAMRRKGVTRRSFLKFCSLAATSLGLGAGMTPKIAWALENKPRIPVVWIHGL WP_001058312.1 hydrogenase [Salmonella enterica] Length: 372\nNumber of predicted TMHs: 2\nExp number of AAs in TMHs: 36.51738\nExp number, first 60 AAs: 19.72849\nTotal prob of N-in: 0.96170\nPOSSIBLE N-term signal sequence\ninside 1 19\nTMhelix 20 39\noutside 40 325\nTMhelix 326 348\ninside 349 372

19357 GCF_001242655.1_Salmonella_enterica_CVM_N45953_v1.0 Salmonella enterica subsp. enterica serovar Saintpaul Proteobacteria; Gammaproteobacteria; Enterobacterales; Enterobacteriaceae; Salmonella; Salmonella enterica; Salmonella enterica subsp. enterica MTGDNTLITSHGINRRDFMKLCAALAATMGLSSKAAAEMAESVSNPQRPPVIWIGAQECT

WP_000145429.1 hydrogenase 2 small subunit [Salmonella enterica] Length: 372\nNumber of predicted TMHs: 1\nExp number of AAs in TMHs: 24.51309\nExp number, first 60 AAs: 0.2682\nTotal prob of N-in: 0.14402\noutside 1 331\nTMhelix 332 354\ninside 355 372

19358 GCF_002031465.1_ASM203146v1 Salmonella enterica subsp. enterica serovar Saintpaul Proteobacteria; Gammaproteobacteria; Enterobacterales; Enterobacteriaceae; Salmonella; Salmonella enterica; Salmonella enterica subsp. enterica MTGDNTLITSHGINRRDFMKLCAALAATMGLSSKAAAEMAESVSNPQRPPVIWIGAQECT WP_000145429.1 hydrogenase 2 small subunit [Salmonella enterica] Length: 372\nNumber of predicted TMHs: 1\nExp number of AAs in TMHs: 24.51309\nExp number, first 60 AAs: 0.2682\nTotal prob of N-in: 0.14402\noutside 1 331\nTMhelix 332 354\ninside 355 372

19359 GCF_001833965.1_ASM183396v1 Salmonella enterica subsp. enterica serovar Saintpaul Proteobacteria; Gammaproteobacteria; Enterobacterales; Enterobacteriaceae; Salmonella; Salmonella enterica; Salmonella enterica subsp. enterica MTGDNTLITSHGINRRDFMKLCAALAATMGLSSKAAAEMAESVSNPQRPPVIWIGAQECT WP_000145429.1 hydrogenase 2 small subunit [Salmonella enterica] Length: 372\nNumber of predicted TMHs: 1\nExp number of AAs in TMHs: 24.51309\nExp number, first 60 AAs: 0.2682\nTotal prob of N-in: 0.14402\noutside 1 331\nTMhelix 332 354\ninside 355 372

19360 GCF_001834495.1_ASM183449v1 Salmonella enterica subsp. enterica serovar Saintpaul Proteobacteria; Gammaproteobacteria; Enterobacterales; Enterobacteriaceae; Salmonella; Salmonella enterica; Salmonella enterica subsp. enterica MNNEETFYQAMRRKGVTRRSFLKFCSLAATSLGLGAGMTPKIAWALENKPRIPVVWIHGL WP_001058311.1 hydrogenase [Salmonella enterica] Length: 372\nNumber of predicted TMHs: 2\nExp number of AAs in TMHs: 37.4699899999999\nExp number, first 60 AAs: 19.72699\nTotal prob of N-in: 0.96171\nPOSSIBLE N-term signal sequence\ninside 1 19\nTMhelix 20 39\noutside 40 325\nTMhelix 326 348\ninside 349 372

19361 GCF_001241665.1_Salmonella_enterica_CVM_N45394_v1.0 Salmonella enterica subsp. enterica serovar Saintpaul Proteobacteria; Gammaproteobacteria; Enterobacterales; Enterobacteriaceae; Salmonella; Salmonella enterica; Salmonella enterica subsp. enterica MNNEETFYQAMRRKGVTRRSFLKFCSLAATSLGLGAGMTPKIAWALENKPRIPVVWIHGL WP_001058312.1 hydrogenase [Salmonella enterica] Length: 372\nNumber of predicted TMHs: 2\nExp number of AAs in TMHs: 36.51738\nExp number, first 60 AAs: 19.72849\nTotal prob of N-in: 0.96170\nPOSSIBLE N-term signal sequence\ninside 1 19\nTMhelix 20 39\noutside 40 325\nTMhelix 326 348\ninside 349 372

19362 GCF_001478685.1_Salmonella_enterica_CVM_N29357-SQ_v1.0 Salmonella enterica subsp. enterica serovar Saintpaul Proteobacteria; Gammaproteobacteria; Enterobacterales; Enterobacteriaceae; Salmonella; Salmonella enterica; Salmonella enterica subsp. enterica MTGDNTLITSHGINRRDFMKLCAALAATMGLSSKAAAEMAESVSNPQRPPVIWIGAQECT WP_000145429.1 hydrogenase 2 small subunit [Salmonella enterica] Length: 372\nNumber of predicted TMHs: 1\nExp number of AAs in TMHs: 24.51309\nExp number, first 60 AAs: 0.2682\nTotal prob of N-in: 0.14402\noutside 1 331\nTMhelix 332 354\ninside 355 372

19363 GCF_001833815.1_ASM183381v1 Salmonella enterica subsp. enterica serovar Saintpaul Proteobacteria; Gammaproteobacteria; Enterobacterales; Enterobacteriaceae; Salmonella; Salmonella enterica; Salmonella enterica subsp. enterica MTGDNTLITSHGINRRDFMKLCAALAATMGLSSKAAAEMAESVSNPQRPPVIWIGAQECT WP_000145429.1 hydrogenase 2 small subunit [Salmonella enterica] Length: 372\nNumber of predicted TMHs: 1\nExp number of AAs in TMHs: 24.51309\nExp number, first 60 AAs: 0.2682\nTotal prob of N-in: 0.14402\noutside 1 331\nTMhelix 332 354\ninside 355 372

19364 GCF_002031275.1_ASM203127v1 Salmonella enterica subsp. enterica serovar Saintpaul Proteobacteria; Gammaproteobacteria; Enterobacterales; Enterobacteriaceae; Salmonella; Salmonella enterica; Salmonella enterica subsp. enterica MNNEETFYQAMRRKGVTRRSFLKFCSLAATSLGLGAGMTPKIAWALENKPRIPVVWIHGL WP_001058312.1 hydrogenase [Salmonella enterica] Length: 372\nNumber of predicted TMHs: 2\nExp number of AAs in TMHs: 36.51738\nExp number, first 60 AAs: 19.72849\nTotal prob of N-in: 0.96170\nPOSSIBLE N-term signal sequence\ninside 1 19\nTMhelix 20 39\noutside 40 325\nTMhelix 326 348\ninside 349 372

19365 GCF_001953695.1_ASM195369v1 Salmonella enterica subsp. enterica serovar Saintpaul Proteobacteria; Gammaproteobacteria; Enterobacterales; Enterobacteriaceae; Salmonella; Salmonella enterica; Salmonella enterica subsp. enterica MNNEETFYQAMRRKGVTRRSFLKFCSLAATSLGLGAGMTPKIAWALENKPRIPVVWIHGL WP_001058312.1 hydrogenase [Salmonella enterica] Length: 372\nNumber of predicted TMHs: 2\nExp number of AAs in TMHs: 36.51738\nExp number, first 60 AAs: 19.72849\nTotal prob of N-in: 0.96170\nPOSSIBLE N-term signal sequence\ninside 1 19\nTMhelix 20 39\noutside 40 325\nTMhelix 326 348\ninside 349 372

19366 GCF_001479385.1_Salmonella_enterica_CVM_N32053-SQ_v1.0 Salmonella enterica subsp. enterica serovar Saintpaul Proteobacteria; Gammaproteobacteria; Enterobacterales; Enterobacteriaceae; Salmonella; Salmonella enterica; Salmonella enterica subsp. enterica MTGDNTLITSHGINRRDFMKLCAALAATMGLSSKAAAEMAESVSNPQRPPVIWIGAQECT WP_000145429.1 hydrogenase 2 small subunit [Salmonella enterica] Length: 372\nNumber of predicted TMHs: 1\nExp number of AAs in TMHs: 24.51309\nExp number, first 60 AAs: 0.2682\nTotal prob of N-in: 0.14402\noutside 1 331\nTMhelix 332 354\ninside 355 372

19367 GCF_001479215.1_Salmonella_enterica_CVM_N31593-SQ_v1.0 Salmonella enterica subsp. enterica serovar Saintpaul Proteobacteria; Gammaproteobacteria; Enterobacterales; Enterobacteriaceae; Salmonella; Salmonella enterica; Salmonella enterica subsp. enterica MTGDNTLITSHGINRRDFMKLCAALAATMGLSSKAAAEMAESVSNPQRPPVIWIGAQECT WP_000145429.1 hydrogenase 2 small subunit [Salmonella enterica] Length: 372\nNumber of predicted TMHs: 1\nExp number of AAs in TMHs: 24.51309\nExp number, first 60 AAs: 0.2682\nTotal prob of N-in: 0.14402\noutside 1 331\nTMhelix 332 354\ninside 355 372

19368 GCF_001479385.1_Salmonella_enterica_CVM_N32053-SQ_v1.0 Salmonella enterica subsp. enterica serovar Saintpaul Proteobacteria; Gammaproteobacteria; Enterobacterales; Enterobacteriaceae; Salmonella; Salmonella enterica; Salmonella enterica subsp. enterica MNNEETFYQAMRRKGVTRRSFLKFCSLAATSLGLGAGMTPKIAWALENKPRIPVVWIHGL WP_001058312.1 hydrogenase [Salmonella enterica] Length: 372\nNumber of predicted TMHs: 2\nExp number of AAs in TMHs: 36.51738\nExp number, first 60 AAs: 19.72849\nTotal prob of N-in: 0.96170\nPOSSIBLE N-term signal sequence\ninside 1 19\nTMhelix 20 39\noutside 40 325\nTMhelix 326 348\ninside 349 372

19369 GCF_001832855.1_ASM183285v1 Salmonella enterica subsp. enterica serovar Saintpaul Proteobacteria; Gammaproteobacteria; Enterobacterales; Enterobacteriaceae; Salmonella; Salmonella enterica; Salmonella enterica subsp. enterica MTGDNTLITSHGINRRDFMKLCAALAATMGLSSKAAAEMAESVSNPQRPPVIWIGAQECT WP_000145429.1 hydrogenase 2 small subunit [Salmonella enterica] Length: 372\nNumber of predicted TMHs: 1\nExp number of AAs in TMHs: 24.51309\nExp number, first 60 AAs: 0.2682\nTotal prob of N-in: 0.14402\noutside 1 331\nTMhelix 332 354\ninside 355 372

19370 GCF_001952995.1_ASM195299v1 Salmonella enterica subsp. enterica serovar Saintpaul Proteobacteria; Gammaproteobacteria; Enterobacterales; Enterobacteriaceae; Salmonella; Salmonella enterica; Salmonella enterica subsp. enterica MNNEETFYQAMRRKGVTRRSFLKFCSLAATSLGLGAGMTPKIAWALENKPRIPVVWIHGL WP_001058312.1 hydrogenase [Salmonella enterica] Length: 372\nNumber of predicted TMHs: 2\nExp number of AAs in TMHs: 36.51738\nExp number, first 60 AAs: 19.72849\nTotal prob of N-in: 0.96170\nPOSSIBLE N-term signal sequence\ninside 1 19\nTMhelix 20 39\noutside 40 325\nTMhelix 326 348\ninside 349 372

19371 GCF_001953695.1_ASM195369v1 Salmonella enterica subsp. enterica serovar Saintpaul Proteobacteria; Gammaproteobacteria; Enterobacterales; Enterobacteriaceae; Salmonella; Salmonella enterica; Salmonella enterica subsp. enterica MTGDNTLITSHGINRRDFMKLCAALAATMGLSSKAAAEMAESVSNPQRPPVIWIGAQECT WP_000145429.1 hydrogenase 2 small subunit [Salmonella enterica] Length: 372\nNumber of predicted TMHs: 1\nExp number of AAs in TMHs: 24.51309\nExp number, first 60 AAs: 0.2682\nTotal prob of N-in: 0.14402\noutside 1 331\nTMhelix 332 354\ninside 355 372

19372 GCF_001478225.1_Salmonella_enterica_CVM_N32750-SQ_v1.0 Salmonella enterica subsp. enterica serovar Saintpaul Proteobacteria; Gammaproteobacteria; Enterobacterales; Enterobacteriaceae; Salmonella; Salmonella enterica; Salmonella enterica subsp. enterica MTGDNTLITSHGINRRDFMKLCAALAATMGLSSKAAAEMAESVSNPQRPPVIWIGAQECT WP_000145429.1 hydrogenase 2 small subunit [Salmonella enterica] Length: 372\nNumber of predicted TMHs: 1\nExp number of AAs in TMHs: 24.51309\nExp number, first 60 AAs: 0.2682\nTotal prob of N-in: 0.14402\noutside 1 331\nTMhelix 332 354\ninside 355 372

19373 GCF_001478225.1_Salmonella_enterica_CVM_N32750-SQ_v1.0 Salmonella enterica subsp. enterica serovar Saintpaul Proteobacteria; Gammaproteobacteria; Enterobacterales; Enterobacteriaceae; Salmonella; Salmonella enterica; Salmonella enterica subsp. enterica MNNEETFYQAMRRKGVTRRSFLKFCSLAATSLGLGAGMTPKIAWALENKPRIPVVWIHGL WP_001058312.1 hydrogenase [Salmonella enterica] Length: 372\nNumber of predicted TMHs: 2\nExp number of AAs in TMHs: 36.51738\nExp number, first 60 AAs: 19.72849\nTotal prob of N-in: 0.96170\nPOSSIBLE N-term signal sequence\ninside 1 19\nTMhelix 20 39\noutside 40 325\nTMhelix 326 348\ninside 349 372

19374 GCF_001477905.1_Salmonella_enterica_CVM_N29360-SQ_v1.0 Salmonella enterica subsp. enterica serovar Saintpaul Proteobacteria; Gammaproteobacteria; Enterobacterales; Enterobacteriaceae; Salmonella; Salmonella enterica; Salmonella enterica subsp. enterica MTGDNTLITSHGINRRDFMKLCAALAATMGLSSKAAAEMAESVSNPQRPPVIWIGAQECT WP_000145429.1 hydrogenase 2 small subunit [Salmonella enterica] Length: 372\nNumber of

predicted TMHs:  1\nExp number of AAs in TMHs: 24.51309\nExp number, first 60 AAs:  0.2682\nTotal prob of N-in: 0.14402\noutside   1   331\nTMhelix   332   354\ninside   355   372

19375 GCF_001833935.1_ASM183393v1 Salmonella enterica subsp. enterica serovar Saintpaul Proteobacteria; Gammaproteobacteria; Enterobacterales; Enterobacteriaceae; Salmonella; Salmonella enterica; Salmonella enterica subsp. enterica MTGDNTLITSHGINRRDFMKLCAALAATMGLSSKAAAEMAESVSNPQRPPVIWIGAQECT WP_000145429.1 hydrogenase 2 small subunit [Salmonella enterica] Length: 372\nNumber of predicted TMHs:  1\nExp number of AAs in TMHs: 24.51309\nExp number, first 60 AAs:  0.2682\nTotal prob of N-in: 0.14402\noutside   1   331\nTMhelix   332   354\ninside   355   372

19376 GCF_001833855.1_ASM183385v1 Salmonella enterica subsp. enterica serovar Saintpaul Proteobacteria; Gammaproteobacteria; Enterobacterales; Enterobacteriaceae; Salmonella; Salmonella enterica; Salmonella enterica subsp. enterica MTGDNTLITSHGINRRDFMKLCAALAATMGLSSKAAAEMAESVSNPQRPPVIWIGAQECT WP_000145429.1 hydrogenase 2 small subunit [Salmonella enterica] Length: 372\nNumber of predicted TMHs:  1\nExp number of AAs in TMHs: 24.51309\nExp number, first 60 AAs:  0.2682\nTotal prob of N-in: 0.14402\noutside   1   331\nTMhelix   332   354\ninside   355   372

19377 GCF_001479215.1_Salmonella_enterica_CVM_N31593-SQ_v1.0 Salmonella enterica subsp. enterica serovar Saintpaul Proteobacteria; Gammaproteobacteria; Enterobacterales; Enterobacteriaceae; Salmonella; Salmonella enterica; Salmonella enterica subsp. enterica MNNEETFYQAMRRKGVTRRSFLKFCSLAATSLGLGAGMTPKIAWALENKPRIPVVWIHGL WP_001058312.1 hydrogenase [Salmonella enterica] Length: 372\nNumber of predicted TMHs:  2\nExp number of AAs in TMHs: 36.51738\nExp number, first 60 AAs:  19.72849\nTotal prob of N-in:   0.96170\nPOSSIBLE N-term signal sequence\ninside   1   19\nTMhelix   20   39\noutside   40   325\nTMhelix   326   348\ninside   349   372

19378 GCF_001833655.1_ASM183365v1 Salmonella enterica subsp. enterica serovar Saintpaul Proteobacteria; Gammaproteobacteria; Enterobacterales; Enterobacteriaceae; Salmonella; Salmonella enterica; Salmonella enterica subsp. enterica MTGDNTLITSHGINRRDFMKLCAALAATMGLSSKAAAEMAESVSNPQRPPVIWIGAQECT WP_000145429.1 hydrogenase 2 small subunit [Salmonella enterica] Length: 372\nNumber of predicted TMHs:  1\nExp number of AAs in TMHs: 24.51309\nExp number, first 60 AAs:  0.2682\nTotal prob of N-in: 0.14402\noutside   1   331\nTMhelix   332   354\ninside   355   372

19379 GCF_001833315.1_ASM183331v1 Salmonella enterica subsp. enterica serovar Saintpaul Proteobacteria; Gammaproteobacteria; Enterobacterales; Enterobacteriaceae; Salmonella; Salmonella enterica; Salmonella enterica subsp. enterica MNNEETFYQAMRRKGVTRRSFLKFCSLAATSLGLGAGMTPKIAWALENKPRIPVVWIHGL WP_001058311.1 hydrogenase [Salmonella enterica] Length: 372\nNumber of predicted TMHs:  2\nExp number of AAs in TMHs: 37.4699899999999\nExp number, first 60 AAs:  19.72699\nTotal prob of N-in: 0.96171\nPOSSIBLE N-term signal sequence\ninside   1   19\nTMhelix   20   39\noutside   40   325\nTMhelix   326 348\ninside   349   372

19380 GCF_001834525.1_ASM183452v1 Salmonella enterica subsp. enterica serovar Saintpaul Proteobacteria; Gammaproteobacteria; Enterobacterales; Enterobacteriaceae; Salmonella; Salmonella enterica; Salmonella enterica subsp. enterica MNNEETFYQAMRRKGVTRRSFLKFCSLAATSLGLGAGMTPKIAWALENKPRIPVVWIHGL WP_001058311.1 hydrogenase [Salmonella enterica] Length: 372\nNumber of predicted TMHs:  2\nExp number of AAs in TMHs: 37.4699899999999\nExp number, first 60 AAs:  19.72699\nTotal prob of N-in: 0.96171\nPOSSIBLE N-term signal sequence\ninside   1   19\nTMhelix   20   39\noutside   40   325\nTMhelix   326 348\ninside   349   372

19381 GCF_001833975.1_ASM183397v1 Salmonella enterica subsp. enterica serovar Saintpaul Proteobacteria; Gammaproteobacteria; Enterobacterales; Enterobacteriaceae; Salmonella; Salmonella enterica; Salmonella enterica subsp. enterica MNNEETFYQAMRRKGVTRRSFLKFCSLAATSLGLGAGMTPKIAWALENKPRIPVVWIHGL WP_001058311.1 hydrogenase [Salmonella enterica] Length: 372\nNumber of predicted TMHs:  2\nExp number of AAs in TMHs: 37.4699899999999\nExp number, first 60 AAs:  19.72699\nTotal prob of N-in: 0.96171\nPOSSIBLE N-term signal sequence\ninside   1   19\nTMhelix   20   39\noutside   40   325\nTMhelix   326 348\ninside   349   372

19382 GCF_001477905.1_Salmonella_enterica_CVM_N29360-SQ_v1.0 Salmonella enterica subsp. enterica serovar Saintpaul Proteobacteria; Gammaproteobacteria; Enterobacterales; Enterobacteriaceae; Salmonella; Salmonella enterica; Salmonella enterica subsp. enterica MNNEETFYQAMRRKGVTRRSFLKFCSLAATSLGLGAGMTPKIAWALENKPRIPVVWIHGL WP_001058312.1 hydrogenase [Salmonella enterica] Length: 372\nNumber of predicted TMHs:  2\nExp number of AAs in TMHs: 36.51738\nExp number, first 60 AAs:  19.72849\nTotal prob of N-in:   0.96170\nPOSSIBLE N-term signal sequence\ninside   1   19\nTMhelix   20   39\noutside   40   325\nTMhelix   326   348\ninside   349   372

19383 GCF_002032665.1_ASM203266v1 Salmonella enterica subsp. enterica serovar Saintpaul Proteobacteria; Gammaproteobacteria; Enterobacterales; Enterobacteriaceae; Salmonella; Salmonella enterica; Salmonella enterica subsp. enterica MNNEETFYQAMRRKGVTRRSFLKFCSLAATSLGLGAGMTPKIAWALENKPRIPVVWIHGL WP_001058312.1 hydrogenase [Salmonella enterica] Length: 372\nNumber of predicted TMHs: 2\nExp number of AAs in TMHs: 36.51738\nExp number, first 60 AAs: 19.72849\nTotal prob of N-in: 0.96170\nPOSSIBLE N-term signal sequence\ninside 1 19\nTMhelix 20 39\noutside 40 325\nTMhelix 326 348\ninside 349 372

19384 GCF_002032665.1_ASM203266v1 Salmonella enterica subsp. enterica serovar Saintpaul Proteobacteria; Gammaproteobacteria; Enterobacterales; Enterobacteriaceae; Salmonella; Salmonella enterica; Salmonella enterica subsp. enterica MTGDNTLITSHGINRRDFMKLCAALAATMGLSSKAAAEMAESVSNPQRPPVIWIGAQECT WP_000145429.1 hydrogenase 2 small subunit [Salmonella enterica] Length: 372\nNumber of predicted TMHs: 1\nExp number of AAs in TMHs: 24.51309\nExp number, first 60 AAs: 0.2682\nTotal prob of N-in: 0.14402\noutside 1 331\nTMhelix 332 354\ninside 355 372

19385 GCF_002032025.1_ASM203202v1 Salmonella enterica subsp. enterica serovar Saintpaul Proteobacteria; Gammaproteobacteria; Enterobacterales; Enterobacteriaceae; Salmonella; Salmonella enterica; Salmonella enterica subsp. enterica MNNEETFYQAMRRKGVTRRSFLKFCSLAATSLGLGAGMTPKIAWALENKPRIPVVWIHGL WP_001058312.1 hydrogenase [Salmonella enterica] Length: 372\nNumber of predicted TMHs: 2\nExp number of AAs in TMHs: 36.51738\nExp number, first 60 AAs: 19.72849\nTotal prob of N-in: 0.96170\nPOSSIBLE N-term signal sequence\ninside 1 19\nTMhelix 20 39\noutside 40 325\nTMhelix 326 348\ninside 349 372

19386 GCF_001834565.1_ASM183456v1 Salmonella enterica subsp. enterica serovar Saintpaul Proteobacteria; Gammaproteobacteria; Enterobacterales; Enterobacteriaceae; Salmonella; Salmonella enterica; Salmonella enterica subsp. enterica MNNEETFYQAMRRKGVTRRSFLKFCSLAATSLGLGAGMTPKIAWALENKPRIPVVWIHGL WP_001058311.1 hydrogenase [Salmonella enterica] Length: 372\nNumber of predicted TMHs: 2\nExp number of AAs in TMHs: 37.4699899999999\nExp number, first 60 AAs: 19.72699\nTotal prob of N-in: 0.96171\nPOSSIBLE N-term signal sequence\ninside 1 19\nTMhelix 20 39\noutside 40 325\nTMhelix 326 348\ninside 349 372

19387 GCF_001834605.1_ASM183460v1 Salmonella enterica subsp. enterica serovar Saintpaul Proteobacteria; Gammaproteobacteria; Enterobacterales; Enterobacteriaceae; Salmonella; Salmonella enterica; Salmonella enterica subsp. enterica MNNEETFYQAMRRKGVTRRSFLKFCSLAATSLGLGAGMTPKIAWALENKPRIPVVWIHGL WP_001058311.1 hydrogenase [Salmonella enterica] Length: 372\nNumber of predicted TMHs: 2\nExp number of AAs in TMHs: 37.4699899999999\nExp number, first 60 AAs: 19.72699\nTotal prob of N-in: 0.96171\nPOSSIBLE N-term signal sequence\ninside 1 19\nTMhelix 20 39\noutside 40 325\nTMhelix 326 348\ninside 349 372

19388 GCF_001833975.1_ASM183397v1 Salmonella enterica subsp. enterica serovar Saintpaul Proteobacteria; Gammaproteobacteria; Enterobacterales; Enterobacteriaceae; Salmonella; Salmonella enterica; Salmonella enterica subsp. enterica MTGDNTLITSHGINRRDFMKLCAALAATMGLSSKAAAEMAESVSNPQRPPVIWIGAQECT WP_000145429.1 hydrogenase 2 small subunit [Salmonella enterica] Length: 372\nNumber of predicted TMHs: 1\nExp number of AAs in TMHs: 24.51309\nExp number, first 60 AAs: 0.2682\nTotal prob of N-in: 0.14402\noutside 1 331\nTMhelix 332 354\ninside 355 372

19389 GCF_001834445.1_ASM183444v1 Salmonella enterica subsp. enterica serovar Saintpaul Proteobacteria; Gammaproteobacteria; Enterobacterales; Enterobacteriaceae; Salmonella; Salmonella enterica; Salmonella enterica subsp. enterica MNNEETFYQAMRRKGVTRRSFLKFCSLAATSLGLGAGMTPKIAWALENKPRIPVVWIHGL WP_001058311.1 hydrogenase [Salmonella enterica] Length: 372\nNumber of predicted TMHs: 2\nExp number of AAs in TMHs: 37.4699899999999\nExp number, first 60 AAs: 19.72699\nTotal prob of N-in: 0.96171\nPOSSIBLE N-term signal sequence\ninside 1 19\nTMhelix 20 39\noutside 40 325\nTMhelix 326 348\ninside 349 372

19390 GCF_002032025.1_ASM203202v1 Salmonella enterica subsp. enterica serovar Saintpaul Proteobacteria; Gammaproteobacteria; Enterobacterales; Enterobacteriaceae; Salmonella; Salmonella enterica; Salmonella enterica subsp. enterica MTGDNTLITSHGINRRDFMKLCAALAATMGLSSKAAAEMAESVSNPQRPPVIWIGAQECT WP_000145429.1 hydrogenase 2 small subunit [Salmonella enterica] Length: 372\nNumber of predicted TMHs: 1\nExp number of AAs in TMHs: 24.51309\nExp number, first 60 AAs: 0.2682\nTotal prob of N-in: 0.14402\noutside 1 331\nTMhelix 332 354\ninside 355 372

19391 GCF_001832595.1_ASM183259v1 Salmonella enterica subsp. enterica serovar Saintpaul Proteobacteria; Gammaproteobacteria; Enterobacterales; Enterobacteriaceae; Salmonella; Salmonella enterica;

Salmonella enterica subsp. enterica MNNEETFYQAMRRKGVTRRSFLKFCSLAATSLGLGAGMTPKIAWALENKPRIPVVWIHGL WP_001058311.1 hydrogenase [Salmonella enterica] Length: 372\nNumber of predicted TMHs: 2\nExp number of AAs in TMHs: 37.4699899999999\nExp number, first 60 AAs: 19.72699\nTotal prob of N-in: 0.96171\nPOSSIBLE N-term signal sequence\ninside 1 19\nTMhelix 20 39\noutside 40 325\nTMhelix 326 348\ninside 349 372

19392 GCF_001834005.1_ASM183400v1 Salmonella enterica subsp. enterica serovar Saintpaul Proteobacteria; Gammaproteobacteria; Enterobacterales; Enterobacteriaceae; Salmonella; Salmonella enterica; Salmonella enterica subsp. enterica MNNEETFYQAMRRKGVTRRSFLKFCSLAATSLGLGAGMTPKIAWALENKPRIPVVWIHGL WP_001058311.1 hydrogenase [Salmonella enterica] Length: 372\nNumber of predicted TMHs: 2\nExp number of AAs in TMHs: 37.4699899999999\nExp number, first 60 AAs: 19.72699\nTotal prob of N-in: 0.96171\nPOSSIBLE N-term signal sequence\ninside 1 19\nTMhelix 20 39\noutside 40 325\nTMhelix 326 348\ninside 349 372

19393 GCF_001834305.1_ASM183430v1 Salmonella enterica subsp. enterica serovar Saintpaul Proteobacteria; Gammaproteobacteria; Enterobacterales; Enterobacteriaceae; Salmonella; Salmonella enterica; Salmonella enterica subsp. enterica MNNEETFYQAMRRKGVTRRSFLKFCSLAATSLGLGAGMTPKIAWALENKPRIPVVWIHGL WP_001058311.1 hydrogenase [Salmonella enterica] Length: 372\nNumber of predicted TMHs: 2\nExp number of AAs in TMHs: 37.4699899999999\nExp number, first 60 AAs: 19.72699\nTotal prob of N-in: 0.96171\nPOSSIBLE N-term signal sequence\ninside 1 19\nTMhelix 20 39\noutside 40 325\nTMhelix 326 348\ninside 349 372

19394 GCF_001833935.1_ASM183393v1 Salmonella enterica subsp. enterica serovar Saintpaul Proteobacteria; Gammaproteobacteria; Enterobacterales; Enterobacteriaceae; Salmonella; Salmonella enterica; Salmonella enterica subsp. enterica MNNEETFYQAMRRKGVTRRSFLKFCSLAATSLGLGAGMTPKIAWALENKPRIPVVWIHGL WP_001058311.1 hydrogenase [Salmonella enterica] Length: 372\nNumber of predicted TMHs: 2\nExp number of AAs in TMHs: 37.4699899999999\nExp number, first 60 AAs: 19.72699\nTotal prob of N-in: 0.96171\nPOSSIBLE N-term signal sequence\ninside 1 19\nTMhelix 20 39\noutside 40 325\nTMhelix 326 348\ninside 349 372

19395 GCF_001833465.1_ASM183346v1 Salmonella enterica subsp. enterica serovar Saintpaul Proteobacteria; Gammaproteobacteria; Enterobacterales; Enterobacteriaceae; Salmonella; Salmonella enterica; Salmonella enterica subsp. enterica MNNEETFYQAMRRKGVTRRSFLKFCSLAATSLGLGAGMTPKIAWALENKPRIPVVWIHGL WP_001058311.1 hydrogenase [Salmonella enterica] Length: 372\nNumber of predicted TMHs: 2\nExp number of AAs in TMHs: 37.4699899999999\nExp number, first 60 AAs: 19.72699\nTotal prob of N-in: 0.96171\nPOSSIBLE N-term signal sequence\ninside 1 19\nTMhelix 20 39\noutside 40 325\nTMhelix 326 348\ninside 349 372

19396 GCF_001834485.1_ASM183448v1 Salmonella enterica subsp. enterica serovar Saintpaul Proteobacteria; Gammaproteobacteria; Enterobacterales; Enterobacteriaceae; Salmonella; Salmonella enterica; Salmonella enterica subsp. enterica MNNEETFYQAMRRKGVTRRSFLKFCSLAATSLGLGAGMTPKIAWALENKPRIPVVWIHGL WP_001058311.1 hydrogenase [Salmonella enterica] Length: 372\nNumber of predicted TMHs: 2\nExp number of AAs in TMHs: 37.4699899999999\nExp number, first 60 AAs: 19.72699\nTotal prob of N-in: 0.96171\nPOSSIBLE N-term signal sequence\ninside 1 19\nTMhelix 20 39\noutside 40 325\nTMhelix 326 348\ninside 349 372

19397 GCF_001478745.1_Salmonella_enterica_CVM_N29369-R_v1.0 Salmonella enterica subsp. enterica serovar Saintpaul Proteobacteria; Gammaproteobacteria; Enterobacterales; Enterobacteriaceae; Salmonella; Salmonella enterica; Salmonella enterica subsp. enterica MTGDNTLITSHGINRRDFMKLCAALAATMGLSSKAAAEMAESVSNPQRPPVIWIGAQECT WP_000145429.1 hydrogenase 2 small subunit [Salmonella enterica] Length: 372\nNumber of predicted TMHs: 1\nExp number of AAs in TMHs: 24.51309\nExp number, first 60 AAs: 0.2682\nTotal prob of N-in: 0.14402\noutside 1 331\nTMhelix 332 354\ninside 355 372

19398 GCF_001478745.1_Salmonella_enterica_CVM_N29369-R_v1.0 Salmonella enterica subsp. enterica serovar Saintpaul Proteobacteria; Gammaproteobacteria; Enterobacterales; Enterobacteriaceae; Salmonella; Salmonella enterica; Salmonella enterica subsp. enterica MNNEETFYQAMRRKGVTRRSFLKFCSLAATSLGLGAGMTPKIAWALENKPRIPVVWIHGL WP_001058312.1 hydrogenase [Salmonella enterica] Length: 372\nNumber of predicted TMHs: 2\nExp number of AAs in TMHs: 36.51738\nExp number, first 60 AAs: 19.72849\nTotal prob of N-in: 0.96170\nPOSSIBLE N-term signal sequence\ninside 1 19\nTMhelix 20 39\noutside 40 325\nTMhelix 326 348\ninside 349 372

19399 GCF_002063965.1_ASM206396v1 Salmonella enterica subsp. enterica serovar Sandiego Proteobacteria; Gammaproteobacteria; Enterobacterales; Enterobacteriaceae; Salmonella; Salmonella enterica; Salmonella enterica subsp. enterica MNNEETFYQAMRRKGVTRRSFLKFCSLAATSLGLGAGITPKIAWALENKPRIPVVWIHGL WP_080210458.1 [Ni/Fe] hydrogenase small subunit [Salmonella enterica] Length: 372\nNumber of predicted TMHs: 2\nExp number of AAs in TMHs: 37.06689\nExp number, first 60 AAs: 20.27828\nTotal prob of N-in: 0.97175\nPOSSIBLE N-term signal sequence\ninside 1 20\nTMhelix 21 43\noutside 44 325\nTMhelix 326 348\ninside 349 372

19400 GCF_002063965.1_ASM206396v1 Salmonella enterica subsp. enterica serovar Sandiego Proteobacteria; Gammaproteobacteria; Enterobacterales; Enterobacteriaceae; Salmonella; Salmonella enterica; Salmonella enterica subsp. enterica MTGDNTLITSHGINRRDFMKLCAALAATMGLSSKAAAEMAESVSNPQRPPVIWIGAQECT WP_000145429.1 hydrogenase 2 small subunit [Salmonella enterica] Length: 372\nNumber of predicted TMHs: 1\nExp number of AAs in TMHs: 24.51309\nExp number, first 60 AAs: 0.2682\nTotal prob of N-in: 0.14402\noutside 1 331\nTMhelix 332 354\ninside 355 372

19401 GCF_002108105.1_ASM210810v1 Salmonella enterica subsp. enterica serovar Saphra Proteobacteria; Gammaproteobacteria; Enterobacterales; Enterobacteriaceae; Salmonella; Salmonella enterica; Salmonella enterica subsp. enterica MTGDNTLITSHGINRRDFMKLCAALAATMGLSSKAAAEMAESVSNPQRPPVIWIGAQECT WP_000145429.1 hydrogenase 2 small subunit [Salmonella enterica] Length: 372\nNumber of predicted TMHs: 1\nExp number of AAs in TMHs: 24.51309\nExp number, first 60 AAs: 0.2682\nTotal prob of N-in: 0.14402\noutside 1 331\nTMhelix 332 354\ninside 355 372

19402 GCF_002108105.1_ASM210810v1 Salmonella enterica subsp. enterica serovar Saphra Proteobacteria; Gammaproteobacteria; Enterobacterales; Enterobacteriaceae; Salmonella; Salmonella enterica; Salmonella enterica subsp. enterica MNNEETFYQAMRRKGVTRRSFLKFCSLAATSLGLGAGMTPKIAWALENKPRIPVVWIHGL WP_085353658.1 [Ni/Fe] hydrogenase small subunit [Salmonella enterica] Length: 372\nNumber of predicted TMHs: 2\nExp number of AAs in TMHs: 36.5180799999999\nExp number, first 60 AAs: 19.72844\nTotal prob of N-in: 0.96170\nPOSSIBLE N-term signal sequence\ninside 1 19\nTMhelix 20 39\noutside 40 325\nTMhelix 326 348\ninside 349 372

19403 GCF_002034855.1_ASM203485v1 Salmonella enterica subsp. enterica serovar Saphra Proteobacteria; Gammaproteobacteria; Enterobacterales; Enterobacteriaceae; Salmonella; Salmonella enterica; Salmonella enterica subsp. enterica MNNEETFYQAMRRKGVTRRSFLKFCSLAATSLGLGAGMTPKIAWALENKPRIPVVWIHGL WP_001058312.1 hydrogenase [Salmonella enterica] Length: 372\nNumber of predicted TMHs: 2\nExp number of AAs in TMHs: 36.51738\nExp number, first 60 AAs: 19.72849\nTotal prob of N-in: 0.96170\nPOSSIBLE N-term signal sequence\ninside 1 19\nTMhelix 20 39\noutside 40 325\nTMhelix 326 348\ninside 349 372

19404 GCF_002034855.1_ASM203485v1 Salmonella enterica subsp. enterica serovar Saphra Proteobacteria; Gammaproteobacteria; Enterobacterales; Enterobacteriaceae; Salmonella; Salmonella enterica; Salmonella enterica subsp. enterica MTGDNTLITSHGINRRDFMKLCAALAATMGLSSKAAAEMAESVSNPQRPPVIWIGAQECT WP_000145429.1 hydrogenase 2 small subunit [Salmonella enterica] Length: 372\nNumber of predicted TMHs: 1\nExp number of AAs in TMHs: 24.51309\nExp number, first 60 AAs: 0.2682\nTotal prob of N-in: 0.14402\noutside 1 331\nTMhelix 332 354\ninside 355 372

19405 GCF_002062905.1_ASM206290v1 Salmonella enterica subsp. enterica serovar Schwardzengrund Proteobacteria; Gammaproteobacteria; Enterobacterales; Enterobacteriaceae; Salmonella; Salmonella enterica; Salmonella enterica subsp. enterica MNNEETFYQAMRRKGVTRRSFLKFCSLAATSLGLGAGMTPKIAWALENKPRIPVVWIHGL WP_001058312.1 hydrogenase [Salmonella enterica] Length: 372\nNumber of predicted TMHs: 2\nExp number of AAs in TMHs: 36.51738\nExp number, first 60 AAs: 19.72849\nTotal prob of N-in: 0.96170\nPOSSIBLE N-term signal sequence\ninside 1 19\nTMhelix 20 39\noutside 40 325\nTMhelix 326 348\ninside 349 372

19406 GCF_002062905.1_ASM206290v1 Salmonella enterica subsp. enterica serovar Schwardzengrund Proteobacteria; Gammaproteobacteria; Enterobacterales; Enterobacteriaceae; Salmonella; Salmonella enterica; Salmonella enterica subsp. enterica MTGDNTLITSHGINRRDFMKLCAALAATMGLSSKAAAEMAESVSNPQRPPVIWIGAQECT WP_000145431.1 hydrogenase 2 small subunit [Salmonella enterica] Length: 372\nNumber of predicted TMHs: 1\nExp number of AAs in TMHs: 24.51505\nExp number, first 60 AAs: 0.2682\nTotal prob of N-in: 0.14406\noutside 1 331\nTMhelix 332 354\ninside 355 372

19407 GCF_002063915.1_ASM206391v1 Salmonella enterica subsp. enterica serovar Schwarzengrund Proteobacteria; Gammaproteobacteria; Enterobacterales; Enterobacteriaceae; Salmonella; Salmonella enterica; Salmonella enterica subsp. enterica MTGDNTLITSHGINRRDFMKLCAALAATMGLSSKAAAEMAESVSNPQRPPVIWIGAQECT

WP_000145429.1 hydrogenase 2 small subunit [Salmonella enterica] Length: 372\nNumber of predicted TMHs: 1\nExp number of AAs in TMHs: 24.51309\nExp number, first 60 AAs: 0.2682\nTotal prob of N-in: 0.14402\noutside 1 331\nTMhelix 332 354\ninside 355 372

19408 GCF_002063915.1_ASM206391v1 Salmonella enterica subsp. enterica serovar Schwarzengrund Proteobacteria; Gammaproteobacteria; Enterobacterales; Enterobacteriaceae; Salmonella; Salmonella enterica; Salmonella enterica subsp. enterica MNNEETFYQAMRRKGVTRRSFLKFCSLAATSLGLGAGMTPKIAWALENKPRIPVVWIHGL WP_001058312.1 hydrogenase [Salmonella enterica] Length: 372\nNumber of predicted TMHs: 2\nExp number of AAs in TMHs: 36.51738\nExp number, first 60 AAs: 19.72849\nTotal prob of N-in: 0.96170\nPOSSIBLE N-term signal sequence\ninside 1 19\nTMhelix 20 39\noutside 40 325\nTMhelix 326 348\ninside 349 372

19409 GCF_002063955.1_ASM206395v1 Salmonella enterica subsp. enterica serovar Schwarzengrund Proteobacteria; Gammaproteobacteria; Enterobacterales; Enterobacteriaceae; Salmonella; Salmonella enterica; Salmonella enterica subsp. enterica MTGDNTLITSHGINRRDFMKLCAALAATMGLSSKAAAEMAESVSNPQRPPVIWIGAQECT WP_000145431.1 hydrogenase 2 small subunit [Salmonella enterica] Length: 372\nNumber of predicted TMHs: 1\nExp number of AAs in TMHs: 24.51505\nExp number, first 60 AAs: 0.2682\nTotal prob of N-in: 0.14406\noutside 1 331\nTMhelix 332 354\ninside 355 372

19410 GCF_002063955.1_ASM206395v1 Salmonella enterica subsp. enterica serovar Schwarzengrund Proteobacteria; Gammaproteobacteria; Enterobacterales; Enterobacteriaceae; Salmonella; Salmonella enterica; Salmonella enterica subsp. enterica MNNEETFYQAMRRKGVTRRSFLKFCSLAATSLGLGAGMTPKIAWALENKPRIPVVWIHGL WP_001058312.1 hydrogenase [Salmonella enterica] Length: 372\nNumber of predicted TMHs: 2\nExp number of AAs in TMHs: 36.51738\nExp number, first 60 AAs: 19.72849\nTotal prob of N-in: 0.96170\nPOSSIBLE N-term signal sequence\ninside 1 19\nTMhelix 20 39\noutside 40 325\nTMhelix 326 348\ninside 349 372

19411 GCF_002205805.1_ASM220580v1 Salmonella enterica subsp. enterica serovar Schwarzengrund Proteobacteria; Gammaproteobacteria; Enterobacterales; Enterobacteriaceae; Salmonella; Salmonella enterica; Salmonella enterica subsp. enterica MTGDNTLITSHGINRRDFMKLCAALAATMGLSSKAAAEMAESVSNPQRPPVIWIGAQECT WP_000145431.1 hydrogenase 2 small subunit [Salmonella enterica] Length: 372\nNumber of predicted TMHs: 1\nExp number of AAs in TMHs: 24.51505\nExp number, first 60 AAs: 0.2682\nTotal prob of N-in: 0.14406\noutside 1 331\nTMhelix 332 354\ninside 355 372

19412 GCF_002205805.1_ASM220580v1 Salmonella enterica subsp. enterica serovar Schwarzengrund Proteobacteria; Gammaproteobacteria; Enterobacterales; Enterobacteriaceae; Salmonella; Salmonella enterica; Salmonella enterica subsp. enterica MNNEETFYQAMRRKGVTRRSFLKFCSLAATSLGLGAGMTPKIAWALENKPRIPVVWIHGL WP_001058312.1 hydrogenase [Salmonella enterica] Length: 372\nNumber of predicted TMHs: 2\nExp number of AAs in TMHs: 36.51738\nExp number, first 60 AAs: 19.72849\nTotal prob of N-in: 0.96170\nPOSSIBLE N-term signal sequence\ninside 1 19\nTMhelix 20 39\noutside 40 325\nTMhelix 326 348\ninside 349 372

19413 GCF_002065105.1_ASM206510v1 Salmonella enterica subsp. enterica serovar Schwarzengrund Proteobacteria; Gammaproteobacteria; Enterobacterales; Enterobacteriaceae; Salmonella; Salmonella enterica; Salmonella enterica subsp. enterica MTGDNTLITSHGINRRDFMKLCAALAATMGLSSKAAAEMAESVSNPQRPPVIWIGAQECT WP_000145431.1 hydrogenase 2 small subunit [Salmonella enterica] Length: 372\nNumber of predicted TMHs: 1\nExp number of AAs in TMHs: 24.51505\nExp number, first 60 AAs: 0.2682\nTotal prob of N-in: 0.14406\noutside 1 331\nTMhelix 332 354\ninside 355 372

19414 GCF_002065105.1_ASM206510v1 Salmonella enterica subsp. enterica serovar Schwarzengrund Proteobacteria; Gammaproteobacteria; Enterobacterales; Enterobacteriaceae; Salmonella; Salmonella enterica; Salmonella enterica subsp. enterica MNNEETFYQAMRRKGVTRRSFLKFCSLAATSLGLGAGMTPKIAWALENKPRIPVVWIHGL WP_001058312.1 hydrogenase [Salmonella enterica] Length: 372\nNumber of predicted TMHs: 2\nExp number of AAs in TMHs: 36.51738\nExp number, first 60 AAs: 19.72849\nTotal prob of N-in: 0.96170\nPOSSIBLE N-term signal sequence\ninside 1 19\nTMhelix 20 39\noutside 40 325\nTMhelix 326 348\ninside 349 372

19415 GCF_001240625.1_Salmonella_enterica_CVM_N43479_v1.0 Salmonella enterica subsp. enterica serovar Schwarzengrund Proteobacteria; Gammaproteobacteria; Enterobacterales; Enterobacteriaceae; Salmonella; Salmonella enterica; Salmonella enterica subsp. enterica MTGDNTLITSHGINRRDFMKLCAALAATMGLSSKAAAEMAESVSNPQRPPVIWIGAQECT WP_000145431.1 hydrogenase 2 small subunit [Salmonella enterica] Length: 372\nNumber of predicted TMHs: 1\nExp number of AAs in TMHs: 24.51505\nExp number, first 60 AAs: 0.2682\nTotal prob of N-in: 0.14406\noutside 1 331\nTMhelix 332 354\ninside 355 372

19416 GCF_001246985.1_Salmonella_enterica_CVM_N51289_v1.0 Salmonella enterica subsp. enterica serovar Schwarzengrund Proteobacteria; Gammaproteobacteria; Enterobacterales; Enterobacteriaceae; Salmonella; Salmonella enterica; Salmonella enterica subsp. enterica MTGDNTLITSHGINRRDFMKLCAALAATMGLSSKAAAEMAESVSNPQRPPVIWIGAQECT WP_000145431.1 hydrogenase 2 small subunit [Salmonella enterica] Length: 372\nNumber of predicted TMHs: 1\nExp number of AAs in TMHs: 24.51505\nExp number, first 60 AAs: 0.2682\nTotal prob of N-in: 0.14406\noutside 1 331\nTMhelix 332 354\ninside 355 372

19417 GCF_002033085.1_ASM203308v1 Salmonella enterica subsp. enterica serovar Schwarzengrund Proteobacteria; Gammaproteobacteria; Enterobacterales; Enterobacteriaceae; Salmonella; Salmonella enterica; Salmonella enterica subsp. enterica MTGDNTLITSHGINRRDFMKLCAALAATMGLSSKAAAEMAESVSNPQRPPVIWIGAQECT WP_000145431.1 hydrogenase 2 small subunit [Salmonella enterica] Length: 372\nNumber of predicted TMHs: 1\nExp number of AAs in TMHs: 24.51505\nExp number, first 60 AAs: 0.2682\nTotal prob of N-in: 0.14406\noutside 1 331\nTMhelix 332 354\ninside 355 372

19418 GCF_002033125.1_ASM203312v1 Salmonella enterica subsp. enterica serovar Schwarzengrund Proteobacteria; Gammaproteobacteria; Enterobacterales; Enterobacteriaceae; Salmonella; Salmonella enterica; Salmonella enterica subsp. enterica MNNEETFYQAMRRKGVTRRSFLKFCSLAATSLGLGAGMTPKIAWALENKPRIPVVWIHGL WP_001058312.1 hydrogenase [Salmonella enterica] Length: 372\nNumber of predicted TMHs: 2\nExp number of AAs in TMHs: 36.51738\nExp number, first 60 AAs: 19.72849\nTotal prob of N-in: 0.96170\nPOSSIBLE N-term signal sequence\ninside 1 19\nTMhelix 20 39\noutside 40 325\nTMhelix 326 348\ninside 349 372

19419 GCF_001246555.1_Salmonella_enterica_CVM_N51267_v1.0 Salmonella enterica subsp. enterica serovar Schwarzengrund Proteobacteria; Gammaproteobacteria; Enterobacterales; Enterobacteriaceae; Salmonella; Salmonella enterica; Salmonella enterica subsp. enterica MTGDNTLITSHGINRRDFMKLCAALAATMGLSSKAAAEMAESVSNPQRPPVIWIGAQECT WP_000145431.1 hydrogenase 2 small subunit [Salmonella enterica] Length: 372\nNumber of predicted TMHs: 1\nExp number of AAs in TMHs: 24.51505\nExp number, first 60 AAs: 0.2682\nTotal prob of N-in: 0.14406\noutside 1 331\nTMhelix 332 354\ninside 355 372

19420 GCF_001246305.1_Salmonella_enterica_CVM_N51257_v1.0 Salmonella enterica subsp. enterica serovar Schwarzengrund Proteobacteria; Gammaproteobacteria; Enterobacterales; Enterobacteriaceae; Salmonella; Salmonella enterica; Salmonella enterica subsp. enterica MNNEETFYQAMRRKGVTRRSFLKFCSLAATSLGLGAGMTPKIAWALENKPRIPVVWIHGL WP_001058312.1 hydrogenase [Salmonella enterica] Length: 372\nNumber of predicted TMHs: 2\nExp number of AAs in TMHs: 36.51738\nExp number, first 60 AAs: 19.72849\nTotal prob of N-in: 0.96170\nPOSSIBLE N-term signal sequence\ninside 1 19\nTMhelix 20 39\noutside 40 325\nTMhelix 326 348\ninside 349 372

19421 GCF_001247455.1_Salmonella_enterica_CVM_N51311_v1.0 Salmonella enterica subsp. enterica serovar Schwarzengrund Proteobacteria; Gammaproteobacteria; Enterobacterales; Enterobacteriaceae; Salmonella; Salmonella enterica; Salmonella enterica subsp. enterica MTGDNTLITSHGINRRDFMKLCAALAATMGLSSKAAAEMAESVSNPQRPPVIWIGAQECT WP_000145431.1 hydrogenase 2 small subunit [Salmonella enterica] Length: 372\nNumber of predicted TMHs: 1\nExp number of AAs in TMHs: 24.51505\nExp number, first 60 AAs: 0.2682\nTotal prob of N-in: 0.14406\noutside 1 331\nTMhelix 332 354\ninside 355 372

19422 GCF_001245545.1_Salmonella_enterica_CVM_N50434_v1.0 Salmonella enterica subsp. enterica serovar Schwarzengrund Proteobacteria; Gammaproteobacteria; Enterobacterales; Enterobacteriaceae; Salmonella; Salmonella enterica; Salmonella enterica subsp. enterica MTGDNTLITSHGINRRDFMKLCAALAATMGLSSKAAAEMAESVSNPQRPPVIWIGAQECT WP_000145431.1 hydrogenase 2 small subunit [Salmonella enterica] Length: 372\nNumber of predicted TMHs: 1\nExp number of AAs in TMHs: 24.51505\nExp number, first 60 AAs: 0.2682\nTotal prob of N-in: 0.14406\noutside 1 331\nTMhelix 332 354\ninside 355 372

19423 GCF_001240625.1_Salmonella_enterica_CVM_N43479_v1.0 Salmonella enterica subsp. enterica serovar Schwarzengrund Proteobacteria; Gammaproteobacteria; Enterobacterales; Enterobacteriaceae; Salmonella; Salmonella enterica; Salmonella enterica subsp. enterica MNNEETFYQAMRRKGVTRRSFLKFCSLAATSLGLGAGMTPKIAWALENKPRIPVVWIHGL WP_001058312.1 hydrogenase [Salmonella enterica] Length: 372\nNumber of predicted TMHs: 2\nExp number of AAs in TMHs: 36.51738\nExp number, first 60 AAs: 19.72849\nTotal prob of N-in: 0.96170\nPOSSIBLE N-term signal sequence\ninside 1 19\nTMhelix 20 39\noutside 40 325\nTMhelix 326 348\ninside 349 372

19424 GCF_001243745.1_Salmonella_enterica_CVM_N47711_v1.0 Salmonella enterica subsp. enterica serovar Schwarzengrund Proteobacteria; Gammaproteobacteria; Enterobacterales; Enterobacteriaceae; Salmonella; Salmonella enterica; Salmonella enterica subsp. enterica MNNEETFYQAMRRKGVTRRSFLKFCSLAATSLGLGAGMTPKIAWALENKPRIPVVWIHGL WP_001058312.1 hydrogenase [Salmonella enterica] Length: 372\nNumber of predicted TMHs: 2\nExp number of AAs in TMHs: 36.51738\nExp number, first 60 AAs: 19.72849\nTotal prob of N-in: 0.96170\nPOSSIBLE N-term signal sequence\ninside 1 19\nTMhelix 20 39\noutside 40 325\nTMhelix 326 348\ninside 349 372

19425 GCF_001241265.1_Salmonella_enterica_CVM_N44700-R_v1.0 Salmonella enterica subsp. enterica serovar Schwarzengrund Proteobacteria; Gammaproteobacteria; Enterobacterales; Enterobacteriaceae; Salmonella; Salmonella enterica; Salmonella enterica subsp. enterica MNNEETFYQAMRRKGVTRRSFLKFCSLAATSLGLGAGMTPKIAWALENKPRIPVVWIHGL WP_001058312.1 hydrogenase [Salmonella enterica] Length: 372\nNumber of predicted TMHs: 2\nExp number of AAs in TMHs: 36.51738\nExp number, first 60 AAs: 19.72849\nTotal prob of N-in: 0.96170\nPOSSIBLE N-term signal sequence\ninside 1 19\nTMhelix 20 39\noutside 40 325\nTMhelix 326 348\ninside 349 372

19426 GCF_001242665.1_Salmonella_enterica_CVM_N45952_v1.0 Salmonella enterica subsp. enterica serovar Schwarzengrund Proteobacteria; Gammaproteobacteria; Enterobacterales; Enterobacteriaceae; Salmonella; Salmonella enterica; Salmonella enterica subsp. enterica MTGDNTLITSHGINRRDFMKLCAALAATMGLSSKAAAEMAESVSNPQRPPVIWIGAQECT WP_000145431.1 hydrogenase 2 small subunit [Salmonella enterica] Length: 372\nNumber of predicted TMHs: 1\nExp number of AAs in TMHs: 24.51505\nExp number, first 60 AAs: 0.2682\nTotal prob of N-in: 0.14406\noutside 1 331\nTMhelix 332 354\ninside 355 372

19427 GCF_001241265.1_Salmonella_enterica_CVM_N44700-R_v1.0 Salmonella enterica subsp. enterica serovar Schwarzengrund Proteobacteria; Gammaproteobacteria; Enterobacterales; Enterobacteriaceae; Salmonella; Salmonella enterica; Salmonella enterica subsp. enterica MTGDNTLITSHGINRRDFMKLCAALAATMGLSSKAAAEMAESVSNPQRPPVIWIGAQECT WP_000145431.1 hydrogenase 2 small subunit [Salmonella enterica] Length: 372\nNumber of predicted TMHs: 1\nExp number of AAs in TMHs: 24.51505\nExp number, first 60 AAs: 0.2682\nTotal prob of N-in: 0.14406\noutside 1 331\nTMhelix 332 354\ninside 355 372

19428 GCF_001243745.1_Salmonella_enterica_CVM_N47711_v1.0 Salmonella enterica subsp. enterica serovar Schwarzengrund Proteobacteria; Gammaproteobacteria; Enterobacterales; Enterobacteriaceae; Salmonella; Salmonella enterica; Salmonella enterica subsp. enterica MTGDNTLITSHGINRRDFMKLCAALAATMGLSSKAAAEMAESVSNPQRPPVIWIGAQECT WP_000145431.1 hydrogenase 2 small subunit [Salmonella enterica] Length: 372\nNumber of predicted TMHs: 1\nExp number of AAs in TMHs: 24.51505\nExp number, first 60 AAs: 0.2682\nTotal prob of N-in: 0.14406\noutside 1 331\nTMhelix 332 354\ninside 355 372

19429 GCF_001246375.1_Salmonella_enterica_CVM_N51259_v1.0 Salmonella enterica subsp. enterica serovar Schwarzengrund Proteobacteria; Gammaproteobacteria; Enterobacterales; Enterobacteriaceae; Salmonella; Salmonella enterica; Salmonella enterica subsp. enterica MTGDNTLITSHGINRRDFMKLCAALAATMGLSSKAAAEMAESVSNPQRPPVIWIGAQECT WP_000145431.1 hydrogenase 2 small subunit [Salmonella enterica] Length: 372\nNumber of predicted TMHs: 1\nExp number of AAs in TMHs: 24.51505\nExp number, first 60 AAs: 0.2682\nTotal prob of N-in: 0.14406\noutside 1 331\nTMhelix 332 354\ninside 355 372

19430 GCF_001240275.1_Salmonella_enterica_CVM_N43459_v1.0 Salmonella enterica subsp. enterica serovar Schwarzengrund Proteobacteria; Gammaproteobacteria; Enterobacterales; Enterobacteriaceae; Salmonella; Salmonella enterica; Salmonella enterica subsp. enterica MNNEETFYQAMRRKGVTRRSFLKFCSLAATSLGLGAGMTPKIAWALENKPRIPVVWIHGL WP_001058312.1 hydrogenase [Salmonella enterica] Length: 372\nNumber of predicted TMHs: 2\nExp number of AAs in TMHs: 36.51738\nExp number, first 60 AAs: 19.72849\nTotal prob of N-in: 0.96170\nPOSSIBLE N-term signal sequence\ninside 1 19\nTMhelix 20 39\noutside 40 325\nTMhelix 326 348\ninside 349 372

19431 GCF_001246375.1_Salmonella_enterica_CVM_N51259_v1.0 Salmonella enterica subsp. enterica serovar Schwarzengrund Proteobacteria; Gammaproteobacteria; Enterobacterales; Enterobacteriaceae; Salmonella; Salmonella enterica; Salmonella enterica subsp. enterica MNNEETFYQAMRRKGVTRRSFLKFCSLAATSLGLGAGMTPKIAWALENKPRIPVVWIHGL WP_001058312.1 hydrogenase [Salmonella enterica] Length: 372\nNumber of predicted TMHs: 2\nExp number of AAs in

TMHs: 36.51738\nExp number, first 60 AAs: 19.72849\nTotal prob of N-in: 0.96170\nPOSSIBLE N-term signal sequence\ninside 1 19\nTMhelix 20 39\noutside 40 325\nTMhelix 326 348\ninside 349 372

19432 GCF_001242265.1_Salmonella_enterica_CVM_N45932_v1.0 Salmonella enterica subsp. enterica serovar Schwarzengrund Proteobacteria; Gammaproteobacteria; Enterobacterales; Enterobacteriaceae; Salmonella; Salmonella enterica; Salmonella enterica subsp. enterica MTGDNTLITSHGINRRDFMKLCAALAATMGLSSKAAAEMAESVSNPQRPPVIWIGAQECT WP_000145431.1 hydrogenase 2 small subunit [Salmonella enterica] Length: 372\nNumber of predicted TMHs: 1\nExp number of AAs in TMHs: 24.51505\nExp number, first 60 AAs: 0.2682\nTotal prob of N-in: 0.14406\noutside 1 331\nTMhelix 332 354\ninside 355 372

19433 GCF_001242265.1_Salmonella_enterica_CVM_N45932_v1.0 Salmonella enterica subsp. enterica serovar Schwarzengrund Proteobacteria; Gammaproteobacteria; Enterobacterales; Enterobacteriaceae; Salmonella; Salmonella enterica; Salmonella enterica subsp. enterica MNNEETFYQAMRRKGVTRRSFLKFCSLAATSLGLGAGMTPKIAWALENKPRIPVVWIHGL WP_001058312.1 hydrogenase [Salmonella enterica] Length: 372\nNumber of predicted TMHs: 2\nExp number of AAs in TMHs: 36.51738\nExp number, first 60 AAs: 19.72849\nTotal prob of N-in: 0.96170\nPOSSIBLE N-term signal sequence\ninside 1 19\nTMhelix 20 39\noutside 40 325\nTMhelix 326 348\ninside 349 372

19434 GCF_001244375.1_Salmonella_enterica_CVM_N48682_v1.0 Salmonella enterica subsp. enterica serovar Schwarzengrund Proteobacteria; Gammaproteobacteria; Enterobacterales; Enterobacteriaceae; Salmonella; Salmonella enterica; Salmonella enterica subsp. enterica MTGDNTLITSHGINRRDFMKLCAALAATMGLSSKAAAEMAESVSNPQRPPVIWIGAQECT WP_000145431.1 hydrogenase 2 small subunit [Salmonella enterica] Length: 372\nNumber of predicted TMHs: 1\nExp number of AAs in TMHs: 24.51505\nExp number, first 60 AAs: 0.2682\nTotal prob of N-in: 0.14406\noutside 1 331\nTMhelix 332 354\ninside 355 372

19435 GCF_001240275.1_Salmonella_enterica_CVM_N43459_v1.0 Salmonella enterica subsp. enterica serovar Schwarzengrund Proteobacteria; Gammaproteobacteria; Enterobacterales; Enterobacteriaceae; Salmonella; Salmonella enterica; Salmonella enterica subsp. enterica MTGDNTLITSHGINRRDFMKLCAALAATMGLSSKAAAEMAESVSNPQRPPVIWIGAQECT WP_000145431.1 hydrogenase 2 small subunit [Salmonella enterica] Length: 372\nNumber of predicted TMHs: 1\nExp number of AAs in TMHs: 24.51505\nExp number, first 60 AAs: 0.2682\nTotal prob of N-in: 0.14406\noutside 1 331\nTMhelix 332 354\ninside 355 372

19436 GCF_001271815.1_Salmonella_enterica_CVM_N47712_v1.0 Salmonella enterica subsp. enterica serovar Schwarzengrund Proteobacteria; Gammaproteobacteria; Enterobacterales; Enterobacteriaceae; Salmonella; Salmonella enterica; Salmonella enterica subsp. enterica MTGDNTLITSHGINRRDFMKLCAALAATMGLSSKAAAEMAESVSNPQRPPVIWIGAQECT WP_000145431.1 hydrogenase 2 small subunit [Salmonella enterica] Length: 372\nNumber of predicted TMHs: 1\nExp number of AAs in TMHs: 24.51505\nExp number, first 60 AAs: 0.2682\nTotal prob of N-in: 0.14406\noutside 1 331\nTMhelix 332 354\ninside 355 372

19437 GCF_001244375.1_Salmonella_enterica_CVM_N48682_v1.0 Salmonella enterica subsp. enterica serovar Schwarzengrund Proteobacteria; Gammaproteobacteria; Enterobacterales; Enterobacteriaceae; Salmonella; Salmonella enterica; Salmonella enterica subsp. enterica MNNEETFYQAMRRKGVTRRSFLKFCSLAATSLGLGAGMTPKIAWALENKPRIPVVWIHGL WP_001058312.1 hydrogenase [Salmonella enterica] Length: 372\nNumber of predicted TMHs: 2\nExp number of AAs in TMHs: 36.51738\nExp number, first 60 AAs: 19.72849\nTotal prob of N-in: 0.96170\nPOSSIBLE N-term signal sequence\ninside 1 19\nTMhelix 20 39\noutside 40 325\nTMhelix 326 348\ninside 349 372

19438 GCF_001271815.1_Salmonella_enterica_CVM_N47712_v1.0 Salmonella enterica subsp. enterica serovar Schwarzengrund Proteobacteria; Gammaproteobacteria; Enterobacterales; Enterobacteriaceae; Salmonella; Salmonella enterica; Salmonella enterica subsp. enterica MNNEETFYQAMRRKGVTRRSFLKFCSLAATSLGLGAGMTPKIAWALENKPRIPVVWIHGL WP_001058312.1 hydrogenase [Salmonella enterica] Length: 372\nNumber of predicted TMHs: 2\nExp number of AAs in TMHs: 36.51738\nExp number, first 60 AAs: 19.72849\nTotal prob of N-in: 0.96170\nPOSSIBLE N-term signal sequence\ninside 1 19\nTMhelix 20 39\noutside 40 325\nTMhelix 326 348\ninside 349 372

19439 GCF_001242665.1_Salmonella_enterica_CVM_N45952_v1.0 Salmonella enterica subsp. enterica serovar Schwarzengrund Proteobacteria; Gammaproteobacteria; Enterobacterales; Enterobacteriaceae; Salmonella; Salmonella

enterica; Salmonella enterica subsp. enterica
MNNEETFYQAMRRKGVTRRSFLKFCSLAATSLGLGAGMTPKIAWALENKPRIPVVWIHGL WP_001058312.1 hydrogenase [Salmonella enterica] Length: 372\nNumber of predicted TMHs: 2\nExp number of AAs in TMHs: 36.51738\nExp number, first 60 AAs: 19.72849\nTotal prob of N-in: 0.96170\nPOSSIBLE N-term signal sequence\ninside 1 19\nTMhelix 20 39\noutside 40 325\nTMhelix 326 348\ninside 349 372

19440 GCF_002031705.1_ASM203170v1 Salmonella enterica subsp. enterica serovar Schwarzengrund Proteobacteria; Gammaproteobacteria; Enterobacterales; Enterobacteriaceae; Salmonella; Salmonella enterica; Salmonella enterica subsp. enterica MNNEETFYQAMRRKGVTRRSFLKFCSLAATSLGLGAGMTPKIAWALENKPRIPVVWIHGL WP_001058312.1 hydrogenase [Salmonella enterica] Length: 372\nNumber of predicted TMHs: 2\nExp number of AAs in TMHs: 36.51738\nExp number, first 60 AAs: 19.72849\nTotal prob of N-in: 0.96170\nPOSSIBLE N-term signal sequence\ninside 1 19\nTMhelix 20 39\noutside 40 325\nTMhelix 326 348\ninside 349 372

19441 GCF_002031705.1_ASM203170v1 Salmonella enterica subsp. enterica serovar Schwarzengrund Proteobacteria; Gammaproteobacteria; Enterobacterales; Enterobacteriaceae; Salmonella; Salmonella enterica; Salmonella enterica subsp. enterica MTGDNTLITSHGINRRDFMKLCAALAATMGLSSKAAAEMAESVSNPQRPPVIWIGAQECT WP_000145431.1 hydrogenase 2 small subunit [Salmonella enterica] Length: 372\nNumber of predicted TMHs: 1\nExp number of AAs in TMHs: 24.51505\nExp number, first 60 AAs: 0.2682\nTotal prob of N-in: 0.14406\noutside 1 331\nTMhelix 332 354\ninside 355 372

19442 GCF_002063925.1_ASM206392v1 Salmonella enterica subsp. enterica serovar Schwarzengrund Proteobacteria; Gammaproteobacteria; Enterobacterales; Enterobacteriaceae; Salmonella; Salmonella enterica; Salmonella enterica subsp. enterica MNNEETFYQAMRRKGVTRRSFLKFCSLAATSLGLGAGMTPKIAWALENKPRIPVVWIHGL WP_001058312.1 hydrogenase [Salmonella enterica] Length: 372\nNumber of predicted TMHs: 2\nExp number of AAs in TMHs: 36.51738\nExp number, first 60 AAs: 19.72849\nTotal prob of N-in: 0.96170\nPOSSIBLE N-term signal sequence\ninside 1 19\nTMhelix 20 39\noutside 40 325\nTMhelix 326 348\ninside 349 372

19443 GCF_002031905.1_ASM203190v1 Salmonella enterica subsp. enterica serovar Schwarzengrund Proteobacteria; Gammaproteobacteria; Enterobacterales; Enterobacteriaceae; Salmonella; Salmonella enterica; Salmonella enterica subsp. enterica MTGDNTLITSHGINRRDFMKLCAALAATMGLSSKAAAEMAESVSNPQRPPVIWIGAQECT WP_000145431.1 hydrogenase 2 small subunit [Salmonella enterica] Length: 372\nNumber of predicted TMHs: 1\nExp number of AAs in TMHs: 24.51505\nExp number, first 60 AAs: 0.2682\nTotal prob of N-in: 0.14406\noutside 1 331\nTMhelix 332 354\ninside 355 372

19444 GCF_002032925.1_ASM203292v1 Salmonella enterica subsp. enterica serovar Schwarzengrund Proteobacteria; Gammaproteobacteria; Enterobacterales; Enterobacteriaceae; Salmonella; Salmonella enterica; Salmonella enterica subsp. enterica MTGDNTLITSHGINRRDFMKLCAALAATMGLSSKAAAEMAESVSNPQRPPVIWIGAQECT WP_000145431.1 hydrogenase 2 small subunit [Salmonella enterica] Length: 372\nNumber of predicted TMHs: 1\nExp number of AAs in TMHs: 24.51505\nExp number, first 60 AAs: 0.2682\nTotal prob of N-in: 0.14406\noutside 1 331\nTMhelix 332 354\ninside 355 372

19445 GCF_002063925.1_ASM206392v1 Salmonella enterica subsp. enterica serovar Schwarzengrund Proteobacteria; Gammaproteobacteria; Enterobacterales; Enterobacteriaceae; Salmonella; Salmonella enterica; Salmonella enterica subsp. enterica MTGDNTLITSHGINRRDFMKLCAALAATMGLSSKAAAEMAESVSNPQRPPVIWIGAQECT WP_000145429.1 hydrogenase 2 small subunit [Salmonella enterica] Length: 372\nNumber of predicted TMHs: 1\nExp number of AAs in TMHs: 24.51309\nExp number, first 60 AAs: 0.2682\nTotal prob of N-in: 0.14402\noutside 1 331\nTMhelix 332 354\ninside 355 372

19446 GCF_002030415.1_ASM203041v1 Salmonella enterica subsp. enterica serovar Schwarzengrund Proteobacteria; Gammaproteobacteria; Enterobacterales; Enterobacteriaceae; Salmonella; Salmonella enterica; Salmonella enterica subsp. enterica MTGDNTLITSHGINRRDFMKLCAALAATMGLSSKAAAEMAESVSNPQRPPVIWIGAQECT WP_000145431.1 hydrogenase 2 small subunit [Salmonella enterica] Length: 372\nNumber of predicted TMHs: 1\nExp number of AAs in TMHs: 24.51505\nExp number, first 60 AAs: 0.2682\nTotal prob of N-in: 0.14406\noutside 1 331\nTMhelix 332 354\ninside 355 372

19447 GCF_001480325.1_Salmonella_enterica_CVM_N41734-SQ_v1.0 Salmonella enterica subsp. enterica serovar Schwarzengrund Proteobacteria; Gammaproteobacteria; Enterobacterales; Enterobacteriaceae; Salmonella; Salmonella enterica; Salmonella enterica subsp. enterica
MTGDNTLITSHGINRRDFMKLCAALAATMGLSSKAAAEMAESVSNPQRPPVIWIGAQECT WP_000145431.1 hydrogenase 2 small subunit [Salmonella enterica] Length: 372\nNumber of predicted TMHs: 1\nExp

number of AAs in TMHs: 24.51505\nExp number, first 60 AAs: 0.2682\nTotal prob of N-in: 0.14406\noutside 1 331\nTMhelix 332 354\ninside 355 372

19448 GCF_001480535.1_Salmonella_enterica_CVM_N38910-SQ_v1.0 Salmonella enterica subsp. enterica serovar Schwarzengrund Proteobacteria; Gammaproteobacteria; Enterobacterales; Enterobacteriaceae; Salmonella; Salmonella enterica; Salmonella enterica subsp. enterica MTGDNTLITSHGINRRDFMKLCAALAATMGLSSKAAAEMAESVSNPQRPPVIWIGAQECT WP_000145431.1 hydrogenase 2 small subunit [Salmonella enterica] Length: 372\nNumber of predicted TMHs: 1\nExp number of AAs in TMHs: 24.51505\nExp number, first 60 AAs: 0.2682\nTotal prob of N-in: 0.14406\noutside 1 331\nTMhelix 332 354\ninside 355 372

19449 GCF_001245545.1_Salmonella_enterica_CVM_N50434_v1.0 Salmonella enterica subsp. enterica serovar Schwarzengrund Proteobacteria; Gammaproteobacteria; Enterobacterales; Enterobacteriaceae; Salmonella; Salmonella enterica; Salmonella enterica subsp. enterica MNNEETFYQAMRRKGVTRRSFLKFCSLAATSLGLGAGMTPKIAWALENKPRIPVVWIHGL WP_001058312.1 hydrogenase [Salmonella enterica] Length: 372\nNumber of predicted TMHs: 2\nExp number of AAs in TMHs: 36.51738\nExp number, first 60 AAs: 19.72849\nTotal prob of N-in: 0.96170\nPOSSIBLE N-term signal sequence\ninside 1 19\nTMhelix 20 39\noutside 40 325\nTMhelix 326 348\ninside 349 372

19450 GCF_001246555.1_Salmonella_enterica_CVM_N51267_v1.0 Salmonella enterica subsp. enterica serovar Schwarzengrund Proteobacteria; Gammaproteobacteria; Enterobacterales; Enterobacteriaceae; Salmonella; Salmonella enterica; Salmonella enterica subsp. enterica MNNEETFYQAMRRKGVTRRSFLKFCSLAATSLGLGAGMTPKIAWALENKPRIPVVWIHGL WP_001058312.1 hydrogenase [Salmonella enterica] Length: 372\nNumber of predicted TMHs: 2\nExp number of AAs in TMHs: 36.51738\nExp number, first 60 AAs: 19.72849\nTotal prob of N-in: 0.96170\nPOSSIBLE N-term signal sequence\ninside 1 19\nTMhelix 20 39\noutside 40 325\nTMhelix 326 348\ninside 349 372

19451 GCF_001247455.1_Salmonella_enterica_CVM_N51311_v1.0 Salmonella enterica subsp. enterica serovar Schwarzengrund Proteobacteria; Gammaproteobacteria; Enterobacterales; Enterobacteriaceae; Salmonella; Salmonella enterica; Salmonella enterica subsp. enterica MNNEETFYQAMRRKGVTRRSFLKFCSLAATSLGLGAGMTPKIAWALENKPRIPVVWIHGL WP_001058312.1 hydrogenase [Salmonella enterica] Length: 372\nNumber of predicted TMHs: 2\nExp number of AAs in TMHs: 36.51738\nExp number, first 60 AAs: 19.72849\nTotal prob of N-in: 0.96170\nPOSSIBLE N-term signal sequence\ninside 1 19\nTMhelix 20 39\noutside 40 325\nTMhelix 326 348\ninside 349 372

19452 GCF_001614045.1_ASM161404v1 Salmonella enterica subsp. enterica serovar Schwarzengrund Proteobacteria; Gammaproteobacteria; Enterobacterales; Enterobacteriaceae; Salmonella; Salmonella enterica; Salmonella enterica subsp. enterica MNNEETFYQAMRRKGVTRRSFLKFCSLAATSLGLGAGMTPKIAWALENKPRIPVVWIHGL WP_001058312.1 hydrogenase [Salmonella enterica] Length: 372\nNumber of predicted TMHs: 2\nExp number of AAs in TMHs: 36.51738\nExp number, first 60 AAs: 19.72849\nTotal prob of N-in: 0.96170\nPOSSIBLE N-term signal sequence\ninside 1 19\nTMhelix 20 39\noutside 40 325\nTMhelix 326 348\ninside 349 372

19453 GCF_002033125.1_ASM203312v1 Salmonella enterica subsp. enterica serovar Schwarzengrund Proteobacteria; Gammaproteobacteria; Enterobacterales; Enterobacteriaceae; Salmonella; Salmonella enterica; Salmonella enterica subsp. enterica MTGDNTLITSHGINRRDFMKLCAALAATMGLSSKAAAEMAESVSNPQRPPVIWIGAQECT WP_000145431.1 hydrogenase 2 small subunit [Salmonella enterica] Length: 372\nNumber of predicted TMHs: 1\nExp number of AAs in TMHs: 24.51505\nExp number, first 60 AAs: 0.2682\nTotal prob of N-in: 0.14406\noutside 1 331\nTMhelix 332 354\ninside 355 372

19454 GCF_001478525.1_Salmonella_enterica_CVM_N41900-SQ_v1.0 Salmonella enterica subsp. enterica serovar Schwarzengrund Proteobacteria; Gammaproteobacteria; Enterobacterales; Enterobacteriaceae; Salmonella; Salmonella enterica; Salmonella enterica subsp. enterica MTGDNTLITSHGINRRDFMKLCAALAATMGLSSKAAAEMAESVSNPQRPPVIWIGAQECT WP_000145431.1 hydrogenase 2 small subunit [Salmonella enterica] Length: 372\nNumber of predicted TMHs: 1\nExp number of AAs in TMHs: 24.51505\nExp number, first 60 AAs: 0.2682\nTotal prob of N-in: 0.14406\noutside 1 331\nTMhelix 332 354\ninside 355 372

19455 GCF_001614045.1_ASM161404v1 Salmonella enterica subsp. enterica serovar Schwarzengrund Proteobacteria; Gammaproteobacteria; Enterobacterales; Enterobacteriaceae; Salmonella; Salmonella enterica; Salmonella enterica subsp. enterica MTGDNTLITSHGINRRDFMKLCAALAATMGLSSKAAAEMAESVSNPQRPPVIWIGAQECT WP_000145431.1 hydrogenase 2 small subunit [Salmonella enterica] Length: 372\nNumber of

predicted TMHs:  1\nExp number of AAs in TMHs: 24.51505\nExp number, first 60 AAs:  0.2682\nTotal prob of N-in: 0.14406\noutside   1   331\nTMhelix   332   354\ninside   355   372

19456 GCF_001478525.1_Salmonella_enterica_CVM_N41900-SQ_v1.0 Salmonella enterica subsp. enterica serovar Schwarzengrund Proteobacteria; Gammaproteobacteria; Enterobacterales; Enterobacteriaceae; Salmonella; Salmonella enterica; Salmonella enterica subsp. enterica MNNEETFYQAMRRKGVTRRSFLKFCSLAATSLGLGAGMTPKIAWALENKPRIPVVWIHGL WP_001058312.1 hydrogenase [Salmonella enterica] Length: 372\nNumber of predicted TMHs:  2\nExp number of AAs in TMHs: 36.51738\nExp number, first 60 AAs:  19.72849\nTotal prob of N-in:   0.96170\nPOSSIBLE N-term signal sequence\ninside   1   19\nTMhelix   20   39\noutside   40   325\nTMhelix   326   348\ninside   349   372

19457 GCF_001246305.1_Salmonella_enterica_CVM_N51257_v1.0 Salmonella enterica subsp. enterica serovar Schwarzengrund Proteobacteria; Gammaproteobacteria; Enterobacterales; Enterobacteriaceae; Salmonella; Salmonella enterica; Salmonella enterica subsp. enterica MTGDNTLITSHGINRRDFMKLCAALAATMGLSSKAAAEMAESVSNPQRPPVIWIGAQECT WP_000145431.1 hydrogenase 2 small subunit [Salmonella enterica] Length: 372\nNumber of predicted TMHs:  1\nExp number of AAs in TMHs: 24.51505\nExp number, first 60 AAs:  0.2682\nTotal prob of N-in:   0.14406\noutside   1   331\nTMhelix   332   354\ninside   355   372

19458 GCF_001480535.1_Salmonella_enterica_CVM_N38910-SQ_v1.0 Salmonella enterica subsp. enterica serovar Schwarzengrund Proteobacteria; Gammaproteobacteria; Enterobacterales; Enterobacteriaceae; Salmonella; Salmonella enterica; Salmonella enterica subsp. enterica MNNEETFYQAMRRKGVTRRSFLKFCSLAATSLGLGAGMTPKIAWALENKPRIPVVWIHGL WP_001058312.1 hydrogenase [Salmonella enterica] Length: 372\nNumber of predicted TMHs:  2\nExp number of AAs in TMHs: 36.51738\nExp number, first 60 AAs:  19.72849\nTotal prob of N-in:   0.96170\nPOSSIBLE N-term signal sequence\ninside   1   19\nTMhelix   20   39\noutside   40   325\nTMhelix   326   348\ninside   349   372

19459 GCF_001480325.1_Salmonella_enterica_CVM_N41734-SQ_v1.0 Salmonella enterica subsp. enterica serovar Schwarzengrund Proteobacteria; Gammaproteobacteria; Enterobacterales; Enterobacteriaceae; Salmonella; Salmonella enterica; Salmonella enterica subsp. enterica MNNEETFYQAMRRKGVTRRSFLKFCSLAATSLGLGAGMTPKIAWALENKPRIPVVWIHGL WP_001058312.1 hydrogenase [Salmonella enterica] Length: 372\nNumber of predicted TMHs:  2\nExp number of AAs in TMHs: 36.51738\nExp number, first 60 AAs:  19.72849\nTotal prob of N-in:   0.96170\nPOSSIBLE N-term signal sequence\ninside   1   19\nTMhelix   20   39\noutside   40   325\nTMhelix   326   348\ninside   349   372

19460 GCF_001614095.1_ASM161409v1 Salmonella enterica subsp. enterica serovar Schwarzengrund Proteobacteria; Gammaproteobacteria; Enterobacterales; Enterobacteriaceae; Salmonella; Salmonella enterica; Salmonella enterica subsp. enterica MNNEETFYQAMRRKGVTRRSFLKFCSLAATSLGLGAGMTPKIAWALENKPRIPVVWIHGL WP_001058312.1 hydrogenase [Salmonella enterica] Length: 372\nNumber of predicted TMHs:  2\nExp number of AAs in TMHs: 36.51738\nExp number, first 60 AAs:  19.72849\nTotal prob of N-in:   0.96170\nPOSSIBLE N-term signal sequence\ninside   1   19\nTMhelix   20   39\noutside   40   325\nTMhelix   326   348\ninside   349   372

19461 GCF_002033085.1_ASM203308v1 Salmonella enterica subsp. enterica serovar Schwarzengrund Proteobacteria; Gammaproteobacteria; Enterobacterales; Enterobacteriaceae; Salmonella; Salmonella enterica; Salmonella enterica subsp. enterica MNNEETFYQAMRRKGVTRRSFLKFCSLAATSLGLGAGMTPKIAWALENKPRIPVVWIHGL WP_001058312.1 hydrogenase [Salmonella enterica] Length: 372\nNumber of predicted TMHs:  2\nExp number of AAs in TMHs: 36.51738\nExp number, first 60 AAs:  19.72849\nTotal prob of N-in:   0.96170\nPOSSIBLE N-term signal sequence\ninside   1   19\nTMhelix   20   39\noutside   40   325\nTMhelix   326   348\ninside   349   372

19462 GCF_001614095.1_ASM161409v1 Salmonella enterica subsp. enterica serovar Schwarzengrund Proteobacteria; Gammaproteobacteria; Enterobacterales; Enterobacteriaceae; Salmonella; Salmonella enterica; Salmonella enterica subsp. enterica MTGDNTLITSHGINRRDFMKLCAALAATMGLSSKAAAEMAESVSNPQRPPVIWIGAQECT WP_000145431.1 hydrogenase 2 small subunit [Salmonella enterica] Length: 372\nNumber of predicted TMHs:  1\nExp number of AAs in TMHs: 24.51505\nExp number, first 60 AAs:  0.2682\nTotal prob of N-in: 0.14406\noutside   1   331\nTMhelix   332   354\ninside   355   372

19463 GCF_001246985.1_Salmonella_enterica_CVM_N51289_v1.0 Salmonella enterica subsp. enterica serovar Schwarzengrund Proteobacteria; Gammaproteobacteria; Enterobacterales; Enterobacteriaceae; Salmonella; Salmonella enterica; Salmonella enterica subsp. enterica MNNEETFYQAMRRKGVTRRSFLKFCSLAATSLGLGAGMTPKIAWALENKPRIPVVWIHGL WP_001058312.1 hydrogenase [Salmonella enterica] Length: 372\nNumber of predicted TMHs:  2\nExp number of AAs in

TMHs: 36.51738\nExp number, first 60 AAs: 19.72849\nTotal prob of N-in: 0.96170\nPOSSIBLE N-term signal sequence\ninside 1 19\nTMhelix 20 39\noutside 40 325\nTMhelix 326 348\ninside 349 372

19464 GCF_002031125.1_ASM203112v1 Salmonella enterica subsp. enterica serovar Schwarzengrund Proteobacteria; Gammaproteobacteria; Enterobacterales; Enterobacteriaceae; Salmonella; Salmonella enterica; Salmonella enterica subsp. enterica MTGDNTLITSHGINRRDFMKLCAALAATMGLSSKAAAEMAESVSNPQRPPVIWIGAQECT WP_000145431.1 hydrogenase 2 small subunit [Salmonella enterica] Length: 372\nNumber of predicted TMHs: 1\nExp number of AAs in TMHs: 24.51505\nExp number, first 60 AAs: 0.2682\nTotal prob of N-in: 0.14406\noutside 1 331\nTMhelix 332 354\ninside 355 372

19465 GCF_001479805.1_Salmonella_enterica_CVM_N38851-SQ_v1.0 Salmonella enterica subsp. enterica serovar Schwarzengrund Proteobacteria; Gammaproteobacteria; Enterobacterales; Enterobacteriaceae; Salmonella; Salmonella enterica; Salmonella enterica subsp. enterica MTGDNTLITSHGINRRDFMKLCAALAATMGLSSKAAAEMAESVSNPQRPPVIWIGAQECT WP_000145431.1 hydrogenase 2 small subunit [Salmonella enterica] Length: 372\nNumber of predicted TMHs: 1\nExp number of AAs in TMHs: 24.51505\nExp number, first 60 AAs: 0.2682\nTotal prob of N-in: 0.14406\noutside 1 331\nTMhelix 332 354\ninside 355 372

19466 GCF_001479805.1_Salmonella_enterica_CVM_N38851-SQ_v1.0 Salmonella enterica subsp. enterica serovar Schwarzengrund Proteobacteria; Gammaproteobacteria; Enterobacterales; Enterobacteriaceae; Salmonella; Salmonella enterica; Salmonella enterica subsp. enterica MNNEETFYQAMRRKGVTRRSFLKFCSLAATSLGLGAGMTPKIAWALENKPRIPVVWIHGL WP_001058312.1 hydrogenase [Salmonella enterica] Length: 372\nNumber of predicted TMHs: 2\nExp number of AAs in TMHs: 36.51738\nExp number, first 60 AAs: 19.72849\nTotal prob of N-in: 0.96170\nPOSSIBLE N-term signal sequence\ninside 1 19\nTMhelix 20 39\noutside 40 325\nTMhelix 326 348\ninside 349 372

19467 GCF_001614065.1_ASM161406v1 Salmonella enterica subsp. enterica serovar Schwarzengrund Proteobacteria; Gammaproteobacteria; Enterobacterales; Enterobacteriaceae; Salmonella; Salmonella enterica; Salmonella enterica subsp. enterica MTGDNTLITSHGINRRDFMKLCAALAATMGLSSKAAAEMAESVSNPQRPPVIWIGAQECT WP_000145431.1 hydrogenase 2 small subunit [Salmonella enterica] Length: 372\nNumber of predicted TMHs: 1\nExp number of AAs in TMHs: 24.51505\nExp number, first 60 AAs: 0.2682\nTotal prob of N-in: 0.14406\noutside 1 331\nTMhelix 332 354\ninside 355 372

19468 GCF_002032925.1_ASM203292v1 Salmonella enterica subsp. enterica serovar Schwarzengrund Proteobacteria; Gammaproteobacteria; Enterobacterales; Enterobacteriaceae; Salmonella; Salmonella enterica; Salmonella enterica subsp. enterica MNNEETFYQAMRRKGVTRRSFLKFCSLAATSLGLGAGMTPKIAWALENKPRIPVVWIHGL WP_001058312.1 hydrogenase [Salmonella enterica] Length: 372\nNumber of predicted TMHs: 2\nExp number of AAs in TMHs: 36.51738\nExp number, first 60 AAs: 19.72849\nTotal prob of N-in: 0.96170\nPOSSIBLE N-term signal sequence\ninside 1 19\nTMhelix 20 39\noutside 40 325\nTMhelix 326 348\ninside 349 372

19469 GCF_001614105.1_ASM161410v1 Salmonella enterica subsp. enterica serovar Schwarzengrund Proteobacteria; Gammaproteobacteria; Enterobacterales; Enterobacteriaceae; Salmonella; Salmonella enterica; Salmonella enterica subsp. enterica MTGDNTLITSHGINRRDFMKLCAALAATMGLSSKAAAEMAESVSNPQRPPVIWIGAQECT WP_000145431.1 hydrogenase 2 small subunit [Salmonella enterica] Length: 372\nNumber of predicted TMHs: 1\nExp number of AAs in TMHs: 24.51505\nExp number, first 60 AAs: 0.2682\nTotal prob of N-in: 0.14406\noutside 1 331\nTMhelix 332 354\ninside 355 372

19470 GCF_001480895.1_Salmonella_enterica_CVM_N39866-SQ_v1.0 Salmonella enterica subsp. enterica serovar Schwarzengrund Proteobacteria; Gammaproteobacteria; Enterobacterales; Enterobacteriaceae; Salmonella; Salmonella enterica; Salmonella enterica subsp. enterica MNNEETFYQAMRRKGVTRRSFLKFCSLAATSLGLGAGMTPKIAWALENKPRIPVVWIHGL WP_001058312.1 hydrogenase [Salmonella enterica] Length: 372\nNumber of predicted TMHs: 2\nExp number of AAs in TMHs: 36.51738\nExp number, first 60 AAs: 19.72849\nTotal prob of N-in: 0.96170\nPOSSIBLE N-term signal sequence\ninside 1 19\nTMhelix 20 39\noutside 40 325\nTMhelix 326 348\ninside 349 372

19471 GCF_002030415.1_ASM203041v1 Salmonella enterica subsp. enterica serovar Schwarzengrund Proteobacteria; Gammaproteobacteria; Enterobacterales; Enterobacteriaceae; Salmonella; Salmonella enterica; Salmonella enterica subsp. enterica MNNEETFYQAMRRKGVTRRSFLKFCSLAATSLGLGAGMTPKIAWALENKPRIPVVWIHGL WP_001058312.1 hydrogenase [Salmonella enterica] Length: 372\nNumber of predicted TMHs: 2\nExp number of AAs in TMHs: 36.51738\nExp number, first 60 AAs: 19.72849\nTotal prob of N-in: 0.96170\nPOSSIBLE N-term signal sequence\ninside 1 19\nTMhelix 20 39\noutside 40 325\nTMhelix 326 348\ninside 349 372

19472 GCF_002031905.1_ASM203190v1 Salmonella enterica subsp. enterica serovar Schwarzengrund Proteobacteria; Gammaproteobacteria; Enterobacterales; Enterobacteriaceae; Salmonella; Salmonella enterica; Salmonella enterica subsp. enterica MNNEETFYQAMRRKGVTRRSFLKFCSLAATSLGLGAGMTPKIAWALENKPRIPVVWIHGL WP_001058312.1 hydrogenase [Salmonella enterica] Length: 372\nNumber of predicted TMHs: 2\nExp number of AAs in TMHs: 36.51738\nExp number, first 60 AAs: 19.72849\nTotal prob of N-in: 0.96170\nPOSSIBLE N-term signal sequence\ninside 1 19\nTMhelix 20 39\noutside 40 325\nTMhelix 326 348\ninside 349 372

19473 GCF_002031125.1_ASM203112v1 Salmonella enterica subsp. enterica serovar Schwarzengrund Proteobacteria; Gammaproteobacteria; Enterobacterales; Enterobacteriaceae; Salmonella; Salmonella enterica; Salmonella enterica subsp. enterica MNNEETFYQAMRRKGVTRRSFLKFCSLAATSLGLGAGMTPKIAWALENKPRIPVVWIHGL WP_001058312.1 hydrogenase [Salmonella enterica] Length: 372\nNumber of predicted TMHs: 2\nExp number of AAs in TMHs: 36.51738\nExp number, first 60 AAs: 19.72849\nTotal prob of N-in: 0.96170\nPOSSIBLE N-term signal sequence\ninside 1 19\nTMhelix 20 39\noutside 40 325\nTMhelix 326 348\ninside 349 372

19474 GCF_001614065.1_ASM161406v1 Salmonella enterica subsp. enterica serovar Schwarzengrund Proteobacteria; Gammaproteobacteria; Enterobacterales; Enterobacteriaceae; Salmonella; Salmonella enterica; Salmonella enterica subsp. enterica MNNEETFYQAMRRKGVTRRSFLKFCSLAATSLGLGAGMTPKIAWALENKPRIPVVWIHGL WP_001058312.1 hydrogenase [Salmonella enterica] Length: 372\nNumber of predicted TMHs: 2\nExp number of AAs in TMHs: 36.51738\nExp number, first 60 AAs: 19.72849\nTotal prob of N-in: 0.96170\nPOSSIBLE N-term signal sequence\ninside 1 19\nTMhelix 20 39\noutside 40 325\nTMhelix 326 348\ninside 349 372

19475 GCF_001614105.1_ASM161410v1 Salmonella enterica subsp. enterica serovar Schwarzengrund Proteobacteria; Gammaproteobacteria; Enterobacterales; Enterobacteriaceae; Salmonella; Salmonella enterica; Salmonella enterica subsp. enterica MNNEETFYQAMRRKGVTRRSFLKFCSLAATSLGLGAGMTPKIAWALENKPRIPVVWIHGL WP_001058312.1 hydrogenase [Salmonella enterica] Length: 372\nNumber of predicted TMHs: 2\nExp number of AAs in TMHs: 36.51738\nExp number, first 60 AAs: 19.72849\nTotal prob of N-in: 0.96170\nPOSSIBLE N-term signal sequence\ninside 1 19\nTMhelix 20 39\noutside 40 325\nTMhelix 326 348\ninside 349 372

19476 GCF_001480895.1_Salmonella_enterica_CVM_N39866-SQ_v1.0 Salmonella enterica subsp. enterica serovar Schwarzengrund Proteobacteria; Gammaproteobacteria; Enterobacterales; Enterobacteriaceae; Salmonella; Salmonella enterica; Salmonella enterica subsp. enterica MTGDNTLITSHGINRRDFMKLCAALAATMGLSSKAAAEMAESVSNPQRPPVIWIGAQECT WP_000145431.1 hydrogenase 2 small subunit [Salmonella enterica] Length: 372\nNumber of predicted TMHs: 1\nExp number of AAs in TMHs: 24.51505\nExp number, first 60 AAs: 0.2682\nTotal prob of N-in: 0.14406\noutside 1 331\nTMhelix 332 354\ninside 355 372

19477 GCF_002106675.1_ASM210667v1 Salmonella enterica subsp. enterica serovar Seftenburg Proteobacteria; Gammaproteobacteria; Enterobacterales; Enterobacteriaceae; Salmonella; Salmonella enterica; Salmonella enterica subsp. enterica MNNEETFYQAMRRKGVTRRSFLKFCSLAATSLGLGAGMTPKIAWALENKPRIPVVWIHGL WP_001058312.1 hydrogenase [Salmonella enterica] Length: 372\nNumber of predicted TMHs: 2\nExp number of AAs in TMHs: 36.51738\nExp number, first 60 AAs: 19.72849\nTotal prob of N-in: 0.96170\nPOSSIBLE N-term signal sequence\ninside 1 19\nTMhelix 20 39\noutside 40 325\nTMhelix 326 348\ninside 349 372

19478 GCF_002106675.1_ASM210667v1 Salmonella enterica subsp. enterica serovar Seftenburg Proteobacteria; Gammaproteobacteria; Enterobacterales; Enterobacteriaceae; Salmonella; Salmonella enterica; Salmonella enterica subsp. enterica MTGDNTLITSHGINRRDFMKLCAALAATMGLSSKAAAEMAESVSNPQRPPVIWIGAQECT WP_000145429.1 hydrogenase 2 small subunit [Salmonella enterica] Length: 372\nNumber of predicted TMHs: 1\nExp number of AAs in TMHs: 24.51309\nExp number, first 60 AAs: 0.2682\nTotal prob of N-in: 0.14402\noutside 1 331\nTMhelix 332 354\ninside 355 372

19479 GCF_002062745.1_ASM206274v1 Salmonella enterica subsp. enterica serovar Senftenberg Proteobacteria; Gammaproteobacteria; Enterobacterales; Enterobacteriaceae; Salmonella; Salmonella enterica; Salmonella enterica subsp. enterica MTGDNTLITSHGINRRDFMKLCAALAATMGLSSKAAAEMAESVSNPQRPPVIWIGAQECT WP_000145429.1 hydrogenase 2 small subunit [Salmonella enterica] Length: 372\nNumber of predicted TMHs: 1\nExp number of AAs in TMHs: 24.51309\nExp number, first 60 AAs: 0.2682\nTotal prob of N-in: 0.14402\noutside 1 331\nTMhelix 332 354\ninside 355 372

19480 GCF_002062745.1_ASM206274v1 Salmonella enterica subsp. enterica serovar Senftenberg Proteobacteria; Gammaproteobacteria; Enterobacterales; Enterobacteriaceae; Salmonella; Salmonella enterica; Salmonella enterica subsp. enterica MNNEETFYQAMRRKGVTRRSFLKFCSLAATSLGLGAGMTPKIAWALENKPRIPVVWIHGL WP_001058307.1 hydrogenase [Salmonella enterica] Length: 372\nNumber of predicted TMHs: 2\nExp

number of AAs in TMHs: 36.51738\nExp number, first 60 AAs: 19.72849\nTotal prob of N-in: 0.96170\nPOSSIBLE N-term signal sequence\ninside 1 19\nTMhelix 20 39\noutside 40 325\nTMhelix 326 348\ninside 349 372

19481 GCF_001247445.1_Salmonella_enterica_CVM_N51312_v1.0 Salmonella enterica subsp. enterica serovar Senftenberg Proteobacteria; Gammaproteobacteria; Enterobacterales; Enterobacteriaceae; Salmonella; Salmonella enterica; Salmonella enterica subsp. enterica MTGDNTLITSHGINRRDFMKLCAALAATMGLSSKAAAEMAESVSNPQRPPVIWIGAQECT WP_000145429.1 hydrogenase 2 small subunit [Salmonella enterica] Length: 372\nNumber of predicted TMHs: 1\nExp number of AAs in TMHs: 24.51309\nExp number, first 60 AAs: 0.2682\nTotal prob of N-in: 0.14402\noutside 1 331\nTMhelix 332 354\ninside 355 372

19482 GCF_001276765.1_ASM127676v1 Salmonella enterica subsp. enterica serovar Senftenberg Proteobacteria; Gammaproteobacteria; Enterobacterales; Enterobacteriaceae; Salmonella; Salmonella enterica; Salmonella enterica subsp. enterica MTGDNTLITSHGINRRDFMKLCAALAATMGLSSKAAAEMAESVSNPQRPPVIWIGAQECT WP_000145429.1 hydrogenase 2 small subunit [Salmonella enterica] Length: 372\nNumber of predicted TMHs: 1\nExp number of AAs in TMHs: 24.51309\nExp number, first 60 AAs: 0.2682\nTotal prob of N-in: 0.14402\noutside 1 331\nTMhelix 332 354\ninside 355 372

19483 GCF_001247445.1_Salmonella_enterica_CVM_N51312_v1.0 Salmonella enterica subsp. enterica serovar Senftenberg Proteobacteria; Gammaproteobacteria; Enterobacterales; Enterobacteriaceae; Salmonella; Salmonella enterica; Salmonella enterica subsp. enterica MNNEETFYQAMRRKGVTRRSFLKFCSLAATSLGLGAGMTPKIAWALENKPRIPVVWIHGL WP_001058312.1 hydrogenase [Salmonella enterica] Length: 372\nNumber of predicted TMHs: 2\nExp number of AAs in TMHs: 36.51738\nExp number, first 60 AAs: 19.72849\nTotal prob of N-in: 0.96170\nPOSSIBLE N-term signal sequence\ninside 1 19\nTMhelix 20 39\noutside 40 325\nTMhelix 326 348\ninside 349 372

19484 GCF_001276765.1_ASM127676v1 Salmonella enterica subsp. enterica serovar Senftenberg Proteobacteria; Gammaproteobacteria; Enterobacterales; Enterobacteriaceae; Salmonella; Salmonella enterica; Salmonella enterica subsp. enterica MNNEETFYQAMRRKGVTRRSFLKFCSLAATSLGLGAGMTPKIAWALENKPRIPVVWIHGL WP_001058312.1 hydrogenase [Salmonella enterica] Length: 372\nNumber of predicted TMHs: 2\nExp number of AAs in TMHs: 36.51738\nExp number, first 60 AAs: 19.72849\nTotal prob of N-in: 0.96170\nPOSSIBLE N-term signal sequence\ninside 1 19\nTMhelix 20 39\noutside 40 325\nTMhelix 326 348\ninside 349 372

19485 GCF_001276775.1_ASM127677v1 Salmonella enterica subsp. enterica serovar Senftenberg Proteobacteria; Gammaproteobacteria; Enterobacterales; Enterobacteriaceae; Salmonella; Salmonella enterica; Salmonella enterica subsp. enterica MTGDNTLITSHGINRRDFMKLCAALAATMGLSSKAAAEMAESVSNPQRPPVIWIGAQECT WP_000145429.1 hydrogenase 2 small subunit [Salmonella enterica] Length: 372\nNumber of predicted TMHs: 1\nExp number of AAs in TMHs: 24.51309\nExp number, first 60 AAs: 0.2682\nTotal prob of N-in: 0.14402\noutside 1 331\nTMhelix 332 354\ninside 355 372

19486 GCF_001246655.1_Salmonella_enterica_CVM_N51274_v1.0 Salmonella enterica subsp. enterica serovar Senftenberg Proteobacteria; Gammaproteobacteria; Enterobacterales; Enterobacteriaceae; Salmonella; Salmonella enterica; Salmonella enterica subsp. enterica MTGDNTLITSHGINRRDFMKLCAALAATMGLSSKAAAEMAESVSNPQRPPVIWIGAQECT WP_000145429.1 hydrogenase 2 small subunit [Salmonella enterica] Length: 372\nNumber of predicted TMHs: 1\nExp number of AAs in TMHs: 24.51309\nExp number, first 60 AAs: 0.2682\nTotal prob of N-in: 0.14402\noutside 1 331\nTMhelix 332 354\ninside 355 372

19487 GCF_002030925.1_ASM203092v1 Salmonella enterica subsp. enterica serovar Senftenberg Proteobacteria; Gammaproteobacteria; Enterobacterales; Enterobacteriaceae; Salmonella; Salmonella enterica; Salmonella enterica subsp. enterica MNNEETFYQAMRRKGVTRRSFLKFCSLAATSLGLGAGMTPKIAWALENKPRIPVVWIHGL WP_001058312.1 hydrogenase [Salmonella enterica] Length: 372\nNumber of predicted TMHs: 2\nExp number of AAs in TMHs: 36.51738\nExp number, first 60 AAs: 19.72849\nTotal prob of N-in: 0.96170\nPOSSIBLE N-term signal sequence\ninside 1 19\nTMhelix 20 39\noutside 40 325\nTMhelix 326 348\ninside 349 372

19488 GCF_001477795.1_Salmonella_enterica_CVM_N29321-SQ_v1.0 Salmonella enterica subsp. enterica serovar Senftenberg Proteobacteria; Gammaproteobacteria; Enterobacterales; Enterobacteriaceae; Salmonella; Salmonella enterica; Salmonella enterica subsp. enterica MTGDNTLITSHGINRRDFMKLCAALAATMGLSSKAAAEMAESVSNPQRPPVIWIGAQECT WP_000145429.1 hydrogenase 2 small subunit [Salmonella enterica] Length: 372\nNumber of predicted TMHs: 1\nExp

number of AAs in TMHs: 24.51309\nExp number, first 60 AAs: 0.2682\nTotal prob of N-in: 0.14402\noutside 1 331\nTMhelix 332 354\ninside 355 372

19489 GCF_001479095.1_Salmonella_enterica_CVM_N31402-SQ_v1.0 Salmonella enterica subsp. enterica serovar Senftenberg Proteobacteria; Gammaproteobacteria; Enterobacterales; Enterobacteriaceae; Salmonella; Salmonella enterica; Salmonella enterica subsp. enterica MNNEETFYQAMRRKGVTRRSFLKFCSLAATSLGLGAGMTPKIAWALENKPRIPVVWIHGL WP_001058312.1 hydrogenase [Salmonella enterica] Length: 372\nNumber of predicted TMHs: 2\nExp number of AAs in TMHs: 36.51738\nExp number, first 60 AAs: 19.72849\nTotal prob of N-in: 0.96170\nPOSSIBLE N-term signal sequence\ninside 1 19\nTMhelix 20 39\noutside 40 325\nTMhelix 326 348\ninside 349 372

19490 GCF_001479525.1_Salmonella_enterica_CVM_N32779-R_v1.0 Salmonella enterica subsp. enterica serovar Senftenberg Proteobacteria; Gammaproteobacteria; Enterobacterales; Enterobacteriaceae; Salmonella; Salmonella enterica; Salmonella enterica subsp. enterica MTGDNTLITSHGINRRDFMKLCAALAATMGLSSKAAAEMAESVSNPQRPPVIWIGAQECT WP_000145429.1 hydrogenase 2 small subunit [Salmonella enterica] Length: 372\nNumber of predicted TMHs: 1\nExp number of AAs in TMHs: 24.51309\nExp number, first 60 AAs: 0.2682\nTotal prob of N-in: 0.14402\noutside 1 331\nTMhelix 332 354\ninside 355 372

19491 GCF_001969425.1_ASM196942v1 Salmonella enterica subsp. enterica serovar Senftenberg Proteobacteria; Gammaproteobacteria; Enterobacterales; Enterobacteriaceae; Salmonella; Salmonella enterica; Salmonella enterica subsp. enterica MTGDNTLITSHGINRRDFMKLCAALAATMGLSSKAAAEMAESVSNPQRPPVIWIGAQECT WP_000145429.1 hydrogenase 2 small subunit [Salmonella enterica] Length: 372\nNumber of predicted TMHs: 1\nExp number of AAs in TMHs: 24.51309\nExp number, first 60 AAs: 0.2682\nTotal prob of N-in: 0.14402\noutside 1 331\nTMhelix 332 354\ninside 355 372

19492 GCF_001276925.1_ASM127692v1 Salmonella enterica subsp. enterica serovar Senftenberg Proteobacteria; Gammaproteobacteria; Enterobacterales; Enterobacteriaceae; Salmonella; Salmonella enterica; Salmonella enterica subsp. enterica MTGDNTLITSHGINRRDFMKLCAALAATMGLSSKAAAEMAESVSNPQRPPVIWIGAQECT WP_000145429.1 hydrogenase 2 small subunit [Salmonella enterica] Length: 372\nNumber of predicted TMHs: 1\nExp number of AAs in TMHs: 24.51309\nExp number, first 60 AAs: 0.2682\nTotal prob of N-in: 0.14402\noutside 1 331\nTMhelix 332 354\ninside 355 372

19493 GCF_001479465.1_Salmonella_enterica_CVM_N32755-R_v1.0 Salmonella enterica subsp. enterica serovar Senftenberg Proteobacteria; Gammaproteobacteria; Enterobacterales; Enterobacteriaceae; Salmonella; Salmonella enterica; Salmonella enterica subsp. enterica MNNEETFYQAMRRKGVTRRSFLKFCSLAATSLGLGAGMTPKIAWALENKPRIPVVWIHGL WP_001058312.1 hydrogenase [Salmonella enterica] Length: 372\nNumber of predicted TMHs: 2\nExp number of AAs in TMHs: 36.51738\nExp number, first 60 AAs: 19.72849\nTotal prob of N-in: 0.96170\nPOSSIBLE N-term signal sequence\ninside 1 19\nTMhelix 20 39\noutside 40 325\nTMhelix 326 348\ninside 349 372

19494 GCF_001479005.1_Salmonella_enterica_CVM_N30685-SQ_v1.0 Salmonella enterica subsp. enterica serovar Senftenberg Proteobacteria; Gammaproteobacteria; Enterobacterales; Enterobacteriaceae; Salmonella; Salmonella enterica; Salmonella enterica subsp. enterica MNNEETFYQAMRRKGVTRRSFLKFCSLAATSLGLGAGMTPKIAWALENKPRIPVVWIHGL WP_001058312.1 hydrogenase [Salmonella enterica] Length: 372\nNumber of predicted TMHs: 2\nExp number of AAs in TMHs: 36.51738\nExp number, first 60 AAs: 19.72849\nTotal prob of N-in: 0.96170\nPOSSIBLE N-term signal sequence\ninside 1 19\nTMhelix 20 39\noutside 40 325\nTMhelix 326 348\ninside 349 372

19495 GCF_001479465.1_Salmonella_enterica_CVM_N32755-R_v1.0 Salmonella enterica subsp. enterica serovar Senftenberg Proteobacteria; Gammaproteobacteria; Enterobacterales; Enterobacteriaceae; Salmonella; Salmonella enterica; Salmonella enterica subsp. enterica MTGDNTLITSHGINRRDFMKLCAALAATMGLSSKAAAEMAESVSNPQRPPVIWIGAQECT WP_000145429.1 hydrogenase 2 small subunit [Salmonella enterica] Length: 372\nNumber of predicted TMHs: 1\nExp number of AAs in TMHs: 24.51309\nExp number, first 60 AAs: 0.2682\nTotal prob of N-in: 0.14402\noutside 1 331\nTMhelix 332 354\ninside 355 372

19496 GCF_001478345.1_Salmonella_enterica_CVM_N38864-R_v1.0 Salmonella enterica subsp. enterica serovar Senftenberg Proteobacteria; Gammaproteobacteria; Enterobacterales; Enterobacteriaceae; Salmonella; Salmonella enterica; Salmonella enterica subsp. enterica MNNEETFYQAMRRKGVTRRSFLKFCSLAATSLGLGAGMTPKIAWALENKPRIPVVWIHGL WP_001058312.1

hydrogenase [Salmonella enterica]	Length: 372\nNumber of predicted TMHs:  2\nExp number of AAs in TMHs: 36.51738\nExp number, first 60 AAs:  19.72849\nTotal prob of N-in:   0.96170\nPOSSIBLE N-term signal sequence\ninside   1   19\nTMhelix   20   39\noutside   40   325\nTMhelix   326   348\ninside   349   372

19497	GCF_001481035.1_Salmonella_enterica_CVM_N42232_v1.0	Salmonella enterica subsp. enterica serovar Senftenberg	Proteobacteria; Gammaproteobacteria; Enterobacterales; Enterobacteriaceae; Salmonella; Salmonella enterica; Salmonella enterica subsp. enterica	MNNEETFYQAMRRKGVTRRSFLKFCSLAATSLGLGAGMTPKIAWALENKPRIPVVWIHGL	WP_001058312.1	hydrogenase [Salmonella enterica]	Length: 372\nNumber of predicted TMHs:  2\nExp number of AAs in TMHs: 36.51738\nExp number, first 60 AAs:  19.72849\nTotal prob of N-in:   0.96170\nPOSSIBLE N-term signal sequence\ninside   1   19\nTMhelix   20   39\noutside   40   325\nTMhelix   326   348\ninside   349   372

19498	GCF_002031755.1_ASM203175v1	Salmonella enterica subsp. enterica serovar Senftenberg	Proteobacteria; Gammaproteobacteria; Enterobacterales; Enterobacteriaceae; Salmonella; Salmonella enterica; Salmonella enterica subsp. enterica	MNNEETFYQAMRRKGVTRRSFLKFCSLAATSLGLGAGMTPKIAWALENKPRIPVVWIHGL	WP_001058312.1	hydrogenase [Salmonella enterica]	Length: 372\nNumber of predicted TMHs:  2\nExp number of AAs in TMHs: 36.51738\nExp number, first 60 AAs:  19.72849\nTotal prob of N-in:   0.96170\nPOSSIBLE N-term signal sequence\ninside   1   19\nTMhelix   20   39\noutside   40   325\nTMhelix   326   348\ninside   349   372

19499	GCF_001457675.1_NCTC10384	Salmonella enterica subsp. enterica serovar Senftenberg	Proteobacteria; Gammaproteobacteria; Enterobacterales; Enterobacteriaceae; Salmonella; Salmonella enterica; Salmonella enterica subsp. enterica	MNNEETFYQAMRRKGVTRRSFLKFCSLAATSLGLGAGMTPKIAWALENKPRIPVVWIHGL	WP_001058312.1	hydrogenase [Salmonella enterica]	Length: 372\nNumber of predicted TMHs:  2\nExp number of AAs in TMHs: 36.51738\nExp number, first 60 AAs:  19.72849\nTotal prob of N-in:   0.96170\nPOSSIBLE N-term signal sequence\ninside   1   19\nTMhelix   20   39\noutside   40   325\nTMhelix   326   348\ninside   349   372

19500	GCF_001481035.1_Salmonella_enterica_CVM_N42232_v1.0	Salmonella enterica subsp. enterica serovar Senftenberg	Proteobacteria; Gammaproteobacteria; Enterobacterales; Enterobacteriaceae; Salmonella; Salmonella enterica; Salmonella enterica subsp. enterica	MTGDNTLITSHGINRRDFMKLCAALAATMGLSSKAAAEMAESVSNPQRPPVIWIGAQECT	WP_000145429.1	hydrogenase 2 small subunit [Salmonella enterica]	Length: 372\nNumber of predicted TMHs:  1\nExp number of AAs in TMHs: 24.51309\nExp number, first 60 AAs:  0.2682\nTotal prob of N-in:   0.14402\noutside   1   331\nTMhelix   332   354\ninside   355   372

19501	GCF_001478295.1_Salmonella_enterica_CVM_N37931-SQ_v1.0	Salmonella enterica subsp. enterica serovar Senftenberg	Proteobacteria; Gammaproteobacteria; Enterobacterales; Enterobacteriaceae; Salmonella; Salmonella enterica; Salmonella enterica subsp. enterica	MTGDNTLITSHGINRRDFMKLCAALAATMGLSSKAAAEMAESVSNPQRPPVIWIGAQECT	WP_000145429.1	hydrogenase 2 small subunit [Salmonella enterica]	Length: 372\nNumber of predicted TMHs:  1\nExp number of AAs in TMHs: 24.51309\nExp number, first 60 AAs:  0.2682\nTotal prob of N-in:   0.14402\noutside   1   331\nTMhelix   332   354\ninside   355   372

19502	GCF_001478295.1_Salmonella_enterica_CVM_N37931-SQ_v1.0	Salmonella enterica subsp. enterica serovar Senftenberg	Proteobacteria; Gammaproteobacteria; Enterobacterales; Enterobacteriaceae; Salmonella; Salmonella enterica; Salmonella enterica subsp. enterica	MNNEETFYQAMRRKGVTRRSFLKFCSLAATSLGLGAGMTPKIAWALENKPRIPVVWIHGL	WP_001058312.1	hydrogenase [Salmonella enterica]	Length: 372\nNumber of predicted TMHs:  2\nExp number of AAs in TMHs: 36.51738\nExp number, first 60 AAs:  19.72849\nTotal prob of N-in:   0.96170\nPOSSIBLE N-term signal sequence\ninside   1   19\nTMhelix   20   39\noutside   40   325\nTMhelix   326   348\ninside   349   372

19503	GCF_001479005.1_Salmonella_enterica_CVM_N30685-SQ_v1.0	Salmonella enterica subsp. enterica serovar Senftenberg	Proteobacteria; Gammaproteobacteria; Enterobacterales; Enterobacteriaceae; Salmonella; Salmonella enterica; Salmonella enterica subsp. enterica	MTGDNTLITSHGINRRDFMKLCAALAATMGLSSKAAAEMAESVSNPQRPPVIWIGAQECT	WP_000145429.1	hydrogenase 2 small subunit [Salmonella enterica]	Length: 372\nNumber of predicted TMHs:  1\nExp number of AAs in TMHs: 24.51309\nExp number, first 60 AAs:  0.2682\nTotal prob of N-in:   0.14402\noutside   1   331\nTMhelix   332   354\ninside   355   372

19504	GCF_001457675.1_NCTC10384	Salmonella enterica subsp. enterica serovar Senftenberg	Proteobacteria; Gammaproteobacteria; Enterobacterales; Enterobacteriaceae; Salmonella; Salmonella enterica; Salmonella enterica subsp. enterica	MTGDNTLITSHGINRRDFMKLCAALAATMGLSSKAAAEMAESVSNPQRPPVIWIGAQECT

WP_000145429.1 hydrogenase 2 small subunit [Salmonella enterica] Length: 372\nNumber of predicted TMHs: 1\nExp number of AAs in TMHs: 24.51309\nExp number, first 60 AAs: 0.2682\nTotal prob of N-in: 0.14402\noutside 1 331\nTMhelix 332 354\ninside 355 372

19505 GCF_001478345.1_Salmonella_enterica_CVM_N38864-R_v1.0 Salmonella enterica subsp. enterica serovar Senftenberg Proteobacteria; Gammaproteobacteria; Enterobacterales; Enterobacteriaceae; Salmonella; Salmonella enterica; Salmonella enterica subsp. enterica MTGDNTLITSHGINRRDFMKLCAALAATMGLSSKAAAEMAESVSNPQRPPVIWIGAQECT WP_000145429.1 hydrogenase 2 small subunit [Salmonella enterica] Length: 372\nNumber of predicted TMHs: 1\nExp number of AAs in TMHs: 24.51309\nExp number, first 60 AAs: 0.2682\nTotal prob of N-in: 0.14402\noutside 1 331\nTMhelix 332 354\ninside 355 372

19506 GCF_001276925.1_ASM127692v1 Salmonella enterica subsp. enterica serovar Senftenberg Proteobacteria; Gammaproteobacteria; Enterobacterales; Enterobacteriaceae; Salmonella; Salmonella enterica; Salmonella enterica subsp. enterica MNNEETFYQAMRRKGVTRRSFLKFCSLAATSLGLGAGMTPKIAWALENKPRIPVVWIHGL WP_001058312.1 hydrogenase [Salmonella enterica] Length: 372\nNumber of predicted TMHs: 2\nExp number of AAs in TMHs: 36.51738\nExp number, first 60 AAs: 19.72849\nTotal prob of N-in: 0.96170\nPOSSIBLE N-term signal sequence\ninside 1 19\nTMhelix 20 39\noutside 40 325\nTMhelix 326 348\ninside 349 372

19507 GCF_001276745.1_ASM127674v1 Salmonella enterica subsp. enterica serovar Senftenberg Proteobacteria; Gammaproteobacteria; Enterobacterales; Enterobacteriaceae; Salmonella; Salmonella enterica; Salmonella enterica subsp. enterica MTGDNTLITSHGINRRDFMKLCAALAATMGLSSKAAAEMAESVSNPQRPPVIWIGAQECT WP_000145429.1 hydrogenase 2 small subunit [Salmonella enterica] Length: 372\nNumber of predicted TMHs: 1\nExp number of AAs in TMHs: 24.51309\nExp number, first 60 AAs: 0.2682\nTotal prob of N-in: 0.14402\noutside 1 331\nTMhelix 332 354\ninside 355 372

19508 GCF_002031375.1_ASM203137v1 Salmonella enterica subsp. enterica serovar Senftenberg Proteobacteria; Gammaproteobacteria; Enterobacterales; Enterobacteriaceae; Salmonella; Salmonella enterica; Salmonella enterica subsp. enterica MTGDNTLITSHGINRRDFMKLCAALAATMGLSSKAAAEMAESVSNPQRPPVIWIGAQECT WP_000145429.1 hydrogenase 2 small subunit [Salmonella enterica] Length: 372\nNumber of predicted TMHs: 1\nExp number of AAs in TMHs: 24.51309\nExp number, first 60 AAs: 0.2682\nTotal prob of N-in: 0.14402\noutside 1 331\nTMhelix 332 354\ninside 355 372

19509 GCF_001479095.1_Salmonella_enterica_CVM_N31402-SQ_v1.0 Salmonella enterica subsp. enterica serovar Senftenberg Proteobacteria; Gammaproteobacteria; Enterobacterales; Enterobacteriaceae; Salmonella; Salmonella enterica; Salmonella enterica subsp. enterica MTGDNTLITSHGINRRDFMKLCAALAATMGLSSKAAAEMAESVSNPQRPPVIWIGAQECT WP_000145429.1 hydrogenase 2 small subunit [Salmonella enterica] Length: 372\nNumber of predicted TMHs: 1\nExp number of AAs in TMHs: 24.51309\nExp number, first 60 AAs: 0.2682\nTotal prob of N-in: 0.14402\noutside 1 331\nTMhelix 332 354\ninside 355 372

19510 GCF_001246655.1_Salmonella_enterica_CVM_N51274_v1.0 Salmonella enterica subsp. enterica serovar Senftenberg Proteobacteria; Gammaproteobacteria; Enterobacterales; Enterobacteriaceae; Salmonella; Salmonella enterica; Salmonella enterica subsp. enterica MNNEETFYQAMRRKGVTRRSFLKFCSLAATSLGLGAGMTPKIAWALENKPRIPVVWIHGL WP_001058312.1 hydrogenase [Salmonella enterica] Length: 372\nNumber of predicted TMHs: 2\nExp number of AAs in TMHs: 36.51738\nExp number, first 60 AAs: 19.72849\nTotal prob of N-in: 0.96170\nPOSSIBLE N-term signal sequence\ninside 1 19\nTMhelix 20 39\noutside 40 325\nTMhelix 326 348\ninside 349 372

19511 GCF_002031375.1_ASM203137v1 Salmonella enterica subsp. enterica serovar Senftenberg Proteobacteria; Gammaproteobacteria; Enterobacterales; Enterobacteriaceae; Salmonella; Salmonella enterica; Salmonella enterica subsp. enterica MNNEETFYQAMRRKGVTRRSFLKFCSLAATSLGLGAGMTPKIAWALENKPRIPVVWIHGL WP_001058312.1 hydrogenase [Salmonella enterica] Length: 372\nNumber of predicted TMHs: 2\nExp number of AAs in TMHs: 36.51738\nExp number, first 60 AAs: 19.72849\nTotal prob of N-in: 0.96170\nPOSSIBLE N-term signal sequence\ninside 1 19\nTMhelix 20 39\noutside 40 325\nTMhelix 326 348\ninside 349 372

19512 GCF_001479525.1_Salmonella_enterica_CVM_N32779-R_v1.0 Salmonella enterica subsp. enterica serovar Senftenberg Proteobacteria; Gammaproteobacteria; Enterobacterales; Enterobacteriaceae; Salmonella; Salmonella enterica; Salmonella enterica subsp. enterica MNNEETFYQAMRRKGVTRRSFLKFCSLAATSLGLGAGMTPKIAWALENKPRIPVVWIHGL WP_001058312.1 hydrogenase [Salmonella enterica] Length: 372\nNumber of predicted TMHs: 2\nExp number of AAs in

TMHs: 36.51738\nExp number, first 60 AAs:  19.72849\nTotal prob of N-in:   0.96170\nPOSSIBLE N-term signal sequence\ninside   1   19\nTMhelix   20   39\noutside   40   325\nTMhelix   326   348\ninside   349   372

19513 GCF_001477795.1_Salmonella_enterica_CVM_N29321-SQ_v1.0 Salmonella enterica subsp. enterica serovar Senftenberg Proteobacteria; Gammaproteobacteria; Enterobacterales; Enterobacteriaceae; Salmonella; Salmonella enterica; Salmonella enterica subsp. enterica MNNEETFYQAMRRKGVTRRSFLKFCSLAATSLGLGAGMTPKIAWALENKPRIPVVWIHGL WP_001058312.1 hydrogenase [Salmonella enterica] Length: 372\nNumber of predicted TMHs:  2\nExp number of AAs in TMHs: 36.51738\nExp number, first 60 AAs:  19.72849\nTotal prob of N-in:   0.96170\nPOSSIBLE N-term signal sequence\ninside   1   19\nTMhelix   20   39\noutside   40   325\nTMhelix   326   348\ninside   349   372

19514 GCF_001478905.1_Salmonella_enterica_CVM_N30663-SQ_v1.0 Salmonella enterica subsp. enterica serovar Senftenberg Proteobacteria; Gammaproteobacteria; Enterobacterales; Enterobacteriaceae; Salmonella; Salmonella enterica; Salmonella enterica subsp. enterica MTGDNTLITSHGINRRDFMKLCAALAATMGLSSKAAAEMAESVSNPQRPPVIWIGAQECT WP_000145429.1 hydrogenase 2 small subunit [Salmonella enterica] Length: 372\nNumber of predicted TMHs:  1\nExp number of AAs in TMHs: 24.51309\nExp number, first 60 AAs:  0.2682\nTotal prob of N-in:   0.14402\noutside   1   331\nTMhelix   332   354\ninside   355   372

19515 GCF_001240395.1_Salmonella_enterica_CVM_N43462_v1.0 Salmonella enterica subsp. enterica serovar Senftenberg Proteobacteria; Gammaproteobacteria; Enterobacterales; Enterobacteriaceae; Salmonella; Salmonella enterica; Salmonella enterica subsp. enterica MTGDNTLITSHGINRRDFMKLCAALAATMGLSSKAAAEMAESVSNPQRPPVIWIGAQECT WP_000145429.1 hydrogenase 2 small subunit [Salmonella enterica] Length: 372\nNumber of predicted TMHs:  1\nExp number of AAs in TMHs: 24.51309\nExp number, first 60 AAs:  0.2682\nTotal prob of N-in:   0.14402\noutside   1   331\nTMhelix   332   354\ninside   355   372

19516 GCF_001478905.1_Salmonella_enterica_CVM_N30663-SQ_v1.0 Salmonella enterica subsp. enterica serovar Senftenberg Proteobacteria; Gammaproteobacteria; Enterobacterales; Enterobacteriaceae; Salmonella; Salmonella enterica; Salmonella enterica subsp. enterica MNNEETFYQAMRRKGVTRRSFLKFCSLAATSLGLGAGMTPKIAWALENKPRIPVVWIHGL WP_001058312.1 hydrogenase [Salmonella enterica] Length: 372\nNumber of predicted TMHs:  2\nExp number of AAs in TMHs: 36.51738\nExp number, first 60 AAs:  19.72849\nTotal prob of N-in:   0.96170\nPOSSIBLE N-term signal sequence\ninside   1   19\nTMhelix   20   39\noutside   40   325\nTMhelix   326   348\ninside   349   372

19517 GCF_001276775.1_ASM127677v1 Salmonella enterica subsp. enterica serovar Senftenberg Proteobacteria; Gammaproteobacteria; Enterobacterales; Enterobacteriaceae; Salmonella; Salmonella enterica; Salmonella enterica subsp. enterica MNNEETFYQAMRRKGVTRRSFLKFCSLAATSLGLGAGMTPKIAWALENKPRIPVVWIHGL WP_001058312.1 hydrogenase [Salmonella enterica] Length: 372\nNumber of predicted TMHs:  2\nExp number of AAs in TMHs: 36.51738\nExp number, first 60 AAs:  19.72849\nTotal prob of N-in:   0.96170\nPOSSIBLE N-term signal sequence\ninside   1   19\nTMhelix   20   39\noutside   40   325\nTMhelix   326   348\ninside   349   372

19518 GCF_001276745.1_ASM127674v1 Salmonella enterica subsp. enterica serovar Senftenberg Proteobacteria; Gammaproteobacteria; Enterobacterales; Enterobacteriaceae; Salmonella; Salmonella enterica; Salmonella enterica subsp. enterica MNNEETFYQAMRRKGVTRRSFLKFCSLAATSLGLGAGMTPKIAWALENKPRIPVVWIHGL WP_001058312.1 hydrogenase [Salmonella enterica] Length: 372\nNumber of predicted TMHs:  2\nExp number of AAs in TMHs: 36.51738\nExp number, first 60 AAs:  19.72849\nTotal prob of N-in:   0.96170\nPOSSIBLE N-term signal sequence\ninside   1   19\nTMhelix   20   39\noutside   40   325\nTMhelix   326   348\ninside   349   372

19519 GCF_002030925.1_ASM203092v1 Salmonella enterica subsp. enterica serovar Senftenberg Proteobacteria; Gammaproteobacteria; Enterobacterales; Enterobacteriaceae; Salmonella; Salmonella enterica; Salmonella enterica subsp. enterica MTGDNTLITSHGINRRDFMKLCAALAATMGLSSKAAAEMAESVSNPQRPPVIWIGAQECT WP_000145429.1 hydrogenase 2 small subunit [Salmonella enterica] Length: 372\nNumber of predicted TMHs:  1\nExp number of AAs in TMHs: 24.51309\nExp number, first 60 AAs:  0.2682\nTotal prob of N-in:   0.14402\noutside   1   331\nTMhelix   332   354\ninside   355   372

19520 GCF_001240395.1_Salmonella_enterica_CVM_N43462_v1.0 Salmonella enterica subsp. enterica serovar Senftenberg Proteobacteria; Gammaproteobacteria; Enterobacterales; Enterobacteriaceae; Salmonella; Salmonella enterica; Salmonella enterica subsp. enterica MNNEETFYQAMRRKGVTRRSFLKFCSLAATSLGLGAGMTPKIAWALENKPRIPVVWIHGL WP_001058312.1 hydrogenase [Salmonella enterica] Length: 372\nNumber of predicted TMHs:  2\nExp number of AAs in

TMHs: 36.51738\nExp number, first 60 AAs: 19.72849\nTotal prob of N-in: 0.96170\nPOSSIBLE N-term signal sequence\ninside 1 19\nTMhelix 20 39\noutside 40 325\nTMhelix 326 348\ninside 349 372

19521 GCF_001969425.1_ASM196942v1 Salmonella enterica subsp. enterica serovar Senftenberg Proteobacteria; Gammaproteobacteria; Enterobacterales; Enterobacteriaceae; Salmonella; Salmonella enterica; Salmonella enterica subsp. enterica MNNEETFYQAMRRKGVTRRSFLKFCSLAATSLGLGAGMTPKIAWALENKPRIPVVWIHGL WP_001058311.1 hydrogenase [Salmonella enterica] Length: 372\nNumber of predicted TMHs: 2\nExp number of AAs in TMHs: 37.4699899999999\nExp number, first 60 AAs: 19.72699\nTotal prob of N-in: 0.96171\nPOSSIBLE N-term signal sequence\ninside 1 19\nTMhelix 20 39\noutside 40 325\nTMhelix 326 348\ninside 349 372

19522 GCF_002031755.1_ASM203175v1 Salmonella enterica subsp. enterica serovar Senftenberg Proteobacteria; Gammaproteobacteria; Enterobacterales; Enterobacteriaceae; Salmonella; Salmonella enterica; Salmonella enterica subsp. enterica MTGDNTLITSHGINRRDFMKLCAALAATMGLSSKAAAEMAESVSNPQRPPVIWIGAQECT WP_000145429.1 hydrogenase 2 small subunit [Salmonella enterica] Length: 372\nNumber of predicted TMHs: 1\nExp number of AAs in TMHs: 24.51309\nExp number, first 60 AAs: 0.2682\nTotal prob of N-in: 0.14402\noutside 1 331\nTMhelix 332 354\ninside 355 372

19523 GCF_002062125.1_ASM206212v1 Salmonella enterica subsp. enterica serovar Senftenberg Proteobacteria; Gammaproteobacteria; Enterobacterales; Enterobacteriaceae; Salmonella; Salmonella enterica; Salmonella enterica subsp. enterica MTGDNTLITSHGINRRDFMKLCAALAATMGLSSKAAAEMAESVSNPQRPPVIWIGAQECT WP_000145429.1 hydrogenase 2 small subunit [Salmonella enterica] Length: 372\nNumber of predicted TMHs: 1\nExp number of AAs in TMHs: 24.51309\nExp number, first 60 AAs: 0.2682\nTotal prob of N-in: 0.14402\noutside 1 331\nTMhelix 332 354\ninside 355 372

19524 GCF_002062125.1_ASM206212v1 Salmonella enterica subsp. enterica serovar Senftenberg Proteobacteria; Gammaproteobacteria; Enterobacterales; Enterobacteriaceae; Salmonella; Salmonella enterica; Salmonella enterica subsp. enterica MNNEETFYQAMRRKGVTRRSFLKFCSLAATSLGLGAGMTPKIAWALENKPRIPVVWIHGL WP_001058312.1 hydrogenase [Salmonella enterica] Length: 372\nNumber of predicted TMHs: 2\nExp number of AAs in TMHs: 36.51738\nExp number, first 60 AAs: 19.72849\nTotal prob of N-in: 0.96170\nPOSSIBLE N-term signal sequence\ninside 1 19\nTMhelix 20 39\noutside 40 325\nTMhelix 326 348\ninside 349 372

19525 GCF_002062575.1_ASM206257v1 Salmonella enterica subsp. enterica serovar Singapore Proteobacteria; Gammaproteobacteria; Enterobacterales; Enterobacteriaceae; Salmonella; Salmonella enterica; Salmonella enterica subsp. enterica MNNEETFYQAMRRKGVTRRSFLKFCSLAATSLGLGAGMTPKIAWALENKPRIPVVWIHGL WP_001058312.1 hydrogenase [Salmonella enterica] Length: 372\nNumber of predicted TMHs: 2\nExp number of AAs in TMHs: 36.51738\nExp number, first 60 AAs: 19.72849\nTotal prob of N-in: 0.96170\nPOSSIBLE N-term signal sequence\ninside 1 19\nTMhelix 20 39\noutside 40 325\nTMhelix 326 348\ninside 349 372

19526 GCF_002062575.1_ASM206257v1 Salmonella enterica subsp. enterica serovar Singapore Proteobacteria; Gammaproteobacteria; Enterobacterales; Enterobacteriaceae; Salmonella; Salmonella enterica; Salmonella enterica subsp. enterica MTGDNTLITSHGINRRDFMKLCAALAATMGLSSKAAAEMAESVSNPQRPPVIWIGAQECT WP_000145429.1 hydrogenase 2 small subunit [Salmonella enterica] Length: 372\nNumber of predicted TMHs: 1\nExp number of AAs in TMHs: 24.51309\nExp number, first 60 AAs: 0.2682\nTotal prob of N-in: 0.14402\noutside 1 331\nTMhelix 332 354\ninside 355 372

19527 GCF_002063875.1_ASM206387v1 Salmonella enterica subsp. enterica serovar Singapore Proteobacteria; Gammaproteobacteria; Enterobacterales; Enterobacteriaceae; Salmonella; Salmonella enterica; Salmonella enterica subsp. enterica MTGDNTLITSHGINRRDFMKLCAALAATMGLSSKAAAEMAESVSNPQRPPVIWIGAQECT WP_000145429.1 hydrogenase 2 small subunit [Salmonella enterica] Length: 372\nNumber of predicted TMHs: 1\nExp number of AAs in TMHs: 24.51309\nExp number, first 60 AAs: 0.2682\nTotal prob of N-in: 0.14402\noutside 1 331\nTMhelix 332 354\ninside 355 372

19528 GCF_002063805.1_ASM206380v1 Salmonella enterica subsp. enterica serovar Stanley Proteobacteria; Gammaproteobacteria; Enterobacterales; Enterobacteriaceae; Salmonella; Salmonella enterica; Salmonella enterica subsp. enterica MNNEETFYQAMRRKGVTRRSFLKFCSLAATSLGLGAGMTPKIAWALENKPRIPVVWIHGL WP_001058310.1 hydrogenase [Salmonella enterica] Length: 372\nNumber of predicted TMHs: 2\nExp number of AAs in TMHs: 37.4567299999999\nExp number, first 60 AAs: 19.72694\nTotal prob of N-in: 0.96171\nPOSSIBLE N-term signal sequence\ninside 1 19\nTMhelix 20 39\noutside 40 325\nTMhelix 326 348\ninside 349 372

19529	GCF_002063805.1_ASM206380v1	Salmonella enterica subsp. enterica serovar Stanley
	Proteobacteria; Gammaproteobacteria; Enterobacterales; Enterobacteriaceae; Salmonella; Salmonella enterica; Salmonella enterica subsp. enterica	MTGDNTLITSHGINRRDFMKLCAALAATMGLSSKAAAEMAESVSNPQRPPVIWIGAQECT
	WP_000145429.1	hydrogenase 2 small subunit [Salmonella enterica]	Length: 372\nNumber of predicted TMHs:  1\nExp number of AAs in TMHs: 24.51309\nExp number, first 60 AAs:  0.2682\nTotal prob of N-in: 0.14402\noutside   1   331\nTMhelix   332   354\ninside   355   372

19530	GCF_002063845.1_ASM206384v1	Salmonella enterica subsp. enterica serovar Stanley
	Proteobacteria; Gammaproteobacteria; Enterobacterales; Enterobacteriaceae; Salmonella; Salmonella enterica; Salmonella enterica subsp. enterica	MTGDNTLITSHGINRRDFMKLCAALAATMGLSSKAAAEMAESVSNPQRPPVIWIGAQECT
	WP_000145429.1	hydrogenase 2 small subunit [Salmonella enterica]	Length: 372\nNumber of predicted TMHs:  1\nExp number of AAs in TMHs: 24.51309\nExp number, first 60 AAs:  0.2682\nTotal prob of N-in: 0.14402\noutside   1   331\nTMhelix   332   354\ninside   355   372

19531	GCF_002063845.1_ASM206384v1	Salmonella enterica subsp. enterica serovar Stanley
	Proteobacteria; Gammaproteobacteria; Enterobacterales; Enterobacteriaceae; Salmonella; Salmonella enterica; Salmonella enterica subsp. enterica	MNNEETFYQAMRRKGVTRRSFLKFCSLAATSLGLGAGMTPKIAWALENKPRIPVVWIHGL
	WP_001058310.1	hydrogenase [Salmonella enterica]	Length: 372\nNumber of predicted TMHs:  2\nExp number of AAs in TMHs: 37.4567299999999\nExp number, first 60 AAs:  19.72694\nTotal prob of N-in: 0.96171\nPOSSIBLE N-term signal sequence\ninside   1   19\nTMhelix   20   39\noutside   40   325\nTMhelix   326   348\ninside   349   372

19532	GCF_002065125.1_ASM206512v1	Salmonella enterica subsp. enterica serovar Stanley
	Proteobacteria; Gammaproteobacteria; Enterobacterales; Enterobacteriaceae; Salmonella; Salmonella enterica; Salmonella enterica subsp. enterica	MTGDNTLITSHGINRRDFMKLCAALAATMGLSSKAAAEMAESVSNPQRPPVIWIGAQECT
	WP_000145429.1	hydrogenase 2 small subunit [Salmonella enterica]	Length: 372\nNumber of predicted TMHs:  1\nExp number of AAs in TMHs: 24.51309\nExp number, first 60 AAs:  0.2682\nTotal prob of N-in: 0.14402\noutside   1   331\nTMhelix   332   354\ninside   355   372

19533	GCF_002065125.1_ASM206512v1	Salmonella enterica subsp. enterica serovar Stanley
	Proteobacteria; Gammaproteobacteria; Enterobacterales; Enterobacteriaceae; Salmonella; Salmonella enterica; Salmonella enterica subsp. enterica	MNNEETFYQAMRRKGVTRRSFLKFCSLAATSLGLGAGMTPKIAWALENKPRIPVVWIHGL
	WP_001058310.1	hydrogenase [Salmonella enterica]	Length: 372\nNumber of predicted TMHs:  2\nExp number of AAs in TMHs: 37.4567299999999\nExp number, first 60 AAs:  19.72694\nTotal prob of N-in: 0.96171\nPOSSIBLE N-term signal sequence\ninside   1   19\nTMhelix   20   39\noutside   40   325\nTMhelix   326   348\ninside   349   372

19534	GCF_002033845.1_ASM203384v1	Salmonella enterica subsp. enterica serovar Stanley
	Proteobacteria; Gammaproteobacteria; Enterobacterales; Enterobacteriaceae; Salmonella; Salmonella enterica; Salmonella enterica subsp. enterica	MNNEETFYQAMRRKGVTRRSFLKFCSLAATSLGLGAGMTPKIAWALENKPRIPVVWIHGL
	WP_001058310.1	hydrogenase [Salmonella enterica]	Length: 372\nNumber of predicted TMHs:  2\nExp number of AAs in TMHs: 37.4567299999999\nExp number, first 60 AAs:  19.72694\nTotal prob of N-in: 0.96171\nPOSSIBLE N-term signal sequence\ninside   1   19\nTMhelix   20   39\noutside   40   325\nTMhelix   326   348\ninside   349   372

19535	GCF_002033845.1_ASM203384v1	Salmonella enterica subsp. enterica serovar Stanley
	Proteobacteria; Gammaproteobacteria; Enterobacterales; Enterobacteriaceae; Salmonella; Salmonella enterica; Salmonella enterica subsp. enterica	MTGDNTLITSHGINRRDFMKLCAALAATMGLSSKAAAEMAESVSNPQRPPVIWIGAQECT
	WP_000145429.1	hydrogenase 2 small subunit [Salmonella enterica]	Length: 372\nNumber of predicted TMHs:  1\nExp number of AAs in TMHs: 24.51309\nExp number, first 60 AAs:  0.2682\nTotal prob of N-in: 0.14402\noutside   1   331\nTMhelix   332   354\ninside   355   372

19536	GCF_002063835.1_ASM206383v1	Salmonella enterica subsp. enterica serovar Tarshyne
	Proteobacteria; Gammaproteobacteria; Enterobacterales; Enterobacteriaceae; Salmonella; Salmonella enterica; Salmonella enterica subsp. enterica	MNNEETFYQAMRRKGVTRRSFLKFCSLAATSLGLGAGMTPKIAWALENKPRIPVVWIHGL
	WP_001058312.1	hydrogenase [Salmonella enterica]	Length: 372\nNumber of predicted TMHs:  2\nExp number of AAs in TMHs: 36.51738\nExp number, first 60 AAs:  19.72849\nTotal prob of N-in:   0.96170\nPOSSIBLE N-term signal sequence\ninside   1   19\nTMhelix   20   39\noutside   40   325\nTMhelix   326   348\ninside   349   372

19537	GCF_002063835.1_ASM206383v1	Salmonella enterica subsp. enterica serovar Tarshyne
	Proteobacteria; Gammaproteobacteria; Enterobacterales; Enterobacteriaceae; Salmonella; Salmonella enterica;

Salmonella enterica subsp. enterica MTGDNTLITSHGINRRDFMKLCAALAATMGLSSKAAAEMAESVSNPQRPPVIWIGAQECT WP_000145429.1 hydrogenase 2 small subunit [Salmonella enterica] Length: 372\nNumber of predicted TMHs: 1\nExp number of AAs in TMHs: 24.51309\nExp number, first 60 AAs: 0.2682\nTotal prob of N-in: 0.14402\noutside 1 331\nTMhelix 332 354\ninside 355 372

19538 GCF_002063835.1_ASM206383v1 Salmonella enterica subsp. enterica serovar Tarshyne Proteobacteria; Gammaproteobacteria; Enterobacterales; Enterobacteriaceae; Salmonella; Salmonella enterica; Salmonella enterica subsp. enterica MLRFRTIFLKKIFTLKNSILARRGAVIVIVSAVFTSIMFFAHSWASDKEVAMTLSVLNSN WP_080151465.1 hypothetical protein [Salmonella enterica] Length: 204\nNumber of predicted TMHs: 2\nExp number of AAs in TMHs: 44.99538\nExp number, first 60 AAs: 22.34317\nTotal prob of N-in: 0.89744\nPOSSIBLE N-term signal sequence\ninside 1 19\nTMhelix 20 42\noutside 43 171\nTMhelix 172 194\ninside 195 204

19539 GCF_002047275.1_ASM204727v1 Salmonella enterica subsp. enterica serovar Telelkebir Proteobacteria; Gammaproteobacteria; Enterobacterales; Enterobacteriaceae; Salmonella; Salmonella enterica; Salmonella enterica subsp. enterica MTGDNTLITSHGINRRDFMKLCAALAATMGLSSKAAAEMAESVSNPQRPPVIWIGAQECT WP_000145429.1 hydrogenase 2 small subunit [Salmonella enterica] Length: 372\nNumber of predicted TMHs: 1\nExp number of AAs in TMHs: 24.51309\nExp number, first 60 AAs: 0.2682\nTotal prob of N-in: 0.14402\noutside 1 331\nTMhelix 332 354\ninside 355 372

19540 GCF_002048465.1_ASM204846v1 Salmonella enterica subsp. enterica serovar Telelkebir Proteobacteria; Gammaproteobacteria; Enterobacterales; Enterobacteriaceae; Salmonella; Salmonella enterica; Salmonella enterica subsp. enterica MTGDNTLITSHGINRRDFMKLCAALAATMGLSSKAAAEMAESVSNPQRPPVIWIGAQECT WP_000145429.1 hydrogenase 2 small subunit [Salmonella enterica] Length: 372\nNumber of predicted TMHs: 1\nExp number of AAs in TMHs: 24.51309\nExp number, first 60 AAs: 0.2682\nTotal prob of N-in: 0.14402\noutside 1 331\nTMhelix 332 354\ninside 355 372

19541 GCF_002047435.1_ASM204743v1 Salmonella enterica subsp. enterica serovar Telelkebir Proteobacteria; Gammaproteobacteria; Enterobacterales; Enterobacteriaceae; Salmonella; Salmonella enterica; Salmonella enterica subsp. enterica MTGDNTLITSHGINRRDFMKLCAALAATMGLSSKAAAEMAESVSNPQRPPVIWIGAQECT WP_000145429.1 hydrogenase 2 small subunit [Salmonella enterica] Length: 372\nNumber of predicted TMHs: 1\nExp number of AAs in TMHs: 24.51309\nExp number, first 60 AAs: 0.2682\nTotal prob of N-in: 0.14402\noutside 1 331\nTMhelix 332 354\ninside 355 372

19542 GCF_002047205.1_ASM204720v1 Salmonella enterica subsp. enterica serovar Telelkebir Proteobacteria; Gammaproteobacteria; Enterobacterales; Enterobacteriaceae; Salmonella; Salmonella enterica; Salmonella enterica subsp. enterica MTGDNTLITSHGINRRDFMKLCAALAATMGLSSKAAAEMAESVSNPQRPPVIWIGAQECT WP_000145429.1 hydrogenase 2 small subunit [Salmonella enterica] Length: 372\nNumber of predicted TMHs: 1\nExp number of AAs in TMHs: 24.51309\nExp number, first 60 AAs: 0.2682\nTotal prob of N-in: 0.14402\noutside 1 331\nTMhelix 332 354\ninside 355 372

19543 GCF_002048425.1_ASM204842v1 Salmonella enterica subsp. enterica serovar Telelkebir Proteobacteria; Gammaproteobacteria; Enterobacterales; Enterobacteriaceae; Salmonella; Salmonella enterica; Salmonella enterica subsp. enterica MTGDNTLITSHGINRRDFMKLCAALAATMGLSSKAAAEMAESVSNPQRPPVIWIGAQECT WP_000145429.1 hydrogenase 2 small subunit [Salmonella enterica] Length: 372\nNumber of predicted TMHs: 1\nExp number of AAs in TMHs: 24.51309\nExp number, first 60 AAs: 0.2682\nTotal prob of N-in: 0.14402\noutside 1 331\nTMhelix 332 354\ninside 355 372

19544 GCF_002047365.1_ASM204736v1 Salmonella enterica subsp. enterica serovar Telelkebir Proteobacteria; Gammaproteobacteria; Enterobacterales; Enterobacteriaceae; Salmonella; Salmonella enterica; Salmonella enterica subsp. enterica MTGDNTLITSHGINRRDFMKLCAALAATMGLSSKAAAEMAESVSNPQRPPVIWIGAQECT WP_000145429.1 hydrogenase 2 small subunit [Salmonella enterica] Length: 372\nNumber of predicted TMHs: 1\nExp number of AAs in TMHs: 24.51309\nExp number, first 60 AAs: 0.2682\nTotal prob of N-in: 0.14402\noutside 1 331\nTMhelix 332 354\ninside 355 372

19545 GCF_001620665.1_ASM162066v1 Salmonella enterica subsp. enterica serovar Tennessee Proteobacteria; Gammaproteobacteria; Enterobacterales; Enterobacteriaceae; Salmonella; Salmonella enterica; Salmonella enterica subsp. enterica MNNEETFYQAMRRKGVTRRSFLKFCSLAATSLGLGAGMTPKIAWALENKPRIPVVWIHGL WP_001058312.1 hydrogenase [Salmonella enterica] Length: 372\nNumber of predicted TMHs: 2\nExp number of AAs in TMHs: 36.51738\nExp number, first 60 AAs: 19.72849\nTotal prob of N-in: 0.96170\nPOSSIBLE N-term signal sequence\ninside 1 19\nTMhelix 20 39\noutside 40 325\nTMhelix 326 348\ninside 349 372

19546 GCF_001620605.1_ASM162060v1 Salmonella enterica subsp. enterica serovar Tennessee Proteobacteria; Gammaproteobacteria; Enterobacterales; Enterobacteriaceae; Salmonella; Salmonella enterica; Salmonella enterica subsp. enterica MTGDNTLITSHGINRRDFMKLCAALAATMGLSSKAAAEMAESVSNPQRPPVIWIGAQECT WP_000145429.1 hydrogenase 2 small subunit [Salmonella enterica] Length: 372\nNumber of predicted TMHs: 1\nExp number of AAs in TMHs: 24.51309\nExp number, first 60 AAs: 0.2682\nTotal prob of N-in: 0.14402\noutside 1 331\nTMhelix 332 354\ninside 355 372

19547 GCF_001620495.1_ASM162049v1 Salmonella enterica subsp. enterica serovar Tennessee Proteobacteria; Gammaproteobacteria; Enterobacterales; Enterobacteriaceae; Salmonella; Salmonella enterica; Salmonella enterica subsp. enterica MNNEETFYQAMRRKGVTRRSFLKFCSLAATSLGLGAGMTPKIAWALENKPRIPVVWIHGL WP_001058312.1 hydrogenase [Salmonella enterica] Length: 372\nNumber of predicted TMHs: 2\nExp number of AAs in TMHs: 36.51738\nExp number, first 60 AAs: 19.72849\nTotal prob of N-in: 0.96170\nPOSSIBLE N-term signal sequence\ninside 1 19\nTMhelix 20 39\noutside 40 325\nTMhelix 326 348\ninside 349 372

19548 GCF_001620665.1_ASM162066v1 Salmonella enterica subsp. enterica serovar Tennessee Proteobacteria; Gammaproteobacteria; Enterobacterales; Enterobacteriaceae; Salmonella; Salmonella enterica; Salmonella enterica subsp. enterica MTGDNTLITSHGINRRDFMKLCAALAATMGLSSKAAAEMAESVSNPQRPPVIWIGAQECT WP_000145429.1 hydrogenase 2 small subunit [Salmonella enterica] Length: 372\nNumber of predicted TMHs: 1\nExp number of AAs in TMHs: 24.51309\nExp number, first 60 AAs: 0.2682\nTotal prob of N-in: 0.14402\noutside 1 331\nTMhelix 332 354\ninside 355 372

19549 GCF_001974765.1_ASM197476v1 Salmonella enterica subsp. enterica serovar Tennessee Proteobacteria; Gammaproteobacteria; Enterobacterales; Enterobacteriaceae; Salmonella; Salmonella enterica; Salmonella enterica subsp. enterica MNNEETFYQAMRRKGVTRRSFLKFCSLAATSLGLGAGMTPKIAWALENKPRIPVVWIHGL WP_001058312.1 hydrogenase [Salmonella enterica] Length: 372\nNumber of predicted TMHs: 2\nExp number of AAs in TMHs: 36.51738\nExp number, first 60 AAs: 19.72849\nTotal prob of N-in: 0.96170\nPOSSIBLE N-term signal sequence\ninside 1 19\nTMhelix 20 39\noutside 40 325\nTMhelix 326 348\ninside 349 372

19550 GCF_001620585.1_ASM162058v1 Salmonella enterica subsp. enterica serovar Tennessee Proteobacteria; Gammaproteobacteria; Enterobacterales; Enterobacteriaceae; Salmonella; Salmonella enterica; Salmonella enterica subsp. enterica MTGDNTLITSHGINRRDFMKLCAALAATMGLSSKAAAEMAESVSNPQRPPVIWIGAQECT WP_000145429.1 hydrogenase 2 small subunit [Salmonella enterica] Length: 372\nNumber of predicted TMHs: 1\nExp number of AAs in TMHs: 24.51309\nExp number, first 60 AAs: 0.2682\nTotal prob of N-in: 0.14402\noutside 1 331\nTMhelix 332 354\ninside 355 372

19551 GCF_001630295.1_ASM163029v1 Salmonella enterica subsp. enterica serovar Tennessee Proteobacteria; Gammaproteobacteria; Enterobacterales; Enterobacteriaceae; Salmonella; Salmonella enterica; Salmonella enterica subsp. enterica MTGDNTLITSHGINRRDFMKLCAALAATMGLSSKAAAEMAESVSNPQRPPVIWIGAQECT WP_000145429.1 hydrogenase 2 small subunit [Salmonella enterica] Length: 372\nNumber of predicted TMHs: 1\nExp number of AAs in TMHs: 24.51309\nExp number, first 60 AAs: 0.2682\nTotal prob of N-in: 0.14402\noutside 1 331\nTMhelix 332 354\ninside 355 372

19552 GCF_001620605.1_ASM162060v1 Salmonella enterica subsp. enterica serovar Tennessee Proteobacteria; Gammaproteobacteria; Enterobacterales; Enterobacteriaceae; Salmonella; Salmonella enterica; Salmonella enterica subsp. enterica MNNEETFYQAMRRKGVTRRSFLKFCSLAATSLGLGAGMTPKIAWALENKPRIPVVWIHGL WP_001058312.1 hydrogenase [Salmonella enterica] Length: 372\nNumber of predicted TMHs: 2\nExp number of AAs in TMHs: 36.51738\nExp number, first 60 AAs: 19.72849\nTotal prob of N-in: 0.96170\nPOSSIBLE N-term signal sequence\ninside 1 19\nTMhelix 20 39\noutside 40 325\nTMhelix 326 348\ninside 349 372

19553 GCF_001630135.1_ASM163013v1 Salmonella enterica subsp. enterica serovar Tennessee Proteobacteria; Gammaproteobacteria; Enterobacterales; Enterobacteriaceae; Salmonella; Salmonella enterica; Salmonella enterica subsp. enterica MTGDNTLITSHGINRRDFMKLCAALAATMGLSSKAAAEMAESVSNPQRPPVIWIGAQECT WP_000145429.1 hydrogenase 2 small subunit [Salmonella enterica] Length: 372\nNumber of predicted TMHs: 1\nExp number of AAs in TMHs: 24.51309\nExp number, first 60 AAs: 0.2682\nTotal prob of N-in: 0.14402\noutside 1 331\nTMhelix 332 354\ninside 355 372

19554 GCF_001630135.1_ASM163013v1 Salmonella enterica subsp. enterica serovar Tennessee Proteobacteria; Gammaproteobacteria; Enterobacterales; Enterobacteriaceae; Salmonella; Salmonella enterica; Salmonella enterica subsp. enterica MNNEETFYQAMRRKGVTRRSFLKFCSLAATSLGLGAGMTPKIAWALENKPRIPVVWIHGL WP_001058312.1 hydrogenase [Salmonella enterica] Length: 372\nNumber of predicted TMHs: 2\nExp

number of AAs in TMHs: 36.51738\nExp number, first 60 AAs: 19.72849\nTotal prob of N-in: 0.96170\nPOSSIBLE N-term signal sequence\ninside 1 19\nTMhelix 20 39\noutside 40 325\nTMhelix 326 348\ninside 349 372

19555 GCF_001630235.1_ASM163023v1 Salmonella enterica subsp. enterica serovar Tennessee Proteobacteria; Gammaproteobacteria; Enterobacterales; Enterobacteriaceae; Salmonella; Salmonella enterica; Salmonella enterica subsp. enterica MTGDNTLITSHGINRRDFMKLCAALAATMGLSSKAAAEMAESVSNPQRPPVIWIGAQECT WP_000145429.1 hydrogenase 2 small subunit [Salmonella enterica] Length: 372\nNumber of predicted TMHs: 1\nExp number of AAs in TMHs: 24.51309\nExp number, first 60 AAs: 0.2682\nTotal prob of N-in: 0.14402\noutside 1 331\nTMhelix 332 354\ninside 355 372

19556 GCF_001620505.1_ASM162050v1 Salmonella enterica subsp. enterica serovar Tennessee Proteobacteria; Gammaproteobacteria; Enterobacterales; Enterobacteriaceae; Salmonella; Salmonella enterica; Salmonella enterica subsp. enterica MTGDNTLITSHGINRRDFMKLCAALAATMGLSSKAAAEMAESVSNPQRPPVIWIGAQECT WP_000145429.1 hydrogenase 2 small subunit [Salmonella enterica] Length: 372\nNumber of predicted TMHs: 1\nExp number of AAs in TMHs: 24.51309\nExp number, first 60 AAs: 0.2682\nTotal prob of N-in: 0.14402\noutside 1 331\nTMhelix 332 354\ninside 355 372

19557 GCF_001620565.1_ASM162056v1 Salmonella enterica subsp. enterica serovar Tennessee Proteobacteria; Gammaproteobacteria; Enterobacterales; Enterobacteriaceae; Salmonella; Salmonella enterica; Salmonella enterica subsp. enterica MTGDNTLITSHGINRRDFMKLCAALAATMGLSSKAAAEMAESVSNPQRPPVIWIGAQECT WP_000145429.1 hydrogenase 2 small subunit [Salmonella enterica] Length: 372\nNumber of predicted TMHs: 1\nExp number of AAs in TMHs: 24.51309\nExp number, first 60 AAs: 0.2682\nTotal prob of N-in: 0.14402\noutside 1 331\nTMhelix 332 354\ninside 355 372

19558 GCF_001620425.1_ASM162042v1 Salmonella enterica subsp. enterica serovar Tennessee Proteobacteria; Gammaproteobacteria; Enterobacterales; Enterobacteriaceae; Salmonella; Salmonella enterica; Salmonella enterica subsp. enterica MNNEETFYQAMRRKGVTRRSFLKFCSLAATSLGLGAGMTPKIAWALENKPRIPVVWIHGL WP_001058312.1 hydrogenase [Salmonella enterica] Length: 372\nNumber of predicted TMHs: 2\nExp number of AAs in TMHs: 36.51738\nExp number, first 60 AAs: 19.72849\nTotal prob of N-in: 0.96170\nPOSSIBLE N-term signal sequence\ninside 1 19\nTMhelix 20 39\noutside 40 325\nTMhelix 326 348\ninside 349 372

19559 GCF_001620815.1_ASM162081v1 Salmonella enterica subsp. enterica serovar Tennessee Proteobacteria; Gammaproteobacteria; Enterobacterales; Enterobacteriaceae; Salmonella; Salmonella enterica; Salmonella enterica subsp. enterica MTGDNTLITSHGINRRDFMKLCAALAATMGLSSKAAAEMAESVSNPQRPPVIWIGAQECT WP_000145429.1 hydrogenase 2 small subunit [Salmonella enterica] Length: 372\nNumber of predicted TMHs: 1\nExp number of AAs in TMHs: 24.51309\nExp number, first 60 AAs: 0.2682\nTotal prob of N-in: 0.14402\noutside 1 331\nTMhelix 332 354\ninside 355 372

19560 GCF_001620755.2_ASM162075v2 Salmonella enterica subsp. enterica serovar Tennessee Proteobacteria; Gammaproteobacteria; Enterobacterales; Enterobacteriaceae; Salmonella; Salmonella enterica; Salmonella enterica subsp. enterica MNNEETFYQAMRRKGVTRRSFLKFCSLAATSLGLGAGMTPKIAWALENKPRIPVVWIHGL WP_001058312.1 hydrogenase [Salmonella enterica] Length: 372\nNumber of predicted TMHs: 2\nExp number of AAs in TMHs: 36.51738\nExp number, first 60 AAs: 19.72849\nTotal prob of N-in: 0.96170\nPOSSIBLE N-term signal sequence\ninside 1 19\nTMhelix 20 39\noutside 40 325\nTMhelix 326 348\ninside 349 372

19561 GCF_001620405.1_ASM162040v1 Salmonella enterica subsp. enterica serovar Tennessee Proteobacteria; Gammaproteobacteria; Enterobacterales; Enterobacteriaceae; Salmonella; Salmonella enterica; Salmonella enterica subsp. enterica MNNEETFYQAMRRKGVTRRSFLKFCSLAATSLGLGAGMTPKIAWALENKPRIPVVWIHGL WP_001058312.1 hydrogenase [Salmonella enterica] Length: 372\nNumber of predicted TMHs: 2\nExp number of AAs in TMHs: 36.51738\nExp number, first 60 AAs: 19.72849\nTotal prob of N-in: 0.96170\nPOSSIBLE N-term signal sequence\ninside 1 19\nTMhelix 20 39\noutside 40 325\nTMhelix 326 348\ninside 349 372

19562 GCF_001620485.1_ASM162048v1 Salmonella enterica subsp. enterica serovar Tennessee Proteobacteria; Gammaproteobacteria; Enterobacterales; Enterobacteriaceae; Salmonella; Salmonella enterica; Salmonella enterica subsp. enterica MNNEETFYQAMRRKGVTRRSFLKFCSLAATSLGLGAGMTPKIAWALENKPRIPVVWIHGL WP_001058312.1 hydrogenase [Salmonella enterica] Length: 372\nNumber of predicted TMHs: 2\nExp number of AAs in TMHs: 36.51738\nExp number, first 60 AAs: 19.72849\nTotal prob of N-in: 0.96170\nPOSSIBLE N-term signal sequence\ninside 1 19\nTMhelix 20 39\noutside 40 325\nTMhelix 326 348\ninside 349 372

19563 GCF_001535415.1_ASM153541v1 Salmonella enterica subsp. enterica serovar Tennessee Proteobacteria; Gammaproteobacteria; Enterobacterales; Enterobacteriaceae; Salmonella; Salmonella enterica;

Salmonella enterica subsp. enterica	MTGDNTLITSHGINRRDFMKLCAALAATMGLSSKAAAEMAESVSNPQRPPVIWIGAQECT	WP_000145429.1	hydrogenase 2 small subunit [Salmonella enterica]	Length: 372\nNumber of predicted TMHs:  1\nExp number of AAs in TMHs: 24.51309\nExp number, first 60 AAs:  0.2682\nTotal prob of N-in:  0.14402\noutside   1   331\nTMhelix   332   354\ninside   355   372

19564	GCF_001630165.1_ASM163016v1	Salmonella enterica subsp. enterica serovar Tennessee	Proteobacteria; Gammaproteobacteria; Enterobacterales; Enterobacteriaceae; Salmonella; Salmonella enterica; Salmonella enterica subsp. enterica	MNNEETFYQAMRRKGVTRRSFLKFCSLAATSLGLGAGMTPKIAWALENKPRIPVVWIHGL	WP_001058312.1	hydrogenase [Salmonella enterica]	Length: 372\nNumber of predicted TMHs:  2\nExp number of AAs in TMHs: 36.51738\nExp number, first 60 AAs:  19.72849\nTotal prob of N-in:   0.96170\nPOSSIBLE N-term signal sequence\ninside   1   19\nTMhelix   20   39\noutside   40   325\nTMhelix   326   348\ninside   349   372

19565	GCF_001535315.2_ASM153531v2	Salmonella enterica subsp. enterica serovar Tennessee	Proteobacteria; Gammaproteobacteria; Enterobacterales; Enterobacteriaceae; Salmonella; Salmonella enterica; Salmonella enterica subsp. enterica	MNNEETFYQAMRRKGVTRRSFLKFCSLAATSLGLGAGMTPKIAWALENKPRIPVVWIHGL	WP_001058312.1	hydrogenase [Salmonella enterica]	Length: 372\nNumber of predicted TMHs:  2\nExp number of AAs in TMHs: 36.51738\nExp number, first 60 AAs:  19.72849\nTotal prob of N-in:   0.96170\nPOSSIBLE N-term signal sequence\ninside   1   19\nTMhelix   20   39\noutside   40   325\nTMhelix   326   348\ninside   349   372

19566	GCF_001535315.2_ASM153531v2	Salmonella enterica subsp. enterica serovar Tennessee	Proteobacteria; Gammaproteobacteria; Enterobacterales; Enterobacteriaceae; Salmonella; Salmonella enterica; Salmonella enterica subsp. enterica	MTGDNTLITSHGINRRDFMKLCAALAATMGLSSKAAAEMAESVSNPQRPPVIWIGAQECT	WP_000145429.1	hydrogenase 2 small subunit [Salmonella enterica]	Length: 372\nNumber of predicted TMHs:  1\nExp number of AAs in TMHs: 24.51309\nExp number, first 60 AAs:  0.2682\nTotal prob of N-in:  0.14402\noutside   1   331\nTMhelix   332   354\ninside   355   372

19567	GCF_001535085.1_ASM153508v1	Salmonella enterica subsp. enterica serovar Tennessee	Proteobacteria; Gammaproteobacteria; Enterobacterales; Enterobacteriaceae; Salmonella; Salmonella enterica; Salmonella enterica subsp. enterica	MNNEETFYQAMRRKGVTRRSFLKFCSLAATSLGLGAGMTPKIAWALENKPRIPVVWIHGL	WP_001058312.1	hydrogenase [Salmonella enterica]	Length: 372\nNumber of predicted TMHs:  2\nExp number of AAs in TMHs: 36.51738\nExp number, first 60 AAs:  19.72849\nTotal prob of N-in:   0.96170\nPOSSIBLE N-term signal sequence\ninside   1   19\nTMhelix   20   39\noutside   40   325\nTMhelix   326   348\ninside   349   372

19568	GCF_001630315.1_ASM163031v1	Salmonella enterica subsp. enterica serovar Tennessee	Proteobacteria; Gammaproteobacteria; Enterobacterales; Enterobacteriaceae; Salmonella; Salmonella enterica; Salmonella enterica subsp. enterica	MNNEETFYQAMRRKGVTRRSFLKFCSLAATSLGLGAGMTPKIAWALENKPRIPVVWIHGL	WP_001058312.1	hydrogenase [Salmonella enterica]	Length: 372\nNumber of predicted TMHs:  2\nExp number of AAs in TMHs: 36.51738\nExp number, first 60 AAs:  19.72849\nTotal prob of N-in:   0.96170\nPOSSIBLE N-term signal sequence\ninside   1   19\nTMhelix   20   39\noutside   40   325\nTMhelix   326   348\ninside   349   372

19569	GCF_001535455.1_ASM153545v1	Salmonella enterica subsp. enterica serovar Tennessee	Proteobacteria; Gammaproteobacteria; Enterobacterales; Enterobacteriaceae; Salmonella; Salmonella enterica; Salmonella enterica subsp. enterica	MTGDNTLITSHGINRRDFMKLCAALAATMGLSSKAAAEMAESVSNPQRPPVIWIGAQECT	WP_000145429.1	hydrogenase 2 small subunit [Salmonella enterica]	Length: 372\nNumber of predicted TMHs:  1\nExp number of AAs in TMHs: 24.51309\nExp number, first 60 AAs:  0.2682\nTotal prob of N-in:  0.14402\noutside   1   331\nTMhelix   332   354\ninside   355   372

19570	GCF_001620815.1_ASM162081v1	Salmonella enterica subsp. enterica serovar Tennessee	Proteobacteria; Gammaproteobacteria; Enterobacterales; Enterobacteriaceae; Salmonella; Salmonella enterica; Salmonella enterica subsp. enterica	MNNEETFYQAMRRKGVTRRSFLKFCSLAATSLGLGAGMTPKIAWALENKPRIPVVWIHGL	WP_001058312.1	hydrogenase [Salmonella enterica]	Length: 372\nNumber of predicted TMHs:  2\nExp number of AAs in TMHs: 36.51738\nExp number, first 60 AAs:  19.72849\nTotal prob of N-in:   0.96170\nPOSSIBLE N-term signal sequence\ninside   1   19\nTMhelix   20   39\noutside   40   325\nTMhelix   326   348\ninside   349   372

19571	GCF_001276845.1_ASM127684v1	Salmonella enterica subsp. enterica serovar Tennessee	Proteobacteria; Gammaproteobacteria; Enterobacterales; Enterobacteriaceae; Salmonella; Salmonella enterica; Salmonella enterica subsp. enterica	MNNEETFYQAMRRKGVTRRSFLKFCSLAATSLGLGAGMTPKIAWALENKPRIPVVWIHGL	WP_001058312.1	hydrogenase [Salmonella enterica]	Length: 372\nNumber of predicted TMHs:  2\nExp number of AAs in TMHs: 36.51738\nExp number, first 60 AAs:  19.72849\nTotal prob of N-in:   0.96170\nPOSSIBLE N-term signal sequence\ninside   1   19\nTMhelix   20   39\noutside   40   325\nTMhelix   326   348\ninside   349   372

19572 GCF_001620425.1_ASM162042v1 Salmonella enterica subsp. enterica serovar Tennessee Proteobacteria; Gammaproteobacteria; Enterobacterales; Enterobacteriaceae; Salmonella; Salmonella enterica; Salmonella enterica subsp. enterica MTGDNTLITSHGINRRDFMKLCAALAATMGLSSKAAAEMAESVSNPQRPPVIWIGAQECT WP_000145429.1 hydrogenase 2 small subunit [Salmonella enterica] Length: 372\nNumber of predicted TMHs: 1\nExp number of AAs in TMHs: 24.51309\nExp number, first 60 AAs: 0.2682\nTotal prob of N-in: 0.14402\noutside 1 331\nTMhelix 332 354\ninside 355 372

19573 GCF_001630255.1_ASM163025v1 Salmonella enterica subsp. enterica serovar Tennessee Proteobacteria; Gammaproteobacteria; Enterobacterales; Enterobacteriaceae; Salmonella; Salmonella enterica; Salmonella enterica subsp. enterica MTGDNTLITSHGINRRDFMKLCAALAATMGLSSKAAAEMAESVSNPQRPPVIWIGAQECT WP_000145429.1 hydrogenase 2 small subunit [Salmonella enterica] Length: 372\nNumber of predicted TMHs: 1\nExp number of AAs in TMHs: 24.51309\nExp number, first 60 AAs: 0.2682\nTotal prob of N-in: 0.14402\noutside 1 331\nTMhelix 332 354\ninside 355 372

19574 GCF_001620485.1_ASM162048v1 Salmonella enterica subsp. enterica serovar Tennessee Proteobacteria; Gammaproteobacteria; Enterobacterales; Enterobacteriaceae; Salmonella; Salmonella enterica; Salmonella enterica subsp. enterica MTGDNTLITSHGINRRDFMKLCAALAATMGLSSKAAAEMAESVSNPQRPPVIWIGAQECT WP_000145429.1 hydrogenase 2 small subunit [Salmonella enterica] Length: 372\nNumber of predicted TMHs: 1\nExp number of AAs in TMHs: 24.51309\nExp number, first 60 AAs: 0.2682\nTotal prob of N-in: 0.14402\noutside 1 331\nTMhelix 332 354\ninside 355 372

19575 GCF_001620865.1_ASM162086v1 Salmonella enterica subsp. enterica serovar Tennessee Proteobacteria; Gammaproteobacteria; Enterobacterales; Enterobacteriaceae; Salmonella; Salmonella enterica; Salmonella enterica subsp. enterica MTGDNTLITSHGINRRDFMKLCAALAATMGLSSKAAAEMAESVSNPQRPPVIWIGAQECT WP_000145429.1 hydrogenase 2 small subunit [Salmonella enterica] Length: 372\nNumber of predicted TMHs: 1\nExp number of AAs in TMHs: 24.51309\nExp number, first 60 AAs: 0.2682\nTotal prob of N-in: 0.14402\noutside 1 331\nTMhelix 332 354\ninside 355 372

19576 GCF_001535175.1_ASM153517v1 Salmonella enterica subsp. enterica serovar Tennessee Proteobacteria; Gammaproteobacteria; Enterobacterales; Enterobacteriaceae; Salmonella; Salmonella enterica; Salmonella enterica subsp. enterica MTGDNTLITSHGINRRDFMKLCAALAATMGLSSKAAAEMAESVSNPQRPPVIWIGAQECT WP_000145429.1 hydrogenase 2 small subunit [Salmonella enterica] Length: 372\nNumber of predicted TMHs: 1\nExp number of AAs in TMHs: 24.51309\nExp number, first 60 AAs: 0.2682\nTotal prob of N-in: 0.14402\noutside 1 331\nTMhelix 332 354\ninside 355 372

19577 GCF_001535415.1_ASM153541v1 Salmonella enterica subsp. enterica serovar Tennessee Proteobacteria; Gammaproteobacteria; Enterobacterales; Enterobacteriaceae; Salmonella; Salmonella enterica; Salmonella enterica subsp. enterica MNNEETFYQAMRRKGVTRRSFLKFCSLAATSLGLGAGMTPKIAWALENKPRIPVVWIHGL WP_001058312.1 hydrogenase [Salmonella enterica] Length: 372\nNumber of predicted TMHs: 2\nExp number of AAs in TMHs: 36.51738\nExp number, first 60 AAs: 19.72849\nTotal prob of N-in: 0.96170\nPOSSIBLE N-term signal sequence\ninside 1 19\nTMhelix 20 39\noutside 40 325\nTMhelix 326 348\ninside 349 372

19578 GCF_001276675.1_ASM127667v1 Salmonella enterica subsp. enterica serovar Tennessee Proteobacteria; Gammaproteobacteria; Enterobacterales; Enterobacteriaceae; Salmonella; Salmonella enterica; Salmonella enterica subsp. enterica MNNEETFYQAMRRKGVTRRSFLKFCSLAATSLGLGAGMTPKIAWALENKPRIPVVWIHGL WP_001058312.1 hydrogenase [Salmonella enterica] Length: 372\nNumber of predicted TMHs: 2\nExp number of AAs in TMHs: 36.51738\nExp number, first 60 AAs: 19.72849\nTotal prob of N-in: 0.96170\nPOSSIBLE N-term signal sequence\ninside 1 19\nTMhelix 20 39\noutside 40 325\nTMhelix 326 348\ninside 349 372

19579 GCF_001620405.1_ASM162040v1 Salmonella enterica subsp. enterica serovar Tennessee Proteobacteria; Gammaproteobacteria; Enterobacterales; Enterobacteriaceae; Salmonella; Salmonella enterica; Salmonella enterica subsp. enterica MTGDNTLITSHGINRRDFMKLCAALAATMGLSSKAAAEMAESVSNPQRPPVIWIGAQECT WP_000145429.1 hydrogenase 2 small subunit [Salmonella enterica] Length: 372\nNumber of predicted TMHs: 1\nExp number of AAs in TMHs: 24.51309\nExp number, first 60 AAs: 0.2682\nTotal prob of N-in: 0.14402\noutside 1 331\nTMhelix 332 354\ninside 355 372

19580 GCF_001535085.1_ASM153508v1 Salmonella enterica subsp. enterica serovar Tennessee Proteobacteria; Gammaproteobacteria; Enterobacterales; Enterobacteriaceae; Salmonella; Salmonella enterica; Salmonella enterica subsp. enterica MTGDNTLITSHGINRRDFMKLCAALAATMGLSSKAAAEMAESVSNPQRPPVIWIGAQECT WP_000145429.1 hydrogenase 2 small subunit [Salmonella enterica] Length: 372\nNumber of

predicted TMHs:  1\nExp number of AAs in TMHs: 24.51309\nExp number, first 60 AAs:  0.2682\nTotal prob of N-in: 0.14402\noutside   1   331\nTMhelix   332   354\ninside   355   372

19581 GCF_001535255.1_ASM153525v1 Salmonella enterica subsp. enterica serovar Tennessee Proteobacteria; Gammaproteobacteria; Enterobacterales; Enterobacteriaceae; Salmonella; Salmonella enterica; Salmonella enterica subsp. enterica MTGDNTLITSHGINRRDFMKLCAALAATMGLSSKAAAEMAESVSNPQRPPVIWIGAQECT WP_000145429.1 hydrogenase 2 small subunit [Salmonella enterica] Length: 372\nNumber of predicted TMHs:  1\nExp number of AAs in TMHs: 24.51309\nExp number, first 60 AAs:  0.2682\nTotal prob of N-in: 0.14402\noutside   1   331\nTMhelix   332   354\ninside   355   372

19582 GCF_001620785.1_ASM162078v1 Salmonella enterica subsp. enterica serovar Tennessee Proteobacteria; Gammaproteobacteria; Enterobacterales; Enterobacteriaceae; Salmonella; Salmonella enterica; Salmonella enterica subsp. enterica MTGDNTLITSHGINRRDFMKLCAALAATMGLSSKAAAEMAESVSNPQRPPVIWIGAQECT WP_000145429.1 hydrogenase 2 small subunit [Salmonella enterica] Length: 372\nNumber of predicted TMHs:  1\nExp number of AAs in TMHs: 24.51309\nExp number, first 60 AAs:  0.2682\nTotal prob of N-in: 0.14402\noutside   1   331\nTMhelix   332   354\ninside   355   372

19583 GCF_001620645.1_ASM162064v1 Salmonella enterica subsp. enterica serovar Tennessee Proteobacteria; Gammaproteobacteria; Enterobacterales; Enterobacteriaceae; Salmonella; Salmonella enterica; Salmonella enterica subsp. enterica MNNEETFYQAMRRKGVTRRSFLKFCSLAATSLGLGAGMTPKIAWALENKPRIPVVWIHGL WP_001058312.1 hydrogenase [Salmonella enterica] Length: 372\nNumber of predicted TMHs:  2\nExp number of AAs in TMHs: 36.51738\nExp number, first 60 AAs:  19.72849\nTotal prob of N-in:   0.96170\nPOSSIBLE N-term signal sequence\ninside   1   19\nTMhelix   20   39\noutside   40   325\nTMhelix   326   348\ninside   349   372

19584 GCF_001620505.1_ASM162050v1 Salmonella enterica subsp. enterica serovar Tennessee Proteobacteria; Gammaproteobacteria; Enterobacterales; Enterobacteriaceae; Salmonella; Salmonella enterica; Salmonella enterica subsp. enterica MNNEETFYQAMRRKGVTRRSFLKFCSLAATSLGLGAGMTPKIAWALENKPRIPVVWIHGL WP_001058312.1 hydrogenase [Salmonella enterica] Length: 372\nNumber of predicted TMHs:  2\nExp number of AAs in TMHs: 36.51738\nExp number, first 60 AAs:  19.72849\nTotal prob of N-in:   0.96170\nPOSSIBLE N-term signal sequence\ninside   1   19\nTMhelix   20   39\noutside   40   325\nTMhelix   326   348\ninside   349   372

19585 GCF_001620565.1_ASM162056v1 Salmonella enterica subsp. enterica serovar Tennessee Proteobacteria; Gammaproteobacteria; Enterobacterales; Enterobacteriaceae; Salmonella; Salmonella enterica; Salmonella enterica subsp. enterica MNNEETFYQAMRRKGVTRRSFLKFCSLAATSLGLGAGMTPKIAWALENKPRIPVVWIHGL WP_001058312.1 hydrogenase [Salmonella enterica] Length: 372\nNumber of predicted TMHs:  2\nExp number of AAs in TMHs: 36.51738\nExp number, first 60 AAs:  19.72849\nTotal prob of N-in:   0.96170\nPOSSIBLE N-term signal sequence\ninside   1   19\nTMhelix   20   39\noutside   40   325\nTMhelix   326   348\ninside   349   372

19586 GCF_001630225.1_ASM163022v1 Salmonella enterica subsp. enterica serovar Tennessee Proteobacteria; Gammaproteobacteria; Enterobacterales; Enterobacteriaceae; Salmonella; Salmonella enterica; Salmonella enterica subsp. enterica MNNEETFYQAMRRKGVTRRSFLKFCSLAATSLGLGAGMTPKIAWALENKPRIPVVWIHGL WP_001058312.1 hydrogenase [Salmonella enterica] Length: 372\nNumber of predicted TMHs:  2\nExp number of AAs in TMHs: 36.51738\nExp number, first 60 AAs:  19.72849\nTotal prob of N-in:   0.96170\nPOSSIBLE N-term signal sequence\ninside   1   19\nTMhelix   20   39\noutside   40   325\nTMhelix   326   348\ninside   349   372

19587 GCF_001620725.1_ASM162072v1 Salmonella enterica subsp. enterica serovar Tennessee Proteobacteria; Gammaproteobacteria; Enterobacterales; Enterobacteriaceae; Salmonella; Salmonella enterica; Salmonella enterica subsp. enterica MNNEETFYQAMRRKGVTRRSFLKFCSLAATSLGLGAGMTPKIAWALENKPRIPVVWIHGL WP_001058312.1 hydrogenase [Salmonella enterica] Length: 372\nNumber of predicted TMHs:  2\nExp number of AAs in TMHs: 36.51738\nExp number, first 60 AAs:  19.72849\nTotal prob of N-in:   0.96170\nPOSSIBLE N-term signal sequence\ninside   1   19\nTMhelix   20   39\noutside   40   325\nTMhelix   326   348\ninside   349   372

19588 GCF_001620685.1_ASM162068v1 Salmonella enterica subsp. enterica serovar Tennessee Proteobacteria; Gammaproteobacteria; Enterobacterales; Enterobacteriaceae; Salmonella; Salmonella enterica; Salmonella enterica subsp. enterica MNNEETFYQAMRRKGVTRRSFLKFCSLAATSLGLGAGMTPKIAWALENKPRIPVVWIHGL WP_001058312.1 hydrogenase [Salmonella enterica] Length: 372\nNumber of predicted TMHs:  2\nExp number of AAs in TMHs: 36.51738\nExp number, first 60 AAs:  19.72849\nTotal prob of N-in:   0.96170\nPOSSIBLE N-term signal sequence\ninside   1   19\nTMhelix   20   39\noutside   40   325\nTMhelix   326   348\ninside   349   372

19589 GCF_001535255.1_ASM153525v1 Salmonella enterica subsp. enterica serovar Tennessee Proteobacteria; Gammaproteobacteria; Enterobacterales; Enterobacteriaceae; Salmonella; Salmonella enterica;

Salmonella enterica subsp. enterica MNNEETFYQAMRRKGVTRRSFLKFCSLAATSLGLGAGMTPKIAWALENKPRIPVVWIHGL WP_001058312.1 hydrogenase [Salmonella enterica] Length: 372\nNumber of predicted TMHs: 2\nExp number of AAs in TMHs: 36.51738\nExp number, first 60 AAs: 19.72849\nTotal prob of N-in: 0.96170\nPOSSIBLE N-term signal sequence\ninside 1 19\nTMhelix 20 39\noutside 40 325\nTMhelix 326 348\ninside 349 372

19590 GCF_001535145.1_ASM153514v1 Salmonella enterica subsp. enterica serovar Tennessee Proteobacteria; Gammaproteobacteria; Enterobacterales; Enterobacteriaceae; Salmonella; Salmonella enterica; Salmonella enterica subsp. enterica MTGDNTLITSHGINRRDFMKLCAALAATMGLSSKAAAEMAESVSNPQRPPVIWIGAQECT WP_000145429.1 hydrogenase 2 small subunit [Salmonella enterica] Length: 372\nNumber of predicted TMHs: 1\nExp number of AAs in TMHs: 24.51309\nExp number, first 60 AAs: 0.2682\nTotal prob of N-in: 0.14402\noutside 1 331\nTMhelix 332 354\ninside 355 372

19591 GCF_001630145.1_ASM163014v1 Salmonella enterica subsp. enterica serovar Tennessee Proteobacteria; Gammaproteobacteria; Enterobacterales; Enterobacteriaceae; Salmonella; Salmonella enterica; Salmonella enterica subsp. enterica MNNEETFYQAMRRKGVTRRSFLKFCSLAATSLGLGAGMTPKIAWALENKPRIPVVWIHGL WP_001058312.1 hydrogenase [Salmonella enterica] Length: 372\nNumber of predicted TMHs: 2\nExp number of AAs in TMHs: 36.51738\nExp number, first 60 AAs: 19.72849\nTotal prob of N-in: 0.96170\nPOSSIBLE N-term signal sequence\ninside 1 19\nTMhelix 20 39\noutside 40 325\nTMhelix 326 348\ninside 349 372

19592 GCF_001276845.1_ASM127684v1 Salmonella enterica subsp. enterica serovar Tennessee Proteobacteria; Gammaproteobacteria; Enterobacterales; Enterobacteriaceae; Salmonella; Salmonella enterica; Salmonella enterica subsp. enterica MTGDNTLITSHGINRRDFMKLCAALAATMGLSSKAAAEMAESVSNPQRPPVIWIGAQECT WP_000145429.1 hydrogenase 2 small subunit [Salmonella enterica] Length: 372\nNumber of predicted TMHs: 1\nExp number of AAs in TMHs: 24.51309\nExp number, first 60 AAs: 0.2682\nTotal prob of N-in: 0.14402\noutside 1 331\nTMhelix 332 354\ninside 355 372

19593 GCF_001620865.1_ASM162086v1 Salmonella enterica subsp. enterica serovar Tennessee Proteobacteria; Gammaproteobacteria; Enterobacterales; Enterobacteriaceae; Salmonella; Salmonella enterica; Salmonella enterica subsp. enterica MNNEETFYQAMRRKGVTRRSFLKFCSLAATSLGLGAGMTPKIAWALENKPRIPVVWIHGL WP_001058312.1 hydrogenase [Salmonella enterica] Length: 372\nNumber of predicted TMHs: 2\nExp number of AAs in TMHs: 36.51738\nExp number, first 60 AAs: 19.72849\nTotal prob of N-in: 0.96170\nPOSSIBLE N-term signal sequence\ninside 1 19\nTMhelix 20 39\noutside 40 325\nTMhelix 326 348\ninside 349 372

19594 GCF_001534985.1_ASM153498v1 Salmonella enterica subsp. enterica serovar Tennessee Proteobacteria; Gammaproteobacteria; Enterobacterales; Enterobacteriaceae; Salmonella; Salmonella enterica; Salmonella enterica subsp. enterica MTGDNTLITSHGINRRDFMKLCAALAATMGLSSKAAAEMAESVSNPQRPPVIWIGAQECT WP_000145429.1 hydrogenase 2 small subunit [Salmonella enterica] Length: 372\nNumber of predicted TMHs: 1\nExp number of AAs in TMHs: 24.51309\nExp number, first 60 AAs: 0.2682\nTotal prob of N-in: 0.14402\noutside 1 331\nTMhelix 332 354\ninside 355 372

19595 GCF_001630145.1_ASM163014v1 Salmonella enterica subsp. enterica serovar Tennessee Proteobacteria; Gammaproteobacteria; Enterobacterales; Enterobacteriaceae; Salmonella; Salmonella enterica; Salmonella enterica subsp. enterica MTGDNTLITSHGINRRDFMKLCAALAATMGLSSKAAAEMAESVSNPQRPPVIWIGAQECT WP_000145429.1 hydrogenase 2 small subunit [Salmonella enterica] Length: 372\nNumber of predicted TMHs: 1\nExp number of AAs in TMHs: 24.51309\nExp number, first 60 AAs: 0.2682\nTotal prob of N-in: 0.14402\noutside 1 331\nTMhelix 332 354\ninside 355 372

19596 GCF_001630165.1_ASM163016v1 Salmonella enterica subsp. enterica serovar Tennessee Proteobacteria; Gammaproteobacteria; Enterobacterales; Enterobacteriaceae; Salmonella; Salmonella enterica; Salmonella enterica subsp. enterica MTGDNTLITSHGINRRDFMKLCAALAATMGLSSKAAAEMAESVSNPQRPPVIWIGAQECT WP_000145429.1 hydrogenase 2 small subunit [Salmonella enterica] Length: 372\nNumber of predicted TMHs: 1\nExp number of AAs in TMHs: 24.51309\nExp number, first 60 AAs: 0.2682\nTotal prob of N-in: 0.14402\noutside 1 331\nTMhelix 332 354\ninside 355 372

19597 GCF_001276665.1_ASM127666v1 Salmonella enterica subsp. enterica serovar Tennessee Proteobacteria; Gammaproteobacteria; Enterobacterales; Enterobacteriaceae; Salmonella; Salmonella enterica; Salmonella enterica subsp. enterica MTGDNTLITSHGINRRDFMKLCAALAATMGLSSKAAAEMAESVSNPQRPPVIWIGAQECT WP_000145429.1 hydrogenase 2 small subunit [Salmonella enterica] Length: 372\nNumber of predicted TMHs: 1\nExp number of AAs in TMHs: 24.51309\nExp number, first 60 AAs: 0.2682\nTotal prob of N-in: 0.14402\noutside 1 331\nTMhelix 332 354\ninside 355 372

19598	GCF_001276665.1_ASM127666v1	Salmonella enterica subsp. enterica serovar Tennessee	Proteobacteria; Gammaproteobacteria; Enterobacterales; Enterobacteriaceae; Salmonella; Salmonella enterica; Salmonella enterica subsp. enterica	MNNEETFYQAMRRKGVTRRSFLKFCSLAATSLGLGAGMTPKIAWALENKPRIPVVWIHGL	WP_001058312.1	hydrogenase [Salmonella enterica]	Length: 372\nNumber of predicted TMHs:  2\nExp number of AAs in TMHs: 36.51738\nExp number, first 60 AAs:  19.72849\nTotal prob of N-in:   0.96170\nPOSSIBLE N-term signal sequence\ninside   1   19\nTMhelix   20   39\noutside   40   325\nTMhelix   326   348\ninside   349   372

19599	GCF_001620705.1_ASM162070v1	Salmonella enterica subsp. enterica serovar Tennessee	Proteobacteria; Gammaproteobacteria; Enterobacterales; Enterobacteriaceae; Salmonella; Salmonella enterica; Salmonella enterica subsp. enterica	MNNEETFYQAMRRKGVTRRSFLKFCSLAATSLGLGAGMTPKIAWALENKPRIPVVWIHGL	WP_001058312.1	hydrogenase [Salmonella enterica]	Length: 372\nNumber of predicted TMHs:  2\nExp number of AAs in TMHs: 36.51738\nExp number, first 60 AAs:  19.72849\nTotal prob of N-in:   0.96170\nPOSSIBLE N-term signal sequence\ninside   1   19\nTMhelix   20   39\noutside   40   325\nTMhelix   326   348\ninside   349   372

19600	GCF_001620795.1_ASM162079v1	Salmonella enterica subsp. enterica serovar Tennessee	Proteobacteria; Gammaproteobacteria; Enterobacterales; Enterobacteriaceae; Salmonella; Salmonella enterica; Salmonella enterica subsp. enterica	MNNEETFYQAMRRKGVTRRSFLKFCSLAATSLGLGAGMTPKIAWALENKPRIPVVWIHGL	WP_001058312.1	hydrogenase [Salmonella enterica]	Length: 372\nNumber of predicted TMHs:  2\nExp number of AAs in TMHs: 36.51738\nExp number, first 60 AAs:  19.72849\nTotal prob of N-in:   0.96170\nPOSSIBLE N-term signal sequence\ninside   1   19\nTMhelix   20   39\noutside   40   325\nTMhelix   326   348\ninside   349   372

19601	GCF_001620735.1_ASM162073v1	Salmonella enterica subsp. enterica serovar Tennessee	Proteobacteria; Gammaproteobacteria; Enterobacterales; Enterobacteriaceae; Salmonella; Salmonella enterica; Salmonella enterica subsp. enterica	MTGDNTLITSHGINRRDFMKLCAALAATMGLSSKAAAEMAESVSNPQRPPVIWIGAQECT	WP_000145429.1	hydrogenase 2 small subunit [Salmonella enterica]	Length: 372\nNumber of predicted TMHs:  1\nExp number of AAs in TMHs: 24.51309\nExp number, first 60 AAs:  0.2682\nTotal prob of N-in:   0.14402\noutside   1   331\nTMhelix   332   354\ninside   355   372

19602	GCF_001620705.1_ASM162070v1	Salmonella enterica subsp. enterica serovar Tennessee	Proteobacteria; Gammaproteobacteria; Enterobacterales; Enterobacteriaceae; Salmonella; Salmonella enterica; Salmonella enterica subsp. enterica	MTGDNTLITSHGINRRDFMKLCAALAATMGLSSKAAAEMAESVSNPQRPPVIWIGAQECT	WP_000145429.1	hydrogenase 2 small subunit [Salmonella enterica]	Length: 372\nNumber of predicted TMHs:  1\nExp number of AAs in TMHs: 24.51309\nExp number, first 60 AAs:  0.2682\nTotal prob of N-in:   0.14402\noutside   1   331\nTMhelix   332   354\ninside   355   372

19603	GCF_001535485.1_ASM153548v1	Salmonella enterica subsp. enterica serovar Tennessee	Proteobacteria; Gammaproteobacteria; Enterobacterales; Enterobacteriaceae; Salmonella; Salmonella enterica; Salmonella enterica subsp. enterica	MNNEETFYQAMRRKGVTRRSFLKFCSLAATSLGLGAGMTPKIAWALENKPRIPVVWIHGL	WP_001058312.1	hydrogenase [Salmonella enterica]	Length: 372\nNumber of predicted TMHs:  2\nExp number of AAs in TMHs: 36.51738\nExp number, first 60 AAs:  19.72849\nTotal prob of N-in:   0.96170\nPOSSIBLE N-term signal sequence\ninside   1   19\nTMhelix   20   39\noutside   40   325\nTMhelix   326   348\ninside   349   372

19604	GCF_001620795.1_ASM162079v1	Salmonella enterica subsp. enterica serovar Tennessee	Proteobacteria; Gammaproteobacteria; Enterobacterales; Enterobacteriaceae; Salmonella; Salmonella enterica; Salmonella enterica subsp. enterica	MTGDNTLITSHGINRRDFMKLCAALAATMGLSSKAAAEMAESVSNPQRPPVIWIGAQECT	WP_000145429.1	hydrogenase 2 small subunit [Salmonella enterica]	Length: 372\nNumber of predicted TMHs:  1\nExp number of AAs in TMHs: 24.51309\nExp number, first 60 AAs:  0.2682\nTotal prob of N-in:   0.14402\noutside   1   331\nTMhelix   332   354\ninside   355   372

19605	GCF_001630155.1_ASM163015v1	Salmonella enterica subsp. enterica serovar Tennessee	Proteobacteria; Gammaproteobacteria; Enterobacterales; Enterobacteriaceae; Salmonella; Salmonella enterica; Salmonella enterica subsp. enterica	MTGDNTLITSHGINRRDFMKLCAALAATMGLSSKAAAEMAESVSNPQRPPVIWIGAQECT	WP_000145429.1	hydrogenase 2 small subunit [Salmonella enterica]	Length: 372\nNumber of predicted TMHs:  1\nExp number of AAs in TMHs: 24.51309\nExp number, first 60 AAs:  0.2682\nTotal prob of N-in:   0.14402\noutside   1   331\nTMhelix   332   354\ninside   355   372

19606	GCF_001630215.1_ASM163021v1	Salmonella enterica subsp. enterica serovar Tennessee	Proteobacteria; Gammaproteobacteria; Enterobacterales; Enterobacteriaceae; Salmonella; Salmonella enterica; Salmonella enterica subsp. enterica	MTGDNTLITSHGINRRDFMKLCAALAATMGLSSKAAAEMAESVSNPQRPPVIWIGAQECT	WP_000145429.1	hydrogenase 2 small subunit [Salmonella enterica]	Length: 372\nNumber of

predicted TMHs: 1\nExp number of AAs in TMHs: 24.51309\nExp number, first 60 AAs: 0.2682\nTotal prob of N-in: 0.14402\noutside 1 331\nTMhelix 332 354\ninside 355 372

19607 GCF_001630155.1_ASM163015v1 Salmonella enterica subsp. enterica serovar Tennessee Proteobacteria; Gammaproteobacteria; Enterobacterales; Enterobacteriaceae; Salmonella; Salmonella enterica; Salmonella enterica subsp. enterica MNNEETFYQAMRRKGVTRRSFLKFCSLAATSLGLGAGMTPKIAWALENKPRIPVVWIHGL WP_001058312.1 hydrogenase [Salmonella enterica] Length: 372\nNumber of predicted TMHs: 2\nExp number of AAs in TMHs: 36.51738\nExp number, first 60 AAs: 19.72849\nTotal prob of N-in: 0.96170\nPOSSIBLE N-term signal sequence\ninside 1 19\nTMhelix 20 39\noutside 40 325\nTMhelix 326 348\ninside 349 372

19608 GCF_001630215.1_ASM163021v1 Salmonella enterica subsp. enterica serovar Tennessee Proteobacteria; Gammaproteobacteria; Enterobacterales; Enterobacteriaceae; Salmonella; Salmonella enterica; Salmonella enterica subsp. enterica MNNEETFYQAMRRKGVTRRSFLKFCSLAATSLGLGAGMTPKIAWALENKPRIPVVWIHGL WP_001058312.1 hydrogenase [Salmonella enterica] Length: 372\nNumber of predicted TMHs: 2\nExp number of AAs in TMHs: 36.51738\nExp number, first 60 AAs: 19.72849\nTotal prob of N-in: 0.96170\nPOSSIBLE N-term signal sequence\ninside 1 19\nTMhelix 20 39\noutside 40 325\nTMhelix 326 348\ninside 349 372

19609 GCF_001630235.1_ASM163023v1 Salmonella enterica subsp. enterica serovar Tennessee Proteobacteria; Gammaproteobacteria; Enterobacterales; Enterobacteriaceae; Salmonella; Salmonella enterica; Salmonella enterica subsp. enterica MNNEETFYQAMRRKGVTRRSFLKFCSLAATSLGLGAGMTPKIAWALENKPRIPVVWIHGL WP_001058312.1 hydrogenase [Salmonella enterica] Length: 372\nNumber of predicted TMHs: 2\nExp number of AAs in TMHs: 36.51738\nExp number, first 60 AAs: 19.72849\nTotal prob of N-in: 0.96170\nPOSSIBLE N-term signal sequence\ninside 1 19\nTMhelix 20 39\noutside 40 325\nTMhelix 326 348\ninside 349 372

19610 GCF_001620435.1_ASM162043v1 Salmonella enterica subsp. enterica serovar Tennessee Proteobacteria; Gammaproteobacteria; Enterobacterales; Enterobacteriaceae; Salmonella; Salmonella enterica; Salmonella enterica subsp. enterica MNNEETFYQAMRRKGVTRRSFLKFCSLAATSLGLGAGMTPKIAWALENKPRIPVVWIHGL WP_001058312.1 hydrogenase [Salmonella enterica] Length: 372\nNumber of predicted TMHs: 2\nExp number of AAs in TMHs: 36.51738\nExp number, first 60 AAs: 19.72849\nTotal prob of N-in: 0.96170\nPOSSIBLE N-term signal sequence\ninside 1 19\nTMhelix 20 39\noutside 40 325\nTMhelix 326 348\ninside 349 372

19611 GCF_001620495.1_ASM162049v1 Salmonella enterica subsp. enterica serovar Tennessee Proteobacteria; Gammaproteobacteria; Enterobacterales; Enterobacteriaceae; Salmonella; Salmonella enterica; Salmonella enterica subsp. enterica MTGDNTLITSHGINRRDFMKLCAALAATMGLSSKAAAEMAESVSNPQRPPVIWIGAQECT WP_000145429.1 hydrogenase 2 small subunit [Salmonella enterica] Length: 372\nNumber of predicted TMHs: 1\nExp number of AAs in TMHs: 24.51309\nExp number, first 60 AAs: 0.2682\nTotal prob of N-in: 0.14402\noutside 1 331\nTMhelix 332 354\ninside 355 372

19612 GCF_001630295.1_ASM163029v1 Salmonella enterica subsp. enterica serovar Tennessee Proteobacteria; Gammaproteobacteria; Enterobacterales; Enterobacteriaceae; Salmonella; Salmonella enterica; Salmonella enterica subsp. enterica MNNEETFYQAMRRKGVTRRSFLKFCSLAATSLGLGAGMTPKIAWALENKPRIPVVWIHGL WP_001058312.1 hydrogenase [Salmonella enterica] Length: 372\nNumber of predicted TMHs: 2\nExp number of AAs in TMHs: 36.51738\nExp number, first 60 AAs: 19.72849\nTotal prob of N-in: 0.96170\nPOSSIBLE N-term signal sequence\ninside 1 19\nTMhelix 20 39\noutside 40 325\nTMhelix 326 348\ninside 349 372

19613 GCF_001620435.1_ASM162043v1 Salmonella enterica subsp. enterica serovar Tennessee Proteobacteria; Gammaproteobacteria; Enterobacterales; Enterobacteriaceae; Salmonella; Salmonella enterica; Salmonella enterica subsp. enterica MTGDNTLITSHGINRRDFMKLCAALAATMGLSSKAAAEMAESVSNPQRPPVIWIGAQECT WP_000145429.1 hydrogenase 2 small subunit [Salmonella enterica] Length: 372\nNumber of predicted TMHs: 1\nExp number of AAs in TMHs: 24.51309\nExp number, first 60 AAs: 0.2682\nTotal prob of N-in: 0.14402\noutside 1 331\nTMhelix 332 354\ninside 355 372

19614 GCF_001620415.1_ASM162041v1 Salmonella enterica subsp. enterica serovar Tennessee Proteobacteria; Gammaproteobacteria; Enterobacterales; Enterobacteriaceae; Salmonella; Salmonella enterica; Salmonella enterica subsp. enterica MNNEETFYQAMRRKGVTRRSFLKFCSLAATSLGLGAGMTPKIAWALENKPRIPVVWIHGL WP_001058312.1 hydrogenase [Salmonella enterica] Length: 372\nNumber of predicted TMHs: 2\nExp number of AAs in TMHs: 36.51738\nExp number, first 60 AAs: 19.72849\nTotal prob of N-in: 0.96170\nPOSSIBLE N-term signal sequence\ninside 1 19\nTMhelix 20 39\noutside 40 325\nTMhelix 326 348\ninside 349 372

19615 GCF_001620415.1_ASM162041v1 Salmonella enterica subsp. enterica serovar Tennessee Proteobacteria; Gammaproteobacteria; Enterobacterales; Enterobacteriaceae; Salmonella; Salmonella enterica;

Salmonella enterica subsp. enterica	MTGDNTLITSHGINRRDFMKLCAALAATMGLSSKAAAEMAESVSNPQRPPVIWIGAQECT	WP_000145429.1	hydrogenase 2 small subunit [Salmonella enterica]	Length: 372\nNumber of predicted TMHs:  1\nExp number of AAs in TMHs: 24.51309\nExp number, first 60 AAs:  0.2682\nTotal prob of N-in:  0.14402\noutside   1   331\nTMhelix   332   354\ninside   355   372

19616	GCF_001974765.1_ASM197476v1	Salmonella enterica subsp. enterica serovar Tennessee	Proteobacteria; Gammaproteobacteria; Enterobacterales; Enterobacteriaceae; Salmonella; Salmonella enterica; Salmonella enterica subsp. enterica	MTGDNTLITSHGINRRDFMKLCAALAATMGLSSKAAAEMAESVSNPQRPPVIWIGAQECT	WP_000145429.1	hydrogenase 2 small subunit [Salmonella enterica]	Length: 372\nNumber of predicted TMHs:  1\nExp number of AAs in TMHs: 24.51309\nExp number, first 60 AAs:  0.2682\nTotal prob of N-in:  0.14402\noutside   1   331\nTMhelix   332   354\ninside   355   372

19617	GCF_001620515.1_ASM162051v1	Salmonella enterica subsp. enterica serovar Tennessee	Proteobacteria; Gammaproteobacteria; Enterobacterales; Enterobacteriaceae; Salmonella; Salmonella enterica; Salmonella enterica subsp. enterica	MTGDNTLITSHGINRRDFMKLCAALAATMGLSSKAAAEMAESVSNPQRPPVIWIGAQECT	WP_000145429.1	hydrogenase 2 small subunit [Salmonella enterica]	Length: 372\nNumber of predicted TMHs:  1\nExp number of AAs in TMHs: 24.51309\nExp number, first 60 AAs:  0.2682\nTotal prob of N-in:  0.14402\noutside   1   331\nTMhelix   332   354\ninside   355   372

19618	GCF_001535125.1_ASM153512v1	Salmonella enterica subsp. enterica serovar Tennessee	Proteobacteria; Gammaproteobacteria; Enterobacterales; Enterobacteriaceae; Salmonella; Salmonella enterica; Salmonella enterica subsp. enterica	MTGDNTLITSHGINRRDFMKLCAALAATMGLSSKAAAEMAESVSNPQRPPVIWIGAQECT	WP_000145429.1	hydrogenase 2 small subunit [Salmonella enterica]	Length: 372\nNumber of predicted TMHs:  1\nExp number of AAs in TMHs: 24.51309\nExp number, first 60 AAs:  0.2682\nTotal prob of N-in:  0.14402\noutside   1   331\nTMhelix   332   354\ninside   355   372

19619	GCF_001535125.1_ASM153512v1	Salmonella enterica subsp. enterica serovar Tennessee	Proteobacteria; Gammaproteobacteria; Enterobacterales; Enterobacteriaceae; Salmonella; Salmonella enterica; Salmonella enterica subsp. enterica	MNNEETFYQAMRRKGVTRRSFLKFCSLAATSLGLGAGMTPKIAWALENKPRIPVVWIHGL	WP_001058312.1	hydrogenase [Salmonella enterica]	Length: 372\nNumber of predicted TMHs:  2\nExp number of AAs in TMHs: 36.51738\nExp number, first 60 AAs:  19.72849\nTotal prob of N-in:   0.96170\nPOSSIBLE N-term signal sequence\ninside   1   19\nTMhelix   20   39\noutside   40   325\nTMhelix   326   348\ninside   349   372

19620	GCF_001535445.1_ASM153544v1	Salmonella enterica subsp. enterica serovar Tennessee	Proteobacteria; Gammaproteobacteria; Enterobacterales; Enterobacteriaceae; Salmonella; Salmonella enterica; Salmonella enterica subsp. enterica	MTGDNTLITSHGINRRDFMKLCAALAATMGLSSKAAAEMAESVSNPQRPPVIWIGAQECT	WP_000145429.1	hydrogenase 2 small subunit [Salmonella enterica]	Length: 372\nNumber of predicted TMHs:  1\nExp number of AAs in TMHs: 24.51309\nExp number, first 60 AAs:  0.2682\nTotal prob of N-in:  0.14402\noutside   1   331\nTMhelix   332   354\ninside   355   372

19621	GCF_001535485.1_ASM153548v1	Salmonella enterica subsp. enterica serovar Tennessee	Proteobacteria; Gammaproteobacteria; Enterobacterales; Enterobacteriaceae; Salmonella; Salmonella enterica; Salmonella enterica subsp. enterica	MTGDNTLITSHGINRRDFMKLCAALAATMGLSSKAAAEMAESVSNPQRPPVIWIGAQECT	WP_000145429.1	hydrogenase 2 small subunit [Salmonella enterica]	Length: 372\nNumber of predicted TMHs:  1\nExp number of AAs in TMHs: 24.51309\nExp number, first 60 AAs:  0.2682\nTotal prob of N-in:  0.14402\noutside   1   331\nTMhelix   332   354\ninside   355   372

19622	GCF_001535405.1_ASM153540v1	Salmonella enterica subsp. enterica serovar Tennessee	Proteobacteria; Gammaproteobacteria; Enterobacterales; Enterobacteriaceae; Salmonella; Salmonella enterica; Salmonella enterica subsp. enterica	MNNEETFYQAMRRKGVTRRSFLKFCSLAATSLGLGAGMTPKIAWALENKPRIPVVWIHGL	WP_001058312.1	hydrogenase [Salmonella enterica]	Length: 372\nNumber of predicted TMHs:  2\nExp number of AAs in TMHs: 36.51738\nExp number, first 60 AAs:  19.72849\nTotal prob of N-in:   0.96170\nPOSSIBLE N-term signal sequence\ninside   1   19\nTMhelix   20   39\noutside   40   325\nTMhelix   326   348\ninside   349   372

19623	GCF_001535405.1_ASM153540v1	Salmonella enterica subsp. enterica serovar Tennessee	Proteobacteria; Gammaproteobacteria; Enterobacterales; Enterobacteriaceae; Salmonella; Salmonella enterica; Salmonella enterica subsp. enterica	MTGDNTLITSHGINRRDFMKLCAALAATMGLSSKAAAEMAESVSNPQRPPVIWIGAQECT	WP_000145429.1	hydrogenase 2 small subunit [Salmonella enterica]	Length: 372\nNumber of predicted TMHs:  1\nExp number of AAs in TMHs: 24.51309\nExp number, first 60 AAs:  0.2682\nTotal prob of N-in:  0.14402\noutside   1   331\nTMhelix   332   354\ninside   355   372

19624 GCF_001535445.1_ASM153544v1 Salmonella enterica subsp. enterica serovar Tennessee Proteobacteria; Gammaproteobacteria; Enterobacterales; Enterobacteriaceae; Salmonella; Salmonella enterica; Salmonella enterica subsp. enterica MNNEETFYQAMRRKGVTRRSFLKFCSLAATSLGLGAGMTPKIAWALENKPRIPVVWIHGL WP_001058312.1 hydrogenase [Salmonella enterica] Length: 372\nNumber of predicted TMHs: 2\nExp number of AAs in TMHs: 36.51738\nExp number, first 60 AAs: 19.72849\nTotal prob of N-in: 0.96170\nPOSSIBLE N-term signal sequence\ninside 1 19\nTMhelix 20 39\noutside 40 325\nTMhelix 326 348\ninside 349 372

19625 GCF_001534905.1_ASM153490v1 Salmonella enterica subsp. enterica serovar Tennessee Proteobacteria; Gammaproteobacteria; Enterobacterales; Enterobacteriaceae; Salmonella; Salmonella enterica; Salmonella enterica subsp. enterica MTGDNTLITSHGINRRDFMKLCAALAATMGLSSKAAAEMAESVSNPQRPPVIWIGAQECT WP_000145429.1 hydrogenase 2 small subunit [Salmonella enterica] Length: 372\nNumber of predicted TMHs: 1\nExp number of AAs in TMHs: 24.51309\nExp number, first 60 AAs: 0.2682\nTotal prob of N-in: 0.14402\noutside 1 331\nTMhelix 332 354\ninside 355 372

19626 GCF_001534835.1_ASM153483v1 Salmonella enterica subsp. enterica serovar Tennessee Proteobacteria; Gammaproteobacteria; Enterobacterales; Enterobacteriaceae; Salmonella; Salmonella enterica; Salmonella enterica subsp. enterica MTGDNTLITSHGINRRDFMKLCAALAATMGLSSKAAAEMAESVSNPQRPPVIWIGAQECT WP_000145429.1 hydrogenase 2 small subunit [Salmonella enterica] Length: 372\nNumber of predicted TMHs: 1\nExp number of AAs in TMHs: 24.51309\nExp number, first 60 AAs: 0.2682\nTotal prob of N-in: 0.14402\noutside 1 331\nTMhelix 332 354\ninside 355 372

19627 GCF_001535165.1_ASM153516v1 Salmonella enterica subsp. enterica serovar Tennessee Proteobacteria; Gammaproteobacteria; Enterobacterales; Enterobacteriaceae; Salmonella; Salmonella enterica; Salmonella enterica subsp. enterica MNNEETFYQAMRRKGVTRRSFLKFCSLAATSLGLGAGMTPKIAWALENKPRIPVVWIHGL WP_001058312.1 hydrogenase [Salmonella enterica] Length: 372\nNumber of predicted TMHs: 2\nExp number of AAs in TMHs: 36.51738\nExp number, first 60 AAs: 19.72849\nTotal prob of N-in: 0.96170\nPOSSIBLE N-term signal sequence\ninside 1 19\nTMhelix 20 39\noutside 40 325\nTMhelix 326 348\ninside 349 372

19628 GCF_001535285.1_ASM153528v1 Salmonella enterica subsp. enterica serovar Tennessee Proteobacteria; Gammaproteobacteria; Enterobacterales; Enterobacteriaceae; Salmonella; Salmonella enterica; Salmonella enterica subsp. enterica MTGDNTLITSHGINRRDFMKLCAALAATMGLSSKAAAEMAESVSNPQRPPVIWIGAQECT WP_000145429.1 hydrogenase 2 small subunit [Salmonella enterica] Length: 372\nNumber of predicted TMHs: 1\nExp number of AAs in TMHs: 24.51309\nExp number, first 60 AAs: 0.2682\nTotal prob of N-in: 0.14402\noutside 1 331\nTMhelix 332 354\ninside 355 372

19629 GCF_001535195.1_ASM153519v1 Salmonella enterica subsp. enterica serovar Tennessee Proteobacteria; Gammaproteobacteria; Enterobacterales; Enterobacteriaceae; Salmonella; Salmonella enterica; Salmonella enterica subsp. enterica MTGDNTLITSHGINRRDFMKLCAALAATMGLSSKAAAEMAESVSNPQRPPVIWIGAQECT WP_000145429.1 hydrogenase 2 small subunit [Salmonella enterica] Length: 372\nNumber of predicted TMHs: 1\nExp number of AAs in TMHs: 24.51309\nExp number, first 60 AAs: 0.2682\nTotal prob of N-in: 0.14402\noutside 1 331\nTMhelix 332 354\ninside 355 372

19630 GCF_001535165.1_ASM153516v1 Salmonella enterica subsp. enterica serovar Tennessee Proteobacteria; Gammaproteobacteria; Enterobacterales; Enterobacteriaceae; Salmonella; Salmonella enterica; Salmonella enterica subsp. enterica MTGDNTLITSHGINRRDFMKLCAALAATMGLSSKAAAEMAESVSNPQRPPVIWIGAQECT WP_000145429.1 hydrogenase 2 small subunit [Salmonella enterica] Length: 372\nNumber of predicted TMHs: 1\nExp number of AAs in TMHs: 24.51309\nExp number, first 60 AAs: 0.2682\nTotal prob of N-in: 0.14402\noutside 1 331\nTMhelix 332 354\ninside 355 372

19631 GCF_001534905.1_ASM153490v1 Salmonella enterica subsp. enterica serovar Tennessee Proteobacteria; Gammaproteobacteria; Enterobacterales; Enterobacteriaceae; Salmonella; Salmonella enterica; Salmonella enterica subsp. enterica MNNEETFYQAMRRKGVTRRSFLKFCSLAATSLGLGAGMTPKIAWALENKPRIPVVWIHGL WP_001058312.1 hydrogenase [Salmonella enterica] Length: 372\nNumber of predicted TMHs: 2\nExp number of AAs in TMHs: 36.51738\nExp number, first 60 AAs: 19.72849\nTotal prob of N-in: 0.96170\nPOSSIBLE N-term signal sequence\ninside 1 19\nTMhelix 20 39\noutside 40 325\nTMhelix 326 348\ninside 349 372

19632 GCF_001535025.1_ASM153502v1 Salmonella enterica subsp. enterica serovar Tennessee Proteobacteria; Gammaproteobacteria; Enterobacterales; Enterobacteriaceae; Salmonella; Salmonella enterica; Salmonella enterica subsp. enterica MNNEETFYQAMRRKGVTRRSFLKFCSLAATSLGLGAGMTPKIAWALENKPRIPVVWIHGL WP_001058312.1 hydrogenase [Salmonella enterica] Length: 372\nNumber of predicted TMHs: 2\nExp

number of AAs in TMHs: 36.51738\nExp number, first 60 AAs: 19.72849\nTotal prob of N-in: 0.96170\nPOSSIBLE N-term signal sequence\ninside 1 19\nTMhelix 20 39\noutside 40 325\nTMhelix 326 348\ninside 349 372

19633 GCF_001535095.1_ASM153509v1 Salmonella enterica subsp. enterica serovar Tennessee Proteobacteria; Gammaproteobacteria; Enterobacterales; Enterobacteriaceae; Salmonella; Salmonella enterica; Salmonella enterica subsp. enterica MNNEETFYQAMRRKGVTRRSFLKFCSLAATSLGLGAGMTPKIAWALENKPRIPVVWIHGL WP_001058312.1 hydrogenase [Salmonella enterica] Length: 372\nNumber of predicted TMHs: 2\nExp number of AAs in TMHs: 36.51738\nExp number, first 60 AAs: 19.72849\nTotal prob of N-in: 0.96170\nPOSSIBLE N-term signal sequence\ninside 1 19\nTMhelix 20 39\noutside 40 325\nTMhelix 326 348\ninside 349 372

19634 GCF_001534835.1_ASM153483v1 Salmonella enterica subsp. enterica serovar Tennessee Proteobacteria; Gammaproteobacteria; Enterobacterales; Enterobacteriaceae; Salmonella; Salmonella enterica; Salmonella enterica subsp. enterica MNNEETFYQAMRRKGVTRRSFLKFCSLAATSLGLGAGMTPKIAWALENKPRIPVVWIHGL WP_001058312.1 hydrogenase [Salmonella enterica] Length: 372\nNumber of predicted TMHs: 2\nExp number of AAs in TMHs: 36.51738\nExp number, first 60 AAs: 19.72849\nTotal prob of N-in: 0.96170\nPOSSIBLE N-term signal sequence\ninside 1 19\nTMhelix 20 39\noutside 40 325\nTMhelix 326 348\ninside 349 372

19635 GCF_001535025.1_ASM153502v1 Salmonella enterica subsp. enterica serovar Tennessee Proteobacteria; Gammaproteobacteria; Enterobacterales; Enterobacteriaceae; Salmonella; Salmonella enterica; Salmonella enterica subsp. enterica MTGDNTLITSHGINRRDFMKLCAALAATMGLSSKAAAEMAESVSNPQRPPVIWIGAQECT WP_000145429.1 hydrogenase 2 small subunit [Salmonella enterica] Length: 372\nNumber of predicted TMHs: 1\nExp number of AAs in TMHs: 24.51309\nExp number, first 60 AAs: 0.2682\nTotal prob of N-in: 0.14402\noutside 1 331\nTMhelix 332 354\ninside 355 372

19636 GCF_001535095.1_ASM153509v1 Salmonella enterica subsp. enterica serovar Tennessee Proteobacteria; Gammaproteobacteria; Enterobacterales; Enterobacteriaceae; Salmonella; Salmonella enterica; Salmonella enterica subsp. enterica MTGDNTLITSHGINRRDFMKLCAALAATMGLSSKAAAEMAESVSNPQRPPVIWIGAQECT WP_000145429.1 hydrogenase 2 small subunit [Salmonella enterica] Length: 372\nNumber of predicted TMHs: 1\nExp number of AAs in TMHs: 24.51309\nExp number, first 60 AAs: 0.2682\nTotal prob of N-in: 0.14402\noutside 1 331\nTMhelix 332 354\ninside 355 372

19637 GCF_001620835.1_ASM162083v1 Salmonella enterica subsp. enterica serovar Tennessee Proteobacteria; Gammaproteobacteria; Enterobacterales; Enterobacteriaceae; Salmonella; Salmonella enterica; Salmonella enterica subsp. enterica MTGDNTLITSHGINRRDFMKLCAALAATMGLSSKAAAEMAESVSNPQRPPVIWIGAQECT WP_000145429.1 hydrogenase 2 small subunit [Salmonella enterica] Length: 372\nNumber of predicted TMHs: 1\nExp number of AAs in TMHs: 24.51309\nExp number, first 60 AAs: 0.2682\nTotal prob of N-in: 0.14402\noutside 1 331\nTMhelix 332 354\ninside 355 372

19638 GCF_001620835.1_ASM162083v1 Salmonella enterica subsp. enterica serovar Tennessee Proteobacteria; Gammaproteobacteria; Enterobacterales; Enterobacteriaceae; Salmonella; Salmonella enterica; Salmonella enterica subsp. enterica MNNEETFYQAMRRKGVTRRSFLKFCSLAATSLGLGAGMTPKIAWALENKPRIPVVWIHGL WP_001058312.1 hydrogenase [Salmonella enterica] Length: 372\nNumber of predicted TMHs: 2\nExp number of AAs in TMHs: 36.51738\nExp number, first 60 AAs: 19.72849\nTotal prob of N-in: 0.96170\nPOSSIBLE N-term signal sequence\ninside 1 19\nTMhelix 20 39\noutside 40 325\nTMhelix 326 348\ninside 349 372

19639 GCF_001620585.1_ASM162058v1 Salmonella enterica subsp. enterica serovar Tennessee Proteobacteria; Gammaproteobacteria; Enterobacterales; Enterobacteriaceae; Salmonella; Salmonella enterica; Salmonella enterica subsp. enterica MNNEETFYQAMRRKGVTRRSFLKFCSLAATSLGLGAGMTPKIAWALENKPRIPVVWIHGL WP_001058312.1 hydrogenase [Salmonella enterica] Length: 372\nNumber of predicted TMHs: 2\nExp number of AAs in TMHs: 36.51738\nExp number, first 60 AAs: 19.72849\nTotal prob of N-in: 0.96170\nPOSSIBLE N-term signal sequence\ninside 1 19\nTMhelix 20 39\noutside 40 325\nTMhelix 326 348\ninside 349 372

19640 GCF_001620515.1_ASM162051v1 Salmonella enterica subsp. enterica serovar Tennessee Proteobacteria; Gammaproteobacteria; Enterobacterales; Enterobacteriaceae; Salmonella; Salmonella enterica; Salmonella enterica subsp. enterica MNNEETFYQAMRRKGVTRRSFLKFCSLAATSLGLGAGMTPKIAWALENKPRIPVVWIHGL WP_001058312.1 hydrogenase [Salmonella enterica] Length: 372\nNumber of predicted TMHs: 2\nExp number of AAs in TMHs: 36.51738\nExp number, first 60 AAs: 19.72849\nTotal prob of N-in: 0.96170\nPOSSIBLE N-term signal sequence\ninside 1 19\nTMhelix 20 39\noutside 40 325\nTMhelix 326 348\ninside 349 372

19641 GCF_001620735.1_ASM162073v1 Salmonella enterica subsp. enterica serovar Tennessee Proteobacteria; Gammaproteobacteria; Enterobacterales; Enterobacteriaceae; Salmonella; Salmonella enterica;

Salmonella enterica subsp. enterica MNNEETFYQAMRRKGVTRRSFLKFCSLAATSLGLGAGMTPKIAWALENKPRIPVVWIHGL WP_001058312.1 hydrogenase [Salmonella enterica] Length: 372\nNumber of predicted TMHs: 2\nExp number of AAs in TMHs: 36.51738\nExp number, first 60 AAs: 19.72849\nTotal prob of N-in: 0.96170\nPOSSIBLE N-term signal sequence\ninside 1 19\nTMhelix 20 39\noutside 40 325\nTMhelix 326 348\ninside 349 372

19642 GCF_001620595.1_ASM162059v1 Salmonella enterica subsp. enterica serovar Tennessee Proteobacteria; Gammaproteobacteria; Enterobacterales; Enterobacteriaceae; Salmonella; Salmonella enterica; Salmonella enterica subsp. enterica MTGDNTLITSHGINRRDFMKLCAALAATMGLSSKAAAEMAESVSNPQRPPVIWIGAQECT WP_000145429.1 hydrogenase 2 small subunit [Salmonella enterica] Length: 372\nNumber of predicted TMHs: 1\nExp number of AAs in TMHs: 24.51309\nExp number, first 60 AAs: 0.2682\nTotal prob of N-in: 0.14402\noutside 1 331\nTMhelix 332 354\ninside 355 372

19643 GCF_001276675.1_ASM127667v1 Salmonella enterica subsp. enterica serovar Tennessee Proteobacteria; Gammaproteobacteria; Enterobacterales; Enterobacteriaceae; Salmonella; Salmonella enterica; Salmonella enterica subsp. enterica MTGDNTLITSHGINRRDFMKLCAALAATMGLSSKAAAEMAESVSNPQRPPVIWIGAQECT WP_000145429.1 hydrogenase 2 small subunit [Salmonella enterica] Length: 372\nNumber of predicted TMHs: 1\nExp number of AAs in TMHs: 24.51309\nExp number, first 60 AAs: 0.2682\nTotal prob of N-in: 0.14402\noutside 1 331\nTMhelix 332 354\ninside 355 372

19644 GCF_001620645.1_ASM162064v1 Salmonella enterica subsp. enterica serovar Tennessee Proteobacteria; Gammaproteobacteria; Enterobacterales; Enterobacteriaceae; Salmonella; Salmonella enterica; Salmonella enterica subsp. enterica MTGDNTLITSHGINRRDFMKLCAALAATMGLSSKAAAEMAESVSNPQRPPVIWIGAQECT WP_000145429.1 hydrogenase 2 small subunit [Salmonella enterica] Length: 372\nNumber of predicted TMHs: 1\nExp number of AAs in TMHs: 24.51309\nExp number, first 60 AAs: 0.2682\nTotal prob of N-in: 0.14402\noutside 1 331\nTMhelix 332 354\ninside 355 372

19645 GCF_001620725.1_ASM162072v1 Salmonella enterica subsp. enterica serovar Tennessee Proteobacteria; Gammaproteobacteria; Enterobacterales; Enterobacteriaceae; Salmonella; Salmonella enterica; Salmonella enterica subsp. enterica MTGDNTLITSHGINRRDFMKLCAALAATMGLSSKAAAEMAESVSNPQRPPVIWIGAQECT WP_000145429.1 hydrogenase 2 small subunit [Salmonella enterica] Length: 372\nNumber of predicted TMHs: 1\nExp number of AAs in TMHs: 24.51309\nExp number, first 60 AAs: 0.2682\nTotal prob of N-in: 0.14402\noutside 1 331\nTMhelix 332 354\ninside 355 372

19646 GCF_001630255.1_ASM163025v1 Salmonella enterica subsp. enterica serovar Tennessee Proteobacteria; Gammaproteobacteria; Enterobacterales; Enterobacteriaceae; Salmonella; Salmonella enterica; Salmonella enterica subsp. enterica MNNEETFYQAMRRKGVTRRSFLKFCSLAATSLGLGAGMTPKIAWALENKPRIPVVWIHGL WP_001058312.1 hydrogenase [Salmonella enterica] Length: 372\nNumber of predicted TMHs: 2\nExp number of AAs in TMHs: 36.51738\nExp number, first 60 AAs: 19.72849\nTotal prob of N-in: 0.96170\nPOSSIBLE N-term signal sequence\ninside 1 19\nTMhelix 20 39\noutside 40 325\nTMhelix 326 348\ninside 349 372

19647 GCF_001620595.1_ASM162059v1 Salmonella enterica subsp. enterica serovar Tennessee Proteobacteria; Gammaproteobacteria; Enterobacterales; Enterobacteriaceae; Salmonella; Salmonella enterica; Salmonella enterica subsp. enterica MNNEETFYQAMRRKGVTRRSFLKFCSLAATSLGLGAGMTPKIAWALENKPRIPVVWIHGL WP_001058312.1 hydrogenase [Salmonella enterica] Length: 372\nNumber of predicted TMHs: 2\nExp number of AAs in TMHs: 36.51738\nExp number, first 60 AAs: 19.72849\nTotal prob of N-in: 0.96170\nPOSSIBLE N-term signal sequence\ninside 1 19\nTMhelix 20 39\noutside 40 325\nTMhelix 326 348\ninside 349 372

19648 GCF_001535455.1_ASM153545v1 Salmonella enterica subsp. enterica serovar Tennessee Proteobacteria; Gammaproteobacteria; Enterobacterales; Enterobacteriaceae; Salmonella; Salmonella enterica; Salmonella enterica subsp. enterica MNNEETFYQAMRRKGVTRRSFLKFCSLAATSLGLGAGMTPKIAWALENKPRIPVVWIHGL WP_001058312.1 hydrogenase [Salmonella enterica] Length: 372\nNumber of predicted TMHs: 2\nExp number of AAs in TMHs: 36.51738\nExp number, first 60 AAs: 19.72849\nTotal prob of N-in: 0.96170\nPOSSIBLE N-term signal sequence\ninside 1 19\nTMhelix 20 39\noutside 40 325\nTMhelix 326 348\ninside 349 372

19649 GCF_001620755.2_ASM162075v2 Salmonella enterica subsp. enterica serovar Tennessee Proteobacteria; Gammaproteobacteria; Enterobacterales; Enterobacteriaceae; Salmonella; Salmonella enterica; Salmonella enterica subsp. enterica MTGDNTLITSHGINRRDFMKLCAALAATMGLSSKAAAEMAESVSNPQRPPVIWIGAQECT WP_000145429.1 hydrogenase 2 small subunit [Salmonella enterica] Length: 372\nNumber of predicted TMHs: 1\nExp number of AAs in TMHs: 24.51309\nExp number, first 60 AAs: 0.2682\nTotal prob of N-in: 0.14402\noutside 1 331\nTMhelix 332 354\ninside 355 372

19650 GCF_001630315.1_ASM163031v1 Salmonella enterica subsp. enterica serovar Tennessee Proteobacteria; Gammaproteobacteria; Enterobacterales; Enterobacteriaceae; Salmonella; Salmonella enterica; Salmonella enterica subsp. enterica MTGDNTLITSHGINRRDFMKLCAALAATMGLSSKAAAEMAESVSNPQRPPVIWIGAQECT WP_000145429.1 hydrogenase 2 small subunit [Salmonella enterica] Length: 372\nNumber of predicted TMHs: 1\nExp number of AAs in TMHs: 24.51309\nExp number, first 60 AAs: 0.2682\nTotal prob of N-in: 0.14402\noutside 1 331\nTMhelix 332 354\ninside 355 372

19651 GCF_001630225.1_ASM163022v1 Salmonella enterica subsp. enterica serovar Tennessee Proteobacteria; Gammaproteobacteria; Enterobacterales; Enterobacteriaceae; Salmonella; Salmonella enterica; Salmonella enterica subsp. enterica MTGDNTLITSHGINRRDFMKLCAALAATMGLSSKAAAEMAESVSNPQRPPVIWIGAQECT WP_000145429.1 hydrogenase 2 small subunit [Salmonella enterica] Length: 372\nNumber of predicted TMHs: 1\nExp number of AAs in TMHs: 24.51309\nExp number, first 60 AAs: 0.2682\nTotal prob of N-in: 0.14402\noutside 1 331\nTMhelix 332 354\ninside 355 372

19652 GCF_001535385.1_ASM153538v1 Salmonella enterica subsp. enterica serovar Tennessee Proteobacteria; Gammaproteobacteria; Enterobacterales; Enterobacteriaceae; Salmonella; Salmonella enterica; Salmonella enterica subsp. enterica MNNEETFYQAMRRKGVTRRSFLKFCSLAATSLGLGAGMTPKIAWALENKPRIPVVWIHGL WP_001058312.1 hydrogenase [Salmonella enterica] Length: 372\nNumber of predicted TMHs: 2\nExp number of AAs in TMHs: 36.51738\nExp number, first 60 AAs: 19.72849\nTotal prob of N-in: 0.96170\nPOSSIBLE N-term signal sequence\ninside 1 19\nTMhelix 20 39\noutside 40 325\nTMhelix 326 348\ninside 349 372

19653 GCF_001534845.1_ASM153484v1 Salmonella enterica subsp. enterica serovar Tennessee Proteobacteria; Gammaproteobacteria; Enterobacterales; Enterobacteriaceae; Salmonella; Salmonella enterica; Salmonella enterica subsp. enterica MTGDNTLITSHGINRRDFMKLCAALAATMGLSSKAAAEMAESVSNPQRPPVIWIGAQECT WP_000145429.1 hydrogenase 2 small subunit [Salmonella enterica] Length: 372\nNumber of predicted TMHs: 1\nExp number of AAs in TMHs: 24.51309\nExp number, first 60 AAs: 0.2682\nTotal prob of N-in: 0.14402\noutside 1 331\nTMhelix 332 354\ninside 355 372

19654 GCF_001534845.1_ASM153484v1 Salmonella enterica subsp. enterica serovar Tennessee Proteobacteria; Gammaproteobacteria; Enterobacterales; Enterobacteriaceae; Salmonella; Salmonella enterica; Salmonella enterica subsp. enterica MNNEETFYQAMRRKGVTRRSFLKFCSLAATSLGLGAGMTPKIAWALENKPRIPVVWIHGL WP_001058312.1 hydrogenase [Salmonella enterica] Length: 372\nNumber of predicted TMHs: 2\nExp number of AAs in TMHs: 36.51738\nExp number, first 60 AAs: 19.72849\nTotal prob of N-in: 0.96170\nPOSSIBLE N-term signal sequence\ninside 1 19\nTMhelix 20 39\noutside 40 325\nTMhelix 326 348\ninside 349 372

19655 GCF_001534985.1_ASM153498v1 Salmonella enterica subsp. enterica serovar Tennessee Proteobacteria; Gammaproteobacteria; Enterobacterales; Enterobacteriaceae; Salmonella; Salmonella enterica; Salmonella enterica subsp. enterica MNNEETFYQAMRRKGVTRRSFLKFCSLAATSLGLGAGMTPKIAWALENKPRIPVVWIHGL WP_001058312.1 hydrogenase [Salmonella enterica] Length: 372\nNumber of predicted TMHs: 2\nExp number of AAs in TMHs: 36.51738\nExp number, first 60 AAs: 19.72849\nTotal prob of N-in: 0.96170\nPOSSIBLE N-term signal sequence\ninside 1 19\nTMhelix 20 39\noutside 40 325\nTMhelix 326 348\ninside 349 372

19656 GCF_001535175.1_ASM153517v1 Salmonella enterica subsp. enterica serovar Tennessee Proteobacteria; Gammaproteobacteria; Enterobacterales; Enterobacteriaceae; Salmonella; Salmonella enterica; Salmonella enterica subsp. enterica MNNEETFYQAMRRKGVTRRSFLKFCSLAATSLGLGAGMTPKIAWALENKPRIPVVWIHGL WP_001058312.1 hydrogenase [Salmonella enterica] Length: 372\nNumber of predicted TMHs: 2\nExp number of AAs in TMHs: 36.51738\nExp number, first 60 AAs: 19.72849\nTotal prob of N-in: 0.96170\nPOSSIBLE N-term signal sequence\ninside 1 19\nTMhelix 20 39\noutside 40 325\nTMhelix 326 348\ninside 349 372

19657 GCF_001535145.1_ASM153514v1 Salmonella enterica subsp. enterica serovar Tennessee Proteobacteria; Gammaproteobacteria; Enterobacterales; Enterobacteriaceae; Salmonella; Salmonella enterica; Salmonella enterica subsp. enterica MNNEETFYQAMRRKGVTRRSFLKFCSLAATSLGLGAGMTPKIAWALENKPRIPVVWIHGL WP_001058312.1 hydrogenase [Salmonella enterica] Length: 372\nNumber of predicted TMHs: 2\nExp number of AAs in TMHs: 36.51738\nExp number, first 60 AAs: 19.72849\nTotal prob of N-in: 0.96170\nPOSSIBLE N-term signal sequence\ninside 1 19\nTMhelix 20 39\noutside 40 325\nTMhelix 326 348\ninside 349 372

19658 GCF_001620685.1_ASM162068v1 Salmonella enterica subsp. enterica serovar Tennessee Proteobacteria; Gammaproteobacteria; Enterobacterales; Enterobacteriaceae; Salmonella; Salmonella enterica; Salmonella enterica subsp. enterica MTGDNTLITSHGINRRDFMKLCAALAATMGLSSKAAAEMAESVSNPQRPPVIWIGAQECT WP_000145429.1 hydrogenase 2 small subunit [Salmonella enterica] Length: 372\nNumber of

predicted TMHs: 1\nExp number of AAs in TMHs: 24.51309\nExp number, first 60 AAs: 0.2682\nTotal prob of N-in: 0.14402\noutside 1 331\nTMhelix 332 354\ninside 355 372

19659 GCF_001242585.1_Salmonella_enterica_CVM_N45945_v1.0 Salmonella enterica subsp. enterica serovar Thompson Proteobacteria; Gammaproteobacteria; Enterobacterales; Enterobacteriaceae; Salmonella; Salmonella enterica; Salmonella enterica subsp. enterica MTGDNTLITSHGINRRDFMKLCAALAATMGLSSKAAAEMAESVSNPQRPPVIWIGAQECT WP_000145429.1 hydrogenase 2 small subunit [Salmonella enterica] Length: 372\nNumber of predicted TMHs: 1\nExp number of AAs in TMHs: 24.51309\nExp number, first 60 AAs: 0.2682\nTotal prob of N-in: 0.14402\noutside 1 331\nTMhelix 332 354\ninside 355 372

19660 GCF_001242585.1_Salmonella_enterica_CVM_N45945_v1.0 Salmonella enterica subsp. enterica serovar Thompson Proteobacteria; Gammaproteobacteria; Enterobacterales; Enterobacteriaceae; Salmonella; Salmonella enterica; Salmonella enterica subsp. enterica MNNEETFYQAMRRKGVTRRSFLKFCSLAATSLGLGAGMTPKIAWALENKPRIPVVWIHGL WP_001058310.1 hydrogenase [Salmonella enterica] Length: 372\nNumber of predicted TMHs: 2\nExp number of AAs in TMHs: 37.4567299999999\nExp number, first 60 AAs: 19.72694\nTotal prob of N-in: 0.96171\nPOSSIBLE N-term signal sequence\ninside 1 19\nTMhelix 20 39\noutside 40 325\nTMhelix 326 348\ninside 349 372

19661 GCF_002106065.1_ASM210606v1 Salmonella enterica subsp. enterica serovar Thompson Proteobacteria; Gammaproteobacteria; Enterobacterales; Enterobacteriaceae; Salmonella; Salmonella enterica; Salmonella enterica subsp. enterica MTGDNTLITSHGINRRDFMKLCAALAATMGLSSKAAAEMAESVSNPQRPPVIWIGAQECT WP_000145429.1 hydrogenase 2 small subunit [Salmonella enterica] Length: 372\nNumber of predicted TMHs: 1\nExp number of AAs in TMHs: 24.51309\nExp number, first 60 AAs: 0.2682\nTotal prob of N-in: 0.14402\noutside 1 331\nTMhelix 332 354\ninside 355 372

19662 GCF_002106375.1_ASM210637v1 Salmonella enterica subsp. enterica serovar Thompson Proteobacteria; Gammaproteobacteria; Enterobacterales; Enterobacteriaceae; Salmonella; Salmonella enterica; Salmonella enterica subsp. enterica MNNEETFYQAMRRKGVTRRSFLKFCSLAATSLGLGAGMTPKIAWALENKPRIPVVWIHGL WP_001058310.1 hydrogenase [Salmonella enterica] Length: 372\nNumber of predicted TMHs: 2\nExp number of AAs in TMHs: 37.4567299999999\nExp number, first 60 AAs: 19.72694\nTotal prob of N-in: 0.96171\nPOSSIBLE N-term signal sequence\ninside 1 19\nTMhelix 20 39\noutside 40 325\nTMhelix 326 348\ninside 349 372

19663 GCF_002106795.1_ASM210679v1 Salmonella enterica subsp. enterica serovar Thompson Proteobacteria; Gammaproteobacteria; Enterobacterales; Enterobacteriaceae; Salmonella; Salmonella enterica; Salmonella enterica subsp. enterica MTGDNTLITSHGINRRDFMKLCAALAATMGLSSKAAAEMAESVSNPQRPPVIWIGAQECT WP_000145429.1 hydrogenase 2 small subunit [Salmonella enterica] Length: 372\nNumber of predicted TMHs: 1\nExp number of AAs in TMHs: 24.51309\nExp number, first 60 AAs: 0.2682\nTotal prob of N-in: 0.14402\noutside 1 331\nTMhelix 332 354\ninside 355 372

19664 GCF_002107385.1_ASM210738v1 Salmonella enterica subsp. enterica serovar Thompson Proteobacteria; Gammaproteobacteria; Enterobacterales; Enterobacteriaceae; Salmonella; Salmonella enterica; Salmonella enterica subsp. enterica MNNEETFYQAMRRKGVTRRSFLKFCSLAATSLGLGAGMTPKIAWALENKPRIPVVWIHGL WP_001058310.1 hydrogenase [Salmonella enterica] Length: 372\nNumber of predicted TMHs: 2\nExp number of AAs in TMHs: 37.4567299999999\nExp number, first 60 AAs: 19.72694\nTotal prob of N-in: 0.96171\nPOSSIBLE N-term signal sequence\ninside 1 19\nTMhelix 20 39\noutside 40 325\nTMhelix 326 348\ninside 349 372

19665 GCF_002107615.1_ASM210761v1 Salmonella enterica subsp. enterica serovar Thompson Proteobacteria; Gammaproteobacteria; Enterobacterales; Enterobacteriaceae; Salmonella; Salmonella enterica; Salmonella enterica subsp. enterica MNNEETFYQAMRRKGVTRRSFLKFCSLAATSLGLGAGMTPKIAWALENKPRIPVVWIHGL WP_001058310.1 hydrogenase [Salmonella enterica] Length: 372\nNumber of predicted TMHs: 2\nExp number of AAs in TMHs: 37.4567299999999\nExp number, first 60 AAs: 19.72694\nTotal prob of N-in: 0.96171\nPOSSIBLE N-term signal sequence\ninside 1 19\nTMhelix 20 39\noutside 40 325\nTMhelix 326 348\ninside 349 372

19666 GCF_000973955.1_ABBSB1050-2.id13_v1.0 Salmonella enterica subsp. enterica serovar Thompson Proteobacteria; Gammaproteobacteria; Enterobacterales; Enterobacteriaceae; Salmonella; Salmonella enterica; Salmonella enterica subsp. enterica MTGDNTLITSHGINRRDFMKLCAALAATMGLSSKAAAEMAESVSNPQRPPVIWIGAQECT WP_000145429.1 hydrogenase 2 small subunit [Salmonella enterica] Length: 372\nNumber of

predicted TMHs:  1\nExp number of AAs in TMHs: 24.51309\nExp number, first 60 AAs:  0.2682\nTotal prob of N-in: 0.14402\noutside   1   331\nTMhelix   332   354\ninside   355   372

19667 GCF_002106375.1_ASM210637v1 Salmonella enterica subsp. enterica serovar Thompson Proteobacteria; Gammaproteobacteria; Enterobacterales; Enterobacteriaceae; Salmonella; Salmonella enterica; Salmonella enterica subsp. enterica MTGDNTLITSHGINRRDFMKLCAALAATMGLSSKAAAEMAESVSNPQRPPVIWIGAQECT WP_000145429.1 hydrogenase 2 small subunit [Salmonella enterica] Length: 372\nNumber of predicted TMHs:  1\nExp number of AAs in TMHs: 24.51309\nExp number, first 60 AAs:  0.2682\nTotal prob of N-in: 0.14402\noutside   1   331\nTMhelix   332   354\ninside   355   372

19668 GCF_002106065.1_ASM210606v1 Salmonella enterica subsp. enterica serovar Thompson Proteobacteria; Gammaproteobacteria; Enterobacterales; Enterobacteriaceae; Salmonella; Salmonella enterica; Salmonella enterica subsp. enterica MNNEETFYQAMRRKGVTRRSFLKFCSLAATSLGLGAGMTPKIAWALENKPRIPVVWIHGL WP_001058310.1 hydrogenase [Salmonella enterica] Length: 372\nNumber of predicted TMHs:  2\nExp number of AAs in TMHs: 37.4567299999999\nExp number, first 60 AAs:  19.72694\nTotal prob of N-in: 0.96171\nPOSSIBLE N-term signal sequence\ninside   1   19\nTMhelix   20   39\noutside   40   325\nTMhelix   326 348\ninside   349   372

19669 GCF_001478265.1_Salmonella_enterica_CVM_N37925-SQ_v1.0 Salmonella enterica subsp. enterica serovar Thompson Proteobacteria; Gammaproteobacteria; Enterobacterales; Enterobacteriaceae; Salmonella; Salmonella enterica; Salmonella enterica subsp. enterica MNNEETFYQAMRRKGVTRRSFLKFCSLAATSLGLGAGMTPKIAWALENKPRIPVVWIHGL WP_001058310.1 hydrogenase [Salmonella enterica] Length: 372\nNumber of predicted TMHs:  2\nExp number of AAs in TMHs: 37.4567299999999\nExp number, first 60 AAs:  19.72694\nTotal prob of N-in:   0.96171\nPOSSIBLE N-term signal sequence\ninside   1   19\nTMhelix   20   39\noutside   40   325\nTMhelix   326   348\ninside   349   372

19670 GCF_002061345.1_ASM206134v1 Salmonella enterica subsp. enterica serovar Thompson Proteobacteria; Gammaproteobacteria; Enterobacterales; Enterobacteriaceae; Salmonella; Salmonella enterica; Salmonella enterica subsp. enterica MTGDNTLITSHGINRRDFMKLCAALAATMGLSSKAAAEMAESVSNPQRPPVIWIGAQECT WP_000145429.1 hydrogenase 2 small subunit [Salmonella enterica] Length: 372\nNumber of predicted TMHs:  1\nExp number of AAs in TMHs: 24.51309\nExp number, first 60 AAs:  0.2682\nTotal prob of N-in: 0.14402\noutside   1   331\nTMhelix   332   354\ninside   355   372

19671 GCF_001478265.1_Salmonella_enterica_CVM_N37925-SQ_v1.0 Salmonella enterica subsp. enterica serovar Thompson Proteobacteria; Gammaproteobacteria; Enterobacterales; Enterobacteriaceae; Salmonella; Salmonella enterica; Salmonella enterica subsp. enterica MTGDNTLITSHGINRRDFMKLCAALAATMGLSSKAAAEMAESVSNPQRPPVIWIGAQECT WP_000145429.1 hydrogenase 2 small subunit [Salmonella enterica] Length: 372\nNumber of predicted TMHs:  1\nExp number of AAs in TMHs: 24.51309\nExp number, first 60 AAs:  0.2682\nTotal prob of N-in:   0.14402\noutside   1 331\nTMhelix   332   354\ninside   355   372

19672 GCF_001479545.1_Salmonella_enterica_CVM_N32781-SQ_v1.0 Salmonella enterica subsp. enterica serovar Thompson Proteobacteria; Gammaproteobacteria; Enterobacterales; Enterobacteriaceae; Salmonella; Salmonella enterica; Salmonella enterica subsp. enterica MTGDNTLITSHGINRRDFMKLCAALAATMGLSSKAAAEMAESVSNPQRPPVIWIGAQECT WP_000145429.1 hydrogenase 2 small subunit [Salmonella enterica] Length: 372\nNumber of predicted TMHs:  1\nExp number of AAs in TMHs: 24.51309\nExp number, first 60 AAs:  0.2682\nTotal prob of N-in:   0.14402\noutside   1 331\nTMhelix   332   354\ninside   355   372

19673 GCF_001479545.1_Salmonella_enterica_CVM_N32781-SQ_v1.0 Salmonella enterica subsp. enterica serovar Thompson Proteobacteria; Gammaproteobacteria; Enterobacterales; Enterobacteriaceae; Salmonella; Salmonella enterica; Salmonella enterica subsp. enterica MNNEETFYQAMRRKGVTRRSFLKFCSLAATSLGLGAGMTPKIAWALENKPRIPVVWIHGL WP_001058310.1 hydrogenase [Salmonella enterica] Length: 372\nNumber of predicted TMHs:  2\nExp number of AAs in TMHs: 37.4567299999999\nExp number, first 60 AAs:  19.72694\nTotal prob of N-in:   0.96171\nPOSSIBLE N-term signal sequence\ninside   1   19\nTMhelix   20   39\noutside   40   325\nTMhelix   326   348\ninside   349   372

19674 GCF_002059915.1_ASM205991v1 Salmonella enterica subsp. enterica serovar Thompson Proteobacteria; Gammaproteobacteria; Enterobacterales; Enterobacteriaceae; Salmonella; Salmonella enterica; Salmonella enterica subsp. enterica MNNEETFYQAMRRKGVTRRSFLKFCSLAATSLGLGAGMTPKIAWALENKPRIPVVWIHGL WP_001058311.1 hydrogenase [Salmonella enterica] Length: 372\nNumber of predicted TMHs:  2\nExp

number of AAs in TMHs: 37.4699899999999\nExp number, first 60 AAs: 19.72699\nTotal prob of N-in: 0.96171\nPOSSIBLE N-term signal sequence\ninside 1 19\nTMhelix 20 39\noutside 40 325\nTMhelix 326 348\ninside 349 372

19675 GCF_002061205.1_ASM206120v1 Salmonella enterica subsp. enterica serovar Thompson Proteobacteria; Gammaproteobacteria; Enterobacterales; Enterobacteriaceae; Salmonella; Salmonella enterica; Salmonella enterica subsp. enterica MTGDNTLITSHGINRRDFMKLCAALAATMGLSSKAAAEMAESVSNPQRPPVIWIGAQECT WP_000145429.1 hydrogenase 2 small subunit [Salmonella enterica] Length: 372\nNumber of predicted TMHs: 1\nExp number of AAs in TMHs: 24.51309\nExp number, first 60 AAs: 0.2682\nTotal prob of N-in: 0.14402\noutside 1 331\nTMhelix 332 354\ninside 355 372

19676 GCF_002059885.1_ASM205988v1 Salmonella enterica subsp. enterica serovar Thompson Proteobacteria; Gammaproteobacteria; Enterobacterales; Enterobacteriaceae; Salmonella; Salmonella enterica; Salmonella enterica subsp. enterica MNNEETFYQAMRRKGVTRRSFLKFCSLAATSLGLGAGMTPKIAWALENKPRIPVVWIHGL WP_001058312.1 hydrogenase [Salmonella enterica] Length: 372\nNumber of predicted TMHs: 2\nExp number of AAs in TMHs: 36.51738\nExp number, first 60 AAs: 19.72849\nTotal prob of N-in: 0.96170\nPOSSIBLE N-term signal sequence\ninside 1 19\nTMhelix 20 39\noutside 40 325\nTMhelix 326 348\ninside 349 372

19677 GCF_001691965.1_ASM169196v1 Salmonella enterica subsp. enterica serovar Thompson Proteobacteria; Gammaproteobacteria; Enterobacterales; Enterobacteriaceae; Salmonella; Salmonella enterica; Salmonella enterica subsp. enterica MNNEETFYQAMRRKGVTRRSFLKFCSLAATSLGLGAGMTPKIAWALENKPRIPVVWIHGL WP_001058310.1 hydrogenase [Salmonella enterica] Length: 372\nNumber of predicted TMHs: 2\nExp number of AAs in TMHs: 37.4567299999999\nExp number, first 60 AAs: 19.72694\nTotal prob of N-in: 0.96171\nPOSSIBLE N-term signal sequence\ninside 1 19\nTMhelix 20 39\noutside 40 325\nTMhelix 326 348\ninside 349 372

19678 GCF_001305835.1_ASM130583v1 Salmonella enterica subsp. enterica serovar Thompson Proteobacteria; Gammaproteobacteria; Enterobacterales; Enterobacteriaceae; Salmonella; Salmonella enterica; Salmonella enterica subsp. enterica MNNEETFYQAMRRKGVTRRSFLKFCSLAATSLGLGAGMTPKIAWALENKPRIPVVWIHGL WP_001058310.1 hydrogenase [Salmonella enterica] Length: 372\nNumber of predicted TMHs: 2\nExp number of AAs in TMHs: 37.4567299999999\nExp number, first 60 AAs: 19.72694\nTotal prob of N-in: 0.96171\nPOSSIBLE N-term signal sequence\ninside 1 19\nTMhelix 20 39\noutside 40 325\nTMhelix 326 348\ninside 349 372

19679 GCF_001953015.1_ASM195301v1 Salmonella enterica subsp. enterica serovar Thompson Proteobacteria; Gammaproteobacteria; Enterobacterales; Enterobacteriaceae; Salmonella; Salmonella enterica; Salmonella enterica subsp. enterica MNNEETFYQAMRRKGVTRRSFLKFCSLAATSLGLGAGMTPKIAWALENKPRIPVVWIHGL WP_001058310.1 hydrogenase [Salmonella enterica] Length: 372\nNumber of predicted TMHs: 2\nExp number of AAs in TMHs: 37.4567299999999\nExp number, first 60 AAs: 19.72694\nTotal prob of N-in: 0.96171\nPOSSIBLE N-term signal sequence\ninside 1 19\nTMhelix 20 39\noutside 40 325\nTMhelix 326 348\ninside 349 372

19680 GCF_002059885.1_ASM205988v1 Salmonella enterica subsp. enterica serovar Thompson Proteobacteria; Gammaproteobacteria; Enterobacterales; Enterobacteriaceae; Salmonella; Salmonella enterica; Salmonella enterica subsp. enterica MTGDNTLITSHGINRRDFMKLCAALAATMGLSSNAAAEMAESVSNPQRPPVIWIGAQECT WP_080214612.1 hydrogenase 2 small subunit [Salmonella enterica] Length: 372\nNumber of predicted TMHs: 1\nExp number of AAs in TMHs: 25.2552\nExp number, first 60 AAs: 1.05097\nTotal prob of N-in: 0.17767\noutside 1 331\nTMhelix 332 354\ninside 355 372

19681 GCF_001691965.1_ASM169196v1 Salmonella enterica subsp. enterica serovar Thompson Proteobacteria; Gammaproteobacteria; Enterobacterales; Enterobacteriaceae; Salmonella; Salmonella enterica; Salmonella enterica subsp. enterica MTGDNTLITSHGINRRDFMKLCAALAATMGLSSKAAAEMAESVSNPQRPPVIWIGAQECT WP_000145429.1 hydrogenase 2 small subunit [Salmonella enterica] Length: 372\nNumber of predicted TMHs: 1\nExp number of AAs in TMHs: 24.51309\nExp number, first 60 AAs: 0.2682\nTotal prob of N-in: 0.14402\noutside 1 331\nTMhelix 332 354\ninside 355 372

19682 GCF_001305835.1_ASM130583v1 Salmonella enterica subsp. enterica serovar Thompson Proteobacteria; Gammaproteobacteria; Enterobacterales; Enterobacteriaceae; Salmonella; Salmonella enterica; Salmonella enterica subsp. enterica MTGDNTLITSHGINRRDFMKLCAALAATMGLSSKAAAEMAESVSNPQRPPVIWIGAQECT WP_000145429.1 hydrogenase 2 small subunit [Salmonella enterica] Length: 372\nNumber of

predicted TMHs:  1\nExp number of AAs in TMHs: 24.51309\nExp number, first 60 AAs:  0.2682\nTotal prob of N-in: 0.14402\noutside   1   331\nTMhelix   332   354\ninside   355   372

19683 GCF_002106665.1_ASM210666v1 Salmonella enterica subsp. enterica serovar Thompson Proteobacteria; Gammaproteobacteria; Enterobacterales; Enterobacteriaceae; Salmonella; Salmonella enterica; Salmonella enterica subsp. enterica MTGDNTLITSHGINRRDFMKLCAALAATMGLSSKAAAEMAESVSNPQRPPVIWIGAQECT WP_000145429.1 hydrogenase 2 small subunit [Salmonella enterica] Length: 372\nNumber of predicted TMHs:  1\nExp number of AAs in TMHs: 24.51309\nExp number, first 60 AAs:  0.2682\nTotal prob of N-in: 0.14402\noutside   1   331\nTMhelix   332   354\ninside   355   372

19684 GCF_002108215.1_ASM210821v1 Salmonella enterica subsp. enterica serovar Thompson Proteobacteria; Gammaproteobacteria; Enterobacterales; Enterobacteriaceae; Salmonella; Salmonella enterica; Salmonella enterica subsp. enterica MNNEETFYQAMRRKGVTRRSFLKFCSLAATSLGLGAGMTPKIAWALENKPRIPVVWIHGL WP_001058310.1 hydrogenase [Salmonella enterica] Length: 372\nNumber of predicted TMHs:  2\nExp number of AAs in TMHs: 37.4567299999999\nExp number, first 60 AAs:  19.72694\nTotal prob of N-in: 0.96171\nPOSSIBLE N-term signal sequence\ninside   1   19\nTMhelix   20   39\noutside   40   325\nTMhelix   326   348\ninside   349   372

19685 GCF_002108215.1_ASM210821v1 Salmonella enterica subsp. enterica serovar Thompson Proteobacteria; Gammaproteobacteria; Enterobacterales; Enterobacteriaceae; Salmonella; Salmonella enterica; Salmonella enterica subsp. enterica MTGDNTLITSHGINRRDFMKLCAALAATMGLSSKAAAEMAESVSNPQRPPVIWIGAQECT WP_000145429.1 hydrogenase 2 small subunit [Salmonella enterica] Length: 372\nNumber of predicted TMHs:  1\nExp number of AAs in TMHs: 24.51309\nExp number, first 60 AAs:  0.2682\nTotal prob of N-in: 0.14402\noutside   1   331\nTMhelix   332   354\ninside   355   372

19686 GCF_002106635.1_ASM210663v1 Salmonella enterica subsp. enterica serovar Thompson Proteobacteria; Gammaproteobacteria; Enterobacterales; Enterobacteriaceae; Salmonella; Salmonella enterica; Salmonella enterica subsp. enterica MNNEETFYQAMRRKGVTRRSFLKFCSLAATSLGLGAGMTPKIAWALENKPRIPVVWIHGL WP_001058310.1 hydrogenase [Salmonella enterica] Length: 372\nNumber of predicted TMHs:  2\nExp number of AAs in TMHs: 37.4567299999999\nExp number, first 60 AAs:  19.72694\nTotal prob of N-in: 0.96171\nPOSSIBLE N-term signal sequence\ninside   1   19\nTMhelix   20   39\noutside   40   325\nTMhelix   326   348\ninside   349   372

19687 GCF_002106665.1_ASM210666v1 Salmonella enterica subsp. enterica serovar Thompson Proteobacteria; Gammaproteobacteria; Enterobacterales; Enterobacteriaceae; Salmonella; Salmonella enterica; Salmonella enterica subsp. enterica MNNEETFYQAMRRKGVTRRSFLKFCSLAATSLGLGAGMTPKIAWALENKPRIPVVWIHGL WP_001058310.1 hydrogenase [Salmonella enterica] Length: 372\nNumber of predicted TMHs:  2\nExp number of AAs in TMHs: 37.4567299999999\nExp number, first 60 AAs:  19.72694\nTotal prob of N-in: 0.96171\nPOSSIBLE N-term signal sequence\ninside   1   19\nTMhelix   20   39\noutside   40   325\nTMhelix   326   348\ninside   349   372

19688 GCF_002059565.1_ASM205956v1 Salmonella enterica subsp. enterica serovar Thompson Proteobacteria; Gammaproteobacteria; Enterobacterales; Enterobacteriaceae; Salmonella; Salmonella enterica; Salmonella enterica subsp. enterica MNNEETFYQAMRRKGVTRRSFLKFCSLAATSLGLGAGMTPKIAWALENKPRIPVVWIHGL WP_001058310.1 hydrogenase [Salmonella enterica] Length: 372\nNumber of predicted TMHs:  2\nExp number of AAs in TMHs: 37.4567299999999\nExp number, first 60 AAs:  19.72694\nTotal prob of N-in: 0.96171\nPOSSIBLE N-term signal sequence\ninside   1   19\nTMhelix   20   39\noutside   40   325\nTMhelix   326   348\ninside   349   372

19689 GCF_002105905.1_ASM210590v1 Salmonella enterica subsp. enterica serovar Thompson Proteobacteria; Gammaproteobacteria; Enterobacterales; Enterobacteriaceae; Salmonella; Salmonella enterica; Salmonella enterica subsp. enterica MNNEETFYQAMRRKGVTRRSFLKFCSLAATSLGLGAGMTPKIAWALENKPRIPVVWIHGL WP_001058310.1 hydrogenase [Salmonella enterica] Length: 372\nNumber of predicted TMHs:  2\nExp number of AAs in TMHs: 37.4567299999999\nExp number, first 60 AAs:  19.72694\nTotal prob of N-in: 0.96171\nPOSSIBLE N-term signal sequence\ninside   1   19\nTMhelix   20   39\noutside   40   325\nTMhelix   326   348\ninside   349   372

19690 GCF_002059915.1_ASM205991v1 Salmonella enterica subsp. enterica serovar Thompson Proteobacteria; Gammaproteobacteria; Enterobacterales; Enterobacteriaceae; Salmonella; Salmonella enterica; Salmonella enterica subsp. enterica MTGDNTLITSHGINRRDFMKLCAALAATMGLSSKAAAEMAESVSNPQRPPVIWIGAQECT WP_000145429.1 hydrogenase 2 small subunit [Salmonella enterica] Length: 372\nNumber of

predicted TMHs:  1\nExp number of AAs in TMHs: 24.51309\nExp number, first 60 AAs:  0.2682\nTotal prob of N-in: 0.14402\noutside   1   331\nTMhelix   332   354\ninside   355   372

19691 GCF_002059565.1_ASM205956v1 Salmonella enterica subsp. enterica serovar Thompson Proteobacteria; Gammaproteobacteria; Enterobacterales; Enterobacteriaceae; Salmonella; Salmonella enterica; Salmonella enterica subsp. enterica MTGDNTLITSHGINRRDFMKLCAALAATMGLSSKAAAEMAESVSNPQRPPVIWIGAQECT WP_000145429.1 hydrogenase 2 small subunit [Salmonella enterica] Length: 372\nNumber of predicted TMHs:  1\nExp number of AAs in TMHs: 24.51309\nExp number, first 60 AAs:  0.2682\nTotal prob of N-in: 0.14402\noutside   1   331\nTMhelix   332   354\ninside   355   372

19692 GCF_002061345.1_ASM206134v1 Salmonella enterica subsp. enterica serovar Thompson Proteobacteria; Gammaproteobacteria; Enterobacterales; Enterobacteriaceae; Salmonella; Salmonella enterica; Salmonella enterica subsp. enterica MNNEETFYQAMRRKGVTRRSFLKFCSLAATSLGLGAGMTPKIAWALENKPRIPVVWIHGL WP_001058310.1 hydrogenase [Salmonella enterica] Length: 372\nNumber of predicted TMHs:  2\nExp number of AAs in TMHs: 37.4567299999999\nExp number, first 60 AAs:  19.72694\nTotal prob of N-in: 0.96171\nPOSSIBLE N-term signal sequence\ninside   1   19\nTMhelix   20   39\noutside   40   325\nTMhelix   326 348\ninside   349   372

19693 GCF_002105955.1_ASM210595v1 Salmonella enterica subsp. enterica serovar Thompson Proteobacteria; Gammaproteobacteria; Enterobacterales; Enterobacteriaceae; Salmonella; Salmonella enterica; Salmonella enterica subsp. enterica MNNEETFYQAMRRKGVTRRSFLKFCSLAATSLGLGAGMTPKIAWALENKPRIPVVWIHGL WP_001058310.1 hydrogenase [Salmonella enterica] Length: 372\nNumber of predicted TMHs:  2\nExp number of AAs in TMHs: 37.4567299999999\nExp number, first 60 AAs:  19.72694\nTotal prob of N-in: 0.96171\nPOSSIBLE N-term signal sequence\ninside   1   19\nTMhelix   20   39\noutside   40   325\nTMhelix   326 348\ninside   349   372

19694 GCF_002059875.1_ASM205987v1 Salmonella enterica subsp. enterica serovar Thompson Proteobacteria; Gammaproteobacteria; Enterobacterales; Enterobacteriaceae; Salmonella; Salmonella enterica; Salmonella enterica subsp. enterica MTGDNTLITSHGINRRDFMKLCAALAATMGLSSNAAAEMAESVSNPQRPPVIWIGAQECT WP_080214612.1 hydrogenase 2 small subunit [Salmonella enterica] Length: 372\nNumber of predicted TMHs:  1\nExp number of AAs in TMHs: 25.2552\nExp number, first 60 AAs:  1.05097\nTotal prob of N-in: 0.17767\noutside   1   331\nTMhelix   332   354\ninside   355   372

19695 GCF_002105955.1_ASM210595v1 Salmonella enterica subsp. enterica serovar Thompson Proteobacteria; Gammaproteobacteria; Enterobacterales; Enterobacteriaceae; Salmonella; Salmonella enterica; Salmonella enterica subsp. enterica MTGDNTLITSHGINRRDFMKLCAALAATMGLSSKAAAEMAESVSNPQRPPVIWIGAQECT WP_000145429.1 hydrogenase 2 small subunit [Salmonella enterica] Length: 372\nNumber of predicted TMHs:  1\nExp number of AAs in TMHs: 24.51309\nExp number, first 60 AAs:  0.2682\nTotal prob of N-in: 0.14402\noutside   1   331\nTMhelix   332   354\ninside   355   372

19696 GCF_002106235.1_ASM210623v1 Salmonella enterica subsp. enterica serovar Thompson Proteobacteria; Gammaproteobacteria; Enterobacterales; Enterobacteriaceae; Salmonella; Salmonella enterica; Salmonella enterica subsp. enterica MTGDNTLITSHGINRRDFMKLCAALAATMGLSSKAAAEMAESVSNPQRPPVIWIGAQECT WP_000145429.1 hydrogenase 2 small subunit [Salmonella enterica] Length: 372\nNumber of predicted TMHs:  1\nExp number of AAs in TMHs: 24.51309\nExp number, first 60 AAs:  0.2682\nTotal prob of N-in: 0.14402\noutside   1   331\nTMhelix   332   354\ninside   355   372

19697 GCF_002062295.1_ASM206229v1 Salmonella enterica subsp. enterica serovar Thompson Proteobacteria; Gammaproteobacteria; Enterobacterales; Enterobacteriaceae; Salmonella; Salmonella enterica; Salmonella enterica subsp. enterica MTGDNTLITSHGINRRDFMKLCAALAATMGLSSKAAAEMAESVSNPQRPPVIWIGAQECT WP_000145429.1 hydrogenase 2 small subunit [Salmonella enterica] Length: 372\nNumber of predicted TMHs:  1\nExp number of AAs in TMHs: 24.51309\nExp number, first 60 AAs:  0.2682\nTotal prob of N-in: 0.14402\noutside   1   331\nTMhelix   332   354\ninside   355   372

19698 GCF_002061205.1_ASM206120v1 Salmonella enterica subsp. enterica serovar Thompson Proteobacteria; Gammaproteobacteria; Enterobacterales; Enterobacteriaceae; Salmonella; Salmonella enterica; Salmonella enterica subsp. enterica MNNEETFYQAMRRKGVTRRSFLKFCSLAATSLGLGAGMTPKIAWALENKPRIPVVWIHGL WP_001058310.1 hydrogenase [Salmonella enterica] Length: 372\nNumber of predicted TMHs:  2\nExp number of AAs in TMHs: 37.4567299999999\nExp number, first 60 AAs:  19.72694\nTotal prob of N-in: 0.96171\nPOSSIBLE N-term signal sequence\ninside   1   19\nTMhelix   20   39\noutside   40   325\nTMhelix   326 348\ninside   349   372

19699 GCF_002105905.1_ASM210590v1 Salmonella enterica subsp. enterica serovar Thompson Proteobacteria; Gammaproteobacteria; Enterobacterales; Enterobacteriaceae; Salmonella; Salmonella enterica; Salmonella enterica subsp. enterica MTGDNTLITSHGINRRDFMKLCAALAATMGLSSKAAAEMAESVSNPQRPPVIWIGAQECT WP_000145429.1 hydrogenase 2 small subunit [Salmonella enterica] Length: 372\nNumber of predicted TMHs: 1\nExp number of AAs in TMHs: 24.51309\nExp number, first 60 AAs: 0.2682\nTotal prob of N-in: 0.14402\noutside 1 331\nTMhelix 332 354\ninside 355 372

19700 GCF_002062295.1_ASM206229v1 Salmonella enterica subsp. enterica serovar Thompson Proteobacteria; Gammaproteobacteria; Enterobacterales; Enterobacteriaceae; Salmonella; Salmonella enterica; Salmonella enterica subsp. enterica MNNEETFYQAMRRKGVTRRSFLKFCSLAATSLGLGAGMTPKIAWALENKPRIPVVWIHGL WP_001058310.1 hydrogenase [Salmonella enterica] Length: 372\nNumber of predicted TMHs: 2\nExp number of AAs in TMHs: 37.4567299999999\nExp number, first 60 AAs: 19.72694\nTotal prob of N-in: 0.96171\nPOSSIBLE N-term signal sequence\ninside 1 19\nTMhelix 20 39\noutside 40 325\nTMhelix 326 348\ninside 349 372

19701 GCF_002059875.1_ASM205987v1 Salmonella enterica subsp. enterica serovar Thompson Proteobacteria; Gammaproteobacteria; Enterobacterales; Enterobacteriaceae; Salmonella; Salmonella enterica; Salmonella enterica subsp. enterica MNNEETFYQAMRRKGVTRRSFLKFCSLAATSLGLGAGMTPKIAWALENKPRIPVVWIHGL WP_001058312.1 hydrogenase [Salmonella enterica] Length: 372\nNumber of predicted TMHs: 2\nExp number of AAs in TMHs: 36.51738\nExp number, first 60 AAs: 19.72849\nTotal prob of N-in: 0.96170\nPOSSIBLE N-term signal sequence\ninside 1 19\nTMhelix 20 39\noutside 40 325\nTMhelix 326 348\ninside 349 372

19702 GCF_002106635.1_ASM210663v1 Salmonella enterica subsp. enterica serovar Thompson Proteobacteria; Gammaproteobacteria; Enterobacterales; Enterobacteriaceae; Salmonella; Salmonella enterica; Salmonella enterica subsp. enterica MTGDNTLITSHGINRRDFMKLCAALAATMGLSSKAAAEMAESVSNPQRPPVIWIGAQECT WP_000145429.1 hydrogenase 2 small subunit [Salmonella enterica] Length: 372\nNumber of predicted TMHs: 1\nExp number of AAs in TMHs: 24.51309\nExp number, first 60 AAs: 0.2682\nTotal prob of N-in: 0.14402\noutside 1 331\nTMhelix 332 354\ninside 355 372

19703 GCF_002106235.1_ASM210623v1 Salmonella enterica subsp. enterica serovar Thompson Proteobacteria; Gammaproteobacteria; Enterobacterales; Enterobacteriaceae; Salmonella; Salmonella enterica; Salmonella enterica subsp. enterica MNNEETFYQAMRRKGVTRRSFLKFCSLAATSLGLGAGMTPKIAWALENKPRIPVVWIHGL WP_001058310.1 hydrogenase [Salmonella enterica] Length: 372\nNumber of predicted TMHs: 2\nExp number of AAs in TMHs: 37.4567299999999\nExp number, first 60 AAs: 19.72694\nTotal prob of N-in: 0.96171\nPOSSIBLE N-term signal sequence\ninside 1 19\nTMhelix 20 39\noutside 40 325\nTMhelix 326 348\ninside 349 372

19704 GCF_001240735.1_Salmonella_enterica_CVM_N43821_v1.0 Salmonella enterica subsp. enterica serovar Thompson Proteobacteria; Gammaproteobacteria; Enterobacterales; Enterobacteriaceae; Salmonella; Salmonella enterica; Salmonella enterica subsp. enterica MTGDNTLITSHGINRRDFMKLCAALAATMGLSSKAAAEMAESVSNPQRPPVIWIGAQECT WP_000145429.1 hydrogenase 2 small subunit [Salmonella enterica] Length: 372\nNumber of predicted TMHs: 1\nExp number of AAs in TMHs: 24.51309\nExp number, first 60 AAs: 0.2682\nTotal prob of N-in: 0.14402\noutside 1 331\nTMhelix 332 354\ninside 355 372

19705 GCF_001240555.1_Salmonella_enterica_CVM_N43474_v1.0 Salmonella enterica subsp. enterica serovar Thompson Proteobacteria; Gammaproteobacteria; Enterobacterales; Enterobacteriaceae; Salmonella; Salmonella enterica; Salmonella enterica subsp. enterica MTGDNTLITSHGINRRDFMKLCAALAATMGLSSKAAAEMAESVSNPQRPPVIWIGAQECT WP_000145429.1 hydrogenase 2 small subunit [Salmonella enterica] Length: 372\nNumber of predicted TMHs: 1\nExp number of AAs in TMHs: 24.51309\nExp number, first 60 AAs: 0.2682\nTotal prob of N-in: 0.14402\noutside 1 331\nTMhelix 332 354\ninside 355 372

19706 GCF_001240525.1_Salmonella_enterica_CVM_N43472_v1.0 Salmonella enterica subsp. enterica serovar Thompson Proteobacteria; Gammaproteobacteria; Enterobacterales; Enterobacteriaceae; Salmonella; Salmonella enterica; Salmonella enterica subsp. enterica MTGDNTLITSHGINRRDFMKLCAALAATMGLSSKAAAEMAESVSNPQRPPVIWIGAQECT WP_000145429.1 hydrogenase 2 small subunit [Salmonella enterica] Length: 372\nNumber of predicted TMHs: 1\nExp number of AAs in TMHs: 24.51309\nExp number, first 60 AAs: 0.2682\nTotal prob of N-in: 0.14402\noutside 1 331\nTMhelix 332 354\ninside 355 372

19707 GCF_001305815.1_ASM130581v1 Salmonella enterica subsp. enterica serovar Thompson Proteobacteria; Gammaproteobacteria; Enterobacterales; Enterobacteriaceae; Salmonella; Salmonella enterica; Salmonella enterica subsp. enterica MTGDNTLITSHGINRRDFMKLCAALAATMGLSSKAAAEMAESVSNPQRPPVIWIGAQECT WP_000145429.1 hydrogenase 2 small subunit [Salmonella enterica] Length: 372\nNumber of predicted TMHs: 1\nExp number of AAs in TMHs: 24.51309\nExp number, first 60 AAs: 0.2682\nTotal prob of N-in: 0.14402\noutside 1 331\nTMhelix 332 354\ninside 355 372

19708 GCF_001305815.1_ASM130581v1 Salmonella enterica subsp. enterica serovar Thompson Proteobacteria; Gammaproteobacteria; Enterobacterales; Enterobacteriaceae; Salmonella; Salmonella enterica; Salmonella enterica subsp. enterica MNNEETFYQAMRRKGVTRRSFLKFCSLAATSLGLGAGMTPKIAWALENKPRIPVVWIHGL WP_001058310.1 hydrogenase [Salmonella enterica] Length: 372\nNumber of predicted TMHs: 2\nExp number of AAs in TMHs: 37.4567299999999\nExp number, first 60 AAs: 19.72694\nTotal prob of N-in: 0.96171\nPOSSIBLE N-term signal sequence\ninside 1 19\nTMhelix 20 39\noutside 40 325\nTMhelix 326 348\ninside 349 372

19709 GCF_000973955.1_ABBSB1050-2.id13_v1.0 Salmonella enterica subsp. enterica serovar Thompson Proteobacteria; Gammaproteobacteria; Enterobacterales; Enterobacteriaceae; Salmonella; Salmonella enterica; Salmonella enterica subsp. enterica MNNEETFYQAMRRKGVTRRSFLKFCSLAATSLGLGAGMTPKIAWALENKPRIPVVWIHGL WP_001058310.1 hydrogenase [Salmonella enterica] Length: 372\nNumber of predicted TMHs: 2\nExp number of AAs in TMHs: 37.4567299999999\nExp number, first 60 AAs: 19.72694\nTotal prob of N-in: 0.96171\nPOSSIBLE N-term signal sequence\ninside 1 19\nTMhelix 20 39\noutside 40 325\nTMhelix 326 348\ninside 349 372

19710 GCF_001240735.1_Salmonella_enterica_CVM_N43821_v1.0 Salmonella enterica subsp. enterica serovar Thompson Proteobacteria; Gammaproteobacteria; Enterobacterales; Enterobacteriaceae; Salmonella; Salmonella enterica; Salmonella enterica subsp. enterica MNNEETFYQAMRRKGVTRRSFLKFCSLAATSLGLGAGMTPKIAWALENKPRIPVVWIHGL WP_001058310.1 hydrogenase [Salmonella enterica] Length: 372\nNumber of predicted TMHs: 2\nExp number of AAs in TMHs: 37.4567299999999\nExp number, first 60 AAs: 19.72694\nTotal prob of N-in: 0.96171\nPOSSIBLE N-term signal sequence\ninside 1 19\nTMhelix 20 39\noutside 40 325\nTMhelix 326 348\ninside 349 372

19711 GCF_001240555.1_Salmonella_enterica_CVM_N43474_v1.0 Salmonella enterica subsp. enterica serovar Thompson Proteobacteria; Gammaproteobacteria; Enterobacterales; Enterobacteriaceae; Salmonella; Salmonella enterica; Salmonella enterica subsp. enterica MNNEETFYQAMRRKGVTRRSFLKFCSLAATSLGLGAGMTPKIAWALENKPRIPVVWIHGL WP_001058310.1 hydrogenase [Salmonella enterica] Length: 372\nNumber of predicted TMHs: 2\nExp number of AAs in TMHs: 37.4567299999999\nExp number, first 60 AAs: 19.72694\nTotal prob of N-in: 0.96171\nPOSSIBLE N-term signal sequence\ninside 1 19\nTMhelix 20 39\noutside 40 325\nTMhelix 326 348\ninside 349 372

19712 GCF_001240525.1_Salmonella_enterica_CVM_N43472_v1.0 Salmonella enterica subsp. enterica serovar Thompson Proteobacteria; Gammaproteobacteria; Enterobacterales; Enterobacteriaceae; Salmonella; Salmonella enterica; Salmonella enterica subsp. enterica MNNEETFYQAMRRKGVTRRSFLKFCSLAATSLGLGAGMTPKIAWALENKPRIPVVWIHGL WP_001058310.1 hydrogenase [Salmonella enterica] Length: 372\nNumber of predicted TMHs: 2\nExp number of AAs in TMHs: 37.4567299999999\nExp number, first 60 AAs: 19.72694\nTotal prob of N-in: 0.96171\nPOSSIBLE N-term signal sequence\ninside 1 19\nTMhelix 20 39\noutside 40 325\nTMhelix 326 348\ninside 349 372

19713 GCF_002106145.1_ASM210614v1 Salmonella enterica subsp. enterica serovar Thompson Proteobacteria; Gammaproteobacteria; Enterobacterales; Enterobacteriaceae; Salmonella; Salmonella enterica; Salmonella enterica subsp. enterica MNNEETFYQAMRRKGVTRRSFLKFCSLAATSLGLGAGMTPKIAWALENKPRIPVVWIHGL WP_001058310.1 hydrogenase [Salmonella enterica] Length: 372\nNumber of predicted TMHs: 2\nExp number of AAs in TMHs: 37.4567299999999\nExp number, first 60 AAs: 19.72694\nTotal prob of N-in: 0.96171\nPOSSIBLE N-term signal sequence\ninside 1 19\nTMhelix 20 39\noutside 40 325\nTMhelix 326 348\ninside 349 372

19714 GCF_002106795.1_ASM210679v1 Salmonella enterica subsp. enterica serovar Thompson Proteobacteria; Gammaproteobacteria; Enterobacterales; Enterobacteriaceae; Salmonella; Salmonella enterica; Salmonella enterica subsp. enterica MNNEETFYQAMRRKGVTRRSFLKFCSLAATSLGLGAGMTPKIAWALENKPRIPVVWIHGL WP_001058310.1 hydrogenase [Salmonella enterica] Length: 372\nNumber of predicted TMHs: 2\nExp number of AAs in TMHs: 37.4567299999999\nExp number, first 60 AAs: 19.72694\nTotal prob of N-in:

0.96171\nPOSSIBLE N-term signal sequence\ninside 1 19\nTMhelix 20 39\noutside 40 325\nTMhelix 326 348\ninside 349 372

19715 GCF_002107385.1_ASM210738v1 Salmonella enterica subsp. enterica serovar Thompson Proteobacteria; Gammaproteobacteria; Enterobacterales; Enterobacteriaceae; Salmonella; Salmonella enterica; Salmonella enterica subsp. enterica MTGDNTLITSHGINRRDFMKLCAALAATMGLSSKAAAEMAESVSNPQRPPVIWIGAQECT WP_000145429.1 hydrogenase 2 small subunit [Salmonella enterica] Length: 372\nNumber of predicted TMHs: 1\nExp number of AAs in TMHs: 24.51309\nExp number, first 60 AAs: 0.2682\nTotal prob of N-in: 0.14402\noutside 1 331\nTMhelix 332 354\ninside 355 372

19716 GCF_002063795.1_ASM206379v1 Salmonella enterica subsp. enterica serovar Thompson Proteobacteria; Gammaproteobacteria; Enterobacterales; Enterobacteriaceae; Salmonella; Salmonella enterica; Salmonella enterica subsp. enterica MTGDNTLITSHGINRRDFMKLCAALAATMGLSSKAAAEMAESVSNPQRPPVIWIGAQECT WP_000145429.1 hydrogenase 2 small subunit [Salmonella enterica] Length: 372\nNumber of predicted TMHs: 1\nExp number of AAs in TMHs: 24.51309\nExp number, first 60 AAs: 0.2682\nTotal prob of N-in: 0.14402\noutside 1 331\nTMhelix 332 354\ninside 355 372

19717 GCF_002063795.1_ASM206379v1 Salmonella enterica subsp. enterica serovar Thompson Proteobacteria; Gammaproteobacteria; Enterobacterales; Enterobacteriaceae; Salmonella; Salmonella enterica; Salmonella enterica subsp. enterica MNNEETFYQAMRRKGVTRRSFLKFCSLAATSLGLGAGMTPKIAWALENKPRIPVVWIHGL WP_001058310.1 hydrogenase [Salmonella enterica] Length: 372\nNumber of predicted TMHs: 2\nExp number of AAs in TMHs: 37.4567299999999\nExp number, first 60 AAs: 19.72694\nTotal prob of N-in: 0.96171\nPOSSIBLE N-term signal sequence\ninside 1 19\nTMhelix 20 39\noutside 40 325\nTMhelix 326 348\ninside 349 372

19718 GCF_002107305.1_ASM210730v1 Salmonella enterica subsp. enterica serovar Thompson Proteobacteria; Gammaproteobacteria; Enterobacterales; Enterobacteriaceae; Salmonella; Salmonella enterica; Salmonella enterica subsp. enterica MNNEETFYQAMRRKGVTRRSFLKFCSLAATSLGLGAGMTPKIAWALENKPRIPVVWIHGL WP_001058310.1 hydrogenase [Salmonella enterica] Length: 372\nNumber of predicted TMHs: 2\nExp number of AAs in TMHs: 37.4567299999999\nExp number, first 60 AAs: 19.72694\nTotal prob of N-in: 0.96171\nPOSSIBLE N-term signal sequence\ninside 1 19\nTMhelix 20 39\noutside 40 325\nTMhelix 326 348\ninside 349 372

19719 GCF_002107615.1_ASM210761v1 Salmonella enterica subsp. enterica serovar Thompson Proteobacteria; Gammaproteobacteria; Enterobacterales; Enterobacteriaceae; Salmonella; Salmonella enterica; Salmonella enterica subsp. enterica MTGDNTLITSHGINRRDFMKLCAALAATMGLSSKAAAEMAESVSNPQRPPVIWIGAQECT WP_000145429.1 hydrogenase 2 small subunit [Salmonella enterica] Length: 372\nNumber of predicted TMHs: 1\nExp number of AAs in TMHs: 24.51309\nExp number, first 60 AAs: 0.2682\nTotal prob of N-in: 0.14402\noutside 1 331\nTMhelix 332 354\ninside 355 372

19720 GCF_002106145.1_ASM210614v1 Salmonella enterica subsp. enterica serovar Thompson Proteobacteria; Gammaproteobacteria; Enterobacterales; Enterobacteriaceae; Salmonella; Salmonella enterica; Salmonella enterica subsp. enterica MTGDNTLITSHGINRRDFMKLCAALAATMGLSSKAAAEMAESVSNPQRPPVIWIGAQECT WP_000145429.1 hydrogenase 2 small subunit [Salmonella enterica] Length: 372\nNumber of predicted TMHs: 1\nExp number of AAs in TMHs: 24.51309\nExp number, first 60 AAs: 0.2682\nTotal prob of N-in: 0.14402\noutside 1 331\nTMhelix 332 354\ninside 355 372

19721 GCF_002107305.1_ASM210730v1 Salmonella enterica subsp. enterica serovar Thompson Proteobacteria; Gammaproteobacteria; Enterobacterales; Enterobacteriaceae; Salmonella; Salmonella enterica; Salmonella enterica subsp. enterica MTGDNTLITSHGINRRDFMKLCAALAATMGLSSKAAAEMAESVSNPQRPPVIWIGAQECT WP_000145429.1 hydrogenase 2 small subunit [Salmonella enterica] Length: 372\nNumber of predicted TMHs: 1\nExp number of AAs in TMHs: 24.51309\nExp number, first 60 AAs: 0.2682\nTotal prob of N-in: 0.14402\noutside 1 331\nTMhelix 332 354\ninside 355 372

19722 GCF_001692105.1_ASM169210v1 Salmonella enterica subsp. enterica serovar Thompson Proteobacteria; Gammaproteobacteria; Enterobacterales; Enterobacteriaceae; Salmonella; Salmonella enterica; Salmonella enterica subsp. enterica MTGDNTLITSHGINRRDFMKLCAALAATMGLSSKAAAEMAESVSNPQRPPVIWIGAQECT WP_000145429.1 hydrogenase 2 small subunit [Salmonella enterica] Length: 372\nNumber of predicted TMHs: 1\nExp number of AAs in TMHs: 24.51309\nExp number, first 60 AAs: 0.2682\nTotal prob of N-in: 0.14402\noutside 1 331\nTMhelix 332 354\ninside 355 372

19723	GCF_001692105.1_ASM169210v1	Salmonella enterica subsp. enterica serovar Thompson	Proteobacteria; Gammaproteobacteria; Enterobacterales; Enterobacteriaceae; Salmonella; Salmonella enterica; Salmonella enterica subsp. enterica	MNNEETFYQAMRRKGVTRRSFLKFCSLAATSLGLGAGMTPKIAWALENKPRIPVVWIHGL	WP_001058310.1	hydrogenase [Salmonella enterica]	Length: 372\nNumber of predicted TMHs:  2\nExp number of AAs in TMHs: 37.4567299999999\nExp number, first 60 AAs:  19.72694\nTotal prob of N-in: 0.96171\nPOSSIBLE N-term signal sequence\ninside   1   19\nTMhelix   20   39\noutside   40   325\nTMhelix   326   348\ninside   349   372

19724	GCF_001953015.1_ASM195301v1	Salmonella enterica subsp. enterica serovar Thompson	Proteobacteria; Gammaproteobacteria; Enterobacterales; Enterobacteriaceae; Salmonella; Salmonella enterica; Salmonella enterica subsp. enterica	MTGDNTLITSHGINRRDFMKLCAALAATMGLSSKAAAEMAESVSNPQRPPVIWIGAQECT	WP_000145429.1	hydrogenase 2 small subunit [Salmonella enterica]	Length: 372\nNumber of predicted TMHs:  1\nExp number of AAs in TMHs: 24.51309\nExp number, first 60 AAs:  0.2682\nTotal prob of N-in: 0.14402\noutside   1   331\nTMhelix   332   354\ninside   355   372

19725	GCF_002034395.1_ASM203439v1	Salmonella enterica subsp. enterica serovar Tornow	Proteobacteria; Gammaproteobacteria; Enterobacterales; Enterobacteriaceae; Salmonella; Salmonella enterica; Salmonella enterica subsp. enterica	MNNEETFYQAMRRKGVTRRSFLKFCSLAATSLGLGAGMTPKIAWALENKPRIPVVWIHGL	WP_079841295.1	[Ni/Fe] hydrogenase small subunit [Salmonella enterica]	Length: 372\nNumber of predicted TMHs:  2\nExp number of AAs in TMHs: 36.54746\nExp number, first 60 AAs:  19.72849\nTotal prob of N-in: 0.96170\nPOSSIBLE N-term signal sequence\ninside   1   19\nTMhelix   20   39\noutside   40   325\nTMhelix   326   348\ninside   349   372

19726	GCF_002034395.1_ASM203439v1	Salmonella enterica subsp. enterica serovar Tornow	Proteobacteria; Gammaproteobacteria; Enterobacterales; Enterobacteriaceae; Salmonella; Salmonella enterica; Salmonella enterica subsp. enterica	MTGDNTLITSHGINRRDFMKLCAALAATMGLSSKAAAEMAESVSNPQRPPVIWIGAQECT	WP_000145429.1	hydrogenase 2 small subunit [Salmonella enterica]	Length: 372\nNumber of predicted TMHs:  1\nExp number of AAs in TMHs: 24.51309\nExp number, first 60 AAs:  0.2682\nTotal prob of N-in: 0.14402\noutside   1   331\nTMhelix   332   354\ninside   355   372

19727	GCF_001367455.1_10607_2_30	Salmonella enterica subsp. enterica serovar Typhi	Proteobacteria; Gammaproteobacteria; Enterobacterales; Enterobacteriaceae; Salmonella; Salmonella enterica; Salmonella enterica subsp. enterica	MTGDNTLITSHGINRRDFMKLCAALAATMGLSSKAATEMAESVSNPQRPPVIWIGAQECT	WP_001782304.1	hydrogenase 2 small subunit [Salmonella enterica]	Length: 372\nNumber of predicted TMHs:  1\nExp number of AAs in TMHs: 24.41173\nExp number, first 60 AAs:  0.16124\nTotal prob of N-in: 0.13947\noutside   1   331\nTMhelix   332   354\ninside   355   372

19728	GCF_001362555.1_10562_2_6	Salmonella enterica subsp. enterica serovar Typhi	Proteobacteria; Gammaproteobacteria; Enterobacterales; Enterobacteriaceae; Salmonella; Salmonella enterica; Salmonella enterica subsp. enterica	MTGDNTLITSHGINRRDFMKLCAALAATMGLSSKAAAEMAESVSNPQRPPVIWIGAQECT	WP_000145429.1	hydrogenase 2 small subunit [Salmonella enterica]	Length: 372\nNumber of predicted TMHs:  1\nExp number of AAs in TMHs: 24.51309\nExp number, first 60 AAs:  0.2682\nTotal prob of N-in: 0.14402\noutside   1   331\nTMhelix   332   354\ninside   355   372

19729	GCF_001362515.1_10540_1_27	Salmonella enterica subsp. enterica serovar Typhi	Proteobacteria; Gammaproteobacteria; Enterobacterales; Enterobacteriaceae; Salmonella; Salmonella enterica; Salmonella enterica subsp. enterica	MTGDNTLITSHGINRRDFMKLCAALAATMGLSSKAAAEMAESVSNPQRPPVIWIGAQECT	WP_000145429.1	hydrogenase 2 small subunit [Salmonella enterica]	Length: 372\nNumber of predicted TMHs:  1\nExp number of AAs in TMHs: 24.51309\nExp number, first 60 AAs:  0.2682\nTotal prob of N-in: 0.14402\noutside   1   331\nTMhelix   332   354\ninside   355   372

19730	GCF_001362595.1_10607_1_62	Salmonella enterica subsp. enterica serovar Typhi	Proteobacteria; Gammaproteobacteria; Enterobacterales; Enterobacteriaceae; Salmonella; Salmonella enterica; Salmonella enterica subsp. enterica	MTGDNTLITSHGINRRDFMKLCAALAATMGLSSKAAAEMAESVSNPQRPPVIWIGAQECT	WP_000145429.1	hydrogenase 2 small subunit [Salmonella enterica]	Length: 372\nNumber of predicted TMHs:  1\nExp number of AAs in TMHs: 24.51309\nExp number, first 60 AAs:  0.2682\nTotal prob of N-in: 0.14402\noutside   1   331\nTMhelix   332   354\ninside   355   372

19731	GCF_001362475.1_10540_1_48	Salmonella enterica subsp. enterica serovar Typhi	Proteobacteria; Gammaproteobacteria; Enterobacterales; Enterobacteriaceae; Salmonella; Salmonella enterica; Salmonella enterica subsp. enterica	MTGDNTLITSHGINRRDFMKLCAALAATMGLSSKAAAEMAESVSNPQRPPVIWIGAQECT

WP_000145429.1 hydrogenase 2 small subunit [Salmonella enterica] Length: 372\nNumber of predicted TMHs: 1\nExp number of AAs in TMHs: 24.51309\nExp number, first 60 AAs: 0.2682\nTotal prob of N-in: 0.14402\noutside 1 331\nTMhelix 332 354\ninside 355 372

19732 GCF_001362635.1_10540_1_42 Salmonella enterica subsp. enterica serovar Typhi Proteobacteria; Gammaproteobacteria; Enterobacterales; Enterobacteriaceae; Salmonella; Salmonella enterica; Salmonella enterica subsp. enterica MTGDNTLITSHGINRRDFMKLCAALAATMGLSSKAAAEMAESVSNPQRPPVIWIGAQECT WP_000145429.1 hydrogenase 2 small subunit [Salmonella enterica] Length: 372\nNumber of predicted TMHs: 1\nExp number of AAs in TMHs: 24.51309\nExp number, first 60 AAs: 0.2682\nTotal prob of N-in: 0.14402\noutside 1 331\nTMhelix 332 354\ninside 355 372

19733 GCF_001362435.1_10561_2_88 Salmonella enterica subsp. enterica serovar Typhi Proteobacteria; Gammaproteobacteria; Enterobacterales; Enterobacteriaceae; Salmonella; Salmonella enterica; Salmonella enterica subsp. enterica MTGDNTLITSHGINRRDFMKLCAALAATMGLSSKAAAEMAESVSNPQRPPVIWIGAQECT WP_000145429.1 hydrogenase 2 small subunit [Salmonella enterica] Length: 372\nNumber of predicted TMHs: 1\nExp number of AAs in TMHs: 24.51309\nExp number, first 60 AAs: 0.2682\nTotal prob of N-in: 0.14402\noutside 1 331\nTMhelix 332 354\ninside 355 372

19734 GCF_001362395.1_10426_1_1 Salmonella enterica subsp. enterica serovar Typhi Proteobacteria; Gammaproteobacteria; Enterobacterales; Enterobacteriaceae; Salmonella; Salmonella enterica; Salmonella enterica subsp. enterica MTGDNTLITSHGINRRDFMKLCAALAATMGLSSKAAAEMAESVSNPQRPPVIWIGAQECT WP_000145429.1 hydrogenase 2 small subunit [Salmonella enterica] Length: 372\nNumber of predicted TMHs: 1\nExp number of AAs in TMHs: 24.51309\nExp number, first 60 AAs: 0.2682\nTotal prob of N-in: 0.14402\noutside 1 331\nTMhelix 332 354\ninside 355 372

19735 GCF_001362675.1_10562_2_83 Salmonella enterica subsp. enterica serovar Typhi Proteobacteria; Gammaproteobacteria; Enterobacterales; Enterobacteriaceae; Salmonella; Salmonella enterica; Salmonella enterica subsp. enterica MTGDNTLITSHGINRRDFMKLCAALAATMGLSSKAAAEMAESVSNPQRPPVIWIGAQECT WP_000145429.1 hydrogenase 2 small subunit [Salmonella enterica] Length: 372\nNumber of predicted TMHs: 1\nExp number of AAs in TMHs: 24.51309\nExp number, first 60 AAs: 0.2682\nTotal prob of N-in: 0.14402\noutside 1 331\nTMhelix 332 354\ninside 355 372

19736 GCF_001362715.1_10540_1_28 Salmonella enterica subsp. enterica serovar Typhi Proteobacteria; Gammaproteobacteria; Enterobacterales; Enterobacteriaceae; Salmonella; Salmonella enterica; Salmonella enterica subsp. enterica MTGDNTLITSHGINRRDFMKLCAALAATMGLSSKAAAEMAESVSNPQRPPVIWIGAQECT WP_000145429.1 hydrogenase 2 small subunit [Salmonella enterica] Length: 372\nNumber of predicted TMHs: 1\nExp number of AAs in TMHs: 24.51309\nExp number, first 60 AAs: 0.2682\nTotal prob of N-in: 0.14402\noutside 1 331\nTMhelix 332 354\ninside 355 372

19737 GCF_001362755.1_10426_1_10 Salmonella enterica subsp. enterica serovar Typhi Proteobacteria; Gammaproteobacteria; Enterobacterales; Enterobacteriaceae; Salmonella; Salmonella enterica; Salmonella enterica subsp. enterica MTGDNTLITSHGINRRDFMKLCAALAATMGLSSKAAAEMAESVSNPQRPPVIWIGAQECT WP_000145429.1 hydrogenase 2 small subunit [Salmonella enterica] Length: 372\nNumber of predicted TMHs: 1\nExp number of AAs in TMHs: 24.51309\nExp number, first 60 AAs: 0.2682\nTotal prob of N-in: 0.14402\noutside 1 331\nTMhelix 332 354\ninside 355 372

19738 GCF_001362355.1_10349_1_87 Salmonella enterica subsp. enterica serovar Typhi Proteobacteria; Gammaproteobacteria; Enterobacterales; Enterobacteriaceae; Salmonella; Salmonella enterica; Salmonella enterica subsp. enterica MTGDNTLITSHGINRRDFMKLCAALAATMGLSSKAATEMAESVSNPQRPPVIWIGAQECT WP_001782304.1 hydrogenase 2 small subunit [Salmonella enterica] Length: 372\nNumber of predicted TMHs: 1\nExp number of AAs in TMHs: 24.41173\nExp number, first 60 AAs: 0.16124\nTotal prob of N-in: 0.13947\noutside 1 331\nTMhelix 332 354\ninside 355 372

19739 GCF_001362795.1_10426_1_28 Salmonella enterica subsp. enterica serovar Typhi Proteobacteria; Gammaproteobacteria; Enterobacterales; Enterobacteriaceae; Salmonella; Salmonella enterica; Salmonella enterica subsp. enterica MTGDNTLITSHGINRRDFMKLCAALAATMGLSSKAAAEMAESVSNPQRPPVIWIGAQECT WP_000145429.1 hydrogenase 2 small subunit [Salmonella enterica] Length: 372\nNumber of predicted TMHs: 1\nExp number of AAs in TMHs: 24.51309\nExp number, first 60 AAs: 0.2682\nTotal prob of N-in: 0.14402\noutside 1 331\nTMhelix 332 354\ninside 355 372

19740 GCF_001362835.1_10561_2_82 Salmonella enterica subsp. enterica serovar Typhi Proteobacteria; Gammaproteobacteria; Enterobacterales; Enterobacteriaceae; Salmonella; Salmonella enterica; Salmonella enterica subsp. enterica MTGDNTLITSHGINRRDFMKLCAALAATMGLSSKAAAEMAESVSNPQRPPVIWIGAQECT WP_000145429.1 hydrogenase 2 small subunit [Salmonella enterica] Length: 372\nNumber of predicted TMHs: 1\nExp number of AAs in TMHs: 24.51309\nExp number, first 60 AAs: 0.2682\nTotal prob of N-in: 0.14402\noutside 1 331\nTMhelix 332 354\ninside 355 372

19741 GCF_001362875.1_10540_1_12 Salmonella enterica subsp. enterica serovar Typhi Proteobacteria; Gammaproteobacteria; Enterobacterales; Enterobacteriaceae; Salmonella; Salmonella enterica; Salmonella enterica subsp. enterica MTGDNTLITSHGINRRDFMKLCAALAATMGLSSKAAAEMAESVSNPQRPPVIWIGAQECT WP_000145429.1 hydrogenase 2 small subunit [Salmonella enterica] Length: 372\nNumber of predicted TMHs: 1\nExp number of AAs in TMHs: 24.51309\nExp number, first 60 AAs: 0.2682\nTotal prob of N-in: 0.14402\noutside 1 331\nTMhelix 332 354\ninside 355 372

19742 GCF_001362895.1_10540_1_70 Salmonella enterica subsp. enterica serovar Typhi Proteobacteria; Gammaproteobacteria; Enterobacterales; Enterobacteriaceae; Salmonella; Salmonella enterica; Salmonella enterica subsp. enterica MTGDNTLITSHGINRRDFMKLCAALAATMGLSSKAAAEMAESVSNPQRPPVIWIGAQECT WP_000145429.1 hydrogenase 2 small subunit [Salmonella enterica] Length: 372\nNumber of predicted TMHs: 1\nExp number of AAs in TMHs: 24.51309\nExp number, first 60 AAs: 0.2682\nTotal prob of N-in: 0.14402\noutside 1 331\nTMhelix 332 354\ninside 355 372

19743 GCF_001362935.1_10493_1_69 Salmonella enterica subsp. enterica serovar Typhi Proteobacteria; Gammaproteobacteria; Enterobacterales; Enterobacteriaceae; Salmonella; Salmonella enterica; Salmonella enterica subsp. enterica MTGDNTLITSHGINRRDFMKLCAALAATMGLSSKAAAEMAESVSNPQRPPVIWIGAQECT WP_000145429.1 hydrogenase 2 small subunit [Salmonella enterica] Length: 372\nNumber of predicted TMHs: 1\nExp number of AAs in TMHs: 24.51309\nExp number, first 60 AAs: 0.2682\nTotal prob of N-in: 0.14402\noutside 1 331\nTMhelix 332 354\ninside 355 372

19744 GCF_001362975.1_10608_2_67 Salmonella enterica subsp. enterica serovar Typhi Proteobacteria; Gammaproteobacteria; Enterobacterales; Enterobacteriaceae; Salmonella; Salmonella enterica; Salmonella enterica subsp. enterica MTGDNTLITSHGINRRDFMKLCAALAATMGLSSKAAAEMAESVSNPQRPPVIWIGAQECT WP_000145429.1 hydrogenase 2 small subunit [Salmonella enterica] Length: 372\nNumber of predicted TMHs: 1\nExp number of AAs in TMHs: 24.51309\nExp number, first 60 AAs: 0.2682\nTotal prob of N-in: 0.14402\noutside 1 331\nTMhelix 332 354\ninside 355 372

19745 GCF_001363015.1_10561_2_19 Salmonella enterica subsp. enterica serovar Typhi Proteobacteria; Gammaproteobacteria; Enterobacterales; Enterobacteriaceae; Salmonella; Salmonella enterica; Salmonella enterica subsp. enterica MTGDNTLITSHGINRRDFMKLCAALAATMGLSSKAAAEMAESVSNPQRPPVIWIGAQECT WP_000145429.1 hydrogenase 2 small subunit [Salmonella enterica] Length: 372\nNumber of predicted TMHs: 1\nExp number of AAs in TMHs: 24.51309\nExp number, first 60 AAs: 0.2682\nTotal prob of N-in: 0.14402\noutside 1 331\nTMhelix 332 354\ninside 355 372

19746 GCF_001363095.1_10608_2_12 Salmonella enterica subsp. enterica serovar Typhi Proteobacteria; Gammaproteobacteria; Enterobacterales; Enterobacteriaceae; Salmonella; Salmonella enterica; Salmonella enterica subsp. enterica MTGDNTLITSHGINRRDFMKLCAALAATMGLSSKAAAEMAESVSNPQRPPVIWIGAQECT WP_000145429.1 hydrogenase 2 small subunit [Salmonella enterica] Length: 372\nNumber of predicted TMHs: 1\nExp number of AAs in TMHs: 24.51309\nExp number, first 60 AAs: 0.2682\nTotal prob of N-in: 0.14402\noutside 1 331\nTMhelix 332 354\ninside 355 372

19747 GCF_001363055.1_10540_1_35 Salmonella enterica subsp. enterica serovar Typhi Proteobacteria; Gammaproteobacteria; Enterobacterales; Enterobacteriaceae; Salmonella; Salmonella enterica; Salmonella enterica subsp. enterica MTGDNTLITSHGINRRDFMKLCAALAATMGLSSKAAAEMAESVSNPQRPPVIWIGAQECT WP_000145429.1 hydrogenase 2 small subunit [Salmonella enterica] Length: 372\nNumber of predicted TMHs: 1\nExp number of AAs in TMHs: 24.51309\nExp number, first 60 AAs: 0.2682\nTotal prob of N-in: 0.14402\noutside 1 331\nTMhelix 332 354\ninside 355 372

19748 GCF_001363135.1_10562_2_33 Salmonella enterica subsp. enterica serovar Typhi Proteobacteria; Gammaproteobacteria; Enterobacterales; Enterobacteriaceae; Salmonella; Salmonella enterica; Salmonella enterica subsp. enterica MTGDNTLITSHGINRRDFMKLCAALAATMGLSSKAAAEMAESVSNPQRPPVIWIGAQECT WP_000145429.1 hydrogenase 2 small subunit [Salmonella enterica] Length: 372\nNumber of

predicted TMHs:  1\nExp number of AAs in TMHs: 24.51309\nExp number, first 60 AAs:  0.2682\nTotal prob of N-in: 0.14402\noutside   1   331\nTMhelix   332   354\ninside   355   372

19749 GCF_001363175.1_10426_1_95 Salmonella enterica subsp. enterica serovar Typhi Proteobacteria; Gammaproteobacteria; Enterobacterales; Enterobacteriaceae; Salmonella; Salmonella enterica; Salmonella enterica subsp. enterica MTGDNTLITSHGINRRDFMKLCAALAATMGLSSKAAAEMAESVSNPQRPPVIWIGAQECT WP_000145429.1 hydrogenase 2 small subunit [Salmonella enterica] Length: 372\nNumber of predicted TMHs:  1\nExp number of AAs in TMHs: 24.51309\nExp number, first 60 AAs:  0.2682\nTotal prob of N-in: 0.14402\noutside   1   331\nTMhelix   332   354\ninside   355   372

19750 GCF_001363215.1_10426_1_52 Salmonella enterica subsp. enterica serovar Typhi Proteobacteria; Gammaproteobacteria; Enterobacterales; Enterobacteriaceae; Salmonella; Salmonella enterica; Salmonella enterica subsp. enterica MTGDNTLITSHGINRRDFMKLCAALAATMGLSSKAAAEMAESVSNPQRPPVIWIGAQECT WP_000145429.1 hydrogenase 2 small subunit [Salmonella enterica] Length: 372\nNumber of predicted TMHs:  1\nExp number of AAs in TMHs: 24.51309\nExp number, first 60 AAs:  0.2682\nTotal prob of N-in: 0.14402\noutside   1   331\nTMhelix   332   354\ninside   355   372

19751 GCF_001363255.1_10607_1_45 Salmonella enterica subsp. enterica serovar Typhi Proteobacteria; Gammaproteobacteria; Enterobacterales; Enterobacteriaceae; Salmonella; Salmonella enterica; Salmonella enterica subsp. enterica MTGDNTLITSHGINRRDFMKLCAALAATMGLSSKAAAEMAESVSNPQRPPVIWIGAQECT WP_000145429.1 hydrogenase 2 small subunit [Salmonella enterica] Length: 372\nNumber of predicted TMHs:  1\nExp number of AAs in TMHs: 24.51309\nExp number, first 60 AAs:  0.2682\nTotal prob of N-in: 0.14402\noutside   1   331\nTMhelix   332   354\ninside   355   372

19752 GCF_001363295.1_10540_1_54 Salmonella enterica subsp. enterica serovar Typhi Proteobacteria; Gammaproteobacteria; Enterobacterales; Enterobacteriaceae; Salmonella; Salmonella enterica; Salmonella enterica subsp. enterica MTGDNTLITSHGINRRDFMKLCAALAATMGLSSKAAAEMAESVSNPQRPPVIWIGAQECT WP_000145429.1 hydrogenase 2 small subunit [Salmonella enterica] Length: 372\nNumber of predicted TMHs:  1\nExp number of AAs in TMHs: 24.51309\nExp number, first 60 AAs:  0.2682\nTotal prob of N-in: 0.14402\noutside   1   331\nTMhelix   332   354\ninside   355   372

19753 GCF_001363335.1_10540_1_19 Salmonella enterica subsp. enterica serovar Typhi Proteobacteria; Gammaproteobacteria; Enterobacterales; Enterobacteriaceae; Salmonella; Salmonella enterica; Salmonella enterica subsp. enterica MTGDNTLITSHGINRRDFMKLCAALAATMGLSSKAAAEMAESVSNPQRPPVIWIGAQECT WP_000145429.1 hydrogenase 2 small subunit [Salmonella enterica] Length: 372\nNumber of predicted TMHs:  1\nExp number of AAs in TMHs: 24.51309\nExp number, first 60 AAs:  0.2682\nTotal prob of N-in: 0.14402\noutside   1   331\nTMhelix   332   354\ninside   355   372

19754 GCF_001363375.1_10608_2_18 Salmonella enterica subsp. enterica serovar Typhi Proteobacteria; Gammaproteobacteria; Enterobacterales; Enterobacteriaceae; Salmonella; Salmonella enterica; Salmonella enterica subsp. enterica MTGDNTLITSHGINRRDFMKLCAALAATMGLSSKAAAEMAESVSNPQRPPVIWIGAQECT WP_000145429.1 hydrogenase 2 small subunit [Salmonella enterica] Length: 372\nNumber of predicted TMHs:  1\nExp number of AAs in TMHs: 24.51309\nExp number, first 60 AAs:  0.2682\nTotal prob of N-in: 0.14402\noutside   1   331\nTMhelix   332   354\ninside   355   372

19755 GCF_001363415.1_10561_2_76 Salmonella enterica subsp. enterica serovar Typhi Proteobacteria; Gammaproteobacteria; Enterobacterales; Enterobacteriaceae; Salmonella; Salmonella enterica; Salmonella enterica subsp. enterica MTGDNTLITSHGINRRDFMKLCAALAATMGLSSKAAAEMAESVSNPQRPPVIWIGAQECT WP_000145429.1 hydrogenase 2 small subunit [Salmonella enterica] Length: 372\nNumber of predicted TMHs:  1\nExp number of AAs in TMHs: 24.51309\nExp number, first 60 AAs:  0.2682\nTotal prob of N-in: 0.14402\noutside   1   331\nTMhelix   332   354\ninside   355   372

19756 GCF_001363595.1_10492_1_30 Salmonella enterica subsp. enterica serovar Typhi Proteobacteria; Gammaproteobacteria; Enterobacterales; Enterobacteriaceae; Salmonella; Salmonella enterica; Salmonella enterica subsp. enterica MTGDNTLITSHGINRRDFMKLCAALAATMGLSSKAAAEMAESVSNPQRPPVIWIGAQECT WP_000145429.1 hydrogenase 2 small subunit [Salmonella enterica] Length: 372\nNumber of predicted TMHs:  1\nExp number of AAs in TMHs: 24.51309\nExp number, first 60 AAs:  0.2682\nTotal prob of N-in: 0.14402\noutside   1   331\nTMhelix   332   354\ninside   355   372

19757 GCF_001363535.1_10561_2_47 Salmonella enterica subsp. enterica serovar Typhi Proteobacteria; Gammaproteobacteria; Enterobacterales; Enterobacteriaceae; Salmonella; Salmonella enterica;

Salmonella enterica subsp. enterica	MTGDNTLITSHGINRRDFMKLCAALAATMGLSSKAAAEMAESVSNPQRPPVIWIGAQECT
WP_000145429.1	hydrogenase 2 small subunit [Salmonella enterica]	Length: 372\nNumber of predicted TMHs: 1\nExp number of AAs in TMHs: 24.51309\nExp number, first 60 AAs: 0.2682\nTotal prob of N-in: 0.14402\noutside 1 331\nTMhelix 332 354\ninside 355 372

19758	GCF_001363495.1_10540_1_94	Salmonella enterica subsp. enterica serovar Typhi
Proteobacteria; Gammaproteobacteria; Enterobacterales; Enterobacteriaceae; Salmonella; Salmonella enterica; Salmonella enterica subsp. enterica	MTGDNTLITSHGINRRDFMKLCAALAATMGLSSKAAAEMAESVSNPQRPPVIWIGAQECT
WP_000145429.1	hydrogenase 2 small subunit [Salmonella enterica]	Length: 372\nNumber of predicted TMHs: 1\nExp number of AAs in TMHs: 24.51309\nExp number, first 60 AAs: 0.2682\nTotal prob of N-in: 0.14402\noutside 1 331\nTMhelix 332 354\ninside 355 372

19759	GCF_001363455.1_10426_1_69	Salmonella enterica subsp. enterica serovar Typhi
Proteobacteria; Gammaproteobacteria; Enterobacterales; Enterobacteriaceae; Salmonella; Salmonella enterica; Salmonella enterica subsp. enterica	MTGDNTLITSHGINRRDFMKLCAALAATMGLSSKAAAEMAESVSNPQRPPVIWIGAQECT
WP_000145429.1	hydrogenase 2 small subunit [Salmonella enterica]	Length: 372\nNumber of predicted TMHs: 1\nExp number of AAs in TMHs: 24.51309\nExp number, first 60 AAs: 0.2682\nTotal prob of N-in: 0.14402\noutside 1 331\nTMhelix 332 354\ninside 355 372

19760	GCF_001363575.1_10426_1_88	Salmonella enterica subsp. enterica serovar Typhi
Proteobacteria; Gammaproteobacteria; Enterobacterales; Enterobacteriaceae; Salmonella; Salmonella enterica; Salmonella enterica subsp. enterica	MTGDNTLITSHGINRRDFMKLCAALAATMGLSSKAAAEMAESVSNPQRPPVIWIGAQECT
WP_000145429.1	hydrogenase 2 small subunit [Salmonella enterica]	Length: 372\nNumber of predicted TMHs: 1\nExp number of AAs in TMHs: 24.51309\nExp number, first 60 AAs: 0.2682\nTotal prob of N-in: 0.14402\noutside 1 331\nTMhelix 332 354\ninside 355 372

19761	GCF_001363635.1_10562_2_43	Salmonella enterica subsp. enterica serovar Typhi
Proteobacteria; Gammaproteobacteria; Enterobacterales; Enterobacteriaceae; Salmonella; Salmonella enterica; Salmonella enterica subsp. enterica	MTGDNTLITSHGINRRDFMKLCAALAATMGLSSKAAAEMAESVSNPQRPPVIWIGAQECT
WP_000145429.1	hydrogenase 2 small subunit [Salmonella enterica]	Length: 372\nNumber of predicted TMHs: 1\nExp number of AAs in TMHs: 24.51309\nExp number, first 60 AAs: 0.2682\nTotal prob of N-in: 0.14402\noutside 1 331\nTMhelix 332 354\ninside 355 372

19762	GCF_001363675.1_10541_2_38	Salmonella enterica subsp. enterica serovar Typhi
Proteobacteria; Gammaproteobacteria; Enterobacterales; Enterobacteriaceae; Salmonella; Salmonella enterica; Salmonella enterica subsp. enterica	MTGDNTLITSHGINRRDFMKLCAALAATMGLSSKAAAEMAESVSNPQRPPVIWIGAQECT
WP_000145429.1	hydrogenase 2 small subunit [Salmonella enterica]	Length: 372\nNumber of predicted TMHs: 1\nExp number of AAs in TMHs: 24.51309\nExp number, first 60 AAs: 0.2682\nTotal prob of N-in: 0.14402\noutside 1 331\nTMhelix 332 354\ninside 355 372

19763	GCF_001363715.1_10540_1_49	Salmonella enterica subsp. enterica serovar Typhi
Proteobacteria; Gammaproteobacteria; Enterobacterales; Enterobacteriaceae; Salmonella; Salmonella enterica; Salmonella enterica subsp. enterica	MTGDNTLITSHGINRRDFMKLCAALAATMGLSSKAAAEMAESVSNPQRPPVIWIGAQECT
WP_000145429.1	hydrogenase 2 small subunit [Salmonella enterica]	Length: 372\nNumber of predicted TMHs: 1\nExp number of AAs in TMHs: 24.51309\nExp number, first 60 AAs: 0.2682\nTotal prob of N-in: 0.14402\noutside 1 331\nTMhelix 332 354\ninside 355 372

19764	GCF_001363755.1_10561_2_90	Salmonella enterica subsp. enterica serovar Typhi
Proteobacteria; Gammaproteobacteria; Enterobacterales; Enterobacteriaceae; Salmonella; Salmonella enterica; Salmonella enterica subsp. enterica	MTGDNTLITSHGINRRDFMKLCAALAATMGLSSKAAAEMAESVSNPQRPPVIWIGAQECT
WP_000145429.1	hydrogenase 2 small subunit [Salmonella enterica]	Length: 372\nNumber of predicted TMHs: 1\nExp number of AAs in TMHs: 24.51309\nExp number, first 60 AAs: 0.2682\nTotal prob of N-in: 0.14402\noutside 1 331\nTMhelix 332 354\ninside 355 372

19765	GCF_001363795.1_10561_2_86	Salmonella enterica subsp. enterica serovar Typhi
Proteobacteria; Gammaproteobacteria; Enterobacterales; Enterobacteriaceae; Salmonella; Salmonella enterica; Salmonella enterica subsp. enterica	MTGDNTLITSHGINRRDFMKLCAALAATMGLSSKAAAEMAESVSNPQRPPVIWIGAQECT
WP_000145429.1	hydrogenase 2 small subunit [Salmonella enterica]	Length: 372\nNumber of predicted TMHs: 1\nExp number of AAs in TMHs: 24.51309\nExp number, first 60 AAs: 0.2682\nTotal prob of N-in: 0.14402\noutside 1 331\nTMhelix 332 354\ninside 355 372

19766 GCF_001363835.1_10607_2_20 Salmonella enterica subsp. enterica serovar Typhi Proteobacteria; Gammaproteobacteria; Enterobacterales; Enterobacteriaceae; Salmonella; Salmonella enterica; Salmonella enterica subsp. enterica MTGDNTLITSHGINRRDFMKLCAALAATMGLSSKAAAEMAESVSNPQRPPVIWIGAQECT WP_000145429.1 hydrogenase 2 small subunit [Salmonella enterica] Length: 372\nNumber of predicted TMHs: 1\nExp number of AAs in TMHs: 24.51309\nExp number, first 60 AAs: 0.2682\nTotal prob of N-in: 0.14402\noutside 1 331\nTMhelix 332 354\ninside 355 372

19767 GCF_001363915.1_10562_2_49 Salmonella enterica subsp. enterica serovar Typhi Proteobacteria; Gammaproteobacteria; Enterobacterales; Enterobacteriaceae; Salmonella; Salmonella enterica; Salmonella enterica subsp. enterica MTGDNTLITSHGINRRDFMKLCAALAATMGLSSKAAAEMAESVSNPQRPPVIWIGAQECT WP_000145429.1 hydrogenase 2 small subunit [Salmonella enterica] Length: 372\nNumber of predicted TMHs: 1\nExp number of AAs in TMHs: 24.51309\nExp number, first 60 AAs: 0.2682\nTotal prob of N-in: 0.14402\noutside 1 331\nTMhelix 332 354\ninside 355 372

19768 GCF_001363875.1_10540_1_76 Salmonella enterica subsp. enterica serovar Typhi Proteobacteria; Gammaproteobacteria; Enterobacterales; Enterobacteriaceae; Salmonella; Salmonella enterica; Salmonella enterica subsp. enterica MTGDNTLITSHGINRRDFMKLCAALAATMGLSSKAAAEMAESVSNPQRPPVIWIGAQECT WP_000145429.1 hydrogenase 2 small subunit [Salmonella enterica] Length: 372\nNumber of predicted TMHs: 1\nExp number of AAs in TMHs: 24.51309\nExp number, first 60 AAs: 0.2682\nTotal prob of N-in: 0.14402\noutside 1 331\nTMhelix 332 354\ninside 355 372

19769 GCF_001363955.1_10426_1_38 Salmonella enterica subsp. enterica serovar Typhi Proteobacteria; Gammaproteobacteria; Enterobacterales; Enterobacteriaceae; Salmonella; Salmonella enterica; Salmonella enterica subsp. enterica MTGDNTLITSHGINRRDFMKLCAALAATMGLSSKAAAEMAESVSNPQRPPVIWIGAQECT WP_000145429.1 hydrogenase 2 small subunit [Salmonella enterica] Length: 372\nNumber of predicted TMHs: 1\nExp number of AAs in TMHs: 24.51309\nExp number, first 60 AAs: 0.2682\nTotal prob of N-in: 0.14402\noutside 1 331\nTMhelix 332 354\ninside 355 372

19770 GCF_001363995.1_10608_2_36 Salmonella enterica subsp. enterica serovar Typhi Proteobacteria; Gammaproteobacteria; Enterobacterales; Enterobacteriaceae; Salmonella; Salmonella enterica; Salmonella enterica subsp. enterica MTGDNTLITSHGINRRDFMKLCAALAATMGLSSKAAAEMAESVSNPQRPPVIWIGAQECT WP_000145429.1 hydrogenase 2 small subunit [Salmonella enterica] Length: 372\nNumber of predicted TMHs: 1\nExp number of AAs in TMHs: 24.51309\nExp number, first 60 AAs: 0.2682\nTotal prob of N-in: 0.14402\noutside 1 331\nTMhelix 332 354\ninside 355 372

19771 GCF_001364035.1_10607_2_28 Salmonella enterica subsp. enterica serovar Typhi Proteobacteria; Gammaproteobacteria; Enterobacterales; Enterobacteriaceae; Salmonella; Salmonella enterica; Salmonella enterica subsp. enterica MTGDNTLITSHGINRRDFMKLCAALAATMGLSSKAAAEMAESVSNPQRPPVIWIGAQECT WP_000145429.1 hydrogenase 2 small subunit [Salmonella enterica] Length: 372\nNumber of predicted TMHs: 1\nExp number of AAs in TMHs: 24.51309\nExp number, first 60 AAs: 0.2682\nTotal prob of N-in: 0.14402\noutside 1 331\nTMhelix 332 354\ninside 355 372

19772 GCF_001364075.1_10493_1_38 Salmonella enterica subsp. enterica serovar Typhi Proteobacteria; Gammaproteobacteria; Enterobacterales; Enterobacteriaceae; Salmonella; Salmonella enterica; Salmonella enterica subsp. enterica MTGDNTLITSHGINRRDFMKLCAALAATMGLSSKAAAEMAESVSNPQRPPVIWIGAQECT WP_000145429.1 hydrogenase 2 small subunit [Salmonella enterica] Length: 372\nNumber of predicted TMHs: 1\nExp number of AAs in TMHs: 24.51309\nExp number, first 60 AAs: 0.2682\nTotal prob of N-in: 0.14402\noutside 1 331\nTMhelix 332 354\ninside 355 372

19773 GCF_001364115.1_10426_1_74 Salmonella enterica subsp. enterica serovar Typhi Proteobacteria; Gammaproteobacteria; Enterobacterales; Enterobacteriaceae; Salmonella; Salmonella enterica; Salmonella enterica subsp. enterica MTGDNTLITSHGINRRDFMKLCAALAATMGLSSKAAAEMAESVSNPQRPPVIWIGAQECT WP_000145429.1 hydrogenase 2 small subunit [Salmonella enterica] Length: 372\nNumber of predicted TMHs: 1\nExp number of AAs in TMHs: 24.51309\nExp number, first 60 AAs: 0.2682\nTotal prob of N-in: 0.14402\noutside 1 331\nTMhelix 332 354\ninside 355 372

19774 GCF_001364155.1_10540_1_96 Salmonella enterica subsp. enterica serovar Typhi Proteobacteria; Gammaproteobacteria; Enterobacterales; Enterobacteriaceae; Salmonella; Salmonella enterica; Salmonella enterica subsp. enterica MTGDNTLITSHGINRRDFMKLCAALAATMGLSSKAAAEMAESVSNPQRPPVIWIGAQECT WP_000145429.1 hydrogenase 2 small subunit [Salmonella enterica] Length: 372\nNumber of

predicted TMHs:  1\nExp number of AAs in TMHs: 24.51309\nExp number, first 60 AAs:  0.2682\nTotal prob of N-in: 0.14402\noutside   1   331\nTMhelix   332   354\ninside   355   372

19775 GCF_001364195.1_10493_1_52 Salmonella enterica subsp. enterica serovar Typhi Proteobacteria; Gammaproteobacteria; Enterobacterales; Enterobacteriaceae; Salmonella; Salmonella enterica; Salmonella enterica subsp. enterica MTGDNTLITSHGINRRDFMKLCAALAATMGLSSKAAAEMAESVSNPQRPPVIWIGAQECT WP_000145429.1 hydrogenase 2 small subunit [Salmonella enterica] Length: 372\nNumber of predicted TMHs:  1\nExp number of AAs in TMHs: 24.51309\nExp number, first 60 AAs:  0.2682\nTotal prob of N-in: 0.14402\noutside   1   331\nTMhelix   332   354\ninside   355   372

19776 GCF_001364235.1_10607_1_75 Salmonella enterica subsp. enterica serovar Typhi Proteobacteria; Gammaproteobacteria; Enterobacterales; Enterobacteriaceae; Salmonella; Salmonella enterica; Salmonella enterica subsp. enterica MTGDNTLITSHGINRRDFMKLCAALAATMGLSSKAAAEMAESVSNPQRPPVIWIGAQECT WP_000145429.1 hydrogenase 2 small subunit [Salmonella enterica] Length: 372\nNumber of predicted TMHs:  1\nExp number of AAs in TMHs: 24.51309\nExp number, first 60 AAs:  0.2682\nTotal prob of N-in: 0.14402\noutside   1   331\nTMhelix   332   354\ninside   355   372

19777 GCF_001364275.1_10608_2_5 Salmonella enterica subsp. enterica serovar Typhi Proteobacteria; Gammaproteobacteria; Enterobacterales; Enterobacteriaceae; Salmonella; Salmonella enterica; Salmonella enterica subsp. enterica MTGDNTLITSHGINRRDFMKLCAALAATMGLSSKAAAEMAESVSNPQRPPVIWIGAQECT WP_000145429.1 hydrogenase 2 small subunit [Salmonella enterica] Length: 372\nNumber of predicted TMHs:  1\nExp number of AAs in TMHs: 24.51309\nExp number, first 60 AAs:  0.2682\nTotal prob of N-in: 0.14402\noutside   1   331\nTMhelix   332   354\ninside   355   372

19778 GCF_001364295.1_10540_1_8 Salmonella enterica subsp. enterica serovar Typhi Proteobacteria; Gammaproteobacteria; Enterobacterales; Enterobacteriaceae; Salmonella; Salmonella enterica; Salmonella enterica subsp. enterica MTGDNTLITSHGINRRDFMKLCAALAATMGLSSKAAAEMAESVSNPQRPPVIWIGAQECT WP_000145429.1 hydrogenase 2 small subunit [Salmonella enterica] Length: 372\nNumber of predicted TMHs:  1\nExp number of AAs in TMHs: 24.51309\nExp number, first 60 AAs:  0.2682\nTotal prob of N-in: 0.14402\noutside   1   331\nTMhelix   332   354\ninside   355   372

19779 GCF_001364335.1_10492_1_48 Salmonella enterica subsp. enterica serovar Typhi Proteobacteria; Gammaproteobacteria; Enterobacterales; Enterobacteriaceae; Salmonella; Salmonella enterica; Salmonella enterica subsp. enterica MTGDNTLITSHGINRRDFMKLCAALAATMGLSSKAAAEMAESVSNPQRPPVIWIGAQECT WP_000145429.1 hydrogenase 2 small subunit [Salmonella enterica] Length: 372\nNumber of predicted TMHs:  1\nExp number of AAs in TMHs: 24.51309\nExp number, first 60 AAs:  0.2682\nTotal prob of N-in: 0.14402\noutside   1   331\nTMhelix   332   354\ninside   355   372

19780 GCF_001364415.1_10425_1_73 Salmonella enterica subsp. enterica serovar Typhi Proteobacteria; Gammaproteobacteria; Enterobacterales; Enterobacteriaceae; Salmonella; Salmonella enterica; Salmonella enterica subsp. enterica MTGDNTLITSHGINRRDFMKLCAALAATMGLSSKAAAEMAESVSNPQRPPVIWIGAQECT WP_000145429.1 hydrogenase 2 small subunit [Salmonella enterica] Length: 372\nNumber of predicted TMHs:  1\nExp number of AAs in TMHs: 24.51309\nExp number, first 60 AAs:  0.2682\nTotal prob of N-in: 0.14402\noutside   1   331\nTMhelix   332   354\ninside   355   372

19781 GCF_001364375.1_10608_2_26 Salmonella enterica subsp. enterica serovar Typhi Proteobacteria; Gammaproteobacteria; Enterobacterales; Enterobacteriaceae; Salmonella; Salmonella enterica; Salmonella enterica subsp. enterica MTGDNTLITSHGINRRDFMKLCAALAATMGLSSKAAAEMAESVSNPQRPPVIWIGAQECT WP_000145429.1 hydrogenase 2 small subunit [Salmonella enterica] Length: 372\nNumber of predicted TMHs:  1\nExp number of AAs in TMHs: 24.51309\nExp number, first 60 AAs:  0.2682\nTotal prob of N-in: 0.14402\noutside   1   331\nTMhelix   332   354\ninside   355   372

19782 GCF_001364455.1_10426_1_2 Salmonella enterica subsp. enterica serovar Typhi Proteobacteria; Gammaproteobacteria; Enterobacterales; Enterobacteriaceae; Salmonella; Salmonella enterica; Salmonella enterica subsp. enterica MTGDNTLITSHGINRRDFMKLCAALAATMGLSSKAAAEMAESVSNPQRPPVIWIGAQECT WP_000145429.1 hydrogenase 2 small subunit [Salmonella enterica] Length: 372\nNumber of predicted TMHs:  1\nExp number of AAs in TMHs: 24.51309\nExp number, first 60 AAs:  0.2682\nTotal prob of N-in: 0.14402\noutside   1   331\nTMhelix   332   354\ninside   355   372

19783 GCF_001364495.1_10540_1_58 Salmonella enterica subsp. enterica serovar Typhi Proteobacteria; Gammaproteobacteria; Enterobacterales; Enterobacteriaceae; Salmonella; Salmonella enterica;

Salmonella enterica subsp. enterica MTGDNTLITSHGINRRDFMKLCAALAATMGLSSKAAAEMAESVSNPQRPPVIWIGAQECT WP_000145429.1 hydrogenase 2 small subunit [Salmonella enterica] Length: 372\nNumber of predicted TMHs: 1\nExp number of AAs in TMHs: 24.51309\nExp number, first 60 AAs: 0.2682\nTotal prob of N-in: 0.14402\noutside 1 331\nTMhelix 332 354\ninside 355 372

19784 GCF_001364535.1_10562_2_8 Salmonella enterica subsp. enterica serovar Typhi Proteobacteria; Gammaproteobacteria; Enterobacterales; Enterobacteriaceae; Salmonella; Salmonella enterica; Salmonella enterica subsp. enterica MTGDNTLITSHGINRRDFMKLCAALAATMGLSSKAAAEMAESVSNPQRPPVIWIGAQECT WP_000145429.1 hydrogenase 2 small subunit [Salmonella enterica] Length: 372\nNumber of predicted TMHs: 1\nExp number of AAs in TMHs: 24.51309\nExp number, first 60 AAs: 0.2682\nTotal prob of N-in: 0.14402\noutside 1 331\nTMhelix 332 354\ninside 355 372

19785 GCF_001364575.1_10492_1_54 Salmonella enterica subsp. enterica serovar Typhi Proteobacteria; Gammaproteobacteria; Enterobacterales; Enterobacteriaceae; Salmonella; Salmonella enterica; Salmonella enterica subsp. enterica MTGDNTLITSHGINRRDFMKLCAALAATMGLSSKAAAEMAESVSNPQRPPVIWIGAQECT WP_000145429.1 hydrogenase 2 small subunit [Salmonella enterica] Length: 372\nNumber of predicted TMHs: 1\nExp number of AAs in TMHs: 24.51309\nExp number, first 60 AAs: 0.2682\nTotal prob of N-in: 0.14402\noutside 1 331\nTMhelix 332 354\ninside 355 372

19786 GCF_001364615.1_10561_2_8 Salmonella enterica subsp. enterica serovar Typhi Proteobacteria; Gammaproteobacteria; Enterobacterales; Enterobacteriaceae; Salmonella; Salmonella enterica; Salmonella enterica subsp. enterica MTGDNTLITSHGINRRDFMKLCAALAATMGLSSKAAAEMAESVSNPQRPPVIWIGAQECT WP_000145429.1 hydrogenase 2 small subunit [Salmonella enterica] Length: 372\nNumber of predicted TMHs: 1\nExp number of AAs in TMHs: 24.51309\nExp number, first 60 AAs: 0.2682\nTotal prob of N-in: 0.14402\noutside 1 331\nTMhelix 332 354\ninside 355 372

19787 GCF_001364655.1_10540_1_47 Salmonella enterica subsp. enterica serovar Typhi Proteobacteria; Gammaproteobacteria; Enterobacterales; Enterobacteriaceae; Salmonella; Salmonella enterica; Salmonella enterica subsp. enterica MTGDNTLITSHGINRRDFMKLCAALAATMGLSSKAAAEMAESVSNPQRPPVIWIGAQECT WP_000145429.1 hydrogenase 2 small subunit [Salmonella enterica] Length: 372\nNumber of predicted TMHs: 1\nExp number of AAs in TMHs: 24.51309\nExp number, first 60 AAs: 0.2682\nTotal prob of N-in: 0.14402\noutside 1 331\nTMhelix 332 354\ninside 355 372

19788 GCF_001364695.1_10540_1_53 Salmonella enterica subsp. enterica serovar Typhi Proteobacteria; Gammaproteobacteria; Enterobacterales; Enterobacteriaceae; Salmonella; Salmonella enterica; Salmonella enterica subsp. enterica MTGDNTLITSHGINRRDFMKLCAALAATMGLSSKAAAEMAESVSNPQRPPVIWIGAQECT WP_000145429.1 hydrogenase 2 small subunit [Salmonella enterica] Length: 372\nNumber of predicted TMHs: 1\nExp number of AAs in TMHs: 24.51309\nExp number, first 60 AAs: 0.2682\nTotal prob of N-in: 0.14402\noutside 1 331\nTMhelix 332 354\ninside 355 372

19789 GCF_001364735.1_10561_2_9 Salmonella enterica subsp. enterica serovar Typhi Proteobacteria; Gammaproteobacteria; Enterobacterales; Enterobacteriaceae; Salmonella; Salmonella enterica; Salmonella enterica subsp. enterica MTGDNTLITSHGINRRDFMKLCAALAATMGLSSKAAAEMAESVSNPQRPPVIWIGAQECT WP_000145429.1 hydrogenase 2 small subunit [Salmonella enterica] Length: 372\nNumber of predicted TMHs: 1\nExp number of AAs in TMHs: 24.51309\nExp number, first 60 AAs: 0.2682\nTotal prob of N-in: 0.14402\noutside 1 331\nTMhelix 332 354\ninside 355 372

19790 GCF_001364775.1_10492_1_85 Salmonella enterica subsp. enterica serovar Typhi Proteobacteria; Gammaproteobacteria; Enterobacterales; Enterobacteriaceae; Salmonella; Salmonella enterica; Salmonella enterica subsp. enterica MTGDNTLITSHGINRRDFMKLCAALAATMGLSSKAAAEMAESVSNPQRPPVIWIGAQECT WP_000145429.1 hydrogenase 2 small subunit [Salmonella enterica] Length: 372\nNumber of predicted TMHs: 1\nExp number of AAs in TMHs: 24.51309\nExp number, first 60 AAs: 0.2682\nTotal prob of N-in: 0.14402\noutside 1 331\nTMhelix 332 354\ninside 355 372

19791 GCF_001364815.1_10540_1_72 Salmonella enterica subsp. enterica serovar Typhi Proteobacteria; Gammaproteobacteria; Enterobacterales; Enterobacteriaceae; Salmonella; Salmonella enterica; Salmonella enterica subsp. enterica MTGDNTLITSHGINRRDFMKLCAALAATMGLSSKAAAEMAESVSNPQRPPVIWIGAQECT WP_000145429.1 hydrogenase 2 small subunit [Salmonella enterica] Length: 372\nNumber of predicted TMHs: 1\nExp number of AAs in TMHs: 24.51309\nExp number, first 60 AAs: 0.2682\nTotal prob of N-in: 0.14402\noutside 1 331\nTMhelix 332 354\ninside 355 372

19792 GCF_001364855.1_10540_1_40 Salmonella enterica subsp. enterica serovar Typhi Proteobacteria; Gammaproteobacteria; Enterobacterales; Enterobacteriaceae; Salmonella; Salmonella enterica; Salmonella enterica subsp. enterica MTGDNTLITSHGINRRDFMKLCAALAATMGLSSKAAAEMAESVSNPQRPPVIWIGAQECT WP_000145429.1 hydrogenase 2 small subunit [Salmonella enterica] Length: 372\nNumber of predicted TMHs: 1\nExp number of AAs in TMHs: 24.51309\nExp number, first 60 AAs: 0.2682\nTotal prob of N-in: 0.14402\noutside 1 331\nTMhelix 332 354\ninside 355 372

19793 GCF_001364975.1_10493_1_7 Salmonella enterica subsp. enterica serovar Typhi Proteobacteria; Gammaproteobacteria; Enterobacterales; Enterobacteriaceae; Salmonella; Salmonella enterica; Salmonella enterica subsp. enterica MTGDNTLITSHGINRRDFMKLCAALAATMGLSSKAAAEMAESVSNPQRPPVIWIGAQECT WP_000145429.1 hydrogenase 2 small subunit [Salmonella enterica] Length: 372\nNumber of predicted TMHs: 1\nExp number of AAs in TMHs: 24.51309\nExp number, first 60 AAs: 0.2682\nTotal prob of N-in: 0.14402\noutside 1 331\nTMhelix 332 354\ninside 355 372

19794 GCF_001364895.1_10426_1_27 Salmonella enterica subsp. enterica serovar Typhi Proteobacteria; Gammaproteobacteria; Enterobacterales; Enterobacteriaceae; Salmonella; Salmonella enterica; Salmonella enterica subsp. enterica MTGDNTLITSHGINRRDFMKLCAALAATMGLSSKAAAEMAESVSNPQRPPVIWIGAQECT WP_000145429.1 hydrogenase 2 small subunit [Salmonella enterica] Length: 372\nNumber of predicted TMHs: 1\nExp number of AAs in TMHs: 24.51309\nExp number, first 60 AAs: 0.2682\nTotal prob of N-in: 0.14402\noutside 1 331\nTMhelix 332 354\ninside 355 372

19795 GCF_001364935.1_10562_2_39 Salmonella enterica subsp. enterica serovar Typhi Proteobacteria; Gammaproteobacteria; Enterobacterales; Enterobacteriaceae; Salmonella; Salmonella enterica; Salmonella enterica subsp. enterica MTGDNTLITSHGINRRDFMKLCAALAATMGLSSKAAAEMAESVSNPQRPPVIWIGAQECT WP_000145429.1 hydrogenase 2 small subunit [Salmonella enterica] Length: 372\nNumber of predicted TMHs: 1\nExp number of AAs in TMHs: 24.51309\nExp number, first 60 AAs: 0.2682\nTotal prob of N-in: 0.14402\noutside 1 331\nTMhelix 332 354\ninside 355 372

19796 GCF_001365075.1_10561_2_77 Salmonella enterica subsp. enterica serovar Typhi Proteobacteria; Gammaproteobacteria; Enterobacterales; Enterobacteriaceae; Salmonella; Salmonella enterica; Salmonella enterica subsp. enterica MTGDNTLITSHGINRRDFMKLCAALAATMGLSSKAAAEMAESVSNPQRPPVIWIGAQECT WP_000145429.1 hydrogenase 2 small subunit [Salmonella enterica] Length: 372\nNumber of predicted TMHs: 1\nExp number of AAs in TMHs: 24.51309\nExp number, first 60 AAs: 0.2682\nTotal prob of N-in: 0.14402\noutside 1 331\nTMhelix 332 354\ninside 355 372

19797 GCF_001364995.1_10540_1_57 Salmonella enterica subsp. enterica serovar Typhi Proteobacteria; Gammaproteobacteria; Enterobacterales; Enterobacteriaceae; Salmonella; Salmonella enterica; Salmonella enterica subsp. enterica MTGDNTLITSHGINRRDFMKLCAALAATMGLSSKAAAEMAESVSNPQRPPVIWIGAQECT WP_000145429.1 hydrogenase 2 small subunit [Salmonella enterica] Length: 372\nNumber of predicted TMHs: 1\nExp number of AAs in TMHs: 24.51309\nExp number, first 60 AAs: 0.2682\nTotal prob of N-in: 0.14402\noutside 1 331\nTMhelix 332 354\ninside 355 372

19798 GCF_001365035.1_10425_1_43 Salmonella enterica subsp. enterica serovar Typhi Proteobacteria; Gammaproteobacteria; Enterobacterales; Enterobacteriaceae; Salmonella; Salmonella enterica; Salmonella enterica subsp. enterica MTGDNTLITSHGINRRDFMKLCAALAATMGLSSKAAAEMAESVSNPQRPPVIWIGAQECT WP_000145429.1 hydrogenase 2 small subunit [Salmonella enterica] Length: 372\nNumber of predicted TMHs: 1\nExp number of AAs in TMHs: 24.51309\nExp number, first 60 AAs: 0.2682\nTotal prob of N-in: 0.14402\noutside 1 331\nTMhelix 332 354\ninside 355 372

19799 GCF_001365115.1_10608_2_22 Salmonella enterica subsp. enterica serovar Typhi Proteobacteria; Gammaproteobacteria; Enterobacterales; Enterobacteriaceae; Salmonella; Salmonella enterica; Salmonella enterica subsp. enterica MTGDNTLITSHGINRRDFMKLCAALAATMGLSSKAAAEMAESVSNPQRPPVIWIGAQECT WP_000145429.1 hydrogenase 2 small subunit [Salmonella enterica] Length: 372\nNumber of predicted TMHs: 1\nExp number of AAs in TMHs: 24.51309\nExp number, first 60 AAs: 0.2682\nTotal prob of N-in: 0.14402\noutside 1 331\nTMhelix 332 354\ninside 355 372

19800 GCF_001365155.1_10540_1_74 Salmonella enterica subsp. enterica serovar Typhi Proteobacteria; Gammaproteobacteria; Enterobacterales; Enterobacteriaceae; Salmonella; Salmonella enterica; Salmonella enterica subsp. enterica MTGDNTLITSHGINRRDFMKLCAALAATMGLSSKAAAEMAESVSNPQRPPVIWIGAQECT WP_000145429.1 hydrogenase 2 small subunit [Salmonella enterica] Length: 372\nNumber of

predicted TMHs:  1\nExp number of AAs in TMHs: 24.51309\nExp number, first 60 AAs:  0.2682\nTotal prob of N-in: 0.14402\noutside   1   331\nTMhelix   332   354\ninside   355   372

19801 GCF_001365195.1_10607_2_23 Salmonella enterica subsp. enterica serovar Typhi Proteobacteria; Gammaproteobacteria; Enterobacterales; Enterobacteriaceae; Salmonella; Salmonella enterica; Salmonella enterica subsp. enterica MTGDNTLITSHGINRRDFMKLCAALAATMGLSSKAAAEMAESVSNPQRPPVIWIGAQECT WP_000145429.1 hydrogenase 2 small subunit [Salmonella enterica] Length: 372\nNumber of predicted TMHs:  1\nExp number of AAs in TMHs: 24.51309\nExp number, first 60 AAs:  0.2682\nTotal prob of N-in: 0.14402\noutside   1   331\nTMhelix   332   354\ninside   355   372

19802 GCF_001365235.1_10561_2_43 Salmonella enterica subsp. enterica serovar Typhi Proteobacteria; Gammaproteobacteria; Enterobacterales; Enterobacteriaceae; Salmonella; Salmonella enterica; Salmonella enterica subsp. enterica MTGDNTLITSHGINRRDFMKLCAALAATMGLSSKAAAEMAESVSNPQRPPVIWIGAQECT WP_000145429.1 hydrogenase 2 small subunit [Salmonella enterica] Length: 372\nNumber of predicted TMHs:  1\nExp number of AAs in TMHs: 24.51309\nExp number, first 60 AAs:  0.2682\nTotal prob of N-in: 0.14402\noutside   1   331\nTMhelix   332   354\ninside   355   372

19803 GCF_001365315.1_10607_1_47 Salmonella enterica subsp. enterica serovar Typhi Proteobacteria; Gammaproteobacteria; Enterobacterales; Enterobacteriaceae; Salmonella; Salmonella enterica; Salmonella enterica subsp. enterica MTGDNTLITSHGINRRDFMKLCAALAATMGLSSKAAAEMAESVSNPQRPPVIWIGAQECT WP_000145429.1 hydrogenase 2 small subunit [Salmonella enterica] Length: 372\nNumber of predicted TMHs:  1\nExp number of AAs in TMHs: 24.51309\nExp number, first 60 AAs:  0.2682\nTotal prob of N-in: 0.14402\noutside   1   331\nTMhelix   332   354\ninside   355   372

19804 GCF_001365275.1_10607_2_56 Salmonella enterica subsp. enterica serovar Typhi Proteobacteria; Gammaproteobacteria; Enterobacterales; Enterobacteriaceae; Salmonella; Salmonella enterica; Salmonella enterica subsp. enterica MTGDNTLITSHGINRRDFMKLCAALAATMGLSSKAAAEMAESVSNPQRPPVIWIGAQECT WP_000145429.1 hydrogenase 2 small subunit [Salmonella enterica] Length: 372\nNumber of predicted TMHs:  1\nExp number of AAs in TMHs: 24.51309\nExp number, first 60 AAs:  0.2682\nTotal prob of N-in: 0.14402\noutside   1   331\nTMhelix   332   354\ninside   355   372

19805 GCF_001365355.1_10492_1_13 Salmonella enterica subsp. enterica serovar Typhi Proteobacteria; Gammaproteobacteria; Enterobacterales; Enterobacteriaceae; Salmonella; Salmonella enterica; Salmonella enterica subsp. enterica MTGDNTLITSHGINRRDFMKLCAALAATMGLSSKAAAEMAESVSNPQRPPVIWIGAQECT WP_000145429.1 hydrogenase 2 small subunit [Salmonella enterica] Length: 372\nNumber of predicted TMHs:  1\nExp number of AAs in TMHs: 24.51309\nExp number, first 60 AAs:  0.2682\nTotal prob of N-in: 0.14402\noutside   1   331\nTMhelix   332   354\ninside   355   372

19806 GCF_001365395.1_10561_2_50 Salmonella enterica subsp. enterica serovar Typhi Proteobacteria; Gammaproteobacteria; Enterobacterales; Enterobacteriaceae; Salmonella; Salmonella enterica; Salmonella enterica subsp. enterica MTGDNTLITSHGINRRDFMKLCAALAATMGLSSKAAAEMAESVSNPQRPPVIWIGAQECT WP_000145429.1 hydrogenase 2 small subunit [Salmonella enterica] Length: 372\nNumber of predicted TMHs:  1\nExp number of AAs in TMHs: 24.51309\nExp number, first 60 AAs:  0.2682\nTotal prob of N-in: 0.14402\noutside   1   331\nTMhelix   332   354\ninside   355   372

19807 GCF_001365435.1_10561_2_29 Salmonella enterica subsp. enterica serovar Typhi Proteobacteria; Gammaproteobacteria; Enterobacterales; Enterobacteriaceae; Salmonella; Salmonella enterica; Salmonella enterica subsp. enterica MTGDNTLITSHGINRRDFMKLCAALAATMGLSSKAAAEMAESVSNPQRPPVIWIGAQECT WP_000145429.1 hydrogenase 2 small subunit [Salmonella enterica] Length: 372\nNumber of predicted TMHs:  1\nExp number of AAs in TMHs: 24.51309\nExp number, first 60 AAs:  0.2682\nTotal prob of N-in: 0.14402\noutside   1   331\nTMhelix   332   354\ninside   355   372

19808 GCF_001365475.1_10561_2_13 Salmonella enterica subsp. enterica serovar Typhi Proteobacteria; Gammaproteobacteria; Enterobacterales; Enterobacteriaceae; Salmonella; Salmonella enterica; Salmonella enterica subsp. enterica MTGDNTLITSHGINRRDFMKLCAALAATMGLSSKAAAEMAESVSNPQRPPVIWIGAQECT WP_000145429.1 hydrogenase 2 small subunit [Salmonella enterica] Length: 372\nNumber of predicted TMHs:  1\nExp number of AAs in TMHs: 24.51309\nExp number, first 60 AAs:  0.2682\nTotal prob of N-in: 0.14402\noutside   1   331\nTMhelix   332   354\ninside   355   372

19809 GCF_001365515.1_10608_2_51 Salmonella enterica subsp. enterica serovar Typhi Proteobacteria; Gammaproteobacteria; Enterobacterales; Enterobacteriaceae; Salmonella; Salmonella enterica;

Salmonella enterica subsp. enterica MTGDNTLITSHGINRRDFMKLCAALAATMGLSSKAAAEMAESVSNPQRPPVIWIGAQECT WP_000145429.1 hydrogenase 2 small subunit [Salmonella enterica] Length: 372\nNumber of predicted TMHs: 1\nExp number of AAs in TMHs: 24.51309\nExp number, first 60 AAs: 0.2682\nTotal prob of N-in: 0.14402\noutside 1 331\nTMhelix 332 354\ninside 355 372

19810 GCF_001365555.1_10492_1_51 Salmonella enterica subsp. enterica serovar Typhi Proteobacteria; Gammaproteobacteria; Enterobacterales; Enterobacteriaceae; Salmonella; Salmonella enterica; Salmonella enterica subsp. enterica MTGDNTLITSHGINRRDFMKLCAALAATMGLSSKAAAEMAESVSNPQRPPVIWIGAQECT WP_000145429.1 hydrogenase 2 small subunit [Salmonella enterica] Length: 372\nNumber of predicted TMHs: 1\nExp number of AAs in TMHs: 24.51309\nExp number, first 60 AAs: 0.2682\nTotal prob of N-in: 0.14402\noutside 1 331\nTMhelix 332 354\ninside 355 372

19811 GCF_001365595.1_10607_2_7 Salmonella enterica subsp. enterica serovar Typhi Proteobacteria; Gammaproteobacteria; Enterobacterales; Enterobacteriaceae; Salmonella; Salmonella enterica; Salmonella enterica subsp. enterica MTGDNTLITSHGINRRDFMKLCAALAATMGLSSKAAAEMAESVSNPQRPPVIWIGAQECT WP_000145429.1 hydrogenase 2 small subunit [Salmonella enterica] Length: 372\nNumber of predicted TMHs: 1\nExp number of AAs in TMHs: 24.51309\nExp number, first 60 AAs: 0.2682\nTotal prob of N-in: 0.14402\noutside 1 331\nTMhelix 332 354\ninside 355 372

19812 GCF_001365635.1_10607_1_76 Salmonella enterica subsp. enterica serovar Typhi Proteobacteria; Gammaproteobacteria; Enterobacterales; Enterobacteriaceae; Salmonella; Salmonella enterica; Salmonella enterica subsp. enterica MTGDNTLITSHGINRRDFMKLCAALAATMGLSSKAAAEMAESVSNPQRPPVIWIGAQECT WP_000145429.1 hydrogenase 2 small subunit [Salmonella enterica] Length: 372\nNumber of predicted TMHs: 1\nExp number of AAs in TMHs: 24.51309\nExp number, first 60 AAs: 0.2682\nTotal prob of N-in: 0.14402\noutside 1 331\nTMhelix 332 354\ninside 355 372

19813 GCF_001365675.1_10426_1_61 Salmonella enterica subsp. enterica serovar Typhi Proteobacteria; Gammaproteobacteria; Enterobacterales; Enterobacteriaceae; Salmonella; Salmonella enterica; Salmonella enterica subsp. enterica MTGDNTLITSHGINRRDFMKLCAALAATMGLSSKAAAEMAESVSNPQRPPVIWIGAQECT WP_000145429.1 hydrogenase 2 small subunit [Salmonella enterica] Length: 372\nNumber of predicted TMHs: 1\nExp number of AAs in TMHs: 24.51309\nExp number, first 60 AAs: 0.2682\nTotal prob of N-in: 0.14402\noutside 1 331\nTMhelix 332 354\ninside 355 372

19814 GCF_001365695.1_10493_1_14 Salmonella enterica subsp. enterica serovar Typhi Proteobacteria; Gammaproteobacteria; Enterobacterales; Enterobacteriaceae; Salmonella; Salmonella enterica; Salmonella enterica subsp. enterica MTGDNTLITSHGINRRDFMKLCAALAATMGLSSKAAAEMAESVSNPQRPPVIWIGAQECT WP_000145429.1 hydrogenase 2 small subunit [Salmonella enterica] Length: 372\nNumber of predicted TMHs: 1\nExp number of AAs in TMHs: 24.51309\nExp number, first 60 AAs: 0.2682\nTotal prob of N-in: 0.14402\noutside 1 331\nTMhelix 332 354\ninside 355 372

19815 GCF_001362315.1_10349_1_95 Salmonella enterica subsp. enterica serovar Typhi Proteobacteria; Gammaproteobacteria; Enterobacterales; Enterobacteriaceae; Salmonella; Salmonella enterica; Salmonella enterica subsp. enterica MTGDNTLITSHGINRRDFMKLCAALAATMGLSSKAAAEMAESVSNPQRPPVIWIGAQECT WP_000145429.1 hydrogenase 2 small subunit [Salmonella enterica] Length: 372\nNumber of predicted TMHs: 1\nExp number of AAs in TMHs: 24.51309\nExp number, first 60 AAs: 0.2682\nTotal prob of N-in: 0.14402\noutside 1 331\nTMhelix 332 354\ninside 355 372

19816 GCF_001365735.1_10540_1_61 Salmonella enterica subsp. enterica serovar Typhi Proteobacteria; Gammaproteobacteria; Enterobacterales; Enterobacteriaceae; Salmonella; Salmonella enterica; Salmonella enterica subsp. enterica MTGDNTLITSHGINRRDFMKLCAALAATMGLSSKAAAEMAESVSNPQRPPVIWIGAQECT WP_000145429.1 hydrogenase 2 small subunit [Salmonella enterica] Length: 372\nNumber of predicted TMHs: 1\nExp number of AAs in TMHs: 24.51309\nExp number, first 60 AAs: 0.2682\nTotal prob of N-in: 0.14402\noutside 1 331\nTMhelix 332 354\ninside 355 372

19817 GCF_001365775.1_10493_1_57 Salmonella enterica subsp. enterica serovar Typhi Proteobacteria; Gammaproteobacteria; Enterobacterales; Enterobacteriaceae; Salmonella; Salmonella enterica; Salmonella enterica subsp. enterica MTGDNTLITSHGINRRDFMKLCAALAATMGLSSKAAAEMAESVSNPQRPPVIWIGAQECT WP_000145429.1 hydrogenase 2 small subunit [Salmonella enterica] Length: 372\nNumber of predicted TMHs: 1\nExp number of AAs in TMHs: 24.51309\nExp number, first 60 AAs: 0.2682\nTotal prob of N-in: 0.14402\noutside 1 331\nTMhelix 332 354\ninside 355 372

19818 GCF_001365855.1_10561_2_70 Salmonella enterica subsp. enterica serovar Typhi Proteobacteria; Gammaproteobacteria; Enterobacterales; Enterobacteriaceae; Salmonella; Salmonella enterica; Salmonella enterica subsp. enterica MTGDNTLITSHGINRRDFMKLCAALAATMGLSSKAAAEMAESVSNPQRPPVIWIGAQECT WP_000145429.1 hydrogenase 2 small subunit [Salmonella enterica] Length: 372\nNumber of predicted TMHs: 1\nExp number of AAs in TMHs: 24.51309\nExp number, first 60 AAs: 0.2682\nTotal prob of N-in: 0.14402\noutside 1 331\nTMhelix 332 354\ninside 355 372

19819 GCF_001365815.1_10561_2_48 Salmonella enterica subsp. enterica serovar Typhi Proteobacteria; Gammaproteobacteria; Enterobacterales; Enterobacteriaceae; Salmonella; Salmonella enterica; Salmonella enterica subsp. enterica MTGDNTLITSHGINRRDFMKLCAALAATMGLSSKAAAEMAESVSNPQRPPVIWIGAQECT WP_000145429.1 hydrogenase 2 small subunit [Salmonella enterica] Length: 372\nNumber of predicted TMHs: 1\nExp number of AAs in TMHs: 24.51309\nExp number, first 60 AAs: 0.2682\nTotal prob of N-in: 0.14402\noutside 1 331\nTMhelix 332 354\ninside 355 372

19820 GCF_001365895.1_10608_2_37 Salmonella enterica subsp. enterica serovar Typhi Proteobacteria; Gammaproteobacteria; Enterobacterales; Enterobacteriaceae; Salmonella; Salmonella enterica; Salmonella enterica subsp. enterica MTGDNTLITSHGINRRDFMKLCAALAATMGLSSKAAAEMAESVSNPQRPPVIWIGAQECT WP_000145429.1 hydrogenase 2 small subunit [Salmonella enterica] Length: 372\nNumber of predicted TMHs: 1\nExp number of AAs in TMHs: 24.51309\nExp number, first 60 AAs: 0.2682\nTotal prob of N-in: 0.14402\noutside 1 331\nTMhelix 332 354\ninside 355 372

19821 GCF_001365935.1_10540_1_69 Salmonella enterica subsp. enterica serovar Typhi Proteobacteria; Gammaproteobacteria; Enterobacterales; Enterobacteriaceae; Salmonella; Salmonella enterica; Salmonella enterica subsp. enterica MTGDNTLITSHGINRRDFMKLCAALAATMGLSSKAAAEMAESVSNPQRPPVIWIGAQECT WP_000145429.1 hydrogenase 2 small subunit [Salmonella enterica] Length: 372\nNumber of predicted TMHs: 1\nExp number of AAs in TMHs: 24.51309\nExp number, first 60 AAs: 0.2682\nTotal prob of N-in: 0.14402\noutside 1 331\nTMhelix 332 354\ninside 355 372

19822 GCF_001365975.1_10426_1_30 Salmonella enterica subsp. enterica serovar Typhi Proteobacteria; Gammaproteobacteria; Enterobacterales; Enterobacteriaceae; Salmonella; Salmonella enterica; Salmonella enterica subsp. enterica MTGDNTLITSHGINRRDFMKLCAALAATMGLSSKAAAEMAESVSNPQRPPVIWIGAQECT WP_000145429.1 hydrogenase 2 small subunit [Salmonella enterica] Length: 372\nNumber of predicted TMHs: 1\nExp number of AAs in TMHs: 24.51309\nExp number, first 60 AAs: 0.2682\nTotal prob of N-in: 0.14402\noutside 1 331\nTMhelix 332 354\ninside 355 372

19823 GCF_001366015.1_10492_1_76 Salmonella enterica subsp. enterica serovar Typhi Proteobacteria; Gammaproteobacteria; Enterobacterales; Enterobacteriaceae; Salmonella; Salmonella enterica; Salmonella enterica subsp. enterica MTGDNTLITSHGINRRDFMKLCAALAATMGLSSKAAAEMAESVSNPQRPPVIWIGAQECT WP_000145429.1 hydrogenase 2 small subunit [Salmonella enterica] Length: 372\nNumber of predicted TMHs: 1\nExp number of AAs in TMHs: 24.51309\nExp number, first 60 AAs: 0.2682\nTotal prob of N-in: 0.14402\noutside 1 331\nTMhelix 332 354\ninside 355 372

19824 GCF_001366035.1_10608_2_44 Salmonella enterica subsp. enterica serovar Typhi Proteobacteria; Gammaproteobacteria; Enterobacterales; Enterobacteriaceae; Salmonella; Salmonella enterica; Salmonella enterica subsp. enterica MTGDNTLITSHGINRRDFMKLCAALAATMGLSSKAAAEMAESVSNPQRPPVIWIGAQECT WP_000145429.1 hydrogenase 2 small subunit [Salmonella enterica] Length: 372\nNumber of predicted TMHs: 1\nExp number of AAs in TMHs: 24.51309\nExp number, first 60 AAs: 0.2682\nTotal prob of N-in: 0.14402\noutside 1 331\nTMhelix 332 354\ninside 355 372

19825 GCF_001366055.1_10561_2_24 Salmonella enterica subsp. enterica serovar Typhi Proteobacteria; Gammaproteobacteria; Enterobacterales; Enterobacteriaceae; Salmonella; Salmonella enterica; Salmonella enterica subsp. enterica MTGDNTLITSHGINRRDFMKLCAALAATMGLSSKAAAEMAESVSNPQRPPVIWIGAQECT WP_000145429.1 hydrogenase 2 small subunit [Salmonella enterica] Length: 372\nNumber of predicted TMHs: 1\nExp number of AAs in TMHs: 24.51309\nExp number, first 60 AAs: 0.2682\nTotal prob of N-in: 0.14402\noutside 1 331\nTMhelix 332 354\ninside 355 372

19826 GCF_001366075.1_10608_1_1 Salmonella enterica subsp. enterica serovar Typhi Proteobacteria; Gammaproteobacteria; Enterobacterales; Enterobacteriaceae; Salmonella; Salmonella enterica; Salmonella enterica subsp. enterica MTGDNTLITSHGINRRDFMKLCAALAATMGLSSKAAAEMAESVSNPQRPPVIWIGAQECT WP_000145429.1 hydrogenase 2 small subunit [Salmonella enterica] Length: 372\nNumber of

predicted TMHs: 1\nExp number of AAs in TMHs: 24.51309\nExp number, first 60 AAs: 0.2682\nTotal prob of N-in: 0.14402\noutside 1 331\nTMhelix 332 354\ninside 355 372

19827 GCF_001366095.1_10540_1_68 Salmonella enterica subsp. enterica serovar Typhi Proteobacteria; Gammaproteobacteria; Enterobacterales; Enterobacteriaceae; Salmonella; Salmonella enterica; Salmonella enterica subsp. enterica MTGDNTLITSHGINRRDFMKLCAALAATMGLSSKAAAEMAESVSNPQRPPVIWIGAQECT WP_000145429.1 hydrogenase 2 small subunit [Salmonella enterica] Length: 372\nNumber of predicted TMHs: 1\nExp number of AAs in TMHs: 24.51309\nExp number, first 60 AAs: 0.2682\nTotal prob of N-in: 0.14402\noutside 1 331\nTMhelix 332 354\ninside 355 372

19828 GCF_001366115.1_10425_1_23 Salmonella enterica subsp. enterica serovar Typhi Proteobacteria; Gammaproteobacteria; Enterobacterales; Enterobacteriaceae; Salmonella; Salmonella enterica; Salmonella enterica subsp. enterica MTGDNTLITSHGINRRDFMKLCAALAATMGLSSKAAAEMAESVSNPQRPPVIWIGAQECT WP_000145429.1 hydrogenase 2 small subunit [Salmonella enterica] Length: 372\nNumber of predicted TMHs: 1\nExp number of AAs in TMHs: 24.51309\nExp number, first 60 AAs: 0.2682\nTotal prob of N-in: 0.14402\noutside 1 331\nTMhelix 332 354\ninside 355 372

19829 GCF_001362275.1_10349_1_29 Salmonella enterica subsp. enterica serovar Typhi Proteobacteria; Gammaproteobacteria; Enterobacterales; Enterobacteriaceae; Salmonella; Salmonella enterica; Salmonella enterica subsp. enterica MTGDNTLITSHGINRRDFMKLCAALAATMGLSSKAAAEMAESVSNPQRPPVIWIGAQECT WP_000145429.1 hydrogenase 2 small subunit [Salmonella enterica] Length: 372\nNumber of predicted TMHs: 1\nExp number of AAs in TMHs: 24.51309\nExp number, first 60 AAs: 0.2682\nTotal prob of N-in: 0.14402\noutside 1 331\nTMhelix 332 354\ninside 355 372

19830 GCF_001366135.1_10540_1_79 Salmonella enterica subsp. enterica serovar Typhi Proteobacteria; Gammaproteobacteria; Enterobacterales; Enterobacteriaceae; Salmonella; Salmonella enterica; Salmonella enterica subsp. enterica MTGDNTLITSHGINRRDFMKLCAALAATMGLSSKAAAEMAESVSNPQRPPVIWIGAQECT WP_000145429.1 hydrogenase 2 small subunit [Salmonella enterica] Length: 372\nNumber of predicted TMHs: 1\nExp number of AAs in TMHs: 24.51309\nExp number, first 60 AAs: 0.2682\nTotal prob of N-in: 0.14402\noutside 1 331\nTMhelix 332 354\ninside 355 372

19831 GCF_001366155.1_10493_1_85 Salmonella enterica subsp. enterica serovar Typhi Proteobacteria; Gammaproteobacteria; Enterobacterales; Enterobacteriaceae; Salmonella; Salmonella enterica; Salmonella enterica subsp. enterica MTGDNTLITSHGINRRDFMKLCAALAATMGLSSKAAAEMAESVSNPQRPPVIWIGAQECT WP_000145429.1 hydrogenase 2 small subunit [Salmonella enterica] Length: 372\nNumber of predicted TMHs: 1\nExp number of AAs in TMHs: 24.51309\nExp number, first 60 AAs: 0.2682\nTotal prob of N-in: 0.14402\noutside 1 331\nTMhelix 332 354\ninside 355 372

19832 GCF_001366175.1_10540_1_11 Salmonella enterica subsp. enterica serovar Typhi Proteobacteria; Gammaproteobacteria; Enterobacterales; Enterobacteriaceae; Salmonella; Salmonella enterica; Salmonella enterica subsp. enterica MTGDNTLITSHGINRRDFMKLCAALAATMGLSSKAAAEMAESVSNPQRPPVIWIGAQECT WP_000145429.1 hydrogenase 2 small subunit [Salmonella enterica] Length: 372\nNumber of predicted TMHs: 1\nExp number of AAs in TMHs: 24.51309\nExp number, first 60 AAs: 0.2682\nTotal prob of N-in: 0.14402\noutside 1 331\nTMhelix 332 354\ninside 355 372

19833 GCF_001366215.1_10540_1_32 Salmonella enterica subsp. enterica serovar Typhi Proteobacteria; Gammaproteobacteria; Enterobacterales; Enterobacteriaceae; Salmonella; Salmonella enterica; Salmonella enterica subsp. enterica MTGDNTLITSHGINRRDFMKLCAALAATMGLSSKAAAEMAESVSNPQRPPVIWIGAQECT WP_000145429.1 hydrogenase 2 small subunit [Salmonella enterica] Length: 372\nNumber of predicted TMHs: 1\nExp number of AAs in TMHs: 24.51309\nExp number, first 60 AAs: 0.2682\nTotal prob of N-in: 0.14402\noutside 1 331\nTMhelix 332 354\ninside 355 372

19834 GCF_001366195.1_10608_2_21 Salmonella enterica subsp. enterica serovar Typhi Proteobacteria; Gammaproteobacteria; Enterobacterales; Enterobacteriaceae; Salmonella; Salmonella enterica; Salmonella enterica subsp. enterica MTGDNTLITSHGINRRDFMKLCAALAATMGLSSKAAAEMAESVSNPQRPPVIWIGAQECT WP_000145429.1 hydrogenase 2 small subunit [Salmonella enterica] Length: 372\nNumber of predicted TMHs: 1\nExp number of AAs in TMHs: 24.51309\nExp number, first 60 AAs: 0.2682\nTotal prob of N-in: 0.14402\noutside 1 331\nTMhelix 332 354\ninside 355 372

19835 GCF_001362195.1_10349_1_80 Salmonella enterica subsp. enterica serovar Typhi Proteobacteria; Gammaproteobacteria; Enterobacterales; Enterobacteriaceae; Salmonella; Salmonella enterica;

Salmonella enterica subsp. enterica MTGDNTLITSHGINRRDFMKLCAALAATMGLSSKAAAEMAESVSNPQRPPVIWIGAQECT WP_000145429.1 hydrogenase 2 small subunit [Salmonella enterica] Length: 372\nNumber of predicted TMHs: 1\nExp number of AAs in TMHs: 24.51309\nExp number, first 60 AAs: 0.2682\nTotal prob of N-in: 0.14402\noutside 1 331\nTMhelix 332 354\ninside 355 372

19836 GCF_001362175.1_10349_1_79 Salmonella enterica subsp. enterica serovar Typhi Proteobacteria; Gammaproteobacteria; Enterobacterales; Enterobacteriaceae; Salmonella; Salmonella enterica; Salmonella enterica subsp. enterica MTGDNTLITSHGINRRDFMKLCAALAATMGLSSKAAAEMAESVSNPQRPPVIWIGAQECT WP_000145429.1 hydrogenase 2 small subunit [Salmonella enterica] Length: 372\nNumber of predicted TMHs: 1\nExp number of AAs in TMHs: 24.51309\nExp number, first 60 AAs: 0.2682\nTotal prob of N-in: 0.14402\noutside 1 331\nTMhelix 332 354\ninside 355 372

19837 GCF_001362235.1_10349_1_48 Salmonella enterica subsp. enterica serovar Typhi Proteobacteria; Gammaproteobacteria; Enterobacterales; Enterobacteriaceae; Salmonella; Salmonella enterica; Salmonella enterica subsp. enterica MTGDNTLITSHGINRRDFMKLCAALAATMGLSSKAAAEMAESVSNPQRPPVIWIGAQECT WP_000145429.1 hydrogenase 2 small subunit [Salmonella enterica] Length: 372\nNumber of predicted TMHs: 1\nExp number of AAs in TMHs: 24.51309\nExp number, first 60 AAs: 0.2682\nTotal prob of N-in: 0.14402\noutside 1 331\nTMhelix 332 354\ninside 355 372

19838 GCF_001362095.1_10349_1_12 Salmonella enterica subsp. enterica serovar Typhi Proteobacteria; Gammaproteobacteria; Enterobacterales; Enterobacteriaceae; Salmonella; Salmonella enterica; Salmonella enterica subsp. enterica MTGDNTLITSHGINRRDFMKLCAALAATMGLSSKAAAEMAESVSNPQRPPVIWIGAQECT WP_000145429.1 hydrogenase 2 small subunit [Salmonella enterica] Length: 372\nNumber of predicted TMHs: 1\nExp number of AAs in TMHs: 24.51309\nExp number, first 60 AAs: 0.2682\nTotal prob of N-in: 0.14402\noutside 1 331\nTMhelix 332 354\ninside 355 372

19839 GCF_001362055.1_10349_1_10 Salmonella enterica subsp. enterica serovar Typhi Proteobacteria; Gammaproteobacteria; Enterobacterales; Enterobacteriaceae; Salmonella; Salmonella enterica; Salmonella enterica subsp. enterica MTGDNTLITSHGINRRDFMKLCAALAATMGLSSKAAAEMAESVSNPQRPPVIWIGAQECT WP_000145429.1 hydrogenase 2 small subunit [Salmonella enterica] Length: 372\nNumber of predicted TMHs: 1\nExp number of AAs in TMHs: 24.51309\nExp number, first 60 AAs: 0.2682\nTotal prob of N-in: 0.14402\noutside 1 331\nTMhelix 332 354\ninside 355 372

19840 GCF_001362135.1_10349_1_73 Salmonella enterica subsp. enterica serovar Typhi Proteobacteria; Gammaproteobacteria; Enterobacterales; Enterobacteriaceae; Salmonella; Salmonella enterica; Salmonella enterica subsp. enterica MTGDNTLITSHGINRRDFMKLCAALAATMGLSSKAAAEMAESVSNPQRPPVIWIGAQECT WP_000145429.1 hydrogenase 2 small subunit [Salmonella enterica] Length: 372\nNumber of predicted TMHs: 1\nExp number of AAs in TMHs: 24.51309\nExp number, first 60 AAs: 0.2682\nTotal prob of N-in: 0.14402\noutside 1 331\nTMhelix 332 354\ninside 355 372

19841 GCF_001366255.1_10492_1_58 Salmonella enterica subsp. enterica serovar Typhi Proteobacteria; Gammaproteobacteria; Enterobacterales; Enterobacteriaceae; Salmonella; Salmonella enterica; Salmonella enterica subsp. enterica MTGDNTLITSHGINRRDFMKLCAALAATMGLSSKAAAEMAESVSNPQRPPVIWIGAQECT WP_000145429.1 hydrogenase 2 small subunit [Salmonella enterica] Length: 372\nNumber of predicted TMHs: 1\nExp number of AAs in TMHs: 24.51309\nExp number, first 60 AAs: 0.2682\nTotal prob of N-in: 0.14402\noutside 1 331\nTMhelix 332 354\ninside 355 372

19842 GCF_001362015.1_10209_5_42 Salmonella enterica subsp. enterica serovar Typhi Proteobacteria; Gammaproteobacteria; Enterobacterales; Enterobacteriaceae; Salmonella; Salmonella enterica; Salmonella enterica subsp. enterica MTGDNTLITSHGINRRDFMKLCAALAATMGLSSKAAAEMAESVSNPQRPPVIWIGAQECT WP_000145429.1 hydrogenase 2 small subunit [Salmonella enterica] Length: 372\nNumber of predicted TMHs: 1\nExp number of AAs in TMHs: 24.51309\nExp number, first 60 AAs: 0.2682\nTotal prob of N-in: 0.14402\noutside 1 331\nTMhelix 332 354\ninside 355 372

19843 GCF_001361975.1_10209_5_21 Salmonella enterica subsp. enterica serovar Typhi Proteobacteria; Gammaproteobacteria; Enterobacterales; Enterobacteriaceae; Salmonella; Salmonella enterica; Salmonella enterica subsp. enterica MTGDNTLITSHGINRRDFMKLCAALAATMGLSSKAAAEMAESVSNPQRPPVIWIGAQECT WP_000145429.1 hydrogenase 2 small subunit [Salmonella enterica] Length: 372\nNumber of predicted TMHs: 1\nExp number of AAs in TMHs: 24.51309\nExp number, first 60 AAs: 0.2682\nTotal prob of N-in: 0.14402\noutside 1 331\nTMhelix 332 354\ninside 355 372

19844	GCF_001361935.1_10209_5_3	Salmonella enterica subsp. enterica serovar Typhi	Proteobacteria; Gammaproteobacteria; Enterobacterales; Enterobacteriaceae; Salmonella; Salmonella enterica; Salmonella enterica subsp. enterica	MTGDNTLITSHGINRRDFMKLCAALAATMGLSSKAAAEMAESVSNPQRPPVIWIGAQECT	WP_000145429.1	hydrogenase 2 small subunit [Salmonella enterica]	Length: 372\nNumber of predicted TMHs: 1\nExp number of AAs in TMHs: 24.51309\nExp number, first 60 AAs: 0.2682\nTotal prob of N-in: 0.14402\noutside 1 331\nTMhelix 332 354\ninside 355 372

19845	GCF_001361895.1_10071_8_53	Salmonella enterica subsp. enterica serovar Typhi	Proteobacteria; Gammaproteobacteria; Enterobacterales; Enterobacteriaceae; Salmonella; Salmonella enterica; Salmonella enterica subsp. enterica	MTGDNTLITSHGINRRDFMKLCAALAATMGLSSKAAAEMAESVSNPQRPPVIWIGAQECT	WP_000145429.1	hydrogenase 2 small subunit [Salmonella enterica]	Length: 372\nNumber of predicted TMHs: 1\nExp number of AAs in TMHs: 24.51309\nExp number, first 60 AAs: 0.2682\nTotal prob of N-in: 0.14402\noutside 1 331\nTMhelix 332 354\ninside 355 372

19846	GCF_001361855.1_10071_8_2	Salmonella enterica subsp. enterica serovar Typhi	Proteobacteria; Gammaproteobacteria; Enterobacterales; Enterobacteriaceae; Salmonella; Salmonella enterica; Salmonella enterica subsp. enterica	MTGDNTLITSHGINRRDFMKLCAALAATMGLSSKAAAEMAESVSNPQRPPVIWIGAQECT	WP_000145429.1	hydrogenase 2 small subunit [Salmonella enterica]	Length: 372\nNumber of predicted TMHs: 1\nExp number of AAs in TMHs: 24.51309\nExp number, first 60 AAs: 0.2682\nTotal prob of N-in: 0.14402\noutside 1 331\nTMhelix 332 354\ninside 355 372

19847	GCF_001361815.1_10209_5_26	Salmonella enterica subsp. enterica serovar Typhi	Proteobacteria; Gammaproteobacteria; Enterobacterales; Enterobacteriaceae; Salmonella; Salmonella enterica; Salmonella enterica subsp. enterica	MTGDNTLITSHGINRRDFMKLCAALAATMGLSSKAAAEMAESVSNPQRPPVIWIGAQECT	WP_000145429.1	hydrogenase 2 small subunit [Salmonella enterica]	Length: 372\nNumber of predicted TMHs: 1\nExp number of AAs in TMHs: 24.51309\nExp number, first 60 AAs: 0.2682\nTotal prob of N-in: 0.14402\noutside 1 331\nTMhelix 332 354\ninside 355 372

19848	GCF_001361775.1_10071_8_79	Salmonella enterica subsp. enterica serovar Typhi	Proteobacteria; Gammaproteobacteria; Enterobacterales; Enterobacteriaceae; Salmonella; Salmonella enterica; Salmonella enterica subsp. enterica	MTGDNTLITSHGINRRDFMKLCAALAATMGLSSKAAAEMAESVSNPQRPPVIWIGAQECT	WP_000145429.1	hydrogenase 2 small subunit [Salmonella enterica]	Length: 372\nNumber of predicted TMHs: 1\nExp number of AAs in TMHs: 24.51309\nExp number, first 60 AAs: 0.2682\nTotal prob of N-in: 0.14402\noutside 1 331\nTMhelix 332 354\ninside 355 372

19849	GCF_001361735.1_10209_5_61	Salmonella enterica subsp. enterica serovar Typhi	Proteobacteria; Gammaproteobacteria; Enterobacterales; Enterobacteriaceae; Salmonella; Salmonella enterica; Salmonella enterica subsp. enterica	MTGDNTLITSHGINRRDFMKLCAALAATMGLSSKAAAEMAESVSNPQRPPVIWIGAQECT	WP_000145429.1	hydrogenase 2 small subunit [Salmonella enterica]	Length: 372\nNumber of predicted TMHs: 1\nExp number of AAs in TMHs: 24.51309\nExp number, first 60 AAs: 0.2682\nTotal prob of N-in: 0.14402\noutside 1 331\nTMhelix 332 354\ninside 355 372

19850	GCF_001366235.1_10426_1_63	Salmonella enterica subsp. enterica serovar Typhi	Proteobacteria; Gammaproteobacteria; Enterobacterales; Enterobacteriaceae; Salmonella; Salmonella enterica; Salmonella enterica subsp. enterica	MTGDNTLITSHGINRRDFMKLCAALAATMGLSSKAAAEMAESVSNPQRPPVIWIGAQECT	WP_000145429.1	hydrogenase 2 small subunit [Salmonella enterica]	Length: 372\nNumber of predicted TMHs: 1\nExp number of AAs in TMHs: 24.51309\nExp number, first 60 AAs: 0.2682\nTotal prob of N-in: 0.14402\noutside 1 331\nTMhelix 332 354\ninside 355 372

19851	GCF_001366275.1_10426_1_42	Salmonella enterica subsp. enterica serovar Typhi	Proteobacteria; Gammaproteobacteria; Enterobacterales; Enterobacteriaceae; Salmonella; Salmonella enterica; Salmonella enterica subsp. enterica	MTGDNTLITSHGINRRDFMKLCAALAATMGLSSKAAAEMAESVSNPQRPPVIWIGAQECT	WP_000145429.1	hydrogenase 2 small subunit [Salmonella enterica]	Length: 372\nNumber of predicted TMHs: 1\nExp number of AAs in TMHs: 24.51309\nExp number, first 60 AAs: 0.2682\nTotal prob of N-in: 0.14402\noutside 1 331\nTMhelix 332 354\ninside 355 372

19852	GCF_001366295.1_10608_2_30	Salmonella enterica subsp. enterica serovar Typhi	Proteobacteria; Gammaproteobacteria; Enterobacterales; Enterobacteriaceae; Salmonella; Salmonella enterica; Salmonella enterica subsp. enterica	MTGDNTLITSHGINRRDFMKLCAALAATMGLSSKAAAEMAESVSNPQRPPVIWIGAQECT	WP_000145429.1	hydrogenase 2 small subunit [Salmonella enterica]	Length: 372\nNumber of

predicted TMHs: 1\nExp number of AAs in TMHs: 24.51309\nExp number, first 60 AAs: 0.2682\nTotal prob of N-in: 0.14402\noutside 1 331\nTMhelix 332 354\ninside 355 372

19853 GCF_001366335.1_10426_1_48 Salmonella enterica subsp. enterica serovar Typhi Proteobacteria; Gammaproteobacteria; Enterobacterales; Enterobacteriaceae; Salmonella; Salmonella enterica; Salmonella enterica subsp. enterica MTGDNTLITSHGINRRDFMKLCAALAATMGLSSKAAAEMAESVSNPQRPPVIWIGAQECT WP_000145429.1 hydrogenase 2 small subunit [Salmonella enterica] Length: 372\nNumber of predicted TMHs: 1\nExp number of AAs in TMHs: 24.51309\nExp number, first 60 AAs: 0.2682\nTotal prob of N-in: 0.14402\noutside 1 331\nTMhelix 332 354\ninside 355 372

19854 GCF_001366315.1_10608_1_2 Salmonella enterica subsp. enterica serovar Typhi Proteobacteria; Gammaproteobacteria; Enterobacterales; Enterobacteriaceae; Salmonella; Salmonella enterica; Salmonella enterica subsp. enterica MTGDNTLITSHGINRRDFMKLCAALAATMGLSSKAAAEMAESVSNPQRPPVIWIGAQECT WP_000145429.1 hydrogenase 2 small subunit [Salmonella enterica] Length: 372\nNumber of predicted TMHs: 1\nExp number of AAs in TMHs: 24.51309\nExp number, first 60 AAs: 0.2682\nTotal prob of N-in: 0.14402\noutside 1 331\nTMhelix 332 354\ninside 355 372

19855 GCF_001366355.1_10607_2_4 Salmonella enterica subsp. enterica serovar Typhi Proteobacteria; Gammaproteobacteria; Enterobacterales; Enterobacteriaceae; Salmonella; Salmonella enterica; Salmonella enterica subsp. enterica MTGDNTLITSHGINRRDFMKLCAALAATMGLSSKAAAEMAESVSNPQRPPVIWIGAQECT WP_000145429.1 hydrogenase 2 small subunit [Salmonella enterica] Length: 372\nNumber of predicted TMHs: 1\nExp number of AAs in TMHs: 24.51309\nExp number, first 60 AAs: 0.2682\nTotal prob of N-in: 0.14402\noutside 1 331\nTMhelix 332 354\ninside 355 372

19856 GCF_001366375.1_10426_1_17 Salmonella enterica subsp. enterica serovar Typhi Proteobacteria; Gammaproteobacteria; Enterobacterales; Enterobacteriaceae; Salmonella; Salmonella enterica; Salmonella enterica subsp. enterica MTGDNTLITSHGINRRDFMKLCAALAATMGLSSKAAAEMAESVSNPQRPPVIWIGAQECT WP_000145429.1 hydrogenase 2 small subunit [Salmonella enterica] Length: 372\nNumber of predicted TMHs: 1\nExp number of AAs in TMHs: 24.51309\nExp number, first 60 AAs: 0.2682\nTotal prob of N-in: 0.14402\noutside 1 331\nTMhelix 332 354\ninside 355 372

19857 GCF_001366395.1_10493_1_29 Salmonella enterica subsp. enterica serovar Typhi Proteobacteria; Gammaproteobacteria; Enterobacterales; Enterobacteriaceae; Salmonella; Salmonella enterica; Salmonella enterica subsp. enterica MTGDNTLITSHGINRRDFMKLCAALAATMGLSSKAAAEMAESVSNPQRPPVIWIGAQECT WP_000145429.1 hydrogenase 2 small subunit [Salmonella enterica] Length: 372\nNumber of predicted TMHs: 1\nExp number of AAs in TMHs: 24.51309\nExp number, first 60 AAs: 0.2682\nTotal prob of N-in: 0.14402\noutside 1 331\nTMhelix 332 354\ninside 355 372

19858 GCF_001366435.1_10493_1_44 Salmonella enterica subsp. enterica serovar Typhi Proteobacteria; Gammaproteobacteria; Enterobacterales; Enterobacteriaceae; Salmonella; Salmonella enterica; Salmonella enterica subsp. enterica MTGDNTLITSHGINRRDFMKLCAALAATMGLSSKAAAEMAESVSNPQRPPVIWIGAQECT WP_000145429.1 hydrogenase 2 small subunit [Salmonella enterica] Length: 372\nNumber of predicted TMHs: 1\nExp number of AAs in TMHs: 24.51309\nExp number, first 60 AAs: 0.2682\nTotal prob of N-in: 0.14402\noutside 1 331\nTMhelix 332 354\ninside 355 372

19859 GCF_001366455.1_10540_1_90 Salmonella enterica subsp. enterica serovar Typhi Proteobacteria; Gammaproteobacteria; Enterobacterales; Enterobacteriaceae; Salmonella; Salmonella enterica; Salmonella enterica subsp. enterica MTGDNTLITSHGINRRDFMKLCAALAATMGLSSKAAAEMAESVSNPQRPPVIWIGAQECT WP_000145429.1 hydrogenase 2 small subunit [Salmonella enterica] Length: 372\nNumber of predicted TMHs: 1\nExp number of AAs in TMHs: 24.51309\nExp number, first 60 AAs: 0.2682\nTotal prob of N-in: 0.14402\noutside 1 331\nTMhelix 332 354\ninside 355 372

19860 GCF_001366475.1_10561_2_45 Salmonella enterica subsp. enterica serovar Typhi Proteobacteria; Gammaproteobacteria; Enterobacterales; Enterobacteriaceae; Salmonella; Salmonella enterica; Salmonella enterica subsp. enterica MTGDNTLITSHGINRRDFMKLCAALAATMGLSSKAAAEMAESVSNPQRPPVIWIGAQECT WP_000145429.1 hydrogenase 2 small subunit [Salmonella enterica] Length: 372\nNumber of predicted TMHs: 1\nExp number of AAs in TMHs: 24.51309\nExp number, first 60 AAs: 0.2682\nTotal prob of N-in: 0.14402\noutside 1 331\nTMhelix 332 354\ninside 355 372

19861 GCF_001366495.1_10540_1_51 Salmonella enterica subsp. enterica serovar Typhi Proteobacteria; Gammaproteobacteria; Enterobacterales; Enterobacteriaceae; Salmonella; Salmonella enterica;

Salmonella enterica subsp. enterica MTGDNTLITSHGINRRDFMKLCAALAATMGLSSKAAAEMAESVSNPQRPPVIWIGAQECT WP_000145429.1 hydrogenase 2 small subunit [Salmonella enterica] Length: 372\nNumber of predicted TMHs: 1\nExp number of AAs in TMHs: 24.51309\nExp number, first 60 AAs: 0.2682\nTotal prob of N-in: 0.14402\noutside 1 331\nTMhelix 332 354\ninside 355 372

19862 GCF_001366515.1_10608_2_60 Salmonella enterica subsp. enterica serovar Typhi Proteobacteria; Gammaproteobacteria; Enterobacterales; Enterobacteriaceae; Salmonella; Salmonella enterica; Salmonella enterica subsp. enterica MTGDNTLITSHGINRRDFMKLCAALAATMGLSSKAAAEMAESVSNPQRPPVIWIGAQECT WP_000145429.1 hydrogenase 2 small subunit [Salmonella enterica] Length: 372\nNumber of predicted TMHs: 1\nExp number of AAs in TMHs: 24.51309\nExp number, first 60 AAs: 0.2682\nTotal prob of N-in: 0.14402\noutside 1 331\nTMhelix 332 354\ninside 355 372

19863 GCF_001366535.1_10540_1_46 Salmonella enterica subsp. enterica serovar Typhi Proteobacteria; Gammaproteobacteria; Enterobacterales; Enterobacteriaceae; Salmonella; Salmonella enterica; Salmonella enterica subsp. enterica MTGDNTLITSHGINRRDFMKLCAALAATMGLSSKAAAEMAESVSNPQRPPVIWIGAQECT WP_000145429.1 hydrogenase 2 small subunit [Salmonella enterica] Length: 372\nNumber of predicted TMHs: 1\nExp number of AAs in TMHs: 24.51309\nExp number, first 60 AAs: 0.2682\nTotal prob of N-in: 0.14402\noutside 1 331\nTMhelix 332 354\ninside 355 372

19864 GCF_001366555.1_10607_1_50 Salmonella enterica subsp. enterica serovar Typhi Proteobacteria; Gammaproteobacteria; Enterobacterales; Enterobacteriaceae; Salmonella; Salmonella enterica; Salmonella enterica subsp. enterica MTGDNTLITSHGINRRDFMKLCAALAATMGLSSKAAAEMAESVSNPQRPPVIWIGAQECT WP_000145429.1 hydrogenase 2 small subunit [Salmonella enterica] Length: 372\nNumber of predicted TMHs: 1\nExp number of AAs in TMHs: 24.51309\nExp number, first 60 AAs: 0.2682\nTotal prob of N-in: 0.14402\noutside 1 331\nTMhelix 332 354\ninside 355 372

19865 GCF_001366575.1_10426_1_21 Salmonella enterica subsp. enterica serovar Typhi Proteobacteria; Gammaproteobacteria; Enterobacterales; Enterobacteriaceae; Salmonella; Salmonella enterica; Salmonella enterica subsp. enterica MTGDNTLITSHGINRRDFMKLCAALAATMGLSSKAAAEMAESVSNPQRPPVIWIGAQECT WP_000145429.1 hydrogenase 2 small subunit [Salmonella enterica] Length: 372\nNumber of predicted TMHs: 1\nExp number of AAs in TMHs: 24.51309\nExp number, first 60 AAs: 0.2682\nTotal prob of N-in: 0.14402\noutside 1 331\nTMhelix 332 354\ninside 355 372

19866 GCF_001366595.1_10607_1_90 Salmonella enterica subsp. enterica serovar Typhi Proteobacteria; Gammaproteobacteria; Enterobacterales; Enterobacteriaceae; Salmonella; Salmonella enterica; Salmonella enterica subsp. enterica MTGDNTLITSHGINRRDFMKLCAALAATMGLSSKAAAEMAESVSNPQRPPVIWIGAQECT WP_000145429.1 hydrogenase 2 small subunit [Salmonella enterica] Length: 372\nNumber of predicted TMHs: 1\nExp number of AAs in TMHs: 24.51309\nExp number, first 60 AAs: 0.2682\nTotal prob of N-in: 0.14402\noutside 1 331\nTMhelix 332 354\ninside 355 372

19867 GCF_001367235.1_10426_1_25 Salmonella enterica subsp. enterica serovar Typhi Proteobacteria; Gammaproteobacteria; Enterobacterales; Enterobacteriaceae; Salmonella; Salmonella enterica; Salmonella enterica subsp. enterica MTGDNTLITSHGINRRDFMKLCAALAATMGLSSKAAAEMAESVSNPQRPPVIWIGAQECT WP_000145429.1 hydrogenase 2 small subunit [Salmonella enterica] Length: 372\nNumber of predicted TMHs: 1\nExp number of AAs in TMHs: 24.51309\nExp number, first 60 AAs: 0.2682\nTotal prob of N-in: 0.14402\noutside 1 331\nTMhelix 332 354\ninside 355 372

19868 GCF_001368695.1_10541_1_58 Salmonella enterica subsp. enterica serovar Typhi Proteobacteria; Gammaproteobacteria; Enterobacterales; Enterobacteriaceae; Salmonella; Salmonella enterica; Salmonella enterica subsp. enterica MTGDNTLITSHGINRRDFMKLCAALAATMGLSSKAAAEMAESVSNPQRPPVIWIGAQECT WP_000145429.1 hydrogenase 2 small subunit [Salmonella enterica] Length: 372\nNumber of predicted TMHs: 1\nExp number of AAs in TMHs: 24.51309\nExp number, first 60 AAs: 0.2682\nTotal prob of N-in: 0.14402\noutside 1 331\nTMhelix 332 354\ninside 355 372

19869 GCF_001368675.1_10541_1_74 Salmonella enterica subsp. enterica serovar Typhi Proteobacteria; Gammaproteobacteria; Enterobacterales; Enterobacteriaceae; Salmonella; Salmonella enterica; Salmonella enterica subsp. enterica MTGDNTLITSHGINRRDFMKLCAALAATMGLSSKAAAEMAESVSNPQRPPVIWIGAQECT WP_000145429.1 hydrogenase 2 small subunit [Salmonella enterica] Length: 372\nNumber of predicted TMHs: 1\nExp number of AAs in TMHs: 24.51309\nExp number, first 60 AAs: 0.2682\nTotal prob of N-in: 0.14402\noutside 1 331\nTMhelix 332 354\ninside 355 372

19870 GCF_001368655.1_10541_1_71 Salmonella enterica subsp. enterica serovar Typhi Proteobacteria; Gammaproteobacteria; Enterobacterales; Enterobacteriaceae; Salmonella; Salmonella enterica; Salmonella enterica subsp. enterica MTGDNTLITSHGINRRDFMKLCAALAATMGLSSKAAAEMAESVSNPQRPPVIWIGAQECT WP_000145429.1 hydrogenase 2 small subunit [Salmonella enterica] Length: 372\nNumber of predicted TMHs: 1\nExp number of AAs in TMHs: 24.51309\nExp number, first 60 AAs: 0.2682\nTotal prob of N-in: 0.14402\noutside 1 331\nTMhelix 332 354\ninside 355 372

19871 GCF_001368635.1_10541_1_64 Salmonella enterica subsp. enterica serovar Typhi Proteobacteria; Gammaproteobacteria; Enterobacterales; Enterobacteriaceae; Salmonella; Salmonella enterica; Salmonella enterica subsp. enterica MTGDNTLITSHGINRRDFMKLCAALAATMGLSSKAAAEMAESVSNPQRPPVIWIGAQECT WP_000145429.1 hydrogenase 2 small subunit [Salmonella enterica] Length: 372\nNumber of predicted TMHs: 1\nExp number of AAs in TMHs: 24.51309\nExp number, first 60 AAs: 0.2682\nTotal prob of N-in: 0.14402\noutside 1 331\nTMhelix 332 354\ninside 355 372

19872 GCF_001368615.1_10562_2_78 Salmonella enterica subsp. enterica serovar Typhi Proteobacteria; Gammaproteobacteria; Enterobacterales; Enterobacteriaceae; Salmonella; Salmonella enterica; Salmonella enterica subsp. enterica MTGDNTLITSHGINRRDFMKLCAALAATMGLSSKAAAEMAESVSNPQRPPVIWIGAQECT WP_000145429.1 hydrogenase 2 small subunit [Salmonella enterica] Length: 372\nNumber of predicted TMHs: 1\nExp number of AAs in TMHs: 24.51309\nExp number, first 60 AAs: 0.2682\nTotal prob of N-in: 0.14402\noutside 1 331\nTMhelix 332 354\ninside 355 372

19873 GCF_001368575.1_10608_2_59 Salmonella enterica subsp. enterica serovar Typhi Proteobacteria; Gammaproteobacteria; Enterobacterales; Enterobacteriaceae; Salmonella; Salmonella enterica; Salmonella enterica subsp. enterica MTGDNTLITSHGINRRDFMKLCAALAATMGLSSKAAAEMAESVSNPQRPPVIWIGAQECT WP_000145429.1 hydrogenase 2 small subunit [Salmonella enterica] Length: 372\nNumber of predicted TMHs: 1\nExp number of AAs in TMHs: 24.51309\nExp number, first 60 AAs: 0.2682\nTotal prob of N-in: 0.14402\noutside 1 331\nTMhelix 332 354\ninside 355 372

19874 GCF_001368595.1_10426_1_41 Salmonella enterica subsp. enterica serovar Typhi Proteobacteria; Gammaproteobacteria; Enterobacterales; Enterobacteriaceae; Salmonella; Salmonella enterica; Salmonella enterica subsp. enterica MTGDNTLITSHGINRRDFMKLCAALAATMGLSSKAAAEMAESVSNPQRPPVIWIGAQECT WP_000145429.1 hydrogenase 2 small subunit [Salmonella enterica] Length: 372\nNumber of predicted TMHs: 1\nExp number of AAs in TMHs: 24.51309\nExp number, first 60 AAs: 0.2682\nTotal prob of N-in: 0.14402\noutside 1 331\nTMhelix 332 354\ninside 355 372

19875 GCF_001368555.1_10493_1_62 Salmonella enterica subsp. enterica serovar Typhi Proteobacteria; Gammaproteobacteria; Enterobacterales; Enterobacteriaceae; Salmonella; Salmonella enterica; Salmonella enterica subsp. enterica MTGDNTLITSHGINRRDFMKLCAALAATMGLSSKAAAEMAESVSNPQRPPVIWIGAQECT WP_000145429.1 hydrogenase 2 small subunit [Salmonella enterica] Length: 372\nNumber of predicted TMHs: 1\nExp number of AAs in TMHs: 24.51309\nExp number, first 60 AAs: 0.2682\nTotal prob of N-in: 0.14402\noutside 1 331\nTMhelix 332 354\ninside 355 372

19876 GCF_001368535.1_10426_1_12 Salmonella enterica subsp. enterica serovar Typhi Proteobacteria; Gammaproteobacteria; Enterobacterales; Enterobacteriaceae; Salmonella; Salmonella enterica; Salmonella enterica subsp. enterica MTGDNTLITSHGINRRDFMKLCAALAATMGLSSKAAAEMAESVSNPQRPPVIWIGAQECT WP_000145429.1 hydrogenase 2 small subunit [Salmonella enterica] Length: 372\nNumber of predicted TMHs: 1\nExp number of AAs in TMHs: 24.51309\nExp number, first 60 AAs: 0.2682\nTotal prob of N-in: 0.14402\noutside 1 331\nTMhelix 332 354\ninside 355 372

19877 GCF_001368495.1_10540_1_14 Salmonella enterica subsp. enterica serovar Typhi Proteobacteria; Gammaproteobacteria; Enterobacterales; Enterobacteriaceae; Salmonella; Salmonella enterica; Salmonella enterica subsp. enterica MTGDNTLITSHGINRRDFMKLCAALAATMGLSSKAAAEMAESVSNPQRPPVIWIGAQECT WP_000145429.1 hydrogenase 2 small subunit [Salmonella enterica] Length: 372\nNumber of predicted TMHs: 1\nExp number of AAs in TMHs: 24.51309\nExp number, first 60 AAs: 0.2682\nTotal prob of N-in: 0.14402\noutside 1 331\nTMhelix 332 354\ninside 355 372

19878 GCF_001368515.1_10607_2_41 Salmonella enterica subsp. enterica serovar Typhi Proteobacteria; Gammaproteobacteria; Enterobacterales; Enterobacteriaceae; Salmonella; Salmonella enterica; Salmonella enterica subsp. enterica MTGDNTLITSHGINRRDFMKLCAALAATMGLSSKAAAEMAESVSNPQRPPVIWIGAQECT WP_000145429.1 hydrogenase 2 small subunit [Salmonella enterica] Length: 372\nNumber of

predicted TMHs:  1\nExp number of AAs in TMHs: 24.51309\nExp number, first 60 AAs:  0.2682\nTotal prob of N-in: 0.14402\noutside   1   331\nTMhelix   332   354\ninside   355   372

19879 GCF_001368475.1_10561_2_68 Salmonella enterica subsp. enterica serovar Typhi Proteobacteria; Gammaproteobacteria; Enterobacterales; Enterobacteriaceae; Salmonella; Salmonella enterica; Salmonella enterica subsp. enterica MTGDNTLITSHGINRRDFMKLCAALAATMGLSSKAAAEMAESVSNPQRPPVIWIGAQECT WP_000145429.1 hydrogenase 2 small subunit [Salmonella enterica] Length: 372\nNumber of predicted TMHs:  1\nExp number of AAs in TMHs: 24.51309\nExp number, first 60 AAs:  0.2682\nTotal prob of N-in: 0.14402\noutside   1   331\nTMhelix   332   354\ninside   355   372

19880 GCF_001368455.1_10426_1_82 Salmonella enterica subsp. enterica serovar Typhi Proteobacteria; Gammaproteobacteria; Enterobacterales; Enterobacteriaceae; Salmonella; Salmonella enterica; Salmonella enterica subsp. enterica MTGDNTLITSHGINRRDFMKLCAALAATMGLSSKAAAEMAESVSNPQRPPVIWIGAQECT WP_000145429.1 hydrogenase 2 small subunit [Salmonella enterica] Length: 372\nNumber of predicted TMHs:  1\nExp number of AAs in TMHs: 24.51309\nExp number, first 60 AAs:  0.2682\nTotal prob of N-in: 0.14402\noutside   1   331\nTMhelix   332   354\ninside   355   372

19881 GCF_001368435.1_10426_1_81 Salmonella enterica subsp. enterica serovar Typhi Proteobacteria; Gammaproteobacteria; Enterobacterales; Enterobacteriaceae; Salmonella; Salmonella enterica; Salmonella enterica subsp. enterica MTGDNTLITSHGINRRDFMKLCAALAATMGLSSKAAAEMAESVSNPQRPPVIWIGAQECT WP_000145429.1 hydrogenase 2 small subunit [Salmonella enterica] Length: 372\nNumber of predicted TMHs:  1\nExp number of AAs in TMHs: 24.51309\nExp number, first 60 AAs:  0.2682\nTotal prob of N-in: 0.14402\noutside   1   331\nTMhelix   332   354\ninside   355   372

19882 GCF_001368415.1_10562_2_81 Salmonella enterica subsp. enterica serovar Typhi Proteobacteria; Gammaproteobacteria; Enterobacterales; Enterobacteriaceae; Salmonella; Salmonella enterica; Salmonella enterica subsp. enterica MTGDNTLITSHGINRRDFMKLCAALAATMGLSSKAAAEMAESVSNPQRPPVIWIGAQECT WP_000145429.1 hydrogenase 2 small subunit [Salmonella enterica] Length: 372\nNumber of predicted TMHs:  1\nExp number of AAs in TMHs: 24.51309\nExp number, first 60 AAs:  0.2682\nTotal prob of N-in: 0.14402\noutside   1   331\nTMhelix   332   354\ninside   355   372

19883 GCF_001368395.1_10607_1_83 Salmonella enterica subsp. enterica serovar Typhi Proteobacteria; Gammaproteobacteria; Enterobacterales; Enterobacteriaceae; Salmonella; Salmonella enterica; Salmonella enterica subsp. enterica MTGDNTLITSHGINRRDFMKLCAALAATMGLSSKAAAEMAESVSNPQRPPVIWIGAQECT WP_000145429.1 hydrogenase 2 small subunit [Salmonella enterica] Length: 372\nNumber of predicted TMHs:  1\nExp number of AAs in TMHs: 24.51309\nExp number, first 60 AAs:  0.2682\nTotal prob of N-in: 0.14402\noutside   1   331\nTMhelix   332   354\ninside   355   372

19884 GCF_001368375.1_10607_1_48 Salmonella enterica subsp. enterica serovar Typhi Proteobacteria; Gammaproteobacteria; Enterobacterales; Enterobacteriaceae; Salmonella; Salmonella enterica; Salmonella enterica subsp. enterica MTGDNTLITSHGINRRDFMKLCAALAATMGLSSKAAAEMAESVSNPQRPPVIWIGAQECT WP_000145429.1 hydrogenase 2 small subunit [Salmonella enterica] Length: 372\nNumber of predicted TMHs:  1\nExp number of AAs in TMHs: 24.51309\nExp number, first 60 AAs:  0.2682\nTotal prob of N-in: 0.14402\noutside   1   331\nTMhelix   332   354\ninside   355   372

19885 GCF_001368335.1_10540_1_73 Salmonella enterica subsp. enterica serovar Typhi Proteobacteria; Gammaproteobacteria; Enterobacterales; Enterobacteriaceae; Salmonella; Salmonella enterica; Salmonella enterica subsp. enterica MTGDNTLITSHGINRRDFMKLCAALAATMGLSSKAAAEMAESVSNPQRPPVIWIGAQECT WP_000145429.1 hydrogenase 2 small subunit [Salmonella enterica] Length: 372\nNumber of predicted TMHs:  1\nExp number of AAs in TMHs: 24.51309\nExp number, first 60 AAs:  0.2682\nTotal prob of N-in: 0.14402\noutside   1   331\nTMhelix   332   354\ninside   355   372

19886 GCF_001368315.1_10426_1_71 Salmonella enterica subsp. enterica serovar Typhi Proteobacteria; Gammaproteobacteria; Enterobacterales; Enterobacteriaceae; Salmonella; Salmonella enterica; Salmonella enterica subsp. enterica MTGDNTLITSHGINRRDFMKLCAALAATMGLSSKAAAEMAESVSNPQRPPVIWIGAQECT WP_000145429.1 hydrogenase 2 small subunit [Salmonella enterica] Length: 372\nNumber of predicted TMHs:  1\nExp number of AAs in TMHs: 24.51309\nExp number, first 60 AAs:  0.2682\nTotal prob of N-in: 0.14402\noutside   1   331\nTMhelix   332   354\ninside   355   372

19887 GCF_001368355.1_10541_2_50 Salmonella enterica subsp. enterica serovar Typhi Proteobacteria; Gammaproteobacteria; Enterobacterales; Enterobacteriaceae; Salmonella; Salmonella enterica;

Salmonella enterica subsp. enterica	MTGDNTLITSHGINRRDFMKLCAALAATMGLSSKAAAEMAESVSNPQRPPVIWIGAQECT
	WP_000145429.1	hydrogenase 2 small subunit [Salmonella enterica]	Length: 372\nNumber of predicted TMHs:  1\nExp number of AAs in TMHs: 24.51309\nExp number, first 60 AAs:  0.2682\nTotal prob of N-in: 0.14402\noutside   1   331\nTMhelix   332   354\ninside   355   372

19888	GCF_001368295.1_10426_1_78	Salmonella enterica subsp. enterica serovar Typhi
	Proteobacteria; Gammaproteobacteria; Enterobacterales; Enterobacteriaceae; Salmonella; Salmonella enterica; Salmonella enterica subsp. enterica	MTGDNTLITSHGINRRDFMKLCAALAATMGLSSKAAAEMAESVSNPQRPPVIWIGAQECT
	WP_000145429.1	hydrogenase 2 small subunit [Salmonella enterica]	Length: 372\nNumber of predicted TMHs:  1\nExp number of AAs in TMHs: 24.51309\nExp number, first 60 AAs:  0.2682\nTotal prob of N-in: 0.14402\noutside   1   331\nTMhelix   332   354\ninside   355   372

19889	GCF_001368275.1_10540_1_77	Salmonella enterica subsp. enterica serovar Typhi
	Proteobacteria; Gammaproteobacteria; Enterobacterales; Enterobacteriaceae; Salmonella; Salmonella enterica; Salmonella enterica subsp. enterica	MTGDNTLITSHGINRRDFMKLCAALAATMGLSSKAAAEMAESVSNPQRPPVIWIGAQECT
	WP_000145429.1	hydrogenase 2 small subunit [Salmonella enterica]	Length: 372\nNumber of predicted TMHs:  1\nExp number of AAs in TMHs: 24.51309\nExp number, first 60 AAs:  0.2682\nTotal prob of N-in: 0.14402\noutside   1   331\nTMhelix   332   354\ninside   355   372

19890	GCF_001368235.1_10540_1_30	Salmonella enterica subsp. enterica serovar Typhi
	Proteobacteria; Gammaproteobacteria; Enterobacterales; Enterobacteriaceae; Salmonella; Salmonella enterica; Salmonella enterica subsp. enterica	MTGDNTLITSHGINRRDFMKLCAALAATMGLSSKAAAEMAESVSNPQRPPVIWIGAQECT
	WP_000145429.1	hydrogenase 2 small subunit [Salmonella enterica]	Length: 372\nNumber of predicted TMHs:  1\nExp number of AAs in TMHs: 24.51309\nExp number, first 60 AAs:  0.2682\nTotal prob of N-in: 0.14402\noutside   1   331\nTMhelix   332   354\ninside   355   372

19891	GCF_001368255.1_10540_1_2	Salmonella enterica subsp. enterica serovar Typhi
	Proteobacteria; Gammaproteobacteria; Enterobacterales; Enterobacteriaceae; Salmonella; Salmonella enterica; Salmonella enterica subsp. enterica	MTGDNTLITSHGINRRDFMKLCAALAATMGLSSKAAAEMAESVSNPQRPPVIWIGAQECT
	WP_000145429.1	hydrogenase 2 small subunit [Salmonella enterica]	Length: 372\nNumber of predicted TMHs:  1\nExp number of AAs in TMHs: 24.51309\nExp number, first 60 AAs:  0.2682\nTotal prob of N-in: 0.14402\noutside   1   331\nTMhelix   332   354\ninside   355   372

19892	GCF_001368215.1_10492_1_29	Salmonella enterica subsp. enterica serovar Typhi
	Proteobacteria; Gammaproteobacteria; Enterobacterales; Enterobacteriaceae; Salmonella; Salmonella enterica; Salmonella enterica subsp. enterica	MTGDNTLITSHGINRRDFMKLCAALAATMGLSSKAAAEMAESVSNPQRPPVIWIGAQECT
	WP_000145429.1	hydrogenase 2 small subunit [Salmonella enterica]	Length: 372\nNumber of predicted TMHs:  1\nExp number of AAs in TMHs: 24.51309\nExp number, first 60 AAs:  0.2682\nTotal prob of N-in: 0.14402\noutside   1   331\nTMhelix   332   354\ninside   355   372

19893	GCF_001368195.1_10561_2_10	Salmonella enterica subsp. enterica serovar Typhi
	Proteobacteria; Gammaproteobacteria; Enterobacterales; Enterobacteriaceae; Salmonella; Salmonella enterica; Salmonella enterica subsp. enterica	MTGDNTLITSHGINRRDFMKLCAALAATMGLSSKAAAEMAESVSNPQRPPVIWIGAQECT
	WP_000145429.1	hydrogenase 2 small subunit [Salmonella enterica]	Length: 372\nNumber of predicted TMHs:  1\nExp number of AAs in TMHs: 24.51309\nExp number, first 60 AAs:  0.2682\nTotal prob of N-in: 0.14402\noutside   1   331\nTMhelix   332   354\ninside   355   372

19894	GCF_001368175.1_10426_1_62	Salmonella enterica subsp. enterica serovar Typhi
	Proteobacteria; Gammaproteobacteria; Enterobacterales; Enterobacteriaceae; Salmonella; Salmonella enterica; Salmonella enterica subsp. enterica	MTGDNTLITSHGINRRDFMKLCAALAATMGLSSKAAAEMAESVSNPQRPPVIWIGAQECT
	WP_000145429.1	hydrogenase 2 small subunit [Salmonella enterica]	Length: 372\nNumber of predicted TMHs:  1\nExp number of AAs in TMHs: 24.51309\nExp number, first 60 AAs:  0.2682\nTotal prob of N-in: 0.14402\noutside   1   331\nTMhelix   332   354\ninside   355   372

19895	GCF_001366615.1_10540_1_41	Salmonella enterica subsp. enterica serovar Typhi
	Proteobacteria; Gammaproteobacteria; Enterobacterales; Enterobacteriaceae; Salmonella; Salmonella enterica; Salmonella enterica subsp. enterica	MTGDNTLITSHGINRRDFMKLCAALAATMGLSSKAAAEMAESVSNPQRPPVIWIGAQECT
	WP_000145429.1	hydrogenase 2 small subunit [Salmonella enterica]	Length: 372\nNumber of predicted TMHs:  1\nExp number of AAs in TMHs: 24.51309\nExp number, first 60 AAs:  0.2682\nTotal prob of N-in: 0.14402\noutside   1   331\nTMhelix   332   354\ninside   355   372

19896 GCF_001368155.1_10562_2_82 Salmonella enterica subsp. enterica serovar Typhi Proteobacteria; Gammaproteobacteria; Enterobacterales; Enterobacteriaceae; Salmonella; Salmonella enterica; Salmonella enterica subsp. enterica MTGDNTLITSHGINRRDFMKLCAALAATMGLSSKAAAEMAESVSNPQRPPVIWIGAQECT WP_000145429.1 hydrogenase 2 small subunit [Salmonella enterica] Length: 372\nNumber of predicted TMHs: 1\nExp number of AAs in TMHs: 24.51309\nExp number, first 60 AAs: 0.2682\nTotal prob of N-in: 0.14402\noutside 1 331\nTMhelix 332 354\ninside 355 372

19897 GCF_001368135.1_10426_1_72 Salmonella enterica subsp. enterica serovar Typhi Proteobacteria; Gammaproteobacteria; Enterobacterales; Enterobacteriaceae; Salmonella; Salmonella enterica; Salmonella enterica subsp. enterica MTGDNTLITSHGINRRDFMKLCAALAATMGLSSKAAAEMAESVSNPQRPPVIWIGAQECT WP_000145429.1 hydrogenase 2 small subunit [Salmonella enterica] Length: 372\nNumber of predicted TMHs: 1\nExp number of AAs in TMHs: 24.51309\nExp number, first 60 AAs: 0.2682\nTotal prob of N-in: 0.14402\noutside 1 331\nTMhelix 332 354\ninside 355 372

19898 GCF_001366655.1_10540_1_52 Salmonella enterica subsp. enterica serovar Typhi Proteobacteria; Gammaproteobacteria; Enterobacterales; Enterobacteriaceae; Salmonella; Salmonella enterica; Salmonella enterica subsp. enterica MTGDNTLITSHGINRRDFMKLCAALAATMGLSSKAAAEMAESVSNPQRPPVIWIGAQECT WP_000145429.1 hydrogenase 2 small subunit [Salmonella enterica] Length: 372\nNumber of predicted TMHs: 1\nExp number of AAs in TMHs: 24.51309\nExp number, first 60 AAs: 0.2682\nTotal prob of N-in: 0.14402\noutside 1 331\nTMhelix 332 354\ninside 355 372

19899 GCF_001366635.1_10607_1_80 Salmonella enterica subsp. enterica serovar Typhi Proteobacteria; Gammaproteobacteria; Enterobacterales; Enterobacteriaceae; Salmonella; Salmonella enterica; Salmonella enterica subsp. enterica MTGDNTLITSHGINRRDFMKLCAALAATMGLSSKAAAEMAESVSNPQRPPVIWIGAQECT WP_000145429.1 hydrogenase 2 small subunit [Salmonella enterica] Length: 372\nNumber of predicted TMHs: 1\nExp number of AAs in TMHs: 24.51309\nExp number, first 60 AAs: 0.2682\nTotal prob of N-in: 0.14402\noutside 1 331\nTMhelix 332 354\ninside 355 372

19900 GCF_001366675.1_10607_2_55 Salmonella enterica subsp. enterica serovar Typhi Proteobacteria; Gammaproteobacteria; Enterobacterales; Enterobacteriaceae; Salmonella; Salmonella enterica; Salmonella enterica subsp. enterica MTGDNTLITSHGINRRDFMKLCAALAATMGLSSKAAAEMAESVSNPQRPPVIWIGAQECT WP_000145429.1 hydrogenase 2 small subunit [Salmonella enterica] Length: 372\nNumber of predicted TMHs: 1\nExp number of AAs in TMHs: 24.51309\nExp number, first 60 AAs: 0.2682\nTotal prob of N-in: 0.14402\noutside 1 331\nTMhelix 332 354\ninside 355 372

19901 GCF_001366695.1_10608_2_7 Salmonella enterica subsp. enterica serovar Typhi Proteobacteria; Gammaproteobacteria; Enterobacterales; Enterobacteriaceae; Salmonella; Salmonella enterica; Salmonella enterica subsp. enterica MTGDNTLITSHGINRRDFMKLCAALAATMGLSSKAAAEMAESVSNPQRPPVIWIGAQECT WP_000145429.1 hydrogenase 2 small subunit [Salmonella enterica] Length: 372\nNumber of predicted TMHs: 1\nExp number of AAs in TMHs: 24.51309\nExp number, first 60 AAs: 0.2682\nTotal prob of N-in: 0.14402\noutside 1 331\nTMhelix 332 354\ninside 355 372

19902 GCF_001366715.1_10540_1_55 Salmonella enterica subsp. enterica serovar Typhi Proteobacteria; Gammaproteobacteria; Enterobacterales; Enterobacteriaceae; Salmonella; Salmonella enterica; Salmonella enterica subsp. enterica MTGDNTLITSHGINRRDFMKLCAALAATMGLSSKAAAEMAESVSNPQRPPVIWIGAQECT WP_000145429.1 hydrogenase 2 small subunit [Salmonella enterica] Length: 372\nNumber of predicted TMHs: 1\nExp number of AAs in TMHs: 24.51309\nExp number, first 60 AAs: 0.2682\nTotal prob of N-in: 0.14402\noutside 1 331\nTMhelix 332 354\ninside 355 372

19903 GCF_001366735.1_10607_1_93 Salmonella enterica subsp. enterica serovar Typhi Proteobacteria; Gammaproteobacteria; Enterobacterales; Enterobacteriaceae; Salmonella; Salmonella enterica; Salmonella enterica subsp. enterica MTGDNTLITSHGINRRDFMKLCAALAATMGLSSKAAAEMAESVSNPQRPPVIWIGAQECT WP_000145429.1 hydrogenase 2 small subunit [Salmonella enterica] Length: 372\nNumber of predicted TMHs: 1\nExp number of AAs in TMHs: 24.51309\nExp number, first 60 AAs: 0.2682\nTotal prob of N-in: 0.14402\noutside 1 331\nTMhelix 332 354\ninside 355 372

19904 GCF_001366755.1_10426_1_24 Salmonella enterica subsp. enterica serovar Typhi Proteobacteria; Gammaproteobacteria; Enterobacterales; Enterobacteriaceae; Salmonella; Salmonella enterica; Salmonella enterica subsp. enterica MTGDNTLITSHGINRRDFMKLCAALAATMGLSSKAAAEMAESVSNPQRPPVIWIGAQECT WP_000145429.1 hydrogenase 2 small subunit [Salmonella enterica] Length: 372\nNumber of

predicted TMHs: 1\nExp number of AAs in TMHs: 24.51309\nExp number, first 60 AAs: 0.2682\nTotal prob of N-in: 0.14402\noutside 1 331\nTMhelix 332 354\ninside 355 372

19905 GCF_001366775.1_10561_2_4 Salmonella enterica subsp. enterica serovar Typhi Proteobacteria; Gammaproteobacteria; Enterobacterales; Enterobacteriaceae; Salmonella; Salmonella enterica; Salmonella enterica subsp. enterica MTGDNTLITSHGINRRDFMKLCAALAATMGLSSKAAAEMAESVSNPQRPPVIWIGAQECT WP_000145429.1 hydrogenase 2 small subunit [Salmonella enterica] Length: 372\nNumber of predicted TMHs: 1\nExp number of AAs in TMHs: 24.51309\nExp number, first 60 AAs: 0.2682\nTotal prob of N-in: 0.14402\noutside 1 331\nTMhelix 332 354\ninside 355 372

19906 GCF_001368075.1_10426_1_34 Salmonella enterica subsp. enterica serovar Typhi Proteobacteria; Gammaproteobacteria; Enterobacterales; Enterobacteriaceae; Salmonella; Salmonella enterica; Salmonella enterica subsp. enterica MTGDNTLITSHGINRRDFMKLCAALAATMGLSSKAAAEMAESVSNPQRPPVIWIGAQECT WP_000145429.1 hydrogenase 2 small subunit [Salmonella enterica] Length: 372\nNumber of predicted TMHs: 1\nExp number of AAs in TMHs: 24.51309\nExp number, first 60 AAs: 0.2682\nTotal prob of N-in: 0.14402\noutside 1 331\nTMhelix 332 354\ninside 355 372

19907 GCF_001368115.1_10607_1_78 Salmonella enterica subsp. enterica serovar Typhi Proteobacteria; Gammaproteobacteria; Enterobacterales; Enterobacteriaceae; Salmonella; Salmonella enterica; Salmonella enterica subsp. enterica MTGDNTLITSHGINRRDFMKLCAALAATMGLSSKAAAEMAESVSNPQRPPVIWIGAQECT WP_000145429.1 hydrogenase 2 small subunit [Salmonella enterica] Length: 372\nNumber of predicted TMHs: 1\nExp number of AAs in TMHs: 24.51309\nExp number, first 60 AAs: 0.2682\nTotal prob of N-in: 0.14402\noutside 1 331\nTMhelix 332 354\ninside 355 372

19908 GCF_001366795.1_10426_1_70 Salmonella enterica subsp. enterica serovar Typhi Proteobacteria; Gammaproteobacteria; Enterobacterales; Enterobacteriaceae; Salmonella; Salmonella enterica; Salmonella enterica subsp. enterica MTGDNTLITSHGINRRDFMKLCAALAATMGLSSKAAAEMAESVSNPQRPPVIWIGAQECT WP_000145429.1 hydrogenase 2 small subunit [Salmonella enterica] Length: 372\nNumber of predicted TMHs: 1\nExp number of AAs in TMHs: 24.51309\nExp number, first 60 AAs: 0.2682\nTotal prob of N-in: 0.14402\noutside 1 331\nTMhelix 332 354\ninside 355 372

19909 GCF_001366815.1_10540_1_89 Salmonella enterica subsp. enterica serovar Typhi Proteobacteria; Gammaproteobacteria; Enterobacterales; Enterobacteriaceae; Salmonella; Salmonella enterica; Salmonella enterica subsp. enterica MTGDNTLITSHGINRRDFMKLCAALAATMGLSSKAAAEMAESVSNPQRPPVIWIGAQECT WP_000145429.1 hydrogenase 2 small subunit [Salmonella enterica] Length: 372\nNumber of predicted TMHs: 1\nExp number of AAs in TMHs: 24.51309\nExp number, first 60 AAs: 0.2682\nTotal prob of N-in: 0.14402\noutside 1 331\nTMhelix 332 354\ninside 355 372

19910 GCF_001368095.1_10426_1_54 Salmonella enterica subsp. enterica serovar Typhi Proteobacteria; Gammaproteobacteria; Enterobacterales; Enterobacteriaceae; Salmonella; Salmonella enterica; Salmonella enterica subsp. enterica MTGDNTLITSHGINRRDFMKLCAALAATMGLSSKAAAEMAESVSNPQRPPVIWIGAQECT WP_000145429.1 hydrogenase 2 small subunit [Salmonella enterica] Length: 372\nNumber of predicted TMHs: 1\nExp number of AAs in TMHs: 24.51309\nExp number, first 60 AAs: 0.2682\nTotal prob of N-in: 0.14402\noutside 1 331\nTMhelix 332 354\ninside 355 372

19911 GCF_001366835.1_10492_1_5 Salmonella enterica subsp. enterica serovar Typhi Proteobacteria; Gammaproteobacteria; Enterobacterales; Enterobacteriaceae; Salmonella; Salmonella enterica; Salmonella enterica subsp. enterica MTGDNTLITSHGINRRDFMKLCAALAATMGLSSKAAAEMAESVSNPQRPPVIWIGAQECT WP_000145429.1 hydrogenase 2 small subunit [Salmonella enterica] Length: 372\nNumber of predicted TMHs: 1\nExp number of AAs in TMHs: 24.51309\nExp number, first 60 AAs: 0.2682\nTotal prob of N-in: 0.14402\noutside 1 331\nTMhelix 332 354\ninside 355 372

19912 GCF_001368055.1_10540_1_38 Salmonella enterica subsp. enterica serovar Typhi Proteobacteria; Gammaproteobacteria; Enterobacterales; Enterobacteriaceae; Salmonella; Salmonella enterica; Salmonella enterica subsp. enterica MTGDNTLITSHGINRRDFMKLCAALAATMGLSSKAAAEMAESVSNPQRPPVIWIGAQECT WP_000145429.1 hydrogenase 2 small subunit [Salmonella enterica] Length: 372\nNumber of predicted TMHs: 1\nExp number of AAs in TMHs: 24.51309\nExp number, first 60 AAs: 0.2682\nTotal prob of N-in: 0.14402\noutside 1 331\nTMhelix 332 354\ninside 355 372

19913 GCF_001368035.1_10561_2_42 Salmonella enterica subsp. enterica serovar Typhi Proteobacteria; Gammaproteobacteria; Enterobacterales; Enterobacteriaceae; Salmonella; Salmonella enterica;

Salmonella enterica subsp. enterica	MTGDNTLITSHGINRRDFMKLCAALAATMGLSSKAAAEMAESVSNPQRPPVIWIGAQECT
WP_000145429.1	hydrogenase 2 small subunit [Salmonella enterica]	Length: 372\nNumber of predicted TMHs:  1\nExp number of AAs in TMHs: 24.51309\nExp number, first 60 AAs:  0.2682\nTotal prob of N-in: 0.14402\noutside   1   331\nTMhelix   332   354\ninside   355   372

19914	GCF_001366875.1_10541_2_8	Salmonella enterica subsp. enterica serovar Typhi
Proteobacteria; Gammaproteobacteria; Enterobacterales; Enterobacteriaceae; Salmonella; Salmonella enterica; Salmonella enterica subsp. enterica	MTGDNTLITSHGINRRDFMKLCAALAATMGLSSKAAAEMAESVSNPQRPPVIWIGAQECT
WP_000145429.1	hydrogenase 2 small subunit [Salmonella enterica]	Length: 372\nNumber of predicted TMHs:  1\nExp number of AAs in TMHs: 24.51309\nExp number, first 60 AAs:  0.2682\nTotal prob of N-in: 0.14402\noutside   1   331\nTMhelix   332   354\ninside   355   372

19915	GCF_001368015.1_10561_2_18	Salmonella enterica subsp. enterica serovar Typhi
Proteobacteria; Gammaproteobacteria; Enterobacterales; Enterobacteriaceae; Salmonella; Salmonella enterica; Salmonella enterica subsp. enterica	MTGDNTLITSHGINRRDFMKLCAALAATMGLSSKAAAEMAESVSNPQRPPVIWIGAQECT
WP_000145429.1	hydrogenase 2 small subunit [Salmonella enterica]	Length: 372\nNumber of predicted TMHs:  1\nExp number of AAs in TMHs: 24.51309\nExp number, first 60 AAs:  0.2682\nTotal prob of N-in: 0.14402\noutside   1   331\nTMhelix   332   354\ninside   355   372

19916	GCF_001367995.1_10607_1_44	Salmonella enterica subsp. enterica serovar Typhi
Proteobacteria; Gammaproteobacteria; Enterobacterales; Enterobacteriaceae; Salmonella; Salmonella enterica; Salmonella enterica subsp. enterica	MTGDNTLITSHGINRRDFMKLCAALAATMGLSSKAAAEMAESVSNPQRPPVIWIGAQECT
WP_000145429.1	hydrogenase 2 small subunit [Salmonella enterica]	Length: 372\nNumber of predicted TMHs:  1\nExp number of AAs in TMHs: 24.51309\nExp number, first 60 AAs:  0.2682\nTotal prob of N-in: 0.14402\noutside   1   331\nTMhelix   332   354\ninside   355   372

19917	GCF_001367975.1_10426_1_15	Salmonella enterica subsp. enterica serovar Typhi
Proteobacteria; Gammaproteobacteria; Enterobacterales; Enterobacteriaceae; Salmonella; Salmonella enterica; Salmonella enterica subsp. enterica	MTGDNTLITSHGINRRDFMKLCAALAATMGLSSKAAAEMAESVSNPQRPPVIWIGAQECT
WP_000145429.1	hydrogenase 2 small subunit [Salmonella enterica]	Length: 372\nNumber of predicted TMHs:  1\nExp number of AAs in TMHs: 24.51309\nExp number, first 60 AAs:  0.2682\nTotal prob of N-in: 0.14402\noutside   1   331\nTMhelix   332   354\ninside   355   372

19918	GCF_001367795.1_10425_1_5	Salmonella enterica subsp. enterica serovar Typhi
Proteobacteria; Gammaproteobacteria; Enterobacterales; Enterobacteriaceae; Salmonella; Salmonella enterica; Salmonella enterica subsp. enterica	MTGDNTLITSHGINRRDFMKLCAALAATMGLSSKAATEMAESVSNPQRPPVIWIGAQECT
WP_001782304.1	hydrogenase 2 small subunit [Salmonella enterica]	Length: 372\nNumber of predicted TMHs:  1\nExp number of AAs in TMHs: 24.41173\nExp number, first 60 AAs:  0.16124\nTotal prob of N-in: 0.13947\noutside   1   331\nTMhelix   332   354\ninside   355   372

19919	GCF_001367955.1_10426_1_67	Salmonella enterica subsp. enterica serovar Typhi
Proteobacteria; Gammaproteobacteria; Enterobacterales; Enterobacteriaceae; Salmonella; Salmonella enterica; Salmonella enterica subsp. enterica	MTGDNTLITSHGINRRDFMKLCAALAATMGLSSKAAAEMAESVSNPQRPPVIWIGAQECT
WP_000145429.1	hydrogenase 2 small subunit [Salmonella enterica]	Length: 372\nNumber of predicted TMHs:  1\nExp number of AAs in TMHs: 24.51309\nExp number, first 60 AAs:  0.2682\nTotal prob of N-in: 0.14402\noutside   1   331\nTMhelix   332   354\ninside   355   372

19920	GCF_001367935.1_10561_2_84	Salmonella enterica subsp. enterica serovar Typhi
Proteobacteria; Gammaproteobacteria; Enterobacterales; Enterobacteriaceae; Salmonella; Salmonella enterica; Salmonella enterica subsp. enterica	MTGDNTLITSHGINRRDFMKLCAALAATMGLSSKAAAEMAESVSNPQRPPVIWIGAQECT
WP_000145429.1	hydrogenase 2 small subunit [Salmonella enterica]	Length: 372\nNumber of predicted TMHs:  1\nExp number of AAs in TMHs: 24.51309\nExp number, first 60 AAs:  0.2682\nTotal prob of N-in: 0.14402\noutside   1   331\nTMhelix   332   354\ninside   355   372

19921	GCF_001367895.1_10492_1_86	Salmonella enterica subsp. enterica serovar Typhi
Proteobacteria; Gammaproteobacteria; Enterobacterales; Enterobacteriaceae; Salmonella; Salmonella enterica; Salmonella enterica subsp. enterica	MTGDNTLITSHGINRRDFMKLCAALAATMGLSSKAAAEMAESVSNPQRPPVIWIGAQECT
WP_000145429.1	hydrogenase 2 small subunit [Salmonella enterica]	Length: 372\nNumber of predicted TMHs:  1\nExp number of AAs in TMHs: 24.51309\nExp number, first 60 AAs:  0.2682\nTotal prob of N-in: 0.14402\noutside   1   331\nTMhelix   332   354\ninside   355   372

19922 GCF_001367915.1_10493_1_6 Salmonella enterica subsp. enterica serovar Typhi Proteobacteria; Gammaproteobacteria; Enterobacterales; Enterobacteriaceae; Salmonella; Salmonella enterica; Salmonella enterica subsp. enterica MTGDNTLITSHGINRRDFMKLCAALAATMGLSSKAAAEMAESVSNPQRPPVIWIGAQECT WP_000145429.1 hydrogenase 2 small subunit [Salmonella enterica] Length: 372\nNumber of predicted TMHs: 1\nExp number of AAs in TMHs: 24.51309\nExp number, first 60 AAs: 0.2682\nTotal prob of N-in: 0.14402\noutside 1 331\nTMhelix 332 354\ninside 355 372

19923 GCF_001367855.1_10608_2_28 Salmonella enterica subsp. enterica serovar Typhi Proteobacteria; Gammaproteobacteria; Enterobacterales; Enterobacteriaceae; Salmonella; Salmonella enterica; Salmonella enterica subsp. enterica MTGDNTLITSHGINRRDFMKLCAALAATMGLSSKAAAEMAESVSNPQRPPVIWIGAQECT WP_000145429.1 hydrogenase 2 small subunit [Salmonella enterica] Length: 372\nNumber of predicted TMHs: 1\nExp number of AAs in TMHs: 24.51309\nExp number, first 60 AAs: 0.2682\nTotal prob of N-in: 0.14402\noutside 1 331\nTMhelix 332 354\ninside 355 372

19924 GCF_001367835.1_10540_1_62 Salmonella enterica subsp. enterica serovar Typhi Proteobacteria; Gammaproteobacteria; Enterobacterales; Enterobacteriaceae; Salmonella; Salmonella enterica; Salmonella enterica subsp. enterica MTGDNTLITSHGINRRDFMKLCAALAATMGLSSKAAAEMAESVSNPQRPPVIWIGAQECT WP_000145429.1 hydrogenase 2 small subunit [Salmonella enterica] Length: 372\nNumber of predicted TMHs: 1\nExp number of AAs in TMHs: 24.51309\nExp number, first 60 AAs: 0.2682\nTotal prob of N-in: 0.14402\noutside 1 331\nTMhelix 332 354\ninside 355 372

19925 GCF_001367875.1_10540_1_43 Salmonella enterica subsp. enterica serovar Typhi Proteobacteria; Gammaproteobacteria; Enterobacterales; Enterobacteriaceae; Salmonella; Salmonella enterica; Salmonella enterica subsp. enterica MTGDNTLITSHGINRRDFMKLCAALAATMGLSSKAAAEMAESVSNPQRPPVIWIGAQECT WP_000145429.1 hydrogenase 2 small subunit [Salmonella enterica] Length: 372\nNumber of predicted TMHs: 1\nExp number of AAs in TMHs: 24.51309\nExp number, first 60 AAs: 0.2682\nTotal prob of N-in: 0.14402\noutside 1 331\nTMhelix 332 354\ninside 355 372

19926 GCF_001367775.1_10607_2_13 Salmonella enterica subsp. enterica serovar Typhi Proteobacteria; Gammaproteobacteria; Enterobacterales; Enterobacteriaceae; Salmonella; Salmonella enterica; Salmonella enterica subsp. enterica MTGDNTLITSHGINRRDFMKLCAALAATMGLSSKAAAEMAESVSNPQRPPVIWIGAQECT WP_000145429.1 hydrogenase 2 small subunit [Salmonella enterica] Length: 372\nNumber of predicted TMHs: 1\nExp number of AAs in TMHs: 24.51309\nExp number, first 60 AAs: 0.2682\nTotal prob of N-in: 0.14402\noutside 1 331\nTMhelix 332 354\ninside 355 372

19927 GCF_001367815.1_10540_1_91 Salmonella enterica subsp. enterica serovar Typhi Proteobacteria; Gammaproteobacteria; Enterobacterales; Enterobacteriaceae; Salmonella; Salmonella enterica; Salmonella enterica subsp. enterica MTGDNTLITSHGINRRDFMKLCAALAATMGLSSKAAAEMAESVSNPQRPPVIWIGAQECT WP_000145429.1 hydrogenase 2 small subunit [Salmonella enterica] Length: 372\nNumber of predicted TMHs: 1\nExp number of AAs in TMHs: 24.51309\nExp number, first 60 AAs: 0.2682\nTotal prob of N-in: 0.14402\noutside 1 331\nTMhelix 332 354\ninside 355 372

19928 GCF_001367735.1_10540_1_17 Salmonella enterica subsp. enterica serovar Typhi Proteobacteria; Gammaproteobacteria; Enterobacterales; Enterobacteriaceae; Salmonella; Salmonella enterica; Salmonella enterica subsp. enterica MTGDNTLITSHGINRRDFMKLCAALAATMGLSSKAAAEMAESVSNPQRPPVIWIGAQECT WP_000145429.1 hydrogenase 2 small subunit [Salmonella enterica] Length: 372\nNumber of predicted TMHs: 1\nExp number of AAs in TMHs: 24.51309\nExp number, first 60 AAs: 0.2682\nTotal prob of N-in: 0.14402\noutside 1 331\nTMhelix 332 354\ninside 355 372

19929 GCF_001367755.1_10541_2_12 Salmonella enterica subsp. enterica serovar Typhi Proteobacteria; Gammaproteobacteria; Enterobacterales; Enterobacteriaceae; Salmonella; Salmonella enterica; Salmonella enterica subsp. enterica MTGDNTLITSHGINRRDFMKLCAALAATMGLSSKAAAEMAESVSNPQRPPVIWIGAQECT WP_000145429.1 hydrogenase 2 small subunit [Salmonella enterica] Length: 372\nNumber of predicted TMHs: 1\nExp number of AAs in TMHs: 24.51309\nExp number, first 60 AAs: 0.2682\nTotal prob of N-in: 0.14402\noutside 1 331\nTMhelix 332 354\ninside 355 372

19930 GCF_001367715.1_10426_1_92 Salmonella enterica subsp. enterica serovar Typhi Proteobacteria; Gammaproteobacteria; Enterobacterales; Enterobacteriaceae; Salmonella; Salmonella enterica; Salmonella enterica subsp. enterica MTGDNTLITSHGINRRDFMKLCAALAATMGLSSKAAAEMAESVSNPQRPPVIWIGAQECT WP_000145429.1 hydrogenase 2 small subunit [Salmonella enterica] Length: 372\nNumber of

predicted TMHs:  1\nExp number of AAs in TMHs: 24.51309\nExp number, first 60 AAs:  0.2682\nTotal prob of N-in: 0.14402\noutside   1   331\nTMhelix   332   354\ninside   355   372

19931 GCF_001367675.1_10607_2_3 Salmonella enterica subsp. enterica serovar Typhi Proteobacteria; Gammaproteobacteria; Enterobacterales; Enterobacteriaceae; Salmonella; Salmonella enterica; Salmonella enterica subsp. enterica MTGDNTLITSHGINRRDFMKLCAALAATMGLSSKAAAEMAESVSNPQRPPVIWIGAQECT WP_000145429.1 hydrogenase 2 small subunit [Salmonella enterica] Length: 372\nNumber of predicted TMHs:  1\nExp number of AAs in TMHs: 24.51309\nExp number, first 60 AAs:  0.2682\nTotal prob of N-in: 0.14402\noutside   1   331\nTMhelix   332   354\ninside   355   372

19932 GCF_001367695.1_10608_2_42 Salmonella enterica subsp. enterica serovar Typhi Proteobacteria; Gammaproteobacteria; Enterobacterales; Enterobacteriaceae; Salmonella; Salmonella enterica; Salmonella enterica subsp. enterica MTGDNTLITSHGINRRDFMKLCAALAATMGLSSKAAAEMAESVSNPQRPPVIWIGAQECT WP_000145429.1 hydrogenase 2 small subunit [Salmonella enterica] Length: 372\nNumber of predicted TMHs:  1\nExp number of AAs in TMHs: 24.51309\nExp number, first 60 AAs:  0.2682\nTotal prob of N-in: 0.14402\noutside   1   331\nTMhelix   332   354\ninside   355   372

19933 GCF_001367655.1_10540_1_24 Salmonella enterica subsp. enterica serovar Typhi Proteobacteria; Gammaproteobacteria; Enterobacterales; Enterobacteriaceae; Salmonella; Salmonella enterica; Salmonella enterica subsp. enterica MTGDNTLITSHGINRRDFMKLCAALAATMGLSSKAAAEMAESVSNPQRPPVIWIGAQECT WP_000145429.1 hydrogenase 2 small subunit [Salmonella enterica] Length: 372\nNumber of predicted TMHs:  1\nExp number of AAs in TMHs: 24.51309\nExp number, first 60 AAs:  0.2682\nTotal prob of N-in: 0.14402\noutside   1   331\nTMhelix   332   354\ninside   355   372

19934 GCF_001367635.1_10493_1_47 Salmonella enterica subsp. enterica serovar Typhi Proteobacteria; Gammaproteobacteria; Enterobacterales; Enterobacteriaceae; Salmonella; Salmonella enterica; Salmonella enterica subsp. enterica MTGDNTLITSHGINRRDFMKLCAALAATMGLSSKAAAEMAESVSNPQRPPVIWIGAQECT WP_000145429.1 hydrogenase 2 small subunit [Salmonella enterica] Length: 372\nNumber of predicted TMHs:  1\nExp number of AAs in TMHs: 24.51309\nExp number, first 60 AAs:  0.2682\nTotal prob of N-in: 0.14402\noutside   1   331\nTMhelix   332   354\ninside   355   372

19935 GCF_001367595.1_10561_2_69 Salmonella enterica subsp. enterica serovar Typhi Proteobacteria; Gammaproteobacteria; Enterobacterales; Enterobacteriaceae; Salmonella; Salmonella enterica; Salmonella enterica subsp. enterica MTGDNTLITSHGINRRDFMKLCAALAATMGLSSKAAAEMAESVSNPQRPPVIWIGAQECT WP_000145429.1 hydrogenase 2 small subunit [Salmonella enterica] Length: 372\nNumber of predicted TMHs:  1\nExp number of AAs in TMHs: 24.51309\nExp number, first 60 AAs:  0.2682\nTotal prob of N-in: 0.14402\noutside   1   331\nTMhelix   332   354\ninside   355   372

19936 GCF_001367615.1_10608_2_61 Salmonella enterica subsp. enterica serovar Typhi Proteobacteria; Gammaproteobacteria; Enterobacterales; Enterobacteriaceae; Salmonella; Salmonella enterica; Salmonella enterica subsp. enterica MTGDNTLITSHGINRRDFMKLCAALAATMGLSSKAAAEMAESVSNPQRPPVIWIGAQECT WP_000145429.1 hydrogenase 2 small subunit [Salmonella enterica] Length: 372\nNumber of predicted TMHs:  1\nExp number of AAs in TMHs: 24.51309\nExp number, first 60 AAs:  0.2682\nTotal prob of N-in: 0.14402\noutside   1   331\nTMhelix   332   354\ninside   355   372

19937 GCF_001367575.1_10540_1_21 Salmonella enterica subsp. enterica serovar Typhi Proteobacteria; Gammaproteobacteria; Enterobacterales; Enterobacteriaceae; Salmonella; Salmonella enterica; Salmonella enterica subsp. enterica MTGDNTLITSHGINRRDFMKLCAALAATMGLSSKAAAEMAESVSNPQRPPVIWIGAQECT WP_000145429.1 hydrogenase 2 small subunit [Salmonella enterica] Length: 372\nNumber of predicted TMHs:  1\nExp number of AAs in TMHs: 24.51309\nExp number, first 60 AAs:  0.2682\nTotal prob of N-in: 0.14402\noutside   1   331\nTMhelix   332   354\ninside   355   372

19938 GCF_001367555.1_10607_2_36 Salmonella enterica subsp. enterica serovar Typhi Proteobacteria; Gammaproteobacteria; Enterobacterales; Enterobacteriaceae; Salmonella; Salmonella enterica; Salmonella enterica subsp. enterica MTGDNTLITSHGINRRDFMKLCAALAATMGLSSKAAAEMAESVSNPQRPPVIWIGAQECT WP_000145429.1 hydrogenase 2 small subunit [Salmonella enterica] Length: 372\nNumber of predicted TMHs:  1\nExp number of AAs in TMHs: 24.51309\nExp number, first 60 AAs:  0.2682\nTotal prob of N-in: 0.14402\noutside   1   331\nTMhelix   332   354\ninside   355   372

19939 GCF_001367535.1_10426_1_29 Salmonella enterica subsp. enterica serovar Typhi Proteobacteria; Gammaproteobacteria; Enterobacterales; Enterobacteriaceae; Salmonella; Salmonella enterica;

Salmonella enterica subsp. enterica	MTGDNTLITSHGINRRDFMKLCAALAATMGLSSKAAAEMAESVSNPQRPPVIWIGAQECT
	WP_000145429.1	hydrogenase 2 small subunit [Salmonella enterica]	Length: 372\nNumber of predicted TMHs:  1\nExp number of AAs in TMHs: 24.51309\nExp number, first 60 AAs:  0.2682\nTotal prob of N-in: 0.14402\noutside	1	331\nTMhelix	332	354\ninside	355	372

19940	GCF_001367515.1_10607_2_22	Salmonella enterica subsp. enterica serovar Typhi
	Proteobacteria; Gammaproteobacteria; Enterobacterales; Enterobacteriaceae; Salmonella; Salmonella enterica; Salmonella enterica subsp. enterica	MTGDNTLITSHGINRRDFMKLCAALAATMGLSSKAAAEMAESVSNPQRPPVIWIGAQECT
	WP_000145429.1	hydrogenase 2 small subunit [Salmonella enterica]	Length: 372\nNumber of predicted TMHs:  1\nExp number of AAs in TMHs: 24.51309\nExp number, first 60 AAs:  0.2682\nTotal prob of N-in: 0.14402\noutside	1	331\nTMhelix	332	354\ninside	355	372

19941	GCF_001367495.1_10607_1_82	Salmonella enterica subsp. enterica serovar Typhi
	Proteobacteria; Gammaproteobacteria; Enterobacterales; Enterobacteriaceae; Salmonella; Salmonella enterica; Salmonella enterica subsp. enterica	MTGDNTLITSHGINRRDFMKLCAALAATMGLSSKAAAEMAESVSNPQRPPVIWIGAQECT
	WP_000145429.1	hydrogenase 2 small subunit [Salmonella enterica]	Length: 372\nNumber of predicted TMHs:  1\nExp number of AAs in TMHs: 24.51309\nExp number, first 60 AAs:  0.2682\nTotal prob of N-in: 0.14402\noutside	1	331\nTMhelix	332	354\ninside	355	372

19942	GCF_001367475.1_10608_2_32	Salmonella enterica subsp. enterica serovar Typhi
	Proteobacteria; Gammaproteobacteria; Enterobacterales; Enterobacteriaceae; Salmonella; Salmonella enterica; Salmonella enterica subsp. enterica	MTGDNTLITSHGINRRDFMKLCAALAATMGLSSKAAAEMAESVSNPQRPPVIWIGAQECT
	WP_000145429.1	hydrogenase 2 small subunit [Salmonella enterica]	Length: 372\nNumber of predicted TMHs:  1\nExp number of AAs in TMHs: 24.51309\nExp number, first 60 AAs:  0.2682\nTotal prob of N-in: 0.14402\noutside	1	331\nTMhelix	332	354\ninside	355	372

19943	GCF_001367435.1_10607_1_79	Salmonella enterica subsp. enterica serovar Typhi
	Proteobacteria; Gammaproteobacteria; Enterobacterales; Enterobacteriaceae; Salmonella; Salmonella enterica; Salmonella enterica subsp. enterica	MTGDNTLITSHGINRRDFMKLCAALAATMGLSSKAAAEMAESVSNPQRPPVIWIGAQECT
	WP_000145429.1	hydrogenase 2 small subunit [Salmonella enterica]	Length: 372\nNumber of predicted TMHs:  1\nExp number of AAs in TMHs: 24.51309\nExp number, first 60 AAs:  0.2682\nTotal prob of N-in: 0.14402\noutside	1	331\nTMhelix	332	354\ninside	355	372

19944	GCF_001367395.1_10493_1_10	Salmonella enterica subsp. enterica serovar Typhi
	Proteobacteria; Gammaproteobacteria; Enterobacterales; Enterobacteriaceae; Salmonella; Salmonella enterica; Salmonella enterica subsp. enterica	MTGDNTLITSHGINRRDFMKLCAALAATMGLSSKAAAEMAESVSNPQRPPVIWIGAQECT
	WP_000145429.1	hydrogenase 2 small subunit [Salmonella enterica]	Length: 372\nNumber of predicted TMHs:  1\nExp number of AAs in TMHs: 24.51309\nExp number, first 60 AAs:  0.2682\nTotal prob of N-in: 0.14402\noutside	1	331\nTMhelix	332	354\ninside	355	372

19945	GCF_001367415.1_10426_1_80	Salmonella enterica subsp. enterica serovar Typhi
	Proteobacteria; Gammaproteobacteria; Enterobacterales; Enterobacteriaceae; Salmonella; Salmonella enterica; Salmonella enterica subsp. enterica	MTGDNTLITSHGINRRDFMKLCAALAATMGLSSKAAAEMAESVSNPQRPPVIWIGAQECT
	WP_000145429.1	hydrogenase 2 small subunit [Salmonella enterica]	Length: 372\nNumber of predicted TMHs:  1\nExp number of AAs in TMHs: 24.51309\nExp number, first 60 AAs:  0.2682\nTotal prob of N-in: 0.14402\noutside	1	331\nTMhelix	332	354\ninside	355	372

19946	GCF_001367375.1_10607_1_81	Salmonella enterica subsp. enterica serovar Typhi
	Proteobacteria; Gammaproteobacteria; Enterobacterales; Enterobacteriaceae; Salmonella; Salmonella enterica; Salmonella enterica subsp. enterica	MTGDNTLITSHGINRRDFMKLCAALAATMGLSSKAAAEMAESVSNPQRPPVIWIGAQECT
	WP_000145429.1	hydrogenase 2 small subunit [Salmonella enterica]	Length: 372\nNumber of predicted TMHs:  1\nExp number of AAs in TMHs: 24.51309\nExp number, first 60 AAs:  0.2682\nTotal prob of N-in: 0.14402\noutside	1	331\nTMhelix	332	354\ninside	355	372

19947	GCF_001367355.1_10425_1_2	Salmonella enterica subsp. enterica serovar Typhi
	Proteobacteria; Gammaproteobacteria; Enterobacterales; Enterobacteriaceae; Salmonella; Salmonella enterica; Salmonella enterica subsp. enterica	MTGDNTLITSHGINRRDFMKLCAALAATMGLSSKAAAEMAESVSNPQRPPVIWIGAQECT
	WP_000145429.1	hydrogenase 2 small subunit [Salmonella enterica]	Length: 372\nNumber of predicted TMHs:  1\nExp number of AAs in TMHs: 24.51309\nExp number, first 60 AAs:  0.2682\nTotal prob of N-in: 0.14402\noutside	1	331\nTMhelix	332	354\ninside	355	372

19948 GCF_001367335.1_10540_1_82 Salmonella enterica subsp. enterica serovar Typhi Proteobacteria; Gammaproteobacteria; Enterobacterales; Enterobacteriaceae; Salmonella; Salmonella enterica; Salmonella enterica subsp. enterica MTGDNTLITSHGINRRDFMKLCAALAATMGLSSKAAAEMAESVSNPQRPPVIWIGAQECT WP_000145429.1 hydrogenase 2 small subunit [Salmonella enterica] Length: 372\nNumber of predicted TMHs: 1\nExp number of AAs in TMHs: 24.51309\nExp number, first 60 AAs: 0.2682\nTotal prob of N-in: 0.14402\noutside 1 331\nTMhelix 332 354\ninside 355 372

19949 GCF_001367315.1_10607_1_84 Salmonella enterica subsp. enterica serovar Typhi Proteobacteria; Gammaproteobacteria; Enterobacterales; Enterobacteriaceae; Salmonella; Salmonella enterica; Salmonella enterica subsp. enterica MTGDNTLITSHGINRRDFMKLCAALAATMGLSSKAAAEMAESVSNPQRPPVIWIGAQECT WP_000145429.1 hydrogenase 2 small subunit [Salmonella enterica] Length: 372\nNumber of predicted TMHs: 1\nExp number of AAs in TMHs: 24.51309\nExp number, first 60 AAs: 0.2682\nTotal prob of N-in: 0.14402\noutside 1 331\nTMhelix 332 354\ninside 355 372

19950 GCF_001367295.1_10607_2_25 Salmonella enterica subsp. enterica serovar Typhi Proteobacteria; Gammaproteobacteria; Enterobacterales; Enterobacteriaceae; Salmonella; Salmonella enterica; Salmonella enterica subsp. enterica MTGDNTLITSHGINRRDFMKLCAALAATMGLSSKAAAEMAESVSNPQRPPVIWIGAQECT WP_000145429.1 hydrogenase 2 small subunit [Salmonella enterica] Length: 372\nNumber of predicted TMHs: 1\nExp number of AAs in TMHs: 24.51309\nExp number, first 60 AAs: 0.2682\nTotal prob of N-in: 0.14402\noutside 1 331\nTMhelix 332 354\ninside 355 372

19951 GCF_001367275.1_10492_1_19 Salmonella enterica subsp. enterica serovar Typhi Proteobacteria; Gammaproteobacteria; Enterobacterales; Enterobacteriaceae; Salmonella; Salmonella enterica; Salmonella enterica subsp. enterica MTGDNTLITSHGINRRDFMKLCAALAATMGLSSKAAAEMAESVSNPQRPPVIWIGAQECT WP_000145429.1 hydrogenase 2 small subunit [Salmonella enterica] Length: 372\nNumber of predicted TMHs: 1\nExp number of AAs in TMHs: 24.51309\nExp number, first 60 AAs: 0.2682\nTotal prob of N-in: 0.14402\noutside 1 331\nTMhelix 332 354\ninside 355 372

19952 GCF_001366855.1_10607_2_11 Salmonella enterica subsp. enterica serovar Typhi Proteobacteria; Gammaproteobacteria; Enterobacterales; Enterobacteriaceae; Salmonella; Salmonella enterica; Salmonella enterica subsp. enterica MTGDNTLITSHGINRRDFMKLCAALAATMGLSSKAAAEMAESVSNPQRPPVIWIGAQECT WP_000145429.1 hydrogenase 2 small subunit [Salmonella enterica] Length: 372\nNumber of predicted TMHs: 1\nExp number of AAs in TMHs: 24.51309\nExp number, first 60 AAs: 0.2682\nTotal prob of N-in: 0.14402\noutside 1 331\nTMhelix 332 354\ninside 355 372

19953 GCF_001366915.1_10426_1_96 Salmonella enterica subsp. enterica serovar Typhi Proteobacteria; Gammaproteobacteria; Enterobacterales; Enterobacteriaceae; Salmonella; Salmonella enterica; Salmonella enterica subsp. enterica MTGDNTLITSHGINRRDFMKLCAALAATMGLSSKAAAEMAESVSNPQRPPVIWIGAQECT WP_000145429.1 hydrogenase 2 small subunit [Salmonella enterica] Length: 372\nNumber of predicted TMHs: 1\nExp number of AAs in TMHs: 24.51309\nExp number, first 60 AAs: 0.2682\nTotal prob of N-in: 0.14402\noutside 1 331\nTMhelix 332 354\ninside 355 372

19954 GCF_001366895.1_10540_1_20 Salmonella enterica subsp. enterica serovar Typhi Proteobacteria; Gammaproteobacteria; Enterobacterales; Enterobacteriaceae; Salmonella; Salmonella enterica; Salmonella enterica subsp. enterica MTGDNTLITSHGINRRDFMKLCAALAATMGLSSKAAAEMAESVSNPQRPPVIWIGAQECT WP_000145429.1 hydrogenase 2 small subunit [Salmonella enterica] Length: 372\nNumber of predicted TMHs: 1\nExp number of AAs in TMHs: 24.51309\nExp number, first 60 AAs: 0.2682\nTotal prob of N-in: 0.14402\noutside 1 331\nTMhelix 332 354\ninside 355 372

19955 GCF_001366935.1_10493_1_36 Salmonella enterica subsp. enterica serovar Typhi Proteobacteria; Gammaproteobacteria; Enterobacterales; Enterobacteriaceae; Salmonella; Salmonella enterica; Salmonella enterica subsp. enterica MTGDNTLITSHGINRRDFMKLCAALAATMGLSSKAAAEMAESVSNPQRPPVIWIGAQECT WP_000145429.1 hydrogenase 2 small subunit [Salmonella enterica] Length: 372\nNumber of predicted TMHs: 1\nExp number of AAs in TMHs: 24.51309\nExp number, first 60 AAs: 0.2682\nTotal prob of N-in: 0.14402\noutside 1 331\nTMhelix 332 354\ninside 355 372

19956 GCF_001366955.1_10426_1_37 Salmonella enterica subsp. enterica serovar Typhi Proteobacteria; Gammaproteobacteria; Enterobacterales; Enterobacteriaceae; Salmonella; Salmonella enterica; Salmonella enterica subsp. enterica MTGDNTLITSHGINRRDFMKLCAALAATMGLSSKAAAEMAESVSNPQRPPVIWIGAQECT WP_000145429.1 hydrogenase 2 small subunit [Salmonella enterica] Length: 372\nNumber of

predicted TMHs:  1\nExp number of AAs in TMHs: 24.51309\nExp number, first 60 AAs:  0.2682\nTotal prob of N-in: 0.14402\noutside   1   331\nTMhelix   332   354\ninside   355   372

19957 GCF_001366975.1_10540_1_44 Salmonella enterica subsp. enterica serovar Typhi Proteobacteria; Gammaproteobacteria; Enterobacterales; Enterobacteriaceae; Salmonella; Salmonella enterica; Salmonella enterica subsp. enterica MTGDNTLITSHGINRRDFMKLCAALAATMGLSSKAAAEMAESVSNPQRPPVIWIGAQECT WP_000145429.1 hydrogenase 2 small subunit [Salmonella enterica] Length: 372\nNumber of predicted TMHs:  1\nExp number of AAs in TMHs: 24.51309\nExp number, first 60 AAs:  0.2682\nTotal prob of N-in: 0.14402\noutside   1   331\nTMhelix   332   354\ninside   355   372

19958 GCF_001366995.1_10426_1_50 Salmonella enterica subsp. enterica serovar Typhi Proteobacteria; Gammaproteobacteria; Enterobacterales; Enterobacteriaceae; Salmonella; Salmonella enterica; Salmonella enterica subsp. enterica MTGDNTLITSHGINRRDFMKLCAALAATMGLSSKAAAEMAESVSNPQRPPVIWIGAQECT WP_000145429.1 hydrogenase 2 small subunit [Salmonella enterica] Length: 372\nNumber of predicted TMHs:  1\nExp number of AAs in TMHs: 24.51309\nExp number, first 60 AAs:  0.2682\nTotal prob of N-in: 0.14402\noutside   1   331\nTMhelix   332   354\ninside   355   372

19959 GCF_001367015.1_10607_2_24 Salmonella enterica subsp. enterica serovar Typhi Proteobacteria; Gammaproteobacteria; Enterobacterales; Enterobacteriaceae; Salmonella; Salmonella enterica; Salmonella enterica subsp. enterica MTGDNTLITSHGINRRDFMKLCAALAATMGLSSKAAAEMAESVSNPQRPPVIWIGAQECT WP_000145429.1 hydrogenase 2 small subunit [Salmonella enterica] Length: 372\nNumber of predicted TMHs:  1\nExp number of AAs in TMHs: 24.51309\nExp number, first 60 AAs:  0.2682\nTotal prob of N-in: 0.14402\noutside   1   331\nTMhelix   332   354\ninside   355   372

19960 GCF_001367055.1_10540_1_39 Salmonella enterica subsp. enterica serovar Typhi Proteobacteria; Gammaproteobacteria; Enterobacterales; Enterobacteriaceae; Salmonella; Salmonella enterica; Salmonella enterica subsp. enterica MTGDNTLITSHGINRRDFMKLCAALAATMGLSSKAAAEMAESVSNPQRPPVIWIGAQECT WP_000145429.1 hydrogenase 2 small subunit [Salmonella enterica] Length: 372\nNumber of predicted TMHs:  1\nExp number of AAs in TMHs: 24.51309\nExp number, first 60 AAs:  0.2682\nTotal prob of N-in: 0.14402\noutside   1   331\nTMhelix   332   354\ninside   355   372

19961 GCF_001367035.1_10493_1_66 Salmonella enterica subsp. enterica serovar Typhi Proteobacteria; Gammaproteobacteria; Enterobacterales; Enterobacteriaceae; Salmonella; Salmonella enterica; Salmonella enterica subsp. enterica MTGDNTLITSHGINRRDFMKLCAALAATMGLSSKAAAEMAESVSNPQRPPVIWIGAQECT WP_000145429.1 hydrogenase 2 small subunit [Salmonella enterica] Length: 372\nNumber of predicted TMHs:  1\nExp number of AAs in TMHs: 24.51309\nExp number, first 60 AAs:  0.2682\nTotal prob of N-in: 0.14402\noutside   1   331\nTMhelix   332   354\ninside   355   372

19962 GCF_001367075.1_10607_1_58 Salmonella enterica subsp. enterica serovar Typhi Proteobacteria; Gammaproteobacteria; Enterobacterales; Enterobacteriaceae; Salmonella; Salmonella enterica; Salmonella enterica subsp. enterica MTGDNTLITSHGINRRDFMKLCAALAATMGLSSKAAAEMAESVSNPQRPPVIWIGAQECT WP_000145429.1 hydrogenase 2 small subunit [Salmonella enterica] Length: 372\nNumber of predicted TMHs:  1\nExp number of AAs in TMHs: 24.51309\nExp number, first 60 AAs:  0.2682\nTotal prob of N-in: 0.14402\noutside   1   331\nTMhelix   332   354\ninside   355   372

19963 GCF_001367095.1_10607_1_42 Salmonella enterica subsp. enterica serovar Typhi Proteobacteria; Gammaproteobacteria; Enterobacterales; Enterobacteriaceae; Salmonella; Salmonella enterica; Salmonella enterica subsp. enterica MTGDNTLITSHGINRRDFMKLCAALAATMGLSSKAAAEMAESVSNPQRPPVIWIGAQECT WP_000145429.1 hydrogenase 2 small subunit [Salmonella enterica] Length: 372\nNumber of predicted TMHs:  1\nExp number of AAs in TMHs: 24.51309\nExp number, first 60 AAs:  0.2682\nTotal prob of N-in: 0.14402\noutside   1   331\nTMhelix   332   354\ninside   355   372

19964 GCF_001367115.1_10540_1_67 Salmonella enterica subsp. enterica serovar Typhi Proteobacteria; Gammaproteobacteria; Enterobacterales; Enterobacteriaceae; Salmonella; Salmonella enterica; Salmonella enterica subsp. enterica MTGDNTLITSHGINRRDFMKLCAALAATMGLSSKAAAEMAESVSNPQRPPVIWIGAQECT WP_000145429.1 hydrogenase 2 small subunit [Salmonella enterica] Length: 372\nNumber of predicted TMHs:  1\nExp number of AAs in TMHs: 24.51309\nExp number, first 60 AAs:  0.2682\nTotal prob of N-in: 0.14402\noutside   1   331\nTMhelix   332   354\ninside   355   372

19965 GCF_001367135.1_10540_1_26 Salmonella enterica subsp. enterica serovar Typhi Proteobacteria; Gammaproteobacteria; Enterobacterales; Enterobacteriaceae; Salmonella; Salmonella enterica;

Salmonella enterica subsp. enterica MTGDNTLITSHGINRRDFMKLCAALAATMGLSSKAAAEMAESVSNPQRPPVIWIGAQECT WP_000145429.1 hydrogenase 2 small subunit [Salmonella enterica] Length: 372\nNumber of predicted TMHs: 1\nExp number of AAs in TMHs: 24.51309\nExp number, first 60 AAs: 0.2682\nTotal prob of N-in: 0.14402\noutside 1 331\nTMhelix 332 354\ninside 355 372

19966 GCF_001367255.1_10540_1_31 Salmonella enterica subsp. enterica serovar Typhi Proteobacteria; Gammaproteobacteria; Enterobacterales; Enterobacteriaceae; Salmonella; Salmonella enterica; Salmonella enterica subsp. enterica MTGDNTLITSHGINRRDFMKLCAALAATMGLSSKAAAEMAESVSNPQRPPVIWIGAQECT WP_000145429.1 hydrogenase 2 small subunit [Salmonella enterica] Length: 372\nNumber of predicted TMHs: 1\nExp number of AAs in TMHs: 24.51309\nExp number, first 60 AAs: 0.2682\nTotal prob of N-in: 0.14402\noutside 1 331\nTMhelix 332 354\ninside 355 372

19967 GCF_001367175.1_10426_1_84 Salmonella enterica subsp. enterica serovar Typhi Proteobacteria; Gammaproteobacteria; Enterobacterales; Enterobacteriaceae; Salmonella; Salmonella enterica; Salmonella enterica subsp. enterica MTGDNTLITSHGINRRDFMKLCAALAATMGLSSKAAAEMAESVSNPQRPPVIWIGAQECT WP_000145429.1 hydrogenase 2 small subunit [Salmonella enterica] Length: 372\nNumber of predicted TMHs: 1\nExp number of AAs in TMHs: 24.51309\nExp number, first 60 AAs: 0.2682\nTotal prob of N-in: 0.14402\noutside 1 331\nTMhelix 332 354\ninside 355 372

19968 GCF_001367155.1_10608_2_19 Salmonella enterica subsp. enterica serovar Typhi Proteobacteria; Gammaproteobacteria; Enterobacterales; Enterobacteriaceae; Salmonella; Salmonella enterica; Salmonella enterica subsp. enterica MTGDNTLITSHGINRRDFMKLCAALAATMGLSSKAAAEMAESVSNPQRPPVIWIGAQECT WP_000145429.1 hydrogenase 2 small subunit [Salmonella enterica] Length: 372\nNumber of predicted TMHs: 1\nExp number of AAs in TMHs: 24.51309\nExp number, first 60 AAs: 0.2682\nTotal prob of N-in: 0.14402\noutside 1 331\nTMhelix 332 354\ninside 355 372

19969 GCF_001367195.1_10540_1_59 Salmonella enterica subsp. enterica serovar Typhi Proteobacteria; Gammaproteobacteria; Enterobacterales; Enterobacteriaceae; Salmonella; Salmonella enterica; Salmonella enterica subsp. enterica MTGDNTLITSHGINRRDFMKLCAALAATMGLSSKAAAEMAESVSNPQRPPVIWIGAQECT WP_000145429.1 hydrogenase 2 small subunit [Salmonella enterica] Length: 372\nNumber of predicted TMHs: 1\nExp number of AAs in TMHs: 24.51309\nExp number, first 60 AAs: 0.2682\nTotal prob of N-in: 0.14402\noutside 1 331\nTMhelix 332 354\ninside 355 372

19970 GCF_001367215.1_10425_1_32 Salmonella enterica subsp. enterica serovar Typhi Proteobacteria; Gammaproteobacteria; Enterobacterales; Enterobacteriaceae; Salmonella; Salmonella enterica; Salmonella enterica subsp. enterica MTGDNTLITSHGINRRDFMKLCAALAATMGLSSKAAAEMAESVSNPQRPPVIWIGAQECT WP_000145429.1 hydrogenase 2 small subunit [Salmonella enterica] Length: 372\nNumber of predicted TMHs: 1\nExp number of AAs in TMHs: 24.51309\nExp number, first 60 AAs: 0.2682\nTotal prob of N-in: 0.14402\noutside 1 331\nTMhelix 332 354\ninside 355 372

19971 GCF_001114745.1_9953_5_84 Salmonella enterica subsp. enterica serovar Typhi Proteobacteria; Gammaproteobacteria; Enterobacterales; Enterobacteriaceae; Salmonella; Salmonella enterica; Salmonella enterica subsp. enterica MTGDNTLITSHGINRRDFMKLCAALAATMGLSSKAAAEMAESVSNPQRPPVIWIGAQECT WP_000145429.1 hydrogenase 2 small subunit [Salmonella enterica] Length: 372\nNumber of predicted TMHs: 1\nExp number of AAs in TMHs: 24.51309\nExp number, first 60 AAs: 0.2682\nTotal prob of N-in: 0.14402\noutside 1 331\nTMhelix 332 354\ninside 355 372

19972 GCF_001048055.1_10492_1_84 Salmonella enterica subsp. enterica serovar Typhi Proteobacteria; Gammaproteobacteria; Enterobacterales; Enterobacteriaceae; Salmonella; Salmonella enterica; Salmonella enterica subsp. enterica MTGDNTLITSHGINRRDFMKLCAALAATMGLSSKAAAEMAESVSNPQRPPVIWIGAQECT WP_000145429.1 hydrogenase 2 small subunit [Salmonella enterica] Length: 372\nNumber of predicted TMHs: 1\nExp number of AAs in TMHs: 24.51309\nExp number, first 60 AAs: 0.2682\nTotal prob of N-in: 0.14402\noutside 1 331\nTMhelix 332 354\ninside 355 372

19973 GCF_001048135.1_10493_1_63 Salmonella enterica subsp. enterica serovar Typhi Proteobacteria; Gammaproteobacteria; Enterobacterales; Enterobacteriaceae; Salmonella; Salmonella enterica; Salmonella enterica subsp. enterica MTGDNTLITSHGINRRDFMKLCAALAATMGLSSKAAAEMAESVSNPQRPPVIWIGAQECT WP_000145429.1 hydrogenase 2 small subunit [Salmonella enterica] Length: 372\nNumber of predicted TMHs: 1\nExp number of AAs in TMHs: 24.51309\nExp number, first 60 AAs: 0.2682\nTotal prob of N-in: 0.14402\noutside 1 331\nTMhelix 332 354\ninside 355 372

19974 GCF_001048095.1_10493_1_17 Salmonella enterica subsp. enterica serovar Typhi Proteobacteria; Gammaproteobacteria; Enterobacterales; Enterobacteriaceae; Salmonella; Salmonella enterica; Salmonella enterica subsp. enterica MTGDNTLITSHGINRRDFMKLCAALAATMGLSSKAAAEMAESVSNPQRPPVIWIGAQECT WP_000145429.1 hydrogenase 2 small subunit [Salmonella enterica] Length: 372\nNumber of predicted TMHs: 1\nExp number of AAs in TMHs: 24.51309\nExp number, first 60 AAs: 0.2682\nTotal prob of N-in: 0.14402\noutside 1 331\nTMhelix 332 354\ninside 355 372

19975 GCF_001048175.1_10493_1_50 Salmonella enterica subsp. enterica serovar Typhi Proteobacteria; Gammaproteobacteria; Enterobacterales; Enterobacteriaceae; Salmonella; Salmonella enterica; Salmonella enterica subsp. enterica MTGDNTLITSHGINRRDFMKLCAALAATMGLSSKAAAEMAESVSNPQRPPVIWIGAQECT WP_000145429.1 hydrogenase 2 small subunit [Salmonella enterica] Length: 372\nNumber of predicted TMHs: 1\nExp number of AAs in TMHs: 24.51309\nExp number, first 60 AAs: 0.2682\nTotal prob of N-in: 0.14402\noutside 1 331\nTMhelix 332 354\ninside 355 372

19976 GCF_001048215.1_10492_1_23 Salmonella enterica subsp. enterica serovar Typhi Proteobacteria; Gammaproteobacteria; Enterobacterales; Enterobacteriaceae; Salmonella; Salmonella enterica; Salmonella enterica subsp. enterica MTGDNTLITSHGINRRDFMKLCAALAATMGLSSKAAAEMAESVSNPQRPPVIWIGAQECT WP_000145429.1 hydrogenase 2 small subunit [Salmonella enterica] Length: 372\nNumber of predicted TMHs: 1\nExp number of AAs in TMHs: 24.51309\nExp number, first 60 AAs: 0.2682\nTotal prob of N-in: 0.14402\noutside 1 331\nTMhelix 332 354\ninside 355 372

19977 GCF_001048255.1_10493_1_86 Salmonella enterica subsp. enterica serovar Typhi Proteobacteria; Gammaproteobacteria; Enterobacterales; Enterobacteriaceae; Salmonella; Salmonella enterica; Salmonella enterica subsp. enterica MTGDNTLITSHGINRRDFMKLCAALAATMGLSSKAAAEMAESVSNPQRPPVIWIGAQECT WP_000145429.1 hydrogenase 2 small subunit [Salmonella enterica] Length: 372\nNumber of predicted TMHs: 1\nExp number of AAs in TMHs: 24.51309\nExp number, first 60 AAs: 0.2682\nTotal prob of N-in: 0.14402\noutside 1 331\nTMhelix 332 354\ninside 355 372

19978 GCF_001078925.1_10493_1_24 Salmonella enterica subsp. enterica serovar Typhi Proteobacteria; Gammaproteobacteria; Enterobacterales; Enterobacteriaceae; Salmonella; Salmonella enterica; Salmonella enterica subsp. enterica MTGDNTLITSHGINRRDFMKLCAALAATMGLSSKAAAEMAESVSNPQRPPVIWIGAQECT WP_000145429.1 hydrogenase 2 small subunit [Salmonella enterica] Length: 372\nNumber of predicted TMHs: 1\nExp number of AAs in TMHs: 24.51309\nExp number, first 60 AAs: 0.2682\nTotal prob of N-in: 0.14402\noutside 1 331\nTMhelix 332 354\ninside 355 372

19979 GCF_001078965.1_10071_3_27 Salmonella enterica subsp. enterica serovar Typhi Proteobacteria; Gammaproteobacteria; Enterobacterales; Enterobacteriaceae; Salmonella; Salmonella enterica; Salmonella enterica subsp. enterica MTGDNTLITSHGINRRDFMKLCAALAATMGLSSKAAAEMAESVSNPQRPPVIWIGAQECT WP_000145429.1 hydrogenase 2 small subunit [Salmonella enterica] Length: 372\nNumber of predicted TMHs: 1\nExp number of AAs in TMHs: 24.51309\nExp number, first 60 AAs: 0.2682\nTotal prob of N-in: 0.14402\noutside 1 331\nTMhelix 332 354\ninside 355 372

19980 GCF_001079005.1_10071_3_52 Salmonella enterica subsp. enterica serovar Typhi Proteobacteria; Gammaproteobacteria; Enterobacterales; Enterobacteriaceae; Salmonella; Salmonella enterica; Salmonella enterica subsp. enterica MTGDNTLITSHGINRRDFMKLCAALAATMGLSSKAAAEMAESVSNPQRPPVIWIGAQECT WP_000145429.1 hydrogenase 2 small subunit [Salmonella enterica] Length: 372\nNumber of predicted TMHs: 1\nExp number of AAs in TMHs: 24.51309\nExp number, first 60 AAs: 0.2682\nTotal prob of N-in: 0.14402\noutside 1 331\nTMhelix 332 354\ninside 355 372

19981 GCF_001079045.1_10071_8_13 Salmonella enterica subsp. enterica serovar Typhi Proteobacteria; Gammaproteobacteria; Enterobacterales; Enterobacteriaceae; Salmonella; Salmonella enterica; Salmonella enterica subsp. enterica MTGDNTLITSHGINRRDFMKLCAALAATMGLSSKAAAEMAESVSNPQRPPVIWIGAQECT WP_000145429.1 hydrogenase 2 small subunit [Salmonella enterica] Length: 372\nNumber of predicted TMHs: 1\nExp number of AAs in TMHs: 24.51309\nExp number, first 60 AAs: 0.2682\nTotal prob of N-in: 0.14402\noutside 1 331\nTMhelix 332 354\ninside 355 372

19982 GCF_001079065.1_10060_6_27 Salmonella enterica subsp. enterica serovar Typhi Proteobacteria; Gammaproteobacteria; Enterobacterales; Enterobacteriaceae; Salmonella; Salmonella enterica; Salmonella enterica subsp. enterica MTGDNTLITSHGINRRDFMKLCAALAATMGLSSKAAAEMAESVSNPQRPPVIWIGAQECT WP_000145429.1 hydrogenase 2 small subunit [Salmonella enterica] Length: 372\nNumber of

predicted TMHs: 1\nExp number of AAs in TMHs: 24.51309\nExp number, first 60 AAs: 0.2682\nTotal prob of N-in: 0.14402\noutside 1 331\nTMhelix 332 354\ninside 355 372

19983 GCF_001079105.1_10060_6_87 Salmonella enterica subsp. enterica serovar Typhi Proteobacteria; Gammaproteobacteria; Enterobacterales; Enterobacteriaceae; Salmonella; Salmonella enterica; Salmonella enterica subsp. enterica MTGDNTLITSHGINRRDFMKLCAALAATMGLSSKAAAEMAESVSNPQRPPVIWIGAQECT WP_000145429.1 hydrogenase 2 small subunit [Salmonella enterica] Length: 372\nNumber of predicted TMHs: 1\nExp number of AAs in TMHs: 24.51309\nExp number, first 60 AAs: 0.2682\nTotal prob of N-in: 0.14402\noutside 1 331\nTMhelix 332 354\ninside 355 372

19984 GCF_001079705.1_10060_6_47 Salmonella enterica subsp. enterica serovar Typhi Proteobacteria; Gammaproteobacteria; Enterobacterales; Enterobacteriaceae; Salmonella; Salmonella enterica; Salmonella enterica subsp. enterica MTGDNTLITSHGINRRDFMKLCAALAATMGLSSKAAAEMAESVSNPQRPPVIWIGAQECT WP_000145429.1 hydrogenase 2 small subunit [Salmonella enterica] Length: 372\nNumber of predicted TMHs: 1\nExp number of AAs in TMHs: 24.51309\nExp number, first 60 AAs: 0.2682\nTotal prob of N-in: 0.14402\noutside 1 331\nTMhelix 332 354\ninside 355 372

19985 GCF_001079785.1_10071_3_60 Salmonella enterica subsp. enterica serovar Typhi Proteobacteria; Gammaproteobacteria; Enterobacterales; Enterobacteriaceae; Salmonella; Salmonella enterica; Salmonella enterica subsp. enterica MTGDNTLITSHGINRRDFMKLCAALAATMGLSSKAAAEMAESVSNPQRPPVIWIGAQECT WP_000145429.1 hydrogenase 2 small subunit [Salmonella enterica] Length: 372\nNumber of predicted TMHs: 1\nExp number of AAs in TMHs: 24.51309\nExp number, first 60 AAs: 0.2682\nTotal prob of N-in: 0.14402\noutside 1 331\nTMhelix 332 354\ninside 355 372

19986 GCF_001079745.1_10060_6_38 Salmonella enterica subsp. enterica serovar Typhi Proteobacteria; Gammaproteobacteria; Enterobacterales; Enterobacteriaceae; Salmonella; Salmonella enterica; Salmonella enterica subsp. enterica MTGDNTLITSHGINRRDFMKLCAALAATMGLSSKAAAEMAESVSNPQRPPVIWIGAQECT WP_000145429.1 hydrogenase 2 small subunit [Salmonella enterica] Length: 372\nNumber of predicted TMHs: 1\nExp number of AAs in TMHs: 24.51309\nExp number, first 60 AAs: 0.2682\nTotal prob of N-in: 0.14402\noutside 1 331\nTMhelix 332 354\ninside 355 372

19987 GCF_001079825.1_10060_6_93 Salmonella enterica subsp. enterica serovar Typhi Proteobacteria; Gammaproteobacteria; Enterobacterales; Enterobacteriaceae; Salmonella; Salmonella enterica; Salmonella enterica subsp. enterica MTGDNTLITSHGINRRDFMKLCAALAATMGLSSKAAAEMAESVSNPQRPPVIWIGAQECT WP_000145429.1 hydrogenase 2 small subunit [Salmonella enterica] Length: 372\nNumber of predicted TMHs: 1\nExp number of AAs in TMHs: 24.51309\nExp number, first 60 AAs: 0.2682\nTotal prob of N-in: 0.14402\noutside 1 331\nTMhelix 332 354\ninside 355 372

19988 GCF_001079905.1_10071_8_47 Salmonella enterica subsp. enterica serovar Typhi Proteobacteria; Gammaproteobacteria; Enterobacterales; Enterobacteriaceae; Salmonella; Salmonella enterica; Salmonella enterica subsp. enterica MTGDNTLITSHGINRRDFMKLCAALAATMGLSSKAAAEMAESVSNPQRPPVIWIGAQECT WP_000145429.1 hydrogenase 2 small subunit [Salmonella enterica] Length: 372\nNumber of predicted TMHs: 1\nExp number of AAs in TMHs: 24.51309\nExp number, first 60 AAs: 0.2682\nTotal prob of N-in: 0.14402\noutside 1 331\nTMhelix 332 354\ninside 355 372

19989 GCF_001079865.1_10060_5_84 Salmonella enterica subsp. enterica serovar Typhi Proteobacteria; Gammaproteobacteria; Enterobacterales; Enterobacteriaceae; Salmonella; Salmonella enterica; Salmonella enterica subsp. enterica MTGDNTLITSHGINRRDFMKLCAALAATMGLSSKAAAEMAESVSNPQRPPVIWIGAQECT WP_000145429.1 hydrogenase 2 small subunit [Salmonella enterica] Length: 372\nNumber of predicted TMHs: 1\nExp number of AAs in TMHs: 24.51309\nExp number, first 60 AAs: 0.2682\nTotal prob of N-in: 0.14402\noutside 1 331\nTMhelix 332 354\ninside 355 372

19990 GCF_001080485.1_10071_8_17 Salmonella enterica subsp. enterica serovar Typhi Proteobacteria; Gammaproteobacteria; Enterobacterales; Enterobacteriaceae; Salmonella; Salmonella enterica; Salmonella enterica subsp. enterica MTGDNTLITSHGINRRDFMKLCAALAATMGLSSKAAAEMAESVSNPQRPPVIWIGAQECT WP_000145429.1 hydrogenase 2 small subunit [Salmonella enterica] Length: 372\nNumber of predicted TMHs: 1\nExp number of AAs in TMHs: 24.51309\nExp number, first 60 AAs: 0.2682\nTotal prob of N-in: 0.14402\noutside 1 331\nTMhelix 332 354\ninside 355 372

19991 GCF_001080245.1_10071_3_51 Salmonella enterica subsp. enterica serovar Typhi Proteobacteria; Gammaproteobacteria; Enterobacterales; Enterobacteriaceae; Salmonella; Salmonella enterica;

Salmonella enterica subsp. enterica	MTGDNTLITSHGINRRDFMKLCAALAATMGLSSKAATEMAESVSNPQRPPVIWIGAQECT	WP_001782304.1	hydrogenase 2 small subunit [Salmonella enterica]	Length: 372\nNumber of predicted TMHs:  1\nExp number of AAs in TMHs: 24.41173\nExp number, first 60 AAs:  0.16124\nTotal prob of N-in: 0.13947\noutside   1   331\nTMhelix   332   354\ninside   355   372

19992	GCF_001080525.1_10071_3_53	Salmonella enterica subsp. enterica serovar Typhi	Proteobacteria; Gammaproteobacteria; Enterobacterales; Enterobacteriaceae; Salmonella; Salmonella enterica; Salmonella enterica subsp. enterica	MTGDNTLITSHGINRRDFMKLCAALAATMGLSSKAAAEMAESVSNPQRPPVIWIGAQECT	WP_000145429.1	hydrogenase 2 small subunit [Salmonella enterica]	Length: 372\nNumber of predicted TMHs:  1\nExp number of AAs in TMHs: 24.51309\nExp number, first 60 AAs:  0.2682\nTotal prob of N-in: 0.14402\noutside   1   331\nTMhelix   332   354\ninside   355   372

19993	GCF_001080365.1_10071_3_61	Salmonella enterica subsp. enterica serovar Typhi	Proteobacteria; Gammaproteobacteria; Enterobacterales; Enterobacteriaceae; Salmonella; Salmonella enterica; Salmonella enterica subsp. enterica	MTGDNTLITSHGINRRDFMKLCAALAATMGLSSKAATEMAESVSNPQRPPVIWIGAQECT	WP_001782304.1	hydrogenase 2 small subunit [Salmonella enterica]	Length: 372\nNumber of predicted TMHs:  1\nExp number of AAs in TMHs: 24.41173\nExp number, first 60 AAs:  0.16124\nTotal prob of N-in: 0.13947\noutside   1   331\nTMhelix   332   354\ninside   355   372

19994	GCF_001080565.1_10071_3_48	Salmonella enterica subsp. enterica serovar Typhi	Proteobacteria; Gammaproteobacteria; Enterobacterales; Enterobacteriaceae; Salmonella; Salmonella enterica; Salmonella enterica subsp. enterica	MTGDNTLITSHGINRRDFMKLCAALAATMGLSSKAAAEMAESVSNPQRPPVIWIGAQECT	WP_000145429.1	hydrogenase 2 small subunit [Salmonella enterica]	Length: 372\nNumber of predicted TMHs:  1\nExp number of AAs in TMHs: 24.51309\nExp number, first 60 AAs:  0.2682\nTotal prob of N-in: 0.14402\noutside   1   331\nTMhelix   332   354\ninside   355   372

19995	GCF_001080605.1_10060_6_68	Salmonella enterica subsp. enterica serovar Typhi	Proteobacteria; Gammaproteobacteria; Enterobacterales; Enterobacteriaceae; Salmonella; Salmonella enterica; Salmonella enterica subsp. enterica	MTGDNTLITSHGINRRDFMKLCAALAATMGLSSKAAAEMAESVSNPQRPPVIWIGAQECT	WP_000145429.1	hydrogenase 2 small subunit [Salmonella enterica]	Length: 372\nNumber of predicted TMHs:  1\nExp number of AAs in TMHs: 24.51309\nExp number, first 60 AAs:  0.2682\nTotal prob of N-in: 0.14402\noutside   1   331\nTMhelix   332   354\ninside   355   372

19996	GCF_001080945.1_10071_8_5	Salmonella enterica subsp. enterica serovar Typhi	Proteobacteria; Gammaproteobacteria; Enterobacterales; Enterobacteriaceae; Salmonella; Salmonella enterica; Salmonella enterica subsp. enterica	MTGDNTLITSHGINRRDFMKLCAALAATMGLSSKAAAEMAESVSNPQRPPVIWIGAQECT	WP_000145429.1	hydrogenase 2 small subunit [Salmonella enterica]	Length: 372\nNumber of predicted TMHs:  1\nExp number of AAs in TMHs: 24.51309\nExp number, first 60 AAs:  0.2682\nTotal prob of N-in: 0.14402\noutside   1   331\nTMhelix   332   354\ninside   355   372

19997	GCF_001080985.1_10071_3_40	Salmonella enterica subsp. enterica serovar Typhi	Proteobacteria; Gammaproteobacteria; Enterobacterales; Enterobacteriaceae; Salmonella; Salmonella enterica; Salmonella enterica subsp. enterica	MTGDNTLITSHGINRRDFMKLCAALAATMGLSSKAAAEMAESVSNPQRPPVIWIGAQECT	WP_000145429.1	hydrogenase 2 small subunit [Salmonella enterica]	Length: 372\nNumber of predicted TMHs:  1\nExp number of AAs in TMHs: 24.51309\nExp number, first 60 AAs:  0.2682\nTotal prob of N-in: 0.14402\noutside   1   331\nTMhelix   332   354\ninside   355   372

19998	GCF_001081025.1_10060_6_45	Salmonella enterica subsp. enterica serovar Typhi	Proteobacteria; Gammaproteobacteria; Enterobacterales; Enterobacteriaceae; Salmonella; Salmonella enterica; Salmonella enterica subsp. enterica	MTGDNTLITSHGINRRDFMKLCAALAATMGLSSKAAAEMAESVSNPQRPPVIWIGAQECT	WP_000145429.1	hydrogenase 2 small subunit [Salmonella enterica]	Length: 372\nNumber of predicted TMHs:  1\nExp number of AAs in TMHs: 24.51309\nExp number, first 60 AAs:  0.2682\nTotal prob of N-in: 0.14402\noutside   1   331\nTMhelix   332   354\ninside   355   372

19999	GCF_001081065.1_10071_8_68	Salmonella enterica subsp. enterica serovar Typhi	Proteobacteria; Gammaproteobacteria; Enterobacterales; Enterobacteriaceae; Salmonella; Salmonella enterica; Salmonella enterica subsp. enterica	MTGDNTLITSHGINRRDFMKLCAALAATMGLSSKAAAEMAESVSNPQRPPVIWIGAQECT	WP_000145429.1	hydrogenase 2 small subunit [Salmonella enterica]	Length: 372\nNumber of predicted TMHs:  1\nExp number of AAs in TMHs: 24.51309\nExp number, first 60 AAs:  0.2682\nTotal prob of N-in: 0.14402\noutside   1   331\nTMhelix   332   354\ninside   355   372

20000	GCF_001081105.1_10071_8_59	Salmonella enterica subsp. enterica serovar Typhi	Proteobacteria; Gammaproteobacteria; Enterobacterales; Enterobacteriaceae; Salmonella; Salmonella enterica; Salmonella enterica subsp. enterica	MTGDNTLITSHGINRRDFMKLCAALAATMGLSSKAAAEMAESVSNPQRPPVIWIGAQECT	WP_000145429.1	hydrogenase 2 small subunit [Salmonella enterica]	Length: 372\nNumber of predicted TMHs: 1\nExp number of AAs in TMHs: 24.51309\nExp number, first 60 AAs: 0.2682\nTotal prob of N-in: 0.14402\noutside 1 331\nTMhelix 332 354\ninside 355 372

20001	GCF_001081185.1_10071_3_64	Salmonella enterica subsp. enterica serovar Typhi	Proteobacteria; Gammaproteobacteria; Enterobacterales; Enterobacteriaceae; Salmonella; Salmonella enterica; Salmonella enterica subsp. enterica	MTGDNTLITSHGINRRDFMKLCAALAATMGLSSKAAAEMAESVSNPQRPPVIWIGAQECT	WP_000145429.1	hydrogenase 2 small subunit [Salmonella enterica]	Length: 372\nNumber of predicted TMHs: 1\nExp number of AAs in TMHs: 24.51309\nExp number, first 60 AAs: 0.2682\nTotal prob of N-in: 0.14402\noutside 1 331\nTMhelix 332 354\ninside 355 372

20002	GCF_001081685.1_10071_8_83	Salmonella enterica subsp. enterica serovar Typhi	Proteobacteria; Gammaproteobacteria; Enterobacterales; Enterobacteriaceae; Salmonella; Salmonella enterica; Salmonella enterica subsp. enterica	MTGDNTLITSHGINRRDFMKLCAALAATMGLSSKAAAEMAESVSNPQRPPVIWIGAQECT	WP_000145429.1	hydrogenase 2 small subunit [Salmonella enterica]	Length: 372\nNumber of predicted TMHs: 1\nExp number of AAs in TMHs: 24.51309\nExp number, first 60 AAs: 0.2682\nTotal prob of N-in: 0.14402\noutside 1 331\nTMhelix 332 354\ninside 355 372

20003	GCF_001081725.1_10071_8_64	Salmonella enterica subsp. enterica serovar Typhi	Proteobacteria; Gammaproteobacteria; Enterobacterales; Enterobacteriaceae; Salmonella; Salmonella enterica; Salmonella enterica subsp. enterica	MTGDNTLITSHGINRRDFMKLCAALAATMGLSSKAAAEMAESVSNPQRPPVIWIGAQECT	WP_000145429.1	hydrogenase 2 small subunit [Salmonella enterica]	Length: 372\nNumber of predicted TMHs: 1\nExp number of AAs in TMHs: 24.51309\nExp number, first 60 AAs: 0.2682\nTotal prob of N-in: 0.14402\noutside 1 331\nTMhelix 332 354\ninside 355 372

20004	GCF_001081805.1_10060_5_79	Salmonella enterica subsp. enterica serovar Typhi	Proteobacteria; Gammaproteobacteria; Enterobacterales; Enterobacteriaceae; Salmonella; Salmonella enterica; Salmonella enterica subsp. enterica	MTGDNTLITSHGINRRDFMKLCAALAATMGLSSKAAAEMAESVSNPQRPPVIWIGAQECT	WP_000145429.1	hydrogenase 2 small subunit [Salmonella enterica]	Length: 372\nNumber of predicted TMHs: 1\nExp number of AAs in TMHs: 24.51309\nExp number, first 60 AAs: 0.2682\nTotal prob of N-in: 0.14402\noutside 1 331\nTMhelix 332 354\ninside 355 372

20005	GCF_001081445.1_10071_3_9	Salmonella enterica subsp. enterica serovar Typhi	Proteobacteria; Gammaproteobacteria; Enterobacterales; Enterobacteriaceae; Salmonella; Salmonella enterica; Salmonella enterica subsp. enterica	MTGDNTLITSHGINRRDFMKLCAALAATMGLSSKAATEMAESVSNPQRPPVIWIGAQECT	WP_001782304.1	hydrogenase 2 small subunit [Salmonella enterica]	Length: 372\nNumber of predicted TMHs: 1\nExp number of AAs in TMHs: 24.41173\nExp number, first 60 AAs: 0.16124\nTotal prob of N-in: 0.13947\noutside 1 331\nTMhelix 332 354\ninside 355 372

20006	GCF_001081845.1_10071_8_14	Salmonella enterica subsp. enterica serovar Typhi	Proteobacteria; Gammaproteobacteria; Enterobacterales; Enterobacteriaceae; Salmonella; Salmonella enterica; Salmonella enterica subsp. enterica	MTGDNTLITSHGINRRDFMKLCAALAATMGLSSKAAAEMAESVSNPQRPPVIWIGAQECT	WP_000145429.1	hydrogenase 2 small subunit [Salmonella enterica]	Length: 372\nNumber of predicted TMHs: 1\nExp number of AAs in TMHs: 24.51309\nExp number, first 60 AAs: 0.2682\nTotal prob of N-in: 0.14402\noutside 1 331\nTMhelix 332 354\ninside 355 372

20007	GCF_001081925.1_10060_6_1	Salmonella enterica subsp. enterica serovar Typhi	Proteobacteria; Gammaproteobacteria; Enterobacterales; Enterobacteriaceae; Salmonella; Salmonella enterica; Salmonella enterica subsp. enterica	MTGDNTLITSHGINRRDFMKLCAALAATMGLSSKAAAEMAESVSNPQRPPVIWIGAQECT	WP_000145429.1	hydrogenase 2 small subunit [Salmonella enterica]	Length: 372\nNumber of predicted TMHs: 1\nExp number of AAs in TMHs: 24.51309\nExp number, first 60 AAs: 0.2682\nTotal prob of N-in: 0.14402\noutside 1 331\nTMhelix 332 354\ninside 355 372

20008	GCF_001081885.1_10060_5_21	Salmonella enterica subsp. enterica serovar Typhi	Proteobacteria; Gammaproteobacteria; Enterobacterales; Enterobacteriaceae; Salmonella; Salmonella enterica; Salmonella enterica subsp. enterica	MTGDNTLITSHGINRRDFMKLCAALAATMGLSSKAAAEMAESVSNPQRPPVIWIGAQECT	WP_000145429.1	hydrogenase 2 small subunit [Salmonella enterica]	Length: 372\nNumber of

predicted TMHs: 1\nExp number of AAs in TMHs: 24.51309\nExp number, first 60 AAs: 0.2682\nTotal prob of N-in: 0.14402\noutside 1 331\nTMhelix 332 354\ninside 355 372

20009 GCF_001082505.1_10071_8_51 Salmonella enterica subsp. enterica serovar Typhi Proteobacteria; Gammaproteobacteria; Enterobacterales; Enterobacteriaceae; Salmonella; Salmonella enterica; Salmonella enterica subsp. enterica MTGDNTLITSHGINRRDFMKLCAALAATMGLSSKAAAEMAESVSNPQRPPVIWIGAQECT WP_000145429.1 hydrogenase 2 small subunit [Salmonella enterica] Length: 372\nNumber of predicted TMHs: 1\nExp number of AAs in TMHs: 24.51309\nExp number, first 60 AAs: 0.2682\nTotal prob of N-in: 0.14402\noutside 1 331\nTMhelix 332 354\ninside 355 372

20010 GCF_001082545.1_10071_8_30 Salmonella enterica subsp. enterica serovar Typhi Proteobacteria; Gammaproteobacteria; Enterobacterales; Enterobacteriaceae; Salmonella; Salmonella enterica; Salmonella enterica subsp. enterica MTGDNTLITSHGINRRDFMKLCAALAATMGLSSKAAAEMAESVSNPQRPPVIWIGAQECT WP_000145429.1 hydrogenase 2 small subunit [Salmonella enterica] Length: 372\nNumber of predicted TMHs: 1\nExp number of AAs in TMHs: 24.51309\nExp number, first 60 AAs: 0.2682\nTotal prob of N-in: 0.14402\noutside 1 331\nTMhelix 332 354\ninside 355 372

20011 GCF_001082585.1_10071_3_92 Salmonella enterica subsp. enterica serovar Typhi Proteobacteria; Gammaproteobacteria; Enterobacterales; Enterobacteriaceae; Salmonella; Salmonella enterica; Salmonella enterica subsp. enterica MTGDNTLITSHGINRRDFMKLCAALAATMGLSSKAAAEMAESVSNPQRPPVIWIGAQECT WP_000145429.1 hydrogenase 2 small subunit [Salmonella enterica] Length: 372\nNumber of predicted TMHs: 1\nExp number of AAs in TMHs: 24.51309\nExp number, first 60 AAs: 0.2682\nTotal prob of N-in: 0.14402\noutside 1 331\nTMhelix 332 354\ninside 355 372

20012 GCF_001082625.1_10071_3_2 Salmonella enterica subsp. enterica serovar Typhi Proteobacteria; Gammaproteobacteria; Enterobacterales; Enterobacteriaceae; Salmonella; Salmonella enterica; Salmonella enterica subsp. enterica MTGDNTLITSHGINRRDFMKLCAALAATMGLSSKAAAEMAESVSNPQRPPVIWIGAQECT WP_000145429.1 hydrogenase 2 small subunit [Salmonella enterica] Length: 372\nNumber of predicted TMHs: 1\nExp number of AAs in TMHs: 24.51309\nExp number, first 60 AAs: 0.2682\nTotal prob of N-in: 0.14402\noutside 1 331\nTMhelix 332 354\ninside 355 372

20013 GCF_001082665.1_10071_8_67 Salmonella enterica subsp. enterica serovar Typhi Proteobacteria; Gammaproteobacteria; Enterobacterales; Enterobacteriaceae; Salmonella; Salmonella enterica; Salmonella enterica subsp. enterica MTGDNTLITSHGINRRDFMKLCAALAATMGLSSKAAAEMAESVSNPQRPPVIWIGAQECT WP_000145429.1 hydrogenase 2 small subunit [Salmonella enterica] Length: 372\nNumber of predicted TMHs: 1\nExp number of AAs in TMHs: 24.51309\nExp number, first 60 AAs: 0.2682\nTotal prob of N-in: 0.14402\noutside 1 331\nTMhelix 332 354\ninside 355 372

20014 GCF_001082705.1_10060_6_53 Salmonella enterica subsp. enterica serovar Typhi Proteobacteria; Gammaproteobacteria; Enterobacterales; Enterobacteriaceae; Salmonella; Salmonella enterica; Salmonella enterica subsp. enterica MTGDNTLITSHGINRRDFMKLCAALAATMGLSSKAAAEMAESVSNPQRPPVIWIGAQECT WP_000145429.1 hydrogenase 2 small subunit [Salmonella enterica] Length: 372\nNumber of predicted TMHs: 1\nExp number of AAs in TMHs: 24.51309\nExp number, first 60 AAs: 0.2682\nTotal prob of N-in: 0.14402\noutside 1 331\nTMhelix 332 354\ninside 355 372

20015 GCF_001084965.1_10493_1_80 Salmonella enterica subsp. enterica serovar Typhi Proteobacteria; Gammaproteobacteria; Enterobacterales; Enterobacteriaceae; Salmonella; Salmonella enterica; Salmonella enterica subsp. enterica MTGDNTLITSHGINRRDFMKLCAALAATMGLSSKAAAEMAESVSNPQRPPVIWIGAQECT WP_000145429.1 hydrogenase 2 small subunit [Salmonella enterica] Length: 372\nNumber of predicted TMHs: 1\nExp number of AAs in TMHs: 24.51309\nExp number, first 60 AAs: 0.2682\nTotal prob of N-in: 0.14402\noutside 1 331\nTMhelix 332 354\ninside 355 372

20016 GCF_001085025.1_10562_2_40 Salmonella enterica subsp. enterica serovar Typhi Proteobacteria; Gammaproteobacteria; Enterobacterales; Enterobacteriaceae; Salmonella; Salmonella enterica; Salmonella enterica subsp. enterica MTGDNTLITSHGINRRDFMKLCAALAATMGLSSKAAAEMAESVSNPQRPPVIWIGAQECT WP_000145429.1 hydrogenase 2 small subunit [Salmonella enterica] Length: 372\nNumber of predicted TMHs: 1\nExp number of AAs in TMHs: 24.51309\nExp number, first 60 AAs: 0.2682\nTotal prob of N-in: 0.14402\noutside 1 331\nTMhelix 332 354\ninside 355 372

20017 GCF_001085225.1_9953_5_62 Salmonella enterica subsp. enterica serovar Typhi Proteobacteria; Gammaproteobacteria; Enterobacterales; Enterobacteriaceae; Salmonella; Salmonella enterica;

Salmonella enterica subsp. enterica MTGDNTLITSHGINRRDFMKLCAALAATMGLSSKAAAEMAESVSNPQRPPVIWIGAQECT WP_000145429.1 hydrogenase 2 small subunit [Salmonella enterica] Length: 372\nNumber of predicted TMHs: 1\nExp number of AAs in TMHs: 24.51309\nExp number, first 60 AAs: 0.2682\nTotal prob of N-in: 0.14402\noutside 1 331\nTMhelix 332 354\ninside 355 372

20018 GCF_001085185.1_10349_1_8 Salmonella enterica subsp. enterica serovar Typhi Proteobacteria; Gammaproteobacteria; Enterobacterales; Enterobacteriaceae; Salmonella; Salmonella enterica; Salmonella enterica subsp. enterica MTGDNTLITSHGINRRDFMKLCAALAATMGLSSKAAAEMAESVSNPQRPPVIWIGAQECT WP_000145429.1 hydrogenase 2 small subunit [Salmonella enterica] Length: 372\nNumber of predicted TMHs: 1\nExp number of AAs in TMHs: 24.51309\nExp number, first 60 AAs: 0.2682\nTotal prob of N-in: 0.14402\noutside 1 331\nTMhelix 332 354\ninside 355 372

20019 GCF_001085365.1_9870_8_76 Salmonella enterica subsp. enterica serovar Typhi Proteobacteria; Gammaproteobacteria; Enterobacterales; Enterobacteriaceae; Salmonella; Salmonella enterica; Salmonella enterica subsp. enterica MTGDNTLITSHGINRRDFMKLCAALAATMGLSSKAAAEMAESVSNPQRPPVIWIGAQECT WP_000145429.1 hydrogenase 2 small subunit [Salmonella enterica] Length: 372\nNumber of predicted TMHs: 1\nExp number of AAs in TMHs: 24.51309\nExp number, first 60 AAs: 0.2682\nTotal prob of N-in: 0.14402\noutside 1 331\nTMhelix 332 354\ninside 355 372

20020 GCF_001085525.1_10592_2_70 Salmonella enterica subsp. enterica serovar Typhi Proteobacteria; Gammaproteobacteria; Enterobacterales; Enterobacteriaceae; Salmonella; Salmonella enterica; Salmonella enterica subsp. enterica MTGDNTLITSHGINRRDFMKLCAALAATMGLSSKAAAEMAESVSNPQRPPVIWIGAQECT WP_000145429.1 hydrogenase 2 small subunit [Salmonella enterica] Length: 372\nNumber of predicted TMHs: 1\nExp number of AAs in TMHs: 24.51309\nExp number, first 60 AAs: 0.2682\nTotal prob of N-in: 0.14402\noutside 1 331\nTMhelix 332 354\ninside 355 372

20021 GCF_001085625.1_10349_1_25 Salmonella enterica subsp. enterica serovar Typhi Proteobacteria; Gammaproteobacteria; Enterobacterales; Enterobacteriaceae; Salmonella; Salmonella enterica; Salmonella enterica subsp. enterica MTGDNTLITSHGINRRDFMKLCAALAATMGLSSKAAAEMAESVSNPQRPPVIWIGAQECT WP_000145429.1 hydrogenase 2 small subunit [Salmonella enterica] Length: 372\nNumber of predicted TMHs: 1\nExp number of AAs in TMHs: 24.51309\nExp number, first 60 AAs: 0.2682\nTotal prob of N-in: 0.14402\noutside 1 331\nTMhelix 332 354\ninside 355 372

20022 GCF_001089065.1_10607_1_63 Salmonella enterica subsp. enterica serovar Typhi Proteobacteria; Gammaproteobacteria; Enterobacterales; Enterobacteriaceae; Salmonella; Salmonella enterica; Salmonella enterica subsp. enterica MTGDNTLITSHGINRRDFMKLCAALAATMGLSSKAAAEMAESVSNPQRPPVIWIGAQECT WP_000145429.1 hydrogenase 2 small subunit [Salmonella enterica] Length: 372\nNumber of predicted TMHs: 1\nExp number of AAs in TMHs: 24.51309\nExp number, first 60 AAs: 0.2682\nTotal prob of N-in: 0.14402\noutside 1 331\nTMhelix 332 354\ninside 355 372

20023 GCF_001089545.1_10349_1_57 Salmonella enterica subsp. enterica serovar Typhi Proteobacteria; Gammaproteobacteria; Enterobacterales; Enterobacteriaceae; Salmonella; Salmonella enterica; Salmonella enterica subsp. enterica MTGDNTLITSHGINRRDFMKLCAALAATMGLSSKAAAEMAESVSNPQRPPVIWIGAQECT WP_000145429.1 hydrogenase 2 small subunit [Salmonella enterica] Length: 372\nNumber of predicted TMHs: 1\nExp number of AAs in TMHs: 24.51309\nExp number, first 60 AAs: 0.2682\nTotal prob of N-in: 0.14402\noutside 1 331\nTMhelix 332 354\ninside 355 372

20024 GCF_001089625.1_10592_2_40 Salmonella enterica subsp. enterica serovar Typhi Proteobacteria; Gammaproteobacteria; Enterobacterales; Enterobacteriaceae; Salmonella; Salmonella enterica; Salmonella enterica subsp. enterica MTGDNTLITSHGINRRDFMKLCAALAATMGLSSKAAAEMAESVSNPQRPPVIWIGAQECT WP_000145429.1 hydrogenase 2 small subunit [Salmonella enterica] Length: 372\nNumber of predicted TMHs: 1\nExp number of AAs in TMHs: 24.51309\nExp number, first 60 AAs: 0.2682\nTotal prob of N-in: 0.14402\noutside 1 331\nTMhelix 332 354\ninside 355 372

20025 GCF_001089985.1_10561_2_34 Salmonella enterica subsp. enterica serovar Typhi Proteobacteria; Gammaproteobacteria; Enterobacterales; Enterobacteriaceae; Salmonella; Salmonella enterica; Salmonella enterica subsp. enterica MTGDNTLITSHGINRRDFMKLCAALAATMGLSSKAAAEMAESVSNPQRPPVIWIGAQECT WP_000145429.1 hydrogenase 2 small subunit [Salmonella enterica] Length: 372\nNumber of predicted TMHs: 1\nExp number of AAs in TMHs: 24.51309\nExp number, first 60 AAs: 0.2682\nTotal prob of N-in: 0.14402\noutside 1 331\nTMhelix 332 354\ninside 355 372

20026 GCF_001089865.1_9475_6_9Salmonella enterica subsp. enterica serovar Typhi Proteobacteria; Gammaproteobacteria; Enterobacterales; Enterobacteriaceae; Salmonella; Salmonella enterica; Salmonella enterica subsp. enterica MTGDNTLITSHGINRRDFMKLCAALAATMGLSSKAAAEMAESVSNPQRPPVIWIGAQECT WP_000145429.1 hydrogenase 2 small subunit [Salmonella enterica] Length: 372\nNumber of predicted TMHs: 1\nExp number of AAs in TMHs: 24.51309\nExp number, first 60 AAs: 0.2682\nTotal prob of N-in: 0.14402\noutside 1 331\nTMhelix 332 354\ninside 355 372

20027 GCF_001090265.1_8616_4_51 Salmonella enterica subsp. enterica serovar Typhi Proteobacteria; Gammaproteobacteria; Enterobacterales; Enterobacteriaceae; Salmonella; Salmonella enterica; Salmonella enterica subsp. enterica MTGDNTLITSHGINRRDFMKLCAALAATMGLSSKAAAEMAESVSNPQRPPVIWIGAQECT WP_000145429.1 hydrogenase 2 small subunit [Salmonella enterica] Length: 372\nNumber of predicted TMHs: 1\nExp number of AAs in TMHs: 24.51309\nExp number, first 60 AAs: 0.2682\nTotal prob of N-in: 0.14402\noutside 1 331\nTMhelix 332 354\ninside 355 372

20028 GCF_001093025.1_8447_8_31 Salmonella enterica subsp. enterica serovar Typhi Proteobacteria; Gammaproteobacteria; Enterobacterales; Enterobacteriaceae; Salmonella; Salmonella enterica; Salmonella enterica subsp. enterica MTGDNTLITSHGINRRDFMKLCAALAATMGLSSKAAAEMAESVSNPQRPPVIWIGAQECT WP_000145429.1 hydrogenase 2 small subunit [Salmonella enterica] Length: 372\nNumber of predicted TMHs: 1\nExp number of AAs in TMHs: 24.51309\nExp number, first 60 AAs: 0.2682\nTotal prob of N-in: 0.14402\noutside 1 331\nTMhelix 332 354\ninside 355 372

20029 GCF_001093265.1_10561_2_57 Salmonella enterica subsp. enterica serovar Typhi Proteobacteria; Gammaproteobacteria; Enterobacterales; Enterobacteriaceae; Salmonella; Salmonella enterica; Salmonella enterica subsp. enterica MTGDNTLITSHGINRRDFMKLCAALAATMGLSSKAAAEMAESVSNPQRPPVIWIGAQECT WP_000145429.1 hydrogenase 2 small subunit [Salmonella enterica] Length: 372\nNumber of predicted TMHs: 1\nExp number of AAs in TMHs: 24.51309\nExp number, first 60 AAs: 0.2682\nTotal prob of N-in: 0.14402\noutside 1 331\nTMhelix 332 354\ninside 355 372

20030 GCF_001093505.1_10608_2_23 Salmonella enterica subsp. enterica serovar Typhi Proteobacteria; Gammaproteobacteria; Enterobacterales; Enterobacteriaceae; Salmonella; Salmonella enterica; Salmonella enterica subsp. enterica MTGDNTLITSHGINRRDFMKLCAALAATMGLSSKAAAEMAESVSNPQRPPVIWIGAQECT WP_000145429.1 hydrogenase 2 small subunit [Salmonella enterica] Length: 372\nNumber of predicted TMHs: 1\nExp number of AAs in TMHs: 24.51309\nExp number, first 60 AAs: 0.2682\nTotal prob of N-in: 0.14402\noutside 1 331\nTMhelix 332 354\ninside 355 372

20031 GCF_001093585.1_10492_1_7 Salmonella enterica subsp. enterica serovar Typhi Proteobacteria; Gammaproteobacteria; Enterobacterales; Enterobacteriaceae; Salmonella; Salmonella enterica; Salmonella enterica subsp. enterica MTGDNTLITSHGINRRDFMKLCAALAATMGLSSKAAAEMAESVSNPQRPPVIWIGAQECT WP_000145429.1 hydrogenase 2 small subunit [Salmonella enterica] Length: 372\nNumber of predicted TMHs: 1\nExp number of AAs in TMHs: 24.51309\nExp number, first 60 AAs: 0.2682\nTotal prob of N-in: 0.14402\noutside 1 331\nTMhelix 332 354\ninside 355 372

20032 GCF_001093825.1_10349_1_66 Salmonella enterica subsp. enterica serovar Typhi Proteobacteria; Gammaproteobacteria; Enterobacterales; Enterobacteriaceae; Salmonella; Salmonella enterica; Salmonella enterica subsp. enterica MTGDNTLITSHGINRRDFMKLCAALAATMGLSSKAAAEMAESVSNPQRPPVIWIGAQECT WP_000145429.1 hydrogenase 2 small subunit [Salmonella enterica] Length: 372\nNumber of predicted TMHs: 1\nExp number of AAs in TMHs: 24.51309\nExp number, first 60 AAs: 0.2682\nTotal prob of N-in: 0.14402\noutside 1 331\nTMhelix 332 354\ninside 355 372

20033 GCF_001094025.1_10562_2_42 Salmonella enterica subsp. enterica serovar Typhi Proteobacteria; Gammaproteobacteria; Enterobacterales; Enterobacteriaceae; Salmonella; Salmonella enterica; Salmonella enterica subsp. enterica MTGDNTLITSHGINRRDFMKLCAALAATMGLSSKAAAEMAESVSNPQRPPVIWIGAQECT WP_000145429.1 hydrogenase 2 small subunit [Salmonella enterica] Length: 372\nNumber of predicted TMHs: 1\nExp number of AAs in TMHs: 24.51309\nExp number, first 60 AAs: 0.2682\nTotal prob of N-in: 0.14402\noutside 1 331\nTMhelix 332 354\ninside 355 372

20034 GCF_001094105.1_10541_2_53 Salmonella enterica subsp. enterica serovar Typhi Proteobacteria; Gammaproteobacteria; Enterobacterales; Enterobacteriaceae; Salmonella; Salmonella enterica; Salmonella enterica subsp. enterica MTGDNTLITSHGINRRDFMKLCAALAATMGLSSKAAAEMAESVSNPQRPPVIWIGAQECT WP_000145429.1 hydrogenase 2 small subunit [Salmonella enterica] Length: 372\nNumber of

predicted TMHs:  1\nExp number of AAs in TMHs: 24.51309\nExp number, first 60 AAs:  0.2682\nTotal prob of N-in: 0.14402\noutside   1   331\nTMhelix   332   354\ninside   355   372

20035 GCF_001096745.1_10209_5_59 Salmonella enterica subsp. enterica serovar Typhi Proteobacteria; Gammaproteobacteria; Enterobacterales; Enterobacteriaceae; Salmonella; Salmonella enterica; Salmonella enterica subsp. enterica MTGDNTLITSHGINRRDFMKLCAALAATMGLSSKAAAEMAESVSNPQRPPVIWIGAQECT WP_000145429.1 hydrogenase 2 small subunit [Salmonella enterica] Length: 372\nNumber of predicted TMHs:  1\nExp number of AAs in TMHs: 24.51309\nExp number, first 60 AAs:  0.2682\nTotal prob of N-in: 0.14402\noutside   1   331\nTMhelix   332   354\ninside   355   372

20036 GCF_001096965.1_9870_8_89 Salmonella enterica subsp. enterica serovar Typhi Proteobacteria; Gammaproteobacteria; Enterobacterales; Enterobacteriaceae; Salmonella; Salmonella enterica; Salmonella enterica subsp. enterica MTGDNTLITSHGINRRDFMKLCAALAATMGLSSKAAAEMAESVSNPQRPPVIWIGAQECT WP_000145429.1 hydrogenase 2 small subunit [Salmonella enterica] Length: 372\nNumber of predicted TMHs:  1\nExp number of AAs in TMHs: 24.51309\nExp number, first 60 AAs:  0.2682\nTotal prob of N-in: 0.14402\noutside   1   331\nTMhelix   332   354\ninside   355   372

20037 GCF_001097065.1_10562_2_52 Salmonella enterica subsp. enterica serovar Typhi Proteobacteria; Gammaproteobacteria; Enterobacterales; Enterobacteriaceae; Salmonella; Salmonella enterica; Salmonella enterica subsp. enterica MTGDNTLITSHGINRRDFMKLCAALAATMGLSSKAAAEMAESVSNPQRPPVIWIGAQECT WP_000145429.1 hydrogenase 2 small subunit [Salmonella enterica] Length: 372\nNumber of predicted TMHs:  1\nExp number of AAs in TMHs: 24.51309\nExp number, first 60 AAs:  0.2682\nTotal prob of N-in: 0.14402\noutside   1   331\nTMhelix   332   354\ninside   355   372

20038 GCF_001097005.1_9953_5_23 Salmonella enterica subsp. enterica serovar Typhi Proteobacteria; Gammaproteobacteria; Enterobacterales; Enterobacteriaceae; Salmonella; Salmonella enterica; Salmonella enterica subsp. enterica MTGDNTLITSHGINRRDFMKLCAALAATMGLSSKAAAEMAESVSNPQRPPVIWIGAQECT WP_000145429.1 hydrogenase 2 small subunit [Salmonella enterica] Length: 372\nNumber of predicted TMHs:  1\nExp number of AAs in TMHs: 24.51309\nExp number, first 60 AAs:  0.2682\nTotal prob of N-in: 0.14402\noutside   1   331\nTMhelix   332   354\ninside   355   372

20039 GCF_001097165.1_10593_2_83 Salmonella enterica subsp. enterica serovar Typhi Proteobacteria; Gammaproteobacteria; Enterobacterales; Enterobacteriaceae; Salmonella; Salmonella enterica; Salmonella enterica subsp. enterica MTGDNTLITSHGINRRDFMKLCAALAATMGLSSKAAAEMAESVSNPQRPPVIWIGAQECT WP_000145429.1 hydrogenase 2 small subunit [Salmonella enterica] Length: 372\nNumber of predicted TMHs:  1\nExp number of AAs in TMHs: 24.51309\nExp number, first 60 AAs:  0.2682\nTotal prob of N-in: 0.14402\noutside   1   331\nTMhelix   332   354\ninside   355   372

20040 GCF_001097525.1_10562_2_1 Salmonella enterica subsp. enterica serovar Typhi Proteobacteria; Gammaproteobacteria; Enterobacterales; Enterobacteriaceae; Salmonella; Salmonella enterica; Salmonella enterica subsp. enterica MTGDNTLITSHGINRRDFMKLCAALAATMGLSSKAAAEMAESVSNPQRPPVIWIGAQECT WP_000145429.1 hydrogenase 2 small subunit [Salmonella enterica] Length: 372\nNumber of predicted TMHs:  1\nExp number of AAs in TMHs: 24.51309\nExp number, first 60 AAs:  0.2682\nTotal prob of N-in: 0.14402\noutside   1   331\nTMhelix   332   354\ninside   355   372

20041 GCF_001097625.1_10607_2_38 Salmonella enterica subsp. enterica serovar Typhi Proteobacteria; Gammaproteobacteria; Enterobacterales; Enterobacteriaceae; Salmonella; Salmonella enterica; Salmonella enterica subsp. enterica MTGDNTLITSHGINRRDFMKLCAALAATMGLSSKAAAEMAESVSNPQRPPVIWIGAQECT WP_000145429.1 hydrogenase 2 small subunit [Salmonella enterica] Length: 372\nNumber of predicted TMHs:  1\nExp number of AAs in TMHs: 24.51309\nExp number, first 60 AAs:  0.2682\nTotal prob of N-in: 0.14402\noutside   1   331\nTMhelix   332   354\ninside   355   372

20042 GCF_001100465.1_10540_1_5 Salmonella enterica subsp. enterica serovar Typhi Proteobacteria; Gammaproteobacteria; Enterobacterales; Enterobacteriaceae; Salmonella; Salmonella enterica; Salmonella enterica subsp. enterica MTGDNTLITSHGINRRDFMKLCAALAATMGLSSKAAAEMAESVSNPQRPPVIWIGAQECT WP_000145429.1 hydrogenase 2 small subunit [Salmonella enterica] Length: 372\nNumber of predicted TMHs:  1\nExp number of AAs in TMHs: 24.51309\nExp number, first 60 AAs:  0.2682\nTotal prob of N-in: 0.14402\noutside   1   331\nTMhelix   332   354\ninside   355   372

20043 GCF_001101505.1_10562_2_54 Salmonella enterica subsp. enterica serovar Typhi Proteobacteria; Gammaproteobacteria; Enterobacterales; Enterobacteriaceae; Salmonella; Salmonella enterica;

Salmonella enterica subsp. enterica MTGDNTLITSHGINRRDFMKLCAALAATMGLSSKAAAEMAESVSNPQRPPVIWIGAQECT WP_000145429.1 hydrogenase 2 small subunit [Salmonella enterica] Length: 372\nNumber of predicted TMHs: 1\nExp number of AAs in TMHs: 24.51309\nExp number, first 60 AAs: 0.2682\nTotal prob of N-in: 0.14402\noutside 1 331\nTMhelix 332 354\ninside 355 372

20044 GCF_001101625.1_10562_2_66 Salmonella enterica subsp. enterica serovar Typhi Proteobacteria; Gammaproteobacteria; Enterobacterales; Enterobacteriaceae; Salmonella; Salmonella enterica; Salmonella enterica subsp. enterica MTGDNTLITSHGINRRDFMKLCAALAATMGLSSKAAAEMAESVSNPQRPPVIWIGAQECT WP_000145429.1 hydrogenase 2 small subunit [Salmonella enterica] Length: 372\nNumber of predicted TMHs: 1\nExp number of AAs in TMHs: 24.51309\nExp number, first 60 AAs: 0.2682\nTotal prob of N-in: 0.14402\noutside 1 331\nTMhelix 332 354\ninside 355 372

20045 GCF_001101825.1_8490_6_39 Salmonella enterica subsp. enterica serovar Typhi Proteobacteria; Gammaproteobacteria; Enterobacterales; Enterobacteriaceae; Salmonella; Salmonella enterica; Salmonella enterica subsp. enterica MTGDNTLITSHGINRRDFMKLCAALAATMGLSSKAAAEMAESVSNPQRPPVIWIGAQECT WP_000145429.1 hydrogenase 2 small subunit [Salmonella enterica] Length: 372\nNumber of predicted TMHs: 1\nExp number of AAs in TMHs: 24.51309\nExp number, first 60 AAs: 0.2682\nTotal prob of N-in: 0.14402\noutside 1 331\nTMhelix 332 354\ninside 355 372

20046 GCF_001102465.1_10562_2_84 Salmonella enterica subsp. enterica serovar Typhi Proteobacteria; Gammaproteobacteria; Enterobacterales; Enterobacteriaceae; Salmonella; Salmonella enterica; Salmonella enterica subsp. enterica MTGDNTLITSHGINRRDFMKLCAALAATMGLSSKAAAEMAESVSNPQRPPVIWIGAQECT WP_000145429.1 hydrogenase 2 small subunit [Salmonella enterica] Length: 372\nNumber of predicted TMHs: 1\nExp number of AAs in TMHs: 24.51309\nExp number, first 60 AAs: 0.2682\nTotal prob of N-in: 0.14402\noutside 1 331\nTMhelix 332 354\ninside 355 372

20047 GCF_001102325.1_10608_2_65 Salmonella enterica subsp. enterica serovar Typhi Proteobacteria; Gammaproteobacteria; Enterobacterales; Enterobacteriaceae; Salmonella; Salmonella enterica; Salmonella enterica subsp. enterica MTGDNTLITSHGINRRDFMKLCAALAATMGLSSKAAAEMAESVSNPQRPPVIWIGAQECT WP_000145429.1 hydrogenase 2 small subunit [Salmonella enterica] Length: 372\nNumber of predicted TMHs: 1\nExp number of AAs in TMHs: 24.51309\nExp number, first 60 AAs: 0.2682\nTotal prob of N-in: 0.14402\noutside 1 331\nTMhelix 332 354\ninside 355 372

20048 GCF_001101365.1_7468_7_85 Salmonella enterica subsp. enterica serovar Typhi Proteobacteria; Gammaproteobacteria; Enterobacterales; Enterobacteriaceae; Salmonella; Salmonella enterica; Salmonella enterica subsp. enterica MTGDNTLITSHGINRRDFMKLCAALAATMGLSSKAATEMAESVSNPQRPPVIWIGAQECT WP_001782304.1 hydrogenase 2 small subunit [Salmonella enterica] Length: 372\nNumber of predicted TMHs: 1\nExp number of AAs in TMHs: 24.41173\nExp number, first 60 AAs: 0.16124\nTotal prob of N-in: 0.13947\noutside 1 331\nTMhelix 332 354\ninside 355 372

20049 GCF_001103385.1_8490_5_4Salmonella enterica subsp. enterica serovar Typhi Proteobacteria; Gammaproteobacteria; Enterobacterales; Enterobacteriaceae; Salmonella; Salmonella enterica; Salmonella enterica subsp. enterica MTGDNTLITSHGINRRDFMKLCAALAATMGLSSKAAAEMAESVSNPQRPPVIWIGAQECT WP_000145429.1 hydrogenase 2 small subunit [Salmonella enterica] Length: 372\nNumber of predicted TMHs: 1\nExp number of AAs in TMHs: 24.51309\nExp number, first 60 AAs: 0.2682\nTotal prob of N-in: 0.14402\noutside 1 331\nTMhelix 332 354\ninside 355 372

20050 GCF_001103505.1_8447_8_20 Salmonella enterica subsp. enterica serovar Typhi Proteobacteria; Gammaproteobacteria; Enterobacterales; Enterobacteriaceae; Salmonella; Salmonella enterica; Salmonella enterica subsp. enterica MTGDNTLITSHGINRRDFMKLCAALAATMGLSSKAAAEMAESVSNPQRPPVIWIGAQECT WP_000145429.1 hydrogenase 2 small subunit [Salmonella enterica] Length: 372\nNumber of predicted TMHs: 1\nExp number of AAs in TMHs: 24.51309\nExp number, first 60 AAs: 0.2682\nTotal prob of N-in: 0.14402\noutside 1 331\nTMhelix 332 354\ninside 355 372

20051 GCF_001103885.1_10607_2_8 Salmonella enterica subsp. enterica serovar Typhi Proteobacteria; Gammaproteobacteria; Enterobacterales; Enterobacteriaceae; Salmonella; Salmonella enterica; Salmonella enterica subsp. enterica MTGDNTLITSHGINRRDFMKLCAALAATMGLSSKAAAEMAESVSNPQRPPVIWIGAQECT WP_000145429.1 hydrogenase 2 small subunit [Salmonella enterica] Length: 372\nNumber of predicted TMHs: 1\nExp number of AAs in TMHs: 24.51309\nExp number, first 60 AAs: 0.2682\nTotal prob of N-in: 0.14402\noutside 1 331\nTMhelix 332 354\ninside 355 372

20052 GCF_001103945.1_8490_5_28 Salmonella enterica subsp. enterica serovar Typhi Proteobacteria; Gammaproteobacteria; Enterobacterales; Enterobacteriaceae; Salmonella; Salmonella enterica; Salmonella enterica subsp. enterica MTGDNTLITSHGINRRDFMKLCAALAATMGLSSKAAAEMAESVSNPQRPPVIWIGAQECT WP_000145429.1 hydrogenase 2 small subunit [Salmonella enterica] Length: 372\nNumber of predicted TMHs: 1\nExp number of AAs in TMHs: 24.51309\nExp number, first 60 AAs: 0.2682\nTotal prob of N-in: 0.14402\noutside 1 331\nTMhelix 332 354\ninside 355 372

20053 GCF_001104065.1_10607_1_43 Salmonella enterica subsp. enterica serovar Typhi Proteobacteria; Gammaproteobacteria; Enterobacterales; Enterobacteriaceae; Salmonella; Salmonella enterica; Salmonella enterica subsp. enterica MTGDNTLITSHGINRRDFMKLCAALAATMGLSSKAAAEMAESVSNPQRPPVIWIGAQECT WP_000145429.1 hydrogenase 2 small subunit [Salmonella enterica] Length: 372\nNumber of predicted TMHs: 1\nExp number of AAs in TMHs: 24.51309\nExp number, first 60 AAs: 0.2682\nTotal prob of N-in: 0.14402\noutside 1 331\nTMhelix 332 354\ninside 355 372

20054 GCF_001104205.1_8490_6_60 Salmonella enterica subsp. enterica serovar Typhi Proteobacteria; Gammaproteobacteria; Enterobacterales; Enterobacteriaceae; Salmonella; Salmonella enterica; Salmonella enterica subsp. enterica MTGDNTLITSHGINRRDFMKLCAALAATMGLSSKAAAEMAESVSNPQRPPVIWIGAQECT WP_000145429.1 hydrogenase 2 small subunit [Salmonella enterica] Length: 372\nNumber of predicted TMHs: 1\nExp number of AAs in TMHs: 24.51309\nExp number, first 60 AAs: 0.2682\nTotal prob of N-in: 0.14402\noutside 1 331\nTMhelix 332 354\ninside 355 372

20055 GCF_001104545.1_10493_1_54 Salmonella enterica subsp. enterica serovar Typhi Proteobacteria; Gammaproteobacteria; Enterobacterales; Enterobacteriaceae; Salmonella; Salmonella enterica; Salmonella enterica subsp. enterica MTGDNTLITSHGINRRDFMKLCAALAATMGLSSKAAAEMAESVSNPQRPPVIWIGAQECT WP_000145429.1 hydrogenase 2 small subunit [Salmonella enterica] Length: 372\nNumber of predicted TMHs: 1\nExp number of AAs in TMHs: 24.51309\nExp number, first 60 AAs: 0.2682\nTotal prob of N-in: 0.14402\noutside 1 331\nTMhelix 332 354\ninside 355 372

20056 GCF_001106785.1_8490_5_11 Salmonella enterica subsp. enterica serovar Typhi Proteobacteria; Gammaproteobacteria; Enterobacterales; Enterobacteriaceae; Salmonella; Salmonella enterica; Salmonella enterica subsp. enterica MTGDNTLITSHGINRRDFMKLCAALAATMGLSSKAAAEMAESVSNPQRPPVIWIGAQECT WP_000145429.1 hydrogenase 2 small subunit [Salmonella enterica] Length: 372\nNumber of predicted TMHs: 1\nExp number of AAs in TMHs: 24.51309\nExp number, first 60 AAs: 0.2682\nTotal prob of N-in: 0.14402\noutside 1 331\nTMhelix 332 354\ninside 355 372

20057 GCF_001106665.1_10492_1_65 Salmonella enterica subsp. enterica serovar Typhi Proteobacteria; Gammaproteobacteria; Enterobacterales; Enterobacteriaceae; Salmonella; Salmonella enterica; Salmonella enterica subsp. enterica MTGDNTLITSHGINRRDFMKLCAALAATMGLSSKAAAEMAESVSNPQRPPVIWIGAQECT WP_000145429.1 hydrogenase 2 small subunit [Salmonella enterica] Length: 372\nNumber of predicted TMHs: 1\nExp number of AAs in TMHs: 24.51309\nExp number, first 60 AAs: 0.2682\nTotal prob of N-in: 0.14402\noutside 1 331\nTMhelix 332 354\ninside 355 372

20058 GCF_001107045.1_8490_6_72 Salmonella enterica subsp. enterica serovar Typhi Proteobacteria; Gammaproteobacteria; Enterobacterales; Enterobacteriaceae; Salmonella; Salmonella enterica; Salmonella enterica subsp. enterica MTGDNTLITSHGINRRDFMKLCAALAATMGLSSKAAAEMAESVSNPQRPPVIWIGAQECT WP_000145429.1 hydrogenase 2 small subunit [Salmonella enterica] Length: 372\nNumber of predicted TMHs: 1\nExp number of AAs in TMHs: 24.51309\nExp number, first 60 AAs: 0.2682\nTotal prob of N-in: 0.14402\noutside 1 331\nTMhelix 332 354\ninside 355 372

20059 GCF_001107225.1_10561_2_17 Salmonella enterica subsp. enterica serovar Typhi Proteobacteria; Gammaproteobacteria; Enterobacterales; Enterobacteriaceae; Salmonella; Salmonella enterica; Salmonella enterica subsp. enterica MTGDNTLITSHGINRRDFMKLCAALAATMGLSSKAAAEMAESVSNPQRPPVIWIGAQECT WP_000145429.1 hydrogenase 2 small subunit [Salmonella enterica] Length: 372\nNumber of predicted TMHs: 1\nExp number of AAs in TMHs: 24.51309\nExp number, first 60 AAs: 0.2682\nTotal prob of N-in: 0.14402\noutside 1 331\nTMhelix 332 354\ninside 355 372

20060 GCF_001107405.1_10492_1_40 Salmonella enterica subsp. enterica serovar Typhi Proteobacteria; Gammaproteobacteria; Enterobacterales; Enterobacteriaceae; Salmonella; Salmonella enterica; Salmonella enterica subsp. enterica MTGDNTLITSHGINRRDFMKLCAALAATMGLSSKAAAEMAESVSNPQRPPVIWIGAQECT WP_000145429.1 hydrogenase 2 small subunit [Salmonella enterica] Length: 372\nNumber of

predicted TMHs: 1\nExp number of AAs in TMHs: 24.51309\nExp number, first 60 AAs: 0.2682\nTotal prob of N-in: 0.14402\noutside 1 331\nTMhelix 332 354\ninside 355 372

20061 GCF_001107665.1_10493_1_15 Salmonella enterica subsp. enterica serovar Typhi Proteobacteria; Gammaproteobacteria; Enterobacterales; Enterobacteriaceae; Salmonella; Salmonella enterica; Salmonella enterica subsp. enterica MTGDNTLITSHGINRRDFMKLCAALAATMGLSSKAAAEMAESVSNPQRPPVIWIGAQECT WP_000145429.1 hydrogenase 2 small subunit [Salmonella enterica] Length: 372\nNumber of predicted TMHs: 1\nExp number of AAs in TMHs: 24.51309\nExp number, first 60 AAs: 0.2682\nTotal prob of N-in: 0.14402\noutside 1 331\nTMhelix 332 354\ninside 355 372

20062 GCF_001107745.1_10071_8_91 Salmonella enterica subsp. enterica serovar Typhi Proteobacteria; Gammaproteobacteria; Enterobacterales; Enterobacteriaceae; Salmonella; Salmonella enterica; Salmonella enterica subsp. enterica MTGDNTLITSHGINRRDFMKLCAALAATMGLSSKAAAEMAESVSNPQRPPVIWIGAQECT WP_000145429.1 hydrogenase 2 small subunit [Salmonella enterica] Length: 372\nNumber of predicted TMHs: 1\nExp number of AAs in TMHs: 24.51309\nExp number, first 60 AAs: 0.2682\nTotal prob of N-in: 0.14402\noutside 1 331\nTMhelix 332 354\ninside 355 372

20063 GCF_001110685.1_10492_1_62 Salmonella enterica subsp. enterica serovar Typhi Proteobacteria; Gammaproteobacteria; Enterobacterales; Enterobacteriaceae; Salmonella; Salmonella enterica; Salmonella enterica subsp. enterica MTGDNTLITSHGINRRDFMKLCAALAATMGLSSKAAAEMAESVSNPQRPPVIWIGAQECT WP_000145429.1 hydrogenase 2 small subunit [Salmonella enterica] Length: 372\nNumber of predicted TMHs: 1\nExp number of AAs in TMHs: 24.51309\nExp number, first 60 AAs: 0.2682\nTotal prob of N-in: 0.14402\noutside 1 331\nTMhelix 332 354\ninside 355 372

20064 GCF_001111085.1_10561_2_52 Salmonella enterica subsp. enterica serovar Typhi Proteobacteria; Gammaproteobacteria; Enterobacterales; Enterobacteriaceae; Salmonella; Salmonella enterica; Salmonella enterica subsp. enterica MTGDNTLITSHGINRRDFMKLCAALAATMGLSSKAAAEMAESVSNPQRPPVIWIGAQECT WP_000145429.1 hydrogenase 2 small subunit [Salmonella enterica] Length: 372\nNumber of predicted TMHs: 1\nExp number of AAs in TMHs: 24.51309\nExp number, first 60 AAs: 0.2682\nTotal prob of N-in: 0.14402\noutside 1 331\nTMhelix 332 354\ninside 355 372

20065 GCF_001111225.1_8447_8_19 Salmonella enterica subsp. enterica serovar Typhi Proteobacteria; Gammaproteobacteria; Enterobacterales; Enterobacteriaceae; Salmonella; Salmonella enterica; Salmonella enterica subsp. enterica MTGDNTLITSHGINRRDFMKLCAALAATMGLSSKAAAEMAESVSNPQRPPVIWIGAQECT WP_000145429.1 hydrogenase 2 small subunit [Salmonella enterica] Length: 372\nNumber of predicted TMHs: 1\nExp number of AAs in TMHs: 24.51309\nExp number, first 60 AAs: 0.2682\nTotal prob of N-in: 0.14402\noutside 1 331\nTMhelix 332 354\ninside 355 372

20066 GCF_001111385.1_10593_2_22 Salmonella enterica subsp. enterica serovar Typhi Proteobacteria; Gammaproteobacteria; Enterobacterales; Enterobacteriaceae; Salmonella; Salmonella enterica; Salmonella enterica subsp. enterica MTGDNTLITSHGINRRDFMKLCAALAATMGLSSKAAAEMAESVSNPQRPPVIWIGAQECT WP_000145429.1 hydrogenase 2 small subunit [Salmonella enterica] Length: 372\nNumber of predicted TMHs: 1\nExp number of AAs in TMHs: 24.51309\nExp number, first 60 AAs: 0.2682\nTotal prob of N-in: 0.14402\noutside 1 331\nTMhelix 332 354\ninside 355 372

20067 GCF_001111425.1_8490_5_9Salmonella enterica subsp. enterica serovar Typhi Proteobacteria; Gammaproteobacteria; Enterobacterales; Enterobacteriaceae; Salmonella; Salmonella enterica; Salmonella enterica subsp. enterica MTGDNTLITSHGINRRDFMKLCAALAATMGLSSKAAAEMAESVSNPQRPPVIWIGAQECT WP_000145429.1 hydrogenase 2 small subunit [Salmonella enterica] Length: 372\nNumber of predicted TMHs: 1\nExp number of AAs in TMHs: 24.51309\nExp number, first 60 AAs: 0.2682\nTotal prob of N-in: 0.14402\noutside 1 331\nTMhelix 332 354\ninside 355 372

20068 GCF_001111685.1_9953_5_91 Salmonella enterica subsp. enterica serovar Typhi Proteobacteria; Gammaproteobacteria; Enterobacterales; Enterobacteriaceae; Salmonella; Salmonella enterica; Salmonella enterica subsp. enterica MTGDNTLITSHGINRRDFMKLCAALAATMGLSSKAAAEMAESVSNPQRPPVIWIGAQECT WP_000145429.1 hydrogenase 2 small subunit [Salmonella enterica] Length: 372\nNumber of predicted TMHs: 1\nExp number of AAs in TMHs: 24.51309\nExp number, first 60 AAs: 0.2682\nTotal prob of N-in: 0.14402\noutside 1 331\nTMhelix 332 354\ninside 355 372

20069 GCF_001114305.1_8525_3_34 Salmonella enterica subsp. enterica serovar Typhi Proteobacteria; Gammaproteobacteria; Enterobacterales; Enterobacteriaceae; Salmonella; Salmonella enterica;

Salmonella enterica subsp. enterica MTGDNTLITSHGINRRDFMKLCAALAATMGLSSKAAAEMAESVSNPQRPPVIWIGAQECT WP_000145429.1 hydrogenase 2 small subunit [Salmonella enterica] Length: 372\nNumber of predicted TMHs: 1\nExp number of AAs in TMHs: 24.51309\nExp number, first 60 AAs: 0.2682\nTotal prob of N-in: 0.14402\noutside 1 331\nTMhelix 332 354\ninside 355 372

20070 GCF_001114385.1_10607_2_18 Salmonella enterica subsp. enterica serovar Typhi Proteobacteria; Gammaproteobacteria; Enterobacterales; Enterobacteriaceae; Salmonella; Salmonella enterica; Salmonella enterica subsp. enterica MTGDNTLITSHGINRRDFMKLCAALAATMGLSSKAAAEMAESVSNPQRPPVIWIGAQECT WP_000145429.1 hydrogenase 2 small subunit [Salmonella enterica] Length: 372\nNumber of predicted TMHs: 1\nExp number of AAs in TMHs: 24.51309\nExp number, first 60 AAs: 0.2682\nTotal prob of N-in: 0.14402\noutside 1 331\nTMhelix 332 354\ninside 355 372

20071 GCF_001114565.1_10541_2_46 Salmonella enterica subsp. enterica serovar Typhi Proteobacteria; Gammaproteobacteria; Enterobacterales; Enterobacteriaceae; Salmonella; Salmonella enterica; Salmonella enterica subsp. enterica MTGDNTLITSHGINRRDFMKLCAALAATMGLSSKAAAEMAESVSNPQRPPVIWIGAQECT WP_000145429.1 hydrogenase 2 small subunit [Salmonella enterica] Length: 372\nNumber of predicted TMHs: 1\nExp number of AAs in TMHs: 24.51309\nExp number, first 60 AAs: 0.2682\nTotal prob of N-in: 0.14402\noutside 1 331\nTMhelix 332 354\ninside 355 372

20072 GCF_001113925.1_10492_1_37 Salmonella enterica subsp. enterica serovar Typhi Proteobacteria; Gammaproteobacteria; Enterobacterales; Enterobacteriaceae; Salmonella; Salmonella enterica; Salmonella enterica subsp. enterica MTGDNTLITSHGINRRDFMKLCAALAATMGLSSKAATEMAESVSNPQRPPVIWIGAQECT WP_001782304.1 hydrogenase 2 small subunit [Salmonella enterica] Length: 372\nNumber of predicted TMHs: 1\nExp number of AAs in TMHs: 24.41173\nExp number, first 60 AAs: 0.16124\nTotal prob of N-in: 0.13947\noutside 1 331\nTMhelix 332 354\ninside 355 372

20073 GCF_001118425.1_10607_2_2 Salmonella enterica subsp. enterica serovar Typhi Proteobacteria; Gammaproteobacteria; Enterobacterales; Enterobacteriaceae; Salmonella; Salmonella enterica; Salmonella enterica subsp. enterica MTGDNTLITSHGINRRDFMKLCAALAATMGLSSKAAAEMAESVSNPQRPPVIWIGAQECT WP_000145429.1 hydrogenase 2 small subunit [Salmonella enterica] Length: 372\nNumber of predicted TMHs: 1\nExp number of AAs in TMHs: 24.51309\nExp number, first 60 AAs: 0.2682\nTotal prob of N-in: 0.14402\noutside 1 331\nTMhelix 332 354\ninside 355 372

20074 GCF_001118745.1_10561_2_55 Salmonella enterica subsp. enterica serovar Typhi Proteobacteria; Gammaproteobacteria; Enterobacterales; Enterobacteriaceae; Salmonella; Salmonella enterica; Salmonella enterica subsp. enterica MTGDNTLITSHGINRRDFMKLCAALAATMGLSSKAAAEMAESVSNPQRPPVIWIGAQECT WP_000145429.1 hydrogenase 2 small subunit [Salmonella enterica] Length: 372\nNumber of predicted TMHs: 1\nExp number of AAs in TMHs: 24.51309\nExp number, first 60 AAs: 0.2682\nTotal prob of N-in: 0.14402\noutside 1 331\nTMhelix 332 354\ninside 355 372

20075 GCF_001118905.1_10541_2_30 Salmonella enterica subsp. enterica serovar Typhi Proteobacteria; Gammaproteobacteria; Enterobacterales; Enterobacteriaceae; Salmonella; Salmonella enterica; Salmonella enterica subsp. enterica MTGDNTLITSHGINRRDFMKLCAALAATMGLSSKAAAEMAESVSNPQRPPVIWIGAQECT WP_000145429.1 hydrogenase 2 small subunit [Salmonella enterica] Length: 372\nNumber of predicted TMHs: 1\nExp number of AAs in TMHs: 24.51309\nExp number, first 60 AAs: 0.2682\nTotal prob of N-in: 0.14402\noutside 1 331\nTMhelix 332 354\ninside 355 372

20076 GCF_001119225.1_10493_1_81 Salmonella enterica subsp. enterica serovar Typhi Proteobacteria; Gammaproteobacteria; Enterobacterales; Enterobacteriaceae; Salmonella; Salmonella enterica; Salmonella enterica subsp. enterica MTGDNTLITSHGINRRDFMKLCAALAATMGLSSKAAAEMAESVSNPQRPPVIWIGAQECT WP_000145429.1 hydrogenase 2 small subunit [Salmonella enterica] Length: 372\nNumber of predicted TMHs: 1\nExp number of AAs in TMHs: 24.51309\nExp number, first 60 AAs: 0.2682\nTotal prob of N-in: 0.14402\noutside 1 331\nTMhelix 332 354\ninside 355 372

20077 GCF_001119445.1_9870_8_38 Salmonella enterica subsp. enterica serovar Typhi Proteobacteria; Gammaproteobacteria; Enterobacterales; Enterobacteriaceae; Salmonella; Salmonella enterica; Salmonella enterica subsp. enterica MTGDNTLITSHGINRRDFMKLCAALAATMGLSSKAAAEMAESVSNPQRPPVIWIGAQECT WP_000145429.1 hydrogenase 2 small subunit [Salmonella enterica] Length: 372\nNumber of predicted TMHs: 1\nExp number of AAs in TMHs: 24.51309\nExp number, first 60 AAs: 0.2682\nTotal prob of N-in: 0.14402\noutside 1 331\nTMhelix 332 354\ninside 355 372

20078 GCF_001119745.1_9475_6_26 Salmonella enterica subsp. enterica serovar Typhi Proteobacteria; Gammaproteobacteria; Enterobacterales; Enterobacteriaceae; Salmonella; Salmonella enterica; Salmonella enterica subsp. enterica MTGDNTLITSHGINRRDFMKLCAALAATMGLSSKAAAEMAESVSNPQRPPVIWIGAQECT WP_000145429.1 hydrogenase 2 small subunit [Salmonella enterica] Length: 372\nNumber of predicted TMHs: 1\nExp number of AAs in TMHs: 24.51309\nExp number, first 60 AAs: 0.2682\nTotal prob of N-in: 0.14402\noutside 1 331\nTMhelix 332 354\ninside 355 372

20079 GCF_001122625.1_10492_1_47 Salmonella enterica subsp. enterica serovar Typhi Proteobacteria; Gammaproteobacteria; Enterobacterales; Enterobacteriaceae; Salmonella; Salmonella enterica; Salmonella enterica subsp. enterica MTGDNTLITSHGINRRDFMKLCAALAATMGLSSKAAAEMAESVSNPQRPPVIWIGAQECT WP_000145429.1 hydrogenase 2 small subunit [Salmonella enterica] Length: 372\nNumber of predicted TMHs: 1\nExp number of AAs in TMHs: 24.51309\nExp number, first 60 AAs: 0.2682\nTotal prob of N-in: 0.14402\noutside 1 331\nTMhelix 332 354\ninside 355 372

20080 GCF_001122805.1_10562_2_36 Salmonella enterica subsp. enterica serovar Typhi Proteobacteria; Gammaproteobacteria; Enterobacterales; Enterobacteriaceae; Salmonella; Salmonella enterica; Salmonella enterica subsp. enterica MTGDNTLITSHGINRRDFMKLCAALAATMGLSSKAAAEMAESVSNPQRPPVIWIGAQECT WP_000145429.1 hydrogenase 2 small subunit [Salmonella enterica] Length: 372\nNumber of predicted TMHs: 1\nExp number of AAs in TMHs: 24.51309\nExp number, first 60 AAs: 0.2682\nTotal prob of N-in: 0.14402\noutside 1 331\nTMhelix 332 354\ninside 355 372

20081 GCF_001123045.1_10608_2_45 Salmonella enterica subsp. enterica serovar Typhi Proteobacteria; Gammaproteobacteria; Enterobacterales; Enterobacteriaceae; Salmonella; Salmonella enterica; Salmonella enterica subsp. enterica MTGDNTLITSHGINRRDFMKLCAALAATMGLSSKAAAEMAESVSNPQRPPVIWIGAQECT WP_000145429.1 hydrogenase 2 small subunit [Salmonella enterica] Length: 372\nNumber of predicted TMHs: 1\nExp number of AAs in TMHs: 24.51309\nExp number, first 60 AAs: 0.2682\nTotal prob of N-in: 0.14402\noutside 1 331\nTMhelix 332 354\ninside 355 372

20082 GCF_001123365.1_7468_7_66 Salmonella enterica subsp. enterica serovar Typhi Proteobacteria; Gammaproteobacteria; Enterobacterales; Enterobacteriaceae; Salmonella; Salmonella enterica; Salmonella enterica subsp. enterica MTGDNTLITSHGINRRDFMKLCAALAATMGLSSKAAAEMAESVSNPQRPPVIWIGAQECT WP_000145429.1 hydrogenase 2 small subunit [Salmonella enterica] Length: 372\nNumber of predicted TMHs: 1\nExp number of AAs in TMHs: 24.51309\nExp number, first 60 AAs: 0.2682\nTotal prob of N-in: 0.14402\noutside 1 331\nTMhelix 332 354\ninside 355 372

20083 GCF_001123465.1_10562_2_9 Salmonella enterica subsp. enterica serovar Typhi Proteobacteria; Gammaproteobacteria; Enterobacterales; Enterobacteriaceae; Salmonella; Salmonella enterica; Salmonella enterica subsp. enterica MTGDNTLITSHGINRRDFMKLCAALAATMGLSSKAAAEMAESVSNPQRPPVIWIGAQECT WP_000145429.1 hydrogenase 2 small subunit [Salmonella enterica] Length: 372\nNumber of predicted TMHs: 1\nExp number of AAs in TMHs: 24.51309\nExp number, first 60 AAs: 0.2682\nTotal prob of N-in: 0.14402\noutside 1 331\nTMhelix 332 354\ninside 355 372

20084 GCF_001123625.1_10608_2_43 Salmonella enterica subsp. enterica serovar Typhi Proteobacteria; Gammaproteobacteria; Enterobacterales; Enterobacteriaceae; Salmonella; Salmonella enterica; Salmonella enterica subsp. enterica MTGDNTLITSHGINRRDFMKLCAALAATMGLSSKAAAEMAESVSNPQRPPVIWIGAQECT WP_000145429.1 hydrogenase 2 small subunit [Salmonella enterica] Length: 372\nNumber of predicted TMHs: 1\nExp number of AAs in TMHs: 24.51309\nExp number, first 60 AAs: 0.2682\nTotal prob of N-in: 0.14402\noutside 1 331\nTMhelix 332 354\ninside 355 372

20085 GCF_001123685.1_7468_7_86 Salmonella enterica subsp. enterica serovar Typhi Proteobacteria; Gammaproteobacteria; Enterobacterales; Enterobacteriaceae; Salmonella; Salmonella enterica; Salmonella enterica subsp. enterica MTGDNTLITSHGINRRDFMKLCAALAATMGLSSKAAAEMAESVSNPQRPPVIWIGAQECT WP_000145429.1 hydrogenase 2 small subunit [Salmonella enterica] Length: 372\nNumber of predicted TMHs: 1\nExp number of AAs in TMHs: 24.51309\nExp number, first 60 AAs: 0.2682\nTotal prob of N-in: 0.14402\noutside 1 331\nTMhelix 332 354\ninside 355 372

20086 GCF_001126405.1_10607_2_17 Salmonella enterica subsp. enterica serovar Typhi Proteobacteria; Gammaproteobacteria; Enterobacterales; Enterobacteriaceae; Salmonella; Salmonella enterica; Salmonella enterica subsp. enterica MTGDNTLITSHGINRRDFMKLCAALAATMGLSSKAAAEMAESVSNPQRPPVIWIGAQECT WP_000145429.1 hydrogenase 2 small subunit [Salmonella enterica] Length: 372\nNumber of

predicted TMHs:  1\nExp number of AAs in TMHs: 24.51309\nExp number, first 60 AAs:  0.2682\nTotal prob of N-in: 0.14402\noutside   1   331\nTMhelix   332   354\ninside   355   372

20087 GCF_001126745.1_8616_4_37 Salmonella enterica subsp. enterica serovar Typhi Proteobacteria; Gammaproteobacteria; Enterobacterales; Enterobacteriaceae; Salmonella; Salmonella enterica; Salmonella enterica subsp. enterica MTGDNTLITSHGINRRDFMKLCAALAATMGLSSKAAAEMAESVSNPQRPPVIWIGAQECT WP_000145429.1 hydrogenase 2 small subunit [Salmonella enterica] Length: 372\nNumber of predicted TMHs:  1\nExp number of AAs in TMHs: 24.51309\nExp number, first 60 AAs:  0.2682\nTotal prob of N-in: 0.14402\noutside   1   331\nTMhelix   332   354\ninside   355   372

20088 GCF_001126885.1_10349_1_38 Salmonella enterica subsp. enterica serovar Typhi Proteobacteria; Gammaproteobacteria; Enterobacterales; Enterobacteriaceae; Salmonella; Salmonella enterica; Salmonella enterica subsp. enterica MTGDNTLITSHGINRRDFMKLCAALAATMGLSSKAAAEMAESVSNPQRPPVIWIGAQECT WP_000145429.1 hydrogenase 2 small subunit [Salmonella enterica] Length: 372\nNumber of predicted TMHs:  1\nExp number of AAs in TMHs: 24.51309\nExp number, first 60 AAs:  0.2682\nTotal prob of N-in: 0.14402\noutside   1   331\nTMhelix   332   354\ninside   355   372

20089 GCF_001127185.1_10492_1_77 Salmonella enterica subsp. enterica serovar Typhi Proteobacteria; Gammaproteobacteria; Enterobacterales; Enterobacteriaceae; Salmonella; Salmonella enterica; Salmonella enterica subsp. enterica MTGDNTLITSHGINRRDFMKLCAALAATMGLSSKAAAEMAESVSNPQRPPVIWIGAQECT WP_000145429.1 hydrogenase 2 small subunit [Salmonella enterica] Length: 372\nNumber of predicted TMHs:  1\nExp number of AAs in TMHs: 24.51309\nExp number, first 60 AAs:  0.2682\nTotal prob of N-in: 0.14402\noutside   1   331\nTMhelix   332   354\ninside   355   372

20090 GCF_001128025.1_10349_1_47 Salmonella enterica subsp. enterica serovar Typhi Proteobacteria; Gammaproteobacteria; Enterobacterales; Enterobacteriaceae; Salmonella; Salmonella enterica; Salmonella enterica subsp. enterica MTGDNTLITSHGINRRDFMKLCAALAATMGLSSKAAAEMAESVSNPQRPPVIWIGAQECT WP_000145429.1 hydrogenase 2 small subunit [Salmonella enterica] Length: 372\nNumber of predicted TMHs:  1\nExp number of AAs in TMHs: 24.51309\nExp number, first 60 AAs:  0.2682\nTotal prob of N-in: 0.14402\noutside   1   331\nTMhelix   332   354\ninside   355   372

20091 GCF_001128065.1_8447_8_11 Salmonella enterica subsp. enterica serovar Typhi Proteobacteria; Gammaproteobacteria; Enterobacterales; Enterobacteriaceae; Salmonella; Salmonella enterica; Salmonella enterica subsp. enterica MTGDNTLITSHGINRRDFMKLCAALAATMGLSSKAAAEMAESVSNPQRPPVIWIGAQECT WP_000145429.1 hydrogenase 2 small subunit [Salmonella enterica] Length: 372\nNumber of predicted TMHs:  1\nExp number of AAs in TMHs: 24.51309\nExp number, first 60 AAs:  0.2682\nTotal prob of N-in: 0.14402\noutside   1   331\nTMhelix   332   354\ninside   355   372

20092 GCF_001131605.1_8525_3_59 Salmonella enterica subsp. enterica serovar Typhi Proteobacteria; Gammaproteobacteria; Enterobacterales; Enterobacteriaceae; Salmonella; Salmonella enterica; Salmonella enterica subsp. enterica MTGDNTLITSHGINRRDFMKLCAALAATMGLSSKAAAEMAESVSNPQRPPVIWIGAQECT WP_000145429.1 hydrogenase 2 small subunit [Salmonella enterica] Length: 372\nNumber of predicted TMHs:  1\nExp number of AAs in TMHs: 24.51309\nExp number, first 60 AAs:  0.2682\nTotal prob of N-in: 0.14402\noutside   1   331\nTMhelix   332   354\ninside   355   372

20093 GCF_001131845.1_10492_1_27 Salmonella enterica subsp. enterica serovar Typhi Proteobacteria; Gammaproteobacteria; Enterobacterales; Enterobacteriaceae; Salmonella; Salmonella enterica; Salmonella enterica subsp. enterica MTGDNTLITSHGINRRDFMKLCAALAATMGLSSKAAAEMAESVSNPQRPPVIWIGAQECT WP_000145429.1 hydrogenase 2 small subunit [Salmonella enterica] Length: 372\nNumber of predicted TMHs:  1\nExp number of AAs in TMHs: 24.51309\nExp number, first 60 AAs:  0.2682\nTotal prob of N-in: 0.14402\noutside   1   331\nTMhelix   332   354\ninside   355   372

20094 GCF_001132065.1_8490_5_1Salmonella enterica subsp. enterica serovar Typhi Proteobacteria; Gammaproteobacteria; Enterobacterales; Enterobacteriaceae; Salmonella; Salmonella enterica; Salmonella enterica subsp. enterica MTGDNTLITSHGINRRDFMKLCAALAATMGLSSKAAAEMAESVSNPQRPPVIWIGAQECT WP_000145429.1 hydrogenase 2 small subunit [Salmonella enterica] Length: 372\nNumber of predicted TMHs:  1\nExp number of AAs in TMHs: 24.51309\nExp number, first 60 AAs:  0.2682\nTotal prob of N-in:   0.14402\noutside   1 331\nTMhelix   332   354\ninside   355   372

20095 GCF_001132285.1_7468_7_67 Salmonella enterica subsp. enterica serovar Typhi Proteobacteria; Gammaproteobacteria; Enterobacterales; Enterobacteriaceae; Salmonella; Salmonella enterica;

Salmonella enterica subsp. enterica MTGDNTLITSHGINRRDFMKLCAALAATMGLSSKAAAEMAESVSNPQRPPVIWIGAQECT WP_000145429.1 hydrogenase 2 small subunit [Salmonella enterica] Length: 372\nNumber of predicted TMHs: 1\nExp number of AAs in TMHs: 24.51309\nExp number, first 60 AAs: 0.2682\nTotal prob of N-in: 0.14402\noutside 1 331\nTMhelix 332 354\ninside 355 372

20096 GCF_001132445.1_10593_2_4 Salmonella enterica subsp. enterica serovar Typhi Proteobacteria; Gammaproteobacteria; Enterobacterales; Enterobacteriaceae; Salmonella; Salmonella enterica; Salmonella enterica subsp. enterica MTGDNTLITSHGINRRDFMKLCAALAATMGLSSKAAAEMAESVSNPQRPPVIWIGAQECT WP_000145429.1 hydrogenase 2 small subunit [Salmonella enterica] Length: 372\nNumber of predicted TMHs: 1\nExp number of AAs in TMHs: 24.51309\nExp number, first 60 AAs: 0.2682\nTotal prob of N-in: 0.14402\noutside 1 331\nTMhelix 332 354\ninside 355 372

20097 GCF_001132705.1_10562_2_44 Salmonella enterica subsp. enterica serovar Typhi Proteobacteria; Gammaproteobacteria; Enterobacterales; Enterobacteriaceae; Salmonella; Salmonella enterica; Salmonella enterica subsp. enterica MTGDNTLITSHGINRRDFMKLCAALAATMGLSSKAAAEMAESVSNPQRPPVIWIGAQECT WP_000145429.1 hydrogenase 2 small subunit [Salmonella enterica] Length: 372\nNumber of predicted TMHs: 1\nExp number of AAs in TMHs: 24.51309\nExp number, first 60 AAs: 0.2682\nTotal prob of N-in: 0.14402\noutside 1 331\nTMhelix 332 354\ninside 355 372

20098 GCF_001132785.1_10541_2_56 Salmonella enterica subsp. enterica serovar Typhi Proteobacteria; Gammaproteobacteria; Enterobacterales; Enterobacteriaceae; Salmonella; Salmonella enterica; Salmonella enterica subsp. enterica MTGDNTLITSHGINRRDFMKLCAALAATMGLSSKAAAEMAESVSNPQRPPVIWIGAQECT WP_000145429.1 hydrogenase 2 small subunit [Salmonella enterica] Length: 372\nNumber of predicted TMHs: 1\nExp number of AAs in TMHs: 24.51309\nExp number, first 60 AAs: 0.2682\nTotal prob of N-in: 0.14402\noutside 1 331\nTMhelix 332 354\ninside 355 372

20099 GCF_001135805.1_10349_1_31 Salmonella enterica subsp. enterica serovar Typhi Proteobacteria; Gammaproteobacteria; Enterobacterales; Enterobacteriaceae; Salmonella; Salmonella enterica; Salmonella enterica subsp. enterica MTGDNTLITSHGINRRDFMKLCAALAATMGLSSKAAAEMAESVSNPQRPPVIWIGAQECT WP_000145429.1 hydrogenase 2 small subunit [Salmonella enterica] Length: 372\nNumber of predicted TMHs: 1\nExp number of AAs in TMHs: 24.51309\nExp number, first 60 AAs: 0.2682\nTotal prob of N-in: 0.14402\noutside 1 331\nTMhelix 332 354\ninside 355 372

20100 GCF_001136005.1_10493_1_58 Salmonella enterica subsp. enterica serovar Typhi Proteobacteria; Gammaproteobacteria; Enterobacterales; Enterobacteriaceae; Salmonella; Salmonella enterica; Salmonella enterica subsp. enterica MTGDNTLITSHGINRRDFMKLCAALAATMGLSSKAAAEMAESVSNPQRPPVIWIGAQECT WP_000145429.1 hydrogenase 2 small subunit [Salmonella enterica] Length: 372\nNumber of predicted TMHs: 1\nExp number of AAs in TMHs: 24.51309\nExp number, first 60 AAs: 0.2682\nTotal prob of N-in: 0.14402\noutside 1 331\nTMhelix 332 354\ninside 355 372

20101 GCF_001136605.1_10593_2_65 Salmonella enterica subsp. enterica serovar Typhi Proteobacteria; Gammaproteobacteria; Enterobacterales; Enterobacteriaceae; Salmonella; Salmonella enterica; Salmonella enterica subsp. enterica MTGDNTLITSHGINRRDFMKLCAALAATMGLSSKAAAEMAESVSNPQRPPVIWIGAQECT WP_000145429.1 hydrogenase 2 small subunit [Salmonella enterica] Length: 372\nNumber of predicted TMHs: 1\nExp number of AAs in TMHs: 24.51309\nExp number, first 60 AAs: 0.2682\nTotal prob of N-in: 0.14402\noutside 1 331\nTMhelix 332 354\ninside 355 372

20102 GCF_001137065.1_10593_2_46 Salmonella enterica subsp. enterica serovar Typhi Proteobacteria; Gammaproteobacteria; Enterobacterales; Enterobacteriaceae; Salmonella; Salmonella enterica; Salmonella enterica subsp. enterica MTGDNTLITSHGINRRDFMKLCAALAATMGLSSKAAAEMAESVSNPQRPPVIWIGAQECT WP_000145429.1 hydrogenase 2 small subunit [Salmonella enterica] Length: 372\nNumber of predicted TMHs: 1\nExp number of AAs in TMHs: 24.51309\nExp number, first 60 AAs: 0.2682\nTotal prob of N-in: 0.14402\noutside 1 331\nTMhelix 332 354\ninside 355 372

20103 GCF_001137305.1_9953_5_38 Salmonella enterica subsp. enterica serovar Typhi Proteobacteria; Gammaproteobacteria; Enterobacterales; Enterobacteriaceae; Salmonella; Salmonella enterica; Salmonella enterica subsp. enterica MTGDNTLITSHGINRRDFMKLCAALAATMGLSSKAAAEMAESVSNPQRPPVIWIGAQECT WP_000145429.1 hydrogenase 2 small subunit [Salmonella enterica] Length: 372\nNumber of predicted TMHs: 1\nExp number of AAs in TMHs: 24.51309\nExp number, first 60 AAs: 0.2682\nTotal prob of N-in: 0.14402\noutside 1 331\nTMhelix 332 354\ninside 355 372

20104 GCF_001137545.1_9870_8_3Salmonella enterica subsp. enterica serovar Typhi Proteobacteria; Gammaproteobacteria; Enterobacterales; Enterobacteriaceae; Salmonella; Salmonella enterica; Salmonella enterica subsp. enterica MTGDNTLITSHGINRRDFMKLCAALAATMGLSSKAAAEMAESVSNPQRPPVIWIGAQECT WP_000145429.1 hydrogenase 2 small subunit [Salmonella enterica] Length: 372\nNumber of predicted TMHs: 1\nExp number of AAs in TMHs: 24.51309\nExp number, first 60 AAs: 0.2682\nTotal prob of N-in: 0.14402\noutside 1 331\nTMhelix 332 354\ninside 355 372

20105 GCF_001137705.1_9870_8_66 Salmonella enterica subsp. enterica serovar Typhi Proteobacteria; Gammaproteobacteria; Enterobacterales; Enterobacteriaceae; Salmonella; Salmonella enterica; Salmonella enterica subsp. enterica MTGDNTLITSHGINRRDFMKLCAALAATMGLSSKAAAEMAESVSNPQRPPVIWIGAQECT WP_000145429.1 hydrogenase 2 small subunit [Salmonella enterica] Length: 372\nNumber of predicted TMHs: 1\nExp number of AAs in TMHs: 24.51309\nExp number, first 60 AAs: 0.2682\nTotal prob of N-in: 0.14402\noutside 1 331\nTMhelix 332 354\ninside 355 372

20106 GCF_001141085.1_10493_1_34 Salmonella enterica subsp. enterica serovar Typhi Proteobacteria; Gammaproteobacteria; Enterobacterales; Enterobacteriaceae; Salmonella; Salmonella enterica; Salmonella enterica subsp. enterica MTGDNTLITSHGINRRDFMKLCAALAATMGLSSKAAAEMAESVSNPQRPPVIWIGAQECT WP_000145429.1 hydrogenase 2 small subunit [Salmonella enterica] Length: 372\nNumber of predicted TMHs: 1\nExp number of AAs in TMHs: 24.51309\nExp number, first 60 AAs: 0.2682\nTotal prob of N-in: 0.14402\noutside 1 331\nTMhelix 332 354\ninside 355 372

20107 GCF_001141125.1_10592_2_4 Salmonella enterica subsp. enterica serovar Typhi Proteobacteria; Gammaproteobacteria; Enterobacterales; Enterobacteriaceae; Salmonella; Salmonella enterica; Salmonella enterica subsp. enterica MTGDNTLITSHGINRRDFMKLCAALAATMGLSSKAAAEMAESVSNPQRPPVIWIGAQECT WP_000145429.1 hydrogenase 2 small subunit [Salmonella enterica] Length: 372\nNumber of predicted TMHs: 1\nExp number of AAs in TMHs: 24.51309\nExp number, first 60 AAs: 0.2682\nTotal prob of N-in: 0.14402\noutside 1 331\nTMhelix 332 354\ninside 355 372

20108 GCF_001141205.1_10541_2_14 Salmonella enterica subsp. enterica serovar Typhi Proteobacteria; Gammaproteobacteria; Enterobacterales; Enterobacteriaceae; Salmonella; Salmonella enterica; Salmonella enterica subsp. enterica MTGDNTLITSHGINRRDFMKLCAALAATMGLSSKAAAEMAESVSNPQRPPVIWIGAQECT WP_000145429.1 hydrogenase 2 small subunit [Salmonella enterica] Length: 372\nNumber of predicted TMHs: 1\nExp number of AAs in TMHs: 24.51309\nExp number, first 60 AAs: 0.2682\nTotal prob of N-in: 0.14402\noutside 1 331\nTMhelix 332 354\ninside 355 372

20109 GCF_001141445.1_10562_2_73 Salmonella enterica subsp. enterica serovar Typhi Proteobacteria; Gammaproteobacteria; Enterobacterales; Enterobacteriaceae; Salmonella; Salmonella enterica; Salmonella enterica subsp. enterica MTGDNTLITSHGINRRDFMKLCAALAATMGLSSKAAAEMAESVSNPQRPPVIWIGAQECT WP_000145429.1 hydrogenase 2 small subunit [Salmonella enterica] Length: 372\nNumber of predicted TMHs: 1\nExp number of AAs in TMHs: 24.51309\nExp number, first 60 AAs: 0.2682\nTotal prob of N-in: 0.14402\noutside 1 331\nTMhelix 332 354\ninside 355 372

20110 GCF_001141905.1_10492_1_3 Salmonella enterica subsp. enterica serovar Typhi Proteobacteria; Gammaproteobacteria; Enterobacterales; Enterobacteriaceae; Salmonella; Salmonella enterica; Salmonella enterica subsp. enterica MTGDNTLITSHGINRRDFMKLCAALAATMGLSSKAAAEMAESVSNPQRPPVIWIGAQECT WP_000145429.1 hydrogenase 2 small subunit [Salmonella enterica] Length: 372\nNumber of predicted TMHs: 1\nExp number of AAs in TMHs: 24.51309\nExp number, first 60 AAs: 0.2682\nTotal prob of N-in: 0.14402\noutside 1 331\nTMhelix 332 354\ninside 355 372

20111 GCF_001140345.1_10562_2_71 Salmonella enterica subsp. enterica serovar Typhi Proteobacteria; Gammaproteobacteria; Enterobacterales; Enterobacteriaceae; Salmonella; Salmonella enterica; Salmonella enterica subsp. enterica MTGDNTLITSHGINRRDFMKLCAALAATMGLSSKAATEMAESVSNPQRPPVIWIGAQECT WP_001782304.1 hydrogenase 2 small subunit [Salmonella enterica] Length: 372\nNumber of predicted TMHs: 1\nExp number of AAs in TMHs: 24.41173\nExp number, first 60 AAs: 0.16124\nTotal prob of N-in: 0.13947\noutside 1 331\nTMhelix 332 354\ninside 355 372

20112 GCF_001142085.1_10562_2_45 Salmonella enterica subsp. enterica serovar Typhi Proteobacteria; Gammaproteobacteria; Enterobacterales; Enterobacteriaceae; Salmonella; Salmonella enterica; Salmonella enterica subsp. enterica MTGDNTLITSHGINRRDFMKLCAALAATMGLSSKAAAEMAESVSNPQRPPVIWIGAQECT WP_000145429.1 hydrogenase 2 small subunit [Salmonella enterica] Length: 372\nNumber of

predicted TMHs: 1\nExp number of AAs in TMHs: 24.51309\nExp number, first 60 AAs: 0.2682\nTotal prob of N-in: 0.14402\noutside 1 331\nTMhelix 332 354\ninside 355 372

20113 GCF_001145385.1_10561_2_23 Salmonella enterica subsp. enterica serovar Typhi Proteobacteria; Gammaproteobacteria; Enterobacterales; Enterobacteriaceae; Salmonella; Salmonella enterica; Salmonella enterica subsp. enterica MTGDNTLITSHGINRRDFMKLCAALAATMGLSSKAAAEMAESVSNPQRPPVIWIGAQECT WP_000145429.1 hydrogenase 2 small subunit [Salmonella enterica] Length: 372\nNumber of predicted TMHs: 1\nExp number of AAs in TMHs: 24.51309\nExp number, first 60 AAs: 0.2682\nTotal prob of N-in: 0.14402\noutside 1 331\nTMhelix 332 354\ninside 355 372

20114 GCF_001145565.1_10541_2_36 Salmonella enterica subsp. enterica serovar Typhi Proteobacteria; Gammaproteobacteria; Enterobacterales; Enterobacteriaceae; Salmonella; Salmonella enterica; Salmonella enterica subsp. enterica MTGDNTLITSHGINRRDFMKLCAALAATMGLSSKAAAEMAESVSNPQRPPVIWIGAQECT WP_000145429.1 hydrogenase 2 small subunit [Salmonella enterica] Length: 372\nNumber of predicted TMHs: 1\nExp number of AAs in TMHs: 24.51309\nExp number, first 60 AAs: 0.2682\nTotal prob of N-in: 0.14402\noutside 1 331\nTMhelix 332 354\ninside 355 372

20115 GCF_001145945.1_10541_2_21 Salmonella enterica subsp. enterica serovar Typhi Proteobacteria; Gammaproteobacteria; Enterobacterales; Enterobacteriaceae; Salmonella; Salmonella enterica; Salmonella enterica subsp. enterica MTGDNTLITSHGINRRDFMKLCAALAATMGLSSKAAAEMAESVSNPQRPPVIWIGAQECT WP_000145429.1 hydrogenase 2 small subunit [Salmonella enterica] Length: 372\nNumber of predicted TMHs: 1\nExp number of AAs in TMHs: 24.51309\nExp number, first 60 AAs: 0.2682\nTotal prob of N-in: 0.14402\noutside 1 331\nTMhelix 332 354\ninside 355 372

20116 GCF_001145105.1_10541_2_4 Salmonella enterica subsp. enterica serovar Typhi Proteobacteria; Gammaproteobacteria; Enterobacterales; Enterobacteriaceae; Salmonella; Salmonella enterica; Salmonella enterica subsp. enterica MTGDNTLITSHGINRRDFMKLCAALAATMGLSSKAATEMAESVSNPQRPPVIWIGAQECT WP_001782304.1 hydrogenase 2 small subunit [Salmonella enterica] Length: 372\nNumber of predicted TMHs: 1\nExp number of AAs in TMHs: 24.41173\nExp number, first 60 AAs: 0.16124\nTotal prob of N-in: 0.13947\noutside 1 331\nTMhelix 332 354\ninside 355 372

20117 GCF_001146045.1_10592_2_12 Salmonella enterica subsp. enterica serovar Typhi Proteobacteria; Gammaproteobacteria; Enterobacterales; Enterobacteriaceae; Salmonella; Salmonella enterica; Salmonella enterica subsp. enterica MTGDNTLITSHGINRRDFMKLCAALAATMGLSSKAAAEMAESVSNPQRPPVIWIGAQECT WP_000145429.1 hydrogenase 2 small subunit [Salmonella enterica] Length: 372\nNumber of predicted TMHs: 1\nExp number of AAs in TMHs: 24.51309\nExp number, first 60 AAs: 0.2682\nTotal prob of N-in: 0.14402\noutside 1 331\nTMhelix 332 354\ninside 355 372

20118 GCF_001148345.1_9953_5_81 Salmonella enterica subsp. enterica serovar Typhi Proteobacteria; Gammaproteobacteria; Enterobacterales; Enterobacteriaceae; Salmonella; Salmonella enterica; Salmonella enterica subsp. enterica MTGDNTLITSHGINRRDFMKLCAALAATMGLSSKAAAEMAESVSNPQRPPVIWIGAQECT WP_000145429.1 hydrogenase 2 small subunit [Salmonella enterica] Length: 372\nNumber of predicted TMHs: 1\nExp number of AAs in TMHs: 24.51309\nExp number, first 60 AAs: 0.2682\nTotal prob of N-in: 0.14402\noutside 1 331\nTMhelix 332 354\ninside 355 372

20119 GCF_001148665.1_10493_1_77 Salmonella enterica subsp. enterica serovar Typhi Proteobacteria; Gammaproteobacteria; Enterobacterales; Enterobacteriaceae; Salmonella; Salmonella enterica; Salmonella enterica subsp. enterica MTGDNTLITSHGINRRDFMKLCAALAATMGLSSKAAAEMAESVSNPQRPPVIWIGAQECT WP_000145429.1 hydrogenase 2 small subunit [Salmonella enterica] Length: 372\nNumber of predicted TMHs: 1\nExp number of AAs in TMHs: 24.51309\nExp number, first 60 AAs: 0.2682\nTotal prob of N-in: 0.14402\noutside 1 331\nTMhelix 332 354\ninside 355 372

20120 GCF_001148825.1_9870_8_17 Salmonella enterica subsp. enterica serovar Typhi Proteobacteria; Gammaproteobacteria; Enterobacterales; Enterobacteriaceae; Salmonella; Salmonella enterica; Salmonella enterica subsp. enterica MTGDNTLITSHGINRRDFMKLCAALAATMGLSSKAAAEMAESVSNPQRPPVIWIGAQECT WP_000145429.1 hydrogenase 2 small subunit [Salmonella enterica] Length: 372\nNumber of predicted TMHs: 1\nExp number of AAs in TMHs: 24.51309\nExp number, first 60 AAs: 0.2682\nTotal prob of N-in: 0.14402\noutside 1 331\nTMhelix 332 354\ninside 355 372

20121 GCF_001148945.1_10349_1_55 Salmonella enterica subsp. enterica serovar Typhi Proteobacteria; Gammaproteobacteria; Enterobacterales; Enterobacteriaceae; Salmonella; Salmonella enterica;

Salmonella enterica subsp. enterica MTGDNTLITSHGINRRDFMKLCAALAATMGLSSKAAAEMAESVSNPQRPPVIWIGAQECT WP_000145429.1 hydrogenase 2 small subunit [Salmonella enterica] Length: 372\nNumber of predicted TMHs: 1\nExp number of AAs in TMHs: 24.51309\nExp number, first 60 AAs: 0.2682\nTotal prob of N-in: 0.14402\noutside 1 331\nTMhelix 332 354\ninside 355 372

20122 GCF_001149145.1_9870_8_9Salmonella enterica subsp. enterica serovar Typhi Proteobacteria; Gammaproteobacteria; Enterobacterales; Enterobacteriaceae; Salmonella; Salmonella enterica; Salmonella enterica subsp. enterica MTGDNTLITSHGINRRDFMKLCAALAATMGLSSKAAAEMAESVSNPQRPPVIWIGAQECT WP_000145429.1 hydrogenase 2 small subunit [Salmonella enterica] Length: 372\nNumber of predicted TMHs: 1\nExp number of AAs in TMHs: 24.51309\nExp number, first 60 AAs: 0.2682\nTotal prob of N-in: 0.14402\noutside 1 331\nTMhelix 332 354\ninside 355 372

20123 GCF_001151725.1_7468_7_72 Salmonella enterica subsp. enterica serovar Typhi Proteobacteria; Gammaproteobacteria; Enterobacterales; Enterobacteriaceae; Salmonella; Salmonella enterica; Salmonella enterica subsp. enterica MTGDNTLITSHGINRRDFMKLCAALAATMGLSSKAATEMAESVSNPQRPPVIWIGAQECT WP_001782304.1 hydrogenase 2 small subunit [Salmonella enterica] Length: 372\nNumber of predicted TMHs: 1\nExp number of AAs in TMHs: 24.41173\nExp number, first 60 AAs: 0.16124\nTotal prob of N-in: 0.13947\noutside 1 331\nTMhelix 332 354\ninside 355 372

20124 GCF_001153265.1_10561_2_28 Salmonella enterica subsp. enterica serovar Typhi Proteobacteria; Gammaproteobacteria; Enterobacterales; Enterobacteriaceae; Salmonella; Salmonella enterica; Salmonella enterica subsp. enterica MTGDNTLITSHGINRRDFMKLCAALAATMGLSSKAAAEMAESVSNPQRPPVIWIGAQECT WP_000145429.1 hydrogenase 2 small subunit [Salmonella enterica] Length: 372\nNumber of predicted TMHs: 1\nExp number of AAs in TMHs: 24.51309\nExp number, first 60 AAs: 0.2682\nTotal prob of N-in: 0.14402\noutside 1 331\nTMhelix 332 354\ninside 355 372

20125 GCF_001153465.1_8490_6_42 Salmonella enterica subsp. enterica serovar Typhi Proteobacteria; Gammaproteobacteria; Enterobacterales; Enterobacteriaceae; Salmonella; Salmonella enterica; Salmonella enterica subsp. enterica MTGDNTLITSHGINRRDFMKLCAALAATMGLSSKAAAEMAESVSNPQRPPVIWIGAQECT WP_000145429.1 hydrogenase 2 small subunit [Salmonella enterica] Length: 372\nNumber of predicted TMHs: 1\nExp number of AAs in TMHs: 24.51309\nExp number, first 60 AAs: 0.2682\nTotal prob of N-in: 0.14402\noutside 1 331\nTMhelix 332 354\ninside 355 372

20126 GCF_001153605.1_10592_2_21 Salmonella enterica subsp. enterica serovar Typhi Proteobacteria; Gammaproteobacteria; Enterobacterales; Enterobacteriaceae; Salmonella; Salmonella enterica; Salmonella enterica subsp. enterica MTGDNTLITSHGINRRDFMKLCAALAATMGLSSKAAAEMAESVSNPQRPPVIWIGAQECT WP_000145429.1 hydrogenase 2 small subunit [Salmonella enterica] Length: 372\nNumber of predicted TMHs: 1\nExp number of AAs in TMHs: 24.51309\nExp number, first 60 AAs: 0.2682\nTotal prob of N-in: 0.14402\noutside 1 331\nTMhelix 332 354\ninside 355 372

20127 GCF_001153825.1_8615_4_29 Salmonella enterica subsp. enterica serovar Typhi Proteobacteria; Gammaproteobacteria; Enterobacterales; Enterobacteriaceae; Salmonella; Salmonella enterica; Salmonella enterica subsp. enterica MTGDNTLITSHGINRRDFMKLCAALAATMGLSSKAAAEMAESVSNPQRPPVIWIGAQECT WP_000145429.1 hydrogenase 2 small subunit [Salmonella enterica] Length: 372\nNumber of predicted TMHs: 1\nExp number of AAs in TMHs: 24.51309\nExp number, first 60 AAs: 0.2682\nTotal prob of N-in: 0.14402\noutside 1 331\nTMhelix 332 354\ninside 355 372

20128 GCF_001154205.1_9953_5_63 Salmonella enterica subsp. enterica serovar Typhi Proteobacteria; Gammaproteobacteria; Enterobacterales; Enterobacteriaceae; Salmonella; Salmonella enterica; Salmonella enterica subsp. enterica MTGDNTLITSHGINRRDFMKLCAALAATMGLSSKAAAEMAESVSNPQRPPVIWIGAQECT WP_000145429.1 hydrogenase 2 small subunit [Salmonella enterica] Length: 372\nNumber of predicted TMHs: 1\nExp number of AAs in TMHs: 24.51309\nExp number, first 60 AAs: 0.2682\nTotal prob of N-in: 0.14402\noutside 1 331\nTMhelix 332 354\ninside 355 372

20129 GCF_001154045.1_10608_2_9 Salmonella enterica subsp. enterica serovar Typhi Proteobacteria; Gammaproteobacteria; Enterobacterales; Enterobacteriaceae; Salmonella; Salmonella enterica; Salmonella enterica subsp. enterica MTGDNTLITSHGINRRDFMKLCAALAATMGLSSKAAAEMAESVSNPQRPPVIWIGAQECT WP_000145429.1 hydrogenase 2 small subunit [Salmonella enterica] Length: 372\nNumber of predicted TMHs: 1\nExp number of AAs in TMHs: 24.51309\nExp number, first 60 AAs: 0.2682\nTotal prob of N-in: 0.14402\noutside 1 331\nTMhelix 332 354\ninside 355 372

20130 GCF_001157305.1_10541_2_18 Salmonella enterica subsp. enterica serovar Typhi Proteobacteria; Gammaproteobacteria; Enterobacterales; Enterobacteriaceae; Salmonella; Salmonella enterica; Salmonella enterica subsp. enterica MTGDNTLITSHGINRRDFMKLCAALAATMGLSSKAAAEMAESVSNPQRPPVIWIGAQECT WP_000145429.1 hydrogenase 2 small subunit [Salmonella enterica] Length: 372\nNumber of predicted TMHs: 1\nExp number of AAs in TMHs: 24.51309\nExp number, first 60 AAs: 0.2682\nTotal prob of N-in: 0.14402\noutside 1 331\nTMhelix 332 354\ninside 355 372

20131 GCF_001157925.1_10608_2_50 Salmonella enterica subsp. enterica serovar Typhi Proteobacteria; Gammaproteobacteria; Enterobacterales; Enterobacteriaceae; Salmonella; Salmonella enterica; Salmonella enterica subsp. enterica MTGDNTLITSHGINRRDFMKLCAALAATMGLSSKAAAEMAESVSNPQRPPVIWIGAQECT WP_000145429.1 hydrogenase 2 small subunit [Salmonella enterica] Length: 372\nNumber of predicted TMHs: 1\nExp number of AAs in TMHs: 24.51309\nExp number, first 60 AAs: 0.2682\nTotal prob of N-in: 0.14402\noutside 1 331\nTMhelix 332 354\ninside 355 372

20132 GCF_001157765.1_10209_5_38 Salmonella enterica subsp. enterica serovar Typhi Proteobacteria; Gammaproteobacteria; Enterobacterales; Enterobacteriaceae; Salmonella; Salmonella enterica; Salmonella enterica subsp. enterica MTGDNTLITSHGINRRDFMKLCAALAATMGLSSKAAAEMAESVSNPQRPPVIWIGAQECT WP_000145429.1 hydrogenase 2 small subunit [Salmonella enterica] Length: 372\nNumber of predicted TMHs: 1\nExp number of AAs in TMHs: 24.51309\nExp number, first 60 AAs: 0.2682\nTotal prob of N-in: 0.14402\noutside 1 331\nTMhelix 332 354\ninside 355 372

20133 GCF_001156265.1_10592_2_42 Salmonella enterica subsp. enterica serovar Typhi Proteobacteria; Gammaproteobacteria; Enterobacterales; Enterobacteriaceae; Salmonella; Salmonella enterica; Salmonella enterica subsp. enterica MTGDNTLITSHGINRRDFMKLCAALAATMGLSSKAATEMAESVSNPQRPPVIWIGAQECT WP_001782304.1 hydrogenase 2 small subunit [Salmonella enterica] Length: 372\nNumber of predicted TMHs: 1\nExp number of AAs in TMHs: 24.41173\nExp number, first 60 AAs: 0.16124\nTotal prob of N-in: 0.13947\noutside 1 331\nTMhelix 332 354\ninside 355 372

20134 GCF_001158025.1_10593_2_57 Salmonella enterica subsp. enterica serovar Typhi Proteobacteria; Gammaproteobacteria; Enterobacterales; Enterobacteriaceae; Salmonella; Salmonella enterica; Salmonella enterica subsp. enterica MTGDNTLITSHGINRRDFMKLCAALAATMGLSSKAAAEMAESVSNPQRPPVIWIGAQECT WP_000145429.1 hydrogenase 2 small subunit [Salmonella enterica] Length: 372\nNumber of predicted TMHs: 1\nExp number of AAs in TMHs: 24.51309\nExp number, first 60 AAs: 0.2682\nTotal prob of N-in: 0.14402\noutside 1 331\nTMhelix 332 354\ninside 355 372

20135 GCF_001158125.1_8490_5_33 Salmonella enterica subsp. enterica serovar Typhi Proteobacteria; Gammaproteobacteria; Enterobacterales; Enterobacteriaceae; Salmonella; Salmonella enterica; Salmonella enterica subsp. enterica MTGDNTLITSHGINRRDFMKLCAALAATMGLSSKAAAEMAESVSNPQRPPVIWIGAQECT WP_000145429.1 hydrogenase 2 small subunit [Salmonella enterica] Length: 372\nNumber of predicted TMHs: 1\nExp number of AAs in TMHs: 24.51309\nExp number, first 60 AAs: 0.2682\nTotal prob of N-in: 0.14402\noutside 1 331\nTMhelix 332 354\ninside 355 372

20136 GCF_001158585.1_10592_2_3 Salmonella enterica subsp. enterica serovar Typhi Proteobacteria; Gammaproteobacteria; Enterobacterales; Enterobacteriaceae; Salmonella; Salmonella enterica; Salmonella enterica subsp. enterica MTGDNTLITSHGINRRDFMKLCAALAATMGLSSKAAAEMAESVSNPQRPPVIWIGAQECT WP_000145429.1 hydrogenase 2 small subunit [Salmonella enterica] Length: 372\nNumber of predicted TMHs: 1\nExp number of AAs in TMHs: 24.51309\nExp number, first 60 AAs: 0.2682\nTotal prob of N-in: 0.14402\noutside 1 331\nTMhelix 332 354\ninside 355 372

20137 GCF_001161005.1_10492_1_6 Salmonella enterica subsp. enterica serovar Typhi Proteobacteria; Gammaproteobacteria; Enterobacterales; Enterobacteriaceae; Salmonella; Salmonella enterica; Salmonella enterica subsp. enterica MTGDNTLITSHGINRRDFMKLCAALAATMGLSSKAAAEMAESVSNPQRPPVIWIGAQECT WP_000145429.1 hydrogenase 2 small subunit [Salmonella enterica] Length: 372\nNumber of predicted TMHs: 1\nExp number of AAs in TMHs: 24.51309\nExp number, first 60 AAs: 0.2682\nTotal prob of N-in: 0.14402\noutside 1 331\nTMhelix 332 354\ninside 355 372

20138 GCF_001161385.1_8490_6_53 Salmonella enterica subsp. enterica serovar Typhi Proteobacteria; Gammaproteobacteria; Enterobacterales; Enterobacteriaceae; Salmonella; Salmonella enterica; Salmonella enterica subsp. enterica MTGDNTLITSHGINRRDFMKLCAALAATMGLSSKAAAEMAESVSNPQRPPVIWIGAQECT WP_000145429.1 hydrogenase 2 small subunit [Salmonella enterica] Length: 372\nNumber of

predicted TMHs: 1\nExp number of AAs in TMHs: 24.51309\nExp number, first 60 AAs: 0.2682\nTotal prob of N-in: 0.14402\noutside 1 331\nTMhelix 332 354\ninside 355 372

20139 GCF_001161525.1_10492_1_17 Salmonella enterica subsp. enterica serovar Typhi Proteobacteria; Gammaproteobacteria; Enterobacterales; Enterobacteriaceae; Salmonella; Salmonella enterica; Salmonella enterica subsp. enterica MTGDNTLITSHGINRRDFMKLCAALAATMGLSSKAAAEMAESVSNPQRPPVIWIGAQECT WP_000145429.1 hydrogenase 2 small subunit [Salmonella enterica] Length: 372\nNumber of predicted TMHs: 1\nExp number of AAs in TMHs: 24.51309\nExp number, first 60 AAs: 0.2682\nTotal prob of N-in: 0.14402\noutside 1 331\nTMhelix 332 354\ninside 355 372

20140 GCF_001161765.1_10492_1_21 Salmonella enterica subsp. enterica serovar Typhi Proteobacteria; Gammaproteobacteria; Enterobacterales; Enterobacteriaceae; Salmonella; Salmonella enterica; Salmonella enterica subsp. enterica MTGDNTLITSHGINRRDFMKLCAALAATMGLSSKAAAEMAESVSNPQRPPVIWIGAQECT WP_000145429.1 hydrogenase 2 small subunit [Salmonella enterica] Length: 372\nNumber of predicted TMHs: 1\nExp number of AAs in TMHs: 24.51309\nExp number, first 60 AAs: 0.2682\nTotal prob of N-in: 0.14402\noutside 1 331\nTMhelix 332 354\ninside 355 372

20141 GCF_001162345.1_10349_1_30 Salmonella enterica subsp. enterica serovar Typhi Proteobacteria; Gammaproteobacteria; Enterobacterales; Enterobacteriaceae; Salmonella; Salmonella enterica; Salmonella enterica subsp. enterica MTGDNTLITSHGINRRDFMKLCAALAATMGLSSKAAAEMAESVSNPQRPPVIWIGAQECT WP_000145429.1 hydrogenase 2 small subunit [Salmonella enterica] Length: 372\nNumber of predicted TMHs: 1\nExp number of AAs in TMHs: 24.51309\nExp number, first 60 AAs: 0.2682\nTotal prob of N-in: 0.14402\noutside 1 331\nTMhelix 332 354\ninside 355 372

20142 GCF_001162085.1_10209_5_41 Salmonella enterica subsp. enterica serovar Typhi Proteobacteria; Gammaproteobacteria; Enterobacterales; Enterobacteriaceae; Salmonella; Salmonella enterica; Salmonella enterica subsp. enterica MTGDNTLITSHGINRRDFMKLCAALAATMGLSSKAAAEMAESVSNPQRPPVIWIGAQECT WP_000145429.1 hydrogenase 2 small subunit [Salmonella enterica] Length: 372\nNumber of predicted TMHs: 1\nExp number of AAs in TMHs: 24.51309\nExp number, first 60 AAs: 0.2682\nTotal prob of N-in: 0.14402\noutside 1 331\nTMhelix 332 354\ninside 355 372

20143 GCF_001162605.1_9475_6_50 Salmonella enterica subsp. enterica serovar Typhi Proteobacteria; Gammaproteobacteria; Enterobacterales; Enterobacteriaceae; Salmonella; Salmonella enterica; Salmonella enterica subsp. enterica MTGDNTLITSHGINRRDFMKLCAALAATMGLSSKAAAEMAESVSNPQRPPVIWIGAQECT WP_000145429.1 hydrogenase 2 small subunit [Salmonella enterica] Length: 372\nNumber of predicted TMHs: 1\nExp number of AAs in TMHs: 24.51309\nExp number, first 60 AAs: 0.2682\nTotal prob of N-in: 0.14402\noutside 1 331\nTMhelix 332 354\ninside 355 372

20144 GCF_001164525.1_10607_2_54 Salmonella enterica subsp. enterica serovar Typhi Proteobacteria; Gammaproteobacteria; Enterobacterales; Enterobacteriaceae; Salmonella; Salmonella enterica; Salmonella enterica subsp. enterica MTGDNTLITSHGINRRDFMKLCAALAATMGLSSKAAAEMAESVSNPQRPPVIWIGAQECT WP_000145429.1 hydrogenase 2 small subunit [Salmonella enterica] Length: 372\nNumber of predicted TMHs: 1\nExp number of AAs in TMHs: 24.51309\nExp number, first 60 AAs: 0.2682\nTotal prob of N-in: 0.14402\noutside 1 331\nTMhelix 332 354\ninside 355 372

20145 GCF_001165785.1_10349_1_7 Salmonella enterica subsp. enterica serovar Typhi Proteobacteria; Gammaproteobacteria; Enterobacterales; Enterobacteriaceae; Salmonella; Salmonella enterica; Salmonella enterica subsp. enterica MTGDNTLITSHGINRRDFMKLCAALAATMGLSSKAAAEMAESVSNPQRPPVIWIGAQECT WP_000145429.1 hydrogenase 2 small subunit [Salmonella enterica] Length: 372\nNumber of predicted TMHs: 1\nExp number of AAs in TMHs: 24.51309\nExp number, first 60 AAs: 0.2682\nTotal prob of N-in: 0.14402\noutside 1 331\nTMhelix 332 354\ninside 355 372

20146 GCF_001165305.1_10592_2_2 Salmonella enterica subsp. enterica serovar Typhi Proteobacteria; Gammaproteobacteria; Enterobacterales; Enterobacteriaceae; Salmonella; Salmonella enterica; Salmonella enterica subsp. enterica MTGDNTLITSHGINRRDFMKLCAALAATMGLSSKAAAEMAESVSNPQRPPVIWIGAQECT WP_000145429.1 hydrogenase 2 small subunit [Salmonella enterica] Length: 372\nNumber of predicted TMHs: 1\nExp number of AAs in TMHs: 24.51309\nExp number, first 60 AAs: 0.2682\nTotal prob of N-in: 0.14402\noutside 1 331\nTMhelix 332 354\ninside 355 372

20147 GCF_001165665.1_9953_5_73 Salmonella enterica subsp. enterica serovar Typhi Proteobacteria; Gammaproteobacteria; Enterobacterales; Enterobacteriaceae; Salmonella; Salmonella enterica;

Salmonella enterica subsp. enterica	MTGDNTLITSHGINRRDFMKLCAALAATMGLSSKAAAEMAESVSNPQRPPVIWIGAQECT
	WP_000145429.1	hydrogenase 2 small subunit [Salmonella enterica]	Length: 372\nNumber of predicted TMHs: 1\nExp number of AAs in TMHs: 24.51309\nExp number, first 60 AAs: 0.2682\nTotal prob of N-in: 0.14402\noutside 1 331\nTMhelix 332 354\ninside 355 372

20148	GCF_001165605.1_9953_5_35	Salmonella enterica subsp. enterica serovar Typhi
	Proteobacteria; Gammaproteobacteria; Enterobacterales; Enterobacteriaceae; Salmonella; Salmonella enterica; Salmonella enterica subsp. enterica	MTGDNTLITSHGINRRDFMKLCAALAATMGLSSKAAAEMAESVSNPQRPPVIWIGAQECT
	WP_000145429.1	hydrogenase 2 small subunit [Salmonella enterica]	Length: 372\nNumber of predicted TMHs: 1\nExp number of AAs in TMHs: 24.51309\nExp number, first 60 AAs: 0.2682\nTotal prob of N-in: 0.14402\noutside 1 331\nTMhelix 332 354\ninside 355 372

20149	GCF_001166125.1_9953_5_89	Salmonella enterica subsp. enterica serovar Typhi
	Proteobacteria; Gammaproteobacteria; Enterobacterales; Enterobacteriaceae; Salmonella; Salmonella enterica; Salmonella enterica subsp. enterica	MTGDNTLITSHGINRRDFMKLCAALAATMGLSSKAAAEMAESVSNPQRPPVIWIGAQECT
	WP_000145429.1	hydrogenase 2 small subunit [Salmonella enterica]	Length: 372\nNumber of predicted TMHs: 1\nExp number of AAs in TMHs: 24.51309\nExp number, first 60 AAs: 0.2682\nTotal prob of N-in: 0.14402\noutside 1 331\nTMhelix 332 354\ninside 355 372

20150	GCF_001168985.1_10562_2_34	Salmonella enterica subsp. enterica serovar Typhi
	Proteobacteria; Gammaproteobacteria; Enterobacterales; Enterobacteriaceae; Salmonella; Salmonella enterica; Salmonella enterica subsp. enterica	MTGDNTLITSHGINRRDFMKLCAALAATMGLSSKAAAEMAESVSNPQRPPVIWIGAQECT
	WP_000145429.1	hydrogenase 2 small subunit [Salmonella enterica]	Length: 372\nNumber of predicted TMHs: 1\nExp number of AAs in TMHs: 24.51309\nExp number, first 60 AAs: 0.2682\nTotal prob of N-in: 0.14402\noutside 1 331\nTMhelix 332 354\ninside 355 372

20151	GCF_001170145.1_10209_5_33	Salmonella enterica subsp. enterica serovar Typhi
	Proteobacteria; Gammaproteobacteria; Enterobacterales; Enterobacteriaceae; Salmonella; Salmonella enterica; Salmonella enterica subsp. enterica	MTGDNTLITSHGINRRDFMKLCAALAATMGLSSKAAAEMAESVSNPQRPPVIWIGAQECT
	WP_000145429.1	hydrogenase 2 small subunit [Salmonella enterica]	Length: 372\nNumber of predicted TMHs: 1\nExp number of AAs in TMHs: 24.51309\nExp number, first 60 AAs: 0.2682\nTotal prob of N-in: 0.14402\noutside 1 331\nTMhelix 332 354\ninside 355 372

20152	GCF_001169845.1_10561_2_27	Salmonella enterica subsp. enterica serovar Typhi
	Proteobacteria; Gammaproteobacteria; Enterobacterales; Enterobacteriaceae; Salmonella; Salmonella enterica; Salmonella enterica subsp. enterica	MTGDNTLITSHGINRRDFMKLCAALAATMGLSSKAAAEMAESVSNPQRPPVIWIGAQECT
	WP_000145429.1	hydrogenase 2 small subunit [Salmonella enterica]	Length: 372\nNumber of predicted TMHs: 1\nExp number of AAs in TMHs: 24.51309\nExp number, first 60 AAs: 0.2682\nTotal prob of N-in: 0.14402\noutside 1 331\nTMhelix 332 354\ninside 355 372

20153	GCF_001170245.1_9870_8_93	Salmonella enterica subsp. enterica serovar Typhi
	Proteobacteria; Gammaproteobacteria; Enterobacterales; Enterobacteriaceae; Salmonella; Salmonella enterica; Salmonella enterica subsp. enterica	MTGDNTLITSHGINRRDFMKLCAALAATMGLSSKAAAEMAESVSNPQRPPVIWIGAQECT
	WP_000145429.1	hydrogenase 2 small subunit [Salmonella enterica]	Length: 372\nNumber of predicted TMHs: 1\nExp number of AAs in TMHs: 24.51309\nExp number, first 60 AAs: 0.2682\nTotal prob of N-in: 0.14402\noutside 1 331\nTMhelix 332 354\ninside 355 372

20154	GCF_001170345.1_10541_1_75	Salmonella enterica subsp. enterica serovar Typhi
	Proteobacteria; Gammaproteobacteria; Enterobacterales; Enterobacteriaceae; Salmonella; Salmonella enterica; Salmonella enterica subsp. enterica	MTGDNTLITSHGINRRDFMKLCAALAATMGLSSKAAAEMAESVSNPQRPPVIWIGAQECT
	WP_000145429.1	hydrogenase 2 small subunit [Salmonella enterica]	Length: 372\nNumber of predicted TMHs: 1\nExp number of AAs in TMHs: 24.51309\nExp number, first 60 AAs: 0.2682\nTotal prob of N-in: 0.14402\noutside 1 331\nTMhelix 332 354\ninside 355 372

20155	GCF_001170925.1_10607_2_34	Salmonella enterica subsp. enterica serovar Typhi
	Proteobacteria; Gammaproteobacteria; Enterobacterales; Enterobacteriaceae; Salmonella; Salmonella enterica; Salmonella enterica subsp. enterica	MTGDNTLITSHGINRRDFMKLCAALAATMGLSSKAAAEMAESVSNPQRPPVIWIGAQECT
	WP_000145429.1	hydrogenase 2 small subunit [Salmonella enterica]	Length: 372\nNumber of predicted TMHs: 1\nExp number of AAs in TMHs: 24.51309\nExp number, first 60 AAs: 0.2682\nTotal prob of N-in: 0.14402\noutside 1 331\nTMhelix 332 354\ninside 355 372

20156 GCF_001171105.1_9870_8_60 Salmonella enterica subsp. enterica serovar Typhi Proteobacteria; Gammaproteobacteria; Enterobacterales; Enterobacteriaceae; Salmonella; Salmonella enterica; Salmonella enterica subsp. enterica MTGDNTLITSHGINRRDFMKLCAALAATMGLSSKAAAEMAESVSNPQRPPVIWIGAQECT WP_000145429.1 hydrogenase 2 small subunit [Salmonella enterica] Length: 372\nNumber of predicted TMHs: 1\nExp number of AAs in TMHs: 24.51309\nExp number, first 60 AAs: 0.2682\nTotal prob of N-in: 0.14402\noutside 1 331\nTMhelix 332 354\ninside 355 372

20157 GCF_001174425.1_10493_1_41 Salmonella enterica subsp. enterica serovar Typhi Proteobacteria; Gammaproteobacteria; Enterobacterales; Enterobacteriaceae; Salmonella; Salmonella enterica; Salmonella enterica subsp. enterica MTGDNTLITSHGINRRDFMKLCAALAATMGLSSKAAAEMAESVSNPQRPPVIWIGAQECT WP_000145429.1 hydrogenase 2 small subunit [Salmonella enterica] Length: 372\nNumber of predicted TMHs: 1\nExp number of AAs in TMHs: 24.51309\nExp number, first 60 AAs: 0.2682\nTotal prob of N-in: 0.14402\noutside 1 331\nTMhelix 332 354\ninside 355 372

20158 GCF_001174645.1_9870_8_39 Salmonella enterica subsp. enterica serovar Typhi Proteobacteria; Gammaproteobacteria; Enterobacterales; Enterobacteriaceae; Salmonella; Salmonella enterica; Salmonella enterica subsp. enterica MTGDNTLITSHGINRRDFMKLCAALAATMGLSSKAAAEMAESVSNPQRPPVIWIGAQECT WP_000145429.1 hydrogenase 2 small subunit [Salmonella enterica] Length: 372\nNumber of predicted TMHs: 1\nExp number of AAs in TMHs: 24.51309\nExp number, first 60 AAs: 0.2682\nTotal prob of N-in: 0.14402\noutside 1 331\nTMhelix 332 354\ninside 355 372

20159 GCF_001216505.1_10349_1_2 Salmonella enterica subsp. enterica serovar Typhi Proteobacteria; Gammaproteobacteria; Enterobacterales; Enterobacteriaceae; Salmonella; Salmonella enterica; Salmonella enterica subsp. enterica MTGDNTLITSHGINRRDFMKLCAALAATMGLSSKAAAEMAESVSNPQRPPVIWIGAQECT WP_000145429.1 hydrogenase 2 small subunit [Salmonella enterica] Length: 372\nNumber of predicted TMHs: 1\nExp number of AAs in TMHs: 24.51309\nExp number, first 60 AAs: 0.2682\nTotal prob of N-in: 0.14402\noutside 1 331\nTMhelix 332 354\ninside 355 372

20160 GCF_001239985.1_10425_1_7 Salmonella enterica subsp. enterica serovar Typhi Proteobacteria; Gammaproteobacteria; Enterobacterales; Enterobacteriaceae; Salmonella; Salmonella enterica; Salmonella enterica subsp. enterica MTGDNTLITSHGINRRDFMKLCAALAATMGLSSKAAAEMAESVSNPQRPPVIWIGAQECT WP_000145429.1 hydrogenase 2 small subunit [Salmonella enterica] Length: 372\nNumber of predicted TMHs: 1\nExp number of AAs in TMHs: 24.51309\nExp number, first 60 AAs: 0.2682\nTotal prob of N-in: 0.14402\noutside 1 331\nTMhelix 332 354\ninside 355 372

20161 GCF_001240325.1_10425_1_30 Salmonella enterica subsp. enterica serovar Typhi Proteobacteria; Gammaproteobacteria; Enterobacterales; Enterobacteriaceae; Salmonella; Salmonella enterica; Salmonella enterica subsp. enterica MTGDNTLITSHGINRRDFMKLCAALAATMGLSSKAAAEMAESVSNPQRPPVIWIGAQECT WP_000145429.1 hydrogenase 2 small subunit [Salmonella enterica] Length: 372\nNumber of predicted TMHs: 1\nExp number of AAs in TMHs: 24.51309\nExp number, first 60 AAs: 0.2682\nTotal prob of N-in: 0.14402\noutside 1 331\nTMhelix 332 354\ninside 355 372

20162 GCF_001240755.1_10425_1_75 Salmonella enterica subsp. enterica serovar Typhi Proteobacteria; Gammaproteobacteria; Enterobacterales; Enterobacteriaceae; Salmonella; Salmonella enterica; Salmonella enterica subsp. enterica MTGDNTLITSHGINRRDFMKLCAALAATMGLSSKAAAEMAESVSNPQRPPVIWIGAQECT WP_000145429.1 hydrogenase 2 small subunit [Salmonella enterica] Length: 372\nNumber of predicted TMHs: 1\nExp number of AAs in TMHs: 24.51309\nExp number, first 60 AAs: 0.2682\nTotal prob of N-in: 0.14402\noutside 1 331\nTMhelix 332 354\ninside 355 372

20163 GCF_001241235.1_10425_1_27 Salmonella enterica subsp. enterica serovar Typhi Proteobacteria; Gammaproteobacteria; Enterobacterales; Enterobacteriaceae; Salmonella; Salmonella enterica; Salmonella enterica subsp. enterica MTGDNTLITSHGINRRDFMKLCAALAATMGLSSKAAAEMAESVSNPQRPPVIWIGAQECT WP_000145429.1 hydrogenase 2 small subunit [Salmonella enterica] Length: 372\nNumber of predicted TMHs: 1\nExp number of AAs in TMHs: 24.51309\nExp number, first 60 AAs: 0.2682\nTotal prob of N-in: 0.14402\noutside 1 331\nTMhelix 332 354\ninside 355 372

20164 GCF_001241705.1_10425_1_81 Salmonella enterica subsp. enterica serovar Typhi Proteobacteria; Gammaproteobacteria; Enterobacterales; Enterobacteriaceae; Salmonella; Salmonella enterica; Salmonella enterica subsp. enterica MTGDNTLITSHGINRRDFMKLCAALAATMGLSSKAAAEMAESVSNPQRPPVIWIGAQECT WP_000145429.1 hydrogenase 2 small subunit [Salmonella enterica] Length: 372\nNumber of

predicted TMHs:  1\nExp number of AAs in TMHs: 24.51309\nExp number, first 60 AAs:  0.2682\nTotal prob of N-in: 0.14402\noutside   1   331\nTMhelix   332   354\ninside   355   372

20165 GCF_001241805.1_10425_1_46 Salmonella enterica subsp. enterica serovar Typhi Proteobacteria; Gammaproteobacteria; Enterobacterales; Enterobacteriaceae; Salmonella; Salmonella enterica; Salmonella enterica subsp. enterica MTGDNTLITSHGINRRDFMKLCAALAATMGLSSKAAAEMAESVSNPQRPPVIWIGAQECT WP_000145429.1 hydrogenase 2 small subunit [Salmonella enterica] Length: 372\nNumber of predicted TMHs:  1\nExp number of AAs in TMHs: 24.51309\nExp number, first 60 AAs:  0.2682\nTotal prob of N-in: 0.14402\noutside   1   331\nTMhelix   332   354\ninside   355   372

20166 GCF_001283125.1_10593_2_80 Salmonella enterica subsp. enterica serovar Typhi Proteobacteria; Gammaproteobacteria; Enterobacterales; Enterobacteriaceae; Salmonella; Salmonella enterica; Salmonella enterica subsp. enterica MTGDNTLITSHGINRRDFMKLCAALAATMGLSSKAAAEMAESVSNPQRPPVIWIGAQECT WP_000145429.1 hydrogenase 2 small subunit [Salmonella enterica] Length: 372\nNumber of predicted TMHs:  1\nExp number of AAs in TMHs: 24.51309\nExp number, first 60 AAs:  0.2682\nTotal prob of N-in: 0.14402\noutside   1   331\nTMhelix   332   354\ninside   355   372

20167 GCF_001317645.1_9475_4_55 Salmonella enterica subsp. enterica serovar Typhi Proteobacteria; Gammaproteobacteria; Enterobacterales; Enterobacteriaceae; Salmonella; Salmonella enterica; Salmonella enterica subsp. enterica MTGDNTLITSHGINRRDFMKLCAALAATMGLSSKAAAEMAESVSNPQRPPVIWIGAQECT WP_000145429.1 hydrogenase 2 small subunit [Salmonella enterica] Length: 372\nNumber of predicted TMHs:  1\nExp number of AAs in TMHs: 24.51309\nExp number, first 60 AAs:  0.2682\nTotal prob of N-in: 0.14402\noutside   1   331\nTMhelix   332   354\ninside   355   372

20168 GCF_001317685.1_9472_4_78 Salmonella enterica subsp. enterica serovar Typhi Proteobacteria; Gammaproteobacteria; Enterobacterales; Enterobacteriaceae; Salmonella; Salmonella enterica; Salmonella enterica subsp. enterica MTGDNTLITSHGINRRDFMKLCAALAATMGLSSKAAAEMAESVSNPQRPPVIWIGAQECT WP_000145429.1 hydrogenase 2 small subunit [Salmonella enterica] Length: 372\nNumber of predicted TMHs:  1\nExp number of AAs in TMHs: 24.51309\nExp number, first 60 AAs:  0.2682\nTotal prob of N-in: 0.14402\noutside   1   331\nTMhelix   332   354\ninside   355   372

20169 GCF_001350295.1_8616_4_56 Salmonella enterica subsp. enterica serovar Typhi Proteobacteria; Gammaproteobacteria; Enterobacterales; Enterobacteriaceae; Salmonella; Salmonella enterica; Salmonella enterica subsp. enterica MTGDNTLITSHGINRRDFMKLCAALAATMGLSSKAAAEMAESVSNPQRPPVIWIGAQECT WP_000145429.1 hydrogenase 2 small subunit [Salmonella enterica] Length: 372\nNumber of predicted TMHs:  1\nExp number of AAs in TMHs: 24.51309\nExp number, first 60 AAs:  0.2682\nTotal prob of N-in: 0.14402\noutside   1   331\nTMhelix   332   354\ninside   355   372

20170 GCF_001350335.1_8616_4_54 Salmonella enterica subsp. enterica serovar Typhi Proteobacteria; Gammaproteobacteria; Enterobacterales; Enterobacteriaceae; Salmonella; Salmonella enterica; Salmonella enterica subsp. enterica MTGDNTLITSHGINRRDFMKLCAALAATMGLSSKAAAEMAESVSNPQRPPVIWIGAQECT WP_000145429.1 hydrogenase 2 small subunit [Salmonella enterica] Length: 372\nNumber of predicted TMHs:  1\nExp number of AAs in TMHs: 24.51309\nExp number, first 60 AAs:  0.2682\nTotal prob of N-in: 0.14402\noutside   1   331\nTMhelix   332   354\ninside   355   372

20171 GCF_001350375.1_8616_4_53 Salmonella enterica subsp. enterica serovar Typhi Proteobacteria; Gammaproteobacteria; Enterobacterales; Enterobacteriaceae; Salmonella; Salmonella enterica; Salmonella enterica subsp. enterica MTGDNTLITSHGINRRDFMKLCAALAATMGLSSKAAAEMAESVSNPQRPPVIWIGAQECT WP_000145429.1 hydrogenase 2 small subunit [Salmonella enterica] Length: 372\nNumber of predicted TMHs:  1\nExp number of AAs in TMHs: 24.51309\nExp number, first 60 AAs:  0.2682\nTotal prob of N-in: 0.14402\noutside   1   331\nTMhelix   332   354\ninside   355   372

20172 GCF_001328875.1_7468_7_53 Salmonella enterica subsp. enterica serovar Typhi Proteobacteria; Gammaproteobacteria; Enterobacterales; Enterobacteriaceae; Salmonella; Salmonella enterica; Salmonella enterica subsp. enterica MTGDNTLITSHGINRRDFMKLCAALAATMGLSSKAATEMAESVSNPQRPPVIWIGAQECT WP_001782304.1 hydrogenase 2 small subunit [Salmonella enterica] Length: 372\nNumber of predicted TMHs:  1\nExp number of AAs in TMHs: 24.41173\nExp number, first 60 AAs:  0.16124\nTotal prob of N-in: 0.13947\noutside   1   331\nTMhelix   332   354\ninside   355   372

20173 GCF_001355615.1_8447_8_28 Salmonella enterica subsp. enterica serovar Typhi Proteobacteria; Gammaproteobacteria; Enterobacterales; Enterobacteriaceae; Salmonella; Salmonella enterica;

Salmonella enterica subsp. enterica MTGDNTLITSHGINRRDFMKLCAALAATMGLSSKAAAEMAESVSNPQRPPVIWIGAQECT WP_000145429.1 hydrogenase 2 small subunit [Salmonella enterica] Length: 372\nNumber of predicted TMHs: 1\nExp number of AAs in TMHs: 24.51309\nExp number, first 60 AAs: 0.2682\nTotal prob of N-in: 0.14402\noutside 1 331\nTMhelix 332 354\ninside 355 372

20174 GCF_001355575.1_8525_3_33 Salmonella enterica subsp. enterica serovar Typhi Proteobacteria; Gammaproteobacteria; Enterobacterales; Enterobacteriaceae; Salmonella; Salmonella enterica; Salmonella enterica subsp. enterica MTGDNTLITSHGINRRDFMKLCAALAATMGLSSKAAAEMAESVSNPQRPPVIWIGAQECT WP_000145429.1 hydrogenase 2 small subunit [Salmonella enterica] Length: 372\nNumber of predicted TMHs: 1\nExp number of AAs in TMHs: 24.51309\nExp number, first 60 AAs: 0.2682\nTotal prob of N-in: 0.14402\noutside 1 331\nTMhelix 332 354\ninside 355 372

20175 GCF_001355535.1_8525_3_71 Salmonella enterica subsp. enterica serovar Typhi Proteobacteria; Gammaproteobacteria; Enterobacterales; Enterobacteriaceae; Salmonella; Salmonella enterica; Salmonella enterica subsp. enterica MTGDNTLITSHGINRRDFMKLCAALAATMGLSSKAAAEMAESVSNPQRPPVIWIGAQECT WP_000145429.1 hydrogenase 2 small subunit [Salmonella enterica] Length: 372\nNumber of predicted TMHs: 1\nExp number of AAs in TMHs: 24.51309\nExp number, first 60 AAs: 0.2682\nTotal prob of N-in: 0.14402\noutside 1 331\nTMhelix 332 354\ninside 355 372

20176 GCF_001355655.1_8490_5_22 Salmonella enterica subsp. enterica serovar Typhi Proteobacteria; Gammaproteobacteria; Enterobacterales; Enterobacteriaceae; Salmonella; Salmonella enterica; Salmonella enterica subsp. enterica MTGDNTLITSHGINRRDFMKLCAALAATMGLSSKAAAEMAESVSNPQRPPVIWIGAQECT WP_000145429.1 hydrogenase 2 small subunit [Salmonella enterica] Length: 372\nNumber of predicted TMHs: 1\nExp number of AAs in TMHs: 24.51309\nExp number, first 60 AAs: 0.2682\nTotal prob of N-in: 0.14402\noutside 1 331\nTMhelix 332 354\ninside 355 372

20177 GCF_001355695.1_8447_8_27 Salmonella enterica subsp. enterica serovar Typhi Proteobacteria; Gammaproteobacteria; Enterobacterales; Enterobacteriaceae; Salmonella; Salmonella enterica; Salmonella enterica subsp. enterica MTGDNTLITSHGINRRDFMKLCAALAATMGLSSKAAAEMAESVSNPQRPPVIWIGAQECT WP_000145429.1 hydrogenase 2 small subunit [Salmonella enterica] Length: 372\nNumber of predicted TMHs: 1\nExp number of AAs in TMHs: 24.51309\nExp number, first 60 AAs: 0.2682\nTotal prob of N-in: 0.14402\noutside 1 331\nTMhelix 332 354\ninside 355 372

20178 GCF_001355735.1_8490_5_14 Salmonella enterica subsp. enterica serovar Typhi Proteobacteria; Gammaproteobacteria; Enterobacterales; Enterobacteriaceae; Salmonella; Salmonella enterica; Salmonella enterica subsp. enterica MTGDNTLITSHGINRRDFMKLCAALAATMGLSSKAAAEMAESVSNPQRPPVIWIGAQECT WP_000145429.1 hydrogenase 2 small subunit [Salmonella enterica] Length: 372\nNumber of predicted TMHs: 1\nExp number of AAs in TMHs: 24.51309\nExp number, first 60 AAs: 0.2682\nTotal prob of N-in: 0.14402\noutside 1 331\nTMhelix 332 354\ninside 355 372

20179 GCF_001355775.1_8447_8_22 Salmonella enterica subsp. enterica serovar Typhi Proteobacteria; Gammaproteobacteria; Enterobacterales; Enterobacteriaceae; Salmonella; Salmonella enterica; Salmonella enterica subsp. enterica MTGDNTLITSHGINRRDFMKLCAALAATMGLSSKAAAEMAESVSNPQRPPVIWIGAQECT WP_000145429.1 hydrogenase 2 small subunit [Salmonella enterica] Length: 372\nNumber of predicted TMHs: 1\nExp number of AAs in TMHs: 24.51309\nExp number, first 60 AAs: 0.2682\nTotal prob of N-in: 0.14402\noutside 1 331\nTMhelix 332 354\ninside 355 372

20180 GCF_001356615.1_9475_6_54 Salmonella enterica subsp. enterica serovar Typhi Proteobacteria; Gammaproteobacteria; Enterobacterales; Enterobacteriaceae; Salmonella; Salmonella enterica; Salmonella enterica subsp. enterica MTGDNTLITSHGINRRDFMKLCAALAATMGLSSKAAAEMAESVSNPQRPPVIWIGAQECT WP_000145429.1 hydrogenase 2 small subunit [Salmonella enterica] Length: 372\nNumber of predicted TMHs: 1\nExp number of AAs in TMHs: 24.51309\nExp number, first 60 AAs: 0.2682\nTotal prob of N-in: 0.14402\noutside 1 331\nTMhelix 332 354\ninside 355 372

20181 GCF_001356655.1_9475_6_73 Salmonella enterica subsp. enterica serovar Typhi Proteobacteria; Gammaproteobacteria; Enterobacterales; Enterobacteriaceae; Salmonella; Salmonella enterica; Salmonella enterica subsp. enterica MTGDNTLITSHGINRRDFMKLCAALAATMGLSSKAAAEMAESVSNPQRPPVIWIGAQECT WP_000145429.1 hydrogenase 2 small subunit [Salmonella enterica] Length: 372\nNumber of predicted TMHs: 1\nExp number of AAs in TMHs: 24.51309\nExp number, first 60 AAs: 0.2682\nTotal prob of N-in: 0.14402\noutside 1 331\nTMhelix 332 354\ninside 355 372

20182 GCF_001356695.1_9475_6_80 Salmonella enterica subsp. enterica serovar Typhi Proteobacteria; Gammaproteobacteria; Enterobacterales; Enterobacteriaceae; Salmonella; Salmonella enterica; Salmonella enterica subsp. enterica MTGDNTLITSHGINRRDFMKLCAALAATMGLSSKAAAEMAESVSNPQRPPVIWIGAQECT WP_000145429.1 hydrogenase 2 small subunit [Salmonella enterica] Length: 372\nNumber of predicted TMHs: 1\nExp number of AAs in TMHs: 24.51309\nExp number, first 60 AAs: 0.2682\nTotal prob of N-in: 0.14402\noutside 1 331\nTMhelix 332 354\ninside 355 372

20183 GCF_001356735.1_10592_2_29 Salmonella enterica subsp. enterica serovar Typhi Proteobacteria; Gammaproteobacteria; Enterobacterales; Enterobacteriaceae; Salmonella; Salmonella enterica; Salmonella enterica subsp. enterica MTGDNTLITSHGINRRDFMKLCAALAATMGLSSKAAAEMAESVSNPQRPPVIWIGAQECT WP_000145429.1 hydrogenase 2 small subunit [Salmonella enterica] Length: 372\nNumber of predicted TMHs: 1\nExp number of AAs in TMHs: 24.51309\nExp number, first 60 AAs: 0.2682\nTotal prob of N-in: 0.14402\noutside 1 331\nTMhelix 332 354\ninside 355 372

20184 GCF_001356815.1_10592_2_34 Salmonella enterica subsp. enterica serovar Typhi Proteobacteria; Gammaproteobacteria; Enterobacterales; Enterobacteriaceae; Salmonella; Salmonella enterica; Salmonella enterica subsp. enterica MTGDNTLITSHGINRRDFMKLCAALAATMGLSSKAAAEMAESVSNPQRPPVIWIGAQECT WP_000145429.1 hydrogenase 2 small subunit [Salmonella enterica] Length: 372\nNumber of predicted TMHs: 1\nExp number of AAs in TMHs: 24.51309\nExp number, first 60 AAs: 0.2682\nTotal prob of N-in: 0.14402\noutside 1 331\nTMhelix 332 354\ninside 355 372

20185 GCF_001356855.1_10592_2_5 Salmonella enterica subsp. enterica serovar Typhi Proteobacteria; Gammaproteobacteria; Enterobacterales; Enterobacteriaceae; Salmonella; Salmonella enterica; Salmonella enterica subsp. enterica MTGDNTLITSHGINRRDFMKLCAALAATMGLSSKAAAEMAESVSNPQRPPVIWIGAQECT WP_000145429.1 hydrogenase 2 small subunit [Salmonella enterica] Length: 372\nNumber of predicted TMHs: 1\nExp number of AAs in TMHs: 24.51309\nExp number, first 60 AAs: 0.2682\nTotal prob of N-in: 0.14402\noutside 1 331\nTMhelix 332 354\ninside 355 372

20186 GCF_001357095.1_10592_2_62 Salmonella enterica subsp. enterica serovar Typhi Proteobacteria; Gammaproteobacteria; Enterobacterales; Enterobacteriaceae; Salmonella; Salmonella enterica; Salmonella enterica subsp. enterica MTGDNTLITSHGINRRDFMKLCAALAATMGLSSKAAAEMAESVSNPQRPPVIWIGAQECT WP_000145429.1 hydrogenase 2 small subunit [Salmonella enterica] Length: 372\nNumber of predicted TMHs: 1\nExp number of AAs in TMHs: 24.51309\nExp number, first 60 AAs: 0.2682\nTotal prob of N-in: 0.14402\noutside 1 331\nTMhelix 332 354\ninside 355 372

20187 GCF_001357135.1_10593_2_5 Salmonella enterica subsp. enterica serovar Typhi Proteobacteria; Gammaproteobacteria; Enterobacterales; Enterobacteriaceae; Salmonella; Salmonella enterica; Salmonella enterica subsp. enterica MTGDNTLITSHGINRRDFMKLCAALAATMGLSSKAAAEMAESVSNPQRPPVIWIGAQECT WP_000145429.1 hydrogenase 2 small subunit [Salmonella enterica] Length: 372\nNumber of predicted TMHs: 1\nExp number of AAs in TMHs: 24.51309\nExp number, first 60 AAs: 0.2682\nTotal prob of N-in: 0.14402\noutside 1 331\nTMhelix 332 354\ninside 355 372

20188 GCF_001357255.1_10593_2_81 Salmonella enterica subsp. enterica serovar Typhi Proteobacteria; Gammaproteobacteria; Enterobacterales; Enterobacteriaceae; Salmonella; Salmonella enterica; Salmonella enterica subsp. enterica MTGDNTLITSHGINRRDFMKLCAALAATMGLSSKAAAEMAESVSNPQRPPVIWIGAQECT WP_000145429.1 hydrogenase 2 small subunit [Salmonella enterica] Length: 372\nNumber of predicted TMHs: 1\nExp number of AAs in TMHs: 24.51309\nExp number, first 60 AAs: 0.2682\nTotal prob of N-in: 0.14402\noutside 1 331\nTMhelix 332 354\ninside 355 372

20189 GCF_001357315.1_10593_2_13 Salmonella enterica subsp. enterica serovar Typhi Proteobacteria; Gammaproteobacteria; Enterobacterales; Enterobacteriaceae; Salmonella; Salmonella enterica; Salmonella enterica subsp. enterica MTGDNTLITSHGINRRDFMKLCAALAATMGLSSKAAAEMAESVSNPQRPPVIWIGAQECT WP_000145429.1 hydrogenase 2 small subunit [Salmonella enterica] Length: 372\nNumber of predicted TMHs: 1\nExp number of AAs in TMHs: 24.51309\nExp number, first 60 AAs: 0.2682\nTotal prob of N-in: 0.14402\noutside 1 331\nTMhelix 332 354\ninside 355 372

20190 GCF_001357355.1_10593_2_39 Salmonella enterica subsp. enterica serovar Typhi Proteobacteria; Gammaproteobacteria; Enterobacterales; Enterobacteriaceae; Salmonella; Salmonella enterica; Salmonella enterica subsp. enterica MTGDNTLITSHGINRRDFMKLCAALAATMGLSSKAAAEMAESVSNPQRPPVIWIGAQECT WP_000145429.1 hydrogenase 2 small subunit [Salmonella enterica] Length: 372\nNumber of

predicted TMHs:  1\nExp number of AAs in TMHs: 24.51309\nExp number, first 60 AAs:  0.2682\nTotal prob of N-in: 0.14402\noutside   1   331\nTMhelix   332   354\ninside   355   372

20191 GCF_001357635.1_10593_2_64 Salmonella enterica subsp. enterica serovar Typhi Proteobacteria; Gammaproteobacteria; Enterobacterales; Enterobacteriaceae; Salmonella; Salmonella enterica; Salmonella enterica subsp. enterica MTGDNTLITSHGINRRDFMKLCAALAATMGLSSKAAAEMAESVSNPQRPPVIWIGAQECT WP_000145429.1 hydrogenase 2 small subunit [Salmonella enterica] Length: 372\nNumber of predicted TMHs:  1\nExp number of AAs in TMHs: 24.51309\nExp number, first 60 AAs:  0.2682\nTotal prob of N-in: 0.14402\noutside   1   331\nTMhelix   332   354\ninside   355   372

20192 GCF_001357675.1_10593_2_32 Salmonella enterica subsp. enterica serovar Typhi Proteobacteria; Gammaproteobacteria; Enterobacterales; Enterobacteriaceae; Salmonella; Salmonella enterica; Salmonella enterica subsp. enterica MTGDNTLITSHGINRRDFMKLCAALAATMGLSSKAAAEMAESVSNPQRPPVIWIGAQECT WP_000145429.1 hydrogenase 2 small subunit [Salmonella enterica] Length: 372\nNumber of predicted TMHs:  1\nExp number of AAs in TMHs: 24.51309\nExp number, first 60 AAs:  0.2682\nTotal prob of N-in: 0.14402\noutside   1   331\nTMhelix   332   354\ninside   355   372

20193 GCF_001357715.1_10593_2_23 Salmonella enterica subsp. enterica serovar Typhi Proteobacteria; Gammaproteobacteria; Enterobacterales; Enterobacteriaceae; Salmonella; Salmonella enterica; Salmonella enterica subsp. enterica MTGDNTLITSHGINRRDFMKLCAALAATMGLSSKAAAEMAESVSNPQRPPVIWIGAQECT WP_000145429.1 hydrogenase 2 small subunit [Salmonella enterica] Length: 372\nNumber of predicted TMHs:  1\nExp number of AAs in TMHs: 24.51309\nExp number, first 60 AAs:  0.2682\nTotal prob of N-in: 0.14402\noutside   1   331\nTMhelix   332   354\ninside   355   372

20194 GCF_001357755.1_10593_2_34 Salmonella enterica subsp. enterica serovar Typhi Proteobacteria; Gammaproteobacteria; Enterobacterales; Enterobacteriaceae; Salmonella; Salmonella enterica; Salmonella enterica subsp. enterica MTGDNTLITSHGINRRDFMKLCAALAATMGLSSKAAAEMAESVSNPQRPPVIWIGAQECT WP_000145429.1 hydrogenase 2 small subunit [Salmonella enterica] Length: 372\nNumber of predicted TMHs:  1\nExp number of AAs in TMHs: 24.51309\nExp number, first 60 AAs:  0.2682\nTotal prob of N-in: 0.14402\noutside   1   331\nTMhelix   332   354\ninside   355   372

20195 GCF_001357795.1_10592_2_76 Salmonella enterica subsp. enterica serovar Typhi Proteobacteria; Gammaproteobacteria; Enterobacterales; Enterobacteriaceae; Salmonella; Salmonella enterica; Salmonella enterica subsp. enterica MTGDNTLITSHGINRRDFMKLCAALAATMGLSSKAAAEMAESVSNPQRPPVIWIGAQECT WP_000145429.1 hydrogenase 2 small subunit [Salmonella enterica] Length: 372\nNumber of predicted TMHs:  1\nExp number of AAs in TMHs: 24.51309\nExp number, first 60 AAs:  0.2682\nTotal prob of N-in: 0.14402\noutside   1   331\nTMhelix   332   354\ninside   355   372

20196 GCF_001357835.1_10593_2_47 Salmonella enterica subsp. enterica serovar Typhi Proteobacteria; Gammaproteobacteria; Enterobacterales; Enterobacteriaceae; Salmonella; Salmonella enterica; Salmonella enterica subsp. enterica MTGDNTLITSHGINRRDFMKLCAALAATMGLSSKAAAEMAESVSNPQRPPVIWIGAQECT WP_000145429.1 hydrogenase 2 small subunit [Salmonella enterica] Length: 372\nNumber of predicted TMHs:  1\nExp number of AAs in TMHs: 24.51309\nExp number, first 60 AAs:  0.2682\nTotal prob of N-in: 0.14402\noutside   1   331\nTMhelix   332   354\ninside   355   372

20197 GCF_001357875.1_10593_2_50 Salmonella enterica subsp. enterica serovar Typhi Proteobacteria; Gammaproteobacteria; Enterobacterales; Enterobacteriaceae; Salmonella; Salmonella enterica; Salmonella enterica subsp. enterica MTGDNTLITSHGINRRDFMKLCAALAATMGLSSKAAAEMAESVSNPQRPPVIWIGAQECT WP_000145429.1 hydrogenase 2 small subunit [Salmonella enterica] Length: 372\nNumber of predicted TMHs:  1\nExp number of AAs in TMHs: 24.51309\nExp number, first 60 AAs:  0.2682\nTotal prob of N-in: 0.14402\noutside   1   331\nTMhelix   332   354\ninside   355   372

20198 GCF_001358415.1_10060_6_78 Salmonella enterica subsp. enterica serovar Typhi Proteobacteria; Gammaproteobacteria; Enterobacterales; Enterobacteriaceae; Salmonella; Salmonella enterica; Salmonella enterica subsp. enterica MTGDNTLITSHGINRRDFMKLCAALAATMGLSSKAAAEMAESVSNPQRPPVIWIGAQECT WP_000145429.1 hydrogenase 2 small subunit [Salmonella enterica] Length: 372\nNumber of predicted TMHs:  1\nExp number of AAs in TMHs: 24.51309\nExp number, first 60 AAs:  0.2682\nTotal prob of N-in: 0.14402\noutside   1   331\nTMhelix   332   354\ninside   355   372

20199 GCF_001358495.1_10540_1_9 Salmonella enterica subsp. enterica serovar Typhi Proteobacteria; Gammaproteobacteria; Enterobacterales; Enterobacteriaceae; Salmonella; Salmonella enterica;

Salmonella enterica subsp. enterica MTGDNTLITSHGINRRDFMKLCAALAATMGLSSKAAAEMAESVSNPQRPPVIWIGAQECT WP_000145429.1 hydrogenase 2 small subunit [Salmonella enterica] Length: 372\nNumber of predicted TMHs: 1\nExp number of AAs in TMHs: 24.51309\nExp number, first 60 AAs: 0.2682\nTotal prob of N-in: 0.14402\noutside 1 331\nTMhelix 332 354\ninside 355 372

20200 GCF_001358535.1_10060_6_86 Salmonella enterica subsp. enterica serovar Typhi Proteobacteria; Gammaproteobacteria; Enterobacterales; Enterobacteriaceae; Salmonella; Salmonella enterica; Salmonella enterica subsp. enterica MTGDNTLITSHGINRRDFMKLCAALAATMGLSSKAAAEMAESVSNPQRPPVIWIGAQECT WP_000145429.1 hydrogenase 2 small subunit [Salmonella enterica] Length: 372\nNumber of predicted TMHs: 1\nExp number of AAs in TMHs: 24.51309\nExp number, first 60 AAs: 0.2682\nTotal prob of N-in: 0.14402\noutside 1 331\nTMhelix 332 354\ninside 355 372

20201 GCF_001358615.1_10426_1_9 Salmonella enterica subsp. enterica serovar Typhi Proteobacteria; Gammaproteobacteria; Enterobacterales; Enterobacteriaceae; Salmonella; Salmonella enterica; Salmonella enterica subsp. enterica MTGDNTLITSHGINRRDFMKLCAALAATMGLSSKAAAEMAESVSNPQRPPVIWIGAQECT WP_000145429.1 hydrogenase 2 small subunit [Salmonella enterica] Length: 372\nNumber of predicted TMHs: 1\nExp number of AAs in TMHs: 24.51309\nExp number, first 60 AAs: 0.2682\nTotal prob of N-in: 0.14402\noutside 1 331\nTMhelix 332 354\ninside 355 372

20202 GCF_001358575.1_10607_2_1 Salmonella enterica subsp. enterica serovar Typhi Proteobacteria; Gammaproteobacteria; Enterobacterales; Enterobacteriaceae; Salmonella; Salmonella enterica; Salmonella enterica subsp. enterica MTGDNTLITSHGINRRDFMKLCAALAATMGLSSKAAAEMAESVSNPQRPPVIWIGAQECT WP_000145429.1 hydrogenase 2 small subunit [Salmonella enterica] Length: 372\nNumber of predicted TMHs: 1\nExp number of AAs in TMHs: 24.51309\nExp number, first 60 AAs: 0.2682\nTotal prob of N-in: 0.14402\noutside 1 331\nTMhelix 332 354\ninside 355 372

20203 GCF_001358655.1_10426_1_3 Salmonella enterica subsp. enterica serovar Typhi Proteobacteria; Gammaproteobacteria; Enterobacterales; Enterobacteriaceae; Salmonella; Salmonella enterica; Salmonella enterica subsp. enterica MTGDNTLITSHGINRRDFMKLCAALAATMGLSSKAAAEMAESVSNPQRPPVIWIGAQECT WP_000145429.1 hydrogenase 2 small subunit [Salmonella enterica] Length: 372\nNumber of predicted TMHs: 1\nExp number of AAs in TMHs: 24.51309\nExp number, first 60 AAs: 0.2682\nTotal prob of N-in: 0.14402\noutside 1 331\nTMhelix 332 354\ninside 355 372

20204 GCF_001359575.1_9870_8_23 Salmonella enterica subsp. enterica serovar Typhi Proteobacteria; Gammaproteobacteria; Enterobacterales; Enterobacteriaceae; Salmonella; Salmonella enterica; Salmonella enterica subsp. enterica MTGDNTLITSHGINRRDFMKLCAALAATMGLSSKAAAEMAESVSNPQRPPVIWIGAQECT WP_000145429.1 hydrogenase 2 small subunit [Salmonella enterica] Length: 372\nNumber of predicted TMHs: 1\nExp number of AAs in TMHs: 24.51309\nExp number, first 60 AAs: 0.2682\nTotal prob of N-in: 0.14402\noutside 1 331\nTMhelix 332 354\ninside 355 372

20205 GCF_001359615.1_9953_5_60 Salmonella enterica subsp. enterica serovar Typhi Proteobacteria; Gammaproteobacteria; Enterobacterales; Enterobacteriaceae; Salmonella; Salmonella enterica; Salmonella enterica subsp. enterica MTGDNTLITSHGINRRDFMKLCAALAATMGLSSKAAAEMAESVSNPQRPPVIWIGAQECT WP_000145429.1 hydrogenase 2 small subunit [Salmonella enterica] Length: 372\nNumber of predicted TMHs: 1\nExp number of AAs in TMHs: 24.51309\nExp number, first 60 AAs: 0.2682\nTotal prob of N-in: 0.14402\noutside 1 331\nTMhelix 332 354\ninside 355 372

20206 GCF_001359655.1_10060_6_3 Salmonella enterica subsp. enterica serovar Typhi Proteobacteria; Gammaproteobacteria; Enterobacterales; Enterobacteriaceae; Salmonella; Salmonella enterica; Salmonella enterica subsp. enterica MTGDNTLITSHGINRRDFMKLCAALAATMGLSSKAAAEMAESVSNPQRPPVIWIGAQECT WP_000145429.1 hydrogenase 2 small subunit [Salmonella enterica] Length: 372\nNumber of predicted TMHs: 1\nExp number of AAs in TMHs: 24.51309\nExp number, first 60 AAs: 0.2682\nTotal prob of N-in: 0.14402\noutside 1 331\nTMhelix 332 354\ninside 355 372

20207 GCF_001359695.1_10060_6_15 Salmonella enterica subsp. enterica serovar Typhi Proteobacteria; Gammaproteobacteria; Enterobacterales; Enterobacteriaceae; Salmonella; Salmonella enterica; Salmonella enterica subsp. enterica MTGDNTLITSHGINRRDFMKLCAALAATMGLSSKAAAEMAESVSNPQRPPVIWIGAQECT WP_000145429.1 hydrogenase 2 small subunit [Salmonella enterica] Length: 372\nNumber of predicted TMHs: 1\nExp number of AAs in TMHs: 24.51309\nExp number, first 60 AAs: 0.2682\nTotal prob of N-in: 0.14402\noutside 1 331\nTMhelix 332 354\ninside 355 372

20208 GCF_001359455.1_10071_3_15 Salmonella enterica subsp. enterica serovar Typhi Proteobacteria; Gammaproteobacteria; Enterobacterales; Enterobacteriaceae; Salmonella; Salmonella enterica; Salmonella enterica subsp. enterica MTGDNTLITSHGINRRDFMKLCAALAATMGLSSKAATEMAESVSNPQRPPVIWIGAQECT WP_001782304.1 hydrogenase 2 small subunit [Salmonella enterica] Length: 372\nNumber of predicted TMHs: 1\nExp number of AAs in TMHs: 24.41173\nExp number, first 60 AAs: 0.16124\nTotal prob of N-in: 0.13947\noutside 1 331\nTMhelix 332 354\ninside 355 372

20209 GCF_001359495.1_10071_3_54 Salmonella enterica subsp. enterica serovar Typhi Proteobacteria; Gammaproteobacteria; Enterobacterales; Enterobacteriaceae; Salmonella; Salmonella enterica; Salmonella enterica subsp. enterica MTGDNTLITSHGINRRDFMKLCAALAATMGLSSKAATEMAESVSNPQRPPVIWIGAQECT WP_001782304.1 hydrogenase 2 small subunit [Salmonella enterica] Length: 372\nNumber of predicted TMHs: 1\nExp number of AAs in TMHs: 24.41173\nExp number, first 60 AAs: 0.16124\nTotal prob of N-in: 0.13947\noutside 1 331\nTMhelix 332 354\ninside 355 372

20210 GCF_001360515.1_9870_8_13 Salmonella enterica subsp. enterica serovar Typhi Proteobacteria; Gammaproteobacteria; Enterobacterales; Enterobacteriaceae; Salmonella; Salmonella enterica; Salmonella enterica subsp. enterica MTGDNTLITSHGINRRDFMKLCAALAATMGLSSKAAAEMAESVSNPQRPPVIWIGAQECT WP_000145429.1 hydrogenase 2 small subunit [Salmonella enterica] Length: 372\nNumber of predicted TMHs: 1\nExp number of AAs in TMHs: 24.51309\nExp number, first 60 AAs: 0.2682\nTotal prob of N-in: 0.14402\noutside 1 331\nTMhelix 332 354\ninside 355 372

20211 GCF_001360595.1_9870_8_55 Salmonella enterica subsp. enterica serovar Typhi Proteobacteria; Gammaproteobacteria; Enterobacterales; Enterobacteriaceae; Salmonella; Salmonella enterica; Salmonella enterica subsp. enterica MTGDNTLITSHGINRRDFMKLCAALAATMGLSSKAAAEMAESVSNPQRPPVIWIGAQECT WP_000145429.1 hydrogenase 2 small subunit [Salmonella enterica] Length: 372\nNumber of predicted TMHs: 1\nExp number of AAs in TMHs: 24.51309\nExp number, first 60 AAs: 0.2682\nTotal prob of N-in: 0.14402\noutside 1 331\nTMhelix 332 354\ninside 355 372

20212 GCF_001360635.1_9870_8_67 Salmonella enterica subsp. enterica serovar Typhi Proteobacteria; Gammaproteobacteria; Enterobacterales; Enterobacteriaceae; Salmonella; Salmonella enterica; Salmonella enterica subsp. enterica MTGDNTLITSHGINRRDFMKLCAALAATMGLSSKAAAEMAESVSNPQRPPVIWIGAQECT WP_000145429.1 hydrogenase 2 small subunit [Salmonella enterica] Length: 372\nNumber of predicted TMHs: 1\nExp number of AAs in TMHs: 24.51309\nExp number, first 60 AAs: 0.2682\nTotal prob of N-in: 0.14402\noutside 1 331\nTMhelix 332 354\ninside 355 372

20213 GCF_001360715.1_10060_5_12 Salmonella enterica subsp. enterica serovar Typhi Proteobacteria; Gammaproteobacteria; Enterobacterales; Enterobacteriaceae; Salmonella; Salmonella enterica; Salmonella enterica subsp. enterica MTGDNTLITSHGINRRDFMKLCAALAATMGLSSKAAAEMAESVSNPQRPPVIWIGAQECT WP_000145429.1 hydrogenase 2 small subunit [Salmonella enterica] Length: 372\nNumber of predicted TMHs: 1\nExp number of AAs in TMHs: 24.51309\nExp number, first 60 AAs: 0.2682\nTotal prob of N-in: 0.14402\noutside 1 331\nTMhelix 332 354\ninside 355 372

20214 GCF_001360675.1_9870_8_91 Salmonella enterica subsp. enterica serovar Typhi Proteobacteria; Gammaproteobacteria; Enterobacterales; Enterobacteriaceae; Salmonella; Salmonella enterica; Salmonella enterica subsp. enterica MTGDNTLITSHGINRRDFMKLCAALAATMGLSSKAAAEMAESVSNPQRPPVIWIGAQECT WP_000145429.1 hydrogenase 2 small subunit [Salmonella enterica] Length: 372\nNumber of predicted TMHs: 1\nExp number of AAs in TMHs: 24.51309\nExp number, first 60 AAs: 0.2682\nTotal prob of N-in: 0.14402\noutside 1 331\nTMhelix 332 354\ninside 355 372

20215 GCF_001360755.1_10060_5_18 Salmonella enterica subsp. enterica serovar Typhi Proteobacteria; Gammaproteobacteria; Enterobacterales; Enterobacteriaceae; Salmonella; Salmonella enterica; Salmonella enterica subsp. enterica MTGDNTLITSHGINRRDFMKLCAALAATMGLSSKAAAEMAESVSNPQRPPVIWIGAQECT WP_000145429.1 hydrogenase 2 small subunit [Salmonella enterica] Length: 372\nNumber of predicted TMHs: 1\nExp number of AAs in TMHs: 24.51309\nExp number, first 60 AAs: 0.2682\nTotal prob of N-in: 0.14402\noutside 1 331\nTMhelix 332 354\ninside 355 372

20216 GCF_001361135.1_10071_3_43 Salmonella enterica subsp. enterica serovar Typhi Proteobacteria; Gammaproteobacteria; Enterobacterales; Enterobacteriaceae; Salmonella; Salmonella enterica; Salmonella enterica subsp. enterica MTGDNTLITSHGINRRDFMKLCAALAATMGLSSKAAAEMAESVSNPQRPPVIWIGAQECT WP_000145429.1 hydrogenase 2 small subunit [Salmonella enterica] Length: 372\nNumber of

predicted TMHs: 1\nExp number of AAs in TMHs: 24.51309\nExp number, first 60 AAs: 0.2682\nTotal prob of N-in: 0.14402\noutside 1 331\nTMhelix 332 354\ninside 355 372

20217 GCF_001361175.1_9953_5_46 Salmonella enterica subsp. enterica serovar Typhi Proteobacteria; Gammaproteobacteria; Enterobacterales; Enterobacteriaceae; Salmonella; Salmonella enterica; Salmonella enterica subsp. enterica MTGDNTLITSHGINRRDFMKLCAALAATMGLSSKAAAEMAESVSNPQRPPVIWIGAQECT WP_000145429.1 hydrogenase 2 small subunit [Salmonella enterica] Length: 372\nNumber of predicted TMHs: 1\nExp number of AAs in TMHs: 24.51309\nExp number, first 60 AAs: 0.2682\nTotal prob of N-in: 0.14402\noutside 1 331\nTMhelix 332 354\ninside 355 372

20218 GCF_001361015.1_10071_3_7 Salmonella enterica subsp. enterica serovar Typhi Proteobacteria; Gammaproteobacteria; Enterobacterales; Enterobacteriaceae; Salmonella; Salmonella enterica; Salmonella enterica subsp. enterica MTGDNTLITSHGINRRDFMKLCAALAATMGLSSKAATEMAESVSNPQRPPVIWIGAQECT WP_001782304.1 hydrogenase 2 small subunit [Salmonella enterica] Length: 372\nNumber of predicted TMHs: 1\nExp number of AAs in TMHs: 24.41173\nExp number, first 60 AAs: 0.16124\nTotal prob of N-in: 0.13947\noutside 1 331\nTMhelix 332 354\ninside 355 372

20219 GCF_001361215.1_9953_5_58 Salmonella enterica subsp. enterica serovar Typhi Proteobacteria; Gammaproteobacteria; Enterobacterales; Enterobacteriaceae; Salmonella; Salmonella enterica; Salmonella enterica subsp. enterica MTGDNTLITSHGINRRDFMKLCAALAATMGLSSKAAAEMAESVSNPQRPPVIWIGAQECT WP_000145429.1 hydrogenase 2 small subunit [Salmonella enterica] Length: 372\nNumber of predicted TMHs: 1\nExp number of AAs in TMHs: 24.51309\nExp number, first 60 AAs: 0.2682\nTotal prob of N-in: 0.14402\noutside 1 331\nTMhelix 332 354\ninside 355 372

20220 GCF_001361255.1_9953_5_82 Salmonella enterica subsp. enterica serovar Typhi Proteobacteria; Gammaproteobacteria; Enterobacterales; Enterobacteriaceae; Salmonella; Salmonella enterica; Salmonella enterica subsp. enterica MTGDNTLITSHGINRRDFMKLCAALAATMGLSSKAAAEMAESVSNPQRPPVIWIGAQECT WP_000145429.1 hydrogenase 2 small subunit [Salmonella enterica] Length: 372\nNumber of predicted TMHs: 1\nExp number of AAs in TMHs: 24.51309\nExp number, first 60 AAs: 0.2682\nTotal prob of N-in: 0.14402\noutside 1 331\nTMhelix 332 354\ninside 355 372

20221 GCF_001361295.1_10060_6_4 Salmonella enterica subsp. enterica serovar Typhi Proteobacteria; Gammaproteobacteria; Enterobacterales; Enterobacteriaceae; Salmonella; Salmonella enterica; Salmonella enterica subsp. enterica MTGDNTLITSHGINRRDFMKLCAALAATMGLSSKAAAEMAESVSNPQRPPVIWIGAQECT WP_000145429.1 hydrogenase 2 small subunit [Salmonella enterica] Length: 372\nNumber of predicted TMHs: 1\nExp number of AAs in TMHs: 24.51309\nExp number, first 60 AAs: 0.2682\nTotal prob of N-in: 0.14402\noutside 1 331\nTMhelix 332 354\ninside 355 372

20222 GCF_001361335.1_10060_6_10 Salmonella enterica subsp. enterica serovar Typhi Proteobacteria; Gammaproteobacteria; Enterobacterales; Enterobacteriaceae; Salmonella; Salmonella enterica; Salmonella enterica subsp. enterica MTGDNTLITSHGINRRDFMKLCAALAATMGLSSKAAAEMAESVSNPQRPPVIWIGAQECT WP_000145429.1 hydrogenase 2 small subunit [Salmonella enterica] Length: 372\nNumber of predicted TMHs: 1\nExp number of AAs in TMHs: 24.51309\nExp number, first 60 AAs: 0.2682\nTotal prob of N-in: 0.14402\noutside 1 331\nTMhelix 332 354\ninside 355 372

20223 GCF_001361875.1_10071_8_38 Salmonella enterica subsp. enterica serovar Typhi Proteobacteria; Gammaproteobacteria; Enterobacterales; Enterobacteriaceae; Salmonella; Salmonella enterica; Salmonella enterica subsp. enterica MTGDNTLITSHGINRRDFMKLCAALAATMGLSSKAAAEMAESVSNPQRPPVIWIGAQECT WP_000145429.1 hydrogenase 2 small subunit [Salmonella enterica] Length: 372\nNumber of predicted TMHs: 1\nExp number of AAs in TMHs: 24.51309\nExp number, first 60 AAs: 0.2682\nTotal prob of N-in: 0.14402\noutside 1 331\nTMhelix 332 354\ninside 355 372

20224 GCF_001361915.1_10071_8_56 Salmonella enterica subsp. enterica serovar Typhi Proteobacteria; Gammaproteobacteria; Enterobacterales; Enterobacteriaceae; Salmonella; Salmonella enterica; Salmonella enterica subsp. enterica MTGDNTLITSHGINRRDFMKLCAALAATMGLSSKAAAEMAESVSNPQRPPVIWIGAQECT WP_000145429.1 hydrogenase 2 small subunit [Salmonella enterica] Length: 372\nNumber of predicted TMHs: 1\nExp number of AAs in TMHs: 24.51309\nExp number, first 60 AAs: 0.2682\nTotal prob of N-in: 0.14402\noutside 1 331\nTMhelix 332 354\ninside 355 372

20225 GCF_001361955.1_10209_5_18 Salmonella enterica subsp. enterica serovar Typhi Proteobacteria; Gammaproteobacteria; Enterobacterales; Enterobacteriaceae; Salmonella; Salmonella enterica;

Salmonella enterica subsp. enterica MTGDNTLITSHGINRRDFMKLCAALAATMGLSSKAAAEMAESVSNPQRPPVIWIGAQECT WP_000145429.1 hydrogenase 2 small subunit [Salmonella enterica] Length: 372\nNumber of predicted TMHs: 1\nExp number of AAs in TMHs: 24.51309\nExp number, first 60 AAs: 0.2682\nTotal prob of N-in: 0.14402\noutside 1 331\nTMhelix 332 354\ninside 355 372

20226 GCF_001361995.1_10209_5_27 Salmonella enterica subsp. enterica serovar Typhi Proteobacteria; Gammaproteobacteria; Enterobacterales; Enterobacteriaceae; Salmonella; Salmonella enterica; Salmonella enterica subsp. enterica MTGDNTLITSHGINRRDFMKLCAALAATMGLSSKAAAEMAESVSNPQRPPVIWIGAQECT WP_000145429.1 hydrogenase 2 small subunit [Salmonella enterica] Length: 372\nNumber of predicted TMHs: 1\nExp number of AAs in TMHs: 24.51309\nExp number, first 60 AAs: 0.2682\nTotal prob of N-in: 0.14402\noutside 1 331\nTMhelix 332 354\ninside 355 372

20227 GCF_001362035.1_10209_5_54 Salmonella enterica subsp. enterica serovar Typhi Proteobacteria; Gammaproteobacteria; Enterobacterales; Enterobacteriaceae; Salmonella; Salmonella enterica; Salmonella enterica subsp. enterica MTGDNTLITSHGINRRDFMKLCAALAATMGLSSKAAAEMAESVSNPQRPPVIWIGAQECT WP_000145429.1 hydrogenase 2 small subunit [Salmonella enterica] Length: 372\nNumber of predicted TMHs: 1\nExp number of AAs in TMHs: 24.51309\nExp number, first 60 AAs: 0.2682\nTotal prob of N-in: 0.14402\noutside 1 331\nTMhelix 332 354\ninside 355 372

20228 GCF_001362075.1_10349_1_59 Salmonella enterica subsp. enterica serovar Typhi Proteobacteria; Gammaproteobacteria; Enterobacterales; Enterobacteriaceae; Salmonella; Salmonella enterica; Salmonella enterica subsp. enterica MTGDNTLITSHGINRRDFMKLCAALAATMGLSSKAAAEMAESVSNPQRPPVIWIGAQECT WP_000145429.1 hydrogenase 2 small subunit [Salmonella enterica] Length: 372\nNumber of predicted TMHs: 1\nExp number of AAs in TMHs: 24.51309\nExp number, first 60 AAs: 0.2682\nTotal prob of N-in: 0.14402\noutside 1 331\nTMhelix 332 354\ninside 355 372

20229 GCF_001362115.1_10349_1_71 Salmonella enterica subsp. enterica serovar Typhi Proteobacteria; Gammaproteobacteria; Enterobacterales; Enterobacteriaceae; Salmonella; Salmonella enterica; Salmonella enterica subsp. enterica MTGDNTLITSHGINRRDFMKLCAALAATMGLSSKAAAEMAESVSNPQRPPVIWIGAQECT WP_000145429.1 hydrogenase 2 small subunit [Salmonella enterica] Length: 372\nNumber of predicted TMHs: 1\nExp number of AAs in TMHs: 24.51309\nExp number, first 60 AAs: 0.2682\nTotal prob of N-in: 0.14402\noutside 1 331\nTMhelix 332 354\ninside 355 372

20230 GCF_001362695.1_10540_1_22 Salmonella enterica subsp. enterica serovar Typhi Proteobacteria; Gammaproteobacteria; Enterobacterales; Enterobacteriaceae; Salmonella; Salmonella enterica; Salmonella enterica subsp. enterica MTGDNTLITSHGINRRDFMKLCAALAATMGLSSKAAAEMAESVSNPQRPPVIWIGAQECT WP_000145429.1 hydrogenase 2 small subunit [Salmonella enterica] Length: 372\nNumber of predicted TMHs: 1\nExp number of AAs in TMHs: 24.51309\nExp number, first 60 AAs: 0.2682\nTotal prob of N-in: 0.14402\noutside 1 331\nTMhelix 332 354\ninside 355 372

20231 GCF_001362735.1_10562_2_79 Salmonella enterica subsp. enterica serovar Typhi Proteobacteria; Gammaproteobacteria; Enterobacterales; Enterobacteriaceae; Salmonella; Salmonella enterica; Salmonella enterica subsp. enterica MTGDNTLITSHGINRRDFMKLCAALAATMGLSSKAAAEMAESVSNPQRPPVIWIGAQECT WP_000145429.1 hydrogenase 2 small subunit [Salmonella enterica] Length: 372\nNumber of predicted TMHs: 1\nExp number of AAs in TMHs: 24.51309\nExp number, first 60 AAs: 0.2682\nTotal prob of N-in: 0.14402\noutside 1 331\nTMhelix 332 354\ninside 355 372

20232 GCF_001362775.1_10426_1_40 Salmonella enterica subsp. enterica serovar Typhi Proteobacteria; Gammaproteobacteria; Enterobacterales; Enterobacteriaceae; Salmonella; Salmonella enterica; Salmonella enterica subsp. enterica MTGDNTLITSHGINRRDFMKLCAALAATMGLSSKAAAEMAESVSNPQRPPVIWIGAQECT WP_000145429.1 hydrogenase 2 small subunit [Salmonella enterica] Length: 372\nNumber of predicted TMHs: 1\nExp number of AAs in TMHs: 24.51309\nExp number, first 60 AAs: 0.2682\nTotal prob of N-in: 0.14402\noutside 1 331\nTMhelix 332 354\ninside 355 372

20233 GCF_001362815.1_10492_1_69 Salmonella enterica subsp. enterica serovar Typhi Proteobacteria; Gammaproteobacteria; Enterobacterales; Enterobacteriaceae; Salmonella; Salmonella enterica; Salmonella enterica subsp. enterica MTGDNTLITSHGINRRDFMKLCAALAATMGLSSKAAAEMAESVSNPQRPPVIWIGAQECT WP_000145429.1 hydrogenase 2 small subunit [Salmonella enterica] Length: 372\nNumber of predicted TMHs: 1\nExp number of AAs in TMHs: 24.51309\nExp number, first 60 AAs: 0.2682\nTotal prob of N-in: 0.14402\noutside 1 331\nTMhelix 332 354\ninside 355 372

20234 GCF_001362855.1_10426_1_49 Salmonella enterica subsp. enterica serovar Typhi Proteobacteria; Gammaproteobacteria; Enterobacterales; Enterobacteriaceae; Salmonella; Salmonella enterica; Salmonella enterica subsp. enterica MTGDNTLITSHGINRRDFMKLCAALAATMGLSSKAAAEMAESVSNPQRPPVIWIGAQECT WP_000145429.1 hydrogenase 2 small subunit [Salmonella enterica] Length: 372\nNumber of predicted TMHs: 1\nExp number of AAs in TMHs: 24.51309\nExp number, first 60 AAs: 0.2682\nTotal prob of N-in: 0.14402\noutside 1 331\nTMhelix 332 354\ninside 355 372

20235 GCF_001362915.1_10561_2_71 Salmonella enterica subsp. enterica serovar Typhi Proteobacteria; Gammaproteobacteria; Enterobacterales; Enterobacteriaceae; Salmonella; Salmonella enterica; Salmonella enterica subsp. enterica MTGDNTLITSHGINRRDFMKLCAALAATMGLSSKAAAEMAESVSNPQRPPVIWIGAQECT WP_000145429.1 hydrogenase 2 small subunit [Salmonella enterica] Length: 372\nNumber of predicted TMHs: 1\nExp number of AAs in TMHs: 24.51309\nExp number, first 60 AAs: 0.2682\nTotal prob of N-in: 0.14402\noutside 1 331\nTMhelix 332 354\ninside 355 372

20236 GCF_001363475.1_10608_2_63 Salmonella enterica subsp. enterica serovar Typhi Proteobacteria; Gammaproteobacteria; Enterobacterales; Enterobacteriaceae; Salmonella; Salmonella enterica; Salmonella enterica subsp. enterica MTGDNTLITSHGINRRDFMKLCAALAATMGLSSKAAAEMAESVSNPQRPPVIWIGAQECT WP_000145429.1 hydrogenase 2 small subunit [Salmonella enterica] Length: 372\nNumber of predicted TMHs: 1\nExp number of AAs in TMHs: 24.51309\nExp number, first 60 AAs: 0.2682\nTotal prob of N-in: 0.14402\noutside 1 331\nTMhelix 332 354\ninside 355 372

20237 GCF_001363555.1_10608_2_11 Salmonella enterica subsp. enterica serovar Typhi Proteobacteria; Gammaproteobacteria; Enterobacterales; Enterobacteriaceae; Salmonella; Salmonella enterica; Salmonella enterica subsp. enterica MTGDNTLITSHGINRRDFMKLCAALAATMGLSSKAAAEMAESVSNPQRPPVIWIGAQECT WP_000145429.1 hydrogenase 2 small subunit [Salmonella enterica] Length: 372\nNumber of predicted TMHs: 1\nExp number of AAs in TMHs: 24.51309\nExp number, first 60 AAs: 0.2682\nTotal prob of N-in: 0.14402\noutside 1 331\nTMhelix 332 354\ninside 355 372

20238 GCF_001363515.1_10493_1_16 Salmonella enterica subsp. enterica serovar Typhi Proteobacteria; Gammaproteobacteria; Enterobacterales; Enterobacteriaceae; Salmonella; Salmonella enterica; Salmonella enterica subsp. enterica MTGDNTLITSHGINRRDFMKLCAALAATMGLSSKAAAEMAESVSNPQRPPVIWIGAQECT WP_000145429.1 hydrogenase 2 small subunit [Salmonella enterica] Length: 372\nNumber of predicted TMHs: 1\nExp number of AAs in TMHs: 24.51309\nExp number, first 60 AAs: 0.2682\nTotal prob of N-in: 0.14402\noutside 1 331\nTMhelix 332 354\ninside 355 372

20239 GCF_001363355.1_10562_2_15 Salmonella enterica subsp. enterica serovar Typhi Proteobacteria; Gammaproteobacteria; Enterobacterales; Enterobacteriaceae; Salmonella; Salmonella enterica; Salmonella enterica subsp. enterica MTGDNTLITSHGINRRDFMKLCAALAATMGLSSKAATEMAESVSNPQRPPVIWIGAQECT WP_001782304.1 hydrogenase 2 small subunit [Salmonella enterica] Length: 372\nNumber of predicted TMHs: 1\nExp number of AAs in TMHs: 24.41173\nExp number, first 60 AAs: 0.16124\nTotal prob of N-in: 0.13947\noutside 1 331\nTMhelix 332 354\ninside 355 372

20240 GCF_001363615.1_10540_1_60 Salmonella enterica subsp. enterica serovar Typhi Proteobacteria; Gammaproteobacteria; Enterobacterales; Enterobacteriaceae; Salmonella; Salmonella enterica; Salmonella enterica subsp. enterica MTGDNTLITSHGINRRDFMKLCAALAATMGLSSKAAAEMAESVSNPQRPPVIWIGAQECT WP_000145429.1 hydrogenase 2 small subunit [Salmonella enterica] Length: 372\nNumber of predicted TMHs: 1\nExp number of AAs in TMHs: 24.51309\nExp number, first 60 AAs: 0.2682\nTotal prob of N-in: 0.14402\noutside 1 331\nTMhelix 332 354\ninside 355 372

20241 GCF_001363655.1_10426_1_19 Salmonella enterica subsp. enterica serovar Typhi Proteobacteria; Gammaproteobacteria; Enterobacterales; Enterobacteriaceae; Salmonella; Salmonella enterica; Salmonella enterica subsp. enterica MTGDNTLITSHGINRRDFMKLCAALAATMGLSSKAAAEMAESVSNPQRPPVIWIGAQECT WP_000145429.1 hydrogenase 2 small subunit [Salmonella enterica] Length: 372\nNumber of predicted TMHs: 1\nExp number of AAs in TMHs: 24.51309\nExp number, first 60 AAs: 0.2682\nTotal prob of N-in: 0.14402\noutside 1 331\nTMhelix 332 354\ninside 355 372

20242 GCF_001364255.1_10607_1_51 Salmonella enterica subsp. enterica serovar Typhi Proteobacteria; Gammaproteobacteria; Enterobacterales; Enterobacteriaceae; Salmonella; Salmonella enterica; Salmonella enterica subsp. enterica MTGDNTLITSHGINRRDFMKLCAALAATMGLSSKAAAEMAESVSNPQRPPVIWIGAQECT WP_000145429.1 hydrogenase 2 small subunit [Salmonella enterica] Length: 372\nNumber of

predicted TMHs:  1\nExp number of AAs in TMHs: 24.51309\nExp number, first 60 AAs:  0.2682\nTotal prob of N-in: 0.14402\noutside   1   331\nTMhelix   332   354\ninside   355   372

20243 GCF_001364315.1_10562_2_64 Salmonella enterica subsp. enterica serovar Typhi Proteobacteria; Gammaproteobacteria; Enterobacterales; Enterobacteriaceae; Salmonella; Salmonella enterica; Salmonella enterica subsp. enterica MTGDNTLITSHGINRRDFMKLCAALAATMGLSSKAAAEMAESVSNPQRPPVIWIGAQECT WP_000145429.1 hydrogenase 2 small subunit [Salmonella enterica] Length: 372\nNumber of predicted TMHs:  1\nExp number of AAs in TMHs: 24.51309\nExp number, first 60 AAs:  0.2682\nTotal prob of N-in: 0.14402\noutside   1   331\nTMhelix   332   354\ninside   355   372

20244 GCF_001364355.1_10561_2_72 Salmonella enterica subsp. enterica serovar Typhi Proteobacteria; Gammaproteobacteria; Enterobacterales; Enterobacteriaceae; Salmonella; Salmonella enterica; Salmonella enterica subsp. enterica MTGDNTLITSHGINRRDFMKLCAALAATMGLSSKAAAEMAESVSNPQRPPVIWIGAQECT WP_000145429.1 hydrogenase 2 small subunit [Salmonella enterica] Length: 372\nNumber of predicted TMHs:  1\nExp number of AAs in TMHs: 24.51309\nExp number, first 60 AAs:  0.2682\nTotal prob of N-in: 0.14402\noutside   1   331\nTMhelix   332   354\ninside   355   372

20245 GCF_001364395.1_10561_2_49 Salmonella enterica subsp. enterica serovar Typhi Proteobacteria; Gammaproteobacteria; Enterobacterales; Enterobacteriaceae; Salmonella; Salmonella enterica; Salmonella enterica subsp. enterica MTGDNTLITSHGINRRDFMKLCAALAATMGLSSKAAAEMAESVSNPQRPPVIWIGAQECT WP_000145429.1 hydrogenase 2 small subunit [Salmonella enterica] Length: 372\nNumber of predicted TMHs:  1\nExp number of AAs in TMHs: 24.51309\nExp number, first 60 AAs:  0.2682\nTotal prob of N-in: 0.14402\noutside   1   331\nTMhelix   332   354\ninside   355   372

20246 GCF_001364435.1_10493_1_35 Salmonella enterica subsp. enterica serovar Typhi Proteobacteria; Gammaproteobacteria; Enterobacterales; Enterobacteriaceae; Salmonella; Salmonella enterica; Salmonella enterica subsp. enterica MTGDNTLITSHGINRRDFMKLCAALAATMGLSSKAAAEMAESVSNPQRPPVIWIGAQECT WP_000145429.1 hydrogenase 2 small subunit [Salmonella enterica] Length: 372\nNumber of predicted TMHs:  1\nExp number of AAs in TMHs: 24.51309\nExp number, first 60 AAs:  0.2682\nTotal prob of N-in: 0.14402\noutside   1   331\nTMhelix   332   354\ninside   355   372

20247 GCF_001364755.1_10540_1_93 Salmonella enterica subsp. enterica serovar Typhi Proteobacteria; Gammaproteobacteria; Enterobacterales; Enterobacteriaceae; Salmonella; Salmonella enterica; Salmonella enterica subsp. enterica MTGDNTLITSHGINRRDFMKLCAALAATMGLSSKAAAEMAESVSNPQRPPVIWIGAQECT WP_000145429.1 hydrogenase 2 small subunit [Salmonella enterica] Length: 372\nNumber of predicted TMHs:  1\nExp number of AAs in TMHs: 24.51309\nExp number, first 60 AAs:  0.2682\nTotal prob of N-in: 0.14402\noutside   1   331\nTMhelix   332   354\ninside   355   372

20248 GCF_001364795.1_10607_1_86 Salmonella enterica subsp. enterica serovar Typhi Proteobacteria; Gammaproteobacteria; Enterobacterales; Enterobacteriaceae; Salmonella; Salmonella enterica; Salmonella enterica subsp. enterica MTGDNTLITSHGINRRDFMKLCAALAATMGLSSKAAAEMAESVSNPQRPPVIWIGAQECT WP_000145429.1 hydrogenase 2 small subunit [Salmonella enterica] Length: 372\nNumber of predicted TMHs:  1\nExp number of AAs in TMHs: 24.51309\nExp number, first 60 AAs:  0.2682\nTotal prob of N-in: 0.14402\noutside   1   331\nTMhelix   332   354\ninside   355   372

20249 GCF_001364835.1_10562_2_12 Salmonella enterica subsp. enterica serovar Typhi Proteobacteria; Gammaproteobacteria; Enterobacterales; Enterobacteriaceae; Salmonella; Salmonella enterica; Salmonella enterica subsp. enterica MTGDNTLITSHGINRRDFMKLCAALAATMGLSSKAAAEMAESVSNPQRPPVIWIGAQECT WP_000145429.1 hydrogenase 2 small subunit [Salmonella enterica] Length: 372\nNumber of predicted TMHs:  1\nExp number of AAs in TMHs: 24.51309\nExp number, first 60 AAs:  0.2682\nTotal prob of N-in: 0.14402\noutside   1   331\nTMhelix   332   354\ninside   355   372

20250 GCF_001364875.1_10561_2_92 Salmonella enterica subsp. enterica serovar Typhi Proteobacteria; Gammaproteobacteria; Enterobacterales; Enterobacteriaceae; Salmonella; Salmonella enterica; Salmonella enterica subsp. enterica MTGDNTLITSHGINRRDFMKLCAALAATMGLSSKAAAEMAESVSNPQRPPVIWIGAQECT WP_000145429.1 hydrogenase 2 small subunit [Salmonella enterica] Length: 372\nNumber of predicted TMHs:  1\nExp number of AAs in TMHs: 24.51309\nExp number, first 60 AAs:  0.2682\nTotal prob of N-in: 0.14402\noutside   1   331\nTMhelix   332   354\ninside   355   372

20251 GCF_001364915.1_10426_1_51 Salmonella enterica subsp. enterica serovar Typhi Proteobacteria; Gammaproteobacteria; Enterobacterales; Enterobacteriaceae; Salmonella; Salmonella enterica;

Salmonella enterica subsp. enterica MTGDNTLITSHGINRRDFMKLCAALAATMGLSSKAAAEMAESVSNPQRPPVIWIGAQECT WP_000145429.1 hydrogenase 2 small subunit [Salmonella enterica] Length: 372\nNumber of predicted TMHs: 1\nExp number of AAs in TMHs: 24.51309\nExp number, first 60 AAs: 0.2682\nTotal prob of N-in: 0.14402\noutside 1 331\nTMhelix 332 354\ninside 355 372

20252 GCF_001364955.1_10426_1_13 Salmonella enterica subsp. enterica serovar Typhi Proteobacteria; Gammaproteobacteria; Enterobacterales; Enterobacteriaceae; Salmonella; Salmonella enterica; Salmonella enterica subsp. enterica MTGDNTLITSHGINRRDFMKLCAALAATMGLSSKAAAEMAESVSNPQRPPVIWIGAQECT WP_000145429.1 hydrogenase 2 small subunit [Salmonella enterica] Length: 372\nNumber of predicted TMHs: 1\nExp number of AAs in TMHs: 24.51309\nExp number, first 60 AAs: 0.2682\nTotal prob of N-in: 0.14402\noutside 1 331\nTMhelix 332 354\ninside 355 372

20253 GCF_001365535.1_10540_1_15 Salmonella enterica subsp. enterica serovar Typhi Proteobacteria; Gammaproteobacteria; Enterobacterales; Enterobacteriaceae; Salmonella; Salmonella enterica; Salmonella enterica subsp. enterica MTGDNTLITSHGINRRDFMKLCAALAATMGLSSKAAAEMAESVSNPQRPPVIWIGAQECT WP_000145429.1 hydrogenase 2 small subunit [Salmonella enterica] Length: 372\nNumber of predicted TMHs: 1\nExp number of AAs in TMHs: 24.51309\nExp number, first 60 AAs: 0.2682\nTotal prob of N-in: 0.14402\noutside 1 331\nTMhelix 332 354\ninside 355 372

20254 GCF_001365615.1_10493_1_45 Salmonella enterica subsp. enterica serovar Typhi Proteobacteria; Gammaproteobacteria; Enterobacterales; Enterobacteriaceae; Salmonella; Salmonella enterica; Salmonella enterica subsp. enterica MTGDNTLITSHGINRRDFMKLCAALAATMGLSSKAAAEMAESVSNPQRPPVIWIGAQECT WP_000145429.1 hydrogenase 2 small subunit [Salmonella enterica] Length: 372\nNumber of predicted TMHs: 1\nExp number of AAs in TMHs: 24.51309\nExp number, first 60 AAs: 0.2682\nTotal prob of N-in: 0.14402\noutside 1 331\nTMhelix 332 354\ninside 355 372

20255 GCF_001365575.1_10426_1_79 Salmonella enterica subsp. enterica serovar Typhi Proteobacteria; Gammaproteobacteria; Enterobacterales; Enterobacteriaceae; Salmonella; Salmonella enterica; Salmonella enterica subsp. enterica MTGDNTLITSHGINRRDFMKLCAALAATMGLSSKAAAEMAESVSNPQRPPVIWIGAQECT WP_000145429.1 hydrogenase 2 small subunit [Salmonella enterica] Length: 372\nNumber of predicted TMHs: 1\nExp number of AAs in TMHs: 24.51309\nExp number, first 60 AAs: 0.2682\nTotal prob of N-in: 0.14402\noutside 1 331\nTMhelix 332 354\ninside 355 372

20256 GCF_001365655.1_10561_2_21 Salmonella enterica subsp. enterica serovar Typhi Proteobacteria; Gammaproteobacteria; Enterobacterales; Enterobacteriaceae; Salmonella; Salmonella enterica; Salmonella enterica subsp. enterica MTGDNTLITSHGINRRDFMKLCAALAATMGLSSKAAAEMAESVSNPQRPPVIWIGAQECT WP_000145429.1 hydrogenase 2 small subunit [Salmonella enterica] Length: 372\nNumber of predicted TMHs: 1\nExp number of AAs in TMHs: 24.51309\nExp number, first 60 AAs: 0.2682\nTotal prob of N-in: 0.14402\noutside 1 331\nTMhelix 332 354\ninside 355 372

20257 GCF_001365995.1_10561_2_80 Salmonella enterica subsp. enterica serovar Typhi Proteobacteria; Gammaproteobacteria; Enterobacterales; Enterobacteriaceae; Salmonella; Salmonella enterica; Salmonella enterica subsp. enterica MTGDNTLITSHGINRRDFMKLCAALAATMGLSSKAAAEMAESVSNPQRPPVIWIGAQECT WP_000145429.1 hydrogenase 2 small subunit [Salmonella enterica] Length: 372\nNumber of predicted TMHs: 1\nExp number of AAs in TMHs: 24.51309\nExp number, first 60 AAs: 0.2682\nTotal prob of N-in: 0.14402\noutside 1 331\nTMhelix 332 354\ninside 355 372

20258 GCF_001365715.1_10561_2_25 Salmonella enterica subsp. enterica serovar Typhi Proteobacteria; Gammaproteobacteria; Enterobacterales; Enterobacteriaceae; Salmonella; Salmonella enterica; Salmonella enterica subsp. enterica MTGDNTLITSHGINRRDFMKLCAALAATMGLSSKAATEMAESVSNPQRPPVIWIGAQECT WP_001782304.1 hydrogenase 2 small subunit [Salmonella enterica] Length: 372\nNumber of predicted TMHs: 1\nExp number of AAs in TMHs: 24.41173\nExp number, first 60 AAs: 0.16124\nTotal prob of N-in: 0.13947\noutside 1 331\nTMhelix 332 354\ninside 355 372

20259 GCF_001566195.1_10426_1_83 Salmonella enterica subsp. enterica serovar Typhi Proteobacteria; Gammaproteobacteria; Enterobacterales; Enterobacteriaceae; Salmonella; Salmonella enterica; Salmonella enterica subsp. enterica MTGDNTLITSHGINRRDFMKLCAALAATMGLSSKAAAEMAESVSNPQRPPVIWIGAQECT WP_000145429.1 hydrogenase 2 small subunit [Salmonella enterica] Length: 372\nNumber of predicted TMHs: 1\nExp number of AAs in TMHs: 24.51309\nExp number, first 60 AAs: 0.2682\nTotal prob of N-in: 0.14402\noutside 1 331\nTMhelix 332 354\ninside 355 372

20260	GCF_900033735.1_14555_2_66	Salmonella enterica subsp. enterica serovar Typhi	Proteobacteria; Gammaproteobacteria; Enterobacterales; Enterobacteriaceae; Salmonella; Salmonella enterica; Salmonella enterica subsp. enterica	MTGDNTLITSHGINRRDFMKLCAALAATMGLSSKAAAEMAESVSNPQRPPVIWIGAQECT	WP_000145429.1	hydrogenase 2 small subunit [Salmonella enterica]	Length: 372\nNumber of predicted TMHs: 1\nExp number of AAs in TMHs: 24.51309\nExp number, first 60 AAs: 0.2682\nTotal prob of N-in: 0.14402\noutside 1 331\nTMhelix 332 354\ninside 355 372

20261	GCF_900035575.1_14555_2_53	Salmonella enterica subsp. enterica serovar Typhi	Proteobacteria; Gammaproteobacteria; Enterobacterales; Enterobacteriaceae; Salmonella; Salmonella enterica; Salmonella enterica subsp. enterica	MTGDNTLITSHGINRRDFMKLCAALAATMGLSSKAAAEMAESVSNPQRPPVIWIGAQECT	WP_000145429.1	hydrogenase 2 small subunit [Salmonella enterica]	Length: 372\nNumber of predicted TMHs: 1\nExp number of AAs in TMHs: 24.51309\nExp number, first 60 AAs: 0.2682\nTotal prob of N-in: 0.14402\noutside 1 331\nTMhelix 332 354\ninside 355 372

20262	GCF_900036295.1_13566_1_3	Salmonella enterica subsp. enterica serovar Typhi	Proteobacteria; Gammaproteobacteria; Enterobacterales; Enterobacteriaceae; Salmonella; Salmonella enterica; Salmonella enterica subsp. enterica	MTGDNTLITSHGINRRDFMKLCAALAATMGLSSKAAAEMAESVSNPQRPPVIWIGAQECT	WP_000145429.1	hydrogenase 2 small subunit [Salmonella enterica]	Length: 372\nNumber of predicted TMHs: 1\nExp number of AAs in TMHs: 24.51309\nExp number, first 60 AAs: 0.2682\nTotal prob of N-in: 0.14402\noutside 1 331\nTMhelix 332 354\ninside 355 372

20263	GCF_900036315.1_13566_1_83	Salmonella enterica subsp. enterica serovar Typhi	Proteobacteria; Gammaproteobacteria; Enterobacterales; Enterobacteriaceae; Salmonella; Salmonella enterica; Salmonella enterica subsp. enterica	MTGDNTLITSHGINRRDFMKLCAALAATMGLSSKAAAEMAESVSNPQRPPVIWIGAQECT	WP_000145429.1	hydrogenase 2 small subunit [Salmonella enterica]	Length: 372\nNumber of predicted TMHs: 1\nExp number of AAs in TMHs: 24.51309\nExp number, first 60 AAs: 0.2682\nTotal prob of N-in: 0.14402\noutside 1 331\nTMhelix 332 354\ninside 355 372

20264	GCF_900047125.1_14555_2_55	Salmonella enterica subsp. enterica serovar Typhi	Proteobacteria; Gammaproteobacteria; Enterobacterales; Enterobacteriaceae; Salmonella; Salmonella enterica; Salmonella enterica subsp. enterica	MTGDNTLITSHGINRRDFMKLCAALAATMGLSSKAAAEMAESVSNPQRPPVIWIGAQECT	WP_000145429.1	hydrogenase 2 small subunit [Salmonella enterica]	Length: 372\nNumber of predicted TMHs: 1\nExp number of AAs in TMHs: 24.51309\nExp number, first 60 AAs: 0.2682\nTotal prob of N-in: 0.14402\noutside 1 331\nTMhelix 332 354\ninside 355 372

20265	GCF_900018645.1_14555_2_62	Salmonella enterica subsp. enterica serovar Typhi	Proteobacteria; Gammaproteobacteria; Enterobacterales; Enterobacteriaceae; Salmonella; Salmonella enterica; Salmonella enterica subsp. enterica	MTGDNTLITSHGINRRDFMKLCAALAATMGLSSKAAAEMAESVSNPQRPPVIWIGAQECT	WP_000145429.1	hydrogenase 2 small subunit [Salmonella enterica]	Length: 372\nNumber of predicted TMHs: 1\nExp number of AAs in TMHs: 24.51309\nExp number, first 60 AAs: 0.2682\nTotal prob of N-in: 0.14402\noutside 1 331\nTMhelix 332 354\ninside 355 372

20266	GCF_900018665.1_14555_2_91	Salmonella enterica subsp. enterica serovar Typhi	Proteobacteria; Gammaproteobacteria; Enterobacterales; Enterobacteriaceae; Salmonella; Salmonella enterica; Salmonella enterica subsp. enterica	MTGDNTLITSHGINRRDFMKLCAALAATMGLSSKAAAEMAESVSNPQRPPVIWIGAQECT	WP_000145429.1	hydrogenase 2 small subunit [Salmonella enterica]	Length: 372\nNumber of predicted TMHs: 1\nExp number of AAs in TMHs: 24.51309\nExp number, first 60 AAs: 0.2682\nTotal prob of N-in: 0.14402\noutside 1 331\nTMhelix 332 354\ninside 355 372

20267	GCF_900022565.1_14555_2_47	Salmonella enterica subsp. enterica serovar Typhi	Proteobacteria; Gammaproteobacteria; Enterobacterales; Enterobacteriaceae; Salmonella; Salmonella enterica; Salmonella enterica subsp. enterica	MTGDNTLITSHGINRRDFMKLCAALAATMGLSSKAAAEMAESVSNPQRPPVIWIGAQECT	WP_000145429.1	hydrogenase 2 small subunit [Salmonella enterica]	Length: 372\nNumber of predicted TMHs: 1\nExp number of AAs in TMHs: 24.51309\nExp number, first 60 AAs: 0.2682\nTotal prob of N-in: 0.14402\noutside 1 331\nTMhelix 332 354\ninside 355 372

20268	GCF_900024945.1_14555_2_48	Salmonella enterica subsp. enterica serovar Typhi	Proteobacteria; Gammaproteobacteria; Enterobacterales; Enterobacteriaceae; Salmonella; Salmonella enterica; Salmonella enterica subsp. enterica	MTGDNTLITSHGINRRDFMKLCAALAATMGLSSKAAAEMAESVSNPQRPPVIWIGAQECT	WP_000145429.1	hydrogenase 2 small subunit [Salmonella enterica]	Length: 372\nNumber of

predicted TMHs:  1\nExp number of AAs in TMHs: 24.51309\nExp number, first 60 AAs:  0.2682\nTotal prob of N-in:  0.14402\noutside   1   331\nTMhelix   332   354\ninside   355   372

20269 GCF_900029285.1_13097_1_74 Salmonella enterica subsp. enterica serovar Typhi Proteobacteria; Gammaproteobacteria; Enterobacterales; Enterobacteriaceae; Salmonella; Salmonella enterica; Salmonella enterica subsp. enterica MTGDNTLITSHGINRRDFMKLCAALAATMGLSSKAAAEMAESVSNPQRPPVIWIGAQECT WP_000145429.1 hydrogenase 2 small subunit [Salmonella enterica] Length: 372\nNumber of predicted TMHs:  1\nExp number of AAs in TMHs: 24.51309\nExp number, first 60 AAs:  0.2682\nTotal prob of N-in:  0.14402\noutside   1   331\nTMhelix   332   354\ninside   355   372

20270 GCF_900029305.1_13103_1_12 Salmonella enterica subsp. enterica serovar Typhi Proteobacteria; Gammaproteobacteria; Enterobacterales; Enterobacteriaceae; Salmonella; Salmonella enterica; Salmonella enterica subsp. enterica MTGDNTLITSHGINRRDFMKLCAALAATMGLSSKAAAEMAESVSNPQRPPVIWIGAQECT WP_000145429.1 hydrogenase 2 small subunit [Salmonella enterica] Length: 372\nNumber of predicted TMHs:  1\nExp number of AAs in TMHs: 24.51309\nExp number, first 60 AAs:  0.2682\nTotal prob of N-in:  0.14402\noutside   1   331\nTMhelix   332   354\ninside   355   372

20271 GCF_900029325.1_13134_1_21 Salmonella enterica subsp. enterica serovar Typhi Proteobacteria; Gammaproteobacteria; Enterobacterales; Enterobacteriaceae; Salmonella; Salmonella enterica; Salmonella enterica subsp. enterica MTGDNTLITSHGINRRDFMKLCAALAATMGLSSKAAAEMAESVSNPQRPPVIWIGAQECT WP_000145429.1 hydrogenase 2 small subunit [Salmonella enterica] Length: 372\nNumber of predicted TMHs:  1\nExp number of AAs in TMHs: 24.51309\nExp number, first 60 AAs:  0.2682\nTotal prob of N-in:  0.14402\noutside   1   331\nTMhelix   332   354\ninside   355   372

20272 GCF_900033715.1_14555_2_32 Salmonella enterica subsp. enterica serovar Typhi Proteobacteria; Gammaproteobacteria; Enterobacterales; Enterobacteriaceae; Salmonella; Salmonella enterica; Salmonella enterica subsp. enterica MTGDNTLITSHGINRRDFMKLCAALAATMGLSSKAAAEMAESVSNPQRPPVIWIGAQECT WP_000145429.1 hydrogenase 2 small subunit [Salmonella enterica] Length: 372\nNumber of predicted TMHs:  1\nExp number of AAs in TMHs: 24.51309\nExp number, first 60 AAs:  0.2682\nTotal prob of N-in:  0.14402\noutside   1   331\nTMhelix   332   354\ninside   355   372

20273 GCF_900036335.1_14555_2_28 Salmonella enterica subsp. enterica serovar Typhi Proteobacteria; Gammaproteobacteria; Enterobacterales; Enterobacteriaceae; Salmonella; Salmonella enterica; Salmonella enterica subsp. enterica MTGDNTLITSHGINRRDFMKLCAALAATMGLSSKAAAEMAESVSNPQRPPVIWIGAQECT WP_000145429.1 hydrogenase 2 small subunit [Salmonella enterica] Length: 372\nNumber of predicted TMHs:  1\nExp number of AAs in TMHs: 24.51309\nExp number, first 60 AAs:  0.2682\nTotal prob of N-in:  0.14402\noutside   1   331\nTMhelix   332   354\ninside   355   372

20274 GCF_900036865.1_14555_2_90 Salmonella enterica subsp. enterica serovar Typhi Proteobacteria; Gammaproteobacteria; Enterobacterales; Enterobacteriaceae; Salmonella; Salmonella enterica; Salmonella enterica subsp. enterica MTGDNTLITSHGINRRDFMKLCAALAATMGLSSKAAAEMAESVSNPQRPPVIWIGAQECT WP_000145429.1 hydrogenase 2 small subunit [Salmonella enterica] Length: 372\nNumber of predicted TMHs:  1\nExp number of AAs in TMHs: 24.51309\nExp number, first 60 AAs:  0.2682\nTotal prob of N-in:  0.14402\noutside   1   331\nTMhelix   332   354\ninside   355   372

20275 GCF_900037575.1_14555_2_15 Salmonella enterica subsp. enterica serovar Typhi Proteobacteria; Gammaproteobacteria; Enterobacterales; Enterobacteriaceae; Salmonella; Salmonella enterica; Salmonella enterica subsp. enterica MTGDNTLITSHGINRRDFMKLCAALAATMGLSSKAAAEMAESVSNPQRPPVIWIGAQECT WP_000145429.1 hydrogenase 2 small subunit [Salmonella enterica] Length: 372\nNumber of predicted TMHs:  1\nExp number of AAs in TMHs: 24.51309\nExp number, first 60 AAs:  0.2682\nTotal prob of N-in:  0.14402\noutside   1   331\nTMhelix   332   354\ninside   355   372

20276 GCF_900038085.1_14555_2_14 Salmonella enterica subsp. enterica serovar Typhi Proteobacteria; Gammaproteobacteria; Enterobacterales; Enterobacteriaceae; Salmonella; Salmonella enterica; Salmonella enterica subsp. enterica MTGDNTLITSHGINRRDFMKLCAALAATMGLSSKAAAEMAESVSNPQRPPVIWIGAQECT WP_000145429.1 hydrogenase 2 small subunit [Salmonella enterica] Length: 372\nNumber of predicted TMHs:  1\nExp number of AAs in TMHs: 24.51309\nExp number, first 60 AAs:  0.2682\nTotal prob of N-in:  0.14402\noutside   1   331\nTMhelix   332   354\ninside   355   372

20277 GCF_900039625.1_13566_1_80 Salmonella enterica subsp. enterica serovar Typhi Proteobacteria; Gammaproteobacteria; Enterobacterales; Enterobacteriaceae; Salmonella; Salmonella enterica;

Salmonella enterica subsp. enterica MTGDNTLITSHGINRRDFMKLCAALAATMGLSSKAAAEMAESVSNPQRPPVIWIGAQECT WP_000145429.1 hydrogenase 2 small subunit [Salmonella enterica] Length: 372\nNumber of predicted TMHs: 1\nExp number of AAs in TMHs: 24.51309\nExp number, first 60 AAs: 0.2682\nTotal prob of N-in: 0.14402\noutside 1 331\nTMhelix 332 354\ninside 355 372

20278 GCF_900040045.1_14555_2_52 Salmonella enterica subsp. enterica serovar Typhi Proteobacteria; Gammaproteobacteria; Enterobacterales; Enterobacteriaceae; Salmonella; Salmonella enterica; Salmonella enterica subsp. enterica MTGDNTLITSHGINRRDFMKLCAALAATMGLSSKAAAEMAESVSNPQRPPVIWIGAQECT WP_000145429.1 hydrogenase 2 small subunit [Salmonella enterica] Length: 372\nNumber of predicted TMHs: 1\nExp number of AAs in TMHs: 24.51309\nExp number, first 60 AAs: 0.2682\nTotal prob of N-in: 0.14402\noutside 1 331\nTMhelix 332 354\ninside 355 372

20279 GCF_900002625.1_JX2005-56 Salmonella enterica subsp. enterica serovar Typhi Proteobacteria; Gammaproteobacteria; Enterobacterales; Enterobacteriaceae; Salmonella; Salmonella enterica; Salmonella enterica subsp. enterica MTGDNTLITSHGINRRDFMKLCAALAATMGLSSKAAAEMAESVSNPQRPPVIWIGAQECT WP_000145429.1 hydrogenase 2 small subunit [Salmonella enterica] Length: 372\nNumber of predicted TMHs: 1\nExp number of AAs in TMHs: 24.51309\nExp number, first 60 AAs: 0.2682\nTotal prob of N-in: 0.14402\noutside 1 331\nTMhelix 332 354\ninside 355 372

20280 GCF_900040685.1_13566_1_69 Salmonella enterica subsp. enterica serovar Typhi Proteobacteria; Gammaproteobacteria; Enterobacterales; Enterobacteriaceae; Salmonella; Salmonella enterica; Salmonella enterica subsp. enterica MTGDNTLITSHGINRRDFMKLCAALAATMGLSSKAAAEMAESVSNPQRPPVIWIGAQECT WP_000145429.1 hydrogenase 2 small subunit [Salmonella enterica] Length: 372\nNumber of predicted TMHs: 1\nExp number of AAs in TMHs: 24.51309\nExp number, first 60 AAs: 0.2682\nTotal prob of N-in: 0.14402\noutside 1 331\nTMhelix 332 354\ninside 355 372

20281 GCF_900041445.1_13097_1_80 Salmonella enterica subsp. enterica serovar Typhi Proteobacteria; Gammaproteobacteria; Enterobacterales; Enterobacteriaceae; Salmonella; Salmonella enterica; Salmonella enterica subsp. enterica MTGDNTLITSHGINRRDFMKLCAALAATMGLSSKAAAEMAESVSNPQRPPVIWIGAQECT WP_000145429.1 hydrogenase 2 small subunit [Salmonella enterica] Length: 372\nNumber of predicted TMHs: 1\nExp number of AAs in TMHs: 24.51309\nExp number, first 60 AAs: 0.2682\nTotal prob of N-in: 0.14402\noutside 1 331\nTMhelix 332 354\ninside 355 372

20282 GCF_900041595.1_14555_2_30 Salmonella enterica subsp. enterica serovar Typhi Proteobacteria; Gammaproteobacteria; Enterobacterales; Enterobacteriaceae; Salmonella; Salmonella enterica; Salmonella enterica subsp. enterica MTGDNTLITSHGINRRDFMKLCAALAATMGLSSKAAAEMAESVSNPQRPPVIWIGAQECT WP_000145429.1 hydrogenase 2 small subunit [Salmonella enterica] Length: 372\nNumber of predicted TMHs: 1\nExp number of AAs in TMHs: 24.51309\nExp number, first 60 AAs: 0.2682\nTotal prob of N-in: 0.14402\noutside 1 331\nTMhelix 332 354\ninside 355 372

20283 GCF_900041615.1_14555_2_41 Salmonella enterica subsp. enterica serovar Typhi Proteobacteria; Gammaproteobacteria; Enterobacterales; Enterobacteriaceae; Salmonella; Salmonella enterica; Salmonella enterica subsp. enterica MTGDNTLITSHGINRRDFMKLCAALAATMGLSSKAAAEMAESVSNPQRPPVIWIGAQECT WP_000145429.1 hydrogenase 2 small subunit [Salmonella enterica] Length: 372\nNumber of predicted TMHs: 1\nExp number of AAs in TMHs: 24.51309\nExp number, first 60 AAs: 0.2682\nTotal prob of N-in: 0.14402\noutside 1 331\nTMhelix 332 354\ninside 355 372

20284 GCF_900041645.1_14555_2_58 Salmonella enterica subsp. enterica serovar Typhi Proteobacteria; Gammaproteobacteria; Enterobacterales; Enterobacteriaceae; Salmonella; Salmonella enterica; Salmonella enterica subsp. enterica MTGDNTLITSHGINRRDFMKLCAALAATMGLSSKAAAEMAESVSNPQRPPVIWIGAQECT WP_000145429.1 hydrogenase 2 small subunit [Salmonella enterica] Length: 372\nNumber of predicted TMHs: 1\nExp number of AAs in TMHs: 24.51309\nExp number, first 60 AAs: 0.2682\nTotal prob of N-in: 0.14402\noutside 1 331\nTMhelix 332 354\ninside 355 372

20285 GCF_900042405.1_13071_1_91 Salmonella enterica subsp. enterica serovar Typhi Proteobacteria; Gammaproteobacteria; Enterobacterales; Enterobacteriaceae; Salmonella; Salmonella enterica; Salmonella enterica subsp. enterica MTGDNTLITSHGINRRDFMKLCAALAATMGLSSKAAAEMAESVSNPQRPPVIWIGAQECT WP_000145429.1 hydrogenase 2 small subunit [Salmonella enterica] Length: 372\nNumber of predicted TMHs: 1\nExp number of AAs in TMHs: 24.51309\nExp number, first 60 AAs: 0.2682\nTotal prob of N-in: 0.14402\noutside 1 331\nTMhelix 332 354\ninside 355 372

20286	GCF_900042425.1_13097_1_79	Salmonella enterica subsp. enterica serovar Typhi	Proteobacteria; Gammaproteobacteria; Enterobacterales; Enterobacteriaceae; Salmonella; Salmonella enterica; Salmonella enterica subsp. enterica	MTGDNTLITSHGINRRDFMKLCAALAATMGLSSKAAAEMAESVSNPQRPPVIWIGAQECT	WP_000145429.1	hydrogenase 2 small subunit [Salmonella enterica]	Length: 372\nNumber of predicted TMHs:  1\nExp number of AAs in TMHs: 24.51309\nExp number, first 60 AAs:  0.2682\nTotal prob of N-in:  0.14402\noutside   1   331\nTMhelix   332   354\ninside   355   372

20287	GCF_900042445.1_13097_1_83	Salmonella enterica subsp. enterica serovar Typhi	Proteobacteria; Gammaproteobacteria; Enterobacterales; Enterobacteriaceae; Salmonella; Salmonella enterica; Salmonella enterica subsp. enterica	MTGDNTLITSHGINRRDFMKLCAALAATMGLSSKAAAEMAESVSNPQRPPVIWIGAQECT	WP_000145429.1	hydrogenase 2 small subunit [Salmonella enterica]	Length: 372\nNumber of predicted TMHs:  1\nExp number of AAs in TMHs: 24.51309\nExp number, first 60 AAs:  0.2682\nTotal prob of N-in:  0.14402\noutside   1   331\nTMhelix   332   354\ninside   355   372

20288	GCF_900002615.1_St1428-XJ2010	Salmonella enterica subsp. enterica serovar Typhi	Proteobacteria; Gammaproteobacteria; Enterobacterales; Enterobacteriaceae; Salmonella; Salmonella enterica; Salmonella enterica subsp. enterica	MTGDNTLITSHGINRRDFMKLCAALAATMGLSSKAAAEMAESVSNPQRPPVIWIGAQECT	WP_000145429.1	hydrogenase 2 small subunit [Salmonella enterica]	Length: 372\nNumber of predicted TMHs:  1\nExp number of AAs in TMHs: 24.51309\nExp number, first 60 AAs:  0.2682\nTotal prob of N-in:  0.14402\noutside   1   331\nTMhelix   332   354\ninside   355   372

20289	GCF_900043515.1_14555_2_65	Salmonella enterica subsp. enterica serovar Typhi	Proteobacteria; Gammaproteobacteria; Enterobacterales; Enterobacteriaceae; Salmonella; Salmonella enterica; Salmonella enterica subsp. enterica	MTGDNTLITSHGINRRDFMKLCAALAATMGLSSKAAAEMAESVSNPQRPPVIWIGAQECT	WP_000145429.1	hydrogenase 2 small subunit [Salmonella enterica]	Length: 372\nNumber of predicted TMHs:  1\nExp number of AAs in TMHs: 24.51309\nExp number, first 60 AAs:  0.2682\nTotal prob of N-in:  0.14402\noutside   1   331\nTMhelix   332   354\ninside   355   372

20290	GCF_900045465.1_14555_2_89	Salmonella enterica subsp. enterica serovar Typhi	Proteobacteria; Gammaproteobacteria; Enterobacterales; Enterobacteriaceae; Salmonella; Salmonella enterica; Salmonella enterica subsp. enterica	MTGDNTLITSHGINRRDFMKLCAALAATMGLSSKAAAEMAESVSNPQRPPVIWIGAQECT	WP_000145429.1	hydrogenase 2 small subunit [Salmonella enterica]	Length: 372\nNumber of predicted TMHs:  1\nExp number of AAs in TMHs: 24.51309\nExp number, first 60 AAs:  0.2682\nTotal prob of N-in:  0.14402\noutside   1   331\nTMhelix   332   354\ninside   355   372

20291	GCF_900046755.1_13103_1_9	Salmonella enterica subsp. enterica serovar Typhi	Proteobacteria; Gammaproteobacteria; Enterobacterales; Enterobacteriaceae; Salmonella; Salmonella enterica; Salmonella enterica subsp. enterica	MTGDNTLITSHGINRRDFMKLCAALAATMGLSSKAAAEMAESVSNPQRPPVIWIGAQECT	WP_000145429.1	hydrogenase 2 small subunit [Salmonella enterica]	Length: 372\nNumber of predicted TMHs:  1\nExp number of AAs in TMHs: 24.51309\nExp number, first 60 AAs:  0.2682\nTotal prob of N-in:  0.14402\noutside   1   331\nTMhelix   332   354\ninside   355   372

20292	GCF_900047075.1_13097_1_77	Salmonella enterica subsp. enterica serovar Typhi	Proteobacteria; Gammaproteobacteria; Enterobacterales; Enterobacteriaceae; Salmonella; Salmonella enterica; Salmonella enterica subsp. enterica	MTGDNTLITSHGINRRDFMKLCAALAATMGLSSKAAAEMAESVSNPQRPPVIWIGAQECT	WP_000145429.1	hydrogenase 2 small subunit [Salmonella enterica]	Length: 372\nNumber of predicted TMHs:  1\nExp number of AAs in TMHs: 24.51309\nExp number, first 60 AAs:  0.2682\nTotal prob of N-in:  0.14402\noutside   1   331\nTMhelix   332   354\ninside   355   372

20293	GCF_900047095.1_13103_1_4	Salmonella enterica subsp. enterica serovar Typhi	Proteobacteria; Gammaproteobacteria; Enterobacterales; Enterobacteriaceae; Salmonella; Salmonella enterica; Salmonella enterica subsp. enterica	MTGDNTLITSHGINRRDFMKLCAALAATMGLSSKAAAEMAESVSNPQRPPVIWIGAQECT	WP_000145429.1	hydrogenase 2 small subunit [Salmonella enterica]	Length: 372\nNumber of predicted TMHs:  1\nExp number of AAs in TMHs: 24.51309\nExp number, first 60 AAs:  0.2682\nTotal prob of N-in:  0.14402\noutside   1   331\nTMhelix   332   354\ninside   355   372

20294	GCF_001131365.1_10209_5_30	Salmonella enterica subsp. enterica serovar Typhi	Proteobacteria; Gammaproteobacteria; Enterobacterales; Enterobacteriaceae; Salmonella; Salmonella enterica; Salmonella enterica subsp. enterica	MTGDNTLITSHGINRRDFMKLCAALAATMGLSSKAAAEMAESVSNPQRPPVIWIGAQECT	WP_000145429.1	hydrogenase 2 small subunit [Salmonella enterica]	Length: 372\nNumber of

predicted TMHs: 1\nExp number of AAs in TMHs: 24.51309\nExp number, first 60 AAs: 0.2682\nTotal prob of N-in: 0.14402\noutside 1 331\nTMhelix 332 354\ninside 355 372

20295 GCF_001130905.1_10608_2_62 Salmonella enterica subsp. enterica serovar Typhi Proteobacteria; Gammaproteobacteria; Enterobacterales; Enterobacteriaceae; Salmonella; Salmonella enterica; Salmonella enterica subsp. enterica MTGDNTLITSHGINRRDFMKLCAALAATMGLSSKAAAEMAESVSNPQRPPVIWIGAQECT WP_000145429.1 hydrogenase 2 small subunit [Salmonella enterica] Length: 372\nNumber of predicted TMHs: 1\nExp number of AAs in TMHs: 24.51309\nExp number, first 60 AAs: 0.2682\nTotal prob of N-in: 0.14402\noutside 1 331\nTMhelix 332 354\ninside 355 372

20296 GCF_001130345.1_8447_8_9Salmonella enterica subsp. enterica serovar Typhi Proteobacteria; Gammaproteobacteria; Enterobacterales; Enterobacteriaceae; Salmonella; Salmonella enterica; Salmonella enterica subsp. enterica MTGDNTLITSHGINRRDFMKLCAALAATMGLSSKAAAEMAESVSNPQRPPVIWIGAQECT WP_000145429.1 hydrogenase 2 small subunit [Salmonella enterica] Length: 372\nNumber of predicted TMHs: 1\nExp number of AAs in TMHs: 24.51309\nExp number, first 60 AAs: 0.2682\nTotal prob of N-in: 0.14402\noutside 1 331\nTMhelix 332 354\ninside 355 372

20297 GCF_001130285.1_8525_3_45 Salmonella enterica subsp. enterica serovar Typhi Proteobacteria; Gammaproteobacteria; Enterobacterales; Enterobacteriaceae; Salmonella; Salmonella enterica; Salmonella enterica subsp. enterica MTGDNTLITSHGINRRDFMKLCAALAATMGLSSKAAAEMAESVSNPQRPPVIWIGAQECT WP_000145429.1 hydrogenase 2 small subunit [Salmonella enterica] Length: 372\nNumber of predicted TMHs: 1\nExp number of AAs in TMHs: 24.51309\nExp number, first 60 AAs: 0.2682\nTotal prob of N-in: 0.14402\noutside 1 331\nTMhelix 332 354\ninside 355 372

20298 GCF_001129345.1_10541_2_45 Salmonella enterica subsp. enterica serovar Typhi Proteobacteria; Gammaproteobacteria; Enterobacterales; Enterobacteriaceae; Salmonella; Salmonella enterica; Salmonella enterica subsp. enterica MTGDNTLITSHGINRRDFMKLCAALAATMGLSSKAAAEMAESVSNPQRPPVIWIGAQECT WP_000145429.1 hydrogenase 2 small subunit [Salmonella enterica] Length: 372\nNumber of predicted TMHs: 1\nExp number of AAs in TMHs: 24.51309\nExp number, first 60 AAs: 0.2682\nTotal prob of N-in: 0.14402\noutside 1 331\nTMhelix 332 354\ninside 355 372

20299 GCF_001125205.1_10541_2_43 Salmonella enterica subsp. enterica serovar Typhi Proteobacteria; Gammaproteobacteria; Enterobacterales; Enterobacteriaceae; Salmonella; Salmonella enterica; Salmonella enterica subsp. enterica MTGDNTLITSHGINRRDFMKLCAALAATMGLSSKAAAEMAESVSNPQRPPVIWIGAQECT WP_000145429.1 hydrogenase 2 small subunit [Salmonella enterica] Length: 372\nNumber of predicted TMHs: 1\nExp number of AAs in TMHs: 24.51309\nExp number, first 60 AAs: 0.2682\nTotal prob of N-in: 0.14402\noutside 1 331\nTMhelix 332 354\ninside 355 372

20300 GCF_001124105.1_10492_1_70 Salmonella enterica subsp. enterica serovar Typhi Proteobacteria; Gammaproteobacteria; Enterobacterales; Enterobacteriaceae; Salmonella; Salmonella enterica; Salmonella enterica subsp. enterica MTGDNTLITSHGINRRDFMKLCAALAATMGLSSKAATEMAESVSNPQRPPVIWIGAQECT WP_001782304.1 hydrogenase 2 small subunit [Salmonella enterica] Length: 372\nNumber of predicted TMHs: 1\nExp number of AAs in TMHs: 24.41173\nExp number, first 60 AAs: 0.16124\nTotal prob of N-in: 0.13947\noutside 1 331\nTMhelix 332 354\ninside 355 372

20301 GCF_001124945.1_10562_2_4 Salmonella enterica subsp. enterica serovar Typhi Proteobacteria; Gammaproteobacteria; Enterobacterales; Enterobacteriaceae; Salmonella; Salmonella enterica; Salmonella enterica subsp. enterica MTGDNTLITSHGINRRDFMKLCAALAATMGLSSKAAAEMAESVSNPQRPPVIWIGAQECT WP_000145429.1 hydrogenase 2 small subunit [Salmonella enterica] Length: 372\nNumber of predicted TMHs: 1\nExp number of AAs in TMHs: 24.51309\nExp number, first 60 AAs: 0.2682\nTotal prob of N-in: 0.14402\noutside 1 331\nTMhelix 332 354\ninside 355 372

20302 GCF_001124685.1_9953_5_17 Salmonella enterica subsp. enterica serovar Typhi Proteobacteria; Gammaproteobacteria; Enterobacterales; Enterobacteriaceae; Salmonella; Salmonella enterica; Salmonella enterica subsp. enterica MTGDNTLITSHGINRRDFMKLCAALAATMGLSSKAAAEMAESVSNPQRPPVIWIGAQECT WP_000145429.1 hydrogenase 2 small subunit [Salmonella enterica] Length: 372\nNumber of predicted TMHs: 1\nExp number of AAs in TMHs: 24.51309\nExp number, first 60 AAs: 0.2682\nTotal prob of N-in: 0.14402\noutside 1 331\nTMhelix 332 354\ninside 355 372

20303 GCF_001124805.1_10562_2_27 Salmonella enterica subsp. enterica serovar Typhi Proteobacteria; Gammaproteobacteria; Enterobacterales; Enterobacteriaceae; Salmonella; Salmonella enterica;

Salmonella enterica subsp. enterica MTGDNTLITSHGINRRDFMKLCAALAATMGLSSKAAAEMAESVSNPQRPPVIWIGAQECT WP_000145429.1 hydrogenase 2 small subunit [Salmonella enterica] Length: 372\nNumber of predicted TMHs: 1\nExp number of AAs in TMHs: 24.51309\nExp number, first 60 AAs: 0.2682\nTotal prob of N-in: 0.14402\noutside 1 331\nTMhelix 332 354\ninside 355 372

20304 GCF_001124405.1_10541_2_26 Salmonella enterica subsp. enterica serovar Typhi Proteobacteria; Gammaproteobacteria; Enterobacterales; Enterobacteriaceae; Salmonella; Salmonella enterica; Salmonella enterica subsp. enterica MTGDNTLITSHGINRRDFMKLCAALAATMGLSSKAAAEMAESVSNPQRPPVIWIGAQECT WP_000145429.1 hydrogenase 2 small subunit [Salmonella enterica] Length: 372\nNumber of predicted TMHs: 1\nExp number of AAs in TMHs: 24.51309\nExp number, first 60 AAs: 0.2682\nTotal prob of N-in: 0.14402\noutside 1 331\nTMhelix 332 354\ninside 355 372

20305 GCF_001124365.1_10209_5_19 Salmonella enterica subsp. enterica serovar Typhi Proteobacteria; Gammaproteobacteria; Enterobacterales; Enterobacteriaceae; Salmonella; Salmonella enterica; Salmonella enterica subsp. enterica MTGDNTLITSHGINRRDFMKLCAALAATMGLSSKAAAEMAESVSNPQRPPVIWIGAQECT WP_000145429.1 hydrogenase 2 small subunit [Salmonella enterica] Length: 372\nNumber of predicted TMHs: 1\nExp number of AAs in TMHs: 24.51309\nExp number, first 60 AAs: 0.2682\nTotal prob of N-in: 0.14402\noutside 1 331\nTMhelix 332 354\ninside 355 372

20306 GCF_001120845.1_10592_2_35 Salmonella enterica subsp. enterica serovar Typhi Proteobacteria; Gammaproteobacteria; Enterobacterales; Enterobacteriaceae; Salmonella; Salmonella enterica; Salmonella enterica subsp. enterica MTGDNTLITSHGINRRDFMKLCAALAATMGLSSKAAAEMAESVSNPQRPPVIWIGAQECT WP_000145429.1 hydrogenase 2 small subunit [Salmonella enterica] Length: 372\nNumber of predicted TMHs: 1\nExp number of AAs in TMHs: 24.51309\nExp number, first 60 AAs: 0.2682\nTotal prob of N-in: 0.14402\noutside 1 331\nTMhelix 332 354\ninside 355 372

20307 GCF_001121045.1_10540_1_4 Salmonella enterica subsp. enterica serovar Typhi Proteobacteria; Gammaproteobacteria; Enterobacterales; Enterobacteriaceae; Salmonella; Salmonella enterica; Salmonella enterica subsp. enterica MTGDNTLITSHGINRRDFMKLCAALAATMGLSSKAAAEMAESVSNPQRPPVIWIGAQECT WP_000145429.1 hydrogenase 2 small subunit [Salmonella enterica] Length: 372\nNumber of predicted TMHs: 1\nExp number of AAs in TMHs: 24.51309\nExp number, first 60 AAs: 0.2682\nTotal prob of N-in: 0.14402\noutside 1 331\nTMhelix 332 354\ninside 355 372

20308 GCF_001151125.1_10492_1_18 Salmonella enterica subsp. enterica serovar Typhi Proteobacteria; Gammaproteobacteria; Enterobacterales; Enterobacteriaceae; Salmonella; Salmonella enterica; Salmonella enterica subsp. enterica MTGDNTLITSHGINRRDFMKLCAALAATMGLSSKAAAEMAESVSNPQRPPVIWIGAQECT WP_000145429.1 hydrogenase 2 small subunit [Salmonella enterica] Length: 372\nNumber of predicted TMHs: 1\nExp number of AAs in TMHs: 24.51309\nExp number, first 60 AAs: 0.2682\nTotal prob of N-in: 0.14402\noutside 1 331\nTMhelix 332 354\ninside 355 372

20309 GCF_001120645.1_9870_8_43 Salmonella enterica subsp. enterica serovar Typhi Proteobacteria; Gammaproteobacteria; Enterobacterales; Enterobacteriaceae; Salmonella; Salmonella enterica; Salmonella enterica subsp. enterica MTGDNTLITSHGINRRDFMKLCAALAATMGLSSKAAAEMAESVSNPQRPPVIWIGAQECT WP_000145429.1 hydrogenase 2 small subunit [Salmonella enterica] Length: 372\nNumber of predicted TMHs: 1\nExp number of AAs in TMHs: 24.51309\nExp number, first 60 AAs: 0.2682\nTotal prob of N-in: 0.14402\noutside 1 331\nTMhelix 332 354\ninside 355 372

20310 GCF_001120605.1_9475_6_68 Salmonella enterica subsp. enterica serovar Typhi Proteobacteria; Gammaproteobacteria; Enterobacterales; Enterobacteriaceae; Salmonella; Salmonella enterica; Salmonella enterica subsp. enterica MTGDNTLITSHGINRRDFMKLCAALAATMGLSSKAAAEMAESVSNPQRPPVIWIGAQECT WP_000145429.1 hydrogenase 2 small subunit [Salmonella enterica] Length: 372\nNumber of predicted TMHs: 1\nExp number of AAs in TMHs: 24.51309\nExp number, first 60 AAs: 0.2682\nTotal prob of N-in: 0.14402\noutside 1 331\nTMhelix 332 354\ninside 355 372

20311 GCF_001120405.1_10492_1_61 Salmonella enterica subsp. enterica serovar Typhi Proteobacteria; Gammaproteobacteria; Enterobacterales; Enterobacteriaceae; Salmonella; Salmonella enterica; Salmonella enterica subsp. enterica MTGDNTLITSHGINRRDFMKLCAALAATMGLSSKAAAEMAESVSNPQRPPVIWIGAQECT WP_000145429.1 hydrogenase 2 small subunit [Salmonella enterica] Length: 372\nNumber of predicted TMHs: 1\nExp number of AAs in TMHs: 24.51309\nExp number, first 60 AAs: 0.2682\nTotal prob of N-in: 0.14402\noutside 1 331\nTMhelix 332 354\ninside 355 372

20312	GCF_001328675.1_7468_7_89	Salmonella enterica subsp. enterica serovar Typhi	Proteobacteria; Gammaproteobacteria; Enterobacterales; Enterobacteriaceae; Salmonella; Salmonella enterica; Salmonella enterica subsp. enterica	MTGDNTLITSHGINRRDFMKLCAALAATMGLSSKAAAEMAESVSNPQRPPVIWIGAQECT	WP_000145429.1	hydrogenase 2 small subunit [Salmonella enterica]	Length: 372\nNumber of predicted TMHs:  1\nExp number of AAs in TMHs: 24.51309\nExp number, first 60 AAs:  0.2682\nTotal prob of N-in: 0.14402\noutside	1	331\nTMhelix	332	354\ninside	355	372

20313	GCF_001120085.1_9870_8_85	Salmonella enterica subsp. enterica serovar Typhi	Proteobacteria; Gammaproteobacteria; Enterobacterales; Enterobacteriaceae; Salmonella; Salmonella enterica; Salmonella enterica subsp. enterica	MTGDNTLITSHGINRRDFMKLCAALAATMGLSSKAAAEMAESVSNPQRPPVIWIGAQECT	WP_000145429.1	hydrogenase 2 small subunit [Salmonella enterica]	Length: 372\nNumber of predicted TMHs:  1\nExp number of AAs in TMHs: 24.51309\nExp number, first 60 AAs:  0.2682\nTotal prob of N-in: 0.14402\noutside	1	331\nTMhelix	332	354\ninside	355	372

20314	GCF_001120005.1_10592_2_58	Salmonella enterica subsp. enterica serovar Typhi	Proteobacteria; Gammaproteobacteria; Enterobacterales; Enterobacteriaceae; Salmonella; Salmonella enterica; Salmonella enterica subsp. enterica	MTGDNTLITSHGINRRDFMKLCAALAATMGLSSKAAAEMAESVSNPQRPPVIWIGAQECT	WP_000145429.1	hydrogenase 2 small subunit [Salmonella enterica]	Length: 372\nNumber of predicted TMHs:  1\nExp number of AAs in TMHs: 24.51309\nExp number, first 60 AAs:  0.2682\nTotal prob of N-in: 0.14402\noutside	1	331\nTMhelix	332	354\ninside	355	372

20315	GCF_001151265.1_10608_2_34	Salmonella enterica subsp. enterica serovar Typhi	Proteobacteria; Gammaproteobacteria; Enterobacterales; Enterobacteriaceae; Salmonella; Salmonella enterica; Salmonella enterica subsp. enterica	MTGDNTLITSHGINRRDFMKLCAALAATMGLSSKAAAEMAESVSNPQRPPVIWIGAQECT	WP_000145429.1	hydrogenase 2 small subunit [Salmonella enterica]	Length: 372\nNumber of predicted TMHs:  1\nExp number of AAs in TMHs: 24.51309\nExp number, first 60 AAs:  0.2682\nTotal prob of N-in: 0.14402\noutside	1	331\nTMhelix	332	354\ninside	355	372

20316	GCF_001117125.1_9870_8_20	Salmonella enterica subsp. enterica serovar Typhi	Proteobacteria; Gammaproteobacteria; Enterobacterales; Enterobacteriaceae; Salmonella; Salmonella enterica; Salmonella enterica subsp. enterica	MTGDNTLITSHGINRRDFMKLCAALAATMGLSSKAAAEMAESVSNPQRPPVIWIGAQECT	WP_000145429.1	hydrogenase 2 small subunit [Salmonella enterica]	Length: 372\nNumber of predicted TMHs:  1\nExp number of AAs in TMHs: 24.51309\nExp number, first 60 AAs:  0.2682\nTotal prob of N-in: 0.14402\noutside	1	331\nTMhelix	332	354\ninside	355	372

20317	GCF_001154765.1_9475_6_10	Salmonella enterica subsp. enterica serovar Typhi	Proteobacteria; Gammaproteobacteria; Enterobacterales; Enterobacteriaceae; Salmonella; Salmonella enterica; Salmonella enterica subsp. enterica	MTGDNTLITSHGINRRDFMKLCAALAATMGLSSKAAAEMAESVSNPQRPPVIWIGAQECT	WP_000145429.1	hydrogenase 2 small subunit [Salmonella enterica]	Length: 372\nNumber of predicted TMHs:  1\nExp number of AAs in TMHs: 24.51309\nExp number, first 60 AAs:  0.2682\nTotal prob of N-in: 0.14402\noutside	1	331\nTMhelix	332	354\ninside	355	372

20318	GCF_001116685.1_8490_6_40	Salmonella enterica subsp. enterica serovar Typhi	Proteobacteria; Gammaproteobacteria; Enterobacterales; Enterobacteriaceae; Salmonella; Salmonella enterica; Salmonella enterica subsp. enterica	MTGDNTLITSHGINRRDFMKLCAALAATMGLSSKAAAEMAESVSNPQRPPVIWIGAQECT	WP_000145429.1	hydrogenase 2 small subunit [Salmonella enterica]	Length: 372\nNumber of predicted TMHs:  1\nExp number of AAs in TMHs: 24.51309\nExp number, first 60 AAs:  0.2682\nTotal prob of N-in: 0.14402\noutside	1	331\nTMhelix	332	354\ninside	355	372

20319	GCF_001116345.1_10593_2_77	Salmonella enterica subsp. enterica serovar Typhi	Proteobacteria; Gammaproteobacteria; Enterobacterales; Enterobacteriaceae; Salmonella; Salmonella enterica; Salmonella enterica subsp. enterica	MTGDNTLITSHGINRRDFMKLCAALAATMGLSSKAAAEMAESVSNPQRPPVIWIGAQECT	WP_000145429.1	hydrogenase 2 small subunit [Salmonella enterica]	Length: 372\nNumber of predicted TMHs:  1\nExp number of AAs in TMHs: 24.51309\nExp number, first 60 AAs:  0.2682\nTotal prob of N-in: 0.14402\noutside	1	331\nTMhelix	332	354\ninside	355	372

20320	GCF_001116145.1_10349_1_26	Salmonella enterica subsp. enterica serovar Typhi	Proteobacteria; Gammaproteobacteria; Enterobacterales; Enterobacteriaceae; Salmonella; Salmonella enterica; Salmonella enterica subsp. enterica	MTGDNTLITSHGINRRDFMKLCAALAATMGLSSKAAAEMAESVSNPQRPPVIWIGAQECT	WP_000145429.1	hydrogenase 2 small subunit [Salmonella enterica]	Length: 372\nNumber of

predicted TMHs:  1\nExp number of AAs in TMHs: 24.51309\nExp number, first 60 AAs:  0.2682\nTotal prob of N-in: 0.14402\noutside   1   331\nTMhelix   332   354\ninside   355   372

20321 GCF_001154865.1_10541_2_37 Salmonella enterica subsp. enterica serovar Typhi Proteobacteria; Gammaproteobacteria; Enterobacterales; Enterobacteriaceae; Salmonella; Salmonella enterica; Salmonella enterica subsp. enterica MTGDNTLITSHGINRRDFMKLCAALAATMGLSSKAAAEMAESVSNPQRPPVIWIGAQECT WP_000145429.1 hydrogenase 2 small subunit [Salmonella enterica] Length: 372\nNumber of predicted TMHs:  1\nExp number of AAs in TMHs: 24.51309\nExp number, first 60 AAs:  0.2682\nTotal prob of N-in: 0.14402\noutside   1   331\nTMhelix   332   354\ninside   355   372

20322 GCF_001115785.1_9953_5_31 Salmonella enterica subsp. enterica serovar Typhi Proteobacteria; Gammaproteobacteria; Enterobacterales; Enterobacteriaceae; Salmonella; Salmonella enterica; Salmonella enterica subsp. enterica MTGDNTLITSHGINRRDFMKLCAALAATMGLSSKAAAEMAESVSNPQRPPVIWIGAQECT WP_000145429.1 hydrogenase 2 small subunit [Salmonella enterica] Length: 372\nNumber of predicted TMHs:  1\nExp number of AAs in TMHs: 24.51309\nExp number, first 60 AAs:  0.2682\nTotal prob of N-in: 0.14402\noutside   1   331\nTMhelix   332   354\ninside   355   372

20323 GCF_001114985.1_9475_6_57 Salmonella enterica subsp. enterica serovar Typhi Proteobacteria; Gammaproteobacteria; Enterobacterales; Enterobacteriaceae; Salmonella; Salmonella enterica; Salmonella enterica subsp. enterica MTGDNTLITSHGINRRDFMKLCAALAATMGLSSKAAAEMAESVSNPQRPPVIWIGAQECT WP_000145429.1 hydrogenase 2 small subunit [Salmonella enterica] Length: 372\nNumber of predicted TMHs:  1\nExp number of AAs in TMHs: 24.51309\nExp number, first 60 AAs:  0.2682\nTotal prob of N-in: 0.14402\noutside   1   331\nTMhelix   332   354\ninside   355   372

20324 GCF_001115105.1_9870_8_74 Salmonella enterica subsp. enterica serovar Typhi Proteobacteria; Gammaproteobacteria; Enterobacterales; Enterobacteriaceae; Salmonella; Salmonella enterica; Salmonella enterica subsp. enterica MTGDNTLITSHGINRRDFMKLCAALAATMGLSSKAAAEMAESVSNPQRPPVIWIGAQECT WP_000145429.1 hydrogenase 2 small subunit [Salmonella enterica] Length: 372\nNumber of predicted TMHs:  1\nExp number of AAs in TMHs: 24.51309\nExp number, first 60 AAs:  0.2682\nTotal prob of N-in: 0.14402\noutside   1   331\nTMhelix   332   354\ninside   355   372

20325 GCF_001155045.1_8525_3_37 Salmonella enterica subsp. enterica serovar Typhi Proteobacteria; Gammaproteobacteria; Enterobacterales; Enterobacteriaceae; Salmonella; Salmonella enterica; Salmonella enterica subsp. enterica MTGDNTLITSHGINRRDFMKLCAALAATMGLSSKAAAEMAESVSNPQRPPVIWIGAQECT WP_000145429.1 hydrogenase 2 small subunit [Salmonella enterica] Length: 372\nNumber of predicted TMHs:  1\nExp number of AAs in TMHs: 24.51309\nExp number, first 60 AAs:  0.2682\nTotal prob of N-in: 0.14402\noutside   1   331\nTMhelix   332   354\ninside   355   372

20326 GCF_001112345.1_7468_7_94 Salmonella enterica subsp. enterica serovar Typhi Proteobacteria; Gammaproteobacteria; Enterobacterales; Enterobacteriaceae; Salmonella; Salmonella enterica; Salmonella enterica subsp. enterica MTGDNTLITSHGINRRDFMKLCAALAATMGLSSKAAAEMAESVSNPQRPPVIWIGAQECT WP_000145429.1 hydrogenase 2 small subunit [Salmonella enterica] Length: 372\nNumber of predicted TMHs:  1\nExp number of AAs in TMHs: 24.51309\nExp number, first 60 AAs:  0.2682\nTotal prob of N-in: 0.14402\noutside   1   331\nTMhelix   332   354\ninside   355   372

20327 GCF_001155405.1_10561_2_61 Salmonella enterica subsp. enterica serovar Typhi Proteobacteria; Gammaproteobacteria; Enterobacterales; Enterobacteriaceae; Salmonella; Salmonella enterica; Salmonella enterica subsp. enterica MTGDNTLITSHGINRRDFMKLCAALAATMGLSSKAAAEMAESVSNPQRPPVIWIGAQECT WP_000145429.1 hydrogenase 2 small subunit [Salmonella enterica] Length: 372\nNumber of predicted TMHs:  1\nExp number of AAs in TMHs: 24.51309\nExp number, first 60 AAs:  0.2682\nTotal prob of N-in: 0.14402\noutside   1   331\nTMhelix   332   354\ninside   355   372

20328 GCF_001155545.1_10209_5_44 Salmonella enterica subsp. enterica serovar Typhi Proteobacteria; Gammaproteobacteria; Enterobacterales; Enterobacteriaceae; Salmonella; Salmonella enterica; Salmonella enterica subsp. enterica MTGDNTLITSHGINRRDFMKLCAALAATMGLSSKAAAEMAESVSNPQRPPVIWIGAQECT WP_000145429.1 hydrogenase 2 small subunit [Salmonella enterica] Length: 372\nNumber of predicted TMHs:  1\nExp number of AAs in TMHs: 24.51309\nExp number, first 60 AAs:  0.2682\nTotal prob of N-in: 0.14402\noutside   1   331\nTMhelix   332   354\ninside   355   372

20329 GCF_001158985.1_10561_2_53 Salmonella enterica subsp. enterica serovar Typhi Proteobacteria; Gammaproteobacteria; Enterobacterales; Enterobacteriaceae; Salmonella; Salmonella enterica;

Salmonella enterica subsp. enterica	MTGDNTLITSHGINRRDFMKLCAALAATMGLSSKAAAEMAESVSNPQRPPVIWIGAQECT	WP_000145429.1	hydrogenase 2 small subunit [Salmonella enterica]	Length: 372\nNumber of predicted TMHs:  1\nExp number of AAs in TMHs: 24.51309\nExp number, first 60 AAs:  0.2682\nTotal prob of N-in:  0.14402\noutside   1   331\nTMhelix   332   354\ninside   355   372

20330	GCF_001112225.1_9870_8_25	Salmonella enterica subsp. enterica serovar Typhi	Proteobacteria; Gammaproteobacteria; Enterobacterales; Enterobacteriaceae; Salmonella; Salmonella enterica; Salmonella enterica subsp. enterica	MTGDNTLITSHGINRRDFMKLCAALAATMGLSSKAAAEMAESVSNPQRPPVIWIGAQECT	WP_000145429.1	hydrogenase 2 small subunit [Salmonella enterica]	Length: 372\nNumber of predicted TMHs:  1\nExp number of AAs in TMHs: 24.51309\nExp number, first 60 AAs:  0.2682\nTotal prob of N-in:  0.14402\noutside   1   331\nTMhelix   332   354\ninside   355   372

20331	GCF_001158805.1_10562_2_55	Salmonella enterica subsp. enterica serovar Typhi	Proteobacteria; Gammaproteobacteria; Enterobacterales; Enterobacteriaceae; Salmonella; Salmonella enterica; Salmonella enterica subsp. enterica	MTGDNTLITSHGINRRDFMKLCAALAATMGLSSKAAAEMAESVSNPQRPPVIWIGAQECT	WP_000145429.1	hydrogenase 2 small subunit [Salmonella enterica]	Length: 372\nNumber of predicted TMHs:  1\nExp number of AAs in TMHs: 24.51309\nExp number, first 60 AAs:  0.2682\nTotal prob of N-in:  0.14402\noutside   1   331\nTMhelix   332   354\ninside   355   372

20332	GCF_001112085.1_10607_1_74	Salmonella enterica subsp. enterica serovar Typhi	Proteobacteria; Gammaproteobacteria; Enterobacterales; Enterobacteriaceae; Salmonella; Salmonella enterica; Salmonella enterica subsp. enterica	MTGDNTLITSHGINRRDFMKLCAALAATMGLSSKAAAEMAESVSNPQRPPVIWIGAQECT	WP_000145429.1	hydrogenase 2 small subunit [Salmonella enterica]	Length: 372\nNumber of predicted TMHs:  1\nExp number of AAs in TMHs: 24.51309\nExp number, first 60 AAs:  0.2682\nTotal prob of N-in:  0.14402\noutside   1   331\nTMhelix   332   354\ninside   355   372

20333	GCF_001159465.1_10592_2_15	Salmonella enterica subsp. enterica serovar Typhi	Proteobacteria; Gammaproteobacteria; Enterobacterales; Enterobacteriaceae; Salmonella; Salmonella enterica; Salmonella enterica subsp. enterica	MTGDNTLITSHGINRRDFMKLCAALAATMGLSSKAAAEMAESVSNPQRPPVIWIGAQECT	WP_000145429.1	hydrogenase 2 small subunit [Salmonella enterica]	Length: 372\nNumber of predicted TMHs:  1\nExp number of AAs in TMHs: 24.51309\nExp number, first 60 AAs:  0.2682\nTotal prob of N-in:  0.14402\noutside   1   331\nTMhelix   332   354\ninside   355   372

20334	GCF_001159605.1_9475_4_53	Salmonella enterica subsp. enterica serovar Typhi	Proteobacteria; Gammaproteobacteria; Enterobacterales; Enterobacteriaceae; Salmonella; Salmonella enterica; Salmonella enterica subsp. enterica	MTGDNTLITSHGINRRDFMKLCAALAATMGLSSKAAAEMAESVSNPQRPPVIWIGAQECT	WP_000145429.1	hydrogenase 2 small subunit [Salmonella enterica]	Length: 372\nNumber of predicted TMHs:  1\nExp number of AAs in TMHs: 24.51309\nExp number, first 60 AAs:  0.2682\nTotal prob of N-in:  0.14402\noutside   1   331\nTMhelix   332   354\ninside   355   372

20335	GCF_001110925.1_10562_2_86	Salmonella enterica subsp. enterica serovar Typhi	Proteobacteria; Gammaproteobacteria; Enterobacterales; Enterobacteriaceae; Salmonella; Salmonella enterica; Salmonella enterica subsp. enterica	MTGDNTLITSHGINRRDFMKLCAALAATMGLSSKAATEMAESVSNPQRPPVIWIGAQECT	WP_001782304.1	hydrogenase 2 small subunit [Salmonella enterica]	Length: 372\nNumber of predicted TMHs:  1\nExp number of AAs in TMHs: 24.41173\nExp number, first 60 AAs:  0.16124\nTotal prob of N-in:  0.13947\noutside   1   331\nTMhelix   332   354\ninside   355   372

20336	GCF_001111965.1_9953_5_76	Salmonella enterica subsp. enterica serovar Typhi	Proteobacteria; Gammaproteobacteria; Enterobacterales; Enterobacteriaceae; Salmonella; Salmonella enterica; Salmonella enterica subsp. enterica	MTGDNTLITSHGINRRDFMKLCAALAATMGLSSKAAAEMAESVSNPQRPPVIWIGAQECT	WP_000145429.1	hydrogenase 2 small subunit [Salmonella enterica]	Length: 372\nNumber of predicted TMHs:  1\nExp number of AAs in TMHs: 24.51309\nExp number, first 60 AAs:  0.2682\nTotal prob of N-in:  0.14402\noutside   1   331\nTMhelix   332   354\ninside   355   372

20337	GCF_001111865.1_10493_1_27	Salmonella enterica subsp. enterica serovar Typhi	Proteobacteria; Gammaproteobacteria; Enterobacterales; Enterobacteriaceae; Salmonella; Salmonella enterica; Salmonella enterica subsp. enterica	MTGDNTLITSHGINRRDFMKLCAALAATMGLSSKAAAEMAESVSNPQRPPVIWIGAQECT	WP_000145429.1	hydrogenase 2 small subunit [Salmonella enterica]	Length: 372\nNumber of predicted TMHs:  1\nExp number of AAs in TMHs: 24.51309\nExp number, first 60 AAs:  0.2682\nTotal prob of N-in:  0.14402\noutside   1   331\nTMhelix   332   354\ninside   355   372

20338 GCF_001108905.1_9870_8_11 Salmonella enterica subsp. enterica serovar Typhi Proteobacteria; Gammaproteobacteria; Enterobacterales; Enterobacteriaceae; Salmonella; Salmonella enterica; Salmonella enterica subsp. enterica MTGDNTLITSHGINRRDFMKLCAALAATMGLSSKAAAEMAESVSNPQRPPVIWIGAQECT WP_000145429.1 hydrogenase 2 small subunit [Salmonella enterica] Length: 372\nNumber of predicted TMHs: 1\nExp number of AAs in TMHs: 24.51309\nExp number, first 60 AAs: 0.2682\nTotal prob of N-in: 0.14402\noutside 1 331\nTMhelix 332 354\ninside 355 372

20339 GCF_001109205.1_8490_6_67 Salmonella enterica subsp. enterica serovar Typhi Proteobacteria; Gammaproteobacteria; Enterobacterales; Enterobacteriaceae; Salmonella; Salmonella enterica; Salmonella enterica subsp. enterica MTGDNTLITSHGINRRDFMKLCAALAATMGLSSKAAAEMAESVSNPQRPPVIWIGAQECT WP_000145429.1 hydrogenase 2 small subunit [Salmonella enterica] Length: 372\nNumber of predicted TMHs: 1\nExp number of AAs in TMHs: 24.51309\nExp number, first 60 AAs: 0.2682\nTotal prob of N-in: 0.14402\noutside 1 331\nTMhelix 332 354\ninside 355 372

20340 GCF_001108725.1_9953_5_33 Salmonella enterica subsp. enterica serovar Typhi Proteobacteria; Gammaproteobacteria; Enterobacterales; Enterobacteriaceae; Salmonella; Salmonella enterica; Salmonella enterica subsp. enterica MTGDNTLITSHGINRRDFMKLCAALAATMGLSSKAAAEMAESVSNPQRPPVIWIGAQECT WP_000145429.1 hydrogenase 2 small subunit [Salmonella enterica] Length: 372\nNumber of predicted TMHs: 1\nExp number of AAs in TMHs: 24.51309\nExp number, first 60 AAs: 0.2682\nTotal prob of N-in: 0.14402\noutside 1 331\nTMhelix 332 354\ninside 355 372

20341 GCF_001108325.1_10426_1_5 Salmonella enterica subsp. enterica serovar Typhi Proteobacteria; Gammaproteobacteria; Enterobacterales; Enterobacteriaceae; Salmonella; Salmonella enterica; Salmonella enterica subsp. enterica MTGDNTLITSHGINRRDFMKLCAALAATMGLSSKAAAEMAESVSNPQRPPVIWIGAQECT WP_000145429.1 hydrogenase 2 small subunit [Salmonella enterica] Length: 372\nNumber of predicted TMHs: 1\nExp number of AAs in TMHs: 24.51309\nExp number, first 60 AAs: 0.2682\nTotal prob of N-in: 0.14402\noutside 1 331\nTMhelix 332 354\ninside 355 372

20342 GCF_001108125.1_8490_6_47 Salmonella enterica subsp. enterica serovar Typhi Proteobacteria; Gammaproteobacteria; Enterobacterales; Enterobacteriaceae; Salmonella; Salmonella enterica; Salmonella enterica subsp. enterica MTGDNTLITSHGINRRDFMKLCAALAATMGLSSKAAAEMAESVSNPQRPPVIWIGAQECT WP_000145429.1 hydrogenase 2 small subunit [Salmonella enterica] Length: 372\nNumber of predicted TMHs: 1\nExp number of AAs in TMHs: 24.51309\nExp number, first 60 AAs: 0.2682\nTotal prob of N-in: 0.14402\noutside 1 331\nTMhelix 332 354\ninside 355 372

20343 GCF_001162845.1_10561_2_66 Salmonella enterica subsp. enterica serovar Typhi Proteobacteria; Gammaproteobacteria; Enterobacterales; Enterobacteriaceae; Salmonella; Salmonella enterica; Salmonella enterica subsp. enterica MTGDNTLITSHGINRRDFMKLCAALAATMGLSSKAAAEMAESVSNPQRPPVIWIGAQECT WP_000145429.1 hydrogenase 2 small subunit [Salmonella enterica] Length: 372\nNumber of predicted TMHs: 1\nExp number of AAs in TMHs: 24.51309\nExp number, first 60 AAs: 0.2682\nTotal prob of N-in: 0.14402\noutside 1 331\nTMhelix 332 354\ninside 355 372

20344 GCF_001163005.1_8490_5_6Salmonella enterica subsp. enterica serovar Typhi Proteobacteria; Gammaproteobacteria; Enterobacterales; Enterobacteriaceae; Salmonella; Salmonella enterica; Salmonella enterica subsp. enterica MTGDNTLITSHGINRRDFMKLCAALAATMGLSSKAAAEMAESVSNPQRPPVIWIGAQECT WP_000145429.1 hydrogenase 2 small subunit [Salmonella enterica] Length: 372\nNumber of predicted TMHs: 1\nExp number of AAs in TMHs: 24.51309\nExp number, first 60 AAs: 0.2682\nTotal prob of N-in: 0.14402\noutside 1 331\nTMhelix 332 354\ninside 355 372

20345 GCF_001163165.1_10593_2_29 Salmonella enterica subsp. enterica serovar Typhi Proteobacteria; Gammaproteobacteria; Enterobacterales; Enterobacteriaceae; Salmonella; Salmonella enterica; Salmonella enterica subsp. enterica MTGDNTLITSHGINRRDFMKLCAALAATMGLSSKAAAEMAESVSNPQRPPVIWIGAQECT WP_000145429.1 hydrogenase 2 small subunit [Salmonella enterica] Length: 372\nNumber of predicted TMHs: 1\nExp number of AAs in TMHs: 24.51309\nExp number, first 60 AAs: 0.2682\nTotal prob of N-in: 0.14402\noutside 1 331\nTMhelix 332 354\ninside 355 372

20346 GCF_001107945.1_9953_5_80 Salmonella enterica subsp. enterica serovar Typhi Proteobacteria; Gammaproteobacteria; Enterobacterales; Enterobacteriaceae; Salmonella; Salmonella enterica; Salmonella enterica subsp. enterica MTGDNTLITSHGINRRDFMKLCAALAATMGLSSKAAAEMAESVSNPQRPPVIWIGAQECT WP_000145429.1 hydrogenase 2 small subunit [Salmonella enterica] Length: 372\nNumber of

predicted TMHs:  1\nExp number of AAs in TMHs: 24.51309\nExp number, first 60 AAs:  0.2682\nTotal prob of N-in: 0.14402\noutside   1   331\nTMhelix   332   354\ninside   355   372

20347 GCF_001107845.1_7468_7_87 Salmonella enterica subsp. enterica serovar Typhi Proteobacteria; Gammaproteobacteria; Enterobacterales; Enterobacteriaceae; Salmonella; Salmonella enterica; Salmonella enterica subsp. enterica MTGDNTLITSHGINRRDFMKLCAALAATMGLSSKAAAEMAESVSNPQRPPVIWIGAQECT WP_000145429.1 hydrogenase 2 small subunit [Salmonella enterica] Length: 372\nNumber of predicted TMHs:  1\nExp number of AAs in TMHs: 24.51309\nExp number, first 60 AAs:  0.2682\nTotal prob of N-in: 0.14402\noutside   1   331\nTMhelix   332   354\ninside   355   372

20348 GCF_001105205.1_8490_5_31 Salmonella enterica subsp. enterica serovar Typhi Proteobacteria; Gammaproteobacteria; Enterobacterales; Enterobacteriaceae; Salmonella; Salmonella enterica; Salmonella enterica subsp. enterica MTGDNTLITSHGINRRDFMKLCAALAATMGLSSKAAAEMAESVSNPQRPPVIWIGAQECT WP_000145429.1 hydrogenase 2 small subunit [Salmonella enterica] Length: 372\nNumber of predicted TMHs:  1\nExp number of AAs in TMHs: 24.51309\nExp number, first 60 AAs:  0.2682\nTotal prob of N-in: 0.14402\noutside   1   331\nTMhelix   332   354\ninside   355   372

20349 GCF_001105025.1_8490_6_38 Salmonella enterica subsp. enterica serovar Typhi Proteobacteria; Gammaproteobacteria; Enterobacterales; Enterobacteriaceae; Salmonella; Salmonella enterica; Salmonella enterica subsp. enterica MTGDNTLITSHGINRRDFMKLCAALAATMGLSSKAAAEMAESVSNPQRPPVIWIGAQECT WP_000145429.1 hydrogenase 2 small subunit [Salmonella enterica] Length: 372\nNumber of predicted TMHs:  1\nExp number of AAs in TMHs: 24.51309\nExp number, first 60 AAs:  0.2682\nTotal prob of N-in: 0.14402\noutside   1   331\nTMhelix   332   354\ninside   355   372

20350 GCF_001104925.1_10492_1_31 Salmonella enterica subsp. enterica serovar Typhi Proteobacteria; Gammaproteobacteria; Enterobacterales; Enterobacteriaceae; Salmonella; Salmonella enterica; Salmonella enterica subsp. enterica MTGDNTLITSHGINRRDFMKLCAALAATMGLSSKAAAEMAESVSNPQRPPVIWIGAQECT WP_000145429.1 hydrogenase 2 small subunit [Salmonella enterica] Length: 372\nNumber of predicted TMHs:  1\nExp number of AAs in TMHs: 24.51309\nExp number, first 60 AAs:  0.2682\nTotal prob of N-in: 0.14402\noutside   1   331\nTMhelix   332   354\ninside   355   372

20351 GCF_001104685.1_10493_1_68 Salmonella enterica subsp. enterica serovar Typhi Proteobacteria; Gammaproteobacteria; Enterobacterales; Enterobacteriaceae; Salmonella; Salmonella enterica; Salmonella enterica subsp. enterica MTGDNTLITSHGINRRDFMKLCAALAATMGLSSKAAAEMAESVSNPQRPPVIWIGAQECT WP_000145429.1 hydrogenase 2 small subunit [Salmonella enterica] Length: 372\nNumber of predicted TMHs:  1\nExp number of AAs in TMHs: 24.51309\nExp number, first 60 AAs:  0.2682\nTotal prob of N-in: 0.14402\noutside   1   331\nTMhelix   332   354\ninside   355   372

20352 GCF_001163605.1_8525_2_54 Salmonella enterica subsp. enterica serovar Typhi Proteobacteria; Gammaproteobacteria; Enterobacterales; Enterobacteriaceae; Salmonella; Salmonella enterica; Salmonella enterica subsp. enterica MTGDNTLITSHGINRRDFMKLCAALAATMGLSSKAAAEMAESVSNPQRPPVIWIGAQECT WP_000145429.1 hydrogenase 2 small subunit [Salmonella enterica] Length: 372\nNumber of predicted TMHs:  1\nExp number of AAs in TMHs: 24.51309\nExp number, first 60 AAs:  0.2682\nTotal prob of N-in: 0.14402\noutside   1   331\nTMhelix   332   354\ninside   355   372

20353 GCF_001163925.1_9475_6_59 Salmonella enterica subsp. enterica serovar Typhi Proteobacteria; Gammaproteobacteria; Enterobacterales; Enterobacteriaceae; Salmonella; Salmonella enterica; Salmonella enterica subsp. enterica MTGDNTLITSHGINRRDFMKLCAALAATMGLSSKAAAEMAESVSNPQRPPVIWIGAQECT WP_000145429.1 hydrogenase 2 small subunit [Salmonella enterica] Length: 372\nNumber of predicted TMHs:  1\nExp number of AAs in TMHs: 24.51309\nExp number, first 60 AAs:  0.2682\nTotal prob of N-in: 0.14402\noutside   1   331\nTMhelix   332   354\ninside   355   372

20354 GCF_001164325.1_10593_2_1 Salmonella enterica subsp. enterica serovar Typhi Proteobacteria; Gammaproteobacteria; Enterobacterales; Enterobacteriaceae; Salmonella; Salmonella enterica; Salmonella enterica subsp. enterica MTGDNTLITSHGINRRDFMKLCAALAATMGLSSKAAAEMAESVSNPQRPPVIWIGAQECT WP_000145429.1 hydrogenase 2 small subunit [Salmonella enterica] Length: 372\nNumber of predicted TMHs:  1\nExp number of AAs in TMHs: 24.51309\nExp number, first 60 AAs:  0.2682\nTotal prob of N-in: 0.14402\noutside   1   331\nTMhelix   332   354\ninside   355   372

20355 GCF_001104885.1_10349_1_89 Salmonella enterica subsp. enterica serovar Typhi Proteobacteria; Gammaproteobacteria; Enterobacterales; Enterobacteriaceae; Salmonella; Salmonella enterica;

Salmonella enterica subsp. enterica MTGDNTLITSHGINRRDFMKLCAALAATMGLSSKAAAEMAESVSNPQRPPVIWIGAQECT WP_000145429.1 hydrogenase 2 small subunit [Salmonella enterica] Length: 372\nNumber of predicted TMHs: 1\nExp number of AAs in TMHs: 24.51309\nExp number, first 60 AAs: 0.2682\nTotal prob of N-in: 0.14402\noutside 1 331\nTMhelix 332 354\ninside 355 372

20356 GCF_001104365.1_8490_5_32 Salmonella enterica subsp. enterica serovar Typhi Proteobacteria; Gammaproteobacteria; Enterobacterales; Enterobacteriaceae; Salmonella; Salmonella enterica; Salmonella enterica subsp. enterica MTGDNTLITSHGINRRDFMKLCAALAATMGLSSKAAAEMAESVSNPQRPPVIWIGAQECT WP_000145429.1 hydrogenase 2 small subunit [Salmonella enterica] Length: 372\nNumber of predicted TMHs: 1\nExp number of AAs in TMHs: 24.51309\nExp number, first 60 AAs: 0.2682\nTotal prob of N-in: 0.14402\noutside 1 331\nTMhelix 332 354\ninside 355 372

20357 GCF_001167605.1_10492_1_74 Salmonella enterica subsp. enterica serovar Typhi Proteobacteria; Gammaproteobacteria; Enterobacterales; Enterobacteriaceae; Salmonella; Salmonella enterica; Salmonella enterica subsp. enterica MTGDNTLITSHGINRRDFMKLCAALAATMGLSSKAAAEMAESVSNPQRPPVIWIGAQECT WP_000145429.1 hydrogenase 2 small subunit [Salmonella enterica] Length: 372\nNumber of predicted TMHs: 1\nExp number of AAs in TMHs: 24.51309\nExp number, first 60 AAs: 0.2682\nTotal prob of N-in: 0.14402\noutside 1 331\nTMhelix 332 354\ninside 355 372

20358 GCF_001167865.1_9870_8_94 Salmonella enterica subsp. enterica serovar Typhi Proteobacteria; Gammaproteobacteria; Enterobacterales; Enterobacteriaceae; Salmonella; Salmonella enterica; Salmonella enterica subsp. enterica MTGDNTLITSHGINRRDFMKLCAALAATMGLSSKAAAEMAESVSNPQRPPVIWIGAQECT WP_000145429.1 hydrogenase 2 small subunit [Salmonella enterica] Length: 372\nNumber of predicted TMHs: 1\nExp number of AAs in TMHs: 24.51309\nExp number, first 60 AAs: 0.2682\nTotal prob of N-in: 0.14402\noutside 1 331\nTMhelix 332 354\ninside 355 372

20359 GCF_001168125.1_10592_2_50 Salmonella enterica subsp. enterica serovar Typhi Proteobacteria; Gammaproteobacteria; Enterobacterales; Enterobacteriaceae; Salmonella; Salmonella enterica; Salmonella enterica subsp. enterica MTGDNTLITSHGINRRDFMKLCAALAATMGLSSKAAAEMAESVSNPQRPPVIWIGAQECT WP_000145429.1 hydrogenase 2 small subunit [Salmonella enterica] Length: 372\nNumber of predicted TMHs: 1\nExp number of AAs in TMHs: 24.51309\nExp number, first 60 AAs: 0.2682\nTotal prob of N-in: 0.14402\noutside 1 331\nTMhelix 332 354\ninside 355 372

20360 GCF_001168305.1_8525_3_32 Salmonella enterica subsp. enterica serovar Typhi Proteobacteria; Gammaproteobacteria; Enterobacterales; Enterobacteriaceae; Salmonella; Salmonella enterica; Salmonella enterica subsp. enterica MTGDNTLITSHGINRRDFMKLCAALAATMGLSSKAAAEMAESVSNPQRPPVIWIGAQECT WP_000145429.1 hydrogenase 2 small subunit [Salmonella enterica] Length: 372\nNumber of predicted TMHs: 1\nExp number of AAs in TMHs: 24.51309\nExp number, first 60 AAs: 0.2682\nTotal prob of N-in: 0.14402\noutside 1 331\nTMhelix 332 354\ninside 355 372

20361 GCF_001100225.1_9953_5_26 Salmonella enterica subsp. enterica serovar Typhi Proteobacteria; Gammaproteobacteria; Enterobacterales; Enterobacteriaceae; Salmonella; Salmonella enterica; Salmonella enterica subsp. enterica MTGDNTLITSHGINRRDFMKLCAALAATMGLSSKAAAEMAESVSNPQRPPVIWIGAQECT WP_000145429.1 hydrogenase 2 small subunit [Salmonella enterica] Length: 372\nNumber of predicted TMHs: 1\nExp number of AAs in TMHs: 24.51309\nExp number, first 60 AAs: 0.2682\nTotal prob of N-in: 0.14402\noutside 1 331\nTMhelix 332 354\ninside 355 372

20362 GCF_001168525.1_10562_2_61 Salmonella enterica subsp. enterica serovar Typhi Proteobacteria; Gammaproteobacteria; Enterobacterales; Enterobacteriaceae; Salmonella; Salmonella enterica; Salmonella enterica subsp. enterica MTGDNTLITSHGINRRDFMKLCAALAATMGLSSKAAAEMAESVSNPQRPPVIWIGAQECT WP_000145429.1 hydrogenase 2 small subunit [Salmonella enterica] Length: 372\nNumber of predicted TMHs: 1\nExp number of AAs in TMHs: 24.51309\nExp number, first 60 AAs: 0.2682\nTotal prob of N-in: 0.14402\noutside 1 331\nTMhelix 332 354\ninside 355 372

20363 GCF_001100065.1_10607_1_52 Salmonella enterica subsp. enterica serovar Typhi Proteobacteria; Gammaproteobacteria; Enterobacterales; Enterobacteriaceae; Salmonella; Salmonella enterica; Salmonella enterica subsp. enterica MTGDNTLITSHGINRRDFMKLCAALAATMGLSSKAAAEMAESVSNPQRPPVIWIGAQECT WP_000145429.1 hydrogenase 2 small subunit [Salmonella enterica] Length: 372\nNumber of predicted TMHs: 1\nExp number of AAs in TMHs: 24.51309\nExp number, first 60 AAs: 0.2682\nTotal prob of N-in: 0.14402\noutside 1 331\nTMhelix 332 354\ninside 355 372

20364 GCF_001168785.1_10209_5_50 Salmonella enterica subsp. enterica serovar Typhi Proteobacteria; Gammaproteobacteria; Enterobacterales; Enterobacteriaceae; Salmonella; Salmonella enterica; Salmonella enterica subsp. enterica MTGDNTLITSHGINRRDFMKLCAALAATMGLSSKAAAEMAESVSNPQRPPVIWIGAQECT WP_000145429.1 hydrogenase 2 small subunit [Salmonella enterica] Length: 372\nNumber of predicted TMHs: 1\nExp number of AAs in TMHs: 24.51309\nExp number, first 60 AAs: 0.2682\nTotal prob of N-in: 0.14402\noutside 1 331\nTMhelix 332 354\ninside 355 372

20365 GCF_001173245.1_8490_5_15 Salmonella enterica subsp. enterica serovar Typhi Proteobacteria; Gammaproteobacteria; Enterobacterales; Enterobacteriaceae; Salmonella; Salmonella enterica; Salmonella enterica subsp. enterica MTGDNTLITSHGINRRDFMKLCAALAATMGLSSKAAAEMAESVSNPQRPPVIWIGAQECT WP_000145429.1 hydrogenase 2 small subunit [Salmonella enterica] Length: 372\nNumber of predicted TMHs: 1\nExp number of AAs in TMHs: 24.51309\nExp number, first 60 AAs: 0.2682\nTotal prob of N-in: 0.14402\noutside 1 331\nTMhelix 332 354\ninside 355 372

20366 GCF_001173525.1_10493_1_75 Salmonella enterica subsp. enterica serovar Typhi Proteobacteria; Gammaproteobacteria; Enterobacterales; Enterobacteriaceae; Salmonella; Salmonella enterica; Salmonella enterica subsp. enterica MTGDNTLITSHGINRRDFMKLCAALAATMGLSSKAAAEMAESVSNPQRPPVIWIGAQECT WP_000145429.1 hydrogenase 2 small subunit [Salmonella enterica] Length: 372\nNumber of predicted TMHs: 1\nExp number of AAs in TMHs: 24.51309\nExp number, first 60 AAs: 0.2682\nTotal prob of N-in: 0.14402\noutside 1 331\nTMhelix 332 354\ninside 355 372

20367 GCF_001173745.1_10608_2_48 Salmonella enterica subsp. enterica serovar Typhi Proteobacteria; Gammaproteobacteria; Enterobacterales; Enterobacteriaceae; Salmonella; Salmonella enterica; Salmonella enterica subsp. enterica MTGDNTLITSHGINRRDFMKLCAALAATMGLSSKAAAEMAESVSNPQRPPVIWIGAQECT WP_000145429.1 hydrogenase 2 small subunit [Salmonella enterica] Length: 372\nNumber of predicted TMHs: 1\nExp number of AAs in TMHs: 24.51309\nExp number, first 60 AAs: 0.2682\nTotal prob of N-in: 0.14402\noutside 1 331\nTMhelix 332 354\ninside 355 372

20368 GCF_001099985.1_9870_8_40 Salmonella enterica subsp. enterica serovar Typhi Proteobacteria; Gammaproteobacteria; Enterobacterales; Enterobacteriaceae; Salmonella; Salmonella enterica; Salmonella enterica subsp. enterica MTGDNTLITSHGINRRDFMKLCAALAATMGLSSKAAAEMAESVSNPQRPPVIWIGAQECT WP_000145429.1 hydrogenase 2 small subunit [Salmonella enterica] Length: 372\nNumber of predicted TMHs: 1\nExp number of AAs in TMHs: 24.51309\nExp number, first 60 AAs: 0.2682\nTotal prob of N-in: 0.14402\noutside 1 331\nTMhelix 332 354\ninside 355 372

20369 GCF_001099785.1_10608_2_49 Salmonella enterica subsp. enterica serovar Typhi Proteobacteria; Gammaproteobacteria; Enterobacterales; Enterobacteriaceae; Salmonella; Salmonella enterica; Salmonella enterica subsp. enterica MTGDNTLITSHGINRRDFMKLCAALAATMGLSSKAAAEMAESVSNPQRPPVIWIGAQECT WP_000145429.1 hydrogenase 2 small subunit [Salmonella enterica] Length: 372\nNumber of predicted TMHs: 1\nExp number of AAs in TMHs: 24.51309\nExp number, first 60 AAs: 0.2682\nTotal prob of N-in: 0.14402\noutside 1 331\nTMhelix 332 354\ninside 355 372

20370 GCF_001099705.1_9475_6_78 Salmonella enterica subsp. enterica serovar Typhi Proteobacteria; Gammaproteobacteria; Enterobacterales; Enterobacteriaceae; Salmonella; Salmonella enterica; Salmonella enterica subsp. enterica MTGDNTLITSHGINRRDFMKLCAALAATMGLSSKAAAEMAESVSNPQRPPVIWIGAQECT WP_000145429.1 hydrogenase 2 small subunit [Salmonella enterica] Length: 372\nNumber of predicted TMHs: 1\nExp number of AAs in TMHs: 24.51309\nExp number, first 60 AAs: 0.2682\nTotal prob of N-in: 0.14402\noutside 1 331\nTMhelix 332 354\ninside 355 372

20371 GCF_001174065.1_10562_2_53 Salmonella enterica subsp. enterica serovar Typhi Proteobacteria; Gammaproteobacteria; Enterobacterales; Enterobacteriaceae; Salmonella; Salmonella enterica; Salmonella enterica subsp. enterica MTGDNTLITSHGINRRDFMKLCAALAATMGLSSKAAAEMAESVSNPQRPPVIWIGAQECT WP_000145429.1 hydrogenase 2 small subunit [Salmonella enterica] Length: 372\nNumber of predicted TMHs: 1\nExp number of AAs in TMHs: 24.51309\nExp number, first 60 AAs: 0.2682\nTotal prob of N-in: 0.14402\noutside 1 331\nTMhelix 332 354\ninside 355 372

20372 GCF_001099585.1_10349_1_53 Salmonella enterica subsp. enterica serovar Typhi Proteobacteria; Gammaproteobacteria; Enterobacterales; Enterobacteriaceae; Salmonella; Salmonella enterica; Salmonella enterica subsp. enterica MTGDNTLITSHGINRRDFMKLCAALAATMGLSSKAAAEMAESVSNPQRPPVIWIGAQECT WP_000145429.1 hydrogenase 2 small subunit [Salmonella enterica] Length: 372\nNumber of

predicted TMHs: 1\nExp number of AAs in TMHs: 24.51309\nExp number, first 60 AAs: 0.2682\nTotal prob of N-in: 0.14402\noutside 1 331\nTMhelix 332 354\ninside 355 372

20373 GCF_001099405.1_8525_3_56 Salmonella enterica subsp. enterica serovar Typhi Proteobacteria; Gammaproteobacteria; Enterobacterales; Enterobacteriaceae; Salmonella; Salmonella enterica; Salmonella enterica subsp. enterica MTGDNTLITSHGINRRDFMKLCAALAATMGLSSKAAAEMAESVSNPQRPPVIWIGAQECT WP_000145429.1 hydrogenase 2 small subunit [Salmonella enterica] Length: 372\nNumber of predicted TMHs: 1\nExp number of AAs in TMHs: 24.51309\nExp number, first 60 AAs: 0.2682\nTotal prob of N-in: 0.14402\noutside 1 331\nTMhelix 332 354\ninside 355 372

20374 GCF_001096665.1_10492_1_73 Salmonella enterica subsp. enterica serovar Typhi Proteobacteria; Gammaproteobacteria; Enterobacterales; Enterobacteriaceae; Salmonella; Salmonella enterica; Salmonella enterica subsp. enterica MTGDNTLITSHGINRRDFMKLCAALAATMGLSSKAAAEMAESVSNPQRPPVIWIGAQECT WP_000145429.1 hydrogenase 2 small subunit [Salmonella enterica] Length: 372\nNumber of predicted TMHs: 1\nExp number of AAs in TMHs: 24.51309\nExp number, first 60 AAs: 0.2682\nTotal prob of N-in: 0.14402\noutside 1 331\nTMhelix 332 354\ninside 355 372

20375 GCF_001174285.1_10592_2_24 Salmonella enterica subsp. enterica serovar Typhi Proteobacteria; Gammaproteobacteria; Enterobacterales; Enterobacteriaceae; Salmonella; Salmonella enterica; Salmonella enterica subsp. enterica MTGDNTLITSHGINRRDFMKLCAALAATMGLSSKAAAEMAESVSNPQRPPVIWIGAQECT WP_000145429.1 hydrogenase 2 small subunit [Salmonella enterica] Length: 372\nNumber of predicted TMHs: 1\nExp number of AAs in TMHs: 24.51309\nExp number, first 60 AAs: 0.2682\nTotal prob of N-in: 0.14402\noutside 1 331\nTMhelix 332 354\ninside 355 372

20376 GCF_001096485.1_9475_6_69 Salmonella enterica subsp. enterica serovar Typhi Proteobacteria; Gammaproteobacteria; Enterobacterales; Enterobacteriaceae; Salmonella; Salmonella enterica; Salmonella enterica subsp. enterica MTGDNTLITSHGINRRDFMKLCAALAATMGLSSKAAAEMAESVSNPQRPPVIWIGAQECT WP_000145429.1 hydrogenase 2 small subunit [Salmonella enterica] Length: 372\nNumber of predicted TMHs: 1\nExp number of AAs in TMHs: 24.51309\nExp number, first 60 AAs: 0.2682\nTotal prob of N-in: 0.14402\noutside 1 331\nTMhelix 332 354\ninside 355 372

20377 GCF_002035945.1_ASM203594v1 Salmonella enterica subsp. enterica serovar Typhi Proteobacteria; Gammaproteobacteria; Enterobacterales; Enterobacteriaceae; Salmonella; Salmonella enterica; Salmonella enterica subsp. enterica MTGDNTLITSHGINRRDFMKLCAALAATMGLSSKAAAEMAESVSNPQRPPVIWIGAQECT WP_000145429.1 hydrogenase 2 small subunit [Salmonella enterica] Length: 372\nNumber of predicted TMHs: 1\nExp number of AAs in TMHs: 24.51309\nExp number, first 60 AAs: 0.2682\nTotal prob of N-in: 0.14402\noutside 1 331\nTMhelix 332 354\ninside 355 372

20378 GCF_001096345.1_10541_2_40 Salmonella enterica subsp. enterica serovar Typhi Proteobacteria; Gammaproteobacteria; Enterobacterales; Enterobacteriaceae; Salmonella; Salmonella enterica; Salmonella enterica subsp. enterica MTGDNTLITSHGINRRDFMKLCAALAATMGLSSKAAAEMAESVSNPQRPPVIWIGAQECT WP_000145429.1 hydrogenase 2 small subunit [Salmonella enterica] Length: 372\nNumber of predicted TMHs: 1\nExp number of AAs in TMHs: 24.51309\nExp number, first 60 AAs: 0.2682\nTotal prob of N-in: 0.14402\noutside 1 331\nTMhelix 332 354\ninside 355 372

20379 GCF_001096105.1_10562_2_21 Salmonella enterica subsp. enterica serovar Typhi Proteobacteria; Gammaproteobacteria; Enterobacterales; Enterobacteriaceae; Salmonella; Salmonella enterica; Salmonella enterica subsp. enterica MTGDNTLITSHGINRRDFMKLCAALAATMGLSSKAAAEMAESVSNPQRPPVIWIGAQECT WP_000145429.1 hydrogenase 2 small subunit [Salmonella enterica] Length: 372\nNumber of predicted TMHs: 1\nExp number of AAs in TMHs: 24.51309\nExp number, first 60 AAs: 0.2682\nTotal prob of N-in: 0.14402\noutside 1 331\nTMhelix 332 354\ninside 355 372

20380 GCF_001095705.1_7468_7_80 Salmonella enterica subsp. enterica serovar Typhi Proteobacteria; Gammaproteobacteria; Enterobacterales; Enterobacteriaceae; Salmonella; Salmonella enterica; Salmonella enterica subsp. enterica MTGDNTLITSHGINRRDFMKLCAALAATMGLSSKAAAEMAESVSNPQRPPVIWIGAQECT WP_000145429.1 hydrogenase 2 small subunit [Salmonella enterica] Length: 372\nNumber of predicted TMHs: 1\nExp number of AAs in TMHs: 24.51309\nExp number, first 60 AAs: 0.2682\nTotal prob of N-in: 0.14402\noutside 1 331\nTMhelix 332 354\ninside 355 372

20381 GCF_001092785.1_10607_1_87 Salmonella enterica subsp. enterica serovar Typhi Proteobacteria; Gammaproteobacteria; Enterobacterales; Enterobacteriaceae; Salmonella; Salmonella enterica;

Salmonella enterica subsp. enterica MTGDNTLITSHGINRRDFMKLCAALAATMGLSSKAAAEMAESVSNPQRPPVIWIGAQECT WP_000145429.1 hydrogenase 2 small subunit [Salmonella enterica] Length: 372\nNumber of predicted TMHs: 1\nExp number of AAs in TMHs: 24.51309\nExp number, first 60 AAs: 0.2682\nTotal prob of N-in: 0.14402\noutside 1 331\nTMhelix 332 354\ninside 355 372

20382 GCF_001092745.1_10349_1_75 Salmonella enterica subsp. enterica serovar Typhi Proteobacteria; Gammaproteobacteria; Enterobacterales; Enterobacteriaceae; Salmonella; Salmonella enterica; Salmonella enterica subsp. enterica MTGDNTLITSHGINRRDFMKLCAALAATMGLSSKAAAEMAESVSNPQRPPVIWIGAQECT WP_000145429.1 hydrogenase 2 small subunit [Salmonella enterica] Length: 372\nNumber of predicted TMHs: 1\nExp number of AAs in TMHs: 24.51309\nExp number, first 60 AAs: 0.2682\nTotal prob of N-in: 0.14402\noutside 1 331\nTMhelix 332 354\ninside 355 372

20383 GCF_001092485.1_10562_2_77 Salmonella enterica subsp. enterica serovar Typhi Proteobacteria; Gammaproteobacteria; Enterobacterales; Enterobacteriaceae; Salmonella; Salmonella enterica; Salmonella enterica subsp. enterica MTGDNTLITSHGINRRDFMKLCAALAATMGLSSKAAAEMAESVSNPQRPPVIWIGAQECT WP_000145429.1 hydrogenase 2 small subunit [Salmonella enterica] Length: 372\nNumber of predicted TMHs: 1\nExp number of AAs in TMHs: 24.51309\nExp number, first 60 AAs: 0.2682\nTotal prob of N-in: 0.14402\noutside 1 331\nTMhelix 332 354\ninside 355 372

20384 GCF_001092385.1_7468_7_64 Salmonella enterica subsp. enterica serovar Typhi Proteobacteria; Gammaproteobacteria; Enterobacterales; Enterobacteriaceae; Salmonella; Salmonella enterica; Salmonella enterica subsp. enterica MTGDNTLITSHGINRRDFMKLCAALAATMGLSSKAAAEMAESVSNPQRPPVIWIGAQECT WP_000145429.1 hydrogenase 2 small subunit [Salmonella enterica] Length: 372\nNumber of predicted TMHs: 1\nExp number of AAs in TMHs: 24.51309\nExp number, first 60 AAs: 0.2682\nTotal prob of N-in: 0.14402\noutside 1 331\nTMhelix 332 354\ninside 355 372

20385 GCF_001092205.1_10492_1_1 Salmonella enterica subsp. enterica serovar Typhi Proteobacteria; Gammaproteobacteria; Enterobacterales; Enterobacteriaceae; Salmonella; Salmonella enterica; Salmonella enterica subsp. enterica MTGDNTLITSHGINRRDFMKLCAALAATMGLSSKAAAEMAESVSNPQRPPVIWIGAQECT WP_000145429.1 hydrogenase 2 small subunit [Salmonella enterica] Length: 372\nNumber of predicted TMHs: 1\nExp number of AAs in TMHs: 24.51309\nExp number, first 60 AAs: 0.2682\nTotal prob of N-in: 0.14402\noutside 1 331\nTMhelix 332 354\ninside 355 372

20386 GCF_001091845.1_9475_6_4Salmonella enterica subsp. enterica serovar Typhi Proteobacteria; Gammaproteobacteria; Enterobacterales; Enterobacteriaceae; Salmonella; Salmonella enterica; Salmonella enterica subsp. enterica MTGDNTLITSHGINRRDFMKLCAALAATMGLSSKAAAEMAESVSNPQRPPVIWIGAQECT WP_000145429.1 hydrogenase 2 small subunit [Salmonella enterica] Length: 372\nNumber of predicted TMHs: 1\nExp number of AAs in TMHs: 24.51309\nExp number, first 60 AAs: 0.2682\nTotal prob of N-in: 0.14402\noutside 1 331\nTMhelix 332 354\ninside 355 372

20387 GCF_001091725.1_9870_8_86 Salmonella enterica subsp. enterica serovar Typhi Proteobacteria; Gammaproteobacteria; Enterobacterales; Enterobacteriaceae; Salmonella; Salmonella enterica; Salmonella enterica subsp. enterica MTGDNTLITSHGINRRDFMKLCAALAATMGLSSKAAAEMAESVSNPQRPPVIWIGAQECT WP_000145429.1 hydrogenase 2 small subunit [Salmonella enterica] Length: 372\nNumber of predicted TMHs: 1\nExp number of AAs in TMHs: 24.51309\nExp number, first 60 AAs: 0.2682\nTotal prob of N-in: 0.14402\noutside 1 331\nTMhelix 332 354\ninside 355 372

20388 GCF_001088765.1_10593_2_11 Salmonella enterica subsp. enterica serovar Typhi Proteobacteria; Gammaproteobacteria; Enterobacterales; Enterobacteriaceae; Salmonella; Salmonella enterica; Salmonella enterica subsp. enterica MTGDNTLITSHGINRRDFMKLCAALAATMGLSSKAAAEMAESVSNPQRPPVIWIGAQECT WP_000145429.1 hydrogenase 2 small subunit [Salmonella enterica] Length: 372\nNumber of predicted TMHs: 1\nExp number of AAs in TMHs: 24.51309\nExp number, first 60 AAs: 0.2682\nTotal prob of N-in: 0.14402\noutside 1 331\nTMhelix 332 354\ninside 355 372

20389 GCF_001089345.1_10349_1_50 Salmonella enterica subsp. enterica serovar Typhi Proteobacteria; Gammaproteobacteria; Enterobacterales; Enterobacteriaceae; Salmonella; Salmonella enterica; Salmonella enterica subsp. enterica MTGDNTLITSHGINRRDFMKLCAALAATMGLSSKAAAEMAESVSNPQRPPVIWIGAQECT WP_000145429.1 hydrogenase 2 small subunit [Salmonella enterica] Length: 372\nNumber of predicted TMHs: 1\nExp number of AAs in TMHs: 24.51309\nExp number, first 60 AAs: 0.2682\nTotal prob of N-in: 0.14402\noutside 1 331\nTMhelix 332 354\ninside 355 372

20390 GCF_001088345.1_10349_1_21 Salmonella enterica subsp. enterica serovar Typhi Proteobacteria; Gammaproteobacteria; Enterobacterales; Enterobacteriaceae; Salmonella; Salmonella enterica; Salmonella enterica subsp. enterica MTGDNTLITSHGINRRDFMKLCAALAATMGLSSKAAAEMAESVSNPQRPPVIWIGAQECT WP_000145429.1 hydrogenase 2 small subunit [Salmonella enterica] Length: 372\nNumber of predicted TMHs: 1\nExp number of AAs in TMHs: 24.51309\nExp number, first 60 AAs: 0.2682\nTotal prob of N-in: 0.14402\noutside 1 331\nTMhelix 332 354\ninside 355 372

20391 GCF_001088085.1_10493_1_83 Salmonella enterica subsp. enterica serovar Typhi Proteobacteria; Gammaproteobacteria; Enterobacterales; Enterobacteriaceae; Salmonella; Salmonella enterica; Salmonella enterica subsp. enterica MTGDNTLITSHGINRRDFMKLCAALAATMGLSSKAAAEMAESVSNPQRPPVIWIGAQECT WP_000145429.1 hydrogenase 2 small subunit [Salmonella enterica] Length: 372\nNumber of predicted TMHs: 1\nExp number of AAs in TMHs: 24.51309\nExp number, first 60 AAs: 0.2682\nTotal prob of N-in: 0.14402\noutside 1 331\nTMhelix 332 354\ninside 355 372

20392 GCF_001087985.1_9870_8_63 Salmonella enterica subsp. enterica serovar Typhi Proteobacteria; Gammaproteobacteria; Enterobacterales; Enterobacteriaceae; Salmonella; Salmonella enterica; Salmonella enterica subsp. enterica MTGDNTLITSHGINRRDFMKLCAALAATMGLSSKAAAEMAESVSNPQRPPVIWIGAQECT WP_000145429.1 hydrogenase 2 small subunit [Salmonella enterica] Length: 372\nNumber of predicted TMHs: 1\nExp number of AAs in TMHs: 24.51309\nExp number, first 60 AAs: 0.2682\nTotal prob of N-in: 0.14402\noutside 1 331\nTMhelix 332 354\ninside 355 372

20393 GCF_001087625.1_10541_2_24 Salmonella enterica subsp. enterica serovar Typhi Proteobacteria; Gammaproteobacteria; Enterobacterales; Enterobacteriaceae; Salmonella; Salmonella enterica; Salmonella enterica subsp. enterica MTGDNTLITSHGINRRDFMKLCAALAATMGLSSKAAAEMAESVSNPQRPPVIWIGAQECT WP_000145429.1 hydrogenase 2 small subunit [Salmonella enterica] Length: 372\nNumber of predicted TMHs: 1\nExp number of AAs in TMHs: 24.51309\nExp number, first 60 AAs: 0.2682\nTotal prob of N-in: 0.14402\noutside 1 331\nTMhelix 332 354\ninside 355 372

20394 GCF_001084585.1_10608_2_35 Salmonella enterica subsp. enterica serovar Typhi Proteobacteria; Gammaproteobacteria; Enterobacterales; Enterobacteriaceae; Salmonella; Salmonella enterica; Salmonella enterica subsp. enterica MTGDNTLITSHGINRRDFMKLCAALAATMGLSSKAAAEMAESVSNPQRPPVIWIGAQECT WP_000145429.1 hydrogenase 2 small subunit [Salmonella enterica] Length: 372\nNumber of predicted TMHs: 1\nExp number of AAs in TMHs: 24.51309\nExp number, first 60 AAs: 0.2682\nTotal prob of N-in: 0.14402\noutside 1 331\nTMhelix 332 354\ninside 355 372

20395 GCF_001084325.1_10561_2_59 Salmonella enterica subsp. enterica serovar Typhi Proteobacteria; Gammaproteobacteria; Enterobacterales; Enterobacteriaceae; Salmonella; Salmonella enterica; Salmonella enterica subsp. enterica MTGDNTLITSHGINRRDFMKLCAALAATMGLSSKAAAEMAESVSNPQRPPVIWIGAQECT WP_000145429.1 hydrogenase 2 small subunit [Salmonella enterica] Length: 372\nNumber of predicted TMHs: 1\nExp number of AAs in TMHs: 24.51309\nExp number, first 60 AAs: 0.2682\nTotal prob of N-in: 0.14402\noutside 1 331\nTMhelix 332 354\ninside 355 372

20396 GCF_001084205.1_10607_1_49 Salmonella enterica subsp. enterica serovar Typhi Proteobacteria; Gammaproteobacteria; Enterobacterales; Enterobacteriaceae; Salmonella; Salmonella enterica; Salmonella enterica subsp. enterica MTGDNTLITSHGINRRDFMKLCAALAATMGLSSKAAAEMAESVSNPQRPPVIWIGAQECT WP_000145429.1 hydrogenase 2 small subunit [Salmonella enterica] Length: 372\nNumber of predicted TMHs: 1\nExp number of AAs in TMHs: 24.51309\nExp number, first 60 AAs: 0.2682\nTotal prob of N-in: 0.14402\noutside 1 331\nTMhelix 332 354\ninside 355 372

20397 GCF_001084005.1_9953_5_8Salmonella enterica subsp. enterica serovar Typhi Proteobacteria; Gammaproteobacteria; Enterobacterales; Enterobacteriaceae; Salmonella; Salmonella enterica; Salmonella enterica subsp. enterica MTGDNTLITSHGINRRDFMKLCAALAATMGLSSKAAAEMAESVSNPQRPPVIWIGAQECT WP_000145429.1 hydrogenase 2 small subunit [Salmonella enterica] Length: 372\nNumber of predicted TMHs: 1\nExp number of AAs in TMHs: 24.51309\nExp number, first 60 AAs: 0.2682\nTotal prob of N-in: 0.14402\noutside 1 331\nTMhelix 332 354\ninside 355 372

20398 GCF_001083605.1_10562_2_65 Salmonella enterica subsp. enterica serovar Typhi Proteobacteria; Gammaproteobacteria; Enterobacterales; Enterobacteriaceae; Salmonella; Salmonella enterica; Salmonella enterica subsp. enterica MTGDNTLITSHGINRRDFMKLCAALAATMGLSSKAAAEMAESVSNPQRPPVIWIGAQECT WP_000145429.1 hydrogenase 2 small subunit [Salmonella enterica] Length: 372\nNumber of

predicted TMHs: 1\nExp number of AAs in TMHs: 24.51309\nExp number, first 60 AAs: 0.2682\nTotal prob of N-in: 0.14402\noutside 1 331\nTMhelix 332 354\ninside 355 372

20399 GCF_001083185.1_10608_2_17 Salmonella enterica subsp. enterica serovar Typhi Proteobacteria; Gammaproteobacteria; Enterobacterales; Enterobacteriaceae; Salmonella; Salmonella enterica; Salmonella enterica subsp. enterica MTGDNTLITSHGINRRDFMKLCAALAATMGLSSKAAAEMAESVSNPQRPPVIWIGAQECT WP_000145429.1 hydrogenase 2 small subunit [Salmonella enterica] Length: 372\nNumber of predicted TMHs: 1\nExp number of AAs in TMHs: 24.51309\nExp number, first 60 AAs: 0.2682\nTotal prob of N-in: 0.14402\noutside 1 331\nTMhelix 332 354\ninside 355 372

20400 GCF_001328795.1_7468_7_54 Salmonella enterica subsp. enterica serovar Typhi Proteobacteria; Gammaproteobacteria; Enterobacterales; Enterobacteriaceae; Salmonella; Salmonella enterica; Salmonella enterica subsp. enterica MTGDNTLITSHGINRRDFMKLCAALAATMGLSSKAATEMAESVSNPQRPPVIWIGAQECT WP_001782304.1 hydrogenase 2 small subunit [Salmonella enterica] Length: 372\nNumber of predicted TMHs: 1\nExp number of AAs in TMHs: 24.41173\nExp number, first 60 AAs: 0.16124\nTotal prob of N-in: 0.13947\noutside 1 331\nTMhelix 332 354\ninside 355 372

20401 GCF_001083065.1_10492_1_38 Salmonella enterica subsp. enterica serovar Typhi Proteobacteria; Gammaproteobacteria; Enterobacterales; Enterobacteriaceae; Salmonella; Salmonella enterica; Salmonella enterica subsp. enterica MTGDNTLITSHGINRRDFMKLCAALAATMGLSSKAAAEMAESVSNPQRPPVIWIGAQECT WP_000145429.1 hydrogenase 2 small subunit [Salmonella enterica] Length: 372\nNumber of predicted TMHs: 1\nExp number of AAs in TMHs: 24.51309\nExp number, first 60 AAs: 0.2682\nTotal prob of N-in: 0.14402\noutside 1 331\nTMhelix 332 354\ninside 355 372

20402 GCF_001082465.1_10060_5_23 Salmonella enterica subsp. enterica serovar Typhi Proteobacteria; Gammaproteobacteria; Enterobacterales; Enterobacteriaceae; Salmonella; Salmonella enterica; Salmonella enterica subsp. enterica MTGDNTLITSHGINRRDFMKLCAALAATMGLSSKAAAEMAESVSNPQRPPVIWIGAQECT WP_000145429.1 hydrogenase 2 small subunit [Salmonella enterica] Length: 372\nNumber of predicted TMHs: 1\nExp number of AAs in TMHs: 24.51309\nExp number, first 60 AAs: 0.2682\nTotal prob of N-in: 0.14402\noutside 1 331\nTMhelix 332 354\ninside 355 372

20403 GCF_001240165.1_10425_1_82 Salmonella enterica subsp. enterica serovar Typhi Proteobacteria; Gammaproteobacteria; Enterobacterales; Enterobacteriaceae; Salmonella; Salmonella enterica; Salmonella enterica subsp. enterica MTGDNTLITSHGINRRDFMKLCAALAATMGLSSKAAAEMAESVSNPQRPPVIWIGAQECT WP_000145429.1 hydrogenase 2 small subunit [Salmonella enterica] Length: 372\nNumber of predicted TMHs: 1\nExp number of AAs in TMHs: 24.51309\nExp number, first 60 AAs: 0.2682\nTotal prob of N-in: 0.14402\noutside 1 331\nTMhelix 332 354\ninside 355 372

20404 GCF_001082425.1_10071_8_46 Salmonella enterica subsp. enterica serovar Typhi Proteobacteria; Gammaproteobacteria; Enterobacterales; Enterobacteriaceae; Salmonella; Salmonella enterica; Salmonella enterica subsp. enterica MTGDNTLITSHGINRRDFMKLCAALAATMGLSSKAAAEMAESVSNPQRPPVIWIGAQECT WP_000145429.1 hydrogenase 2 small subunit [Salmonella enterica] Length: 372\nNumber of predicted TMHs: 1\nExp number of AAs in TMHs: 24.51309\nExp number, first 60 AAs: 0.2682\nTotal prob of N-in: 0.14402\noutside 1 331\nTMhelix 332 354\ninside 355 372

20405 GCF_001082345.1_10060_6_76 Salmonella enterica subsp. enterica serovar Typhi Proteobacteria; Gammaproteobacteria; Enterobacterales; Enterobacteriaceae; Salmonella; Salmonella enterica; Salmonella enterica subsp. enterica MTGDNTLITSHGINRRDFMKLCAALAATMGLSSKAAAEMAESVSNPQRPPVIWIGAQECT WP_000145429.1 hydrogenase 2 small subunit [Salmonella enterica] Length: 372\nNumber of predicted TMHs: 1\nExp number of AAs in TMHs: 24.51309\nExp number, first 60 AAs: 0.2682\nTotal prob of N-in: 0.14402\noutside 1 331\nTMhelix 332 354\ninside 355 372

20406 GCF_001082105.1_10071_3_5 Salmonella enterica subsp. enterica serovar Typhi Proteobacteria; Gammaproteobacteria; Enterobacterales; Enterobacteriaceae; Salmonella; Salmonella enterica; Salmonella enterica subsp. enterica MTGDNTLITSHGINRRDFMKLCAALAATMGLSSKAATEMAESVSNPQRPPVIWIGAQECT WP_001782304.1 hydrogenase 2 small subunit [Salmonella enterica] Length: 372\nNumber of predicted TMHs: 1\nExp number of AAs in TMHs: 24.41173\nExp number, first 60 AAs: 0.16124\nTotal prob of N-in: 0.13947\noutside 1 331\nTMhelix 332 354\ninside 355 372

20407 GCF_001082385.1_10071_8_20 Salmonella enterica subsp. enterica serovar Typhi Proteobacteria; Gammaproteobacteria; Enterobacterales; Enterobacteriaceae; Salmonella; Salmonella enterica;

Salmonella enterica subsp. enterica MTGDNTLITSHGINRRDFMKLCAALAATMGLSSKAAAEMAESVSNPQRPPVIWIGAQECT WP_000145429.1 hydrogenase 2 small subunit [Salmonella enterica] Length: 372\nNumber of predicted TMHs: 1\nExp number of AAs in TMHs: 24.51309\nExp number, first 60 AAs: 0.2682\nTotal prob of N-in: 0.14402\noutside 1 331\nTMhelix 332 354\ninside 355 372

20408 GCF_001239745.1_10425_1_21 Salmonella enterica subsp. enterica serovar Typhi Proteobacteria; Gammaproteobacteria; Enterobacterales; Enterobacteriaceae; Salmonella; Salmonella enterica; Salmonella enterica subsp. enterica MTGDNTLITSHGINRRDFMKLCAALAATMGLSSKAAAEMAESVSNPQRPPVIWIGAQECT WP_000145429.1 hydrogenase 2 small subunit [Salmonella enterica] Length: 372\nNumber of predicted TMHs: 1\nExp number of AAs in TMHs: 24.51309\nExp number, first 60 AAs: 0.2682\nTotal prob of N-in: 0.14402\noutside 1 331\nTMhelix 332 354\ninside 355 372

20409 GCF_001239765.1_10425_1_18 Salmonella enterica subsp. enterica serovar Typhi Proteobacteria; Gammaproteobacteria; Enterobacterales; Enterobacteriaceae; Salmonella; Salmonella enterica; Salmonella enterica subsp. enterica MTGDNTLITSHGINRRDFMKLCAALAATMGLSSKAAAEMAESVSNPQRPPVIWIGAQECT WP_000145429.1 hydrogenase 2 small subunit [Salmonella enterica] Length: 372\nNumber of predicted TMHs: 1\nExp number of AAs in TMHs: 24.51309\nExp number, first 60 AAs: 0.2682\nTotal prob of N-in: 0.14402\noutside 1 331\nTMhelix 332 354\ninside 355 372

20410 GCF_001082305.1_10071_8_60 Salmonella enterica subsp. enterica serovar Typhi Proteobacteria; Gammaproteobacteria; Enterobacterales; Enterobacteriaceae; Salmonella; Salmonella enterica; Salmonella enterica subsp. enterica MTGDNTLITSHGINRRDFMKLCAALAATMGLSSKAAAEMAESVSNPQRPPVIWIGAQECT WP_000145429.1 hydrogenase 2 small subunit [Salmonella enterica] Length: 372\nNumber of predicted TMHs: 1\nExp number of AAs in TMHs: 24.51309\nExp number, first 60 AAs: 0.2682\nTotal prob of N-in: 0.14402\noutside 1 331\nTMhelix 332 354\ninside 355 372

20411 GCF_001082265.1_10071_8_89 Salmonella enterica subsp. enterica serovar Typhi Proteobacteria; Gammaproteobacteria; Enterobacterales; Enterobacteriaceae; Salmonella; Salmonella enterica; Salmonella enterica subsp. enterica MTGDNTLITSHGINRRDFMKLCAALAATMGLSSKAAAEMAESVSNPQRPPVIWIGAQECT WP_000145429.1 hydrogenase 2 small subunit [Salmonella enterica] Length: 372\nNumber of predicted TMHs: 1\nExp number of AAs in TMHs: 24.51309\nExp number, first 60 AAs: 0.2682\nTotal prob of N-in: 0.14402\noutside 1 331\nTMhelix 332 354\ninside 355 372

20412 GCF_001081765.1_10071_8_48 Salmonella enterica subsp. enterica serovar Typhi Proteobacteria; Gammaproteobacteria; Enterobacterales; Enterobacteriaceae; Salmonella; Salmonella enterica; Salmonella enterica subsp. enterica MTGDNTLITSHGINRRDFMKLCAALAATMGLSSKAAAEMAESVSNPQRPPVIWIGAQECT WP_000145429.1 hydrogenase 2 small subunit [Salmonella enterica] Length: 372\nNumber of predicted TMHs: 1\nExp number of AAs in TMHs: 24.51309\nExp number, first 60 AAs: 0.2682\nTotal prob of N-in: 0.14402\noutside 1 331\nTMhelix 332 354\ninside 355 372

20413 GCF_001081645.1_10071_3_14 Salmonella enterica subsp. enterica serovar Typhi Proteobacteria; Gammaproteobacteria; Enterobacterales; Enterobacteriaceae; Salmonella; Salmonella enterica; Salmonella enterica subsp. enterica MTGDNTLITSHGINRRDFMKLCAALAATMGLSSKAAAEMAESVSNPQRPPVIWIGAQECT WP_000145429.1 hydrogenase 2 small subunit [Salmonella enterica] Length: 372\nNumber of predicted TMHs: 1\nExp number of AAs in TMHs: 24.51309\nExp number, first 60 AAs: 0.2682\nTotal prob of N-in: 0.14402\noutside 1 331\nTMhelix 332 354\ninside 355 372

20414 GCF_001081565.1_10060_6_85 Salmonella enterica subsp. enterica serovar Typhi Proteobacteria; Gammaproteobacteria; Enterobacterales; Enterobacteriaceae; Salmonella; Salmonella enterica; Salmonella enterica subsp. enterica MTGDNTLITSHGINRRDFMKLCAALAATMGLSSKAAAEMAESVSNPQRPPVIWIGAQECT WP_000145429.1 hydrogenase 2 small subunit [Salmonella enterica] Length: 372\nNumber of predicted TMHs: 1\nExp number of AAs in TMHs: 24.51309\nExp number, first 60 AAs: 0.2682\nTotal prob of N-in: 0.14402\noutside 1 331\nTMhelix 332 354\ninside 355 372

20415 GCF_001081525.1_10060_6_36 Salmonella enterica subsp. enterica serovar Typhi Proteobacteria; Gammaproteobacteria; Enterobacterales; Enterobacteriaceae; Salmonella; Salmonella enterica; Salmonella enterica subsp. enterica MTGDNTLITSHGINRRDFMKLCAALAATMGLSSKAAAEMAESVSNPQRPPVIWIGAQECT WP_000145429.1 hydrogenase 2 small subunit [Salmonella enterica] Length: 372\nNumber of predicted TMHs: 1\nExp number of AAs in TMHs: 24.51309\nExp number, first 60 AAs: 0.2682\nTotal prob of N-in: 0.14402\noutside 1 331\nTMhelix 332 354\ninside 355 372

20416 GCF_001081605.1_10071_8_19 Salmonella enterica subsp. enterica serovar Typhi
Proteobacteria; Gammaproteobacteria; Enterobacterales; Enterobacteriaceae; Salmonella; Salmonella enterica; Salmonella enterica subsp. enterica MTGDNTLITSHGINRRDFMKLCAALAATMGLSSKAAAEMAESVSNPQRPPVIWIGAQECT WP_000145429.1 hydrogenase 2 small subunit [Salmonella enterica] Length: 372\nNumber of predicted TMHs: 1\nExp number of AAs in TMHs: 24.51309\nExp number, first 60 AAs: 0.2682\nTotal prob of N-in: 0.14402\noutside 1 331\nTMhelix 332 354\ninside 355 372

20417 GCF_001081485.1_10060_6_44 Salmonella enterica subsp. enterica serovar Typhi
Proteobacteria; Gammaproteobacteria; Enterobacterales; Enterobacteriaceae; Salmonella; Salmonella enterica; Salmonella enterica subsp. enterica MTGDNTLITSHGINRRDFMKLCAALAATMGLSSKAAAEMAESVSNPQRPPVIWIGAQECT WP_000145429.1 hydrogenase 2 small subunit [Salmonella enterica] Length: 372\nNumber of predicted TMHs: 1\nExp number of AAs in TMHs: 24.51309\nExp number, first 60 AAs: 0.2682\nTotal prob of N-in: 0.14402\noutside 1 331\nTMhelix 332 354\ninside 355 372

20418 GCF_001081405.1_10071_3_37 Salmonella enterica subsp. enterica serovar Typhi
Proteobacteria; Gammaproteobacteria; Enterobacterales; Enterobacteriaceae; Salmonella; Salmonella enterica; Salmonella enterica subsp. enterica MTGDNTLITSHGINRRDFMKLCAALAATMGLSSKAAAEMAESVSNPQRPPVIWIGAQECT WP_000145429.1 hydrogenase 2 small subunit [Salmonella enterica] Length: 372\nNumber of predicted TMHs: 1\nExp number of AAs in TMHs: 24.51309\nExp number, first 60 AAs: 0.2682\nTotal prob of N-in: 0.14402\noutside 1 331\nTMhelix 332 354\ninside 355 372

20419 GCF_001080905.1_10071_8_16 Salmonella enterica subsp. enterica serovar Typhi
Proteobacteria; Gammaproteobacteria; Enterobacterales; Enterobacteriaceae; Salmonella; Salmonella enterica; Salmonella enterica subsp. enterica MTGDNTLITSHGINRRDFMKLCAALAATMGLSSKAAAEMAESVSNPQRPPVIWIGAQECT WP_000145429.1 hydrogenase 2 small subunit [Salmonella enterica] Length: 372\nNumber of predicted TMHs: 1\nExp number of AAs in TMHs: 24.51309\nExp number, first 60 AAs: 0.2682\nTotal prob of N-in: 0.14402\noutside 1 331\nTMhelix 332 354\ninside 355 372

20420 GCF_001080865.1_10060_6_33 Salmonella enterica subsp. enterica serovar Typhi
Proteobacteria; Gammaproteobacteria; Enterobacterales; Enterobacteriaceae; Salmonella; Salmonella enterica; Salmonella enterica subsp. enterica MTGDNTLITSHGINRRDFMKLCAALAATMGLSSKAAAEMAESVSNPQRPPVIWIGAQECT WP_000145429.1 hydrogenase 2 small subunit [Salmonella enterica] Length: 372\nNumber of predicted TMHs: 1\nExp number of AAs in TMHs: 24.51309\nExp number, first 60 AAs: 0.2682\nTotal prob of N-in: 0.14402\noutside 1 331\nTMhelix 332 354\ninside 355 372

20421 GCF_001080785.1_10071_8_66 Salmonella enterica subsp. enterica serovar Typhi
Proteobacteria; Gammaproteobacteria; Enterobacterales; Enterobacteriaceae; Salmonella; Salmonella enterica; Salmonella enterica subsp. enterica MTGDNTLITSHGINRRDFMKLCAALAATMGLSSKAAAEMAESVSNPQRPPVIWIGAQECT WP_000145429.1 hydrogenase 2 small subunit [Salmonella enterica] Length: 372\nNumber of predicted TMHs: 1\nExp number of AAs in TMHs: 24.51309\nExp number, first 60 AAs: 0.2682\nTotal prob of N-in: 0.14402\noutside 1 331\nTMhelix 332 354\ninside 355 372

20422 GCF_001080705.1_10071_3_10 Salmonella enterica subsp. enterica serovar Typhi
Proteobacteria; Gammaproteobacteria; Enterobacterales; Enterobacteriaceae; Salmonella; Salmonella enterica; Salmonella enterica subsp. enterica MTGDNTLITSHGINRRDFMKLCAALAATMGLSSKAAAEMAESVSNPQRPPVIWIGAQECT WP_000145429.1 hydrogenase 2 small subunit [Salmonella enterica] Length: 372\nNumber of predicted TMHs: 1\nExp number of AAs in TMHs: 24.51309\nExp number, first 60 AAs: 0.2682\nTotal prob of N-in: 0.14402\noutside 1 331\nTMhelix 332 354\ninside 355 372

20423 GCF_001080745.1_10071_3_89 Salmonella enterica subsp. enterica serovar Typhi
Proteobacteria; Gammaproteobacteria; Enterobacterales; Enterobacteriaceae; Salmonella; Salmonella enterica; Salmonella enterica subsp. enterica MTGDNTLITSHGINRRDFMKLCAALAATMGLSSKAAAEMAESVSNPQRPPVIWIGAQECT WP_000145429.1 hydrogenase 2 small subunit [Salmonella enterica] Length: 372\nNumber of predicted TMHs: 1\nExp number of AAs in TMHs: 24.51309\nExp number, first 60 AAs: 0.2682\nTotal prob of N-in: 0.14402\noutside 1 331\nTMhelix 332 354\ninside 355 372

20424 GCF_001080825.1_10071_8_34 Salmonella enterica subsp. enterica serovar Typhi
Proteobacteria; Gammaproteobacteria; Enterobacterales; Enterobacteriaceae; Salmonella; Salmonella enterica; Salmonella enterica subsp. enterica MTGDNTLITSHGINRRDFMKLCAALAATMGLSSKAAAEMAESVSNPQRPPVIWIGAQECT WP_000145429.1 hydrogenase 2 small subunit [Salmonella enterica] Length: 372\nNumber of

predicted TMHs:  1\nExp number of AAs in TMHs: 24.51309\nExp number, first 60 AAs:  0.2682\nTotal prob of N-in: 0.14402\noutside   1   331\nTMhelix   332   354\ninside   355   372

20425 GCF_001080665.1_10071_8_55 Salmonella enterica subsp. enterica serovar Typhi Proteobacteria; Gammaproteobacteria; Enterobacterales; Enterobacteriaceae; Salmonella; Salmonella enterica; Salmonella enterica subsp. enterica MTGDNTLITSHGINRRDFMKLCAALAATMGLSSKAAAEMAESVSNPQRPPVIWIGAQECT WP_000145429.1 hydrogenase 2 small subunit [Salmonella enterica] Length: 372\nNumber of predicted TMHs:  1\nExp number of AAs in TMHs: 24.51309\nExp number, first 60 AAs:  0.2682\nTotal prob of N-in: 0.14402\noutside   1   331\nTMhelix   332   354\ninside   355   372

20426 GCF_001080125.1_10071_8_40 Salmonella enterica subsp. enterica serovar Typhi Proteobacteria; Gammaproteobacteria; Enterobacterales; Enterobacteriaceae; Salmonella; Salmonella enterica; Salmonella enterica subsp. enterica MTGDNTLITSHGINRRDFMKLCAALAATMGLSSKAAAEMAESVSNPQRPPVIWIGAQECT WP_000145429.1 hydrogenase 2 small subunit [Salmonella enterica] Length: 372\nNumber of predicted TMHs:  1\nExp number of AAs in TMHs: 24.51309\nExp number, first 60 AAs:  0.2682\nTotal prob of N-in: 0.14402\noutside   1   331\nTMhelix   332   354\ninside   355   372

20427 GCF_001080085.1_10071_8_12 Salmonella enterica subsp. enterica serovar Typhi Proteobacteria; Gammaproteobacteria; Enterobacterales; Enterobacteriaceae; Salmonella; Salmonella enterica; Salmonella enterica subsp. enterica MTGDNTLITSHGINRRDFMKLCAALAATMGLSSKAAAEMAESVSNPQRPPVIWIGAQECT WP_000145429.1 hydrogenase 2 small subunit [Salmonella enterica] Length: 372\nNumber of predicted TMHs:  1\nExp number of AAs in TMHs: 24.51309\nExp number, first 60 AAs:  0.2682\nTotal prob of N-in: 0.14402\noutside   1   331\nTMhelix   332   354\ninside   355   372

20428 GCF_001080045.1_10071_8_39 Salmonella enterica subsp. enterica serovar Typhi Proteobacteria; Gammaproteobacteria; Enterobacterales; Enterobacteriaceae; Salmonella; Salmonella enterica; Salmonella enterica subsp. enterica MTGDNTLITSHGINRRDFMKLCAALAATMGLSSKAAAEMAESVSNPQRPPVIWIGAQECT WP_000145429.1 hydrogenase 2 small subunit [Salmonella enterica] Length: 372\nNumber of predicted TMHs:  1\nExp number of AAs in TMHs: 24.51309\nExp number, first 60 AAs:  0.2682\nTotal prob of N-in: 0.14402\noutside   1   331\nTMhelix   332   354\ninside   355   372

20429 GCF_001080005.1_10071_8_28 Salmonella enterica subsp. enterica serovar Typhi Proteobacteria; Gammaproteobacteria; Enterobacterales; Enterobacteriaceae; Salmonella; Salmonella enterica; Salmonella enterica subsp. enterica MTGDNTLITSHGINRRDFMKLCAALAATMGLSSKAAAEMAESVSNPQRPPVIWIGAQECT WP_000145429.1 hydrogenase 2 small subunit [Salmonella enterica] Length: 372\nNumber of predicted TMHs:  1\nExp number of AAs in TMHs: 24.51309\nExp number, first 60 AAs:  0.2682\nTotal prob of N-in: 0.14402\noutside   1   331\nTMhelix   332   354\ninside   355   372

20430 GCF_001079625.1_10071_8_84 Salmonella enterica subsp. enterica serovar Typhi Proteobacteria; Gammaproteobacteria; Enterobacterales; Enterobacteriaceae; Salmonella; Salmonella enterica; Salmonella enterica subsp. enterica MTGDNTLITSHGINRRDFMKLCAALAATMGLSSKAATEMAESVSNPQRPPVIWIGAQECT WP_001782304.1 hydrogenase 2 small subunit [Salmonella enterica] Length: 372\nNumber of predicted TMHs:  1\nExp number of AAs in TMHs: 24.41173\nExp number, first 60 AAs:  0.16124\nTotal prob of N-in: 0.13947\noutside   1   331\nTMhelix   332   354\ninside   355   372

20431 GCF_001079965.1_10071_8_3 Salmonella enterica subsp. enterica serovar Typhi Proteobacteria; Gammaproteobacteria; Enterobacterales; Enterobacteriaceae; Salmonella; Salmonella enterica; Salmonella enterica subsp. enterica MTGDNTLITSHGINRRDFMKLCAALAATMGLSSKAAAEMAESVSNPQRPPVIWIGAQECT WP_000145429.1 hydrogenase 2 small subunit [Salmonella enterica] Length: 372\nNumber of predicted TMHs:  1\nExp number of AAs in TMHs: 24.51309\nExp number, first 60 AAs:  0.2682\nTotal prob of N-in: 0.14402\noutside   1   331\nTMhelix   332   354\ninside   355   372

20432 GCF_001079385.1_10071_3_22 Salmonella enterica subsp. enterica serovar Typhi Proteobacteria; Gammaproteobacteria; Enterobacterales; Enterobacteriaceae; Salmonella; Salmonella enterica; Salmonella enterica subsp. enterica MTGDNTLITSHGINRRDFMKLCAALAATMGLSSKAAAEMAESVSNPQRPPVIWIGAQECT WP_000145429.1 hydrogenase 2 small subunit [Salmonella enterica] Length: 372\nNumber of predicted TMHs:  1\nExp number of AAs in TMHs: 24.51309\nExp number, first 60 AAs:  0.2682\nTotal prob of N-in: 0.14402\noutside   1   331\nTMhelix   332   354\ninside   355   372

20433 GCF_001597665.1_ASM159766v1 Salmonella enterica subsp. enterica serovar Typhi Proteobacteria; Gammaproteobacteria; Enterobacterales; Enterobacteriaceae; Salmonella; Salmonella enterica;

Salmonella enterica subsp. enterica MTGDNTLITSHGINRRDFMKLCAALAATMGLSSKAAAEMAESVSNPQRPPVIWIGAQECT WP_000145429.1 hydrogenase 2 small subunit [Salmonella enterica] Length: 372\nNumber of predicted TMHs: 1\nExp number of AAs in TMHs: 24.51309\nExp number, first 60 AAs: 0.2682\nTotal prob of N-in: 0.14402\noutside 1 331\nTMhelix 332 354\ninside 355 372

20434 GCF_001079345.1_10071_3_74 Salmonella enterica subsp. enterica serovar Typhi Proteobacteria; Gammaproteobacteria; Enterobacterales; Enterobacteriaceae; Salmonella; Salmonella enterica; Salmonella enterica subsp. enterica MTGDNTLITSHGINRRDFMKLCAALAATMGLSSKAAAEMAESVSNPQRPPVIWIGAQECT WP_000145429.1 hydrogenase 2 small subunit [Salmonella enterica] Length: 372\nNumber of predicted TMHs: 1\nExp number of AAs in TMHs: 24.51309\nExp number, first 60 AAs: 0.2682\nTotal prob of N-in: 0.14402\noutside 1 331\nTMhelix 332 354\ninside 355 372

20435 GCF_001079305.1_10071_3_21 Salmonella enterica subsp. enterica serovar Typhi Proteobacteria; Gammaproteobacteria; Enterobacterales; Enterobacteriaceae; Salmonella; Salmonella enterica; Salmonella enterica subsp. enterica MTGDNTLITSHGINRRDFMKLCAALAATMGLSSKAAAEMAESVSNPQRPPVIWIGAQECT WP_000145429.1 hydrogenase 2 small subunit [Salmonella enterica] Length: 372\nNumber of predicted TMHs: 1\nExp number of AAs in TMHs: 24.51309\nExp number, first 60 AAs: 0.2682\nTotal prob of N-in: 0.14402\noutside 1 331\nTMhelix 332 354\ninside 355 372

20436 GCF_001079265.1_10071_8_88 Salmonella enterica subsp. enterica serovar Typhi Proteobacteria; Gammaproteobacteria; Enterobacterales; Enterobacteriaceae; Salmonella; Salmonella enterica; Salmonella enterica subsp. enterica MTGDNTLITSHGINRRDFMKLCAALAATMGLSSKAAAEMAESVSNPQRPPVIWIGAQECT WP_000145429.1 hydrogenase 2 small subunit [Salmonella enterica] Length: 372\nNumber of predicted TMHs: 1\nExp number of AAs in TMHs: 24.51309\nExp number, first 60 AAs: 0.2682\nTotal prob of N-in: 0.14402\noutside 1 331\nTMhelix 332 354\ninside 355 372

20437 GCF_001079185.1_10060_5_36 Salmonella enterica subsp. enterica serovar Typhi Proteobacteria; Gammaproteobacteria; Enterobacterales; Enterobacteriaceae; Salmonella; Salmonella enterica; Salmonella enterica subsp. enterica MTGDNTLITSHGINRRDFMKLCAALAATMGLSSKAAAEMAESVSNPQRPPVIWIGAQECT WP_000145429.1 hydrogenase 2 small subunit [Salmonella enterica] Length: 372\nNumber of predicted TMHs: 1\nExp number of AAs in TMHs: 24.51309\nExp number, first 60 AAs: 0.2682\nTotal prob of N-in: 0.14402\noutside 1 331\nTMhelix 332 354\ninside 355 372

20438 GCF_001079145.1_10060_6_56 Salmonella enterica subsp. enterica serovar Typhi Proteobacteria; Gammaproteobacteria; Enterobacterales; Enterobacteriaceae; Salmonella; Salmonella enterica; Salmonella enterica subsp. enterica MTGDNTLITSHGINRRDFMKLCAALAATMGLSSKAAAEMAESVSNPQRPPVIWIGAQECT WP_000145429.1 hydrogenase 2 small subunit [Salmonella enterica] Length: 372\nNumber of predicted TMHs: 1\nExp number of AAs in TMHs: 24.51309\nExp number, first 60 AAs: 0.2682\nTotal prob of N-in: 0.14402\noutside 1 331\nTMhelix 332 354\ninside 355 372

20439 GCF_001239825.1_10425_1_70 Salmonella enterica subsp. enterica serovar Typhi Proteobacteria; Gammaproteobacteria; Enterobacterales; Enterobacteriaceae; Salmonella; Salmonella enterica; Salmonella enterica subsp. enterica MTGDNTLITSHGINRRDFMKLCAALAATMGLSSKAAAEMAESVSNPQRPPVIWIGAQECT WP_000145429.1 hydrogenase 2 small subunit [Salmonella enterica] Length: 372\nNumber of predicted TMHs: 1\nExp number of AAs in TMHs: 24.51309\nExp number, first 60 AAs: 0.2682\nTotal prob of N-in: 0.14402\noutside 1 331\nTMhelix 332 354\ninside 355 372

20440 GCF_001047995.1_10349_1_3 Salmonella enterica subsp. enterica serovar Typhi Proteobacteria; Gammaproteobacteria; Enterobacterales; Enterobacteriaceae; Salmonella; Salmonella enterica; Salmonella enterica subsp. enterica MTGDNTLITSHGINRRDFMKLCAALAATMGLSSKAAAEMAESVSNPQRPPVIWIGAQECT WP_000145429.1 hydrogenase 2 small subunit [Salmonella enterica] Length: 372\nNumber of predicted TMHs: 1\nExp number of AAs in TMHs: 24.51309\nExp number, first 60 AAs: 0.2682\nTotal prob of N-in: 0.14402\noutside 1 331\nTMhelix 332 354\ninside 355 372

20441 GCF_001047955.1_10349_1_22 Salmonella enterica subsp. enterica serovar Typhi Proteobacteria; Gammaproteobacteria; Enterobacterales; Enterobacteriaceae; Salmonella; Salmonella enterica; Salmonella enterica subsp. enterica MTGDNTLITSHGINRRDFMKLCAALAATMGLSSKAAAEMAESVSNPQRPPVIWIGAQECT WP_000145429.1 hydrogenase 2 small subunit [Salmonella enterica] Length: 372\nNumber of predicted TMHs: 1\nExp number of AAs in TMHs: 24.51309\nExp number, first 60 AAs: 0.2682\nTotal prob of N-in: 0.14402\noutside 1 331\nTMhelix 332 354\ninside 355 372

20442 GCF_001239905.1_10425_1_93 Salmonella enterica subsp. enterica serovar Typhi Proteobacteria; Gammaproteobacteria; Enterobacterales; Enterobacteriaceae; Salmonella; Salmonella enterica; Salmonella enterica subsp. enterica MTGDNTLITSHGINRRDFMKLCAALAATMGLSSKAAAEMAESVSNPQRPPVIWIGAQECT WP_000145429.1 hydrogenase 2 small subunit [Salmonella enterica] Length: 372\nNumber of predicted TMHs: 1\nExp number of AAs in TMHs: 24.51309\nExp number, first 60 AAs: 0.2682\nTotal prob of N-in: 0.14402\noutside 1 331\nTMhelix 332 354\ninside 355 372

20443 GCF_001240665.1_10425_1_15 Salmonella enterica subsp. enterica serovar Typhi Proteobacteria; Gammaproteobacteria; Enterobacterales; Enterobacteriaceae; Salmonella; Salmonella enterica; Salmonella enterica subsp. enterica MTGDNTLITSHGINRRDFMKLCAALAATMGLSSKAAAEMAESVSNPQRPPVIWIGAQECT WP_000145429.1 hydrogenase 2 small subunit [Salmonella enterica] Length: 372\nNumber of predicted TMHs: 1\nExp number of AAs in TMHs: 24.51309\nExp number, first 60 AAs: 0.2682\nTotal prob of N-in: 0.14402\noutside 1 331\nTMhelix 332 354\ninside 355 372

20444 GCF_001047915.1_10593_2_12 Salmonella enterica subsp. enterica serovar Typhi Proteobacteria; Gammaproteobacteria; Enterobacterales; Enterobacteriaceae; Salmonella; Salmonella enterica; Salmonella enterica subsp. enterica MTGDNTLITSHGINRRDFMKLCAALAATMGLSSKAAAEMAESVSNPQRPPVIWIGAQECT WP_000145429.1 hydrogenase 2 small subunit [Salmonella enterica] Length: 372\nNumber of predicted TMHs: 1\nExp number of AAs in TMHs: 24.51309\nExp number, first 60 AAs: 0.2682\nTotal prob of N-in: 0.14402\noutside 1 331\nTMhelix 332 354\ninside 355 372

20445 GCF_001240985.1_10425_1_54 Salmonella enterica subsp. enterica serovar Typhi Proteobacteria; Gammaproteobacteria; Enterobacterales; Enterobacteriaceae; Salmonella; Salmonella enterica; Salmonella enterica subsp. enterica MTGDNTLITSHGINRRDFMKLCAALAATMGLSSKAAAEMAESVSNPQRPPVIWIGAQECT WP_000145429.1 hydrogenase 2 small subunit [Salmonella enterica] Length: 372\nNumber of predicted TMHs: 1\nExp number of AAs in TMHs: 24.51309\nExp number, first 60 AAs: 0.2682\nTotal prob of N-in: 0.14402\noutside 1 331\nTMhelix 332 354\ninside 355 372

20446 GCF_001241045.1_10425_1_19 Salmonella enterica subsp. enterica serovar Typhi Proteobacteria; Gammaproteobacteria; Enterobacterales; Enterobacteriaceae; Salmonella; Salmonella enterica; Salmonella enterica subsp. enterica MTGDNTLITSHGINRRDFMKLCAALAATMGLSSKAAAEMAESVSNPQRPPVIWIGAQECT WP_000145429.1 hydrogenase 2 small subunit [Salmonella enterica] Length: 372\nNumber of predicted TMHs: 1\nExp number of AAs in TMHs: 24.51309\nExp number, first 60 AAs: 0.2682\nTotal prob of N-in: 0.14402\noutside 1 331\nTMhelix 332 354\ninside 355 372

20447 GCF_001241615.1_10425_1_29 Salmonella enterica subsp. enterica serovar Typhi Proteobacteria; Gammaproteobacteria; Enterobacterales; Enterobacteriaceae; Salmonella; Salmonella enterica; Salmonella enterica subsp. enterica MTGDNTLITSHGINRRDFMKLCAALAATMGLSSKAAAEMAESVSNPQRPPVIWIGAQECT WP_000145429.1 hydrogenase 2 small subunit [Salmonella enterica] Length: 372\nNumber of predicted TMHs: 1\nExp number of AAs in TMHs: 24.51309\nExp number, first 60 AAs: 0.2682\nTotal prob of N-in: 0.14402\noutside 1 331\nTMhelix 332 354\ninside 355 372

20448 GCF_001242105.1_10425_1_62 Salmonella enterica subsp. enterica serovar Typhi Proteobacteria; Gammaproteobacteria; Enterobacterales; Enterobacteriaceae; Salmonella; Salmonella enterica; Salmonella enterica subsp. enterica MTGDNTLITSHGINRRDFMKLCAALAATMGLSSKAAAEMAESVSNPQRPPVIWIGAQECT WP_000145429.1 hydrogenase 2 small subunit [Salmonella enterica] Length: 372\nNumber of predicted TMHs: 1\nExp number of AAs in TMHs: 24.51309\nExp number, first 60 AAs: 0.2682\nTotal prob of N-in: 0.14402\noutside 1 331\nTMhelix 332 354\ninside 355 372

20449 GCF_001365495.1_10607_2_21 Salmonella enterica subsp. enterica serovar Typhi Proteobacteria; Gammaproteobacteria; Enterobacterales; Enterobacteriaceae; Salmonella; Salmonella enterica; Salmonella enterica subsp. enterica MTGDNTLITSHGINRRDFMKLCAALAATMGLSSKAAAEMAESVSNPQRPPVIWIGAQECT WP_000145429.1 hydrogenase 2 small subunit [Salmonella enterica] Length: 372\nNumber of predicted TMHs: 1\nExp number of AAs in TMHs: 24.51309\nExp number, first 60 AAs: 0.2682\nTotal prob of N-in: 0.14402\noutside 1 331\nTMhelix 332 354\ninside 355 372

20450 GCF_001365455.1_10562_2_47 Salmonella enterica subsp. enterica serovar Typhi Proteobacteria; Gammaproteobacteria; Enterobacterales; Enterobacteriaceae; Salmonella; Salmonella enterica; Salmonella enterica subsp. enterica MTGDNTLITSHGINRRDFMKLCAALAATMGLSSKAAAEMAESVSNPQRPPVIWIGAQECT WP_000145429.1 hydrogenase 2 small subunit [Salmonella enterica] Length: 372\nNumber of

predicted TMHs:  1\nExp number of AAs in TMHs: 24.51309\nExp number, first 60 AAs:  0.2682\nTotal prob of N-in: 0.14402\noutside   1   331\nTMhelix   332   354\ninside   355   372

20451 GCF_001365415.1_10426_1_11 Salmonella enterica subsp. enterica serovar Typhi Proteobacteria; Gammaproteobacteria; Enterobacterales; Enterobacteriaceae; Salmonella; Salmonella enterica; Salmonella enterica subsp. enterica MTGDNTLITSHGINRRDFMKLCAALAATMGLSSKAAAEMAESVSNPQRPPVIWIGAQECT WP_000145429.1 hydrogenase 2 small subunit [Salmonella enterica] Length: 372\nNumber of predicted TMHs:  1\nExp number of AAs in TMHs: 24.51309\nExp number, first 60 AAs:  0.2682\nTotal prob of N-in: 0.14402\noutside   1   331\nTMhelix   332   354\ninside   355   372

20452 GCF_001365375.1_10608_2_16 Salmonella enterica subsp. enterica serovar Typhi Proteobacteria; Gammaproteobacteria; Enterobacterales; Enterobacteriaceae; Salmonella; Salmonella enterica; Salmonella enterica subsp. enterica MTGDNTLITSHGINRRDFMKLCAALAATMGLSSKAAAEMAESVSNPQRPPVIWIGAQECT WP_000145429.1 hydrogenase 2 small subunit [Salmonella enterica] Length: 372\nNumber of predicted TMHs:  1\nExp number of AAs in TMHs: 24.51309\nExp number, first 60 AAs:  0.2682\nTotal prob of N-in: 0.14402\noutside   1   331\nTMhelix   332   354\ninside   355   372

20453 GCF_001365295.1_10540_1_65 Salmonella enterica subsp. enterica serovar Typhi Proteobacteria; Gammaproteobacteria; Enterobacterales; Enterobacteriaceae; Salmonella; Salmonella enterica; Salmonella enterica subsp. enterica MTGDNTLITSHGINRRDFMKLCAALAATMGLSSKAAAEMAESVSNPQRPPVIWIGAQECT WP_000145429.1 hydrogenase 2 small subunit [Salmonella enterica] Length: 372\nNumber of predicted TMHs:  1\nExp number of AAs in TMHs: 24.51309\nExp number, first 60 AAs:  0.2682\nTotal prob of N-in: 0.14402\noutside   1   331\nTMhelix   332   354\ninside   355   372

20454 GCF_001365335.1_10493_1_61 Salmonella enterica subsp. enterica serovar Typhi Proteobacteria; Gammaproteobacteria; Enterobacterales; Enterobacteriaceae; Salmonella; Salmonella enterica; Salmonella enterica subsp. enterica MTGDNTLITSHGINRRDFMKLCAALAATMGLSSKAAAEMAESVSNPQRPPVIWIGAQECT WP_000145429.1 hydrogenase 2 small subunit [Salmonella enterica] Length: 372\nNumber of predicted TMHs:  1\nExp number of AAs in TMHs: 24.51309\nExp number, first 60 AAs:  0.2682\nTotal prob of N-in: 0.14402\noutside   1   331\nTMhelix   332   354\ninside   355   372

20455 GCF_001365255.1_10426_1_58 Salmonella enterica subsp. enterica serovar Typhi Proteobacteria; Gammaproteobacteria; Enterobacterales; Enterobacteriaceae; Salmonella; Salmonella enterica; Salmonella enterica subsp. enterica MTGDNTLITSHGINRRDFMKLCAALAATMGLSSKAAAEMAESVSNPQRPPVIWIGAQECT WP_000145429.1 hydrogenase 2 small subunit [Salmonella enterica] Length: 372\nNumber of predicted TMHs:  1\nExp number of AAs in TMHs: 24.51309\nExp number, first 60 AAs:  0.2682\nTotal prob of N-in: 0.14402\noutside   1   331\nTMhelix   332   354\ninside   355   372

20456 GCF_001364715.1_10540_1_88 Salmonella enterica subsp. enterica serovar Typhi Proteobacteria; Gammaproteobacteria; Enterobacterales; Enterobacteriaceae; Salmonella; Salmonella enterica; Salmonella enterica subsp. enterica MTGDNTLITSHGINRRDFMKLCAALAATMGLSSKAAAEMAESVSNPQRPPVIWIGAQECT WP_000145429.1 hydrogenase 2 small subunit [Salmonella enterica] Length: 372\nNumber of predicted TMHs:  1\nExp number of AAs in TMHs: 24.51309\nExp number, first 60 AAs:  0.2682\nTotal prob of N-in: 0.14402\noutside   1   331\nTMhelix   332   354\ninside   355   372

20457 GCF_001364675.1_10540_1_18 Salmonella enterica subsp. enterica serovar Typhi Proteobacteria; Gammaproteobacteria; Enterobacterales; Enterobacteriaceae; Salmonella; Salmonella enterica; Salmonella enterica subsp. enterica MTGDNTLITSHGINRRDFMKLCAALAATMGLSSKAAAEMAESVSNPQRPPVIWIGAQECT WP_000145429.1 hydrogenase 2 small subunit [Salmonella enterica] Length: 372\nNumber of predicted TMHs:  1\nExp number of AAs in TMHs: 24.51309\nExp number, first 60 AAs:  0.2682\nTotal prob of N-in: 0.14402\noutside   1   331\nTMhelix   332   354\ninside   355   372

20458 GCF_001364635.1_10426_1_94 Salmonella enterica subsp. enterica serovar Typhi Proteobacteria; Gammaproteobacteria; Enterobacterales; Enterobacteriaceae; Salmonella; Salmonella enterica; Salmonella enterica subsp. enterica MTGDNTLITSHGINRRDFMKLCAALAATMGLSSKAAAEMAESVSNPQRPPVIWIGAQECT WP_000145429.1 hydrogenase 2 small subunit [Salmonella enterica] Length: 372\nNumber of predicted TMHs:  1\nExp number of AAs in TMHs: 24.51309\nExp number, first 60 AAs:  0.2682\nTotal prob of N-in: 0.14402\noutside   1   331\nTMhelix   332   354\ninside   355   372

20459 GCF_001364595.1_10540_1_92 Salmonella enterica subsp. enterica serovar Typhi Proteobacteria; Gammaproteobacteria; Enterobacterales; Enterobacteriaceae; Salmonella; Salmonella enterica;

Salmonella enterica subsp. enterica MTGDNTLITSHGINRRDFMKLCAALAATMGLSSKAAAEMAESVSNPQRPPVIWIGAQECT WP_000145429.1 hydrogenase 2 small subunit [Salmonella enterica] Length: 372\nNumber of predicted TMHs: 1\nExp number of AAs in TMHs: 24.51309\nExp number, first 60 AAs: 0.2682\nTotal prob of N-in: 0.14402\noutside 1 331\nTMhelix 332 354\ninside 355 372

20460 GCF_001364555.1_10540_1_66 Salmonella enterica subsp. enterica serovar Typhi Proteobacteria; Gammaproteobacteria; Enterobacterales; Enterobacteriaceae; Salmonella; Salmonella enterica; Salmonella enterica subsp. enterica MTGDNTLITSHGINRRDFMKLCAALAATMGLSSKAAAEMAESVSNPQRPPVIWIGAQECT WP_000145429.1 hydrogenase 2 small subunit [Salmonella enterica] Length: 372\nNumber of predicted TMHs: 1\nExp number of AAs in TMHs: 24.51309\nExp number, first 60 AAs: 0.2682\nTotal prob of N-in: 0.14402\noutside 1 331\nTMhelix 332 354\ninside 355 372

20461 GCF_001364515.1_10608_2_24 Salmonella enterica subsp. enterica serovar Typhi Proteobacteria; Gammaproteobacteria; Enterobacterales; Enterobacteriaceae; Salmonella; Salmonella enterica; Salmonella enterica subsp. enterica MTGDNTLITSHGINRRDFMKLCAALAATMGLSSKAAAEMAESVSNPQRPPVIWIGAQECT WP_000145429.1 hydrogenase 2 small subunit [Salmonella enterica] Length: 372\nNumber of predicted TMHs: 1\nExp number of AAs in TMHs: 24.51309\nExp number, first 60 AAs: 0.2682\nTotal prob of N-in: 0.14402\noutside 1 331\nTMhelix 332 354\ninside 355 372

20462 GCF_001364475.1_10541_2_15 Salmonella enterica subsp. enterica serovar Typhi Proteobacteria; Gammaproteobacteria; Enterobacterales; Enterobacteriaceae; Salmonella; Salmonella enterica; Salmonella enterica subsp. enterica MTGDNTLITSHGINRRDFMKLCAALAATMGLSSKAAAEMAESVSNPQRPPVIWIGAQECT WP_000145429.1 hydrogenase 2 small subunit [Salmonella enterica] Length: 372\nNumber of predicted TMHs: 1\nExp number of AAs in TMHs: 24.51309\nExp number, first 60 AAs: 0.2682\nTotal prob of N-in: 0.14402\noutside 1 331\nTMhelix 332 354\ninside 355 372

20463 GCF_001363935.1_10540_1_86 Salmonella enterica subsp. enterica serovar Typhi Proteobacteria; Gammaproteobacteria; Enterobacterales; Enterobacteriaceae; Salmonella; Salmonella enterica; Salmonella enterica subsp. enterica MTGDNTLITSHGINRRDFMKLCAALAATMGLSSKAAAEMAESVSNPQRPPVIWIGAQECT WP_000145429.1 hydrogenase 2 small subunit [Salmonella enterica] Length: 372\nNumber of predicted TMHs: 1\nExp number of AAs in TMHs: 24.51309\nExp number, first 60 AAs: 0.2682\nTotal prob of N-in: 0.14402\noutside 1 331\nTMhelix 332 354\ninside 355 372

20464 GCF_000982755.1_10492_1_10 Salmonella enterica subsp. enterica serovar Typhi Proteobacteria; Gammaproteobacteria; Enterobacterales; Enterobacteriaceae; Salmonella; Salmonella enterica; Salmonella enterica subsp. enterica MTGDNTLITSHGINRRDFMKLCAALAATMGLSSKAATEMAESVSNPQRPPVIWIGAQECT WP_001782304.1 hydrogenase 2 small subunit [Salmonella enterica] Length: 372\nNumber of predicted TMHs: 1\nExp number of AAs in TMHs: 24.41173\nExp number, first 60 AAs: 0.16124\nTotal prob of N-in: 0.13947\noutside 1 331\nTMhelix 332 354\ninside 355 372

20465 GCF_001363895.1_10540_1_78 Salmonella enterica subsp. enterica serovar Typhi Proteobacteria; Gammaproteobacteria; Enterobacterales; Enterobacteriaceae; Salmonella; Salmonella enterica; Salmonella enterica subsp. enterica MTGDNTLITSHGINRRDFMKLCAALAATMGLSSKAAAEMAESVSNPQRPPVIWIGAQECT WP_000145429.1 hydrogenase 2 small subunit [Salmonella enterica] Length: 372\nNumber of predicted TMHs: 1\nExp number of AAs in TMHs: 24.51309\nExp number, first 60 AAs: 0.2682\nTotal prob of N-in: 0.14402\noutside 1 331\nTMhelix 332 354\ninside 355 372

20466 GCF_001363855.1_10426_1_14 Salmonella enterica subsp. enterica serovar Typhi Proteobacteria; Gammaproteobacteria; Enterobacterales; Enterobacteriaceae; Salmonella; Salmonella enterica; Salmonella enterica subsp. enterica MTGDNTLITSHGINRRDFMKLCAALAATMGLSSKAAAEMAESVSNPQRPPVIWIGAQECT WP_000145429.1 hydrogenase 2 small subunit [Salmonella enterica] Length: 372\nNumber of predicted TMHs: 1\nExp number of AAs in TMHs: 24.51309\nExp number, first 60 AAs: 0.2682\nTotal prob of N-in: 0.14402\noutside 1 331\nTMhelix 332 354\ninside 355 372

20467 GCF_001363815.1_10540_1_45 Salmonella enterica subsp. enterica serovar Typhi Proteobacteria; Gammaproteobacteria; Enterobacterales; Enterobacteriaceae; Salmonella; Salmonella enterica; Salmonella enterica subsp. enterica MTGDNTLITSHGINRRDFMKLCAALAATMGLSSKAAAEMAESVSNPQRPPVIWIGAQECT WP_000145429.1 hydrogenase 2 small subunit [Salmonella enterica] Length: 372\nNumber of predicted TMHs: 1\nExp number of AAs in TMHs: 24.51309\nExp number, first 60 AAs: 0.2682\nTotal prob of N-in: 0.14402\noutside 1 331\nTMhelix 332 354\ninside 355 372

20468 GCF_001363775.1_10426_1_23 Salmonella enterica subsp. enterica serovar Typhi Proteobacteria; Gammaproteobacteria; Enterobacterales; Enterobacteriaceae; Salmonella; Salmonella enterica; Salmonella enterica subsp. enterica MTGDNTLITSHGINRRDFMKLCAALAATMGLSSKAAAEMAESVSNPQRPPVIWIGAQECT WP_000145429.1 hydrogenase 2 small subunit [Salmonella enterica] Length: 372\nNumber of predicted TMHs: 1\nExp number of AAs in TMHs: 24.51309\nExp number, first 60 AAs: 0.2682\nTotal prob of N-in: 0.14402\noutside 1 331\nTMhelix 332 354\ninside 355 372

20469 GCF_001363735.1_10426_1_85 Salmonella enterica subsp. enterica serovar Typhi Proteobacteria; Gammaproteobacteria; Enterobacterales; Enterobacteriaceae; Salmonella; Salmonella enterica; Salmonella enterica subsp. enterica MTGDNTLITSHGINRRDFMKLCAALAATMGLSSKAAAEMAESVSNPQRPPVIWIGAQECT WP_000145429.1 hydrogenase 2 small subunit [Salmonella enterica] Length: 372\nNumber of predicted TMHs: 1\nExp number of AAs in TMHs: 24.51309\nExp number, first 60 AAs: 0.2682\nTotal prob of N-in: 0.14402\noutside 1 331\nTMhelix 332 354\ninside 355 372

20470 GCF_001363695.1_10561_2_35 Salmonella enterica subsp. enterica serovar Typhi Proteobacteria; Gammaproteobacteria; Enterobacterales; Enterobacteriaceae; Salmonella; Salmonella enterica; Salmonella enterica subsp. enterica MTGDNTLITSHGINRRDFMKLCAALAATMGLSSKAAAEMAESVSNPQRPPVIWIGAQECT WP_000145429.1 hydrogenase 2 small subunit [Salmonella enterica] Length: 372\nNumber of predicted TMHs: 1\nExp number of AAs in TMHs: 24.51309\nExp number, first 60 AAs: 0.2682\nTotal prob of N-in: 0.14402\noutside 1 331\nTMhelix 332 354\ninside 355 372

20471 GCF_001363155.1_10540_1_29 Salmonella enterica subsp. enterica serovar Typhi Proteobacteria; Gammaproteobacteria; Enterobacterales; Enterobacteriaceae; Salmonella; Salmonella enterica; Salmonella enterica subsp. enterica MTGDNTLITSHGINRRDFMKLCAALAATMGLSSKAAAEMAESVSNPQRPPVIWIGAQECT WP_000145429.1 hydrogenase 2 small subunit [Salmonella enterica] Length: 372\nNumber of predicted TMHs: 1\nExp number of AAs in TMHs: 24.51309\nExp number, first 60 AAs: 0.2682\nTotal prob of N-in: 0.14402\noutside 1 331\nTMhelix 332 354\ninside 355 372

20472 GCF_001363115.1_10540_1_34 Salmonella enterica subsp. enterica serovar Typhi Proteobacteria; Gammaproteobacteria; Enterobacterales; Enterobacteriaceae; Salmonella; Salmonella enterica; Salmonella enterica subsp. enterica MTGDNTLITSHGINRRDFMKLCAALAATMGLSSKAAAEMAESVSNPQRPPVIWIGAQECT WP_000145429.1 hydrogenase 2 small subunit [Salmonella enterica] Length: 372\nNumber of predicted TMHs: 1\nExp number of AAs in TMHs: 24.51309\nExp number, first 60 AAs: 0.2682\nTotal prob of N-in: 0.14402\noutside 1 331\nTMhelix 332 354\ninside 355 372

20473 GCF_001363075.1_10426_1_60 Salmonella enterica subsp. enterica serovar Typhi Proteobacteria; Gammaproteobacteria; Enterobacterales; Enterobacteriaceae; Salmonella; Salmonella enterica; Salmonella enterica subsp. enterica MTGDNTLITSHGINRRDFMKLCAALAATMGLSSKAAAEMAESVSNPQRPPVIWIGAQECT WP_000145429.1 hydrogenase 2 small subunit [Salmonella enterica] Length: 372\nNumber of predicted TMHs: 1\nExp number of AAs in TMHs: 24.51309\nExp number, first 60 AAs: 0.2682\nTotal prob of N-in: 0.14402\noutside 1 331\nTMhelix 332 354\ninside 355 372

20474 GCF_001363035.1_10541_2_52 Salmonella enterica subsp. enterica serovar Typhi Proteobacteria; Gammaproteobacteria; Enterobacterales; Enterobacteriaceae; Salmonella; Salmonella enterica; Salmonella enterica subsp. enterica MTGDNTLITSHGINRRDFMKLCAALAATMGLSSKAAAEMAESVSNPQRPPVIWIGAQECT WP_000145429.1 hydrogenase 2 small subunit [Salmonella enterica] Length: 372\nNumber of predicted TMHs: 1\nExp number of AAs in TMHs: 24.51309\nExp number, first 60 AAs: 0.2682\nTotal prob of N-in: 0.14402\noutside 1 331\nTMhelix 332 354\ninside 355 372

20475 GCF_001362995.1_10540_1_85 Salmonella enterica subsp. enterica serovar Typhi Proteobacteria; Gammaproteobacteria; Enterobacterales; Enterobacteriaceae; Salmonella; Salmonella enterica; Salmonella enterica subsp. enterica MTGDNTLITSHGINRRDFMKLCAALAATMGLSSKAAAEMAESVSNPQRPPVIWIGAQECT WP_000145429.1 hydrogenase 2 small subunit [Salmonella enterica] Length: 372\nNumber of predicted TMHs: 1\nExp number of AAs in TMHs: 24.51309\nExp number, first 60 AAs: 0.2682\nTotal prob of N-in: 0.14402\noutside 1 331\nTMhelix 332 354\ninside 355 372

20476 GCF_001362955.1_10541_2_47 Salmonella enterica subsp. enterica serovar Typhi Proteobacteria; Gammaproteobacteria; Enterobacterales; Enterobacteriaceae; Salmonella; Salmonella enterica; Salmonella enterica subsp. enterica MTGDNTLITSHGINRRDFMKLCAALAATMGLSSKAAAEMAESVSNPQRPPVIWIGAQECT WP_000145429.1 hydrogenase 2 small subunit [Salmonella enterica] Length: 372\nNumber of

predicted TMHs:  1\nExp number of AAs in TMHs: 24.51309\nExp number, first 60 AAs:  0.2682\nTotal prob of N-in: 0.14402\noutside   1   331\nTMhelix   332   354\ninside   355   372

20477 GCF_001362415.1_10426_1_55 Salmonella enterica subsp. enterica serovar Typhi Proteobacteria; Gammaproteobacteria; Enterobacterales; Enterobacteriaceae; Salmonella; Salmonella enterica; Salmonella enterica subsp. enterica MTGDNTLITSHGINRRDFMKLCAALAATMGLSSKAAAEMAESVSNPQRPPVIWIGAQECT WP_000145429.1 hydrogenase 2 small subunit [Salmonella enterica] Length: 372\nNumber of predicted TMHs:  1\nExp number of AAs in TMHs: 24.51309\nExp number, first 60 AAs:  0.2682\nTotal prob of N-in: 0.14402\noutside   1   331\nTMhelix   332   354\ninside   355   372

20478 GCF_001362375.1_10607_2_29 Salmonella enterica subsp. enterica serovar Typhi Proteobacteria; Gammaproteobacteria; Enterobacterales; Enterobacteriaceae; Salmonella; Salmonella enterica; Salmonella enterica subsp. enterica MTGDNTLITSHGINRRDFMKLCAALAATMGLSSKAAAEMAESVSNPQRPPVIWIGAQECT WP_000145429.1 hydrogenase 2 small subunit [Salmonella enterica] Length: 372\nNumber of predicted TMHs:  1\nExp number of AAs in TMHs: 24.51309\nExp number, first 60 AAs:  0.2682\nTotal prob of N-in: 0.14402\noutside   1   331\nTMhelix   332   354\ninside   355   372

20479 GCF_001362335.1_10349_1_84 Salmonella enterica subsp. enterica serovar Typhi Proteobacteria; Gammaproteobacteria; Enterobacterales; Enterobacteriaceae; Salmonella; Salmonella enterica; Salmonella enterica subsp. enterica MTGDNTLITSHGINRRDFMKLCAALAATMGLSSKAAAEMAESVSNPQRPPVIWIGAQECT WP_000145429.1 hydrogenase 2 small subunit [Salmonella enterica] Length: 372\nNumber of predicted TMHs:  1\nExp number of AAs in TMHs: 24.51309\nExp number, first 60 AAs:  0.2682\nTotal prob of N-in: 0.14402\noutside   1   331\nTMhelix   332   354\ninside   355   372

20480 GCF_001362295.1_10349_1_44 Salmonella enterica subsp. enterica serovar Typhi Proteobacteria; Gammaproteobacteria; Enterobacterales; Enterobacteriaceae; Salmonella; Salmonella enterica; Salmonella enterica subsp. enterica MTGDNTLITSHGINRRDFMKLCAALAATMGLSSKAAAEMAESVSNPQRPPVIWIGAQECT WP_000145429.1 hydrogenase 2 small subunit [Salmonella enterica] Length: 372\nNumber of predicted TMHs:  1\nExp number of AAs in TMHs: 24.51309\nExp number, first 60 AAs:  0.2682\nTotal prob of N-in: 0.14402\noutside   1   331\nTMhelix   332   354\ninside   355   372

20481 GCF_001362255.1_10349_1_14 Salmonella enterica subsp. enterica serovar Typhi Proteobacteria; Gammaproteobacteria; Enterobacterales; Enterobacteriaceae; Salmonella; Salmonella enterica; Salmonella enterica subsp. enterica MTGDNTLITSHGINRRDFMKLCAALAATMGLSSKAAAEMAESVSNPQRPPVIWIGAQECT WP_000145429.1 hydrogenase 2 small subunit [Salmonella enterica] Length: 372\nNumber of predicted TMHs:  1\nExp number of AAs in TMHs: 24.51309\nExp number, first 60 AAs:  0.2682\nTotal prob of N-in: 0.14402\noutside   1   331\nTMhelix   332   354\ninside   355   372

20482 GCF_001362215.1_10349_1_86 Salmonella enterica subsp. enterica serovar Typhi Proteobacteria; Gammaproteobacteria; Enterobacterales; Enterobacteriaceae; Salmonella; Salmonella enterica; Salmonella enterica subsp. enterica MTGDNTLITSHGINRRDFMKLCAALAATMGLSSKAAAEMAESVSNPQRPPVIWIGAQECT WP_000145429.1 hydrogenase 2 small subunit [Salmonella enterica] Length: 372\nNumber of predicted TMHs:  1\nExp number of AAs in TMHs: 24.51309\nExp number, first 60 AAs:  0.2682\nTotal prob of N-in: 0.14402\noutside   1   331\nTMhelix   332   354\ninside   355   372

20483 GCF_001362155.1_10349_1_77 Salmonella enterica subsp. enterica serovar Typhi Proteobacteria; Gammaproteobacteria; Enterobacterales; Enterobacteriaceae; Salmonella; Salmonella enterica; Salmonella enterica subsp. enterica MTGDNTLITSHGINRRDFMKLCAALAATMGLSSKAAAEMAESVSNPQRPPVIWIGAQECT WP_000145429.1 hydrogenase 2 small subunit [Salmonella enterica] Length: 372\nNumber of predicted TMHs:  1\nExp number of AAs in TMHs: 24.51309\nExp number, first 60 AAs:  0.2682\nTotal prob of N-in: 0.14402\noutside   1   331\nTMhelix   332   354\ninside   355   372

20484 GCF_001361555.1_10060_6_64 Salmonella enterica subsp. enterica serovar Typhi Proteobacteria; Gammaproteobacteria; Enterobacterales; Enterobacteriaceae; Salmonella; Salmonella enterica; Salmonella enterica subsp. enterica MTGDNTLITSHGINRRDFMKLCAALAATMGLSSKAAAEMAESVSNPQRPPVIWIGAQECT WP_000145429.1 hydrogenase 2 small subunit [Salmonella enterica] Length: 372\nNumber of predicted TMHs:  1\nExp number of AAs in TMHs: 24.51309\nExp number, first 60 AAs:  0.2682\nTotal prob of N-in: 0.14402\noutside   1   331\nTMhelix   332   354\ninside   355   372

20485 GCF_001361515.1_10060_6_58 Salmonella enterica subsp. enterica serovar Typhi Proteobacteria; Gammaproteobacteria; Enterobacterales; Enterobacteriaceae; Salmonella; Salmonella enterica;

Salmonella enterica subsp. enterica	MTGDNTLITSHGINRRDFMKLCAALAATMGLSSKAAAEMAESVSNPQRPPVIWIGAQECT	WP_000145429.1	hydrogenase 2 small subunit [Salmonella enterica]	Length: 372\nNumber of predicted TMHs:  1\nExp number of AAs in TMHs: 24.51309\nExp number, first 60 AAs:  0.2682\nTotal prob of N-in:  0.14402\noutside   1   331\nTMhelix   332   354\ninside   355   372

20486	GCF_001361455.1_10060_6_43	Salmonella enterica subsp. enterica serovar Typhi	Proteobacteria; Gammaproteobacteria; Enterobacterales; Enterobacteriaceae; Salmonella; Salmonella enterica; Salmonella enterica subsp. enterica	MTGDNTLITSHGINRRDFMKLCAALAATMGLSSKAAAEMAESVSNPQRPPVIWIGAQECT	WP_000145429.1	hydrogenase 2 small subunit [Salmonella enterica]	Length: 372\nNumber of predicted TMHs:  1\nExp number of AAs in TMHs: 24.51309\nExp number, first 60 AAs:  0.2682\nTotal prob of N-in:  0.14402\noutside   1   331\nTMhelix   332   354\ninside   355   372

20487	GCF_001361415.1_10060_6_31	Salmonella enterica subsp. enterica serovar Typhi	Proteobacteria; Gammaproteobacteria; Enterobacterales; Enterobacteriaceae; Salmonella; Salmonella enterica; Salmonella enterica subsp. enterica	MTGDNTLITSHGINRRDFMKLCAALAATMGLSSKAAAEMAESVSNPQRPPVIWIGAQECT	WP_000145429.1	hydrogenase 2 small subunit [Salmonella enterica]	Length: 372\nNumber of predicted TMHs:  1\nExp number of AAs in TMHs: 24.51309\nExp number, first 60 AAs:  0.2682\nTotal prob of N-in:  0.14402\noutside   1   331\nTMhelix   332   354\ninside   355   372

20488	GCF_001361375.1_10060_6_19	Salmonella enterica subsp. enterica serovar Typhi	Proteobacteria; Gammaproteobacteria; Enterobacterales; Enterobacteriaceae; Salmonella; Salmonella enterica; Salmonella enterica subsp. enterica	MTGDNTLITSHGINRRDFMKLCAALAATMGLSSKAAAEMAESVSNPQRPPVIWIGAQECT	WP_000145429.1	hydrogenase 2 small subunit [Salmonella enterica]	Length: 372\nNumber of predicted TMHs:  1\nExp number of AAs in TMHs: 24.51309\nExp number, first 60 AAs:  0.2682\nTotal prob of N-in:  0.14402\noutside   1   331\nTMhelix   332   354\ninside   355   372

20489	GCF_001361055.1_10071_3_16	Salmonella enterica subsp. enterica serovar Typhi	Proteobacteria; Gammaproteobacteria; Enterobacterales; Enterobacteriaceae; Salmonella; Salmonella enterica; Salmonella enterica subsp. enterica	MTGDNTLITSHGINRRDFMKLCAALAATMGLSSKAATEMAESVSNPQRPPVIWIGAQECT	WP_001782304.1	hydrogenase 2 small subunit [Salmonella enterica]	Length: 372\nNumber of predicted TMHs:  1\nExp number of AAs in TMHs: 24.41173\nExp number, first 60 AAs:  0.16124\nTotal prob of N-in:  0.13947\noutside   1   331\nTMhelix   332   354\ninside   355   372

20490	GCF_001360555.1_9870_8_31	Salmonella enterica subsp. enterica serovar Typhi	Proteobacteria; Gammaproteobacteria; Enterobacterales; Enterobacteriaceae; Salmonella; Salmonella enterica; Salmonella enterica subsp. enterica	MTGDNTLITSHGINRRDFMKLCAALAATMGLSSKAAAEMAESVSNPQRPPVIWIGAQECT	WP_000145429.1	hydrogenase 2 small subunit [Salmonella enterica]	Length: 372\nNumber of predicted TMHs:  1\nExp number of AAs in TMHs: 24.51309\nExp number, first 60 AAs:  0.2682\nTotal prob of N-in:  0.14402\noutside   1   331\nTMhelix   332   354\ninside   355   372

20491	GCF_001360475.1_9870_8_7Salmonella enterica subsp. enterica serovar Typhi	Proteobacteria; Gammaproteobacteria; Enterobacterales; Enterobacteriaceae; Salmonella; Salmonella enterica; Salmonella enterica subsp. enterica	MTGDNTLITSHGINRRDFMKLCAALAATMGLSSKAAAEMAESVSNPQRPPVIWIGAQECT	WP_000145429.1	hydrogenase 2 small subunit [Salmonella enterica]	Length: 372\nNumber of predicted TMHs:  1\nExp number of AAs in TMHs: 24.51309\nExp number, first 60 AAs:  0.2682\nTotal prob of N-in:  0.14402\noutside   1   331\nTMhelix   332   354\ninside   355   372

20492	GCF_001360435.1_9953_5_77	Salmonella enterica subsp. enterica serovar Typhi	Proteobacteria; Gammaproteobacteria; Enterobacterales; Enterobacteriaceae; Salmonella; Salmonella enterica; Salmonella enterica subsp. enterica	MTGDNTLITSHGINRRDFMKLCAALAATMGLSSKAAAEMAESVSNPQRPPVIWIGAQECT	WP_000145429.1	hydrogenase 2 small subunit [Salmonella enterica]	Length: 372\nNumber of predicted TMHs:  1\nExp number of AAs in TMHs: 24.51309\nExp number, first 60 AAs:  0.2682\nTotal prob of N-in:  0.14402\noutside   1   331\nTMhelix   332   354\ninside   355   372

20493	GCF_001360395.1_9953_5_53	Salmonella enterica subsp. enterica serovar Typhi	Proteobacteria; Gammaproteobacteria; Enterobacterales; Enterobacteriaceae; Salmonella; Salmonella enterica; Salmonella enterica subsp. enterica	MTGDNTLITSHGINRRDFMKLCAALAATMGLSSKAAAEMAESVSNPQRPPVIWIGAQECT	WP_000145429.1	hydrogenase 2 small subunit [Salmonella enterica]	Length: 372\nNumber of predicted TMHs:  1\nExp number of AAs in TMHs: 24.51309\nExp number, first 60 AAs:  0.2682\nTotal prob of N-in:  0.14402\noutside   1   331\nTMhelix   332   354\ninside   355   372

20494	GCF_001360355.1_9953_5_41	Salmonella enterica subsp. enterica serovar Typhi	Proteobacteria; Gammaproteobacteria; Enterobacterales; Enterobacteriaceae; Salmonella; Salmonella enterica; Salmonella enterica subsp. enterica	MTGDNTLITSHGINRRDFMKLCAALAATMGLSSKAAAEMAESVSNPQRPPVIWIGAQECT	WP_000145429.1	hydrogenase 2 small subunit [Salmonella enterica]	Length: 372\nNumber of predicted TMHs:  1\nExp number of AAs in TMHs: 24.51309\nExp number, first 60 AAs:  0.2682\nTotal prob of N-in:  0.14402\noutside   1   331\nTMhelix   332   354\ninside   355   372

20495	GCF_001360275.1_10071_3_23	Salmonella enterica subsp. enterica serovar Typhi	Proteobacteria; Gammaproteobacteria; Enterobacterales; Enterobacteriaceae; Salmonella; Salmonella enterica; Salmonella enterica subsp. enterica	MTGDNTLITSHGINRRDFMKLCAALAATMGLSSKAAAEMAESVSNPQRPPVIWIGAQECT	WP_000145429.1	hydrogenase 2 small subunit [Salmonella enterica]	Length: 372\nNumber of predicted TMHs:  1\nExp number of AAs in TMHs: 24.51309\nExp number, first 60 AAs:  0.2682\nTotal prob of N-in:  0.14402\noutside   1   331\nTMhelix   332   354\ninside   355   372

20496	GCF_001360315.1_10071_3_71	Salmonella enterica subsp. enterica serovar Typhi	Proteobacteria; Gammaproteobacteria; Enterobacterales; Enterobacteriaceae; Salmonella; Salmonella enterica; Salmonella enterica subsp. enterica	MTGDNTLITSHGINRRDFMKLCAALAATMGLSSKAAAEMAESVSNPQRPPVIWIGAQECT	WP_000145429.1	hydrogenase 2 small subunit [Salmonella enterica]	Length: 372\nNumber of predicted TMHs:  1\nExp number of AAs in TMHs: 24.51309\nExp number, first 60 AAs:  0.2682\nTotal prob of N-in:  0.14402\noutside   1   331\nTMhelix   332   354\ninside   355   372

20497	GCF_001359975.1_10060_5_16	Salmonella enterica subsp. enterica serovar Typhi	Proteobacteria; Gammaproteobacteria; Enterobacterales; Enterobacteriaceae; Salmonella; Salmonella enterica; Salmonella enterica subsp. enterica	MTGDNTLITSHGINRRDFMKLCAALAATMGLSSKAAAEMAESVSNPQRPPVIWIGAQECT	WP_000145429.1	hydrogenase 2 small subunit [Salmonella enterica]	Length: 372\nNumber of predicted TMHs:  1\nExp number of AAs in TMHs: 24.51309\nExp number, first 60 AAs:  0.2682\nTotal prob of N-in:  0.14402\noutside   1   331\nTMhelix   332   354\ninside   355   372

20498	GCF_001359935.1_10060_6_69	Salmonella enterica subsp. enterica serovar Typhi	Proteobacteria; Gammaproteobacteria; Enterobacterales; Enterobacteriaceae; Salmonella; Salmonella enterica; Salmonella enterica subsp. enterica	MTGDNTLITSHGINRRDFMKLCAALAATMGLSSKAAAEMAESVSNPQRPPVIWIGAQECT	WP_000145429.1	hydrogenase 2 small subunit [Salmonella enterica]	Length: 372\nNumber of predicted TMHs:  1\nExp number of AAs in TMHs: 24.51309\nExp number, first 60 AAs:  0.2682\nTotal prob of N-in:  0.14402\noutside   1   331\nTMhelix   332   354\ninside   355   372

20499	GCF_001359895.1_10060_6_51	Salmonella enterica subsp. enterica serovar Typhi	Proteobacteria; Gammaproteobacteria; Enterobacterales; Enterobacteriaceae; Salmonella; Salmonella enterica; Salmonella enterica subsp. enterica	MTGDNTLITSHGINRRDFMKLCAALAATMGLSSKAAAEMAESVSNPQRPPVIWIGAQECT	WP_000145429.1	hydrogenase 2 small subunit [Salmonella enterica]	Length: 372\nNumber of predicted TMHs:  1\nExp number of AAs in TMHs: 24.51309\nExp number, first 60 AAs:  0.2682\nTotal prob of N-in:  0.14402\noutside   1   331\nTMhelix   332   354\ninside   355   372

20500	GCF_001359855.1_10060_6_30	Salmonella enterica subsp. enterica serovar Typhi	Proteobacteria; Gammaproteobacteria; Enterobacterales; Enterobacteriaceae; Salmonella; Salmonella enterica; Salmonella enterica subsp. enterica	MTGDNTLITSHGINRRDFMKLCAALAATMGLSSKAAAEMAESVSNPQRPPVIWIGAQECT	WP_000145429.1	hydrogenase 2 small subunit [Salmonella enterica]	Length: 372\nNumber of predicted TMHs:  1\nExp number of AAs in TMHs: 24.51309\nExp number, first 60 AAs:  0.2682\nTotal prob of N-in:  0.14402\noutside   1   331\nTMhelix   332   354\ninside   355   372

20501	GCF_001359815.1_9870_8_80	Salmonella enterica subsp. enterica serovar Typhi	Proteobacteria; Gammaproteobacteria; Enterobacterales; Enterobacteriaceae; Salmonella; Salmonella enterica; Salmonella enterica subsp. enterica	MTGDNTLITSHGINRRDFMKLCAALAATMGLSSKAAAEMAESVSNPQRPPVIWIGAQECT	WP_000145429.1	hydrogenase 2 small subunit [Salmonella enterica]	Length: 372\nNumber of predicted TMHs:  1\nExp number of AAs in TMHs: 24.51309\nExp number, first 60 AAs:  0.2682\nTotal prob of N-in:  0.14402\noutside   1   331\nTMhelix   332   354\ninside   355   372

20502	GCF_001359775.1_9870_8_71	Salmonella enterica subsp. enterica serovar Typhi	Proteobacteria; Gammaproteobacteria; Enterobacterales; Enterobacteriaceae; Salmonella; Salmonella enterica; Salmonella enterica subsp. enterica	MTGDNTLITSHGINRRDFMKLCAALAATMGLSSKAAAEMAESVSNPQRPPVIWIGAQECT	WP_000145429.1	hydrogenase 2 small subunit [Salmonella enterica]	Length: 372\nNumber of

predicted TMHs: 1\nExp number of AAs in TMHs: 24.51309\nExp number, first 60 AAs: 0.2682\nTotal prob of N-in: 0.14402\noutside 1 331\nTMhelix 332 354\ninside 355 372

20503 GCF_001359735.1_9870_8_53 Salmonella enterica subsp. enterica serovar Typhi Proteobacteria; Gammaproteobacteria; Enterobacterales; Enterobacteriaceae; Salmonella; Salmonella enterica; Salmonella enterica subsp. enterica MTGDNTLITSHGINRRDFMKLCAALAATMGLSSKAAAEMAESVSNPQRPPVIWIGAQECT WP_000145429.1 hydrogenase 2 small subunit [Salmonella enterica] Length: 372\nNumber of predicted TMHs: 1\nExp number of AAs in TMHs: 24.51309\nExp number, first 60 AAs: 0.2682\nTotal prob of N-in: 0.14402\noutside 1 331\nTMhelix 332 354\ninside 355 372

20504 GCF_001359275.1_10060_5_50 Salmonella enterica subsp. enterica serovar Typhi Proteobacteria; Gammaproteobacteria; Enterobacterales; Enterobacteriaceae; Salmonella; Salmonella enterica; Salmonella enterica subsp. enterica MTGDNTLITSHGINRRDFMKLCAALAATMGLSSKAAAEMAESVSNPQRPPVIWIGAQECT WP_000145429.1 hydrogenase 2 small subunit [Salmonella enterica] Length: 372\nNumber of predicted TMHs: 1\nExp number of AAs in TMHs: 24.51309\nExp number, first 60 AAs: 0.2682\nTotal prob of N-in: 0.14402\noutside 1 331\nTMhelix 332 354\ninside 355 372

20505 GCF_001359195.1_10060_5_14 Salmonella enterica subsp. enterica serovar Typhi Proteobacteria; Gammaproteobacteria; Enterobacterales; Enterobacteriaceae; Salmonella; Salmonella enterica; Salmonella enterica subsp. enterica MTGDNTLITSHGINRRDFMKLCAALAATMGLSSKAAAEMAESVSNPQRPPVIWIGAQECT WP_000145429.1 hydrogenase 2 small subunit [Salmonella enterica] Length: 372\nNumber of predicted TMHs: 1\nExp number of AAs in TMHs: 24.51309\nExp number, first 60 AAs: 0.2682\nTotal prob of N-in: 0.14402\noutside 1 331\nTMhelix 332 354\ninside 355 372

20506 GCF_001359155.1_9870_8_69 Salmonella enterica subsp. enterica serovar Typhi Proteobacteria; Gammaproteobacteria; Enterobacterales; Enterobacteriaceae; Salmonella; Salmonella enterica; Salmonella enterica subsp. enterica MTGDNTLITSHGINRRDFMKLCAALAATMGLSSKAAAEMAESVSNPQRPPVIWIGAQECT WP_000145429.1 hydrogenase 2 small subunit [Salmonella enterica] Length: 372\nNumber of predicted TMHs: 1\nExp number of AAs in TMHs: 24.51309\nExp number, first 60 AAs: 0.2682\nTotal prob of N-in: 0.14402\noutside 1 331\nTMhelix 332 354\ninside 355 372

20507 GCF_001359115.1_9870_8_42 Salmonella enterica subsp. enterica serovar Typhi Proteobacteria; Gammaproteobacteria; Enterobacterales; Enterobacteriaceae; Salmonella; Salmonella enterica; Salmonella enterica subsp. enterica MTGDNTLITSHGINRRDFMKLCAALAATMGLSSKAAAEMAESVSNPQRPPVIWIGAQECT WP_000145429.1 hydrogenase 2 small subunit [Salmonella enterica] Length: 372\nNumber of predicted TMHs: 1\nExp number of AAs in TMHs: 24.51309\nExp number, first 60 AAs: 0.2682\nTotal prob of N-in: 0.14402\noutside 1 331\nTMhelix 332 354\ninside 355 372

20508 GCF_001359075.1_9870_8_21 Salmonella enterica subsp. enterica serovar Typhi Proteobacteria; Gammaproteobacteria; Enterobacterales; Enterobacteriaceae; Salmonella; Salmonella enterica; Salmonella enterica subsp. enterica MTGDNTLITSHGINRRDFMKLCAALAATMGLSSKAAAEMAESVSNPQRPPVIWIGAQECT WP_000145429.1 hydrogenase 2 small subunit [Salmonella enterica] Length: 372\nNumber of predicted TMHs: 1\nExp number of AAs in TMHs: 24.51309\nExp number, first 60 AAs: 0.2682\nTotal prob of N-in: 0.14402\noutside 1 331\nTMhelix 332 354\ninside 355 372

20509 GCF_001359035.1_10060_6_89 Salmonella enterica subsp. enterica serovar Typhi Proteobacteria; Gammaproteobacteria; Enterobacterales; Enterobacteriaceae; Salmonella; Salmonella enterica; Salmonella enterica subsp. enterica MTGDNTLITSHGINRRDFMKLCAALAATMGLSSKAAAEMAESVSNPQRPPVIWIGAQECT WP_000145429.1 hydrogenase 2 small subunit [Salmonella enterica] Length: 372\nNumber of predicted TMHs: 1\nExp number of AAs in TMHs: 24.51309\nExp number, first 60 AAs: 0.2682\nTotal prob of N-in: 0.14402\noutside 1 331\nTMhelix 332 354\ninside 355 372

20510 GCF_001358955.1_10060_6_62 Salmonella enterica subsp. enterica serovar Typhi Proteobacteria; Gammaproteobacteria; Enterobacterales; Enterobacteriaceae; Salmonella; Salmonella enterica; Salmonella enterica subsp. enterica MTGDNTLITSHGINRRDFMKLCAALAATMGLSSKAAAEMAESVSNPQRPPVIWIGAQECT WP_000145429.1 hydrogenase 2 small subunit [Salmonella enterica] Length: 372\nNumber of predicted TMHs: 1\nExp number of AAs in TMHs: 24.51309\nExp number, first 60 AAs: 0.2682\nTotal prob of N-in: 0.14402\noutside 1 331\nTMhelix 332 354\ninside 355 372

20511 GCF_001358455.1_10607_2_6 Salmonella enterica subsp. enterica serovar Typhi Proteobacteria; Gammaproteobacteria; Enterobacterales; Enterobacteriaceae; Salmonella; Salmonella enterica;

Salmonella enterica subsp. enterica MTGDNTLITSHGINRRDFMKLCAALAATMGLSSKAAAEMAESVSNPQRPPVIWIGAQECT WP_000145429.1 hydrogenase 2 small subunit [Salmonella enterica] Length: 372\nNumber of predicted TMHs: 1\nExp number of AAs in TMHs: 24.51309\nExp number, first 60 AAs: 0.2682\nTotal prob of N-in: 0.14402\noutside 1 331\nTMhelix 332 354\ninside 355 372

20512 GCF_001358375.1_10060_5_31 Salmonella enterica subsp. enterica serovar Typhi Proteobacteria; Gammaproteobacteria; Enterobacterales; Enterobacteriaceae; Salmonella; Salmonella enterica; Salmonella enterica subsp. enterica MTGDNTLITSHGINRRDFMKLCAALAATMGLSSKAAAEMAESVSNPQRPPVIWIGAQECT WP_000145429.1 hydrogenase 2 small subunit [Salmonella enterica] Length: 372\nNumber of predicted TMHs: 1\nExp number of AAs in TMHs: 24.51309\nExp number, first 60 AAs: 0.2682\nTotal prob of N-in: 0.14402\noutside 1 331\nTMhelix 332 354\ninside 355 372

20513 GCF_001358335.1_9953_5_85 Salmonella enterica subsp. enterica serovar Typhi Proteobacteria; Gammaproteobacteria; Enterobacterales; Enterobacteriaceae; Salmonella; Salmonella enterica; Salmonella enterica subsp. enterica MTGDNTLITSHGINRRDFMKLCAALAATMGLSSKAAAEMAESVSNPQRPPVIWIGAQECT WP_000145429.1 hydrogenase 2 small subunit [Salmonella enterica] Length: 372\nNumber of predicted TMHs: 1\nExp number of AAs in TMHs: 24.51309\nExp number, first 60 AAs: 0.2682\nTotal prob of N-in: 0.14402\noutside 1 331\nTMhelix 332 354\ninside 355 372

20514 GCF_001358295.1_10541_2_1 Salmonella enterica subsp. enterica serovar Typhi Proteobacteria; Gammaproteobacteria; Enterobacterales; Enterobacteriaceae; Salmonella; Salmonella enterica; Salmonella enterica subsp. enterica MTGDNTLITSHGINRRDFMKLCAALAATMGLSSKAAAEMAESVSNPQRPPVIWIGAQECT WP_000145429.1 hydrogenase 2 small subunit [Salmonella enterica] Length: 372\nNumber of predicted TMHs: 1\nExp number of AAs in TMHs: 24.51309\nExp number, first 60 AAs: 0.2682\nTotal prob of N-in: 0.14402\noutside 1 331\nTMhelix 332 354\ninside 355 372

20515 GCF_001358255.1_10060_6_82 Salmonella enterica subsp. enterica serovar Typhi Proteobacteria; Gammaproteobacteria; Enterobacterales; Enterobacteriaceae; Salmonella; Salmonella enterica; Salmonella enterica subsp. enterica MTGDNTLITSHGINRRDFMKLCAALAATMGLSSKAAAEMAESVSNPQRPPVIWIGAQECT WP_000145429.1 hydrogenase 2 small subunit [Salmonella enterica] Length: 372\nNumber of predicted TMHs: 1\nExp number of AAs in TMHs: 24.51309\nExp number, first 60 AAs: 0.2682\nTotal prob of N-in: 0.14402\noutside 1 331\nTMhelix 332 354\ninside 355 372

20516 GCF_001358215.1_10607_1_59 Salmonella enterica subsp. enterica serovar Typhi Proteobacteria; Gammaproteobacteria; Enterobacterales; Enterobacteriaceae; Salmonella; Salmonella enterica; Salmonella enterica subsp. enterica MTGDNTLITSHGINRRDFMKLCAALAATMGLSSKAAAEMAESVSNPQRPPVIWIGAQECT WP_000145429.1 hydrogenase 2 small subunit [Salmonella enterica] Length: 372\nNumber of predicted TMHs: 1\nExp number of AAs in TMHs: 24.51309\nExp number, first 60 AAs: 0.2682\nTotal prob of N-in: 0.14402\noutside 1 331\nTMhelix 332 354\ninside 355 372

20517 GCF_001357055.1_10592_2_59 Salmonella enterica subsp. enterica serovar Typhi Proteobacteria; Gammaproteobacteria; Enterobacterales; Enterobacteriaceae; Salmonella; Salmonella enterica; Salmonella enterica subsp. enterica MTGDNTLITSHGINRRDFMKLCAALAATMGLSSKAAAEMAESVSNPQRPPVIWIGAQECT WP_000145429.1 hydrogenase 2 small subunit [Salmonella enterica] Length: 372\nNumber of predicted TMHs: 1\nExp number of AAs in TMHs: 24.51309\nExp number, first 60 AAs: 0.2682\nTotal prob of N-in: 0.14402\noutside 1 331\nTMhelix 332 354\ninside 355 372

20518 GCF_001357175.1_10593_2_79 Salmonella enterica subsp. enterica serovar Typhi Proteobacteria; Gammaproteobacteria; Enterobacterales; Enterobacteriaceae; Salmonella; Salmonella enterica; Salmonella enterica subsp. enterica MTGDNTLITSHGINRRDFMKLCAALAATMGLSSKAAAEMAESVSNPQRPPVIWIGAQECT WP_000145429.1 hydrogenase 2 small subunit [Salmonella enterica] Length: 372\nNumber of predicted TMHs: 1\nExp number of AAs in TMHs: 24.51309\nExp number, first 60 AAs: 0.2682\nTotal prob of N-in: 0.14402\noutside 1 331\nTMhelix 332 354\ninside 355 372

20519 GCF_001357015.1_10592_2_51 Salmonella enterica subsp. enterica serovar Typhi Proteobacteria; Gammaproteobacteria; Enterobacterales; Enterobacteriaceae; Salmonella; Salmonella enterica; Salmonella enterica subsp. enterica MTGDNTLITSHGINRRDFMKLCAALAATMGLSSKAAAEMAESVSNPQRPPVIWIGAQECT WP_000145429.1 hydrogenase 2 small subunit [Salmonella enterica] Length: 372\nNumber of predicted TMHs: 1\nExp number of AAs in TMHs: 24.51309\nExp number, first 60 AAs: 0.2682\nTotal prob of N-in: 0.14402\noutside 1 331\nTMhelix 332 354\ninside 355 372

20520	GCF_001356775.1_10592_2_16	Salmonella enterica subsp. enterica serovar Typhi	Proteobacteria; Gammaproteobacteria; Enterobacterales; Enterobacteriaceae; Salmonella; Salmonella enterica; Salmonella enterica subsp. enterica	MTGDNTLITSHGINRRDFMKLCAALAATMGLSSKAATEMAESVSNPQRPPVIWIGAQECT	WP_001782304.1	hydrogenase 2 small subunit [Salmonella enterica]	Length: 372\nNumber of predicted TMHs:  1\nExp number of AAs in TMHs: 24.41173\nExp number, first 60 AAs:  0.16124\nTotal prob of N-in: 0.13947\noutside	1	331\nTMhelix	332	354\ninside	355	372

20521	GCF_001356975.1_10592_2_38	Salmonella enterica subsp. enterica serovar Typhi	Proteobacteria; Gammaproteobacteria; Enterobacterales; Enterobacteriaceae; Salmonella; Salmonella enterica; Salmonella enterica subsp. enterica	MTGDNTLITSHGINRRDFMKLCAALAATMGLSSKAAAEMAESVSNPQRPPVIWIGAQECT	WP_000145429.1	hydrogenase 2 small subunit [Salmonella enterica]	Length: 372\nNumber of predicted TMHs:  1\nExp number of AAs in TMHs: 24.51309\nExp number, first 60 AAs:  0.2682\nTotal prob of N-in: 0.14402\noutside	1	331\nTMhelix	332	354\ninside	355	372

20522	GCF_001356935.1_10592_2_48	Salmonella enterica subsp. enterica serovar Typhi	Proteobacteria; Gammaproteobacteria; Enterobacterales; Enterobacteriaceae; Salmonella; Salmonella enterica; Salmonella enterica subsp. enterica	MTGDNTLITSHGINRRDFMKLCAALAATMGLSSKAAAEMAESVSNPQRPPVIWIGAQECT	WP_000145429.1	hydrogenase 2 small subunit [Salmonella enterica]	Length: 372\nNumber of predicted TMHs:  1\nExp number of AAs in TMHs: 24.51309\nExp number, first 60 AAs:  0.2682\nTotal prob of N-in: 0.14402\noutside	1	331\nTMhelix	332	354\ninside	355	372

20523	GCF_001356895.1_10592_2_26	Salmonella enterica subsp. enterica serovar Typhi	Proteobacteria; Gammaproteobacteria; Enterobacterales; Enterobacteriaceae; Salmonella; Salmonella enterica; Salmonella enterica subsp. enterica	MTGDNTLITSHGINRRDFMKLCAALAATMGLSSKAAAEMAESVSNPQRPPVIWIGAQECT	WP_000145429.1	hydrogenase 2 small subunit [Salmonella enterica]	Length: 372\nNumber of predicted TMHs:  1\nExp number of AAs in TMHs: 24.51309\nExp number, first 60 AAs:  0.2682\nTotal prob of N-in: 0.14402\noutside	1	331\nTMhelix	332	354\ninside	355	372

20524	GCF_001356555.1_9475_6_49	Salmonella enterica subsp. enterica serovar Typhi	Proteobacteria; Gammaproteobacteria; Enterobacterales; Enterobacteriaceae; Salmonella; Salmonella enterica; Salmonella enterica subsp. enterica	MTGDNTLITSHGINRRDFMKLCAALAATMGLSSKAAAEMAESVSNPQRPPVIWIGAQECT	WP_000145429.1	hydrogenase 2 small subunit [Salmonella enterica]	Length: 372\nNumber of predicted TMHs:  1\nExp number of AAs in TMHs: 24.51309\nExp number, first 60 AAs:  0.2682\nTotal prob of N-in: 0.14402\noutside	1	331\nTMhelix	332	354\ninside	355	372

20525	GCF_001356515.1_9475_6_33	Salmonella enterica subsp. enterica serovar Typhi	Proteobacteria; Gammaproteobacteria; Enterobacterales; Enterobacteriaceae; Salmonella; Salmonella enterica; Salmonella enterica subsp. enterica	MTGDNTLITSHGINRRDFMKLCAALAATMGLSSKAAAEMAESVSNPQRPPVIWIGAQECT	WP_000145429.1	hydrogenase 2 small subunit [Salmonella enterica]	Length: 372\nNumber of predicted TMHs:  1\nExp number of AAs in TMHs: 24.51309\nExp number, first 60 AAs:  0.2682\nTotal prob of N-in: 0.14402\noutside	1	331\nTMhelix	332	354\ninside	355	372

20526	GCF_001356475.1_9475_6_24	Salmonella enterica subsp. enterica serovar Typhi	Proteobacteria; Gammaproteobacteria; Enterobacterales; Enterobacteriaceae; Salmonella; Salmonella enterica; Salmonella enterica subsp. enterica	MTGDNTLITSHGINRRDFMKLCAALAATMGLSSKAAAEMAESVSNPQRPPVIWIGAQECT	WP_000145429.1	hydrogenase 2 small subunit [Salmonella enterica]	Length: 372\nNumber of predicted TMHs:  1\nExp number of AAs in TMHs: 24.51309\nExp number, first 60 AAs:  0.2682\nTotal prob of N-in: 0.14402\noutside	1	331\nTMhelix	332	354\ninside	355	372

20527	GCF_001356435.1_9475_6_17	Salmonella enterica subsp. enterica serovar Typhi	Proteobacteria; Gammaproteobacteria; Enterobacterales; Enterobacteriaceae; Salmonella; Salmonella enterica; Salmonella enterica subsp. enterica	MTGDNTLITSHGINRRDFMKLCAALAATMGLSSKAAAEMAESVSNPQRPPVIWIGAQECT	WP_000145429.1	hydrogenase 2 small subunit [Salmonella enterica]	Length: 372\nNumber of predicted TMHs:  1\nExp number of AAs in TMHs: 24.51309\nExp number, first 60 AAs:  0.2682\nTotal prob of N-in: 0.14402\noutside	1	331\nTMhelix	332	354\ninside	355	372

20528	GCF_001356395.1_9475_6_14	Salmonella enterica subsp. enterica serovar Typhi	Proteobacteria; Gammaproteobacteria; Enterobacterales; Enterobacteriaceae; Salmonella; Salmonella enterica; Salmonella enterica subsp. enterica	MTGDNTLITSHGINRRDFMKLCAALAATMGLSSKAAAEMAESVSNPQRPPVIWIGAQECT	WP_000145429.1	hydrogenase 2 small subunit [Salmonella enterica]	Length: 372\nNumber of

predicted TMHs: 1\nExp number of AAs in TMHs: 24.51309\nExp number, first 60 AAs: 0.2682\nTotal prob of N-in: 0.14402\noutside 1 331\nTMhelix 332 354\ninside 355 372

20529 GCF_001356355.1_9475_6_3Salmonella enterica subsp. enterica serovar Typhi Proteobacteria; Gammaproteobacteria; Enterobacterales; Enterobacteriaceae; Salmonella; Salmonella enterica; Salmonella enterica subsp. enterica MTGDNTLITSHGINRRDFMKLCAALAATMGLSSKAAAEMAESVSNPQRPPVIWIGAQECT WP_000145429.1 hydrogenase 2 small subunit [Salmonella enterica] Length: 372\nNumber of predicted TMHs: 1\nExp number of AAs in TMHs: 24.51309\nExp number, first 60 AAs: 0.2682\nTotal prob of N-in: 0.14402\noutside 1 331\nTMhelix 332 354\ninside 355 372

20530 GCF_001350415.1_8616_4_57 Salmonella enterica subsp. enterica serovar Typhi Proteobacteria; Gammaproteobacteria; Enterobacterales; Enterobacteriaceae; Salmonella; Salmonella enterica; Salmonella enterica subsp. enterica MTGDNTLITSHGINRRDFMKLCAALAATMGLSSKAATEMAESVSNPQRPPVIWIGAQECT WP_001782304.1 hydrogenase 2 small subunit [Salmonella enterica] Length: 372\nNumber of predicted TMHs: 1\nExp number of AAs in TMHs: 24.41173\nExp number, first 60 AAs: 0.16124\nTotal prob of N-in: 0.13947\noutside 1 331\nTMhelix 332 354\ninside 355 372

20531 GCF_001355495.1_8525_3_58 Salmonella enterica subsp. enterica serovar Typhi Proteobacteria; Gammaproteobacteria; Enterobacterales; Enterobacteriaceae; Salmonella; Salmonella enterica; Salmonella enterica subsp. enterica MTGDNTLITSHGINRRDFMKLCAALAATMGLSSKAAAEMAESVSNPQRPPVIWIGAQECT WP_000145429.1 hydrogenase 2 small subunit [Salmonella enterica] Length: 372\nNumber of predicted TMHs: 1\nExp number of AAs in TMHs: 24.51309\nExp number, first 60 AAs: 0.2682\nTotal prob of N-in: 0.14402\noutside 1 331\nTMhelix 332 354\ninside 355 372

20532 GCF_001355455.1_8525_3_61 Salmonella enterica subsp. enterica serovar Typhi Proteobacteria; Gammaproteobacteria; Enterobacterales; Enterobacteriaceae; Salmonella; Salmonella enterica; Salmonella enterica subsp. enterica MTGDNTLITSHGINRRDFMKLCAALAATMGLSSKAAAEMAESVSNPQRPPVIWIGAQECT WP_000145429.1 hydrogenase 2 small subunit [Salmonella enterica] Length: 372\nNumber of predicted TMHs: 1\nExp number of AAs in TMHs: 24.51309\nExp number, first 60 AAs: 0.2682\nTotal prob of N-in: 0.14402\noutside 1 331\nTMhelix 332 354\ninside 355 372

20533 GCF_001355415.1_8525_3_48 Salmonella enterica subsp. enterica serovar Typhi Proteobacteria; Gammaproteobacteria; Enterobacterales; Enterobacteriaceae; Salmonella; Salmonella enterica; Salmonella enterica subsp. enterica MTGDNTLITSHGINRRDFMKLCAALAATMGLSSKAAAEMAESVSNPQRPPVIWIGAQECT WP_000145429.1 hydrogenase 2 small subunit [Salmonella enterica] Length: 372\nNumber of predicted TMHs: 1\nExp number of AAs in TMHs: 24.51309\nExp number, first 60 AAs: 0.2682\nTotal prob of N-in: 0.14402\noutside 1 331\nTMhelix 332 354\ninside 355 372

20534 GCF_001350455.1_8616_4_52 Salmonella enterica subsp. enterica serovar Typhi Proteobacteria; Gammaproteobacteria; Enterobacterales; Enterobacteriaceae; Salmonella; Salmonella enterica; Salmonella enterica subsp. enterica MTGDNTLITSHGINRRDFMKLCAALAATMGLSSKAAAEMAESVSNPQRPPVIWIGAQECT WP_000145429.1 hydrogenase 2 small subunit [Salmonella enterica] Length: 372\nNumber of predicted TMHs: 1\nExp number of AAs in TMHs: 24.51309\nExp number, first 60 AAs: 0.2682\nTotal prob of N-in: 0.14402\noutside 1 331\nTMhelix 332 354\ninside 355 372

20535 GCF_001328835.1_7468_7_95 Salmonella enterica subsp. enterica serovar Typhi Proteobacteria; Gammaproteobacteria; Enterobacterales; Enterobacteriaceae; Salmonella; Salmonella enterica; Salmonella enterica subsp. enterica MTGDNTLITSHGINRRDFMKLCAALAATMGLSSKAAAEMAESVSNPQRPPVIWIGAQECT WP_000145429.1 hydrogenase 2 small subunit [Salmonella enterica] Length: 372\nNumber of predicted TMHs: 1\nExp number of AAs in TMHs: 24.51309\nExp number, first 60 AAs: 0.2682\nTotal prob of N-in: 0.14402\noutside 1 331\nTMhelix 332 354\ninside 355 372

20536 GCF_001328755.1_7468_7_92 Salmonella enterica subsp. enterica serovar Typhi Proteobacteria; Gammaproteobacteria; Enterobacterales; Enterobacteriaceae; Salmonella; Salmonella enterica; Salmonella enterica subsp. enterica MTGDNTLITSHGINRRDFMKLCAALAATMGLSSKAAAEMAESVSNPQRPPVIWIGAQECT WP_000145429.1 hydrogenase 2 small subunit [Salmonella enterica] Length: 372\nNumber of predicted TMHs: 1\nExp number of AAs in TMHs: 24.51309\nExp number, first 60 AAs: 0.2682\nTotal prob of N-in: 0.14402\noutside 1 331\nTMhelix 332 354\ninside 355 372

20537 GCF_001328715.1_7468_7_88 Salmonella enterica subsp. enterica serovar Typhi Proteobacteria; Gammaproteobacteria; Enterobacterales; Enterobacteriaceae; Salmonella; Salmonella enterica;

Salmonella enterica subsp. enterica	MTGDNTLITSHGINRRDFMKLCAALAATMGLSSKAAAEMAESVSNPQRPPVIWIGAQECT	WP_000145429.1	hydrogenase 2 small subunit [Salmonella enterica]	Length: 372\nNumber of predicted TMHs: 1\nExp number of AAs in TMHs: 24.51309\nExp number, first 60 AAs: 0.2682\nTotal prob of N-in: 0.14402\noutside 1 331\nTMhelix 332 354\ninside 355 372

20538	GCF_001150125.1_9870_8_34	Salmonella enterica subsp. enterica serovar Typhi	Proteobacteria; Gammaproteobacteria; Enterobacterales; Enterobacteriaceae; Salmonella; Salmonella enterica; Salmonella enterica subsp. enterica	MTGDNTLITSHGINRRDFMKLCAALAATMGLSSKAAAEMAESVSNPQRPPVIWIGAQECT	WP_000145429.1	hydrogenase 2 small subunit [Salmonella enterica]	Length: 372\nNumber of predicted TMHs: 1\nExp number of AAs in TMHs: 24.51309\nExp number, first 60 AAs: 0.2682\nTotal prob of N-in: 0.14402\noutside 1 331\nTMhelix 332 354\ninside 355 372

20539	GCF_001150805.1_10493_1_76	Salmonella enterica subsp. enterica serovar Typhi	Proteobacteria; Gammaproteobacteria; Enterobacterales; Enterobacteriaceae; Salmonella; Salmonella enterica; Salmonella enterica subsp. enterica	MTGDNTLITSHGINRRDFMKLCAALAATMGLSSKAAAEMAESVSNPQRPPVIWIGAQECT	WP_000145429.1	hydrogenase 2 small subunit [Salmonella enterica]	Length: 372\nNumber of predicted TMHs: 1\nExp number of AAs in TMHs: 24.51309\nExp number, first 60 AAs: 0.2682\nTotal prob of N-in: 0.14402\noutside 1 331\nTMhelix 332 354\ninside 355 372

20540	GCF_001149905.1_10493_1_43	Salmonella enterica subsp. enterica serovar Typhi	Proteobacteria; Gammaproteobacteria; Enterobacterales; Enterobacteriaceae; Salmonella; Salmonella enterica; Salmonella enterica subsp. enterica	MTGDNTLITSHGINRRDFMKLCAALAATMGLSSKAAAEMAESVSNPQRPPVIWIGAQECT	WP_000145429.1	hydrogenase 2 small subunit [Salmonella enterica]	Length: 372\nNumber of predicted TMHs: 1\nExp number of AAs in TMHs: 24.51309\nExp number, first 60 AAs: 0.2682\nTotal prob of N-in: 0.14402\noutside 1 331\nTMhelix 332 354\ninside 355 372

20541	GCF_001149745.1_10562_2_11	Salmonella enterica subsp. enterica serovar Typhi	Proteobacteria; Gammaproteobacteria; Enterobacterales; Enterobacteriaceae; Salmonella; Salmonella enterica; Salmonella enterica subsp. enterica	MTGDNTLITSHGINRRDFMKLCAALAATMGLSSKAAAEMAESVSNPQRPPVIWIGAQECT	WP_000145429.1	hydrogenase 2 small subunit [Salmonella enterica]	Length: 372\nNumber of predicted TMHs: 1\nExp number of AAs in TMHs: 24.51309\nExp number, first 60 AAs: 0.2682\nTotal prob of N-in: 0.14402\noutside 1 331\nTMhelix 332 354\ninside 355 372

20542	GCF_001148045.1_10592_2_43	Salmonella enterica subsp. enterica serovar Typhi	Proteobacteria; Gammaproteobacteria; Enterobacterales; Enterobacteriaceae; Salmonella; Salmonella enterica; Salmonella enterica subsp. enterica	MTGDNTLITSHGINRRDFMKLCAALAATMGLSSKAAAEMAESVSNPQRPPVIWIGAQECT	WP_000145429.1	hydrogenase 2 small subunit [Salmonella enterica]	Length: 372\nNumber of predicted TMHs: 1\nExp number of AAs in TMHs: 24.51309\nExp number, first 60 AAs: 0.2682\nTotal prob of N-in: 0.14402\noutside 1 331\nTMhelix 332 354\ninside 355 372

20543	GCF_001149445.1_9870_8_36	Salmonella enterica subsp. enterica serovar Typhi	Proteobacteria; Gammaproteobacteria; Enterobacterales; Enterobacteriaceae; Salmonella; Salmonella enterica; Salmonella enterica subsp. enterica	MTGDNTLITSHGINRRDFMKLCAALAATMGLSSKAAAEMAESVSNPQRPPVIWIGAQECT	WP_000145429.1	hydrogenase 2 small subunit [Salmonella enterica]	Length: 372\nNumber of predicted TMHs: 1\nExp number of AAs in TMHs: 24.51309\nExp number, first 60 AAs: 0.2682\nTotal prob of N-in: 0.14402\noutside 1 331\nTMhelix 332 354\ninside 355 372

20544	GCF_001147885.1_9870_8_8Salmonella enterica subsp. enterica serovar Typhi	Proteobacteria; Gammaproteobacteria; Enterobacterales; Enterobacteriaceae; Salmonella; Salmonella enterica; Salmonella enterica subsp. enterica	MTGDNTLITSHGINRRDFMKLCAALAATMGLSSKAAAEMAESVSNPQRPPVIWIGAQECT	WP_000145429.1	hydrogenase 2 small subunit [Salmonella enterica]	Length: 372\nNumber of predicted TMHs: 1\nExp number of AAs in TMHs: 24.51309\nExp number, first 60 AAs: 0.2682\nTotal prob of N-in: 0.14402\noutside 1 331\nTMhelix 332 354\ninside 355 372

20545	GCF_001146985.1_10607_2_33	Salmonella enterica subsp. enterica serovar Typhi	Proteobacteria; Gammaproteobacteria; Enterobacterales; Enterobacteriaceae; Salmonella; Salmonella enterica; Salmonella enterica subsp. enterica	MTGDNTLITSHGINRRDFMKLCAALAATMGLSSKAAAEMAESVSNPQRPPVIWIGAQECT	WP_000145429.1	hydrogenase 2 small subunit [Salmonella enterica]	Length: 372\nNumber of predicted TMHs: 1\nExp number of AAs in TMHs: 24.51309\nExp number, first 60 AAs: 0.2682\nTotal prob of N-in: 0.14402\noutside 1 331\nTMhelix 332 354\ninside 355 372

20546 GCF_001146645.1_9870_8_19 Salmonella enterica subsp. enterica serovar Typhi Proteobacteria; Gammaproteobacteria; Enterobacterales; Enterobacteriaceae; Salmonella; Salmonella enterica; Salmonella enterica subsp. enterica MTGDNTLITSHGINRRDFMKLCAALAATMGLSSKAAAEMAESVSNPQRPPVIWIGAQECT WP_000145429.1 hydrogenase 2 small subunit [Salmonella enterica] Length: 372\nNumber of predicted TMHs: 1\nExp number of AAs in TMHs: 24.51309\nExp number, first 60 AAs: 0.2682\nTotal prob of N-in: 0.14402\noutside 1 331\nTMhelix 332 354\ninside 355 372

20547 GCF_001146745.1_10593_2_40 Salmonella enterica subsp. enterica serovar Typhi Proteobacteria; Gammaproteobacteria; Enterobacterales; Enterobacteriaceae; Salmonella; Salmonella enterica; Salmonella enterica subsp. enterica MTGDNTLITSHGINRRDFMKLCAALAATMGLSSKAAAEMAESVSNPQRPPVIWIGAQECT WP_000145429.1 hydrogenase 2 small subunit [Salmonella enterica] Length: 372\nNumber of predicted TMHs: 1\nExp number of AAs in TMHs: 24.51309\nExp number, first 60 AAs: 0.2682\nTotal prob of N-in: 0.14402\noutside 1 331\nTMhelix 332 354\ninside 355 372

20548 GCF_001146305.1_9870_8_16 Salmonella enterica subsp. enterica serovar Typhi Proteobacteria; Gammaproteobacteria; Enterobacterales; Enterobacteriaceae; Salmonella; Salmonella enterica; Salmonella enterica subsp. enterica MTGDNTLITSHGINRRDFMKLCAALAATMGLSSKAAAEMAESVSNPQRPPVIWIGAQECT WP_000145429.1 hydrogenase 2 small subunit [Salmonella enterica] Length: 372\nNumber of predicted TMHs: 1\nExp number of AAs in TMHs: 24.51309\nExp number, first 60 AAs: 0.2682\nTotal prob of N-in: 0.14402\noutside 1 331\nTMhelix 332 354\ninside 355 372

20549 GCF_001146145.1_9475_6_81 Salmonella enterica subsp. enterica serovar Typhi Proteobacteria; Gammaproteobacteria; Enterobacterales; Enterobacteriaceae; Salmonella; Salmonella enterica; Salmonella enterica subsp. enterica MTGDNTLITSHGINRRDFMKLCAALAATMGLSSKAAAEMAESVSNPQRPPVIWIGAQECT WP_000145429.1 hydrogenase 2 small subunit [Salmonella enterica] Length: 372\nNumber of predicted TMHs: 1\nExp number of AAs in TMHs: 24.51309\nExp number, first 60 AAs: 0.2682\nTotal prob of N-in: 0.14402\noutside 1 331\nTMhelix 332 354\ninside 355 372

20550 GCF_001143065.1_10209_5_34 Salmonella enterica subsp. enterica serovar Typhi Proteobacteria; Gammaproteobacteria; Enterobacterales; Enterobacteriaceae; Salmonella; Salmonella enterica; Salmonella enterica subsp. enterica MTGDNTLITSHGINRRDFMKLCAALAATMGLSSKAAAEMAESVSNPQRPPVIWIGAQECT WP_000145429.1 hydrogenase 2 small subunit [Salmonella enterica] Length: 372\nNumber of predicted TMHs: 1\nExp number of AAs in TMHs: 24.51309\nExp number, first 60 AAs: 0.2682\nTotal prob of N-in: 0.14402\noutside 1 331\nTMhelix 332 354\ninside 355 372

20551 GCF_001143025.1_10541_2_17 Salmonella enterica subsp. enterica serovar Typhi Proteobacteria; Gammaproteobacteria; Enterobacterales; Enterobacteriaceae; Salmonella; Salmonella enterica; Salmonella enterica subsp. enterica MTGDNTLITSHGINRRDFMKLCAALAATMGLSSKAAAEMAESVSNPQRPPVIWIGAQECT WP_000145429.1 hydrogenase 2 small subunit [Salmonella enterica] Length: 372\nNumber of predicted TMHs: 1\nExp number of AAs in TMHs: 24.51309\nExp number, first 60 AAs: 0.2682\nTotal prob of N-in: 0.14402\noutside 1 331\nTMhelix 332 354\ninside 355 372

20552 GCF_001142885.1_10541_1_57 Salmonella enterica subsp. enterica serovar Typhi Proteobacteria; Gammaproteobacteria; Enterobacterales; Enterobacteriaceae; Salmonella; Salmonella enterica; Salmonella enterica subsp. enterica MTGDNTLITSHGINRRDFMKLCAALAATMGLSSKAAAEMAESVSNPQRPPVIWIGAQECT WP_000145429.1 hydrogenase 2 small subunit [Salmonella enterica] Length: 372\nNumber of predicted TMHs: 1\nExp number of AAs in TMHs: 24.51309\nExp number, first 60 AAs: 0.2682\nTotal prob of N-in: 0.14402\noutside 1 331\nTMhelix 332 354\ninside 355 372

20553 GCF_001142625.1_10492_1_12 Salmonella enterica subsp. enterica serovar Typhi Proteobacteria; Gammaproteobacteria; Enterobacterales; Enterobacteriaceae; Salmonella; Salmonella enterica; Salmonella enterica subsp. enterica MTGDNTLITSHGINRRDFMKLCAALAATMGLSSKAAAEMAESVSNPQRPPVIWIGAQECT WP_000145429.1 hydrogenase 2 small subunit [Salmonella enterica] Length: 372\nNumber of predicted TMHs: 1\nExp number of AAs in TMHs: 24.51309\nExp number, first 60 AAs: 0.2682\nTotal prob of N-in: 0.14402\noutside 1 331\nTMhelix 332 354\ninside 355 372

20554 GCF_001142525.1_9953_5_3Salmonella enterica subsp. enterica serovar Typhi Proteobacteria; Gammaproteobacteria; Enterobacterales; Enterobacteriaceae; Salmonella; Salmonella enterica; Salmonella enterica subsp. enterica MTGDNTLITSHGINRRDFMKLCAALAATMGLSSKAAAEMAESVSNPQRPPVIWIGAQECT WP_000145429.1 hydrogenase 2 small subunit [Salmonella enterica] Length: 372\nNumber of predicted TMHs: 1\nExp

number of AAs in TMHs: 24.51309\nExp number, first 60 AAs:  0.2682\nTotal prob of N-in:   0.14402\noutside   1   331\nTMhelix   332   354\ninside   355   372

20555 GCF_001142325.1_9870_8_83 Salmonella enterica subsp. enterica serovar Typhi Proteobacteria; Gammaproteobacteria; Enterobacterales; Enterobacteriaceae; Salmonella; Salmonella enterica; Salmonella enterica subsp. enterica MTGDNTLITSHGINRRDFMKLCAALAATMGLSSKAAAEMAESVSNPQRPPVIWIGAQECT WP_000145429.1 hydrogenase 2 small subunit [Salmonella enterica] Length: 372\nNumber of predicted TMHs:  1\nExp number of AAs in TMHs: 24.51309\nExp number, first 60 AAs:  0.2682\nTotal prob of N-in: 0.14402\noutside   1   331\nTMhelix   332   354\ninside   355   372

20556 GCF_001138905.1_10492_1_56 Salmonella enterica subsp. enterica serovar Typhi Proteobacteria; Gammaproteobacteria; Enterobacterales; Enterobacteriaceae; Salmonella; Salmonella enterica; Salmonella enterica subsp. enterica MTGDNTLITSHGINRRDFMKLCAALAATMGLSSKAAAEMAESVSNPQRPPVIWIGAQECT WP_000145429.1 hydrogenase 2 small subunit [Salmonella enterica] Length: 372\nNumber of predicted TMHs:  1\nExp number of AAs in TMHs: 24.51309\nExp number, first 60 AAs:  0.2682\nTotal prob of N-in: 0.14402\noutside   1   331\nTMhelix   332   354\ninside   355   372

20557 GCF_001138465.1_10209_5_47 Salmonella enterica subsp. enterica serovar Typhi Proteobacteria; Gammaproteobacteria; Enterobacterales; Enterobacteriaceae; Salmonella; Salmonella enterica; Salmonella enterica subsp. enterica MTGDNTLITSHGINRRDFMKLCAALAATMGLSSKAAAEMAESVSNPQRPPVIWIGAQECT WP_000145429.1 hydrogenase 2 small subunit [Salmonella enterica] Length: 372\nNumber of predicted TMHs:  1\nExp number of AAs in TMHs: 24.51309\nExp number, first 60 AAs:  0.2682\nTotal prob of N-in: 0.14402\noutside   1   331\nTMhelix   332   354\ninside   355   372

20558 GCF_001138725.1_8490_6_65 Salmonella enterica subsp. enterica serovar Typhi Proteobacteria; Gammaproteobacteria; Enterobacterales; Enterobacteriaceae; Salmonella; Salmonella enterica; Salmonella enterica subsp. enterica MTGDNTLITSHGINRRDFMKLCAALAATMGLSSKAAAEMAESVSNPQRPPVIWIGAQECT WP_000145429.1 hydrogenase 2 small subunit [Salmonella enterica] Length: 372\nNumber of predicted TMHs:  1\nExp number of AAs in TMHs: 24.51309\nExp number, first 60 AAs:  0.2682\nTotal prob of N-in: 0.14402\noutside   1   331\nTMhelix   332   354\ninside   355   372

20559 GCF_001138345.1_10209_5_62 Salmonella enterica subsp. enterica serovar Typhi Proteobacteria; Gammaproteobacteria; Enterobacterales; Enterobacteriaceae; Salmonella; Salmonella enterica; Salmonella enterica subsp. enterica MTGDNTLITSHGINRRDFMKLCAALAATMGLSSKAAAEMAESVSNPQRPPVIWIGAQECT WP_000145429.1 hydrogenase 2 small subunit [Salmonella enterica] Length: 372\nNumber of predicted TMHs:  1\nExp number of AAs in TMHs: 24.51309\nExp number, first 60 AAs:  0.2682\nTotal prob of N-in: 0.14402\noutside   1   331\nTMhelix   332   354\ninside   355   372

20560 GCF_001138305.1_10541_2_16 Salmonella enterica subsp. enterica serovar Typhi Proteobacteria; Gammaproteobacteria; Enterobacterales; Enterobacteriaceae; Salmonella; Salmonella enterica; Salmonella enterica subsp. enterica MTGDNTLITSHGINRRDFMKLCAALAATMGLSSKAAAEMAESVSNPQRPPVIWIGAQECT WP_000145429.1 hydrogenase 2 small subunit [Salmonella enterica] Length: 372\nNumber of predicted TMHs:  1\nExp number of AAs in TMHs: 24.51309\nExp number, first 60 AAs:  0.2682\nTotal prob of N-in: 0.14402\noutside   1   331\nTMhelix   332   354\ninside   355   372

20561 GCF_001138145.1_10349_1_72 Salmonella enterica subsp. enterica serovar Typhi Proteobacteria; Gammaproteobacteria; Enterobacterales; Enterobacteriaceae; Salmonella; Salmonella enterica; Salmonella enterica subsp. enterica MTGDNTLITSHGINRRDFMKLCAALAATMGLSSKAAAEMAESVSNPQRPPVIWIGAQECT WP_000145429.1 hydrogenase 2 small subunit [Salmonella enterica] Length: 372\nNumber of predicted TMHs:  1\nExp number of AAs in TMHs: 24.51309\nExp number, first 60 AAs:  0.2682\nTotal prob of N-in: 0.14402\noutside   1   331\nTMhelix   332   354\ninside   355   372

20562 GCF_001134005.1_9953_5_86 Salmonella enterica subsp. enterica serovar Typhi Proteobacteria; Gammaproteobacteria; Enterobacterales; Enterobacteriaceae; Salmonella; Salmonella enterica; Salmonella enterica subsp. enterica MTGDNTLITSHGINRRDFMKLCAALAATMGLSSKAAAEMAESVSNPQRPPVIWIGAQECT WP_000145429.1 hydrogenase 2 small subunit [Salmonella enterica] Length: 372\nNumber of predicted TMHs:  1\nExp number of AAs in TMHs: 24.51309\nExp number, first 60 AAs:  0.2682\nTotal prob of N-in: 0.14402\noutside   1   331\nTMhelix   332   354\ninside   355   372

20563 GCF_001134125.1_10349_1_62 Salmonella enterica subsp. enterica serovar Typhi Proteobacteria; Gammaproteobacteria; Enterobacterales; Enterobacteriaceae; Salmonella; Salmonella enterica;

Salmonella enterica subsp. enterica	MTGDNTLITSHGINRRDFMKLCAALAATMGLSSKAAAEMAESVSNPQRPPVIWIGAQECT
	WP_000145429.1	hydrogenase 2 small subunit [Salmonella enterica]	Length: 372\nNumber of predicted TMHs: 1\nExp number of AAs in TMHs: 24.51309\nExp number, first 60 AAs: 0.2682\nTotal prob of N-in: 0.14402\noutside 1 331\nTMhelix 332 354\ninside 355 372

20564	GCF_001133745.1_8616_4_58	Salmonella enterica subsp. enterica serovar Typhi
	Proteobacteria; Gammaproteobacteria; Enterobacterales; Enterobacteriaceae; Salmonella; Salmonella enterica; Salmonella enterica subsp. enterica	MTGDNTLITSHGINRRDFMKLCAALAATMGLSSKAAAEMAESVSNPQRPPVIWIGAQECT
	WP_000145429.1	hydrogenase 2 small subunit [Salmonella enterica]	Length: 372\nNumber of predicted TMHs: 1\nExp number of AAs in TMHs: 24.51309\nExp number, first 60 AAs: 0.2682\nTotal prob of N-in: 0.14402\noutside 1 331\nTMhelix 332 354\ninside 355 372

20565	GCF_001133585.1_7468_7_55	Salmonella enterica subsp. enterica serovar Typhi
	Proteobacteria; Gammaproteobacteria; Enterobacterales; Enterobacteriaceae; Salmonella; Salmonella enterica; Salmonella enterica subsp. enterica	MTGDNTLITSHGINRRDFMKLCAALAATMGLSSKAAAEMAESVSNPQRPPVIWIGAQECT
	WP_000145429.1	hydrogenase 2 small subunit [Salmonella enterica]	Length: 372\nNumber of predicted TMHs: 1\nExp number of AAs in TMHs: 24.51309\nExp number, first 60 AAs: 0.2682\nTotal prob of N-in: 0.14402\noutside 1 331\nTMhelix 332 354\ninside 355 372

20566	GCF_001133065.1_10541_2_35	Salmonella enterica subsp. enterica serovar Typhi
	Proteobacteria; Gammaproteobacteria; Enterobacterales; Enterobacteriaceae; Salmonella; Salmonella enterica; Salmonella enterica subsp. enterica	MTGDNTLITSHGINRRDFMKLCAALAATMGLSSKAAAEMAESVSNPQRPPVIWIGAQECT
	WP_000145429.1	hydrogenase 2 small subunit [Salmonella enterica]	Length: 372\nNumber of predicted TMHs: 1\nExp number of AAs in TMHs: 24.51309\nExp number, first 60 AAs: 0.2682\nTotal prob of N-in: 0.14402\noutside 1 331\nTMhelix 332 354\ninside 355 372

20567	GCF_001133225.1_10349_1_49	Salmonella enterica subsp. enterica serovar Typhi
	Proteobacteria; Gammaproteobacteria; Enterobacterales; Enterobacteriaceae; Salmonella; Salmonella enterica; Salmonella enterica subsp. enterica	MTGDNTLITSHGINRRDFMKLCAALAATMGLSSKAAAEMAESVSNPQRPPVIWIGAQECT
	WP_000145429.1	hydrogenase 2 small subunit [Salmonella enterica]	Length: 372\nNumber of predicted TMHs: 1\nExp number of AAs in TMHs: 24.51309\nExp number, first 60 AAs: 0.2682\nTotal prob of N-in: 0.14402\noutside 1 331\nTMhelix 332 354\ninside 355 372

20568	GCF_001132925.1_10541_2_49	Salmonella enterica subsp. enterica serovar Typhi
	Proteobacteria; Gammaproteobacteria; Enterobacterales; Enterobacteriaceae; Salmonella; Salmonella enterica; Salmonella enterica subsp. enterica	MTGDNTLITSHGINRRDFMKLCAALAATMGLSSKAAAEMAESVSNPQRPPVIWIGAQECT
	WP_000145429.1	hydrogenase 2 small subunit [Salmonella enterica]	Length: 372\nNumber of predicted TMHs: 1\nExp number of AAs in TMHs: 24.51309\nExp number, first 60 AAs: 0.2682\nTotal prob of N-in: 0.14402\noutside 1 331\nTMhelix 332 354\ninside 355 372

20569	GCF_001131505.1_10561_2_36	Salmonella enterica subsp. enterica serovar Typhi
	Proteobacteria; Gammaproteobacteria; Enterobacterales; Enterobacteriaceae; Salmonella; Salmonella enterica; Salmonella enterica subsp. enterica	MTGDNTLITSHGINRRDFMKLCAALAATMGLSSKAAAEMAESVSNPQRPPVIWIGAQECT
	WP_000145429.1	hydrogenase 2 small subunit [Salmonella enterica]	Length: 372\nNumber of predicted TMHs: 1\nExp number of AAs in TMHs: 24.51309\nExp number, first 60 AAs: 0.2682\nTotal prob of N-in: 0.14402\noutside 1 331\nTMhelix 332 354\ninside 355 372

20570	GCF_001160145.1_8447_8_23	Salmonella enterica subsp. enterica serovar Typhi
	Proteobacteria; Gammaproteobacteria; Enterobacterales; Enterobacteriaceae; Salmonella; Salmonella enterica; Salmonella enterica subsp. enterica	MTGDNTLITSHGINRRDFMKLCAALAATMGLSSKAAAEMAESVSNPQRPPVIWIGAQECT
	WP_000145429.1	hydrogenase 2 small subunit [Salmonella enterica]	Length: 372\nNumber of predicted TMHs: 1\nExp number of AAs in TMHs: 24.51309\nExp number, first 60 AAs: 0.2682\nTotal prob of N-in: 0.14402\noutside 1 331\nTMhelix 332 354\ninside 355 372

20571	GCF_001160365.1_10209_5_58	Salmonella enterica subsp. enterica serovar Typhi
	Proteobacteria; Gammaproteobacteria; Enterobacterales; Enterobacteriaceae; Salmonella; Salmonella enterica; Salmonella enterica subsp. enterica	MTGDNTLITSHGINRRDFMKLCAALAATMGLSSKAAAEMAESVSNPQRPPVIWIGAQECT
	WP_000145429.1	hydrogenase 2 small subunit [Salmonella enterica]	Length: 372\nNumber of predicted TMHs: 1\nExp number of AAs in TMHs: 24.51309\nExp number, first 60 AAs: 0.2682\nTotal prob of N-in: 0.14402\noutside 1 331\nTMhelix 332 354\ninside 355 372

20572 GCF_001160585.1_10493_1_20 Salmonella enterica subsp. enterica serovar Typhi Proteobacteria; Gammaproteobacteria; Enterobacterales; Enterobacteriaceae; Salmonella; Salmonella enterica; Salmonella enterica subsp. enterica MTGDNTLITSHGINRRDFMKLCAALAATMGLSSKAAAEMAESVSNPQRPPVIWIGAQECT WP_000145429.1 hydrogenase 2 small subunit [Salmonella enterica] Length: 372\nNumber of predicted TMHs: 1\nExp number of AAs in TMHs: 24.51309\nExp number, first 60 AAs: 0.2682\nTotal prob of N-in: 0.14402\noutside 1 331\nTMhelix 332 354\ninside 355 372

20573 GCF_001160945.1_10541_2_39 Salmonella enterica subsp. enterica serovar Typhi Proteobacteria; Gammaproteobacteria; Enterobacterales; Enterobacteriaceae; Salmonella; Salmonella enterica; Salmonella enterica subsp. enterica MTGDNTLITSHGINRRDFMKLCAALAATMGLSSKAAAEMAESVSNPQRPPVIWIGAQECT WP_000145429.1 hydrogenase 2 small subunit [Salmonella enterica] Length: 372\nNumber of predicted TMHs: 1\nExp number of AAs in TMHs: 24.51309\nExp number, first 60 AAs: 0.2682\nTotal prob of N-in: 0.14402\noutside 1 331\nTMhelix 332 354\ninside 355 372

20574 GCF_001161125.1_10607_1_92 Salmonella enterica subsp. enterica serovar Typhi Proteobacteria; Gammaproteobacteria; Enterobacterales; Enterobacteriaceae; Salmonella; Salmonella enterica; Salmonella enterica subsp. enterica MTGDNTLITSHGINRRDFMKLCAALAATMGLSSKAAAEMAESVSNPQRPPVIWIGAQECT WP_000145429.1 hydrogenase 2 small subunit [Salmonella enterica] Length: 372\nNumber of predicted TMHs: 1\nExp number of AAs in TMHs: 24.51309\nExp number, first 60 AAs: 0.2682\nTotal prob of N-in: 0.14402\noutside 1 331\nTMhelix 332 354\ninside 355 372

20575 GCF_001161545.1_8616_4_42 Salmonella enterica subsp. enterica serovar Typhi Proteobacteria; Gammaproteobacteria; Enterobacterales; Enterobacteriaceae; Salmonella; Salmonella enterica; Salmonella enterica subsp. enterica MTGDNTLITSHGINRRDFMKLCAALAATMGLSSKAAAEMAESVSNPQRPPVIWIGAQECT WP_000145429.1 hydrogenase 2 small subunit [Salmonella enterica] Length: 372\nNumber of predicted TMHs: 1\nExp number of AAs in TMHs: 24.51309\nExp number, first 60 AAs: 0.2682\nTotal prob of N-in: 0.14402\noutside 1 331\nTMhelix 332 354\ninside 355 372

20576 GCF_001162325.1_9475_4_48 Salmonella enterica subsp. enterica serovar Typhi Proteobacteria; Gammaproteobacteria; Enterobacterales; Enterobacteriaceae; Salmonella; Salmonella enterica; Salmonella enterica subsp. enterica MTGDNTLITSHGINRRDFMKLCAALAATMGLSSKAAAEMAESVSNPQRPPVIWIGAQECT WP_000145429.1 hydrogenase 2 small subunit [Salmonella enterica] Length: 372\nNumber of predicted TMHs: 1\nExp number of AAs in TMHs: 24.51309\nExp number, first 60 AAs: 0.2682\nTotal prob of N-in: 0.14402\noutside 1 331\nTMhelix 332 354\ninside 355 372

20577 GCF_001160705.1_10592_2_6 Salmonella enterica subsp. enterica serovar Typhi Proteobacteria; Gammaproteobacteria; Enterobacterales; Enterobacteriaceae; Salmonella; Salmonella enterica; Salmonella enterica subsp. enterica MTGDNTLITSHGINRRDFMKLCAALAATMGLSSKAATEMAESVSNPQRPPVIWIGAQECT WP_001782304.1 hydrogenase 2 small subunit [Salmonella enterica] Length: 372\nNumber of predicted TMHs: 1\nExp number of AAs in TMHs: 24.41173\nExp number, first 60 AAs: 0.16124\nTotal prob of N-in: 0.13947\noutside 1 331\nTMhelix 332 354\ninside 355 372

20578 GCF_001162745.1_10541_2_6 Salmonella enterica subsp. enterica serovar Typhi Proteobacteria; Gammaproteobacteria; Enterobacterales; Enterobacteriaceae; Salmonella; Salmonella enterica; Salmonella enterica subsp. enterica MTGDNTLITSHGINRRDFMKLCAALAATMGLSSKAAAEMAESVSNPQRPPVIWIGAQECT WP_000145429.1 hydrogenase 2 small subunit [Salmonella enterica] Length: 372\nNumber of predicted TMHs: 1\nExp number of AAs in TMHs: 24.51309\nExp number, first 60 AAs: 0.2682\nTotal prob of N-in: 0.14402\noutside 1 331\nTMhelix 332 354\ninside 355 372

20579 GCF_001162945.1_10593_2_19 Salmonella enterica subsp. enterica serovar Typhi Proteobacteria; Gammaproteobacteria; Enterobacterales; Enterobacteriaceae; Salmonella; Salmonella enterica; Salmonella enterica subsp. enterica MTGDNTLITSHGINRRDFMKLCAALAATMGLSSKAAAEMAESVSNPQRPPVIWIGAQECT WP_000145429.1 hydrogenase 2 small subunit [Salmonella enterica] Length: 372\nNumber of predicted TMHs: 1\nExp number of AAs in TMHs: 24.51309\nExp number, first 60 AAs: 0.2682\nTotal prob of N-in: 0.14402\noutside 1 331\nTMhelix 332 354\ninside 355 372

20580 GCF_001163385.1_9953_5_7Salmonella enterica subsp. enterica serovar Typhi Proteobacteria; Gammaproteobacteria; Enterobacterales; Enterobacteriaceae; Salmonella; Salmonella enterica; Salmonella enterica subsp. enterica MTGDNTLITSHGINRRDFMKLCAALAATMGLSSKAAAEMAESVSNPQRPPVIWIGAQECT WP_000145429.1 hydrogenase 2 small subunit [Salmonella enterica] Length: 372\nNumber of predicted TMHs: 1\nExp

number of AAs in TMHs: 24.51309\nExp number, first 60 AAs: 0.2682\nTotal prob of N-in: 0.14402\noutside 1 331\nTMhelix 332 354\ninside 355 372

20581 GCF_001161505.1_10493_1_13 Salmonella enterica subsp. enterica serovar Typhi Proteobacteria; Gammaproteobacteria; Enterobacterales; Enterobacteriaceae; Salmonella; Salmonella enterica; Salmonella enterica subsp. enterica MTGDNTLITSHGINRRDFMKLCAALAATMGLSSKAATEMAESVSNPQRPPVIWIGAQECT WP_001782304.1 hydrogenase 2 small subunit [Salmonella enterica] Length: 372\nNumber of predicted TMHs: 1\nExp number of AAs in TMHs: 24.41173\nExp number, first 60 AAs: 0.16124\nTotal prob of N-in: 0.13947\noutside 1 331\nTMhelix 332 354\ninside 355 372

20582 GCF_001163445.1_10541_1_73 Salmonella enterica subsp. enterica serovar Typhi Proteobacteria; Gammaproteobacteria; Enterobacterales; Enterobacteriaceae; Salmonella; Salmonella enterica; Salmonella enterica subsp. enterica MTGDNTLITSHGINRRDFMKLCAALAATMGLSSKAAAEMAESVSNPQRPPVIWIGAQECT WP_000145429.1 hydrogenase 2 small subunit [Salmonella enterica] Length: 372\nNumber of predicted TMHs: 1\nExp number of AAs in TMHs: 24.51309\nExp number, first 60 AAs: 0.2682\nTotal prob of N-in: 0.14402\noutside 1 331\nTMhelix 332 354\ninside 355 372

20583 GCF_001163665.1_10349_1_36 Salmonella enterica subsp. enterica serovar Typhi Proteobacteria; Gammaproteobacteria; Enterobacterales; Enterobacteriaceae; Salmonella; Salmonella enterica; Salmonella enterica subsp. enterica MTGDNTLITSHGINRRDFMKLCAALAATMGLSSKAAAEMAESVSNPQRPPVIWIGAQECT WP_000145429.1 hydrogenase 2 small subunit [Salmonella enterica] Length: 372\nNumber of predicted TMHs: 1\nExp number of AAs in TMHs: 24.51309\nExp number, first 60 AAs: 0.2682\nTotal prob of N-in: 0.14402\noutside 1 331\nTMhelix 332 354\ninside 355 372

20584 GCF_001163985.1_10562_2_38 Salmonella enterica subsp. enterica serovar Typhi Proteobacteria; Gammaproteobacteria; Enterobacterales; Enterobacteriaceae; Salmonella; Salmonella enterica; Salmonella enterica subsp. enterica MTGDNTLITSHGINRRDFMKLCAALAATMGLSSKAAAEMAESVSNPQRPPVIWIGAQECT WP_000145429.1 hydrogenase 2 small subunit [Salmonella enterica] Length: 372\nNumber of predicted TMHs: 1\nExp number of AAs in TMHs: 24.51309\nExp number, first 60 AAs: 0.2682\nTotal prob of N-in: 0.14402\noutside 1 331\nTMhelix 332 354\ninside 355 372

20585 GCF_001164165.1_10541_2_41 Salmonella enterica subsp. enterica serovar Typhi Proteobacteria; Gammaproteobacteria; Enterobacterales; Enterobacteriaceae; Salmonella; Salmonella enterica; Salmonella enterica subsp. enterica MTGDNTLITSHGINRRDFMKLCAALAATMGLSSKAAAEMAESVSNPQRPPVIWIGAQECT WP_000145429.1 hydrogenase 2 small subunit [Salmonella enterica] Length: 372\nNumber of predicted TMHs: 1\nExp number of AAs in TMHs: 24.51309\nExp number, first 60 AAs: 0.2682\nTotal prob of N-in: 0.14402\noutside 1 331\nTMhelix 332 354\ninside 355 372

20586 GCF_001164345.1_10492_1_24 Salmonella enterica subsp. enterica serovar Typhi Proteobacteria; Gammaproteobacteria; Enterobacterales; Enterobacteriaceae; Salmonella; Salmonella enterica; Salmonella enterica subsp. enterica MTGDNTLITSHGINRRDFMKLCAALAATMGLSSKAAAEMAESVSNPQRPPVIWIGAQECT WP_000145429.1 hydrogenase 2 small subunit [Salmonella enterica] Length: 372\nNumber of predicted TMHs: 1\nExp number of AAs in TMHs: 24.51309\nExp number, first 60 AAs: 0.2682\nTotal prob of N-in: 0.14402\noutside 1 331\nTMhelix 332 354\ninside 355 372

20587 GCF_001163025.1_10349_1_82 Salmonella enterica subsp. enterica serovar Typhi Proteobacteria; Gammaproteobacteria; Enterobacterales; Enterobacteriaceae; Salmonella; Salmonella enterica; Salmonella enterica subsp. enterica MTGDNTLITSHGINRRDFMKLCAALAATMGLSSKAATEMAESVSNPQRPPVIWIGAQECT WP_001782304.1 hydrogenase 2 small subunit [Salmonella enterica] Length: 372\nNumber of predicted TMHs: 1\nExp number of AAs in TMHs: 24.41173\nExp number, first 60 AAs: 0.16124\nTotal prob of N-in: 0.13947\noutside 1 331\nTMhelix 332 354\ninside 355 372

20588 GCF_001164605.1_8490_6_68 Salmonella enterica subsp. enterica serovar Typhi Proteobacteria; Gammaproteobacteria; Enterobacterales; Enterobacteriaceae; Salmonella; Salmonella enterica; Salmonella enterica subsp. enterica MTGDNTLITSHGINRRDFMKLCAALAATMGLSSKAAAEMAESVSNPQRPPVIWIGAQECT WP_000145429.1 hydrogenase 2 small subunit [Salmonella enterica] Length: 372\nNumber of predicted TMHs: 1\nExp number of AAs in TMHs: 24.51309\nExp number, first 60 AAs: 0.2682\nTotal prob of N-in: 0.14402\noutside 1 331\nTMhelix 332 354\ninside 355 372

20589 GCF_001165325.1_9475_6_6Salmonella enterica subsp. enterica serovar Typhi Proteobacteria; Gammaproteobacteria; Enterobacterales; Enterobacteriaceae; Salmonella; Salmonella enterica; Salmonella enterica subsp.

enterica MTGDNTLITSHGINRRDFMKLCAALAATMGLSSKAAAEMAESVSNPQRPPVIWIGAQECT WP_000145429.1 hydrogenase 2 small subunit [Salmonella enterica] Length: 372\nNumber of predicted TMHs: 1\nExp number of AAs in TMHs: 24.51309\nExp number, first 60 AAs: 0.2682\nTotal prob of N-in: 0.14402\noutside 1 331\nTMhelix 332 354\ninside 355 372

20590 GCF_001165625.1_8447_8_10 Salmonella enterica subsp. enterica serovar Typhi Proteobacteria; Gammaproteobacteria; Enterobacterales; Enterobacteriaceae; Salmonella; Salmonella enterica; Salmonella enterica subsp. enterica MTGDNTLITSHGINRRDFMKLCAALAATMGLSSKAAAEMAESVSNPQRPPVIWIGAQECT WP_000145429.1 hydrogenase 2 small subunit [Salmonella enterica] Length: 372\nNumber of predicted TMHs: 1\nExp number of AAs in TMHs: 24.51309\nExp number, first 60 AAs: 0.2682\nTotal prob of N-in: 0.14402\noutside 1 331\nTMhelix 332 354\ninside 355 372

20591 GCF_001165745.1_10209_5_51 Salmonella enterica subsp. enterica serovar Typhi Proteobacteria; Gammaproteobacteria; Enterobacterales; Enterobacteriaceae; Salmonella; Salmonella enterica; Salmonella enterica subsp. enterica MTGDNTLITSHGINRRDFMKLCAALAATMGLSSKAAAEMAESVSNPQRPPVIWIGAQECT WP_000145429.1 hydrogenase 2 small subunit [Salmonella enterica] Length: 372\nNumber of predicted TMHs: 1\nExp number of AAs in TMHs: 24.51309\nExp number, first 60 AAs: 0.2682\nTotal prob of N-in: 0.14402\noutside 1 331\nTMhelix 332 354\ninside 355 372

20592 GCF_001166045.1_9475_6_62 Salmonella enterica subsp. enterica serovar Typhi Proteobacteria; Gammaproteobacteria; Enterobacterales; Enterobacteriaceae; Salmonella; Salmonella enterica; Salmonella enterica subsp. enterica MTGDNTLITSHGINRRDFMKLCAALAATMGLSSKAAAEMAESVSNPQRPPVIWIGAQECT WP_000145429.1 hydrogenase 2 small subunit [Salmonella enterica] Length: 372\nNumber of predicted TMHs: 1\nExp number of AAs in TMHs: 24.51309\nExp number, first 60 AAs: 0.2682\nTotal prob of N-in: 0.14402\noutside 1 331\nTMhelix 332 354\ninside 355 372

20593 GCF_001166205.1_7468_7_70 Salmonella enterica subsp. enterica serovar Typhi Proteobacteria; Gammaproteobacteria; Enterobacterales; Enterobacteriaceae; Salmonella; Salmonella enterica; Salmonella enterica subsp. enterica MTGDNTLITSHGINRRDFMKLCAALAATMGLSSKAAAEMAESVSNPQRPPVIWIGAQECT WP_000145429.1 hydrogenase 2 small subunit [Salmonella enterica] Length: 372\nNumber of predicted TMHs: 1\nExp number of AAs in TMHs: 24.51309\nExp number, first 60 AAs: 0.2682\nTotal prob of N-in: 0.14402\noutside 1 331\nTMhelix 332 354\ninside 355 372

20594 GCF_001166405.1_8525_3_42 Salmonella enterica subsp. enterica serovar Typhi Proteobacteria; Gammaproteobacteria; Enterobacterales; Enterobacteriaceae; Salmonella; Salmonella enterica; Salmonella enterica subsp. enterica MTGDNTLITSHGINRRDFMKLCAALAATMGLSSKAAAEMAESVSNPQRPPVIWIGAQECT WP_000145429.1 hydrogenase 2 small subunit [Salmonella enterica] Length: 372\nNumber of predicted TMHs: 1\nExp number of AAs in TMHs: 24.51309\nExp number, first 60 AAs: 0.2682\nTotal prob of N-in: 0.14402\noutside 1 331\nTMhelix 332 354\ninside 355 372

20595 GCF_001166565.1_10349_1_28 Salmonella enterica subsp. enterica serovar Typhi Proteobacteria; Gammaproteobacteria; Enterobacterales; Enterobacteriaceae; Salmonella; Salmonella enterica; Salmonella enterica subsp. enterica MTGDNTLITSHGINRRDFMKLCAALAATMGLSSKAAAEMAESVSNPQRPPVIWIGAQECT WP_000145429.1 hydrogenase 2 small subunit [Salmonella enterica] Length: 372\nNumber of predicted TMHs: 1\nExp number of AAs in TMHs: 24.51309\nExp number, first 60 AAs: 0.2682\nTotal prob of N-in: 0.14402\noutside 1 331\nTMhelix 332 354\ninside 355 372

20596 GCF_001166665.1_10593_2_69 Salmonella enterica subsp. enterica serovar Typhi Proteobacteria; Gammaproteobacteria; Enterobacterales; Enterobacteriaceae; Salmonella; Salmonella enterica; Salmonella enterica subsp. enterica MTGDNTLITSHGINRRDFMKLCAALAATMGLSSKAAAEMAESVSNPQRPPVIWIGAQECT WP_000145429.1 hydrogenase 2 small subunit [Salmonella enterica] Length: 372\nNumber of predicted TMHs: 1\nExp number of AAs in TMHs: 24.51309\nExp number, first 60 AAs: 0.2682\nTotal prob of N-in: 0.14402\noutside 1 331\nTMhelix 332 354\ninside 355 372

20597 GCF_001167165.1_10607_1_91 Salmonella enterica subsp. enterica serovar Typhi Proteobacteria; Gammaproteobacteria; Enterobacterales; Enterobacteriaceae; Salmonella; Salmonella enterica; Salmonella enterica subsp. enterica MTGDNTLITSHGINRRDFMKLCAALAATMGLSSKAAAEMAESVSNPQRPPVIWIGAQECT WP_000145429.1 hydrogenase 2 small subunit [Salmonella enterica] Length: 372\nNumber of predicted TMHs: 1\nExp number of AAs in TMHs: 24.51309\nExp number, first 60 AAs: 0.2682\nTotal prob of N-in: 0.14402\noutside 1 331\nTMhelix 332 354\ninside 355 372

20598 GCF_001167045.1_10561_2_39 Salmonella enterica subsp. enterica serovar Typhi Proteobacteria; Gammaproteobacteria; Enterobacterales; Enterobacteriaceae; Salmonella; Salmonella enterica; Salmonella enterica subsp. enterica MTGDNTLITSHGINRRDFMKLCAALAATMGLSSKAAAEMAESVSNPQRPPVIWIGAQECT WP_000145429.1 hydrogenase 2 small subunit [Salmonella enterica] Length: 372\nNumber of predicted TMHs: 1\nExp number of AAs in TMHs: 24.51309\nExp number, first 60 AAs: 0.2682\nTotal prob of N-in: 0.14402\noutside 1 331\nTMhelix 332 354\ninside 355 372

20599 GCF_001167345.1_10492_1_50 Salmonella enterica subsp. enterica serovar Typhi Proteobacteria; Gammaproteobacteria; Enterobacterales; Enterobacteriaceae; Salmonella; Salmonella enterica; Salmonella enterica subsp. enterica MTGDNTLITSHGINRRDFMKLCAALAATMGLSSKAAAEMAESVSNPQRPPVIWIGAQECT WP_000145429.1 hydrogenase 2 small subunit [Salmonella enterica] Length: 372\nNumber of predicted TMHs: 1\nExp number of AAs in TMHs: 24.51309\nExp number, first 60 AAs: 0.2682\nTotal prob of N-in: 0.14402\noutside 1 331\nTMhelix 332 354\ninside 355 372

20600 GCF_001167405.1_10562_2_41 Salmonella enterica subsp. enterica serovar Typhi Proteobacteria; Gammaproteobacteria; Enterobacterales; Enterobacteriaceae; Salmonella; Salmonella enterica; Salmonella enterica subsp. enterica MTGDNTLITSHGINRRDFMKLCAALAATMGLSSKAAAEMAESVSNPQRPPVIWIGAQECT WP_000145429.1 hydrogenase 2 small subunit [Salmonella enterica] Length: 372\nNumber of predicted TMHs: 1\nExp number of AAs in TMHs: 24.51309\nExp number, first 60 AAs: 0.2682\nTotal prob of N-in: 0.14402\noutside 1 331\nTMhelix 332 354\ninside 355 372

20601 GCF_001167765.1_9870_8_59 Salmonella enterica subsp. enterica serovar Typhi Proteobacteria; Gammaproteobacteria; Enterobacterales; Enterobacteriaceae; Salmonella; Salmonella enterica; Salmonella enterica subsp. enterica MTGDNTLITSHGINRRDFMKLCAALAATMGLSSKAAAEMAESVSNPQRPPVIWIGAQECT WP_000145429.1 hydrogenase 2 small subunit [Salmonella enterica] Length: 372\nNumber of predicted TMHs: 1\nExp number of AAs in TMHs: 24.51309\nExp number, first 60 AAs: 0.2682\nTotal prob of N-in: 0.14402\noutside 1 331\nTMhelix 332 354\ninside 355 372

20602 GCF_001168005.1_9475_6_27 Salmonella enterica subsp. enterica serovar Typhi Proteobacteria; Gammaproteobacteria; Enterobacterales; Enterobacteriaceae; Salmonella; Salmonella enterica; Salmonella enterica subsp. enterica MTGDNTLITSHGINRRDFMKLCAALAATMGLSSKAAAEMAESVSNPQRPPVIWIGAQECT WP_000145429.1 hydrogenase 2 small subunit [Salmonella enterica] Length: 372\nNumber of predicted TMHs: 1\nExp number of AAs in TMHs: 24.51309\nExp number, first 60 AAs: 0.2682\nTotal prob of N-in: 0.14402\noutside 1 331\nTMhelix 332 354\ninside 355 372

20603 GCF_001168365.1_10592_2_55 Salmonella enterica subsp. enterica serovar Typhi Proteobacteria; Gammaproteobacteria; Enterobacterales; Enterobacteriaceae; Salmonella; Salmonella enterica; Salmonella enterica subsp. enterica MTGDNTLITSHGINRRDFMKLCAALAATMGLSSKAAAEMAESVSNPQRPPVIWIGAQECT WP_000145429.1 hydrogenase 2 small subunit [Salmonella enterica] Length: 372\nNumber of predicted TMHs: 1\nExp number of AAs in TMHs: 24.51309\nExp number, first 60 AAs: 0.2682\nTotal prob of N-in: 0.14402\noutside 1 331\nTMhelix 332 354\ninside 355 372

20604 GCF_001168285.1_10607_2_26 Salmonella enterica subsp. enterica serovar Typhi Proteobacteria; Gammaproteobacteria; Enterobacterales; Enterobacteriaceae; Salmonella; Salmonella enterica; Salmonella enterica subsp. enterica MTGDNTLITSHGINRRDFMKLCAALAATMGLSSKAAAEMAESVSNPQRPPVIWIGAQECT WP_000145429.1 hydrogenase 2 small subunit [Salmonella enterica] Length: 372\nNumber of predicted TMHs: 1\nExp number of AAs in TMHs: 24.51309\nExp number, first 60 AAs: 0.2682\nTotal prob of N-in: 0.14402\noutside 1 331\nTMhelix 332 354\ninside 355 372

20605 GCF_001168685.1_9475_6_47 Salmonella enterica subsp. enterica serovar Typhi Proteobacteria; Gammaproteobacteria; Enterobacterales; Enterobacteriaceae; Salmonella; Salmonella enterica; Salmonella enterica subsp. enterica MTGDNTLITSHGINRRDFMKLCAALAATMGLSSKAAAEMAESVSNPQRPPVIWIGAQECT WP_000145429.1 hydrogenase 2 small subunit [Salmonella enterica] Length: 372\nNumber of predicted TMHs: 1\nExp number of AAs in TMHs: 24.51309\nExp number, first 60 AAs: 0.2682\nTotal prob of N-in: 0.14402\noutside 1 331\nTMhelix 332 354\ninside 355 372

20606 GCF_001168825.1_9953_5_92 Salmonella enterica subsp. enterica serovar Typhi Proteobacteria; Gammaproteobacteria; Enterobacterales; Enterobacteriaceae; Salmonella; Salmonella enterica; Salmonella enterica subsp. enterica MTGDNTLITSHGINRRDFMKLCAALAATMGLSSKAAAEMAESVSNPQRPPVIWIGAQECT WP_000145429.1 hydrogenase 2 small subunit [Salmonella enterica] Length: 372\nNumber of

predicted TMHs: 1\nExp number of AAs in TMHs: 24.51309\nExp number, first 60 AAs: 0.2682\nTotal prob of N-in: 0.14402\noutside 1 331\nTMhelix 332 354\ninside 355 372

20607 GCF_001169045.1_10349_1_54 Salmonella enterica subsp. enterica serovar Typhi Proteobacteria; Gammaproteobacteria; Enterobacterales; Enterobacteriaceae; Salmonella; Salmonella enterica; Salmonella enterica subsp. enterica MTGDNTLITSHGINRRDFMKLCAALAATMGLSSKAAAEMAESVSNPQRPPVIWIGAQECT WP_000145429.1 hydrogenase 2 small subunit [Salmonella enterica] Length: 372\nNumber of predicted TMHs: 1\nExp number of AAs in TMHs: 24.51309\nExp number, first 60 AAs: 0.2682\nTotal prob of N-in: 0.14402\noutside 1 331\nTMhelix 332 354\ninside 355 372

20608 GCF_001169645.1_9475_6_58 Salmonella enterica subsp. enterica serovar Typhi Proteobacteria; Gammaproteobacteria; Enterobacterales; Enterobacteriaceae; Salmonella; Salmonella enterica; Salmonella enterica subsp. enterica MTGDNTLITSHGINRRDFMKLCAALAATMGLSSKAAAEMAESVSNPQRPPVIWIGAQECT WP_000145429.1 hydrogenase 2 small subunit [Salmonella enterica] Length: 372\nNumber of predicted TMHs: 1\nExp number of AAs in TMHs: 24.51309\nExp number, first 60 AAs: 0.2682\nTotal prob of N-in: 0.14402\noutside 1 331\nTMhelix 332 354\ninside 355 372

20609 GCF_001170105.1_9953_5_36 Salmonella enterica subsp. enterica serovar Typhi Proteobacteria; Gammaproteobacteria; Enterobacterales; Enterobacteriaceae; Salmonella; Salmonella enterica; Salmonella enterica subsp. enterica MTGDNTLITSHGINRRDFMKLCAALAATMGLSSKAAAEMAESVSNPQRPPVIWIGAQECT WP_000145429.1 hydrogenase 2 small subunit [Salmonella enterica] Length: 372\nNumber of predicted TMHs: 1\nExp number of AAs in TMHs: 24.51309\nExp number, first 60 AAs: 0.2682\nTotal prob of N-in: 0.14402\noutside 1 331\nTMhelix 332 354\ninside 355 372

20610 GCF_001170165.1_10349_1_64 Salmonella enterica subsp. enterica serovar Typhi Proteobacteria; Gammaproteobacteria; Enterobacterales; Enterobacteriaceae; Salmonella; Salmonella enterica; Salmonella enterica subsp. enterica MTGDNTLITSHGINRRDFMKLCAALAATMGLSSKAAAEMAESVSNPQRPPVIWIGAQECT WP_000145429.1 hydrogenase 2 small subunit [Salmonella enterica] Length: 372\nNumber of predicted TMHs: 1\nExp number of AAs in TMHs: 24.51309\nExp number, first 60 AAs: 0.2682\nTotal prob of N-in: 0.14402\noutside 1 331\nTMhelix 332 354\ninside 355 372

20611 GCF_001170285.1_10349_1_68 Salmonella enterica subsp. enterica serovar Typhi Proteobacteria; Gammaproteobacteria; Enterobacterales; Enterobacteriaceae; Salmonella; Salmonella enterica; Salmonella enterica subsp. enterica MTGDNTLITSHGINRRDFMKLCAALAATMGLSSKAAAEMAESVSNPQRPPVIWIGAQECT WP_000145429.1 hydrogenase 2 small subunit [Salmonella enterica] Length: 372\nNumber of predicted TMHs: 1\nExp number of AAs in TMHs: 24.51309\nExp number, first 60 AAs: 0.2682\nTotal prob of N-in: 0.14402\noutside 1 331\nTMhelix 332 354\ninside 355 372

20612 GCF_001170905.1_8490_5_21 Salmonella enterica subsp. enterica serovar Typhi Proteobacteria; Gammaproteobacteria; Enterobacterales; Enterobacteriaceae; Salmonella; Salmonella enterica; Salmonella enterica subsp. enterica MTGDNTLITSHGINRRDFMKLCAALAATMGLSSKAAAEMAESVSNPQRPPVIWIGAQECT WP_000145429.1 hydrogenase 2 small subunit [Salmonella enterica] Length: 372\nNumber of predicted TMHs: 1\nExp number of AAs in TMHs: 24.51309\nExp number, first 60 AAs: 0.2682\nTotal prob of N-in: 0.14402\noutside 1 331\nTMhelix 332 354\ninside 355 372

20613 GCF_001171245.1_9475_4_49 Salmonella enterica subsp. enterica serovar Typhi Proteobacteria; Gammaproteobacteria; Enterobacterales; Enterobacteriaceae; Salmonella; Salmonella enterica; Salmonella enterica subsp. enterica MTGDNTLITSHGINRRDFMKLCAALAATMGLSSKAAAEMAESVSNPQRPPVIWIGAQECT WP_000145429.1 hydrogenase 2 small subunit [Salmonella enterica] Length: 372\nNumber of predicted TMHs: 1\nExp number of AAs in TMHs: 24.51309\nExp number, first 60 AAs: 0.2682\nTotal prob of N-in: 0.14402\noutside 1 331\nTMhelix 332 354\ninside 355 372

20614 GCF_001171125.1_10592_2_47 Salmonella enterica subsp. enterica serovar Typhi Proteobacteria; Gammaproteobacteria; Enterobacterales; Enterobacteriaceae; Salmonella; Salmonella enterica; Salmonella enterica subsp. enterica MTGDNTLITSHGINRRDFMKLCAALAATMGLSSKAAAEMAESVSNPQRPPVIWIGAQECT WP_000145429.1 hydrogenase 2 small subunit [Salmonella enterica] Length: 372\nNumber of predicted TMHs: 1\nExp number of AAs in TMHs: 24.51309\nExp number, first 60 AAs: 0.2682\nTotal prob of N-in: 0.14402\noutside 1 331\nTMhelix 332 354\ninside 355 372

20615 GCF_001172385.1_10209_5_35 Salmonella enterica subsp. enterica serovar Typhi Proteobacteria; Gammaproteobacteria; Enterobacterales; Enterobacteriaceae; Salmonella; Salmonella enterica;

Salmonella enterica subsp. enterica MTGDNTLITSHGINRRDFMKLCAALAATMGLSSKAAAEMAESVSNPQRPPVIWIGAQECT WP_000145429.1 hydrogenase 2 small subunit [Salmonella enterica] Length: 372\nNumber of predicted TMHs: 1\nExp number of AAs in TMHs: 24.51309\nExp number, first 60 AAs: 0.2682\nTotal prob of N-in: 0.14402\noutside 1 331\nTMhelix 332 354\ninside 355 372

20616 GCF_001170985.1_10541_2_2 Salmonella enterica subsp. enterica serovar Typhi Proteobacteria; Gammaproteobacteria; Enterobacterales; Enterobacteriaceae; Salmonella; Salmonella enterica; Salmonella enterica subsp. enterica MTGDNTLITSHGINRRDFMKLCAALAATMGLSSKAATEMAESVSNPQRPPVIWIGAQECT WP_001782304.1 hydrogenase 2 small subunit [Salmonella enterica] Length: 372\nNumber of predicted TMHs: 1\nExp number of AAs in TMHs: 24.41173\nExp number, first 60 AAs: 0.16124\nTotal prob of N-in: 0.13947\noutside 1 331\nTMhelix 332 354\ninside 355 372

20617 GCF_001172105.1_10492_1_71 Salmonella enterica subsp. enterica serovar Typhi Proteobacteria; Gammaproteobacteria; Enterobacterales; Enterobacteriaceae; Salmonella; Salmonella enterica; Salmonella enterica subsp. enterica MTGDNTLITSHGINRRDFMKLCAALAATMGLSSKAATEMAESVSNPQRPPVIWIGAQECT WP_001782304.1 hydrogenase 2 small subunit [Salmonella enterica] Length: 372\nNumber of predicted TMHs: 1\nExp number of AAs in TMHs: 24.41173\nExp number, first 60 AAs: 0.16124\nTotal prob of N-in: 0.13947\noutside 1 331\nTMhelix 332 354\ninside 355 372

20618 GCF_001173685.1_9953_5_83 Salmonella enterica subsp. enterica serovar Typhi Proteobacteria; Gammaproteobacteria; Enterobacterales; Enterobacteriaceae; Salmonella; Salmonella enterica; Salmonella enterica subsp. enterica MTGDNTLITSHGINRRDFMKLCAALAATMGLSSKAAAEMAESVSNPQRPPVIWIGAQECT WP_000145429.1 hydrogenase 2 small subunit [Salmonella enterica] Length: 372\nNumber of predicted TMHs: 1\nExp number of AAs in TMHs: 24.51309\nExp number, first 60 AAs: 0.2682\nTotal prob of N-in: 0.14402\noutside 1 331\nTMhelix 332 354\ninside 355 372

20619 GCF_001172485.1_10492_1_64 Salmonella enterica subsp. enterica serovar Typhi Proteobacteria; Gammaproteobacteria; Enterobacterales; Enterobacteriaceae; Salmonella; Salmonella enterica; Salmonella enterica subsp. enterica MTGDNTLITSHGINRRDFMKLCAALAATMGLSSKAAAEMAESVSNPQRPPVIWIGAQECT WP_000145429.1 hydrogenase 2 small subunit [Salmonella enterica] Length: 372\nNumber of predicted TMHs: 1\nExp number of AAs in TMHs: 24.51309\nExp number, first 60 AAs: 0.2682\nTotal prob of N-in: 0.14402\noutside 1 331\nTMhelix 332 354\ninside 355 372

20620 GCF_001174365.1_8447_8_30 Salmonella enterica subsp. enterica serovar Typhi Proteobacteria; Gammaproteobacteria; Enterobacterales; Enterobacteriaceae; Salmonella; Salmonella enterica; Salmonella enterica subsp. enterica MTGDNTLITSHGINRRDFMKLCAALAATMGLSSKAAAEMAESVSNPQRPPVIWIGAQECT WP_000145429.1 hydrogenase 2 small subunit [Salmonella enterica] Length: 372\nNumber of predicted TMHs: 1\nExp number of AAs in TMHs: 24.51309\nExp number, first 60 AAs: 0.2682\nTotal prob of N-in: 0.14402\noutside 1 331\nTMhelix 332 354\ninside 355 372

20621 GCF_001172705.1_10592_2_39 Salmonella enterica subsp. enterica serovar Typhi Proteobacteria; Gammaproteobacteria; Enterobacterales; Enterobacteriaceae; Salmonella; Salmonella enterica; Salmonella enterica subsp. enterica MTGDNTLITSHGINRRDFMKLCAALAATMGLSSKAAAEMAESVSNPQRPPVIWIGAQECT WP_000145429.1 hydrogenase 2 small subunit [Salmonella enterica] Length: 372\nNumber of predicted TMHs: 1\nExp number of AAs in TMHs: 24.51309\nExp number, first 60 AAs: 0.2682\nTotal prob of N-in: 0.14402\noutside 1 331\nTMhelix 332 354\ninside 355 372

20622 GCF_001173505.1_10607_2_35 Salmonella enterica subsp. enterica serovar Typhi Proteobacteria; Gammaproteobacteria; Enterobacterales; Enterobacteriaceae; Salmonella; Salmonella enterica; Salmonella enterica subsp. enterica MTGDNTLITSHGINRRDFMKLCAALAATMGLSSKAAAEMAESVSNPQRPPVIWIGAQECT WP_000145429.1 hydrogenase 2 small subunit [Salmonella enterica] Length: 372\nNumber of predicted TMHs: 1\nExp number of AAs in TMHs: 24.51309\nExp number, first 60 AAs: 0.2682\nTotal prob of N-in: 0.14402\noutside 1 331\nTMhelix 332 354\ninside 355 372

20623 GCF_001173945.1_10608_2_58 Salmonella enterica subsp. enterica serovar Typhi Proteobacteria; Gammaproteobacteria; Enterobacterales; Enterobacteriaceae; Salmonella; Salmonella enterica; Salmonella enterica subsp. enterica MTGDNTLITSHGINRRDFMKLCAALAATMGLSSKAAAEMAESVSNPQRPPVIWIGAQECT WP_000145429.1 hydrogenase 2 small subunit [Salmonella enterica] Length: 372\nNumber of predicted TMHs: 1\nExp number of AAs in TMHs: 24.51309\nExp number, first 60 AAs: 0.2682\nTotal prob of N-in: 0.14402\noutside 1 331\nTMhelix 332 354\ninside 355 372

20624 GCF_001174705.1_8490_6_63 Salmonella enterica subsp. enterica serovar Typhi Proteobacteria; Gammaproteobacteria; Enterobacterales; Enterobacteriaceae; Salmonella; Salmonella enterica; Salmonella enterica subsp. enterica MTGDNTLITSHGINRRDFMKLCAALAATMGLSSKAAAEMAESVSNPQRPPVIWIGAQECT WP_000145429.1 hydrogenase 2 small subunit [Salmonella enterica] Length: 372\nNumber of predicted TMHs: 1\nExp number of AAs in TMHs: 24.51309\nExp number, first 60 AAs: 0.2682\nTotal prob of N-in: 0.14402\noutside 1 331\nTMhelix 332 354\ninside 355 372

20625 GCF_001174625.1_10607_1_66 Salmonella enterica subsp. enterica serovar Typhi Proteobacteria; Gammaproteobacteria; Enterobacterales; Enterobacteriaceae; Salmonella; Salmonella enterica; Salmonella enterica subsp. enterica MTGDNTLITSHGINRRDFMKLCAALAATMGLSSKAAAEMAESVSNPQRPPVIWIGAQECT WP_000145429.1 hydrogenase 2 small subunit [Salmonella enterica] Length: 372\nNumber of predicted TMHs: 1\nExp number of AAs in TMHs: 24.51309\nExp number, first 60 AAs: 0.2682\nTotal prob of N-in: 0.14402\noutside 1 331\nTMhelix 332 354\ninside 355 372

20626 GCF_001174245.1_10492_1_45 Salmonella enterica subsp. enterica serovar Typhi Proteobacteria; Gammaproteobacteria; Enterobacterales; Enterobacteriaceae; Salmonella; Salmonella enterica; Salmonella enterica subsp. enterica MTGDNTLITSHGINRRDFMKLCAALAATMGLSSKAATEMAESVSNPQRPPVIWIGAQECT WP_001782304.1 hydrogenase 2 small subunit [Salmonella enterica] Length: 372\nNumber of predicted TMHs: 1\nExp number of AAs in TMHs: 24.41173\nExp number, first 60 AAs: 0.16124\nTotal prob of N-in: 0.13947\noutside 1 331\nTMhelix 332 354\ninside 355 372

20627 GCF_001355715.1_8490_6_48 Salmonella enterica subsp. enterica serovar Typhi Proteobacteria; Gammaproteobacteria; Enterobacterales; Enterobacteriaceae; Salmonella; Salmonella enterica; Salmonella enterica subsp. enterica MTGDNTLITSHGINRRDFMKLCAALAATMGLSSKAAAEMAESVSNPQRPPVIWIGAQECT WP_000145429.1 hydrogenase 2 small subunit [Salmonella enterica] Length: 372\nNumber of predicted TMHs: 1\nExp number of AAs in TMHs: 24.51309\nExp number, first 60 AAs: 0.2682\nTotal prob of N-in: 0.14402\noutside 1 331\nTMhelix 332 354\ninside 355 372

20628 GCF_001355675.1_8490_5_3Salmonella enterica subsp. enterica serovar Typhi Proteobacteria; Gammaproteobacteria; Enterobacterales; Enterobacteriaceae; Salmonella; Salmonella enterica; Salmonella enterica subsp. enterica MTGDNTLITSHGINRRDFMKLCAALAATMGLSSKAAAEMAESVSNPQRPPVIWIGAQECT WP_000145429.1 hydrogenase 2 small subunit [Salmonella enterica] Length: 372\nNumber of predicted TMHs: 1\nExp number of AAs in TMHs: 24.51309\nExp number, first 60 AAs: 0.2682\nTotal prob of N-in: 0.14402\noutside 1 331\nTMhelix 332 354\ninside 355 372

20629 GCF_001083745.1_10561_2_2 Salmonella enterica subsp. enterica serovar Typhi Proteobacteria; Gammaproteobacteria; Enterobacterales; Enterobacteriaceae; Salmonella; Salmonella enterica; Salmonella enterica subsp. enterica MTGDNTLITSHGINRRDFMKLCAALAATMGLSSKAATEMAESVSNPQRPPVIWIGAQECT WP_001782304.1 hydrogenase 2 small subunit [Salmonella enterica] Length: 372\nNumber of predicted TMHs: 1\nExp number of AAs in TMHs: 24.41173\nExp number, first 60 AAs: 0.16124\nTotal prob of N-in: 0.13947\noutside 1 331\nTMhelix 332 354\ninside 355 372

20630 GCF_001047895.1_10592_2_52 Salmonella enterica subsp. enterica serovar Typhi Proteobacteria; Gammaproteobacteria; Enterobacterales; Enterobacteriaceae; Salmonella; Salmonella enterica; Salmonella enterica subsp. enterica MTGDNTLITSHGINRRDFMKLCAALAATMGLSSKAAAEMAESVSNPQRPPVIWIGAQECT WP_000145429.1 hydrogenase 2 small subunit [Salmonella enterica] Length: 372\nNumber of predicted TMHs: 1\nExp number of AAs in TMHs: 24.51309\nExp number, first 60 AAs: 0.2682\nTotal prob of N-in: 0.14402\noutside 1 331\nTMhelix 332 354\ninside 355 372

20631 GCF_001047935.1_10492_1_52 Salmonella enterica subsp. enterica serovar Typhi Proteobacteria; Gammaproteobacteria; Enterobacterales; Enterobacteriaceae; Salmonella; Salmonella enterica; Salmonella enterica subsp. enterica MTGDNTLITSHGINRRDFMKLCAALAATMGLSSKAAAEMAESVSNPQRPPVIWIGAQECT WP_000145429.1 hydrogenase 2 small subunit [Salmonella enterica] Length: 372\nNumber of predicted TMHs: 1\nExp number of AAs in TMHs: 24.51309\nExp number, first 60 AAs: 0.2682\nTotal prob of N-in: 0.14402\noutside 1 331\nTMhelix 332 354\ninside 355 372

20632 GCF_001047975.1_10349_1_70 Salmonella enterica subsp. enterica serovar Typhi Proteobacteria; Gammaproteobacteria; Enterobacterales; Enterobacteriaceae; Salmonella; Salmonella enterica; Salmonella enterica subsp. enterica MTGDNTLITSHGINRRDFMKLCAALAATMGLSSKAAAEMAESVSNPQRPPVIWIGAQECT WP_000145429.1 hydrogenase 2 small subunit [Salmonella enterica] Length: 372\nNumber of

predicted TMHs:  1\nExp number of AAs in TMHs: 24.51309\nExp number, first 60 AAs:  0.2682\nTotal prob of N-in: 0.14402\noutside   1   331\nTMhelix   332   354\ninside   355   372

20633 GCF_001048035.1_10349_1_60 Salmonella enterica subsp. enterica serovar Typhi Proteobacteria; Gammaproteobacteria; Enterobacterales; Enterobacteriaceae; Salmonella; Salmonella enterica; Salmonella enterica subsp. enterica MTGDNTLITSHGINRRDFMKLCAALAATMGLSSKAAAEMAESVSNPQRPPVIWIGAQECT WP_000145429.1 hydrogenase 2 small subunit [Salmonella enterica] Length: 372\nNumber of predicted TMHs:  1\nExp number of AAs in TMHs: 24.51309\nExp number, first 60 AAs:  0.2682\nTotal prob of N-in: 0.14402\noutside   1   331\nTMhelix   332   354\ninside   355   372

20634 GCF_001048015.1_10349_1_45 Salmonella enterica subsp. enterica serovar Typhi Proteobacteria; Gammaproteobacteria; Enterobacterales; Enterobacteriaceae; Salmonella; Salmonella enterica; Salmonella enterica subsp. enterica MTGDNTLITSHGINRRDFMKLCAALAATMGLSSKAAAEMAESVSNPQRPPVIWIGAQECT WP_000145429.1 hydrogenase 2 small subunit [Salmonella enterica] Length: 372\nNumber of predicted TMHs:  1\nExp number of AAs in TMHs: 24.51309\nExp number, first 60 AAs:  0.2682\nTotal prob of N-in: 0.14402\noutside   1   331\nTMhelix   332   354\ninside   355   372

20635 GCF_001048115.1_10493_1_49 Salmonella enterica subsp. enterica serovar Typhi Proteobacteria; Gammaproteobacteria; Enterobacterales; Enterobacteriaceae; Salmonella; Salmonella enterica; Salmonella enterica subsp. enterica MTGDNTLITSHGINRRDFMKLCAALAATMGLSSKAAAEMAESVSNPQRPPVIWIGAQECT WP_000145429.1 hydrogenase 2 small subunit [Salmonella enterica] Length: 372\nNumber of predicted TMHs:  1\nExp number of AAs in TMHs: 24.51309\nExp number, first 60 AAs:  0.2682\nTotal prob of N-in: 0.14402\noutside   1   331\nTMhelix   332   354\ninside   355   372

20636 GCF_001048155.1_10561_2_40 Salmonella enterica subsp. enterica serovar Typhi Proteobacteria; Gammaproteobacteria; Enterobacterales; Enterobacteriaceae; Salmonella; Salmonella enterica; Salmonella enterica subsp. enterica MTGDNTLITSHGINRRDFMKLCAALAATMGLSSKAAAEMAESVSNPQRPPVIWIGAQECT WP_000145429.1 hydrogenase 2 small subunit [Salmonella enterica] Length: 372\nNumber of predicted TMHs:  1\nExp number of AAs in TMHs: 24.51309\nExp number, first 60 AAs:  0.2682\nTotal prob of N-in: 0.14402\noutside   1   331\nTMhelix   332   354\ninside   355   372

20637 GCF_001048195.1_10562_2_18 Salmonella enterica subsp. enterica serovar Typhi Proteobacteria; Gammaproteobacteria; Enterobacterales; Enterobacteriaceae; Salmonella; Salmonella enterica; Salmonella enterica subsp. enterica MTGDNTLITSHGINRRDFMKLCAALAATMGLSSKAAAEMAESVSNPQRPPVIWIGAQECT WP_000145429.1 hydrogenase 2 small subunit [Salmonella enterica] Length: 372\nNumber of predicted TMHs:  1\nExp number of AAs in TMHs: 24.51309\nExp number, first 60 AAs:  0.2682\nTotal prob of N-in: 0.14402\noutside   1   331\nTMhelix   332   354\ninside   355   372

20638 GCF_001048075.1_10493_1_11 Salmonella enterica subsp. enterica serovar Typhi Proteobacteria; Gammaproteobacteria; Enterobacterales; Enterobacteriaceae; Salmonella; Salmonella enterica; Salmonella enterica subsp. enterica MTGDNTLITSHGINRRDFMKLCAALAATMGLSSKAATEMAESVSNPQRPPVIWIGAQECT WP_001782304.1 hydrogenase 2 small subunit [Salmonella enterica] Length: 372\nNumber of predicted TMHs:  1\nExp number of AAs in TMHs: 24.41173\nExp number, first 60 AAs:  0.16124\nTotal prob of N-in: 0.13947\noutside   1   331\nTMhelix   332   354\ninside   355   372

20639 GCF_001048235.1_10492_1_33 Salmonella enterica subsp. enterica serovar Typhi Proteobacteria; Gammaproteobacteria; Enterobacterales; Enterobacteriaceae; Salmonella; Salmonella enterica; Salmonella enterica subsp. enterica MTGDNTLITSHGINRRDFMKLCAALAATMGLSSKAAAEMAESVSNPQRPPVIWIGAQECT WP_000145429.1 hydrogenase 2 small subunit [Salmonella enterica] Length: 372\nNumber of predicted TMHs:  1\nExp number of AAs in TMHs: 24.51309\nExp number, first 60 AAs:  0.2682\nTotal prob of N-in: 0.14402\noutside   1   331\nTMhelix   332   354\ninside   355   372

20640 GCF_001048275.1_10493_1_22 Salmonella enterica subsp. enterica serovar Typhi Proteobacteria; Gammaproteobacteria; Enterobacterales; Enterobacteriaceae; Salmonella; Salmonella enterica; Salmonella enterica subsp. enterica MTGDNTLITSHGINRRDFMKLCAALAATMGLSSKAAAEMAESVSNPQRPPVIWIGAQECT WP_000145429.1 hydrogenase 2 small subunit [Salmonella enterica] Length: 372\nNumber of predicted TMHs:  1\nExp number of AAs in TMHs: 24.51309\nExp number, first 60 AAs:  0.2682\nTotal prob of N-in: 0.14402\noutside   1   331\nTMhelix   332   354\ninside   355   372

20641 GCF_001048395.1_10541_2_10 Salmonella enterica subsp. enterica serovar Typhi Proteobacteria; Gammaproteobacteria; Enterobacterales; Enterobacteriaceae; Salmonella; Salmonella enterica;

Salmonella enterica subsp. enterica	MTGDNTLITSHGINRRDFMKLCAALAATMGLSSKAAAEMAESVSNPQRPPVIWIGAQECT	WP_000145429.1	hydrogenase 2 small subunit [Salmonella enterica]	Length: 372\nNumber of predicted TMHs:  1\nExp number of AAs in TMHs: 24.51309\nExp number, first 60 AAs:  0.2682\nTotal prob of N-in: 0.14402\noutside   1   331\nTMhelix   332   354\ninside   355   372

20642	GCF_001048355.1_10562_2_35	Salmonella enterica subsp. enterica serovar Typhi	Proteobacteria; Gammaproteobacteria; Enterobacterales; Enterobacteriaceae; Salmonella; Salmonella enterica; Salmonella enterica subsp. enterica	MTGDNTLITSHGINRRDFMKLCAALAATMGLSSKAAAEMAESVSNPQRPPVIWIGAQECT	WP_000145429.1	hydrogenase 2 small subunit [Salmonella enterica]	Length: 372\nNumber of predicted TMHs:  1\nExp number of AAs in TMHs: 24.51309\nExp number, first 60 AAs:  0.2682\nTotal prob of N-in: 0.14402\noutside   1   331\nTMhelix   332   354\ninside   355   372

20643	GCF_001048435.1_10562_2_10	Salmonella enterica subsp. enterica serovar Typhi	Proteobacteria; Gammaproteobacteria; Enterobacterales; Enterobacteriaceae; Salmonella; Salmonella enterica; Salmonella enterica subsp. enterica	MTGDNTLITSHGINRRDFMKLCAALAATMGLSSKAAAEMAESVSNPQRPPVIWIGAQECT	WP_000145429.1	hydrogenase 2 small subunit [Salmonella enterica]	Length: 372\nNumber of predicted TMHs:  1\nExp number of AAs in TMHs: 24.51309\nExp number, first 60 AAs:  0.2682\nTotal prob of N-in: 0.14402\noutside   1   331\nTMhelix   332   354\ninside   355   372

20644	GCF_001048315.1_10562_2_17	Salmonella enterica subsp. enterica serovar Typhi	Proteobacteria; Gammaproteobacteria; Enterobacterales; Enterobacteriaceae; Salmonella; Salmonella enterica; Salmonella enterica subsp. enterica	MTGDNTLITSHGINRRDFMKLCAALAATMGLSSKAATEMAESVSNPQRPPVIWIGAQECT	WP_001782304.1	hydrogenase 2 small subunit [Salmonella enterica]	Length: 372\nNumber of predicted TMHs:  1\nExp number of AAs in TMHs: 24.41173\nExp number, first 60 AAs:  0.16124\nTotal prob of N-in: 0.13947\noutside   1   331\nTMhelix   332   354\ninside   355   372

20645	GCF_001079025.1_10071_8_65	Salmonella enterica subsp. enterica serovar Typhi	Proteobacteria; Gammaproteobacteria; Enterobacterales; Enterobacteriaceae; Salmonella; Salmonella enterica; Salmonella enterica subsp. enterica	MTGDNTLITSHGINRRDFMKLCAALAATMGLSSKAAAEMAESVSNPQRPPVIWIGAQECT	WP_000145429.1	hydrogenase 2 small subunit [Salmonella enterica]	Length: 372\nNumber of predicted TMHs:  1\nExp number of AAs in TMHs: 24.51309\nExp number, first 60 AAs:  0.2682\nTotal prob of N-in: 0.14402\noutside   1   331\nTMhelix   332   354\ninside   355   372

20646	GCF_001078725.1_10541_1_60	Salmonella enterica subsp. enterica serovar Typhi	Proteobacteria; Gammaproteobacteria; Enterobacterales; Enterobacteriaceae; Salmonella; Salmonella enterica; Salmonella enterica subsp. enterica	MTGDNTLITSHGINRRDFMKLCAALAATMGLSSKAAAEMAESVSNPQRPPVIWIGAQECT	WP_000145429.1	hydrogenase 2 small subunit [Salmonella enterica]	Length: 372\nNumber of predicted TMHs:  1\nExp number of AAs in TMHs: 24.51309\nExp number, first 60 AAs:  0.2682\nTotal prob of N-in: 0.14402\noutside   1   331\nTMhelix   332   354\ninside   355   372

20647	GCF_001078765.1_10562_2_46	Salmonella enterica subsp. enterica serovar Typhi	Proteobacteria; Gammaproteobacteria; Enterobacterales; Enterobacteriaceae; Salmonella; Salmonella enterica; Salmonella enterica subsp. enterica	MTGDNTLITSHGINRRDFMKLCAALAATMGLSSKAAAEMAESVSNPQRPPVIWIGAQECT	WP_000145429.1	hydrogenase 2 small subunit [Salmonella enterica]	Length: 372\nNumber of predicted TMHs:  1\nExp number of AAs in TMHs: 24.51309\nExp number, first 60 AAs:  0.2682\nTotal prob of N-in: 0.14402\noutside   1   331\nTMhelix   332   354\ninside   355   372

20648	GCF_001078825.1_10541_2_25	Salmonella enterica subsp. enterica serovar Typhi	Proteobacteria; Gammaproteobacteria; Enterobacterales; Enterobacteriaceae; Salmonella; Salmonella enterica; Salmonella enterica subsp. enterica	MTGDNTLITSHGINRRDFMKLCAALAATMGLSSKAAAEMAESVSNPQRPPVIWIGAQECT	WP_000145429.1	hydrogenase 2 small subunit [Salmonella enterica]	Length: 372\nNumber of predicted TMHs:  1\nExp number of AAs in TMHs: 24.51309\nExp number, first 60 AAs:  0.2682\nTotal prob of N-in: 0.14402\noutside   1   331\nTMhelix   332   354\ninside   355   372

20649	GCF_001078865.1_10561_2_64	Salmonella enterica subsp. enterica serovar Typhi	Proteobacteria; Gammaproteobacteria; Enterobacterales; Enterobacteriaceae; Salmonella; Salmonella enterica; Salmonella enterica subsp. enterica	MTGDNTLITSHGINRRDFMKLCAALAATMGLSSKAAAEMAESVSNPQRPPVIWIGAQECT	WP_000145429.1	hydrogenase 2 small subunit [Salmonella enterica]	Length: 372\nNumber of predicted TMHs:  1\nExp number of AAs in TMHs: 24.51309\nExp number, first 60 AAs:  0.2682\nTotal prob of N-in: 0.14402\noutside   1   331\nTMhelix   332   354\ninside   355   372

20650 GCF_001078905.1_10493_1_70 Salmonella enterica subsp. enterica serovar Typhi Proteobacteria; Gammaproteobacteria; Enterobacterales; Enterobacteriaceae; Salmonella; Salmonella enterica; Salmonella enterica subsp. enterica MTGDNTLITSHGINRRDFMKLCAALAATMGLSSKAAAEMAESVSNPQRPPVIWIGAQECT WP_000145429.1 hydrogenase 2 small subunit [Salmonella enterica] Length: 372\nNumber of predicted TMHs: 1\nExp number of AAs in TMHs: 24.51309\nExp number, first 60 AAs: 0.2682\nTotal prob of N-in: 0.14402\noutside 1 331\nTMhelix 332 354\ninside 355 372

20651 GCF_001078945.1_10060_6_22 Salmonella enterica subsp. enterica serovar Typhi Proteobacteria; Gammaproteobacteria; Enterobacterales; Enterobacteriaceae; Salmonella; Salmonella enterica; Salmonella enterica subsp. enterica MTGDNTLITSHGINRRDFMKLCAALAATMGLSSKAAAEMAESVSNPQRPPVIWIGAQECT WP_000145429.1 hydrogenase 2 small subunit [Salmonella enterica] Length: 372\nNumber of predicted TMHs: 1\nExp number of AAs in TMHs: 24.51309\nExp number, first 60 AAs: 0.2682\nTotal prob of N-in: 0.14402\noutside 1 331\nTMhelix 332 354\ninside 355 372

20652 GCF_001078985.1_10060_6_81 Salmonella enterica subsp. enterica serovar Typhi Proteobacteria; Gammaproteobacteria; Enterobacterales; Enterobacteriaceae; Salmonella; Salmonella enterica; Salmonella enterica subsp. enterica MTGDNTLITSHGINRRDFMKLCAALAATMGLSSKAAAEMAESVSNPQRPPVIWIGAQECT WP_000145429.1 hydrogenase 2 small subunit [Salmonella enterica] Length: 372\nNumber of predicted TMHs: 1\nExp number of AAs in TMHs: 24.51309\nExp number, first 60 AAs: 0.2682\nTotal prob of N-in: 0.14402\noutside 1 331\nTMhelix 332 354\ninside 355 372

20653 GCF_001079085.1_10060_6_91 Salmonella enterica subsp. enterica serovar Typhi Proteobacteria; Gammaproteobacteria; Enterobacterales; Enterobacteriaceae; Salmonella; Salmonella enterica; Salmonella enterica subsp. enterica MTGDNTLITSHGINRRDFMKLCAALAATMGLSSKAAAEMAESVSNPQRPPVIWIGAQECT WP_000145429.1 hydrogenase 2 small subunit [Salmonella enterica] Length: 372\nNumber of predicted TMHs: 1\nExp number of AAs in TMHs: 24.51309\nExp number, first 60 AAs: 0.2682\nTotal prob of N-in: 0.14402\noutside 1 331\nTMhelix 332 354\ninside 355 372

20654 GCF_001079125.1_10071_3_59 Salmonella enterica subsp. enterica serovar Typhi Proteobacteria; Gammaproteobacteria; Enterobacterales; Enterobacteriaceae; Salmonella; Salmonella enterica; Salmonella enterica subsp. enterica MTGDNTLITSHGINRRDFMKLCAALAATMGLSSKAAAEMAESVSNPQRPPVIWIGAQECT WP_000145429.1 hydrogenase 2 small subunit [Salmonella enterica] Length: 372\nNumber of predicted TMHs: 1\nExp number of AAs in TMHs: 24.51309\nExp number, first 60 AAs: 0.2682\nTotal prob of N-in: 0.14402\noutside 1 331\nTMhelix 332 354\ninside 355 372

20655 GCF_001079165.1_10060_6_46 Salmonella enterica subsp. enterica serovar Typhi Proteobacteria; Gammaproteobacteria; Enterobacterales; Enterobacteriaceae; Salmonella; Salmonella enterica; Salmonella enterica subsp. enterica MTGDNTLITSHGINRRDFMKLCAALAATMGLSSKAAAEMAESVSNPQRPPVIWIGAQECT WP_000145429.1 hydrogenase 2 small subunit [Salmonella enterica] Length: 372\nNumber of predicted TMHs: 1\nExp number of AAs in TMHs: 24.51309\nExp number, first 60 AAs: 0.2682\nTotal prob of N-in: 0.14402\noutside 1 331\nTMhelix 332 354\ninside 355 372

20656 GCF_001079205.1_10071_8_15 Salmonella enterica subsp. enterica serovar Typhi Proteobacteria; Gammaproteobacteria; Enterobacterales; Enterobacteriaceae; Salmonella; Salmonella enterica; Salmonella enterica subsp. enterica MTGDNTLITSHGINRRDFMKLCAALAATMGLSSKAAAEMAESVSNPQRPPVIWIGAQECT WP_000145429.1 hydrogenase 2 small subunit [Salmonella enterica] Length: 372\nNumber of predicted TMHs: 1\nExp number of AAs in TMHs: 24.51309\nExp number, first 60 AAs: 0.2682\nTotal prob of N-in: 0.14402\noutside 1 331\nTMhelix 332 354\ninside 355 372

20657 GCF_001079225.1_10071_3_47 Salmonella enterica subsp. enterica serovar Typhi Proteobacteria; Gammaproteobacteria; Enterobacterales; Enterobacteriaceae; Salmonella; Salmonella enterica; Salmonella enterica subsp. enterica MTGDNTLITSHGINRRDFMKLCAALAATMGLSSKAAAEMAESVSNPQRPPVIWIGAQECT WP_000145429.1 hydrogenase 2 small subunit [Salmonella enterica] Length: 372\nNumber of predicted TMHs: 1\nExp number of AAs in TMHs: 24.51309\nExp number, first 60 AAs: 0.2682\nTotal prob of N-in: 0.14402\noutside 1 331\nTMhelix 332 354\ninside 355 372

20658 GCF_001079245.1_10071_8_78 Salmonella enterica subsp. enterica serovar Typhi Proteobacteria; Gammaproteobacteria; Enterobacterales; Enterobacteriaceae; Salmonella; Salmonella enterica; Salmonella enterica subsp. enterica MTGDNTLITSHGINRRDFMKLCAALAATMGLSSKAAAEMAESVSNPQRPPVIWIGAQECT WP_000145429.1 hydrogenase 2 small subunit [Salmonella enterica] Length: 372\nNumber of

predicted TMHs:  1\nExp number of AAs in TMHs: 24.51309\nExp number, first 60 AAs:  0.2682\nTotal prob of N-in: 0.14402\noutside   1   331\nTMhelix   332   354\ninside   355   372

20659 GCF_001079285.1_10060_5_77 Salmonella enterica subsp. enterica serovar Typhi Proteobacteria; Gammaproteobacteria; Enterobacterales; Enterobacteriaceae; Salmonella; Salmonella enterica; Salmonella enterica subsp. enterica MTGDNTLITSHGINRRDFMKLCAALAATMGLSSKAAAEMAESVSNPQRPPVIWIGAQECT WP_000145429.1 hydrogenase 2 small subunit [Salmonella enterica] Length: 372\nNumber of predicted TMHs:  1\nExp number of AAs in TMHs: 24.51309\nExp number, first 60 AAs:  0.2682\nTotal prob of N-in: 0.14402\noutside   1   331\nTMhelix   332   354\ninside   355   372

20660 GCF_001079325.1_10071_8_10 Salmonella enterica subsp. enterica serovar Typhi Proteobacteria; Gammaproteobacteria; Enterobacterales; Enterobacteriaceae; Salmonella; Salmonella enterica; Salmonella enterica subsp. enterica MTGDNTLITSHGINRRDFMKLCAALAATMGLSSKAAAEMAESVSNPQRPPVIWIGAQECT WP_000145429.1 hydrogenase 2 small subunit [Salmonella enterica] Length: 372\nNumber of predicted TMHs:  1\nExp number of AAs in TMHs: 24.51309\nExp number, first 60 AAs:  0.2682\nTotal prob of N-in: 0.14402\noutside   1   331\nTMhelix   332   354\ninside   355   372

20661 GCF_001079365.1_10060_5_28 Salmonella enterica subsp. enterica serovar Typhi Proteobacteria; Gammaproteobacteria; Enterobacterales; Enterobacteriaceae; Salmonella; Salmonella enterica; Salmonella enterica subsp. enterica MTGDNTLITSHGINRRDFMKLCAALAATMGLSSKAAAEMAESVSNPQRPPVIWIGAQECT WP_000145429.1 hydrogenase 2 small subunit [Salmonella enterica] Length: 372\nNumber of predicted TMHs:  1\nExp number of AAs in TMHs: 24.51309\nExp number, first 60 AAs:  0.2682\nTotal prob of N-in: 0.14402\noutside   1   331\nTMhelix   332   354\ninside   355   372

20662 GCF_001079405.1_10071_8_27 Salmonella enterica subsp. enterica serovar Typhi Proteobacteria; Gammaproteobacteria; Enterobacterales; Enterobacteriaceae; Salmonella; Salmonella enterica; Salmonella enterica subsp. enterica MTGDNTLITSHGINRRDFMKLCAALAATMGLSSKAAAEMAESVSNPQRPPVIWIGAQECT WP_000145429.1 hydrogenase 2 small subunit [Salmonella enterica] Length: 372\nNumber of predicted TMHs:  1\nExp number of AAs in TMHs: 24.51309\nExp number, first 60 AAs:  0.2682\nTotal prob of N-in: 0.14402\noutside   1   331\nTMhelix   332   354\ninside   355   372

20663 GCF_001079445.1_10060_6_63 Salmonella enterica subsp. enterica serovar Typhi Proteobacteria; Gammaproteobacteria; Enterobacterales; Enterobacteriaceae; Salmonella; Salmonella enterica; Salmonella enterica subsp. enterica MTGDNTLITSHGINRRDFMKLCAALAATMGLSSKAAAEMAESVSNPQRPPVIWIGAQECT WP_000145429.1 hydrogenase 2 small subunit [Salmonella enterica] Length: 372\nNumber of predicted TMHs:  1\nExp number of AAs in TMHs: 24.51309\nExp number, first 60 AAs:  0.2682\nTotal prob of N-in: 0.14402\noutside   1   331\nTMhelix   332   354\ninside   355   372

20664 GCF_001079525.1_10060_6_52 Salmonella enterica subsp. enterica serovar Typhi Proteobacteria; Gammaproteobacteria; Enterobacterales; Enterobacteriaceae; Salmonella; Salmonella enterica; Salmonella enterica subsp. enterica MTGDNTLITSHGINRRDFMKLCAALAATMGLSSKAAAEMAESVSNPQRPPVIWIGAQECT WP_000145429.1 hydrogenase 2 small subunit [Salmonella enterica] Length: 372\nNumber of predicted TMHs:  1\nExp number of AAs in TMHs: 24.51309\nExp number, first 60 AAs:  0.2682\nTotal prob of N-in: 0.14402\noutside   1   331\nTMhelix   332   354\ninside   355   372

20665 GCF_001079485.1_10060_6_29 Salmonella enterica subsp. enterica serovar Typhi Proteobacteria; Gammaproteobacteria; Enterobacterales; Enterobacteriaceae; Salmonella; Salmonella enterica; Salmonella enterica subsp. enterica MTGDNTLITSHGINRRDFMKLCAALAATMGLSSKAAAEMAESVSNPQRPPVIWIGAQECT WP_000145429.1 hydrogenase 2 small subunit [Salmonella enterica] Length: 372\nNumber of predicted TMHs:  1\nExp number of AAs in TMHs: 24.51309\nExp number, first 60 AAs:  0.2682\nTotal prob of N-in: 0.14402\noutside   1   331\nTMhelix   332   354\ninside   355   372

20666 GCF_001079565.1_10071_8_31 Salmonella enterica subsp. enterica serovar Typhi Proteobacteria; Gammaproteobacteria; Enterobacterales; Enterobacteriaceae; Salmonella; Salmonella enterica; Salmonella enterica subsp. enterica MTGDNTLITSHGINRRDFMKLCAALAATMGLSSKAAAEMAESVSNPQRPPVIWIGAQECT WP_000145429.1 hydrogenase 2 small subunit [Salmonella enterica] Length: 372\nNumber of predicted TMHs:  1\nExp number of AAs in TMHs: 24.51309\nExp number, first 60 AAs:  0.2682\nTotal prob of N-in: 0.14402\noutside   1   331\nTMhelix   332   354\ninside   355   372

20667 GCF_001079605.1_10060_5_4 Salmonella enterica subsp. enterica serovar Typhi Proteobacteria; Gammaproteobacteria; Enterobacterales; Enterobacteriaceae; Salmonella; Salmonella enterica;

Salmonella enterica subsp. enterica	MTGDNTLITSHGINRRDFMKLCAALAATMGLSSKAAAEMAESVSNPQRPPVIWIGAQECT
WP_000145429.1	hydrogenase 2 small subunit [Salmonella enterica]	Length: 372\nNumber of predicted TMHs: 1\nExp number of AAs in TMHs: 24.51309\nExp number, first 60 AAs: 0.2682\nTotal prob of N-in: 0.14402\noutside 1 331\nTMhelix 332 354\ninside 355 372

20668	GCF_001079645.1_10071_3_44	Salmonella enterica subsp. enterica serovar Typhi
Proteobacteria; Gammaproteobacteria; Enterobacterales; Enterobacteriaceae; Salmonella; Salmonella enterica; Salmonella enterica subsp. enterica	MTGDNTLITSHGINRRDFMKLCAALAATMGLSSKAAAEMAESVSNPQRPPVIWIGAQECT
WP_000145429.1	hydrogenase 2 small subunit [Salmonella enterica]	Length: 372\nNumber of predicted TMHs: 1\nExp number of AAs in TMHs: 24.51309\nExp number, first 60 AAs: 0.2682\nTotal prob of N-in: 0.14402\noutside 1 331\nTMhelix 332 354\ninside 355 372

20669	GCF_001079725.1_10060_6_79	Salmonella enterica subsp. enterica serovar Typhi
Proteobacteria; Gammaproteobacteria; Enterobacterales; Enterobacteriaceae; Salmonella; Salmonella enterica; Salmonella enterica subsp. enterica	MTGDNTLITSHGINRRDFMKLCAALAATMGLSSKAAAEMAESVSNPQRPPVIWIGAQECT
WP_000145429.1	hydrogenase 2 small subunit [Salmonella enterica]	Length: 372\nNumber of predicted TMHs: 1\nExp number of AAs in TMHs: 24.51309\nExp number, first 60 AAs: 0.2682\nTotal prob of N-in: 0.14402\noutside 1 331\nTMhelix 332 354\ninside 355 372

20670	GCF_001079685.1_10071_3_31	Salmonella enterica subsp. enterica serovar Typhi
Proteobacteria; Gammaproteobacteria; Enterobacterales; Enterobacteriaceae; Salmonella; Salmonella enterica; Salmonella enterica subsp. enterica	MTGDNTLITSHGINRRDFMKLCAALAATMGLSSKAAAEMAESVSNPQRPPVIWIGAQECT
WP_000145429.1	hydrogenase 2 small subunit [Salmonella enterica]	Length: 372\nNumber of predicted TMHs: 1\nExp number of AAs in TMHs: 24.51309\nExp number, first 60 AAs: 0.2682\nTotal prob of N-in: 0.14402\noutside 1 331\nTMhelix 332 354\ninside 355 372

20671	GCF_001079765.1_10060_6_9	Salmonella enterica subsp. enterica serovar Typhi
Proteobacteria; Gammaproteobacteria; Enterobacterales; Enterobacteriaceae; Salmonella; Salmonella enterica; Salmonella enterica subsp. enterica	MTGDNTLITSHGINRRDFMKLCAALAATMGLSSKAAAEMAESVSNPQRPPVIWIGAQECT
WP_000145429.1	hydrogenase 2 small subunit [Salmonella enterica]	Length: 372\nNumber of predicted TMHs: 1\nExp number of AAs in TMHs: 24.51309\nExp number, first 60 AAs: 0.2682\nTotal prob of N-in: 0.14402\noutside 1 331\nTMhelix 332 354\ninside 355 372

20672	GCF_001079805.1_10060_6_34	Salmonella enterica subsp. enterica serovar Typhi
Proteobacteria; Gammaproteobacteria; Enterobacterales; Enterobacteriaceae; Salmonella; Salmonella enterica; Salmonella enterica subsp. enterica	MTGDNTLITSHGINRRDFMKLCAALAATMGLSSKAAAEMAESVSNPQRPPVIWIGAQECT
WP_000145429.1	hydrogenase 2 small subunit [Salmonella enterica]	Length: 372\nNumber of predicted TMHs: 1\nExp number of AAs in TMHs: 24.51309\nExp number, first 60 AAs: 0.2682\nTotal prob of N-in: 0.14402\noutside 1 331\nTMhelix 332 354\ninside 355 372

20673	GCF_001079845.1_10071_3_93	Salmonella enterica subsp. enterica serovar Typhi
Proteobacteria; Gammaproteobacteria; Enterobacterales; Enterobacteriaceae; Salmonella; Salmonella enterica; Salmonella enterica subsp. enterica	MTGDNTLITSHGINRRDFMKLCAALAATMGLSSKAAAEMAESVSNPQRPPVIWIGAQECT
WP_000145429.1	hydrogenase 2 small subunit [Salmonella enterica]	Length: 372\nNumber of predicted TMHs: 1\nExp number of AAs in TMHs: 24.51309\nExp number, first 60 AAs: 0.2682\nTotal prob of N-in: 0.14402\noutside 1 331\nTMhelix 332 354\ninside 355 372

20674	GCF_001079885.1_10060_6_12	Salmonella enterica subsp. enterica serovar Typhi
Proteobacteria; Gammaproteobacteria; Enterobacterales; Enterobacteriaceae; Salmonella; Salmonella enterica; Salmonella enterica subsp. enterica	MTGDNTLITSHGINRRDFMKLCAALAATMGLSSKAAAEMAESVSNPQRPPVIWIGAQECT
WP_000145429.1	hydrogenase 2 small subunit [Salmonella enterica]	Length: 372\nNumber of predicted TMHs: 1\nExp number of AAs in TMHs: 24.51309\nExp number, first 60 AAs: 0.2682\nTotal prob of N-in: 0.14402\noutside 1 331\nTMhelix 332 354\ninside 355 372

20675	GCF_001079925.1_10060_6_18	Salmonella enterica subsp. enterica serovar Typhi
Proteobacteria; Gammaproteobacteria; Enterobacterales; Enterobacteriaceae; Salmonella; Salmonella enterica; Salmonella enterica subsp. enterica	MTGDNTLITSHGINRRDFMKLCAALAATMGLSSKAAAEMAESVSNPQRPPVIWIGAQECT
WP_000145429.1	hydrogenase 2 small subunit [Salmonella enterica]	Length: 372\nNumber of predicted TMHs: 1\nExp number of AAs in TMHs: 24.51309\nExp number, first 60 AAs: 0.2682\nTotal prob of N-in: 0.14402\noutside 1 331\nTMhelix 332 354\ninside 355 372

20676 GCF_001079945.1_10071_3_66 Salmonella enterica subsp. enterica serovar Typhi
Proteobacteria; Gammaproteobacteria; Enterobacterales; Enterobacteriaceae; Salmonella; Salmonella enterica; Salmonella enterica subsp. enterica MTGDNTLITSHGINRRDFMKLCAALAATMGLSSKAAAEMAESVSNPQRPPVIWIGAQECT
WP_000145429.1 hydrogenase 2 small subunit [Salmonella enterica] Length: 372\nNumber of predicted TMHs: 1\nExp number of AAs in TMHs: 24.51309\nExp number, first 60 AAs: 0.2682\nTotal prob of N-in: 0.14402\noutside 1 331\nTMhelix 332 354\ninside 355 372

20677 GCF_001079985.1_10071_8_71 Salmonella enterica subsp. enterica serovar Typhi
Proteobacteria; Gammaproteobacteria; Enterobacterales; Enterobacteriaceae; Salmonella; Salmonella enterica; Salmonella enterica subsp. enterica MTGDNTLITSHGINRRDFMKLCAALAATMGLSSKAAAEMAESVSNPQRPPVIWIGAQECT
WP_000145429.1 hydrogenase 2 small subunit [Salmonella enterica] Length: 372\nNumber of predicted TMHs: 1\nExp number of AAs in TMHs: 24.51309\nExp number, first 60 AAs: 0.2682\nTotal prob of N-in: 0.14402\noutside 1 331\nTMhelix 332 354\ninside 355 372

20678 GCF_001080065.1_10493_1_19 Salmonella enterica subsp. enterica serovar Typhi
Proteobacteria; Gammaproteobacteria; Enterobacterales; Enterobacteriaceae; Salmonella; Salmonella enterica; Salmonella enterica subsp. enterica MTGDNTLITSHGINRRDFMKLCAALAATMGLSSKAAAEMAESVSNPQRPPVIWIGAQECT
WP_000145429.1 hydrogenase 2 small subunit [Salmonella enterica] Length: 372\nNumber of predicted TMHs: 1\nExp number of AAs in TMHs: 24.51309\nExp number, first 60 AAs: 0.2682\nTotal prob of N-in: 0.14402\noutside 1 331\nTMhelix 332 354\ninside 355 372

20679 GCF_001080105.1_10071_8_25 Salmonella enterica subsp. enterica serovar Typhi
Proteobacteria; Gammaproteobacteria; Enterobacterales; Enterobacteriaceae; Salmonella; Salmonella enterica; Salmonella enterica subsp. enterica MTGDNTLITSHGINRRDFMKLCAALAATMGLSSKAAAEMAESVSNPQRPPVIWIGAQECT
WP_000145429.1 hydrogenase 2 small subunit [Salmonella enterica] Length: 372\nNumber of predicted TMHs: 1\nExp number of AAs in TMHs: 24.51309\nExp number, first 60 AAs: 0.2682\nTotal prob of N-in: 0.14402\noutside 1 331\nTMhelix 332 354\ninside 355 372

20680 GCF_001080145.1_10071_3_33 Salmonella enterica subsp. enterica serovar Typhi
Proteobacteria; Gammaproteobacteria; Enterobacterales; Enterobacteriaceae; Salmonella; Salmonella enterica; Salmonella enterica subsp. enterica MTGDNTLITSHGINRRDFMKLCAALAATMGLSSKAAAEMAESVSNPQRPPVIWIGAQECT
WP_000145429.1 hydrogenase 2 small subunit [Salmonella enterica] Length: 372\nNumber of predicted TMHs: 1\nExp number of AAs in TMHs: 24.51309\nExp number, first 60 AAs: 0.2682\nTotal prob of N-in: 0.14402\noutside 1 331\nTMhelix 332 354\ninside 355 372

20681 GCF_001080185.1_10071_3_34 Salmonella enterica subsp. enterica serovar Typhi
Proteobacteria; Gammaproteobacteria; Enterobacterales; Enterobacteriaceae; Salmonella; Salmonella enterica; Salmonella enterica subsp. enterica MTGDNTLITSHGINRRDFMKLCAALAATMGLSSKAAAEMAESVSNPQRPPVIWIGAQECT
WP_000145429.1 hydrogenase 2 small subunit [Salmonella enterica] Length: 372\nNumber of predicted TMHs: 1\nExp number of AAs in TMHs: 24.51309\nExp number, first 60 AAs: 0.2682\nTotal prob of N-in: 0.14402\noutside 1 331\nTMhelix 332 354\ninside 355 372

20682 GCF_001080225.1_10071_8_8 Salmonella enterica subsp. enterica serovar Typhi
Proteobacteria; Gammaproteobacteria; Enterobacterales; Enterobacteriaceae; Salmonella; Salmonella enterica; Salmonella enterica subsp. enterica MTGDNTLITSHGINRRDFMKLCAALAATMGLSSKAAAEMAESVSNPQRPPVIWIGAQECT
WP_000145429.1 hydrogenase 2 small subunit [Salmonella enterica] Length: 372\nNumber of predicted TMHs: 1\nExp number of AAs in TMHs: 24.51309\nExp number, first 60 AAs: 0.2682\nTotal prob of N-in: 0.14402\noutside 1 331\nTMhelix 332 354\ninside 355 372

20683 GCF_001080265.1_10060_5_3 Salmonella enterica subsp. enterica serovar Typhi
Proteobacteria; Gammaproteobacteria; Enterobacterales; Enterobacteriaceae; Salmonella; Salmonella enterica; Salmonella enterica subsp. enterica MTGDNTLITSHGINRRDFMKLCAALAATMGLSSKAAAEMAESVSNPQRPPVIWIGAQECT
WP_000145429.1 hydrogenase 2 small subunit [Salmonella enterica] Length: 372\nNumber of predicted TMHs: 1\nExp number of AAs in TMHs: 24.51309\nExp number, first 60 AAs: 0.2682\nTotal prob of N-in: 0.14402\noutside 1 331\nTMhelix 332 354\ninside 355 372

20684 GCF_001080305.1_10060_6_73 Salmonella enterica subsp. enterica serovar Typhi
Proteobacteria; Gammaproteobacteria; Enterobacterales; Enterobacteriaceae; Salmonella; Salmonella enterica; Salmonella enterica subsp. enterica MTGDNTLITSHGINRRDFMKLCAALAATMGLSSKAAAEMAESVSNPQRPPVIWIGAQECT
WP_000145429.1 hydrogenase 2 small subunit [Salmonella enterica] Length: 372\nNumber of

predicted TMHs:  1\nExp number of AAs in TMHs: 24.51309\nExp number, first 60 AAs:  0.2682\nTotal prob of N-in: 0.14402\noutside   1   331\nTMhelix   332   354\ninside   355   372

20685 GCF_001080025.1_10071_3_12 Salmonella enterica subsp. enterica serovar Typhi Proteobacteria; Gammaproteobacteria; Enterobacterales; Enterobacteriaceae; Salmonella; Salmonella enterica; Salmonella enterica subsp. enterica MTGDNTLITSHGINRRDFMKLCAALAATMGLSSKAATEMAESVSNPQRPPVIWIGAQECT WP_001782304.1 hydrogenase 2 small subunit [Salmonella enterica] Length: 372\nNumber of predicted TMHs:  1\nExp number of AAs in TMHs: 24.41173\nExp number, first 60 AAs:  0.16124\nTotal prob of N-in: 0.13947\noutside   1   331\nTMhelix   332   354\ninside   355   372

20686 GCF_001080345.1_10071_8_50 Salmonella enterica subsp. enterica serovar Typhi Proteobacteria; Gammaproteobacteria; Enterobacterales; Enterobacteriaceae; Salmonella; Salmonella enterica; Salmonella enterica subsp. enterica MTGDNTLITSHGINRRDFMKLCAALAATMGLSSKAAAEMAESVSNPQRPPVIWIGAQECT WP_000145429.1 hydrogenase 2 small subunit [Salmonella enterica] Length: 372\nNumber of predicted TMHs:  1\nExp number of AAs in TMHs: 24.51309\nExp number, first 60 AAs:  0.2682\nTotal prob of N-in: 0.14402\noutside   1   331\nTMhelix   332   354\ninside   355   372

20687 GCF_001080385.1_10060_6_37 Salmonella enterica subsp. enterica serovar Typhi Proteobacteria; Gammaproteobacteria; Enterobacterales; Enterobacteriaceae; Salmonella; Salmonella enterica; Salmonella enterica subsp. enterica MTGDNTLITSHGINRRDFMKLCAALAATMGLSSKAAAEMAESVSNPQRPPVIWIGAQECT WP_000145429.1 hydrogenase 2 small subunit [Salmonella enterica] Length: 372\nNumber of predicted TMHs:  1\nExp number of AAs in TMHs: 24.51309\nExp number, first 60 AAs:  0.2682\nTotal prob of N-in: 0.14402\noutside   1   331\nTMhelix   332   354\ninside   355   372

20688 GCF_001080465.1_10071_8_37 Salmonella enterica subsp. enterica serovar Typhi Proteobacteria; Gammaproteobacteria; Enterobacterales; Enterobacteriaceae; Salmonella; Salmonella enterica; Salmonella enterica subsp. enterica MTGDNTLITSHGINRRDFMKLCAALAATMGLSSKAAAEMAESVSNPQRPPVIWIGAQECT WP_000145429.1 hydrogenase 2 small subunit [Salmonella enterica] Length: 372\nNumber of predicted TMHs:  1\nExp number of AAs in TMHs: 24.51309\nExp number, first 60 AAs:  0.2682\nTotal prob of N-in: 0.14402\noutside   1   331\nTMhelix   332   354\ninside   355   372

20689 GCF_001080425.1_10071_3_36 Salmonella enterica subsp. enterica serovar Typhi Proteobacteria; Gammaproteobacteria; Enterobacterales; Enterobacteriaceae; Salmonella; Salmonella enterica; Salmonella enterica subsp. enterica MTGDNTLITSHGINRRDFMKLCAALAATMGLSSKAAAEMAESVSNPQRPPVIWIGAQECT WP_000145429.1 hydrogenase 2 small subunit [Salmonella enterica] Length: 372\nNumber of predicted TMHs:  1\nExp number of AAs in TMHs: 24.51309\nExp number, first 60 AAs:  0.2682\nTotal prob of N-in: 0.14402\noutside   1   331\nTMhelix   332   354\ninside   355   372

20690 GCF_001080505.1_10060_5_87 Salmonella enterica subsp. enterica serovar Typhi Proteobacteria; Gammaproteobacteria; Enterobacterales; Enterobacteriaceae; Salmonella; Salmonella enterica; Salmonella enterica subsp. enterica MTGDNTLITSHGINRRDFMKLCAALAATMGLSSKAAAEMAESVSNPQRPPVIWIGAQECT WP_000145429.1 hydrogenase 2 small subunit [Salmonella enterica] Length: 372\nNumber of predicted TMHs:  1\nExp number of AAs in TMHs: 24.51309\nExp number, first 60 AAs:  0.2682\nTotal prob of N-in: 0.14402\noutside   1   331\nTMhelix   332   354\ninside   355   372

20691 GCF_001080545.1_10071_8_76 Salmonella enterica subsp. enterica serovar Typhi Proteobacteria; Gammaproteobacteria; Enterobacterales; Enterobacteriaceae; Salmonella; Salmonella enterica; Salmonella enterica subsp. enterica MTGDNTLITSHGINRRDFMKLCAALAATMGLSSKAAAEMAESVSNPQRPPVIWIGAQECT WP_000145429.1 hydrogenase 2 small subunit [Salmonella enterica] Length: 372\nNumber of predicted TMHs:  1\nExp number of AAs in TMHs: 24.51309\nExp number, first 60 AAs:  0.2682\nTotal prob of N-in: 0.14402\noutside   1   331\nTMhelix   332   354\ninside   355   372

20692 GCF_001080645.1_10071_8_44 Salmonella enterica subsp. enterica serovar Typhi Proteobacteria; Gammaproteobacteria; Enterobacterales; Enterobacteriaceae; Salmonella; Salmonella enterica; Salmonella enterica subsp. enterica MTGDNTLITSHGINRRDFMKLCAALAATMGLSSKAAAEMAESVSNPQRPPVIWIGAQECT WP_000145429.1 hydrogenase 2 small subunit [Salmonella enterica] Length: 372\nNumber of predicted TMHs:  1\nExp number of AAs in TMHs: 24.51309\nExp number, first 60 AAs:  0.2682\nTotal prob of N-in: 0.14402\noutside   1   331\nTMhelix   332   354\ninside   355   372

20693 GCF_001080585.1_10060_5_66 Salmonella enterica subsp. enterica serovar Typhi Proteobacteria; Gammaproteobacteria; Enterobacterales; Enterobacteriaceae; Salmonella; Salmonella enterica;

Salmonella enterica subsp. enterica	MTGDNTLITSHGINRRDFMKLCAALAATMGLSSKAAAEMAESVSNPQRPPVIWIGAQECT	WP_000145429.1	hydrogenase 2 small subunit [Salmonella enterica]	Length: 372\nNumber of predicted TMHs:  1\nExp number of AAs in TMHs: 24.51309\nExp number, first 60 AAs:  0.2682\nTotal prob of N-in: 0.14402\noutside   1   331\nTMhelix   332   354\ninside   355   372

20694	GCF_001080625.1_10071_8_95	Salmonella enterica subsp. enterica serovar Typhi	Proteobacteria; Gammaproteobacteria; Enterobacterales; Enterobacteriaceae; Salmonella; Salmonella enterica; Salmonella enterica subsp. enterica	MTGDNTLITSHGINRRDFMKLCAALAATMGLSSKAAAEMAESVSNPQRPPVIWIGAQECT	WP_000145429.1	hydrogenase 2 small subunit [Salmonella enterica]	Length: 372\nNumber of predicted TMHs:  1\nExp number of AAs in TMHs: 24.51309\nExp number, first 60 AAs:  0.2682\nTotal prob of N-in: 0.14402\noutside   1   331\nTMhelix   332   354\ninside   355   372

20695	GCF_001080685.1_10060_5_11	Salmonella enterica subsp. enterica serovar Typhi	Proteobacteria; Gammaproteobacteria; Enterobacterales; Enterobacteriaceae; Salmonella; Salmonella enterica; Salmonella enterica subsp. enterica	MTGDNTLITSHGINRRDFMKLCAALAATMGLSSKAAAEMAESVSNPQRPPVIWIGAQECT	WP_000145429.1	hydrogenase 2 small subunit [Salmonella enterica]	Length: 372\nNumber of predicted TMHs:  1\nExp number of AAs in TMHs: 24.51309\nExp number, first 60 AAs:  0.2682\nTotal prob of N-in: 0.14402\noutside   1   331\nTMhelix   332   354\ninside   355   372

20696	GCF_001080725.1_10060_5_70	Salmonella enterica subsp. enterica serovar Typhi	Proteobacteria; Gammaproteobacteria; Enterobacterales; Enterobacteriaceae; Salmonella; Salmonella enterica; Salmonella enterica subsp. enterica	MTGDNTLITSHGINRRDFMKLCAALAATMGLSSKAAAEMAESVSNPQRPPVIWIGAQECT	WP_000145429.1	hydrogenase 2 small subunit [Salmonella enterica]	Length: 372\nNumber of predicted TMHs:  1\nExp number of AAs in TMHs: 24.51309\nExp number, first 60 AAs:  0.2682\nTotal prob of N-in: 0.14402\noutside   1   331\nTMhelix   332   354\ninside   355   372

20697	GCF_001080765.1_10060_5_40	Salmonella enterica subsp. enterica serovar Typhi	Proteobacteria; Gammaproteobacteria; Enterobacterales; Enterobacteriaceae; Salmonella; Salmonella enterica; Salmonella enterica subsp. enterica	MTGDNTLITSHGINRRDFMKLCAALAATMGLSSKAAAEMAESVSNPQRPPVIWIGAQECT	WP_000145429.1	hydrogenase 2 small subunit [Salmonella enterica]	Length: 372\nNumber of predicted TMHs:  1\nExp number of AAs in TMHs: 24.51309\nExp number, first 60 AAs:  0.2682\nTotal prob of N-in: 0.14402\noutside   1   331\nTMhelix   332   354\ninside   355   372

20698	GCF_001080845.1_10071_8_58	Salmonella enterica subsp. enterica serovar Typhi	Proteobacteria; Gammaproteobacteria; Enterobacterales; Enterobacteriaceae; Salmonella; Salmonella enterica; Salmonella enterica subsp. enterica	MTGDNTLITSHGINRRDFMKLCAALAATMGLSSKAAAEMAESVSNPQRPPVIWIGAQECT	WP_000145429.1	hydrogenase 2 small subunit [Salmonella enterica]	Length: 372\nNumber of predicted TMHs:  1\nExp number of AAs in TMHs: 24.51309\nExp number, first 60 AAs:  0.2682\nTotal prob of N-in: 0.14402\noutside   1   331\nTMhelix   332   354\ninside   355   372

20699	GCF_001080805.1_10071_3_65	Salmonella enterica subsp. enterica serovar Typhi	Proteobacteria; Gammaproteobacteria; Enterobacterales; Enterobacteriaceae; Salmonella; Salmonella enterica; Salmonella enterica subsp. enterica	MTGDNTLITSHGINRRDFMKLCAALAATMGLSSKAAAEMAESVSNPQRPPVIWIGAQECT	WP_000145429.1	hydrogenase 2 small subunit [Salmonella enterica]	Length: 372\nNumber of predicted TMHs:  1\nExp number of AAs in TMHs: 24.51309\nExp number, first 60 AAs:  0.2682\nTotal prob of N-in: 0.14402\noutside   1   331\nTMhelix   332   354\ninside   355   372

20700	GCF_001080885.1_10060_6_21	Salmonella enterica subsp. enterica serovar Typhi	Proteobacteria; Gammaproteobacteria; Enterobacterales; Enterobacteriaceae; Salmonella; Salmonella enterica; Salmonella enterica subsp. enterica	MTGDNTLITSHGINRRDFMKLCAALAATMGLSSKAAAEMAESVSNPQRPPVIWIGAQECT	WP_000145429.1	hydrogenase 2 small subunit [Salmonella enterica]	Length: 372\nNumber of predicted TMHs:  1\nExp number of AAs in TMHs: 24.51309\nExp number, first 60 AAs:  0.2682\nTotal prob of N-in: 0.14402\noutside   1   331\nTMhelix   332   354\ninside   355   372

20701	GCF_001080925.1_10071_3_56	Salmonella enterica subsp. enterica serovar Typhi	Proteobacteria; Gammaproteobacteria; Enterobacterales; Enterobacteriaceae; Salmonella; Salmonella enterica; Salmonella enterica subsp. enterica	MTGDNTLITSHGINRRDFMKLCAALAATMGLSSKAAAEMAESVSNPQRPPVIWIGAQECT	WP_000145429.1	hydrogenase 2 small subunit [Salmonella enterica]	Length: 372\nNumber of predicted TMHs:  1\nExp number of AAs in TMHs: 24.51309\nExp number, first 60 AAs:  0.2682\nTotal prob of N-in: 0.14402\noutside   1   331\nTMhelix   332   354\ninside   355   372

20702 GCF_001080965.1_10071_3_67 Salmonella enterica subsp. enterica serovar Typhi Proteobacteria; Gammaproteobacteria; Enterobacterales; Enterobacteriaceae; Salmonella; Salmonella enterica; Salmonella enterica subsp. enterica MTGDNTLITSHGINRRDFMKLCAALAATMGLSSKAAAEMAESVSNPQRPPVIWIGAQECT WP_000145429.1 hydrogenase 2 small subunit [Salmonella enterica] Length: 372\nNumber of predicted TMHs: 1\nExp number of AAs in TMHs: 24.51309\nExp number, first 60 AAs: 0.2682\nTotal prob of N-in: 0.14402\noutside 1 331\nTMhelix 332 354\ninside 355 372

20703 GCF_001081005.1_10071_8_43 Salmonella enterica subsp. enterica serovar Typhi Proteobacteria; Gammaproteobacteria; Enterobacterales; Enterobacteriaceae; Salmonella; Salmonella enterica; Salmonella enterica subsp. enterica MTGDNTLITSHGINRRDFMKLCAALAATMGLSSKAAAEMAESVSNPQRPPVIWIGAQECT WP_000145429.1 hydrogenase 2 small subunit [Salmonella enterica] Length: 372\nNumber of predicted TMHs: 1\nExp number of AAs in TMHs: 24.51309\nExp number, first 60 AAs: 0.2682\nTotal prob of N-in: 0.14402\noutside 1 331\nTMhelix 332 354\ninside 355 372

20704 GCF_001081045.1_10071_3_88 Salmonella enterica subsp. enterica serovar Typhi Proteobacteria; Gammaproteobacteria; Enterobacterales; Enterobacteriaceae; Salmonella; Salmonella enterica; Salmonella enterica subsp. enterica MTGDNTLITSHGINRRDFMKLCAALAATMGLSSKAAAEMAESVSNPQRPPVIWIGAQECT WP_000145429.1 hydrogenase 2 small subunit [Salmonella enterica] Length: 372\nNumber of predicted TMHs: 1\nExp number of AAs in TMHs: 24.51309\nExp number, first 60 AAs: 0.2682\nTotal prob of N-in: 0.14402\noutside 1 331\nTMhelix 332 354\ninside 355 372

20705 GCF_001081125.1_10071_8_7 Salmonella enterica subsp. enterica serovar Typhi Proteobacteria; Gammaproteobacteria; Enterobacterales; Enterobacteriaceae; Salmonella; Salmonella enterica; Salmonella enterica subsp. enterica MTGDNTLITSHGINRRDFMKLCAALAATMGLSSKAAAEMAESVSNPQRPPVIWIGAQECT WP_000145429.1 hydrogenase 2 small subunit [Salmonella enterica] Length: 372\nNumber of predicted TMHs: 1\nExp number of AAs in TMHs: 24.51309\nExp number, first 60 AAs: 0.2682\nTotal prob of N-in: 0.14402\noutside 1 331\nTMhelix 332 354\ninside 355 372

20706 GCF_001081085.1_10060_5_5 Salmonella enterica subsp. enterica serovar Typhi Proteobacteria; Gammaproteobacteria; Enterobacterales; Enterobacteriaceae; Salmonella; Salmonella enterica; Salmonella enterica subsp. enterica MTGDNTLITSHGINRRDFMKLCAALAATMGLSSKAAAEMAESVSNPQRPPVIWIGAQECT WP_000145429.1 hydrogenase 2 small subunit [Salmonella enterica] Length: 372\nNumber of predicted TMHs: 1\nExp number of AAs in TMHs: 24.51309\nExp number, first 60 AAs: 0.2682\nTotal prob of N-in: 0.14402\noutside 1 331\nTMhelix 332 354\ninside 355 372

20707 GCF_001081165.1_10071_3_91 Salmonella enterica subsp. enterica serovar Typhi Proteobacteria; Gammaproteobacteria; Enterobacterales; Enterobacteriaceae; Salmonella; Salmonella enterica; Salmonella enterica subsp. enterica MTGDNTLITSHGINRRDFMKLCAALAATMGLSSKAAAEMAESVSNPQRPPVIWIGAQECT WP_000145429.1 hydrogenase 2 small subunit [Salmonella enterica] Length: 372\nNumber of predicted TMHs: 1\nExp number of AAs in TMHs: 24.51309\nExp number, first 60 AAs: 0.2682\nTotal prob of N-in: 0.14402\noutside 1 331\nTMhelix 332 354\ninside 355 372

20708 GCF_001081245.1_10071_8_54 Salmonella enterica subsp. enterica serovar Typhi Proteobacteria; Gammaproteobacteria; Enterobacterales; Enterobacteriaceae; Salmonella; Salmonella enterica; Salmonella enterica subsp. enterica MTGDNTLITSHGINRRDFMKLCAALAATMGLSSKAAAEMAESVSNPQRPPVIWIGAQECT WP_000145429.1 hydrogenase 2 small subunit [Salmonella enterica] Length: 372\nNumber of predicted TMHs: 1\nExp number of AAs in TMHs: 24.51309\nExp number, first 60 AAs: 0.2682\nTotal prob of N-in: 0.14402\noutside 1 331\nTMhelix 332 354\ninside 355 372

20709 GCF_001081205.1_10071_3_32 Salmonella enterica subsp. enterica serovar Typhi Proteobacteria; Gammaproteobacteria; Enterobacterales; Enterobacteriaceae; Salmonella; Salmonella enterica; Salmonella enterica subsp. enterica MTGDNTLITSHGINRRDFMKLCAALAATMGLSSKAAAEMAESVSNPQRPPVIWIGAQECT WP_000145429.1 hydrogenase 2 small subunit [Salmonella enterica] Length: 372\nNumber of predicted TMHs: 1\nExp number of AAs in TMHs: 24.51309\nExp number, first 60 AAs: 0.2682\nTotal prob of N-in: 0.14402\noutside 1 331\nTMhelix 332 354\ninside 355 372

20710 GCF_001081285.1_10060_5_94 Salmonella enterica subsp. enterica serovar Typhi Proteobacteria; Gammaproteobacteria; Enterobacterales; Enterobacteriaceae; Salmonella; Salmonella enterica; Salmonella enterica subsp. enterica MTGDNTLITSHGINRRDFMKLCAALAATMGLSSKAAAEMAESVSNPQRPPVIWIGAQECT WP_000145429.1 hydrogenase 2 small subunit [Salmonella enterica] Length: 372\nNumber of

predicted TMHs: 1\nExp number of AAs in TMHs: 24.51309\nExp number, first 60 AAs: 0.2682\nTotal prob of N-in: 0.14402\noutside 1 331\nTMhelix 332 354\ninside 355 372

20711 GCF_001081325.1_10060_5_54 Salmonella enterica subsp. enterica serovar Typhi Proteobacteria; Gammaproteobacteria; Enterobacterales; Enterobacteriaceae; Salmonella; Salmonella enterica; Salmonella enterica subsp. enterica MTGDNTLITSHGINRRDFMKLCAALAATMGLSSKAAAEMAESVSNPQRPPVIWIGAQECT WP_000145429.1 hydrogenase 2 small subunit [Salmonella enterica] Length: 372\nNumber of predicted TMHs: 1\nExp number of AAs in TMHs: 24.51309\nExp number, first 60 AAs: 0.2682\nTotal prob of N-in: 0.14402\noutside 1 331\nTMhelix 332 354\ninside 355 372

20712 GCF_001081345.1_10060_5_72 Salmonella enterica subsp. enterica serovar Typhi Proteobacteria; Gammaproteobacteria; Enterobacterales; Enterobacteriaceae; Salmonella; Salmonella enterica; Salmonella enterica subsp. enterica MTGDNTLITSHGINRRDFMKLCAALAATMGLSSKAAAEMAESVSNPQRPPVIWIGAQECT WP_000145429.1 hydrogenase 2 small subunit [Salmonella enterica] Length: 372\nNumber of predicted TMHs: 1\nExp number of AAs in TMHs: 24.51309\nExp number, first 60 AAs: 0.2682\nTotal prob of N-in: 0.14402\noutside 1 331\nTMhelix 332 354\ninside 355 372

20713 GCF_001081385.1_10060_5_57 Salmonella enterica subsp. enterica serovar Typhi Proteobacteria; Gammaproteobacteria; Enterobacterales; Enterobacteriaceae; Salmonella; Salmonella enterica; Salmonella enterica subsp. enterica MTGDNTLITSHGINRRDFMKLCAALAATMGLSSKAAAEMAESVSNPQRPPVIWIGAQECT WP_000145429.1 hydrogenase 2 small subunit [Salmonella enterica] Length: 372\nNumber of predicted TMHs: 1\nExp number of AAs in TMHs: 24.51309\nExp number, first 60 AAs: 0.2682\nTotal prob of N-in: 0.14402\noutside 1 331\nTMhelix 332 354\ninside 355 372

20714 GCF_001081425.1_10060_5_67 Salmonella enterica subsp. enterica serovar Typhi Proteobacteria; Gammaproteobacteria; Enterobacterales; Enterobacteriaceae; Salmonella; Salmonella enterica; Salmonella enterica subsp. enterica MTGDNTLITSHGINRRDFMKLCAALAATMGLSSKAAAEMAESVSNPQRPPVIWIGAQECT WP_000145429.1 hydrogenase 2 small subunit [Salmonella enterica] Length: 372\nNumber of predicted TMHs: 1\nExp number of AAs in TMHs: 24.51309\nExp number, first 60 AAs: 0.2682\nTotal prob of N-in: 0.14402\noutside 1 331\nTMhelix 332 354\ninside 355 372

20715 GCF_001081465.1_10060_5_45 Salmonella enterica subsp. enterica serovar Typhi Proteobacteria; Gammaproteobacteria; Enterobacterales; Enterobacteriaceae; Salmonella; Salmonella enterica; Salmonella enterica subsp. enterica MTGDNTLITSHGINRRDFMKLCAALAATMGLSSKAAAEMAESVSNPQRPPVIWIGAQECT WP_000145429.1 hydrogenase 2 small subunit [Salmonella enterica] Length: 372\nNumber of predicted TMHs: 1\nExp number of AAs in TMHs: 24.51309\nExp number, first 60 AAs: 0.2682\nTotal prob of N-in: 0.14402\noutside 1 331\nTMhelix 332 354\ninside 355 372

20716 GCF_001081585.1_10071_8_33 Salmonella enterica subsp. enterica serovar Typhi Proteobacteria; Gammaproteobacteria; Enterobacterales; Enterobacteriaceae; Salmonella; Salmonella enterica; Salmonella enterica subsp. enterica MTGDNTLITSHGINRRDFMKLCAALAATMGLSSKAAAEMAESVSNPQRPPVIWIGAQECT WP_000145429.1 hydrogenase 2 small subunit [Salmonella enterica] Length: 372\nNumber of predicted TMHs: 1\nExp number of AAs in TMHs: 24.51309\nExp number, first 60 AAs: 0.2682\nTotal prob of N-in: 0.14402\noutside 1 331\nTMhelix 332 354\ninside 355 372

20717 GCF_001081505.1_10060_6_39 Salmonella enterica subsp. enterica serovar Typhi Proteobacteria; Gammaproteobacteria; Enterobacterales; Enterobacteriaceae; Salmonella; Salmonella enterica; Salmonella enterica subsp. enterica MTGDNTLITSHGINRRDFMKLCAALAATMGLSSKAAAEMAESVSNPQRPPVIWIGAQECT WP_000145429.1 hydrogenase 2 small subunit [Salmonella enterica] Length: 372\nNumber of predicted TMHs: 1\nExp number of AAs in TMHs: 24.51309\nExp number, first 60 AAs: 0.2682\nTotal prob of N-in: 0.14402\noutside 1 331\nTMhelix 332 354\ninside 355 372

20718 GCF_001081545.1_10071_8_62 Salmonella enterica subsp. enterica serovar Typhi Proteobacteria; Gammaproteobacteria; Enterobacterales; Enterobacteriaceae; Salmonella; Salmonella enterica; Salmonella enterica subsp. enterica MTGDNTLITSHGINRRDFMKLCAALAATMGLSSKAAAEMAESVSNPQRPPVIWIGAQECT WP_000145429.1 hydrogenase 2 small subunit [Salmonella enterica] Length: 372\nNumber of predicted TMHs: 1\nExp number of AAs in TMHs: 24.51309\nExp number, first 60 AAs: 0.2682\nTotal prob of N-in: 0.14402\noutside 1 331\nTMhelix 332 354\ninside 355 372

20719 GCF_001081625.1_10071_8_35 Salmonella enterica subsp. enterica serovar Typhi Proteobacteria; Gammaproteobacteria; Enterobacterales; Enterobacteriaceae; Salmonella; Salmonella enterica;

Salmonella enterica subsp. enterica	MTGDNTLITSHGINRRDFMKLCAALAATMGLSSKAAAEMAESVSNPQRPPVIWIGAQECT	WP_000145429.1	hydrogenase 2 small subunit [Salmonella enterica]	Length: 372\nNumber of predicted TMHs: 1\nExp number of AAs in TMHs: 24.51309\nExp number, first 60 AAs: 0.2682\nTotal prob of N-in: 0.14402\noutside 1 331\nTMhelix 332 354\ninside 355 372

20720	GCF_001081665.1_10060_6_5	Salmonella enterica subsp. enterica serovar Typhi	Proteobacteria; Gammaproteobacteria; Enterobacterales; Enterobacteriaceae; Salmonella; Salmonella enterica; Salmonella enterica subsp. enterica	MTGDNTLITSHGINRRDFMKLCAALAATMGLSSKAAAEMAESVSNPQRPPVIWIGAQECT	WP_000145429.1	hydrogenase 2 small subunit [Salmonella enterica]	Length: 372\nNumber of predicted TMHs: 1\nExp number of AAs in TMHs: 24.51309\nExp number, first 60 AAs: 0.2682\nTotal prob of N-in: 0.14402\noutside 1 331\nTMhelix 332 354\ninside 355 372

20721	GCF_001081705.1_10060_6_59	Salmonella enterica subsp. enterica serovar Typhi	Proteobacteria; Gammaproteobacteria; Enterobacterales; Enterobacteriaceae; Salmonella; Salmonella enterica; Salmonella enterica subsp. enterica	MTGDNTLITSHGINRRDFMKLCAALAATMGLSSKAAAEMAESVSNPQRPPVIWIGAQECT	WP_000145429.1	hydrogenase 2 small subunit [Salmonella enterica]	Length: 372\nNumber of predicted TMHs: 1\nExp number of AAs in TMHs: 24.51309\nExp number, first 60 AAs: 0.2682\nTotal prob of N-in: 0.14402\noutside 1 331\nTMhelix 332 354\ninside 355 372

20722	GCF_001081745.1_10060_5_13	Salmonella enterica subsp. enterica serovar Typhi	Proteobacteria; Gammaproteobacteria; Enterobacterales; Enterobacteriaceae; Salmonella; Salmonella enterica; Salmonella enterica subsp. enterica	MTGDNTLITSHGINRRDFMKLCAALAATMGLSSKAAAEMAESVSNPQRPPVIWIGAQECT	WP_000145429.1	hydrogenase 2 small subunit [Salmonella enterica]	Length: 372\nNumber of predicted TMHs: 1\nExp number of AAs in TMHs: 24.51309\nExp number, first 60 AAs: 0.2682\nTotal prob of N-in: 0.14402\noutside 1 331\nTMhelix 332 354\ninside 355 372

20723	GCF_001081785.1_10071_8_29	Salmonella enterica subsp. enterica serovar Typhi	Proteobacteria; Gammaproteobacteria; Enterobacterales; Enterobacteriaceae; Salmonella; Salmonella enterica; Salmonella enterica subsp. enterica	MTGDNTLITSHGINRRDFMKLCAALAATMGLSSKAAAEMAESVSNPQRPPVIWIGAQECT	WP_000145429.1	hydrogenase 2 small subunit [Salmonella enterica]	Length: 372\nNumber of predicted TMHs: 1\nExp number of AAs in TMHs: 24.51309\nExp number, first 60 AAs: 0.2682\nTotal prob of N-in: 0.14402\noutside 1 331\nTMhelix 332 354\ninside 355 372

20724	GCF_001081825.1_10060_6_16	Salmonella enterica subsp. enterica serovar Typhi	Proteobacteria; Gammaproteobacteria; Enterobacterales; Enterobacteriaceae; Salmonella; Salmonella enterica; Salmonella enterica subsp. enterica	MTGDNTLITSHGINRRDFMKLCAALAATMGLSSKAAAEMAESVSNPQRPPVIWIGAQECT	WP_000145429.1	hydrogenase 2 small subunit [Salmonella enterica]	Length: 372\nNumber of predicted TMHs: 1\nExp number of AAs in TMHs: 24.51309\nExp number, first 60 AAs: 0.2682\nTotal prob of N-in: 0.14402\noutside 1 331\nTMhelix 332 354\ninside 355 372

20725	GCF_001081865.1_10071_8_82	Salmonella enterica subsp. enterica serovar Typhi	Proteobacteria; Gammaproteobacteria; Enterobacterales; Enterobacteriaceae; Salmonella; Salmonella enterica; Salmonella enterica subsp. enterica	MTGDNTLITSHGINRRDFMKLCAALAATMGLSSKAAAEMAESVSNPQRPPVIWIGAQECT	WP_000145429.1	hydrogenase 2 small subunit [Salmonella enterica]	Length: 372\nNumber of predicted TMHs: 1\nExp number of AAs in TMHs: 24.51309\nExp number, first 60 AAs: 0.2682\nTotal prob of N-in: 0.14402\noutside 1 331\nTMhelix 332 354\ninside 355 372

20726	GCF_001081905.1_10060_5_85	Salmonella enterica subsp. enterica serovar Typhi	Proteobacteria; Gammaproteobacteria; Enterobacterales; Enterobacteriaceae; Salmonella; Salmonella enterica; Salmonella enterica subsp. enterica	MTGDNTLITSHGINRRDFMKLCAALAATMGLSSKAAAEMAESVSNPQRPPVIWIGAQECT	WP_000145429.1	hydrogenase 2 small subunit [Salmonella enterica]	Length: 372\nNumber of predicted TMHs: 1\nExp number of AAs in TMHs: 24.51309\nExp number, first 60 AAs: 0.2682\nTotal prob of N-in: 0.14402\noutside 1 331\nTMhelix 332 354\ninside 355 372

20727	GCF_001081945.1_10071_8_6	Salmonella enterica subsp. enterica serovar Typhi	Proteobacteria; Gammaproteobacteria; Enterobacterales; Enterobacteriaceae; Salmonella; Salmonella enterica; Salmonella enterica subsp. enterica	MTGDNTLITSHGINRRDFMKLCAALAATMGLSSKAAAEMAESVSNPQRPPVIWIGAQECT	WP_000145429.1	hydrogenase 2 small subunit [Salmonella enterica]	Length: 372\nNumber of predicted TMHs: 1\nExp number of AAs in TMHs: 24.51309\nExp number, first 60 AAs: 0.2682\nTotal prob of N-in: 0.14402\noutside 1 331\nTMhelix 332 354\ninside 355 372

20728 GCF_001081985.1_10071_8_21 Salmonella enterica subsp. enterica serovar Typhi Proteobacteria; Gammaproteobacteria; Enterobacterales; Enterobacteriaceae; Salmonella; Salmonella enterica; Salmonella enterica subsp. enterica MTGDNTLITSHGINRRDFMKLCAALAATMGLSSKAAAEMAESVSNPQRPPVIWIGAQECT WP_000145429.1 hydrogenase 2 small subunit [Salmonella enterica] Length: 372\nNumber of predicted TMHs: 1\nExp number of AAs in TMHs: 24.51309\nExp number, first 60 AAs: 0.2682\nTotal prob of N-in: 0.14402\noutside 1 331\nTMhelix 332 354\ninside 355 372

20729 GCF_001082045.1_10071_8_77 Salmonella enterica subsp. enterica serovar Typhi Proteobacteria; Gammaproteobacteria; Enterobacterales; Enterobacteriaceae; Salmonella; Salmonella enterica; Salmonella enterica subsp. enterica MTGDNTLITSHGINRRDFMKLCAALAATMGLSSKAAAEMAESVSNPQRPPVIWIGAQECT WP_000145429.1 hydrogenase 2 small subunit [Salmonella enterica] Length: 372\nNumber of predicted TMHs: 1\nExp number of AAs in TMHs: 24.51309\nExp number, first 60 AAs: 0.2682\nTotal prob of N-in: 0.14402\noutside 1 331\nTMhelix 332 354\ninside 355 372

20730 GCF_001082025.1_10071_3_76 Salmonella enterica subsp. enterica serovar Typhi Proteobacteria; Gammaproteobacteria; Enterobacterales; Enterobacteriaceae; Salmonella; Salmonella enterica; Salmonella enterica subsp. enterica MTGDNTLITSHGINRRDFMKLCAALAATMGLSSKAAAEMAESVSNPQRPPVIWIGAQECT WP_000145429.1 hydrogenase 2 small subunit [Salmonella enterica] Length: 372\nNumber of predicted TMHs: 1\nExp number of AAs in TMHs: 24.51309\nExp number, first 60 AAs: 0.2682\nTotal prob of N-in: 0.14402\noutside 1 331\nTMhelix 332 354\ninside 355 372

20731 GCF_001082085.1_10060_5_38 Salmonella enterica subsp. enterica serovar Typhi Proteobacteria; Gammaproteobacteria; Enterobacterales; Enterobacteriaceae; Salmonella; Salmonella enterica; Salmonella enterica subsp. enterica MTGDNTLITSHGINRRDFMKLCAALAATMGLSSKAAAEMAESVSNPQRPPVIWIGAQECT WP_000145429.1 hydrogenase 2 small subunit [Salmonella enterica] Length: 372\nNumber of predicted TMHs: 1\nExp number of AAs in TMHs: 24.51309\nExp number, first 60 AAs: 0.2682\nTotal prob of N-in: 0.14402\noutside 1 331\nTMhelix 332 354\ninside 355 372

20732 GCF_001082125.1_10071_8_45 Salmonella enterica subsp. enterica serovar Typhi Proteobacteria; Gammaproteobacteria; Enterobacterales; Enterobacteriaceae; Salmonella; Salmonella enterica; Salmonella enterica subsp. enterica MTGDNTLITSHGINRRDFMKLCAALAATMGLSSKAAAEMAESVSNPQRPPVIWIGAQECT WP_000145429.1 hydrogenase 2 small subunit [Salmonella enterica] Length: 372\nNumber of predicted TMHs: 1\nExp number of AAs in TMHs: 24.51309\nExp number, first 60 AAs: 0.2682\nTotal prob of N-in: 0.14402\noutside 1 331\nTMhelix 332 354\ninside 355 372

20733 GCF_001082165.1_10071_3_35 Salmonella enterica subsp. enterica serovar Typhi Proteobacteria; Gammaproteobacteria; Enterobacterales; Enterobacteriaceae; Salmonella; Salmonella enterica; Salmonella enterica subsp. enterica MTGDNTLITSHGINRRDFMKLCAALAATMGLSSKAAAEMAESVSNPQRPPVIWIGAQECT WP_000145429.1 hydrogenase 2 small subunit [Salmonella enterica] Length: 372\nNumber of predicted TMHs: 1\nExp number of AAs in TMHs: 24.51309\nExp number, first 60 AAs: 0.2682\nTotal prob of N-in: 0.14402\noutside 1 331\nTMhelix 332 354\ninside 355 372

20734 GCF_001082205.1_10071_3_84 Salmonella enterica subsp. enterica serovar Typhi Proteobacteria; Gammaproteobacteria; Enterobacterales; Enterobacteriaceae; Salmonella; Salmonella enterica; Salmonella enterica subsp. enterica MTGDNTLITSHGINRRDFMKLCAALAATMGLSSKAAAEMAESVSNPQRPPVIWIGAQECT WP_000145429.1 hydrogenase 2 small subunit [Salmonella enterica] Length: 372\nNumber of predicted TMHs: 1\nExp number of AAs in TMHs: 24.51309\nExp number, first 60 AAs: 0.2682\nTotal prob of N-in: 0.14402\noutside 1 331\nTMhelix 332 354\ninside 355 372

20735 GCF_001082285.1_10071_8_32 Salmonella enterica subsp. enterica serovar Typhi Proteobacteria; Gammaproteobacteria; Enterobacterales; Enterobacteriaceae; Salmonella; Salmonella enterica; Salmonella enterica subsp. enterica MTGDNTLITSHGINRRDFMKLCAALAATMGLSSKAAAEMAESVSNPQRPPVIWIGAQECT WP_000145429.1 hydrogenase 2 small subunit [Salmonella enterica] Length: 372\nNumber of predicted TMHs: 1\nExp number of AAs in TMHs: 24.51309\nExp number, first 60 AAs: 0.2682\nTotal prob of N-in: 0.14402\noutside 1 331\nTMhelix 332 354\ninside 355 372

20736 GCF_001082245.1_10071_3_70 Salmonella enterica subsp. enterica serovar Typhi Proteobacteria; Gammaproteobacteria; Enterobacterales; Enterobacteriaceae; Salmonella; Salmonella enterica; Salmonella enterica subsp. enterica MTGDNTLITSHGINRRDFMKLCAALAATMGLSSKAAAEMAESVSNPQRPPVIWIGAQECT WP_000145429.1 hydrogenase 2 small subunit [Salmonella enterica] Length: 372\nNumber of

predicted TMHs:  1\nExp number of AAs in TMHs: 24.51309\nExp number, first 60 AAs:  0.2682\nTotal prob of N-in: 0.14402\noutside   1   331\nTMhelix   332   354\ninside   355   372

20737 GCF_001082325.1_10060_6_11 Salmonella enterica subsp. enterica serovar Typhi Proteobacteria; Gammaproteobacteria; Enterobacterales; Enterobacteriaceae; Salmonella; Salmonella enterica; Salmonella enterica subsp. enterica MTGDNTLITSHGINRRDFMKLCAALAATMGLSSKAAAEMAESVSNPQRPPVIWIGAQECT WP_000145429.1 hydrogenase 2 small subunit [Salmonella enterica] Length: 372\nNumber of predicted TMHs:  1\nExp number of AAs in TMHs: 24.51309\nExp number, first 60 AAs:  0.2682\nTotal prob of N-in: 0.14402\noutside   1   331\nTMhelix   332   354\ninside   355   372

20738 GCF_001082365.1_10071_3_11 Salmonella enterica subsp. enterica serovar Typhi Proteobacteria; Gammaproteobacteria; Enterobacterales; Enterobacteriaceae; Salmonella; Salmonella enterica; Salmonella enterica subsp. enterica MTGDNTLITSHGINRRDFMKLCAALAATMGLSSKAAAEMAESVSNPQRPPVIWIGAQECT WP_000145429.1 hydrogenase 2 small subunit [Salmonella enterica] Length: 372\nNumber of predicted TMHs:  1\nExp number of AAs in TMHs: 24.51309\nExp number, first 60 AAs:  0.2682\nTotal prob of N-in: 0.14402\noutside   1   331\nTMhelix   332   354\ninside   355   372

20739 GCF_001082405.1_10071_3_85 Salmonella enterica subsp. enterica serovar Typhi Proteobacteria; Gammaproteobacteria; Enterobacterales; Enterobacteriaceae; Salmonella; Salmonella enterica; Salmonella enterica subsp. enterica MTGDNTLITSHGINRRDFMKLCAALAATMGLSSKAAAEMAESVSNPQRPPVIWIGAQECT WP_000145429.1 hydrogenase 2 small subunit [Salmonella enterica] Length: 372\nNumber of predicted TMHs:  1\nExp number of AAs in TMHs: 24.51309\nExp number, first 60 AAs:  0.2682\nTotal prob of N-in: 0.14402\noutside   1   331\nTMhelix   332   354\ninside   355   372

20740 GCF_001082445.1_10060_5_65 Salmonella enterica subsp. enterica serovar Typhi Proteobacteria; Gammaproteobacteria; Enterobacterales; Enterobacteriaceae; Salmonella; Salmonella enterica; Salmonella enterica subsp. enterica MTGDNTLITSHGINRRDFMKLCAALAATMGLSSKAAAEMAESVSNPQRPPVIWIGAQECT WP_000145429.1 hydrogenase 2 small subunit [Salmonella enterica] Length: 372\nNumber of predicted TMHs:  1\nExp number of AAs in TMHs: 24.51309\nExp number, first 60 AAs:  0.2682\nTotal prob of N-in: 0.14402\noutside   1   331\nTMhelix   332   354\ninside   355   372

20741 GCF_001082485.1_10060_5_29 Salmonella enterica subsp. enterica serovar Typhi Proteobacteria; Gammaproteobacteria; Enterobacterales; Enterobacteriaceae; Salmonella; Salmonella enterica; Salmonella enterica subsp. enterica MTGDNTLITSHGINRRDFMKLCAALAATMGLSSKAAAEMAESVSNPQRPPVIWIGAQECT WP_000145429.1 hydrogenase 2 small subunit [Salmonella enterica] Length: 372\nNumber of predicted TMHs:  1\nExp number of AAs in TMHs: 24.51309\nExp number, first 60 AAs:  0.2682\nTotal prob of N-in: 0.14402\noutside   1   331\nTMhelix   332   354\ninside   355   372

20742 GCF_001082525.1_10071_3_20 Salmonella enterica subsp. enterica serovar Typhi Proteobacteria; Gammaproteobacteria; Enterobacterales; Enterobacteriaceae; Salmonella; Salmonella enterica; Salmonella enterica subsp. enterica MTGDNTLITSHGINRRDFMKLCAALAATMGLSSKAAAEMAESVSNPQRPPVIWIGAQECT WP_000145429.1 hydrogenase 2 small subunit [Salmonella enterica] Length: 372\nNumber of predicted TMHs:  1\nExp number of AAs in TMHs: 24.51309\nExp number, first 60 AAs:  0.2682\nTotal prob of N-in: 0.14402\noutside   1   331\nTMhelix   332   354\ninside   355   372

20743 GCF_001082605.1_10071_3_29 Salmonella enterica subsp. enterica serovar Typhi Proteobacteria; Gammaproteobacteria; Enterobacterales; Enterobacteriaceae; Salmonella; Salmonella enterica; Salmonella enterica subsp. enterica MTGDNTLITSHGINRRDFMKLCAALAATMGLSSKAAAEMAESVSNPQRPPVIWIGAQECT WP_000145429.1 hydrogenase 2 small subunit [Salmonella enterica] Length: 372\nNumber of predicted TMHs:  1\nExp number of AAs in TMHs: 24.51309\nExp number, first 60 AAs:  0.2682\nTotal prob of N-in: 0.14402\noutside   1   331\nTMhelix   332   354\ninside   355   372

20744 GCF_001082645.1_10060_6_75 Salmonella enterica subsp. enterica serovar Typhi Proteobacteria; Gammaproteobacteria; Enterobacterales; Enterobacteriaceae; Salmonella; Salmonella enterica; Salmonella enterica subsp. enterica MTGDNTLITSHGINRRDFMKLCAALAATMGLSSKAAAEMAESVSNPQRPPVIWIGAQECT WP_000145429.1 hydrogenase 2 small subunit [Salmonella enterica] Length: 372\nNumber of predicted TMHs:  1\nExp number of AAs in TMHs: 24.51309\nExp number, first 60 AAs:  0.2682\nTotal prob of N-in: 0.14402\noutside   1   331\nTMhelix   332   354\ninside   355   372

20745 GCF_001082685.1_10060_6_26 Salmonella enterica subsp. enterica serovar Typhi Proteobacteria; Gammaproteobacteria; Enterobacterales; Enterobacteriaceae; Salmonella; Salmonella enterica;

Salmonella enterica subsp. enterica MTGDNTLITSHGINRRDFMKLCAALAATMGLSSKAAAEMAESVSNPQRPPVIWIGAQECT WP_000145429.1 hydrogenase 2 small subunit [Salmonella enterica] Length: 372\nNumber of predicted TMHs: 1\nExp number of AAs in TMHs: 24.51309\nExp number, first 60 AAs: 0.2682\nTotal prob of N-in: 0.14402\noutside 1 331\nTMhelix 332 354\ninside 355 372

20746 GCF_001082725.1_10060_5_51 Salmonella enterica subsp. enterica serovar Typhi Proteobacteria; Gammaproteobacteria; Enterobacterales; Enterobacteriaceae; Salmonella; Salmonella enterica; Salmonella enterica subsp. enterica MTGDNTLITSHGINRRDFMKLCAALAATMGLSSKAAAEMAESVSNPQRPPVIWIGAQECT WP_000145429.1 hydrogenase 2 small subunit [Salmonella enterica] Length: 372\nNumber of predicted TMHs: 1\nExp number of AAs in TMHs: 24.51309\nExp number, first 60 AAs: 0.2682\nTotal prob of N-in: 0.14402\noutside 1 331\nTMhelix 332 354\ninside 355 372

20747 GCF_001082745.1_10071_8_70 Salmonella enterica subsp. enterica serovar Typhi Proteobacteria; Gammaproteobacteria; Enterobacterales; Enterobacteriaceae; Salmonella; Salmonella enterica; Salmonella enterica subsp. enterica MTGDNTLITSHGINRRDFMKLCAALAATMGLSSKAAAEMAESVSNPQRPPVIWIGAQECT WP_000145429.1 hydrogenase 2 small subunit [Salmonella enterica] Length: 372\nNumber of predicted TMHs: 1\nExp number of AAs in TMHs: 24.51309\nExp number, first 60 AAs: 0.2682\nTotal prob of N-in: 0.14402\noutside 1 331\nTMhelix 332 354\ninside 355 372

20748 GCF_001082785.1_10060_5_53 Salmonella enterica subsp. enterica serovar Typhi Proteobacteria; Gammaproteobacteria; Enterobacterales; Enterobacteriaceae; Salmonella; Salmonella enterica; Salmonella enterica subsp. enterica MTGDNTLITSHGINRRDFMKLCAALAATMGLSSKAAAEMAESVSNPQRPPVIWIGAQECT WP_000145429.1 hydrogenase 2 small subunit [Salmonella enterica] Length: 372\nNumber of predicted TMHs: 1\nExp number of AAs in TMHs: 24.51309\nExp number, first 60 AAs: 0.2682\nTotal prob of N-in: 0.14402\noutside 1 331\nTMhelix 332 354\ninside 355 372

20749 GCF_001082565.1_10071_3_3 Salmonella enterica subsp. enterica serovar Typhi Proteobacteria; Gammaproteobacteria; Enterobacterales; Enterobacteriaceae; Salmonella; Salmonella enterica; Salmonella enterica subsp. enterica MTGDNTLITSHGINRRDFMKLCAALAATMGLSSKAATEMAESVSNPQRPPVIWIGAQECT WP_001782304.1 hydrogenase 2 small subunit [Salmonella enterica] Length: 372\nNumber of predicted TMHs: 1\nExp number of AAs in TMHs: 24.41173\nExp number, first 60 AAs: 0.16124\nTotal prob of N-in: 0.13947\noutside 1 331\nTMhelix 332 354\ninside 355 372

20750 GCF_001082825.1_10060_6_66 Salmonella enterica subsp. enterica serovar Typhi Proteobacteria; Gammaproteobacteria; Enterobacterales; Enterobacteriaceae; Salmonella; Salmonella enterica; Salmonella enterica subsp. enterica MTGDNTLITSHGINRRDFMKLCAALAATMGLSSKAAAEMAESVSNPQRPPVIWIGAQECT WP_000145429.1 hydrogenase 2 small subunit [Salmonella enterica] Length: 372\nNumber of predicted TMHs: 1\nExp number of AAs in TMHs: 24.51309\nExp number, first 60 AAs: 0.2682\nTotal prob of N-in: 0.14402\noutside 1 331\nTMhelix 332 354\ninside 355 372

20751 GCF_001082865.1_10071_8_11 Salmonella enterica subsp. enterica serovar Typhi Proteobacteria; Gammaproteobacteria; Enterobacterales; Enterobacteriaceae; Salmonella; Salmonella enterica; Salmonella enterica subsp. enterica MTGDNTLITSHGINRRDFMKLCAALAATMGLSSKAAAEMAESVSNPQRPPVIWIGAQECT WP_000145429.1 hydrogenase 2 small subunit [Salmonella enterica] Length: 372\nNumber of predicted TMHs: 1\nExp number of AAs in TMHs: 24.51309\nExp number, first 60 AAs: 0.2682\nTotal prob of N-in: 0.14402\noutside 1 331\nTMhelix 332 354\ninside 355 372

20752 GCF_001082905.1_10071_3_8 Salmonella enterica subsp. enterica serovar Typhi Proteobacteria; Gammaproteobacteria; Enterobacterales; Enterobacteriaceae; Salmonella; Salmonella enterica; Salmonella enterica subsp. enterica MTGDNTLITSHGINRRDFMKLCAALAATMGLSSKAAAEMAESVSNPQRPPVIWIGAQECT WP_000145429.1 hydrogenase 2 small subunit [Salmonella enterica] Length: 372\nNumber of predicted TMHs: 1\nExp number of AAs in TMHs: 24.51309\nExp number, first 60 AAs: 0.2682\nTotal prob of N-in: 0.14402\noutside 1 331\nTMhelix 332 354\ninside 355 372

20753 GCF_001083005.1_10592_2_65 Salmonella enterica subsp. enterica serovar Typhi Proteobacteria; Gammaproteobacteria; Enterobacterales; Enterobacteriaceae; Salmonella; Salmonella enterica; Salmonella enterica subsp. enterica MTGDNTLITSHGINRRDFMKLCAALAATMGLSSKAAAEMAESVSNPQRPPVIWIGAQECT WP_000145429.1 hydrogenase 2 small subunit [Salmonella enterica] Length: 372\nNumber of predicted TMHs: 1\nExp number of AAs in TMHs: 24.51309\nExp number, first 60 AAs: 0.2682\nTotal prob of N-in: 0.14402\noutside 1 331\nTMhelix 332 354\ninside 355 372

20754 GCF_001082945.1_10060_5_39 Salmonella enterica subsp. enterica serovar Typhi Proteobacteria; Gammaproteobacteria; Enterobacterales; Enterobacteriaceae; Salmonella; Salmonella enterica; Salmonella enterica subsp. enterica MTGDNTLITSHGINRRDFMKLCAALAATMGLSSKAAAEMAESVSNPQRPPVIWIGAQECT WP_000145429.1 hydrogenase 2 small subunit [Salmonella enterica] Length: 372\nNumber of predicted TMHs: 1\nExp number of AAs in TMHs: 24.51309\nExp number, first 60 AAs: 0.2682\nTotal prob of N-in: 0.14402\noutside 1 331\nTMhelix 332 354\ninside 355 372

20755 GCF_001083165.1_10608_2_68 Salmonella enterica subsp. enterica serovar Typhi Proteobacteria; Gammaproteobacteria; Enterobacterales; Enterobacteriaceae; Salmonella; Salmonella enterica; Salmonella enterica subsp. enterica MTGDNTLITSHGINRRDFMKLCAALAATMGLSSKAAAEMAESVSNPQRPPVIWIGAQECT WP_000145429.1 hydrogenase 2 small subunit [Salmonella enterica] Length: 372\nNumber of predicted TMHs: 1\nExp number of AAs in TMHs: 24.51309\nExp number, first 60 AAs: 0.2682\nTotal prob of N-in: 0.14402\noutside 1 331\nTMhelix 332 354\ninside 355 372

20756 GCF_001083325.1_10209_5_36 Salmonella enterica subsp. enterica serovar Typhi Proteobacteria; Gammaproteobacteria; Enterobacterales; Enterobacteriaceae; Salmonella; Salmonella enterica; Salmonella enterica subsp. enterica MTGDNTLITSHGINRRDFMKLCAALAATMGLSSKAAAEMAESVSNPQRPPVIWIGAQECT WP_000145429.1 hydrogenase 2 small subunit [Salmonella enterica] Length: 372\nNumber of predicted TMHs: 1\nExp number of AAs in TMHs: 24.51309\nExp number, first 60 AAs: 0.2682\nTotal prob of N-in: 0.14402\noutside 1 331\nTMhelix 332 354\ninside 355 372

20757 GCF_001084165.1_10492_1_36 Salmonella enterica subsp. enterica serovar Typhi Proteobacteria; Gammaproteobacteria; Enterobacterales; Enterobacteriaceae; Salmonella; Salmonella enterica; Salmonella enterica subsp. enterica MTGDNTLITSHGINRRDFMKLCAALAATMGLSSKAAAEMAESVSNPQRPPVIWIGAQECT WP_000145429.1 hydrogenase 2 small subunit [Salmonella enterica] Length: 372\nNumber of predicted TMHs: 1\nExp number of AAs in TMHs: 24.51309\nExp number, first 60 AAs: 0.2682\nTotal prob of N-in: 0.14402\noutside 1 331\nTMhelix 332 354\ninside 355 372

20758 GCF_001084245.1_10561_2_58 Salmonella enterica subsp. enterica serovar Typhi Proteobacteria; Gammaproteobacteria; Enterobacterales; Enterobacteriaceae; Salmonella; Salmonella enterica; Salmonella enterica subsp. enterica MTGDNTLITSHGINRRDFMKLCAALAATMGLSSKAAAEMAESVSNPQRPPVIWIGAQECT WP_000145429.1 hydrogenase 2 small subunit [Salmonella enterica] Length: 372\nNumber of predicted TMHs: 1\nExp number of AAs in TMHs: 24.51309\nExp number, first 60 AAs: 0.2682\nTotal prob of N-in: 0.14402\noutside 1 331\nTMhelix 332 354\ninside 355 372

20759 GCF_001084525.1_10593_2_26 Salmonella enterica subsp. enterica serovar Typhi Proteobacteria; Gammaproteobacteria; Enterobacterales; Enterobacteriaceae; Salmonella; Salmonella enterica; Salmonella enterica subsp. enterica MTGDNTLITSHGINRRDFMKLCAALAATMGLSSKAAAEMAESVSNPQRPPVIWIGAQECT WP_000145429.1 hydrogenase 2 small subunit [Salmonella enterica] Length: 372\nNumber of predicted TMHs: 1\nExp number of AAs in TMHs: 24.51309\nExp number, first 60 AAs: 0.2682\nTotal prob of N-in: 0.14402\noutside 1 331\nTMhelix 332 354\ninside 355 372

20760 GCF_001084945.1_10492_1_26 Salmonella enterica subsp. enterica serovar Typhi Proteobacteria; Gammaproteobacteria; Enterobacterales; Enterobacteriaceae; Salmonella; Salmonella enterica; Salmonella enterica subsp. enterica MTGDNTLITSHGINRRDFMKLCAALAATMGLSSKAAAEMAESVSNPQRPPVIWIGAQECT WP_000145429.1 hydrogenase 2 small subunit [Salmonella enterica] Length: 372\nNumber of predicted TMHs: 1\nExp number of AAs in TMHs: 24.51309\nExp number, first 60 AAs: 0.2682\nTotal prob of N-in: 0.14402\noutside 1 331\nTMhelix 332 354\ninside 355 372

20761 GCF_001084985.1_10593_2_18 Salmonella enterica subsp. enterica serovar Typhi Proteobacteria; Gammaproteobacteria; Enterobacterales; Enterobacteriaceae; Salmonella; Salmonella enterica; Salmonella enterica subsp. enterica MTGDNTLITSHGINRRDFMKLCAALAATMGLSSKAAAEMAESVSNPQRPPVIWIGAQECT WP_000145429.1 hydrogenase 2 small subunit [Salmonella enterica] Length: 372\nNumber of predicted TMHs: 1\nExp number of AAs in TMHs: 24.51309\nExp number, first 60 AAs: 0.2682\nTotal prob of N-in: 0.14402\noutside 1 331\nTMhelix 332 354\ninside 355 372

20762 GCF_001085205.1_10492_1_83 Salmonella enterica subsp. enterica serovar Typhi Proteobacteria; Gammaproteobacteria; Enterobacterales; Enterobacteriaceae; Salmonella; Salmonella enterica; Salmonella enterica subsp. enterica MTGDNTLITSHGINRRDFMKLCAALAATMGLSSKAAAEMAESVSNPQRPPVIWIGAQECT WP_000145429.1 hydrogenase 2 small subunit [Salmonella enterica] Length: 372\nNumber of

predicted TMHs:  1\nExp number of AAs in TMHs: 24.51309\nExp number, first 60 AAs:  0.2682\nTotal prob of N-in: 0.14402\noutside   1   331\nTMhelix   332   354\ninside   355   372

20763 GCF_001085245.1_9870_8_65 Salmonella enterica subsp. enterica serovar Typhi Proteobacteria; Gammaproteobacteria; Enterobacterales; Enterobacteriaceae; Salmonella; Salmonella enterica; Salmonella enterica subsp. enterica MTGDNTLITSHGINRRDFMKLCAALAATMGLSSKAAAEMAESVSNPQRPPVIWIGAQECT WP_000145429.1 hydrogenase 2 small subunit [Salmonella enterica] Length: 372\nNumber of predicted TMHs:  1\nExp number of AAs in TMHs: 24.51309\nExp number, first 60 AAs:  0.2682\nTotal prob of N-in: 0.14402\noutside   1   331\nTMhelix   332   354\ninside   355   372

20764 GCF_001085325.1_10593_2_44 Salmonella enterica subsp. enterica serovar Typhi Proteobacteria; Gammaproteobacteria; Enterobacterales; Enterobacteriaceae; Salmonella; Salmonella enterica; Salmonella enterica subsp. enterica MTGDNTLITSHGINRRDFMKLCAALAATMGLSSKAAAEMAESVSNPQRPPVIWIGAQECT WP_000145429.1 hydrogenase 2 small subunit [Salmonella enterica] Length: 372\nNumber of predicted TMHs:  1\nExp number of AAs in TMHs: 24.51309\nExp number, first 60 AAs:  0.2682\nTotal prob of N-in: 0.14402\noutside   1   331\nTMhelix   332   354\ninside   355   372

20765 GCF_001085485.1_10209_5_10 Salmonella enterica subsp. enterica serovar Typhi Proteobacteria; Gammaproteobacteria; Enterobacterales; Enterobacteriaceae; Salmonella; Salmonella enterica; Salmonella enterica subsp. enterica MTGDNTLITSHGINRRDFMKLCAALAATMGLSSKAAAEMAESVSNPQRPPVIWIGAQECT WP_000145429.1 hydrogenase 2 small subunit [Salmonella enterica] Length: 372\nNumber of predicted TMHs:  1\nExp number of AAs in TMHs: 24.51309\nExp number, first 60 AAs:  0.2682\nTotal prob of N-in: 0.14402\noutside   1   331\nTMhelix   332   354\ninside   355   372

20766 GCF_001085605.1_10592_2_14 Salmonella enterica subsp. enterica serovar Typhi Proteobacteria; Gammaproteobacteria; Enterobacterales; Enterobacteriaceae; Salmonella; Salmonella enterica; Salmonella enterica subsp. enterica MTGDNTLITSHGINRRDFMKLCAALAATMGLSSKAAAEMAESVSNPQRPPVIWIGAQECT WP_000145429.1 hydrogenase 2 small subunit [Salmonella enterica] Length: 372\nNumber of predicted TMHs:  1\nExp number of AAs in TMHs: 24.51309\nExp number, first 60 AAs:  0.2682\nTotal prob of N-in: 0.14402\noutside   1   331\nTMhelix   332   354\ninside   355   372

20767 GCF_001085905.1_10541_2_34 Salmonella enterica subsp. enterica serovar Typhi Proteobacteria; Gammaproteobacteria; Enterobacterales; Enterobacteriaceae; Salmonella; Salmonella enterica; Salmonella enterica subsp. enterica MTGDNTLITSHGINRRDFMKLCAALAATMGLSSKAAAEMAESVSNPQRPPVIWIGAQECT WP_000145429.1 hydrogenase 2 small subunit [Salmonella enterica] Length: 372\nNumber of predicted TMHs:  1\nExp number of AAs in TMHs: 24.51309\nExp number, first 60 AAs:  0.2682\nTotal prob of N-in: 0.14402\noutside   1   331\nTMhelix   332   354\ninside   355   372

20768 GCF_001086425.1_8525_3_55 Salmonella enterica subsp. enterica serovar Typhi Proteobacteria; Gammaproteobacteria; Enterobacterales; Enterobacteriaceae; Salmonella; Salmonella enterica; Salmonella enterica subsp. enterica MTGDNTLITSHGINRRDFMKLCAALAATMGLSSKAAAEMAESVSNPQRPPVIWIGAQECT WP_000145429.1 hydrogenase 2 small subunit [Salmonella enterica] Length: 372\nNumber of predicted TMHs:  1\nExp number of AAs in TMHs: 24.51309\nExp number, first 60 AAs:  0.2682\nTotal prob of N-in: 0.14402\noutside   1   331\nTMhelix   332   354\ninside   355   372

20769 GCF_001086265.1_10561_2_54 Salmonella enterica subsp. enterica serovar Typhi Proteobacteria; Gammaproteobacteria; Enterobacterales; Enterobacteriaceae; Salmonella; Salmonella enterica; Salmonella enterica subsp. enterica MTGDNTLITSHGINRRDFMKLCAALAATMGLSSKAAAEMAESVSNPQRPPVIWIGAQECT WP_000145429.1 hydrogenase 2 small subunit [Salmonella enterica] Length: 372\nNumber of predicted TMHs:  1\nExp number of AAs in TMHs: 24.51309\nExp number, first 60 AAs:  0.2682\nTotal prob of N-in: 0.14402\noutside   1   331\nTMhelix   332   354\ninside   355   372

20770 GCF_001086945.1_10209_5_60 Salmonella enterica subsp. enterica serovar Typhi Proteobacteria; Gammaproteobacteria; Enterobacterales; Enterobacteriaceae; Salmonella; Salmonella enterica; Salmonella enterica subsp. enterica MTGDNTLITSHGINRRDFMKLCAALAATMGLSSKAAAEMAESVSNPQRPPVIWIGAQECT WP_000145429.1 hydrogenase 2 small subunit [Salmonella enterica] Length: 372\nNumber of predicted TMHs:  1\nExp number of AAs in TMHs: 24.51309\nExp number, first 60 AAs:  0.2682\nTotal prob of N-in: 0.14402\noutside   1   331\nTMhelix   332   354\ninside   355   372

20771 GCF_001087065.1_8490_5_12 Salmonella enterica subsp. enterica serovar Typhi Proteobacteria; Gammaproteobacteria; Enterobacterales; Enterobacteriaceae; Salmonella; Salmonella enterica;

Salmonella enterica subsp. enterica MTGDNTLITSHGINRRDFMKLCAALAATMGLSSKAAAEMAESVSNPQRPPVIWIGAQECT WP_000145429.1 hydrogenase 2 small subunit [Salmonella enterica] Length: 372\nNumber of predicted TMHs: 1\nExp number of AAs in TMHs: 24.51309\nExp number, first 60 AAs: 0.2682\nTotal prob of N-in: 0.14402\noutside 1 331\nTMhelix 332 354\ninside 355 372

20772 GCF_001087165.1_8525_3_53 Salmonella enterica subsp. enterica serovar Typhi Proteobacteria; Gammaproteobacteria; Enterobacterales; Enterobacteriaceae; Salmonella; Salmonella enterica; Salmonella enterica subsp. enterica MTGDNTLITSHGINRRDFMKLCAALAATMGLSSKAAAEMAESVSNPQRPPVIWIGAQECT WP_000145429.1 hydrogenase 2 small subunit [Salmonella enterica] Length: 372\nNumber of predicted TMHs: 1\nExp number of AAs in TMHs: 24.51309\nExp number, first 60 AAs: 0.2682\nTotal prob of N-in: 0.14402\noutside 1 331\nTMhelix 332 354\ninside 355 372

20773 GCF_001087545.1_10608_2_57 Salmonella enterica subsp. enterica serovar Typhi Proteobacteria; Gammaproteobacteria; Enterobacterales; Enterobacteriaceae; Salmonella; Salmonella enterica; Salmonella enterica subsp. enterica MTGDNTLITSHGINRRDFMKLCAALAATMGLSSKAAAEMAESVSNPQRPPVIWIGAQECT WP_000145429.1 hydrogenase 2 small subunit [Salmonella enterica] Length: 372\nNumber of predicted TMHs: 1\nExp number of AAs in TMHs: 24.51309\nExp number, first 60 AAs: 0.2682\nTotal prob of N-in: 0.14402\noutside 1 331\nTMhelix 332 354\ninside 355 372

20774 GCF_001087725.1_9953_5_67 Salmonella enterica subsp. enterica serovar Typhi Proteobacteria; Gammaproteobacteria; Enterobacterales; Enterobacteriaceae; Salmonella; Salmonella enterica; Salmonella enterica subsp. enterica MTGDNTLITSHGINRRDFMKLCAALAATMGLSSKAAAEMAESVSNPQRPPVIWIGAQECT WP_000145429.1 hydrogenase 2 small subunit [Salmonella enterica] Length: 372\nNumber of predicted TMHs: 1\nExp number of AAs in TMHs: 24.51309\nExp number, first 60 AAs: 0.2682\nTotal prob of N-in: 0.14402\noutside 1 331\nTMhelix 332 354\ninside 355 372

20775 GCF_001088065.1_10493_1_31 Salmonella enterica subsp. enterica serovar Typhi Proteobacteria; Gammaproteobacteria; Enterobacterales; Enterobacteriaceae; Salmonella; Salmonella enterica; Salmonella enterica subsp. enterica MTGDNTLITSHGINRRDFMKLCAALAATMGLSSKAAAEMAESVSNPQRPPVIWIGAQECT WP_000145429.1 hydrogenase 2 small subunit [Salmonella enterica] Length: 372\nNumber of predicted TMHs: 1\nExp number of AAs in TMHs: 24.51309\nExp number, first 60 AAs: 0.2682\nTotal prob of N-in: 0.14402\noutside 1 331\nTMhelix 332 354\ninside 355 372

20776 GCF_001088305.1_9475_6_29 Salmonella enterica subsp. enterica serovar Typhi Proteobacteria; Gammaproteobacteria; Enterobacterales; Enterobacteriaceae; Salmonella; Salmonella enterica; Salmonella enterica subsp. enterica MTGDNTLITSHGINRRDFMKLCAALAATMGLSSKAAAEMAESVSNPQRPPVIWIGAQECT WP_000145429.1 hydrogenase 2 small subunit [Salmonella enterica] Length: 372\nNumber of predicted TMHs: 1\nExp number of AAs in TMHs: 24.51309\nExp number, first 60 AAs: 0.2682\nTotal prob of N-in: 0.14402\noutside 1 331\nTMhelix 332 354\ninside 355 372

20777 GCF_001088605.1_10592_2_46 Salmonella enterica subsp. enterica serovar Typhi Proteobacteria; Gammaproteobacteria; Enterobacterales; Enterobacteriaceae; Salmonella; Salmonella enterica; Salmonella enterica subsp. enterica MTGDNTLITSHGINRRDFMKLCAALAATMGLSSKAAAEMAESVSNPQRPPVIWIGAQECT WP_000145429.1 hydrogenase 2 small subunit [Salmonella enterica] Length: 372\nNumber of predicted TMHs: 1\nExp number of AAs in TMHs: 24.51309\nExp number, first 60 AAs: 0.2682\nTotal prob of N-in: 0.14402\noutside 1 331\nTMhelix 332 354\ninside 355 372

20778 GCF_001088965.1_10493_1_37 Salmonella enterica subsp. enterica serovar Typhi Proteobacteria; Gammaproteobacteria; Enterobacterales; Enterobacteriaceae; Salmonella; Salmonella enterica; Salmonella enterica subsp. enterica MTGDNTLITSHGINRRDFMKLCAALAATMGLSSKAAAEMAESVSNPQRPPVIWIGAQECT WP_000145429.1 hydrogenase 2 small subunit [Salmonella enterica] Length: 372\nNumber of predicted TMHs: 1\nExp number of AAs in TMHs: 24.51309\nExp number, first 60 AAs: 0.2682\nTotal prob of N-in: 0.14402\noutside 1 331\nTMhelix 332 354\ninside 355 372

20779 GCF_001089365.1_10493_1_55 Salmonella enterica subsp. enterica serovar Typhi Proteobacteria; Gammaproteobacteria; Enterobacterales; Enterobacteriaceae; Salmonella; Salmonella enterica; Salmonella enterica subsp. enterica MTGDNTLITSHGINRRDFMKLCAALAATMGLSSKAAAEMAESVSNPQRPPVIWIGAQECT WP_000145429.1 hydrogenase 2 small subunit [Salmonella enterica] Length: 372\nNumber of predicted TMHs: 1\nExp number of AAs in TMHs: 24.51309\nExp number, first 60 AAs: 0.2682\nTotal prob of N-in: 0.14402\noutside 1 331\nTMhelix 332 354\ninside 355 372

20780 GCF_001089225.1_10349_1_39 Salmonella enterica subsp. enterica serovar Typhi Proteobacteria; Gammaproteobacteria; Enterobacterales; Enterobacteriaceae; Salmonella; Salmonella enterica; Salmonella enterica subsp. enterica MTGDNTLITSHGINRRDFMKLCAALAATMGLSSKAAAEMAESVSNPQRPPVIWIGAQECT WP_000145429.1 hydrogenase 2 small subunit [Salmonella enterica] Length: 372\nNumber of predicted TMHs: 1\nExp number of AAs in TMHs: 24.51309\nExp number, first 60 AAs: 0.2682\nTotal prob of N-in: 0.14402\noutside 1 331\nTMhelix 332 354\ninside 355 372

20781 GCF_001089465.1_9953_5_57 Salmonella enterica subsp. enterica serovar Typhi Proteobacteria; Gammaproteobacteria; Enterobacterales; Enterobacteriaceae; Salmonella; Salmonella enterica; Salmonella enterica subsp. enterica MTGDNTLITSHGINRRDFMKLCAALAATMGLSSKAAAEMAESVSNPQRPPVIWIGAQECT WP_000145429.1 hydrogenase 2 small subunit [Salmonella enterica] Length: 372\nNumber of predicted TMHs: 1\nExp number of AAs in TMHs: 24.51309\nExp number, first 60 AAs: 0.2682\nTotal prob of N-in: 0.14402\noutside 1 331\nTMhelix 332 354\ninside 355 372

20782 GCF_001089565.1_10492_1_16 Salmonella enterica subsp. enterica serovar Typhi Proteobacteria; Gammaproteobacteria; Enterobacterales; Enterobacteriaceae; Salmonella; Salmonella enterica; Salmonella enterica subsp. enterica MTGDNTLITSHGINRRDFMKLCAALAATMGLSSKAAAEMAESVSNPQRPPVIWIGAQECT WP_000145429.1 hydrogenase 2 small subunit [Salmonella enterica] Length: 372\nNumber of predicted TMHs: 1\nExp number of AAs in TMHs: 24.51309\nExp number, first 60 AAs: 0.2682\nTotal prob of N-in: 0.14402\noutside 1 331\nTMhelix 332 354\ninside 355 372

20783 GCF_001089805.1_9475_6_1Salmonella enterica subsp. enterica serovar Typhi Proteobacteria; Gammaproteobacteria; Enterobacterales; Enterobacteriaceae; Salmonella; Salmonella enterica; Salmonella enterica subsp. enterica MTGDNTLITSHGINRRDFMKLCAALAATMGLSSKAAAEMAESVSNPQRPPVIWIGAQECT WP_000145429.1 hydrogenase 2 small subunit [Salmonella enterica] Length: 372\nNumber of predicted TMHs: 1\nExp number of AAs in TMHs: 24.51309\nExp number, first 60 AAs: 0.2682\nTotal prob of N-in: 0.14402\noutside 1 331\nTMhelix 332 354\ninside 355 372

20784 GCF_001090105.1_8447_8_4Salmonella enterica subsp. enterica serovar Typhi Proteobacteria; Gammaproteobacteria; Enterobacterales; Enterobacteriaceae; Salmonella; Salmonella enterica; Salmonella enterica subsp. enterica MTGDNTLITSHGINRRDFMKLCAALAATMGLSSKAAAEMAESVSNPQRPPVIWIGAQECT WP_000145429.1 hydrogenase 2 small subunit [Salmonella enterica] Length: 372\nNumber of predicted TMHs: 1\nExp number of AAs in TMHs: 24.51309\nExp number, first 60 AAs: 0.2682\nTotal prob of N-in: 0.14402\noutside 1 331\nTMhelix 332 354\ninside 355 372

20785 GCF_001089925.1_10209_5_28 Salmonella enterica subsp. enterica serovar Typhi Proteobacteria; Gammaproteobacteria; Enterobacterales; Enterobacteriaceae; Salmonella; Salmonella enterica; Salmonella enterica subsp. enterica MTGDNTLITSHGINRRDFMKLCAALAATMGLSSKAAAEMAESVSNPQRPPVIWIGAQECT WP_000145429.1 hydrogenase 2 small subunit [Salmonella enterica] Length: 372\nNumber of predicted TMHs: 1\nExp number of AAs in TMHs: 24.51309\nExp number, first 60 AAs: 0.2682\nTotal prob of N-in: 0.14402\noutside 1 331\nTMhelix 332 354\ninside 355 372

20786 GCF_001090325.1_10493_1_51 Salmonella enterica subsp. enterica serovar Typhi Proteobacteria; Gammaproteobacteria; Enterobacterales; Enterobacteriaceae; Salmonella; Salmonella enterica; Salmonella enterica subsp. enterica MTGDNTLITSHGINRRDFMKLCAALAATMGLSSKAAAEMAESVSNPQRPPVIWIGAQECT WP_000145429.1 hydrogenase 2 small subunit [Salmonella enterica] Length: 372\nNumber of predicted TMHs: 1\nExp number of AAs in TMHs: 24.51309\nExp number, first 60 AAs: 0.2682\nTotal prob of N-in: 0.14402\noutside 1 331\nTMhelix 332 354\ninside 355 372

20787 GCF_001090485.1_10607_2_10 Salmonella enterica subsp. enterica serovar Typhi Proteobacteria; Gammaproteobacteria; Enterobacterales; Enterobacteriaceae; Salmonella; Salmonella enterica; Salmonella enterica subsp. enterica MTGDNTLITSHGINRRDFMKLCAALAATMGLSSKAAAEMAESVSNPQRPPVIWIGAQECT WP_000145429.1 hydrogenase 2 small subunit [Salmonella enterica] Length: 372\nNumber of predicted TMHs: 1\nExp number of AAs in TMHs: 24.51309\nExp number, first 60 AAs: 0.2682\nTotal prob of N-in: 0.14402\noutside 1 331\nTMhelix 332 354\ninside 355 372

20788 GCF_001090585.1_10349_1_46 Salmonella enterica subsp. enterica serovar Typhi Proteobacteria; Gammaproteobacteria; Enterobacterales; Enterobacteriaceae; Salmonella; Salmonella enterica; Salmonella enterica subsp. enterica MTGDNTLITSHGINRRDFMKLCAALAATMGLSSKAAAEMAESVSNPQRPPVIWIGAQECT WP_000145429.1 hydrogenase 2 small subunit [Salmonella enterica] Length: 372\nNumber of

predicted TMHs: 1\nExp number of AAs in TMHs: 24.51309\nExp number, first 60 AAs: 0.2682\nTotal prob of N-in: 0.14402\noutside 1 331\nTMhelix 332 354\ninside 355 372

20789 GCF_001090925.1_9870_8_48 Salmonella enterica subsp. enterica serovar Typhi Proteobacteria; Gammaproteobacteria; Enterobacterales; Enterobacteriaceae; Salmonella; Salmonella enterica; Salmonella enterica subsp. enterica MTGDNTLITSHGINRRDFMKLCAALAATMGLSSKAAAEMAESVSNPQRPPVIWIGAQECT WP_000145429.1 hydrogenase 2 small subunit [Salmonella enterica] Length: 372\nNumber of predicted TMHs: 1\nExp number of AAs in TMHs: 24.51309\nExp number, first 60 AAs: 0.2682\nTotal prob of N-in: 0.14402\noutside 1 331\nTMhelix 332 354\ninside 355 372

20790 GCF_001091105.1_10593_2_60 Salmonella enterica subsp. enterica serovar Typhi Proteobacteria; Gammaproteobacteria; Enterobacterales; Enterobacteriaceae; Salmonella; Salmonella enterica; Salmonella enterica subsp. enterica MTGDNTLITSHGINRRDFMKLCAALAATMGLSSKAAAEMAESVSNPQRPPVIWIGAQECT WP_000145429.1 hydrogenase 2 small subunit [Salmonella enterica] Length: 372\nNumber of predicted TMHs: 1\nExp number of AAs in TMHs: 24.51309\nExp number, first 60 AAs: 0.2682\nTotal prob of N-in: 0.14402\noutside 1 331\nTMhelix 332 354\ninside 355 372

20791 GCF_001091385.1_8447_8_7Salmonella enterica subsp. enterica serovar Typhi Proteobacteria; Gammaproteobacteria; Enterobacterales; Enterobacteriaceae; Salmonella; Salmonella enterica; Salmonella enterica subsp. enterica MTGDNTLITSHGINRRDFMKLCAALAATMGLSSKAAAEMAESVSNPQRPPVIWIGAQECT WP_000145429.1 hydrogenase 2 small subunit [Salmonella enterica] Length: 372\nNumber of predicted TMHs: 1\nExp number of AAs in TMHs: 24.51309\nExp number, first 60 AAs: 0.2682\nTotal prob of N-in: 0.14402\noutside 1 331\nTMhelix 332 354\ninside 355 372

20792 GCF_001091665.1_9953_5_2Salmonella enterica subsp. enterica serovar Typhi Proteobacteria; Gammaproteobacteria; Enterobacterales; Enterobacteriaceae; Salmonella; Salmonella enterica; Salmonella enterica subsp. enterica MTGDNTLITSHGINRRDFMKLCAALAATMGLSSKAAAEMAESVSNPQRPPVIWIGAQECT WP_000145429.1 hydrogenase 2 small subunit [Salmonella enterica] Length: 372\nNumber of predicted TMHs: 1\nExp number of AAs in TMHs: 24.51309\nExp number, first 60 AAs: 0.2682\nTotal prob of N-in: 0.14402\noutside 1 331\nTMhelix 332 354\ninside 355 372

20793 GCF_001092145.1_10592_2_71 Salmonella enterica subsp. enterica serovar Typhi Proteobacteria; Gammaproteobacteria; Enterobacterales; Enterobacteriaceae; Salmonella; Salmonella enterica; Salmonella enterica subsp. enterica MTGDNTLITSHGINRRDFMKLCAALAATMGLSSKAAAEMAESVSNPQRPPVIWIGAQECT WP_000145429.1 hydrogenase 2 small subunit [Salmonella enterica] Length: 372\nNumber of predicted TMHs: 1\nExp number of AAs in TMHs: 24.51309\nExp number, first 60 AAs: 0.2682\nTotal prob of N-in: 0.14402\noutside 1 331\nTMhelix 332 354\ninside 355 372

20794 GCF_001091785.1_10608_2_13 Salmonella enterica subsp. enterica serovar Typhi Proteobacteria; Gammaproteobacteria; Enterobacterales; Enterobacteriaceae; Salmonella; Salmonella enterica; Salmonella enterica subsp. enterica MTGDNTLITSHGINRRDFMKLCAALAATMGLSSKAAAEMAESVSNPQRPPVIWIGAQECT WP_000145429.1 hydrogenase 2 small subunit [Salmonella enterica] Length: 372\nNumber of predicted TMHs: 1\nExp number of AAs in TMHs: 24.51309\nExp number, first 60 AAs: 0.2682\nTotal prob of N-in: 0.14402\noutside 1 331\nTMhelix 332 354\ninside 355 372

20795 GCF_001092245.1_9475_6_25 Salmonella enterica subsp. enterica serovar Typhi Proteobacteria; Gammaproteobacteria; Enterobacterales; Enterobacteriaceae; Salmonella; Salmonella enterica; Salmonella enterica subsp. enterica MTGDNTLITSHGINRRDFMKLCAALAATMGLSSKAAAEMAESVSNPQRPPVIWIGAQECT WP_000145429.1 hydrogenase 2 small subunit [Salmonella enterica] Length: 372\nNumber of predicted TMHs: 1\nExp number of AAs in TMHs: 24.51309\nExp number, first 60 AAs: 0.2682\nTotal prob of N-in: 0.14402\noutside 1 331\nTMhelix 332 354\ninside 355 372

20796 GCF_001092405.1_10209_5_4 Salmonella enterica subsp. enterica serovar Typhi Proteobacteria; Gammaproteobacteria; Enterobacterales; Enterobacteriaceae; Salmonella; Salmonella enterica; Salmonella enterica subsp. enterica MTGDNTLITSHGINRRDFMKLCAALAATMGLSSKAAAEMAESVSNPQRPPVIWIGAQECT WP_000145429.1 hydrogenase 2 small subunit [Salmonella enterica] Length: 372\nNumber of predicted TMHs: 1\nExp number of AAs in TMHs: 24.51309\nExp number, first 60 AAs: 0.2682\nTotal prob of N-in: 0.14402\noutside 1 331\nTMhelix 332 354\ninside 355 372

20797 GCF_001092585.1_10593_2_20 Salmonella enterica subsp. enterica serovar Typhi Proteobacteria; Gammaproteobacteria; Enterobacterales; Enterobacteriaceae; Salmonella; Salmonella enterica;

Salmonella enterica subsp. enterica	MTGDNTLITSHGINRRDFMKLCAALAATMGLSSKAAAEMAESVSNPQRPPVIWIGAQECT	WP_000145429.1	hydrogenase 2 small subunit [Salmonella enterica]	Length: 372\nNumber of predicted TMHs: 1\nExp number of AAs in TMHs: 24.51309\nExp number, first 60 AAs: 0.2682\nTotal prob of N-in: 0.14402\noutside 1 331\nTMhelix 332 354\ninside 355 372

20798	GCF_001092625.1_9953_5_22	Salmonella enterica subsp. enterica serovar Typhi	Proteobacteria; Gammaproteobacteria; Enterobacterales; Enterobacteriaceae; Salmonella; Salmonella enterica; Salmonella enterica subsp. enterica	MTGDNTLITSHGINRRDFMKLCAALAATMGLSSKAAAEMAESVSNPQRPPVIWIGAQECT	WP_000145429.1	hydrogenase 2 small subunit [Salmonella enterica]	Length: 372\nNumber of predicted TMHs: 1\nExp number of AAs in TMHs: 24.51309\nExp number, first 60 AAs: 0.2682\nTotal prob of N-in: 0.14402\noutside 1 331\nTMhelix 332 354\ninside 355 372

20799	GCF_001092765.1_10209_5_29	Salmonella enterica subsp. enterica serovar Typhi	Proteobacteria; Gammaproteobacteria; Enterobacterales; Enterobacteriaceae; Salmonella; Salmonella enterica; Salmonella enterica subsp. enterica	MTGDNTLITSHGINRRDFMKLCAALAATMGLSSKAAAEMAESVSNPQRPPVIWIGAQECT	WP_000145429.1	hydrogenase 2 small subunit [Salmonella enterica]	Length: 372\nNumber of predicted TMHs: 1\nExp number of AAs in TMHs: 24.51309\nExp number, first 60 AAs: 0.2682\nTotal prob of N-in: 0.14402\noutside 1 331\nTMhelix 332 354\ninside 355 372

20800	GCF_001092865.1_10493_1_84	Salmonella enterica subsp. enterica serovar Typhi	Proteobacteria; Gammaproteobacteria; Enterobacterales; Enterobacteriaceae; Salmonella; Salmonella enterica; Salmonella enterica subsp. enterica	MTGDNTLITSHGINRRDFMKLCAALAATMGLSSKAAAEMAESVSNPQRPPVIWIGAQECT	WP_000145429.1	hydrogenase 2 small subunit [Salmonella enterica]	Length: 372\nNumber of predicted TMHs: 1\nExp number of AAs in TMHs: 24.51309\nExp number, first 60 AAs: 0.2682\nTotal prob of N-in: 0.14402\noutside 1 331\nTMhelix 332 354\ninside 355 372

20801	GCF_001093245.1_10541_2_11	Salmonella enterica subsp. enterica serovar Typhi	Proteobacteria; Gammaproteobacteria; Enterobacterales; Enterobacteriaceae; Salmonella; Salmonella enterica; Salmonella enterica subsp. enterica	MTGDNTLITSHGINRRDFMKLCAALAATMGLSSKAAAEMAESVSNPQRPPVIWIGAQECT	WP_000145429.1	hydrogenase 2 small subunit [Salmonella enterica]	Length: 372\nNumber of predicted TMHs: 1\nExp number of AAs in TMHs: 24.51309\nExp number, first 60 AAs: 0.2682\nTotal prob of N-in: 0.14402\noutside 1 331\nTMhelix 332 354\ninside 355 372

20802	GCF_001093385.1_9953_5_45	Salmonella enterica subsp. enterica serovar Typhi	Proteobacteria; Gammaproteobacteria; Enterobacterales; Enterobacteriaceae; Salmonella; Salmonella enterica; Salmonella enterica subsp. enterica	MTGDNTLITSHGINRRDFMKLCAALAATMGLSSKAAAEMAESVSNPQRPPVIWIGAQECT	WP_000145429.1	hydrogenase 2 small subunit [Salmonella enterica]	Length: 372\nNumber of predicted TMHs: 1\nExp number of AAs in TMHs: 24.51309\nExp number, first 60 AAs: 0.2682\nTotal prob of N-in: 0.14402\noutside 1 331\nTMhelix 332 354\ninside 355 372

20803	GCF_001093525.1_10349_1_41	Salmonella enterica subsp. enterica serovar Typhi	Proteobacteria; Gammaproteobacteria; Enterobacterales; Enterobacteriaceae; Salmonella; Salmonella enterica; Salmonella enterica subsp. enterica	MTGDNTLITSHGINRRDFMKLCAALAATMGLSSKAAAEMAESVSNPQRPPVIWIGAQECT	WP_000145429.1	hydrogenase 2 small subunit [Salmonella enterica]	Length: 372\nNumber of predicted TMHs: 1\nExp number of AAs in TMHs: 24.51309\nExp number, first 60 AAs: 0.2682\nTotal prob of N-in: 0.14402\noutside 1 331\nTMhelix 332 354\ninside 355 372

20804	GCF_001093885.1_10349_1_33	Salmonella enterica subsp. enterica serovar Typhi	Proteobacteria; Gammaproteobacteria; Enterobacterales; Enterobacteriaceae; Salmonella; Salmonella enterica; Salmonella enterica subsp. enterica	MTGDNTLITSHGINRRDFMKLCAALAATMGLSSKAAAEMAESVSNPQRPPVIWIGAQECT	WP_000145429.1	hydrogenase 2 small subunit [Salmonella enterica]	Length: 372\nNumber of predicted TMHs: 1\nExp number of AAs in TMHs: 24.51309\nExp number, first 60 AAs: 0.2682\nTotal prob of N-in: 0.14402\noutside 1 331\nTMhelix 332 354\ninside 355 372

20805	GCF_001094085.1_9953_5_65	Salmonella enterica subsp. enterica serovar Typhi	Proteobacteria; Gammaproteobacteria; Enterobacterales; Enterobacteriaceae; Salmonella; Salmonella enterica; Salmonella enterica subsp. enterica	MTGDNTLITSHGINRRDFMKLCAALAATMGLSSKAAAEMAESVSNPQRPPVIWIGAQECT	WP_000145429.1	hydrogenase 2 small subunit [Salmonella enterica]	Length: 372\nNumber of predicted TMHs: 1\nExp number of AAs in TMHs: 24.51309\nExp number, first 60 AAs: 0.2682\nTotal prob of N-in: 0.14402\noutside 1 331\nTMhelix 332 354\ninside 355 372

20806	GCF_001094245.1_10492_1_60	Salmonella enterica subsp. enterica serovar Typhi	Proteobacteria; Gammaproteobacteria; Enterobacterales; Enterobacteriaceae; Salmonella; Salmonella enterica; Salmonella enterica subsp. enterica	MTGDNTLITSHGINRRDFMKLCAALAATMGLSSKAAAEMAESVSNPQRPPVIWIGAQECT	WP_000145429.1	hydrogenase 2 small subunit [Salmonella enterica]	Length: 372\nNumber of predicted TMHs:  1\nExp number of AAs in TMHs: 24.51309\nExp number, first 60 AAs:  0.2682\nTotal prob of N-in:  0.14402\noutside	1	331\nTMhelix	332	354\ninside	355	372

20807	GCF_001094465.1_10492_1_44	Salmonella enterica subsp. enterica serovar Typhi	Proteobacteria; Gammaproteobacteria; Enterobacterales; Enterobacteriaceae; Salmonella; Salmonella enterica; Salmonella enterica subsp. enterica	MTGDNTLITSHGINRRDFMKLCAALAATMGLSSKAAAEMAESVSNPQRPPVIWIGAQECT	WP_000145429.1	hydrogenase 2 small subunit [Salmonella enterica]	Length: 372\nNumber of predicted TMHs:  1\nExp number of AAs in TMHs: 24.51309\nExp number, first 60 AAs:  0.2682\nTotal prob of N-in:  0.14402\noutside	1	331\nTMhelix	332	354\ninside	355	372

20808	GCF_001094625.1_10492_1_80	Salmonella enterica subsp. enterica serovar Typhi	Proteobacteria; Gammaproteobacteria; Enterobacterales; Enterobacteriaceae; Salmonella; Salmonella enterica; Salmonella enterica subsp. enterica	MTGDNTLITSHGINRRDFMKLCAALAATMGLSSKAAAEMAESVSNPQRPPVIWIGAQECT	WP_000145429.1	hydrogenase 2 small subunit [Salmonella enterica]	Length: 372\nNumber of predicted TMHs:  1\nExp number of AAs in TMHs: 24.51309\nExp number, first 60 AAs:  0.2682\nTotal prob of N-in:  0.14402\noutside	1	331\nTMhelix	332	354\ninside	355	372

20809	GCF_001094745.1_9475_6_94	Salmonella enterica subsp. enterica serovar Typhi	Proteobacteria; Gammaproteobacteria; Enterobacterales; Enterobacteriaceae; Salmonella; Salmonella enterica; Salmonella enterica subsp. enterica	MTGDNTLITSHGINRRDFMKLCAALAATMGLSSKAAAEMAESVSNPQRPPVIWIGAQECT	WP_000145429.1	hydrogenase 2 small subunit [Salmonella enterica]	Length: 372\nNumber of predicted TMHs:  1\nExp number of AAs in TMHs: 24.51309\nExp number, first 60 AAs:  0.2682\nTotal prob of N-in:  0.14402\noutside	1	331\nTMhelix	332	354\ninside	355	372

20810	GCF_001094925.1_10493_1_5	Salmonella enterica subsp. enterica serovar Typhi	Proteobacteria; Gammaproteobacteria; Enterobacterales; Enterobacteriaceae; Salmonella; Salmonella enterica; Salmonella enterica subsp. enterica	MTGDNTLITSHGINRRDFMKLCAALAATMGLSSKAAAEMAESVSNPQRPPVIWIGAQECT	WP_000145429.1	hydrogenase 2 small subunit [Salmonella enterica]	Length: 372\nNumber of predicted TMHs:  1\nExp number of AAs in TMHs: 24.51309\nExp number, first 60 AAs:  0.2682\nTotal prob of N-in:  0.14402\noutside	1	331\nTMhelix	332	354\ninside	355	372

20811	GCF_001095145.1_10492_1_9	Salmonella enterica subsp. enterica serovar Typhi	Proteobacteria; Gammaproteobacteria; Enterobacterales; Enterobacteriaceae; Salmonella; Salmonella enterica; Salmonella enterica subsp. enterica	MTGDNTLITSHGINRRDFMKLCAALAATMGLSSKAAAEMAESVSNPQRPPVIWIGAQECT	WP_000145429.1	hydrogenase 2 small subunit [Salmonella enterica]	Length: 372\nNumber of predicted TMHs:  1\nExp number of AAs in TMHs: 24.51309\nExp number, first 60 AAs:  0.2682\nTotal prob of N-in:  0.14402\noutside	1	331\nTMhelix	332	354\ninside	355	372

20812	GCF_001095325.1_9953_5_6Salmonella enterica subsp. enterica serovar Typhi	Proteobacteria; Gammaproteobacteria; Enterobacterales; Enterobacteriaceae; Salmonella; Salmonella enterica; Salmonella enterica subsp. enterica	MTGDNTLITSHGINRRDFMKLCAALAATMGLSSKAAAEMAESVSNPQRPPVIWIGAQECT	WP_000145429.1	hydrogenase 2 small subunit [Salmonella enterica]	Length: 372\nNumber of predicted TMHs:  1\nExp number of AAs in TMHs: 24.51309\nExp number, first 60 AAs:  0.2682\nTotal prob of N-in:  0.14402\noutside	1	331\nTMhelix	332	354\ninside	355	372

20813	GCF_001095585.1_10349_1_69	Salmonella enterica subsp. enterica serovar Typhi	Proteobacteria; Gammaproteobacteria; Enterobacterales; Enterobacteriaceae; Salmonella; Salmonella enterica; Salmonella enterica subsp. enterica	MTGDNTLITSHGINRRDFMKLCAALAATMGLSSKAAAEMAESVSNPQRPPVIWIGAQECT	WP_000145429.1	hydrogenase 2 small subunit [Salmonella enterica]	Length: 372\nNumber of predicted TMHs:  1\nExp number of AAs in TMHs: 24.51309\nExp number, first 60 AAs:  0.2682\nTotal prob of N-in:  0.14402\noutside	1	331\nTMhelix	332	354\ninside	355	372

20814	GCF_001095885.1_9870_8_61	Salmonella enterica subsp. enterica serovar Typhi	Proteobacteria; Gammaproteobacteria; Enterobacterales; Enterobacteriaceae; Salmonella; Salmonella enterica; Salmonella enterica subsp. enterica	MTGDNTLITSHGINRRDFMKLCAALAATMGLSSKAAAEMAESVSNPQRPPVIWIGAQECT	WP_000145429.1	hydrogenase 2 small subunit [Salmonella enterica]	Length: 372\nNumber of

predicted TMHs: 1\nExp number of AAs in TMHs: 24.51309\nExp number, first 60 AAs: 0.2682\nTotal prob of N-in: 0.14402\noutside 1 331\nTMhelix 332 354\ninside 355 372

20815 GCF_001095965.1_10592_2_18 Salmonella enterica subsp. enterica serovar Typhi Proteobacteria; Gammaproteobacteria; Enterobacterales; Enterobacteriaceae; Salmonella; Salmonella enterica; Salmonella enterica subsp. enterica MTGDNTLITSHGINRRDFMKLCAALAATMGLSSKAAAEMAESVSNPQRPPVIWIGAQECT WP_000145429.1 hydrogenase 2 small subunit [Salmonella enterica] Length: 372\nNumber of predicted TMHs: 1\nExp number of AAs in TMHs: 24.51309\nExp number, first 60 AAs: 0.2682\nTotal prob of N-in: 0.14402\noutside 1 331\nTMhelix 332 354\ninside 355 372

20816 GCF_001096385.1_10593_2_49 Salmonella enterica subsp. enterica serovar Typhi Proteobacteria; Gammaproteobacteria; Enterobacterales; Enterobacteriaceae; Salmonella; Salmonella enterica; Salmonella enterica subsp. enterica MTGDNTLITSHGINRRDFMKLCAALAATMGLSSKAAAEMAESVSNPQRPPVIWIGAQECT WP_000145429.1 hydrogenase 2 small subunit [Salmonella enterica] Length: 372\nNumber of predicted TMHs: 1\nExp number of AAs in TMHs: 24.51309\nExp number, first 60 AAs: 0.2682\nTotal prob of N-in: 0.14402\noutside 1 331\nTMhelix 332 354\ninside 355 372

20817 GCF_001096285.1_10562_2_29 Salmonella enterica subsp. enterica serovar Typhi Proteobacteria; Gammaproteobacteria; Enterobacterales; Enterobacteriaceae; Salmonella; Salmonella enterica; Salmonella enterica subsp. enterica MTGDNTLITSHGINRRDFMKLCAALAATMGLSSKAAAEMAESVSNPQRPPVIWIGAQECT WP_000145429.1 hydrogenase 2 small subunit [Salmonella enterica] Length: 372\nNumber of predicted TMHs: 1\nExp number of AAs in TMHs: 24.51309\nExp number, first 60 AAs: 0.2682\nTotal prob of N-in: 0.14402\noutside 1 331\nTMhelix 332 354\ninside 355 372

20818 GCF_001096505.1_10608_2_52 Salmonella enterica subsp. enterica serovar Typhi Proteobacteria; Gammaproteobacteria; Enterobacterales; Enterobacteriaceae; Salmonella; Salmonella enterica; Salmonella enterica subsp. enterica MTGDNTLITSHGINRRDFMKLCAALAATMGLSSKAAAEMAESVSNPQRPPVIWIGAQECT WP_000145429.1 hydrogenase 2 small subunit [Salmonella enterica] Length: 372\nNumber of predicted TMHs: 1\nExp number of AAs in TMHs: 24.51309\nExp number, first 60 AAs: 0.2682\nTotal prob of N-in: 0.14402\noutside 1 331\nTMhelix 332 354\ninside 355 372

20819 GCF_001096905.1_9475_6_2Salmonella enterica subsp. enterica serovar Typhi Proteobacteria; Gammaproteobacteria; Enterobacterales; Enterobacteriaceae; Salmonella; Salmonella enterica; Salmonella enterica subsp. enterica MTGDNTLITSHGINRRDFMKLCAALAATMGLSSKAAAEMAESVSNPQRPPVIWIGAQECT WP_000145429.1 hydrogenase 2 small subunit [Salmonella enterica] Length: 372\nNumber of predicted TMHs: 1\nExp number of AAs in TMHs: 24.51309\nExp number, first 60 AAs: 0.2682\nTotal prob of N-in: 0.14402\noutside 1 331\nTMhelix 332 354\ninside 355 372

20820 GCF_001096985.1_10425_1_3 Salmonella enterica subsp. enterica serovar Typhi Proteobacteria; Gammaproteobacteria; Enterobacterales; Enterobacteriaceae; Salmonella; Salmonella enterica; Salmonella enterica subsp. enterica MTGDNTLITSHGINRRDFMKLCAALAATMGLSSKAAAEMAESVSNPQRPPVIWIGAQECT WP_000145429.1 hydrogenase 2 small subunit [Salmonella enterica] Length: 372\nNumber of predicted TMHs: 1\nExp number of AAs in TMHs: 24.51309\nExp number, first 60 AAs: 0.2682\nTotal prob of N-in: 0.14402\noutside 1 331\nTMhelix 332 354\ninside 355 372

20821 GCF_001097145.1_10209_5_23 Salmonella enterica subsp. enterica serovar Typhi Proteobacteria; Gammaproteobacteria; Enterobacterales; Enterobacteriaceae; Salmonella; Salmonella enterica; Salmonella enterica subsp. enterica MTGDNTLITSHGINRRDFMKLCAALAATMGLSSKAAAEMAESVSNPQRPPVIWIGAQECT WP_000145429.1 hydrogenase 2 small subunit [Salmonella enterica] Length: 372\nNumber of predicted TMHs: 1\nExp number of AAs in TMHs: 24.51309\nExp number, first 60 AAs: 0.2682\nTotal prob of N-in: 0.14402\noutside 1 331\nTMhelix 332 354\ninside 355 372

20822 GCF_001097285.1_10492_1_79 Salmonella enterica subsp. enterica serovar Typhi Proteobacteria; Gammaproteobacteria; Enterobacterales; Enterobacteriaceae; Salmonella; Salmonella enterica; Salmonella enterica subsp. enterica MTGDNTLITSHGINRRDFMKLCAALAATMGLSSKAAAEMAESVSNPQRPPVIWIGAQECT WP_000145429.1 hydrogenase 2 small subunit [Salmonella enterica] Length: 372\nNumber of predicted TMHs: 1\nExp number of AAs in TMHs: 24.51309\nExp number, first 60 AAs: 0.2682\nTotal prob of N-in: 0.14402\noutside 1 331\nTMhelix 332 354\ninside 355 372

20823 GCF_001097585.1_10349_1_78 Salmonella enterica subsp. enterica serovar Typhi Proteobacteria; Gammaproteobacteria; Enterobacterales; Enterobacteriaceae; Salmonella; Salmonella enterica;

Salmonella enterica subsp. enterica MTGDNTLITSHGINRRDFMKLCAALAATMGLSSKAAAEMAESVSNPQRPPVIWIGAQECT WP_000145429.1 hydrogenase 2 small subunit [Salmonella enterica] Length: 372\nNumber of predicted TMHs: 1\nExp number of AAs in TMHs: 24.51309\nExp number, first 60 AAs: 0.2682\nTotal prob of N-in: 0.14402\noutside 1 331\nTMhelix 332 354\ninside 355 372

20824 GCF_001097825.1_10592_2_30 Salmonella enterica subsp. enterica serovar Typhi Proteobacteria; Gammaproteobacteria; Enterobacterales; Enterobacteriaceae; Salmonella; Salmonella enterica; Salmonella enterica subsp. enterica MTGDNTLITSHGINRRDFMKLCAALAATMGLSSKAAAEMAESVSNPQRPPVIWIGAQECT WP_000145429.1 hydrogenase 2 small subunit [Salmonella enterica] Length: 372\nNumber of predicted TMHs: 1\nExp number of AAs in TMHs: 24.51309\nExp number, first 60 AAs: 0.2682\nTotal prob of N-in: 0.14402\noutside 1 331\nTMhelix 332 354\ninside 355 372

20825 GCF_001098225.1_8447_8_15 Salmonella enterica subsp. enterica serovar Typhi Proteobacteria; Gammaproteobacteria; Enterobacterales; Enterobacteriaceae; Salmonella; Salmonella enterica; Salmonella enterica subsp. enterica MTGDNTLITSHGINRRDFMKLCAALAATMGLSSKAAAEMAESVSNPQRPPVIWIGAQECT WP_000145429.1 hydrogenase 2 small subunit [Salmonella enterica] Length: 372\nNumber of predicted TMHs: 1\nExp number of AAs in TMHs: 24.51309\nExp number, first 60 AAs: 0.2682\nTotal prob of N-in: 0.14402\noutside 1 331\nTMhelix 332 354\ninside 355 372

20826 GCF_001098645.1_10493_1_3 Salmonella enterica subsp. enterica serovar Typhi Proteobacteria; Gammaproteobacteria; Enterobacterales; Enterobacteriaceae; Salmonella; Salmonella enterica; Salmonella enterica subsp. enterica MTGDNTLITSHGINRRDFMKLCAALAATMGLSSKAAAEMAESVSNPQRPPVIWIGAQECT WP_000145429.1 hydrogenase 2 small subunit [Salmonella enterica] Length: 372\nNumber of predicted TMHs: 1\nExp number of AAs in TMHs: 24.51309\nExp number, first 60 AAs: 0.2682\nTotal prob of N-in: 0.14402\noutside 1 331\nTMhelix 332 354\ninside 355 372

20827 GCF_001098685.1_9870_8_32 Salmonella enterica subsp. enterica serovar Typhi Proteobacteria; Gammaproteobacteria; Enterobacterales; Enterobacteriaceae; Salmonella; Salmonella enterica; Salmonella enterica subsp. enterica MTGDNTLITSHGINRRDFMKLCAALAATMGLSSKAAAEMAESVSNPQRPPVIWIGAQECT WP_000145429.1 hydrogenase 2 small subunit [Salmonella enterica] Length: 372\nNumber of predicted TMHs: 1\nExp number of AAs in TMHs: 24.51309\nExp number, first 60 AAs: 0.2682\nTotal prob of N-in: 0.14402\noutside 1 331\nTMhelix 332 354\ninside 355 372

20828 GCF_001099045.1_9475_6_65 Salmonella enterica subsp. enterica serovar Typhi Proteobacteria; Gammaproteobacteria; Enterobacterales; Enterobacteriaceae; Salmonella; Salmonella enterica; Salmonella enterica subsp. enterica MTGDNTLITSHGINRRDFMKLCAALAATMGLSSKAAAEMAESVSNPQRPPVIWIGAQECT WP_000145429.1 hydrogenase 2 small subunit [Salmonella enterica] Length: 372\nNumber of predicted TMHs: 1\nExp number of AAs in TMHs: 24.51309\nExp number, first 60 AAs: 0.2682\nTotal prob of N-in: 0.14402\noutside 1 331\nTMhelix 332 354\ninside 355 372

20829 GCF_001099145.1_10593_2_14 Salmonella enterica subsp. enterica serovar Typhi Proteobacteria; Gammaproteobacteria; Enterobacterales; Enterobacteriaceae; Salmonella; Salmonella enterica; Salmonella enterica subsp. enterica MTGDNTLITSHGINRRDFMKLCAALAATMGLSSKAAAEMAESVSNPQRPPVIWIGAQECT WP_000145429.1 hydrogenase 2 small subunit [Salmonella enterica] Length: 372\nNumber of predicted TMHs: 1\nExp number of AAs in TMHs: 24.51309\nExp number, first 60 AAs: 0.2682\nTotal prob of N-in: 0.14402\noutside 1 331\nTMhelix 332 354\ninside 355 372

20830 GCF_001099265.1_10209_5_53 Salmonella enterica subsp. enterica serovar Typhi Proteobacteria; Gammaproteobacteria; Enterobacterales; Enterobacteriaceae; Salmonella; Salmonella enterica; Salmonella enterica subsp. enterica MTGDNTLITSHGINRRDFMKLCAALAATMGLSSKAAAEMAESVSNPQRPPVIWIGAQECT WP_000145429.1 hydrogenase 2 small subunit [Salmonella enterica] Length: 372\nNumber of predicted TMHs: 1\nExp number of AAs in TMHs: 24.51309\nExp number, first 60 AAs: 0.2682\nTotal prob of N-in: 0.14402\noutside 1 331\nTMhelix 332 354\ninside 355 372

20831 GCF_001099285.1_9475_6_48 Salmonella enterica subsp. enterica serovar Typhi Proteobacteria; Gammaproteobacteria; Enterobacterales; Enterobacteriaceae; Salmonella; Salmonella enterica; Salmonella enterica subsp. enterica MTGDNTLITSHGINRRDFMKLCAALAATMGLSSKAAAEMAESVSNPQRPPVIWIGAQECT WP_000145429.1 hydrogenase 2 small subunit [Salmonella enterica] Length: 372\nNumber of predicted TMHs: 1\nExp number of AAs in TMHs: 24.51309\nExp number, first 60 AAs: 0.2682\nTotal prob of N-in: 0.14402\noutside 1 331\nTMhelix 332 354\ninside 355 372

20832 GCF_001099365.1_10493_1_72 Salmonella enterica subsp. enterica serovar Typhi Proteobacteria; Gammaproteobacteria; Enterobacterales; Enterobacteriaceae; Salmonella; Salmonella enterica; Salmonella enterica subsp. enterica MTGDNTLITSHGINRRDFMKLCAALAATMGLSSKAAAEMAESVSNPQRPPVIWIGAQECT WP_000145429.1 hydrogenase 2 small subunit [Salmonella enterica] Length: 372\nNumber of predicted TMHs: 1\nExp number of AAs in TMHs: 24.51309\nExp number, first 60 AAs: 0.2682\nTotal prob of N-in: 0.14402\noutside 1 331\nTMhelix 332 354\ninside 355 372

20833 GCF_001099465.1_10562_2_31 Salmonella enterica subsp. enterica serovar Typhi Proteobacteria; Gammaproteobacteria; Enterobacterales; Enterobacteriaceae; Salmonella; Salmonella enterica; Salmonella enterica subsp. enterica MTGDNTLITSHGINRRDFMKLCAALAATMGLSSKAAAEMAESVSNPQRPPVIWIGAQECT WP_000145429.1 hydrogenase 2 small subunit [Salmonella enterica] Length: 372\nNumber of predicted TMHs: 1\nExp number of AAs in TMHs: 24.51309\nExp number, first 60 AAs: 0.2682\nTotal prob of N-in: 0.14402\noutside 1 331\nTMhelix 332 354\ninside 355 372

20834 GCF_001099645.1_7468_7_59 Salmonella enterica subsp. enterica serovar Typhi Proteobacteria; Gammaproteobacteria; Enterobacterales; Enterobacteriaceae; Salmonella; Salmonella enterica; Salmonella enterica subsp. enterica MTGDNTLITSHGINRRDFMKLCAALAATMGLSSKAAAEMAESVSNPQRPPVIWIGAQECT WP_000145429.1 hydrogenase 2 small subunit [Salmonella enterica] Length: 372\nNumber of predicted TMHs: 1\nExp number of AAs in TMHs: 24.51309\nExp number, first 60 AAs: 0.2682\nTotal prob of N-in: 0.14402\noutside 1 331\nTMhelix 332 354\ninside 355 372

20835 GCF_001099925.1_9953_5_78 Salmonella enterica subsp. enterica serovar Typhi Proteobacteria; Gammaproteobacteria; Enterobacterales; Enterobacteriaceae; Salmonella; Salmonella enterica; Salmonella enterica subsp. enterica MTGDNTLITSHGINRRDFMKLCAALAATMGLSSKAAAEMAESVSNPQRPPVIWIGAQECT WP_000145429.1 hydrogenase 2 small subunit [Salmonella enterica] Length: 372\nNumber of predicted TMHs: 1\nExp number of AAs in TMHs: 24.51309\nExp number, first 60 AAs: 0.2682\nTotal prob of N-in: 0.14402\noutside 1 331\nTMhelix 332 354\ninside 355 372

20836 GCF_001100005.1_10561_2_60 Salmonella enterica subsp. enterica serovar Typhi Proteobacteria; Gammaproteobacteria; Enterobacterales; Enterobacteriaceae; Salmonella; Salmonella enterica; Salmonella enterica subsp. enterica MTGDNTLITSHGINRRDFMKLCAALAATMGLSSKAAAEMAESVSNPQRPPVIWIGAQECT WP_000145429.1 hydrogenase 2 small subunit [Salmonella enterica] Length: 372\nNumber of predicted TMHs: 1\nExp number of AAs in TMHs: 24.51309\nExp number, first 60 AAs: 0.2682\nTotal prob of N-in: 0.14402\noutside 1 331\nTMhelix 332 354\ninside 355 372

20837 GCF_001100105.1_10349_1_37 Salmonella enterica subsp. enterica serovar Typhi Proteobacteria; Gammaproteobacteria; Enterobacterales; Enterobacteriaceae; Salmonella; Salmonella enterica; Salmonella enterica subsp. enterica MTGDNTLITSHGINRRDFMKLCAALAATMGLSSKAAAEMAESVSNPQRPPVIWIGAQECT WP_000145429.1 hydrogenase 2 small subunit [Salmonella enterica] Length: 372\nNumber of predicted TMHs: 1\nExp number of AAs in TMHs: 24.51309\nExp number, first 60 AAs: 0.2682\nTotal prob of N-in: 0.14402\noutside 1 331\nTMhelix 332 354\ninside 355 372

20838 GCF_001100425.1_10492_1_35 Salmonella enterica subsp. enterica serovar Typhi Proteobacteria; Gammaproteobacteria; Enterobacterales; Enterobacteriaceae; Salmonella; Salmonella enterica; Salmonella enterica subsp. enterica MTGDNTLITSHGINRRDFMKLCAALAATMGLSSKAAAEMAESVSNPQRPPVIWIGAQECT WP_000145429.1 hydrogenase 2 small subunit [Salmonella enterica] Length: 372\nNumber of predicted TMHs: 1\nExp number of AAs in TMHs: 24.51309\nExp number, first 60 AAs: 0.2682\nTotal prob of N-in: 0.14402\noutside 1 331\nTMhelix 332 354\ninside 355 372

20839 GCF_001101305.1_10562_2_48 Salmonella enterica subsp. enterica serovar Typhi Proteobacteria; Gammaproteobacteria; Enterobacterales; Enterobacteriaceae; Salmonella; Salmonella enterica; Salmonella enterica subsp. enterica MTGDNTLITSHGINRRDFMKLCAALAATMGLSSKAAAEMAESVSNPQRPPVIWIGAQECT WP_000145429.1 hydrogenase 2 small subunit [Salmonella enterica] Length: 372\nNumber of predicted TMHs: 1\nExp number of AAs in TMHs: 24.51309\nExp number, first 60 AAs: 0.2682\nTotal prob of N-in: 0.14402\noutside 1 331\nTMhelix 332 354\ninside 355 372

20840 GCF_001101445.1_10492_1_46 Salmonella enterica subsp. enterica serovar Typhi Proteobacteria; Gammaproteobacteria; Enterobacterales; Enterobacteriaceae; Salmonella; Salmonella enterica; Salmonella enterica subsp. enterica MTGDNTLITSHGINRRDFMKLCAALAATMGLSSKAAAEMAESVSNPQRPPVIWIGAQECT WP_000145429.1 hydrogenase 2 small subunit [Salmonella enterica] Length: 372\nNumber of

predicted TMHs: 1\nExp number of AAs in TMHs: 24.51309\nExp number, first 60 AAs: 0.2682\nTotal prob of N-in: 0.14402\noutside 1 331\nTMhelix 332 354\ninside 355 372

20841 GCF_001099765.1_7468_7_78 Salmonella enterica subsp. enterica serovar Typhi Proteobacteria; Gammaproteobacteria; Enterobacterales; Enterobacteriaceae; Salmonella; Salmonella enterica; Salmonella enterica subsp. enterica MTGDNTLITSHGINRRDFMKLCAALAATMGLSSKAATEMAESVSNPQRPPVIWIGAQECT WP_001782304.1 hydrogenase 2 small subunit [Salmonella enterica] Length: 372\nNumber of predicted TMHs: 1\nExp number of AAs in TMHs: 24.41173\nExp number, first 60 AAs: 0.16124\nTotal prob of N-in: 0.13947\noutside 1 331\nTMhelix 332 354\ninside 355 372

20842 GCF_001101805.1_10349_1_67 Salmonella enterica subsp. enterica serovar Typhi Proteobacteria; Gammaproteobacteria; Enterobacterales; Enterobacteriaceae; Salmonella; Salmonella enterica; Salmonella enterica subsp. enterica MTGDNTLITSHGINRRDFMKLCAALAATMGLSSKAAAEMAESVSNPQRPPVIWIGAQECT WP_000145429.1 hydrogenase 2 small subunit [Salmonella enterica] Length: 372\nNumber of predicted TMHs: 1\nExp number of AAs in TMHs: 24.51309\nExp number, first 60 AAs: 0.2682\nTotal prob of N-in: 0.14402\noutside 1 331\nTMhelix 332 354\ninside 355 372

20843 GCF_001102245.1_9870_8_62 Salmonella enterica subsp. enterica serovar Typhi Proteobacteria; Gammaproteobacteria; Enterobacterales; Enterobacteriaceae; Salmonella; Salmonella enterica; Salmonella enterica subsp. enterica MTGDNTLITSHGINRRDFMKLCAALAATMGLSSKAAAEMAESVSNPQRPPVIWIGAQECT WP_000145429.1 hydrogenase 2 small subunit [Salmonella enterica] Length: 372\nNumber of predicted TMHs: 1\nExp number of AAs in TMHs: 24.51309\nExp number, first 60 AAs: 0.2682\nTotal prob of N-in: 0.14402\noutside 1 331\nTMhelix 332 354\ninside 355 372

20844 GCF_001102385.1_10561_2_51 Salmonella enterica subsp. enterica serovar Typhi Proteobacteria; Gammaproteobacteria; Enterobacterales; Enterobacteriaceae; Salmonella; Salmonella enterica; Salmonella enterica subsp. enterica MTGDNTLITSHGINRRDFMKLCAALAATMGLSSKAAAEMAESVSNPQRPPVIWIGAQECT WP_000145429.1 hydrogenase 2 small subunit [Salmonella enterica] Length: 372\nNumber of predicted TMHs: 1\nExp number of AAs in TMHs: 24.51309\nExp number, first 60 AAs: 0.2682\nTotal prob of N-in: 0.14402\noutside 1 331\nTMhelix 332 354\ninside 355 372

20845 GCF_001102485.1_8490_6_66 Salmonella enterica subsp. enterica serovar Typhi Proteobacteria; Gammaproteobacteria; Enterobacterales; Enterobacteriaceae; Salmonella; Salmonella enterica; Salmonella enterica subsp. enterica MTGDNTLITSHGINRRDFMKLCAALAATMGLSSKAAAEMAESVSNPQRPPVIWIGAQECT WP_000145429.1 hydrogenase 2 small subunit [Salmonella enterica] Length: 372\nNumber of predicted TMHs: 1\nExp number of AAs in TMHs: 24.51309\nExp number, first 60 AAs: 0.2682\nTotal prob of N-in: 0.14402\noutside 1 331\nTMhelix 332 354\ninside 355 372

20846 GCF_001102845.1_9870_8_47 Salmonella enterica subsp. enterica serovar Typhi Proteobacteria; Gammaproteobacteria; Enterobacterales; Enterobacteriaceae; Salmonella; Salmonella enterica; Salmonella enterica subsp. enterica MTGDNTLITSHGINRRDFMKLCAALAATMGLSSKAAAEMAESVSNPQRPPVIWIGAQECT WP_000145429.1 hydrogenase 2 small subunit [Salmonella enterica] Length: 372\nNumber of predicted TMHs: 1\nExp number of AAs in TMHs: 24.51309\nExp number, first 60 AAs: 0.2682\nTotal prob of N-in: 0.14402\noutside 1 331\nTMhelix 332 354\ninside 355 372

20847 GCF_001103005.1_10349_1_63 Salmonella enterica subsp. enterica serovar Typhi Proteobacteria; Gammaproteobacteria; Enterobacterales; Enterobacteriaceae; Salmonella; Salmonella enterica; Salmonella enterica subsp. enterica MTGDNTLITSHGINRRDFMKLCAALAATMGLSSKAAAEMAESVSNPQRPPVIWIGAQECT WP_000145429.1 hydrogenase 2 small subunit [Salmonella enterica] Length: 372\nNumber of predicted TMHs: 1\nExp number of AAs in TMHs: 24.51309\nExp number, first 60 AAs: 0.2682\nTotal prob of N-in: 0.14402\noutside 1 331\nTMhelix 332 354\ninside 355 372

20848 GCF_001101585.1_9475_6_43 Salmonella enterica subsp. enterica serovar Typhi Proteobacteria; Gammaproteobacteria; Enterobacterales; Enterobacteriaceae; Salmonella; Salmonella enterica; Salmonella enterica subsp. enterica MTGDNTLITSHGINRRDFMKLCAALAATMGLSSKAATEMAESVSNPQRPPVIWIGAQECT WP_001782304.1 hydrogenase 2 small subunit [Salmonella enterica] Length: 372\nNumber of predicted TMHs: 1\nExp number of AAs in TMHs: 24.41173\nExp number, first 60 AAs: 0.16124\nTotal prob of N-in: 0.13947\noutside 1 331\nTMhelix 332 354\ninside 355 372

20849 GCF_001103165.1_8525_3_30 Salmonella enterica subsp. enterica serovar Typhi Proteobacteria; Gammaproteobacteria; Enterobacterales; Enterobacteriaceae; Salmonella; Salmonella enterica;

Salmonella enterica subsp. enterica MTGDNTLITSHGINRRDFMKLCAALAATMGLSSKAAAEMAESVSNPQRPPVIWIGAQECT WP_000145429.1 hydrogenase 2 small subunit [Salmonella enterica] Length: 372\nNumber of predicted TMHs: 1\nExp number of AAs in TMHs: 24.51309\nExp number, first 60 AAs: 0.2682\nTotal prob of N-in: 0.14402\noutside 1 331\nTMhelix 332 354\ninside 355 372

20850 GCF_001103305.1_10561_2_16 Salmonella enterica subsp. enterica serovar Typhi Proteobacteria; Gammaproteobacteria; Enterobacterales; Enterobacteriaceae; Salmonella; Salmonella enterica; Salmonella enterica subsp. enterica MTGDNTLITSHGINRRDFMKLCAALAATMGLSSKAAAEMAESVSNPQRPPVIWIGAQECT WP_000145429.1 hydrogenase 2 small subunit [Salmonella enterica] Length: 372\nNumber of predicted TMHs: 1\nExp number of AAs in TMHs: 24.51309\nExp number, first 60 AAs: 0.2682\nTotal prob of N-in: 0.14402\noutside 1 331\nTMhelix 332 354\ninside 355 372

20851 GCF_001103345.1_10593_2_82 Salmonella enterica subsp. enterica serovar Typhi Proteobacteria; Gammaproteobacteria; Enterobacterales; Enterobacteriaceae; Salmonella; Salmonella enterica; Salmonella enterica subsp. enterica MTGDNTLITSHGINRRDFMKLCAALAATMGLSSKAAAEMAESVSNPQRPPVIWIGAQECT WP_000145429.1 hydrogenase 2 small subunit [Salmonella enterica] Length: 372\nNumber of predicted TMHs: 1\nExp number of AAs in TMHs: 24.51309\nExp number, first 60 AAs: 0.2682\nTotal prob of N-in: 0.14402\noutside 1 331\nTMhelix 332 354\ninside 355 372

20852 GCF_001103485.1_10592_2_19 Salmonella enterica subsp. enterica serovar Typhi Proteobacteria; Gammaproteobacteria; Enterobacterales; Enterobacteriaceae; Salmonella; Salmonella enterica; Salmonella enterica subsp. enterica MTGDNTLITSHGINRRDFMKLCAALAATMGLSSKAAAEMAESVSNPQRPPVIWIGAQECT WP_000145429.1 hydrogenase 2 small subunit [Salmonella enterica] Length: 372\nNumber of predicted TMHs: 1\nExp number of AAs in TMHs: 24.51309\nExp number, first 60 AAs: 0.2682\nTotal prob of N-in: 0.14402\noutside 1 331\nTMhelix 332 354\ninside 355 372

20853 GCF_001103665.1_10593_2_33 Salmonella enterica subsp. enterica serovar Typhi Proteobacteria; Gammaproteobacteria; Enterobacterales; Enterobacteriaceae; Salmonella; Salmonella enterica; Salmonella enterica subsp. enterica MTGDNTLITSHGINRRDFMKLCAALAATMGLSSKAAAEMAESVSNPQRPPVIWIGAQECT WP_000145429.1 hydrogenase 2 small subunit [Salmonella enterica] Length: 372\nNumber of predicted TMHs: 1\nExp number of AAs in TMHs: 24.51309\nExp number, first 60 AAs: 0.2682\nTotal prob of N-in: 0.14402\noutside 1 331\nTMhelix 332 354\ninside 355 372

20854 GCF_001103905.1_10541_2_55 Salmonella enterica subsp. enterica serovar Typhi Proteobacteria; Gammaproteobacteria; Enterobacterales; Enterobacteriaceae; Salmonella; Salmonella enterica; Salmonella enterica subsp. enterica MTGDNTLITSHGINRRDFMKLCAALAATMGLSSKAAAEMAESVSNPQRPPVIWIGAQECT WP_000145429.1 hydrogenase 2 small subunit [Salmonella enterica] Length: 372\nNumber of predicted TMHs: 1\nExp number of AAs in TMHs: 24.51309\nExp number, first 60 AAs: 0.2682\nTotal prob of N-in: 0.14402\noutside 1 331\nTMhelix 332 354\ninside 355 372

20855 GCF_001104005.1_10209_5_45 Salmonella enterica subsp. enterica serovar Typhi Proteobacteria; Gammaproteobacteria; Enterobacterales; Enterobacteriaceae; Salmonella; Salmonella enterica; Salmonella enterica subsp. enterica MTGDNTLITSHGINRRDFMKLCAALAATMGLSSKAAAEMAESVSNPQRPPVIWIGAQECT WP_000145429.1 hydrogenase 2 small subunit [Salmonella enterica] Length: 372\nNumber of predicted TMHs: 1\nExp number of AAs in TMHs: 24.51309\nExp number, first 60 AAs: 0.2682\nTotal prob of N-in: 0.14402\noutside 1 331\nTMhelix 332 354\ninside 355 372

20856 GCF_001104165.1_10349_1_88 Salmonella enterica subsp. enterica serovar Typhi Proteobacteria; Gammaproteobacteria; Enterobacterales; Enterobacteriaceae; Salmonella; Salmonella enterica; Salmonella enterica subsp. enterica MTGDNTLITSHGINRRDFMKLCAALAATMGLSSKAAAEMAESVSNPQRPPVIWIGAQECT WP_000145429.1 hydrogenase 2 small subunit [Salmonella enterica] Length: 372\nNumber of predicted TMHs: 1\nExp number of AAs in TMHs: 24.51309\nExp number, first 60 AAs: 0.2682\nTotal prob of N-in: 0.14402\noutside 1 331\nTMhelix 332 354\ninside 355 372

20857 GCF_001104345.1_10592_2_73 Salmonella enterica subsp. enterica serovar Typhi Proteobacteria; Gammaproteobacteria; Enterobacterales; Enterobacteriaceae; Salmonella; Salmonella enterica; Salmonella enterica subsp. enterica MTGDNTLITSHGINRRDFMKLCAALAATMGLSSKAAAEMAESVSNPQRPPVIWIGAQECT WP_000145429.1 hydrogenase 2 small subunit [Salmonella enterica] Length: 372\nNumber of predicted TMHs: 1\nExp number of AAs in TMHs: 24.51309\nExp number, first 60 AAs: 0.2682\nTotal prob of N-in: 0.14402\noutside 1 331\nTMhelix 332 354\ninside 355 372

20858 GCF_001104465.1_10607_1_85 Salmonella enterica subsp. enterica serovar Typhi Proteobacteria; Gammaproteobacteria; Enterobacterales; Enterobacteriaceae; Salmonella; Salmonella enterica; Salmonella enterica subsp. enterica MTGDNTLITSHGINRRDFMKLCAALAATMGLSSKAAAEMAESVSNPQRPPVIWIGAQECT WP_000145429.1 hydrogenase 2 small subunit [Salmonella enterica] Length: 372\nNumber of predicted TMHs: 1\nExp number of AAs in TMHs: 24.51309\nExp number, first 60 AAs: 0.2682\nTotal prob of N-in: 0.14402\noutside 1 331\nTMhelix 332 354\ninside 355 372

20859 GCF_001104565.1_10493_1_2 Salmonella enterica subsp. enterica serovar Typhi Proteobacteria; Gammaproteobacteria; Enterobacterales; Enterobacteriaceae; Salmonella; Salmonella enterica; Salmonella enterica subsp. enterica MTGDNTLITSHGINRRDFMKLCAALAATMGLSSKAAAEMAESVSNPQRPPVIWIGAQECT WP_000145429.1 hydrogenase 2 small subunit [Salmonella enterica] Length: 372\nNumber of predicted TMHs: 1\nExp number of AAs in TMHs: 24.51309\nExp number, first 60 AAs: 0.2682\nTotal prob of N-in: 0.14402\noutside 1 331\nTMhelix 332 354\ninside 355 372

20860 GCF_001104705.1_10562_2_20 Salmonella enterica subsp. enterica serovar Typhi Proteobacteria; Gammaproteobacteria; Enterobacterales; Enterobacteriaceae; Salmonella; Salmonella enterica; Salmonella enterica subsp. enterica MTGDNTLITSHGINRRDFMKLCAALAATMGLSSKAAAEMAESVSNPQRPPVIWIGAQECT WP_000145429.1 hydrogenase 2 small subunit [Salmonella enterica] Length: 372\nNumber of predicted TMHs: 1\nExp number of AAs in TMHs: 24.51309\nExp number, first 60 AAs: 0.2682\nTotal prob of N-in: 0.14402\noutside 1 331\nTMhelix 332 354\ninside 355 372

20861 GCF_001104905.1_9870_8_26 Salmonella enterica subsp. enterica serovar Typhi Proteobacteria; Gammaproteobacteria; Enterobacterales; Enterobacteriaceae; Salmonella; Salmonella enterica; Salmonella enterica subsp. enterica MTGDNTLITSHGINRRDFMKLCAALAATMGLSSKAAAEMAESVSNPQRPPVIWIGAQECT WP_000145429.1 hydrogenase 2 small subunit [Salmonella enterica] Length: 372\nNumber of predicted TMHs: 1\nExp number of AAs in TMHs: 24.51309\nExp number, first 60 AAs: 0.2682\nTotal prob of N-in: 0.14402\noutside 1 331\nTMhelix 332 354\ninside 355 372

20862 GCF_001104965.1_10209_5_31 Salmonella enterica subsp. enterica serovar Typhi Proteobacteria; Gammaproteobacteria; Enterobacterales; Enterobacteriaceae; Salmonella; Salmonella enterica; Salmonella enterica subsp. enterica MTGDNTLITSHGINRRDFMKLCAALAATMGLSSKAAAEMAESVSNPQRPPVIWIGAQECT WP_000145429.1 hydrogenase 2 small subunit [Salmonella enterica] Length: 372\nNumber of predicted TMHs: 1\nExp number of AAs in TMHs: 24.51309\nExp number, first 60 AAs: 0.2682\nTotal prob of N-in: 0.14402\noutside 1 331\nTMhelix 332 354\ninside 355 372

20863 GCF_001105225.1_10607_2_15 Salmonella enterica subsp. enterica serovar Typhi Proteobacteria; Gammaproteobacteria; Enterobacterales; Enterobacteriaceae; Salmonella; Salmonella enterica; Salmonella enterica subsp. enterica MTGDNTLITSHGINRRDFMKLCAALAATMGLSSKAAAEMAESVSNPQRPPVIWIGAQECT WP_000145429.1 hydrogenase 2 small subunit [Salmonella enterica] Length: 372\nNumber of predicted TMHs: 1\nExp number of AAs in TMHs: 24.51309\nExp number, first 60 AAs: 0.2682\nTotal prob of N-in: 0.14402\noutside 1 331\nTMhelix 332 354\ninside 355 372

20864 GCF_001105525.1_10349_1_5 Salmonella enterica subsp. enterica serovar Typhi Proteobacteria; Gammaproteobacteria; Enterobacterales; Enterobacteriaceae; Salmonella; Salmonella enterica; Salmonella enterica subsp. enterica MTGDNTLITSHGINRRDFMKLCAALAATMGLSSKAAAEMAESVSNPQRPPVIWIGAQECT WP_000145429.1 hydrogenase 2 small subunit [Salmonella enterica] Length: 372\nNumber of predicted TMHs: 1\nExp number of AAs in TMHs: 24.51309\nExp number, first 60 AAs: 0.2682\nTotal prob of N-in: 0.14402\noutside 1 331\nTMhelix 332 354\ninside 355 372

20865 GCF_001105645.1_10608_2_25 Salmonella enterica subsp. enterica serovar Typhi Proteobacteria; Gammaproteobacteria; Enterobacterales; Enterobacteriaceae; Salmonella; Salmonella enterica; Salmonella enterica subsp. enterica MTGDNTLITSHGINRRDFMKLCAALAATMGLSSKAAAEMAESVSNPQRPPVIWIGAQECT WP_000145429.1 hydrogenase 2 small subunit [Salmonella enterica] Length: 372\nNumber of predicted TMHs: 1\nExp number of AAs in TMHs: 24.51309\nExp number, first 60 AAs: 0.2682\nTotal prob of N-in: 0.14402\noutside 1 331\nTMhelix 332 354\ninside 355 372

20866 GCF_001105825.1_10607_1_89 Salmonella enterica subsp. enterica serovar Typhi Proteobacteria; Gammaproteobacteria; Enterobacterales; Enterobacteriaceae; Salmonella; Salmonella enterica; Salmonella enterica subsp. enterica MTGDNTLITSHGINRRDFMKLCAALAATMGLSSKAAAEMAESVSNPQRPPVIWIGAQECT WP_000145429.1 hydrogenase 2 small subunit [Salmonella enterica] Length: 372\nNumber of

predicted TMHs: 1\nExp number of AAs in TMHs: 24.51309\nExp number, first 60 AAs: 0.2682\nTotal prob of N-in: 0.14402\noutside 1 331\nTMhelix 332 354\ninside 355 372

20867 GCF_001106065.1_10561_2_33 Salmonella enterica subsp. enterica serovar Typhi Proteobacteria; Gammaproteobacteria; Enterobacterales; Enterobacteriaceae; Salmonella; Salmonella enterica; Salmonella enterica subsp. enterica MTGDNTLITSHGINRRDFMKLCAALAATMGLSSKAAAEMAESVSNPQRPPVIWIGAQECT WP_000145429.1 hydrogenase 2 small subunit [Salmonella enterica] Length: 372\nNumber of predicted TMHs: 1\nExp number of AAs in TMHs: 24.51309\nExp number, first 60 AAs: 0.2682\nTotal prob of N-in: 0.14402\noutside 1 331\nTMhelix 332 354\ninside 355 372

20868 GCF_001106305.1_10607_2_27 Salmonella enterica subsp. enterica serovar Typhi Proteobacteria; Gammaproteobacteria; Enterobacterales; Enterobacteriaceae; Salmonella; Salmonella enterica; Salmonella enterica subsp. enterica MTGDNTLITSHGINRRDFMKLCAALAATMGLSSKAAAEMAESVSNPQRPPVIWIGAQECT WP_000145429.1 hydrogenase 2 small subunit [Salmonella enterica] Length: 372\nNumber of predicted TMHs: 1\nExp number of AAs in TMHs: 24.51309\nExp number, first 60 AAs: 0.2682\nTotal prob of N-in: 0.14402\noutside 1 331\nTMhelix 332 354\ninside 355 372

20869 GCF_001106745.1_10209_5_39 Salmonella enterica subsp. enterica serovar Typhi Proteobacteria; Gammaproteobacteria; Enterobacterales; Enterobacteriaceae; Salmonella; Salmonella enterica; Salmonella enterica subsp. enterica MTGDNTLITSHGINRRDFMKLCAALAATMGLSSKAAAEMAESVSNPQRPPVIWIGAQECT WP_000145429.1 hydrogenase 2 small subunit [Salmonella enterica] Length: 372\nNumber of predicted TMHs: 1\nExp number of AAs in TMHs: 24.51309\nExp number, first 60 AAs: 0.2682\nTotal prob of N-in: 0.14402\noutside 1 331\nTMhelix 332 354\ninside 355 372

20870 GCF_001106645.1_9870_8_84 Salmonella enterica subsp. enterica serovar Typhi Proteobacteria; Gammaproteobacteria; Enterobacterales; Enterobacteriaceae; Salmonella; Salmonella enterica; Salmonella enterica subsp. enterica MTGDNTLITSHGINRRDFMKLCAALAATMGLSSKAAAEMAESVSNPQRPPVIWIGAQECT WP_000145429.1 hydrogenase 2 small subunit [Salmonella enterica] Length: 372\nNumber of predicted TMHs: 1\nExp number of AAs in TMHs: 24.51309\nExp number, first 60 AAs: 0.2682\nTotal prob of N-in: 0.14402\noutside 1 331\nTMhelix 332 354\ninside 355 372

20871 GCF_001106825.1_9953_5_27 Salmonella enterica subsp. enterica serovar Typhi Proteobacteria; Gammaproteobacteria; Enterobacterales; Enterobacteriaceae; Salmonella; Salmonella enterica; Salmonella enterica subsp. enterica MTGDNTLITSHGINRRDFMKLCAALAATMGLSSKAAAEMAESVSNPQRPPVIWIGAQECT WP_000145429.1 hydrogenase 2 small subunit [Salmonella enterica] Length: 372\nNumber of predicted TMHs: 1\nExp number of AAs in TMHs: 24.51309\nExp number, first 60 AAs: 0.2682\nTotal prob of N-in: 0.14402\noutside 1 331\nTMhelix 332 354\ninside 355 372

20872 GCF_001107065.1_10592_2_36 Salmonella enterica subsp. enterica serovar Typhi Proteobacteria; Gammaproteobacteria; Enterobacterales; Enterobacteriaceae; Salmonella; Salmonella enterica; Salmonella enterica subsp. enterica MTGDNTLITSHGINRRDFMKLCAALAATMGLSSKAAAEMAESVSNPQRPPVIWIGAQECT WP_000145429.1 hydrogenase 2 small subunit [Salmonella enterica] Length: 372\nNumber of predicted TMHs: 1\nExp number of AAs in TMHs: 24.51309\nExp number, first 60 AAs: 0.2682\nTotal prob of N-in: 0.14402\noutside 1 331\nTMhelix 332 354\ninside 355 372

20873 GCF_001105925.1_7468_7_81 Salmonella enterica subsp. enterica serovar Typhi Proteobacteria; Gammaproteobacteria; Enterobacterales; Enterobacteriaceae; Salmonella; Salmonella enterica; Salmonella enterica subsp. enterica MTGDNTLITSHGINRRDFMKLCAALAATMGLSSKAATEMAESVSNPQRPPVIWIGAQECT WP_001782304.1 hydrogenase 2 small subunit [Salmonella enterica] Length: 372\nNumber of predicted TMHs: 1\nExp number of AAs in TMHs: 24.41173\nExp number, first 60 AAs: 0.16124\nTotal prob of N-in: 0.13947\noutside 1 331\nTMhelix 332 354\ninside 355 372

20874 GCF_001107265.1_10562_2_51 Salmonella enterica subsp. enterica serovar Typhi Proteobacteria; Gammaproteobacteria; Enterobacterales; Enterobacteriaceae; Salmonella; Salmonella enterica; Salmonella enterica subsp. enterica MTGDNTLITSHGINRRDFMKLCAALAATMGLSSKAAAEMAESVSNPQRPPVIWIGAQECT WP_000145429.1 hydrogenase 2 small subunit [Salmonella enterica] Length: 372\nNumber of predicted TMHs: 1\nExp number of AAs in TMHs: 24.51309\nExp number, first 60 AAs: 0.2682\nTotal prob of N-in: 0.14402\noutside 1 331\nTMhelix 332 354\ninside 355 372

20875 GCF_001107585.1_9953_5_59 Salmonella enterica subsp. enterica serovar Typhi Proteobacteria; Gammaproteobacteria; Enterobacterales; Enterobacteriaceae; Salmonella; Salmonella enterica;

Salmonella enterica subsp. enterica	MTGDNTLITSHGINRRDFMKLCAALAATMGLSSKAAAEMAESVSNPQRPPVIWIGAQECT
WP_000145429.1	hydrogenase 2 small subunit [Salmonella enterica]	Length: 372\nNumber of predicted TMHs:  1\nExp number of AAs in TMHs: 24.51309\nExp number, first 60 AAs:  0.2682\nTotal prob of N-in: 0.14402\noutside   1   331\nTMhelix   332   354\ninside   355   372

20876	GCF_001107705.1_8616_4_44	Salmonella enterica subsp. enterica serovar Typhi
Proteobacteria; Gammaproteobacteria; Enterobacterales; Enterobacteriaceae; Salmonella; Salmonella enterica; Salmonella enterica subsp. enterica	MTGDNTLITSHGINRRDFMKLCAALAATMGLSSKAAAEMAESVSNPQRPPVIWIGAQECT
WP_000145429.1	hydrogenase 2 small subunit [Salmonella enterica]	Length: 372\nNumber of predicted TMHs:  1\nExp number of AAs in TMHs: 24.51309\nExp number, first 60 AAs:  0.2682\nTotal prob of N-in: 0.14402\noutside   1   331\nTMhelix   332   354\ninside   355   372

20877	GCF_001107805.1_10209_5_17	Salmonella enterica subsp. enterica serovar Typhi
Proteobacteria; Gammaproteobacteria; Enterobacterales; Enterobacteriaceae; Salmonella; Salmonella enterica; Salmonella enterica subsp. enterica	MTGDNTLITSHGINRRDFMKLCAALAATMGLSSKAAAEMAESVSNPQRPPVIWIGAQECT
WP_000145429.1	hydrogenase 2 small subunit [Salmonella enterica]	Length: 372\nNumber of predicted TMHs:  1\nExp number of AAs in TMHs: 24.51309\nExp number, first 60 AAs:  0.2682\nTotal prob of N-in: 0.14402\noutside   1   331\nTMhelix   332   354\ninside   355   372

20878	GCF_001107885.1_10562_2_85	Salmonella enterica subsp. enterica serovar Typhi
Proteobacteria; Gammaproteobacteria; Enterobacterales; Enterobacteriaceae; Salmonella; Salmonella enterica; Salmonella enterica subsp. enterica	MTGDNTLITSHGINRRDFMKLCAALAATMGLSSKAAAEMAESVSNPQRPPVIWIGAQECT
WP_000145429.1	hydrogenase 2 small subunit [Salmonella enterica]	Length: 372\nNumber of predicted TMHs:  1\nExp number of AAs in TMHs: 24.51309\nExp number, first 60 AAs:  0.2682\nTotal prob of N-in: 0.14402\noutside   1   331\nTMhelix   332   354\ninside   355   372

20879	GCF_001108025.1_9475_6_31	Salmonella enterica subsp. enterica serovar Typhi
Proteobacteria; Gammaproteobacteria; Enterobacterales; Enterobacteriaceae; Salmonella; Salmonella enterica; Salmonella enterica subsp. enterica	MTGDNTLITSHGINRRDFMKLCAALAATMGLSSKAAAEMAESVSNPQRPPVIWIGAQECT
WP_000145429.1	hydrogenase 2 small subunit [Salmonella enterica]	Length: 372\nNumber of predicted TMHs:  1\nExp number of AAs in TMHs: 24.51309\nExp number, first 60 AAs:  0.2682\nTotal prob of N-in: 0.14402\noutside   1   331\nTMhelix   332   354\ninside   355   372

20880	GCF_001108525.1_8490_5_35	Salmonella enterica subsp. enterica serovar Typhi
Proteobacteria; Gammaproteobacteria; Enterobacterales; Enterobacteriaceae; Salmonella; Salmonella enterica; Salmonella enterica subsp. enterica	MTGDNTLITSHGINRRDFMKLCAALAATMGLSSKAAAEMAESVSNPQRPPVIWIGAQECT
WP_000145429.1	hydrogenase 2 small subunit [Salmonella enterica]	Length: 372\nNumber of predicted TMHs:  1\nExp number of AAs in TMHs: 24.51309\nExp number, first 60 AAs:  0.2682\nTotal prob of N-in: 0.14402\noutside   1   331\nTMhelix   332   354\ninside   355   372

20881	GCF_001108265.1_9870_8_81	Salmonella enterica subsp. enterica serovar Typhi
Proteobacteria; Gammaproteobacteria; Enterobacterales; Enterobacteriaceae; Salmonella; Salmonella enterica; Salmonella enterica subsp. enterica	MTGDNTLITSHGINRRDFMKLCAALAATMGLSSKAAAEMAESVSNPQRPPVIWIGAQECT
WP_000145429.1	hydrogenase 2 small subunit [Salmonella enterica]	Length: 372\nNumber of predicted TMHs:  1\nExp number of AAs in TMHs: 24.51309\nExp number, first 60 AAs:  0.2682\nTotal prob of N-in: 0.14402\noutside   1   331\nTMhelix   332   354\ninside   355   372

20882	GCF_001108685.1_10593_2_68	Salmonella enterica subsp. enterica serovar Typhi
Proteobacteria; Gammaproteobacteria; Enterobacterales; Enterobacteriaceae; Salmonella; Salmonella enterica; Salmonella enterica subsp. enterica	MTGDNTLITSHGINRRDFMKLCAALAATMGLSSKAAAEMAESVSNPQRPPVIWIGAQECT
WP_000145429.1	hydrogenase 2 small subunit [Salmonella enterica]	Length: 372\nNumber of predicted TMHs:  1\nExp number of AAs in TMHs: 24.51309\nExp number, first 60 AAs:  0.2682\nTotal prob of N-in: 0.14402\noutside   1   331\nTMhelix   332   354\ninside   355   372

20883	GCF_001108885.1_9953_5_20	Salmonella enterica subsp. enterica serovar Typhi
Proteobacteria; Gammaproteobacteria; Enterobacterales; Enterobacteriaceae; Salmonella; Salmonella enterica; Salmonella enterica subsp. enterica	MTGDNTLITSHGINRRDFMKLCAALAATMGLSSKAAAEMAESVSNPQRPPVIWIGAQECT
WP_000145429.1	hydrogenase 2 small subunit [Salmonella enterica]	Length: 372\nNumber of predicted TMHs:  1\nExp number of AAs in TMHs: 24.51309\nExp number, first 60 AAs:  0.2682\nTotal prob of N-in: 0.14402\noutside   1   331\nTMhelix   332   354\ninside   355   372

20884 GCF_001109025.1_10608_2_39 Salmonella enterica subsp. enterica serovar Typhi Proteobacteria; Gammaproteobacteria; Enterobacterales; Enterobacteriaceae; Salmonella; Salmonella enterica; Salmonella enterica subsp. enterica MTGDNTLITSHGINRRDFMKLCAALAATMGLSSKAAAEMAESVSNPQRPPVIWIGAQECT WP_000145429.1 hydrogenase 2 small subunit [Salmonella enterica] Length: 372\nNumber of predicted TMHs: 1\nExp number of AAs in TMHs: 24.51309\nExp number, first 60 AAs: 0.2682\nTotal prob of N-in: 0.14402\noutside 1 331\nTMhelix 332 354\ninside 355 372

20885 GCF_001109225.1_10592_2_45 Salmonella enterica subsp. enterica serovar Typhi Proteobacteria; Gammaproteobacteria; Enterobacterales; Enterobacteriaceae; Salmonella; Salmonella enterica; Salmonella enterica subsp. enterica MTGDNTLITSHGINRRDFMKLCAALAATMGLSSKAAAEMAESVSNPQRPPVIWIGAQECT WP_000145429.1 hydrogenase 2 small subunit [Salmonella enterica] Length: 372\nNumber of predicted TMHs: 1\nExp number of AAs in TMHs: 24.51309\nExp number, first 60 AAs: 0.2682\nTotal prob of N-in: 0.14402\noutside 1 331\nTMhelix 332 354\ninside 355 372

20886 GCF_001109325.1_9953_5_40 Salmonella enterica subsp. enterica serovar Typhi Proteobacteria; Gammaproteobacteria; Enterobacterales; Enterobacteriaceae; Salmonella; Salmonella enterica; Salmonella enterica subsp. enterica MTGDNTLITSHGINRRDFMKLCAALAATMGLSSKAAAEMAESVSNPQRPPVIWIGAQECT WP_000145429.1 hydrogenase 2 small subunit [Salmonella enterica] Length: 372\nNumber of predicted TMHs: 1\nExp number of AAs in TMHs: 24.51309\nExp number, first 60 AAs: 0.2682\nTotal prob of N-in: 0.14402\noutside 1 331\nTMhelix 332 354\ninside 355 372

20887 GCF_001109525.1_10541_1_62 Salmonella enterica subsp. enterica serovar Typhi Proteobacteria; Gammaproteobacteria; Enterobacterales; Enterobacteriaceae; Salmonella; Salmonella enterica; Salmonella enterica subsp. enterica MTGDNTLITSHGINRRDFMKLCAALAATMGLSSKAAAEMAESVSNPQRPPVIWIGAQECT WP_000145429.1 hydrogenase 2 small subunit [Salmonella enterica] Length: 372\nNumber of predicted TMHs: 1\nExp number of AAs in TMHs: 24.51309\nExp number, first 60 AAs: 0.2682\nTotal prob of N-in: 0.14402\noutside 1 331\nTMhelix 332 354\ninside 355 372

20888 GCF_001109625.1_10492_1_42 Salmonella enterica subsp. enterica serovar Typhi Proteobacteria; Gammaproteobacteria; Enterobacterales; Enterobacteriaceae; Salmonella; Salmonella enterica; Salmonella enterica subsp. enterica MTGDNTLITSHGINRRDFMKLCAALAATMGLSSKAAAEMAESVSNPQRPPVIWIGAQECT WP_000145429.1 hydrogenase 2 small subunit [Salmonella enterica] Length: 372\nNumber of predicted TMHs: 1\nExp number of AAs in TMHs: 24.51309\nExp number, first 60 AAs: 0.2682\nTotal prob of N-in: 0.14402\noutside 1 331\nTMhelix 332 354\ninside 355 372

20889 GCF_001109745.1_10349_1_23 Salmonella enterica subsp. enterica serovar Typhi Proteobacteria; Gammaproteobacteria; Enterobacterales; Enterobacteriaceae; Salmonella; Salmonella enterica; Salmonella enterica subsp. enterica MTGDNTLITSHGINRRDFMKLCAALAATMGLSSKAAAEMAESVSNPQRPPVIWIGAQECT WP_000145429.1 hydrogenase 2 small subunit [Salmonella enterica] Length: 372\nNumber of predicted TMHs: 1\nExp number of AAs in TMHs: 24.51309\nExp number, first 60 AAs: 0.2682\nTotal prob of N-in: 0.14402\noutside 1 331\nTMhelix 332 354\ninside 355 372

20890 GCF_001110185.1_8447_8_3Salmonella enterica subsp. enterica serovar Typhi Proteobacteria; Gammaproteobacteria; Enterobacterales; Enterobacteriaceae; Salmonella; Salmonella enterica; Salmonella enterica subsp. enterica MTGDNTLITSHGINRRDFMKLCAALAATMGLSSKAAAEMAESVSNPQRPPVIWIGAQECT WP_000145429.1 hydrogenase 2 small subunit [Salmonella enterica] Length: 372\nNumber of predicted TMHs: 1\nExp number of AAs in TMHs: 24.51309\nExp number, first 60 AAs: 0.2682\nTotal prob of N-in: 0.14402\noutside 1 331\nTMhelix 332 354\ninside 355 372

20891 GCF_001110445.1_10541_2_48 Salmonella enterica subsp. enterica serovar Typhi Proteobacteria; Gammaproteobacteria; Enterobacterales; Enterobacteriaceae; Salmonella; Salmonella enterica; Salmonella enterica subsp. enterica MTGDNTLITSHGINRRDFMKLCAALAATMGLSSKAAAEMAESVSNPQRPPVIWIGAQECT WP_000145429.1 hydrogenase 2 small subunit [Salmonella enterica] Length: 372\nNumber of predicted TMHs: 1\nExp number of AAs in TMHs: 24.51309\nExp number, first 60 AAs: 0.2682\nTotal prob of N-in: 0.14402\noutside 1 331\nTMhelix 332 354\ninside 355 372

20892 GCF_001110645.1_10593_2_52 Salmonella enterica subsp. enterica serovar Typhi Proteobacteria; Gammaproteobacteria; Enterobacterales; Enterobacteriaceae; Salmonella; Salmonella enterica; Salmonella enterica subsp. enterica MTGDNTLITSHGINRRDFMKLCAALAATMGLSSKAAAEMAESVSNPQRPPVIWIGAQECT WP_000145429.1 hydrogenase 2 small subunit [Salmonella enterica] Length: 372\nNumber of

predicted TMHs: 1\nExp number of AAs in TMHs: 24.51309\nExp number, first 60 AAs: 0.2682\nTotal prob of N-in: 0.14402\noutside 1 331\nTMhelix 332 354\ninside 355 372

20893 GCF_001111005.1_9953_5_11 Salmonella enterica subsp. enterica serovar Typhi Proteobacteria; Gammaproteobacteria; Enterobacterales; Enterobacteriaceae; Salmonella; Salmonella enterica; Salmonella enterica subsp. enterica MTGDNTLITSHGINRRDFMKLCAALAATMGLSSKAAAEMAESVSNPQRPPVIWIGAQECT WP_000145429.1 hydrogenase 2 small subunit [Salmonella enterica] Length: 372\nNumber of predicted TMHs: 1\nExp number of AAs in TMHs: 24.51309\nExp number, first 60 AAs: 0.2682\nTotal prob of N-in: 0.14402\noutside 1 331\nTMhelix 332 354\ninside 355 372

20894 GCF_001111185.1_9870_8_30 Salmonella enterica subsp. enterica serovar Typhi Proteobacteria; Gammaproteobacteria; Enterobacterales; Enterobacteriaceae; Salmonella; Salmonella enterica; Salmonella enterica subsp. enterica MTGDNTLITSHGINRRDFMKLCAALAATMGLSSKAAAEMAESVSNPQRPPVIWIGAQECT WP_000145429.1 hydrogenase 2 small subunit [Salmonella enterica] Length: 372\nNumber of predicted TMHs: 1\nExp number of AAs in TMHs: 24.51309\nExp number, first 60 AAs: 0.2682\nTotal prob of N-in: 0.14402\noutside 1 331\nTMhelix 332 354\ninside 355 372

20895 GCF_001111405.1_9953_5_29 Salmonella enterica subsp. enterica serovar Typhi Proteobacteria; Gammaproteobacteria; Enterobacterales; Enterobacteriaceae; Salmonella; Salmonella enterica; Salmonella enterica subsp. enterica MTGDNTLITSHGINRRDFMKLCAALAATMGLSSKAAAEMAESVSNPQRPPVIWIGAQECT WP_000145429.1 hydrogenase 2 small subunit [Salmonella enterica] Length: 372\nNumber of predicted TMHs: 1\nExp number of AAs in TMHs: 24.51309\nExp number, first 60 AAs: 0.2682\nTotal prob of N-in: 0.14402\noutside 1 331\nTMhelix 332 354\ninside 355 372

20896 GCF_001111305.1_9870_8_15 Salmonella enterica subsp. enterica serovar Typhi Proteobacteria; Gammaproteobacteria; Enterobacterales; Enterobacteriaceae; Salmonella; Salmonella enterica; Salmonella enterica subsp. enterica MTGDNTLITSHGINRRDFMKLCAALAATMGLSSKAAAEMAESVSNPQRPPVIWIGAQECT WP_000145429.1 hydrogenase 2 small subunit [Salmonella enterica] Length: 372\nNumber of predicted TMHs: 1\nExp number of AAs in TMHs: 24.51309\nExp number, first 60 AAs: 0.2682\nTotal prob of N-in: 0.14402\noutside 1 331\nTMhelix 332 354\ninside 355 372

20897 GCF_001111545.1_8490_5_13 Salmonella enterica subsp. enterica serovar Typhi Proteobacteria; Gammaproteobacteria; Enterobacterales; Enterobacteriaceae; Salmonella; Salmonella enterica; Salmonella enterica subsp. enterica MTGDNTLITSHGINRRDFMKLCAALAATMGLSSKAAAEMAESVSNPQRPPVIWIGAQECT WP_000145429.1 hydrogenase 2 small subunit [Salmonella enterica] Length: 372\nNumber of predicted TMHs: 1\nExp number of AAs in TMHs: 24.51309\nExp number, first 60 AAs: 0.2682\nTotal prob of N-in: 0.14402\noutside 1 331\nTMhelix 332 354\ninside 355 372

20898 GCF_001111825.1_10209_5_15 Salmonella enterica subsp. enterica serovar Typhi Proteobacteria; Gammaproteobacteria; Enterobacterales; Enterobacteriaceae; Salmonella; Salmonella enterica; Salmonella enterica subsp. enterica MTGDNTLITSHGINRRDFMKLCAALAATMGLSSKAAAEMAESVSNPQRPPVIWIGAQECT WP_000145429.1 hydrogenase 2 small subunit [Salmonella enterica] Length: 372\nNumber of predicted TMHs: 1\nExp number of AAs in TMHs: 24.51309\nExp number, first 60 AAs: 0.2682\nTotal prob of N-in: 0.14402\noutside 1 331\nTMhelix 332 354\ninside 355 372

20899 GCF_001111725.1_9870_8_18 Salmonella enterica subsp. enterica serovar Typhi Proteobacteria; Gammaproteobacteria; Enterobacterales; Enterobacteriaceae; Salmonella; Salmonella enterica; Salmonella enterica subsp. enterica MTGDNTLITSHGINRRDFMKLCAALAATMGLSSKAAAEMAESVSNPQRPPVIWIGAQECT WP_000145429.1 hydrogenase 2 small subunit [Salmonella enterica] Length: 372\nNumber of predicted TMHs: 1\nExp number of AAs in TMHs: 24.51309\nExp number, first 60 AAs: 0.2682\nTotal prob of N-in: 0.14402\noutside 1 331\nTMhelix 332 354\ninside 355 372

20900 GCF_001112065.1_9475_6_42 Salmonella enterica subsp. enterica serovar Typhi Proteobacteria; Gammaproteobacteria; Enterobacterales; Enterobacteriaceae; Salmonella; Salmonella enterica; Salmonella enterica subsp. enterica MTGDNTLITSHGINRRDFMKLCAALAATMGLSSKAAAEMAESVSNPQRPPVIWIGAQECT WP_000145429.1 hydrogenase 2 small subunit [Salmonella enterica] Length: 372\nNumber of predicted TMHs: 1\nExp number of AAs in TMHs: 24.51309\nExp number, first 60 AAs: 0.2682\nTotal prob of N-in: 0.14402\noutside 1 331\nTMhelix 332 354\ninside 355 372

20901 GCF_001112185.1_10493_1_1 Salmonella enterica subsp. enterica serovar Typhi Proteobacteria; Gammaproteobacteria; Enterobacterales; Enterobacteriaceae; Salmonella; Salmonella enterica;

Salmonella enterica subsp. enterica	MTGDNTLITSHGINRRDFMKLCAALAATMGLSSKAAAEMAESVSNPQRPPVIWIGAQECT
WP_000145429.1	hydrogenase 2 small subunit [Salmonella enterica]	Length: 372\nNumber of predicted TMHs: 1\nExp number of AAs in TMHs: 24.51309\nExp number, first 60 AAs: 0.2682\nTotal prob of N-in: 0.14402\noutside 1 331\nTMhelix 332 354\ninside 355 372

20902	GCF_001112305.1_10209_5_12	Salmonella enterica subsp. enterica serovar Typhi
Proteobacteria; Gammaproteobacteria; Enterobacterales; Enterobacteriaceae; Salmonella; Salmonella enterica; Salmonella enterica subsp. enterica	MTGDNTLITSHGINRRDFMKLCAALAATMGLSSKAAAEMAESVSNPQRPPVIWIGAQECT
WP_000145429.1	hydrogenase 2 small subunit [Salmonella enterica]	Length: 372\nNumber of predicted TMHs: 1\nExp number of AAs in TMHs: 24.51309\nExp number, first 60 AAs: 0.2682\nTotal prob of N-in: 0.14402\noutside 1 331\nTMhelix 332 354\ninside 355 372

20903	GCF_001112445.1_9953_5_90	Salmonella enterica subsp. enterica serovar Typhi
Proteobacteria; Gammaproteobacteria; Enterobacterales; Enterobacteriaceae; Salmonella; Salmonella enterica; Salmonella enterica subsp. enterica	MTGDNTLITSHGINRRDFMKLCAALAATMGLSSKAAAEMAESVSNPQRPPVIWIGAQECT
WP_000145429.1	hydrogenase 2 small subunit [Salmonella enterica]	Length: 372\nNumber of predicted TMHs: 1\nExp number of AAs in TMHs: 24.51309\nExp number, first 60 AAs: 0.2682\nTotal prob of N-in: 0.14402\noutside 1 331\nTMhelix 332 354\ninside 355 372

20904	GCF_001112665.1_9870_8_24	Salmonella enterica subsp. enterica serovar Typhi
Proteobacteria; Gammaproteobacteria; Enterobacterales; Enterobacteriaceae; Salmonella; Salmonella enterica; Salmonella enterica subsp. enterica	MTGDNTLITSHGINRRDFMKLCAALAATMGLSSKAAAEMAESVSNPQRPPVIWIGAQECT
WP_000145429.1	hydrogenase 2 small subunit [Salmonella enterica]	Length: 372\nNumber of predicted TMHs: 1\nExp number of AAs in TMHs: 24.51309\nExp number, first 60 AAs: 0.2682\nTotal prob of N-in: 0.14402\noutside 1 331\nTMhelix 332 354\ninside 355 372

20905	GCF_001112885.1_8447_8_24	Salmonella enterica subsp. enterica serovar Typhi
Proteobacteria; Gammaproteobacteria; Enterobacterales; Enterobacteriaceae; Salmonella; Salmonella enterica; Salmonella enterica subsp. enterica	MTGDNTLITSHGINRRDFMKLCAALAATMGLSSKAAAEMAESVSNPQRPPVIWIGAQECT
WP_000145429.1	hydrogenase 2 small subunit [Salmonella enterica]	Length: 372\nNumber of predicted TMHs: 1\nExp number of AAs in TMHs: 24.51309\nExp number, first 60 AAs: 0.2682\nTotal prob of N-in: 0.14402\noutside 1 331\nTMhelix 332 354\ninside 355 372

20906	GCF_001113065.1_9475_6_64	Salmonella enterica subsp. enterica serovar Typhi
Proteobacteria; Gammaproteobacteria; Enterobacterales; Enterobacteriaceae; Salmonella; Salmonella enterica; Salmonella enterica subsp. enterica	MTGDNTLITSHGINRRDFMKLCAALAATMGLSSKAAAEMAESVSNPQRPPVIWIGAQECT
WP_000145429.1	hydrogenase 2 small subunit [Salmonella enterica]	Length: 372\nNumber of predicted TMHs: 1\nExp number of AAs in TMHs: 24.51309\nExp number, first 60 AAs: 0.2682\nTotal prob of N-in: 0.14402\noutside 1 331\nTMhelix 332 354\ninside 355 372

20907	GCF_001113145.1_9953_5_48	Salmonella enterica subsp. enterica serovar Typhi
Proteobacteria; Gammaproteobacteria; Enterobacterales; Enterobacteriaceae; Salmonella; Salmonella enterica; Salmonella enterica subsp. enterica	MTGDNTLITSHGINRRDFMKLCAALAATMGLSSKAAAEMAESVSNPQRPPVIWIGAQECT
WP_000145429.1	hydrogenase 2 small subunit [Salmonella enterica]	Length: 372\nNumber of predicted TMHs: 1\nExp number of AAs in TMHs: 24.51309\nExp number, first 60 AAs: 0.2682\nTotal prob of N-in: 0.14402\noutside 1 331\nTMhelix 332 354\ninside 355 372

20908	GCF_001113485.1_10561_2_14	Salmonella enterica subsp. enterica serovar Typhi
Proteobacteria; Gammaproteobacteria; Enterobacterales; Enterobacteriaceae; Salmonella; Salmonella enterica; Salmonella enterica subsp. enterica	MTGDNTLITSHGINRRDFMKLCAALAATMGLSSKAAAEMAESVSNPQRPPVIWIGAQECT
WP_000145429.1	hydrogenase 2 small subunit [Salmonella enterica]	Length: 372\nNumber of predicted TMHs: 1\nExp number of AAs in TMHs: 24.51309\nExp number, first 60 AAs: 0.2682\nTotal prob of N-in: 0.14402\noutside 1 331\nTMhelix 332 354\ninside 355 372

20909	GCF_001113605.1_10349_1_76	Salmonella enterica subsp. enterica serovar Typhi
Proteobacteria; Gammaproteobacteria; Enterobacterales; Enterobacteriaceae; Salmonella; Salmonella enterica; Salmonella enterica subsp. enterica	MTGDNTLITSHGINRRDFMKLCAALAATMGLSSKAAAEMAESVSNPQRPPVIWIGAQECT
WP_000145429.1	hydrogenase 2 small subunit [Salmonella enterica]	Length: 372\nNumber of predicted TMHs: 1\nExp number of AAs in TMHs: 24.51309\nExp number, first 60 AAs: 0.2682\nTotal prob of N-in: 0.14402\noutside 1 331\nTMhelix 332 354\ninside 355 372

20910 GCF_001113765.1_10209_5_57 Salmonella enterica subsp. enterica serovar Typhi Proteobacteria; Gammaproteobacteria; Enterobacterales; Enterobacteriaceae; Salmonella; Salmonella enterica; Salmonella enterica subsp. enterica MTGDNTLITSHGINRRDFMKLCAALAATMGLSSKAAAEMAESVSNPQRPPVIWIGAQECT WP_000145429.1 hydrogenase 2 small subunit [Salmonella enterica] Length: 372\nNumber of predicted TMHs: 1\nExp number of AAs in TMHs: 24.51309\nExp number, first 60 AAs: 0.2682\nTotal prob of N-in: 0.14402\noutside 1 331\nTMhelix 332 354\ninside 355 372

20911 GCF_001113945.1_10607_1_64 Salmonella enterica subsp. enterica serovar Typhi Proteobacteria; Gammaproteobacteria; Enterobacterales; Enterobacteriaceae; Salmonella; Salmonella enterica; Salmonella enterica subsp. enterica MTGDNTLITSHGINRRDFMKLCAALAATMGLSSKAAAEMAESVSNPQRPPVIWIGAQECT WP_000145429.1 hydrogenase 2 small subunit [Salmonella enterica] Length: 372\nNumber of predicted TMHs: 1\nExp number of AAs in TMHs: 24.51309\nExp number, first 60 AAs: 0.2682\nTotal prob of N-in: 0.14402\noutside 1 331\nTMhelix 332 354\ninside 355 372

20912 GCF_001114365.1_10562_2_16 Salmonella enterica subsp. enterica serovar Typhi Proteobacteria; Gammaproteobacteria; Enterobacterales; Enterobacteriaceae; Salmonella; Salmonella enterica; Salmonella enterica subsp. enterica MTGDNTLITSHGINRRDFMKLCAALAATMGLSSKAAAEMAESVSNPQRPPVIWIGAQECT WP_000145429.1 hydrogenase 2 small subunit [Salmonella enterica] Length: 372\nNumber of predicted TMHs: 1\nExp number of AAs in TMHs: 24.51309\nExp number, first 60 AAs: 0.2682\nTotal prob of N-in: 0.14402\noutside 1 331\nTMhelix 332 354\ninside 355 372

20913 GCF_001114445.1_9870_8_54 Salmonella enterica subsp. enterica serovar Typhi Proteobacteria; Gammaproteobacteria; Enterobacterales; Enterobacteriaceae; Salmonella; Salmonella enterica; Salmonella enterica subsp. enterica MTGDNTLITSHGINRRDFMKLCAALAATMGLSSKAAAEMAESVSNPQRPPVIWIGAQECT WP_000145429.1 hydrogenase 2 small subunit [Salmonella enterica] Length: 372\nNumber of predicted TMHs: 1\nExp number of AAs in TMHs: 24.51309\nExp number, first 60 AAs: 0.2682\nTotal prob of N-in: 0.14402\noutside 1 331\nTMhelix 332 354\ninside 355 372

20914 GCF_001114665.1_10492_1_49 Salmonella enterica subsp. enterica serovar Typhi Proteobacteria; Gammaproteobacteria; Enterobacterales; Enterobacteriaceae; Salmonella; Salmonella enterica; Salmonella enterica subsp. enterica MTGDNTLITSHGINRRDFMKLCAALAATMGLSSKAAAEMAESVSNPQRPPVIWIGAQECT WP_000145429.1 hydrogenase 2 small subunit [Salmonella enterica] Length: 372\nNumber of predicted TMHs: 1\nExp number of AAs in TMHs: 24.51309\nExp number, first 60 AAs: 0.2682\nTotal prob of N-in: 0.14402\noutside 1 331\nTMhelix 332 354\ninside 355 372

20915 GCF_001114845.1_10592_2_13 Salmonella enterica subsp. enterica serovar Typhi Proteobacteria; Gammaproteobacteria; Enterobacterales; Enterobacteriaceae; Salmonella; Salmonella enterica; Salmonella enterica subsp. enterica MTGDNTLITSHGINRRDFMKLCAALAATMGLSSKAAAEMAESVSNPQRPPVIWIGAQECT WP_000145429.1 hydrogenase 2 small subunit [Salmonella enterica] Length: 372\nNumber of predicted TMHs: 1\nExp number of AAs in TMHs: 24.51309\nExp number, first 60 AAs: 0.2682\nTotal prob of N-in: 0.14402\noutside 1 331\nTMhelix 332 354\ninside 355 372

20916 GCF_001114945.1_10592_2_1 Salmonella enterica subsp. enterica serovar Typhi Proteobacteria; Gammaproteobacteria; Enterobacterales; Enterobacteriaceae; Salmonella; Salmonella enterica; Salmonella enterica subsp. enterica MTGDNTLITSHGINRRDFMKLCAALAATMGLSSKAAAEMAESVSNPQRPPVIWIGAQECT WP_000145429.1 hydrogenase 2 small subunit [Salmonella enterica] Length: 372\nNumber of predicted TMHs: 1\nExp number of AAs in TMHs: 24.51309\nExp number, first 60 AAs: 0.2682\nTotal prob of N-in: 0.14402\noutside 1 331\nTMhelix 332 354\ninside 355 372

20917 GCF_001115025.1_9475_6_63 Salmonella enterica subsp. enterica serovar Typhi Proteobacteria; Gammaproteobacteria; Enterobacterales; Enterobacteriaceae; Salmonella; Salmonella enterica; Salmonella enterica subsp. enterica MTGDNTLITSHGINRRDFMKLCAALAATMGLSSKAAAEMAESVSNPQRPPVIWIGAQECT WP_000145429.1 hydrogenase 2 small subunit [Salmonella enterica] Length: 372\nNumber of predicted TMHs: 1\nExp number of AAs in TMHs: 24.51309\nExp number, first 60 AAs: 0.2682\nTotal prob of N-in: 0.14402\noutside 1 331\nTMhelix 332 354\ninside 355 372

20918 GCF_001115685.1_10561_2_15 Salmonella enterica subsp. enterica serovar Typhi Proteobacteria; Gammaproteobacteria; Enterobacterales; Enterobacteriaceae; Salmonella; Salmonella enterica; Salmonella enterica subsp. enterica MTGDNTLITSHGINRRDFMKLCAALAATMGLSSKAAAEMAESVSNPQRPPVIWIGAQECT WP_000145429.1 hydrogenase 2 small subunit [Salmonella enterica] Length: 372\nNumber of

predicted TMHs:  1\nExp number of AAs in TMHs: 24.51309\nExp number, first 60 AAs:  0.2682\nTotal prob of N-in: 0.14402\noutside   1   331\nTMhelix   332   354\ninside   355   372

20919 GCF_001115965.1_9870_8_12 Salmonella enterica subsp. enterica serovar Typhi Proteobacteria; Gammaproteobacteria; Enterobacterales; Enterobacteriaceae; Salmonella; Salmonella enterica; Salmonella enterica subsp. enterica MTGDNTLITSHGINRRDFMKLCAALAATMGLSSKAAAEMAESVSNPQRPPVIWIGAQECT WP_000145429.1 hydrogenase 2 small subunit [Salmonella enterica] Length: 372\nNumber of predicted TMHs:  1\nExp number of AAs in TMHs: 24.51309\nExp number, first 60 AAs:  0.2682\nTotal prob of N-in: 0.14402\noutside   1   331\nTMhelix   332   354\ninside   355   372

20920 GCF_001116565.1_10493_1_4 Salmonella enterica subsp. enterica serovar Typhi Proteobacteria; Gammaproteobacteria; Enterobacterales; Enterobacteriaceae; Salmonella; Salmonella enterica; Salmonella enterica subsp. enterica MTGDNTLITSHGINRRDFMKLCAALAATMGLSSKAAAEMAESVSNPQRPPVIWIGAQECT WP_000145429.1 hydrogenase 2 small subunit [Salmonella enterica] Length: 372\nNumber of predicted TMHs:  1\nExp number of AAs in TMHs: 24.51309\nExp number, first 60 AAs:  0.2682\nTotal prob of N-in: 0.14402\noutside   1   331\nTMhelix   332   354\ninside   355   372

20921 GCF_001116225.1_10492_1_59 Salmonella enterica subsp. enterica serovar Typhi Proteobacteria; Gammaproteobacteria; Enterobacterales; Enterobacteriaceae; Salmonella; Salmonella enterica; Salmonella enterica subsp. enterica MTGDNTLITSHGINRRDFMKLCAALAATMGLSSKAAAEMAESVSNPQRPPVIWIGAQECT WP_000145429.1 hydrogenase 2 small subunit [Salmonella enterica] Length: 372\nNumber of predicted TMHs:  1\nExp number of AAs in TMHs: 24.51309\nExp number, first 60 AAs:  0.2682\nTotal prob of N-in: 0.14402\noutside   1   331\nTMhelix   332   354\ninside   355   372

20922 GCF_001117145.1_10593_2_2 Salmonella enterica subsp. enterica serovar Typhi Proteobacteria; Gammaproteobacteria; Enterobacterales; Enterobacteriaceae; Salmonella; Salmonella enterica; Salmonella enterica subsp. enterica MTGDNTLITSHGINRRDFMKLCAALAATMGLSSKAAAEMAESVSNPQRPPVIWIGAQECT WP_000145429.1 hydrogenase 2 small subunit [Salmonella enterica] Length: 372\nNumber of predicted TMHs:  1\nExp number of AAs in TMHs: 24.51309\nExp number, first 60 AAs:  0.2682\nTotal prob of N-in: 0.14402\noutside   1   331\nTMhelix   332   354\ninside   355   372

20923 GCF_001117685.1_10592_2_8 Salmonella enterica subsp. enterica serovar Typhi Proteobacteria; Gammaproteobacteria; Enterobacterales; Enterobacteriaceae; Salmonella; Salmonella enterica; Salmonella enterica subsp. enterica MTGDNTLITSHGINRRDFMKLCAALAATMGLSSKAAAEMAESVSNPQRPPVIWIGAQECT WP_000145429.1 hydrogenase 2 small subunit [Salmonella enterica] Length: 372\nNumber of predicted TMHs:  1\nExp number of AAs in TMHs: 24.51309\nExp number, first 60 AAs:  0.2682\nTotal prob of N-in: 0.14402\noutside   1   331\nTMhelix   332   354\ninside   355   372

20924 GCF_001117465.1_10492_1_43 Salmonella enterica subsp. enterica serovar Typhi Proteobacteria; Gammaproteobacteria; Enterobacterales; Enterobacteriaceae; Salmonella; Salmonella enterica; Salmonella enterica subsp. enterica MTGDNTLITSHGINRRDFMKLCAALAATMGLSSKAAAEMAESVSNPQRPPVIWIGAQECT WP_000145429.1 hydrogenase 2 small subunit [Salmonella enterica] Length: 372\nNumber of predicted TMHs:  1\nExp number of AAs in TMHs: 24.51309\nExp number, first 60 AAs:  0.2682\nTotal prob of N-in: 0.14402\noutside   1   331\nTMhelix   332   354\ninside   355   372

20925 GCF_001117965.1_10607_1_70 Salmonella enterica subsp. enterica serovar Typhi Proteobacteria; Gammaproteobacteria; Enterobacterales; Enterobacteriaceae; Salmonella; Salmonella enterica; Salmonella enterica subsp. enterica MTGDNTLITSHGINRRDFMKLCAALAATMGLSSKAAAEMAESVSNPQRPPVIWIGAQECT WP_000145429.1 hydrogenase 2 small subunit [Salmonella enterica] Length: 372\nNumber of predicted TMHs:  1\nExp number of AAs in TMHs: 24.51309\nExp number, first 60 AAs:  0.2682\nTotal prob of N-in: 0.14402\noutside   1   331\nTMhelix   332   354\ninside   355   372

20926 GCF_001118185.1_10561_2_30 Salmonella enterica subsp. enterica serovar Typhi Proteobacteria; Gammaproteobacteria; Enterobacterales; Enterobacteriaceae; Salmonella; Salmonella enterica; Salmonella enterica subsp. enterica MTGDNTLITSHGINRRDFMKLCAALAATMGLSSKAAAEMAESVSNPQRPPVIWIGAQECT WP_000145429.1 hydrogenase 2 small subunit [Salmonella enterica] Length: 372\nNumber of predicted TMHs:  1\nExp number of AAs in TMHs: 24.51309\nExp number, first 60 AAs:  0.2682\nTotal prob of N-in: 0.14402\noutside   1   331\nTMhelix   332   354\ninside   355   372

20927 GCF_001118125.1_10492_1_2 Salmonella enterica subsp. enterica serovar Typhi Proteobacteria; Gammaproteobacteria; Enterobacterales; Enterobacteriaceae; Salmonella; Salmonella enterica;

Salmonella enterica subsp. enterica MTGDNTLITSHGINRRDFMKLCAALAATMGLSSKAAAEMAESVSNPQRPPVIWIGAQECT WP_000145429.1 hydrogenase 2 small subunit [Salmonella enterica] Length: 372\nNumber of predicted TMHs: 1\nExp number of AAs in TMHs: 24.51309\nExp number, first 60 AAs: 0.2682\nTotal prob of N-in: 0.14402\noutside 1 331\nTMhelix 332 354\ninside 355 372

20928 GCF_001118345.1_9475_6_16 Salmonella enterica subsp. enterica serovar Typhi Proteobacteria; Gammaproteobacteria; Enterobacterales; Enterobacteriaceae; Salmonella; Salmonella enterica; Salmonella enterica subsp. enterica MTGDNTLITSHGINRRDFMKLCAALAATMGLSSKAAAEMAESVSNPQRPPVIWIGAQECT WP_000145429.1 hydrogenase 2 small subunit [Salmonella enterica] Length: 372\nNumber of predicted TMHs: 1\nExp number of AAs in TMHs: 24.51309\nExp number, first 60 AAs: 0.2682\nTotal prob of N-in: 0.14402\noutside 1 331\nTMhelix 332 354\ninside 355 372

20929 GCF_001118665.1_10492_1_41 Salmonella enterica subsp. enterica serovar Typhi Proteobacteria; Gammaproteobacteria; Enterobacterales; Enterobacteriaceae; Salmonella; Salmonella enterica; Salmonella enterica subsp. enterica MTGDNTLITSHGINRRDFMKLCAALAATMGLSSKAAAEMAESVSNPQRPPVIWIGAQECT WP_000145429.1 hydrogenase 2 small subunit [Salmonella enterica] Length: 372\nNumber of predicted TMHs: 1\nExp number of AAs in TMHs: 24.51309\nExp number, first 60 AAs: 0.2682\nTotal prob of N-in: 0.14402\noutside 1 331\nTMhelix 332 354\ninside 355 372

20930 GCF_001117285.1_10562_2_22 Salmonella enterica subsp. enterica serovar Typhi Proteobacteria; Gammaproteobacteria; Enterobacterales; Enterobacteriaceae; Salmonella; Salmonella enterica; Salmonella enterica subsp. enterica MTGDNTLITSHGINRRDFMKLCAALAATMGLSSKAATEMAESVSNPQRPPVIWIGAQECT WP_001782304.1 hydrogenase 2 small subunit [Salmonella enterica] Length: 372\nNumber of predicted TMHs: 1\nExp number of AAs in TMHs: 24.41173\nExp number, first 60 AAs: 0.16124\nTotal prob of N-in: 0.13947\noutside 1 331\nTMhelix 332 354\ninside 355 372

20931 GCF_001118805.1_9953_5_16 Salmonella enterica subsp. enterica serovar Typhi Proteobacteria; Gammaproteobacteria; Enterobacterales; Enterobacteriaceae; Salmonella; Salmonella enterica; Salmonella enterica subsp. enterica MTGDNTLITSHGINRRDFMKLCAALAATMGLSSKAAAEMAESVSNPQRPPVIWIGAQECT WP_000145429.1 hydrogenase 2 small subunit [Salmonella enterica] Length: 372\nNumber of predicted TMHs: 1\nExp number of AAs in TMHs: 24.51309\nExp number, first 60 AAs: 0.2682\nTotal prob of N-in: 0.14402\noutside 1 331\nTMhelix 332 354\ninside 355 372

20932 GCF_001118965.1_10349_1_32 Salmonella enterica subsp. enterica serovar Typhi Proteobacteria; Gammaproteobacteria; Enterobacterales; Enterobacteriaceae; Salmonella; Salmonella enterica; Salmonella enterica subsp. enterica MTGDNTLITSHGINRRDFMKLCAALAATMGLSSKAAAEMAESVSNPQRPPVIWIGAQECT WP_000145429.1 hydrogenase 2 small subunit [Salmonella enterica] Length: 372\nNumber of predicted TMHs: 1\nExp number of AAs in TMHs: 24.51309\nExp number, first 60 AAs: 0.2682\nTotal prob of N-in: 0.14402\noutside 1 331\nTMhelix 332 354\ninside 355 372

20933 GCF_001119125.1_9475_6_18 Salmonella enterica subsp. enterica serovar Typhi Proteobacteria; Gammaproteobacteria; Enterobacterales; Enterobacteriaceae; Salmonella; Salmonella enterica; Salmonella enterica subsp. enterica MTGDNTLITSHGINRRDFMKLCAALAATMGLSSKAAAEMAESVSNPQRPPVIWIGAQECT WP_000145429.1 hydrogenase 2 small subunit [Salmonella enterica] Length: 372\nNumber of predicted TMHs: 1\nExp number of AAs in TMHs: 24.51309\nExp number, first 60 AAs: 0.2682\nTotal prob of N-in: 0.14402\noutside 1 331\nTMhelix 332 354\ninside 355 372

20934 GCF_001119485.1_10349_1_18 Salmonella enterica subsp. enterica serovar Typhi Proteobacteria; Gammaproteobacteria; Enterobacterales; Enterobacteriaceae; Salmonella; Salmonella enterica; Salmonella enterica subsp. enterica MTGDNTLITSHGINRRDFMKLCAALAATMGLSSKAAAEMAESVSNPQRPPVIWIGAQECT WP_000145429.1 hydrogenase 2 small subunit [Salmonella enterica] Length: 372\nNumber of predicted TMHs: 1\nExp number of AAs in TMHs: 24.51309\nExp number, first 60 AAs: 0.2682\nTotal prob of N-in: 0.14402\noutside 1 331\nTMhelix 332 354\ninside 355 372

20935 GCF_001119245.1_10592_2_63 Salmonella enterica subsp. enterica serovar Typhi Proteobacteria; Gammaproteobacteria; Enterobacterales; Enterobacteriaceae; Salmonella; Salmonella enterica; Salmonella enterica subsp. enterica MTGDNTLITSHGINRRDFMKLCAALAATMGLSSKAAAEMAESVSNPQRPPVIWIGAQECT WP_000145429.1 hydrogenase 2 small subunit [Salmonella enterica] Length: 372\nNumber of predicted TMHs: 1\nExp number of AAs in TMHs: 24.51309\nExp number, first 60 AAs: 0.2682\nTotal prob of N-in: 0.14402\noutside 1 331\nTMhelix 332 354\ninside 355 372

20936 GCF_001119705.1_10492_1_68 Salmonella enterica subsp. enterica serovar Typhi Proteobacteria; Gammaproteobacteria; Enterobacterales; Enterobacteriaceae; Salmonella; Salmonella enterica; Salmonella enterica subsp. enterica MTGDNTLITSHGINRRDFMKLCAALAATMGLSSKAAAEMAESVSNPQRPPVIWIGAQECT WP_000145429.1 hydrogenase 2 small subunit [Salmonella enterica] Length: 372\nNumber of predicted TMHs: 1\nExp number of AAs in TMHs: 24.51309\nExp number, first 60 AAs: 0.2682\nTotal prob of N-in: 0.14402\noutside 1 331\nTMhelix 332 354\ninside 355 372

20937 GCF_001119805.1_10593_2_30 Salmonella enterica subsp. enterica serovar Typhi Proteobacteria; Gammaproteobacteria; Enterobacterales; Enterobacteriaceae; Salmonella; Salmonella enterica; Salmonella enterica subsp. enterica MTGDNTLITSHGINRRDFMKLCAALAATMGLSSKAAAEMAESVSNPQRPPVIWIGAQECT WP_000145429.1 hydrogenase 2 small subunit [Salmonella enterica] Length: 372\nNumber of predicted TMHs: 1\nExp number of AAs in TMHs: 24.51309\nExp number, first 60 AAs: 0.2682\nTotal prob of N-in: 0.14402\noutside 1 331\nTMhelix 332 354\ninside 355 372

20938 GCF_001123485.1_8490_5_27 Salmonella enterica subsp. enterica serovar Typhi Proteobacteria; Gammaproteobacteria; Enterobacterales; Enterobacteriaceae; Salmonella; Salmonella enterica; Salmonella enterica subsp. enterica MTGDNTLITSHGINRRDFMKLCAALAATMGLSSKAAAEMAESVSNPQRPPVIWIGAQECT WP_000145429.1 hydrogenase 2 small subunit [Salmonella enterica] Length: 372\nNumber of predicted TMHs: 1\nExp number of AAs in TMHs: 24.51309\nExp number, first 60 AAs: 0.2682\nTotal prob of N-in: 0.14402\noutside 1 331\nTMhelix 332 354\ninside 355 372

20939 GCF_001120445.1_10209_5_25 Salmonella enterica subsp. enterica serovar Typhi Proteobacteria; Gammaproteobacteria; Enterobacterales; Enterobacteriaceae; Salmonella; Salmonella enterica; Salmonella enterica subsp. enterica MTGDNTLITSHGINRRDFMKLCAALAATMGLSSKAAAEMAESVSNPQRPPVIWIGAQECT WP_000145429.1 hydrogenase 2 small subunit [Salmonella enterica] Length: 372\nNumber of predicted TMHs: 1\nExp number of AAs in TMHs: 24.51309\nExp number, first 60 AAs: 0.2682\nTotal prob of N-in: 0.14402\noutside 1 331\nTMhelix 332 354\ninside 355 372

20940 GCF_001120625.1_10541_2_19 Salmonella enterica subsp. enterica serovar Typhi Proteobacteria; Gammaproteobacteria; Enterobacterales; Enterobacteriaceae; Salmonella; Salmonella enterica; Salmonella enterica subsp. enterica MTGDNTLITSHGINRRDFMKLCAALAATMGLSSKAAAEMAESVSNPQRPPVIWIGAQECT WP_000145429.1 hydrogenase 2 small subunit [Salmonella enterica] Length: 372\nNumber of predicted TMHs: 1\nExp number of AAs in TMHs: 24.51309\nExp number, first 60 AAs: 0.2682\nTotal prob of N-in: 0.14402\noutside 1 331\nTMhelix 332 354\ninside 355 372

20941 GCF_001121125.1_10541_2_5 Salmonella enterica subsp. enterica serovar Typhi Proteobacteria; Gammaproteobacteria; Enterobacterales; Enterobacteriaceae; Salmonella; Salmonella enterica; Salmonella enterica subsp. enterica MTGDNTLITSHGINRRDFMKLCAALAATMGLSSKAAAEMAESVSNPQRPPVIWIGAQECT WP_000145429.1 hydrogenase 2 small subunit [Salmonella enterica] Length: 372\nNumber of predicted TMHs: 1\nExp number of AAs in TMHs: 24.51309\nExp number, first 60 AAs: 0.2682\nTotal prob of N-in: 0.14402\noutside 1 331\nTMhelix 332 354\ninside 355 372

20942 GCF_001121605.1_8616_4_39 Salmonella enterica subsp. enterica serovar Typhi Proteobacteria; Gammaproteobacteria; Enterobacterales; Enterobacteriaceae; Salmonella; Salmonella enterica; Salmonella enterica subsp. enterica MTGDNTLITSHGINRRDFMKLCAALAATMGLSSKAAAEMAESVSNPQRPPVIWIGAQECT WP_000145429.1 hydrogenase 2 small subunit [Salmonella enterica] Length: 372\nNumber of predicted TMHs: 1\nExp number of AAs in TMHs: 24.51309\nExp number, first 60 AAs: 0.2682\nTotal prob of N-in: 0.14402\noutside 1 331\nTMhelix 332 354\ninside 355 372

20943 GCF_001121865.1_10349_1_90 Salmonella enterica subsp. enterica serovar Typhi Proteobacteria; Gammaproteobacteria; Enterobacterales; Enterobacteriaceae; Salmonella; Salmonella enterica; Salmonella enterica subsp. enterica MTGDNTLITSHGINRRDFMKLCAALAATMGLSSKAAAEMAESVSNPQRPPVIWIGAQECT WP_000145429.1 hydrogenase 2 small subunit [Salmonella enterica] Length: 372\nNumber of predicted TMHs: 1\nExp number of AAs in TMHs: 24.51309\nExp number, first 60 AAs: 0.2682\nTotal prob of N-in: 0.14402\noutside 1 331\nTMhelix 332 354\ninside 355 372

20944 GCF_001122025.1_10493_1_82 Salmonella enterica subsp. enterica serovar Typhi Proteobacteria; Gammaproteobacteria; Enterobacterales; Enterobacteriaceae; Salmonella; Salmonella enterica; Salmonella enterica subsp. enterica MTGDNTLITSHGINRRDFMKLCAALAATMGLSSKAAAEMAESVSNPQRPPVIWIGAQECT WP_000145429.1 hydrogenase 2 small subunit [Salmonella enterica] Length: 372\nNumber of

predicted TMHs:  1\nExp number of AAs in TMHs: 24.51309\nExp number, first 60 AAs:  0.2682\nTotal prob of N-in: 0.14402\noutside   1   331\nTMhelix   332   354\ninside   355   372

20945 GCF_001121005.1_7468_7_56 Salmonella enterica subsp. enterica serovar Typhi Proteobacteria; Gammaproteobacteria; Enterobacterales; Enterobacteriaceae; Salmonella; Salmonella enterica; Salmonella enterica subsp. enterica MTGDNTLITSHGINRRDFMKLCAALAATMGLSSKAATEMAESVSNPQRPPVIWIGAQECT WP_001782304.1 hydrogenase 2 small subunit [Salmonella enterica] Length: 372\nNumber of predicted TMHs:  1\nExp number of AAs in TMHs: 24.41173\nExp number, first 60 AAs:  0.16124\nTotal prob of N-in: 0.13947\noutside   1   331\nTMhelix   332   354\ninside   355   372

20946 GCF_001122105.1_8525_3_49 Salmonella enterica subsp. enterica serovar Typhi Proteobacteria; Gammaproteobacteria; Enterobacterales; Enterobacteriaceae; Salmonella; Salmonella enterica; Salmonella enterica subsp. enterica MTGDNTLITSHGINRRDFMKLCAALAATMGLSSKAAAEMAESVSNPQRPPVIWIGAQECT WP_000145429.1 hydrogenase 2 small subunit [Salmonella enterica] Length: 372\nNumber of predicted TMHs:  1\nExp number of AAs in TMHs: 24.51309\nExp number, first 60 AAs:  0.2682\nTotal prob of N-in: 0.14402\noutside   1   331\nTMhelix   332   354\ninside   355   372

20947 GCF_001122265.1_10562_2_62 Salmonella enterica subsp. enterica serovar Typhi Proteobacteria; Gammaproteobacteria; Enterobacterales; Enterobacteriaceae; Salmonella; Salmonella enterica; Salmonella enterica subsp. enterica MTGDNTLITSHGINRRDFMKLCAALAATMGLSSKAAAEMAESVSNPQRPPVIWIGAQECT WP_000145429.1 hydrogenase 2 small subunit [Salmonella enterica] Length: 372\nNumber of predicted TMHs:  1\nExp number of AAs in TMHs: 24.51309\nExp number, first 60 AAs:  0.2682\nTotal prob of N-in: 0.14402\noutside   1   331\nTMhelix   332   354\ninside   355   372

20948 GCF_001122365.1_10562_2_37 Salmonella enterica subsp. enterica serovar Typhi Proteobacteria; Gammaproteobacteria; Enterobacterales; Enterobacteriaceae; Salmonella; Salmonella enterica; Salmonella enterica subsp. enterica MTGDNTLITSHGINRRDFMKLCAALAATMGLSSKAAAEMAESVSNPQRPPVIWIGAQECT WP_000145429.1 hydrogenase 2 small subunit [Salmonella enterica] Length: 372\nNumber of predicted TMHs:  1\nExp number of AAs in TMHs: 24.51309\nExp number, first 60 AAs:  0.2682\nTotal prob of N-in: 0.14402\noutside   1   331\nTMhelix   332   354\ninside   355   372

20949 GCF_001122465.1_10426_1_6 Salmonella enterica subsp. enterica serovar Typhi Proteobacteria; Gammaproteobacteria; Enterobacterales; Enterobacteriaceae; Salmonella; Salmonella enterica; Salmonella enterica subsp. enterica MTGDNTLITSHGINRRDFMKLCAALAATMGLSSKAAAEMAESVSNPQRPPVIWIGAQECT WP_000145429.1 hydrogenase 2 small subunit [Salmonella enterica] Length: 372\nNumber of predicted TMHs:  1\nExp number of AAs in TMHs: 24.51309\nExp number, first 60 AAs:  0.2682\nTotal prob of N-in: 0.14402\noutside   1   331\nTMhelix   332   354\ninside   355   372

20950 GCF_001122765.1_10561_2_56 Salmonella enterica subsp. enterica serovar Typhi Proteobacteria; Gammaproteobacteria; Enterobacterales; Enterobacteriaceae; Salmonella; Salmonella enterica; Salmonella enterica subsp. enterica MTGDNTLITSHGINRRDFMKLCAALAATMGLSSKAAAEMAESVSNPQRPPVIWIGAQECT WP_000145429.1 hydrogenase 2 small subunit [Salmonella enterica] Length: 372\nNumber of predicted TMHs:  1\nExp number of AAs in TMHs: 24.51309\nExp number, first 60 AAs:  0.2682\nTotal prob of N-in: 0.14402\noutside   1   331\nTMhelix   332   354\ninside   355   372

20951 GCF_001122965.1_9475_6_41 Salmonella enterica subsp. enterica serovar Typhi Proteobacteria; Gammaproteobacteria; Enterobacterales; Enterobacteriaceae; Salmonella; Salmonella enterica; Salmonella enterica subsp. enterica MTGDNTLITSHGINRRDFMKLCAALAATMGLSSKAAAEMAESVSNPQRPPVIWIGAQECT WP_000145429.1 hydrogenase 2 small subunit [Salmonella enterica] Length: 372\nNumber of predicted TMHs:  1\nExp number of AAs in TMHs: 24.51309\nExp number, first 60 AAs:  0.2682\nTotal prob of N-in: 0.14402\noutside   1   331\nTMhelix   332   354\ninside   355   372

20952 GCF_001123065.1_8490_6_54 Salmonella enterica subsp. enterica serovar Typhi Proteobacteria; Gammaproteobacteria; Enterobacterales; Enterobacteriaceae; Salmonella; Salmonella enterica; Salmonella enterica subsp. enterica MTGDNTLITSHGINRRDFMKLCAALAATMGLSSKAAAEMAESVSNPQRPPVIWIGAQECT WP_000145429.1 hydrogenase 2 small subunit [Salmonella enterica] Length: 372\nNumber of predicted TMHs:  1\nExp number of AAs in TMHs: 24.51309\nExp number, first 60 AAs:  0.2682\nTotal prob of N-in: 0.14402\noutside   1   331\nTMhelix   332   354\ninside   355   372

20953 GCF_001123445.1_10562_2_76 Salmonella enterica subsp. enterica serovar Typhi Proteobacteria; Gammaproteobacteria; Enterobacterales; Enterobacteriaceae; Salmonella; Salmonella enterica;

Salmonella enterica subsp. enterica	MTGDNTLITSHGINRRDFMKLCAALAATMGLSSKAAAEMAESVSNPQRPPVIWIGAQECT	WP_000145429.1	hydrogenase 2 small subunit [Salmonella enterica]	Length: 372\nNumber of predicted TMHs:  1\nExp number of AAs in TMHs: 24.51309\nExp number, first 60 AAs:  0.2682\nTotal prob of N-in: 0.14402\noutside	1	331\nTMhelix	332	354\ninside	355	372

20954	GCF_001123645.1_10608_2_66	Salmonella enterica subsp. enterica serovar Typhi	Proteobacteria; Gammaproteobacteria; Enterobacterales; Enterobacteriaceae; Salmonella; Salmonella enterica; Salmonella enterica subsp. enterica	MTGDNTLITSHGINRRDFMKLCAALAATMGLSSKAAAEMAESVSNPQRPPVIWIGAQECT	WP_000145429.1	hydrogenase 2 small subunit [Salmonella enterica]	Length: 372\nNumber of predicted TMHs:  1\nExp number of AAs in TMHs: 24.51309\nExp number, first 60 AAs:  0.2682\nTotal prob of N-in: 0.14402\noutside	1	331\nTMhelix	332	354\ninside	355	372

20955	GCF_001124025.1_9953_5_44	Salmonella enterica subsp. enterica serovar Typhi	Proteobacteria; Gammaproteobacteria; Enterobacterales; Enterobacteriaceae; Salmonella; Salmonella enterica; Salmonella enterica subsp. enterica	MTGDNTLITSHGINRRDFMKLCAALAATMGLSSKAAAEMAESVSNPQRPPVIWIGAQECT	WP_000145429.1	hydrogenase 2 small subunit [Salmonella enterica]	Length: 372\nNumber of predicted TMHs:  1\nExp number of AAs in TMHs: 24.51309\nExp number, first 60 AAs:  0.2682\nTotal prob of N-in: 0.14402\noutside	1	331\nTMhelix	332	354\ninside	355	372

20956	GCF_001124385.1_10607_1_46	Salmonella enterica subsp. enterica serovar Typhi	Proteobacteria; Gammaproteobacteria; Enterobacterales; Enterobacteriaceae; Salmonella; Salmonella enterica; Salmonella enterica subsp. enterica	MTGDNTLITSHGINRRDFMKLCAALAATMGLSSKAAAEMAESVSNPQRPPVIWIGAQECT	WP_000145429.1	hydrogenase 2 small subunit [Salmonella enterica]	Length: 372\nNumber of predicted TMHs:  1\nExp number of AAs in TMHs: 24.51309\nExp number, first 60 AAs:  0.2682\nTotal prob of N-in: 0.14402\noutside	1	331\nTMhelix	332	354\ninside	355	372

20957	GCF_001124525.1_10592_2_10	Salmonella enterica subsp. enterica serovar Typhi	Proteobacteria; Gammaproteobacteria; Enterobacterales; Enterobacteriaceae; Salmonella; Salmonella enterica; Salmonella enterica subsp. enterica	MTGDNTLITSHGINRRDFMKLCAALAATMGLSSKAAAEMAESVSNPQRPPVIWIGAQECT	WP_000145429.1	hydrogenase 2 small subunit [Salmonella enterica]	Length: 372\nNumber of predicted TMHs:  1\nExp number of AAs in TMHs: 24.51309\nExp number, first 60 AAs:  0.2682\nTotal prob of N-in: 0.14402\noutside	1	331\nTMhelix	332	354\ninside	355	372

20958	GCF_001124725.1_9953_5_10	Salmonella enterica subsp. enterica serovar Typhi	Proteobacteria; Gammaproteobacteria; Enterobacterales; Enterobacteriaceae; Salmonella; Salmonella enterica; Salmonella enterica subsp. enterica	MTGDNTLITSHGINRRDFMKLCAALAATMGLSSKAAAEMAESVSNPQRPPVIWIGAQECT	WP_000145429.1	hydrogenase 2 small subunit [Salmonella enterica]	Length: 372\nNumber of predicted TMHs:  1\nExp number of AAs in TMHs: 24.51309\nExp number, first 60 AAs:  0.2682\nTotal prob of N-in: 0.14402\noutside	1	331\nTMhelix	332	354\ninside	355	372

20959	GCF_001124885.1_9953_5_72	Salmonella enterica subsp. enterica serovar Typhi	Proteobacteria; Gammaproteobacteria; Enterobacterales; Enterobacteriaceae; Salmonella; Salmonella enterica; Salmonella enterica subsp. enterica	MTGDNTLITSHGINRRDFMKLCAALAATMGLSSKAAAEMAESVSNPQRPPVIWIGAQECT	WP_000145429.1	hydrogenase 2 small subunit [Salmonella enterica]	Length: 372\nNumber of predicted TMHs:  1\nExp number of AAs in TMHs: 24.51309\nExp number, first 60 AAs:  0.2682\nTotal prob of N-in: 0.14402\noutside	1	331\nTMhelix	332	354\ninside	355	372

20960	GCF_001124985.1_9953_5_75	Salmonella enterica subsp. enterica serovar Typhi	Proteobacteria; Gammaproteobacteria; Enterobacterales; Enterobacteriaceae; Salmonella; Salmonella enterica; Salmonella enterica subsp. enterica	MTGDNTLITSHGINRRDFMKLCAALAATMGLSSKAAAEMAESVSNPQRPPVIWIGAQECT	WP_000145429.1	hydrogenase 2 small subunit [Salmonella enterica]	Length: 372\nNumber of predicted TMHs:  1\nExp number of AAs in TMHs: 24.51309\nExp number, first 60 AAs:  0.2682\nTotal prob of N-in: 0.14402\noutside	1	331\nTMhelix	332	354\ninside	355	372

20961	GCF_001124285.1_7468_7_82	Salmonella enterica subsp. enterica serovar Typhi	Proteobacteria; Gammaproteobacteria; Enterobacterales; Enterobacteriaceae; Salmonella; Salmonella enterica; Salmonella enterica subsp. enterica	MTGDNTLITSHGINRRDFMKLCAALAATMGLSSKAATEMAESVSNPQRPPVIWIGAQECT	WP_001782304.1	hydrogenase 2 small subunit [Salmonella enterica]	Length: 372\nNumber of predicted TMHs:  1\nExp number of AAs in TMHs: 24.41173\nExp number, first 60 AAs:  0.16124\nTotal prob of N-in: 0.13947\noutside	1	331\nTMhelix	332	354\ninside	355	372

20962 GCF_001125085.1_10561_2_6 Salmonella enterica subsp. enterica serovar Typhi Proteobacteria; Gammaproteobacteria; Enterobacterales; Enterobacteriaceae; Salmonella; Salmonella enterica; Salmonella enterica subsp. enterica MTGDNTLITSHGINRRDFMKLCAALAATMGLSSKAAAEMAESVSNPQRPPVIWIGAQECT WP_000145429.1 hydrogenase 2 small subunit [Salmonella enterica] Length: 372\nNumber of predicted TMHs: 1\nExp number of AAs in TMHs: 24.51309\nExp number, first 60 AAs: 0.2682\nTotal prob of N-in: 0.14402\noutside 1 331\nTMhelix 332 354\ninside 355 372

20963 GCF_001125245.1_10561_2_12 Salmonella enterica subsp. enterica serovar Typhi Proteobacteria; Gammaproteobacteria; Enterobacterales; Enterobacteriaceae; Salmonella; Salmonella enterica; Salmonella enterica subsp. enterica MTGDNTLITSHGINRRDFMKLCAALAATMGLSSKAAAEMAESVSNPQRPPVIWIGAQECT WP_000145429.1 hydrogenase 2 small subunit [Salmonella enterica] Length: 372\nNumber of predicted TMHs: 1\nExp number of AAs in TMHs: 24.51309\nExp number, first 60 AAs: 0.2682\nTotal prob of N-in: 0.14402\noutside 1 331\nTMhelix 332 354\ninside 355 372

20964 GCF_001125465.1_10426_1_4 Salmonella enterica subsp. enterica serovar Typhi Proteobacteria; Gammaproteobacteria; Enterobacterales; Enterobacteriaceae; Salmonella; Salmonella enterica; Salmonella enterica subsp. enterica MTGDNTLITSHGINRRDFMKLCAALAATMGLSSKAAAEMAESVSNPQRPPVIWIGAQECT WP_000145429.1 hydrogenase 2 small subunit [Salmonella enterica] Length: 372\nNumber of predicted TMHs: 1\nExp number of AAs in TMHs: 24.51309\nExp number, first 60 AAs: 0.2682\nTotal prob of N-in: 0.14402\noutside 1 331\nTMhelix 332 354\ninside 355 372

20965 GCF_001125845.1_10209_5_5 Salmonella enterica subsp. enterica serovar Typhi Proteobacteria; Gammaproteobacteria; Enterobacterales; Enterobacteriaceae; Salmonella; Salmonella enterica; Salmonella enterica subsp. enterica MTGDNTLITSHGINRRDFMKLCAALAATMGLSSKAAAEMAESVSNPQRPPVIWIGAQECT WP_000145429.1 hydrogenase 2 small subunit [Salmonella enterica] Length: 372\nNumber of predicted TMHs: 1\nExp number of AAs in TMHs: 24.51309\nExp number, first 60 AAs: 0.2682\nTotal prob of N-in: 0.14402\noutside 1 331\nTMhelix 332 354\ninside 355 372

20966 GCF_001125965.1_10541_2_13 Salmonella enterica subsp. enterica serovar Typhi Proteobacteria; Gammaproteobacteria; Enterobacterales; Enterobacteriaceae; Salmonella; Salmonella enterica; Salmonella enterica subsp. enterica MTGDNTLITSHGINRRDFMKLCAALAATMGLSSKAAAEMAESVSNPQRPPVIWIGAQECT WP_000145429.1 hydrogenase 2 small subunit [Salmonella enterica] Length: 372\nNumber of predicted TMHs: 1\nExp number of AAs in TMHs: 24.51309\nExp number, first 60 AAs: 0.2682\nTotal prob of N-in: 0.14402\noutside 1 331\nTMhelix 332 354\ninside 355 372

20967 GCF_001126165.1_9475_6_34 Salmonella enterica subsp. enterica serovar Typhi Proteobacteria; Gammaproteobacteria; Enterobacterales; Enterobacteriaceae; Salmonella; Salmonella enterica; Salmonella enterica subsp. enterica MTGDNTLITSHGINRRDFMKLCAALAATMGLSSKAAAEMAESVSNPQRPPVIWIGAQECT WP_000145429.1 hydrogenase 2 small subunit [Salmonella enterica] Length: 372\nNumber of predicted TMHs: 1\nExp number of AAs in TMHs: 24.51309\nExp number, first 60 AAs: 0.2682\nTotal prob of N-in: 0.14402\noutside 1 331\nTMhelix 332 354\ninside 355 372

20968 GCF_001126385.1_10349_1_6 Salmonella enterica subsp. enterica serovar Typhi Proteobacteria; Gammaproteobacteria; Enterobacterales; Enterobacteriaceae; Salmonella; Salmonella enterica; Salmonella enterica subsp. enterica MTGDNTLITSHGINRRDFMKLCAALAATMGLSSKAAAEMAESVSNPQRPPVIWIGAQECT WP_000145429.1 hydrogenase 2 small subunit [Salmonella enterica] Length: 372\nNumber of predicted TMHs: 1\nExp number of AAs in TMHs: 24.51309\nExp number, first 60 AAs: 0.2682\nTotal prob of N-in: 0.14402\noutside 1 331\nTMhelix 332 354\ninside 355 372

20969 GCF_001126705.1_10592_2_27 Salmonella enterica subsp. enterica serovar Typhi Proteobacteria; Gammaproteobacteria; Enterobacterales; Enterobacteriaceae; Salmonella; Salmonella enterica; Salmonella enterica subsp. enterica MTGDNTLITSHGINRRDFMKLCAALAATMGLSSKAAAEMAESVSNPQRPPVIWIGAQECT WP_000145429.1 hydrogenase 2 small subunit [Salmonella enterica] Length: 372\nNumber of predicted TMHs: 1\nExp number of AAs in TMHs: 24.51309\nExp number, first 60 AAs: 0.2682\nTotal prob of N-in: 0.14402\noutside 1 331\nTMhelix 332 354\ninside 355 372

20970 GCF_001126865.1_8490_5_16 Salmonella enterica subsp. enterica serovar Typhi Proteobacteria; Gammaproteobacteria; Enterobacterales; Enterobacteriaceae; Salmonella; Salmonella enterica; Salmonella enterica subsp. enterica MTGDNTLITSHGINRRDFMKLCAALAATMGLSSKAAAEMAESVSNPQRPPVIWIGAQECT WP_000145429.1 hydrogenase 2 small subunit [Salmonella enterica] Length: 372\nNumber of

predicted TMHs:  1\nExp number of AAs in TMHs: 24.51309\nExp number, first 60 AAs:  0.2682\nTotal prob of N-in: 0.14402\noutside   1   331\nTMhelix   332   354\ninside   355   372

20971 GCF_001127105.1_10209_5_48 Salmonella enterica subsp. enterica serovar Typhi Proteobacteria; Gammaproteobacteria; Enterobacterales; Enterobacteriaceae; Salmonella; Salmonella enterica; Salmonella enterica subsp. enterica MTGDNTLITSHGINRRDFMKLCAALAATMGLSSKAAAEMAESVSNPQRPPVIWIGAQECT WP_000145429.1 hydrogenase 2 small subunit [Salmonella enterica] Length: 372\nNumber of predicted TMHs:  1\nExp number of AAs in TMHs: 24.51309\nExp number, first 60 AAs:  0.2682\nTotal prob of N-in: 0.14402\noutside   1   331\nTMhelix   332   354\ninside   355   372

20972 GCF_001127385.1_10562_2_69 Salmonella enterica subsp. enterica serovar Typhi Proteobacteria; Gammaproteobacteria; Enterobacterales; Enterobacteriaceae; Salmonella; Salmonella enterica; Salmonella enterica subsp. enterica MTGDNTLITSHGINRRDFMKLCAALAATMGLSSKAAAEMAESVSNPQRPPVIWIGAQECT WP_000145429.1 hydrogenase 2 small subunit [Salmonella enterica] Length: 372\nNumber of predicted TMHs:  1\nExp number of AAs in TMHs: 24.51309\nExp number, first 60 AAs:  0.2682\nTotal prob of N-in: 0.14402\noutside   1   331\nTMhelix   332   354\ninside   355   372

20973 GCF_001127505.1_10541_2_28 Salmonella enterica subsp. enterica serovar Typhi Proteobacteria; Gammaproteobacteria; Enterobacterales; Enterobacteriaceae; Salmonella; Salmonella enterica; Salmonella enterica subsp. enterica MTGDNTLITSHGINRRDFMKLCAALAATMGLSSKAAAEMAESVSNPQRPPVIWIGAQECT WP_000145429.1 hydrogenase 2 small subunit [Salmonella enterica] Length: 372\nNumber of predicted TMHs:  1\nExp number of AAs in TMHs: 24.51309\nExp number, first 60 AAs:  0.2682\nTotal prob of N-in: 0.14402\noutside   1   331\nTMhelix   332   354\ninside   355   372

20974 GCF_001127685.1_9953_5_9Salmonella enterica subsp. enterica serovar Typhi Proteobacteria; Gammaproteobacteria; Enterobacterales; Enterobacteriaceae; Salmonella; Salmonella enterica; Salmonella enterica subsp. enterica MTGDNTLITSHGINRRDFMKLCAALAATMGLSSKAAAEMAESVSNPQRPPVIWIGAQECT WP_000145429.1 hydrogenase 2 small subunit [Salmonella enterica] Length: 372\nNumber of predicted TMHs:  1\nExp number of AAs in TMHs: 24.51309\nExp number, first 60 AAs:  0.2682\nTotal prob of N-in:   0.14402\noutside   1 331\nTMhelix   332   354\ninside   355   372

20975 GCF_001128305.1_10561_2_32 Salmonella enterica subsp. enterica serovar Typhi Proteobacteria; Gammaproteobacteria; Enterobacterales; Enterobacteriaceae; Salmonella; Salmonella enterica; Salmonella enterica subsp. enterica MTGDNTLITSHGINRRDFMKLCAALAATMGLSSKAAAEMAESVSNPQRPPVIWIGAQECT WP_000145429.1 hydrogenase 2 small subunit [Salmonella enterica] Length: 372\nNumber of predicted TMHs:  1\nExp number of AAs in TMHs: 24.51309\nExp number, first 60 AAs:  0.2682\nTotal prob of N-in: 0.14402\noutside   1   331\nTMhelix   332   354\ninside   355   372

20976 GCF_001128545.1_9953_5_56 Salmonella enterica subsp. enterica serovar Typhi Proteobacteria; Gammaproteobacteria; Enterobacterales; Enterobacteriaceae; Salmonella; Salmonella enterica; Salmonella enterica subsp. enterica MTGDNTLITSHGINRRDFMKLCAALAATMGLSSKAAAEMAESVSNPQRPPVIWIGAQECT WP_000145429.1 hydrogenase 2 small subunit [Salmonella enterica] Length: 372\nNumber of predicted TMHs:  1\nExp number of AAs in TMHs: 24.51309\nExp number, first 60 AAs:  0.2682\nTotal prob of N-in: 0.14402\noutside   1   331\nTMhelix   332   354\ninside   355   372

20977 GCF_001129145.1_10349_1_58 Salmonella enterica subsp. enterica serovar Typhi Proteobacteria; Gammaproteobacteria; Enterobacterales; Enterobacteriaceae; Salmonella; Salmonella enterica; Salmonella enterica subsp. enterica MTGDNTLITSHGINRRDFMKLCAALAATMGLSSKAAAEMAESVSNPQRPPVIWIGAQECT WP_000145429.1 hydrogenase 2 small subunit [Salmonella enterica] Length: 372\nNumber of predicted TMHs:  1\nExp number of AAs in TMHs: 24.51309\nExp number, first 60 AAs:  0.2682\nTotal prob of N-in: 0.14402\noutside   1   331\nTMhelix   332   354\ninside   355   372

20978 GCF_001128685.1_10209_5_16 Salmonella enterica subsp. enterica serovar Typhi Proteobacteria; Gammaproteobacteria; Enterobacterales; Enterobacteriaceae; Salmonella; Salmonella enterica; Salmonella enterica subsp. enterica MTGDNTLITSHGINRRDFMKLCAALAATMGLSSKAAAEMAESVSNPQRPPVIWIGAQECT WP_000145429.1 hydrogenase 2 small subunit [Salmonella enterica] Length: 372\nNumber of predicted TMHs:  1\nExp number of AAs in TMHs: 24.51309\nExp number, first 60 AAs:  0.2682\nTotal prob of N-in: 0.14402\noutside   1   331\nTMhelix   332   354\ninside   355   372

20979 GCF_001129245.1_9953_5_34 Salmonella enterica subsp. enterica serovar Typhi Proteobacteria; Gammaproteobacteria; Enterobacterales; Enterobacteriaceae; Salmonella; Salmonella enterica;

Salmonella enterica subsp. enterica MTGDNTLITSHGINRRDFMKLCAALAATMGLSSKAAAEMAESVSNPQRPPVIWIGAQECT WP_000145429.1 hydrogenase 2 small subunit [Salmonella enterica] Length: 372\nNumber of predicted TMHs: 1\nExp number of AAs in TMHs: 24.51309\nExp number, first 60 AAs: 0.2682\nTotal prob of N-in: 0.14402\noutside 1 331\nTMhelix 332 354\ninside 355 372

20980 GCF_001129325.1_10562_2_60 Salmonella enterica subsp. enterica serovar Typhi Proteobacteria; Gammaproteobacteria; Enterobacterales; Enterobacteriaceae; Salmonella; Salmonella enterica; Salmonella enterica subsp. enterica MTGDNTLITSHGINRRDFMKLCAALAATMGLSSKAAAEMAESVSNPQRPPVIWIGAQECT WP_000145429.1 hydrogenase 2 small subunit [Salmonella enterica] Length: 372\nNumber of predicted TMHs: 1\nExp number of AAs in TMHs: 24.51309\nExp number, first 60 AAs: 0.2682\nTotal prob of N-in: 0.14402\noutside 1 331\nTMhelix 332 354\ninside 355 372

20981 GCF_001129985.1_9475_6_13 Salmonella enterica subsp. enterica serovar Typhi Proteobacteria; Gammaproteobacteria; Enterobacterales; Enterobacteriaceae; Salmonella; Salmonella enterica; Salmonella enterica subsp. enterica MTGDNTLITSHGINRRDFMKLCAALAATMGLSSKAAAEMAESVSNPQRPPVIWIGAQECT WP_000145429.1 hydrogenase 2 small subunit [Salmonella enterica] Length: 372\nNumber of predicted TMHs: 1\nExp number of AAs in TMHs: 24.51309\nExp number, first 60 AAs: 0.2682\nTotal prob of N-in: 0.14402\noutside 1 331\nTMhelix 332 354\ninside 355 372

20982 GCF_001130645.1_10592_2_9 Salmonella enterica subsp. enterica serovar Typhi Proteobacteria; Gammaproteobacteria; Enterobacterales; Enterobacteriaceae; Salmonella; Salmonella enterica; Salmonella enterica subsp. enterica MTGDNTLITSHGINRRDFMKLCAALAATMGLSSKAAAEMAESVSNPQRPPVIWIGAQECT WP_000145429.1 hydrogenase 2 small subunit [Salmonella enterica] Length: 372\nNumber of predicted TMHs: 1\nExp number of AAs in TMHs: 24.51309\nExp number, first 60 AAs: 0.2682\nTotal prob of N-in: 0.14402\noutside 1 331\nTMhelix 332 354\ninside 355 372

20983 GCF_001131425.1_10592_2_22 Salmonella enterica subsp. enterica serovar Typhi Proteobacteria; Gammaproteobacteria; Enterobacterales; Enterobacteriaceae; Salmonella; Salmonella enterica; Salmonella enterica subsp. enterica MTGDNTLITSHGINRRDFMKLCAALAATMGLSSKAAAEMAESVSNPQRPPVIWIGAQECT WP_000145429.1 hydrogenase 2 small subunit [Salmonella enterica] Length: 372\nNumber of predicted TMHs: 1\nExp number of AAs in TMHs: 24.51309\nExp number, first 60 AAs: 0.2682\nTotal prob of N-in: 0.14402\noutside 1 331\nTMhelix 332 354\ninside 355 372

20984 GCF_001131765.1_9475_6_55 Salmonella enterica subsp. enterica serovar Typhi Proteobacteria; Gammaproteobacteria; Enterobacterales; Enterobacteriaceae; Salmonella; Salmonella enterica; Salmonella enterica subsp. enterica MTGDNTLITSHGINRRDFMKLCAALAATMGLSSKAAAEMAESVSNPQRPPVIWIGAQECT WP_000145429.1 hydrogenase 2 small subunit [Salmonella enterica] Length: 372\nNumber of predicted TMHs: 1\nExp number of AAs in TMHs: 24.51309\nExp number, first 60 AAs: 0.2682\nTotal prob of N-in: 0.14402\noutside 1 331\nTMhelix 332 354\ninside 355 372

20985 GCF_001132005.1_9870_8_33 Salmonella enterica subsp. enterica serovar Typhi Proteobacteria; Gammaproteobacteria; Enterobacterales; Enterobacteriaceae; Salmonella; Salmonella enterica; Salmonella enterica subsp. enterica MTGDNTLITSHGINRRDFMKLCAALAATMGLSSKAAAEMAESVSNPQRPPVIWIGAQECT WP_000145429.1 hydrogenase 2 small subunit [Salmonella enterica] Length: 372\nNumber of predicted TMHs: 1\nExp number of AAs in TMHs: 24.51309\nExp number, first 60 AAs: 0.2682\nTotal prob of N-in: 0.14402\noutside 1 331\nTMhelix 332 354\ninside 355 372

20986 GCF_001132265.1_9475_6_79 Salmonella enterica subsp. enterica serovar Typhi Proteobacteria; Gammaproteobacteria; Enterobacterales; Enterobacteriaceae; Salmonella; Salmonella enterica; Salmonella enterica subsp. enterica MTGDNTLITSHGINRRDFMKLCAALAATMGLSSKAAAEMAESVSNPQRPPVIWIGAQECT WP_000145429.1 hydrogenase 2 small subunit [Salmonella enterica] Length: 372\nNumber of predicted TMHs: 1\nExp number of AAs in TMHs: 24.51309\nExp number, first 60 AAs: 0.2682\nTotal prob of N-in: 0.14402\noutside 1 331\nTMhelix 332 354\ninside 355 372

20987 GCF_001132425.1_9953_5_43 Salmonella enterica subsp. enterica serovar Typhi Proteobacteria; Gammaproteobacteria; Enterobacterales; Enterobacteriaceae; Salmonella; Salmonella enterica; Salmonella enterica subsp. enterica MTGDNTLITSHGINRRDFMKLCAALAATMGLSSKAAAEMAESVSNPQRPPVIWIGAQECT WP_000145429.1 hydrogenase 2 small subunit [Salmonella enterica] Length: 372\nNumber of predicted TMHs: 1\nExp number of AAs in TMHs: 24.51309\nExp number, first 60 AAs: 0.2682\nTotal prob of N-in: 0.14402\noutside 1 331\nTMhelix 332 354\ninside 355 372

20988 GCF_001132605.1_9953_5_71 Salmonella enterica subsp. enterica serovar Typhi Proteobacteria; Gammaproteobacteria; Enterobacterales; Enterobacteriaceae; Salmonella; Salmonella enterica; Salmonella enterica subsp. enterica MTGDNTLITSHGINRRDFMKLCAALAATMGLSSKAAAEMAESVSNPQRPPVIWIGAQECT WP_000145429.1 hydrogenase 2 small subunit [Salmonella enterica] Length: 372\nNumber of predicted TMHs: 1\nExp number of AAs in TMHs: 24.51309\nExp number, first 60 AAs: 0.2682\nTotal prob of N-in: 0.14402\noutside 1 331\nTMhelix 332 354\ninside 355 372

20989 GCF_001132765.1_9870_8_5Salmonella enterica subsp. enterica serovar Typhi Proteobacteria; Gammaproteobacteria; Enterobacterales; Enterobacteriaceae; Salmonella; Salmonella enterica; Salmonella enterica subsp. enterica MTGDNTLITSHGINRRDFMKLCAALAATMGLSSKAAAEMAESVSNPQRPPVIWIGAQECT WP_000145429.1 hydrogenase 2 small subunit [Salmonella enterica] Length: 372\nNumber of predicted TMHs: 1\nExp number of AAs in TMHs: 24.51309\nExp number, first 60 AAs: 0.2682\nTotal prob of N-in: 0.14402\noutside 1 331\nTMhelix 332 354\ninside 355 372

20990 GCF_001132905.1_8490_6_62 Salmonella enterica subsp. enterica serovar Typhi Proteobacteria; Gammaproteobacteria; Enterobacterales; Enterobacteriaceae; Salmonella; Salmonella enterica; Salmonella enterica subsp. enterica MTGDNTLITSHGINRRDFMKLCAALAATMGLSSKAAAEMAESVSNPQRPPVIWIGAQECT WP_000145429.1 hydrogenase 2 small subunit [Salmonella enterica] Length: 372\nNumber of predicted TMHs: 1\nExp number of AAs in TMHs: 24.51309\nExp number, first 60 AAs: 0.2682\nTotal prob of N-in: 0.14402\noutside 1 331\nTMhelix 332 354\ninside 355 372

20991 GCF_001133025.1_10592_2_56 Salmonella enterica subsp. enterica serovar Typhi Proteobacteria; Gammaproteobacteria; Enterobacterales; Enterobacteriaceae; Salmonella; Salmonella enterica; Salmonella enterica subsp. enterica MTGDNTLITSHGINRRDFMKLCAALAATMGLSSKAAAEMAESVSNPQRPPVIWIGAQECT WP_000145429.1 hydrogenase 2 small subunit [Salmonella enterica] Length: 372\nNumber of predicted TMHs: 1\nExp number of AAs in TMHs: 24.51309\nExp number, first 60 AAs: 0.2682\nTotal prob of N-in: 0.14402\noutside 1 331\nTMhelix 332 354\ninside 355 372

20992 GCF_001133105.1_10492_1_20 Salmonella enterica subsp. enterica serovar Typhi Proteobacteria; Gammaproteobacteria; Enterobacterales; Enterobacteriaceae; Salmonella; Salmonella enterica; Salmonella enterica subsp. enterica MTGDNTLITSHGINRRDFMKLCAALAATMGLSSKAAAEMAESVSNPQRPPVIWIGAQECT WP_000145429.1 hydrogenase 2 small subunit [Salmonella enterica] Length: 372\nNumber of predicted TMHs: 1\nExp number of AAs in TMHs: 24.51309\nExp number, first 60 AAs: 0.2682\nTotal prob of N-in: 0.14402\noutside 1 331\nTMhelix 332 354\ninside 355 372

20993 GCF_001133305.1_10492_1_66 Salmonella enterica subsp. enterica serovar Typhi Proteobacteria; Gammaproteobacteria; Enterobacterales; Enterobacteriaceae; Salmonella; Salmonella enterica; Salmonella enterica subsp. enterica MTGDNTLITSHGINRRDFMKLCAALAATMGLSSKAAAEMAESVSNPQRPPVIWIGAQECT WP_000145429.1 hydrogenase 2 small subunit [Salmonella enterica] Length: 372\nNumber of predicted TMHs: 1\nExp number of AAs in TMHs: 24.51309\nExp number, first 60 AAs: 0.2682\nTotal prob of N-in: 0.14402\noutside 1 331\nTMhelix 332 354\ninside 355 372

20994 GCF_001133645.1_10492_1_55 Salmonella enterica subsp. enterica serovar Typhi Proteobacteria; Gammaproteobacteria; Enterobacterales; Enterobacteriaceae; Salmonella; Salmonella enterica; Salmonella enterica subsp. enterica MTGDNTLITSHGINRRDFMKLCAALAATMGLSSKAAAEMAESVSNPQRPPVIWIGAQECT WP_000145429.1 hydrogenase 2 small subunit [Salmonella enterica] Length: 372\nNumber of predicted TMHs: 1\nExp number of AAs in TMHs: 24.51309\nExp number, first 60 AAs: 0.2682\nTotal prob of N-in: 0.14402\noutside 1 331\nTMhelix 332 354\ninside 355 372

20995 GCF_001133765.1_10541_2_9 Salmonella enterica subsp. enterica serovar Typhi Proteobacteria; Gammaproteobacteria; Enterobacterales; Enterobacteriaceae; Salmonella; Salmonella enterica; Salmonella enterica subsp. enterica MTGDNTLITSHGINRRDFMKLCAALAATMGLSSKAAAEMAESVSNPQRPPVIWIGAQECT WP_000145429.1 hydrogenase 2 small subunit [Salmonella enterica] Length: 372\nNumber of predicted TMHs: 1\nExp number of AAs in TMHs: 24.51309\nExp number, first 60 AAs: 0.2682\nTotal prob of N-in: 0.14402\noutside 1 331\nTMhelix 332 354\ninside 355 372

20996 GCF_001133925.1_10492_1_81 Salmonella enterica subsp. enterica serovar Typhi Proteobacteria; Gammaproteobacteria; Enterobacterales; Enterobacteriaceae; Salmonella; Salmonella enterica; Salmonella enterica subsp. enterica MTGDNTLITSHGINRRDFMKLCAALAATMGLSSKAAAEMAESVSNPQRPPVIWIGAQECT WP_000145429.1 hydrogenase 2 small subunit [Salmonella enterica] Length: 372\nNumber of

predicted TMHs: 1\nExp number of AAs in TMHs: 24.51309\nExp number, first 60 AAs: 0.2682\nTotal prob of N-in: 0.14402\noutside 1 331\nTMhelix 332 354\ninside 355 372

20997 GCF_001134065.1_10541_2_42 Salmonella enterica subsp. enterica serovar Typhi Proteobacteria; Gammaproteobacteria; Enterobacterales; Enterobacteriaceae; Salmonella; Salmonella enterica; Salmonella enterica subsp. enterica MTGDNTLITSHGINRRDFMKLCAALAATMGLSSKAAAEMAESVSNPQRPPVIWIGAQECT WP_000145429.1 hydrogenase 2 small subunit [Salmonella enterica] Length: 372\nNumber of predicted TMHs: 1\nExp number of AAs in TMHs: 24.51309\nExp number, first 60 AAs: 0.2682\nTotal prob of N-in: 0.14402\noutside 1 331\nTMhelix 332 354\ninside 355 372

20998 GCF_001134165.1_10493_1_8 Salmonella enterica subsp. enterica serovar Typhi Proteobacteria; Gammaproteobacteria; Enterobacterales; Enterobacteriaceae; Salmonella; Salmonella enterica; Salmonella enterica subsp. enterica MTGDNTLITSHGINRRDFMKLCAALAATMGLSSKAAAEMAESVSNPQRPPVIWIGAQECT WP_000145429.1 hydrogenase 2 small subunit [Salmonella enterica] Length: 372\nNumber of predicted TMHs: 1\nExp number of AAs in TMHs: 24.51309\nExp number, first 60 AAs: 0.2682\nTotal prob of N-in: 0.14402\noutside 1 331\nTMhelix 332 354\ninside 355 372

20999 GCF_001134225.1_10492_1_53 Salmonella enterica subsp. enterica serovar Typhi Proteobacteria; Gammaproteobacteria; Enterobacterales; Enterobacteriaceae; Salmonella; Salmonella enterica; Salmonella enterica subsp. enterica MTGDNTLITSHGINRRDFMKLCAALAATMGLSSKAAAEMAESVSNPQRPPVIWIGAQECT WP_000145429.1 hydrogenase 2 small subunit [Salmonella enterica] Length: 372\nNumber of predicted TMHs: 1\nExp number of AAs in TMHs: 24.51309\nExp number, first 60 AAs: 0.2682\nTotal prob of N-in: 0.14402\noutside 1 331\nTMhelix 332 354\ninside 355 372

21000 GCF_001134365.1_9953_5_69 Salmonella enterica subsp. enterica serovar Typhi Proteobacteria; Gammaproteobacteria; Enterobacterales; Enterobacteriaceae; Salmonella; Salmonella enterica; Salmonella enterica subsp. enterica MTGDNTLITSHGINRRDFMKLCAALAATMGLSSKAAAEMAESVSNPQRPPVIWIGAQECT WP_000145429.1 hydrogenase 2 small subunit [Salmonella enterica] Length: 372\nNumber of predicted TMHs: 1\nExp number of AAs in TMHs: 24.51309\nExp number, first 60 AAs: 0.2682\nTotal prob of N-in: 0.14402\noutside 1 331\nTMhelix 332 354\ninside 355 372

21001 GCF_001134485.1_10608_2_31 Salmonella enterica subsp. enterica serovar Typhi Proteobacteria; Gammaproteobacteria; Enterobacterales; Enterobacteriaceae; Salmonella; Salmonella enterica; Salmonella enterica subsp. enterica MTGDNTLITSHGINRRDFMKLCAALAATMGLSSKAAAEMAESVSNPQRPPVIWIGAQECT WP_000145429.1 hydrogenase 2 small subunit [Salmonella enterica] Length: 372\nNumber of predicted TMHs: 1\nExp number of AAs in TMHs: 24.51309\nExp number, first 60 AAs: 0.2682\nTotal prob of N-in: 0.14402\noutside 1 331\nTMhelix 332 354\ninside 355 372

21002 GCF_001134985.1_9870_8_4Salmonella enterica subsp. enterica serovar Typhi Proteobacteria; Gammaproteobacteria; Enterobacterales; Enterobacteriaceae; Salmonella; Salmonella enterica; Salmonella enterica subsp. enterica MTGDNTLITSHGINRRDFMKLCAALAATMGLSSKAAAEMAESVSNPQRPPVIWIGAQECT WP_000145429.1 hydrogenase 2 small subunit [Salmonella enterica] Length: 372\nNumber of predicted TMHs: 1\nExp number of AAs in TMHs: 24.51309\nExp number, first 60 AAs: 0.2682\nTotal prob of N-in: 0.14402\noutside 1 331\nTMhelix 332 354\ninside 355 372

21003 GCF_001135105.1_9870_8_87 Salmonella enterica subsp. enterica serovar Typhi Proteobacteria; Gammaproteobacteria; Enterobacterales; Enterobacteriaceae; Salmonella; Salmonella enterica; Salmonella enterica subsp. enterica MTGDNTLITSHGINRRDFMKLCAALAATMGLSSKAAAEMAESVSNPQRPPVIWIGAQECT WP_000145429.1 hydrogenase 2 small subunit [Salmonella enterica] Length: 372\nNumber of predicted TMHs: 1\nExp number of AAs in TMHs: 24.51309\nExp number, first 60 AAs: 0.2682\nTotal prob of N-in: 0.14402\noutside 1 331\nTMhelix 332 354\ninside 355 372

21004 GCF_001135425.1_10493_1_73 Salmonella enterica subsp. enterica serovar Typhi Proteobacteria; Gammaproteobacteria; Enterobacterales; Enterobacteriaceae; Salmonella; Salmonella enterica; Salmonella enterica subsp. enterica MTGDNTLITSHGINRRDFMKLCAALAATMGLSSKAAAEMAESVSNPQRPPVIWIGAQECT WP_000145429.1 hydrogenase 2 small subunit [Salmonella enterica] Length: 372\nNumber of predicted TMHs: 1\nExp number of AAs in TMHs: 24.51309\nExp number, first 60 AAs: 0.2682\nTotal prob of N-in: 0.14402\noutside 1 331\nTMhelix 332 354\ninside 355 372

21005 GCF_001135905.1_10071_8_94 Salmonella enterica subsp. enterica serovar Typhi Proteobacteria; Gammaproteobacteria; Enterobacterales; Enterobacteriaceae; Salmonella; Salmonella enterica;

Salmonella enterica subsp. enterica MTGDNTLITSHGINRRDFMKLCAALAATMGLSSKAAAEMAESVSNPQRPPVIWIGAQECT WP_000145429.1 hydrogenase 2 small subunit [Salmonella enterica] Length: 372\nNumber of predicted TMHs: 1\nExp number of AAs in TMHs: 24.51309\nExp number, first 60 AAs: 0.2682\nTotal prob of N-in: 0.14402\noutside 1 331\nTMhelix 332 354\ninside 355 372

21006 GCF_001136385.1_9475_4_51 Salmonella enterica subsp. enterica serovar Typhi Proteobacteria; Gammaproteobacteria; Enterobacterales; Enterobacteriaceae; Salmonella; Salmonella enterica; Salmonella enterica subsp. enterica MTGDNTLITSHGINRRDFMKLCAALAATMGLSSKAAAEMAESVSNPQRPPVIWIGAQECT WP_000145429.1 hydrogenase 2 small subunit [Salmonella enterica] Length: 372\nNumber of predicted TMHs: 1\nExp number of AAs in TMHs: 24.51309\nExp number, first 60 AAs: 0.2682\nTotal prob of N-in: 0.14402\noutside 1 331\nTMhelix 332 354\ninside 355 372

21007 GCF_001137405.1_10349_1_35 Salmonella enterica subsp. enterica serovar Typhi Proteobacteria; Gammaproteobacteria; Enterobacterales; Enterobacteriaceae; Salmonella; Salmonella enterica; Salmonella enterica subsp. enterica MTGDNTLITSHGINRRDFMKLCAALAATMGLSSKAAAEMAESVSNPQRPPVIWIGAQECT WP_000145429.1 hydrogenase 2 small subunit [Salmonella enterica] Length: 372\nNumber of predicted TMHs: 1\nExp number of AAs in TMHs: 24.51309\nExp number, first 60 AAs: 0.2682\nTotal prob of N-in: 0.14402\noutside 1 331\nTMhelix 332 354\ninside 355 372

21008 GCF_001136805.1_7468_7_90 Salmonella enterica subsp. enterica serovar Typhi Proteobacteria; Gammaproteobacteria; Enterobacterales; Enterobacteriaceae; Salmonella; Salmonella enterica; Salmonella enterica subsp. enterica MTGDNTLITSHGINRRDFMKLCAALAATMGLSSKAAAEMAESVSNPQRPPVIWIGAQECT WP_000145429.1 hydrogenase 2 small subunit [Salmonella enterica] Length: 372\nNumber of predicted TMHs: 1\nExp number of AAs in TMHs: 24.51309\nExp number, first 60 AAs: 0.2682\nTotal prob of N-in: 0.14402\noutside 1 331\nTMhelix 332 354\ninside 355 372

21009 GCF_001137205.1_10492_1_28 Salmonella enterica subsp. enterica serovar Typhi Proteobacteria; Gammaproteobacteria; Enterobacterales; Enterobacteriaceae; Salmonella; Salmonella enterica; Salmonella enterica subsp. enterica MTGDNTLITSHGINRRDFMKLCAALAATMGLSSKAAAEMAESVSNPQRPPVIWIGAQECT WP_000145429.1 hydrogenase 2 small subunit [Salmonella enterica] Length: 372\nNumber of predicted TMHs: 1\nExp number of AAs in TMHs: 24.51309\nExp number, first 60 AAs: 0.2682\nTotal prob of N-in: 0.14402\noutside 1 331\nTMhelix 332 354\ninside 355 372

21010 GCF_001137605.1_10493_1_74 Salmonella enterica subsp. enterica serovar Typhi Proteobacteria; Gammaproteobacteria; Enterobacterales; Enterobacteriaceae; Salmonella; Salmonella enterica; Salmonella enterica subsp. enterica MTGDNTLITSHGINRRDFMKLCAALAATMGLSSKAAAEMAESVSNPQRPPVIWIGAQECT WP_000145429.1 hydrogenase 2 small subunit [Salmonella enterica] Length: 372\nNumber of predicted TMHs: 1\nExp number of AAs in TMHs: 24.51309\nExp number, first 60 AAs: 0.2682\nTotal prob of N-in: 0.14402\noutside 1 331\nTMhelix 332 354\ninside 355 372

21011 GCF_001138085.1_10561_2_46 Salmonella enterica subsp. enterica serovar Typhi Proteobacteria; Gammaproteobacteria; Enterobacterales; Enterobacteriaceae; Salmonella; Salmonella enterica; Salmonella enterica subsp. enterica MTGDNTLITSHGINRRDFMKLCAALAATMGLSSKAAAEMAESVSNPQRPPVIWIGAQECT WP_000145429.1 hydrogenase 2 small subunit [Salmonella enterica] Length: 372\nNumber of predicted TMHs: 1\nExp number of AAs in TMHs: 24.51309\nExp number, first 60 AAs: 0.2682\nTotal prob of N-in: 0.14402\noutside 1 331\nTMhelix 332 354\ninside 355 372

21012 GCF_001137745.1_9953_5_68 Salmonella enterica subsp. enterica serovar Typhi Proteobacteria; Gammaproteobacteria; Enterobacterales; Enterobacteriaceae; Salmonella; Salmonella enterica; Salmonella enterica subsp. enterica MTGDNTLITSHGINRRDFMKLCAALAATMGLSSKAAAEMAESVSNPQRPPVIWIGAQECT WP_000145429.1 hydrogenase 2 small subunit [Salmonella enterica] Length: 372\nNumber of predicted TMHs: 1\nExp number of AAs in TMHs: 24.51309\nExp number, first 60 AAs: 0.2682\nTotal prob of N-in: 0.14402\noutside 1 331\nTMhelix 332 354\ninside 355 372

21013 GCF_001138105.1_9953_5_19 Salmonella enterica subsp. enterica serovar Typhi Proteobacteria; Gammaproteobacteria; Enterobacterales; Enterobacteriaceae; Salmonella; Salmonella enterica; Salmonella enterica subsp. enterica MTGDNTLITSHGINRRDFMKLCAALAATMGLSSKAAAEMAESVSNPQRPPVIWIGAQECT WP_000145429.1 hydrogenase 2 small subunit [Salmonella enterica] Length: 372\nNumber of predicted TMHs: 1\nExp number of AAs in TMHs: 24.51309\nExp number, first 60 AAs: 0.2682\nTotal prob of N-in: 0.14402\noutside 1 331\nTMhelix 332 354\ninside 355 372

21014 GCF_001138245.1_9475_6_52 Salmonella enterica subsp. enterica serovar Typhi Proteobacteria; Gammaproteobacteria; Enterobacterales; Enterobacteriaceae; Salmonella; Salmonella enterica; Salmonella enterica subsp. enterica MTGDNTLITSHGINRRDFMKLCAALAATMGLSSKAAAEMAESVSNPQRPPVIWIGAQECT WP_000145429.1 hydrogenase 2 small subunit [Salmonella enterica] Length: 372\nNumber of predicted TMHs: 1\nExp number of AAs in TMHs: 24.51309\nExp number, first 60 AAs: 0.2682\nTotal prob of N-in: 0.14402\noutside 1 331\nTMhelix 332 354\ninside 355 372

21015 GCF_001138325.1_8525_3_54 Salmonella enterica subsp. enterica serovar Typhi Proteobacteria; Gammaproteobacteria; Enterobacterales; Enterobacteriaceae; Salmonella; Salmonella enterica; Salmonella enterica subsp. enterica MTGDNTLITSHGINRRDFMKLCAALAATMGLSSKAAAEMAESVSNPQRPPVIWIGAQECT WP_000145429.1 hydrogenase 2 small subunit [Salmonella enterica] Length: 372\nNumber of predicted TMHs: 1\nExp number of AAs in TMHs: 24.51309\nExp number, first 60 AAs: 0.2682\nTotal prob of N-in: 0.14402\noutside 1 331\nTMhelix 332 354\ninside 355 372

21016 GCF_001138365.1_10492_1_32 Salmonella enterica subsp. enterica serovar Typhi Proteobacteria; Gammaproteobacteria; Enterobacterales; Enterobacteriaceae; Salmonella; Salmonella enterica; Salmonella enterica subsp. enterica MTGDNTLITSHGINRRDFMKLCAALAATMGLSSKAAAEMAESVSNPQRPPVIWIGAQECT WP_000145429.1 hydrogenase 2 small subunit [Salmonella enterica] Length: 372\nNumber of predicted TMHs: 1\nExp number of AAs in TMHs: 24.51309\nExp number, first 60 AAs: 0.2682\nTotal prob of N-in: 0.14402\noutside 1 331\nTMhelix 332 354\ninside 355 372

21017 GCF_001138705.1_10608_1_3 Salmonella enterica subsp. enterica serovar Typhi Proteobacteria; Gammaproteobacteria; Enterobacterales; Enterobacteriaceae; Salmonella; Salmonella enterica; Salmonella enterica subsp. enterica MTGDNTLITSHGINRRDFMKLCAALAATMGLSSKAAAEMAESVSNPQRPPVIWIGAQECT WP_000145429.1 hydrogenase 2 small subunit [Salmonella enterica] Length: 372\nNumber of predicted TMHs: 1\nExp number of AAs in TMHs: 24.51309\nExp number, first 60 AAs: 0.2682\nTotal prob of N-in: 0.14402\noutside 1 331\nTMhelix 332 354\ninside 355 372

21018 GCF_001138885.1_10493_1_78 Salmonella enterica subsp. enterica serovar Typhi Proteobacteria; Gammaproteobacteria; Enterobacterales; Enterobacteriaceae; Salmonella; Salmonella enterica; Salmonella enterica subsp. enterica MTGDNTLITSHGINRRDFMKLCAALAATMGLSSKAAAEMAESVSNPQRPPVIWIGAQECT WP_000145429.1 hydrogenase 2 small subunit [Salmonella enterica] Length: 372\nNumber of predicted TMHs: 1\nExp number of AAs in TMHs: 24.51309\nExp number, first 60 AAs: 0.2682\nTotal prob of N-in: 0.14402\noutside 1 331\nTMhelix 332 354\ninside 355 372

21019 GCF_001139045.1_10593_2_3 Salmonella enterica subsp. enterica serovar Typhi Proteobacteria; Gammaproteobacteria; Enterobacterales; Enterobacteriaceae; Salmonella; Salmonella enterica; Salmonella enterica subsp. enterica MTGDNTLITSHGINRRDFMKLCAALAATMGLSSKAAAEMAESVSNPQRPPVIWIGAQECT WP_000145429.1 hydrogenase 2 small subunit [Salmonella enterica] Length: 372\nNumber of predicted TMHs: 1\nExp number of AAs in TMHs: 24.51309\nExp number, first 60 AAs: 0.2682\nTotal prob of N-in: 0.14402\noutside 1 331\nTMhelix 332 354\ninside 355 372

21020 GCF_001139705.1_9870_8_64 Salmonella enterica subsp. enterica serovar Typhi Proteobacteria; Gammaproteobacteria; Enterobacterales; Enterobacteriaceae; Salmonella; Salmonella enterica; Salmonella enterica subsp. enterica MTGDNTLITSHGINRRDFMKLCAALAATMGLSSKAAAEMAESVSNPQRPPVIWIGAQECT WP_000145429.1 hydrogenase 2 small subunit [Salmonella enterica] Length: 372\nNumber of predicted TMHs: 1\nExp number of AAs in TMHs: 24.51309\nExp number, first 60 AAs: 0.2682\nTotal prob of N-in: 0.14402\noutside 1 331\nTMhelix 332 354\ninside 355 372

21021 GCF_001140305.1_10541_2_54 Salmonella enterica subsp. enterica serovar Typhi Proteobacteria; Gammaproteobacteria; Enterobacterales; Enterobacteriaceae; Salmonella; Salmonella enterica; Salmonella enterica subsp. enterica MTGDNTLITSHGINRRDFMKLCAALAATMGLSSKAAAEMAESVSNPQRPPVIWIGAQECT WP_000145429.1 hydrogenase 2 small subunit [Salmonella enterica] Length: 372\nNumber of predicted TMHs: 1\nExp number of AAs in TMHs: 24.51309\nExp number, first 60 AAs: 0.2682\nTotal prob of N-in: 0.14402\noutside 1 331\nTMhelix 332 354\ninside 355 372

21022 GCF_001140605.1_9953_5_37 Salmonella enterica subsp. enterica serovar Typhi Proteobacteria; Gammaproteobacteria; Enterobacterales; Enterobacteriaceae; Salmonella; Salmonella enterica; Salmonella enterica subsp. enterica MTGDNTLITSHGINRRDFMKLCAALAATMGLSSKAAAEMAESVSNPQRPPVIWIGAQECT WP_000145429.1 hydrogenase 2 small subunit [Salmonella enterica] Length: 372\nNumber of

predicted TMHs:  1\nExp number of AAs in TMHs: 24.51309\nExp number, first 60 AAs:  0.2682\nTotal prob of N-in: 0.14402\noutside   1   331\nTMhelix   332   354\ninside   355   372

21023 GCF_001140985.1_10349_1_51 Salmonella enterica subsp. enterica serovar Typhi Proteobacteria; Gammaproteobacteria; Enterobacterales; Enterobacteriaceae; Salmonella; Salmonella enterica; Salmonella enterica subsp. enterica MTGDNTLITSHGINRRDFMKLCAALAATMGLSSKAAAEMAESVSNPQRPPVIWIGAQECT WP_000145429.1 hydrogenase 2 small subunit [Salmonella enterica] Length: 372\nNumber of predicted TMHs:  1\nExp number of AAs in TMHs: 24.51309\nExp number, first 60 AAs:  0.2682\nTotal prob of N-in: 0.14402\noutside   1   331\nTMhelix   332   354\ninside   355   372

21024 GCF_001140485.1_10541_2_44 Salmonella enterica subsp. enterica serovar Typhi Proteobacteria; Gammaproteobacteria; Enterobacterales; Enterobacteriaceae; Salmonella; Salmonella enterica; Salmonella enterica subsp. enterica MTGDNTLITSHGINRRDFMKLCAALAATMGLSSKAAAEMAESVSNPQRPPVIWIGAQECT WP_000145429.1 hydrogenase 2 small subunit [Salmonella enterica] Length: 372\nNumber of predicted TMHs:  1\nExp number of AAs in TMHs: 24.51309\nExp number, first 60 AAs:  0.2682\nTotal prob of N-in: 0.14402\noutside   1   331\nTMhelix   332   354\ninside   355   372

21025 GCF_001141065.1_10592_2_32 Salmonella enterica subsp. enterica serovar Typhi Proteobacteria; Gammaproteobacteria; Enterobacterales; Enterobacteriaceae; Salmonella; Salmonella enterica; Salmonella enterica subsp. enterica MTGDNTLITSHGINRRDFMKLCAALAATMGLSSKAAAEMAESVSNPQRPPVIWIGAQECT WP_000145429.1 hydrogenase 2 small subunit [Salmonella enterica] Length: 372\nNumber of predicted TMHs:  1\nExp number of AAs in TMHs: 24.51309\nExp number, first 60 AAs:  0.2682\nTotal prob of N-in: 0.14402\noutside   1   331\nTMhelix   332   354\ninside   355   372

21026 GCF_001141105.1_10349_1_24 Salmonella enterica subsp. enterica serovar Typhi Proteobacteria; Gammaproteobacteria; Enterobacterales; Enterobacteriaceae; Salmonella; Salmonella enterica; Salmonella enterica subsp. enterica MTGDNTLITSHGINRRDFMKLCAALAATMGLSSKAAAEMAESVSNPQRPPVIWIGAQECT WP_000145429.1 hydrogenase 2 small subunit [Salmonella enterica] Length: 372\nNumber of predicted TMHs:  1\nExp number of AAs in TMHs: 24.51309\nExp number, first 60 AAs:  0.2682\nTotal prob of N-in: 0.14402\noutside   1   331\nTMhelix   332   354\ninside   355   372

21027 GCF_001141165.1_10593_2_51 Salmonella enterica subsp. enterica serovar Typhi Proteobacteria; Gammaproteobacteria; Enterobacterales; Enterobacteriaceae; Salmonella; Salmonella enterica; Salmonella enterica subsp. enterica MTGDNTLITSHGINRRDFMKLCAALAATMGLSSKAAAEMAESVSNPQRPPVIWIGAQECT WP_000145429.1 hydrogenase 2 small subunit [Salmonella enterica] Length: 372\nNumber of predicted TMHs:  1\nExp number of AAs in TMHs: 24.51309\nExp number, first 60 AAs:  0.2682\nTotal prob of N-in: 0.14402\noutside   1   331\nTMhelix   332   354\ninside   355   372

21028 GCF_001141225.1_10425_1_6 Salmonella enterica subsp. enterica serovar Typhi Proteobacteria; Gammaproteobacteria; Enterobacterales; Enterobacteriaceae; Salmonella; Salmonella enterica; Salmonella enterica subsp. enterica MTGDNTLITSHGINRRDFMKLCAALAATMGLSSKAAAEMAESVSNPQRPPVIWIGAQECT WP_000145429.1 hydrogenase 2 small subunit [Salmonella enterica] Length: 372\nNumber of predicted TMHs:  1\nExp number of AAs in TMHs: 24.51309\nExp number, first 60 AAs:  0.2682\nTotal prob of N-in: 0.14402\noutside   1   331\nTMhelix   332   354\ninside   355   372

21029 GCF_001141645.1_10607_1_65 Salmonella enterica subsp. enterica serovar Typhi Proteobacteria; Gammaproteobacteria; Enterobacterales; Enterobacteriaceae; Salmonella; Salmonella enterica; Salmonella enterica subsp. enterica MTGDNTLITSHGINRRDFMKLCAALAATMGLSSKAAAEMAESVSNPQRPPVIWIGAQECT WP_000145429.1 hydrogenase 2 small subunit [Salmonella enterica] Length: 372\nNumber of predicted TMHs:  1\nExp number of AAs in TMHs: 24.51309\nExp number, first 60 AAs:  0.2682\nTotal prob of N-in: 0.14402\noutside   1   331\nTMhelix   332   354\ninside   355   372

21030 GCF_001141825.1_9475_6_45 Salmonella enterica subsp. enterica serovar Typhi Proteobacteria; Gammaproteobacteria; Enterobacterales; Enterobacteriaceae; Salmonella; Salmonella enterica; Salmonella enterica subsp. enterica MTGDNTLITSHGINRRDFMKLCAALAATMGLSSKAAAEMAESVSNPQRPPVIWIGAQECT WP_000145429.1 hydrogenase 2 small subunit [Salmonella enterica] Length: 372\nNumber of predicted TMHs:  1\nExp number of AAs in TMHs: 24.51309\nExp number, first 60 AAs:  0.2682\nTotal prob of N-in: 0.14402\noutside   1   331\nTMhelix   332   354\ninside   355   372

21031 GCF_001141945.1_10349_1_43 Salmonella enterica subsp. enterica serovar Typhi Proteobacteria; Gammaproteobacteria; Enterobacterales; Enterobacteriaceae; Salmonella; Salmonella enterica;

Salmonella enterica subsp. enterica MTGDNTLITSHGINRRDFMKLCAALAATMGLSSKAAAEMAESVSNPQRPPVIWIGAQECT WP_000145429.1 hydrogenase 2 small subunit [Salmonella enterica] Length: 372\nNumber of predicted TMHs: 1\nExp number of AAs in TMHs: 24.51309\nExp number, first 60 AAs: 0.2682\nTotal prob of N-in: 0.14402\noutside 1 331\nTMhelix 332 354\ninside 355 372

21032 GCF_001142285.1_10592_2_23 Salmonella enterica subsp. enterica serovar Typhi Proteobacteria; Gammaproteobacteria; Enterobacterales; Enterobacteriaceae; Salmonella; Salmonella enterica; Salmonella enterica subsp. enterica MTGDNTLITSHGINRRDFMKLCAALAATMGLSSKAAAEMAESVSNPQRPPVIWIGAQECT WP_000145429.1 hydrogenase 2 small subunit [Salmonella enterica] Length: 372\nNumber of predicted TMHs: 1\nExp number of AAs in TMHs: 24.51309\nExp number, first 60 AAs: 0.2682\nTotal prob of N-in: 0.14402\noutside 1 331\nTMhelix 332 354\ninside 355 372

21033 GCF_001142345.1_10562_2_75 Salmonella enterica subsp. enterica serovar Typhi Proteobacteria; Gammaproteobacteria; Enterobacterales; Enterobacteriaceae; Salmonella; Salmonella enterica; Salmonella enterica subsp. enterica MTGDNTLITSHGINRRDFMKLCAALAATMGLSSKAAAEMAESVSNPQRPPVIWIGAQECT WP_000145429.1 hydrogenase 2 small subunit [Salmonella enterica] Length: 372\nNumber of predicted TMHs: 1\nExp number of AAs in TMHs: 24.51309\nExp number, first 60 AAs: 0.2682\nTotal prob of N-in: 0.14402\noutside 1 331\nTMhelix 332 354\ninside 355 372

21034 GCF_001142565.1_10492_1_72 Salmonella enterica subsp. enterica serovar Typhi Proteobacteria; Gammaproteobacteria; Enterobacterales; Enterobacteriaceae; Salmonella; Salmonella enterica; Salmonella enterica subsp. enterica MTGDNTLITSHGINRRDFMKLCAALAATMGLSSKAAAEMAESVSNPQRPPVIWIGAQECT WP_000145429.1 hydrogenase 2 small subunit [Salmonella enterica] Length: 372\nNumber of predicted TMHs: 1\nExp number of AAs in TMHs: 24.51309\nExp number, first 60 AAs: 0.2682\nTotal prob of N-in: 0.14402\noutside 1 331\nTMhelix 332 354\ninside 355 372

21035 GCF_001142665.1_10562_2_23 Salmonella enterica subsp. enterica serovar Typhi Proteobacteria; Gammaproteobacteria; Enterobacterales; Enterobacteriaceae; Salmonella; Salmonella enterica; Salmonella enterica subsp. enterica MTGDNTLITSHGINRRDFMKLCAALAATMGLSSKAAAEMAESVSNPQRPPVIWIGAQECT WP_000145429.1 hydrogenase 2 small subunit [Salmonella enterica] Length: 372\nNumber of predicted TMHs: 1\nExp number of AAs in TMHs: 24.51309\nExp number, first 60 AAs: 0.2682\nTotal prob of N-in: 0.14402\noutside 1 331\nTMhelix 332 354\ninside 355 372

21036 GCF_001142805.1_10493_1_9 Salmonella enterica subsp. enterica serovar Typhi Proteobacteria; Gammaproteobacteria; Enterobacterales; Enterobacteriaceae; Salmonella; Salmonella enterica; Salmonella enterica subsp. enterica MTGDNTLITSHGINRRDFMKLCAALAATMGLSSKAAAEMAESVSNPQRPPVIWIGAQECT WP_000145429.1 hydrogenase 2 small subunit [Salmonella enterica] Length: 372\nNumber of predicted TMHs: 1\nExp number of AAs in TMHs: 24.51309\nExp number, first 60 AAs: 0.2682\nTotal prob of N-in: 0.14402\noutside 1 331\nTMhelix 332 354\ninside 355 372

21037 GCF_001142965.1_9953_5_87 Salmonella enterica subsp. enterica serovar Typhi Proteobacteria; Gammaproteobacteria; Enterobacterales; Enterobacteriaceae; Salmonella; Salmonella enterica; Salmonella enterica subsp. enterica MTGDNTLITSHGINRRDFMKLCAALAATMGLSSKAAAEMAESVSNPQRPPVIWIGAQECT WP_000145429.1 hydrogenase 2 small subunit [Salmonella enterica] Length: 372\nNumber of predicted TMHs: 1\nExp number of AAs in TMHs: 24.51309\nExp number, first 60 AAs: 0.2682\nTotal prob of N-in: 0.14402\noutside 1 331\nTMhelix 332 354\ninside 355 372

21038 GCF_001143045.1_10493_1_79 Salmonella enterica subsp. enterica serovar Typhi Proteobacteria; Gammaproteobacteria; Enterobacterales; Enterobacteriaceae; Salmonella; Salmonella enterica; Salmonella enterica subsp. enterica MTGDNTLITSHGINRRDFMKLCAALAATMGLSSKAAAEMAESVSNPQRPPVIWIGAQECT WP_000145429.1 hydrogenase 2 small subunit [Salmonella enterica] Length: 372\nNumber of predicted TMHs: 1\nExp number of AAs in TMHs: 24.51309\nExp number, first 60 AAs: 0.2682\nTotal prob of N-in: 0.14402\noutside 1 331\nTMhelix 332 354\ninside 355 372

21039 GCF_001143105.1_10593_2_45 Salmonella enterica subsp. enterica serovar Typhi Proteobacteria; Gammaproteobacteria; Enterobacterales; Enterobacteriaceae; Salmonella; Salmonella enterica; Salmonella enterica subsp. enterica MTGDNTLITSHGINRRDFMKLCAALAATMGLSSKAAAEMAESVSNPQRPPVIWIGAQECT WP_000145429.1 hydrogenase 2 small subunit [Salmonella enterica] Length: 372\nNumber of predicted TMHs: 1\nExp number of AAs in TMHs: 24.51309\nExp number, first 60 AAs: 0.2682\nTotal prob of N-in: 0.14402\noutside 1 331\nTMhelix 332 354\ninside 355 372

21040 GCF_001143485.1_9475_4_56 Salmonella enterica subsp. enterica serovar Typhi Proteobacteria; Gammaproteobacteria; Enterobacterales; Enterobacteriaceae; Salmonella; Salmonella enterica; Salmonella enterica subsp. enterica MTGDNTLITSHGINRRDFMKLCAALAATMGLSSKAAAEMAESVSNPQRPPVIWIGAQECT WP_000145429.1 hydrogenase 2 small subunit [Salmonella enterica] Length: 372\nNumber of predicted TMHs: 1\nExp number of AAs in TMHs: 24.51309\nExp number, first 60 AAs: 0.2682\nTotal prob of N-in: 0.14402\noutside 1 331\nTMhelix 332 354\ninside 355 372

21041 GCF_001143965.1_10209_5_49 Salmonella enterica subsp. enterica serovar Typhi Proteobacteria; Gammaproteobacteria; Enterobacterales; Enterobacteriaceae; Salmonella; Salmonella enterica; Salmonella enterica subsp. enterica MTGDNTLITSHGINRRDFMKLCAALAATMGLSSKAAAEMAESVSNPQRPPVIWIGAQECT WP_000145429.1 hydrogenase 2 small subunit [Salmonella enterica] Length: 372\nNumber of predicted TMHs: 1\nExp number of AAs in TMHs: 24.51309\nExp number, first 60 AAs: 0.2682\nTotal prob of N-in: 0.14402\noutside 1 331\nTMhelix 332 354\ninside 355 372

21042 GCF_001144045.1_9475_6_67 Salmonella enterica subsp. enterica serovar Typhi Proteobacteria; Gammaproteobacteria; Enterobacterales; Enterobacteriaceae; Salmonella; Salmonella enterica; Salmonella enterica subsp. enterica MTGDNTLITSHGINRRDFMKLCAALAATMGLSSKAAAEMAESVSNPQRPPVIWIGAQECT WP_000145429.1 hydrogenase 2 small subunit [Salmonella enterica] Length: 372\nNumber of predicted TMHs: 1\nExp number of AAs in TMHs: 24.51309\nExp number, first 60 AAs: 0.2682\nTotal prob of N-in: 0.14402\noutside 1 331\nTMhelix 332 354\ninside 355 372

21043 GCF_001144345.1_10607_2_37 Salmonella enterica subsp. enterica serovar Typhi Proteobacteria; Gammaproteobacteria; Enterobacterales; Enterobacteriaceae; Salmonella; Salmonella enterica; Salmonella enterica subsp. enterica MTGDNTLITSHGINRRDFMKLCAALAATMGLSSKAAAEMAESVSNPQRPPVIWIGAQECT WP_000145429.1 hydrogenase 2 small subunit [Salmonella enterica] Length: 372\nNumber of predicted TMHs: 1\nExp number of AAs in TMHs: 24.51309\nExp number, first 60 AAs: 0.2682\nTotal prob of N-in: 0.14402\noutside 1 331\nTMhelix 332 354\ninside 355 372

21044 GCF_001144645.1_9475_6_66 Salmonella enterica subsp. enterica serovar Typhi Proteobacteria; Gammaproteobacteria; Enterobacterales; Enterobacteriaceae; Salmonella; Salmonella enterica; Salmonella enterica subsp. enterica MTGDNTLITSHGINRRDFMKLCAALAATMGLSSKAAAEMAESVSNPQRPPVIWIGAQECT WP_000145429.1 hydrogenase 2 small subunit [Salmonella enterica] Length: 372\nNumber of predicted TMHs: 1\nExp number of AAs in TMHs: 24.51309\nExp number, first 60 AAs: 0.2682\nTotal prob of N-in: 0.14402\noutside 1 331\nTMhelix 332 354\ninside 355 372

21045 GCF_001145065.1_8525_3_31 Salmonella enterica subsp. enterica serovar Typhi Proteobacteria; Gammaproteobacteria; Enterobacterales; Enterobacteriaceae; Salmonella; Salmonella enterica; Salmonella enterica subsp. enterica MTGDNTLITSHGINRRDFMKLCAALAATMGLSSKAAAEMAESVSNPQRPPVIWIGAQECT WP_000145429.1 hydrogenase 2 small subunit [Salmonella enterica] Length: 372\nNumber of predicted TMHs: 1\nExp number of AAs in TMHs: 24.51309\nExp number, first 60 AAs: 0.2682\nTotal prob of N-in: 0.14402\noutside 1 331\nTMhelix 332 354\ninside 355 372

21046 GCF_001145205.1_10493_1_59 Salmonella enterica subsp. enterica serovar Typhi Proteobacteria; Gammaproteobacteria; Enterobacterales; Enterobacteriaceae; Salmonella; Salmonella enterica; Salmonella enterica subsp. enterica MTGDNTLITSHGINRRDFMKLCAALAATMGLSSKAAAEMAESVSNPQRPPVIWIGAQECT WP_000145429.1 hydrogenase 2 small subunit [Salmonella enterica] Length: 372\nNumber of predicted TMHs: 1\nExp number of AAs in TMHs: 24.51309\nExp number, first 60 AAs: 0.2682\nTotal prob of N-in: 0.14402\noutside 1 331\nTMhelix 332 354\ninside 355 372

21047 GCF_001145485.1_10592_2_11 Salmonella enterica subsp. enterica serovar Typhi Proteobacteria; Gammaproteobacteria; Enterobacterales; Enterobacteriaceae; Salmonella; Salmonella enterica; Salmonella enterica subsp. enterica MTGDNTLITSHGINRRDFMKLCAALAATMGLSSKAAAEMAESVSNPQRPPVIWIGAQECT WP_000145429.1 hydrogenase 2 small subunit [Salmonella enterica] Length: 372\nNumber of predicted TMHs: 1\nExp number of AAs in TMHs: 24.51309\nExp number, first 60 AAs: 0.2682\nTotal prob of N-in: 0.14402\noutside 1 331\nTMhelix 332 354\ninside 355 372

21048 GCF_001145545.1_10492_1_25 Salmonella enterica subsp. enterica serovar Typhi Proteobacteria; Gammaproteobacteria; Enterobacterales; Enterobacteriaceae; Salmonella; Salmonella enterica; Salmonella enterica subsp. enterica MTGDNTLITSHGINRRDFMKLCAALAATMGLSSKAAAEMAESVSNPQRPPVIWIGAQECT WP_000145429.1 hydrogenase 2 small subunit [Salmonella enterica] Length: 372\nNumber of

predicted TMHs:  1\nExp number of AAs in TMHs: 24.51309\nExp number, first 60 AAs:  0.2682\nTotal prob of N-in: 0.14402\noutside   1   331\nTMhelix   332   354\ninside   355   372

21049 GCF_001145585.1_8490_6_59 Salmonella enterica subsp. enterica serovar Typhi Proteobacteria; Gammaproteobacteria; Enterobacterales; Enterobacteriaceae; Salmonella; Salmonella enterica; Salmonella enterica subsp. enterica MTGDNTLITSHGINRRDFMKLCAALAATMGLSSKAAAEMAESVSNPQRPPVIWIGAQECT WP_000145429.1 hydrogenase 2 small subunit [Salmonella enterica] Length: 372\nNumber of predicted TMHs:  1\nExp number of AAs in TMHs: 24.51309\nExp number, first 60 AAs:  0.2682\nTotal prob of N-in: 0.14402\noutside   1   331\nTMhelix   332   354\ninside   355   372

21050 GCF_001146025.1_10349_1_19 Salmonella enterica subsp. enterica serovar Typhi Proteobacteria; Gammaproteobacteria; Enterobacterales; Enterobacteriaceae; Salmonella; Salmonella enterica; Salmonella enterica subsp. enterica MTGDNTLITSHGINRRDFMKLCAALAATMGLSSKAAAEMAESVSNPQRPPVIWIGAQECT WP_000145429.1 hydrogenase 2 small subunit [Salmonella enterica] Length: 372\nNumber of predicted TMHs:  1\nExp number of AAs in TMHs: 24.51309\nExp number, first 60 AAs:  0.2682\nTotal prob of N-in: 0.14402\noutside   1   331\nTMhelix   332   354\ninside   355   372

21051 GCF_001146125.1_10492_1_57 Salmonella enterica subsp. enterica serovar Typhi Proteobacteria; Gammaproteobacteria; Enterobacterales; Enterobacteriaceae; Salmonella; Salmonella enterica; Salmonella enterica subsp. enterica MTGDNTLITSHGINRRDFMKLCAALAATMGLSSKAAAEMAESVSNPQRPPVIWIGAQECT WP_000145429.1 hydrogenase 2 small subunit [Salmonella enterica] Length: 372\nNumber of predicted TMHs:  1\nExp number of AAs in TMHs: 24.51309\nExp number, first 60 AAs:  0.2682\nTotal prob of N-in: 0.14402\noutside   1   331\nTMhelix   332   354\ninside   355   372

21052 GCF_001146205.1_10492_1_11 Salmonella enterica subsp. enterica serovar Typhi Proteobacteria; Gammaproteobacteria; Enterobacterales; Enterobacteriaceae; Salmonella; Salmonella enterica; Salmonella enterica subsp. enterica MTGDNTLITSHGINRRDFMKLCAALAATMGLSSKAAAEMAESVSNPQRPPVIWIGAQECT WP_000145429.1 hydrogenase 2 small subunit [Salmonella enterica] Length: 372\nNumber of predicted TMHs:  1\nExp number of AAs in TMHs: 24.51309\nExp number, first 60 AAs:  0.2682\nTotal prob of N-in: 0.14402\noutside   1   331\nTMhelix   332   354\ninside   355   372

21053 GCF_001146725.1_10607_2_9 Salmonella enterica subsp. enterica serovar Typhi Proteobacteria; Gammaproteobacteria; Enterobacterales; Enterobacteriaceae; Salmonella; Salmonella enterica; Salmonella enterica subsp. enterica MTGDNTLITSHGINRRDFMKLCAALAATMGLSSKAAAEMAESVSNPQRPPVIWIGAQECT WP_000145429.1 hydrogenase 2 small subunit [Salmonella enterica] Length: 372\nNumber of predicted TMHs:  1\nExp number of AAs in TMHs: 24.51309\nExp number, first 60 AAs:  0.2682\nTotal prob of N-in: 0.14402\noutside   1   331\nTMhelix   332   354\ninside   355   372

21054 GCF_001146825.1_10608_2_4 Salmonella enterica subsp. enterica serovar Typhi Proteobacteria; Gammaproteobacteria; Enterobacterales; Enterobacteriaceae; Salmonella; Salmonella enterica; Salmonella enterica subsp. enterica MTGDNTLITSHGINRRDFMKLCAALAATMGLSSKAAAEMAESVSNPQRPPVIWIGAQECT WP_000145429.1 hydrogenase 2 small subunit [Salmonella enterica] Length: 372\nNumber of predicted TMHs:  1\nExp number of AAs in TMHs: 24.51309\nExp number, first 60 AAs:  0.2682\nTotal prob of N-in: 0.14402\noutside   1   331\nTMhelix   332   354\ninside   355   372

21055 GCF_001147085.1_10493_1_33 Salmonella enterica subsp. enterica serovar Typhi Proteobacteria; Gammaproteobacteria; Enterobacterales; Enterobacteriaceae; Salmonella; Salmonella enterica; Salmonella enterica subsp. enterica MTGDNTLITSHGINRRDFMKLCAALAATMGLSSKAAAEMAESVSNPQRPPVIWIGAQECT WP_000145429.1 hydrogenase 2 small subunit [Salmonella enterica] Length: 372\nNumber of predicted TMHs:  1\nExp number of AAs in TMHs: 24.51309\nExp number, first 60 AAs:  0.2682\nTotal prob of N-in: 0.14402\noutside   1   331\nTMhelix   332   354\ninside   355   372

21056 GCF_001147765.1_10492_1_63 Salmonella enterica subsp. enterica serovar Typhi Proteobacteria; Gammaproteobacteria; Enterobacterales; Enterobacteriaceae; Salmonella; Salmonella enterica; Salmonella enterica subsp. enterica MTGDNTLITSHGINRRDFMKLCAALAATMGLSSKAAAEMAESVSNPQRPPVIWIGAQECT WP_000145429.1 hydrogenase 2 small subunit [Salmonella enterica] Length: 372\nNumber of predicted TMHs:  1\nExp number of AAs in TMHs: 24.51309\nExp number, first 60 AAs:  0.2682\nTotal prob of N-in: 0.14402\noutside   1   331\nTMhelix   332   354\ninside   355   372

21057 GCF_001148005.1_9475_6_39 Salmonella enterica subsp. enterica serovar Typhi Proteobacteria; Gammaproteobacteria; Enterobacterales; Enterobacteriaceae; Salmonella; Salmonella enterica;

Salmonella enterica subsp. enterica MTGDNTLITSHGINRRDFMKLCAALAATMGLSSKAAAEMAESVSNPQRPPVIWIGAQECT WP_000145429.1 hydrogenase 2 small subunit [Salmonella enterica] Length: 372\nNumber of predicted TMHs: 1\nExp number of AAs in TMHs: 24.51309\nExp number, first 60 AAs: 0.2682\nTotal prob of N-in: 0.14402\noutside 1 331\nTMhelix 332 354\ninside 355 372

21058 GCF_001148265.1_9475_6_85 Salmonella enterica subsp. enterica serovar Typhi Proteobacteria; Gammaproteobacteria; Enterobacterales; Enterobacteriaceae; Salmonella; Salmonella enterica; Salmonella enterica subsp. enterica MTGDNTLITSHGINRRDFMKLCAALAATMGLSSKAAAEMAESVSNPQRPPVIWIGAQECT WP_000145429.1 hydrogenase 2 small subunit [Salmonella enterica] Length: 372\nNumber of predicted TMHs: 1\nExp number of AAs in TMHs: 24.51309\nExp number, first 60 AAs: 0.2682\nTotal prob of N-in: 0.14402\noutside 1 331\nTMhelix 332 354\ninside 355 372

21059 GCF_001148125.1_10349_1_4 Salmonella enterica subsp. enterica serovar Typhi Proteobacteria; Gammaproteobacteria; Enterobacterales; Enterobacteriaceae; Salmonella; Salmonella enterica; Salmonella enterica subsp. enterica MTGDNTLITSHGINRRDFMKLCAALAATMGLSSKAAAEMAESVSNPQRPPVIWIGAQECT WP_000145429.1 hydrogenase 2 small subunit [Salmonella enterica] Length: 372\nNumber of predicted TMHs: 1\nExp number of AAs in TMHs: 24.51309\nExp number, first 60 AAs: 0.2682\nTotal prob of N-in: 0.14402\noutside 1 331\nTMhelix 332 354\ninside 355 372

21060 GCF_001148325.1_10492_1_34 Salmonella enterica subsp. enterica serovar Typhi Proteobacteria; Gammaproteobacteria; Enterobacterales; Enterobacteriaceae; Salmonella; Salmonella enterica; Salmonella enterica subsp. enterica MTGDNTLITSHGINRRDFMKLCAALAATMGLSSKAAAEMAESVSNPQRPPVIWIGAQECT WP_000145429.1 hydrogenase 2 small subunit [Salmonella enterica] Length: 372\nNumber of predicted TMHs: 1\nExp number of AAs in TMHs: 24.51309\nExp number, first 60 AAs: 0.2682\nTotal prob of N-in: 0.14402\noutside 1 331\nTMhelix 332 354\ninside 355 372

21061 GCF_001148625.1_10561_2_63 Salmonella enterica subsp. enterica serovar Typhi Proteobacteria; Gammaproteobacteria; Enterobacterales; Enterobacteriaceae; Salmonella; Salmonella enterica; Salmonella enterica subsp. enterica MTGDNTLITSHGINRRDFMKLCAALAATMGLSSKAAAEMAESVSNPQRPPVIWIGAQECT WP_000145429.1 hydrogenase 2 small subunit [Salmonella enterica] Length: 372\nNumber of predicted TMHs: 1\nExp number of AAs in TMHs: 24.51309\nExp number, first 60 AAs: 0.2682\nTotal prob of N-in: 0.14402\noutside 1 331\nTMhelix 332 354\ninside 355 372

21062 GCF_001148805.1_9870_8_1Salmonella enterica subsp. enterica serovar Typhi Proteobacteria; Gammaproteobacteria; Enterobacterales; Enterobacteriaceae; Salmonella; Salmonella enterica; Salmonella enterica subsp. enterica MTGDNTLITSHGINRRDFMKLCAALAATMGLSSKAAAEMAESVSNPQRPPVIWIGAQECT WP_000145429.1 hydrogenase 2 small subunit [Salmonella enterica] Length: 372\nNumber of predicted TMHs: 1\nExp number of AAs in TMHs: 24.51309\nExp number, first 60 AAs: 0.2682\nTotal prob of N-in: 0.14402\noutside 1 331\nTMhelix 332 354\ninside 355 372

21063 GCF_001148845.1_10209_5_7 Salmonella enterica subsp. enterica serovar Typhi Proteobacteria; Gammaproteobacteria; Enterobacterales; Enterobacteriaceae; Salmonella; Salmonella enterica; Salmonella enterica subsp. enterica MTGDNTLITSHGINRRDFMKLCAALAATMGLSSKAAAEMAESVSNPQRPPVIWIGAQECT WP_000145429.1 hydrogenase 2 small subunit [Salmonella enterica] Length: 372\nNumber of predicted TMHs: 1\nExp number of AAs in TMHs: 24.51309\nExp number, first 60 AAs: 0.2682\nTotal prob of N-in: 0.14402\noutside 1 331\nTMhelix 332 354\ninside 355 372

21064 GCF_001148885.1_10562_2_59 Salmonella enterica subsp. enterica serovar Typhi Proteobacteria; Gammaproteobacteria; Enterobacterales; Enterobacteriaceae; Salmonella; Salmonella enterica; Salmonella enterica subsp. enterica MTGDNTLITSHGINRRDFMKLCAALAATMGLSSKAAAEMAESVSNPQRPPVIWIGAQECT WP_000145429.1 hydrogenase 2 small subunit [Salmonella enterica] Length: 372\nNumber of predicted TMHs: 1\nExp number of AAs in TMHs: 24.51309\nExp number, first 60 AAs: 0.2682\nTotal prob of N-in: 0.14402\noutside 1 331\nTMhelix 332 354\ninside 355 372

21065 GCF_001148085.1_10562_2_7 Salmonella enterica subsp. enterica serovar Typhi Proteobacteria; Gammaproteobacteria; Enterobacterales; Enterobacteriaceae; Salmonella; Salmonella enterica; Salmonella enterica subsp. enterica MTGDNTLITSHGINRRDFMKLCAALAATMGLSSKAATEMAESVSNPQRPPVIWIGAQECT WP_001782304.1 hydrogenase 2 small subunit [Salmonella enterica] Length: 372\nNumber of predicted TMHs: 1\nExp number of AAs in TMHs: 24.41173\nExp number, first 60 AAs: 0.16124\nTotal prob of N-in: 0.13947\noutside 1 331\nTMhelix 332 354\ninside 355 372

21066 GCF_001149025.1_10592_2_7 Salmonella enterica subsp. enterica serovar Typhi Proteobacteria; Gammaproteobacteria; Enterobacterales; Enterobacteriaceae; Salmonella; Salmonella enterica; Salmonella enterica subsp. enterica MTGDNTLITSHGINRRDFMKLCAALAATMGLSSKAAAEMAESVSNPQRPPVIWIGAQECT WP_000145429.1 hydrogenase 2 small subunit [Salmonella enterica] Length: 372\nNumber of predicted TMHs: 1\nExp number of AAs in TMHs: 24.51309\nExp number, first 60 AAs: 0.2682\nTotal prob of N-in: 0.14402\noutside 1 331\nTMhelix 332 354\ninside 355 372

21067 GCF_001149165.1_10492_1_4 Salmonella enterica subsp. enterica serovar Typhi Proteobacteria; Gammaproteobacteria; Enterobacterales; Enterobacteriaceae; Salmonella; Salmonella enterica; Salmonella enterica subsp. enterica MTGDNTLITSHGINRRDFMKLCAALAATMGLSSKAAAEMAESVSNPQRPPVIWIGAQECT WP_000145429.1 hydrogenase 2 small subunit [Salmonella enterica] Length: 372\nNumber of predicted TMHs: 1\nExp number of AAs in TMHs: 24.51309\nExp number, first 60 AAs: 0.2682\nTotal prob of N-in: 0.14402\noutside 1 331\nTMhelix 332 354\ninside 355 372

21068 GCF_001149685.1_10607_2_14 Salmonella enterica subsp. enterica serovar Typhi Proteobacteria; Gammaproteobacteria; Enterobacterales; Enterobacteriaceae; Salmonella; Salmonella enterica; Salmonella enterica subsp. enterica MTGDNTLITSHGINRRDFMKLCAALAATMGLSSKAAAEMAESVSNPQRPPVIWIGAQECT WP_000145429.1 hydrogenase 2 small subunit [Salmonella enterica] Length: 372\nNumber of predicted TMHs: 1\nExp number of AAs in TMHs: 24.51309\nExp number, first 60 AAs: 0.2682\nTotal prob of N-in: 0.14402\noutside 1 331\nTMhelix 332 354\ninside 355 372

21069 GCF_001149885.1_9475_6_61 Salmonella enterica subsp. enterica serovar Typhi Proteobacteria; Gammaproteobacteria; Enterobacterales; Enterobacteriaceae; Salmonella; Salmonella enterica; Salmonella enterica subsp. enterica MTGDNTLITSHGINRRDFMKLCAALAATMGLSSKAAAEMAESVSNPQRPPVIWIGAQECT WP_000145429.1 hydrogenase 2 small subunit [Salmonella enterica] Length: 372\nNumber of predicted TMHs: 1\nExp number of AAs in TMHs: 24.51309\nExp number, first 60 AAs: 0.2682\nTotal prob of N-in: 0.14402\noutside 1 331\nTMhelix 332 354\ninside 355 372

21070 GCF_001150065.1_8490_5_20 Salmonella enterica subsp. enterica serovar Typhi Proteobacteria; Gammaproteobacteria; Enterobacterales; Enterobacteriaceae; Salmonella; Salmonella enterica; Salmonella enterica subsp. enterica MTGDNTLITSHGINRRDFMKLCAALAATMGLSSKAAAEMAESVSNPQRPPVIWIGAQECT WP_000145429.1 hydrogenase 2 small subunit [Salmonella enterica] Length: 372\nNumber of predicted TMHs: 1\nExp number of AAs in TMHs: 24.51309\nExp number, first 60 AAs: 0.2682\nTotal prob of N-in: 0.14402\noutside 1 331\nTMhelix 332 354\ninside 355 372

21071 GCF_001151045.1_10493_1_64 Salmonella enterica subsp. enterica serovar Typhi Proteobacteria; Gammaproteobacteria; Enterobacterales; Enterobacteriaceae; Salmonella; Salmonella enterica; Salmonella enterica subsp. enterica MTGDNTLITSHGINRRDFMKLCAALAATMGLSSKAAAEMAESVSNPQRPPVIWIGAQECT WP_000145429.1 hydrogenase 2 small subunit [Salmonella enterica] Length: 372\nNumber of predicted TMHs: 1\nExp number of AAs in TMHs: 24.51309\nExp number, first 60 AAs: 0.2682\nTotal prob of N-in: 0.14402\noutside 1 331\nTMhelix 332 354\ninside 355 372

21072 GCF_001150445.1_10607_1_96 Salmonella enterica subsp. enterica serovar Typhi Proteobacteria; Gammaproteobacteria; Enterobacterales; Enterobacteriaceae; Salmonella; Salmonella enterica; Salmonella enterica subsp. enterica MTGDNTLITSHGINRRDFMKLCAALAATMGLSSKAAAEMAESVSNPQRPPVIWIGAQECT WP_000145429.1 hydrogenase 2 small subunit [Salmonella enterica] Length: 372\nNumber of predicted TMHs: 1\nExp number of AAs in TMHs: 24.51309\nExp number, first 60 AAs: 0.2682\nTotal prob of N-in: 0.14402\noutside 1 331\nTMhelix 332 354\ninside 355 372

21073 GCF_001151225.1_10561_2_44 Salmonella enterica subsp. enterica serovar Typhi Proteobacteria; Gammaproteobacteria; Enterobacterales; Enterobacteriaceae; Salmonella; Salmonella enterica; Salmonella enterica subsp. enterica MTGDNTLITSHGINRRDFMKLCAALAATMGLSSKAAAEMAESVSNPQRPPVIWIGAQECT WP_000145429.1 hydrogenase 2 small subunit [Salmonella enterica] Length: 372\nNumber of predicted TMHs: 1\nExp number of AAs in TMHs: 24.51309\nExp number, first 60 AAs: 0.2682\nTotal prob of N-in: 0.14402\noutside 1 331\nTMhelix 332 354\ninside 355 372

21074 GCF_001151325.1_9953_5_51 Salmonella enterica subsp. enterica serovar Typhi Proteobacteria; Gammaproteobacteria; Enterobacterales; Enterobacteriaceae; Salmonella; Salmonella enterica; Salmonella enterica subsp. enterica MTGDNTLITSHGINRRDFMKLCAALAATMGLSSKAAAEMAESVSNPQRPPVIWIGAQECT WP_000145429.1 hydrogenase 2 small subunit [Salmonella enterica] Length: 372\nNumber of

predicted TMHs: 1\nExp number of AAs in TMHs: 24.51309\nExp number, first 60 AAs: 0.2682\nTotal prob of N-in: 0.14402\noutside 1 331\nTMhelix 332 354\ninside 355 372

21075 GCF_001151665.1_10209_5_1 Salmonella enterica subsp. enterica serovar Typhi Proteobacteria; Gammaproteobacteria; Enterobacterales; Enterobacteriaceae; Salmonella; Salmonella enterica; Salmonella enterica subsp. enterica MTGDNTLITSHGINRRDFMKLCAALAATMGLSSKAAAEMAESVSNPQRPPVIWIGAQECT WP_000145429.1 hydrogenase 2 small subunit [Salmonella enterica] Length: 372\nNumber of predicted TMHs: 1\nExp number of AAs in TMHs: 24.51309\nExp number, first 60 AAs: 0.2682\nTotal prob of N-in: 0.14402\noutside 1 331\nTMhelix 332 354\ninside 355 372

21076 GCF_001151825.1_8525_3_46 Salmonella enterica subsp. enterica serovar Typhi Proteobacteria; Gammaproteobacteria; Enterobacterales; Enterobacteriaceae; Salmonella; Salmonella enterica; Salmonella enterica subsp. enterica MTGDNTLITSHGINRRDFMKLCAALAATMGLSSKAAAEMAESVSNPQRPPVIWIGAQECT WP_000145429.1 hydrogenase 2 small subunit [Salmonella enterica] Length: 372\nNumber of predicted TMHs: 1\nExp number of AAs in TMHs: 24.51309\nExp number, first 60 AAs: 0.2682\nTotal prob of N-in: 0.14402\noutside 1 331\nTMhelix 332 354\ninside 355 372

21077 GCF_001151985.1_10593_2_67 Salmonella enterica subsp. enterica serovar Typhi Proteobacteria; Gammaproteobacteria; Enterobacterales; Enterobacteriaceae; Salmonella; Salmonella enterica; Salmonella enterica subsp. enterica MTGDNTLITSHGINRRDFMKLCAALAATMGLSSKAAAEMAESVSNPQRPPVIWIGAQECT WP_000145429.1 hydrogenase 2 small subunit [Salmonella enterica] Length: 372\nNumber of predicted TMHs: 1\nExp number of AAs in TMHs: 24.51309\nExp number, first 60 AAs: 0.2682\nTotal prob of N-in: 0.14402\noutside 1 331\nTMhelix 332 354\ninside 355 372

21078 GCF_001152145.1_9870_8_37 Salmonella enterica subsp. enterica serovar Typhi Proteobacteria; Gammaproteobacteria; Enterobacterales; Enterobacteriaceae; Salmonella; Salmonella enterica; Salmonella enterica subsp. enterica MTGDNTLITSHGINRRDFMKLCAALAATMGLSSKAAAEMAESVSNPQRPPVIWIGAQECT WP_000145429.1 hydrogenase 2 small subunit [Salmonella enterica] Length: 372\nNumber of predicted TMHs: 1\nExp number of AAs in TMHs: 24.51309\nExp number, first 60 AAs: 0.2682\nTotal prob of N-in: 0.14402\noutside 1 331\nTMhelix 332 354\ninside 355 372

21079 GCF_001152345.1_10209_5_9 Salmonella enterica subsp. enterica serovar Typhi Proteobacteria; Gammaproteobacteria; Enterobacterales; Enterobacteriaceae; Salmonella; Salmonella enterica; Salmonella enterica subsp. enterica MTGDNTLITSHGINRRDFMKLCAALAATMGLSSKAAAEMAESVSNPQRPPVIWIGAQECT WP_000145429.1 hydrogenase 2 small subunit [Salmonella enterica] Length: 372\nNumber of predicted TMHs: 1\nExp number of AAs in TMHs: 24.51309\nExp number, first 60 AAs: 0.2682\nTotal prob of N-in: 0.14402\noutside 1 331\nTMhelix 332 354\ninside 355 372

21080 GCF_001152465.1_9953_5_5Salmonella enterica subsp. enterica serovar Typhi Proteobacteria; Gammaproteobacteria; Enterobacterales; Enterobacteriaceae; Salmonella; Salmonella enterica; Salmonella enterica subsp. enterica MTGDNTLITSHGINRRDFMKLCAALAATMGLSSKAAAEMAESVSNPQRPPVIWIGAQECT WP_000145429.1 hydrogenase 2 small subunit [Salmonella enterica] Length: 372\nNumber of predicted TMHs: 1\nExp number of AAs in TMHs: 24.51309\nExp number, first 60 AAs: 0.2682\nTotal prob of N-in: 0.14402\noutside 1 331\nTMhelix 332 354\ninside 355 372

21081 GCF_001152705.1_10562_2_50 Salmonella enterica subsp. enterica serovar Typhi Proteobacteria; Gammaproteobacteria; Enterobacterales; Enterobacteriaceae; Salmonella; Salmonella enterica; Salmonella enterica subsp. enterica MTGDNTLITSHGINRRDFMKLCAALAATMGLSSKAAAEMAESVSNPQRPPVIWIGAQECT WP_000145429.1 hydrogenase 2 small subunit [Salmonella enterica] Length: 372\nNumber of predicted TMHs: 1\nExp number of AAs in TMHs: 24.51309\nExp number, first 60 AAs: 0.2682\nTotal prob of N-in: 0.14402\noutside 1 331\nTMhelix 332 354\ninside 355 372

21082 GCF_001153245.1_9953_5_66 Salmonella enterica subsp. enterica serovar Typhi Proteobacteria; Gammaproteobacteria; Enterobacterales; Enterobacteriaceae; Salmonella; Salmonella enterica; Salmonella enterica subsp. enterica MTGDNTLITSHGINRRDFMKLCAALAATMGLSSKAAAEMAESVSNPQRPPVIWIGAQECT WP_000145429.1 hydrogenase 2 small subunit [Salmonella enterica] Length: 372\nNumber of predicted TMHs: 1\nExp number of AAs in TMHs: 24.51309\nExp number, first 60 AAs: 0.2682\nTotal prob of N-in: 0.14402\noutside 1 331\nTMhelix 332 354\ninside 355 372

21083 GCF_001153405.1_10541_2_27 Salmonella enterica subsp. enterica serovar Typhi Proteobacteria; Gammaproteobacteria; Enterobacterales; Enterobacteriaceae; Salmonella; Salmonella enterica;

Salmonella enterica subsp. enterica	MTGDNTLITSHGINRRDFMKLCAALAATMGLSSKAAAEMAESVSNPQRPPVIWIGAQECT	WP_000145429.1	hydrogenase 2 small subunit [Salmonella enterica]	Length: 372\nNumber of predicted TMHs:  1\nExp number of AAs in TMHs: 24.51309\nExp number, first 60 AAs:  0.2682\nTotal prob of N-in: 0.14402\noutside   1   331\nTMhelix   332   354\ninside   355   372

21084	GCF_001153665.1_10493_1_40	Salmonella enterica subsp. enterica serovar Typhi	Proteobacteria; Gammaproteobacteria; Enterobacterales; Enterobacteriaceae; Salmonella; Salmonella enterica; Salmonella enterica subsp. enterica	MTGDNTLITSHGINRRDFMKLCAALAATMGLSSKAAAEMAESVSNPQRPPVIWIGAQECT	WP_000145429.1	hydrogenase 2 small subunit [Salmonella enterica]	Length: 372\nNumber of predicted TMHs:  1\nExp number of AAs in TMHs: 24.51309\nExp number, first 60 AAs:  0.2682\nTotal prob of N-in: 0.14402\noutside   1   331\nTMhelix   332   354\ninside   355   372

21085	GCF_001154025.1_10209_5_43	Salmonella enterica subsp. enterica serovar Typhi	Proteobacteria; Gammaproteobacteria; Enterobacterales; Enterobacteriaceae; Salmonella; Salmonella enterica; Salmonella enterica subsp. enterica	MTGDNTLITSHGINRRDFMKLCAALAATMGLSSKAAAEMAESVSNPQRPPVIWIGAQECT	WP_000145429.1	hydrogenase 2 small subunit [Salmonella enterica]	Length: 372\nNumber of predicted TMHs:  1\nExp number of AAs in TMHs: 24.51309\nExp number, first 60 AAs:  0.2682\nTotal prob of N-in: 0.14402\noutside   1   331\nTMhelix   332   354\ninside   355   372

21086	GCF_001154125.1_10607_1_69	Salmonella enterica subsp. enterica serovar Typhi	Proteobacteria; Gammaproteobacteria; Enterobacterales; Enterobacteriaceae; Salmonella; Salmonella enterica; Salmonella enterica subsp. enterica	MTGDNTLITSHGINRRDFMKLCAALAATMGLSSKAAAEMAESVSNPQRPPVIWIGAQECT	WP_000145429.1	hydrogenase 2 small subunit [Salmonella enterica]	Length: 372\nNumber of predicted TMHs:  1\nExp number of AAs in TMHs: 24.51309\nExp number, first 60 AAs:  0.2682\nTotal prob of N-in: 0.14402\noutside   1   331\nTMhelix   332   354\ninside   355   372

21087	GCF_001154225.1_10561_2_31	Salmonella enterica subsp. enterica serovar Typhi	Proteobacteria; Gammaproteobacteria; Enterobacterales; Enterobacteriaceae; Salmonella; Salmonella enterica; Salmonella enterica subsp. enterica	MTGDNTLITSHGINRRDFMKLCAALAATMGLSSKAAAEMAESVSNPQRPPVIWIGAQECT	WP_000145429.1	hydrogenase 2 small subunit [Salmonella enterica]	Length: 372\nNumber of predicted TMHs:  1\nExp number of AAs in TMHs: 24.51309\nExp number, first 60 AAs:  0.2682\nTotal prob of N-in: 0.14402\noutside   1   331\nTMhelix   332   354\ninside   355   372

21088	GCF_001154425.1_9953_5_15	Salmonella enterica subsp. enterica serovar Typhi	Proteobacteria; Gammaproteobacteria; Enterobacterales; Enterobacteriaceae; Salmonella; Salmonella enterica; Salmonella enterica subsp. enterica	MTGDNTLITSHGINRRDFMKLCAALAATMGLSSKAAAEMAESVSNPQRPPVIWIGAQECT	WP_000145429.1	hydrogenase 2 small subunit [Salmonella enterica]	Length: 372\nNumber of predicted TMHs:  1\nExp number of AAs in TMHs: 24.51309\nExp number, first 60 AAs:  0.2682\nTotal prob of N-in: 0.14402\noutside   1   331\nTMhelix   332   354\ninside   355   372

21089	GCF_001153585.1_10071_8_93	Salmonella enterica subsp. enterica serovar Typhi	Proteobacteria; Gammaproteobacteria; Enterobacterales; Enterobacteriaceae; Salmonella; Salmonella enterica; Salmonella enterica subsp. enterica	MTGDNTLITSHGINRRDFMKLCAALAATMGLSSKAATEMAESVSNPQRPPVIWIGAQECT	WP_001782304.1	hydrogenase 2 small subunit [Salmonella enterica]	Length: 372\nNumber of predicted TMHs:  1\nExp number of AAs in TMHs: 24.41173\nExp number, first 60 AAs:  0.16124\nTotal prob of N-in: 0.13947\noutside   1   331\nTMhelix   332   354\ninside   355   372

21090	GCF_001155025.1_9953_5_42	Salmonella enterica subsp. enterica serovar Typhi	Proteobacteria; Gammaproteobacteria; Enterobacterales; Enterobacteriaceae; Salmonella; Salmonella enterica; Salmonella enterica subsp. enterica	MTGDNTLITSHGINRRDFMKLCAALAATMGLSSKAAAEMAESVSNPQRPPVIWIGAQECT	WP_000145429.1	hydrogenase 2 small subunit [Salmonella enterica]	Length: 372\nNumber of predicted TMHs:  1\nExp number of AAs in TMHs: 24.51309\nExp number, first 60 AAs:  0.2682\nTotal prob of N-in: 0.14402\noutside   1   331\nTMhelix   332   354\ninside   355   372

21091	GCF_001155445.1_10492_1_8	Salmonella enterica subsp. enterica serovar Typhi	Proteobacteria; Gammaproteobacteria; Enterobacterales; Enterobacteriaceae; Salmonella; Salmonella enterica; Salmonella enterica subsp. enterica	MTGDNTLITSHGINRRDFMKLCAALAATMGLSSKAAAEMAESVSNPQRPPVIWIGAQECT	WP_000145429.1	hydrogenase 2 small subunit [Salmonella enterica]	Length: 372\nNumber of predicted TMHs:  1\nExp number of AAs in TMHs: 24.51309\nExp number, first 60 AAs:  0.2682\nTotal prob of N-in: 0.14402\noutside   1   331\nTMhelix   332   354\ninside   355   372

21092 GCF_001155365.1_8490_5_36 Salmonella enterica subsp. enterica serovar Typhi Proteobacteria; Gammaproteobacteria; Enterobacterales; Enterobacteriaceae; Salmonella; Salmonella enterica; Salmonella enterica subsp. enterica MTGDNTLITSHGINRRDFMKLCAALAATMGLSSKAAAEMAESVSNPQRPPVIWIGAQECT WP_000145429.1 hydrogenase 2 small subunit [Salmonella enterica] Length: 372\nNumber of predicted TMHs: 1\nExp number of AAs in TMHs: 24.51309\nExp number, first 60 AAs: 0.2682\nTotal prob of N-in: 0.14402\noutside 1 331\nTMhelix 332 354\ninside 355 372

21093 GCF_001155725.1_10209_5_14 Salmonella enterica subsp. enterica serovar Typhi Proteobacteria; Gammaproteobacteria; Enterobacterales; Enterobacteriaceae; Salmonella; Salmonella enterica; Salmonella enterica subsp. enterica MTGDNTLITSHGINRRDFMKLCAALAATMGLSSKAAAEMAESVSNPQRPPVIWIGAQECT WP_000145429.1 hydrogenase 2 small subunit [Salmonella enterica] Length: 372\nNumber of predicted TMHs: 1\nExp number of AAs in TMHs: 24.51309\nExp number, first 60 AAs: 0.2682\nTotal prob of N-in: 0.14402\noutside 1 331\nTMhelix 332 354\ninside 355 372

21094 GCF_001155745.1_9870_8_46 Salmonella enterica subsp. enterica serovar Typhi Proteobacteria; Gammaproteobacteria; Enterobacterales; Enterobacteriaceae; Salmonella; Salmonella enterica; Salmonella enterica subsp. enterica MTGDNTLITSHGINRRDFMKLCAALAATMGLSSKAAAEMAESVSNPQRPPVIWIGAQECT WP_000145429.1 hydrogenase 2 small subunit [Salmonella enterica] Length: 372\nNumber of predicted TMHs: 1\nExp number of AAs in TMHs: 24.51309\nExp number, first 60 AAs: 0.2682\nTotal prob of N-in: 0.14402\noutside 1 331\nTMhelix 332 354\ninside 355 372

21095 GCF_001156285.1_10541_1_68 Salmonella enterica subsp. enterica serovar Typhi Proteobacteria; Gammaproteobacteria; Enterobacterales; Enterobacteriaceae; Salmonella; Salmonella enterica; Salmonella enterica subsp. enterica MTGDNTLITSHGINRRDFMKLCAALAATMGLSSKAAAEMAESVSNPQRPPVIWIGAQECT WP_000145429.1 hydrogenase 2 small subunit [Salmonella enterica] Length: 372\nNumber of predicted TMHs: 1\nExp number of AAs in TMHs: 24.51309\nExp number, first 60 AAs: 0.2682\nTotal prob of N-in: 0.14402\noutside 1 331\nTMhelix 332 354\ninside 355 372

21096 GCF_001156385.1_10562_2_26 Salmonella enterica subsp. enterica serovar Typhi Proteobacteria; Gammaproteobacteria; Enterobacterales; Enterobacteriaceae; Salmonella; Salmonella enterica; Salmonella enterica subsp. enterica MTGDNTLITSHGINRRDFMKLCAALAATMGLSSKAAAEMAESVSNPQRPPVIWIGAQECT WP_000145429.1 hydrogenase 2 small subunit [Salmonella enterica] Length: 372\nNumber of predicted TMHs: 1\nExp number of AAs in TMHs: 24.51309\nExp number, first 60 AAs: 0.2682\nTotal prob of N-in: 0.14402\noutside 1 331\nTMhelix 332 354\ninside 355 372

21097 GCF_001157165.1_10562_2_5 Salmonella enterica subsp. enterica serovar Typhi Proteobacteria; Gammaproteobacteria; Enterobacterales; Enterobacteriaceae; Salmonella; Salmonella enterica; Salmonella enterica subsp. enterica MTGDNTLITSHGINRRDFMKLCAALAATMGLSSKAAAEMAESVSNPQRPPVIWIGAQECT WP_000145429.1 hydrogenase 2 small subunit [Salmonella enterica] Length: 372\nNumber of predicted TMHs: 1\nExp number of AAs in TMHs: 24.51309\nExp number, first 60 AAs: 0.2682\nTotal prob of N-in: 0.14402\noutside 1 331\nTMhelix 332 354\ninside 355 372

21098 GCF_001156465.1_8490_6_49 Salmonella enterica subsp. enterica serovar Typhi Proteobacteria; Gammaproteobacteria; Enterobacterales; Enterobacteriaceae; Salmonella; Salmonella enterica; Salmonella enterica subsp. enterica MTGDNTLITSHGINRRDFMKLCAALAATMGLSSKAAAEMAESVSNPQRPPVIWIGAQECT WP_000145429.1 hydrogenase 2 small subunit [Salmonella enterica] Length: 372\nNumber of predicted TMHs: 1\nExp number of AAs in TMHs: 24.51309\nExp number, first 60 AAs: 0.2682\nTotal prob of N-in: 0.14402\noutside 1 331\nTMhelix 332 354\ninside 355 372

21099 GCF_001156885.1_10592_2_79 Salmonella enterica subsp. enterica serovar Typhi Proteobacteria; Gammaproteobacteria; Enterobacterales; Enterobacteriaceae; Salmonella; Salmonella enterica; Salmonella enterica subsp. enterica MTGDNTLITSHGINRRDFMKLCAALAATMGLSSKAAAEMAESVSNPQRPPVIWIGAQECT WP_000145429.1 hydrogenase 2 small subunit [Salmonella enterica] Length: 372\nNumber of predicted TMHs: 1\nExp number of AAs in TMHs: 24.51309\nExp number, first 60 AAs: 0.2682\nTotal prob of N-in: 0.14402\noutside 1 331\nTMhelix 332 354\ninside 355 372

21100 GCF_001157245.2_ERL12148Salmonella enterica subsp. enterica serovar Typhi Proteobacteria; Gammaproteobacteria; Enterobacterales; Enterobacteriaceae; Salmonella; Salmonella enterica; Salmonella enterica subsp. enterica MTGDNTLITSHGINRRDFMKLCAALAATMGLSSKAAAEMAESVSNPQRPPVIWIGAQECT WP_000145429.1 hydrogenase 2 small subunit [Salmonella enterica] Length: 372\nNumber of predicted TMHs: 1\nExp

number of AAs in TMHs: 24.51309\nExp number, first 60 AAs: 0.2682\nTotal prob of N-in: 0.14402\noutside 1 331\nTMhelix 332 354\ninside 355 372

21101 GCF_001157565.1_10425_1_9 Salmonella enterica subsp. enterica serovar Typhi Proteobacteria; Gammaproteobacteria; Enterobacterales; Enterobacteriaceae; Salmonella; Salmonella enterica; Salmonella enterica subsp. enterica MTGDNTLITSHGINRRDFMKLCAALAATMGLSSKAAAEMAESVSNPQRPPVIWIGAQECT WP_000145429.1 hydrogenase 2 small subunit [Salmonella enterica] Length: 372\nNumber of predicted TMHs: 1\nExp number of AAs in TMHs: 24.51309\nExp number, first 60 AAs: 0.2682\nTotal prob of N-in: 0.14402\noutside 1 331\nTMhelix 332 354\ninside 355 372

21102 GCF_001157725.1_10561_2_5 Salmonella enterica subsp. enterica serovar Typhi Proteobacteria; Gammaproteobacteria; Enterobacterales; Enterobacteriaceae; Salmonella; Salmonella enterica; Salmonella enterica subsp. enterica MTGDNTLITSHGINRRDFMKLCAALAATMGLSSKAAAEMAESVSNPQRPPVIWIGAQECT WP_000145429.1 hydrogenase 2 small subunit [Salmonella enterica] Length: 372\nNumber of predicted TMHs: 1\nExp number of AAs in TMHs: 24.51309\nExp number, first 60 AAs: 0.2682\nTotal prob of N-in: 0.14402\noutside 1 331\nTMhelix 332 354\ninside 355 372

21103 GCF_001157865.1_10592_2_54 Salmonella enterica subsp. enterica serovar Typhi Proteobacteria; Gammaproteobacteria; Enterobacterales; Enterobacteriaceae; Salmonella; Salmonella enterica; Salmonella enterica subsp. enterica MTGDNTLITSHGINRRDFMKLCAALAATMGLSSKAAAEMAESVSNPQRPPVIWIGAQECT WP_000145429.1 hydrogenase 2 small subunit [Salmonella enterica] Length: 372\nNumber of predicted TMHs: 1\nExp number of AAs in TMHs: 24.51309\nExp number, first 60 AAs: 0.2682\nTotal prob of N-in: 0.14402\noutside 1 331\nTMhelix 332 354\ninside 355 372

21104 GCF_001158005.1_9953_5_49 Salmonella enterica subsp. enterica serovar Typhi Proteobacteria; Gammaproteobacteria; Enterobacterales; Enterobacteriaceae; Salmonella; Salmonella enterica; Salmonella enterica subsp. enterica MTGDNTLITSHGINRRDFMKLCAALAATMGLSSKAAAEMAESVSNPQRPPVIWIGAQECT WP_000145429.1 hydrogenase 2 small subunit [Salmonella enterica] Length: 372\nNumber of predicted TMHs: 1\nExp number of AAs in TMHs: 24.51309\nExp number, first 60 AAs: 0.2682\nTotal prob of N-in: 0.14402\noutside 1 331\nTMhelix 332 354\ninside 355 372

21105 GCF_001158085.1_10608_2_15 Salmonella enterica subsp. enterica serovar Typhi Proteobacteria; Gammaproteobacteria; Enterobacterales; Enterobacteriaceae; Salmonella; Salmonella enterica; Salmonella enterica subsp. enterica MTGDNTLITSHGINRRDFMKLCAALAATMGLSSKAAAEMAESVSNPQRPPVIWIGAQECT WP_000145429.1 hydrogenase 2 small subunit [Salmonella enterica] Length: 372\nNumber of predicted TMHs: 1\nExp number of AAs in TMHs: 24.51309\nExp number, first 60 AAs: 0.2682\nTotal prob of N-in: 0.14402\noutside 1 331\nTMhelix 332 354\ninside 355 372

21106 GCF_001158165.1_10349_1_52 Salmonella enterica subsp. enterica serovar Typhi Proteobacteria; Gammaproteobacteria; Enterobacterales; Enterobacteriaceae; Salmonella; Salmonella enterica; Salmonella enterica subsp. enterica MTGDNTLITSHGINRRDFMKLCAALAATMGLSSKAAAEMAESVSNPQRPPVIWIGAQECT WP_000145429.1 hydrogenase 2 small subunit [Salmonella enterica] Length: 372\nNumber of predicted TMHs: 1\nExp number of AAs in TMHs: 24.51309\nExp number, first 60 AAs: 0.2682\nTotal prob of N-in: 0.14402\noutside 1 331\nTMhelix 332 354\ninside 355 372

21107 GCF_001158745.1_10562_2_70 Salmonella enterica subsp. enterica serovar Typhi Proteobacteria; Gammaproteobacteria; Enterobacterales; Enterobacteriaceae; Salmonella; Salmonella enterica; Salmonella enterica subsp. enterica MTGDNTLITSHGINRRDFMKLCAALAATMGLSSKAAAEMAESVSNPQRPPVIWIGAQECT WP_000145429.1 hydrogenase 2 small subunit [Salmonella enterica] Length: 372\nNumber of predicted TMHs: 1\nExp number of AAs in TMHs: 24.51309\nExp number, first 60 AAs: 0.2682\nTotal prob of N-in: 0.14402\noutside 1 331\nTMhelix 332 354\ninside 355 372

21108 GCF_001158905.1_10349_1_15 Salmonella enterica subsp. enterica serovar Typhi Proteobacteria; Gammaproteobacteria; Enterobacterales; Enterobacteriaceae; Salmonella; Salmonella enterica; Salmonella enterica subsp. enterica MTGDNTLITSHGINRRDFMKLCAALAATMGLSSKAAAEMAESVSNPQRPPVIWIGAQECT WP_000145429.1 hydrogenase 2 small subunit [Salmonella enterica] Length: 372\nNumber of predicted TMHs: 1\nExp number of AAs in TMHs: 24.51309\nExp number, first 60 AAs: 0.2682\nTotal prob of N-in: 0.14402\noutside 1 331\nTMhelix 332 354\ninside 355 372

21109 GCF_001159385.1_10562_2_72 Salmonella enterica subsp. enterica serovar Typhi Proteobacteria; Gammaproteobacteria; Enterobacterales; Enterobacteriaceae; Salmonella; Salmonella enterica;

Salmonella enterica subsp. enterica	MTGDNTLITSHGINRRDFMKLCAALAATMGLSSKAAAEMAESVSNPQRPPVIWIGAQECT	WP_000145429.1	hydrogenase 2 small subunit [Salmonella enterica]	Length: 372\nNumber of predicted TMHs: 1\nExp number of AAs in TMHs: 24.51309\nExp number, first 60 AAs: 0.2682\nTotal prob of N-in: 0.14402\noutside 1 331\nTMhelix 332 354\ninside 355 372

21110	GCF_001159425.1_10592_2_49	Salmonella enterica subsp. enterica serovar Typhi	Proteobacteria; Gammaproteobacteria; Enterobacterales; Enterobacteriaceae; Salmonella; Salmonella enterica; Salmonella enterica subsp. enterica	MTGDNTLITSHGINRRDFMKLCAALAATMGLSSKAAAEMAESVSNPQRPPVIWIGAQECT	WP_000145429.1	hydrogenase 2 small subunit [Salmonella enterica]	Length: 372\nNumber of predicted TMHs: 1\nExp number of AAs in TMHs: 24.51309\nExp number, first 60 AAs: 0.2682\nTotal prob of N-in: 0.14402\noutside 1 331\nTMhelix 332 354\ninside 355 372

21111	GCF_001159525.1_10349_1_27	Salmonella enterica subsp. enterica serovar Typhi	Proteobacteria; Gammaproteobacteria; Enterobacterales; Enterobacteriaceae; Salmonella; Salmonella enterica; Salmonella enterica subsp. enterica	MTGDNTLITSHGINRRDFMKLCAALAATMGLSSKAAAEMAESVSNPQRPPVIWIGAQECT	WP_000145429.1	hydrogenase 2 small subunit [Salmonella enterica]	Length: 372\nNumber of predicted TMHs: 1\nExp number of AAs in TMHs: 24.51309\nExp number, first 60 AAs: 0.2682\nTotal prob of N-in: 0.14402\noutside 1 331\nTMhelix 332 354\ninside 355 372

21112	GCF_001159545.1_9870_8_14	Salmonella enterica subsp. enterica serovar Typhi	Proteobacteria; Gammaproteobacteria; Enterobacterales; Enterobacteriaceae; Salmonella; Salmonella enterica; Salmonella enterica subsp. enterica	MTGDNTLITSHGINRRDFMKLCAALAATMGLSSKAAAEMAESVSNPQRPPVIWIGAQECT	WP_000145429.1	hydrogenase 2 small subunit [Salmonella enterica]	Length: 372\nNumber of predicted TMHs: 1\nExp number of AAs in TMHs: 24.51309\nExp number, first 60 AAs: 0.2682\nTotal prob of N-in: 0.14402\noutside 1 331\nTMhelix 332 354\ninside 355 372

21113	GCF_001159785.1_10593_2_28	Salmonella enterica subsp. enterica serovar Typhi	Proteobacteria; Gammaproteobacteria; Enterobacterales; Enterobacteriaceae; Salmonella; Salmonella enterica; Salmonella enterica subsp. enterica	MTGDNTLITSHGINRRDFMKLCAALAATMGLSSKAAAEMAESVSNPQRPPVIWIGAQECT	WP_000145429.1	hydrogenase 2 small subunit [Salmonella enterica]	Length: 372\nNumber of predicted TMHs: 1\nExp number of AAs in TMHs: 24.51309\nExp number, first 60 AAs: 0.2682\nTotal prob of N-in: 0.14402\noutside 1 331\nTMhelix 332 354\ninside 355 372

21114	GCF_001160065.1_10493_1_42	Salmonella enterica subsp. enterica serovar Typhi	Proteobacteria; Gammaproteobacteria; Enterobacterales; Enterobacteriaceae; Salmonella; Salmonella enterica; Salmonella enterica subsp. enterica	MTGDNTLITSHGINRRDFMKLCAALAATMGLSSKAAAEMAESVSNPQRPPVIWIGAQECT	WP_000145429.1	hydrogenase 2 small subunit [Salmonella enterica]	Length: 372\nNumber of predicted TMHs: 1\nExp number of AAs in TMHs: 24.51309\nExp number, first 60 AAs: 0.2682\nTotal prob of N-in: 0.14402\noutside 1 331\nTMhelix 332 354\ninside 355 372

21115	GCF_001355755.1_8490_5_2Salmonella enterica subsp. enterica serovar Typhi	Proteobacteria; Gammaproteobacteria; Enterobacterales; Enterobacteriaceae; Salmonella; Salmonella enterica; Salmonella enterica subsp. enterica	MTGDNTLITSHGINRRDFMKLCAALAATMGLSSKAAAEMAESVSNPQRPPVIWIGAQECT	WP_000145429.1	hydrogenase 2 small subunit [Salmonella enterica]	Length: 372\nNumber of predicted TMHs: 1\nExp number of AAs in TMHs: 24.51309\nExp number, first 60 AAs: 0.2682\nTotal prob of N-in: 0.14402\noutside 1 331\nTMhelix 332 354\ninside 355 372

21116	GCF_001355795.1_8447_8_12	Salmonella enterica subsp. enterica serovar Typhi	Proteobacteria; Gammaproteobacteria; Enterobacterales; Enterobacteriaceae; Salmonella; Salmonella enterica; Salmonella enterica subsp. enterica	MTGDNTLITSHGINRRDFMKLCAALAATMGLSSKAAAEMAESVSNPQRPPVIWIGAQECT	WP_000145429.1	hydrogenase 2 small subunit [Salmonella enterica]	Length: 372\nNumber of predicted TMHs: 1\nExp number of AAs in TMHs: 24.51309\nExp number, first 60 AAs: 0.2682\nTotal prob of N-in: 0.14402\noutside 1 331\nTMhelix 332 354\ninside 355 372

21117	GCF_001355835.1_8490_5_29	Salmonella enterica subsp. enterica serovar Typhi	Proteobacteria; Gammaproteobacteria; Enterobacterales; Enterobacteriaceae; Salmonella; Salmonella enterica; Salmonella enterica subsp. enterica	MTGDNTLITSHGINRRDFMKLCAALAATMGLSSKAAAEMAESVSNPQRPPVIWIGAQECT	WP_000145429.1	hydrogenase 2 small subunit [Salmonella enterica]	Length: 372\nNumber of predicted TMHs: 1\nExp number of AAs in TMHs: 24.51309\nExp number, first 60 AAs: 0.2682\nTotal prob of N-in: 0.14402\noutside 1 331\nTMhelix 332 354\ninside 355 372

21118 GCF_001239885.1_10425_1_8 Salmonella enterica subsp. enterica serovar Typhi Proteobacteria; Gammaproteobacteria; Enterobacterales; Enterobacteriaceae; Salmonella; Salmonella enterica; Salmonella enterica subsp. enterica MTGDNTLITSHGINRRDFMKLCAALAATMGLSSKAAAEMAESVSNPQRPPVIWIGAQECT WP_000145429.1 hydrogenase 2 small subunit [Salmonella enterica] Length: 372\nNumber of predicted TMHs: 1\nExp number of AAs in TMHs: 24.51309\nExp number, first 60 AAs: 0.2682\nTotal prob of N-in: 0.14402\noutside 1 331\nTMhelix 332 354\ninside 355 372

21119 GCF_001355875.1_8490_5_24 Salmonella enterica subsp. enterica serovar Typhi Proteobacteria; Gammaproteobacteria; Enterobacterales; Enterobacteriaceae; Salmonella; Salmonella enterica; Salmonella enterica subsp. enterica MTGDNTLITSHGINRRDFMKLCAALAATMGLSSKAAAEMAESVSNPQRPPVIWIGAQECT WP_000145429.1 hydrogenase 2 small subunit [Salmonella enterica] Length: 372\nNumber of predicted TMHs: 1\nExp number of AAs in TMHs: 24.51309\nExp number, first 60 AAs: 0.2682\nTotal prob of N-in: 0.14402\noutside 1 331\nTMhelix 332 354\ninside 355 372

21120 GCF_001355935.1_8490_6_55 Salmonella enterica subsp. enterica serovar Typhi Proteobacteria; Gammaproteobacteria; Enterobacterales; Enterobacteriaceae; Salmonella; Salmonella enterica; Salmonella enterica subsp. enterica MTGDNTLITSHGINRRDFMKLCAALAATMGLSSKAAAEMAESVSNPQRPPVIWIGAQECT WP_000145429.1 hydrogenase 2 small subunit [Salmonella enterica] Length: 372\nNumber of predicted TMHs: 1\nExp number of AAs in TMHs: 24.51309\nExp number, first 60 AAs: 0.2682\nTotal prob of N-in: 0.14402\noutside 1 331\nTMhelix 332 354\ninside 355 372

21121 GCF_001355895.1_8447_8_13 Salmonella enterica subsp. enterica serovar Typhi Proteobacteria; Gammaproteobacteria; Enterobacterales; Enterobacteriaceae; Salmonella; Salmonella enterica; Salmonella enterica subsp. enterica MTGDNTLITSHGINRRDFMKLCAALAATMGLSSKAAAEMAESVSNPQRPPVIWIGAQECT WP_000145429.1 hydrogenase 2 small subunit [Salmonella enterica] Length: 372\nNumber of predicted TMHs: 1\nExp number of AAs in TMHs: 24.51309\nExp number, first 60 AAs: 0.2682\nTotal prob of N-in: 0.14402\noutside 1 331\nTMhelix 332 354\ninside 355 372

21122 GCF_001328695.1_7468_7_60 Salmonella enterica subsp. enterica serovar Typhi Proteobacteria; Gammaproteobacteria; Enterobacterales; Enterobacteriaceae; Salmonella; Salmonella enterica; Salmonella enterica subsp. enterica MTGDNTLITSHGINRRDFMKLCAALAATMGLSSKAAAEMAESVSNPQRPPVIWIGAQECT WP_000145429.1 hydrogenase 2 small subunit [Salmonella enterica] Length: 372\nNumber of predicted TMHs: 1\nExp number of AAs in TMHs: 24.51309\nExp number, first 60 AAs: 0.2682\nTotal prob of N-in: 0.14402\noutside 1 331\nTMhelix 332 354\ninside 355 372

21123 GCF_001239785.1_10425_1_57 Salmonella enterica subsp. enterica serovar Typhi Proteobacteria; Gammaproteobacteria; Enterobacterales; Enterobacteriaceae; Salmonella; Salmonella enterica; Salmonella enterica subsp. enterica MTGDNTLITSHGINRRDFMKLCAALAATMGLSSKAAAEMAESVSNPQRPPVIWIGAQECT WP_000145429.1 hydrogenase 2 small subunit [Salmonella enterica] Length: 372\nNumber of predicted TMHs: 1\nExp number of AAs in TMHs: 24.51309\nExp number, first 60 AAs: 0.2682\nTotal prob of N-in: 0.14402\noutside 1 331\nTMhelix 332 354\ninside 355 372

21124 GCF_001355975.1_8490_6_37 Salmonella enterica subsp. enterica serovar Typhi Proteobacteria; Gammaproteobacteria; Enterobacterales; Enterobacteriaceae; Salmonella; Salmonella enterica; Salmonella enterica subsp. enterica MTGDNTLITSHGINRRDFMKLCAALAATMGLSSKAAAEMAESVSNPQRPPVIWIGAQECT WP_000145429.1 hydrogenase 2 small subunit [Salmonella enterica] Length: 372\nNumber of predicted TMHs: 1\nExp number of AAs in TMHs: 24.51309\nExp number, first 60 AAs: 0.2682\nTotal prob of N-in: 0.14402\noutside 1 331\nTMhelix 332 354\ninside 355 372

21125 GCF_001356015.1_8490_6_46 Salmonella enterica subsp. enterica serovar Typhi Proteobacteria; Gammaproteobacteria; Enterobacterales; Enterobacteriaceae; Salmonella; Salmonella enterica; Salmonella enterica subsp. enterica MTGDNTLITSHGINRRDFMKLCAALAATMGLSSKAAAEMAESVSNPQRPPVIWIGAQECT WP_000145429.1 hydrogenase 2 small subunit [Salmonella enterica] Length: 372\nNumber of predicted TMHs: 1\nExp number of AAs in TMHs: 24.51309\nExp number, first 60 AAs: 0.2682\nTotal prob of N-in: 0.14402\noutside 1 331\nTMhelix 332 354\ninside 355 372

21126 GCF_001356055.1_8447_8_1Salmonella enterica subsp. enterica serovar Typhi Proteobacteria; Gammaproteobacteria; Enterobacterales; Enterobacteriaceae; Salmonella; Salmonella enterica; Salmonella enterica subsp. enterica MTGDNTLITSHGINRRDFMKLCAALAATMGLSSKAAAEMAESVSNPQRPPVIWIGAQECT WP_000145429.1 hydrogenase 2 small subunit [Salmonella enterica] Length: 372\nNumber of predicted TMHs: 1\nExp

number of AAs in TMHs: 24.51309\nExp number, first 60 AAs: 0.2682\nTotal prob of N-in: 0.14402\noutside 1 331\nTMhelix 332 354\ninside 355 372

21127 GCF_001356095.1_8490_5_7Salmonella enterica subsp. enterica serovar Typhi Proteobacteria; Gammaproteobacteria; Enterobacterales; Enterobacteriaceae; Salmonella; Salmonella enterica; Salmonella enterica subsp. enterica MTGDNTLITSHGINRRDFMKLCAALAATMGLSSKAAAEMAESVSNPQRPPVIWIGAQECT WP_000145429.1 hydrogenase 2 small subunit [Salmonella enterica] Length: 372\nNumber of predicted TMHs: 1\nExp number of AAs in TMHs: 24.51309\nExp number, first 60 AAs: 0.2682\nTotal prob of N-in: 0.14402\noutside 1 331\nTMhelix 332 354\ninside 355 372

21128 GCF_001239845.1_10425_1_58 Salmonella enterica subsp. enterica serovar Typhi Proteobacteria; Gammaproteobacteria; Enterobacterales; Enterobacteriaceae; Salmonella; Salmonella enterica; Salmonella enterica subsp. enterica MTGDNTLITSHGINRRDFMKLCAALAATMGLSSKAAAEMAESVSNPQRPPVIWIGAQECT WP_000145429.1 hydrogenase 2 small subunit [Salmonella enterica] Length: 372\nNumber of predicted TMHs: 1\nExp number of AAs in TMHs: 24.51309\nExp number, first 60 AAs: 0.2682\nTotal prob of N-in: 0.14402\noutside 1 331\nTMhelix 332 354\ninside 355 372

21129 GCF_001239925.1_10425_1_26 Salmonella enterica subsp. enterica serovar Typhi Proteobacteria; Gammaproteobacteria; Enterobacterales; Enterobacteriaceae; Salmonella; Salmonella enterica; Salmonella enterica subsp. enterica MTGDNTLITSHGINRRDFMKLCAALAATMGLSSKAAAEMAESVSNPQRPPVIWIGAQECT WP_000145429.1 hydrogenase 2 small subunit [Salmonella enterica] Length: 372\nNumber of predicted TMHs: 1\nExp number of AAs in TMHs: 24.51309\nExp number, first 60 AAs: 0.2682\nTotal prob of N-in: 0.14402\noutside 1 331\nTMhelix 332 354\ninside 355 372

21130 GCF_001239945.1_10425_1_42 Salmonella enterica subsp. enterica serovar Typhi Proteobacteria; Gammaproteobacteria; Enterobacterales; Enterobacteriaceae; Salmonella; Salmonella enterica; Salmonella enterica subsp. enterica MTGDNTLITSHGINRRDFMKLCAALAATMGLSSKAAAEMAESVSNPQRPPVIWIGAQECT WP_000145429.1 hydrogenase 2 small subunit [Salmonella enterica] Length: 372\nNumber of predicted TMHs: 1\nExp number of AAs in TMHs: 24.51309\nExp number, first 60 AAs: 0.2682\nTotal prob of N-in: 0.14402\noutside 1 331\nTMhelix 332 354\ninside 355 372

21131 GCF_001239965.1_10425_1_33 Salmonella enterica subsp. enterica serovar Typhi Proteobacteria; Gammaproteobacteria; Enterobacterales; Enterobacteriaceae; Salmonella; Salmonella enterica; Salmonella enterica subsp. enterica MTGDNTLITSHGINRRDFMKLCAALAATMGLSSKAAAEMAESVSNPQRPPVIWIGAQECT WP_000145429.1 hydrogenase 2 small subunit [Salmonella enterica] Length: 372\nNumber of predicted TMHs: 1\nExp number of AAs in TMHs: 24.51309\nExp number, first 60 AAs: 0.2682\nTotal prob of N-in: 0.14402\noutside 1 331\nTMhelix 332 354\ninside 355 372

21132 GCF_001240005.1_10425_1_76 Salmonella enterica subsp. enterica serovar Typhi Proteobacteria; Gammaproteobacteria; Enterobacterales; Enterobacteriaceae; Salmonella; Salmonella enterica; Salmonella enterica subsp. enterica MTGDNTLITSHGINRRDFMKLCAALAATMGLSSKAAAEMAESVSNPQRPPVIWIGAQECT WP_000145429.1 hydrogenase 2 small subunit [Salmonella enterica] Length: 372\nNumber of predicted TMHs: 1\nExp number of AAs in TMHs: 24.51309\nExp number, first 60 AAs: 0.2682\nTotal prob of N-in: 0.14402\noutside 1 331\nTMhelix 332 354\ninside 355 372

21133 GCF_001240045.1_10425_1_51 Salmonella enterica subsp. enterica serovar Typhi Proteobacteria; Gammaproteobacteria; Enterobacterales; Enterobacteriaceae; Salmonella; Salmonella enterica; Salmonella enterica subsp. enterica MTGDNTLITSHGINRRDFMKLCAALAATMGLSSKAAAEMAESVSNPQRPPVIWIGAQECT WP_000145429.1 hydrogenase 2 small subunit [Salmonella enterica] Length: 372\nNumber of predicted TMHs: 1\nExp number of AAs in TMHs: 24.51309\nExp number, first 60 AAs: 0.2682\nTotal prob of N-in: 0.14402\noutside 1 331\nTMhelix 332 354\ninside 355 372

21134 GCF_001240085.1_10425_1_85 Salmonella enterica subsp. enterica serovar Typhi Proteobacteria; Gammaproteobacteria; Enterobacterales; Enterobacteriaceae; Salmonella; Salmonella enterica; Salmonella enterica subsp. enterica MTGDNTLITSHGINRRDFMKLCAALAATMGLSSKAAAEMAESVSNPQRPPVIWIGAQECT WP_000145429.1 hydrogenase 2 small subunit [Salmonella enterica] Length: 372\nNumber of predicted TMHs: 1\nExp number of AAs in TMHs: 24.51309\nExp number, first 60 AAs: 0.2682\nTotal prob of N-in: 0.14402\noutside 1 331\nTMhelix 332 354\ninside 355 372

21135 GCF_001241145.1_10425_1_37 Salmonella enterica subsp. enterica serovar Typhi Proteobacteria; Gammaproteobacteria; Enterobacterales; Enterobacteriaceae; Salmonella; Salmonella enterica;

Salmonella enterica subsp. enterica MTGDNTLITSHGINRRDFMKLCAALAATMGLSSKAAAEMAESVSNPQRPPVIWIGAQECT WP_000145429.1 hydrogenase 2 small subunit [Salmonella enterica] Length: 372\nNumber of predicted TMHs: 1\nExp number of AAs in TMHs: 24.51309\nExp number, first 60 AAs: 0.2682\nTotal prob of N-in: 0.14402\noutside 1 331\nTMhelix 332 354\ninside 355 372

21136 GCF_001240865.1_10425_1_48 Salmonella enterica subsp. enterica serovar Typhi Proteobacteria; Gammaproteobacteria; Enterobacterales; Enterobacteriaceae; Salmonella; Salmonella enterica; Salmonella enterica subsp. enterica MTGDNTLITSHGINRRDFMKLCAALAATMGLSSKAAAEMAESVSNPQRPPVIWIGAQECT WP_000145429.1 hydrogenase 2 small subunit [Salmonella enterica] Length: 372\nNumber of predicted TMHs: 1\nExp number of AAs in TMHs: 24.51309\nExp number, first 60 AAs: 0.2682\nTotal prob of N-in: 0.14402\noutside 1 331\nTMhelix 332 354\ninside 355 372

21137 GCF_001241445.1_10425_1_40 Salmonella enterica subsp. enterica serovar Typhi Proteobacteria; Gammaproteobacteria; Enterobacterales; Enterobacteriaceae; Salmonella; Salmonella enterica; Salmonella enterica subsp. enterica MTGDNTLITSHGINRRDFMKLCAALAATMGLSSKAAAEMAESVSNPQRPPVIWIGAQECT WP_000145429.1 hydrogenase 2 small subunit [Salmonella enterica] Length: 372\nNumber of predicted TMHs: 1\nExp number of AAs in TMHs: 24.51309\nExp number, first 60 AAs: 0.2682\nTotal prob of N-in: 0.14402\noutside 1 331\nTMhelix 332 354\ninside 355 372

21138 GCF_001241885.1_10425_1_1 Salmonella enterica subsp. enterica serovar Typhi Proteobacteria; Gammaproteobacteria; Enterobacterales; Enterobacteriaceae; Salmonella; Salmonella enterica; Salmonella enterica subsp. enterica MTGDNTLITSHGINRRDFMKLCAALAATMGLSSKAAAEMAESVSNPQRPPVIWIGAQECT WP_000145429.1 hydrogenase 2 small subunit [Salmonella enterica] Length: 372\nNumber of predicted TMHs: 1\nExp number of AAs in TMHs: 24.51309\nExp number, first 60 AAs: 0.2682\nTotal prob of N-in: 0.14402\noutside 1 331\nTMhelix 332 354\ninside 355 372

21139 GCF_001242025.1_10425_1_67 Salmonella enterica subsp. enterica serovar Typhi Proteobacteria; Gammaproteobacteria; Enterobacterales; Enterobacteriaceae; Salmonella; Salmonella enterica; Salmonella enterica subsp. enterica MTGDNTLITSHGINRRDFMKLCAALAATMGLSSKAAAEMAESVSNPQRPPVIWIGAQECT WP_000145429.1 hydrogenase 2 small subunit [Salmonella enterica] Length: 372\nNumber of predicted TMHs: 1\nExp number of AAs in TMHs: 24.51309\nExp number, first 60 AAs: 0.2682\nTotal prob of N-in: 0.14402\noutside 1 331\nTMhelix 332 354\ninside 355 372

21140 GCF_001242455.1_10425_1_31 Salmonella enterica subsp. enterica serovar Typhi Proteobacteria; Gammaproteobacteria; Enterobacterales; Enterobacteriaceae; Salmonella; Salmonella enterica; Salmonella enterica subsp. enterica MTGDNTLITSHGINRRDFMKLCAALAATMGLSSKAAAEMAESVSNPQRPPVIWIGAQECT WP_000145429.1 hydrogenase 2 small subunit [Salmonella enterica] Length: 372\nNumber of predicted TMHs: 1\nExp number of AAs in TMHs: 24.51309\nExp number, first 60 AAs: 0.2682\nTotal prob of N-in: 0.14402\noutside 1 331\nTMhelix 332 354\ninside 355 372

21141 GCF_001242595.1_10425_1_11 Salmonella enterica subsp. enterica serovar Typhi Proteobacteria; Gammaproteobacteria; Enterobacterales; Enterobacteriaceae; Salmonella; Salmonella enterica; Salmonella enterica subsp. enterica MTGDNTLITSHGINRRDFMKLCAALAATMGLSSKAAAEMAESVSNPQRPPVIWIGAQECT WP_000145429.1 hydrogenase 2 small subunit [Salmonella enterica] Length: 372\nNumber of predicted TMHs: 1\nExp number of AAs in TMHs: 24.51309\nExp number, first 60 AAs: 0.2682\nTotal prob of N-in: 0.14402\noutside 1 331\nTMhelix 332 354\ninside 355 372

21142 GCF_001355595.1_8490_6_41 Salmonella enterica subsp. enterica serovar Typhi Proteobacteria; Gammaproteobacteria; Enterobacterales; Enterobacteriaceae; Salmonella; Salmonella enterica; Salmonella enterica subsp. enterica MTGDNTLITSHGINRRDFMKLCAALAATMGLSSKAAAEMAESVSNPQRPPVIWIGAQECT WP_000145429.1 hydrogenase 2 small subunit [Salmonella enterica] Length: 372\nNumber of predicted TMHs: 1\nExp number of AAs in TMHs: 24.51309\nExp number, first 60 AAs: 0.2682\nTotal prob of N-in: 0.14402\noutside 1 331\nTMhelix 332 354\ninside 355 372

21143 GCF_001355635.1_8490_6_61 Salmonella enterica subsp. enterica serovar Typhi Proteobacteria; Gammaproteobacteria; Enterobacterales; Enterobacteriaceae; Salmonella; Salmonella enterica; Salmonella enterica subsp. enterica MTGDNTLITSHGINRRDFMKLCAALAATMGLSSKAAAEMAESVSNPQRPPVIWIGAQECT WP_000145429.1 hydrogenase 2 small subunit [Salmonella enterica] Length: 372\nNumber of predicted TMHs: 1\nExp number of AAs in TMHs: 24.51309\nExp number, first 60 AAs: 0.2682\nTotal prob of N-in: 0.14402\noutside 1 331\nTMhelix 332 354\ninside 355 372

21144 GCF_001302605.1_ASM130260v1 Salmonella enterica subsp. enterica serovar Typhi Proteobacteria; Gammaproteobacteria; Enterobacterales; Enterobacteriaceae; Salmonella; Salmonella enterica; Salmonella enterica subsp. enterica MTGDNTLITSHGINRRDFMKLCAALAATMGLSSKAAAEMAESVSNPQRPPVIWIGAQECT WP_000145429.1 hydrogenase 2 small subunit [Salmonella enterica] Length: 372\nNumber of predicted TMHs: 1\nExp number of AAs in TMHs: 24.51309\nExp number, first 60 AAs: 0.2682\nTotal prob of N-in: 0.14402\noutside 1 331\nTMhelix 332 354\ninside 355 372

21145 GCF_001328655.1_7468_7_71 Salmonella enterica subsp. enterica serovar Typhi Proteobacteria; Gammaproteobacteria; Enterobacterales; Enterobacteriaceae; Salmonella; Salmonella enterica; Salmonella enterica subsp. enterica MTGDNTLITSHGINRRDFMKLCAALAATMGLSSKAAAEMAESVSNPQRPPVIWIGAQECT WP_000145429.1 hydrogenase 2 small subunit [Salmonella enterica] Length: 372\nNumber of predicted TMHs: 1\nExp number of AAs in TMHs: 24.51309\nExp number, first 60 AAs: 0.2682\nTotal prob of N-in: 0.14402\noutside 1 331\nTMhelix 332 354\ninside 355 372

21146 GCF_001317585.1_9475_4_50 Salmonella enterica subsp. enterica serovar Typhi Proteobacteria; Gammaproteobacteria; Enterobacterales; Enterobacteriaceae; Salmonella; Salmonella enterica; Salmonella enterica subsp. enterica MTGDNTLITSHGINRRDFMKLCAALAATMGLSSKAAAEMAESVSNPQRPPVIWIGAQECT WP_000145429.1 hydrogenase 2 small subunit [Salmonella enterica] Length: 372\nNumber of predicted TMHs: 1\nExp number of AAs in TMHs: 24.51309\nExp number, first 60 AAs: 0.2682\nTotal prob of N-in: 0.14402\noutside 1 331\nTMhelix 332 354\ninside 355 372

21147 GCF_001317625.1_9475_4_52 Salmonella enterica subsp. enterica serovar Typhi Proteobacteria; Gammaproteobacteria; Enterobacterales; Enterobacteriaceae; Salmonella; Salmonella enterica; Salmonella enterica subsp. enterica MTGDNTLITSHGINRRDFMKLCAALAATMGLSSKAAAEMAESVSNPQRPPVIWIGAQECT WP_000145429.1 hydrogenase 2 small subunit [Salmonella enterica] Length: 372\nNumber of predicted TMHs: 1\nExp number of AAs in TMHs: 24.51309\nExp number, first 60 AAs: 0.2682\nTotal prob of N-in: 0.14402\noutside 1 331\nTMhelix 332 354\ninside 355 372

21148 GCF_001317665.1_9475_4_57 Salmonella enterica subsp. enterica serovar Typhi Proteobacteria; Gammaproteobacteria; Enterobacterales; Enterobacteriaceae; Salmonella; Salmonella enterica; Salmonella enterica subsp. enterica MTGDNTLITSHGINRRDFMKLCAALAATMGLSSKAAAEMAESVSNPQRPPVIWIGAQECT WP_000145429.1 hydrogenase 2 small subunit [Salmonella enterica] Length: 372\nNumber of predicted TMHs: 1\nExp number of AAs in TMHs: 24.51309\nExp number, first 60 AAs: 0.2682\nTotal prob of N-in: 0.14402\noutside 1 331\nTMhelix 332 354\ninside 355 372

21149 GCF_001328735.1_7468_7_79 Salmonella enterica subsp. enterica serovar Typhi Proteobacteria; Gammaproteobacteria; Enterobacterales; Enterobacteriaceae; Salmonella; Salmonella enterica; Salmonella enterica subsp. enterica MTGDNTLITSHGINRRDFMKLCAALAATMGLSSKAAAEMAESVSNPQRPPVIWIGAQECT WP_000145429.1 hydrogenase 2 small subunit [Salmonella enterica] Length: 372\nNumber of predicted TMHs: 1\nExp number of AAs in TMHs: 24.51309\nExp number, first 60 AAs: 0.2682\nTotal prob of N-in: 0.14402\noutside 1 331\nTMhelix 332 354\ninside 355 372

21150 GCF_001328775.1_7468_7_50 Salmonella enterica subsp. enterica serovar Typhi Proteobacteria; Gammaproteobacteria; Enterobacterales; Enterobacteriaceae; Salmonella; Salmonella enterica; Salmonella enterica subsp. enterica MTGDNTLITSHGINRRDFMKLCAALAATMGLSSKAAAEMAESVSNPQRPPVIWIGAQECT WP_000145429.1 hydrogenase 2 small subunit [Salmonella enterica] Length: 372\nNumber of predicted TMHs: 1\nExp number of AAs in TMHs: 24.51309\nExp number, first 60 AAs: 0.2682\nTotal prob of N-in: 0.14402\noutside 1 331\nTMhelix 332 354\ninside 355 372

21151 GCF_001328815.1_7468_7_77 Salmonella enterica subsp. enterica serovar Typhi Proteobacteria; Gammaproteobacteria; Enterobacterales; Enterobacteriaceae; Salmonella; Salmonella enterica; Salmonella enterica subsp. enterica MTGDNTLITSHGINRRDFMKLCAALAATMGLSSKAAAEMAESVSNPQRPPVIWIGAQECT WP_000145429.1 hydrogenase 2 small subunit [Salmonella enterica] Length: 372\nNumber of predicted TMHs: 1\nExp number of AAs in TMHs: 24.51309\nExp number, first 60 AAs: 0.2682\nTotal prob of N-in: 0.14402\noutside 1 331\nTMhelix 332 354\ninside 355 372

21152 GCF_001328615.1_7468_7_91 Salmonella enterica subsp. enterica serovar Typhi Proteobacteria; Gammaproteobacteria; Enterobacterales; Enterobacteriaceae; Salmonella; Salmonella enterica; Salmonella enterica subsp. enterica MTGDNTLITSHGINRRDFMKLCAALAATMGLSSKAATEMAESVSNPQRPPVIWIGAQECT WP_001782304.1 hydrogenase 2 small subunit [Salmonella enterica] Length: 372\nNumber of

predicted TMHs:  1\nExp number of AAs in TMHs: 24.41173\nExp number, first 60 AAs:  0.16124\nTotal prob of N-in: 0.13947\noutside   1   331\nTMhelix   332   354\ninside   355   372

21153 GCF_001328855.1_7468_7_69 Salmonella enterica subsp. enterica serovar Typhi Proteobacteria; Gammaproteobacteria; Enterobacterales; Enterobacteriaceae; Salmonella; Salmonella enterica; Salmonella enterica subsp. enterica MTGDNTLITSHGINRRDFMKLCAALAATMGLSSKAAAEMAESVSNPQRPPVIWIGAQECT WP_000145429.1 hydrogenase 2 small subunit [Salmonella enterica] Length: 372\nNumber of predicted TMHs:  1\nExp number of AAs in TMHs: 24.51309\nExp number, first 60 AAs:  0.2682\nTotal prob of N-in: 0.14402\noutside   1   331\nTMhelix   332   354\ninside   355   372

21154 GCF_001350315.1_8616_4_48 Salmonella enterica subsp. enterica serovar Typhi Proteobacteria; Gammaproteobacteria; Enterobacterales; Enterobacteriaceae; Salmonella; Salmonella enterica; Salmonella enterica subsp. enterica MTGDNTLITSHGINRRDFMKLCAALAATMGLSSKAAAEMAESVSNPQRPPVIWIGAQECT WP_000145429.1 hydrogenase 2 small subunit [Salmonella enterica] Length: 372\nNumber of predicted TMHs:  1\nExp number of AAs in TMHs: 24.51309\nExp number, first 60 AAs:  0.2682\nTotal prob of N-in: 0.14402\noutside   1   331\nTMhelix   332   354\ninside   355   372

21155 GCF_001350355.1_8616_4_40 Salmonella enterica subsp. enterica serovar Typhi Proteobacteria; Gammaproteobacteria; Enterobacterales; Enterobacteriaceae; Salmonella; Salmonella enterica; Salmonella enterica subsp. enterica MTGDNTLITSHGINRRDFMKLCAALAATMGLSSKAAAEMAESVSNPQRPPVIWIGAQECT WP_000145429.1 hydrogenase 2 small subunit [Salmonella enterica] Length: 372\nNumber of predicted TMHs:  1\nExp number of AAs in TMHs: 24.51309\nExp number, first 60 AAs:  0.2682\nTotal prob of N-in: 0.14402\noutside   1   331\nTMhelix   332   354\ninside   355   372

21156 GCF_001350395.1_8616_4_41 Salmonella enterica subsp. enterica serovar Typhi Proteobacteria; Gammaproteobacteria; Enterobacterales; Enterobacteriaceae; Salmonella; Salmonella enterica; Salmonella enterica subsp. enterica MTGDNTLITSHGINRRDFMKLCAALAATMGLSSKAAAEMAESVSNPQRPPVIWIGAQECT WP_000145429.1 hydrogenase 2 small subunit [Salmonella enterica] Length: 372\nNumber of predicted TMHs:  1\nExp number of AAs in TMHs: 24.51309\nExp number, first 60 AAs:  0.2682\nTotal prob of N-in: 0.14402\noutside   1   331\nTMhelix   332   354\ninside   355   372

21157 GCF_001350435.1_8616_4_50 Salmonella enterica subsp. enterica serovar Typhi Proteobacteria; Gammaproteobacteria; Enterobacterales; Enterobacteriaceae; Salmonella; Salmonella enterica; Salmonella enterica subsp. enterica MTGDNTLITSHGINRRDFMKLCAALAATMGLSSKAAAEMAESVSNPQRPPVIWIGAQECT WP_000145429.1 hydrogenase 2 small subunit [Salmonella enterica] Length: 372\nNumber of predicted TMHs:  1\nExp number of AAs in TMHs: 24.51309\nExp number, first 60 AAs:  0.2682\nTotal prob of N-in: 0.14402\noutside   1   331\nTMhelix   332   354\ninside   355   372

21158 GCF_001355395.1_8525_3_41 Salmonella enterica subsp. enterica serovar Typhi Proteobacteria; Gammaproteobacteria; Enterobacterales; Enterobacteriaceae; Salmonella; Salmonella enterica; Salmonella enterica subsp. enterica MTGDNTLITSHGINRRDFMKLCAALAATMGLSSKAAAEMAESVSNPQRPPVIWIGAQECT WP_000145429.1 hydrogenase 2 small subunit [Salmonella enterica] Length: 372\nNumber of predicted TMHs:  1\nExp number of AAs in TMHs: 24.51309\nExp number, first 60 AAs:  0.2682\nTotal prob of N-in: 0.14402\noutside   1   331\nTMhelix   332   354\ninside   355   372

21159 GCF_001355515.1_8525_3_51 Salmonella enterica subsp. enterica serovar Typhi Proteobacteria; Gammaproteobacteria; Enterobacterales; Enterobacteriaceae; Salmonella; Salmonella enterica; Salmonella enterica subsp. enterica MTGDNTLITSHGINRRDFMKLCAALAATMGLSSKAAAEMAESVSNPQRPPVIWIGAQECT WP_000145429.1 hydrogenase 2 small subunit [Salmonella enterica] Length: 372\nNumber of predicted TMHs:  1\nExp number of AAs in TMHs: 24.51309\nExp number, first 60 AAs:  0.2682\nTotal prob of N-in: 0.14402\noutside   1   331\nTMhelix   332   354\ninside   355   372

21160 GCF_001355475.1_8525_3_40 Salmonella enterica subsp. enterica serovar Typhi Proteobacteria; Gammaproteobacteria; Enterobacterales; Enterobacteriaceae; Salmonella; Salmonella enterica; Salmonella enterica subsp. enterica MTGDNTLITSHGINRRDFMKLCAALAATMGLSSKAAAEMAESVSNPQRPPVIWIGAQECT WP_000145429.1 hydrogenase 2 small subunit [Salmonella enterica] Length: 372\nNumber of predicted TMHs:  1\nExp number of AAs in TMHs: 24.51309\nExp number, first 60 AAs:  0.2682\nTotal prob of N-in: 0.14402\noutside   1   331\nTMhelix   332   354\ninside   355   372

21161 GCF_001355435.1_8525_3_60 Salmonella enterica subsp. enterica serovar Typhi Proteobacteria; Gammaproteobacteria; Enterobacterales; Enterobacteriaceae; Salmonella; Salmonella enterica;

Salmonella enterica subsp. enterica MTGDNTLITSHGINRRDFMKLCAALAATMGLSSKAAAEMAESVSNPQRPPVIWIGAQECT WP_000145429.1 hydrogenase 2 small subunit [Salmonella enterica] Length: 372\nNumber of predicted TMHs: 1\nExp number of AAs in TMHs: 24.51309\nExp number, first 60 AAs: 0.2682\nTotal prob of N-in: 0.14402\noutside 1 331\nTMhelix 332 354\ninside 355 372

21162 GCF_001355555.1_8525_3_47 Salmonella enterica subsp. enterica serovar Typhi Proteobacteria; Gammaproteobacteria; Enterobacterales; Enterobacteriaceae; Salmonella; Salmonella enterica; Salmonella enterica subsp. enterica MTGDNTLITSHGINRRDFMKLCAALAATMGLSSKAAAEMAESVSNPQRPPVIWIGAQECT WP_000145429.1 hydrogenase 2 small subunit [Salmonella enterica] Length: 372\nNumber of predicted TMHs: 1\nExp number of AAs in TMHs: 24.51309\nExp number, first 60 AAs: 0.2682\nTotal prob of N-in: 0.14402\noutside 1 331\nTMhelix 332 354\ninside 355 372

21163 GCF_001356135.1_8490_5_17 Salmonella enterica subsp. enterica serovar Typhi Proteobacteria; Gammaproteobacteria; Enterobacterales; Enterobacteriaceae; Salmonella; Salmonella enterica; Salmonella enterica subsp. enterica MTGDNTLITSHGINRRDFMKLCAALAATMGLSSKAAAEMAESVSNPQRPPVIWIGAQECT WP_000145429.1 hydrogenase 2 small subunit [Salmonella enterica] Length: 372\nNumber of predicted TMHs: 1\nExp number of AAs in TMHs: 24.51309\nExp number, first 60 AAs: 0.2682\nTotal prob of N-in: 0.14402\noutside 1 331\nTMhelix 332 354\ninside 355 372

21164 GCF_001356175.1_8490_5_19 Salmonella enterica subsp. enterica serovar Typhi Proteobacteria; Gammaproteobacteria; Enterobacterales; Enterobacteriaceae; Salmonella; Salmonella enterica; Salmonella enterica subsp. enterica MTGDNTLITSHGINRRDFMKLCAALAATMGLSSKAAAEMAESVSNPQRPPVIWIGAQECT WP_000145429.1 hydrogenase 2 small subunit [Salmonella enterica] Length: 372\nNumber of predicted TMHs: 1\nExp number of AAs in TMHs: 24.51309\nExp number, first 60 AAs: 0.2682\nTotal prob of N-in: 0.14402\noutside 1 331\nTMhelix 332 354\ninside 355 372

21165 GCF_001356215.1_8490_6_44 Salmonella enterica subsp. enterica serovar Typhi Proteobacteria; Gammaproteobacteria; Enterobacterales; Enterobacteriaceae; Salmonella; Salmonella enterica; Salmonella enterica subsp. enterica MTGDNTLITSHGINRRDFMKLCAALAATMGLSSKAAAEMAESVSNPQRPPVIWIGAQECT WP_000145429.1 hydrogenase 2 small subunit [Salmonella enterica] Length: 372\nNumber of predicted TMHs: 1\nExp number of AAs in TMHs: 24.51309\nExp number, first 60 AAs: 0.2682\nTotal prob of N-in: 0.14402\noutside 1 331\nTMhelix 332 354\ninside 355 372

21166 GCF_001356255.1_8447_8_17 Salmonella enterica subsp. enterica serovar Typhi Proteobacteria; Gammaproteobacteria; Enterobacterales; Enterobacteriaceae; Salmonella; Salmonella enterica; Salmonella enterica subsp. enterica MTGDNTLITSHGINRRDFMKLCAALAATMGLSSKAAAEMAESVSNPQRPPVIWIGAQECT WP_000145429.1 hydrogenase 2 small subunit [Salmonella enterica] Length: 372\nNumber of predicted TMHs: 1\nExp number of AAs in TMHs: 24.51309\nExp number, first 60 AAs: 0.2682\nTotal prob of N-in: 0.14402\noutside 1 331\nTMhelix 332 354\ninside 355 372

21167 GCF_001356295.1_8490_6_70 Salmonella enterica subsp. enterica serovar Typhi Proteobacteria; Gammaproteobacteria; Enterobacterales; Enterobacteriaceae; Salmonella; Salmonella enterica; Salmonella enterica subsp. enterica MTGDNTLITSHGINRRDFMKLCAALAATMGLSSKAAAEMAESVSNPQRPPVIWIGAQECT WP_000145429.1 hydrogenase 2 small subunit [Salmonella enterica] Length: 372\nNumber of predicted TMHs: 1\nExp number of AAs in TMHs: 24.51309\nExp number, first 60 AAs: 0.2682\nTotal prob of N-in: 0.14402\noutside 1 331\nTMhelix 332 354\ninside 355 372

21168 GCF_001356335.1_8447_8_18 Salmonella enterica subsp. enterica serovar Typhi Proteobacteria; Gammaproteobacteria; Enterobacterales; Enterobacteriaceae; Salmonella; Salmonella enterica; Salmonella enterica subsp. enterica MTGDNTLITSHGINRRDFMKLCAALAATMGLSSKAAAEMAESVSNPQRPPVIWIGAQECT WP_000145429.1 hydrogenase 2 small subunit [Salmonella enterica] Length: 372\nNumber of predicted TMHs: 1\nExp number of AAs in TMHs: 24.51309\nExp number, first 60 AAs: 0.2682\nTotal prob of N-in: 0.14402\noutside 1 331\nTMhelix 332 354\ninside 355 372

21169 GCF_001356375.1_9475_6_8Salmonella enterica subsp. enterica serovar Typhi Proteobacteria; Gammaproteobacteria; Enterobacterales; Enterobacteriaceae; Salmonella; Salmonella enterica; Salmonella enterica subsp. enterica MTGDNTLITSHGINRRDFMKLCAALAATMGLSSKAAAEMAESVSNPQRPPVIWIGAQECT WP_000145429.1 hydrogenase 2 small subunit [Salmonella enterica] Length: 372\nNumber of predicted TMHs: 1\nExp number of AAs in TMHs: 24.51309\nExp number, first 60 AAs: 0.2682\nTotal prob of N-in: 0.14402\noutside 1 331\nTMhelix 332 354\ninside 355 372

21170	GCF_001356415.1_9475_6_20	Salmonella enterica subsp. enterica serovar Typhi	Proteobacteria; Gammaproteobacteria; Enterobacterales; Enterobacteriaceae; Salmonella; Salmonella enterica; Salmonella enterica subsp. enterica	MTGDNTLITSHGINRRDFMKLCAALAATMGLSSKAAAEMAESVSNPQRPPVIWIGAQECT	WP_000145429.1	hydrogenase 2 small subunit [Salmonella enterica]	Length: 372\nNumber of predicted TMHs:  1\nExp number of AAs in TMHs: 24.51309\nExp number, first 60 AAs:  0.2682\nTotal prob of N-in: 0.14402\noutside	1	331\nTMhelix	332	354\ninside	355	372

21171	GCF_001356495.1_9475_6_32	Salmonella enterica subsp. enterica serovar Typhi	Proteobacteria; Gammaproteobacteria; Enterobacterales; Enterobacteriaceae; Salmonella; Salmonella enterica; Salmonella enterica subsp. enterica	MTGDNTLITSHGINRRDFMKLCAALAATMGLSSKAAAEMAESVSNPQRPPVIWIGAQECT	WP_000145429.1	hydrogenase 2 small subunit [Salmonella enterica]	Length: 372\nNumber of predicted TMHs:  1\nExp number of AAs in TMHs: 24.51309\nExp number, first 60 AAs:  0.2682\nTotal prob of N-in: 0.14402\noutside	1	331\nTMhelix	332	354\ninside	355	372

21172	GCF_001356535.1_9475_6_37	Salmonella enterica subsp. enterica serovar Typhi	Proteobacteria; Gammaproteobacteria; Enterobacterales; Enterobacteriaceae; Salmonella; Salmonella enterica; Salmonella enterica subsp. enterica	MTGDNTLITSHGINRRDFMKLCAALAATMGLSSKAAAEMAESVSNPQRPPVIWIGAQECT	WP_000145429.1	hydrogenase 2 small subunit [Salmonella enterica]	Length: 372\nNumber of predicted TMHs:  1\nExp number of AAs in TMHs: 24.51309\nExp number, first 60 AAs:  0.2682\nTotal prob of N-in: 0.14402\noutside	1	331\nTMhelix	332	354\ninside	355	372

21173	GCF_001356455.1_9475_6_19	Salmonella enterica subsp. enterica serovar Typhi	Proteobacteria; Gammaproteobacteria; Enterobacterales; Enterobacteriaceae; Salmonella; Salmonella enterica; Salmonella enterica subsp. enterica	MTGDNTLITSHGINRRDFMKLCAALAATMGLSSKAAAEMAESVSNPQRPPVIWIGAQECT	WP_000145429.1	hydrogenase 2 small subunit [Salmonella enterica]	Length: 372\nNumber of predicted TMHs:  1\nExp number of AAs in TMHs: 24.51309\nExp number, first 60 AAs:  0.2682\nTotal prob of N-in: 0.14402\noutside	1	331\nTMhelix	332	354\ninside	355	372

21174	GCF_001356575.1_9475_6_53	Salmonella enterica subsp. enterica serovar Typhi	Proteobacteria; Gammaproteobacteria; Enterobacterales; Enterobacteriaceae; Salmonella; Salmonella enterica; Salmonella enterica subsp. enterica	MTGDNTLITSHGINRRDFMKLCAALAATMGLSSKAAAEMAESVSNPQRPPVIWIGAQECT	WP_000145429.1	hydrogenase 2 small subunit [Salmonella enterica]	Length: 372\nNumber of predicted TMHs:  1\nExp number of AAs in TMHs: 24.51309\nExp number, first 60 AAs:  0.2682\nTotal prob of N-in: 0.14402\noutside	1	331\nTMhelix	332	354\ninside	355	372

21175	GCF_001356635.1_9475_6_72	Salmonella enterica subsp. enterica serovar Typhi	Proteobacteria; Gammaproteobacteria; Enterobacterales; Enterobacteriaceae; Salmonella; Salmonella enterica; Salmonella enterica subsp. enterica	MTGDNTLITSHGINRRDFMKLCAALAATMGLSSKAAAEMAESVSNPQRPPVIWIGAQECT	WP_000145429.1	hydrogenase 2 small subunit [Salmonella enterica]	Length: 372\nNumber of predicted TMHs:  1\nExp number of AAs in TMHs: 24.51309\nExp number, first 60 AAs:  0.2682\nTotal prob of N-in: 0.14402\noutside	1	331\nTMhelix	332	354\ninside	355	372

21176	GCF_001356595.1_9475_6_51	Salmonella enterica subsp. enterica serovar Typhi	Proteobacteria; Gammaproteobacteria; Enterobacterales; Enterobacteriaceae; Salmonella; Salmonella enterica; Salmonella enterica subsp. enterica	MTGDNTLITSHGINRRDFMKLCAALAATMGLSSKAAAEMAESVSNPQRPPVIWIGAQECT	WP_000145429.1	hydrogenase 2 small subunit [Salmonella enterica]	Length: 372\nNumber of predicted TMHs:  1\nExp number of AAs in TMHs: 24.51309\nExp number, first 60 AAs:  0.2682\nTotal prob of N-in: 0.14402\noutside	1	331\nTMhelix	332	354\ninside	355	372

21177	GCF_001356675.1_9475_6_76	Salmonella enterica subsp. enterica serovar Typhi	Proteobacteria; Gammaproteobacteria; Enterobacterales; Enterobacteriaceae; Salmonella; Salmonella enterica; Salmonella enterica subsp. enterica	MTGDNTLITSHGINRRDFMKLCAALAATMGLSSKAAAEMAESVSNPQRPPVIWIGAQECT	WP_000145429.1	hydrogenase 2 small subunit [Salmonella enterica]	Length: 372\nNumber of predicted TMHs:  1\nExp number of AAs in TMHs: 24.51309\nExp number, first 60 AAs:  0.2682\nTotal prob of N-in: 0.14402\noutside	1	331\nTMhelix	332	354\ninside	355	372

21178	GCF_001356715.1_10592_2_20	Salmonella enterica subsp. enterica serovar Typhi	Proteobacteria; Gammaproteobacteria; Enterobacterales; Enterobacteriaceae; Salmonella; Salmonella enterica; Salmonella enterica subsp. enterica	MTGDNTLITSHGINRRDFMKLCAALAATMGLSSKAAAEMAESVSNPQRPPVIWIGAQECT	WP_000145429.1	hydrogenase 2 small subunit [Salmonella enterica]	Length: 372\nNumber of

predicted TMHs: 1\nExp number of AAs in TMHs: 24.51309\nExp number, first 60 AAs: 0.2682\nTotal prob of N-in: 0.14402\noutside 1 331\nTMhelix 332 354\ninside 355 372

21179 GCF_001356755.1_10592_2_28 Salmonella enterica subsp. enterica serovar Typhi Proteobacteria; Gammaproteobacteria; Enterobacterales; Enterobacteriaceae; Salmonella; Salmonella enterica; Salmonella enterica subsp. enterica MTGDNTLITSHGINRRDFMKLCAALAATMGLSSKAAAEMAESVSNPQRPPVIWIGAQECT WP_000145429.1 hydrogenase 2 small subunit [Salmonella enterica] Length: 372\nNumber of predicted TMHs: 1\nExp number of AAs in TMHs: 24.51309\nExp number, first 60 AAs: 0.2682\nTotal prob of N-in: 0.14402\noutside 1 331\nTMhelix 332 354\ninside 355 372

21180 GCF_001356795.1_10592_2_33 Salmonella enterica subsp. enterica serovar Typhi Proteobacteria; Gammaproteobacteria; Enterobacterales; Enterobacteriaceae; Salmonella; Salmonella enterica; Salmonella enterica subsp. enterica MTGDNTLITSHGINRRDFMKLCAALAATMGLSSKAAAEMAESVSNPQRPPVIWIGAQECT WP_000145429.1 hydrogenase 2 small subunit [Salmonella enterica] Length: 372\nNumber of predicted TMHs: 1\nExp number of AAs in TMHs: 24.51309\nExp number, first 60 AAs: 0.2682\nTotal prob of N-in: 0.14402\noutside 1 331\nTMhelix 332 354\ninside 355 372

21181 GCF_001356875.1_10592_2_25 Salmonella enterica subsp. enterica serovar Typhi Proteobacteria; Gammaproteobacteria; Enterobacterales; Enterobacteriaceae; Salmonella; Salmonella enterica; Salmonella enterica subsp. enterica MTGDNTLITSHGINRRDFMKLCAALAATMGLSSKAAAEMAESVSNPQRPPVIWIGAQECT WP_000145429.1 hydrogenase 2 small subunit [Salmonella enterica] Length: 372\nNumber of predicted TMHs: 1\nExp number of AAs in TMHs: 24.51309\nExp number, first 60 AAs: 0.2682\nTotal prob of N-in: 0.14402\noutside 1 331\nTMhelix 332 354\ninside 355 372

21182 GCF_001356915.1_10592_2_66 Salmonella enterica subsp. enterica serovar Typhi Proteobacteria; Gammaproteobacteria; Enterobacterales; Enterobacteriaceae; Salmonella; Salmonella enterica; Salmonella enterica subsp. enterica MTGDNTLITSHGINRRDFMKLCAALAATMGLSSKAAAEMAESVSNPQRPPVIWIGAQECT WP_000145429.1 hydrogenase 2 small subunit [Salmonella enterica] Length: 372\nNumber of predicted TMHs: 1\nExp number of AAs in TMHs: 24.51309\nExp number, first 60 AAs: 0.2682\nTotal prob of N-in: 0.14402\noutside 1 331\nTMhelix 332 354\ninside 355 372

21183 GCF_001357035.1_10593_2_43 Salmonella enterica subsp. enterica serovar Typhi Proteobacteria; Gammaproteobacteria; Enterobacterales; Enterobacteriaceae; Salmonella; Salmonella enterica; Salmonella enterica subsp. enterica MTGDNTLITSHGINRRDFMKLCAALAATMGLSSKAAAEMAESVSNPQRPPVIWIGAQECT WP_000145429.1 hydrogenase 2 small subunit [Salmonella enterica] Length: 372\nNumber of predicted TMHs: 1\nExp number of AAs in TMHs: 24.51309\nExp number, first 60 AAs: 0.2682\nTotal prob of N-in: 0.14402\noutside 1 331\nTMhelix 332 354\ninside 355 372

21184 GCF_001356955.1_10592_2_41 Salmonella enterica subsp. enterica serovar Typhi Proteobacteria; Gammaproteobacteria; Enterobacterales; Enterobacteriaceae; Salmonella; Salmonella enterica; Salmonella enterica subsp. enterica MTGDNTLITSHGINRRDFMKLCAALAATMGLSSKAAAEMAESVSNPQRPPVIWIGAQECT WP_000145429.1 hydrogenase 2 small subunit [Salmonella enterica] Length: 372\nNumber of predicted TMHs: 1\nExp number of AAs in TMHs: 24.51309\nExp number, first 60 AAs: 0.2682\nTotal prob of N-in: 0.14402\noutside 1 331\nTMhelix 332 354\ninside 355 372

21185 GCF_001356995.1_10592_2_68 Salmonella enterica subsp. enterica serovar Typhi Proteobacteria; Gammaproteobacteria; Enterobacterales; Enterobacteriaceae; Salmonella; Salmonella enterica; Salmonella enterica subsp. enterica MTGDNTLITSHGINRRDFMKLCAALAATMGLSSKAAAEMAESVSNPQRPPVIWIGAQECT WP_000145429.1 hydrogenase 2 small subunit [Salmonella enterica] Length: 372\nNumber of predicted TMHs: 1\nExp number of AAs in TMHs: 24.51309\nExp number, first 60 AAs: 0.2682\nTotal prob of N-in: 0.14402\noutside 1 331\nTMhelix 332 354\ninside 355 372

21186 GCF_001357115.1_10592_2_44 Salmonella enterica subsp. enterica serovar Typhi Proteobacteria; Gammaproteobacteria; Enterobacterales; Enterobacteriaceae; Salmonella; Salmonella enterica; Salmonella enterica subsp. enterica MTGDNTLITSHGINRRDFMKLCAALAATMGLSSKAAAEMAESVSNPQRPPVIWIGAQECT WP_000145429.1 hydrogenase 2 small subunit [Salmonella enterica] Length: 372\nNumber of predicted TMHs: 1\nExp number of AAs in TMHs: 24.51309\nExp number, first 60 AAs: 0.2682\nTotal prob of N-in: 0.14402\noutside 1 331\nTMhelix 332 354\ninside 355 372

21187 GCF_001357075.1_10592_2_74 Salmonella enterica subsp. enterica serovar Typhi Proteobacteria; Gammaproteobacteria; Enterobacterales; Enterobacteriaceae; Salmonella; Salmonella enterica;

Salmonella enterica subsp. enterica MTGDNTLITSHGINRRDFMKLCAALAATMGLSSKAAAEMAESVSNPQRPPVIWIGAQECT WP_000145429.1 hydrogenase 2 small subunit [Salmonella enterica] Length: 372\nNumber of predicted TMHs: 1\nExp number of AAs in TMHs: 24.51309\nExp number, first 60 AAs: 0.2682\nTotal prob of N-in: 0.14402\noutside 1 331\nTMhelix 332 354\ninside 355 372

21188 GCF_001357155.1_10593_2_59 Salmonella enterica subsp. enterica serovar Typhi Proteobacteria; Gammaproteobacteria; Enterobacterales; Enterobacteriaceae; Salmonella; Salmonella enterica; Salmonella enterica subsp. enterica MTGDNTLITSHGINRRDFMKLCAALAATMGLSSKAAAEMAESVSNPQRPPVIWIGAQECT WP_000145429.1 hydrogenase 2 small subunit [Salmonella enterica] Length: 372\nNumber of predicted TMHs: 1\nExp number of AAs in TMHs: 24.51309\nExp number, first 60 AAs: 0.2682\nTotal prob of N-in: 0.14402\noutside 1 331\nTMhelix 332 354\ninside 355 372

21189 GCF_001357195.1_10592_2_69 Salmonella enterica subsp. enterica serovar Typhi Proteobacteria; Gammaproteobacteria; Enterobacterales; Enterobacteriaceae; Salmonella; Salmonella enterica; Salmonella enterica subsp. enterica MTGDNTLITSHGINRRDFMKLCAALAATMGLSSKAAAEMAESVSNPQRPPVIWIGAQECT WP_000145429.1 hydrogenase 2 small subunit [Salmonella enterica] Length: 372\nNumber of predicted TMHs: 1\nExp number of AAs in TMHs: 24.51309\nExp number, first 60 AAs: 0.2682\nTotal prob of N-in: 0.14402\noutside 1 331\nTMhelix 332 354\ninside 355 372

21190 GCF_001356835.1_10592_2_17 Salmonella enterica subsp. enterica serovar Typhi Proteobacteria; Gammaproteobacteria; Enterobacterales; Enterobacteriaceae; Salmonella; Salmonella enterica; Salmonella enterica subsp. enterica MTGDNTLITSHGINRRDFMKLCAALAATMGLSSKAATEMAESVSNPQRPPVIWIGAQECT WP_001782304.1 hydrogenase 2 small subunit [Salmonella enterica] Length: 372\nNumber of predicted TMHs: 1\nExp number of AAs in TMHs: 24.41173\nExp number, first 60 AAs: 0.16124\nTotal prob of N-in: 0.13947\noutside 1 331\nTMhelix 332 354\ninside 355 372

21191 GCF_001357275.1_10593_2_55 Salmonella enterica subsp. enterica serovar Typhi Proteobacteria; Gammaproteobacteria; Enterobacterales; Enterobacteriaceae; Salmonella; Salmonella enterica; Salmonella enterica subsp. enterica MTGDNTLITSHGINRRDFMKLCAALAATMGLSSKAAAEMAESVSNPQRPPVIWIGAQECT WP_000145429.1 hydrogenase 2 small subunit [Salmonella enterica] Length: 372\nNumber of predicted TMHs: 1\nExp number of AAs in TMHs: 24.51309\nExp number, first 60 AAs: 0.2682\nTotal prob of N-in: 0.14402\noutside 1 331\nTMhelix 332 354\ninside 355 372

21192 GCF_001357335.1_10593_2_78 Salmonella enterica subsp. enterica serovar Typhi Proteobacteria; Gammaproteobacteria; Enterobacterales; Enterobacteriaceae; Salmonella; Salmonella enterica; Salmonella enterica subsp. enterica MTGDNTLITSHGINRRDFMKLCAALAATMGLSSKAAAEMAESVSNPQRPPVIWIGAQECT WP_000145429.1 hydrogenase 2 small subunit [Salmonella enterica] Length: 372\nNumber of predicted TMHs: 1\nExp number of AAs in TMHs: 24.51309\nExp number, first 60 AAs: 0.2682\nTotal prob of N-in: 0.14402\noutside 1 331\nTMhelix 332 354\ninside 355 372

21193 GCF_001357295.1_10593_2_24 Salmonella enterica subsp. enterica serovar Typhi Proteobacteria; Gammaproteobacteria; Enterobacterales; Enterobacteriaceae; Salmonella; Salmonella enterica; Salmonella enterica subsp. enterica MTGDNTLITSHGINRRDFMKLCAALAATMGLSSKAAAEMAESVSNPQRPPVIWIGAQECT WP_000145429.1 hydrogenase 2 small subunit [Salmonella enterica] Length: 372\nNumber of predicted TMHs: 1\nExp number of AAs in TMHs: 24.51309\nExp number, first 60 AAs: 0.2682\nTotal prob of N-in: 0.14402\noutside 1 331\nTMhelix 332 354\ninside 355 372

21194 GCF_001357375.1_10593_2_10 Salmonella enterica subsp. enterica serovar Typhi Proteobacteria; Gammaproteobacteria; Enterobacterales; Enterobacteriaceae; Salmonella; Salmonella enterica; Salmonella enterica subsp. enterica MTGDNTLITSHGINRRDFMKLCAALAATMGLSSKAAAEMAESVSNPQRPPVIWIGAQECT WP_000145429.1 hydrogenase 2 small subunit [Salmonella enterica] Length: 372\nNumber of predicted TMHs: 1\nExp number of AAs in TMHs: 24.51309\nExp number, first 60 AAs: 0.2682\nTotal prob of N-in: 0.14402\noutside 1 331\nTMhelix 332 354\ninside 355 372

21195 GCF_001357495.1_10593_2_66 Salmonella enterica subsp. enterica serovar Typhi Proteobacteria; Gammaproteobacteria; Enterobacterales; Enterobacteriaceae; Salmonella; Salmonella enterica; Salmonella enterica subsp. enterica MTGDNTLITSHGINRRDFMKLCAALAATMGLSSKAAAEMAESVSNPQRPPVIWIGAQECT WP_000145429.1 hydrogenase 2 small subunit [Salmonella enterica] Length: 372\nNumber of predicted TMHs: 1\nExp number of AAs in TMHs: 24.51309\nExp number, first 60 AAs: 0.2682\nTotal prob of N-in: 0.14402\noutside 1 331\nTMhelix 332 354\ninside 355 372

21196 GCF_001357415.1_10593_2_58 Salmonella enterica subsp. enterica serovar Typhi Proteobacteria; Gammaproteobacteria; Enterobacterales; Enterobacteriaceae; Salmonella; Salmonella enterica; Salmonella enterica subsp. enterica MTGDNTLITSHGINRRDFMKLCAALAATMGLSSKAAAEMAESVSNPQRPPVIWIGAQECT WP_000145429.1 hydrogenase 2 small subunit [Salmonella enterica] Length: 372\nNumber of predicted TMHs: 1\nExp number of AAs in TMHs: 24.51309\nExp number, first 60 AAs: 0.2682\nTotal prob of N-in: 0.14402\noutside 1 331\nTMhelix 332 354\ninside 355 372

21197 GCF_001357455.1_10593_2_27 Salmonella enterica subsp. enterica serovar Typhi Proteobacteria; Gammaproteobacteria; Enterobacterales; Enterobacteriaceae; Salmonella; Salmonella enterica; Salmonella enterica subsp. enterica MTGDNTLITSHGINRRDFMKLCAALAATMGLSSKAAAEMAESVSNPQRPPVIWIGAQECT WP_000145429.1 hydrogenase 2 small subunit [Salmonella enterica] Length: 372\nNumber of predicted TMHs: 1\nExp number of AAs in TMHs: 24.51309\nExp number, first 60 AAs: 0.2682\nTotal prob of N-in: 0.14402\noutside 1 331\nTMhelix 332 354\ninside 355 372

21198 GCF_001357535.1_10593_2_53 Salmonella enterica subsp. enterica serovar Typhi Proteobacteria; Gammaproteobacteria; Enterobacterales; Enterobacteriaceae; Salmonella; Salmonella enterica; Salmonella enterica subsp. enterica MTGDNTLITSHGINRRDFMKLCAALAATMGLSSKAAAEMAESVSNPQRPPVIWIGAQECT WP_000145429.1 hydrogenase 2 small subunit [Salmonella enterica] Length: 372\nNumber of predicted TMHs: 1\nExp number of AAs in TMHs: 24.51309\nExp number, first 60 AAs: 0.2682\nTotal prob of N-in: 0.14402\noutside 1 331\nTMhelix 332 354\ninside 355 372

21199 GCF_001357575.1_10593_2_15 Salmonella enterica subsp. enterica serovar Typhi Proteobacteria; Gammaproteobacteria; Enterobacterales; Enterobacteriaceae; Salmonella; Salmonella enterica; Salmonella enterica subsp. enterica MTGDNTLITSHGINRRDFMKLCAALAATMGLSSKAAAEMAESVSNPQRPPVIWIGAQECT WP_000145429.1 hydrogenase 2 small subunit [Salmonella enterica] Length: 372\nNumber of predicted TMHs: 1\nExp number of AAs in TMHs: 24.51309\nExp number, first 60 AAs: 0.2682\nTotal prob of N-in: 0.14402\noutside 1 331\nTMhelix 332 354\ninside 355 372

21200 GCF_001357615.1_10593_2_73 Salmonella enterica subsp. enterica serovar Typhi Proteobacteria; Gammaproteobacteria; Enterobacterales; Enterobacteriaceae; Salmonella; Salmonella enterica; Salmonella enterica subsp. enterica MTGDNTLITSHGINRRDFMKLCAALAATMGLSSKAAAEMAESVSNPQRPPVIWIGAQECT WP_000145429.1 hydrogenase 2 small subunit [Salmonella enterica] Length: 372\nNumber of predicted TMHs: 1\nExp number of AAs in TMHs: 24.51309\nExp number, first 60 AAs: 0.2682\nTotal prob of N-in: 0.14402\noutside 1 331\nTMhelix 332 354\ninside 355 372

21201 GCF_001357235.1_10592_2_67 Salmonella enterica subsp. enterica serovar Typhi Proteobacteria; Gammaproteobacteria; Enterobacterales; Enterobacteriaceae; Salmonella; Salmonella enterica; Salmonella enterica subsp. enterica MTGDNTLITSHGINRRDFMKLCAALAATMGLSSKAATEMAESVSNPQRPPVIWIGAQECT WP_001782304.1 hydrogenase 2 small subunit [Salmonella enterica] Length: 372\nNumber of predicted TMHs: 1\nExp number of AAs in TMHs: 24.41173\nExp number, first 60 AAs: 0.16124\nTotal prob of N-in: 0.13947\noutside 1 331\nTMhelix 332 354\ninside 355 372

21202 GCF_001357655.1_10593_2_74 Salmonella enterica subsp. enterica serovar Typhi Proteobacteria; Gammaproteobacteria; Enterobacterales; Enterobacteriaceae; Salmonella; Salmonella enterica; Salmonella enterica subsp. enterica MTGDNTLITSHGINRRDFMKLCAALAATMGLSSKAAAEMAESVSNPQRPPVIWIGAQECT WP_000145429.1 hydrogenase 2 small subunit [Salmonella enterica] Length: 372\nNumber of predicted TMHs: 1\nExp number of AAs in TMHs: 24.51309\nExp number, first 60 AAs: 0.2682\nTotal prob of N-in: 0.14402\noutside 1 331\nTMhelix 332 354\ninside 355 372

21203 GCF_001357695.1_10593_2_42 Salmonella enterica subsp. enterica serovar Typhi Proteobacteria; Gammaproteobacteria; Enterobacterales; Enterobacteriaceae; Salmonella; Salmonella enterica; Salmonella enterica subsp. enterica MTGDNTLITSHGINRRDFMKLCAALAATMGLSSKAAAEMAESVSNPQRPPVIWIGAQECT WP_000145429.1 hydrogenase 2 small subunit [Salmonella enterica] Length: 372\nNumber of predicted TMHs: 1\nExp number of AAs in TMHs: 24.51309\nExp number, first 60 AAs: 0.2682\nTotal prob of N-in: 0.14402\noutside 1 331\nTMhelix 332 354\ninside 355 372

21204 GCF_001357735.1_10592_2_72 Salmonella enterica subsp. enterica serovar Typhi Proteobacteria; Gammaproteobacteria; Enterobacterales; Enterobacteriaceae; Salmonella; Salmonella enterica; Salmonella enterica subsp. enterica MTGDNTLITSHGINRRDFMKLCAALAATMGLSSKAAAEMAESVSNPQRPPVIWIGAQECT WP_000145429.1 hydrogenase 2 small subunit [Salmonella enterica] Length: 372\nNumber of

predicted TMHs:  1\nExp number of AAs in TMHs: 24.51309\nExp number, first 60 AAs:  0.2682\nTotal prob of N-in: 0.14402\noutside   1   331\nTMhelix   332   354\ninside   355   372

21205 GCF_001357775.1_10593_2_35 Salmonella enterica subsp. enterica serovar Typhi Proteobacteria; Gammaproteobacteria; Enterobacterales; Enterobacteriaceae; Salmonella; Salmonella enterica; Salmonella enterica subsp. enterica MTGDNTLITSHGINRRDFMKLCAALAATMGLSSKAAAEMAESVSNPQRPPVIWIGAQECT WP_000145429.1 hydrogenase 2 small subunit [Salmonella enterica] Length: 372\nNumber of predicted TMHs:  1\nExp number of AAs in TMHs: 24.51309\nExp number, first 60 AAs:  0.2682\nTotal prob of N-in: 0.14402\noutside   1   331\nTMhelix   332   354\ninside   355   372

21206 GCF_001357815.1_10593_2_61 Salmonella enterica subsp. enterica serovar Typhi Proteobacteria; Gammaproteobacteria; Enterobacterales; Enterobacteriaceae; Salmonella; Salmonella enterica; Salmonella enterica subsp. enterica MTGDNTLITSHGINRRDFMKLCAALAATMGLSSKAAAEMAESVSNPQRPPVIWIGAQECT WP_000145429.1 hydrogenase 2 small subunit [Salmonella enterica] Length: 372\nNumber of predicted TMHs:  1\nExp number of AAs in TMHs: 24.51309\nExp number, first 60 AAs:  0.2682\nTotal prob of N-in: 0.14402\noutside   1   331\nTMhelix   332   354\ninside   355   372

21207 GCF_001357855.1_10593_2_76 Salmonella enterica subsp. enterica serovar Typhi Proteobacteria; Gammaproteobacteria; Enterobacterales; Enterobacteriaceae; Salmonella; Salmonella enterica; Salmonella enterica subsp. enterica MTGDNTLITSHGINRRDFMKLCAALAATMGLSSKAAAEMAESVSNPQRPPVIWIGAQECT WP_000145429.1 hydrogenase 2 small subunit [Salmonella enterica] Length: 372\nNumber of predicted TMHs:  1\nExp number of AAs in TMHs: 24.51309\nExp number, first 60 AAs:  0.2682\nTotal prob of N-in: 0.14402\noutside   1   331\nTMhelix   332   354\ninside   355   372

21208 GCF_001357895.1_10593_2_72 Salmonella enterica subsp. enterica serovar Typhi Proteobacteria; Gammaproteobacteria; Enterobacterales; Enterobacteriaceae; Salmonella; Salmonella enterica; Salmonella enterica subsp. enterica MTGDNTLITSHGINRRDFMKLCAALAATMGLSSKAAAEMAESVSNPQRPPVIWIGAQECT WP_000145429.1 hydrogenase 2 small subunit [Salmonella enterica] Length: 372\nNumber of predicted TMHs:  1\nExp number of AAs in TMHs: 24.51309\nExp number, first 60 AAs:  0.2682\nTotal prob of N-in: 0.14402\noutside   1   331\nTMhelix   332   354\ninside   355   372

21209 GCF_001357975.1_10592_2_82 Salmonella enterica subsp. enterica serovar Typhi Proteobacteria; Gammaproteobacteria; Enterobacterales; Enterobacteriaceae; Salmonella; Salmonella enterica; Salmonella enterica subsp. enterica MTGDNTLITSHGINRRDFMKLCAALAATMGLSSKAAAEMAESVSNPQRPPVIWIGAQECT WP_000145429.1 hydrogenase 2 small subunit [Salmonella enterica] Length: 372\nNumber of predicted TMHs:  1\nExp number of AAs in TMHs: 24.51309\nExp number, first 60 AAs:  0.2682\nTotal prob of N-in: 0.14402\noutside   1   331\nTMhelix   332   354\ninside   355   372

21210 GCF_001357935.1_10592_2_61 Salmonella enterica subsp. enterica serovar Typhi Proteobacteria; Gammaproteobacteria; Enterobacterales; Enterobacteriaceae; Salmonella; Salmonella enterica; Salmonella enterica subsp. enterica MTGDNTLITSHGINRRDFMKLCAALAATMGLSSKAAAEMAESVSNPQRPPVIWIGAQECT WP_000145429.1 hydrogenase 2 small subunit [Salmonella enterica] Length: 372\nNumber of predicted TMHs:  1\nExp number of AAs in TMHs: 24.51309\nExp number, first 60 AAs:  0.2682\nTotal prob of N-in: 0.14402\noutside   1   331\nTMhelix   332   354\ninside   355   372

21211 GCF_001357995.1_10592_2_81 Salmonella enterica subsp. enterica serovar Typhi Proteobacteria; Gammaproteobacteria; Enterobacterales; Enterobacteriaceae; Salmonella; Salmonella enterica; Salmonella enterica subsp. enterica MTGDNTLITSHGINRRDFMKLCAALAATMGLSSKAAAEMAESVSNPQRPPVIWIGAQECT WP_000145429.1 hydrogenase 2 small subunit [Salmonella enterica] Length: 372\nNumber of predicted TMHs:  1\nExp number of AAs in TMHs: 24.51309\nExp number, first 60 AAs:  0.2682\nTotal prob of N-in: 0.14402\noutside   1   331\nTMhelix   332   354\ninside   355   372

21212 GCF_001358035.1_10593_2_63 Salmonella enterica subsp. enterica serovar Typhi Proteobacteria; Gammaproteobacteria; Enterobacterales; Enterobacteriaceae; Salmonella; Salmonella enterica; Salmonella enterica subsp. enterica MTGDNTLITSHGINRRDFMKLCAALAATMGLSSKAAAEMAESVSNPQRPPVIWIGAQECT WP_000145429.1 hydrogenase 2 small subunit [Salmonella enterica] Length: 372\nNumber of predicted TMHs:  1\nExp number of AAs in TMHs: 24.51309\nExp number, first 60 AAs:  0.2682\nTotal prob of N-in: 0.14402\noutside   1   331\nTMhelix   332   354\ninside   355   372

21213 GCF_001358075.1_10593_2_48 Salmonella enterica subsp. enterica serovar Typhi Proteobacteria; Gammaproteobacteria; Enterobacterales; Enterobacteriaceae; Salmonella; Salmonella enterica;

Salmonella enterica subsp. enterica	MTGDNTLITSHGINRRDFMKLCAALAATMGLSSKAAAEMAESVSNPQRPPVIWIGAQECT
WP_000145429.1	hydrogenase 2 small subunit [Salmonella enterica]	Length: 372\nNumber of predicted TMHs: 1\nExp number of AAs in TMHs: 24.51309\nExp number, first 60 AAs: 0.2682\nTotal prob of N-in: 0.14402\noutside 1 331\nTMhelix 332 354\ninside 355 372

21214	GCF_001358115.1_10593_2_21	Salmonella enterica subsp. enterica serovar Typhi
Proteobacteria; Gammaproteobacteria; Enterobacterales; Enterobacteriaceae; Salmonella; Salmonella enterica; Salmonella enterica subsp. enterica	MTGDNTLITSHGINRRDFMKLCAALAATMGLSSKAAAEMAESVSNPQRPPVIWIGAQECT
WP_000145429.1	hydrogenase 2 small subunit [Salmonella enterica]	Length: 372\nNumber of predicted TMHs: 1\nExp number of AAs in TMHs: 24.51309\nExp number, first 60 AAs: 0.2682\nTotal prob of N-in: 0.14402\noutside 1 331\nTMhelix 332 354\ninside 355 372

21215	GCF_001358155.1_10593_2_16	Salmonella enterica subsp. enterica serovar Typhi
Proteobacteria; Gammaproteobacteria; Enterobacterales; Enterobacteriaceae; Salmonella; Salmonella enterica; Salmonella enterica subsp. enterica	MTGDNTLITSHGINRRDFMKLCAALAATMGLSSKAAAEMAESVSNPQRPPVIWIGAQECT
WP_000145429.1	hydrogenase 2 small subunit [Salmonella enterica]	Length: 372\nNumber of predicted TMHs: 1\nExp number of AAs in TMHs: 24.51309\nExp number, first 60 AAs: 0.2682\nTotal prob of N-in: 0.14402\noutside 1 331\nTMhelix 332 354\ninside 355 372

21216	GCF_001358195.1_10060_5_27	Salmonella enterica subsp. enterica serovar Typhi
Proteobacteria; Gammaproteobacteria; Enterobacterales; Enterobacteriaceae; Salmonella; Salmonella enterica; Salmonella enterica subsp. enterica	MTGDNTLITSHGINRRDFMKLCAALAATMGLSSKAAAEMAESVSNPQRPPVIWIGAQECT
WP_000145429.1	hydrogenase 2 small subunit [Salmonella enterica]	Length: 372\nNumber of predicted TMHs: 1\nExp number of AAs in TMHs: 24.51309\nExp number, first 60 AAs: 0.2682\nTotal prob of N-in: 0.14402\noutside 1 331\nTMhelix 332 354\ninside 355 372

21217	GCF_001358235.1_10071_3_75	Salmonella enterica subsp. enterica serovar Typhi
Proteobacteria; Gammaproteobacteria; Enterobacterales; Enterobacteriaceae; Salmonella; Salmonella enterica; Salmonella enterica subsp. enterica	MTGDNTLITSHGINRRDFMKLCAALAATMGLSSKAAAEMAESVSNPQRPPVIWIGAQECT
WP_000145429.1	hydrogenase 2 small subunit [Salmonella enterica]	Length: 372\nNumber of predicted TMHs: 1\nExp number of AAs in TMHs: 24.51309\nExp number, first 60 AAs: 0.2682\nTotal prob of N-in: 0.14402\noutside 1 331\nTMhelix 332 354\ninside 355 372

21218	GCF_001358275.1_10540_1_63	Salmonella enterica subsp. enterica serovar Typhi
Proteobacteria; Gammaproteobacteria; Enterobacterales; Enterobacteriaceae; Salmonella; Salmonella enterica; Salmonella enterica subsp. enterica	MTGDNTLITSHGINRRDFMKLCAALAATMGLSSKAAAEMAESVSNPQRPPVIWIGAQECT
WP_000145429.1	hydrogenase 2 small subunit [Salmonella enterica]	Length: 372\nNumber of predicted TMHs: 1\nExp number of AAs in TMHs: 24.51309\nExp number, first 60 AAs: 0.2682\nTotal prob of N-in: 0.14402\noutside 1 331\nTMhelix 332 354\ninside 355 372

21219	GCF_001358355.1_10561_2_26	Salmonella enterica subsp. enterica serovar Typhi
Proteobacteria; Gammaproteobacteria; Enterobacterales; Enterobacteriaceae; Salmonella; Salmonella enterica; Salmonella enterica subsp. enterica	MTGDNTLITSHGINRRDFMKLCAALAATMGLSSKAAAEMAESVSNPQRPPVIWIGAQECT
WP_000145429.1	hydrogenase 2 small subunit [Salmonella enterica]	Length: 372\nNumber of predicted TMHs: 1\nExp number of AAs in TMHs: 24.51309\nExp number, first 60 AAs: 0.2682\nTotal prob of N-in: 0.14402\noutside 1 331\nTMhelix 332 354\ninside 355 372

21220	GCF_001358315.1_10562_2_74	Salmonella enterica subsp. enterica serovar Typhi
Proteobacteria; Gammaproteobacteria; Enterobacterales; Enterobacteriaceae; Salmonella; Salmonella enterica; Salmonella enterica subsp. enterica	MTGDNTLITSHGINRRDFMKLCAALAATMGLSSKAAAEMAESVSNPQRPPVIWIGAQECT
WP_000145429.1	hydrogenase 2 small subunit [Salmonella enterica]	Length: 372\nNumber of predicted TMHs: 1\nExp number of AAs in TMHs: 24.51309\nExp number, first 60 AAs: 0.2682\nTotal prob of N-in: 0.14402\noutside 1 331\nTMhelix 332 354\ninside 355 372

21221	GCF_001358395.1_10426_1_16	Salmonella enterica subsp. enterica serovar Typhi
Proteobacteria; Gammaproteobacteria; Enterobacterales; Enterobacteriaceae; Salmonella; Salmonella enterica; Salmonella enterica subsp. enterica	MTGDNTLITSHGINRRDFMKLCAALAATMGLSSKAAAEMAESVSNPQRPPVIWIGAQECT
WP_000145429.1	hydrogenase 2 small subunit [Salmonella enterica]	Length: 372\nNumber of predicted TMHs: 1\nExp number of AAs in TMHs: 24.51309\nExp number, first 60 AAs: 0.2682\nTotal prob of N-in: 0.14402\noutside 1 331\nTMhelix 332 354\ninside 355 372

21222 GCF_001358475.1_10349_1_1 Salmonella enterica subsp. enterica serovar Typhi Proteobacteria; Gammaproteobacteria; Enterobacterales; Enterobacteriaceae; Salmonella; Salmonella enterica; Salmonella enterica subsp. enterica MTGDNTLITSHGINRRDFMKLCAALAATMGLSSKAAAEMAESVSNPQRPPVIWIGAQECT WP_000145429.1 hydrogenase 2 small subunit [Salmonella enterica] Length: 372\nNumber of predicted TMHs: 1\nExp number of AAs in TMHs: 24.51309\nExp number, first 60 AAs: 0.2682\nTotal prob of N-in: 0.14402\noutside 1 331\nTMhelix 332 354\ninside 355 372

21223 GCF_001358435.1_10071_3_50 Salmonella enterica subsp. enterica serovar Typhi Proteobacteria; Gammaproteobacteria; Enterobacterales; Enterobacteriaceae; Salmonella; Salmonella enterica; Salmonella enterica subsp. enterica MTGDNTLITSHGINRRDFMKLCAALAATMGLSSKAAAEMAESVSNPQRPPVIWIGAQECT WP_000145429.1 hydrogenase 2 small subunit [Salmonella enterica] Length: 372\nNumber of predicted TMHs: 1\nExp number of AAs in TMHs: 24.51309\nExp number, first 60 AAs: 0.2682\nTotal prob of N-in: 0.14402\noutside 1 331\nTMhelix 332 354\ninside 355 372

21224 GCF_001358515.1_10071_3_82 Salmonella enterica subsp. enterica serovar Typhi Proteobacteria; Gammaproteobacteria; Enterobacterales; Enterobacteriaceae; Salmonella; Salmonella enterica; Salmonella enterica subsp. enterica MTGDNTLITSHGINRRDFMKLCAALAATMGLSSKAAAEMAESVSNPQRPPVIWIGAQECT WP_000145429.1 hydrogenase 2 small subunit [Salmonella enterica] Length: 372\nNumber of predicted TMHs: 1\nExp number of AAs in TMHs: 24.51309\nExp number, first 60 AAs: 0.2682\nTotal prob of N-in: 0.14402\noutside 1 331\nTMhelix 332 354\ninside 355 372

21225 GCF_001358555.1_10071_3_58 Salmonella enterica subsp. enterica serovar Typhi Proteobacteria; Gammaproteobacteria; Enterobacterales; Enterobacteriaceae; Salmonella; Salmonella enterica; Salmonella enterica subsp. enterica MTGDNTLITSHGINRRDFMKLCAALAATMGLSSKAAAEMAESVSNPQRPPVIWIGAQECT WP_000145429.1 hydrogenase 2 small subunit [Salmonella enterica] Length: 372\nNumber of predicted TMHs: 1\nExp number of AAs in TMHs: 24.51309\nExp number, first 60 AAs: 0.2682\nTotal prob of N-in: 0.14402\noutside 1 331\nTMhelix 332 354\ninside 355 372

21226 GCF_001358595.1_10607_2_19 Salmonella enterica subsp. enterica serovar Typhi Proteobacteria; Gammaproteobacteria; Enterobacterales; Enterobacteriaceae; Salmonella; Salmonella enterica; Salmonella enterica subsp. enterica MTGDNTLITSHGINRRDFMKLCAALAATMGLSSKAAAEMAESVSNPQRPPVIWIGAQECT WP_000145429.1 hydrogenase 2 small subunit [Salmonella enterica] Length: 372\nNumber of predicted TMHs: 1\nExp number of AAs in TMHs: 24.51309\nExp number, first 60 AAs: 0.2682\nTotal prob of N-in: 0.14402\noutside 1 331\nTMhelix 332 354\ninside 355 372

21227 GCF_001358635.1_10607_2_53 Salmonella enterica subsp. enterica serovar Typhi Proteobacteria; Gammaproteobacteria; Enterobacterales; Enterobacteriaceae; Salmonella; Salmonella enterica; Salmonella enterica subsp. enterica MTGDNTLITSHGINRRDFMKLCAALAATMGLSSKAAAEMAESVSNPQRPPVIWIGAQECT WP_000145429.1 hydrogenase 2 small subunit [Salmonella enterica] Length: 372\nNumber of predicted TMHs: 1\nExp number of AAs in TMHs: 24.51309\nExp number, first 60 AAs: 0.2682\nTotal prob of N-in: 0.14402\noutside 1 331\nTMhelix 332 354\ninside 355 372

21228 GCF_001358675.1_10493_1_56 Salmonella enterica subsp. enterica serovar Typhi Proteobacteria; Gammaproteobacteria; Enterobacterales; Enterobacteriaceae; Salmonella; Salmonella enterica; Salmonella enterica subsp. enterica MTGDNTLITSHGINRRDFMKLCAALAATMGLSSKAAAEMAESVSNPQRPPVIWIGAQECT WP_000145429.1 hydrogenase 2 small subunit [Salmonella enterica] Length: 372\nNumber of predicted TMHs: 1\nExp number of AAs in TMHs: 24.51309\nExp number, first 60 AAs: 0.2682\nTotal prob of N-in: 0.14402\noutside 1 331\nTMhelix 332 354\ninside 355 372

21229 GCF_001358695.1_10071_8_4 Salmonella enterica subsp. enterica serovar Typhi Proteobacteria; Gammaproteobacteria; Enterobacterales; Enterobacteriaceae; Salmonella; Salmonella enterica; Salmonella enterica subsp. enterica MTGDNTLITSHGINRRDFMKLCAALAATMGLSSKAAAEMAESVSNPQRPPVIWIGAQECT WP_000145429.1 hydrogenase 2 small subunit [Salmonella enterica] Length: 372\nNumber of predicted TMHs: 1\nExp number of AAs in TMHs: 24.51309\nExp number, first 60 AAs: 0.2682\nTotal prob of N-in: 0.14402\noutside 1 331\nTMhelix 332 354\ninside 355 372

21230 GCF_001358855.1_10060_6_17 Salmonella enterica subsp. enterica serovar Typhi Proteobacteria; Gammaproteobacteria; Enterobacterales; Enterobacteriaceae; Salmonella; Salmonella enterica; Salmonella enterica subsp. enterica MTGDNTLITSHGINRRDFMKLCAALAATMGLSSKAAAEMAESVSNPQRPPVIWIGAQECT WP_000145429.1 hydrogenase 2 small subunit [Salmonella enterica] Length: 372\nNumber of

predicted TMHs:  1\nExp number of AAs in TMHs: 24.51309\nExp number, first 60 AAs:  0.2682\nTotal prob of N-in: 0.14402\noutside   1   331\nTMhelix   332   354\ninside   355   372

21231 GCF_001358935.1_10060_6_50 Salmonella enterica subsp. enterica serovar Typhi Proteobacteria; Gammaproteobacteria; Enterobacterales; Enterobacteriaceae; Salmonella; Salmonella enterica; Salmonella enterica subsp. enterica MTGDNTLITSHGINRRDFMKLCAALAATMGLSSKAAAEMAESVSNPQRPPVIWIGAQECT WP_000145429.1 hydrogenase 2 small subunit [Salmonella enterica] Length: 372\nNumber of predicted TMHs:  1\nExp number of AAs in TMHs: 24.51309\nExp number, first 60 AAs:  0.2682\nTotal prob of N-in: 0.14402\noutside   1   331\nTMhelix   332   354\ninside   355   372

21232 GCF_001358895.1_10060_6_32 Salmonella enterica subsp. enterica serovar Typhi Proteobacteria; Gammaproteobacteria; Enterobacterales; Enterobacteriaceae; Salmonella; Salmonella enterica; Salmonella enterica subsp. enterica MTGDNTLITSHGINRRDFMKLCAALAATMGLSSKAAAEMAESVSNPQRPPVIWIGAQECT WP_000145429.1 hydrogenase 2 small subunit [Salmonella enterica] Length: 372\nNumber of predicted TMHs:  1\nExp number of AAs in TMHs: 24.51309\nExp number, first 60 AAs:  0.2682\nTotal prob of N-in: 0.14402\noutside   1   331\nTMhelix   332   354\ninside   355   372

21233 GCF_001358775.1_10426_1_57 Salmonella enterica subsp. enterica serovar Typhi Proteobacteria; Gammaproteobacteria; Enterobacterales; Enterobacteriaceae; Salmonella; Salmonella enterica; Salmonella enterica subsp. enterica MTGDNTLITSHGINRRDFMKLCAALAATMGLSSKAAAEMAESVSNPQRPPVIWIGAQECT WP_000145429.1 hydrogenase 2 small subunit [Salmonella enterica] Length: 372\nNumber of predicted TMHs:  1\nExp number of AAs in TMHs: 24.51309\nExp number, first 60 AAs:  0.2682\nTotal prob of N-in: 0.14402\noutside   1   331\nTMhelix   332   354\ninside   355   372

21234 GCF_001358815.1_10426_1_64 Salmonella enterica subsp. enterica serovar Typhi Proteobacteria; Gammaproteobacteria; Enterobacterales; Enterobacteriaceae; Salmonella; Salmonella enterica; Salmonella enterica subsp. enterica MTGDNTLITSHGINRRDFMKLCAALAATMGLSSKAAAEMAESVSNPQRPPVIWIGAQECT WP_000145429.1 hydrogenase 2 small subunit [Salmonella enterica] Length: 372\nNumber of predicted TMHs:  1\nExp number of AAs in TMHs: 24.51309\nExp number, first 60 AAs:  0.2682\nTotal prob of N-in: 0.14402\noutside   1   331\nTMhelix   332   354\ninside   355   372

21235 GCF_001358735.1_10060_5_26 Salmonella enterica subsp. enterica serovar Typhi Proteobacteria; Gammaproteobacteria; Enterobacterales; Enterobacteriaceae; Salmonella; Salmonella enterica; Salmonella enterica subsp. enterica MTGDNTLITSHGINRRDFMKLCAALAATMGLSSKAAAEMAESVSNPQRPPVIWIGAQECT WP_000145429.1 hydrogenase 2 small subunit [Salmonella enterica] Length: 372\nNumber of predicted TMHs:  1\nExp number of AAs in TMHs: 24.51309\nExp number, first 60 AAs:  0.2682\nTotal prob of N-in: 0.14402\noutside   1   331\nTMhelix   332   354\ninside   355   372

21236 GCF_001358975.1_10060_6_71 Salmonella enterica subsp. enterica serovar Typhi Proteobacteria; Gammaproteobacteria; Enterobacterales; Enterobacteriaceae; Salmonella; Salmonella enterica; Salmonella enterica subsp. enterica MTGDNTLITSHGINRRDFMKLCAALAATMGLSSKAAAEMAESVSNPQRPPVIWIGAQECT WP_000145429.1 hydrogenase 2 small subunit [Salmonella enterica] Length: 372\nNumber of predicted TMHs:  1\nExp number of AAs in TMHs: 24.51309\nExp number, first 60 AAs:  0.2682\nTotal prob of N-in: 0.14402\noutside   1   331\nTMhelix   332   354\ninside   355   372

21237 GCF_001359015.1_10060_6_83 Salmonella enterica subsp. enterica serovar Typhi Proteobacteria; Gammaproteobacteria; Enterobacterales; Enterobacteriaceae; Salmonella; Salmonella enterica; Salmonella enterica subsp. enterica MTGDNTLITSHGINRRDFMKLCAALAATMGLSSKAAAEMAESVSNPQRPPVIWIGAQECT WP_000145429.1 hydrogenase 2 small subunit [Salmonella enterica] Length: 372\nNumber of predicted TMHs:  1\nExp number of AAs in TMHs: 24.51309\nExp number, first 60 AAs:  0.2682\nTotal prob of N-in: 0.14402\noutside   1   331\nTMhelix   332   354\ninside   355   372

21238 GCF_001359055.1_10060_6_92 Salmonella enterica subsp. enterica serovar Typhi Proteobacteria; Gammaproteobacteria; Enterobacterales; Enterobacteriaceae; Salmonella; Salmonella enterica; Salmonella enterica subsp. enterica MTGDNTLITSHGINRRDFMKLCAALAATMGLSSKAAAEMAESVSNPQRPPVIWIGAQECT WP_000145429.1 hydrogenase 2 small subunit [Salmonella enterica] Length: 372\nNumber of predicted TMHs:  1\nExp number of AAs in TMHs: 24.51309\nExp number, first 60 AAs:  0.2682\nTotal prob of N-in: 0.14402\noutside   1   331\nTMhelix   332   354\ninside   355   372

21239 GCF_001359095.1_9870_8_27 Salmonella enterica subsp. enterica serovar Typhi Proteobacteria; Gammaproteobacteria; Enterobacterales; Enterobacteriaceae; Salmonella; Salmonella enterica;

Salmonella enterica subsp. enterica MTGDNTLITSHGINRRDFMKLCAALAATMGLSSKAAAEMAESVSNPQRPPVIWIGAQECT WP_000145429.1 hydrogenase 2 small subunit [Salmonella enterica] Length: 372\nNumber of predicted TMHs: 1\nExp number of AAs in TMHs: 24.51309\nExp number, first 60 AAs: 0.2682\nTotal prob of N-in: 0.14402\noutside 1 331\nTMhelix 332 354\ninside 355 372

21240 GCF_001359135.1_9870_8_45 Salmonella enterica subsp. enterica serovar Typhi Proteobacteria; Gammaproteobacteria; Enterobacterales; Enterobacteriaceae; Salmonella; Salmonella enterica; Salmonella enterica subsp. enterica MTGDNTLITSHGINRRDFMKLCAALAATMGLSSKAAAEMAESVSNPQRPPVIWIGAQECT WP_000145429.1 hydrogenase 2 small subunit [Salmonella enterica] Length: 372\nNumber of predicted TMHs: 1\nExp number of AAs in TMHs: 24.51309\nExp number, first 60 AAs: 0.2682\nTotal prob of N-in: 0.14402\noutside 1 331\nTMhelix 332 354\ninside 355 372

21241 GCF_001359175.1_10060_5_2 Salmonella enterica subsp. enterica serovar Typhi Proteobacteria; Gammaproteobacteria; Enterobacterales; Enterobacteriaceae; Salmonella; Salmonella enterica; Salmonella enterica subsp. enterica MTGDNTLITSHGINRRDFMKLCAALAATMGLSSKAAAEMAESVSNPQRPPVIWIGAQECT WP_000145429.1 hydrogenase 2 small subunit [Salmonella enterica] Length: 372\nNumber of predicted TMHs: 1\nExp number of AAs in TMHs: 24.51309\nExp number, first 60 AAs: 0.2682\nTotal prob of N-in: 0.14402\noutside 1 331\nTMhelix 332 354\ninside 355 372

21242 GCF_001359215.1_10060_5_41 Salmonella enterica subsp. enterica serovar Typhi Proteobacteria; Gammaproteobacteria; Enterobacterales; Enterobacteriaceae; Salmonella; Salmonella enterica; Salmonella enterica subsp. enterica MTGDNTLITSHGINRRDFMKLCAALAATMGLSSKAAAEMAESVSNPQRPPVIWIGAQECT WP_000145429.1 hydrogenase 2 small subunit [Salmonella enterica] Length: 372\nNumber of predicted TMHs: 1\nExp number of AAs in TMHs: 24.51309\nExp number, first 60 AAs: 0.2682\nTotal prob of N-in: 0.14402\noutside 1 331\nTMhelix 332 354\ninside 355 372

21243 GCF_001359295.1_10060_5_59 Salmonella enterica subsp. enterica serovar Typhi Proteobacteria; Gammaproteobacteria; Enterobacterales; Enterobacteriaceae; Salmonella; Salmonella enterica; Salmonella enterica subsp. enterica MTGDNTLITSHGINRRDFMKLCAALAATMGLSSKAAAEMAESVSNPQRPPVIWIGAQECT WP_000145429.1 hydrogenase 2 small subunit [Salmonella enterica] Length: 372\nNumber of predicted TMHs: 1\nExp number of AAs in TMHs: 24.51309\nExp number, first 60 AAs: 0.2682\nTotal prob of N-in: 0.14402\noutside 1 331\nTMhelix 332 354\ninside 355 372

21244 GCF_001359255.1_10060_5_47 Salmonella enterica subsp. enterica serovar Typhi Proteobacteria; Gammaproteobacteria; Enterobacterales; Enterobacteriaceae; Salmonella; Salmonella enterica; Salmonella enterica subsp. enterica MTGDNTLITSHGINRRDFMKLCAALAATMGLSSKAAAEMAESVSNPQRPPVIWIGAQECT WP_000145429.1 hydrogenase 2 small subunit [Salmonella enterica] Length: 372\nNumber of predicted TMHs: 1\nExp number of AAs in TMHs: 24.51309\nExp number, first 60 AAs: 0.2682\nTotal prob of N-in: 0.14402\noutside 1 331\nTMhelix 332 354\ninside 355 372

21245 GCF_001359375.1_10060_5_80 Salmonella enterica subsp. enterica serovar Typhi Proteobacteria; Gammaproteobacteria; Enterobacterales; Enterobacteriaceae; Salmonella; Salmonella enterica; Salmonella enterica subsp. enterica MTGDNTLITSHGINRRDFMKLCAALAATMGLSSKAAAEMAESVSNPQRPPVIWIGAQECT WP_000145429.1 hydrogenase 2 small subunit [Salmonella enterica] Length: 372\nNumber of predicted TMHs: 1\nExp number of AAs in TMHs: 24.51309\nExp number, first 60 AAs: 0.2682\nTotal prob of N-in: 0.14402\noutside 1 331\nTMhelix 332 354\ninside 355 372

21246 GCF_001359335.1_10060_5_68 Salmonella enterica subsp. enterica serovar Typhi Proteobacteria; Gammaproteobacteria; Enterobacterales; Enterobacteriaceae; Salmonella; Salmonella enterica; Salmonella enterica subsp. enterica MTGDNTLITSHGINRRDFMKLCAALAATMGLSSKAAAEMAESVSNPQRPPVIWIGAQECT WP_000145429.1 hydrogenase 2 small subunit [Salmonella enterica] Length: 372\nNumber of predicted TMHs: 1\nExp number of AAs in TMHs: 24.51309\nExp number, first 60 AAs: 0.2682\nTotal prob of N-in: 0.14402\noutside 1 331\nTMhelix 332 354\ninside 355 372

21247 GCF_001359395.1_10060_5_86 Salmonella enterica subsp. enterica serovar Typhi Proteobacteria; Gammaproteobacteria; Enterobacterales; Enterobacteriaceae; Salmonella; Salmonella enterica; Salmonella enterica subsp. enterica MTGDNTLITSHGINRRDFMKLCAALAATMGLSSKAAAEMAESVSNPQRPPVIWIGAQECT WP_000145429.1 hydrogenase 2 small subunit [Salmonella enterica] Length: 372\nNumber of predicted TMHs: 1\nExp number of AAs in TMHs: 24.51309\nExp number, first 60 AAs: 0.2682\nTotal prob of N-in: 0.14402\noutside 1 331\nTMhelix 332 354\ninside 355 372

21248 GCF_001359475.1_10071_3_30 Salmonella enterica subsp. enterica serovar Typhi Proteobacteria; Gammaproteobacteria; Enterobacterales; Enterobacteriaceae; Salmonella; Salmonella enterica; Salmonella enterica subsp. enterica MTGDNTLITSHGINRRDFMKLCAALAATMGLSSKAAAEMAESVSNPQRPPVIWIGAQECT WP_000145429.1 hydrogenase 2 small subunit [Salmonella enterica] Length: 372\nNumber of predicted TMHs: 1\nExp number of AAs in TMHs: 24.51309\nExp number, first 60 AAs: 0.2682\nTotal prob of N-in: 0.14402\noutside 1 331\nTMhelix 332 354\ninside 355 372

21249 GCF_001359435.1_10071_3_6 Salmonella enterica subsp. enterica serovar Typhi Proteobacteria; Gammaproteobacteria; Enterobacterales; Enterobacteriaceae; Salmonella; Salmonella enterica; Salmonella enterica subsp. enterica MTGDNTLITSHGINRRDFMKLCAALAATMGLSSKAATEMAESVSNPQRPPVIWIGAQECT WP_001782304.1 hydrogenase 2 small subunit [Salmonella enterica] Length: 372\nNumber of predicted TMHs: 1\nExp number of AAs in TMHs: 24.41173\nExp number, first 60 AAs: 0.16124\nTotal prob of N-in: 0.13947\noutside 1 331\nTMhelix 332 354\ninside 355 372

21250 GCF_001359555.1_9870_8_2Salmonella enterica subsp. enterica serovar Typhi Proteobacteria; Gammaproteobacteria; Enterobacterales; Enterobacteriaceae; Salmonella; Salmonella enterica; Salmonella enterica subsp. enterica MTGDNTLITSHGINRRDFMKLCAALAATMGLSSKAAAEMAESVSNPQRPPVIWIGAQECT WP_000145429.1 hydrogenase 2 small subunit [Salmonella enterica] Length: 372\nNumber of predicted TMHs: 1\nExp number of AAs in TMHs: 24.51309\nExp number, first 60 AAs: 0.2682\nTotal prob of N-in: 0.14402\noutside 1 331\nTMhelix 332 354\ninside 355 372

21251 GCF_001359595.1_9870_8_35 Salmonella enterica subsp. enterica serovar Typhi Proteobacteria; Gammaproteobacteria; Enterobacterales; Enterobacteriaceae; Salmonella; Salmonella enterica; Salmonella enterica subsp. enterica MTGDNTLITSHGINRRDFMKLCAALAATMGLSSKAAAEMAESVSNPQRPPVIWIGAQECT WP_000145429.1 hydrogenase 2 small subunit [Salmonella enterica] Length: 372\nNumber of predicted TMHs: 1\nExp number of AAs in TMHs: 24.51309\nExp number, first 60 AAs: 0.2682\nTotal prob of N-in: 0.14402\noutside 1 331\nTMhelix 332 354\ninside 355 372

21252 GCF_001359635.1_9953_5_93 Salmonella enterica subsp. enterica serovar Typhi Proteobacteria; Gammaproteobacteria; Enterobacterales; Enterobacteriaceae; Salmonella; Salmonella enterica; Salmonella enterica subsp. enterica MTGDNTLITSHGINRRDFMKLCAALAATMGLSSKAAAEMAESVSNPQRPPVIWIGAQECT WP_000145429.1 hydrogenase 2 small subunit [Salmonella enterica] Length: 372\nNumber of predicted TMHs: 1\nExp number of AAs in TMHs: 24.51309\nExp number, first 60 AAs: 0.2682\nTotal prob of N-in: 0.14402\noutside 1 331\nTMhelix 332 354\ninside 355 372

21253 GCF_001359675.1_10060_6_6 Salmonella enterica subsp. enterica serovar Typhi Proteobacteria; Gammaproteobacteria; Enterobacterales; Enterobacteriaceae; Salmonella; Salmonella enterica; Salmonella enterica subsp. enterica MTGDNTLITSHGINRRDFMKLCAALAATMGLSSKAAAEMAESVSNPQRPPVIWIGAQECT WP_000145429.1 hydrogenase 2 small subunit [Salmonella enterica] Length: 372\nNumber of predicted TMHs: 1\nExp number of AAs in TMHs: 24.51309\nExp number, first 60 AAs: 0.2682\nTotal prob of N-in: 0.14402\noutside 1 331\nTMhelix 332 354\ninside 355 372

21254 GCF_001359715.1_9870_8_50 Salmonella enterica subsp. enterica serovar Typhi Proteobacteria; Gammaproteobacteria; Enterobacterales; Enterobacteriaceae; Salmonella; Salmonella enterica; Salmonella enterica subsp. enterica MTGDNTLITSHGINRRDFMKLCAALAATMGLSSKAAAEMAESVSNPQRPPVIWIGAQECT WP_000145429.1 hydrogenase 2 small subunit [Salmonella enterica] Length: 372\nNumber of predicted TMHs: 1\nExp number of AAs in TMHs: 24.51309\nExp number, first 60 AAs: 0.2682\nTotal prob of N-in: 0.14402\noutside 1 331\nTMhelix 332 354\ninside 355 372

21255 GCF_001359795.1_9870_8_77 Salmonella enterica subsp. enterica serovar Typhi Proteobacteria; Gammaproteobacteria; Enterobacterales; Enterobacteriaceae; Salmonella; Salmonella enterica; Salmonella enterica subsp. enterica MTGDNTLITSHGINRRDFMKLCAALAATMGLSSKAAAEMAESVSNPQRPPVIWIGAQECT WP_000145429.1 hydrogenase 2 small subunit [Salmonella enterica] Length: 372\nNumber of predicted TMHs: 1\nExp number of AAs in TMHs: 24.51309\nExp number, first 60 AAs: 0.2682\nTotal prob of N-in: 0.14402\noutside 1 331\nTMhelix 332 354\ninside 355 372

21256 GCF_001359755.1_9870_8_56 Salmonella enterica subsp. enterica serovar Typhi Proteobacteria; Gammaproteobacteria; Enterobacterales; Enterobacteriaceae; Salmonella; Salmonella enterica; Salmonella enterica subsp. enterica MTGDNTLITSHGINRRDFMKLCAALAATMGLSSKAAAEMAESVSNPQRPPVIWIGAQECT WP_000145429.1 hydrogenase 2 small subunit [Salmonella enterica] Length: 372\nNumber of

predicted TMHs: 1\nExp number of AAs in TMHs: 24.51309\nExp number, first 60 AAs: 0.2682\nTotal prob of N-in: 0.14402\noutside 1 331\nTMhelix 332 354\ninside 355 372

21257 GCF_001359835.1_10060_6_24 Salmonella enterica subsp. enterica serovar Typhi Proteobacteria; Gammaproteobacteria; Enterobacterales; Enterobacteriaceae; Salmonella; Salmonella enterica; Salmonella enterica subsp. enterica MTGDNTLITSHGINRRDFMKLCAALAATMGLSSKAAAEMAESVSNPQRPPVIWIGAQECT WP_000145429.1 hydrogenase 2 small subunit [Salmonella enterica] Length: 372\nNumber of predicted TMHs: 1\nExp number of AAs in TMHs: 24.51309\nExp number, first 60 AAs: 0.2682\nTotal prob of N-in: 0.14402\noutside 1 331\nTMhelix 332 354\ninside 355 372

21258 GCF_001359875.1_10060_6_48 Salmonella enterica subsp. enterica serovar Typhi Proteobacteria; Gammaproteobacteria; Enterobacterales; Enterobacteriaceae; Salmonella; Salmonella enterica; Salmonella enterica subsp. enterica MTGDNTLITSHGINRRDFMKLCAALAATMGLSSKAAAEMAESVSNPQRPPVIWIGAQECT WP_000145429.1 hydrogenase 2 small subunit [Salmonella enterica] Length: 372\nNumber of predicted TMHs: 1\nExp number of AAs in TMHs: 24.51309\nExp number, first 60 AAs: 0.2682\nTotal prob of N-in: 0.14402\noutside 1 331\nTMhelix 332 354\ninside 355 372

21259 GCF_001359515.1_10071_3_63 Salmonella enterica subsp. enterica serovar Typhi Proteobacteria; Gammaproteobacteria; Enterobacterales; Enterobacteriaceae; Salmonella; Salmonella enterica; Salmonella enterica subsp. enterica MTGDNTLITSHGINRRDFMKLCAALAATMGLSSKAATEMAESVSNPQRPPVIWIGAQECT WP_001782304.1 hydrogenase 2 small subunit [Salmonella enterica] Length: 372\nNumber of predicted TMHs: 1\nExp number of AAs in TMHs: 24.41173\nExp number, first 60 AAs: 0.16124\nTotal prob of N-in: 0.13947\noutside 1 331\nTMhelix 332 354\ninside 355 372

21260 GCF_001359915.1_10060_6_60 Salmonella enterica subsp. enterica serovar Typhi Proteobacteria; Gammaproteobacteria; Enterobacterales; Enterobacteriaceae; Salmonella; Salmonella enterica; Salmonella enterica subsp. enterica MTGDNTLITSHGINRRDFMKLCAALAATMGLSSKAAAEMAESVSNPQRPPVIWIGAQECT WP_000145429.1 hydrogenase 2 small subunit [Salmonella enterica] Length: 372\nNumber of predicted TMHs: 1\nExp number of AAs in TMHs: 24.51309\nExp number, first 60 AAs: 0.2682\nTotal prob of N-in: 0.14402\noutside 1 331\nTMhelix 332 354\ninside 355 372

21261 GCF_001359955.1_10060_6_72 Salmonella enterica subsp. enterica serovar Typhi Proteobacteria; Gammaproteobacteria; Enterobacterales; Enterobacteriaceae; Salmonella; Salmonella enterica; Salmonella enterica subsp. enterica MTGDNTLITSHGINRRDFMKLCAALAATMGLSSKAAAEMAESVSNPQRPPVIWIGAQECT WP_000145429.1 hydrogenase 2 small subunit [Salmonella enterica] Length: 372\nNumber of predicted TMHs: 1\nExp number of AAs in TMHs: 24.51309\nExp number, first 60 AAs: 0.2682\nTotal prob of N-in: 0.14402\noutside 1 331\nTMhelix 332 354\ninside 355 372

21262 GCF_001359995.1_10060_5_22 Salmonella enterica subsp. enterica serovar Typhi Proteobacteria; Gammaproteobacteria; Enterobacterales; Enterobacteriaceae; Salmonella; Salmonella enterica; Salmonella enterica subsp. enterica MTGDNTLITSHGINRRDFMKLCAALAATMGLSSKAAAEMAESVSNPQRPPVIWIGAQECT WP_000145429.1 hydrogenase 2 small subunit [Salmonella enterica] Length: 372\nNumber of predicted TMHs: 1\nExp number of AAs in TMHs: 24.51309\nExp number, first 60 AAs: 0.2682\nTotal prob of N-in: 0.14402\noutside 1 331\nTMhelix 332 354\ninside 355 372

21263 GCF_001360035.1_10060_5_37 Salmonella enterica subsp. enterica serovar Typhi Proteobacteria; Gammaproteobacteria; Enterobacterales; Enterobacteriaceae; Salmonella; Salmonella enterica; Salmonella enterica subsp. enterica MTGDNTLITSHGINRRDFMKLCAALAATMGLSSKAAAEMAESVSNPQRPPVIWIGAQECT WP_000145429.1 hydrogenase 2 small subunit [Salmonella enterica] Length: 372\nNumber of predicted TMHs: 1\nExp number of AAs in TMHs: 24.51309\nExp number, first 60 AAs: 0.2682\nTotal prob of N-in: 0.14402\noutside 1 331\nTMhelix 332 354\ninside 355 372

21264 GCF_001360075.1_10060_5_58 Salmonella enterica subsp. enterica serovar Typhi Proteobacteria; Gammaproteobacteria; Enterobacterales; Enterobacteriaceae; Salmonella; Salmonella enterica; Salmonella enterica subsp. enterica MTGDNTLITSHGINRRDFMKLCAALAATMGLSSKAAAEMAESVSNPQRPPVIWIGAQECT WP_000145429.1 hydrogenase 2 small subunit [Salmonella enterica] Length: 372\nNumber of predicted TMHs: 1\nExp number of AAs in TMHs: 24.51309\nExp number, first 60 AAs: 0.2682\nTotal prob of N-in: 0.14402\noutside 1 331\nTMhelix 332 354\ninside 355 372

21265 GCF_001360095.1_10060_5_61 Salmonella enterica subsp. enterica serovar Typhi Proteobacteria; Gammaproteobacteria; Enterobacterales; Enterobacteriaceae; Salmonella; Salmonella enterica;

Salmonella enterica subsp. enterica	MTGDNTLITSHGINRRDFMKLCAALAATMGLSSKAAAEMAESVSNPQRPPVIWIGAQECT
WP_000145429.1	hydrogenase 2 small subunit [Salmonella enterica]	Length: 372\nNumber of predicted TMHs: 1\nExp number of AAs in TMHs: 24.51309\nExp number, first 60 AAs: 0.2682\nTotal prob of N-in: 0.14402\noutside 1 331\nTMhelix 332 354\ninside 355 372

21266	GCF_001360135.1_10060_6_90	Salmonella enterica subsp. enterica serovar Typhi
Proteobacteria; Gammaproteobacteria; Enterobacterales; Enterobacteriaceae; Salmonella; Salmonella enterica; Salmonella enterica subsp. enterica	MTGDNTLITSHGINRRDFMKLCAALAATMGLSSKAAAEMAESVSNPQRPPVIWIGAQECT
WP_000145429.1	hydrogenase 2 small subunit [Salmonella enterica]	Length: 372\nNumber of predicted TMHs: 1\nExp number of AAs in TMHs: 24.51309\nExp number, first 60 AAs: 0.2682\nTotal prob of N-in: 0.14402\noutside 1 331\nTMhelix 332 354\ninside 355 372

21267	GCF_001360215.1_10060_5_88	Salmonella enterica subsp. enterica serovar Typhi
Proteobacteria; Gammaproteobacteria; Enterobacterales; Enterobacteriaceae; Salmonella; Salmonella enterica; Salmonella enterica subsp. enterica	MTGDNTLITSHGINRRDFMKLCAALAATMGLSSKAAAEMAESVSNPQRPPVIWIGAQECT
WP_000145429.1	hydrogenase 2 small subunit [Salmonella enterica]	Length: 372\nNumber of predicted TMHs: 1\nExp number of AAs in TMHs: 24.51309\nExp number, first 60 AAs: 0.2682\nTotal prob of N-in: 0.14402\noutside 1 331\nTMhelix 332 354\ninside 355 372

21268	GCF_001360175.1_10060_5_73	Salmonella enterica subsp. enterica serovar Typhi
Proteobacteria; Gammaproteobacteria; Enterobacterales; Enterobacteriaceae; Salmonella; Salmonella enterica; Salmonella enterica subsp. enterica	MTGDNTLITSHGINRRDFMKLCAALAATMGLSSKAAAEMAESVSNPQRPPVIWIGAQECT
WP_000145429.1	hydrogenase 2 small subunit [Salmonella enterica]	Length: 372\nNumber of predicted TMHs: 1\nExp number of AAs in TMHs: 24.51309\nExp number, first 60 AAs: 0.2682\nTotal prob of N-in: 0.14402\noutside 1 331\nTMhelix 332 354\ninside 355 372

21269	GCF_001360255.1_10071_3_17	Salmonella enterica subsp. enterica serovar Typhi
Proteobacteria; Gammaproteobacteria; Enterobacterales; Enterobacteriaceae; Salmonella; Salmonella enterica; Salmonella enterica subsp. enterica	MTGDNTLITSHGINRRDFMKLCAALAATMGLSSKAAAEMAESVSNPQRPPVIWIGAQECT
WP_000145429.1	hydrogenase 2 small subunit [Salmonella enterica]	Length: 372\nNumber of predicted TMHs: 1\nExp number of AAs in TMHs: 24.51309\nExp number, first 60 AAs: 0.2682\nTotal prob of N-in: 0.14402\noutside 1 331\nTMhelix 332 354\ninside 355 372

21270	GCF_001360295.1_10071_3_26	Salmonella enterica subsp. enterica serovar Typhi
Proteobacteria; Gammaproteobacteria; Enterobacterales; Enterobacteriaceae; Salmonella; Salmonella enterica; Salmonella enterica subsp. enterica	MTGDNTLITSHGINRRDFMKLCAALAATMGLSSKAAAEMAESVSNPQRPPVIWIGAQECT
WP_000145429.1	hydrogenase 2 small subunit [Salmonella enterica]	Length: 372\nNumber of predicted TMHs: 1\nExp number of AAs in TMHs: 24.51309\nExp number, first 60 AAs: 0.2682\nTotal prob of N-in: 0.14402\noutside 1 331\nTMhelix 332 354\ninside 355 372

21271	GCF_001360335.1_10071_3_86	Salmonella enterica subsp. enterica serovar Typhi
Proteobacteria; Gammaproteobacteria; Enterobacterales; Enterobacteriaceae; Salmonella; Salmonella enterica; Salmonella enterica subsp. enterica	MTGDNTLITSHGINRRDFMKLCAALAATMGLSSKAAAEMAESVSNPQRPPVIWIGAQECT
WP_000145429.1	hydrogenase 2 small subunit [Salmonella enterica]	Length: 372\nNumber of predicted TMHs: 1\nExp number of AAs in TMHs: 24.51309\nExp number, first 60 AAs: 0.2682\nTotal prob of N-in: 0.14402\noutside 1 331\nTMhelix 332 354\ninside 355 372

21272	GCF_001360375.1_9953_5_50	Salmonella enterica subsp. enterica serovar Typhi
Proteobacteria; Gammaproteobacteria; Enterobacterales; Enterobacteriaceae; Salmonella; Salmonella enterica; Salmonella enterica subsp. enterica	MTGDNTLITSHGINRRDFMKLCAALAATMGLSSKAAAEMAESVSNPQRPPVIWIGAQECT
WP_000145429.1	hydrogenase 2 small subunit [Salmonella enterica]	Length: 372\nNumber of predicted TMHs: 1\nExp number of AAs in TMHs: 24.51309\nExp number, first 60 AAs: 0.2682\nTotal prob of N-in: 0.14402\noutside 1 331\nTMhelix 332 354\ninside 355 372

21273	GCF_001360415.1_9953_5_74	Salmonella enterica subsp. enterica serovar Typhi
Proteobacteria; Gammaproteobacteria; Enterobacterales; Enterobacteriaceae; Salmonella; Salmonella enterica; Salmonella enterica subsp. enterica	MTGDNTLITSHGINRRDFMKLCAALAATMGLSSKAAAEMAESVSNPQRPPVIWIGAQECT
WP_000145429.1	hydrogenase 2 small subunit [Salmonella enterica]	Length: 372\nNumber of predicted TMHs: 1\nExp number of AAs in TMHs: 24.51309\nExp number, first 60 AAs: 0.2682\nTotal prob of N-in: 0.14402\noutside 1 331\nTMhelix 332 354\ninside 355 372

21274	GCF_001360455.1_10060_6_8	Salmonella enterica subsp. enterica serovar Typhi
Proteobacteria; Gammaproteobacteria; Enterobacterales; Enterobacteriaceae; Salmonella; Salmonella enterica; Salmonella enterica subsp. enterica	MTGDNTLITSHGINRRDFMKLCAALAATMGLSSKAAAEMAESVSNPQRPPVIWIGAQECT	WP_000145429.1	hydrogenase 2 small subunit [Salmonella enterica]	Length: 372\nNumber of predicted TMHs:  1\nExp number of AAs in TMHs: 24.51309\nExp number, first 60 AAs:  0.2682\nTotal prob of N-in: 0.14402\noutside	1	331\nTMhelix	332	354\ninside	355	372

21275	GCF_001360495.1_9870_8_10	Salmonella enterica subsp. enterica serovar Typhi
Proteobacteria; Gammaproteobacteria; Enterobacterales; Enterobacteriaceae; Salmonella; Salmonella enterica; Salmonella enterica subsp. enterica	MTGDNTLITSHGINRRDFMKLCAALAATMGLSSKAAAEMAESVSNPQRPPVIWIGAQECT	WP_000145429.1	hydrogenase 2 small subunit [Salmonella enterica]	Length: 372\nNumber of predicted TMHs:  1\nExp number of AAs in TMHs: 24.51309\nExp number, first 60 AAs:  0.2682\nTotal prob of N-in: 0.14402\noutside	1	331\nTMhelix	332	354\ninside	355	372

21276	GCF_001360535.1_9870_8_22	Salmonella enterica subsp. enterica serovar Typhi
Proteobacteria; Gammaproteobacteria; Enterobacterales; Enterobacteriaceae; Salmonella; Salmonella enterica; Salmonella enterica subsp. enterica	MTGDNTLITSHGINRRDFMKLCAALAATMGLSSKAAAEMAESVSNPQRPPVIWIGAQECT	WP_000145429.1	hydrogenase 2 small subunit [Salmonella enterica]	Length: 372\nNumber of predicted TMHs:  1\nExp number of AAs in TMHs: 24.51309\nExp number, first 60 AAs:  0.2682\nTotal prob of N-in: 0.14402\noutside	1	331\nTMhelix	332	354\ninside	355	372

21277	GCF_001360575.1_9870_8_49	Salmonella enterica subsp. enterica serovar Typhi
Proteobacteria; Gammaproteobacteria; Enterobacterales; Enterobacteriaceae; Salmonella; Salmonella enterica; Salmonella enterica subsp. enterica	MTGDNTLITSHGINRRDFMKLCAALAATMGLSSKAAAEMAESVSNPQRPPVIWIGAQECT	WP_000145429.1	hydrogenase 2 small subunit [Salmonella enterica]	Length: 372\nNumber of predicted TMHs:  1\nExp number of AAs in TMHs: 24.51309\nExp number, first 60 AAs:  0.2682\nTotal prob of N-in: 0.14402\noutside	1	331\nTMhelix	332	354\ninside	355	372

21278	GCF_001360615.1_9870_8_58	Salmonella enterica subsp. enterica serovar Typhi
Proteobacteria; Gammaproteobacteria; Enterobacterales; Enterobacteriaceae; Salmonella; Salmonella enterica; Salmonella enterica subsp. enterica	MTGDNTLITSHGINRRDFMKLCAALAATMGLSSKAAAEMAESVSNPQRPPVIWIGAQECT	WP_000145429.1	hydrogenase 2 small subunit [Salmonella enterica]	Length: 372\nNumber of predicted TMHs:  1\nExp number of AAs in TMHs: 24.51309\nExp number, first 60 AAs:  0.2682\nTotal prob of N-in: 0.14402\noutside	1	331\nTMhelix	332	354\ninside	355	372

21279	GCF_001360655.1_9870_8_73	Salmonella enterica subsp. enterica serovar Typhi
Proteobacteria; Gammaproteobacteria; Enterobacterales; Enterobacteriaceae; Salmonella; Salmonella enterica; Salmonella enterica subsp. enterica	MTGDNTLITSHGINRRDFMKLCAALAATMGLSSKAAAEMAESVSNPQRPPVIWIGAQECT	WP_000145429.1	hydrogenase 2 small subunit [Salmonella enterica]	Length: 372\nNumber of predicted TMHs:  1\nExp number of AAs in TMHs: 24.51309\nExp number, first 60 AAs:  0.2682\nTotal prob of N-in: 0.14402\noutside	1	331\nTMhelix	332	354\ninside	355	372

21280	GCF_001360695.1_10060_5_9	Salmonella enterica subsp. enterica serovar Typhi
Proteobacteria; Gammaproteobacteria; Enterobacterales; Enterobacteriaceae; Salmonella; Salmonella enterica; Salmonella enterica subsp. enterica	MTGDNTLITSHGINRRDFMKLCAALAATMGLSSKAAAEMAESVSNPQRPPVIWIGAQECT	WP_000145429.1	hydrogenase 2 small subunit [Salmonella enterica]	Length: 372\nNumber of predicted TMHs:  1\nExp number of AAs in TMHs: 24.51309\nExp number, first 60 AAs:  0.2682\nTotal prob of N-in: 0.14402\noutside	1	331\nTMhelix	332	354\ninside	355	372

21281	GCF_001360795.1_10060_5_30	Salmonella enterica subsp. enterica serovar Typhi
Proteobacteria; Gammaproteobacteria; Enterobacterales; Enterobacteriaceae; Salmonella; Salmonella enterica; Salmonella enterica subsp. enterica	MTGDNTLITSHGINRRDFMKLCAALAATMGLSSKAAAEMAESVSNPQRPPVIWIGAQECT	WP_000145429.1	hydrogenase 2 small subunit [Salmonella enterica]	Length: 372\nNumber of predicted TMHs:  1\nExp number of AAs in TMHs: 24.51309\nExp number, first 60 AAs:  0.2682\nTotal prob of N-in: 0.14402\noutside	1	331\nTMhelix	332	354\ninside	355	372

21282	GCF_001360735.1_10060_5_15	Salmonella enterica subsp. enterica serovar Typhi
Proteobacteria; Gammaproteobacteria; Enterobacterales; Enterobacteriaceae; Salmonella; Salmonella enterica; Salmonella enterica subsp. enterica	MTGDNTLITSHGINRRDFMKLCAALAATMGLSSKAAAEMAESVSNPQRPPVIWIGAQECT	WP_000145429.1	hydrogenase 2 small subunit [Salmonella enterica]	Length: 372\nNumber of

predicted TMHs: 1\nExp number of AAs in TMHs: 24.51309\nExp number, first 60 AAs: 0.2682\nTotal prob of N-in: 0.14402\noutside 1 331\nTMhelix 332 354\ninside 355 372

21283 GCF_001360775.1_10060_5_24 Salmonella enterica subsp. enterica serovar Typhi Proteobacteria; Gammaproteobacteria; Enterobacterales; Enterobacteriaceae; Salmonella; Salmonella enterica; Salmonella enterica subsp. enterica MTGDNTLITSHGINRRDFMKLCAALAATMGLSSKAAAEMAESVSNPQRPPVIWIGAQECT WP_000145429.1 hydrogenase 2 small subunit [Salmonella enterica] Length: 372\nNumber of predicted TMHs: 1\nExp number of AAs in TMHs: 24.51309\nExp number, first 60 AAs: 0.2682\nTotal prob of N-in: 0.14402\noutside 1 331\nTMhelix 332 354\ninside 355 372

21284 GCF_001360835.1_10060_5_42 Salmonella enterica subsp. enterica serovar Typhi Proteobacteria; Gammaproteobacteria; Enterobacterales; Enterobacteriaceae; Salmonella; Salmonella enterica; Salmonella enterica subsp. enterica MTGDNTLITSHGINRRDFMKLCAALAATMGLSSKAAAEMAESVSNPQRPPVIWIGAQECT WP_000145429.1 hydrogenase 2 small subunit [Salmonella enterica] Length: 372\nNumber of predicted TMHs: 1\nExp number of AAs in TMHs: 24.51309\nExp number, first 60 AAs: 0.2682\nTotal prob of N-in: 0.14402\noutside 1 331\nTMhelix 332 354\ninside 355 372

21285 GCF_001360875.1_10060_5_60 Salmonella enterica subsp. enterica serovar Typhi Proteobacteria; Gammaproteobacteria; Enterobacterales; Enterobacteriaceae; Salmonella; Salmonella enterica; Salmonella enterica subsp. enterica MTGDNTLITSHGINRRDFMKLCAALAATMGLSSKAAAEMAESVSNPQRPPVIWIGAQECT WP_000145429.1 hydrogenase 2 small subunit [Salmonella enterica] Length: 372\nNumber of predicted TMHs: 1\nExp number of AAs in TMHs: 24.51309\nExp number, first 60 AAs: 0.2682\nTotal prob of N-in: 0.14402\noutside 1 331\nTMhelix 332 354\ninside 355 372

21286 GCF_001360915.1_10060_5_78 Salmonella enterica subsp. enterica serovar Typhi Proteobacteria; Gammaproteobacteria; Enterobacterales; Enterobacteriaceae; Salmonella; Salmonella enterica; Salmonella enterica subsp. enterica MTGDNTLITSHGINRRDFMKLCAALAATMGLSSKAAAEMAESVSNPQRPPVIWIGAQECT WP_000145429.1 hydrogenase 2 small subunit [Salmonella enterica] Length: 372\nNumber of predicted TMHs: 1\nExp number of AAs in TMHs: 24.51309\nExp number, first 60 AAs: 0.2682\nTotal prob of N-in: 0.14402\noutside 1 331\nTMhelix 332 354\ninside 355 372

21287 GCF_001360955.1_10060_5_90 Salmonella enterica subsp. enterica serovar Typhi Proteobacteria; Gammaproteobacteria; Enterobacterales; Enterobacteriaceae; Salmonella; Salmonella enterica; Salmonella enterica subsp. enterica MTGDNTLITSHGINRRDFMKLCAALAATMGLSSKAAAEMAESVSNPQRPPVIWIGAQECT WP_000145429.1 hydrogenase 2 small subunit [Salmonella enterica] Length: 372\nNumber of predicted TMHs: 1\nExp number of AAs in TMHs: 24.51309\nExp number, first 60 AAs: 0.2682\nTotal prob of N-in: 0.14402\noutside 1 331\nTMhelix 332 354\ninside 355 372

21288 GCF_001360995.1_10071_3_1 Salmonella enterica subsp. enterica serovar Typhi Proteobacteria; Gammaproteobacteria; Enterobacterales; Enterobacteriaceae; Salmonella; Salmonella enterica; Salmonella enterica subsp. enterica MTGDNTLITSHGINRRDFMKLCAALAATMGLSSKAAAEMAESVSNPQRPPVIWIGAQECT WP_000145429.1 hydrogenase 2 small subunit [Salmonella enterica] Length: 372\nNumber of predicted TMHs: 1\nExp number of AAs in TMHs: 24.51309\nExp number, first 60 AAs: 0.2682\nTotal prob of N-in: 0.14402\noutside 1 331\nTMhelix 332 354\ninside 355 372

21289 GCF_001361035.1_10071_3_13 Salmonella enterica subsp. enterica serovar Typhi Proteobacteria; Gammaproteobacteria; Enterobacterales; Enterobacteriaceae; Salmonella; Salmonella enterica; Salmonella enterica subsp. enterica MTGDNTLITSHGINRRDFMKLCAALAATMGLSSKAAAEMAESVSNPQRPPVIWIGAQECT WP_000145429.1 hydrogenase 2 small subunit [Salmonella enterica] Length: 372\nNumber of predicted TMHs: 1\nExp number of AAs in TMHs: 24.51309\nExp number, first 60 AAs: 0.2682\nTotal prob of N-in: 0.14402\noutside 1 331\nTMhelix 332 354\ninside 355 372

21290 GCF_001361075.1_10071_3_19 Salmonella enterica subsp. enterica serovar Typhi Proteobacteria; Gammaproteobacteria; Enterobacterales; Enterobacteriaceae; Salmonella; Salmonella enterica; Salmonella enterica subsp. enterica MTGDNTLITSHGINRRDFMKLCAALAATMGLSSKAAAEMAESVSNPQRPPVIWIGAQECT WP_000145429.1 hydrogenase 2 small subunit [Salmonella enterica] Length: 372\nNumber of predicted TMHs: 1\nExp number of AAs in TMHs: 24.51309\nExp number, first 60 AAs: 0.2682\nTotal prob of N-in: 0.14402\noutside 1 331\nTMhelix 332 354\ninside 355 372

21291 GCF_001361195.1_9953_5_52 Salmonella enterica subsp. enterica serovar Typhi Proteobacteria; Gammaproteobacteria; Enterobacterales; Enterobacteriaceae; Salmonella; Salmonella enterica;

Salmonella enterica subsp. enterica MTGDNTLITSHGINRRDFMKLCAALAATMGLSSKAAAEMAESVSNPQRPPVIWIGAQECT
WP_000145429.1 hydrogenase 2 small subunit [Salmonella enterica] Length: 372\nNumber of predicted TMHs: 1\nExp number of AAs in TMHs: 24.51309\nExp number, first 60 AAs: 0.2682\nTotal prob of N-in: 0.14402\noutside 1 331\nTMhelix 332 354\ninside 355 372

21292 GCF_001361115.1_10071_3_28 Salmonella enterica subsp. enterica serovar Typhi
Proteobacteria; Gammaproteobacteria; Enterobacterales; Enterobacteriaceae; Salmonella; Salmonella enterica; Salmonella enterica subsp. enterica MTGDNTLITSHGINRRDFMKLCAALAATMGLSSKAAAEMAESVSNPQRPPVIWIGAQECT
WP_000145429.1 hydrogenase 2 small subunit [Salmonella enterica] Length: 372\nNumber of predicted TMHs: 1\nExp number of AAs in TMHs: 24.51309\nExp number, first 60 AAs: 0.2682\nTotal prob of N-in: 0.14402\noutside 1 331\nTMhelix 332 354\ninside 355 372

21293 GCF_001361235.1_9953_5_70 Salmonella enterica subsp. enterica serovar Typhi
Proteobacteria; Gammaproteobacteria; Enterobacterales; Enterobacteriaceae; Salmonella; Salmonella enterica; Salmonella enterica subsp. enterica MTGDNTLITSHGINRRDFMKLCAALAATMGLSSKAAAEMAESVSNPQRPPVIWIGAQECT
WP_000145429.1 hydrogenase 2 small subunit [Salmonella enterica] Length: 372\nNumber of predicted TMHs: 1\nExp number of AAs in TMHs: 24.51309\nExp number, first 60 AAs: 0.2682\nTotal prob of N-in: 0.14402\noutside 1 331\nTMhelix 332 354\ninside 355 372

21294 GCF_001361275.1_9953_5_88 Salmonella enterica subsp. enterica serovar Typhi
Proteobacteria; Gammaproteobacteria; Enterobacterales; Enterobacteriaceae; Salmonella; Salmonella enterica; Salmonella enterica subsp. enterica MTGDNTLITSHGINRRDFMKLCAALAATMGLSSKAAAEMAESVSNPQRPPVIWIGAQECT
WP_000145429.1 hydrogenase 2 small subunit [Salmonella enterica] Length: 372\nNumber of predicted TMHs: 1\nExp number of AAs in TMHs: 24.51309\nExp number, first 60 AAs: 0.2682\nTotal prob of N-in: 0.14402\noutside 1 331\nTMhelix 332 354\ninside 355 372

21295 GCF_001361355.1_10060_6_13 Salmonella enterica subsp. enterica serovar Typhi
Proteobacteria; Gammaproteobacteria; Enterobacterales; Enterobacteriaceae; Salmonella; Salmonella enterica; Salmonella enterica subsp. enterica MTGDNTLITSHGINRRDFMKLCAALAATMGLSSKAAAEMAESVSNPQRPPVIWIGAQECT
WP_000145429.1 hydrogenase 2 small subunit [Salmonella enterica] Length: 372\nNumber of predicted TMHs: 1\nExp number of AAs in TMHs: 24.51309\nExp number, first 60 AAs: 0.2682\nTotal prob of N-in: 0.14402\noutside 1 331\nTMhelix 332 354\ninside 355 372

21296 GCF_001361315.1_10060_6_7 Salmonella enterica subsp. enterica serovar Typhi
Proteobacteria; Gammaproteobacteria; Enterobacterales; Enterobacteriaceae; Salmonella; Salmonella enterica; Salmonella enterica subsp. enterica MTGDNTLITSHGINRRDFMKLCAALAATMGLSSKAAAEMAESVSNPQRPPVIWIGAQECT
WP_000145429.1 hydrogenase 2 small subunit [Salmonella enterica] Length: 372\nNumber of predicted TMHs: 1\nExp number of AAs in TMHs: 24.51309\nExp number, first 60 AAs: 0.2682\nTotal prob of N-in: 0.14402\noutside 1 331\nTMhelix 332 354\ninside 355 372

21297 GCF_001361395.1_10060_6_25 Salmonella enterica subsp. enterica serovar Typhi
Proteobacteria; Gammaproteobacteria; Enterobacterales; Enterobacteriaceae; Salmonella; Salmonella enterica; Salmonella enterica subsp. enterica MTGDNTLITSHGINRRDFMKLCAALAATMGLSSKAAAEMAESVSNPQRPPVIWIGAQECT
WP_000145429.1 hydrogenase 2 small subunit [Salmonella enterica] Length: 372\nNumber of predicted TMHs: 1\nExp number of AAs in TMHs: 24.51309\nExp number, first 60 AAs: 0.2682\nTotal prob of N-in: 0.14402\noutside 1 331\nTMhelix 332 354\ninside 355 372

21298 GCF_001361155.1_10071_3_55 Salmonella enterica subsp. enterica serovar Typhi
Proteobacteria; Gammaproteobacteria; Enterobacterales; Enterobacteriaceae; Salmonella; Salmonella enterica; Salmonella enterica subsp. enterica MTGDNTLITSHGINRRDFMKLCAALAATMGLSSKAATEMAESVSNPQRPPVIWIGAQECT
WP_001782304.1 hydrogenase 2 small subunit [Salmonella enterica] Length: 372\nNumber of predicted TMHs: 1\nExp number of AAs in TMHs: 24.41173\nExp number, first 60 AAs: 0.16124\nTotal prob of N-in: 0.13947\noutside 1 331\nTMhelix 332 354\ninside 355 372

21299 GCF_001361435.1_10060_6_40 Salmonella enterica subsp. enterica serovar Typhi
Proteobacteria; Gammaproteobacteria; Enterobacterales; Enterobacteriaceae; Salmonella; Salmonella enterica; Salmonella enterica subsp. enterica MTGDNTLITSHGINRRDFMKLCAALAATMGLSSKAAAEMAESVSNPQRPPVIWIGAQECT
WP_000145429.1 hydrogenase 2 small subunit [Salmonella enterica] Length: 372\nNumber of predicted TMHs: 1\nExp number of AAs in TMHs: 24.51309\nExp number, first 60 AAs: 0.2682\nTotal prob of N-in: 0.14402\noutside 1 331\nTMhelix 332 354\ninside 355 372

21300 GCF_001361475.1_10060_6_49 Salmonella enterica subsp. enterica serovar Typhi Proteobacteria; Gammaproteobacteria; Enterobacterales; Enterobacteriaceae; Salmonella; Salmonella enterica; Salmonella enterica subsp. enterica MTGDNTLITSHGINRRDFMKLCAALAATMGLSSKAAAEMAESVSNPQRPPVIWIGAQECT WP_000145429.1 hydrogenase 2 small subunit [Salmonella enterica] Length: 372\nNumber of predicted TMHs: 1\nExp number of AAs in TMHs: 24.51309\nExp number, first 60 AAs: 0.2682\nTotal prob of N-in: 0.14402\noutside 1 331\nTMhelix 332 354\ninside 355 372

21301 GCF_001361495.1_10060_6_55 Salmonella enterica subsp. enterica serovar Typhi Proteobacteria; Gammaproteobacteria; Enterobacterales; Enterobacteriaceae; Salmonella; Salmonella enterica; Salmonella enterica subsp. enterica MTGDNTLITSHGINRRDFMKLCAALAATMGLSSKAAAEMAESVSNPQRPPVIWIGAQECT WP_000145429.1 hydrogenase 2 small subunit [Salmonella enterica] Length: 372\nNumber of predicted TMHs: 1\nExp number of AAs in TMHs: 24.51309\nExp number, first 60 AAs: 0.2682\nTotal prob of N-in: 0.14402\noutside 1 331\nTMhelix 332 354\ninside 355 372

21302 GCF_001361535.1_10060_6_61 Salmonella enterica subsp. enterica serovar Typhi Proteobacteria; Gammaproteobacteria; Enterobacterales; Enterobacteriaceae; Salmonella; Salmonella enterica; Salmonella enterica subsp. enterica MTGDNTLITSHGINRRDFMKLCAALAATMGLSSKAAAEMAESVSNPQRPPVIWIGAQECT WP_000145429.1 hydrogenase 2 small subunit [Salmonella enterica] Length: 372\nNumber of predicted TMHs: 1\nExp number of AAs in TMHs: 24.51309\nExp number, first 60 AAs: 0.2682\nTotal prob of N-in: 0.14402\noutside 1 331\nTMhelix 332 354\ninside 355 372

21303 GCF_001361615.1_10071_8_18 Salmonella enterica subsp. enterica serovar Typhi Proteobacteria; Gammaproteobacteria; Enterobacterales; Enterobacteriaceae; Salmonella; Salmonella enterica; Salmonella enterica subsp. enterica MTGDNTLITSHGINRRDFMKLCAALAATMGLSSKAAAEMAESVSNPQRPPVIWIGAQECT WP_000145429.1 hydrogenase 2 small subunit [Salmonella enterica] Length: 372\nNumber of predicted TMHs: 1\nExp number of AAs in TMHs: 24.51309\nExp number, first 60 AAs: 0.2682\nTotal prob of N-in: 0.14402\noutside 1 331\nTMhelix 332 354\ninside 355 372

21304 GCF_001361575.1_10060_6_67 Salmonella enterica subsp. enterica serovar Typhi Proteobacteria; Gammaproteobacteria; Enterobacterales; Enterobacteriaceae; Salmonella; Salmonella enterica; Salmonella enterica subsp. enterica MTGDNTLITSHGINRRDFMKLCAALAATMGLSSKAAAEMAESVSNPQRPPVIWIGAQECT WP_000145429.1 hydrogenase 2 small subunit [Salmonella enterica] Length: 372\nNumber of predicted TMHs: 1\nExp number of AAs in TMHs: 24.51309\nExp number, first 60 AAs: 0.2682\nTotal prob of N-in: 0.14402\noutside 1 331\nTMhelix 332 354\ninside 355 372

21305 GCF_001361655.1_10071_8_42 Salmonella enterica subsp. enterica serovar Typhi Proteobacteria; Gammaproteobacteria; Enterobacterales; Enterobacteriaceae; Salmonella; Salmonella enterica; Salmonella enterica subsp. enterica MTGDNTLITSHGINRRDFMKLCAALAATMGLSSKAAAEMAESVSNPQRPPVIWIGAQECT WP_000145429.1 hydrogenase 2 small subunit [Salmonella enterica] Length: 372\nNumber of predicted TMHs: 1\nExp number of AAs in TMHs: 24.51309\nExp number, first 60 AAs: 0.2682\nTotal prob of N-in: 0.14402\noutside 1 331\nTMhelix 332 354\ninside 355 372

21306 GCF_001361695.1_10209_5_46 Salmonella enterica subsp. enterica serovar Typhi Proteobacteria; Gammaproteobacteria; Enterobacterales; Enterobacteriaceae; Salmonella; Salmonella enterica; Salmonella enterica subsp. enterica MTGDNTLITSHGINRRDFMKLCAALAATMGLSSKAAAEMAESVSNPQRPPVIWIGAQECT WP_000145429.1 hydrogenase 2 small subunit [Salmonella enterica] Length: 372\nNumber of predicted TMHs: 1\nExp number of AAs in TMHs: 24.51309\nExp number, first 60 AAs: 0.2682\nTotal prob of N-in: 0.14402\noutside 1 331\nTMhelix 332 354\ninside 355 372

21307 GCF_900036805.1_13566_1_66 Salmonella enterica subsp. enterica serovar Typhi Proteobacteria; Gammaproteobacteria; Enterobacterales; Enterobacteriaceae; Salmonella; Salmonella enterica; Salmonella enterica subsp. enterica MTGDNTLITSHGINRRDFMKLCAALAATMGLSSKAAAEMAESVSNPQRPPVIWIGAQECT WP_000145429.1 hydrogenase 2 small subunit [Salmonella enterica] Length: 372\nNumber of predicted TMHs: 1\nExp number of AAs in TMHs: 24.51309\nExp number, first 60 AAs: 0.2682\nTotal prob of N-in: 0.14402\noutside 1 331\nTMhelix 332 354\ninside 355 372

21308 GCF_900036845.1_14555_2_37 Salmonella enterica subsp. enterica serovar Typhi Proteobacteria; Gammaproteobacteria; Enterobacterales; Enterobacteriaceae; Salmonella; Salmonella enterica; Salmonella enterica subsp. enterica MTGDNTLITSHGINRRDFMKLCAALAATMGLSSKAAAEMAESVSNPQRPPVIWIGAQECT WP_000145429.1 hydrogenase 2 small subunit [Salmonella enterica] Length: 372\nNumber of

predicted TMHs:  1\nExp number of AAs in TMHs: 24.51309\nExp number, first 60 AAs:  0.2682\nTotal prob of N-in: 0.14402\noutside   1   331\nTMhelix   332   354\ninside   355   372

21309 GCF_900041465.1_13103_1_8 Salmonella enterica subsp. enterica serovar Typhi Proteobacteria; Gammaproteobacteria; Enterobacterales; Enterobacteriaceae; Salmonella; Salmonella enterica; Salmonella enterica subsp. enterica MTGDNTLITSHGINRRDFMKLCAALAATMGLSSKAAAEMAESVSNPQRPPVIWIGAQECT WP_000145429.1 hydrogenase 2 small subunit [Salmonella enterica] Length: 372\nNumber of predicted TMHs:  1\nExp number of AAs in TMHs: 24.51309\nExp number, first 60 AAs:  0.2682\nTotal prob of N-in: 0.14402\noutside   1   331\nTMhelix   332   354\ninside   355   372

21310 GCF_900041485.1_13134_1_25 Salmonella enterica subsp. enterica serovar Typhi Proteobacteria; Gammaproteobacteria; Enterobacterales; Enterobacteriaceae; Salmonella; Salmonella enterica; Salmonella enterica subsp. enterica MTGDNTLITSHGINRRDFMKLCAALAATMGLSSKAAAEMAESVSNPQRPPVIWIGAQECT WP_000145429.1 hydrogenase 2 small subunit [Salmonella enterica] Length: 372\nNumber of predicted TMHs:  1\nExp number of AAs in TMHs: 24.51309\nExp number, first 60 AAs:  0.2682\nTotal prob of N-in: 0.14402\noutside   1   331\nTMhelix   332   354\ninside   355   372

21311 GCF_900041505.1_13566_1_45 Salmonella enterica subsp. enterica serovar Typhi Proteobacteria; Gammaproteobacteria; Enterobacterales; Enterobacteriaceae; Salmonella; Salmonella enterica; Salmonella enterica subsp. enterica MTGDNTLITSHGINRRDFMKLCAALAATMGLSSKAAAEMAESVSNPQRPPVIWIGAQECT WP_000145429.1 hydrogenase 2 small subunit [Salmonella enterica] Length: 372\nNumber of predicted TMHs:  1\nExp number of AAs in TMHs: 24.51309\nExp number, first 60 AAs:  0.2682\nTotal prob of N-in: 0.14402\noutside   1   331\nTMhelix   332   354\ninside   355   372

21312 GCF_900041525.1_13566_1_47 Salmonella enterica subsp. enterica serovar Typhi Proteobacteria; Gammaproteobacteria; Enterobacterales; Enterobacteriaceae; Salmonella; Salmonella enterica; Salmonella enterica subsp. enterica MTGDNTLITSHGINRRDFMKLCAALAATMGLSSKAAAEMAESVSNPQRPPVIWIGAQECT WP_000145429.1 hydrogenase 2 small subunit [Salmonella enterica] Length: 372\nNumber of predicted TMHs:  1\nExp number of AAs in TMHs: 24.51309\nExp number, first 60 AAs:  0.2682\nTotal prob of N-in: 0.14402\noutside   1   331\nTMhelix   332   354\ninside   355   372

21313 GCF_900041545.1_14555_2_9 Salmonella enterica subsp. enterica serovar Typhi Proteobacteria; Gammaproteobacteria; Enterobacterales; Enterobacteriaceae; Salmonella; Salmonella enterica; Salmonella enterica subsp. enterica MTGDNTLITSHGINRRDFMKLCAALAATMGLSSKAAAEMAESVSNPQRPPVIWIGAQECT WP_000145429.1 hydrogenase 2 small subunit [Salmonella enterica] Length: 372\nNumber of predicted TMHs:  1\nExp number of AAs in TMHs: 24.51309\nExp number, first 60 AAs:  0.2682\nTotal prob of N-in: 0.14402\noutside   1   331\nTMhelix   332   354\ninside   355   372

21314 GCF_900041565.1_14555_2_21 Salmonella enterica subsp. enterica serovar Typhi Proteobacteria; Gammaproteobacteria; Enterobacterales; Enterobacteriaceae; Salmonella; Salmonella enterica; Salmonella enterica subsp. enterica MTGDNTLITSHGINRRDFMKLCAALAATMGLSSKAAAEMAESVSNPQRPPVIWIGAQECT WP_000145429.1 hydrogenase 2 small subunit [Salmonella enterica] Length: 372\nNumber of predicted TMHs:  1\nExp number of AAs in TMHs: 24.51309\nExp number, first 60 AAs:  0.2682\nTotal prob of N-in: 0.14402\noutside   1   331\nTMhelix   332   354\ninside   355   372

21315 GCF_900041585.1_14555_2_25 Salmonella enterica subsp. enterica serovar Typhi Proteobacteria; Gammaproteobacteria; Enterobacterales; Enterobacteriaceae; Salmonella; Salmonella enterica; Salmonella enterica subsp. enterica MTGDNTLITSHGINRRDFMKLCAALAATMGLSSKAAAEMAESVSNPQRPPVIWIGAQECT WP_000145429.1 hydrogenase 2 small subunit [Salmonella enterica] Length: 372\nNumber of predicted TMHs:  1\nExp number of AAs in TMHs: 24.51309\nExp number, first 60 AAs:  0.2682\nTotal prob of N-in: 0.14402\noutside   1   331\nTMhelix   332   354\ninside   355   372

21316 GCF_900018615.1_13566_1_33 Salmonella enterica subsp. enterica serovar Typhi Proteobacteria; Gammaproteobacteria; Enterobacterales; Enterobacteriaceae; Salmonella; Salmonella enterica; Salmonella enterica subsp. enterica MTGDNTLITSHGINRRDFMKLCAALAATMGLSSKAAAEMAESVSNPQRPPVIWIGAQECT WP_000145429.1 hydrogenase 2 small subunit [Salmonella enterica] Length: 372\nNumber of predicted TMHs:  1\nExp number of AAs in TMHs: 24.51309\nExp number, first 60 AAs:  0.2682\nTotal prob of N-in: 0.14402\noutside   1   331\nTMhelix   332   354\ninside   355   372

21317 GCF_900046775.1_14555_2_2 Salmonella enterica subsp. enterica serovar Typhi Proteobacteria; Gammaproteobacteria; Enterobacterales; Enterobacteriaceae; Salmonella; Salmonella enterica;

Salmonella enterica subsp. enterica	MTGDNTLITSHGINRRDFMKLCAALAATMGLSSKAATEMAESVSNPQRPPVIWIGAQECT	WP_001782304.1	hydrogenase 2 small subunit [Salmonella enterica]	Length: 372\nNumber of predicted TMHs: 1\nExp number of AAs in TMHs: 24.41173\nExp number, first 60 AAs: 0.16124\nTotal prob of N-in: 0.13947\noutside 1 331\nTMhelix 332 354\ninside 355 372

21318	GCF_900044305.1_14555_2_60	Salmonella enterica subsp. enterica serovar Typhi	Proteobacteria; Gammaproteobacteria; Enterobacterales; Enterobacteriaceae; Salmonella; Salmonella enterica; Salmonella enterica subsp. enterica	MTGDNTLITSHGINRRDFMKLCAALAATMGLSSKAAAEMAESVSNPQRPPVIWIGAQECT	WP_000145429.1	hydrogenase 2 small subunit [Salmonella enterica]	Length: 372\nNumber of predicted TMHs: 1\nExp number of AAs in TMHs: 24.51309\nExp number, first 60 AAs: 0.2682\nTotal prob of N-in: 0.14402\noutside 1 331\nTMhelix 332 354\ninside 355 372

21319	GCF_900018635.1_13566_1_82	Salmonella enterica subsp. enterica serovar Typhi	Proteobacteria; Gammaproteobacteria; Enterobacterales; Enterobacteriaceae; Salmonella; Salmonella enterica; Salmonella enterica subsp. enterica	MTGDNTLITSHGINRRDFMKLCAALAATMGLSSKAAAEMAESVSNPQRPPVIWIGAQECT	WP_000145429.1	hydrogenase 2 small subunit [Salmonella enterica]	Length: 372\nNumber of predicted TMHs: 1\nExp number of AAs in TMHs: 24.51309\nExp number, first 60 AAs: 0.2682\nTotal prob of N-in: 0.14402\noutside 1 331\nTMhelix 332 354\ninside 355 372

21320	GCF_900043535.1_14555_2_92	Salmonella enterica subsp. enterica serovar Typhi	Proteobacteria; Gammaproteobacteria; Enterobacterales; Enterobacteriaceae; Salmonella; Salmonella enterica; Salmonella enterica subsp. enterica	MTGDNTLITSHGINRRDFMKLCAALAATMGLSSKAAAEMAESVSNPQRPPVIWIGAQECT	WP_000145429.1	hydrogenase 2 small subunit [Salmonella enterica]	Length: 372\nNumber of predicted TMHs: 1\nExp number of AAs in TMHs: 24.51309\nExp number, first 60 AAs: 0.2682\nTotal prob of N-in: 0.14402\noutside 1 331\nTMhelix 332 354\ninside 355 372

21321	GCF_900185485.1_BL60006	Salmonella enterica subsp. enterica serovar Typhi	Proteobacteria; Gammaproteobacteria; Enterobacterales; Enterobacteriaceae; Salmonella; Salmonella enterica; Salmonella enterica subsp. enterica	MTGDNTLITSHGINRRDFMKLCAALAATMGLSSKAAAEMAESVSNPQRPPVIWIGAQECT	WP_000145429.1	hydrogenase 2 small subunit [Salmonella enterica]	Length: 372\nNumber of predicted TMHs: 1\nExp number of AAs in TMHs: 24.51309\nExp number, first 60 AAs: 0.2682\nTotal prob of N-in: 0.14402\noutside 1 331\nTMhelix 332 354\ninside 355 372

21322	GCF_900048075.1_14555_2_35	Salmonella enterica subsp. enterica serovar Typhi	Proteobacteria; Gammaproteobacteria; Enterobacterales; Enterobacteriaceae; Salmonella; Salmonella enterica; Salmonella enterica subsp. enterica	MTGDNTLITSHGINRRDFMKLCAALAATMGLSSKAAAEMAESVSNPQRPPVIWIGAQECT	WP_000145429.1	hydrogenase 2 small subunit [Salmonella enterica]	Length: 372\nNumber of predicted TMHs: 1\nExp number of AAs in TMHs: 24.51309\nExp number, first 60 AAs: 0.2682\nTotal prob of N-in: 0.14402\noutside 1 331\nTMhelix 332 354\ninside 355 372

21323	GCF_900044765.1_14555_2_83	Salmonella enterica subsp. enterica serovar Typhi	Proteobacteria; Gammaproteobacteria; Enterobacterales; Enterobacteriaceae; Salmonella; Salmonella enterica; Salmonella enterica subsp. enterica	MTGDNTLITSHGINRRDFMKLCAALAATMGLSSKAAAEMAESVSNPQRPPVIWIGAQECT	WP_000145429.1	hydrogenase 2 small subunit [Salmonella enterica]	Length: 372\nNumber of predicted TMHs: 1\nExp number of AAs in TMHs: 24.51309\nExp number, first 60 AAs: 0.2682\nTotal prob of N-in: 0.14402\noutside 1 331\nTMhelix 332 354\ninside 355 372

21324	GCF_900044745.1_13134_1_24	Salmonella enterica subsp. enterica serovar Typhi	Proteobacteria; Gammaproteobacteria; Enterobacterales; Enterobacteriaceae; Salmonella; Salmonella enterica; Salmonella enterica subsp. enterica	MTGDNTLITSHGINRRDFMKLCAALAATMGLSSKAAAEMAESVSNPQRPPVIWIGAQECT	WP_000145429.1	hydrogenase 2 small subunit [Salmonella enterica]	Length: 372\nNumber of predicted TMHs: 1\nExp number of AAs in TMHs: 24.51309\nExp number, first 60 AAs: 0.2682\nTotal prob of N-in: 0.14402\noutside 1 331\nTMhelix 332 354\ninside 355 372

21325	GCF_900045455.1_14555_2_87	Salmonella enterica subsp. enterica serovar Typhi	Proteobacteria; Gammaproteobacteria; Enterobacterales; Enterobacteriaceae; Salmonella; Salmonella enterica; Salmonella enterica subsp. enterica	MTGDNTLITSHGINRRDFMKLCAALAATMGLSSKAAAEMAESVSNPQRPPVIWIGAQECT	WP_000145429.1	hydrogenase 2 small subunit [Salmonella enterica]	Length: 372\nNumber of predicted TMHs: 1\nExp number of AAs in TMHs: 24.51309\nExp number, first 60 AAs: 0.2682\nTotal prob of N-in: 0.14402\noutside 1 331\nTMhelix 332 354\ninside 355 372

21326 GCF_900045435.1_13566_1_49 Salmonella enterica subsp. enterica serovar Typhi Proteobacteria; Gammaproteobacteria; Enterobacterales; Enterobacteriaceae; Salmonella; Salmonella enterica; Salmonella enterica subsp. enterica MTGDNTLITSHGINRRDFMKLCAALAATMGLSSKAAAEMAESVSNPQRPPVIWIGAQECT WP_000145429.1 hydrogenase 2 small subunit [Salmonella enterica] Length: 372\nNumber of predicted TMHs: 1\nExp number of AAs in TMHs: 24.51309\nExp number, first 60 AAs: 0.2682\nTotal prob of N-in: 0.14402\noutside 1 331\nTMhelix 332 354\ninside 355 372

21327 GCF_900002855.1_St382-GX1995 Salmonella enterica subsp. enterica serovar Typhi Proteobacteria; Gammaproteobacteria; Enterobacterales; Enterobacteriaceae; Salmonella; Salmonella enterica; Salmonella enterica subsp. enterica MTGDNTLITSHGINRRDFMKLCAALAATMGLSSKAAAEMAESVSNPQRPPVIWIGAQECT WP_000145429.1 hydrogenase 2 small subunit [Salmonella enterica] Length: 372\nNumber of predicted TMHs: 1\nExp number of AAs in TMHs: 24.51309\nExp number, first 60 AAs: 0.2682\nTotal prob of N-in: 0.14402\noutside 1 331\nTMhelix 332 354\ninside 355 372

21328 GCF_900002875.1_St383-GX1995 Salmonella enterica subsp. enterica serovar Typhi Proteobacteria; Gammaproteobacteria; Enterobacterales; Enterobacteriaceae; Salmonella; Salmonella enterica; Salmonella enterica subsp. enterica MTGDNTLITSHGINRRDFMKLCAALAATMGLSSKAAAEMAESVSNPQRPPVIWIGAQECT WP_000145429.1 hydrogenase 2 small subunit [Salmonella enterica] Length: 372\nNumber of predicted TMHs: 1\nExp number of AAs in TMHs: 24.51309\nExp number, first 60 AAs: 0.2682\nTotal prob of N-in: 0.14402\noutside 1 331\nTMhelix 332 354\ninside 355 372

21329 GCF_900045415.1_13566_1_11 Salmonella enterica subsp. enterica serovar Typhi Proteobacteria; Gammaproteobacteria; Enterobacterales; Enterobacteriaceae; Salmonella; Salmonella enterica; Salmonella enterica subsp. enterica MTGDNTLITSHGINRRDFMKLCAALAATMGLSSKAAAEMAESVSNPQRPPVIWIGAQECT WP_000145429.1 hydrogenase 2 small subunit [Salmonella enterica] Length: 372\nNumber of predicted TMHs: 1\nExp number of AAs in TMHs: 24.51309\nExp number, first 60 AAs: 0.2682\nTotal prob of N-in: 0.14402\noutside 1 331\nTMhelix 332 354\ninside 355 372

21330 GCF_900016635.1_ZJ98-11 Salmonella enterica subsp. enterica serovar Typhi Proteobacteria; Gammaproteobacteria; Enterobacterales; Enterobacteriaceae; Salmonella; Salmonella enterica; Salmonella enterica subsp. enterica MTGDNTLITSHGINRRDFMKLCAALAATMGLSSKAAAEMAESVSNPQRPPVIWIGAQECT WP_000145429.1 hydrogenase 2 small subunit [Salmonella enterica] Length: 372\nNumber of predicted TMHs: 1\nExp number of AAs in TMHs: 24.51309\nExp number, first 60 AAs: 0.2682\nTotal prob of N-in: 0.14402\noutside 1 331\nTMhelix 332 354\ninside 355 372

21331 GCF_900017245.1_13134_1_20 Salmonella enterica subsp. enterica serovar Typhi Proteobacteria; Gammaproteobacteria; Enterobacterales; Enterobacteriaceae; Salmonella; Salmonella enterica; Salmonella enterica subsp. enterica MTGDNTLITSHGINRRDFMKLCAALAATMGLSSKAAAEMAESVSNPQRPPVIWIGAQECT WP_000145429.1 hydrogenase 2 small subunit [Salmonella enterica] Length: 372\nNumber of predicted TMHs: 1\nExp number of AAs in TMHs: 24.51309\nExp number, first 60 AAs: 0.2682\nTotal prob of N-in: 0.14402\noutside 1 331\nTMhelix 332 354\ninside 355 372

21332 GCF_900031585.1_13566_1_67 Salmonella enterica subsp. enterica serovar Typhi Proteobacteria; Gammaproteobacteria; Enterobacterales; Enterobacteriaceae; Salmonella; Salmonella enterica; Salmonella enterica subsp. enterica MTGDNTLITSHGINRRDFMKLCAALAATMGLSSKAAAEMAESVSNPQRPPVIWIGAQECT WP_000145429.1 hydrogenase 2 small subunit [Salmonella enterica] Length: 372\nNumber of predicted TMHs: 1\nExp number of AAs in TMHs: 24.51309\nExp number, first 60 AAs: 0.2682\nTotal prob of N-in: 0.14402\noutside 1 331\nTMhelix 332 354\ninside 355 372

21333 GCF_900031595.1_14555_2_72 Salmonella enterica subsp. enterica serovar Typhi Proteobacteria; Gammaproteobacteria; Enterobacterales; Enterobacteriaceae; Salmonella; Salmonella enterica; Salmonella enterica subsp. enterica MTGDNTLITSHGINRRDFMKLCAALAATMGLSSKAAAEMAESVSNPQRPPVIWIGAQECT WP_000145429.1 hydrogenase 2 small subunit [Salmonella enterica] Length: 372\nNumber of predicted TMHs: 1\nExp number of AAs in TMHs: 24.51309\nExp number, first 60 AAs: 0.2682\nTotal prob of N-in: 0.14402\noutside 1 331\nTMhelix 332 354\ninside 355 372

21334 GCF_900018595.1_13566_1_26 Salmonella enterica subsp. enterica serovar Typhi Proteobacteria; Gammaproteobacteria; Enterobacterales; Enterobacteriaceae; Salmonella; Salmonella enterica; Salmonella enterica subsp. enterica MTGDNTLITSHGINRRDFMKLCAALAATMGLSSKAAAEMAESVSNPQRPPVIWIGAQECT WP_000145429.1 hydrogenase 2 small subunit [Salmonella enterica] Length: 372\nNumber of

predicted TMHs:  1\nExp number of AAs in TMHs: 24.51309\nExp number, first 60 AAs:  0.2682\nTotal prob of N-in: 0.14402\noutside   1   331\nTMhelix   332   354\ninside   355   372

21335 GCF_900031615.1_14555_2_75 Salmonella enterica subsp. enterica serovar Typhi Proteobacteria; Gammaproteobacteria; Enterobacterales; Enterobacteriaceae; Salmonella; Salmonella enterica; Salmonella enterica subsp. enterica MTGDNTLITSHGINRRDFMKLCAALAATMGLSSKAAAEMAESVSNPQRPPVIWIGAQECT WP_000145429.1 hydrogenase 2 small subunit [Salmonella enterica] Length: 372\nNumber of predicted TMHs:  1\nExp number of AAs in TMHs: 24.51309\nExp number, first 60 AAs:  0.2682\nTotal prob of N-in: 0.14402\noutside   1   331\nTMhelix   332   354\ninside   355   372

21336 GCF_900031635.1_14555_2_81 Salmonella enterica subsp. enterica serovar Typhi Proteobacteria; Gammaproteobacteria; Enterobacterales; Enterobacteriaceae; Salmonella; Salmonella enterica; Salmonella enterica subsp. enterica MTGDNTLITSHGINRRDFMKLCAALAATMGLSSKAAAEMAESVSNPQRPPVIWIGAQECT WP_000145429.1 hydrogenase 2 small subunit [Salmonella enterica] Length: 372\nNumber of predicted TMHs:  1\nExp number of AAs in TMHs: 24.51309\nExp number, first 60 AAs:  0.2682\nTotal prob of N-in: 0.14402\noutside   1   331\nTMhelix   332   354\ninside   355   372

21337 GCF_900036355.1_14555_2_31 Salmonella enterica subsp. enterica serovar Typhi Proteobacteria; Gammaproteobacteria; Enterobacterales; Enterobacteriaceae; Salmonella; Salmonella enterica; Salmonella enterica subsp. enterica MTGDNTLITSHGINRRDFMKLCAALAATMGLSSKAAAEMAESVSNPQRPPVIWIGAQECT WP_000145429.1 hydrogenase 2 small subunit [Salmonella enterica] Length: 372\nNumber of predicted TMHs:  1\nExp number of AAs in TMHs: 24.51309\nExp number, first 60 AAs:  0.2682\nTotal prob of N-in: 0.14402\noutside   1   331\nTMhelix   332   354\ninside   355   372

21338 GCF_900036375.1_14555_2_43 Salmonella enterica subsp. enterica serovar Typhi Proteobacteria; Gammaproteobacteria; Enterobacterales; Enterobacteriaceae; Salmonella; Salmonella enterica; Salmonella enterica subsp. enterica MTGDNTLITSHGINRRDFMKLCAALAATMGLSSKAAAEMAESVSNPQRPPVIWIGAQECT WP_000145429.1 hydrogenase 2 small subunit [Salmonella enterica] Length: 372\nNumber of predicted TMHs:  1\nExp number of AAs in TMHs: 24.51309\nExp number, first 60 AAs:  0.2682\nTotal prob of N-in: 0.14402\noutside   1   331\nTMhelix   332   354\ninside   355   372

21339 GCF_900036745.1_13103_1_6 Salmonella enterica subsp. enterica serovar Typhi Proteobacteria; Gammaproteobacteria; Enterobacterales; Enterobacteriaceae; Salmonella; Salmonella enterica; Salmonella enterica subsp. enterica MTGDNTLITSHGINRRDFMKLCAALAATMGLSSKAAAEMAESVSNPQRPPVIWIGAQECT WP_000145429.1 hydrogenase 2 small subunit [Salmonella enterica] Length: 372\nNumber of predicted TMHs:  1\nExp number of AAs in TMHs: 24.51309\nExp number, first 60 AAs:  0.2682\nTotal prob of N-in: 0.14402\noutside   1   331\nTMhelix   332   354\ninside   355   372

21340 GCF_900036765.1_13566_1_53 Salmonella enterica subsp. enterica serovar Typhi Proteobacteria; Gammaproteobacteria; Enterobacterales; Enterobacteriaceae; Salmonella; Salmonella enterica; Salmonella enterica subsp. enterica MTGDNTLITSHGINRRDFMKLCAALAATMGLSSKAAAEMAESVSNPQRPPVIWIGAQECT WP_000145429.1 hydrogenase 2 small subunit [Salmonella enterica] Length: 372\nNumber of predicted TMHs:  1\nExp number of AAs in TMHs: 24.51309\nExp number, first 60 AAs:  0.2682\nTotal prob of N-in: 0.14402\noutside   1   331\nTMhelix   332   354\ninside   355   372

21341 GCF_900036785.1_13566_1_60 Salmonella enterica subsp. enterica serovar Typhi Proteobacteria; Gammaproteobacteria; Enterobacterales; Enterobacteriaceae; Salmonella; Salmonella enterica; Salmonella enterica subsp. enterica MTGDNTLITSHGINRRDFMKLCAALAATMGLSSKAAAEMAESVSNPQRPPVIWIGAQECT WP_000145429.1 hydrogenase 2 small subunit [Salmonella enterica] Length: 372\nNumber of predicted TMHs:  1\nExp number of AAs in TMHs: 24.51309\nExp number, first 60 AAs:  0.2682\nTotal prob of N-in: 0.14402\noutside   1   331\nTMhelix   332   354\ninside   355   372

21342 GCF_900036755.1_13566_1_17 Salmonella enterica subsp. enterica serovar Typhi Proteobacteria; Gammaproteobacteria; Enterobacterales; Enterobacteriaceae; Salmonella; Salmonella enterica; Salmonella enterica subsp. enterica MTGDNTLITSHGINRRDFMKLCAALAATMGLSSKAAAEMAESVSNPQRPPVIWIGAQECT WP_000145429.1 hydrogenase 2 small subunit [Salmonella enterica] Length: 372\nNumber of predicted TMHs:  1\nExp number of AAs in TMHs: 24.51309\nExp number, first 60 AAs:  0.2682\nTotal prob of N-in: 0.14402\noutside   1   331\nTMhelix   332   354\ninside   355   372

21343 GCF_900036385.1_14555_2_44 Salmonella enterica subsp. enterica serovar Typhi Proteobacteria; Gammaproteobacteria; Enterobacterales; Enterobacteriaceae; Salmonella; Salmonella enterica;

Salmonella enterica subsp. enterica MTGDNTLITSHGINRRDFMKLCAALAATMGLSSKAAAEMAESVSNPQRPPVIWIGAQECT WP_000145429.1 hydrogenase 2 small subunit [Salmonella enterica] Length: 372\nNumber of predicted TMHs: 1\nExp number of AAs in TMHs: 24.51309\nExp number, first 60 AAs: 0.2682\nTotal prob of N-in: 0.14402\noutside 1 331\nTMhelix 332 354\ninside 355 372

21344 GCF_900043525.1_14555_2_71 Salmonella enterica subsp. enterica serovar Typhi Proteobacteria; Gammaproteobacteria; Enterobacterales; Enterobacteriaceae; Salmonella; Salmonella enterica; Salmonella enterica subsp. enterica MTGDNTLITSHGINRRDFMKLCAALAATMGLSSKAAAEMAESVSNPQRPPVIWIGAQECT WP_000145429.1 hydrogenase 2 small subunit [Salmonella enterica] Length: 372\nNumber of predicted TMHs: 1\nExp number of AAs in TMHs: 24.51309\nExp number, first 60 AAs: 0.2682\nTotal prob of N-in: 0.14402\noutside 1 331\nTMhelix 332 354\ninside 355 372

21345 GCF_900036365.1_14555_2_39 Salmonella enterica subsp. enterica serovar Typhi Proteobacteria; Gammaproteobacteria; Enterobacterales; Enterobacteriaceae; Salmonella; Salmonella enterica; Salmonella enterica subsp. enterica MTGDNTLITSHGINRRDFMKLCAALAATMGLSSKAAAEMAESVSNPQRPPVIWIGAQECT WP_000145429.1 hydrogenase 2 small subunit [Salmonella enterica] Length: 372\nNumber of predicted TMHs: 1\nExp number of AAs in TMHs: 24.51309\nExp number, first 60 AAs: 0.2682\nTotal prob of N-in: 0.14402\noutside 1 331\nTMhelix 332 354\ninside 355 372

21346 GCF_900044085.1_14555_2_78 Salmonella enterica subsp. enterica serovar Typhi Proteobacteria; Gammaproteobacteria; Enterobacterales; Enterobacteriaceae; Salmonella; Salmonella enterica; Salmonella enterica subsp. enterica MTGDNTLITSHGINRRDFMKLCAALAATMGLSSKAAAEMAESVSNPQRPPVIWIGAQECT WP_000145429.1 hydrogenase 2 small subunit [Salmonella enterica] Length: 372\nNumber of predicted TMHs: 1\nExp number of AAs in TMHs: 24.51309\nExp number, first 60 AAs: 0.2682\nTotal prob of N-in: 0.14402\noutside 1 331\nTMhelix 332 354\ninside 355 372

21347 GCF_900044315.1_14555_2_61 Salmonella enterica subsp. enterica serovar Typhi Proteobacteria; Gammaproteobacteria; Enterobacterales; Enterobacteriaceae; Salmonella; Salmonella enterica; Salmonella enterica subsp. enterica MTGDNTLITSHGINRRDFMKLCAALAATMGLSSKAAAEMAESVSNPQRPPVIWIGAQECT WP_000145429.1 hydrogenase 2 small subunit [Salmonella enterica] Length: 372\nNumber of predicted TMHs: 1\nExp number of AAs in TMHs: 24.51309\nExp number, first 60 AAs: 0.2682\nTotal prob of N-in: 0.14402\noutside 1 331\nTMhelix 332 354\ninside 355 372

21348 GCF_900045405.1_13566_1_10 Salmonella enterica subsp. enterica serovar Typhi Proteobacteria; Gammaproteobacteria; Enterobacterales; Enterobacteriaceae; Salmonella; Salmonella enterica; Salmonella enterica subsp. enterica MTGDNTLITSHGINRRDFMKLCAALAATMGLSSKAAAEMAESVSNPQRPPVIWIGAQECT WP_000145429.1 hydrogenase 2 small subunit [Salmonella enterica] Length: 372\nNumber of predicted TMHs: 1\nExp number of AAs in TMHs: 24.51309\nExp number, first 60 AAs: 0.2682\nTotal prob of N-in: 0.14402\noutside 1 331\nTMhelix 332 354\ninside 355 372

21349 GCF_900036345.1_14555_2_29 Salmonella enterica subsp. enterica serovar Typhi Proteobacteria; Gammaproteobacteria; Enterobacterales; Enterobacteriaceae; Salmonella; Salmonella enterica; Salmonella enterica subsp. enterica MTGDNTLITSHGINRRDFMKLCAALAATMGLSSKAAAEMAESVSNPQRPPVIWIGAQECT WP_000145429.1 hydrogenase 2 small subunit [Salmonella enterica] Length: 372\nNumber of predicted TMHs: 1\nExp number of AAs in TMHs: 24.51309\nExp number, first 60 AAs: 0.2682\nTotal prob of N-in: 0.14402\noutside 1 331\nTMhelix 332 354\ninside 355 372

21350 GCF_900036325.1_14555_2_27 Salmonella enterica subsp. enterica serovar Typhi Proteobacteria; Gammaproteobacteria; Enterobacterales; Enterobacteriaceae; Salmonella; Salmonella enterica; Salmonella enterica subsp. enterica MTGDNTLITSHGINRRDFMKLCAALAATMGLSSKAAAEMAESVSNPQRPPVIWIGAQECT WP_000145429.1 hydrogenase 2 small subunit [Salmonella enterica] Length: 372\nNumber of predicted TMHs: 1\nExp number of AAs in TMHs: 24.51309\nExp number, first 60 AAs: 0.2682\nTotal prob of N-in: 0.14402\noutside 1 331\nTMhelix 332 354\ninside 355 372

21351 GCF_900036305.1_13566_1_21 Salmonella enterica subsp. enterica serovar Typhi Proteobacteria; Gammaproteobacteria; Enterobacterales; Enterobacteriaceae; Salmonella; Salmonella enterica; Salmonella enterica subsp. enterica MTGDNTLITSHGINRRDFMKLCAALAATMGLSSKAAAEMAESVSNPQRPPVIWIGAQECT WP_000145429.1 hydrogenase 2 small subunit [Salmonella enterica] Length: 372\nNumber of predicted TMHs: 1\nExp number of AAs in TMHs: 24.51309\nExp number, first 60 AAs: 0.2682\nTotal prob of N-in: 0.14402\noutside 1 331\nTMhelix 332 354\ninside 355 372

21352	GCF_900036285.1_13134_1_16	Salmonella enterica subsp. enterica serovar Typhi	Proteobacteria; Gammaproteobacteria; Enterobacterales; Enterobacteriaceae; Salmonella; Salmonella enterica; Salmonella enterica subsp. enterica	MTGDNTLITSHGINRRDFMKLCAALAATMGLSSKAAAEMAESVSNPQRPPVIWIGAQECT	WP_000145429.1	hydrogenase 2 small subunit [Salmonella enterica]	Length: 372\nNumber of predicted TMHs:  1\nExp number of AAs in TMHs: 24.51309\nExp number, first 60 AAs:  0.2682\nTotal prob of N-in: 0.14402\noutside	1	331\nTMhelix	332	354\ninside	355	372

21353	GCF_900044755.1_13566_1_42	Salmonella enterica subsp. enterica serovar Typhi	Proteobacteria; Gammaproteobacteria; Enterobacterales; Enterobacteriaceae; Salmonella; Salmonella enterica; Salmonella enterica subsp. enterica	MTGDNTLITSHGINRRDFMKLCAALAATMGLSSKAAAEMAESVSNPQRPPVIWIGAQECT	WP_000145429.1	hydrogenase 2 small subunit [Salmonella enterica]	Length: 372\nNumber of predicted TMHs:  1\nExp number of AAs in TMHs: 24.51309\nExp number, first 60 AAs:  0.2682\nTotal prob of N-in: 0.14402\noutside	1	331\nTMhelix	332	354\ninside	355	372

21354	GCF_900045445.1_13566_1_52	Salmonella enterica subsp. enterica serovar Typhi	Proteobacteria; Gammaproteobacteria; Enterobacterales; Enterobacteriaceae; Salmonella; Salmonella enterica; Salmonella enterica subsp. enterica	MTGDNTLITSHGINRRDFMKLCAALAATMGLSSKAAAEMAESVSNPQRPPVIWIGAQECT	WP_000145429.1	hydrogenase 2 small subunit [Salmonella enterica]	Length: 372\nNumber of predicted TMHs:  1\nExp number of AAs in TMHs: 24.51309\nExp number, first 60 AAs:  0.2682\nTotal prob of N-in: 0.14402\noutside	1	331\nTMhelix	332	354\ninside	355	372

21355	GCF_900041535.1_14555_2_8	Salmonella enterica subsp. enterica serovar Typhi	Proteobacteria; Gammaproteobacteria; Enterobacterales; Enterobacteriaceae; Salmonella; Salmonella enterica; Salmonella enterica subsp. enterica	MTGDNTLITSHGINRRDFMKLCAALAATMGLSSKAAAEMAESVSNPQRPPVIWIGAQECT	WP_000145429.1	hydrogenase 2 small subunit [Salmonella enterica]	Length: 372\nNumber of predicted TMHs:  1\nExp number of AAs in TMHs: 24.51309\nExp number, first 60 AAs:  0.2682\nTotal prob of N-in: 0.14402\noutside	1	331\nTMhelix	332	354\ninside	355	372

21356	GCF_900045425.1_13566_1_6	Salmonella enterica subsp. enterica serovar Typhi	Proteobacteria; Gammaproteobacteria; Enterobacterales; Enterobacteriaceae; Salmonella; Salmonella enterica; Salmonella enterica subsp. enterica	MTGDNTLITSHGINRRDFMKLCAALAATMGLSSKAAAEMAESVSNPQRPPVIWIGAQECT	WP_000145429.1	hydrogenase 2 small subunit [Salmonella enterica]	Length: 372\nNumber of predicted TMHs:  1\nExp number of AAs in TMHs: 24.51309\nExp number, first 60 AAs:  0.2682\nTotal prob of N-in: 0.14402\noutside	1	331\nTMhelix	332	354\ninside	355	372

21357	GCF_900046745.1_13103_1_7	Salmonella enterica subsp. enterica serovar Typhi	Proteobacteria; Gammaproteobacteria; Enterobacterales; Enterobacteriaceae; Salmonella; Salmonella enterica; Salmonella enterica subsp. enterica	MTGDNTLITSHGINRRDFMKLCAALAATMGLSSKAAAEMAESVSNPQRPPVIWIGAQECT	WP_000145429.1	hydrogenase 2 small subunit [Salmonella enterica]	Length: 372\nNumber of predicted TMHs:  1\nExp number of AAs in TMHs: 24.51309\nExp number, first 60 AAs:  0.2682\nTotal prob of N-in: 0.14402\noutside	1	331\nTMhelix	332	354\ninside	355	372

21358	GCF_900035565.1_14555_2_51	Salmonella enterica subsp. enterica serovar Typhi	Proteobacteria; Gammaproteobacteria; Enterobacterales; Enterobacteriaceae; Salmonella; Salmonella enterica; Salmonella enterica subsp. enterica	MTGDNTLITSHGINRRDFMKLCAALAATMGLSSKAAAEMAESVSNPQRPPVIWIGAQECT	WP_000145429.1	hydrogenase 2 small subunit [Salmonella enterica]	Length: 372\nNumber of predicted TMHs:  1\nExp number of AAs in TMHs: 24.51309\nExp number, first 60 AAs:  0.2682\nTotal prob of N-in: 0.14402\noutside	1	331\nTMhelix	332	354\ninside	355	372

21359	GCF_900046765.1_13566_1_2	Salmonella enterica subsp. enterica serovar Typhi	Proteobacteria; Gammaproteobacteria; Enterobacterales; Enterobacteriaceae; Salmonella; Salmonella enterica; Salmonella enterica subsp. enterica	MTGDNTLITSHGINRRDFMKLCAALAATMGLSSKAAAEMAESVSNPQRPPVIWIGAQECT	WP_000145429.1	hydrogenase 2 small subunit [Salmonella enterica]	Length: 372\nNumber of predicted TMHs:  1\nExp number of AAs in TMHs: 24.51309\nExp number, first 60 AAs:  0.2682\nTotal prob of N-in: 0.14402\noutside	1	331\nTMhelix	332	354\ninside	355	372

21360	GCF_900041515.1_13566_1_46	Salmonella enterica subsp. enterica serovar Typhi	Proteobacteria; Gammaproteobacteria; Enterobacterales; Enterobacteriaceae; Salmonella; Salmonella enterica; Salmonella enterica subsp. enterica	MTGDNTLITSHGINRRDFMKLCAALAATMGLSSKAAAEMAESVSNPQRPPVIWIGAQECT	WP_000145429.1	hydrogenase 2 small subunit [Salmonella enterica]	Length: 372\nNumber of

predicted TMHs: 1\nExp number of AAs in TMHs: 24.51309\nExp number, first 60 AAs: 0.2682\nTotal prob of N-in: 0.14402\noutside 1 331\nTMhelix 332 354\ninside 355 372

21361 GCF_900046785.1_14555_2_3 Salmonella enterica subsp. enterica serovar Typhi Proteobacteria; Gammaproteobacteria; Enterobacterales; Enterobacteriaceae; Salmonella; Salmonella enterica; Salmonella enterica subsp. enterica MTGDNTLITSHGINRRDFMKLCAALAATMGLSSKAAAEMAESVSNPQRPPVIWIGAQECT WP_000145429.1 hydrogenase 2 small subunit [Salmonella enterica] Length: 372\nNumber of predicted TMHs: 1\nExp number of AAs in TMHs: 24.51309\nExp number, first 60 AAs: 0.2682\nTotal prob of N-in: 0.14402\noutside 1 331\nTMhelix 332 354\ninside 355 372

21362 GCF_900041575.1_14555_2_24 Salmonella enterica subsp. enterica serovar Typhi Proteobacteria; Gammaproteobacteria; Enterobacterales; Enterobacteriaceae; Salmonella; Salmonella enterica; Salmonella enterica subsp. enterica MTGDNTLITSHGINRRDFMKLCAALAATMGLSSKAAAEMAESVSNPQRPPVIWIGAQECT WP_000145429.1 hydrogenase 2 small subunit [Salmonella enterica] Length: 372\nNumber of predicted TMHs: 1\nExp number of AAs in TMHs: 24.51309\nExp number, first 60 AAs: 0.2682\nTotal prob of N-in: 0.14402\noutside 1 331\nTMhelix 332 354\ninside 355 372

21363 GCF_900041495.1_13566_1_25 Salmonella enterica subsp. enterica serovar Typhi Proteobacteria; Gammaproteobacteria; Enterobacterales; Enterobacteriaceae; Salmonella; Salmonella enterica; Salmonella enterica subsp. enterica MTGDNTLITSHGINRRDFMKLCAALAATMGLSSKAAAEMAESVSNPQRPPVIWIGAQECT WP_000145429.1 hydrogenase 2 small subunit [Salmonella enterica] Length: 372\nNumber of predicted TMHs: 1\nExp number of AAs in TMHs: 24.51309\nExp number, first 60 AAs: 0.2682\nTotal prob of N-in: 0.14402\noutside 1 331\nTMhelix 332 354\ninside 355 372

21364 GCF_900041475.1_13134_1_23 Salmonella enterica subsp. enterica serovar Typhi Proteobacteria; Gammaproteobacteria; Enterobacterales; Enterobacteriaceae; Salmonella; Salmonella enterica; Salmonella enterica subsp. enterica MTGDNTLITSHGINRRDFMKLCAALAATMGLSSKAAAEMAESVSNPQRPPVIWIGAQECT WP_000145429.1 hydrogenase 2 small subunit [Salmonella enterica] Length: 372\nNumber of predicted TMHs: 1\nExp number of AAs in TMHs: 24.51309\nExp number, first 60 AAs: 0.2682\nTotal prob of N-in: 0.14402\noutside 1 331\nTMhelix 332 354\ninside 355 372

21365 GCF_900041435.1_13097_1_78 Salmonella enterica subsp. enterica serovar Typhi Proteobacteria; Gammaproteobacteria; Enterobacterales; Enterobacteriaceae; Salmonella; Salmonella enterica; Salmonella enterica subsp. enterica MTGDNTLITSHGINRRDFMKLCAALAATMGLSSKAAAEMAESVSNPQRPPVIWIGAQECT WP_000145429.1 hydrogenase 2 small subunit [Salmonella enterica] Length: 372\nNumber of predicted TMHs: 1\nExp number of AAs in TMHs: 24.51309\nExp number, first 60 AAs: 0.2682\nTotal prob of N-in: 0.14402\noutside 1 331\nTMhelix 332 354\ninside 355 372

21366 GCF_900041605.1_14555_2_36 Salmonella enterica subsp. enterica serovar Typhi Proteobacteria; Gammaproteobacteria; Enterobacterales; Enterobacteriaceae; Salmonella; Salmonella enterica; Salmonella enterica subsp. enterica MTGDNTLITSHGINRRDFMKLCAALAATMGLSSKAAAEMAESVSNPQRPPVIWIGAQECT WP_000145429.1 hydrogenase 2 small subunit [Salmonella enterica] Length: 372\nNumber of predicted TMHs: 1\nExp number of AAs in TMHs: 24.51309\nExp number, first 60 AAs: 0.2682\nTotal prob of N-in: 0.14402\noutside 1 331\nTMhelix 332 354\ninside 355 372

21367 GCF_900041625.1_14555_2_54 Salmonella enterica subsp. enterica serovar Typhi Proteobacteria; Gammaproteobacteria; Enterobacterales; Enterobacteriaceae; Salmonella; Salmonella enterica; Salmonella enterica subsp. enterica MTGDNTLITSHGINRRDFMKLCAALAATMGLSSKAAAEMAESVSNPQRPPVIWIGAQECT WP_000145429.1 hydrogenase 2 small subunit [Salmonella enterica] Length: 372\nNumber of predicted TMHs: 1\nExp number of AAs in TMHs: 24.51309\nExp number, first 60 AAs: 0.2682\nTotal prob of N-in: 0.14402\noutside 1 331\nTMhelix 332 354\ninside 355 372

21368 GCF_900033725.1_14555_2_64 Salmonella enterica subsp. enterica serovar Typhi Proteobacteria; Gammaproteobacteria; Enterobacterales; Enterobacteriaceae; Salmonella; Salmonella enterica; Salmonella enterica subsp. enterica MTGDNTLITSHGINRRDFMKLCAALAATMGLSSKAAAEMAESVSNPQRPPVIWIGAQECT WP_000145429.1 hydrogenase 2 small subunit [Salmonella enterica] Length: 372\nNumber of predicted TMHs: 1\nExp number of AAs in TMHs: 24.51309\nExp number, first 60 AAs: 0.2682\nTotal prob of N-in: 0.14402\noutside 1 331\nTMhelix 332 354\ninside 355 372

21369 GCF_900041455.1_13097_1_85 Salmonella enterica subsp. enterica serovar Typhi Proteobacteria; Gammaproteobacteria; Enterobacterales; Enterobacteriaceae; Salmonella; Salmonella enterica;

Salmonella enterica subsp. enterica	MTGDNTLITSHGINRRDFMKLCAALAATMGLSSKAAAEMAESVSNPQRPPVIWIGAQECT
	WP_000145429.1	hydrogenase 2 small subunit [Salmonella enterica]	Length: 372\nNumber of predicted TMHs:  1\nExp number of AAs in TMHs: 24.51309\nExp number, first 60 AAs:  0.2682\nTotal prob of N-in: 0.14402\noutside   1   331\nTMhelix   332   354\ninside   355   372

21370	GCF_900033705.1_13103_1_13	Salmonella enterica subsp. enterica serovar Typhi
	Proteobacteria; Gammaproteobacteria; Enterobacterales; Enterobacteriaceae; Salmonella; Salmonella enterica; Salmonella enterica subsp. enterica	MTGDNTLITSHGINRRDFMKLCAALAATMGLSSKAAAEMAESVSNPQRPPVIWIGAQECT
	WP_000145429.1	hydrogenase 2 small subunit [Salmonella enterica]	Length: 372\nNumber of predicted TMHs:  1\nExp number of AAs in TMHs: 24.51309\nExp number, first 60 AAs:  0.2682\nTotal prob of N-in: 0.14402\noutside   1   331\nTMhelix   332   354\ninside   355   372

21371	GCF_900040675.1_13566_1_40	Salmonella enterica subsp. enterica serovar Typhi
	Proteobacteria; Gammaproteobacteria; Enterobacterales; Enterobacteriaceae; Salmonella; Salmonella enterica; Salmonella enterica subsp. enterica	MTGDNTLITSHGINRRDFMKLCAALAATMGLSSKAAAEMAESVSNPQRPPVIWIGAQECT
	WP_000145429.1	hydrogenase 2 small subunit [Salmonella enterica]	Length: 372\nNumber of predicted TMHs:  1\nExp number of AAs in TMHs: 24.51309\nExp number, first 60 AAs:  0.2682\nTotal prob of N-in: 0.14402\noutside   1   331\nTMhelix   332   354\ninside   355   372

21372	GCF_900031625.1_14555_2_79	Salmonella enterica subsp. enterica serovar Typhi
	Proteobacteria; Gammaproteobacteria; Enterobacterales; Enterobacteriaceae; Salmonella; Salmonella enterica; Salmonella enterica subsp. enterica	MTGDNTLITSHGINRRDFMKLCAALAATMGLSSKAAAEMAESVSNPQRPPVIWIGAQECT
	WP_000145429.1	hydrogenase 2 small subunit [Salmonella enterica]	Length: 372\nNumber of predicted TMHs:  1\nExp number of AAs in TMHs: 24.51309\nExp number, first 60 AAs:  0.2682\nTotal prob of N-in: 0.14402\noutside   1   331\nTMhelix   332   354\ninside   355   372

21373	GCF_900041635.1_14555_2_56	Salmonella enterica subsp. enterica serovar Typhi
	Proteobacteria; Gammaproteobacteria; Enterobacterales; Enterobacteriaceae; Salmonella; Salmonella enterica; Salmonella enterica subsp. enterica	MTGDNTLITSHGINRRDFMKLCAALAATMGLSSKAAAEMAESVSNPQRPPVIWIGAQECT
	WP_000145429.1	hydrogenase 2 small subunit [Salmonella enterica]	Length: 372\nNumber of predicted TMHs:  1\nExp number of AAs in TMHs: 24.51309\nExp number, first 60 AAs:  0.2682\nTotal prob of N-in: 0.14402\noutside   1   331\nTMhelix   332   354\ninside   355   372

21374	GCF_900041655.1_14555_2_63	Salmonella enterica subsp. enterica serovar Typhi
	Proteobacteria; Gammaproteobacteria; Enterobacterales; Enterobacteriaceae; Salmonella; Salmonella enterica; Salmonella enterica subsp. enterica	MTGDNTLITSHGINRRDFMKLCAALAATMGLSSKAAAEMAESVSNPQRPPVIWIGAQECT
	WP_000145429.1	hydrogenase 2 small subunit [Salmonella enterica]	Length: 372\nNumber of predicted TMHs:  1\nExp number of AAs in TMHs: 24.51309\nExp number, first 60 AAs:  0.2682\nTotal prob of N-in: 0.14402\noutside   1   331\nTMhelix   332   354\ninside   355   372

21375	GCF_900031605.1_14555_2_73	Salmonella enterica subsp. enterica serovar Typhi
	Proteobacteria; Gammaproteobacteria; Enterobacterales; Enterobacteriaceae; Salmonella; Salmonella enterica; Salmonella enterica subsp. enterica	MTGDNTLITSHGINRRDFMKLCAALAATMGLSSKAAAEMAESVSNPQRPPVIWIGAQECT
	WP_000145429.1	hydrogenase 2 small subunit [Salmonella enterica]	Length: 372\nNumber of predicted TMHs:  1\nExp number of AAs in TMHs: 24.51309\nExp number, first 60 AAs:  0.2682\nTotal prob of N-in: 0.14402\noutside   1   331\nTMhelix   332   354\ninside   355   372

21376	GCF_900031575.1_13566_1_35	Salmonella enterica subsp. enterica serovar Typhi
	Proteobacteria; Gammaproteobacteria; Enterobacterales; Enterobacteriaceae; Salmonella; Salmonella enterica; Salmonella enterica subsp. enterica	MTGDNTLITSHGINRRDFMKLCAALAATMGLSSKAAAEMAESVSNPQRPPVIWIGAQECT
	WP_000145429.1	hydrogenase 2 small subunit [Salmonella enterica]	Length: 372\nNumber of predicted TMHs:  1\nExp number of AAs in TMHs: 24.51309\nExp number, first 60 AAs:  0.2682\nTotal prob of N-in: 0.14402\noutside   1   331\nTMhelix   332   354\ninside   355   372

21377	GCF_900031565.1_13566_1_14	Salmonella enterica subsp. enterica serovar Typhi
	Proteobacteria; Gammaproteobacteria; Enterobacterales; Enterobacteriaceae; Salmonella; Salmonella enterica; Salmonella enterica subsp. enterica	MTGDNTLITSHGINRRDFMKLCAALAATMGLSSKAAAEMAESVSNPQRPPVIWIGAQECT
	WP_000145429.1	hydrogenase 2 small subunit [Salmonella enterica]	Length: 372\nNumber of predicted TMHs:  1\nExp number of AAs in TMHs: 24.51309\nExp number, first 60 AAs:  0.2682\nTotal prob of N-in: 0.14402\noutside   1   331\nTMhelix   332   354\ninside   355   372

21378	GCF_900040035.1_14555_2_20	Salmonella enterica subsp. enterica serovar Typhi	Proteobacteria; Gammaproteobacteria; Enterobacterales; Enterobacteriaceae; Salmonella; Salmonella enterica; Salmonella enterica subsp. enterica	MTGDNTLITSHGINRRDFMKLCAALAATMGLSSKAAAEMAESVSNPQRPPVIWIGAQECT	WP_000145429.1	hydrogenase 2 small subunit [Salmonella enterica]	Length: 372\nNumber of predicted TMHs: 1\nExp number of AAs in TMHs: 24.51309\nExp number, first 60 AAs: 0.2682\nTotal prob of N-in: 0.14402\noutside 1 331\nTMhelix 332 354\ninside 355 372

21379	GCF_900031555.1_13566_1_4	Salmonella enterica subsp. enterica serovar Typhi	Proteobacteria; Gammaproteobacteria; Enterobacterales; Enterobacteriaceae; Salmonella; Salmonella enterica; Salmonella enterica subsp. enterica	MTGDNTLITSHGINRRDFMKLCAALAATMGLSSKAAAEMAESVSNPQRPPVIWIGAQECT	WP_000145429.1	hydrogenase 2 small subunit [Salmonella enterica]	Length: 372\nNumber of predicted TMHs: 1\nExp number of AAs in TMHs: 24.51309\nExp number, first 60 AAs: 0.2682\nTotal prob of N-in: 0.14402\noutside 1 331\nTMhelix 332 354\ninside 355 372

21380	GCF_900029315.1_13134_1_18	Salmonella enterica subsp. enterica serovar Typhi	Proteobacteria; Gammaproteobacteria; Enterobacterales; Enterobacteriaceae; Salmonella; Salmonella enterica; Salmonella enterica subsp. enterica	MTGDNTLITSHGINRRDFMKLCAALAATMGLSSKAAAEMAESVSNPQRPPVIWIGAQECT	WP_000145429.1	hydrogenase 2 small subunit [Salmonella enterica]	Length: 372\nNumber of predicted TMHs: 1\nExp number of AAs in TMHs: 24.51309\nExp number, first 60 AAs: 0.2682\nTotal prob of N-in: 0.14402\noutside 1 331\nTMhelix 332 354\ninside 355 372

21381	GCF_900042415.1_13071_1_93	Salmonella enterica subsp. enterica serovar Typhi	Proteobacteria; Gammaproteobacteria; Enterobacterales; Enterobacteriaceae; Salmonella; Salmonella enterica; Salmonella enterica subsp. enterica	MTGDNTLITSHGINRRDFMKLCAALAATMGLSSKAAAEMAESVSNPQRPPVIWIGAQECT	WP_000145429.1	hydrogenase 2 small subunit [Salmonella enterica]	Length: 372\nNumber of predicted TMHs: 1\nExp number of AAs in TMHs: 24.51309\nExp number, first 60 AAs: 0.2682\nTotal prob of N-in: 0.14402\noutside 1 331\nTMhelix 332 354\ninside 355 372

21382	GCF_900039615.1_13566_1_81	Salmonella enterica subsp. enterica serovar Typhi	Proteobacteria; Gammaproteobacteria; Enterobacterales; Enterobacteriaceae; Salmonella; Salmonella enterica; Salmonella enterica subsp. enterica	MTGDNTLITSHGINRRDFMKLCAALAATMGLSSKAAAEMAESVSNPQRPPVIWIGAQECT	WP_000145429.1	hydrogenase 2 small subunit [Salmonella enterica]	Length: 372\nNumber of predicted TMHs: 1\nExp number of AAs in TMHs: 24.51309\nExp number, first 60 AAs: 0.2682\nTotal prob of N-in: 0.14402\noutside 1 331\nTMhelix 332 354\ninside 355 372

21383	GCF_900029295.1_13103_1_2	Salmonella enterica subsp. enterica serovar Typhi	Proteobacteria; Gammaproteobacteria; Enterobacterales; Enterobacteriaceae; Salmonella; Salmonella enterica; Salmonella enterica subsp. enterica	MTGDNTLITSHGINRRDFMKLCAALAATMGLSSKAAAEMAESVSNPQRPPVIWIGAQECT	WP_000145429.1	hydrogenase 2 small subunit [Salmonella enterica]	Length: 372\nNumber of predicted TMHs: 1\nExp number of AAs in TMHs: 24.51309\nExp number, first 60 AAs: 0.2682\nTotal prob of N-in: 0.14402\noutside 1 331\nTMhelix 332 354\ninside 355 372

21384	GCF_900037585.1_14555_2_16	Salmonella enterica subsp. enterica serovar Typhi	Proteobacteria; Gammaproteobacteria; Enterobacterales; Enterobacteriaceae; Salmonella; Salmonella enterica; Salmonella enterica subsp. enterica	MTGDNTLITSHGINRRDFMKLCAALAATMGLSSKAAAEMAESVSNPQRPPVIWIGAQECT	WP_000145429.1	hydrogenase 2 small subunit [Salmonella enterica]	Length: 372\nNumber of predicted TMHs: 1\nExp number of AAs in TMHs: 24.51309\nExp number, first 60 AAs: 0.2682\nTotal prob of N-in: 0.14402\noutside 1 331\nTMhelix 332 354\ninside 355 372

21385	GCF_900024935.1_13566_1_39	Salmonella enterica subsp. enterica serovar Typhi	Proteobacteria; Gammaproteobacteria; Enterobacterales; Enterobacteriaceae; Salmonella; Salmonella enterica; Salmonella enterica subsp. enterica	MTGDNTLITSHGINRRDFMKLCAALAATMGLSSKAAAEMAESVSNPQRPPVIWIGAQECT	WP_000145429.1	hydrogenase 2 small subunit [Salmonella enterica]	Length: 372\nNumber of predicted TMHs: 1\nExp number of AAs in TMHs: 24.51309\nExp number, first 60 AAs: 0.2682\nTotal prob of N-in: 0.14402\noutside 1 331\nTMhelix 332 354\ninside 355 372

21386	GCF_900024955.1_14555_2_49	Salmonella enterica subsp. enterica serovar Typhi	Proteobacteria; Gammaproteobacteria; Enterobacterales; Enterobacteriaceae; Salmonella; Salmonella enterica; Salmonella enterica subsp. enterica	MTGDNTLITSHGINRRDFMKLCAALAATMGLSSKAAAEMAESVSNPQRPPVIWIGAQECT	WP_000145429.1	hydrogenase 2 small subunit [Salmonella enterica]	Length: 372\nNumber of

predicted TMHs:  1\nExp number of AAs in TMHs: 24.51309\nExp number, first 60 AAs:  0.2682\nTotal prob of N-in:  0.14402\noutside   1   331\nTMhelix   332   354\ninside   355   372

21387 GCF_900022555.1_13566_1_13 Salmonella enterica subsp. enterica serovar Typhi Proteobacteria; Gammaproteobacteria; Enterobacterales; Enterobacteriaceae; Salmonella; Salmonella enterica; Salmonella enterica subsp. enterica MTGDNTLITSHGINRRDFMKLCAALAATMGLSSKAAAEMAESVSNPQRPPVIWIGAQECT WP_000145429.1 hydrogenase 2 small subunit [Salmonella enterica] Length: 372\nNumber of predicted TMHs:  1\nExp number of AAs in TMHs: 24.51309\nExp number, first 60 AAs:  0.2682\nTotal prob of N-in:  0.14402\noutside   1   331\nTMhelix   332   354\ninside   355   372

21388 GCF_900018655.1_14555_2_70 Salmonella enterica subsp. enterica serovar Typhi Proteobacteria; Gammaproteobacteria; Enterobacterales; Enterobacteriaceae; Salmonella; Salmonella enterica; Salmonella enterica subsp. enterica MTGDNTLITSHGINRRDFMKLCAALAATMGLSSKAAAEMAESVSNPQRPPVIWIGAQECT WP_000145429.1 hydrogenase 2 small subunit [Salmonella enterica] Length: 372\nNumber of predicted TMHs:  1\nExp number of AAs in TMHs: 24.51309\nExp number, first 60 AAs:  0.2682\nTotal prob of N-in:  0.14402\noutside   1   331\nTMhelix   332   354\ninside   355   372

21389 GCF_900018625.1_13566_1_59 Salmonella enterica subsp. enterica serovar Typhi Proteobacteria; Gammaproteobacteria; Enterobacterales; Enterobacteriaceae; Salmonella; Salmonella enterica; Salmonella enterica subsp. enterica MTGDNTLITSHGINRRDFMKLCAALAATMGLSSKAAAEMAESVSNPQRPPVIWIGAQECT WP_000145429.1 hydrogenase 2 small subunit [Salmonella enterica] Length: 372\nNumber of predicted TMHs:  1\nExp number of AAs in TMHs: 24.51309\nExp number, first 60 AAs:  0.2682\nTotal prob of N-in:  0.14402\noutside   1   331\nTMhelix   332   354\ninside   355   372

21390 GCF_900018605.1_13566_1_28 Salmonella enterica subsp. enterica serovar Typhi Proteobacteria; Gammaproteobacteria; Enterobacterales; Enterobacteriaceae; Salmonella; Salmonella enterica; Salmonella enterica subsp. enterica MTGDNTLITSHGINRRDFMKLCAALAATMGLSSKAAAEMAESVSNPQRPPVIWIGAQECT WP_000145429.1 hydrogenase 2 small subunit [Salmonella enterica] Length: 372\nNumber of predicted TMHs:  1\nExp number of AAs in TMHs: 24.51309\nExp number, first 60 AAs:  0.2682\nTotal prob of N-in:  0.14402\noutside   1   331\nTMhelix   332   354\ninside   355   372

21391 GCF_900017255.1_13566_1_55 Salmonella enterica subsp. enterica serovar Typhi Proteobacteria; Gammaproteobacteria; Enterobacterales; Enterobacteriaceae; Salmonella; Salmonella enterica; Salmonella enterica subsp. enterica MTGDNTLITSHGINRRDFMKLCAALAATMGLSSKAAAEMAESVSNPQRPPVIWIGAQECT WP_000145429.1 hydrogenase 2 small subunit [Salmonella enterica] Length: 372\nNumber of predicted TMHs:  1\nExp number of AAs in TMHs: 24.51309\nExp number, first 60 AAs:  0.2682\nTotal prob of N-in:  0.14402\noutside   1   331\nTMhelix   332   354\ninside   355   372

21392 GCF_900017235.1_13134_1_14 Salmonella enterica subsp. enterica serovar Typhi Proteobacteria; Gammaproteobacteria; Enterobacterales; Enterobacteriaceae; Salmonella; Salmonella enterica; Salmonella enterica subsp. enterica MTGDNTLITSHGINRRDFMKLCAALAATMGLSSKAAAEMAESVSNPQRPPVIWIGAQECT WP_000145429.1 hydrogenase 2 small subunit [Salmonella enterica] Length: 372\nNumber of predicted TMHs:  1\nExp number of AAs in TMHs: 24.51309\nExp number, first 60 AAs:  0.2682\nTotal prob of N-in:  0.14402\noutside   1   331\nTMhelix   332   354\ninside   355   372

21393 GCF_900002885.1_St1481-XJ2012 Salmonella enterica subsp. enterica serovar Typhi Proteobacteria; Gammaproteobacteria; Enterobacterales; Enterobacteriaceae; Salmonella; Salmonella enterica; Salmonella enterica subsp. enterica MTGDNTLITSHGINRRDFMKLCAALAATMGLSSKAAAEMAESVSNPQRPPVIWIGAQECT WP_000145429.1 hydrogenase 2 small subunit [Salmonella enterica] Length: 372\nNumber of predicted TMHs:  1\nExp number of AAs in TMHs: 24.51309\nExp number, first 60 AAs:  0.2682\nTotal prob of N-in:  0.14402\noutside   1   331\nTMhelix   332   354\ninside   355   372

21394 GCF_900002865.1_GX-St669 Salmonella enterica subsp. enterica serovar Typhi Proteobacteria; Gammaproteobacteria; Enterobacterales; Enterobacteriaceae; Salmonella; Salmonella enterica; Salmonella enterica subsp. enterica MTGDNTLITSHGINRRDFMKLCAALAATMGLSSKAAAEMAESVSNPQRPPVIWIGAQECT WP_000145429.1 hydrogenase 2 small subunit [Salmonella enterica] Length: 372\nNumber of predicted TMHs:  1\nExp number of AAs in TMHs: 24.51309\nExp number, first 60 AAs:  0.2682\nTotal prob of N-in:   0.14402\noutside   1   331\nTMhelix   332   354\ninside   355   372

21395 GCF_900002845.1_St512-GX1998 Salmonella enterica subsp. enterica serovar Typhi Proteobacteria; Gammaproteobacteria; Enterobacterales; Enterobacteriaceae; Salmonella; Salmonella enterica;

Salmonella enterica subsp. enterica MTGDNTLITSHGINRRDFMKLCAALAATMGLSSKAAAEMAESVSNPQRPPVIWIGAQECT WP_000145429.1 hydrogenase 2 small subunit [Salmonella enterica] Length: 372\nNumber of predicted TMHs: 1\nExp number of AAs in TMHs: 24.51309\nExp number, first 60 AAs: 0.2682\nTotal prob of N-in: 0.14402\noutside 1 331\nTMhelix 332 354\ninside 355 372

21396 GCF_900002605.1_St1332-GZ2011 Salmonella enterica subsp. enterica serovar Typhi Proteobacteria; Gammaproteobacteria; Enterobacterales; Enterobacteriaceae; Salmonella; Salmonella enterica; Salmonella enterica subsp. enterica MTGDNTLITSHGINRRDFMKLCAALAATMGLSSKAAAEMAESVSNPQRPPVIWIGAQECT WP_000145429.1 hydrogenase 2 small subunit [Salmonella enterica] Length: 372\nNumber of predicted TMHs: 1\nExp number of AAs in TMHs: 24.51309\nExp number, first 60 AAs: 0.2682\nTotal prob of N-in: 0.14402\noutside 1 331\nTMhelix 332 354\ninside 355 372

21397 GCF_900002595.1_GZ-18 Salmonella enterica subsp. enterica serovar Typhi Proteobacteria; Gammaproteobacteria; Enterobacterales; Enterobacteriaceae; Salmonella; Salmonella enterica; Salmonella enterica subsp. enterica MTGDNTLITSHGINRRDFMKLCAALAATMGLSSKAAAEMAESVSNPQRPPVIWIGAQECT WP_000145429.1 hydrogenase 2 small subunit [Salmonella enterica] Length: 372\nNumber of predicted TMHs: 1\nExp number of AAs in TMHs: 24.51309\nExp number, first 60 AAs: 0.2682\nTotal prob of N-in: 0.14402\noutside 1 331\nTMhelix 332 354\ninside 355 372

21398 GCF_900002585.1_St508-GX1999 Salmonella enterica subsp. enterica serovar Typhi Proteobacteria; Gammaproteobacteria; Enterobacterales; Enterobacteriaceae; Salmonella; Salmonella enterica; Salmonella enterica subsp. enterica MTGDNTLITSHGINRRDFMKLCAALAATMGLSSKAAAEMAESVSNPQRPPVIWIGAQECT WP_000145429.1 hydrogenase 2 small subunit [Salmonella enterica] Length: 372\nNumber of predicted TMHs: 1\nExp number of AAs in TMHs: 24.51309\nExp number, first 60 AAs: 0.2682\nTotal prob of N-in: 0.14402\noutside 1 331\nTMhelix 332 354\ninside 355 372

21399 GCF_900047085.1_13097_1_84 Salmonella enterica subsp. enterica serovar Typhi Proteobacteria; Gammaproteobacteria; Enterobacterales; Enterobacteriaceae; Salmonella; Salmonella enterica; Salmonella enterica subsp. enterica MTGDNTLITSHGINRRDFMKLCAALAATMGLSSKAAAEMAESVSNPQRPPVIWIGAQECT WP_000145429.1 hydrogenase 2 small subunit [Salmonella enterica] Length: 372\nNumber of predicted TMHs: 1\nExp number of AAs in TMHs: 24.51309\nExp number, first 60 AAs: 0.2682\nTotal prob of N-in: 0.14402\noutside 1 331\nTMhelix 332 354\ninside 355 372

21400 GCF_900047105.1_13103_1_11 Salmonella enterica subsp. enterica serovar Typhi Proteobacteria; Gammaproteobacteria; Enterobacterales; Enterobacteriaceae; Salmonella; Salmonella enterica; Salmonella enterica subsp. enterica MTGDNTLITSHGINRRDFMKLCAALAATMGLSSKAAAEMAESVSNPQRPPVIWIGAQECT WP_000145429.1 hydrogenase 2 small subunit [Salmonella enterica] Length: 372\nNumber of predicted TMHs: 1\nExp number of AAs in TMHs: 24.51309\nExp number, first 60 AAs: 0.2682\nTotal prob of N-in: 0.14402\noutside 1 331\nTMhelix 332 354\ninside 355 372

21401 GCF_900036875.1_14555_2_95 Salmonella enterica subsp. enterica serovar Typhi Proteobacteria; Gammaproteobacteria; Enterobacterales; Enterobacteriaceae; Salmonella; Salmonella enterica; Salmonella enterica subsp. enterica MTGDNTLITSHGINRRDFMKLCAALAATMGLSSKAAAEMAESVSNPQRPPVIWIGAQECT WP_000145429.1 hydrogenase 2 small subunit [Salmonella enterica] Length: 372\nNumber of predicted TMHs: 1\nExp number of AAs in TMHs: 24.51309\nExp number, first 60 AAs: 0.2682\nTotal prob of N-in: 0.14402\noutside 1 331\nTMhelix 332 354\ninside 355 372

21402 GCF_900047115.1_14555_2_17 Salmonella enterica subsp. enterica serovar Typhi Proteobacteria; Gammaproteobacteria; Enterobacterales; Enterobacteriaceae; Salmonella; Salmonella enterica; Salmonella enterica subsp. enterica MTGDNTLITSHGINRRDFMKLCAALAATMGLSSKAAAEMAESVSNPQRPPVIWIGAQECT WP_000145429.1 hydrogenase 2 small subunit [Salmonella enterica] Length: 372\nNumber of predicted TMHs: 1\nExp number of AAs in TMHs: 24.51309\nExp number, first 60 AAs: 0.2682\nTotal prob of N-in: 0.14402\noutside 1 331\nTMhelix 332 354\ninside 355 372

21403 GCF_900036855.1_14555_2_67 Salmonella enterica subsp. enterica serovar Typhi Proteobacteria; Gammaproteobacteria; Enterobacterales; Enterobacteriaceae; Salmonella; Salmonella enterica; Salmonella enterica subsp. enterica MTGDNTLITSHGINRRDFMKLCAALAATMGLSSKAAAEMAESVSNPQRPPVIWIGAQECT WP_000145429.1 hydrogenase 2 small subunit [Salmonella enterica] Length: 372\nNumber of predicted TMHs: 1\nExp number of AAs in TMHs: 24.51309\nExp number, first 60 AAs: 0.2682\nTotal prob of N-in: 0.14402\noutside 1 331\nTMhelix 332 354\ninside 355 372

21404 GCF_900048065.1_13566_1_56 Salmonella enterica subsp. enterica serovar Typhi Proteobacteria; Gammaproteobacteria; Enterobacterales; Enterobacteriaceae; Salmonella; Salmonella enterica; Salmonella enterica subsp. enterica MTGDNTLITSHGINRRDFMKLCAALAATMGLSSKAAAEMAESVSNPQRPPVIWIGAQECT WP_000145429.1 hydrogenase 2 small subunit [Salmonella enterica] Length: 372\nNumber of predicted TMHs: 1\nExp number of AAs in TMHs: 24.51309\nExp number, first 60 AAs: 0.2682\nTotal prob of N-in: 0.14402\noutside 1 331\nTMhelix 332 354\ninside 355 372

21405 GCF_900041555.1_14555_2_18 Salmonella enterica subsp. enterica serovar Typhi Proteobacteria; Gammaproteobacteria; Enterobacterales; Enterobacteriaceae; Salmonella; Salmonella enterica; Salmonella enterica subsp. enterica MTGDNTLITSHGINRRDFMKLCAALAATMGLSSKAAAEMAESVSNPQRPPVIWIGAQECT WP_000145429.1 hydrogenase 2 small subunit [Salmonella enterica] Length: 372\nNumber of predicted TMHs: 1\nExp number of AAs in TMHs: 24.51309\nExp number, first 60 AAs: 0.2682\nTotal prob of N-in: 0.14402\noutside 1 331\nTMhelix 332 354\ninside 355 372

21406 GCF_900042435.1_13097_1_81 Salmonella enterica subsp. enterica serovar Typhi Proteobacteria; Gammaproteobacteria; Enterobacterales; Enterobacteriaceae; Salmonella; Salmonella enterica; Salmonella enterica subsp. enterica MTGDNTLITSHGINRRDFMKLCAALAATMGLSSKAAAEMAESVSNPQRPPVIWIGAQECT WP_000145429.1 hydrogenase 2 small subunit [Salmonella enterica] Length: 372\nNumber of predicted TMHs: 1\nExp number of AAs in TMHs: 24.51309\nExp number, first 60 AAs: 0.2682\nTotal prob of N-in: 0.14402\noutside 1 331\nTMhelix 332 354\ninside 355 372

21407 GCF_900036835.1_14555_2_12 Salmonella enterica subsp. enterica serovar Typhi Proteobacteria; Gammaproteobacteria; Enterobacterales; Enterobacteriaceae; Salmonella; Salmonella enterica; Salmonella enterica subsp. enterica MTGDNTLITSHGINRRDFMKLCAALAATMGLSSKAAAEMAESVSNPQRPPVIWIGAQECT WP_000145429.1 hydrogenase 2 small subunit [Salmonella enterica] Length: 372\nNumber of predicted TMHs: 1\nExp number of AAs in TMHs: 24.51309\nExp number, first 60 AAs: 0.2682\nTotal prob of N-in: 0.14402\noutside 1 331\nTMhelix 332 354\ninside 355 372

21408 GCF_900036825.1_13566_1_76 Salmonella enterica subsp. enterica serovar Typhi Proteobacteria; Gammaproteobacteria; Enterobacterales; Enterobacteriaceae; Salmonella; Salmonella enterica; Salmonella enterica subsp. enterica MTGDNTLITSHGINRRDFMKLCAALAATMGLSSKAAAEMAESVSNPQRPPVIWIGAQECT WP_000145429.1 hydrogenase 2 small subunit [Salmonella enterica] Length: 372\nNumber of predicted TMHs: 1\nExp number of AAs in TMHs: 24.51309\nExp number, first 60 AAs: 0.2682\nTotal prob of N-in: 0.14402\noutside 1 331\nTMhelix 332 354\ninside 355 372

21409 GCF_900042455.1_14555_2_45 Salmonella enterica subsp. enterica serovar Typhi Proteobacteria; Gammaproteobacteria; Enterobacterales; Enterobacteriaceae; Salmonella; Salmonella enterica; Salmonella enterica subsp. enterica MTGDNTLITSHGINRRDFMKLCAALAATMGLSSKAAAEMAESVSNPQRPPVIWIGAQECT WP_000145429.1 hydrogenase 2 small subunit [Salmonella enterica] Length: 372\nNumber of predicted TMHs: 1\nExp number of AAs in TMHs: 24.51309\nExp number, first 60 AAs: 0.2682\nTotal prob of N-in: 0.14402\noutside 1 331\nTMhelix 332 354\ninside 355 372

21410 GCF_900036815.1_13566_1_74 Salmonella enterica subsp. enterica serovar Typhi Proteobacteria; Gammaproteobacteria; Enterobacterales; Enterobacteriaceae; Salmonella; Salmonella enterica; Salmonella enterica subsp. enterica MTGDNTLITSHGINRRDFMKLCAALAATMGLSSKAAAEMAESVSNPQRPPVIWIGAQECT WP_000145429.1 hydrogenase 2 small subunit [Salmonella enterica] Length: 372\nNumber of predicted TMHs: 1\nExp number of AAs in TMHs: 24.51309\nExp number, first 60 AAs: 0.2682\nTotal prob of N-in: 0.14402\noutside 1 331\nTMhelix 332 354\ninside 355 372

21411 GCF_900036795.1_13566_1_61 Salmonella enterica subsp. enterica serovar Typhi Proteobacteria; Gammaproteobacteria; Enterobacterales; Enterobacteriaceae; Salmonella; Salmonella enterica; Salmonella enterica subsp. enterica MTGDNTLITSHGINRRDFMKLCAALAATMGLSSKAAAEMAESVSNPQRPPVIWIGAQECT WP_000145429.1 hydrogenase 2 small subunit [Salmonella enterica] Length: 372\nNumber of predicted TMHs: 1\nExp number of AAs in TMHs: 24.51309\nExp number, first 60 AAs: 0.2682\nTotal prob of N-in: 0.14402\noutside 1 331\nTMhelix 332 354\ninside 355 372

21412 GCF_900036775.1_13566_1_54 Salmonella enterica subsp. enterica serovar Typhi Proteobacteria; Gammaproteobacteria; Enterobacterales; Enterobacteriaceae; Salmonella; Salmonella enterica; Salmonella enterica subsp. enterica MTGDNTLITSHGINRRDFMKLCAALAATMGLSSKAAAEMAESVSNPQRPPVIWIGAQECT WP_000145429.1 hydrogenase 2 small subunit [Salmonella enterica] Length: 372\nNumber of

predicted TMHs:  1\nExp number of AAs in TMHs: 24.51309\nExp number, first 60 AAs:  0.2682\nTotal prob of N-in:  0.14402\noutside   1   331\nTMhelix   332   354\ninside   355   372

21413 GCF_001156405.1_10209_5_22 Salmonella enterica subsp. enterica serovar Typhi Proteobacteria; Gammaproteobacteria; Enterobacterales; Enterobacteriaceae; Salmonella; Salmonella enterica; Salmonella enterica subsp. enterica MTGDNTLITSHGINRRDFMKLCAALAATMGLSSKAAAEMAESVSNPQRPPVIWIGAQECT WP_000145429.1 hydrogenase 2 small subunit [Salmonella enterica] Length: 372\nNumber of predicted TMHs:  1\nExp number of AAs in TMHs: 24.51309\nExp number, first 60 AAs:  0.2682\nTotal prob of N-in:  0.14402\noutside   1   331\nTMhelix   332   354\ninside   355   372

21414 GCF_001156325.1_10349_1_81 Salmonella enterica subsp. enterica serovar Typhi Proteobacteria; Gammaproteobacteria; Enterobacterales; Enterobacteriaceae; Salmonella; Salmonella enterica; Salmonella enterica subsp. enterica MTGDNTLITSHGINRRDFMKLCAALAATMGLSSKAAAEMAESVSNPQRPPVIWIGAQECT WP_000145429.1 hydrogenase 2 small subunit [Salmonella enterica] Length: 372\nNumber of predicted TMHs:  1\nExp number of AAs in TMHs: 24.51309\nExp number, first 60 AAs:  0.2682\nTotal prob of N-in:  0.14402\noutside   1   331\nTMhelix   332   354\ninside   355   372

21415 GCF_001155705.1_9475_6_71 Salmonella enterica subsp. enterica serovar Typhi Proteobacteria; Gammaproteobacteria; Enterobacterales; Enterobacteriaceae; Salmonella; Salmonella enterica; Salmonella enterica subsp. enterica MTGDNTLITSHGINRRDFMKLCAALAATMGLSSKAAAEMAESVSNPQRPPVIWIGAQECT WP_000145429.1 hydrogenase 2 small subunit [Salmonella enterica] Length: 372\nNumber of predicted TMHs:  1\nExp number of AAs in TMHs: 24.51309\nExp number, first 60 AAs:  0.2682\nTotal prob of N-in:  0.14402\noutside   1   331\nTMhelix   332   354\ninside   355   372

21416 GCF_001153125.1_9953_5_39 Salmonella enterica subsp. enterica serovar Typhi Proteobacteria; Gammaproteobacteria; Enterobacterales; Enterobacteriaceae; Salmonella; Salmonella enterica; Salmonella enterica subsp. enterica MTGDNTLITSHGINRRDFMKLCAALAATMGLSSKAAAEMAESVSNPQRPPVIWIGAQECT WP_000145429.1 hydrogenase 2 small subunit [Salmonella enterica] Length: 372\nNumber of predicted TMHs:  1\nExp number of AAs in TMHs: 24.51309\nExp number, first 60 AAs:  0.2682\nTotal prob of N-in:  0.14402\noutside   1   331\nTMhelix   332   354\ninside   355   372

21417 GCF_001152845.1_10492_1_14 Salmonella enterica subsp. enterica serovar Typhi Proteobacteria; Gammaproteobacteria; Enterobacterales; Enterobacteriaceae; Salmonella; Salmonella enterica; Salmonella enterica subsp. enterica MTGDNTLITSHGINRRDFMKLCAALAATMGLSSKAAAEMAESVSNPQRPPVIWIGAQECT WP_000145429.1 hydrogenase 2 small subunit [Salmonella enterica] Length: 372\nNumber of predicted TMHs:  1\nExp number of AAs in TMHs: 24.51309\nExp number, first 60 AAs:  0.2682\nTotal prob of N-in:  0.14402\noutside   1   331\nTMhelix   332   354\ninside   355   372

21418 GCF_001152445.1_10562_2_56 Salmonella enterica subsp. enterica serovar Typhi Proteobacteria; Gammaproteobacteria; Enterobacterales; Enterobacteriaceae; Salmonella; Salmonella enterica; Salmonella enterica subsp. enterica MTGDNTLITSHGINRRDFMKLCAALAATMGLSSKAAAEMAESVSNPQRPPVIWIGAQECT WP_000145429.1 hydrogenase 2 small subunit [Salmonella enterica] Length: 372\nNumber of predicted TMHs:  1\nExp number of AAs in TMHs: 24.51309\nExp number, first 60 AAs:  0.2682\nTotal prob of N-in:  0.14402\noutside   1   331\nTMhelix   332   354\ninside   355   372

21419 GCF_001152505.1_10209_5_24 Salmonella enterica subsp. enterica serovar Typhi Proteobacteria; Gammaproteobacteria; Enterobacterales; Enterobacteriaceae; Salmonella; Salmonella enterica; Salmonella enterica subsp. enterica MTGDNTLITSHGINRRDFMKLCAALAATMGLSSKAAAEMAESVSNPQRPPVIWIGAQECT WP_000145429.1 hydrogenase 2 small subunit [Salmonella enterica] Length: 372\nNumber of predicted TMHs:  1\nExp number of AAs in TMHs: 24.51309\nExp number, first 60 AAs:  0.2682\nTotal prob of N-in:  0.14402\noutside   1   331\nTMhelix   332   354\ninside   355   372

21420 GCF_001152105.1_8447_8_14 Salmonella enterica subsp. enterica serovar Typhi Proteobacteria; Gammaproteobacteria; Enterobacterales; Enterobacteriaceae; Salmonella; Salmonella enterica; Salmonella enterica subsp. enterica MTGDNTLITSHGINRRDFMKLCAALAATMGLSSKAAAEMAESVSNPQRPPVIWIGAQECT WP_000145429.1 hydrogenase 2 small subunit [Salmonella enterica] Length: 372\nNumber of predicted TMHs:  1\nExp number of AAs in TMHs: 24.51309\nExp number, first 60 AAs:  0.2682\nTotal prob of N-in:  0.14402\noutside   1   331\nTMhelix   332   354\ninside   355   372

21421 GCF_001151945.1_9870_8_75 Salmonella enterica subsp. enterica serovar Typhi Proteobacteria; Gammaproteobacteria; Enterobacterales; Enterobacteriaceae; Salmonella; Salmonella enterica;

Salmonella enterica subsp. enterica MTGDNTLITSHGINRRDFMKLCAALAATMGLSSKAAAEMAESVSNPQRPPVIWIGAQECT WP_000145429.1 hydrogenase 2 small subunit [Salmonella enterica] Length: 372\nNumber of predicted TMHs: 1\nExp number of AAs in TMHs: 24.51309\nExp number, first 60 AAs: 0.2682\nTotal prob of N-in: 0.14402\noutside 1 331\nTMhelix 332 354\ninside 355 372

21422 GCF_001151505.1_10425_1_13 Salmonella enterica subsp. enterica serovar Typhi Proteobacteria; Gammaproteobacteria; Enterobacterales; Enterobacteriaceae; Salmonella; Salmonella enterica; Salmonella enterica subsp. enterica MTGDNTLITSHGINRRDFMKLCAALAATMGLSSKAAAEMAESVSNPQRPPVIWIGAQECT WP_000145429.1 hydrogenase 2 small subunit [Salmonella enterica] Length: 372\nNumber of predicted TMHs: 1\nExp number of AAs in TMHs: 24.51309\nExp number, first 60 AAs: 0.2682\nTotal prob of N-in: 0.14402\noutside 1 331\nTMhelix 332 354\ninside 355 372

21423 GCF_001148305.1_10349_1_34 Salmonella enterica subsp. enterica serovar Typhi Proteobacteria; Gammaproteobacteria; Enterobacterales; Enterobacteriaceae; Salmonella; Salmonella enterica; Salmonella enterica subsp. enterica MTGDNTLITSHGINRRDFMKLCAALAATMGLSSKAAAEMAESVSNPQRPPVIWIGAQECT WP_000145429.1 hydrogenase 2 small subunit [Salmonella enterica] Length: 372\nNumber of predicted TMHs: 1\nExp number of AAs in TMHs: 24.51309\nExp number, first 60 AAs: 0.2682\nTotal prob of N-in: 0.14402\noutside 1 331\nTMhelix 332 354\ninside 355 372

21424 GCF_001148225.1_8490_5_18 Salmonella enterica subsp. enterica serovar Typhi Proteobacteria; Gammaproteobacteria; Enterobacterales; Enterobacteriaceae; Salmonella; Salmonella enterica; Salmonella enterica subsp. enterica MTGDNTLITSHGINRRDFMKLCAALAATMGLSSKAAAEMAESVSNPQRPPVIWIGAQECT WP_000145429.1 hydrogenase 2 small subunit [Salmonella enterica] Length: 372\nNumber of predicted TMHs: 1\nExp number of AAs in TMHs: 24.51309\nExp number, first 60 AAs: 0.2682\nTotal prob of N-in: 0.14402\noutside 1 331\nTMhelix 332 354\ninside 355 372

21425 GCF_001147545.1_7468_7_73 Salmonella enterica subsp. enterica serovar Typhi Proteobacteria; Gammaproteobacteria; Enterobacterales; Enterobacteriaceae; Salmonella; Salmonella enterica; Salmonella enterica subsp. enterica MTGDNTLITSHGINRRDFMKLCAALAATMGLSSKAATEMAESVSNPQRPPVIWIGAQECT WP_001782304.1 hydrogenase 2 small subunit [Salmonella enterica] Length: 372\nNumber of predicted TMHs: 1\nExp number of AAs in TMHs: 24.41173\nExp number, first 60 AAs: 0.16124\nTotal prob of N-in: 0.13947\noutside 1 331\nTMhelix 332 354\ninside 355 372

21426 GCF_001148105.1_10541_2_51 Salmonella enterica subsp. enterica serovar Typhi Proteobacteria; Gammaproteobacteria; Enterobacterales; Enterobacteriaceae; Salmonella; Salmonella enterica; Salmonella enterica subsp. enterica MTGDNTLITSHGINRRDFMKLCAALAATMGLSSKAAAEMAESVSNPQRPPVIWIGAQECT WP_000145429.1 hydrogenase 2 small subunit [Salmonella enterica] Length: 372\nNumber of predicted TMHs: 1\nExp number of AAs in TMHs: 24.51309\nExp number, first 60 AAs: 0.2682\nTotal prob of N-in: 0.14402\noutside 1 331\nTMhelix 332 354\ninside 355 372

21427 GCF_001142785.1_10561_2_37 Salmonella enterica subsp. enterica serovar Typhi Proteobacteria; Gammaproteobacteria; Enterobacterales; Enterobacteriaceae; Salmonella; Salmonella enterica; Salmonella enterica subsp. enterica MTGDNTLITSHGINRRDFMKLCAALAATMGLSSKAATEMAESVSNPQRPPVIWIGAQECT WP_001782304.1 hydrogenase 2 small subunit [Salmonella enterica] Length: 372\nNumber of predicted TMHs: 1\nExp number of AAs in TMHs: 24.41173\nExp number, first 60 AAs: 0.16124\nTotal prob of N-in: 0.13947\noutside 1 331\nTMhelix 332 354\ninside 355 372

21428 GCF_001144665.1_10541_2_22 Salmonella enterica subsp. enterica serovar Typhi Proteobacteria; Gammaproteobacteria; Enterobacterales; Enterobacteriaceae; Salmonella; Salmonella enterica; Salmonella enterica subsp. enterica MTGDNTLITSHGINRRDFMKLCAALAATMGLSSKAAAEMAESVSNPQRPPVIWIGAQECT WP_000145429.1 hydrogenase 2 small subunit [Salmonella enterica] Length: 372\nNumber of predicted TMHs: 1\nExp number of AAs in TMHs: 24.51309\nExp number, first 60 AAs: 0.2682\nTotal prob of N-in: 0.14402\noutside 1 331\nTMhelix 332 354\ninside 355 372

21429 GCF_001144465.1_10562_2_32 Salmonella enterica subsp. enterica serovar Typhi Proteobacteria; Gammaproteobacteria; Enterobacterales; Enterobacteriaceae; Salmonella; Salmonella enterica; Salmonella enterica subsp. enterica MTGDNTLITSHGINRRDFMKLCAALAATMGLSSKAAAEMAESVSNPQRPPVIWIGAQECT WP_000145429.1 hydrogenase 2 small subunit [Salmonella enterica] Length: 372\nNumber of predicted TMHs: 1\nExp number of AAs in TMHs: 24.51309\nExp number, first 60 AAs: 0.2682\nTotal prob of N-in: 0.14402\noutside 1 331\nTMhelix 332 354\ninside 355 372

21430	GCF_001144245.1_10593_2_6	Salmonella enterica subsp. enterica serovar Typhi	Proteobacteria; Gammaproteobacteria; Enterobacterales; Enterobacteriaceae; Salmonella; Salmonella enterica; Salmonella enterica subsp. enterica	MTGDNTLITSHGINRRDFMKLCAALAATMGLSSKAAAEMAESVSNPQRPPVIWIGAQECT	WP_000145429.1	hydrogenase 2 small subunit [Salmonella enterica]	Length: 372\nNumber of predicted TMHs:  1\nExp number of AAs in TMHs: 24.51309\nExp number, first 60 AAs:  0.2682\nTotal prob of N-in: 0.14402\noutside   1   331\nTMhelix   332   354\ninside   355   372

21431	GCF_001143985.1_9475_6_90	Salmonella enterica subsp. enterica serovar Typhi	Proteobacteria; Gammaproteobacteria; Enterobacterales; Enterobacteriaceae; Salmonella; Salmonella enterica; Salmonella enterica subsp. enterica	MTGDNTLITSHGINRRDFMKLCAALAATMGLSSKAAAEMAESVSNPQRPPVIWIGAQECT	WP_000145429.1	hydrogenase 2 small subunit [Salmonella enterica]	Length: 372\nNumber of predicted TMHs:  1\nExp number of AAs in TMHs: 24.51309\nExp number, first 60 AAs:  0.2682\nTotal prob of N-in: 0.14402\noutside   1   331\nTMhelix   332   354\ninside   355   372

21432	GCF_001143625.1_10562_2_25	Salmonella enterica subsp. enterica serovar Typhi	Proteobacteria; Gammaproteobacteria; Enterobacterales; Enterobacteriaceae; Salmonella; Salmonella enterica; Salmonella enterica subsp. enterica	MTGDNTLITSHGINRRDFMKLCAALAATMGLSSKAAAEMAESVSNPQRPPVIWIGAQECT	WP_000145429.1	hydrogenase 2 small subunit [Salmonella enterica]	Length: 372\nNumber of predicted TMHs:  1\nExp number of AAs in TMHs: 24.51309\nExp number, first 60 AAs:  0.2682\nTotal prob of N-in: 0.14402\noutside   1   331\nTMhelix   332   354\ninside   355   372

21433	GCF_001143405.1_10493_1_48	Salmonella enterica subsp. enterica serovar Typhi	Proteobacteria; Gammaproteobacteria; Enterobacterales; Enterobacteriaceae; Salmonella; Salmonella enterica; Salmonella enterica subsp. enterica	MTGDNTLITSHGINRRDFMKLCAALAATMGLSSKAAAEMAESVSNPQRPPVIWIGAQECT	WP_000145429.1	hydrogenase 2 small subunit [Salmonella enterica]	Length: 372\nNumber of predicted TMHs:  1\nExp number of AAs in TMHs: 24.51309\nExp number, first 60 AAs:  0.2682\nTotal prob of N-in: 0.14402\noutside   1   331\nTMhelix   332   354\ninside   355   372

21434	GCF_001141045.1_10493_1_71	Salmonella enterica subsp. enterica serovar Typhi	Proteobacteria; Gammaproteobacteria; Enterobacterales; Enterobacteriaceae; Salmonella; Salmonella enterica; Salmonella enterica subsp. enterica	MTGDNTLITSHGINRRDFMKLCAALAATMGLSSKAAAEMAESVSNPQRPPVIWIGAQECT	WP_000145429.1	hydrogenase 2 small subunit [Salmonella enterica]	Length: 372\nNumber of predicted TMHs:  1\nExp number of AAs in TMHs: 24.51309\nExp number, first 60 AAs:  0.2682\nTotal prob of N-in: 0.14402\noutside   1   331\nTMhelix   332   354\ninside   355   372

21435	GCF_001139385.1_10562_2_2	Salmonella enterica subsp. enterica serovar Typhi	Proteobacteria; Gammaproteobacteria; Enterobacterales; Enterobacteriaceae; Salmonella; Salmonella enterica; Salmonella enterica subsp. enterica	MTGDNTLITSHGINRRDFMKLCAALAATMGLSSKAATEMAESVSNPQRPPVIWIGAQECT	WP_001782304.1	hydrogenase 2 small subunit [Salmonella enterica]	Length: 372\nNumber of predicted TMHs:  1\nExp number of AAs in TMHs: 24.41173\nExp number, first 60 AAs:  0.16124\nTotal prob of N-in: 0.13947\noutside   1   331\nTMhelix   332   354\ninside   355   372

21436	GCF_001140585.1_10396_8_20	Salmonella enterica subsp. enterica serovar Typhi	Proteobacteria; Gammaproteobacteria; Enterobacterales; Enterobacteriaceae; Salmonella; Salmonella enterica; Salmonella enterica subsp. enterica	MTGDNTLITSHGINRRDFMKLCAALAATMGLSSKAAAEMAESVSNPQRPPVIWIGAQECT	WP_000145429.1	hydrogenase 2 small subunit [Salmonella enterica]	Length: 372\nNumber of predicted TMHs:  1\nExp number of AAs in TMHs: 24.51309\nExp number, first 60 AAs:  0.2682\nTotal prob of N-in: 0.14402\noutside   1   331\nTMhelix   332   354\ninside   355   372

21437	GCF_001139745.1_10493_1_53	Salmonella enterica subsp. enterica serovar Typhi	Proteobacteria; Gammaproteobacteria; Enterobacterales; Enterobacteriaceae; Salmonella; Salmonella enterica; Salmonella enterica subsp. enterica	MTGDNTLITSHGINRRDFMKLCAALAATMGLSSKAAAEMAESVSNPQRPPVIWIGAQECT	WP_000145429.1	hydrogenase 2 small subunit [Salmonella enterica]	Length: 372\nNumber of predicted TMHs:  1\nExp number of AAs in TMHs: 24.51309\nExp number, first 60 AAs:  0.2682\nTotal prob of N-in: 0.14402\noutside   1   331\nTMhelix   332   354\ninside   355   372

21438	GCF_001139245.1_10540_1_7	Salmonella enterica subsp. enterica serovar Typhi	Proteobacteria; Gammaproteobacteria; Enterobacterales; Enterobacteriaceae; Salmonella; Salmonella enterica; Salmonella enterica subsp. enterica	MTGDNTLITSHGINRRDFMKLCAALAATMGLSSKAAAEMAESVSNPQRPPVIWIGAQECT	WP_000145429.1	hydrogenase 2 small subunit [Salmonella enterica]	Length: 372\nNumber of

predicted TMHs:  1\nExp number of AAs in TMHs: 24.51309\nExp number, first 60 AAs:  0.2682\nTotal prob of N-in: 0.14402\noutside   1   331\nTMhelix   332   354\ninside   355   372

21439 GCF_001135305.1_8447_8_16 Salmonella enterica subsp. enterica serovar Typhi Proteobacteria; Gammaproteobacteria; Enterobacterales; Enterobacteriaceae; Salmonella; Salmonella enterica; Salmonella enterica subsp. enterica MTGDNTLITSHGINRRDFMKLCAALAATMGLSSKAAAEMAESVSNPQRPPVIWIGAQECT WP_000145429.1 hydrogenase 2 small subunit [Salmonella enterica] Length: 372\nNumber of predicted TMHs:  1\nExp number of AAs in TMHs: 24.51309\nExp number, first 60 AAs:  0.2682\nTotal prob of N-in: 0.14402\noutside   1   331\nTMhelix   332   354\ninside   355   372

21440 GCF_001135825.1_8447_8_26 Salmonella enterica subsp. enterica serovar Typhi Proteobacteria; Gammaproteobacteria; Enterobacterales; Enterobacteriaceae; Salmonella; Salmonella enterica; Salmonella enterica subsp. enterica MTGDNTLITSHGINRRDFMKLCAALAATMGLSSKAAAEMAESVSNPQRPPVIWIGAQECT WP_000145429.1 hydrogenase 2 small subunit [Salmonella enterica] Length: 372\nNumber of predicted TMHs:  1\nExp number of AAs in TMHs: 24.51309\nExp number, first 60 AAs:  0.2682\nTotal prob of N-in: 0.14402\noutside   1   331\nTMhelix   332   354\ninside   355   372

21441 GCF_001135005.1_10493_1_23 Salmonella enterica subsp. enterica serovar Typhi Proteobacteria; Gammaproteobacteria; Enterobacterales; Enterobacteriaceae; Salmonella; Salmonella enterica; Salmonella enterica subsp. enterica MTGDNTLITSHGINRRDFMKLCAALAATMGLSSKAAAEMAESVSNPQRPPVIWIGAQECT WP_000145429.1 hydrogenase 2 small subunit [Salmonella enterica] Length: 372\nNumber of predicted TMHs:  1\nExp number of AAs in TMHs: 24.51309\nExp number, first 60 AAs:  0.2682\nTotal prob of N-in: 0.14402\noutside   1   331\nTMhelix   332   354\ninside   355   372

21442 GCF_001134605.1_10541_2_31 Salmonella enterica subsp. enterica serovar Typhi Proteobacteria; Gammaproteobacteria; Enterobacterales; Enterobacteriaceae; Salmonella; Salmonella enterica; Salmonella enterica subsp. enterica MTGDNTLITSHGINRRDFMKLCAALAATMGLSSKAAAEMAESVSNPQRPPVIWIGAQECT WP_000145429.1 hydrogenase 2 small subunit [Salmonella enterica] Length: 372\nNumber of predicted TMHs:  1\nExp number of AAs in TMHs: 24.51309\nExp number, first 60 AAs:  0.2682\nTotal prob of N-in: 0.14402\noutside   1   331\nTMhelix   332   354\ninside   355   372

21443 GCF_001134405.1_9870_8_88 Salmonella enterica subsp. enterica serovar Typhi Proteobacteria; Gammaproteobacteria; Enterobacterales; Enterobacteriaceae; Salmonella; Salmonella enterica; Salmonella enterica subsp. enterica MTGDNTLITSHGINRRDFMKLCAALAATMGLSSKAAAEMAESVSNPQRPPVIWIGAQECT WP_000145429.1 hydrogenase 2 small subunit [Salmonella enterica] Length: 372\nNumber of predicted TMHs:  1\nExp number of AAs in TMHs: 24.51309\nExp number, first 60 AAs:  0.2682\nTotal prob of N-in: 0.14402\noutside   1   331\nTMhelix   332   354\ninside   355   372

21444 GCF_001134345.1_10561_2_1 Salmonella enterica subsp. enterica serovar Typhi Proteobacteria; Gammaproteobacteria; Enterobacterales; Enterobacteriaceae; Salmonella; Salmonella enterica; Salmonella enterica subsp. enterica MTGDNTLITSHGINRRDFMKLCAALAATMGLSSKAAAEMAESVSNPQRPPVIWIGAQECT WP_000145429.1 hydrogenase 2 small subunit [Salmonella enterica] Length: 372\nNumber of predicted TMHs:  1\nExp number of AAs in TMHs: 24.51309\nExp number, first 60 AAs:  0.2682\nTotal prob of N-in: 0.14402\noutside   1   331\nTMhelix   332   354\ninside   355   372

21445 GCF_001134205.1_10349_1_56 Salmonella enterica subsp. enterica serovar Typhi Proteobacteria; Gammaproteobacteria; Enterobacterales; Enterobacteriaceae; Salmonella; Salmonella enterica; Salmonella enterica subsp. enterica MTGDNTLITSHGINRRDFMKLCAALAATMGLSSKAAAEMAESVSNPQRPPVIWIGAQECT WP_000145429.1 hydrogenase 2 small subunit [Salmonella enterica] Length: 372\nNumber of predicted TMHs:  1\nExp number of AAs in TMHs: 24.51309\nExp number, first 60 AAs:  0.2682\nTotal prob of N-in: 0.14402\noutside   1   331\nTMhelix   332   354\ninside   355   372

21446 GCF_001127645.1_10562_2_19 Salmonella enterica subsp. enterica serovar Typhi Proteobacteria; Gammaproteobacteria; Enterobacterales; Enterobacteriaceae; Salmonella; Salmonella enterica; Salmonella enterica subsp. enterica MTGDNTLITSHGINRRDFMKLCAALAATMGLSSKAATEMAESVSNPQRPPVIWIGAQECT WP_001782304.1 hydrogenase 2 small subunit [Salmonella enterica] Length: 372\nNumber of predicted TMHs:  1\nExp number of AAs in TMHs: 24.41173\nExp number, first 60 AAs:  0.16124\nTotal prob of N-in: 0.13947\noutside   1   331\nTMhelix   332   354\ninside   355   372

21447 GCF_001127485.1_10349_1_16 Salmonella enterica subsp. enterica serovar Typhi Proteobacteria; Gammaproteobacteria; Enterobacterales; Enterobacteriaceae; Salmonella; Salmonella enterica;

Salmonella enterica subsp. enterica	MTGDNTLITSHGINRRDFMKLCAALAATMGLSSKAATEMAESVSNPQRPPVIWIGAQECT	WP_001782304.1	hydrogenase 2 small subunit [Salmonella enterica]	Length: 372\nNumber of predicted TMHs: 1\nExp number of AAs in TMHs: 24.41173\nExp number, first 60 AAs: 0.16124\nTotal prob of N-in: 0.13947\noutside 1 331\nTMhelix 332 354\ninside 355 372

21448	GCF_001129305.1_9475_6_21	Salmonella enterica subsp. enterica serovar Typhi	Proteobacteria; Gammaproteobacteria; Enterobacterales; Enterobacteriaceae; Salmonella; Salmonella enterica; Salmonella enterica subsp. enterica	MTGDNTLITSHGINRRDFMKLCAALAATMGLSSKAAAEMAESVSNPQRPPVIWIGAQECT	WP_000145429.1	hydrogenase 2 small subunit [Salmonella enterica]	Length: 372\nNumber of predicted TMHs: 1\nExp number of AAs in TMHs: 24.51309\nExp number, first 60 AAs: 0.2682\nTotal prob of N-in: 0.14402\noutside 1 331\nTMhelix 332 354\ninside 355 372

21449	GCF_001129225.1_10593_2_37	Salmonella enterica subsp. enterica serovar Typhi	Proteobacteria; Gammaproteobacteria; Enterobacterales; Enterobacteriaceae; Salmonella; Salmonella enterica; Salmonella enterica subsp. enterica	MTGDNTLITSHGINRRDFMKLCAALAATMGLSSKAAAEMAESVSNPQRPPVIWIGAQECT	WP_000145429.1	hydrogenase 2 small subunit [Salmonella enterica]	Length: 372\nNumber of predicted TMHs: 1\nExp number of AAs in TMHs: 24.51309\nExp number, first 60 AAs: 0.2682\nTotal prob of N-in: 0.14402\noutside 1 331\nTMhelix 332 354\ninside 355 372

21450	GCF_001129105.1_9953_5_14	Salmonella enterica subsp. enterica serovar Typhi	Proteobacteria; Gammaproteobacteria; Enterobacterales; Enterobacteriaceae; Salmonella; Salmonella enterica; Salmonella enterica subsp. enterica	MTGDNTLITSHGINRRDFMKLCAALAATMGLSSKAAAEMAESVSNPQRPPVIWIGAQECT	WP_000145429.1	hydrogenase 2 small subunit [Salmonella enterica]	Length: 372\nNumber of predicted TMHs: 1\nExp number of AAs in TMHs: 24.51309\nExp number, first 60 AAs: 0.2682\nTotal prob of N-in: 0.14402\noutside 1 331\nTMhelix 332 354\ninside 355 372

21451	GCF_001128625.1_10349_1_61	Salmonella enterica subsp. enterica serovar Typhi	Proteobacteria; Gammaproteobacteria; Enterobacterales; Enterobacteriaceae; Salmonella; Salmonella enterica; Salmonella enterica subsp. enterica	MTGDNTLITSHGINRRDFMKLCAALAATMGLSSKAAAEMAESVSNPQRPPVIWIGAQECT	WP_000145429.1	hydrogenase 2 small subunit [Salmonella enterica]	Length: 372\nNumber of predicted TMHs: 1\nExp number of AAs in TMHs: 24.51309\nExp number, first 60 AAs: 0.2682\nTotal prob of N-in: 0.14402\noutside 1 331\nTMhelix 332 354\ninside 355 372

21452	GCF_001128425.1_8490_6_58	Salmonella enterica subsp. enterica serovar Typhi	Proteobacteria; Gammaproteobacteria; Enterobacterales; Enterobacteriaceae; Salmonella; Salmonella enterica; Salmonella enterica subsp. enterica	MTGDNTLITSHGINRRDFMKLCAALAATMGLSSKAAAEMAESVSNPQRPPVIWIGAQECT	WP_000145429.1	hydrogenase 2 small subunit [Salmonella enterica]	Length: 372\nNumber of predicted TMHs: 1\nExp number of AAs in TMHs: 24.51309\nExp number, first 60 AAs: 0.2682\nTotal prob of N-in: 0.14402\noutside 1 331\nTMhelix 332 354\ninside 355 372

21453	GCF_001126185.1_10541_1_66	Salmonella enterica subsp. enterica serovar Typhi	Proteobacteria; Gammaproteobacteria; Enterobacterales; Enterobacteriaceae; Salmonella; Salmonella enterica; Salmonella enterica subsp. enterica	MTGDNTLITSHGINRRDFMKLCAALAATMGLSSKAAAEMAESVSNPQRPPVIWIGAQECT	WP_000145429.1	hydrogenase 2 small subunit [Salmonella enterica]	Length: 372\nNumber of predicted TMHs: 1\nExp number of AAs in TMHs: 24.51309\nExp number, first 60 AAs: 0.2682\nTotal prob of N-in: 0.14402\noutside 1 331\nTMhelix 332 354\ninside 355 372

21454	GCF_001126125.1_10541_2_32	Salmonella enterica subsp. enterica serovar Typhi	Proteobacteria; Gammaproteobacteria; Enterobacterales; Enterobacteriaceae; Salmonella; Salmonella enterica; Salmonella enterica subsp. enterica	MTGDNTLITSHGINRRDFMKLCAALAATMGLSSKAAAEMAESVSNPQRPPVIWIGAQECT	WP_000145429.1	hydrogenase 2 small subunit [Salmonella enterica]	Length: 372\nNumber of predicted TMHs: 1\nExp number of AAs in TMHs: 24.51309\nExp number, first 60 AAs: 0.2682\nTotal prob of N-in: 0.14402\noutside 1 331\nTMhelix 332 354\ninside 355 372

21455	GCF_001125885.1_8490_6_43	Salmonella enterica subsp. enterica serovar Typhi	Proteobacteria; Gammaproteobacteria; Enterobacterales; Enterobacteriaceae; Salmonella; Salmonella enterica; Salmonella enterica subsp. enterica	MTGDNTLITSHGINRRDFMKLCAALAATMGLSSKAAAEMAESVSNPQRPPVIWIGAQECT	WP_000145429.1	hydrogenase 2 small subunit [Salmonella enterica]	Length: 372\nNumber of predicted TMHs: 1\nExp number of AAs in TMHs: 24.51309\nExp number, first 60 AAs: 0.2682\nTotal prob of N-in: 0.14402\noutside 1 331\nTMhelix 332 354\ninside 355 372

21456 GCF_001125485.1_9475_6_56 Salmonella enterica subsp. enterica serovar Typhi Proteobacteria; Gammaproteobacteria; Enterobacterales; Enterobacteriaceae; Salmonella; Salmonella enterica; Salmonella enterica subsp. enterica MTGDNTLITSHGINRRDFMKLCAALAATMGLSSKAAAEMAESVSNPQRPPVIWIGAQECT WP_000145429.1 hydrogenase 2 small subunit [Salmonella enterica] Length: 372\nNumber of predicted TMHs: 1\nExp number of AAs in TMHs: 24.51309\nExp number, first 60 AAs: 0.2682\nTotal prob of N-in: 0.14402\noutside 1 331\nTMhelix 332 354\ninside 355 372

21457 GCF_001125345.1_10541_2_29 Salmonella enterica subsp. enterica serovar Typhi Proteobacteria; Gammaproteobacteria; Enterobacterales; Enterobacteriaceae; Salmonella; Salmonella enterica; Salmonella enterica subsp. enterica MTGDNTLITSHGINRRDFMKLCAALAATMGLSSKAAAEMAESVSNPQRPPVIWIGAQECT WP_000145429.1 hydrogenase 2 small subunit [Salmonella enterica] Length: 372\nNumber of predicted TMHs: 1\nExp number of AAs in TMHs: 24.51309\nExp number, first 60 AAs: 0.2682\nTotal prob of N-in: 0.14402\noutside 1 331\nTMhelix 332 354\ninside 355 372

21458 GCF_001125005.1_10493_1_32 Salmonella enterica subsp. enterica serovar Typhi Proteobacteria; Gammaproteobacteria; Enterobacterales; Enterobacteriaceae; Salmonella; Salmonella enterica; Salmonella enterica subsp. enterica MTGDNTLITSHGINRRDFMKLCAALAATMGLSSKAAAEMAESVSNPQRPPVIWIGAQECT WP_000145429.1 hydrogenase 2 small subunit [Salmonella enterica] Length: 372\nNumber of predicted TMHs: 1\nExp number of AAs in TMHs: 24.51309\nExp number, first 60 AAs: 0.2682\nTotal prob of N-in: 0.14402\noutside 1 331\nTMhelix 332 354\ninside 355 372

21459 GCF_001122325.1_10209_5_37 Salmonella enterica subsp. enterica serovar Typhi Proteobacteria; Gammaproteobacteria; Enterobacterales; Enterobacteriaceae; Salmonella; Salmonella enterica; Salmonella enterica subsp. enterica MTGDNTLITSHGINRRDFMKLCAALAATMGLSSKAAAEMAESVSNPQRPPVIWIGAQECT WP_000145429.1 hydrogenase 2 small subunit [Salmonella enterica] Length: 372\nNumber of predicted TMHs: 1\nExp number of AAs in TMHs: 24.51309\nExp number, first 60 AAs: 0.2682\nTotal prob of N-in: 0.14402\noutside 1 331\nTMhelix 332 354\ninside 355 372

21460 GCF_001122165.1_8490_5_23 Salmonella enterica subsp. enterica serovar Typhi Proteobacteria; Gammaproteobacteria; Enterobacterales; Enterobacteriaceae; Salmonella; Salmonella enterica; Salmonella enterica subsp. enterica MTGDNTLITSHGINRRDFMKLCAALAATMGLSSKAAAEMAESVSNPQRPPVIWIGAQECT WP_000145429.1 hydrogenase 2 small subunit [Salmonella enterica] Length: 372\nNumber of predicted TMHs: 1\nExp number of AAs in TMHs: 24.51309\nExp number, first 60 AAs: 0.2682\nTotal prob of N-in: 0.14402\noutside 1 331\nTMhelix 332 354\ninside 355 372

21461 GCF_001122085.1_9953_5_28 Salmonella enterica subsp. enterica serovar Typhi Proteobacteria; Gammaproteobacteria; Enterobacterales; Enterobacteriaceae; Salmonella; Salmonella enterica; Salmonella enterica subsp. enterica MTGDNTLITSHGINRRDFMKLCAALAATMGLSSKAAAEMAESVSNPQRPPVIWIGAQECT WP_000145429.1 hydrogenase 2 small subunit [Salmonella enterica] Length: 372\nNumber of predicted TMHs: 1\nExp number of AAs in TMHs: 24.51309\nExp number, first 60 AAs: 0.2682\nTotal prob of N-in: 0.14402\noutside 1 331\nTMhelix 332 354\ninside 355 372

21462 GCF_001121925.1_10492_1_22 Salmonella enterica subsp. enterica serovar Typhi Proteobacteria; Gammaproteobacteria; Enterobacterales; Enterobacteriaceae; Salmonella; Salmonella enterica; Salmonella enterica subsp. enterica MTGDNTLITSHGINRRDFMKLCAALAATMGLSSKAAAEMAESVSNPQRPPVIWIGAQECT WP_000145429.1 hydrogenase 2 small subunit [Salmonella enterica] Length: 372\nNumber of predicted TMHs: 1\nExp number of AAs in TMHs: 24.51309\nExp number, first 60 AAs: 0.2682\nTotal prob of N-in: 0.14402\noutside 1 331\nTMhelix 332 354\ninside 355 372

21463 GCF_001121805.1_10493_1_60 Salmonella enterica subsp. enterica serovar Typhi Proteobacteria; Gammaproteobacteria; Enterobacterales; Enterobacteriaceae; Salmonella; Salmonella enterica; Salmonella enterica subsp. enterica MTGDNTLITSHGINRRDFMKLCAALAATMGLSSKAAAEMAESVSNPQRPPVIWIGAQECT WP_000145429.1 hydrogenase 2 small subunit [Salmonella enterica] Length: 372\nNumber of predicted TMHs: 1\nExp number of AAs in TMHs: 24.51309\nExp number, first 60 AAs: 0.2682\nTotal prob of N-in: 0.14402\noutside 1 331\nTMhelix 332 354\ninside 355 372

21464 GCF_001119685.1_10561_2_20 Salmonella enterica subsp. enterica serovar Typhi Proteobacteria; Gammaproteobacteria; Enterobacterales; Enterobacteriaceae; Salmonella; Salmonella enterica; Salmonella enterica subsp. enterica MTGDNTLITSHGINRRDFMKLCAALAATMGLSSKAATEMAESVSNPQRPPVIWIGAQECT WP_001782304.1 hydrogenase 2 small subunit [Salmonella enterica] Length: 372\nNumber of

predicted TMHs: 1\nExp number of AAs in TMHs: 24.41173\nExp number, first 60 AAs: 0.16124\nTotal prob of N-in: 0.13947\noutside 1 331\nTMhelix 332 354\ninside 355 372

21465 GCF_001121185.1_10562_2_3 Salmonella enterica subsp. enterica serovar Typhi Proteobacteria; Gammaproteobacteria; Enterobacterales; Enterobacteriaceae; Salmonella; Salmonella enterica; Salmonella enterica subsp. enterica MTGDNTLITSHGINRRDFMKLCAALAATMGLSSKAAAEMAESVSNPQRPPVIWIGAQECT WP_000145429.1 hydrogenase 2 small subunit [Salmonella enterica] Length: 372\nNumber of predicted TMHs: 1\nExp number of AAs in TMHs: 24.51309\nExp number, first 60 AAs: 0.2682\nTotal prob of N-in: 0.14402\noutside 1 331\nTMhelix 332 354\ninside 355 372

21466 GCF_000944835.1_H125160566_illumina_only.fasta Salmonella enterica subsp. enterica serovar Typhi Proteobacteria; Gammaproteobacteria; Enterobacterales; Enterobacteriaceae; Salmonella; Salmonella enterica; Salmonella enterica subsp. enterica MTGDNTLITSHGINRRDFMKLCAALAATMGLSSKAAAEMAESVSNPQRPPVIWIGAQECT WP_000145429.1 hydrogenase 2 small subunit [Salmonella enterica] Length: 372\nNumber of predicted TMHs: 1\nExp number of AAs in TMHs: 24.51309\nExp number, first 60 AAs: 0.2682\nTotal prob of N-in: 0.14402\noutside 1 331\nTMhelix 332 354\ninside 355 372

21467 GCF_001118305.1_10607_1_67 Salmonella enterica subsp. enterica serovar Typhi Proteobacteria; Gammaproteobacteria; Enterobacterales; Enterobacteriaceae; Salmonella; Salmonella enterica; Salmonella enterica subsp. enterica MTGDNTLITSHGINRRDFMKLCAALAATMGLSSKAAAEMAESVSNPQRPPVIWIGAQECT WP_000145429.1 hydrogenase 2 small subunit [Salmonella enterica] Length: 372\nNumber of predicted TMHs: 1\nExp number of AAs in TMHs: 24.51309\nExp number, first 60 AAs: 0.2682\nTotal prob of N-in: 0.14402\noutside 1 331\nTMhelix 332 354\ninside 355 372

21468 GCF_001118165.1_8616_4_45 Salmonella enterica subsp. enterica serovar Typhi Proteobacteria; Gammaproteobacteria; Enterobacterales; Enterobacteriaceae; Salmonella; Salmonella enterica; Salmonella enterica subsp. enterica MTGDNTLITSHGINRRDFMKLCAALAATMGLSSKAAAEMAESVSNPQRPPVIWIGAQECT WP_000145429.1 hydrogenase 2 small subunit [Salmonella enterica] Length: 372\nNumber of predicted TMHs: 1\nExp number of AAs in TMHs: 24.51309\nExp number, first 60 AAs: 0.2682\nTotal prob of N-in: 0.14402\noutside 1 331\nTMhelix 332 354\ninside 355 372

21469 GCF_001117985.1_10562_2_63 Salmonella enterica subsp. enterica serovar Typhi Proteobacteria; Gammaproteobacteria; Enterobacterales; Enterobacteriaceae; Salmonella; Salmonella enterica; Salmonella enterica subsp. enterica MTGDNTLITSHGINRRDFMKLCAALAATMGLSSKAAAEMAESVSNPQRPPVIWIGAQECT WP_000145429.1 hydrogenase 2 small subunit [Salmonella enterica] Length: 372\nNumber of predicted TMHs: 1\nExp number of AAs in TMHs: 24.51309\nExp number, first 60 AAs: 0.2682\nTotal prob of N-in: 0.14402\noutside 1 331\nTMhelix 332 354\ninside 355 372

21470 GCF_001117925.1_10349_1_9 Salmonella enterica subsp. enterica serovar Typhi Proteobacteria; Gammaproteobacteria; Enterobacterales; Enterobacteriaceae; Salmonella; Salmonella enterica; Salmonella enterica subsp. enterica MTGDNTLITSHGINRRDFMKLCAALAATMGLSSKAAAEMAESVSNPQRPPVIWIGAQECT WP_000145429.1 hydrogenase 2 small subunit [Salmonella enterica] Length: 372\nNumber of predicted TMHs: 1\nExp number of AAs in TMHs: 24.51309\nExp number, first 60 AAs: 0.2682\nTotal prob of N-in: 0.14402\noutside 1 331\nTMhelix 332 354\ninside 355 372

21471 GCF_001117485.1_9870_8_68 Salmonella enterica subsp. enterica serovar Typhi Proteobacteria; Gammaproteobacteria; Enterobacterales; Enterobacteriaceae; Salmonella; Salmonella enterica; Salmonella enterica subsp. enterica MTGDNTLITSHGINRRDFMKLCAALAATMGLSSKAAAEMAESVSNPQRPPVIWIGAQECT WP_000145429.1 hydrogenase 2 small subunit [Salmonella enterica] Length: 372\nNumber of predicted TMHs: 1\nExp number of AAs in TMHs: 24.51309\nExp number, first 60 AAs: 0.2682\nTotal prob of N-in: 0.14402\noutside 1 331\nTMhelix 332 354\ninside 355 372

21472 GCF_001117365.1_10561_2_38 Salmonella enterica subsp. enterica serovar Typhi Proteobacteria; Gammaproteobacteria; Enterobacterales; Enterobacteriaceae; Salmonella; Salmonella enterica; Salmonella enterica subsp. enterica MTGDNTLITSHGINRRDFMKLCAALAATMGLSSKAAAEMAESVSNPQRPPVIWIGAQECT WP_000145429.1 hydrogenase 2 small subunit [Salmonella enterica] Length: 372\nNumber of predicted TMHs: 1\nExp number of AAs in TMHs: 24.51309\nExp number, first 60 AAs: 0.2682\nTotal prob of N-in: 0.14402\noutside 1 331\nTMhelix 332 354\ninside 355 372

21473 GCF_001117225.1_8447_8_32 Salmonella enterica subsp. enterica serovar Typhi Proteobacteria; Gammaproteobacteria; Enterobacterales; Enterobacteriaceae; Salmonella; Salmonella enterica;

Salmonella enterica subsp. enterica MTGDNTLITSHGINRRDFMKLCAALAATMGLSSKAAAEMAESVSNPQRPPVIWIGAQECT WP_000145429.1 hydrogenase 2 small subunit [Salmonella enterica] Length: 372\nNumber of predicted TMHs: 1\nExp number of AAs in TMHs: 24.51309\nExp number, first 60 AAs: 0.2682\nTotal prob of N-in: 0.14402\noutside 1 331\nTMhelix 332 354\ninside 355 372

21474 GCF_001113665.1_9475_6_92 Salmonella enterica subsp. enterica serovar Typhi Proteobacteria; Gammaproteobacteria; Enterobacterales; Enterobacteriaceae; Salmonella; Salmonella enterica; Salmonella enterica subsp. enterica MTGDNTLITSHGINRRDFMKLCAALAATMGLSSKAAAEMAESVSNPQRPPVIWIGAQECT WP_000145429.1 hydrogenase 2 small subunit [Salmonella enterica] Length: 372\nNumber of predicted TMHs: 1\nExp number of AAs in TMHs: 24.51309\nExp number, first 60 AAs: 0.2682\nTotal prob of N-in: 0.14402\noutside 1 331\nTMhelix 332 354\ninside 355 372

21475 GCF_001113525.1_8490_5_8Salmonella enterica subsp. enterica serovar Typhi Proteobacteria; Gammaproteobacteria; Enterobacterales; Enterobacteriaceae; Salmonella; Salmonella enterica; Salmonella enterica subsp. enterica MTGDNTLITSHGINRRDFMKLCAALAATMGLSSKAAAEMAESVSNPQRPPVIWIGAQECT WP_000145429.1 hydrogenase 2 small subunit [Salmonella enterica] Length: 372\nNumber of predicted TMHs: 1\nExp number of AAs in TMHs: 24.51309\nExp number, first 60 AAs: 0.2682\nTotal prob of N-in: 0.14402\noutside 1 331\nTMhelix 332 354\ninside 355 372

21476 GCF_001113325.1_9870_8_82 Salmonella enterica subsp. enterica serovar Typhi Proteobacteria; Gammaproteobacteria; Enterobacterales; Enterobacteriaceae; Salmonella; Salmonella enterica; Salmonella enterica subsp. enterica MTGDNTLITSHGINRRDFMKLCAALAATMGLSSKAAAEMAESVSNPQRPPVIWIGAQECT WP_000145429.1 hydrogenase 2 small subunit [Salmonella enterica] Length: 372\nNumber of predicted TMHs: 1\nExp number of AAs in TMHs: 24.51309\nExp number, first 60 AAs: 0.2682\nTotal prob of N-in: 0.14402\noutside 1 331\nTMhelix 332 354\ninside 355 372

21477 GCF_001113105.1_8490_6_56 Salmonella enterica subsp. enterica serovar Typhi Proteobacteria; Gammaproteobacteria; Enterobacterales; Enterobacteriaceae; Salmonella; Salmonella enterica; Salmonella enterica subsp. enterica MTGDNTLITSHGINRRDFMKLCAALAATMGLSSKAAAEMAESVSNPQRPPVIWIGAQECT WP_000145429.1 hydrogenase 2 small subunit [Salmonella enterica] Length: 372\nNumber of predicted TMHs: 1\nExp number of AAs in TMHs: 24.51309\nExp number, first 60 AAs: 0.2682\nTotal prob of N-in: 0.14402\noutside 1 331\nTMhelix 332 354\ninside 355 372

21478 GCF_001112705.1_10562_2_28 Salmonella enterica subsp. enterica serovar Typhi Proteobacteria; Gammaproteobacteria; Enterobacterales; Enterobacteriaceae; Salmonella; Salmonella enterica; Salmonella enterica subsp. enterica MTGDNTLITSHGINRRDFMKLCAALAATMGLSSKAAAEMAESVSNPQRPPVIWIGAQECT WP_000145429.1 hydrogenase 2 small subunit [Salmonella enterica] Length: 372\nNumber of predicted TMHs: 1\nExp number of AAs in TMHs: 24.51309\nExp number, first 60 AAs: 0.2682\nTotal prob of N-in: 0.14402\noutside 1 331\nTMhelix 332 354\ninside 355 372

21479 GCF_001112965.1_9475_6_35 Salmonella enterica subsp. enterica serovar Typhi Proteobacteria; Gammaproteobacteria; Enterobacterales; Enterobacteriaceae; Salmonella; Salmonella enterica; Salmonella enterica subsp. enterica MTGDNTLITSHGINRRDFMKLCAALAATMGLSSKAAAEMAESVSNPQRPPVIWIGAQECT WP_000145429.1 hydrogenase 2 small subunit [Salmonella enterica] Length: 372\nNumber of predicted TMHs: 1\nExp number of AAs in TMHs: 24.51309\nExp number, first 60 AAs: 0.2682\nTotal prob of N-in: 0.14402\noutside 1 331\nTMhelix 332 354\ninside 355 372

21480 GCF_001112465.1_10493_1_46 Salmonella enterica subsp. enterica serovar Typhi Proteobacteria; Gammaproteobacteria; Enterobacterales; Enterobacteriaceae; Salmonella; Salmonella enterica; Salmonella enterica subsp. enterica MTGDNTLITSHGINRRDFMKLCAALAATMGLSSKAAAEMAESVSNPQRPPVIWIGAQECT WP_000145429.1 hydrogenase 2 small subunit [Salmonella enterica] Length: 372\nNumber of predicted TMHs: 1\nExp number of AAs in TMHs: 24.51309\nExp number, first 60 AAs: 0.2682\nTotal prob of N-in: 0.14402\noutside 1 331\nTMhelix 332 354\ninside 355 372

21481 GCF_001110605.1_9475_6_46 Salmonella enterica subsp. enterica serovar Typhi Proteobacteria; Gammaproteobacteria; Enterobacterales; Enterobacteriaceae; Salmonella; Salmonella enterica; Salmonella enterica subsp. enterica MTGDNTLITSHGINRRDFMKLCAALAATMGLSSKAAAEMAESVSNPQRPPVIWIGAQECT WP_000145429.1 hydrogenase 2 small subunit [Salmonella enterica] Length: 372\nNumber of predicted TMHs: 1\nExp number of AAs in TMHs: 24.51309\nExp number, first 60 AAs: 0.2682\nTotal prob of N-in: 0.14402\noutside 1 331\nTMhelix 332 354\ninside 355 372

21482 GCF_001110425.1_10593_2_8 Salmonella enterica subsp. enterica serovar Typhi Proteobacteria; Gammaproteobacteria; Enterobacterales; Enterobacteriaceae; Salmonella; Salmonella enterica; Salmonella enterica subsp. enterica MTGDNTLITSHGINRRDFMKLCAALAATMGLSSKAAAEMAESVSNPQRPPVIWIGAQECT WP_000145429.1 hydrogenase 2 small subunit [Salmonella enterica] Length: 372\nNumber of predicted TMHs: 1\nExp number of AAs in TMHs: 24.51309\nExp number, first 60 AAs: 0.2682\nTotal prob of N-in: 0.14402\noutside 1 331\nTMhelix 332 354\ninside 355 372

21483 GCF_001109985.1_10349_1_17 Salmonella enterica subsp. enterica serovar Typhi Proteobacteria; Gammaproteobacteria; Enterobacterales; Enterobacteriaceae; Salmonella; Salmonella enterica; Salmonella enterica subsp. enterica MTGDNTLITSHGINRRDFMKLCAALAATMGLSSKAAAEMAESVSNPQRPPVIWIGAQECT WP_000145429.1 hydrogenase 2 small subunit [Salmonella enterica] Length: 372\nNumber of predicted TMHs: 1\nExp number of AAs in TMHs: 24.51309\nExp number, first 60 AAs: 0.2682\nTotal prob of N-in: 0.14402\noutside 1 331\nTMhelix 332 354\ninside 355 372

21484 GCF_001109645.1_9870_8_41 Salmonella enterica subsp. enterica serovar Typhi Proteobacteria; Gammaproteobacteria; Enterobacterales; Enterobacteriaceae; Salmonella; Salmonella enterica; Salmonella enterica subsp. enterica MTGDNTLITSHGINRRDFMKLCAALAATMGLSSKAAAEMAESVSNPQRPPVIWIGAQECT WP_000145429.1 hydrogenase 2 small subunit [Salmonella enterica] Length: 372\nNumber of predicted TMHs: 1\nExp number of AAs in TMHs: 24.51309\nExp number, first 60 AAs: 0.2682\nTotal prob of N-in: 0.14402\noutside 1 331\nTMhelix 332 354\ninside 355 372

21485 GCF_001109605.1_10492_1_82 Salmonella enterica subsp. enterica serovar Typhi Proteobacteria; Gammaproteobacteria; Enterobacterales; Enterobacteriaceae; Salmonella; Salmonella enterica; Salmonella enterica subsp. enterica MTGDNTLITSHGINRRDFMKLCAALAATMGLSSKAAAEMAESVSNPQRPPVIWIGAQECT WP_000145429.1 hydrogenase 2 small subunit [Salmonella enterica] Length: 372\nNumber of predicted TMHs: 1\nExp number of AAs in TMHs: 24.51309\nExp number, first 60 AAs: 0.2682\nTotal prob of N-in: 0.14402\noutside 1 331\nTMhelix 332 354\ninside 355 372

21486 GCF_001109485.1_9870_8_44 Salmonella enterica subsp. enterica serovar Typhi Proteobacteria; Gammaproteobacteria; Enterobacterales; Enterobacteriaceae; Salmonella; Salmonella enterica; Salmonella enterica subsp. enterica MTGDNTLITSHGINRRDFMKLCAALAATMGLSSKAAAEMAESVSNPQRPPVIWIGAQECT WP_000145429.1 hydrogenase 2 small subunit [Salmonella enterica] Length: 372\nNumber of predicted TMHs: 1\nExp number of AAs in TMHs: 24.51309\nExp number, first 60 AAs: 0.2682\nTotal prob of N-in: 0.14402\noutside 1 331\nTMhelix 332 354\ninside 355 372

21487 GCF_001109285.1_10492_1_78 Salmonella enterica subsp. enterica serovar Typhi Proteobacteria; Gammaproteobacteria; Enterobacterales; Enterobacteriaceae; Salmonella; Salmonella enterica; Salmonella enterica subsp. enterica MTGDNTLITSHGINRRDFMKLCAALAATMGLSSKAAAEMAESVSNPQRPPVIWIGAQECT WP_000145429.1 hydrogenase 2 small subunit [Salmonella enterica] Length: 372\nNumber of predicted TMHs: 1\nExp number of AAs in TMHs: 24.51309\nExp number, first 60 AAs: 0.2682\nTotal prob of N-in: 0.14402\noutside 1 331\nTMhelix 332 354\ninside 355 372

21488 GCF_001106625.1_10071_8_92 Salmonella enterica subsp. enterica serovar Typhi Proteobacteria; Gammaproteobacteria; Enterobacterales; Enterobacteriaceae; Salmonella; Salmonella enterica; Salmonella enterica subsp. enterica MTGDNTLITSHGINRRDFMKLCAALAATMGLSSKAAAEMAESVSNPQRPPVIWIGAQECT WP_000145429.1 hydrogenase 2 small subunit [Salmonella enterica] Length: 372\nNumber of predicted TMHs: 1\nExp number of AAs in TMHs: 24.51309\nExp number, first 60 AAs: 0.2682\nTotal prob of N-in: 0.14402\noutside 1 331\nTMhelix 332 354\ninside 355 372

21489 GCF_001048295.1_10493_1_28 Salmonella enterica subsp. enterica serovar Typhi Proteobacteria; Gammaproteobacteria; Enterobacterales; Enterobacteriaceae; Salmonella; Salmonella enterica; Salmonella enterica subsp. enterica MTGDNTLITSHGINRRDFMKLCAALAATMGLSSKAAAEMAESVSNPQRPPVIWIGAQECT WP_000145429.1 hydrogenase 2 small subunit [Salmonella enterica] Length: 372\nNumber of predicted TMHs: 1\nExp number of AAs in TMHs: 24.51309\nExp number, first 60 AAs: 0.2682\nTotal prob of N-in: 0.14402\noutside 1 331\nTMhelix 332 354\ninside 355 372

21490 GCF_001048335.1_10492_1_15 Salmonella enterica subsp. enterica serovar Typhi Proteobacteria; Gammaproteobacteria; Enterobacterales; Enterobacteriaceae; Salmonella; Salmonella enterica; Salmonella enterica subsp. enterica MTGDNTLITSHGINRRDFMKLCAALAATMGLSSKAAAEMAESVSNPQRPPVIWIGAQECT WP_000145429.1 hydrogenase 2 small subunit [Salmonella enterica] Length: 372\nNumber of

predicted TMHs: 1\nExp number of AAs in TMHs: 24.51309\nExp number, first 60 AAs: 0.2682\nTotal prob of N-in: 0.14402\noutside 1 331\nTMhelix 332 354\ninside 355 372

21491 GCF_001048375.1_10425_1_10 Salmonella enterica subsp. enterica serovar Typhi Proteobacteria; Gammaproteobacteria; Enterobacterales; Enterobacteriaceae; Salmonella; Salmonella enterica; Salmonella enterica subsp. enterica MTGDNTLITSHGINRRDFMKLCAALAATMGLSSKAAAEMAESVSNPQRPPVIWIGAQECT WP_000145429.1 hydrogenase 2 small subunit [Salmonella enterica] Length: 372\nNumber of predicted TMHs: 1\nExp number of AAs in TMHs: 24.51309\nExp number, first 60 AAs: 0.2682\nTotal prob of N-in: 0.14402\noutside 1 331\nTMhelix 332 354\ninside 355 372

21492 GCF_001048415.1_10492_1_39 Salmonella enterica subsp. enterica serovar Typhi Proteobacteria; Gammaproteobacteria; Enterobacterales; Enterobacteriaceae; Salmonella; Salmonella enterica; Salmonella enterica subsp. enterica MTGDNTLITSHGINRRDFMKLCAALAATMGLSSKAAAEMAESVSNPQRPPVIWIGAQECT WP_000145429.1 hydrogenase 2 small subunit [Salmonella enterica] Length: 372\nNumber of predicted TMHs: 1\nExp number of AAs in TMHs: 24.51309\nExp number, first 60 AAs: 0.2682\nTotal prob of N-in: 0.14402\noutside 1 331\nTMhelix 332 354\ninside 355 372

21493 GCF_001048455.1_10562_2_14 Salmonella enterica subsp. enterica serovar Typhi Proteobacteria; Gammaproteobacteria; Enterobacterales; Enterobacteriaceae; Salmonella; Salmonella enterica; Salmonella enterica subsp. enterica MTGDNTLITSHGINRRDFMKLCAALAATMGLSSKAAAEMAESVSNPQRPPVIWIGAQECT WP_000145429.1 hydrogenase 2 small subunit [Salmonella enterica] Length: 372\nNumber of predicted TMHs: 1\nExp number of AAs in TMHs: 24.51309\nExp number, first 60 AAs: 0.2682\nTotal prob of N-in: 0.14402\noutside 1 331\nTMhelix 332 354\ninside 355 372

21494 GCF_001106085.1_10593_2_75 Salmonella enterica subsp. enterica serovar Typhi Proteobacteria; Gammaproteobacteria; Enterobacterales; Enterobacteriaceae; Salmonella; Salmonella enterica; Salmonella enterica subsp. enterica MTGDNTLITSHGINRRDFMKLCAALAATMGLSSKAAAEMAESVSNPQRPPVIWIGAQECT WP_000145429.1 hydrogenase 2 small subunit [Salmonella enterica] Length: 372\nNumber of predicted TMHs: 1\nExp number of AAs in TMHs: 24.51309\nExp number, first 60 AAs: 0.2682\nTotal prob of N-in: 0.14402\noutside 1 331\nTMhelix 332 354\ninside 355 372

21495 GCF_001051385.1_ASM105138v1 Salmonella enterica subsp. enterica serovar Typhi Proteobacteria; Gammaproteobacteria; Enterobacterales; Enterobacteriaceae; Salmonella; Salmonella enterica; Salmonella enterica subsp. enterica MTGDNTLITSHGINRRDFMKLCAALAATMGLSSKAAAEMAESVSNPQRPPVIWIGAQECT WP_000145429.1 hydrogenase 2 small subunit [Salmonella enterica] Length: 372\nNumber of predicted TMHs: 1\nExp number of AAs in TMHs: 24.51309\nExp number, first 60 AAs: 0.2682\nTotal prob of N-in: 0.14402\noutside 1 331\nTMhelix 332 354\ninside 355 372

21496 GCF_001078745.1_10492_1_67 Salmonella enterica subsp. enterica serovar Typhi Proteobacteria; Gammaproteobacteria; Enterobacterales; Enterobacteriaceae; Salmonella; Salmonella enterica; Salmonella enterica subsp. enterica MTGDNTLITSHGINRRDFMKLCAALAATMGLSSKAAAEMAESVSNPQRPPVIWIGAQECT WP_000145429.1 hydrogenase 2 small subunit [Salmonella enterica] Length: 372\nNumber of predicted TMHs: 1\nExp number of AAs in TMHs: 24.51309\nExp number, first 60 AAs: 0.2682\nTotal prob of N-in: 0.14402\noutside 1 331\nTMhelix 332 354\ninside 355 372

21497 GCF_001078785.1_10493_1_39 Salmonella enterica subsp. enterica serovar Typhi Proteobacteria; Gammaproteobacteria; Enterobacterales; Enterobacteriaceae; Salmonella; Salmonella enterica; Salmonella enterica subsp. enterica MTGDNTLITSHGINRRDFMKLCAALAATMGLSSKAAAEMAESVSNPQRPPVIWIGAQECT WP_000145429.1 hydrogenase 2 small subunit [Salmonella enterica] Length: 372\nNumber of predicted TMHs: 1\nExp number of AAs in TMHs: 24.51309\nExp number, first 60 AAs: 0.2682\nTotal prob of N-in: 0.14402\noutside 1 331\nTMhelix 332 354\ninside 355 372

21498 GCF_001078845.1_10493_1_21 Salmonella enterica subsp. enterica serovar Typhi Proteobacteria; Gammaproteobacteria; Enterobacterales; Enterobacteriaceae; Salmonella; Salmonella enterica; Salmonella enterica subsp. enterica MTGDNTLITSHGINRRDFMKLCAALAATMGLSSKAAAEMAESVSNPQRPPVIWIGAQECT WP_000145429.1 hydrogenase 2 small subunit [Salmonella enterica] Length: 372\nNumber of predicted TMHs: 1\nExp number of AAs in TMHs: 24.51309\nExp number, first 60 AAs: 0.2682\nTotal prob of N-in: 0.14402\noutside 1 331\nTMhelix 332 354\ninside 355 372

21499 GCF_001106045.1_9953_5_64 Salmonella enterica subsp. enterica serovar Typhi Proteobacteria; Gammaproteobacteria; Enterobacterales; Enterobacteriaceae; Salmonella; Salmonella enterica;

Salmonella enterica subsp. enterica	MTGDNTLITSHGINRRDFMKLCAALAATMGLSSKAAAEMAESVSNPQRPPVIWIGAQECT
WP_000145429.1	hydrogenase 2 small subunit [Salmonella enterica]	Length: 372\nNumber of predicted TMHs: 1\nExp number of AAs in TMHs: 24.51309\nExp number, first 60 AAs: 0.2682\nTotal prob of N-in: 0.14402\noutside 1 331\nTMhelix 332 354\ninside 355 372

21500	GCF_001078885.1_10562_2_13	Salmonella enterica subsp. enterica serovar Typhi
Proteobacteria; Gammaproteobacteria; Enterobacterales; Enterobacteriaceae; Salmonella; Salmonella enterica; Salmonella enterica subsp. enterica	MTGDNTLITSHGINRRDFMKLCAALAATMGLSSKAAAEMAESVSNPQRPPVIWIGAQECT
WP_000145429.1	hydrogenase 2 small subunit [Salmonella enterica]	Length: 372\nNumber of predicted TMHs: 1\nExp number of AAs in TMHs: 24.51309\nExp number, first 60 AAs: 0.2682\nTotal prob of N-in: 0.14402\noutside 1 331\nTMhelix 332 354\ninside 355 372

21501	GCF_001105845.1_8490_6_51	Salmonella enterica subsp. enterica serovar Typhi
Proteobacteria; Gammaproteobacteria; Enterobacterales; Enterobacteriaceae; Salmonella; Salmonella enterica; Salmonella enterica subsp. enterica	MTGDNTLITSHGINRRDFMKLCAALAATMGLSSKAAAEMAESVSNPQRPPVIWIGAQECT
WP_000145429.1	hydrogenase 2 small subunit [Salmonella enterica]	Length: 372\nNumber of predicted TMHs: 1\nExp number of AAs in TMHs: 24.51309\nExp number, first 60 AAs: 0.2682\nTotal prob of N-in: 0.14402\noutside 1 331\nTMhelix 332 354\ninside 355 372

21502	GCF_001079465.1_10060_5_91	Salmonella enterica subsp. enterica serovar Typhi
Proteobacteria; Gammaproteobacteria; Enterobacterales; Enterobacteriaceae; Salmonella; Salmonella enterica; Salmonella enterica subsp. enterica	MTGDNTLITSHGINRRDFMKLCAALAATMGLSSKAAAEMAESVSNPQRPPVIWIGAQECT
WP_000145429.1	hydrogenase 2 small subunit [Salmonella enterica]	Length: 372\nNumber of predicted TMHs: 1\nExp number of AAs in TMHs: 24.51309\nExp number, first 60 AAs: 0.2682\nTotal prob of N-in: 0.14402\noutside 1 331\nTMhelix 332 354\ninside 355 372

21503	GCF_001079425.1_10071_8_90	Salmonella enterica subsp. enterica serovar Typhi
Proteobacteria; Gammaproteobacteria; Enterobacterales; Enterobacteriaceae; Salmonella; Salmonella enterica; Salmonella enterica subsp. enterica	MTGDNTLITSHGINRRDFMKLCAALAATMGLSSKAAAEMAESVSNPQRPPVIWIGAQECT
WP_000145429.1	hydrogenase 2 small subunit [Salmonella enterica]	Length: 372\nNumber of predicted TMHs: 1\nExp number of AAs in TMHs: 24.51309\nExp number, first 60 AAs: 0.2682\nTotal prob of N-in: 0.14402\noutside 1 331\nTMhelix 332 354\ninside 355 372

21504	GCF_001105805.1_10493_1_18	Salmonella enterica subsp. enterica serovar Typhi
Proteobacteria; Gammaproteobacteria; Enterobacterales; Enterobacteriaceae; Salmonella; Salmonella enterica; Salmonella enterica subsp. enterica	MTGDNTLITSHGINRRDFMKLCAALAATMGLSSKAAAEMAESVSNPQRPPVIWIGAQECT
WP_000145429.1	hydrogenase 2 small subunit [Salmonella enterica]	Length: 372\nNumber of predicted TMHs: 1\nExp number of AAs in TMHs: 24.51309\nExp number, first 60 AAs: 0.2682\nTotal prob of N-in: 0.14402\noutside 1 331\nTMhelix 332 354\ninside 355 372

21505	GCF_001079505.1_10060_5_17	Salmonella enterica subsp. enterica serovar Typhi
Proteobacteria; Gammaproteobacteria; Enterobacterales; Enterobacteriaceae; Salmonella; Salmonella enterica; Salmonella enterica subsp. enterica	MTGDNTLITSHGINRRDFMKLCAALAATMGLSSKAAAEMAESVSNPQRPPVIWIGAQECT
WP_000145429.1	hydrogenase 2 small subunit [Salmonella enterica]	Length: 372\nNumber of predicted TMHs: 1\nExp number of AAs in TMHs: 24.51309\nExp number, first 60 AAs: 0.2682\nTotal prob of N-in: 0.14402\noutside 1 331\nTMhelix 332 354\ninside 355 372

21506	GCF_001079545.1_10071_8_9	Salmonella enterica subsp. enterica serovar Typhi
Proteobacteria; Gammaproteobacteria; Enterobacterales; Enterobacteriaceae; Salmonella; Salmonella enterica; Salmonella enterica subsp. enterica	MTGDNTLITSHGINRRDFMKLCAALAATMGLSSKAAAEMAESVSNPQRPPVIWIGAQECT
WP_000145429.1	hydrogenase 2 small subunit [Salmonella enterica]	Length: 372\nNumber of predicted TMHs: 1\nExp number of AAs in TMHs: 24.51309\nExp number, first 60 AAs: 0.2682\nTotal prob of N-in: 0.14402\noutside 1 331\nTMhelix 332 354\ninside 355 372

21507	GCF_001079585.1_10071_8_22	Salmonella enterica subsp. enterica serovar Typhi
Proteobacteria; Gammaproteobacteria; Enterobacterales; Enterobacteriaceae; Salmonella; Salmonella enterica; Salmonella enterica subsp. enterica	MTGDNTLITSHGINRRDFMKLCAALAATMGLSSKAAAEMAESVSNPQRPPVIWIGAQECT
WP_000145429.1	hydrogenase 2 small subunit [Salmonella enterica]	Length: 372\nNumber of predicted TMHs: 1\nExp number of AAs in TMHs: 24.51309\nExp number, first 60 AAs: 0.2682\nTotal prob of N-in: 0.14402\noutside 1 331\nTMhelix 332 354\ninside 355 372

21508 GCF_001079665.1_10071_8_81 Salmonella enterica subsp. enterica serovar Typhi Proteobacteria; Gammaproteobacteria; Enterobacterales; Enterobacteriaceae; Salmonella; Salmonella enterica; Salmonella enterica subsp. enterica MTGDNTLITSHGINRRDFMKLCAALAATMGLSSKAAAEMAESVSNPQRPPVIWIGAQECT WP_000145429.1 hydrogenase 2 small subunit [Salmonella enterica] Length: 372\nNumber of predicted TMHs: 1\nExp number of AAs in TMHs: 24.51309\nExp number, first 60 AAs: 0.2682\nTotal prob of N-in: 0.14402\noutside 1 331\nTMhelix 332 354\ninside 355 372

21509 GCF_001080165.1_10060_6_20 Salmonella enterica subsp. enterica serovar Typhi Proteobacteria; Gammaproteobacteria; Enterobacterales; Enterobacteriaceae; Salmonella; Salmonella enterica; Salmonella enterica subsp. enterica MTGDNTLITSHGINRRDFMKLCAALAATMGLSSKAAAEMAESVSNPQRPPVIWIGAQECT WP_000145429.1 hydrogenase 2 small subunit [Salmonella enterica] Length: 372\nNumber of predicted TMHs: 1\nExp number of AAs in TMHs: 24.51309\nExp number, first 60 AAs: 0.2682\nTotal prob of N-in: 0.14402\noutside 1 331\nTMhelix 332 354\ninside 355 372

21510 GCF_001080205.1_10060_6_42 Salmonella enterica subsp. enterica serovar Typhi Proteobacteria; Gammaproteobacteria; Enterobacterales; Enterobacteriaceae; Salmonella; Salmonella enterica; Salmonella enterica subsp. enterica MTGDNTLITSHGINRRDFMKLCAALAATMGLSSKAAAEMAESVSNPQRPPVIWIGAQECT WP_000145429.1 hydrogenase 2 small subunit [Salmonella enterica] Length: 372\nNumber of predicted TMHs: 1\nExp number of AAs in TMHs: 24.51309\nExp number, first 60 AAs: 0.2682\nTotal prob of N-in: 0.14402\noutside 1 331\nTMhelix 332 354\ninside 355 372

21511 GCF_001080285.1_10071_8_74 Salmonella enterica subsp. enterica serovar Typhi Proteobacteria; Gammaproteobacteria; Enterobacterales; Enterobacteriaceae; Salmonella; Salmonella enterica; Salmonella enterica subsp. enterica MTGDNTLITSHGINRRDFMKLCAALAATMGLSSKAAAEMAESVSNPQRPPVIWIGAQECT WP_000145429.1 hydrogenase 2 small subunit [Salmonella enterica] Length: 372\nNumber of predicted TMHs: 1\nExp number of AAs in TMHs: 24.51309\nExp number, first 60 AAs: 0.2682\nTotal prob of N-in: 0.14402\noutside 1 331\nTMhelix 332 354\ninside 355 372

21512 GCF_001080325.1_10060_6_28 Salmonella enterica subsp. enterica serovar Typhi Proteobacteria; Gammaproteobacteria; Enterobacterales; Enterobacteriaceae; Salmonella; Salmonella enterica; Salmonella enterica subsp. enterica MTGDNTLITSHGINRRDFMKLCAALAATMGLSSKAAAEMAESVSNPQRPPVIWIGAQECT WP_000145429.1 hydrogenase 2 small subunit [Salmonella enterica] Length: 372\nNumber of predicted TMHs: 1\nExp number of AAs in TMHs: 24.51309\nExp number, first 60 AAs: 0.2682\nTotal prob of N-in: 0.14402\noutside 1 331\nTMhelix 332 354\ninside 355 372

21513 GCF_001080405.1_10060_6_57 Salmonella enterica subsp. enterica serovar Typhi Proteobacteria; Gammaproteobacteria; Enterobacterales; Enterobacteriaceae; Salmonella; Salmonella enterica; Salmonella enterica subsp. enterica MTGDNTLITSHGINRRDFMKLCAALAATMGLSSKAAAEMAESVSNPQRPPVIWIGAQECT WP_000145429.1 hydrogenase 2 small subunit [Salmonella enterica] Length: 372\nNumber of predicted TMHs: 1\nExp number of AAs in TMHs: 24.51309\nExp number, first 60 AAs: 0.2682\nTotal prob of N-in: 0.14402\noutside 1 331\nTMhelix 332 354\ninside 355 372

21514 GCF_001080445.1_10071_8_63 Salmonella enterica subsp. enterica serovar Typhi Proteobacteria; Gammaproteobacteria; Enterobacterales; Enterobacteriaceae; Salmonella; Salmonella enterica; Salmonella enterica subsp. enterica MTGDNTLITSHGINRRDFMKLCAALAATMGLSSKAAAEMAESVSNPQRPPVIWIGAQECT WP_000145429.1 hydrogenase 2 small subunit [Salmonella enterica] Length: 372\nNumber of predicted TMHs: 1\nExp number of AAs in TMHs: 24.51309\nExp number, first 60 AAs: 0.2682\nTotal prob of N-in: 0.14402\noutside 1 331\nTMhelix 332 354\ninside 355 372

21515 GCF_001105625.1_10562_2_68 Salmonella enterica subsp. enterica serovar Typhi Proteobacteria; Gammaproteobacteria; Enterobacterales; Enterobacteriaceae; Salmonella; Salmonella enterica; Salmonella enterica subsp. enterica MTGDNTLITSHGINRRDFMKLCAALAATMGLSSKAAAEMAESVSNPQRPPVIWIGAQECT WP_000145429.1 hydrogenase 2 small subunit [Salmonella enterica] Length: 372\nNumber of predicted TMHs: 1\nExp number of AAs in TMHs: 24.51309\nExp number, first 60 AAs: 0.2682\nTotal prob of N-in: 0.14402\noutside 1 331\nTMhelix 332 354\ninside 355 372

21516 GCF_001081145.1_10060_6_80 Salmonella enterica subsp. enterica serovar Typhi Proteobacteria; Gammaproteobacteria; Enterobacterales; Enterobacteriaceae; Salmonella; Salmonella enterica; Salmonella enterica subsp. enterica MTGDNTLITSHGINRRDFMKLCAALAATMGLSSKAAAEMAESVSNPQRPPVIWIGAQECT WP_000145429.1 hydrogenase 2 small subunit [Salmonella enterica] Length: 372\nNumber of

predicted TMHs:  1\nExp number of AAs in TMHs: 24.51309\nExp number, first 60 AAs:  0.2682\nTotal prob of N-in: 0.14402\noutside   1   331\nTMhelix   332   354\ninside   355   372

21517 GCF_001081265.1_10071_8_23 Salmonella enterica subsp. enterica serovar Typhi Proteobacteria; Gammaproteobacteria; Enterobacterales; Enterobacteriaceae; Salmonella; Salmonella enterica; Salmonella enterica subsp. enterica MTGDNTLITSHGINRRDFMKLCAALAATMGLSSKAAAEMAESVSNPQRPPVIWIGAQECT WP_000145429.1 hydrogenase 2 small subunit [Salmonella enterica] Length: 372\nNumber of predicted TMHs:  1\nExp number of AAs in TMHs: 24.51309\nExp number, first 60 AAs:  0.2682\nTotal prob of N-in: 0.14402\noutside   1   331\nTMhelix   332   354\ninside   355   372

21518 GCF_001081225.1_10060_6_54 Salmonella enterica subsp. enterica serovar Typhi Proteobacteria; Gammaproteobacteria; Enterobacterales; Enterobacteriaceae; Salmonella; Salmonella enterica; Salmonella enterica subsp. enterica MTGDNTLITSHGINRRDFMKLCAALAATMGLSSKAAAEMAESVSNPQRPPVIWIGAQECT WP_000145429.1 hydrogenase 2 small subunit [Salmonella enterica] Length: 372\nNumber of predicted TMHs:  1\nExp number of AAs in TMHs: 24.51309\nExp number, first 60 AAs:  0.2682\nTotal prob of N-in: 0.14402\noutside   1   331\nTMhelix   332   354\ninside   355   372

21519 GCF_001103325.1_8616_4_38 Salmonella enterica subsp. enterica serovar Typhi Proteobacteria; Gammaproteobacteria; Enterobacterales; Enterobacteriaceae; Salmonella; Salmonella enterica; Salmonella enterica subsp. enterica MTGDNTLITSHGINRRDFMKLCAALAATMGLSSKAAAEMAESVSNPQRPPVIWIGAQECT WP_000145429.1 hydrogenase 2 small subunit [Salmonella enterica] Length: 372\nNumber of predicted TMHs:  1\nExp number of AAs in TMHs: 24.51309\nExp number, first 60 AAs:  0.2682\nTotal prob of N-in: 0.14402\noutside   1   331\nTMhelix   332   354\ninside   355   372

21520 GCF_001081305.1_10071_8_80 Salmonella enterica subsp. enterica serovar Typhi Proteobacteria; Gammaproteobacteria; Enterobacterales; Enterobacteriaceae; Salmonella; Salmonella enterica; Salmonella enterica subsp. enterica MTGDNTLITSHGINRRDFMKLCAALAATMGLSSKAAAEMAESVSNPQRPPVIWIGAQECT WP_000145429.1 hydrogenase 2 small subunit [Salmonella enterica] Length: 372\nNumber of predicted TMHs:  1\nExp number of AAs in TMHs: 24.51309\nExp number, first 60 AAs:  0.2682\nTotal prob of N-in: 0.14402\noutside   1   331\nTMhelix   332   354\ninside   355   372

21521 GCF_001081365.1_10071_8_72 Salmonella enterica subsp. enterica serovar Typhi Proteobacteria; Gammaproteobacteria; Enterobacterales; Enterobacteriaceae; Salmonella; Salmonella enterica; Salmonella enterica subsp. enterica MTGDNTLITSHGINRRDFMKLCAALAATMGLSSKAAAEMAESVSNPQRPPVIWIGAQECT WP_000145429.1 hydrogenase 2 small subunit [Salmonella enterica] Length: 372\nNumber of predicted TMHs:  1\nExp number of AAs in TMHs: 24.51309\nExp number, first 60 AAs:  0.2682\nTotal prob of N-in: 0.14402\noutside   1   331\nTMhelix   332   354\ninside   355   372

21522 GCF_001103265.1_9870_8_72 Salmonella enterica subsp. enterica serovar Typhi Proteobacteria; Gammaproteobacteria; Enterobacterales; Enterobacteriaceae; Salmonella; Salmonella enterica; Salmonella enterica subsp. enterica MTGDNTLITSHGINRRDFMKLCAALAATMGLSSKAAAEMAESVSNPQRPPVIWIGAQECT WP_000145429.1 hydrogenase 2 small subunit [Salmonella enterica] Length: 372\nNumber of predicted TMHs:  1\nExp number of AAs in TMHs: 24.51309\nExp number, first 60 AAs:  0.2682\nTotal prob of N-in: 0.14402\noutside   1   331\nTMhelix   332   354\ninside   355   372

21523 GCF_001081965.1_10071_8_26 Salmonella enterica subsp. enterica serovar Typhi Proteobacteria; Gammaproteobacteria; Enterobacterales; Enterobacteriaceae; Salmonella; Salmonella enterica; Salmonella enterica subsp. enterica MTGDNTLITSHGINRRDFMKLCAALAATMGLSSKAAAEMAESVSNPQRPPVIWIGAQECT WP_000145429.1 hydrogenase 2 small subunit [Salmonella enterica] Length: 372\nNumber of predicted TMHs:  1\nExp number of AAs in TMHs: 24.51309\nExp number, first 60 AAs:  0.2682\nTotal prob of N-in: 0.14402\noutside   1   331\nTMhelix   332   354\ninside   355   372

21524 GCF_001082005.1_10060_5_35 Salmonella enterica subsp. enterica serovar Typhi Proteobacteria; Gammaproteobacteria; Enterobacterales; Enterobacteriaceae; Salmonella; Salmonella enterica; Salmonella enterica subsp. enterica MTGDNTLITSHGINRRDFMKLCAALAATMGLSSKAAAEMAESVSNPQRPPVIWIGAQECT WP_000145429.1 hydrogenase 2 small subunit [Salmonella enterica] Length: 372\nNumber of predicted TMHs:  1\nExp number of AAs in TMHs: 24.51309\nExp number, first 60 AAs:  0.2682\nTotal prob of N-in: 0.14402\noutside   1   331\nTMhelix   332   354\ninside   355   372

21525 GCF_001082065.1_10060_5_75 Salmonella enterica subsp. enterica serovar Typhi Proteobacteria; Gammaproteobacteria; Enterobacterales; Enterobacteriaceae; Salmonella; Salmonella enterica;

Salmonella enterica subsp. enterica MTGDNTLITSHGINRRDFMKLCAALAATMGLSSKAAAEMAESVSNPQRPPVIWIGAQECT WP_000145429.1 hydrogenase 2 small subunit [Salmonella enterica] Length: 372\nNumber of predicted TMHs: 1\nExp number of AAs in TMHs: 24.51309\nExp number, first 60 AAs: 0.2682\nTotal prob of N-in: 0.14402\noutside 1 331\nTMhelix 332 354\ninside 355 372

21526 GCF_001082185.1_10071_8_36 Salmonella enterica subsp. enterica serovar Typhi Proteobacteria; Gammaproteobacteria; Enterobacterales; Enterobacteriaceae; Salmonella; Salmonella enterica; Salmonella enterica subsp. enterica MTGDNTLITSHGINRRDFMKLCAALAATMGLSSKAAAEMAESVSNPQRPPVIWIGAQECT WP_000145429.1 hydrogenase 2 small subunit [Salmonella enterica] Length: 372\nNumber of predicted TMHs: 1\nExp number of AAs in TMHs: 24.51309\nExp number, first 60 AAs: 0.2682\nTotal prob of N-in: 0.14402\noutside 1 331\nTMhelix 332 354\ninside 355 372

21527 GCF_001082145.1_10071_8_49 Salmonella enterica subsp. enterica serovar Typhi Proteobacteria; Gammaproteobacteria; Enterobacterales; Enterobacteriaceae; Salmonella; Salmonella enterica; Salmonella enterica subsp. enterica MTGDNTLITSHGINRRDFMKLCAALAATMGLSSKAAAEMAESVSNPQRPPVIWIGAQECT WP_000145429.1 hydrogenase 2 small subunit [Salmonella enterica] Length: 372\nNumber of predicted TMHs: 1\nExp number of AAs in TMHs: 24.51309\nExp number, first 60 AAs: 0.2682\nTotal prob of N-in: 0.14402\noutside 1 331\nTMhelix 332 354\ninside 355 372

21528 GCF_001102885.1_8490_6_69 Salmonella enterica subsp. enterica serovar Typhi Proteobacteria; Gammaproteobacteria; Enterobacterales; Enterobacteriaceae; Salmonella; Salmonella enterica; Salmonella enterica subsp. enterica MTGDNTLITSHGINRRDFMKLCAALAATMGLSSKAAAEMAESVSNPQRPPVIWIGAQECT WP_000145429.1 hydrogenase 2 small subunit [Salmonella enterica] Length: 372\nNumber of predicted TMHs: 1\nExp number of AAs in TMHs: 24.51309\nExp number, first 60 AAs: 0.2682\nTotal prob of N-in: 0.14402\noutside 1 331\nTMhelix 332 354\ninside 355 372

21529 GCF_001082225.1_10071_8_41 Salmonella enterica subsp. enterica serovar Typhi Proteobacteria; Gammaproteobacteria; Enterobacterales; Enterobacteriaceae; Salmonella; Salmonella enterica; Salmonella enterica subsp. enterica MTGDNTLITSHGINRRDFMKLCAALAATMGLSSKAAAEMAESVSNPQRPPVIWIGAQECT WP_000145429.1 hydrogenase 2 small subunit [Salmonella enterica] Length: 372\nNumber of predicted TMHs: 1\nExp number of AAs in TMHs: 24.51309\nExp number, first 60 AAs: 0.2682\nTotal prob of N-in: 0.14402\noutside 1 331\nTMhelix 332 354\ninside 355 372

21530 GCF_001082765.1_10071_3_73 Salmonella enterica subsp. enterica serovar Typhi Proteobacteria; Gammaproteobacteria; Enterobacterales; Enterobacteriaceae; Salmonella; Salmonella enterica; Salmonella enterica subsp. enterica MTGDNTLITSHGINRRDFMKLCAALAATMGLSSKAAAEMAESVSNPQRPPVIWIGAQECT WP_000145429.1 hydrogenase 2 small subunit [Salmonella enterica] Length: 372\nNumber of predicted TMHs: 1\nExp number of AAs in TMHs: 24.51309\nExp number, first 60 AAs: 0.2682\nTotal prob of N-in: 0.14402\noutside 1 331\nTMhelix 332 354\ninside 355 372

21531 GCF_001082805.1_10060_5_55 Salmonella enterica subsp. enterica serovar Typhi Proteobacteria; Gammaproteobacteria; Enterobacterales; Enterobacteriaceae; Salmonella; Salmonella enterica; Salmonella enterica subsp. enterica MTGDNTLITSHGINRRDFMKLCAALAATMGLSSKAAAEMAESVSNPQRPPVIWIGAQECT WP_000145429.1 hydrogenase 2 small subunit [Salmonella enterica] Length: 372\nNumber of predicted TMHs: 1\nExp number of AAs in TMHs: 24.51309\nExp number, first 60 AAs: 0.2682\nTotal prob of N-in: 0.14402\noutside 1 331\nTMhelix 332 354\ninside 355 372

21532 GCF_001103145.1_10493_1_67 Salmonella enterica subsp. enterica serovar Typhi Proteobacteria; Gammaproteobacteria; Enterobacterales; Enterobacteriaceae; Salmonella; Salmonella enterica; Salmonella enterica subsp. enterica MTGDNTLITSHGINRRDFMKLCAALAATMGLSSKAAAEMAESVSNPQRPPVIWIGAQECT WP_000145429.1 hydrogenase 2 small subunit [Salmonella enterica] Length: 372\nNumber of predicted TMHs: 1\nExp number of AAs in TMHs: 24.51309\nExp number, first 60 AAs: 0.2682\nTotal prob of N-in: 0.14402\noutside 1 331\nTMhelix 332 354\ninside 355 372

21533 GCF_001082845.1_10071_3_49 Salmonella enterica subsp. enterica serovar Typhi Proteobacteria; Gammaproteobacteria; Enterobacterales; Enterobacteriaceae; Salmonella; Salmonella enterica; Salmonella enterica subsp. enterica MTGDNTLITSHGINRRDFMKLCAALAATMGLSSKAAAEMAESVSNPQRPPVIWIGAQECT WP_000145429.1 hydrogenase 2 small subunit [Salmonella enterica] Length: 372\nNumber of predicted TMHs: 1\nExp number of AAs in TMHs: 24.51309\nExp number, first 60 AAs: 0.2682\nTotal prob of N-in: 0.14402\noutside 1 331\nTMhelix 332 354\ninside 355 372

21534 GCF_001082885.1_10060_6_77 Salmonella enterica subsp. enterica serovar Typhi Proteobacteria; Gammaproteobacteria; Enterobacterales; Enterobacteriaceae; Salmonella; Salmonella enterica; Salmonella enterica subsp. enterica MTGDNTLITSHGINRRDFMKLCAALAATMGLSSKAAAEMAESVSNPQRPPVIWIGAQECT WP_000145429.1 hydrogenase 2 small subunit [Salmonella enterica] Length: 372\nNumber of predicted TMHs: 1\nExp number of AAs in TMHs: 24.51309\nExp number, first 60 AAs: 0.2682\nTotal prob of N-in: 0.14402\noutside 1 331\nTMhelix 332 354\ninside 355 372

21535 GCF_001082925.1_10060_5_76 Salmonella enterica subsp. enterica serovar Typhi Proteobacteria; Gammaproteobacteria; Enterobacterales; Enterobacteriaceae; Salmonella; Salmonella enterica; Salmonella enterica subsp. enterica MTGDNTLITSHGINRRDFMKLCAALAATMGLSSKAAAEMAESVSNPQRPPVIWIGAQECT WP_000145429.1 hydrogenase 2 small subunit [Salmonella enterica] Length: 372\nNumber of predicted TMHs: 1\nExp number of AAs in TMHs: 24.51309\nExp number, first 60 AAs: 0.2682\nTotal prob of N-in: 0.14402\noutside 1 331\nTMhelix 332 354\ninside 355 372

21536 GCF_001082965.1_10060_5_46 Salmonella enterica subsp. enterica serovar Typhi Proteobacteria; Gammaproteobacteria; Enterobacterales; Enterobacteriaceae; Salmonella; Salmonella enterica; Salmonella enterica subsp. enterica MTGDNTLITSHGINRRDFMKLCAALAATMGLSSKAAAEMAESVSNPQRPPVIWIGAQECT WP_000145429.1 hydrogenase 2 small subunit [Salmonella enterica] Length: 372\nNumber of predicted TMHs: 1\nExp number of AAs in TMHs: 24.51309\nExp number, first 60 AAs: 0.2682\nTotal prob of N-in: 0.14402\noutside 1 331\nTMhelix 332 354\ninside 355 372

21537 GCF_001085745.1_9475_6_28 Salmonella enterica subsp. enterica serovar Typhi Proteobacteria; Gammaproteobacteria; Enterobacterales; Enterobacteriaceae; Salmonella; Salmonella enterica; Salmonella enterica subsp. enterica MTGDNTLITSHGINRRDFMKLCAALAATMGLSSKAAAEMAESVSNPQRPPVIWIGAQECT WP_000145429.1 hydrogenase 2 small subunit [Salmonella enterica] Length: 372\nNumber of predicted TMHs: 1\nExp number of AAs in TMHs: 24.51309\nExp number, first 60 AAs: 0.2682\nTotal prob of N-in: 0.14402\noutside 1 331\nTMhelix 332 354\ninside 355 372

21538 GCF_001085925.1_9953_5_12 Salmonella enterica subsp. enterica serovar Typhi Proteobacteria; Gammaproteobacteria; Enterobacterales; Enterobacteriaceae; Salmonella; Salmonella enterica; Salmonella enterica subsp. enterica MTGDNTLITSHGINRRDFMKLCAALAATMGLSSKAAAEMAESVSNPQRPPVIWIGAQECT WP_000145429.1 hydrogenase 2 small subunit [Salmonella enterica] Length: 372\nNumber of predicted TMHs: 1\nExp number of AAs in TMHs: 24.51309\nExp number, first 60 AAs: 0.2682\nTotal prob of N-in: 0.14402\noutside 1 331\nTMhelix 332 354\ninside 355 372

21539 GCF_001086325.1_10349_1_91 Salmonella enterica subsp. enterica serovar Typhi Proteobacteria; Gammaproteobacteria; Enterobacterales; Enterobacteriaceae; Salmonella; Salmonella enterica; Salmonella enterica subsp. enterica MTGDNTLITSHGINRRDFMKLCAALAATMGLSSKAAAEMAESVSNPQRPPVIWIGAQECT WP_000145429.1 hydrogenase 2 small subunit [Salmonella enterica] Length: 372\nNumber of predicted TMHs: 1\nExp number of AAs in TMHs: 24.51309\nExp number, first 60 AAs: 0.2682\nTotal prob of N-in: 0.14402\noutside 1 331\nTMhelix 332 354\ninside 355 372

21540 GCF_001086485.1_10561_2_62 Salmonella enterica subsp. enterica serovar Typhi Proteobacteria; Gammaproteobacteria; Enterobacterales; Enterobacteriaceae; Salmonella; Salmonella enterica; Salmonella enterica subsp. enterica MTGDNTLITSHGINRRDFMKLCAALAATMGLSSKAAAEMAESVSNPQRPPVIWIGAQECT WP_000145429.1 hydrogenase 2 small subunit [Salmonella enterica] Length: 372\nNumber of predicted TMHs: 1\nExp number of AAs in TMHs: 24.51309\nExp number, first 60 AAs: 0.2682\nTotal prob of N-in: 0.14402\noutside 1 331\nTMhelix 332 354\ninside 355 372

21541 GCF_001087005.1_10209_5_11 Salmonella enterica subsp. enterica serovar Typhi Proteobacteria; Gammaproteobacteria; Enterobacterales; Enterobacteriaceae; Salmonella; Salmonella enterica; Salmonella enterica subsp. enterica MTGDNTLITSHGINRRDFMKLCAALAATMGLSSKAAAEMAESVSNPQRPPVIWIGAQECT WP_000145429.1 hydrogenase 2 small subunit [Salmonella enterica] Length: 372\nNumber of predicted TMHs: 1\nExp number of AAs in TMHs: 24.51309\nExp number, first 60 AAs: 0.2682\nTotal prob of N-in: 0.14402\noutside 1 331\nTMhelix 332 354\ninside 355 372

21542 GCF_001087105.1_10349_1_65 Salmonella enterica subsp. enterica serovar Typhi Proteobacteria; Gammaproteobacteria; Enterobacterales; Enterobacteriaceae; Salmonella; Salmonella enterica; Salmonella enterica subsp. enterica MTGDNTLITSHGINRRDFMKLCAALAATMGLSSKAAAEMAESVSNPQRPPVIWIGAQECT WP_000145429.1 hydrogenase 2 small subunit [Salmonella enterica] Length: 372\nNumber of

predicted TMHs:  1\nExp number of AAs in TMHs: 24.51309\nExp number, first 60 AAs:  0.2682\nTotal prob of N-in: 0.14402\noutside   1   331\nTMhelix   332   354\ninside   355   372

21543 GCF_001087345.1_7468_7_68 Salmonella enterica subsp. enterica serovar Typhi Proteobacteria; Gammaproteobacteria; Enterobacterales; Enterobacteriaceae; Salmonella; Salmonella enterica; Salmonella enterica subsp. enterica MTGDNTLITSHGINRRDFMKLCAALAATMGLSSKAAAEMAESVSNPQRPPVIWIGAQECT WP_000145429.1 hydrogenase 2 small subunit [Salmonella enterica] Length: 372\nNumber of predicted TMHs:  1\nExp number of AAs in TMHs: 24.51309\nExp number, first 60 AAs:  0.2682\nTotal prob of N-in: 0.14402\noutside   1   331\nTMhelix   332   354\ninside   355   372

21544 GCF_001090905.1_10493_1_12 Salmonella enterica subsp. enterica serovar Typhi Proteobacteria; Gammaproteobacteria; Enterobacterales; Enterobacteriaceae; Salmonella; Salmonella enterica; Salmonella enterica subsp. enterica MTGDNTLITSHGINRRDFMKLCAALAATMGLSSKAAAEMAESVSNPQRPPVIWIGAQECT WP_000145429.1 hydrogenase 2 small subunit [Salmonella enterica] Length: 372\nNumber of predicted TMHs:  1\nExp number of AAs in TMHs: 24.51309\nExp number, first 60 AAs:  0.2682\nTotal prob of N-in: 0.14402\noutside   1   331\nTMhelix   332   354\ninside   355   372

21545 GCF_001090505.1_10607_2_32 Salmonella enterica subsp. enterica serovar Typhi Proteobacteria; Gammaproteobacteria; Enterobacterales; Enterobacteriaceae; Salmonella; Salmonella enterica; Salmonella enterica subsp. enterica MTGDNTLITSHGINRRDFMKLCAALAATMGLSSKAAAEMAESVSNPQRPPVIWIGAQECT WP_000145429.1 hydrogenase 2 small subunit [Salmonella enterica] Length: 372\nNumber of predicted TMHs:  1\nExp number of AAs in TMHs: 24.51309\nExp number, first 60 AAs:  0.2682\nTotal prob of N-in: 0.14402\noutside   1   331\nTMhelix   332   354\ninside   355   372

21546 GCF_001090385.1_9870_8_79 Salmonella enterica subsp. enterica serovar Typhi Proteobacteria; Gammaproteobacteria; Enterobacterales; Enterobacteriaceae; Salmonella; Salmonella enterica; Salmonella enterica subsp. enterica MTGDNTLITSHGINRRDFMKLCAALAATMGLSSKAAAEMAESVSNPQRPPVIWIGAQECT WP_000145429.1 hydrogenase 2 small subunit [Salmonella enterica] Length: 372\nNumber of predicted TMHs:  1\nExp number of AAs in TMHs: 24.51309\nExp number, first 60 AAs:  0.2682\nTotal prob of N-in: 0.14402\noutside   1   331\nTMhelix   332   354\ninside   355   372

21547 GCF_001090965.1_9870_8_51 Salmonella enterica subsp. enterica serovar Typhi Proteobacteria; Gammaproteobacteria; Enterobacterales; Enterobacteriaceae; Salmonella; Salmonella enterica; Salmonella enterica subsp. enterica MTGDNTLITSHGINRRDFMKLCAALAATMGLSSKAAAEMAESVSNPQRPPVIWIGAQECT WP_000145429.1 hydrogenase 2 small subunit [Salmonella enterica] Length: 372\nNumber of predicted TMHs:  1\nExp number of AAs in TMHs: 24.51309\nExp number, first 60 AAs:  0.2682\nTotal prob of N-in: 0.14402\noutside   1   331\nTMhelix   332   354\ninside   355   372

21548 GCF_001091245.1_8616_4_49 Salmonella enterica subsp. enterica serovar Typhi Proteobacteria; Gammaproteobacteria; Enterobacterales; Enterobacteriaceae; Salmonella; Salmonella enterica; Salmonella enterica subsp. enterica MTGDNTLITSHGINRRDFMKLCAALAATMGLSSKAAAEMAESVSNPQRPPVIWIGAQECT WP_000145429.1 hydrogenase 2 small subunit [Salmonella enterica] Length: 372\nNumber of predicted TMHs:  1\nExp number of AAs in TMHs: 24.51309\nExp number, first 60 AAs:  0.2682\nTotal prob of N-in: 0.14402\noutside   1   331\nTMhelix   332   354\ninside   355   372

21549 GCF_001091425.1_9870_8_52 Salmonella enterica subsp. enterica serovar Typhi Proteobacteria; Gammaproteobacteria; Enterobacterales; Enterobacteriaceae; Salmonella; Salmonella enterica; Salmonella enterica subsp. enterica MTGDNTLITSHGINRRDFMKLCAALAATMGLSSKAAAEMAESVSNPQRPPVIWIGAQECT WP_000145429.1 hydrogenase 2 small subunit [Salmonella enterica] Length: 372\nNumber of predicted TMHs:  1\nExp number of AAs in TMHs: 24.51309\nExp number, first 60 AAs:  0.2682\nTotal prob of N-in: 0.14402\noutside   1   331\nTMhelix   332   354\ninside   355   372

21550 GCF_001091525.1_8490_5_25 Salmonella enterica subsp. enterica serovar Typhi Proteobacteria; Gammaproteobacteria; Enterobacterales; Enterobacteriaceae; Salmonella; Salmonella enterica; Salmonella enterica subsp. enterica MTGDNTLITSHGINRRDFMKLCAALAATMGLSSKAAAEMAESVSNPQRPPVIWIGAQECT WP_000145429.1 hydrogenase 2 small subunit [Salmonella enterica] Length: 372\nNumber of predicted TMHs:  1\nExp number of AAs in TMHs: 24.51309\nExp number, first 60 AAs:  0.2682\nTotal prob of N-in: 0.14402\noutside   1   331\nTMhelix   332   354\ninside   355   372

21551 GCF_001094425.1_10607_1_60 Salmonella enterica subsp. enterica serovar Typhi Proteobacteria; Gammaproteobacteria; Enterobacterales; Enterobacteriaceae; Salmonella; Salmonella enterica;

Salmonella enterica subsp. enterica MTGDNTLITSHGINRRDFMKLCAALAATMGLSSKAAAEMAESVSNPQRPPVIWIGAQECT WP_000145429.1 hydrogenase 2 small subunit [Salmonella enterica] Length: 372\nNumber of predicted TMHs: 1\nExp number of AAs in TMHs: 24.51309\nExp number, first 60 AAs: 0.2682\nTotal prob of N-in: 0.14402\noutside 1 331\nTMhelix 332 354\ninside 355 372

21552 GCF_001094525.1_9475_6_44 Salmonella enterica subsp. enterica serovar Typhi Proteobacteria; Gammaproteobacteria; Enterobacterales; Enterobacteriaceae; Salmonella; Salmonella enterica; Salmonella enterica subsp. enterica MTGDNTLITSHGINRRDFMKLCAALAATMGLSSKAAAEMAESVSNPQRPPVIWIGAQECT WP_000145429.1 hydrogenase 2 small subunit [Salmonella enterica] Length: 372\nNumber of predicted TMHs: 1\nExp number of AAs in TMHs: 24.51309\nExp number, first 60 AAs: 0.2682\nTotal prob of N-in: 0.14402\noutside 1 331\nTMhelix 332 354\ninside 355 372

21553 GCF_001094645.1_8616_4_47 Salmonella enterica subsp. enterica serovar Typhi Proteobacteria; Gammaproteobacteria; Enterobacterales; Enterobacteriaceae; Salmonella; Salmonella enterica; Salmonella enterica subsp. enterica MTGDNTLITSHGINRRDFMKLCAALAATMGLSSKAAAEMAESVSNPQRPPVIWIGAQECT WP_000145429.1 hydrogenase 2 small subunit [Salmonella enterica] Length: 372\nNumber of predicted TMHs: 1\nExp number of AAs in TMHs: 24.51309\nExp number, first 60 AAs: 0.2682\nTotal prob of N-in: 0.14402\noutside 1 331\nTMhelix 332 354\ninside 355 372

21554 GCF_001094765.1_9870_8_28 Salmonella enterica subsp. enterica serovar Typhi Proteobacteria; Gammaproteobacteria; Enterobacterales; Enterobacteriaceae; Salmonella; Salmonella enterica; Salmonella enterica subsp. enterica MTGDNTLITSHGINRRDFMKLCAALAATMGLSSKAAAEMAESVSNPQRPPVIWIGAQECT WP_000145429.1 hydrogenase 2 small subunit [Salmonella enterica] Length: 372\nNumber of predicted TMHs: 1\nExp number of AAs in TMHs: 24.51309\nExp number, first 60 AAs: 0.2682\nTotal prob of N-in: 0.14402\noutside 1 331\nTMhelix 332 354\ninside 355 372

21555 GCF_001095125.1_9475_6_93 Salmonella enterica subsp. enterica serovar Typhi Proteobacteria; Gammaproteobacteria; Enterobacterales; Enterobacteriaceae; Salmonella; Salmonella enterica; Salmonella enterica subsp. enterica MTGDNTLITSHGINRRDFMKLCAALAATMGLSSKAAAEMAESVSNPQRPPVIWIGAQECT WP_000145429.1 hydrogenase 2 small subunit [Salmonella enterica] Length: 372\nNumber of predicted TMHs: 1\nExp number of AAs in TMHs: 24.51309\nExp number, first 60 AAs: 0.2682\nTotal prob of N-in: 0.14402\noutside 1 331\nTMhelix 332 354\ninside 355 372

21556 GCF_001095305.1_10592_2_31 Salmonella enterica subsp. enterica serovar Typhi Proteobacteria; Gammaproteobacteria; Enterobacterales; Enterobacteriaceae; Salmonella; Salmonella enterica; Salmonella enterica subsp. enterica MTGDNTLITSHGINRRDFMKLCAALAATMGLSSKAAAEMAESVSNPQRPPVIWIGAQECT WP_000145429.1 hydrogenase 2 small subunit [Salmonella enterica] Length: 372\nNumber of predicted TMHs: 1\nExp number of AAs in TMHs: 24.51309\nExp number, first 60 AAs: 0.2682\nTotal prob of N-in: 0.14402\noutside 1 331\nTMhelix 332 354\ninside 355 372

21557 GCF_001102745.1_10607_1_95 Salmonella enterica subsp. enterica serovar Typhi Proteobacteria; Gammaproteobacteria; Enterobacterales; Enterobacteriaceae; Salmonella; Salmonella enterica; Salmonella enterica subsp. enterica MTGDNTLITSHGINRRDFMKLCAALAATMGLSSKAAAEMAESVSNPQRPPVIWIGAQECT WP_000145429.1 hydrogenase 2 small subunit [Salmonella enterica] Length: 372\nNumber of predicted TMHs: 1\nExp number of AAs in TMHs: 24.51309\nExp number, first 60 AAs: 0.2682\nTotal prob of N-in: 0.14402\noutside 1 331\nTMhelix 332 354\ninside 355 372

21558 GCF_001098065.1_9870_8_57 Salmonella enterica subsp. enterica serovar Typhi Proteobacteria; Gammaproteobacteria; Enterobacterales; Enterobacteriaceae; Salmonella; Salmonella enterica; Salmonella enterica subsp. enterica MTGDNTLITSHGINRRDFMKLCAALAATMGLSSKAAAEMAESVSNPQRPPVIWIGAQECT WP_000145429.1 hydrogenase 2 small subunit [Salmonella enterica] Length: 372\nNumber of predicted TMHs: 1\nExp number of AAs in TMHs: 24.51309\nExp number, first 60 AAs: 0.2682\nTotal prob of N-in: 0.14402\noutside 1 331\nTMhelix 332 354\ninside 355 372

21559 GCF_001098485.1_10541_2_7 Salmonella enterica subsp. enterica serovar Typhi Proteobacteria; Gammaproteobacteria; Enterobacterales; Enterobacteriaceae; Salmonella; Salmonella enterica; Salmonella enterica subsp. enterica MTGDNTLITSHGINRRDFMKLCAALAATMGLSSKAAAEMAESVSNPQRPPVIWIGAQECT WP_000145429.1 hydrogenase 2 small subunit [Salmonella enterica] Length: 372\nNumber of predicted TMHs: 1\nExp number of AAs in TMHs: 24.51309\nExp number, first 60 AAs: 0.2682\nTotal prob of N-in: 0.14402\noutside 1 331\nTMhelix 332 354\ninside 355 372

21560 GCF_001098665.1_10593_2_38 Salmonella enterica subsp. enterica serovar Typhi Proteobacteria; Gammaproteobacteria; Enterobacterales; Enterobacteriaceae; Salmonella; Salmonella enterica; Salmonella enterica subsp. enterica MTGDNTLITSHGINRRDFMKLCAALAATMGLSSKAAAEMAESVSNPQRPPVIWIGAQECT WP_000145429.1 hydrogenase 2 small subunit [Salmonella enterica] Length: 372\nNumber of predicted TMHs: 1\nExp number of AAs in TMHs: 24.51309\nExp number, first 60 AAs: 0.2682\nTotal prob of N-in: 0.14402\noutside 1 331\nTMhelix 332 354\ninside 355 372

21561 GCF_001098945.1_9475_6_77 Salmonella enterica subsp. enterica serovar Typhi Proteobacteria; Gammaproteobacteria; Enterobacterales; Enterobacteriaceae; Salmonella; Salmonella enterica; Salmonella enterica subsp. enterica MTGDNTLITSHGINRRDFMKLCAALAATMGLSSKAAAEMAESVSNPQRPPVIWIGAQECT WP_000145429.1 hydrogenase 2 small subunit [Salmonella enterica] Length: 372\nNumber of predicted TMHs: 1\nExp number of AAs in TMHs: 24.51309\nExp number, first 60 AAs: 0.2682\nTotal prob of N-in: 0.14402\noutside 1 331\nTMhelix 332 354\ninside 355 372

21562 GCF_001365875.1_10492_1_75 Salmonella enterica subsp. enterica serovar Typhi Proteobacteria; Gammaproteobacteria; Enterobacterales; Enterobacteriaceae; Salmonella; Salmonella enterica; Salmonella enterica subsp. enterica MTGDNTLITSHGINRRDFMKLCAALAATMGLSSKAAAEMAESVSNPQRPPVIWIGAQECT WP_000145429.1 hydrogenase 2 small subunit [Salmonella enterica] Length: 372\nNumber of predicted TMHs: 1\nExp number of AAs in TMHs: 24.51309\nExp number, first 60 AAs: 0.2682\nTotal prob of N-in: 0.14402\noutside 1 331\nTMhelix 332 354\ninside 355 372

21563 GCF_001365915.1_10426_1_36 Salmonella enterica subsp. enterica serovar Typhi Proteobacteria; Gammaproteobacteria; Enterobacterales; Enterobacteriaceae; Salmonella; Salmonella enterica; Salmonella enterica subsp. enterica MTGDNTLITSHGINRRDFMKLCAALAATMGLSSKAAAEMAESVSNPQRPPVIWIGAQECT WP_000145429.1 hydrogenase 2 small subunit [Salmonella enterica] Length: 372\nNumber of predicted TMHs: 1\nExp number of AAs in TMHs: 24.51309\nExp number, first 60 AAs: 0.2682\nTotal prob of N-in: 0.14402\noutside 1 331\nTMhelix 332 354\ninside 355 372

21564 GCF_001365835.1_10493_1_26 Salmonella enterica subsp. enterica serovar Typhi Proteobacteria; Gammaproteobacteria; Enterobacterales; Enterobacteriaceae; Salmonella; Salmonella enterica; Salmonella enterica subsp. enterica MTGDNTLITSHGINRRDFMKLCAALAATMGLSSKAAAEMAESVSNPQRPPVIWIGAQECT WP_000145429.1 hydrogenase 2 small subunit [Salmonella enterica] Length: 372\nNumber of predicted TMHs: 1\nExp number of AAs in TMHs: 24.51309\nExp number, first 60 AAs: 0.2682\nTotal prob of N-in: 0.14402\noutside 1 331\nTMhelix 332 354\ninside 355 372

21565 GCF_001365955.1_10561_2_79 Salmonella enterica subsp. enterica serovar Typhi Proteobacteria; Gammaproteobacteria; Enterobacterales; Enterobacteriaceae; Salmonella; Salmonella enterica; Salmonella enterica subsp. enterica MTGDNTLITSHGINRRDFMKLCAALAATMGLSSKAAAEMAESVSNPQRPPVIWIGAQECT WP_000145429.1 hydrogenase 2 small subunit [Salmonella enterica] Length: 372\nNumber of predicted TMHs: 1\nExp number of AAs in TMHs: 24.51309\nExp number, first 60 AAs: 0.2682\nTotal prob of N-in: 0.14402\noutside 1 331\nTMhelix 332 354\ninside 355 372

21566 GCF_001365795.1_10561_2_74 Salmonella enterica subsp. enterica serovar Typhi Proteobacteria; Gammaproteobacteria; Enterobacterales; Enterobacteriaceae; Salmonella; Salmonella enterica; Salmonella enterica subsp. enterica MTGDNTLITSHGINRRDFMKLCAALAATMGLSSKAAAEMAESVSNPQRPPVIWIGAQECT WP_000145429.1 hydrogenase 2 small subunit [Salmonella enterica] Length: 372\nNumber of predicted TMHs: 1\nExp number of AAs in TMHs: 24.51309\nExp number, first 60 AAs: 0.2682\nTotal prob of N-in: 0.14402\noutside 1 331\nTMhelix 332 354\ninside 355 372

21567 GCF_001365755.1_10562_2_80 Salmonella enterica subsp. enterica serovar Typhi Proteobacteria; Gammaproteobacteria; Enterobacterales; Enterobacteriaceae; Salmonella; Salmonella enterica; Salmonella enterica subsp. enterica MTGDNTLITSHGINRRDFMKLCAALAATMGLSSKAAAEMAESVSNPQRPPVIWIGAQECT WP_000145429.1 hydrogenase 2 small subunit [Salmonella enterica] Length: 372\nNumber of predicted TMHs: 1\nExp number of AAs in TMHs: 24.51309\nExp number, first 60 AAs: 0.2682\nTotal prob of N-in: 0.14402\noutside 1 331\nTMhelix 332 354\ninside 355 372

21568 GCF_001365215.1_10607_2_12 Salmonella enterica subsp. enterica serovar Typhi Proteobacteria; Gammaproteobacteria; Enterobacterales; Enterobacteriaceae; Salmonella; Salmonella enterica; Salmonella enterica subsp. enterica MTGDNTLITSHGINRRDFMKLCAALAATMGLSSKAAAEMAESVSNPQRPPVIWIGAQECT WP_000145429.1 hydrogenase 2 small subunit [Salmonella enterica] Length: 372\nNumber of

predicted TMHs: 1\nExp number of AAs in TMHs: 24.51309\nExp number, first 60 AAs: 0.2682\nTotal prob of N-in: 0.14402\noutside 1 331\nTMhelix 332 354\ninside 355 372

21569 GCF_001365175.1_10561_2_67 Salmonella enterica subsp. enterica serovar Typhi Proteobacteria; Gammaproteobacteria; Enterobacterales; Enterobacteriaceae; Salmonella; Salmonella enterica; Salmonella enterica subsp. enterica MTGDNTLITSHGINRRDFMKLCAALAATMGLSSKAAAEMAESVSNPQRPPVIWIGAQECT WP_000145429.1 hydrogenase 2 small subunit [Salmonella enterica] Length: 372\nNumber of predicted TMHs: 1\nExp number of AAs in TMHs: 24.51309\nExp number, first 60 AAs: 0.2682\nTotal prob of N-in: 0.14402\noutside 1 331\nTMhelix 332 354\ninside 355 372

21570 GCF_001365135.1_10540_1_36 Salmonella enterica subsp. enterica serovar Typhi Proteobacteria; Gammaproteobacteria; Enterobacterales; Enterobacteriaceae; Salmonella; Salmonella enterica; Salmonella enterica subsp. enterica MTGDNTLITSHGINRRDFMKLCAALAATMGLSSKAAAEMAESVSNPQRPPVIWIGAQECT WP_000145429.1 hydrogenase 2 small subunit [Salmonella enterica] Length: 372\nNumber of predicted TMHs: 1\nExp number of AAs in TMHs: 24.51309\nExp number, first 60 AAs: 0.2682\nTotal prob of N-in: 0.14402\noutside 1 331\nTMhelix 332 354\ninside 355 372

21571 GCF_001365095.1_10561_2_78 Salmonella enterica subsp. enterica serovar Typhi Proteobacteria; Gammaproteobacteria; Enterobacterales; Enterobacteriaceae; Salmonella; Salmonella enterica; Salmonella enterica subsp. enterica MTGDNTLITSHGINRRDFMKLCAALAATMGLSSKAAAEMAESVSNPQRPPVIWIGAQECT WP_000145429.1 hydrogenase 2 small subunit [Salmonella enterica] Length: 372\nNumber of predicted TMHs: 1\nExp number of AAs in TMHs: 24.51309\nExp number, first 60 AAs: 0.2682\nTotal prob of N-in: 0.14402\noutside 1 331\nTMhelix 332 354\ninside 355 372

21572 GCF_001365055.1_10493_1_65 Salmonella enterica subsp. enterica serovar Typhi Proteobacteria; Gammaproteobacteria; Enterobacterales; Enterobacteriaceae; Salmonella; Salmonella enterica; Salmonella enterica subsp. enterica MTGDNTLITSHGINRRDFMKLCAALAATMGLSSKAAAEMAESVSNPQRPPVIWIGAQECT WP_000145429.1 hydrogenase 2 small subunit [Salmonella enterica] Length: 372\nNumber of predicted TMHs: 1\nExp number of AAs in TMHs: 24.51309\nExp number, first 60 AAs: 0.2682\nTotal prob of N-in: 0.14402\noutside 1 331\nTMhelix 332 354\ninside 355 372

21573 GCF_001365015.1_10426_1_47 Salmonella enterica subsp. enterica serovar Typhi Proteobacteria; Gammaproteobacteria; Enterobacterales; Enterobacteriaceae; Salmonella; Salmonella enterica; Salmonella enterica subsp. enterica MTGDNTLITSHGINRRDFMKLCAALAATMGLSSKAAAEMAESVSNPQRPPVIWIGAQECT WP_000145429.1 hydrogenase 2 small subunit [Salmonella enterica] Length: 372\nNumber of predicted TMHs: 1\nExp number of AAs in TMHs: 24.51309\nExp number, first 60 AAs: 0.2682\nTotal prob of N-in: 0.14402\noutside 1 331\nTMhelix 332 354\ninside 355 372

21574 GCF_001364215.1_10425_1_24 Salmonella enterica subsp. enterica serovar Typhi Proteobacteria; Gammaproteobacteria; Enterobacterales; Enterobacteriaceae; Salmonella; Salmonella enterica; Salmonella enterica subsp. enterica MTGDNTLITSHGINRRDFMKLCAALAATMGLSSKAAAEMAESVSNPQRPPVIWIGAQECT WP_000145429.1 hydrogenase 2 small subunit [Salmonella enterica] Length: 372\nNumber of predicted TMHs: 1\nExp number of AAs in TMHs: 24.51309\nExp number, first 60 AAs: 0.2682\nTotal prob of N-in: 0.14402\noutside 1 331\nTMhelix 332 354\ninside 355 372

21575 GCF_001364175.1_10540_1_50 Salmonella enterica subsp. enterica serovar Typhi Proteobacteria; Gammaproteobacteria; Enterobacterales; Enterobacteriaceae; Salmonella; Salmonella enterica; Salmonella enterica subsp. enterica MTGDNTLITSHGINRRDFMKLCAALAATMGLSSKAAAEMAESVSNPQRPPVIWIGAQECT WP_000145429.1 hydrogenase 2 small subunit [Salmonella enterica] Length: 372\nNumber of predicted TMHs: 1\nExp number of AAs in TMHs: 24.51309\nExp number, first 60 AAs: 0.2682\nTotal prob of N-in: 0.14402\noutside 1 331\nTMhelix 332 354\ninside 355 372

21576 GCF_001364135.1_10426_1_26 Salmonella enterica subsp. enterica serovar Typhi Proteobacteria; Gammaproteobacteria; Enterobacterales; Enterobacteriaceae; Salmonella; Salmonella enterica; Salmonella enterica subsp. enterica MTGDNTLITSHGINRRDFMKLCAALAATMGLSSKAAAEMAESVSNPQRPPVIWIGAQECT WP_000145429.1 hydrogenase 2 small subunit [Salmonella enterica] Length: 372\nNumber of predicted TMHs: 1\nExp number of AAs in TMHs: 24.51309\nExp number, first 60 AAs: 0.2682\nTotal prob of N-in: 0.14402\noutside 1 331\nTMhelix 332 354\ninside 355 372

21577 GCF_001364095.1_10540_1_23 Salmonella enterica subsp. enterica serovar Typhi Proteobacteria; Gammaproteobacteria; Enterobacterales; Enterobacteriaceae; Salmonella; Salmonella enterica;

Salmonella enterica subsp. enterica MTGDNTLITSHGINRRDFMKLCAALAATMGLSSKAAAEMAESVSNPQRPPVIWIGAQECT WP_000145429.1 hydrogenase 2 small subunit [Salmonella enterica] Length: 372\nNumber of predicted TMHs: 1\nExp number of AAs in TMHs: 24.51309\nExp number, first 60 AAs: 0.2682\nTotal prob of N-in: 0.14402\noutside 1 331\nTMhelix 332 354\ninside 355 372

21578 GCF_001364055.1_10540_1_83 Salmonella enterica subsp. enterica serovar Typhi Proteobacteria; Gammaproteobacteria; Enterobacterales; Enterobacteriaceae; Salmonella; Salmonella enterica; Salmonella enterica subsp. enterica MTGDNTLITSHGINRRDFMKLCAALAATMGLSSKAAAEMAESVSNPQRPPVIWIGAQECT WP_000145429.1 hydrogenase 2 small subunit [Salmonella enterica] Length: 372\nNumber of predicted TMHs: 1\nExp number of AAs in TMHs: 24.51309\nExp number, first 60 AAs: 0.2682\nTotal prob of N-in: 0.14402\noutside 1 331\nTMhelix 332 354\ninside 355 372

21579 GCF_001364015.1_10426_1_35 Salmonella enterica subsp. enterica serovar Typhi Proteobacteria; Gammaproteobacteria; Enterobacterales; Enterobacteriaceae; Salmonella; Salmonella enterica; Salmonella enterica subsp. enterica MTGDNTLITSHGINRRDFMKLCAALAATMGLSSKAAAEMAESVSNPQRPPVIWIGAQECT WP_000145429.1 hydrogenase 2 small subunit [Salmonella enterica] Length: 372\nNumber of predicted TMHs: 1\nExp number of AAs in TMHs: 24.51309\nExp number, first 60 AAs: 0.2682\nTotal prob of N-in: 0.14402\noutside 1 331\nTMhelix 332 354\ninside 355 372

21580 GCF_001363975.1_10540_1_16 Salmonella enterica subsp. enterica serovar Typhi Proteobacteria; Gammaproteobacteria; Enterobacterales; Enterobacteriaceae; Salmonella; Salmonella enterica; Salmonella enterica subsp. enterica MTGDNTLITSHGINRRDFMKLCAALAATMGLSSKAAAEMAESVSNPQRPPVIWIGAQECT WP_000145429.1 hydrogenase 2 small subunit [Salmonella enterica] Length: 372\nNumber of predicted TMHs: 1\nExp number of AAs in TMHs: 24.51309\nExp number, first 60 AAs: 0.2682\nTotal prob of N-in: 0.14402\noutside 1 331\nTMhelix 332 354\ninside 355 372

21581 GCF_001363435.1_10607_1_73 Salmonella enterica subsp. enterica serovar Typhi Proteobacteria; Gammaproteobacteria; Enterobacterales; Enterobacteriaceae; Salmonella; Salmonella enterica; Salmonella enterica subsp. enterica MTGDNTLITSHGINRRDFMKLCAALAATMGLSSKAAAEMAESVSNPQRPPVIWIGAQECT WP_000145429.1 hydrogenase 2 small subunit [Salmonella enterica] Length: 372\nNumber of predicted TMHs: 1\nExp number of AAs in TMHs: 24.51309\nExp number, first 60 AAs: 0.2682\nTotal prob of N-in: 0.14402\noutside 1 331\nTMhelix 332 354\ninside 355 372

21582 GCF_001363395.1_10541_2_3 Salmonella enterica subsp. enterica serovar Typhi Proteobacteria; Gammaproteobacteria; Enterobacterales; Enterobacteriaceae; Salmonella; Salmonella enterica; Salmonella enterica subsp. enterica MTGDNTLITSHGINRRDFMKLCAALAATMGLSSKAAAEMAESVSNPQRPPVIWIGAQECT WP_000145429.1 hydrogenase 2 small subunit [Salmonella enterica] Length: 372\nNumber of predicted TMHs: 1\nExp number of AAs in TMHs: 24.51309\nExp number, first 60 AAs: 0.2682\nTotal prob of N-in: 0.14402\noutside 1 331\nTMhelix 332 354\ninside 355 372

21583 GCF_001363315.1_10426_1_43 Salmonella enterica subsp. enterica serovar Typhi Proteobacteria; Gammaproteobacteria; Enterobacterales; Enterobacteriaceae; Salmonella; Salmonella enterica; Salmonella enterica subsp. enterica MTGDNTLITSHGINRRDFMKLCAALAATMGLSSKAAAEMAESVSNPQRPPVIWIGAQECT WP_000145429.1 hydrogenase 2 small subunit [Salmonella enterica] Length: 372\nNumber of predicted TMHs: 1\nExp number of AAs in TMHs: 24.51309\nExp number, first 60 AAs: 0.2682\nTotal prob of N-in: 0.14402\noutside 1 331\nTMhelix 332 354\ninside 355 372

21584 GCF_001363275.1_10426_1_31 Salmonella enterica subsp. enterica serovar Typhi Proteobacteria; Gammaproteobacteria; Enterobacterales; Enterobacteriaceae; Salmonella; Salmonella enterica; Salmonella enterica subsp. enterica MTGDNTLITSHGINRRDFMKLCAALAATMGLSSKAAAEMAESVSNPQRPPVIWIGAQECT WP_000145429.1 hydrogenase 2 small subunit [Salmonella enterica] Length: 372\nNumber of predicted TMHs: 1\nExp number of AAs in TMHs: 24.51309\nExp number, first 60 AAs: 0.2682\nTotal prob of N-in: 0.14402\noutside 1 331\nTMhelix 332 354\ninside 355 372

21585 GCF_001363235.1_10561_2_3 Salmonella enterica subsp. enterica serovar Typhi Proteobacteria; Gammaproteobacteria; Enterobacterales; Enterobacteriaceae; Salmonella; Salmonella enterica; Salmonella enterica subsp. enterica MTGDNTLITSHGINRRDFMKLCAALAATMGLSSKAAAEMAESVSNPQRPPVIWIGAQECT WP_000145429.1 hydrogenase 2 small subunit [Salmonella enterica] Length: 372\nNumber of predicted TMHs: 1\nExp number of AAs in TMHs: 24.51309\nExp number, first 60 AAs: 0.2682\nTotal prob of N-in: 0.14402\noutside 1 331\nTMhelix 332 354\ninside 355 372

21586 GCF_001363195.1_10426_1_59 Salmonella enterica subsp. enterica serovar Typhi Proteobacteria; Gammaproteobacteria; Enterobacterales; Enterobacteriaceae; Salmonella; Salmonella enterica; Salmonella enterica subsp. enterica MTGDNTLITSHGINRRDFMKLCAALAATMGLSSKAAAEMAESVSNPQRPPVIWIGAQECT WP_000145429.1 hydrogenase 2 small subunit [Salmonella enterica] Length: 372\nNumber of predicted TMHs: 1\nExp number of AAs in TMHs: 24.51309\nExp number, first 60 AAs: 0.2682\nTotal prob of N-in: 0.14402\noutside 1 331\nTMhelix 332 354\ninside 355 372

21587 GCF_001362655.1_10607_1_77 Salmonella enterica subsp. enterica serovar Typhi Proteobacteria; Gammaproteobacteria; Enterobacterales; Enterobacteriaceae; Salmonella; Salmonella enterica; Salmonella enterica subsp. enterica MTGDNTLITSHGINRRDFMKLCAALAATMGLSSKAAAEMAESVSNPQRPPVIWIGAQECT WP_000145429.1 hydrogenase 2 small subunit [Salmonella enterica] Length: 372\nNumber of predicted TMHs: 1\nExp number of AAs in TMHs: 24.51309\nExp number, first 60 AAs: 0.2682\nTotal prob of N-in: 0.14402\noutside 1 331\nTMhelix 332 354\ninside 355 372

21588 GCF_001362575.1_10607_2_5 Salmonella enterica subsp. enterica serovar Typhi Proteobacteria; Gammaproteobacteria; Enterobacterales; Enterobacteriaceae; Salmonella; Salmonella enterica; Salmonella enterica subsp. enterica MTGDNTLITSHGINRRDFMKLCAALAATMGLSSKAAAEMAESVSNPQRPPVIWIGAQECT WP_000145429.1 hydrogenase 2 small subunit [Salmonella enterica] Length: 372\nNumber of predicted TMHs: 1\nExp number of AAs in TMHs: 24.51309\nExp number, first 60 AAs: 0.2682\nTotal prob of N-in: 0.14402\noutside 1 331\nTMhelix 332 354\ninside 355 372

21589 GCF_001362615.1_10562_2_67 Salmonella enterica subsp. enterica serovar Typhi Proteobacteria; Gammaproteobacteria; Enterobacterales; Enterobacteriaceae; Salmonella; Salmonella enterica; Salmonella enterica subsp. enterica MTGDNTLITSHGINRRDFMKLCAALAATMGLSSKAAAEMAESVSNPQRPPVIWIGAQECT WP_000145429.1 hydrogenase 2 small subunit [Salmonella enterica] Length: 372\nNumber of predicted TMHs: 1\nExp number of AAs in TMHs: 24.51309\nExp number, first 60 AAs: 0.2682\nTotal prob of N-in: 0.14402\noutside 1 331\nTMhelix 332 354\ninside 355 372

21590 GCF_001362535.1_10540_1_10 Salmonella enterica subsp. enterica serovar Typhi Proteobacteria; Gammaproteobacteria; Enterobacterales; Enterobacteriaceae; Salmonella; Salmonella enterica; Salmonella enterica subsp. enterica MTGDNTLITSHGINRRDFMKLCAALAATMGLSSKAAAEMAESVSNPQRPPVIWIGAQECT WP_000145429.1 hydrogenase 2 small subunit [Salmonella enterica] Length: 372\nNumber of predicted TMHs: 1\nExp number of AAs in TMHs: 24.51309\nExp number, first 60 AAs: 0.2682\nTotal prob of N-in: 0.14402\noutside 1 331\nTMhelix 332 354\ninside 355 372

21591 GCF_001362495.1_10540_1_64 Salmonella enterica subsp. enterica serovar Typhi Proteobacteria; Gammaproteobacteria; Enterobacterales; Enterobacteriaceae; Salmonella; Salmonella enterica; Salmonella enterica subsp. enterica MTGDNTLITSHGINRRDFMKLCAALAATMGLSSKAAAEMAESVSNPQRPPVIWIGAQECT WP_000145429.1 hydrogenase 2 small subunit [Salmonella enterica] Length: 372\nNumber of predicted TMHs: 1\nExp number of AAs in TMHs: 24.51309\nExp number, first 60 AAs: 0.2682\nTotal prob of N-in: 0.14402\noutside 1 331\nTMhelix 332 354\ninside 355 372

21592 GCF_001362455.1_10561_2_73 Salmonella enterica subsp. enterica serovar Typhi Proteobacteria; Gammaproteobacteria; Enterobacterales; Enterobacteriaceae; Salmonella; Salmonella enterica; Salmonella enterica subsp. enterica MTGDNTLITSHGINRRDFMKLCAALAATMGLSSKAAAEMAESVSNPQRPPVIWIGAQECT WP_000145429.1 hydrogenase 2 small subunit [Salmonella enterica] Length: 372\nNumber of predicted TMHs: 1\nExp number of AAs in TMHs: 24.51309\nExp number, first 60 AAs: 0.2682\nTotal prob of N-in: 0.14402\noutside 1 331\nTMhelix 332 354\ninside 355 372

21593 GCF_001361835.1_10209_5_32 Salmonella enterica subsp. enterica serovar Typhi Proteobacteria; Gammaproteobacteria; Enterobacterales; Enterobacteriaceae; Salmonella; Salmonella enterica; Salmonella enterica subsp. enterica MTGDNTLITSHGINRRDFMKLCAALAATMGLSSKAAAEMAESVSNPQRPPVIWIGAQECT WP_000145429.1 hydrogenase 2 small subunit [Salmonella enterica] Length: 372\nNumber of predicted TMHs: 1\nExp number of AAs in TMHs: 24.51309\nExp number, first 60 AAs: 0.2682\nTotal prob of N-in: 0.14402\noutside 1 331\nTMhelix 332 354\ninside 355 372

21594 GCF_001361795.1_10209_5_2 Salmonella enterica subsp. enterica serovar Typhi Proteobacteria; Gammaproteobacteria; Enterobacterales; Enterobacteriaceae; Salmonella; Salmonella enterica; Salmonella enterica subsp. enterica MTGDNTLITSHGINRRDFMKLCAALAATMGLSSKAAAEMAESVSNPQRPPVIWIGAQECT WP_000145429.1 hydrogenase 2 small subunit [Salmonella enterica] Length: 372\nNumber of

predicted TMHs:  1\nExp number of AAs in TMHs: 24.51309\nExp number, first 60 AAs:  0.2682\nTotal prob of N-in: 0.14402\noutside   1   331\nTMhelix   332   354\ninside   355   372

21595 GCF_001361715.1_10209_5_52 Salmonella enterica subsp. enterica serovar Typhi Proteobacteria; Gammaproteobacteria; Enterobacterales; Enterobacteriaceae; Salmonella; Salmonella enterica; Salmonella enterica subsp. enterica MTGDNTLITSHGINRRDFMKLCAALAATMGLSSKAAAEMAESVSNPQRPPVIWIGAQECT WP_000145429.1 hydrogenase 2 small subunit [Salmonella enterica] Length: 372\nNumber of predicted TMHs:  1\nExp number of AAs in TMHs: 24.51309\nExp number, first 60 AAs:  0.2682\nTotal prob of N-in: 0.14402\noutside   1   331\nTMhelix   332   354\ninside   355   372

21596 GCF_001361755.1_10071_8_1 Salmonella enterica subsp. enterica serovar Typhi Proteobacteria; Gammaproteobacteria; Enterobacterales; Enterobacteriaceae; Salmonella; Salmonella enterica; Salmonella enterica subsp. enterica MTGDNTLITSHGINRRDFMKLCAALAATMGLSSKAAAEMAESVSNPQRPPVIWIGAQECT WP_000145429.1 hydrogenase 2 small subunit [Salmonella enterica] Length: 372\nNumber of predicted TMHs:  1\nExp number of AAs in TMHs: 24.51309\nExp number, first 60 AAs:  0.2682\nTotal prob of N-in: 0.14402\noutside   1   331\nTMhelix   332   354\ninside   355   372

21597 GCF_001361675.1_10071_8_69 Salmonella enterica subsp. enterica serovar Typhi Proteobacteria; Gammaproteobacteria; Enterobacterales; Enterobacteriaceae; Salmonella; Salmonella enterica; Salmonella enterica subsp. enterica MTGDNTLITSHGINRRDFMKLCAALAATMGLSSKAAAEMAESVSNPQRPPVIWIGAQECT WP_000145429.1 hydrogenase 2 small subunit [Salmonella enterica] Length: 372\nNumber of predicted TMHs:  1\nExp number of AAs in TMHs: 24.51309\nExp number, first 60 AAs:  0.2682\nTotal prob of N-in: 0.14402\noutside   1   331\nTMhelix   332   354\ninside   355   372

21598 GCF_001361635.1_10071_8_24 Salmonella enterica subsp. enterica serovar Typhi Proteobacteria; Gammaproteobacteria; Enterobacterales; Enterobacteriaceae; Salmonella; Salmonella enterica; Salmonella enterica subsp. enterica MTGDNTLITSHGINRRDFMKLCAALAATMGLSSKAAAEMAESVSNPQRPPVIWIGAQECT WP_000145429.1 hydrogenase 2 small subunit [Salmonella enterica] Length: 372\nNumber of predicted TMHs:  1\nExp number of AAs in TMHs: 24.51309\nExp number, first 60 AAs:  0.2682\nTotal prob of N-in: 0.14402\noutside   1   331\nTMhelix   332   354\ninside   355   372

21599 GCF_001361595.1_10060_6_88 Salmonella enterica subsp. enterica serovar Typhi Proteobacteria; Gammaproteobacteria; Enterobacterales; Enterobacteriaceae; Salmonella; Salmonella enterica; Salmonella enterica subsp. enterica MTGDNTLITSHGINRRDFMKLCAALAATMGLSSKAAAEMAESVSNPQRPPVIWIGAQECT WP_000145429.1 hydrogenase 2 small subunit [Salmonella enterica] Length: 372\nNumber of predicted TMHs:  1\nExp number of AAs in TMHs: 24.51309\nExp number, first 60 AAs:  0.2682\nTotal prob of N-in: 0.14402\noutside   1   331\nTMhelix   332   354\ninside   355   372

21600 GCF_001361095.1_10071_3_25 Salmonella enterica subsp. enterica serovar Typhi Proteobacteria; Gammaproteobacteria; Enterobacterales; Enterobacteriaceae; Salmonella; Salmonella enterica; Salmonella enterica subsp. enterica MTGDNTLITSHGINRRDFMKLCAALAATMGLSSKAAAEMAESVSNPQRPPVIWIGAQECT WP_000145429.1 hydrogenase 2 small subunit [Salmonella enterica] Length: 372\nNumber of predicted TMHs:  1\nExp number of AAs in TMHs: 24.51309\nExp number, first 60 AAs:  0.2682\nTotal prob of N-in: 0.14402\noutside   1   331\nTMhelix   332   354\ninside   355   372

21601 GCF_001360975.1_10060_5_92 Salmonella enterica subsp. enterica serovar Typhi Proteobacteria; Gammaproteobacteria; Enterobacterales; Enterobacteriaceae; Salmonella; Salmonella enterica; Salmonella enterica subsp. enterica MTGDNTLITSHGINRRDFMKLCAALAATMGLSSKAAAEMAESVSNPQRPPVIWIGAQECT WP_000145429.1 hydrogenase 2 small subunit [Salmonella enterica] Length: 372\nNumber of predicted TMHs:  1\nExp number of AAs in TMHs: 24.51309\nExp number, first 60 AAs:  0.2682\nTotal prob of N-in: 0.14402\noutside   1   331\nTMhelix   332   354\ninside   355   372

21602 GCF_001360935.1_10060_5_81 Salmonella enterica subsp. enterica serovar Typhi Proteobacteria; Gammaproteobacteria; Enterobacterales; Enterobacteriaceae; Salmonella; Salmonella enterica; Salmonella enterica subsp. enterica MTGDNTLITSHGINRRDFMKLCAALAATMGLSSKAAAEMAESVSNPQRPPVIWIGAQECT WP_000145429.1 hydrogenase 2 small subunit [Salmonella enterica] Length: 372\nNumber of predicted TMHs:  1\nExp number of AAs in TMHs: 24.51309\nExp number, first 60 AAs:  0.2682\nTotal prob of N-in: 0.14402\noutside   1   331\nTMhelix   332   354\ninside   355   372

21603 GCF_001360895.1_10060_5_63 Salmonella enterica subsp. enterica serovar Typhi Proteobacteria; Gammaproteobacteria; Enterobacterales; Enterobacteriaceae; Salmonella; Salmonella enterica;

Salmonella enterica subsp. enterica	MTGDNTLITSHGINRRDFMKLCAALAATMGLSSKAAAEMAESVSNPQRPPVIWIGAQECT	WP_000145429.1	hydrogenase 2 small subunit [Salmonella enterica]	Length: 372\nNumber of predicted TMHs:  1\nExp number of AAs in TMHs: 24.51309\nExp number, first 60 AAs:  0.2682\nTotal prob of N-in: 0.14402\noutside   1   331\nTMhelix   332   354\ninside   355   372

21604	GCF_001360855.1_10060_5_48	Salmonella enterica subsp. enterica serovar Typhi	Proteobacteria; Gammaproteobacteria; Enterobacterales; Enterobacteriaceae; Salmonella; Salmonella enterica; Salmonella enterica subsp. enterica	MTGDNTLITSHGINRRDFMKLCAALAATMGLSSKAAAEMAESVSNPQRPPVIWIGAQECT	WP_000145429.1	hydrogenase 2 small subunit [Salmonella enterica]	Length: 372\nNumber of predicted TMHs:  1\nExp number of AAs in TMHs: 24.51309\nExp number, first 60 AAs:  0.2682\nTotal prob of N-in: 0.14402\noutside   1   331\nTMhelix   332   354\ninside   355   372

21605	GCF_001360815.1_10060_5_33	Salmonella enterica subsp. enterica serovar Typhi	Proteobacteria; Gammaproteobacteria; Enterobacterales; Enterobacteriaceae; Salmonella; Salmonella enterica; Salmonella enterica subsp. enterica	MTGDNTLITSHGINRRDFMKLCAALAATMGLSSKAAAEMAESVSNPQRPPVIWIGAQECT	WP_000145429.1	hydrogenase 2 small subunit [Salmonella enterica]	Length: 372\nNumber of predicted TMHs:  1\nExp number of AAs in TMHs: 24.51309\nExp number, first 60 AAs:  0.2682\nTotal prob of N-in: 0.14402\noutside   1   331\nTMhelix   332   354\ninside   355   372

21606	GCF_001360235.1_10060_5_93	Salmonella enterica subsp. enterica serovar Typhi	Proteobacteria; Gammaproteobacteria; Enterobacterales; Enterobacteriaceae; Salmonella; Salmonella enterica; Salmonella enterica subsp. enterica	MTGDNTLITSHGINRRDFMKLCAALAATMGLSSKAAAEMAESVSNPQRPPVIWIGAQECT	WP_000145429.1	hydrogenase 2 small subunit [Salmonella enterica]	Length: 372\nNumber of predicted TMHs:  1\nExp number of AAs in TMHs: 24.51309\nExp number, first 60 AAs:  0.2682\nTotal prob of N-in: 0.14402\noutside   1   331\nTMhelix   332   354\ninside   355   372

21607	GCF_001360195.1_10060_5_82	Salmonella enterica subsp. enterica serovar Typhi	Proteobacteria; Gammaproteobacteria; Enterobacterales; Enterobacteriaceae; Salmonella; Salmonella enterica; Salmonella enterica subsp. enterica	MTGDNTLITSHGINRRDFMKLCAALAATMGLSSKAAAEMAESVSNPQRPPVIWIGAQECT	WP_000145429.1	hydrogenase 2 small subunit [Salmonella enterica]	Length: 372\nNumber of predicted TMHs:  1\nExp number of AAs in TMHs: 24.51309\nExp number, first 60 AAs:  0.2682\nTotal prob of N-in: 0.14402\noutside   1   331\nTMhelix   332   354\ninside   355   372

21608	GCF_001360155.1_10060_5_64	Salmonella enterica subsp. enterica serovar Typhi	Proteobacteria; Gammaproteobacteria; Enterobacterales; Enterobacteriaceae; Salmonella; Salmonella enterica; Salmonella enterica subsp. enterica	MTGDNTLITSHGINRRDFMKLCAALAATMGLSSKAAAEMAESVSNPQRPPVIWIGAQECT	WP_000145429.1	hydrogenase 2 small subunit [Salmonella enterica]	Length: 372\nNumber of predicted TMHs:  1\nExp number of AAs in TMHs: 24.51309\nExp number, first 60 AAs:  0.2682\nTotal prob of N-in: 0.14402\noutside   1   331\nTMhelix   332   354\ninside   355   372

21609	GCF_001360115.1_10060_6_84	Salmonella enterica subsp. enterica serovar Typhi	Proteobacteria; Gammaproteobacteria; Enterobacterales; Enterobacteriaceae; Salmonella; Salmonella enterica; Salmonella enterica subsp. enterica	MTGDNTLITSHGINRRDFMKLCAALAATMGLSSKAAAEMAESVSNPQRPPVIWIGAQECT	WP_000145429.1	hydrogenase 2 small subunit [Salmonella enterica]	Length: 372\nNumber of predicted TMHs:  1\nExp number of AAs in TMHs: 24.51309\nExp number, first 60 AAs:  0.2682\nTotal prob of N-in: 0.14402\noutside   1   331\nTMhelix   332   354\ninside   355   372

21610	GCF_001360055.1_10060_5_49	Salmonella enterica subsp. enterica serovar Typhi	Proteobacteria; Gammaproteobacteria; Enterobacterales; Enterobacteriaceae; Salmonella; Salmonella enterica; Salmonella enterica subsp. enterica	MTGDNTLITSHGINRRDFMKLCAALAATMGLSSKAAAEMAESVSNPQRPPVIWIGAQECT	WP_000145429.1	hydrogenase 2 small subunit [Salmonella enterica]	Length: 372\nNumber of predicted TMHs:  1\nExp number of AAs in TMHs: 24.51309\nExp number, first 60 AAs:  0.2682\nTotal prob of N-in: 0.14402\noutside   1   331\nTMhelix   332   354\ninside   355   372

21611	GCF_001360015.1_10060_5_34	Salmonella enterica subsp. enterica serovar Typhi	Proteobacteria; Gammaproteobacteria; Enterobacterales; Enterobacteriaceae; Salmonella; Salmonella enterica; Salmonella enterica subsp. enterica	MTGDNTLITSHGINRRDFMKLCAALAATMGLSSKAAAEMAESVSNPQRPPVIWIGAQECT	WP_000145429.1	hydrogenase 2 small subunit [Salmonella enterica]	Length: 372\nNumber of predicted TMHs:  1\nExp number of AAs in TMHs: 24.51309\nExp number, first 60 AAs:  0.2682\nTotal prob of N-in: 0.14402\noutside   1   331\nTMhelix   332   354\ninside   355   372

21612 GCF_001359535.1_9953_5_18 Salmonella enterica subsp. enterica serovar Typhi Proteobacteria; Gammaproteobacteria; Enterobacterales; Enterobacteriaceae; Salmonella; Salmonella enterica; Salmonella enterica subsp. enterica MTGDNTLITSHGINRRDFMKLCAALAATMGLSSKAAAEMAESVSNPQRPPVIWIGAQECT WP_000145429.1 hydrogenase 2 small subunit [Salmonella enterica] Length: 372\nNumber of predicted TMHs: 1\nExp number of AAs in TMHs: 24.51309\nExp number, first 60 AAs: 0.2682\nTotal prob of N-in: 0.14402\noutside 1 331\nTMhelix 332 354\ninside 355 372

21613 GCF_001359415.1_10060_5_89 Salmonella enterica subsp. enterica serovar Typhi Proteobacteria; Gammaproteobacteria; Enterobacterales; Enterobacteriaceae; Salmonella; Salmonella enterica; Salmonella enterica subsp. enterica MTGDNTLITSHGINRRDFMKLCAALAATMGLSSKAAAEMAESVSNPQRPPVIWIGAQECT WP_000145429.1 hydrogenase 2 small subunit [Salmonella enterica] Length: 372\nNumber of predicted TMHs: 1\nExp number of AAs in TMHs: 24.51309\nExp number, first 60 AAs: 0.2682\nTotal prob of N-in: 0.14402\noutside 1 331\nTMhelix 332 354\ninside 355 372

21614 GCF_001359355.1_10060_5_71 Salmonella enterica subsp. enterica serovar Typhi Proteobacteria; Gammaproteobacteria; Enterobacterales; Enterobacteriaceae; Salmonella; Salmonella enterica; Salmonella enterica subsp. enterica MTGDNTLITSHGINRRDFMKLCAALAATMGLSSKAAAEMAESVSNPQRPPVIWIGAQECT WP_000145429.1 hydrogenase 2 small subunit [Salmonella enterica] Length: 372\nNumber of predicted TMHs: 1\nExp number of AAs in TMHs: 24.51309\nExp number, first 60 AAs: 0.2682\nTotal prob of N-in: 0.14402\noutside 1 331\nTMhelix 332 354\ninside 355 372

21615 GCF_001359315.1_10060_5_62 Salmonella enterica subsp. enterica serovar Typhi Proteobacteria; Gammaproteobacteria; Enterobacterales; Enterobacteriaceae; Salmonella; Salmonella enterica; Salmonella enterica subsp. enterica MTGDNTLITSHGINRRDFMKLCAALAATMGLSSKAAAEMAESVSNPQRPPVIWIGAQECT WP_000145429.1 hydrogenase 2 small subunit [Salmonella enterica] Length: 372\nNumber of predicted TMHs: 1\nExp number of AAs in TMHs: 24.51309\nExp number, first 60 AAs: 0.2682\nTotal prob of N-in: 0.14402\noutside 1 331\nTMhelix 332 354\ninside 355 372

21616 GCF_001359235.1_10060_5_44 Salmonella enterica subsp. enterica serovar Typhi Proteobacteria; Gammaproteobacteria; Enterobacterales; Enterobacteriaceae; Salmonella; Salmonella enterica; Salmonella enterica subsp. enterica MTGDNTLITSHGINRRDFMKLCAALAATMGLSSKAAAEMAESVSNPQRPPVIWIGAQECT WP_000145429.1 hydrogenase 2 small subunit [Salmonella enterica] Length: 372\nNumber of predicted TMHs: 1\nExp number of AAs in TMHs: 24.51309\nExp number, first 60 AAs: 0.2682\nTotal prob of N-in: 0.14402\noutside 1 331\nTMhelix 332 354\ninside 355 372

21617 GCF_001358995.1_10060_6_74 Salmonella enterica subsp. enterica serovar Typhi Proteobacteria; Gammaproteobacteria; Enterobacterales; Enterobacteriaceae; Salmonella; Salmonella enterica; Salmonella enterica subsp. enterica MTGDNTLITSHGINRRDFMKLCAALAATMGLSSKAATEMAESVSNPQRPPVIWIGAQECT WP_001782304.1 hydrogenase 2 small subunit [Salmonella enterica] Length: 372\nNumber of predicted TMHs: 1\nExp number of AAs in TMHs: 24.41173\nExp number, first 60 AAs: 0.16124\nTotal prob of N-in: 0.13947\noutside 1 331\nTMhelix 332 354\ninside 355 372

21618 GCF_001358915.1_10060_6_35 Salmonella enterica subsp. enterica serovar Typhi Proteobacteria; Gammaproteobacteria; Enterobacterales; Enterobacteriaceae; Salmonella; Salmonella enterica; Salmonella enterica subsp. enterica MTGDNTLITSHGINRRDFMKLCAALAATMGLSSKAAAEMAESVSNPQRPPVIWIGAQECT WP_000145429.1 hydrogenase 2 small subunit [Salmonella enterica] Length: 372\nNumber of predicted TMHs: 1\nExp number of AAs in TMHs: 24.51309\nExp number, first 60 AAs: 0.2682\nTotal prob of N-in: 0.14402\noutside 1 331\nTMhelix 332 354\ninside 355 372

21619 GCF_001358835.1_10060_6_14 Salmonella enterica subsp. enterica serovar Typhi Proteobacteria; Gammaproteobacteria; Enterobacterales; Enterobacteriaceae; Salmonella; Salmonella enterica; Salmonella enterica subsp. enterica MTGDNTLITSHGINRRDFMKLCAALAATMGLSSKAAAEMAESVSNPQRPPVIWIGAQECT WP_000145429.1 hydrogenase 2 small subunit [Salmonella enterica] Length: 372\nNumber of predicted TMHs: 1\nExp number of AAs in TMHs: 24.51309\nExp number, first 60 AAs: 0.2682\nTotal prob of N-in: 0.14402\noutside 1 331\nTMhelix 332 354\ninside 355 372

21620 GCF_001358875.1_10060_6_23 Salmonella enterica subsp. enterica serovar Typhi Proteobacteria; Gammaproteobacteria; Enterobacterales; Enterobacteriaceae; Salmonella; Salmonella enterica; Salmonella enterica subsp. enterica MTGDNTLITSHGINRRDFMKLCAALAATMGLSSKAAAEMAESVSNPQRPPVIWIGAQECT WP_000145429.1 hydrogenase 2 small subunit [Salmonella enterica] Length: 372\nNumber of

predicted TMHs:  1\nExp number of AAs in TMHs: 24.51309\nExp number, first 60 AAs:  0.2682\nTotal prob of N-in: 0.14402\noutside   1   331\nTMhelix   332   354\ninside   355   372

21621 GCF_001358795.1_10562_2_58 Salmonella enterica subsp. enterica serovar Typhi Proteobacteria; Gammaproteobacteria; Enterobacterales; Enterobacteriaceae; Salmonella; Salmonella enterica; Salmonella enterica subsp. enterica MTGDNTLITSHGINRRDFMKLCAALAATMGLSSKAAAEMAESVSNPQRPPVIWIGAQECT WP_000145429.1 hydrogenase 2 small subunit [Salmonella enterica] Length: 372\nNumber of predicted TMHs:  1\nExp number of AAs in TMHs: 24.51309\nExp number, first 60 AAs:  0.2682\nTotal prob of N-in: 0.14402\noutside   1   331\nTMhelix   332   354\ninside   355   372

21622 GCF_001358755.1_10071_3_68 Salmonella enterica subsp. enterica serovar Typhi Proteobacteria; Gammaproteobacteria; Enterobacterales; Enterobacteriaceae; Salmonella; Salmonella enterica; Salmonella enterica subsp. enterica MTGDNTLITSHGINRRDFMKLCAALAATMGLSSKAAAEMAESVSNPQRPPVIWIGAQECT WP_000145429.1 hydrogenase 2 small subunit [Salmonella enterica] Length: 372\nNumber of predicted TMHs:  1\nExp number of AAs in TMHs: 24.51309\nExp number, first 60 AAs:  0.2682\nTotal prob of N-in: 0.14402\noutside   1   331\nTMhelix   332   354\ninside   355   372

21623 GCF_001358715.1_10607_2_40 Salmonella enterica subsp. enterica serovar Typhi Proteobacteria; Gammaproteobacteria; Enterobacterales; Enterobacteriaceae; Salmonella; Salmonella enterica; Salmonella enterica subsp. enterica MTGDNTLITSHGINRRDFMKLCAALAATMGLSSKAAAEMAESVSNPQRPPVIWIGAQECT WP_000145429.1 hydrogenase 2 small subunit [Salmonella enterica] Length: 372\nNumber of predicted TMHs:  1\nExp number of AAs in TMHs: 24.51309\nExp number, first 60 AAs:  0.2682\nTotal prob of N-in: 0.14402\noutside   1   331\nTMhelix   332   354\ninside   355   372

21624 GCF_001358175.1_10426_1_76 Salmonella enterica subsp. enterica serovar Typhi Proteobacteria; Gammaproteobacteria; Enterobacterales; Enterobacteriaceae; Salmonella; Salmonella enterica; Salmonella enterica subsp. enterica MTGDNTLITSHGINRRDFMKLCAALAATMGLSSKAAAEMAESVSNPQRPPVIWIGAQECT WP_000145429.1 hydrogenase 2 small subunit [Salmonella enterica] Length: 372\nNumber of predicted TMHs:  1\nExp number of AAs in TMHs: 24.51309\nExp number, first 60 AAs:  0.2682\nTotal prob of N-in: 0.14402\noutside   1   331\nTMhelix   332   354\ninside   355   372

21625 GCF_001358135.1_10592_2_80 Salmonella enterica subsp. enterica serovar Typhi Proteobacteria; Gammaproteobacteria; Enterobacterales; Enterobacteriaceae; Salmonella; Salmonella enterica; Salmonella enterica subsp. enterica MTGDNTLITSHGINRRDFMKLCAALAATMGLSSKAAAEMAESVSNPQRPPVIWIGAQECT WP_000145429.1 hydrogenase 2 small subunit [Salmonella enterica] Length: 372\nNumber of predicted TMHs:  1\nExp number of AAs in TMHs: 24.51309\nExp number, first 60 AAs:  0.2682\nTotal prob of N-in: 0.14402\noutside   1   331\nTMhelix   332   354\ninside   355   372

21626 GCF_001358095.1_10593_2_36 Salmonella enterica subsp. enterica serovar Typhi Proteobacteria; Gammaproteobacteria; Enterobacterales; Enterobacteriaceae; Salmonella; Salmonella enterica; Salmonella enterica subsp. enterica MTGDNTLITSHGINRRDFMKLCAALAATMGLSSKAAAEMAESVSNPQRPPVIWIGAQECT WP_000145429.1 hydrogenase 2 small subunit [Salmonella enterica] Length: 372\nNumber of predicted TMHs:  1\nExp number of AAs in TMHs: 24.51309\nExp number, first 60 AAs:  0.2682\nTotal prob of N-in: 0.14402\noutside   1   331\nTMhelix   332   354\ninside   355   372

21627 GCF_001358055.1_10593_2_31 Salmonella enterica subsp. enterica serovar Typhi Proteobacteria; Gammaproteobacteria; Enterobacterales; Enterobacteriaceae; Salmonella; Salmonella enterica; Salmonella enterica subsp. enterica MTGDNTLITSHGINRRDFMKLCAALAATMGLSSKAAAEMAESVSNPQRPPVIWIGAQECT WP_000145429.1 hydrogenase 2 small subunit [Salmonella enterica] Length: 372\nNumber of predicted TMHs:  1\nExp number of AAs in TMHs: 24.51309\nExp number, first 60 AAs:  0.2682\nTotal prob of N-in: 0.14402\noutside   1   331\nTMhelix   332   354\ninside   355   372

21628 GCF_001358015.1_10592_2_53 Salmonella enterica subsp. enterica serovar Typhi Proteobacteria; Gammaproteobacteria; Enterobacterales; Enterobacteriaceae; Salmonella; Salmonella enterica; Salmonella enterica subsp. enterica MTGDNTLITSHGINRRDFMKLCAALAATMGLSSKAAAEMAESVSNPQRPPVIWIGAQECT WP_000145429.1 hydrogenase 2 small subunit [Salmonella enterica] Length: 372\nNumber of predicted TMHs:  1\nExp number of AAs in TMHs: 24.51309\nExp number, first 60 AAs:  0.2682\nTotal prob of N-in: 0.14402\noutside   1   331\nTMhelix   332   354\ninside   355   372

21629 GCF_001357915.1_10592_2_37 Salmonella enterica subsp. enterica serovar Typhi Proteobacteria; Gammaproteobacteria; Enterobacterales; Enterobacteriaceae; Salmonella; Salmonella enterica;

Salmonella enterica subsp. enterica MTGDNTLITSHGINRRDFMKLCAALAATMGLSSKAAAEMAESVSNPQRPPVIWIGAQECT WP_000145429.1 hydrogenase 2 small subunit [Salmonella enterica] Length: 372\nNumber of predicted TMHs: 1\nExp number of AAs in TMHs: 24.51309\nExp number, first 60 AAs: 0.2682\nTotal prob of N-in: 0.14402\noutside 1 331\nTMhelix 332 354\ninside 355 372

21630 GCF_001357955.1_10592_2_75 Salmonella enterica subsp. enterica serovar Typhi Proteobacteria; Gammaproteobacteria; Enterobacterales; Enterobacteriaceae; Salmonella; Salmonella enterica; Salmonella enterica subsp. enterica MTGDNTLITSHGINRRDFMKLCAALAATMGLSSKAAAEMAESVSNPQRPPVIWIGAQECT WP_000145429.1 hydrogenase 2 small subunit [Salmonella enterica] Length: 372\nNumber of predicted TMHs: 1\nExp number of AAs in TMHs: 24.51309\nExp number, first 60 AAs: 0.2682\nTotal prob of N-in: 0.14402\noutside 1 331\nTMhelix 332 354\ninside 355 372

21631 GCF_001357595.1_10592_2_78 Salmonella enterica subsp. enterica serovar Typhi Proteobacteria; Gammaproteobacteria; Enterobacterales; Enterobacteriaceae; Salmonella; Salmonella enterica; Salmonella enterica subsp. enterica MTGDNTLITSHGINRRDFMKLCAALAATMGLSSKAAAEMAESVSNPQRPPVIWIGAQECT WP_000145429.1 hydrogenase 2 small subunit [Salmonella enterica] Length: 372\nNumber of predicted TMHs: 1\nExp number of AAs in TMHs: 24.51309\nExp number, first 60 AAs: 0.2682\nTotal prob of N-in: 0.14402\noutside 1 331\nTMhelix 332 354\ninside 355 372

21632 GCF_001357555.1_10593_2_56 Salmonella enterica subsp. enterica serovar Typhi Proteobacteria; Gammaproteobacteria; Enterobacterales; Enterobacteriaceae; Salmonella; Salmonella enterica; Salmonella enterica subsp. enterica MTGDNTLITSHGINRRDFMKLCAALAATMGLSSKAAAEMAESVSNPQRPPVIWIGAQECT WP_000145429.1 hydrogenase 2 small subunit [Salmonella enterica] Length: 372\nNumber of predicted TMHs: 1\nExp number of AAs in TMHs: 24.51309\nExp number, first 60 AAs: 0.2682\nTotal prob of N-in: 0.14402\noutside 1 331\nTMhelix 332 354\ninside 355 372

21633 GCF_001357515.1_10592_2_77 Salmonella enterica subsp. enterica serovar Typhi Proteobacteria; Gammaproteobacteria; Enterobacterales; Enterobacteriaceae; Salmonella; Salmonella enterica; Salmonella enterica subsp. enterica MTGDNTLITSHGINRRDFMKLCAALAATMGLSSKAAAEMAESVSNPQRPPVIWIGAQECT WP_000145429.1 hydrogenase 2 small subunit [Salmonella enterica] Length: 372\nNumber of predicted TMHs: 1\nExp number of AAs in TMHs: 24.51309\nExp number, first 60 AAs: 0.2682\nTotal prob of N-in: 0.14402\noutside 1 331\nTMhelix 332 354\ninside 355 372

21634 GCF_001357475.1_10593_2_62 Salmonella enterica subsp. enterica serovar Typhi Proteobacteria; Gammaproteobacteria; Enterobacterales; Enterobacteriaceae; Salmonella; Salmonella enterica; Salmonella enterica subsp. enterica MTGDNTLITSHGINRRDFMKLCAALAATMGLSSKAAAEMAESVSNPQRPPVIWIGAQECT WP_000145429.1 hydrogenase 2 small subunit [Salmonella enterica] Length: 372\nNumber of predicted TMHs: 1\nExp number of AAs in TMHs: 24.51309\nExp number, first 60 AAs: 0.2682\nTotal prob of N-in: 0.14402\noutside 1 331\nTMhelix 332 354\ninside 355 372

21635 GCF_001099125.1_9953_5_30 Salmonella enterica subsp. enterica serovar Typhi Proteobacteria; Gammaproteobacteria; Enterobacterales; Enterobacteriaceae; Salmonella; Salmonella enterica; Salmonella enterica subsp. enterica MTGDNTLITSHGINRRDFMKLCAALAATMGLSSKAAAEMAESVSNPQRPPVIWIGAQECT WP_000145429.1 hydrogenase 2 small subunit [Salmonella enterica] Length: 372\nNumber of predicted TMHs: 1\nExp number of AAs in TMHs: 24.51309\nExp number, first 60 AAs: 0.2682\nTotal prob of N-in: 0.14402\noutside 1 331\nTMhelix 332 354\ninside 355 372

21636 GCF_001099205.1_8490_5_30 Salmonella enterica subsp. enterica serovar Typhi Proteobacteria; Gammaproteobacteria; Enterobacterales; Enterobacteriaceae; Salmonella; Salmonella enterica; Salmonella enterica subsp. enterica MTGDNTLITSHGINRRDFMKLCAALAATMGLSSKAAAEMAESVSNPQRPPVIWIGAQECT WP_000145429.1 hydrogenase 2 small subunit [Salmonella enterica] Length: 372\nNumber of predicted TMHs: 1\nExp number of AAs in TMHs: 24.51309\nExp number, first 60 AAs: 0.2682\nTotal prob of N-in: 0.14402\noutside 1 331\nTMhelix 332 354\ninside 355 372

21637 GCF_001102545.1_9475_6_5Salmonella enterica subsp. enterica serovar Typhi Proteobacteria; Gammaproteobacteria; Enterobacterales; Enterobacteriaceae; Salmonella; Salmonella enterica; Salmonella enterica subsp. enterica MTGDNTLITSHGINRRDFMKLCAALAATMGLSSKAAAEMAESVSNPQRPPVIWIGAQECT WP_000145429.1 hydrogenase 2 small subunit [Salmonella enterica] Length: 372\nNumber of predicted TMHs: 1\nExp number of AAs in TMHs: 24.51309\nExp number, first 60 AAs: 0.2682\nTotal prob of N-in: 0.14402\noutside 1 331\nTMhelix 332 354\ninside 355 372

21638 GCF_001357435.1_10593_2_17 Salmonella enterica subsp. enterica serovar Typhi Proteobacteria; Gammaproteobacteria; Enterobacterales; Enterobacteriaceae; Salmonella; Salmonella enterica; Salmonella enterica subsp. enterica MTGDNTLITSHGINRRDFMKLCAALAATMGLSSKAAAEMAESVSNPQRPPVIWIGAQECT WP_000145429.1 hydrogenase 2 small subunit [Salmonella enterica] Length: 372\nNumber of predicted TMHs: 1\nExp number of AAs in TMHs: 24.51309\nExp number, first 60 AAs: 0.2682\nTotal prob of N-in: 0.14402\noutside 1 331\nTMhelix 332 354\ninside 355 372

21639 GCF_001357395.1_10593_2_7 Salmonella enterica subsp. enterica serovar Typhi Proteobacteria; Gammaproteobacteria; Enterobacterales; Enterobacteriaceae; Salmonella; Salmonella enterica; Salmonella enterica subsp. enterica MTGDNTLITSHGINRRDFMKLCAALAATMGLSSKAAAEMAESVSNPQRPPVIWIGAQECT WP_000145429.1 hydrogenase 2 small subunit [Salmonella enterica] Length: 372\nNumber of predicted TMHs: 1\nExp number of AAs in TMHs: 24.51309\nExp number, first 60 AAs: 0.2682\nTotal prob of N-in: 0.14402\noutside 1 331\nTMhelix 332 354\ninside 355 372

21640 GCF_001099305.1_10592_2_57 Salmonella enterica subsp. enterica serovar Typhi Proteobacteria; Gammaproteobacteria; Enterobacterales; Enterobacteriaceae; Salmonella; Salmonella enterica; Salmonella enterica subsp. enterica MTGDNTLITSHGINRRDFMKLCAALAATMGLSSKAAAEMAESVSNPQRPPVIWIGAQECT WP_000145429.1 hydrogenase 2 small subunit [Salmonella enterica] Length: 372\nNumber of predicted TMHs: 1\nExp number of AAs in TMHs: 24.51309\nExp number, first 60 AAs: 0.2682\nTotal prob of N-in: 0.14402\noutside 1 331\nTMhelix 332 354\ninside 355 372

21641 GCF_001357215.1_10593_2_41 Salmonella enterica subsp. enterica serovar Typhi Proteobacteria; Gammaproteobacteria; Enterobacterales; Enterobacteriaceae; Salmonella; Salmonella enterica; Salmonella enterica subsp. enterica MTGDNTLITSHGINRRDFMKLCAALAATMGLSSKAATEMAESVSNPQRPPVIWIGAQECT WP_001782304.1 hydrogenase 2 small subunit [Salmonella enterica] Length: 372\nNumber of predicted TMHs: 1\nExp number of AAs in TMHs: 24.41173\nExp number, first 60 AAs: 0.16124\nTotal prob of N-in: 0.13947\noutside 1 331\nTMhelix 332 354\ninside 355 372

21642 GCF_001356315.1_8447_8_8Salmonella enterica subsp. enterica serovar Typhi Proteobacteria; Gammaproteobacteria; Enterobacterales; Enterobacteriaceae; Salmonella; Salmonella enterica; Salmonella enterica subsp. enterica MTGDNTLITSHGINRRDFMKLCAALAATMGLSSKAAAEMAESVSNPQRPPVIWIGAQECT WP_000145429.1 hydrogenase 2 small subunit [Salmonella enterica] Length: 372\nNumber of predicted TMHs: 1\nExp number of AAs in TMHs: 24.51309\nExp number, first 60 AAs: 0.2682\nTotal prob of N-in: 0.14402\noutside 1 331\nTMhelix 332 354\ninside 355 372

21643 GCF_001356275.1_8490_5_34 Salmonella enterica subsp. enterica serovar Typhi Proteobacteria; Gammaproteobacteria; Enterobacterales; Enterobacteriaceae; Salmonella; Salmonella enterica; Salmonella enterica subsp. enterica MTGDNTLITSHGINRRDFMKLCAALAATMGLSSKAAAEMAESVSNPQRPPVIWIGAQECT WP_000145429.1 hydrogenase 2 small subunit [Salmonella enterica] Length: 372\nNumber of predicted TMHs: 1\nExp number of AAs in TMHs: 24.51309\nExp number, first 60 AAs: 0.2682\nTotal prob of N-in: 0.14402\noutside 1 331\nTMhelix 332 354\ninside 355 372

21644 GCF_001356235.1_8490_6_57 Salmonella enterica subsp. enterica serovar Typhi Proteobacteria; Gammaproteobacteria; Enterobacterales; Enterobacteriaceae; Salmonella; Salmonella enterica; Salmonella enterica subsp. enterica MTGDNTLITSHGINRRDFMKLCAALAATMGLSSKAAAEMAESVSNPQRPPVIWIGAQECT WP_000145429.1 hydrogenase 2 small subunit [Salmonella enterica] Length: 372\nNumber of predicted TMHs: 1\nExp number of AAs in TMHs: 24.51309\nExp number, first 60 AAs: 0.2682\nTotal prob of N-in: 0.14402\noutside 1 331\nTMhelix 332 354\ninside 355 372

21645 GCF_001356195.1_8490_6_71 Salmonella enterica subsp. enterica serovar Typhi Proteobacteria; Gammaproteobacteria; Enterobacterales; Enterobacteriaceae; Salmonella; Salmonella enterica; Salmonella enterica subsp. enterica MTGDNTLITSHGINRRDFMKLCAALAATMGLSSKAAAEMAESVSNPQRPPVIWIGAQECT WP_000145429.1 hydrogenase 2 small subunit [Salmonella enterica] Length: 372\nNumber of predicted TMHs: 1\nExp number of AAs in TMHs: 24.51309\nExp number, first 60 AAs: 0.2682\nTotal prob of N-in: 0.14402\noutside 1 331\nTMhelix 332 354\ninside 355 372

21646 GCF_001356155.1_8490_6_45 Salmonella enterica subsp. enterica serovar Typhi Proteobacteria; Gammaproteobacteria; Enterobacterales; Enterobacteriaceae; Salmonella; Salmonella enterica; Salmonella enterica subsp. enterica MTGDNTLITSHGINRRDFMKLCAALAATMGLSSKAAAEMAESVSNPQRPPVIWIGAQECT WP_000145429.1 hydrogenase 2 small subunit [Salmonella enterica] Length: 372\nNumber of

predicted TMHs: 1\nExp number of AAs in TMHs: 24.51309\nExp number, first 60 AAs: 0.2682\nTotal prob of N-in: 0.14402\noutside 1 331\nTMhelix 332 354\ninside 355 372

21647 GCF_001356075.1_8490_6_52 Salmonella enterica subsp. enterica serovar Typhi Proteobacteria; Gammaproteobacteria; Enterobacterales; Enterobacteriaceae; Salmonella; Salmonella enterica; Salmonella enterica subsp. enterica MTGDNTLITSHGINRRDFMKLCAALAATMGLSSKAAAEMAESVSNPQRPPVIWIGAQECT WP_000145429.1 hydrogenase 2 small subunit [Salmonella enterica] Length: 372\nNumber of predicted TMHs: 1\nExp number of AAs in TMHs: 24.51309\nExp number, first 60 AAs: 0.2682\nTotal prob of N-in: 0.14402\noutside 1 331\nTMhelix 332 354\ninside 355 372

21648 GCF_001356115.1_8447_8_5Salmonella enterica subsp. enterica serovar Typhi Proteobacteria; Gammaproteobacteria; Enterobacterales; Enterobacteriaceae; Salmonella; Salmonella enterica; Salmonella enterica subsp. enterica MTGDNTLITSHGINRRDFMKLCAALAATMGLSSKAAAEMAESVSNPQRPPVIWIGAQECT WP_000145429.1 hydrogenase 2 small subunit [Salmonella enterica] Length: 372\nNumber of predicted TMHs: 1\nExp number of AAs in TMHs: 24.51309\nExp number, first 60 AAs: 0.2682\nTotal prob of N-in: 0.14402\noutside 1 331\nTMhelix 332 354\ninside 355 372

21649 GCF_001356035.1_8447_8_29 Salmonella enterica subsp. enterica serovar Typhi Proteobacteria; Gammaproteobacteria; Enterobacterales; Enterobacteriaceae; Salmonella; Salmonella enterica; Salmonella enterica subsp. enterica MTGDNTLITSHGINRRDFMKLCAALAATMGLSSKAAAEMAESVSNPQRPPVIWIGAQECT WP_000145429.1 hydrogenase 2 small subunit [Salmonella enterica] Length: 372\nNumber of predicted TMHs: 1\nExp number of AAs in TMHs: 24.51309\nExp number, first 60 AAs: 0.2682\nTotal prob of N-in: 0.14402\noutside 1 331\nTMhelix 332 354\ninside 355 372

21650 GCF_001355995.1_8447_8_33 Salmonella enterica subsp. enterica serovar Typhi Proteobacteria; Gammaproteobacteria; Enterobacterales; Enterobacteriaceae; Salmonella; Salmonella enterica; Salmonella enterica subsp. enterica MTGDNTLITSHGINRRDFMKLCAALAATMGLSSKAAAEMAESVSNPQRPPVIWIGAQECT WP_000145429.1 hydrogenase 2 small subunit [Salmonella enterica] Length: 372\nNumber of predicted TMHs: 1\nExp number of AAs in TMHs: 24.51309\nExp number, first 60 AAs: 0.2682\nTotal prob of N-in: 0.14402\noutside 1 331\nTMhelix 332 354\ninside 355 372

21651 GCF_001355955.1_8490_5_26 Salmonella enterica subsp. enterica serovar Typhi Proteobacteria; Gammaproteobacteria; Enterobacterales; Enterobacteriaceae; Salmonella; Salmonella enterica; Salmonella enterica subsp. enterica MTGDNTLITSHGINRRDFMKLCAALAATMGLSSKAAAEMAESVSNPQRPPVIWIGAQECT WP_000145429.1 hydrogenase 2 small subunit [Salmonella enterica] Length: 372\nNumber of predicted TMHs: 1\nExp number of AAs in TMHs: 24.51309\nExp number, first 60 AAs: 0.2682\nTotal prob of N-in: 0.14402\noutside 1 331\nTMhelix 332 354\ninside 355 372

21652 GCF_001355915.1_8447_8_2Salmonella enterica subsp. enterica serovar Typhi Proteobacteria; Gammaproteobacteria; Enterobacterales; Enterobacteriaceae; Salmonella; Salmonella enterica; Salmonella enterica subsp. enterica MTGDNTLITSHGINRRDFMKLCAALAATMGLSSKAAAEMAESVSNPQRPPVIWIGAQECT WP_000145429.1 hydrogenase 2 small subunit [Salmonella enterica] Length: 372\nNumber of predicted TMHs: 1\nExp number of AAs in TMHs: 24.51309\nExp number, first 60 AAs: 0.2682\nTotal prob of N-in: 0.14402\noutside 1 331\nTMhelix 332 354\ninside 355 372

21653 GCF_001355855.1_8490_5_10 Salmonella enterica subsp. enterica serovar Typhi Proteobacteria; Gammaproteobacteria; Enterobacterales; Enterobacteriaceae; Salmonella; Salmonella enterica; Salmonella enterica subsp. enterica MTGDNTLITSHGINRRDFMKLCAALAATMGLSSKAAAEMAESVSNPQRPPVIWIGAQECT WP_000145429.1 hydrogenase 2 small subunit [Salmonella enterica] Length: 372\nNumber of predicted TMHs: 1\nExp number of AAs in TMHs: 24.51309\nExp number, first 60 AAs: 0.2682\nTotal prob of N-in: 0.14402\noutside 1 331\nTMhelix 332 354\ninside 355 372

21654 GCF_001355815.1_8490_6_64 Salmonella enterica subsp. enterica serovar Typhi Proteobacteria; Gammaproteobacteria; Enterobacterales; Enterobacteriaceae; Salmonella; Salmonella enterica; Salmonella enterica subsp. enterica MTGDNTLITSHGINRRDFMKLCAALAATMGLSSKAAAEMAESVSNPQRPPVIWIGAQECT WP_000145429.1 hydrogenase 2 small subunit [Salmonella enterica] Length: 372\nNumber of predicted TMHs: 1\nExp number of AAs in TMHs: 24.51309\nExp number, first 60 AAs: 0.2682\nTotal prob of N-in: 0.14402\noutside 1 331\nTMhelix 332 354\ninside 355 372

21655 GCF_001328635.1_7468_7_57 Salmonella enterica subsp. enterica serovar Typhi Proteobacteria; Gammaproteobacteria; Enterobacterales; Enterobacteriaceae; Salmonella; Salmonella enterica;

Salmonella enterica subsp. enterica MTGDNTLITSHGINRRDFMKLCAALAATMGLSSKAAAEMAESVSNPQRPPVIWIGAQECT WP_000145429.1 hydrogenase 2 small subunit [Salmonella enterica] Length: 372\nNumber of predicted TMHs: 1\nExp number of AAs in TMHs: 24.51309\nExp number, first 60 AAs: 0.2682\nTotal prob of N-in: 0.14402\noutside 1 331\nTMhelix 332 354\ninside 355 372

21656 GCF_001328575.1_7468_7_61 Salmonella enterica subsp. enterica serovar Typhi Proteobacteria; Gammaproteobacteria; Enterobacterales; Enterobacteriaceae; Salmonella; Salmonella enterica; Salmonella enterica subsp. enterica MTGDNTLITSHGINRRDFMKLCAALAATMGLSSKAAAEMAESVSNPQRPPVIWIGAQECT WP_000145429.1 hydrogenase 2 small subunit [Salmonella enterica] Length: 372\nNumber of predicted TMHs: 1\nExp number of AAs in TMHs: 24.51309\nExp number, first 60 AAs: 0.2682\nTotal prob of N-in: 0.14402\noutside 1 331\nTMhelix 332 354\ninside 355 372

21657 GCF_001328595.1_7468_7_65 Salmonella enterica subsp. enterica serovar Typhi Proteobacteria; Gammaproteobacteria; Enterobacterales; Enterobacteriaceae; Salmonella; Salmonella enterica; Salmonella enterica subsp. enterica MTGDNTLITSHGINRRDFMKLCAALAATMGLSSKAAAEMAESVSNPQRPPVIWIGAQECT WP_000145429.1 hydrogenase 2 small subunit [Salmonella enterica] Length: 372\nNumber of predicted TMHs: 1\nExp number of AAs in TMHs: 24.51309\nExp number, first 60 AAs: 0.2682\nTotal prob of N-in: 0.14402\noutside 1 331\nTMhelix 332 354\ninside 355 372

21658 GCF_001317605.1_9475_4_54 Salmonella enterica subsp. enterica serovar Typhi Proteobacteria; Gammaproteobacteria; Enterobacterales; Enterobacteriaceae; Salmonella; Salmonella enterica; Salmonella enterica subsp. enterica MTGDNTLITSHGINRRDFMKLCAALAATMGLSSKAAAEMAESVSNPQRPPVIWIGAQECT WP_000145429.1 hydrogenase 2 small subunit [Salmonella enterica] Length: 372\nNumber of predicted TMHs: 1\nExp number of AAs in TMHs: 24.51309\nExp number, first 60 AAs: 0.2682\nTotal prob of N-in: 0.14402\noutside 1 331\nTMhelix 332 354\ninside 355 372

21659 GCF_001242295.1_10425_1_34 Salmonella enterica subsp. enterica serovar Typhi Proteobacteria; Gammaproteobacteria; Enterobacterales; Enterobacteriaceae; Salmonella; Salmonella enterica; Salmonella enterica subsp. enterica MTGDNTLITSHGINRRDFMKLCAALAATMGLSSKAAAEMAESVSNPQRPPVIWIGAQECT WP_000145429.1 hydrogenase 2 small subunit [Salmonella enterica] Length: 372\nNumber of predicted TMHs: 1\nExp number of AAs in TMHs: 24.51309\nExp number, first 60 AAs: 0.2682\nTotal prob of N-in: 0.14402\noutside 1 331\nTMhelix 332 354\ninside 355 372

21660 GCF_001242365.1_10425_1_60 Salmonella enterica subsp. enterica serovar Typhi Proteobacteria; Gammaproteobacteria; Enterobacterales; Enterobacteriaceae; Salmonella; Salmonella enterica; Salmonella enterica subsp. enterica MTGDNTLITSHGINRRDFMKLCAALAATMGLSSKAAAEMAESVSNPQRPPVIWIGAQECT WP_000145429.1 hydrogenase 2 small subunit [Salmonella enterica] Length: 372\nNumber of predicted TMHs: 1\nExp number of AAs in TMHs: 24.51309\nExp number, first 60 AAs: 0.2682\nTotal prob of N-in: 0.14402\noutside 1 331\nTMhelix 332 354\ninside 355 372

21661 GCF_001242225.1_10425_1_96 Salmonella enterica subsp. enterica serovar Typhi Proteobacteria; Gammaproteobacteria; Enterobacterales; Enterobacteriaceae; Salmonella; Salmonella enterica; Salmonella enterica subsp. enterica MTGDNTLITSHGINRRDFMKLCAALAATMGLSSKAAAEMAESVSNPQRPPVIWIGAQECT WP_000145429.1 hydrogenase 2 small subunit [Salmonella enterica] Length: 372\nNumber of predicted TMHs: 1\nExp number of AAs in TMHs: 24.51309\nExp number, first 60 AAs: 0.2682\nTotal prob of N-in: 0.14402\noutside 1 331\nTMhelix 332 354\ninside 355 372

21662 GCF_001241545.1_10425_1_68 Salmonella enterica subsp. enterica serovar Typhi Proteobacteria; Gammaproteobacteria; Enterobacterales; Enterobacteriaceae; Salmonella; Salmonella enterica; Salmonella enterica subsp. enterica MTGDNTLITSHGINRRDFMKLCAALAATMGLSSKAAAEMAESVSNPQRPPVIWIGAQECT WP_000145429.1 hydrogenase 2 small subunit [Salmonella enterica] Length: 372\nNumber of predicted TMHs: 1\nExp number of AAs in TMHs: 24.51309\nExp number, first 60 AAs: 0.2682\nTotal prob of N-in: 0.14402\noutside 1 331\nTMhelix 332 354\ninside 355 372

21663 GCF_001241345.1_10425_1_72 Salmonella enterica subsp. enterica serovar Typhi Proteobacteria; Gammaproteobacteria; Enterobacterales; Enterobacteriaceae; Salmonella; Salmonella enterica; Salmonella enterica subsp. enterica MTGDNTLITSHGINRRDFMKLCAALAATMGLSSKAAAEMAESVSNPQRPPVIWIGAQECT WP_000145429.1 hydrogenase 2 small subunit [Salmonella enterica] Length: 372\nNumber of predicted TMHs: 1\nExp number of AAs in TMHs: 24.51309\nExp number, first 60 AAs: 0.2682\nTotal prob of N-in: 0.14402\noutside 1 331\nTMhelix 332 354\ninside 355 372

21664	GCF_001240535.1_10425_1_16	Salmonella enterica subsp. enterica serovar Typhi	Proteobacteria; Gammaproteobacteria; Enterobacterales; Enterobacteriaceae; Salmonella; Salmonella enterica; Salmonella enterica subsp. enterica	MTGDNTLITSHGINRRDFMKLCAALAATMGLSSKAAAEMAESVSNPQRPPVIWIGAQECT	WP_000145429.1	hydrogenase 2 small subunit [Salmonella enterica]	Length: 372\nNumber of predicted TMHs:  1\nExp number of AAs in TMHs: 24.51309\nExp number, first 60 AAs:  0.2682\nTotal prob of N-in: 0.14402\noutside	1	331\nTMhelix	332	354\ninside	355	372

21665	GCF_001240435.1_10425_1_4	Salmonella enterica subsp. enterica serovar Typhi	Proteobacteria; Gammaproteobacteria; Enterobacterales; Enterobacteriaceae; Salmonella; Salmonella enterica; Salmonella enterica subsp. enterica	MTGDNTLITSHGINRRDFMKLCAALAATMGLSSKAAAEMAESVSNPQRPPVIWIGAQECT	WP_000145429.1	hydrogenase 2 small subunit [Salmonella enterica]	Length: 372\nNumber of predicted TMHs:  1\nExp number of AAs in TMHs: 24.51309\nExp number, first 60 AAs:  0.2682\nTotal prob of N-in: 0.14402\noutside	1	331\nTMhelix	332	354\ninside	355	372

21666	GCF_001240245.1_10425_1_20	Salmonella enterica subsp. enterica serovar Typhi	Proteobacteria; Gammaproteobacteria; Enterobacterales; Enterobacteriaceae; Salmonella; Salmonella enterica; Salmonella enterica subsp. enterica	MTGDNTLITSHGINRRDFMKLCAALAATMGLSSKAAAEMAESVSNPQRPPVIWIGAQECT	WP_000145429.1	hydrogenase 2 small subunit [Salmonella enterica]	Length: 372\nNumber of predicted TMHs:  1\nExp number of AAs in TMHs: 24.51309\nExp number, first 60 AAs:  0.2682\nTotal prob of N-in: 0.14402\noutside	1	331\nTMhelix	332	354\ninside	355	372

21667	GCF_001240105.1_10425_1_94	Salmonella enterica subsp. enterica serovar Typhi	Proteobacteria; Gammaproteobacteria; Enterobacterales; Enterobacteriaceae; Salmonella; Salmonella enterica; Salmonella enterica subsp. enterica	MTGDNTLITSHGINRRDFMKLCAALAATMGLSSKAAAEMAESVSNPQRPPVIWIGAQECT	WP_000145429.1	hydrogenase 2 small subunit [Salmonella enterica]	Length: 372\nNumber of predicted TMHs:  1\nExp number of AAs in TMHs: 24.51309\nExp number, first 60 AAs:  0.2682\nTotal prob of N-in: 0.14402\noutside	1	331\nTMhelix	332	354\ninside	355	372

21668	GCF_001240065.1_10425_1_55	Salmonella enterica subsp. enterica serovar Typhi	Proteobacteria; Gammaproteobacteria; Enterobacterales; Enterobacteriaceae; Salmonella; Salmonella enterica; Salmonella enterica subsp. enterica	MTGDNTLITSHGINRRDFMKLCAALAATMGLSSKAAAEMAESVSNPQRPPVIWIGAQECT	WP_000145429.1	hydrogenase 2 small subunit [Salmonella enterica]	Length: 372\nNumber of predicted TMHs:  1\nExp number of AAs in TMHs: 24.51309\nExp number, first 60 AAs:  0.2682\nTotal prob of N-in: 0.14402\noutside	1	331\nTMhelix	332	354\ninside	355	372

21669	GCF_001240025.1_10425_1_17	Salmonella enterica subsp. enterica serovar Typhi	Proteobacteria; Gammaproteobacteria; Enterobacterales; Enterobacteriaceae; Salmonella; Salmonella enterica; Salmonella enterica subsp. enterica	MTGDNTLITSHGINRRDFMKLCAALAATMGLSSKAAAEMAESVSNPQRPPVIWIGAQECT	WP_000145429.1	hydrogenase 2 small subunit [Salmonella enterica]	Length: 372\nNumber of predicted TMHs:  1\nExp number of AAs in TMHs: 24.51309\nExp number, first 60 AAs:  0.2682\nTotal prob of N-in: 0.14402\noutside	1	331\nTMhelix	332	354\ninside	355	372

21670	GCF_001239865.1_10425_1_45	Salmonella enterica subsp. enterica serovar Typhi	Proteobacteria; Gammaproteobacteria; Enterobacterales; Enterobacteriaceae; Salmonella; Salmonella enterica; Salmonella enterica subsp. enterica	MTGDNTLITSHGINRRDFMKLCAALAATMGLSSKAAAEMAESVSNPQRPPVIWIGAQECT	WP_000145429.1	hydrogenase 2 small subunit [Salmonella enterica]	Length: 372\nNumber of predicted TMHs:  1\nExp number of AAs in TMHs: 24.51309\nExp number, first 60 AAs:  0.2682\nTotal prob of N-in: 0.14402\noutside	1	331\nTMhelix	332	354\ninside	355	372

21671	GCF_001172465.1_9870_8_70	Salmonella enterica subsp. enterica serovar Typhi	Proteobacteria; Gammaproteobacteria; Enterobacterales; Enterobacteriaceae; Salmonella; Salmonella enterica; Salmonella enterica subsp. enterica	MTGDNTLITSHGINRRDFMKLCAALAATMGLSSKAAAEMAESVSNPQRPPVIWIGAQECT	WP_000145429.1	hydrogenase 2 small subunit [Salmonella enterica]	Length: 372\nNumber of predicted TMHs:  1\nExp number of AAs in TMHs: 24.51309\nExp number, first 60 AAs:  0.2682\nTotal prob of N-in: 0.14402\noutside	1	331\nTMhelix	332	354\ninside	355	372

21672	GCF_001172125.1_10593_2_9	Salmonella enterica subsp. enterica serovar Typhi	Proteobacteria; Gammaproteobacteria; Enterobacterales; Enterobacteriaceae; Salmonella; Salmonella enterica; Salmonella enterica subsp. enterica	MTGDNTLITSHGINRRDFMKLCAALAATMGLSSKAAAEMAESVSNPQRPPVIWIGAQECT	WP_000145429.1	hydrogenase 2 small subunit [Salmonella enterica]	Length: 372\nNumber of

predicted TMHs: 1\nExp number of AAs in TMHs: 24.51309\nExp number, first 60 AAs: 0.2682\nTotal prob of N-in: 0.14402\noutside 1 331\nTMhelix 332 354\ninside 355 372

21673 GCF_001171345.1_10562_2_30 Salmonella enterica subsp. enterica serovar Typhi Proteobacteria; Gammaproteobacteria; Enterobacterales; Enterobacteriaceae; Salmonella; Salmonella enterica; Salmonella enterica subsp. enterica MTGDNTLITSHGINRRDFMKLCAALAATMGLSSKAAAEMAESVSNPQRPPVIWIGAQECT WP_000145429.1 hydrogenase 2 small subunit [Salmonella enterica] Length: 372\nNumber of predicted TMHs: 1\nExp number of AAs in TMHs: 24.51309\nExp number, first 60 AAs: 0.2682\nTotal prob of N-in: 0.14402\noutside 1 331\nTMhelix 332 354\ninside 355 372

21674 GCF_001171165.1_9953_5_55 Salmonella enterica subsp. enterica serovar Typhi Proteobacteria; Gammaproteobacteria; Enterobacterales; Enterobacteriaceae; Salmonella; Salmonella enterica; Salmonella enterica subsp. enterica MTGDNTLITSHGINRRDFMKLCAALAATMGLSSKAAAEMAESVSNPQRPPVIWIGAQECT WP_000145429.1 hydrogenase 2 small subunit [Salmonella enterica] Length: 372\nNumber of predicted TMHs: 1\nExp number of AAs in TMHs: 24.51309\nExp number, first 60 AAs: 0.2682\nTotal prob of N-in: 0.14402\noutside 1 331\nTMhelix 332 354\ninside 355 372

21675 GCF_001169625.1_7468_7_93 Salmonella enterica subsp. enterica serovar Typhi Proteobacteria; Gammaproteobacteria; Enterobacterales; Enterobacteriaceae; Salmonella; Salmonella enterica; Salmonella enterica subsp. enterica MTGDNTLITSHGINRRDFMKLCAALAATMGLSSKAATEMAESVSNPQRPPVIWIGAQECT WP_001782304.1 hydrogenase 2 small subunit [Salmonella enterica] Length: 372\nNumber of predicted TMHs: 1\nExp number of AAs in TMHs: 24.41173\nExp number, first 60 AAs: 0.16124\nTotal prob of N-in: 0.13947\noutside 1 331\nTMhelix 332 354\ninside 355 372

21676 GCF_001167305.1_8490_5_5Salmonella enterica subsp. enterica serovar Typhi Proteobacteria; Gammaproteobacteria; Enterobacterales; Enterobacteriaceae; Salmonella; Salmonella enterica; Salmonella enterica subsp. enterica MTGDNTLITSHGINRRDFMKLCAALAATMGLSSKAAAEMAESVSNPQRPPVIWIGAQECT WP_000145429.1 hydrogenase 2 small subunit [Salmonella enterica] Length: 372\nNumber of predicted TMHs: 1\nExp number of AAs in TMHs: 24.51309\nExp number, first 60 AAs: 0.2682\nTotal prob of N-in: 0.14402\noutside 1 331\nTMhelix 332 354\ninside 355 372

21677 GCF_001167105.1_8447_8_21 Salmonella enterica subsp. enterica serovar Typhi Proteobacteria; Gammaproteobacteria; Enterobacterales; Enterobacteriaceae; Salmonella; Salmonella enterica; Salmonella enterica subsp. enterica MTGDNTLITSHGINRRDFMKLCAALAATMGLSSKAAAEMAESVSNPQRPPVIWIGAQECT WP_000145429.1 hydrogenase 2 small subunit [Salmonella enterica] Length: 372\nNumber of predicted TMHs: 1\nExp number of AAs in TMHs: 24.51309\nExp number, first 60 AAs: 0.2682\nTotal prob of N-in: 0.14402\noutside 1 331\nTMhelix 332 354\ninside 355 372

21678 GCF_001166905.1_9953_5_54 Salmonella enterica subsp. enterica serovar Typhi Proteobacteria; Gammaproteobacteria; Enterobacterales; Enterobacteriaceae; Salmonella; Salmonella enterica; Salmonella enterica subsp. enterica MTGDNTLITSHGINRRDFMKLCAALAATMGLSSKAAAEMAESVSNPQRPPVIWIGAQECT WP_000145429.1 hydrogenase 2 small subunit [Salmonella enterica] Length: 372\nNumber of predicted TMHs: 1\nExp number of AAs in TMHs: 24.51309\nExp number, first 60 AAs: 0.2682\nTotal prob of N-in: 0.14402\noutside 1 331\nTMhelix 332 354\ninside 355 372

21679 GCF_001166585.1_9475_6_36 Salmonella enterica subsp. enterica serovar Typhi Proteobacteria; Gammaproteobacteria; Enterobacterales; Enterobacteriaceae; Salmonella; Salmonella enterica; Salmonella enterica subsp. enterica MTGDNTLITSHGINRRDFMKLCAALAATMGLSSKAAAEMAESVSNPQRPPVIWIGAQECT WP_000145429.1 hydrogenase 2 small subunit [Salmonella enterica] Length: 372\nNumber of predicted TMHs: 1\nExp number of AAs in TMHs: 24.51309\nExp number, first 60 AAs: 0.2682\nTotal prob of N-in: 0.14402\noutside 1 331\nTMhelix 332 354\ninside 355 372

21680 GCF_001166545.1_10541_2_20 Salmonella enterica subsp. enterica serovar Typhi Proteobacteria; Gammaproteobacteria; Enterobacterales; Enterobacteriaceae; Salmonella; Salmonella enterica; Salmonella enterica subsp. enterica MTGDNTLITSHGINRRDFMKLCAALAATMGLSSKAAAEMAESVSNPQRPPVIWIGAQECT WP_000145429.1 hydrogenase 2 small subunit [Salmonella enterica] Length: 372\nNumber of predicted TMHs: 1\nExp number of AAs in TMHs: 24.51309\nExp number, first 60 AAs: 0.2682\nTotal prob of N-in: 0.14402\noutside 1 331\nTMhelix 332 354\ninside 355 372

21681 GCF_001165145.1_10562_2_24 Salmonella enterica subsp. enterica serovar Typhi Proteobacteria; Gammaproteobacteria; Enterobacterales; Enterobacteriaceae; Salmonella; Salmonella enterica;

Salmonella enterica subsp. enterica MTGDNTLITSHGINRRDFMKLCAALAATMGLSSKAATEMAESVSNPQRPPVIWIGAQECT WP_001782304.1 hydrogenase 2 small subunit [Salmonella enterica] Length: 372\nNumber of predicted TMHs: 1\nExp number of AAs in TMHs: 24.41173\nExp number, first 60 AAs: 0.16124\nTotal prob of N-in: 0.13947\noutside 1 331\nTMhelix 332 354\ninside 355 372

21682 GCF_001166345.1_9475_6_12 Salmonella enterica subsp. enterica serovar Typhi Proteobacteria; Gammaproteobacteria; Enterobacterales; Enterobacteriaceae; Salmonella; Salmonella enterica; Salmonella enterica subsp. enterica MTGDNTLITSHGINRRDFMKLCAALAATMGLSSKAAAEMAESVSNPQRPPVIWIGAQECT WP_000145429.1 hydrogenase 2 small subunit [Salmonella enterica] Length: 372\nNumber of predicted TMHs: 1\nExp number of AAs in TMHs: 24.51309\nExp number, first 60 AAs: 0.2682\nTotal prob of N-in: 0.14402\noutside 1 331\nTMhelix 332 354\ninside 355 372

21683 GCF_001160865.1_9953_5_61 Salmonella enterica subsp. enterica serovar Typhi Proteobacteria; Gammaproteobacteria; Enterobacterales; Enterobacteriaceae; Salmonella; Salmonella enterica; Salmonella enterica subsp. enterica MTGDNTLITSHGINRRDFMKLCAALAATMGLSSKAAAEMAESVSNPQRPPVIWIGAQECT WP_000145429.1 hydrogenase 2 small subunit [Salmonella enterica] Length: 372\nNumber of predicted TMHs: 1\nExp number of AAs in TMHs: 24.51309\nExp number, first 60 AAs: 0.2682\nTotal prob of N-in: 0.14402\noutside 1 331\nTMhelix 332 354\ninside 355 372

21684 GCF_001160665.1_8490_6_50 Salmonella enterica subsp. enterica serovar Typhi Proteobacteria; Gammaproteobacteria; Enterobacterales; Enterobacteriaceae; Salmonella; Salmonella enterica; Salmonella enterica subsp. enterica MTGDNTLITSHGINRRDFMKLCAALAATMGLSSKAAAEMAESVSNPQRPPVIWIGAQECT WP_000145429.1 hydrogenase 2 small subunit [Salmonella enterica] Length: 372\nNumber of predicted TMHs: 1\nExp number of AAs in TMHs: 24.51309\nExp number, first 60 AAs: 0.2682\nTotal prob of N-in: 0.14402\noutside 1 331\nTMhelix 332 354\ninside 355 372

21685 GCF_001160385.1_10349_1_40 Salmonella enterica subsp. enterica serovar Typhi Proteobacteria; Gammaproteobacteria; Enterobacterales; Enterobacteriaceae; Salmonella; Salmonella enterica; Salmonella enterica subsp. enterica MTGDNTLITSHGINRRDFMKLCAALAATMGLSSKAAAEMAESVSNPQRPPVIWIGAQECT WP_000145429.1 hydrogenase 2 small subunit [Salmonella enterica] Length: 372\nNumber of predicted TMHs: 1\nExp number of AAs in TMHs: 24.51309\nExp number, first 60 AAs: 0.2682\nTotal prob of N-in: 0.14402\noutside 1 331\nTMhelix 332 354\ninside 355 372

21686 GCF_001160285.1_8447_8_6Salmonella enterica subsp. enterica serovar Typhi Proteobacteria; Gammaproteobacteria; Enterobacterales; Enterobacteriaceae; Salmonella; Salmonella enterica; Salmonella enterica subsp. enterica MTGDNTLITSHGINRRDFMKLCAALAATMGLSSKAAAEMAESVSNPQRPPVIWIGAQECT WP_000145429.1 hydrogenase 2 small subunit [Salmonella enterica] Length: 372\nNumber of predicted TMHs: 1\nExp number of AAs in TMHs: 24.51309\nExp number, first 60 AAs: 0.2682\nTotal prob of N-in: 0.14402\noutside 1 331\nTMhelix 332 354\ninside 355 372

21687 GCF_001160125.1_10493_1_30 Salmonella enterica subsp. enterica serovar Typhi Proteobacteria; Gammaproteobacteria; Enterobacterales; Enterobacteriaceae; Salmonella; Salmonella enterica; Salmonella enterica subsp. enterica MTGDNTLITSHGINRRDFMKLCAALAATMGLSSKAAAEMAESVSNPQRPPVIWIGAQECT WP_000145429.1 hydrogenase 2 small subunit [Salmonella enterica] Length: 372\nNumber of predicted TMHs: 1\nExp number of AAs in TMHs: 24.51309\nExp number, first 60 AAs: 0.2682\nTotal prob of N-in: 0.14402\noutside 1 331\nTMhelix 332 354\ninside 355 372

21688 GCF_001159985.1_8616_4_46 Salmonella enterica subsp. enterica serovar Typhi Proteobacteria; Gammaproteobacteria; Enterobacterales; Enterobacteriaceae; Salmonella; Salmonella enterica; Salmonella enterica subsp. enterica MTGDNTLITSHGINRRDFMKLCAALAATMGLSSKAAAEMAESVSNPQRPPVIWIGAQECT WP_000145429.1 hydrogenase 2 small subunit [Salmonella enterica] Length: 372\nNumber of predicted TMHs: 1\nExp number of AAs in TMHs: 24.51309\nExp number, first 60 AAs: 0.2682\nTotal prob of N-in: 0.14402\noutside 1 331\nTMhelix 332 354\ninside 355 372

21689 GCF_001157685.1_10593_2_54 Salmonella enterica subsp. enterica serovar Typhi Proteobacteria; Gammaproteobacteria; Enterobacterales; Enterobacteriaceae; Salmonella; Salmonella enterica; Salmonella enterica subsp. enterica MTGDNTLITSHGINRRDFMKLCAALAATMGLSSKAAAEMAESVSNPQRPPVIWIGAQECT WP_000145429.1 hydrogenase 2 small subunit [Salmonella enterica] Length: 372\nNumber of predicted TMHs: 1\nExp number of AAs in TMHs: 24.51309\nExp number, first 60 AAs: 0.2682\nTotal prob of N-in: 0.14402\noutside 1 331\nTMhelix 332 354\ninside 355 372

21690 GCF_001157185.1_10209_5_13 Salmonella enterica subsp. enterica serovar Typhi Proteobacteria; Gammaproteobacteria; Enterobacterales; Enterobacteriaceae; Salmonella; Salmonella enterica; Salmonella enterica subsp. enterica MTGDNTLITSHGINRRDFMKLCAALAATMGLSSKAAAEMAESVSNPQRPPVIWIGAQECT WP_000145429.1 hydrogenase 2 small subunit [Salmonella enterica] Length: 372\nNumber of predicted TMHs: 1\nExp number of AAs in TMHs: 24.51309\nExp number, first 60 AAs: 0.2682\nTotal prob of N-in: 0.14402\noutside 1 331\nTMhelix 332 354\ninside 355 372

21691 GCF_001157105.1_9870_8_78 Salmonella enterica subsp. enterica serovar Typhi Proteobacteria; Gammaproteobacteria; Enterobacterales; Enterobacteriaceae; Salmonella; Salmonella enterica; Salmonella enterica subsp. enterica MTGDNTLITSHGINRRDFMKLCAALAATMGLSSKAAAEMAESVSNPQRPPVIWIGAQECT WP_000145429.1 hydrogenase 2 small subunit [Salmonella enterica] Length: 372\nNumber of predicted TMHs: 1\nExp number of AAs in TMHs: 24.51309\nExp number, first 60 AAs: 0.2682\nTotal prob of N-in: 0.14402\noutside 1 331\nTMhelix 332 354\ninside 355 372

21692 GCF_001156625.1_7468_7_49 Salmonella enterica subsp. enterica serovar Typhi Proteobacteria; Gammaproteobacteria; Enterobacterales; Enterobacteriaceae; Salmonella; Salmonella enterica; Salmonella enterica subsp. enterica MTGDNTLITSHGINRRDFMKLCAALAATMGLSSKAAAEMAESVSNPQRPPVIWIGAQECT WP_000145429.1 hydrogenase 2 small subunit [Salmonella enterica] Length: 372\nNumber of predicted TMHs: 1\nExp number of AAs in TMHs: 24.51309\nExp number, first 60 AAs: 0.2682\nTotal prob of N-in: 0.14402\noutside 1 331\nTMhelix 332 354\ninside 355 372

21693 GCF_001588355.1_CFSAN033899_01.0Salmonella enterica subsp. enterica serovar Typhimurium Proteobacteria; Gammaproteobacteria; Enterobacterales; Enterobacteriaceae; Salmonella; Salmonella enterica; Salmonella enterica subsp. enterica MNNEETFYQAMRRKGVTRRSFLKFCSLAATSLGLGAGMTPKIAWALENKPRIPVVWIHGL WP_001058312.1 hydrogenase [Salmonella enterica] Length: 372\nNumber of predicted TMHs: 2\nExp number of AAs in TMHs: 36.51738\nExp number, first 60 AAs: 19.72849\nTotal prob of N-in: 0.96170\nPOSSIBLE N-term signal sequence\ninside 1 19\nTMhelix 20 39\noutside 40 325\nTMhelix 326 348\ninside 349 372

21694 GCF_000784235.1_ASM78423v1 Salmonella enterica subsp. enterica serovar Typhimurium Proteobacteria; Gammaproteobacteria; Enterobacterales; Enterobacteriaceae; Salmonella; Salmonella enterica; Salmonella enterica subsp. enterica MNNEETFYQAMRRKGVTRRSFLKFCSLAATSLGLGAGMTPKIAWALENKPRIPVVWIHGL WP_001058312.1 hydrogenase [Salmonella enterica] Length: 372\nNumber of predicted TMHs: 2\nExp number of AAs in TMHs: 36.51738\nExp number, first 60 AAs: 19.72849\nTotal prob of N-in: 0.96170\nPOSSIBLE N-term signal sequence\ninside 1 19\nTMhelix 20 39\noutside 40 325\nTMhelix 326 348\ninside 349 372

21695 GCF_000784295.1_ASM78429v1 Salmonella enterica subsp. enterica serovar Typhimurium Proteobacteria; Gammaproteobacteria; Enterobacterales; Enterobacteriaceae; Salmonella; Salmonella enterica; Salmonella enterica subsp. enterica MNNEETFYQAMRRKGVTRRSFLKFCSLAATSLGLGAGMTPKIAWALENKPRIPVVWIHGL WP_001058312.1 hydrogenase [Salmonella enterica] Length: 372\nNumber of predicted TMHs: 2\nExp number of AAs in TMHs: 36.51738\nExp number, first 60 AAs: 19.72849\nTotal prob of N-in: 0.96170\nPOSSIBLE N-term signal sequence\ninside 1 19\nTMhelix 20 39\noutside 40 325\nTMhelix 326 348\ninside 349 372

21696 GCF_000784215.1_ASM78421v1 Salmonella enterica subsp. enterica serovar Typhimurium Proteobacteria; Gammaproteobacteria; Enterobacterales; Enterobacteriaceae; Salmonella; Salmonella enterica; Salmonella enterica subsp. enterica MNNEETFYQAMRRKGVTRRSFLKFCSLAATSLGLGAGMTPKIAWALENKPRIPVVWIHGL WP_001058312.1 hydrogenase [Salmonella enterica] Length: 372\nNumber of predicted TMHs: 2\nExp number of AAs in TMHs: 36.51738\nExp number, first 60 AAs: 19.72849\nTotal prob of N-in: 0.96170\nPOSSIBLE N-term signal sequence\ninside 1 19\nTMhelix 20 39\noutside 40 325\nTMhelix 326 348\ninside 349 372

21697 GCF_000973845.1_SALF-276-1.id8_v1.0 Salmonella enterica subsp. enterica serovar Typhimurium Proteobacteria; Gammaproteobacteria; Enterobacterales; Enterobacteriaceae; Salmonella; Salmonella enterica; Salmonella enterica subsp. enterica MTGDNTLITSHGINRRDFMKLCAALAATMGLSSKAAAEMAESVSNPQRPPVIWIGAQECT WP_000145429.1 hydrogenase 2 small subunit [Salmonella enterica] Length: 372\nNumber of predicted TMHs: 1\nExp number of AAs in TMHs: 24.51309\nExp number, first 60 AAs: 0.2682\nTotal prob of N-in: 0.14402\noutside 1 331\nTMhelix 332 354\ninside 355 372

21698 GCF_000974125.1_ABBSB1116-2.id10_v1.0 Salmonella enterica subsp. enterica serovar Typhimurium Proteobacteria; Gammaproteobacteria; Enterobacterales; Enterobacteriaceae; Salmonella; Salmonella enterica; Salmonella enterica subsp. enterica MTGDNTLITSHGINRRDFMKLCAALAATMGLSSKAAAEMAESVSNPQRPPVIWIGAQECT WP_000145429.1 hydrogenase 2 small subunit [Salmonella enterica] Length: 372\nNumber of

predicted TMHs:  1\nExp number of AAs in TMHs: 24.51309\nExp number, first 60 AAs:  0.2682\nTotal prob of N-in: 0.14402\noutside   1   331\nTMhelix   332   354\ninside   355   372

21699 GCF_000974215.1_SALF-297-3.id2_v1.0 Salmonella enterica subsp. enterica serovar Typhimurium Proteobacteria; Gammaproteobacteria; Enterobacterales; Enterobacteriaceae; Salmonella; Salmonella enterica; Salmonella enterica subsp. enterica MTGDNTLITSHGINRRDFMKLCAALAATMGLSSKAAAEMAESVSNPQRPPVIWIGAQECT WP_000145429.1 hydrogenase 2 small subunit [Salmonella enterica] Length: 372\nNumber of predicted TMHs:  1\nExp number of AAs in TMHs: 24.51309\nExp number, first 60 AAs:  0.2682\nTotal prob of N-in: 0.14402\noutside   1   331\nTMhelix   332   354\ninside   355   372

21700 GCF_000974125.1_ABBSB1116-2.id10_v1.0 Salmonella enterica subsp. enterica serovar Typhimurium Proteobacteria; Gammaproteobacteria; Enterobacterales; Enterobacteriaceae; Salmonella; Salmonella enterica; Salmonella enterica subsp. enterica MNNEETFYQAMRRKGVTRRSFLKFCSLAATSLGLGAGMTPKIAWALENKPRIPVVWIHGL WP_001058312.1 hydrogenase [Salmonella enterica] Length: 372\nNumber of predicted TMHs:  2\nExp number of AAs in TMHs: 36.51738\nExp number, first 60 AAs:  19.72849\nTotal prob of N-in:   0.96170\nPOSSIBLE N-term signal sequence\ninside   1   19\nTMhelix   20   39\noutside   40   325\nTMhelix   326   348\ninside   349   372

21701 GCF_000974215.1_SALF-297-3.id2_v1.0 Salmonella enterica subsp. enterica serovar Typhimurium Proteobacteria; Gammaproteobacteria; Enterobacterales; Enterobacteriaceae; Salmonella; Salmonella enterica; Salmonella enterica subsp. enterica MNNEETFYQAMRRKGVTRRSFLKFCSLAATSLGLGAGMTPKIAWALENKPRIPVVWIHGL WP_001058312.1 hydrogenase [Salmonella enterica] Length: 372\nNumber of predicted TMHs:  2\nExp number of AAs in TMHs: 36.51738\nExp number, first 60 AAs:  19.72849\nTotal prob of N-in:   0.96170\nPOSSIBLE N-term signal sequence\ninside   1   19\nTMhelix   20   39\noutside   40   325\nTMhelix   326   348\ninside   349   372

21702 GCF_001295705.1_ASM129570v1 Salmonella enterica subsp. enterica serovar Typhimurium Proteobacteria; Gammaproteobacteria; Enterobacterales; Enterobacteriaceae; Salmonella; Salmonella enterica; Salmonella enterica subsp. enterica MTGDNTLITSHGINRRDFMKLCAALAATMGLSSKAAAEMAESVSNPQRPPVIWIGAQECT WP_000145429.1 hydrogenase 2 small subunit [Salmonella enterica] Length: 372\nNumber of predicted TMHs:  1\nExp number of AAs in TMHs: 24.51309\nExp number, first 60 AAs:  0.2682\nTotal prob of N-in: 0.14402\noutside   1   331\nTMhelix   332   354\ninside   355   372

21703 GCF_001240925.1_Salmonella_enterica_CVM_N43829-R_v1.0 Salmonella enterica subsp. enterica serovar Typhimurium Proteobacteria; Gammaproteobacteria; Enterobacterales; Enterobacteriaceae; Salmonella; Salmonella enterica; Salmonella enterica subsp. enterica MTGDNTLITSHGINRRDFMKLCAALAATMGLSSKAAAEMAESVSNPQRPPVIWIGAQECT WP_000145429.1 hydrogenase 2 small subunit [Salmonella enterica] Length: 372\nNumber of predicted TMHs:  1\nExp number of AAs in TMHs: 24.51309\nExp number, first 60 AAs:  0.2682\nTotal prob of N-in:   0.14402\noutside   1   331\nTMhelix   332   354\ninside   355   372

21704 GCF_001245645.1_Salmonella_enterica_CVM_N50438_v1.0 Salmonella enterica subsp. enterica serovar Typhimurium Proteobacteria; Gammaproteobacteria; Enterobacterales; Enterobacteriaceae; Salmonella; Salmonella enterica; Salmonella enterica subsp. enterica MTGDNTLITSHGINRRDFMKLCAALAATMGLSSKAAAEMAESVSNPQRPPVIWIGAQECT WP_000145429.1 hydrogenase 2 small subunit [Salmonella enterica] Length: 372\nNumber of predicted TMHs:  1\nExp number of AAs in TMHs: 24.51309\nExp number, first 60 AAs:  0.2682\nTotal prob of N-in:   0.14402\noutside   1   331\nTMhelix   332   354\ninside   355   372

21705 GCF_001245645.1_Salmonella_enterica_CVM_N50438_v1.0 Salmonella enterica subsp. enterica serovar Typhimurium Proteobacteria; Gammaproteobacteria; Enterobacterales; Enterobacteriaceae; Salmonella; Salmonella enterica; Salmonella enterica subsp. enterica MNNEETFYQAMRRKGVTRRSFLKFCSLAATSLGLGAGMTPKIAWALENKPRIPVVWIHGL WP_001058312.1 hydrogenase [Salmonella enterica] Length: 372\nNumber of predicted TMHs:  2\nExp number of AAs in TMHs: 36.51738\nExp number, first 60 AAs:  19.72849\nTotal prob of N-in:   0.96170\nPOSSIBLE N-term signal sequence\ninside   1   19\nTMhelix   20   39\noutside   40   325\nTMhelix   326   348\ninside   349   372

21706 GCF_001246225.1_Salmonella_enterica_CVM_N51254_v1.0 Salmonella enterica subsp. enterica serovar Typhimurium Proteobacteria; Gammaproteobacteria; Enterobacterales; Enterobacteriaceae; Salmonella; Salmonella enterica; Salmonella enterica subsp. enterica MTGDNTLITSHGINRRDFMKLCAALAATMGLSSKAAAEMAESVSNPQRPPVIWIGAQECT WP_000145429.1 hydrogenase 2 small subunit [Salmonella enterica] Length: 372\nNumber of predicted TMHs:  1\nExp

number of AAs in TMHs: 24.51309\nExp number, first 60 AAs: 0.2682\nTotal prob of N-in: 0.14402\noutside 1 331\nTMhelix 332 354\ninside 355 372

21707 GCF_001272415.1_Salmonella_enterica_CVM_N51295_v1.0 Salmonella enterica subsp. enterica serovar Typhimurium Proteobacteria; Gammaproteobacteria; Enterobacterales; Enterobacteriaceae; Salmonella; Salmonella enterica; Salmonella enterica subsp. enterica MNNEETFYQAMRRKGVTRRSFLKFCSLAATSLGLGAGMTPKIAWALENKPRIPVVWIHGL WP_001058312.1 hydrogenase [Salmonella enterica] Length: 372\nNumber of predicted TMHs: 2\nExp number of AAs in TMHs: 36.51738\nExp number, first 60 AAs: 19.72849\nTotal prob of N-in: 0.96170\nPOSSIBLE N-term signal sequence\ninside 1 19\nTMhelix 20 39\noutside 40 325\nTMhelix 326 348\ninside 349 372

21708 GCF_001295385.1_ASM129538v1 Salmonella enterica subsp. enterica serovar Typhimurium Proteobacteria; Gammaproteobacteria; Enterobacterales; Enterobacteriaceae; Salmonella; Salmonella enterica; Salmonella enterica subsp. enterica MTGDNTLITSHGINRRDFMKLCAALAATMGLSSKAAAEMAESVSNPQRPPVIWIGAQECT WP_000145429.1 hydrogenase 2 small subunit [Salmonella enterica] Length: 372\nNumber of predicted TMHs: 1\nExp number of AAs in TMHs: 24.51309\nExp number, first 60 AAs: 0.2682\nTotal prob of N-in: 0.14402\noutside 1 331\nTMhelix 332 354\ninside 355 372

21709 GCF_001295385.1_ASM129538v1 Salmonella enterica subsp. enterica serovar Typhimurium Proteobacteria; Gammaproteobacteria; Enterobacterales; Enterobacteriaceae; Salmonella; Salmonella enterica; Salmonella enterica subsp. enterica MNNEETFYQAMRRKGVTRRSFLKFCSLAATSLGLGAGMTPKIAWALENKPRIPVVWIHGL WP_001058312.1 hydrogenase [Salmonella enterica] Length: 372\nNumber of predicted TMHs: 2\nExp number of AAs in TMHs: 36.51738\nExp number, first 60 AAs: 19.72849\nTotal prob of N-in: 0.96170\nPOSSIBLE N-term signal sequence\ninside 1 19\nTMhelix 20 39\noutside 40 325\nTMhelix 326 348\ninside 349 372

21710 GCF_001295595.1_ASM129559v1 Salmonella enterica subsp. enterica serovar Typhimurium Proteobacteria; Gammaproteobacteria; Enterobacterales; Enterobacteriaceae; Salmonella; Salmonella enterica; Salmonella enterica subsp. enterica MNNEETFYQAMRRKGVTRRSFLKFCSLAATSLGLGAGMTPKIAWALENKPRIPVVWIHGL WP_001058312.1 hydrogenase [Salmonella enterica] Length: 372\nNumber of predicted TMHs: 2\nExp number of AAs in TMHs: 36.51738\nExp number, first 60 AAs: 19.72849\nTotal prob of N-in: 0.96170\nPOSSIBLE N-term signal sequence\ninside 1 19\nTMhelix 20 39\noutside 40 325\nTMhelix 326 348\ninside 349 372

21711 GCF_001295705.1_ASM129570v1 Salmonella enterica subsp. enterica serovar Typhimurium Proteobacteria; Gammaproteobacteria; Enterobacterales; Enterobacteriaceae; Salmonella; Salmonella enterica; Salmonella enterica subsp. enterica MNNEETFYQAMRRKGVTRRSFLKFCSLAATSLGLGAGMTPKIAWALENKPRIPVVWIHGL WP_001058312.1 hydrogenase [Salmonella enterica] Length: 372\nNumber of predicted TMHs: 2\nExp number of AAs in TMHs: 36.51738\nExp number, first 60 AAs: 19.72849\nTotal prob of N-in: 0.96170\nPOSSIBLE N-term signal sequence\ninside 1 19\nTMhelix 20 39\noutside 40 325\nTMhelix 326 348\ninside 349 372

21712 GCF_001478125.1_Salmonella_enterica_CVM_N31395-SQ_v1.0 Salmonella enterica subsp. enterica serovar Typhimurium Proteobacteria; Gammaproteobacteria; Enterobacterales; Enterobacteriaceae; Salmonella; Salmonella enterica; Salmonella enterica subsp. enterica MTGDNTLITSHGINRRDFMKLCAALAATMGLSSKAAAEMAESVSNPQRPPVIWIGAQECT WP_000145429.1 hydrogenase 2 small subunit [Salmonella enterica] Length: 372\nNumber of predicted TMHs: 1\nExp number of AAs in TMHs: 24.51309\nExp number, first 60 AAs: 0.2682\nTotal prob of N-in: 0.14402\noutside 1 331\nTMhelix 332 354\ninside 355 372

21713 GCF_001480115.1_Salmonella_enterica_CVM_N38929-SQ_v1.0 Salmonella enterica subsp. enterica serovar Typhimurium Proteobacteria; Gammaproteobacteria; Enterobacterales; Enterobacteriaceae; Salmonella; Salmonella enterica; Salmonella enterica subsp. enterica MNNEETFYQAMRRKGVTRRSFLKFCSLAATSLGLGAGMTPKIAWALENKPRIPVVWIHGL WP_001058312.1 hydrogenase [Salmonella enterica] Length: 372\nNumber of predicted TMHs: 2\nExp number of AAs in TMHs: 36.51738\nExp number, first 60 AAs: 19.72849\nTotal prob of N-in: 0.96170\nPOSSIBLE N-term signal sequence\ninside 1 19\nTMhelix 20 39\noutside 40 325\nTMhelix 326 348\ninside 349 372

21714 GCF_001480485.1_Salmonella_enterica_CVM_N42450-SQ_v1.0 Salmonella enterica subsp. enterica serovar Typhimurium Proteobacteria; Gammaproteobacteria; Enterobacterales; Enterobacteriaceae; Salmonella; Salmonella enterica; Salmonella enterica subsp. enterica MTGDNTLITSHGINRRDFMKLCAALAATMGLSSKAAAEMAESVSNPQRPPVIWIGAQECT WP_000145429.1 hydrogenase 2 small subunit [Salmonella enterica] Length: 372\nNumber of predicted TMHs: 1\nExp

number of AAs in TMHs: 24.51309\nExp number, first 60 AAs:  0.2682\nTotal prob of N-in:   0.14402\noutside   1   331\nTMhelix   332   354\ninside   355   372

21715 GCF_001540845.1_SO4698_09 Salmonella enterica subsp. enterica serovar Typhimurium Proteobacteria; Gammaproteobacteria; Enterobacterales; Enterobacteriaceae; Salmonella; Salmonella enterica; Salmonella enterica subsp. enterica MTGDNTLITSHGINRRDFMKLCAALAATMGLSSKAAAEMAESVSNPQRPPVIWIGAQECT WP_000145429.1 hydrogenase 2 small subunit [Salmonella enterica] Length: 372\nNumber of predicted TMHs:  1\nExp number of AAs in TMHs: 24.51309\nExp number, first 60 AAs:  0.2682\nTotal prob of N-in:   0.14402\noutside   1   331\nTMhelix   332   354\ninside   355   372

21716 GCF_001587755.1_CFSAN033938_01.0Salmonella enterica subsp. enterica serovar Typhimurium Proteobacteria; Gammaproteobacteria; Enterobacterales; Enterobacteriaceae; Salmonella; Salmonella enterica; Salmonella enterica subsp. enterica MNNEETFYQAMRRKGVTRRSFLKFCSLAATSLGLGAGMTPKIAWALENKPRIPVVWIHGL WP_001058312.1 hydrogenase [Salmonella enterica] Length: 372\nNumber of predicted TMHs:  2\nExp number of AAs in TMHs: 36.51738\nExp number, first 60 AAs:  19.72849\nTotal prob of N-in:   0.96170\nPOSSIBLE N-term signal sequence\ninside   1   19\nTMhelix   20   39\noutside   40   325\nTMhelix   326   348\ninside   349   372

21717 GCF_001587965.1_CFSAN033925_01.0Salmonella enterica subsp. enterica serovar Typhimurium Proteobacteria; Gammaproteobacteria; Enterobacterales; Enterobacteriaceae; Salmonella; Salmonella enterica; Salmonella enterica subsp. enterica MTGDNTLITSHGINRRDFMKLCAALAATMGLSSKAAAEMAESVSNPQRPPVIWIGAQECT WP_000145429.1 hydrogenase 2 small subunit [Salmonella enterica] Length: 372\nNumber of predicted TMHs:  1\nExp number of AAs in TMHs: 24.51309\nExp number, first 60 AAs:  0.2682\nTotal prob of N-in:   0.14402\noutside   1   331\nTMhelix   332   354\ninside   355   372

21718 GCF_001588015.1_CFSAN033921_01.0Salmonella enterica subsp. enterica serovar Typhimurium Proteobacteria; Gammaproteobacteria; Enterobacterales; Enterobacteriaceae; Salmonella; Salmonella enterica; Salmonella enterica subsp. enterica MTGDNTLITSHGINRRDFMKLCAALAATMGLSSKAAAEMAESVSNPQRPPVIWIGAQECT WP_000145429.1 hydrogenase 2 small subunit [Salmonella enterica] Length: 372\nNumber of predicted TMHs:  1\nExp number of AAs in TMHs: 24.51309\nExp number, first 60 AAs:  0.2682\nTotal prob of N-in:   0.14402\noutside   1   331\nTMhelix   332   354\ninside   355   372

21719 GCF_001588045.1_CFSAN033918_01.0Salmonella enterica subsp. enterica serovar Typhimurium Proteobacteria; Gammaproteobacteria; Enterobacterales; Enterobacteriaceae; Salmonella; Salmonella enterica; Salmonella enterica subsp. enterica MTGDNTLITSHGINRRDFMKLCAALAATMGLSSKAAAEMAESVSNPQRPPVIWIGAQECT WP_000145429.1 hydrogenase 2 small subunit [Salmonella enterica] Length: 372\nNumber of predicted TMHs:  1\nExp number of AAs in TMHs: 24.51309\nExp number, first 60 AAs:  0.2682\nTotal prob of N-in:   0.14402\noutside   1   331\nTMhelix   332   354\ninside   355   372

21720 GCF_001588075.1_CFSAN033917_01.0Salmonella enterica subsp. enterica serovar Typhimurium Proteobacteria; Gammaproteobacteria; Enterobacterales; Enterobacteriaceae; Salmonella; Salmonella enterica; Salmonella enterica subsp. enterica MTGDNTLITSHGINRRDFMKLCAALAATMGLSSKAAAEMAESVSNPQRPPVIWIGAQECT WP_000145429.1 hydrogenase 2 small subunit [Salmonella enterica] Length: 372\nNumber of predicted TMHs:  1\nExp number of AAs in TMHs: 24.51309\nExp number, first 60 AAs:  0.2682\nTotal prob of N-in:   0.14402\noutside   1   331\nTMhelix   332   354\ninside   355   372

21721 GCF_001587965.1_CFSAN033925_01.0Salmonella enterica subsp. enterica serovar Typhimurium Proteobacteria; Gammaproteobacteria; Enterobacterales; Enterobacteriaceae; Salmonella; Salmonella enterica; Salmonella enterica subsp. enterica MNNEETFYQAMRRKGVTRRSFLKFCSLAATSLGLGAGMTPKIAWALENKPRIPVVWIHGL WP_001058312.1 hydrogenase [Salmonella enterica] Length: 372\nNumber of predicted TMHs:  2\nExp number of AAs in TMHs: 36.51738\nExp number, first 60 AAs:  19.72849\nTotal prob of N-in:   0.96170\nPOSSIBLE N-term signal sequence\ninside   1   19\nTMhelix   20   39\noutside   40   325\nTMhelix   326   348\ninside   349   372

21722 GCF_001588515.1_CFSAN033878_01.0Salmonella enterica subsp. enterica serovar Typhimurium Proteobacteria; Gammaproteobacteria; Enterobacterales; Enterobacteriaceae; Salmonella; Salmonella enterica; Salmonella enterica subsp. enterica MNNEETFYQAMRRKGVTRRSFLKFCSLAATSLGLGAGMTPKIAWALENKPRIPVVWIHGL WP_001058312.1 hydrogenase [Salmonella enterica] Length: 372\nNumber of predicted TMHs:  2\nExp number of AAs in TMHs: 36.51738\nExp number, first 60 AAs:  19.72849\nTotal prob of N-in:   0.96170\nPOSSIBLE N-term signal sequence\ninside   1   19\nTMhelix   20   39\noutside   40   325\nTMhelix   326   348\ninside   349   372

21723 GCF_001588545.1_CFSAN033868_01.0Salmonella enterica subsp. enterica serovar Typhimurium Proteobacteria; Gammaproteobacteria; Enterobacterales; Enterobacteriaceae; Salmonella; Salmonella enterica;

Salmonella enterica subsp. enterica MNNEETFYQAMRRKGVTRRSFLKFCSLAATSLGLGAGMTPKIAWALENKPRIPVVWIHGL WP_001058312.1 hydrogenase [Salmonella enterica] Length: 372\nNumber of predicted TMHs: 2\nExp number of AAs in TMHs: 36.51738\nExp number, first 60 AAs: 19.72849\nTotal prob of N-in: 0.96170\nPOSSIBLE N-term signal sequence\ninside 1 19\nTMhelix 20 39\noutside 40 325\nTMhelix 326 348\ninside 349 372

21724 GCF_001588275.1_CFSAN033904_01.0Salmonella enterica subsp. enterica serovar Typhimurium Proteobacteria; Gammaproteobacteria; Enterobacterales; Enterobacteriaceae; Salmonella; Salmonella enterica; Salmonella enterica subsp. enterica MNNEETFYQAMRRKGVTRRSFLKFCSLAATSLGLGAGMTPKIAWALENKPRIPVVWIHGL WP_001058312.1 hydrogenase [Salmonella enterica] Length: 372\nNumber of predicted TMHs: 2\nExp number of AAs in TMHs: 36.51738\nExp number, first 60 AAs: 19.72849\nTotal prob of N-in: 0.96170\nPOSSIBLE N-term signal sequence\ninside 1 19\nTMhelix 20 39\noutside 40 325\nTMhelix 326 348\ninside 349 372

21725 GCF_001588235.1_CFSAN033906_01.0Salmonella enterica subsp. enterica serovar Typhimurium Proteobacteria; Gammaproteobacteria; Enterobacterales; Enterobacteriaceae; Salmonella; Salmonella enterica; Salmonella enterica subsp. enterica MNNEETFYQAMRRKGVTRRSFLKFCSLAATSLGLGAGMTPKIAWALENKPRIPVVWIHGL WP_001058312.1 hydrogenase [Salmonella enterica] Length: 372\nNumber of predicted TMHs: 2\nExp number of AAs in TMHs: 36.51738\nExp number, first 60 AAs: 19.72849\nTotal prob of N-in: 0.96170\nPOSSIBLE N-term signal sequence\ninside 1 19\nTMhelix 20 39\noutside 40 325\nTMhelix 326 348\ninside 349 372

21726 GCF_001588315.1_CFSAN033902_01.0Salmonella enterica subsp. enterica serovar Typhimurium Proteobacteria; Gammaproteobacteria; Enterobacterales; Enterobacteriaceae; Salmonella; Salmonella enterica; Salmonella enterica subsp. enterica MNNEETFYQAMRRKGVTRRSFLKFCSLAATSLGLGAGMTPKIAWALENKPRIPVVWIHGL WP_001058312.1 hydrogenase [Salmonella enterica] Length: 372\nNumber of predicted TMHs: 2\nExp number of AAs in TMHs: 36.51738\nExp number, first 60 AAs: 19.72849\nTotal prob of N-in: 0.96170\nPOSSIBLE N-term signal sequence\ninside 1 19\nTMhelix 20 39\noutside 40 325\nTMhelix 326 348\ninside 349 372

21727 GCF_001594715.1_CFSAN033888_01.0Salmonella enterica subsp. enterica serovar Typhimurium Proteobacteria; Gammaproteobacteria; Enterobacterales; Enterobacteriaceae; Salmonella; Salmonella enterica; Salmonella enterica subsp. enterica MTGDNTLITSHGINRRDFMKLCAALAATMGLSSKAAAEMAESVSNPQRPPVIWIGAQECT WP_000145429.1 hydrogenase 2 small subunit [Salmonella enterica] Length: 372\nNumber of predicted TMHs: 1\nExp number of AAs in TMHs: 24.51309\nExp number, first 60 AAs: 0.2682\nTotal prob of N-in: 0.14402\noutside 1 331\nTMhelix 332 354\ninside 355 372

21728 GCF_001594765.1_CFSAN033883_01.0Salmonella enterica subsp. enterica serovar Typhimurium Proteobacteria; Gammaproteobacteria; Enterobacterales; Enterobacteriaceae; Salmonella; Salmonella enterica; Salmonella enterica subsp. enterica MTGDNTLITSHGINRRDFMKLCAALAATMGLSSKAAAEMAESVSNPQRPPVIWIGAQECT WP_000145429.1 hydrogenase 2 small subunit [Salmonella enterica] Length: 372\nNumber of predicted TMHs: 1\nExp number of AAs in TMHs: 24.51309\nExp number, first 60 AAs: 0.2682\nTotal prob of N-in: 0.14402\noutside 1 331\nTMhelix 332 354\ninside 355 372

21729 GCF_001594715.1_CFSAN033888_01.0Salmonella enterica subsp. enterica serovar Typhimurium Proteobacteria; Gammaproteobacteria; Enterobacterales; Enterobacteriaceae; Salmonella; Salmonella enterica; Salmonella enterica subsp. enterica MNNEETFYQAMRRKGVTRRSFLKFCSLAATSLGLGAGMTPKIAWALENKPRIPVVWIHGL WP_001058312.1 hydrogenase [Salmonella enterica] Length: 372\nNumber of predicted TMHs: 2\nExp number of AAs in TMHs: 36.51738\nExp number, first 60 AAs: 19.72849\nTotal prob of N-in: 0.96170\nPOSSIBLE N-term signal sequence\ninside 1 19\nTMhelix 20 39\noutside 40 325\nTMhelix 326 348\ninside 349 372

21730 GCF_001594765.1_CFSAN033883_01.0Salmonella enterica subsp. enterica serovar Typhimurium Proteobacteria; Gammaproteobacteria; Enterobacterales; Enterobacteriaceae; Salmonella; Salmonella enterica; Salmonella enterica subsp. enterica MNNEETFYQAMRRKGVTRRSFLKFCSLAATSLGLGAGMTPKIAWALENKPRIPVVWIHGL WP_001058312.1 hydrogenase [Salmonella enterica] Length: 372\nNumber of predicted TMHs: 2\nExp number of AAs in TMHs: 36.51738\nExp number, first 60 AAs: 19.72849\nTotal prob of N-in: 0.96170\nPOSSIBLE N-term signal sequence\ninside 1 19\nTMhelix 20 39\noutside 40 325\nTMhelix 326 348\ninside 349 372

21731 GCF_001594515.1_CFSAN033920_01.0Salmonella enterica subsp. enterica serovar Typhimurium Proteobacteria; Gammaproteobacteria; Enterobacterales; Enterobacteriaceae; Salmonella; Salmonella enterica; Salmonella enterica subsp. enterica MNNEETFYQAMRRKGVTRRSFLKFCSLAATSLGLGAGMTPKIAWALENKPRIPVVWIHGL WP_001058312.1 hydrogenase [Salmonella enterica] Length: 372\nNumber of predicted TMHs: 2\nExp number of AAs in TMHs: 36.51738\nExp number, first 60 AAs: 19.72849\nTotal prob of N-in: 0.96170\nPOSSIBLE N-term signal sequence\ninside 1 19\nTMhelix 20 39\noutside 40 325\nTMhelix 326 348\ninside 349 372

21732 GCF_001594595.1_CFSAN033901_01.0Salmonella enterica subsp. enterica serovar Typhimurium Proteobacteria; Gammaproteobacteria; Enterobacterales; Enterobacteriaceae; Salmonella; Salmonella enterica; Salmonella enterica subsp. enterica MNNEETFYQAMRRKGVTRRSFLKFCSLAATSLGLGAGMTPKIAWALENKPRIPVVWIHGL WP_001058312.1 hydrogenase [Salmonella enterica] Length: 372\nNumber of predicted TMHs: 2\nExp number of AAs in TMHs: 36.51738\nExp number, first 60 AAs: 19.72849\nTotal prob of N-in: 0.96170\nPOSSIBLE N-term signal sequence\ninside 1 19\nTMhelix 20 39\noutside 40 325\nTMhelix 326 348\ninside 349 372

21733 GCF_001595145.1_CFSAN033861_01.0Salmonella enterica subsp. enterica serovar Typhimurium Proteobacteria; Gammaproteobacteria; Enterobacterales; Enterobacteriaceae; Salmonella; Salmonella enterica; Salmonella enterica subsp. enterica MTGDNTLITSHGINRRDFMKLCAALAATMGLSSKAAAEMAESVSNPQRPPVIWIGAQECT WP_000145429.1 hydrogenase 2 small subunit [Salmonella enterica] Length: 372\nNumber of predicted TMHs: 1\nExp number of AAs in TMHs: 24.51309\nExp number, first 60 AAs: 0.2682\nTotal prob of N-in: 0.14402\noutside 1 331\nTMhelix 332 354\ninside 355 372

21734 GCF_001594905.1_CFSAN033874_01.0Salmonella enterica subsp. enterica serovar Typhimurium Proteobacteria; Gammaproteobacteria; Enterobacterales; Enterobacteriaceae; Salmonella; Salmonella enterica; Salmonella enterica subsp. enterica MNNEETFYQAMRRKGVTRRSFLKFCSLAATSLGLGAGMTPKIAWALENKPRIPVVWIHGL WP_001058312.1 hydrogenase [Salmonella enterica] Length: 372\nNumber of predicted TMHs: 2\nExp number of AAs in TMHs: 36.51738\nExp number, first 60 AAs: 19.72849\nTotal prob of N-in: 0.96170\nPOSSIBLE N-term signal sequence\ninside 1 19\nTMhelix 20 39\noutside 40 325\nTMhelix 326 348\ninside 349 372

21735 GCF_001595185.1_CFSAN033858_01.0Salmonella enterica subsp. enterica serovar Typhimurium Proteobacteria; Gammaproteobacteria; Enterobacterales; Enterobacteriaceae; Salmonella; Salmonella enterica; Salmonella enterica subsp. enterica MTGDNTLITSHGINRRDFMKLCAALAATMGLSSKAAAEMAESVSNPQRPPVIWIGAQECT WP_000145429.1 hydrogenase 2 small subunit [Salmonella enterica] Length: 372\nNumber of predicted TMHs: 1\nExp number of AAs in TMHs: 24.51309\nExp number, first 60 AAs: 0.2682\nTotal prob of N-in: 0.14402\noutside 1 331\nTMhelix 332 354\ninside 355 372

21736 GCF_001595225.1_CFSAN033856_01.0Salmonella enterica subsp. enterica serovar Typhimurium Proteobacteria; Gammaproteobacteria; Enterobacterales; Enterobacteriaceae; Salmonella; Salmonella enterica; Salmonella enterica subsp. enterica MTGDNTLITSHGINRRDFMKLCAALAATMGLSSKAAAEMAESVSNPQRPPVIWIGAQECT WP_000145429.1 hydrogenase 2 small subunit [Salmonella enterica] Length: 372\nNumber of predicted TMHs: 1\nExp number of AAs in TMHs: 24.51309\nExp number, first 60 AAs: 0.2682\nTotal prob of N-in: 0.14402\noutside 1 331\nTMhelix 332 354\ninside 355 372

21737 GCF_001595265.1_CFSAN033853_01.0Salmonella enterica subsp. enterica serovar Typhimurium Proteobacteria; Gammaproteobacteria; Enterobacterales; Enterobacteriaceae; Salmonella; Salmonella enterica; Salmonella enterica subsp. enterica MTGDNTLITSHGINRRDFMKLCAALAATMGLSSKAAAEMAESVSNPQRPPVIWIGAQECT WP_000145429.1 hydrogenase 2 small subunit [Salmonella enterica] Length: 372\nNumber of predicted TMHs: 1\nExp number of AAs in TMHs: 24.51309\nExp number, first 60 AAs: 0.2682\nTotal prob of N-in: 0.14402\noutside 1 331\nTMhelix 332 354\ninside 355 372

21738 GCF_001595185.1_CFSAN033858_01.0Salmonella enterica subsp. enterica serovar Typhimurium Proteobacteria; Gammaproteobacteria; Enterobacterales; Enterobacteriaceae; Salmonella; Salmonella enterica; Salmonella enterica subsp. enterica MNNEETFYQAMRRKGVTRRSFLKFCSLAATSLGLGAGMTPKIAWALENKPRIPVVWIHGL WP_001058312.1 hydrogenase [Salmonella enterica] Length: 372\nNumber of predicted TMHs: 2\nExp number of AAs in TMHs: 36.51738\nExp number, first 60 AAs: 19.72849\nTotal prob of N-in: 0.96170\nPOSSIBLE N-term signal sequence\ninside 1 19\nTMhelix 20 39\noutside 40 325\nTMhelix 326 348\ninside 349 372

21739 GCF_001595225.1_CFSAN033856_01.0Salmonella enterica subsp. enterica serovar Typhimurium Proteobacteria; Gammaproteobacteria; Enterobacterales; Enterobacteriaceae; Salmonella; Salmonella enterica; Salmonella enterica subsp. enterica MNNEETFYQAMRRKGVTRRSFLKFCSLAATSLGLGAGMTPKIAWALENKPRIPVVWIHGL WP_001058312.1 hydrogenase [Salmonella enterica] Length: 372\nNumber of predicted TMHs: 2\nExp number of AAs in TMHs: 36.51738\nExp number, first 60 AAs: 19.72849\nTotal prob of N-in: 0.96170\nPOSSIBLE N-term signal sequence\ninside 1 19\nTMhelix 20 39\noutside 40 325\nTMhelix 326 348\ninside 349 372

21740 GCF_001595345.1_CFSAN033849_01.0Salmonella enterica subsp. enterica serovar Typhimurium Proteobacteria; Gammaproteobacteria; Enterobacterales; Enterobacteriaceae; Salmonella; Salmonella enterica; Salmonella enterica subsp. enterica MNNEETFYQAMRRKGVTRRSFLKFCSLAATSLGLGAGMTPKIAWALENKPRIPVVWIHGL WP_001058312.1 hydrogenase [Salmonella enterica] Length: 372\nNumber of predicted TMHs: 2\nExp

number of AAs in TMHs: 36.51738\nExp number, first 60 AAs: 19.72849\nTotal prob of N-in: 0.96170\nPOSSIBLE N-term signal sequence\ninside 1 19\nTMhelix 20 39\noutside 40 325\nTMhelix 326 348\ninside 349 372

21741 GCF_001705365.1_ASM170536v1 Salmonella enterica subsp. enterica serovar Typhimurium Proteobacteria; Gammaproteobacteria; Enterobacterales; Enterobacteriaceae; Salmonella; Salmonella enterica; Salmonella enterica subsp. enterica MTGDNTLITSHGINRRDFMKLCAALAATMGLSSKAAAEMAESVSNPQRPPVIWIGAQECT WP_000145429.1 hydrogenase 2 small subunit [Salmonella enterica] Length: 372\nNumber of predicted TMHs: 1\nExp number of AAs in TMHs: 24.51309\nExp number, first 60 AAs: 0.2682\nTotal prob of N-in: 0.14402\noutside 1 331\nTMhelix 332 354\ninside 355 372

21742 GCF_001705365.1_ASM170536v1 Salmonella enterica subsp. enterica serovar Typhimurium Proteobacteria; Gammaproteobacteria; Enterobacterales; Enterobacteriaceae; Salmonella; Salmonella enterica; Salmonella enterica subsp. enterica MNNEETFYQAMRRKGVTRRSFLKFCSLAATSLGLGAGMTPKIAWALENKPRIPVVWIHGL WP_001058312.1 hydrogenase [Salmonella enterica] Length: 372\nNumber of predicted TMHs: 2\nExp number of AAs in TMHs: 36.51738\nExp number, first 60 AAs: 19.72849\nTotal prob of N-in: 0.96170\nPOSSIBLE N-term signal sequence\ninside 1 19\nTMhelix 20 39\noutside 40 325\nTMhelix 326 348\ninside 349 372

21743 GCF_001755865.1_CFSAN008084 Salmonella enterica subsp. enterica serovar Typhimurium Proteobacteria; Gammaproteobacteria; Enterobacterales; Enterobacteriaceae; Salmonella; Salmonella enterica; Salmonella enterica subsp. enterica MTGDNTLITSHGINRRDFMKLCAALAATMGLSSKAAAEMAESVSNPQRPPVIWIGAQECT WP_000145429.1 hydrogenase 2 small subunit [Salmonella enterica] Length: 372\nNumber of predicted TMHs: 1\nExp number of AAs in TMHs: 24.51309\nExp number, first 60 AAs: 0.2682\nTotal prob of N-in: 0.14402\noutside 1 331\nTMhelix 332 354\ninside 355 372

21744 GCF_001729225.1_ASM172922v1 Salmonella enterica subsp. enterica serovar Typhimurium Proteobacteria; Gammaproteobacteria; Enterobacterales; Enterobacteriaceae; Salmonella; Salmonella enterica; Salmonella enterica subsp. enterica MTGDNTLITSHGINRRDFMKLCAALAATMGLSSKAAAEMAESVSNPQRPPVIWIGAQECT WP_000145429.1 hydrogenase 2 small subunit [Salmonella enterica] Length: 372\nNumber of predicted TMHs: 1\nExp number of AAs in TMHs: 24.51309\nExp number, first 60 AAs: 0.2682\nTotal prob of N-in: 0.14402\noutside 1 331\nTMhelix 332 354\ninside 355 372

21745 GCF_001755885.1_CFSAN008089 Salmonella enterica subsp. enterica serovar Typhimurium Proteobacteria; Gammaproteobacteria; Enterobacterales; Enterobacteriaceae; Salmonella; Salmonella enterica; Salmonella enterica subsp. enterica MTGDNTLITSHGINRRDFMKLCAALAATMGLSSKAAAEMAESVSNPQRPPVIWIGAQECT WP_000145429.1 hydrogenase 2 small subunit [Salmonella enterica] Length: 372\nNumber of predicted TMHs: 1\nExp number of AAs in TMHs: 24.51309\nExp number, first 60 AAs: 0.2682\nTotal prob of N-in: 0.14402\noutside 1 331\nTMhelix 332 354\ninside 355 372

21746 GCF_001755945.1_CFSAN008079 Salmonella enterica subsp. enterica serovar Typhimurium Proteobacteria; Gammaproteobacteria; Enterobacterales; Enterobacteriaceae; Salmonella; Salmonella enterica; Salmonella enterica subsp. enterica MTGDNTLITSHGINRRDFMKLCAALAATMGLSSKAAAEMAESVSNPQRPPVIWIGAQECT WP_000145429.1 hydrogenase 2 small subunit [Salmonella enterica] Length: 372\nNumber of predicted TMHs: 1\nExp number of AAs in TMHs: 24.51309\nExp number, first 60 AAs: 0.2682\nTotal prob of N-in: 0.14402\noutside 1 331\nTMhelix 332 354\ninside 355 372

21747 GCF_001756025.1_CFSAN008081 Salmonella enterica subsp. enterica serovar Typhimurium Proteobacteria; Gammaproteobacteria; Enterobacterales; Enterobacteriaceae; Salmonella; Salmonella enterica; Salmonella enterica subsp. enterica MTGDNTLITSHGINRRDFMKLCAALAATMGLSSKAAAEMAESVSNPQRPPVIWIGAQECT WP_000145429.1 hydrogenase 2 small subunit [Salmonella enterica] Length: 372\nNumber of predicted TMHs: 1\nExp number of AAs in TMHs: 24.51309\nExp number, first 60 AAs: 0.2682\nTotal prob of N-in: 0.14402\noutside 1 331\nTMhelix 332 354\ninside 355 372

21748 GCF_001756025.1_CFSAN008081 Salmonella enterica subsp. enterica serovar Typhimurium Proteobacteria; Gammaproteobacteria; Enterobacterales; Enterobacteriaceae; Salmonella; Salmonella enterica; Salmonella enterica subsp. enterica MNNEETFYQAMRRKGVTRRSFLKFCSLAATSLGLGAGMTPKIAWAPENKPRIPVVWIHGL WP_070130171.1 hydrogenase [Salmonella enterica] Length: 372\nNumber of predicted TMHs: 2\nExp number of AAs in TMHs: 36.1294\nExp number, first 60 AAs: 19.29582\nTotal prob of N-in: 0.94342\nPOSSIBLE N-term signal sequence\ninside 1 19\nTMhelix 20 39\noutside 40 325\nTMhelix 326 348\ninside 349 372

21749 GCF_001922185.1_ASM192218v1 Salmonella enterica subsp. enterica serovar Typhimurium Proteobacteria; Gammaproteobacteria; Enterobacterales; Enterobacteriaceae; Salmonella; Salmonella enterica;

Salmonella enterica subsp. enterica MTGDNTLITSHGINRRDFMKLCAALAATMGLSSKAAAEMAESVSNPQRPPVIWIGAQECT WP_000145429.1 hydrogenase 2 small subunit [Salmonella enterica] Length: 372\nNumber of predicted TMHs: 1\nExp number of AAs in TMHs: 24.51309\nExp number, first 60 AAs: 0.2682\nTotal prob of N-in: 0.14402\noutside 1 331\nTMhelix 332 354\ninside 355 372

21750 GCF_001922185.1_ASM192218v1 Salmonella enterica subsp. enterica serovar Typhimurium Proteobacteria; Gammaproteobacteria; Enterobacterales; Enterobacteriaceae; Salmonella; Salmonella enterica; Salmonella enterica subsp. enterica MNNEETFYQAMRRKGVTRRSFLKFCSLAATSLGLGAGMTPKIAWALENKPRIPVVWIHGL WP_001058312.1 hydrogenase [Salmonella enterica] Length: 372\nNumber of predicted TMHs: 2\nExp number of AAs in TMHs: 36.51738\nExp number, first 60 AAs: 19.72849\nTotal prob of N-in: 0.96170\nPOSSIBLE N-term signal sequence\ninside 1 19\nTMhelix 20 39\noutside 40 325\nTMhelix 326 348\ninside 349 372

21751 GCF_001955315.1_ASM195531v1 Salmonella enterica subsp. enterica serovar Typhimurium Proteobacteria; Gammaproteobacteria; Enterobacterales; Enterobacteriaceae; Salmonella; Salmonella enterica; Salmonella enterica subsp. enterica MNNEETFYQAMRRKGVTRRSFLKFCSLAATSLGLGAGMTPKIAWALENKPRIPVVWIHGL WP_001058312.1 hydrogenase [Salmonella enterica] Length: 372\nNumber of predicted TMHs: 2\nExp number of AAs in TMHs: 36.51738\nExp number, first 60 AAs: 19.72849\nTotal prob of N-in: 0.96170\nPOSSIBLE N-term signal sequence\ninside 1 19\nTMhelix 20 39\noutside 40 325\nTMhelix 326 348\ninside 349 372

21752 GCF_001955295.1_ASM195529v1 Salmonella enterica subsp. enterica serovar Typhimurium Proteobacteria; Gammaproteobacteria; Enterobacterales; Enterobacteriaceae; Salmonella; Salmonella enterica; Salmonella enterica subsp. enterica MNNEETFYQAMRRKGVTRRSFLKFCSLAATSLGLGAGMTPKIAWALENKPRIPVVWIHGL WP_001058312.1 hydrogenase [Salmonella enterica] Length: 372\nNumber of predicted TMHs: 2\nExp number of AAs in TMHs: 36.51738\nExp number, first 60 AAs: 19.72849\nTotal prob of N-in: 0.96170\nPOSSIBLE N-term signal sequence\ninside 1 19\nTMhelix 20 39\noutside 40 325\nTMhelix 326 348\ninside 349 372

21753 GCF_002009155.1_ASM200915v1 Salmonella enterica subsp. enterica serovar Typhimurium Proteobacteria; Gammaproteobacteria; Enterobacterales; Enterobacteriaceae; Salmonella; Salmonella enterica; Salmonella enterica subsp. enterica MTGDNTLITSHGINRRDFMKLCAALAATMGLSSKAAAEMAESVSNPQRPPVIWIGAQECT WP_000145429.1 hydrogenase 2 small subunit [Salmonella enterica] Length: 372\nNumber of predicted TMHs: 1\nExp number of AAs in TMHs: 24.51309\nExp number, first 60 AAs: 0.2682\nTotal prob of N-in: 0.14402\noutside 1 331\nTMhelix 332 354\ninside 355 372

21754 GCF_002018355.1_ASM201835v1 Salmonella enterica subsp. enterica serovar Typhimurium Proteobacteria; Gammaproteobacteria; Enterobacterales; Enterobacteriaceae; Salmonella; Salmonella enterica; Salmonella enterica subsp. enterica MTGDNTLITSHGINRRDFMKLCAALAATMGLSSKAAAEMAESVSNPQRPPVIWIGAQECT WP_000145429.1 hydrogenase 2 small subunit [Salmonella enterica] Length: 372\nNumber of predicted TMHs: 1\nExp number of AAs in TMHs: 24.51309\nExp number, first 60 AAs: 0.2682\nTotal prob of N-in: 0.14402\noutside 1 331\nTMhelix 332 354\ninside 355 372

21755 GCF_002018355.1_ASM201835v1 Salmonella enterica subsp. enterica serovar Typhimurium Proteobacteria; Gammaproteobacteria; Enterobacterales; Enterobacteriaceae; Salmonella; Salmonella enterica; Salmonella enterica subsp. enterica MNNEETFYQAMRRKGVTRRSFLKFCSLAATSLGLGAGMTPKIAWALENKPRIPVVWIHGL WP_001058312.1 hydrogenase [Salmonella enterica] Length: 372\nNumber of predicted TMHs: 2\nExp number of AAs in TMHs: 36.51738\nExp number, first 60 AAs: 19.72849\nTotal prob of N-in: 0.96170\nPOSSIBLE N-term signal sequence\ninside 1 19\nTMhelix 20 39\noutside 40 325\nTMhelix 326 348\ninside 349 372

21756 GCF_001989635.1_ASM198963v1 Salmonella enterica subsp. enterica serovar Typhimurium Proteobacteria; Gammaproteobacteria; Enterobacterales; Enterobacteriaceae; Salmonella; Salmonella enterica; Salmonella enterica subsp. enterica MNNEETFYQAMRRKGVTRRSFLKFCSLAATSLGLGAGMTPKIAWALENKPRIPVVWIHGL WP_001058312.1 hydrogenase [Salmonella enterica] Length: 372\nNumber of predicted TMHs: 2\nExp number of AAs in TMHs: 36.51738\nExp number, first 60 AAs: 19.72849\nTotal prob of N-in: 0.96170\nPOSSIBLE N-term signal sequence\ninside 1 19\nTMhelix 20 39\noutside 40 325\nTMhelix 326 348\ninside 349 372

21757 GCF_002029675.1_ASM202967v1 Salmonella enterica subsp. enterica serovar Typhimurium Proteobacteria; Gammaproteobacteria; Enterobacterales; Enterobacteriaceae; Salmonella; Salmonella enterica; Salmonella enterica subsp. enterica MNNEETFYQAMRRKGVTRRSFLKFCSLAATSLGLGAGMTPKIAWALENKPRIPVVWIHGL WP_001058312.1 hydrogenase [Salmonella enterica] Length: 372\nNumber of predicted TMHs: 2\nExp number of AAs in TMHs: 36.51738\nExp number, first 60 AAs: 19.72849\nTotal prob of N-in: 0.96170\nPOSSIBLE N-term signal sequence\ninside 1 19\nTMhelix 20 39\noutside 40 325\nTMhelix 326 348\ninside 349 372

21758 GCF_002028635.1_ASM202863v1 Salmonella enterica subsp. enterica serovar Typhimurium Proteobacteria; Gammaproteobacteria; Enterobacterales; Enterobacteriaceae; Salmonella; Salmonella enterica; Salmonella enterica subsp. enterica MNNEETFYQAMRRKGVTRRSFLKFCSLAATSLGLGAGMTPKIAWALENKPRIPVVWIHGL WP_001058312.1 hydrogenase [Salmonella enterica] Length: 372\nNumber of predicted TMHs: 2\nExp number of AAs in TMHs: 36.51738\nExp number, first 60 AAs: 19.72849\nTotal prob of N-in: 0.96170\nPOSSIBLE N-term signal sequence\ninside 1 19\nTMhelix 20 39\noutside 40 325\nTMhelix 326 348\ninside 349 372

21759 GCF_002029765.1_ASM202976v1 Salmonella enterica subsp. enterica serovar Typhimurium Proteobacteria; Gammaproteobacteria; Enterobacterales; Enterobacteriaceae; Salmonella; Salmonella enterica; Salmonella enterica subsp. enterica MTGDNTLITSHGINRRDFMKLCAALAATMGLSSKAAAEMAESVSNPQRPPVIWIGAQECT WP_000145429.1 hydrogenase 2 small subunit [Salmonella enterica] Length: 372\nNumber of predicted TMHs: 1\nExp number of AAs in TMHs: 24.51309\nExp number, first 60 AAs: 0.2682\nTotal prob of N-in: 0.14402\noutside 1 331\nTMhelix 332 354\ninside 355 372

21760 GCF_002029795.1_ASM202979v1 Salmonella enterica subsp. enterica serovar Typhimurium Proteobacteria; Gammaproteobacteria; Enterobacterales; Enterobacteriaceae; Salmonella; Salmonella enterica; Salmonella enterica subsp. enterica MTGDNTLITSHGINRRDFMKLCAALAATMGLSSKAAAEMAESVSNPQRPPVIWIGAQECT WP_000145429.1 hydrogenase 2 small subunit [Salmonella enterica] Length: 372\nNumber of predicted TMHs: 1\nExp number of AAs in TMHs: 24.51309\nExp number, first 60 AAs: 0.2682\nTotal prob of N-in: 0.14402\noutside 1 331\nTMhelix 332 354\ninside 355 372

21761 GCF_002029835.1_ASM202983v1 Salmonella enterica subsp. enterica serovar Typhimurium Proteobacteria; Gammaproteobacteria; Enterobacterales; Enterobacteriaceae; Salmonella; Salmonella enterica; Salmonella enterica subsp. enterica MTGDNTLITSHGINRRDFMKLCAALAATMGLSSKAAAEMAESVSNPQRPPVIWIGAQECT WP_000145429.1 hydrogenase 2 small subunit [Salmonella enterica] Length: 372\nNumber of predicted TMHs: 1\nExp number of AAs in TMHs: 24.51309\nExp number, first 60 AAs: 0.2682\nTotal prob of N-in: 0.14402\noutside 1 331\nTMhelix 332 354\ninside 355 372

21762 GCF_002029765.1_ASM202976v1 Salmonella enterica subsp. enterica serovar Typhimurium Proteobacteria; Gammaproteobacteria; Enterobacterales; Enterobacteriaceae; Salmonella; Salmonella enterica; Salmonella enterica subsp. enterica MNNEETFYQAMRRKGVTRRSFLKFCSLAATSLGLGAGMTPKIAWALENKPRIPVVWIHGL WP_001058312.1 hydrogenase [Salmonella enterica] Length: 372\nNumber of predicted TMHs: 2\nExp number of AAs in TMHs: 36.51738\nExp number, first 60 AAs: 19.72849\nTotal prob of N-in: 0.96170\nPOSSIBLE N-term signal sequence\ninside 1 19\nTMhelix 20 39\noutside 40 325\nTMhelix 326 348\ninside 349 372

21763 GCF_002029795.1_ASM202979v1 Salmonella enterica subsp. enterica serovar Typhimurium Proteobacteria; Gammaproteobacteria; Enterobacterales; Enterobacteriaceae; Salmonella; Salmonella enterica; Salmonella enterica subsp. enterica MNNEETFYQAMRRKGVTRRSFLKFCSLAATSLGLGAGMTPKIAWALENKPRIPVVWIHGL WP_001058312.1 hydrogenase [Salmonella enterica] Length: 372\nNumber of predicted TMHs: 2\nExp number of AAs in TMHs: 36.51738\nExp number, first 60 AAs: 19.72849\nTotal prob of N-in: 0.96170\nPOSSIBLE N-term signal sequence\ninside 1 19\nTMhelix 20 39\noutside 40 325\nTMhelix 326 348\ninside 349 372

21764 GCF_002033825.1_ASM203382v1 Salmonella enterica subsp. enterica serovar Typhimurium Proteobacteria; Gammaproteobacteria; Enterobacterales; Enterobacteriaceae; Salmonella; Salmonella enterica; Salmonella enterica subsp. enterica MTGDNTLITSHGINRRDFMKLCAALAATMGLSSKAAAEMAESVSNPQRPPVIWIGAQECT WP_000145429.1 hydrogenase 2 small subunit [Salmonella enterica] Length: 372\nNumber of predicted TMHs: 1\nExp number of AAs in TMHs: 24.51309\nExp number, first 60 AAs: 0.2682\nTotal prob of N-in: 0.14402\noutside 1 331\nTMhelix 332 354\ninside 355 372

21765 GCF_002035135.1_ASM203513v1 Salmonella enterica subsp. enterica serovar Typhimurium Proteobacteria; Gammaproteobacteria; Enterobacterales; Enterobacteriaceae; Salmonella; Salmonella enterica; Salmonella enterica subsp. enterica MNNEETFYQAMRRKGVTRRSFLKFCSLAATSLGLGAGMTPKIAWALENKPRIPVVWIHGL WP_001058312.1 hydrogenase [Salmonella enterica] Length: 372\nNumber of predicted TMHs: 2\nExp number of AAs in TMHs: 36.51738\nExp number, first 60 AAs: 19.72849\nTotal prob of N-in: 0.96170\nPOSSIBLE N-term signal sequence\ninside 1 19\nTMhelix 20 39\noutside 40 325\nTMhelix 326 348\ninside 349 372

21766 GCF_002035445.1_ASM203544v1 Salmonella enterica subsp. enterica serovar Typhimurium Proteobacteria; Gammaproteobacteria; Enterobacterales; Enterobacteriaceae; Salmonella; Salmonella enterica; Salmonella enterica subsp. enterica MTGDNTLITSHGINRRDFMKLCAALAATMGLSSKAAAEMAESVSNPQRPPVIWIGAQECT WP_000145429.1 hydrogenase 2 small subunit [Salmonella enterica] Length: 372\nNumber of

predicted TMHs: 1\nExp number of AAs in TMHs: 24.51309\nExp number, first 60 AAs: 0.2682\nTotal prob of N-in: 0.14402\noutside 1 331\nTMhelix 332 354\ninside 355 372

21767 GCF_002035445.1_ASM203544v1 Salmonella enterica subsp. enterica serovar Typhimurium Proteobacteria; Gammaproteobacteria; Enterobacterales; Enterobacteriaceae; Salmonella; Salmonella enterica; Salmonella enterica subsp. enterica MNNEETFYQAMRRKGVTRRSFLKFCSLAATSLGLGAGMTPKIAWALENKPRIPVVWIHGL WP_001058312.1 hydrogenase [Salmonella enterica] Length: 372\nNumber of predicted TMHs: 2\nExp number of AAs in TMHs: 36.51738\nExp number, first 60 AAs: 19.72849\nTotal prob of N-in: 0.96170\nPOSSIBLE N-term signal sequence\ninside 1 19\nTMhelix 20 39\noutside 40 325\nTMhelix 326 348\ninside 349 372

21768 GCF_002036205.1_ASM203620v1 Salmonella enterica subsp. enterica serovar Typhimurium Proteobacteria; Gammaproteobacteria; Enterobacterales; Enterobacteriaceae; Salmonella; Salmonella enterica; Salmonella enterica subsp. enterica MTGDNTLITSHGINRRDFMKLCAALAATMGLSSKAAAEMAESVSNPQRPPVIWIGAQECT WP_000145429.1 hydrogenase 2 small subunit [Salmonella enterica] Length: 372\nNumber of predicted TMHs: 1\nExp number of AAs in TMHs: 24.51309\nExp number, first 60 AAs: 0.2682\nTotal prob of N-in: 0.14402\noutside 1 331\nTMhelix 332 354\ninside 355 372

21769 GCF_002036315.1_ASM203631v1 Salmonella enterica subsp. enterica serovar Typhimurium Proteobacteria; Gammaproteobacteria; Enterobacterales; Enterobacteriaceae; Salmonella; Salmonella enterica; Salmonella enterica subsp. enterica MTGDNTLITSHGINRRDFMKLCAALAATMGLSSKAAAEMAESVSNPQRPPVIWIGAQECT WP_000145429.1 hydrogenase 2 small subunit [Salmonella enterica] Length: 372\nNumber of predicted TMHs: 1\nExp number of AAs in TMHs: 24.51309\nExp number, first 60 AAs: 0.2682\nTotal prob of N-in: 0.14402\noutside 1 331\nTMhelix 332 354\ninside 355 372

21770 GCF_002057985.1_ASM205798v1 Salmonella enterica subsp. enterica serovar Typhimurium Proteobacteria; Gammaproteobacteria; Enterobacterales; Enterobacteriaceae; Salmonella; Salmonella enterica; Salmonella enterica subsp. enterica MTGDNTLITSHGINRRDFMKLCAALAATMGLSSKAAAEMAESVSNPQRPPVIWIGAQECT WP_000145429.1 hydrogenase 2 small subunit [Salmonella enterica] Length: 372\nNumber of predicted TMHs: 1\nExp number of AAs in TMHs: 24.51309\nExp number, first 60 AAs: 0.2682\nTotal prob of N-in: 0.14402\noutside 1 331\nTMhelix 332 354\ninside 355 372

21771 GCF_002057985.1_ASM205798v1 Salmonella enterica subsp. enterica serovar Typhimurium Proteobacteria; Gammaproteobacteria; Enterobacterales; Enterobacteriaceae; Salmonella; Salmonella enterica; Salmonella enterica subsp. enterica MNNEETFYQAMRRKGVTRRSFLKFCSLAATSLGLGAGMTPKIAWALENKPRIPVVWIHGL WP_001058312.1 hydrogenase [Salmonella enterica] Length: 372\nNumber of predicted TMHs: 2\nExp number of AAs in TMHs: 36.51738\nExp number, first 60 AAs: 19.72849\nTotal prob of N-in: 0.96170\nPOSSIBLE N-term signal sequence\ninside 1 19\nTMhelix 20 39\noutside 40 325\nTMhelix 326 348\ninside 349 372

21772 GCF_002059555.1_ASM205955v1 Salmonella enterica subsp. enterica serovar Typhimurium Proteobacteria; Gammaproteobacteria; Enterobacterales; Enterobacteriaceae; Salmonella; Salmonella enterica; Salmonella enterica subsp. enterica MTGDNTLITSHGINRRDFMKLCAALAATMGLSSKAAAEMAESVSNPQRPPVIWIGAQECT WP_000145429.1 hydrogenase 2 small subunit [Salmonella enterica] Length: 372\nNumber of predicted TMHs: 1\nExp number of AAs in TMHs: 24.51309\nExp number, first 60 AAs: 0.2682\nTotal prob of N-in: 0.14402\noutside 1 331\nTMhelix 332 354\ninside 355 372

21773 GCF_002059555.1_ASM205955v1 Salmonella enterica subsp. enterica serovar Typhimurium Proteobacteria; Gammaproteobacteria; Enterobacterales; Enterobacteriaceae; Salmonella; Salmonella enterica; Salmonella enterica subsp. enterica MNNEETFYQAMRRKGVTRRSFLKFCSLAATSLGLGAGMTPKIAWALENKPRIPVVWIHGL WP_001058312.1 hydrogenase [Salmonella enterica] Length: 372\nNumber of predicted TMHs: 2\nExp number of AAs in TMHs: 36.51738\nExp number, first 60 AAs: 19.72849\nTotal prob of N-in: 0.96170\nPOSSIBLE N-term signal sequence\ninside 1 19\nTMhelix 20 39\noutside 40 325\nTMhelix 326 348\ninside 349 372

21774 GCF_900184385.1_STMU2UK Salmonella enterica subsp. enterica serovar Typhimurium Proteobacteria; Gammaproteobacteria; Enterobacterales; Enterobacteriaceae; Salmonella; Salmonella enterica; Salmonella enterica subsp. enterica MTGDNTLITSHGINRRDFMKLCAALAATMGLSSKAAAEMAESVSNPQRPPVIWIGAQECT WP_000145429.1 hydrogenase 2 small subunit [Salmonella enterica] Length: 372\nNumber of predicted TMHs: 1\nExp number of AAs in TMHs: 24.51309\nExp number, first 60 AAs: 0.2682\nTotal prob of N-in: 0.14402\noutside 1 331\nTMhelix 332 354\ninside 355 372

21775 GCF_002074335.1_ASM207433v1 Salmonella enterica subsp. enterica serovar Typhimurium Proteobacteria; Gammaproteobacteria; Enterobacterales; Enterobacteriaceae; Salmonella; Salmonella enterica;

Salmonella enterica subsp. enterica MNNEETFYQAMRRKGVTRRSFLKFCSLAATSLGLGAGMTPKIAWALENKPRIPVVWIHGL WP_001058312.1 hydrogenase [Salmonella enterica] Length: 372\nNumber of predicted TMHs: 2\nExp number of AAs in TMHs: 36.51738\nExp number, first 60 AAs: 19.72849\nTotal prob of N-in: 0.96170\nPOSSIBLE N-term signal sequence\ninside 1 19\nTMhelix 20 39\noutside 40 325\nTMhelix 326 348\ninside 349 372

21776 GCF_002074435.1_ASM207443v1 Salmonella enterica subsp. enterica serovar Typhimurium Proteobacteria; Gammaproteobacteria; Enterobacterales; Enterobacteriaceae; Salmonella; Salmonella enterica; Salmonella enterica subsp. enterica MNNEETFYQAMRRKGVTRRSFLKFCSLAATSLGLGAGMTPKIAWALENKPRIPVVWIHGL WP_001058312.1 hydrogenase [Salmonella enterica] Length: 372\nNumber of predicted TMHs: 2\nExp number of AAs in TMHs: 36.51738\nExp number, first 60 AAs: 19.72849\nTotal prob of N-in: 0.96170\nPOSSIBLE N-term signal sequence\ninside 1 19\nTMhelix 20 39\noutside 40 325\nTMhelix 326 348\ninside 349 372

21777 GCF_002074455.1_ASM207445v1 Salmonella enterica subsp. enterica serovar Typhimurium Proteobacteria; Gammaproteobacteria; Enterobacterales; Enterobacteriaceae; Salmonella; Salmonella enterica; Salmonella enterica subsp. enterica MNNEETFYQAMRRKGVTRRSFLKFCSLAATSLGLGAGMTPKIAWALENKPRIPVVWIHGL WP_001058312.1 hydrogenase [Salmonella enterica] Length: 372\nNumber of predicted TMHs: 2\nExp number of AAs in TMHs: 36.51738\nExp number, first 60 AAs: 19.72849\nTotal prob of N-in: 0.96170\nPOSSIBLE N-term signal sequence\ninside 1 19\nTMhelix 20 39\noutside 40 325\nTMhelix 326 348\ninside 349 372

21778 GCF_002074515.1_ASM207451v1 Salmonella enterica subsp. enterica serovar Typhimurium Proteobacteria; Gammaproteobacteria; Enterobacterales; Enterobacteriaceae; Salmonella; Salmonella enterica; Salmonella enterica subsp. enterica MNNEETFYQAMRRKGVTRRSFLKFCSLAATSLGLGAGMTPKIAWALENKPRIPVVWIHGL WP_001058312.1 hydrogenase [Salmonella enterica] Length: 372\nNumber of predicted TMHs: 2\nExp number of AAs in TMHs: 36.51738\nExp number, first 60 AAs: 19.72849\nTotal prob of N-in: 0.96170\nPOSSIBLE N-term signal sequence\ninside 1 19\nTMhelix 20 39\noutside 40 325\nTMhelix 326 348\ninside 349 372

21779 GCF_002074545.1_ASM207454v1 Salmonella enterica subsp. enterica serovar Typhimurium Proteobacteria; Gammaproteobacteria; Enterobacterales; Enterobacteriaceae; Salmonella; Salmonella enterica; Salmonella enterica subsp. enterica MNNEETFYQAMRRKGVTRRSFLKFCSLAATSLGLGAGMTPKIAWALENKPRIPVVWIHGL WP_001058312.1 hydrogenase [Salmonella enterica] Length: 372\nNumber of predicted TMHs: 2\nExp number of AAs in TMHs: 36.51738\nExp number, first 60 AAs: 19.72849\nTotal prob of N-in: 0.96170\nPOSSIBLE N-term signal sequence\ninside 1 19\nTMhelix 20 39\noutside 40 325\nTMhelix 326 348\ninside 349 372

21780 GCF_002074595.1_ASM207459v1 Salmonella enterica subsp. enterica serovar Typhimurium Proteobacteria; Gammaproteobacteria; Enterobacterales; Enterobacteriaceae; Salmonella; Salmonella enterica; Salmonella enterica subsp. enterica MTGDNTLITSHGINRRDFMKLCAALAATMGLSSKAAAEMAESVSNPQRPPVIWIGAQECT WP_000145429.1 hydrogenase 2 small subunit [Salmonella enterica] Length: 372\nNumber of predicted TMHs: 1\nExp number of AAs in TMHs: 24.51309\nExp number, first 60 AAs: 0.2682\nTotal prob of N-in: 0.14402\noutside 1 331\nTMhelix 332 354\ninside 355 372

21781 GCF_002074635.1_ASM207463v1 Salmonella enterica subsp. enterica serovar Typhimurium Proteobacteria; Gammaproteobacteria; Enterobacterales; Enterobacteriaceae; Salmonella; Salmonella enterica; Salmonella enterica subsp. enterica MTGDNTLITSHGINRRDFMKLCAALAATMGLSSKAAAEMAESVSNPQRPPVIWIGAQECT WP_000145429.1 hydrogenase 2 small subunit [Salmonella enterica] Length: 372\nNumber of predicted TMHs: 1\nExp number of AAs in TMHs: 24.51309\nExp number, first 60 AAs: 0.2682\nTotal prob of N-in: 0.14402\noutside 1 331\nTMhelix 332 354\ninside 355 372

21782 GCF_002074675.1_ASM207467v1 Salmonella enterica subsp. enterica serovar Typhimurium Proteobacteria; Gammaproteobacteria; Enterobacterales; Enterobacteriaceae; Salmonella; Salmonella enterica; Salmonella enterica subsp. enterica MTGDNTLITSHGINRRDFMKLCAALAATMGLSSKAAAEMAESVSNPQRPPVIWIGAQECT WP_000145429.1 hydrogenase 2 small subunit [Salmonella enterica] Length: 372\nNumber of predicted TMHs: 1\nExp number of AAs in TMHs: 24.51309\nExp number, first 60 AAs: 0.2682\nTotal prob of N-in: 0.14402\noutside 1 331\nTMhelix 332 354\ninside 355 372

21783 GCF_002074685.1_ASM207468v1 Salmonella enterica subsp. enterica serovar Typhimurium Proteobacteria; Gammaproteobacteria; Enterobacterales; Enterobacteriaceae; Salmonella; Salmonella enterica; Salmonella enterica subsp. enterica MTGDNTLITSHGINRRDFMKLCAALAATMGLSSKAAAEMAESVSNPQRPPVIWIGAQECT WP_000145429.1 hydrogenase 2 small subunit [Salmonella enterica] Length: 372\nNumber of predicted TMHs: 1\nExp number of AAs in TMHs: 24.51309\nExp number, first 60 AAs: 0.2682\nTotal prob of N-in: 0.14402\noutside 1 331\nTMhelix 332 354\ninside 355 372

21784 GCF_002074715.1_ASM207471v1 Salmonella enterica subsp. enterica serovar Typhimurium Proteobacteria; Gammaproteobacteria; Enterobacterales; Enterobacteriaceae; Salmonella; Salmonella enterica; Salmonella enterica subsp. enterica MTGDNTLITSHGINRRDFMKLCAALAATMGLSSKAAAEMAESVSNPQRPPVIWIGAQECT WP_000145429.1 hydrogenase 2 small subunit [Salmonella enterica] Length: 372\nNumber of predicted TMHs: 1\nExp number of AAs in TMHs: 24.51309\nExp number, first 60 AAs: 0.2682\nTotal prob of N-in: 0.14402\noutside 1 331\nTMhelix 332 354\ninside 355 372

21785 GCF_002074745.1_ASM207474v1 Salmonella enterica subsp. enterica serovar Typhimurium Proteobacteria; Gammaproteobacteria; Enterobacterales; Enterobacteriaceae; Salmonella; Salmonella enterica; Salmonella enterica subsp. enterica MTGDNTLITSHGINRRDFMKLCAALAATMGLSSKAAAEMAESVSNPQRPPVIWIGAQECT WP_000145429.1 hydrogenase 2 small subunit [Salmonella enterica] Length: 372\nNumber of predicted TMHs: 1\nExp number of AAs in TMHs: 24.51309\nExp number, first 60 AAs: 0.2682\nTotal prob of N-in: 0.14402\noutside 1 331\nTMhelix 332 354\ninside 355 372

21786 GCF_002074795.1_ASM207479v1 Salmonella enterica subsp. enterica serovar Typhimurium Proteobacteria; Gammaproteobacteria; Enterobacterales; Enterobacteriaceae; Salmonella; Salmonella enterica; Salmonella enterica subsp. enterica MTGDNTLITSHGINRRDFMKLCAALAATMGLSSKAAAEMAESVSNPQRPPVIWIGAQECT WP_000145429.1 hydrogenase 2 small subunit [Salmonella enterica] Length: 372\nNumber of predicted TMHs: 1\nExp number of AAs in TMHs: 24.51309\nExp number, first 60 AAs: 0.2682\nTotal prob of N-in: 0.14402\noutside 1 331\nTMhelix 332 354\ninside 355 372

21787 GCF_002074795.1_ASM207479v1 Salmonella enterica subsp. enterica serovar Typhimurium Proteobacteria; Gammaproteobacteria; Enterobacterales; Enterobacteriaceae; Salmonella; Salmonella enterica; Salmonella enterica subsp. enterica MNNEETFYQAMRRKGVTRRSFLKFCSLAATSLGLGAGMTPKIAWALENKPRIPVVWIHGL WP_001058312.1 hydrogenase [Salmonella enterica] Length: 372\nNumber of predicted TMHs: 2\nExp number of AAs in TMHs: 36.51738\nExp number, first 60 AAs: 19.72849\nTotal prob of N-in: 0.96170\nPOSSIBLE N-term signal sequence\ninside 1 19\nTMhelix 20 39\noutside 40 325\nTMhelix 326 348\ninside 349 372

21788 GCF_002074595.1_ASM207459v1 Salmonella enterica subsp. enterica serovar Typhimurium Proteobacteria; Gammaproteobacteria; Enterobacterales; Enterobacteriaceae; Salmonella; Salmonella enterica; Salmonella enterica subsp. enterica MNNEETFYQAMRRKGVTRRSFLKFCSLAATSLGLGAGMTPKIAWALENKPRIPVVWIHGL WP_001058312.1 hydrogenase [Salmonella enterica] Length: 372\nNumber of predicted TMHs: 2\nExp number of AAs in TMHs: 36.51738\nExp number, first 60 AAs: 19.72849\nTotal prob of N-in: 0.96170\nPOSSIBLE N-term signal sequence\ninside 1 19\nTMhelix 20 39\noutside 40 325\nTMhelix 326 348\ninside 349 372

21789 GCF_002066815.1_ASM206681v1 Salmonella enterica subsp. enterica serovar Typhimurium Proteobacteria; Gammaproteobacteria; Enterobacterales; Enterobacteriaceae; Salmonella; Salmonella enterica; Salmonella enterica subsp. enterica MNNEETFYQAMRRKGVTRRSFLKFCSLAATSLGLGAGMTPKIAWALENKPRIPVVWIHGL WP_001058312.1 hydrogenase [Salmonella enterica] Length: 372\nNumber of predicted TMHs: 2\nExp number of AAs in TMHs: 36.51738\nExp number, first 60 AAs: 19.72849\nTotal prob of N-in: 0.96170\nPOSSIBLE N-term signal sequence\ninside 1 19\nTMhelix 20 39\noutside 40 325\nTMhelix 326 348\ninside 349 372

21790 GCF_002074635.1_ASM207463v1 Salmonella enterica subsp. enterica serovar Typhimurium Proteobacteria; Gammaproteobacteria; Enterobacterales; Enterobacteriaceae; Salmonella; Salmonella enterica; Salmonella enterica subsp. enterica MNNEETFYQAMRRKGVTRRSFLKFCSLAATSLGLGAGMTPKIAWALENKPRIPVVWIHGL WP_001058312.1 hydrogenase [Salmonella enterica] Length: 372\nNumber of predicted TMHs: 2\nExp number of AAs in TMHs: 36.51738\nExp number, first 60 AAs: 19.72849\nTotal prob of N-in: 0.96170\nPOSSIBLE N-term signal sequence\ninside 1 19\nTMhelix 20 39\noutside 40 325\nTMhelix 326 348\ninside 349 372

21791 GCF_002074675.1_ASM207467v1 Salmonella enterica subsp. enterica serovar Typhimurium Proteobacteria; Gammaproteobacteria; Enterobacterales; Enterobacteriaceae; Salmonella; Salmonella enterica; Salmonella enterica subsp. enterica MNNEETFYQAMRRKGVTRRSFLKFCSLAATSLGLGAGMTPKIAWALENKPRIPVVWIHGL WP_001058312.1 hydrogenase [Salmonella enterica] Length: 372\nNumber of predicted TMHs: 2\nExp number of AAs in TMHs: 36.51738\nExp number, first 60 AAs: 19.72849\nTotal prob of N-in: 0.96170\nPOSSIBLE N-term signal sequence\ninside 1 19\nTMhelix 20 39\noutside 40 325\nTMhelix 326 348\ninside 349 372

21792 GCF_002065035.1_ASM206503v1 Salmonella enterica subsp. enterica serovar Typhimurium Proteobacteria; Gammaproteobacteria; Enterobacterales; Enterobacteriaceae; Salmonella; Salmonella enterica; Salmonella enterica subsp. enterica MNNEETFYQAMRRKGVTRRSFLKFCSLAATSLGLGAGMTPKIAWALENKPRIPVVWIHGL WP_001058312.1 hydrogenase [Salmonella enterica] Length: 372\nNumber of predicted TMHs: 2\nExp

number of AAs in TMHs: 36.51738\nExp number, first 60 AAs: 19.72849\nTotal prob of N-in: 0.96170\nPOSSIBLE N-term signal sequence\ninside 1 19\nTMhelix 20 39\noutside 40 325\nTMhelix 326 348\ninside 349 372

21793 GCF_002065035.1_ASM206503v1 Salmonella enterica subsp. enterica serovar Typhimurium Proteobacteria; Gammaproteobacteria; Enterobacterales; Enterobacteriaceae; Salmonella; Salmonella enterica; Salmonella enterica subsp. enterica MTGDNTLITSHGINRRDFMKLCAALAATMGLSSKAAAEMAESVSNPQRPPVIWIGAQECT WP_000145429.1 hydrogenase 2 small subunit [Salmonella enterica] Length: 372\nNumber of predicted TMHs: 1\nExp number of AAs in TMHs: 24.51309\nExp number, first 60 AAs: 0.2682\nTotal prob of N-in: 0.14402\noutside 1 331\nTMhelix 332 354\ninside 355 372

21794 GCF_002074685.1_ASM207468v1 Salmonella enterica subsp. enterica serovar Typhimurium Proteobacteria; Gammaproteobacteria; Enterobacterales; Enterobacteriaceae; Salmonella; Salmonella enterica; Salmonella enterica subsp. enterica MNNEETFYQAMRRKGVTRRSFLKFCSLAATSLGLGAGMTPKIAWALENKPRIPVVWIHGL WP_001058312.1 hydrogenase [Salmonella enterica] Length: 372\nNumber of predicted TMHs: 2\nExp number of AAs in TMHs: 36.51738\nExp number, first 60 AAs: 19.72849\nTotal prob of N-in: 0.96170\nPOSSIBLE N-term signal sequence\ninside 1 19\nTMhelix 20 39\noutside 40 325\nTMhelix 326 348\ninside 349 372

21795 GCF_002074715.1_ASM207471v1 Salmonella enterica subsp. enterica serovar Typhimurium Proteobacteria; Gammaproteobacteria; Enterobacterales; Enterobacteriaceae; Salmonella; Salmonella enterica; Salmonella enterica subsp. enterica MNNEETFYQAMRRKGVTRRSFLKFCSLAATSLGLGAGMTPKIAWALENKPRIPVVWIHGL WP_001058312.1 hydrogenase [Salmonella enterica] Length: 372\nNumber of predicted TMHs: 2\nExp number of AAs in TMHs: 36.51738\nExp number, first 60 AAs: 19.72849\nTotal prob of N-in: 0.96170\nPOSSIBLE N-term signal sequence\ninside 1 19\nTMhelix 20 39\noutside 40 325\nTMhelix 326 348\ninside 349 372

21796 GCF_002063725.1_ASM206372v1 Salmonella enterica subsp. enterica serovar Typhimurium Proteobacteria; Gammaproteobacteria; Enterobacterales; Enterobacteriaceae; Salmonella; Salmonella enterica; Salmonella enterica subsp. enterica MNNEETFYQAMRRKGVTRRSFLKFCSLAATSLGLGAGMTPKIAWALENKPRIPVVWIHGL WP_001058312.1 hydrogenase [Salmonella enterica] Length: 372\nNumber of predicted TMHs: 2\nExp number of AAs in TMHs: 36.51738\nExp number, first 60 AAs: 19.72849\nTotal prob of N-in: 0.96170\nPOSSIBLE N-term signal sequence\ninside 1 19\nTMhelix 20 39\noutside 40 325\nTMhelix 326 348\ninside 349 372

21797 GCF_002074745.1_ASM207474v1 Salmonella enterica subsp. enterica serovar Typhimurium Proteobacteria; Gammaproteobacteria; Enterobacterales; Enterobacteriaceae; Salmonella; Salmonella enterica; Salmonella enterica subsp. enterica MNNEETFYQAMRRKGVTRRSFLKFCSLAATSLGLGAGMTPKIAWALENKPRIPVVWIHGL WP_001058312.1 hydrogenase [Salmonella enterica] Length: 372\nNumber of predicted TMHs: 2\nExp number of AAs in TMHs: 36.51738\nExp number, first 60 AAs: 19.72849\nTotal prob of N-in: 0.96170\nPOSSIBLE N-term signal sequence\ninside 1 19\nTMhelix 20 39\noutside 40 325\nTMhelix 326 348\ninside 349 372

21798 GCF_002063665.1_ASM206366v1 Salmonella enterica subsp. enterica serovar Typhimurium Proteobacteria; Gammaproteobacteria; Enterobacterales; Enterobacteriaceae; Salmonella; Salmonella enterica; Salmonella enterica subsp. enterica MNNEETFYQAMRRKGVTRRSFLKFCSLAATSLGLGAGMTPKIAWALENKPRIPVVWIHGL WP_001058312.1 hydrogenase [Salmonella enterica] Length: 372\nNumber of predicted TMHs: 2\nExp number of AAs in TMHs: 36.51738\nExp number, first 60 AAs: 19.72849\nTotal prob of N-in: 0.96170\nPOSSIBLE N-term signal sequence\ninside 1 19\nTMhelix 20 39\noutside 40 325\nTMhelix 326 348\ninside 349 372

21799 GCF_002063725.1_ASM206372v1 Salmonella enterica subsp. enterica serovar Typhimurium Proteobacteria; Gammaproteobacteria; Enterobacterales; Enterobacteriaceae; Salmonella; Salmonella enterica; Salmonella enterica subsp. enterica MTGDNTLITSHGINRRDFMKLCAALAATMGLSSKAAAEMAESVSNPQRPPVIWIGAQECT WP_000145429.1 hydrogenase 2 small subunit [Salmonella enterica] Length: 372\nNumber of predicted TMHs: 1\nExp number of AAs in TMHs: 24.51309\nExp number, first 60 AAs: 0.2682\nTotal prob of N-in: 0.14402\noutside 1 331\nTMhelix 332 354\ninside 355 372

21800 GCF_002063665.1_ASM206366v1 Salmonella enterica subsp. enterica serovar Typhimurium Proteobacteria; Gammaproteobacteria; Enterobacterales; Enterobacteriaceae; Salmonella; Salmonella enterica; Salmonella enterica subsp. enterica MTGDNTLITSHGINRRDFMKLCAALAATMGLSSKAAAEMAESVSNPQRPPVIWIGAQECT WP_000145429.1 hydrogenase 2 small subunit [Salmonella enterica] Length: 372\nNumber of predicted TMHs: 1\nExp number of AAs in TMHs: 24.51309\nExp number, first 60 AAs: 0.2682\nTotal prob of N-in: 0.14402\noutside 1 331\nTMhelix 332 354\ninside 355 372

21801 GCF_002063625.1_ASM206362v1 Salmonella enterica subsp. enterica serovar Typhimurium Proteobacteria; Gammaproteobacteria; Enterobacterales; Enterobacteriaceae; Salmonella; Salmonella enterica;

Salmonella enterica subsp. enterica MTGDNTLITSHGINRRDFMKLCAALAATMGLSSKAAAEMAESVSNPQRPPVIWIGAQECT WP_000145429.1 hydrogenase 2 small subunit [Salmonella enterica] Length: 372\nNumber of predicted TMHs: 1\nExp number of AAs in TMHs: 24.51309\nExp number, first 60 AAs: 0.2682\nTotal prob of N-in: 0.14402\noutside 1 331\nTMhelix 332 354\ninside 355 372

21802 GCF_002062935.1_ASM206293v1 Salmonella enterica subsp. enterica serovar Typhimurium Proteobacteria; Gammaproteobacteria; Enterobacterales; Enterobacteriaceae; Salmonella; Salmonella enterica; Salmonella enterica subsp. enterica MNNEETFYQAMRRKGVTRRSFLKFCSLAATSLGLGAGMTPKIAWALENKPRIPVVWIHGL WP_001058312.1 hydrogenase [Salmonella enterica] Length: 372\nNumber of predicted TMHs: 2\nExp number of AAs in TMHs: 36.51738\nExp number, first 60 AAs: 19.72849\nTotal prob of N-in: 0.96170\nPOSSIBLE N-term signal sequence\ninside 1 19\nTMhelix 20 39\noutside 40 325\nTMhelix 326 348\ninside 349 372

21803 GCF_002090935.1_ASM209093v1 Salmonella enterica subsp. enterica serovar Typhimurium Proteobacteria; Gammaproteobacteria; Enterobacterales; Enterobacteriaceae; Salmonella; Salmonella enterica; Salmonella enterica subsp. enterica MTGDNTLITSHGINRRDFMKLCAALAATMGLSSKAAAEMAESVSNPQRPPVIWIGAQECT WP_000145429.1 hydrogenase 2 small subunit [Salmonella enterica] Length: 372\nNumber of predicted TMHs: 1\nExp number of AAs in TMHs: 24.51309\nExp number, first 60 AAs: 0.2682\nTotal prob of N-in: 0.14402\noutside 1 331\nTMhelix 332 354\ninside 355 372

21804 GCF_002062935.1_ASM206293v1 Salmonella enterica subsp. enterica serovar Typhimurium Proteobacteria; Gammaproteobacteria; Enterobacterales; Enterobacteriaceae; Salmonella; Salmonella enterica; Salmonella enterica subsp. enterica MTGDNTLITSHGINRRDFMKLCAALAATMGLSSKAAAEMAESVSNPQRPPVIWIGAQECT WP_000145429.1 hydrogenase 2 small subunit [Salmonella enterica] Length: 372\nNumber of predicted TMHs: 1\nExp number of AAs in TMHs: 24.51309\nExp number, first 60 AAs: 0.2682\nTotal prob of N-in: 0.14402\noutside 1 331\nTMhelix 332 354\ninside 355 372

21805 GCF_002090935.1_ASM209093v1 Salmonella enterica subsp. enterica serovar Typhimurium Proteobacteria; Gammaproteobacteria; Enterobacterales; Enterobacteriaceae; Salmonella; Salmonella enterica; Salmonella enterica subsp. enterica MNNEETFYQAMRRKGVTRRSFLKFCSLAATSLGLGAGMTPKIAWALENKPRIPVVWIHGL WP_001058312.1 hydrogenase [Salmonella enterica] Length: 372\nNumber of predicted TMHs: 2\nExp number of AAs in TMHs: 36.51738\nExp number, first 60 AAs: 19.72849\nTotal prob of N-in: 0.96170\nPOSSIBLE N-term signal sequence\ninside 1 19\nTMhelix 20 39\noutside 40 325\nTMhelix 326 348\ninside 349 372

21806 GCF_002091015.1_ASM209101v1 Salmonella enterica subsp. enterica serovar Typhimurium Proteobacteria; Gammaproteobacteria; Enterobacterales; Enterobacteriaceae; Salmonella; Salmonella enterica; Salmonella enterica subsp. enterica MTGDNTLITSHGINRRDFMKLCAALAATMGLSSKAAAEMAESVSNPQRPPVIWIGAQECT WP_000145429.1 hydrogenase 2 small subunit [Salmonella enterica] Length: 372\nNumber of predicted TMHs: 1\nExp number of AAs in TMHs: 24.51309\nExp number, first 60 AAs: 0.2682\nTotal prob of N-in: 0.14402\noutside 1 331\nTMhelix 332 354\ninside 355 372

21807 GCF_002090955.1_ASM209095v1 Salmonella enterica subsp. enterica serovar Typhimurium Proteobacteria; Gammaproteobacteria; Enterobacterales; Enterobacteriaceae; Salmonella; Salmonella enterica; Salmonella enterica subsp. enterica MTGDNTLITSHGINRRDFMKLCAALAATMGLSSKAAAEMAESVSNPQRPPVIWIGAQECT WP_000145429.1 hydrogenase 2 small subunit [Salmonella enterica] Length: 372\nNumber of predicted TMHs: 1\nExp number of AAs in TMHs: 24.51309\nExp number, first 60 AAs: 0.2682\nTotal prob of N-in: 0.14402\noutside 1 331\nTMhelix 332 354\ninside 355 372

21808 GCF_002091045.1_ASM209104v1 Salmonella enterica subsp. enterica serovar Typhimurium Proteobacteria; Gammaproteobacteria; Enterobacterales; Enterobacteriaceae; Salmonella; Salmonella enterica; Salmonella enterica subsp. enterica MTGDNTLITSHGINRRDFMKLCAALAATMGLSSKAAAEMAESVSNPQRPPVIWIGAQECT WP_000145429.1 hydrogenase 2 small subunit [Salmonella enterica] Length: 372\nNumber of predicted TMHs: 1\nExp number of AAs in TMHs: 24.51309\nExp number, first 60 AAs: 0.2682\nTotal prob of N-in: 0.14402\noutside 1 331\nTMhelix 332 354\ninside 355 372

21809 GCF_002091095.1_ASM209109v1 Salmonella enterica subsp. enterica serovar Typhimurium Proteobacteria; Gammaproteobacteria; Enterobacterales; Enterobacteriaceae; Salmonella; Salmonella enterica; Salmonella enterica subsp. enterica MTGDNTLITSHGINRRDFMKLCAALAATMGLSSKAAAEMAESVSNPQRPPVIWIGAQECT WP_000145429.1 hydrogenase 2 small subunit [Salmonella enterica] Length: 372\nNumber of predicted TMHs: 1\nExp number of AAs in TMHs: 24.51309\nExp number, first 60 AAs: 0.2682\nTotal prob of N-in: 0.14402\noutside 1 331\nTMhelix 332 354\ninside 355 372

21810 GCF_002091125.1_ASM209112v1 Salmonella enterica subsp. enterica serovar Typhimurium Proteobacteria; Gammaproteobacteria; Enterobacterales; Enterobacteriaceae; Salmonella; Salmonella enterica; Salmonella enterica subsp. enterica MTGDNTLITSHGINRRDFMKLCAALAATMGLSSKAAAEMAESVSNPQRPPVIWIGAQECT WP_000145429.1 hydrogenase 2 small subunit [Salmonella enterica] Length: 372\nNumber of predicted TMHs: 1\nExp number of AAs in TMHs: 24.51309\nExp number, first 60 AAs: 0.2682\nTotal prob of N-in: 0.14402\noutside 1 331\nTMhelix 332 354\ninside 355 372

21811 GCF_002091015.1_ASM209101v1 Salmonella enterica subsp. enterica serovar Typhimurium Proteobacteria; Gammaproteobacteria; Enterobacterales; Enterobacteriaceae; Salmonella; Salmonella enterica; Salmonella enterica subsp. enterica MNNEETFYQAMRRKGVTRRSFLKFCSLAATSLGLGAGMTPKIAWALENKPRIPVVWIHGL WP_001058312.1 hydrogenase [Salmonella enterica] Length: 372\nNumber of predicted TMHs: 2\nExp number of AAs in TMHs: 36.51738\nExp number, first 60 AAs: 19.72849\nTotal prob of N-in: 0.96170\nPOSSIBLE N-term signal sequence\ninside 1 19\nTMhelix 20 39\noutside 40 325\nTMhelix 326 348\ninside 349 372

21812 GCF_002091215.1_ASM209121v1 Salmonella enterica subsp. enterica serovar Typhimurium Proteobacteria; Gammaproteobacteria; Enterobacterales; Enterobacteriaceae; Salmonella; Salmonella enterica; Salmonella enterica subsp. enterica MTGDNTLITSHGINRRDFMKLCAALAATMGLSSKAAAEMAESVSNPQRPPVIWIGAQECT WP_000145429.1 hydrogenase 2 small subunit [Salmonella enterica] Length: 372\nNumber of predicted TMHs: 1\nExp number of AAs in TMHs: 24.51309\nExp number, first 60 AAs: 0.2682\nTotal prob of N-in: 0.14402\noutside 1 331\nTMhelix 332 354\ninside 355 372

21813 GCF_002091215.1_ASM209121v1 Salmonella enterica subsp. enterica serovar Typhimurium Proteobacteria; Gammaproteobacteria; Enterobacterales; Enterobacteriaceae; Salmonella; Salmonella enterica; Salmonella enterica subsp. enterica MNNEETFYQAMRRKGVTRRSFLKFCSLAATSLGLGAGMTPKIAWALENKPRIPVVWIHGL WP_001058312.1 hydrogenase [Salmonella enterica] Length: 372\nNumber of predicted TMHs: 2\nExp number of AAs in TMHs: 36.51738\nExp number, first 60 AAs: 19.72849\nTotal prob of N-in: 0.96170\nPOSSIBLE N-term signal sequence\ninside 1 19\nTMhelix 20 39\noutside 40 325\nTMhelix 326 348\ninside 349 372

21814 GCF_002090955.1_ASM209095v1 Salmonella enterica subsp. enterica serovar Typhimurium Proteobacteria; Gammaproteobacteria; Enterobacterales; Enterobacteriaceae; Salmonella; Salmonella enterica; Salmonella enterica subsp. enterica MNNEETFYQAMRRKGVTRRSFLKFCSLAATSLGLGAGMTPKIAWALENKPRIPVVWIHGL WP_001058312.1 hydrogenase [Salmonella enterica] Length: 372\nNumber of predicted TMHs: 2\nExp number of AAs in TMHs: 36.51738\nExp number, first 60 AAs: 19.72849\nTotal prob of N-in: 0.96170\nPOSSIBLE N-term signal sequence\ninside 1 19\nTMhelix 20 39\noutside 40 325\nTMhelix 326 348\ninside 349 372

21815 GCF_002091335.1_ASM209133v1 Salmonella enterica subsp. enterica serovar Typhimurium Proteobacteria; Gammaproteobacteria; Enterobacterales; Enterobacteriaceae; Salmonella; Salmonella enterica; Salmonella enterica subsp. enterica MTGDNTLITSHGINRRDFMKLCAALAATMGLSSKAAAEMAESVSNPQRPPVIWIGAQECT WP_000145429.1 hydrogenase 2 small subunit [Salmonella enterica] Length: 372\nNumber of predicted TMHs: 1\nExp number of AAs in TMHs: 24.51309\nExp number, first 60 AAs: 0.2682\nTotal prob of N-in: 0.14402\noutside 1 331\nTMhelix 332 354\ninside 355 372

21816 GCF_002091255.1_ASM209125v1 Salmonella enterica subsp. enterica serovar Typhimurium Proteobacteria; Gammaproteobacteria; Enterobacterales; Enterobacteriaceae; Salmonella; Salmonella enterica; Salmonella enterica subsp. enterica MTGDNTLITSHGINRRDFMKLCAALAATMGLSSKAAAEMAESVSNPQRPPVIWIGAQECT WP_000145429.1 hydrogenase 2 small subunit [Salmonella enterica] Length: 372\nNumber of predicted TMHs: 1\nExp number of AAs in TMHs: 24.51309\nExp number, first 60 AAs: 0.2682\nTotal prob of N-in: 0.14402\noutside 1 331\nTMhelix 332 354\ninside 355 372

21817 GCF_002091335.1_ASM209133v1 Salmonella enterica subsp. enterica serovar Typhimurium Proteobacteria; Gammaproteobacteria; Enterobacterales; Enterobacteriaceae; Salmonella; Salmonella enterica; Salmonella enterica subsp. enterica MNNEETFYQAMRRKGVTRRSFLKFCSLAATSLGLGAGMTPKIAWALENKPRIPVVWIHGL WP_001058312.1 hydrogenase [Salmonella enterica] Length: 372\nNumber of predicted TMHs: 2\nExp number of AAs in TMHs: 36.51738\nExp number, first 60 AAs: 19.72849\nTotal prob of N-in: 0.96170\nPOSSIBLE N-term signal sequence\ninside 1 19\nTMhelix 20 39\noutside 40 325\nTMhelix 326 348\ninside 349 372

21818 GCF_002091255.1_ASM209125v1 Salmonella enterica subsp. enterica serovar Typhimurium Proteobacteria; Gammaproteobacteria; Enterobacterales; Enterobacteriaceae; Salmonella; Salmonella enterica; Salmonella enterica subsp. enterica MNNEETFYQAMRRKGVTRRSFLKFCSLAATSLGLGAGMTPKIAWALENKPRIPVVWIHGL WP_001058312.1 hydrogenase [Salmonella enterica] Length: 372\nNumber of predicted TMHs: 2\nExp

number of AAs in TMHs: 36.51738\nExp number, first 60 AAs: 19.72849\nTotal prob of N-in: 0.96170\nPOSSIBLE N-term signal sequence\ninside 1 19\nTMhelix 20 39\noutside 40 325\nTMhelix 326 348\ninside 349 372

21819 GCF_002091045.1_ASM209104v1 Salmonella enterica subsp. enterica serovar Typhimurium Proteobacteria; Gammaproteobacteria; Enterobacterales; Enterobacteriaceae; Salmonella; Salmonella enterica; Salmonella enterica subsp. enterica MNNEETFYQAMRRKGVTRRSFLKFCSLAATSLGLGAGMTPKIAWALENKPRIPVVWIHGL WP_001058312.1 hydrogenase [Salmonella enterica] Length: 372\nNumber of predicted TMHs: 2\nExp number of AAs in TMHs: 36.51738\nExp number, first 60 AAs: 19.72849\nTotal prob of N-in: 0.96170\nPOSSIBLE N-term signal sequence\ninside 1 19\nTMhelix 20 39\noutside 40 325\nTMhelix 326 348\ninside 349 372

21820 GCF_002091285.1_ASM209128v1 Salmonella enterica subsp. enterica serovar Typhimurium Proteobacteria; Gammaproteobacteria; Enterobacterales; Enterobacteriaceae; Salmonella; Salmonella enterica; Salmonella enterica subsp. enterica MTGDNTLITSHGINRRDFMKLCAALAATMGLSSKAAAEMAESVSNPQRPPVIWIGAQECT WP_000145429.1 hydrogenase 2 small subunit [Salmonella enterica] Length: 372\nNumber of predicted TMHs: 1\nExp number of AAs in TMHs: 24.51309\nExp number, first 60 AAs: 0.2682\nTotal prob of N-in: 0.14402\noutside 1 331\nTMhelix 332 354\ninside 355 372

21821 GCF_002091285.1_ASM209128v1 Salmonella enterica subsp. enterica serovar Typhimurium Proteobacteria; Gammaproteobacteria; Enterobacterales; Enterobacteriaceae; Salmonella; Salmonella enterica; Salmonella enterica subsp. enterica MNNEETFYQAMRRKGVTRRSFLKFCSLAATSLGLGAGMTPKIAWALENKPRIPVVWIHGL WP_001058312.1 hydrogenase [Salmonella enterica] Length: 372\nNumber of predicted TMHs: 2\nExp number of AAs in TMHs: 36.51738\nExp number, first 60 AAs: 19.72849\nTotal prob of N-in: 0.96170\nPOSSIBLE N-term signal sequence\ninside 1 19\nTMhelix 20 39\noutside 40 325\nTMhelix 326 348\ninside 349 372

21822 GCF_002091095.1_ASM209109v1 Salmonella enterica subsp. enterica serovar Typhimurium Proteobacteria; Gammaproteobacteria; Enterobacterales; Enterobacteriaceae; Salmonella; Salmonella enterica; Salmonella enterica subsp. enterica MNNEETFYQAMRRKGVTRRSFLKFCSLAATSLGLGAGMTPKIAWALENKPRIPVVWIHGL WP_001058312.1 hydrogenase [Salmonella enterica] Length: 372\nNumber of predicted TMHs: 2\nExp number of AAs in TMHs: 36.51738\nExp number, first 60 AAs: 19.72849\nTotal prob of N-in: 0.96170\nPOSSIBLE N-term signal sequence\ninside 1 19\nTMhelix 20 39\noutside 40 325\nTMhelix 326 348\ninside 349 372

21823 GCF_002091125.1_ASM209112v1 Salmonella enterica subsp. enterica serovar Typhimurium Proteobacteria; Gammaproteobacteria; Enterobacterales; Enterobacteriaceae; Salmonella; Salmonella enterica; Salmonella enterica subsp. enterica MNNEETFYQAMRRKGVTRRSFLKFCSLAATSLGLGAGMTPKIAWALENKPRIPVVWIHGL WP_001058312.1 hydrogenase [Salmonella enterica] Length: 372\nNumber of predicted TMHs: 2\nExp number of AAs in TMHs: 36.51738\nExp number, first 60 AAs: 19.72849\nTotal prob of N-in: 0.96170\nPOSSIBLE N-term signal sequence\ninside 1 19\nTMhelix 20 39\noutside 40 325\nTMhelix 326 348\ninside 349 372

21824 GCF_002061615.1_ASM206161v1 Salmonella enterica subsp. enterica serovar Typhimurium Proteobacteria; Gammaproteobacteria; Enterobacterales; Enterobacteriaceae; Salmonella; Salmonella enterica; Salmonella enterica subsp. enterica MTGDNTLITSHGINRRDFMKLCAALAATMGLSSKAAAEMAESVSNPQRPPVIWIGAQECT WP_000145431.1 hydrogenase 2 small subunit [Salmonella enterica] Length: 372\nNumber of predicted TMHs: 1\nExp number of AAs in TMHs: 24.51505\nExp number, first 60 AAs: 0.2682\nTotal prob of N-in: 0.14406\noutside 1 331\nTMhelix 332 354\ninside 355 372

21825 GCF_002106175.1_ASM210617v1 Salmonella enterica subsp. enterica serovar Typhimurium Proteobacteria; Gammaproteobacteria; Enterobacterales; Enterobacteriaceae; Salmonella; Salmonella enterica; Salmonella enterica subsp. enterica MTGDNTLITSHGINRRDFMKLCAALAATMGLSSKAAAEMAESVSNPQRPPVIWIGAQECT WP_000145429.1 hydrogenase 2 small subunit [Salmonella enterica] Length: 372\nNumber of predicted TMHs: 1\nExp number of AAs in TMHs: 24.51309\nExp number, first 60 AAs: 0.2682\nTotal prob of N-in: 0.14402\noutside 1 331\nTMhelix 332 354\ninside 355 372

21826 GCF_002106175.1_ASM210617v1 Salmonella enterica subsp. enterica serovar Typhimurium Proteobacteria; Gammaproteobacteria; Enterobacterales; Enterobacteriaceae; Salmonella; Salmonella enterica; Salmonella enterica subsp. enterica MNNEETFYQAMRRKGVTRRSFLKFCSLAATSLGLGAGMTPKIAWALENKPRIPVVWIHGL WP_001058312.1 hydrogenase [Salmonella enterica] Length: 372\nNumber of predicted TMHs: 2\nExp number of AAs in TMHs: 36.51738\nExp number, first 60 AAs: 19.72849\nTotal prob of N-in: 0.96170\nPOSSIBLE N-term signal sequence\ninside 1 19\nTMhelix 20 39\noutside 40 325\nTMhelix 326 348\ninside 349 372

21827 GCF_002074175.1_ASM207417v1 Salmonella enterica subsp. enterica serovar Typhimurium Proteobacteria; Gammaproteobacteria; Enterobacterales; Enterobacteriaceae; Salmonella; Salmonella enterica;

Salmonella enterica subsp. enterica MTGDNTLITSHGINRRDFMKLCAALAATMGLSSKAAAEMAESVSNPQRPPVIWIGAQECT WP_000145429.1 hydrogenase 2 small subunit [Salmonella enterica] Length: 372\nNumber of predicted TMHs: 1\nExp number of AAs in TMHs: 24.51309\nExp number, first 60 AAs: 0.2682\nTotal prob of N-in: 0.14402\noutside 1 331\nTMhelix 332 354\ninside 355 372

21828 GCF_001691985.1_ASM169198v1 Salmonella enterica subsp. enterica serovar Typhimurium Proteobacteria; Gammaproteobacteria; Enterobacterales; Enterobacteriaceae; Salmonella; Salmonella enterica; Salmonella enterica subsp. enterica MTGDNTLITSHGINRRDFMKLCAALAATMGLSSKAAAEMAESVSNPQRPPVIWIGAQECT WP_000145429.1 hydrogenase 2 small subunit [Salmonella enterica] Length: 372\nNumber of predicted TMHs: 1\nExp number of AAs in TMHs: 24.51309\nExp number, first 60 AAs: 0.2682\nTotal prob of N-in: 0.14402\noutside 1 331\nTMhelix 332 354\ninside 355 372

21829 GCF_001691985.1_ASM169198v1 Salmonella enterica subsp. enterica serovar Typhimurium Proteobacteria; Gammaproteobacteria; Enterobacterales; Enterobacteriaceae; Salmonella; Salmonella enterica; Salmonella enterica subsp. enterica MNNEETFYQAMRRKGVTRRSFLKFCSLAATSLGLGAGMTPKIAWALENKPRIPVVWIHGL WP_001058312.1 hydrogenase [Salmonella enterica] Length: 372\nNumber of predicted TMHs: 2\nExp number of AAs in TMHs: 36.51738\nExp number, first 60 AAs: 19.72849\nTotal prob of N-in: 0.96170\nPOSSIBLE N-term signal sequence\ninside 1 19\nTMhelix 20 39\noutside 40 325\nTMhelix 326 348\ninside 349 372

21830 GCF_001692145.1_ASM169214v1 Salmonella enterica subsp. enterica serovar Typhimurium Proteobacteria; Gammaproteobacteria; Enterobacterales; Enterobacteriaceae; Salmonella; Salmonella enterica; Salmonella enterica subsp. enterica MTGDNTLITSHGINRRDFMKLCAALAATMGLSSKAAAEMAESVSNPQRPPVIWIGAQECT WP_000145429.1 hydrogenase 2 small subunit [Salmonella enterica] Length: 372\nNumber of predicted TMHs: 1\nExp number of AAs in TMHs: 24.51309\nExp number, first 60 AAs: 0.2682\nTotal prob of N-in: 0.14402\noutside 1 331\nTMhelix 332 354\ninside 355 372

21831 GCF_001729035.1_ASM172903v1 Salmonella enterica subsp. enterica serovar Typhimurium Proteobacteria; Gammaproteobacteria; Enterobacterales; Enterobacteriaceae; Salmonella; Salmonella enterica; Salmonella enterica subsp. enterica MTGDNTLITSHGINRRDFMKLCAALAATMGLSSKAAAEMAESVSNPQRPPVIWIGAQECT WP_000145429.1 hydrogenase 2 small subunit [Salmonella enterica] Length: 372\nNumber of predicted TMHs: 1\nExp number of AAs in TMHs: 24.51309\nExp number, first 60 AAs: 0.2682\nTotal prob of N-in: 0.14402\noutside 1 331\nTMhelix 332 354\ninside 355 372

21832 GCF_002066815.1_ASM206681v1 Salmonella enterica subsp. enterica serovar Typhimurium Proteobacteria; Gammaproteobacteria; Enterobacterales; Enterobacteriaceae; Salmonella; Salmonella enterica; Salmonella enterica subsp. enterica MTGDNTLITSHGINRRDFMKLCAALAATMGLSSKAAAEMAESVSNPQRPPVIWIGAQECT WP_080248566.1 hydrogenase 2 small subunit [Salmonella enterica] Length: 372\nNumber of predicted TMHs: 1\nExp number of AAs in TMHs: 24.51398\nExp number, first 60 AAs: 0.26828\nTotal prob of N-in: 0.14404\noutside 1 331\nTMhelix 332 354\ninside 355 372

21833 GCF_002074215.1_ASM207421v1 Salmonella enterica subsp. enterica serovar Typhimurium Proteobacteria; Gammaproteobacteria; Enterobacterales; Enterobacteriaceae; Salmonella; Salmonella enterica; Salmonella enterica subsp. enterica MTGDNTLITSHGINRRDFMKLCAALAATMGLSSKAAAEMAESVSNPQRPPVIWIGAQECT WP_000145429.1 hydrogenase 2 small subunit [Salmonella enterica] Length: 372\nNumber of predicted TMHs: 1\nExp number of AAs in TMHs: 24.51309\nExp number, first 60 AAs: 0.2682\nTotal prob of N-in: 0.14402\noutside 1 331\nTMhelix 332 354\ninside 355 372

21834 GCF_002035045.1_ASM203504v1 Salmonella enterica subsp. enterica serovar Typhimurium Proteobacteria; Gammaproteobacteria; Enterobacterales; Enterobacteriaceae; Salmonella; Salmonella enterica; Salmonella enterica subsp. enterica MNNEETFYQAMRRKGVTRRSFLKFCSLAATSLGLGAGMTPKIAWALENKPRIPVVWIHGL WP_001058312.1 hydrogenase [Salmonella enterica] Length: 372\nNumber of predicted TMHs: 2\nExp number of AAs in TMHs: 36.51738\nExp number, first 60 AAs: 19.72849\nTotal prob of N-in: 0.96170\nPOSSIBLE N-term signal sequence\ninside 1 19\nTMhelix 20 39\noutside 40 325\nTMhelix 326 348\ninside 349 372

21835 GCF_001729045.1_ASM172904v1 Salmonella enterica subsp. enterica serovar Typhimurium Proteobacteria; Gammaproteobacteria; Enterobacterales; Enterobacteriaceae; Salmonella; Salmonella enterica; Salmonella enterica subsp. enterica MTGDNTLITSHGINRRDFMKLCAALAATMGLSSKAAAEMAESVSNPQRPPVIWIGAQECT WP_000145429.1 hydrogenase 2 small subunit [Salmonella enterica] Length: 372\nNumber of predicted TMHs: 1\nExp number of AAs in TMHs: 24.51309\nExp number, first 60 AAs: 0.2682\nTotal prob of N-in: 0.14402\noutside 1 331\nTMhelix 332 354\ninside 355 372

21836 GCF_001756075.1_CFSAN008077 Salmonella enterica subsp. enterica serovar Typhimurium Proteobacteria; Gammaproteobacteria; Enterobacterales; Enterobacteriaceae; Salmonella; Salmonella enterica; Salmonella enterica subsp. enterica MTGDNTLITSHGINRRDFMKLCAALAATMGLSSKAAAEMAESVSNPQRPPVIWIGAQECT WP_000145429.1 hydrogenase 2 small subunit [Salmonella enterica] Length: 372\nNumber of predicted TMHs: 1\nExp number of AAs in TMHs: 24.51309\nExp number, first 60 AAs: 0.2682\nTotal prob of N-in: 0.14402\noutside 1 331\nTMhelix 332 354\ninside 355 372

21837 GCF_002034985.1_ASM203498v1 Salmonella enterica subsp. enterica serovar Typhimurium Proteobacteria; Gammaproteobacteria; Enterobacterales; Enterobacteriaceae; Salmonella; Salmonella enterica; Salmonella enterica subsp. enterica MTGDNTLITSHGINRRDFMKLCAALAATMGLSSKAAAEMAESVSNPQRPPVIWIGAQECT WP_000145429.1 hydrogenase 2 small subunit [Salmonella enterica] Length: 372\nNumber of predicted TMHs: 1\nExp number of AAs in TMHs: 24.51309\nExp number, first 60 AAs: 0.2682\nTotal prob of N-in: 0.14402\noutside 1 331\nTMhelix 332 354\ninside 355 372

21838 GCF_001755955.1_CFSAN008080 Salmonella enterica subsp. enterica serovar Typhimurium Proteobacteria; Gammaproteobacteria; Enterobacterales; Enterobacteriaceae; Salmonella; Salmonella enterica; Salmonella enterica subsp. enterica MTGDNTLITSHGINRRDFMKLCAALAATMGLSSKAAAEMAESVSNPQRPPVIWIGAQECT WP_000145429.1 hydrogenase 2 small subunit [Salmonella enterica] Length: 372\nNumber of predicted TMHs: 1\nExp number of AAs in TMHs: 24.51309\nExp number, first 60 AAs: 0.2682\nTotal prob of N-in: 0.14402\noutside 1 331\nTMhelix 332 354\ninside 355 372

21839 GCF_002034225.1_ASM203422v1 Salmonella enterica subsp. enterica serovar Typhimurium Proteobacteria; Gammaproteobacteria; Enterobacterales; Enterobacteriaceae; Salmonella; Salmonella enterica; Salmonella enterica subsp. enterica MNNEETFYQAMRRKGVTRRSFLKFCSLAATSLGLGAGMTPKIAWALENKPRIPVVWIHGL WP_001058312.1 hydrogenase [Salmonella enterica] Length: 372\nNumber of predicted TMHs: 2\nExp number of AAs in TMHs: 36.51738\nExp number, first 60 AAs: 19.72849\nTotal prob of N-in: 0.96170\nPOSSIBLE N-term signal sequence\ninside 1 19\nTMhelix 20 39\noutside 40 325\nTMhelix 326 348\ninside 349 372

21840 GCF_001756035.1_CFSAN008086 Salmonella enterica subsp. enterica serovar Typhimurium Proteobacteria; Gammaproteobacteria; Enterobacterales; Enterobacteriaceae; Salmonella; Salmonella enterica; Salmonella enterica subsp. enterica MNNEETFYQAMRRKGVTRRSFLKFCSLAATSLGLGAGMTPKIAWALENKPRIPVVWIHGL WP_001058312.1 hydrogenase [Salmonella enterica] Length: 372\nNumber of predicted TMHs: 2\nExp number of AAs in TMHs: 36.51738\nExp number, first 60 AAs: 19.72849\nTotal prob of N-in: 0.96170\nPOSSIBLE N-term signal sequence\ninside 1 19\nTMhelix 20 39\noutside 40 325\nTMhelix 326 348\ninside 349 372

21841 GCF_002034225.1_ASM203422v1 Salmonella enterica subsp. enterica serovar Typhimurium Proteobacteria; Gammaproteobacteria; Enterobacterales; Enterobacteriaceae; Salmonella; Salmonella enterica; Salmonella enterica subsp. enterica MTGDNTLITSHGINRRDFMKLCAALAATMGLSSKAAAEMAESVSNPQRPPVIWIGAQECT WP_000145429.1 hydrogenase 2 small subunit [Salmonella enterica] Length: 372\nNumber of predicted TMHs: 1\nExp number of AAs in TMHs: 24.51309\nExp number, first 60 AAs: 0.2682\nTotal prob of N-in: 0.14402\noutside 1 331\nTMhelix 332 354\ninside 355 372

21842 GCF_001755955.1_CFSAN008080 Salmonella enterica subsp. enterica serovar Typhimurium Proteobacteria; Gammaproteobacteria; Enterobacterales; Enterobacteriaceae; Salmonella; Salmonella enterica; Salmonella enterica subsp. enterica MNNEETFYQAMRRKGVTRRSFLKFCSLAATSLGLGAGMTPKIAWALENKPRIPVVWIHGL WP_001058312.1 hydrogenase [Salmonella enterica] Length: 372\nNumber of predicted TMHs: 2\nExp number of AAs in TMHs: 36.51738\nExp number, first 60 AAs: 19.72849\nTotal prob of N-in: 0.96170\nPOSSIBLE N-term signal sequence\ninside 1 19\nTMhelix 20 39\noutside 40 325\nTMhelix 326 348\ninside 349 372

21843 GCF_002074235.1_ASM207423v1 Salmonella enterica subsp. enterica serovar Typhimurium Proteobacteria; Gammaproteobacteria; Enterobacterales; Enterobacteriaceae; Salmonella; Salmonella enterica; Salmonella enterica subsp. enterica MTGDNTLITSHGINRRDFMKLCAALAATMGLSSKAAAEMAESVSNPQRPPVIWIGAQECT WP_000145429.1 hydrogenase 2 small subunit [Salmonella enterica] Length: 372\nNumber of predicted TMHs: 1\nExp number of AAs in TMHs: 24.51309\nExp number, first 60 AAs: 0.2682\nTotal prob of N-in: 0.14402\noutside 1 331\nTMhelix 332 354\ninside 355 372

21844 GCF_900184385.1_STMU2UK Salmonella enterica subsp. enterica serovar Typhimurium Proteobacteria; Gammaproteobacteria; Enterobacterales; Enterobacteriaceae; Salmonella; Salmonella enterica; Salmonella enterica subsp. enterica MNNEETFYQAMRRKGVTRRSFLKFCSLAATSLGLGAGMTPKIAWALENKPRIPVVWIHGL WP_001058312.1 hydrogenase [Salmonella enterica] Length: 372\nNumber of predicted TMHs: 2\nExp

number of AAs in TMHs: 36.51738\nExp number, first 60 AAs: 19.72849\nTotal prob of N-in: 0.96170\nPOSSIBLE N-term signal sequence\ninside 1 19\nTMhelix 20 39\noutside 40 325\nTMhelix 326 348\ninside 349 372

21845 GCF_002074295.1_ASM207429v1 Salmonella enterica subsp. enterica serovar Typhimurium Proteobacteria; Gammaproteobacteria; Enterobacterales; Enterobacteriaceae; Salmonella; Salmonella enterica; Salmonella enterica subsp. enterica MTGDNTLITSHGINRRDFMKLCAALAATMGLSSKAAAEMAESVSNPQRPPVIWIGAQECT WP_000145429.1 hydrogenase 2 small subunit [Salmonella enterica] Length: 372\nNumber of predicted TMHs: 1\nExp number of AAs in TMHs: 24.51309\nExp number, first 60 AAs: 0.2682\nTotal prob of N-in: 0.14402\noutside 1 331\nTMhelix 332 354\ninside 355 372

21846 GCF_002074335.1_ASM207433v1 Salmonella enterica subsp. enterica serovar Typhimurium Proteobacteria; Gammaproteobacteria; Enterobacterales; Enterobacteriaceae; Salmonella; Salmonella enterica; Salmonella enterica subsp. enterica MTGDNTLITSHGINRRDFMKLCAALAATMGLSSKAAAEMAESVSNPQRPPVIWIGAQECT WP_000145429.1 hydrogenase 2 small subunit [Salmonella enterica] Length: 372\nNumber of predicted TMHs: 1\nExp number of AAs in TMHs: 24.51309\nExp number, first 60 AAs: 0.2682\nTotal prob of N-in: 0.14402\noutside 1 331\nTMhelix 332 354\ninside 355 372

21847 GCF_002032975.1_ASM203297v1 Salmonella enterica subsp. enterica serovar Typhimurium Proteobacteria; Gammaproteobacteria; Enterobacterales; Enterobacteriaceae; Salmonella; Salmonella enterica; Salmonella enterica subsp. enterica MTGDNTLITSHGINRRDFMKLCAALAATMGLSSKAAAEMAESVSNPQRPPVIWIGAQECT WP_000145429.1 hydrogenase 2 small subunit [Salmonella enterica] Length: 372\nNumber of predicted TMHs: 1\nExp number of AAs in TMHs: 24.51309\nExp number, first 60 AAs: 0.2682\nTotal prob of N-in: 0.14402\noutside 1 331\nTMhelix 332 354\ninside 355 372

21848 GCF_001756075.1_CFSAN008077 Salmonella enterica subsp. enterica serovar Typhimurium Proteobacteria; Gammaproteobacteria; Enterobacterales; Enterobacteriaceae; Salmonella; Salmonella enterica; Salmonella enterica subsp. enterica MNNEETFYQAMRRKGVTRRSFLKFCSLAATSLGLGAGMTPKIAWALENKPRIPVVWIHGL WP_001058312.1 hydrogenase [Salmonella enterica] Length: 372\nNumber of predicted TMHs: 2\nExp number of AAs in TMHs: 36.51738\nExp number, first 60 AAs: 19.72849\nTotal prob of N-in: 0.96170\nPOSSIBLE N-term signal sequence\ninside 1 19\nTMhelix 20 39\noutside 40 325\nTMhelix 326 348\ninside 349 372

21849 GCF_002074215.1_ASM207421v1 Salmonella enterica subsp. enterica serovar Typhimurium Proteobacteria; Gammaproteobacteria; Enterobacterales; Enterobacteriaceae; Salmonella; Salmonella enterica; Salmonella enterica subsp. enterica MNNEETFYQAMRRKGVTRRSFLKFCSLAATSLGLGAGMTPKIAWALENKPRIPVVWIHGL WP_001058312.1 hydrogenase [Salmonella enterica] Length: 372\nNumber of predicted TMHs: 2\nExp number of AAs in TMHs: 36.51738\nExp number, first 60 AAs: 19.72849\nTotal prob of N-in: 0.96170\nPOSSIBLE N-term signal sequence\ninside 1 19\nTMhelix 20 39\noutside 40 325\nTMhelix 326 348\ninside 349 372

21850 GCF_002074235.1_ASM207423v1 Salmonella enterica subsp. enterica serovar Typhimurium Proteobacteria; Gammaproteobacteria; Enterobacterales; Enterobacteriaceae; Salmonella; Salmonella enterica; Salmonella enterica subsp. enterica MNNEETFYQAMRRKGVTRRSFLKFCSLAATSLGLGAGMTPKIAWALENKPRIPVVWIHGL WP_001058312.1 hydrogenase [Salmonella enterica] Length: 372\nNumber of predicted TMHs: 2\nExp number of AAs in TMHs: 36.51738\nExp number, first 60 AAs: 19.72849\nTotal prob of N-in: 0.96170\nPOSSIBLE N-term signal sequence\ninside 1 19\nTMhelix 20 39\noutside 40 325\nTMhelix 326 348\ninside 349 372

21851 GCF_002074175.1_ASM207417v1 Salmonella enterica subsp. enterica serovar Typhimurium Proteobacteria; Gammaproteobacteria; Enterobacterales; Enterobacteriaceae; Salmonella; Salmonella enterica; Salmonella enterica subsp. enterica MNNEETFYQAMRRKGVTRRSFLKFCSLAATSLGLGAGMTPKIAWALENKPRIPVVWIHGL WP_001058312.1 hydrogenase [Salmonella enterica] Length: 372\nNumber of predicted TMHs: 2\nExp number of AAs in TMHs: 36.51738\nExp number, first 60 AAs: 19.72849\nTotal prob of N-in: 0.96170\nPOSSIBLE N-term signal sequence\ninside 1 19\nTMhelix 20 39\noutside 40 325\nTMhelix 326 348\ninside 349 372

21852 GCF_002074385.1_ASM207438v1 Salmonella enterica subsp. enterica serovar Typhimurium Proteobacteria; Gammaproteobacteria; Enterobacterales; Enterobacteriaceae; Salmonella; Salmonella enterica; Salmonella enterica subsp. enterica MTGDNTLITSHGINRRDFMKLCAALAATMGLSSKAAAEMAESVSNPQRPPVIWIGAQECT WP_000145429.1 hydrogenase 2 small subunit [Salmonella enterica] Length: 372\nNumber of predicted TMHs: 1\nExp number of AAs in TMHs: 24.51309\nExp number, first 60 AAs: 0.2682\nTotal prob of N-in: 0.14402\noutside 1 331\nTMhelix 332 354\ninside 355 372

21853 GCF_002074435.1_ASM207443v1 Salmonella enterica subsp. enterica serovar Typhimurium Proteobacteria; Gammaproteobacteria; Enterobacterales; Enterobacteriaceae; Salmonella; Salmonella enterica;

Salmonella enterica subsp. enterica MTGDNTLITSHGINRRDFMKLCAALAATMGLSSKAAAEMAESVSNPQRPPVIWIGAQECT WP_000145429.1 hydrogenase 2 small subunit [Salmonella enterica] Length: 372\nNumber of predicted TMHs: 1\nExp number of AAs in TMHs: 24.51309\nExp number, first 60 AAs: 0.2682\nTotal prob of N-in: 0.14402\noutside 1 331\nTMhelix 332 354\ninside 355 372

21854 GCF_002074455.1_ASM207445v1 Salmonella enterica subsp. enterica serovar Typhimurium Proteobacteria; Gammaproteobacteria; Enterobacterales; Enterobacteriaceae; Salmonella; Salmonella enterica; Salmonella enterica subsp. enterica MTGDNTLITSHGINRRDFMKLCAALAATMGLSSKAAAEMAESVSNPQRPPVIWIGAQECT WP_000145429.1 hydrogenase 2 small subunit [Salmonella enterica] Length: 372\nNumber of predicted TMHs: 1\nExp number of AAs in TMHs: 24.51309\nExp number, first 60 AAs: 0.2682\nTotal prob of N-in: 0.14402\noutside 1 331\nTMhelix 332 354\ninside 355 372

21855 GCF_002074515.1_ASM207451v1 Salmonella enterica subsp. enterica serovar Typhimurium Proteobacteria; Gammaproteobacteria; Enterobacterales; Enterobacteriaceae; Salmonella; Salmonella enterica; Salmonella enterica subsp. enterica MTGDNTLITSHGINRRDFMKLCAALAATMGLSSKAAAEMAESVSNPQRPPVIWIGAQECT WP_000145429.1 hydrogenase 2 small subunit [Salmonella enterica] Length: 372\nNumber of predicted TMHs: 1\nExp number of AAs in TMHs: 24.51309\nExp number, first 60 AAs: 0.2682\nTotal prob of N-in: 0.14402\noutside 1 331\nTMhelix 332 354\ninside 355 372

21856 GCF_002074545.1_ASM207454v1 Salmonella enterica subsp. enterica serovar Typhimurium Proteobacteria; Gammaproteobacteria; Enterobacterales; Enterobacteriaceae; Salmonella; Salmonella enterica; Salmonella enterica subsp. enterica MTGDNTLITSHGINRRDFMKLCAALAATMGLSSKAAAEMAESVSNPQRPPVIWIGAQECT WP_000145429.1 hydrogenase 2 small subunit [Salmonella enterica] Length: 372\nNumber of predicted TMHs: 1\nExp number of AAs in TMHs: 24.51309\nExp number, first 60 AAs: 0.2682\nTotal prob of N-in: 0.14402\noutside 1 331\nTMhelix 332 354\ninside 355 372

21857 GCF_002030885.1_ASM203088v1 Salmonella enterica subsp. enterica serovar Typhimurium Proteobacteria; Gammaproteobacteria; Enterobacterales; Enterobacteriaceae; Salmonella; Salmonella enterica; Salmonella enterica subsp. enterica MTGDNTLITSHGINRRDFMKLCAALAATMGLSSKAAAEMAESVSNPQRPPVIWIGAQECT WP_000145429.1 hydrogenase 2 small subunit [Salmonella enterica] Length: 372\nNumber of predicted TMHs: 1\nExp number of AAs in TMHs: 24.51309\nExp number, first 60 AAs: 0.2682\nTotal prob of N-in: 0.14402\noutside 1 331\nTMhelix 332 354\ninside 355 372

21858 GCF_002074385.1_ASM207438v1 Salmonella enterica subsp. enterica serovar Typhimurium Proteobacteria; Gammaproteobacteria; Enterobacterales; Enterobacteriaceae; Salmonella; Salmonella enterica; Salmonella enterica subsp. enterica MNNEETFYQAMRRKGVTRRSFLKFCSLAATSLGLGAGMTPKIAWALENKPRIPVVWIHGL WP_001058312.1 hydrogenase [Salmonella enterica] Length: 372\nNumber of predicted TMHs: 2\nExp number of AAs in TMHs: 36.51738\nExp number, first 60 AAs: 19.72849\nTotal prob of N-in: 0.96170\nPOSSIBLE N-term signal sequence\ninside 1 19\nTMhelix 20 39\noutside 40 325\nTMhelix 326 348\ninside 349 372

21859 GCF_002074295.1_ASM207429v1 Salmonella enterica subsp. enterica serovar Typhimurium Proteobacteria; Gammaproteobacteria; Enterobacterales; Enterobacteriaceae; Salmonella; Salmonella enterica; Salmonella enterica subsp. enterica MNNEETFYQAMRRKGVTRRSFLKFCSLAATSLGLGAGMTPKIAWALENKPRIPVVWIHGL WP_001058312.1 hydrogenase [Salmonella enterica] Length: 372\nNumber of predicted TMHs: 2\nExp number of AAs in TMHs: 36.51738\nExp number, first 60 AAs: 19.72849\nTotal prob of N-in: 0.96170\nPOSSIBLE N-term signal sequence\ninside 1 19\nTMhelix 20 39\noutside 40 325\nTMhelix 326 348\ninside 349 372

21860 GCF_002029825.1_ASM202982v1 Salmonella enterica subsp. enterica serovar Typhimurium Proteobacteria; Gammaproteobacteria; Enterobacterales; Enterobacteriaceae; Salmonella; Salmonella enterica; Salmonella enterica subsp. enterica MTGDNTLITSHGINRRDFMKLCAALAATMGLSSKAAAEMAESVSNPQRPPVIWIGAQECT WP_000145429.1 hydrogenase 2 small subunit [Salmonella enterica] Length: 372\nNumber of predicted TMHs: 1\nExp number of AAs in TMHs: 24.51309\nExp number, first 60 AAs: 0.2682\nTotal prob of N-in: 0.14402\noutside 1 331\nTMhelix 332 354\ninside 355 372

21861 GCF_002029395.1_ASM202939v1 Salmonella enterica subsp. enterica serovar Typhimurium Proteobacteria; Gammaproteobacteria; Enterobacterales; Enterobacteriaceae; Salmonella; Salmonella enterica; Salmonella enterica subsp. enterica MNNEETFYQAMRRKGVTRRSFLKFCSLAATSLGLGAGMTPKIAWALENKPRIPVVWIHGL WP_001058312.1 hydrogenase [Salmonella enterica] Length: 372\nNumber of predicted TMHs: 2\nExp number of AAs in TMHs: 36.51738\nExp number, first 60 AAs: 19.72849\nTotal prob of N-in: 0.96170\nPOSSIBLE N-term signal sequence\ninside 1 19\nTMhelix 20 39\noutside 40 325\nTMhelix 326 348\ninside 349 372

21862 GCF_002029785.1_ASM202978v1 Salmonella enterica subsp. enterica serovar Typhimurium Proteobacteria; Gammaproteobacteria; Enterobacterales; Enterobacteriaceae; Salmonella; Salmonella enterica; Salmonella enterica subsp. enterica MTGDNTLITSHGINRRDFMKLCAALAATMGLSSKAAAEMAESVSNPQRPPVIWIGAQECT WP_000145429.1 hydrogenase 2 small subunit [Salmonella enterica] Length: 372\nNumber of predicted TMHs: 1\nExp number of AAs in TMHs: 24.51309\nExp number, first 60 AAs: 0.2682\nTotal prob of N-in: 0.14402\noutside 1 331\nTMhelix 332 354\ninside 355 372

21863 GCF_002029375.1_ASM202937v1 Salmonella enterica subsp. enterica serovar Typhimurium Proteobacteria; Gammaproteobacteria; Enterobacterales; Enterobacteriaceae; Salmonella; Salmonella enterica; Salmonella enterica subsp. enterica MNNEETFYQAMRRKGVTRRSFLKFCSLAATSLGLGAGMTPKIAWALENKPRIPVVWIHGL WP_001058312.1 hydrogenase [Salmonella enterica] Length: 372\nNumber of predicted TMHs: 2\nExp number of AAs in TMHs: 36.51738\nExp number, first 60 AAs: 19.72849\nTotal prob of N-in: 0.96170\nPOSSIBLE N-term signal sequence\ninside 1 19\nTMhelix 20 39\noutside 40 325\nTMhelix 326 348\ninside 349 372

21864 GCF_002029375.1_ASM202937v1 Salmonella enterica subsp. enterica serovar Typhimurium Proteobacteria; Gammaproteobacteria; Enterobacterales; Enterobacteriaceae; Salmonella; Salmonella enterica; Salmonella enterica subsp. enterica MTGDNTLITSHGINRRDFMKLCAALAATMGLSSKAAAEMAESVSNPQRPPVIWIGAQECT WP_000145429.1 hydrogenase 2 small subunit [Salmonella enterica] Length: 372\nNumber of predicted TMHs: 1\nExp number of AAs in TMHs: 24.51309\nExp number, first 60 AAs: 0.2682\nTotal prob of N-in: 0.14402\noutside 1 331\nTMhelix 332 354\ninside 355 372

21865 GCF_001997115.1_ASM199711v1 Salmonella enterica subsp. enterica serovar Typhimurium Proteobacteria; Gammaproteobacteria; Enterobacterales; Enterobacteriaceae; Salmonella; Salmonella enterica; Salmonella enterica subsp. enterica MNNEETFYQAMRRKGVTRRSFLKFCSLAATSLGLGAGMTPKIAWALENKPRIPVVWIHGL WP_001058312.1 hydrogenase [Salmonella enterica] Length: 372\nNumber of predicted TMHs: 2\nExp number of AAs in TMHs: 36.51738\nExp number, first 60 AAs: 19.72849\nTotal prob of N-in: 0.96170\nPOSSIBLE N-term signal sequence\ninside 1 19\nTMhelix 20 39\noutside 40 325\nTMhelix 326 348\ninside 349 372

21866 GCF_002028675.1_ASM202867v1 Salmonella enterica subsp. enterica serovar Typhimurium Proteobacteria; Gammaproteobacteria; Enterobacterales; Enterobacteriaceae; Salmonella; Salmonella enterica; Salmonella enterica subsp. enterica MNNEETFYQAMRRKGVTRRSFLKFCSLAATSLGLGAGMTPKIAWALENKPRIPVVWIHGL WP_001058312.1 hydrogenase [Salmonella enterica] Length: 372\nNumber of predicted TMHs: 2\nExp number of AAs in TMHs: 36.51738\nExp number, first 60 AAs: 19.72849\nTotal prob of N-in: 0.96170\nPOSSIBLE N-term signal sequence\ninside 1 19\nTMhelix 20 39\noutside 40 325\nTMhelix 326 348\ninside 349 372

21867 GCF_002156155.1_ASM215615v1 Salmonella enterica subsp. enterica serovar Typhimurium Proteobacteria; Gammaproteobacteria; Enterobacterales; Enterobacteriaceae; Salmonella; Salmonella enterica; Salmonella enterica subsp. enterica MNNEETFYQAMRRKGVTRRSFLKFCSLAATSLGLGAGMTPKIAWALENKPRIPVVWIHGL WP_001058312.1 hydrogenase [Salmonella enterica] Length: 372\nNumber of predicted TMHs: 2\nExp number of AAs in TMHs: 36.51738\nExp number, first 60 AAs: 19.72849\nTotal prob of N-in: 0.96170\nPOSSIBLE N-term signal sequence\ninside 1 19\nTMhelix 20 39\noutside 40 325\nTMhelix 326 348\ninside 349 372

21868 GCF_002028585.1_ASM202858v1 Salmonella enterica subsp. enterica serovar Typhimurium Proteobacteria; Gammaproteobacteria; Enterobacterales; Enterobacteriaceae; Salmonella; Salmonella enterica; Salmonella enterica subsp. enterica MNNEETFYQAMRRKGVTRRSFLKFCSLAATSLGLGAGMTPKIAWALENKPRIPVVWIHGL WP_001058312.1 hydrogenase [Salmonella enterica] Length: 372\nNumber of predicted TMHs: 2\nExp number of AAs in TMHs: 36.51738\nExp number, first 60 AAs: 19.72849\nTotal prob of N-in: 0.96170\nPOSSIBLE N-term signal sequence\ninside 1 19\nTMhelix 20 39\noutside 40 325\nTMhelix 326 348\ninside 349 372

21869 GCF_002029365.1_ASM202936v1 Salmonella enterica subsp. enterica serovar Typhimurium Proteobacteria; Gammaproteobacteria; Enterobacterales; Enterobacteriaceae; Salmonella; Salmonella enterica; Salmonella enterica subsp. enterica MTGDNTLITSHGINRRDFMKLCAALAATMGLSSKAAAEMAESVSNPQRPPVIWIGAQECT WP_000145429.1 hydrogenase 2 small subunit [Salmonella enterica] Length: 372\nNumber of predicted TMHs: 1\nExp number of AAs in TMHs: 24.51309\nExp number, first 60 AAs: 0.2682\nTotal prob of N-in: 0.14402\noutside 1 331\nTMhelix 332 354\ninside 355 372

21870 GCF_002028625.1_ASM202862v1 Salmonella enterica subsp. enterica serovar Typhimurium Proteobacteria; Gammaproteobacteria; Enterobacterales; Enterobacteriaceae; Salmonella; Salmonella enterica; Salmonella enterica subsp. enterica MTGDNTLITSHGINRRDFMKLCAALAATMGLSSKAAAEMAESVSNPQRPPVIWIGAQECT WP_000145429.1 hydrogenase 2 small subunit [Salmonella enterica] Length: 372\nNumber of

predicted TMHs:  1\nExp number of AAs in TMHs: 24.51309\nExp number, first 60 AAs:  0.2682\nTotal prob of N-in: 0.14402\noutside   1   331\nTMhelix   332   354\ninside   355   372

21871 GCF_002193535.1_ASM219353v1 Salmonella enterica subsp. enterica serovar Typhimurium Proteobacteria; Gammaproteobacteria; Enterobacterales; Enterobacteriaceae; Salmonella; Salmonella enterica; Salmonella enterica subsp. enterica MTGDNTLITSHGINRRDFMKLCAALAATMGLSSKAAAEMAESVSNPQRPPVIWIGAQECT WP_000145429.1 hydrogenase 2 small subunit [Salmonella enterica] Length: 372\nNumber of predicted TMHs:  1\nExp number of AAs in TMHs: 24.51309\nExp number, first 60 AAs:  0.2682\nTotal prob of N-in: 0.14402\noutside   1   331\nTMhelix   332   354\ninside   355   372

21872 GCF_002193595.1_ASM219359v1 Salmonella enterica subsp. enterica serovar Typhimurium Proteobacteria; Gammaproteobacteria; Enterobacterales; Enterobacteriaceae; Salmonella; Salmonella enterica; Salmonella enterica subsp. enterica MTGDNTLITSHGINRRDFMKLCAALAATMGLSSKAAAEMAESVSNPQRPPVIWIGAQECT WP_000145429.1 hydrogenase 2 small subunit [Salmonella enterica] Length: 372\nNumber of predicted TMHs:  1\nExp number of AAs in TMHs: 24.51309\nExp number, first 60 AAs:  0.2682\nTotal prob of N-in: 0.14402\noutside   1   331\nTMhelix   332   354\ninside   355   372

21873 GCF_002193595.1_ASM219359v1 Salmonella enterica subsp. enterica serovar Typhimurium Proteobacteria; Gammaproteobacteria; Enterobacterales; Enterobacteriaceae; Salmonella; Salmonella enterica; Salmonella enterica subsp. enterica MNNEETFYQAMRRKGVTRRSFLKFCSLAATSLGLGAGMTPKIAWALENKPRIPVVWIHGL WP_001058312.1 hydrogenase [Salmonella enterica] Length: 372\nNumber of predicted TMHs:  2\nExp number of AAs in TMHs: 36.51738\nExp number, first 60 AAs:  19.72849\nTotal prob of N-in:   0.96170\nPOSSIBLE N-term signal sequence\ninside   1   19\nTMhelix   20   39\noutside   40   325\nTMhelix   326   348\ninside   349   372

21874 GCF_002025495.1_ASM202549v1 Salmonella enterica subsp. enterica serovar Typhimurium Proteobacteria; Gammaproteobacteria; Enterobacterales; Enterobacteriaceae; Salmonella; Salmonella enterica; Salmonella enterica subsp. enterica MTGDNTLITSHGINRRDFMKLCAALAATMGLSSKAAAEMAESVSNPQRPPVIWIGAQECT WP_000145429.1 hydrogenase 2 small subunit [Salmonella enterica] Length: 372\nNumber of predicted TMHs:  1\nExp number of AAs in TMHs: 24.51309\nExp number, first 60 AAs:  0.2682\nTotal prob of N-in: 0.14402\noutside   1   331\nTMhelix   332   354\ninside   355   372

21875 GCF_001997115.1_ASM199711v1 Salmonella enterica subsp. enterica serovar Typhimurium Proteobacteria; Gammaproteobacteria; Enterobacterales; Enterobacteriaceae; Salmonella; Salmonella enterica; Salmonella enterica subsp. enterica MTGDNTLITSHGINRRDFMKLCAALAATMGLSSKAAAEMAESVSNPQRPPVIWIGAQECT WP_000145429.1 hydrogenase 2 small subunit [Salmonella enterica] Length: 372\nNumber of predicted TMHs:  1\nExp number of AAs in TMHs: 24.51309\nExp number, first 60 AAs:  0.2682\nTotal prob of N-in: 0.14402\noutside   1   331\nTMhelix   332   354\ninside   355   372

21876 GCF_002193615.1_ASM219361v1 Salmonella enterica subsp. enterica serovar Typhimurium Proteobacteria; Gammaproteobacteria; Enterobacterales; Enterobacteriaceae; Salmonella; Salmonella enterica; Salmonella enterica subsp. enterica MTGDNTLITSHGINRRDFMKLCAALAATMGLSSKAAAEMAESVSNPQRPPVIWIGAQECT WP_000145429.1 hydrogenase 2 small subunit [Salmonella enterica] Length: 372\nNumber of predicted TMHs:  1\nExp number of AAs in TMHs: 24.51309\nExp number, first 60 AAs:  0.2682\nTotal prob of N-in: 0.14402\noutside   1   331\nTMhelix   332   354\ninside   355   372

21877 GCF_002193615.1_ASM219361v1 Salmonella enterica subsp. enterica serovar Typhimurium Proteobacteria; Gammaproteobacteria; Enterobacterales; Enterobacteriaceae; Salmonella; Salmonella enterica; Salmonella enterica subsp. enterica MNNEETFYQAMRRKGVTRRSFLKFCSLAATSLGLGAGMTPKIAWALENKPRIPVVWIHGL WP_001058312.1 hydrogenase [Salmonella enterica] Length: 372\nNumber of predicted TMHs:  2\nExp number of AAs in TMHs: 36.51738\nExp number, first 60 AAs:  19.72849\nTotal prob of N-in:   0.96170\nPOSSIBLE N-term signal sequence\ninside   1   19\nTMhelix   20   39\noutside   40   325\nTMhelix   326   348\ninside   349   372

21878 GCF_002193535.1_ASM219353v1 Salmonella enterica subsp. enterica serovar Typhimurium Proteobacteria; Gammaproteobacteria; Enterobacterales; Enterobacteriaceae; Salmonella; Salmonella enterica; Salmonella enterica subsp. enterica MNNEETFYQAMRRKGVTRRSFLKFCSLAATSLGLGAGMTPKIAWALENKPRIPVVWIHGL WP_001058312.1 hydrogenase [Salmonella enterica] Length: 372\nNumber of predicted TMHs:  2\nExp number of AAs in TMHs: 36.51738\nExp number, first 60 AAs:  19.72849\nTotal prob of N-in:   0.96170\nPOSSIBLE N-term signal sequence\ninside   1   19\nTMhelix   20   39\noutside   40   325\nTMhelix   326   348\ninside   349   372

21879 GCF_001955195.1_ASM195519v1 Salmonella enterica subsp. enterica serovar Typhimurium Proteobacteria; Gammaproteobacteria; Enterobacterales; Enterobacteriaceae; Salmonella; Salmonella enterica;

Salmonella enterica subsp. enterica MNNEETFYQAMRRKGVTRRSFLKFCSLAATSLGLGAGMTPKIAWALENKPRIPVVWIHGL WP_001058312.1 hydrogenase [Salmonella enterica] Length: 372\nNumber of predicted TMHs: 2\nExp number of AAs in TMHs: 36.51738\nExp number, first 60 AAs: 19.72849\nTotal prob of N-in: 0.96170\nPOSSIBLE N-term signal sequence\ninside 1 19\nTMhelix 20 39\noutside 40 325\nTMhelix 326 348\ninside 349 372

21880 GCF_001955415.1_ASM195541v1 Salmonella enterica subsp. enterica serovar Typhimurium Proteobacteria; Gammaproteobacteria; Enterobacterales; Enterobacteriaceae; Salmonella; Salmonella enterica; Salmonella enterica subsp. enterica MNNEETFYQAMRRKGVTRRSFLKFCSLAATSLGLGAGMTPKIAWALENKPRIPVVWIHGL WP_001058312.1 hydrogenase [Salmonella enterica] Length: 372\nNumber of predicted TMHs: 2\nExp number of AAs in TMHs: 36.51738\nExp number, first 60 AAs: 19.72849\nTotal prob of N-in: 0.96170\nPOSSIBLE N-term signal sequence\ninside 1 19\nTMhelix 20 39\noutside 40 325\nTMhelix 326 348\ninside 349 372

21881 GCF_001955385.1_ASM195538v1 Salmonella enterica subsp. enterica serovar Typhimurium Proteobacteria; Gammaproteobacteria; Enterobacterales; Enterobacteriaceae; Salmonella; Salmonella enterica; Salmonella enterica subsp. enterica MNNEETFYQAMRRKGVTRRSFLKFCSLAATSLGLGAGMTPKIAWALENKPRIPVVWIHGL WP_001058312.1 hydrogenase [Salmonella enterica] Length: 372\nNumber of predicted TMHs: 2\nExp number of AAs in TMHs: 36.51738\nExp number, first 60 AAs: 19.72849\nTotal prob of N-in: 0.96170\nPOSSIBLE N-term signal sequence\ninside 1 19\nTMhelix 20 39\noutside 40 325\nTMhelix 326 348\ninside 349 372

21882 GCF_001955345.1_ASM195534v1 Salmonella enterica subsp. enterica serovar Typhimurium Proteobacteria; Gammaproteobacteria; Enterobacterales; Enterobacteriaceae; Salmonella; Salmonella enterica; Salmonella enterica subsp. enterica MNNEETFYQAMRRKGVTRRSFLKFCSLAATSLGLGAGMTPKIAWALENKPRIPVVWIHGL WP_001058312.1 hydrogenase [Salmonella enterica] Length: 372\nNumber of predicted TMHs: 2\nExp number of AAs in TMHs: 36.51738\nExp number, first 60 AAs: 19.72849\nTotal prob of N-in: 0.96170\nPOSSIBLE N-term signal sequence\ninside 1 19\nTMhelix 20 39\noutside 40 325\nTMhelix 326 348\ninside 349 372

21883 GCF_001954895.1_ASM195489v1 Salmonella enterica subsp. enterica serovar Typhimurium Proteobacteria; Gammaproteobacteria; Enterobacterales; Enterobacteriaceae; Salmonella; Salmonella enterica; Salmonella enterica subsp. enterica MTGDNTLITSHGINRRDFMKLCAALAATMGLSSKAAAEMAESVSNPQRPPVIWIGAQECT WP_000145429.1 hydrogenase 2 small subunit [Salmonella enterica] Length: 372\nNumber of predicted TMHs: 1\nExp number of AAs in TMHs: 24.51309\nExp number, first 60 AAs: 0.2682\nTotal prob of N-in: 0.14402\noutside 1 331\nTMhelix 332 354\ninside 355 372

21884 GCF_001887015.1_ASM188701v1 Salmonella enterica subsp. enterica serovar Typhimurium Proteobacteria; Gammaproteobacteria; Enterobacterales; Enterobacteriaceae; Salmonella; Salmonella enterica; Salmonella enterica subsp. enterica MTGDNTLITSHGINRRDFMKLCAALAATMGLSSKAAAEMAESVSNPQRPPVIWIGAQECT WP_000145429.1 hydrogenase 2 small subunit [Salmonella enterica] Length: 372\nNumber of predicted TMHs: 1\nExp number of AAs in TMHs: 24.51309\nExp number, first 60 AAs: 0.2682\nTotal prob of N-in: 0.14402\noutside 1 331\nTMhelix 332 354\ninside 355 372

21885 GCF_001758525.1_ASM175852v1 Salmonella enterica subsp. enterica serovar Typhimurium Proteobacteria; Gammaproteobacteria; Enterobacterales; Enterobacteriaceae; Salmonella; Salmonella enterica; Salmonella enterica subsp. enterica MNNEETFYQAMRRKGVTRRSFLKFCSLAATSLGLGAGMTPKIAWALENKPRIPVVWIHGL WP_001058312.1 hydrogenase [Salmonella enterica] Length: 372\nNumber of predicted TMHs: 2\nExp number of AAs in TMHs: 36.51738\nExp number, first 60 AAs: 19.72849\nTotal prob of N-in: 0.96170\nPOSSIBLE N-term signal sequence\ninside 1 19\nTMhelix 20 39\noutside 40 325\nTMhelix 326 348\ninside 349 372

21886 GCF_001756245.1_CFSAN008082 Salmonella enterica subsp. enterica serovar Typhimurium Proteobacteria; Gammaproteobacteria; Enterobacterales; Enterobacteriaceae; Salmonella; Salmonella enterica; Salmonella enterica subsp. enterica MNNEETFYQAMRRKGVTRRSFLKFCSLAATSLGLGAGMTPKIAWALENKPRIPVVWIHGL WP_001058312.1 hydrogenase [Salmonella enterica] Length: 372\nNumber of predicted TMHs: 2\nExp number of AAs in TMHs: 36.51738\nExp number, first 60 AAs: 19.72849\nTotal prob of N-in: 0.96170\nPOSSIBLE N-term signal sequence\ninside 1 19\nTMhelix 20 39\noutside 40 325\nTMhelix 326 348\ninside 349 372

21887 GCF_001756155.1_CFSAN008090 Salmonella enterica subsp. enterica serovar Typhimurium Proteobacteria; Gammaproteobacteria; Enterobacterales; Enterobacteriaceae; Salmonella; Salmonella enterica; Salmonella enterica subsp. enterica MTGDNTLITSHGINRRDFMKLCAALAATMGLSSKAAAEMAESVSNPQRPPVIWIGAQECT WP_000145429.1 hydrogenase 2 small subunit [Salmonella enterica] Length: 372\nNumber of predicted TMHs: 1\nExp number of AAs in TMHs: 24.51309\nExp number, first 60 AAs: 0.2682\nTotal prob of N-in: 0.14402\noutside 1 331\nTMhelix 332 354\ninside 355 372

21888 GCF_000715155.1_ASM71515v2 Salmonella enterica subsp. enterica serovar Typhimurium Proteobacteria; Gammaproteobacteria; Enterobacterales; Enterobacteriaceae; Salmonella; Salmonella enterica; Salmonella enterica subsp. enterica MTGDNTLITSHGINRRDFMKLCAALAATMGLSSKAAAEMAESVSNPQRPPVIWIGAQECT WP_000145429.1 hydrogenase 2 small subunit [Salmonella enterica] Length: 372\nNumber of predicted TMHs: 1\nExp number of AAs in TMHs: 24.51309\nExp number, first 60 AAs: 0.2682\nTotal prob of N-in: 0.14402\noutside 1 331\nTMhelix 332 354\ninside 355 372

21889 GCF_000974095.1_ABBSB1113-1.id1_v1.0 Salmonella enterica subsp. enterica serovar Typhimurium Proteobacteria; Gammaproteobacteria; Enterobacterales; Enterobacteriaceae; Salmonella; Salmonella enterica; Salmonella enterica subsp. enterica MTGDNTLITSHGINRRDFMKLCAALAATMGLSSKAAAEMAESVSNPQRPPVIWIGAQECT WP_000145429.1 hydrogenase 2 small subunit [Salmonella enterica] Length: 372\nNumber of predicted TMHs: 1\nExp number of AAs in TMHs: 24.51309\nExp number, first 60 AAs: 0.2682\nTotal prob of N-in: 0.14402\noutside 1 331\nTMhelix 332 354\ninside 355 372

21890 GCF_000974095.1_ABBSB1113-1.id1_v1.0 Salmonella enterica subsp. enterica serovar Typhimurium Proteobacteria; Gammaproteobacteria; Enterobacterales; Enterobacteriaceae; Salmonella; Salmonella enterica; Salmonella enterica subsp. enterica MNNEETFYQAMRRKGVTRRSFLKFCSLAATSLGLGAGMTPKIAWALENKPRIPVVWIHGL WP_001058312.1 hydrogenase [Salmonella enterica] Length: 372\nNumber of predicted TMHs: 2\nExp number of AAs in TMHs: 36.51738\nExp number, first 60 AAs: 19.72849\nTotal prob of N-in: 0.96170\nPOSSIBLE N-term signal sequence\ninside 1 19\nTMhelix 20 39\noutside 40 325\nTMhelix 326 348\ninside 349 372

21891 GCF_000973915.1_ABB1162-2.id9_v1.0 Salmonella enterica subsp. enterica serovar Typhimurium Proteobacteria; Gammaproteobacteria; Enterobacterales; Enterobacteriaceae; Salmonella; Salmonella enterica; Salmonella enterica subsp. enterica MNNEETFYQAMRRKGVTRRSFLKFCSLAATSLGLGAGMTPKIAWALENKPRIPVVWIHGL WP_001058312.1 hydrogenase [Salmonella enterica] Length: 372\nNumber of predicted TMHs: 2\nExp number of AAs in TMHs: 36.51738\nExp number, first 60 AAs: 19.72849\nTotal prob of N-in: 0.96170\nPOSSIBLE N-term signal sequence\ninside 1 19\nTMhelix 20 39\noutside 40 325\nTMhelix 326 348\ninside 349 372

21892 GCF_002091345.1_ASM209134v1 Salmonella enterica subsp. enterica serovar Typhimurium Proteobacteria; Gammaproteobacteria; Enterobacterales; Enterobacteriaceae; Salmonella; Salmonella enterica; Salmonella enterica subsp. enterica MNNEETFYQAMRRKGVTRRSFLKFCSLAATSLGLGAGMTPKIAWALENKPRIPVVWIHGL WP_001058312.1 hydrogenase [Salmonella enterica] Length: 372\nNumber of predicted TMHs: 2\nExp number of AAs in TMHs: 36.51738\nExp number, first 60 AAs: 19.72849\nTotal prob of N-in: 0.96170\nPOSSIBLE N-term signal sequence\ninside 1 19\nTMhelix 20 39\noutside 40 325\nTMhelix 326 348\ninside 349 372

21893 GCF_002091185.1_ASM209118v1 Salmonella enterica subsp. enterica serovar Typhimurium Proteobacteria; Gammaproteobacteria; Enterobacterales; Enterobacteriaceae; Salmonella; Salmonella enterica; Salmonella enterica subsp. enterica MNNEETFYQAMRRKGVTRRSFLKFCSLAATSLGLGAGMTPKIAWALENKPRIPVVWIHGL WP_001058312.1 hydrogenase [Salmonella enterica] Length: 372\nNumber of predicted TMHs: 2\nExp number of AAs in TMHs: 36.51738\nExp number, first 60 AAs: 19.72849\nTotal prob of N-in: 0.96170\nPOSSIBLE N-term signal sequence\ninside 1 19\nTMhelix 20 39\noutside 40 325\nTMhelix 326 348\ninside 349 372

21894 GCF_002091235.1_ASM209123v1 Salmonella enterica subsp. enterica serovar Typhimurium Proteobacteria; Gammaproteobacteria; Enterobacterales; Enterobacteriaceae; Salmonella; Salmonella enterica; Salmonella enterica subsp. enterica MNNEETFYQAMRRKGVTRRSFLKFCSLAATSLGLGAGMTPKIAWALENKPRIPVVWIHGL WP_001058312.1 hydrogenase [Salmonella enterica] Length: 372\nNumber of predicted TMHs: 2\nExp number of AAs in TMHs: 36.51738\nExp number, first 60 AAs: 19.72849\nTotal prob of N-in: 0.96170\nPOSSIBLE N-term signal sequence\ninside 1 19\nTMhelix 20 39\noutside 40 325\nTMhelix 326 348\ninside 349 372

21895 GCF_002193545.1_ASM219354v1 Salmonella enterica subsp. enterica serovar Typhimurium Proteobacteria; Gammaproteobacteria; Enterobacterales; Enterobacteriaceae; Salmonella; Salmonella enterica; Salmonella enterica subsp. enterica MTGDNTLITSHGINRRDFMKLCAALAATMGLSSKAAAEMAESVSNPQRPPVIWIGAQECT WP_000145429.1 hydrogenase 2 small subunit [Salmonella enterica] Length: 372\nNumber of predicted TMHs: 1\nExp number of AAs in TMHs: 24.51309\nExp number, first 60 AAs: 0.2682\nTotal prob of N-in: 0.14402\noutside 1 331\nTMhelix 332 354\ninside 355 372

21896 GCF_002193545.1_ASM219354v1 Salmonella enterica subsp. enterica serovar Typhimurium Proteobacteria; Gammaproteobacteria; Enterobacterales; Enterobacteriaceae; Salmonella; Salmonella enterica; Salmonella enterica subsp. enterica MNNEETFYQAMRRKGVTRRSFLKFCSLAATSLGLGAGMTPKIAWALENKPRIPVVWIHGL WP_001058312.1 hydrogenase [Salmonella enterica] Length: 372\nNumber of predicted TMHs: 2\nExp

number of AAs in TMHs: 36.51738\nExp number, first 60 AAs: 19.72849\nTotal prob of N-in: 0.96170\nPOSSIBLE N-term signal sequence\ninside 1 19\nTMhelix 20 39\noutside 40 325\nTMhelix 326 348\ninside 349 372

21897 GCF_002193605.1_ASM219360v1 Salmonella enterica subsp. enterica serovar Typhimurium Proteobacteria; Gammaproteobacteria; Enterobacterales; Enterobacteriaceae; Salmonella; Salmonella enterica; Salmonella enterica subsp. enterica MTGDNTLITSHGINRRDFMKLCAALAATMGLSSKAAAEMAESVSNPQRPPVIWIGAQECT WP_000145429.1 hydrogenase 2 small subunit [Salmonella enterica] Length: 372\nNumber of predicted TMHs: 1\nExp number of AAs in TMHs: 24.51309\nExp number, first 60 AAs: 0.2682\nTotal prob of N-in: 0.14402\noutside 1 331\nTMhelix 332 354\ninside 355 372

21898 GCF_002091115.1_ASM209111v1 Salmonella enterica subsp. enterica serovar Typhimurium Proteobacteria; Gammaproteobacteria; Enterobacterales; Enterobacteriaceae; Salmonella; Salmonella enterica; Salmonella enterica subsp. enterica MTGDNTLITSHGINRRDFMKLCAALAATMGLSSKAAAEMAESVSNPQRPPVIWIGAQECT WP_000145429.1 hydrogenase 2 small subunit [Salmonella enterica] Length: 372\nNumber of predicted TMHs: 1\nExp number of AAs in TMHs: 24.51309\nExp number, first 60 AAs: 0.2682\nTotal prob of N-in: 0.14402\noutside 1 331\nTMhelix 332 354\ninside 355 372

21899 GCF_002091275.1_ASM209127v1 Salmonella enterica subsp. enterica serovar Typhimurium Proteobacteria; Gammaproteobacteria; Enterobacterales; Enterobacteriaceae; Salmonella; Salmonella enterica; Salmonella enterica subsp. enterica MNNEETFYQAMRRKGVTRRSFLKFCSLAATSLGLGAGMTPKIAWALENKPRIPVVWIHGL WP_001058312.1 hydrogenase [Salmonella enterica] Length: 372\nNumber of predicted TMHs: 2\nExp number of AAs in TMHs: 36.51738\nExp number, first 60 AAs: 19.72849\nTotal prob of N-in: 0.96170\nPOSSIBLE N-term signal sequence\ninside 1 19\nTMhelix 20 39\noutside 40 325\nTMhelix 326 348\ninside 349 372

21900 GCF_002193525.1_ASM219352v1 Salmonella enterica subsp. enterica serovar Typhimurium Proteobacteria; Gammaproteobacteria; Enterobacterales; Enterobacteriaceae; Salmonella; Salmonella enterica; Salmonella enterica subsp. enterica MNNEETFYQAMRRKGVTRRSFLKFCSLAATSLGLGAGMTPKIAWALENKPRIPVVWIHGL WP_001058312.1 hydrogenase [Salmonella enterica] Length: 372\nNumber of predicted TMHs: 2\nExp number of AAs in TMHs: 36.51738\nExp number, first 60 AAs: 19.72849\nTotal prob of N-in: 0.96170\nPOSSIBLE N-term signal sequence\ninside 1 19\nTMhelix 20 39\noutside 40 325\nTMhelix 326 348\ninside 349 372

21901 GCF_002091075.1_ASM209107v1 Salmonella enterica subsp. enterica serovar Typhimurium Proteobacteria; Gammaproteobacteria; Enterobacterales; Enterobacteriaceae; Salmonella; Salmonella enterica; Salmonella enterica subsp. enterica MNNEETFYQAMRRKGVTRRSFLKFCSLAATSLGLGAGMTPKIAWALENKPRIPVVWIHGL WP_001058312.1 hydrogenase [Salmonella enterica] Length: 372\nNumber of predicted TMHs: 2\nExp number of AAs in TMHs: 36.51738\nExp number, first 60 AAs: 19.72849\nTotal prob of N-in: 0.96170\nPOSSIBLE N-term signal sequence\ninside 1 19\nTMhelix 20 39\noutside 40 325\nTMhelix 326 348\ninside 349 372

21902 GCF_000784315.1_ASM78431v1 Salmonella enterica subsp. enterica serovar Typhimurium Proteobacteria; Gammaproteobacteria; Enterobacterales; Enterobacteriaceae; Salmonella; Salmonella enterica; Salmonella enterica subsp. enterica MTGDNTLITSHGINRRDFMKLCAALAATMGLSSKAAAEMAESVSNPQRPPVIWIGAQECT WP_000145429.1 hydrogenase 2 small subunit [Salmonella enterica] Length: 372\nNumber of predicted TMHs: 1\nExp number of AAs in TMHs: 24.51309\nExp number, first 60 AAs: 0.2682\nTotal prob of N-in: 0.14402\noutside 1 331\nTMhelix 332 354\ninside 355 372

21903 GCF_002091185.1_ASM209118v1 Salmonella enterica subsp. enterica serovar Typhimurium Proteobacteria; Gammaproteobacteria; Enterobacterales; Enterobacteriaceae; Salmonella; Salmonella enterica; Salmonella enterica subsp. enterica MTGDNTLITSHGINRRDFMKLCAALAATMGLSSKAAAEMAESVSNPQRPPVIWIGAQECT WP_000145429.1 hydrogenase 2 small subunit [Salmonella enterica] Length: 372\nNumber of predicted TMHs: 1\nExp number of AAs in TMHs: 24.51309\nExp number, first 60 AAs: 0.2682\nTotal prob of N-in: 0.14402\noutside 1 331\nTMhelix 332 354\ninside 355 372

21904 GCF_002091235.1_ASM209123v1 Salmonella enterica subsp. enterica serovar Typhimurium Proteobacteria; Gammaproteobacteria; Enterobacterales; Enterobacteriaceae; Salmonella; Salmonella enterica; Salmonella enterica subsp. enterica MTGDNTLITSHGINRRDFMKLCAALAATMGLSSKAAAEMAESVSNPQRPPVIWIGAQECT WP_000145429.1 hydrogenase 2 small subunit [Salmonella enterica] Length: 372\nNumber of predicted TMHs: 1\nExp number of AAs in TMHs: 24.51309\nExp number, first 60 AAs: 0.2682\nTotal prob of N-in: 0.14402\noutside 1 331\nTMhelix 332 354\ninside 355 372

21905 GCF_002091275.1_ASM209127v1 Salmonella enterica subsp. enterica serovar Typhimurium Proteobacteria; Gammaproteobacteria; Enterobacterales; Enterobacteriaceae; Salmonella; Salmonella enterica;

Salmonella enterica subsp. enterica MTGDNTLITSHGINRRDFMKLCAALAATMGLSSKAAAEMAESVSNPQRPPVIWIGAQECT WP_000145429.1 hydrogenase 2 small subunit [Salmonella enterica] Length: 372\nNumber of predicted TMHs: 1\nExp number of AAs in TMHs: 24.51309\nExp number, first 60 AAs: 0.2682\nTotal prob of N-in: 0.14402\noutside 1 331\nTMhelix 332 354\ninside 355 372

21906 GCF_002193525.1_ASM219352v1 Salmonella enterica subsp. enterica serovar Typhimurium Proteobacteria; Gammaproteobacteria; Enterobacterales; Enterobacteriaceae; Salmonella; Salmonella enterica; Salmonella enterica subsp. enterica MTGDNTLITSHGINRRDFMKLCAALAATMGLSSKAAAEMAESVSNPQRPPVIWIGAQECT WP_000145429.1 hydrogenase 2 small subunit [Salmonella enterica] Length: 372\nNumber of predicted TMHs: 1\nExp number of AAs in TMHs: 24.51309\nExp number, first 60 AAs: 0.2682\nTotal prob of N-in: 0.14402\noutside 1 331\nTMhelix 332 354\ninside 355 372

21907 GCF_001595195.1_CFSAN033857_01.0Salmonella enterica subsp. enterica serovar Typhimurium Proteobacteria; Gammaproteobacteria; Enterobacterales; Enterobacteriaceae; Salmonella; Salmonella enterica; Salmonella enterica subsp. enterica MTGDNTLITSHGINRRDFMKLCAALAATMGLSSKAAAEMAESVSNPQRPPVIWIGAQECT WP_000145429.1 hydrogenase 2 small subunit [Salmonella enterica] Length: 372\nNumber of predicted TMHs: 1\nExp number of AAs in TMHs: 24.51309\nExp number, first 60 AAs: 0.2682\nTotal prob of N-in: 0.14402\noutside 1 331\nTMhelix 332 354\ninside 355 372

21908 GCF_001454965.1_ASM145496v1 Salmonella enterica subsp. enterica serovar Typhimurium Proteobacteria; Gammaproteobacteria; Enterobacterales; Enterobacteriaceae; Salmonella; Salmonella enterica; Salmonella enterica subsp. enterica MTGDNTLITSHGINRRDFMKLCAALAATMGLSSKAAAEMAESVSNPQRPPVIWIGAQECT WP_000145429.1 hydrogenase 2 small subunit [Salmonella enterica] Length: 372\nNumber of predicted TMHs: 1\nExp number of AAs in TMHs: 24.51309\nExp number, first 60 AAs: 0.2682\nTotal prob of N-in: 0.14402\noutside 1 331\nTMhelix 332 354\ninside 355 372

21909 GCF_001457365.1_ASM145736v1 Salmonella enterica subsp. enterica serovar Typhimurium Proteobacteria; Gammaproteobacteria; Enterobacterales; Enterobacteriaceae; Salmonella; Salmonella enterica; Salmonella enterica subsp. enterica MTGDNTLITSHGINRRDFMKLCAALAATMGLSSKAAAEMAESVSNPQRPPVIWIGAQECT WP_000145429.1 hydrogenase 2 small subunit [Salmonella enterica] Length: 372\nNumber of predicted TMHs: 1\nExp number of AAs in TMHs: 24.51309\nExp number, first 60 AAs: 0.2682\nTotal prob of N-in: 0.14402\noutside 1 331\nTMhelix 332 354\ninside 355 372

21910 GCF_001454965.1_ASM145496v1 Salmonella enterica subsp. enterica serovar Typhimurium Proteobacteria; Gammaproteobacteria; Enterobacterales; Enterobacteriaceae; Salmonella; Salmonella enterica; Salmonella enterica subsp. enterica MNNEETFYQAMRRKGVTRRSFLKFCSLAATSLGLGAGMTPKIAWALENKPRIPVVWIHGL WP_001058312.1 hydrogenase [Salmonella enterica] Length: 372\nNumber of predicted TMHs: 2\nExp number of AAs in TMHs: 36.51738\nExp number, first 60 AAs: 19.72849\nTotal prob of N-in: 0.96170\nPOSSIBLE N-term signal sequence\ninside 1 19\nTMhelix 20 39\noutside 40 325\nTMhelix 326 348\ninside 349 372

21911 GCF_001457365.1_ASM145736v1 Salmonella enterica subsp. enterica serovar Typhimurium Proteobacteria; Gammaproteobacteria; Enterobacterales; Enterobacteriaceae; Salmonella; Salmonella enterica; Salmonella enterica subsp. enterica MNNEETFYQAMRRKGVTRRSFLKFCSLAATSLGLGAGMTPKIAWALENKPRIPVVWIHGL WP_001058312.1 hydrogenase [Salmonella enterica] Length: 372\nNumber of predicted TMHs: 2\nExp number of AAs in TMHs: 36.51738\nExp number, first 60 AAs: 19.72849\nTotal prob of N-in: 0.96170\nPOSSIBLE N-term signal sequence\ninside 1 19\nTMhelix 20 39\noutside 40 325\nTMhelix 326 348\ninside 349 372

21912 GCF_001466375.1_ASM146637v1 Salmonella enterica subsp. enterica serovar Typhimurium Proteobacteria; Gammaproteobacteria; Enterobacterales; Enterobacteriaceae; Salmonella; Salmonella enterica; Salmonella enterica subsp. enterica MTGDNTLITSHGINRRDFMKLCAALAATMGLSSKAAAEMAESVSNPQRPPVIWIGAQECT WP_000145429.1 hydrogenase 2 small subunit [Salmonella enterica] Length: 372\nNumber of predicted TMHs: 1\nExp number of AAs in TMHs: 24.51309\nExp number, first 60 AAs: 0.2682\nTotal prob of N-in: 0.14402\noutside 1 331\nTMhelix 332 354\ninside 355 372

21913 GCF_001466375.1_ASM146637v1 Salmonella enterica subsp. enterica serovar Typhimurium Proteobacteria; Gammaproteobacteria; Enterobacterales; Enterobacteriaceae; Salmonella; Salmonella enterica; Salmonella enterica subsp. enterica MNNEETFYQAMRRKGVTRRSFLKFCSLAATSLGLGAGMTPKIAWALENKPRIPVVWIHGL WP_001058312.1 hydrogenase [Salmonella enterica] Length: 372\nNumber of predicted TMHs: 2\nExp number of AAs in TMHs: 36.51738\nExp number, first 60 AAs: 19.72849\nTotal prob of N-in: 0.96170\nPOSSIBLE N-term signal sequence\ninside 1 19\nTMhelix 20 39\noutside 40 325\nTMhelix 326 348\ninside 349 372

21914 GCF_001477735.1_Salmonella_enterica_CVM_N29313-SQ_v1.0 Salmonella enterica subsp. enterica serovar Typhimurium Proteobacteria; Gammaproteobacteria; Enterobacterales; Enterobacteriaceae; Salmonella; Salmonella enterica; Salmonella enterica subsp. enterica
MTGDNTLITSHGINRRDFMKLCAALAATMGLSSKAAAEMAESVSNPQRPPVIWIGAQECT WP_000145429.1
hydrogenase 2 small subunit [Salmonella enterica] Length: 372\nNumber of predicted TMHs: 1\nExp number of AAs in TMHs: 24.51309\nExp number, first 60 AAs: 0.2682\nTotal prob of N-in: 0.14402\noutside 1 331\nTMhelix 332 354\ninside 355 372

21915 GCF_001477805.1_Salmonella_enterica_CVM_N29338-SQ_v1.0 Salmonella enterica subsp. enterica serovar Typhimurium Proteobacteria; Gammaproteobacteria; Enterobacterales; Enterobacteriaceae; Salmonella; Salmonella enterica; Salmonella enterica subsp. enterica
MTGDNTLITSHGINRRDFMKLCAALAATMGLSSKAAAEMAESVSNPQRPPVIWIGAQECT WP_000145429.1
hydrogenase 2 small subunit [Salmonella enterica] Length: 372\nNumber of predicted TMHs: 1\nExp number of AAs in TMHs: 24.51309\nExp number, first 60 AAs: 0.2682\nTotal prob of N-in: 0.14402\noutside 1 331\nTMhelix 332 354\ninside 355 372

21916 GCF_001477735.1_Salmonella_enterica_CVM_N29313-SQ_v1.0 Salmonella enterica subsp. enterica serovar Typhimurium Proteobacteria; Gammaproteobacteria; Enterobacterales; Enterobacteriaceae; Salmonella; Salmonella enterica; Salmonella enterica subsp. enterica
MNNEETFYQAMRRKGVTRRSFLKFCSLAATSLGLGAGMTPKIAWALENKPRIPVVWIHGL WP_001058312.1
hydrogenase [Salmonella enterica] Length: 372\nNumber of predicted TMHs: 2\nExp number of AAs in TMHs: 36.51738\nExp number, first 60 AAs: 19.72849\nTotal prob of N-in: 0.96170\nPOSSIBLE N-term signal sequence\ninside 1 19\nTMhelix 20 39\noutside 40 325\nTMhelix 326 348\ninside 349 372

21917 GCF_001477805.1_Salmonella_enterica_CVM_N29338-SQ_v1.0 Salmonella enterica subsp. enterica serovar Typhimurium Proteobacteria; Gammaproteobacteria; Enterobacterales; Enterobacteriaceae; Salmonella; Salmonella enterica; Salmonella enterica subsp. enterica
MNNEETFYQAMRRKGVTRRSFLKFCSLAATSLGLGAGMTPKIAWALENKPRIPVVWIHGL WP_001058312.1
hydrogenase [Salmonella enterica] Length: 372\nNumber of predicted TMHs: 2\nExp number of AAs in TMHs: 36.51738\nExp number, first 60 AAs: 19.72849\nTotal prob of N-in: 0.96170\nPOSSIBLE N-term signal sequence\ninside 1 19\nTMhelix 20 39\noutside 40 325\nTMhelix 326 348\ninside 349 372

21918 GCF_001478065.1_Salmonella_enterica_CVM_N30696-SQ_v1.0 Salmonella enterica subsp. enterica serovar Typhimurium Proteobacteria; Gammaproteobacteria; Enterobacterales; Enterobacteriaceae; Salmonella; Salmonella enterica; Salmonella enterica subsp. enterica
MTGDNTLITSHGINRRDFMKLCAALAATMGLSSKAAAEMAESVSNPQRPPVIWIGAQECT WP_000145429.1
hydrogenase 2 small subunit [Salmonella enterica] Length: 372\nNumber of predicted TMHs: 1\nExp number of AAs in TMHs: 24.51309\nExp number, first 60 AAs: 0.2682\nTotal prob of N-in: 0.14402\noutside 1 331\nTMhelix 332 354\ninside 355 372

21919 GCF_001478065.1_Salmonella_enterica_CVM_N30696-SQ_v1.0 Salmonella enterica subsp. enterica serovar Typhimurium Proteobacteria; Gammaproteobacteria; Enterobacterales; Enterobacteriaceae; Salmonella; Salmonella enterica; Salmonella enterica subsp. enterica
MNNEETFYQAMRRKGVTRRSFLKFCSLAATSLGLGAGMTPKIAWALENKPRIPVVWIHGL WP_001058312.1
hydrogenase [Salmonella enterica] Length: 372\nNumber of predicted TMHs: 2\nExp number of AAs in TMHs: 36.51738\nExp number, first 60 AAs: 19.72849\nTotal prob of N-in: 0.96170\nPOSSIBLE N-term signal sequence\ninside 1 19\nTMhelix 20 39\noutside 40 325\nTMhelix 326 348\ninside 349 372

21920 GCF_001478425.1_Salmonella_enterica_CVM_N38925-SQ_v1.0 Salmonella enterica subsp. enterica serovar Typhimurium Proteobacteria; Gammaproteobacteria; Enterobacterales; Enterobacteriaceae; Salmonella; Salmonella enterica; Salmonella enterica subsp. enterica
MTGDNTLITSHGINRRDFMKLCAALAATMGLSSKAAAEMAESVSNPQRPPVIWIGAQECT WP_000145429.1
hydrogenase 2 small subunit [Salmonella enterica] Length: 372\nNumber of predicted TMHs: 1\nExp number of AAs in TMHs: 24.51309\nExp number, first 60 AAs: 0.2682\nTotal prob of N-in: 0.14402\noutside 1 331\nTMhelix 332 354\ninside 355 372

21921 GCF_001478425.1_Salmonella_enterica_CVM_N38925-SQ_v1.0 Salmonella enterica subsp. enterica serovar Typhimurium Proteobacteria; Gammaproteobacteria; Enterobacterales; Enterobacteriaceae; Salmonella; Salmonella enterica; Salmonella enterica subsp. enterica
MNNEETFYQAMRRKGVTRRSFLKFCSLAATSLGLGAGMTPKIAWALENKPRIPVVWIHGL WP_001058312.1
hydrogenase [Salmonella enterica] Length: 372\nNumber of predicted TMHs: 2\nExp number of AAs in

TMHs: 36.51738\nExp number, first 60 AAs: 19.72849\nTotal prob of N-in: 0.96170\nPOSSIBLE N-term signal sequence\ninside 1 19\nTMhelix 20 39\noutside 40 325\nTMhelix 326 348\ninside 349 372

21922 GCF_001479125.1_Salmonella_enterica_CVM_N31409-SQ_v1.0 Salmonella enterica subsp. enterica serovar Typhimurium Proteobacteria; Gammaproteobacteria; Enterobacterales; Enterobacteriaceae; Salmonella; Salmonella enterica; Salmonella enterica subsp. enterica MTGDNTLITSHGINRRDFMKLCAALAATMGLSSKAAAEMAESVSNPQRPPVIWIGAQECT WP_000145429.1 hydrogenase 2 small subunit [Salmonella enterica] Length: 372\nNumber of predicted TMHs: 1\nExp number of AAs in TMHs: 24.51309\nExp number, first 60 AAs: 0.2682\nTotal prob of N-in: 0.14402\noutside 1 331\nTMhelix 332 354\ninside 355 372

21923 GCF_001479125.1_Salmonella_enterica_CVM_N31409-SQ_v1.0 Salmonella enterica subsp. enterica serovar Typhimurium Proteobacteria; Gammaproteobacteria; Enterobacterales; Enterobacteriaceae; Salmonella; Salmonella enterica; Salmonella enterica subsp. enterica MNNEETFYQAMRRKGVTRRSFLKFCSLAATSLGLGAGMTPKIAWALENKPRIPVVWIHGL WP_001058312.1 hydrogenase [Salmonella enterica] Length: 372\nNumber of predicted TMHs: 2\nExp number of AAs in TMHs: 36.51738\nExp number, first 60 AAs: 19.72849\nTotal prob of N-in: 0.96170\nPOSSIBLE N-term signal sequence\ninside 1 19\nTMhelix 20 39\noutside 40 325\nTMhelix 326 348\ninside 349 372

21924 GCF_001479335.1_Salmonella_enterica_CVM_N32048-SQ_v1.0 Salmonella enterica subsp. enterica serovar Typhimurium Proteobacteria; Gammaproteobacteria; Enterobacterales; Enterobacteriaceae; Salmonella; Salmonella enterica; Salmonella enterica subsp. enterica MTGDNTLITSHGINRRDFMKLCAALAATMGLSSKAAAEMAESVSNPQRPPVIWIGAQECT WP_000145429.1 hydrogenase 2 small subunit [Salmonella enterica] Length: 372\nNumber of predicted TMHs: 1\nExp number of AAs in TMHs: 24.51309\nExp number, first 60 AAs: 0.2682\nTotal prob of N-in: 0.14402\noutside 1 331\nTMhelix 332 354\ninside 355 372

21925 GCF_001479335.1_Salmonella_enterica_CVM_N32048-SQ_v1.0 Salmonella enterica subsp. enterica serovar Typhimurium Proteobacteria; Gammaproteobacteria; Enterobacterales; Enterobacteriaceae; Salmonella; Salmonella enterica; Salmonella enterica subsp. enterica MNNEETFYQAMRRKGVTRRSFLKFCSLAATSLGLGAGMTPKIAWALENKPRIPVVWIHGL WP_001058312.1 hydrogenase [Salmonella enterica] Length: 372\nNumber of predicted TMHs: 2\nExp number of AAs in TMHs: 36.51738\nExp number, first 60 AAs: 19.72849\nTotal prob of N-in: 0.96170\nPOSSIBLE N-term signal sequence\ninside 1 19\nTMhelix 20 39\noutside 40 325\nTMhelix 326 348\ninside 349 372

21926 GCF_001480055.1_Salmonella_enterica_CVM_N38917-SQ_v1.0 Salmonella enterica subsp. enterica serovar Typhimurium Proteobacteria; Gammaproteobacteria; Enterobacterales; Enterobacteriaceae; Salmonella; Salmonella enterica; Salmonella enterica subsp. enterica MTGDNTLITSHGINRRDFMKLCAALAATMGLSSKAAAEMAESVSNPQRPPVIWIGAQECT WP_000145429.1 hydrogenase 2 small subunit [Salmonella enterica] Length: 372\nNumber of predicted TMHs: 1\nExp number of AAs in TMHs: 24.51309\nExp number, first 60 AAs: 0.2682\nTotal prob of N-in: 0.14402\noutside 1 331\nTMhelix 332 354\ninside 355 372

21927 GCF_001480055.1_Salmonella_enterica_CVM_N38917-SQ_v1.0 Salmonella enterica subsp. enterica serovar Typhimurium Proteobacteria; Gammaproteobacteria; Enterobacterales; Enterobacteriaceae; Salmonella; Salmonella enterica; Salmonella enterica subsp. enterica MNNEETFYQAMRRKGVTRRSFLKFCSLAATSLGLGAGMTPKIAWALENKPRIPVVWIHGL WP_001058312.1 hydrogenase [Salmonella enterica] Length: 372\nNumber of predicted TMHs: 2\nExp number of AAs in TMHs: 36.51738\nExp number, first 60 AAs: 19.72849\nTotal prob of N-in: 0.96170\nPOSSIBLE N-term signal sequence\ninside 1 19\nTMhelix 20 39\noutside 40 325\nTMhelix 326 348\ninside 349 372

21928 GCF_001480855.1_Salmonella_enterica_CVM_N38914-SQ_v1.0 Salmonella enterica subsp. enterica serovar Typhimurium Proteobacteria; Gammaproteobacteria; Enterobacterales; Enterobacteriaceae; Salmonella; Salmonella enterica; Salmonella enterica subsp. enterica MTGDNTLITSHGINRRDFMKLCAALAATMGLSSKAAAEMAESVSNPQRPPVIWIGAQECT WP_000145429.1 hydrogenase 2 small subunit [Salmonella enterica] Length: 372\nNumber of predicted TMHs: 1\nExp number of AAs in TMHs: 24.51309\nExp number, first 60 AAs: 0.2682\nTotal prob of N-in: 0.14402\noutside 1 331\nTMhelix 332 354\ninside 355 372

21929 GCF_001480855.1_Salmonella_enterica_CVM_N38914-SQ_v1.0 Salmonella enterica subsp. enterica serovar Typhimurium Proteobacteria; Gammaproteobacteria; Enterobacterales; Enterobacteriaceae; Salmonella; Salmonella

enterica; Salmonella enterica subsp. enterica
MNNEETFYQAMRRKGVTRRSFLKFCSLAATSLGLGAGMTPKIAWALENKPRIPVVWIHGL WP_001058312.1 hydrogenase [Salmonella enterica] Length: 372\nNumber of predicted TMHs: 2\nExp number of AAs in TMHs: 36.51738\nExp number, first 60 AAs: 19.72849\nTotal prob of N-in: 0.96170\nPOSSIBLE N-term signal sequence\ninside 1 19\nTMhelix 20 39\noutside 40 325\nTMhelix 326 348\ninside 349 372

21930 GCF_001481075.1_Salmonella_enterica_CVM_N42467_v1.0 Salmonella enterica subsp. enterica serovar Typhimurium Proteobacteria; Gammaproteobacteria; Enterobacterales; Enterobacteriaceae; Salmonella; Salmonella enterica; Salmonella enterica subsp. enterica
MTGDNTLITSHGINRRDFMKLCAALAATMGLSSKAAAEMAESVSNPQRPPVIWIGAQECT WP_000145429.1 hydrogenase 2 small subunit [Salmonella enterica] Length: 372\nNumber of predicted TMHs: 1\nExp number of AAs in TMHs: 24.51309\nExp number, first 60 AAs: 0.2682\nTotal prob of N-in: 0.14402\noutside 1 331\nTMhelix 332 354\ninside 355 372

21931 GCF_001481075.1_Salmonella_enterica_CVM_N42467_v1.0 Salmonella enterica subsp. enterica serovar Typhimurium Proteobacteria; Gammaproteobacteria; Enterobacterales; Enterobacteriaceae; Salmonella; Salmonella enterica; Salmonella enterica subsp. enterica
MNNEETFYQAMRRKGVTRRSFLKFCSLAATSLGLGAGMTPKIAWALENKPRIPVVWIHGL WP_001058312.1 hydrogenase [Salmonella enterica] Length: 372\nNumber of predicted TMHs: 2\nExp number of AAs in TMHs: 36.51738\nExp number, first 60 AAs: 19.72849\nTotal prob of N-in: 0.96170\nPOSSIBLE N-term signal sequence\ninside 1 19\nTMhelix 20 39\noutside 40 325\nTMhelix 326 348\ninside 349 372

21932 GCF_001576255.1_ASM157625v1 Salmonella enterica subsp. enterica serovar Typhimurium Proteobacteria; Gammaproteobacteria; Enterobacterales; Enterobacteriaceae; Salmonella; Salmonella enterica; Salmonella enterica subsp. enterica MTGDNTLITSHGINRRDFMKLCAALAATMGLSSKAAAEMAESVSNPQRPPVIWIGAQECT WP_000145429.1 hydrogenase 2 small subunit [Salmonella enterica] Length: 372\nNumber of predicted TMHs: 1\nExp number of AAs in TMHs: 24.51309\nExp number, first 60 AAs: 0.2682\nTotal prob of N-in: 0.14402\noutside 1 331\nTMhelix 332 354\ninside 355 372

21933 GCF_001576255.1_ASM157625v1 Salmonella enterica subsp. enterica serovar Typhimurium Proteobacteria; Gammaproteobacteria; Enterobacterales; Enterobacteriaceae; Salmonella; Salmonella enterica; Salmonella enterica subsp. enterica MNNEETFYQAMRRKGVTRRSFLKFCSLAATSLGLGAGMTPKIAWALENKPRIPVVWIHGL WP_001058312.1 hydrogenase [Salmonella enterica] Length: 372\nNumber of predicted TMHs: 2\nExp number of AAs in TMHs: 36.51738\nExp number, first 60 AAs: 19.72849\nTotal prob of N-in: 0.96170\nPOSSIBLE N-term signal sequence\ninside 1 19\nTMhelix 20 39\noutside 40 325\nTMhelix 326 348\ninside 349 372

21934 GCF_001577505.1_ASM157750v1 Salmonella enterica subsp. enterica serovar Typhimurium Proteobacteria; Gammaproteobacteria; Enterobacterales; Enterobacteriaceae; Salmonella; Salmonella enterica; Salmonella enterica subsp. enterica MTGDNTLITSHGINRRDFMKLCAALAATMGLSSKAAAEMAESVSNPQRPPVIWIGAQECT WP_000145429.1 hydrogenase 2 small subunit [Salmonella enterica] Length: 372\nNumber of predicted TMHs: 1\nExp number of AAs in TMHs: 24.51309\nExp number, first 60 AAs: 0.2682\nTotal prob of N-in: 0.14402\noutside 1 331\nTMhelix 332 354\ninside 355 372

21935 GCF_001577505.1_ASM157750v1 Salmonella enterica subsp. enterica serovar Typhimurium Proteobacteria; Gammaproteobacteria; Enterobacterales; Enterobacteriaceae; Salmonella; Salmonella enterica; Salmonella enterica subsp. enterica MNNEETFYQAMRRKGVTRRSFLKFCSLAATSLGLGAGMTPKIAWALENKPRIPVVWIHGL WP_001058312.1 hydrogenase [Salmonella enterica] Length: 372\nNumber of predicted TMHs: 2\nExp number of AAs in TMHs: 36.51738\nExp number, first 60 AAs: 19.72849\nTotal prob of N-in: 0.96170\nPOSSIBLE N-term signal sequence\ninside 1 19\nTMhelix 20 39\noutside 40 325\nTMhelix 326 348\ninside 349 372

21936 GCF_001587055.1_ASM158705v1 Salmonella enterica subsp. enterica serovar Typhimurium Proteobacteria; Gammaproteobacteria; Enterobacterales; Enterobacteriaceae; Salmonella; Salmonella enterica; Salmonella enterica subsp. enterica MTGDNTLITSHGINRRDFMKLCAALAATMGLSSKAAAEMAESVSNPQRPPVIWIGAQECT WP_000145429.1 hydrogenase 2 small subunit [Salmonella enterica] Length: 372\nNumber of predicted TMHs: 1\nExp number of AAs in TMHs: 24.51309\nExp number, first 60 AAs: 0.2682\nTotal prob of N-in: 0.14402\noutside 1 331\nTMhelix 332 354\ninside 355 372

21937 GCF_001587765.1_CFSAN033935_01.0Salmonella enterica subsp. enterica serovar Typhimurium Proteobacteria; Gammaproteobacteria; Enterobacterales; Enterobacteriaceae; Salmonella; Salmonella enterica; Salmonella enterica subsp. enterica MTGDNTLITSHGINRRDFMKLCAALAATMGLSSKAAAEMAESVSNPQRPPVIWIGAQECT WP_000145429.1 hydrogenase 2 small subunit [Salmonella enterica] Length: 372\nNumber of

predicted TMHs: 1\nExp number of AAs in TMHs: 24.51309\nExp number, first 60 AAs: 0.2682\nTotal prob of N-in: 0.14402\noutside 1 331\nTMhelix 332 354\ninside 355 372

21938 GCF_001587785.1_CFSAN033937_01.0Salmonella enterica subsp. enterica serovar Typhimurium Proteobacteria; Gammaproteobacteria; Enterobacterales; Enterobacteriaceae; Salmonella; Salmonella enterica; Salmonella enterica subsp. enterica MTGDNTLITSHGINRRDFMKLCAALAATMGLSSKAAAEMAESVSNPQRPPVIWIGAQECT WP_000145429.1 hydrogenase 2 small subunit [Salmonella enterica] Length: 372\nNumber of predicted TMHs: 1\nExp number of AAs in TMHs: 24.51309\nExp number, first 60 AAs: 0.2682\nTotal prob of N-in: 0.14402\noutside 1 331\nTMhelix 332 354\ninside 355 372

21939 GCF_001587845.1_CFSAN033933_01.0Salmonella enterica subsp. enterica serovar Typhimurium Proteobacteria; Gammaproteobacteria; Enterobacterales; Enterobacteriaceae; Salmonella; Salmonella enterica; Salmonella enterica subsp. enterica MTGDNTLITSHGINRRDFMKLCAALAATMGLSSKAAAEMAESVSNPQRPPVIWIGAQECT WP_000145429.1 hydrogenase 2 small subunit [Salmonella enterica] Length: 372\nNumber of predicted TMHs: 1\nExp number of AAs in TMHs: 24.51309\nExp number, first 60 AAs: 0.2682\nTotal prob of N-in: 0.14402\noutside 1 331\nTMhelix 332 354\ninside 355 372

21940 GCF_001587055.1_ASM158705v1 Salmonella enterica subsp. enterica serovar Typhimurium Proteobacteria; Gammaproteobacteria; Enterobacterales; Enterobacteriaceae; Salmonella; Salmonella enterica; Salmonella enterica subsp. enterica MNNEETFYQAMRRKGVTRRSFLKFCSLAATSLGLGAGMTPKIAWALENKPRIPVVWIHGL WP_001058312.1 hydrogenase [Salmonella enterica] Length: 372\nNumber of predicted TMHs: 2\nExp number of AAs in TMHs: 36.51738\nExp number, first 60 AAs: 19.72849\nTotal prob of N-in: 0.96170\nPOSSIBLE N-term signal sequence\ninside 1 19\nTMhelix 20 39\noutside 40 325\nTMhelix 326 348\ninside 349 372

21941 GCF_001587925.1_CFSAN033927_01.0Salmonella enterica subsp. enterica serovar Typhimurium Proteobacteria; Gammaproteobacteria; Enterobacterales; Enterobacteriaceae; Salmonella; Salmonella enterica; Salmonella enterica subsp. enterica MTGDNTLITSHGINRRDFMKLCAALAATMGLSSKAAAEMAESVSNPQRPPVIWIGAQECT WP_000145429.1 hydrogenase 2 small subunit [Salmonella enterica] Length: 372\nNumber of predicted TMHs: 1\nExp number of AAs in TMHs: 24.51309\nExp number, first 60 AAs: 0.2682\nTotal prob of N-in: 0.14402\noutside 1 331\nTMhelix 332 354\ninside 355 372

21942 GCF_001587885.1_CFSAN033930_01.0Salmonella enterica subsp. enterica serovar Typhimurium Proteobacteria; Gammaproteobacteria; Enterobacterales; Enterobacteriaceae; Salmonella; Salmonella enterica; Salmonella enterica subsp. enterica MTGDNTLITSHGINRRDFMKLCAALAATMGLSSKAAAEMAESVSNPQRPPVIWIGAQECT WP_000145429.1 hydrogenase 2 small subunit [Salmonella enterica] Length: 372\nNumber of predicted TMHs: 1\nExp number of AAs in TMHs: 24.51309\nExp number, first 60 AAs: 0.2682\nTotal prob of N-in: 0.14402\noutside 1 331\nTMhelix 332 354\ninside 355 372

21943 GCF_001587925.1_CFSAN033927_01.0Salmonella enterica subsp. enterica serovar Typhimurium Proteobacteria; Gammaproteobacteria; Enterobacterales; Enterobacteriaceae; Salmonella; Salmonella enterica; Salmonella enterica subsp. enterica MNNEETFYQAMRRKGVTRRSFLKFCSLAATSLGLGAGMTPKIAWALENKPRIPVVWIHGL WP_001058312.1 hydrogenase [Salmonella enterica] Length: 372\nNumber of predicted TMHs: 2\nExp number of AAs in TMHs: 36.51738\nExp number, first 60 AAs: 19.72849\nTotal prob of N-in: 0.96170\nPOSSIBLE N-term signal sequence\ninside 1 19\nTMhelix 20 39\noutside 40 325\nTMhelix 326 348\ninside 349 372

21944 GCF_001587885.1_CFSAN033930_01.0Salmonella enterica subsp. enterica serovar Typhimurium Proteobacteria; Gammaproteobacteria; Enterobacterales; Enterobacteriaceae; Salmonella; Salmonella enterica; Salmonella enterica subsp. enterica MNNEETFYQAMRRKGVTRRSFLKFCSLAATSLGLGAGMTPKIAWALENKPRIPVVWIHGL WP_001058312.1 hydrogenase [Salmonella enterica] Length: 372\nNumber of predicted TMHs: 2\nExp number of AAs in TMHs: 36.51738\nExp number, first 60 AAs: 19.72849\nTotal prob of N-in: 0.96170\nPOSSIBLE N-term signal sequence\ninside 1 19\nTMhelix 20 39\noutside 40 325\nTMhelix 326 348\ninside 349 372

21945 GCF_001587785.1_CFSAN033937_01.0Salmonella enterica subsp. enterica serovar Typhimurium Proteobacteria; Gammaproteobacteria; Enterobacterales; Enterobacteriaceae; Salmonella; Salmonella enterica; Salmonella enterica subsp. enterica MNNEETFYQAMRRKGVTRRSFLKFCSLAATSLGLGAGMTPKIAWALENKPRIPVVWIHGL WP_001058312.1 hydrogenase [Salmonella enterica] Length: 372\nNumber of predicted TMHs: 2\nExp number of AAs in TMHs: 36.51738\nExp number, first 60 AAs: 19.72849\nTotal prob of N-in: 0.96170\nPOSSIBLE N-term signal sequence\ninside 1 19\nTMhelix 20 39\noutside 40 325\nTMhelix 326 348\ninside 349 372

21946 GCF_001587945.1_CFSAN033926_01.0Salmonella enterica subsp. enterica serovar Typhimurium Proteobacteria; Gammaproteobacteria; Enterobacterales; Enterobacteriaceae; Salmonella; Salmonella enterica;

Salmonella enterica subsp. enterica MTGDNTLITSHGINRRDFMKLCAALAATMGLSSKAAAEMAESVSNPQRPPVIWIGAQECT WP_000145429.1 hydrogenase 2 small subunit [Salmonella enterica] Length: 372\nNumber of predicted TMHs: 1\nExp number of AAs in TMHs: 24.51309\nExp number, first 60 AAs: 0.2682\nTotal prob of N-in: 0.14402\noutside 1 331\nTMhelix 332 354\ninside 355 372

21947 GCF_001587945.1_CFSAN033926_01.0Salmonella enterica subsp. enterica serovar Typhimurium Proteobacteria; Gammaproteobacteria; Enterobacterales; Enterobacteriaceae; Salmonella; Salmonella enterica; Salmonella enterica subsp. enterica MNNEETFYQAMRRKGVTRRSFLKFCSLAATSLGLGAGMTPKIAWALENKPRIPVVWIHGL WP_001058312.1 hydrogenase [Salmonella enterica] Length: 372\nNumber of predicted TMHs: 2\nExp number of AAs in TMHs: 36.51738\nExp number, first 60 AAs: 19.72849\nTotal prob of N-in: 0.96170\nPOSSIBLE N-term signal sequence\ninside 1 19\nTMhelix 20 39\noutside 40 325\nTMhelix 326 348\ninside 349 372

21948 GCF_001587845.1_CFSAN033933_01.0Salmonella enterica subsp. enterica serovar Typhimurium Proteobacteria; Gammaproteobacteria; Enterobacterales; Enterobacteriaceae; Salmonella; Salmonella enterica; Salmonella enterica subsp. enterica MNNEETFYQAMRRKGVTRRSFLKFCSLAATSLGLGAGMTPKIAWALENKPRIPVVWIHGL WP_001058312.1 hydrogenase [Salmonella enterica] Length: 372\nNumber of predicted TMHs: 2\nExp number of AAs in TMHs: 36.51738\nExp number, first 60 AAs: 19.72849\nTotal prob of N-in: 0.96170\nPOSSIBLE N-term signal sequence\ninside 1 19\nTMhelix 20 39\noutside 40 325\nTMhelix 326 348\ninside 349 372

21949 GCF_001587765.1_CFSAN033935_01.0Salmonella enterica subsp. enterica serovar Typhimurium Proteobacteria; Gammaproteobacteria; Enterobacterales; Enterobacteriaceae; Salmonella; Salmonella enterica; Salmonella enterica subsp. enterica MNNEETFYQAMRRKGVTRRSFLKFCSLAATSLGLGAGMTPKIAWALENKPRIPVVWIHGL WP_001058312.1 hydrogenase [Salmonella enterica] Length: 372\nNumber of predicted TMHs: 2\nExp number of AAs in TMHs: 36.51738\nExp number, first 60 AAs: 19.72849\nTotal prob of N-in: 0.96170\nPOSSIBLE N-term signal sequence\ninside 1 19\nTMhelix 20 39\noutside 40 325\nTMhelix 326 348\ninside 349 372

21950 GCF_001587995.1_CFSAN033922_01.0Salmonella enterica subsp. enterica serovar Typhimurium Proteobacteria; Gammaproteobacteria; Enterobacterales; Enterobacteriaceae; Salmonella; Salmonella enterica; Salmonella enterica subsp. enterica MTGDNTLITSHGINRRDFMKLCAALAATMGLSSKAAAEMAESVSNPQRPPVIWIGAQECT WP_000145429.1 hydrogenase 2 small subunit [Salmonella enterica] Length: 372\nNumber of predicted TMHs: 1\nExp number of AAs in TMHs: 24.51309\nExp number, first 60 AAs: 0.2682\nTotal prob of N-in: 0.14402\noutside 1 331\nTMhelix 332 354\ninside 355 372

21951 GCF_001588025.1_CFSAN033919_01.0Salmonella enterica subsp. enterica serovar Typhimurium Proteobacteria; Gammaproteobacteria; Enterobacterales; Enterobacteriaceae; Salmonella; Salmonella enterica; Salmonella enterica subsp. enterica MTGDNTLITSHGINRRDFMKLCAALAATMGLSSKAAAEMAESVSNPQRPPVIWIGAQECT WP_000145429.1 hydrogenase 2 small subunit [Salmonella enterica] Length: 372\nNumber of predicted TMHs: 1\nExp number of AAs in TMHs: 24.51309\nExp number, first 60 AAs: 0.2682\nTotal prob of N-in: 0.14402\noutside 1 331\nTMhelix 332 354\ninside 355 372

21952 GCF_001588125.1_CFSAN033913_01.0Salmonella enterica subsp. enterica serovar Typhimurium Proteobacteria; Gammaproteobacteria; Enterobacterales; Enterobacteriaceae; Salmonella; Salmonella enterica; Salmonella enterica subsp. enterica MTGDNTLITSHGINRRDFMKLCAALAATMGLSSKAAAEMAESVSNPQRPPVIWIGAQECT WP_000145429.1 hydrogenase 2 small subunit [Salmonella enterica] Length: 372\nNumber of predicted TMHs: 1\nExp number of AAs in TMHs: 24.51309\nExp number, first 60 AAs: 0.2682\nTotal prob of N-in: 0.14402\noutside 1 331\nTMhelix 332 354\ninside 355 372

21953 GCF_001588095.1_CFSAN033915_01.0Salmonella enterica subsp. enterica serovar Typhimurium Proteobacteria; Gammaproteobacteria; Enterobacterales; Enterobacteriaceae; Salmonella; Salmonella enterica; Salmonella enterica subsp. enterica MTGDNTLITSHGINRRDFMKLCAALAATMGLSSKAAAEMAESVSNPQRPPVIWIGAQECT WP_000145429.1 hydrogenase 2 small subunit [Salmonella enterica] Length: 372\nNumber of predicted TMHs: 1\nExp number of AAs in TMHs: 24.51309\nExp number, first 60 AAs: 0.2682\nTotal prob of N-in: 0.14402\noutside 1 331\nTMhelix 332 354\ninside 355 372

21954 GCF_001588165.1_CFSAN033910_01.0Salmonella enterica subsp. enterica serovar Typhimurium Proteobacteria; Gammaproteobacteria; Enterobacterales; Enterobacteriaceae; Salmonella; Salmonella enterica; Salmonella enterica subsp. enterica MTGDNTLITSHGINRRDFMKLCAALAATMGLSSKAAAEMAESVSNPQRPPVIWIGAQECT WP_000145429.1 hydrogenase 2 small subunit [Salmonella enterica] Length: 372\nNumber of predicted TMHs: 1\nExp number of AAs in TMHs: 24.51309\nExp number, first 60 AAs: 0.2682\nTotal prob of N-in: 0.14402\noutside 1 331\nTMhelix 332 354\ninside 355 372

21955 GCF_001588215.1_CFSAN033908_01.0Salmonella enterica subsp. enterica serovar Typhimurium Proteobacteria; Gammaproteobacteria; Enterobacterales; Enterobacteriaceae; Salmonella; Salmonella enterica; Salmonella enterica subsp. enterica MTGDNTLITSHGINRRDFMKLCAALAATMGLSSKAAAEMAESVSNPQRPPVIWIGAQECT WP_000145429.1 hydrogenase 2 small subunit [Salmonella enterica] Length: 372\nNumber of predicted TMHs: 1\nExp number of AAs in TMHs: 24.51309\nExp number, first 60 AAs: 0.2682\nTotal prob of N-in: 0.14402\noutside 1 331\nTMhelix 332 354\ninside 355 372

21956 GCF_001587995.1_CFSAN033922_01.0Salmonella enterica subsp. enterica serovar Typhimurium Proteobacteria; Gammaproteobacteria; Enterobacterales; Enterobacteriaceae; Salmonella; Salmonella enterica; Salmonella enterica subsp. enterica MNNEETFYQAMRRKGVTRRSFLKFCSLAATSLGLGAGMTPKIAWALENKPRIPVVWIHGL WP_001058312.1 hydrogenase [Salmonella enterica] Length: 372\nNumber of predicted TMHs: 2\nExp number of AAs in TMHs: 36.51738\nExp number, first 60 AAs: 19.72849\nTotal prob of N-in: 0.96170\nPOSSIBLE N-term signal sequence\ninside 1 19\nTMhelix 20 39\noutside 40 325\nTMhelix 326 348\ninside 349 372

21957 GCF_001588025.1_CFSAN033919_01.0Salmonella enterica subsp. enterica serovar Typhimurium Proteobacteria; Gammaproteobacteria; Enterobacterales; Enterobacteriaceae; Salmonella; Salmonella enterica; Salmonella enterica subsp. enterica MNNEETFYQAMRRKGVTRRSFLKFCSLAATSLGLGAGMTPKIAWALENKPRIPVVWIHGL WP_001058312.1 hydrogenase [Salmonella enterica] Length: 372\nNumber of predicted TMHs: 2\nExp number of AAs in TMHs: 36.51738\nExp number, first 60 AAs: 19.72849\nTotal prob of N-in: 0.96170\nPOSSIBLE N-term signal sequence\ninside 1 19\nTMhelix 20 39\noutside 40 325\nTMhelix 326 348\ninside 349 372

21958 GCF_001588125.1_CFSAN033913_01.0Salmonella enterica subsp. enterica serovar Typhimurium Proteobacteria; Gammaproteobacteria; Enterobacterales; Enterobacteriaceae; Salmonella; Salmonella enterica; Salmonella enterica subsp. enterica MNNEETFYQAMRRKGVTRRSFLKFCSLAATSLGLGAGMTPKIAWALENKPRIPVVWIHGL WP_001058312.1 hydrogenase [Salmonella enterica] Length: 372\nNumber of predicted TMHs: 2\nExp number of AAs in TMHs: 36.51738\nExp number, first 60 AAs: 19.72849\nTotal prob of N-in: 0.96170\nPOSSIBLE N-term signal sequence\ninside 1 19\nTMhelix 20 39\noutside 40 325\nTMhelix 326 348\ninside 349 372

21959 GCF_001588095.1_CFSAN033915_01.0Salmonella enterica subsp. enterica serovar Typhimurium Proteobacteria; Gammaproteobacteria; Enterobacterales; Enterobacteriaceae; Salmonella; Salmonella enterica; Salmonella enterica subsp. enterica MNNEETFYQAMRRKGVTRRSFLKFCSLAATSLGLGAGMTPKIAWALENKPRIPVVWIHGL WP_001058312.1 hydrogenase [Salmonella enterica] Length: 372\nNumber of predicted TMHs: 2\nExp number of AAs in TMHs: 36.51738\nExp number, first 60 AAs: 19.72849\nTotal prob of N-in: 0.96170\nPOSSIBLE N-term signal sequence\ninside 1 19\nTMhelix 20 39\noutside 40 325\nTMhelix 326 348\ninside 349 372

21960 GCF_001588255.1_CFSAN033905_01.0Salmonella enterica subsp. enterica serovar Typhimurium Proteobacteria; Gammaproteobacteria; Enterobacterales; Enterobacteriaceae; Salmonella; Salmonella enterica; Salmonella enterica subsp. enterica MTGDNTLITSHGINRRDFMKLCAALAATMGLSSKAAAEMAESVSNPQRPPVIWIGAQECT WP_000145429.1 hydrogenase 2 small subunit [Salmonella enterica] Length: 372\nNumber of predicted TMHs: 1\nExp number of AAs in TMHs: 24.51309\nExp number, first 60 AAs: 0.2682\nTotal prob of N-in: 0.14402\noutside 1 331\nTMhelix 332 354\ninside 355 372

21961 GCF_001588255.1_CFSAN033905_01.0Salmonella enterica subsp. enterica serovar Typhimurium Proteobacteria; Gammaproteobacteria; Enterobacterales; Enterobacteriaceae; Salmonella; Salmonella enterica; Salmonella enterica subsp. enterica MNNEETFYQAMRRKGVTRRSFLKFCSLAATSLGLGAGMTPKIAWALENKPRIPVVWIHGL WP_001058312.1 hydrogenase [Salmonella enterica] Length: 372\nNumber of predicted TMHs: 2\nExp number of AAs in TMHs: 36.51738\nExp number, first 60 AAs: 19.72849\nTotal prob of N-in: 0.96170\nPOSSIBLE N-term signal sequence\ninside 1 19\nTMhelix 20 39\noutside 40 325\nTMhelix 326 348\ninside 349 372

21962 GCF_001588165.1_CFSAN033910_01.0Salmonella enterica subsp. enterica serovar Typhimurium Proteobacteria; Gammaproteobacteria; Enterobacterales; Enterobacteriaceae; Salmonella; Salmonella enterica; Salmonella enterica subsp. enterica MNNEETFYQAMRRKGVTRRSFLKFCSLAATSLGLGAGMTPKIAWALENKPRIPVVWIHGL WP_001058312.1 hydrogenase [Salmonella enterica] Length: 372\nNumber of predicted TMHs: 2\nExp number of AAs in TMHs: 36.51738\nExp number, first 60 AAs: 19.72849\nTotal prob of N-in: 0.96170\nPOSSIBLE N-term signal sequence\ninside 1 19\nTMhelix 20 39\noutside 40 325\nTMhelix 326 348\ninside 349 372

21963 GCF_001588285.1_CFSAN033903_01.0Salmonella enterica subsp. enterica serovar Typhimurium Proteobacteria; Gammaproteobacteria; Enterobacterales; Enterobacteriaceae; Salmonella; Salmonella enterica; Salmonella enterica subsp. enterica MTGDNTLITSHGINRRDFMKLCAALAATMGLSSKAAAEMAESVSNPQRPPVIWIGAQECT WP_000145429.1 hydrogenase 2 small subunit [Salmonella enterica] Length: 372\nNumber of

predicted TMHs:  1\nExp number of AAs in TMHs: 24.51309\nExp number, first 60 AAs:  0.2682\nTotal prob of N-in:  0.14402\noutside   1   331\nTMhelix   332   354\ninside   355   372

21964 GCF_001588335.1_CFSAN033900_01.0Salmonella enterica subsp. enterica serovar Typhimurium Proteobacteria; Gammaproteobacteria; Enterobacterales; Enterobacteriaceae; Salmonella; Salmonella enterica; Salmonella enterica subsp. enterica MTGDNTLITSHGINRRDFMKLCAALAATMGLSSKAAAEMAESVSNPQRPPVIWIGAQECT WP_000145429.1 hydrogenase 2 small subunit [Salmonella enterica] Length: 372\nNumber of predicted TMHs:  1\nExp number of AAs in TMHs: 24.51309\nExp number, first 60 AAs:  0.2682\nTotal prob of N-in:  0.14402\noutside   1   331\nTMhelix   332   354\ninside   355   372

21965 GCF_001588285.1_CFSAN033903_01.0Salmonella enterica subsp. enterica serovar Typhimurium Proteobacteria; Gammaproteobacteria; Enterobacterales; Enterobacteriaceae; Salmonella; Salmonella enterica; Salmonella enterica subsp. enterica MNNEETFYQAMRRKGVTRRSFLKFCSLAATSLGLGAGMTPKIAWALENKPRIPVVWIHGL WP_001058312.1 hydrogenase [Salmonella enterica] Length: 372\nNumber of predicted TMHs:  2\nExp number of AAs in TMHs: 36.51738\nExp number, first 60 AAs:  19.72849\nTotal prob of N-in:   0.96170\nPOSSIBLE N-term signal sequence\ninside   1   19\nTMhelix   20   39\noutside   40   325\nTMhelix   326   348\ninside   349   372

21966 GCF_001588335.1_CFSAN033900_01.0Salmonella enterica subsp. enterica serovar Typhimurium Proteobacteria; Gammaproteobacteria; Enterobacterales; Enterobacteriaceae; Salmonella; Salmonella enterica; Salmonella enterica subsp. enterica MNNEETFYQAMRRKGVTRRSFLKFCSLAATSLGLGAGMTPKIAWALENKPRIPVVWIHGL WP_001058312.1 hydrogenase [Salmonella enterica] Length: 372\nNumber of predicted TMHs:  2\nExp number of AAs in TMHs: 36.51738\nExp number, first 60 AAs:  19.72849\nTotal prob of N-in:   0.96170\nPOSSIBLE N-term signal sequence\ninside   1   19\nTMhelix   20   39\noutside   40   325\nTMhelix   326   348\ninside   349   372

21967 GCF_001588215.1_CFSAN033908_01.0Salmonella enterica subsp. enterica serovar Typhimurium Proteobacteria; Gammaproteobacteria; Enterobacterales; Enterobacteriaceae; Salmonella; Salmonella enterica; Salmonella enterica subsp. enterica MNNEETFYQAMRRKGVTRRSFLKFCSLAATSLGLGAGMTPKIAWALENKPRIPVVWIHGL WP_001058312.1 hydrogenase [Salmonella enterica] Length: 372\nNumber of predicted TMHs:  2\nExp number of AAs in TMHs: 36.51738\nExp number, first 60 AAs:  19.72849\nTotal prob of N-in:   0.96170\nPOSSIBLE N-term signal sequence\ninside   1   19\nTMhelix   20   39\noutside   40   325\nTMhelix   326   348\ninside   349   372

21968 GCF_001588365.1_CFSAN033898_01.0Salmonella enterica subsp. enterica serovar Typhimurium Proteobacteria; Gammaproteobacteria; Enterobacterales; Enterobacteriaceae; Salmonella; Salmonella enterica; Salmonella enterica subsp. enterica MTGDNTLITSHGINRRDFMKLCAALAATMGLSSKAAAEMAESVSNPQRPPVIWIGAQECT WP_000145429.1 hydrogenase 2 small subunit [Salmonella enterica] Length: 372\nNumber of predicted TMHs:  1\nExp number of AAs in TMHs: 24.51309\nExp number, first 60 AAs:  0.2682\nTotal prob of N-in:  0.14402\noutside   1   331\nTMhelix   332   354\ninside   355   372

21969 GCF_001588495.1_CFSAN033885_01.0Salmonella enterica subsp. enterica serovar Typhimurium Proteobacteria; Gammaproteobacteria; Enterobacterales; Enterobacteriaceae; Salmonella; Salmonella enterica; Salmonella enterica subsp. enterica MTGDNTLITSHGINRRDFMKLCAALAATMGLSSKAAAEMAESVSNPQRPPVIWIGAQECT WP_000145429.1 hydrogenase 2 small subunit [Salmonella enterica] Length: 372\nNumber of predicted TMHs:  1\nExp number of AAs in TMHs: 24.51309\nExp number, first 60 AAs:  0.2682\nTotal prob of N-in:  0.14402\noutside   1   331\nTMhelix   332   354\ninside   355   372

21970 GCF_001588535.1_CFSAN033875_01.0Salmonella enterica subsp. enterica serovar Typhimurium Proteobacteria; Gammaproteobacteria; Enterobacterales; Enterobacteriaceae; Salmonella; Salmonella enterica; Salmonella enterica subsp. enterica MTGDNTLITSHGINRRDFMKLCAALAATMGLSSKAAAEMAESVSNPQRPPVIWIGAQECT WP_000145429.1 hydrogenase 2 small subunit [Salmonella enterica] Length: 372\nNumber of predicted TMHs:  1\nExp number of AAs in TMHs: 24.51309\nExp number, first 60 AAs:  0.2682\nTotal prob of N-in:  0.14402\noutside   1   331\nTMhelix   332   354\ninside   355   372

21971 GCF_001588365.1_CFSAN033898_01.0Salmonella enterica subsp. enterica serovar Typhimurium Proteobacteria; Gammaproteobacteria; Enterobacterales; Enterobacteriaceae; Salmonella; Salmonella enterica; Salmonella enterica subsp. enterica MNNEETFYQAMRRKGVTRRSFLKFCSLAATSLGLGAGMTPKIAWALENKPRIPVVWIHGL WP_001058312.1 hydrogenase [Salmonella enterica] Length: 372\nNumber of predicted TMHs:  2\nExp number of AAs in TMHs: 36.51738\nExp number, first 60 AAs:  19.72849\nTotal prob of N-in:   0.96170\nPOSSIBLE N-term signal sequence\ninside   1   19\nTMhelix   20   39\noutside   40   325\nTMhelix   326   348\ninside   349   372

21972 GCF_001588415.1_CFSAN033894_01.0Salmonella enterica subsp. enterica serovar Typhimurium Proteobacteria; Gammaproteobacteria; Enterobacterales; Enterobacteriaceae; Salmonella; Salmonella enterica;

Salmonella enterica subsp. enterica MNNEETFYQAMRRKGVTRRSFLKFCSLAATSLGLGAGMTPKIAWALENKPRIPVVWIHGL WP_001058312.1 hydrogenase [Salmonella enterica] Length: 372\nNumber of predicted TMHs: 2\nExp number of AAs in TMHs: 36.51738\nExp number, first 60 AAs: 19.72849\nTotal prob of N-in: 0.96170\nPOSSIBLE N-term signal sequence\ninside 1 19\nTMhelix 20 39\noutside 40 325\nTMhelix 326 348\ninside 349 372

21973 GCF_001588565.1_CFSAN033859_01.0Salmonella enterica subsp. enterica serovar Typhimurium Proteobacteria; Gammaproteobacteria; Enterobacterales; Enterobacteriaceae; Salmonella; Salmonella enterica; Salmonella enterica subsp. enterica MTGDNTLITSHGINRRDFMKLCAALAATMGLSSKAAAEMAESVSNPQRPPVIWIGAQECT WP_000145429.1 hydrogenase 2 small subunit [Salmonella enterica] Length: 372\nNumber of predicted TMHs: 1\nExp number of AAs in TMHs: 24.51309\nExp number, first 60 AAs: 0.2682\nTotal prob of N-in: 0.14402\noutside 1 331\nTMhelix 332 354\ninside 355 372

21974 GCF_001588455.1_CFSAN033887_01.0Salmonella enterica subsp. enterica serovar Typhimurium Proteobacteria; Gammaproteobacteria; Enterobacterales; Enterobacteriaceae; Salmonella; Salmonella enterica; Salmonella enterica subsp. enterica MNNEETFYQAMRRKGVTRRSFLKFCSLAATSLGLGAGMTPKIAWALENKPRIPVVWIHGL WP_001058312.1 hydrogenase [Salmonella enterica] Length: 372\nNumber of predicted TMHs: 2\nExp number of AAs in TMHs: 36.51738\nExp number, first 60 AAs: 19.72849\nTotal prob of N-in: 0.96170\nPOSSIBLE N-term signal sequence\ninside 1 19\nTMhelix 20 39\noutside 40 325\nTMhelix 326 348\ninside 349 372

21975 GCF_001594425.1_CFSAN033939_02.0Salmonella enterica subsp. enterica serovar Typhimurium Proteobacteria; Gammaproteobacteria; Enterobacterales; Enterobacteriaceae; Salmonella; Salmonella enterica; Salmonella enterica subsp. enterica MTGDNTLITSHGINRRDFMKLCAALAATMGLSSKAAAEMAESVSNPQRPPVIWIGAQECT WP_000145429.1 hydrogenase 2 small subunit [Salmonella enterica] Length: 372\nNumber of predicted TMHs: 1\nExp number of AAs in TMHs: 24.51309\nExp number, first 60 AAs: 0.2682\nTotal prob of N-in: 0.14402\noutside 1 331\nTMhelix 332 354\ninside 355 372

21976 GCF_001594445.1_CFSAN033931_01.0Salmonella enterica subsp. enterica serovar Typhimurium Proteobacteria; Gammaproteobacteria; Enterobacterales; Enterobacteriaceae; Salmonella; Salmonella enterica; Salmonella enterica subsp. enterica MTGDNTLITSHGINRRDFMKLCAALAATMGLSSKAAAEMAESVSNPQRPPVIWIGAQECT WP_000145429.1 hydrogenase 2 small subunit [Salmonella enterica] Length: 372\nNumber of predicted TMHs: 1\nExp number of AAs in TMHs: 24.51309\nExp number, first 60 AAs: 0.2682\nTotal prob of N-in: 0.14402\noutside 1 331\nTMhelix 332 354\ninside 355 372

21977 GCF_001594505.1_CFSAN033923_01.0Salmonella enterica subsp. enterica serovar Typhimurium Proteobacteria; Gammaproteobacteria; Enterobacterales; Enterobacteriaceae; Salmonella; Salmonella enterica; Salmonella enterica subsp. enterica MTGDNTLITSHGINRRDFMKLCAALAATMGLSSKAAAEMAESVSNPQRPPVIWIGAQECT WP_000145429.1 hydrogenase 2 small subunit [Salmonella enterica] Length: 372\nNumber of predicted TMHs: 1\nExp number of AAs in TMHs: 24.51309\nExp number, first 60 AAs: 0.2682\nTotal prob of N-in: 0.14402\noutside 1 331\nTMhelix 332 354\ninside 355 372

21978 GCF_001588455.1_CFSAN033887_01.0Salmonella enterica subsp. enterica serovar Typhimurium Proteobacteria; Gammaproteobacteria; Enterobacterales; Enterobacteriaceae; Salmonella; Salmonella enterica; Salmonella enterica subsp. enterica MTGDNTLITSHGINRRDFMKLCAALAATMGLSSKAAAEMAESVSNPQRPPVIWIGAQECT WP_061584213.1 hydrogenase 2 small subunit [Salmonella enterica] Length: 372\nNumber of predicted TMHs: 1\nExp number of AAs in TMHs: 24.51809\nExp number, first 60 AAs: 0.26818\nTotal prob of N-in: 0.14414\noutside 1 331\nTMhelix 332 354\ninside 355 372

21979 GCF_001594425.1_CFSAN033939_02.0Salmonella enterica subsp. enterica serovar Typhimurium Proteobacteria; Gammaproteobacteria; Enterobacterales; Enterobacteriaceae; Salmonella; Salmonella enterica; Salmonella enterica subsp. enterica MNNEETFYQAMRRKGVTRRSFLKFCSLAATSLGLGAGMTPKIAWALENKPRIPVVWIHGL WP_001058312.1 hydrogenase [Salmonella enterica] Length: 372\nNumber of predicted TMHs: 2\nExp number of AAs in TMHs: 36.51738\nExp number, first 60 AAs: 19.72849\nTotal prob of N-in: 0.96170\nPOSSIBLE N-term signal sequence\ninside 1 19\nTMhelix 20 39\noutside 40 325\nTMhelix 326 348\ninside 349 372

21980 GCF_001594445.1_CFSAN033931_01.0Salmonella enterica subsp. enterica serovar Typhimurium Proteobacteria; Gammaproteobacteria; Enterobacterales; Enterobacteriaceae; Salmonella; Salmonella enterica; Salmonella enterica subsp. enterica MNNEETFYQAMRRKGVTRRSFLKFCSLAATSLGLGAGMTPKIAWALENKPRIPVVWIHGL WP_001058312.1 hydrogenase [Salmonella enterica] Length: 372\nNumber of predicted TMHs: 2\nExp number of AAs in TMHs: 36.51738\nExp number, first 60 AAs: 19.72849\nTotal prob of N-in: 0.96170\nPOSSIBLE N-term signal sequence\ninside 1 19\nTMhelix 20 39\noutside 40 325\nTMhelix 326 348\ninside 349 372

21981 GCF_001588495.1_CFSAN033885_01.0Salmonella enterica subsp. enterica serovar Typhimurium Proteobacteria; Gammaproteobacteria; Enterobacterales; Enterobacteriaceae; Salmonella; Salmonella enterica; Salmonella enterica subsp. enterica MNNEETFYQAMRRKGVTRRSFLKFCSLAATSLGLGAGMTPKIAWALENKPRIPVVWIHGL WP_001058312.1 hydrogenase [Salmonella enterica] Length: 372\nNumber of predicted TMHs: 2\nExp number of AAs in TMHs: 36.51738\nExp number, first 60 AAs: 19.72849\nTotal prob of N-in: 0.96170\nPOSSIBLE N-term signal sequence\ninside 1 19\nTMhelix 20 39\noutside 40 325\nTMhelix 326 348\ninside 349 372

21982 GCF_001588415.1_CFSAN033894_01.0Salmonella enterica subsp. enterica serovar Typhimurium Proteobacteria; Gammaproteobacteria; Enterobacterales; Enterobacteriaceae; Salmonella; Salmonella enterica; Salmonella enterica subsp. enterica MTGDNTLITSHGINRRDFMKLCAALAATMGLSSKAAAEMAESVSNPQRPPVIWIGAQECT WP_061584213.1 hydrogenase 2 small subunit [Salmonella enterica] Length: 372\nNumber of predicted TMHs: 1\nExp number of AAs in TMHs: 24.51809\nExp number, first 60 AAs: 0.26818\nTotal prob of N-in: 0.14414\noutside 1 331\nTMhelix 332 354\ninside 355 372

21983 GCF_001588535.1_CFSAN033875_01.0Salmonella enterica subsp. enterica serovar Typhimurium Proteobacteria; Gammaproteobacteria; Enterobacterales; Enterobacteriaceae; Salmonella; Salmonella enterica; Salmonella enterica subsp. enterica MNNEETFYQAMRRKGVTRRSFLKFCSLAATSLGLGAGMTPKIAWALENKPRIPVVWIHGL WP_001058312.1 hydrogenase [Salmonella enterica] Length: 372\nNumber of predicted TMHs: 2\nExp number of AAs in TMHs: 36.51738\nExp number, first 60 AAs: 19.72849\nTotal prob of N-in: 0.96170\nPOSSIBLE N-term signal sequence\ninside 1 19\nTMhelix 20 39\noutside 40 325\nTMhelix 326 348\ninside 349 372

21984 GCF_001588565.1_CFSAN033859_01.0Salmonella enterica subsp. enterica serovar Typhimurium Proteobacteria; Gammaproteobacteria; Enterobacterales; Enterobacteriaceae; Salmonella; Salmonella enterica; Salmonella enterica subsp. enterica MNNEETFYQAMRRKGVTRRSFLKFCSLAATSLGLGAGMTPKIAWALENKPRIPVVWIHGL WP_001058312.1 hydrogenase [Salmonella enterica] Length: 372\nNumber of predicted TMHs: 2\nExp number of AAs in TMHs: 36.51738\nExp number, first 60 AAs: 19.72849\nTotal prob of N-in: 0.96170\nPOSSIBLE N-term signal sequence\ninside 1 19\nTMhelix 20 39\noutside 40 325\nTMhelix 326 348\ninside 349 372

21985 GCF_001594525.1_CFSAN033916_01.0Salmonella enterica subsp. enterica serovar Typhimurium Proteobacteria; Gammaproteobacteria; Enterobacterales; Enterobacteriaceae; Salmonella; Salmonella enterica; Salmonella enterica subsp. enterica MTGDNTLITSHGINRRDFMKLCAALAATMGLSSKAAAEMAESVSNPQRPPVIWIGAQECT WP_000145429.1 hydrogenase 2 small subunit [Salmonella enterica] Length: 372\nNumber of predicted TMHs: 1\nExp number of AAs in TMHs: 24.51309\nExp number, first 60 AAs: 0.2682\nTotal prob of N-in: 0.14402\noutside 1 331\nTMhelix 332 354\ninside 355 372

21986 GCF_001594585.1_CFSAN033907_01.0Salmonella enterica subsp. enterica serovar Typhimurium Proteobacteria; Gammaproteobacteria; Enterobacterales; Enterobacteriaceae; Salmonella; Salmonella enterica; Salmonella enterica subsp. enterica MTGDNTLITSHGINRRDFMKLCAALAATMGLSSKAAAEMAESVSNPQRPPVIWIGAQECT WP_000145429.1 hydrogenase 2 small subunit [Salmonella enterica] Length: 372\nNumber of predicted TMHs: 1\nExp number of AAs in TMHs: 24.51309\nExp number, first 60 AAs: 0.2682\nTotal prob of N-in: 0.14402\noutside 1 331\nTMhelix 332 354\ninside 355 372

21987 GCF_001594525.1_CFSAN033916_01.0Salmonella enterica subsp. enterica serovar Typhimurium Proteobacteria; Gammaproteobacteria; Enterobacterales; Enterobacteriaceae; Salmonella; Salmonella enterica; Salmonella enterica subsp. enterica MNNEETFYQAMRRKGVTRRSFLKFCSLAATSLGLGAGMTPKIAWALENKPRIPVVWIHGL WP_001058312.1 hydrogenase [Salmonella enterica] Length: 372\nNumber of predicted TMHs: 2\nExp number of AAs in TMHs: 36.51738\nExp number, first 60 AAs: 19.72849\nTotal prob of N-in: 0.96170\nPOSSIBLE N-term signal sequence\ninside 1 19\nTMhelix 20 39\noutside 40 325\nTMhelix 326 348\ninside 349 372

21988 GCF_001594585.1_CFSAN033907_01.0Salmonella enterica subsp. enterica serovar Typhimurium Proteobacteria; Gammaproteobacteria; Enterobacterales; Enterobacteriaceae; Salmonella; Salmonella enterica; Salmonella enterica subsp. enterica MNNEETFYQAMRRKGVTRRSFLKFCSLAATSLGLGAGMTPKIAWALENKPRIPVVWIHGL WP_001058312.1 hydrogenase [Salmonella enterica] Length: 372\nNumber of predicted TMHs: 2\nExp number of AAs in TMHs: 36.51738\nExp number, first 60 AAs: 19.72849\nTotal prob of N-in: 0.96170\nPOSSIBLE N-term signal sequence\ninside 1 19\nTMhelix 20 39\noutside 40 325\nTMhelix 326 348\ninside 349 372

21989 GCF_001594505.1_CFSAN033923_01.0Salmonella enterica subsp. enterica serovar Typhimurium Proteobacteria; Gammaproteobacteria; Enterobacterales; Enterobacteriaceae; Salmonella; Salmonella enterica; Salmonella enterica subsp. enterica MNNEETFYQAMRRKGVTRRSFLKFCSLAATSLGLGAGMTPKIAWALENKPRIPVVWIHGL WP_001058312.1 hydrogenase [Salmonella enterica] Length: 372\nNumber of predicted TMHs: 2\nExp

number of AAs in TMHs: 36.51738\nExp number, first 60 AAs: 19.72849\nTotal prob of N-in: 0.96170\nPOSSIBLE N-term signal sequence\ninside 1 19\nTMhelix 20 39\noutside 40 325\nTMhelix 326 348\ninside 349 372

21990 GCF_001594615.1_CFSAN033897_01.0Salmonella enterica subsp. enterica serovar Typhimurium Proteobacteria; Gammaproteobacteria; Enterobacterales; Enterobacteriaceae; Salmonella; Salmonella enterica; Salmonella enterica subsp. enterica MTGDNTLITSHGINRRDFMKLCAALAATMGLSSKAAAEMAESVSNPQRPPVIWIGAQECT WP_000145429.1 hydrogenase 2 small subunit [Salmonella enterica] Length: 372\nNumber of predicted TMHs: 1\nExp number of AAs in TMHs: 24.51309\nExp number, first 60 AAs: 0.2682\nTotal prob of N-in: 0.14402\noutside 1 331\nTMhelix 332 354\ninside 355 372

21991 GCF_001594665.1_CFSAN033893_01.0Salmonella enterica subsp. enterica serovar Typhimurium Proteobacteria; Gammaproteobacteria; Enterobacterales; Enterobacteriaceae; Salmonella; Salmonella enterica; Salmonella enterica subsp. enterica MTGDNTLITSHGINRRDFMKLCAALAATMGLSSKAAAEMAESVSNPQRPPVIWIGAQECT WP_000145429.1 hydrogenase 2 small subunit [Salmonella enterica] Length: 372\nNumber of predicted TMHs: 1\nExp number of AAs in TMHs: 24.51309\nExp number, first 60 AAs: 0.2682\nTotal prob of N-in: 0.14402\noutside 1 331\nTMhelix 332 354\ninside 355 372

21992 GCF_001594695.1_CFSAN033890_01.0Salmonella enterica subsp. enterica serovar Typhimurium Proteobacteria; Gammaproteobacteria; Enterobacterales; Enterobacteriaceae; Salmonella; Salmonella enterica; Salmonella enterica subsp. enterica MTGDNTLITSHGINRRDFMKLCAALAATMGLSSKAAAEMAESVSNPQRPPVIWIGAQECT WP_000145429.1 hydrogenase 2 small subunit [Salmonella enterica] Length: 372\nNumber of predicted TMHs: 1\nExp number of AAs in TMHs: 24.51309\nExp number, first 60 AAs: 0.2682\nTotal prob of N-in: 0.14402\noutside 1 331\nTMhelix 332 354\ninside 355 372

21993 GCF_001594615.1_CFSAN033897_01.0Salmonella enterica subsp. enterica serovar Typhimurium Proteobacteria; Gammaproteobacteria; Enterobacterales; Enterobacteriaceae; Salmonella; Salmonella enterica; Salmonella enterica subsp. enterica MNNEETFYQAMRRKGVTRRSFLKFCSLAATSLGLGAGMTPKIAWALENKPRIPVVWIHGL WP_001058312.1 hydrogenase [Salmonella enterica] Length: 372\nNumber of predicted TMHs: 2\nExp number of AAs in TMHs: 36.51738\nExp number, first 60 AAs: 19.72849\nTotal prob of N-in: 0.96170\nPOSSIBLE N-term signal sequence\ninside 1 19\nTMhelix 20 39\noutside 40 325\nTMhelix 326 348\ninside 349 372

21994 GCF_001594825.1_CFSAN033880_01.0Salmonella enterica subsp. enterica serovar Typhimurium Proteobacteria; Gammaproteobacteria; Enterobacterales; Enterobacteriaceae; Salmonella; Salmonella enterica; Salmonella enterica subsp. enterica MTGDNTLITSHGINRRDFMKLCAALAATMGLSSKAAAEMAESVSNPQRPPVIWIGAQECT WP_000145429.1 hydrogenase 2 small subunit [Salmonella enterica] Length: 372\nNumber of predicted TMHs: 1\nExp number of AAs in TMHs: 24.51309\nExp number, first 60 AAs: 0.2682\nTotal prob of N-in: 0.14402\noutside 1 331\nTMhelix 332 354\ninside 355 372

21995 GCF_001594665.1_CFSAN033893_01.0Salmonella enterica subsp. enterica serovar Typhimurium Proteobacteria; Gammaproteobacteria; Enterobacterales; Enterobacteriaceae; Salmonella; Salmonella enterica; Salmonella enterica subsp. enterica MNNEETFYQAMRRKGVTRRSFLKFCSLAATSLGLGAGMTPKIAWALENKPRIPVVWIHGL WP_001058312.1 hydrogenase [Salmonella enterica] Length: 372\nNumber of predicted TMHs: 2\nExp number of AAs in TMHs: 36.51738\nExp number, first 60 AAs: 19.72849\nTotal prob of N-in: 0.96170\nPOSSIBLE N-term signal sequence\ninside 1 19\nTMhelix 20 39\noutside 40 325\nTMhelix 326 348\ninside 349 372

21996 GCF_001594955.1_CFSAN033871_01.0Salmonella enterica subsp. enterica serovar Typhimurium Proteobacteria; Gammaproteobacteria; Enterobacterales; Enterobacteriaceae; Salmonella; Salmonella enterica; Salmonella enterica subsp. enterica MTGDNTLITSHGINRRDFMKLCAALAATMGLSSKAAAEMAESVSNPQRPPVIWIGAQECT WP_000145429.1 hydrogenase 2 small subunit [Salmonella enterica] Length: 372\nNumber of predicted TMHs: 1\nExp number of AAs in TMHs: 24.51309\nExp number, first 60 AAs: 0.2682\nTotal prob of N-in: 0.14402\noutside 1 331\nTMhelix 332 354\ninside 355 372

21997 GCF_001594915.1_CFSAN033873_01.0Salmonella enterica subsp. enterica serovar Typhimurium Proteobacteria; Gammaproteobacteria; Enterobacterales; Enterobacteriaceae; Salmonella; Salmonella enterica; Salmonella enterica subsp. enterica MTGDNTLITSHGINRRDFMKLCAALAATMGLSSKAAAEMAESVSNPQRPPVIWIGAQECT WP_000145429.1 hydrogenase 2 small subunit [Salmonella enterica] Length: 372\nNumber of predicted TMHs: 1\nExp number of AAs in TMHs: 24.51309\nExp number, first 60 AAs: 0.2682\nTotal prob of N-in: 0.14402\noutside 1 331\nTMhelix 332 354\ninside 355 372

21998 GCF_001594745.1_CFSAN033884_01.0Salmonella enterica subsp. enterica serovar Typhimurium Proteobacteria; Gammaproteobacteria; Enterobacterales; Enterobacteriaceae; Salmonella; Salmonella enterica;

Salmonella enterica subsp. enterica MNNEETFYQAMRRKGVTRRSFLKFCSLAATSLGLGAGMTPKIAWALENKPRIPVVWIHGL WP_001058312.1 hydrogenase [Salmonella enterica] Length: 372\nNumber of predicted TMHs: 2\nExp number of AAs in TMHs: 36.51738\nExp number, first 60 AAs: 19.72849\nTotal prob of N-in: 0.96170\nPOSSIBLE N-term signal sequence\ninside 1 19\nTMhelix 20 39\noutside 40 325\nTMhelix 326 348\ninside 349 372

21999 GCF_001594915.1_CFSAN033873_01.0Salmonella enterica subsp. enterica serovar Typhimurium Proteobacteria; Gammaproteobacteria; Enterobacterales; Enterobacteriaceae; Salmonella; Salmonella enterica; Salmonella enterica subsp. enterica MNNEETFYQAMRRKGVTRRSFLKFCSLAATSLGLGAGMTPKIAWALENKPRIPVVWIHGL WP_001058312.1 hydrogenase [Salmonella enterica] Length: 372\nNumber of predicted TMHs: 2\nExp number of AAs in TMHs: 36.51738\nExp number, first 60 AAs: 19.72849\nTotal prob of N-in: 0.96170\nPOSSIBLE N-term signal sequence\ninside 1 19\nTMhelix 20 39\noutside 40 325\nTMhelix 326 348\ninside 349 372

22000 GCF_001594955.1_CFSAN033871_01.0Salmonella enterica subsp. enterica serovar Typhimurium Proteobacteria; Gammaproteobacteria; Enterobacterales; Enterobacteriaceae; Salmonella; Salmonella enterica; Salmonella enterica subsp. enterica MNNEETFYQAMRRKGVTRRSFLKFCSLAATSLGLGAGMTPKIAWALENKPRIPVVWIHGL WP_001058312.1 hydrogenase [Salmonella enterica] Length: 372\nNumber of predicted TMHs: 2\nExp number of AAs in TMHs: 36.51738\nExp number, first 60 AAs: 19.72849\nTotal prob of N-in: 0.96170\nPOSSIBLE N-term signal sequence\ninside 1 19\nTMhelix 20 39\noutside 40 325\nTMhelix 326 348\ninside 349 372

22001 GCF_001594745.1_CFSAN033884_01.0Salmonella enterica subsp. enterica serovar Typhimurium Proteobacteria; Gammaproteobacteria; Enterobacterales; Enterobacteriaceae; Salmonella; Salmonella enterica; Salmonella enterica subsp. enterica MTGDNTLITSHGINRRDFMKLCAALAATMGLSSKAAAEMAESVSNPQRPPVIWIGAQECT WP_061584213.1 hydrogenase 2 small subunit [Salmonella enterica] Length: 372\nNumber of predicted TMHs: 1\nExp number of AAs in TMHs: 24.51809\nExp number, first 60 AAs: 0.26818\nTotal prob of N-in: 0.14414\noutside 1 331\nTMhelix 332 354\ninside 355 372

22002 GCF_001594695.1_CFSAN033890_01.0Salmonella enterica subsp. enterica serovar Typhimurium Proteobacteria; Gammaproteobacteria; Enterobacterales; Enterobacteriaceae; Salmonella; Salmonella enterica; Salmonella enterica subsp. enterica MNNEETFYQAMRRKGVTRRSFLKFCSLAATSLGLGAGMTPKIAWALENKPRIPVVWIHGL WP_001058312.1 hydrogenase [Salmonella enterica] Length: 372\nNumber of predicted TMHs: 2\nExp number of AAs in TMHs: 36.51738\nExp number, first 60 AAs: 19.72849\nTotal prob of N-in: 0.96170\nPOSSIBLE N-term signal sequence\ninside 1 19\nTMhelix 20 39\noutside 40 325\nTMhelix 326 348\ninside 349 372

22003 GCF_001594775.1_CFSAN033882_01.0Salmonella enterica subsp. enterica serovar Typhimurium Proteobacteria; Gammaproteobacteria; Enterobacterales; Enterobacteriaceae; Salmonella; Salmonella enterica; Salmonella enterica subsp. enterica MNNEETFYQAMRRKGVTRRSFLKFCSLAATSLGLGAGMTPKIAWALENKPRIPVVWIHGL WP_001058312.1 hydrogenase [Salmonella enterica] Length: 372\nNumber of predicted TMHs: 2\nExp number of AAs in TMHs: 36.51738\nExp number, first 60 AAs: 19.72849\nTotal prob of N-in: 0.96170\nPOSSIBLE N-term signal sequence\ninside 1 19\nTMhelix 20 39\noutside 40 325\nTMhelix 326 348\ninside 349 372

22004 GCF_001594995.1_CFSAN033869_01.0Salmonella enterica subsp. enterica serovar Typhimurium Proteobacteria; Gammaproteobacteria; Enterobacterales; Enterobacteriaceae; Salmonella; Salmonella enterica; Salmonella enterica subsp. enterica MTGDNTLITSHGINRRDFMKLCAALAATMGLSSKAAAEMAESVSNPQRPPVIWIGAQECT WP_000145429.1 hydrogenase 2 small subunit [Salmonella enterica] Length: 372\nNumber of predicted TMHs: 1\nExp number of AAs in TMHs: 24.51309\nExp number, first 60 AAs: 0.2682\nTotal prob of N-in: 0.14402\noutside 1 331\nTMhelix 332 354\ninside 355 372

22005 GCF_001594875.1_CFSAN033876_01.0Salmonella enterica subsp. enterica serovar Typhimurium Proteobacteria; Gammaproteobacteria; Enterobacterales; Enterobacteriaceae; Salmonella; Salmonella enterica; Salmonella enterica subsp. enterica MNNEETFYQAMRRKGVTRRSFLKFCSLAATSLGLGAGMTPKIAWALENKPRIPVVWIHGL WP_001058312.1 hydrogenase [Salmonella enterica] Length: 372\nNumber of predicted TMHs: 2\nExp number of AAs in TMHs: 36.51738\nExp number, first 60 AAs: 19.72849\nTotal prob of N-in: 0.96170\nPOSSIBLE N-term signal sequence\ninside 1 19\nTMhelix 20 39\noutside 40 325\nTMhelix 326 348\ninside 349 372

22006 GCF_001594775.1_CFSAN033882_01.0Salmonella enterica subsp. enterica serovar Typhimurium Proteobacteria; Gammaproteobacteria; Enterobacterales; Enterobacteriaceae; Salmonella; Salmonella enterica; Salmonella enterica subsp. enterica MTGDNTLITSHGINRRDFMKLCAALAATMGLSSKAAAEMAESVSNPQRPPVIWIGAQECT WP_061584213.1 hydrogenase 2 small subunit [Salmonella enterica] Length: 372\nNumber of predicted TMHs: 1\nExp number of AAs in TMHs: 24.51809\nExp number, first 60 AAs: 0.26818\nTotal prob of N-in: 0.14414\noutside 1 331\nTMhelix 332 354\ninside 355 372

22007 GCF_001594995.1_CFSAN033869_01.0Salmonella enterica subsp. enterica serovar Typhimurium Proteobacteria; Gammaproteobacteria; Enterobacterales; Enterobacteriaceae; Salmonella; Salmonella enterica; Salmonella enterica subsp. enterica MNNEETFYQAMRRKGVTRRSFLKFCSLAATSLGLGAGMTPKIAWALENKPRIPVVWIHGL WP_001058312.1 hydrogenase [Salmonella enterica] Length: 372\nNumber of predicted TMHs: 2\nExp number of AAs in TMHs: 36.51738\nExp number, first 60 AAs: 19.72849\nTotal prob of N-in: 0.96170\nPOSSIBLE N-term signal sequence\ninside 1 19\nTMhelix 20 39\noutside 40 325\nTMhelix 326 348\ninside 349 372

22008 GCF_001594875.1_CFSAN033876_01.0Salmonella enterica subsp. enterica serovar Typhimurium Proteobacteria; Gammaproteobacteria; Enterobacterales; Enterobacteriaceae; Salmonella; Salmonella enterica; Salmonella enterica subsp. enterica MTGDNTLITSHGINRRDFMKLCAALAATMGLSSKAAAEMAESVSNPQRPPVIWIGAQECT WP_061584213.1 hydrogenase 2 small subunit [Salmonella enterica] Length: 372\nNumber of predicted TMHs: 1\nExp number of AAs in TMHs: 24.51809\nExp number, first 60 AAs: 0.26818\nTotal prob of N-in: 0.14414\noutside 1 331\nTMhelix 332 354\ninside 355 372

22009 GCF_001594825.1_CFSAN033880_01.0Salmonella enterica subsp. enterica serovar Typhimurium Proteobacteria; Gammaproteobacteria; Enterobacterales; Enterobacteriaceae; Salmonella; Salmonella enterica; Salmonella enterica subsp. enterica MNNEETFYQAMRRKGVTRRSFLKFCSLAATSLGLGAGMTPKIAWALENKPRIPVVWIHGL WP_001058312.1 hydrogenase [Salmonella enterica] Length: 372\nNumber of predicted TMHs: 2\nExp number of AAs in TMHs: 36.51738\nExp number, first 60 AAs: 19.72849\nTotal prob of N-in: 0.96170\nPOSSIBLE N-term signal sequence\ninside 1 19\nTMhelix 20 39\noutside 40 325\nTMhelix 326 348\ninside 349 372

22010 GCF_001595015.1_CFSAN033867_01.0Salmonella enterica subsp. enterica serovar Typhimurium Proteobacteria; Gammaproteobacteria; Enterobacterales; Enterobacteriaceae; Salmonella; Salmonella enterica; Salmonella enterica subsp. enterica MTGDNTLITSHGINRRDFMKLCAALAATMGLSSKAAAEMAESVSNPQRPPVIWIGAQECT WP_000145429.1 hydrogenase 2 small subunit [Salmonella enterica] Length: 372\nNumber of predicted TMHs: 1\nExp number of AAs in TMHs: 24.51309\nExp number, first 60 AAs: 0.2682\nTotal prob of N-in: 0.14402\noutside 1 331\nTMhelix 332 354\ninside 355 372

22011 GCF_001595035.1_CFSAN033866_01.0Salmonella enterica subsp. enterica serovar Typhimurium Proteobacteria; Gammaproteobacteria; Enterobacterales; Enterobacteriaceae; Salmonella; Salmonella enterica; Salmonella enterica subsp. enterica MTGDNTLITSHGINRRDFMKLCAALAATMGLSSKAAAEMAESVSNPQRPPVIWIGAQECT WP_000145429.1 hydrogenase 2 small subunit [Salmonella enterica] Length: 372\nNumber of predicted TMHs: 1\nExp number of AAs in TMHs: 24.51309\nExp number, first 60 AAs: 0.2682\nTotal prob of N-in: 0.14402\noutside 1 331\nTMhelix 332 354\ninside 355 372

22012 GCF_001595075.1_CFSAN033864_01.0Salmonella enterica subsp. enterica serovar Typhimurium Proteobacteria; Gammaproteobacteria; Enterobacterales; Enterobacteriaceae; Salmonella; Salmonella enterica; Salmonella enterica subsp. enterica MTGDNTLITSHGINRRDFMKLCAALAATMGLSSKAAAEMAESVSNPQRPPVIWIGAQECT WP_000145429.1 hydrogenase 2 small subunit [Salmonella enterica] Length: 372\nNumber of predicted TMHs: 1\nExp number of AAs in TMHs: 24.51309\nExp number, first 60 AAs: 0.2682\nTotal prob of N-in: 0.14402\noutside 1 331\nTMhelix 332 354\ninside 355 372

22013 GCF_001595115.1_CFSAN033862_01.0Salmonella enterica subsp. enterica serovar Typhimurium Proteobacteria; Gammaproteobacteria; Enterobacterales; Enterobacteriaceae; Salmonella; Salmonella enterica; Salmonella enterica subsp. enterica MTGDNTLITSHGINRRDFMKLCAALAATMGLSSKAAAEMAESVSNPQRPPVIWIGAQECT WP_000145429.1 hydrogenase 2 small subunit [Salmonella enterica] Length: 372\nNumber of predicted TMHs: 1\nExp number of AAs in TMHs: 24.51309\nExp number, first 60 AAs: 0.2682\nTotal prob of N-in: 0.14402\noutside 1 331\nTMhelix 332 354\ninside 355 372

22014 GCF_001595155.1_CFSAN033860_01.0Salmonella enterica subsp. enterica serovar Typhimurium Proteobacteria; Gammaproteobacteria; Enterobacterales; Enterobacteriaceae; Salmonella; Salmonella enterica; Salmonella enterica subsp. enterica MTGDNTLITSHGINRRDFMKLCAALAATMGLSSKAAAEMAESVSNPQRPPVIWIGAQECT WP_000145429.1 hydrogenase 2 small subunit [Salmonella enterica] Length: 372\nNumber of predicted TMHs: 1\nExp number of AAs in TMHs: 24.51309\nExp number, first 60 AAs: 0.2682\nTotal prob of N-in: 0.14402\noutside 1 331\nTMhelix 332 354\ninside 355 372

22015 GCF_001595035.1_CFSAN033866_01.0Salmonella enterica subsp. enterica serovar Typhimurium Proteobacteria; Gammaproteobacteria; Enterobacterales; Enterobacteriaceae; Salmonella; Salmonella enterica; Salmonella enterica subsp. enterica MNNEETFYQAMRRKGVTRRSFLKFCSLAATSLGLGAGMTPKIAWALENKPRIPVVWIHGL WP_001058312.1 hydrogenase [Salmonella enterica] Length: 372\nNumber of predicted TMHs: 2\nExp

number of AAs in TMHs: 36.51738\nExp number, first 60 AAs:  19.72849\nTotal prob of N-in:   0.96170\nPOSSIBLE N-term signal sequence\ninside   1   19\nTMhelix   20   39\noutside   40   325\nTMhelix   326   348\ninside   349   372

22016 GCF_001595015.1_CFSAN033867_01.0Salmonella enterica subsp. enterica serovar Typhimurium Proteobacteria; Gammaproteobacteria; Enterobacterales; Enterobacteriaceae; Salmonella; Salmonella enterica; Salmonella enterica subsp. enterica MNNEETFYQAMRRKGVTRRSFLKFCSLAATSLGLGAGMTPKIAWALENKPRIPVVWIHGL WP_001058312.1 hydrogenase [Salmonella enterica] Length: 372\nNumber of predicted TMHs:  2\nExp number of AAs in TMHs: 36.51738\nExp number, first 60 AAs:  19.72849\nTotal prob of N-in:   0.96170\nPOSSIBLE N-term signal sequence\ninside   1   19\nTMhelix   20   39\noutside   40   325\nTMhelix   326   348\ninside   349   372

22017 GCF_001595235.1_CFSAN033854_01.0Salmonella enterica subsp. enterica serovar Typhimurium Proteobacteria; Gammaproteobacteria; Enterobacterales; Enterobacteriaceae; Salmonella; Salmonella enterica; Salmonella enterica subsp. enterica MTGDNTLITSHGINRRDFMKLCAALAATMGLSSKAAAEMAESVSNPQRPPVIWIGAQECT WP_000145429.1 hydrogenase 2 small subunit [Salmonella enterica] Length: 372\nNumber of predicted TMHs:  1\nExp number of AAs in TMHs: 24.51309\nExp number, first 60 AAs:  0.2682\nTotal prob of N-in: 0.14402\noutside   1   331\nTMhelix   332   354\ninside   355   372

22018 GCF_001595275.1_CFSAN033852_01.0Salmonella enterica subsp. enterica serovar Typhimurium Proteobacteria; Gammaproteobacteria; Enterobacterales; Enterobacteriaceae; Salmonella; Salmonella enterica; Salmonella enterica subsp. enterica MTGDNTLITSHGINRRDFMKLCAALAATMGLSSKAAAEMAESVSNPQRPPVIWIGAQECT WP_000145429.1 hydrogenase 2 small subunit [Salmonella enterica] Length: 372\nNumber of predicted TMHs:  1\nExp number of AAs in TMHs: 24.51309\nExp number, first 60 AAs:  0.2682\nTotal prob of N-in: 0.14402\noutside   1   331\nTMhelix   332   354\ninside   355   372

22019 GCF_001595235.1_CFSAN033854_01.0Salmonella enterica subsp. enterica serovar Typhimurium Proteobacteria; Gammaproteobacteria; Enterobacterales; Enterobacteriaceae; Salmonella; Salmonella enterica; Salmonella enterica subsp. enterica MNNEETFYQAMRRKGVTRRSFLKFCSLAATSLGLGAGMTPKIAWALENKPRIPVVWIHGL WP_001058312.1 hydrogenase [Salmonella enterica] Length: 372\nNumber of predicted TMHs:  2\nExp number of AAs in TMHs: 36.51738\nExp number, first 60 AAs:  19.72849\nTotal prob of N-in:   0.96170\nPOSSIBLE N-term signal sequence\ninside   1   19\nTMhelix   20   39\noutside   40   325\nTMhelix   326   348\ninside   349   372

22020 GCF_001595275.1_CFSAN033852_01.0Salmonella enterica subsp. enterica serovar Typhimurium Proteobacteria; Gammaproteobacteria; Enterobacterales; Enterobacteriaceae; Salmonella; Salmonella enterica; Salmonella enterica subsp. enterica MNNEETFYQAMRRKGVTRRSFLKFCSLAATSLGLGAGMTPKIAWALENKPRIPVVWIHGL WP_001058312.1 hydrogenase [Salmonella enterica] Length: 372\nNumber of predicted TMHs:  2\nExp number of AAs in TMHs: 36.51738\nExp number, first 60 AAs:  19.72849\nTotal prob of N-in:   0.96170\nPOSSIBLE N-term signal sequence\ninside   1   19\nTMhelix   20   39\noutside   40   325\nTMhelix   326   348\ninside   349   372

22021 GCF_001595075.1_CFSAN033864_01.0Salmonella enterica subsp. enterica serovar Typhimurium Proteobacteria; Gammaproteobacteria; Enterobacterales; Enterobacteriaceae; Salmonella; Salmonella enterica; Salmonella enterica subsp. enterica MNNEETFYQAMRRKGVTRRSFLKFCSLAATSLGLGAGMTPKIAWALENKPRIPVVWIHGL WP_001058312.1 hydrogenase [Salmonella enterica] Length: 372\nNumber of predicted TMHs:  2\nExp number of AAs in TMHs: 36.51738\nExp number, first 60 AAs:  19.72849\nTotal prob of N-in:   0.96170\nPOSSIBLE N-term signal sequence\ninside   1   19\nTMhelix   20   39\noutside   40   325\nTMhelix   326   348\ninside   349   372

22022 GCF_001595155.1_CFSAN033860_01.0Salmonella enterica subsp. enterica serovar Typhimurium Proteobacteria; Gammaproteobacteria; Enterobacterales; Enterobacteriaceae; Salmonella; Salmonella enterica; Salmonella enterica subsp. enterica MNNEETFYQAMRRKGVTRRSFLKFCSLAATSLGLGAGMTPKIAWALENKPRIPVVWIHGL WP_001058312.1 hydrogenase [Salmonella enterica] Length: 372\nNumber of predicted TMHs:  2\nExp number of AAs in TMHs: 36.51738\nExp number, first 60 AAs:  19.72849\nTotal prob of N-in:   0.96170\nPOSSIBLE N-term signal sequence\ninside   1   19\nTMhelix   20   39\noutside   40   325\nTMhelix   326   348\ninside   349   372

22023 GCF_001595315.1_CFSAN033850_01.0Salmonella enterica subsp. enterica serovar Typhimurium Proteobacteria; Gammaproteobacteria; Enterobacterales; Enterobacteriaceae; Salmonella; Salmonella enterica; Salmonella enterica subsp. enterica MTGDNTLITSHGINRRDFMKLCAALAATMGLSSKAAAEMAESVSNPQRPPVIWIGAQECT WP_000145429.1 hydrogenase 2 small subunit [Salmonella enterica] Length: 372\nNumber of predicted TMHs:  1\nExp number of AAs in TMHs: 24.51309\nExp number, first 60 AAs:  0.2682\nTotal prob of N-in: 0.14402\noutside   1   331\nTMhelix   332   354\ninside   355   372

22024 GCF_001595315.1_CFSAN033850_01.0Salmonella enterica subsp. enterica serovar Typhimurium Proteobacteria; Gammaproteobacteria; Enterobacterales; Enterobacteriaceae; Salmonella; Salmonella enterica;

Salmonella enterica subsp. enterica MNNEETFYQAMRRKGVTRRSFLKFCSLAATSLGLGAGMTPKIAWALENKPRIPVVWIHGL WP_001058312.1 hydrogenase [Salmonella enterica] Length: 372\nNumber of predicted TMHs: 2\nExp number of AAs in TMHs: 36.51738\nExp number, first 60 AAs: 19.72849\nTotal prob of N-in: 0.96170\nPOSSIBLE N-term signal sequence\ninside 1 19\nTMhelix 20 39\noutside 40 325\nTMhelix 326 348\ninside 349 372

22025 GCF_001595115.1_CFSAN033862_01.0Salmonella enterica subsp. enterica serovar Typhimurium Proteobacteria; Gammaproteobacteria; Enterobacterales; Enterobacteriaceae; Salmonella; Salmonella enterica; Salmonella enterica subsp. enterica MNNEETFYQAMRRKGVTRRSFLKFCSLAATSLGLGAGMTPKIAWALENKPRIPVVWIHGL WP_001058312.1 hydrogenase [Salmonella enterica] Length: 372\nNumber of predicted TMHs: 2\nExp number of AAs in TMHs: 36.51738\nExp number, first 60 AAs: 19.72849\nTotal prob of N-in: 0.96170\nPOSSIBLE N-term signal sequence\ninside 1 19\nTMhelix 20 39\noutside 40 325\nTMhelix 326 348\ninside 349 372

22026 GCF_001595195.1_CFSAN033857_01.0Salmonella enterica subsp. enterica serovar Typhimurium Proteobacteria; Gammaproteobacteria; Enterobacterales; Enterobacteriaceae; Salmonella; Salmonella enterica; Salmonella enterica subsp. enterica MNNEETFYQAMRRKGVTRRSFLKFCSLAATSLGLGAGMTPKIAWALENKPRIPVVWIHGL WP_001058312.1 hydrogenase [Salmonella enterica] Length: 372\nNumber of predicted TMHs: 2\nExp number of AAs in TMHs: 36.51738\nExp number, first 60 AAs: 19.72849\nTotal prob of N-in: 0.96170\nPOSSIBLE N-term signal sequence\ninside 1 19\nTMhelix 20 39\noutside 40 325\nTMhelix 326 348\ninside 349 372

22027 GCF_001595355.1_CFSAN033848_01.0Salmonella enterica subsp. enterica serovar Typhimurium Proteobacteria; Gammaproteobacteria; Enterobacterales; Enterobacteriaceae; Salmonella; Salmonella enterica; Salmonella enterica subsp. enterica MTGDNTLITSHGINRRDFMKLCAALAATMGLSSKAAAEMAESVSNPQRPPVIWIGAQECT WP_000145429.1 hydrogenase 2 small subunit [Salmonella enterica] Length: 372\nNumber of predicted TMHs: 1\nExp number of AAs in TMHs: 24.51309\nExp number, first 60 AAs: 0.2682\nTotal prob of N-in: 0.14402\noutside 1 331\nTMhelix 332 354\ninside 355 372

22028 GCF_001595355.1_CFSAN033848_01.0Salmonella enterica subsp. enterica serovar Typhimurium Proteobacteria; Gammaproteobacteria; Enterobacterales; Enterobacteriaceae; Salmonella; Salmonella enterica; Salmonella enterica subsp. enterica MNNEETFYQAMRRKGVTRRSFLKFCSLAATSLGLGAGMTPKIAWALENKPRIPVVWIHGL WP_001058312.1 hydrogenase [Salmonella enterica] Length: 372\nNumber of predicted TMHs: 2\nExp number of AAs in TMHs: 36.51738\nExp number, first 60 AAs: 19.72849\nTotal prob of N-in: 0.96170\nPOSSIBLE N-term signal sequence\ninside 1 19\nTMhelix 20 39\noutside 40 325\nTMhelix 326 348\ninside 349 372

22029 GCF_001614025.1_ASM161402v1 Salmonella enterica subsp. enterica serovar Typhimurium Proteobacteria; Gammaproteobacteria; Enterobacterales; Enterobacteriaceae; Salmonella; Salmonella enterica; Salmonella enterica subsp. enterica MTGDNTLITSHGINRRDFMKLCAALAATMGLSSKAAAEMAESVSNPQRPPVIWIGAQECT WP_000145429.1 hydrogenase 2 small subunit [Salmonella enterica] Length: 372\nNumber of predicted TMHs: 1\nExp number of AAs in TMHs: 24.51309\nExp number, first 60 AAs: 0.2682\nTotal prob of N-in: 0.14402\noutside 1 331\nTMhelix 332 354\ninside 355 372

22030 GCF_001617585.1_ASM161758v1 Salmonella enterica subsp. enterica serovar Typhimurium Proteobacteria; Gammaproteobacteria; Enterobacterales; Enterobacteriaceae; Salmonella; Salmonella enterica; Salmonella enterica subsp. enterica MTGDNTLITSHGINRRDFMKLCAALAATMGLSSKAAAEMAESVSNPQRPPVIWIGAQECT WP_000145429.1 hydrogenase 2 small subunit [Salmonella enterica] Length: 372\nNumber of predicted TMHs: 1\nExp number of AAs in TMHs: 24.51309\nExp number, first 60 AAs: 0.2682\nTotal prob of N-in: 0.14402\noutside 1 331\nTMhelix 332 354\ninside 355 372

22031 GCF_001614025.1_ASM161402v1 Salmonella enterica subsp. enterica serovar Typhimurium Proteobacteria; Gammaproteobacteria; Enterobacterales; Enterobacteriaceae; Salmonella; Salmonella enterica; Salmonella enterica subsp. enterica MNNEETFYQAMRRKGVTRRSFLKFCSLAATSLGLGAGMTPKIAWALENKPRIPVVWIHGL WP_001058312.1 hydrogenase [Salmonella enterica] Length: 372\nNumber of predicted TMHs: 2\nExp number of AAs in TMHs: 36.51738\nExp number, first 60 AAs: 19.72849\nTotal prob of N-in: 0.96170\nPOSSIBLE N-term signal sequence\ninside 1 19\nTMhelix 20 39\noutside 40 325\nTMhelix 326 348\ninside 349 372

22032 GCF_001617585.1_ASM161758v1 Salmonella enterica subsp. enterica serovar Typhimurium Proteobacteria; Gammaproteobacteria; Enterobacterales; Enterobacteriaceae; Salmonella; Salmonella enterica; Salmonella enterica subsp. enterica MNNEETFYQAMRRKGVTRRSFLKFCSLAATSLGLGAGMTPKIAWALENKPRIPVVWIHGL WP_001058312.1 hydrogenase [Salmonella enterica] Length: 372\nNumber of predicted TMHs: 2\nExp number of AAs in TMHs: 36.51738\nExp number, first 60 AAs: 19.72849\nTotal prob of N-in: 0.96170\nPOSSIBLE N-term signal sequence\ninside 1 19\nTMhelix 20 39\noutside 40 325\nTMhelix 326 348\ninside 349 372

22033 GCF_001692145.1_ASM169214v1 Salmonella enterica subsp. enterica serovar Typhimurium Proteobacteria; Gammaproteobacteria; Enterobacterales; Enterobacteriaceae; Salmonella; Salmonella enterica; Salmonella enterica subsp. enterica MNNEETFYQAMRRKGVTRRSFLKFCSLAATSLGLGAGMTPKIAWALENKPRIPVVWIHGL WP_001058312.1 hydrogenase [Salmonella enterica] Length: 372\nNumber of predicted TMHs: 2\nExp number of AAs in TMHs: 36.51738\nExp number, first 60 AAs: 19.72849\nTotal prob of N-in: 0.96170\nPOSSIBLE N-term signal sequence\ninside 1 19\nTMhelix 20 39\noutside 40 325\nTMhelix 326 348\ninside 349 372

22034 GCF_001755875.1_CFSAN008071 Salmonella enterica subsp. enterica serovar Typhimurium Proteobacteria; Gammaproteobacteria; Enterobacterales; Enterobacteriaceae; Salmonella; Salmonella enterica; Salmonella enterica subsp. enterica MTGDNTLITSHGINRRDFMKLCAALAATMGLSSKAAAEMAESVSNPQRPPVIWIGAQECT WP_000145429.1 hydrogenase 2 small subunit [Salmonella enterica] Length: 372\nNumber of predicted TMHs: 1\nExp number of AAs in TMHs: 24.51309\nExp number, first 60 AAs: 0.2682\nTotal prob of N-in: 0.14402\noutside 1 331\nTMhelix 332 354\ninside 355 372

22035 GCF_001729035.1_ASM172903v1 Salmonella enterica subsp. enterica serovar Typhimurium Proteobacteria; Gammaproteobacteria; Enterobacterales; Enterobacteriaceae; Salmonella; Salmonella enterica; Salmonella enterica subsp. enterica MNNEETFYQAMRRKGVTRRSFLKFCSLAATSLGLGAGMTPKIAWALENKPRIPVVWIHGL WP_001058312.1 hydrogenase [Salmonella enterica] Length: 372\nNumber of predicted TMHs: 2\nExp number of AAs in TMHs: 36.51738\nExp number, first 60 AAs: 19.72849\nTotal prob of N-in: 0.96170\nPOSSIBLE N-term signal sequence\ninside 1 19\nTMhelix 20 39\noutside 40 325\nTMhelix 326 348\ninside 349 372

22036 GCF_001755895.1_CFSAN008074 Salmonella enterica subsp. enterica serovar Typhimurium Proteobacteria; Gammaproteobacteria; Enterobacterales; Enterobacteriaceae; Salmonella; Salmonella enterica; Salmonella enterica subsp. enterica MTGDNTLITSHGINRRDFMKLCAALAATMGLSSKAAAEMAESVSNPQRPPVIWIGAQECT WP_000145429.1 hydrogenase 2 small subunit [Salmonella enterica] Length: 372\nNumber of predicted TMHs: 1\nExp number of AAs in TMHs: 24.51309\nExp number, first 60 AAs: 0.2682\nTotal prob of N-in: 0.14402\noutside 1 331\nTMhelix 332 354\ninside 355 372

22037 GCF_001755995.1_CFSAN008083 Salmonella enterica subsp. enterica serovar Typhimurium Proteobacteria; Gammaproteobacteria; Enterobacterales; Enterobacteriaceae; Salmonella; Salmonella enterica; Salmonella enterica subsp. enterica MTGDNTLITSHGINRRDFMKLCAALAATMGLSSKAAAEMAESVSNPQRPPVIWIGAQECT WP_000145429.1 hydrogenase 2 small subunit [Salmonella enterica] Length: 372\nNumber of predicted TMHs: 1\nExp number of AAs in TMHs: 24.51309\nExp number, first 60 AAs: 0.2682\nTotal prob of N-in: 0.14402\noutside 1 331\nTMhelix 332 354\ninside 355 372

22038 GCF_001755895.1_CFSAN008074 Salmonella enterica subsp. enterica serovar Typhimurium Proteobacteria; Gammaproteobacteria; Enterobacterales; Enterobacteriaceae; Salmonella; Salmonella enterica; Salmonella enterica subsp. enterica MNNEETFYQAMRRKGVTRRSFLKFCSLAATSLGLGAGMTPKIAWALENKPRIPVVWIHGL WP_001058312.1 hydrogenase [Salmonella enterica] Length: 372\nNumber of predicted TMHs: 2\nExp number of AAs in TMHs: 36.51738\nExp number, first 60 AAs: 19.72849\nTotal prob of N-in: 0.96170\nPOSSIBLE N-term signal sequence\ninside 1 19\nTMhelix 20 39\noutside 40 325\nTMhelix 326 348\ninside 349 372

22039 GCF_001729045.1_ASM172904v1 Salmonella enterica subsp. enterica serovar Typhimurium Proteobacteria; Gammaproteobacteria; Enterobacterales; Enterobacteriaceae; Salmonella; Salmonella enterica; Salmonella enterica subsp. enterica MNNEETFYQAMRRKGVTRRSFLKFCSLAATSLGLGAGMTPKIAWALENKPRIPVVWIHGL WP_001058312.1 hydrogenase [Salmonella enterica] Length: 372\nNumber of predicted TMHs: 2\nExp number of AAs in TMHs: 36.51738\nExp number, first 60 AAs: 19.72849\nTotal prob of N-in: 0.96170\nPOSSIBLE N-term signal sequence\ninside 1 19\nTMhelix 20 39\noutside 40 325\nTMhelix 326 348\ninside 349 372

22040 GCF_001756035.1_CFSAN008086 Salmonella enterica subsp. enterica serovar Typhimurium Proteobacteria; Gammaproteobacteria; Enterobacterales; Enterobacteriaceae; Salmonella; Salmonella enterica; Salmonella enterica subsp. enterica MTGDNTLITSHGINRRDFMKLCAALAATMGLSSKAAAEMAESVSNPQRPPVIWIGAQECT WP_000145429.1 hydrogenase 2 small subunit [Salmonella enterica] Length: 372\nNumber of predicted TMHs: 1\nExp number of AAs in TMHs: 24.51309\nExp number, first 60 AAs: 0.2682\nTotal prob of N-in: 0.14402\noutside 1 331\nTMhelix 332 354\ninside 355 372

22041 GCF_001756125.1_CFSAN008075 Salmonella enterica subsp. enterica serovar Typhimurium Proteobacteria; Gammaproteobacteria; Enterobacterales; Enterobacteriaceae; Salmonella; Salmonella enterica; Salmonella enterica subsp. enterica MTGDNTLITSHGINRRDFMKLCAALAATMGLSSKAAAEMAESVSNPQRPPVIWIGAQECT WP_000145429.1 hydrogenase 2 small subunit [Salmonella enterica] Length: 372\nNumber of

predicted TMHs: 1\nExp number of AAs in TMHs: 24.51309\nExp number, first 60 AAs: 0.2682\nTotal prob of N-in: 0.14402\noutside 1 331\nTMhelix 332 354\ninside 355 372

22042 GCF_001756125.1_CFSAN008075 Salmonella enterica subsp. enterica serovar Typhimurium Proteobacteria; Gammaproteobacteria; Enterobacterales; Enterobacteriaceae; Salmonella; Salmonella enterica; Salmonella enterica subsp. enterica MNNEETFYQAMRRKGVTRRSFLKFCSLAATSLGLGAGMTPKIAWALENKPRIPVVWIHGL WP_001058312.1 hydrogenase [Salmonella enterica] Length: 372\nNumber of predicted TMHs: 2\nExp number of AAs in TMHs: 36.51738\nExp number, first 60 AAs: 19.72849\nTotal prob of N-in: 0.96170\nPOSSIBLE N-term signal sequence\ninside 1 19\nTMhelix 20 39\noutside 40 325\nTMhelix 326 348\ninside 349 372

22043 GCF_001756155.1_CFSAN008090 Salmonella enterica subsp. enterica serovar Typhimurium Proteobacteria; Gammaproteobacteria; Enterobacterales; Enterobacteriaceae; Salmonella; Salmonella enterica; Salmonella enterica subsp. enterica MNNEETFYQAMRRKGVTRRSFLKFCSLAATSLGLGAGMTPKIAWALENKPRIPVVWIHGL WP_023198237.1 hydrogenase 1, small subunit [Salmonella enterica] Length: 372\nNumber of predicted TMHs: 2\nExp number of AAs in TMHs: 36.39423\nExp number, first 60 AAs: 19.72877\nTotal prob of N-in: 0.96162\nPOSSIBLE N-term signal sequence\ninside 1 19\nTMhelix 20 39\noutside 40 325\nTMhelix 326 348\ninside 349 372

22044 GCF_001755875.1_CFSAN008071 Salmonella enterica subsp. enterica serovar Typhimurium Proteobacteria; Gammaproteobacteria; Enterobacterales; Enterobacteriaceae; Salmonella; Salmonella enterica; Salmonella enterica subsp. enterica MNNEETFYQAMRRKGVTRRSFLKFCSLAATSLGLGAGMTPKIAWALENKPRIPVVWIHGL WP_001058312.1 hydrogenase [Salmonella enterica] Length: 372\nNumber of predicted TMHs: 2\nExp number of AAs in TMHs: 36.51738\nExp number, first 60 AAs: 19.72849\nTotal prob of N-in: 0.96170\nPOSSIBLE N-term signal sequence\ninside 1 19\nTMhelix 20 39\noutside 40 325\nTMhelix 326 348\ninside 349 372

22045 GCF_001756205.1_CFSAN008087 Salmonella enterica subsp. enterica serovar Typhimurium Proteobacteria; Gammaproteobacteria; Enterobacterales; Enterobacteriaceae; Salmonella; Salmonella enterica; Salmonella enterica subsp. enterica MTGDNTLITSHGINRRDFMKLCAALAATMGLSSKAAAEMAESVSNPQRPPVIWIGAQECT WP_000145429.1 hydrogenase 2 small subunit [Salmonella enterica] Length: 372\nNumber of predicted TMHs: 1\nExp number of AAs in TMHs: 24.51309\nExp number, first 60 AAs: 0.2682\nTotal prob of N-in: 0.14402\noutside 1 331\nTMhelix 332 354\ninside 355 372

22046 GCF_001756205.1_CFSAN008087 Salmonella enterica subsp. enterica serovar Typhimurium Proteobacteria; Gammaproteobacteria; Enterobacterales; Enterobacteriaceae; Salmonella; Salmonella enterica; Salmonella enterica subsp. enterica MNNEETFYQAMRRKGVTRRSFLKFCSLAATSLGLGAGMTPKIAWALENKPRIPVVWIHGL WP_001058312.1 hydrogenase [Salmonella enterica] Length: 372\nNumber of predicted TMHs: 2\nExp number of AAs in TMHs: 36.51738\nExp number, first 60 AAs: 19.72849\nTotal prob of N-in: 0.96170\nPOSSIBLE N-term signal sequence\ninside 1 19\nTMhelix 20 39\noutside 40 325\nTMhelix 326 348\ninside 349 372

22047 GCF_001756245.1_CFSAN008082 Salmonella enterica subsp. enterica serovar Typhimurium Proteobacteria; Gammaproteobacteria; Enterobacterales; Enterobacteriaceae; Salmonella; Salmonella enterica; Salmonella enterica subsp. enterica MTGDNTLITSHGINRRDFMKLCAALAATMGLSSKAAAEMAESVSNPQRPPVIWIGAQECT WP_000145429.1 hydrogenase 2 small subunit [Salmonella enterica] Length: 372\nNumber of predicted TMHs: 1\nExp number of AAs in TMHs: 24.51309\nExp number, first 60 AAs: 0.2682\nTotal prob of N-in: 0.14402\noutside 1 331\nTMhelix 332 354\ninside 355 372

22048 GCF_001887015.1_ASM188701v1 Salmonella enterica subsp. enterica serovar Typhimurium Proteobacteria; Gammaproteobacteria; Enterobacterales; Enterobacteriaceae; Salmonella; Salmonella enterica; Salmonella enterica subsp. enterica MNNEETFYQAMRRKGVTRRSFLKFCSLAATSLGLGAGMTPKIAWALENKPRIPVVWIHGL WP_001058312.1 hydrogenase [Salmonella enterica] Length: 372\nNumber of predicted TMHs: 2\nExp number of AAs in TMHs: 36.51738\nExp number, first 60 AAs: 19.72849\nTotal prob of N-in: 0.96170\nPOSSIBLE N-term signal sequence\ninside 1 19\nTMhelix 20 39\noutside 40 325\nTMhelix 326 348\ninside 349 372

22049 GCF_001884505.1_ASM188450v1 Salmonella enterica subsp. enterica serovar Typhimurium Proteobacteria; Gammaproteobacteria; Enterobacterales; Enterobacteriaceae; Salmonella; Salmonella enterica; Salmonella enterica subsp. enterica MTGDNTLITSHGINRRDFMKLCAALAATMGLSSKAAAEMAESVSNPQRPPVIWIGAQECT WP_000145429.1 hydrogenase 2 small subunit [Salmonella enterica] Length: 372\nNumber of predicted TMHs: 1\nExp number of AAs in TMHs: 24.51309\nExp number, first 60 AAs: 0.2682\nTotal prob of N-in: 0.14402\noutside 1 331\nTMhelix 332 354\ninside 355 372

22050 GCF_001884505.1_ASM188450v1 Salmonella enterica subsp. enterica serovar Typhimurium Proteobacteria; Gammaproteobacteria; Enterobacterales; Enterobacteriaceae; Salmonella; Salmonella enterica; Salmonella enterica subsp. enterica MNNEETFYQAMRRKGVTRRSFLKFCSLAATSLGLGAGMTPKIAWALENKPRIPVVWIHGL WP_001058312.1 hydrogenase [Salmonella enterica] Length: 372\nNumber of predicted TMHs: 2\nExp number of AAs in TMHs: 36.51738\nExp number, first 60 AAs: 19.72849\nTotal prob of N-in: 0.96170\nPOSSIBLE N-term signal sequence\ninside 1 19\nTMhelix 20 39\noutside 40 325\nTMhelix 326 348\ninside 349 372

22051 GCF_001954895.1_ASM195489v1 Salmonella enterica subsp. enterica serovar Typhimurium Proteobacteria; Gammaproteobacteria; Enterobacterales; Enterobacteriaceae; Salmonella; Salmonella enterica; Salmonella enterica subsp. enterica MNNEETFYQAMRRKGVTRRSFLKFCSLAATSLGLGAGMTPKIAWALENKPRIPVVWIHGL WP_001058312.1 hydrogenase [Salmonella enterica] Length: 372\nNumber of predicted TMHs: 2\nExp number of AAs in TMHs: 36.51738\nExp number, first 60 AAs: 19.72849\nTotal prob of N-in: 0.96170\nPOSSIBLE N-term signal sequence\ninside 1 19\nTMhelix 20 39\noutside 40 325\nTMhelix 326 348\ninside 349 372

22052 GCF_001955195.1_ASM195519v1 Salmonella enterica subsp. enterica serovar Typhimurium Proteobacteria; Gammaproteobacteria; Enterobacterales; Enterobacteriaceae; Salmonella; Salmonella enterica; Salmonella enterica subsp. enterica MTGDNTLITSHGINRRDFMKLCAALAATMGLSSKAAAEMAESVSNPQRPPVIWIGAQECT WP_000145429.1 hydrogenase 2 small subunit [Salmonella enterica] Length: 372\nNumber of predicted TMHs: 1\nExp number of AAs in TMHs: 24.51309\nExp number, first 60 AAs: 0.2682\nTotal prob of N-in: 0.14402\noutside 1 331\nTMhelix 332 354\ninside 355 372

22053 GCF_001955225.1_ASM195522v1 Salmonella enterica subsp. enterica serovar Typhimurium Proteobacteria; Gammaproteobacteria; Enterobacterales; Enterobacteriaceae; Salmonella; Salmonella enterica; Salmonella enterica subsp. enterica MTGDNTLITSHGINRRDFMKLCAALAATMGLSSKAAAEMAESVSNPQRPPVIWIGAQECT WP_000145429.1 hydrogenase 2 small subunit [Salmonella enterica] Length: 372\nNumber of predicted TMHs: 1\nExp number of AAs in TMHs: 24.51309\nExp number, first 60 AAs: 0.2682\nTotal prob of N-in: 0.14402\noutside 1 331\nTMhelix 332 354\ninside 355 372

22054 GCF_001955265.1_ASM195526v1 Salmonella enterica subsp. enterica serovar Typhimurium Proteobacteria; Gammaproteobacteria; Enterobacterales; Enterobacteriaceae; Salmonella; Salmonella enterica; Salmonella enterica subsp. enterica MTGDNTLITSHGINRRDFMKLCAALAATMGLSSKAAAEMAESVSNPQRPPVIWIGAQECT WP_000145429.1 hydrogenase 2 small subunit [Salmonella enterica] Length: 372\nNumber of predicted TMHs: 1\nExp number of AAs in TMHs: 24.51309\nExp number, first 60 AAs: 0.2682\nTotal prob of N-in: 0.14402\noutside 1 331\nTMhelix 332 354\ninside 355 372

22055 GCF_001955305.1_ASM195530v1 Salmonella enterica subsp. enterica serovar Typhimurium Proteobacteria; Gammaproteobacteria; Enterobacterales; Enterobacteriaceae; Salmonella; Salmonella enterica; Salmonella enterica subsp. enterica MTGDNTLITSHGINRRDFMKLCAALAATMGLSSKAAAEMAESVSNPQRPPVIWIGAQECT WP_000145429.1 hydrogenase 2 small subunit [Salmonella enterica] Length: 372\nNumber of predicted TMHs: 1\nExp number of AAs in TMHs: 24.51309\nExp number, first 60 AAs: 0.2682\nTotal prob of N-in: 0.14402\noutside 1 331\nTMhelix 332 354\ninside 355 372

22056 GCF_001955345.1_ASM195534v1 Salmonella enterica subsp. enterica serovar Typhimurium Proteobacteria; Gammaproteobacteria; Enterobacterales; Enterobacteriaceae; Salmonella; Salmonella enterica; Salmonella enterica subsp. enterica MTGDNTLITSHGINRRDFMKLCAALAATMGLSSKAAAEMAESVSNPQRPPVIWIGAQECT WP_000145429.1 hydrogenase 2 small subunit [Salmonella enterica] Length: 372\nNumber of predicted TMHs: 1\nExp number of AAs in TMHs: 24.51309\nExp number, first 60 AAs: 0.2682\nTotal prob of N-in: 0.14402\noutside 1 331\nTMhelix 332 354\ninside 355 372

22057 GCF_001955415.1_ASM195541v1 Salmonella enterica subsp. enterica serovar Typhimurium Proteobacteria; Gammaproteobacteria; Enterobacterales; Enterobacteriaceae; Salmonella; Salmonella enterica; Salmonella enterica subsp. enterica MTGDNTLITSHGINRRDFMKLCAALAATMGLSSKAAAEMAESVSNPQRPPVIWIGAQECT WP_000145429.1 hydrogenase 2 small subunit [Salmonella enterica] Length: 372\nNumber of predicted TMHs: 1\nExp number of AAs in TMHs: 24.51309\nExp number, first 60 AAs: 0.2682\nTotal prob of N-in: 0.14402\noutside 1 331\nTMhelix 332 354\ninside 355 372

22058 GCF_001955385.1_ASM195538v1 Salmonella enterica subsp. enterica serovar Typhimurium Proteobacteria; Gammaproteobacteria; Enterobacterales; Enterobacteriaceae; Salmonella; Salmonella enterica; Salmonella enterica subsp. enterica MTGDNTLITSHGINRRDFMKLCAALAATMGLSSKAAAEMAESVSNPQRPPVIWIGAQECT WP_000145429.1 hydrogenase 2 small subunit [Salmonella enterica] Length: 372\nNumber of

predicted TMHs: 1\nExp number of AAs in TMHs: 24.51309\nExp number, first 60 AAs: 0.2682\nTotal prob of N-in: 0.14402\noutside 1 331\nTMhelix 332 354\ninside 355 372

22059 GCF_001955225.1_ASM195522v1 Salmonella enterica subsp. enterica serovar Typhimurium Proteobacteria; Gammaproteobacteria; Enterobacterales; Enterobacteriaceae; Salmonella; Salmonella enterica; Salmonella enterica subsp. enterica MNNEETFYQAMRRKGVTRRSFLKFCSLAATSLGLGAGMTPKIAWALENKPRIPVVWIHGL WP_001058312.1 hydrogenase [Salmonella enterica] Length: 372\nNumber of predicted TMHs: 2\nExp number of AAs in TMHs: 36.51738\nExp number, first 60 AAs: 19.72849\nTotal prob of N-in: 0.96170\nPOSSIBLE N-term signal sequence\ninside 1 19\nTMhelix 20 39\noutside 40 325\nTMhelix 326 348\ninside 349 372

22060 GCF_001955265.1_ASM195526v1 Salmonella enterica subsp. enterica serovar Typhimurium Proteobacteria; Gammaproteobacteria; Enterobacterales; Enterobacteriaceae; Salmonella; Salmonella enterica; Salmonella enterica subsp. enterica MNNEETFYQAMRRKGVTRRSFLKFCSLAATSLGLGAGMTPKIAWALENKPRIPVVWIHGL WP_001058312.1 hydrogenase [Salmonella enterica] Length: 372\nNumber of predicted TMHs: 2\nExp number of AAs in TMHs: 36.51738\nExp number, first 60 AAs: 19.72849\nTotal prob of N-in: 0.96170\nPOSSIBLE N-term signal sequence\ninside 1 19\nTMhelix 20 39\noutside 40 325\nTMhelix 326 348\ninside 349 372

22061 GCF_001955305.1_ASM195530v1 Salmonella enterica subsp. enterica serovar Typhimurium Proteobacteria; Gammaproteobacteria; Enterobacterales; Enterobacteriaceae; Salmonella; Salmonella enterica; Salmonella enterica subsp. enterica MNNEETFYQAMRRKGVTRRSFLKFCSLAATSLGLGAGMTPKIAWALENKPRIPVVWIHGL WP_001058312.1 hydrogenase [Salmonella enterica] Length: 372\nNumber of predicted TMHs: 2\nExp number of AAs in TMHs: 36.51738\nExp number, first 60 AAs: 19.72849\nTotal prob of N-in: 0.96170\nPOSSIBLE N-term signal sequence\ninside 1 19\nTMhelix 20 39\noutside 40 325\nTMhelix 326 348\ninside 349 372

22062 GCF_001955465.1_ASM195546v1 Salmonella enterica subsp. enterica serovar Typhimurium Proteobacteria; Gammaproteobacteria; Enterobacterales; Enterobacteriaceae; Salmonella; Salmonella enterica; Salmonella enterica subsp. enterica MTGDNTLITSHGINRRDFMKLCAALAATMGLSSKAAAEMAESVSNPQRPPVIWIGAQECT WP_000145429.1 hydrogenase 2 small subunit [Salmonella enterica] Length: 372\nNumber of predicted TMHs: 1\nExp number of AAs in TMHs: 24.51309\nExp number, first 60 AAs: 0.2682\nTotal prob of N-in: 0.14402\noutside 1 331\nTMhelix 332 354\ninside 355 372

22063 GCF_001955465.1_ASM195546v1 Salmonella enterica subsp. enterica serovar Typhimurium Proteobacteria; Gammaproteobacteria; Enterobacterales; Enterobacteriaceae; Salmonella; Salmonella enterica; Salmonella enterica subsp. enterica MNNEETFYQAMRRKGVTRRSFLKFCSLAATSLGLGAGMTPKIAWALENKPRIPVVWIHGL WP_001058312.1 hydrogenase [Salmonella enterica] Length: 372\nNumber of predicted TMHs: 2\nExp number of AAs in TMHs: 36.51738\nExp number, first 60 AAs: 19.72849\nTotal prob of N-in: 0.96170\nPOSSIBLE N-term signal sequence\ninside 1 19\nTMhelix 20 39\noutside 40 325\nTMhelix 326 348\ninside 349 372

22064 GCF_002025495.1_ASM202549v1 Salmonella enterica subsp. enterica serovar Typhimurium Proteobacteria; Gammaproteobacteria; Enterobacterales; Enterobacteriaceae; Salmonella; Salmonella enterica; Salmonella enterica subsp. enterica MNNEETFYQAMRRKGVTRRSFLKFCSLAATSLGLGAGMTPKIAWALENKPRIPVVWIHGL WP_001058312.1 hydrogenase [Salmonella enterica] Length: 372\nNumber of predicted TMHs: 2\nExp number of AAs in TMHs: 36.51738\nExp number, first 60 AAs: 19.72849\nTotal prob of N-in: 0.96170\nPOSSIBLE N-term signal sequence\ninside 1 19\nTMhelix 20 39\noutside 40 325\nTMhelix 326 348\ninside 349 372

22065 GCF_002018325.1_ASM201832v1 Salmonella enterica subsp. enterica serovar Typhimurium Proteobacteria; Gammaproteobacteria; Enterobacterales; Enterobacteriaceae; Salmonella; Salmonella enterica; Salmonella enterica subsp. enterica MTGDNTLITSHGINRRDFMKLCAALAATMGLSSKAAAEMAESVSNPQRPPVIWIGAQECT WP_000145429.1 hydrogenase 2 small subunit [Salmonella enterica] Length: 372\nNumber of predicted TMHs: 1\nExp number of AAs in TMHs: 24.51309\nExp number, first 60 AAs: 0.2682\nTotal prob of N-in: 0.14402\noutside 1 331\nTMhelix 332 354\ninside 355 372

22066 GCF_002018325.1_ASM201832v1 Salmonella enterica subsp. enterica serovar Typhimurium Proteobacteria; Gammaproteobacteria; Enterobacterales; Enterobacteriaceae; Salmonella; Salmonella enterica; Salmonella enterica subsp. enterica MNNEETFYQAMRRKGVTRRSFLKFCSLAATSLGLGAGMTPKIAWALENKPRIPVVWIHGL WP_001058312.1 hydrogenase [Salmonella enterica] Length: 372\nNumber of predicted TMHs: 2\nExp number of AAs in TMHs: 36.51738\nExp number, first 60 AAs: 19.72849\nTotal prob of N-in: 0.96170\nPOSSIBLE N-term signal sequence\ninside 1 19\nTMhelix 20 39\noutside 40 325\nTMhelix 326 348\ninside 349 372

22067 GCF_002028585.1_ASM202858v1 Salmonella enterica subsp. enterica serovar Typhimurium Proteobacteria; Gammaproteobacteria; Enterobacterales; Enterobacteriaceae; Salmonella; Salmonella enterica;

Salmonella enterica subsp. enterica	MTGDNTLITSHGINRRDFMKLCAALAATMGLSSKAAAEMAESVSNPQRPPVIWIGAQECT
WP_000145429.1	hydrogenase 2 small subunit [Salmonella enterica]	Length: 372\nNumber of predicted TMHs: 1\nExp number of AAs in TMHs: 24.51309\nExp number, first 60 AAs: 0.2682\nTotal prob of N-in: 0.14402\noutside 1 331\nTMhelix 332 354\ninside 355 372

22068	GCF_002028675.1_ASM202867v1	Salmonella enterica subsp. enterica serovar Typhimurium
Proteobacteria; Gammaproteobacteria; Enterobacterales; Enterobacteriaceae; Salmonella; Salmonella enterica; Salmonella enterica subsp. enterica	MTGDNTLITSHGINRRDFMKLCAALAATMGLSSKAAAEMAESVSNPQRPPVIWIGAQECT
WP_000145429.1	hydrogenase 2 small subunit [Salmonella enterica]	Length: 372\nNumber of predicted TMHs: 1\nExp number of AAs in TMHs: 24.51309\nExp number, first 60 AAs: 0.2682\nTotal prob of N-in: 0.14402\noutside 1 331\nTMhelix 332 354\ninside 355 372

22069	GCF_002028625.1_ASM202862v1	Salmonella enterica subsp. enterica serovar Typhimurium
Proteobacteria; Gammaproteobacteria; Enterobacterales; Enterobacteriaceae; Salmonella; Salmonella enterica; Salmonella enterica subsp. enterica	MNNEETFYQAMRRKGVTRRSFLKFCSLAATSLGLGAGMTPKIAWALENKPRIPVVWIHGL
WP_001058312.1	hydrogenase [Salmonella enterica]	Length: 372\nNumber of predicted TMHs: 2\nExp number of AAs in TMHs: 36.51738\nExp number, first 60 AAs: 19.72849\nTotal prob of N-in: 0.96170\nPOSSIBLE N-term signal sequence\ninside 1 19\nTMhelix 20 39\noutside 40 325\nTMhelix 326 348\ninside 349 372

22070	GCF_002029365.1_ASM202936v1	Salmonella enterica subsp. enterica serovar Typhimurium
Proteobacteria; Gammaproteobacteria; Enterobacterales; Enterobacteriaceae; Salmonella; Salmonella enterica; Salmonella enterica subsp. enterica	MNNEETFYQAMRRKGVTRRSFLKFCSLAATSLGLGAGMTPKIAWALENKPRIPVVWIHGL
WP_001058312.1	hydrogenase [Salmonella enterica]	Length: 372\nNumber of predicted TMHs: 2\nExp number of AAs in TMHs: 36.51738\nExp number, first 60 AAs: 19.72849\nTotal prob of N-in: 0.96170\nPOSSIBLE N-term signal sequence\ninside 1 19\nTMhelix 20 39\noutside 40 325\nTMhelix 326 348\ninside 349 372

22071	GCF_002029395.1_ASM202939v1	Salmonella enterica subsp. enterica serovar Typhimurium
Proteobacteria; Gammaproteobacteria; Enterobacterales; Enterobacteriaceae; Salmonella; Salmonella enterica; Salmonella enterica subsp. enterica	MTGDNTLITSHGINRRDFMKLCAALAATMGLSSKAAAEMAESVSNPQRPPVIWIGAQECT
WP_000145429.1	hydrogenase 2 small subunit [Salmonella enterica]	Length: 372\nNumber of predicted TMHs: 1\nExp number of AAs in TMHs: 24.51309\nExp number, first 60 AAs: 0.2682\nTotal prob of N-in: 0.14402\noutside 1 331\nTMhelix 332 354\ninside 355 372

22072	GCF_002029455.1_ASM202945v1	Salmonella enterica subsp. enterica serovar Typhimurium
Proteobacteria; Gammaproteobacteria; Enterobacterales; Enterobacteriaceae; Salmonella; Salmonella enterica; Salmonella enterica subsp. enterica	MTGDNTLITSHGINRRDFMKLCAALAATMGLSSKAAAEMAESVSNPQRPPVIWIGAQECT
WP_000145429.1	hydrogenase 2 small subunit [Salmonella enterica]	Length: 372\nNumber of predicted TMHs: 1\nExp number of AAs in TMHs: 24.51309\nExp number, first 60 AAs: 0.2682\nTotal prob of N-in: 0.14402\noutside 1 331\nTMhelix 332 354\ninside 355 372

22073	GCF_002029665.1_ASM202966v1	Salmonella enterica subsp. enterica serovar Typhimurium
Proteobacteria; Gammaproteobacteria; Enterobacterales; Enterobacteriaceae; Salmonella; Salmonella enterica; Salmonella enterica subsp. enterica	MTGDNTLITSHGINRRDFMKLCAALAATMGLSSKAAAEMAESVSNPQRPPVIWIGAQECT
WP_000145429.1	hydrogenase 2 small subunit [Salmonella enterica]	Length: 372\nNumber of predicted TMHs: 1\nExp number of AAs in TMHs: 24.51309\nExp number, first 60 AAs: 0.2682\nTotal prob of N-in: 0.14402\noutside 1 331\nTMhelix 332 354\ninside 355 372

22074	GCF_002029705.1_ASM202970v1	Salmonella enterica subsp. enterica serovar Typhimurium
Proteobacteria; Gammaproteobacteria; Enterobacterales; Enterobacteriaceae; Salmonella; Salmonella enterica; Salmonella enterica subsp. enterica	MTGDNTLITSHGINRRDFMKLCAALAATMGLSSKAAAEMAESVSNPQRPPVIWIGAQECT
WP_000145429.1	hydrogenase 2 small subunit [Salmonella enterica]	Length: 372\nNumber of predicted TMHs: 1\nExp number of AAs in TMHs: 24.51309\nExp number, first 60 AAs: 0.2682\nTotal prob of N-in: 0.14402\noutside 1 331\nTMhelix 332 354\ninside 355 372

22075	GCF_002029745.1_ASM202974v1	Salmonella enterica subsp. enterica serovar Typhimurium
Proteobacteria; Gammaproteobacteria; Enterobacterales; Enterobacteriaceae; Salmonella; Salmonella enterica; Salmonella enterica subsp. enterica	MTGDNTLITSHGINRRDFMKLCAALAATMGLSSKAAAEMAESVSNPQRPPVIWIGAQECT
WP_000145429.1	hydrogenase 2 small subunit [Salmonella enterica]	Length: 372\nNumber of predicted TMHs: 1\nExp number of AAs in TMHs: 24.51309\nExp number, first 60 AAs: 0.2682\nTotal prob of N-in: 0.14402\noutside 1 331\nTMhelix 332 354\ninside 355 372

22076 GCF_002029825.1_ASM202982v1 Salmonella enterica subsp. enterica serovar Typhimurium Proteobacteria; Gammaproteobacteria; Enterobacterales; Enterobacteriaceae; Salmonella; Salmonella enterica; Salmonella enterica subsp. enterica MNNEETFYQAMRRKGVTRRSFLKFCSLAATSLGLGAGMTPKIAWALENKPRIPVVWIHGL WP_001058312.1 hydrogenase [Salmonella enterica] Length: 372\nNumber of predicted TMHs: 2\nExp number of AAs in TMHs: 36.51738\nExp number, first 60 AAs: 19.72849\nTotal prob of N-in: 0.96170\nPOSSIBLE N-term signal sequence\ninside 1 19\nTMhelix 20 39\noutside 40 325\nTMhelix 326 348\ninside 349 372

22077 GCF_002029455.1_ASM202945v1 Salmonella enterica subsp. enterica serovar Typhimurium Proteobacteria; Gammaproteobacteria; Enterobacterales; Enterobacteriaceae; Salmonella; Salmonella enterica; Salmonella enterica subsp. enterica MNNEETFYQAMRRKGVTRRSFLKFCSLAATSLGLGAGMTPKIAWALENKPRIPVVWIHGL WP_001058312.1 hydrogenase [Salmonella enterica] Length: 372\nNumber of predicted TMHs: 2\nExp number of AAs in TMHs: 36.51738\nExp number, first 60 AAs: 19.72849\nTotal prob of N-in: 0.96170\nPOSSIBLE N-term signal sequence\ninside 1 19\nTMhelix 20 39\noutside 40 325\nTMhelix 326 348\ninside 349 372

22078 GCF_002029665.1_ASM202966v1 Salmonella enterica subsp. enterica serovar Typhimurium Proteobacteria; Gammaproteobacteria; Enterobacterales; Enterobacteriaceae; Salmonella; Salmonella enterica; Salmonella enterica subsp. enterica MNNEETFYQAMRRKGVTRRSFLKFCSLAATSLGLGAGMTPKIAWALENKPRIPVVWIHGL WP_001058312.1 hydrogenase [Salmonella enterica] Length: 372\nNumber of predicted TMHs: 2\nExp number of AAs in TMHs: 36.51738\nExp number, first 60 AAs: 19.72849\nTotal prob of N-in: 0.96170\nPOSSIBLE N-term signal sequence\ninside 1 19\nTMhelix 20 39\noutside 40 325\nTMhelix 326 348\ninside 349 372

22079 GCF_002029705.1_ASM202970v1 Salmonella enterica subsp. enterica serovar Typhimurium Proteobacteria; Gammaproteobacteria; Enterobacterales; Enterobacteriaceae; Salmonella; Salmonella enterica; Salmonella enterica subsp. enterica MNNEETFYQAMRRKGVTRRSFLKFCSLAATSLGLGAGMTPKIAWALENKPRIPVVWIHGL WP_001058312.1 hydrogenase [Salmonella enterica] Length: 372\nNumber of predicted TMHs: 2\nExp number of AAs in TMHs: 36.51738\nExp number, first 60 AAs: 19.72849\nTotal prob of N-in: 0.96170\nPOSSIBLE N-term signal sequence\ninside 1 19\nTMhelix 20 39\noutside 40 325\nTMhelix 326 348\ninside 349 372

22080 GCF_002029745.1_ASM202974v1 Salmonella enterica subsp. enterica serovar Typhimurium Proteobacteria; Gammaproteobacteria; Enterobacterales; Enterobacteriaceae; Salmonella; Salmonella enterica; Salmonella enterica subsp. enterica MNNEETFYQAMRRKGVTRRSFLKFCSLAATSLGLGAGMTPKIAWALENKPRIPVVWIHGL WP_001058312.1 hydrogenase [Salmonella enterica] Length: 372\nNumber of predicted TMHs: 2\nExp number of AAs in TMHs: 36.51738\nExp number, first 60 AAs: 19.72849\nTotal prob of N-in: 0.96170\nPOSSIBLE N-term signal sequence\ninside 1 19\nTMhelix 20 39\noutside 40 325\nTMhelix 326 348\ninside 349 372

22081 GCF_002029785.1_ASM202978v1 Salmonella enterica subsp. enterica serovar Typhimurium Proteobacteria; Gammaproteobacteria; Enterobacterales; Enterobacteriaceae; Salmonella; Salmonella enterica; Salmonella enterica subsp. enterica MNNEETFYQAMRRKGVTRRSFLKFCSLAATSLGLGAGMTPKIAWALENKPRIPVVWIHGL WP_001058312.1 hydrogenase [Salmonella enterica] Length: 372\nNumber of predicted TMHs: 2\nExp number of AAs in TMHs: 36.51738\nExp number, first 60 AAs: 19.72849\nTotal prob of N-in: 0.96170\nPOSSIBLE N-term signal sequence\ninside 1 19\nTMhelix 20 39\noutside 40 325\nTMhelix 326 348\ninside 349 372

22082 GCF_002030885.1_ASM203088v1 Salmonella enterica subsp. enterica serovar Typhimurium Proteobacteria; Gammaproteobacteria; Enterobacterales; Enterobacteriaceae; Salmonella; Salmonella enterica; Salmonella enterica subsp. enterica MNNEETFYQAMRRKGVTRRSFLKFCSLAATSLGLGAGMTPKIAWALENKPRIPVVWIHGL WP_001058312.1 hydrogenase [Salmonella enterica] Length: 372\nNumber of predicted TMHs: 2\nExp number of AAs in TMHs: 36.51738\nExp number, first 60 AAs: 19.72849\nTotal prob of N-in: 0.96170\nPOSSIBLE N-term signal sequence\ninside 1 19\nTMhelix 20 39\noutside 40 325\nTMhelix 326 348\ninside 349 372

22083 GCF_002032975.1_ASM203297v1 Salmonella enterica subsp. enterica serovar Typhimurium Proteobacteria; Gammaproteobacteria; Enterobacterales; Enterobacteriaceae; Salmonella; Salmonella enterica; Salmonella enterica subsp. enterica MNNEETFYQAMRRKGVTRRSFLKFCSLAATSLGLGAGMTPKIAWALENKPRIPVVWIHGL WP_001058312.1 hydrogenase [Salmonella enterica] Length: 372\nNumber of predicted TMHs: 2\nExp number of AAs in TMHs: 36.51738\nExp number, first 60 AAs: 19.72849\nTotal prob of N-in: 0.96170\nPOSSIBLE N-term signal sequence\ninside 1 19\nTMhelix 20 39\noutside 40 325\nTMhelix 326 348\ninside 349 372

22084 GCF_002035025.1_ASM203502v1 Salmonella enterica subsp. enterica serovar Typhimurium Proteobacteria; Gammaproteobacteria; Enterobacterales; Enterobacteriaceae; Salmonella; Salmonella enterica; Salmonella enterica subsp. enterica MTGDNTLITSHGINRRDFMKLCAALAATMGLSSKAAAEMAESVSNPQRPPVIWIGAQECT WP_000145429.1 hydrogenase 2 small subunit [Salmonella enterica] Length: 372\nNumber of

predicted TMHs: 1\nExp number of AAs in TMHs: 24.51309\nExp number, first 60 AAs: 0.2682\nTotal prob of N-in: 0.14402\noutside 1 331\nTMhelix 332 354\ninside 355 372

22085 GCF_002035045.1_ASM203504v1 Salmonella enterica subsp. enterica serovar Typhimurium Proteobacteria; Gammaproteobacteria; Enterobacterales; Enterobacteriaceae; Salmonella; Salmonella enterica; Salmonella enterica subsp. enterica MTGDNTLITSHGINRRDFMKLCAALAATMGLSSKAAAEMAESVSNPQRPPVIWIGAQECT WP_000145429.1 hydrogenase 2 small subunit [Salmonella enterica] Length: 372\nNumber of predicted TMHs: 1\nExp number of AAs in TMHs: 24.51309\nExp number, first 60 AAs: 0.2682\nTotal prob of N-in: 0.14402\noutside 1 331\nTMhelix 332 354\ninside 355 372

22086 GCF_002035105.1_ASM203510v1 Salmonella enterica subsp. enterica serovar Typhimurium Proteobacteria; Gammaproteobacteria; Enterobacterales; Enterobacteriaceae; Salmonella; Salmonella enterica; Salmonella enterica subsp. enterica MTGDNTLITSHGINRRDFMKLCAALAATMGLSSKAAAEMAESVSNPQRPPVIWIGAQECT WP_000145429.1 hydrogenase 2 small subunit [Salmonella enterica] Length: 372\nNumber of predicted TMHs: 1\nExp number of AAs in TMHs: 24.51309\nExp number, first 60 AAs: 0.2682\nTotal prob of N-in: 0.14402\noutside 1 331\nTMhelix 332 354\ninside 355 372

22087 GCF_002034985.1_ASM203498v1 Salmonella enterica subsp. enterica serovar Typhimurium Proteobacteria; Gammaproteobacteria; Enterobacterales; Enterobacteriaceae; Salmonella; Salmonella enterica; Salmonella enterica subsp. enterica MNNEETFYQAMRRKGVTRRSFLKFCSLAATSLGLGAGMTPKIAWALENKPRIPVVWIHGL WP_001058312.1 hydrogenase [Salmonella enterica] Length: 372\nNumber of predicted TMHs: 2\nExp number of AAs in TMHs: 36.51738\nExp number, first 60 AAs: 19.72849\nTotal prob of N-in: 0.96170\nPOSSIBLE N-term signal sequence\ninside 1 19\nTMhelix 20 39\noutside 40 325\nTMhelix 326 348\ninside 349 372

22088 GCF_002035025.1_ASM203502v1 Salmonella enterica subsp. enterica serovar Typhimurium Proteobacteria; Gammaproteobacteria; Enterobacterales; Enterobacteriaceae; Salmonella; Salmonella enterica; Salmonella enterica subsp. enterica MNNEETFYQAMRRKGVTRRSFLKFCSLAATSLGLGAGMTPKIAWALENKPRIPVVWIHGL WP_001058312.1 hydrogenase [Salmonella enterica] Length: 372\nNumber of predicted TMHs: 2\nExp number of AAs in TMHs: 36.51738\nExp number, first 60 AAs: 19.72849\nTotal prob of N-in: 0.96170\nPOSSIBLE N-term signal sequence\ninside 1 19\nTMhelix 20 39\noutside 40 325\nTMhelix 326 348\ninside 349 372

22089 GCF_002035105.1_ASM203510v1 Salmonella enterica subsp. enterica serovar Typhimurium Proteobacteria; Gammaproteobacteria; Enterobacterales; Enterobacteriaceae; Salmonella; Salmonella enterica; Salmonella enterica subsp. enterica MNNEETFYQAMRRKGVTRRSFLKFCSLAATSLGLGAGMTPKIAWALENKPRIPVVWIHGL WP_001058312.1 hydrogenase [Salmonella enterica] Length: 372\nNumber of predicted TMHs: 2\nExp number of AAs in TMHs: 36.51738\nExp number, first 60 AAs: 19.72849\nTotal prob of N-in: 0.96170\nPOSSIBLE N-term signal sequence\ninside 1 19\nTMhelix 20 39\noutside 40 325\nTMhelix 326 348\ninside 349 372

22090 GCF_002035415.1_ASM203541v1 Salmonella enterica subsp. enterica serovar Typhimurium Proteobacteria; Gammaproteobacteria; Enterobacterales; Enterobacteriaceae; Salmonella; Salmonella enterica; Salmonella enterica subsp. enterica MTGDNTLITSHGINRRDFMKLCAALAATMGLSSKAAAEMAESVSNPQRPPVIWIGAQECT WP_000145429.1 hydrogenase 2 small subunit [Salmonella enterica] Length: 372\nNumber of predicted TMHs: 1\nExp number of AAs in TMHs: 24.51309\nExp number, first 60 AAs: 0.2682\nTotal prob of N-in: 0.14402\noutside 1 331\nTMhelix 332 354\ninside 355 372

22091 GCF_002035415.1_ASM203541v1 Salmonella enterica subsp. enterica serovar Typhimurium Proteobacteria; Gammaproteobacteria; Enterobacterales; Enterobacteriaceae; Salmonella; Salmonella enterica; Salmonella enterica subsp. enterica MNNEETFYQAMRRKGVTRRSFLKFCSLAATSLGLGAGMTPKIAWALENKPRIPVVWIHGL WP_001058312.1 hydrogenase [Salmonella enterica] Length: 372\nNumber of predicted TMHs: 2\nExp number of AAs in TMHs: 36.51738\nExp number, first 60 AAs: 19.72849\nTotal prob of N-in: 0.96170\nPOSSIBLE N-term signal sequence\ninside 1 19\nTMhelix 20 39\noutside 40 325\nTMhelix 326 348\ninside 349 372

22092 GCF_002058205.1_ASM205820v1 Salmonella enterica subsp. enterica serovar Typhimurium Proteobacteria; Gammaproteobacteria; Enterobacterales; Enterobacteriaceae; Salmonella; Salmonella enterica; Salmonella enterica subsp. enterica MTGDNTLITSHGINRRDFMKLCAALAATMGLSSKAAAEMAESVSNPQRPPVIWIGAQECT WP_000145429.1 hydrogenase 2 small subunit [Salmonella enterica] Length: 372\nNumber of predicted TMHs: 1\nExp number of AAs in TMHs: 24.51309\nExp number, first 60 AAs: 0.2682\nTotal prob of N-in: 0.14402\noutside 1 331\nTMhelix 332 354\ninside 355 372

22093 GCF_002058205.1_ASM205820v1 Salmonella enterica subsp. enterica serovar Typhimurium Proteobacteria; Gammaproteobacteria; Enterobacterales; Enterobacteriaceae; Salmonella; Salmonella enterica;

Salmonella enterica subsp. enterica MNNEETFYQAMRRKGVTRRSFLKFCSLAATSLGLGAGMTPKIAWALENKPRIPVVWIHGL WP_001058312.1 hydrogenase [Salmonella enterica] Length: 372\nNumber of predicted TMHs: 2\nExp number of AAs in TMHs: 36.51738\nExp number, first 60 AAs: 19.72849\nTotal prob of N-in: 0.96170\nPOSSIBLE N-term signal sequence\ninside 1 19\nTMhelix 20 39\noutside 40 325\nTMhelix 326 348\ninside 349 372

22094 GCF_002061615.1_ASM206161v1 Salmonella enterica subsp. enterica serovar Typhimurium Proteobacteria; Gammaproteobacteria; Enterobacterales; Enterobacteriaceae; Salmonella; Salmonella enterica; Salmonella enterica subsp. enterica MNNEETFYQAMRRKGVTRRSFLKFCSLAATSLGLGAGMTPKIAWALENKPRIPVVWIHGL WP_001058312.1 hydrogenase [Salmonella enterica] Length: 372\nNumber of predicted TMHs: 2\nExp number of AAs in TMHs: 36.51738\nExp number, first 60 AAs: 19.72849\nTotal prob of N-in: 0.96170\nPOSSIBLE N-term signal sequence\ninside 1 19\nTMhelix 20 39\noutside 40 325\nTMhelix 326 348\ninside 349 372

22095 GCF_001587855.1_CFSAN033932_01.0 Salmonella enterica subsp. enterica serovar Typhimurium Proteobacteria; Gammaproteobacteria; Enterobacterales; Enterobacteriaceae; Salmonella; Salmonella enterica; Salmonella enterica subsp. enterica MTGDNTLITSHGINRRDFMKLCAALAATMGLSSKAAAEMAESVSNPQRPPVIWIGAQECT WP_000145429.1 hydrogenase 2 small subunit [Salmonella enterica] Length: 372\nNumber of predicted TMHs: 1\nExp number of AAs in TMHs: 24.51309\nExp number, first 60 AAs: 0.2682\nTotal prob of N-in: 0.14402\noutside 1 331\nTMhelix 332 354\ninside 355 372

22096 GCF_002030385.1_ASM203038v1 Salmonella enterica subsp. enterica serovar Typhimurium Proteobacteria; Gammaproteobacteria; Enterobacterales; Enterobacteriaceae; Salmonella; Salmonella enterica; Salmonella enterica subsp. enterica MNNEETFYQAMRRKGVTRRSFLKFCSLAATSLGLGAGMTPKIAWALENKPRIPVVWIHGL WP_001058312.1 hydrogenase [Salmonella enterica] Length: 372\nNumber of predicted TMHs: 2\nExp number of AAs in TMHs: 36.51738\nExp number, first 60 AAs: 19.72849\nTotal prob of N-in: 0.96170\nPOSSIBLE N-term signal sequence\ninside 1 19\nTMhelix 20 39\noutside 40 325\nTMhelix 326 348\ninside 349 372

22097 GCF_002029715.1_ASM202971v1 Salmonella enterica subsp. enterica serovar Typhimurium Proteobacteria; Gammaproteobacteria; Enterobacterales; Enterobacteriaceae; Salmonella; Salmonella enterica; Salmonella enterica subsp. enterica MNNEETFYQAMRRKGVTRRSFLKFCSLAATSLGLGAGMTPKIAWALENKPRIPVVWIHGL WP_001058312.1 hydrogenase [Salmonella enterica] Length: 372\nNumber of predicted TMHs: 2\nExp number of AAs in TMHs: 36.51738\nExp number, first 60 AAs: 19.72849\nTotal prob of N-in: 0.96170\nPOSSIBLE N-term signal sequence\ninside 1 19\nTMhelix 20 39\noutside 40 325\nTMhelix 326 348\ninside 349 372

22098 GCF_002029445.1_ASM202944v1 Salmonella enterica subsp. enterica serovar Typhimurium Proteobacteria; Gammaproteobacteria; Enterobacterales; Enterobacteriaceae; Salmonella; Salmonella enterica; Salmonella enterica subsp. enterica MNNEETFYQAMRRKGVTRRSFLKFCSLAATSLGLGAGMTPKIAWALENKPRIPVVWIHGL WP_001058312.1 hydrogenase [Salmonella enterica] Length: 372\nNumber of predicted TMHs: 2\nExp number of AAs in TMHs: 36.51738\nExp number, first 60 AAs: 19.72849\nTotal prob of N-in: 0.96170\nPOSSIBLE N-term signal sequence\ninside 1 19\nTMhelix 20 39\noutside 40 325\nTMhelix 326 348\ninside 349 372

22099 GCF_002029835.1_ASM202983v1 Salmonella enterica subsp. enterica serovar Typhimurium Proteobacteria; Gammaproteobacteria; Enterobacterales; Enterobacteriaceae; Salmonella; Salmonella enterica; Salmonella enterica subsp. enterica MNNEETFYQAMRRKGVTRRSFLKFCSLAATSLGLGAGMTPKIAWALENKPRIPVVWIHGL WP_001058312.1 hydrogenase [Salmonella enterica] Length: 372\nNumber of predicted TMHs: 2\nExp number of AAs in TMHs: 36.51738\nExp number, first 60 AAs: 19.72849\nTotal prob of N-in: 0.96170\nPOSSIBLE N-term signal sequence\ninside 1 19\nTMhelix 20 39\noutside 40 325\nTMhelix 326 348\ninside 349 372

22100 GCF_001614175.1_ASM161417v1 Salmonella enterica subsp. enterica serovar Typhimurium Proteobacteria; Gammaproteobacteria; Enterobacterales; Enterobacteriaceae; Salmonella; Salmonella enterica; Salmonella enterica subsp. enterica MTGDNTLITSHGINRRDFMKLCAALAATMGLSSKAAAEMAESVSNPQRPPVIWIGAQECT WP_000145429.1 hydrogenase 2 small subunit [Salmonella enterica] Length: 372\nNumber of predicted TMHs: 1\nExp number of AAs in TMHs: 24.51309\nExp number, first 60 AAs: 0.2682\nTotal prob of N-in: 0.14402\noutside 1 331\nTMhelix 332 354\ninside 355 372

22101 GCF_002029385.1_ASM202938v1 Salmonella enterica subsp. enterica serovar Typhimurium Proteobacteria; Gammaproteobacteria; Enterobacterales; Enterobacteriaceae; Salmonella; Salmonella enterica; Salmonella enterica subsp. enterica MTGDNTLITSHGINRRDFMKLCAALAATMGLSSKAAAEMAESVSNPQRPPVIWIGAQECT WP_000145429.1 hydrogenase 2 small subunit [Salmonella enterica] Length: 372\nNumber of predicted TMHs: 1\nExp number of AAs in TMHs: 24.51309\nExp number, first 60 AAs: 0.2682\nTotal prob of N-in: 0.14402\noutside 1 331\nTMhelix 332 354\ninside 355 372

22102 GCF_000743055.1_ASM74305v1 Salmonella enterica subsp. enterica serovar Typhimurium Proteobacteria; Gammaproteobacteria; Enterobacterales; Enterobacteriaceae; Salmonella; Salmonella enterica; Salmonella enterica subsp. enterica MNNEETFYQAMRRKGVTRRSFLKFCSLAATSLGLGAGMTPKIAWALENKPRIPVVWIHGL WP_001058312.1 hydrogenase [Salmonella enterica] Length: 372\nNumber of predicted TMHs: 2\nExp number of AAs in TMHs: 36.51738\nExp number, first 60 AAs: 19.72849\nTotal prob of N-in: 0.96170\nPOSSIBLE N-term signal sequence\ninside 1 19\nTMhelix 20 39\noutside 40 325\nTMhelix 326 348\ninside 349 372

22103 GCF_002033825.1_ASM203382v1 Salmonella enterica subsp. enterica serovar Typhimurium Proteobacteria; Gammaproteobacteria; Enterobacterales; Enterobacteriaceae; Salmonella; Salmonella enterica; Salmonella enterica subsp. enterica MNNEETFYQAMRRKGVTRRSFLKFCSLAATSLGLGAGMTPKIAWALENKPRIPVVWIHGL WP_001058312.1 hydrogenase [Salmonella enterica] Length: 372\nNumber of predicted TMHs: 2\nExp number of AAs in TMHs: 36.51738\nExp number, first 60 AAs: 19.72849\nTotal prob of N-in: 0.96170\nPOSSIBLE N-term signal sequence\ninside 1 19\nTMhelix 20 39\noutside 40 325\nTMhelix 326 348\ninside 349 372

22104 GCF_002028595.1_ASM202859v1 Salmonella enterica subsp. enterica serovar Typhimurium Proteobacteria; Gammaproteobacteria; Enterobacterales; Enterobacteriaceae; Salmonella; Salmonella enterica; Salmonella enterica subsp. enterica MNNEETFYQAMRRKGVTRRSFLKFCSLAATSLGLGAGMTPKIAWALENKPRIPVVWIHGL WP_001058312.1 hydrogenase [Salmonella enterica] Length: 372\nNumber of predicted TMHs: 2\nExp number of AAs in TMHs: 36.51738\nExp number, first 60 AAs: 19.72849\nTotal prob of N-in: 0.96170\nPOSSIBLE N-term signal sequence\ninside 1 19\nTMhelix 20 39\noutside 40 325\nTMhelix 326 348\ninside 349 372

22105 GCF_001886995.1_ASM188699v1 Salmonella enterica subsp. enterica serovar Typhimurium Proteobacteria; Gammaproteobacteria; Enterobacterales; Enterobacteriaceae; Salmonella; Salmonella enterica; Salmonella enterica subsp. enterica MTGDNTLITSHGINRRDFMKLCAALAATMGLSSKAAAEMAESVSNPQRPPVIWIGAQECT WP_000145429.1 hydrogenase 2 small subunit [Salmonella enterica] Length: 372\nNumber of predicted TMHs: 1\nExp number of AAs in TMHs: 24.51309\nExp number, first 60 AAs: 0.2682\nTotal prob of N-in: 0.14402\noutside 1 331\nTMhelix 332 354\ninside 355 372

22106 GCF_001246225.1_Salmonella_enterica_CVM_N51254_v1.0 Salmonella enterica subsp. enterica serovar Typhimurium Proteobacteria; Gammaproteobacteria; Enterobacterales; Enterobacteriaceae; Salmonella; Salmonella enterica; Salmonella enterica subsp. enterica MNNEETFYQAMRRKGVTRRSFLKFCSLAATSLGLGAGMTPKIAWALENKPRIPVVWIHGL WP_001058312.1 hydrogenase [Salmonella enterica] Length: 372\nNumber of predicted TMHs: 2\nExp number of AAs in TMHs: 36.51738\nExp number, first 60 AAs: 19.72849\nTotal prob of N-in: 0.96170\nPOSSIBLE N-term signal sequence\ninside 1 19\nTMhelix 20 39\noutside 40 325\nTMhelix 326 348\ninside 349 372

22107 GCF_002028635.1_ASM202863v1 Salmonella enterica subsp. enterica serovar Typhimurium Proteobacteria; Gammaproteobacteria; Enterobacterales; Enterobacteriaceae; Salmonella; Salmonella enterica; Salmonella enterica subsp. enterica MTGDNTLITSHGINRRDFMKLCAALAATMGLSSKAAAEMAESVSNPQRPPVIWIGAQECT WP_000145429.1 hydrogenase 2 small subunit [Salmonella enterica] Length: 372\nNumber of predicted TMHs: 1\nExp number of AAs in TMHs: 24.51309\nExp number, first 60 AAs: 0.2682\nTotal prob of N-in: 0.14402\noutside 1 331\nTMhelix 332 354\ninside 355 372

22108 GCF_002028525.1_ASM202852v1 Salmonella enterica subsp. enterica serovar Typhimurium Proteobacteria; Gammaproteobacteria; Enterobacterales; Enterobacteriaceae; Salmonella; Salmonella enterica; Salmonella enterica subsp. enterica MNNEETFYQAMRRKGVTRRSFLKFCSLAATSLGLGAGMTPKIAWALENKPRIPVVWIHGL WP_001058312.1 hydrogenase [Salmonella enterica] Length: 372\nNumber of predicted TMHs: 2\nExp number of AAs in TMHs: 36.51738\nExp number, first 60 AAs: 19.72849\nTotal prob of N-in: 0.96170\nPOSSIBLE N-term signal sequence\ninside 1 19\nTMhelix 20 39\noutside 40 325\nTMhelix 326 348\ninside 349 372

22109 GCF_002028595.1_ASM202859v1 Salmonella enterica subsp. enterica serovar Typhimurium Proteobacteria; Gammaproteobacteria; Enterobacterales; Enterobacteriaceae; Salmonella; Salmonella enterica; Salmonella enterica subsp. enterica MTGDNTLITSHGINRRDFMKLCAALAATMGLSSKAAAEMAESVSNPQRPPVIWIGAQECT WP_000145429.1 hydrogenase 2 small subunit [Salmonella enterica] Length: 372\nNumber of predicted TMHs: 1\nExp number of AAs in TMHs: 24.51309\nExp number, first 60 AAs: 0.2682\nTotal prob of N-in: 0.14402\noutside 1 331\nTMhelix 332 354\ninside 355 372

22110 GCF_001594855.1_CFSAN033877_01.0Salmonella enterica subsp. enterica serovar Typhimurium Proteobacteria; Gammaproteobacteria; Enterobacterales; Enterobacteriaceae; Salmonella; Salmonella enterica; Salmonella enterica subsp. enterica MTGDNTLITSHGINRRDFMKLCAALAATMGLSSKAAAEMAESVSNPQRPPVIWIGAQECT WP_061584213.1 hydrogenase 2 small subunit [Salmonella enterica] Length: 372\nNumber of

predicted TMHs:  1\nExp number of AAs in TMHs: 24.51809\nExp number, first 60 AAs:  0.26818\nTotal prob of N-in: 0.14414\noutside   1   331\nTMhelix   332   354\ninside   355   372

22111 GCF_001595105.1_CFSAN033863_01.0Salmonella enterica subsp. enterica serovar Typhimurium Proteobacteria; Gammaproteobacteria; Enterobacterales; Enterobacteriaceae; Salmonella; Salmonella enterica; Salmonella enterica subsp. enterica MNNEETFYQAMRRKGVTRRSFLKFCSLAATSLGLGAGMTPKIAWALENKPRIPVVWIHGL WP_001058312.1 hydrogenase [Salmonella enterica] Length: 372\nNumber of predicted TMHs:  2\nExp number of AAs in TMHs: 36.51738\nExp number, first 60 AAs:  19.72849\nTotal prob of N-in:   0.96170\nPOSSIBLE N-term signal sequence\ninside   1   19\nTMhelix   20   39\noutside   40   325\nTMhelix   326   348\ninside   349   372

22112 GCF_002036315.1_ASM203631v1 Salmonella enterica subsp. enterica serovar Typhimurium Proteobacteria; Gammaproteobacteria; Enterobacterales; Enterobacteriaceae; Salmonella; Salmonella enterica; Salmonella enterica subsp. enterica MNNEETFYQAMRRKGVTRRSFLKFCSLAATSLGLGAGMTPKIAWALENKPRIPVVWIHGL WP_001058312.1 hydrogenase [Salmonella enterica] Length: 372\nNumber of predicted TMHs:  2\nExp number of AAs in TMHs: 36.51738\nExp number, first 60 AAs:  19.72849\nTotal prob of N-in:   0.96170\nPOSSIBLE N-term signal sequence\ninside   1   19\nTMhelix   20   39\noutside   40   325\nTMhelix   326   348\ninside   349   372

22113 GCF_001594985.1_CFSAN033870_01.0Salmonella enterica subsp. enterica serovar Typhimurium Proteobacteria; Gammaproteobacteria; Enterobacterales; Enterobacteriaceae; Salmonella; Salmonella enterica; Salmonella enterica subsp. enterica MNNEETFYQAMRRKGVTRRSFLKFCSLAATSLGLGAGMTPKIAWALENKPRIPVVWIHGL WP_001058312.1 hydrogenase [Salmonella enterica] Length: 372\nNumber of predicted TMHs:  2\nExp number of AAs in TMHs: 36.51738\nExp number, first 60 AAs:  19.72849\nTotal prob of N-in:   0.96170\nPOSSIBLE N-term signal sequence\ninside   1   19\nTMhelix   20   39\noutside   40   325\nTMhelix   326   348\ninside   349   372

22114 GCF_001595065.1_CFSAN033865_01.0Salmonella enterica subsp. enterica serovar Typhimurium Proteobacteria; Gammaproteobacteria; Enterobacterales; Enterobacteriaceae; Salmonella; Salmonella enterica; Salmonella enterica subsp. enterica MNNEETFYQAMRRKGVTRRSFLKFCSLAATSLGLGAGMTPKIAWALENKPRIPVVWIHGL WP_001058312.1 hydrogenase [Salmonella enterica] Length: 372\nNumber of predicted TMHs:  2\nExp number of AAs in TMHs: 36.51738\nExp number, first 60 AAs:  19.72849\nTotal prob of N-in:   0.96170\nPOSSIBLE N-term signal sequence\ninside   1   19\nTMhelix   20   39\noutside   40   325\nTMhelix   326   348\ninside   349   372

22115 GCF_001594935.1_CFSAN033872_01.0Salmonella enterica subsp. enterica serovar Typhimurium Proteobacteria; Gammaproteobacteria; Enterobacterales; Enterobacteriaceae; Salmonella; Salmonella enterica; Salmonella enterica subsp. enterica MNNEETFYQAMRRKGVTRRSFLKFCSLAATSLGLGAGMTPKIAWALENKPRIPVVWIHGL WP_001058312.1 hydrogenase [Salmonella enterica] Length: 372\nNumber of predicted TMHs:  2\nExp number of AAs in TMHs: 36.51738\nExp number, first 60 AAs:  19.72849\nTotal prob of N-in:   0.96170\nPOSSIBLE N-term signal sequence\ninside   1   19\nTMhelix   20   39\noutside   40   325\nTMhelix   326   348\ninside   349   372

22116 GCF_000784295.1_ASM78429v1 Salmonella enterica subsp. enterica serovar Typhimurium Proteobacteria; Gammaproteobacteria; Enterobacterales; Enterobacteriaceae; Salmonella; Salmonella enterica; Salmonella enterica subsp. enterica MTGDNTLITSHGINRRDFMKLCAALAATMGLSSKAAAEMAESVSNPQRPPVIWIGAQECT WP_000145429.1 hydrogenase 2 small subunit [Salmonella enterica] Length: 372\nNumber of predicted TMHs:  1\nExp number of AAs in TMHs: 24.51309\nExp number, first 60 AAs:  0.2682\nTotal prob of N-in: 0.14402\noutside   1   331\nTMhelix   332   354\ninside   355   372

22117 GCF_000784215.1_ASM78421v1 Salmonella enterica subsp. enterica serovar Typhimurium Proteobacteria; Gammaproteobacteria; Enterobacterales; Enterobacteriaceae; Salmonella; Salmonella enterica; Salmonella enterica subsp. enterica MTGDNTLITSHGINRRDFMKLCAALAATMGLSSKAAAEMAESVSNPQRPPVIWIGAQECT WP_000145429.1 hydrogenase 2 small subunit [Salmonella enterica] Length: 372\nNumber of predicted TMHs:  1\nExp number of AAs in TMHs: 24.51309\nExp number, first 60 AAs:  0.2682\nTotal prob of N-in: 0.14402\noutside   1   331\nTMhelix   332   354\ninside   355   372

22118 GCF_000784235.1_ASM78423v1 Salmonella enterica subsp. enterica serovar Typhimurium Proteobacteria; Gammaproteobacteria; Enterobacterales; Enterobacteriaceae; Salmonella; Salmonella enterica; Salmonella enterica subsp. enterica MTGDNTLITSHGINRRDFMKLCAALAATMGLSSKAAAEMAESVSNPQRPPVIWIGAQECT WP_000145429.1 hydrogenase 2 small subunit [Salmonella enterica] Length: 372\nNumber of predicted TMHs:  1\nExp number of AAs in TMHs: 24.51309\nExp number, first 60 AAs:  0.2682\nTotal prob of N-in: 0.14402\noutside   1   331\nTMhelix   332   354\ninside   355   372

22119 GCF_000973845.1_SALF-276-1.id8_v1.0 Salmonella enterica subsp. enterica serovar Typhimurium Proteobacteria; Gammaproteobacteria; Enterobacterales; Enterobacteriaceae; Salmonella; Salmonella enterica;

Salmonella enterica subsp. enterica MNNEETFYQAMRRKGVTRRSFLKFCSLAATSLGLGAGMTPKIAWALENKPRIPVVWIHGL WP_001058312.1 hydrogenase [Salmonella enterica] Length: 372\nNumber of predicted TMHs: 2\nExp number of AAs in TMHs: 36.51738\nExp number, first 60 AAs: 19.72849\nTotal prob of N-in: 0.96170\nPOSSIBLE N-term signal sequence\ninside 1 19\nTMhelix 20 39\noutside 40 325\nTMhelix 326 348\ninside 349 372

22120 GCF_001595265.1_CFSAN033853_01.0Salmonella enterica subsp. enterica serovar Typhimurium Proteobacteria; Gammaproteobacteria; Enterobacterales; Enterobacteriaceae; Salmonella; Salmonella enterica; Salmonella enterica subsp. enterica MNNEETFYQAMRRKGVTRRSFLKFCSLAATSLGLGAGMTPKIAWALENKPRIPVVWIHGL WP_001058312.1 hydrogenase [Salmonella enterica] Length: 372\nNumber of predicted TMHs: 2\nExp number of AAs in TMHs: 36.51738\nExp number, first 60 AAs: 19.72849\nTotal prob of N-in: 0.96170\nPOSSIBLE N-term signal sequence\ninside 1 19\nTMhelix 20 39\noutside 40 325\nTMhelix 326 348\ninside 349 372

22121 GCF_001594905.1_CFSAN033874_01.0Salmonella enterica subsp. enterica serovar Typhimurium Proteobacteria; Gammaproteobacteria; Enterobacterales; Enterobacteriaceae; Salmonella; Salmonella enterica; Salmonella enterica subsp. enterica MTGDNTLITSHGINRRDFMKLCAALAATMGLSSKAAAEMAESVSNPQRPPVIWIGAQECT WP_000145429.1 hydrogenase 2 small subunit [Salmonella enterica] Length: 372\nNumber of predicted TMHs: 1\nExp number of AAs in TMHs: 24.51309\nExp number, first 60 AAs: 0.2682\nTotal prob of N-in: 0.14402\noutside 1 331\nTMhelix 332 354\ninside 355 372

22122 GCF_001594685.1_CFSAN033891_01.0Salmonella enterica subsp. enterica serovar Typhimurium Proteobacteria; Gammaproteobacteria; Enterobacterales; Enterobacteriaceae; Salmonella; Salmonella enterica; Salmonella enterica subsp. enterica MTGDNTLITSHGINRRDFMKLCAALAATMGLSSKAAAEMAESVSNPQRPPVIWIGAQECT WP_061584213.1 hydrogenase 2 small subunit [Salmonella enterica] Length: 372\nNumber of predicted TMHs: 1\nExp number of AAs in TMHs: 24.51809\nExp number, first 60 AAs: 0.26818\nTotal prob of N-in: 0.14414\noutside 1 331\nTMhelix 332 354\ninside 355 372

22123 GCF_001594845.1_CFSAN033879_01.0Salmonella enterica subsp. enterica serovar Typhimurium Proteobacteria; Gammaproteobacteria; Enterobacterales; Enterobacteriaceae; Salmonella; Salmonella enterica; Salmonella enterica subsp. enterica MNNEETFYQAMRRKGVTRRSFLKFCSLAATSLGLGAGMTPKIAWALENKPRIPVVWIHGL WP_001058312.1 hydrogenase [Salmonella enterica] Length: 372\nNumber of predicted TMHs: 2\nExp number of AAs in TMHs: 36.51738\nExp number, first 60 AAs: 19.72849\nTotal prob of N-in: 0.96170\nPOSSIBLE N-term signal sequence\ninside 1 19\nTMhelix 20 39\noutside 40 325\nTMhelix 326 348\ninside 349 372

22124 GCF_001594795.1_CFSAN033881_01.0Salmonella enterica subsp. enterica serovar Typhimurium Proteobacteria; Gammaproteobacteria; Enterobacterales; Enterobacteriaceae; Salmonella; Salmonella enterica; Salmonella enterica subsp. enterica MNNEETFYQAMRRKGVTRRSFLKFCSLAATSLGLGAGMTPKIAWALENKPRIPVVWIHGL WP_001058312.1 hydrogenase [Salmonella enterica] Length: 372\nNumber of predicted TMHs: 2\nExp number of AAs in TMHs: 36.51738\nExp number, first 60 AAs: 19.72849\nTotal prob of N-in: 0.96170\nPOSSIBLE N-term signal sequence\ninside 1 19\nTMhelix 20 39\noutside 40 325\nTMhelix 326 348\ninside 349 372

22125 GCF_001594685.1_CFSAN033891_01.0Salmonella enterica subsp. enterica serovar Typhimurium Proteobacteria; Gammaproteobacteria; Enterobacterales; Enterobacteriaceae; Salmonella; Salmonella enterica; Salmonella enterica subsp. enterica MNNEETFYQAMRRKGVTRRSFLKFCSLAATSLGLGAGMTPKIAWALENKPRIPVVWIHGL WP_001058312.1 hydrogenase [Salmonella enterica] Length: 372\nNumber of predicted TMHs: 2\nExp number of AAs in TMHs: 36.51738\nExp number, first 60 AAs: 19.72849\nTotal prob of N-in: 0.96170\nPOSSIBLE N-term signal sequence\ninside 1 19\nTMhelix 20 39\noutside 40 325\nTMhelix 326 348\ninside 349 372

22126 GCF_001594845.1_CFSAN033879_01.0Salmonella enterica subsp. enterica serovar Typhimurium Proteobacteria; Gammaproteobacteria; Enterobacterales; Enterobacteriaceae; Salmonella; Salmonella enterica; Salmonella enterica subsp. enterica MTGDNTLITSHGINRRDFMKLCAALAATMGLSSKAAAEMAESVSNPQRPPVIWIGAQECT WP_000145429.1 hydrogenase 2 small subunit [Salmonella enterica] Length: 372\nNumber of predicted TMHs: 1\nExp number of AAs in TMHs: 24.51309\nExp number, first 60 AAs: 0.2682\nTotal prob of N-in: 0.14402\noutside 1 331\nTMhelix 332 354\ninside 355 372

22127 GCF_001594625.1_CFSAN033896_01.0Salmonella enterica subsp. enterica serovar Typhimurium Proteobacteria; Gammaproteobacteria; Enterobacterales; Enterobacteriaceae; Salmonella; Salmonella enterica; Salmonella enterica subsp. enterica MNNEETFYQAMRRKGVTRRSFLKFCSLAATSLGLGAGMTPKIAWALENKPRIPVVWIHGL WP_001058312.1 hydrogenase [Salmonella enterica] Length: 372\nNumber of predicted TMHs: 2\nExp number of AAs in TMHs: 36.51738\nExp number, first 60 AAs: 19.72849\nTotal prob of N-in: 0.96170\nPOSSIBLE N-term signal sequence\ninside 1 19\nTMhelix 20 39\noutside 40 325\nTMhelix 326 348\ninside 349 372

22128 GCF_001594455.1_CFSAN033928_01.0Salmonella enterica subsp. enterica serovar Typhimurium Proteobacteria; Gammaproteobacteria; Enterobacterales; Enterobacteriaceae; Salmonella; Salmonella enterica; Salmonella enterica subsp. enterica MTGDNTLITSHGINRRDFMKLCAALAATMGLSSKAAAEMAESVSNPQRPPVIWIGAQECT WP_000145429.1 hydrogenase 2 small subunit [Salmonella enterica] Length: 372\nNumber of predicted TMHs: 1\nExp number of AAs in TMHs: 24.51309\nExp number, first 60 AAs: 0.2682\nTotal prob of N-in: 0.14402\noutside 1 331\nTMhelix 332 354\ninside 355 372

22129 GCF_001594535.1_CFSAN033912_01.0Salmonella enterica subsp. enterica serovar Typhimurium Proteobacteria; Gammaproteobacteria; Enterobacterales; Enterobacteriaceae; Salmonella; Salmonella enterica; Salmonella enterica subsp. enterica MTGDNTLITSHGINRRDFMKLCAALAATMGLSSKAAAEMAESVSNPQRPPVIWIGAQECT WP_000145429.1 hydrogenase 2 small subunit [Salmonella enterica] Length: 372\nNumber of predicted TMHs: 1\nExp number of AAs in TMHs: 24.51309\nExp number, first 60 AAs: 0.2682\nTotal prob of N-in: 0.14402\noutside 1 331\nTMhelix 332 354\ninside 355 372

22130 GCF_001588475.1_CFSAN033886_01.0Salmonella enterica subsp. enterica serovar Typhimurium Proteobacteria; Gammaproteobacteria; Enterobacterales; Enterobacteriaceae; Salmonella; Salmonella enterica; Salmonella enterica subsp. enterica MTGDNTLITSHGINRRDFMKLCAALAATMGLSSKAAAEMAESVSNPQRPPVIWIGAQECT WP_061584213.1 hydrogenase 2 small subunit [Salmonella enterica] Length: 372\nNumber of predicted TMHs: 1\nExp number of AAs in TMHs: 24.51809\nExp number, first 60 AAs: 0.26818\nTotal prob of N-in: 0.14414\noutside 1 331\nTMhelix 332 354\ninside 355 372

22131 GCF_001594435.1_CFSAN033924_01.0Salmonella enterica subsp. enterica serovar Typhimurium Proteobacteria; Gammaproteobacteria; Enterobacterales; Enterobacteriaceae; Salmonella; Salmonella enterica; Salmonella enterica subsp. enterica MTGDNTLITSHGINRRDFMKLCAALAATMGLSSKAAAEMAESVSNPQRPPVIWIGAQECT WP_000145429.1 hydrogenase 2 small subunit [Salmonella enterica] Length: 372\nNumber of predicted TMHs: 1\nExp number of AAs in TMHs: 24.51309\nExp number, first 60 AAs: 0.2682\nTotal prob of N-in: 0.14402\noutside 1 331\nTMhelix 332 354\ninside 355 372

22132 GCF_001714645.1_ASM171464v1 Salmonella enterica subsp. enterica serovar Typhimurium Proteobacteria; Gammaproteobacteria; Enterobacterales; Enterobacteriaceae; Salmonella; Salmonella enterica; Salmonella enterica subsp. enterica MTGDNTLITSHGINRRDFMKLCAALAATMGLSSKAAAEMAESVSNPQRPPVIWIGAQECT WP_000145429.1 hydrogenase 2 small subunit [Salmonella enterica] Length: 372\nNumber of predicted TMHs: 1\nExp number of AAs in TMHs: 24.51309\nExp number, first 60 AAs: 0.2682\nTotal prob of N-in: 0.14402\noutside 1 331\nTMhelix 332 354\ninside 355 372

22133 GCF_000743055.1_ASM74305v1 Salmonella enterica subsp. enterica serovar Typhimurium Proteobacteria; Gammaproteobacteria; Enterobacterales; Enterobacteriaceae; Salmonella; Salmonella enterica; Salmonella enterica subsp. enterica MTGDNTLITSHGINRRDFMKLCAALAATMGLSSKAAAEMAESVSNPQRPPVIWIGAQECT WP_000145429.1 hydrogenase 2 small subunit [Salmonella enterica] Length: 372\nNumber of predicted TMHs: 1\nExp number of AAs in TMHs: 24.51309\nExp number, first 60 AAs: 0.2682\nTotal prob of N-in: 0.14402\noutside 1 331\nTMhelix 332 354\ninside 355 372

22134 GCF_002028525.1_ASM202852v1 Salmonella enterica subsp. enterica serovar Typhimurium Proteobacteria; Gammaproteobacteria; Enterobacterales; Enterobacteriaceae; Salmonella; Salmonella enterica; Salmonella enterica subsp. enterica MTGDNTLITSHGINRRDFMKLCAALAATMGLSSKAAAEMAESVSNPQRPPVIWIGAQECT WP_000145429.1 hydrogenase 2 small subunit [Salmonella enterica] Length: 372\nNumber of predicted TMHs: 1\nExp number of AAs in TMHs: 24.51309\nExp number, first 60 AAs: 0.2682\nTotal prob of N-in: 0.14402\noutside 1 331\nTMhelix 332 354\ninside 355 372

22135 GCF_001588475.1_CFSAN033886_01.0Salmonella enterica subsp. enterica serovar Typhimurium Proteobacteria; Gammaproteobacteria; Enterobacterales; Enterobacteriaceae; Salmonella; Salmonella enterica; Salmonella enterica subsp. enterica MNNEETFYQAMRRKGVTRRSFLKFCSLAATSLGLGAGMTPKIAWALENKPRIPVVWIHGL WP_001058312.1 hydrogenase [Salmonella enterica] Length: 372\nNumber of predicted TMHs: 2\nExp number of AAs in TMHs: 36.51738\nExp number, first 60 AAs: 19.72849\nTotal prob of N-in: 0.96170\nPOSSIBLE N-term signal sequence\ninside 1 19\nTMhelix 20 39\noutside 40 325\nTMhelix 326 348\ninside 349 372

22136 GCF_001588395.1_CFSAN033895_01.0Salmonella enterica subsp. enterica serovar Typhimurium Proteobacteria; Gammaproteobacteria; Enterobacterales; Enterobacteriaceae; Salmonella; Salmonella enterica; Salmonella enterica subsp. enterica MTGDNTLITSHGINRRDFMKLCAALAATMGLSSKAAAEMAESVSNPQRPPVIWIGAQECT WP_000145429.1 hydrogenase 2 small subunit [Salmonella enterica] Length: 372\nNumber of

predicted TMHs: 1\nExp number of AAs in TMHs: 24.51309\nExp number, first 60 AAs: 0.2682\nTotal prob of N-in: 0.14402\noutside 1 331\nTMhelix 332 354\ninside 355 372

22137 GCF_001588315.1_CFSAN033902_01.0Salmonella enterica subsp. enterica serovar Typhimurium Proteobacteria; Gammaproteobacteria; Enterobacterales; Enterobacteriaceae; Salmonella; Salmonella enterica; Salmonella enterica subsp. enterica MTGDNTLITSHGINRRDFMKLCAALAATMGLSSKAAAEMAESVSNPQRPPVIWIGAQECT WP_000145429.1 hydrogenase 2 small subunit [Salmonella enterica] Length: 372\nNumber of predicted TMHs: 1\nExp number of AAs in TMHs: 24.51309\nExp number, first 60 AAs: 0.2682\nTotal prob of N-in: 0.14402\noutside 1 331\nTMhelix 332 354\ninside 355 372

22138 GCF_001479325.1_Salmonella_enterica_CVM_N32045-SQ_v1.0 Salmonella enterica subsp. enterica serovar Typhimurium Proteobacteria; Gammaproteobacteria; Enterobacterales; Enterobacteriaceae; Salmonella; Salmonella enterica; Salmonella enterica subsp. enterica MTGDNTLITSHGINRRDFMKLCAALAATMGLSSKAAAEMAESVSNPQRPPVIWIGAQECT WP_000145429.1 hydrogenase 2 small subunit [Salmonella enterica] Length: 372\nNumber of predicted TMHs: 1\nExp number of AAs in TMHs: 24.51309\nExp number, first 60 AAs: 0.2682\nTotal prob of N-in: 0.14402\noutside 1 331\nTMhelix 332 354\ninside 355 372

22139 GCF_001588355.1_CFSAN033899_01.0Salmonella enterica subsp. enterica serovar Typhimurium Proteobacteria; Gammaproteobacteria; Enterobacterales; Enterobacteriaceae; Salmonella; Salmonella enterica; Salmonella enterica subsp. enterica MTGDNTLITSHGINRRDFMKLCAALAATMGLSSKAAAEMAESVSNPQRPPVIWIGAQECT WP_000145429.1 hydrogenase 2 small subunit [Salmonella enterica] Length: 372\nNumber of predicted TMHs: 1\nExp number of AAs in TMHs: 24.51309\nExp number, first 60 AAs: 0.2682\nTotal prob of N-in: 0.14402\noutside 1 331\nTMhelix 332 354\ninside 355 372

22140 GCF_001588275.1_CFSAN033904_01.0Salmonella enterica subsp. enterica serovar Typhimurium Proteobacteria; Gammaproteobacteria; Enterobacterales; Enterobacteriaceae; Salmonella; Salmonella enterica; Salmonella enterica subsp. enterica MTGDNTLITSHGINRRDFMKLCAALAATMGLSSKAAAEMAESVSNPQRPPVIWIGAQECT WP_000145429.1 hydrogenase 2 small subunit [Salmonella enterica] Length: 372\nNumber of predicted TMHs: 1\nExp number of AAs in TMHs: 24.51309\nExp number, first 60 AAs: 0.2682\nTotal prob of N-in: 0.14402\noutside 1 331\nTMhelix 332 354\ninside 355 372

22141 GCF_001480115.1_Salmonella_enterica_CVM_N38929-SQ_v1.0 Salmonella enterica subsp. enterica serovar Typhimurium Proteobacteria; Gammaproteobacteria; Enterobacterales; Enterobacteriaceae; Salmonella; Salmonella enterica; Salmonella enterica subsp. enterica MTGDNTLITSHGINRRDFMKLCAALAATMGLSSKAAAEMAESVSNPQRPPVIWIGAQECT WP_000145429.1 hydrogenase 2 small subunit [Salmonella enterica] Length: 372\nNumber of predicted TMHs: 1\nExp number of AAs in TMHs: 24.51309\nExp number, first 60 AAs: 0.2682\nTotal prob of N-in: 0.14402\noutside 1 331\nTMhelix 332 354\ninside 355 372

22142 GCF_001955215.1_ASM195521v1 Salmonella enterica subsp. enterica serovar Typhimurium Proteobacteria; Gammaproteobacteria; Enterobacterales; Enterobacteriaceae; Salmonella; Salmonella enterica; Salmonella enterica subsp. enterica MTGDNTLITSHGINRRDFMKLCAALAATMGLSSKAAAEMAESVSNPQRPPVIWIGAQECT WP_000145429.1 hydrogenase 2 small subunit [Salmonella enterica] Length: 372\nNumber of predicted TMHs: 1\nExp number of AAs in TMHs: 24.51309\nExp number, first 60 AAs: 0.2682\nTotal prob of N-in: 0.14402\noutside 1 331\nTMhelix 332 354\ninside 355 372

22143 GCF_001955215.1_ASM195521v1 Salmonella enterica subsp. enterica serovar Typhimurium Proteobacteria; Gammaproteobacteria; Enterobacterales; Enterobacteriaceae; Salmonella; Salmonella enterica; Salmonella enterica subsp. enterica MNNEETFYQAMRRKGVTRRSFLKFCSLAATSLGLGAGMTPKIAWALENKPRIPVVWIHGL WP_001058312.1 hydrogenase [Salmonella enterica] Length: 372\nNumber of predicted TMHs: 2\nExp number of AAs in TMHs: 36.51738\nExp number, first 60 AAs: 19.72849\nTotal prob of N-in: 0.96170\nPOSSIBLE N-term signal sequence\ninside 1 19\nTMhelix 20 39\noutside 40 325\nTMhelix 326 348\ninside 349 372

22144 GCF_001955235.1_ASM195523v1 Salmonella enterica subsp. enterica serovar Typhimurium Proteobacteria; Gammaproteobacteria; Enterobacterales; Enterobacteriaceae; Salmonella; Salmonella enterica; Salmonella enterica subsp. enterica MTGDNTLITSHGINRRDFMKLCAALAATMGLSSKAAAEMAESVSNPQRPPVIWIGAQECT WP_000145429.1 hydrogenase 2 small subunit [Salmonella enterica] Length: 372\nNumber of predicted TMHs: 1\nExp number of AAs in TMHs: 24.51309\nExp number, first 60 AAs: 0.2682\nTotal prob of N-in: 0.14402\noutside 1 331\nTMhelix 332 354\ninside 355 372

22145 GCF_001588235.1_CFSAN033906_01.0 Salmonella enterica subsp. enterica serovar Typhimurium Proteobacteria; Gammaproteobacteria; Enterobacterales; Enterobacteriaceae; Salmonella; Salmonella enterica; Salmonella enterica subsp. enterica MTGDNTLITSHGINRRDFMKLCAALAATMGLSSKAAAEMAESVSNPQRPPVIWIGAQECT WP_000145429.1 hydrogenase 2 small subunit [Salmonella enterica] Length: 372\nNumber of predicted TMHs: 1\nExp number of AAs in TMHs: 24.51309\nExp number, first 60 AAs: 0.2682\nTotal prob of N-in: 0.14402\noutside 1 331\nTMhelix 332 354\ninside 355 372

22146 GCF_001758445.1_ASM175844v1 Salmonella enterica subsp. enterica serovar Typhimurium Proteobacteria; Gammaproteobacteria; Enterobacterales; Enterobacteriaceae; Salmonella; Salmonella enterica; Salmonella enterica subsp. enterica MNNEETFYQAMRRKGVTRRSFLKFCSLAATSLGLGAGMTPKIAWALENKPRIPVVWIHGL WP_001058312.1 hydrogenase [Salmonella enterica] Length: 372\nNumber of predicted TMHs: 2\nExp number of AAs in TMHs: 36.51738\nExp number, first 60 AAs: 19.72849\nTotal prob of N-in: 0.96170\nPOSSIBLE N-term signal sequence\ninside 1 19\nTMhelix 20 39\noutside 40 325\nTMhelix 326 348\ninside 349 372

22147 GCF_001756215.1_CFSAN008078 Salmonella enterica subsp. enterica serovar Typhimurium Proteobacteria; Gammaproteobacteria; Enterobacterales; Enterobacteriaceae; Salmonella; Salmonella enterica; Salmonella enterica subsp. enterica MNNEETFYQAMRRKGVTRRSFLKFCSLAATSLGLGAGMTPKIAWALENKPRIPVVWIHGL WP_001058312.1 hydrogenase [Salmonella enterica] Length: 372\nNumber of predicted TMHs: 2\nExp number of AAs in TMHs: 36.51738\nExp number, first 60 AAs: 19.72849\nTotal prob of N-in: 0.96170\nPOSSIBLE N-term signal sequence\ninside 1 19\nTMhelix 20 39\noutside 40 325\nTMhelix 326 348\ninside 349 372

22148 GCF_001758445.1_ASM175844v1 Salmonella enterica subsp. enterica serovar Typhimurium Proteobacteria; Gammaproteobacteria; Enterobacterales; Enterobacteriaceae; Salmonella; Salmonella enterica; Salmonella enterica subsp. enterica MTGDNTLITSHGINRRDFMKLCAALAATMGLSSKAAAEMAESVSNPQRPPVIWIGAQECT WP_000145429.1 hydrogenase 2 small subunit [Salmonella enterica] Length: 372\nNumber of predicted TMHs: 1\nExp number of AAs in TMHs: 24.51309\nExp number, first 60 AAs: 0.2682\nTotal prob of N-in: 0.14402\noutside 1 331\nTMhelix 332 354\ninside 355 372

22149 GCF_001756215.1_CFSAN008078 Salmonella enterica subsp. enterica serovar Typhimurium Proteobacteria; Gammaproteobacteria; Enterobacterales; Enterobacteriaceae; Salmonella; Salmonella enterica; Salmonella enterica subsp. enterica MTGDNTLITSHGINRRDFMKLCAALAATMGLSSKAAAEMAESVSNPQRPPVIWIGAQECT WP_000145429.1 hydrogenase 2 small subunit [Salmonella enterica] Length: 372\nNumber of predicted TMHs: 1\nExp number of AAs in TMHs: 24.51309\nExp number, first 60 AAs: 0.2682\nTotal prob of N-in: 0.14402\noutside 1 331\nTMhelix 332 354\ninside 355 372

22150 GCF_001755885.1_CFSAN008089 Salmonella enterica subsp. enterica serovar Typhimurium Proteobacteria; Gammaproteobacteria; Enterobacterales; Enterobacteriaceae; Salmonella; Salmonella enterica; Salmonella enterica subsp. enterica MNNEETFYQAMRRKGVTRRSFLKFCSLAATSLGLGAGMTPKIAWALENKPRIPVVWIHGL WP_001058312.1 hydrogenase [Salmonella enterica] Length: 372\nNumber of predicted TMHs: 2\nExp number of AAs in TMHs: 36.51738\nExp number, first 60 AAs: 19.72849\nTotal prob of N-in: 0.96170\nPOSSIBLE N-term signal sequence\ninside 1 19\nTMhelix 20 39\noutside 40 325\nTMhelix 326 348\ninside 349 372

22151 GCF_001756185.1_CFSAN008072 Salmonella enterica subsp. enterica serovar Typhimurium Proteobacteria; Gammaproteobacteria; Enterobacterales; Enterobacteriaceae; Salmonella; Salmonella enterica; Salmonella enterica subsp. enterica MNNEETFYQAMRRKGVTRRSFLKFCSLAATSLGLGAGMTPKIAWALENKPRIPVVWIHGL WP_001058312.1 hydrogenase [Salmonella enterica] Length: 372\nNumber of predicted TMHs: 2\nExp number of AAs in TMHs: 36.51738\nExp number, first 60 AAs: 19.72849\nTotal prob of N-in: 0.96170\nPOSSIBLE N-term signal sequence\ninside 1 19\nTMhelix 20 39\noutside 40 325\nTMhelix 326 348\ninside 349 372

22152 GCF_001756105.1_CFSAN008085 Salmonella enterica subsp. enterica serovar Typhimurium Proteobacteria; Gammaproteobacteria; Enterobacterales; Enterobacteriaceae; Salmonella; Salmonella enterica; Salmonella enterica subsp. enterica MTGDNTLITSHGINRRDFMKLCAALAATMGLSSKAAAEMAESVSNPQRPPVIWIGAQECT WP_000145429.1 hydrogenase 2 small subunit [Salmonella enterica] Length: 372\nNumber of predicted TMHs: 1\nExp number of AAs in TMHs: 24.51309\nExp number, first 60 AAs: 0.2682\nTotal prob of N-in: 0.14402\noutside 1 331\nTMhelix 332 354\ninside 355 372

22153 GCF_001955295.1_ASM195529v1 Salmonella enterica subsp. enterica serovar Typhimurium Proteobacteria; Gammaproteobacteria; Enterobacterales; Enterobacteriaceae; Salmonella; Salmonella enterica; Salmonella enterica subsp. enterica MTGDNTLITSHGINRRDFMKLCAALAATMGLSSKAAAEMAESVSNPQRPPVIWIGAQECT WP_000145429.1 hydrogenase 2 small subunit [Salmonella enterica] Length: 372\nNumber of

predicted TMHs: 1\nExp number of AAs in TMHs: 24.51309\nExp number, first 60 AAs: 0.2682\nTotal prob of N-in: 0.14402\noutside 1 331\nTMhelix 332 354\ninside 355 372

22154 GCF_001756065.1_CFSAN008076 Salmonella enterica subsp. enterica serovar Typhimurium Proteobacteria; Gammaproteobacteria; Enterobacterales; Enterobacteriaceae; Salmonella; Salmonella enterica; Salmonella enterica subsp. enterica MNNEETFYQAMRRKGVTRRSFLKFCSLAATSLGLGAGMTPKIAWALENKPRIPVVWIHGL WP_001058312.1 hydrogenase [Salmonella enterica] Length: 372\nNumber of predicted TMHs: 2\nExp number of AAs in TMHs: 36.51738\nExp number, first 60 AAs: 19.72849\nTotal prob of N-in: 0.96170\nPOSSIBLE N-term signal sequence\ninside 1 19\nTMhelix 20 39\noutside 40 325\nTMhelix 326 348\ninside 349 372

22155 GCF_001755985.1_CFSAN008073 Salmonella enterica subsp. enterica serovar Typhimurium Proteobacteria; Gammaproteobacteria; Enterobacterales; Enterobacteriaceae; Salmonella; Salmonella enterica; Salmonella enterica subsp. enterica MNNEETFYQAMRRKGVTRRSFLKFCSLAATSLGLGAGMTPKIAWALENKPRIPVVWIHGL WP_001058312.1 hydrogenase [Salmonella enterica] Length: 372\nNumber of predicted TMHs: 2\nExp number of AAs in TMHs: 36.51738\nExp number, first 60 AAs: 19.72849\nTotal prob of N-in: 0.96170\nPOSSIBLE N-term signal sequence\ninside 1 19\nTMhelix 20 39\noutside 40 325\nTMhelix 326 348\ninside 349 372

22156 GCF_001755945.1_CFSAN008079 Salmonella enterica subsp. enterica serovar Typhimurium Proteobacteria; Gammaproteobacteria; Enterobacterales; Enterobacteriaceae; Salmonella; Salmonella enterica; Salmonella enterica subsp. enterica MNNEETFYQAMRRKGVTRRSFLKFCSLAATSLGLGAGMTPKIAWALENKPRIPVVWIHGL WP_001058312.1 hydrogenase [Salmonella enterica] Length: 372\nNumber of predicted TMHs: 2\nExp number of AAs in TMHs: 36.51738\nExp number, first 60 AAs: 19.72849\nTotal prob of N-in: 0.96170\nPOSSIBLE N-term signal sequence\ninside 1 19\nTMhelix 20 39\noutside 40 325\nTMhelix 326 348\ninside 349 372

22157 GCF_001756065.1_CFSAN008076 Salmonella enterica subsp. enterica serovar Typhimurium Proteobacteria; Gammaproteobacteria; Enterobacterales; Enterobacteriaceae; Salmonella; Salmonella enterica; Salmonella enterica subsp. enterica MTGDNTLITSHGINRRDFMKLCAALAATMGLSSKAAAEMAESVSNPQRPPVIWIGAQECT WP_000145429.1 hydrogenase 2 small subunit [Salmonella enterica] Length: 372\nNumber of predicted TMHs: 1\nExp number of AAs in TMHs: 24.51309\nExp number, first 60 AAs: 0.2682\nTotal prob of N-in: 0.14402\noutside 1 331\nTMhelix 332 354\ninside 355 372

22158 GCF_001755985.1_CFSAN008073 Salmonella enterica subsp. enterica serovar Typhimurium Proteobacteria; Gammaproteobacteria; Enterobacterales; Enterobacteriaceae; Salmonella; Salmonella enterica; Salmonella enterica subsp. enterica MTGDNTLITSHGINRRDFMKLCAALAATMGLSSKAAAEMAESVSNPQRPPVIWIGAQECT WP_000145429.1 hydrogenase 2 small subunit [Salmonella enterica] Length: 372\nNumber of predicted TMHs: 1\nExp number of AAs in TMHs: 24.51309\nExp number, first 60 AAs: 0.2682\nTotal prob of N-in: 0.14402\noutside 1 331\nTMhelix 332 354\ninside 355 372

22159 GCF_001955375.1_ASM195537v1 Salmonella enterica subsp. enterica serovar Typhimurium Proteobacteria; Gammaproteobacteria; Enterobacterales; Enterobacteriaceae; Salmonella; Salmonella enterica; Salmonella enterica subsp. enterica MNNEETFYQAMRRKGVTRRSFLKFCSLAATSLGLGAGMTPKIAWALENKPRIPVVWIHGL WP_001058312.1 hydrogenase [Salmonella enterica] Length: 372\nNumber of predicted TMHs: 2\nExp number of AAs in TMHs: 36.51738\nExp number, first 60 AAs: 19.72849\nTotal prob of N-in: 0.96170\nPOSSIBLE N-term signal sequence\ninside 1 19\nTMhelix 20 39\noutside 40 325\nTMhelix 326 348\ninside 349 372

22160 GCF_002035055.1_ASM203505v1 Salmonella enterica subsp. enterica serovar Typhimurium Proteobacteria; Gammaproteobacteria; Enterobacterales; Enterobacteriaceae; Salmonella; Salmonella enterica; Salmonella enterica subsp. enterica MNNEETFYQAMRRKGVTRRSFLKFCSLAATSLGLGAGMTPKIAWALENKPRIPVVWIHGL WP_001058312.1 hydrogenase [Salmonella enterica] Length: 372\nNumber of predicted TMHs: 2\nExp number of AAs in TMHs: 36.51738\nExp number, first 60 AAs: 19.72849\nTotal prob of N-in: 0.96170\nPOSSIBLE N-term signal sequence\ninside 1 19\nTMhelix 20 39\noutside 40 325\nTMhelix 326 348\ninside 349 372

22161 GCF_001955395.1_ASM195539v1 Salmonella enterica subsp. enterica serovar Typhimurium Proteobacteria; Gammaproteobacteria; Enterobacterales; Enterobacteriaceae; Salmonella; Salmonella enterica; Salmonella enterica subsp. enterica MNNEETFYQAMRRKGVTRRSFLKFCSLAATSLGLGAGMTPKIAWALENKPRIPVVWIHGL WP_001058312.1 hydrogenase [Salmonella enterica] Length: 372\nNumber of predicted TMHs: 2\nExp number of AAs in TMHs: 36.51738\nExp number, first 60 AAs: 19.72849\nTotal prob of N-in: 0.96170\nPOSSIBLE N-term signal sequence\ninside 1 19\nTMhelix 20 39\noutside 40 325\nTMhelix 326 348\ninside 349 372

22162 GCF_001477895.1_Salmonella_enterica_CVM_N29323-SQ_v1.0 Salmonella enterica subsp. enterica serovar Typhimurium Proteobacteria; Gammaproteobacteria; Enterobacterales; Enterobacteriaceae; Salmonella; Salmonella

enterica; Salmonella enterica subsp. enterica
MTGDNTLITSHGINRRDFMKLCAALAATMGLSSKAAAEMAESVSNPQRPPVIWIGAQECT WP_000145429.1 hydrogenase 2 small subunit [Salmonella enterica] Length: 372\nNumber of predicted TMHs: 1\nExp number of AAs in TMHs: 24.51309\nExp number, first 60 AAs: 0.2682\nTotal prob of N-in: 0.14402\noutside 1 331\nTMhelix 332 354\ninside 355 372

22163 GCF_002035135.1_ASM203513v1 Salmonella enterica subsp. enterica serovar Typhimurium Proteobacteria; Gammaproteobacteria; Enterobacterales; Enterobacteriaceae; Salmonella; Salmonella enterica; Salmonella enterica subsp. enterica MTGDNTLITSHGINRRDFMKLCAALAATMGLSSKAAAEMAESVSNPQRPPVIWIGAQECT WP_000145429.1 hydrogenase 2 small subunit [Salmonella enterica] Length: 372\nNumber of predicted TMHs: 1\nExp number of AAs in TMHs: 24.51309\nExp number, first 60 AAs: 0.2682\nTotal prob of N-in: 0.14402\noutside 1 331\nTMhelix 332 354\ninside 355 372

22164 GCF_001588195.1_CFSAN033909_01.0Salmonella enterica subsp. enterica serovar Typhimurium Proteobacteria; Gammaproteobacteria; Enterobacterales; Enterobacteriaceae; Salmonella; Salmonella enterica; Salmonella enterica subsp. enterica MTGDNTLITSHGINRRDFMKLCAALAATMGLSSKAAAEMAESVSNPQRPPVIWIGAQECT WP_000145429.1 hydrogenase 2 small subunit [Salmonella enterica] Length: 372\nNumber of predicted TMHs: 1\nExp number of AAs in TMHs: 24.51309\nExp number, first 60 AAs: 0.2682\nTotal prob of N-in: 0.14402\noutside 1 331\nTMhelix 332 354\ninside 355 372

22165 GCF_001479325.1_Salmonella_enterica_CVM_N32045-SQ_v1.0 Salmonella enterica subsp. enterica serovar Typhimurium Proteobacteria; Gammaproteobacteria; Enterobacterales; Enterobacteriaceae; Salmonella; Salmonella enterica; Salmonella enterica subsp. enterica
MNNEETFYQAMRRKGVTRRSFLKFCSLAATSLGLGAGMTPKIAWALENKPRIPVVWIHGL WP_001058312.1 hydrogenase [Salmonella enterica] Length: 372\nNumber of predicted TMHs: 2\nExp number of AAs in TMHs: 36.51738\nExp number, first 60 AAs: 19.72849\nTotal prob of N-in: 0.96170\nPOSSIBLE N-term signal sequence\ninside 1 19\nTMhelix 20 39\noutside 40 325\nTMhelix 326 348\ninside 349 372

22166 GCF_001955475.1_ASM195547v1 Salmonella enterica subsp. enterica serovar Typhimurium Proteobacteria; Gammaproteobacteria; Enterobacterales; Enterobacteriaceae; Salmonella; Salmonella enterica; Salmonella enterica subsp. enterica MNNEETFYQAMRRKGVTRRSFLKFCSLAATSLGLGAGMTPKIAWALENKPRIPVVWIHGL WP_001058312.1 hydrogenase [Salmonella enterica] Length: 372\nNumber of predicted TMHs: 2\nExp number of AAs in TMHs: 36.51738\nExp number, first 60 AAs: 19.72849\nTotal prob of N-in: 0.96170\nPOSSIBLE N-term signal sequence\ninside 1 19\nTMhelix 20 39\noutside 40 325\nTMhelix 326 348\ninside 349 372

22167 GCF_001955455.1_ASM195545v1 Salmonella enterica subsp. enterica serovar Typhimurium Proteobacteria; Gammaproteobacteria; Enterobacterales; Enterobacteriaceae; Salmonella; Salmonella enterica; Salmonella enterica subsp. enterica MNNEETFYQAMRRKGVTRRSFLKFCSLAATSLGLGAGMTPKIAWALENKPRIPVVWIHGL WP_001058312.1 hydrogenase [Salmonella enterica] Length: 372\nNumber of predicted TMHs: 2\nExp number of AAs in TMHs: 36.51738\nExp number, first 60 AAs: 19.72849\nTotal prob of N-in: 0.96170\nPOSSIBLE N-term signal sequence\ninside 1 19\nTMhelix 20 39\noutside 40 325\nTMhelix 326 348\ninside 349 372

22168 GCF_001295415.1_ASM129541v1 Salmonella enterica subsp. enterica serovar Typhimurium Proteobacteria; Gammaproteobacteria; Enterobacterales; Enterobacteriaceae; Salmonella; Salmonella enterica; Salmonella enterica subsp. enterica MTGDNTLITSHGINRRDFMKLCAALAATMGLSSKAAAEMAESVSNPQRPPVIWIGAQECT WP_000145429.1 hydrogenase 2 small subunit [Salmonella enterica] Length: 372\nNumber of predicted TMHs: 1\nExp number of AAs in TMHs: 24.51309\nExp number, first 60 AAs: 0.2682\nTotal prob of N-in: 0.14402\noutside 1 331\nTMhelix 332 354\ninside 355 372

22169 GCF_001295465.1_ASM129546v1 Salmonella enterica subsp. enterica serovar Typhimurium Proteobacteria; Gammaproteobacteria; Enterobacterales; Enterobacteriaceae; Salmonella; Salmonella enterica; Salmonella enterica subsp. enterica MTGDNTLITSHGINRRDFMKLCAALAATMGLSSKAAAEMAESVSNPQRPPVIWIGAQECT WP_000145429.1 hydrogenase 2 small subunit [Salmonella enterica] Length: 372\nNumber of predicted TMHs: 1\nExp number of AAs in TMHs: 24.51309\nExp number, first 60 AAs: 0.2682\nTotal prob of N-in: 0.14402\noutside 1 331\nTMhelix 332 354\ninside 355 372

22170 GCF_001588155.1_CFSAN033911_01.0Salmonella enterica subsp. enterica serovar Typhimurium Proteobacteria; Gammaproteobacteria; Enterobacterales; Enterobacteriaceae; Salmonella; Salmonella enterica; Salmonella enterica subsp. enterica MTGDNTLITSHGINRRDFMKLCAALAATMGLSSKAAAEMAESVSNPQRPPVIWIGAQECT WP_000145429.1 hydrogenase 2 small subunit [Salmonella enterica] Length: 372\nNumber of

predicted TMHs:  1\nExp number of AAs in TMHs: 24.51309\nExp number, first 60 AAs:  0.2682\nTotal prob of N-in: 0.14402\noutside   1   331\nTMhelix   332   354\ninside   355   372

22171 GCF_001587915.1_CFSAN033929_01.0Salmonella enterica subsp. enterica serovar Typhimurium Proteobacteria; Gammaproteobacteria; Enterobacterales; Enterobacteriaceae; Salmonella; Salmonella enterica; Salmonella enterica subsp. enterica MNNEETFYQAMRRKGVTRRSFLKFCSLAATSLGLGAGMTPKIAWALENKPRIPVVWIHGL WP_001058312.1 hydrogenase [Salmonella enterica] Length: 372\nNumber of predicted TMHs:  2\nExp number of AAs in TMHs: 36.51738\nExp number, first 60 AAs:  19.72849\nTotal prob of N-in:   0.96170\nPOSSIBLE N-term signal sequence\ninside   1   19\nTMhelix   20   39\noutside   40   325\nTMhelix   326   348\ninside   349   372

22172 GCF_001295545.1_ASM129554v1 Salmonella enterica subsp. enterica serovar Typhimurium Proteobacteria; Gammaproteobacteria; Enterobacterales; Enterobacteriaceae; Salmonella; Salmonella enterica; Salmonella enterica subsp. enterica MTGDNTLITSHGINRRDFMKLCAALAATMGLSSKAAAEMAESVSNPQRPPVIWIGAQECT WP_000145429.1 hydrogenase 2 small subunit [Salmonella enterica] Length: 372\nNumber of predicted TMHs:  1\nExp number of AAs in TMHs: 24.51309\nExp number, first 60 AAs:  0.2682\nTotal prob of N-in: 0.14402\noutside   1   331\nTMhelix   332   354\ninside   355   372

22173 GCF_001587835.1_CFSAN033892_01.0Salmonella enterica subsp. enterica serovar Typhimurium Proteobacteria; Gammaproteobacteria; Enterobacterales; Enterobacteriaceae; Salmonella; Salmonella enterica; Salmonella enterica subsp. enterica MNNEETFYQAMRRKGVTRRSFLKFCSLAATSLGLGAGMTPKIAWALENKPRIPVVWIHGL WP_001058312.1 hydrogenase [Salmonella enterica] Length: 372\nNumber of predicted TMHs:  2\nExp number of AAs in TMHs: 36.51738\nExp number, first 60 AAs:  19.72849\nTotal prob of N-in:   0.96170\nPOSSIBLE N-term signal sequence\ninside   1   19\nTMhelix   20   39\noutside   40   325\nTMhelix   326   348\ninside   349   372

22174 GCF_001295665.1_ASM129566v1 Salmonella enterica subsp. enterica serovar Typhimurium Proteobacteria; Gammaproteobacteria; Enterobacterales; Enterobacteriaceae; Salmonella; Salmonella enterica; Salmonella enterica subsp. enterica MNNEETFYQAMRRKGVTRRSFLKFCSLAATSLGLGAGMTPKIAWALENKPRIPVVWIHGL WP_001058312.1 hydrogenase [Salmonella enterica] Length: 372\nNumber of predicted TMHs:  2\nExp number of AAs in TMHs: 36.51738\nExp number, first 60 AAs:  19.72849\nTotal prob of N-in:   0.96170\nPOSSIBLE N-term signal sequence\ninside   1   19\nTMhelix   20   39\noutside   40   325\nTMhelix   326   348\ninside   349   372

22175 GCF_001587775.1_CFSAN033934_01.0Salmonella enterica subsp. enterica serovar Typhimurium Proteobacteria; Gammaproteobacteria; Enterobacterales; Enterobacteriaceae; Salmonella; Salmonella enterica; Salmonella enterica subsp. enterica MNNEETFYQAMRRKGVTRRSFLKFCSLAATSLGLGAGMTPKIAWALENKPRIPVVWIHGL WP_001058312.1 hydrogenase [Salmonella enterica] Length: 372\nNumber of predicted TMHs:  2\nExp number of AAs in TMHs: 36.51738\nExp number, first 60 AAs:  19.72849\nTotal prob of N-in:   0.96170\nPOSSIBLE N-term signal sequence\ninside   1   19\nTMhelix   20   39\noutside   40   325\nTMhelix   326   348\ninside   349   372

22176 GCF_001587915.1_CFSAN033929_01.0Salmonella enterica subsp. enterica serovar Typhimurium Proteobacteria; Gammaproteobacteria; Enterobacterales; Enterobacteriaceae; Salmonella; Salmonella enterica; Salmonella enterica subsp. enterica MTGDNTLITSHGINRRDFMKLCAALAATMGLSSKAAAEMAESVSNPQRPPVIWIGAQECT WP_000145429.1 hydrogenase 2 small subunit [Salmonella enterica] Length: 372\nNumber of predicted TMHs:  1\nExp number of AAs in TMHs: 24.51309\nExp number, first 60 AAs:  0.2682\nTotal prob of N-in: 0.14402\noutside   1   331\nTMhelix   332   354\ninside   355   372

22177 GCF_001576275.1_ASM157627v1 Salmonella enterica subsp. enterica serovar Typhimurium Proteobacteria; Gammaproteobacteria; Enterobacterales; Enterobacteriaceae; Salmonella; Salmonella enterica; Salmonella enterica subsp. enterica MTGDNTLITSHGINRRDFMKLCAALAATMGLSSKAAAEMAESVSNPQRPPVIWIGAQECT WP_000145429.1 hydrogenase 2 small subunit [Salmonella enterica] Length: 372\nNumber of predicted TMHs:  1\nExp number of AAs in TMHs: 24.51309\nExp number, first 60 AAs:  0.2682\nTotal prob of N-in: 0.14402\noutside   1   331\nTMhelix   332   354\ninside   355   372

22178 GCF_001480085.1_Salmonella_enterica_CVM_N38921-SQ_v1.0 Salmonella enterica subsp. enterica serovar Typhimurium Proteobacteria; Gammaproteobacteria; Enterobacterales; Enterobacteriaceae; Salmonella; Salmonella enterica; Salmonella enterica subsp. enterica MTGDNTLITSHGINRRDFMKLCAALAATMGLSSKAAAEMAESVSNPQRPPVIWIGAQECT WP_000145429.1 hydrogenase 2 small subunit [Salmonella enterica] Length: 372\nNumber of predicted TMHs:  1\nExp number of AAs in TMHs: 24.51309\nExp number, first 60 AAs:  0.2682\nTotal prob of N-in:   0.14402\noutside   1 331\nTMhelix   332   354\ninside   355   372

22179 GCF_001714645.1_ASM171464v1 Salmonella enterica subsp. enterica serovar Typhimurium Proteobacteria; Gammaproteobacteria; Enterobacterales; Enterobacteriaceae; Salmonella; Salmonella enterica; Salmonella enterica subsp. enterica MNNEETFYQAMRRKGVTRRSFLKFCSLAATSLGLGAGMTPKIAWALENKPRIPVVWIHGL WP_001058312.1 hydrogenase [Salmonella enterica] Length: 372\nNumber of predicted TMHs: 2\nExp number of AAs in TMHs: 36.51738\nExp number, first 60 AAs: 19.72849\nTotal prob of N-in: 0.96170\nPOSSIBLE N-term signal sequence\ninside 1 19\nTMhelix 20 39\noutside 40 325\nTMhelix 326 348\ninside 349 372

22180 GCF_001587775.1_CFSAN033934_01.0Salmonella enterica subsp. enterica serovar Typhimurium Proteobacteria; Gammaproteobacteria; Enterobacterales; Enterobacteriaceae; Salmonella; Salmonella enterica; Salmonella enterica subsp. enterica MTGDNTLITSHGINRRDFMKLCAALAATMGLSSKAAAEMAESVSNPQRPPVIWIGAQECT WP_000145429.1 hydrogenase 2 small subunit [Salmonella enterica] Length: 372\nNumber of predicted TMHs: 1\nExp number of AAs in TMHs: 24.51309\nExp number, first 60 AAs: 0.2682\nTotal prob of N-in: 0.14402\noutside 1 331\nTMhelix 332 354\ninside 355 372

22181 GCF_001587835.1_CFSAN033892_01.0Salmonella enterica subsp. enterica serovar Typhimurium Proteobacteria; Gammaproteobacteria; Enterobacterales; Enterobacteriaceae; Salmonella; Salmonella enterica; Salmonella enterica subsp. enterica MTGDNTLITSHGINRRDFMKLCAALAATMGLSSKAAAEMAESVSNPQRPPVIWIGAQECT WP_000145429.1 hydrogenase 2 small subunit [Salmonella enterica] Length: 372\nNumber of predicted TMHs: 1\nExp number of AAs in TMHs: 24.51309\nExp number, first 60 AAs: 0.2682\nTotal prob of N-in: 0.14402\noutside 1 331\nTMhelix 332 354\ninside 355 372

22182 GCF_002074395.1_ASM207439v1 Salmonella enterica subsp. enterica serovar Typhimurium Proteobacteria; Gammaproteobacteria; Enterobacterales; Enterobacteriaceae; Salmonella; Salmonella enterica; Salmonella enterica subsp. enterica MTGDNTLITSHGINRRDFMKLCAALAATMGLSSKAAAEMAESVSNPQRPPVIWIGAQECT WP_000145429.1 hydrogenase 2 small subunit [Salmonella enterica] Length: 372\nNumber of predicted TMHs: 1\nExp number of AAs in TMHs: 24.51309\nExp number, first 60 AAs: 0.2682\nTotal prob of N-in: 0.14402\noutside 1 331\nTMhelix 332 354\ninside 355 372

22183 GCF_002074245.1_ASM207424v1 Salmonella enterica subsp. enterica serovar Typhimurium Proteobacteria; Gammaproteobacteria; Enterobacterales; Enterobacteriaceae; Salmonella; Salmonella enterica; Salmonella enterica subsp. enterica MNNEETFYQAMRRKGVTRRSFLKFCSLAATSLGLGAGMTPKIAWALENKPRIPVVWIHGL WP_001058312.1 hydrogenase [Salmonella enterica] Length: 372\nNumber of predicted TMHs: 2\nExp number of AAs in TMHs: 36.51738\nExp number, first 60 AAs: 19.72849\nTotal prob of N-in: 0.96170\nPOSSIBLE N-term signal sequence\ninside 1 19\nTMhelix 20 39\noutside 40 325\nTMhelix 326 348\ninside 349 372

22184 GCF_002074375.1_ASM207437v1 Salmonella enterica subsp. enterica serovar Typhimurium Proteobacteria; Gammaproteobacteria; Enterobacterales; Enterobacteriaceae; Salmonella; Salmonella enterica; Salmonella enterica subsp. enterica MTGDNTLITSHGINRRDFMKLCAALAATMGLSSKAAAEMAESVSNPQRPPVIWIGAQECT WP_000145429.1 hydrogenase 2 small subunit [Salmonella enterica] Length: 372\nNumber of predicted TMHs: 1\nExp number of AAs in TMHs: 24.51309\nExp number, first 60 AAs: 0.2682\nTotal prob of N-in: 0.14402\noutside 1 331\nTMhelix 332 354\ninside 355 372

22185 GCF_002074305.1_ASM207430v1 Salmonella enterica subsp. enterica serovar Typhimurium Proteobacteria; Gammaproteobacteria; Enterobacterales; Enterobacteriaceae; Salmonella; Salmonella enterica; Salmonella enterica subsp. enterica MNNEETFYQAMRRKGVTRRSFLKFCSLAATSLGLGAGMTPKIAWALENKPRIPVVWIHGL WP_001058312.1 hydrogenase [Salmonella enterica] Length: 372\nNumber of predicted TMHs: 2\nExp number of AAs in TMHs: 36.51738\nExp number, first 60 AAs: 19.72849\nTotal prob of N-in: 0.96170\nPOSSIBLE N-term signal sequence\ninside 1 19\nTMhelix 20 39\noutside 40 325\nTMhelix 326 348\ninside 349 372

22186 GCF_002074345.1_ASM207434v1 Salmonella enterica subsp. enterica serovar Typhimurium Proteobacteria; Gammaproteobacteria; Enterobacterales; Enterobacteriaceae; Salmonella; Salmonella enterica; Salmonella enterica subsp. enterica MTGDNTLITSHGINRRDFMKLCAALAATMGLSSKAAAEMAESVSNPQRPPVIWIGAQECT WP_000145429.1 hydrogenase 2 small subunit [Salmonella enterica] Length: 372\nNumber of predicted TMHs: 1\nExp number of AAs in TMHs: 24.51309\nExp number, first 60 AAs: 0.2682\nTotal prob of N-in: 0.14402\noutside 1 331\nTMhelix 332 354\ninside 355 372

22187 GCF_002074345.1_ASM207434v1 Salmonella enterica subsp. enterica serovar Typhimurium Proteobacteria; Gammaproteobacteria; Enterobacterales; Enterobacteriaceae; Salmonella; Salmonella enterica; Salmonella enterica subsp. enterica MNNEETFYQAMRRKGVTRRSFLKFCSLAATSLGLGAGMTPKIAWALENKPRIPVVWIHGL WP_001058312.1 hydrogenase [Salmonella enterica] Length: 372\nNumber of predicted TMHs: 2\nExp

number of AAs in TMHs: 36.51738\nExp number, first 60 AAs: 19.72849\nTotal prob of N-in: 0.96170\nPOSSIBLE N-term signal sequence\ninside 1 19\nTMhelix 20 39\noutside 40 325\nTMhelix 326 348\ninside 349 372

22188 GCF_002074305.1_ASM207430v1 Salmonella enterica subsp. enterica serovar Typhimurium Proteobacteria; Gammaproteobacteria; Enterobacterales; Enterobacteriaceae; Salmonella; Salmonella enterica; Salmonella enterica subsp. enterica MTGDNTLITSHGINRRDFMKLCAALAATMGLSSKAAAEMAESVSNPQRPPVIWIGAQECT WP_000145429.1 hydrogenase 2 small subunit [Salmonella enterica] Length: 372\nNumber of predicted TMHs: 1\nExp number of AAs in TMHs: 24.51309\nExp number, first 60 AAs: 0.2682\nTotal prob of N-in: 0.14402\noutside 1 331\nTMhelix 332 354\ninside 355 372

22189 GCF_002074245.1_ASM207424v1 Salmonella enterica subsp. enterica serovar Typhimurium Proteobacteria; Gammaproteobacteria; Enterobacterales; Enterobacteriaceae; Salmonella; Salmonella enterica; Salmonella enterica subsp. enterica MTGDNTLITSHGINRRDFMKLCAALAATMGLSSKAAAEMAESVSNPQRPPVIWIGAQECT WP_000145429.1 hydrogenase 2 small subunit [Salmonella enterica] Length: 372\nNumber of predicted TMHs: 1\nExp number of AAs in TMHs: 24.51309\nExp number, first 60 AAs: 0.2682\nTotal prob of N-in: 0.14402\noutside 1 331\nTMhelix 332 354\ninside 355 372

22190 GCF_002074225.1_ASM207422v1 Salmonella enterica subsp. enterica serovar Typhimurium Proteobacteria; Gammaproteobacteria; Enterobacterales; Enterobacteriaceae; Salmonella; Salmonella enterica; Salmonella enterica subsp. enterica MNNEETFYQAMRRKGVTRRSFLKFCSLAATSLGLGAGMTPKIAWALENKPRIPVVWIHGL WP_001058312.1 hydrogenase [Salmonella enterica] Length: 372\nNumber of predicted TMHs: 2\nExp number of AAs in TMHs: 36.51738\nExp number, first 60 AAs: 19.72849\nTotal prob of N-in: 0.96170\nPOSSIBLE N-term signal sequence\ninside 1 19\nTMhelix 20 39\noutside 40 325\nTMhelix 326 348\ninside 349 372

22191 GCF_002074445.1_ASM207444v1 Salmonella enterica subsp. enterica serovar Typhimurium Proteobacteria; Gammaproteobacteria; Enterobacterales; Enterobacteriaceae; Salmonella; Salmonella enterica; Salmonella enterica subsp. enterica MTGDNTLITSHGINRRDFMKLCAALAATMGLSSKAAAEMAESVSNPQRPPVIWIGAQECT WP_000145429.1 hydrogenase 2 small subunit [Salmonella enterica] Length: 372\nNumber of predicted TMHs: 1\nExp number of AAs in TMHs: 24.51309\nExp number, first 60 AAs: 0.2682\nTotal prob of N-in: 0.14402\noutside 1 331\nTMhelix 332 354\ninside 355 372

22192 GCF_002074445.1_ASM207444v1 Salmonella enterica subsp. enterica serovar Typhimurium Proteobacteria; Gammaproteobacteria; Enterobacterales; Enterobacteriaceae; Salmonella; Salmonella enterica; Salmonella enterica subsp. enterica MNNEETFYQAMRRKGVTRRSFLKFCSLAATSLGLGAGMTPKIAWALENKPRIPVVWIHGL WP_001058312.1 hydrogenase [Salmonella enterica] Length: 372\nNumber of predicted TMHs: 2\nExp number of AAs in TMHs: 36.51738\nExp number, first 60 AAs: 19.72849\nTotal prob of N-in: 0.96170\nPOSSIBLE N-term signal sequence\ninside 1 19\nTMhelix 20 39\noutside 40 325\nTMhelix 326 348\ninside 349 372

22193 GCF_002074375.1_ASM207437v1 Salmonella enterica subsp. enterica serovar Typhimurium Proteobacteria; Gammaproteobacteria; Enterobacterales; Enterobacteriaceae; Salmonella; Salmonella enterica; Salmonella enterica subsp. enterica MNNEETFYQAMRRKGVTRRSFLKFCSLAATSLGLGAGMTPKIAWALENKPRIPVVWIHGL WP_001058312.1 hydrogenase [Salmonella enterica] Length: 372\nNumber of predicted TMHs: 2\nExp number of AAs in TMHs: 36.51738\nExp number, first 60 AAs: 19.72849\nTotal prob of N-in: 0.96170\nPOSSIBLE N-term signal sequence\ninside 1 19\nTMhelix 20 39\noutside 40 325\nTMhelix 326 348\ninside 349 372

22194 GCF_002074395.1_ASM207439v1 Salmonella enterica subsp. enterica serovar Typhimurium Proteobacteria; Gammaproteobacteria; Enterobacterales; Enterobacteriaceae; Salmonella; Salmonella enterica; Salmonella enterica subsp. enterica MNNEETFYQAMRRKGVTRRSFLKFCSLAATSLGLGAGMTPKIAWALENKPRIPVVWIHGL WP_001058312.1 hydrogenase [Salmonella enterica] Length: 372\nNumber of predicted TMHs: 2\nExp number of AAs in TMHs: 36.51738\nExp number, first 60 AAs: 19.72849\nTotal prob of N-in: 0.96170\nPOSSIBLE N-term signal sequence\ninside 1 19\nTMhelix 20 39\noutside 40 325\nTMhelix 326 348\ninside 349 372

22195 GCF_002074465.1_ASM207446v1 Salmonella enterica subsp. enterica serovar Typhimurium Proteobacteria; Gammaproteobacteria; Enterobacterales; Enterobacteriaceae; Salmonella; Salmonella enterica; Salmonella enterica subsp. enterica MTGDNTLITSHGINRRDFMKLCAALAATMGLSSKAAAEMAESVSNPQRPPVIWIGAQECT WP_000145429.1 hydrogenase 2 small subunit [Salmonella enterica] Length: 372\nNumber of predicted TMHs: 1\nExp number of AAs in TMHs: 24.51309\nExp number, first 60 AAs: 0.2682\nTotal prob of N-in: 0.14402\noutside 1 331\nTMhelix 332 354\ninside 355 372

22196 GCF_002074555.1_ASM207455v1 Salmonella enterica subsp. enterica serovar Typhimurium Proteobacteria; Gammaproteobacteria; Enterobacterales; Enterobacteriaceae; Salmonella; Salmonella enterica;

Salmonella enterica subsp. enterica MTGDNTLITSHGINRRDFMKLCAALAATMGLSSKAAAEMAESVSNPQRPPVIWIGAQECT WP_000145429.1 hydrogenase 2 small subunit [Salmonella enterica] Length: 372\nNumber of predicted TMHs: 1\nExp number of AAs in TMHs: 24.51309\nExp number, first 60 AAs: 0.2682\nTotal prob of N-in: 0.14402\noutside 1 331\nTMhelix 332 354\ninside 355 372

22197 GCF_002074185.1_ASM207418v1 Salmonella enterica subsp. enterica serovar Typhimurium Proteobacteria; Gammaproteobacteria; Enterobacterales; Enterobacteriaceae; Salmonella; Salmonella enterica; Salmonella enterica subsp. enterica MNNEETFYQAMRRKGVTRRSFLKFCSLAATSLGLGAGMTPKIAWALENKPRIPVVWIHGL WP_001058312.1 hydrogenase [Salmonella enterica] Length: 372\nNumber of predicted TMHs: 2\nExp number of AAs in TMHs: 36.51738\nExp number, first 60 AAs: 19.72849\nTotal prob of N-in: 0.96170\nPOSSIBLE N-term signal sequence\ninside 1 19\nTMhelix 20 39\noutside 40 325\nTMhelix 326 348\ninside 349 372

22198 GCF_002074525.1_ASM207452v1 Salmonella enterica subsp. enterica serovar Typhimurium Proteobacteria; Gammaproteobacteria; Enterobacterales; Enterobacteriaceae; Salmonella; Salmonella enterica; Salmonella enterica subsp. enterica MTGDNTLITSHGINRRDFMKLCAALAATMGLSSKAAAEMAESVSNPQRPPVIWIGAQECT WP_000145429.1 hydrogenase 2 small subunit [Salmonella enterica] Length: 372\nNumber of predicted TMHs: 1\nExp number of AAs in TMHs: 24.51309\nExp number, first 60 AAs: 0.2682\nTotal prob of N-in: 0.14402\noutside 1 331\nTMhelix 332 354\ninside 355 372

22199 GCF_002074605.1_ASM207460v1 Salmonella enterica subsp. enterica serovar Typhimurium Proteobacteria; Gammaproteobacteria; Enterobacterales; Enterobacteriaceae; Salmonella; Salmonella enterica; Salmonella enterica subsp. enterica MTGDNTLITSHGINRRDFMKLCAALAATMGLSSKAAAEMAESVSNPQRPPVIWIGAQECT WP_000145429.1 hydrogenase 2 small subunit [Salmonella enterica] Length: 372\nNumber of predicted TMHs: 1\nExp number of AAs in TMHs: 24.51309\nExp number, first 60 AAs: 0.2682\nTotal prob of N-in: 0.14402\noutside 1 331\nTMhelix 332 354\ninside 355 372

22200 GCF_002074655.1_ASM207465v1 Salmonella enterica subsp. enterica serovar Typhimurium Proteobacteria; Gammaproteobacteria; Enterobacterales; Enterobacteriaceae; Salmonella; Salmonella enterica; Salmonella enterica subsp. enterica MTGDNTLITSHGINRRDFMKLCAALAATMGLSSKAAAEMAESVSNPQRPPVIWIGAQECT WP_000145429.1 hydrogenase 2 small subunit [Salmonella enterica] Length: 372\nNumber of predicted TMHs: 1\nExp number of AAs in TMHs: 24.51309\nExp number, first 60 AAs: 0.2682\nTotal prob of N-in: 0.14402\noutside 1 331\nTMhelix 332 354\ninside 355 372

22201 GCF_002074465.1_ASM207446v1 Salmonella enterica subsp. enterica serovar Typhimurium Proteobacteria; Gammaproteobacteria; Enterobacterales; Enterobacteriaceae; Salmonella; Salmonella enterica; Salmonella enterica subsp. enterica MNNEETFYQAMRRKGVTRRSFLKFCSLAATSLGLGAGMTPKIAWALENKPRIPVVWIHGL WP_001058312.1 hydrogenase [Salmonella enterica] Length: 372\nNumber of predicted TMHs: 2\nExp number of AAs in TMHs: 36.51738\nExp number, first 60 AAs: 19.72849\nTotal prob of N-in: 0.96170\nPOSSIBLE N-term signal sequence\ninside 1 19\nTMhelix 20 39\noutside 40 325\nTMhelix 326 348\ninside 349 372

22202 GCF_002074225.1_ASM207422v1 Salmonella enterica subsp. enterica serovar Typhimurium Proteobacteria; Gammaproteobacteria; Enterobacterales; Enterobacteriaceae; Salmonella; Salmonella enterica; Salmonella enterica subsp. enterica MTGDNTLITSHGINRRDFMKLCAALAATMGLSSKAAAEMAESVSNPQRPPVIWIGAQECT WP_000145429.1 hydrogenase 2 small subunit [Salmonella enterica] Length: 372\nNumber of predicted TMHs: 1\nExp number of AAs in TMHs: 24.51309\nExp number, first 60 AAs: 0.2682\nTotal prob of N-in: 0.14402\noutside 1 331\nTMhelix 332 354\ninside 355 372

22203 GCF_002074735.1_ASM207473v1 Salmonella enterica subsp. enterica serovar Typhimurium Proteobacteria; Gammaproteobacteria; Enterobacterales; Enterobacteriaceae; Salmonella; Salmonella enterica; Salmonella enterica subsp. enterica MTGDNTLITSHGINRRDFMKLCAALAATMGLSSKAAAEMAESVSNPQRPPVIWIGAQECT WP_000145429.1 hydrogenase 2 small subunit [Salmonella enterica] Length: 372\nNumber of predicted TMHs: 1\nExp number of AAs in TMHs: 24.51309\nExp number, first 60 AAs: 0.2682\nTotal prob of N-in: 0.14402\noutside 1 331\nTMhelix 332 354\ninside 355 372

22204 GCF_002074735.1_ASM207473v1 Salmonella enterica subsp. enterica serovar Typhimurium Proteobacteria; Gammaproteobacteria; Enterobacterales; Enterobacteriaceae; Salmonella; Salmonella enterica; Salmonella enterica subsp. enterica MNNEETFYQAMRRKGVTRRSFLKFCSLAATSLGLGAGMTPKIAWALENKPRIPVVWIHGL WP_001058312.1 hydrogenase [Salmonella enterica] Length: 372\nNumber of predicted TMHs: 2\nExp number of AAs in TMHs: 36.51738\nExp number, first 60 AAs: 19.72849\nTotal prob of N-in: 0.96170\nPOSSIBLE N-term signal sequence\ninside 1 19\nTMhelix 20 39\noutside 40 325\nTMhelix 326 348\ninside 349 372

22205	GCF_002074185.1_ASM207418v1	Salmonella enterica subsp. enterica serovar Typhimurium	Proteobacteria; Gammaproteobacteria; Enterobacterales; Enterobacteriaceae; Salmonella; Salmonella enterica; Salmonella enterica subsp. enterica	MTGDNTLITSHGINRRDFMKLCAALAATMGLSSKAAAEMAESVSNPQRPPVIWIGAQECT	WP_000145429.1	hydrogenase 2 small subunit [Salmonella enterica]	Length: 372\nNumber of predicted TMHs:  1\nExp number of AAs in TMHs: 24.51309\nExp number, first 60 AAs:  0.2682\nTotal prob of N-in:  0.14402\noutside   1   331\nTMhelix   332   354\ninside   355   372

22206	GCF_002074555.1_ASM207455v1	Salmonella enterica subsp. enterica serovar Typhimurium	Proteobacteria; Gammaproteobacteria; Enterobacterales; Enterobacteriaceae; Salmonella; Salmonella enterica; Salmonella enterica subsp. enterica	MNNEETFYQAMRRKGVTRRSFLKFCSLAATSLGLGAGMTPKIAWALENKPRIPVVWIHGL	WP_001058312.1	hydrogenase [Salmonella enterica]	Length: 372\nNumber of predicted TMHs:  2\nExp number of AAs in TMHs: 36.51738\nExp number, first 60 AAs:  19.72849\nTotal prob of N-in:   0.96170\nPOSSIBLE N-term signal sequence\ninside   1   19\nTMhelix   20   39\noutside   40   325\nTMhelix   326   348\ninside   349   372

22207	GCF_002074525.1_ASM207452v1	Salmonella enterica subsp. enterica serovar Typhimurium	Proteobacteria; Gammaproteobacteria; Enterobacterales; Enterobacteriaceae; Salmonella; Salmonella enterica; Salmonella enterica subsp. enterica	MNNEETFYQAMRRKGVTRRSFLKFCSLAATSLGLGAGMTPKIAWALENKPRIPVVWIHGL	WP_001058312.1	hydrogenase [Salmonella enterica]	Length: 372\nNumber of predicted TMHs:  2\nExp number of AAs in TMHs: 36.51738\nExp number, first 60 AAs:  19.72849\nTotal prob of N-in:   0.96170\nPOSSIBLE N-term signal sequence\ninside   1   19\nTMhelix   20   39\noutside   40   325\nTMhelix   326   348\ninside   349   372

22208	GCF_002074775.1_ASM207477v1	Salmonella enterica subsp. enterica serovar Typhimurium	Proteobacteria; Gammaproteobacteria; Enterobacterales; Enterobacteriaceae; Salmonella; Salmonella enterica; Salmonella enterica subsp. enterica	MTGDNTLITSHGINRRDFMKLCAALAATMGLSSKAAAEMAESVSNPQRPPVIWIGAQECT	WP_000145429.1	hydrogenase 2 small subunit [Salmonella enterica]	Length: 372\nNumber of predicted TMHs:  1\nExp number of AAs in TMHs: 24.51309\nExp number, first 60 AAs:  0.2682\nTotal prob of N-in:  0.14402\noutside   1   331\nTMhelix   332   354\ninside   355   372

22209	GCF_002074775.1_ASM207477v1	Salmonella enterica subsp. enterica serovar Typhimurium	Proteobacteria; Gammaproteobacteria; Enterobacterales; Enterobacteriaceae; Salmonella; Salmonella enterica; Salmonella enterica subsp. enterica	MNNEETFYQAMRRKGVTRRSFLKFCSLAATSLGLGAGMTPKIAWALENKPRIPVVWIHGL	WP_001058312.1	hydrogenase [Salmonella enterica]	Length: 372\nNumber of predicted TMHs:  2\nExp number of AAs in TMHs: 36.51738\nExp number, first 60 AAs:  19.72849\nTotal prob of N-in:   0.96170\nPOSSIBLE N-term signal sequence\ninside   1   19\nTMhelix   20   39\noutside   40   325\nTMhelix   326   348\ninside   349   372

22210	GCF_002074605.1_ASM207460v1	Salmonella enterica subsp. enterica serovar Typhimurium	Proteobacteria; Gammaproteobacteria; Enterobacterales; Enterobacteriaceae; Salmonella; Salmonella enterica; Salmonella enterica subsp. enterica	MNNEETFYQAMRRKGVTRRSFLKFCSLAATSLGLGAGMTPKIAWALENKPRIPVVWIHGL	WP_001058312.1	hydrogenase [Salmonella enterica]	Length: 372\nNumber of predicted TMHs:  2\nExp number of AAs in TMHs: 36.51738\nExp number, first 60 AAs:  19.72849\nTotal prob of N-in:   0.96170\nPOSSIBLE N-term signal sequence\ninside   1   19\nTMhelix   20   39\noutside   40   325\nTMhelix   326   348\ninside   349   372

22211	GCF_002066505.1_ASM206650v1	Salmonella enterica subsp. enterica serovar Typhimurium	Proteobacteria; Gammaproteobacteria; Enterobacterales; Enterobacteriaceae; Salmonella; Salmonella enterica; Salmonella enterica subsp. enterica	MNNEETFYQAMRRKGVTRRSFLKFCSLAATSLGLGAGMTPKIAWALENKPRIPVVWIHGL	WP_001058312.1	hydrogenase [Salmonella enterica]	Length: 372\nNumber of predicted TMHs:  2\nExp number of AAs in TMHs: 36.51738\nExp number, first 60 AAs:  19.72849\nTotal prob of N-in:   0.96170\nPOSSIBLE N-term signal sequence\ninside   1   19\nTMhelix   20   39\noutside   40   325\nTMhelix   326   348\ninside   349   372

22212	GCF_000784315.1_ASM78431v1	Salmonella enterica subsp. enterica serovar Typhimurium	Proteobacteria; Gammaproteobacteria; Enterobacterales; Enterobacteriaceae; Salmonella; Salmonella enterica; Salmonella enterica subsp. enterica	MNNEETFYQAMRRKGVTRRSFLKFCSLAATSLGLGAGMTPKIAWALENKPRIPVVWIHGL	WP_001058312.1	hydrogenase [Salmonella enterica]	Length: 372\nNumber of predicted TMHs:  2\nExp number of AAs in TMHs: 36.51738\nExp number, first 60 AAs:  19.72849\nTotal prob of N-in:   0.96170\nPOSSIBLE N-term signal sequence\ninside   1   19\nTMhelix   20   39\noutside   40   325\nTMhelix   326   348\ninside   349   372

22213	GCF_002066505.1_ASM206650v1	Salmonella enterica subsp. enterica serovar Typhimurium	Proteobacteria; Gammaproteobacteria; Enterobacterales; Enterobacteriaceae; Salmonella; Salmonella enterica; Salmonella enterica subsp. enterica	MTGDNTLITSHGINRRDFMKLCAALAATMGLSSKAAAEMAESVSNPQRPPVIWIGAQECT	WP_000145429.1	hydrogenase 2 small subunit [Salmonella enterica]	Length: 372\nNumber of

predicted TMHs: 1\nExp number of AAs in TMHs: 24.51309\nExp number, first 60 AAs: 0.2682\nTotal prob of N-in: 0.14402\noutside 1 331\nTMhelix 332 354\ninside 355 372

22214 GCF_002065055.1_ASM206505v1 Salmonella enterica subsp. enterica serovar Typhimurium Proteobacteria; Gammaproteobacteria; Enterobacterales; Enterobacteriaceae; Salmonella; Salmonella enterica; Salmonella enterica subsp. enterica MNNEETFYQAMRRKGVTRRSFLKFCSLAATSLGLGAGMTPKIAWALENKPRIPVVWIHGL WP_001058312.1 hydrogenase [Salmonella enterica] Length: 372\nNumber of predicted TMHs: 2\nExp number of AAs in TMHs: 36.51738\nExp number, first 60 AAs: 19.72849\nTotal prob of N-in: 0.96170\nPOSSIBLE N-term signal sequence\ninside 1 19\nTMhelix 20 39\noutside 40 325\nTMhelix 326 348\ninside 349 372

22215 GCF_002065055.1_ASM206505v1 Salmonella enterica subsp. enterica serovar Typhimurium Proteobacteria; Gammaproteobacteria; Enterobacterales; Enterobacteriaceae; Salmonella; Salmonella enterica; Salmonella enterica subsp. enterica MTGDNTLITSHGINRRDFMKLCAALAATMGLSSKAAAEMAESVSNPQRPPVIWIGAQECT WP_000145429.1 hydrogenase 2 small subunit [Salmonella enterica] Length: 372\nNumber of predicted TMHs: 1\nExp number of AAs in TMHs: 24.51309\nExp number, first 60 AAs: 0.2682\nTotal prob of N-in: 0.14402\noutside 1 331\nTMhelix 332 354\ninside 355 372

22216 GCF_002063685.1_ASM206368v1 Salmonella enterica subsp. enterica serovar Typhimurium Proteobacteria; Gammaproteobacteria; Enterobacterales; Enterobacteriaceae; Salmonella; Salmonella enterica; Salmonella enterica subsp. enterica MNNEETFYQAMRRKGVTRRSFLKFCSLAATSLGLGAGMTPKIAWALENKPRIPVVWIHGL WP_001058312.1 hydrogenase [Salmonella enterica] Length: 372\nNumber of predicted TMHs: 2\nExp number of AAs in TMHs: 36.51738\nExp number, first 60 AAs: 19.72849\nTotal prob of N-in: 0.96170\nPOSSIBLE N-term signal sequence\ninside 1 19\nTMhelix 20 39\noutside 40 325\nTMhelix 326 348\ninside 349 372

22217 GCF_002063745.1_ASM206374v1 Salmonella enterica subsp. enterica serovar Typhimurium Proteobacteria; Gammaproteobacteria; Enterobacterales; Enterobacteriaceae; Salmonella; Salmonella enterica; Salmonella enterica subsp. enterica MNNEETFYQAMRRKGVTRRSFLKFCSLAATSLGLGAGMTPKIAWALENKPRIPVVWIHGL WP_001058312.1 hydrogenase [Salmonella enterica] Length: 372\nNumber of predicted TMHs: 2\nExp number of AAs in TMHs: 36.51738\nExp number, first 60 AAs: 19.72849\nTotal prob of N-in: 0.96170\nPOSSIBLE N-term signal sequence\ninside 1 19\nTMhelix 20 39\noutside 40 325\nTMhelix 326 348\ninside 349 372

22218 GCF_000973915.1_ABB1162-2.id9_v1.0 Salmonella enterica subsp. enterica serovar Typhimurium Proteobacteria; Gammaproteobacteria; Enterobacterales; Enterobacteriaceae; Salmonella; Salmonella enterica; Salmonella enterica subsp. enterica MTGDNTLITSHGINRRDFMKLCAALAATMGLSSKAAAEMAESVSNPQRPPVIWIGAQECT WP_000145429.1 hydrogenase 2 small subunit [Salmonella enterica] Length: 372\nNumber of predicted TMHs: 1\nExp number of AAs in TMHs: 24.51309\nExp number, first 60 AAs: 0.2682\nTotal prob of N-in: 0.14402\noutside 1 331\nTMhelix 332 354\ninside 355 372

22219 GCF_002063685.1_ASM206368v1 Salmonella enterica subsp. enterica serovar Typhimurium Proteobacteria; Gammaproteobacteria; Enterobacterales; Enterobacteriaceae; Salmonella; Salmonella enterica; Salmonella enterica subsp. enterica MTGDNTLITSHGINRRDFMKLCAALAATMGLSSKAAAEMAESVSNPQRPPVIWIGAQECT WP_000145429.1 hydrogenase 2 small subunit [Salmonella enterica] Length: 372\nNumber of predicted TMHs: 1\nExp number of AAs in TMHs: 24.51309\nExp number, first 60 AAs: 0.2682\nTotal prob of N-in: 0.14402\noutside 1 331\nTMhelix 332 354\ninside 355 372

22220 GCF_002074655.1_ASM207465v1 Salmonella enterica subsp. enterica serovar Typhimurium Proteobacteria; Gammaproteobacteria; Enterobacterales; Enterobacteriaceae; Salmonella; Salmonella enterica; Salmonella enterica subsp. enterica MNNEETFYQAMRRKGVTRRSFLKFCSLAATSLGLGAGMTPKIAWALENKPRIPVVWIHGL WP_001058312.1 hydrogenase [Salmonella enterica] Length: 372\nNumber of predicted TMHs: 2\nExp number of AAs in TMHs: 36.51738\nExp number, first 60 AAs: 19.72849\nTotal prob of N-in: 0.96170\nPOSSIBLE N-term signal sequence\ninside 1 19\nTMhelix 20 39\noutside 40 325\nTMhelix 326 348\ninside 349 372

22221 GCF_002063745.1_ASM206374v1 Salmonella enterica subsp. enterica serovar Typhimurium Proteobacteria; Gammaproteobacteria; Enterobacterales; Enterobacteriaceae; Salmonella; Salmonella enterica; Salmonella enterica subsp. enterica MTGDNTLITSHGINRRDFMKLCAALAATMGLSSKAAAEMAESVSNPQRPPVIWIGAQECT WP_000145429.1 hydrogenase 2 small subunit [Salmonella enterica] Length: 372\nNumber of predicted TMHs: 1\nExp number of AAs in TMHs: 24.51309\nExp number, first 60 AAs: 0.2682\nTotal prob of N-in: 0.14402\noutside 1 331\nTMhelix 332 354\ninside 355 372

22222 GCF_002063655.1_ASM206365v1 Salmonella enterica subsp. enterica serovar Typhimurium Proteobacteria; Gammaproteobacteria; Enterobacterales; Enterobacteriaceae; Salmonella; Salmonella enterica;

Salmonella enterica subsp. enterica	MTGDNTLITSHGINRRDFMKLCAALAATMGLSSKAAAEMAESVSNPQRPPVIWIGAQECT
	WP_000145429.1	hydrogenase 2 small subunit [Salmonella enterica]	Length: 372\nNumber of predicted TMHs:  1\nExp number of AAs in TMHs: 24.51309\nExp number, first 60 AAs:  0.2682\nTotal prob of N-in:  0.14402\noutside   1   331\nTMhelix   332   354\ninside   355   372

22223	GCF_002062795.1_ASM206279v1	Salmonella enterica subsp. enterica serovar Typhimurium	Proteobacteria; Gammaproteobacteria; Enterobacterales; Enterobacteriaceae; Salmonella; Salmonella enterica; Salmonella enterica subsp. enterica	MNNEETFYQAMRRKGVTRRSFLKFCSLAATSLGLGAGMTPKIAWALENKPRIPVVWIHGL
	WP_001058312.1	hydrogenase [Salmonella enterica]	Length: 372\nNumber of predicted TMHs:  2\nExp number of AAs in TMHs: 36.51738\nExp number, first 60 AAs:  19.72849\nTotal prob of N-in:   0.96170\nPOSSIBLE N-term signal sequence\ninside   1   19\nTMhelix   20   39\noutside   40   325\nTMhelix   326   348\ninside   349   372

22224	GCF_002062795.1_ASM206279v1	Salmonella enterica subsp. enterica serovar Typhimurium	Proteobacteria; Gammaproteobacteria; Enterobacterales; Enterobacteriaceae; Salmonella; Salmonella enterica; Salmonella enterica subsp. enterica	MTGDNTLITSHGINRRDFMKLCAALAATMGLSSKAAAEMAESVSNPQRPPVIWIGAQECT
	WP_000145429.1	hydrogenase 2 small subunit [Salmonella enterica]	Length: 372\nNumber of predicted TMHs:  1\nExp number of AAs in TMHs: 24.51309\nExp number, first 60 AAs:  0.2682\nTotal prob of N-in:  0.14402\noutside   1   331\nTMhelix   332   354\ninside   355   372

22225	GCF_002193605.1_ASM219360v1	Salmonella enterica subsp. enterica serovar Typhimurium	Proteobacteria; Gammaproteobacteria; Enterobacterales; Enterobacteriaceae; Salmonella; Salmonella enterica; Salmonella enterica subsp. enterica	MNNEETFYQAMRRKGVTRRSFLKFCSLAATSLGLGAGMTPKIAWALENKPRIPVVWIHGL
	WP_001058312.1	hydrogenase [Salmonella enterica]	Length: 372\nNumber of predicted TMHs:  2\nExp number of AAs in TMHs: 36.51738\nExp number, first 60 AAs:  19.72849\nTotal prob of N-in:   0.96170\nPOSSIBLE N-term signal sequence\ninside   1   19\nTMhelix   20   39\noutside   40   325\nTMhelix   326   348\ninside   349   372

22226	GCF_002091345.1_ASM209134v1	Salmonella enterica subsp. enterica serovar Typhimurium	Proteobacteria; Gammaproteobacteria; Enterobacterales; Enterobacteriaceae; Salmonella; Salmonella enterica; Salmonella enterica subsp. enterica	MTGDNTLITSHGINRRDFMKLCAALAATMGLSSKAAAEMAESVSNPQRPPVIWIGAQECT
	WP_000145429.1	hydrogenase 2 small subunit [Salmonella enterica]	Length: 372\nNumber of predicted TMHs:  1\nExp number of AAs in TMHs: 24.51309\nExp number, first 60 AAs:  0.2682\nTotal prob of N-in:  0.14402\noutside   1   331\nTMhelix   332   354\ninside   355   372

22227	GCF_002091115.1_ASM209111v1	Salmonella enterica subsp. enterica serovar Typhimurium	Proteobacteria; Gammaproteobacteria; Enterobacterales; Enterobacteriaceae; Salmonella; Salmonella enterica; Salmonella enterica subsp. enterica	MNNEETFYQAMRRKGVTRRSFLKFCSLAATSLGLGAGMTPKIAWALENKPRIPVVWIHGL
	WP_001058312.1	hydrogenase [Salmonella enterica]	Length: 372\nNumber of predicted TMHs:  2\nExp number of AAs in TMHs: 36.51738\nExp number, first 60 AAs:  19.72849\nTotal prob of N-in:   0.96170\nPOSSIBLE N-term signal sequence\ninside   1   19\nTMhelix   20   39\noutside   40   325\nTMhelix   326   348\ninside   349   372

22228	GCF_002091315.1_ASM209131v1	Salmonella enterica subsp. enterica serovar Typhimurium	Proteobacteria; Gammaproteobacteria; Enterobacterales; Enterobacteriaceae; Salmonella; Salmonella enterica; Salmonella enterica subsp. enterica	MNNEETFYQAMRRKGVTRRSFLKFCSLAATSLGLGAGMTPKIAWALENKPRIPVVWIHGL
	WP_001058312.1	hydrogenase [Salmonella enterica]	Length: 372\nNumber of predicted TMHs:  2\nExp number of AAs in TMHs: 36.51738\nExp number, first 60 AAs:  19.72849\nTotal prob of N-in:   0.96170\nPOSSIBLE N-term signal sequence\ninside   1   19\nTMhelix   20   39\noutside   40   325\nTMhelix   326   348\ninside   349   372

22229	GCF_002091315.1_ASM209131v1	Salmonella enterica subsp. enterica serovar Typhimurium	Proteobacteria; Gammaproteobacteria; Enterobacterales; Enterobacteriaceae; Salmonella; Salmonella enterica; Salmonella enterica subsp. enterica	MTGDNTLITSHGINRRDFMKLCAALAATMGLSSKAAAEMAESVSNPQRPPVIWIGAQECT
	WP_000145429.1	hydrogenase 2 small subunit [Salmonella enterica]	Length: 372\nNumber of predicted TMHs:  1\nExp number of AAs in TMHs: 24.51309\nExp number, first 60 AAs:  0.2682\nTotal prob of N-in:  0.14402\noutside   1   331\nTMhelix   332   354\ninside   355   372

22230	GCF_002091025.1_ASM209102v1	Salmonella enterica subsp. enterica serovar Typhimurium	Proteobacteria; Gammaproteobacteria; Enterobacterales; Enterobacteriaceae; Salmonella; Salmonella enterica; Salmonella enterica subsp. enterica	MNNEETFYQAMRRKGVTRRSFLKFCSLAATSLGLGAGMTPKIAWALENKPRIPVVWIHGL
	WP_001058312.1	hydrogenase [Salmonella enterica]	Length: 372\nNumber of predicted TMHs:  2\nExp number of AAs in TMHs: 36.51738\nExp number, first 60 AAs:  19.72849\nTotal prob of N-in:   0.96170\nPOSSIBLE N-term signal sequence\ninside   1   19\nTMhelix   20   39\noutside   40   325\nTMhelix   326   348\ninside   349   372

22231 GCF_002091075.1_ASM209107v1 Salmonella enterica subsp. enterica serovar Typhimurium Proteobacteria; Gammaproteobacteria; Enterobacterales; Enterobacteriaceae; Salmonella; Salmonella enterica; Salmonella enterica subsp. enterica MTGDNTLITSHGINRRDFMKLCAALAATMGLSSKAAAEMAESVSNPQRPPVIWIGAQECT WP_000145429.1 hydrogenase 2 small subunit [Salmonella enterica] Length: 372\nNumber of predicted TMHs: 1\nExp number of AAs in TMHs: 24.51309\nExp number, first 60 AAs: 0.2682\nTotal prob of N-in: 0.14402\noutside 1 331\nTMhelix 332 354\ninside 355 372

22232 GCF_002091025.1_ASM209102v1 Salmonella enterica subsp. enterica serovar Typhimurium Proteobacteria; Gammaproteobacteria; Enterobacterales; Enterobacteriaceae; Salmonella; Salmonella enterica; Salmonella enterica subsp. enterica MTGDNTLITSHGINRRDFMKLCAALAATMGLSSKAAAEMAESVSNPQRPPVIWIGAQECT WP_000145429.1 hydrogenase 2 small subunit [Salmonella enterica] Length: 372\nNumber of predicted TMHs: 1\nExp number of AAs in TMHs: 24.51309\nExp number, first 60 AAs: 0.2682\nTotal prob of N-in: 0.14402\noutside 1 331\nTMhelix 332 354\ninside 355 372

22233 GCF_002090945.1_ASM209094v1 Salmonella enterica subsp. enterica serovar Typhimurium Proteobacteria; Gammaproteobacteria; Enterobacterales; Enterobacteriaceae; Salmonella; Salmonella enterica; Salmonella enterica subsp. enterica MNNEETFYQAMRRKGVTRRSFLKFCSLAATSLGLGAGMTPKIAWALENKPRIPVVWIHGL WP_001058312.1 hydrogenase [Salmonella enterica] Length: 372\nNumber of predicted TMHs: 2\nExp number of AAs in TMHs: 36.51738\nExp number, first 60 AAs: 19.72849\nTotal prob of N-in: 0.96170\nPOSSIBLE N-term signal sequence\ninside 1 19\nTMhelix 20 39\noutside 40 325\nTMhelix 326 348\ninside 349 372

22234 GCF_002090995.1_ASM209099v1 Salmonella enterica subsp. enterica serovar Typhimurium Proteobacteria; Gammaproteobacteria; Enterobacterales; Enterobacteriaceae; Salmonella; Salmonella enterica; Salmonella enterica subsp. enterica MNNEETFYQAMRRKGVTRRSFLKFCSLAATSLGLGAGMTPKIAWALENKPRIPVVWIHGL WP_001058312.1 hydrogenase [Salmonella enterica] Length: 372\nNumber of predicted TMHs: 2\nExp number of AAs in TMHs: 36.51738\nExp number, first 60 AAs: 19.72849\nTotal prob of N-in: 0.96170\nPOSSIBLE N-term signal sequence\ninside 1 19\nTMhelix 20 39\noutside 40 325\nTMhelix 326 348\ninside 349 372

22235 GCF_002090995.1_ASM209099v1 Salmonella enterica subsp. enterica serovar Typhimurium Proteobacteria; Gammaproteobacteria; Enterobacterales; Enterobacteriaceae; Salmonella; Salmonella enterica; Salmonella enterica subsp. enterica MTGDNTLITSHGINRRDFMKLCAALAATMGLSSKAAAEMAESVSNPQRPPVIWIGAQECT WP_000145429.1 hydrogenase 2 small subunit [Salmonella enterica] Length: 372\nNumber of predicted TMHs: 1\nExp number of AAs in TMHs: 24.51309\nExp number, first 60 AAs: 0.2682\nTotal prob of N-in: 0.14402\noutside 1 331\nTMhelix 332 354\ninside 355 372

22236 GCF_002090945.1_ASM209094v1 Salmonella enterica subsp. enterica serovar Typhimurium Proteobacteria; Gammaproteobacteria; Enterobacterales; Enterobacteriaceae; Salmonella; Salmonella enterica; Salmonella enterica subsp. enterica MTGDNTLITSHGINRRDFMKLCAALAATMGLSSKAAAEMAESVSNPQRPPVIWIGAQECT WP_000145429.1 hydrogenase 2 small subunit [Salmonella enterica] Length: 372\nNumber of predicted TMHs: 1\nExp number of AAs in TMHs: 24.51309\nExp number, first 60 AAs: 0.2682\nTotal prob of N-in: 0.14402\noutside 1 331\nTMhelix 332 354\ninside 355 372

22237 GCF_001594535.1_CFSAN033912_01.0Salmonella enterica subsp. enterica serovar Typhimurium Proteobacteria; Gammaproteobacteria; Enterobacterales; Enterobacteriaceae; Salmonella; Salmonella enterica; Salmonella enterica subsp. enterica MNNEETFYQAMRRKGVTRRSFLKFCSLAATSLGLGAGMTPKIAWALENKPRIPVVWIHGL WP_001058312.1 hydrogenase [Salmonella enterica] Length: 372\nNumber of predicted TMHs: 2\nExp number of AAs in TMHs: 36.51738\nExp number, first 60 AAs: 19.72849\nTotal prob of N-in: 0.96170\nPOSSIBLE N-term signal sequence\ninside 1 19\nTMhelix 20 39\noutside 40 325\nTMhelix 326 348\ninside 349 372

22238 GCF_000973985.1_Senterica.svTyphimurium.strABBSB1189-1.id14_v1.0 Salmonella enterica subsp. enterica serovar Typhimurium Proteobacteria; Gammaproteobacteria; Enterobacterales; Enterobacteriaceae; Salmonella; Salmonella enterica; Salmonella enterica subsp. enterica MTGDNTLITSHGINRRDFMKLCAALAATMGLSSKAAAEMAESVSNPQRPPVIWIGAQECT WP_000145429.1 hydrogenase 2 small subunit [Salmonella enterica] Length: 372\nNumber of predicted TMHs: 1\nExp number of AAs in TMHs: 24.51309\nExp number, first 60 AAs: 0.2682\nTotal prob of N-in: 0.14402\noutside 1 331\nTMhelix 332 354\ninside 355 372

22239 GCF_000974015.1_SALF-297-2.id5_v1.0 Salmonella enterica subsp. enterica serovar Typhimurium Proteobacteria; Gammaproteobacteria; Enterobacterales; Enterobacteriaceae; Salmonella; Salmonella enterica; Salmonella enterica subsp. enterica MTGDNTLITSHGINRRDFMKLCAALAATMGLSSKAAAEMAESVSNPQRPPVIWIGAQECT WP_000145429.1 hydrogenase 2 small subunit [Salmonella enterica] Length: 372\nNumber of

predicted TMHs: 1\nExp number of AAs in TMHs: 24.51309\nExp number, first 60 AAs: 0.2682\nTotal prob of N-in: 0.14402\noutside 1 331\nTMhelix 332 354\ninside 355 372

22240 GCF_000974015.1_SALF-297-2.id5_v1.0 Salmonella enterica subsp. enterica serovar Typhimurium Proteobacteria; Gammaproteobacteria; Enterobacterales; Enterobacteriaceae; Salmonella; Salmonella enterica; Salmonella enterica subsp. enterica MNNEETFYQAMRRKGVTRRSFLKFCSLAATSLGLGAGMTPKIAWALENKPRIPVVWIHGL WP_001058312.1 hydrogenase [Salmonella enterica] Length: 372\nNumber of predicted TMHs: 2\nExp number of AAs in TMHs: 36.51738\nExp number, first 60 AAs: 19.72849\nTotal prob of N-in: 0.96170\nPOSSIBLE N-term signal sequence\ninside 1 19\nTMhelix 20 39\noutside 40 325\nTMhelix 326 348\ninside 349 372

22241 GCF_000973985.1_Senterica.svTyphimurium.strABBSB1189-1.id14_v1.0 Salmonella enterica subsp. enterica serovar Typhimurium Proteobacteria; Gammaproteobacteria; Enterobacterales; Enterobacteriaceae; Salmonella; Salmonella enterica; Salmonella enterica subsp. enterica MNNEETFYQAMRRKGVTRRSFLKFCSLAATSLGLGAGMTPKIAWALENKPRIPVVWIHGL WP_001058312.1 hydrogenase [Salmonella enterica] Length: 372\nNumber of predicted TMHs: 2\nExp number of AAs in TMHs: 36.51738\nExp number, first 60 AAs: 19.72849\nTotal prob of N-in: 0.96170\nPOSSIBLE N-term signal sequence\ninside 1 19\nTMhelix 20 39\noutside 40 325\nTMhelix 326 348\ninside 349 372

22242 GCF_001295665.1_ASM129566v1 Salmonella enterica subsp. enterica serovar Typhimurium Proteobacteria; Gammaproteobacteria; Enterobacterales; Enterobacteriaceae; Salmonella; Salmonella enterica; Salmonella enterica subsp. enterica MTGDNTLITSHGINRRDFMKLCAALAATMGLSSKAAAEMAESVSNPQRPPVIWIGAQECT WP_000145429.1 hydrogenase 2 small subunit [Salmonella enterica] Length: 372\nNumber of predicted TMHs: 1\nExp number of AAs in TMHs: 24.51309\nExp number, first 60 AAs: 0.2682\nTotal prob of N-in: 0.14402\noutside 1 331\nTMhelix 332 354\ninside 355 372

22243 GCF_001240925.1_Salmonella_enterica_CVM_N43829-R_v1.0 Salmonella enterica subsp. enterica serovar Typhimurium Proteobacteria; Gammaproteobacteria; Enterobacterales; Enterobacteriaceae; Salmonella; Salmonella enterica; Salmonella enterica subsp. enterica MNNEETFYQAMRRKGVTRRSFLKFCSLAATSLGLGAGMTPKIAWALENKPRIPVVWIHGL WP_001058312.1 hydrogenase [Salmonella enterica] Length: 372\nNumber of predicted TMHs: 2\nExp number of AAs in TMHs: 36.51738\nExp number, first 60 AAs: 19.72849\nTotal prob of N-in: 0.96170\nPOSSIBLE N-term signal sequence\ninside 1 19\nTMhelix 20 39\noutside 40 325\nTMhelix 326 348\ninside 349 372

22244 GCF_001272415.1_Salmonella_enterica_CVM_N51295_v1.0 Salmonella enterica subsp. enterica serovar Typhimurium Proteobacteria; Gammaproteobacteria; Enterobacterales; Enterobacteriaceae; Salmonella; Salmonella enterica; Salmonella enterica subsp. enterica MTGDNTLITSHGINRRDFMKLCAALAATMGLSSKAAAEMAESVSNPQRPPVIWIGAQECT WP_000145429.1 hydrogenase 2 small subunit [Salmonella enterica] Length: 372\nNumber of predicted TMHs: 1\nExp number of AAs in TMHs: 24.51309\nExp number, first 60 AAs: 0.2682\nTotal prob of N-in: 0.14402\noutside 1 331\nTMhelix 332 354\ninside 355 372

22245 GCF_001295415.1_ASM129541v1 Salmonella enterica subsp. enterica serovar Typhimurium Proteobacteria; Gammaproteobacteria; Enterobacterales; Enterobacteriaceae; Salmonella; Salmonella enterica; Salmonella enterica subsp. enterica MNNEETFYQAMRRKGVTRRSFLKFCSLAATSLGLGAGMTPKIAWALENKPRIPVVWIHGL WP_001058312.1 hydrogenase [Salmonella enterica] Length: 372\nNumber of predicted TMHs: 2\nExp number of AAs in TMHs: 36.51738\nExp number, first 60 AAs: 19.72849\nTotal prob of N-in: 0.96170\nPOSSIBLE N-term signal sequence\ninside 1 19\nTMhelix 20 39\noutside 40 325\nTMhelix 326 348\ninside 349 372

22246 GCF_001295595.1_ASM129559v1 Salmonella enterica subsp. enterica serovar Typhimurium Proteobacteria; Gammaproteobacteria; Enterobacterales; Enterobacteriaceae; Salmonella; Salmonella enterica; Salmonella enterica subsp. enterica MTGDNTLITSHGINRRDFMKLCAALAATMGLSSKAAAEMAESVSNPQRPPVIWIGAQECT WP_000145429.1 hydrogenase 2 small subunit [Salmonella enterica] Length: 372\nNumber of predicted TMHs: 1\nExp number of AAs in TMHs: 24.51309\nExp number, first 60 AAs: 0.2682\nTotal prob of N-in: 0.14402\noutside 1 331\nTMhelix 332 354\ninside 355 372

22247 GCF_001295465.1_ASM129546v1 Salmonella enterica subsp. enterica serovar Typhimurium Proteobacteria; Gammaproteobacteria; Enterobacterales; Enterobacteriaceae; Salmonella; Salmonella enterica; Salmonella enterica subsp. enterica MNNEETFYQAMRRKGVTRRSFLKFCSLAATSLGLGAGMTPKIAWALENKPRIPVVWIHGL WP_001058312.1 hydrogenase [Salmonella enterica] Length: 372\nNumber of predicted TMHs: 2\nExp number of AAs in TMHs: 36.51738\nExp number, first 60 AAs: 19.72849\nTotal prob of N-in: 0.96170\nPOSSIBLE N-term signal sequence\ninside 1 19\nTMhelix 20 39\noutside 40 325\nTMhelix 326 348\ninside 349 372

22248 GCF_001295745.1_ASM129574v1 Salmonella enterica subsp. enterica serovar Typhimurium Proteobacteria; Gammaproteobacteria; Enterobacterales; Enterobacteriaceae; Salmonella; Salmonella enterica; Salmonella enterica subsp. enterica MTGDNTLITSHGINRRDFMKLCAALAATMGLSSKAAAEMAESVSNPQRPPVIWIGAQECT WP_000145429.1 hydrogenase 2 small subunit [Salmonella enterica] Length: 372\nNumber of predicted TMHs: 1\nExp number of AAs in TMHs: 24.51309\nExp number, first 60 AAs: 0.2682\nTotal prob of N-in: 0.14402\noutside 1 331\nTMhelix 332 354\ninside 355 372

22249 GCF_001295745.1_ASM129574v1 Salmonella enterica subsp. enterica serovar Typhimurium Proteobacteria; Gammaproteobacteria; Enterobacterales; Enterobacteriaceae; Salmonella; Salmonella enterica; Salmonella enterica subsp. enterica MNNEETFYQAMRRKGVTRRSFLKFCSLAATSLGLGAGMTPKIAWALENKPRIPVVWIHGL WP_001058312.1 hydrogenase [Salmonella enterica] Length: 372\nNumber of predicted TMHs: 2\nExp number of AAs in TMHs: 36.51738\nExp number, first 60 AAs: 19.72849\nTotal prob of N-in: 0.96170\nPOSSIBLE N-term signal sequence\ninside 1 19\nTMhelix 20 39\noutside 40 325\nTMhelix 326 348\ninside 349 372

22250 GCF_001477895.1_Salmonella_enterica_CVM_N29323-SQ_v1.0 Salmonella enterica subsp. enterica serovar Typhimurium Proteobacteria; Gammaproteobacteria; Enterobacterales; Enterobacteriaceae; Salmonella; Salmonella enterica; Salmonella enterica subsp. enterica MNNEETFYQAMRRKGVTRRSFLKFCSLAATSLGLGAGMTPKIAWALENKPRIPVVWIHGL WP_001058312.1 hydrogenase [Salmonella enterica] Length: 372\nNumber of predicted TMHs: 2\nExp number of AAs in TMHs: 36.51738\nExp number, first 60 AAs: 19.72849\nTotal prob of N-in: 0.96170\nPOSSIBLE N-term signal sequence\ninside 1 19\nTMhelix 20 39\noutside 40 325\nTMhelix 326 348\ninside 349 372

22251 GCF_001478125.1_Salmonella_enterica_CVM_N31395-SQ_v1.0 Salmonella enterica subsp. enterica serovar Typhimurium Proteobacteria; Gammaproteobacteria; Enterobacterales; Enterobacteriaceae; Salmonella; Salmonella enterica; Salmonella enterica subsp. enterica MNNEETFYQAMRRKGVTRRSFLKFCSLAATSLGLGAGMTPKIAWALENKPRIPVVWIHGL WP_001058312.1 hydrogenase [Salmonella enterica] Length: 372\nNumber of predicted TMHs: 2\nExp number of AAs in TMHs: 36.51738\nExp number, first 60 AAs: 19.72849\nTotal prob of N-in: 0.96170\nPOSSIBLE N-term signal sequence\ninside 1 19\nTMhelix 20 39\noutside 40 325\nTMhelix 326 348\ninside 349 372

22252 GCF_001480085.1_Salmonella_enterica_CVM_N38921-SQ_v1.0 Salmonella enterica subsp. enterica serovar Typhimurium Proteobacteria; Gammaproteobacteria; Enterobacterales; Enterobacteriaceae; Salmonella; Salmonella enterica; Salmonella enterica subsp. enterica MNNEETFYQAMRRKGVTRRSFLKFCSLAATSLGLGAGMTPKIAWALENKPRIPVVWIHGL WP_001058312.1 hydrogenase [Salmonella enterica] Length: 372\nNumber of predicted TMHs: 2\nExp number of AAs in TMHs: 36.51738\nExp number, first 60 AAs: 19.72849\nTotal prob of N-in: 0.96170\nPOSSIBLE N-term signal sequence\ninside 1 19\nTMhelix 20 39\noutside 40 325\nTMhelix 326 348\ninside 349 372

22253 GCF_001480485.1_Salmonella_enterica_CVM_N42450-SQ_v1.0 Salmonella enterica subsp. enterica serovar Typhimurium Proteobacteria; Gammaproteobacteria; Enterobacterales; Enterobacteriaceae; Salmonella; Salmonella enterica; Salmonella enterica subsp. enterica MNNEETFYQAMRRKGVTRRSFLKFCSLAATSLGLGAGMTPKIAWALENKPRIPVVWIHGL WP_001058312.1 hydrogenase [Salmonella enterica] Length: 372\nNumber of predicted TMHs: 2\nExp number of AAs in TMHs: 36.51738\nExp number, first 60 AAs: 19.72849\nTotal prob of N-in: 0.96170\nPOSSIBLE N-term signal sequence\ninside 1 19\nTMhelix 20 39\noutside 40 325\nTMhelix 326 348\ninside 349 372

22254 GCF_001559855.1_ASM155985v1 Salmonella enterica subsp. enterica serovar Typhimurium Proteobacteria; Gammaproteobacteria; Enterobacterales; Enterobacteriaceae; Salmonella; Salmonella enterica; Salmonella enterica subsp. enterica MTGDNTLITSHGINRRDFMKLCAALAATMGLSSKAAAEMAESVSNPQRPPVIWIGAQECT WP_000145429.1 hydrogenase 2 small subunit [Salmonella enterica] Length: 372\nNumber of predicted TMHs: 1\nExp number of AAs in TMHs: 24.51309\nExp number, first 60 AAs: 0.2682\nTotal prob of N-in: 0.14402\noutside 1 331\nTMhelix 332 354\ninside 355 372

22255 GCF_001559855.1_ASM155985v1 Salmonella enterica subsp. enterica serovar Typhimurium Proteobacteria; Gammaproteobacteria; Enterobacterales; Enterobacteriaceae; Salmonella; Salmonella enterica; Salmonella enterica subsp. enterica MNNEETFYQAMRRKGVTRRSFLKFCSLAATSLGLGAGMTPKIAWALENKPRIPVVWIHGL WP_001058312.1 hydrogenase [Salmonella enterica] Length: 372\nNumber of predicted TMHs: 2\nExp number of AAs in TMHs: 36.51738\nExp number, first 60 AAs: 19.72849\nTotal prob of N-in: 0.96170\nPOSSIBLE N-term signal sequence\ninside 1 19\nTMhelix 20 39\noutside 40 325\nTMhelix 326 348\ninside 349 372

22256	GCF_001540845.1_SO4698_09	Salmonella enterica subsp. enterica serovar Typhimurium	Proteobacteria; Gammaproteobacteria; Enterobacterales; Enterobacteriaceae; Salmonella; Salmonella enterica; Salmonella enterica subsp. enterica	MNNEETFYQAMRRKGVTRRSFLKFCSLAATSLGLGAGMTPKIAWALENKPRIPVVWIHGL	WP_001058312.1	hydrogenase [Salmonella enterica]	Length: 372\nNumber of predicted TMHs:  2\nExp number of AAs in TMHs: 36.51738\nExp number, first 60 AAs:  19.72849\nTotal prob of N-in:   0.96170\nPOSSIBLE N-term signal sequence\ninside   1   19\nTMhelix   20   39\noutside   40   325\nTMhelix   326   348\ninside   349   372

22257	GCF_001576275.1_ASM157627v1	Salmonella enterica subsp. enterica serovar Typhimurium	Proteobacteria; Gammaproteobacteria; Enterobacterales; Enterobacteriaceae; Salmonella; Salmonella enterica; Salmonella enterica subsp. enterica	MNNEETFYQAMRRKGVTRRSFLKFCSLAATSLGLGAGMTPKIAWALENKPRIPVVWIHGL	WP_001058312.1	hydrogenase [Salmonella enterica]	Length: 372\nNumber of predicted TMHs:  2\nExp number of AAs in TMHs: 36.51738\nExp number, first 60 AAs:  19.72849\nTotal prob of N-in:   0.96170\nPOSSIBLE N-term signal sequence\ninside   1   19\nTMhelix   20   39\noutside   40   325\nTMhelix   326   348\ninside   349   372

22258	GCF_001587755.1_CFSAN033938_01.0Salmonella enterica subsp. enterica serovar Typhimurium	Proteobacteria; Gammaproteobacteria; Enterobacterales; Enterobacteriaceae; Salmonella; Salmonella enterica; Salmonella enterica subsp. enterica	MTGDNTLITSHGINRRDFMKLCAALAATMGLSSKAAAEMAESVSNPQRPPVIWIGAQECT	WP_000145429.1	hydrogenase 2 small subunit [Salmonella enterica]	Length: 372\nNumber of predicted TMHs:  1\nExp number of AAs in TMHs: 24.51309\nExp number, first 60 AAs:  0.2682\nTotal prob of N-in:   0.14402\noutside   1   331\nTMhelix   332   354\ninside   355   372

22259	GCF_001588015.1_CFSAN033921_01.0Salmonella enterica subsp. enterica serovar Typhimurium	Proteobacteria; Gammaproteobacteria; Enterobacterales; Enterobacteriaceae; Salmonella; Salmonella enterica; Salmonella enterica subsp. enterica	MNNEETFYQAMRRKGVTRRSFLKFCSLAATSLGLGAGMTPKIAWALENKPRIPVVWIHGL	WP_001058312.1	hydrogenase [Salmonella enterica]	Length: 372\nNumber of predicted TMHs:  2\nExp number of AAs in TMHs: 36.51738\nExp number, first 60 AAs:  19.72849\nTotal prob of N-in:   0.96170\nPOSSIBLE N-term signal sequence\ninside   1   19\nTMhelix   20   39\noutside   40   325\nTMhelix   326   348\ninside   349   372

22260	GCF_001587855.1_CFSAN033932_01.0Salmonella enterica subsp. enterica serovar Typhimurium	Proteobacteria; Gammaproteobacteria; Enterobacterales; Enterobacteriaceae; Salmonella; Salmonella enterica; Salmonella enterica subsp. enterica	MNNEETFYQAMRRKGVTRRSFLKFCSLAATSLGLGAGMTPKIAWALENKPRIPVVWIHGL	WP_001058312.1	hydrogenase [Salmonella enterica]	Length: 372\nNumber of predicted TMHs:  2\nExp number of AAs in TMHs: 36.51738\nExp number, first 60 AAs:  19.72849\nTotal prob of N-in:   0.96170\nPOSSIBLE N-term signal sequence\ninside   1   19\nTMhelix   20   39\noutside   40   325\nTMhelix   326   348\ninside   349   372

22261	GCF_001588105.1_CFSAN033914_01.0Salmonella enterica subsp. enterica serovar Typhimurium	Proteobacteria; Gammaproteobacteria; Enterobacterales; Enterobacteriaceae; Salmonella; Salmonella enterica; Salmonella enterica subsp. enterica	MTGDNTLITSHGINRRDFMKLCAALAATMGLSSKAAAEMAESVSNPQRPPVIWIGAQECT	WP_000145429.1	hydrogenase 2 small subunit [Salmonella enterica]	Length: 372\nNumber of predicted TMHs:  1\nExp number of AAs in TMHs: 24.51309\nExp number, first 60 AAs:  0.2682\nTotal prob of N-in:   0.14402\noutside   1   331\nTMhelix   332   354\ninside   355   372

22262	GCF_001588105.1_CFSAN033914_01.0Salmonella enterica subsp. enterica serovar Typhimurium	Proteobacteria; Gammaproteobacteria; Enterobacterales; Enterobacteriaceae; Salmonella; Salmonella enterica; Salmonella enterica subsp. enterica	MNNEETFYQAMRRKGVTRRSFLKFCSLAATSLGLGAGMTPKIAWALENKPRIPVVWIHGL	WP_001058312.1	hydrogenase [Salmonella enterica]	Length: 372\nNumber of predicted TMHs:  2\nExp number of AAs in TMHs: 36.51738\nExp number, first 60 AAs:  19.72849\nTotal prob of N-in:   0.96170\nPOSSIBLE N-term signal sequence\ninside   1   19\nTMhelix   20   39\noutside   40   325\nTMhelix   326   348\ninside   349   372

22263	GCF_001588045.1_CFSAN033918_01.0Salmonella enterica subsp. enterica serovar Typhimurium	Proteobacteria; Gammaproteobacteria; Enterobacterales; Enterobacteriaceae; Salmonella; Salmonella enterica; Salmonella enterica subsp. enterica	MNNEETFYQAMRRKGVTRRSFLKFCSLAATSLGLGAGMTPKIAWALENKPRIPVVWIHGL	WP_001058312.1	hydrogenase [Salmonella enterica]	Length: 372\nNumber of predicted TMHs:  2\nExp number of AAs in TMHs: 36.51738\nExp number, first 60 AAs:  19.72849\nTotal prob of N-in:   0.96170\nPOSSIBLE N-term signal sequence\ninside   1   19\nTMhelix   20   39\noutside   40   325\nTMhelix   326   348\ninside   349   372

22264	GCF_001588075.1_CFSAN033917_01.0Salmonella enterica subsp. enterica serovar Typhimurium	Proteobacteria; Gammaproteobacteria; Enterobacterales; Enterobacteriaceae; Salmonella; Salmonella enterica; Salmonella enterica subsp. enterica	MNNEETFYQAMRRKGVTRRSFLKFCSLAATSLGLGAGMTPKIAWALENKPRIPVVWIHGL	WP_001058312.1	hydrogenase [Salmonella enterica]	Length: 372\nNumber of predicted TMHs:  2\nExp

number of AAs in TMHs: 36.51738\nExp number, first 60 AAs: 19.72849\nTotal prob of N-in: 0.96170\nPOSSIBLE N-term signal sequence\ninside 1 19\nTMhelix 20 39\noutside 40 325\nTMhelix 326 348\ninside 349 372

22265 GCF_001588395.1_CFSAN033895_01.0Salmonella enterica subsp. enterica serovar Typhimurium Proteobacteria; Gammaproteobacteria; Enterobacterales; Enterobacteriaceae; Salmonella; Salmonella enterica; Salmonella enterica subsp. enterica MNNEETFYQAMRRKGVTRRSFLKFCSLAATSLGLGAGMTPKIAWALENKPRIPVVWIHGL WP_001058312.1 hydrogenase [Salmonella enterica] Length: 372\nNumber of predicted TMHs: 2\nExp number of AAs in TMHs: 36.51738\nExp number, first 60 AAs: 19.72849\nTotal prob of N-in: 0.96170\nPOSSIBLE N-term signal sequence\ninside 1 19\nTMhelix 20 39\noutside 40 325\nTMhelix 326 348\ninside 349 372

22266 GCF_001588155.1_CFSAN033911_01.0Salmonella enterica subsp. enterica serovar Typhimurium Proteobacteria; Gammaproteobacteria; Enterobacterales; Enterobacteriaceae; Salmonella; Salmonella enterica; Salmonella enterica subsp. enterica MNNEETFYQAMRRKGVTRRSFLKFCSLAATSLGLGAGMTPKIAWALENKPRIPVVWIHGL WP_001058312.1 hydrogenase [Salmonella enterica] Length: 372\nNumber of predicted TMHs: 2\nExp number of AAs in TMHs: 36.51738\nExp number, first 60 AAs: 19.72849\nTotal prob of N-in: 0.96170\nPOSSIBLE N-term signal sequence\ninside 1 19\nTMhelix 20 39\noutside 40 325\nTMhelix 326 348\ninside 349 372

22267 GCF_001588435.1_CFSAN033889_01.0Salmonella enterica subsp. enterica serovar Typhimurium Proteobacteria; Gammaproteobacteria; Enterobacterales; Enterobacteriaceae; Salmonella; Salmonella enterica; Salmonella enterica subsp. enterica MTGDNTLITSHGINRRDFMKLCAALAATMGLSSKAAAEMAESVSNPQRPPVIWIGAQECT WP_000145429.1 hydrogenase 2 small subunit [Salmonella enterica] Length: 372\nNumber of predicted TMHs: 1\nExp number of AAs in TMHs: 24.51309\nExp number, first 60 AAs: 0.2682\nTotal prob of N-in: 0.14402\noutside 1 331\nTMhelix 332 354\ninside 355 372

22268 GCF_001588435.1_CFSAN033889_01.0Salmonella enterica subsp. enterica serovar Typhimurium Proteobacteria; Gammaproteobacteria; Enterobacterales; Enterobacteriaceae; Salmonella; Salmonella enterica; Salmonella enterica subsp. enterica MNNEETFYQAMRRKGVTRRSFLKFCSLAATSLGLGAGMTPKIAWALENKPRIPVVWIHGL WP_001058312.1 hydrogenase [Salmonella enterica] Length: 372\nNumber of predicted TMHs: 2\nExp number of AAs in TMHs: 36.51738\nExp number, first 60 AAs: 19.72849\nTotal prob of N-in: 0.96170\nPOSSIBLE N-term signal sequence\ninside 1 19\nTMhelix 20 39\noutside 40 325\nTMhelix 326 348\ninside 349 372

22269 GCF_001588195.1_CFSAN033909_01.0Salmonella enterica subsp. enterica serovar Typhimurium Proteobacteria; Gammaproteobacteria; Enterobacterales; Enterobacteriaceae; Salmonella; Salmonella enterica; Salmonella enterica subsp. enterica MNNEETFYQAMRRKGVTRRSFLKFCSLAATSLGLGAGMTPKIAWALENKPRIPVVWIHGL WP_001058312.1 hydrogenase [Salmonella enterica] Length: 372\nNumber of predicted TMHs: 2\nExp number of AAs in TMHs: 36.51738\nExp number, first 60 AAs: 19.72849\nTotal prob of N-in: 0.96170\nPOSSIBLE N-term signal sequence\ninside 1 19\nTMhelix 20 39\noutside 40 325\nTMhelix 326 348\ninside 349 372

22270 GCF_001588545.1_CFSAN033868_01.0Salmonella enterica subsp. enterica serovar Typhimurium Proteobacteria; Gammaproteobacteria; Enterobacterales; Enterobacteriaceae; Salmonella; Salmonella enterica; Salmonella enterica subsp. enterica MTGDNTLITSHGINRRDFMKLCAALAATMGLSSKAAAEMAESVSNPQRPPVIWIGAQECT WP_000145429.1 hydrogenase 2 small subunit [Salmonella enterica] Length: 372\nNumber of predicted TMHs: 1\nExp number of AAs in TMHs: 24.51309\nExp number, first 60 AAs: 0.2682\nTotal prob of N-in: 0.14402\noutside 1 331\nTMhelix 332 354\ninside 355 372

22271 GCF_001588515.1_CFSAN033878_01.0Salmonella enterica subsp. enterica serovar Typhimurium Proteobacteria; Gammaproteobacteria; Enterobacterales; Enterobacteriaceae; Salmonella; Salmonella enterica; Salmonella enterica subsp. enterica MTGDNTLITSHGINRRDFMKLCAALAATMGLSSKAAAEMAESVSNPQRPPVIWIGAQECT WP_000145429.1 hydrogenase 2 small subunit [Salmonella enterica] Length: 372\nNumber of predicted TMHs: 1\nExp number of AAs in TMHs: 24.51309\nExp number, first 60 AAs: 0.2682\nTotal prob of N-in: 0.14402\noutside 1 331\nTMhelix 332 354\ninside 355 372

22272 GCF_001594515.1_CFSAN033920_01.0Salmonella enterica subsp. enterica serovar Typhimurium Proteobacteria; Gammaproteobacteria; Enterobacterales; Enterobacteriaceae; Salmonella; Salmonella enterica; Salmonella enterica subsp. enterica MTGDNTLITSHGINRRDFMKLCAALAATMGLSSKAAAEMAESVSNPQRPPVIWIGAQECT WP_000145429.1 hydrogenase 2 small subunit [Salmonella enterica] Length: 372\nNumber of predicted TMHs: 1\nExp number of AAs in TMHs: 24.51309\nExp number, first 60 AAs: 0.2682\nTotal prob of N-in: 0.14402\noutside 1 331\nTMhelix 332 354\ninside 355 372

22273 GCF_001594595.1_CFSAN033901_01.0Salmonella enterica subsp. enterica serovar Typhimurium Proteobacteria; Gammaproteobacteria; Enterobacterales; Enterobacteriaceae; Salmonella; Salmonella enterica;

Salmonella enterica subsp. enterica MTGDNTLITSHGINRRDFMKLCAALAATMGLSSKAAAEMAESVSNPQRPPVIWIGAQECT WP_000145429.1 hydrogenase 2 small subunit [Salmonella enterica] Length: 372\nNumber of predicted TMHs: 1\nExp number of AAs in TMHs: 24.51309\nExp number, first 60 AAs: 0.2682\nTotal prob of N-in: 0.14402\noutside 1 331\nTMhelix 332 354\ninside 355 372

22274 GCF_001594625.1_CFSAN033896_01.0Salmonella enterica subsp. enterica serovar Typhimurium Proteobacteria; Gammaproteobacteria; Enterobacterales; Enterobacteriaceae; Salmonella; Salmonella enterica; Salmonella enterica subsp. enterica MTGDNTLITSHGINRRDFMKLCAALAATMGLSSKAAAEMAESVSNPQRPPVIWIGAQECT WP_000145429.1 hydrogenase 2 small subunit [Salmonella enterica] Length: 372\nNumber of predicted TMHs: 1\nExp number of AAs in TMHs: 24.51309\nExp number, first 60 AAs: 0.2682\nTotal prob of N-in: 0.14402\noutside 1 331\nTMhelix 332 354\ninside 355 372

22275 GCF_001594455.1_CFSAN033928_01.0Salmonella enterica subsp. enterica serovar Typhimurium Proteobacteria; Gammaproteobacteria; Enterobacterales; Enterobacteriaceae; Salmonella; Salmonella enterica; Salmonella enterica subsp. enterica MNNEETFYQAMRRKGVTRRSFLKFCSLAATSLGLGAGMTPKIAWALENKPRIPVVWIHGL WP_001058312.1 hydrogenase [Salmonella enterica] Length: 372\nNumber of predicted TMHs: 2\nExp number of AAs in TMHs: 36.51738\nExp number, first 60 AAs: 19.72849\nTotal prob of N-in: 0.96170\nPOSSIBLE N-term signal sequence\ninside 1 19\nTMhelix 20 39\noutside 40 325\nTMhelix 326 348\ninside 349 372

22276 GCF_001594435.1_CFSAN033924_01.0Salmonella enterica subsp. enterica serovar Typhimurium Proteobacteria; Gammaproteobacteria; Enterobacterales; Enterobacteriaceae; Salmonella; Salmonella enterica; Salmonella enterica subsp. enterica MNNEETFYQAMRRKGVTRRSFLKFCSLAATSLGLGAGMTPKIAWALENKPRIPVVWIHGL WP_001058312.1 hydrogenase [Salmonella enterica] Length: 372\nNumber of predicted TMHs: 2\nExp number of AAs in TMHs: 36.51738\nExp number, first 60 AAs: 19.72849\nTotal prob of N-in: 0.96170\nPOSSIBLE N-term signal sequence\ninside 1 19\nTMhelix 20 39\noutside 40 325\nTMhelix 326 348\ninside 349 372

22277 GCF_001594795.1_CFSAN033881_01.0Salmonella enterica subsp. enterica serovar Typhimurium Proteobacteria; Gammaproteobacteria; Enterobacterales; Enterobacteriaceae; Salmonella; Salmonella enterica; Salmonella enterica subsp. enterica MTGDNTLITSHGINRRDFMKLCAALAATMGLSSKAAAEMAESVSNPQRPPVIWIGAQECT WP_061584213.1 hydrogenase 2 small subunit [Salmonella enterica] Length: 372\nNumber of predicted TMHs: 1\nExp number of AAs in TMHs: 24.51809\nExp number, first 60 AAs: 0.26818\nTotal prob of N-in: 0.14414\noutside 1 331\nTMhelix 332 354\ninside 355 372

22278 GCF_001594935.1_CFSAN033872_01.0Salmonella enterica subsp. enterica serovar Typhimurium Proteobacteria; Gammaproteobacteria; Enterobacterales; Enterobacteriaceae; Salmonella; Salmonella enterica; Salmonella enterica subsp. enterica MTGDNTLITSHGINRRDFMKLCAALAATMGLSSKAAAEMAESVSNPQRPPVIWIGAQECT WP_000145429.1 hydrogenase 2 small subunit [Salmonella enterica] Length: 372\nNumber of predicted TMHs: 1\nExp number of AAs in TMHs: 24.51309\nExp number, first 60 AAs: 0.2682\nTotal prob of N-in: 0.14402\noutside 1 331\nTMhelix 332 354\ninside 355 372

22279 GCF_001594985.1_CFSAN033870_01.0Salmonella enterica subsp. enterica serovar Typhimurium Proteobacteria; Gammaproteobacteria; Enterobacterales; Enterobacteriaceae; Salmonella; Salmonella enterica; Salmonella enterica subsp. enterica MTGDNTLITSHGINRRDFMKLCAALAATMGLSSKAAAEMAESVSNPQRPPVIWIGAQECT WP_000145429.1 hydrogenase 2 small subunit [Salmonella enterica] Length: 372\nNumber of predicted TMHs: 1\nExp number of AAs in TMHs: 24.51309\nExp number, first 60 AAs: 0.2682\nTotal prob of N-in: 0.14402\noutside 1 331\nTMhelix 332 354\ninside 355 372

22280 GCF_001595065.1_CFSAN033865_01.0Salmonella enterica subsp. enterica serovar Typhimurium Proteobacteria; Gammaproteobacteria; Enterobacterales; Enterobacteriaceae; Salmonella; Salmonella enterica; Salmonella enterica subsp. enterica MTGDNTLITSHGINRRDFMKLCAALAATMGLSSKAAAEMAESVSNPQRPPVIWIGAQECT WP_000145429.1 hydrogenase 2 small subunit [Salmonella enterica] Length: 372\nNumber of predicted TMHs: 1\nExp number of AAs in TMHs: 24.51309\nExp number, first 60 AAs: 0.2682\nTotal prob of N-in: 0.14402\noutside 1 331\nTMhelix 332 354\ninside 355 372

22281 GCF_001595105.1_CFSAN033863_01.0Salmonella enterica subsp. enterica serovar Typhimurium Proteobacteria; Gammaproteobacteria; Enterobacterales; Enterobacteriaceae; Salmonella; Salmonella enterica; Salmonella enterica subsp. enterica MTGDNTLITSHGINRRDFMKLCAALAATMGLSSKAAAEMAESVSNPQRPPVIWIGAQECT WP_000145429.1 hydrogenase 2 small subunit [Salmonella enterica] Length: 372\nNumber of predicted TMHs: 1\nExp number of AAs in TMHs: 24.51309\nExp number, first 60 AAs: 0.2682\nTotal prob of N-in: 0.14402\noutside 1 331\nTMhelix 332 354\ninside 355 372

22282 GCF_001594855.1_CFSAN033877_01.0Salmonella enterica subsp. enterica serovar Typhimurium Proteobacteria; Gammaproteobacteria; Enterobacterales; Enterobacteriaceae; Salmonella; Salmonella enterica; Salmonella enterica subsp. enterica MNNEETFYQAMRRKGVTRRSFLKFCSLAATSLGLGAGMTPKIAWALENKPRIPVVWIHGL WP_001058312.1 hydrogenase [Salmonella enterica] Length: 372\nNumber of predicted TMHs: 2\nExp number of AAs in TMHs: 36.51738\nExp number, first 60 AAs: 19.72849\nTotal prob of N-in: 0.96170\nPOSSIBLE N-term signal sequence\ninside 1 19\nTMhelix 20 39\noutside 40 325\nTMhelix 326 348\ninside 349 372

22283 GCF_001595145.1_CFSAN033861_01.0Salmonella enterica subsp. enterica serovar Typhimurium Proteobacteria; Gammaproteobacteria; Enterobacterales; Enterobacteriaceae; Salmonella; Salmonella enterica; Salmonella enterica subsp. enterica MNNEETFYQAMRRKGVTRRSFLKFCSLAATSLGLGAGMTPKIAWALENKPRIPVVWIHGL WP_001058312.1 hydrogenase [Salmonella enterica] Length: 372\nNumber of predicted TMHs: 2\nExp number of AAs in TMHs: 36.51738\nExp number, first 60 AAs: 19.72849\nTotal prob of N-in: 0.96170\nPOSSIBLE N-term signal sequence\ninside 1 19\nTMhelix 20 39\noutside 40 325\nTMhelix 326 348\ninside 349 372

22284 GCF_001595305.1_CFSAN033851_01.0Salmonella enterica subsp. enterica serovar Typhimurium Proteobacteria; Gammaproteobacteria; Enterobacterales; Enterobacteriaceae; Salmonella; Salmonella enterica; Salmonella enterica subsp. enterica MTGDNTLITSHGINRRDFMKLCAALAATMGLSSKAAAEMAESVSNPQRPPVIWIGAQECT WP_000145429.1 hydrogenase 2 small subunit [Salmonella enterica] Length: 372\nNumber of predicted TMHs: 1\nExp number of AAs in TMHs: 24.51309\nExp number, first 60 AAs: 0.2682\nTotal prob of N-in: 0.14402\noutside 1 331\nTMhelix 332 354\ninside 355 372

22285 GCF_001595345.1_CFSAN033849_01.0Salmonella enterica subsp. enterica serovar Typhimurium Proteobacteria; Gammaproteobacteria; Enterobacterales; Enterobacteriaceae; Salmonella; Salmonella enterica; Salmonella enterica subsp. enterica MTGDNTLITSHGINRRDFMKLCAALAATMGLSSKAAAEMAESVSNPQRPPVIWIGAQECT WP_000145429.1 hydrogenase 2 small subunit [Salmonella enterica] Length: 372\nNumber of predicted TMHs: 1\nExp number of AAs in TMHs: 24.51309\nExp number, first 60 AAs: 0.2682\nTotal prob of N-in: 0.14402\noutside 1 331\nTMhelix 332 354\ninside 355 372

22286 GCF_001595305.1_CFSAN033851_01.0Salmonella enterica subsp. enterica serovar Typhimurium Proteobacteria; Gammaproteobacteria; Enterobacterales; Enterobacteriaceae; Salmonella; Salmonella enterica; Salmonella enterica subsp. enterica MNNEETFYQAMRRKGVTRRSFLKFCSLAATSLGLGAGMTPKIAWALENKPRIPVVWIHGL WP_001058312.1 hydrogenase [Salmonella enterica] Length: 372\nNumber of predicted TMHs: 2\nExp number of AAs in TMHs: 36.51738\nExp number, first 60 AAs: 19.72849\nTotal prob of N-in: 0.96170\nPOSSIBLE N-term signal sequence\ninside 1 19\nTMhelix 20 39\noutside 40 325\nTMhelix 326 348\ninside 349 372

22287 GCF_001614175.1_ASM161417v1 Salmonella enterica subsp. enterica serovar Typhimurium Proteobacteria; Gammaproteobacteria; Enterobacterales; Enterobacteriaceae; Salmonella; Salmonella enterica; Salmonella enterica subsp. enterica MNNEETFYQAMRRKGVTRRSFLKFCSLAATSLGLGAGMTPKIAWALENKPRIPVVWIHGL WP_001058312.1 hydrogenase [Salmonella enterica] Length: 372\nNumber of predicted TMHs: 2\nExp number of AAs in TMHs: 36.51738\nExp number, first 60 AAs: 19.72849\nTotal prob of N-in: 0.96170\nPOSSIBLE N-term signal sequence\ninside 1 19\nTMhelix 20 39\noutside 40 325\nTMhelix 326 348\ninside 349 372

22288 GCF_001692065.1_ASM169206v1 Salmonella enterica subsp. enterica serovar Typhimurium Proteobacteria; Gammaproteobacteria; Enterobacterales; Enterobacteriaceae; Salmonella; Salmonella enterica; Salmonella enterica subsp. enterica MTGDNTLITSHGINRRDFMKLCAALAATMGLSSKAAAEMAESVSNPQRPPVIWIGAQECT WP_000145429.1 hydrogenase 2 small subunit [Salmonella enterica] Length: 372\nNumber of predicted TMHs: 1\nExp number of AAs in TMHs: 24.51309\nExp number, first 60 AAs: 0.2682\nTotal prob of N-in: 0.14402\noutside 1 331\nTMhelix 332 354\ninside 355 372

22289 GCF_001692065.1_ASM169206v1 Salmonella enterica subsp. enterica serovar Typhimurium Proteobacteria; Gammaproteobacteria; Enterobacterales; Enterobacteriaceae; Salmonella; Salmonella enterica; Salmonella enterica subsp. enterica MNNEETFYQAMRRKGVTRRSFLKFCSLAATSLGLGAGMTPKIAWALENKPRIPVVWIHGL WP_001058312.1 hydrogenase [Salmonella enterica] Length: 372\nNumber of predicted TMHs: 2\nExp number of AAs in TMHs: 36.51738\nExp number, first 60 AAs: 19.72849\nTotal prob of N-in: 0.96170\nPOSSIBLE N-term signal sequence\ninside 1 19\nTMhelix 20 39\noutside 40 325\nTMhelix 326 348\ninside 349 372

22290 GCF_001756135.1_CFSAN008088 Salmonella enterica subsp. enterica serovar Typhimurium Proteobacteria; Gammaproteobacteria; Enterobacterales; Enterobacteriaceae; Salmonella; Salmonella enterica; Salmonella enterica subsp. enterica MTGDNTLITSHGINRRDFMKLCAALAATMGLSSKAAAEMAESVSNPQRPPVIWIGAQECT WP_000145429.1 hydrogenase 2 small subunit [Salmonella enterica] Length: 372\nNumber of

predicted TMHs: 1\nExp number of AAs in TMHs: 24.51309\nExp number, first 60 AAs: 0.2682\nTotal prob of N-in: 0.14402\noutside 1 331\nTMhelix 332 354\ninside 355 372

22291 GCF_001756135.1_CFSAN008088 Salmonella enterica subsp. enterica serovar Typhimurium Proteobacteria; Gammaproteobacteria; Enterobacterales; Enterobacteriaceae; Salmonella; Salmonella enterica; Salmonella enterica subsp. enterica MNNEETFYQAMRRKGVTRRSFLKFCSLAATSLGLGAGMTPKIAWALENKPRIPVVWIHGL WP_001058312.1 hydrogenase [Salmonella enterica] Length: 372\nNumber of predicted TMHs: 2\nExp number of AAs in TMHs: 36.51738\nExp number, first 60 AAs: 19.72849\nTotal prob of N-in: 0.96170\nPOSSIBLE N-term signal sequence\ninside 1 19\nTMhelix 20 39\noutside 40 325\nTMhelix 326 348\ninside 349 372

22292 GCF_001756105.1_CFSAN008085 Salmonella enterica subsp. enterica serovar Typhimurium Proteobacteria; Gammaproteobacteria; Enterobacterales; Enterobacteriaceae; Salmonella; Salmonella enterica; Salmonella enterica subsp. enterica MNNEETFYQAMRRKGVTRRSFLKFCSLAATSLGLGAGMTPKIAWALENKPRIPVVWIHGL WP_001058312.1 hydrogenase [Salmonella enterica] Length: 372\nNumber of predicted TMHs: 2\nExp number of AAs in TMHs: 36.51738\nExp number, first 60 AAs: 19.72849\nTotal prob of N-in: 0.96170\nPOSSIBLE N-term signal sequence\ninside 1 19\nTMhelix 20 39\noutside 40 325\nTMhelix 326 348\ninside 349 372

22293 GCF_001755865.1_CFSAN008084 Salmonella enterica subsp. enterica serovar Typhimurium Proteobacteria; Gammaproteobacteria; Enterobacterales; Enterobacteriaceae; Salmonella; Salmonella enterica; Salmonella enterica subsp. enterica MNNEETFYQAMRRKGVTRRSFLKFCSLAATSLGLGAGMTPKIAWALENKPRIPVVWIHGL WP_001058312.1 hydrogenase [Salmonella enterica] Length: 372\nNumber of predicted TMHs: 2\nExp number of AAs in TMHs: 36.51738\nExp number, first 60 AAs: 19.72849\nTotal prob of N-in: 0.96170\nPOSSIBLE N-term signal sequence\ninside 1 19\nTMhelix 20 39\noutside 40 325\nTMhelix 326 348\ninside 349 372

22294 GCF_001729225.1_ASM172922v1 Salmonella enterica subsp. enterica serovar Typhimurium Proteobacteria; Gammaproteobacteria; Enterobacterales; Enterobacteriaceae; Salmonella; Salmonella enterica; Salmonella enterica subsp. enterica MNNEETFYQAMRRKGVTRRSFLKFCSLAATSLGLGAGMTPKIAWALENKPRIPVVWIHGL WP_001058312.1 hydrogenase [Salmonella enterica] Length: 372\nNumber of predicted TMHs: 2\nExp number of AAs in TMHs: 36.51738\nExp number, first 60 AAs: 19.72849\nTotal prob of N-in: 0.96170\nPOSSIBLE N-term signal sequence\ninside 1 19\nTMhelix 20 39\noutside 40 325\nTMhelix 326 348\ninside 349 372

22295 GCF_001756185.1_CFSAN008072 Salmonella enterica subsp. enterica serovar Typhimurium Proteobacteria; Gammaproteobacteria; Enterobacterales; Enterobacteriaceae; Salmonella; Salmonella enterica; Salmonella enterica subsp. enterica MTGDNTLITSHGINRRDFMKLCAALAATMGLSSKAAAEMAESVSNPQRPPVIWIGAQECT WP_000145429.1 hydrogenase 2 small subunit [Salmonella enterica] Length: 372\nNumber of predicted TMHs: 1\nExp number of AAs in TMHs: 24.51309\nExp number, first 60 AAs: 0.2682\nTotal prob of N-in: 0.14402\noutside 1 331\nTMhelix 332 354\ninside 355 372

22296 GCF_001955375.1_ASM195537v1 Salmonella enterica subsp. enterica serovar Typhimurium Proteobacteria; Gammaproteobacteria; Enterobacterales; Enterobacteriaceae; Salmonella; Salmonella enterica; Salmonella enterica subsp. enterica MTGDNTLITSHGINRRDFMKLCAALAATMGLSSKAAAEMAESVSNPQRPPVIWIGAQECT WP_000145429.1 hydrogenase 2 small subunit [Salmonella enterica] Length: 372\nNumber of predicted TMHs: 1\nExp number of AAs in TMHs: 24.51309\nExp number, first 60 AAs: 0.2682\nTotal prob of N-in: 0.14402\noutside 1 331\nTMhelix 332 354\ninside 355 372

22297 GCF_001955315.1_ASM195531v1 Salmonella enterica subsp. enterica serovar Typhimurium Proteobacteria; Gammaproteobacteria; Enterobacterales; Enterobacteriaceae; Salmonella; Salmonella enterica; Salmonella enterica subsp. enterica MTGDNTLITSHGINRRDFMKLCAALAATMGLSSKAAAEMAESVSNPQRPPVIWIGAQECT WP_000145429.1 hydrogenase 2 small subunit [Salmonella enterica] Length: 372\nNumber of predicted TMHs: 1\nExp number of AAs in TMHs: 24.51309\nExp number, first 60 AAs: 0.2682\nTotal prob of N-in: 0.14402\noutside 1 331\nTMhelix 332 354\ninside 355 372

22298 GCF_001955395.1_ASM195539v1 Salmonella enterica subsp. enterica serovar Typhimurium Proteobacteria; Gammaproteobacteria; Enterobacterales; Enterobacteriaceae; Salmonella; Salmonella enterica; Salmonella enterica subsp. enterica MTGDNTLITSHGINRRDFMKLCAALAATMGLSSKAAAEMAESVSNPQRPPVIWIGAQECT WP_000145429.1 hydrogenase 2 small subunit [Salmonella enterica] Length: 372\nNumber of predicted TMHs: 1\nExp number of AAs in TMHs: 24.51309\nExp number, first 60 AAs: 0.2682\nTotal prob of N-in: 0.14402\noutside 1 331\nTMhelix 332 354\ninside 355 372

22299 GCF_001955455.1_ASM195545v1 Salmonella enterica subsp. enterica serovar Typhimurium Proteobacteria; Gammaproteobacteria; Enterobacterales; Enterobacteriaceae; Salmonella; Salmonella enterica;

Salmonella enterica subsp. enterica MTGDNTLITSHGINRRDFMKLCAALAATMGLSSKAAAEMAESVSNPQRPPVIWIGAQECT WP_000145429.1 hydrogenase 2 small subunit [Salmonella enterica] Length: 372\nNumber of predicted TMHs: 1\nExp number of AAs in TMHs: 24.51309\nExp number, first 60 AAs: 0.2682\nTotal prob of N-in: 0.14402\noutside 1 331\nTMhelix 332 354\ninside 355 372

22300 GCF_001955235.1_ASM195523v1 Salmonella enterica subsp. enterica serovar Typhimurium Proteobacteria; Gammaproteobacteria; Enterobacterales; Enterobacteriaceae; Salmonella; Salmonella enterica; Salmonella enterica subsp. enterica MNNEETFYQAMRRKGVTRRSFLKFCSLAATSLGLGAGMTPKIAWALENKPRIPVVWIHGL WP_001058312.1 hydrogenase [Salmonella enterica] Length: 372\nNumber of predicted TMHs: 2\nExp number of AAs in TMHs: 36.51738\nExp number, first 60 AAs: 19.72849\nTotal prob of N-in: 0.96170\nPOSSIBLE N-term signal sequence\ninside 1 19\nTMhelix 20 39\noutside 40 325\nTMhelix 326 348\ninside 349 372

22301 GCF_001955475.1_ASM195547v1 Salmonella enterica subsp. enterica serovar Typhimurium Proteobacteria; Gammaproteobacteria; Enterobacterales; Enterobacteriaceae; Salmonella; Salmonella enterica; Salmonella enterica subsp. enterica MTGDNTLITSHGINRRDFMKLCAALAATMGLSSKAAAEMAESVSNPQRPPVIWIGAQECT WP_000145429.1 hydrogenase 2 small subunit [Salmonella enterica] Length: 372\nNumber of predicted TMHs: 1\nExp number of AAs in TMHs: 24.51309\nExp number, first 60 AAs: 0.2682\nTotal prob of N-in: 0.14402\noutside 1 331\nTMhelix 332 354\ninside 355 372

22302 GCF_001989635.1_ASM198963v1 Salmonella enterica subsp. enterica serovar Typhimurium Proteobacteria; Gammaproteobacteria; Enterobacterales; Enterobacteriaceae; Salmonella; Salmonella enterica; Salmonella enterica subsp. enterica MTGDNTLITSHGINRRDFMKLCAALAATMGLSSKAAAEMAESVSNPQRPPVIWIGAQECT WP_000145429.1 hydrogenase 2 small subunit [Salmonella enterica] Length: 372\nNumber of predicted TMHs: 1\nExp number of AAs in TMHs: 24.51309\nExp number, first 60 AAs: 0.2682\nTotal prob of N-in: 0.14402\noutside 1 331\nTMhelix 332 354\ninside 355 372

22303 GCF_002029385.1_ASM202938v1 Salmonella enterica subsp. enterica serovar Typhimurium Proteobacteria; Gammaproteobacteria; Enterobacterales; Enterobacteriaceae; Salmonella; Salmonella enterica; Salmonella enterica subsp. enterica MNNEETFYQAMRRKGVTRRSFLKFCSLAATSLGLGAGMTPKIAWALENKPRIPVVWIHGL WP_001058312.1 hydrogenase [Salmonella enterica] Length: 372\nNumber of predicted TMHs: 2\nExp number of AAs in TMHs: 36.51738\nExp number, first 60 AAs: 19.72849\nTotal prob of N-in: 0.96170\nPOSSIBLE N-term signal sequence\ninside 1 19\nTMhelix 20 39\noutside 40 325\nTMhelix 326 348\ninside 349 372

22304 GCF_002029485.1_ASM202948v1 Salmonella enterica subsp. enterica serovar Typhimurium Proteobacteria; Gammaproteobacteria; Enterobacterales; Enterobacteriaceae; Salmonella; Salmonella enterica; Salmonella enterica subsp. enterica MTGDNTLITSHGINRRDFMKLCAALAATMGLSSKAAAEMAESVSNPQRPPVIWIGAQECT WP_000145429.1 hydrogenase 2 small subunit [Salmonella enterica] Length: 372\nNumber of predicted TMHs: 1\nExp number of AAs in TMHs: 24.51309\nExp number, first 60 AAs: 0.2682\nTotal prob of N-in: 0.14402\noutside 1 331\nTMhelix 332 354\ninside 355 372

22305 GCF_002029445.1_ASM202944v1 Salmonella enterica subsp. enterica serovar Typhimurium Proteobacteria; Gammaproteobacteria; Enterobacterales; Enterobacteriaceae; Salmonella; Salmonella enterica; Salmonella enterica subsp. enterica MTGDNTLITSHGINRRDFMKLCAALAATMGLSSKAAAEMAESVSNPQRPPVIWIGAQECT WP_000145429.1 hydrogenase 2 small subunit [Salmonella enterica] Length: 372\nNumber of predicted TMHs: 1\nExp number of AAs in TMHs: 24.51309\nExp number, first 60 AAs: 0.2682\nTotal prob of N-in: 0.14402\noutside 1 331\nTMhelix 332 354\ninside 355 372

22306 GCF_002029675.1_ASM202967v1 Salmonella enterica subsp. enterica serovar Typhimurium Proteobacteria; Gammaproteobacteria; Enterobacterales; Enterobacteriaceae; Salmonella; Salmonella enterica; Salmonella enterica subsp. enterica MTGDNTLITSHGINRRDFMKLCAALAATMGLSSKAAAEMAESVSNPQRPPVIWIGAQECT WP_000145429.1 hydrogenase 2 small subunit [Salmonella enterica] Length: 372\nNumber of predicted TMHs: 1\nExp number of AAs in TMHs: 24.51309\nExp number, first 60 AAs: 0.2682\nTotal prob of N-in: 0.14402\noutside 1 331\nTMhelix 332 354\ninside 355 372

22307 GCF_002029715.1_ASM202971v1 Salmonella enterica subsp. enterica serovar Typhimurium Proteobacteria; Gammaproteobacteria; Enterobacterales; Enterobacteriaceae; Salmonella; Salmonella enterica; Salmonella enterica subsp. enterica MTGDNTLITSHGINRRDFMKLCAALAATMGLSSKAAAEMAESVSNPQRPPVIWIGAQECT WP_000145429.1 hydrogenase 2 small subunit [Salmonella enterica] Length: 372\nNumber of predicted TMHs: 1\nExp number of AAs in TMHs: 24.51309\nExp number, first 60 AAs: 0.2682\nTotal prob of N-in: 0.14402\noutside 1 331\nTMhelix 332 354\ninside 355 372

22308 GCF_002029485.1_ASM202948v1 Salmonella enterica subsp. enterica serovar Typhimurium Proteobacteria; Gammaproteobacteria; Enterobacterales; Enterobacteriaceae; Salmonella; Salmonella enterica; Salmonella enterica subsp. enterica MNNEETFYQAMRRKGVTRRSFLKFCSLAATSLGLGAGMTPKIAWALENKPRIPVVWIHGL WP_001058312.1 hydrogenase [Salmonella enterica] Length: 372\nNumber of predicted TMHs: 2\nExp number of AAs in TMHs: 36.51738\nExp number, first 60 AAs: 19.72849\nTotal prob of N-in: 0.96170\nPOSSIBLE N-term signal sequence\ninside 1 19\nTMhelix 20 39\noutside 40 325\nTMhelix 326 348\ninside 349 372

22309 GCF_002030385.1_ASM203038v1 Salmonella enterica subsp. enterica serovar Typhimurium Proteobacteria; Gammaproteobacteria; Enterobacterales; Enterobacteriaceae; Salmonella; Salmonella enterica; Salmonella enterica subsp. enterica MTGDNTLITSHGINRRDFMKLCAALAATMGLSSKAAAEMAESVSNPQRPPVIWIGAQECT WP_000145429.1 hydrogenase 2 small subunit [Salmonella enterica] Length: 372\nNumber of predicted TMHs: 1\nExp number of AAs in TMHs: 24.51309\nExp number, first 60 AAs: 0.2682\nTotal prob of N-in: 0.14402\noutside 1 331\nTMhelix 332 354\ninside 355 372

22310 GCF_002030565.1_ASM203056v1 Salmonella enterica subsp. enterica serovar Typhimurium Proteobacteria; Gammaproteobacteria; Enterobacterales; Enterobacteriaceae; Salmonella; Salmonella enterica; Salmonella enterica subsp. enterica MTGDNTLITSHGINRRDFMKLCAALAATMGLSSKAAAEMAESVSNPQRPPVIWIGAQECT WP_000145429.1 hydrogenase 2 small subunit [Salmonella enterica] Length: 372\nNumber of predicted TMHs: 1\nExp number of AAs in TMHs: 24.51309\nExp number, first 60 AAs: 0.2682\nTotal prob of N-in: 0.14402\noutside 1 331\nTMhelix 332 354\ninside 355 372

22311 GCF_002030565.1_ASM203056v1 Salmonella enterica subsp. enterica serovar Typhimurium Proteobacteria; Gammaproteobacteria; Enterobacterales; Enterobacteriaceae; Salmonella; Salmonella enterica; Salmonella enterica subsp. enterica MNNEETFYQAMRRKGVTRRSFLKFCSLAATSLGLGAGMTPKIAWALENKPRIPVVWIHGL WP_001058312.1 hydrogenase [Salmonella enterica] Length: 372\nNumber of predicted TMHs: 2\nExp number of AAs in TMHs: 36.51738\nExp number, first 60 AAs: 19.72849\nTotal prob of N-in: 0.96170\nPOSSIBLE N-term signal sequence\ninside 1 19\nTMhelix 20 39\noutside 40 325\nTMhelix 326 348\ninside 349 372

22312 GCF_002035115.1_ASM203511v1 Salmonella enterica subsp. enterica serovar Typhimurium Proteobacteria; Gammaproteobacteria; Enterobacterales; Enterobacteriaceae; Salmonella; Salmonella enterica; Salmonella enterica subsp. enterica MTGDNTLITSHGINRRDFMKLCAALAATMGLSSKAAAEMAESVSNPQRPPVIWIGAQECT WP_000145429.1 hydrogenase 2 small subunit [Salmonella enterica] Length: 372\nNumber of predicted TMHs: 1\nExp number of AAs in TMHs: 24.51309\nExp number, first 60 AAs: 0.2682\nTotal prob of N-in: 0.14402\noutside 1 331\nTMhelix 332 354\ninside 355 372

22313 GCF_002035035.1_ASM203503v1 Salmonella enterica subsp. enterica serovar Typhimurium Proteobacteria; Gammaproteobacteria; Enterobacterales; Enterobacteriaceae; Salmonella; Salmonella enterica; Salmonella enterica subsp. enterica MTGDNTLITSHGINRRDFMKLCAALAATMGLSSKAAAEMAESVSNPQRPPVIWIGAQECT WP_000145429.1 hydrogenase 2 small subunit [Salmonella enterica] Length: 372\nNumber of predicted TMHs: 1\nExp number of AAs in TMHs: 24.51309\nExp number, first 60 AAs: 0.2682\nTotal prob of N-in: 0.14402\noutside 1 331\nTMhelix 332 354\ninside 355 372

22314 GCF_002035035.1_ASM203503v1 Salmonella enterica subsp. enterica serovar Typhimurium Proteobacteria; Gammaproteobacteria; Enterobacterales; Enterobacteriaceae; Salmonella; Salmonella enterica; Salmonella enterica subsp. enterica MNNEETFYQAMRRKGVTRRSFLKFCSLAATSLGLGAGMTPKIAWALENKPRIPVVWIHGL WP_001058312.1 hydrogenase [Salmonella enterica] Length: 372\nNumber of predicted TMHs: 2\nExp number of AAs in TMHs: 36.51738\nExp number, first 60 AAs: 19.72849\nTotal prob of N-in: 0.96170\nPOSSIBLE N-term signal sequence\ninside 1 19\nTMhelix 20 39\noutside 40 325\nTMhelix 326 348\ninside 349 372

22315 GCF_002035055.1_ASM203505v1 Salmonella enterica subsp. enterica serovar Typhimurium Proteobacteria; Gammaproteobacteria; Enterobacterales; Enterobacteriaceae; Salmonella; Salmonella enterica; Salmonella enterica subsp. enterica MTGDNTLITSHGINRRDFMKLCAALAATMGLSSKAAAEMAESVSNPQRPPVIWIGAQECT WP_000145429.1 hydrogenase 2 small subunit [Salmonella enterica] Length: 372\nNumber of predicted TMHs: 1\nExp number of AAs in TMHs: 24.51309\nExp number, first 60 AAs: 0.2682\nTotal prob of N-in: 0.14402\noutside 1 331\nTMhelix 332 354\ninside 355 372

22316 GCF_002036205.1_ASM203620v1 Salmonella enterica subsp. enterica serovar Typhimurium Proteobacteria; Gammaproteobacteria; Enterobacterales; Enterobacteriaceae; Salmonella; Salmonella enterica; Salmonella enterica subsp. enterica MNNEETFYQAMRRKGVTRRSFLKFCSLAATSLGLGAGMTPKIAWALENKPRIPVVWIHGL WP_001058312.1 hydrogenase [Salmonella enterica] Length: 372\nNumber of predicted TMHs: 2\nExp

number of AAs in TMHs: 36.51738\nExp number, first 60 AAs: 19.72849\nTotal prob of N-in: 0.96170\nPOSSIBLE N-term signal sequence\ninside 1 19\nTMhelix 20 39\noutside 40 325\nTMhelix 326 348\ninside 349 372

22317 GCF_002056945.1_ASM205694v1 Salmonella enterica subsp. enterica serovar Typhimurium Proteobacteria; Gammaproteobacteria; Enterobacterales; Enterobacteriaceae; Salmonella; Salmonella enterica; Salmonella enterica subsp. enterica MTGDNTLITSHGINRRDFMKLCAALAATMGLSSKAAAEMAESVSNPQRPPVIWIGAQECT WP_000145429.1 hydrogenase 2 small subunit [Salmonella enterica] Length: 372\nNumber of predicted TMHs: 1\nExp number of AAs in TMHs: 24.51309\nExp number, first 60 AAs: 0.2682\nTotal prob of N-in: 0.14402\noutside 1 331\nTMhelix 332 354\ninside 355 372

22318 GCF_002056945.1_ASM205694v1 Salmonella enterica subsp. enterica serovar Typhimurium Proteobacteria; Gammaproteobacteria; Enterobacterales; Enterobacteriaceae; Salmonella; Salmonella enterica; Salmonella enterica subsp. enterica MNNEETFYQAMRRKGVTRRSFLKFCSLAATSLGLGAGMTPKIAWALENKPRIPVVWIHGL WP_001058312.1 hydrogenase [Salmonella enterica] Length: 372\nNumber of predicted TMHs: 2\nExp number of AAs in TMHs: 36.51738\nExp number, first 60 AAs: 19.72849\nTotal prob of N-in: 0.96170\nPOSSIBLE N-term signal sequence\ninside 1 19\nTMhelix 20 39\noutside 40 325\nTMhelix 326 348\ninside 349 372

22319 GCF_000636135.1_ASM63613v1 Salmonella enterica subsp. enterica serovar Typhimurium Proteobacteria; Gammaproteobacteria; Enterobacterales; Enterobacteriaceae; Salmonella; Salmonella enterica; Salmonella enterica subsp. enterica MTGDNTLITSHGINRRDFMKLCAALAATMGLSSKAAAEMAESVSNPQRPPVIWIGAQECT WP_000145429.1 hydrogenase 2 small subunit [Salmonella enterica] Length: 372\nNumber of predicted TMHs: 1\nExp number of AAs in TMHs: 24.51309\nExp number, first 60 AAs: 0.2682\nTotal prob of N-in: 0.14402\noutside 1 331\nTMhelix 332 354\ninside 355 372

22320 GCF_001247185.1_Salmonella_enterica_CVM_N51299_v1.0 Salmonella enterica subsp. enterica serovar Typhimurium Proteobacteria; Gammaproteobacteria; Enterobacterales; Enterobacteriaceae; Salmonella; Salmonella enterica; Salmonella enterica subsp. enterica MNNEETFYQAMRRKGVTRRSFLKFCSLAATSLGLGAGMTPKIAWALENKPRIPVVWIHGL WP_001058312.1 hydrogenase [Salmonella enterica] Length: 372\nNumber of predicted TMHs: 2\nExp number of AAs in TMHs: 36.51738\nExp number, first 60 AAs: 19.72849\nTotal prob of N-in: 0.96170\nPOSSIBLE N-term signal sequence\ninside 1 19\nTMhelix 20 39\noutside 40 325\nTMhelix 326 348\ninside 349 372

22321 GCF_001295635.1_ASM129563v1 Salmonella enterica subsp. enterica serovar Typhimurium Proteobacteria; Gammaproteobacteria; Enterobacterales; Enterobacteriaceae; Salmonella; Salmonella enterica; Salmonella enterica subsp. enterica MNNEETFYQAMRRKGVTRRSFLKFCSLAATSLGLGAGMTPKIAWALENKPRIPVVWIHGL WP_001058312.1 hydrogenase [Salmonella enterica] Length: 372\nNumber of predicted TMHs: 2\nExp number of AAs in TMHs: 36.51738\nExp number, first 60 AAs: 19.72849\nTotal prob of N-in: 0.96170\nPOSSIBLE N-term signal sequence\ninside 1 19\nTMhelix 20 39\noutside 40 325\nTMhelix 326 348\ninside 349 372

22322 GCF_001295675.1_ASM129567v1 Salmonella enterica subsp. enterica serovar Typhimurium Proteobacteria; Gammaproteobacteria; Enterobacterales; Enterobacteriaceae; Salmonella; Salmonella enterica; Salmonella enterica subsp. enterica MTGDNTLITSHGINRRDFMKLCAALAATMGLSSKAAAEMAESVSNPQRPPVIWIGAQECT WP_000145429.1 hydrogenase 2 small subunit [Salmonella enterica] Length: 372\nNumber of predicted TMHs: 1\nExp number of AAs in TMHs: 24.51309\nExp number, first 60 AAs: 0.2682\nTotal prob of N-in: 0.14402\noutside 1 331\nTMhelix 332 354\ninside 355 372

22323 GCF_001295555.1_ASM129555v1 Salmonella enterica subsp. enterica serovar Typhimurium Proteobacteria; Gammaproteobacteria; Enterobacterales; Enterobacteriaceae; Salmonella; Salmonella enterica; Salmonella enterica subsp. enterica MNNEETFYQAMRRKGVTRRSFLKFCSLAATSLGLGAGMTPKIAWALENKPRIPVVWIHGL WP_001058312.1 hydrogenase [Salmonella enterica] Length: 372\nNumber of predicted TMHs: 2\nExp number of AAs in TMHs: 36.51738\nExp number, first 60 AAs: 19.72849\nTotal prob of N-in: 0.96170\nPOSSIBLE N-term signal sequence\ninside 1 19\nTMhelix 20 39\noutside 40 325\nTMhelix 326 348\ninside 349 372

22324 GCF_001295405.1_ASM129540v1 Salmonella enterica subsp. enterica serovar Typhimurium Proteobacteria; Gammaproteobacteria; Enterobacterales; Enterobacteriaceae; Salmonella; Salmonella enterica; Salmonella enterica subsp. enterica MNNEETFYQAMRRKGVTRRSFLKFCSLAATSLGLGAGMTPKIAWALENKPRIPVVWIHGL WP_001058312.1 hydrogenase [Salmonella enterica] Length: 372\nNumber of predicted TMHs: 2\nExp number of AAs in TMHs: 36.51738\nExp number, first 60 AAs: 19.72849\nTotal prob of N-in: 0.96170\nPOSSIBLE N-term signal sequence\ninside 1 19\nTMhelix 20 39\noutside 40 325\nTMhelix 326 348\ninside 349 372

22325 GCF_001295555.1_ASM129555v1 Salmonella enterica subsp. enterica serovar Typhimurium Proteobacteria; Gammaproteobacteria; Enterobacterales; Enterobacteriaceae; Salmonella; Salmonella enterica; Salmonella enterica subsp. enterica MTGDNTLITSHGINRRDFMKLCAALAATMGLSSKAAAEMAESVSNPQRPPVIWIGAQECT WP_000145429.1 hydrogenase 2 small subunit [Salmonella enterica] Length: 372\nNumber of predicted TMHs: 1\nExp number of AAs in TMHs: 24.51309\nExp number, first 60 AAs: 0.2682\nTotal prob of N-in: 0.14402\noutside 1 331\nTMhelix 332 354\ninside 355 372

22326 GCF_001295405.1_ASM129540v1 Salmonella enterica subsp. enterica serovar Typhimurium Proteobacteria; Gammaproteobacteria; Enterobacterales; Enterobacteriaceae; Salmonella; Salmonella enterica; Salmonella enterica subsp. enterica MTGDNTLITSHGINRRDFMKLCAALAATMGLSSKAAAEMAESVSNPQRPPVIWIGAQECT WP_000145429.1 hydrogenase 2 small subunit [Salmonella enterica] Length: 372\nNumber of predicted TMHs: 1\nExp number of AAs in TMHs: 24.51309\nExp number, first 60 AAs: 0.2682\nTotal prob of N-in: 0.14402\noutside 1 331\nTMhelix 332 354\ninside 355 372

22327 GCF_001295725.1_ASM129572v1 Salmonella enterica subsp. enterica serovar Typhimurium Proteobacteria; Gammaproteobacteria; Enterobacterales; Enterobacteriaceae; Salmonella; Salmonella enterica; Salmonella enterica subsp. enterica MTGDNTLITSHGINRRDFMKLCAALAATMGLSSKAAAEMAESVSNPQRPPVIWIGAQECT WP_000145429.1 hydrogenase 2 small subunit [Salmonella enterica] Length: 372\nNumber of predicted TMHs: 1\nExp number of AAs in TMHs: 24.51309\nExp number, first 60 AAs: 0.2682\nTotal prob of N-in: 0.14402\noutside 1 331\nTMhelix 332 354\ninside 355 372

22328 GCF_001271965.1_Salmonella_enterica_CVM_N43825_v1.0 Salmonella enterica subsp. enterica serovar Typhimurium Proteobacteria; Gammaproteobacteria; Enterobacterales; Enterobacteriaceae; Salmonella; Salmonella enterica; Salmonella enterica subsp. enterica MNNEETFYQAMRRKGVTRRSFLKFCSLAATSLGLGAGMTPKIAWALENKPRIPVVWIHGL WP_001058312.1 hydrogenase [Salmonella enterica] Length: 372\nNumber of predicted TMHs: 2\nExp number of AAs in TMHs: 36.51738\nExp number, first 60 AAs: 19.72849\nTotal prob of N-in: 0.96170\nPOSSIBLE N-term signal sequence\ninside 1 19\nTMhelix 20 39\noutside 40 325\nTMhelix 326 348\ninside 349 372

22329 GCF_001272535.1_Salmonella_enterica_CVM_N51314_v1.0 Salmonella enterica subsp. enterica serovar Typhimurium Proteobacteria; Gammaproteobacteria; Enterobacterales; Enterobacteriaceae; Salmonella; Salmonella enterica; Salmonella enterica subsp. enterica MNNEETFYQAMRRKGVTRRSFLKFCSLAATSLGLGAGMTPKIAWALENKPRIPVVWIHGL WP_001058312.1 hydrogenase [Salmonella enterica] Length: 372\nNumber of predicted TMHs: 2\nExp number of AAs in TMHs: 36.51738\nExp number, first 60 AAs: 19.72849\nTotal prob of N-in: 0.96170\nPOSSIBLE N-term signal sequence\ninside 1 19\nTMhelix 20 39\noutside 40 325\nTMhelix 326 348\ninside 349 372

22330 GCF_001271965.1_Salmonella_enterica_CVM_N43825_v1.0 Salmonella enterica subsp. enterica serovar Typhimurium Proteobacteria; Gammaproteobacteria; Enterobacterales; Enterobacteriaceae; Salmonella; Salmonella enterica; Salmonella enterica subsp. enterica MTGDNTLITSHGINRRDFMKLCAALAATMGLSSKAAAEMAESVSNPQRPPVIWIGAQECT WP_000145429.1 hydrogenase 2 small subunit [Salmonella enterica] Length: 372\nNumber of predicted TMHs: 1\nExp number of AAs in TMHs: 24.51309\nExp number, first 60 AAs: 0.2682\nTotal prob of N-in: 0.14402\noutside 1 331\nTMhelix 332 354\ninside 355 372

22331 GCF_001246705.1_Salmonella_enterica_CVM_N51276_v1.0 Salmonella enterica subsp. enterica serovar Typhimurium Proteobacteria; Gammaproteobacteria; Enterobacterales; Enterobacteriaceae; Salmonella; Salmonella enterica; Salmonella enterica subsp. enterica MNNEETFYQAMRRKGVTRRSFLKFCSLAATSLGLGAGMTPKIAWALENKPRIPVVWIHGL WP_001058312.1 hydrogenase [Salmonella enterica] Length: 372\nNumber of predicted TMHs: 2\nExp number of AAs in TMHs: 36.51738\nExp number, first 60 AAs: 19.72849\nTotal prob of N-in: 0.96170\nPOSSIBLE N-term signal sequence\ninside 1 19\nTMhelix 20 39\noutside 40 325\nTMhelix 326 348\ninside 349 372

22332 GCF_001246705.1_Salmonella_enterica_CVM_N51276_v1.0 Salmonella enterica subsp. enterica serovar Typhimurium Proteobacteria; Gammaproteobacteria; Enterobacterales; Enterobacteriaceae; Salmonella; Salmonella enterica; Salmonella enterica subsp. enterica MTGDNTLITSHGINRRDFMKLCAALAATMGLSSKAAAEMAESVSNPQRPPVIWIGAQECT WP_000145429.1 hydrogenase 2 small subunit [Salmonella enterica] Length: 372\nNumber of predicted TMHs: 1\nExp number of AAs in TMHs: 24.51309\nExp number, first 60 AAs: 0.2682\nTotal prob of N-in: 0.14402\noutside 1 331\nTMhelix 332 354\ninside 355 372

22333 GCF_001246875.1_Salmonella_enterica_CVM_N51283_v1.0 Salmonella enterica subsp. enterica serovar Typhimurium Proteobacteria; Gammaproteobacteria; Enterobacterales; Enterobacteriaceae; Salmonella; Salmonella enterica; Salmonella enterica subsp. enterica MNNEETFYQAMRRKGVTRRSFLKFCSLAATSLGLGAGMTPKIAWALENKPRIPVVWIHGL WP_001058312.1 hydrogenase [Salmonella enterica] Length: 372\nNumber of predicted TMHs: 2\nExp number of AAs in TMHs: 36.51738\nExp number, first 60 AAs: 19.72849\nTotal prob of N-in: 0.96170\nPOSSIBLE N-term signal sequence\ninside 1 19\nTMhelix 20 39\noutside 40 325\nTMhelix 326 348\ninside 349 372

22334 GCF_000993725.1_ASM99372v1 Salmonella enterica subsp. enterica serovar Typhimurium Proteobacteria; Gammaproteobacteria; Enterobacterales; Enterobacteriaceae; Salmonella; Salmonella enterica; Salmonella enterica subsp. enterica MNNEETFYQAMRRKGVTRRSFLKFCSLAATSLGLGAGMTPKIAWALENKPRIPVVWIHGL WP_001058311.1 hydrogenase [Salmonella enterica] Length: 372\nNumber of predicted TMHs: 2\nExp number of AAs in TMHs: 37.4699899999999\nExp number, first 60 AAs: 19.72699\nTotal prob of N-in: 0.96171\nPOSSIBLE N-term signal sequence\ninside 1 19\nTMhelix 20 39\noutside 40 325\nTMhelix 326 348\ninside 349 372

22335 GCF_001272535.1_Salmonella_enterica_CVM_N51314_v1.0 Salmonella enterica subsp. enterica serovar Typhimurium Proteobacteria; Gammaproteobacteria; Enterobacterales; Enterobacteriaceae; Salmonella; Salmonella enterica; Salmonella enterica subsp. enterica MTGDNTLITSHGINRRDFMKLCAALAATMGLSSKAAAEMAESVSNPQRPPVIWIGAQECT WP_000145429.1 hydrogenase 2 small subunit [Salmonella enterica] Length: 372\nNumber of predicted TMHs: 1\nExp number of AAs in TMHs: 24.51309\nExp number, first 60 AAs: 0.2682\nTotal prob of N-in: 0.14402\noutside 1 331\nTMhelix 332 354\ninside 355 372

22336 GCF_000993725.1_ASM99372v1 Salmonella enterica subsp. enterica serovar Typhimurium Proteobacteria; Gammaproteobacteria; Enterobacterales; Enterobacteriaceae; Salmonella; Salmonella enterica; Salmonella enterica subsp. enterica MTGDNTLITSHGINRRDFMKLCAALAATMGLSSKAAAEMAESVSNPQRPPVIWIGAQECT WP_000145429.1 hydrogenase 2 small subunit [Salmonella enterica] Length: 372\nNumber of predicted TMHs: 1\nExp number of AAs in TMHs: 24.51309\nExp number, first 60 AAs: 0.2682\nTotal prob of N-in: 0.14402\noutside 1 331\nTMhelix 332 354\ninside 355 372

22337 GCF_000973885.1_SALH-394-2.id3_v1.0 Salmonella enterica subsp. enterica serovar Typhimurium Proteobacteria; Gammaproteobacteria; Enterobacterales; Enterobacteriaceae; Salmonella; Salmonella enterica; Salmonella enterica subsp. enterica MNNEETFYQAMRRKGVTRRSFLKFCSLAATSLGLGAGMTPKIAWALENKPRIPVVWIHGL WP_001058312.1 hydrogenase [Salmonella enterica] Length: 372\nNumber of predicted TMHs: 2\nExp number of AAs in TMHs: 36.51738\nExp number, first 60 AAs: 19.72849\nTotal prob of N-in: 0.96170\nPOSSIBLE N-term signal sequence\ninside 1 19\nTMhelix 20 39\noutside 40 325\nTMhelix 326 348\ninside 349 372

22338 GCF_000973885.1_SALH-394-2.id3_v1.0 Salmonella enterica subsp. enterica serovar Typhimurium Proteobacteria; Gammaproteobacteria; Enterobacterales; Enterobacteriaceae; Salmonella; Salmonella enterica; Salmonella enterica subsp. enterica MTGDNTLITSHGINRRDFMKLCAALAATMGLSSKAAAEMAESVSNPQRPPVIWIGAQECT WP_000145429.1 hydrogenase 2 small subunit [Salmonella enterica] Length: 372\nNumber of predicted TMHs: 1\nExp number of AAs in TMHs: 24.51309\nExp number, first 60 AAs: 0.2682\nTotal prob of N-in: 0.14402\noutside 1 331\nTMhelix 332 354\ninside 355 372

22339 GCF_001271645.1_Salmonella_enterica_CVM_N46810_v1.0 Salmonella enterica subsp. enterica serovar Typhimurium Proteobacteria; Gammaproteobacteria; Enterobacterales; Enterobacteriaceae; Salmonella; Salmonella enterica; Salmonella enterica subsp. enterica MNNEETFYQAMRRKGVTRRSFLKFCSLAATSLGLGAGMTPKIAWALENKPRIPVVWIHGL WP_001058312.1 hydrogenase [Salmonella enterica] Length: 372\nNumber of predicted TMHs: 2\nExp number of AAs in TMHs: 36.51738\nExp number, first 60 AAs: 19.72849\nTotal prob of N-in: 0.96170\nPOSSIBLE N-term signal sequence\ninside 1 19\nTMhelix 20 39\noutside 40 325\nTMhelix 326 348\ninside 349 372

22340 GCF_001271645.1_Salmonella_enterica_CVM_N46810_v1.0 Salmonella enterica subsp. enterica serovar Typhimurium Proteobacteria; Gammaproteobacteria; Enterobacterales; Enterobacteriaceae; Salmonella; Salmonella enterica; Salmonella enterica subsp. enterica MTGDNTLITSHGINRRDFMKLCAALAATMGLSSKAAAEMAESVSNPQRPPVIWIGAQECT WP_000145429.1 hydrogenase 2 small subunit [Salmonella enterica] Length: 372\nNumber of predicted TMHs: 1\nExp number of AAs in TMHs: 24.51309\nExp number, first 60 AAs: 0.2682\nTotal prob of N-in: 0.14402\noutside 1 331\nTMhelix 332 354\ninside 355 372

22341 GCF_000715155.1_ASM71515v2 Salmonella enterica subsp. enterica serovar Typhimurium Proteobacteria; Gammaproteobacteria; Enterobacterales; Enterobacteriaceae; Salmonella; Salmonella enterica; Salmonella enterica subsp. enterica MNNEETFYQAMRRKGVTRRSFLKFCSLAATSLGLGAGMTPKIAWALENKPRIPVVWIHGL WP_001058312.1 hydrogenase [Salmonella enterica] Length: 372\nNumber of predicted TMHs: 2\nExp number of AAs in TMHs: 36.51738\nExp number, first 60 AAs: 19.72849\nTotal prob of N-in: 0.96170\nPOSSIBLE N-term signal sequence\ninside 1 19\nTMhelix 20 39\noutside 40 325\nTMhelix 326 348\ninside 349 372

22342 GCF_001295425.1_ASM129542v1 Salmonella enterica subsp. enterica serovar Typhimurium Proteobacteria; Gammaproteobacteria; Enterobacterales; Enterobacteriaceae; Salmonella; Salmonella enterica; Salmonella enterica subsp. enterica MNNEETFYQAMRRKGVTRRSFLKFCSLAATSLGLGAGMTPKIAWALENKPRIPVVWIHGL WP_001058312.1 hydrogenase [Salmonella enterica] Length: 372\nNumber of predicted TMHs: 2\nExp number of AAs in TMHs: 36.51738\nExp number, first 60 AAs: 19.72849\nTotal prob of N-in: 0.96170\nPOSSIBLE N-term signal sequence\ninside 1 19\nTMhelix 20 39\noutside 40 325\nTMhelix 326 348\ninside 349 372

22343 GCF_001246875.1_Salmonella_enterica_CVM_N51283_v1.0 Salmonella enterica subsp. enterica serovar Typhimurium Proteobacteria; Gammaproteobacteria; Enterobacterales; Enterobacteriaceae; Salmonella; Salmonella enterica; Salmonella enterica subsp. enterica MTGDNTLITSHGINRRDFMKLCAALAATMGLSSKAAAEMAESVSNPQRPPVIWIGAQECT WP_000145429.1 hydrogenase 2 small subunit [Salmonella enterica] Length: 372\nNumber of predicted TMHs: 1\nExp number of AAs in TMHs: 24.51309\nExp number, first 60 AAs: 0.2682\nTotal prob of N-in: 0.14402\noutside 1 331\nTMhelix 332 354\ninside 355 372

22344 GCF_001295635.1_ASM129563v1 Salmonella enterica subsp. enterica serovar Typhimurium Proteobacteria; Gammaproteobacteria; Enterobacterales; Enterobacteriaceae; Salmonella; Salmonella enterica; Salmonella enterica subsp. enterica MTGDNTLITSHGINRRDFMKLCAALAATMGLSSKAAAEMAESVSNPQRPPVIWIGAQECT WP_000145429.1 hydrogenase 2 small subunit [Salmonella enterica] Length: 372\nNumber of predicted TMHs: 1\nExp number of AAs in TMHs: 24.51309\nExp number, first 60 AAs: 0.2682\nTotal prob of N-in: 0.14402\noutside 1 331\nTMhelix 332 354\ninside 355 372

22345 GCF_000784245.1_ASM78424v1 Salmonella enterica subsp. enterica serovar Typhimurium Proteobacteria; Gammaproteobacteria; Enterobacterales; Enterobacteriaceae; Salmonella; Salmonella enterica; Salmonella enterica subsp. enterica MTGDNTLITSHGINRRDFMKLCAALAATMGLSSKAAAEMAESVSNPQRPPVIWIGAQECT WP_000145429.1 hydrogenase 2 small subunit [Salmonella enterica] Length: 372\nNumber of predicted TMHs: 1\nExp number of AAs in TMHs: 24.51309\nExp number, first 60 AAs: 0.2682\nTotal prob of N-in: 0.14402\noutside 1 331\nTMhelix 332 354\ninside 355 372

22346 GCF_000784225.1_ASM78422v1 Salmonella enterica subsp. enterica serovar Typhimurium Proteobacteria; Gammaproteobacteria; Enterobacterales; Enterobacteriaceae; Salmonella; Salmonella enterica; Salmonella enterica subsp. enterica MTGDNTLITSHGINRRDFMKLCAALAATMGLSSKAAAEMAESVSNPQRPPVIWIGAQECT WP_000145429.1 hydrogenase 2 small subunit [Salmonella enterica] Length: 372\nNumber of predicted TMHs: 1\nExp number of AAs in TMHs: 24.51309\nExp number, first 60 AAs: 0.2682\nTotal prob of N-in: 0.14402\noutside 1 331\nTMhelix 332 354\ninside 355 372

22347 GCF_000784225.1_ASM78422v1 Salmonella enterica subsp. enterica serovar Typhimurium Proteobacteria; Gammaproteobacteria; Enterobacterales; Enterobacteriaceae; Salmonella; Salmonella enterica; Salmonella enterica subsp. enterica MNNEETFYQAMRRKGVTRRSFLKFCSLAATSLGLGAGMTPKIAWALENKPRIPVVWIHGL WP_001058312.1 hydrogenase [Salmonella enterica] Length: 372\nNumber of predicted TMHs: 2\nExp number of AAs in TMHs: 36.51738\nExp number, first 60 AAs: 19.72849\nTotal prob of N-in: 0.96170\nPOSSIBLE N-term signal sequence\ninside 1 19\nTMhelix 20 39\noutside 40 325\nTMhelix 326 348\ninside 349 372

22348 GCF_000784245.1_ASM78424v1 Salmonella enterica subsp. enterica serovar Typhimurium Proteobacteria; Gammaproteobacteria; Enterobacterales; Enterobacteriaceae; Salmonella; Salmonella enterica; Salmonella enterica subsp. enterica MNNEETFYQAMRRKGVTRRSFLKFCSLAATSLGLGAGMTPKIAWALENKPRIPVVWIHGL WP_001058312.1 hydrogenase [Salmonella enterica] Length: 372\nNumber of predicted TMHs: 2\nExp number of AAs in TMHs: 36.51738\nExp number, first 60 AAs: 19.72849\nTotal prob of N-in: 0.96170\nPOSSIBLE N-term signal sequence\ninside 1 19\nTMhelix 20 39\noutside 40 325\nTMhelix 326 348\ninside 349 372

22349 GCF_000636135.1_ASM63613v1 Salmonella enterica subsp. enterica serovar Typhimurium Proteobacteria; Gammaproteobacteria; Enterobacterales; Enterobacteriaceae; Salmonella; Salmonella enterica; Salmonella enterica subsp. enterica MNNEETFYQAMRRKGVTRRSFLKFCSLAATSLGLGAGMTPKIAWALENKPRIPVVWIHGL WP_001058312.1 hydrogenase [Salmonella enterica] Length: 372\nNumber of predicted TMHs: 2\nExp

number of AAs in TMHs: 36.51738\nExp number, first 60 AAs: 19.72849\nTotal prob of N-in: 0.96170\nPOSSIBLE N-term signal sequence\ninside 1 19\nTMhelix 20 39\noutside 40 325\nTMhelix 326 348\ninside 349 372

22350 GCF_001247185.1_Salmonella_enterica_CVM_N51299_v1.0 Salmonella enterica subsp. enterica serovar Typhimurium Proteobacteria; Gammaproteobacteria; Enterobacterales; Enterobacteriaceae; Salmonella; Salmonella enterica; Salmonella enterica subsp. enterica MTGDNTLITSHGINRRDFMKLCAALAATMGLSSKAAAEMAESVSNPQRPPVIWIGAQECT WP_000145429.1 hydrogenase 2 small subunit [Salmonella enterica] Length: 372\nNumber of predicted TMHs: 1\nExp number of AAs in TMHs: 24.51309\nExp number, first 60 AAs: 0.2682\nTotal prob of N-in: 0.14402\noutside 1 331\nTMhelix 332 354\ninside 355 372

22351 GCF_001293505.1_ASM129350v1 Salmonella enterica subsp. enterica serovar Typhimurium Proteobacteria; Gammaproteobacteria; Enterobacterales; Enterobacteriaceae; Salmonella; Salmonella enterica; Salmonella enterica subsp. enterica MTGDNTLITSHGINRRDFMKLCAALAATMGLSSKAAAEMAESVSNPQRPPVIWIGAQECT WP_000145429.1 hydrogenase 2 small subunit [Salmonella enterica] Length: 372\nNumber of predicted TMHs: 1\nExp number of AAs in TMHs: 24.51309\nExp number, first 60 AAs: 0.2682\nTotal prob of N-in: 0.14402\noutside 1 331\nTMhelix 332 354\ninside 355 372

22352 GCF_001295425.1_ASM129542v1 Salmonella enterica subsp. enterica serovar Typhimurium Proteobacteria; Gammaproteobacteria; Enterobacterales; Enterobacteriaceae; Salmonella; Salmonella enterica; Salmonella enterica subsp. enterica MTGDNTLITSHGINRRDFMKLCAALAATMGLSSKAAAEMAESVSNPQRPPVIWIGAQECT WP_000145429.1 hydrogenase 2 small subunit [Salmonella enterica] Length: 372\nNumber of predicted TMHs: 1\nExp number of AAs in TMHs: 24.51309\nExp number, first 60 AAs: 0.2682\nTotal prob of N-in: 0.14402\noutside 1 331\nTMhelix 332 354\ninside 355 372

22353 GCF_001293505.1_ASM129350v1 Salmonella enterica subsp. enterica serovar Typhimurium Proteobacteria; Gammaproteobacteria; Enterobacterales; Enterobacteriaceae; Salmonella; Salmonella enterica; Salmonella enterica subsp. enterica MNNEETFYQAMRRKGVTRRSFLKFCSLAATSLGLGAGMTPKIAWALENKPRIPVVWIHGL WP_001058312.1 hydrogenase [Salmonella enterica] Length: 372\nNumber of predicted TMHs: 2\nExp number of AAs in TMHs: 36.51738\nExp number, first 60 AAs: 19.72849\nTotal prob of N-in: 0.96170\nPOSSIBLE N-term signal sequence\ninside 1 19\nTMhelix 20 39\noutside 40 325\nTMhelix 326 348\ninside 349 372

22354 GCF_001295675.1_ASM129567v1 Salmonella enterica subsp. enterica serovar Typhimurium Proteobacteria; Gammaproteobacteria; Enterobacterales; Enterobacteriaceae; Salmonella; Salmonella enterica; Salmonella enterica subsp. enterica MNNEETFYQAMRRKGVTRRSFLKFCSLAATSLGLGAGMTPKIAWALENKPRIPVVWIHGL WP_001058312.1 hydrogenase [Salmonella enterica] Length: 372\nNumber of predicted TMHs: 2\nExp number of AAs in TMHs: 36.51738\nExp number, first 60 AAs: 19.72849\nTotal prob of N-in: 0.96170\nPOSSIBLE N-term signal sequence\ninside 1 19\nTMhelix 20 39\noutside 40 325\nTMhelix 326 348\ninside 349 372

22355 GCF_001295525.1_ASM129552v1 Salmonella enterica subsp. enterica serovar Typhimurium Proteobacteria; Gammaproteobacteria; Enterobacterales; Enterobacteriaceae; Salmonella; Salmonella enterica; Salmonella enterica subsp. enterica MTGDNTLITSHGINRRDFMKLCAALAATMGLSSKAAAEMAESVSNPQRPPVIWIGAQECT WP_000145429.1 hydrogenase 2 small subunit [Salmonella enterica] Length: 372\nNumber of predicted TMHs: 1\nExp number of AAs in TMHs: 24.51309\nExp number, first 60 AAs: 0.2682\nTotal prob of N-in: 0.14402\noutside 1 331\nTMhelix 332 354\ninside 355 372

22356 GCF_001295485.1_ASM129548v1 Salmonella enterica subsp. enterica serovar Typhimurium Proteobacteria; Gammaproteobacteria; Enterobacterales; Enterobacteriaceae; Salmonella; Salmonella enterica; Salmonella enterica subsp. enterica MNNEETFYQAMRRKGVTRRSFLKFCSLAATSLGLGAGMTPKIAWALENKPRIPVVWIHGL WP_001058312.1 hydrogenase [Salmonella enterica] Length: 372\nNumber of predicted TMHs: 2\nExp number of AAs in TMHs: 36.51738\nExp number, first 60 AAs: 19.72849\nTotal prob of N-in: 0.96170\nPOSSIBLE N-term signal sequence\ninside 1 19\nTMhelix 20 39\noutside 40 325\nTMhelix 326 348\ninside 349 372

22357 GCF_001295725.1_ASM129572v1 Salmonella enterica subsp. enterica serovar Typhimurium Proteobacteria; Gammaproteobacteria; Enterobacterales; Enterobacteriaceae; Salmonella; Salmonella enterica; Salmonella enterica subsp. enterica MNNEETFYQAMRRKGVTRRSFLKFCSLAATSLGLGAGMTPKIAWALENKPRIPVVWIHGL WP_001058312.1 hydrogenase [Salmonella enterica] Length: 372\nNumber of predicted TMHs: 2\nExp number of AAs in TMHs: 36.51738\nExp number, first 60 AAs: 19.72849\nTotal prob of N-in: 0.96170\nPOSSIBLE N-term signal sequence\ninside 1 19\nTMhelix 20 39\noutside 40 325\nTMhelix 326 348\ninside 349 372

22358	GCF_001295485.1_ASM129548v1	Salmonella enterica subsp. enterica serovar Typhimurium	Proteobacteria; Gammaproteobacteria; Enterobacterales; Enterobacteriaceae; Salmonella; Salmonella enterica; Salmonella enterica subsp. enterica	MTGDNTLITSHGINRRDFMKLCAALAATMGLSSKAAAEMAESVSNPQRPPVIWIGAQECT	WP_000145429.1	hydrogenase 2 small subunit [Salmonella enterica]	Length: 372\nNumber of predicted TMHs: 1\nExp number of AAs in TMHs: 24.51309\nExp number, first 60 AAs: 0.2682\nTotal prob of N-in: 0.14402\noutside 1 331\nTMhelix 332 354\ninside 355 372

22359	GCF_001295155.1_ASM129515v1	Salmonella enterica subsp. enterica serovar Typhimurium	Proteobacteria; Gammaproteobacteria; Enterobacterales; Enterobacteriaceae; Salmonella; Salmonella enterica; Salmonella enterica subsp. enterica	MNNEETFYQAMRRKGVTRRSFLKFCSLAATSLGLGAGMTPKIAWALENKPRIPVVWIHGL	WP_001058312.1	hydrogenase [Salmonella enterica]	Length: 372\nNumber of predicted TMHs: 2\nExp number of AAs in TMHs: 36.51738\nExp number, first 60 AAs: 19.72849\nTotal prob of N-in: 0.96170\nPOSSIBLE N-term signal sequence\ninside 1 19\nTMhelix 20 39\noutside 40 325\nTMhelix 326 348\ninside 349 372

22360	GCF_001295525.1_ASM129552v1	Salmonella enterica subsp. enterica serovar Typhimurium	Proteobacteria; Gammaproteobacteria; Enterobacterales; Enterobacteriaceae; Salmonella; Salmonella enterica; Salmonella enterica subsp. enterica	MNNEETFYQAMRRKGVTRRSFLKFCSLAATSLGLGAGMTPKIAWALENKPRIPVVWIHGL	WP_001058312.1	hydrogenase [Salmonella enterica]	Length: 372\nNumber of predicted TMHs: 2\nExp number of AAs in TMHs: 36.51738\nExp number, first 60 AAs: 19.72849\nTotal prob of N-in: 0.96170\nPOSSIBLE N-term signal sequence\ninside 1 19\nTMhelix 20 39\noutside 40 325\nTMhelix 326 348\ninside 349 372

22361	GCF_001295155.1_ASM129515v1	Salmonella enterica subsp. enterica serovar Typhimurium	Proteobacteria; Gammaproteobacteria; Enterobacterales; Enterobacteriaceae; Salmonella; Salmonella enterica; Salmonella enterica subsp. enterica	MTGDNTLITSHGINRRDFMKLCAALAATMGLSSKAAAEMAESVSNPQRPPVIWIGAQECT	WP_000145429.1	hydrogenase 2 small subunit [Salmonella enterica]	Length: 372\nNumber of predicted TMHs: 1\nExp number of AAs in TMHs: 24.51309\nExp number, first 60 AAs: 0.2682\nTotal prob of N-in: 0.14402\noutside 1 331\nTMhelix 332 354\ninside 355 372

22362	GCF_002063615.1_ASM206361v1	Salmonella enterica subsp. enterica serovar Uganda	Proteobacteria; Gammaproteobacteria; Enterobacterales; Enterobacteriaceae; Salmonella; Salmonella enterica; Salmonella enterica subsp. enterica	MNNEETFYQAMRRKGVTRRSFLKFCSLAATSLGLGAGMTPKIAWALENKPRIPVVWIHGL	WP_001058311.1	hydrogenase [Salmonella enterica]	Length: 372\nNumber of predicted TMHs: 2\nExp number of AAs in TMHs: 37.4699899999999\nExp number, first 60 AAs: 19.72699\nTotal prob of N-in: 0.96171\nPOSSIBLE N-term signal sequence\ninside 1 19\nTMhelix 20 39\noutside 40 325\nTMhelix 326 348\ninside 349 372

22363	GCF_002063615.1_ASM206361v1	Salmonella enterica subsp. enterica serovar Uganda	Proteobacteria; Gammaproteobacteria; Enterobacterales; Enterobacteriaceae; Salmonella; Salmonella enterica; Salmonella enterica subsp. enterica	MTGDNTLITSHGINRRDFMKLCAALAATMGLSSKVAAEMAESVSNPQRPPVIWIGAQECT	WP_079918295.1	hydrogenase 2 small subunit [Salmonella enterica]	Length: 372\nNumber of predicted TMHs: 1\nExp number of AAs in TMHs: 24.67391\nExp number, first 60 AAs: 0.43696\nTotal prob of N-in: 0.15108\noutside 1 331\nTMhelix 332 354\ninside 355 372

22364	GCF_002047785.1_ASM204778v1	Salmonella enterica subsp. enterica serovar Uganda	Proteobacteria; Gammaproteobacteria; Enterobacterales; Enterobacteriaceae; Salmonella; Salmonella enterica; Salmonella enterica subsp. enterica	MTGDNTLITSHGINRRDFMKLCAALAATMGLSSKAAAEMAESVSNPQRPPVIWIGAQECT	WP_000145429.1	hydrogenase 2 small subunit [Salmonella enterica]	Length: 372\nNumber of predicted TMHs: 1\nExp number of AAs in TMHs: 24.51309\nExp number, first 60 AAs: 0.2682\nTotal prob of N-in: 0.14402\noutside 1 331\nTMhelix 332 354\ninside 355 372

22365	GCF_002030165.1_ASM203016v1	Salmonella enterica subsp. enterica serovar Uganda	Proteobacteria; Gammaproteobacteria; Enterobacterales; Enterobacteriaceae; Salmonella; Salmonella enterica; Salmonella enterica subsp. enterica	MTGDNTLITSHGINRRDFMKLCAALAATMGLSSKAAAEMAESVSNPQRPPVIWIGAQECT	WP_000145429.1	hydrogenase 2 small subunit [Salmonella enterica]	Length: 372\nNumber of predicted TMHs: 1\nExp number of AAs in TMHs: 24.51309\nExp number, first 60 AAs: 0.2682\nTotal prob of N-in: 0.14402\noutside 1 331\nTMhelix 332 354\ninside 355 372

22366	GCF_002047505.1_ASM204750v1	Salmonella enterica subsp. enterica serovar Uganda	Proteobacteria; Gammaproteobacteria; Enterobacterales; Enterobacteriaceae; Salmonella; Salmonella enterica; Salmonella enterica subsp. enterica	MTGDNTLITSHGINRRDFMKLCAALAATMGLSSKVAAEMAESVSNPQRPPVIWIGAQECT	WP_079918295.1	hydrogenase 2 small subunit [Salmonella enterica]	Length: 372\nNumber of

predicted TMHs:  1\nExp number of AAs in TMHs: 24.67391\nExp number, first 60 AAs:  0.43696\nTotal prob of N-in: 0.15108\noutside   1   331\nTMhelix   332   354\ninside   355   372

22367 GCF_002047505.1_ASM204750v1 Salmonella enterica subsp. enterica serovar Uganda Proteobacteria; Gammaproteobacteria; Enterobacterales; Enterobacteriaceae; Salmonella; Salmonella enterica; Salmonella enterica subsp. enterica MNNEETFYQAMRRKGVTRRSFLKFCSLAATSLGLGAGMTPKIAWALENKPRIPVVWIHGL WP_001058311.1 hydrogenase [Salmonella enterica] Length: 372\nNumber of predicted TMHs:  2\nExp number of AAs in TMHs: 37.4699899999999\nExp number, first 60 AAs:  19.72699\nTotal prob of N-in: 0.96171\nPOSSIBLE N-term signal sequence\ninside   1   19\nTMhelix   20   39\noutside   40   325\nTMhelix   326   348\ninside   349   372

22368 GCF_002048685.1_ASM204868v1 Salmonella enterica subsp. enterica serovar Uganda Proteobacteria; Gammaproteobacteria; Enterobacterales; Enterobacteriaceae; Salmonella; Salmonella enterica; Salmonella enterica subsp. enterica MNNEETFYQAMRRKGVTRRSFLKFCSLAATSLGLGAGMTPKIAWALENKPRIPVVWIHGL WP_001058311.1 hydrogenase [Salmonella enterica] Length: 372\nNumber of predicted TMHs:  2\nExp number of AAs in TMHs: 37.4699899999999\nExp number, first 60 AAs:  19.72699\nTotal prob of N-in: 0.96171\nPOSSIBLE N-term signal sequence\ninside   1   19\nTMhelix   20   39\noutside   40   325\nTMhelix   326   348\ninside   349   372

22369 GCF_002047575.1_ASM204757v1 Salmonella enterica subsp. enterica serovar Uganda Proteobacteria; Gammaproteobacteria; Enterobacterales; Enterobacteriaceae; Salmonella; Salmonella enterica; Salmonella enterica subsp. enterica MTGDNTLITSHGINRRDFMKLCAALAATMGLSSKAAAEMAESVSNPQRPPVIWIGAQECT WP_000145429.1 hydrogenase 2 small subunit [Salmonella enterica] Length: 372\nNumber of predicted TMHs:  1\nExp number of AAs in TMHs: 24.51309\nExp number, first 60 AAs:  0.2682\nTotal prob of N-in: 0.14402\noutside   1   331\nTMhelix   332   354\ninside   355   372

22370 GCF_002047785.1_ASM204778v1 Salmonella enterica subsp. enterica serovar Uganda Proteobacteria; Gammaproteobacteria; Enterobacterales; Enterobacteriaceae; Salmonella; Salmonella enterica; Salmonella enterica subsp. enterica MNNEETFYQAMRRKGVTRRSFLKFCSLAATSLGLGAGMTPKIAWALENKPRIPVVWIHGL WP_001058311.1 hydrogenase [Salmonella enterica] Length: 372\nNumber of predicted TMHs:  2\nExp number of AAs in TMHs: 37.4699899999999\nExp number, first 60 AAs:  19.72699\nTotal prob of N-in: 0.96171\nPOSSIBLE N-term signal sequence\ninside   1   19\nTMhelix   20   39\noutside   40   325\nTMhelix   326   348\ninside   349   372

22371 GCF_002047575.1_ASM204757v1 Salmonella enterica subsp. enterica serovar Uganda Proteobacteria; Gammaproteobacteria; Enterobacterales; Enterobacteriaceae; Salmonella; Salmonella enterica; Salmonella enterica subsp. enterica MNNEETFYQAMRRKGVTRRSFLKFCSLAATSLGLGAGMTPKIAWALENKPRIPVVWIHGL WP_001058311.1 hydrogenase [Salmonella enterica] Length: 372\nNumber of predicted TMHs:  2\nExp number of AAs in TMHs: 37.4699899999999\nExp number, first 60 AAs:  19.72699\nTotal prob of N-in: 0.96171\nPOSSIBLE N-term signal sequence\ninside   1   19\nTMhelix   20   39\noutside   40   325\nTMhelix   326   348\ninside   349   372

22372 GCF_002030165.1_ASM203016v1 Salmonella enterica subsp. enterica serovar Uganda Proteobacteria; Gammaproteobacteria; Enterobacterales; Enterobacteriaceae; Salmonella; Salmonella enterica; Salmonella enterica subsp. enterica MNNEETFYQAMRRKGVTRRSFLKFCSLAATSLGLGAGMTPKIAWALENKPRIPVVWIHGL WP_001058311.1 hydrogenase [Salmonella enterica] Length: 372\nNumber of predicted TMHs:  2\nExp number of AAs in TMHs: 37.4699899999999\nExp number, first 60 AAs:  19.72699\nTotal prob of N-in: 0.96171\nPOSSIBLE N-term signal sequence\ninside   1   19\nTMhelix   20   39\noutside   40   325\nTMhelix   326   348\ninside   349   372

22373 GCF_002048685.1_ASM204868v1 Salmonella enterica subsp. enterica serovar Uganda Proteobacteria; Gammaproteobacteria; Enterobacterales; Enterobacteriaceae; Salmonella; Salmonella enterica; Salmonella enterica subsp. enterica MTGDNTLITSHGINRRDFMKLCAALAATMGLSSKAAAEMAESVSNPQRPPVIWIGAQECT WP_000145429.1 hydrogenase 2 small subunit [Salmonella enterica] Length: 372\nNumber of predicted TMHs:  1\nExp number of AAs in TMHs: 24.51309\nExp number, first 60 AAs:  0.2682\nTotal prob of N-in: 0.14402\noutside   1   331\nTMhelix   332   354\ninside   355   372

22374 GCF_002048565.1_ASM204856v1 Salmonella enterica subsp. enterica serovar Urbana Proteobacteria; Gammaproteobacteria; Enterobacterales; Enterobacteriaceae; Salmonella; Salmonella enterica; Salmonella enterica subsp. enterica MTGDNTLITSHGINRRDFMKLCAALAATMGLSSKAAAEMAESVSNPQRPPVIWIGAQECT WP_000145429.1 hydrogenase 2 small subunit [Salmonella enterica] Length: 372\nNumber of

predicted TMHs:  1\nExp number of AAs in TMHs: 24.51309\nExp number, first 60 AAs:  0.2682\nTotal prob of N-in:   0.14402\noutside   1   331\nTMhelix   332   354\ninside   355   372

22375 GCF_002047895.1_ASM204789v1 Salmonella enterica subsp. enterica serovar Urbana Proteobacteria; Gammaproteobacteria; Enterobacterales; Enterobacteriaceae; Salmonella; Salmonella enterica; Salmonella enterica subsp. enterica MNNEETFYQAMRRKGVTRRSFLKFCSLAATSLGLGAGMTPKIAWALENKPRIPVVWIHGL WP_001058312.1 hydrogenase [Salmonella enterica] Length: 372\nNumber of predicted TMHs:  2\nExp number of AAs in TMHs: 36.51738\nExp number, first 60 AAs:  19.72849\nTotal prob of N-in:   0.96170\nPOSSIBLE N-term signal sequence\ninside   1   19\nTMhelix   20   39\noutside   40   325\nTMhelix   326   348\ninside   349   372

22376 GCF_002047735.1_ASM204773v1 Salmonella enterica subsp. enterica serovar Urbana Proteobacteria; Gammaproteobacteria; Enterobacterales; Enterobacteriaceae; Salmonella; Salmonella enterica; Salmonella enterica subsp. enterica MTGDNTLITSHGINRRDFMKLCAALAATMGLSSKAAAEMAESVSNPQRPPVIWIGAQECT WP_000145429.1 hydrogenase 2 small subunit [Salmonella enterica] Length: 372\nNumber of predicted TMHs:  1\nExp number of AAs in TMHs: 24.51309\nExp number, first 60 AAs:  0.2682\nTotal prob of N-in:   0.14402\noutside   1   331\nTMhelix   332   354\ninside   355   372

22377 GCF_002047645.1_ASM204764v1 Salmonella enterica subsp. enterica serovar Urbana Proteobacteria; Gammaproteobacteria; Enterobacterales; Enterobacteriaceae; Salmonella; Salmonella enterica; Salmonella enterica subsp. enterica MTGDNTLITSHGINRRDFMKLCAALAATMGLSSKAAAEMAESVSNPQRPPVIWIGAQECT WP_000145429.1 hydrogenase 2 small subunit [Salmonella enterica] Length: 372\nNumber of predicted TMHs:  1\nExp number of AAs in TMHs: 24.51309\nExp number, first 60 AAs:  0.2682\nTotal prob of N-in:   0.14402\noutside   1   331\nTMhelix   332   354\ninside   355   372

22378 GCF_002048525.1_ASM204852v1 Salmonella enterica subsp. enterica serovar Urbana Proteobacteria; Gammaproteobacteria; Enterobacterales; Enterobacteriaceae; Salmonella; Salmonella enterica; Salmonella enterica subsp. enterica MNNEETFYQAMRRKGVTRRSFLKFCSLAATSLGLGAGMTPKIAWALENKPRIPVVWIHGL WP_001058312.1 hydrogenase [Salmonella enterica] Length: 372\nNumber of predicted TMHs:  2\nExp number of AAs in TMHs: 36.51738\nExp number, first 60 AAs:  19.72849\nTotal prob of N-in:   0.96170\nPOSSIBLE N-term signal sequence\ninside   1   19\nTMhelix   20   39\noutside   40   325\nTMhelix   326   348\ninside   349   372

22379 GCF_002048525.1_ASM204852v1 Salmonella enterica subsp. enterica serovar Urbana Proteobacteria; Gammaproteobacteria; Enterobacterales; Enterobacteriaceae; Salmonella; Salmonella enterica; Salmonella enterica subsp. enterica MTGDNTLITSHGINRRDFMKLCAALAATMGLSSKAAAEMAESVSNPQRPPVIWIGAQECT WP_000145429.1 hydrogenase 2 small subunit [Salmonella enterica] Length: 372\nNumber of predicted TMHs:  1\nExp number of AAs in TMHs: 24.51309\nExp number, first 60 AAs:  0.2682\nTotal prob of N-in:   0.14402\noutside   1   331\nTMhelix   332   354\ninside   355   372

22380 GCF_002035665.1_ASM203566v1 Salmonella enterica subsp. enterica serovar Urbana Proteobacteria; Gammaproteobacteria; Enterobacterales; Enterobacteriaceae; Salmonella; Salmonella enterica; Salmonella enterica subsp. enterica MTGDNTLITSHGINRRDFMKLCAALAATMGLSSKAAAEMAESVSNPQRPPVIWIGAQECT WP_000145429.1 hydrogenase 2 small subunit [Salmonella enterica] Length: 372\nNumber of predicted TMHs:  1\nExp number of AAs in TMHs: 24.51309\nExp number, first 60 AAs:  0.2682\nTotal prob of N-in:   0.14402\noutside   1   331\nTMhelix   332   354\ninside   355   372

22381 GCF_002035665.1_ASM203566v1 Salmonella enterica subsp. enterica serovar Urbana Proteobacteria; Gammaproteobacteria; Enterobacterales; Enterobacteriaceae; Salmonella; Salmonella enterica; Salmonella enterica subsp. enterica MNNEETFYQAMRRKGVTRRSFLKFCSLAATSLGLGAGMTPKIAWALENKPRIPVVWIHGL WP_023198237.1 hydrogenase 1, small subunit [Salmonella enterica] Length: 372\nNumber of predicted TMHs:  2\nExp number of AAs in TMHs: 36.39423\nExp number, first 60 AAs:  19.72877\nTotal prob of N-in:   0.96162\nPOSSIBLE N-term signal sequence\ninside   1   19\nTMhelix   20   39\noutside   40   325\nTMhelix   326   348\ninside   349   372

22382 GCF_002047645.1_ASM204764v1 Salmonella enterica subsp. enterica serovar Urbana Proteobacteria; Gammaproteobacteria; Enterobacterales; Enterobacteriaceae; Salmonella; Salmonella enterica; Salmonella enterica subsp. enterica MNNEETFYQAMRRKGVTRRSFLKFCSLAATSLGLGAGMTPKIAWALENKPRIPVVWIHGL WP_001058312.1 hydrogenase [Salmonella enterica] Length: 372\nNumber of predicted TMHs:  2\nExp number of AAs in TMHs: 36.51738\nExp number, first 60 AAs:  19.72849\nTotal prob of N-in:   0.96170\nPOSSIBLE N-term signal sequence\ninside   1   19\nTMhelix   20   39\noutside   40   325\nTMhelix   326   348\ninside   349   372

22383	GCF_002048565.1_ASM204856v1	Salmonella enterica subsp. enterica serovar Urbana
	Proteobacteria; Gammaproteobacteria; Enterobacterales; Enterobacteriaceae; Salmonella; Salmonella enterica; Salmonella enterica subsp. enterica	MNNEETFYQAMRRKGVTRRSFLKFCSLAATSLGLGAGMTPKIAWALENKPRIPVVWIHGL
	WP_001058312.1	hydrogenase [Salmonella enterica]	Length: 372\nNumber of predicted TMHs: 2\nExp number of AAs in TMHs: 36.51738\nExp number, first 60 AAs: 19.72849\nTotal prob of N-in: 0.96170\nPOSSIBLE N-term signal sequence\ninside 1 19\nTMhelix 20 39\noutside 40 325\nTMhelix 326 348\ninside 349 372

22384	GCF_002047735.1_ASM204773v1	Salmonella enterica subsp. enterica serovar Urbana
	Proteobacteria; Gammaproteobacteria; Enterobacterales; Enterobacteriaceae; Salmonella; Salmonella enterica; Salmonella enterica subsp. enterica	MNNEETFYQAMRRKGVTRRSFLKFCSLAATSLGLGAGMTPKIAWALENKPRIPVVWIHGL
	WP_001058312.1	hydrogenase [Salmonella enterica]	Length: 372\nNumber of predicted TMHs: 2\nExp number of AAs in TMHs: 36.51738\nExp number, first 60 AAs: 19.72849\nTotal prob of N-in: 0.96170\nPOSSIBLE N-term signal sequence\ninside 1 19\nTMhelix 20 39\noutside 40 325\nTMhelix 326 348\ninside 349 372

22385	GCF_002047895.1_ASM204789v1	Salmonella enterica subsp. enterica serovar Urbana
	Proteobacteria; Gammaproteobacteria; Enterobacterales; Enterobacteriaceae; Salmonella; Salmonella enterica; Salmonella enterica subsp. enterica	MTGDNTLITSHGINRRDFMKLCAALAATMGLSSKAAAEMAESVSNPQRPPVIWIGAQECT
	WP_000145429.1	hydrogenase 2 small subunit [Salmonella enterica]	Length: 372\nNumber of predicted TMHs: 1\nExp number of AAs in TMHs: 24.51309\nExp number, first 60 AAs: 0.2682\nTotal prob of N-in: 0.14402\noutside 1 331\nTMhelix 332 354\ninside 355 372

22386	GCF_002063595.1_ASM206359v1	Salmonella enterica subsp. enterica serovar Vinnorrhady
	Proteobacteria; Gammaproteobacteria; Enterobacterales; Enterobacteriaceae; Salmonella; Salmonella enterica; Salmonella enterica subsp. enterica	MNNEETFYQAMRRKGVTRRSFLKFCSLAATSLGLGAGMTPKIAWALENKPRIPVVWIHGL
	WP_001058311.1	hydrogenase [Salmonella enterica]	Length: 372\nNumber of predicted TMHs: 2\nExp number of AAs in TMHs: 37.4699899999999\nExp number, first 60 AAs: 19.72699\nTotal prob of N-in: 0.96171\nPOSSIBLE N-term signal sequence\ninside 1 19\nTMhelix 20 39\noutside 40 325\nTMhelix 326 348\ninside 349 372

22387	GCF_002063595.1_ASM206359v1	Salmonella enterica subsp. enterica serovar Vinnorrhady
	Proteobacteria; Gammaproteobacteria; Enterobacterales; Enterobacteriaceae; Salmonella; Salmonella enterica; Salmonella enterica subsp. enterica	MTGDNTLITSHGINRRDFMKLCAALAATMGLSSKAAAEMAESVSNPQRPPVIWIGAQECT
	WP_080222249.1	hydrogenase 2 small subunit [Salmonella enterica]	Length: 372\nNumber of predicted TMHs: 1\nExp number of AAs in TMHs: 23.92634\nExp number, first 60 AAs: 0.27026\nTotal prob of N-in: 0.16033\noutside 1 331\nTMhelix 332 354\ninside 355 372

22388	GCF_002062615.1_ASM206261v1	Salmonella enterica subsp. enterica serovar Vinobrady
	Proteobacteria; Gammaproteobacteria; Enterobacterales; Enterobacteriaceae; Salmonella; Salmonella enterica; Salmonella enterica subsp. enterica	MNNEETFYQAMRRKGVTRRSFLKFCSLAATSLGLGAGMTPKIAWALENKPRIPVVWIHGL
	WP_001058312.1	hydrogenase [Salmonella enterica]	Length: 372\nNumber of predicted TMHs: 2\nExp number of AAs in TMHs: 36.51738\nExp number, first 60 AAs: 19.72849\nTotal prob of N-in: 0.96170\nPOSSIBLE N-term signal sequence\ninside 1 19\nTMhelix 20 39\noutside 40 325\nTMhelix 326 348\ninside 349 372

22389	GCF_002062615.1_ASM206261v1	Salmonella enterica subsp. enterica serovar Vinobrady
	Proteobacteria; Gammaproteobacteria; Enterobacterales; Enterobacteriaceae; Salmonella; Salmonella enterica; Salmonella enterica subsp. enterica	MTGDNTLITSHGINRRDFMKLCAALAATMGLSSKAAAEMAESVSNPQRPPVIWIGAQECT
	WP_000145429.1	hydrogenase 2 small subunit [Salmonella enterica]	Length: 372\nNumber of predicted TMHs: 1\nExp number of AAs in TMHs: 24.51309\nExp number, first 60 AAs: 0.2682\nTotal prob of N-in: 0.14402\noutside 1 331\nTMhelix 332 354\ninside 355 372

22390	GCF_002062755.1_ASM206275v1	Salmonella enterica subsp. enterica serovar Virchow
	Proteobacteria; Gammaproteobacteria; Enterobacterales; Enterobacteriaceae; Salmonella; Salmonella enterica; Salmonella enterica subsp. enterica	MNNEETFYQAMRRKGVTRRSFLKFCSLAATSLGLGAGMTPKIAWALENKPRIPVVWIHGL
	WP_001058310.1	hydrogenase [Salmonella enterica]	Length: 372\nNumber of predicted TMHs: 2\nExp number of AAs in TMHs: 37.4567299999999\nExp number, first 60 AAs: 19.72694\nTotal prob of N-in: 0.96171\nPOSSIBLE N-term signal sequence\ninside 1 19\nTMhelix 20 39\noutside 40 325\nTMhelix 326 348\ninside 349 372

22391	GCF_002156325.1_ASM215632v1	Salmonella enterica subsp. enterica serovar Virchow
	Proteobacteria; Gammaproteobacteria; Enterobacterales; Enterobacteriaceae; Salmonella; Salmonella enterica; Salmonella enterica subsp. enterica	MTGDNTLITSHGINRRDFMKLCAALAATMGLSSKAAAEMAESVSNPQRPPVIWIGAQECT

WP_000145429.1 hydrogenase 2 small subunit [Salmonella enterica] Length: 372\nNumber of predicted TMHs: 1\nExp number of AAs in TMHs: 24.51309\nExp number, first 60 AAs: 0.2682\nTotal prob of N-in: 0.14402\noutside 1 331\nTMhelix 332 354\ninside 355 372

22392 GCF_002156325.1_ASM215632v1 Salmonella enterica subsp. enterica serovar Virchow Proteobacteria; Gammaproteobacteria; Enterobacterales; Enterobacteriaceae; Salmonella; Salmonella enterica; Salmonella enterica subsp. enterica MNNEETFYQAMRRKGVTRRSFLKFCSLAATSLGLGAGMTPKIAWALENKPRIPVVWIHGL WP_001058310.1 hydrogenase [Salmonella enterica] Length: 372\nNumber of predicted TMHs: 2\nExp number of AAs in TMHs: 37.4567299999999\nExp number, first 60 AAs: 19.72694\nTotal prob of N-in: 0.96171\nPOSSIBLE N-term signal sequence\ninside 1 19\nTMhelix 20 39\noutside 40 325\nTMhelix 326 348\ninside 349 372

22393 GCF_002063545.1_ASM206354v1 Salmonella enterica subsp. enterica serovar Virchow Proteobacteria; Gammaproteobacteria; Enterobacterales; Enterobacteriaceae; Salmonella; Salmonella enterica; Salmonella enterica subsp. enterica MTGDNTLITSHGINRRDFMKLCAALAATMGLSSKAAAEMAESVSNPQRPPVIWIGAQECT WP_000145429.1 hydrogenase 2 small subunit [Salmonella enterica] Length: 372\nNumber of predicted TMHs: 1\nExp number of AAs in TMHs: 24.51309\nExp number, first 60 AAs: 0.2682\nTotal prob of N-in: 0.14402\noutside 1 331\nTMhelix 332 354\ninside 355 372

22394 GCF_002062755.1_ASM206275v1 Salmonella enterica subsp. enterica serovar Virchow Proteobacteria; Gammaproteobacteria; Enterobacterales; Enterobacteriaceae; Salmonella; Salmonella enterica; Salmonella enterica subsp. enterica MTGDNTLITSHGINRRDFMKLCAALAATMGLSSKAAAEMAESVSNPQRPPVIWIGAQECT WP_000145429.1 hydrogenase 2 small subunit [Salmonella enterica] Length: 372\nNumber of predicted TMHs: 1\nExp number of AAs in TMHs: 24.51309\nExp number, first 60 AAs: 0.2682\nTotal prob of N-in: 0.14402\noutside 1 331\nTMhelix 332 354\ninside 355 372

22395 GCF_002063545.1_ASM206354v1 Salmonella enterica subsp. enterica serovar Virchow Proteobacteria; Gammaproteobacteria; Enterobacterales; Enterobacteriaceae; Salmonella; Salmonella enterica; Salmonella enterica subsp. enterica MNNEETFYQAMRRKGVTRRSFLKFCSLAATSLGLGAGMTPKIAWALENKPRIPVVWIHGL WP_001058310.1 hydrogenase [Salmonella enterica] Length: 372\nNumber of predicted TMHs: 2\nExp number of AAs in TMHs: 37.4567299999999\nExp number, first 60 AAs: 19.72694\nTotal prob of N-in: 0.96171\nPOSSIBLE N-term signal sequence\ninside 1 19\nTMhelix 20 39\noutside 40 325\nTMhelix 326 348\ninside 349 372

22396 GCF_002062485.1_ASM206248v1 Salmonella enterica subsp. enterica serovar Virchow Proteobacteria; Gammaproteobacteria; Enterobacterales; Enterobacteriaceae; Salmonella; Salmonella enterica; Salmonella enterica subsp. enterica MNNEETFYQAMRRKGVTRRSFLKFCSLAATSLGLGAGMTPKIAWALENKPRIPVVWIHGL WP_001058310.1 hydrogenase [Salmonella enterica] Length: 372\nNumber of predicted TMHs: 2\nExp number of AAs in TMHs: 37.4567299999999\nExp number, first 60 AAs: 19.72694\nTotal prob of N-in: 0.96171\nPOSSIBLE N-term signal sequence\ninside 1 19\nTMhelix 20 39\noutside 40 325\nTMhelix 326 348\ninside 349 372

22397 GCF_002062485.1_ASM206248v1 Salmonella enterica subsp. enterica serovar Virchow Proteobacteria; Gammaproteobacteria; Enterobacterales; Enterobacteriaceae; Salmonella; Salmonella enterica; Salmonella enterica subsp. enterica MTGDNTLITSHGINRRDFMKLCAALAATMGLSSKAAAEMAESVSNPQRPPVIWIGAQECT WP_000145429.1 hydrogenase 2 small subunit [Salmonella enterica] Length: 372\nNumber of predicted TMHs: 1\nExp number of AAs in TMHs: 24.51309\nExp number, first 60 AAs: 0.2682\nTotal prob of N-in: 0.14402\noutside 1 331\nTMhelix 332 354\ninside 355 372

22398 GCF_002049385.1_ASM204938v1 Salmonella enterica subsp. enterica serovar Virchow Proteobacteria; Gammaproteobacteria; Enterobacterales; Enterobacteriaceae; Salmonella; Salmonella enterica; Salmonella enterica subsp. enterica MNNEETFYQAMRRKGVTRRSFLKFCSLAATSLGLGAGMTPKIAWALENKPRIPVVWIHGL WP_079927855.1 [Ni/Fe] hydrogenase small subunit [Salmonella enterica] Length: 372\nNumber of predicted TMHs: 2\nExp number of AAs in TMHs: 37.4697299999999\nExp number, first 60 AAs: 19.72699\nTotal prob of N-in: 0.96171\nPOSSIBLE N-term signal sequence\ninside 1 19\nTMhelix 20 39\noutside 40 325\nTMhelix 326 348\ninside 349 372

22399 GCF_002049385.1_ASM204938v1 Salmonella enterica subsp. enterica serovar Virchow Proteobacteria; Gammaproteobacteria; Enterobacterales; Enterobacteriaceae; Salmonella; Salmonella enterica; Salmonella enterica subsp. enterica MTGDNTLITSHGINRRDFMKLCAALAATMGLSSKAAAEMAESVSNPQRPPVIWIGAQECT WP_000145429.1 hydrogenase 2 small subunit [Salmonella enterica] Length: 372\nNumber of

predicted TMHs: 1\nExp number of AAs in TMHs: 24.51309\nExp number, first 60 AAs: 0.2682\nTotal prob of N-in: 0.14402\noutside 1 331\nTMhelix 332 354\ninside 355 372

22400 GCF_002049055.1_ASM204905v1 Salmonella enterica subsp. enterica serovar Virchow Proteobacteria; Gammaproteobacteria; Enterobacterales; Enterobacteriaceae; Salmonella; Salmonella enterica; Salmonella enterica subsp. enterica MTGDNTLITSHGINRRDFMKLCAALAATMGLSSKAAAEMAESVSNPQRPPVIWIGAQECT WP_000145429.1 hydrogenase 2 small subunit [Salmonella enterica] Length: 372\nNumber of predicted TMHs: 1\nExp number of AAs in TMHs: 24.51309\nExp number, first 60 AAs: 0.2682\nTotal prob of N-in: 0.14402\noutside 1 331\nTMhelix 332 354\ninside 355 372

22401 GCF_002049055.1_ASM204905v1 Salmonella enterica subsp. enterica serovar Virchow Proteobacteria; Gammaproteobacteria; Enterobacterales; Enterobacteriaceae; Salmonella; Salmonella enterica; Salmonella enterica subsp. enterica MNNEETFYQAMRRKGVTRRSFLKFCSLAATSLGLGAGMTPKIAWALENKPRIPVVWIHGL WP_001058310.1 hydrogenase [Salmonella enterica] Length: 372\nNumber of predicted TMHs: 2\nExp number of AAs in TMHs: 37.4567299999999\nExp number, first 60 AAs: 19.72694\nTotal prob of N-in: 0.96171\nPOSSIBLE N-term signal sequence\ninside 1 19\nTMhelix 20 39\noutside 40 325\nTMhelix 326 348\ninside 349 372

22402 GCF_002063155.1_ASM206315v1 Salmonella enterica subsp. enterica serovar Virchow Proteobacteria; Gammaproteobacteria; Enterobacterales; Enterobacteriaceae; Salmonella; Salmonella enterica; Salmonella enterica subsp. enterica MTGDNTLITSHGINRRDFMKLCAALAATMGLSSKAAAEMAESVSNPQRPPVIWIGAQECT WP_000145429.1 hydrogenase 2 small subunit [Salmonella enterica] Length: 372\nNumber of predicted TMHs: 1\nExp number of AAs in TMHs: 24.51309\nExp number, first 60 AAs: 0.2682\nTotal prob of N-in: 0.14402\noutside 1 331\nTMhelix 332 354\ninside 355 372

22403 GCF_002063155.1_ASM206315v1 Salmonella enterica subsp. enterica serovar Virchow Proteobacteria; Gammaproteobacteria; Enterobacterales; Enterobacteriaceae; Salmonella; Salmonella enterica; Salmonella enterica subsp. enterica MNNEETFYQAMRRKGVTRRSFLKFCSLAATSLGLGAGMTPKIAWALENKPRIPVVWIHGL WP_001058310.1 hydrogenase [Salmonella enterica] Length: 372\nNumber of predicted TMHs: 2\nExp number of AAs in TMHs: 37.4567299999999\nExp number, first 60 AAs: 19.72694\nTotal prob of N-in: 0.96171\nPOSSIBLE N-term signal sequence\ninside 1 19\nTMhelix 20 39\noutside 40 325\nTMhelix 326 348\ninside 349 372

22404 GCF_002063555.1_ASM206355v1 Salmonella enterica subsp. enterica serovar Virchow Proteobacteria; Gammaproteobacteria; Enterobacterales; Enterobacteriaceae; Salmonella; Salmonella enterica; Salmonella enterica subsp. enterica MTGDNTLITSHGINRRDFMKLCAALAATMGLSSKAAAEMAESVSNPQRPPVIWIGAQECT WP_000145429.1 hydrogenase 2 small subunit [Salmonella enterica] Length: 372\nNumber of predicted TMHs: 1\nExp number of AAs in TMHs: 24.51309\nExp number, first 60 AAs: 0.2682\nTotal prob of N-in: 0.14402\noutside 1 331\nTMhelix 332 354\ninside 355 372

22405 GCF_002063525.1_ASM206352v1 Salmonella enterica subsp. enterica serovar Virchow Proteobacteria; Gammaproteobacteria; Enterobacterales; Enterobacteriaceae; Salmonella; Salmonella enterica; Salmonella enterica subsp. enterica MNNEETFYQAMRRKGVTRRSFLKFCSLAATSLGLGAGMTPKIAWALENKPRIPVVWIHGL WP_079927855.1 [Ni/Fe] hydrogenase small subunit [Salmonella enterica] Length: 372\nNumber of predicted TMHs: 2\nExp number of AAs in TMHs: 37.4697299999999\nExp number, first 60 AAs: 19.72699\nTotal prob of N-in: 0.96171\nPOSSIBLE N-term signal sequence\ninside 1 19\nTMhelix 20 39\noutside 40 325\nTMhelix 326 348\ninside 349 372

22406 GCF_002065025.1_ASM206502v1 Salmonella enterica subsp. enterica serovar Virchow Proteobacteria; Gammaproteobacteria; Enterobacterales; Enterobacteriaceae; Salmonella; Salmonella enterica; Salmonella enterica subsp. enterica MTGDNTLITSHGINRRDFMKLCAALAATMGLSSKAAAEMAESVSNPQRPPVIWIGAQECT WP_000145429.1 hydrogenase 2 small subunit [Salmonella enterica] Length: 372\nNumber of predicted TMHs: 1\nExp number of AAs in TMHs: 24.51309\nExp number, first 60 AAs: 0.2682\nTotal prob of N-in: 0.14402\noutside 1 331\nTMhelix 332 354\ninside 355 372

22407 GCF_002063555.1_ASM206355v1 Salmonella enterica subsp. enterica serovar Virchow Proteobacteria; Gammaproteobacteria; Enterobacterales; Enterobacteriaceae; Salmonella; Salmonella enterica; Salmonella enterica subsp. enterica MNNEETFYQAMRRKGVTRRSFLKFCSLAATSLGLGAGMTPKIAWALENKPRIPVVWIHGL WP_001058310.1 hydrogenase [Salmonella enterica] Length: 372\nNumber of predicted TMHs: 2\nExp number of AAs in TMHs: 37.4567299999999\nExp number, first 60 AAs: 19.72694\nTotal prob of N-in:

0.96171\nPOSSIBLE N-term signal sequence\ninside 1 19\nTMhelix 20 39\noutside 40 325\nTMhelix 326 348\ninside 349 372

22408 GCF_002065025.1_ASM206502v1 Salmonella enterica subsp. enterica serovar Virchow Proteobacteria; Gammaproteobacteria; Enterobacterales; Enterobacteriaceae; Salmonella; Salmonella enterica; Salmonella enterica subsp. enterica MNNEETFYQAMRRKGVTRRSFLKFCSLAATSLGLGAGMTPKIAWALENKPRIPVVWIHGL WP_001058310.1 hydrogenase [Salmonella enterica] Length: 372\nNumber of predicted TMHs: 2\nExp number of AAs in TMHs: 37.4567299999999\nExp number, first 60 AAs: 19.72694\nTotal prob of N-in: 0.96171\nPOSSIBLE N-term signal sequence\ninside 1 19\nTMhelix 20 39\noutside 40 325\nTMhelix 326 348\ninside 349 372

22409 GCF_002063525.1_ASM206352v1 Salmonella enterica subsp. enterica serovar Virchow Proteobacteria; Gammaproteobacteria; Enterobacterales; Enterobacteriaceae; Salmonella; Salmonella enterica; Salmonella enterica subsp. enterica MTGDNTLITSHGINRRDFMKLCAALAATMGLSSKAAAEMAESVSNPQRPPVIWIGAQECT WP_000145429.1 hydrogenase 2 small subunit [Salmonella enterica] Length: 372\nNumber of predicted TMHs: 1\nExp number of AAs in TMHs: 24.51309\nExp number, first 60 AAs: 0.2682\nTotal prob of N-in: 0.14402\noutside 1 331\nTMhelix 332 354\ninside 355 372

22410 GCF_002063435.1_ASM206343v1 Salmonella enterica subsp. enterica serovar Weltenvreden Proteobacteria; Gammaproteobacteria; Enterobacterales; Enterobacteriaceae; Salmonella; Salmonella enterica; Salmonella enterica subsp. enterica MNNEETFYQAMRRKGVTRRSFLKFCSLAATSLGLGAGMTPKIAWALENKPRIPVVWIHGL WP_001058307.1 hydrogenase [Salmonella enterica] Length: 372\nNumber of predicted TMHs: 2\nExp number of AAs in TMHs: 36.51738\nExp number, first 60 AAs: 19.72849\nTotal prob of N-in: 0.96170\nPOSSIBLE N-term signal sequence\ninside 1 19\nTMhelix 20 39\noutside 40 325\nTMhelix 326 348\ninside 349 372

22411 GCF_002063435.1_ASM206343v1 Salmonella enterica subsp. enterica serovar Weltenvreden Proteobacteria; Gammaproteobacteria; Enterobacterales; Enterobacteriaceae; Salmonella; Salmonella enterica; Salmonella enterica subsp. enterica MTGDNTLITSHGINRRDFMKLCAALAATMGLSSKAAAEMAESVSNPQRPPVIWIGAQECT WP_000145429.1 hydrogenase 2 small subunit [Salmonella enterica] Length: 372\nNumber of predicted TMHs: 1\nExp number of AAs in TMHs: 24.51309\nExp number, first 60 AAs: 0.2682\nTotal prob of N-in: 0.14402\noutside 1 331\nTMhelix 332 354\ninside 355 372

22412 GCF_002062735.1_ASM206273v1 Salmonella enterica subsp. enterica serovar Weltenvreden Proteobacteria; Gammaproteobacteria; Enterobacterales; Enterobacteriaceae; Salmonella; Salmonella enterica; Salmonella enterica subsp. enterica MTGDNTLITSHGINRRDFMKLCAALAATMGLSSKAAAEMAESVSNPQRPPVIWIGAQECT WP_000145429.1 hydrogenase 2 small subunit [Salmonella enterica] Length: 372\nNumber of predicted TMHs: 1\nExp number of AAs in TMHs: 24.51309\nExp number, first 60 AAs: 0.2682\nTotal prob of N-in: 0.14402\noutside 1 331\nTMhelix 332 354\ninside 355 372

22413 GCF_002062735.1_ASM206273v1 Salmonella enterica subsp. enterica serovar Weltenvreden Proteobacteria; Gammaproteobacteria; Enterobacterales; Enterobacteriaceae; Salmonella; Salmonella enterica; Salmonella enterica subsp. enterica MNNEETFYQAMRRKGVTRRSFLKFCSLAATSLGLGAGMTPKIAWALENKPRIPVVWIHGL WP_001058307.1 hydrogenase [Salmonella enterica] Length: 372\nNumber of predicted TMHs: 2\nExp number of AAs in TMHs: 36.51738\nExp number, first 60 AAs: 19.72849\nTotal prob of N-in: 0.96170\nPOSSIBLE N-term signal sequence\ninside 1 19\nTMhelix 20 39\noutside 40 325\nTMhelix 326 348\ninside 349 372

22414 GCF_002050595.1_ASM205059v1 Salmonella enterica subsp. enterica serovar Weltevreden Proteobacteria; Gammaproteobacteria; Enterobacterales; Enterobacteriaceae; Salmonella; Salmonella enterica; Salmonella enterica subsp. enterica MNNEETFYQAMRRKGVTRRSFLKFCSLAATSLGLGAGMTPKIAWALENKPRIPVVWIHGL WP_001058307.1 hydrogenase [Salmonella enterica] Length: 372\nNumber of predicted TMHs: 2\nExp number of AAs in TMHs: 36.51738\nExp number, first 60 AAs: 19.72849\nTotal prob of N-in: 0.96170\nPOSSIBLE N-term signal sequence\ninside 1 19\nTMhelix 20 39\noutside 40 325\nTMhelix 326 348\ninside 349 372

22415 GCF_001410155.1_12227_3_24 Salmonella enterica subsp. enterica serovar Weltevreden Proteobacteria; Gammaproteobacteria; Enterobacterales; Enterobacteriaceae; Salmonella; Salmonella enterica; Salmonella enterica subsp. enterica MTGDNTLITSHGINRRDFMKLCAALAATMGLSSKAAAEMAESVSNPQRPPVIWIGAQECT WP_000145429.1 hydrogenase 2 small subunit [Salmonella enterica] Length: 372\nNumber of predicted TMHs: 1\nExp number of AAs in TMHs: 24.51309\nExp number, first 60 AAs: 0.2682\nTotal prob of N-in: 0.14402\noutside 1 331\nTMhelix 332 354\ninside 355 372

22416 GCF_002050835.1_ASM205083v1 Salmonella enterica subsp. enterica serovar Weltevreden Proteobacteria; Gammaproteobacteria; Enterobacterales; Enterobacteriaceae; Salmonella; Salmonella enterica; Salmonella enterica subsp. enterica MNNEETFYQAMRRKGVTRRSFLKFCSLAATSLGLGAGMTPKIAWALENKPRIPVVWIHGL WP_001058307.1 hydrogenase [Salmonella enterica] Length: 372\nNumber of predicted TMHs: 2\nExp number of AAs in TMHs: 36.51738\nExp number, first 60 AAs: 19.72849\nTotal prob of N-in: 0.96170\nPOSSIBLE N-term signal sequence\ninside 1 19\nTMhelix 20 39\noutside 40 325\nTMhelix 326 348\ninside 349 372

22417 GCF_001410995.1_12227_3_26 Salmonella enterica subsp. enterica serovar Weltevreden Proteobacteria; Gammaproteobacteria; Enterobacterales; Enterobacteriaceae; Salmonella; Salmonella enterica; Salmonella enterica subsp. enterica MTGDNTLITSHGINRRDFMKLCAALAATMGLSSKAAAEMAESVSNPQRPPVIWIGAQECT WP_000145429.1 hydrogenase 2 small subunit [Salmonella enterica] Length: 372\nNumber of predicted TMHs: 1\nExp number of AAs in TMHs: 24.51309\nExp number, first 60 AAs: 0.2682\nTotal prob of N-in: 0.14402\noutside 1 331\nTMhelix 332 354\ninside 355 372

22418 GCF_001410955.1_10868_1_89 Salmonella enterica subsp. enterica serovar Weltevreden Proteobacteria; Gammaproteobacteria; Enterobacterales; Enterobacteriaceae; Salmonella; Salmonella enterica; Salmonella enterica subsp. enterica MTGDNTLITSHGINRRDFMKLCAALAATMGLSSKAAAEMAESVSNPQRPPVIWIGAQECT WP_000145429.1 hydrogenase 2 small subunit [Salmonella enterica] Length: 372\nNumber of predicted TMHs: 1\nExp number of AAs in TMHs: 24.51309\nExp number, first 60 AAs: 0.2682\nTotal prob of N-in: 0.14402\noutside 1 331\nTMhelix 332 354\ninside 355 372

22419 GCF_001410975.1_12216_4_62 Salmonella enterica subsp. enterica serovar Weltevreden Proteobacteria; Gammaproteobacteria; Enterobacterales; Enterobacteriaceae; Salmonella; Salmonella enterica; Salmonella enterica subsp. enterica MTGDNTLITSHGINRRDFMKLCAALAATMGLSSKAAAEMAESVSNPQRPPVIWIGAQECT WP_000145429.1 hydrogenase 2 small subunit [Salmonella enterica] Length: 372\nNumber of predicted TMHs: 1\nExp number of AAs in TMHs: 24.51309\nExp number, first 60 AAs: 0.2682\nTotal prob of N-in: 0.14402\noutside 1 331\nTMhelix 332 354\ninside 355 372

22420 GCF_001410895.1_12227_3_33 Salmonella enterica subsp. enterica serovar Weltevreden Proteobacteria; Gammaproteobacteria; Enterobacterales; Enterobacteriaceae; Salmonella; Salmonella enterica; Salmonella enterica subsp. enterica MNNEETFYQAMRRKGVTRRSFLKFCSLAATSLGLGAGMTPKIAWALENKPRIPVVWIHGL WP_001058307.1 hydrogenase [Salmonella enterica] Length: 372\nNumber of predicted TMHs: 2\nExp number of AAs in TMHs: 36.51738\nExp number, first 60 AAs: 19.72849\nTotal prob of N-in: 0.96170\nPOSSIBLE N-term signal sequence\ninside 1 19\nTMhelix 20 39\noutside 40 325\nTMhelix 326 348\ninside 349 372

22421 GCF_001410915.1_12216_4_57 Salmonella enterica subsp. enterica serovar Weltevreden Proteobacteria; Gammaproteobacteria; Enterobacterales; Enterobacteriaceae; Salmonella; Salmonella enterica; Salmonella enterica subsp. enterica MNNEETFYQAMRRKGVTRRSFLKFCSLAATSLGLGAGMTPKIAWALENKPRIPVVWIHGL WP_001058307.1 hydrogenase [Salmonella enterica] Length: 372\nNumber of predicted TMHs: 2\nExp number of AAs in TMHs: 36.51738\nExp number, first 60 AAs: 19.72849\nTotal prob of N-in: 0.96170\nPOSSIBLE N-term signal sequence\ninside 1 19\nTMhelix 20 39\noutside 40 325\nTMhelix 326 348\ninside 349 372

22422 GCF_001410875.1_12227_3_34 Salmonella enterica subsp. enterica serovar Weltevreden Proteobacteria; Gammaproteobacteria; Enterobacterales; Enterobacteriaceae; Salmonella; Salmonella enterica; Salmonella enterica subsp. enterica MNNEETFYQAMRRKGVTRRSFLKFCSLAATSLGLGAGMTPKIAWALENKPRIPVVWIHGL WP_001058307.1 hydrogenase [Salmonella enterica] Length: 372\nNumber of predicted TMHs: 2\nExp number of AAs in TMHs: 36.51738\nExp number, first 60 AAs: 19.72849\nTotal prob of N-in: 0.96170\nPOSSIBLE N-term signal sequence\ninside 1 19\nTMhelix 20 39\noutside 40 325\nTMhelix 326 348\ninside 349 372

22423 GCF_001410855.1_12227_3_25 Salmonella enterica subsp. enterica serovar Weltevreden Proteobacteria; Gammaproteobacteria; Enterobacterales; Enterobacteriaceae; Salmonella; Salmonella enterica; Salmonella enterica subsp. enterica MNNEETFYQAMRRKGVTRRSFLKFCSLAATSLGLGAGMTPKIAWALENKPRIPVVWIHGL WP_001058307.1 hydrogenase [Salmonella enterica] Length: 372\nNumber of predicted TMHs: 2\nExp number of AAs in TMHs: 36.51738\nExp number, first 60 AAs: 19.72849\nTotal prob of N-in: 0.96170\nPOSSIBLE N-term signal sequence\ninside 1 19\nTMhelix 20 39\noutside 40 325\nTMhelix 326 348\ninside 349 372

22424 GCF_001410935.1_9472_3_5Salmonella enterica subsp. enterica serovar Weltevreden Proteobacteria; Gammaproteobacteria; Enterobacterales; Enterobacteriaceae; Salmonella; Salmonella enterica; Salmonella enterica subsp. enterica MNNEETFYQAMRRKGVTRRSFLKFCSLAATSLGLGAGMTPKIAWALENKPRIPVVWIHGL WP_001058307.1 hydrogenase [Salmonella enterica] Length: 372\nNumber of predicted TMHs: 2\nExp number of AAs in

TMHs: 36.51738\nExp number, first 60 AAs: 19.72849\nTotal prob of N-in: 0.96170\nPOSSIBLE N-term signal sequence\ninside 1 19\nTMhelix 20 39\noutside 40 325\nTMhelix 326 348\ninside 349 372

22425 GCF_001410935.1_9472_3_5Salmonella enterica subsp. enterica serovar Weltevreden Proteobacteria; Gammaproteobacteria; Enterobacterales; Enterobacteriaceae; Salmonella; Salmonella enterica; Salmonella enterica subsp. enterica MTGDNTLITSHGINRRDFMKLCAALAATMGLSSKAAAEMAESVSNPQRPPVIWIGAQECT WP_000145429.1 hydrogenase 2 small subunit [Salmonella enterica] Length: 372\nNumber of predicted TMHs: 1\nExp number of AAs in TMHs: 24.51309\nExp number, first 60 AAs: 0.2682\nTotal prob of N-in: 0.14402\noutside 1 331\nTMhelix 332 354\ninside 355 372

22426 GCF_001410815.1_10868_1_91 Salmonella enterica subsp. enterica serovar Weltevreden Proteobacteria; Gammaproteobacteria; Enterobacterales; Enterobacteriaceae; Salmonella; Salmonella enterica; Salmonella enterica subsp. enterica MNNEETFYQAMRRKGVTRRSFLKFCSLAATSLGLGAGMTPKIAWALENKPRIPVVWIHGL WP_001058307.1 hydrogenase [Salmonella enterica] Length: 372\nNumber of predicted TMHs: 2\nExp number of AAs in TMHs: 36.51738\nExp number, first 60 AAs: 19.72849\nTotal prob of N-in: 0.96170\nPOSSIBLE N-term signal sequence\ninside 1 19\nTMhelix 20 39\noutside 40 325\nTMhelix 326 348\ninside 349 372

22427 GCF_001410835.1_9472_3_1Salmonella enterica subsp. enterica serovar Weltevreden Proteobacteria; Gammaproteobacteria; Enterobacterales; Enterobacteriaceae; Salmonella; Salmonella enterica; Salmonella enterica subsp. enterica MNNEETFYQAMRRKGVTRRSFLKFCSLAATSLGLGAGMTPKIAWALENKPRIPVVWIHGL WP_001058307.1 hydrogenase [Salmonella enterica] Length: 372\nNumber of predicted TMHs: 2\nExp number of AAs in TMHs: 36.51738\nExp number, first 60 AAs: 19.72849\nTotal prob of N-in: 0.96170\nPOSSIBLE N-term signal sequence\ninside 1 19\nTMhelix 20 39\noutside 40 325\nTMhelix 326 348\ninside 349 372

22428 GCF_001410795.1_9472_3_33 Salmonella enterica subsp. enterica serovar Weltevreden Proteobacteria; Gammaproteobacteria; Enterobacterales; Enterobacteriaceae; Salmonella; Salmonella enterica; Salmonella enterica subsp. enterica MNNEETFYQAMRRKGVTRRSFLKFCSLAATSLGLGAGMTPKIAWALENKPRIPVVWIHGL WP_001058307.1 hydrogenase [Salmonella enterica] Length: 372\nNumber of predicted TMHs: 2\nExp number of AAs in TMHs: 36.51738\nExp number, first 60 AAs: 19.72849\nTotal prob of N-in: 0.96170\nPOSSIBLE N-term signal sequence\ninside 1 19\nTMhelix 20 39\noutside 40 325\nTMhelix 326 348\ninside 349 372

22429 GCF_001410895.1_12227_3_33 Salmonella enterica subsp. enterica serovar Weltevreden Proteobacteria; Gammaproteobacteria; Enterobacterales; Enterobacteriaceae; Salmonella; Salmonella enterica; Salmonella enterica subsp. enterica MTGDNTLITSHGINRRDFMKLCAALAATMGLSSKAAAEMAESVSNPQRPPVIWIGAQECT WP_000145429.1 hydrogenase 2 small subunit [Salmonella enterica] Length: 372\nNumber of predicted TMHs: 1\nExp number of AAs in TMHs: 24.51309\nExp number, first 60 AAs: 0.2682\nTotal prob of N-in: 0.14402\noutside 1 331\nTMhelix 332 354\ninside 355 372

22430 GCF_001410915.1_12216_4_57 Salmonella enterica subsp. enterica serovar Weltevreden Proteobacteria; Gammaproteobacteria; Enterobacterales; Enterobacteriaceae; Salmonella; Salmonella enterica; Salmonella enterica subsp. enterica MTGDNTLITSHGINRRDFMKLCAALAATMGLSSKAAAEMAESVSNPQRPPVIWIGAQECT WP_000145429.1 hydrogenase 2 small subunit [Salmonella enterica] Length: 372\nNumber of predicted TMHs: 1\nExp number of AAs in TMHs: 24.51309\nExp number, first 60 AAs: 0.2682\nTotal prob of N-in: 0.14402\noutside 1 331\nTMhelix 332 354\ninside 355 372

22431 GCF_001410875.1_12227_3_34 Salmonella enterica subsp. enterica serovar Weltevreden Proteobacteria; Gammaproteobacteria; Enterobacterales; Enterobacteriaceae; Salmonella; Salmonella enterica; Salmonella enterica subsp. enterica MTGDNTLITSHGINRRDFMKLCAALAATMGLSSKAAAEMAESVSNPQRPPVIWIGAQECT WP_000145429.1 hydrogenase 2 small subunit [Salmonella enterica] Length: 372\nNumber of predicted TMHs: 1\nExp number of AAs in TMHs: 24.51309\nExp number, first 60 AAs: 0.2682\nTotal prob of N-in: 0.14402\noutside 1 331\nTMhelix 332 354\ninside 355 372

22432 GCF_001410855.1_12227_3_25 Salmonella enterica subsp. enterica serovar Weltevreden Proteobacteria; Gammaproteobacteria; Enterobacterales; Enterobacteriaceae; Salmonella; Salmonella enterica; Salmonella enterica subsp. enterica MTGDNTLITSHGINRRDFMKLCAALAATMGLSSKAAAEMAESVSNPQRPPVIWIGAQECT WP_000145429.1 hydrogenase 2 small subunit [Salmonella enterica] Length: 372\nNumber of predicted TMHs: 1\nExp number of AAs in TMHs: 24.51309\nExp number, first 60 AAs: 0.2682\nTotal prob of N-in: 0.14402\noutside 1 331\nTMhelix 332 354\ninside 355 372

22433 GCF_001410815.1_10868_1_91 Salmonella enterica subsp. enterica serovar Weltevreden Proteobacteria; Gammaproteobacteria; Enterobacterales; Enterobacteriaceae; Salmonella; Salmonella enterica;

Salmonella enterica subsp. enterica MTGDNTLITSHGINRRDFMKLCAALAATMGLSSKAAAEMAESVSNPQRPPVIWIGAQECT WP_000145429.1 hydrogenase 2 small subunit [Salmonella enterica] Length: 372\nNumber of predicted TMHs: 1\nExp number of AAs in TMHs: 24.51309\nExp number, first 60 AAs: 0.2682\nTotal prob of N-in: 0.14402\noutside 1 331\nTMhelix 332 354\ninside 355 372

22434 GCF_001410835.1_9472_3_1Salmonella enterica subsp. enterica serovar Weltevreden Proteobacteria; Gammaproteobacteria; Enterobacterales; Enterobacteriaceae; Salmonella; Salmonella enterica; Salmonella enterica subsp. enterica MTGDNTLITSHGINRRDFMKLCAALAATMGLSSKAAAEMAESVSNPQRPPVIWIGAQECT WP_000145429.1 hydrogenase 2 small subunit [Salmonella enterica] Length: 372\nNumber of predicted TMHs: 1\nExp number of AAs in TMHs: 24.51309\nExp number, first 60 AAs: 0.2682\nTotal prob of N-in: 0.14402\noutside 1 331\nTMhelix 332 354\ninside 355 372

22435 GCF_001410735.1_12227_3_32 Salmonella enterica subsp. enterica serovar Weltevreden Proteobacteria; Gammaproteobacteria; Enterobacterales; Enterobacteriaceae; Salmonella; Salmonella enterica; Salmonella enterica subsp. enterica MNNEETFYQAMRRKGVTRRSFLKFCSLAATSLGLGAGMTPKIAWALENKPRIPVVWIHGL WP_001058307.1 hydrogenase [Salmonella enterica] Length: 372\nNumber of predicted TMHs: 2\nExp number of AAs in TMHs: 36.51738\nExp number, first 60 AAs: 19.72849\nTotal prob of N-in: 0.96170\nPOSSIBLE N-term signal sequence\ninside 1 19\nTMhelix 20 39\noutside 40 325\nTMhelix 326 348\ninside 349 372

22436 GCF_001410755.1_12216_4_54 Salmonella enterica subsp. enterica serovar Weltevreden Proteobacteria; Gammaproteobacteria; Enterobacterales; Enterobacteriaceae; Salmonella; Salmonella enterica; Salmonella enterica subsp. enterica MNNEETFYQAMRRKGVTRRSFLKFCSLAATSLGLGAGMTPKIAWALENKPRIPVVWIHGL WP_001058307.1 hydrogenase [Salmonella enterica] Length: 372\nNumber of predicted TMHs: 2\nExp number of AAs in TMHs: 36.51738\nExp number, first 60 AAs: 19.72849\nTotal prob of N-in: 0.96170\nPOSSIBLE N-term signal sequence\ninside 1 19\nTMhelix 20 39\noutside 40 325\nTMhelix 326 348\ninside 349 372

22437 GCF_001410715.1_9472_3_27 Salmonella enterica subsp. enterica serovar Weltevreden Proteobacteria; Gammaproteobacteria; Enterobacterales; Enterobacteriaceae; Salmonella; Salmonella enterica; Salmonella enterica subsp. enterica MNNEETFYQAMRRKGVTRRSFLKFCSLAATSLGLGAGMTPKIAWALENKPRIPVVWIHGL WP_001058307.1 hydrogenase [Salmonella enterica] Length: 372\nNumber of predicted TMHs: 2\nExp number of AAs in TMHs: 36.51738\nExp number, first 60 AAs: 19.72849\nTotal prob of N-in: 0.96170\nPOSSIBLE N-term signal sequence\ninside 1 19\nTMhelix 20 39\noutside 40 325\nTMhelix 326 348\ninside 349 372

22438 GCF_001410795.1_9472_3_33 Salmonella enterica subsp. enterica serovar Weltevreden Proteobacteria; Gammaproteobacteria; Enterobacterales; Enterobacteriaceae; Salmonella; Salmonella enterica; Salmonella enterica subsp. enterica MTGDNTLITSHGINRRDFMKLCAALAATMGLSSKAAAEMAESVSNPQRPPVIWIGAQECT WP_000145429.1 hydrogenase 2 small subunit [Salmonella enterica] Length: 372\nNumber of predicted TMHs: 1\nExp number of AAs in TMHs: 24.51309\nExp number, first 60 AAs: 0.2682\nTotal prob of N-in: 0.14402\noutside 1 331\nTMhelix 332 354\ninside 355 372

22439 GCF_001410775.1_12216_4_46 Salmonella enterica subsp. enterica serovar Weltevreden Proteobacteria; Gammaproteobacteria; Enterobacterales; Enterobacteriaceae; Salmonella; Salmonella enterica; Salmonella enterica subsp. enterica MTGDNTLITSHGINRRDFMKLCAALAATMGLSSKAAAEMAESVSNPQRPPVIWIGAQECT WP_000145429.1 hydrogenase 2 small subunit [Salmonella enterica] Length: 372\nNumber of predicted TMHs: 1\nExp number of AAs in TMHs: 24.51309\nExp number, first 60 AAs: 0.2682\nTotal prob of N-in: 0.14402\noutside 1 331\nTMhelix 332 354\ninside 355 372

22440 GCF_001410755.1_12216_4_54 Salmonella enterica subsp. enterica serovar Weltevreden Proteobacteria; Gammaproteobacteria; Enterobacterales; Enterobacteriaceae; Salmonella; Salmonella enterica; Salmonella enterica subsp. enterica MTGDNTLITSHGINRRDFMKLCAALAATMGLSSKAAAEMAESVSNPQRPPVIWIGAQECT WP_000145429.1 hydrogenase 2 small subunit [Salmonella enterica] Length: 372\nNumber of predicted TMHs: 1\nExp number of AAs in TMHs: 24.51309\nExp number, first 60 AAs: 0.2682\nTotal prob of N-in: 0.14402\noutside 1 331\nTMhelix 332 354\ninside 355 372

22441 GCF_001410735.1_12227_3_32 Salmonella enterica subsp. enterica serovar Weltevreden Proteobacteria; Gammaproteobacteria; Enterobacterales; Enterobacteriaceae; Salmonella; Salmonella enterica; Salmonella enterica subsp. enterica MTGDNTLITSHGINRRDFMKLCAALAATMGLSSKAAAEMAESVSNPQRPPVIWIGAQECT WP_000145429.1 hydrogenase 2 small subunit [Salmonella enterica] Length: 372\nNumber of predicted TMHs: 1\nExp number of AAs in TMHs: 24.51309\nExp number, first 60 AAs: 0.2682\nTotal prob of N-in: 0.14402\noutside 1 331\nTMhelix 332 354\ninside 355 372

22442 GCF_001410715.1_9472_3_27 Salmonella enterica subsp. enterica serovar Weltevreden Proteobacteria; Gammaproteobacteria; Enterobacterales; Enterobacteriaceae; Salmonella; Salmonella enterica; Salmonella enterica subsp. enterica MTGDNTLITSHGINRRDFMKLCAALAATMGLSSKAAAEMAESVSNPQRPPVIWIGAQECT WP_000145429.1 hydrogenase 2 small subunit [Salmonella enterica] Length: 372\nNumber of predicted TMHs: 1\nExp number of AAs in TMHs: 24.51309\nExp number, first 60 AAs: 0.2682\nTotal prob of N-in: 0.14402\noutside 1 331\nTMhelix 332 354\ninside 355 372

22443 GCF_001410675.1_12227_3_19 Salmonella enterica subsp. enterica serovar Weltevreden Proteobacteria; Gammaproteobacteria; Enterobacterales; Enterobacteriaceae; Salmonella; Salmonella enterica; Salmonella enterica subsp. enterica MNNEETFYQAMRRKGVTRRSFLKFCSLAATSLGLGAGMTPKIAWALENKPRIPVVWIHGL WP_001058307.1 hydrogenase [Salmonella enterica] Length: 372\nNumber of predicted TMHs: 2\nExp number of AAs in TMHs: 36.51738\nExp number, first 60 AAs: 19.72849\nTotal prob of N-in: 0.96170\nPOSSIBLE N-term signal sequence\ninside 1 19\nTMhelix 20 39\noutside 40 325\nTMhelix 326 348\ninside 349 372

22444 GCF_001410655.1_12227_3_41 Salmonella enterica subsp. enterica serovar Weltevreden Proteobacteria; Gammaproteobacteria; Enterobacterales; Enterobacteriaceae; Salmonella; Salmonella enterica; Salmonella enterica subsp. enterica MNNEETFYQAMRRKGVTRRSFLKFCSLAATSLGLGAGMTPKIAWALENKPRIPVVWIHGL WP_001058307.1 hydrogenase [Salmonella enterica] Length: 372\nNumber of predicted TMHs: 2\nExp number of AAs in TMHs: 36.51738\nExp number, first 60 AAs: 19.72849\nTotal prob of N-in: 0.96170\nPOSSIBLE N-term signal sequence\ninside 1 19\nTMhelix 20 39\noutside 40 325\nTMhelix 326 348\ninside 349 372

22445 GCF_001410635.1_12216_4_44 Salmonella enterica subsp. enterica serovar Weltevreden Proteobacteria; Gammaproteobacteria; Enterobacterales; Enterobacteriaceae; Salmonella; Salmonella enterica; Salmonella enterica subsp. enterica MNNEETFYQAMRRKGVTRRSFLKFCSLAATSLGLGAGMTPKIAWALENKPRIPVVWIHGL WP_001058307.1 hydrogenase [Salmonella enterica] Length: 372\nNumber of predicted TMHs: 2\nExp number of AAs in TMHs: 36.51738\nExp number, first 60 AAs: 19.72849\nTotal prob of N-in: 0.96170\nPOSSIBLE N-term signal sequence\ninside 1 19\nTMhelix 20 39\noutside 40 325\nTMhelix 326 348\ninside 349 372

22446 GCF_001410595.1_12227_3_49 Salmonella enterica subsp. enterica serovar Weltevreden Proteobacteria; Gammaproteobacteria; Enterobacterales; Enterobacteriaceae; Salmonella; Salmonella enterica; Salmonella enterica subsp. enterica MNNEETFYQAMRRKGVTRRSFLKFCSLAATSLGLGAGMTPKIAWALENKPRIPVVWIHGL WP_001058307.1 hydrogenase [Salmonella enterica] Length: 372\nNumber of predicted TMHs: 2\nExp number of AAs in TMHs: 36.51738\nExp number, first 60 AAs: 19.72849\nTotal prob of N-in: 0.96170\nPOSSIBLE N-term signal sequence\ninside 1 19\nTMhelix 20 39\noutside 40 325\nTMhelix 326 348\ninside 349 372

22447 GCF_001410615.1_12216_4_51 Salmonella enterica subsp. enterica serovar Weltevreden Proteobacteria; Gammaproteobacteria; Enterobacterales; Enterobacteriaceae; Salmonella; Salmonella enterica; Salmonella enterica subsp. enterica MNNEETFYQAMRRKGVTRRSFLKFCSLAATSLGLGAGMTPKIAWALENKPRIPVVWIHGL WP_001058307.1 hydrogenase [Salmonella enterica] Length: 372\nNumber of predicted TMHs: 2\nExp number of AAs in TMHs: 36.51738\nExp number, first 60 AAs: 19.72849\nTotal prob of N-in: 0.96170\nPOSSIBLE N-term signal sequence\ninside 1 19\nTMhelix 20 39\noutside 40 325\nTMhelix 326 348\ninside 349 372

22448 GCF_001410695.1_12227_3_46 Salmonella enterica subsp. enterica serovar Weltevreden Proteobacteria; Gammaproteobacteria; Enterobacterales; Enterobacteriaceae; Salmonella; Salmonella enterica; Salmonella enterica subsp. enterica MNNEETFYQAMRRKGVTRRSFLKFCSLAATSLGLGAGMTPKIAWALENKPRIPVVWIHGL WP_001058307.1 hydrogenase [Salmonella enterica] Length: 372\nNumber of predicted TMHs: 2\nExp number of AAs in TMHs: 36.51738\nExp number, first 60 AAs: 19.72849\nTotal prob of N-in: 0.96170\nPOSSIBLE N-term signal sequence\ninside 1 19\nTMhelix 20 39\noutside 40 325\nTMhelix 326 348\ninside 349 372

22449 GCF_001410695.1_12227_3_46 Salmonella enterica subsp. enterica serovar Weltevreden Proteobacteria; Gammaproteobacteria; Enterobacterales; Enterobacteriaceae; Salmonella; Salmonella enterica; Salmonella enterica subsp. enterica MTGDNTLITSHGINRRDFMKLCAALAATMGLSSKAAAEMAESVSNPQRPPVIWIGAQECT WP_000145429.1 hydrogenase 2 small subunit [Salmonella enterica] Length: 372\nNumber of predicted TMHs: 1\nExp number of AAs in TMHs: 24.51309\nExp number, first 60 AAs: 0.2682\nTotal prob of N-in: 0.14402\noutside 1 331\nTMhelix 332 354\ninside 355 372

22450 GCF_001410675.1_12227_3_19 Salmonella enterica subsp. enterica serovar Weltevreden Proteobacteria; Gammaproteobacteria; Enterobacterales; Enterobacteriaceae; Salmonella; Salmonella enterica; Salmonella enterica subsp. enterica MTGDNTLITSHGINRRDFMKLCAALAATMGLSSKAAAEMAESVSNPQRPPVIWIGAQECT WP_000145429.1 hydrogenase 2 small subunit [Salmonella enterica] Length: 372\nNumber of

predicted TMHs:  1\nExp number of AAs in TMHs: 24.51309\nExp number, first 60 AAs:  0.2682\nTotal prob of N-in:  0.14402\noutside  1  331\nTMhelix  332  354\ninside  355  372

22451 GCF_001410655.1_12227_3_41 Salmonella enterica subsp. enterica serovar Weltevreden Proteobacteria; Gammaproteobacteria; Enterobacterales; Enterobacteriaceae; Salmonella; Salmonella enterica; Salmonella enterica subsp. enterica MTGDNTLITSHGINRRDFMKLCAALAATMGLSSKAAAEMAESVSNPQRPPVIWIGAQECT WP_000145429.1 hydrogenase 2 small subunit [Salmonella enterica] Length: 372\nNumber of predicted TMHs:  1\nExp number of AAs in TMHs: 24.51309\nExp number, first 60 AAs:  0.2682\nTotal prob of N-in:  0.14402\noutside  1  331\nTMhelix  332  354\ninside  355  372

22452 GCF_001410635.1_12216_4_44 Salmonella enterica subsp. enterica serovar Weltevreden Proteobacteria; Gammaproteobacteria; Enterobacterales; Enterobacteriaceae; Salmonella; Salmonella enterica; Salmonella enterica subsp. enterica MTGDNTLITSHGINRRDFMKLCAALAATMGLSSKAAAEMAESVSNPQRPPVIWIGAQECT WP_000145429.1 hydrogenase 2 small subunit [Salmonella enterica] Length: 372\nNumber of predicted TMHs:  1\nExp number of AAs in TMHs: 24.51309\nExp number, first 60 AAs:  0.2682\nTotal prob of N-in:  0.14402\noutside  1  331\nTMhelix  332  354\ninside  355  372

22453 GCF_001410615.1_12216_4_51 Salmonella enterica subsp. enterica serovar Weltevreden Proteobacteria; Gammaproteobacteria; Enterobacterales; Enterobacteriaceae; Salmonella; Salmonella enterica; Salmonella enterica subsp. enterica MTGDNTLITSHGINRRDFMKLCAALAATMGLSSKAAAEMAESVSNPQRPPVIWIGAQECT WP_000145429.1 hydrogenase 2 small subunit [Salmonella enterica] Length: 372\nNumber of predicted TMHs:  1\nExp number of AAs in TMHs: 24.51309\nExp number, first 60 AAs:  0.2682\nTotal prob of N-in:  0.14402\noutside  1  331\nTMhelix  332  354\ninside  355  372

22454 GCF_001410555.1_9472_3_18 Salmonella enterica subsp. enterica serovar Weltevreden Proteobacteria; Gammaproteobacteria; Enterobacterales; Enterobacteriaceae; Salmonella; Salmonella enterica; Salmonella enterica subsp. enterica MNNEETFYQAMRRKGVTRRSFLKFCSLAATSLGLGAGMTPKIAWALENKPRIPVVWIHGL WP_001058307.1 hydrogenase [Salmonella enterica] Length: 372\nNumber of predicted TMHs:  2\nExp number of AAs in TMHs: 36.51738\nExp number, first 60 AAs:  19.72849\nTotal prob of N-in:  0.96170\nPOSSIBLE N-term signal sequence\ninside  1  19\nTMhelix  20  39\noutside  40  325\nTMhelix  326  348\ninside  349  372

22455 GCF_001410535.1_12227_3_2 Salmonella enterica subsp. enterica serovar Weltevreden Proteobacteria; Gammaproteobacteria; Enterobacterales; Enterobacteriaceae; Salmonella; Salmonella enterica; Salmonella enterica subsp. enterica MNNEETFYQAMRRKGVTRRSFLKFCSLAATSLGLGAGMTPKIAWALENKPRIPVVWIHGL WP_001058307.1 hydrogenase [Salmonella enterica] Length: 372\nNumber of predicted TMHs:  2\nExp number of AAs in TMHs: 36.51738\nExp number, first 60 AAs:  19.72849\nTotal prob of N-in:  0.96170\nPOSSIBLE N-term signal sequence\ninside  1  19\nTMhelix  20  39\noutside  40  325\nTMhelix  326  348\ninside  349  372

22456 GCF_001410575.1_12227_3_35 Salmonella enterica subsp. enterica serovar Weltevreden Proteobacteria; Gammaproteobacteria; Enterobacterales; Enterobacteriaceae; Salmonella; Salmonella enterica; Salmonella enterica subsp. enterica MNNEETFYQAMRRKGVTRRSFLKFCSLAATSLGLGAGMTPKIAWALENKPRIPVVWIHGL WP_001058307.1 hydrogenase [Salmonella enterica] Length: 372\nNumber of predicted TMHs:  2\nExp number of AAs in TMHs: 36.51738\nExp number, first 60 AAs:  19.72849\nTotal prob of N-in:  0.96170\nPOSSIBLE N-term signal sequence\ninside  1  19\nTMhelix  20  39\noutside  40  325\nTMhelix  326  348\ninside  349  372

22457 GCF_001410575.1_12227_3_35 Salmonella enterica subsp. enterica serovar Weltevreden Proteobacteria; Gammaproteobacteria; Enterobacterales; Enterobacteriaceae; Salmonella; Salmonella enterica; Salmonella enterica subsp. enterica MTGDNTLITSHGINRRDFMKLCAALAATMGLSSKAAAEMAESVSNPQRPPVIWIGAQECT WP_000145429.1 hydrogenase 2 small subunit [Salmonella enterica] Length: 372\nNumber of predicted TMHs:  1\nExp number of AAs in TMHs: 24.51309\nExp number, first 60 AAs:  0.2682\nTotal prob of N-in:  0.14402\noutside  1  331\nTMhelix  332  354\ninside  355  372

22458 GCF_001410475.1_10900_1_32 Salmonella enterica subsp. enterica serovar Weltevreden Proteobacteria; Gammaproteobacteria; Enterobacterales; Enterobacteriaceae; Salmonella; Salmonella enterica; Salmonella enterica subsp. enterica MNNEETFYQAMRRKGVTRRSFLKFCSLAATSLGLGAGMTPKIAWALENKPRIPVVWIHGL WP_057521139.1 hydrogenase [Salmonella enterica] Length: 372\nNumber of predicted TMHs:  2\nExp number of AAs in TMHs: 36.76338\nExp number, first 60 AAs:  19.73298\nTotal prob of N-in:  0.96170\nPOSSIBLE N-term signal sequence\ninside  1  19\nTMhelix  20  39\noutside  40  325\nTMhelix  326  348\ninside  349  372

22459 GCF_001410515.1_12227_3_28 Salmonella enterica subsp. enterica serovar Weltevreden Proteobacteria; Gammaproteobacteria; Enterobacterales; Enterobacteriaceae; Salmonella; Salmonella enterica;

Salmonella enterica subsp. enterica	MNNEETFYQAMRRKGVTRRSFLKFCSLAATSLGLGAGMTPKIAWALENKPRIPVVWIHGL	WP_001058307.1	hydrogenase [Salmonella enterica]	Length: 372\nNumber of predicted TMHs:  2\nExp number of AAs in TMHs: 36.51738\nExp number, first 60 AAs:  19.72849\nTotal prob of N-in:   0.96170\nPOSSIBLE N-term signal sequence\ninside   1   19\nTMhelix   20   39\noutside   40   325\nTMhelix   326   348\ninside   349   372

22460	GCF_001410495.1_10868_1_90	Salmonella enterica subsp. enterica serovar Weltevreden	Proteobacteria; Gammaproteobacteria; Enterobacterales; Enterobacteriaceae; Salmonella; Salmonella enterica; Salmonella enterica subsp. enterica	MNNEETFYQAMRRKGVTRRSFLKFCSLAATSLGLGAGMTPKIAWALENKPRIPVVWIHGL	WP_001058307.1	hydrogenase [Salmonella enterica]	Length: 372\nNumber of predicted TMHs:  2\nExp number of AAs in TMHs: 36.51738\nExp number, first 60 AAs:  19.72849\nTotal prob of N-in:   0.96170\nPOSSIBLE N-term signal sequence\ninside   1   19\nTMhelix   20   39\noutside   40   325\nTMhelix   326   348\ninside   349   372

22461	GCF_001410555.1_9472_3_18	Salmonella enterica subsp. enterica serovar Weltevreden	Proteobacteria; Gammaproteobacteria; Enterobacterales; Enterobacteriaceae; Salmonella; Salmonella enterica; Salmonella enterica subsp. enterica	MTGDNTLITSHGINRRDFMKLCAALAATMGLSSKAAAEMAESVSNPQRPPVIWIGAQECT	WP_000145429.1	hydrogenase 2 small subunit [Salmonella enterica]	Length: 372\nNumber of predicted TMHs:  1\nExp number of AAs in TMHs: 24.51309\nExp number, first 60 AAs:  0.2682\nTotal prob of N-in: 0.14402\noutside   1   331\nTMhelix   332   354\ninside   355   372

22462	GCF_001410535.1_12227_3_2	Salmonella enterica subsp. enterica serovar Weltevreden	Proteobacteria; Gammaproteobacteria; Enterobacterales; Enterobacteriaceae; Salmonella; Salmonella enterica; Salmonella enterica subsp. enterica	MTGDNTLITSHGINRRDFMKLCAALAATMGLSSKAAAEMAESVSNPQRPPVIWIGAQECT	WP_000145429.1	hydrogenase 2 small subunit [Salmonella enterica]	Length: 372\nNumber of predicted TMHs:  1\nExp number of AAs in TMHs: 24.51309\nExp number, first 60 AAs:  0.2682\nTotal prob of N-in: 0.14402\noutside   1   331\nTMhelix   332   354\ninside   355   372

22463	GCF_001410515.1_12227_3_28	Salmonella enterica subsp. enterica serovar Weltevreden	Proteobacteria; Gammaproteobacteria; Enterobacterales; Enterobacteriaceae; Salmonella; Salmonella enterica; Salmonella enterica subsp. enterica	MTGDNTLITSHGINRRDFMKLCAALAATMGLSSKAAAEMAESVSNPQRPPVIWIGAQECT	WP_000145429.1	hydrogenase 2 small subunit [Salmonella enterica]	Length: 372\nNumber of predicted TMHs:  1\nExp number of AAs in TMHs: 24.51309\nExp number, first 60 AAs:  0.2682\nTotal prob of N-in: 0.14402\noutside   1   331\nTMhelix   332   354\ninside   355   372

22464	GCF_001410495.1_10868_1_90	Salmonella enterica subsp. enterica serovar Weltevreden	Proteobacteria; Gammaproteobacteria; Enterobacterales; Enterobacteriaceae; Salmonella; Salmonella enterica; Salmonella enterica subsp. enterica	MTGDNTLITSHGINRRDFMKLCAALAATMGLSSKAAAEMAESVSNPQRPPVIWIGAQECT	WP_000145429.1	hydrogenase 2 small subunit [Salmonella enterica]	Length: 372\nNumber of predicted TMHs:  1\nExp number of AAs in TMHs: 24.51309\nExp number, first 60 AAs:  0.2682\nTotal prob of N-in: 0.14402\noutside   1   331\nTMhelix   332   354\ninside   355   372

22465	GCF_001410455.1_9472_3_34	Salmonella enterica subsp. enterica serovar Weltevreden	Proteobacteria; Gammaproteobacteria; Enterobacterales; Enterobacteriaceae; Salmonella; Salmonella enterica; Salmonella enterica subsp. enterica	MNNEETFYQAMRRKGVTRRSFLKFCSLAATSLGLGAGMTPKIAWALENKPRIPVVWIHGL	WP_001058307.1	hydrogenase [Salmonella enterica]	Length: 372\nNumber of predicted TMHs:  2\nExp number of AAs in TMHs: 36.51738\nExp number, first 60 AAs:  19.72849\nTotal prob of N-in:   0.96170\nPOSSIBLE N-term signal sequence\ninside   1   19\nTMhelix   20   39\noutside   40   325\nTMhelix   326   348\ninside   349   372

22466	GCF_001410435.1_9472_3_25	Salmonella enterica subsp. enterica serovar Weltevreden	Proteobacteria; Gammaproteobacteria; Enterobacterales; Enterobacteriaceae; Salmonella; Salmonella enterica; Salmonella enterica subsp. enterica	MNNEETFYQAMRRKGVTRRSFLKFCSLAATSLGLGAGMTPKIAWALENKPRIPVVWIHGL	WP_001058307.1	hydrogenase [Salmonella enterica]	Length: 372\nNumber of predicted TMHs:  2\nExp number of AAs in TMHs: 36.51738\nExp number, first 60 AAs:  19.72849\nTotal prob of N-in:   0.96170\nPOSSIBLE N-term signal sequence\ninside   1   19\nTMhelix   20   39\noutside   40   325\nTMhelix   326   348\ninside   349   372

22467	GCF_001410595.1_12227_3_49	Salmonella enterica subsp. enterica serovar Weltevreden	Proteobacteria; Gammaproteobacteria; Enterobacterales; Enterobacteriaceae; Salmonella; Salmonella enterica; Salmonella enterica subsp. enterica	MTGDNTLITSHGINRRDFMKLCAALAATMGLSSKAAAEMAESVSNPQRPPVIWIGAQECT	WP_000145429.1	hydrogenase 2 small subunit [Salmonella enterica]	Length: 372\nNumber of predicted TMHs:  1\nExp number of AAs in TMHs: 24.51309\nExp number, first 60 AAs:  0.2682\nTotal prob of N-in: 0.14402\noutside   1   331\nTMhelix   332   354\ninside   355   372

22468 GCF_001410415.1_12227_3_44 Salmonella enterica subsp. enterica serovar Weltevreden Proteobacteria; Gammaproteobacteria; Enterobacterales; Enterobacteriaceae; Salmonella; Salmonella enterica; Salmonella enterica subsp. enterica MNNEETFYQAMRRKGVTRRSFLKFCSLAATSLGLGAGMTPKIAWALENKPRIPVVWIHGL WP_001058307.1 hydrogenase [Salmonella enterica] Length: 372\nNumber of predicted TMHs: 2\nExp number of AAs in TMHs: 36.51738\nExp number, first 60 AAs: 19.72849\nTotal prob of N-in: 0.96170\nPOSSIBLE N-term signal sequence\ninside 1 19\nTMhelix 20 39\noutside 40 325\nTMhelix 326 348\ninside 349 372

22469 GCF_001410395.1_12227_3_10 Salmonella enterica subsp. enterica serovar Weltevreden Proteobacteria; Gammaproteobacteria; Enterobacterales; Enterobacteriaceae; Salmonella; Salmonella enterica; Salmonella enterica subsp. enterica MNNEETFYQAMRRKGVTRRSFLKFCSLAATSLGLGAGMTPKIAWALENKPRIPVVWIHGL WP_001058307.1 hydrogenase [Salmonella enterica] Length: 372\nNumber of predicted TMHs: 2\nExp number of AAs in TMHs: 36.51738\nExp number, first 60 AAs: 19.72849\nTotal prob of N-in: 0.96170\nPOSSIBLE N-term signal sequence\ninside 1 19\nTMhelix 20 39\noutside 40 325\nTMhelix 326 348\ninside 349 372

22470 GCF_001410335.1_12227_3_57 Salmonella enterica subsp. enterica serovar Weltevreden Proteobacteria; Gammaproteobacteria; Enterobacterales; Enterobacteriaceae; Salmonella; Salmonella enterica; Salmonella enterica subsp. enterica MNNEETFYQAMRRKGVTRRSFLKFCSLAATSLGLGAGMTPKIAWALENKPRIPVVWIHGL WP_001058307.1 hydrogenase [Salmonella enterica] Length: 372\nNumber of predicted TMHs: 2\nExp number of AAs in TMHs: 36.51738\nExp number, first 60 AAs: 19.72849\nTotal prob of N-in: 0.96170\nPOSSIBLE N-term signal sequence\ninside 1 19\nTMhelix 20 39\noutside 40 325\nTMhelix 326 348\ninside 349 372

22471 GCF_001410355.1_12227_3_37 Salmonella enterica subsp. enterica serovar Weltevreden Proteobacteria; Gammaproteobacteria; Enterobacterales; Enterobacteriaceae; Salmonella; Salmonella enterica; Salmonella enterica subsp. enterica MNNEETFYQAMRRKGVTRRSFLKFCSLAATSLGLGAGMTPKIAWALENKPRIPVVWIHGL WP_001058307.1 hydrogenase [Salmonella enterica] Length: 372\nNumber of predicted TMHs: 2\nExp number of AAs in TMHs: 36.51738\nExp number, first 60 AAs: 19.72849\nTotal prob of N-in: 0.96170\nPOSSIBLE N-term signal sequence\ninside 1 19\nTMhelix 20 39\noutside 40 325\nTMhelix 326 348\ninside 349 372

22472 GCF_001410475.1_10900_1_32 Salmonella enterica subsp. enterica serovar Weltevreden Proteobacteria; Gammaproteobacteria; Enterobacterales; Enterobacteriaceae; Salmonella; Salmonella enterica; Salmonella enterica subsp. enterica MTGDNTLITSHGINRRDFMKLCAALAATMGLSSKAAAEMAESVSNPQRPPVIWIGAQECT WP_000145429.1 hydrogenase 2 small subunit [Salmonella enterica] Length: 372\nNumber of predicted TMHs: 1\nExp number of AAs in TMHs: 24.51309\nExp number, first 60 AAs: 0.2682\nTotal prob of N-in: 0.14402\noutside 1 331\nTMhelix 332 354\ninside 355 372

22473 GCF_001410455.1_9472_3_34 Salmonella enterica subsp. enterica serovar Weltevreden Proteobacteria; Gammaproteobacteria; Enterobacterales; Enterobacteriaceae; Salmonella; Salmonella enterica; Salmonella enterica subsp. enterica MTGDNTLITSHGINRRDFMKLCAALAATMGLSSKAAAEMAESVSNPQRPPVIWIGAQECT WP_000145429.1 hydrogenase 2 small subunit [Salmonella enterica] Length: 372\nNumber of predicted TMHs: 1\nExp number of AAs in TMHs: 24.51309\nExp number, first 60 AAs: 0.2682\nTotal prob of N-in: 0.14402\noutside 1 331\nTMhelix 332 354\ninside 355 372

22474 GCF_001410415.1_12227_3_44 Salmonella enterica subsp. enterica serovar Weltevreden Proteobacteria; Gammaproteobacteria; Enterobacterales; Enterobacteriaceae; Salmonella; Salmonella enterica; Salmonella enterica subsp. enterica MTGDNTLITSHGINRRDFMKLCAALAATMGLSSKAAAEMAESVSNPQRPPVIWIGAQECT WP_000145429.1 hydrogenase 2 small subunit [Salmonella enterica] Length: 372\nNumber of predicted TMHs: 1\nExp number of AAs in TMHs: 24.51309\nExp number, first 60 AAs: 0.2682\nTotal prob of N-in: 0.14402\noutside 1 331\nTMhelix 332 354\ninside 355 372

22475 GCF_001410395.1_12227_3_10 Salmonella enterica subsp. enterica serovar Weltevreden Proteobacteria; Gammaproteobacteria; Enterobacterales; Enterobacteriaceae; Salmonella; Salmonella enterica; Salmonella enterica subsp. enterica MTGDNTLITSHGINRRDFMKLCAALAATMGLSSKAAAEMAESVSNPQRPPVIWIGAQECT WP_000145429.1 hydrogenase 2 small subunit [Salmonella enterica] Length: 372\nNumber of predicted TMHs: 1\nExp number of AAs in TMHs: 24.51309\nExp number, first 60 AAs: 0.2682\nTotal prob of N-in: 0.14402\noutside 1 331\nTMhelix 332 354\ninside 355 372

22476 GCF_001410375.1_12227_3_22 Salmonella enterica subsp. enterica serovar Weltevreden Proteobacteria; Gammaproteobacteria; Enterobacterales; Enterobacteriaceae; Salmonella; Salmonella enterica; Salmonella enterica subsp. enterica MTGDNTLITSHGINRRDFMKLCAALAATMGLSSKAAAEMAESVSNPQRPPVIWIGAQECT WP_000145429.1 hydrogenase 2 small subunit [Salmonella enterica] Length: 372\nNumber of

predicted TMHs: 1\nExp number of AAs in TMHs: 24.51309\nExp number, first 60 AAs: 0.2682\nTotal prob of N-in: 0.14402\noutside 1 331\nTMhelix 332 354\ninside 355 372

22477 GCF_001410355.1_12227_3_37 Salmonella enterica subsp. enterica serovar Weltevreden Proteobacteria; Gammaproteobacteria; Enterobacterales; Enterobacteriaceae; Salmonella; Salmonella enterica; Salmonella enterica subsp. enterica MTGDNTLITSHGINRRDFMKLCAALAATMGLSSKAAAEMAESVSNPQRPPVIWIGAQECT WP_000145429.1 hydrogenase 2 small subunit [Salmonella enterica] Length: 372\nNumber of predicted TMHs: 1\nExp number of AAs in TMHs: 24.51309\nExp number, first 60 AAs: 0.2682\nTotal prob of N-in: 0.14402\noutside 1 331\nTMhelix 332 354\ninside 355 372

22478 GCF_001410335.1_12227_3_57 Salmonella enterica subsp. enterica serovar Weltevreden Proteobacteria; Gammaproteobacteria; Enterobacterales; Enterobacteriaceae; Salmonella; Salmonella enterica; Salmonella enterica subsp. enterica MTGDNTLITSHGINRRDFMKLCAALAATMGLSSKAAAEMAESVSNPQRPPVIWIGAQECT WP_000145429.1 hydrogenase 2 small subunit [Salmonella enterica] Length: 372\nNumber of predicted TMHs: 1\nExp number of AAs in TMHs: 24.51309\nExp number, first 60 AAs: 0.2682\nTotal prob of N-in: 0.14402\noutside 1 331\nTMhelix 332 354\ninside 355 372

22479 GCF_001410295.1_12216_4_49 Salmonella enterica subsp. enterica serovar Weltevreden Proteobacteria; Gammaproteobacteria; Enterobacterales; Enterobacteriaceae; Salmonella; Salmonella enterica; Salmonella enterica subsp. enterica MNNEETFYQAMRRKGVTRRSFLKFCSLAATSLGLGAGMTPKIAWALENKPRIPVVWIHGL WP_001058307.1 hydrogenase [Salmonella enterica] Length: 372\nNumber of predicted TMHs: 2\nExp number of AAs in TMHs: 36.51738\nExp number, first 60 AAs: 19.72849\nTotal prob of N-in: 0.96170\nPOSSIBLE N-term signal sequence\ninside 1 19\nTMhelix 20 39\noutside 40 325\nTMhelix 326 348\ninside 349 372

22480 GCF_001410315.1_12227_3_6 Salmonella enterica subsp. enterica serovar Weltevreden Proteobacteria; Gammaproteobacteria; Enterobacterales; Enterobacteriaceae; Salmonella; Salmonella enterica; Salmonella enterica subsp. enterica MNNEETFYQAMRRKGVTRRSFLKFCSLAATSLGLGAGMTPKIAWALENKPRIPVVWIHGL WP_001058307.1 hydrogenase [Salmonella enterica] Length: 372\nNumber of predicted TMHs: 2\nExp number of AAs in TMHs: 36.51738\nExp number, first 60 AAs: 19.72849\nTotal prob of N-in: 0.96170\nPOSSIBLE N-term signal sequence\ninside 1 19\nTMhelix 20 39\noutside 40 325\nTMhelix 326 348\ninside 349 372

22481 GCF_001410295.1_12216_4_49 Salmonella enterica subsp. enterica serovar Weltevreden Proteobacteria; Gammaproteobacteria; Enterobacterales; Enterobacteriaceae; Salmonella; Salmonella enterica; Salmonella enterica subsp. enterica MTGDNTLITSHGINRRDFMKLCAALAATMGLSSKAAAEMAESVSNPQRPPVIWIGAQECT WP_000145429.1 hydrogenase 2 small subunit [Salmonella enterica] Length: 372\nNumber of predicted TMHs: 1\nExp number of AAs in TMHs: 24.51309\nExp number, first 60 AAs: 0.2682\nTotal prob of N-in: 0.14402\noutside 1 331\nTMhelix 332 354\ninside 355 372

22482 GCF_001410315.1_12227_3_6 Salmonella enterica subsp. enterica serovar Weltevreden Proteobacteria; Gammaproteobacteria; Enterobacterales; Enterobacteriaceae; Salmonella; Salmonella enterica; Salmonella enterica subsp. enterica MTGDNTLITSHGINRRDFMKLCAALAATMGLSSKAAAEMAESVSNPQRPPVIWIGAQECT WP_000145429.1 hydrogenase 2 small subunit [Salmonella enterica] Length: 372\nNumber of predicted TMHs: 1\nExp number of AAs in TMHs: 24.51309\nExp number, first 60 AAs: 0.2682\nTotal prob of N-in: 0.14402\noutside 1 331\nTMhelix 332 354\ninside 355 372

22483 GCF_001410135.1_12216_4_50 Salmonella enterica subsp. enterica serovar Weltevreden Proteobacteria; Gammaproteobacteria; Enterobacterales; Enterobacteriaceae; Salmonella; Salmonella enterica; Salmonella enterica subsp. enterica MNNEETFYQAMRRKGVTRRSFLKFCSLAATSLGLGAGMTPKIAWALENKPRIPVVWIHGL WP_057519867.1 hydrogenase [Salmonella enterica] Length: 372\nNumber of predicted TMHs: 2\nExp number of AAs in TMHs: 36.51816\nExp number, first 60 AAs: 19.72874\nTotal prob of N-in: 0.96170\nPOSSIBLE N-term signal sequence\ninside 1 19\nTMhelix 20 39\noutside 40 325\nTMhelix 326 348\ninside 349 372

22484 GCF_001410275.1_12227_3_47 Salmonella enterica subsp. enterica serovar Weltevreden Proteobacteria; Gammaproteobacteria; Enterobacterales; Enterobacteriaceae; Salmonella; Salmonella enterica; Salmonella enterica subsp. enterica MNNEETFYQAMRRKGVTRRSFLKFCSLAATSLGLGAGMTPKIAWALENKPRIPVVWIHGL WP_001058307.1 hydrogenase [Salmonella enterica] Length: 372\nNumber of predicted TMHs: 2\nExp number of AAs in TMHs: 36.51738\nExp number, first 60 AAs: 19.72849\nTotal prob of N-in: 0.96170\nPOSSIBLE N-term signal sequence\ninside 1 19\nTMhelix 20 39\noutside 40 325\nTMhelix 326 348\ninside 349 372

22485 GCF_001410275.1_12227_3_47 Salmonella enterica subsp. enterica serovar Weltevreden Proteobacteria; Gammaproteobacteria; Enterobacterales; Enterobacteriaceae; Salmonella; Salmonella enterica;

Salmonella enterica subsp. enterica MTGDNTLITSHGINRRDFMKLCAALAATMGLSSKAAAEMAESVSNPQRPPVIWIGAQECT WP_000145429.1 hydrogenase 2 small subunit [Salmonella enterica] Length: 372\nNumber of predicted TMHs: 1\nExp number of AAs in TMHs: 24.51309\nExp number, first 60 AAs: 0.2682\nTotal prob of N-in: 0.14402\noutside 1 331\nTMhelix 332 354\ninside 355 372

22486 GCF_001410175.1_12227_3_14 Salmonella enterica subsp. enterica serovar Weltevreden Proteobacteria; Gammaproteobacteria; Enterobacterales; Enterobacteriaceae; Salmonella; Salmonella enterica; Salmonella enterica subsp. enterica MNNEETFYQAMRRKGVTRRSFLKFCSLAATSLGLGAGMTPKIAWALENKPRIPVVWIHGL WP_001058307.1 hydrogenase [Salmonella enterica] Length: 372\nNumber of predicted TMHs: 2\nExp number of AAs in TMHs: 36.51738\nExp number, first 60 AAs: 19.72849\nTotal prob of N-in: 0.96170\nPOSSIBLE N-term signal sequence\ninside 1 19\nTMhelix 20 39\noutside 40 325\nTMhelix 326 348\ninside 349 372

22487 GCF_001410215.1_9472_3_7Salmonella enterica subsp. enterica serovar Weltevreden Proteobacteria; Gammaproteobacteria; Enterobacterales; Enterobacteriaceae; Salmonella; Salmonella enterica; Salmonella enterica subsp. enterica MNNEETFYQAMRRKGVTRRSFLKFCSLAATSLGLGAGMTPKIAWALENKPRIPVVWIHGL WP_001058307.1 hydrogenase [Salmonella enterica] Length: 372\nNumber of predicted TMHs: 2\nExp number of AAs in TMHs: 36.51738\nExp number, first 60 AAs: 19.72849\nTotal prob of N-in: 0.96170\nPOSSIBLE N-term signal sequence\ninside 1 19\nTMhelix 20 39\noutside 40 325\nTMhelix 326 348\ninside 349 372

22488 GCF_001410435.1_9472_3_25 Salmonella enterica subsp. enterica serovar Weltevreden Proteobacteria; Gammaproteobacteria; Enterobacterales; Enterobacteriaceae; Salmonella; Salmonella enterica; Salmonella enterica subsp. enterica MTGDNTLITSHGINRRDFMKLCAALAATMGLSSKAAAEMAESVSNPQRPPVIWIGAQECT WP_000145429.1 hydrogenase 2 small subunit [Salmonella enterica] Length: 372\nNumber of predicted TMHs: 1\nExp number of AAs in TMHs: 24.51309\nExp number, first 60 AAs: 0.2682\nTotal prob of N-in: 0.14402\noutside 1 331\nTMhelix 332 354\ninside 355 372

22489 GCF_001410255.1_12227_3_38 Salmonella enterica subsp. enterica serovar Weltevreden Proteobacteria; Gammaproteobacteria; Enterobacterales; Enterobacteriaceae; Salmonella; Salmonella enterica; Salmonella enterica subsp. enterica MNNEETFYQAMRRKGVTRRSFLKFCSLAATSLGLGAGMTPKIAWALENKPRIPVVWIHGL WP_001058307.1 hydrogenase [Salmonella enterica] Length: 372\nNumber of predicted TMHs: 2\nExp number of AAs in TMHs: 36.51738\nExp number, first 60 AAs: 19.72849\nTotal prob of N-in: 0.96170\nPOSSIBLE N-term signal sequence\ninside 1 19\nTMhelix 20 39\noutside 40 325\nTMhelix 326 348\ninside 349 372

22490 GCF_001410255.1_12227_3_38 Salmonella enterica subsp. enterica serovar Weltevreden Proteobacteria; Gammaproteobacteria; Enterobacterales; Enterobacteriaceae; Salmonella; Salmonella enterica; Salmonella enterica subsp. enterica MTGDNTLITSHGINRRDFMKLCAALAATMGLSSKAAAEMAESVSNPQRPPVIWIGAQECT WP_000145429.1 hydrogenase 2 small subunit [Salmonella enterica] Length: 372\nNumber of predicted TMHs: 1\nExp number of AAs in TMHs: 24.51309\nExp number, first 60 AAs: 0.2682\nTotal prob of N-in: 0.14402\noutside 1 331\nTMhelix 332 354\ninside 355 372

22491 GCF_001410115.1_9472_3_12 Salmonella enterica subsp. enterica serovar Weltevreden Proteobacteria; Gammaproteobacteria; Enterobacterales; Enterobacteriaceae; Salmonella; Salmonella enterica; Salmonella enterica subsp. enterica MNNEETFYQAMRRKGVTRRSFLKFCSLAATSLGLGAGMTPKIAWALENKPRIPVVWIHGL WP_001058307.1 hydrogenase [Salmonella enterica] Length: 372\nNumber of predicted TMHs: 2\nExp number of AAs in TMHs: 36.51738\nExp number, first 60 AAs: 19.72849\nTotal prob of N-in: 0.96170\nPOSSIBLE N-term signal sequence\ninside 1 19\nTMhelix 20 39\noutside 40 325\nTMhelix 326 348\ninside 349 372

22492 GCF_001410075.1_12227_3_7 Salmonella enterica subsp. enterica serovar Weltevreden Proteobacteria; Gammaproteobacteria; Enterobacterales; Enterobacteriaceae; Salmonella; Salmonella enterica; Salmonella enterica subsp. enterica MGLIYITSHGINRRDFMKLCAALAATMGLSSKAAAEMAESVSNPQRPPVIWIGAQECTGC WP_057516266.1 hydrogenase 2 small subunit [Salmonella enterica] Length: 370\nNumber of predicted TMHs: 1\nExp number of AAs in TMHs: 22.98319\nExp number, first 60 AAs: 0.05008\nTotal prob of N-in: 0.05279\noutside 1 329\nTMhelix 330 352\ninside 353 370

22493 GCF_001410195.1_12227_3_40 Salmonella enterica subsp. enterica serovar Weltevreden Proteobacteria; Gammaproteobacteria; Enterobacterales; Enterobacteriaceae; Salmonella; Salmonella enterica; Salmonella enterica subsp. enterica MNNEETFYQAMRRKGVTRRSFLKFCSLAATSLGLGAGMTPKIAWALENKPRIPVVWIHGL WP_001058307.1 hydrogenase [Salmonella enterica] Length: 372\nNumber of predicted TMHs: 2\nExp number of AAs in TMHs: 36.51738\nExp number, first 60 AAs: 19.72849\nTotal prob of N-in: 0.96170\nPOSSIBLE N-term signal sequence\ninside 1 19\nTMhelix 20 39\noutside 40 325\nTMhelix 326 348\ninside 349 372

22494	GCF_001410235.1_12227_3_29	Salmonella enterica subsp. enterica serovar Weltevreden	Proteobacteria; Gammaproteobacteria; Enterobacterales; Enterobacteriaceae; Salmonella; Salmonella enterica; Salmonella enterica subsp. enterica	MNNEETFYQAMRRKGVTRRSFLKFCSLAATSLGLGAGMTPKIAWALENKPRIPVVWIHGL	WP_001058307.1	hydrogenase [Salmonella enterica]	Length: 372\nNumber of predicted TMHs:  2\nExp number of AAs in TMHs: 36.51738\nExp number, first 60 AAs:  19.72849\nTotal prob of N-in:   0.96170\nPOSSIBLE N-term signal sequence\ninside   1   19\nTMhelix   20   39\noutside   40   325\nTMhelix   326   348\ninside   349   372

22495	GCF_001410195.1_12227_3_40	Salmonella enterica subsp. enterica serovar Weltevreden	Proteobacteria; Gammaproteobacteria; Enterobacterales; Enterobacteriaceae; Salmonella; Salmonella enterica; Salmonella enterica subsp. enterica	MTGDNTLITSHGINRRDFMKLCAALAATMGLSSKAAAEMAESVSNPQRPPVIWIGAQECT	WP_000145429.1	hydrogenase 2 small subunit [Salmonella enterica]	Length: 372\nNumber of predicted TMHs:  1\nExp number of AAs in TMHs: 24.51309\nExp number, first 60 AAs:  0.2682\nTotal prob of N-in:   0.14402\noutside   1   331\nTMhelix   332   354\ninside   355   372

22496	GCF_001410235.1_12227_3_29	Salmonella enterica subsp. enterica serovar Weltevreden	Proteobacteria; Gammaproteobacteria; Enterobacterales; Enterobacteriaceae; Salmonella; Salmonella enterica; Salmonella enterica subsp. enterica	MTGDNTLITSHGINRRDFMKLCAALAATMGLSSKAAAEMAESVSNPQRPPVIWIGAQECT	WP_000145429.1	hydrogenase 2 small subunit [Salmonella enterica]	Length: 372\nNumber of predicted TMHs:  1\nExp number of AAs in TMHs: 24.51309\nExp number, first 60 AAs:  0.2682\nTotal prob of N-in:   0.14402\noutside   1   331\nTMhelix   332   354\ninside   355   372

22497	GCF_001410175.1_12227_3_14	Salmonella enterica subsp. enterica serovar Weltevreden	Proteobacteria; Gammaproteobacteria; Enterobacterales; Enterobacteriaceae; Salmonella; Salmonella enterica; Salmonella enterica subsp. enterica	MTGDNTLITSHGINRRDFMKLCAALAATMGLSSKAAAEMAESVSNPQRPPVIWIGAQECT	WP_000145429.1	hydrogenase 2 small subunit [Salmonella enterica]	Length: 372\nNumber of predicted TMHs:  1\nExp number of AAs in TMHs: 24.51309\nExp number, first 60 AAs:  0.2682\nTotal prob of N-in:   0.14402\noutside   1   331\nTMhelix   332   354\ninside   355   372

22498	GCF_001410215.1_9472_3_7Salmonella enterica subsp. enterica serovar Weltevreden	Proteobacteria; Gammaproteobacteria; Enterobacterales; Enterobacteriaceae; Salmonella; Salmonella enterica; Salmonella enterica subsp. enterica	MTGDNTLITSHGINRRDFMKLCAALAATMGLSSKAAAEMAESVSNPQRPPVIWIGAQECT	WP_000145429.1	hydrogenase 2 small subunit [Salmonella enterica]	Length: 372\nNumber of predicted TMHs:  1\nExp number of AAs in TMHs: 24.51309\nExp number, first 60 AAs:  0.2682\nTotal prob of N-in:   0.14402\noutside   1   331\nTMhelix   332   354\ninside   355   372

22499	GCF_001410135.1_12216_4_50	Salmonella enterica subsp. enterica serovar Weltevreden	Proteobacteria; Gammaproteobacteria; Enterobacterales; Enterobacteriaceae; Salmonella; Salmonella enterica; Salmonella enterica subsp. enterica	MTGDNTLITSHGINRRDFMKLCAALAATMGLSSKAAAEMAESVSNPQRPPVIWIGAQECT	WP_000145429.1	hydrogenase 2 small subunit [Salmonella enterica]	Length: 372\nNumber of predicted TMHs:  1\nExp number of AAs in TMHs: 24.51309\nExp number, first 60 AAs:  0.2682\nTotal prob of N-in:   0.14402\noutside   1   331\nTMhelix   332   354\ninside   355   372

22500	GCF_001410075.1_12227_3_7	Salmonella enterica subsp. enterica serovar Weltevreden	Proteobacteria; Gammaproteobacteria; Enterobacterales; Enterobacteriaceae; Salmonella; Salmonella enterica; Salmonella enterica subsp. enterica	MNNEETFYQAMRRKGVTRRSFLKFCSLAATSLGLGAGMTPKIAWALENKPRIPVVWIHGL	WP_001058307.1	hydrogenase [Salmonella enterica]	Length: 372\nNumber of predicted TMHs:  2\nExp number of AAs in TMHs: 36.51738\nExp number, first 60 AAs:  19.72849\nTotal prob of N-in:   0.96170\nPOSSIBLE N-term signal sequence\ninside   1   19\nTMhelix   20   39\noutside   40   325\nTMhelix   326   348\ninside   349   372

22501	GCF_001410115.1_9472_3_12	Salmonella enterica subsp. enterica serovar Weltevreden	Proteobacteria; Gammaproteobacteria; Enterobacterales; Enterobacteriaceae; Salmonella; Salmonella enterica; Salmonella enterica subsp. enterica	MTGDNTLITSHGINRRDFMKLCAALAATMGLSSKAAAEMAESVSNPQRPPVIWIGAQECT	WP_000145429.1	hydrogenase 2 small subunit [Salmonella enterica]	Length: 372\nNumber of predicted TMHs:  1\nExp number of AAs in TMHs: 24.51309\nExp number, first 60 AAs:  0.2682\nTotal prob of N-in:   0.14402\noutside   1   331\nTMhelix   332   354\ninside   355   372

22502	GCF_001410015.1_9472_3_32	Salmonella enterica subsp. enterica serovar Weltevreden	Proteobacteria; Gammaproteobacteria; Enterobacterales; Enterobacteriaceae; Salmonella; Salmonella enterica; Salmonella enterica subsp. enterica	MNNEETFYQAMRRKGVTRRSFLKFCSLAATSLGLGAGMTPKIAWALENKPRIPVVWIHGL	WP_001058307.1	hydrogenase [Salmonella enterica]	Length: 372\nNumber of predicted TMHs:  2\nExp

number of AAs in TMHs: 36.51738\nExp number, first 60 AAs:  19.72849\nTotal prob of N-in:   0.96170\nPOSSIBLE N-term signal sequence\ninside   1   19\nTMhelix   20   39\noutside   40   325\nTMhelix   326   348\ninside   349   372

22503 GCF_001410095.1_9472_3_31 Salmonella enterica subsp. enterica serovar Weltevreden Proteobacteria; Gammaproteobacteria; Enterobacterales; Enterobacteriaceae; Salmonella; Salmonella enterica; Salmonella enterica subsp. enterica MNNEETFYQAMRRKGVTRRSFLKFCSLAATSLGLGAGMTPKIAWALENKPRIPVVWIHGL WP_001058307.1 hydrogenase [Salmonella enterica] Length: 372\nNumber of predicted TMHs:  2\nExp number of AAs in TMHs: 36.51738\nExp number, first 60 AAs:  19.72849\nTotal prob of N-in:   0.96170\nPOSSIBLE N-term signal sequence\ninside   1   19\nTMhelix   20   39\noutside   40   325\nTMhelix   326   348\ninside   349   372

22504 GCF_001410055.1_12227_3_20 Salmonella enterica subsp. enterica serovar Weltevreden Proteobacteria; Gammaproteobacteria; Enterobacterales; Enterobacteriaceae; Salmonella; Salmonella enterica; Salmonella enterica subsp. enterica MNNEETFYQAMRRKGVTRRSFLKFCSLAATSLGLGAGMTPKIAWALENKPRIPVVWIHGL WP_001058307.1 hydrogenase [Salmonella enterica] Length: 372\nNumber of predicted TMHs:  2\nExp number of AAs in TMHs: 36.51738\nExp number, first 60 AAs:  19.72849\nTotal prob of N-in:   0.96170\nPOSSIBLE N-term signal sequence\ninside   1   19\nTMhelix   20   39\noutside   40   325\nTMhelix   326   348\ninside   349   372

22505 GCF_001410035.1_9472_3_4Salmonella enterica subsp. enterica serovar Weltevreden Proteobacteria; Gammaproteobacteria; Enterobacterales; Enterobacteriaceae; Salmonella; Salmonella enterica; Salmonella enterica subsp. enterica MNNEETFYQAMRRKGVTRRSFLKFCSLAATSLGLGAGMTPKIAWALENKPRIPVVWIHGL WP_001058307.1 hydrogenase [Salmonella enterica] Length: 372\nNumber of predicted TMHs:  2\nExp number of AAs in TMHs: 36.51738\nExp number, first 60 AAs:  19.72849\nTotal prob of N-in:   0.96170\nPOSSIBLE N-term signal sequence\ninside   1   19\nTMhelix   20   39\noutside   40   325\nTMhelix   326   348\ninside   349   372

22506 GCF_001410095.1_9472_3_31 Salmonella enterica subsp. enterica serovar Weltevreden Proteobacteria; Gammaproteobacteria; Enterobacterales; Enterobacteriaceae; Salmonella; Salmonella enterica; Salmonella enterica subsp. enterica MTGDNTLITSHGINRRDFMKLCAALAATMGLSSKAAAEMAESVSNPQRPPVIWIGAQECT WP_000145429.1 hydrogenase 2 small subunit [Salmonella enterica] Length: 372\nNumber of predicted TMHs:  1\nExp number of AAs in TMHs: 24.51309\nExp number, first 60 AAs:  0.2682\nTotal prob of N-in: 0.14402\noutside   1   331\nTMhelix   332   354\ninside   355   372

22507 GCF_001410055.1_12227_3_20 Salmonella enterica subsp. enterica serovar Weltevreden Proteobacteria; Gammaproteobacteria; Enterobacterales; Enterobacteriaceae; Salmonella; Salmonella enterica; Salmonella enterica subsp. enterica MTGDNTLITSHGINRRDFMKLCAALAATMGLSSKAAAEMAESVSNPQRPPVIWIGAQECT WP_000145429.1 hydrogenase 2 small subunit [Salmonella enterica] Length: 372\nNumber of predicted TMHs:  1\nExp number of AAs in TMHs: 24.51309\nExp number, first 60 AAs:  0.2682\nTotal prob of N-in: 0.14402\noutside   1   331\nTMhelix   332   354\ninside   355   372

22508 GCF_001410035.1_9472_3_4Salmonella enterica subsp. enterica serovar Weltevreden Proteobacteria; Gammaproteobacteria; Enterobacterales; Enterobacteriaceae; Salmonella; Salmonella enterica; Salmonella enterica subsp. enterica MTGDNTLITSHGINRRDFMKLCAALAATMGLSSKAAAEMAESVSNPQRPPVIWIGAQECT WP_000145429.1 hydrogenase 2 small subunit [Salmonella enterica] Length: 372\nNumber of predicted TMHs:  1\nExp number of AAs in TMHs: 24.51309\nExp number, first 60 AAs:  0.2682\nTotal prob of N-in:   0.14402\noutside   1 331\nTMhelix   332   354\ninside   355   372

22509 GCF_001409955.1_12216_4_45 Salmonella enterica subsp. enterica serovar Weltevreden Proteobacteria; Gammaproteobacteria; Enterobacterales; Enterobacteriaceae; Salmonella; Salmonella enterica; Salmonella enterica subsp. enterica MNNEETFYQAMRRKGVTRRSFLKFCSLAATSLGLGAGMTPKIAWALENKPRIPVVWIHGL WP_001058307.1 hydrogenase [Salmonella enterica] Length: 372\nNumber of predicted TMHs:  2\nExp number of AAs in TMHs: 36.51738\nExp number, first 60 AAs:  19.72849\nTotal prob of N-in:   0.96170\nPOSSIBLE N-term signal sequence\ninside   1   19\nTMhelix   20   39\noutside   40   325\nTMhelix   326   348\ninside   349   372

22510 GCF_001409935.1_12227_3_17 Salmonella enterica subsp. enterica serovar Weltevreden Proteobacteria; Gammaproteobacteria; Enterobacterales; Enterobacteriaceae; Salmonella; Salmonella enterica; Salmonella enterica subsp. enterica MNNEETFYQAMRRKGVTRRSFLKFCSLAATSLGLGAGMTPKIAWALENKPRIPVVWIHGL WP_001058307.1 hydrogenase [Salmonella enterica] Length: 372\nNumber of predicted TMHs:  2\nExp number of AAs in TMHs: 36.51738\nExp number, first 60 AAs:  19.72849\nTotal prob of N-in:   0.96170\nPOSSIBLE N-term signal sequence\ninside   1   19\nTMhelix   20   39\noutside   40   325\nTMhelix   326   348\ninside   349   372

22511 GCF_001410015.1_9472_3_32 Salmonella enterica subsp. enterica serovar Weltevreden Proteobacteria; Gammaproteobacteria; Enterobacterales; Enterobacteriaceae; Salmonella; Salmonella enterica;

Salmonella enterica subsp. enterica	MTGDNTLITSHGINRRDFMKLCAALAATMGLSSKAAAEMAESVSNPQRPPVIWIGAQECT	WP_000145429.1	hydrogenase 2 small subunit [Salmonella enterica]	Length: 372\nNumber of predicted TMHs:  1\nExp number of AAs in TMHs: 24.51309\nExp number, first 60 AAs:  0.2682\nTotal prob of N-in:  0.14402\noutside   1   331\nTMhelix   332   354\ninside   355   372

22512	GCF_001409995.1_12216_4_47	Salmonella enterica subsp. enterica serovar Weltevreden	Proteobacteria; Gammaproteobacteria; Enterobacterales; Enterobacteriaceae; Salmonella; Salmonella enterica; Salmonella enterica subsp. enterica	MNNEETFYQAMRRKGVTRRSFLKFCSLAATSLGLGAGMTPKIAWALENKPRIPVVWIHGL	WP_001058307.1	hydrogenase [Salmonella enterica]	Length: 372\nNumber of predicted TMHs:  2\nExp number of AAs in TMHs: 36.51738\nExp number, first 60 AAs:  19.72849\nTotal prob of N-in:   0.96170\nPOSSIBLE N-term signal sequence\ninside   1   19\nTMhelix   20   39\noutside   40   325\nTMhelix   326   348\ninside   349   372

22513	GCF_001409955.1_12216_4_45	Salmonella enterica subsp. enterica serovar Weltevreden	Proteobacteria; Gammaproteobacteria; Enterobacterales; Enterobacteriaceae; Salmonella; Salmonella enterica; Salmonella enterica subsp. enterica	MTGDNTLITSHGINRRDFMKLCAALAATMGLSSKAAAEMAESVSNPQRPPVIWIGAQECT	WP_000145429.1	hydrogenase 2 small subunit [Salmonella enterica]	Length: 372\nNumber of predicted TMHs:  1\nExp number of AAs in TMHs: 24.51309\nExp number, first 60 AAs:  0.2682\nTotal prob of N-in:  0.14402\noutside   1   331\nTMhelix   332   354\ninside   355   372

22514	GCF_001409935.1_12227_3_17	Salmonella enterica subsp. enterica serovar Weltevreden	Proteobacteria; Gammaproteobacteria; Enterobacterales; Enterobacteriaceae; Salmonella; Salmonella enterica; Salmonella enterica subsp. enterica	MTGDNTLITSHGINRRDFMKLCAALAATMGLSSKAAAEMAESVSNPQRPPVIWIGAQECT	WP_000145429.1	hydrogenase 2 small subunit [Salmonella enterica]	Length: 372\nNumber of predicted TMHs:  1\nExp number of AAs in TMHs: 24.51309\nExp number, first 60 AAs:  0.2682\nTotal prob of N-in:  0.14402\noutside   1   331\nTMhelix   332   354\ninside   355   372

22515	GCF_001409995.1_12216_4_47	Salmonella enterica subsp. enterica serovar Weltevreden	Proteobacteria; Gammaproteobacteria; Enterobacterales; Enterobacteriaceae; Salmonella; Salmonella enterica; Salmonella enterica subsp. enterica	MTGDNTLITSHGINRRDFMKLCAALAATMGLSSKAAAEMAESVSNPQRPPVIWIGAQECT	WP_000145429.1	hydrogenase 2 small subunit [Salmonella enterica]	Length: 372\nNumber of predicted TMHs:  1\nExp number of AAs in TMHs: 24.51309\nExp number, first 60 AAs:  0.2682\nTotal prob of N-in:  0.14402\noutside   1   331\nTMhelix   332   354\ninside   355   372

22516	GCF_001409875.1_9472_3_11	Salmonella enterica subsp. enterica serovar Weltevreden	Proteobacteria; Gammaproteobacteria; Enterobacterales; Enterobacteriaceae; Salmonella; Salmonella enterica; Salmonella enterica subsp. enterica	MNNEETFYQAMRRKGVTRRSFLKFCSLAATSLGLGAGMTPKIAWALENKPRIPVVWIHGL	WP_001058307.1	hydrogenase [Salmonella enterica]	Length: 372\nNumber of predicted TMHs:  2\nExp number of AAs in TMHs: 36.51738\nExp number, first 60 AAs:  19.72849\nTotal prob of N-in:   0.96170\nPOSSIBLE N-term signal sequence\ninside   1   19\nTMhelix   20   39\noutside   40   325\nTMhelix   326   348\ninside   349   372

22517	GCF_001409915.1_12216_4_60	Salmonella enterica subsp. enterica serovar Weltevreden	Proteobacteria; Gammaproteobacteria; Enterobacterales; Enterobacteriaceae; Salmonella; Salmonella enterica; Salmonella enterica subsp. enterica	MNNEETFYQAMRRKGVTRRSFLKFCSLAATSLGLGAGMTPKIAWALENKPRIPVVWIHGL	WP_001058307.1	hydrogenase [Salmonella enterica]	Length: 372\nNumber of predicted TMHs:  2\nExp number of AAs in TMHs: 36.51738\nExp number, first 60 AAs:  19.72849\nTotal prob of N-in:   0.96170\nPOSSIBLE N-term signal sequence\ninside   1   19\nTMhelix   20   39\noutside   40   325\nTMhelix   326   348\ninside   349   372

22518	GCF_001409915.1_12216_4_60	Salmonella enterica subsp. enterica serovar Weltevreden	Proteobacteria; Gammaproteobacteria; Enterobacterales; Enterobacteriaceae; Salmonella; Salmonella enterica; Salmonella enterica subsp. enterica	MTGDNTLITSHGINRRDFMKLCAALAATMGLSSKAAAEMAESVSNPQRPPVIWIGAQECT	WP_000145429.1	hydrogenase 2 small subunit [Salmonella enterica]	Length: 372\nNumber of predicted TMHs:  1\nExp number of AAs in TMHs: 24.51309\nExp number, first 60 AAs:  0.2682\nTotal prob of N-in:  0.14402\noutside   1   331\nTMhelix   332   354\ninside   355   372

22519	GCF_001409835.1_12227_3_36	Salmonella enterica subsp. enterica serovar Weltevreden	Proteobacteria; Gammaproteobacteria; Enterobacterales; Enterobacteriaceae; Salmonella; Salmonella enterica; Salmonella enterica subsp. enterica	MNNEETFYQAMRRKGVTRRSFLKFCSLAATSLGLGAGMTPKIAWALENKPRIPVVWIHGL	WP_001058307.1	hydrogenase [Salmonella enterica]	Length: 372\nNumber of predicted TMHs:  2\nExp number of AAs in TMHs: 36.51738\nExp number, first 60 AAs:  19.72849\nTotal prob of N-in:   0.96170\nPOSSIBLE N-term signal sequence\ninside   1   19\nTMhelix   20   39\noutside   40   325\nTMhelix   326   348\ninside   349   372

22520 GCF_001409975.1_12227_3_4 Salmonella enterica subsp. enterica serovar Weltevreden Proteobacteria; Gammaproteobacteria; Enterobacterales; Enterobacteriaceae; Salmonella; Salmonella enterica; Salmonella enterica subsp. enterica MTGDNTLITSHGINRRDFMKLCAALAATMGLSSKAAAEMAESVSNPQRPPVIWIGAQECT WP_000145429.1 hydrogenase 2 small subunit [Salmonella enterica] Length: 372\nNumber of predicted TMHs: 1\nExp number of AAs in TMHs: 24.51309\nExp number, first 60 AAs: 0.2682\nTotal prob of N-in: 0.14402\noutside 1 331\nTMhelix 332 354\ninside 355 372

22521 GCF_001409855.1_9472_3_2Salmonella enterica subsp. enterica serovar Weltevreden Proteobacteria; Gammaproteobacteria; Enterobacterales; Enterobacteriaceae; Salmonella; Salmonella enterica; Salmonella enterica subsp. enterica MNNEETFYQAMRRKGVTRRSFLKFCSLAATSLGLGAGMTPKIAWALENKPRIPVVWIHGL WP_001058307.1 hydrogenase [Salmonella enterica] Length: 372\nNumber of predicted TMHs: 2\nExp number of AAs in TMHs: 36.51738\nExp number, first 60 AAs: 19.72849\nTotal prob of N-in: 0.96170\nPOSSIBLE N-term signal sequence\ninside 1 19\nTMhelix 20 39\noutside 40 325\nTMhelix 326 348\ninside 349 372

22522 GCF_001409895.1_12216_4_59 Salmonella enterica subsp. enterica serovar Weltevreden Proteobacteria; Gammaproteobacteria; Enterobacterales; Enterobacteriaceae; Salmonella; Salmonella enterica; Salmonella enterica subsp. enterica MNNEETFYQAMRRKGVTRRSFLKFCSLAATSLGLGAGMTPKIAWALENKPRIPVVWIHGL WP_001058307.1 hydrogenase [Salmonella enterica] Length: 372\nNumber of predicted TMHs: 2\nExp number of AAs in TMHs: 36.51738\nExp number, first 60 AAs: 19.72849\nTotal prob of N-in: 0.96170\nPOSSIBLE N-term signal sequence\ninside 1 19\nTMhelix 20 39\noutside 40 325\nTMhelix 326 348\ninside 349 372

22523 GCF_001409895.1_12216_4_59 Salmonella enterica subsp. enterica serovar Weltevreden Proteobacteria; Gammaproteobacteria; Enterobacterales; Enterobacteriaceae; Salmonella; Salmonella enterica; Salmonella enterica subsp. enterica MTGDNTLITSHGINRRDFMKLCAALAATMGLSSKAAAEMAESVSNPQRPPVIWIGAQECT WP_000145429.1 hydrogenase 2 small subunit [Salmonella enterica] Length: 372\nNumber of predicted TMHs: 1\nExp number of AAs in TMHs: 24.51309\nExp number, first 60 AAs: 0.2682\nTotal prob of N-in: 0.14402\noutside 1 331\nTMhelix 332 354\ninside 355 372

22524 GCF_001409815.1_12216_4_61 Salmonella enterica subsp. enterica serovar Weltevreden Proteobacteria; Gammaproteobacteria; Enterobacterales; Enterobacteriaceae; Salmonella; Salmonella enterica; Salmonella enterica subsp. enterica MNNEETFYQAMRRKGVTRRSFLKFCSLAATSLGLGAGMTPKIAWALENKPRIPVVWIHGL WP_001058307.1 hydrogenase [Salmonella enterica] Length: 372\nNumber of predicted TMHs: 2\nExp number of AAs in TMHs: 36.51738\nExp number, first 60 AAs: 19.72849\nTotal prob of N-in: 0.96170\nPOSSIBLE N-term signal sequence\ninside 1 19\nTMhelix 20 39\noutside 40 325\nTMhelix 326 348\ninside 349 372

22525 GCF_001409795.1_12227_3_56 Salmonella enterica subsp. enterica serovar Weltevreden Proteobacteria; Gammaproteobacteria; Enterobacterales; Enterobacteriaceae; Salmonella; Salmonella enterica; Salmonella enterica subsp. enterica MNNEETFYQAMRRKGVTRRSFLKFCSLAATSLGLGAGMTPKIAWALENKPRIPVVWIHGL WP_001058307.1 hydrogenase [Salmonella enterica] Length: 372\nNumber of predicted TMHs: 2\nExp number of AAs in TMHs: 36.51738\nExp number, first 60 AAs: 19.72849\nTotal prob of N-in: 0.96170\nPOSSIBLE N-term signal sequence\ninside 1 19\nTMhelix 20 39\noutside 40 325\nTMhelix 326 348\ninside 349 372

22526 GCF_001409875.1_9472_3_11 Salmonella enterica subsp. enterica serovar Weltevreden Proteobacteria; Gammaproteobacteria; Enterobacterales; Enterobacteriaceae; Salmonella; Salmonella enterica; Salmonella enterica subsp. enterica MTGDNTLITSHGINRRDFMKLCAALAATMGLSSKAAAEMAESVSNPQRPPVIWIGAQECT WP_000145429.1 hydrogenase 2 small subunit [Salmonella enterica] Length: 372\nNumber of predicted TMHs: 1\nExp number of AAs in TMHs: 24.51309\nExp number, first 60 AAs: 0.2682\nTotal prob of N-in: 0.14402\noutside 1 331\nTMhelix 332 354\ninside 355 372

22527 GCF_001409855.1_9472_3_2Salmonella enterica subsp. enterica serovar Weltevreden Proteobacteria; Gammaproteobacteria; Enterobacterales; Enterobacteriaceae; Salmonella; Salmonella enterica; Salmonella enterica subsp. enterica MTGDNTLITSHGINRRDFMKLCAALAATMGLSSKAAAEMAESVSNPQRPPVIWIGAQECT WP_000145429.1 hydrogenase 2 small subunit [Salmonella enterica] Length: 372\nNumber of predicted TMHs: 1\nExp number of AAs in TMHs: 24.51309\nExp number, first 60 AAs: 0.2682\nTotal prob of N-in: 0.14402\noutside 1 331\nTMhelix 332 354\ninside 355 372

22528 GCF_001409815.1_12216_4_61 Salmonella enterica subsp. enterica serovar Weltevreden Proteobacteria; Gammaproteobacteria; Enterobacterales; Enterobacteriaceae; Salmonella; Salmonella enterica; Salmonella enterica subsp. enterica MTGDNTLITSHGINRRDFMKLCAALAATMGLSSKAAAEMAESVSNPQRPPVIWIGAQECT WP_000145429.1 hydrogenase 2 small subunit [Salmonella enterica] Length: 372\nNumber of

predicted TMHs: 1\nExp number of AAs in TMHs: 24.51309\nExp number, first 60 AAs: 0.2682\nTotal prob of N-in: 0.14402\noutside 1 331\nTMhelix 332 354\ninside 355 372

22529 GCF_001409795.1_12227_3_56 Salmonella enterica subsp. enterica serovar Weltevreden Proteobacteria; Gammaproteobacteria; Enterobacterales; Enterobacteriaceae; Salmonella; Salmonella enterica; Salmonella enterica subsp. enterica MTGDNTLITSHGINRRDFMKLCAALAATMGLSSKAAAEMAESVSNPQRPPVIWIGAQECT WP_000145429.1 hydrogenase 2 small subunit [Salmonella enterica] Length: 372\nNumber of predicted TMHs: 1\nExp number of AAs in TMHs: 24.51309\nExp number, first 60 AAs: 0.2682\nTotal prob of N-in: 0.14402\noutside 1 331\nTMhelix 332 354\ninside 355 372

22530 GCF_001409715.1_9472_3_6Salmonella enterica subsp. enterica serovar Weltevreden Proteobacteria; Gammaproteobacteria; Enterobacterales; Enterobacteriaceae; Salmonella; Salmonella enterica; Salmonella enterica subsp. enterica MNNEETFYQAMRRKGVTRRSFLKFCSLAATSLGLGAGMTPKIAWALENKPRIPVVWIHGL WP_001058307.1 hydrogenase [Salmonella enterica] Length: 372\nNumber of predicted TMHs: 2\nExp number of AAs in TMHs: 36.51738\nExp number, first 60 AAs: 19.72849\nTotal prob of N-in: 0.96170\nPOSSIBLE N-term signal sequence\ninside 1 19\nTMhelix 20 39\noutside 40 325\nTMhelix 326 348\ninside 349 372

22531 GCF_001409695.1_12227_3_11 Salmonella enterica subsp. enterica serovar Weltevreden Proteobacteria; Gammaproteobacteria; Enterobacterales; Enterobacteriaceae; Salmonella; Salmonella enterica; Salmonella enterica subsp. enterica MNNEETFYQAMRRKGVTRRSFLKFCSLAATSLGLGAGMTPKIAWALENKPRIPVVWIHGL WP_001058307.1 hydrogenase [Salmonella enterica] Length: 372\nNumber of predicted TMHs: 2\nExp number of AAs in TMHs: 36.51738\nExp number, first 60 AAs: 19.72849\nTotal prob of N-in: 0.96170\nPOSSIBLE N-term signal sequence\ninside 1 19\nTMhelix 20 39\noutside 40 325\nTMhelix 326 348\ninside 349 372

22532 GCF_001409735.1_12227_3_54 Salmonella enterica subsp. enterica serovar Weltevreden Proteobacteria; Gammaproteobacteria; Enterobacterales; Enterobacteriaceae; Salmonella; Salmonella enterica; Salmonella enterica subsp. enterica MNNEETFYQAMRRKGVTRRSFLKFCSLAATSLGLGAGMTPKIAWALENKPRIPVVWIHGL WP_001058307.1 hydrogenase [Salmonella enterica] Length: 372\nNumber of predicted TMHs: 2\nExp number of AAs in TMHs: 36.51738\nExp number, first 60 AAs: 19.72849\nTotal prob of N-in: 0.96170\nPOSSIBLE N-term signal sequence\ninside 1 19\nTMhelix 20 39\noutside 40 325\nTMhelix 326 348\ninside 349 372

22533 GCF_001409675.1_12227_3_45 Salmonella enterica subsp. enterica serovar Weltevreden Proteobacteria; Gammaproteobacteria; Enterobacterales; Enterobacteriaceae; Salmonella; Salmonella enterica; Salmonella enterica subsp. enterica MNNEETFYQAMRRKGVTRRSFLKFCSLAATSLGLGAGMTPKIAWALENKPRIPVVWIHGL WP_001058307.1 hydrogenase [Salmonella enterica] Length: 372\nNumber of predicted TMHs: 2\nExp number of AAs in TMHs: 36.51738\nExp number, first 60 AAs: 19.72849\nTotal prob of N-in: 0.96170\nPOSSIBLE N-term signal sequence\ninside 1 19\nTMhelix 20 39\noutside 40 325\nTMhelix 326 348\ninside 349 372

22534 GCF_001409755.1_12227_3_51 Salmonella enterica subsp. enterica serovar Weltevreden Proteobacteria; Gammaproteobacteria; Enterobacterales; Enterobacteriaceae; Salmonella; Salmonella enterica; Salmonella enterica subsp. enterica MNNEETFYQAMRRKGVTRRSFLKFCSLAATSLGLGAGMTPKIAWALENKPRIPVVWIHGL WP_001058307.1 hydrogenase [Salmonella enterica] Length: 372\nNumber of predicted TMHs: 2\nExp number of AAs in TMHs: 36.51738\nExp number, first 60 AAs: 19.72849\nTotal prob of N-in: 0.96170\nPOSSIBLE N-term signal sequence\ninside 1 19\nTMhelix 20 39\noutside 40 325\nTMhelix 326 348\ninside 349 372

22535 GCF_001409775.1_12216_4_58 Salmonella enterica subsp. enterica serovar Weltevreden Proteobacteria; Gammaproteobacteria; Enterobacterales; Enterobacteriaceae; Salmonella; Salmonella enterica; Salmonella enterica subsp. enterica MNNEETFYQAMRRKGVTRRSFLKFCSLAATSLGLGAGMTPKIAWALENKPRIPVVWIHGL WP_001058307.1 hydrogenase [Salmonella enterica] Length: 372\nNumber of predicted TMHs: 2\nExp number of AAs in TMHs: 36.51738\nExp number, first 60 AAs: 19.72849\nTotal prob of N-in: 0.96170\nPOSSIBLE N-term signal sequence\ninside 1 19\nTMhelix 20 39\noutside 40 325\nTMhelix 326 348\ninside 349 372

22536 GCF_001409775.1_12216_4_58 Salmonella enterica subsp. enterica serovar Weltevreden Proteobacteria; Gammaproteobacteria; Enterobacterales; Enterobacteriaceae; Salmonella; Salmonella enterica; Salmonella enterica subsp. enterica MTGDNTLITSHGINRRDFMKLCAALAATMGLSSKAAAEMAESVSNPQRPPVIWIGAQECT WP_000145429.1 hydrogenase 2 small subunit [Salmonella enterica] Length: 372\nNumber of predicted TMHs: 1\nExp number of AAs in TMHs: 24.51309\nExp number, first 60 AAs: 0.2682\nTotal prob of N-in: 0.14402\noutside 1 331\nTMhelix 332 354\ninside 355 372

22537 GCF_001409755.1_12227_3_51 Salmonella enterica subsp. enterica serovar Weltevreden Proteobacteria; Gammaproteobacteria; Enterobacterales; Enterobacteriaceae; Salmonella; Salmonella enterica;

Salmonella enterica subsp. enterica MTGDNTLITSHGINRRDFMKLCAALAATMGLSSKAAAEMAESVSNPQRPPVIWIGAQECT WP_000145429.1 hydrogenase 2 small subunit [Salmonella enterica] Length: 372\nNumber of predicted TMHs: 1\nExp number of AAs in TMHs: 24.51309\nExp number, first 60 AAs: 0.2682\nTotal prob of N-in: 0.14402\noutside 1 331\nTMhelix 332 354\ninside 355 372

22538 GCF_001409635.1_9472_3_22 Salmonella enterica subsp. enterica serovar Weltevreden Proteobacteria; Gammaproteobacteria; Enterobacterales; Enterobacteriaceae; Salmonella; Salmonella enterica; Salmonella enterica subsp. enterica MNNEETFYQAMRRKGVTRRSFLKFCSLAATSLGLGAGMTPKIAWALENKPRIPVVWIHGL WP_001058307.1 hydrogenase [Salmonella enterica] Length: 372\nNumber of predicted TMHs: 2\nExp number of AAs in TMHs: 36.51738\nExp number, first 60 AAs: 19.72849\nTotal prob of N-in: 0.96170\nPOSSIBLE N-term signal sequence\ninside 1 19\nTMhelix 20 39\noutside 40 325\nTMhelix 326 348\ninside 349 372

22539 GCF_001409655.1_12227_3_52 Salmonella enterica subsp. enterica serovar Weltevreden Proteobacteria; Gammaproteobacteria; Enterobacterales; Enterobacteriaceae; Salmonella; Salmonella enterica; Salmonella enterica subsp. enterica MNNEETFYQAMRRKGVTRRSFLKFCSLAATSLGLGAGMTPKIAWALENKPRIPVVWIHGL WP_001058307.1 hydrogenase [Salmonella enterica] Length: 372\nNumber of predicted TMHs: 2\nExp number of AAs in TMHs: 36.51738\nExp number, first 60 AAs: 19.72849\nTotal prob of N-in: 0.96170\nPOSSIBLE N-term signal sequence\ninside 1 19\nTMhelix 20 39\noutside 40 325\nTMhelix 326 348\ninside 349 372

22540 GCF_001409715.1_9472_3_6Salmonella enterica subsp. enterica serovar Weltevreden Proteobacteria; Gammaproteobacteria; Enterobacterales; Enterobacteriaceae; Salmonella; Salmonella enterica; Salmonella enterica subsp. enterica MTGDNTLITSHGINRRDFMKLCAALAATMGLSSKAAAEMAESVSNPQRPPVIWIGAQECT WP_000145429.1 hydrogenase 2 small subunit [Salmonella enterica] Length: 372\nNumber of predicted TMHs: 1\nExp number of AAs in TMHs: 24.51309\nExp number, first 60 AAs: 0.2682\nTotal prob of N-in: 0.14402\noutside 1 331\nTMhelix 332 354\ninside 355 372

22541 GCF_001409735.1_12227_3_54 Salmonella enterica subsp. enterica serovar Weltevreden Proteobacteria; Gammaproteobacteria; Enterobacterales; Enterobacteriaceae; Salmonella; Salmonella enterica; Salmonella enterica subsp. enterica MTGDNTLITSHGINRRDFMKLCAALAATMGLSSKAAAEMAESVSNPQRPPVIWIGAQECT WP_000145429.1 hydrogenase 2 small subunit [Salmonella enterica] Length: 372\nNumber of predicted TMHs: 1\nExp number of AAs in TMHs: 24.51309\nExp number, first 60 AAs: 0.2682\nTotal prob of N-in: 0.14402\noutside 1 331\nTMhelix 332 354\ninside 355 372

22542 GCF_001409695.1_12227_3_11 Salmonella enterica subsp. enterica serovar Weltevreden Proteobacteria; Gammaproteobacteria; Enterobacterales; Enterobacteriaceae; Salmonella; Salmonella enterica; Salmonella enterica subsp. enterica MTGDNTLITSHGINRRDFMKLCAALAATMGLSSKAAAEMAESVSNPQRPPVIWIGAQECT WP_000145429.1 hydrogenase 2 small subunit [Salmonella enterica] Length: 372\nNumber of predicted TMHs: 1\nExp number of AAs in TMHs: 24.51309\nExp number, first 60 AAs: 0.2682\nTotal prob of N-in: 0.14402\noutside 1 331\nTMhelix 332 354\ninside 355 372

22543 GCF_001409655.1_12227_3_52 Salmonella enterica subsp. enterica serovar Weltevreden Proteobacteria; Gammaproteobacteria; Enterobacterales; Enterobacteriaceae; Salmonella; Salmonella enterica; Salmonella enterica subsp. enterica MTGDNTLITSHGINRRDFMKLCAALAATMGLSSKAAAEMAESVSNPQRPPVIWIGAQECT WP_000145429.1 hydrogenase 2 small subunit [Salmonella enterica] Length: 372\nNumber of predicted TMHs: 1\nExp number of AAs in TMHs: 24.51309\nExp number, first 60 AAs: 0.2682\nTotal prob of N-in: 0.14402\noutside 1 331\nTMhelix 332 354\ninside 355 372

22544 GCF_001409675.1_12227_3_45 Salmonella enterica subsp. enterica serovar Weltevreden Proteobacteria; Gammaproteobacteria; Enterobacterales; Enterobacteriaceae; Salmonella; Salmonella enterica; Salmonella enterica subsp. enterica MTGDNTLITSHGINRRDFMKLCAALAATMGLSSKAAAEMAESVSNPQRPPVIWIGAQECT WP_000145429.1 hydrogenase 2 small subunit [Salmonella enterica] Length: 372\nNumber of predicted TMHs: 1\nExp number of AAs in TMHs: 24.51309\nExp number, first 60 AAs: 0.2682\nTotal prob of N-in: 0.14402\noutside 1 331\nTMhelix 332 354\ninside 355 372

22545 GCF_001409595.1_12227_3_3 Salmonella enterica subsp. enterica serovar Weltevreden Proteobacteria; Gammaproteobacteria; Enterobacterales; Enterobacteriaceae; Salmonella; Salmonella enterica; Salmonella enterica subsp. enterica MNNEETFYQAMRRKGVTRRSFLKFCSLAATSLGLGAGMTPKIAWALENKPRIPVVWIHGL WP_001058307.1 hydrogenase [Salmonella enterica] Length: 372\nNumber of predicted TMHs: 2\nExp number of AAs in TMHs: 36.51738\nExp number, first 60 AAs: 19.72849\nTotal prob of N-in: 0.96170\nPOSSIBLE N-term signal sequence\ninside 1 19\nTMhelix 20 39\noutside 40 325\nTMhelix 326 348\ninside 349 372

22546	GCF_001409575.1_12227_3_55	Salmonella enterica subsp. enterica serovar Weltevreden	Proteobacteria; Gammaproteobacteria; Enterobacterales; Enterobacteriaceae; Salmonella; Salmonella enterica; Salmonella enterica subsp. enterica	MNNEETFYQAMRRKGVTRRSFLKFCSLAATSLGLGAGMTPKIAWALENKPRIPVVWIHGL	WP_001058307.1	hydrogenase [Salmonella enterica]	Length: 372\nNumber of predicted TMHs:  2\nExp number of AAs in TMHs: 36.51738\nExp number, first 60 AAs:  19.72849\nTotal prob of N-in:   0.96170\nPOSSIBLE N-term signal sequence\ninside   1   19\nTMhelix   20   39\noutside   40   325\nTMhelix   326   348\ninside   349   372

22547	GCF_001409635.1_9472_3_22	Salmonella enterica subsp. enterica serovar Weltevreden	Proteobacteria; Gammaproteobacteria; Enterobacterales; Enterobacteriaceae; Salmonella; Salmonella enterica; Salmonella enterica subsp. enterica	MTGDNTLITSHGINRRDFMKLCAALAATMGLSSKAAAEMAESVSNPQRPPVIWIGAQECT	WP_000145429.1	hydrogenase 2 small subunit [Salmonella enterica]	Length: 372\nNumber of predicted TMHs:  1\nExp number of AAs in TMHs: 24.51309\nExp number, first 60 AAs:  0.2682\nTotal prob of N-in:   0.14402\noutside   1   331\nTMhelix   332   354\ninside   355   372

22548	GCF_001409615.1_9472_3_15	Salmonella enterica subsp. enterica serovar Weltevreden	Proteobacteria; Gammaproteobacteria; Enterobacterales; Enterobacteriaceae; Salmonella; Salmonella enterica; Salmonella enterica subsp. enterica	MTGDNTLITSHGINRRDFMKLCAALAATMGLSSKAAAEMAESVSNPQRPPVIWIGAQECT	WP_000145429.1	hydrogenase 2 small subunit [Salmonella enterica]	Length: 372\nNumber of predicted TMHs:  1\nExp number of AAs in TMHs: 24.51309\nExp number, first 60 AAs:  0.2682\nTotal prob of N-in:   0.14402\noutside   1   331\nTMhelix   332   354\ninside   355   372

22549	GCF_001409595.1_12227_3_3	Salmonella enterica subsp. enterica serovar Weltevreden	Proteobacteria; Gammaproteobacteria; Enterobacterales; Enterobacteriaceae; Salmonella; Salmonella enterica; Salmonella enterica subsp. enterica	MTGDNTLITSHGINRRDFMKLCAALAATMGLSSKAAAEMAESVSNPQRPPVIWIGAQECT	WP_000145429.1	hydrogenase 2 small subunit [Salmonella enterica]	Length: 372\nNumber of predicted TMHs:  1\nExp number of AAs in TMHs: 24.51309\nExp number, first 60 AAs:  0.2682\nTotal prob of N-in:   0.14402\noutside   1   331\nTMhelix   332   354\ninside   355   372

22550	GCF_001409575.1_12227_3_55	Salmonella enterica subsp. enterica serovar Weltevreden	Proteobacteria; Gammaproteobacteria; Enterobacterales; Enterobacteriaceae; Salmonella; Salmonella enterica; Salmonella enterica subsp. enterica	MTGDNTLITSHGINRRDFMKLCAALAATMGLSSKAAAEMAESVSNPQRPPVIWIGAQECT	WP_000145429.1	hydrogenase 2 small subunit [Salmonella enterica]	Length: 372\nNumber of predicted TMHs:  1\nExp number of AAs in TMHs: 24.51309\nExp number, first 60 AAs:  0.2682\nTotal prob of N-in:   0.14402\noutside   1   331\nTMhelix   332   354\ninside   355   372

22551	GCF_001409555.1_12227_3_16	Salmonella enterica subsp. enterica serovar Weltevreden	Proteobacteria; Gammaproteobacteria; Enterobacterales; Enterobacteriaceae; Salmonella; Salmonella enterica; Salmonella enterica subsp. enterica	MNNEETFYQAMRRKGVTRRSFLKFCSLAATSLGLGAGMTPKIAWALENKPRIPVVWIHGL	WP_001058307.1	hydrogenase [Salmonella enterica]	Length: 372\nNumber of predicted TMHs:  2\nExp number of AAs in TMHs: 36.51738\nExp number, first 60 AAs:  19.72849\nTotal prob of N-in:   0.96170\nPOSSIBLE N-term signal sequence\ninside   1   19\nTMhelix   20   39\noutside   40   325\nTMhelix   326   348\ninside   349   372

22552	GCF_001409535.1_12227_3_9	Salmonella enterica subsp. enterica serovar Weltevreden	Proteobacteria; Gammaproteobacteria; Enterobacterales; Enterobacteriaceae; Salmonella; Salmonella enterica; Salmonella enterica subsp. enterica	MNNEETFYQAMRRKGVTRRSFLKFCSLAATSLGLGAGMTPKIAWALENKPRIPVVWIHGL	WP_001058307.1	hydrogenase [Salmonella enterica]	Length: 372\nNumber of predicted TMHs:  2\nExp number of AAs in TMHs: 36.51738\nExp number, first 60 AAs:  19.72849\nTotal prob of N-in:   0.96170\nPOSSIBLE N-term signal sequence\ninside   1   19\nTMhelix   20   39\noutside   40   325\nTMhelix   326   348\ninside   349   372

22553	GCF_001409835.1_12227_3_36	Salmonella enterica subsp. enterica serovar Weltevreden	Proteobacteria; Gammaproteobacteria; Enterobacterales; Enterobacteriaceae; Salmonella; Salmonella enterica; Salmonella enterica subsp. enterica	MTGDNTLITSHGINRRDFMKLCAALAATMGLSSKAAAEMAESVSNPQRPPVIWIGAQECT	WP_000145429.1	hydrogenase 2 small subunit [Salmonella enterica]	Length: 372\nNumber of predicted TMHs:  1\nExp number of AAs in TMHs: 24.51309\nExp number, first 60 AAs:  0.2682\nTotal prob of N-in:   0.14402\noutside   1   331\nTMhelix   332   354\ninside   355   372

22554	GCF_001409475.1_12227_3_48	Salmonella enterica subsp. enterica serovar Weltevreden	Proteobacteria; Gammaproteobacteria; Enterobacterales; Enterobacteriaceae; Salmonella; Salmonella enterica; Salmonella enterica subsp. enterica	MNNEETFYQAMRRKGVTRRSFLKFCSLAATSLGLGAGMTPKIAWALENKPRIPVVWIHGL	WP_001058307.1	hydrogenase [Salmonella enterica]	Length: 372\nNumber of predicted TMHs:  2\nExp

number of AAs in TMHs: 36.51738\nExp number, first 60 AAs: 19.72849\nTotal prob of N-in: 0.96170\nPOSSIBLE N-term signal sequence\ninside 1 19\nTMhelix 20 39\noutside 40 325\nTMhelix 326 348\ninside 349 372

22555 GCF_001409495.1_10900_1_31 Salmonella enterica subsp. enterica serovar Weltevreden Proteobacteria; Gammaproteobacteria; Enterobacterales; Enterobacteriaceae; Salmonella; Salmonella enterica; Salmonella enterica subsp. enterica MNNEETFYQAMRRKGVTRRSFLKFCSLAATSLGLGAGMTPKIAWALENKPRIPVVWIHGL WP_001058307.1 hydrogenase [Salmonella enterica] Length: 372\nNumber of predicted TMHs: 2\nExp number of AAs in TMHs: 36.51738\nExp number, first 60 AAs: 19.72849\nTotal prob of N-in: 0.96170\nPOSSIBLE N-term signal sequence\ninside 1 19\nTMhelix 20 39\noutside 40 325\nTMhelix 326 348\ninside 349 372

22556 GCF_001409415.1_12216_4_40 Salmonella enterica subsp. enterica serovar Weltevreden Proteobacteria; Gammaproteobacteria; Enterobacterales; Enterobacteriaceae; Salmonella; Salmonella enterica; Salmonella enterica subsp. enterica MNNEETFYQAMRRKGVTRRSFLKFCSLAATSLGLGAGMTPKIAWALENKPRIPVVWIHGL WP_001058307.1 hydrogenase [Salmonella enterica] Length: 372\nNumber of predicted TMHs: 2\nExp number of AAs in TMHs: 36.51738\nExp number, first 60 AAs: 19.72849\nTotal prob of N-in: 0.96170\nPOSSIBLE N-term signal sequence\ninside 1 19\nTMhelix 20 39\noutside 40 325\nTMhelix 326 348\ninside 349 372

22557 GCF_001409515.1_9472_3_24 Salmonella enterica subsp. enterica serovar Weltevreden Proteobacteria; Gammaproteobacteria; Enterobacterales; Enterobacteriaceae; Salmonella; Salmonella enterica; Salmonella enterica subsp. enterica MNNEETFYQAMRRKGVTRRSFLKFCSLAATSLGLGAGMTPKIAWALENKPRIPVVWIHGL WP_001058307.1 hydrogenase [Salmonella enterica] Length: 372\nNumber of predicted TMHs: 2\nExp number of AAs in TMHs: 36.51738\nExp number, first 60 AAs: 19.72849\nTotal prob of N-in: 0.96170\nPOSSIBLE N-term signal sequence\ninside 1 19\nTMhelix 20 39\noutside 40 325\nTMhelix 326 348\ninside 349 372

22558 GCF_001409555.1_12227_3_16 Salmonella enterica subsp. enterica serovar Weltevreden Proteobacteria; Gammaproteobacteria; Enterobacterales; Enterobacteriaceae; Salmonella; Salmonella enterica; Salmonella enterica subsp. enterica MTGDNTLITSHGINRRDFMKLCAALAATMGLSSKAAAEMAESVSNPQRPPVIWIGAQECT WP_000145429.1 hydrogenase 2 small subunit [Salmonella enterica] Length: 372\nNumber of predicted TMHs: 1\nExp number of AAs in TMHs: 24.51309\nExp number, first 60 AAs: 0.2682\nTotal prob of N-in: 0.14402\noutside 1 331\nTMhelix 332 354\ninside 355 372

22559 GCF_001409515.1_9472_3_24 Salmonella enterica subsp. enterica serovar Weltevreden Proteobacteria; Gammaproteobacteria; Enterobacterales; Enterobacteriaceae; Salmonella; Salmonella enterica; Salmonella enterica subsp. enterica MTGDNTLITSHGINRRDFMKLCAALAATMGLSSKAAAEMAESVSNPQRPPVIWIGAQECT WP_000145429.1 hydrogenase 2 small subunit [Salmonella enterica] Length: 372\nNumber of predicted TMHs: 1\nExp number of AAs in TMHs: 24.51309\nExp number, first 60 AAs: 0.2682\nTotal prob of N-in: 0.14402\noutside 1 331\nTMhelix 332 354\ninside 355 372

22560 GCF_001409495.1_10900_1_31 Salmonella enterica subsp. enterica serovar Weltevreden Proteobacteria; Gammaproteobacteria; Enterobacterales; Enterobacteriaceae; Salmonella; Salmonella enterica; Salmonella enterica subsp. enterica MTGDNTLITSHGINRRDFMKLCAALAATMGLSSKAAAEMAESVSNPQRPPVIWIGAQECT WP_000145429.1 hydrogenase 2 small subunit [Salmonella enterica] Length: 372\nNumber of predicted TMHs: 1\nExp number of AAs in TMHs: 24.51309\nExp number, first 60 AAs: 0.2682\nTotal prob of N-in: 0.14402\noutside 1 331\nTMhelix 332 354\ninside 355 372

22561 GCF_001409535.1_12227_3_9 Salmonella enterica subsp. enterica serovar Weltevreden Proteobacteria; Gammaproteobacteria; Enterobacterales; Enterobacteriaceae; Salmonella; Salmonella enterica; Salmonella enterica subsp. enterica MTGDNTLITSHGINRRDFMKLCAALAATMGLSSKAAAEMAESVSNPQRPPVIWIGAQECT WP_000145429.1 hydrogenase 2 small subunit [Salmonella enterica] Length: 372\nNumber of predicted TMHs: 1\nExp number of AAs in TMHs: 24.51309\nExp number, first 60 AAs: 0.2682\nTotal prob of N-in: 0.14402\noutside 1 331\nTMhelix 332 354\ninside 355 372

22562 GCF_001409475.1_12227_3_48 Salmonella enterica subsp. enterica serovar Weltevreden Proteobacteria; Gammaproteobacteria; Enterobacterales; Enterobacteriaceae; Salmonella; Salmonella enterica; Salmonella enterica subsp. enterica MTGDNTLITSHGINRRDFMKLCAALAATMGLSSKAAAEMAESVSNPQRPPVIWIGAQECT WP_000145429.1 hydrogenase 2 small subunit [Salmonella enterica] Length: 372\nNumber of predicted TMHs: 1\nExp number of AAs in TMHs: 24.51309\nExp number, first 60 AAs: 0.2682\nTotal prob of N-in: 0.14402\noutside 1 331\nTMhelix 332 354\ninside 355 372

22563 GCF_001409375.1_12227_3_23 Salmonella enterica subsp. enterica serovar Weltevreden Proteobacteria; Gammaproteobacteria; Enterobacterales; Enterobacteriaceae; Salmonella; Salmonella enterica;

Salmonella enterica subsp. enterica	MNNEETFYQAMRRKGVTRRSFLKFCSLAATSLGLGAGMTPKIAWALENKPRIPVVWIHGL	WP_001058307.1	hydrogenase [Salmonella enterica]	Length: 372\nNumber of predicted TMHs:  2\nExp number of AAs in TMHs: 36.51738\nExp number, first 60 AAs:  19.72849\nTotal prob of N-in:   0.96170\nPOSSIBLE N-term signal sequence\ninside   1   19\nTMhelix   20   39\noutside   40   325\nTMhelix   326   348\ninside   349   372

22564	GCF_001409395.1_9472_3_17	Salmonella enterica subsp. enterica serovar Weltevreden	Proteobacteria; Gammaproteobacteria; Enterobacterales; Enterobacteriaceae; Salmonella; Salmonella enterica; Salmonella enterica subsp. enterica	MNNEETFYQAMRRKGVTRRSFLKFCSLAATSLGLGAGMTPKIAWALENKPRIPVVWIHGL	WP_001058307.1	hydrogenase [Salmonella enterica]	Length: 372\nNumber of predicted TMHs:  2\nExp number of AAs in TMHs: 36.51738\nExp number, first 60 AAs:  19.72849\nTotal prob of N-in:   0.96170\nPOSSIBLE N-term signal sequence\ninside   1   19\nTMhelix   20   39\noutside   40   325\nTMhelix   326   348\ninside   349   372

22565	GCF_001409455.1_12227_3_53	Salmonella enterica subsp. enterica serovar Weltevreden	Proteobacteria; Gammaproteobacteria; Enterobacterales; Enterobacteriaceae; Salmonella; Salmonella enterica; Salmonella enterica subsp. enterica	MNNEETFYQAMRRKGVTRRSFLKFCSLAATSLGLGAGMTPKIAWALENKPRIPVVWIHGL	WP_001058307.1	hydrogenase [Salmonella enterica]	Length: 372\nNumber of predicted TMHs:  2\nExp number of AAs in TMHs: 36.51738\nExp number, first 60 AAs:  19.72849\nTotal prob of N-in:   0.96170\nPOSSIBLE N-term signal sequence\ninside   1   19\nTMhelix   20   39\noutside   40   325\nTMhelix   326   348\ninside   349   372

22566	GCF_001409435.1_9472_3_14	Salmonella enterica subsp. enterica serovar Weltevreden	Proteobacteria; Gammaproteobacteria; Enterobacterales; Enterobacteriaceae; Salmonella; Salmonella enterica; Salmonella enterica subsp. enterica	MNNEETFYQAMRRKGVTRRSFLKFCSLAATSLGLGAGMTPKIAWALENKPRIPVVWIHGL	WP_001058307.1	hydrogenase [Salmonella enterica]	Length: 372\nNumber of predicted TMHs:  2\nExp number of AAs in TMHs: 36.51738\nExp number, first 60 AAs:  19.72849\nTotal prob of N-in:   0.96170\nPOSSIBLE N-term signal sequence\ninside   1   19\nTMhelix   20   39\noutside   40   325\nTMhelix   326   348\ninside   349   372

22567	GCF_001409455.1_12227_3_53	Salmonella enterica subsp. enterica serovar Weltevreden	Proteobacteria; Gammaproteobacteria; Enterobacterales; Enterobacteriaceae; Salmonella; Salmonella enterica; Salmonella enterica subsp. enterica	MTGDNTLITSHGINRRDFMKLCAALAATMGLSSKAAAEMAESVSNPQRPPVIWIGAQECT	WP_000145429.1	hydrogenase 2 small subunit [Salmonella enterica]	Length: 372\nNumber of predicted TMHs:  1\nExp number of AAs in TMHs: 24.51309\nExp number, first 60 AAs:  0.2682\nTotal prob of N-in: 0.14402\noutside   1   331\nTMhelix   332   354\ninside   355   372

22568	GCF_001409435.1_9472_3_14	Salmonella enterica subsp. enterica serovar Weltevreden	Proteobacteria; Gammaproteobacteria; Enterobacterales; Enterobacteriaceae; Salmonella; Salmonella enterica; Salmonella enterica subsp. enterica	MTGDNTLITSHGINRRDFMKLCAALAATMGLSSKAAAEMAESVSNPQRPPVIWIGAQECT	WP_000145429.1	hydrogenase 2 small subunit [Salmonella enterica]	Length: 372\nNumber of predicted TMHs:  1\nExp number of AAs in TMHs: 24.51309\nExp number, first 60 AAs:  0.2682\nTotal prob of N-in: 0.14402\noutside   1   331\nTMhelix   332   354\ninside   355   372

22569	GCF_001409415.1_12216_4_40	Salmonella enterica subsp. enterica serovar Weltevreden	Proteobacteria; Gammaproteobacteria; Enterobacterales; Enterobacteriaceae; Salmonella; Salmonella enterica; Salmonella enterica subsp. enterica	MTGDNTLITSHGINRRDFMKLCAALAATMGLSSKAAAEMAESVSNPQRPPVIWIGAQECT	WP_000145429.1	hydrogenase 2 small subunit [Salmonella enterica]	Length: 372\nNumber of predicted TMHs:  1\nExp number of AAs in TMHs: 24.51309\nExp number, first 60 AAs:  0.2682\nTotal prob of N-in: 0.14402\noutside   1   331\nTMhelix   332   354\ninside   355   372

22570	GCF_001409335.1_9472_3_16	Salmonella enterica subsp. enterica serovar Weltevreden	Proteobacteria; Gammaproteobacteria; Enterobacterales; Enterobacteriaceae; Salmonella; Salmonella enterica; Salmonella enterica subsp. enterica	MNNEETFYQAMRRKGVTRRSFLKFCSLAATSLGLGAGMTPKIAWALENKPRIPVVWIHGL	WP_001058307.1	hydrogenase [Salmonella enterica]	Length: 372\nNumber of predicted TMHs:  2\nExp number of AAs in TMHs: 36.51738\nExp number, first 60 AAs:  19.72849\nTotal prob of N-in:   0.96170\nPOSSIBLE N-term signal sequence\ninside   1   19\nTMhelix   20   39\noutside   40   325\nTMhelix   326   348\ninside   349   372

22571	GCF_001409395.1_9472_3_17	Salmonella enterica subsp. enterica serovar Weltevreden	Proteobacteria; Gammaproteobacteria; Enterobacterales; Enterobacteriaceae; Salmonella; Salmonella enterica; Salmonella enterica subsp. enterica	MTGDNTLITSHGINRRDFMKLCAALAATMGLSSKAAAEMAESVSNPQRPPVIWIGAQECT	WP_000145429.1	hydrogenase 2 small subunit [Salmonella enterica]	Length: 372\nNumber of predicted TMHs:  1\nExp number of AAs in TMHs: 24.51309\nExp number, first 60 AAs:  0.2682\nTotal prob of N-in: 0.14402\noutside   1   331\nTMhelix   332   354\ninside   355   372

22572 GCF_001409375.1_12227_3_23 Salmonella enterica subsp. enterica serovar Weltevreden Proteobacteria; Gammaproteobacteria; Enterobacterales; Enterobacteriaceae; Salmonella; Salmonella enterica; Salmonella enterica subsp. enterica MTGDNTLITSHGINRRDFMKLCAALAATMGLSSKAAAEMAESVSNPQRPPVIWIGAQECT WP_000145429.1 hydrogenase 2 small subunit [Salmonella enterica] Length: 372\nNumber of predicted TMHs: 1\nExp number of AAs in TMHs: 24.51309\nExp number, first 60 AAs: 0.2682\nTotal prob of N-in: 0.14402\noutside 1 331\nTMhelix 332 354\ninside 355 372

22573 GCF_001409355.1_12227_3_15 Salmonella enterica subsp. enterica serovar Weltevreden Proteobacteria; Gammaproteobacteria; Enterobacterales; Enterobacteriaceae; Salmonella; Salmonella enterica; Salmonella enterica subsp. enterica MTGDNTLITSHGINRRDFMKLCAALAATMGLSSKAAAEMAESVSNPQRPPVIWIGAQECT WP_000145429.1 hydrogenase 2 small subunit [Salmonella enterica] Length: 372\nNumber of predicted TMHs: 1\nExp number of AAs in TMHs: 24.51309\nExp number, first 60 AAs: 0.2682\nTotal prob of N-in: 0.14402\noutside 1 331\nTMhelix 332 354\ninside 355 372

22574 GCF_001409335.1_9472_3_16 Salmonella enterica subsp. enterica serovar Weltevreden Proteobacteria; Gammaproteobacteria; Enterobacterales; Enterobacteriaceae; Salmonella; Salmonella enterica; Salmonella enterica subsp. enterica MTGDNTLITSHGINRRDFMKLCAALAATMGLSSKAAAEMAESVSNPQRPPVIWIGAQECT WP_000145429.1 hydrogenase 2 small subunit [Salmonella enterica] Length: 372\nNumber of predicted TMHs: 1\nExp number of AAs in TMHs: 24.51309\nExp number, first 60 AAs: 0.2682\nTotal prob of N-in: 0.14402\noutside 1 331\nTMhelix 332 354\ninside 355 372

22575 GCF_001409295.1_9472_3_30 Salmonella enterica subsp. enterica serovar Weltevreden Proteobacteria; Gammaproteobacteria; Enterobacterales; Enterobacteriaceae; Salmonella; Salmonella enterica; Salmonella enterica subsp. enterica MNNEETFYQAMRRKGVTRRSFLKFCSLAATSLGLGAGMTPKIAWALENKPRIPVVWIHGL WP_001058307.1 hydrogenase [Salmonella enterica] Length: 372\nNumber of predicted TMHs: 2\nExp number of AAs in TMHs: 36.51738\nExp number, first 60 AAs: 19.72849\nTotal prob of N-in: 0.96170\nPOSSIBLE N-term signal sequence\ninside 1 19\nTMhelix 20 39\noutside 40 325\nTMhelix 326 348\ninside 349 372

22576 GCF_001409195.1_C2346 Salmonella enterica subsp. enterica serovar Weltevreden Proteobacteria; Gammaproteobacteria; Enterobacterales; Enterobacteriaceae; Salmonella; Salmonella enterica; Salmonella enterica subsp. enterica MNNEETFYQAMRRKGVTRRSFLKFCSLAATSLGLGAGMTPKIAWALENKPRIPVVWIHGL WP_001058307.1 hydrogenase [Salmonella enterica] Length: 372\nNumber of predicted TMHs: 2\nExp number of AAs in TMHs: 36.51738\nExp number, first 60 AAs: 19.72849\nTotal prob of N-in: 0.96170\nPOSSIBLE N-term signal sequence\ninside 1 19\nTMhelix 20 39\noutside 40 325\nTMhelix 326 348\ninside 349 372

22577 GCF_001409175.1_99_3134 Salmonella enterica subsp. enterica serovar Weltevreden Proteobacteria; Gammaproteobacteria; Enterobacterales; Enterobacteriaceae; Salmonella; Salmonella enterica; Salmonella enterica subsp. enterica MNNEETFYQAMRRKGVTRRSFLKFCSLAATSLGLGAGMTPKIAWALENKPRIPVVWIHGL WP_001058307.1 hydrogenase [Salmonella enterica] Length: 372\nNumber of predicted TMHs: 2\nExp number of AAs in TMHs: 36.51738\nExp number, first 60 AAs: 19.72849\nTotal prob of N-in: 0.96170\nPOSSIBLE N-term signal sequence\ninside 1 19\nTMhelix 20 39\noutside 40 325\nTMhelix 326 348\ninside 349 372

22578 GCF_001409155.1_98_11262Salmonella enterica subsp. enterica serovar Weltevreden Proteobacteria; Gammaproteobacteria; Enterobacterales; Enterobacteriaceae; Salmonella; Salmonella enterica; Salmonella enterica subsp. enterica MNNEETFYQAMRRKGVTRRSFLKFCSLAATSLGLGAGMTPKIAWALENKPRIPVVWIHGL WP_001058307.1 hydrogenase [Salmonella enterica] Length: 372\nNumber of predicted TMHs: 2\nExp number of AAs in TMHs: 36.51738\nExp number, first 60 AAs: 19.72849\nTotal prob of N-in: 0.96170\nPOSSIBLE N-term signal sequence\ninside 1 19\nTMhelix 20 39\noutside 40 325\nTMhelix 326 348\ninside 349 372

22579 GCF_001409275.1_12216_4_53 Salmonella enterica subsp. enterica serovar Weltevreden Proteobacteria; Gammaproteobacteria; Enterobacterales; Enterobacteriaceae; Salmonella; Salmonella enterica; Salmonella enterica subsp. enterica MNNEETFYQAMRRKGVTRRSFLKFCSLAATSLGLGAGMTPKIAWALENKPRIPVVWIHGL WP_001058307.1 hydrogenase [Salmonella enterica] Length: 372\nNumber of predicted TMHs: 2\nExp number of AAs in TMHs: 36.51738\nExp number, first 60 AAs: 19.72849\nTotal prob of N-in: 0.96170\nPOSSIBLE N-term signal sequence\ninside 1 19\nTMhelix 20 39\noutside 40 325\nTMhelix 326 348\ninside 349 372

22580 GCF_001409315.1_12216_4_55 Salmonella enterica subsp. enterica serovar Weltevreden Proteobacteria; Gammaproteobacteria; Enterobacterales; Enterobacteriaceae; Salmonella; Salmonella enterica; Salmonella enterica subsp. enterica MNNEETFYQAMRRKGVTRRSFLKFCSLAATSLGLGAGMTPKIAWALENKPRIPVVWIHGL WP_001058307.1 hydrogenase [Salmonella enterica] Length: 372\nNumber of predicted TMHs: 2\nExp

number of AAs in TMHs: 36.51738\nExp number, first 60 AAs: 19.72849\nTotal prob of N-in: 0.96170\nPOSSIBLE N-term signal sequence\ninside 1 19\nTMhelix 20 39\noutside 40 325\nTMhelix 326 348\ninside 349 372

22581 GCF_001409295.1_9472_3_30 Salmonella enterica subsp. enterica serovar Weltevreden Proteobacteria; Gammaproteobacteria; Enterobacterales; Enterobacteriaceae; Salmonella; Salmonella enterica; Salmonella enterica subsp. enterica MTGDNTLITSHGINRRDFMKLCAALAATMGLSSKAAAEMAESVSNPQRPPVIWIGAQECT WP_000145429.1 hydrogenase 2 small subunit [Salmonella enterica] Length: 372\nNumber of predicted TMHs: 1\nExp number of AAs in TMHs: 24.51309\nExp number, first 60 AAs: 0.2682\nTotal prob of N-in: 0.14402\noutside 1 331\nTMhelix 332 354\ninside 355 372

22582 GCF_001409275.1_12216_4_53 Salmonella enterica subsp. enterica serovar Weltevreden Proteobacteria; Gammaproteobacteria; Enterobacterales; Enterobacteriaceae; Salmonella; Salmonella enterica; Salmonella enterica subsp. enterica MTGDNTLITSHGINRRDFMKLCAALAATMGLSSKAAAEMAESVSNPQRPPVIWIGAQECT WP_000145429.1 hydrogenase 2 small subunit [Salmonella enterica] Length: 372\nNumber of predicted TMHs: 1\nExp number of AAs in TMHs: 24.51309\nExp number, first 60 AAs: 0.2682\nTotal prob of N-in: 0.14402\noutside 1 331\nTMhelix 332 354\ninside 355 372

22583 GCF_001409315.1_12216_4_55 Salmonella enterica subsp. enterica serovar Weltevreden Proteobacteria; Gammaproteobacteria; Enterobacterales; Enterobacteriaceae; Salmonella; Salmonella enterica; Salmonella enterica subsp. enterica MTGDNTLITSHGINRRDFMKLCAALAATMGLSSKAAAEMAESVSNPQRPPVIWIGAQECT WP_000145429.1 hydrogenase 2 small subunit [Salmonella enterica] Length: 372\nNumber of predicted TMHs: 1\nExp number of AAs in TMHs: 24.51309\nExp number, first 60 AAs: 0.2682\nTotal prob of N-in: 0.14402\noutside 1 331\nTMhelix 332 354\ninside 355 372

22584 GCF_001409255.1_12227_3_1 Salmonella enterica subsp. enterica serovar Weltevreden Proteobacteria; Gammaproteobacteria; Enterobacterales; Enterobacteriaceae; Salmonella; Salmonella enterica; Salmonella enterica subsp. enterica MTGDNTLITSHGINRRDFMKLCAALAATMGLSSKAAAEMAESVSNPQRPPVIWIGAQECT WP_000145429.1 hydrogenase 2 small subunit [Salmonella enterica] Length: 372\nNumber of predicted TMHs: 1\nExp number of AAs in TMHs: 24.51309\nExp number, first 60 AAs: 0.2682\nTotal prob of N-in: 0.14402\noutside 1 331\nTMhelix 332 354\ninside 355 372

22585 GCF_001409135.1_10259 Salmonella enterica subsp. enterica serovar Weltevreden Proteobacteria; Gammaproteobacteria; Enterobacterales; Enterobacteriaceae; Salmonella; Salmonella enterica; Salmonella enterica subsp. enterica MNNEETFYQAMRRKGVTRRSFLKFCSLAATSLGLGAGMTPKIAWALENKPRIPVVWIHGL WP_001058307.1 hydrogenase [Salmonella enterica] Length: 372\nNumber of predicted TMHs: 2\nExp number of AAs in TMHs: 36.51738\nExp number, first 60 AAs: 19.72849\nTotal prob of N-in: 0.96170\nPOSSIBLE N-term signal sequence\ninside 1 19\nTMhelix 20 39\noutside 40 325\nTMhelix 326 348\ninside 349 372

22586 GCF_001409235.1_12227_3_30 Salmonella enterica subsp. enterica serovar Weltevreden Proteobacteria; Gammaproteobacteria; Enterobacterales; Enterobacteriaceae; Salmonella; Salmonella enterica; Salmonella enterica subsp. enterica MNNEETFYQAMRRKGVTRRSFLKFCSLAATSLGLGAGMTPKIAWALENKPRIPVVWIHGL WP_001058307.1 hydrogenase [Salmonella enterica] Length: 372\nNumber of predicted TMHs: 2\nExp number of AAs in TMHs: 36.51738\nExp number, first 60 AAs: 19.72849\nTotal prob of N-in: 0.96170\nPOSSIBLE N-term signal sequence\ninside 1 19\nTMhelix 20 39\noutside 40 325\nTMhelix 326 348\ninside 349 372

22587 GCF_001409215.1_12227_3_8 Salmonella enterica subsp. enterica serovar Weltevreden Proteobacteria; Gammaproteobacteria; Enterobacterales; Enterobacteriaceae; Salmonella; Salmonella enterica; Salmonella enterica subsp. enterica MNNEETFYQAMRRKGVTRRSFLKFCSLAATSLGLGAGMTPKIAWALENKPRIPVVWIHGL WP_001058307.1 hydrogenase [Salmonella enterica] Length: 372\nNumber of predicted TMHs: 2\nExp number of AAs in TMHs: 36.51738\nExp number, first 60 AAs: 19.72849\nTotal prob of N-in: 0.96170\nPOSSIBLE N-term signal sequence\ninside 1 19\nTMhelix 20 39\noutside 40 325\nTMhelix 326 348\ninside 349 372

22588 GCF_001409235.1_12227_3_30 Salmonella enterica subsp. enterica serovar Weltevreden Proteobacteria; Gammaproteobacteria; Enterobacterales; Enterobacteriaceae; Salmonella; Salmonella enterica; Salmonella enterica subsp. enterica MTGDNTLITSHGINRRDFMKLCAALAATMGLSSKAAAEMAESVSNPQRPPVIWIGAQECT WP_000145429.1 hydrogenase 2 small subunit [Salmonella enterica] Length: 372\nNumber of predicted TMHs: 1\nExp number of AAs in TMHs: 24.51309\nExp number, first 60 AAs: 0.2682\nTotal prob of N-in: 0.14402\noutside 1 331\nTMhelix 332 354\ninside 355 372

22589 GCF_001409215.1_12227_3_8 Salmonella enterica subsp. enterica serovar Weltevreden Proteobacteria; Gammaproteobacteria; Enterobacterales; Enterobacteriaceae; Salmonella; Salmonella enterica;

Salmonella enterica subsp. enterica MTGDNTLITSHGINRRDFMKLCAALAATMGLSSKAAAEMAESVSNPQRPPVIWIGAQECT WP_000145429.1 hydrogenase 2 small subunit [Salmonella enterica] Length: 372\nNumber of predicted TMHs: 1\nExp number of AAs in TMHs: 24.51309\nExp number, first 60 AAs: 0.2682\nTotal prob of N-in: 0.14402\noutside 1 331\nTMhelix 332 354\ninside 355 372

22590 GCF_001409195.1_C2346 Salmonella enterica subsp. enterica serovar Weltevreden Proteobacteria; Gammaproteobacteria; Enterobacterales; Enterobacteriaceae; Salmonella; Salmonella enterica; Salmonella enterica subsp. enterica MTGDNTLITSHGINRRDFMKLCAALAATMGLSSKAAAEMAESVSNPQRPPVIWIGAQECT WP_000145429.1 hydrogenase 2 small subunit [Salmonella enterica] Length: 372\nNumber of predicted TMHs: 1\nExp number of AAs in TMHs: 24.51309\nExp number, first 60 AAs: 0.2682\nTotal prob of N-in: 0.14402\noutside 1 331\nTMhelix 332 354\ninside 355 372

22591 GCF_001409175.1_99_3134 Salmonella enterica subsp. enterica serovar Weltevreden Proteobacteria; Gammaproteobacteria; Enterobacterales; Enterobacteriaceae; Salmonella; Salmonella enterica; Salmonella enterica subsp. enterica MTGDNTLITSHGINRRDFMKLCAALAATMGLSSKAAAEMAESVSNPQRPPVIWIGAQECT WP_000145429.1 hydrogenase 2 small subunit [Salmonella enterica] Length: 372\nNumber of predicted TMHs: 1\nExp number of AAs in TMHs: 24.51309\nExp number, first 60 AAs: 0.2682\nTotal prob of N-in: 0.14402\noutside 1 331\nTMhelix 332 354\ninside 355 372

22592 GCF_001409155.1_98_11262Salmonella enterica subsp. enterica serovar Weltevreden Proteobacteria; Gammaproteobacteria; Enterobacterales; Enterobacteriaceae; Salmonella; Salmonella enterica; Salmonella enterica subsp. enterica MTGDNTLITSHGINRRDFMKLCAALAATMGLSSKAAAEMAESVSNPQRPPVIWIGAQECT WP_000145429.1 hydrogenase 2 small subunit [Salmonella enterica] Length: 372\nNumber of predicted TMHs: 1\nExp number of AAs in TMHs: 24.51309\nExp number, first 60 AAs: 0.2682\nTotal prob of N-in: 0.14402\noutside 1 331\nTMhelix 332 354\ninside 355 372

22593 GCF_001409135.1_10259 Salmonella enterica subsp. enterica serovar Weltevreden Proteobacteria; Gammaproteobacteria; Enterobacterales; Enterobacteriaceae; Salmonella; Salmonella enterica; Salmonella enterica subsp. enterica MTGDNTLITSHGINRRDFMKLCAALAATMGLSSKAAAEMAESVSNPQRPPVIWIGAQECT WP_000145429.1 hydrogenase 2 small subunit [Salmonella enterica] Length: 372\nNumber of predicted TMHs: 1\nExp number of AAs in TMHs: 24.51309\nExp number, first 60 AAs: 0.2682\nTotal prob of N-in: 0.14402\noutside 1 331\nTMhelix 332 354\ninside 355 372

22594 GCF_001411455.1_9472_3_20 Salmonella enterica subsp. enterica serovar Weltevreden Proteobacteria; Gammaproteobacteria; Enterobacterales; Enterobacteriaceae; Salmonella; Salmonella enterica; Salmonella enterica subsp. enterica MNNEETFYQAMRRKGVTRRSFLKFCSLAATSLGLGAGMTPKIAWALENKPRIPVVWIHGL WP_001058307.1 hydrogenase [Salmonella enterica] Length: 372\nNumber of predicted TMHs: 2\nExp number of AAs in TMHs: 36.51738\nExp number, first 60 AAs: 19.72849\nTotal prob of N-in: 0.96170\nPOSSIBLE N-term signal sequence\ninside 1 19\nTMhelix 20 39\noutside 40 325\nTMhelix 326 348\ninside 349 372

22595 GCF_001411435.1_12227_3_43 Salmonella enterica subsp. enterica serovar Weltevreden Proteobacteria; Gammaproteobacteria; Enterobacterales; Enterobacteriaceae; Salmonella; Salmonella enterica; Salmonella enterica subsp. enterica MNNEETFYQAMRRKGVTRRSFLKFCSLAATSLGLGAGMTPKIAWALENKPRIPVVWIHGL WP_001058307.1 hydrogenase [Salmonella enterica] Length: 372\nNumber of predicted TMHs: 2\nExp number of AAs in TMHs: 36.51738\nExp number, first 60 AAs: 19.72849\nTotal prob of N-in: 0.96170\nPOSSIBLE N-term signal sequence\ninside 1 19\nTMhelix 20 39\noutside 40 325\nTMhelix 326 348\ninside 349 372

22596 GCF_001411395.1_12227_3_31 Salmonella enterica subsp. enterica serovar Weltevreden Proteobacteria; Gammaproteobacteria; Enterobacterales; Enterobacteriaceae; Salmonella; Salmonella enterica; Salmonella enterica subsp. enterica MNNEETFYQAMRRKGVTRRSFLKFCSLAATSLGLGAGMTPKIAWALENKPRIPVVWIHGL WP_001058307.1 hydrogenase [Salmonella enterica] Length: 372\nNumber of predicted TMHs: 2\nExp number of AAs in TMHs: 36.51738\nExp number, first 60 AAs: 19.72849\nTotal prob of N-in: 0.96170\nPOSSIBLE N-term signal sequence\ninside 1 19\nTMhelix 20 39\noutside 40 325\nTMhelix 326 348\ninside 349 372

22597 GCF_001411455.1_9472_3_20 Salmonella enterica subsp. enterica serovar Weltevreden Proteobacteria; Gammaproteobacteria; Enterobacterales; Enterobacteriaceae; Salmonella; Salmonella enterica; Salmonella enterica subsp. enterica MTGDNTLITSHGINRRDFMKLCAALAATMGLSSKAAAEMAESVSNPQRPPVIWIGAQECT WP_000145429.1 hydrogenase 2 small subunit [Salmonella enterica] Length: 372\nNumber of predicted TMHs: 1\nExp number of AAs in TMHs: 24.51309\nExp number, first 60 AAs: 0.2682\nTotal prob of N-in: 0.14402\noutside 1 331\nTMhelix 332 354\ninside 355 372

22598	GCF_001411435.1_12227_3_43	Salmonella enterica subsp. enterica serovar Weltevreden
Proteobacteria; Gammaproteobacteria; Enterobacterales; Enterobacteriaceae; Salmonella; Salmonella enterica; Salmonella enterica subsp. enterica	MTGDNTLITSHGINRRDFMKLCAALAATMGLSSKAAAEMAESVSNPQRPPVIWIGAQECT
WP_000145429.1	hydrogenase 2 small subunit [Salmonella enterica]	Length: 372\nNumber of predicted TMHs: 1\nExp number of AAs in TMHs: 24.51309\nExp number, first 60 AAs: 0.2682\nTotal prob of N-in: 0.14402\noutside 1 331\nTMhelix 332 354\ninside 355 372

22599	GCF_001411395.1_12227_3_31	Salmonella enterica subsp. enterica serovar Weltevreden
Proteobacteria; Gammaproteobacteria; Enterobacterales; Enterobacteriaceae; Salmonella; Salmonella enterica; Salmonella enterica subsp. enterica	MTGDNTLITSHGINRRDFMKLCAALAATMGLSSKAAAEMAESVSNPQRPPVIWIGAQECT
WP_000145429.1	hydrogenase 2 small subunit [Salmonella enterica]	Length: 372\nNumber of predicted TMHs: 1\nExp number of AAs in TMHs: 24.51309\nExp number, first 60 AAs: 0.2682\nTotal prob of N-in: 0.14402\noutside 1 331\nTMhelix 332 354\ninside 355 372

22600	GCF_001411355.1_12227_3_18	Salmonella enterica subsp. enterica serovar Weltevreden
Proteobacteria; Gammaproteobacteria; Enterobacterales; Enterobacteriaceae; Salmonella; Salmonella enterica; Salmonella enterica subsp. enterica	MNNEETFYQAMRRKGVTRRSFLKFCSLAATSLGLGAGMTPKIAWALENKPRIPVVWIHGL
WP_001058307.1	hydrogenase [Salmonella enterica]	Length: 372\nNumber of predicted TMHs: 2\nExp number of AAs in TMHs: 36.51738\nExp number, first 60 AAs: 19.72849\nTotal prob of N-in: 0.96170\nPOSSIBLE N-term signal sequence\ninside 1 19\nTMhelix 20 39\noutside 40 325\nTMhelix 326 348\ninside 349 372

22601	GCF_001411375.1_12227_3_21	Salmonella enterica subsp. enterica serovar Weltevreden
Proteobacteria; Gammaproteobacteria; Enterobacterales; Enterobacteriaceae; Salmonella; Salmonella enterica; Salmonella enterica subsp. enterica	MNNEETFYQAMRRKGVTRRSFLKFCSLAATSLGLGAGMTPKIAWALENKPRIPVVWIHGL
WP_001058307.1	hydrogenase [Salmonella enterica]	Length: 372\nNumber of predicted TMHs: 2\nExp number of AAs in TMHs: 36.51738\nExp number, first 60 AAs: 19.72849\nTotal prob of N-in: 0.96170\nPOSSIBLE N-term signal sequence\ninside 1 19\nTMhelix 20 39\noutside 40 325\nTMhelix 326 348\ninside 349 372

22602	GCF_001411375.1_12227_3_21	Salmonella enterica subsp. enterica serovar Weltevreden
Proteobacteria; Gammaproteobacteria; Enterobacterales; Enterobacteriaceae; Salmonella; Salmonella enterica; Salmonella enterica subsp. enterica	MTGDNTLITSHGINRRDFMKLCAALAATMGLSSKAAAEMAESVSNPQRPPVIWIGAQECT
WP_000145429.1	hydrogenase 2 small subunit [Salmonella enterica]	Length: 372\nNumber of predicted TMHs: 1\nExp number of AAs in TMHs: 24.51309\nExp number, first 60 AAs: 0.2682\nTotal prob of N-in: 0.14402\noutside 1 331\nTMhelix 332 354\ninside 355 372

22603	GCF_001411335.1_12227_3_12	Salmonella enterica subsp. enterica serovar Weltevreden
Proteobacteria; Gammaproteobacteria; Enterobacterales; Enterobacteriaceae; Salmonella; Salmonella enterica; Salmonella enterica subsp. enterica	MNNEETFYQAMRRKGVTRRSFLKFCSLAATSLGLGAGMTPKIAWALENKPRIPVVWIHGL
WP_001058307.1	hydrogenase [Salmonella enterica]	Length: 372\nNumber of predicted TMHs: 2\nExp number of AAs in TMHs: 36.51738\nExp number, first 60 AAs: 19.72849\nTotal prob of N-in: 0.96170\nPOSSIBLE N-term signal sequence\ninside 1 19\nTMhelix 20 39\noutside 40 325\nTMhelix 326 348\ninside 349 372

22604	GCF_001411295.1_9472_3_21	Salmonella enterica subsp. enterica serovar Weltevreden
Proteobacteria; Gammaproteobacteria; Enterobacterales; Enterobacteriaceae; Salmonella; Salmonella enterica; Salmonella enterica subsp. enterica	MNNEETFYQAMRRKGVTRRSFLKFCSLAATSLGLGAGMTPKIAWALENKPRIPVVWIHGL
WP_001058307.1	hydrogenase [Salmonella enterica]	Length: 372\nNumber of predicted TMHs: 2\nExp number of AAs in TMHs: 36.51738\nExp number, first 60 AAs: 19.72849\nTotal prob of N-in: 0.96170\nPOSSIBLE N-term signal sequence\ninside 1 19\nTMhelix 20 39\noutside 40 325\nTMhelix 326 348\ninside 349 372

22605	GCF_001411315.1_12216_4_41	Salmonella enterica subsp. enterica serovar Weltevreden
Proteobacteria; Gammaproteobacteria; Enterobacterales; Enterobacteriaceae; Salmonella; Salmonella enterica; Salmonella enterica subsp. enterica	MNNEETFYQAMRRKGVTRRSFLKFCSLAATSLGLGAGMTPKIAWALENKPRIPVVWIHGL
WP_001058307.1	hydrogenase [Salmonella enterica]	Length: 372\nNumber of predicted TMHs: 2\nExp number of AAs in TMHs: 36.51738\nExp number, first 60 AAs: 19.72849\nTotal prob of N-in: 0.96170\nPOSSIBLE N-term signal sequence\ninside 1 19\nTMhelix 20 39\noutside 40 325\nTMhelix 326 348\ninside 349 372

22606	GCF_001411275.1_9472_3_23	Salmonella enterica subsp. enterica serovar Weltevreden
Proteobacteria; Gammaproteobacteria; Enterobacterales; Enterobacteriaceae; Salmonella; Salmonella enterica; Salmonella enterica subsp. enterica	MNNEETFYQAMRRKGVTRRSFLKFCSLAATSLGLGAGMTPKIAWALENKPRIPVVWIHGL
WP_001058307.1	hydrogenase [Salmonella enterica]	Length: 372\nNumber of predicted TMHs: 2\nExp

number of AAs in TMHs: 36.51738\nExp number, first 60 AAs: 19.72849\nTotal prob of N-in: 0.96170\nPOSSIBLE N-term signal sequence\ninside 1 19\nTMhelix 20 39\noutside 40 325\nTMhelix 326 348\ninside 349 372

22607 GCF_001411255.1_12227_3_13 Salmonella enterica subsp. enterica serovar Weltevreden Proteobacteria; Gammaproteobacteria; Enterobacterales; Enterobacteriaceae; Salmonella; Salmonella enterica; Salmonella enterica subsp. enterica MNNEETFYQAMRRKGVTRRSFLKFCSLAATSLGLGAGMTPKIAWALENKPRIPVVWIHGL WP_001058307.1 hydrogenase [Salmonella enterica] Length: 372\nNumber of predicted TMHs: 2\nExp number of AAs in TMHs: 36.51738\nExp number, first 60 AAs: 19.72849\nTotal prob of N-in: 0.96170\nPOSSIBLE N-term signal sequence\ninside 1 19\nTMhelix 20 39\noutside 40 325\nTMhelix 326 348\ninside 349 372

22608 GCF_001411355.1_12227_3_18 Salmonella enterica subsp. enterica serovar Weltevreden Proteobacteria; Gammaproteobacteria; Enterobacterales; Enterobacteriaceae; Salmonella; Salmonella enterica; Salmonella enterica subsp. enterica MTGDNTLITSHGINRRDFMKLCAALAATMGLSSKAAAEMAESVSNPQRPPVIWIGAQECT WP_000145429.1 hydrogenase 2 small subunit [Salmonella enterica] Length: 372\nNumber of predicted TMHs: 1\nExp number of AAs in TMHs: 24.51309\nExp number, first 60 AAs: 0.2682\nTotal prob of N-in: 0.14402\noutside 1 331\nTMhelix 332 354\ninside 355 372

22609 GCF_001411335.1_12227_3_12 Salmonella enterica subsp. enterica serovar Weltevreden Proteobacteria; Gammaproteobacteria; Enterobacterales; Enterobacteriaceae; Salmonella; Salmonella enterica; Salmonella enterica subsp. enterica MTGDNTLITSHGINRRDFMKLCAALAATMGLSSKAAAEMAESVSNPQRPPVIWIGAQECT WP_000145429.1 hydrogenase 2 small subunit [Salmonella enterica] Length: 372\nNumber of predicted TMHs: 1\nExp number of AAs in TMHs: 24.51309\nExp number, first 60 AAs: 0.2682\nTotal prob of N-in: 0.14402\noutside 1 331\nTMhelix 332 354\ninside 355 372

22610 GCF_001411295.1_9472_3_21 Salmonella enterica subsp. enterica serovar Weltevreden Proteobacteria; Gammaproteobacteria; Enterobacterales; Enterobacteriaceae; Salmonella; Salmonella enterica; Salmonella enterica subsp. enterica MTGDNTLITSHGINRRDFMKLCAALAATMGLSSKAAAEMAESVSNPQRPPVIWIGAQECT WP_000145429.1 hydrogenase 2 small subunit [Salmonella enterica] Length: 372\nNumber of predicted TMHs: 1\nExp number of AAs in TMHs: 24.51309\nExp number, first 60 AAs: 0.2682\nTotal prob of N-in: 0.14402\noutside 1 331\nTMhelix 332 354\ninside 355 372

22611 GCF_001411315.1_12216_4_41 Salmonella enterica subsp. enterica serovar Weltevreden Proteobacteria; Gammaproteobacteria; Enterobacterales; Enterobacteriaceae; Salmonella; Salmonella enterica; Salmonella enterica subsp. enterica MTGDNTLITSHGINRRDFMKLCAALAATMGLSSKAAAEMAESVSNPQRPPVIWIGAQECT WP_000145429.1 hydrogenase 2 small subunit [Salmonella enterica] Length: 372\nNumber of predicted TMHs: 1\nExp number of AAs in TMHs: 24.51309\nExp number, first 60 AAs: 0.2682\nTotal prob of N-in: 0.14402\noutside 1 331\nTMhelix 332 354\ninside 355 372

22612 GCF_001411275.1_9472_3_23 Salmonella enterica subsp. enterica serovar Weltevreden Proteobacteria; Gammaproteobacteria; Enterobacterales; Enterobacteriaceae; Salmonella; Salmonella enterica; Salmonella enterica subsp. enterica MTGDNTLITSHGINRRDFMKLCAALAATMGLSSKAAAEMAESVSNPQRPPVIWIGAQECT WP_000145429.1 hydrogenase 2 small subunit [Salmonella enterica] Length: 372\nNumber of predicted TMHs: 1\nExp number of AAs in TMHs: 24.51309\nExp number, first 60 AAs: 0.2682\nTotal prob of N-in: 0.14402\noutside 1 331\nTMhelix 332 354\ninside 355 372

22613 GCF_001411215.1_12227_3_5 Salmonella enterica subsp. enterica serovar Weltevreden Proteobacteria; Gammaproteobacteria; Enterobacterales; Enterobacteriaceae; Salmonella; Salmonella enterica; Salmonella enterica subsp. enterica MNNEETFYQAMRRKGVTRRSFLKFCSLAATSLGLGAGMTPKIAWALENKPRIPVVWIHGL WP_001058307.1 hydrogenase [Salmonella enterica] Length: 372\nNumber of predicted TMHs: 2\nExp number of AAs in TMHs: 36.51738\nExp number, first 60 AAs: 19.72849\nTotal prob of N-in: 0.96170\nPOSSIBLE N-term signal sequence\ninside 1 19\nTMhelix 20 39\noutside 40 325\nTMhelix 326 348\ninside 349 372

22614 GCF_001411195.1_9472_3_9Salmonella enterica subsp. enterica serovar Weltevreden Proteobacteria; Gammaproteobacteria; Enterobacterales; Enterobacteriaceae; Salmonella; Salmonella enterica; Salmonella enterica subsp. enterica MNNEETFYQAMRRKGVTRRSFLKFCSLAATSLGLGAGMTPKIAWALENKPRIPVVWIHGL WP_001058307.1 hydrogenase [Salmonella enterica] Length: 372\nNumber of predicted TMHs: 2\nExp number of AAs in TMHs: 36.51738\nExp number, first 60 AAs: 19.72849\nTotal prob of N-in: 0.96170\nPOSSIBLE N-term signal sequence\ninside 1 19\nTMhelix 20 39\noutside 40 325\nTMhelix 326 348\ninside 349 372

22615 GCF_001411175.1_9472_3_13 Salmonella enterica subsp. enterica serovar Weltevreden Proteobacteria; Gammaproteobacteria; Enterobacterales; Enterobacteriaceae; Salmonella; Salmonella enterica;

Salmonella enterica subsp. enterica MNNEETFYQAMRRKGVTRRSFLKFCSLAATSLGLGAGMTPKIAWALENKPRIPVVWIHGL WP_001058307.1 hydrogenase [Salmonella enterica] Length: 372\nNumber of predicted TMHs: 2\nExp number of AAs in TMHs: 36.51738\nExp number, first 60 AAs: 19.72849\nTotal prob of N-in: 0.96170\nPOSSIBLE N-term signal sequence\ninside 1 19\nTMhelix 20 39\noutside 40 325\nTMhelix 326 348\ninside 349 372

22616 GCF_001411255.1_12227_3_13 Salmonella enterica subsp. enterica serovar Weltevreden Proteobacteria; Gammaproteobacteria; Enterobacterales; Enterobacteriaceae; Salmonella; Salmonella enterica; Salmonella enterica subsp. enterica MTGDNTLITSHGINRRDFMKLCAALAATMGLSSKAAAEMAESVSNPQRPPVIWIGAQECT WP_000145429.1 hydrogenase 2 small subunit [Salmonella enterica] Length: 372\nNumber of predicted TMHs: 1\nExp number of AAs in TMHs: 24.51309\nExp number, first 60 AAs: 0.2682\nTotal prob of N-in: 0.14402\noutside 1 331\nTMhelix 332 354\ninside 355 372

22617 GCF_001411235.1_12227_3_39 Salmonella enterica subsp. enterica serovar Weltevreden Proteobacteria; Gammaproteobacteria; Enterobacterales; Enterobacteriaceae; Salmonella; Salmonella enterica; Salmonella enterica subsp. enterica MTGDNTLITSHGINRRDFMKLCAALAATMGLSSKAAAEMAESVSNPQRPPVIWIGAQECT WP_000145429.1 hydrogenase 2 small subunit [Salmonella enterica] Length: 372\nNumber of predicted TMHs: 1\nExp number of AAs in TMHs: 24.51309\nExp number, first 60 AAs: 0.2682\nTotal prob of N-in: 0.14402\noutside 1 331\nTMhelix 332 354\ninside 355 372

22618 GCF_001411215.1_12227_3_5 Salmonella enterica subsp. enterica serovar Weltevreden Proteobacteria; Gammaproteobacteria; Enterobacterales; Enterobacteriaceae; Salmonella; Salmonella enterica; Salmonella enterica subsp. enterica MTGDNTLITSHGINRRDFMKLCAALAATMGLSSKAAAEMAESVSNPQRPPVIWIGAQECT WP_000145429.1 hydrogenase 2 small subunit [Salmonella enterica] Length: 372\nNumber of predicted TMHs: 1\nExp number of AAs in TMHs: 24.51309\nExp number, first 60 AAs: 0.2682\nTotal prob of N-in: 0.14402\noutside 1 331\nTMhelix 332 354\ninside 355 372

22619 GCF_001411195.1_9472_3_9Salmonella enterica subsp. enterica serovar Weltevreden Proteobacteria; Gammaproteobacteria; Enterobacterales; Enterobacteriaceae; Salmonella; Salmonella enterica; Salmonella enterica subsp. enterica MTGDNTLITSHGINRRDFMKLCAALAATMGLSSKAAAEMAESVSNPQRPPVIWIGAQECT WP_000145429.1 hydrogenase 2 small subunit [Salmonella enterica] Length: 372\nNumber of predicted TMHs: 1\nExp number of AAs in TMHs: 24.51309\nExp number, first 60 AAs: 0.2682\nTotal prob of N-in: 0.14402\noutside 1 331\nTMhelix 332 354\ninside 355 372

22620 GCF_001411175.1_9472_3_13 Salmonella enterica subsp. enterica serovar Weltevreden Proteobacteria; Gammaproteobacteria; Enterobacterales; Enterobacteriaceae; Salmonella; Salmonella enterica; Salmonella enterica subsp. enterica MTGDNTLITSHGINRRDFMKLCAALAATMGLSSKAAAEMAESVSNPQRPPVIWIGAQECT WP_000145429.1 hydrogenase 2 small subunit [Salmonella enterica] Length: 372\nNumber of predicted TMHs: 1\nExp number of AAs in TMHs: 24.51309\nExp number, first 60 AAs: 0.2682\nTotal prob of N-in: 0.14402\noutside 1 331\nTMhelix 332 354\ninside 355 372

22621 GCF_001411135.1_12227_3_27 Salmonella enterica subsp. enterica serovar Weltevreden Proteobacteria; Gammaproteobacteria; Enterobacterales; Enterobacteriaceae; Salmonella; Salmonella enterica; Salmonella enterica subsp. enterica MNNEETFYQAMRRKGVTRRSFLKFCSLAATSLGLGAGMTPKIAWALENKPRIPVVWIHGL WP_001058307.1 hydrogenase [Salmonella enterica] Length: 372\nNumber of predicted TMHs: 2\nExp number of AAs in TMHs: 36.51738\nExp number, first 60 AAs: 19.72849\nTotal prob of N-in: 0.96170\nPOSSIBLE N-term signal sequence\ninside 1 19\nTMhelix 20 39\noutside 40 325\nTMhelix 326 348\ninside 349 372

22622 GCF_001411115.1_9472_3_10 Salmonella enterica subsp. enterica serovar Weltevreden Proteobacteria; Gammaproteobacteria; Enterobacterales; Enterobacteriaceae; Salmonella; Salmonella enterica; Salmonella enterica subsp. enterica MNNEETFYQAMRRKGVTRRSFLKFCSLAATSLGLGAGMTPKIAWALENKPRIPVVWIHGL WP_001058307.1 hydrogenase [Salmonella enterica] Length: 372\nNumber of predicted TMHs: 2\nExp number of AAs in TMHs: 36.51738\nExp number, first 60 AAs: 19.72849\nTotal prob of N-in: 0.96170\nPOSSIBLE N-term signal sequence\ninside 1 19\nTMhelix 20 39\noutside 40 325\nTMhelix 326 348\ninside 349 372

22623 GCF_001411075.1_12227_3_50 Salmonella enterica subsp. enterica serovar Weltevreden Proteobacteria; Gammaproteobacteria; Enterobacterales; Enterobacteriaceae; Salmonella; Salmonella enterica; Salmonella enterica subsp. enterica MNNEETFYQAMRRKGVTRRSFLKFCSLAATSLGLGAGMTPKIAWALENKPRIPVVWIHGL WP_001058307.1 hydrogenase [Salmonella enterica] Length: 372\nNumber of predicted TMHs: 2\nExp number of AAs in TMHs: 36.51738\nExp number, first 60 AAs: 19.72849\nTotal prob of N-in: 0.96170\nPOSSIBLE N-term signal sequence\ninside 1 19\nTMhelix 20 39\noutside 40 325\nTMhelix 326 348\ninside 349 372

22624 GCF_001411155.1_12216_4_56 Salmonella enterica subsp. enterica serovar Weltevreden Proteobacteria; Gammaproteobacteria; Enterobacterales; Enterobacteriaceae; Salmonella; Salmonella enterica; Salmonella enterica subsp. enterica MNNEETFYQAMRRKGVTRRSFLKFCSLAATSLGLGAGMTPKIAWALENKPRIPVVWIHGL WP_001058307.1 hydrogenase [Salmonella enterica] Length: 372\nNumber of predicted TMHs: 2\nExp number of AAs in TMHs: 36.51738\nExp number, first 60 AAs: 19.72849\nTotal prob of N-in: 0.96170\nPOSSIBLE N-term signal sequence\ninside 1 19\nTMhelix 20 39\noutside 40 325\nTMhelix 326 348\ninside 349 372

22625 GCF_001411155.1_12216_4_56 Salmonella enterica subsp. enterica serovar Weltevreden Proteobacteria; Gammaproteobacteria; Enterobacterales; Enterobacteriaceae; Salmonella; Salmonella enterica; Salmonella enterica subsp. enterica MTGDNTLITSHGINRRDFMKLCAALAATMGLSSKAAAEMAESVSNPQRPPVIWIGAQECT WP_000145429.1 hydrogenase 2 small subunit [Salmonella enterica] Length: 372\nNumber of predicted TMHs: 1\nExp number of AAs in TMHs: 24.51309\nExp number, first 60 AAs: 0.2682\nTotal prob of N-in: 0.14402\noutside 1 331\nTMhelix 332 354\ninside 355 372

22626 GCF_001411095.1_12227_3_42 Salmonella enterica subsp. enterica serovar Weltevreden Proteobacteria; Gammaproteobacteria; Enterobacterales; Enterobacteriaceae; Salmonella; Salmonella enterica; Salmonella enterica subsp. enterica MNNEETFYQAMRRKGVTRRSFLKFCSLAATSLGLGAGMTPKIAWALENKPRIPVVWIHGL WP_001058307.1 hydrogenase [Salmonella enterica] Length: 372\nNumber of predicted TMHs: 2\nExp number of AAs in TMHs: 36.51738\nExp number, first 60 AAs: 19.72849\nTotal prob of N-in: 0.96170\nPOSSIBLE N-term signal sequence\ninside 1 19\nTMhelix 20 39\noutside 40 325\nTMhelix 326 348\ninside 349 372

22627 GCF_001411055.1_9472_3_8Salmonella enterica subsp. enterica serovar Weltevreden Proteobacteria; Gammaproteobacteria; Enterobacterales; Enterobacteriaceae; Salmonella; Salmonella enterica; Salmonella enterica subsp. enterica MNNEETFYQAMRRKGVTRRSFLKFCSLAATSLGLGAGMTPKIAWALENKPRIPVVWIHGL WP_001058307.1 hydrogenase [Salmonella enterica] Length: 372\nNumber of predicted TMHs: 2\nExp number of AAs in TMHs: 36.51738\nExp number, first 60 AAs: 19.72849\nTotal prob of N-in: 0.96170\nPOSSIBLE N-term signal sequence\ninside 1 19\nTMhelix 20 39\noutside 40 325\nTMhelix 326 348\ninside 349 372

22628 GCF_001411035.1_12216_4_48 Salmonella enterica subsp. enterica serovar Weltevreden Proteobacteria; Gammaproteobacteria; Enterobacterales; Enterobacteriaceae; Salmonella; Salmonella enterica; Salmonella enterica subsp. enterica MNNEETFYQAMRRKGVTRRSFLKFCSLAATSLGLGAGMTPKIAWALENKPRIPVVWIHGL WP_001058307.1 hydrogenase [Salmonella enterica] Length: 372\nNumber of predicted TMHs: 2\nExp number of AAs in TMHs: 36.51738\nExp number, first 60 AAs: 19.72849\nTotal prob of N-in: 0.96170\nPOSSIBLE N-term signal sequence\ninside 1 19\nTMhelix 20 39\noutside 40 325\nTMhelix 326 348\ninside 349 372

22629 GCF_001411015.1_9472_3_19 Salmonella enterica subsp. enterica serovar Weltevreden Proteobacteria; Gammaproteobacteria; Enterobacterales; Enterobacteriaceae; Salmonella; Salmonella enterica; Salmonella enterica subsp. enterica MNNEETFYQAMRRKGVTRRSFLKFCSLAATSLGLGAGMTPKIAWALENKPRIPVVWIHGL WP_001058307.1 hydrogenase [Salmonella enterica] Length: 372\nNumber of predicted TMHs: 2\nExp number of AAs in TMHs: 36.51738\nExp number, first 60 AAs: 19.72849\nTotal prob of N-in: 0.96170\nPOSSIBLE N-term signal sequence\ninside 1 19\nTMhelix 20 39\noutside 40 325\nTMhelix 326 348\ninside 349 372

22630 GCF_001411135.1_12227_3_27 Salmonella enterica subsp. enterica serovar Weltevreden Proteobacteria; Gammaproteobacteria; Enterobacterales; Enterobacteriaceae; Salmonella; Salmonella enterica; Salmonella enterica subsp. enterica MTGDNTLITSHGINRRDFMKLCAALAATMGLSSKAAAEMAESVSNPQRPPVIWIGAQECT WP_000145429.1 hydrogenase 2 small subunit [Salmonella enterica] Length: 372\nNumber of predicted TMHs: 1\nExp number of AAs in TMHs: 24.51309\nExp number, first 60 AAs: 0.2682\nTotal prob of N-in: 0.14402\noutside 1 331\nTMhelix 332 354\ninside 355 372

22631 GCF_001411095.1_12227_3_42 Salmonella enterica subsp. enterica serovar Weltevreden Proteobacteria; Gammaproteobacteria; Enterobacterales; Enterobacteriaceae; Salmonella; Salmonella enterica; Salmonella enterica subsp. enterica MTGDNTLITSHGINRRDFMKLCAALAATMGLSSKAAAEMAESVSNPQRPPVIWIGAQECT WP_000145429.1 hydrogenase 2 small subunit [Salmonella enterica] Length: 372\nNumber of predicted TMHs: 1\nExp number of AAs in TMHs: 24.51309\nExp number, first 60 AAs: 0.2682\nTotal prob of N-in: 0.14402\noutside 1 331\nTMhelix 332 354\ninside 355 372

22632 GCF_001411115.1_9472_3_10 Salmonella enterica subsp. enterica serovar Weltevreden Proteobacteria; Gammaproteobacteria; Enterobacterales; Enterobacteriaceae; Salmonella; Salmonella enterica; Salmonella enterica subsp. enterica MTGDNTLITSHGINRRDFMKLCAALAATMGLSSKAAAEMAESVSNPQRPPVIWIGAQECT WP_000145429.1 hydrogenase 2 small subunit [Salmonella enterica] Length: 372\nNumber of

predicted TMHs:  1\nExp number of AAs in TMHs: 24.51309\nExp number, first 60 AAs:  0.2682\nTotal prob of N-in: 0.14402\noutside   1   331\nTMhelix   332   354\ninside   355   372

22633 GCF_001411075.1_12227_3_50 Salmonella enterica subsp. enterica serovar Weltevreden Proteobacteria; Gammaproteobacteria; Enterobacterales; Enterobacteriaceae; Salmonella; Salmonella enterica; Salmonella enterica subsp. enterica MTGDNTLITSHGINRRDFMKLCAALAATMGLSSKAAAEMAESVSNPQRPPVIWIGAQECT WP_000145429.1 hydrogenase 2 small subunit [Salmonella enterica] Length: 372\nNumber of predicted TMHs:  1\nExp number of AAs in TMHs: 24.51309\nExp number, first 60 AAs:  0.2682\nTotal prob of N-in: 0.14402\noutside   1   331\nTMhelix   332   354\ninside   355   372

22634 GCF_001411055.1_9472_3_8Salmonella enterica subsp. enterica serovar Weltevreden Proteobacteria; Gammaproteobacteria; Enterobacterales; Enterobacteriaceae; Salmonella; Salmonella enterica; Salmonella enterica subsp. enterica MTGDNTLITSHGINRRDFMKLCAALAATMGLSSKAAAEMAESVSNPQRPPVIWIGAQECT WP_000145429.1 hydrogenase 2 small subunit [Salmonella enterica] Length: 372\nNumber of predicted TMHs:  1\nExp number of AAs in TMHs: 24.51309\nExp number, first 60 AAs:  0.2682\nTotal prob of N-in:   0.14402\noutside   1 331\nTMhelix   332   354\ninside   355   372

22635 GCF_001411035.1_12216_4_48 Salmonella enterica subsp. enterica serovar Weltevreden Proteobacteria; Gammaproteobacteria; Enterobacterales; Enterobacteriaceae; Salmonella; Salmonella enterica; Salmonella enterica subsp. enterica MTGDNTLITSHGINRRDFMKLCAALAATMGLSSKAAAEMAESVSNPQRPPVIWIGAQECT WP_000145429.1 hydrogenase 2 small subunit [Salmonella enterica] Length: 372\nNumber of predicted TMHs:  1\nExp number of AAs in TMHs: 24.51309\nExp number, first 60 AAs:  0.2682\nTotal prob of N-in: 0.14402\noutside   1   331\nTMhelix   332   354\ninside   355   372

22636 GCF_001410995.1_12227_3_26 Salmonella enterica subsp. enterica serovar Weltevreden Proteobacteria; Gammaproteobacteria; Enterobacterales; Enterobacteriaceae; Salmonella; Salmonella enterica; Salmonella enterica subsp. enterica MNNEETFYQAMRRKGVTRRSFLKFCSLAATSLGLGAGMTPKIAWALENKPRIPVVWIHGL WP_001058307.1 hydrogenase [Salmonella enterica] Length: 372\nNumber of predicted TMHs:  2\nExp number of AAs in TMHs: 36.51738\nExp number, first 60 AAs:  19.72849\nTotal prob of N-in:   0.96170\nPOSSIBLE N-term signal sequence\ninside   1   19\nTMhelix   20   39\noutside   40   325\nTMhelix   326   348\ninside   349   372

22637 GCF_001410955.1_10868_1_89 Salmonella enterica subsp. enterica serovar Weltevreden Proteobacteria; Gammaproteobacteria; Enterobacterales; Enterobacteriaceae; Salmonella; Salmonella enterica; Salmonella enterica subsp. enterica MNNEETFYQAMRRKGVTRRSFLKFCSLAATSLGLGAGMTPKIAWALENKPRIPVVWIHGL WP_001058307.1 hydrogenase [Salmonella enterica] Length: 372\nNumber of predicted TMHs:  2\nExp number of AAs in TMHs: 36.51738\nExp number, first 60 AAs:  19.72849\nTotal prob of N-in:   0.96170\nPOSSIBLE N-term signal sequence\ninside   1   19\nTMhelix   20   39\noutside   40   325\nTMhelix   326   348\ninside   349   372

22638 GCF_002050595.1_ASM205059v1 Salmonella enterica subsp. enterica serovar Weltevreden Proteobacteria; Gammaproteobacteria; Enterobacterales; Enterobacteriaceae; Salmonella; Salmonella enterica; Salmonella enterica subsp. enterica MTGDNTLITSHGINRRDFMKLCAALAATMGLSSKAAAEMAESVSNPQRPPVIWIGAQECT WP_000145429.1 hydrogenase 2 small subunit [Salmonella enterica] Length: 372\nNumber of predicted TMHs:  1\nExp number of AAs in TMHs: 24.51309\nExp number, first 60 AAs:  0.2682\nTotal prob of N-in: 0.14402\noutside   1   331\nTMhelix   332   354\ninside   355   372

22639 GCF_001411015.1_9472_3_19 Salmonella enterica subsp. enterica serovar Weltevreden Proteobacteria; Gammaproteobacteria; Enterobacterales; Enterobacteriaceae; Salmonella; Salmonella enterica; Salmonella enterica subsp. enterica MTGDNTLITSHGINRRDFMKLCAALAATMGLSSKAAAEMAESVSNPQRPPVIWIGAQECT WP_000145429.1 hydrogenase 2 small subunit [Salmonella enterica] Length: 372\nNumber of predicted TMHs:  1\nExp number of AAs in TMHs: 24.51309\nExp number, first 60 AAs:  0.2682\nTotal prob of N-in: 0.14402\noutside   1   331\nTMhelix   332   354\ninside   355   372

22640 GCF_001410975.1_12216_4_62 Salmonella enterica subsp. enterica serovar Weltevreden Proteobacteria; Gammaproteobacteria; Enterobacterales; Enterobacteriaceae; Salmonella; Salmonella enterica; Salmonella enterica subsp. enterica MNNEETFYQAMRRKGVTRRSFLKFCSLAATSLGLGAGMTPKIAWALENKPRIPVVWIHGL WP_001058307.1 hydrogenase [Salmonella enterica] Length: 372\nNumber of predicted TMHs:  2\nExp number of AAs in TMHs: 36.51738\nExp number, first 60 AAs:  19.72849\nTotal prob of N-in:   0.96170\nPOSSIBLE N-term signal sequence\ninside   1   19\nTMhelix   20   39\noutside   40   325\nTMhelix   326   348\ninside   349   372

22641 GCF_002050835.1_ASM205083v1 Salmonella enterica subsp. enterica serovar Weltevreden Proteobacteria; Gammaproteobacteria; Enterobacterales; Enterobacteriaceae; Salmonella; Salmonella enterica;

Salmonella enterica subsp. enterica MTGDNTLITSHGINRRDFMKLCAALAATMGLSSKAAAEMAESVSNPQRPPVIWIGAQECT WP_000145429.1 hydrogenase 2 small subunit [Salmonella enterica] Length: 372\nNumber of predicted TMHs: 1\nExp number of AAs in TMHs: 24.51309\nExp number, first 60 AAs: 0.2682\nTotal prob of N-in: 0.14402\noutside 1 331\nTMhelix 332 354\ninside 355 372

22642 GCF_002050715.1_ASM205071v1 Salmonella enterica subsp. enterica serovar Weltevreden Proteobacteria; Gammaproteobacteria; Enterobacterales; Enterobacteriaceae; Salmonella; Salmonella enterica; Salmonella enterica subsp. enterica MTGDNTLITSHGINRRDFMKLCAALAATMGLSSKAAAEMAESVSNPQRPPVIWIGAQECT WP_000145429.1 hydrogenase 2 small subunit [Salmonella enterica] Length: 372\nNumber of predicted TMHs: 1\nExp number of AAs in TMHs: 24.51309\nExp number, first 60 AAs: 0.2682\nTotal prob of N-in: 0.14402\noutside 1 331\nTMhelix 332 354\ninside 355 372

22643 GCF_002050715.1_ASM205071v1 Salmonella enterica subsp. enterica serovar Weltevreden Proteobacteria; Gammaproteobacteria; Enterobacterales; Enterobacteriaceae; Salmonella; Salmonella enterica; Salmonella enterica subsp. enterica MNNEETFYQAMRRKGVTRRSFLKFCSLAATSLGLGAGMTPKIAWALENKPRIPVVWIHGL WP_001058307.1 hydrogenase [Salmonella enterica] Length: 372\nNumber of predicted TMHs: 2\nExp number of AAs in TMHs: 36.51738\nExp number, first 60 AAs: 19.72849\nTotal prob of N-in: 0.96170\nPOSSIBLE N-term signal sequence\ninside 1 19\nTMhelix 20 39\noutside 40 325\nTMhelix 326 348\ninside 349 372

22644 GCF_002049345.1_ASM204934v1 Salmonella enterica subsp. enterica serovar Weltevreden Proteobacteria; Gammaproteobacteria; Enterobacterales; Enterobacteriaceae; Salmonella; Salmonella enterica; Salmonella enterica subsp. enterica MTGDNTLITSHGINRRDFMKLCAALAATMGLSSKAAAEMAESVSNPQRPPVIWIGAQECT WP_000145429.1 hydrogenase 2 small subunit [Salmonella enterica] Length: 372\nNumber of predicted TMHs: 1\nExp number of AAs in TMHs: 24.51309\nExp number, first 60 AAs: 0.2682\nTotal prob of N-in: 0.14402\noutside 1 331\nTMhelix 332 354\ninside 355 372

22645 GCF_002049345.1_ASM204934v1 Salmonella enterica subsp. enterica serovar Weltevreden Proteobacteria; Gammaproteobacteria; Enterobacterales; Enterobacteriaceae; Salmonella; Salmonella enterica; Salmonella enterica subsp. enterica MNNEETFYQAMRRKGVTRRSFLKFCSLAATSLGLGAGMTPKIAWALENKPRIPVVWIHGL WP_001058307.1 hydrogenase [Salmonella enterica] Length: 372\nNumber of predicted TMHs: 2\nExp number of AAs in TMHs: 36.51738\nExp number, first 60 AAs: 19.72849\nTotal prob of N-in: 0.96170\nPOSSIBLE N-term signal sequence\ninside 1 19\nTMhelix 20 39\noutside 40 325\nTMhelix 326 348\ninside 349 372

22646 GCF_002071635.1_ASM207163v1 Salmonella enterica subsp. enterica serovar Westthampton Proteobacteria; Gammaproteobacteria; Enterobacterales; Enterobacteriaceae; Salmonella; Salmonella enterica; Salmonella enterica subsp. enterica MNNEETFYQAMRRKGVTRRSFLKFCSLAATSLGLGAGMTPKIAWALENKPRIPVVWIHGL WP_001058311.1 hydrogenase [Salmonella enterica] Length: 372\nNumber of predicted TMHs: 2\nExp number of AAs in TMHs: 37.4699899999999\nExp number, first 60 AAs: 19.72699\nTotal prob of N-in: 0.96171\nPOSSIBLE N-term signal sequence\ninside 1 19\nTMhelix 20 39\noutside 40 325\nTMhelix 326 348\ninside 349 372

22647 GCF_002071635.1_ASM207163v1 Salmonella enterica subsp. enterica serovar Westthampton Proteobacteria; Gammaproteobacteria; Enterobacterales; Enterobacteriaceae; Salmonella; Salmonella enterica; Salmonella enterica subsp. enterica MTGDNTLITSHGINRRDFMKLCAALAATMGLSSKAAAEMAESVSNPQRPPVIWIGAQECT WP_000145429.1 hydrogenase 2 small subunit [Salmonella enterica] Length: 372\nNumber of predicted TMHs: 1\nExp number of AAs in TMHs: 24.51309\nExp number, first 60 AAs: 0.2682\nTotal prob of N-in: 0.14402\noutside 1 331\nTMhelix 332 354\ninside 355 372

22648 GCF_001643815.1_CRJJGF_00058_v1.0Salmonella enterica subsp. enterica serovar Widemarsh Proteobacteria; Gammaproteobacteria; Enterobacterales; Enterobacteriaceae; Salmonella; Salmonella enterica; Salmonella enterica subsp. enterica MNNEETFYQAMRRKGVTRRSFLKFCSLAATSLGLGAGMTPKIAWALENKPRIPVVWIHGL WP_001058311.1 hydrogenase [Salmonella enterica] Length: 372\nNumber of predicted TMHs: 2\nExp number of AAs in TMHs: 37.4699899999999\nExp number, first 60 AAs: 19.72699\nTotal prob of N-in: 0.96171\nPOSSIBLE N-term signal sequence\ninside 1 19\nTMhelix 20 39\noutside 40 325\nTMhelix 326 348\ninside 349 372

22649 GCF_001643815.1_CRJJGF_00058_v1.0Salmonella enterica subsp. enterica serovar Widemarsh Proteobacteria; Gammaproteobacteria; Enterobacterales; Enterobacteriaceae; Salmonella; Salmonella enterica; Salmonella enterica subsp. enterica MTGDNTLITSHGINRRDFMKLCAALAATMGLSSKAAAEMAESVSNPQRPPVIWIGAQECT WP_000145429.1 hydrogenase 2 small subunit [Salmonella enterica] Length: 372\nNumber of

predicted TMHs: 1\nExp number of AAs in TMHs: 24.51309\nExp number, first 60 AAs: 0.2682\nTotal prob of N-in: 0.14402\noutside 1 331\nTMhelix 332 354\ninside 355 372

22650 GCF_002062955.1_ASM206295v1 Salmonella enterica subsp. enterica serovar Wilhemsburg Proteobacteria; Gammaproteobacteria; Enterobacterales; Enterobacteriaceae; Salmonella; Salmonella enterica; Salmonella enterica subsp. enterica MNNEETFYQAMRRKGVTRRSFLKFCSLAATSLGLGAGMTPKIAWALENKPRIPVVWIHGL WP_001058311.1 hydrogenase [Salmonella enterica] Length: 372\nNumber of predicted TMHs: 2\nExp number of AAs in TMHs: 37.4699899999999\nExp number, first 60 AAs: 19.72699\nTotal prob of N-in: 0.96171\nPOSSIBLE N-term signal sequence\ninside 1 19\nTMhelix 20 39\noutside 40 325\nTMhelix 326 348\ninside 349 372

22651 GCF_002062955.1_ASM206295v1 Salmonella enterica subsp. enterica serovar Wilhemsburg Proteobacteria; Gammaproteobacteria; Enterobacterales; Enterobacteriaceae; Salmonella; Salmonella enterica; Salmonella enterica subsp. enterica MTGDNTLITSHGINRRDFMKLCAALAATMGLSSKAAAEMAESVSNPQRPPVIWIGAQECT WP_000145429.1 hydrogenase 2 small subunit [Salmonella enterica] Length: 372\nNumber of predicted TMHs: 1\nExp number of AAs in TMHs: 24.51309\nExp number, first 60 AAs: 0.2682\nTotal prob of N-in: 0.14402\noutside 1 331\nTMhelix 332 354\ninside 355 372

22652 GCF_002066965.1_ASM206696v1 Salmonella enterica subsp. enterica serovar Wippra Proteobacteria; Gammaproteobacteria; Enterobacterales; Enterobacteriaceae; Salmonella; Salmonella enterica; Salmonella enterica subsp. enterica MNNEETFYQAMRRKGVTRRSFLKFCSLAATSLGLGAGMTPKIAWALENKPRIPVVWIHGL WP_001058312.1 hydrogenase [Salmonella enterica] Length: 372\nNumber of predicted TMHs: 2\nExp number of AAs in TMHs: 36.51738\nExp number, first 60 AAs: 19.72849\nTotal prob of N-in: 0.96170\nPOSSIBLE N-term signal sequence\ninside 1 19\nTMhelix 20 39\noutside 40 325\nTMhelix 326 348\ninside 349 372

22653 GCF_002066965.1_ASM206696v1 Salmonella enterica subsp. enterica serovar Wippra Proteobacteria; Gammaproteobacteria; Enterobacterales; Enterobacteriaceae; Salmonella; Salmonella enterica; Salmonella enterica subsp. enterica MTGDNTLITSHGINRRDFMKLCAALAATMGLSSKAAAEMAESVSNPQRPPVIWIGAQECT WP_000145431.1 hydrogenase 2 small subunit [Salmonella enterica] Length: 372\nNumber of predicted TMHs: 1\nExp number of AAs in TMHs: 24.51505\nExp number, first 60 AAs: 0.2682\nTotal prob of N-in: 0.14406\noutside 1 331\nTMhelix 332 354\ninside 355 372

22654 GCF_002063445.1_ASM206344v1 Salmonella enterica subsp. enterica serovar Worthington Proteobacteria; Gammaproteobacteria; Enterobacterales; Enterobacteriaceae; Salmonella; Salmonella enterica; Salmonella enterica subsp. enterica MTGDNTLITSHGINRRDFMKLCAALAATMGLSSKAAAEMAESVSNPQRPPVIWIGAQECT WP_000145429.1 hydrogenase 2 small subunit [Salmonella enterica] Length: 372\nNumber of predicted TMHs: 1\nExp number of AAs in TMHs: 24.51309\nExp number, first 60 AAs: 0.2682\nTotal prob of N-in: 0.14402\noutside 1 331\nTMhelix 332 354\ninside 355 372

22655 GCF_002048255.1_ASM204825v1 Salmonella enterica subsp. enterica serovar Worthington Proteobacteria; Gammaproteobacteria; Enterobacterales; Enterobacteriaceae; Salmonella; Salmonella enterica; Salmonella enterica subsp. enterica MTGDNTLITSHGINRRDFMKLCAALAATMGLSSKAAAEMAESVSNPQRPPVIWIGAQECT WP_000145429.1 hydrogenase 2 small subunit [Salmonella enterica] Length: 372\nNumber of predicted TMHs: 1\nExp number of AAs in TMHs: 24.51309\nExp number, first 60 AAs: 0.2682\nTotal prob of N-in: 0.14402\noutside 1 331\nTMhelix 332 354\ninside 355 372

22656 GCF_002048255.1_ASM204825v1 Salmonella enterica subsp. enterica serovar Worthington Proteobacteria; Gammaproteobacteria; Enterobacterales; Enterobacteriaceae; Salmonella; Salmonella enterica; Salmonella enterica subsp. enterica MNNEETFYQAMRRKGVTRRSFLKFCSLAATSLGLGAGMTPKIAWALENKPRIPVVWIHGL WP_001058312.1 hydrogenase [Salmonella enterica] Length: 372\nNumber of predicted TMHs: 2\nExp number of AAs in TMHs: 36.51738\nExp number, first 60 AAs: 19.72849\nTotal prob of N-in: 0.96170\nPOSSIBLE N-term signal sequence\ninside 1 19\nTMhelix 20 39\noutside 40 325\nTMhelix 326 348\ninside 349 372

22657 GCF_002063055.1_ASM206305v1 Salmonella enterica subsp. enterica serovar Worthington Proteobacteria; Gammaproteobacteria; Enterobacterales; Enterobacteriaceae; Salmonella; Salmonella enterica; Salmonella enterica subsp. enterica MTGDNTLITSHGINRRDFMKLCAALAATMGLSSKAAAEMAESVSNPQRPPVIWIGAQECT WP_000145429.1 hydrogenase 2 small subunit [Salmonella enterica] Length: 372\nNumber of predicted TMHs: 1\nExp number of AAs in TMHs: 24.51309\nExp number, first 60 AAs: 0.2682\nTotal prob of N-in: 0.14402\noutside 1 331\nTMhelix 332 354\ninside 355 372

22658	GCF_002063055.1_ASM206305v1	Salmonella enterica subsp. enterica serovar Worthington	Proteobacteria; Gammaproteobacteria; Enterobacterales; Enterobacteriaceae; Salmonella; Salmonella enterica; Salmonella enterica subsp. enterica	MNNEETFYQAMRRKGVTRRSFLKFCSLAATSLGLGAGMTPKIAWALENKPRIPVVWIHGL	WP_001058312.1	hydrogenase [Salmonella enterica]	Length: 372\nNumber of predicted TMHs:  2\nExp number of AAs in TMHs: 36.51738\nExp number, first 60 AAs:  19.72849\nTotal prob of N-in:   0.96170\nPOSSIBLE N-term signal sequence\ninside   1   19\nTMhelix   20   39\noutside   40   325\nTMhelix   326   348\ninside   349   372

22659	GCF_002063445.1_ASM206344v1	Salmonella enterica subsp. enterica serovar Worthington	Proteobacteria; Gammaproteobacteria; Enterobacterales; Enterobacteriaceae; Salmonella; Salmonella enterica; Salmonella enterica subsp. enterica	MNNEETFYQAMRRKGVTRRSFLKFCSLAATSLGLGAGMTPKIAWALENKPRIPVVWIHGL	WP_001058312.1	hydrogenase [Salmonella enterica]	Length: 372\nNumber of predicted TMHs:  2\nExp number of AAs in TMHs: 36.51738\nExp number, first 60 AAs:  19.72849\nTotal prob of N-in:   0.96170\nPOSSIBLE N-term signal sequence\ninside   1   19\nTMhelix   20   39\noutside   40   325\nTMhelix   326   348\ninside   349   372

22660	GCF_001242245.1_Salmonella_enterica_CVM_N45931_v1.0	Salmonella enterica subsp. enterica serovar Worthington	Proteobacteria; Gammaproteobacteria; Enterobacterales; Enterobacteriaceae; Salmonella; Salmonella enterica; Salmonella enterica subsp. enterica	MTGDNTLITSHGINRRDFMKLCAALAATMGLSSKAAAEMAESVSNPQRPPVIWIGAQECT	WP_000145429.1	hydrogenase 2 small subunit [Salmonella enterica]	Length: 372\nNumber of predicted TMHs:  1\nExp number of AAs in TMHs: 24.51309\nExp number, first 60 AAs:  0.2682\nTotal prob of N-in:   0.14402\noutside   1   331\nTMhelix   332   354\ninside   355   372

22661	GCF_002046845.1_ASM204684v1	Salmonella enterica subsp. enterica serovar Worthington	Proteobacteria; Gammaproteobacteria; Enterobacterales; Enterobacteriaceae; Salmonella; Salmonella enterica; Salmonella enterica subsp. enterica	MNNEETFYQAMRRKGVTRRSFLKFCSLAATSLGLGAGMTPKIAWALENKPRIPVVWIHGL	WP_001058312.1	hydrogenase [Salmonella enterica]	Length: 372\nNumber of predicted TMHs:  2\nExp number of AAs in TMHs: 36.51738\nExp number, first 60 AAs:  19.72849\nTotal prob of N-in:   0.96170\nPOSSIBLE N-term signal sequence\ninside   1   19\nTMhelix   20   39\noutside   40   325\nTMhelix   326   348\ninside   349   372

22662	GCF_001997835.1_ASM199783v1	Salmonella enterica subsp. enterica serovar Worthington	Proteobacteria; Gammaproteobacteria; Enterobacterales; Enterobacteriaceae; Salmonella; Salmonella enterica; Salmonella enterica subsp. enterica	MNNEETFYQAMRRKGVTRRSFLKFCSLAATSLGLGAGMTPKIAWALENKPRIPVVWIHGL	WP_001058312.1	hydrogenase [Salmonella enterica]	Length: 372\nNumber of predicted TMHs:  2\nExp number of AAs in TMHs: 36.51738\nExp number, first 60 AAs:  19.72849\nTotal prob of N-in:   0.96170\nPOSSIBLE N-term signal sequence\ninside   1   19\nTMhelix   20   39\noutside   40   325\nTMhelix   326   348\ninside   349   372

22663	GCF_001242245.1_Salmonella_enterica_CVM_N45931_v1.0	Salmonella enterica subsp. enterica serovar Worthington	Proteobacteria; Gammaproteobacteria; Enterobacterales; Enterobacteriaceae; Salmonella; Salmonella enterica; Salmonella enterica subsp. enterica	MNNEETFYQAMRRKGVTRRSFLKFCSLAATSLGLGAGMTPKIAWALENKPRIPVVWIHGL	WP_001058312.1	hydrogenase [Salmonella enterica]	Length: 372\nNumber of predicted TMHs:  2\nExp number of AAs in TMHs: 36.51738\nExp number, first 60 AAs:  19.72849\nTotal prob of N-in:   0.96170\nPOSSIBLE N-term signal sequence\ninside   1   19\nTMhelix   20   39\noutside   40   325\nTMhelix   326   348\ninside   349   372

22664	GCF_001247285.1_Salmonella_enterica_CVM_N51304_v1.0	Salmonella enterica subsp. enterica serovar Worthington	Proteobacteria; Gammaproteobacteria; Enterobacterales; Enterobacteriaceae; Salmonella; Salmonella enterica; Salmonella enterica subsp. enterica	MTGDNTLITSHGINRRDFMKLCAALAATMGLSSKAAAEMAESVSNPQRPPVIWIGAQECT	WP_000145429.1	hydrogenase 2 small subunit [Salmonella enterica]	Length: 372\nNumber of predicted TMHs:  1\nExp number of AAs in TMHs: 24.51309\nExp number, first 60 AAs:  0.2682\nTotal prob of N-in:   0.14402\noutside   1   331\nTMhelix   332   354\ninside   355   372

22665	GCF_001480165.1_Salmonella_enterica_CVM_N40374-SQ_v1.0	Salmonella enterica subsp. enterica serovar Worthington	Proteobacteria; Gammaproteobacteria; Enterobacterales; Enterobacteriaceae; Salmonella; Salmonella enterica; Salmonella enterica subsp. enterica	MNNEETFYQAMRRKGVTRRSFLKFCSLAATSLGLGAGMTPKIAWALENKPRIPVVWIHGL	WP_001058312.1	hydrogenase [Salmonella enterica]	Length: 372\nNumber of predicted TMHs:  2\nExp number of AAs in TMHs: 36.51738\nExp number, first 60 AAs:  19.72849\nTotal prob of N-in:   0.96170\nPOSSIBLE N-term signal sequence\ninside   1   19\nTMhelix   20   39\noutside   40   325\nTMhelix   326   348\ninside   349   372

22666 GCF_001997835.1_ASM199783v1 Salmonella enterica subsp. enterica serovar Worthington Proteobacteria; Gammaproteobacteria; Enterobacterales; Enterobacteriaceae; Salmonella; Salmonella enterica; Salmonella enterica subsp. enterica MTGDNTLITSHGINRRDFMKLCAALAATMGLSSKAAAEMAESVSNPQRPPVIWIGAQECT WP_000145429.1 hydrogenase 2 small subunit [Salmonella enterica] Length: 372\nNumber of predicted TMHs: 1\nExp number of AAs in TMHs: 24.51309\nExp number, first 60 AAs: 0.2682\nTotal prob of N-in: 0.14402\noutside 1 331\nTMhelix 332 354\ninside 355 372

22667 GCF_001243275.1_Salmonella_enterica_CVM_N46835_v1.0 Salmonella enterica subsp. enterica serovar Worthington Proteobacteria; Gammaproteobacteria; Enterobacterales; Enterobacteriaceae; Salmonella; Salmonella enterica; Salmonella enterica subsp. enterica MTGDNTLITSHGINRRDFMKLCAALAATMGLSSKAAAEMAESVSNPQRPPVIWIGAQECT WP_000145429.1 hydrogenase 2 small subunit [Salmonella enterica] Length: 372\nNumber of predicted TMHs: 1\nExp number of AAs in TMHs: 24.51309\nExp number, first 60 AAs: 0.2682\nTotal prob of N-in: 0.14402\noutside 1 331\nTMhelix 332 354\ninside 355 372

22668 GCF_001241965.1_Salmonella_enterica_CVM_N45406_v1.0 Salmonella enterica subsp. enterica serovar Worthington Proteobacteria; Gammaproteobacteria; Enterobacterales; Enterobacteriaceae; Salmonella; Salmonella enterica; Salmonella enterica subsp. enterica MTGDNTLITSHGINRRDFMKLCAALAATMGLSSKAAAEMAESVSNPQRPPVIWIGAQECT WP_000145429.1 hydrogenase 2 small subunit [Salmonella enterica] Length: 372\nNumber of predicted TMHs: 1\nExp number of AAs in TMHs: 24.51309\nExp number, first 60 AAs: 0.2682\nTotal prob of N-in: 0.14402\noutside 1 331\nTMhelix 332 354\ninside 355 372

22669 GCF_001241965.1_Salmonella_enterica_CVM_N45406_v1.0 Salmonella enterica subsp. enterica serovar Worthington Proteobacteria; Gammaproteobacteria; Enterobacterales; Enterobacteriaceae; Salmonella; Salmonella enterica; Salmonella enterica subsp. enterica MNNEETFYQAMRRKGVTRRSFLKFCSLAATSLGLGAGMTPKIAWALENKPRIPVVWIHGL WP_001058312.1 hydrogenase [Salmonella enterica] Length: 372\nNumber of predicted TMHs: 2\nExp number of AAs in TMHs: 36.51738\nExp number, first 60 AAs: 19.72849\nTotal prob of N-in: 0.96170\nPOSSIBLE N-term signal sequence\ninside 1 19\nTMhelix 20 39\noutside 40 325\nTMhelix 326 348\ninside 349 372

22670 GCF_001243275.1_Salmonella_enterica_CVM_N46835_v1.0 Salmonella enterica subsp. enterica serovar Worthington Proteobacteria; Gammaproteobacteria; Enterobacterales; Enterobacteriaceae; Salmonella; Salmonella enterica; Salmonella enterica subsp. enterica MNNEETFYQAMRRKGVTRRSFLKFCSLAATSLGLGAGMTPKIAWALENKPRIPVVWIHGL WP_001058312.1 hydrogenase [Salmonella enterica] Length: 372\nNumber of predicted TMHs: 2\nExp number of AAs in TMHs: 36.51738\nExp number, first 60 AAs: 19.72849\nTotal prob of N-in: 0.96170\nPOSSIBLE N-term signal sequence\ninside 1 19\nTMhelix 20 39\noutside 40 325\nTMhelix 326 348\ninside 349 372

22671 GCF_002046845.1_ASM204684v1 Salmonella enterica subsp. enterica serovar Worthington Proteobacteria; Gammaproteobacteria; Enterobacterales; Enterobacteriaceae; Salmonella; Salmonella enterica; Salmonella enterica subsp. enterica MTGDNTLITSHGINRRDFMKLCAALAATMGLSSKAAAEMAESVSNPQRPPVIWIGAQECT WP_000145429.1 hydrogenase 2 small subunit [Salmonella enterica] Length: 372\nNumber of predicted TMHs: 1\nExp number of AAs in TMHs: 24.51309\nExp number, first 60 AAs: 0.2682\nTotal prob of N-in: 0.14402\noutside 1 331\nTMhelix 332 354\ninside 355 372

22672 GCF_001247285.1_Salmonella_enterica_CVM_N51304_v1.0 Salmonella enterica subsp. enterica serovar Worthington Proteobacteria; Gammaproteobacteria; Enterobacterales; Enterobacteriaceae; Salmonella; Salmonella enterica; Salmonella enterica subsp. enterica MNNEETFYQAMRRKGVTRRSFLKFCSLAATSLGLGAGMTPKIAWALENKPRIPVVWIHGL WP_001058312.1 hydrogenase [Salmonella enterica] Length: 372\nNumber of predicted TMHs: 2\nExp number of AAs in TMHs: 36.51738\nExp number, first 60 AAs: 19.72849\nTotal prob of N-in: 0.96170\nPOSSIBLE N-term signal sequence\ninside 1 19\nTMhelix 20 39\noutside 40 325\nTMhelix 326 348\ninside 349 372

22673 GCF_002032305.1_ASM203230v1 Salmonella enterica subsp. enterica serovar Worthington Proteobacteria; Gammaproteobacteria; Enterobacterales; Enterobacteriaceae; Salmonella; Salmonella enterica; Salmonella enterica subsp. enterica MTGDNTLITSHGINRRDFMKLCAALAATMGLSSKAAAEMAESVSNPQRPPVIWIGAQECT WP_000145429.1 hydrogenase 2 small subunit [Salmonella enterica] Length: 372\nNumber of predicted TMHs: 1\nExp number of AAs in TMHs: 24.51309\nExp number, first 60 AAs: 0.2682\nTotal prob of N-in: 0.14402\noutside 1 331\nTMhelix 332 354\ninside 355 372

22674	GCF_002032305.1_ASM203230v1	Salmonella enterica subsp. enterica serovar Worthington	Proteobacteria; Gammaproteobacteria; Enterobacterales; Enterobacteriaceae; Salmonella; Salmonella enterica; Salmonella enterica subsp. enterica	MNNEETFYQAMRRKGVTRRSFLKFCSLAATSLGLGAGMTPKIAWALENKPRIPVVWIHGL	WP_001058312.1	hydrogenase [Salmonella enterica]	Length: 372\nNumber of predicted TMHs: 2\nExp number of AAs in TMHs: 36.51738\nExp number, first 60 AAs: 19.72849\nTotal prob of N-in: 0.96170\nPOSSIBLE N-term signal sequence\ninside 1 19\nTMhelix 20 39\noutside 40 325\nTMhelix 326 348\ninside 349 372

22675	GCF_001480165.1_Salmonella_enterica_CVM_N40374-SQ_v1.0	Salmonella enterica subsp. enterica serovar Worthington	Proteobacteria; Gammaproteobacteria; Enterobacterales; Enterobacteriaceae; Salmonella; Salmonella enterica; Salmonella enterica subsp. enterica	MTGDNTLITSHGINRRDFMKLCAALAATMGLSSKAAAEMAESVSNPQRPPVIWIGAQECT	WP_000145429.1	hydrogenase 2 small subunit [Salmonella enterica]	Length: 372\nNumber of predicted TMHs: 1\nExp number of AAs in TMHs: 24.51309\nExp number, first 60 AAs: 0.2682\nTotal prob of N-in: 0.14402\noutside 1 331\nTMhelix 332 354\ninside 355 372

22676	GCF_002046735.1_ASM204673v1	Salmonella enterica subsp. enterica serovar Worthington	Proteobacteria; Gammaproteobacteria; Enterobacterales; Enterobacteriaceae; Salmonella; Salmonella enterica; Salmonella enterica subsp. enterica	MTGDNTLITSHGINRRDFMKLCAALAATMGLSSKAAAEMAESVSNPQRPPVIWIGAQECT	WP_000145429.1	hydrogenase 2 small subunit [Salmonella enterica]	Length: 372\nNumber of predicted TMHs: 1\nExp number of AAs in TMHs: 24.51309\nExp number, first 60 AAs: 0.2682\nTotal prob of N-in: 0.14402\noutside 1 331\nTMhelix 332 354\ninside 355 372

22677	GCF_002046735.1_ASM204673v1	Salmonella enterica subsp. enterica serovar Worthington	Proteobacteria; Gammaproteobacteria; Enterobacterales; Enterobacteriaceae; Salmonella; Salmonella enterica; Salmonella enterica subsp. enterica	MNNEETFYQAMRRKGVTRRSFLKFCSLAATSLGLGAGMTPKIAWALENKPRIPVVWIHGL	WP_001058312.1	hydrogenase [Salmonella enterica]	Length: 372\nNumber of predicted TMHs: 2\nExp number of AAs in TMHs: 36.51738\nExp number, first 60 AAs: 19.72849\nTotal prob of N-in: 0.96170\nPOSSIBLE N-term signal sequence\ninside 1 19\nTMhelix 20 39\noutside 40 325\nTMhelix 326 348\ninside 349 372

22678	GCF_001479505.1_Salmonella_enterica_CVM_N32773-SQ_v1.0	Salmonella enterica subsp. enterica serovar Worthington	Proteobacteria; Gammaproteobacteria; Enterobacterales; Enterobacteriaceae; Salmonella; Salmonella enterica; Salmonella enterica subsp. enterica	MTGDNTLITSHGINRRDFMKLCAALAATMGLSSKAAAEMAESVSNPQRPPVIWIGAQECT	WP_000145429.1	hydrogenase 2 small subunit [Salmonella enterica]	Length: 372\nNumber of predicted TMHs: 1\nExp number of AAs in TMHs: 24.51309\nExp number, first 60 AAs: 0.2682\nTotal prob of N-in: 0.14402\noutside 1 331\nTMhelix 332 354\ninside 355 372

22679	GCF_002046885.1_ASM204688v1	Salmonella enterica subsp. enterica serovar Worthington	Proteobacteria; Gammaproteobacteria; Enterobacterales; Enterobacteriaceae; Salmonella; Salmonella enterica; Salmonella enterica subsp. enterica	MTGDNTLITSHGINRRDFMKLCAALAATMGLSSKAAAEMAESVSNPQRPPVIWIGAQECT	WP_000145429.1	hydrogenase 2 small subunit [Salmonella enterica]	Length: 372\nNumber of predicted TMHs: 1\nExp number of AAs in TMHs: 24.51309\nExp number, first 60 AAs: 0.2682\nTotal prob of N-in: 0.14402\noutside 1 331\nTMhelix 332 354\ninside 355 372

22680	GCF_002046645.1_ASM204664v1	Salmonella enterica subsp. enterica serovar Worthington	Proteobacteria; Gammaproteobacteria; Enterobacterales; Enterobacteriaceae; Salmonella; Salmonella enterica; Salmonella enterica subsp. enterica	MNNEETFYQAMRRKGVTRRSFLKFCSLAATSLGLGAGMTPKIAWALENKPRIPVVWIHGL	WP_001058312.1	hydrogenase [Salmonella enterica]	Length: 372\nNumber of predicted TMHs: 2\nExp number of AAs in TMHs: 36.51738\nExp number, first 60 AAs: 19.72849\nTotal prob of N-in: 0.96170\nPOSSIBLE N-term signal sequence\ninside 1 19\nTMhelix 20 39\noutside 40 325\nTMhelix 326 348\ninside 349 372

22681	GCF_002046645.1_ASM204664v1	Salmonella enterica subsp. enterica serovar Worthington	Proteobacteria; Gammaproteobacteria; Enterobacterales; Enterobacteriaceae; Salmonella; Salmonella enterica; Salmonella enterica subsp. enterica	MTGDNTLITSHGINRRDFMKLCAALAATMGLSSKAAAEMAESVSNPQRPPVIWIGAQECT	WP_000145429.1	hydrogenase 2 small subunit [Salmonella enterica]	Length: 372\nNumber of predicted TMHs: 1\nExp number of AAs in TMHs: 24.51309\nExp number, first 60 AAs: 0.2682\nTotal prob of N-in: 0.14402\noutside 1 331\nTMhelix 332 354\ninside 355 372

22682	GCF_000335915.1_ASM33591v1	Salmonella enterica subsp. enterica serovar 4,[5],12:i:- str. 08-1700	Proteobacteria; Gammaproteobacteria; Enterobacterales; Enterobacteriaceae; Salmonella; Salmonella enterica; Salmonella enterica subsp. enterica; Salmonella enterica subsp. enterica serovar 4,[5],12

MTGDNTLITSHGINRRDFMKLCAALAATMGLSSKAAAEMAESVSNPQRPPVIWIGAQECT WP_000145429.1 hydrogenase 2 small subunit [Salmonella enterica] Length: 372\nNumber of predicted TMHs: 1\nExp number of AAs in TMHs: 24.51309\nExp number, first 60 AAs: 0.2682\nTotal prob of N-in: 0.14402\noutside 1 331\nTMhelix 332 354\ninside 355 372

22683 GCF_000335915.1_ASM33591v1 Salmonella enterica subsp. enterica serovar 4,[5],12:i:- str. 08-1700 Proteobacteria; Gammaproteobacteria; Enterobacterales; Enterobacteriaceae; Salmonella; Salmonella enterica; Salmonella enterica subsp. enterica; Salmonella enterica subsp. enterica serovar 4,[5],12
MNNEETFYQAMRRKGVTRRSFLKFCSLAATSLGLGAGMTPKIAWALENKPRIPVVWIHGL WP_001058312.1 hydrogenase [Salmonella enterica] Length: 372\nNumber of predicted TMHs: 2\nExp number of AAs in TMHs: 36.51738\nExp number, first 60 AAs: 19.72849\nTotal prob of N-in: 0.96170\nPOSSIBLE N-term signal sequence\ninside 1 19\nTMhelix 20 39\noutside 40 325\nTMhelix 326 348\ninside 349 372

22684 GCF_000430165.1_ASM43016v1 Salmonella enterica subsp. enterica serovar 4,[5],12:i:- str. 08-1736 Proteobacteria; Gammaproteobacteria; Enterobacterales; Enterobacteriaceae; Salmonella; Salmonella enterica; Salmonella enterica subsp. enterica; Salmonella enterica subsp. enterica serovar 4,[5],12
MTGDNTLITSHGINRRDFMKLCAALAATMGLSSKAAAEMAESVSNPQRPPVIWIGAQECT WP_000145429.1 hydrogenase 2 small subunit [Salmonella enterica] Length: 372\nNumber of predicted TMHs: 1\nExp number of AAs in TMHs: 24.51309\nExp number, first 60 AAs: 0.2682\nTotal prob of N-in: 0.14402\noutside 1 331\nTMhelix 332 354\ninside 355 372

22685 GCF_000430165.1_ASM43016v1 Salmonella enterica subsp. enterica serovar 4,[5],12:i:- str. 08-1736 Proteobacteria; Gammaproteobacteria; Enterobacterales; Enterobacteriaceae; Salmonella; Salmonella enterica; Salmonella enterica subsp. enterica; Salmonella enterica subsp. enterica serovar 4,[5],12
MNNEETFYQAMRRKGVTRRSFLKFCSLAATSLGLGAGMTPKIAWALENKPRIPVVWIHGL WP_001058312.1 hydrogenase [Salmonella enterica] Length: 372\nNumber of predicted TMHs: 2\nExp number of AAs in TMHs: 36.51738\nExp number, first 60 AAs: 19.72849\nTotal prob of N-in: 0.96170\nPOSSIBLE N-term signal sequence\ninside 1 19\nTMhelix 20 39\noutside 40 325\nTMhelix 326 348\ninside 349 372

22686 GCF_000335935.1_ASM33593v1 Salmonella enterica subsp. enterica serovar 4,[5],12:i:- str. 08-1739 Proteobacteria; Gammaproteobacteria; Enterobacterales; Enterobacteriaceae; Salmonella; Salmonella enterica; Salmonella enterica subsp. enterica; Salmonella enterica subsp. enterica serovar 4,[5],12
MTGDNTLITSHGINRRDFMKLCAALAATMGLSSKAAAEMAESVSNPQRPPVIWIGAQECT WP_000145429.1 hydrogenase 2 small subunit [Salmonella enterica] Length: 372\nNumber of predicted TMHs: 1\nExp number of AAs in TMHs: 24.51309\nExp number, first 60 AAs: 0.2682\nTotal prob of N-in: 0.14402\noutside 1 331\nTMhelix 332 354\ninside 355 372

22687 GCF_000335935.1_ASM33593v1 Salmonella enterica subsp. enterica serovar 4,[5],12:i:- str. 08-1739 Proteobacteria; Gammaproteobacteria; Enterobacterales; Enterobacteriaceae; Salmonella; Salmonella enterica; Salmonella enterica subsp. enterica; Salmonella enterica subsp. enterica serovar 4,[5],12
MNNEETFYQAMRRKGVTRRSFLKFCSLAATSLGLGAGMTPKIAWALENKPRIPVVWIHGL WP_001058312.1 hydrogenase [Salmonella enterica] Length: 372\nNumber of predicted TMHs: 2\nExp number of AAs in TMHs: 36.51738\nExp number, first 60 AAs: 19.72849\nTotal prob of N-in: 0.96170\nPOSSIBLE N-term signal sequence\ninside 1 19\nTMhelix 20 39\noutside 40 325\nTMhelix 326 348\ninside 349 372

22688 GCF_000170255.1_ASM17025v1 Salmonella enterica subsp. enterica serovar 4,[5],12:i:- str. CVM23701 Proteobacteria; Gammaproteobacteria; Enterobacterales; Enterobacteriaceae; Salmonella; Salmonella enterica; Salmonella enterica subsp. enterica; Salmonella enterica subsp. enterica serovar 4,[5],12
MNNEETFYQAMRRKGVTRRSFLKFCSLAATSLGLGAGMTPKIAWALENKPRIPVVWIHGL WP_001058312.1 hydrogenase [Salmonella enterica] Length: 372\nNumber of predicted TMHs: 2\nExp number of AAs in TMHs: 36.51738\nExp number, first 60 AAs: 19.72849\nTotal prob of N-in: 0.96170\nPOSSIBLE N-term signal sequence\ninside 1 19\nTMhelix 20 39\noutside 40 325\nTMhelix 326 348\ninside 349 372

22689 GCF_000170255.1_ASM17025v1 Salmonella enterica subsp. enterica serovar 4,[5],12:i:- str. CVM23701 Proteobacteria; Gammaproteobacteria; Enterobacterales; Enterobacteriaceae; Salmonella; Salmonella enterica; Salmonella enterica subsp. enterica; Salmonella enterica subsp. enterica serovar 4,[5],12
MTGDNTLITSHGINRRDFMKLCAALAATMGLSSKAAAEMAESVSNPQRPPVIWIGAQECT WP_000145429.1 hydrogenase 2 small subunit [Salmonella enterica] Length: 372\nNumber of predicted TMHs: 1\nExp number of AAs in TMHs: 24.51309\nExp number, first 60 AAs: 0.2682\nTotal prob of N-in: 0.14402\noutside 1 331\nTMhelix 332 354\ninside 355 372

22690 GCF_000648375.2_94293_Ver1 Salmonella enterica subsp. enterica serovar 9,12:l,v:- str. 94293
Proteobacteria; Gammaproteobacteria; Enterobacterales; Enterobacteriaceae; Salmonella; Salmonella enterica; Salmonella enterica subsp. enterica; Salmonella enterica subsp. enterica serovar 9,12
MTGDNTLITSHGINRRDFMKLCAALAATMGLSSKAAAEMAESVSNPQRPPVIWIGAQECT WP_000145429.1
hydrogenase 2 small subunit [Salmonella enterica] Length: 372\nNumber of predicted TMHs: 1\nExp number of AAs in TMHs: 24.51309\nExp number, first 60 AAs: 0.2682\nTotal prob of N-in: 0.14402\noutside 1 331\nTMhelix 332 354\ninside 355 372

22691 GCF_000487915.2_ASM48791v2 Salmonella enterica subsp. enterica serovar Abaetetuba str. ATCC 35640
Proteobacteria; Gammaproteobacteria; Enterobacterales; Enterobacteriaceae; Salmonella; Salmonella enterica; Salmonella enterica subsp. enterica; Salmonella enterica subsp. enterica serovar Abaetetuba
MNNEETFYQAMRRKGVTRRSFLKFCSLAATSLGLGAGMTPKIAWALENKPRIPVVWIHGL WP_001058312.1
hydrogenase [Salmonella enterica] Length: 372\nNumber of predicted TMHs: 2\nExp number of AAs in TMHs: 36.51738\nExp number, first 60 AAs: 19.72849\nTotal prob of N-in: 0.96170\nPOSSIBLE N-term signal sequence\ninside 1 19\nTMhelix 20 39\noutside 40 325\nTMhelix 326 348\ninside 349 372

22692 GCF_000487915.2_ASM48791v2 Salmonella enterica subsp. enterica serovar Abaetetuba str. ATCC 35640
Proteobacteria; Gammaproteobacteria; Enterobacterales; Enterobacteriaceae; Salmonella; Salmonella enterica; Salmonella enterica subsp. enterica; Salmonella enterica subsp. enterica serovar Abaetetuba
MTGDNTLITSHGINRRDFMKLCAALAATMGLSSKAAAEMAESVSNPQRPPVIWIGAQECT WP_000145431.1
hydrogenase 2 small subunit [Salmonella enterica] Length: 372\nNumber of predicted TMHs: 1\nExp number of AAs in TMHs: 24.51505\nExp number, first 60 AAs: 0.2682\nTotal prob of N-in: 0.14406\noutside 1 331\nTMhelix 332 354\ninside 355 372

22693 GCF_000487615.2_ASM48761v2 Salmonella enterica subsp. enterica serovar Abony str. 0014
Proteobacteria; Gammaproteobacteria; Enterobacterales; Enterobacteriaceae; Salmonella; Salmonella enterica; Salmonella enterica subsp. enterica; Salmonella enterica subsp. enterica serovar Abony
MTGDNTLITSHGINRRDFMKLCAALAATMGLSSKAAAEMAESVSNPQRPPVIWIGAQECT WP_000145429.1
hydrogenase 2 small subunit [Salmonella enterica] Length: 372\nNumber of predicted TMHs: 1\nExp number of AAs in TMHs: 24.51309\nExp number, first 60 AAs: 0.2682\nTotal prob of N-in: 0.14402\noutside 1 331\nTMhelix 332 354\ninside 355 372

22694 GCF_000487615.2_ASM48761v2 Salmonella enterica subsp. enterica serovar Abony str. 0014
Proteobacteria; Gammaproteobacteria; Enterobacterales; Enterobacteriaceae; Salmonella; Salmonella enterica; Salmonella enterica subsp. enterica; Salmonella enterica subsp. enterica serovar Abony
MNNEETFYQAMRRKGVTRRSFLKFCSLAATSLGLGAGMTPKIAWALENKPRIPVVWIHGL WP_001058310.1
hydrogenase [Salmonella enterica] Length: 372\nNumber of predicted TMHs: 2\nExp number of AAs in TMHs: 37.4567299999999\nExp number, first 60 AAs: 19.72694\nTotal prob of N-in: 0.96171\nPOSSIBLE N-term signal sequence\ninside 1 19\nTMhelix 20 39\noutside 40 325\nTMhelix 326 348\ninside 349 372

22695 GCF_000388195.1_01.O.05 Salmonella enterica subsp. enterica serovar Agona str. 01.O.05
Proteobacteria; Gammaproteobacteria; Enterobacterales; Enterobacteriaceae; Salmonella; Salmonella enterica; Salmonella enterica subsp. enterica; Salmonella enterica subsp. enterica serovar Agona
MTGDNTLITSHGINRRDFMKLCAALAATMGLSSKAAAEMAESVSNPQRPPVIWIGAQECT WP_000145429.1
hydrogenase 2 small subunit [Salmonella enterica] Length: 372\nNumber of predicted TMHs: 1\nExp number of AAs in TMHs: 24.51309\nExp number, first 60 AAs: 0.2682\nTotal prob of N-in: 0.14402\noutside 1 331\nTMhelix 332 354\ninside 355 372

22696 GCF_000388195.1_01.O.05 Salmonella enterica subsp. enterica serovar Agona str. 01.O.05
Proteobacteria; Gammaproteobacteria; Enterobacterales; Enterobacteriaceae; Salmonella; Salmonella enterica; Salmonella enterica subsp. enterica; Salmonella enterica subsp. enterica serovar Agona
MNNEETFYQAMRRKGVTRRSFLKFCSLAATSLGLSAGMTPKIAWALENKPRIPVVWIHGL WP_001058316.1
hydrogenase [Salmonella enterica] Length: 372\nNumber of predicted TMHs: 2\nExp number of AAs in TMHs: 35.76397\nExp number, first 60 AAs: 18.97201\nTotal prob of N-in: 0.92785\nPOSSIBLE N-term signal sequence\ninside 1 19\nTMhelix 20 39\noutside 40 325\nTMhelix 326 348\ninside 349 372

22697 GCF_000484335.1_ASM48433v1 Salmonella enterica subsp. enterica serovar Agona str. 0292
Proteobacteria; Gammaproteobacteria; Enterobacterales; Enterobacteriaceae; Salmonella; Salmonella enterica; Salmonella enterica subsp. enterica; Salmonella enterica subsp. enterica serovar Agona
MTGDNTLITSHGINRRDFMKLCAALAATMGLSSKAAAEMAESVSNPQRPPVIWIGAQECT WP_000145429.1
hydrogenase 2 small subunit [Salmonella enterica] Length: 372\nNumber of predicted TMHs: 1\nExp

number of AAs in TMHs: 24.51309\nExp number, first 60 AAs: 0.2682\nTotal prob of N-in: 0.14402\noutside 1 331\nTMhelix 332 354\ninside 355 372

22698 GCF_000484335.1_ASM48433v1 Salmonella enterica subsp. enterica serovar Agona str. 0292 Proteobacteria; Gammaproteobacteria; Enterobacterales; Enterobacteriaceae; Salmonella; Salmonella enterica; Salmonella enterica subsp. enterica; Salmonella enterica subsp. enterica serovar Agona MNNEETFYQAMRRKGVTRRSFLKFCSLAATSLGLSAGMTPKIAWALENKPRIPVVWIHGL WP_001058316.1 hydrogenase [Salmonella enterica] Length: 372\nNumber of predicted TMHs: 2\nExp number of AAs in TMHs: 35.76397\nExp number, first 60 AAs: 18.97201\nTotal prob of N-in: 0.92785\nPOSSIBLE N-term signal sequence\ninside 1 19\nTMhelix 20 39\noutside 40 325\nTMhelix 326 348\ninside 349 372

22699 GCF_000388215.1_02.O.05 Salmonella enterica subsp. enterica serovar Agona str. 02.O.05 Proteobacteria; Gammaproteobacteria; Enterobacterales; Enterobacteriaceae; Salmonella; Salmonella enterica; Salmonella enterica subsp. enterica; Salmonella enterica subsp. enterica serovar Agona MTGDNTLITSHGINRRDFMKLCAALAATMGLSSKAAAEMAESVSNPQRPPVIWIGAQECT WP_000145429.1 hydrogenase 2 small subunit [Salmonella enterica] Length: 372\nNumber of predicted TMHs: 1\nExp number of AAs in TMHs: 24.51309\nExp number, first 60 AAs: 0.2682\nTotal prob of N-in: 0.14402\noutside 1 331\nTMhelix 332 354\ninside 355 372

22700 GCF_000388235.1_03.O.05 Salmonella enterica subsp. enterica serovar Agona str. 03.O.05 Proteobacteria; Gammaproteobacteria; Enterobacterales; Enterobacteriaceae; Salmonella; Salmonella enterica; Salmonella enterica subsp. enterica; Salmonella enterica subsp. enterica serovar Agona MTGDNTLITSHGINRRDFMKLCAALAATMGLSSKAAAEMAESVSNPQRPPVIWIGAQECT WP_000145429.1 hydrogenase 2 small subunit [Salmonella enterica] Length: 372\nNumber of predicted TMHs: 1\nExp number of AAs in TMHs: 24.51309\nExp number, first 60 AAs: 0.2682\nTotal prob of N-in: 0.14402\noutside 1 331\nTMhelix 332 354\ninside 355 372

22701 GCF_000388235.1_03.O.05 Salmonella enterica subsp. enterica serovar Agona str. 03.O.05 Proteobacteria; Gammaproteobacteria; Enterobacterales; Enterobacteriaceae; Salmonella; Salmonella enterica; Salmonella enterica subsp. enterica; Salmonella enterica subsp. enterica serovar Agona MNNEETFYQAMRRKGVTRRSFLKFCSLAATSLGLSAGMTPKIAWALENKPRIPVVWIHGL WP_001058316.1 hydrogenase [Salmonella enterica] Length: 372\nNumber of predicted TMHs: 2\nExp number of AAs in TMHs: 35.76397\nExp number, first 60 AAs: 18.97201\nTotal prob of N-in: 0.92785\nPOSSIBLE N-term signal sequence\ninside 1 19\nTMhelix 20 39\noutside 40 325\nTMhelix 326 348\ninside 349 372

22702 GCF_000388255.1_04.O.05 Salmonella enterica subsp. enterica serovar Agona str. 04.O.05 Proteobacteria; Gammaproteobacteria; Enterobacterales; Enterobacteriaceae; Salmonella; Salmonella enterica; Salmonella enterica subsp. enterica; Salmonella enterica subsp. enterica serovar Agona MTGDNTLITSHGINRRDFMKLCAALAATMGLSSKAAAEMAESVSNPQRPPVIWIGAQECT WP_000145429.1 hydrogenase 2 small subunit [Salmonella enterica] Length: 372\nNumber of predicted TMHs: 1\nExp number of AAs in TMHs: 24.51309\nExp number, first 60 AAs: 0.2682\nTotal prob of N-in: 0.14402\noutside 1 331\nTMhelix 332 354\ninside 355 372

22703 GCF_000388275.1_05.O.06 Salmonella enterica subsp. enterica serovar Agona str. 05.O.06 Proteobacteria; Gammaproteobacteria; Enterobacterales; Enterobacteriaceae; Salmonella; Salmonella enterica; Salmonella enterica subsp. enterica; Salmonella enterica subsp. enterica serovar Agona MTGDNTLITSHGINRRDFMKLCAALAATMGLSSKAAAEMAESVSNPQRPPVIWIGAQECT WP_000145429.1 hydrogenase 2 small subunit [Salmonella enterica] Length: 372\nNumber of predicted TMHs: 1\nExp number of AAs in TMHs: 24.51309\nExp number, first 60 AAs: 0.2682\nTotal prob of N-in: 0.14402\noutside 1 331\nTMhelix 332 354\ninside 355 372

22704 GCF_000388295.1_06.O.05 Salmonella enterica subsp. enterica serovar Agona str. 06.O.05 Proteobacteria; Gammaproteobacteria; Enterobacterales; Enterobacteriaceae; Salmonella; Salmonella enterica; Salmonella enterica subsp. enterica; Salmonella enterica subsp. enterica serovar Agona MNNEETFYQAMRRKGVTRRSFLKFCSLAATSLGLSAGMTPKIAWALENKPRIPVVWIHGL WP_001058316.1 hydrogenase [Salmonella enterica] Length: 372\nNumber of predicted TMHs: 2\nExp number of AAs in TMHs: 35.76397\nExp number, first 60 AAs: 18.97201\nTotal prob of N-in: 0.92785\nPOSSIBLE N-term signal sequence\ninside 1 19\nTMhelix 20 39\noutside 40 325\nTMhelix 326 348\ninside 349 372

22705 GCF_000388295.1_06.O.05 Salmonella enterica subsp. enterica serovar Agona str. 06.O.05 Proteobacteria; Gammaproteobacteria; Enterobacterales; Enterobacteriaceae; Salmonella; Salmonella enterica;

Salmonella enterica subsp. enterica; Salmonella enterica subsp. enterica serovar Agona
MTGDNTLITSHGINRRDFMKLCAALAATMGLSSKAAAEMAESVSNPQRPPVIWIGAQECT WP_000145429.1
hydrogenase 2 small subunit [Salmonella enterica] Length: 372\nNumber of predicted TMHs: 1\nExp number of AAs in TMHs: 24.51309\nExp number, first 60 AAs: 0.2682\nTotal prob of N-in: 0.14402\noutside 1 331\nTMhelix 332 354\ninside 355 372

22706 GCF_000388315.1_07.O.05 Salmonella enterica subsp. enterica serovar Agona str. 07.O.05
Proteobacteria; Gammaproteobacteria; Enterobacterales; Enterobacteriaceae; Salmonella; Salmonella enterica; Salmonella enterica subsp. enterica; Salmonella enterica subsp. enterica serovar Agona
MNNEETFYQAMRRKGVTRRSFLKFCSLAATSLGLSAGMTPKIAWALENKPRIPVVWIHGL WP_001058316.1
hydrogenase [Salmonella enterica] Length: 372\nNumber of predicted TMHs: 2\nExp number of AAs in TMHs: 35.76397\nExp number, first 60 AAs: 18.97201\nTotal prob of N-in: 0.92785\nPOSSIBLE N-term signal sequence\ninside 1 19\nTMhelix 20 39\noutside 40 325\nTMhelix 326 348\ninside 349 372

22707 GCF_000388315.1_07.O.05 Salmonella enterica subsp. enterica serovar Agona str. 07.O.05
Proteobacteria; Gammaproteobacteria; Enterobacterales; Enterobacteriaceae; Salmonella; Salmonella enterica; Salmonella enterica subsp. enterica; Salmonella enterica subsp. enterica serovar Agona
MTGDNTLITSHGINRRDFMKLCAALAATMGLSSKAAAEMAESVSNPQRPPVIWIGAQECT WP_000145429.1
hydrogenase 2 small subunit [Salmonella enterica] Length: 372\nNumber of predicted TMHs: 1\nExp number of AAs in TMHs: 24.51309\nExp number, first 60 AAs: 0.2682\nTotal prob of N-in: 0.14402\noutside 1 331\nTMhelix 332 354\ninside 355 372

22708 GCF_000388335.1_08.A.05 Salmonella enterica subsp. enterica serovar Agona str. 08.A.05
Proteobacteria; Gammaproteobacteria; Enterobacterales; Enterobacteriaceae; Salmonella; Salmonella enterica; Salmonella enterica subsp. enterica; Salmonella enterica subsp. enterica serovar Agona
MNNEETFYQAMRRKGVTRRSFLKFCSLAATSLGLSAGMTPKIAWALENKPRIPVVWIHGL WP_001058316.1
hydrogenase [Salmonella enterica] Length: 372\nNumber of predicted TMHs: 2\nExp number of AAs in TMHs: 35.76397\nExp number, first 60 AAs: 18.97201\nTotal prob of N-in: 0.92785\nPOSSIBLE N-term signal sequence\ninside 1 19\nTMhelix 20 39\noutside 40 325\nTMhelix 326 348\ninside 349 372

22709 GCF_000388335.1_08.A.05 Salmonella enterica subsp. enterica serovar Agona str. 08.A.05
Proteobacteria; Gammaproteobacteria; Enterobacterales; Enterobacteriaceae; Salmonella; Salmonella enterica; Salmonella enterica subsp. enterica; Salmonella enterica subsp. enterica serovar Agona
MTGDNTLITSHGINRRDFMKLCAALAATMGLSSKAAAEMAESVSNPQRPPVIWIGAQECT WP_000145429.1
hydrogenase 2 small subunit [Salmonella enterica] Length: 372\nNumber of predicted TMHs: 1\nExp number of AAs in TMHs: 24.51309\nExp number, first 60 AAs: 0.2682\nTotal prob of N-in: 0.14402\noutside 1 331\nTMhelix 332 354\ninside 355 372

22710 GCF_000388355.1_09.F.08 Salmonella enterica subsp. enterica serovar Agona str. 09.F.08
Proteobacteria; Gammaproteobacteria; Enterobacterales; Enterobacteriaceae; Salmonella; Salmonella enterica; Salmonella enterica subsp. enterica; Salmonella enterica subsp. enterica serovar Agona
MNNEETFYQAMRRKGVTRRSFLKFCSLAATSLGLSAGMTPKIAWALENKPRIPVVWIHGL WP_001058316.1
hydrogenase [Salmonella enterica] Length: 372\nNumber of predicted TMHs: 2\nExp number of AAs in TMHs: 35.76397\nExp number, first 60 AAs: 18.97201\nTotal prob of N-in: 0.92785\nPOSSIBLE N-term signal sequence\ninside 1 19\nTMhelix 20 39\noutside 40 325\nTMhelix 326 348\ninside 349 372

22711 GCF_000388355.1_09.F.08 Salmonella enterica subsp. enterica serovar Agona str. 09.F.08
Proteobacteria; Gammaproteobacteria; Enterobacterales; Enterobacteriaceae; Salmonella; Salmonella enterica; Salmonella enterica subsp. enterica; Salmonella enterica subsp. enterica serovar Agona
MTGDNTLITSHGINRRDFMKLCAALAATMGLSSKAAAEMAESVSNPQRPPVIWIGAQECT WP_000145429.1
hydrogenase 2 small subunit [Salmonella enterica] Length: 372\nNumber of predicted TMHs: 1\nExp number of AAs in TMHs: 24.51309\nExp number, first 60 AAs: 0.2682\nTotal prob of N-in: 0.14402\noutside 1 331\nTMhelix 332 354\ninside 355 372

22712 GCF_000388375.1_10.A.05 Salmonella enterica subsp. enterica serovar Agona str. 10.A.05
Proteobacteria; Gammaproteobacteria; Enterobacterales; Enterobacteriaceae; Salmonella; Salmonella enterica; Salmonella enterica subsp. enterica; Salmonella enterica subsp. enterica serovar Agona
MNNEETFYQAMRRKGVTRRSFLKFCSLAATSLGLSAGMTPKIAWALENKPRIPVVWIHGL WP_001058316.1
hydrogenase [Salmonella enterica] Length: 372\nNumber of predicted TMHs: 2\nExp number of AAs in TMHs: 35.76397\nExp number, first 60 AAs: 18.97201\nTotal prob of N-in: 0.92785\nPOSSIBLE N-term signal sequence\ninside 1 19\nTMhelix 20 39\noutside 40 325\nTMhelix 326 348\ninside 349 372

22713 GCF_000388375.1_10.A.05 Salmonella enterica subsp. enterica serovar Agona str. 10.A.05
Proteobacteria; Gammaproteobacteria; Enterobacterales; Enterobacteriaceae; Salmonella; Salmonella enterica; Salmonella enterica subsp. enterica; Salmonella enterica subsp. enterica serovar Agona
MTGDNTLITSHGINRRDFMKLCAALAATMGLSSKAAAEMAESVSNPQRPPVIWIGAQECT WP_000145429.1
hydrogenase 2 small subunit [Salmonella enterica] Length: 372\nNumber of predicted TMHs: 1\nExp number of AAs in TMHs: 24.51309\nExp number, first 60 AAs: 0.2682\nTotal prob of N-in: 0.14402\noutside 1 331\nTMhelix 332 354\ninside 355 372

22714 GCF_000388395.1_11.A.05 Salmonella enterica subsp. enterica serovar Agona str. 11.A.05
Proteobacteria; Gammaproteobacteria; Enterobacterales; Enterobacteriaceae; Salmonella; Salmonella enterica; Salmonella enterica subsp. enterica; Salmonella enterica subsp. enterica serovar Agona
MNNEETFYQAMRRKGVTRRSFLKFCSLAATSLGLSAGMTPKIAWALENKPRIPVVWIHGL WP_001058316.1
hydrogenase [Salmonella enterica] Length: 372\nNumber of predicted TMHs: 2\nExp number of AAs in TMHs: 35.76397\nExp number, first 60 AAs: 18.97201\nTotal prob of N-in: 0.92785\nPOSSIBLE N-term signal sequence\ninside 1 19\nTMhelix 20 39\noutside 40 325\nTMhelix 326 348\ninside 349 372

22715 GCF_000388395.1_11.A.05 Salmonella enterica subsp. enterica serovar Agona str. 11.A.05
Proteobacteria; Gammaproteobacteria; Enterobacterales; Enterobacteriaceae; Salmonella; Salmonella enterica; Salmonella enterica subsp. enterica; Salmonella enterica subsp. enterica serovar Agona
MTGDNTLITSHGINRRDFMKLCAALAATMGLSSKAAAEMAESVSNPQRPPVIWIGAQECT WP_000145429.1
hydrogenase 2 small subunit [Salmonella enterica] Length: 372\nNumber of predicted TMHs: 1\nExp number of AAs in TMHs: 24.51309\nExp number, first 60 AAs: 0.2682\nTotal prob of N-in: 0.14402\noutside 1 331\nTMhelix 332 354\ninside 355 372

22716 GCF_000388415.1_12.A.06 Salmonella enterica subsp. enterica serovar Agona str. 12.A.06
Proteobacteria; Gammaproteobacteria; Enterobacterales; Enterobacteriaceae; Salmonella; Salmonella enterica; Salmonella enterica subsp. enterica; Salmonella enterica subsp. enterica serovar Agona
MTGDNTLITSHGINRRDFMKLCAALAATMGLSSKAAAEMAESVSNPQRPPVIWIGAQECT WP_000145429.1
hydrogenase 2 small subunit [Salmonella enterica] Length: 372\nNumber of predicted TMHs: 1\nExp number of AAs in TMHs: 24.51309\nExp number, first 60 AAs: 0.2682\nTotal prob of N-in: 0.14402\noutside 1 331\nTMhelix 332 354\ninside 355 372

22717 GCF_000388415.1_12.A.06 Salmonella enterica subsp. enterica serovar Agona str. 12.A.06
Proteobacteria; Gammaproteobacteria; Enterobacterales; Enterobacteriaceae; Salmonella; Salmonella enterica; Salmonella enterica subsp. enterica; Salmonella enterica subsp. enterica serovar Agona
MNNEETFYQAMRRKGVTRRSFLKFCSLAATSLGLSAGMTPKIAWALENKPRIPVVWIHGL WP_001058316.1
hydrogenase [Salmonella enterica] Length: 372\nNumber of predicted TMHs: 2\nExp number of AAs in TMHs: 35.76397\nExp number, first 60 AAs: 18.97201\nTotal prob of N-in: 0.92785\nPOSSIBLE N-term signal sequence\ninside 1 19\nTMhelix 20 39\noutside 40 325\nTMhelix 326 348\ninside 349 372

22718 GCF_000388435.1_13.E.05 Salmonella enterica subsp. enterica serovar Agona str. 13.E.05
Proteobacteria; Gammaproteobacteria; Enterobacterales; Enterobacteriaceae; Salmonella; Salmonella enterica; Salmonella enterica subsp. enterica; Salmonella enterica subsp. enterica serovar Agona
MTGDNTLITSHGINRRDFMKLCAALAATMGLSSKAAAEMAESVSNPQRPPVIWIGAQECT WP_000145429.1
hydrogenase 2 small subunit [Salmonella enterica] Length: 372\nNumber of predicted TMHs: 1\nExp number of AAs in TMHs: 24.51309\nExp number, first 60 AAs: 0.2682\nTotal prob of N-in: 0.14402\noutside 1 331\nTMhelix 332 354\ninside 355 372

22719 GCF_000388435.1_13.E.05 Salmonella enterica subsp. enterica serovar Agona str. 13.E.05
Proteobacteria; Gammaproteobacteria; Enterobacterales; Enterobacteriaceae; Salmonella; Salmonella enterica; Salmonella enterica subsp. enterica; Salmonella enterica subsp. enterica serovar Agona
MNNEETFYQAMRRKGVTRRSFLKFCSLAATSLGLSAGMTPKIAWALENKPRIPVVWIHGL WP_001058316.1
hydrogenase [Salmonella enterica] Length: 372\nNumber of predicted TMHs: 2\nExp number of AAs in TMHs: 35.76397\nExp number, first 60 AAs: 18.97201\nTotal prob of N-in: 0.92785\nPOSSIBLE N-term signal sequence\ninside 1 19\nTMhelix 20 39\noutside 40 325\nTMhelix 326 348\ninside 349 372

22720 GCF_000388455.1_14.E.05 Salmonella enterica subsp. enterica serovar Agona str. 14.E.05
Proteobacteria; Gammaproteobacteria; Enterobacterales; Enterobacteriaceae; Salmonella; Salmonella enterica; Salmonella enterica subsp. enterica; Salmonella enterica subsp. enterica serovar Agona
MNNEETFYQAMRRKGVTRRSFLKFCSLAATSLGLSAGMTPKIAWALENKPRIPVVWIHGL WP_001058316.1
hydrogenase [Salmonella enterica] Length: 372\nNumber of predicted TMHs: 2\nExp number of AAs in

TMHs: 35.76397\nExp number, first 60 AAs: 18.97201\nTotal prob of N-in: 0.92785\nPOSSIBLE N-term signal sequence\ninside 1 19\nTMhelix 20 39\noutside 40 325\nTMhelix 326 348\ninside 349 372

22721 GCF_000388455.1_14.E.05 Salmonella enterica subsp. enterica serovar Agona str. 14.E.05 Proteobacteria; Gammaproteobacteria; Enterobacterales; Enterobacteriaceae; Salmonella; Salmonella enterica; Salmonella enterica subsp. enterica; Salmonella enterica subsp. enterica serovar Agona MTGDNTLITSHGINRRDFMKLCAALAATMGLSSKAAAEMAESVSNPQRPPVIWIGAQECT WP_000145429.1 hydrogenase 2 small subunit [Salmonella enterica] Length: 372\nNumber of predicted TMHs: 1\nExp number of AAs in TMHs: 24.51309\nExp number, first 60 AAs: 0.2682\nTotal prob of N-in: 0.14402\noutside 1 331\nTMhelix 332 354\ninside 355 372

22722 GCF_000388475.1_15.H.03 Salmonella enterica subsp. enterica serovar Agona str. 15.H.03 Proteobacteria; Gammaproteobacteria; Enterobacterales; Enterobacteriaceae; Salmonella; Salmonella enterica; Salmonella enterica subsp. enterica; Salmonella enterica subsp. enterica serovar Agona MNNEETFYQAMRRKGVTRRSFLKFCSLAATSLGLSAGMTPKIAWALENKPRIPVVWIHGL WP_001058316.1 hydrogenase [Salmonella enterica] Length: 372\nNumber of predicted TMHs: 2\nExp number of AAs in TMHs: 35.76397\nExp number, first 60 AAs: 18.97201\nTotal prob of N-in: 0.92785\nPOSSIBLE N-term signal sequence\ninside 1 19\nTMhelix 20 39\noutside 40 325\nTMhelix 326 348\ninside 349 372

22723 GCF_000388475.1_15.H.03 Salmonella enterica subsp. enterica serovar Agona str. 15.H.03 Proteobacteria; Gammaproteobacteria; Enterobacterales; Enterobacteriaceae; Salmonella; Salmonella enterica; Salmonella enterica subsp. enterica; Salmonella enterica subsp. enterica serovar Agona MTGDNTLITSHGINRRDFMKLCAALAATMGLSSKAAAEMAESVSNPQRPPVIWIGAQECT WP_000145429.1 hydrogenase 2 small subunit [Salmonella enterica] Length: 372\nNumber of predicted TMHs: 1\nExp number of AAs in TMHs: 24.51309\nExp number, first 60 AAs: 0.2682\nTotal prob of N-in: 0.14402\noutside 1 331\nTMhelix 332 354\ninside 355 372

22724 GCF_001050855.1_ASM105085v1 Salmonella enterica subsp. enterica serovar Agona str. 16.H.08 Proteobacteria; Gammaproteobacteria; Enterobacterales; Enterobacteriaceae; Salmonella; Salmonella enterica; Salmonella enterica subsp. enterica; Salmonella enterica subsp. enterica serovar Agona MTGDNTLITSHGINRRDFMKLCAALAATMGLSSKAAAEMAESVSNPQRPPVIWIGAQECT WP_000145429.1 hydrogenase 2 small subunit [Salmonella enterica] Length: 372\nNumber of predicted TMHs: 1\nExp number of AAs in TMHs: 24.51309\nExp number, first 60 AAs: 0.2682\nTotal prob of N-in: 0.14402\noutside 1 331\nTMhelix 332 354\ninside 355 372

22725 GCF_001050855.1_ASM105085v1 Salmonella enterica subsp. enterica serovar Agona str. 16.H.08 Proteobacteria; Gammaproteobacteria; Enterobacterales; Enterobacteriaceae; Salmonella; Salmonella enterica; Salmonella enterica subsp. enterica; Salmonella enterica subsp. enterica serovar Agona MNNEETFYQAMRRKGVTRRSFLKFCSLAATSLGLSAGMTPKIAWALENKPRIPVVWIHGL WP_001058316.1 hydrogenase [Salmonella enterica] Length: 372\nNumber of predicted TMHs: 2\nExp number of AAs in TMHs: 35.76397\nExp number, first 60 AAs: 18.97201\nTotal prob of N-in: 0.92785\nPOSSIBLE N-term signal sequence\ninside 1 19\nTMhelix 20 39\noutside 40 325\nTMhelix 326 348\ninside 349 372

22726 GCF_000388495.1_17.H.06 Salmonella enterica subsp. enterica serovar Agona str. 17.H.06 Proteobacteria; Gammaproteobacteria; Enterobacterales; Enterobacteriaceae; Salmonella; Salmonella enterica; Salmonella enterica subsp. enterica; Salmonella enterica subsp. enterica serovar Agona MTGDNTLITSHGINRRDFMKLCAALAATMGLSSKAAAEMAESVSNPQRPPVIWIGAQECT WP_000145429.1 hydrogenase 2 small subunit [Salmonella enterica] Length: 372\nNumber of predicted TMHs: 1\nExp number of AAs in TMHs: 24.51309\nExp number, first 60 AAs: 0.2682\nTotal prob of N-in: 0.14402\noutside 1 331\nTMhelix 332 354\ninside 355 372

22727 GCF_000388495.1_17.H.06 Salmonella enterica subsp. enterica serovar Agona str. 17.H.06 Proteobacteria; Gammaproteobacteria; Enterobacterales; Enterobacteriaceae; Salmonella; Salmonella enterica; Salmonella enterica subsp. enterica; Salmonella enterica subsp. enterica serovar Agona MNNEETFYQAMRRKGVTRRSFLKFCSLAATSLGLSAGMTPKIAWALENKPRIPVVWIHGL WP_001058316.1 hydrogenase [Salmonella enterica] Length: 372\nNumber of predicted TMHs: 2\nExp number of AAs in TMHs: 35.76397\nExp number, first 60 AAs: 18.97201\nTotal prob of N-in: 0.92785\nPOSSIBLE N-term signal sequence\ninside 1 19\nTMhelix 20 39\noutside 40 325\nTMhelix 326 348\ninside 349 372

22728 GCF_000388515.1_18.H.07 Salmonella enterica subsp. enterica serovar Agona str. 18.H.07 Proteobacteria; Gammaproteobacteria; Enterobacterales; Enterobacteriaceae; Salmonella; Salmonella enterica;

Salmonella enterica subsp. enterica; Salmonella enterica subsp. enterica serovar Agona
MNNEETFYQAMRRKGVTRRSFLKFCSLAATSLGLSAGMTPKIAWALENKPRIPVVWIHGL WP_001058316.1
hydrogenase [Salmonella enterica] Length: 372\nNumber of predicted TMHs: 2\nExp number of AAs in TMHs: 35.76397\nExp number, first 60 AAs: 18.97201\nTotal prob of N-in: 0.92785\nPOSSIBLE N-term signal sequence\ninside 1 19\nTMhelix 20 39\noutside 40 325\nTMhelix 326 348\ninside 349 372

22729 GCF_000388515.1_18.H.07 Salmonella enterica subsp. enterica serovar Agona str. 18.H.07
Proteobacteria; Gammaproteobacteria; Enterobacterales; Enterobacteriaceae; Salmonella; Salmonella enterica; Salmonella enterica subsp. enterica; Salmonella enterica subsp. enterica serovar Agona
MTGDNTLITSHGINRRDFMKLCAALAATMGLSSKAAAEMAESVSNPQRPPVIWIGAQECT WP_000145429.1
hydrogenase 2 small subunit [Salmonella enterica] Length: 372\nNumber of predicted TMHs: 1\nExp number of AAs in TMHs: 24.51309\nExp number, first 60 AAs: 0.2682\nTotal prob of N-in: 0.14402\noutside 1 331\nTMhelix 332 354\ninside 355 372

22730 GCF_000388535.1_19.F.03 Salmonella enterica subsp. enterica serovar Agona str. 19.F.03
Proteobacteria; Gammaproteobacteria; Enterobacterales; Enterobacteriaceae; Salmonella; Salmonella enterica; Salmonella enterica subsp. enterica; Salmonella enterica subsp. enterica serovar Agona
MTGDNTLITSHGINRRDFMKLCAALAATMGLSSKAAAEMAESVSNPQRPPVIWIGAQECT WP_000145429.1
hydrogenase 2 small subunit [Salmonella enterica] Length: 372\nNumber of predicted TMHs: 1\nExp number of AAs in TMHs: 24.51309\nExp number, first 60 AAs: 0.2682\nTotal prob of N-in: 0.14402\noutside 1 331\nTMhelix 332 354\ninside 355 372

22731 GCF_000388535.1_19.F.03 Salmonella enterica subsp. enterica serovar Agona str. 19.F.03
Proteobacteria; Gammaproteobacteria; Enterobacterales; Enterobacteriaceae; Salmonella; Salmonella enterica; Salmonella enterica subsp. enterica; Salmonella enterica subsp. enterica serovar Agona
MNNEETFYQAMRRKGVTRRSFLKFCSLAATSLGLSAGMTPKIAWALENKPRIPVVWIHGL WP_001058316.1
hydrogenase [Salmonella enterica] Length: 372\nNumber of predicted TMHs: 2\nExp number of AAs in TMHs: 35.76397\nExp number, first 60 AAs: 18.97201\nTotal prob of N-in: 0.92785\nPOSSIBLE N-term signal sequence\ninside 1 19\nTMhelix 20 39\noutside 40 325\nTMhelix 326 348\ninside 349 372

22732 GCF_000388555.1_20.H.06 Salmonella enterica subsp. enterica serovar Agona str. 20.H.06
Proteobacteria; Gammaproteobacteria; Enterobacterales; Enterobacteriaceae; Salmonella; Salmonella enterica; Salmonella enterica subsp. enterica; Salmonella enterica subsp. enterica serovar Agona
MNNEETFYQAMRRKGVTRRSFLKFCSLAATSLGLSAGMTPKIAWALENKPRIPVVWIHGL WP_001058316.1
hydrogenase [Salmonella enterica] Length: 372\nNumber of predicted TMHs: 2\nExp number of AAs in TMHs: 35.76397\nExp number, first 60 AAs: 18.97201\nTotal prob of N-in: 0.92785\nPOSSIBLE N-term signal sequence\ninside 1 19\nTMhelix 20 39\noutside 40 325\nTMhelix 326 348\ninside 349 372

22733 GCF_000388555.1_20.H.06 Salmonella enterica subsp. enterica serovar Agona str. 20.H.06
Proteobacteria; Gammaproteobacteria; Enterobacterales; Enterobacteriaceae; Salmonella; Salmonella enterica; Salmonella enterica subsp. enterica; Salmonella enterica subsp. enterica serovar Agona
MTGDNTLITSHGINRRDFMKLCAALAATMGLSSKAAAEMAESVSNPQRPPVIWIGAQECT WP_000145429.1
hydrogenase 2 small subunit [Salmonella enterica] Length: 372\nNumber of predicted TMHs: 1\nExp number of AAs in TMHs: 24.51309\nExp number, first 60 AAs: 0.2682\nTotal prob of N-in: 0.14402\noutside 1 331\nTMhelix 332 354\ninside 355 372

22734 GCF_000388575.1_21.H.10 Salmonella enterica subsp. enterica serovar Agona str. 21.H.10
Proteobacteria; Gammaproteobacteria; Enterobacterales; Enterobacteriaceae; Salmonella; Salmonella enterica; Salmonella enterica subsp. enterica; Salmonella enterica subsp. enterica serovar Agona
MTGDNTLITSHGINRRDFMKLCAALAATMGLSSKAAAEMAESVSNPQRPPVIWIGAQECT WP_000145429.1
hydrogenase 2 small subunit [Salmonella enterica] Length: 372\nNumber of predicted TMHs: 1\nExp number of AAs in TMHs: 24.51309\nExp number, first 60 AAs: 0.2682\nTotal prob of N-in: 0.14402\noutside 1 331\nTMhelix 332 354\ninside 355 372

22735 GCF_000388575.1_21.H.10 Salmonella enterica subsp. enterica serovar Agona str. 21.H.10
Proteobacteria; Gammaproteobacteria; Enterobacterales; Enterobacteriaceae; Salmonella; Salmonella enterica; Salmonella enterica subsp. enterica; Salmonella enterica subsp. enterica serovar Agona
MNNEETFYQAMRRKGVTRRSFLKFCSLAATSLGLSAGMTPKIAWALENKPRIPVVWIHGL WP_001058316.1
hydrogenase [Salmonella enterica] Length: 372\nNumber of predicted TMHs: 2\nExp number of AAs in TMHs: 35.76397\nExp number, first 60 AAs: 18.97201\nTotal prob of N-in: 0.92785\nPOSSIBLE N-term signal sequence\ninside 1 19\nTMhelix 20 39\noutside 40 325\nTMhelix 326 348\ninside 349 372

22736 GCF_000388595.1_22.H.04 Salmonella enterica subsp. enterica serovar Agona str. 22.H.04 Proteobacteria; Gammaproteobacteria; Enterobacterales; Enterobacteriaceae; Salmonella; Salmonella enterica; Salmonella enterica subsp. enterica; Salmonella enterica subsp. enterica serovar Agona MTGDNTLITSHGINRRDFMKLCAALAATMGLSSKAAAEMAESVSNPQRPPVIWIGAQECT WP_000145429.1 hydrogenase 2 small subunit [Salmonella enterica] Length: 372\nNumber of predicted TMHs: 1\nExp number of AAs in TMHs: 24.51309\nExp number, first 60 AAs: 0.2682\nTotal prob of N-in: 0.14402\noutside 1 331\nTMhelix 332 354\ninside 355 372

22737 GCF_000388595.1_22.H.04 Salmonella enterica subsp. enterica serovar Agona str. 22.H.04 Proteobacteria; Gammaproteobacteria; Enterobacterales; Enterobacteriaceae; Salmonella; Salmonella enterica; Salmonella enterica subsp. enterica; Salmonella enterica subsp. enterica serovar Agona MNNEETFYQAMRRKGVTRRSFLKFCSLAATSLGLSAGMTPKIAWALENKPRIPVVWIHGL WP_001058316.1 hydrogenase [Salmonella enterica] Length: 372\nNumber of predicted TMHs: 2\nExp number of AAs in TMHs: 35.76397\nExp number, first 60 AAs: 18.97201\nTotal prob of N-in: 0.92785\nPOSSIBLE N-term signal sequence\ninside 1 19\nTMhelix 20 39\noutside 40 325\nTMhelix 326 348\ninside 349 372

22738 GCF_000388615.1_23.F.01 Salmonella enterica subsp. enterica serovar Agona str. 23.F.01 Proteobacteria; Gammaproteobacteria; Enterobacterales; Enterobacteriaceae; Salmonella; Salmonella enterica; Salmonella enterica subsp. enterica; Salmonella enterica subsp. enterica serovar Agona MTGDNTLITSHGINRRDFMKLCAALAATMGLSSKAAAEMAESVSNPQRPPVIWIGAQECT WP_000145429.1 hydrogenase 2 small subunit [Salmonella enterica] Length: 372\nNumber of predicted TMHs: 1\nExp number of AAs in TMHs: 24.51309\nExp number, first 60 AAs: 0.2682\nTotal prob of N-in: 0.14402\noutside 1 331\nTMhelix 332 354\ninside 355 372

22739 GCF_000388615.1_23.F.01 Salmonella enterica subsp. enterica serovar Agona str. 23.F.01 Proteobacteria; Gammaproteobacteria; Enterobacterales; Enterobacteriaceae; Salmonella; Salmonella enterica; Salmonella enterica subsp. enterica; Salmonella enterica subsp. enterica serovar Agona MNNEETFYQAMRRKGVTRRSFLKFCSLAATSLGLSAGMTPKIAWALENKPRIPVVWIHGL WP_001058316.1 hydrogenase [Salmonella enterica] Length: 372\nNumber of predicted TMHs: 2\nExp number of AAs in TMHs: 35.76397\nExp number, first 60 AAs: 18.97201\nTotal prob of N-in: 0.92785\nPOSSIBLE N-term signal sequence\ninside 1 19\nTMhelix 20 39\noutside 40 325\nTMhelix 326 348\ninside 349 372

22740 GCF_000494365.1_CFSAN000479_01.0Salmonella enterica subsp. enterica serovar Agona str. 241981 Proteobacteria; Gammaproteobacteria; Enterobacterales; Enterobacteriaceae; Salmonella; Salmonella enterica; Salmonella enterica subsp. enterica; Salmonella enterica subsp. enterica serovar Agona MNNEETFYQAMRRKGVTRRSFLKFCSLAATSLGLSAGMTPKIAWALENKPRIPVVWIHGL WP_001058316.1 hydrogenase [Salmonella enterica] Length: 372\nNumber of predicted TMHs: 2\nExp number of AAs in TMHs: 35.76397\nExp number, first 60 AAs: 18.97201\nTotal prob of N-in: 0.92785\nPOSSIBLE N-term signal sequence\ninside 1 19\nTMhelix 20 39\noutside 40 325\nTMhelix 326 348\ninside 349 372

22741 GCF_000494365.1_CFSAN000479_01.0Salmonella enterica subsp. enterica serovar Agona str. 241981 Proteobacteria; Gammaproteobacteria; Enterobacterales; Enterobacteriaceae; Salmonella; Salmonella enterica; Salmonella enterica subsp. enterica; Salmonella enterica subsp. enterica serovar Agona MTGDNTLITSHGINRRDFMKLCAALAATMGLSSKAAAEMAESVSNPQRPPVIWIGAQECT WP_000145429.1 hydrogenase 2 small subunit [Salmonella enterica] Length: 372\nNumber of predicted TMHs: 1\nExp number of AAs in TMHs: 24.51309\nExp number, first 60 AAs: 0.2682\nTotal prob of N-in: 0.14402\noutside 1 331\nTMhelix 332 354\ninside 355 372

22742 GCF_000503845.1_ASM50384v1 Salmonella enterica subsp. enterica serovar Agona str. 24249 Proteobacteria; Gammaproteobacteria; Enterobacterales; Enterobacteriaceae; Salmonella; Salmonella enterica; Salmonella enterica subsp. enterica; Salmonella enterica subsp. enterica serovar Agona MTGDNTLITSHGINRRDFMKLCAALAATMGLSSKAAAEMAESVSNPQRPPVIWIGAQECT WP_000145429.1 hydrogenase 2 small subunit [Salmonella enterica] Length: 372\nNumber of predicted TMHs: 1\nExp number of AAs in TMHs: 24.51309\nExp number, first 60 AAs: 0.2682\nTotal prob of N-in: 0.14402\noutside 1 331\nTMhelix 332 354\ninside 355 372

22743 GCF_000503845.1_ASM50384v1 Salmonella enterica subsp. enterica serovar Agona str. 24249 Proteobacteria; Gammaproteobacteria; Enterobacterales; Enterobacteriaceae; Salmonella; Salmonella enterica; Salmonella enterica subsp. enterica; Salmonella enterica subsp. enterica serovar Agona MNNEETFYQAMRRKGVTRRSFLKFCSLAATSLGLSAGMTPKIAWALENKPRIPVVWIHGL WP_001058316.1 hydrogenase [Salmonella enterica] Length: 372\nNumber of predicted TMHs: 2\nExp number of AAs in

TMHs: 35.76397\nExp number, first 60 AAs: 18.97201\nTotal prob of N-in: 0.92785\nPOSSIBLE N-term signal sequence\ninside 1 19\nTMhelix 20 39\noutside 40 325\nTMhelix 326 348\ninside 349 372

22744 GCF_000494445.1_CFSAN000484_01.0 Salmonella enterica subsp. enterica serovar Agona str. 246555-3 Proteobacteria; Gammaproteobacteria; Enterobacterales; Enterobacteriaceae; Salmonella; Salmonella enterica; Salmonella enterica subsp. enterica; Salmonella enterica subsp. enterica serovar Agona MTGDNTLITSHGINRRDFMKLCAALAATMGLSSKAAAEMAESVSNPQRPPVIWIGAQECT WP_000145429.1 hydrogenase 2 small subunit [Salmonella enterica] Length: 372\nNumber of predicted TMHs: 1\nExp number of AAs in TMHs: 24.51309\nExp number, first 60 AAs: 0.2682\nTotal prob of N-in: 0.14402\noutside 1 331\nTMhelix 332 354\ninside 355 372

22745 GCF_000494445.1_CFSAN000484_01.0 Salmonella enterica subsp. enterica serovar Agona str. 246555-3 Proteobacteria; Gammaproteobacteria; Enterobacterales; Enterobacteriaceae; Salmonella; Salmonella enterica; Salmonella enterica subsp. enterica; Salmonella enterica subsp. enterica serovar Agona MNNEETFYQAMRRKGVTRRSFLKFCSLAATSLGLSAGMTPKIAWALENKPRIPVVWIHGL WP_001058316.1 hydrogenase [Salmonella enterica] Length: 372\nNumber of predicted TMHs: 2\nExp number of AAs in TMHs: 35.76397\nExp number, first 60 AAs: 18.97201\nTotal prob of N-in: 0.92785\nPOSSIBLE N-term signal sequence\ninside 1 19\nTMhelix 20 39\noutside 40 325\nTMhelix 326 348\ninside 349 372

22746 GCF_000388635.1_24.H.04 Salmonella enterica subsp. enterica serovar Agona str. 24.H.04 Proteobacteria; Gammaproteobacteria; Enterobacterales; Enterobacteriaceae; Salmonella; Salmonella enterica; Salmonella enterica subsp. enterica; Salmonella enterica subsp. enterica serovar Agona MNNEETFYQAMRRKGVTRRSFLKFCSLAATSLGLSAGMTPKIAWALENKPRIPVVWIHGL WP_001058316.1 hydrogenase [Salmonella enterica] Length: 372\nNumber of predicted TMHs: 2\nExp number of AAs in TMHs: 35.76397\nExp number, first 60 AAs: 18.97201\nTotal prob of N-in: 0.92785\nPOSSIBLE N-term signal sequence\ninside 1 19\nTMhelix 20 39\noutside 40 325\nTMhelix 326 348\ninside 349 372

22747 GCF_000388635.1_24.H.04 Salmonella enterica subsp. enterica serovar Agona str. 24.H.04 Proteobacteria; Gammaproteobacteria; Enterobacterales; Enterobacteriaceae; Salmonella; Salmonella enterica; Salmonella enterica subsp. enterica; Salmonella enterica subsp. enterica serovar Agona MTGDNTLITSHGINRRDFMKLCAALAATMGLSSKAAAEMAESVSNPQRPPVIWIGAQECT WP_000145429.1 hydrogenase 2 small subunit [Salmonella enterica] Length: 372\nNumber of predicted TMHs: 1\nExp number of AAs in TMHs: 24.51309\nExp number, first 60 AAs: 0.2682\nTotal prob of N-in: 0.14402\noutside 1 331\nTMhelix 332 354\ninside 355 372

22748 GCF_000388655.1_26.F.98 Salmonella enterica subsp. enterica serovar Agona str. 26.F.98 Proteobacteria; Gammaproteobacteria; Enterobacterales; Enterobacteriaceae; Salmonella; Salmonella enterica; Salmonella enterica subsp. enterica; Salmonella enterica subsp. enterica serovar Agona MNNEETFYQAMRRKGVTRRSFLKFCSLAATSLGLSAGMTPKIAWALENKPRIPVVWIHGL WP_001058316.1 hydrogenase [Salmonella enterica] Length: 372\nNumber of predicted TMHs: 2\nExp number of AAs in TMHs: 35.76397\nExp number, first 60 AAs: 18.97201\nTotal prob of N-in: 0.92785\nPOSSIBLE N-term signal sequence\ninside 1 19\nTMhelix 20 39\noutside 40 325\nTMhelix 326 348\ninside 349 372

22749 GCF_000388655.1_26.F.98 Salmonella enterica subsp. enterica serovar Agona str. 26.F.98 Proteobacteria; Gammaproteobacteria; Enterobacterales; Enterobacteriaceae; Salmonella; Salmonella enterica; Salmonella enterica subsp. enterica; Salmonella enterica subsp. enterica serovar Agona MTGDNTLITSHGINRRDFMKLCAALAATMGLSSKAAAEMAESVSNPQRPPVIWIGAQECT WP_000145429.1 hydrogenase 2 small subunit [Salmonella enterica] Length: 372\nNumber of predicted TMHs: 1\nExp number of AAs in TMHs: 24.51309\nExp number, first 60 AAs: 0.2682\nTotal prob of N-in: 0.14402\noutside 1 331\nTMhelix 332 354\ninside 355 372

22750 GCF_000388675.1_27.O.98 Salmonella enterica subsp. enterica serovar Agona str. 27.O.98 Proteobacteria; Gammaproteobacteria; Enterobacterales; Enterobacteriaceae; Salmonella; Salmonella enterica; Salmonella enterica subsp. enterica; Salmonella enterica subsp. enterica serovar Agona MNNEETFYQAMRRKGVTRRSFLKFCSLAATSLGLSAGMTPKIAWALENKPRIPVVWIHGL WP_001058316.1 hydrogenase [Salmonella enterica] Length: 372\nNumber of predicted TMHs: 2\nExp number of AAs in TMHs: 35.76397\nExp number, first 60 AAs: 18.97201\nTotal prob of N-in: 0.92785\nPOSSIBLE N-term signal sequence\ninside 1 19\nTMhelix 20 39\noutside 40 325\nTMhelix 326 348\ninside 349 372

22751 GCF_000388675.1_27.O.98 Salmonella enterica subsp. enterica serovar Agona str. 27.O.98 Proteobacteria; Gammaproteobacteria; Enterobacterales; Enterobacteriaceae; Salmonella; Salmonella enterica;

Salmonella enterica subsp. enterica; Salmonella enterica subsp. enterica serovar Agona
MTGDNTLITSHGINRRDFMKLCAALAATMGLSSKAAAEMAESVSNPQRPPVIWIGAQECT WP_000145429.1
hydrogenase 2 small subunit [Salmonella enterica] Length: 372\nNumber of predicted TMHs: 1\nExp number of AAs in TMHs: 24.51309\nExp number, first 60 AAs: 0.2682\nTotal prob of N-in: 0.14402\noutside 1 331\nTMhelix 332 354\ninside 355 372

22752 GCF_000388695.1_28.O.08 Salmonella enterica subsp. enterica serovar Agona str. 28.O.08
Proteobacteria; Gammaproteobacteria; Enterobacterales; Enterobacteriaceae; Salmonella; Salmonella enterica; Salmonella enterica subsp. enterica; Salmonella enterica subsp. enterica serovar Agona
MTGDNTLITSHGINRRDFMKLCAALAATMGLSSKAAAEMAESVSNPQRPPVIWIGAQECT WP_000145429.1
hydrogenase 2 small subunit [Salmonella enterica] Length: 372\nNumber of predicted TMHs: 1\nExp number of AAs in TMHs: 24.51309\nExp number, first 60 AAs: 0.2682\nTotal prob of N-in: 0.14402\noutside 1 331\nTMhelix 332 354\ninside 355 372

22753 GCF_000388715.1_29.O.08 Salmonella enterica subsp. enterica serovar Agona str. 29.O.08
Proteobacteria; Gammaproteobacteria; Enterobacterales; Enterobacteriaceae; Salmonella; Salmonella enterica; Salmonella enterica subsp. enterica; Salmonella enterica subsp. enterica serovar Agona
MTGDNTLITSHGINRRDFMKLCAALAATMGLSSKAAAEMAESVSNPQRPPVIWIGAQECT WP_000145429.1
hydrogenase 2 small subunit [Salmonella enterica] Length: 372\nNumber of predicted TMHs: 1\nExp number of AAs in TMHs: 24.51309\nExp number, first 60 AAs: 0.2682\nTotal prob of N-in: 0.14402\noutside 1 331\nTMhelix 332 354\ninside 355 372

22754 GCF_000388735.1_30.H.04 Salmonella enterica subsp. enterica serovar Agona str. 30.H.04
Proteobacteria; Gammaproteobacteria; Enterobacterales; Enterobacteriaceae; Salmonella; Salmonella enterica; Salmonella enterica subsp. enterica; Salmonella enterica subsp. enterica serovar Agona
MTGDNTLITSHGINRRDFMKLCAALAATMGLSSKAAAEMAESVSNPQRPPVIWIGAQECT WP_000145429.1
hydrogenase 2 small subunit [Salmonella enterica] Length: 372\nNumber of predicted TMHs: 1\nExp number of AAs in TMHs: 24.51309\nExp number, first 60 AAs: 0.2682\nTotal prob of N-in: 0.14402\noutside 1 331\nTMhelix 332 354\ninside 355 372

22755 GCF_000388735.1_30.H.04 Salmonella enterica subsp. enterica serovar Agona str. 30.H.04
Proteobacteria; Gammaproteobacteria; Enterobacterales; Enterobacteriaceae; Salmonella; Salmonella enterica; Salmonella enterica subsp. enterica; Salmonella enterica subsp. enterica serovar Agona
MNNEETFYQAMRRKGVTRRSFLKFCSLAATSLGLSAGMTPKIAWALENKPRIPVVWIHGL WP_001058316.1
hydrogenase [Salmonella enterica] Length: 372\nNumber of predicted TMHs: 2\nExp number of AAs in TMHs: 35.76397\nExp number, first 60 AAs: 18.97201\nTotal prob of N-in: 0.92785\nPOSSIBLE N-term signal sequence\ninside 1 19\nTMhelix 20 39\noutside 40 325\nTMhelix 326 348\ninside 349 372

22756 GCF_000484255.1_ASM48425v1 Salmonella enterica subsp. enterica serovar Agona str. 311387-1
Proteobacteria; Gammaproteobacteria; Enterobacterales; Enterobacteriaceae; Salmonella; Salmonella enterica; Salmonella enterica subsp. enterica; Salmonella enterica subsp. enterica serovar Agona
MTGDNTLITSHGINRRDFMKLCAALAATMGLSSKAAAEMAESVSNPQRPPVIWIGAQECT WP_000145429.1
hydrogenase 2 small subunit [Salmonella enterica] Length: 372\nNumber of predicted TMHs: 1\nExp number of AAs in TMHs: 24.51309\nExp number, first 60 AAs: 0.2682\nTotal prob of N-in: 0.14402\noutside 1 331\nTMhelix 332 354\ninside 355 372

22757 GCF_000484255.1_ASM48425v1 Salmonella enterica subsp. enterica serovar Agona str. 311387-1
Proteobacteria; Gammaproteobacteria; Enterobacterales; Enterobacteriaceae; Salmonella; Salmonella enterica; Salmonella enterica subsp. enterica; Salmonella enterica subsp. enterica serovar Agona
MNNEETFYQAMRRKGVTRRSFLKFCSLAATSLGLSAGMTPKIAWALENKPRIPVVWIHGL WP_001058316.1
hydrogenase [Salmonella enterica] Length: 372\nNumber of predicted TMHs: 2\nExp number of AAs in TMHs: 35.76397\nExp number, first 60 AAs: 18.97201\nTotal prob of N-in: 0.92785\nPOSSIBLE N-term signal sequence\ninside 1 19\nTMhelix 20 39\noutside 40 325\nTMhelix 326 348\ninside 349 372

22758 GCF_000388755.1_31.H.09 Salmonella enterica subsp. enterica serovar Agona str. 31.H.09
Proteobacteria; Gammaproteobacteria; Enterobacterales; Enterobacteriaceae; Salmonella; Salmonella enterica; Salmonella enterica subsp. enterica; Salmonella enterica subsp. enterica serovar Agona
MNNEETFYQAMRRKGVTRRSFLKFCSLAATSLGLSAGMTPKIAWALENKPRIPVVWIHGL WP_001058316.1
hydrogenase [Salmonella enterica] Length: 372\nNumber of predicted TMHs: 2\nExp number of AAs in TMHs: 35.76397\nExp number, first 60 AAs: 18.97201\nTotal prob of N-in: 0.92785\nPOSSIBLE N-term signal sequence\ninside 1 19\nTMhelix 20 39\noutside 40 325\nTMhelix 326 348\ninside 349 372

22759 GCF_000388755.1_31.H.09 Salmonella enterica subsp. enterica serovar Agona str. 31.H.09
Proteobacteria; Gammaproteobacteria; Enterobacterales; Enterobacteriaceae; Salmonella; Salmonella enterica; Salmonella enterica subsp. enterica; Salmonella enterica subsp. enterica serovar Agona
MTGDNTLITSHGINRRDFMKLCAALAATMGLSSKAAAEMAESVSNPQRPPVIWIGAQECT WP_000145429.1
hydrogenase 2 small subunit [Salmonella enterica] Length: 372\nNumber of predicted TMHs: 1\nExp number of AAs in TMHs: 24.51309\nExp number, first 60 AAs: 0.2682\nTotal prob of N-in: 0.14402\noutside 1 331\nTMhelix 332 354\ninside 355 372

22760 GCF_000388775.1_32.A.00 Salmonella enterica subsp. enterica serovar Agona str. 32.A.00
Proteobacteria; Gammaproteobacteria; Enterobacterales; Enterobacteriaceae; Salmonella; Salmonella enterica; Salmonella enterica subsp. enterica; Salmonella enterica subsp. enterica serovar Agona
MTGDNTLITSHGINRRDFMKLCAALAATMGLSSKAAAEMAESVSNPQRPPVIWIGAQECT WP_000145429.1
hydrogenase 2 small subunit [Salmonella enterica] Length: 372\nNumber of predicted TMHs: 1\nExp number of AAs in TMHs: 24.51309\nExp number, first 60 AAs: 0.2682\nTotal prob of N-in: 0.14402\noutside 1 331\nTMhelix 332 354\ninside 355 372

22761 GCF_000484135.1_ASM48413v1 Salmonella enterica subsp. enterica serovar Agona str. 339787
Proteobacteria; Gammaproteobacteria; Enterobacterales; Enterobacteriaceae; Salmonella; Salmonella enterica; Salmonella enterica subsp. enterica; Salmonella enterica subsp. enterica serovar Agona
MNNEETFYQAMRRKGVTRRSFLKFCSLAATSLGLSAGMTPKIAWALENKPRIPVVWIHGL WP_001058316.1
hydrogenase [Salmonella enterica] Length: 372\nNumber of predicted TMHs: 2\nExp number of AAs in TMHs: 35.76397\nExp number, first 60 AAs: 18.97201\nTotal prob of N-in: 0.92785\nPOSSIBLE N-term signal sequence\ninside 1 19\nTMhelix 20 39\noutside 40 325\nTMhelix 326 348\ninside 349 372

22762 GCF_000484135.1_ASM48413v1 Salmonella enterica subsp. enterica serovar Agona str. 339787
Proteobacteria; Gammaproteobacteria; Enterobacterales; Enterobacteriaceae; Salmonella; Salmonella enterica; Salmonella enterica subsp. enterica; Salmonella enterica subsp. enterica serovar Agona
MTGDNTLITSHGINRRDFMKLCAALAATMGLSSKAAAEMAESVSNPQRPPVIWIGAQECT WP_000145429.1
hydrogenase 2 small subunit [Salmonella enterica] Length: 372\nNumber of predicted TMHs: 1\nExp number of AAs in TMHs: 24.51309\nExp number, first 60 AAs: 0.2682\nTotal prob of N-in: 0.14402\noutside 1 331\nTMhelix 332 354\ninside 355 372

22763 GCF_000388795.1_33.A.05 Salmonella enterica subsp. enterica serovar Agona str. 33.A.05
Proteobacteria; Gammaproteobacteria; Enterobacterales; Enterobacteriaceae; Salmonella; Salmonella enterica; Salmonella enterica subsp. enterica; Salmonella enterica subsp. enterica serovar Agona
MTGDNTLITSHGINRRDFMKLCAALAATMGLSSKAAAEMAESVSNPQRPPVIWIGAQECT WP_000145429.1
hydrogenase 2 small subunit [Salmonella enterica] Length: 372\nNumber of predicted TMHs: 1\nExp number of AAs in TMHs: 24.51309\nExp number, first 60 AAs: 0.2682\nTotal prob of N-in: 0.14402\noutside 1 331\nTMhelix 332 354\ninside 355 372

22764 GCF_000388795.1_33.A.05 Salmonella enterica subsp. enterica serovar Agona str. 33.A.05
Proteobacteria; Gammaproteobacteria; Enterobacterales; Enterobacteriaceae; Salmonella; Salmonella enterica; Salmonella enterica subsp. enterica; Salmonella enterica subsp. enterica serovar Agona
MNNEETFYQAMRRKGVTRRSFLKFCSLAATSLGLSAGMTPKIAWALENKPRIPVVWIHGL WP_001058316.1
hydrogenase [Salmonella enterica] Length: 372\nNumber of predicted TMHs: 2\nExp number of AAs in TMHs: 35.76397\nExp number, first 60 AAs: 18.97201\nTotal prob of N-in: 0.92785\nPOSSIBLE N-term signal sequence\ninside 1 19\nTMhelix 20 39\noutside 40 325\nTMhelix 326 348\ninside 349 372

22765 GCF_000388815.1_34.H.09 Salmonella enterica subsp. enterica serovar Agona str. 34.H.09
Proteobacteria; Gammaproteobacteria; Enterobacterales; Enterobacteriaceae; Salmonella; Salmonella enterica; Salmonella enterica subsp. enterica; Salmonella enterica subsp. enterica serovar Agona
MTGDNTLITSHGINRRDFMKLCAALAATMGLSSKAAAEMAESVSNPQRPPVIWIGAQECT WP_000145429.1
hydrogenase 2 small subunit [Salmonella enterica] Length: 372\nNumber of predicted TMHs: 1\nExp number of AAs in TMHs: 24.51309\nExp number, first 60 AAs: 0.2682\nTotal prob of N-in: 0.14402\noutside 1 331\nTMhelix 332 354\ninside 355 372

22766 GCF_000388815.1_34.H.09 Salmonella enterica subsp. enterica serovar Agona str. 34.H.09
Proteobacteria; Gammaproteobacteria; Enterobacterales; Enterobacteriaceae; Salmonella; Salmonella enterica; Salmonella enterica subsp. enterica; Salmonella enterica subsp. enterica serovar Agona
MNNEETFYQAMRRKGVTRRSFLKFCSLAATSLGLSAGMTPKIAWALENKPRIPVVWIHGL WP_001058316.1
hydrogenase [Salmonella enterica] Length: 372\nNumber of predicted TMHs: 2\nExp number of AAs in

TMHs: 35.76397\nExp number, first 60 AAs: 18.97201\nTotal prob of N-in: 0.92785\nPOSSIBLE N-term signal sequence\ninside 1 19\nTMhelix 20 39\noutside 40 325\nTMhelix 326 348\ninside 349 372

22767 GCF_000388835.1_35.H.08 Salmonella enterica subsp. enterica serovar Agona str. 35.H.08 Proteobacteria; Gammaproteobacteria; Enterobacterales; Enterobacteriaceae; Salmonella; Salmonella enterica; Salmonella enterica subsp. enterica; Salmonella enterica subsp. enterica serovar Agona MTGDNTLITSHGINRRDFMKLCAALAATMGLSSKAAAEMAESVSNPQRPPVIWIGAQECT WP_000145429.1 hydrogenase 2 small subunit [Salmonella enterica] Length: 372\nNumber of predicted TMHs: 1\nExp number of AAs in TMHs: 24.51309\nExp number, first 60 AAs: 0.2682\nTotal prob of N-in: 0.14402\noutside 1 331\nTMhelix 332 354\ninside 355 372

22768 GCF_000388835.1_35.H.08 Salmonella enterica subsp. enterica serovar Agona str. 35.H.08 Proteobacteria; Gammaproteobacteria; Enterobacterales; Enterobacteriaceae; Salmonella; Salmonella enterica; Salmonella enterica subsp. enterica; Salmonella enterica subsp. enterica serovar Agona MNNEETFYQAMRRKGVTRRSFLKFCSLAATSLGLSAGMTPKIAWALENKPRIPVVWIHGL WP_001058316.1 hydrogenase [Salmonella enterica] Length: 372\nNumber of predicted TMHs: 2\nExp number of AAs in TMHs: 35.76397\nExp number, first 60 AAs: 18.97201\nTotal prob of N-in: 0.92785\nPOSSIBLE N-term signal sequence\ninside 1 19\nTMhelix 20 39\noutside 40 325\nTMhelix 326 348\ninside 349 372

22769 GCF_000388855.1_36.H.00 Salmonella enterica subsp. enterica serovar Agona str. 36.H.00 Proteobacteria; Gammaproteobacteria; Enterobacterales; Enterobacteriaceae; Salmonella; Salmonella enterica; Salmonella enterica subsp. enterica; Salmonella enterica subsp. enterica serovar Agona MTGDNTLITSHGINRRDFMKLCAALAATMGLSSKAAAEMAESVSNPQRPPVIWIGAQECT WP_000145429.1 hydrogenase 2 small subunit [Salmonella enterica] Length: 372\nNumber of predicted TMHs: 1\nExp number of AAs in TMHs: 24.51309\nExp number, first 60 AAs: 0.2682\nTotal prob of N-in: 0.14402\noutside 1 331\nTMhelix 332 354\ninside 355 372

22770 GCF_000388875.1_37.F.02 Salmonella enterica subsp. enterica serovar Agona str. 37.F.02 Proteobacteria; Gammaproteobacteria; Enterobacterales; Enterobacteriaceae; Salmonella; Salmonella enterica; Salmonella enterica subsp. enterica; Salmonella enterica subsp. enterica serovar Agona MNNEETFYQAMRRKGVTRRSFLKFCSLAATSLGLSAGMTPKIAWALENKPRIPVVWIHGL WP_001058316.1 hydrogenase [Salmonella enterica] Length: 372\nNumber of predicted TMHs: 2\nExp number of AAs in TMHs: 35.76397\nExp number, first 60 AAs: 18.97201\nTotal prob of N-in: 0.92785\nPOSSIBLE N-term signal sequence\ninside 1 19\nTMhelix 20 39\noutside 40 325\nTMhelix 326 348\ninside 349 372

22771 GCF_000388875.1_37.F.02 Salmonella enterica subsp. enterica serovar Agona str. 37.F.02 Proteobacteria; Gammaproteobacteria; Enterobacterales; Enterobacteriaceae; Salmonella; Salmonella enterica; Salmonella enterica subsp. enterica; Salmonella enterica subsp. enterica serovar Agona MTGDNTLITSHGINRRDFMKLCAALAATMGLSSKAAAEMAESVSNPQRPPVIWIGAQECT WP_000145429.1 hydrogenase 2 small subunit [Salmonella enterica] Length: 372\nNumber of predicted TMHs: 1\nExp number of AAs in TMHs: 24.51309\nExp number, first 60 AAs: 0.2682\nTotal prob of N-in: 0.14402\noutside 1 331\nTMhelix 332 354\ninside 355 372

22772 GCF_000388895.1_38.O.03 Salmonella enterica subsp. enterica serovar Agona str. 38.O.03 Proteobacteria; Gammaproteobacteria; Enterobacterales; Enterobacteriaceae; Salmonella; Salmonella enterica; Salmonella enterica subsp. enterica; Salmonella enterica subsp. enterica serovar Agona MTGDNTLITSHGINRRDFMKLCAALAATMGLSSKAAAEMAESVSNPQRPPVIWIGAQECT WP_000145429.1 hydrogenase 2 small subunit [Salmonella enterica] Length: 372\nNumber of predicted TMHs: 1\nExp number of AAs in TMHs: 24.51309\nExp number, first 60 AAs: 0.2682\nTotal prob of N-in: 0.14402\noutside 1 331\nTMhelix 332 354\ninside 355 372

22773 GCF_000494465.1_CFSAN000470_01.0Salmonella enterica subsp. enterica serovar Agona str. 392869-1 Proteobacteria; Gammaproteobacteria; Enterobacterales; Enterobacteriaceae; Salmonella; Salmonella enterica; Salmonella enterica subsp. enterica; Salmonella enterica subsp. enterica serovar Agona MNNEETFYQAMRRKGVTRRSFLKFCSLAATSLGLSAGMTPKIAWALENKPRIPVVWIHGL WP_001058316.1 hydrogenase [Salmonella enterica] Length: 372\nNumber of predicted TMHs: 2\nExp number of AAs in TMHs: 35.76397\nExp number, first 60 AAs: 18.97201\nTotal prob of N-in: 0.92785\nPOSSIBLE N-term signal sequence\ninside 1 19\nTMhelix 20 39\noutside 40 325\nTMhelix 326 348\ninside 349 372

22774 GCF_000494465.1_CFSAN000470_01.0Salmonella enterica subsp. enterica serovar Agona str. 392869-1 Proteobacteria; Gammaproteobacteria; Enterobacterales; Enterobacteriaceae; Salmonella; Salmonella enterica;

Salmonella enterica subsp. enterica; Salmonella enterica subsp. enterica serovar Agona
MTGDNTLITSHGINRRDFMKLCAALAATMGLSSKAAAEMAESVSNPQRPPVIWIGAQECT WP_000145429.1
hydrogenase 2 small subunit [Salmonella enterica] Length: 372\nNumber of predicted TMHs: 1\nExp number of AAs in TMHs: 24.51309\nExp number, first 60 AAs: 0.2682\nTotal prob of N-in: 0.14402\noutside 1 331\nTMhelix 332 354\ninside 355 372

22775 GCF_000494385.2_ASM49438v2 Salmonella enterica subsp. enterica serovar Agona str. 392869-2
Proteobacteria; Gammaproteobacteria; Enterobacterales; Enterobacteriaceae; Salmonella; Salmonella enterica; Salmonella enterica subsp. enterica; Salmonella enterica subsp. enterica serovar Agona
MNNEETFYQAMRRKGVTRRSFLKFCSLAATSLGLSAGMTPKIAWALENKPRIPVVWIHGL WP_001058316.1
hydrogenase [Salmonella enterica] Length: 372\nNumber of predicted TMHs: 2\nExp number of AAs in TMHs: 35.76397\nExp number, first 60 AAs: 18.97201\nTotal prob of N-in: 0.92785\nPOSSIBLE N-term signal sequence\ninside 1 19\nTMhelix 20 39\noutside 40 325\nTMhelix 326 348\ninside 349 372

22776 GCF_000494385.2_ASM49438v2 Salmonella enterica subsp. enterica serovar Agona str. 392869-2
Proteobacteria; Gammaproteobacteria; Enterobacterales; Enterobacteriaceae; Salmonella; Salmonella enterica; Salmonella enterica subsp. enterica; Salmonella enterica subsp. enterica serovar Agona
MTGDNTLITSHGINRRDFMKLCAALAATMGLSSKAAAEMAESVSNPQRPPVIWIGAQECT WP_000145429.1
hydrogenase 2 small subunit [Salmonella enterica] Length: 372\nNumber of predicted TMHs: 1\nExp number of AAs in TMHs: 24.51309\nExp number, first 60 AAs: 0.2682\nTotal prob of N-in: 0.14402\noutside 1 331\nTMhelix 332 354\ninside 355 372

22777 GCF_000388915.1_39.O.03 Salmonella enterica subsp. enterica serovar Agona str. 39.O.03
Proteobacteria; Gammaproteobacteria; Enterobacterales; Enterobacteriaceae; Salmonella; Salmonella enterica; Salmonella enterica subsp. enterica; Salmonella enterica subsp. enterica serovar Agona
MTGDNTLITSHGINRRDFMKLCAALAATMGLSSKAAAEMAESVSNPQRPPVIWIGAQECT WP_000145429.1
hydrogenase 2 small subunit [Salmonella enterica] Length: 372\nNumber of predicted TMHs: 1\nExp number of AAs in TMHs: 24.51309\nExp number, first 60 AAs: 0.2682\nTotal prob of N-in: 0.14402\noutside 1 331\nTMhelix 332 354\ninside 355 372

22778 GCF_000494425.1_CFSAN000452_01.0Salmonella enterica subsp. enterica serovar Agona str. 400095 11
Proteobacteria; Gammaproteobacteria; Enterobacterales; Enterobacteriaceae; Salmonella; Salmonella enterica; Salmonella enterica subsp. enterica; Salmonella enterica subsp. enterica serovar Agona
MTGDNTLITSHGINRRDFMKLCAALAATMGLSSKAAAEMAESVSNPQRPPVIWIGAQECT WP_000145429.1
hydrogenase 2 small subunit [Salmonella enterica] Length: 372\nNumber of predicted TMHs: 1\nExp number of AAs in TMHs: 24.51309\nExp number, first 60 AAs: 0.2682\nTotal prob of N-in: 0.14402\noutside 1 331\nTMhelix 332 354\ninside 355 372

22779 GCF_000494425.1_CFSAN000452_01.0Salmonella enterica subsp. enterica serovar Agona str. 400095 11
Proteobacteria; Gammaproteobacteria; Enterobacterales; Enterobacteriaceae; Salmonella; Salmonella enterica; Salmonella enterica subsp. enterica; Salmonella enterica subsp. enterica serovar Agona
MNNEETFYQAMRRKGVTRRSFLKFCSLAATSLGLSAGMTPKIAWALENKPRIPVVWIHGL WP_001058316.1
hydrogenase [Salmonella enterica] Length: 372\nNumber of predicted TMHs: 2\nExp number of AAs in TMHs: 35.76397\nExp number, first 60 AAs: 18.97201\nTotal prob of N-in: 0.92785\nPOSSIBLE N-term signal sequence\ninside 1 19\nTMhelix 20 39\noutside 40 325\nTMhelix 326 348\ninside 349 372

22780 GCF_000494325.1_CFSAN000453_01.0Salmonella enterica subsp. enterica serovar Agona str. 400095 13
Proteobacteria; Gammaproteobacteria; Enterobacterales; Enterobacteriaceae; Salmonella; Salmonella enterica; Salmonella enterica subsp. enterica; Salmonella enterica subsp. enterica serovar Agona
MTGDNTLITSHGINRRDFMKLCAALAATMGLSSKAAAEMAESVSNPQRPPVIWIGAQECT WP_000145429.1
hydrogenase 2 small subunit [Salmonella enterica] Length: 372\nNumber of predicted TMHs: 1\nExp number of AAs in TMHs: 24.51309\nExp number, first 60 AAs: 0.2682\nTotal prob of N-in: 0.14402\noutside 1 331\nTMhelix 332 354\ninside 355 372

22781 GCF_000494325.1_CFSAN000453_01.0Salmonella enterica subsp. enterica serovar Agona str. 400095 13
Proteobacteria; Gammaproteobacteria; Enterobacterales; Enterobacteriaceae; Salmonella; Salmonella enterica; Salmonella enterica subsp. enterica; Salmonella enterica subsp. enterica serovar Agona
MNNEETFYQAMRRKGVTRRSFLKFCSLAATSLGLSAGMTPKIAWALENKPRIPVVWIHGL WP_001058316.1
hydrogenase [Salmonella enterica] Length: 372\nNumber of predicted TMHs: 2\nExp number of AAs in TMHs: 35.76397\nExp number, first 60 AAs: 18.97201\nTotal prob of N-in: 0.92785\nPOSSIBLE N-term signal sequence\ninside 1 19\nTMhelix 20 39\noutside 40 325\nTMhelix 326 348\ninside 349 372

22782 GCF_000494505.1_CFSAN000454_01.0Salmonella enterica subsp. enterica serovar Agona str. 400095 14 Proteobacteria; Gammaproteobacteria; Enterobacterales; Enterobacteriaceae; Salmonella; Salmonella enterica; Salmonella enterica subsp. enterica; Salmonella enterica subsp. enterica serovar Agona MTGDNTLITSHGINRRDFMKLCAALAATMGLSSKAAAEMAESVSNPQRPPVIWIGAQECT WP_000145429.1 hydrogenase 2 small subunit [Salmonella enterica] Length: 372\nNumber of predicted TMHs: 1\nExp number of AAs in TMHs: 24.51309\nExp number, first 60 AAs: 0.2682\nTotal prob of N-in: 0.14402\noutside 1 331\nTMhelix 332 354\ninside 355 372

22783 GCF_000494505.1_CFSAN000454_01.0Salmonella enterica subsp. enterica serovar Agona str. 400095 14 Proteobacteria; Gammaproteobacteria; Enterobacterales; Enterobacteriaceae; Salmonella; Salmonella enterica; Salmonella enterica subsp. enterica; Salmonella enterica subsp. enterica serovar Agona MNNEETFYQAMRRKGVTRRSFLKFCSLAATSLGLSAGMTPKIAWALENKPRIPVVWIHGL WP_001058316.1 hydrogenase [Salmonella enterica] Length: 372\nNumber of predicted TMHs: 2\nExp number of AAs in TMHs: 35.76397\nExp number, first 60 AAs: 18.97201\nTotal prob of N-in: 0.92785\nPOSSIBLE N-term signal sequence\ninside 1 19\nTMhelix 20 39\noutside 40 325\nTMhelix 326 348\ninside 349 372

22784 GCF_000494405.1_CFSAN000455_01.0Salmonella enterica subsp. enterica serovar Agona str. 400095 17 Proteobacteria; Gammaproteobacteria; Enterobacterales; Enterobacteriaceae; Salmonella; Salmonella enterica; Salmonella enterica subsp. enterica; Salmonella enterica subsp. enterica serovar Agona MTGDNTLITSHGINRRDFMKLCAALAATMGLSSKAAAEMAESVSNPQRPPVIWIGAQECT WP_000145429.1 hydrogenase 2 small subunit [Salmonella enterica] Length: 372\nNumber of predicted TMHs: 1\nExp number of AAs in TMHs: 24.51309\nExp number, first 60 AAs: 0.2682\nTotal prob of N-in: 0.14402\noutside 1 331\nTMhelix 332 354\ninside 355 372

22785 GCF_000494405.1_CFSAN000455_01.0Salmonella enterica subsp. enterica serovar Agona str. 400095 17 Proteobacteria; Gammaproteobacteria; Enterobacterales; Enterobacteriaceae; Salmonella; Salmonella enterica; Salmonella enterica subsp. enterica; Salmonella enterica subsp. enterica serovar Agona MNNEETFYQAMRRKGVTRRSFLKFCSLAATSLGLSAGMTPKIAWALENKPRIPVVWIHGL WP_001058316.1 hydrogenase [Salmonella enterica] Length: 372\nNumber of predicted TMHs: 2\nExp number of AAs in TMHs: 35.76397\nExp number, first 60 AAs: 18.97201\nTotal prob of N-in: 0.92785\nPOSSIBLE N-term signal sequence\ninside 1 19\nTMhelix 20 39\noutside 40 325\nTMhelix 326 348\ninside 349 372

22786 GCF_001649455.1_ASM164945v1 Salmonella enterica subsp. enterica serovar Agona str. 400100 30-11 Proteobacteria; Gammaproteobacteria; Enterobacterales; Enterobacteriaceae; Salmonella; Salmonella enterica; Salmonella enterica subsp. enterica; Salmonella enterica subsp. enterica serovar Agona MTGDNTLITSHGINRRDFMKLCAALAATMGLSSKAAAEMAESVSNPQRPPVIWIGAQECT WP_000145429.1 hydrogenase 2 small subunit [Salmonella enterica] Length: 372\nNumber of predicted TMHs: 1\nExp number of AAs in TMHs: 24.51309\nExp number, first 60 AAs: 0.2682\nTotal prob of N-in: 0.14402\noutside 1 331\nTMhelix 332 354\ninside 355 372

22787 GCF_001649455.1_ASM164945v1 Salmonella enterica subsp. enterica serovar Agona str. 400100 30-11 Proteobacteria; Gammaproteobacteria; Enterobacterales; Enterobacteriaceae; Salmonella; Salmonella enterica; Salmonella enterica subsp. enterica; Salmonella enterica subsp. enterica serovar Agona MNNEETFYQAMRRKGVTRRSFLKFCSLAATSLGLSAGMTPKIAWALENKPRIPVVWIHGL WP_001058316.1 hydrogenase [Salmonella enterica] Length: 372\nNumber of predicted TMHs: 2\nExp number of AAs in TMHs: 35.76397\nExp number, first 60 AAs: 18.97201\nTotal prob of N-in: 0.92785\nPOSSIBLE N-term signal sequence\ninside 1 19\nTMhelix 20 39\noutside 40 325\nTMhelix 326 348\ninside 349 372

22788 GCF_000484115.1_ASM48411v1 Salmonella enterica subsp. enterica serovar Agona str. 409753-6 Proteobacteria; Gammaproteobacteria; Enterobacterales; Enterobacteriaceae; Salmonella; Salmonella enterica; Salmonella enterica subsp. enterica; Salmonella enterica subsp. enterica serovar Agona MTGDNTLITSHGINRRDFMKLCAALAATMGLSSKAAAEMAESVSNPQRPPVIWIGAQECT WP_000145429.1 hydrogenase 2 small subunit [Salmonella enterica] Length: 372\nNumber of predicted TMHs: 1\nExp number of AAs in TMHs: 24.51309\nExp number, first 60 AAs: 0.2682\nTotal prob of N-in: 0.14402\noutside 1 331\nTMhelix 332 354\ninside 355 372

22789 GCF_000484115.1_ASM48411v1 Salmonella enterica subsp. enterica serovar Agona str. 409753-6 Proteobacteria; Gammaproteobacteria; Enterobacterales; Enterobacteriaceae; Salmonella; Salmonella enterica; Salmonella enterica subsp. enterica; Salmonella enterica subsp. enterica serovar Agona MNNEETFYQAMRRKGVTRRSFLKFCSLAATSLGLSAGMTPKIAWALENKPRIPVVWIHGL WP_001058316.1 hydrogenase [Salmonella enterica] Length: 372\nNumber of predicted TMHs: 2\nExp number of AAs in

TMHs: 35.76397\nExp number, first 60 AAs: 18.97201\nTotal prob of N-in: 0.92785\nPOSSIBLE N-term signal sequence\ninside 1 19\nTMhelix 20 39\noutside 40 325\nTMhelix 326 348\ninside 349 372

22790 GCF_000388955.1_40.E.08 Salmonella enterica subsp. enterica serovar Agona str. 40.E.08 Proteobacteria; Gammaproteobacteria; Enterobacterales; Enterobacteriaceae; Salmonella; Salmonella enterica; Salmonella enterica subsp. enterica; Salmonella enterica subsp. enterica serovar Agona MNNEETFYQAMRRKGVTRRSFLKFCSLAATSLGLSAGMTPKIAWALENKPRIPVVWIHGL WP_001058316.1 hydrogenase [Salmonella enterica] Length: 372\nNumber of predicted TMHs: 2\nExp number of AAs in TMHs: 35.76397\nExp number, first 60 AAs: 18.97201\nTotal prob of N-in: 0.92785\nPOSSIBLE N-term signal sequence\ninside 1 19\nTMhelix 20 39\noutside 40 325\nTMhelix 326 348\ninside 349 372

22791 GCF_000388955.1_40.E.08 Salmonella enterica subsp. enterica serovar Agona str. 40.E.08 Proteobacteria; Gammaproteobacteria; Enterobacterales; Enterobacteriaceae; Salmonella; Salmonella enterica; Salmonella enterica subsp. enterica; Salmonella enterica subsp. enterica serovar Agona MTGDNTLITSHGINRRDFMKLCAALAATMGLSSKAAAEMAESVSNPQRPPVIWIGAQECT WP_000145429.1 hydrogenase 2 small subunit [Salmonella enterica] Length: 372\nNumber of predicted TMHs: 1\nExp number of AAs in TMHs: 24.51309\nExp number, first 60 AAs: 0.2682\nTotal prob of N-in: 0.14402\noutside 1 331\nTMhelix 332 354\ninside 355 372

22792 GCF_000487495.1_ASM48749v1 Salmonella enterica subsp. enterica serovar Agona str. 419639 2-1 Proteobacteria; Gammaproteobacteria; Enterobacterales; Enterobacteriaceae; Salmonella; Salmonella enterica; Salmonella enterica subsp. enterica; Salmonella enterica subsp. enterica serovar Agona MTGDNTLITSHGINRRDFMKLCAALAATMGLSSKAAAEMAESVSNPQRPPVIWIGAQECT WP_000145429.1 hydrogenase 2 small subunit [Salmonella enterica] Length: 372\nNumber of predicted TMHs: 1\nExp number of AAs in TMHs: 24.51309\nExp number, first 60 AAs: 0.2682\nTotal prob of N-in: 0.14402\noutside 1 331\nTMhelix 332 354\ninside 355 372

22793 GCF_000487495.1_ASM48749v1 Salmonella enterica subsp. enterica serovar Agona str. 419639 2-1 Proteobacteria; Gammaproteobacteria; Enterobacterales; Enterobacteriaceae; Salmonella; Salmonella enterica; Salmonella enterica subsp. enterica; Salmonella enterica subsp. enterica serovar Agona MNNEETFYQAMRRKGVTRRSFLKFCSLAATSLGLSAGMTPKIAWALENKPRIPVVWIHGL WP_001058316.1 hydrogenase [Salmonella enterica] Length: 372\nNumber of predicted TMHs: 2\nExp number of AAs in TMHs: 35.76397\nExp number, first 60 AAs: 18.97201\nTotal prob of N-in: 0.92785\nPOSSIBLE N-term signal sequence\ninside 1 19\nTMhelix 20 39\noutside 40 325\nTMhelix 326 348\ninside 349 372

22794 GCF_000388935.1_41.E.09 Salmonella enterica subsp. enterica serovar Agona str. 41.E.09 Proteobacteria; Gammaproteobacteria; Enterobacterales; Enterobacteriaceae; Salmonella; Salmonella enterica; Salmonella enterica subsp. enterica; Salmonella enterica subsp. enterica serovar Agona MTGDNTLITSHGINRRDFMKLCAALAATMGLSSKAAAEMAESVSNPQRPPVIWIGAQECT WP_000145429.1 hydrogenase 2 small subunit [Salmonella enterica] Length: 372\nNumber of predicted TMHs: 1\nExp number of AAs in TMHs: 24.51309\nExp number, first 60 AAs: 0.2682\nTotal prob of N-in: 0.14402\noutside 1 331\nTMhelix 332 354\ninside 355 372

22795 GCF_000388975.1_42.E.09 Salmonella enterica subsp. enterica serovar Agona str. 42.E.09 Proteobacteria; Gammaproteobacteria; Enterobacterales; Enterobacteriaceae; Salmonella; Salmonella enterica; Salmonella enterica subsp. enterica; Salmonella enterica subsp. enterica serovar Agona MTGDNTLITSHGINRRDFMKLCAALAATMGLSSKAAAEMAESVSNPQRPPVIWIGAQECT WP_000145429.1 hydrogenase 2 small subunit [Salmonella enterica] Length: 372\nNumber of predicted TMHs: 1\nExp number of AAs in TMHs: 24.51309\nExp number, first 60 AAs: 0.2682\nTotal prob of N-in: 0.14402\noutside 1 331\nTMhelix 332 354\ninside 355 372

22796 GCF_000484055.1_ASM48405v1 Salmonella enterica subsp. enterica serovar Agona str. 432613 Proteobacteria; Gammaproteobacteria; Enterobacterales; Enterobacteriaceae; Salmonella; Salmonella enterica; Salmonella enterica subsp. enterica; Salmonella enterica subsp. enterica serovar Agona MTGDNTLITSHGINRRDFMKLCAALAATMGLSSKAAAEMAESVSNPQRPPVIWIGAQECT WP_000145429.1 hydrogenase 2 small subunit [Salmonella enterica] Length: 372\nNumber of predicted TMHs: 1\nExp number of AAs in TMHs: 24.51309\nExp number, first 60 AAs: 0.2682\nTotal prob of N-in: 0.14402\noutside 1 331\nTMhelix 332 354\ninside 355 372

22797 GCF_000484055.1_ASM48405v1 Salmonella enterica subsp. enterica serovar Agona str. 432613 Proteobacteria; Gammaproteobacteria; Enterobacterales; Enterobacteriaceae; Salmonella; Salmonella enterica;

Salmonella enterica subsp. enterica; Salmonella enterica subsp. enterica serovar Agona
MNNEETFYQAMRRKGVTRRSFLKFCSLAATSLGLSAGMTPKIAWALENKPRIPVVWIHGL WP_001058316.1
hydrogenase [Salmonella enterica] Length: 372\nNumber of predicted TMHs: 2\nExp number of AAs in TMHs: 35.76397\nExp number, first 60 AAs: 18.97201\nTotal prob of N-in: 0.92785\nPOSSIBLE N-term signal sequence\ninside 1 19\nTMhelix 20 39\noutside 40 325\nTMhelix 326 348\ninside 349 372

22798 GCF_000388995.1_43.E.09 Salmonella enterica subsp. enterica serovar Agona str. 43.E.09
Proteobacteria; Gammaproteobacteria; Enterobacterales; Enterobacteriaceae; Salmonella; Salmonella enterica; Salmonella enterica subsp. enterica; Salmonella enterica subsp. enterica serovar Agona
MTGDNTLITSHGINRRDFMKLCAALAATMGLSSKAAAEMAESVSNPQRPPVIWIGAQECT WP_000145429.1
hydrogenase 2 small subunit [Salmonella enterica] Length: 372\nNumber of predicted TMHs: 1\nExp number of AAs in TMHs: 24.51309\nExp number, first 60 AAs: 0.2682\nTotal prob of N-in: 0.14402\noutside 1 331\nTMhelix 332 354\ninside 355 372

22799 GCF_000388995.1_43.E.09 Salmonella enterica subsp. enterica serovar Agona str. 43.E.09
Proteobacteria; Gammaproteobacteria; Enterobacterales; Enterobacteriaceae; Salmonella; Salmonella enterica; Salmonella enterica subsp. enterica; Salmonella enterica subsp. enterica serovar Agona
MNNEETFYQAMRRKGVTRRSFLKFCSLAATSLGLSAGMTPKIAWALENKPRIPVVWIHGL WP_001058316.1
hydrogenase [Salmonella enterica] Length: 372\nNumber of predicted TMHs: 2\nExp number of AAs in TMHs: 35.76397\nExp number, first 60 AAs: 18.97201\nTotal prob of N-in: 0.92785\nPOSSIBLE N-term signal sequence\ninside 1 19\nTMhelix 20 39\noutside 40 325\nTMhelix 326 348\ninside 349 372

22800 GCF_000483895.1_ASM48389v1 Salmonella enterica subsp. enterica serovar Agona str. 442692 2-4
Proteobacteria; Gammaproteobacteria; Enterobacterales; Enterobacteriaceae; Salmonella; Salmonella enterica; Salmonella enterica subsp. enterica; Salmonella enterica subsp. enterica serovar Agona
MTGDNTLITSHGINRRDFMKLCAALAATMGLSSKAAAEMAESVSNPQRPPVIWIGAQECT WP_000145429.1
hydrogenase 2 small subunit [Salmonella enterica] Length: 372\nNumber of predicted TMHs: 1\nExp number of AAs in TMHs: 24.51309\nExp number, first 60 AAs: 0.2682\nTotal prob of N-in: 0.14402\noutside 1 331\nTMhelix 332 354\ninside 355 372

22801 GCF_000483895.1_ASM48389v1 Salmonella enterica subsp. enterica serovar Agona str. 442692 2-4
Proteobacteria; Gammaproteobacteria; Enterobacterales; Enterobacteriaceae; Salmonella; Salmonella enterica; Salmonella enterica subsp. enterica; Salmonella enterica subsp. enterica serovar Agona
MNNEETFYQAMRRKGVTRRSFLKFCSLAATSLGLSAGMTPKIAWALENKPRIPVVWIHGL WP_001058316.1
hydrogenase [Salmonella enterica] Length: 372\nNumber of predicted TMHs: 2\nExp number of AAs in TMHs: 35.76397\nExp number, first 60 AAs: 18.97201\nTotal prob of N-in: 0.92785\nPOSSIBLE N-term signal sequence\ninside 1 19\nTMhelix 20 39\noutside 40 325\nTMhelix 326 348\ninside 349 372

22802 GCF_000484075.1_ASM48407v1 Salmonella enterica subsp. enterica serovar Agona str. 442692 2-5
Proteobacteria; Gammaproteobacteria; Enterobacterales; Enterobacteriaceae; Salmonella; Salmonella enterica; Salmonella enterica subsp. enterica; Salmonella enterica subsp. enterica serovar Agona
MTGDNTLITSHGINRRDFMKLCAALAATMGLSSKAAAEMAESVSNPQRPPVIWIGAQECT WP_000145429.1
hydrogenase 2 small subunit [Salmonella enterica] Length: 372\nNumber of predicted TMHs: 1\nExp number of AAs in TMHs: 24.51309\nExp number, first 60 AAs: 0.2682\nTotal prob of N-in: 0.14402\noutside 1 331\nTMhelix 332 354\ninside 355 372

22803 GCF_000484075.1_ASM48407v1 Salmonella enterica subsp. enterica serovar Agona str. 442692 2-5
Proteobacteria; Gammaproteobacteria; Enterobacterales; Enterobacteriaceae; Salmonella; Salmonella enterica; Salmonella enterica subsp. enterica; Salmonella enterica subsp. enterica serovar Agona
MNNEETFYQAMRRKGVTRRSFLKFCSLAATSLGLSAGMTPKIAWALENKPRIPVVWIHGL WP_001058316.1
hydrogenase [Salmonella enterica] Length: 372\nNumber of predicted TMHs: 2\nExp number of AAs in TMHs: 35.76397\nExp number, first 60 AAs: 18.97201\nTotal prob of N-in: 0.92785\nPOSSIBLE N-term signal sequence\ninside 1 19\nTMhelix 20 39\noutside 40 325\nTMhelix 326 348\ninside 349 372

22804 GCF_000484095.1_ASM48409v1 Salmonella enterica subsp. enterica serovar Agona str. 447967 1-1
Proteobacteria; Gammaproteobacteria; Enterobacterales; Enterobacteriaceae; Salmonella; Salmonella enterica; Salmonella enterica subsp. enterica; Salmonella enterica subsp. enterica serovar Agona
MTGDNTLITSHGINRRDFMKLCAALAATMGLSSKAAAEMAESVSNPQRPPVIWIGAQECT WP_000145429.1
hydrogenase 2 small subunit [Salmonella enterica] Length: 372\nNumber of predicted TMHs: 1\nExp number of AAs in TMHs: 24.51309\nExp number, first 60 AAs: 0.2682\nTotal prob of N-in: 0.14402\noutside 1 331\nTMhelix 332 354\ninside 355 372

22805 GCF_000484095.1_ASM48409v1 Salmonella enterica subsp. enterica serovar Agona str. 447967 1-1 Proteobacteria; Gammaproteobacteria; Enterobacterales; Enterobacteriaceae; Salmonella; Salmonella enterica; Salmonella enterica subsp. enterica; Salmonella enterica subsp. enterica serovar Agona MNNEETFYQAMRRKGVTRRSFLKFCSLAATSLGLSAGMTPKIAWALENKPRIPVVWIHGL WP_001058316.1 hydrogenase [Salmonella enterica] Length: 372\nNumber of predicted TMHs: 2\nExp number of AAs in TMHs: 35.76397\nExp number, first 60 AAs: 18.97201\nTotal prob of N-in: 0.92785\nPOSSIBLE N-term signal sequence\ninside 1 19\nTMhelix 20 39\noutside 40 325\nTMhelix 326 348\ninside 349 372

22806 GCF_000484155.1_ASM48415v1 Salmonella enterica subsp. enterica serovar Agona str. 447967 2-1 Proteobacteria; Gammaproteobacteria; Enterobacterales; Enterobacteriaceae; Salmonella; Salmonella enterica; Salmonella enterica subsp. enterica; Salmonella enterica subsp. enterica serovar Agona MNNEETFYQAMRRKGVTRRSFLKFCSLAATSLGLSAGMTPKIAWALENKPRIPVVWIHGL WP_001058316.1 hydrogenase [Salmonella enterica] Length: 372\nNumber of predicted TMHs: 2\nExp number of AAs in TMHs: 35.76397\nExp number, first 60 AAs: 18.97201\nTotal prob of N-in: 0.92785\nPOSSIBLE N-term signal sequence\ninside 1 19\nTMhelix 20 39\noutside 40 325\nTMhelix 326 348\ninside 349 372

22807 GCF_000484155.1_ASM48415v1 Salmonella enterica subsp. enterica serovar Agona str. 447967 2-1 Proteobacteria; Gammaproteobacteria; Enterobacterales; Enterobacteriaceae; Salmonella; Salmonella enterica; Salmonella enterica subsp. enterica; Salmonella enterica subsp. enterica serovar Agona MTGDNTLITSHGINRRDFMKLCAALAATMGLSSKAAAEMAESVSNPQRPPVIWIGAQECT WP_000145429.1 hydrogenase 2 small subunit [Salmonella enterica] Length: 372\nNumber of predicted TMHs: 1\nExp number of AAs in TMHs: 24.51309\nExp number, first 60 AAs: 0.2682\nTotal prob of N-in: 0.14402\noutside 1 331\nTMhelix 332 354\ninside 355 372

22808 GCF_000389015.1_44.E.09 Salmonella enterica subsp. enterica serovar Agona str. 44.E.09 Proteobacteria; Gammaproteobacteria; Enterobacterales; Enterobacteriaceae; Salmonella; Salmonella enterica; Salmonella enterica subsp. enterica; Salmonella enterica subsp. enterica serovar Agona MTGDNTLITSHGINRRDFMKLCAALAATMGLSSKAAAEMAESVSNPQRPPVIWIGAQECT WP_000145429.1 hydrogenase 2 small subunit [Salmonella enterica] Length: 372\nNumber of predicted TMHs: 1\nExp number of AAs in TMHs: 24.51309\nExp number, first 60 AAs: 0.2682\nTotal prob of N-in: 0.14402\noutside 1 331\nTMhelix 332 354\ninside 355 372

22809 GCF_000389035.1_45.E.09 Salmonella enterica subsp. enterica serovar Agona str. 45.E.09 Proteobacteria; Gammaproteobacteria; Enterobacterales; Enterobacteriaceae; Salmonella; Salmonella enterica; Salmonella enterica subsp. enterica; Salmonella enterica subsp. enterica serovar Agona MTGDNTLITSHGINRRDFMKLCAALAATMGLSSKAAAEMAESVSNPQRPPVIWIGAQECT WP_000145429.1 hydrogenase 2 small subunit [Salmonella enterica] Length: 372\nNumber of predicted TMHs: 1\nExp number of AAs in TMHs: 24.51309\nExp number, first 60 AAs: 0.2682\nTotal prob of N-in: 0.14402\noutside 1 331\nTMhelix 332 354\ninside 355 372

22810 GCF_000484175.1_ASM48417v1 Salmonella enterica subsp. enterica serovar Agona str. 460004 1-1 Proteobacteria; Gammaproteobacteria; Enterobacterales; Enterobacteriaceae; Salmonella; Salmonella enterica; Salmonella enterica subsp. enterica; Salmonella enterica subsp. enterica serovar Agona MNNEETFYQAMRRKGVTRRSFLKFCSLAATSLGLSAGMTPKIAWALENKPRIPVVWIHGL WP_001058316.1 hydrogenase [Salmonella enterica] Length: 372\nNumber of predicted TMHs: 2\nExp number of AAs in TMHs: 35.76397\nExp number, first 60 AAs: 18.97201\nTotal prob of N-in: 0.92785\nPOSSIBLE N-term signal sequence\ninside 1 19\nTMhelix 20 39\noutside 40 325\nTMhelix 326 348\ninside 349 372

22811 GCF_000484175.1_ASM48417v1 Salmonella enterica subsp. enterica serovar Agona str. 460004 1-1 Proteobacteria; Gammaproteobacteria; Enterobacterales; Enterobacteriaceae; Salmonella; Salmonella enterica; Salmonella enterica subsp. enterica; Salmonella enterica subsp. enterica serovar Agona MTGDNTLITSHGINRRDFMKLCAALAATMGLSSKAAAEMAESVSNPQRPPVIWIGAQECT WP_000145429.1 hydrogenase 2 small subunit [Salmonella enterica] Length: 372\nNumber of predicted TMHs: 1\nExp number of AAs in TMHs: 24.51309\nExp number, first 60 AAs: 0.2682\nTotal prob of N-in: 0.14402\noutside 1 331\nTMhelix 332 354\ninside 355 372

22812 GCF_000484195.2_ASM48419v2 Salmonella enterica subsp. enterica serovar Agona str. 460004 2-1 Proteobacteria; Gammaproteobacteria; Enterobacterales; Enterobacteriaceae; Salmonella; Salmonella enterica; Salmonella enterica subsp. enterica; Salmonella enterica subsp. enterica serovar Agona MNNEETFYQAMRRKGVTRRSFLKFCSLAATSLGLSAGMTPKIAWALENKPRIPVVWIHGL WP_001058316.1 hydrogenase [Salmonella enterica] Length: 372\nNumber of predicted TMHs: 2\nExp number of AAs in

TMHs: 35.76397\nExp number, first 60 AAs: 18.97201\nTotal prob of N-in: 0.92785\nPOSSIBLE N-term signal sequence\ninside 1 19\nTMhelix 20 39\noutside 40 325\nTMhelix 326 348\ninside 349 372

22813 GCF_000484195.2_ASM48419v2 Salmonella enterica subsp. enterica serovar Agona str. 460004 2-1 Proteobacteria; Gammaproteobacteria; Enterobacterales; Enterobacteriaceae; Salmonella; Salmonella enterica; Salmonella enterica subsp. enterica; Salmonella enterica subsp. enterica serovar Agona MTGDNTLITSHGINRRDFMKLCAALAATMGLSSKAAAEMAESVSNPQRPPVIWIGAQECT WP_000145429.1 hydrogenase 2 small subunit [Salmonella enterica] Length: 372\nNumber of predicted TMHs: 1\nExp number of AAs in TMHs: 24.51309\nExp number, first 60 AAs: 0.2682\nTotal prob of N-in: 0.14402\noutside 1 331\nTMhelix 332 354\ninside 355 372

22814 GCF_000484395.1_ASM48439v1 Salmonella enterica subsp. enterica serovar Agona str. 467481 Proteobacteria; Gammaproteobacteria; Enterobacterales; Enterobacteriaceae; Salmonella; Salmonella enterica; Salmonella enterica subsp. enterica; Salmonella enterica subsp. enterica serovar Agona MNNEETFYQAMRRKGVTRRSFLKFCSLAATSLGLSAGMTPKIAWALENKPRIPVVWIHGL WP_001058316.1 hydrogenase [Salmonella enterica] Length: 372\nNumber of predicted TMHs: 2\nExp number of AAs in TMHs: 35.76397\nExp number, first 60 AAs: 18.97201\nTotal prob of N-in: 0.92785\nPOSSIBLE N-term signal sequence\ninside 1 19\nTMhelix 20 39\noutside 40 325\nTMhelix 326 348\ninside 349 372

22815 GCF_000484395.1_ASM48439v1 Salmonella enterica subsp. enterica serovar Agona str. 467481 Proteobacteria; Gammaproteobacteria; Enterobacterales; Enterobacteriaceae; Salmonella; Salmonella enterica; Salmonella enterica subsp. enterica; Salmonella enterica subsp. enterica serovar Agona MTGDNTLITSHGINRRDFMKLCAALAATMGLSSKAAAEMAESVSNPQRPPVIWIGAQECT WP_000145429.1 hydrogenase 2 small subunit [Salmonella enterica] Length: 372\nNumber of predicted TMHs: 1\nExp number of AAs in TMHs: 24.51309\nExp number, first 60 AAs: 0.2682\nTotal prob of N-in: 0.14402\noutside 1 331\nTMhelix 332 354\ninside 355 372

22816 GCF_000389055.1_46.E.09 Salmonella enterica subsp. enterica serovar Agona str. 46.E.09 Proteobacteria; Gammaproteobacteria; Enterobacterales; Enterobacteriaceae; Salmonella; Salmonella enterica; Salmonella enterica subsp. enterica; Salmonella enterica subsp. enterica serovar Agona MTGDNTLITSHGINRRDFMKLCAALAATMGLSSKAAAEMAESVSNPQRPPVIWIGAQECT WP_000145429.1 hydrogenase 2 small subunit [Salmonella enterica] Length: 372\nNumber of predicted TMHs: 1\nExp number of AAs in TMHs: 24.51309\nExp number, first 60 AAs: 0.2682\nTotal prob of N-in: 0.14402\noutside 1 331\nTMhelix 332 354\ninside 355 372

22817 GCF_000604425.1_47.E.09 Salmonella enterica subsp. enterica serovar Agona str. 47.E.09 Proteobacteria; Gammaproteobacteria; Enterobacterales; Enterobacteriaceae; Salmonella; Salmonella enterica; Salmonella enterica subsp. enterica; Salmonella enterica subsp. enterica serovar Agona MTGDNTLITSHGINRRDFMKLCAALAATMGLSSKAAAEMAESVSNPQRPPVIWIGAQECT WP_000145429.1 hydrogenase 2 small subunit [Salmonella enterica] Length: 372\nNumber of predicted TMHs: 1\nExp number of AAs in TMHs: 24.51309\nExp number, first 60 AAs: 0.2682\nTotal prob of N-in: 0.14402\noutside 1 331\nTMhelix 332 354\ninside 355 372

22818 GCF_000389075.1_48.E.08 Salmonella enterica subsp. enterica serovar Agona str. 48.E.08 Proteobacteria; Gammaproteobacteria; Enterobacterales; Enterobacteriaceae; Salmonella; Salmonella enterica; Salmonella enterica subsp. enterica; Salmonella enterica subsp. enterica serovar Agona MNNEETFYQAMRRKGVTRRSFLKFCSLAATSLGLSAGMTPKIAWALENKPRIPVVWIHGL WP_001058316.1 hydrogenase [Salmonella enterica] Length: 372\nNumber of predicted TMHs: 2\nExp number of AAs in TMHs: 35.76397\nExp number, first 60 AAs: 18.97201\nTotal prob of N-in: 0.92785\nPOSSIBLE N-term signal sequence\ninside 1 19\nTMhelix 20 39\noutside 40 325\nTMhelix 326 348\ninside 349 372

22819 GCF_000389075.1_48.E.08 Salmonella enterica subsp. enterica serovar Agona str. 48.E.08 Proteobacteria; Gammaproteobacteria; Enterobacterales; Enterobacteriaceae; Salmonella; Salmonella enterica; Salmonella enterica subsp. enterica; Salmonella enterica subsp. enterica serovar Agona MTGDNTLITSHGINRRDFMKLCAALAATMGLSSKAAAEMAESVSNPQRPPVIWIGAQECT WP_000145429.1 hydrogenase 2 small subunit [Salmonella enterica] Length: 372\nNumber of predicted TMHs: 1\nExp number of AAs in TMHs: 24.51309\nExp number, first 60 AAs: 0.2682\nTotal prob of N-in: 0.14402\noutside 1 331\nTMhelix 332 354\ninside 355 372

22820 GCF_000389095.1_49.E.09 Salmonella enterica subsp. enterica serovar Agona str. 49.E.09 Proteobacteria; Gammaproteobacteria; Enterobacterales; Enterobacteriaceae; Salmonella; Salmonella enterica;

Salmonella enterica subsp. enterica; Salmonella enterica subsp. enterica serovar Agona
MTGDNTLITSHGINRRDFMKLCAALAATMGLSSKAAAEMAESVSNPQRPPVIWIGAQECT WP_000145429.1
hydrogenase 2 small subunit [Salmonella enterica] Length: 372\nNumber of predicted TMHs: 1\nExp number of AAs in TMHs: 24.51309\nExp number, first 60 AAs: 0.2682\nTotal prob of N-in: 0.14402\noutside 1 331\nTMhelix 332 354\ninside 355 372

22821 GCF_000389115.1_50.E.08 Salmonella enterica subsp. enterica serovar Agona str. 50.E.08
Proteobacteria; Gammaproteobacteria; Enterobacterales; Enterobacteriaceae; Salmonella; Salmonella enterica; Salmonella enterica subsp. enterica; Salmonella enterica subsp. enterica serovar Agona
MNNEETFYQAMRRKGVTRRSFLKFCSLAATSLGLSAGMTPKIAWALENKPRIPVVWIHGL WP_001058316.1
hydrogenase [Salmonella enterica] Length: 372\nNumber of predicted TMHs: 2\nExp number of AAs in TMHs: 35.76397\nExp number, first 60 AAs: 18.97201\nTotal prob of N-in: 0.92785\nPOSSIBLE N-term signal sequence\ninside 1 19\nTMhelix 20 39\noutside 40 325\nTMhelix 326 348\ninside 349 372

22822 GCF_000389115.1_50.E.08 Salmonella enterica subsp. enterica serovar Agona str. 50.E.08
Proteobacteria; Gammaproteobacteria; Enterobacterales; Enterobacteriaceae; Salmonella; Salmonella enterica; Salmonella enterica subsp. enterica; Salmonella enterica subsp. enterica serovar Agona
MTGDNTLITSHGINRRDFMKLCAALAATMGLSSKAAAEMAESVSNPQRPPVIWIGAQECT WP_000145429.1
hydrogenase 2 small subunit [Salmonella enterica] Length: 372\nNumber of predicted TMHs: 1\nExp number of AAs in TMHs: 24.51309\nExp number, first 60 AAs: 0.2682\nTotal prob of N-in: 0.14402\noutside 1 331\nTMhelix 332 354\ninside 355 372

22823 GCF_000389135.1_51.E.09 Salmonella enterica subsp. enterica serovar Agona str. 51.E.09
Proteobacteria; Gammaproteobacteria; Enterobacterales; Enterobacteriaceae; Salmonella; Salmonella enterica; Salmonella enterica subsp. enterica; Salmonella enterica subsp. enterica serovar Agona
MTGDNTLITSHGINRRDFMKLCAALAATMGLSSKAAAEMAESVSNPQRPPVIWIGAQECT WP_000145429.1
hydrogenase 2 small subunit [Salmonella enterica] Length: 372\nNumber of predicted TMHs: 1\nExp number of AAs in TMHs: 24.51309\nExp number, first 60 AAs: 0.2682\nTotal prob of N-in: 0.14402\noutside 1 331\nTMhelix 332 354\ninside 355 372

22824 GCF_000389155.1_52.F.08 Salmonella enterica subsp. enterica serovar Agona str. 52.F.08
Proteobacteria; Gammaproteobacteria; Enterobacterales; Enterobacteriaceae; Salmonella; Salmonella enterica; Salmonella enterica subsp. enterica; Salmonella enterica subsp. enterica serovar Agona
MTGDNTLITSHGINRRDFMKLCAALAATMGLSSKAAAEMAESVSNPQRPPVIWIGAQECT WP_000145429.1
hydrogenase 2 small subunit [Salmonella enterica] Length: 372\nNumber of predicted TMHs: 1\nExp number of AAs in TMHs: 24.51309\nExp number, first 60 AAs: 0.2682\nTotal prob of N-in: 0.14402\noutside 1 331\nTMhelix 332 354\ninside 355 372

22825 GCF_000389155.1_52.F.08 Salmonella enterica subsp. enterica serovar Agona str. 52.F.08
Proteobacteria; Gammaproteobacteria; Enterobacterales; Enterobacteriaceae; Salmonella; Salmonella enterica; Salmonella enterica subsp. enterica; Salmonella enterica subsp. enterica serovar Agona
MNNEETFYQAMRRKGVTRRSFLKFCSLAATSLGLSAGMTPKIAWALENKPRIPVVWIHGL WP_001058316.1
hydrogenase [Salmonella enterica] Length: 372\nNumber of predicted TMHs: 2\nExp number of AAs in TMHs: 35.76397\nExp number, first 60 AAs: 18.97201\nTotal prob of N-in: 0.92785\nPOSSIBLE N-term signal sequence\ninside 1 19\nTMhelix 20 39\noutside 40 325\nTMhelix 326 348\ninside 349 372

22826 GCF_000389175.1_53.F.08 Salmonella enterica subsp. enterica serovar Agona str. 53.F.08
Proteobacteria; Gammaproteobacteria; Enterobacterales; Enterobacteriaceae; Salmonella; Salmonella enterica; Salmonella enterica subsp. enterica; Salmonella enterica subsp. enterica serovar Agona
MNNEETFYQAMRRKGVTRRSFLKFCSLAATSLGLSAGMTPKIAWALENKPRIPVVWIHGL WP_001058316.1
hydrogenase [Salmonella enterica] Length: 372\nNumber of predicted TMHs: 2\nExp number of AAs in TMHs: 35.76397\nExp number, first 60 AAs: 18.97201\nTotal prob of N-in: 0.92785\nPOSSIBLE N-term signal sequence\ninside 1 19\nTMhelix 20 39\noutside 40 325\nTMhelix 326 348\ninside 349 372

22827 GCF_000389175.1_53.F.08 Salmonella enterica subsp. enterica serovar Agona str. 53.F.08
Proteobacteria; Gammaproteobacteria; Enterobacterales; Enterobacteriaceae; Salmonella; Salmonella enterica; Salmonella enterica subsp. enterica; Salmonella enterica subsp. enterica serovar Agona
MTGDNTLITSHGINRRDFMKLCAALAATMGLSSKAAAEMAESVSNPQRPPVIWIGAQECT WP_000145429.1
hydrogenase 2 small subunit [Salmonella enterica] Length: 372\nNumber of predicted TMHs: 1\nExp number of AAs in TMHs: 24.51309\nExp number, first 60 AAs: 0.2682\nTotal prob of N-in: 0.14402\noutside 1 331\nTMhelix 332 354\ninside 355 372

22828 GCF_000389195.1_54.O.08 Salmonella enterica subsp. enterica serovar Agona str. 54.O.08 Proteobacteria; Gammaproteobacteria; Enterobacterales; Enterobacteriaceae; Salmonella; Salmonella enterica; Salmonella enterica subsp. enterica; Salmonella enterica subsp. enterica serovar Agona MTGDNTLITSHGINRRDFMKLCAALAATMGLSSKAAAEMAESVSNPQRPPVIWIGAQECT WP_000145429.1 hydrogenase 2 small subunit [Salmonella enterica] Length: 372\nNumber of predicted TMHs: 1\nExp number of AAs in TMHs: 24.51309\nExp number, first 60 AAs: 0.2682\nTotal prob of N-in: 0.14402\noutside 1 331\nTMhelix 332 354\ninside 355 372

22829 GCF_000389195.1_54.O.08 Salmonella enterica subsp. enterica serovar Agona str. 54.O.08 Proteobacteria; Gammaproteobacteria; Enterobacterales; Enterobacteriaceae; Salmonella; Salmonella enterica; Salmonella enterica subsp. enterica; Salmonella enterica subsp. enterica serovar Agona MNNEETFYQAMRRKGVTRRSFLKFCSLAATSLGLSAGMTPKIAWALENKPRIPVVWIHGL WP_001058316.1 hydrogenase [Salmonella enterica] Length: 372\nNumber of predicted TMHs: 2\nExp number of AAs in TMHs: 35.76397\nExp number, first 60 AAs: 18.97201\nTotal prob of N-in: 0.92785\nPOSSIBLE N-term signal sequence\ninside 1 19\nTMhelix 20 39\noutside 40 325\nTMhelix 326 348\ninside 349 372

22830 GCF_000484375.1_ASM48437v1 Salmonella enterica subsp. enterica serovar Agona str. 557928 Proteobacteria; Gammaproteobacteria; Enterobacterales; Enterobacteriaceae; Salmonella; Salmonella enterica; Salmonella enterica subsp. enterica; Salmonella enterica subsp. enterica serovar Agona MTGDNTLITSHGINRRDFMKLCAALAATMGLSSKAAAEMAESVSNPQRPPVIWIGAQECT WP_000145429.1 hydrogenase 2 small subunit [Salmonella enterica] Length: 372\nNumber of predicted TMHs: 1\nExp number of AAs in TMHs: 24.51309\nExp number, first 60 AAs: 0.2682\nTotal prob of N-in: 0.14402\noutside 1 331\nTMhelix 332 354\ninside 355 372

22831 GCF_000484375.1_ASM48437v1 Salmonella enterica subsp. enterica serovar Agona str. 557928 Proteobacteria; Gammaproteobacteria; Enterobacterales; Enterobacteriaceae; Salmonella; Salmonella enterica; Salmonella enterica subsp. enterica; Salmonella enterica subsp. enterica serovar Agona MNNEETFYQAMRRKGVTRRSFLKFCSLAATSLGLSAGMTPKIAWALENKPRIPVVWIHGL WP_001058316.1 hydrogenase [Salmonella enterica] Length: 372\nNumber of predicted TMHs: 2\nExp number of AAs in TMHs: 35.76397\nExp number, first 60 AAs: 18.97201\nTotal prob of N-in: 0.92785\nPOSSIBLE N-term signal sequence\ninside 1 19\nTMhelix 20 39\noutside 40 325\nTMhelix 326 348\ninside 349 372

22832 GCF_000389215.1_55.U.08 Salmonella enterica subsp. enterica serovar Agona str. 55.U.08 Proteobacteria; Gammaproteobacteria; Enterobacterales; Enterobacteriaceae; Salmonella; Salmonella enterica; Salmonella enterica subsp. enterica; Salmonella enterica subsp. enterica serovar Agona MNNEETFYQAMRRKGVTRRSFLKFCSLAATSLGLSAGMTPKIAWALENKPRIPVVWIHGL WP_001058316.1 hydrogenase [Salmonella enterica] Length: 372\nNumber of predicted TMHs: 2\nExp number of AAs in TMHs: 35.76397\nExp number, first 60 AAs: 18.97201\nTotal prob of N-in: 0.92785\nPOSSIBLE N-term signal sequence\ninside 1 19\nTMhelix 20 39\noutside 40 325\nTMhelix 326 348\ninside 349 372

22833 GCF_000389215.1_55.U.08 Salmonella enterica subsp. enterica serovar Agona str. 55.U.08 Proteobacteria; Gammaproteobacteria; Enterobacterales; Enterobacteriaceae; Salmonella; Salmonella enterica; Salmonella enterica subsp. enterica; Salmonella enterica subsp. enterica serovar Agona MTGDNTLITSHGINRRDFMKLCAALAATMGLSSKAAAEMAESVSNPQRPPVIWIGAQECT WP_000145429.1 hydrogenase 2 small subunit [Salmonella enterica] Length: 372\nNumber of predicted TMHs: 1\nExp number of AAs in TMHs: 24.51309\nExp number, first 60 AAs: 0.2682\nTotal prob of N-in: 0.14402\noutside 1 331\nTMhelix 332 354\ninside 355 372

22834 GCF_000389235.1_56.O.08 Salmonella enterica subsp. enterica serovar Agona str. 56.O.08 Proteobacteria; Gammaproteobacteria; Enterobacterales; Enterobacteriaceae; Salmonella; Salmonella enterica; Salmonella enterica subsp. enterica; Salmonella enterica subsp. enterica serovar Agona MTGDNTLITSHGINRRDFMKLCAALAATMGLSSKAAAEMAESVSNPQRPPVIWIGAQECT WP_000145429.1 hydrogenase 2 small subunit [Salmonella enterica] Length: 372\nNumber of predicted TMHs: 1\nExp number of AAs in TMHs: 24.51309\nExp number, first 60 AAs: 0.2682\nTotal prob of N-in: 0.14402\noutside 1 331\nTMhelix 332 354\ninside 355 372

22835 GCF_000389235.1_56.O.08 Salmonella enterica subsp. enterica serovar Agona str. 56.O.08 Proteobacteria; Gammaproteobacteria; Enterobacterales; Enterobacteriaceae; Salmonella; Salmonella enterica; Salmonella enterica subsp. enterica; Salmonella enterica subsp. enterica serovar Agona MNNEETFYQAMRRKGVTRRSFLKFCSLAATSLGLSAGMTPKIAWALENKPRIPVVWIHGL WP_001058316.1 hydrogenase [Salmonella enterica] Length: 372\nNumber of predicted TMHs: 2\nExp number of AAs in

TMHs: 35.76397\nExp number, first 60 AAs: 18.97201\nTotal prob of N-in: 0.92785\nPOSSIBLE N-term signal sequence\ninside 1 19\nTMhelix 20 39\noutside 40 325\nTMhelix 326 348\ninside 349 372

22836 GCF_000389255.1_57.A.08 Salmonella enterica subsp. enterica serovar Agona str. 57.A.08 Proteobacteria; Gammaproteobacteria; Enterobacterales; Enterobacteriaceae; Salmonella; Salmonella enterica; Salmonella enterica subsp. enterica; Salmonella enterica subsp. enterica serovar Agona MNNEETFYQAMRRKGVTRRSFLKFCSLAATSLGLSAGMTPKIAWALENKPRIPVVWIHGL WP_001058316.1 hydrogenase [Salmonella enterica] Length: 372\nNumber of predicted TMHs: 2\nExp number of AAs in TMHs: 35.76397\nExp number, first 60 AAs: 18.97201\nTotal prob of N-in: 0.92785\nPOSSIBLE N-term signal sequence\ninside 1 19\nTMhelix 20 39\noutside 40 325\nTMhelix 326 348\ninside 349 372

22837 GCF_000389255.1_57.A.08 Salmonella enterica subsp. enterica serovar Agona str. 57.A.08 Proteobacteria; Gammaproteobacteria; Enterobacterales; Enterobacteriaceae; Salmonella; Salmonella enterica; Salmonella enterica subsp. enterica; Salmonella enterica subsp. enterica serovar Agona MTGDNTLITSHGINRRDFMKLCAALAATMGLSSKAAAEMAESVSNPQRPPVIWIGAQECT WP_006895223.1 hydrogenase-2 small chain [Salmonella enterica] Length: 372\nNumber of predicted TMHs: 1\nExp number of AAs in TMHs: 23.96244\nExp number, first 60 AAs: 0.26916\nTotal prob of N-in: 0.12388\noutside 1 331\nTMhelix 332 354\ninside 355 372

22838 GCF_000389275.1_58.E.08 Salmonella enterica subsp. enterica serovar Agona str. 58.E.08 Proteobacteria; Gammaproteobacteria; Enterobacterales; Enterobacteriaceae; Salmonella; Salmonella enterica; Salmonella enterica subsp. enterica; Salmonella enterica subsp. enterica serovar Agona MTGDNTLITSHGINRRDFMKLCAALAATMGLSSKAAAEMAESVSNPQRPPVIWIGAQECT WP_000145429.1 hydrogenase 2 small subunit [Salmonella enterica] Length: 372\nNumber of predicted TMHs: 1\nExp number of AAs in TMHs: 24.51309\nExp number, first 60 AAs: 0.2682\nTotal prob of N-in: 0.14402\noutside 1 331\nTMhelix 332 354\ninside 355 372

22839 GCF_000389275.1_58.E.08 Salmonella enterica subsp. enterica serovar Agona str. 58.E.08 Proteobacteria; Gammaproteobacteria; Enterobacterales; Enterobacteriaceae; Salmonella; Salmonella enterica; Salmonella enterica subsp. enterica; Salmonella enterica subsp. enterica serovar Agona MNNEETFYQAMRRKGVTRRSFLKFCSLAATSLGLSAGMTPKIAWALENKPRIPVVWIHGL WP_001058316.1 hydrogenase [Salmonella enterica] Length: 372\nNumber of predicted TMHs: 2\nExp number of AAs in TMHs: 35.76397\nExp number, first 60 AAs: 18.97201\nTotal prob of N-in: 0.92785\nPOSSIBLE N-term signal sequence\ninside 1 19\nTMhelix 20 39\noutside 40 325\nTMhelix 326 348\ninside 349 372

22840 GCF_000389295.1_59.F.08 Salmonella enterica subsp. enterica serovar Agona str. 59.F.08 Proteobacteria; Gammaproteobacteria; Enterobacterales; Enterobacteriaceae; Salmonella; Salmonella enterica; Salmonella enterica subsp. enterica; Salmonella enterica subsp. enterica serovar Agona MTGDNTLITSHGINRRDFMKLCAALAATMGLSSKAAAEMAESVSNPQRPPVIWIGAQECT WP_000145429.1 hydrogenase 2 small subunit [Salmonella enterica] Length: 372\nNumber of predicted TMHs: 1\nExp number of AAs in TMHs: 24.51309\nExp number, first 60 AAs: 0.2682\nTotal prob of N-in: 0.14402\noutside 1 331\nTMhelix 332 354\ninside 355 372

22841 GCF_000389295.1_59.F.08 Salmonella enterica subsp. enterica serovar Agona str. 59.F.08 Proteobacteria; Gammaproteobacteria; Enterobacterales; Enterobacteriaceae; Salmonella; Salmonella enterica; Salmonella enterica subsp. enterica; Salmonella enterica subsp. enterica serovar Agona MNNEETFYQAMRRKGVTRRSFLKFCSLAATSLGLSAGMTPKIAWALENKPRIPVVWIHGL WP_001058316.1 hydrogenase [Salmonella enterica] Length: 372\nNumber of predicted TMHs: 2\nExp number of AAs in TMHs: 35.76397\nExp number, first 60 AAs: 18.97201\nTotal prob of N-in: 0.92785\nPOSSIBLE N-term signal sequence\ninside 1 19\nTMhelix 20 39\noutside 40 325\nTMhelix 326 348\ninside 349 372

22842 GCF_000389315.1_60.O.08 Salmonella enterica subsp. enterica serovar Agona str. 60.O.08 Proteobacteria; Gammaproteobacteria; Enterobacterales; Enterobacteriaceae; Salmonella; Salmonella enterica; Salmonella enterica subsp. enterica; Salmonella enterica subsp. enterica serovar Agona MTGDNTLITSHGINRRDFMKLCAALAATMGLSSKAAAEMAESVSNPQRPPVIWIGAQECT WP_000145429.1 hydrogenase 2 small subunit [Salmonella enterica] Length: 372\nNumber of predicted TMHs: 1\nExp number of AAs in TMHs: 24.51309\nExp number, first 60 AAs: 0.2682\nTotal prob of N-in: 0.14402\noutside 1 331\nTMhelix 332 354\ninside 355 372

22843 GCF_000389315.1_60.O.08 Salmonella enterica subsp. enterica serovar Agona str. 60.O.08 Proteobacteria; Gammaproteobacteria; Enterobacterales; Enterobacteriaceae; Salmonella; Salmonella enterica;

Salmonella enterica subsp. enterica; Salmonella enterica subsp. enterica serovar Agona
MNNEETFYQAMRRKGVTRRSFLKFCSLAATSLGLSAGMTPKIAWALENKPRIPVVWIHGL WP_001058316.1
hydrogenase [Salmonella enterica] Length: 372\nNumber of predicted TMHs: 2\nExp number of AAs in TMHs: 35.76397\nExp number, first 60 AAs: 18.97201\nTotal prob of N-in: 0.92785\nPOSSIBLE N-term signal sequence\ninside 1 19\nTMhelix 20 39\noutside 40 325\nTMhelix 326 348\ninside 349 372

22844 GCF_000389335.1_61.O.08 Salmonella enterica subsp. enterica serovar Agona str. 61.O.08
Proteobacteria; Gammaproteobacteria; Enterobacterales; Enterobacteriaceae; Salmonella; Salmonella enterica; Salmonella enterica subsp. enterica; Salmonella enterica subsp. enterica serovar Agona
MTGDNTLITSHGINRRDFMKLCAALAATMGLSSKAAAEMAESVSNPQRPPVIWIGAQECT WP_000145429.1
hydrogenase 2 small subunit [Salmonella enterica] Length: 372\nNumber of predicted TMHs: 1\nExp number of AAs in TMHs: 24.51309\nExp number, first 60 AAs: 0.2682\nTotal prob of N-in: 0.14402\noutside 1 331\nTMhelix 332 354\ninside 355 372

22845 GCF_000484415.1_ASM48441v1 Salmonella enterica subsp. enterica serovar Agona str. 620239
Proteobacteria; Gammaproteobacteria; Enterobacterales; Enterobacteriaceae; Salmonella; Salmonella enterica; Salmonella enterica subsp. enterica; Salmonella enterica subsp. enterica serovar Agona
MTGDNTLITSHGINRRDFMKLCAALAATMGLSSKAAAEMAESVSNPQRPPVIWIGAQECT WP_000145429.1
hydrogenase 2 small subunit [Salmonella enterica] Length: 372\nNumber of predicted TMHs: 1\nExp number of AAs in TMHs: 24.51309\nExp number, first 60 AAs: 0.2682\nTotal prob of N-in: 0.14402\noutside 1 331\nTMhelix 332 354\ninside 355 372

22846 GCF_000484415.1_ASM48441v1 Salmonella enterica subsp. enterica serovar Agona str. 620239
Proteobacteria; Gammaproteobacteria; Enterobacterales; Enterobacteriaceae; Salmonella; Salmonella enterica; Salmonella enterica subsp. enterica; Salmonella enterica subsp. enterica serovar Agona
MNNEETFYQAMRRKGVTRRSFLKFCSLAATSLGLSAGMTPKIAWALENKPRIPVVWIHGL WP_001058316.1
hydrogenase [Salmonella enterica] Length: 372\nNumber of predicted TMHs: 2\nExp number of AAs in TMHs: 35.76397\nExp number, first 60 AAs: 18.97201\nTotal prob of N-in: 0.92785\nPOSSIBLE N-term signal sequence\ninside 1 19\nTMhelix 20 39\noutside 40 325\nTMhelix 326 348\ninside 349 372

22847 GCF_000389355.1_62.H.72 Salmonella enterica subsp. enterica serovar Agona str. 62.H.72
Proteobacteria; Gammaproteobacteria; Enterobacterales; Enterobacteriaceae; Salmonella; Salmonella enterica; Salmonella enterica subsp. enterica; Salmonella enterica subsp. enterica serovar Agona
MNNEETFYQAMRRKGVTRRSFLKFCSLAATSLGLSAGMTPKIAWALENKPRIPVVWIHGL WP_001058316.1
hydrogenase [Salmonella enterica] Length: 372\nNumber of predicted TMHs: 2\nExp number of AAs in TMHs: 35.76397\nExp number, first 60 AAs: 18.97201\nTotal prob of N-in: 0.92785\nPOSSIBLE N-term signal sequence\ninside 1 19\nTMhelix 20 39\noutside 40 325\nTMhelix 326 348\ninside 349 372

22848 GCF_000389355.1_62.H.72 Salmonella enterica subsp. enterica serovar Agona str. 62.H.72
Proteobacteria; Gammaproteobacteria; Enterobacterales; Enterobacteriaceae; Salmonella; Salmonella enterica; Salmonella enterica subsp. enterica; Salmonella enterica subsp. enterica serovar Agona
MTGDNTLITSHGINRRDFMKLCAALAATMGLSSKAAAEMAESVSNPQRPPVIWIGAQECT WP_000145429.1
hydrogenase 2 small subunit [Salmonella enterica] Length: 372\nNumber of predicted TMHs: 1\nExp number of AAs in TMHs: 24.51309\nExp number, first 60 AAs: 0.2682\nTotal prob of N-in: 0.14402\noutside 1 331\nTMhelix 332 354\ninside 355 372

22849 GCF_000487555.1_ASM48755v1 Salmonella enterica subsp. enterica serovar Agona str. 632182-2
Proteobacteria; Gammaproteobacteria; Enterobacterales; Enterobacteriaceae; Salmonella; Salmonella enterica; Salmonella enterica subsp. enterica; Salmonella enterica subsp. enterica serovar Agona
MTGDNTLITSHGINRRDFMKLCAALAATMGLSSKAAAEMAESVSNPQRPPVIWIGAQECT WP_000145429.1
hydrogenase 2 small subunit [Salmonella enterica] Length: 372\nNumber of predicted TMHs: 1\nExp number of AAs in TMHs: 24.51309\nExp number, first 60 AAs: 0.2682\nTotal prob of N-in: 0.14402\noutside 1 331\nTMhelix 332 354\ninside 355 372

22850 GCF_000487555.1_ASM48755v1 Salmonella enterica subsp. enterica serovar Agona str. 632182-2
Proteobacteria; Gammaproteobacteria; Enterobacterales; Enterobacteriaceae; Salmonella; Salmonella enterica; Salmonella enterica subsp. enterica; Salmonella enterica subsp. enterica serovar Agona
MNNEETFYQAMRRKGVTRRSFLKFCSLAATSLGLGAGMTPKIAWALENKPRIPVVWIHGL WP_001058311.1
hydrogenase [Salmonella enterica] Length: 372\nNumber of predicted TMHs: 2\nExp number of AAs in TMHs: 37.4699899999999\nExp number, first 60 AAs: 19.72699\nTotal prob of N-in: 0.96171\nPOSSIBLE N-term signal sequence\ninside 1 19\nTMhelix 20 39\noutside 40 325\nTMhelix 326 348\ninside 349 372

22851 GCF_000389375.1_63.H.87 Salmonella enterica subsp. enterica serovar Agona str. 63.H.87 Proteobacteria; Gammaproteobacteria; Enterobacterales; Enterobacteriaceae; Salmonella; Salmonella enterica; Salmonella enterica subsp. enterica; Salmonella enterica subsp. enterica serovar Agona MNNEETFYQAMRRKGVTRRSFLKFCSLAATSLGLSAGMTPKIAWALENKPRIPVVWIHGL WP_001058316.1 hydrogenase [Salmonella enterica] Length: 372\nNumber of predicted TMHs: 2\nExp number of AAs in TMHs: 35.76397\nExp number, first 60 AAs: 18.97201\nTotal prob of N-in: 0.92785\nPOSSIBLE N-term signal sequence\ninside 1 19\nTMhelix 20 39\noutside 40 325\nTMhelix 326 348\ninside 349 372

22852 GCF_000389375.1_63.H.87 Salmonella enterica subsp. enterica serovar Agona str. 63.H.87 Proteobacteria; Gammaproteobacteria; Enterobacterales; Enterobacteriaceae; Salmonella; Salmonella enterica; Salmonella enterica subsp. enterica; Salmonella enterica subsp. enterica serovar Agona MTGDNTLITSHGINRRDFMKLCAALAATMGLSSKAAAEMAESVSNPQRPPVIWIGAQECT WP_000145429.1 hydrogenase 2 small subunit [Salmonella enterica] Length: 372\nNumber of predicted TMHs: 1\nExp number of AAs in TMHs: 24.51309\nExp number, first 60 AAs: 0.2682\nTotal prob of N-in: 0.14402\noutside 1 331\nTMhelix 332 354\ninside 355 372

22853 GCF_000487475.1_ASM48747v1 Salmonella enterica subsp. enterica serovar Agona str. 648586-1 Proteobacteria; Gammaproteobacteria; Enterobacterales; Enterobacteriaceae; Salmonella; Salmonella enterica; Salmonella enterica subsp. enterica; Salmonella enterica subsp. enterica serovar Agona MNNEETFYQAMRRKGVTRRSFLKFCSLAATSLGLSAGMTPKIAWALENKPRIPVVWIHGL WP_001058316.1 hydrogenase [Salmonella enterica] Length: 372\nNumber of predicted TMHs: 2\nExp number of AAs in TMHs: 35.76397\nExp number, first 60 AAs: 18.97201\nTotal prob of N-in: 0.92785\nPOSSIBLE N-term signal sequence\ninside 1 19\nTMhelix 20 39\noutside 40 325\nTMhelix 326 348\ninside 349 372

22854 GCF_000487475.1_ASM48747v1 Salmonella enterica subsp. enterica serovar Agona str. 648586-1 Proteobacteria; Gammaproteobacteria; Enterobacterales; Enterobacteriaceae; Salmonella; Salmonella enterica; Salmonella enterica subsp. enterica; Salmonella enterica subsp. enterica serovar Agona MTGDNTLITSHGINRRDFMKLCAALAATMGLSSKAAAEMAESVSNPQRPPVIWIGAQECT WP_000145429.1 hydrogenase 2 small subunit [Salmonella enterica] Length: 372\nNumber of predicted TMHs: 1\nExp number of AAs in TMHs: 24.51309\nExp number, first 60 AAs: 0.2682\nTotal prob of N-in: 0.14402\noutside 1 331\nTMhelix 332 354\ninside 355 372

22855 GCF_000389395.1_64.H.00 Salmonella enterica subsp. enterica serovar Agona str. 64.H.00 Proteobacteria; Gammaproteobacteria; Enterobacterales; Enterobacteriaceae; Salmonella; Salmonella enterica; Salmonella enterica subsp. enterica; Salmonella enterica subsp. enterica serovar Agona MNNEETFYQAMRRKGVTRRSFLKFCSLAATSLGLSAGMTPKIAWALENKPRIPVVWIHGL WP_001058316.1 hydrogenase [Salmonella enterica] Length: 372\nNumber of predicted TMHs: 2\nExp number of AAs in TMHs: 35.76397\nExp number, first 60 AAs: 18.97201\nTotal prob of N-in: 0.92785\nPOSSIBLE N-term signal sequence\ninside 1 19\nTMhelix 20 39\noutside 40 325\nTMhelix 326 348\ninside 349 372

22856 GCF_000389395.1_64.H.00 Salmonella enterica subsp. enterica serovar Agona str. 64.H.00 Proteobacteria; Gammaproteobacteria; Enterobacterales; Enterobacteriaceae; Salmonella; Salmonella enterica; Salmonella enterica subsp. enterica; Salmonella enterica subsp. enterica serovar Agona MTGDNTLITSHGINRRDFMKLCAALAATMGLSSKAAAEMAESVSNPQRPPVIWIGAQECT WP_000145429.1 hydrogenase 2 small subunit [Salmonella enterica] Length: 372\nNumber of predicted TMHs: 1\nExp number of AAs in TMHs: 24.51309\nExp number, first 60 AAs: 0.2682\nTotal prob of N-in: 0.14402\noutside 1 331\nTMhelix 332 354\ninside 355 372

22857 GCF_000389415.1_65.H.72 Salmonella enterica subsp. enterica serovar Agona str. 65.H.72 Proteobacteria; Gammaproteobacteria; Enterobacterales; Enterobacteriaceae; Salmonella; Salmonella enterica; Salmonella enterica subsp. enterica; Salmonella enterica subsp. enterica serovar Agona MTGDNTLITSHGINRRDFMKLCAALAATMGLSSKAAAEMAESVSNPQRPPVIWIGAQECT WP_000145429.1 hydrogenase 2 small subunit [Salmonella enterica] Length: 372\nNumber of predicted TMHs: 1\nExp number of AAs in TMHs: 24.51309\nExp number, first 60 AAs: 0.2682\nTotal prob of N-in: 0.14402\noutside 1 331\nTMhelix 332 354\ninside 355 372

22858 GCF_000389415.1_65.H.72 Salmonella enterica subsp. enterica serovar Agona str. 65.H.72 Proteobacteria; Gammaproteobacteria; Enterobacterales; Enterobacteriaceae; Salmonella; Salmonella enterica; Salmonella enterica subsp. enterica; Salmonella enterica subsp. enterica serovar Agona MNNEETFYQAMRRKGVTRRSFLKFCSLAATSLGLSAGMTPKIAWALENKPRIPVVWIHGL WP_001058316.1 hydrogenase [Salmonella enterica] Length: 372\nNumber of predicted TMHs: 2\nExp number of AAs in

TMHs: 35.76397\nExp number, first 60 AAs: 18.97201\nTotal prob of N-in: 0.92785\nPOSSIBLE N-term signal sequence\ninside 1 19\nTMhelix 20 39\noutside 40 325\nTMhelix 326 348\ninside 349 372

22859 GCF_000389435.1_66.F.99 Salmonella enterica subsp. enterica serovar Agona str. 66.F.99 Proteobacteria; Gammaproteobacteria; Enterobacterales; Enterobacteriaceae; Salmonella; Salmonella enterica; Salmonella enterica subsp. enterica; Salmonella enterica subsp. enterica serovar Agona MNNEETFYQAMRRKGVTRRSFLKFCSLAATSLGLSAGMTPKIAWALENKPRIPVVWIHGL WP_001058316.1 hydrogenase [Salmonella enterica] Length: 372\nNumber of predicted TMHs: 2\nExp number of AAs in TMHs: 35.76397\nExp number, first 60 AAs: 18.97201\nTotal prob of N-in: 0.92785\nPOSSIBLE N-term signal sequence\ninside 1 19\nTMhelix 20 39\noutside 40 325\nTMhelix 326 348\ninside 349 372

22860 GCF_000389435.1_66.F.99 Salmonella enterica subsp. enterica serovar Agona str. 66.F.99 Proteobacteria; Gammaproteobacteria; Enterobacterales; Enterobacteriaceae; Salmonella; Salmonella enterica; Salmonella enterica subsp. enterica; Salmonella enterica subsp. enterica serovar Agona MTGDNTLITSHGINRRDFMKLCAALAATMGLSSKAAAEMAESVSNPQRPPVIWIGAQECT WP_000145429.1 hydrogenase 2 small subunit [Salmonella enterica] Length: 372\nNumber of predicted TMHs: 1\nExp number of AAs in TMHs: 24.51309\nExp number, first 60 AAs: 0.2682\nTotal prob of N-in: 0.14402\noutside 1 331\nTMhelix 332 354\ninside 355 372

22861 GCF_000389455.1_67.H.09 Salmonella enterica subsp. enterica serovar Agona str. 67.H.09 Proteobacteria; Gammaproteobacteria; Enterobacterales; Enterobacteriaceae; Salmonella; Salmonella enterica; Salmonella enterica subsp. enterica; Salmonella enterica subsp. enterica serovar Agona MTGDNTLITSHGINRRDFMKLCAALAATMGLSSKAAAEMAESVSNPQRPPVIWIGAQECT WP_000145429.1 hydrogenase 2 small subunit [Salmonella enterica] Length: 372\nNumber of predicted TMHs: 1\nExp number of AAs in TMHs: 24.51309\nExp number, first 60 AAs: 0.2682\nTotal prob of N-in: 0.14402\noutside 1 331\nTMhelix 332 354\ninside 355 372

22862 GCF_000389455.1_67.H.09 Salmonella enterica subsp. enterica serovar Agona str. 67.H.09 Proteobacteria; Gammaproteobacteria; Enterobacterales; Enterobacteriaceae; Salmonella; Salmonella enterica; Salmonella enterica subsp. enterica; Salmonella enterica subsp. enterica serovar Agona MNNEETFYQAMRRKGVTRRSFLKFCSLAATSLGLSAGMTPKIAWALENKPRIPVVWIHGL WP_001058316.1 hydrogenase [Salmonella enterica] Length: 372\nNumber of predicted TMHs: 2\nExp number of AAs in TMHs: 35.76397\nExp number, first 60 AAs: 18.97201\nTotal prob of N-in: 0.92785\nPOSSIBLE N-term signal sequence\ninside 1 19\nTMhelix 20 39\noutside 40 325\nTMhelix 326 348\ninside 349 372

22863 GCF_000389475.1_68.U.05 Salmonella enterica subsp. enterica serovar Agona str. 68.U.05 Proteobacteria; Gammaproteobacteria; Enterobacterales; Enterobacteriaceae; Salmonella; Salmonella enterica; Salmonella enterica subsp. enterica; Salmonella enterica subsp. enterica serovar Agona MNNEETFYQAMRRKGVTRRSFLKFCSLAATSLGLSAGMTPKIAWALENKPRIPVVWIHGL WP_001058316.1 hydrogenase [Salmonella enterica] Length: 372\nNumber of predicted TMHs: 2\nExp number of AAs in TMHs: 35.76397\nExp number, first 60 AAs: 18.97201\nTotal prob of N-in: 0.92785\nPOSSIBLE N-term signal sequence\ninside 1 19\nTMhelix 20 39\noutside 40 325\nTMhelix 326 348\ninside 349 372

22864 GCF_000389475.1_68.U.05 Salmonella enterica subsp. enterica serovar Agona str. 68.U.05 Proteobacteria; Gammaproteobacteria; Enterobacterales; Enterobacteriaceae; Salmonella; Salmonella enterica; Salmonella enterica subsp. enterica; Salmonella enterica subsp. enterica serovar Agona MTGDNTLITSHGINRRDFMKLCAALAATMGLSSKAAAEMAESVSNPQRPPVIWIGAQECT WP_000145429.1 hydrogenase 2 small subunit [Salmonella enterica] Length: 372\nNumber of predicted TMHs: 1\nExp number of AAs in TMHs: 24.51309\nExp number, first 60 AAs: 0.2682\nTotal prob of N-in: 0.14402\noutside 1 331\nTMhelix 332 354\ninside 355 372

22865 GCF_000389495.1_69.H.06 Salmonella enterica subsp. enterica serovar Agona str. 69.H.06 Proteobacteria; Gammaproteobacteria; Enterobacterales; Enterobacteriaceae; Salmonella; Salmonella enterica; Salmonella enterica subsp. enterica; Salmonella enterica subsp. enterica serovar Agona MTGDNTLITSHGINRRDFMKLCAALAATMGLSSKAAAEMAESVSNPQRPPVIWIGAQECT WP_000145429.1 hydrogenase 2 small subunit [Salmonella enterica] Length: 372\nNumber of predicted TMHs: 1\nExp number of AAs in TMHs: 24.51309\nExp number, first 60 AAs: 0.2682\nTotal prob of N-in: 0.14402\noutside 1 331\nTMhelix 332 354\ninside 355 372

22866 GCF_000389515.1_70.E.05 Salmonella enterica subsp. enterica serovar Agona str. 70.E.05 Proteobacteria; Gammaproteobacteria; Enterobacterales; Enterobacteriaceae; Salmonella; Salmonella enterica;

Salmonella enterica subsp. enterica; Salmonella enterica subsp. enterica serovar Agona
MNNEETFYQAMRRKGVTRRSFLKFCSLAATSLGLSAGMTPKIAWALENKPRIPVVWIHGL WP_001058316.1
hydrogenase [Salmonella enterica] Length: 372\nNumber of predicted TMHs: 2\nExp number of AAs in TMHs: 35.76397\nExp number, first 60 AAs: 18.97201\nTotal prob of N-in: 0.92785\nPOSSIBLE N-term signal sequence\ninside 1 19\nTMhelix 20 39\noutside 40 325\nTMhelix 326 348\ninside 349 372

22867 GCF_000389515.1_70.E.05 Salmonella enterica subsp. enterica serovar Agona str. 70.E.05
Proteobacteria; Gammaproteobacteria; Enterobacterales; Enterobacteriaceae; Salmonella; Salmonella enterica; Salmonella enterica subsp. enterica; Salmonella enterica subsp. enterica serovar Agona
MTGDNTLITSHGINRRDFMKLCAALAATMGLSSKAAAEMAESVSNPQRPPVIWIGAQECT WP_000145429.1
hydrogenase 2 small subunit [Salmonella enterica] Length: 372\nNumber of predicted TMHs: 1\nExp number of AAs in TMHs: 24.51309\nExp number, first 60 AAs: 0.2682\nTotal prob of N-in: 0.14402\noutside 1 331\nTMhelix 332 354\ninside 355 372

22868 GCF_000389535.1_71.E.05 Salmonella enterica subsp. enterica serovar Agona str. 71.E.05
Proteobacteria; Gammaproteobacteria; Enterobacterales; Enterobacteriaceae; Salmonella; Salmonella enterica; Salmonella enterica subsp. enterica; Salmonella enterica subsp. enterica serovar Agona
MTGDNTLITSHGINRRDFMKLCAALAATMGLSSKAAAEMAESVSNPQRPPVIWIGAQECT WP_000145429.1
hydrogenase 2 small subunit [Salmonella enterica] Length: 372\nNumber of predicted TMHs: 1\nExp number of AAs in TMHs: 24.51309\nExp number, first 60 AAs: 0.2682\nTotal prob of N-in: 0.14402\noutside 1 331\nTMhelix 332 354\ninside 355 372

22869 GCF_000389535.1_71.E.05 Salmonella enterica subsp. enterica serovar Agona str. 71.E.05
Proteobacteria; Gammaproteobacteria; Enterobacterales; Enterobacteriaceae; Salmonella; Salmonella enterica; Salmonella enterica subsp. enterica; Salmonella enterica subsp. enterica serovar Agona
MNNEETFYQAMRRKGVTRRSFLKFCSLAATSLGLSAGMTPKIAWALENKPRIPVVWIHGL WP_001058316.1
hydrogenase [Salmonella enterica] Length: 372\nNumber of predicted TMHs: 2\nExp number of AAs in TMHs: 35.76397\nExp number, first 60 AAs: 18.97201\nTotal prob of N-in: 0.92785\nPOSSIBLE N-term signal sequence\ninside 1 19\nTMhelix 20 39\noutside 40 325\nTMhelix 326 348\ninside 349 372

22870 GCF_000389555.1_72.A.52 Salmonella enterica subsp. enterica serovar Agona str. 72.A.52
Proteobacteria; Gammaproteobacteria; Enterobacterales; Enterobacteriaceae; Salmonella; Salmonella enterica; Salmonella enterica subsp. enterica; Salmonella enterica subsp. enterica serovar Agona
MTGDNTLITSHGINRRDFMKLCAALAATMGLSSKAAAEMAESVSNPQRPPVIWIGAQECT WP_000145429.1
hydrogenase 2 small subunit [Salmonella enterica] Length: 372\nNumber of predicted TMHs: 1\nExp number of AAs in TMHs: 24.51309\nExp number, first 60 AAs: 0.2682\nTotal prob of N-in: 0.14402\noutside 1 331\nTMhelix 332 354\ninside 355 372

22871 GCF_000389555.1_72.A.52 Salmonella enterica subsp. enterica serovar Agona str. 72.A.52
Proteobacteria; Gammaproteobacteria; Enterobacterales; Enterobacteriaceae; Salmonella; Salmonella enterica; Salmonella enterica subsp. enterica; Salmonella enterica subsp. enterica serovar Agona
MNNEETFYQAMRRKGVTRRSFLKFCSLAATSLGLSAGMTPKIAWALENKPRIPVVWIHGL WP_001058316.1
hydrogenase [Salmonella enterica] Length: 372\nNumber of predicted TMHs: 2\nExp number of AAs in TMHs: 35.76397\nExp number, first 60 AAs: 18.97201\nTotal prob of N-in: 0.92785\nPOSSIBLE N-term signal sequence\ninside 1 19\nTMhelix 20 39\noutside 40 325\nTMhelix 326 348\ninside 349 372

22872 GCF_000389575.1_73.H.09 Salmonella enterica subsp. enterica serovar Agona str. 73.H.09
Proteobacteria; Gammaproteobacteria; Enterobacterales; Enterobacteriaceae; Salmonella; Salmonella enterica; Salmonella enterica subsp. enterica; Salmonella enterica subsp. enterica serovar Agona
MTGDNTLITSHGINRRDFMKLCAALAATMGLSSKAAAEMAESVSNPQRPPVIWIGAQECT WP_000145429.1
hydrogenase 2 small subunit [Salmonella enterica] Length: 372\nNumber of predicted TMHs: 1\nExp number of AAs in TMHs: 24.51309\nExp number, first 60 AAs: 0.2682\nTotal prob of N-in: 0.14402\noutside 1 331\nTMhelix 332 354\ninside 355 372

22873 GCF_000389575.1_73.H.09 Salmonella enterica subsp. enterica serovar Agona str. 73.H.09
Proteobacteria; Gammaproteobacteria; Enterobacterales; Enterobacteriaceae; Salmonella; Salmonella enterica; Salmonella enterica subsp. enterica; Salmonella enterica subsp. enterica serovar Agona
MNNEETFYQAMRRKGVTRRSFLKFCSLAATSLGLSAGMTPKIAWALENKPRIPVVWIHGL WP_001058316.1
hydrogenase [Salmonella enterica] Length: 372\nNumber of predicted TMHs: 2\nExp number of AAs in TMHs: 35.76397\nExp number, first 60 AAs: 18.97201\nTotal prob of N-in: 0.92785\nPOSSIBLE N-term signal sequence\ninside 1 19\nTMhelix 20 39\noutside 40 325\nTMhelix 326 348\ninside 349 372

22874 GCF_000487535.1_ASM48753v1 Salmonella enterica subsp. enterica serovar Agona str. ATCC 51957 Proteobacteria; Gammaproteobacteria; Enterobacterales; Enterobacteriaceae; Salmonella; Salmonella enterica; Salmonella enterica subsp. enterica; Salmonella enterica subsp. enterica serovar Agona MTGDNTLITSHGINRRDFMKLCAALAATMGLSSKAAAEMAESVSNPQRPPVIWIGAQECT WP_000145429.1 hydrogenase 2 small subunit [Salmonella enterica] Length: 372\nNumber of predicted TMHs: 1\nExp number of AAs in TMHs: 24.51309\nExp number, first 60 AAs: 0.2682\nTotal prob of N-in: 0.14402\noutside 1 331\nTMhelix 332 354\ninside 355 372

22875 GCF_000487535.1_ASM48753v1 Salmonella enterica subsp. enterica serovar Agona str. ATCC 51957 Proteobacteria; Gammaproteobacteria; Enterobacterales; Enterobacteriaceae; Salmonella; Salmonella enterica; Salmonella enterica subsp. enterica; Salmonella enterica subsp. enterica serovar Agona MNNEETFYQAMRRKGVTRRSFLKFCSLAATSLGLSAGMTPKIAWALENKPRIPVVWIHGL WP_001058316.1 hydrogenase [Salmonella enterica] Length: 372\nNumber of predicted TMHs: 2\nExp number of AAs in TMHs: 35.76397\nExp number, first 60 AAs: 18.97201\nTotal prob of N-in: 0.92785\nPOSSIBLE N-term signal sequence\ninside 1 19\nTMhelix 20 39\noutside 40 325\nTMhelix 326 348\ninside 349 372

22876 GCF_000483935.1_CFSAN000506_01.0Salmonella enterica subsp. enterica serovar Agona str. ATCC BAA-707 Proteobacteria; Gammaproteobacteria; Enterobacterales; Enterobacteriaceae; Salmonella; Salmonella enterica; Salmonella enterica subsp. enterica; Salmonella enterica subsp. enterica serovar Agona MTGDNTLITSHGINRRDFMKLCAALAATMGLSSKAAAEMAESVSNPQRPPVIWIGAQECT WP_000145429.1 hydrogenase 2 small subunit [Salmonella enterica] Length: 372\nNumber of predicted TMHs: 1\nExp number of AAs in TMHs: 24.51309\nExp number, first 60 AAs: 0.2682\nTotal prob of N-in: 0.14402\noutside 1 331\nTMhelix 332 354\ninside 355 372

22877 GCF_000483935.1_CFSAN000506_01.0Salmonella enterica subsp. enterica serovar Agona str. ATCC BAA-707 Proteobacteria; Gammaproteobacteria; Enterobacterales; Enterobacteriaceae; Salmonella; Salmonella enterica; Salmonella enterica subsp. enterica; Salmonella enterica subsp. enterica serovar Agona MNNEETFYQAMRRKGVTRRSFLKFCSLAATSLGLSAGMTPKIAWALENKPRIPVVWIHGL WP_001058316.1 hydrogenase [Salmonella enterica] Length: 372\nNumber of predicted TMHs: 2\nExp number of AAs in TMHs: 35.76397\nExp number, first 60 AAs: 18.97201\nTotal prob of N-in: 0.92785\nPOSSIBLE N-term signal sequence\ninside 1 19\nTMhelix 20 39\noutside 40 325\nTMhelix 326 348\ninside 349 372

22878 GCF_000483815.1_ASM48381v1 Salmonella enterica subsp. enterica serovar Agona str. SA-2 Proteobacteria; Gammaproteobacteria; Enterobacterales; Enterobacteriaceae; Salmonella; Salmonella enterica; Salmonella enterica subsp. enterica; Salmonella enterica subsp. enterica serovar Agona MNNEETFYQAMRRKGVTRRSFLKFCSLAATSLGLSAGMTPKIAWALENKPRIPVVWIHGL WP_001058316.1 hydrogenase [Salmonella enterica] Length: 372\nNumber of predicted TMHs: 2\nExp number of AAs in TMHs: 35.76397\nExp number, first 60 AAs: 18.97201\nTotal prob of N-in: 0.92785\nPOSSIBLE N-term signal sequence\ninside 1 19\nTMhelix 20 39\noutside 40 325\nTMhelix 326 348\ninside 349 372

22879 GCF_000483815.1_ASM48381v1 Salmonella enterica subsp. enterica serovar Agona str. SA-2 Proteobacteria; Gammaproteobacteria; Enterobacterales; Enterobacteriaceae; Salmonella; Salmonella enterica; Salmonella enterica subsp. enterica; Salmonella enterica subsp. enterica serovar Agona MTGDNTLITSHGINRRDFMKLCAALAATMGLSSKAAAEMAESVSNPQRPPVIWIGAQECT WP_000145429.1 hydrogenase 2 small subunit [Salmonella enterica] Length: 372\nNumber of predicted TMHs: 1\nExp number of AAs in TMHs: 24.51309\nExp number, first 60 AAs: 0.2682\nTotal prob of N-in: 0.14402\noutside 1 331\nTMhelix 332 354\ninside 355 372

22880 GCF_000484235.1_ASM48423v1 Salmonella enterica subsp. enterica serovar Agona str. SA-5 Proteobacteria; Gammaproteobacteria; Enterobacterales; Enterobacteriaceae; Salmonella; Salmonella enterica; Salmonella enterica subsp. enterica; Salmonella enterica subsp. enterica serovar Agona MTGDNTLITSHGINRRDFMKLCAALAATMGLSSKAAAEMAESVSNPQRPPVIWIGAQECT WP_000145429.1 hydrogenase 2 small subunit [Salmonella enterica] Length: 372\nNumber of predicted TMHs: 1\nExp number of AAs in TMHs: 24.51309\nExp number, first 60 AAs: 0.2682\nTotal prob of N-in: 0.14402\noutside 1 331\nTMhelix 332 354\ninside 355 372

22881 GCF_000484235.1_ASM48423v1 Salmonella enterica subsp. enterica serovar Agona str. SA-5 Proteobacteria; Gammaproteobacteria; Enterobacterales; Enterobacteriaceae; Salmonella; Salmonella enterica; Salmonella enterica subsp. enterica; Salmonella enterica subsp. enterica serovar Agona MNNEETFYQAMRRKGVTRRSFLKFCSLAATSLGLSAGMTPKIAWALENKPRIPVVWIHGL WP_001058316.1 hydrogenase [Salmonella enterica] Length: 372\nNumber of predicted TMHs: 2\nExp number of AAs in

TMHs: 35.76397\nExp number, first 60 AAs: 18.97201\nTotal prob of N-in: 0.92785\nPOSSIBLE N-term signal sequence\ninside 1 19\nTMhelix 20 39\noutside 40 325\nTMhelix 326 348\ninside 349 372

22882 GCF_000330625.1_ASM33062v1 Salmonella enterica subsp. enterica serovar Agona str. SH08SF124 Proteobacteria; Gammaproteobacteria; Enterobacterales; Enterobacteriaceae; Salmonella; Salmonella enterica; Salmonella enterica subsp. enterica; Salmonella enterica subsp. enterica serovar Agona MNNEETFYQAMRRKGVTRRSFLKFCSLAATSLGLSAGMTPKIAWALENKPRIPVVWIHGL WP_001058316.1 hydrogenase [Salmonella enterica] Length: 372\nNumber of predicted TMHs: 2\nExp number of AAs in TMHs: 35.76397\nExp number, first 60 AAs: 18.97201\nTotal prob of N-in: 0.92785\nPOSSIBLE N-term signal sequence\ninside 1 19\nTMhelix 20 39\noutside 40 325\nTMhelix 326 348\ninside 349 372

22883 GCF_000330625.1_ASM33062v1 Salmonella enterica subsp. enterica serovar Agona str. SH08SF124 Proteobacteria; Gammaproteobacteria; Enterobacterales; Enterobacteriaceae; Salmonella; Salmonella enterica; Salmonella enterica subsp. enterica; Salmonella enterica subsp. enterica serovar Agona MTGDNTLITSHGINRRDFMKLCAALAATMGLSSKAAAEMAESVSNPQRPPVIWIGAQECT WP_000145429.1 hydrogenase 2 small subunit [Salmonella enterica] Length: 372\nNumber of predicted TMHs: 1\nExp number of AAs in TMHs: 24.51309\nExp number, first 60 AAs: 0.2682\nTotal prob of N-in: 0.14402\noutside 1 331\nTMhelix 332 354\ninside 355 372

22884 GCF_000330645.1_ASM33064v1 Salmonella enterica subsp. enterica serovar Agona str. SH10GFN094 Proteobacteria; Gammaproteobacteria; Enterobacterales; Enterobacteriaceae; Salmonella; Salmonella enterica; Salmonella enterica subsp. enterica; Salmonella enterica subsp. enterica serovar Agona MNNEETFYQAMRRKGVTRRSFLKFCSLAATSLGLSAGMTPKIAWALENKPRIPVVWIHGL WP_001058316.1 hydrogenase [Salmonella enterica] Length: 372\nNumber of predicted TMHs: 2\nExp number of AAs in TMHs: 35.76397\nExp number, first 60 AAs: 18.97201\nTotal prob of N-in: 0.92785\nPOSSIBLE N-term signal sequence\ninside 1 19\nTMhelix 20 39\noutside 40 325\nTMhelix 326 348\ninside 349 372

22885 GCF_000330645.1_ASM33064v1 Salmonella enterica subsp. enterica serovar Agona str. SH10GFN094 Proteobacteria; Gammaproteobacteria; Enterobacterales; Enterobacteriaceae; Salmonella; Salmonella enterica; Salmonella enterica subsp. enterica; Salmonella enterica subsp. enterica serovar Agona MTGDNTLITSHGINRRDFMKLCAALAATMGLSSKAAAEMAESVSNPQRPPVIWIGAQECT WP_000145429.1 hydrogenase 2 small subunit [Salmonella enterica] Length: 372\nNumber of predicted TMHs: 1\nExp number of AAs in TMHs: 24.51309\nExp number, first 60 AAs: 0.2682\nTotal prob of N-in: 0.14402\noutside 1 331\nTMhelix 332 354\ninside 355 372

22886 GCF_000330665.1_ASM33066v1 Salmonella enterica subsp. enterica serovar Agona str. SH11G1113 Proteobacteria; Gammaproteobacteria; Enterobacterales; Enterobacteriaceae; Salmonella; Salmonella enterica; Salmonella enterica subsp. enterica; Salmonella enterica subsp. enterica serovar Agona MNNEETFYQAMRRKGVTRRSFLKFCSLAATSLGLSAGMTPKIAWALENKPRIPVVWIHGL WP_001058316.1 hydrogenase [Salmonella enterica] Length: 372\nNumber of predicted TMHs: 2\nExp number of AAs in TMHs: 35.76397\nExp number, first 60 AAs: 18.97201\nTotal prob of N-in: 0.92785\nPOSSIBLE N-term signal sequence\ninside 1 19\nTMhelix 20 39\noutside 40 325\nTMhelix 326 348\ninside 349 372

22887 GCF_000330665.1_ASM33066v1 Salmonella enterica subsp. enterica serovar Agona str. SH11G1113 Proteobacteria; Gammaproteobacteria; Enterobacterales; Enterobacteriaceae; Salmonella; Salmonella enterica; Salmonella enterica subsp. enterica; Salmonella enterica subsp. enterica serovar Agona MTGDNTLITSHGINRRDFMKLCAALAATMGLSSKAAAEMAESVSNPQRPPVIWIGAQECT WP_000145429.1 hydrogenase 2 small subunit [Salmonella enterica] Length: 372\nNumber of predicted TMHs: 1\nExp number of AAs in TMHs: 24.51309\nExp number, first 60 AAs: 0.2682\nTotal prob of N-in: 0.14402\noutside 1 331\nTMhelix 332 354\ninside 355 372

22888 GCF_000020885.1_ASM2088v1 Salmonella enterica subsp. enterica serovar Agona str. SL483 Proteobacteria; Gammaproteobacteria; Enterobacterales; Enterobacteriaceae; Salmonella; Salmonella enterica; Salmonella enterica subsp. enterica; Salmonella enterica subsp. enterica serovar Agona MNNEETFYQAMRRKGVTRRSFLKFCSLAATSLGLSAGMTPKIAWALENKPRIPVVWIHGL WP_001058316.1 hydrogenase [Salmonella enterica] Length: 372\nNumber of predicted TMHs: 2\nExp number of AAs in TMHs: 35.76397\nExp number, first 60 AAs: 18.97201\nTotal prob of N-in: 0.92785\nPOSSIBLE N-term signal sequence\ninside 1 19\nTMhelix 20 39\noutside 40 325\nTMhelix 326 348\ninside 349 372

22889 GCF_000020885.1_ASM2088v1 Salmonella enterica subsp. enterica serovar Agona str. SL483 Proteobacteria; Gammaproteobacteria; Enterobacterales; Enterobacteriaceae; Salmonella; Salmonella enterica;

Salmonella enterica subsp. enterica; Salmonella enterica subsp. enterica serovar Agona
MTGDNTLITSHGINRRDFMKLCAALAATMGLSSKAAAEMAESVSNPQRPPVIWIGAQECT WP_000145429.1
hydrogenase 2 small subunit [Salmonella enterica] Length: 372\nNumber of predicted TMHs: 1\nExp number of AAs in TMHs: 24.51309\nExp number, first 60 AAs: 0.2682\nTotal prob of N-in: 0.14402\noutside 1 331\nTMhelix 332 354\ninside 355 372

22890 GCF_000487515.2_ASM48751v2 Salmonella enterica subsp. enterica serovar Albany str. ATCC 51960
Proteobacteria; Gammaproteobacteria; Enterobacterales; Enterobacteriaceae; Salmonella; Salmonella enterica; Salmonella enterica subsp. enterica; Salmonella enterica subsp. enterica serovar Albany
MTGDNTLITSHGINRRDFMKLCAALAATMGLSSKAAAEMAESVSNPQRPPVIWIGAQECT WP_000145429.1
hydrogenase 2 small subunit [Salmonella enterica] Length: 372\nNumber of predicted TMHs: 1\nExp number of AAs in TMHs: 24.51309\nExp number, first 60 AAs: 0.2682\nTotal prob of N-in: 0.14402\noutside 1 331\nTMhelix 332 354\ninside 355 372

22891 GCF_000487515.2_ASM48751v2 Salmonella enterica subsp. enterica serovar Albany str. ATCC 51960
Proteobacteria; Gammaproteobacteria; Enterobacterales; Enterobacteriaceae; Salmonella; Salmonella enterica; Salmonella enterica subsp. enterica; Salmonella enterica subsp. enterica serovar Albany
MNNEETFYQAMRRKGVTRRSFLKFCSLAATSLGLGAGMTPKIAWALENKPRIPVVWIHGL WP_001058311.1
hydrogenase [Salmonella enterica] Length: 372\nNumber of predicted TMHs: 2\nExp number of AAs in TMHs: 37.4699899999999\nExp number, first 60 AAs: 19.72699\nTotal prob of N-in: 0.96171\nPOSSIBLE N-term signal sequence\ninside 1 19\nTMhelix 20 39\noutside 40 325\nTMhelix 326 348\ninside 349 372

22892 GCF_000412875.1_06-0532Ver1 Salmonella enterica subsp. enterica serovar Anatum str. 06-0532
Proteobacteria; Gammaproteobacteria; Enterobacterales; Enterobacteriaceae; Salmonella; Salmonella enterica; Salmonella enterica subsp. enterica; Salmonella enterica subsp. enterica serovar Anatum
MTGDNTLITSHGINRRDFMKLCAALAATMGLSSKAAAEMAESVSNPQRPPVIWIGAQECT WP_000145429.1
hydrogenase 2 small subunit [Salmonella enterica] Length: 372\nNumber of predicted TMHs: 1\nExp number of AAs in TMHs: 24.51309\nExp number, first 60 AAs: 0.2682\nTotal prob of N-in: 0.14402\noutside 1 331\nTMhelix 332 354\ninside 355 372

22893 GCF_000412875.1_06-0532Ver1 Salmonella enterica subsp. enterica serovar Anatum str. 06-0532
Proteobacteria; Gammaproteobacteria; Enterobacterales; Enterobacteriaceae; Salmonella; Salmonella enterica; Salmonella enterica subsp. enterica; Salmonella enterica subsp. enterica serovar Anatum
MNNEETFYQAMRRKGVTRRSFLKFCSLAATSLGLGAGMTPKIAWALENKPRIPVVWIHGL WP_023243720.1
hydrogenase 1, small subunit [Salmonella enterica] Length: 372\nNumber of predicted TMHs: 2\nExp number of AAs in TMHs: 36.52147\nExp number, first 60 AAs: 19.72838\nTotal prob of N-in: 0.96170\nPOSSIBLE N-term signal sequence\ninside 1 19\nTMhelix 20 39\noutside 40 325\nTMhelix 326 348\ninside 349 372

22894 GCF_000487575.2_ASM48757v2 Salmonella enterica subsp. enterica serovar Anatum str. ATCC BAA-1592
Proteobacteria; Gammaproteobacteria; Enterobacterales; Enterobacteriaceae; Salmonella; Salmonella enterica; Salmonella enterica subsp. enterica; Salmonella enterica subsp. enterica serovar Anatum
MTGDNTLITSHGINRRDFMKLCAALAATMGLSSKAAAEMAESVSNPQRPPVIWIGAQECT WP_000145429.1
hydrogenase 2 small subunit [Salmonella enterica] Length: 372\nNumber of predicted TMHs: 1\nExp number of AAs in TMHs: 24.51309\nExp number, first 60 AAs: 0.2682\nTotal prob of N-in: 0.14402\noutside 1 331\nTMhelix 332 354\ninside 355 372

22895 GCF_000487575.2_ASM48757v2 Salmonella enterica subsp. enterica serovar Anatum str. ATCC BAA-1592
Proteobacteria; Gammaproteobacteria; Enterobacterales; Enterobacteriaceae; Salmonella; Salmonella enterica; Salmonella enterica subsp. enterica; Salmonella enterica subsp. enterica serovar Anatum
MNNEETFYQAMRRKGVTRRSFLKFCSLAATSLGLGAGMTPKIAWALENKPRIPVVWIHGL WP_023243720.1
hydrogenase 1, small subunit [Salmonella enterica] Length: 372\nNumber of predicted TMHs: 2\nExp number of AAs in TMHs: 36.52147\nExp number, first 60 AAs: 19.72838\nTotal prob of N-in: 0.96170\nPOSSIBLE N-term signal sequence\ninside 1 19\nTMhelix 20 39\noutside 40 325\nTMhelix 326 348\ninside 349 372

22896 GCF_000940895.2_ASM94089v2 Salmonella enterica subsp. enterica serovar Anatum str. CDC 06-0532
Proteobacteria; Gammaproteobacteria; Enterobacterales; Enterobacteriaceae; Salmonella; Salmonella enterica; Salmonella enterica subsp. enterica; Salmonella enterica subsp. enterica serovar Anatum
MNNEETFYQAMRRKGVTRRSFLKFCSLAATSLGLGAGMTPKIAWALENKPRIPVVWIHGL WP_023243720.1
hydrogenase 1, small subunit [Salmonella enterica] Length: 372\nNumber of predicted TMHs: 2\nExp number of AAs in TMHs: 36.52147\nExp number, first 60 AAs: 19.72838\nTotal prob of N-in: 0.96170\nPOSSIBLE N-term signal sequence\ninside 1 19\nTMhelix 20 39\noutside 40 325\nTMhelix 326 348\ninside 349 372

22897 GCF_000940895.2_ASM94089v2 Salmonella enterica subsp. enterica serovar Anatum str. CDC 06-0532 Proteobacteria; Gammaproteobacteria; Enterobacterales; Enterobacteriaceae; Salmonella; Salmonella enterica; Salmonella enterica subsp. enterica; Salmonella enterica subsp. enterica serovar Anatum MTGDNTLITSHGINRRDFMKLCAALAATMGLSSKAAAEMAESVSNPQRPPVIWIGAQECT WP_000145429.1 hydrogenase 2 small subunit [Salmonella enterica] Length: 372\nNumber of predicted TMHs: 1\nExp number of AAs in TMHs: 24.51309\nExp number, first 60 AAs: 0.2682\nTotal prob of N-in: 0.14402\noutside 1 331\nTMhelix 332 354\ninside 355 372

22898 GCF_000487595.1_ASM48759v1 Salmonella enterica subsp. enterica serovar Anatum str. USDA 100 Proteobacteria; Gammaproteobacteria; Enterobacterales; Enterobacteriaceae; Salmonella; Salmonella enterica; Salmonella enterica subsp. enterica; Salmonella enterica subsp. enterica serovar Anatum MTGDNTLITSHGINRRDFMKLCAALAATMGLSSKAAAEMAESVSNPQRPPVIWIGAQECT WP_000145429.1 hydrogenase 2 small subunit [Salmonella enterica] Length: 372\nNumber of predicted TMHs: 1\nExp number of AAs in TMHs: 24.51309\nExp number, first 60 AAs: 0.2682\nTotal prob of N-in: 0.14402\noutside 1 331\nTMhelix 332 354\ninside 355 372

22899 GCF_000487595.1_ASM48759v1 Salmonella enterica subsp. enterica serovar Anatum str. USDA 100 Proteobacteria; Gammaproteobacteria; Enterobacterales; Enterobacteriaceae; Salmonella; Salmonella enterica; Salmonella enterica subsp. enterica; Salmonella enterica subsp. enterica serovar Anatum MNNEETFYQAMRRKGVTRRSFLKFCSLAATSLGLGAGMTPKIAWALENKPRIPVVWIHGL WP_023243720.1 hydrogenase 1, small subunit [Salmonella enterica] Length: 372\nNumber of predicted TMHs: 2\nExp number of AAs in TMHs: 36.52147\nExp number, first 60 AAs: 19.72838\nTotal prob of N-in: 0.96170\nPOSSIBLE N-term signal sequence\ninside 1 19\nTMhelix 20 39\noutside 40 325\nTMhelix 326 348\ninside 349 372

22900 GCF_000963535.2_ASM96353v2 Salmonella enterica subsp. enterica serovar Anatum str. USDA-ARS-USMARC-1175 Proteobacteria; Gammaproteobacteria; Enterobacterales; Enterobacteriaceae; Salmonella; Salmonella enterica; Salmonella enterica subsp. enterica; Salmonella enterica subsp. enterica serovar Anatum MTGDNTLITSHGINRRDFMKLCAALAATMGLSSKAAAEMAESVSNPQRPPVIWIGAQECT WP_000145429.1 hydrogenase 2 small subunit [Salmonella enterica] Length: 372\nNumber of predicted TMHs: 1\nExp number of AAs in TMHs: 24.51309\nExp number, first 60 AAs: 0.2682\nTotal prob of N-in: 0.14402\noutside 1 331\nTMhelix 332 354\ninside 355 372

22901 GCF_000963535.2_ASM96353v2 Salmonella enterica subsp. enterica serovar Anatum str. USDA-ARS-USMARC-1175 Proteobacteria; Gammaproteobacteria; Enterobacterales; Enterobacteriaceae; Salmonella; Salmonella enterica; Salmonella enterica subsp. enterica; Salmonella enterica subsp. enterica serovar Anatum MNNEETFYQAMRRKGVTRRSFLKFCSLAATSLGLGAGMTPKIAWALENKPRIPVVWIHGL WP_023243720.1 hydrogenase 1, small subunit [Salmonella enterica] Length: 372\nNumber of predicted TMHs: 2\nExp number of AAs in TMHs: 36.52147\nExp number, first 60 AAs: 19.72838\nTotal prob of N-in: 0.96170\nPOSSIBLE N-term signal sequence\ninside 1 19\nTMhelix 20 39\noutside 40 325\nTMhelix 326 348\ninside 349 372

22902 GCF_001620225.1_ASM162022v1 Salmonella enterica subsp. enterica serovar Anatum str. USDA-ARS-USMARC-1676 Proteobacteria; Gammaproteobacteria; Enterobacterales; Enterobacteriaceae; Salmonella; Salmonella enterica; Salmonella enterica subsp. enterica; Salmonella enterica subsp. enterica serovar Anatum MNNEETFYQAMRRKGVTRRSFLKFCSLAATSLGLGAGMTPKIAWALENKPRIPVVWIHGL WP_023243720.1 hydrogenase 1, small subunit [Salmonella enterica] Length: 372\nNumber of predicted TMHs: 2\nExp number of AAs in TMHs: 36.52147\nExp number, first 60 AAs: 19.72838\nTotal prob of N-in: 0.96170\nPOSSIBLE N-term signal sequence\ninside 1 19\nTMhelix 20 39\noutside 40 325\nTMhelix 326 348\ninside 349 372

22903 GCF_001620225.1_ASM162022v1 Salmonella enterica subsp. enterica serovar Anatum str. USDA-ARS-USMARC-1676 Proteobacteria; Gammaproteobacteria; Enterobacterales; Enterobacteriaceae; Salmonella; Salmonella enterica; Salmonella enterica subsp. enterica; Salmonella enterica subsp. enterica serovar Anatum MTGDNTLITSHGINRRDFMKLCAALAATMGLSSKAAAEMAESVSNPQRPPVIWIGAQECT WP_000145429.1 hydrogenase 2 small subunit [Salmonella enterica] Length: 372\nNumber of predicted TMHs: 1\nExp number of AAs in TMHs: 24.51309\nExp number, first 60 AAs: 0.2682\nTotal prob of N-in: 0.14402\noutside 1 331\nTMhelix 332 354\ninside 355 372

22904 GCF_001623545.2_ASM162354v2 Salmonella enterica subsp. enterica serovar Anatum str. USDA-ARS-USMARC-1677 Proteobacteria; Gammaproteobacteria; Enterobacterales; Enterobacteriaceae; Salmonella; Salmonella enterica; Salmonella enterica subsp. enterica; Salmonella enterica subsp. enterica serovar Anatum MTGDNTLITSHGINRRDFMKLCAALAATMGLSSKAAAEMAESVSNPQRPPVIWIGAQECT WP_000145429.1 hydrogenase 2 small subunit [Salmonella enterica] Length: 372\nNumber of predicted TMHs: 1\nExp

number of AAs in TMHs: 24.51309\nExp number, first 60 AAs: 0.2682\nTotal prob of N-in: 0.14402\noutside 1 331\nTMhelix 332 354\ninside 355 372

22905 GCF_001623545.2_ASM162354v2 Salmonella enterica subsp. enterica serovar Anatum str. USDA-ARS-USMARC-1677 Proteobacteria; Gammaproteobacteria; Enterobacterales; Enterobacteriaceae; Salmonella; Salmonella enterica; Salmonella enterica subsp. enterica; Salmonella enterica subsp. enterica serovar Anatum MNNEETFYQAMRRKGVTRRSFLKFCSLAATSLGLGAGMTPKIAWALENKPRIPVVWIHGL WP_023243720.1 hydrogenase 1, small subunit [Salmonella enterica] Length: 372\nNumber of predicted TMHs: 2\nExp number of AAs in TMHs: 36.52147\nExp number, first 60 AAs: 19.72838\nTotal prob of N-in: 0.96170\nPOSSIBLE N-term signal sequence\ninside 1 19\nTMhelix 20 39\noutside 40 325\nTMhelix 326 348\ninside 349 372

22906 GCF_001620205.2_ASM162020v2 Salmonella enterica subsp. enterica serovar Anatum str. USDA-ARS-USMARC-1727 Proteobacteria; Gammaproteobacteria; Enterobacterales; Enterobacteriaceae; Salmonella; Salmonella enterica; Salmonella enterica subsp. enterica; Salmonella enterica subsp. enterica serovar Anatum MNNEETFYQAMRRKGVTRRSFLKFCSLAATSLGLGAGMTPKIAWALENKPRIPVVWIHGL WP_023243720.1 hydrogenase 1, small subunit [Salmonella enterica] Length: 372\nNumber of predicted TMHs: 2\nExp number of AAs in TMHs: 36.52147\nExp number, first 60 AAs: 19.72838\nTotal prob of N-in: 0.96170\nPOSSIBLE N-term signal sequence\ninside 1 19\nTMhelix 20 39\noutside 40 325\nTMhelix 326 348\ninside 349 372

22907 GCF_001620205.2_ASM162020v2 Salmonella enterica subsp. enterica serovar Anatum str. USDA-ARS-USMARC-1727 Proteobacteria; Gammaproteobacteria; Enterobacterales; Enterobacteriaceae; Salmonella; Salmonella enterica; Salmonella enterica subsp. enterica; Salmonella enterica subsp. enterica serovar Anatum MTGDNTLITSHGINRRDFMKLCAALAATMGLSSKAAAEMAESVSNPQRPPVIWIGAQECT WP_000145429.1 hydrogenase 2 small subunit [Salmonella enterica] Length: 372\nNumber of predicted TMHs: 1\nExp number of AAs in TMHs: 24.51309\nExp number, first 60 AAs: 0.2682\nTotal prob of N-in: 0.14402\noutside 1 331\nTMhelix 332 354\ninside 355 372

22908 GCF_001623585.1_ASM162358v1 Salmonella enterica subsp. enterica serovar Anatum str. USDA-ARS-USMARC-1728 Proteobacteria; Gammaproteobacteria; Enterobacterales; Enterobacteriaceae; Salmonella; Salmonella enterica; Salmonella enterica subsp. enterica; Salmonella enterica subsp. enterica serovar Anatum MNNEETFYQAMRRKGVTRRSFLKFCSLAATSLGLGAGMTPKIAWALENKPRIPVVWIHGL WP_023243720.1 hydrogenase 1, small subunit [Salmonella enterica] Length: 372\nNumber of predicted TMHs: 2\nExp number of AAs in TMHs: 36.52147\nExp number, first 60 AAs: 19.72838\nTotal prob of N-in: 0.96170\nPOSSIBLE N-term signal sequence\ninside 1 19\nTMhelix 20 39\noutside 40 325\nTMhelix 326 348\ninside 349 372

22909 GCF_001623585.1_ASM162358v1 Salmonella enterica subsp. enterica serovar Anatum str. USDA-ARS-USMARC-1728 Proteobacteria; Gammaproteobacteria; Enterobacterales; Enterobacteriaceae; Salmonella; Salmonella enterica; Salmonella enterica subsp. enterica; Salmonella enterica subsp. enterica serovar Anatum MTGDNTLITSHGINRRDFMKLCAALAATMGLSSKAAAEMAESVSNPQRPPVIWIGAQECT WP_000145429.1 hydrogenase 2 small subunit [Salmonella enterica] Length: 372\nNumber of predicted TMHs: 1\nExp number of AAs in TMHs: 24.51309\nExp number, first 60 AAs: 0.2682\nTotal prob of N-in: 0.14402\noutside 1 331\nTMhelix 332 354\ninside 355 372

22910 GCF_000988525.3_ASM98852v3 Salmonella enterica subsp. enterica serovar Anatum str. USDA-ARS-USMARC-1735 Proteobacteria; Gammaproteobacteria; Enterobacterales; Enterobacteriaceae; Salmonella; Salmonella enterica; Salmonella enterica subsp. enterica; Salmonella enterica subsp. enterica serovar Anatum MNNEETFYQAMRRKGVTRRSFLKFCSLAATSLGLGAGMTPKIAWALENKPRIPVVWIHGL WP_023243720.1 hydrogenase 1, small subunit [Salmonella enterica] Length: 372\nNumber of predicted TMHs: 2\nExp number of AAs in TMHs: 36.52147\nExp number, first 60 AAs: 19.72838\nTotal prob of N-in: 0.96170\nPOSSIBLE N-term signal sequence\ninside 1 19\nTMhelix 20 39\noutside 40 325\nTMhelix 326 348\ninside 349 372

22911 GCF_000988525.3_ASM98852v3 Salmonella enterica subsp. enterica serovar Anatum str. USDA-ARS-USMARC-1735 Proteobacteria; Gammaproteobacteria; Enterobacterales; Enterobacteriaceae; Salmonella; Salmonella enterica; Salmonella enterica subsp. enterica; Salmonella enterica subsp. enterica serovar Anatum MTGDNTLITSHGINRRDFMKLCAALAATMGLSSKAAAEMAESVSNPQRPPVIWIGAQECT WP_000145429.1 hydrogenase 2 small subunit [Salmonella enterica] Length: 372\nNumber of predicted TMHs: 1\nExp number of AAs in TMHs: 24.51309\nExp number, first 60 AAs: 0.2682\nTotal prob of N-in: 0.14402\noutside 1 331\nTMhelix 332 354\ninside 355 372

22912 GCF_001620115.1_ASM162011v1 Salmonella enterica subsp. enterica serovar Anatum str. USDA-ARS-USMARC-1736 Proteobacteria; Gammaproteobacteria; Enterobacterales; Enterobacteriaceae; Salmonella; Salmonella

enterica; Salmonella enterica subsp. enterica; Salmonella enterica subsp. enterica serovar Anatum
MTGDNTLITSHGINRRDFMKLCAALAATMGLSSKAAAEMAESVSNPQRPPVIWIGAQECT WP_000145429.1
hydrogenase 2 small subunit [Salmonella enterica] Length: 372\nNumber of predicted TMHs: 1\nExp number of AAs in TMHs: 24.51309\nExp number, first 60 AAs: 0.2682\nTotal prob of N-in: 0.14402\noutside 1 331\nTMhelix 332 354\ninside 355 372

22913 GCF_001620115.1_ASM162011v1 Salmonella enterica subsp. enterica serovar Anatum str. USDA-ARS-USMARC-1736 Proteobacteria; Gammaproteobacteria; Enterobacterales; Enterobacteriaceae; Salmonella; Salmonella enterica; Salmonella enterica subsp. enterica; Salmonella enterica subsp. enterica serovar Anatum
MNNEETFYQAMRRKGVTRRSFLKFCSLAATSLGLGAGMTPKIAWALENKPRIPVVWIHGL WP_023243720.1
hydrogenase 1, small subunit [Salmonella enterica] Length: 372\nNumber of predicted TMHs: 2\nExp number of AAs in TMHs: 36.52147\nExp number, first 60 AAs: 19.72838\nTotal prob of N-in: 0.96170\nPOSSIBLE N-term signal sequence\ninside 1 19\nTMhelix 20 39\noutside 40 325\nTMhelix 326 348\ninside 349 372

22914 GCF_001620135.2_ASM162013v2 Salmonella enterica subsp. enterica serovar Anatum str. USDA-ARS-USMARC-1765 Proteobacteria; Gammaproteobacteria; Enterobacterales; Enterobacteriaceae; Salmonella; Salmonella enterica; Salmonella enterica subsp. enterica; Salmonella enterica subsp. enterica serovar Anatum
MTGDNTLITSHGINRRDFMKLCAALAATMGLSSKAAAEMAESVSNPQRPPVIWIGAQECT WP_000145429.1
hydrogenase 2 small subunit [Salmonella enterica] Length: 372\nNumber of predicted TMHs: 1\nExp number of AAs in TMHs: 24.51309\nExp number, first 60 AAs: 0.2682\nTotal prob of N-in: 0.14402\noutside 1 331\nTMhelix 332 354\ninside 355 372

22915 GCF_001620135.2_ASM162013v2 Salmonella enterica subsp. enterica serovar Anatum str. USDA-ARS-USMARC-1765 Proteobacteria; Gammaproteobacteria; Enterobacterales; Enterobacteriaceae; Salmonella; Salmonella enterica; Salmonella enterica subsp. enterica; Salmonella enterica subsp. enterica serovar Anatum
MNNEETFYQAMRRKGVTRRSFLKFCSLAATSLGLGAGMTPKIAWALENKPRIPVVWIHGL WP_023243720.1
hydrogenase 1, small subunit [Salmonella enterica] Length: 372\nNumber of predicted TMHs: 2\nExp number of AAs in TMHs: 36.52147\nExp number, first 60 AAs: 19.72838\nTotal prob of N-in: 0.96170\nPOSSIBLE N-term signal sequence\ninside 1 19\nTMhelix 20 39\noutside 40 325\nTMhelix 326 348\ninside 349 372

22916 GCF_001623625.1_ASM162362v1 Salmonella enterica subsp. enterica serovar Anatum str. USDA-ARS-USMARC-1766 Proteobacteria; Gammaproteobacteria; Enterobacterales; Enterobacteriaceae; Salmonella; Salmonella enterica; Salmonella enterica subsp. enterica; Salmonella enterica subsp. enterica serovar Anatum
MNNEETFYQAMRRKGVTRRSFLKFCSLAATSLGLGAGMTPKIAWALENKPRIPVVWIHGL WP_023243720.1
hydrogenase 1, small subunit [Salmonella enterica] Length: 372\nNumber of predicted TMHs: 2\nExp number of AAs in TMHs: 36.52147\nExp number, first 60 AAs: 19.72838\nTotal prob of N-in: 0.96170\nPOSSIBLE N-term signal sequence\ninside 1 19\nTMhelix 20 39\noutside 40 325\nTMhelix 326 348\ninside 349 372

22917 GCF_001623625.1_ASM162362v1 Salmonella enterica subsp. enterica serovar Anatum str. USDA-ARS-USMARC-1766 Proteobacteria; Gammaproteobacteria; Enterobacterales; Enterobacteriaceae; Salmonella; Salmonella enterica; Salmonella enterica subsp. enterica; Salmonella enterica subsp. enterica serovar Anatum
MTGDNTLITSHGINRRDFMKLCAALAATMGLSSKAAAEMAESVSNPQRPPVIWIGAQECT WP_000145429.1
hydrogenase 2 small subunit [Salmonella enterica] Length: 372\nNumber of predicted TMHs: 1\nExp number of AAs in TMHs: 24.51309\nExp number, first 60 AAs: 0.2682\nTotal prob of N-in: 0.14402\noutside 1 331\nTMhelix 332 354\ninside 355 372

22918 GCF_001623665.2_ASM162366v2 Salmonella enterica subsp. enterica serovar Anatum str. USDA-ARS-USMARC-1781 Proteobacteria; Gammaproteobacteria; Enterobacterales; Enterobacteriaceae; Salmonella; Salmonella enterica; Salmonella enterica subsp. enterica; Salmonella enterica subsp. enterica serovar Anatum
MTGDNTLITSHGINRRDFMKLCAALAATMGLSSKAAAEMAESVSNPQRPPVIWIGAQECT WP_000145429.1
hydrogenase 2 small subunit [Salmonella enterica] Length: 372\nNumber of predicted TMHs: 1\nExp number of AAs in TMHs: 24.51309\nExp number, first 60 AAs: 0.2682\nTotal prob of N-in: 0.14402\noutside 1 331\nTMhelix 332 354\ninside 355 372

22919 GCF_001623665.2_ASM162366v2 Salmonella enterica subsp. enterica serovar Anatum str. USDA-ARS-USMARC-1781 Proteobacteria; Gammaproteobacteria; Enterobacterales; Enterobacteriaceae; Salmonella; Salmonella enterica; Salmonella enterica subsp. enterica; Salmonella enterica subsp. enterica serovar Anatum
MNNEETFYQAMRRKGVTRRSFLKFCSLAATSLGLGAGMTPKIAWALENKPRIPVVWIHGL WP_023243720.1
hydrogenase 1, small subunit [Salmonella enterica] Length: 372\nNumber of predicted TMHs: 2\nExp number of AAs in TMHs: 36.52147\nExp number, first 60 AAs: 19.72838\nTotal prob of N-in: 0.96170\nPOSSIBLE N-term signal sequence\ninside 1 19\nTMhelix 20 39\noutside 40 325\nTMhelix 326 348\ninside 349 372

22920	GCF_001620155.1_ASM162015v1	Salmonella enterica subsp. enterica serovar Anatum str. USDA-ARS-USMARC-1783	Proteobacteria; Gammaproteobacteria; Enterobacterales; Enterobacteriaceae; Salmonella; Salmonella enterica; Salmonella enterica subsp. enterica; Salmonella enterica subsp. enterica serovar Anatum
	MNNEETFYQAMRRKGVTRRSFLKFCSLAATSLGLGAGMTPKIAWALENKPRIPVVWIHGL	WP_023243720.1
	hydrogenase 1, small subunit [Salmonella enterica]	Length: 372\nNumber of predicted TMHs:  2\nExp number of AAs in TMHs: 36.52147\nExp number, first 60 AAs:  19.72838\nTotal prob of N-in:   0.96170\nPOSSIBLE N-term signal sequence\ninside   1   19\nTMhelix   20   39\noutside   40   325\nTMhelix   326   348\ninside   349   372

22921	GCF_001620155.1_ASM162015v1	Salmonella enterica subsp. enterica serovar Anatum str. USDA-ARS-USMARC-1783	Proteobacteria; Gammaproteobacteria; Enterobacterales; Enterobacteriaceae; Salmonella; Salmonella enterica; Salmonella enterica subsp. enterica; Salmonella enterica subsp. enterica serovar Anatum
	MTGDNTLITSHGINRRDFMKLCAALAATMGLSSKAAAEMAESVSNPQRPPVIWIGAQECT	WP_000145429.1
	hydrogenase 2 small subunit [Salmonella enterica]	Length: 372\nNumber of predicted TMHs:  1\nExp number of AAs in TMHs: 24.51309\nExp number, first 60 AAs:  0.2682\nTotal prob of N-in:   0.14402\noutside   1   331\nTMhelix   332   354\ninside   355   372

22922	GCF_001956735.1_ASM195673v1	Salmonella enterica subsp. enterica serovar Antsalova str. S01-0511
	Proteobacteria; Gammaproteobacteria; Enterobacterales; Enterobacteriaceae; Salmonella; Salmonella enterica; Salmonella enterica subsp. enterica; Salmonella enterica subsp. enterica serovar Antsalova
	MTGDNTLITSHGINRRDFMKLCAALAATMGLSSKAAAEMAESVSNPQRPPVIWIGAQECT	WP_000145431.1
	hydrogenase 2 small subunit [Salmonella enterica]	Length: 372\nNumber of predicted TMHs:  1\nExp number of AAs in TMHs: 24.51505\nExp number, first 60 AAs:  0.2682\nTotal prob of N-in:   0.14406\noutside   1   331\nTMhelix   332   354\ninside   355   372

22923	GCF_001956735.1_ASM195673v1	Salmonella enterica subsp. enterica serovar Antsalova str. S01-0511
	Proteobacteria; Gammaproteobacteria; Enterobacterales; Enterobacteriaceae; Salmonella; Salmonella enterica; Salmonella enterica subsp. enterica; Salmonella enterica subsp. enterica serovar Antsalova
	MNNEETFYQAMRRKGVTRRSFLKFCSLAATSLGLGAGMTPKIAWALENKPRIPVVWIHGL	WP_001058312.1
	hydrogenase [Salmonella enterica]	Length: 372\nNumber of predicted TMHs:  2\nExp number of AAs in TMHs: 36.51738\nExp number, first 60 AAs:  19.72849\nTotal prob of N-in:   0.96170\nPOSSIBLE N-term signal sequence\ninside   1   19\nTMhelix   20   39\noutside   40   325\nTMhelix   326   348\ninside   349   372

22924	GCF_001975285.1_ASM197528v1	Salmonella enterica subsp. enterica serovar Apapa str. SA20060561
	Proteobacteria; Gammaproteobacteria; Enterobacterales; Enterobacteriaceae; Salmonella; Salmonella enterica; Salmonella enterica subsp. enterica; Salmonella enterica subsp. enterica serovar Apapa
	MTGDNTLITSHGINRRDFMKLCAALAATMGLSSKAAAEMAESVSNPQRPPVIWIGAQECT	WP_000145429.1
	hydrogenase 2 small subunit [Salmonella enterica]	Length: 372\nNumber of predicted TMHs:  1\nExp number of AAs in TMHs: 24.51309\nExp number, first 60 AAs:  0.2682\nTotal prob of N-in:   0.14402\noutside   1   331\nTMhelix   332   354\ninside   355   372

22925	GCF_001975285.1_ASM197528v1	Salmonella enterica subsp. enterica serovar Apapa str. SA20060561
	Proteobacteria; Gammaproteobacteria; Enterobacterales; Enterobacteriaceae; Salmonella; Salmonella enterica; Salmonella enterica subsp. enterica; Salmonella enterica subsp. enterica serovar Apapa
	MNNEETFYQAMRRKGVTRRSFLKFCSLAATSLGLGAGMTPKIAWALENKPRIPVVWIHGL	WP_001058311.1
	hydrogenase [Salmonella enterica]	Length: 372\nNumber of predicted TMHs:  2\nExp number of AAs in TMHs: 37.4699899999999\nExp number, first 60 AAs:  19.72699\nTotal prob of N-in:   0.96171\nPOSSIBLE N-term signal sequence\ninside   1   19\nTMhelix   20   39\noutside   40   325\nTMhelix   326   348\ninside   349   372

22926	GCF_000688355.1_Salmonella_Aqua	Salmonella enterica subsp. enterica serovar Aqua str. NVSL2001
	Proteobacteria; Gammaproteobacteria; Enterobacterales; Enterobacteriaceae; Salmonella; Salmonella enterica; Salmonella enterica subsp. enterica; Salmonella enterica subsp. enterica serovar Aqua
	MTGDNTLITSHGINRRDFMKLCAALAATMGLSSKAAAEMAESVSNPQRPPVIWIGAQECT	WP_000145429.1
	hydrogenase 2 small subunit [Salmonella enterica]	Length: 372\nNumber of predicted TMHs:  1\nExp number of AAs in TMHs: 24.51309\nExp number, first 60 AAs:  0.2682\nTotal prob of N-in:   0.14402\noutside   1   331\nTMhelix   332   354\ninside   355   372

22927	GCF_000688355.1_Salmonella_Aqua	Salmonella enterica subsp. enterica serovar Aqua str. NVSL2001
	Proteobacteria; Gammaproteobacteria; Enterobacterales; Enterobacteriaceae; Salmonella; Salmonella enterica; Salmonella enterica subsp. enterica; Salmonella enterica subsp. enterica serovar Aqua
	MNNEETFYQAMRRKGVTRRSFLKFCSLAATSLGLGAGMTPKIAWALENKPRIPVVWIHGL	WP_001058312.1
	hydrogenase [Salmonella enterica]	Length: 372\nNumber of predicted TMHs:  2\nExp number of AAs in

TMHs: 36.51738\nExp number, first 60 AAs: 19.72849\nTotal prob of N-in: 0.96170\nPOSSIBLE N-term signal sequence\ninside 1 19\nTMhelix 20 39\noutside 40 325\nTMhelix 326 348\ninside 349 372

22928 GCF_000293075.1_ASM29307v1 Salmonella enterica subsp. enterica serovar Bareilly str. 06-0784 Proteobacteria; Gammaproteobacteria; Enterobacterales; Enterobacteriaceae; Salmonella; Salmonella enterica; Salmonella enterica subsp. enterica; Salmonella enterica subsp. enterica serovar Bareilly MNNEETFYQAMRRKGVTRRSFLKFCSLAATSLGLGAGMTPKIAWALENKPRIPVVWIHGL WP_001058312.1 hydrogenase [Salmonella enterica] Length: 372\nNumber of predicted TMHs: 2\nExp number of AAs in TMHs: 36.51738\nExp number, first 60 AAs: 19.72849\nTotal prob of N-in: 0.96170\nPOSSIBLE N-term signal sequence\ninside 1 19\nTMhelix 20 39\noutside 40 325\nTMhelix 326 348\ninside 349 372

22929 GCF_000293075.1_ASM29307v1 Salmonella enterica subsp. enterica serovar Bareilly str. 06-0784 Proteobacteria; Gammaproteobacteria; Enterobacterales; Enterobacteriaceae; Salmonella; Salmonella enterica; Salmonella enterica subsp. enterica; Salmonella enterica subsp. enterica serovar Bareilly MTGDNTLITSHGINRRDFMKLCAALAATMGLSSKAAAEMAESVSNPQRPPVIWIGAQECT WP_000145429.1 hydrogenase 2 small subunit [Salmonella enterica] Length: 372\nNumber of predicted TMHs: 1\nExp number of AAs in TMHs: 24.51309\nExp number, first 60 AAs: 0.2682\nTotal prob of N-in: 0.14402\noutside 1 331\nTMhelix 332 354\ninside 355 372

22930 GCF_000412895.1_07-0873_Ver1 Salmonella enterica subsp. enterica serovar Bareilly str. 07-0873 Proteobacteria; Gammaproteobacteria; Enterobacterales; Enterobacteriaceae; Salmonella; Salmonella enterica; Salmonella enterica subsp. enterica; Salmonella enterica subsp. enterica serovar Bareilly MTGDNTLITSHGINRRDFMKLCAALAATMGLSSKAAAEMAESVSNPQRPPVIWIGAQECT WP_000145429.1 hydrogenase 2 small subunit [Salmonella enterica] Length: 372\nNumber of predicted TMHs: 1\nExp number of AAs in TMHs: 24.51309\nExp number, first 60 AAs: 0.2682\nTotal prob of N-in: 0.14402\noutside 1 331\nTMhelix 332 354\ninside 355 372

22931 GCF_000487355.1_ASM48735v1 Salmonella enterica subsp. enterica serovar Bareilly str. ATCC 9115 Proteobacteria; Gammaproteobacteria; Enterobacterales; Enterobacteriaceae; Salmonella; Salmonella enterica; Salmonella enterica subsp. enterica; Salmonella enterica subsp. enterica serovar Bareilly MNNEETFYQAMRRKGVTRRSFLKFCSLAATSLGLGAGMTPKIAWALENKPRIPVVWIHGL WP_001058311.1 hydrogenase [Salmonella enterica] Length: 372\nNumber of predicted TMHs: 2\nExp number of AAs in TMHs: 37.4699899999999\nExp number, first 60 AAs: 19.72699\nTotal prob of N-in: 0.96171\nPOSSIBLE N-term signal sequence\ninside 1 19\nTMhelix 20 39\noutside 40 325\nTMhelix 326 348\ninside 349 372

22932 GCF_000487355.1_ASM48735v1 Salmonella enterica subsp. enterica serovar Bareilly str. ATCC 9115 Proteobacteria; Gammaproteobacteria; Enterobacterales; Enterobacteriaceae; Salmonella; Salmonella enterica; Salmonella enterica subsp. enterica; Salmonella enterica subsp. enterica serovar Bareilly MTGDNTLITSHGINRRDFMKLCAALAATMGLSSKAAAEMAESVSNPQRPPVIWIGAQECT WP_000145429.1 hydrogenase 2 small subunit [Salmonella enterica] Length: 372\nNumber of predicted TMHs: 1\nExp number of AAs in TMHs: 24.51309\nExp number, first 60 AAs: 0.2682\nTotal prob of N-in: 0.14402\noutside 1 331\nTMhelix 332 354\ninside 355 372

22933 GCF_000749455.1_CFSAN000178_01.0Salmonella enterica subsp. enterica serovar Bareilly str. CFSAN000178 Proteobacteria; Gammaproteobacteria; Enterobacterales; Enterobacteriaceae; Salmonella; Salmonella enterica; Salmonella enterica subsp. enterica; Salmonella enterica subsp. enterica serovar Bareilly MNNEETFYQAMRRKGVTRRSFLKFCSLAATSLGLGAGMTPKIAWALENKPRIPVVWIHGL WP_001058311.1 hydrogenase [Salmonella enterica] Length: 372\nNumber of predicted TMHs: 2\nExp number of AAs in TMHs: 37.4699899999999\nExp number, first 60 AAs: 19.72699\nTotal prob of N-in: 0.96171\nPOSSIBLE N-term signal sequence\ninside 1 19\nTMhelix 20 39\noutside 40 325\nTMhelix 326 348\ninside 349 372

22934 GCF_000749455.1_CFSAN000178_01.0Salmonella enterica subsp. enterica serovar Bareilly str. CFSAN000178 Proteobacteria; Gammaproteobacteria; Enterobacterales; Enterobacteriaceae; Salmonella; Salmonella enterica; Salmonella enterica subsp. enterica; Salmonella enterica subsp. enterica serovar Bareilly MTGDNTLITSHGINRRDFMKLCAALAATMGLSSKAAAEMAESVSNPQRPPVIWIGAQECT WP_000145429.1 hydrogenase 2 small subunit [Salmonella enterica] Length: 372\nNumber of predicted TMHs: 1\nExp number of AAs in TMHs: 24.51309\nExp number, first 60 AAs: 0.2682\nTotal prob of N-in: 0.14402\noutside 1 331\nTMhelix 332 354\ninside 355 372

22935 GCF_000698675.1_CFSAN000179_01.0Salmonella enterica subsp. enterica serovar Bareilly str. CFSAN000179 Proteobacteria; Gammaproteobacteria; Enterobacterales; Enterobacteriaceae; Salmonella; Salmonella enterica;

Salmonella enterica subsp. enterica; Salmonella enterica subsp. enterica serovar Bareilly
MTGDNTLITSHGINRRDFMKLCAALAATMGLSSKAAAEMAESVSNPQRPPVIWIGAQECT WP_000145429.1
hydrogenase 2 small subunit [Salmonella enterica] Length: 372\nNumber of predicted TMHs: 1\nExp number of AAs in TMHs: 24.51309\nExp number, first 60 AAs: 0.2682\nTotal prob of N-in: 0.14402\noutside 1 331\nTMhelix 332 354\ninside 355 372

22936 GCF_000698675.1_CFSAN000179_01.0Salmonella enterica subsp. enterica serovar Bareilly str. CFSAN000179
Proteobacteria; Gammaproteobacteria; Enterobacterales; Enterobacteriaceae; Salmonella; Salmonella enterica; Salmonella enterica subsp. enterica; Salmonella enterica subsp. enterica serovar Bareilly
MNNEETFYQAMRRKGVTRRSFLKFCSLAATSLGLGAGMTPKIAWALENKPRIPVVWIHGL WP_001058311.1
hydrogenase [Salmonella enterica] Length: 372\nNumber of predicted TMHs: 2\nExp number of AAs in TMHs: 37.4699899999999\nExp number, first 60 AAs: 19.72699\nTotal prob of N-in: 0.96171\nPOSSIBLE N-term signal sequence\ninside 1 19\nTMhelix 20 39\noutside 40 325\nTMhelix 326 348\ninside 349 372

22937 GCF_000747835.1_CFSAN000180_01.0Salmonella enterica subsp. enterica serovar Bareilly str. CFSAN000180
Proteobacteria; Gammaproteobacteria; Enterobacterales; Enterobacteriaceae; Salmonella; Salmonella enterica; Salmonella enterica subsp. enterica; Salmonella enterica subsp. enterica serovar Bareilly
MTGDNTLITSHGINRRDFMKLCAALAATMGLSSKAAAEMAESVSNPQRPPVIWIGAQECT WP_000145429.1
hydrogenase 2 small subunit [Salmonella enterica] Length: 372\nNumber of predicted TMHs: 1\nExp number of AAs in TMHs: 24.51309\nExp number, first 60 AAs: 0.2682\nTotal prob of N-in: 0.14402\noutside 1 331\nTMhelix 332 354\ninside 355 372

22938 GCF_000747835.1_CFSAN000180_01.0Salmonella enterica subsp. enterica serovar Bareilly str. CFSAN000180
Proteobacteria; Gammaproteobacteria; Enterobacterales; Enterobacteriaceae; Salmonella; Salmonella enterica; Salmonella enterica subsp. enterica; Salmonella enterica subsp. enterica serovar Bareilly
MNNEETFYQAMRRKGVTRRSFLKFCSLAATSLGLGAGMTPKIAWALENKPRIPVVWIHGL WP_001058311.1
hydrogenase [Salmonella enterica] Length: 372\nNumber of predicted TMHs: 2\nExp number of AAs in TMHs: 37.4699899999999\nExp number, first 60 AAs: 19.72699\nTotal prob of N-in: 0.96171\nPOSSIBLE N-term signal sequence\ninside 1 19\nTMhelix 20 39\noutside 40 325\nTMhelix 326 348\ninside 349 372

22939 GCF_000747825.1_CFSAN000181_01.0Salmonella enterica subsp. enterica serovar Bareilly str. CFSAN000181
Proteobacteria; Gammaproteobacteria; Enterobacterales; Enterobacteriaceae; Salmonella; Salmonella enterica; Salmonella enterica subsp. enterica; Salmonella enterica subsp. enterica serovar Bareilly
MTGDNTLITSHGINRRDFMKLCAALAATMGLSSKAAAEMAESVSNPQRPPVIWIGAQECT WP_000145429.1
hydrogenase 2 small subunit [Salmonella enterica] Length: 372\nNumber of predicted TMHs: 1\nExp number of AAs in TMHs: 24.51309\nExp number, first 60 AAs: 0.2682\nTotal prob of N-in: 0.14402\noutside 1 331\nTMhelix 332 354\ninside 355 372

22940 GCF_000747825.1_CFSAN000181_01.0Salmonella enterica subsp. enterica serovar Bareilly str. CFSAN000181
Proteobacteria; Gammaproteobacteria; Enterobacterales; Enterobacteriaceae; Salmonella; Salmonella enterica; Salmonella enterica subsp. enterica; Salmonella enterica subsp. enterica serovar Bareilly
MNNEETFYQAMRRKGVTRRSFLKFCSLAATSLGLGAGMTPKIAWALENKPRIPVVWIHGL WP_001058311.1
hydrogenase [Salmonella enterica] Length: 372\nNumber of predicted TMHs: 2\nExp number of AAs in TMHs: 37.4699899999999\nExp number, first 60 AAs: 19.72699\nTotal prob of N-in: 0.96171\nPOSSIBLE N-term signal sequence\ninside 1 19\nTMhelix 20 39\noutside 40 325\nTMhelix 326 348\ninside 349 372

22941 GCF_000747845.1_CFSAN000182_01.0Salmonella enterica subsp. enterica serovar Bareilly str. CFSAN000182
Proteobacteria; Gammaproteobacteria; Enterobacterales; Enterobacteriaceae; Salmonella; Salmonella enterica; Salmonella enterica subsp. enterica; Salmonella enterica subsp. enterica serovar Bareilly
MNNEETFYQAMRRKGVTRRSFLKFCSLAATSLGLGAGMTPKIAWALENKPRIPVVWIHGL WP_001058311.1
hydrogenase [Salmonella enterica] Length: 372\nNumber of predicted TMHs: 2\nExp number of AAs in TMHs: 37.4699899999999\nExp number, first 60 AAs: 19.72699\nTotal prob of N-in: 0.96171\nPOSSIBLE N-term signal sequence\ninside 1 19\nTMhelix 20 39\noutside 40 325\nTMhelix 326 348\ninside 349 372

22942 GCF_000747845.1_CFSAN000182_01.0Salmonella enterica subsp. enterica serovar Bareilly str. CFSAN000182
Proteobacteria; Gammaproteobacteria; Enterobacterales; Enterobacteriaceae; Salmonella; Salmonella enterica; Salmonella enterica subsp. enterica; Salmonella enterica subsp. enterica serovar Bareilly
MTGDNTLITSHGINRRDFMKLCAALAATMGLSSKAAAEMAESVSNPQRPPVIWIGAQECT WP_000145429.1
hydrogenase 2 small subunit [Salmonella enterica] Length: 372\nNumber of predicted TMHs: 1\nExp number of AAs in TMHs: 24.51309\nExp number, first 60 AAs: 0.2682\nTotal prob of N-in: 0.14402\noutside 1 331\nTMhelix 332 354\ninside 355 372

22943 GCF_000487455.1_ASM48745v1 Salmonella enterica subsp. enterica serovar Bareilly str. CFSAN000183 Proteobacteria; Gammaproteobacteria; Enterobacterales; Enterobacteriaceae; Salmonella; Salmonella enterica; Salmonella enterica subsp. enterica; Salmonella enterica subsp. enterica serovar Bareilly MTGDNTLITSHGINRRDFMKLCAALAATMGLSSKAAAEMAESVSNPQRPPVIWIGAQECT WP_000145429.1 hydrogenase 2 small subunit [Salmonella enterica] Length: 372\nNumber of predicted TMHs: 1\nExp number of AAs in TMHs: 24.51309\nExp number, first 60 AAs: 0.2682\nTotal prob of N-in: 0.14402\noutside 1 331\nTMhelix 332 354\ninside 355 372

22944 GCF_000487455.1_ASM48745v1 Salmonella enterica subsp. enterica serovar Bareilly str. CFSAN000183 Proteobacteria; Gammaproteobacteria; Enterobacterales; Enterobacteriaceae; Salmonella; Salmonella enterica; Salmonella enterica subsp. enterica; Salmonella enterica subsp. enterica serovar Bareilly MNNEETFYQAMRRKGVTRRSFLKFCSLAATSLGLGAGMTPKIAWALENKPRIPVVWIHGL WP_001058311.1 hydrogenase [Salmonella enterica] Length: 372\nNumber of predicted TMHs: 2\nExp number of AAs in TMHs: 37.4699899999999\nExp number, first 60 AAs: 19.72699\nTotal prob of N-in: 0.96171\nPOSSIBLE N-term signal sequence\ninside 1 19\nTMhelix 20 39\noutside 40 325\nTMhelix 326 348\ninside 349 372

22945 GCF_000747885.1_CFSAN000184_01.0Salmonella enterica subsp. enterica serovar Bareilly str. CFSAN000184 Proteobacteria; Gammaproteobacteria; Enterobacterales; Enterobacteriaceae; Salmonella; Salmonella enterica; Salmonella enterica subsp. enterica; Salmonella enterica subsp. enterica serovar Bareilly MNNEETFYQAMRRKGVTRRSFLKFCSLAATSLGLGAGMTPKIAWALENKPRIPVVWIHGL WP_001058311.1 hydrogenase [Salmonella enterica] Length: 372\nNumber of predicted TMHs: 2\nExp number of AAs in TMHs: 37.4699899999999\nExp number, first 60 AAs: 19.72699\nTotal prob of N-in: 0.96171\nPOSSIBLE N-term signal sequence\ninside 1 19\nTMhelix 20 39\noutside 40 325\nTMhelix 326 348\ninside 349 372

22946 GCF_000747885.1_CFSAN000184_01.0Salmonella enterica subsp. enterica serovar Bareilly str. CFSAN000184 Proteobacteria; Gammaproteobacteria; Enterobacterales; Enterobacteriaceae; Salmonella; Salmonella enterica; Salmonella enterica subsp. enterica; Salmonella enterica subsp. enterica serovar Bareilly MTGDNTLITSHGINRRDFMKLCAALAATMGLSSKAAAEMAESVSNPQRPPVIWIGAQECT WP_000145429.1 hydrogenase 2 small subunit [Salmonella enterica] Length: 372\nNumber of predicted TMHs: 1\nExp number of AAs in TMHs: 24.51309\nExp number, first 60 AAs: 0.2682\nTotal prob of N-in: 0.14402\noutside 1 331\nTMhelix 332 354\ninside 355 372

22947 GCF_000749505.1_CFSAN000185_01.0Salmonella enterica subsp. enterica serovar Bareilly str. CFSAN000185 Proteobacteria; Gammaproteobacteria; Enterobacterales; Enterobacteriaceae; Salmonella; Salmonella enterica; Salmonella enterica subsp. enterica; Salmonella enterica subsp. enterica serovar Bareilly MTGDNTLITSHGINRRDFMKLCAALAATMGLSSKAAAEMAESVSNPQRPPVIWIGAQECT WP_000145429.1 hydrogenase 2 small subunit [Salmonella enterica] Length: 372\nNumber of predicted TMHs: 1\nExp number of AAs in TMHs: 24.51309\nExp number, first 60 AAs: 0.2682\nTotal prob of N-in: 0.14402\noutside 1 331\nTMhelix 332 354\ninside 355 372

22948 GCF_000749505.1_CFSAN000185_01.0Salmonella enterica subsp. enterica serovar Bareilly str. CFSAN000185 Proteobacteria; Gammaproteobacteria; Enterobacterales; Enterobacteriaceae; Salmonella; Salmonella enterica; Salmonella enterica subsp. enterica; Salmonella enterica subsp. enterica serovar Bareilly MNNEETFYQAMRRKGVTRRSFLKFCSLAATSLGLGAGMTPKIAWALENKPRIPVVWIHGL WP_001058311.1 hydrogenase [Salmonella enterica] Length: 372\nNumber of predicted TMHs: 2\nExp number of AAs in TMHs: 37.4699899999999\nExp number, first 60 AAs: 19.72699\nTotal prob of N-in: 0.96171\nPOSSIBLE N-term signal sequence\ninside 1 19\nTMhelix 20 39\noutside 40 325\nTMhelix 326 348\ninside 349 372

22949 GCF_000749435.1_CFSAN000186_01.0Salmonella enterica subsp. enterica serovar Bareilly str. CFSAN000186 Proteobacteria; Gammaproteobacteria; Enterobacterales; Enterobacteriaceae; Salmonella; Salmonella enterica; Salmonella enterica subsp. enterica; Salmonella enterica subsp. enterica serovar Bareilly MNNEETFYQAMRRKGVTRRSFLKFCSLAATSLGLGAGMTPKIAWALENKPRIPVVWIHGL WP_001058311.1 hydrogenase [Salmonella enterica] Length: 372\nNumber of predicted TMHs: 2\nExp number of AAs in TMHs: 37.4699899999999\nExp number, first 60 AAs: 19.72699\nTotal prob of N-in: 0.96171\nPOSSIBLE N-term signal sequence\ninside 1 19\nTMhelix 20 39\noutside 40 325\nTMhelix 326 348\ninside 349 372

22950 GCF_000749435.1_CFSAN000186_01.0Salmonella enterica subsp. enterica serovar Bareilly str. CFSAN000186 Proteobacteria; Gammaproteobacteria; Enterobacterales; Enterobacteriaceae; Salmonella; Salmonella enterica; Salmonella enterica subsp. enterica; Salmonella enterica subsp. enterica serovar Bareilly MTGDNTLITSHGINRRDFMKLCAALAATMGLSSKAAAEMAESVSNPQRPPVIWIGAQECT WP_000145429.1 hydrogenase 2 small subunit [Salmonella enterica] Length: 372\nNumber of predicted TMHs: 1\nExp

number of AAs in TMHs: 24.51309\nExp number, first 60 AAs: 0.2682\nTotal prob of N-in: 0.14402\noutside 1 331\nTMhelix 332 354\ninside 355 372

22951 GCF_000749195.1_CFSAN000187_01.0Salmonella enterica subsp. enterica serovar Bareilly str. CFSAN000187 Proteobacteria; Gammaproteobacteria; Enterobacterales; Enterobacteriaceae; Salmonella; Salmonella enterica; Salmonella enterica subsp. enterica; Salmonella enterica subsp. enterica serovar Bareilly MNNEETFYQAMRRKGVTRRSFLKFCSLAATSLGLGAGMTPKIAWALENKPRIPVVWIHGL WP_001058311.1 hydrogenase [Salmonella enterica] Length: 372\nNumber of predicted TMHs: 2\nExp number of AAs in TMHs: 37.4699899999999\nExp number, first 60 AAs: 19.72699\nTotal prob of N-in: 0.96171\nPOSSIBLE N-term signal sequence\ninside 1 19\nTMhelix 20 39\noutside 40 325\nTMhelix 326 348\ninside 349 372

22952 GCF_000749195.1_CFSAN000187_01.0Salmonella enterica subsp. enterica serovar Bareilly str. CFSAN000187 Proteobacteria; Gammaproteobacteria; Enterobacterales; Enterobacteriaceae; Salmonella; Salmonella enterica; Salmonella enterica subsp. enterica; Salmonella enterica subsp. enterica serovar Bareilly MTGDNTLITSHGINRRDFMKLCAALAATMGLSSKAAAEMAESVSNPQRPPVIWIGAQECT WP_000145429.1 hydrogenase 2 small subunit [Salmonella enterica] Length: 372\nNumber of predicted TMHs: 1\nExp number of AAs in TMHs: 24.51309\nExp number, first 60 AAs: 0.2682\nTotal prob of N-in: 0.14402\noutside 1 331\nTMhelix 332 354\ninside 355 372

22953 GCF_000749215.1_CFSAN000188_01.0Salmonella enterica subsp. enterica serovar Bareilly str. CFSAN000188 Proteobacteria; Gammaproteobacteria; Enterobacterales; Enterobacteriaceae; Salmonella; Salmonella enterica; Salmonella enterica subsp. enterica; Salmonella enterica subsp. enterica serovar Bareilly MTGDNTLITSHGINRRDFMKLCAALAATMGLSSKAAAEMAESVSNPQRPPVIWIGAQECT WP_000145429.1 hydrogenase 2 small subunit [Salmonella enterica] Length: 372\nNumber of predicted TMHs: 1\nExp number of AAs in TMHs: 24.51309\nExp number, first 60 AAs: 0.2682\nTotal prob of N-in: 0.14402\noutside 1 331\nTMhelix 332 354\ninside 355 372

22954 GCF_000749215.1_CFSAN000188_01.0Salmonella enterica subsp. enterica serovar Bareilly str. CFSAN000188 Proteobacteria; Gammaproteobacteria; Enterobacterales; Enterobacteriaceae; Salmonella; Salmonella enterica; Salmonella enterica subsp. enterica; Salmonella enterica subsp. enterica serovar Bareilly MNNEETFYQAMRRKGVTRRSFLKFCSLAATSLGLGAGMTPKIAWALENKPRIPVVWIHGL WP_001058311.1 hydrogenase [Salmonella enterica] Length: 372\nNumber of predicted TMHs: 2\nExp number of AAs in TMHs: 37.4699899999999\nExp number, first 60 AAs: 19.72699\nTotal prob of N-in: 0.96171\nPOSSIBLE N-term signal sequence\ninside 1 19\nTMhelix 20 39\noutside 40 325\nTMhelix 326 348\ninside 349 372

22955 GCF_000439415.1_ASM43941v1 Salmonella enterica subsp. enterica serovar Bareilly str. CFSAN000189 Proteobacteria; Gammaproteobacteria; Enterobacterales; Enterobacteriaceae; Salmonella; Salmonella enterica; Salmonella enterica subsp. enterica; Salmonella enterica subsp. enterica serovar Bareilly MNNEETFYQAMRRKGVTRRSFLKFCSLAATSLGLGAGMTPKIAWALENKPRIPVVWIHGL WP_001058311.1 hydrogenase [Salmonella enterica] Length: 372\nNumber of predicted TMHs: 2\nExp number of AAs in TMHs: 37.4699899999999\nExp number, first 60 AAs: 19.72699\nTotal prob of N-in: 0.96171\nPOSSIBLE N-term signal sequence\ninside 1 19\nTMhelix 20 39\noutside 40 325\nTMhelix 326 348\ninside 349 372

22956 GCF_000439415.1_ASM43941v1 Salmonella enterica subsp. enterica serovar Bareilly str. CFSAN000189 Proteobacteria; Gammaproteobacteria; Enterobacterales; Enterobacteriaceae; Salmonella; Salmonella enterica; Salmonella enterica subsp. enterica; Salmonella enterica subsp. enterica serovar Bareilly MTGDNTLITSHGINRRDFMKLCAALAATMGLSSKAAAEMAESVSNPQRPPVIWIGAQECT WP_000145429.1 hydrogenase 2 small subunit [Salmonella enterica] Length: 372\nNumber of predicted TMHs: 1\nExp number of AAs in TMHs: 24.51309\nExp number, first 60 AAs: 0.2682\nTotal prob of N-in: 0.14402\noutside 1 331\nTMhelix 332 354\ninside 355 372

22957 GCF_000749235.1_CFSAN000190_01.0Salmonella enterica subsp. enterica serovar Bareilly str. CFSAN000190 Proteobacteria; Gammaproteobacteria; Enterobacterales; Enterobacteriaceae; Salmonella; Salmonella enterica; Salmonella enterica subsp. enterica; Salmonella enterica subsp. enterica serovar Bareilly MTGDNTLITSHGINRRDFMKLCAALAATMGLSSKAAAEMAESVSNPQRPPVIWIGAQECT WP_000145429.1 hydrogenase 2 small subunit [Salmonella enterica] Length: 372\nNumber of predicted TMHs: 1\nExp number of AAs in TMHs: 24.51309\nExp number, first 60 AAs: 0.2682\nTotal prob of N-in: 0.14402\noutside 1 331\nTMhelix 332 354\ninside 355 372

22958 GCF_000749235.1_CFSAN000190_01.0Salmonella enterica subsp. enterica serovar Bareilly str. CFSAN000190 Proteobacteria; Gammaproteobacteria; Enterobacterales; Enterobacteriaceae; Salmonella; Salmonella enterica;

Salmonella enterica subsp. enterica; Salmonella enterica subsp. enterica serovar Bareilly
MNNEETFYQAMRRKGVTRRSFLKFCSLAATSLGLGAGMTPKIAWALENKPRIPVVWIHGL WP_001058311.1
hydrogenase [Salmonella enterica] Length: 372\nNumber of predicted TMHs: 2\nExp number of AAs in TMHs: 37.4699899999999\nExp number, first 60 AAs: 19.72699\nTotal prob of N-in: 0.96171\nPOSSIBLE N-term signal sequence\ninside 1 19\nTMhelix 20 39\noutside 40 325\nTMhelix 326 348\ninside 349 372

22959 GCF_000698635.1_CFSAN000191_01.0Salmonella enterica subsp. enterica serovar Bareilly str. CFSAN000191
Proteobacteria; Gammaproteobacteria; Enterobacterales; Enterobacteriaceae; Salmonella; Salmonella enterica; Salmonella enterica subsp. enterica; Salmonella enterica subsp. enterica serovar Bareilly
MNNEETFYQAMRRKGVTRRSFLKFCSLAATSLGLGAGMTPKIAWALENKPRIPVVWIHGL WP_001058311.1
hydrogenase [Salmonella enterica] Length: 372\nNumber of predicted TMHs: 2\nExp number of AAs in TMHs: 37.4699899999999\nExp number, first 60 AAs: 19.72699\nTotal prob of N-in: 0.96171\nPOSSIBLE N-term signal sequence\ninside 1 19\nTMhelix 20 39\noutside 40 325\nTMhelix 326 348\ninside 349 372

22960 GCF_000698635.1_CFSAN000191_01.0Salmonella enterica subsp. enterica serovar Bareilly str. CFSAN000191
Proteobacteria; Gammaproteobacteria; Enterobacterales; Enterobacteriaceae; Salmonella; Salmonella enterica; Salmonella enterica subsp. enterica; Salmonella enterica subsp. enterica serovar Bareilly
MTGDNTLITSHGINRRDFMKLCAALAATMGLSSKAAAEMAESVSNPQRPPVIWIGAQECT WP_000145429.1
hydrogenase 2 small subunit [Salmonella enterica] Length: 372\nNumber of predicted TMHs: 1\nExp number of AAs in TMHs: 24.51309\nExp number, first 60 AAs: 0.2682\nTotal prob of N-in: 0.14402\noutside 1 331\nTMhelix 332 354\ninside 355 372

22961 GCF_000749255.1_CFSAN000192_01.0Salmonella enterica subsp. enterica serovar Bareilly str. CFSAN000192
Proteobacteria; Gammaproteobacteria; Enterobacterales; Enterobacteriaceae; Salmonella; Salmonella enterica; Salmonella enterica subsp. enterica; Salmonella enterica subsp. enterica serovar Bareilly
MTGDNTLITSHGINRRDFMKLCAALAATMGLSSKAAAEMAESVSNPQRPPVIWIGAQECT WP_000145429.1
hydrogenase 2 small subunit [Salmonella enterica] Length: 372\nNumber of predicted TMHs: 1\nExp number of AAs in TMHs: 24.51309\nExp number, first 60 AAs: 0.2682\nTotal prob of N-in: 0.14402\noutside 1 331\nTMhelix 332 354\ninside 355 372

22962 GCF_000749255.1_CFSAN000192_01.0Salmonella enterica subsp. enterica serovar Bareilly str. CFSAN000192
Proteobacteria; Gammaproteobacteria; Enterobacterales; Enterobacteriaceae; Salmonella; Salmonella enterica; Salmonella enterica subsp. enterica; Salmonella enterica subsp. enterica serovar Bareilly
MNNEETFYQAMRRKGVTRRSFLKFCSLAATSLGLGAGMTPKIAWALENKPRIPVVWIHGL WP_001058311.1
hydrogenase [Salmonella enterica] Length: 372\nNumber of predicted TMHs: 2\nExp number of AAs in TMHs: 37.4699899999999\nExp number, first 60 AAs: 19.72699\nTotal prob of N-in: 0.96171\nPOSSIBLE N-term signal sequence\ninside 1 19\nTMhelix 20 39\noutside 40 325\nTMhelix 326 348\ninside 349 372

22963 GCF_000749275.1_CFSAN000193_01.0Salmonella enterica subsp. enterica serovar Bareilly str. CFSAN000193
Proteobacteria; Gammaproteobacteria; Enterobacterales; Enterobacteriaceae; Salmonella; Salmonella enterica; Salmonella enterica subsp. enterica; Salmonella enterica subsp. enterica serovar Bareilly
MNNEETFYQAMRRKGVTRRSFLKFCSLAATSLGLGAGMTPKIAWALENKPRIPVVWIHGL WP_001058311.1
hydrogenase [Salmonella enterica] Length: 372\nNumber of predicted TMHs: 2\nExp number of AAs in TMHs: 37.4699899999999\nExp number, first 60 AAs: 19.72699\nTotal prob of N-in: 0.96171\nPOSSIBLE N-term signal sequence\ninside 1 19\nTMhelix 20 39\noutside 40 325\nTMhelix 326 348\ninside 349 372

22964 GCF_000749275.1_CFSAN000193_01.0Salmonella enterica subsp. enterica serovar Bareilly str. CFSAN000193
Proteobacteria; Gammaproteobacteria; Enterobacterales; Enterobacteriaceae; Salmonella; Salmonella enterica; Salmonella enterica subsp. enterica; Salmonella enterica subsp. enterica serovar Bareilly
MTGDNTLITSHGINRRDFMKLCAALAATMGLSSKAAAEMAESVSNPQRPPVIWIGAQECT WP_000145429.1
hydrogenase 2 small subunit [Salmonella enterica] Length: 372\nNumber of predicted TMHs: 1\nExp number of AAs in TMHs: 24.51309\nExp number, first 60 AAs: 0.2682\nTotal prob of N-in: 0.14402\noutside 1 331\nTMhelix 332 354\ninside 355 372

22965 GCF_000748105.1_CFSAN000194_01.0Salmonella enterica subsp. enterica serovar Bareilly str. CFSAN000194
Proteobacteria; Gammaproteobacteria; Enterobacterales; Enterobacteriaceae; Salmonella; Salmonella enterica; Salmonella enterica subsp. enterica; Salmonella enterica subsp. enterica serovar Bareilly
MTGDNTLITSHGINRRDFMKLCAALAATMGLSSKAAAEMAESVSNPQRPPVIWIGAQECT WP_000145429.1
hydrogenase 2 small subunit [Salmonella enterica] Length: 372\nNumber of predicted TMHs: 1\nExp number of AAs in TMHs: 24.51309\nExp number, first 60 AAs: 0.2682\nTotal prob of N-in: 0.14402\noutside 1 331\nTMhelix 332 354\ninside 355 372

22966 GCF_000748105.1_CFSAN000194_01.0Salmonella enterica subsp. enterica serovar Bareilly str. CFSAN000194 Proteobacteria; Gammaproteobacteria; Enterobacterales; Enterobacteriaceae; Salmonella; Salmonella enterica; Salmonella enterica subsp. enterica; Salmonella enterica subsp. enterica serovar Bareilly MNNEETFYQAMRRKGVTRRSFLKFCSLAATSLGLGAGMTPKIAWALENKPRIPVVWIHGL WP_001058311.1 hydrogenase [Salmonella enterica] Length: 372\nNumber of predicted TMHs: 2\nExp number of AAs in TMHs: 37.4699899999999\nExp number, first 60 AAs: 19.72699\nTotal prob of N-in: 0.96171\nPOSSIBLE N-term signal sequence\ninside 1 19\nTMhelix 20 39\noutside 40 325\nTMhelix 326 348\ninside 349 372

22967 GCF_000748125.1_CFSAN000195_01.0Salmonella enterica subsp. enterica serovar Bareilly str. CFSAN000195 Proteobacteria; Gammaproteobacteria; Enterobacterales; Enterobacteriaceae; Salmonella; Salmonella enterica; Salmonella enterica subsp. enterica; Salmonella enterica subsp. enterica serovar Bareilly MNNEETFYQAMRRKGVTRRSFLKFCSLAATSLGLGAGMTPKIAWALENKPRIPVVWIHGL WP_001058311.1 hydrogenase [Salmonella enterica] Length: 372\nNumber of predicted TMHs: 2\nExp number of AAs in TMHs: 37.4699899999999\nExp number, first 60 AAs: 19.72699\nTotal prob of N-in: 0.96171\nPOSSIBLE N-term signal sequence\ninside 1 19\nTMhelix 20 39\noutside 40 325\nTMhelix 326 348\ninside 349 372

22968 GCF_000748125.1_CFSAN000195_01.0Salmonella enterica subsp. enterica serovar Bareilly str. CFSAN000195 Proteobacteria; Gammaproteobacteria; Enterobacterales; Enterobacteriaceae; Salmonella; Salmonella enterica; Salmonella enterica subsp. enterica; Salmonella enterica subsp. enterica serovar Bareilly MTGDNTLITSHGINRRDFMKLCAALAATMGLSSKAAAEMAESVSNPQRPPVIWIGAQECT WP_000145429.1 hydrogenase 2 small subunit [Salmonella enterica] Length: 372\nNumber of predicted TMHs: 1\nExp number of AAs in TMHs: 24.51309\nExp number, first 60 AAs: 0.2682\nTotal prob of N-in: 0.14402\noutside 1 331\nTMhelix 332 354\ninside 355 372

22969 GCF_000748145.1_CFSAN000196_01.0Salmonella enterica subsp. enterica serovar Bareilly str. CFSAN000196 Proteobacteria; Gammaproteobacteria; Enterobacterales; Enterobacteriaceae; Salmonella; Salmonella enterica; Salmonella enterica subsp. enterica; Salmonella enterica subsp. enterica serovar Bareilly MNNEETFYQAMRRKGVTRRSFLKFCSLAATSLGLGAGMTPKIAWALENKPRIPVVWIHGL WP_001058311.1 hydrogenase [Salmonella enterica] Length: 372\nNumber of predicted TMHs: 2\nExp number of AAs in TMHs: 37.4699899999999\nExp number, first 60 AAs: 19.72699\nTotal prob of N-in: 0.96171\nPOSSIBLE N-term signal sequence\ninside 1 19\nTMhelix 20 39\noutside 40 325\nTMhelix 326 348\ninside 349 372

22970 GCF_000748145.1_CFSAN000196_01.0Salmonella enterica subsp. enterica serovar Bareilly str. CFSAN000196 Proteobacteria; Gammaproteobacteria; Enterobacterales; Enterobacteriaceae; Salmonella; Salmonella enterica; Salmonella enterica subsp. enterica; Salmonella enterica subsp. enterica serovar Bareilly MTGDNTLITSHGINRRDFMKLCAALAATMGLSSKAAAEMAESVSNPQRPPVIWIGAQECT WP_000145429.1 hydrogenase 2 small subunit [Salmonella enterica] Length: 372\nNumber of predicted TMHs: 1\nExp number of AAs in TMHs: 24.51309\nExp number, first 60 AAs: 0.2682\nTotal prob of N-in: 0.14402\noutside 1 331\nTMhelix 332 354\ninside 355 372

22971 GCF_000487435.1_ASM48743v1 Salmonella enterica subsp. enterica serovar Bareilly str. CFSAN000197 Proteobacteria; Gammaproteobacteria; Enterobacterales; Enterobacteriaceae; Salmonella; Salmonella enterica; Salmonella enterica subsp. enterica; Salmonella enterica subsp. enterica serovar Bareilly MTGDNTLITSHGINRRDFMKLCAALAATMGLSSKAAAEMAESVSNPQRPPVIWIGAQECT WP_000145429.1 hydrogenase 2 small subunit [Salmonella enterica] Length: 372\nNumber of predicted TMHs: 1\nExp number of AAs in TMHs: 24.51309\nExp number, first 60 AAs: 0.2682\nTotal prob of N-in: 0.14402\noutside 1 331\nTMhelix 332 354\ninside 355 372

22972 GCF_000487435.1_ASM48743v1 Salmonella enterica subsp. enterica serovar Bareilly str. CFSAN000197 Proteobacteria; Gammaproteobacteria; Enterobacterales; Enterobacteriaceae; Salmonella; Salmonella enterica; Salmonella enterica subsp. enterica; Salmonella enterica subsp. enterica serovar Bareilly MNNEETFYQAMRRKGVTRRSFLKFCSLAATSLGLGAGMTPKIAWALENKPRIPVVWIHGL WP_001058312.1 hydrogenase [Salmonella enterica] Length: 372\nNumber of predicted TMHs: 2\nExp number of AAs in TMHs: 36.51738\nExp number, first 60 AAs: 19.72849\nTotal prob of N-in: 0.96170\nPOSSIBLE N-term signal sequence\ninside 1 19\nTMhelix 20 39\noutside 40 325\nTMhelix 326 348\ninside 349 372

22973 GCF_000748165.1_CFSAN000198_01.0Salmonella enterica subsp. enterica serovar Bareilly str. CFSAN000198 Proteobacteria; Gammaproteobacteria; Enterobacterales; Enterobacteriaceae; Salmonella; Salmonella enterica; Salmonella enterica subsp. enterica; Salmonella enterica subsp. enterica serovar Bareilly MTGDNTLITSHGINRRDFMKLCAALAATMGLSSKAAAEMAESVSNPQRPPVIWIGAQECT WP_000145429.1 hydrogenase 2 small subunit [Salmonella enterica] Length: 372\nNumber of predicted TMHs: 1\nExp

number of AAs in TMHs: 24.51309\nExp number, first 60 AAs: 0.2682\nTotal prob of N-in: 0.14402\noutside 1 331\nTMhelix 332 354\ninside 355 372

22974 GCF_000748165.1_CFSAN000198_01.0Salmonella enterica subsp. enterica serovar Bareilly str. CFSAN000198 Proteobacteria; Gammaproteobacteria; Enterobacterales; Enterobacteriaceae; Salmonella; Salmonella enterica; Salmonella enterica subsp. enterica; Salmonella enterica subsp. enterica serovar Bareilly MNNEETFYQAMRRKGVTRRSFLKFCSLAATSLGLGAGMTPKIAWALENKPRIPVVWIHGL WP_001058311.1 hydrogenase [Salmonella enterica] Length: 372\nNumber of predicted TMHs: 2\nExp number of AAs in TMHs: 37.4699899999999\nExp number, first 60 AAs: 19.72699\nTotal prob of N-in: 0.96171\nPOSSIBLE N-term signal sequence\ninside 1 19\nTMhelix 20 39\noutside 40 325\nTMhelix 326 348\ninside 349 372

22975 GCF_000748185.1_CFSAN000199_01.0Salmonella enterica subsp. enterica serovar Bareilly str. CFSAN000199 Proteobacteria; Gammaproteobacteria; Enterobacterales; Enterobacteriaceae; Salmonella; Salmonella enterica; Salmonella enterica subsp. enterica; Salmonella enterica subsp. enterica serovar Bareilly MTGDNTLITSHGINRRDFMKLCAALAATMGLSSKAAAEMAESVSNPQRPPVIWIGAQECT WP_000145429.1 hydrogenase 2 small subunit [Salmonella enterica] Length: 372\nNumber of predicted TMHs: 1\nExp number of AAs in TMHs: 24.51309\nExp number, first 60 AAs: 0.2682\nTotal prob of N-in: 0.14402\noutside 1 331\nTMhelix 332 354\ninside 355 372

22976 GCF_000748185.1_CFSAN000199_01.0Salmonella enterica subsp. enterica serovar Bareilly str. CFSAN000199 Proteobacteria; Gammaproteobacteria; Enterobacterales; Enterobacteriaceae; Salmonella; Salmonella enterica; Salmonella enterica subsp. enterica; Salmonella enterica subsp. enterica serovar Bareilly MNNEETFYQAMRRKGVTRRSFLKFCSLAATSLGLGAGMTPKIAWALENKPRIPVVWIHGL WP_001058311.1 hydrogenase [Salmonella enterica] Length: 372\nNumber of predicted TMHs: 2\nExp number of AAs in TMHs: 37.4699899999999\nExp number, first 60 AAs: 19.72699\nTotal prob of N-in: 0.96171\nPOSSIBLE N-term signal sequence\ninside 1 19\nTMhelix 20 39\noutside 40 325\nTMhelix 326 348\ninside 349 372

22977 GCF_000487415.1_ASM48741v1 Salmonella enterica subsp. enterica serovar Bareilly str. CFSAN000200 Proteobacteria; Gammaproteobacteria; Enterobacterales; Enterobacteriaceae; Salmonella; Salmonella enterica; Salmonella enterica subsp. enterica; Salmonella enterica subsp. enterica serovar Bareilly MTGDNTLITSHGINRRDFMKLCAALAATMGLSSKAAAEMAESVSNPQRPPVIWIGAQECT WP_000145429.1 hydrogenase 2 small subunit [Salmonella enterica] Length: 372\nNumber of predicted TMHs: 1\nExp number of AAs in TMHs: 24.51309\nExp number, first 60 AAs: 0.2682\nTotal prob of N-in: 0.14402\noutside 1 331\nTMhelix 332 354\ninside 355 372

22978 GCF_000487415.1_ASM48741v1 Salmonella enterica subsp. enterica serovar Bareilly str. CFSAN000200 Proteobacteria; Gammaproteobacteria; Enterobacterales; Enterobacteriaceae; Salmonella; Salmonella enterica; Salmonella enterica subsp. enterica; Salmonella enterica subsp. enterica serovar Bareilly MNNEETFYQAMRRKGVTRRSFLKFCSLAATSLGLGAGMTPKIAWALENKPRIPVVWIHGL WP_001058311.1 hydrogenase [Salmonella enterica] Length: 372\nNumber of predicted TMHs: 2\nExp number of AAs in TMHs: 37.4699899999999\nExp number, first 60 AAs: 19.72699\nTotal prob of N-in: 0.96171\nPOSSIBLE N-term signal sequence\ninside 1 19\nTMhelix 20 39\noutside 40 325\nTMhelix 326 348\ninside 349 372

22979 GCF_000748205.1_CFSAN000201.0 Salmonella enterica subsp. enterica serovar Bareilly str. CFSAN000201 Proteobacteria; Gammaproteobacteria; Enterobacterales; Enterobacteriaceae; Salmonella; Salmonella enterica; Salmonella enterica subsp. enterica; Salmonella enterica subsp. enterica serovar Bareilly MNNEETFYQAMRRKGVTRRSFLKFCSLAATSLGLGAGMTPKIAWALENKPRIPVVWIHGL WP_001058311.1 hydrogenase [Salmonella enterica] Length: 372\nNumber of predicted TMHs: 2\nExp number of AAs in TMHs: 37.4699899999999\nExp number, first 60 AAs: 19.72699\nTotal prob of N-in: 0.96171\nPOSSIBLE N-term signal sequence\ninside 1 19\nTMhelix 20 39\noutside 40 325\nTMhelix 326 348\ninside 349 372

22980 GCF_000748205.1_CFSAN000201.0 Salmonella enterica subsp. enterica serovar Bareilly str. CFSAN000201 Proteobacteria; Gammaproteobacteria; Enterobacterales; Enterobacteriaceae; Salmonella; Salmonella enterica; Salmonella enterica subsp. enterica; Salmonella enterica subsp. enterica serovar Bareilly MTGDNTLITSHGINRRDFMKLCAALAATMGLSSKAAAEMAESVSNPQRPPVIWIGAQECT WP_000145429.1 hydrogenase 2 small subunit [Salmonella enterica] Length: 372\nNumber of predicted TMHs: 1\nExp number of AAs in TMHs: 24.51309\nExp number, first 60 AAs: 0.2682\nTotal prob of N-in: 0.14402\noutside 1 331\nTMhelix 332 354\ninside 355 372

22981 GCF_000748225.1_CFSAN000202_01.0Salmonella enterica subsp. enterica serovar Bareilly str. CFSAN000202 Proteobacteria; Gammaproteobacteria; Enterobacterales; Enterobacteriaceae; Salmonella; Salmonella enterica;

Salmonella enterica subsp. enterica; Salmonella enterica subsp. enterica serovar Bareilly
MNNEETFYQAMRRKGVTRRSFLKFCSLAATSLGLGAGMTPKIAWALENKPRIPVVWIHGL WP_001058311.1
hydrogenase [Salmonella enterica] Length: 372\nNumber of predicted TMHs: 2\nExp number of AAs in TMHs: 37.4699899999999\nExp number, first 60 AAs: 19.72699\nTotal prob of N-in: 0.96171\nPOSSIBLE N-term signal sequence\ninside 1 19\nTMhelix 20 39\noutside 40 325\nTMhelix 326 348\ninside 349 372

22982 GCF_000748225.1_CFSAN000202_01.0Salmonella enterica subsp. enterica serovar Bareilly str. CFSAN000202
Proteobacteria; Gammaproteobacteria; Enterobacterales; Enterobacteriaceae; Salmonella; Salmonella enterica; Salmonella enterica subsp. enterica; Salmonella enterica subsp. enterica serovar Bareilly
MTGDNTLITSHGINRRDFMKLCAALAATMGLSSKAAAEMAESVSNPQRPPVIWIGAQECT WP_000145429.1
hydrogenase 2 small subunit [Salmonella enterica] Length: 372\nNumber of predicted TMHs: 1\nExp number of AAs in TMHs: 24.51309\nExp number, first 60 AAs: 0.2682\nTotal prob of N-in: 0.14402\noutside 1 331\nTMhelix 332 354\ninside 355 372

22983 GCF_000698615.1_CFSAN000203_01.0Salmonella enterica subsp. enterica serovar Bareilly str. CFSAN000203
Proteobacteria; Gammaproteobacteria; Enterobacterales; Enterobacteriaceae; Salmonella; Salmonella enterica; Salmonella enterica subsp. enterica; Salmonella enterica subsp. enterica serovar Bareilly
MTGDNTLITSHGINRRDFMKLCAALAATMGLSSKAAAEMAESVSNPQRPPVIWIGAQECT WP_000145429.1
hydrogenase 2 small subunit [Salmonella enterica] Length: 372\nNumber of predicted TMHs: 1\nExp number of AAs in TMHs: 24.51309\nExp number, first 60 AAs: 0.2682\nTotal prob of N-in: 0.14402\noutside 1 331\nTMhelix 332 354\ninside 355 372

22984 GCF_000698615.1_CFSAN000203_01.0Salmonella enterica subsp. enterica serovar Bareilly str. CFSAN000203
Proteobacteria; Gammaproteobacteria; Enterobacterales; Enterobacteriaceae; Salmonella; Salmonella enterica; Salmonella enterica subsp. enterica; Salmonella enterica subsp. enterica serovar Bareilly
MNNEETFYQAMRRKGVTRRSFLKFCSLAATSLGLGAGMTPKIAWALENKPRIPVVWIHGL WP_001058311.1
hydrogenase [Salmonella enterica] Length: 372\nNumber of predicted TMHs: 2\nExp number of AAs in TMHs: 37.4699899999999\nExp number, first 60 AAs: 19.72699\nTotal prob of N-in: 0.96171\nPOSSIBLE N-term signal sequence\ninside 1 19\nTMhelix 20 39\noutside 40 325\nTMhelix 326 348\ninside 349 372

22985 GCF_000748805.1_CFSAN000204_01.0Salmonella enterica subsp. enterica serovar Bareilly str. CFSAN000204
Proteobacteria; Gammaproteobacteria; Enterobacterales; Enterobacteriaceae; Salmonella; Salmonella enterica; Salmonella enterica subsp. enterica; Salmonella enterica subsp. enterica serovar Bareilly
MNNEETFYQAMRRKGVTRRSFLKFCSLAATSLGLGAGMTPKIAWALENKPRIPVVWIHGL WP_001058311.1
hydrogenase [Salmonella enterica] Length: 372\nNumber of predicted TMHs: 2\nExp number of AAs in TMHs: 37.4699899999999\nExp number, first 60 AAs: 19.72699\nTotal prob of N-in: 0.96171\nPOSSIBLE N-term signal sequence\ninside 1 19\nTMhelix 20 39\noutside 40 325\nTMhelix 326 348\ninside 349 372

22986 GCF_000748805.1_CFSAN000204_01.0Salmonella enterica subsp. enterica serovar Bareilly str. CFSAN000204
Proteobacteria; Gammaproteobacteria; Enterobacterales; Enterobacteriaceae; Salmonella; Salmonella enterica; Salmonella enterica subsp. enterica; Salmonella enterica subsp. enterica serovar Bareilly
MTGDNTLITSHGINRRDFMKLCAALAATMGLSSKAAAEMAESVSNPQRPPVIWIGAQECT WP_000145429.1
hydrogenase 2 small subunit [Salmonella enterica] Length: 372\nNumber of predicted TMHs: 1\nExp number of AAs in TMHs: 24.51309\nExp number, first 60 AAs: 0.2682\nTotal prob of N-in: 0.14402\noutside 1 331\nTMhelix 332 354\ninside 355 372

22987 GCF_000748825.1_CFSAN000205_01.0Salmonella enterica subsp. enterica serovar Bareilly str. CFSAN000205
Proteobacteria; Gammaproteobacteria; Enterobacterales; Enterobacteriaceae; Salmonella; Salmonella enterica; Salmonella enterica subsp. enterica; Salmonella enterica subsp. enterica serovar Bareilly
MTGDNTLITSHGINRRDFMKLCAALAATMGLSSKAAAEMAESVSNPQRPPVIWIGAQECT WP_000145429.1
hydrogenase 2 small subunit [Salmonella enterica] Length: 372\nNumber of predicted TMHs: 1\nExp number of AAs in TMHs: 24.51309\nExp number, first 60 AAs: 0.2682\nTotal prob of N-in: 0.14402\noutside 1 331\nTMhelix 332 354\ninside 355 372

22988 GCF_000748825.1_CFSAN000205_01.0Salmonella enterica subsp. enterica serovar Bareilly str. CFSAN000205
Proteobacteria; Gammaproteobacteria; Enterobacterales; Enterobacteriaceae; Salmonella; Salmonella enterica; Salmonella enterica subsp. enterica; Salmonella enterica subsp. enterica serovar Bareilly
MNNEETFYQAMRRKGVTRRSFLKFCSLAATSLGLGAGMTPKIAWALENKPRIPVVWIHGL WP_001058311.1
hydrogenase [Salmonella enterica] Length: 372\nNumber of predicted TMHs: 2\nExp number of AAs in TMHs: 37.4699899999999\nExp number, first 60 AAs: 19.72699\nTotal prob of N-in: 0.96171\nPOSSIBLE N-term signal sequence\ninside 1 19\nTMhelix 20 39\noutside 40 325\nTMhelix 326 348\ninside 349 372

22989 GCF_000748845.1_CFSAN000206_01.0Salmonella enterica subsp. enterica serovar Bareilly str. CFSAN000206 Proteobacteria; Gammaproteobacteria; Enterobacterales; Enterobacteriaceae; Salmonella; Salmonella enterica; Salmonella enterica subsp. enterica; Salmonella enterica subsp. enterica serovar Bareilly MTGDNTLITSHGINRRDFMKLCAALAATMGLSSKAAAEMAESVSNPQRPPVIWIGAQECT WP_000145429.1 hydrogenase 2 small subunit [Salmonella enterica] Length: 372\nNumber of predicted TMHs: 1\nExp number of AAs in TMHs: 24.51309\nExp number, first 60 AAs: 0.2682\nTotal prob of N-in: 0.14402\noutside 1 331\nTMhelix 332 354\ninside 355 372

22990 GCF_000748845.1_CFSAN000206_01.0Salmonella enterica subsp. enterica serovar Bareilly str. CFSAN000206 Proteobacteria; Gammaproteobacteria; Enterobacterales; Enterobacteriaceae; Salmonella; Salmonella enterica; Salmonella enterica subsp. enterica; Salmonella enterica subsp. enterica serovar Bareilly MNNEETFYQAMRRKGVTRRSFLKFCSLAATSLGLGAGMTPKIAWALENKPRIPVVWIHGL WP_001058311.1 hydrogenase [Salmonella enterica] Length: 372\nNumber of predicted TMHs: 2\nExp number of AAs in TMHs: 37.4699899999999\nExp number, first 60 AAs: 19.72699\nTotal prob of N-in: 0.96171\nPOSSIBLE N-term signal sequence\ninside 1 19\nTMhelix 20 39\noutside 40 325\nTMhelix 326 348\ninside 349 372

22991 GCF_000748865.1_CFSAN000207_01.0Salmonella enterica subsp. enterica serovar Bareilly str. CFSAN000207 Proteobacteria; Gammaproteobacteria; Enterobacterales; Enterobacteriaceae; Salmonella; Salmonella enterica; Salmonella enterica subsp. enterica; Salmonella enterica subsp. enterica serovar Bareilly MTGDNTLITSHGINRRDFMKLCAALAATMGLSSKAAAEMAESVSNPQRPPVIWIGAQECT WP_000145429.1 hydrogenase 2 small subunit [Salmonella enterica] Length: 372\nNumber of predicted TMHs: 1\nExp number of AAs in TMHs: 24.51309\nExp number, first 60 AAs: 0.2682\nTotal prob of N-in: 0.14402\noutside 1 331\nTMhelix 332 354\ninside 355 372

22992 GCF_000748865.1_CFSAN000207_01.0Salmonella enterica subsp. enterica serovar Bareilly str. CFSAN000207 Proteobacteria; Gammaproteobacteria; Enterobacterales; Enterobacteriaceae; Salmonella; Salmonella enterica; Salmonella enterica subsp. enterica; Salmonella enterica subsp. enterica serovar Bareilly MNNEETFYQAMRRKGVTRRSFLKFCSLAATSLGLGAGMTPKIAWALENKPRIPVVWIHGL WP_023243720.1 hydrogenase 1, small subunit [Salmonella enterica] Length: 372\nNumber of predicted TMHs: 2\nExp number of AAs in TMHs: 36.52147\nExp number, first 60 AAs: 19.72838\nTotal prob of N-in: 0.96170\nPOSSIBLE N-term signal sequence\ninside 1 19\nTMhelix 20 39\noutside 40 325\nTMhelix 326 348\ninside 349 372

22993 GCF_000748885.1_CFSAN000208_01.0Salmonella enterica subsp. enterica serovar Bareilly str. CFSAN000208 Proteobacteria; Gammaproteobacteria; Enterobacterales; Enterobacteriaceae; Salmonella; Salmonella enterica; Salmonella enterica subsp. enterica; Salmonella enterica subsp. enterica serovar Bareilly MTGDNTLITSHGINRRDFMKLCAALAATMGLSSKAAAEMAESVSNPQRPPVIWIGAQECT WP_000145429.1 hydrogenase 2 small subunit [Salmonella enterica] Length: 372\nNumber of predicted TMHs: 1\nExp number of AAs in TMHs: 24.51309\nExp number, first 60 AAs: 0.2682\nTotal prob of N-in: 0.14402\noutside 1 331\nTMhelix 332 354\ninside 355 372

22994 GCF_000748885.1_CFSAN000208_01.0Salmonella enterica subsp. enterica serovar Bareilly str. CFSAN000208 Proteobacteria; Gammaproteobacteria; Enterobacterales; Enterobacteriaceae; Salmonella; Salmonella enterica; Salmonella enterica subsp. enterica; Salmonella enterica subsp. enterica serovar Bareilly MNNEETFYQAMRRKGVTRRSFLKFCSLAATSLGLGAGMTPKIAWALENKPRIPVVWIHGL WP_001058311.1 hydrogenase [Salmonella enterica] Length: 372\nNumber of predicted TMHs: 2\nExp number of AAs in TMHs: 37.4699899999999\nExp number, first 60 AAs: 19.72699\nTotal prob of N-in: 0.96171\nPOSSIBLE N-term signal sequence\ninside 1 19\nTMhelix 20 39\noutside 40 325\nTMhelix 326 348\ninside 349 372

22995 GCF_000748905.1_CFSAN000209_01.0Salmonella enterica subsp. enterica serovar Bareilly str. CFSAN000209 Proteobacteria; Gammaproteobacteria; Enterobacterales; Enterobacteriaceae; Salmonella; Salmonella enterica; Salmonella enterica subsp. enterica; Salmonella enterica subsp. enterica serovar Bareilly MNNEETFYQAMRRKGVTRRSFLKFCSLAATSLGLGAGMTPKIAWALENKPRIPVVWIHGL WP_001058311.1 hydrogenase [Salmonella enterica] Length: 372\nNumber of predicted TMHs: 2\nExp number of AAs in TMHs: 37.4699899999999\nExp number, first 60 AAs: 19.72699\nTotal prob of N-in: 0.96171\nPOSSIBLE N-term signal sequence\ninside 1 19\nTMhelix 20 39\noutside 40 325\nTMhelix 326 348\ninside 349 372

22996 GCF_000748905.1_CFSAN000209_01.0Salmonella enterica subsp. enterica serovar Bareilly str. CFSAN000209 Proteobacteria; Gammaproteobacteria; Enterobacterales; Enterobacteriaceae; Salmonella; Salmonella enterica; Salmonella enterica subsp. enterica; Salmonella enterica subsp. enterica serovar Bareilly MTGDNTLITSHGINRRDFMKLCAALAATMGLSSKAAAEMAESVSNPQRPPVIWIGAQECT WP_000145429.1 hydrogenase 2 small subunit [Salmonella enterica] Length: 372\nNumber of predicted TMHs: 1\nExp

number of AAs in TMHs: 24.51309\nExp number, first 60 AAs: 0.2682\nTotal prob of N-in: 0.14402\noutside 1 331\nTMhelix 332 354\ninside 355 372

22997 GCF_000748925.1_CFSAN000210_01.0Salmonella enterica subsp. enterica serovar Bareilly str. CFSAN000210 Proteobacteria; Gammaproteobacteria; Enterobacterales; Enterobacteriaceae; Salmonella; Salmonella enterica; Salmonella enterica subsp. enterica; Salmonella enterica subsp. enterica serovar Bareilly MNNEETFYQAMRRKGVTRRSFLKFCSLAATSLGLGAGMTPKIAWALENKPRIPVVWIHGL WP_001058311.1 hydrogenase [Salmonella enterica] Length: 372\nNumber of predicted TMHs: 2\nExp number of AAs in TMHs: 37.4699899999999\nExp number, first 60 AAs: 19.72699\nTotal prob of N-in: 0.96171\nPOSSIBLE N-term signal sequence\ninside 1 19\nTMhelix 20 39\noutside 40 325\nTMhelix 326 348\ninside 349 372

22998 GCF_000748925.1_CFSAN000210_01.0Salmonella enterica subsp. enterica serovar Bareilly str. CFSAN000210 Proteobacteria; Gammaproteobacteria; Enterobacterales; Enterobacteriaceae; Salmonella; Salmonella enterica; Salmonella enterica subsp. enterica; Salmonella enterica subsp. enterica serovar Bareilly MTGDNTLITSHGINRRDFMKLCAALAATMGLSSKAAAEMAESVSNPQRPPVIWIGAQECT WP_000145429.1 hydrogenase 2 small subunit [Salmonella enterica] Length: 372\nNumber of predicted TMHs: 1\nExp number of AAs in TMHs: 24.51309\nExp number, first 60 AAs: 0.2682\nTotal prob of N-in: 0.14402\noutside 1 331\nTMhelix 332 354\ninside 355 372

22999 GCF_000698715.1_CFSAN000211_01.0Salmonella enterica subsp. enterica serovar Bareilly str. CFSAN000211 Proteobacteria; Gammaproteobacteria; Enterobacterales; Enterobacteriaceae; Salmonella; Salmonella enterica; Salmonella enterica subsp. enterica; Salmonella enterica subsp. enterica serovar Bareilly MNNEETFYQAMRRKGVTRRSFLKFCSLAATSLGLGAGMTPKIAWALENKPRIPVVWIHGL WP_001058311.1 hydrogenase [Salmonella enterica] Length: 372\nNumber of predicted TMHs: 2\nExp number of AAs in TMHs: 37.4699899999999\nExp number, first 60 AAs: 19.72699\nTotal prob of N-in: 0.96171\nPOSSIBLE N-term signal sequence\ninside 1 19\nTMhelix 20 39\noutside 40 325\nTMhelix 326 348\ninside 349 372

23000 GCF_000698715.1_CFSAN000211_01.0Salmonella enterica subsp. enterica serovar Bareilly str. CFSAN000211 Proteobacteria; Gammaproteobacteria; Enterobacterales; Enterobacteriaceae; Salmonella; Salmonella enterica; Salmonella enterica subsp. enterica; Salmonella enterica subsp. enterica serovar Bareilly MTGDNTLITSHGINRRDFMKLCAALAATMGLSSKAAAEMAESVSNPQRPPVIWIGAQECT WP_000145429.1 hydrogenase 2 small subunit [Salmonella enterica] Length: 372\nNumber of predicted TMHs: 1\nExp number of AAs in TMHs: 24.51309\nExp number, first 60 AAs: 0.2682\nTotal prob of N-in: 0.14402\noutside 1 331\nTMhelix 332 354\ninside 355 372

23001 GCF_000748245.1_CFSAN000212_01.0Salmonella enterica subsp. enterica serovar Bareilly str. CFSAN000212 Proteobacteria; Gammaproteobacteria; Enterobacterales; Enterobacteriaceae; Salmonella; Salmonella enterica; Salmonella enterica subsp. enterica; Salmonella enterica subsp. enterica serovar Bareilly MNNEETFYQAMRRKGVTRRSFLKFCSLAATSLGLGAGMTPKIAWALENKPRIPVVWIHGL WP_001058311.1 hydrogenase [Salmonella enterica] Length: 372\nNumber of predicted TMHs: 2\nExp number of AAs in TMHs: 37.4699899999999\nExp number, first 60 AAs: 19.72699\nTotal prob of N-in: 0.96171\nPOSSIBLE N-term signal sequence\ninside 1 19\nTMhelix 20 39\noutside 40 325\nTMhelix 326 348\ninside 349 372

23002 GCF_000748245.1_CFSAN000212_01.0Salmonella enterica subsp. enterica serovar Bareilly str. CFSAN000212 Proteobacteria; Gammaproteobacteria; Enterobacterales; Enterobacteriaceae; Salmonella; Salmonella enterica; Salmonella enterica subsp. enterica; Salmonella enterica subsp. enterica serovar Bareilly MTGDNTLITSHGINRRDFMKLCAALAATMGLSSKAAAEMAESVSNPQRPPVIWIGAQECT WP_000145429.1 hydrogenase 2 small subunit [Salmonella enterica] Length: 372\nNumber of predicted TMHs: 1\nExp number of AAs in TMHs: 24.51309\nExp number, first 60 AAs: 0.2682\nTotal prob of N-in: 0.14402\noutside 1 331\nTMhelix 332 354\ninside 355 372

23003 GCF_000748265.1_CFSAN000213_01.0Salmonella enterica subsp. enterica serovar Bareilly str. CFSAN000213 Proteobacteria; Gammaproteobacteria; Enterobacterales; Enterobacteriaceae; Salmonella; Salmonella enterica; Salmonella enterica subsp. enterica; Salmonella enterica subsp. enterica serovar Bareilly MTGDNTLITSHGINRRDFMKLCAALAATMGLSSKAAAEMAESVSNPQRPPVIWIGAQECT WP_000145429.1 hydrogenase 2 small subunit [Salmonella enterica] Length: 372\nNumber of predicted TMHs: 1\nExp number of AAs in TMHs: 24.51309\nExp number, first 60 AAs: 0.2682\nTotal prob of N-in: 0.14402\noutside 1 331\nTMhelix 332 354\ninside 355 372

23004 GCF_000748265.1_CFSAN000213_01.0Salmonella enterica subsp. enterica serovar Bareilly str. CFSAN000213 Proteobacteria; Gammaproteobacteria; Enterobacterales; Enterobacteriaceae; Salmonella; Salmonella enterica;

Salmonella enterica subsp. enterica; Salmonella enterica subsp. enterica serovar Bareilly
MNNEETFYQAMRRKGVTRRSFLKFCSLAATSLGLGAGMTPKIAWALENKPRIPVVWIHGL WP_001058311.1
hydrogenase [Salmonella enterica] Length: 372\nNumber of predicted TMHs: 2\nExp number of AAs in TMHs: 37.4699899999999\nExp number, first 60 AAs: 19.72699\nTotal prob of N-in: 0.96171\nPOSSIBLE N-term signal sequence\ninside 1 19\nTMhelix 20 39\noutside 40 325\nTMhelix 326 348\ninside 349 372

23005 GCF_000748285.1_CFSAN000214_01.0Salmonella enterica subsp. enterica serovar Bareilly str. CFSAN000214
Proteobacteria; Gammaproteobacteria; Enterobacterales; Enterobacteriaceae; Salmonella; Salmonella enterica; Salmonella enterica subsp. enterica; Salmonella enterica subsp. enterica serovar Bareilly
MTGDNTLITSHGINRRDFMKLCAALAATMGLSSKAAAEMAESVSNPQRPPVIWIGAQECT WP_000145429.1
hydrogenase 2 small subunit [Salmonella enterica] Length: 372\nNumber of predicted TMHs: 1\nExp number of AAs in TMHs: 24.51309\nExp number, first 60 AAs: 0.2682\nTotal prob of N-in: 0.14402\noutside 1 331\nTMhelix 332 354\ninside 355 372

23006 GCF_000748285.1_CFSAN000214_01.0Salmonella enterica subsp. enterica serovar Bareilly str. CFSAN000214
Proteobacteria; Gammaproteobacteria; Enterobacterales; Enterobacteriaceae; Salmonella; Salmonella enterica; Salmonella enterica subsp. enterica; Salmonella enterica subsp. enterica serovar Bareilly
MNNEETFYQAMRRKGVTRRSFLKFCSLAATSLGLGAGMTPKIAWALENKPRIPVVWIHGL WP_001058311.1
hydrogenase [Salmonella enterica] Length: 372\nNumber of predicted TMHs: 2\nExp number of AAs in TMHs: 37.4699899999999\nExp number, first 60 AAs: 19.72699\nTotal prob of N-in: 0.96171\nPOSSIBLE N-term signal sequence\ninside 1 19\nTMhelix 20 39\noutside 40 325\nTMhelix 326 348\ninside 349 372

23007 GCF_000748305.1_CFSAN000215_01.0Salmonella enterica subsp. enterica serovar Bareilly str. CFSAN000215
Proteobacteria; Gammaproteobacteria; Enterobacterales; Enterobacteriaceae; Salmonella; Salmonella enterica; Salmonella enterica subsp. enterica; Salmonella enterica subsp. enterica serovar Bareilly
MTGDNTLITSHGINRRDFMKLCAALAATMGLSSKAAAEMAESVSNPQRPPVIWIGAQECT WP_000145429.1
hydrogenase 2 small subunit [Salmonella enterica] Length: 372\nNumber of predicted TMHs: 1\nExp number of AAs in TMHs: 24.51309\nExp number, first 60 AAs: 0.2682\nTotal prob of N-in: 0.14402\noutside 1 331\nTMhelix 332 354\ninside 355 372

23008 GCF_000748305.1_CFSAN000215_01.0Salmonella enterica subsp. enterica serovar Bareilly str. CFSAN000215
Proteobacteria; Gammaproteobacteria; Enterobacterales; Enterobacteriaceae; Salmonella; Salmonella enterica; Salmonella enterica subsp. enterica; Salmonella enterica subsp. enterica serovar Bareilly
MNNEETFYQAMRRKGVTRRSFLKFCSLAATSLGLGAGMTPKIAWALENKPRIPVVWIHGL WP_001058311.1
hydrogenase [Salmonella enterica] Length: 372\nNumber of predicted TMHs: 2\nExp number of AAs in TMHs: 37.4699899999999\nExp number, first 60 AAs: 19.72699\nTotal prob of N-in: 0.96171\nPOSSIBLE N-term signal sequence\ninside 1 19\nTMhelix 20 39\noutside 40 325\nTMhelix 326 348\ninside 349 372

23009 GCF_000748325.1_CFSAN000216_01.0Salmonella enterica subsp. enterica serovar Bareilly str. CFSAN000216
Proteobacteria; Gammaproteobacteria; Enterobacterales; Enterobacteriaceae; Salmonella; Salmonella enterica; Salmonella enterica subsp. enterica; Salmonella enterica subsp. enterica serovar Bareilly
MNNEETFYQAMRRKGVTRRSFLKFCSLAATSLGLGAGMTPKIAWALENKPRIPVVWIHGL WP_001058311.1
hydrogenase [Salmonella enterica] Length: 372\nNumber of predicted TMHs: 2\nExp number of AAs in TMHs: 37.4699899999999\nExp number, first 60 AAs: 19.72699\nTotal prob of N-in: 0.96171\nPOSSIBLE N-term signal sequence\ninside 1 19\nTMhelix 20 39\noutside 40 325\nTMhelix 326 348\ninside 349 372

23010 GCF_000748325.1_CFSAN000216_01.0Salmonella enterica subsp. enterica serovar Bareilly str. CFSAN000216
Proteobacteria; Gammaproteobacteria; Enterobacterales; Enterobacteriaceae; Salmonella; Salmonella enterica; Salmonella enterica subsp. enterica; Salmonella enterica subsp. enterica serovar Bareilly
MTGDNTLITSHGINRRDFMKLCAALAATMGLSSKAAAEMAESVSNPQRPPVIWIGAQECT WP_000145429.1
hydrogenase 2 small subunit [Salmonella enterica] Length: 372\nNumber of predicted TMHs: 1\nExp number of AAs in TMHs: 24.51309\nExp number, first 60 AAs: 0.2682\nTotal prob of N-in: 0.14402\noutside 1 331\nTMhelix 332 354\ninside 355 372

23011 GCF_000748345.1_CFSAN000217_01.0Salmonella enterica subsp. enterica serovar Bareilly str. CFSAN000217
Proteobacteria; Gammaproteobacteria; Enterobacterales; Enterobacteriaceae; Salmonella; Salmonella enterica; Salmonella enterica subsp. enterica; Salmonella enterica subsp. enterica serovar Bareilly
MTGDNTLITSHGINRRDFMKLCAALAATMGLSSKAAAEMAESVSNPQRPPVIWIGAQECT WP_000145429.1
hydrogenase 2 small subunit [Salmonella enterica] Length: 372\nNumber of predicted TMHs: 1\nExp number of AAs in TMHs: 24.51309\nExp number, first 60 AAs: 0.2682\nTotal prob of N-in: 0.14402\noutside 1 331\nTMhelix 332 354\ninside 355 372

23012	GCF_000748345.1_CFSAN000217_01.0Salmonella enterica subsp. enterica serovar Bareilly str. CFSAN000217	Proteobacteria; Gammaproteobacteria; Enterobacterales; Enterobacteriaceae; Salmonella; Salmonella enterica; Salmonella enterica subsp. enterica; Salmonella enterica subsp. enterica serovar Bareilly	MNNEETFYQAMRRKGVTRRSFLKFCSLAATSLGLGAGMTPKIAWALENKPRIPVVWIHGL	WP_001058311.1	hydrogenase [Salmonella enterica]	Length: 372\nNumber of predicted TMHs:  2\nExp number of AAs in TMHs: 37.4699899999999\nExp number, first 60 AAs:  19.72699\nTotal prob of N-in:   0.96171\nPOSSIBLE N-term signal sequence\ninside   1   19\nTMhelix   20   39\noutside   40   325\nTMhelix   326   348\ninside   349   372

23013	GCF_000748365.1_CFSAN000218_01.0Salmonella enterica subsp. enterica serovar Bareilly str. CFSAN000218	Proteobacteria; Gammaproteobacteria; Enterobacterales; Enterobacteriaceae; Salmonella; Salmonella enterica; Salmonella enterica subsp. enterica; Salmonella enterica subsp. enterica serovar Bareilly	MNNEETFYQAMRRKGVTRRSFLKFCSLAATSLGLGAGMTPKIAWALENKPRIPVVWIHGL	WP_001058311.1	hydrogenase [Salmonella enterica]	Length: 372\nNumber of predicted TMHs:  2\nExp number of AAs in TMHs: 37.4699899999999\nExp number, first 60 AAs:  19.72699\nTotal prob of N-in:   0.96171\nPOSSIBLE N-term signal sequence\ninside   1   19\nTMhelix   20   39\noutside   40   325\nTMhelix   326   348\ninside   349   372

23014	GCF_000748365.1_CFSAN000218_01.0Salmonella enterica subsp. enterica serovar Bareilly str. CFSAN000218	Proteobacteria; Gammaproteobacteria; Enterobacterales; Enterobacteriaceae; Salmonella; Salmonella enterica; Salmonella enterica subsp. enterica; Salmonella enterica subsp. enterica serovar Bareilly	MTGDNTLITSHGINRRDFMKLCAALAATMGLSSKAAAEMAESVSNPQRPPVIWIGAQECT	WP_000145429.1	hydrogenase 2 small subunit [Salmonella enterica]	Length: 372\nNumber of predicted TMHs:  1\nExp number of AAs in TMHs: 24.51309\nExp number, first 60 AAs:  0.2682\nTotal prob of N-in:   0.14402\noutside   1   331\nTMhelix   332   354\ninside   355   372

23015	GCF_000747905.1_CFSAN000219_01.0Salmonella enterica subsp. enterica serovar Bareilly str. CFSAN000219	Proteobacteria; Gammaproteobacteria; Enterobacterales; Enterobacteriaceae; Salmonella; Salmonella enterica; Salmonella enterica subsp. enterica; Salmonella enterica subsp. enterica serovar Bareilly	MNNEETFYQAMRRKGVTRRSFLKFCSLAATSLGLGAGMTPKIAWALENKPRIPVVWIHGL	WP_001058311.1	hydrogenase [Salmonella enterica]	Length: 372\nNumber of predicted TMHs:  2\nExp number of AAs in TMHs: 37.4699899999999\nExp number, first 60 AAs:  19.72699\nTotal prob of N-in:   0.96171\nPOSSIBLE N-term signal sequence\ninside   1   19\nTMhelix   20   39\noutside   40   325\nTMhelix   326   348\ninside   349   372

23016	GCF_000747905.1_CFSAN000219_01.0Salmonella enterica subsp. enterica serovar Bareilly str. CFSAN000219	Proteobacteria; Gammaproteobacteria; Enterobacterales; Enterobacteriaceae; Salmonella; Salmonella enterica; Salmonella enterica subsp. enterica; Salmonella enterica subsp. enterica serovar Bareilly	MTGDNTLITSHGINRRDFMKLCAALAATMGLSSKAAAEMAESVSNPQRPPVIWIGAQECT	WP_000145429.1	hydrogenase 2 small subunit [Salmonella enterica]	Length: 372\nNumber of predicted TMHs:  1\nExp number of AAs in TMHs: 24.51309\nExp number, first 60 AAs:  0.2682\nTotal prob of N-in:   0.14402\noutside   1   331\nTMhelix   332   354\ninside   355   372

23017	GCF_000757945.1_CFSAN000220.0	Salmonella enterica subsp. enterica serovar Bareilly str. CFSAN000220	Proteobacteria; Gammaproteobacteria; Enterobacterales; Enterobacteriaceae; Salmonella; Salmonella enterica; Salmonella enterica subsp. enterica; Salmonella enterica subsp. enterica serovar Bareilly	MTGDNTLITSHGINRRDFMKLCAALAATMGLSSKAAAEMAESVSNPQRPPVIWIGAQECT	WP_000145429.1	hydrogenase 2 small subunit [Salmonella enterica]	Length: 372\nNumber of predicted TMHs:  1\nExp number of AAs in TMHs: 24.51309\nExp number, first 60 AAs:  0.2682\nTotal prob of N-in:   0.14402\noutside   1   331\nTMhelix   332   354\ninside   355   372

23018	GCF_000757945.1_CFSAN000220.0	Salmonella enterica subsp. enterica serovar Bareilly str. CFSAN000220	Proteobacteria; Gammaproteobacteria; Enterobacterales; Enterobacteriaceae; Salmonella; Salmonella enterica; Salmonella enterica subsp. enterica; Salmonella enterica subsp. enterica serovar Bareilly	MNNEETFYQAMRRKGVTRRSFLKFCSLAATSLGLGAGMTPKIAWALENKPRIPVVWIHGL	WP_001058311.1	hydrogenase [Salmonella enterica]	Length: 372\nNumber of predicted TMHs:  2\nExp number of AAs in TMHs: 37.4699899999999\nExp number, first 60 AAs:  19.72699\nTotal prob of N-in:   0.96171\nPOSSIBLE N-term signal sequence\ninside   1   19\nTMhelix   20   39\noutside   40   325\nTMhelix   326   348\ninside   349   372

23019	GCF_000747925.1_CFSAN000221_01.0Salmonella enterica subsp. enterica serovar Bareilly str. CFSAN000221	Proteobacteria; Gammaproteobacteria; Enterobacterales; Enterobacteriaceae; Salmonella; Salmonella enterica; Salmonella enterica subsp. enterica; Salmonella enterica subsp. enterica serovar Bareilly	MNNEETFYQAMRRKGVTRRSFLKFCSLAATSLGLGAGMTPKIAWALENKPRIPVVWIHGL	WP_001058311.1	hydrogenase [Salmonella enterica]	Length: 372\nNumber of predicted TMHs:  2\nExp number of AAs in

TMHs: 37.4699899999999\nExp number, first 60 AAs: 19.72699\nTotal prob of N-in: 0.96171\nPOSSIBLE N-term signal sequence\ninside 1 19\nTMhelix 20 39\noutside 40 325\nTMhelix 326 348\ninside 349 372

23020 GCF_000747925.1_CFSAN000221_01.0Salmonella enterica subsp. enterica serovar Bareilly str. CFSAN000221 Proteobacteria; Gammaproteobacteria; Enterobacterales; Enterobacteriaceae; Salmonella; Salmonella enterica; Salmonella enterica subsp. enterica; Salmonella enterica subsp. enterica serovar Bareilly MTGDNTLITSHGINRRDFMKLCAALAATMGLSSKAAAEMAESVSNPQRPPVIWIGAQECT WP_000145429.1 hydrogenase 2 small subunit [Salmonella enterica] Length: 372\nNumber of predicted TMHs: 1\nExp number of AAs in TMHs: 24.51309\nExp number, first 60 AAs: 0.2682\nTotal prob of N-in: 0.14402\noutside 1 331\nTMhelix 332 354\ninside 355 372

23021 GCF_000747945.1_CFSAN000222_01.0Salmonella enterica subsp. enterica serovar Bareilly str. CFSAN000222 Proteobacteria; Gammaproteobacteria; Enterobacterales; Enterobacteriaceae; Salmonella; Salmonella enterica; Salmonella enterica subsp. enterica; Salmonella enterica subsp. enterica serovar Bareilly MTGDNTLITSHGINRRDFMKLCAALAATMGLSSKAAAEMAESVSNPQRPPVIWIGAQECT WP_000145429.1 hydrogenase 2 small subunit [Salmonella enterica] Length: 372\nNumber of predicted TMHs: 1\nExp number of AAs in TMHs: 24.51309\nExp number, first 60 AAs: 0.2682\nTotal prob of N-in: 0.14402\noutside 1 331\nTMhelix 332 354\ninside 355 372

23022 GCF_000747945.1_CFSAN000222_01.0Salmonella enterica subsp. enterica serovar Bareilly str. CFSAN000222 Proteobacteria; Gammaproteobacteria; Enterobacterales; Enterobacteriaceae; Salmonella; Salmonella enterica; Salmonella enterica subsp. enterica; Salmonella enterica subsp. enterica serovar Bareilly MNNEETFYQAMRRKGVTRRSFLKFCSLAATSLGLGAGMTPKIAWALENKPRIPVVWIHGL WP_001058311.1 hydrogenase [Salmonella enterica] Length: 372\nNumber of predicted TMHs: 2\nExp number of AAs in TMHs: 37.4699899999999\nExp number, first 60 AAs: 19.72699\nTotal prob of N-in: 0.96171\nPOSSIBLE N-term signal sequence\ninside 1 19\nTMhelix 20 39\noutside 40 325\nTMhelix 326 348\ninside 349 372

23023 GCF_000747965.1_CFSAN000223_01.0Salmonella enterica subsp. enterica serovar Bareilly str. CFSAN000223 Proteobacteria; Gammaproteobacteria; Enterobacterales; Enterobacteriaceae; Salmonella; Salmonella enterica; Salmonella enterica subsp. enterica; Salmonella enterica subsp. enterica serovar Bareilly MTGDNTLITSHGINRRDFMKLCAALAATMGLSSKAAAEMAESVSNPQRPPVIWIGAQECT WP_000145429.1 hydrogenase 2 small subunit [Salmonella enterica] Length: 372\nNumber of predicted TMHs: 1\nExp number of AAs in TMHs: 24.51309\nExp number, first 60 AAs: 0.2682\nTotal prob of N-in: 0.14402\noutside 1 331\nTMhelix 332 354\ninside 355 372

23024 GCF_000747965.1_CFSAN000223_01.0Salmonella enterica subsp. enterica serovar Bareilly str. CFSAN000223 Proteobacteria; Gammaproteobacteria; Enterobacterales; Enterobacteriaceae; Salmonella; Salmonella enterica; Salmonella enterica subsp. enterica; Salmonella enterica subsp. enterica serovar Bareilly MNNEETFYQAMRRKGVTRRSFLKFCSLAATSLGLGAGMTPKIAWALENKPRIPVVWIHGL WP_001058311.1 hydrogenase [Salmonella enterica] Length: 372\nNumber of predicted TMHs: 2\nExp number of AAs in TMHs: 37.4699899999999\nExp number, first 60 AAs: 19.72699\nTotal prob of N-in: 0.96171\nPOSSIBLE N-term signal sequence\ninside 1 19\nTMhelix 20 39\noutside 40 325\nTMhelix 326 348\ninside 349 372

23025 GCF_000747985.1_CFSAN000224_01.0Salmonella enterica subsp. enterica serovar Bareilly str. CFSAN000224 Proteobacteria; Gammaproteobacteria; Enterobacterales; Enterobacteriaceae; Salmonella; Salmonella enterica; Salmonella enterica subsp. enterica; Salmonella enterica subsp. enterica serovar Bareilly MTGDNTLITSHGINRRDFMKLCAALAATMGLSSKAAAEMAESVSNPQRPPVIWIGAQECT WP_000145429.1 hydrogenase 2 small subunit [Salmonella enterica] Length: 372\nNumber of predicted TMHs: 1\nExp number of AAs in TMHs: 24.51309\nExp number, first 60 AAs: 0.2682\nTotal prob of N-in: 0.14402\noutside 1 331\nTMhelix 332 354\ninside 355 372

23026 GCF_000747985.1_CFSAN000224_01.0Salmonella enterica subsp. enterica serovar Bareilly str. CFSAN000224 Proteobacteria; Gammaproteobacteria; Enterobacterales; Enterobacteriaceae; Salmonella; Salmonella enterica; Salmonella enterica subsp. enterica; Salmonella enterica subsp. enterica serovar Bareilly MNNEETFYQAMRRKGVTRRSFLKFCSLAATSLGLGAGMTPKIAWALENKPRIPVVWIHGL WP_001058311.1 hydrogenase [Salmonella enterica] Length: 372\nNumber of predicted TMHs: 2\nExp number of AAs in TMHs: 37.4699899999999\nExp number, first 60 AAs: 19.72699\nTotal prob of N-in: 0.96171\nPOSSIBLE N-term signal sequence\ninside 1 19\nTMhelix 20 39\noutside 40 325\nTMhelix 326 348\ninside 349 372

23027 GCF_000748005.1_CFSAN000225.0 Salmonella enterica subsp. enterica serovar Bareilly str. CFSAN000225 Proteobacteria; Gammaproteobacteria; Enterobacterales; Enterobacteriaceae; Salmonella; Salmonella enterica;

Salmonella enterica subsp. enterica; Salmonella enterica subsp. enterica serovar Bareilly
MNNEETFYQAMRRKGVTRRSFLKFCSLAATSLGLGAGMTPKIAWALENKPRIPVVWIHGL WP_001058311.1
hydrogenase [Salmonella enterica] Length: 372\nNumber of predicted TMHs: 2\nExp number of AAs in TMHs: 37.4699899999999\nExp number, first 60 AAs: 19.72699\nTotal prob of N-in: 0.96171\nPOSSIBLE N-term signal sequence\ninside 1 19\nTMhelix 20 39\noutside 40 325\nTMhelix 326 348\ninside 349 372

23028 GCF_000748005.1_CFSAN000225.0 Salmonella enterica subsp. enterica serovar Bareilly str. CFSAN000225
Proteobacteria; Gammaproteobacteria; Enterobacterales; Enterobacteriaceae; Salmonella; Salmonella enterica; Salmonella enterica subsp. enterica; Salmonella enterica subsp. enterica serovar Bareilly
MTGDNTLITSHGINRRDFMKLCAALAATMGLSSKAAAEMAESVSNPQRPPVIWIGAQECT WP_000145429.1
hydrogenase 2 small subunit [Salmonella enterica] Length: 372\nNumber of predicted TMHs: 1\nExp number of AAs in TMHs: 24.51309\nExp number, first 60 AAs: 0.2682\nTotal prob of N-in: 0.14402\noutside 1 331\nTMhelix 332 354\ninside 355 372

23029 GCF_000748545.1_CFSAN000227_01.0Salmonella enterica subsp. enterica serovar Bareilly str. CFSAN000227
Proteobacteria; Gammaproteobacteria; Enterobacterales; Enterobacteriaceae; Salmonella; Salmonella enterica; Salmonella enterica subsp. enterica; Salmonella enterica subsp. enterica serovar Bareilly
MNNEETFYQAMRRKGVTRRSFLKFCSLAATSLGLGAGMTPKIAWALENKPRIPVVWIHGL WP_001058311.1
hydrogenase [Salmonella enterica] Length: 372\nNumber of predicted TMHs: 2\nExp number of AAs in TMHs: 37.4699899999999\nExp number, first 60 AAs: 19.72699\nTotal prob of N-in: 0.96171\nPOSSIBLE N-term signal sequence\ninside 1 19\nTMhelix 20 39\noutside 40 325\nTMhelix 326 348\ninside 349 372

23030 GCF_000748545.1_CFSAN000227_01.0Salmonella enterica subsp. enterica serovar Bareilly str. CFSAN000227
Proteobacteria; Gammaproteobacteria; Enterobacterales; Enterobacteriaceae; Salmonella; Salmonella enterica; Salmonella enterica subsp. enterica; Salmonella enterica subsp. enterica serovar Bareilly
MTGDNTLITSHGINRRDFMKLCAALAATMGLSSKAAAEMAESVSNPQRPPVIWIGAQECT WP_000145429.1
hydrogenase 2 small subunit [Salmonella enterica] Length: 372\nNumber of predicted TMHs: 1\nExp number of AAs in TMHs: 24.51309\nExp number, first 60 AAs: 0.2682\nTotal prob of N-in: 0.14402\noutside 1 331\nTMhelix 332 354\ninside 355 372

23031 GCF_000748565.1_CFSAN000228_01.0Salmonella enterica subsp. enterica serovar Bareilly str. CFSAN000228
Proteobacteria; Gammaproteobacteria; Enterobacterales; Enterobacteriaceae; Salmonella; Salmonella enterica; Salmonella enterica subsp. enterica; Salmonella enterica subsp. enterica serovar Bareilly
MTGDNTLITSHGINRRDFMKLCAALAATMGLSSKAAAEMAESVSNPQRPPVIWIGAQECT WP_000145429.1
hydrogenase 2 small subunit [Salmonella enterica] Length: 372\nNumber of predicted TMHs: 1\nExp number of AAs in TMHs: 24.51309\nExp number, first 60 AAs: 0.2682\nTotal prob of N-in: 0.14402\noutside 1 331\nTMhelix 332 354\ninside 355 372

23032 GCF_000748565.1_CFSAN000228_01.0Salmonella enterica subsp. enterica serovar Bareilly str. CFSAN000228
Proteobacteria; Gammaproteobacteria; Enterobacterales; Enterobacteriaceae; Salmonella; Salmonella enterica; Salmonella enterica subsp. enterica; Salmonella enterica subsp. enterica serovar Bareilly
MNNEETFYQAMRRKGVTRRSFLKFCSLAATSLGLGAGMTPKIAWALENKPRIPVVWIHGL WP_001058311.1
hydrogenase [Salmonella enterica] Length: 372\nNumber of predicted TMHs: 2\nExp number of AAs in TMHs: 37.4699899999999\nExp number, first 60 AAs: 19.72699\nTotal prob of N-in: 0.96171\nPOSSIBLE N-term signal sequence\ninside 1 19\nTMhelix 20 39\noutside 40 325\nTMhelix 326 348\ninside 349 372

23033 GCF_000748605.1_CFSAN000231.0 Salmonella enterica subsp. enterica serovar Bareilly str. CFSAN000231
Proteobacteria; Gammaproteobacteria; Enterobacterales; Enterobacteriaceae; Salmonella; Salmonella enterica; Salmonella enterica subsp. enterica; Salmonella enterica subsp. enterica serovar Bareilly
MTGDNTLITSHGINRRDFMKLCAALAATMGLSSKAAAEMAESVSNPQRPPVIWIGAQECT WP_000145429.1
hydrogenase 2 small subunit [Salmonella enterica] Length: 372\nNumber of predicted TMHs: 1\nExp number of AAs in TMHs: 24.51309\nExp number, first 60 AAs: 0.2682\nTotal prob of N-in: 0.14402\noutside 1 331\nTMhelix 332 354\ninside 355 372

23034 GCF_000748945.1_CFSAN000235_01.0Salmonella enterica subsp. enterica serovar Bareilly str. CFSAN000235
Proteobacteria; Gammaproteobacteria; Enterobacterales; Enterobacteriaceae; Salmonella; Salmonella enterica; Salmonella enterica subsp. enterica; Salmonella enterica subsp. enterica serovar Bareilly
MNNEETFYQAMRRKGVTRRSFLKFCSLAATSLGLGAGMTPKIAWALENKPRIPVVWIHGL WP_001058312.1
hydrogenase [Salmonella enterica] Length: 372\nNumber of predicted TMHs: 2\nExp number of AAs in TMHs: 36.51738\nExp number, first 60 AAs: 19.72849\nTotal prob of N-in: 0.96170\nPOSSIBLE N-term signal sequence\ninside 1 19\nTMhelix 20 39\noutside 40 325\nTMhelix 326 348\ninside 349 372

23035 GCF_000748945.1_CFSAN000235_01.0Salmonella enterica subsp. enterica serovar Bareilly str. CFSAN000235 Proteobacteria; Gammaproteobacteria; Enterobacterales; Enterobacteriaceae; Salmonella; Salmonella enterica; Salmonella enterica subsp. enterica; Salmonella enterica subsp. enterica serovar Bareilly MTGDNTLITSHGINRRDFMKLCAALAATMGLSSKAAAEMAESVSNPQRPPVIWIGAQECT WP_000145429.1 hydrogenase 2 small subunit [Salmonella enterica] Length: 372\nNumber of predicted TMHs: 1\nExp number of AAs in TMHs: 24.51309\nExp number, first 60 AAs: 0.2682\nTotal prob of N-in: 0.14402\noutside 1 331\nTMhelix 332 354\ninside 355 372

23036 GCF_000748965.1_CFSAN000236_01.0Salmonella enterica subsp. enterica serovar Bareilly str. CFSAN000236 Proteobacteria; Gammaproteobacteria; Enterobacterales; Enterobacteriaceae; Salmonella; Salmonella enterica; Salmonella enterica subsp. enterica; Salmonella enterica subsp. enterica serovar Bareilly MTGDNTLITSHGINRRDFMKLCAALAATMGLSSKAAAEMAESVSNPQRPPVIWIGAQECT WP_000145429.1 hydrogenase 2 small subunit [Salmonella enterica] Length: 372\nNumber of predicted TMHs: 1\nExp number of AAs in TMHs: 24.51309\nExp number, first 60 AAs: 0.2682\nTotal prob of N-in: 0.14402\noutside 1 331\nTMhelix 332 354\ninside 355 372

23037 GCF_000748965.1_CFSAN000236_01.0Salmonella enterica subsp. enterica serovar Bareilly str. CFSAN000236 Proteobacteria; Gammaproteobacteria; Enterobacterales; Enterobacteriaceae; Salmonella; Salmonella enterica; Salmonella enterica subsp. enterica; Salmonella enterica subsp. enterica serovar Bareilly MNNEETFYQAMRRKGVTRRSFLKFCSLAATSLGLGAGMTPKIAWALENKPRIPVVWIHGL WP_001058312.1 hydrogenase [Salmonella enterica] Length: 372\nNumber of predicted TMHs: 2\nExp number of AAs in TMHs: 36.51738\nExp number, first 60 AAs: 19.72849\nTotal prob of N-in: 0.96170\nPOSSIBLE N-term signal sequence\ninside 1 19\nTMhelix 20 39\noutside 40 325\nTMhelix 326 348\ninside 349 372

23038 GCF_000698515.1_CFSAN000661_01.0Salmonella enterica subsp. enterica serovar Bareilly str. CFSAN000661 Proteobacteria; Gammaproteobacteria; Enterobacterales; Enterobacteriaceae; Salmonella; Salmonella enterica; Salmonella enterica subsp. enterica; Salmonella enterica subsp. enterica serovar Bareilly MNNEETFYQAMRRKGVTRRSFLKFCSLAATSLGLGAGMTPKIAWALENKPRIPVVWIHGL WP_001058311.1 hydrogenase [Salmonella enterica] Length: 372\nNumber of predicted TMHs: 2\nExp number of AAs in TMHs: 37.4699899999999\nExp number, first 60 AAs: 19.72699\nTotal prob of N-in: 0.96171\nPOSSIBLE N-term signal sequence\ninside 1 19\nTMhelix 20 39\noutside 40 325\nTMhelix 326 348\ninside 349 372

23039 GCF_000698515.1_CFSAN000661_01.0Salmonella enterica subsp. enterica serovar Bareilly str. CFSAN000661 Proteobacteria; Gammaproteobacteria; Enterobacterales; Enterobacteriaceae; Salmonella; Salmonella enterica; Salmonella enterica subsp. enterica; Salmonella enterica subsp. enterica serovar Bareilly MTGDNTLITSHGINRRDFMKLCAALAATMGLSSKAAAEMAESVSNPQRPPVIWIGAQECT WP_000145429.1 hydrogenase 2 small subunit [Salmonella enterica] Length: 372\nNumber of predicted TMHs: 1\nExp number of AAs in TMHs: 24.51309\nExp number, first 60 AAs: 0.2682\nTotal prob of N-in: 0.14402\noutside 1 331\nTMhelix 332 354\ninside 355 372

23040 GCF_000748985.1_CFSAN000662_01.0Salmonella enterica subsp. enterica serovar Bareilly str. CFSAN000662 Proteobacteria; Gammaproteobacteria; Enterobacterales; Enterobacteriaceae; Salmonella; Salmonella enterica; Salmonella enterica subsp. enterica; Salmonella enterica subsp. enterica serovar Bareilly MNNEETFYQAMRRKGVTRRSFLKFCSLAATSLGLGAGMTPKIAWALENKPRIPVVWIHGL WP_001058311.1 hydrogenase [Salmonella enterica] Length: 372\nNumber of predicted TMHs: 2\nExp number of AAs in TMHs: 37.4699899999999\nExp number, first 60 AAs: 19.72699\nTotal prob of N-in: 0.96171\nPOSSIBLE N-term signal sequence\ninside 1 19\nTMhelix 20 39\noutside 40 325\nTMhelix 326 348\ninside 349 372

23041 GCF_000748985.1_CFSAN000662_01.0Salmonella enterica subsp. enterica serovar Bareilly str. CFSAN000662 Proteobacteria; Gammaproteobacteria; Enterobacterales; Enterobacteriaceae; Salmonella; Salmonella enterica; Salmonella enterica subsp. enterica; Salmonella enterica subsp. enterica serovar Bareilly MTGDNTLITSHGINRRDFMKLCAALAATMGLSSKAAAEMAESVSNPQRPPVIWIGAQECT WP_000145429.1 hydrogenase 2 small subunit [Salmonella enterica] Length: 372\nNumber of predicted TMHs: 1\nExp number of AAs in TMHs: 24.51309\nExp number, first 60 AAs: 0.2682\nTotal prob of N-in: 0.14402\noutside 1 331\nTMhelix 332 354\ninside 355 372

23042 GCF_000749005.1_CFSAN000669_01.0Salmonella enterica subsp. enterica serovar Bareilly str. CFSAN000669 Proteobacteria; Gammaproteobacteria; Enterobacterales; Enterobacteriaceae; Salmonella; Salmonella enterica; Salmonella enterica subsp. enterica; Salmonella enterica subsp. enterica serovar Bareilly MNNEETFYQAMRRKGVTRRSFLKFCSLAATSLGLGAGMTPKIAWALENKPRIPVVWIHGL WP_001058311.1 hydrogenase [Salmonella enterica] Length: 372\nNumber of predicted TMHs: 2\nExp number of AAs in

TMHs: 37.4699899999999\nExp number, first 60 AAs: 19.72699\nTotal prob of N-in: 0.96171\nPOSSIBLE N-term signal sequence\ninside 1 19\nTMhelix 20 39\noutside 40 325\nTMhelix 326 348\ninside 349 372

23043 GCF_000749005.1_CFSAN000669_01.0Salmonella enterica subsp. enterica serovar Bareilly str. CFSAN000669 Proteobacteria; Gammaproteobacteria; Enterobacterales; Enterobacteriaceae; Salmonella; Salmonella enterica; Salmonella enterica subsp. enterica; Salmonella enterica subsp. enterica serovar Bareilly MTGDNTLITSHGINRRDFMKLCAALAATMGLSSKAAAEMAESVSNPQRPPVIWIGAQECT WP_000145429.1 hydrogenase 2 small subunit [Salmonella enterica] Length: 372\nNumber of predicted TMHs: 1\nExp number of AAs in TMHs: 24.51309\nExp number, first 60 AAs: 0.2682\nTotal prob of N-in: 0.14402\noutside 1 331\nTMhelix 332 354\ninside 355 372

23044 GCF_000749025.1_CFSAN000680_01.0Salmonella enterica subsp. enterica serovar Bareilly str. CFSAN000680 Proteobacteria; Gammaproteobacteria; Enterobacterales; Enterobacteriaceae; Salmonella; Salmonella enterica; Salmonella enterica subsp. enterica; Salmonella enterica subsp. enterica serovar Bareilly MTGDNTLITSHGINRRDFMKLCAALAATMGLSSKAAAEMAESVSNPQRPPVIWIGAQECT WP_000145429.1 hydrogenase 2 small subunit [Salmonella enterica] Length: 372\nNumber of predicted TMHs: 1\nExp number of AAs in TMHs: 24.51309\nExp number, first 60 AAs: 0.2682\nTotal prob of N-in: 0.14402\noutside 1 331\nTMhelix 332 354\ninside 355 372

23045 GCF_000749025.1_CFSAN000680_01.0Salmonella enterica subsp. enterica serovar Bareilly str. CFSAN000680 Proteobacteria; Gammaproteobacteria; Enterobacterales; Enterobacteriaceae; Salmonella; Salmonella enterica; Salmonella enterica subsp. enterica; Salmonella enterica subsp. enterica serovar Bareilly MNNEETFYQAMRRKGVTRRSFLKFCSLAATSLGLGAGMTPKIAWALENKPRIPVVWIHGL WP_001058311.1 hydrogenase [Salmonella enterica] Length: 372\nNumber of predicted TMHs: 2\nExp number of AAs in TMHs: 37.4699899999999\nExp number, first 60 AAs: 19.72699\nTotal prob of N-in: 0.96171\nPOSSIBLE N-term signal sequence\ninside 1 19\nTMhelix 20 39\noutside 40 325\nTMhelix 326 348\ninside 349 372

23046 GCF_000749045.1_CFSAN000700_01.0Salmonella enterica subsp. enterica serovar Bareilly str. CFSAN000700 Proteobacteria; Gammaproteobacteria; Enterobacterales; Enterobacteriaceae; Salmonella; Salmonella enterica; Salmonella enterica subsp. enterica; Salmonella enterica subsp. enterica serovar Bareilly MNNEETFYQAMRRKGVTRRSFLKFCSLAATSLGLGAGMTPKIAWALENKPRIPVVWIHGL WP_001058311.1 hydrogenase [Salmonella enterica] Length: 372\nNumber of predicted TMHs: 2\nExp number of AAs in TMHs: 37.4699899999999\nExp number, first 60 AAs: 19.72699\nTotal prob of N-in: 0.96171\nPOSSIBLE N-term signal sequence\ninside 1 19\nTMhelix 20 39\noutside 40 325\nTMhelix 326 348\ninside 349 372

23047 GCF_000749045.1_CFSAN000700_01.0Salmonella enterica subsp. enterica serovar Bareilly str. CFSAN000700 Proteobacteria; Gammaproteobacteria; Enterobacterales; Enterobacteriaceae; Salmonella; Salmonella enterica; Salmonella enterica subsp. enterica; Salmonella enterica subsp. enterica serovar Bareilly MTGDNTLITSHGINRRDFMKLCAALAATMGLSSKAAAEMAESVSNPQRPPVIWIGAQECT WP_000145429.1 hydrogenase 2 small subunit [Salmonella enterica] Length: 372\nNumber of predicted TMHs: 1\nExp number of AAs in TMHs: 24.51309\nExp number, first 60 AAs: 0.2682\nTotal prob of N-in: 0.14402\noutside 1 331\nTMhelix 332 354\ninside 355 372

23048 GCF_000749065.1_CFSAN000752_01.0Salmonella enterica subsp. enterica serovar Bareilly str. CFSAN000752 Proteobacteria; Gammaproteobacteria; Enterobacterales; Enterobacteriaceae; Salmonella; Salmonella enterica; Salmonella enterica subsp. enterica; Salmonella enterica subsp. enterica serovar Bareilly MNNEETFYQAMRRKGVTRRSFLKFCSLAATSLGLGAGMTPKIAWALENKPRIPVVWIHGL WP_001058311.1 hydrogenase [Salmonella enterica] Length: 372\nNumber of predicted TMHs: 2\nExp number of AAs in TMHs: 37.4699899999999\nExp number, first 60 AAs: 19.72699\nTotal prob of N-in: 0.96171\nPOSSIBLE N-term signal sequence\ninside 1 19\nTMhelix 20 39\noutside 40 325\nTMhelix 326 348\ninside 349 372

23049 GCF_000749065.1_CFSAN000752_01.0Salmonella enterica subsp. enterica serovar Bareilly str. CFSAN000752 Proteobacteria; Gammaproteobacteria; Enterobacterales; Enterobacteriaceae; Salmonella; Salmonella enterica; Salmonella enterica subsp. enterica; Salmonella enterica subsp. enterica serovar Bareilly MTGDNTLITSHGINRRDFMKLCAALAATMGLSSKAAAEMAESVSNPQRPPVIWIGAQECT WP_000145429.1 hydrogenase 2 small subunit [Salmonella enterica] Length: 372\nNumber of predicted TMHs: 1\nExp number of AAs in TMHs: 24.51309\nExp number, first 60 AAs: 0.2682\nTotal prob of N-in: 0.14402\noutside 1 331\nTMhelix 332 354\ninside 355 372

23050 GCF_000749085.1_CFSAN000753_01.0Salmonella enterica subsp. enterica serovar Bareilly str. CFSAN000753 Proteobacteria; Gammaproteobacteria; Enterobacterales; Enterobacteriaceae; Salmonella; Salmonella enterica;

Salmonella enterica subsp. enterica; Salmonella enterica subsp. enterica serovar Bareilly
MTGDNTLITSHGINRRDFMKLCAALAATMGLSSKAAAEMAESVSNPQRPPVIWIGAQECT WP_000145429.1
hydrogenase 2 small subunit [Salmonella enterica] Length: 372\nNumber of predicted TMHs: 1\nExp number of AAs in TMHs: 24.51309\nExp number, first 60 AAs: 0.2682\nTotal prob of N-in: 0.14402\noutside 1 331\nTMhelix 332 354\ninside 355 372

23051 GCF_000749085.1_CFSAN000753_01.0Salmonella enterica subsp. enterica serovar Bareilly str. CFSAN000753
Proteobacteria; Gammaproteobacteria; Enterobacterales; Enterobacteriaceae; Salmonella; Salmonella enterica; Salmonella enterica subsp. enterica; Salmonella enterica subsp. enterica serovar Bareilly
MNNEETFYQAMRRKGVTRRSFLKFCSLAATSLGLGAGMTPKIAWALENKPRIPVVWIHGL WP_001058311.1
hydrogenase [Salmonella enterica] Length: 372\nNumber of predicted TMHs: 2\nExp number of AAs in TMHs: 37.4699899999999\nExp number, first 60 AAs: 19.72699\nTotal prob of N-in: 0.96171\nPOSSIBLE N-term signal sequence\ninside 1 19\nTMhelix 20 39\noutside 40 325\nTMhelix 326 348\ninside 349 372

23052 GCF_000749105.1_CFSAN000754_01.0Salmonella enterica subsp. enterica serovar Bareilly str. CFSAN000754
Proteobacteria; Gammaproteobacteria; Enterobacterales; Enterobacteriaceae; Salmonella; Salmonella enterica; Salmonella enterica subsp. enterica; Salmonella enterica subsp. enterica serovar Bareilly
MNNEETFYQAMRRKGVTRRSFLKFCSLAATSLGLGAGMTPKIAWALENKPRIPVVWIHGL WP_001058312.1
hydrogenase [Salmonella enterica] Length: 372\nNumber of predicted TMHs: 2\nExp number of AAs in TMHs: 36.51738\nExp number, first 60 AAs: 19.72849\nTotal prob of N-in: 0.96170\nPOSSIBLE N-term signal sequence\ninside 1 19\nTMhelix 20 39\noutside 40 325\nTMhelix 326 348\ninside 349 372

23053 GCF_000749105.1_CFSAN000754_01.0Salmonella enterica subsp. enterica serovar Bareilly str. CFSAN000754
Proteobacteria; Gammaproteobacteria; Enterobacterales; Enterobacteriaceae; Salmonella; Salmonella enterica; Salmonella enterica subsp. enterica; Salmonella enterica subsp. enterica serovar Bareilly
MTGDNTLITSHGINRRDFMKLCAALAATMGLSSKAAAEMAESVSNPQRPPVIWIGAQECT WP_000145429.1
hydrogenase 2 small subunit [Salmonella enterica] Length: 372\nNumber of predicted TMHs: 1\nExp number of AAs in TMHs: 24.51309\nExp number, first 60 AAs: 0.2682\nTotal prob of N-in: 0.14402\noutside 1 331\nTMhelix 332 354\ninside 355 372

23054 GCF_000749125.1_CFSAN000755_01.0Salmonella enterica subsp. enterica serovar Bareilly str. CFSAN000755
Proteobacteria; Gammaproteobacteria; Enterobacterales; Enterobacteriaceae; Salmonella; Salmonella enterica; Salmonella enterica subsp. enterica; Salmonella enterica subsp. enterica serovar Bareilly
MNNEETFYQAMRRKGVTRRSFLKFCSLAATSLGLGAGMTPKIAWALENKPRIPVVWIHGL WP_001058311.1
hydrogenase [Salmonella enterica] Length: 372\nNumber of predicted TMHs: 2\nExp number of AAs in TMHs: 37.4699899999999\nExp number, first 60 AAs: 19.72699\nTotal prob of N-in: 0.96171\nPOSSIBLE N-term signal sequence\ninside 1 19\nTMhelix 20 39\noutside 40 325\nTMhelix 326 348\ninside 349 372

23055 GCF_000749125.1_CFSAN000755_01.0Salmonella enterica subsp. enterica serovar Bareilly str. CFSAN000755
Proteobacteria; Gammaproteobacteria; Enterobacterales; Enterobacteriaceae; Salmonella; Salmonella enterica; Salmonella enterica subsp. enterica; Salmonella enterica subsp. enterica serovar Bareilly
MTGDNTLITSHGINRRDFMKLCAALAATMGLSSKAAAEMAESVSNPQRPPVIWIGAQECT WP_000145429.1
hydrogenase 2 small subunit [Salmonella enterica] Length: 372\nNumber of predicted TMHs: 1\nExp number of AAs in TMHs: 24.51309\nExp number, first 60 AAs: 0.2682\nTotal prob of N-in: 0.14402\noutside 1 331\nTMhelix 332 354\ninside 355 372

23056 GCF_000749145.1_CFSAN000951_01.0Salmonella enterica subsp. enterica serovar Bareilly str. CFSAN000951
Proteobacteria; Gammaproteobacteria; Enterobacterales; Enterobacteriaceae; Salmonella; Salmonella enterica; Salmonella enterica subsp. enterica; Salmonella enterica subsp. enterica serovar Bareilly
MTGDNTLITSHGINRRDFMKLCAALAATMGLSSKAAAEMAESVSNPQRPPVIWIGAQECT WP_000145429.1
hydrogenase 2 small subunit [Salmonella enterica] Length: 372\nNumber of predicted TMHs: 1\nExp number of AAs in TMHs: 24.51309\nExp number, first 60 AAs: 0.2682\nTotal prob of N-in: 0.14402\noutside 1 331\nTMhelix 332 354\ninside 355 372

23057 GCF_000749145.1_CFSAN000951_01.0Salmonella enterica subsp. enterica serovar Bareilly str. CFSAN000951
Proteobacteria; Gammaproteobacteria; Enterobacterales; Enterobacteriaceae; Salmonella; Salmonella enterica; Salmonella enterica subsp. enterica; Salmonella enterica subsp. enterica serovar Bareilly
MNNEETFYQAMRRKGVTRRSFLKFCSLAATSLGLGAGMTPKIAWALENKPRIPVVWIHGL WP_001058311.1
hydrogenase [Salmonella enterica] Length: 372\nNumber of predicted TMHs: 2\nExp number of AAs in TMHs: 37.4699899999999\nExp number, first 60 AAs: 19.72699\nTotal prob of N-in: 0.96171\nPOSSIBLE N-term signal sequence\ninside 1 19\nTMhelix 20 39\noutside 40 325\nTMhelix 326 348\ninside 349 372

23058 GCF_000749165.1_CFSAN000952_01.0Salmonella enterica subsp. enterica serovar Bareilly str. CFSAN000952 Proteobacteria; Gammaproteobacteria; Enterobacterales; Enterobacteriaceae; Salmonella; Salmonella enterica; Salmonella enterica subsp. enterica; Salmonella enterica subsp. enterica serovar Bareilly MTGDNTLITSHGINRRDFMKLCAALAATMGLSSKAAAEMAESVSNPQRPPVIWIGAQECT WP_000145429.1 hydrogenase 2 small subunit [Salmonella enterica] Length: 372\nNumber of predicted TMHs: 1\nExp number of AAs in TMHs: 24.51309\nExp number, first 60 AAs: 0.2682\nTotal prob of N-in: 0.14402\noutside 1 331\nTMhelix 332 354\ninside 355 372

23059 GCF_000749165.1_CFSAN000952_01.0Salmonella enterica subsp. enterica serovar Bareilly str. CFSAN000952 Proteobacteria; Gammaproteobacteria; Enterobacterales; Enterobacteriaceae; Salmonella; Salmonella enterica; Salmonella enterica subsp. enterica; Salmonella enterica subsp. enterica serovar Bareilly MNNEETFYQAMRRKGVTRRSFLKFCSLAATSLGLGAGMTPKIAWALENKPRIPVVWIHGL WP_001058311.1 hydrogenase [Salmonella enterica] Length: 372\nNumber of predicted TMHs: 2\nExp number of AAs in TMHs: 37.4699899999999\nExp number, first 60 AAs: 19.72699\nTotal prob of N-in: 0.96171\nPOSSIBLE N-term signal sequence\ninside 1 19\nTMhelix 20 39\noutside 40 325\nTMhelix 326 348\ninside 349 372

23060 GCF_000748385.1_CFSAN000953_01.0Salmonella enterica subsp. enterica serovar Bareilly str. CFSAN000953 Proteobacteria; Gammaproteobacteria; Enterobacterales; Enterobacteriaceae; Salmonella; Salmonella enterica; Salmonella enterica subsp. enterica; Salmonella enterica subsp. enterica serovar Bareilly MTGDNTLITSHGINRRDFMKLCAALAATMGLSSKAAAEMAESVSNPQRPPVIWIGAQECT WP_000145429.1 hydrogenase 2 small subunit [Salmonella enterica] Length: 372\nNumber of predicted TMHs: 1\nExp number of AAs in TMHs: 24.51309\nExp number, first 60 AAs: 0.2682\nTotal prob of N-in: 0.14402\noutside 1 331\nTMhelix 332 354\ninside 355 372

23061 GCF_000748385.1_CFSAN000953_01.0Salmonella enterica subsp. enterica serovar Bareilly str. CFSAN000953 Proteobacteria; Gammaproteobacteria; Enterobacterales; Enterobacteriaceae; Salmonella; Salmonella enterica; Salmonella enterica subsp. enterica; Salmonella enterica subsp. enterica serovar Bareilly MNNEETFYQAMRRKGVTRRSFLKFCSLAATSLGLGAGMTPKIAWALENKPRIPVVWIHGL WP_001058311.1 hydrogenase [Salmonella enterica] Length: 372\nNumber of predicted TMHs: 2\nExp number of AAs in TMHs: 37.4699899999999\nExp number, first 60 AAs: 19.72699\nTotal prob of N-in: 0.96171\nPOSSIBLE N-term signal sequence\ninside 1 19\nTMhelix 20 39\noutside 40 325\nTMhelix 326 348\ninside 349 372

23062 GCF_000748405.1_CFSAN000954_01.0Salmonella enterica subsp. enterica serovar Bareilly str. CFSAN000954 Proteobacteria; Gammaproteobacteria; Enterobacterales; Enterobacteriaceae; Salmonella; Salmonella enterica; Salmonella enterica subsp. enterica; Salmonella enterica subsp. enterica serovar Bareilly MNNEETFYQAMRRKGVTRRSFLKFCSLAATSLGLGAGMTPKIAWALENKPRIPVVWIHGL WP_001058311.1 hydrogenase [Salmonella enterica] Length: 372\nNumber of predicted TMHs: 2\nExp number of AAs in TMHs: 37.4699899999999\nExp number, first 60 AAs: 19.72699\nTotal prob of N-in: 0.96171\nPOSSIBLE N-term signal sequence\ninside 1 19\nTMhelix 20 39\noutside 40 325\nTMhelix 326 348\ninside 349 372

23063 GCF_000748405.1_CFSAN000954_01.0Salmonella enterica subsp. enterica serovar Bareilly str. CFSAN000954 Proteobacteria; Gammaproteobacteria; Enterobacterales; Enterobacteriaceae; Salmonella; Salmonella enterica; Salmonella enterica subsp. enterica; Salmonella enterica subsp. enterica serovar Bareilly MTGDNTLITSHGINRRDFMKLCAALAATMGLSSKAAAEMAESVSNPQRPPVIWIGAQECT WP_000145429.1 hydrogenase 2 small subunit [Salmonella enterica] Length: 372\nNumber of predicted TMHs: 1\nExp number of AAs in TMHs: 24.51309\nExp number, first 60 AAs: 0.2682\nTotal prob of N-in: 0.14402\noutside 1 331\nTMhelix 332 354\ninside 355 372

23064 GCF_000748425.1_CFSAN000955_01.0Salmonella enterica subsp. enterica serovar Bareilly str. CFSAN000955 Proteobacteria; Gammaproteobacteria; Enterobacterales; Enterobacteriaceae; Salmonella; Salmonella enterica; Salmonella enterica subsp. enterica; Salmonella enterica subsp. enterica serovar Bareilly MNNEETFYQAMRRKGVTRRSFLKFCSLAATSLGLGAGMTPKIAWALENKPRIPVVWIHGL WP_001058311.1 hydrogenase [Salmonella enterica] Length: 372\nNumber of predicted TMHs: 2\nExp number of AAs in TMHs: 37.4699899999999\nExp number, first 60 AAs: 19.72699\nTotal prob of N-in: 0.96171\nPOSSIBLE N-term signal sequence\ninside 1 19\nTMhelix 20 39\noutside 40 325\nTMhelix 326 348\ninside 349 372

23065 GCF_000748425.1_CFSAN000955_01.0Salmonella enterica subsp. enterica serovar Bareilly str. CFSAN000955 Proteobacteria; Gammaproteobacteria; Enterobacterales; Enterobacteriaceae; Salmonella; Salmonella enterica; Salmonella enterica subsp. enterica; Salmonella enterica subsp. enterica serovar Bareilly MTGDNTLITSHGINRRDFMKLCAALAATMGLSSKAAAEMAESVSNPQRPPVIWIGAQECT WP_000145429.1 hydrogenase 2 small subunit [Salmonella enterica] Length: 372\nNumber of predicted TMHs: 1\nExp

number of AAs in TMHs: 24.51309\nExp number, first 60 AAs: 0.2682\nTotal prob of N-in: 0.14402\noutside 1 331\nTMhelix 332 354\ninside 355 372

23066 GCF_000757925.1_CFSAN000956_01.0Salmonella enterica subsp. enterica serovar Bareilly str. CFSAN000956 Proteobacteria; Gammaproteobacteria; Enterobacterales; Enterobacteriaceae; Salmonella; Salmonella enterica; Salmonella enterica subsp. enterica; Salmonella enterica subsp. enterica serovar Bareilly MTGDNTLITSHGINRRDFMKLCAALAATMGLSSKAAAEMAESVSNPQRPPVIWIGAQECT WP_000145429.1 hydrogenase 2 small subunit [Salmonella enterica] Length: 372\nNumber of predicted TMHs: 1\nExp number of AAs in TMHs: 24.51309\nExp number, first 60 AAs: 0.2682\nTotal prob of N-in: 0.14402\noutside 1 331\nTMhelix 332 354\ninside 355 372

23067 GCF_000757925.1_CFSAN000956_01.0Salmonella enterica subsp. enterica serovar Bareilly str. CFSAN000956 Proteobacteria; Gammaproteobacteria; Enterobacterales; Enterobacteriaceae; Salmonella; Salmonella enterica; Salmonella enterica subsp. enterica; Salmonella enterica subsp. enterica serovar Bareilly MNNEETFYQAMRRKGVTRRSFLKFCSLAATSLGLGAGMTPKIAWALENKPRIPVVWIHGL WP_001058311.1 hydrogenase [Salmonella enterica] Length: 372\nNumber of predicted TMHs: 2\nExp number of AAs in TMHs: 37.4699899999999\nExp number, first 60 AAs: 19.72699\nTotal prob of N-in: 0.96171\nPOSSIBLE N-term signal sequence\ninside 1 19\nTMhelix 20 39\noutside 40 325\nTMhelix 326 348\ninside 349 372

23068 GCF_000748445.1_CFSAN000957_01.0Salmonella enterica subsp. enterica serovar Bareilly str. CFSAN000957 Proteobacteria; Gammaproteobacteria; Enterobacterales; Enterobacteriaceae; Salmonella; Salmonella enterica; Salmonella enterica subsp. enterica; Salmonella enterica subsp. enterica serovar Bareilly MTGDNTLITSHGINRRDFMKLCAALAATMGLSSKAAAEMAESVSNPQRPPVIWIGAQECT WP_000145429.1 hydrogenase 2 small subunit [Salmonella enterica] Length: 372\nNumber of predicted TMHs: 1\nExp number of AAs in TMHs: 24.51309\nExp number, first 60 AAs: 0.2682\nTotal prob of N-in: 0.14402\noutside 1 331\nTMhelix 332 354\ninside 355 372

23069 GCF_000748445.1_CFSAN000957_01.0Salmonella enterica subsp. enterica serovar Bareilly str. CFSAN000957 Proteobacteria; Gammaproteobacteria; Enterobacterales; Enterobacteriaceae; Salmonella; Salmonella enterica; Salmonella enterica subsp. enterica; Salmonella enterica subsp. enterica serovar Bareilly MNNEETFYQAMRRKGVTRRSFLKFCSLAATSLGLGAGMTPKIAWALENKPRIPVVWIHGL WP_001058311.1 hydrogenase [Salmonella enterica] Length: 372\nNumber of predicted TMHs: 2\nExp number of AAs in TMHs: 37.4699899999999\nExp number, first 60 AAs: 19.72699\nTotal prob of N-in: 0.96171\nPOSSIBLE N-term signal sequence\ninside 1 19\nTMhelix 20 39\noutside 40 325\nTMhelix 326 348\ninside 349 372

23070 GCF_000748465.1_CFSAN000958_01.0Salmonella enterica subsp. enterica serovar Bareilly str. CFSAN000958 Proteobacteria; Gammaproteobacteria; Enterobacterales; Enterobacteriaceae; Salmonella; Salmonella enterica; Salmonella enterica subsp. enterica; Salmonella enterica subsp. enterica serovar Bareilly MNNEETFYQAMRRKGVTRRSFLKFCSLAATSLGLGAGMTPKIAWALENKPRIPVVWIHGL WP_001058311.1 hydrogenase [Salmonella enterica] Length: 372\nNumber of predicted TMHs: 2\nExp number of AAs in TMHs: 37.4699899999999\nExp number, first 60 AAs: 19.72699\nTotal prob of N-in: 0.96171\nPOSSIBLE N-term signal sequence\ninside 1 19\nTMhelix 20 39\noutside 40 325\nTMhelix 326 348\ninside 349 372

23071 GCF_000748465.1_CFSAN000958_01.0Salmonella enterica subsp. enterica serovar Bareilly str. CFSAN000958 Proteobacteria; Gammaproteobacteria; Enterobacterales; Enterobacteriaceae; Salmonella; Salmonella enterica; Salmonella enterica subsp. enterica; Salmonella enterica subsp. enterica serovar Bareilly MTGDNTLITSHGINRRDFMKLCAALAATMGLSSKAAAEMAESVSNPQRPPVIWIGAQECT WP_000145429.1 hydrogenase 2 small subunit [Salmonella enterica] Length: 372\nNumber of predicted TMHs: 1\nExp number of AAs in TMHs: 24.51309\nExp number, first 60 AAs: 0.2682\nTotal prob of N-in: 0.14402\noutside 1 331\nTMhelix 332 354\ninside 355 372

23072 GCF_000748485.1_CFSAN000959_01.0Salmonella enterica subsp. enterica serovar Bareilly str. CFSAN000959 Proteobacteria; Gammaproteobacteria; Enterobacterales; Enterobacteriaceae; Salmonella; Salmonella enterica; Salmonella enterica subsp. enterica; Salmonella enterica subsp. enterica serovar Bareilly MNNEETFYQAMRRKGVTRRSFLKFCSLAATSLGLGAGMTPKIAWALENKPRIPVVWIHGL WP_001058311.1 hydrogenase [Salmonella enterica] Length: 372\nNumber of predicted TMHs: 2\nExp number of AAs in TMHs: 37.4699899999999\nExp number, first 60 AAs: 19.72699\nTotal prob of N-in: 0.96171\nPOSSIBLE N-term signal sequence\ninside 1 19\nTMhelix 20 39\noutside 40 325\nTMhelix 326 348\ninside 349 372

23073 GCF_000748485.1_CFSAN000959_01.0Salmonella enterica subsp. enterica serovar Bareilly str. CFSAN000959 Proteobacteria; Gammaproteobacteria; Enterobacterales; Enterobacteriaceae; Salmonella; Salmonella enterica;

Salmonella enterica subsp. enterica; Salmonella enterica subsp. enterica serovar Bareilly
MTGDNTLITSHGINRRDFMKLCAALAATMGLSSKAAAEMAESVSNPQRPPVIWIGAQECT WP_000145429.1
hydrogenase 2 small subunit [Salmonella enterica] Length: 372\nNumber of predicted TMHs: 1\nExp number of AAs in TMHs: 24.51309\nExp number, first 60 AAs: 0.2682\nTotal prob of N-in: 0.14402\noutside 1 331\nTMhelix 332 354\ninside 355 372

23074 GCF_000748505.1_CFSAN000960_01.0Salmonella enterica subsp. enterica serovar Bareilly str. CFSAN000960
Proteobacteria; Gammaproteobacteria; Enterobacterales; Enterobacteriaceae; Salmonella; Salmonella enterica; Salmonella enterica subsp. enterica; Salmonella enterica subsp. enterica serovar Bareilly
MNNEETFYQAMRRKGVTRRSFLKFCSLAATSLGLGAGMTPKIAWALENKPRIPVVWIHGL WP_001058311.1
hydrogenase [Salmonella enterica] Length: 372\nNumber of predicted TMHs: 2\nExp number of AAs in TMHs: 37.4699899999999\nExp number, first 60 AAs: 19.72699\nTotal prob of N-in: 0.96171\nPOSSIBLE N-term signal sequence\ninside 1 19\nTMhelix 20 39\noutside 40 325\nTMhelix 326 348\ninside 349 372

23075 GCF_000748505.1_CFSAN000960_01.0Salmonella enterica subsp. enterica serovar Bareilly str. CFSAN000960
Proteobacteria; Gammaproteobacteria; Enterobacterales; Enterobacteriaceae; Salmonella; Salmonella enterica; Salmonella enterica subsp. enterica; Salmonella enterica subsp. enterica serovar Bareilly
MTGDNTLITSHGINRRDFMKLCAALAATMGLSSKAAAEMAESVSNPQRPPVIWIGAQECT WP_000145429.1
hydrogenase 2 small subunit [Salmonella enterica] Length: 372\nNumber of predicted TMHs: 1\nExp number of AAs in TMHs: 24.51309\nExp number, first 60 AAs: 0.2682\nTotal prob of N-in: 0.14402\noutside 1 331\nTMhelix 332 354\ninside 355 372

23076 GCF_000748525.1_CFSAN000961_01.0Salmonella enterica subsp. enterica serovar Bareilly str. CFSAN000961
Proteobacteria; Gammaproteobacteria; Enterobacterales; Enterobacteriaceae; Salmonella; Salmonella enterica; Salmonella enterica subsp. enterica; Salmonella enterica subsp. enterica serovar Bareilly
MNNEETFYQAMRRKGVTRRSFLKFCSLAATSLGLGAGMTPKIAWALENKPRIPVVWIHGL WP_001058311.1
hydrogenase [Salmonella enterica] Length: 372\nNumber of predicted TMHs: 2\nExp number of AAs in TMHs: 37.4699899999999\nExp number, first 60 AAs: 19.72699\nTotal prob of N-in: 0.96171\nPOSSIBLE N-term signal sequence\ninside 1 19\nTMhelix 20 39\noutside 40 325\nTMhelix 326 348\ninside 349 372

23077 GCF_000748525.1_CFSAN000961_01.0Salmonella enterica subsp. enterica serovar Bareilly str. CFSAN000961
Proteobacteria; Gammaproteobacteria; Enterobacterales; Enterobacteriaceae; Salmonella; Salmonella enterica; Salmonella enterica subsp. enterica; Salmonella enterica subsp. enterica serovar Bareilly
MTGDNTLITSHGINRRDFMKLCAALAATMGLSSKAAAEMAESVSNPQRPPVIWIGAQECT WP_000145429.1
hydrogenase 2 small subunit [Salmonella enterica] Length: 372\nNumber of predicted TMHs: 1\nExp number of AAs in TMHs: 24.51309\nExp number, first 60 AAs: 0.2682\nTotal prob of N-in: 0.14402\noutside 1 331\nTMhelix 332 354\ninside 355 372

23078 GCF_000757955.1_CFSAN000962_01.0Salmonella enterica subsp. enterica serovar Bareilly str. CFSAN000962
Proteobacteria; Gammaproteobacteria; Enterobacterales; Enterobacteriaceae; Salmonella; Salmonella enterica; Salmonella enterica subsp. enterica; Salmonella enterica subsp. enterica serovar Bareilly
MNNEETFYQAMRRKGVTRRSFLKFCSLAATSLGLGAGMTPKIAWALENKPRIPVVWIHGL WP_001058311.1
hydrogenase [Salmonella enterica] Length: 372\nNumber of predicted TMHs: 2\nExp number of AAs in TMHs: 37.4699899999999\nExp number, first 60 AAs: 19.72699\nTotal prob of N-in: 0.96171\nPOSSIBLE N-term signal sequence\ninside 1 19\nTMhelix 20 39\noutside 40 325\nTMhelix 326 348\ninside 349 372

23079 GCF_000757955.1_CFSAN000962_01.0Salmonella enterica subsp. enterica serovar Bareilly str. CFSAN000962
Proteobacteria; Gammaproteobacteria; Enterobacterales; Enterobacteriaceae; Salmonella; Salmonella enterica; Salmonella enterica subsp. enterica; Salmonella enterica subsp. enterica serovar Bareilly
MTGDNTLITSHGINRRDFMKLCAALAATMGLSSKAAAEMAESVSNPQRPPVIWIGAQECT WP_000145429.1
hydrogenase 2 small subunit [Salmonella enterica] Length: 372\nNumber of predicted TMHs: 1\nExp number of AAs in TMHs: 24.51309\nExp number, first 60 AAs: 0.2682\nTotal prob of N-in: 0.14402\noutside 1 331\nTMhelix 332 354\ninside 355 372

23080 GCF_000749295.1_CFSAN000963_01.0Salmonella enterica subsp. enterica serovar Bareilly str. CFSAN000963
Proteobacteria; Gammaproteobacteria; Enterobacterales; Enterobacteriaceae; Salmonella; Salmonella enterica; Salmonella enterica subsp. enterica; Salmonella enterica subsp. enterica serovar Bareilly
MTGDNTLITSHGINRRDFMKLCAALAATMGLSSKAAAEMAESVSNPQRPPVIWIGAQECT WP_000145429.1
hydrogenase 2 small subunit [Salmonella enterica] Length: 372\nNumber of predicted TMHs: 1\nExp number of AAs in TMHs: 24.51309\nExp number, first 60 AAs: 0.2682\nTotal prob of N-in: 0.14402\noutside 1 331\nTMhelix 332 354\ninside 355 372

23081 GCF_000749295.1_CFSAN000963_01.0Salmonella enterica subsp. enterica serovar Bareilly str. CFSAN000963 Proteobacteria; Gammaproteobacteria; Enterobacterales; Enterobacteriaceae; Salmonella; Salmonella enterica; Salmonella enterica subsp. enterica; Salmonella enterica subsp. enterica serovar Bareilly MNNEETFYQAMRRKGVTRRSFLKFCSLAATSLGLGAGMTPKIAWALENKPRIPVVWIHGL WP_001058311.1 hydrogenase [Salmonella enterica] Length: 372\nNumber of predicted TMHs: 2\nExp number of AAs in TMHs: 37.4699899999999\nExp number, first 60 AAs: 19.72699\nTotal prob of N-in: 0.96171\nPOSSIBLE N-term signal sequence\ninside 1 19\nTMhelix 20 39\noutside 40 325\nTMhelix 326 348\ninside 349 372

23082 GCF_000749315.1_CFSAN000964_01.0Salmonella enterica subsp. enterica serovar Bareilly str. CFSAN000964 Proteobacteria; Gammaproteobacteria; Enterobacterales; Enterobacteriaceae; Salmonella; Salmonella enterica; Salmonella enterica subsp. enterica; Salmonella enterica subsp. enterica serovar Bareilly MTGDNTLITSHGINRRDFMKLCAALAATMGLSSKAAAEMAESVSNPQRPPVIWIGAQECT WP_000145429.1 hydrogenase 2 small subunit [Salmonella enterica] Length: 372\nNumber of predicted TMHs: 1\nExp number of AAs in TMHs: 24.51309\nExp number, first 60 AAs: 0.2682\nTotal prob of N-in: 0.14402\noutside 1 331\nTMhelix 332 354\ninside 355 372

23083 GCF_000749315.1_CFSAN000964_01.0Salmonella enterica subsp. enterica serovar Bareilly str. CFSAN000964 Proteobacteria; Gammaproteobacteria; Enterobacterales; Enterobacteriaceae; Salmonella; Salmonella enterica; Salmonella enterica subsp. enterica; Salmonella enterica subsp. enterica serovar Bareilly MNNEETFYQAMRRKGVTRRSFLKFCSLAATSLGLGAGMTPKIAWALENKPRIPVVWIHGL WP_001058311.1 hydrogenase [Salmonella enterica] Length: 372\nNumber of predicted TMHs: 2\nExp number of AAs in TMHs: 37.4699899999999\nExp number, first 60 AAs: 19.72699\nTotal prob of N-in: 0.96171\nPOSSIBLE N-term signal sequence\ninside 1 19\nTMhelix 20 39\noutside 40 325\nTMhelix 326 348\ninside 349 372

23084 GCF_000749335.1_CFSAN000965_01.0Salmonella enterica subsp. enterica serovar Bareilly str. CFSAN000965 Proteobacteria; Gammaproteobacteria; Enterobacterales; Enterobacteriaceae; Salmonella; Salmonella enterica; Salmonella enterica subsp. enterica; Salmonella enterica subsp. enterica serovar Bareilly MTGDNTLITSHGINRRDFMKLCAALAATMGLSSKAAAEMAESVSNPQRPPVIWIGAQECT WP_000145429.1 hydrogenase 2 small subunit [Salmonella enterica] Length: 372\nNumber of predicted TMHs: 1\nExp number of AAs in TMHs: 24.51309\nExp number, first 60 AAs: 0.2682\nTotal prob of N-in: 0.14402\noutside 1 331\nTMhelix 332 354\ninside 355 372

23085 GCF_000749335.1_CFSAN000965_01.0Salmonella enterica subsp. enterica serovar Bareilly str. CFSAN000965 Proteobacteria; Gammaproteobacteria; Enterobacterales; Enterobacteriaceae; Salmonella; Salmonella enterica; Salmonella enterica subsp. enterica; Salmonella enterica subsp. enterica serovar Bareilly MNNEETFYQAMRRKGVTRRSFLKFCSLAATSLGLGAGMTPKIAWALENKPRIPVVWIHGL WP_001058311.1 hydrogenase [Salmonella enterica] Length: 372\nNumber of predicted TMHs: 2\nExp number of AAs in TMHs: 37.4699899999999\nExp number, first 60 AAs: 19.72699\nTotal prob of N-in: 0.96171\nPOSSIBLE N-term signal sequence\ninside 1 19\nTMhelix 20 39\noutside 40 325\nTMhelix 326 348\ninside 349 372

23086 GCF_000749355.1_CFSAN000966_01.0Salmonella enterica subsp. enterica serovar Bareilly str. CFSAN000966 Proteobacteria; Gammaproteobacteria; Enterobacterales; Enterobacteriaceae; Salmonella; Salmonella enterica; Salmonella enterica subsp. enterica; Salmonella enterica subsp. enterica serovar Bareilly MTGDNTLITSHGINRRDFMKLCAALAATMGLSSKAAAEMAESVSNPQRPPVIWIGAQECT WP_000145429.1 hydrogenase 2 small subunit [Salmonella enterica] Length: 372\nNumber of predicted TMHs: 1\nExp number of AAs in TMHs: 24.51309\nExp number, first 60 AAs: 0.2682\nTotal prob of N-in: 0.14402\noutside 1 331\nTMhelix 332 354\ninside 355 372

23087 GCF_000749355.1_CFSAN000966_01.0Salmonella enterica subsp. enterica serovar Bareilly str. CFSAN000966 Proteobacteria; Gammaproteobacteria; Enterobacterales; Enterobacteriaceae; Salmonella; Salmonella enterica; Salmonella enterica subsp. enterica; Salmonella enterica subsp. enterica serovar Bareilly MNNEETFYQAMRRKGVTRRSFLKFCSLAATSLGLGAGMTPKIAWALENKPRIPVVWIHGL WP_001058311.1 hydrogenase [Salmonella enterica] Length: 372\nNumber of predicted TMHs: 2\nExp number of AAs in TMHs: 37.4699899999999\nExp number, first 60 AAs: 19.72699\nTotal prob of N-in: 0.96171\nPOSSIBLE N-term signal sequence\ninside 1 19\nTMhelix 20 39\noutside 40 325\nTMhelix 326 348\ninside 349 372

23088 GCF_000749375.1_CFSAN000967_01.0Salmonella enterica subsp. enterica serovar Bareilly str. CFSAN000967 Proteobacteria; Gammaproteobacteria; Enterobacterales; Enterobacteriaceae; Salmonella; Salmonella enterica; Salmonella enterica subsp. enterica; Salmonella enterica subsp. enterica serovar Bareilly MTGDNTLITSHGINRRDFMKLCAALAATMGLSSKAAAEMAESVSNPQRPPVIWIGAQECT WP_000145429.1 hydrogenase 2 small subunit [Salmonella enterica] Length: 372\nNumber of predicted TMHs: 1\nExp

number of AAs in TMHs: 24.51309\nExp number, first 60 AAs: 0.2682\nTotal prob of N-in: 0.14402\noutside 1 331\nTMhelix 332 354\ninside 355 372

23089 GCF_000749375.1_CFSAN000967_01.0Salmonella enterica subsp. enterica serovar Bareilly str. CFSAN000967 Proteobacteria; Gammaproteobacteria; Enterobacterales; Enterobacteriaceae; Salmonella; Salmonella enterica; Salmonella enterica subsp. enterica; Salmonella enterica subsp. enterica serovar Bareilly MNNEETFYQAMRRKGVTRRSFLKFCSLAATSLGLGAGMTPKIAWALENKPRIPVVWIHGL WP_001058311.1 hydrogenase [Salmonella enterica] Length: 372\nNumber of predicted TMHs: 2\nExp number of AAs in TMHs: 37.4699899999999\nExp number, first 60 AAs: 19.72699\nTotal prob of N-in: 0.96171\nPOSSIBLE N-term signal sequence\ninside 1 19\nTMhelix 20 39\noutside 40 325\nTMhelix 326 348\ninside 349 372

23090 GCF_000698535.1_CFSAN000968_01.0Salmonella enterica subsp. enterica serovar Bareilly str. CFSAN000968 Proteobacteria; Gammaproteobacteria; Enterobacterales; Enterobacteriaceae; Salmonella; Salmonella enterica; Salmonella enterica subsp. enterica; Salmonella enterica subsp. enterica serovar Bareilly MTGDNTLITSHGINRRDFMKLCAALAATMGLSSKAAAEMAESVSNPQRPPVIWIGAQECT WP_000145429.1 hydrogenase 2 small subunit [Salmonella enterica] Length: 372\nNumber of predicted TMHs: 1\nExp number of AAs in TMHs: 24.51309\nExp number, first 60 AAs: 0.2682\nTotal prob of N-in: 0.14402\noutside 1 331\nTMhelix 332 354\ninside 355 372

23091 GCF_000698535.1_CFSAN000968_01.0Salmonella enterica subsp. enterica serovar Bareilly str. CFSAN000968 Proteobacteria; Gammaproteobacteria; Enterobacterales; Enterobacteriaceae; Salmonella; Salmonella enterica; Salmonella enterica subsp. enterica; Salmonella enterica subsp. enterica serovar Bareilly MNNEETFYQAMRRKGVTRRSFLKFCSLAATSLGLGAGMTPKIAWALENKPRIPVVWIHGL WP_001058311.1 hydrogenase [Salmonella enterica] Length: 372\nNumber of predicted TMHs: 2\nExp number of AAs in TMHs: 37.4699899999999\nExp number, first 60 AAs: 19.72699\nTotal prob of N-in: 0.96171\nPOSSIBLE N-term signal sequence\ninside 1 19\nTMhelix 20 39\noutside 40 325\nTMhelix 326 348\ninside 349 372

23092 GCF_000749395.1_CFSAN000969_01.0Salmonella enterica subsp. enterica serovar Bareilly str. CFSAN000969 Proteobacteria; Gammaproteobacteria; Enterobacterales; Enterobacteriaceae; Salmonella; Salmonella enterica; Salmonella enterica subsp. enterica; Salmonella enterica subsp. enterica serovar Bareilly MNNEETFYQAMRRKGVTRRSFLKFCSLAATSLGLGAGMTPKIAWALENKPRIPVVWIHGL WP_001058311.1 hydrogenase [Salmonella enterica] Length: 372\nNumber of predicted TMHs: 2\nExp number of AAs in TMHs: 37.4699899999999\nExp number, first 60 AAs: 19.72699\nTotal prob of N-in: 0.96171\nPOSSIBLE N-term signal sequence\ninside 1 19\nTMhelix 20 39\noutside 40 325\nTMhelix 326 348\ninside 349 372

23093 GCF_000749395.1_CFSAN000969_01.0Salmonella enterica subsp. enterica serovar Bareilly str. CFSAN000969 Proteobacteria; Gammaproteobacteria; Enterobacterales; Enterobacteriaceae; Salmonella; Salmonella enterica; Salmonella enterica subsp. enterica; Salmonella enterica subsp. enterica serovar Bareilly MTGDNTLITSHGINRRDFMKLCAALAATMGLSSKAAAEMAESVSNPQRPPVIWIGAQECT WP_000145429.1 hydrogenase 2 small subunit [Salmonella enterica] Length: 372\nNumber of predicted TMHs: 1\nExp number of AAs in TMHs: 24.51309\nExp number, first 60 AAs: 0.2682\nTotal prob of N-in: 0.14402\noutside 1 331\nTMhelix 332 354\ninside 355 372

23094 GCF_000749415.1_CFSAN000970_01.0Salmonella enterica subsp. enterica serovar Bareilly str. CFSAN000970 Proteobacteria; Gammaproteobacteria; Enterobacterales; Enterobacteriaceae; Salmonella; Salmonella enterica; Salmonella enterica subsp. enterica; Salmonella enterica subsp. enterica serovar Bareilly MTGDNTLITSHGINRRDFMKLCAALAATMGLSSKAAAEMAESVSNPQRPPVIWIGAQECT WP_000145429.1 hydrogenase 2 small subunit [Salmonella enterica] Length: 372\nNumber of predicted TMHs: 1\nExp number of AAs in TMHs: 24.51309\nExp number, first 60 AAs: 0.2682\nTotal prob of N-in: 0.14402\noutside 1 331\nTMhelix 332 354\ninside 355 372

23095 GCF_000749415.1_CFSAN000970_01.0Salmonella enterica subsp. enterica serovar Bareilly str. CFSAN000970 Proteobacteria; Gammaproteobacteria; Enterobacterales; Enterobacteriaceae; Salmonella; Salmonella enterica; Salmonella enterica subsp. enterica; Salmonella enterica subsp. enterica serovar Bareilly MNNEETFYQAMRRKGVTRRSFLKFCSLAATSLGLGAGMTPKIAWALENKPRIPVVWIHGL WP_001058311.1 hydrogenase [Salmonella enterica] Length: 372\nNumber of predicted TMHs: 2\nExp number of AAs in TMHs: 37.4699899999999\nExp number, first 60 AAs: 19.72699\nTotal prob of N-in: 0.96171\nPOSSIBLE N-term signal sequence\ninside 1 19\nTMhelix 20 39\noutside 40 325\nTMhelix 326 348\ninside 349 372

23096 GCF_000698695.1_CFSAN001087_03.0Salmonella enterica subsp. enterica serovar Bareilly str. CFSAN001087 Proteobacteria; Gammaproteobacteria; Enterobacterales; Enterobacteriaceae; Salmonella; Salmonella enterica;

Salmonella enterica subsp. enterica; Salmonella enterica subsp. enterica serovar Bareilly
MNNEETFYQAMRRKGVTRRSFLKFCSLAATSLGLGAGMTPKIAWALENKPRIPVVWIHGL WP_001058311.1
hydrogenase [Salmonella enterica] Length: 372\nNumber of predicted TMHs: 2\nExp number of AAs in TMHs: 37.4699899999999\nExp number, first 60 AAs: 19.72699\nTotal prob of N-in: 0.96171\nPOSSIBLE N-term signal sequence\ninside 1 19\nTMhelix 20 39\noutside 40 325\nTMhelix 326 348\ninside 349 372

23097 GCF_000698695.1_CFSAN001087_03.0Salmonella enterica subsp. enterica serovar Bareilly str. CFSAN001087
Proteobacteria; Gammaproteobacteria; Enterobacterales; Enterobacteriaceae; Salmonella; Salmonella enterica; Salmonella enterica subsp. enterica; Salmonella enterica subsp. enterica serovar Bareilly
MTGDNTLITSHGINRRDFMKLCAALAATMGLSSKAAAEMAESVSNPQRPPVIWIGAQECT WP_000145429.1
hydrogenase 2 small subunit [Salmonella enterica] Length: 372\nNumber of predicted TMHs: 1\nExp number of AAs in TMHs: 24.51309\nExp number, first 60 AAs: 0.2682\nTotal prob of N-in: 0.14402\noutside 1 331\nTMhelix 332 354\ninside 355 372

23098 GCF_000748665.1_CFSAN001088_01.0Salmonella enterica subsp. enterica serovar Bareilly str. CFSAN001088
Proteobacteria; Gammaproteobacteria; Enterobacterales; Enterobacteriaceae; Salmonella; Salmonella enterica; Salmonella enterica subsp. enterica; Salmonella enterica subsp. enterica serovar Bareilly
MNNEETFYQAMRRKGVTRRSFLKFCSLAATSLGLGAGMTPKIAWALENKPRIPVVWIHGL WP_001058311.1
hydrogenase [Salmonella enterica] Length: 372\nNumber of predicted TMHs: 2\nExp number of AAs in TMHs: 37.4699899999999\nExp number, first 60 AAs: 19.72699\nTotal prob of N-in: 0.96171\nPOSSIBLE N-term signal sequence\ninside 1 19\nTMhelix 20 39\noutside 40 325\nTMhelix 326 348\ninside 349 372

23099 GCF_000748665.1_CFSAN001088_01.0Salmonella enterica subsp. enterica serovar Bareilly str. CFSAN001088
Proteobacteria; Gammaproteobacteria; Enterobacterales; Enterobacteriaceae; Salmonella; Salmonella enterica; Salmonella enterica subsp. enterica; Salmonella enterica subsp. enterica serovar Bareilly
MTGDNTLITSHGINRRDFMKLCAALAATMGLSSKAAAEMAESVSNPQRPPVIWIGAQECT WP_000145429.1
hydrogenase 2 small subunit [Salmonella enterica] Length: 372\nNumber of predicted TMHs: 1\nExp number of AAs in TMHs: 24.51309\nExp number, first 60 AAs: 0.2682\nTotal prob of N-in: 0.14402\noutside 1 331\nTMhelix 332 354\ninside 355 372

23100 GCF_000748685.1_CFSAN001089.0 Salmonella enterica subsp. enterica serovar Bareilly str. CFSAN001089
Proteobacteria; Gammaproteobacteria; Enterobacterales; Enterobacteriaceae; Salmonella; Salmonella enterica; Salmonella enterica subsp. enterica; Salmonella enterica subsp. enterica serovar Bareilly
MNNEETFYQAMRRKGVTRRSFLKFCSLAATSLGLGAGMTPKIAWALENKPRIPVVWIHGL WP_001058311.1
hydrogenase [Salmonella enterica] Length: 372\nNumber of predicted TMHs: 2\nExp number of AAs in TMHs: 37.4699899999999\nExp number, first 60 AAs: 19.72699\nTotal prob of N-in: 0.96171\nPOSSIBLE N-term signal sequence\ninside 1 19\nTMhelix 20 39\noutside 40 325\nTMhelix 326 348\ninside 349 372

23101 GCF_000748685.1_CFSAN001089.0 Salmonella enterica subsp. enterica serovar Bareilly str. CFSAN001089
Proteobacteria; Gammaproteobacteria; Enterobacterales; Enterobacteriaceae; Salmonella; Salmonella enterica; Salmonella enterica subsp. enterica; Salmonella enterica subsp. enterica serovar Bareilly
MTGDNTLITSHGINRRDFMKLCAALAATMGLSSKAAAEMAESVSNPQRPPVIWIGAQECT WP_000145429.1
hydrogenase 2 small subunit [Salmonella enterica] Length: 372\nNumber of predicted TMHs: 1\nExp number of AAs in TMHs: 24.51309\nExp number, first 60 AAs: 0.2682\nTotal prob of N-in: 0.14402\noutside 1 331\nTMhelix 332 354\ninside 355 372

23102 GCF_000758005.1_CFSAN001090_01.0Salmonella enterica subsp. enterica serovar Bareilly str. CFSAN001090
Proteobacteria; Gammaproteobacteria; Enterobacterales; Enterobacteriaceae; Salmonella; Salmonella enterica; Salmonella enterica subsp. enterica; Salmonella enterica subsp. enterica serovar Bareilly
MNNEETFYQAMRRKGVTRRSFLKFCSLAATSLGLGAGMTPKIAWALENKPRIPVVWIHGL WP_001058311.1
hydrogenase [Salmonella enterica] Length: 372\nNumber of predicted TMHs: 2\nExp number of AAs in TMHs: 37.4699899999999\nExp number, first 60 AAs: 19.72699\nTotal prob of N-in: 0.96171\nPOSSIBLE N-term signal sequence\ninside 1 19\nTMhelix 20 39\noutside 40 325\nTMhelix 326 348\ninside 349 372

23103 GCF_000758005.1_CFSAN001090_01.0Salmonella enterica subsp. enterica serovar Bareilly str. CFSAN001090
Proteobacteria; Gammaproteobacteria; Enterobacterales; Enterobacteriaceae; Salmonella; Salmonella enterica; Salmonella enterica subsp. enterica; Salmonella enterica subsp. enterica serovar Bareilly
MTGDNTLITSHGINRRDFMKLCAALAATMGLSSKAAAEMAESVSNPQRPPVIWIGAQECT WP_000145429.1
hydrogenase 2 small subunit [Salmonella enterica] Length: 372\nNumber of predicted TMHs: 1\nExp number of AAs in TMHs: 24.51309\nExp number, first 60 AAs: 0.2682\nTotal prob of N-in: 0.14402\noutside 1 331\nTMhelix 332 354\ninside 355 372

23104 GCF_000748705.1_CFSAN001099_01.0Salmonella enterica subsp. enterica serovar Bareilly str. CFSAN001099 Proteobacteria; Gammaproteobacteria; Enterobacterales; Enterobacteriaceae; Salmonella; Salmonella enterica; Salmonella enterica subsp. enterica; Salmonella enterica subsp. enterica serovar Bareilly MTGDNTLITSHGINRRDFMKLCAALAATMGLSSKAAAEMAESVSNPQRPPVIWIGAQECT WP_000145429.1 hydrogenase 2 small subunit [Salmonella enterica] Length: 372\nNumber of predicted TMHs: 1\nExp number of AAs in TMHs: 24.51309\nExp number, first 60 AAs: 0.2682\nTotal prob of N-in: 0.14402\noutside 1 331\nTMhelix 332 354\ninside 355 372

23105 GCF_000748705.1_CFSAN001099_01.0Salmonella enterica subsp. enterica serovar Bareilly str. CFSAN001099 Proteobacteria; Gammaproteobacteria; Enterobacterales; Enterobacteriaceae; Salmonella; Salmonella enterica; Salmonella enterica subsp. enterica; Salmonella enterica subsp. enterica serovar Bareilly MNNEETFYQAMRRKGVTRRSFLKFCSLAATSLGLGAGMTPKIAWALENKPRIPVVWIHGL WP_001058311.1 hydrogenase [Salmonella enterica] Length: 372\nNumber of predicted TMHs: 2\nExp number of AAs in TMHs: 37.4699899999999\nExp number, first 60 AAs: 19.72699\nTotal prob of N-in: 0.96171\nPOSSIBLE N-term signal sequence\ninside 1 19\nTMhelix 20 39\noutside 40 325\nTMhelix 326 348\ninside 349 372

23106 GCF_000748725.1_CFSAN001102.0 Salmonella enterica subsp. enterica serovar Bareilly str. CFSAN001102 Proteobacteria; Gammaproteobacteria; Enterobacterales; Enterobacteriaceae; Salmonella; Salmonella enterica; Salmonella enterica subsp. enterica; Salmonella enterica subsp. enterica serovar Bareilly MNNEETFYQAMRRKGVTRRSFLKFCSLAATSLGLGAGMTPKIAWALENKPRIPVVWIHGL WP_001058311.1 hydrogenase [Salmonella enterica] Length: 372\nNumber of predicted TMHs: 2\nExp number of AAs in TMHs: 37.4699899999999\nExp number, first 60 AAs: 19.72699\nTotal prob of N-in: 0.96171\nPOSSIBLE N-term signal sequence\ninside 1 19\nTMhelix 20 39\noutside 40 325\nTMhelix 326 348\ninside 349 372

23107 GCF_000748725.1_CFSAN001102.0 Salmonella enterica subsp. enterica serovar Bareilly str. CFSAN001102 Proteobacteria; Gammaproteobacteria; Enterobacterales; Enterobacteriaceae; Salmonella; Salmonella enterica; Salmonella enterica subsp. enterica; Salmonella enterica subsp. enterica serovar Bareilly MTGDNTLITSHGINRRDFMKLCAALAATMGLSSKAAAEMAESVSNPQRPPVIWIGAQECT WP_000145429.1 hydrogenase 2 small subunit [Salmonella enterica] Length: 372\nNumber of predicted TMHs: 1\nExp number of AAs in TMHs: 24.51309\nExp number, first 60 AAs: 0.2682\nTotal prob of N-in: 0.14402\noutside 1 331\nTMhelix 332 354\ninside 355 372

23108 GCF_000748745.1_CFSAN001105_01.0Salmonella enterica subsp. enterica serovar Bareilly str. CFSAN001105 Proteobacteria; Gammaproteobacteria; Enterobacterales; Enterobacteriaceae; Salmonella; Salmonella enterica; Salmonella enterica subsp. enterica; Salmonella enterica subsp. enterica serovar Bareilly MTGDNTLITSHGINRRDFMKLCAALAATMGLSSKAAAEMAESVSNPQRPPVIWIGAQECT WP_000145429.1 hydrogenase 2 small subunit [Salmonella enterica] Length: 372\nNumber of predicted TMHs: 1\nExp number of AAs in TMHs: 24.51309\nExp number, first 60 AAs: 0.2682\nTotal prob of N-in: 0.14402\noutside 1 331\nTMhelix 332 354\ninside 355 372

23109 GCF_000748745.1_CFSAN001105_01.0Salmonella enterica subsp. enterica serovar Bareilly str. CFSAN001105 Proteobacteria; Gammaproteobacteria; Enterobacterales; Enterobacteriaceae; Salmonella; Salmonella enterica; Salmonella enterica subsp. enterica; Salmonella enterica subsp. enterica serovar Bareilly MNNEETFYQAMRRKGVTRRSFLKFCSLAATSLGLGAGMTPKIAWALENKPRIPVVWIHGL WP_001058311.1 hydrogenase [Salmonella enterica] Length: 372\nNumber of predicted TMHs: 2\nExp number of AAs in TMHs: 37.4699899999999\nExp number, first 60 AAs: 19.72699\nTotal prob of N-in: 0.96171\nPOSSIBLE N-term signal sequence\ninside 1 19\nTMhelix 20 39\noutside 40 325\nTMhelix 326 348\ninside 349 372

23110 GCF_000748765.1_CFSAN001108_01.0Salmonella enterica subsp. enterica serovar Bareilly str. CFSAN001108 Proteobacteria; Gammaproteobacteria; Enterobacterales; Enterobacteriaceae; Salmonella; Salmonella enterica; Salmonella enterica subsp. enterica; Salmonella enterica subsp. enterica serovar Bareilly MNNEETFYQAMRRKGVTRRSFLKFCSLAATSLGLGAGMTPKIAWALENKPRIPVVWIHGL WP_001058311.1 hydrogenase [Salmonella enterica] Length: 372\nNumber of predicted TMHs: 2\nExp number of AAs in TMHs: 37.4699899999999\nExp number, first 60 AAs: 19.72699\nTotal prob of N-in: 0.96171\nPOSSIBLE N-term signal sequence\ninside 1 19\nTMhelix 20 39\noutside 40 325\nTMhelix 326 348\ninside 349 372

23111 GCF_000748765.1_CFSAN001108_01.0Salmonella enterica subsp. enterica serovar Bareilly str. CFSAN001108 Proteobacteria; Gammaproteobacteria; Enterobacterales; Enterobacteriaceae; Salmonella; Salmonella enterica; Salmonella enterica subsp. enterica; Salmonella enterica subsp. enterica serovar Bareilly MTGDNTLITSHGINRRDFMKLCAALAATMGLSSKAAAEMAESVSNPQRPPVIWIGAQECT WP_000145429.1 hydrogenase 2 small subunit [Salmonella enterica] Length: 372\nNumber of predicted TMHs: 1\nExp

number of AAs in TMHs: 24.51309\nExp number, first 60 AAs: 0.2682\nTotal prob of N-in: 0.14402\noutside 1 331\nTMhelix 332 354\ninside 355 372

23112 GCF_000748785.1_CFSAN001109_01.0Salmonella enterica subsp. enterica serovar Bareilly str. CFSAN001109 Proteobacteria; Gammaproteobacteria; Enterobacterales; Enterobacteriaceae; Salmonella; Salmonella enterica; Salmonella enterica subsp. enterica; Salmonella enterica subsp. enterica serovar Bareilly MNNEETFYQAMRRKGVTRRSFLKFCSLAATSLGLGAGMTPKIAWALENKPRIPVVWIHGL WP_001058311.1 hydrogenase [Salmonella enterica] Length: 372\nNumber of predicted TMHs: 2\nExp number of AAs in TMHs: 37.4699899999999\nExp number, first 60 AAs: 19.72699\nTotal prob of N-in: 0.96171\nPOSSIBLE N-term signal sequence\ninside 1 19\nTMhelix 20 39\noutside 40 325\nTMhelix 326 348\ninside 349 372

23113 GCF_000748785.1_CFSAN001109_01.0Salmonella enterica subsp. enterica serovar Bareilly str. CFSAN001109 Proteobacteria; Gammaproteobacteria; Enterobacterales; Enterobacteriaceae; Salmonella; Salmonella enterica; Salmonella enterica subsp. enterica; Salmonella enterica subsp. enterica serovar Bareilly MTGDNTLITSHGINRRDFMKLCAALAATMGLSSKAAAEMAESVSNPQRPPVIWIGAQECT WP_000145429.1 hydrogenase 2 small subunit [Salmonella enterica] Length: 372\nNumber of predicted TMHs: 1\nExp number of AAs in TMHs: 24.51309\nExp number, first 60 AAs: 0.2682\nTotal prob of N-in: 0.14402\noutside 1 331\nTMhelix 332 354\ninside 355 372

23114 GCF_001707735.1_ASM170773v1 Salmonella enterica subsp. enterica serovar Bareilly str. CFSAN001111 Proteobacteria; Gammaproteobacteria; Enterobacterales; Enterobacteriaceae; Salmonella; Salmonella enterica; Salmonella enterica subsp. enterica; Salmonella enterica subsp. enterica serovar Bareilly MNNEETFYQAMRRKGVTRRSFLKFCSLAATSLGLGAGMTPKIAWALENKPRIPVVWIHGL WP_001058311.1 hydrogenase [Salmonella enterica] Length: 372\nNumber of predicted TMHs: 2\nExp number of AAs in TMHs: 37.4699899999999\nExp number, first 60 AAs: 19.72699\nTotal prob of N-in: 0.96171\nPOSSIBLE N-term signal sequence\ninside 1 19\nTMhelix 20 39\noutside 40 325\nTMhelix 326 348\ninside 349 372

23115 GCF_001707735.1_ASM170773v1 Salmonella enterica subsp. enterica serovar Bareilly str. CFSAN001111 Proteobacteria; Gammaproteobacteria; Enterobacterales; Enterobacteriaceae; Salmonella; Salmonella enterica; Salmonella enterica subsp. enterica; Salmonella enterica subsp. enterica serovar Bareilly MTGDNTLITSHGINRRDFMKLCAALAATMGLSSKAAAEMAESVSNPQRPPVIWIGAQECT WP_000145429.1 hydrogenase 2 small subunit [Salmonella enterica] Length: 372\nNumber of predicted TMHs: 1\nExp number of AAs in TMHs: 24.51309\nExp number, first 60 AAs: 0.2682\nTotal prob of N-in: 0.14402\noutside 1 331\nTMhelix 332 354\ninside 355 372

23116 GCF_000748025.1_CFSAN001112_01.0Salmonella enterica subsp. enterica serovar Bareilly str. CFSAN001112 Proteobacteria; Gammaproteobacteria; Enterobacterales; Enterobacteriaceae; Salmonella; Salmonella enterica; Salmonella enterica subsp. enterica; Salmonella enterica subsp. enterica serovar Bareilly MNNEETFYQAMRRKGVTRRSFLKFCSLAATSLGLGAGMTPKIAWALENKPRIPVVWIHGL WP_001058311.1 hydrogenase [Salmonella enterica] Length: 372\nNumber of predicted TMHs: 2\nExp number of AAs in TMHs: 37.4699899999999\nExp number, first 60 AAs: 19.72699\nTotal prob of N-in: 0.96171\nPOSSIBLE N-term signal sequence\ninside 1 19\nTMhelix 20 39\noutside 40 325\nTMhelix 326 348\ninside 349 372

23117 GCF_000748025.1_CFSAN001112_01.0Salmonella enterica subsp. enterica serovar Bareilly str. CFSAN001112 Proteobacteria; Gammaproteobacteria; Enterobacterales; Enterobacteriaceae; Salmonella; Salmonella enterica; Salmonella enterica subsp. enterica; Salmonella enterica subsp. enterica serovar Bareilly MTGDNTLITSHGINRRDFMKLCAALAATMGLSSKAAAEMAESVSNPQRPPVIWIGAQECT WP_000145429.1 hydrogenase 2 small subunit [Salmonella enterica] Length: 372\nNumber of predicted TMHs: 1\nExp number of AAs in TMHs: 24.51309\nExp number, first 60 AAs: 0.2682\nTotal prob of N-in: 0.14402\noutside 1 331\nTMhelix 332 354\ninside 355 372

23118 GCF_000748045.1_CFSAN001115_01.0Salmonella enterica subsp. enterica serovar Bareilly str. CFSAN001115 Proteobacteria; Gammaproteobacteria; Enterobacterales; Enterobacteriaceae; Salmonella; Salmonella enterica; Salmonella enterica subsp. enterica; Salmonella enterica subsp. enterica serovar Bareilly MTGDNTLITSHGINRRDFMKLCAALAATMGLSSKAAAEMAESVSNPQRPPVIWIGAQECT WP_000145429.1 hydrogenase 2 small subunit [Salmonella enterica] Length: 372\nNumber of predicted TMHs: 1\nExp number of AAs in TMHs: 24.51309\nExp number, first 60 AAs: 0.2682\nTotal prob of N-in: 0.14402\noutside 1 331\nTMhelix 332 354\ninside 355 372

23119 GCF_000748045.1_CFSAN001115_01.0Salmonella enterica subsp. enterica serovar Bareilly str. CFSAN001115 Proteobacteria; Gammaproteobacteria; Enterobacterales; Enterobacteriaceae; Salmonella; Salmonella enterica;

Salmonella enterica subsp. enterica; Salmonella enterica subsp. enterica serovar Bareilly
MNNEETFYQAMRRKGVTRRSFLKFCSLAATSLGLGAGMTPKIAWALENKPRIPVVWIHGL WP_001058311.1
hydrogenase [Salmonella enterica] Length: 372\nNumber of predicted TMHs: 2\nExp number of AAs in TMHs: 37.4699899999999\nExp number, first 60 AAs: 19.72699\nTotal prob of N-in: 0.96171\nPOSSIBLE N-term signal sequence\ninside 1 19\nTMhelix 20 39\noutside 40 325\nTMhelix 326 348\ninside 349 372

23120 GCF_000748065.1_CFSAN001118_01.0Salmonella enterica subsp. enterica serovar Bareilly str. CFSAN001118
Proteobacteria; Gammaproteobacteria; Enterobacterales; Enterobacteriaceae; Salmonella; Salmonella enterica; Salmonella enterica subsp. enterica; Salmonella enterica subsp. enterica serovar Bareilly
MTGDNTLITSHGINRRDFMKLCAALAATMGLSSKAAAEMAESVSNPQRPPVIWIGAQECT WP_000145429.1
hydrogenase 2 small subunit [Salmonella enterica] Length: 372\nNumber of predicted TMHs: 1\nExp number of AAs in TMHs: 24.51309\nExp number, first 60 AAs: 0.2682\nTotal prob of N-in: 0.14402\noutside 1 331\nTMhelix 332 354\ninside 355 372

23121 GCF_000748065.1_CFSAN001118_01.0Salmonella enterica subsp. enterica serovar Bareilly str. CFSAN001118
Proteobacteria; Gammaproteobacteria; Enterobacterales; Enterobacteriaceae; Salmonella; Salmonella enterica; Salmonella enterica subsp. enterica; Salmonella enterica subsp. enterica serovar Bareilly
MNNEETFYQAMRRKGVTRRSFLKFCSLAATSLGLGAGMTPKIAWALENKPRIPVVWIHGL WP_001058311.1
hydrogenase [Salmonella enterica] Length: 372\nNumber of predicted TMHs: 2\nExp number of AAs in TMHs: 37.4699899999999\nExp number, first 60 AAs: 19.72699\nTotal prob of N-in: 0.96171\nPOSSIBLE N-term signal sequence\ninside 1 19\nTMhelix 20 39\noutside 40 325\nTMhelix 326 348\ninside 349 372

23122 GCF_000748085.1_CFSAN001140_01.0Salmonella enterica subsp. enterica serovar Bareilly str. CFSAN001140
Proteobacteria; Gammaproteobacteria; Enterobacterales; Enterobacteriaceae; Salmonella; Salmonella enterica; Salmonella enterica subsp. enterica; Salmonella enterica subsp. enterica serovar Bareilly
MNNEETFYQAMRRKGVTRRSFLKFCSLAATSLGLGAGMTPKIAWALENKPRIPVVWIHGL WP_001058311.1
hydrogenase [Salmonella enterica] Length: 372\nNumber of predicted TMHs: 2\nExp number of AAs in TMHs: 37.4699899999999\nExp number, first 60 AAs: 19.72699\nTotal prob of N-in: 0.96171\nPOSSIBLE N-term signal sequence\ninside 1 19\nTMhelix 20 39\noutside 40 325\nTMhelix 326 348\ninside 349 372

23123 GCF_000748085.1_CFSAN001140_01.0Salmonella enterica subsp. enterica serovar Bareilly str. CFSAN001140
Proteobacteria; Gammaproteobacteria; Enterobacterales; Enterobacteriaceae; Salmonella; Salmonella enterica; Salmonella enterica subsp. enterica; Salmonella enterica subsp. enterica serovar Bareilly
MTGDNTLITSHGINRRDFMKLCAALAATMGLSSKAAAEMAESVSNPQRPPVIWIGAQECT WP_000145429.1
hydrogenase 2 small subunit [Salmonella enterica] Length: 372\nNumber of predicted TMHs: 1\nExp number of AAs in TMHs: 24.51309\nExp number, first 60 AAs: 0.2682\nTotal prob of N-in: 0.14402\noutside 1 331\nTMhelix 332 354\ninside 355 372

23124 GCF_000832685.1_CFSAN001591_01.0Salmonella enterica subsp. enterica serovar Bareilly str. CFSAN001591
Proteobacteria; Gammaproteobacteria; Enterobacterales; Enterobacteriaceae; Salmonella; Salmonella enterica; Salmonella enterica subsp. enterica; Salmonella enterica subsp. enterica serovar Bareilly
MTGDNTLITSHGINRRDFMKLCAALAATMGLSSKAAAEMAESVSNPQRPPVIWIGAQECT WP_000145429.1
hydrogenase 2 small subunit [Salmonella enterica] Length: 372\nNumber of predicted TMHs: 1\nExp number of AAs in TMHs: 24.51309\nExp number, first 60 AAs: 0.2682\nTotal prob of N-in: 0.14402\noutside 1 331\nTMhelix 332 354\ninside 355 372

23125 GCF_000832685.1_CFSAN001591_01.0Salmonella enterica subsp. enterica serovar Bareilly str. CFSAN001591
Proteobacteria; Gammaproteobacteria; Enterobacterales; Enterobacteriaceae; Salmonella; Salmonella enterica; Salmonella enterica subsp. enterica; Salmonella enterica subsp. enterica serovar Bareilly
MNNEETFYQAMRRKGVTRRSFLKFCSLAATSLGLGAGMTPKIAWALENKPRIPVVWIHGL WP_001058311.1
hydrogenase [Salmonella enterica] Length: 372\nNumber of predicted TMHs: 2\nExp number of AAs in TMHs: 37.4699899999999\nExp number, first 60 AAs: 19.72699\nTotal prob of N-in: 0.96171\nPOSSIBLE N-term signal sequence\ninside 1 19\nTMhelix 20 39\noutside 40 325\nTMhelix 326 348\ninside 349 372

23126 GCF_001975325.1_ASM197532v1 Salmonella enterica subsp. enterica serovar Bergen str. ST350
Proteobacteria; Gammaproteobacteria; Enterobacterales; Enterobacteriaceae; Salmonella; Salmonella enterica; Salmonella enterica subsp. enterica; Salmonella enterica subsp. enterica serovar Bergen
MTGDNTLITSHGINRRDFMKLCAALAATMGLSSKAAAEMAESVSNPQRPPVIWIGAQECT WP_000145429.1
hydrogenase 2 small subunit [Salmonella enterica] Length: 372\nNumber of predicted TMHs: 1\nExp number of AAs in TMHs: 24.51309\nExp number, first 60 AAs: 0.2682\nTotal prob of N-in: 0.14402\noutside 1 331\nTMhelix 332 354\ninside 355 372

23127 GCF_001975325.1_ASM197532v1 Salmonella enterica subsp. enterica serovar Bergen str. ST350 Proteobacteria; Gammaproteobacteria; Enterobacterales; Enterobacteriaceae; Salmonella; Salmonella enterica; Salmonella enterica subsp. enterica; Salmonella enterica subsp. enterica serovar Bergen MNNEETFYQAMRRKGVTRRSFLKFCSLAATSLGLGAGMTPKIAWALENKPRIPVVWIHGL WP_076937197.1 [Ni/Fe] hydrogenase small subunit [Salmonella enterica] Length: 372\nNumber of predicted TMHs: 2\nExp number of AAs in TMHs: 37.4699899999999\nExp number, first 60 AAs: 19.72699\nTotal prob of N-in: 0.96171\nPOSSIBLE N-term signal sequence\ninside 1 19\nTMhelix 20 39\noutside 40 325\nTMhelix 326 348\ninside 349 372

23128 GCF_000964155.1_ASM96415v1 Salmonella enterica subsp. enterica serovar Berta str. ATCC 8392 Proteobacteria; Gammaproteobacteria; Enterobacterales; Enterobacteriaceae; Salmonella; Salmonella enterica; Salmonella enterica subsp. enterica; Salmonella enterica subsp. enterica serovar Berta MNNEETFYQAMRRKGVTRRSFLKFCSLAATSLGLGAGMTPKIAWALENKPRIPVVWIHGL WP_001058311.1 hydrogenase [Salmonella enterica] Length: 372\nNumber of predicted TMHs: 2\nExp number of AAs in TMHs: 37.4699899999999\nExp number, first 60 AAs: 19.72699\nTotal prob of N-in: 0.96171\nPOSSIBLE N-term signal sequence\ninside 1 19\nTMhelix 20 39\noutside 40 325\nTMhelix 326 348\ninside 349 372

23129 GCF_000487375.1_ASM48737v1 Salmonella enterica subsp. enterica serovar Berta str. ATCC 8392 Proteobacteria; Gammaproteobacteria; Enterobacterales; Enterobacteriaceae; Salmonella; Salmonella enterica; Salmonella enterica subsp. enterica; Salmonella enterica subsp. enterica serovar Berta MNNEETFYQAMRRKGVTRRSFLKFCSLAATSLGLGAGMTPKIAWALENKPRIPVVWIHGL WP_001058311.1 hydrogenase [Salmonella enterica] Length: 372\nNumber of predicted TMHs: 2\nExp number of AAs in TMHs: 37.4699899999999\nExp number, first 60 AAs: 19.72699\nTotal prob of N-in: 0.96171\nPOSSIBLE N-term signal sequence\ninside 1 19\nTMhelix 20 39\noutside 40 325\nTMhelix 326 348\ninside 349 372

23130 GCF_000487375.1_ASM48737v1 Salmonella enterica subsp. enterica serovar Berta str. ATCC 8392 Proteobacteria; Gammaproteobacteria; Enterobacterales; Enterobacteriaceae; Salmonella; Salmonella enterica; Salmonella enterica subsp. enterica; Salmonella enterica subsp. enterica serovar Berta MTGDNTLITSHGINRRDFMKLCAALAATMGLSSKAAAEMAESVSNPQRPPVIWIGAQECT WP_000145429.1 hydrogenase 2 small subunit [Salmonella enterica] Length: 372\nNumber of predicted TMHs: 1\nExp number of AAs in TMHs: 24.51309\nExp number, first 60 AAs: 0.2682\nTotal prob of N-in: 0.14402\noutside 1 331\nTMhelix 332 354\ninside 355 372

23131 GCF_000964155.1_ASM96415v1 Salmonella enterica subsp. enterica serovar Berta str. ATCC 8392 Proteobacteria; Gammaproteobacteria; Enterobacterales; Enterobacteriaceae; Salmonella; Salmonella enterica; Salmonella enterica subsp. enterica; Salmonella enterica subsp. enterica serovar Berta MTGDNTLITSHGINRRDFMKLCAALAATMGLSSKAAAEMAESVSNPQRPPVIWIGAQECT WP_000145429.1 hydrogenase 2 small subunit [Salmonella enterica] Length: 372\nNumber of predicted TMHs: 1\nExp number of AAs in TMHs: 24.51309\nExp number, first 60 AAs: 0.2682\nTotal prob of N-in: 0.14402\noutside 1 331\nTMhelix 332 354\ninside 355 372

23132 GCF_001448475.1_ASM144847v1 Salmonella enterica subsp. enterica serovar Berta str. SA20103550 Proteobacteria; Gammaproteobacteria; Enterobacterales; Enterobacteriaceae; Salmonella; Salmonella enterica; Salmonella enterica subsp. enterica; Salmonella enterica subsp. enterica serovar Berta MTGDNTLITSHGINRRDFMKLCAALAATMGLSSKAAAEMAESVSNPQRPPVIWIGAQECT WP_000145429.1 hydrogenase 2 small subunit [Salmonella enterica] Length: 372\nNumber of predicted TMHs: 1\nExp number of AAs in TMHs: 24.51309\nExp number, first 60 AAs: 0.2682\nTotal prob of N-in: 0.14402\noutside 1 331\nTMhelix 332 354\ninside 355 372

23133 GCF_001448475.1_ASM144847v1 Salmonella enterica subsp. enterica serovar Berta str. SA20103550 Proteobacteria; Gammaproteobacteria; Enterobacterales; Enterobacteriaceae; Salmonella; Salmonella enterica; Salmonella enterica subsp. enterica; Salmonella enterica subsp. enterica serovar Berta MNNEETFYQAMRRKGVTRRSFLKFCSLAATSLGLGAGMTPKIAWALENKPRIPVVWIHGL WP_001058311.1 hydrogenase [Salmonella enterica] Length: 372\nNumber of predicted TMHs: 2\nExp number of AAs in TMHs: 37.4699899999999\nExp number, first 60 AAs: 19.72699\nTotal prob of N-in: 0.96171\nPOSSIBLE N-term signal sequence\ninside 1 19\nTMhelix 20 39\noutside 40 325\nTMhelix 326 348\ninside 349 372

23134 GCF_001975265.1_ASM197526v1 Salmonella enterica subsp. enterica serovar Blegdam str. S-1824 Proteobacteria; Gammaproteobacteria; Enterobacterales; Enterobacteriaceae; Salmonella; Salmonella enterica; Salmonella enterica subsp. enterica; Salmonella enterica subsp. enterica serovar Blegdam MNNEETFYQAMRRKGVTRRSFLKFCSLAATSLGLGAGMTPKIAWALENKPRIPVVWIHGL WP_001058307.1

hydrogenase [Salmonella enterica] Length: 372\nNumber of predicted TMHs: 2\nExp number of AAs in TMHs: 36.51738\nExp number, first 60 AAs: 19.72849\nTotal prob of N-in: 0.96170\nPOSSIBLE N-term signal sequence\ninside 1 19\nTMhelix 20 39\noutside 40 325\nTMhelix 326 348\ninside 349 372

23135 GCF_001975265.1_ASM197526v1 Salmonella enterica subsp. enterica serovar Blegdam str. S-1824 Proteobacteria; Gammaproteobacteria; Enterobacterales; Enterobacteriaceae; Salmonella; Salmonella enterica; Salmonella enterica subsp. enterica; Salmonella enterica subsp. enterica serovar Blegdam MTGDNTLITSHGINRRDFMKLCAALAATMGLSSKAAAEMAESVSNPQRPPVIWIGAQECT WP_000145429.1 hydrogenase 2 small subunit [Salmonella enterica] Length: 372\nNumber of predicted TMHs: 1\nExp number of AAs in TMHs: 24.51309\nExp number, first 60 AAs: 0.2682\nTotal prob of N-in: 0.14402\noutside 1 331\nTMhelix 332 354\ninside 355 372

23136 GCF_001975345.1_ASM197534v1 Salmonella enterica subsp. enterica serovar Borreze str. SA20041063 Proteobacteria; Gammaproteobacteria; Enterobacterales; Enterobacteriaceae; Salmonella; Salmonella enterica; Salmonella enterica subsp. enterica; Salmonella enterica subsp. enterica serovar Borreze MNNEETFYQAMRRKGVTRRSFLKFCSLAATSLGLSAGMTPKIAWALENKPRIPVVWIHGL WP_001058316.1 hydrogenase [Salmonella enterica] Length: 372\nNumber of predicted TMHs: 2\nExp number of AAs in TMHs: 35.76397\nExp number, first 60 AAs: 18.97201\nTotal prob of N-in: 0.92785\nPOSSIBLE N-term signal sequence\ninside 1 19\nTMhelix 20 39\noutside 40 325\nTMhelix 326 348\ninside 349 372

23137 GCF_001975345.1_ASM197534v1 Salmonella enterica subsp. enterica serovar Borreze str. SA20041063 Proteobacteria; Gammaproteobacteria; Enterobacterales; Enterobacteriaceae; Salmonella; Salmonella enterica; Salmonella enterica subsp. enterica; Salmonella enterica subsp. enterica serovar Borreze MTGDNTLITSHGINRRDFMKLCAALAATMGLSSKAAAEMAESVSNPQRPPVIWIGAQECT WP_000145429.1 hydrogenase 2 small subunit [Salmonella enterica] Length: 372\nNumber of predicted TMHs: 1\nExp number of AAs in TMHs: 24.51309\nExp number, first 60 AAs: 0.2682\nTotal prob of N-in: 0.14402\noutside 1 331\nTMhelix 332 354\ninside 355 372

23138 GCF_000524415.1_sal185_version2 Salmonella enterica subsp. enterica serovar Bovismorbificans str. Sal185 Proteobacteria; Gammaproteobacteria; Enterobacterales; Enterobacteriaceae; Salmonella; Salmonella enterica; Salmonella enterica subsp. enterica; Salmonella enterica subsp. enterica serovar Bovismorbificans MTGDNTLITSHGINRRDFMKLCAALAATMGLSSKAAAEMAESVSNPQRPPVIWIGAQECT WP_000145429.1 hydrogenase 2 small subunit [Salmonella enterica] Length: 372\nNumber of predicted TMHs: 1\nExp number of AAs in TMHs: 24.51309\nExp number, first 60 AAs: 0.2682\nTotal prob of N-in: 0.14402\noutside 1 331\nTMhelix 332 354\ninside 355 372

23139 GCF_000524415.1_sal185_version2 Salmonella enterica subsp. enterica serovar Bovismorbificans str. Sal185 Proteobacteria; Gammaproteobacteria; Enterobacterales; Enterobacteriaceae; Salmonella; Salmonella enterica; Salmonella enterica subsp. enterica; Salmonella enterica subsp. enterica serovar Bovismorbificans MNNEETFYQAMRRKGVTRRSFLKFCSLAATSLGLGAGMTPKIAWALENKPRIPVVWIHGL WP_001058311.1 hydrogenase [Salmonella enterica] Length: 372\nNumber of predicted TMHs: 2\nExp number of AAs in TMHs: 37.4699899999999\nExp number, first 60 AAs: 19.72699\nTotal prob of N-in: 0.96171\nPOSSIBLE N-term signal sequence\ninside 1 19\nTMhelix 20 39\noutside 40 325\nTMhelix 326 348\ninside 349 372

23140 GCF_000524885.1_sal609 Salmonella enterica subsp. enterica serovar Bovismorbificans str. Sal609 Proteobacteria; Gammaproteobacteria; Enterobacterales; Enterobacteriaceae; Salmonella; Salmonella enterica; Salmonella enterica subsp. enterica; Salmonella enterica subsp. enterica serovar Bovismorbificans MNNEETFYQAMRRKGVTRRSFLKFCSLAATSLGLGAGMTPKIAWALENKPRIPVVWIHGL WP_001058311.1 hydrogenase [Salmonella enterica] Length: 372\nNumber of predicted TMHs: 2\nExp number of AAs in TMHs: 37.4699899999999\nExp number, first 60 AAs: 19.72699\nTotal prob of N-in: 0.96171\nPOSSIBLE N-term signal sequence\ninside 1 19\nTMhelix 20 39\noutside 40 325\nTMhelix 326 348\ninside 349 372

23141 GCF_000524885.1_sal609 Salmonella enterica subsp. enterica serovar Bovismorbificans str. Sal609 Proteobacteria; Gammaproteobacteria; Enterobacterales; Enterobacteriaceae; Salmonella; Salmonella enterica; Salmonella enterica subsp. enterica; Salmonella enterica subsp. enterica serovar Bovismorbificans MTGDNTLITSHGINRRDFMKLCAALAATMGLSSKAAAEMAESVSNPQRPPVIWIGAQECT WP_000145429.1 hydrogenase 2 small subunit [Salmonella enterica] Length: 372\nNumber of predicted TMHs: 1\nExp number of AAs in TMHs: 24.51309\nExp number, first 60 AAs: 0.2682\nTotal prob of N-in: 0.14402\noutside 1 331\nTMhelix 332 354\ninside 355 372

23142 GCF_000524845.1_sal610 Salmonella enterica subsp. enterica serovar Bovismorbificans str. Sal610 Proteobacteria; Gammaproteobacteria; Enterobacterales; Enterobacteriaceae; Salmonella; Salmonella enterica; Salmonella enterica subsp. enterica; Salmonella enterica subsp. enterica serovar Bovismorbificans MTGDNTLITSHGINRRDFMKLCAALAATMGLSSKAAAEMAESVSNPQRPPVIWIGAQECT WP_000145429.1 hydrogenase 2 small subunit [Salmonella enterica] Length: 372\nNumber of predicted TMHs: 1\nExp number of AAs in TMHs: 24.51309\nExp number, first 60 AAs: 0.2682\nTotal prob of N-in: 0.14402\noutside 1 331\nTMhelix 332 354\ninside 355 372

23143 GCF_000524845.1_sal610 Salmonella enterica subsp. enterica serovar Bovismorbificans str. Sal610 Proteobacteria; Gammaproteobacteria; Enterobacterales; Enterobacteriaceae; Salmonella; Salmonella enterica; Salmonella enterica subsp. enterica; Salmonella enterica subsp. enterica serovar Bovismorbificans MNNEETFYQAMRRKGVTRRSFLKFCSLAATSLGLGAGMTPKIAWALENKPRIPVVWIHGL WP_001058311.1 hydrogenase [Salmonella enterica] Length: 372\nNumber of predicted TMHs: 2\nExp number of AAs in TMHs: 37.4699899999999\nExp number, first 60 AAs: 19.72699\nTotal prob of N-in: 0.96171\nPOSSIBLE N-term signal sequence\ninside 1 19\nTMhelix 20 39\noutside 40 325\nTMhelix 326 348\ninside 349 372

23144 GCF_000524865.1_sal615 Salmonella enterica subsp. enterica serovar Bovismorbificans str. Sal615 Proteobacteria; Gammaproteobacteria; Enterobacterales; Enterobacteriaceae; Salmonella; Salmonella enterica; Salmonella enterica subsp. enterica; Salmonella enterica subsp. enterica serovar Bovismorbificans MNNEETFYQAMRRKGVTRRSFLKFCSLAATSLGLGAGMTPKIAWALENKPRIPVVWIHGL WP_001058311.1 hydrogenase [Salmonella enterica] Length: 372\nNumber of predicted TMHs: 2\nExp number of AAs in TMHs: 37.4699899999999\nExp number, first 60 AAs: 19.72699\nTotal prob of N-in: 0.96171\nPOSSIBLE N-term signal sequence\ninside 1 19\nTMhelix 20 39\noutside 40 325\nTMhelix 326 348\ninside 349 372

23145 GCF_000524865.1_sal615 Salmonella enterica subsp. enterica serovar Bovismorbificans str. Sal615 Proteobacteria; Gammaproteobacteria; Enterobacterales; Enterobacteriaceae; Salmonella; Salmonella enterica; Salmonella enterica subsp. enterica; Salmonella enterica subsp. enterica serovar Bovismorbificans MTGDNTLITSHGINRRDFMKLCAALAATMGLSSKAAAEMAESVSNPQRPPVIWIGAQECT WP_000145429.1 hydrogenase 2 small subunit [Salmonella enterica] Length: 372\nNumber of predicted TMHs: 1\nExp number of AAs in TMHs: 24.51309\nExp number, first 60 AAs: 0.2682\nTotal prob of N-in: 0.14402\noutside 1 331\nTMhelix 332 354\ninside 355 372

23146 GCF_000524905.1_sal616 Salmonella enterica subsp. enterica serovar Bovismorbificans str. Sal616 Proteobacteria; Gammaproteobacteria; Enterobacterales; Enterobacteriaceae; Salmonella; Salmonella enterica; Salmonella enterica subsp. enterica; Salmonella enterica subsp. enterica serovar Bovismorbificans MTGDNTLITSHGINRRDFMKLCAALAATMGLSSKAAAEMAESVSNPQRPPVIWIGAQECT WP_000145429.1 hydrogenase 2 small subunit [Salmonella enterica] Length: 372\nNumber of predicted TMHs: 1\nExp number of AAs in TMHs: 24.51309\nExp number, first 60 AAs: 0.2682\nTotal prob of N-in: 0.14402\noutside 1 331\nTMhelix 332 354\ninside 355 372

23147 GCF_000524905.1_sal616 Salmonella enterica subsp. enterica serovar Bovismorbificans str. Sal616 Proteobacteria; Gammaproteobacteria; Enterobacterales; Enterobacteriaceae; Salmonella; Salmonella enterica; Salmonella enterica subsp. enterica; Salmonella enterica subsp. enterica serovar Bovismorbificans MNNEETFYQAMRRKGVTRRSFLKFCSLAATSLGLGAGMTPKIAWALENKPRIPVVWIHGL WP_001058311.1 hydrogenase [Salmonella enterica] Length: 372\nNumber of predicted TMHs: 2\nExp number of AAs in TMHs: 37.4699899999999\nExp number, first 60 AAs: 19.72699\nTotal prob of N-in: 0.96171\nPOSSIBLE N-term signal sequence\ninside 1 19\nTMhelix 20 39\noutside 40 325\nTMhelix 326 348\ninside 349 372

23148 GCF_000524925.1_sal617 Salmonella enterica subsp. enterica serovar Bovismorbificans str. Sal617 Proteobacteria; Gammaproteobacteria; Enterobacterales; Enterobacteriaceae; Salmonella; Salmonella enterica; Salmonella enterica subsp. enterica; Salmonella enterica subsp. enterica serovar Bovismorbificans MTGDNTLITSHGINRRDFMKLCAALAATMGLSSKAAAEMAESVSNPQRPPVIWIGAQECT WP_000145429.1 hydrogenase 2 small subunit [Salmonella enterica] Length: 372\nNumber of predicted TMHs: 1\nExp number of AAs in TMHs: 24.51309\nExp number, first 60 AAs: 0.2682\nTotal prob of N-in: 0.14402\noutside 1 331\nTMhelix 332 354\ninside 355 372

23149 GCF_000524925.1_sal617 Salmonella enterica subsp. enterica serovar Bovismorbificans str. Sal617 Proteobacteria; Gammaproteobacteria; Enterobacterales; Enterobacteriaceae; Salmonella; Salmonella enterica; Salmonella enterica subsp. enterica; Salmonella enterica subsp. enterica serovar Bovismorbificans MNNEETFYQAMRRKGVTRRSFLKFCSLAATSLGLGAGMTPKIAWALENKPRIPVVWIHGL WP_001058311.1 hydrogenase [Salmonella enterica] Length: 372\nNumber of predicted TMHs: 2\nExp number of AAs in

TMHs: 37.4699899999999\nExp number, first 60 AAs: 19.72699\nTotal prob of N-in: 0.96171\nPOSSIBLE N-term signal sequence\ninside 1 19\nTMhelix 20 39\noutside 40 325\nTMhelix 326 348\ninside 349 372

23150 GCF_000524705.1_sal644 Salmonella enterica subsp. enterica serovar Bovismorbificans str. Sal644 Proteobacteria; Gammaproteobacteria; Enterobacterales; Enterobacteriaceae; Salmonella; Salmonella enterica; Salmonella enterica subsp. enterica; Salmonella enterica subsp. enterica serovar Bovismorbificans MNNEETFYQAMRRKGVTRRSFLKFCSLAATSLGLGAGMTPKIAWALENKPRIPVVWIHGL WP_001058311.1 hydrogenase [Salmonella enterica] Length: 372\nNumber of predicted TMHs: 2\nExp number of AAs in TMHs: 37.4699899999999\nExp number, first 60 AAs: 19.72699\nTotal prob of N-in: 0.96171\nPOSSIBLE N-term signal sequence\ninside 1 19\nTMhelix 20 39\noutside 40 325\nTMhelix 326 348\ninside 349 372

23151 GCF_000524705.1_sal644 Salmonella enterica subsp. enterica serovar Bovismorbificans str. Sal644 Proteobacteria; Gammaproteobacteria; Enterobacterales; Enterobacteriaceae; Salmonella; Salmonella enterica; Salmonella enterica subsp. enterica; Salmonella enterica subsp. enterica serovar Bovismorbificans MTGDNTLITSHGINRRDFMKLCAALAATMGLSSKAAAEMAESVSNPQRPPVIWIGAQECT WP_000145429.1 hydrogenase 2 small subunit [Salmonella enterica] Length: 372\nNumber of predicted TMHs: 1\nExp number of AAs in TMHs: 24.51309\nExp number, first 60 AAs: 0.2682\nTotal prob of N-in: 0.14402\noutside 1 331\nTMhelix 332 354\ninside 355 372

23152 GCF_000524805.1_sal676 Salmonella enterica subsp. enterica serovar Bovismorbificans str. Sal676 Proteobacteria; Gammaproteobacteria; Enterobacterales; Enterobacteriaceae; Salmonella; Salmonella enterica; Salmonella enterica subsp. enterica; Salmonella enterica subsp. enterica serovar Bovismorbificans MNNEETFYQAMRRKGVTRRSFLKFCSLAATSLGLGAGMTPKIAWALENKPRIPVVWIHGL WP_001058312.1 hydrogenase [Salmonella enterica] Length: 372\nNumber of predicted TMHs: 2\nExp number of AAs in TMHs: 36.51738\nExp number, first 60 AAs: 19.72849\nTotal prob of N-in: 0.96170\nPOSSIBLE N-term signal sequence\ninside 1 19\nTMhelix 20 39\noutside 40 325\nTMhelix 326 348\ninside 349 372

23153 GCF_000524805.1_sal676 Salmonella enterica subsp. enterica serovar Bovismorbificans str. Sal676 Proteobacteria; Gammaproteobacteria; Enterobacterales; Enterobacteriaceae; Salmonella; Salmonella enterica; Salmonella enterica subsp. enterica; Salmonella enterica subsp. enterica serovar Bovismorbificans MTGDNTLITSHGINRRDFMKLCAALAATMGLSSKAAAEMAESVSNPQRPPVIWIGAQECT WP_000145431.1 hydrogenase 2 small subunit [Salmonella enterica] Length: 372\nNumber of predicted TMHs: 1\nExp number of AAs in TMHs: 24.51505\nExp number, first 60 AAs: 0.2682\nTotal prob of N-in: 0.14406\noutside 1 331\nTMhelix 332 354\ninside 355 372

23154 GCF_000524745.1_sal677 Salmonella enterica subsp. enterica serovar Bovismorbificans str. Sal677 Proteobacteria; Gammaproteobacteria; Enterobacterales; Enterobacteriaceae; Salmonella; Salmonella enterica; Salmonella enterica subsp. enterica; Salmonella enterica subsp. enterica serovar Bovismorbificans MTGDNTLITSHGINRRDFMKLCAALAATMGLSSKAAAEMAESVSNPQRPPVIWIGAQECT WP_000145431.1 hydrogenase 2 small subunit [Salmonella enterica] Length: 372\nNumber of predicted TMHs: 1\nExp number of AAs in TMHs: 24.51505\nExp number, first 60 AAs: 0.2682\nTotal prob of N-in: 0.14406\noutside 1 331\nTMhelix 332 354\ninside 355 372

23155 GCF_000524745.1_sal677 Salmonella enterica subsp. enterica serovar Bovismorbificans str. Sal677 Proteobacteria; Gammaproteobacteria; Enterobacterales; Enterobacteriaceae; Salmonella; Salmonella enterica; Salmonella enterica subsp. enterica; Salmonella enterica subsp. enterica serovar Bovismorbificans MNNEETFYQAMRRKGVTRRSFLKFCSLAATSLGLGAGMTPKIAWALENKPRIPVVWIHGL WP_001058312.1 hydrogenase [Salmonella enterica] Length: 372\nNumber of predicted TMHs: 2\nExp number of AAs in TMHs: 36.51738\nExp number, first 60 AAs: 19.72849\nTotal prob of N-in: 0.96170\nPOSSIBLE N-term signal sequence\ninside 1 19\nTMhelix 20 39\noutside 40 325\nTMhelix 326 348\ninside 349 372

23156 GCF_000524765.1_sal678 Salmonella enterica subsp. enterica serovar Bovismorbificans str. Sal678 Proteobacteria; Gammaproteobacteria; Enterobacterales; Enterobacteriaceae; Salmonella; Salmonella enterica; Salmonella enterica subsp. enterica; Salmonella enterica subsp. enterica serovar Bovismorbificans MTGDNTLITSHGINRRDFMKLCAALAATMGLSSKAAAEMAESVSNPQRPPVIWIGAQECT WP_000145431.1 hydrogenase 2 small subunit [Salmonella enterica] Length: 372\nNumber of predicted TMHs: 1\nExp number of AAs in TMHs: 24.51505\nExp number, first 60 AAs: 0.2682\nTotal prob of N-in: 0.14406\noutside 1 331\nTMhelix 332 354\ninside 355 372

23157 GCF_000524765.1_sal678 Salmonella enterica subsp. enterica serovar Bovismorbificans str. Sal678 Proteobacteria; Gammaproteobacteria; Enterobacterales; Enterobacteriaceae; Salmonella; Salmonella enterica;

Salmonella enterica subsp. enterica; Salmonella enterica subsp. enterica serovar Bovismorbificans
MNNEETFYQAMRRKGVTRRSFLKFCSLAATSLGLGAGMTPKIAWALENKPRIPVVWIHGL WP_001058312.1
hydrogenase [Salmonella enterica] Length: 372\nNumber of predicted TMHs: 2\nExp number of AAs in TMHs: 36.51738\nExp number, first 60 AAs: 19.72849\nTotal prob of N-in: 0.96170\nPOSSIBLE N-term signal sequence\ninside 1 19\nTMhelix 20 39\noutside 40 325\nTMhelix 326 348\ninside 349 372

23158 GCF_000524785.1_sal679 Salmonella enterica subsp. enterica serovar Bovismorbificans str. Sal679
Proteobacteria; Gammaproteobacteria; Enterobacterales; Enterobacteriaceae; Salmonella; Salmonella enterica; Salmonella enterica subsp. enterica; Salmonella enterica subsp. enterica serovar Bovismorbificans
MTGDNTLITSHGINRRDFMKLCAALAATMGLSSKAAAEMAESVSNPQRPPVIWIGAQECT WP_000145431.1
hydrogenase 2 small subunit [Salmonella enterica] Length: 372\nNumber of predicted TMHs: 1\nExp number of AAs in TMHs: 24.51505\nExp number, first 60 AAs: 0.2682\nTotal prob of N-in: 0.14406\noutside 1 331\nTMhelix 332 354\ninside 355 372

23159 GCF_000524785.1_sal679 Salmonella enterica subsp. enterica serovar Bovismorbificans str. Sal679
Proteobacteria; Gammaproteobacteria; Enterobacterales; Enterobacteriaceae; Salmonella; Salmonella enterica; Salmonella enterica subsp. enterica; Salmonella enterica subsp. enterica serovar Bovismorbificans
MNNEETFYQAMRRKGVTRRSFLKFCSLAATSLGLGAGMTPKIAWALENKPRIPVVWIHGL WP_001058312.1
hydrogenase [Salmonella enterica] Length: 372\nNumber of predicted TMHs: 2\nExp number of AAs in TMHs: 36.51738\nExp number, first 60 AAs: 19.72849\nTotal prob of N-in: 0.96170\nPOSSIBLE N-term signal sequence\ninside 1 19\nTMhelix 20 39\noutside 40 325\nTMhelix 326 348\ninside 349 372

23160 GCF_000524825.1_sal680 Salmonella enterica subsp. enterica serovar Bovismorbificans str. Sal680
Proteobacteria; Gammaproteobacteria; Enterobacterales; Enterobacteriaceae; Salmonella; Salmonella enterica; Salmonella enterica subsp. enterica; Salmonella enterica subsp. enterica serovar Bovismorbificans
MTGDNTLITSHGINRRDFMKLCAALAATMGLSSKAAAEMAESVSNPQRPPVIWIGAQECT WP_000145431.1
hydrogenase 2 small subunit [Salmonella enterica] Length: 372\nNumber of predicted TMHs: 1\nExp number of AAs in TMHs: 24.51505\nExp number, first 60 AAs: 0.2682\nTotal prob of N-in: 0.14406\noutside 1 331\nTMhelix 332 354\ninside 355 372

23161 GCF_000524825.1_sal680 Salmonella enterica subsp. enterica serovar Bovismorbificans str. Sal680
Proteobacteria; Gammaproteobacteria; Enterobacterales; Enterobacteriaceae; Salmonella; Salmonella enterica; Salmonella enterica subsp. enterica; Salmonella enterica subsp. enterica serovar Bovismorbificans
MNNEETFYQAMRRKGVTRRSFLKFCSLAATSLGLGAGMTPKIAWALENKPRIPVVWIHGL WP_001058312.1
hydrogenase [Salmonella enterica] Length: 372\nNumber of predicted TMHs: 2\nExp number of AAs in TMHs: 36.51738\nExp number, first 60 AAs: 19.72849\nTotal prob of N-in: 0.96170\nPOSSIBLE N-term signal sequence\ninside 1 19\nTMhelix 20 39\noutside 40 325\nTMhelix 326 348\ninside 349 372

23162 GCF_000524945.1_sal681 Salmonella enterica subsp. enterica serovar Bovismorbificans str. Sal681
Proteobacteria; Gammaproteobacteria; Enterobacterales; Enterobacteriaceae; Salmonella; Salmonella enterica; Salmonella enterica subsp. enterica; Salmonella enterica subsp. enterica serovar Bovismorbificans
MNNEETFYQAMRRKGVTRRSFLKFCSLAATSLGLGAGMTPKIAWALENKPRIPVVWIHGL WP_001058312.1
hydrogenase [Salmonella enterica] Length: 372\nNumber of predicted TMHs: 2\nExp number of AAs in TMHs: 36.51738\nExp number, first 60 AAs: 19.72849\nTotal prob of N-in: 0.96170\nPOSSIBLE N-term signal sequence\ninside 1 19\nTMhelix 20 39\noutside 40 325\nTMhelix 326 348\ninside 349 372

23163 GCF_000524945.1_sal681 Salmonella enterica subsp. enterica serovar Bovismorbificans str. Sal681
Proteobacteria; Gammaproteobacteria; Enterobacterales; Enterobacteriaceae; Salmonella; Salmonella enterica; Salmonella enterica subsp. enterica; Salmonella enterica subsp. enterica serovar Bovismorbificans
MTGDNTLITSHGINRRDFMKLCAALAATMGLSSKAAAEMAESVSNPQRPPVIWIGAQECT WP_000145431.1
hydrogenase 2 small subunit [Salmonella enterica] Length: 372\nNumber of predicted TMHs: 1\nExp number of AAs in TMHs: 24.51505\nExp number, first 60 AAs: 0.2682\nTotal prob of N-in: 0.14406\noutside 1 331\nTMhelix 332 354\ninside 355 372

23164 GCF_000524965.1_sal682v2 Salmonella enterica subsp. enterica serovar Bovismorbificans str. Sal682
Proteobacteria; Gammaproteobacteria; Enterobacterales; Enterobacteriaceae; Salmonella; Salmonella enterica; Salmonella enterica subsp. enterica; Salmonella enterica subsp. enterica serovar Bovismorbificans
MTGDNTLITSHGINRRDFMKLCAALAATMGLSSKAAAEMAESVSNPQRPPVIWIGAQECT WP_000145429.1
hydrogenase 2 small subunit [Salmonella enterica] Length: 372\nNumber of predicted TMHs: 1\nExp number of AAs in TMHs: 24.51309\nExp number, first 60 AAs: 0.2682\nTotal prob of N-in: 0.14402\noutside 1 331\nTMhelix 332 354\ninside 355 372

23165 GCF_000524965.1_sal682v2 Salmonella enterica subsp. enterica serovar Bovismorbificans str. Sal682 Proteobacteria; Gammaproteobacteria; Enterobacterales; Enterobacteriaceae; Salmonella; Salmonella enterica; Salmonella enterica subsp. enterica; Salmonella enterica subsp. enterica serovar Bovismorbificans MNNEETFYQAMRRKGVTRRSFLKFCSLAATSLGLGAGMTPKIAWALENKPRIPVVWIHGL WP_001058311.1 hydrogenase [Salmonella enterica] Length: 372\nNumber of predicted TMHs: 2\nExp number of AAs in TMHs: 37.4699899999999\nExp number, first 60 AAs: 19.72699\nTotal prob of N-in: 0.96171\nPOSSIBLE N-term signal sequence\ninside 1 19\nTMhelix 20 39\noutside 40 325\nTMhelix 326 348\ninside 349 372

23166 GCF_000524725.1_sal683 Salmonella enterica subsp. enterica serovar Bovismorbificans str. Sal683 Proteobacteria; Gammaproteobacteria; Enterobacterales; Enterobacteriaceae; Salmonella; Salmonella enterica; Salmonella enterica subsp. enterica; Salmonella enterica subsp. enterica serovar Bovismorbificans MTGDNTLITSHGINRRDFMKLCAALAATMGLSSKAAAEMAESVSNPQRPPVIWIGAQECT WP_000145431.1 hydrogenase 2 small subunit [Salmonella enterica] Length: 372\nNumber of predicted TMHs: 1\nExp number of AAs in TMHs: 24.51505\nExp number, first 60 AAs: 0.2682\nTotal prob of N-in: 0.14406\noutside 1 331\nTMhelix 332 354\ninside 355 372

23167 GCF_000524725.1_sal683 Salmonella enterica subsp. enterica serovar Bovismorbificans str. Sal683 Proteobacteria; Gammaproteobacteria; Enterobacterales; Enterobacteriaceae; Salmonella; Salmonella enterica; Salmonella enterica subsp. enterica; Salmonella enterica subsp. enterica serovar Bovismorbificans MNNEETFYQAMRRKGVTRRSFLKFCSLAATSLGLGAGMTPKIAWALENKPRIPVVWIHGL WP_001058312.1 hydrogenase [Salmonella enterica] Length: 372\nNumber of predicted TMHs: 2\nExp number of AAs in TMHs: 36.51738\nExp number, first 60 AAs: 19.72849\nTotal prob of N-in: 0.96170\nPOSSIBLE N-term signal sequence\ninside 1 19\nTMhelix 20 39\noutside 40 325\nTMhelix 326 348\ninside 349 372

23168 GCF_000487335.1_ASM48733v1 Salmonella enterica subsp. enterica serovar Braenderup str. ATCC BAA-664 Proteobacteria; Gammaproteobacteria; Enterobacterales; Enterobacteriaceae; Salmonella; Salmonella enterica; Salmonella enterica subsp. enterica; Salmonella enterica subsp. enterica serovar Braenderup MTGDNTLITSHGINRRDFMKLCAALAATMGLSSKAAAEMAESVSNPQRPPVIWIGAQECT WP_000145429.1 hydrogenase 2 small subunit [Salmonella enterica] Length: 372\nNumber of predicted TMHs: 1\nExp number of AAs in TMHs: 24.51309\nExp number, first 60 AAs: 0.2682\nTotal prob of N-in: 0.14402\noutside 1 331\nTMhelix 332 354\ninside 355 372

23169 GCF_000487335.1_ASM48733v1 Salmonella enterica subsp. enterica serovar Braenderup str. ATCC BAA-664 Proteobacteria; Gammaproteobacteria; Enterobacterales; Enterobacteriaceae; Salmonella; Salmonella enterica; Salmonella enterica subsp. enterica; Salmonella enterica subsp. enterica serovar Braenderup MNNEETFYQAMRRKGVTRRSFLKFCSLAATSLGLGAGMTPKIAWALENKPRIPVVWIHGL WP_001058312.1 hydrogenase [Salmonella enterica] Length: 372\nNumber of predicted TMHs: 2\nExp number of AAs in TMHs: 36.51738\nExp number, first 60 AAs: 19.72849\nTotal prob of N-in: 0.96170\nPOSSIBLE N-term signal sequence\ninside 1 19\nTMhelix 20 39\noutside 40 325\nTMhelix 326 348\ninside 349 372

23170 GCF_000487895.1_ASM48789v1 Salmonella enterica subsp. enterica serovar Braenderup str. CFSAN000756 Proteobacteria; Gammaproteobacteria; Enterobacterales; Enterobacteriaceae; Salmonella; Salmonella enterica; Salmonella enterica subsp. enterica; Salmonella enterica subsp. enterica serovar Braenderup MNNEETFYQAMRRKGVTRRSFLKFCSLAATSLGLGAGMTPKIAWALENKPRIPVVWIHGL WP_001058312.1 hydrogenase [Salmonella enterica] Length: 372\nNumber of predicted TMHs: 2\nExp number of AAs in TMHs: 36.51738\nExp number, first 60 AAs: 19.72849\nTotal prob of N-in: 0.96170\nPOSSIBLE N-term signal sequence\ninside 1 19\nTMhelix 20 39\noutside 40 325\nTMhelix 326 348\ninside 349 372

23171 GCF_000487895.1_ASM48789v1 Salmonella enterica subsp. enterica serovar Braenderup str. CFSAN000756 Proteobacteria; Gammaproteobacteria; Enterobacterales; Enterobacteriaceae; Salmonella; Salmonella enterica; Salmonella enterica subsp. enterica; Salmonella enterica subsp. enterica serovar Braenderup MTGDNTLITSHGINRRDFMKLCAALAATMGLSSKAAAEMAESVSNPQRPPVIWIGAQECT WP_000145429.1 hydrogenase 2 small subunit [Salmonella enterica] Length: 372\nNumber of predicted TMHs: 1\nExp number of AAs in TMHs: 24.51309\nExp number, first 60 AAs: 0.2682\nTotal prob of N-in: 0.14402\noutside 1 331\nTMhelix 332 354\ninside 355 372

23172 GCF_000487775.1_ASM48777v2 Salmonella enterica subsp. enterica serovar Bredeney str. CFSAN001080 Proteobacteria; Gammaproteobacteria; Enterobacterales; Enterobacteriaceae; Salmonella; Salmonella enterica; Salmonella enterica subsp. enterica; Salmonella enterica subsp. enterica serovar Bredeney MNNEETFYQAMRRKGVTRRSFLKFCSLAATSLGLGAGMTPKIAWALENKPRIPVVWIHGL WP_001058312.1 hydrogenase [Salmonella enterica] Length: 372\nNumber of predicted TMHs: 2\nExp number of AAs in

TMHs: 36.51738\nExp number, first 60 AAs: 19.72849\nTotal prob of N-in: 0.96170\nPOSSIBLE N-term signal sequence\ninside 1 19\nTMhelix 20 39\noutside 40 325\nTMhelix 326 348\ninside 349 372

23173 GCF_000487775.1_ASM48777v2 Salmonella enterica subsp. enterica serovar Bredeney str. CFSAN001080 Proteobacteria; Gammaproteobacteria; Enterobacterales; Enterobacteriaceae; Salmonella; Salmonella enterica; Salmonella enterica subsp. enterica; Salmonella enterica subsp. enterica serovar Bredeney MTGDNTLITSHGINRRDFMKLCAALAATMGLSSKAAAEMAESVSNPQRPPVIWIGAQECT WP_000145429.1 hydrogenase 2 small subunit [Salmonella enterica] Length: 372\nNumber of predicted TMHs: 1\nExp number of AAs in TMHs: 24.51309\nExp number, first 60 AAs: 0.2682\nTotal prob of N-in: 0.14402\noutside 1 331\nTMhelix 332 354\ninside 355 372

23174 GCF_000487275.1_ASM48727v1 Salmonella enterica subsp. enterica serovar Cerro str. 818 Proteobacteria; Gammaproteobacteria; Enterobacterales; Enterobacteriaceae; Salmonella; Salmonella enterica; Salmonella enterica subsp. enterica; Salmonella enterica subsp. enterica serovar Cerro MTGDNTLITSHGINRRDFMKLCAALAATMGLSSKAAAEMAESVSNPQRPPVIWIGAQECT WP_000145429.1 hydrogenase 2 small subunit [Salmonella enterica] Length: 372\nNumber of predicted TMHs: 1\nExp number of AAs in TMHs: 24.51309\nExp number, first 60 AAs: 0.2682\nTotal prob of N-in: 0.14402\noutside 1 331\nTMhelix 332 354\ninside 355 372

23175 GCF_000487275.1_ASM48727v1 Salmonella enterica subsp. enterica serovar Cerro str. 818 Proteobacteria; Gammaproteobacteria; Enterobacterales; Enterobacteriaceae; Salmonella; Salmonella enterica; Salmonella enterica subsp. enterica; Salmonella enterica subsp. enterica serovar Cerro MNNEETFYQAMRRKGVTRRSFLKFCSLAATSLGLGAGMTPKIAWALENKPRIPVVWIHGL WP_023233644.1 hydrogenase 1, small subunit [Salmonella enterica] Length: 372\nNumber of predicted TMHs: 2\nExp number of AAs in TMHs: 37.4208299999999\nExp number, first 60 AAs: 19.72732\nTotal prob of N-in: 0.96168\nPOSSIBLE N-term signal sequence\ninside 1 19\nTMhelix 20 39\noutside 40 325\nTMhelix 326 348\ninside 349 372

23176 GCF_000505145.1_CFSAN001587_01.0Salmonella enterica subsp. enterica serovar Cerro str. CFSAN001587 Proteobacteria; Gammaproteobacteria; Enterobacterales; Enterobacteriaceae; Salmonella; Salmonella enterica; Salmonella enterica subsp. enterica; Salmonella enterica subsp. enterica serovar Cerro MNNEETFYQAMRRKGVTRRSFLKFCSLAATSLGLGAGMTPKIAWALENKPRIPVVWIHGL WP_023233644.1 hydrogenase 1, small subunit [Salmonella enterica] Length: 372\nNumber of predicted TMHs: 2\nExp number of AAs in TMHs: 37.4208299999999\nExp number, first 60 AAs: 19.72732\nTotal prob of N-in: 0.96168\nPOSSIBLE N-term signal sequence\ninside 1 19\nTMhelix 20 39\noutside 40 325\nTMhelix 326 348\ninside 349 372

23177 GCF_000505145.1_CFSAN001587_01.0Salmonella enterica subsp. enterica serovar Cerro str. CFSAN001587 Proteobacteria; Gammaproteobacteria; Enterobacterales; Enterobacteriaceae; Salmonella; Salmonella enterica; Salmonella enterica subsp. enterica; Salmonella enterica subsp. enterica serovar Cerro MTGDNTLITSHGINRRDFMKLCAALAATMGLSSKAAAEMAESVSNPQRPPVIWIGAQECT WP_000145429.1 hydrogenase 2 small subunit [Salmonella enterica] Length: 372\nNumber of predicted TMHs: 1\nExp number of AAs in TMHs: 24.51309\nExp number, first 60 AAs: 0.2682\nTotal prob of N-in: 0.14402\noutside 1 331\nTMhelix 332 354\ninside 355 372

23178 GCF_000505365.2_ASM50536v2 Salmonella enterica subsp. enterica serovar Cerro str. CFSAN001588 Proteobacteria; Gammaproteobacteria; Enterobacterales; Enterobacteriaceae; Salmonella; Salmonella enterica; Salmonella enterica subsp. enterica; Salmonella enterica subsp. enterica serovar Cerro MTGDNTLITSHGINRRDFMKLCAALAATMGLSSKAAAEMAESVSNPQRPPVIWIGAQECT WP_000145429.1 hydrogenase 2 small subunit [Salmonella enterica] Length: 372\nNumber of predicted TMHs: 1\nExp number of AAs in TMHs: 24.51309\nExp number, first 60 AAs: 0.2682\nTotal prob of N-in: 0.14402\noutside 1 331\nTMhelix 332 354\ninside 355 372

23179 GCF_000505365.2_ASM50536v2 Salmonella enterica subsp. enterica serovar Cerro str. CFSAN001588 Proteobacteria; Gammaproteobacteria; Enterobacterales; Enterobacteriaceae; Salmonella; Salmonella enterica; Salmonella enterica subsp. enterica; Salmonella enterica subsp. enterica serovar Cerro MNNEETFYQAMRRKGVTRRSFLKFCSLAATSLGLGAGMTPKIAWALENKPRIPVVWIHGL WP_023233644.1 hydrogenase 1, small subunit [Salmonella enterica] Length: 372\nNumber of predicted TMHs: 2\nExp number of AAs in TMHs: 37.4208299999999\nExp number, first 60 AAs: 19.72732\nTotal prob of N-in: 0.96168\nPOSSIBLE N-term signal sequence\ninside 1 19\nTMhelix 20 39\noutside 40 325\nTMhelix 326 348\ninside 349 372

23180 GCF_000505385.1_CFSAN001589_01.0Salmonella enterica subsp. enterica serovar Cerro str. CFSAN001589 Proteobacteria; Gammaproteobacteria; Enterobacterales; Enterobacteriaceae; Salmonella; Salmonella enterica; Salmonella enterica subsp. enterica; Salmonella enterica subsp. enterica serovar Cerro MNNEETFYQAMRRKGVTRRSFLKFCSLAATSLGLGAGMTPKIAWALENKPRIPVVWIHGL WP_023233644.1 hydrogenase 1, small subunit [Salmonella enterica] Length: 372\nNumber of predicted TMHs: 2\nExp number of AAs in TMHs: 37.4208299999999\nExp number, first 60 AAs: 19.72732\nTotal prob of N-in: 0.96168\nPOSSIBLE N-term signal sequence\ninside 1 19\nTMhelix 20 39\noutside 40 325\nTMhelix 326 348\ninside 349 372

23181 GCF_000505385.1_CFSAN001589_01.0Salmonella enterica subsp. enterica serovar Cerro str. CFSAN001589 Proteobacteria; Gammaproteobacteria; Enterobacterales; Enterobacteriaceae; Salmonella; Salmonella enterica; Salmonella enterica subsp. enterica; Salmonella enterica subsp. enterica serovar Cerro MTGDNTLITSHGINRRDFMKLCAALAATMGLSSKAAAEMAESVSNPQRPPVIWIGAQECT WP_000145429.1 hydrogenase 2 small subunit [Salmonella enterica] Length: 372\nNumber of predicted TMHs: 1\nExp number of AAs in TMHs: 24.51309\nExp number, first 60 AAs: 0.2682\nTotal prob of N-in: 0.14402\noutside 1 331\nTMhelix 332 354\ninside 355 372

23182 GCF_000505165.1_CFSAN001590_01.0Salmonella enterica subsp. enterica serovar Cerro str. CFSAN001590 Proteobacteria; Gammaproteobacteria; Enterobacterales; Enterobacteriaceae; Salmonella; Salmonella enterica; Salmonella enterica subsp. enterica; Salmonella enterica subsp. enterica serovar Cerro MNNEETFYQAMRRKGVTRRSFLKFCSLAATSLGLGAGMTPKIAWALENKPRIPVVWIHGL WP_023233644.1 hydrogenase 1, small subunit [Salmonella enterica] Length: 372\nNumber of predicted TMHs: 2\nExp number of AAs in TMHs: 37.4208299999999\nExp number, first 60 AAs: 19.72732\nTotal prob of N-in: 0.96168\nPOSSIBLE N-term signal sequence\ninside 1 19\nTMhelix 20 39\noutside 40 325\nTMhelix 326 348\ninside 349 372

23183 GCF_000505165.1_CFSAN001590_01.0Salmonella enterica subsp. enterica serovar Cerro str. CFSAN001590 Proteobacteria; Gammaproteobacteria; Enterobacterales; Enterobacteriaceae; Salmonella; Salmonella enterica; Salmonella enterica subsp. enterica; Salmonella enterica subsp. enterica serovar Cerro MTGDNTLITSHGINRRDFMKLCAALAATMGLSSKAAAEMAESVSNPQRPPVIWIGAQECT WP_000145429.1 hydrogenase 2 small subunit [Salmonella enterica] Length: 372\nNumber of predicted TMHs: 1\nExp number of AAs in TMHs: 24.51309\nExp number, first 60 AAs: 0.2682\nTotal prob of N-in: 0.14402\noutside 1 331\nTMhelix 332 354\ninside 355 372

23184 GCF_000505185.1_CFSAN001669_01.0Salmonella enterica subsp. enterica serovar Cerro str. CFSAN001669 Proteobacteria; Gammaproteobacteria; Enterobacterales; Enterobacteriaceae; Salmonella; Salmonella enterica; Salmonella enterica subsp. enterica; Salmonella enterica subsp. enterica serovar Cerro MNNEETFYQAMRRKGVTRRSFLKFCSLAATSLGLGAGMTPKIAWALENKPRIPVVWIHGL WP_023233644.1 hydrogenase 1, small subunit [Salmonella enterica] Length: 372\nNumber of predicted TMHs: 2\nExp number of AAs in TMHs: 37.4208299999999\nExp number, first 60 AAs: 19.72732\nTotal prob of N-in: 0.96168\nPOSSIBLE N-term signal sequence\ninside 1 19\nTMhelix 20 39\noutside 40 325\nTMhelix 326 348\ninside 349 372

23185 GCF_000505185.1_CFSAN001669_01.0Salmonella enterica subsp. enterica serovar Cerro str. CFSAN001669 Proteobacteria; Gammaproteobacteria; Enterobacterales; Enterobacteriaceae; Salmonella; Salmonella enterica; Salmonella enterica subsp. enterica; Salmonella enterica subsp. enterica serovar Cerro MTGDNTLITSHGINRRDFMKLCAALAATMGLSSKAAAEMAESVSNPQRPPVIWIGAQECT WP_000145429.1 hydrogenase 2 small subunit [Salmonella enterica] Length: 372\nNumber of predicted TMHs: 1\nExp number of AAs in TMHs: 24.51309\nExp number, first 60 AAs: 0.2682\nTotal prob of N-in: 0.14402\noutside 1 331\nTMhelix 332 354\ninside 355 372

23186 GCF_000505405.1_CFSAN001670_01.0Salmonella enterica subsp. enterica serovar Cerro str. CFSAN001670 Proteobacteria; Gammaproteobacteria; Enterobacterales; Enterobacteriaceae; Salmonella; Salmonella enterica; Salmonella enterica subsp. enterica; Salmonella enterica subsp. enterica serovar Cerro MNNEETFYQAMRRKGVTRRSFLKFCSLAATSLGLGAGMTPKIAWALENKPRIPVVWIHGL WP_023233644.1 hydrogenase 1, small subunit [Salmonella enterica] Length: 372\nNumber of predicted TMHs: 2\nExp number of AAs in TMHs: 37.4208299999999\nExp number, first 60 AAs: 19.72732\nTotal prob of N-in: 0.96168\nPOSSIBLE N-term signal sequence\ninside 1 19\nTMhelix 20 39\noutside 40 325\nTMhelix 326 348\ninside 349 372

23187	GCF_000505405.1_CFSAN001670_01.0Salmonella enterica subsp. enterica serovar Cerro str. CFSAN001670
Proteobacteria; Gammaproteobacteria; Enterobacterales; Enterobacteriaceae; Salmonella; Salmonella enterica; Salmonella enterica subsp. enterica; Salmonella enterica subsp. enterica serovar Cerro
MTGDNTLITSHGINRRDFMKLCAALAATMGLSSKAAAEMAESVSNPQRPPVIWIGAQECT	WP_000145429.1
hydrogenase 2 small subunit [Salmonella enterica]	Length: 372\nNumber of predicted TMHs:  1\nExp number of AAs in TMHs: 24.51309\nExp number, first 60 AAs:  0.2682\nTotal prob of N-in:   0.14402\noutside   1 331\nTMhelix   332   354\ninside   355   372

23188	GCF_000505205.1_CFSAN001671_01.0Salmonella enterica subsp. enterica serovar Cerro str. CFSAN001671
Proteobacteria; Gammaproteobacteria; Enterobacterales; Enterobacteriaceae; Salmonella; Salmonella enterica; Salmonella enterica subsp. enterica; Salmonella enterica subsp. enterica serovar Cerro
MTGDNTLITSHGINRRDFMKLCAALAATMGLSSKAAAEMAESVSNPQRPPVIWIGAQECT	WP_000145429.1
hydrogenase 2 small subunit [Salmonella enterica]	Length: 372\nNumber of predicted TMHs:  1\nExp number of AAs in TMHs: 24.51309\nExp number, first 60 AAs:  0.2682\nTotal prob of N-in:   0.14402\noutside   1 331\nTMhelix   332   354\ninside   355   372

23189	GCF_000505205.1_CFSAN001671_01.0Salmonella enterica subsp. enterica serovar Cerro str. CFSAN001671
Proteobacteria; Gammaproteobacteria; Enterobacterales; Enterobacteriaceae; Salmonella; Salmonella enterica; Salmonella enterica subsp. enterica; Salmonella enterica subsp. enterica serovar Cerro
MNNEETFYQAMRRKGVTRRSFLKFCSLAATSLGLGAGMTPKIAWALENKPRIPVVWIHGL	WP_023233644.1
hydrogenase 1, small subunit [Salmonella enterica]	Length: 372\nNumber of predicted TMHs:  2\nExp number of AAs in TMHs: 37.4208299999999\nExp number, first 60 AAs:  19.72732\nTotal prob of N-in: 0.96168\nPOSSIBLE N-term signal sequence\ninside   1   19\nTMhelix   20   39\noutside   40   325\nTMhelix   326 348\ninside   349   372

23190	GCF_000505225.1_CFSAN001673_01.0Salmonella enterica subsp. enterica serovar Cerro str. CFSAN001673
Proteobacteria; Gammaproteobacteria; Enterobacterales; Enterobacteriaceae; Salmonella; Salmonella enterica; Salmonella enterica subsp. enterica; Salmonella enterica subsp. enterica serovar Cerro
MNNEETFYQAMRRKGVTRRSFLKFCSLAATSLGLGAGMTPKIAWALENKPRIPVVWIHGL	WP_023233644.1
hydrogenase 1, small subunit [Salmonella enterica]	Length: 372\nNumber of predicted TMHs:  2\nExp number of AAs in TMHs: 37.4208299999999\nExp number, first 60 AAs:  19.72732\nTotal prob of N-in: 0.96168\nPOSSIBLE N-term signal sequence\ninside   1   19\nTMhelix   20   39\noutside   40   325\nTMhelix   326 348\ninside   349   372

23191	GCF_000505225.1_CFSAN001673_01.0Salmonella enterica subsp. enterica serovar Cerro str. CFSAN001673
Proteobacteria; Gammaproteobacteria; Enterobacterales; Enterobacteriaceae; Salmonella; Salmonella enterica; Salmonella enterica subsp. enterica; Salmonella enterica subsp. enterica serovar Cerro
MTGDNTLITSHGINRRDFMKLCAALAATMGLSSKAAAEMAESVSNPQRPPVIWIGAQECT	WP_000145429.1
hydrogenase 2 small subunit [Salmonella enterica]	Length: 372\nNumber of predicted TMHs:  1\nExp number of AAs in TMHs: 24.51309\nExp number, first 60 AAs:  0.2682\nTotal prob of N-in:   0.14402\noutside   1 331\nTMhelix   332   354\ninside   355   372

23192	GCF_000505425.1_CFSAN001674_01.0Salmonella enterica subsp. enterica serovar Cerro str. CFSAN001674
Proteobacteria; Gammaproteobacteria; Enterobacterales; Enterobacteriaceae; Salmonella; Salmonella enterica; Salmonella enterica subsp. enterica; Salmonella enterica subsp. enterica serovar Cerro
MNNEETFYQAMRRKGVTRRSFLKFCSLAATSLGLGAGMTPKIAWALENKPRIPVVWIHGL	WP_023233644.1
hydrogenase 1, small subunit [Salmonella enterica]	Length: 372\nNumber of predicted TMHs:  2\nExp number of AAs in TMHs: 37.4208299999999\nExp number, first 60 AAs:  19.72732\nTotal prob of N-in: 0.96168\nPOSSIBLE N-term signal sequence\ninside   1   19\nTMhelix   20   39\noutside   40   325\nTMhelix   326 348\ninside   349   372

23193	GCF_000505425.1_CFSAN001674_01.0Salmonella enterica subsp. enterica serovar Cerro str. CFSAN001674
Proteobacteria; Gammaproteobacteria; Enterobacterales; Enterobacteriaceae; Salmonella; Salmonella enterica; Salmonella enterica subsp. enterica; Salmonella enterica subsp. enterica serovar Cerro
MTGDNTLITSHGINRRDFMKLCAALAATMGLSSKAAAEMAESVSNPQRPPVIWIGAQECT	WP_000145429.1
hydrogenase 2 small subunit [Salmonella enterica]	Length: 372\nNumber of predicted TMHs:  1\nExp number of AAs in TMHs: 24.51309\nExp number, first 60 AAs:  0.2682\nTotal prob of N-in:   0.14402\noutside   1 331\nTMhelix   332   354\ninside   355   372

23194	GCF_000505245.1_CFSAN001679_01.0Salmonella enterica subsp. enterica serovar Cerro str. CFSAN001679
Proteobacteria; Gammaproteobacteria; Enterobacterales; Enterobacteriaceae; Salmonella; Salmonella enterica;

Salmonella enterica subsp. enterica; Salmonella enterica subsp. enterica serovar Cerro
MTGDNTLITSHGINRRDFMKLCAALAATMGLSSKAAAEMAESVSNPQRPPVIWIGAQECT WP_000145429.1
hydrogenase 2 small subunit [Salmonella enterica] Length: 372\nNumber of predicted TMHs: 1\nExp number of AAs in TMHs: 24.51309\nExp number, first 60 AAs: 0.2682\nTotal prob of N-in: 0.14402\noutside 1 331\nTMhelix 332 354\ninside 355 372

23195 GCF_000505245.1_CFSAN001679_01.0Salmonella enterica subsp. enterica serovar Cerro str. CFSAN001679
Proteobacteria; Gammaproteobacteria; Enterobacterales; Enterobacteriaceae; Salmonella; Salmonella enterica; Salmonella enterica subsp. enterica; Salmonella enterica subsp. enterica serovar Cerro
MNNEETFYQAMRRKGVTRRSFLKFCSLAATSLGLGAGMTPKIAWALENKPRIPVVWIHGL WP_023233644.1
hydrogenase 1, small subunit [Salmonella enterica] Length: 372\nNumber of predicted TMHs: 2\nExp number of AAs in TMHs: 37.4208299999999\nExp number, first 60 AAs: 19.72732\nTotal prob of N-in: 0.96168\nPOSSIBLE N-term signal sequence\ninside 1 19\nTMhelix 20 39\noutside 40 325\nTMhelix 326 348\ninside 349 372

23196 GCF_000505265.1_CFSAN001680_01.0Salmonella enterica subsp. enterica serovar Cerro str. CFSAN001680
Proteobacteria; Gammaproteobacteria; Enterobacterales; Enterobacteriaceae; Salmonella; Salmonella enterica; Salmonella enterica subsp. enterica; Salmonella enterica subsp. enterica serovar Cerro
MTGDNTLITSHGINRRDFMKLCAALAATMGLSSKAAAEMAESVSNPQRPPVIWIGAQECT WP_000145429.1
hydrogenase 2 small subunit [Salmonella enterica] Length: 372\nNumber of predicted TMHs: 1\nExp number of AAs in TMHs: 24.51309\nExp number, first 60 AAs: 0.2682\nTotal prob of N-in: 0.14402\noutside 1 331\nTMhelix 332 354\ninside 355 372

23197 GCF_000505265.1_CFSAN001680_01.0Salmonella enterica subsp. enterica serovar Cerro str. CFSAN001680
Proteobacteria; Gammaproteobacteria; Enterobacterales; Enterobacteriaceae; Salmonella; Salmonella enterica; Salmonella enterica subsp. enterica; Salmonella enterica subsp. enterica serovar Cerro
MNNEETFYQAMRRKGVTRRSFLKFCSLAATSLGLGAGMTPKIAWALENKPRIPVVWIHGL WP_023233644.1
hydrogenase 1, small subunit [Salmonella enterica] Length: 372\nNumber of predicted TMHs: 2\nExp number of AAs in TMHs: 37.4208299999999\nExp number, first 60 AAs: 19.72732\nTotal prob of N-in: 0.96168\nPOSSIBLE N-term signal sequence\ninside 1 19\nTMhelix 20 39\noutside 40 325\nTMhelix 326 348\ninside 349 372

23198 GCF_000505285.1_CFSAN001681_01.0Salmonella enterica subsp. enterica serovar Cerro str. CFSAN001681
Proteobacteria; Gammaproteobacteria; Enterobacterales; Enterobacteriaceae; Salmonella; Salmonella enterica; Salmonella enterica subsp. enterica; Salmonella enterica subsp. enterica serovar Cerro
MTGDNTLITSHGINRRDFMKLCAALAATMGLSSKAAAEMAESVSNPQRPPVIWIGAQECT WP_000145429.1
hydrogenase 2 small subunit [Salmonella enterica] Length: 372\nNumber of predicted TMHs: 1\nExp number of AAs in TMHs: 24.51309\nExp number, first 60 AAs: 0.2682\nTotal prob of N-in: 0.14402\noutside 1 331\nTMhelix 332 354\ninside 355 372

23199 GCF_000505285.1_CFSAN001681_01.0Salmonella enterica subsp. enterica serovar Cerro str. CFSAN001681
Proteobacteria; Gammaproteobacteria; Enterobacterales; Enterobacteriaceae; Salmonella; Salmonella enterica; Salmonella enterica subsp. enterica; Salmonella enterica subsp. enterica serovar Cerro
MNNEETFYQAMRRKGVTRRSFLKFCSLAATSLGLGAGMTPKIAWALENKPRIPVVWIHGL WP_023233644.1
hydrogenase 1, small subunit [Salmonella enterica] Length: 372\nNumber of predicted TMHs: 2\nExp number of AAs in TMHs: 37.4208299999999\nExp number, first 60 AAs: 19.72732\nTotal prob of N-in: 0.96168\nPOSSIBLE N-term signal sequence\ninside 1 19\nTMhelix 20 39\noutside 40 325\nTMhelix 326 348\ninside 349 372

23200 GCF_000505305.1_CFSAN001690_01.0Salmonella enterica subsp. enterica serovar Cerro str. CFSAN001690
Proteobacteria; Gammaproteobacteria; Enterobacterales; Enterobacteriaceae; Salmonella; Salmonella enterica; Salmonella enterica subsp. enterica; Salmonella enterica subsp. enterica serovar Cerro
MNNEETFYQAMRRKGVTRRSFLKFCSLAATSLGLGAGMTPKIAWALENKPRIPVVWIHGL WP_023233644.1
hydrogenase 1, small subunit [Salmonella enterica] Length: 372\nNumber of predicted TMHs: 2\nExp number of AAs in TMHs: 37.4208299999999\nExp number, first 60 AAs: 19.72732\nTotal prob of N-in: 0.96168\nPOSSIBLE N-term signal sequence\ninside 1 19\nTMhelix 20 39\noutside 40 325\nTMhelix 326 348\ninside 349 372

23201 GCF_000505305.1_CFSAN001690_01.0Salmonella enterica subsp. enterica serovar Cerro str. CFSAN001690
Proteobacteria; Gammaproteobacteria; Enterobacterales; Enterobacteriaceae; Salmonella; Salmonella enterica; Salmonella enterica subsp. enterica; Salmonella enterica subsp. enterica serovar Cerro

MTGDNTLITSHGINRRDFMKLCAALAATMGLSSKAAAEMAESVSNPQRPPVIWIGAQECT WP_000145429.1 hydrogenase 2 small subunit [Salmonella enterica] Length: 372\nNumber of predicted TMHs: 1\nExp number of AAs in TMHs: 24.51309\nExp number, first 60 AAs: 0.2682\nTotal prob of N-in: 0.14402\noutside 1 331\nTMhelix 332 354\ninside 355 372

23202 GCF_000505445.1_CFSAN001691_01.0Salmonella enterica subsp. enterica serovar Cerro str. CFSAN001691 Proteobacteria; Gammaproteobacteria; Enterobacterales; Enterobacteriaceae; Salmonella; Salmonella enterica; Salmonella enterica subsp. enterica; Salmonella enterica subsp. enterica serovar Cerro MTGDNTLITSHGINRRDFMKLCAALAATMGLSSKAAAEMAESVSNPQRPPVIWIGAQECT WP_000145429.1 hydrogenase 2 small subunit [Salmonella enterica] Length: 372\nNumber of predicted TMHs: 1\nExp number of AAs in TMHs: 24.51309\nExp number, first 60 AAs: 0.2682\nTotal prob of N-in: 0.14402\noutside 1 331\nTMhelix 332 354\ninside 355 372

23203 GCF_000505445.1_CFSAN001691_01.0Salmonella enterica subsp. enterica serovar Cerro str. CFSAN001691 Proteobacteria; Gammaproteobacteria; Enterobacterales; Enterobacteriaceae; Salmonella; Salmonella enterica; Salmonella enterica subsp. enterica; Salmonella enterica subsp. enterica serovar Cerro MNNEETFYQAMRRKGVTRRSFLKFCSLAATSLGLGAGMTPKIAWALENKPRIPVVWIHGL WP_023233644.1 hydrogenase 1, small subunit [Salmonella enterica] Length: 372\nNumber of predicted TMHs: 2\nExp number of AAs in TMHs: 37.4208299999999\nExp number, first 60 AAs: 19.72732\nTotal prob of N-in: 0.96168\nPOSSIBLE N-term signal sequence\ninside 1 19\nTMhelix 20 39\noutside 40 325\nTMhelix 326 348\ninside 349 372

23204 GCF_000505325.1_CFSAN001692_01.0Salmonella enterica subsp. enterica serovar Cerro str. CFSAN001692 Proteobacteria; Gammaproteobacteria; Enterobacterales; Enterobacteriaceae; Salmonella; Salmonella enterica; Salmonella enterica subsp. enterica; Salmonella enterica subsp. enterica serovar Cerro MTGDNTLITSHGINRRDFMKLCAALAATMGLSSKAAAEMAESVSNPQRPPVIWIGAQECT WP_000145429.1 hydrogenase 2 small subunit [Salmonella enterica] Length: 372\nNumber of predicted TMHs: 1\nExp number of AAs in TMHs: 24.51309\nExp number, first 60 AAs: 0.2682\nTotal prob of N-in: 0.14402\noutside 1 331\nTMhelix 332 354\ninside 355 372

23205 GCF_000505325.1_CFSAN001692_01.0Salmonella enterica subsp. enterica serovar Cerro str. CFSAN001692 Proteobacteria; Gammaproteobacteria; Enterobacterales; Enterobacteriaceae; Salmonella; Salmonella enterica; Salmonella enterica subsp. enterica; Salmonella enterica subsp. enterica serovar Cerro MNNEETFYQAMRRKGVTRRSFLKFCSLAATSLGLGAGMTPKIAWALENKPRIPVVWIHGL WP_023233644.1 hydrogenase 1, small subunit [Salmonella enterica] Length: 372\nNumber of predicted TMHs: 2\nExp number of AAs in TMHs: 37.4208299999999\nExp number, first 60 AAs: 19.72732\nTotal prob of N-in: 0.96168\nPOSSIBLE N-term signal sequence\ninside 1 19\nTMhelix 20 39\noutside 40 325\nTMhelix 326 348\ninside 349 372

23206 GCF_000505345.1_CFSAN001697_01.0Salmonella enterica subsp. enterica serovar Cerro str. CFSAN001697 Proteobacteria; Gammaproteobacteria; Enterobacterales; Enterobacteriaceae; Salmonella; Salmonella enterica; Salmonella enterica subsp. enterica; Salmonella enterica subsp. enterica serovar Cerro MTGDNTLITSHGINRRDFMKLCAALAATMGLSSKAAAEMAESVSNPQRPPVIWIGAQECT WP_000145429.1 hydrogenase 2 small subunit [Salmonella enterica] Length: 372\nNumber of predicted TMHs: 1\nExp number of AAs in TMHs: 24.51309\nExp number, first 60 AAs: 0.2682\nTotal prob of N-in: 0.14402\noutside 1 331\nTMhelix 332 354\ninside 355 372

23207 GCF_000505345.1_CFSAN001697_01.0Salmonella enterica subsp. enterica serovar Cerro str. CFSAN001697 Proteobacteria; Gammaproteobacteria; Enterobacterales; Enterobacteriaceae; Salmonella; Salmonella enterica; Salmonella enterica subsp. enterica; Salmonella enterica subsp. enterica serovar Cerro MNNEETFYQAMRRKGVTRRSFLKFCSLAATSLGLGAGMTPKIAWALENKPRIPVVWIHGL WP_023233644.1 hydrogenase 1, small subunit [Salmonella enterica] Length: 372\nNumber of predicted TMHs: 2\nExp number of AAs in TMHs: 37.4208299999999\nExp number, first 60 AAs: 19.72732\nTotal prob of N-in: 0.96168\nPOSSIBLE N-term signal sequence\ninside 1 19\nTMhelix 20 39\noutside 40 325\nTMhelix 326 348\ninside 349 372

23208 GCF_000487255.2_ASM48725v2 Salmonella enterica subsp. enterica serovar Chester str. ATCC 11997 Proteobacteria; Gammaproteobacteria; Enterobacterales; Enterobacteriaceae; Salmonella; Salmonella enterica; Salmonella enterica subsp. enterica; Salmonella enterica subsp. enterica serovar Chester MNNEETFYQAMRRKGVTRRSFLKFCSLAATSLGLGAGMTPKIAWALENKPRIPVVWIHGL WP_001058312.1 hydrogenase [Salmonella enterica] Length: 372\nNumber of predicted TMHs: 2\nExp number of AAs in

TMHs: 36.51738\nExp number, first 60 AAs:  19.72849\nTotal prob of N-in:   0.96170\nPOSSIBLE N-term signal sequence\ninside   1   19\nTMhelix   20   39\noutside   40   325\nTMhelix   326   348\ninside   349   372

23209 GCF_000487255.2_ASM48725v2 Salmonella enterica subsp. enterica serovar Chester str. ATCC 11997 Proteobacteria; Gammaproteobacteria; Enterobacterales; Enterobacteriaceae; Salmonella; Salmonella enterica; Salmonella enterica subsp. enterica; Salmonella enterica subsp. enterica serovar Chester MTGDNTLITSHGINRRDFMKLCAALAATMGLSSKAAAEMAESVSNPQRPPVIWIGAQECT WP_000145429.1 hydrogenase 2 small subunit [Salmonella enterica] Length: 372\nNumber of predicted TMHs:  1\nExp number of AAs in TMHs: 24.51309\nExp number, first 60 AAs:  0.2682\nTotal prob of N-in:   0.14402\noutside   1   331\nTMhelix   332   354\ninside   355   372

23210 GCF_000487295.2_ASM48729v2 Salmonella enterica subsp. enterica serovar Choleraesuis str. ATCC 10708 Proteobacteria; Gammaproteobacteria; Enterobacterales; Enterobacteriaceae; Salmonella; Salmonella enterica; Salmonella enterica subsp. enterica; Salmonella enterica subsp. enterica serovar Choleraesuis MTGDNTLITSHGINRRDFMKLCAALAATMGLSSKAAAEMAESVSNPQRPPVIWIGAQECT WP_000145429.1 hydrogenase 2 small subunit [Salmonella enterica] Length: 372\nNumber of predicted TMHs:  1\nExp number of AAs in TMHs: 24.51309\nExp number, first 60 AAs:  0.2682\nTotal prob of N-in:   0.14402\noutside   1   331\nTMhelix   332   354\ninside   355   372

23211 GCF_000487295.2_ASM48729v2 Salmonella enterica subsp. enterica serovar Choleraesuis str. ATCC 10708 Proteobacteria; Gammaproteobacteria; Enterobacterales; Enterobacteriaceae; Salmonella; Salmonella enterica; Salmonella enterica subsp. enterica; Salmonella enterica subsp. enterica serovar Choleraesuis MNNEETFYQAMRRKGVTRRSFLKFCSLAATSLGLGAGMTPKIAWALENKPRIPVVWIHGL WP_001058312.1 hydrogenase [Salmonella enterica] Length: 372\nNumber of predicted TMHs:  2\nExp number of AAs in TMHs: 36.51738\nExp number, first 60 AAs:  19.72849\nTotal prob of N-in:   0.96170\nPOSSIBLE N-term signal sequence\ninside   1   19\nTMhelix   20   39\noutside   40   325\nTMhelix   326   348\ninside   349   372

23212 GCF_000008105.1_ASM810v1 Salmonella enterica subsp. enterica serovar Choleraesuis str. SC-B67 Proteobacteria; Gammaproteobacteria; Enterobacterales; Enterobacteriaceae; Salmonella; Salmonella enterica; Salmonella enterica subsp. enterica; Salmonella enterica subsp. enterica serovar Choleraesuis MNNEETFYQAMRRKGVTRRSFLKFCSLAATSLGLGAGMTPKIAWALENKPRIPVVWIHGL WP_001540183.1 hydrogenase [Salmonella enterica] Length: 372\nNumber of predicted TMHs:  2\nExp number of AAs in TMHs: 37.41804\nExp number, first 60 AAs:  19.72808\nTotal prob of N-in:   0.96169\nPOSSIBLE N-term signal sequence\ninside   1   19\nTMhelix   20   39\noutside   40   325\nTMhelix   326   348\ninside   349   372

23213 GCF_000008105.1_ASM810v1 Salmonella enterica subsp. enterica serovar Choleraesuis str. SC-B67 Proteobacteria; Gammaproteobacteria; Enterobacterales; Enterobacteriaceae; Salmonella; Salmonella enterica; Salmonella enterica subsp. enterica; Salmonella enterica subsp. enterica serovar Choleraesuis MTGDNTLITSHGINRRDFMKLCAALAATMGLSSKAAAEMAESVSNPQRPPVIWIGAQECT WP_000145429.1 hydrogenase 2 small subunit [Salmonella enterica] Length: 372\nNumber of predicted TMHs:  1\nExp number of AAs in TMHs: 24.51309\nExp number, first 60 AAs:  0.2682\nTotal prob of N-in:   0.14402\noutside   1   331\nTMhelix   332   354\ninside   355   372

23214 GCF_000187565.1_ASM18756v1 Salmonella enterica subsp. enterica serovar Choleraesuis str. SCSA50 Proteobacteria; Gammaproteobacteria; Enterobacterales; Enterobacteriaceae; Salmonella; Salmonella enterica; Salmonella enterica subsp. enterica; Salmonella enterica subsp. enterica serovar Choleraesuis MNNEETFYQAMRRKGVTRRSFLKFCSLAATSLGLGAGMTPKIAWALENKPRIPVVWIHGL WP_001540183.1 hydrogenase [Salmonella enterica] Length: 372\nNumber of predicted TMHs:  2\nExp number of AAs in TMHs: 37.41804\nExp number, first 60 AAs:  19.72808\nTotal prob of N-in:   0.96169\nPOSSIBLE N-term signal sequence\ninside   1   19\nTMhelix   20   39\noutside   40   325\nTMhelix   326   348\ninside   349   372

23215 GCF_000187565.1_ASM18756v1 Salmonella enterica subsp. enterica serovar Choleraesuis str. SCSA50 Proteobacteria; Gammaproteobacteria; Enterobacterales; Enterobacteriaceae; Salmonella; Salmonella enterica; Salmonella enterica subsp. enterica; Salmonella enterica subsp. enterica serovar Choleraesuis MTGDNTLITSHGINRRDFMKLCAALAATMGLSSKAAAEMAESVSNPQRPPVIWIGAQECT WP_000145429.1 hydrogenase 2 small subunit [Salmonella enterica] Length: 372\nNumber of predicted TMHs:  1\nExp number of AAs in TMHs: 24.51309\nExp number, first 60 AAs:  0.2682\nTotal prob of N-in:   0.14402\noutside   1   331\nTMhelix   332   354\ninside   355   372

23216 GCF_002045865.1_ASM204586v1 Salmonella enterica subsp. enterica serovar Choleraesuis var. Kunzendorf Proteobacteria; Gammaproteobacteria; Enterobacterales; Enterobacteriaceae; Salmonella; Salmonella enterica;

Salmonella enterica subsp. enterica; Salmonella enterica subsp. enterica serovar Choleraesuis
MTGDNTLITSHGINRRDFMKLCAALAATMGLSSKAAAEMAESVSNPQRPPVIWIGAQECT WP_000145429.1
hydrogenase 2 small subunit [Salmonella enterica] Length: 372\nNumber of predicted TMHs: 1\nExp number of AAs in TMHs: 24.51309\nExp number, first 60 AAs: 0.2682\nTotal prob of N-in: 0.14402\noutside 1 331\nTMhelix 332 354\ninside 355 372

23217 GCF_002045865.1_ASM204586v1 Salmonella enterica subsp. enterica serovar Choleraesuis var. Kunzendorf
Proteobacteria; Gammaproteobacteria; Enterobacterales; Enterobacteriaceae; Salmonella; Salmonella enterica; Salmonella enterica subsp. enterica; Salmonella enterica subsp. enterica serovar Choleraesuis
MNNEETFYQAMRRKGVTRRSFLKFCSLAATSLGLGAGMTPKIAWALENKPRIPVVWIHGL WP_001540183.1
hydrogenase [Salmonella enterica] Length: 372\nNumber of predicted TMHs: 2\nExp number of AAs in TMHs: 37.41804\nExp number, first 60 AAs: 19.72808\nTotal prob of N-in: 0.96169\nPOSSIBLE N-term signal sequence\ninside 1 19\nTMhelix 20 39\noutside 40 325\nTMhelix 326 348\ninside 349 372

23218 GCF_002045165.1_ASM204516v1 Salmonella enterica subsp. enterica serovar Choleraesuis var. Kunzendorf
Proteobacteria; Gammaproteobacteria; Enterobacterales; Enterobacteriaceae; Salmonella; Salmonella enterica; Salmonella enterica subsp. enterica; Salmonella enterica subsp. enterica serovar Choleraesuis
MNNEETFYQAMRRKGVTRRSFLKFCSLAATSLGLGAGMTPKIAWALENKPRIPVVWIHGL WP_023234992.1
hydrogenase 1, small subunit [Salmonella enterica] Length: 372\nNumber of predicted TMHs: 2\nExp number of AAs in TMHs: 37.41309\nExp number, first 60 AAs: 19.72865\nTotal prob of N-in: 0.96169\nPOSSIBLE N-term signal sequence\ninside 1 19\nTMhelix 20 39\noutside 40 325\nTMhelix 326 348\ninside 349 372

23219 GCF_002045165.1_ASM204516v1 Salmonella enterica subsp. enterica serovar Choleraesuis var. Kunzendorf
Proteobacteria; Gammaproteobacteria; Enterobacterales; Enterobacteriaceae; Salmonella; Salmonella enterica; Salmonella enterica subsp. enterica; Salmonella enterica subsp. enterica serovar Choleraesuis
MTGDNTLITSHCINRRDFMKLCAALAATMGLSSKAAAEMAESVSNPQRPPVIWIGAQECT WP_079970246.1
hydrogenase 2 small subunit [Salmonella enterica] Length: 372\nNumber of predicted TMHs: 1\nExp number of AAs in TMHs: 24.58192\nExp number, first 60 AAs: 0.34172\nTotal prob of N-in: 0.14721\noutside 1 331\nTMhelix 332 354\ninside 355 372

23220 GCF_002045915.1_ASM204591v1 Salmonella enterica subsp. enterica serovar Choleraesuis var. Kunzendorf
Proteobacteria; Gammaproteobacteria; Enterobacterales; Enterobacteriaceae; Salmonella; Salmonella enterica; Salmonella enterica subsp. enterica; Salmonella enterica subsp. enterica serovar Choleraesuis
MTGDNTLITSHGINRRDFMKLCAALAATMGLSSKAAAEMAESVSNPQRPPVIWIGAQECT WP_000145429.1
hydrogenase 2 small subunit [Salmonella enterica] Length: 372\nNumber of predicted TMHs: 1\nExp number of AAs in TMHs: 24.51309\nExp number, first 60 AAs: 0.2682\nTotal prob of N-in: 0.14402\noutside 1 331\nTMhelix 332 354\ninside 355 372

23221 GCF_002045915.1_ASM204591v1 Salmonella enterica subsp. enterica serovar Choleraesuis var. Kunzendorf
Proteobacteria; Gammaproteobacteria; Enterobacterales; Enterobacteriaceae; Salmonella; Salmonella enterica; Salmonella enterica subsp. enterica; Salmonella enterica subsp. enterica serovar Choleraesuis
MNNEETFYQAMRRKGVTRRSFLKFCSLAATSLGLGAGMTPKIAWALENKPRIPVVWIHGL WP_001540183.1
hydrogenase [Salmonella enterica] Length: 372\nNumber of predicted TMHs: 2\nExp number of AAs in TMHs: 37.41804\nExp number, first 60 AAs: 19.72808\nTotal prob of N-in: 0.96169\nPOSSIBLE N-term signal sequence\ninside 1 19\nTMhelix 20 39\noutside 40 325\nTMhelix 326 348\ninside 349 372

23222 GCF_001975245.1_ASM197524v1 Salmonella enterica subsp. enterica serovar Crossness str. 1422-74
Proteobacteria; Gammaproteobacteria; Enterobacterales; Enterobacteriaceae; Salmonella; Salmonella enterica; Salmonella enterica subsp. enterica; Salmonella enterica subsp. enterica serovar Crossness
MNNEETFYQAMRRKGVTRRSFLKFCSLAATSLGLGAGMTPKIAWALENKPRIPVVWIHGL WP_001058312.1
hydrogenase [Salmonella enterica] Length: 372\nNumber of predicted TMHs: 2\nExp number of AAs in TMHs: 36.51738\nExp number, first 60 AAs: 19.72849\nTotal prob of N-in: 0.96170\nPOSSIBLE N-term signal sequence\ninside 1 19\nTMhelix 20 39\noutside 40 325\nTMhelix 326 348\ninside 349 372

23223 GCF_001975245.1_ASM197524v1 Salmonella enterica subsp. enterica serovar Crossness str. 1422-74
Proteobacteria; Gammaproteobacteria; Enterobacterales; Enterobacteriaceae; Salmonella; Salmonella enterica; Salmonella enterica subsp. enterica; Salmonella enterica subsp. enterica serovar Crossness
MTGDNTLITSHGINRRDFMKLCAALAATMGLSSKAAAEMAESVSNPQRPPVIWIGAQECT WP_000145429.1
hydrogenase 2 small subunit [Salmonella enterica] Length: 372\nNumber of predicted TMHs: 1\nExp number of AAs in TMHs: 24.51309\nExp number, first 60 AAs: 0.2682\nTotal prob of N-in: 0.14402\noutside 1 331\nTMhelix 332 354\ninside 355 372

23224 GCF_000504705.2_ASM50470v2 Salmonella enterica subsp. enterica serovar Cubana str. 76814 Proteobacteria; Gammaproteobacteria; Enterobacterales; Enterobacteriaceae; Salmonella; Salmonella enterica; Salmonella enterica subsp. enterica; Salmonella enterica subsp. enterica serovar Cubana MNNEETFYQAMRRKGVTRRSFLKFCSLAATSLGLGAGMTPKIAWALENKPRIPVVWIHGL WP_001058311.1 hydrogenase [Salmonella enterica] Length: 372\nNumber of predicted TMHs: 2\nExp number of AAs in TMHs: 37.4699899999999\nExp number, first 60 AAs: 19.72699\nTotal prob of N-in: 0.96171\nPOSSIBLE N-term signal sequence\ninside 1 19\nTMhelix 20 39\noutside 40 325\nTMhelix 326 348\ninside 349 372

23225 GCF_000505845.2_S_enterica_enterica_Cubana_Ver1 Salmonella enterica subsp. enterica serovar Cubana str. 76814 Proteobacteria; Gammaproteobacteria; Enterobacterales; Enterobacteriaceae; Salmonella; Salmonella enterica; Salmonella enterica subsp. enterica; Salmonella enterica subsp. enterica serovar Cubana MTGDNTLITSHGINRRDFMKLCAALAATMGLSSKAAAEMAESVSNPQRPPVIWIGAQECT WP_000145429.1 hydrogenase 2 small subunit [Salmonella enterica] Length: 372\nNumber of predicted TMHs: 1\nExp number of AAs in TMHs: 24.51309\nExp number, first 60 AAs: 0.2682\nTotal prob of N-in: 0.14402\noutside 1 331\nTMhelix 332 354\ninside 355 372

23226 GCF_000504705.2_ASM50470v2 Salmonella enterica subsp. enterica serovar Cubana str. 76814 Proteobacteria; Gammaproteobacteria; Enterobacterales; Enterobacteriaceae; Salmonella; Salmonella enterica; Salmonella enterica subsp. enterica; Salmonella enterica subsp. enterica serovar Cubana MTGDNTLITSHGINRRDFMKLCAALAATMGLSSKAAAEMAESVSNPQRPPVIWIGAQECT WP_000145429.1 hydrogenase 2 small subunit [Salmonella enterica] Length: 372\nNumber of predicted TMHs: 1\nExp number of AAs in TMHs: 24.51309\nExp number, first 60 AAs: 0.2682\nTotal prob of N-in: 0.14402\noutside 1 331\nTMhelix 332 354\ninside 355 372

23227 GCF_000505845.2_S_enterica_enterica_Cubana_Ver1 Salmonella enterica subsp. enterica serovar Cubana str. 76814 Proteobacteria; Gammaproteobacteria; Enterobacterales; Enterobacteriaceae; Salmonella; Salmonella enterica; Salmonella enterica subsp. enterica; Salmonella enterica subsp. enterica serovar Cubana MNNEETFYQAMRRKGVTRRSFLKFCSLAATSLGLGAGMTPKIAWALENKPRIPVVWIHGL WP_001058311.1 hydrogenase [Salmonella enterica] Length: 372\nNumber of predicted TMHs: 2\nExp number of AAs in TMHs: 37.4699899999999\nExp number, first 60 AAs: 19.72699\nTotal prob of N-in: 0.96171\nPOSSIBLE N-term signal sequence\ninside 1 19\nTMhelix 20 39\noutside 40 325\nTMhelix 326 348\ninside 349 372

23228 GCF_000487715.1_ASM48771v1 Salmonella enterica subsp. enterica serovar Cubana str. CFSAN001083 Proteobacteria; Gammaproteobacteria; Enterobacterales; Enterobacteriaceae; Salmonella; Salmonella enterica; Salmonella enterica subsp. enterica; Salmonella enterica subsp. enterica serovar Cubana MTGDNTLITSHGINRRDFMKLCAALAATMGLSSKAAAEMAESVSNPQRPPVIWIGAQECT WP_000145429.1 hydrogenase 2 small subunit [Salmonella enterica] Length: 372\nNumber of predicted TMHs: 1\nExp number of AAs in TMHs: 24.51309\nExp number, first 60 AAs: 0.2682\nTotal prob of N-in: 0.14402\noutside 1 331\nTMhelix 332 354\ninside 355 372

23229 GCF_000430125.1_ASM43012v1 Salmonella enterica subsp. enterica serovar Cubana str. CFSAN002050 Proteobacteria; Gammaproteobacteria; Enterobacterales; Enterobacteriaceae; Salmonella; Salmonella enterica; Salmonella enterica subsp. enterica; Salmonella enterica subsp. enterica serovar Cubana MTGDNTLITSHGINRRDFMKLCAALAATMGLSSKAAAEMAESVSNPQRPPVIWIGAQECT WP_000145429.1 hydrogenase 2 small subunit [Salmonella enterica] Length: 372\nNumber of predicted TMHs: 1\nExp number of AAs in TMHs: 24.51309\nExp number, first 60 AAs: 0.2682\nTotal prob of N-in: 0.14402\noutside 1 331\nTMhelix 332 354\ninside 355 372

23230 GCF_000430125.1_ASM43012v1 Salmonella enterica subsp. enterica serovar Cubana str. CFSAN002050 Proteobacteria; Gammaproteobacteria; Enterobacterales; Enterobacteriaceae; Salmonella; Salmonella enterica; Salmonella enterica subsp. enterica; Salmonella enterica subsp. enterica serovar Cubana MNNEETFYQAMRRKGVTRRSFLKFCSLAATSLGLGAGMTPKIAWALENKPRIPVVWIHGL WP_001058311.1 hydrogenase [Salmonella enterica] Length: 372\nNumber of predicted TMHs: 2\nExp number of AAs in TMHs: 37.4699899999999\nExp number, first 60 AAs: 19.72699\nTotal prob of N-in: 0.96171\nPOSSIBLE N-term signal sequence\ninside 1 19\nTMhelix 20 39\noutside 40 325\nTMhelix 326 348\ninside 349 372

23231 GCF_000500025.1_ASM50002v1 Salmonella enterica subsp. enterica serovar Cubana str. CVM42234 Proteobacteria; Gammaproteobacteria; Enterobacterales; Enterobacteriaceae; Salmonella; Salmonella enterica; Salmonella enterica subsp. enterica; Salmonella enterica subsp. enterica serovar Cubana MTGDNTLITSHGINRRDFMKLCAALAATMGLSSKAAAEMAESVSNPQRPPVIWIGAQECT WP_000145429.1 hydrogenase 2 small subunit [Salmonella enterica] Length: 372\nNumber of predicted TMHs: 1\nExp

number of AAs in TMHs: 24.51309\nExp number, first 60 AAs: 0.2682\nTotal prob of N-in: 0.14402\noutside 1 331\nTMhelix 332 354\ninside 355 372

23232 GCF_000500025.1_ASM50002v1 Salmonella enterica subsp. enterica serovar Cubana str. CVM42234 Proteobacteria; Gammaproteobacteria; Enterobacterales; Enterobacteriaceae; Salmonella; Salmonella enterica; Salmonella enterica subsp. enterica; Salmonella enterica subsp. enterica serovar Cubana MNNEETFYQAMRRKGVTRRSFLKFCSLAATSLGLGAGMTPKIAWALENKPRIPVVWIHGL WP_001058311.1 hydrogenase [Salmonella enterica] Length: 372\nNumber of predicted TMHs: 2\nExp number of AAs in TMHs: 37.4699899999999\nExp number, first 60 AAs: 19.72699\nTotal prob of N-in: 0.96171\nPOSSIBLE N-term signal sequence\ninside 1 19\nTMhelix 20 39\noutside 40 325\nTMhelix 326 348\ninside 349 372

23233 GCF_000487235.1_ASM48723v1 Salmonella enterica subsp. enterica serovar Derby str. 626 Proteobacteria; Gammaproteobacteria; Enterobacterales; Enterobacteriaceae; Salmonella; Salmonella enterica; Salmonella enterica subsp. enterica; Salmonella enterica subsp. enterica serovar Derby MTGDNTLITSHGINRRDFMKLCAALAATMGLSSKAAAEMAESVSNPQRPPVIWIGAQECT WP_000145429.1 hydrogenase 2 small subunit [Salmonella enterica] Length: 372\nNumber of predicted TMHs: 1\nExp number of AAs in TMHs: 24.51309\nExp number, first 60 AAs: 0.2682\nTotal prob of N-in: 0.14402\noutside 1 331\nTMhelix 332 354\ninside 355 372

23234 GCF_000487235.1_ASM48723v1 Salmonella enterica subsp. enterica serovar Derby str. 626 Proteobacteria; Gammaproteobacteria; Enterobacterales; Enterobacteriaceae; Salmonella; Salmonella enterica; Salmonella enterica subsp. enterica; Salmonella enterica subsp. enterica serovar Derby MNNEETFYQAMRRKGVTRRSFLKFCSLAATSLGLGAGMTPKIAWALENKPRIPVVWIHGL WP_001058312.1 hydrogenase [Salmonella enterica] Length: 372\nNumber of predicted TMHs: 2\nExp number of AAs in TMHs: 36.51738\nExp number, first 60 AAs: 19.72849\nTotal prob of N-in: 0.96170\nPOSSIBLE N-term signal sequence\ninside 1 19\nTMhelix 20 39\noutside 40 325\nTMhelix 326 348\ninside 349 372

23235 GCF_001975365.1_ASM197536v1 Salmonella enterica subsp. enterica serovar Djakarta str. S-1087 Proteobacteria; Gammaproteobacteria; Enterobacterales; Enterobacteriaceae; Salmonella; Salmonella enterica; Salmonella enterica subsp. enterica; Salmonella enterica subsp. enterica serovar Djakarta MTGDNTLITSHGINRRDFMKLCAALAATMGLSSKAAAEMAESVSNPQRPPVIWIGAQECT WP_000145429.1 hydrogenase 2 small subunit [Salmonella enterica] Length: 372\nNumber of predicted TMHs: 1\nExp number of AAs in TMHs: 24.51309\nExp number, first 60 AAs: 0.2682\nTotal prob of N-in: 0.14402\noutside 1 331\nTMhelix 332 354\ninside 355 372

23236 GCF_001975365.1_ASM197536v1 Salmonella enterica subsp. enterica serovar Djakarta str. S-1087 Proteobacteria; Gammaproteobacteria; Enterobacterales; Enterobacteriaceae; Salmonella; Salmonella enterica; Salmonella enterica subsp. enterica; Salmonella enterica subsp. enterica serovar Djakarta MNNEETFYQAMRRKGVTRRSFLKFCSLAATSLGLGAGMTPKIAWALENKPRIPVVWIHGL WP_001058312.1 hydrogenase [Salmonella enterica] Length: 372\nNumber of predicted TMHs: 2\nExp number of AAs in TMHs: 36.51738\nExp number, first 60 AAs: 19.72849\nTotal prob of N-in: 0.96170\nPOSSIBLE N-term signal sequence\ninside 1 19\nTMhelix 20 39\noutside 40 325\nTMhelix 326 348\ninside 349 372

23237 GCF_001953035.1_ASM195303v1 Salmonella enterica subsp. enterica serovar Dublin str. ATCC 39184 Proteobacteria; Gammaproteobacteria; Enterobacterales; Enterobacteriaceae; Salmonella; Salmonella enterica; Salmonella enterica subsp. enterica; Salmonella enterica subsp. enterica serovar Dublin MNNEETFYQAMRRKGVTRRSFLKFCSLAATSLGLGAGMTPKIAWALENKPRIPVVWIHGL WP_001058312.1 hydrogenase [Salmonella enterica] Length: 372\nNumber of predicted TMHs: 2\nExp number of AAs in TMHs: 36.51738\nExp number, first 60 AAs: 19.72849\nTotal prob of N-in: 0.96170\nPOSSIBLE N-term signal sequence\ninside 1 19\nTMhelix 20 39\noutside 40 325\nTMhelix 326 348\ninside 349 372

23238 GCF_001953035.1_ASM195303v1 Salmonella enterica subsp. enterica serovar Dublin str. ATCC 39184 Proteobacteria; Gammaproteobacteria; Enterobacterales; Enterobacteriaceae; Salmonella; Salmonella enterica; Salmonella enterica subsp. enterica; Salmonella enterica subsp. enterica serovar Dublin MTGDNTLITSHGINRRDFMKLCAALAATMGLSSKAAAEMAESVSNPQRPPVIWIGAQECT WP_000145428.1 hydrogenase 2 small subunit [Salmonella enterica] Length: 372\nNumber of predicted TMHs: 1\nExp number of AAs in TMHs: 24.50677\nExp number, first 60 AAs: 0.2682\nTotal prob of N-in: 0.14390\noutside 1 331\nTMhelix 332 354\ninside 355 372

23239 GCF_000020925.1_ASM2092v1 Salmonella enterica subsp. enterica serovar Dublin str. CT_02021853 Proteobacteria; Gammaproteobacteria; Enterobacterales; Enterobacteriaceae; Salmonella; Salmonella enterica;

Salmonella enterica subsp. enterica; Salmonella enterica subsp. enterica serovar Dublin
MNNEETFYQAMRRKGVTRRSFLKFCSLAATSLGLGAGMTPKIAWALENKPRIPVVWIHGL WP_001058303.1
hydrogenase [Salmonella enterica] Length: 372\nNumber of predicted TMHs: 2\nExp number of AAs in TMHs: 36.518\nExp number, first 60 AAs: 19.72854\nTotal prob of N-in: 0.96169\nPOSSIBLE N-term signal sequence\ninside 1 19\nTMhelix 20 39\noutside 40 325\nTMhelix 326 348\ninside 349 372

23240 GCF_000020925.1_ASM2092v1 Salmonella enterica subsp. enterica serovar Dublin str. CT_02021853
Proteobacteria; Gammaproteobacteria; Enterobacterales; Enterobacteriaceae; Salmonella; Salmonella enterica; Salmonella enterica subsp. enterica; Salmonella enterica subsp. enterica serovar Dublin
MTGDNTLITSHGINRRDFMKLCAALAATMGLSSKAAAEMAESVSNPQRPPVIWIGAQECT WP_000145428.1
hydrogenase 2 small subunit [Salmonella enterica] Length: 372\nNumber of predicted TMHs: 1\nExp number of AAs in TMHs: 24.50677\nExp number, first 60 AAs: 0.2682\nTotal prob of N-in: 0.14390\noutside 1 331\nTMhelix 332 354\ninside 355 372

23241 GCF_000414745.2_ASM41474v2 Salmonella enterica subsp. enterica serovar Dublin str. DG22
Proteobacteria; Gammaproteobacteria; Enterobacterales; Enterobacteriaceae; Salmonella; Salmonella enterica; Salmonella enterica subsp. enterica; Salmonella enterica subsp. enterica serovar Dublin
MTGDNTLITSHGINRRDFMKLCAALAATMGLSSKAAAEMAESVSNPQRPPVIWIGAQECT WP_000145428.1
hydrogenase 2 small subunit [Salmonella enterica] Length: 372\nNumber of predicted TMHs: 1\nExp number of AAs in TMHs: 24.50677\nExp number, first 60 AAs: 0.2682\nTotal prob of N-in: 0.14390\noutside 1 331\nTMhelix 332 354\ninside 355 372

23242 GCF_000414745.2_ASM41474v2 Salmonella enterica subsp. enterica serovar Dublin str. DG22
Proteobacteria; Gammaproteobacteria; Enterobacterales; Enterobacteriaceae; Salmonella; Salmonella enterica; Salmonella enterica subsp. enterica; Salmonella enterica subsp. enterica serovar Dublin
MNNEETLYQAMRRKGVTRRSFLKFCSLAATSLGLGAGMTPKIAWALENKPRIPVVWIHGL WP_001752445.1
hydrogenase-1 small chain [Salmonella enterica] Length: 372\nNumber of predicted TMHs: 2\nExp number of AAs in TMHs: 36.5123\nExp number, first 60 AAs: 19.72341\nTotal prob of N-in: 0.96144\nPOSSIBLE N-term signal sequence\ninside 1 19\nTMhelix 20 39\noutside 40 325\nTMhelix 326 348\ninside 349 372

23243 GCF_001448615.1_ASM144861v1 Salmonella enterica subsp. enterica serovar Dublin str. SA20093032
Proteobacteria; Gammaproteobacteria; Enterobacterales; Enterobacteriaceae; Salmonella; Salmonella enterica; Salmonella enterica subsp. enterica; Salmonella enterica subsp. enterica serovar Dublin
MNNEETLYQAMRRKGVTRRSFLKFCSLAATSLGLGAGMTPKIAWALENKPRIPVVWIHGL WP_001752445.1
hydrogenase-1 small chain [Salmonella enterica] Length: 372\nNumber of predicted TMHs: 2\nExp number of AAs in TMHs: 36.5123\nExp number, first 60 AAs: 19.72341\nTotal prob of N-in: 0.96144\nPOSSIBLE N-term signal sequence\ninside 1 19\nTMhelix 20 39\noutside 40 325\nTMhelix 326 348\ninside 349 372

23244 GCF_001448615.1_ASM144861v1 Salmonella enterica subsp. enterica serovar Dublin str. SA20093032
Proteobacteria; Gammaproteobacteria; Enterobacterales; Enterobacteriaceae; Salmonella; Salmonella enterica; Salmonella enterica subsp. enterica; Salmonella enterica subsp. enterica serovar Dublin
MTGDNTLITSHGINRRDFMKLCAALAATMGLSSKAAAEMAESVSNPQRPPVIWIGAQECT WP_000145428.1
hydrogenase 2 small subunit [Salmonella enterica] Length: 372\nNumber of predicted TMHs: 1\nExp number of AAs in TMHs: 24.50677\nExp number, first 60 AAs: 0.2682\nTotal prob of N-in: 0.14390\noutside 1 331\nTMhelix 332 354\ninside 355 372

23245 GCF_000192085.1_ASM19208v1 Salmonella enterica subsp. enterica serovar Dublin str. SD3246
Proteobacteria; Gammaproteobacteria; Enterobacterales; Enterobacteriaceae; Salmonella; Salmonella enterica; Salmonella enterica subsp. enterica; Salmonella enterica subsp. enterica serovar Dublin
MNNEETFYQAMRRKGVTRRSFLKFCSLAATSLGLGAGMTPKIAWALENKPRIPVVWIHGL WP_001058312.1
hydrogenase [Salmonella enterica] Length: 372\nNumber of predicted TMHs: 2\nExp number of AAs in TMHs: 36.51738\nExp number, first 60 AAs: 19.72849\nTotal prob of N-in: 0.96170\nPOSSIBLE N-term signal sequence\ninside 1 19\nTMhelix 20 39\noutside 40 325\nTMhelix 326 348\ninside 349 372

23246 GCF_000192085.1_ASM19208v1 Salmonella enterica subsp. enterica serovar Dublin str. SD3246
Proteobacteria; Gammaproteobacteria; Enterobacterales; Enterobacteriaceae; Salmonella; Salmonella enterica; Salmonella enterica subsp. enterica; Salmonella enterica subsp. enterica serovar Dublin
MTGDNTLITSHGINRRDFMKLCAALAATMGLSSKAAAEMAESVSNPQRPPVIWIGAQECT WP_000145428.1
hydrogenase 2 small subunit [Salmonella enterica] Length: 372\nNumber of predicted TMHs: 1\nExp number of AAs in TMHs: 24.50677\nExp number, first 60 AAs: 0.2682\nTotal prob of N-in: 0.14390\noutside 1 331\nTMhelix 332 354\ninside 355 372

23247 GCF_000336015.1_ASM33601v1 Salmonella enterica subsp. enterica serovar Dublin str. SL1438 Proteobacteria; Gammaproteobacteria; Enterobacterales; Enterobacteriaceae; Salmonella; Salmonella enterica; Salmonella enterica subsp. enterica; Salmonella enterica subsp. enterica serovar Dublin MTGDNTLITSHGINRRDFMKLCAALAATMGLSSKAAAEMAESVSNPQRPPVIWIGAQECT WP_000145428.1 hydrogenase 2 small subunit [Salmonella enterica] Length: 372\nNumber of predicted TMHs: 1\nExp number of AAs in TMHs: 24.50677\nExp number, first 60 AAs: 0.2682\nTotal prob of N-in: 0.14390\noutside 1 331\nTMhelix 332 354\ninside 355 372

23248 GCF_000336015.1_ASM33601v1 Salmonella enterica subsp. enterica serovar Dublin str. SL1438 Proteobacteria; Gammaproteobacteria; Enterobacterales; Enterobacteriaceae; Salmonella; Salmonella enterica; Salmonella enterica subsp. enterica; Salmonella enterica subsp. enterica serovar Dublin MNNEETFYQAMRRKGVTRRSFLKFCSLAATSLGLGAGMTPKIAWALENKPRIPVVWIHGL WP_001058312.1 hydrogenase [Salmonella enterica] Length: 372\nNumber of predicted TMHs: 2\nExp number of AAs in TMHs: 36.51738\nExp number, first 60 AAs: 19.72849\nTotal prob of N-in: 0.96170\nPOSSIBLE N-term signal sequence\ninside 1 19\nTMhelix 20 39\noutside 40 325\nTMhelix 326 348\ninside 349 372

23249 GCF_000353305.2_ASM35330v2 Salmonella enterica subsp. enterica serovar Dublin str. UC16 Proteobacteria; Gammaproteobacteria; Enterobacterales; Enterobacteriaceae; Salmonella; Salmonella enterica; Salmonella enterica subsp. enterica; Salmonella enterica subsp. enterica serovar Dublin MNNEETLYQAMRRKGVTRRSFLKFCSLAATSLGLGAGMTPKIAWALENKPRIPVVWIHGL WP_001752445.1 hydrogenase-1 small chain [Salmonella enterica] Length: 372\nNumber of predicted TMHs: 2\nExp number of AAs in TMHs: 36.5123\nExp number, first 60 AAs: 19.72341\nTotal prob of N-in: 0.96144\nPOSSIBLE N-term signal sequence\ninside 1 19\nTMhelix 20 39\noutside 40 325\nTMhelix 326 348\ninside 349 372

23250 GCF_000353305.2_ASM35330v2 Salmonella enterica subsp. enterica serovar Dublin str. UC16 Proteobacteria; Gammaproteobacteria; Enterobacterales; Enterobacteriaceae; Salmonella; Salmonella enterica; Salmonella enterica subsp. enterica; Salmonella enterica subsp. enterica serovar Dublin MTGDNTLITSHGINRRDFMKLCAALAATMGLSSKAAAEMAESVSNPQRPPVIWIGAQECT WP_000145428.1 hydrogenase 2 small subunit [Salmonella enterica] Length: 372\nNumber of predicted TMHs: 1\nExp number of AAs in TMHs: 24.50677\nExp number, first 60 AAs: 0.2682\nTotal prob of N-in: 0.14390\noutside 1 331\nTMhelix 332 354\ninside 355 372

23251 GCF_000487695.1_ASM48769v1 Salmonella enterica subsp. enterica serovar Eastbourne str. CFSAN001084 Proteobacteria; Gammaproteobacteria; Enterobacterales; Enterobacteriaceae; Salmonella; Salmonella enterica; Salmonella enterica subsp. enterica; Salmonella enterica subsp. enterica serovar Eastbourne MTGDNTLITSHGINRRDFMKLCAALAATMGLSSKAAAEMAESVSNPQRPPVIWIGAQECT WP_000145429.1 hydrogenase 2 small subunit [Salmonella enterica] Length: 372\nNumber of predicted TMHs: 1\nExp number of AAs in TMHs: 24.51309\nExp number, first 60 AAs: 0.2682\nTotal prob of N-in: 0.14402\noutside 1 331\nTMhelix 332 354\ninside 355 372

23252 GCF_000487695.1_ASM48769v1 Salmonella enterica subsp. enterica serovar Eastbourne str. CFSAN001084 Proteobacteria; Gammaproteobacteria; Enterobacterales; Enterobacteriaceae; Salmonella; Salmonella enterica; Salmonella enterica subsp. enterica; Salmonella enterica subsp. enterica serovar Eastbourne MNNEETFYQAMRRKGVTRRSFLKFCSLAATSLGLGAGMTPKIAWALENKPRIPVVWIHGL WP_001058312.1 hydrogenase [Salmonella enterica] Length: 372\nNumber of predicted TMHs: 2\nExp number of AAs in TMHs: 36.51738\nExp number, first 60 AAs: 19.72849\nTotal prob of N-in: 0.96170\nPOSSIBLE N-term signal sequence\ninside 1 19\nTMhelix 20 39\noutside 40 325\nTMhelix 326 348\ninside 349 372

23253 GCF_000278505.1_08-0047v1.0 Salmonella enterica subsp. enterica serovar Enteritidis str. 08-0047 Proteobacteria; Gammaproteobacteria; Enterobacterales; Enterobacteriaceae; Salmonella; Salmonella enterica; Salmonella enterica subsp. enterica; Salmonella enterica subsp. enterica serovar Enteritidis MTGDNTLITSHGINRRDFMKLCAALAATMGLSSKAAAEMAESVSNPQRPPVIWIGAQECT WP_000145429.1 hydrogenase 2 small subunit [Salmonella enterica] Length: 372\nNumber of predicted TMHs: 1\nExp number of AAs in TMHs: 24.51309\nExp number, first 60 AAs: 0.2682\nTotal prob of N-in: 0.14402\noutside 1 331\nTMhelix 332 354\ninside 355 372

23254 GCF_000278505.1_08-0047v1.0 Salmonella enterica subsp. enterica serovar Enteritidis str. 08-0047 Proteobacteria; Gammaproteobacteria; Enterobacterales; Enterobacteriaceae; Salmonella; Salmonella enterica; Salmonella enterica subsp. enterica; Salmonella enterica subsp. enterica serovar Enteritidis MNNEETFYQAMRRKGVTRRSFLKFCSLAATSLGLGAGMTPKIAWALENKPRIPVVWIHGL WP_001058307.1 hydrogenase [Salmonella enterica] Length: 372\nNumber of predicted TMHs: 2\nExp number of AAs in

TMHs: 36.51738\nExp number, first 60 AAs: 19.72849\nTotal prob of N-in: 0.96170\nPOSSIBLE N-term signal sequence\ninside 1 19\nTMhelix 20 39\noutside 40 325\nTMhelix 326 348\ninside 349 372

23255 GCF_000278525.1_08-0128v1.0 Salmonella enterica subsp. enterica serovar Enteritidis str. 08-0128 Proteobacteria; Gammaproteobacteria; Enterobacterales; Enterobacteriaceae; Salmonella; Salmonella enterica; Salmonella enterica subsp. enterica; Salmonella enterica subsp. enterica serovar Enteritidis MNNEETFYQAMRRKGVTRRSFLKFCSLAATSLGLGAGMTPKIAWALENKPRIPVVWIHGL WP_001058307.1 hydrogenase [Salmonella enterica] Length: 372\nNumber of predicted TMHs: 2\nExp number of AAs in TMHs: 36.51738\nExp number, first 60 AAs: 19.72849\nTotal prob of N-in: 0.96170\nPOSSIBLE N-term signal sequence\ninside 1 19\nTMhelix 20 39\noutside 40 325\nTMhelix 326 348\ninside 349 372

23256 GCF_000278525.1_08-0128v1.0 Salmonella enterica subsp. enterica serovar Enteritidis str. 08-0128 Proteobacteria; Gammaproteobacteria; Enterobacterales; Enterobacteriaceae; Salmonella; Salmonella enterica; Salmonella enterica subsp. enterica; Salmonella enterica subsp. enterica serovar Enteritidis MTGDNTLITSHGINRRDFMKLCAALAATMGLSSKAAAEMAESVSNPQRPPVIWIGAQECT WP_000145429.1 hydrogenase 2 small subunit [Salmonella enterica] Length: 372\nNumber of predicted TMHs: 1\nExp number of AAs in TMHs: 24.51309\nExp number, first 60 AAs: 0.2682\nTotal prob of N-in: 0.14402\noutside 1 331\nTMhelix 332 354\ninside 355 372

23257 GCF_000278545.1_08-0627v1.0 Salmonella enterica subsp. enterica serovar Enteritidis str. 08-0627 Proteobacteria; Gammaproteobacteria; Enterobacterales; Enterobacteriaceae; Salmonella; Salmonella enterica; Salmonella enterica subsp. enterica; Salmonella enterica subsp. enterica serovar Enteritidis MTGDNTLITSHGINRRDFMKLCAALAATMGLSSKAAAEMAESVSNPQRPPVIWIGAQECT WP_000145429.1 hydrogenase 2 small subunit [Salmonella enterica] Length: 372\nNumber of predicted TMHs: 1\nExp number of AAs in TMHs: 24.51309\nExp number, first 60 AAs: 0.2682\nTotal prob of N-in: 0.14402\noutside 1 331\nTMhelix 332 354\ninside 355 372

23258 GCF_000278545.1_08-0627v1.0 Salmonella enterica subsp. enterica serovar Enteritidis str. 08-0627 Proteobacteria; Gammaproteobacteria; Enterobacterales; Enterobacteriaceae; Salmonella; Salmonella enterica; Salmonella enterica subsp. enterica; Salmonella enterica subsp. enterica serovar Enteritidis MNNEETFYQAMRRKGVTRRSFLKFCSLAATSLGLGAGMTPKIAWALENKPRIPVVWIHGL WP_001058307.1 hydrogenase [Salmonella enterica] Length: 372\nNumber of predicted TMHs: 2\nExp number of AAs in TMHs: 36.51738\nExp number, first 60 AAs: 19.72849\nTotal prob of N-in: 0.96170\nPOSSIBLE N-term signal sequence\ninside 1 19\nTMhelix 20 39\noutside 40 325\nTMhelix 326 348\ninside 349 372

23259 GCF_000414765.2_ASM41476v2 Salmonella enterica subsp. enterica serovar Enteritidis str. 08-1080 Proteobacteria; Gammaproteobacteria; Enterobacterales; Enterobacteriaceae; Salmonella; Salmonella enterica; Salmonella enterica subsp. enterica; Salmonella enterica subsp. enterica serovar Enteritidis MTGDNTLITSHGINRRDFMKLCAALAATMGLSSKAAAEMAESVSNPQRPPVIWIGAQECT WP_000145429.1 hydrogenase 2 small subunit [Salmonella enterica] Length: 372\nNumber of predicted TMHs: 1\nExp number of AAs in TMHs: 24.51309\nExp number, first 60 AAs: 0.2682\nTotal prob of N-in: 0.14402\noutside 1 331\nTMhelix 332 354\ninside 355 372

23260 GCF_000414765.2_ASM41476v2 Salmonella enterica subsp. enterica serovar Enteritidis str. 08-1080 Proteobacteria; Gammaproteobacteria; Enterobacterales; Enterobacteriaceae; Salmonella; Salmonella enterica; Salmonella enterica subsp. enterica; Salmonella enterica subsp. enterica serovar Enteritidis MNNEETFYQAMRRKGVTRRSFLKFCSLAATSLGLGAGMTPKIAWALENKPRIPVVWIHGL WP_001058307.1 hydrogenase [Salmonella enterica] Length: 372\nNumber of predicted TMHs: 2\nExp number of AAs in TMHs: 36.51738\nExp number, first 60 AAs: 19.72849\nTotal prob of N-in: 0.96170\nPOSSIBLE N-term signal sequence\ninside 1 19\nTMhelix 20 39\noutside 40 325\nTMhelix 326 348\ninside 349 372

23261 GCF_000335895.1_ASM33589v1 Salmonella enterica subsp. enterica serovar Enteritidis str. 13-1 Proteobacteria; Gammaproteobacteria; Enterobacterales; Enterobacteriaceae; Salmonella; Salmonella enterica; Salmonella enterica subsp. enterica; Salmonella enterica subsp. enterica serovar Enteritidis MNNEETFYQAMRRKGVTRRSFLKFCSLAATSLGLGAGMTPKIAWALENKPRIPVVWIHGL WP_001058307.1 hydrogenase [Salmonella enterica] Length: 372\nNumber of predicted TMHs: 2\nExp number of AAs in TMHs: 36.51738\nExp number, first 60 AAs: 19.72849\nTotal prob of N-in: 0.96170\nPOSSIBLE N-term signal sequence\ninside 1 19\nTMhelix 20 39\noutside 40 325\nTMhelix 326 348\ninside 349 372

23262 GCF_000335895.1_ASM33589v1 Salmonella enterica subsp. enterica serovar Enteritidis str. 13-1 Proteobacteria; Gammaproteobacteria; Enterobacterales; Enterobacteriaceae; Salmonella; Salmonella enterica;

Salmonella enterica subsp. enterica; Salmonella enterica subsp. enterica serovar Enteritidis
MTGDNTLITSHGINRRDFMKLCAALAATMGLSSKAAAEMAESVSNPQRPPVIWIGAQECT WP_000145429.1 hydrogenase 2 small subunit [Salmonella enterica] Length: 372\nNumber of predicted TMHs: 1\nExp number of AAs in TMHs: 24.51309\nExp number, first 60 AAs: 0.2682\nTotal prob of N-in: 0.14402\noutside 1 331\nTMhelix 332 354\ninside 355 372

23263 GCF_000329985.1_ASM32998v1 Salmonella enterica subsp. enterica serovar Enteritidis str. 17927 Proteobacteria; Gammaproteobacteria; Enterobacterales; Enterobacteriaceae; Salmonella; Salmonella enterica; Salmonella enterica subsp. enterica; Salmonella enterica subsp. enterica serovar Enteritidis
MNNEETFYQAMRRKGVTRRSFLKFCSLAATSLGLGAGMTPKIAWALENKPRIPVVWIHGL WP_001058307.1 hydrogenase [Salmonella enterica] Length: 372\nNumber of predicted TMHs: 2\nExp number of AAs in TMHs: 36.51738\nExp number, first 60 AAs: 19.72849\nTotal prob of N-in: 0.96170\nPOSSIBLE N-term signal sequence\ninside 1 19\nTMhelix 20 39\noutside 40 325\nTMhelix 326 348\ninside 349 372

23264 GCF_000329985.1_ASM32998v1 Salmonella enterica subsp. enterica serovar Enteritidis str. 17927 Proteobacteria; Gammaproteobacteria; Enterobacterales; Enterobacteriaceae; Salmonella; Salmonella enterica; Salmonella enterica subsp. enterica; Salmonella enterica subsp. enterica serovar Enteritidis
MTGDNTLITSHGINRRDFMKLCAALAATMGLSSKAAAEMAESVSNPQRPPVIWIGAQECT WP_000145429.1 hydrogenase 2 small subunit [Salmonella enterica] Length: 372\nNumber of predicted TMHs: 1\nExp number of AAs in TMHs: 24.51309\nExp number, first 60 AAs: 0.2682\nTotal prob of N-in: 0.14402\noutside 1 331\nTMhelix 332 354\ninside 355 372

23265 GCF_000335875.2_ASM33587v2 Salmonella enterica subsp. enterica serovar Enteritidis str. 18569 Proteobacteria; Gammaproteobacteria; Enterobacterales; Enterobacteriaceae; Salmonella; Salmonella enterica; Salmonella enterica subsp. enterica; Salmonella enterica subsp. enterica serovar Enteritidis
MTGDNTLITSHGINRRDFMKLCAALAATMGLSSKAAAEMAESVSNPQRPPVIWIGAQECT WP_000145429.1 hydrogenase 2 small subunit [Salmonella enterica] Length: 372\nNumber of predicted TMHs: 1\nExp number of AAs in TMHs: 24.51309\nExp number, first 60 AAs: 0.2682\nTotal prob of N-in: 0.14402\noutside 1 331\nTMhelix 332 354\ninside 355 372

23266 GCF_000335875.2_ASM33587v2 Salmonella enterica subsp. enterica serovar Enteritidis str. 18569 Proteobacteria; Gammaproteobacteria; Enterobacterales; Enterobacteriaceae; Salmonella; Salmonella enterica; Salmonella enterica subsp. enterica; Salmonella enterica subsp. enterica serovar Enteritidis
MNNEETFYQAMRRKGVTRRSFLKFCSLAATSLGLGAGMTPKIAWALENKPRIPVVWIHGL WP_001058307.1 hydrogenase [Salmonella enterica] Length: 372\nNumber of predicted TMHs: 2\nExp number of AAs in TMHs: 36.51738\nExp number, first 60 AAs: 19.72849\nTotal prob of N-in: 0.96170\nPOSSIBLE N-term signal sequence\ninside 1 19\nTMhelix 20 39\noutside 40 325\nTMhelix 326 348\ninside 349 372

23267 GCF_000336075.1_ASM33607v1 Salmonella enterica subsp. enterica serovar Enteritidis str. 20037 Proteobacteria; Gammaproteobacteria; Enterobacterales; Enterobacteriaceae; Salmonella; Salmonella enterica; Salmonella enterica subsp. enterica; Salmonella enterica subsp. enterica serovar Enteritidis
MTRRSFLKFCSLAATSLGLGAGMTPKIAWALENKPRIPVVWIHGLECTCCTESFIRSSHP WP_001839776.1 hydrogenase 1, small subunit [Salmonella enterica] Length: 357\nNumber of predicted TMHs: 1\nExp number of AAs in TMHs: 19.92233\nExp number, first 60 AAs: 3.08453\nTotal prob of N-in: 0.16445\noutside 1 310\nTMhelix 311 333\ninside 334 357

23268 GCF_000336075.1_ASM33607v1 Salmonella enterica subsp. enterica serovar Enteritidis str. 20037 Proteobacteria; Gammaproteobacteria; Enterobacterales; Enterobacteriaceae; Salmonella; Salmonella enterica; Salmonella enterica subsp. enterica; Salmonella enterica subsp. enterica serovar Enteritidis
MTGDNTLITSHGINRRDFMKLCAALAATMGLSSKAAAEMAESVSNPQRPPVIWIGAQECT WP_000145429.1 hydrogenase 2 small subunit [Salmonella enterica] Length: 372\nNumber of predicted TMHs: 1\nExp number of AAs in TMHs: 24.51309\nExp number, first 60 AAs: 0.2682\nTotal prob of N-in: 0.14402\noutside 1 331\nTMhelix 332 354\ninside 355 372

23269 GCF_000414785.2_ASM41478v2 Salmonella enterica subsp. enterica serovar Enteritidis str. 2009K0958 Proteobacteria; Gammaproteobacteria; Enterobacterales; Enterobacteriaceae; Salmonella; Salmonella enterica; Salmonella enterica subsp. enterica; Salmonella enterica subsp. enterica serovar Enteritidis
MTGDNTLITSHGINRRDFMKLCAALAATMGLSSKAAAEMAESVSNPQRPPVIWIGAQECT WP_000145429.1 hydrogenase 2 small subunit [Salmonella enterica] Length: 372\nNumber of predicted TMHs: 1\nExp number of AAs in TMHs: 24.51309\nExp number, first 60 AAs: 0.2682\nTotal prob of N-in: 0.14402\noutside 1 331\nTMhelix 332 354\ninside 355 372

23270	GCF_000414785.2_ASM41478v2	Salmonella enterica subsp. enterica serovar Enteritidis str. 2009K0958	Proteobacteria; Gammaproteobacteria; Enterobacterales; Enterobacteriaceae; Salmonella; Salmonella enterica; Salmonella enterica subsp. enterica; Salmonella enterica subsp. enterica serovar Enteritidis	MNNEETFYQAMRRKGVTRRSFLKFCSLAATSLGLGAGMTPKIAWALENKPRIPVVWIHGL	WP_001058307.1	hydrogenase [Salmonella enterica]	Length: 372\nNumber of predicted TMHs: 2\nExp number of AAs in TMHs: 36.51738\nExp number, first 60 AAs: 19.72849\nTotal prob of N-in: 0.96170\nPOSSIBLE N-term signal sequence\ninside 1 19\nTMhelix 20 39\noutside 40 325\nTMhelix 326 348\ninside 349 372

23271	GCF_000414805.2_ASM41480v2	Salmonella enterica subsp. enterica serovar Enteritidis str. 2009K1651	Proteobacteria; Gammaproteobacteria; Enterobacterales; Enterobacteriaceae; Salmonella; Salmonella enterica; Salmonella enterica subsp. enterica; Salmonella enterica subsp. enterica serovar Enteritidis	MTGDNTLITSHGINRRDFMKLCAALAATMGLSSKAAAEMAESVSNPQRPPVIWIGAQECT	WP_000145429.1	hydrogenase 2 small subunit [Salmonella enterica]	Length: 372\nNumber of predicted TMHs: 1\nExp number of AAs in TMHs: 24.51309\nExp number, first 60 AAs: 0.2682\nTotal prob of N-in: 0.14402\noutside 1 331\nTMhelix 332 354\ninside 355 372

23272	GCF_000414805.2_ASM41480v2	Salmonella enterica subsp. enterica serovar Enteritidis str. 2009K1651	Proteobacteria; Gammaproteobacteria; Enterobacterales; Enterobacteriaceae; Salmonella; Salmonella enterica; Salmonella enterica subsp. enterica; Salmonella enterica subsp. enterica serovar Enteritidis	MNNEETFYQAMRRKGVTRRSFLKFCSLAATSLGLGAGMTPKIAWALENKPRIPVVWIHGL	WP_001058307.1	hydrogenase [Salmonella enterica]	Length: 372\nNumber of predicted TMHs: 2\nExp number of AAs in TMHs: 36.51738\nExp number, first 60 AAs: 19.72849\nTotal prob of N-in: 0.96170\nPOSSIBLE N-term signal sequence\ninside 1 19\nTMhelix 20 39\noutside 40 325\nTMhelix 326 348\ninside 349 372

23273	GCF_000414825.2_ASM41482v2	Salmonella enterica subsp. enterica serovar Enteritidis str. 2009K1726	Proteobacteria; Gammaproteobacteria; Enterobacterales; Enterobacteriaceae; Salmonella; Salmonella enterica; Salmonella enterica subsp. enterica; Salmonella enterica subsp. enterica serovar Enteritidis	MNNEETFYQAMRRKGVTRRSFLKFCSLAATSLGLGAGMTPKIAWALENKPRIPVVWIHGL	WP_001058307.1	hydrogenase [Salmonella enterica]	Length: 372\nNumber of predicted TMHs: 2\nExp number of AAs in TMHs: 36.51738\nExp number, first 60 AAs: 19.72849\nTotal prob of N-in: 0.96170\nPOSSIBLE N-term signal sequence\ninside 1 19\nTMhelix 20 39\noutside 40 325\nTMhelix 326 348\ninside 349 372

23274	GCF_000414825.2_ASM41482v2	Salmonella enterica subsp. enterica serovar Enteritidis str. 2009K1726	Proteobacteria; Gammaproteobacteria; Enterobacterales; Enterobacteriaceae; Salmonella; Salmonella enterica; Salmonella enterica subsp. enterica; Salmonella enterica subsp. enterica serovar Enteritidis	MTGDNTLITSHGINRRDFMKLCAALAATMGLSSKAAAEMAESVSNPQRPPVIWIGAQECT	WP_000145429.1	hydrogenase 2 small subunit [Salmonella enterica]	Length: 372\nNumber of predicted TMHs: 1\nExp number of AAs in TMHs: 24.51309\nExp number, first 60 AAs: 0.2682\nTotal prob of N-in: 0.14402\noutside 1 331\nTMhelix 332 354\ninside 355 372

23275	GCF_000261985.1_CDC_2010-1237v1.0	Salmonella enterica subsp. enterica serovar Enteritidis str. 2010-1237	Proteobacteria; Gammaproteobacteria; Enterobacterales; Enterobacteriaceae; Salmonella; Salmonella enterica; Salmonella enterica subsp. enterica; Salmonella enterica subsp. enterica serovar Enteritidis	MTGDNTLITSHGINRRDFMKLCAALAATMGLSSKAAAEMAESVSNPQRPPVIWIGAQECT	WP_000145429.1	hydrogenase 2 small subunit [Salmonella enterica]	Length: 372\nNumber of predicted TMHs: 1\nExp number of AAs in TMHs: 24.51309\nExp number, first 60 AAs: 0.2682\nTotal prob of N-in: 0.14402\noutside 1 331\nTMhelix 332 354\ninside 355 372

23276	GCF_000261985.1_CDC_2010-1237v1.0	Salmonella enterica subsp. enterica serovar Enteritidis str. 2010-1237	Proteobacteria; Gammaproteobacteria; Enterobacterales; Enterobacteriaceae; Salmonella; Salmonella enterica; Salmonella enterica subsp. enterica; Salmonella enterica subsp. enterica serovar Enteritidis	MNNEETFYQAMRRKGVTRRSFLKFCSLAATSLGLGAGMTPKIAWALENKPRIPVVWIHGL	WP_001058307.1	hydrogenase [Salmonella enterica]	Length: 372\nNumber of predicted TMHs: 2\nExp number of AAs in TMHs: 36.51738\nExp number, first 60 AAs: 19.72849\nTotal prob of N-in: 0.96170\nPOSSIBLE N-term signal sequence\ninside 1 19\nTMhelix 20 39\noutside 40 325\nTMhelix 326 348\ninside 349 372

23277	GCF_000414845.2_ASM41484v2	Salmonella enterica subsp. enterica serovar Enteritidis str. 2010K-0262	Proteobacteria; Gammaproteobacteria; Enterobacterales; Enterobacteriaceae; Salmonella; Salmonella enterica; Salmonella enterica subsp. enterica; Salmonella enterica subsp. enterica serovar Enteritidis	MNNEETFYQAMRRKGVTRRSFLKFCSLAATSLGLGAGMTPKIAWALENKPRIPVVWIHGL	WP_001058307.1	hydrogenase [Salmonella enterica]	Length: 372\nNumber of predicted TMHs: 2\nExp number of AAs in

TMHs: 36.51738\nExp number, first 60 AAs: 19.72849\nTotal prob of N-in: 0.96170\nPOSSIBLE N-term signal sequence\ninside 1 19\nTMhelix 20 39\noutside 40 325\nTMhelix 326 348\ninside 349 372

23278 GCF_000414845.2_ASM41484v2 Salmonella enterica subsp. enterica serovar Enteritidis str. 2010K-0262 Proteobacteria; Gammaproteobacteria; Enterobacterales; Enterobacteriaceae; Salmonella; Salmonella enterica; Salmonella enterica subsp. enterica; Salmonella enterica subsp. enterica serovar Enteritidis MTGDNTLITSHGINRRDFMKLCAALAATMGLSSKAAAEMAESVSNPQRPPVIWIGAQECT WP_000145429.1 hydrogenase 2 small subunit [Salmonella enterica] Length: 372\nNumber of predicted TMHs: 1\nExp number of AAs in TMHs: 24.51309\nExp number, first 60 AAs: 0.2682\nTotal prob of N-in: 0.14402\noutside 1 331\nTMhelix 332 354\ninside 355 372

23279 GCF_000414865.2_ASM41486v2 Salmonella enterica subsp. enterica serovar Enteritidis str. 2010K-0267 Proteobacteria; Gammaproteobacteria; Enterobacterales; Enterobacteriaceae; Salmonella; Salmonella enterica; Salmonella enterica subsp. enterica; Salmonella enterica subsp. enterica serovar Enteritidis MNNEETFYQAMRRKGVTRRSFLKFCSLAATSLGLGAGMTPKIAWALENKPRIPVVWIHGL WP_001058307.1 hydrogenase [Salmonella enterica] Length: 372\nNumber of predicted TMHs: 2\nExp number of AAs in TMHs: 36.51738\nExp number, first 60 AAs: 19.72849\nTotal prob of N-in: 0.96170\nPOSSIBLE N-term signal sequence\ninside 1 19\nTMhelix 20 39\noutside 40 325\nTMhelix 326 348\ninside 349 372

23280 GCF_000414865.2_ASM41486v2 Salmonella enterica subsp. enterica serovar Enteritidis str. 2010K-0267 Proteobacteria; Gammaproteobacteria; Enterobacterales; Enterobacteriaceae; Salmonella; Salmonella enterica; Salmonella enterica subsp. enterica; Salmonella enterica subsp. enterica serovar Enteritidis MTGDNTLITSHGINRRDFMKLCAALAATMGLSSKAAAEMAESVSNPQRPPVIWIGAQECT WP_000145429.1 hydrogenase 2 small subunit [Salmonella enterica] Length: 372\nNumber of predicted TMHs: 1\nExp number of AAs in TMHs: 24.51309\nExp number, first 60 AAs: 0.2682\nTotal prob of N-in: 0.14402\noutside 1 331\nTMhelix 332 354\ninside 355 372

23281 GCF_000414885.2_ASM41488v2 Salmonella enterica subsp. enterica serovar Enteritidis str. 2010K-0271 Proteobacteria; Gammaproteobacteria; Enterobacterales; Enterobacteriaceae; Salmonella; Salmonella enterica; Salmonella enterica subsp. enterica; Salmonella enterica subsp. enterica serovar Enteritidis MTGDNTLITSHGINRRDFMKLCAALAATMGLSSKAAAEMAESVSNPQRPPVIWIGAQECT WP_000145429.1 hydrogenase 2 small subunit [Salmonella enterica] Length: 372\nNumber of predicted TMHs: 1\nExp number of AAs in TMHs: 24.51309\nExp number, first 60 AAs: 0.2682\nTotal prob of N-in: 0.14402\noutside 1 331\nTMhelix 332 354\ninside 355 372

23282 GCF_000414885.2_ASM41488v2 Salmonella enterica subsp. enterica serovar Enteritidis str. 2010K-0271 Proteobacteria; Gammaproteobacteria; Enterobacterales; Enterobacteriaceae; Salmonella; Salmonella enterica; Salmonella enterica subsp. enterica; Salmonella enterica subsp. enterica serovar Enteritidis MNNEETFYQAMRRKGVTRRSFLKFCSLAATSLGLGAGMTPKIAWALENKPRIPVVWIHGL WP_001058307.1 hydrogenase [Salmonella enterica] Length: 372\nNumber of predicted TMHs: 2\nExp number of AAs in TMHs: 36.51738\nExp number, first 60 AAs: 19.72849\nTotal prob of N-in: 0.96170\nPOSSIBLE N-term signal sequence\ninside 1 19\nTMhelix 20 39\noutside 40 325\nTMhelix 326 348\ninside 349 372

23283 GCF_000414905.2_ASM41490v2 Salmonella enterica subsp. enterica serovar Enteritidis str. 2010K-0284 Proteobacteria; Gammaproteobacteria; Enterobacterales; Enterobacteriaceae; Salmonella; Salmonella enterica; Salmonella enterica subsp. enterica; Salmonella enterica subsp. enterica serovar Enteritidis MNNEETFYQAMRRKGVTRRSFLKFCSLAATSLGLGAGMTPKIAWALENKPRIPVVWIHGL WP_001058307.1 hydrogenase [Salmonella enterica] Length: 372\nNumber of predicted TMHs: 2\nExp number of AAs in TMHs: 36.51738\nExp number, first 60 AAs: 19.72849\nTotal prob of N-in: 0.96170\nPOSSIBLE N-term signal sequence\ninside 1 19\nTMhelix 20 39\noutside 40 325\nTMhelix 326 348\ninside 349 372

23284 GCF_000414905.2_ASM41490v2 Salmonella enterica subsp. enterica serovar Enteritidis str. 2010K-0284 Proteobacteria; Gammaproteobacteria; Enterobacterales; Enterobacteriaceae; Salmonella; Salmonella enterica; Salmonella enterica subsp. enterica; Salmonella enterica subsp. enterica serovar Enteritidis MTGDNTLITSHGINRRDFMKLCAALAATMGLSSKAAAEMAESVSNPQRPPVIWIGAQECT WP_000145429.1 hydrogenase 2 small subunit [Salmonella enterica] Length: 372\nNumber of predicted TMHs: 1\nExp number of AAs in TMHs: 24.51309\nExp number, first 60 AAs: 0.2682\nTotal prob of N-in: 0.14402\noutside 1 331\nTMhelix 332 354\ninside 355 372

23285 GCF_000414925.2_ASM41492v2 Salmonella enterica subsp. enterica serovar Enteritidis str. 2010K-0286 Proteobacteria; Gammaproteobacteria; Enterobacterales; Enterobacteriaceae; Salmonella; Salmonella enterica;

Salmonella enterica subsp. enterica; Salmonella enterica subsp. enterica serovar Enteritidis
MTGDNTLITSHGINRRDFMKLCAALAATMGLSSKAAAEMAESVSNPQRPPVIWIGAQECT WP_000145429.1
hydrogenase 2 small subunit [Salmonella enterica] Length: 372\nNumber of predicted TMHs: 1\nExp number of AAs in TMHs: 24.51309\nExp number, first 60 AAs: 0.2682\nTotal prob of N-in: 0.14402\noutside 1 331\nTMhelix 332 354\ninside 355 372

23286 GCF_000278485.1_CDC-2010K_0669v1.0 Salmonella enterica subsp. enterica serovar Enteritidis str. 2010K-0669 Proteobacteria; Gammaproteobacteria; Enterobacterales; Enterobacteriaceae; Salmonella; Salmonella enterica; Salmonella enterica subsp. enterica; Salmonella enterica subsp. enterica serovar Enteritidis
MTGDNTLITSHGINRRDFMKLCAALAATMGLSSKAAAEMAESVSNPQRPPVIWIGAQECT WP_000145429.1
hydrogenase 2 small subunit [Salmonella enterica] Length: 372\nNumber of predicted TMHs: 1\nExp number of AAs in TMHs: 24.51309\nExp number, first 60 AAs: 0.2682\nTotal prob of N-in: 0.14402\noutside 1 331\nTMhelix 332 354\ninside 355 372

23287 GCF_000278485.1_CDC-2010K_0669v1.0 Salmonella enterica subsp. enterica serovar Enteritidis str. 2010K-0669 Proteobacteria; Gammaproteobacteria; Enterobacterales; Enterobacteriaceae; Salmonella; Salmonella enterica; Salmonella enterica subsp. enterica; Salmonella enterica subsp. enterica serovar Enteritidis
MNNEETFYQAMRRKGVTRRSFLKFCSLAATSLGLGAGMTPKIAWALENKPRIPVVWIHGL WP_001058307.1
hydrogenase [Salmonella enterica] Length: 372\nNumber of predicted TMHs: 2\nExp number of AAs in TMHs: 36.51738\nExp number, first 60 AAs: 19.72849\nTotal prob of N-in: 0.96170\nPOSSIBLE N-term signal sequence\ninside 1 19\nTMhelix 20 39\noutside 40 325\nTMhelix 326 348\ninside 349 372

23288 GCF_000278465.1_CDC_2010K-2599v1.0 Salmonella enterica subsp. enterica serovar Enteritidis str. 2010K-2599 Proteobacteria; Gammaproteobacteria; Enterobacterales; Enterobacteriaceae; Salmonella; Salmonella enterica; Salmonella enterica subsp. enterica; Salmonella enterica subsp. enterica serovar Enteritidis
MTGDNTLITSHGINRRDFMKLCAALAATMGLSSKAAAEMAESVSNPQRPPVIWIGAQECT WP_000145429.1
hydrogenase 2 small subunit [Salmonella enterica] Length: 372\nNumber of predicted TMHs: 1\nExp number of AAs in TMHs: 24.51309\nExp number, first 60 AAs: 0.2682\nTotal prob of N-in: 0.14402\noutside 1 331\nTMhelix 332 354\ninside 355 372

23289 GCF_000278465.1_CDC_2010K-2599v1.0 Salmonella enterica subsp. enterica serovar Enteritidis str. 2010K-2599 Proteobacteria; Gammaproteobacteria; Enterobacterales; Enterobacteriaceae; Salmonella; Salmonella enterica; Salmonella enterica subsp. enterica; Salmonella enterica subsp. enterica serovar Enteritidis
MNNEETFYQAMRRKGVTRRSFLKFCSLAATSLGLGAGMTPKIAWALENKPRIPVVWIHGL WP_001058307.1
hydrogenase [Salmonella enterica] Length: 372\nNumber of predicted TMHs: 2\nExp number of AAs in TMHs: 36.51738\nExp number, first 60 AAs: 19.72849\nTotal prob of N-in: 0.96170\nPOSSIBLE N-term signal sequence\ninside 1 19\nTMhelix 20 39\noutside 40 325\nTMhelix 326 348\ninside 349 372

23290 GCF_000280375.1_ASM28037v1 Salmonella enterica subsp. enterica serovar Enteritidis str. 22510-1 Proteobacteria; Gammaproteobacteria; Enterobacterales; Enterobacteriaceae; Salmonella; Salmonella enterica; Salmonella enterica subsp. enterica; Salmonella enterica subsp. enterica serovar Enteritidis
MNNEETFYQAMRRKGVTRRSFLKFCSLAATSLGLGAGMTPKIAWALENKPRIPVVWIHGL WP_001058307.1
hydrogenase [Salmonella enterica] Length: 372\nNumber of predicted TMHs: 2\nExp number of AAs in TMHs: 36.51738\nExp number, first 60 AAs: 19.72849\nTotal prob of N-in: 0.96170\nPOSSIBLE N-term signal sequence\ninside 1 19\nTMhelix 20 39\noutside 40 325\nTMhelix 326 348\ninside 349 372

23291 GCF_000280375.1_ASM28037v1 Salmonella enterica subsp. enterica serovar Enteritidis str. 22510-1 Proteobacteria; Gammaproteobacteria; Enterobacterales; Enterobacteriaceae; Salmonella; Salmonella enterica; Salmonella enterica subsp. enterica; Salmonella enterica subsp. enterica serovar Enteritidis
MTGDNTLITSHGINRRDFMKLCAALAATMGLSSKAAAEMAESVSNPQRPPVIWIGAQECT WP_000145429.1
hydrogenase 2 small subunit [Salmonella enterica] Length: 372\nNumber of predicted TMHs: 1\nExp number of AAs in TMHs: 24.51309\nExp number, first 60 AAs: 0.2682\nTotal prob of N-in: 0.14402\noutside 1 331\nTMhelix 332 354\ninside 355 372

23292 GCF_000328985.1_ASM32898v1 Salmonella enterica subsp. enterica serovar Enteritidis str. 22704 Proteobacteria; Gammaproteobacteria; Enterobacterales; Enterobacteriaceae; Salmonella; Salmonella enterica; Salmonella enterica subsp. enterica; Salmonella enterica subsp. enterica serovar Enteritidis
MNNEETFYQAMRRKGVTRRSFLKFCSLAATSLGLGAGMTPKIAWALENKPRIPVVWIHGL WP_001058307.1
hydrogenase [Salmonella enterica] Length: 372\nNumber of predicted TMHs: 2\nExp number of AAs in TMHs: 36.51738\nExp number, first 60 AAs: 19.72849\nTotal prob of N-in: 0.96170\nPOSSIBLE N-term signal sequence\ninside 1 19\nTMhelix 20 39\noutside 40 325\nTMhelix 326 348\ninside 349 372

23293 GCF_000328985.1_ASM32898v1 Salmonella enterica subsp. enterica serovar Enteritidis str. 22704 Proteobacteria; Gammaproteobacteria; Enterobacterales; Enterobacteriaceae; Salmonella; Salmonella enterica; Salmonella enterica subsp. enterica; Salmonella enterica subsp. enterica serovar Enteritidis MTGDNTLITSHGINRRDFMKLCAALAATMGLSSKAAAEMAESVSNPQRPPVIWIGAQECT WP_000145429.1 hydrogenase 2 small subunit [Salmonella enterica] Length: 372\nNumber of predicted TMHs: 1\nExp number of AAs in TMHs: 24.51309\nExp number, first 60 AAs: 0.2682\nTotal prob of N-in: 0.14402\noutside 1 331\nTMhelix 332 354\ninside 355 372

23294 GCF_000330425.1_ASM33042v1 Salmonella enterica subsp. enterica serovar Enteritidis str. 33944 Proteobacteria; Gammaproteobacteria; Enterobacterales; Enterobacteriaceae; Salmonella; Salmonella enterica; Salmonella enterica subsp. enterica; Salmonella enterica subsp. enterica serovar Enteritidis MTGDNTLITSHGINRRDFMKLCAALAATMGLSSKAAAEMAESVSNPQRPPVIWIGAQECT WP_000145429.1 hydrogenase 2 small subunit [Salmonella enterica] Length: 372\nNumber of predicted TMHs: 1\nExp number of AAs in TMHs: 24.51309\nExp number, first 60 AAs: 0.2682\nTotal prob of N-in: 0.14402\noutside 1 331\nTMhelix 332 354\ninside 355 372

23295 GCF_000330425.1_ASM33042v1 Salmonella enterica subsp. enterica serovar Enteritidis str. 33944 Proteobacteria; Gammaproteobacteria; Enterobacterales; Enterobacteriaceae; Salmonella; Salmonella enterica; Salmonella enterica subsp. enterica; Salmonella enterica subsp. enterica serovar Enteritidis MNNEETFYQAMRRKGVTRRSFLKFCSLAATSLGLGAGMTPKIAWALENKPRIPVVWIHGL WP_001058307.1 hydrogenase [Salmonella enterica] Length: 372\nNumber of predicted TMHs: 2\nExp number of AAs in TMHs: 36.51738\nExp number, first 60 AAs: 19.72849\nTotal prob of N-in: 0.96170\nPOSSIBLE N-term signal sequence\ninside 1 19\nTMhelix 20 39\noutside 40 325\nTMhelix 326 348\ninside 349 372

23296 GCF_000505105.1_CFSAN001333_01.0Salmonella enterica subsp. enterica serovar Enteritidis str. 3402 Proteobacteria; Gammaproteobacteria; Enterobacterales; Enterobacteriaceae; Salmonella; Salmonella enterica; Salmonella enterica subsp. enterica; Salmonella enterica subsp. enterica serovar Enteritidis MTGDNTLITSHGINRRDFMKLCAALAATMGLSSKAAAEMAESVSNPQRPPVIWIGAQECT WP_000145429.1 hydrogenase 2 small subunit [Salmonella enterica] Length: 372\nNumber of predicted TMHs: 1\nExp number of AAs in TMHs: 24.51309\nExp number, first 60 AAs: 0.2682\nTotal prob of N-in: 0.14402\noutside 1 331\nTMhelix 332 354\ninside 355 372

23297 GCF_000505105.1_CFSAN001333_01.0Salmonella enterica subsp. enterica serovar Enteritidis str. 3402 Proteobacteria; Gammaproteobacteria; Enterobacterales; Enterobacteriaceae; Salmonella; Salmonella enterica; Salmonella enterica subsp. enterica; Salmonella enterica subsp. enterica serovar Enteritidis MNNEETFYQAMRRKGVTRRSFLKFCSLAATSLGLGAGMTPKIAWALENKPRIPVVWIHGL WP_001058312.1 hydrogenase [Salmonella enterica] Length: 372\nNumber of predicted TMHs: 2\nExp number of AAs in TMHs: 36.51738\nExp number, first 60 AAs: 19.72849\nTotal prob of N-in: 0.96170\nPOSSIBLE N-term signal sequence\ninside 1 19\nTMhelix 20 39\noutside 40 325\nTMhelix 326 348\ninside 349 372

23298 GCF_000280175.1_ASM28017v1 Salmonella enterica subsp. enterica serovar Enteritidis str. 485549-17 Proteobacteria; Gammaproteobacteria; Enterobacterales; Enterobacteriaceae; Salmonella; Salmonella enterica; Salmonella enterica subsp. enterica; Salmonella enterica subsp. enterica serovar Enteritidis MNNEETFYQAMRRKGVTRRSFLKFCSLAATSLGLGAGMTPKIAWALENKPRIPVVWIHGL WP_001058307.1 hydrogenase [Salmonella enterica] Length: 372\nNumber of predicted TMHs: 2\nExp number of AAs in TMHs: 36.51738\nExp number, first 60 AAs: 19.72849\nTotal prob of N-in: 0.96170\nPOSSIBLE N-term signal sequence\ninside 1 19\nTMhelix 20 39\noutside 40 325\nTMhelix 326 348\ninside 349 372

23299 GCF_000280175.1_ASM28017v1 Salmonella enterica subsp. enterica serovar Enteritidis str. 485549-17 Proteobacteria; Gammaproteobacteria; Enterobacterales; Enterobacteriaceae; Salmonella; Salmonella enterica; Salmonella enterica subsp. enterica; Salmonella enterica subsp. enterica serovar Enteritidis MTGDNTLITSHGINRRDFMKLCAALAATMGLSSKAAAEMAESVSNPQRPPVIWIGAQECT WP_000145429.1 hydrogenase 2 small subunit [Salmonella enterica] Length: 372\nNumber of predicted TMHs: 1\nExp number of AAs in TMHs: 24.51309\nExp number, first 60 AAs: 0.2682\nTotal prob of N-in: 0.14402\noutside 1 331\nTMhelix 332 354\ninside 355 372

23300 GCF_000280455.1_ASM28045v1 Salmonella enterica subsp. enterica serovar Enteritidis str. 50-3079 Proteobacteria; Gammaproteobacteria; Enterobacterales; Enterobacteriaceae; Salmonella; Salmonella enterica; Salmonella enterica subsp. enterica; Salmonella enterica subsp. enterica serovar Enteritidis MNNEETFYQAMRRKGVTRRSFLKFCSLAATSLGLGAGMTPKIAWALENKPRIPVVWIHGL WP_001058307.1 hydrogenase [Salmonella enterica] Length: 372\nNumber of predicted TMHs: 2\nExp number of AAs in

TMHs: 36.51738\nExp number, first 60 AAs: 19.72849\nTotal prob of N-in: 0.96170\nPOSSIBLE N-term signal sequence\ninside 1 19\nTMhelix 20 39\noutside 40 325\nTMhelix 326 348\ninside 349 372

23301 GCF_000280455.1_ASM28045v1 Salmonella enterica subsp. enterica serovar Enteritidis str. 50-3079 Proteobacteria; Gammaproteobacteria; Enterobacterales; Enterobacteriaceae; Salmonella; Salmonella enterica; Salmonella enterica subsp. enterica; Salmonella enterica subsp. enterica serovar Enteritidis MTGDNTLITSHGINRRDFMKLCAALAATMGLSSKAAAEMAESVSNPQRPPVIWIGAQECT WP_000145429.1 hydrogenase 2 small subunit [Salmonella enterica] Length: 372\nNumber of predicted TMHs: 1\nExp number of AAs in TMHs: 24.51309\nExp number, first 60 AAs: 0.2682\nTotal prob of N-in: 0.14402\noutside 1 331\nTMhelix 332 354\ninside 355 372

23302 GCF_000330585.1_ASM33058v1 Salmonella enterica subsp. enterica serovar Enteritidis str. 53-407 Proteobacteria; Gammaproteobacteria; Enterobacterales; Enterobacteriaceae; Salmonella; Salmonella enterica; Salmonella enterica subsp. enterica; Salmonella enterica subsp. enterica serovar Enteritidis MNNEETFYQAMRRKGVTRRSFLKFCSLAATSLGLGAGMTPKIAWALENKPRIPVVWIHGL WP_001058307.1 hydrogenase [Salmonella enterica] Length: 372\nNumber of predicted TMHs: 2\nExp number of AAs in TMHs: 36.51738\nExp number, first 60 AAs: 19.72849\nTotal prob of N-in: 0.96170\nPOSSIBLE N-term signal sequence\ninside 1 19\nTMhelix 20 39\noutside 40 325\nTMhelix 326 348\ninside 349 372

23303 GCF_000330585.1_ASM33058v1 Salmonella enterica subsp. enterica serovar Enteritidis str. 53-407 Proteobacteria; Gammaproteobacteria; Enterobacterales; Enterobacteriaceae; Salmonella; Salmonella enterica; Salmonella enterica subsp. enterica; Salmonella enterica subsp. enterica serovar Enteritidis MTGDNTLITSHGINRRDFMKLCAALAATMGLSSKAAAEMAESVSNPQRPPVIWIGAQECT WP_000145429.1 hydrogenase 2 small subunit [Salmonella enterica] Length: 372\nNumber of predicted TMHs: 1\nExp number of AAs in TMHs: 24.51309\nExp number, first 60 AAs: 0.2682\nTotal prob of N-in: 0.14402\noutside 1 331\nTMhelix 332 354\ninside 355 372

23304 GCF_000330045.1_ASM33004v1 Salmonella enterica subsp. enterica serovar Enteritidis str. 543463 22-17 Proteobacteria; Gammaproteobacteria; Enterobacterales; Enterobacteriaceae; Salmonella; Salmonella enterica; Salmonella enterica subsp. enterica; Salmonella enterica subsp. enterica serovar Enteritidis MTGDNTLITSHGINRRDFMKLCAALAATMGLSSKAAAEMAESVSNPQRPPVIWIGAQECT WP_000145429.1 hydrogenase 2 small subunit [Salmonella enterica] Length: 372\nNumber of predicted TMHs: 1\nExp number of AAs in TMHs: 24.51309\nExp number, first 60 AAs: 0.2682\nTotal prob of N-in: 0.14402\noutside 1 331\nTMhelix 332 354\ninside 355 372

23305 GCF_000330045.1_ASM33004v1 Salmonella enterica subsp. enterica serovar Enteritidis str. 543463 22-17 Proteobacteria; Gammaproteobacteria; Enterobacterales; Enterobacteriaceae; Salmonella; Salmonella enterica; Salmonella enterica subsp. enterica; Salmonella enterica subsp. enterica serovar Enteritidis MNNEETFYQAMRRKGVTRRSFLKFCSLAATSLGLGAGMTPKIAWALENKPRIPVVWIHGL WP_001058307.1 hydrogenase [Salmonella enterica] Length: 372\nNumber of predicted TMHs: 2\nExp number of AAs in TMHs: 36.51738\nExp number, first 60 AAs: 19.72849\nTotal prob of N-in: 0.96170\nPOSSIBLE N-term signal sequence\ninside 1 19\nTMhelix 20 39\noutside 40 325\nTMhelix 326 348\ninside 349 372

23306 GCF_000330065.1_ASM33006v1 Salmonella enterica subsp. enterica serovar Enteritidis str. 543463 40-18 Proteobacteria; Gammaproteobacteria; Enterobacterales; Enterobacteriaceae; Salmonella; Salmonella enterica; Salmonella enterica subsp. enterica; Salmonella enterica subsp. enterica serovar Enteritidis MNNEETFYQAMRRKGVTRRSFLKFCSLAATSLGLGAGMTPKIAWALENKPRIPVVWIHGL WP_001058307.1 hydrogenase [Salmonella enterica] Length: 372\nNumber of predicted TMHs: 2\nExp number of AAs in TMHs: 36.51738\nExp number, first 60 AAs: 19.72849\nTotal prob of N-in: 0.96170\nPOSSIBLE N-term signal sequence\ninside 1 19\nTMhelix 20 39\noutside 40 325\nTMhelix 326 348\ninside 349 372

23307 GCF_000330065.1_ASM33006v1 Salmonella enterica subsp. enterica serovar Enteritidis str. 543463 40-18 Proteobacteria; Gammaproteobacteria; Enterobacterales; Enterobacteriaceae; Salmonella; Salmonella enterica; Salmonella enterica subsp. enterica; Salmonella enterica subsp. enterica serovar Enteritidis MTGDNTLITSHGINRRDFMKLCAALAATMGLSSKAAAEMAESVSNPQRPPVIWIGAQECT WP_000145429.1 hydrogenase 2 small subunit [Salmonella enterica] Length: 372\nNumber of predicted TMHs: 1\nExp number of AAs in TMHs: 24.51309\nExp number, first 60 AAs: 0.2682\nTotal prob of N-in: 0.14402\noutside 1 331\nTMhelix 332 354\ninside 355 372

23308 GCF_000330365.1_ASM33036v1 Salmonella enterica subsp. enterica serovar Enteritidis str. 543463 42-20 Proteobacteria; Gammaproteobacteria; Enterobacterales; Enterobacteriaceae; Salmonella; Salmonella enterica;

Salmonella enterica subsp. enterica; Salmonella enterica subsp. enterica serovar Enteritidis
MNNEETFYQAMRRKGVTRRSFLKFCSLAATSLGLGAGMTPKIAWALENKPRIPVVWIHGL WP_001058307.1
hydrogenase [Salmonella enterica] Length: 372\nNumber of predicted TMHs: 2\nExp number of AAs in TMHs: 36.51738\nExp number, first 60 AAs: 19.72849\nTotal prob of N-in: 0.96170\nPOSSIBLE N-term signal sequence\ninside 1 19\nTMhelix 20 39\noutside 40 325\nTMhelix 326 348\ninside 349 372

23309 GCF_000330365.1_ASM33036v1 Salmonella enterica subsp. enterica serovar Enteritidis str. 543463 42-20
Proteobacteria; Gammaproteobacteria; Enterobacterales; Enterobacteriaceae; Salmonella; Salmonella enterica; Salmonella enterica subsp. enterica; Salmonella enterica subsp. enterica serovar Enteritidis
MTGDNTLITSHGINRRDFMKLCAALAATMGLSSKAAAEMAESVSNPQRPPVIWIGAQECT WP_000145429.1
hydrogenase 2 small subunit [Salmonella enterica] Length: 372\nNumber of predicted TMHs: 1\nExp number of AAs in TMHs: 24.51309\nExp number, first 60 AAs: 0.2682\nTotal prob of N-in: 0.14402\noutside 1 331\nTMhelix 332 354\ninside 355 372

23310 GCF_000330085.1_ASM33008v1 Salmonella enterica subsp. enterica serovar Enteritidis str. 561362 1-1
Proteobacteria; Gammaproteobacteria; Enterobacterales; Enterobacteriaceae; Salmonella; Salmonella enterica; Salmonella enterica subsp. enterica; Salmonella enterica subsp. enterica serovar Enteritidis
MTGDNTLITSHGINRRDFMKLCAALAATMGLSSKAAAEMAESVSNPQRPPVIWIGAQECT WP_000145429.1
hydrogenase 2 small subunit [Salmonella enterica] Length: 372\nNumber of predicted TMHs: 1\nExp number of AAs in TMHs: 24.51309\nExp number, first 60 AAs: 0.2682\nTotal prob of N-in: 0.14402\noutside 1 331\nTMhelix 332 354\ninside 355 372

23311 GCF_000330085.1_ASM33008v1 Salmonella enterica subsp. enterica serovar Enteritidis str. 561362 1-1
Proteobacteria; Gammaproteobacteria; Enterobacterales; Enterobacteriaceae; Salmonella; Salmonella enterica; Salmonella enterica subsp. enterica; Salmonella enterica subsp. enterica serovar Enteritidis
MNNEETFYQAMRRKGVTRRSFLKFCSLAATSLGLGAGMTPKIAWALENKPRIPVVWIHGL WP_001058307.1
hydrogenase [Salmonella enterica] Length: 372\nNumber of predicted TMHs: 2\nExp number of AAs in TMHs: 36.51738\nExp number, first 60 AAs: 19.72849\nTotal prob of N-in: 0.96170\nPOSSIBLE N-term signal sequence\ninside 1 19\nTMhelix 20 39\noutside 40 325\nTMhelix 326 348\ninside 349 372

23312 GCF_000330345.1_ASM33034v1 Salmonella enterica subsp. enterica serovar Enteritidis str. 561362 9-7
Proteobacteria; Gammaproteobacteria; Enterobacterales; Enterobacteriaceae; Salmonella; Salmonella enterica; Salmonella enterica subsp. enterica; Salmonella enterica subsp. enterica serovar Enteritidis
MTGDNTLITSHGINRRDFMKLCAALAATMGLSSKAAAEMAESVSNPQRPPVIWIGAQECT WP_000145429.1
hydrogenase 2 small subunit [Salmonella enterica] Length: 372\nNumber of predicted TMHs: 1\nExp number of AAs in TMHs: 24.51309\nExp number, first 60 AAs: 0.2682\nTotal prob of N-in: 0.14402\noutside 1 331\nTMhelix 332 354\ninside 355 372

23313 GCF_000330345.1_ASM33034v1 Salmonella enterica subsp. enterica serovar Enteritidis str. 561362 9-7
Proteobacteria; Gammaproteobacteria; Enterobacterales; Enterobacteriaceae; Salmonella; Salmonella enterica; Salmonella enterica subsp. enterica; Salmonella enterica subsp. enterica serovar Enteritidis
MNNEETFYQAMRRKGVTRRSFLKFCSLAATSLGLGAGMTPKIAWALENKPRIPVVWIHGL WP_001058307.1
hydrogenase [Salmonella enterica] Length: 372\nNumber of predicted TMHs: 2\nExp number of AAs in TMHs: 36.51738\nExp number, first 60 AAs: 19.72849\nTotal prob of N-in: 0.96170\nPOSSIBLE N-term signal sequence\ninside 1 19\nTMhelix 20 39\noutside 40 325\nTMhelix 326 348\ninside 349 372

23314 GCF_000329645.1_ASM32964v1 Salmonella enterica subsp. enterica serovar Enteritidis str. 576709
Proteobacteria; Gammaproteobacteria; Enterobacterales; Enterobacteriaceae; Salmonella; Salmonella enterica; Salmonella enterica subsp. enterica; Salmonella enterica subsp. enterica serovar Enteritidis
MNNEETFYQAMRRKGVTRRSFLKFCSLAATSLGLGAGMTPKIAWALENKPRIPVVWIHGL WP_001058307.1
hydrogenase [Salmonella enterica] Length: 372\nNumber of predicted TMHs: 2\nExp number of AAs in TMHs: 36.51738\nExp number, first 60 AAs: 19.72849\nTotal prob of N-in: 0.96170\nPOSSIBLE N-term signal sequence\ninside 1 19\nTMhelix 20 39\noutside 40 325\nTMhelix 326 348\ninside 349 372

23315 GCF_000329645.1_ASM32964v1 Salmonella enterica subsp. enterica serovar Enteritidis str. 576709
Proteobacteria; Gammaproteobacteria; Enterobacterales; Enterobacteriaceae; Salmonella; Salmonella enterica; Salmonella enterica subsp. enterica; Salmonella enterica subsp. enterica serovar Enteritidis
MTGDNTLITSHGINRRDFMKLCAALAATMGLSSKAAAEMAESVSNPQRPPVIWIGAQECT WP_000145429.1
hydrogenase 2 small subunit [Salmonella enterica] Length: 372\nNumber of predicted TMHs: 1\nExp number of AAs in TMHs: 24.51309\nExp number, first 60 AAs: 0.2682\nTotal prob of N-in: 0.14402\noutside 1 331\nTMhelix 332 354\ninside 355 372

23316 GCF_000280475.1_ASM28047v1 Salmonella enterica subsp. enterica serovar Enteritidis str. 58-6482 Proteobacteria; Gammaproteobacteria; Enterobacterales; Enterobacteriaceae; Salmonella; Salmonella enterica; Salmonella enterica subsp. enterica; Salmonella enterica subsp. enterica serovar Enteritidis MNNEETFYQAMRRKGVTRRSFLKFCSLAATSLGLGAGMTPKIAWALENKPRIPVVWIHGL WP_001058307.1 hydrogenase [Salmonella enterica] Length: 372\nNumber of predicted TMHs: 2\nExp number of AAs in TMHs: 36.51738\nExp number, first 60 AAs: 19.72849\nTotal prob of N-in: 0.96170\nPOSSIBLE N-term signal sequence\ninside 1 19\nTMhelix 20 39\noutside 40 325\nTMhelix 326 348\ninside 349 372

23317 GCF_000280475.1_ASM28047v1 Salmonella enterica subsp. enterica serovar Enteritidis str. 58-6482 Proteobacteria; Gammaproteobacteria; Enterobacterales; Enterobacteriaceae; Salmonella; Salmonella enterica; Salmonella enterica subsp. enterica; Salmonella enterica subsp. enterica serovar Enteritidis MTGDNTLITSHGINRRDFMKLCAALAATMGLSSKAAAEMAESVSNPQRPPVIWIGAQECT WP_000145429.1 hydrogenase 2 small subunit [Salmonella enterica] Length: 372\nNumber of predicted TMHs: 1\nExp number of AAs in TMHs: 24.51309\nExp number, first 60 AAs: 0.2682\nTotal prob of N-in: 0.14402\noutside 1 331\nTMhelix 332 354\ninside 355 372

23318 GCF_000280195.1_ASM28019v1 Salmonella enterica subsp. enterica serovar Enteritidis str. 596866-22 Proteobacteria; Gammaproteobacteria; Enterobacterales; Enterobacteriaceae; Salmonella; Salmonella enterica; Salmonella enterica subsp. enterica; Salmonella enterica subsp. enterica serovar Enteritidis MNNEETFYQAMRRKGVTRRSFLKFCSLAATSLGLGAGMTPKIAWALENKPRIPVVWIHGL WP_001058307.1 hydrogenase [Salmonella enterica] Length: 372\nNumber of predicted TMHs: 2\nExp number of AAs in TMHs: 36.51738\nExp number, first 60 AAs: 19.72849\nTotal prob of N-in: 0.96170\nPOSSIBLE N-term signal sequence\ninside 1 19\nTMhelix 20 39\noutside 40 325\nTMhelix 326 348\ninside 349 372

23319 GCF_000280195.1_ASM28019v1 Salmonella enterica subsp. enterica serovar Enteritidis str. 596866-22 Proteobacteria; Gammaproteobacteria; Enterobacterales; Enterobacteriaceae; Salmonella; Salmonella enterica; Salmonella enterica subsp. enterica; Salmonella enterica subsp. enterica serovar Enteritidis MTGDNTLITSHGINRRDFMKLCAALAATMGLSSKAAAEMAESVSNPQRPPVIWIGAQECT WP_000145429.1 hydrogenase 2 small subunit [Salmonella enterica] Length: 372\nNumber of predicted TMHs: 1\nExp number of AAs in TMHs: 24.51309\nExp number, first 60 AAs: 0.2682\nTotal prob of N-in: 0.14402\noutside 1 331\nTMhelix 332 354\ninside 355 372

23320 GCF_000280215.1_ASM28021v1 Salmonella enterica subsp. enterica serovar Enteritidis str. 596866-70 Proteobacteria; Gammaproteobacteria; Enterobacterales; Enterobacteriaceae; Salmonella; Salmonella enterica; Salmonella enterica subsp. enterica; Salmonella enterica subsp. enterica serovar Enteritidis MTGDNTLITSHGINRRDFMKLCAALAATMGLSSKAAAEMAESVSNPQRPPVIWIGAQECT WP_000145429.1 hydrogenase 2 small subunit [Salmonella enterica] Length: 372\nNumber of predicted TMHs: 1\nExp number of AAs in TMHs: 24.51309\nExp number, first 60 AAs: 0.2682\nTotal prob of N-in: 0.14402\noutside 1 331\nTMhelix 332 354\ninside 355 372

23321 GCF_000280215.1_ASM28021v1 Salmonella enterica subsp. enterica serovar Enteritidis str. 596866-70 Proteobacteria; Gammaproteobacteria; Enterobacterales; Enterobacteriaceae; Salmonella; Salmonella enterica; Salmonella enterica subsp. enterica; Salmonella enterica subsp. enterica serovar Enteritidis MNNEETFYQAMRRKGVTRRSFLKFCSLAATSLGLGAGMTPKIAWALENKPRIPVVWIHGL WP_001058307.1 hydrogenase [Salmonella enterica] Length: 372\nNumber of predicted TMHs: 2\nExp number of AAs in TMHs: 36.51738\nExp number, first 60 AAs: 19.72849\nTotal prob of N-in: 0.96170\nPOSSIBLE N-term signal sequence\ninside 1 19\nTMhelix 20 39\noutside 40 325\nTMhelix 326 348\ninside 349 372

23322 GCF_000329725.1_ASM32972v1 Salmonella enterica subsp. enterica serovar Enteritidis str. 607307-2 Proteobacteria; Gammaproteobacteria; Enterobacterales; Enterobacteriaceae; Salmonella; Salmonella enterica; Salmonella enterica subsp. enterica; Salmonella enterica subsp. enterica serovar Enteritidis MNNEETFYQAMRRKGVTRRSFLKFCSLAATSLGLGAGMTPKIAWALENKPRIPVVWIHGL WP_001058307.1 hydrogenase [Salmonella enterica] Length: 372\nNumber of predicted TMHs: 2\nExp number of AAs in TMHs: 36.51738\nExp number, first 60 AAs: 19.72849\nTotal prob of N-in: 0.96170\nPOSSIBLE N-term signal sequence\ninside 1 19\nTMhelix 20 39\noutside 40 325\nTMhelix 326 348\ninside 349 372

23323 GCF_000329725.1_ASM32972v1 Salmonella enterica subsp. enterica serovar Enteritidis str. 607307-2 Proteobacteria; Gammaproteobacteria; Enterobacterales; Enterobacteriaceae; Salmonella; Salmonella enterica; Salmonella enterica subsp. enterica; Salmonella enterica subsp. enterica serovar Enteritidis MTGDNTLITSHGINRRDFMKLCAALAATMGLSSKAAAEMAESVSNPQRPPVIWIGAQECT WP_000145429.1 hydrogenase 2 small subunit [Salmonella enterica] Length: 372\nNumber of predicted TMHs: 1\nExp

number of AAs in TMHs: 24.51309\nExp number, first 60 AAs: 0.2682\nTotal prob of N-in: 0.14402\noutside 1 331\nTMhelix 332 354\ninside 355 372

23324 GCF_000280155.1_ASM28015v1 Salmonella enterica subsp. enterica serovar Enteritidis str. 607307-6 Proteobacteria; Gammaproteobacteria; Enterobacterales; Enterobacteriaceae; Salmonella; Salmonella enterica; Salmonella enterica subsp. enterica; Salmonella enterica subsp. enterica serovar Enteritidis MTGDNTLITSHGINRRDFMKLCAALAATMGLSSKAAAEMAESVSNPQRPPVIWIGAQECT WP_000145429.1 hydrogenase 2 small subunit [Salmonella enterica] Length: 372\nNumber of predicted TMHs: 1\nExp number of AAs in TMHs: 24.51309\nExp number, first 60 AAs: 0.2682\nTotal prob of N-in: 0.14402\noutside 1 331\nTMhelix 332 354\ninside 355 372

23325 GCF_000280155.1_ASM28015v1 Salmonella enterica subsp. enterica serovar Enteritidis str. 607307-6 Proteobacteria; Gammaproteobacteria; Enterobacterales; Enterobacteriaceae; Salmonella; Salmonella enterica; Salmonella enterica subsp. enterica; Salmonella enterica subsp. enterica serovar Enteritidis MNNEETFYQAMRRKGVTRRSFLKFCSLAATSLGLGAGMTPKIAWALENKPRIPVVWIHGL WP_001058307.1 hydrogenase [Salmonella enterica] Length: 372\nNumber of predicted TMHs: 2\nExp number of AAs in TMHs: 36.51738\nExp number, first 60 AAs: 19.72849\nTotal prob of N-in: 0.96170\nPOSSIBLE N-term signal sequence\ninside 1 19\nTMhelix 20 39\noutside 40 325\nTMhelix 326 348\ninside 349 372

23326 GCF_000329685.1_ASM32968v1 Salmonella enterica subsp. enterica serovar Enteritidis str. 607308-16 Proteobacteria; Gammaproteobacteria; Enterobacterales; Enterobacteriaceae; Salmonella; Salmonella enterica; Salmonella enterica subsp. enterica; Salmonella enterica subsp. enterica serovar Enteritidis MNNEETFYQAMRRKGVTRRSFLKFCSLAATSLGLGAGMTPKIAWALENKPRIPVVWIHGL WP_001058307.1 hydrogenase [Salmonella enterica] Length: 372\nNumber of predicted TMHs: 2\nExp number of AAs in TMHs: 36.51738\nExp number, first 60 AAs: 19.72849\nTotal prob of N-in: 0.96170\nPOSSIBLE N-term signal sequence\ninside 1 19\nTMhelix 20 39\noutside 40 325\nTMhelix 326 348\ninside 349 372

23327 GCF_000329685.1_ASM32968v1 Salmonella enterica subsp. enterica serovar Enteritidis str. 607308-16 Proteobacteria; Gammaproteobacteria; Enterobacterales; Enterobacteriaceae; Salmonella; Salmonella enterica; Salmonella enterica subsp. enterica; Salmonella enterica subsp. enterica serovar Enteritidis MTGDNTLITSHGINRRDFMKLCAALAATMGLSSKAAAEMAESVSNPQRPPVIWIGAQECT WP_000145429.1 hydrogenase 2 small subunit [Salmonella enterica] Length: 372\nNumber of predicted TMHs: 1\nExp number of AAs in TMHs: 24.51309\nExp number, first 60 AAs: 0.2682\nTotal prob of N-in: 0.14402\noutside 1 331\nTMhelix 332 354\ninside 355 372

23328 GCF_000329705.1_ASM32970v1 Salmonella enterica subsp. enterica serovar Enteritidis str. 607308-19 Proteobacteria; Gammaproteobacteria; Enterobacterales; Enterobacteriaceae; Salmonella; Salmonella enterica; Salmonella enterica subsp. enterica; Salmonella enterica subsp. enterica serovar Enteritidis MTGDNTLITSHGINRRDFMKLCAALAATMGLSSKAAAEMAESVSNPQRPPVIWIGAQECT WP_000145429.1 hydrogenase 2 small subunit [Salmonella enterica] Length: 372\nNumber of predicted TMHs: 1\nExp number of AAs in TMHs: 24.51309\nExp number, first 60 AAs: 0.2682\nTotal prob of N-in: 0.14402\noutside 1 331\nTMhelix 332 354\ninside 355 372

23329 GCF_000329705.1_ASM32970v1 Salmonella enterica subsp. enterica serovar Enteritidis str. 607308-19 Proteobacteria; Gammaproteobacteria; Enterobacterales; Enterobacteriaceae; Salmonella; Salmonella enterica; Salmonella enterica subsp. enterica; Salmonella enterica subsp. enterica serovar Enteritidis MNNEETFYQAMRRKGVTRRSFLKFCSLAATSLGLGAGMTPKIAWALENKPRIPVVWIHGL WP_001058307.1 hydrogenase [Salmonella enterica] Length: 372\nNumber of predicted TMHs: 2\nExp number of AAs in TMHs: 36.51738\nExp number, first 60 AAs: 19.72849\nTotal prob of N-in: 0.96170\nPOSSIBLE N-term signal sequence\ninside 1 19\nTMhelix 20 39\noutside 40 325\nTMhelix 326 348\ninside 349 372

23330 GCF_000329745.1_ASM32974v1 Salmonella enterica subsp. enterica serovar Enteritidis str. 607308-9 Proteobacteria; Gammaproteobacteria; Enterobacterales; Enterobacteriaceae; Salmonella; Salmonella enterica; Salmonella enterica subsp. enterica; Salmonella enterica subsp. enterica serovar Enteritidis MNNEETFYQAMRRKGVTRRSFLKFCSLAATSLGLGAGMTPKIAWALENKPRIPVVWIHGL WP_001058307.1 hydrogenase [Salmonella enterica] Length: 372\nNumber of predicted TMHs: 2\nExp number of AAs in TMHs: 36.51738\nExp number, first 60 AAs: 19.72849\nTotal prob of N-in: 0.96170\nPOSSIBLE N-term signal sequence\ninside 1 19\nTMhelix 20 39\noutside 40 325\nTMhelix 326 348\ninside 349 372

23331 GCF_000329745.1_ASM32974v1 Salmonella enterica subsp. enterica serovar Enteritidis str. 607308-9 Proteobacteria; Gammaproteobacteria; Enterobacterales; Enterobacteriaceae; Salmonella; Salmonella enterica;

Salmonella enterica subsp. enterica; Salmonella enterica subsp. enterica serovar Enteritidis
MTGDNTLITSHGINRRDFMKLCAALAATMGLSSKAAAEMAESVSNPQRPPVIWIGAQECT WP_000145429.1
hydrogenase 2 small subunit [Salmonella enterica] Length: 372\nNumber of predicted TMHs: 1\nExp number of AAs in TMHs: 24.51309\nExp number, first 60 AAs: 0.2682\nTotal prob of N-in: 0.14402\noutside 1 331\nTMhelix 332 354\ninside 355 372

23332 GCF_000330565.1_ASM33056v1 Salmonella enterica subsp. enterica serovar Enteritidis str. 62-1976
Proteobacteria; Gammaproteobacteria; Enterobacterales; Enterobacteriaceae; Salmonella; Salmonella enterica; Salmonella enterica subsp. enterica; Salmonella enterica subsp. enterica serovar Enteritidis
MTGDNTLITSHGINRRDFMKLCAALAATMGLSSKAAAEMAESVSNPQRPPVIWIGAQECT WP_000145429.1
hydrogenase 2 small subunit [Salmonella enterica] Length: 372\nNumber of predicted TMHs: 1\nExp number of AAs in TMHs: 24.51309\nExp number, first 60 AAs: 0.2682\nTotal prob of N-in: 0.14402\noutside 1 331\nTMhelix 332 354\ninside 355 372

23333 GCF_000330565.1_ASM33056v1 Salmonella enterica subsp. enterica serovar Enteritidis str. 62-1976
Proteobacteria; Gammaproteobacteria; Enterobacterales; Enterobacteriaceae; Salmonella; Salmonella enterica; Salmonella enterica subsp. enterica; Salmonella enterica subsp. enterica serovar Enteritidis
MNNEETFYQAMRRKGVTRRSFLKFCSLAATSLGLGAGMTPKIAWALENKPRIPVVWIHGL WP_001058307.1
hydrogenase [Salmonella enterica] Length: 372\nNumber of predicted TMHs: 2\nExp number of AAs in TMHs: 36.51738\nExp number, first 60 AAs: 19.72849\nTotal prob of N-in: 0.96170\nPOSSIBLE N-term signal sequence\ninside 1 19\nTMhelix 20 39\noutside 40 325\nTMhelix 326 348\ninside 349 372

23334 GCF_000280075.1_ASM28007v1 Salmonella enterica subsp. enterica serovar Enteritidis str. 622731-39
Proteobacteria; Gammaproteobacteria; Enterobacterales; Enterobacteriaceae; Salmonella; Salmonella enterica; Salmonella enterica subsp. enterica; Salmonella enterica subsp. enterica serovar Enteritidis
MNNEETFYQAMRRKGVTRRSFLKFCSLAATSLGLGAGMTPKIAWALENKPRIPVVWIHGL WP_001058307.1
hydrogenase [Salmonella enterica] Length: 372\nNumber of predicted TMHs: 2\nExp number of AAs in TMHs: 36.51738\nExp number, first 60 AAs: 19.72849\nTotal prob of N-in: 0.96170\nPOSSIBLE N-term signal sequence\ninside 1 19\nTMhelix 20 39\noutside 40 325\nTMhelix 326 348\ninside 349 372

23335 GCF_000280075.1_ASM28007v1 Salmonella enterica subsp. enterica serovar Enteritidis str. 622731-39
Proteobacteria; Gammaproteobacteria; Enterobacterales; Enterobacteriaceae; Salmonella; Salmonella enterica; Salmonella enterica subsp. enterica; Salmonella enterica subsp. enterica serovar Enteritidis
MTGDNTLITSHGINRRDFMKLCAALAATMGLSSKAAAEMAESVSNPQRPPVIWIGAQECT WP_000145429.1
hydrogenase 2 small subunit [Salmonella enterica] Length: 372\nNumber of predicted TMHs: 1\nExp number of AAs in TMHs: 24.51309\nExp number, first 60 AAs: 0.2682\nTotal prob of N-in: 0.14402\noutside 1 331\nTMhelix 332 354\ninside 355 372

23336 GCF_000329765.1_ASM32976v1 Salmonella enterica subsp. enterica serovar Enteritidis str. 629163
Proteobacteria; Gammaproteobacteria; Enterobacterales; Enterobacteriaceae; Salmonella; Salmonella enterica; Salmonella enterica subsp. enterica; Salmonella enterica subsp. enterica serovar Enteritidis
MNNEETFYQAMRRKGVTRRSFLKFCSLAATSLGLGAGMTPKIAWALENKPRIPVVWIHGL WP_001058307.1
hydrogenase [Salmonella enterica] Length: 372\nNumber of predicted TMHs: 2\nExp number of AAs in TMHs: 36.51738\nExp number, first 60 AAs: 19.72849\nTotal prob of N-in: 0.96170\nPOSSIBLE N-term signal sequence\ninside 1 19\nTMhelix 20 39\noutside 40 325\nTMhelix 326 348\ninside 349 372

23337 GCF_000329765.1_ASM32976v1 Salmonella enterica subsp. enterica serovar Enteritidis str. 629163
Proteobacteria; Gammaproteobacteria; Enterobacterales; Enterobacteriaceae; Salmonella; Salmonella enterica; Salmonella enterica subsp. enterica; Salmonella enterica subsp. enterica serovar Enteritidis
MTGDNTLITSHGINRRDFMKLCAALAATMGLSSKAAAEMAESVSNPQRPPVIWIGAQECT WP_000145429.1
hydrogenase 2 small subunit [Salmonella enterica] Length: 372\nNumber of predicted TMHs: 1\nExp number of AAs in TMHs: 24.51309\nExp number, first 60 AAs: 0.2682\nTotal prob of N-in: 0.14402\noutside 1 331\nTMhelix 332 354\ninside 355 372

23338 GCF_000280235.1_ASM28023v1 Salmonella enterica subsp. enterica serovar Enteritidis str. 629164-26
Proteobacteria; Gammaproteobacteria; Enterobacterales; Enterobacteriaceae; Salmonella; Salmonella enterica; Salmonella enterica subsp. enterica; Salmonella enterica subsp. enterica serovar Enteritidis
MNNEETFYQAMRRKGVTRRSFLKFCSLAATSLGLGAGMTPKIAWALENKPRIPVVWIHGL WP_001058307.1
hydrogenase [Salmonella enterica] Length: 372\nNumber of predicted TMHs: 2\nExp number of AAs in TMHs: 36.51738\nExp number, first 60 AAs: 19.72849\nTotal prob of N-in: 0.96170\nPOSSIBLE N-term signal sequence\ninside 1 19\nTMhelix 20 39\noutside 40 325\nTMhelix 326 348\ninside 349 372

23339 GCF_000280235.1_ASM28023v1 Salmonella enterica subsp. enterica serovar Enteritidis str. 629164-26 Proteobacteria; Gammaproteobacteria; Enterobacterales; Enterobacteriaceae; Salmonella; Salmonella enterica; Salmonella enterica subsp. enterica; Salmonella enterica subsp. enterica serovar Enteritidis MTGDNTLITSHGINRRDFMKLCAALAATMGLSSKAAAEMAESVSNPQRPPVIWIGAQECT WP_000145429.1 hydrogenase 2 small subunit [Salmonella enterica] Length: 372\nNumber of predicted TMHs: 1\nExp number of AAs in TMHs: 24.51309\nExp number, first 60 AAs: 0.2682\nTotal prob of N-in: 0.14402\noutside 1 331\nTMhelix 332 354\ninside 355 372

23340 GCF_000280255.1_ASM28025v1 Salmonella enterica subsp. enterica serovar Enteritidis str. 629164-37 Proteobacteria; Gammaproteobacteria; Enterobacterales; Enterobacteriaceae; Salmonella; Salmonella enterica; Salmonella enterica subsp. enterica; Salmonella enterica subsp. enterica serovar Enteritidis MNNEETFYQAMRRKGVTRRSFLKFCSLAATSLGLGAGMTPKIAWALENKPRIPVVWIHGL WP_001058307.1 hydrogenase [Salmonella enterica] Length: 372\nNumber of predicted TMHs: 2\nExp number of AAs in TMHs: 36.51738\nExp number, first 60 AAs: 19.72849\nTotal prob of N-in: 0.96170\nPOSSIBLE N-term signal sequence\ninside 1 19\nTMhelix 20 39\noutside 40 325\nTMhelix 326 348\ninside 349 372

23341 GCF_000280255.1_ASM28025v1 Salmonella enterica subsp. enterica serovar Enteritidis str. 629164-37 Proteobacteria; Gammaproteobacteria; Enterobacterales; Enterobacteriaceae; Salmonella; Salmonella enterica; Salmonella enterica subsp. enterica; Salmonella enterica subsp. enterica serovar Enteritidis MTGDNTLITSHGINRRDFMKLCAALAATMGLSSKAAAEMAESVSNPQRPPVIWIGAQECT WP_000145429.1 hydrogenase 2 small subunit [Salmonella enterica] Length: 372\nNumber of predicted TMHs: 1\nExp number of AAs in TMHs: 24.51309\nExp number, first 60 AAs: 0.2682\nTotal prob of N-in: 0.14402\noutside 1 331\nTMhelix 332 354\ninside 355 372

23342 GCF_000329665.1_ASM32966v1 Salmonella enterica subsp. enterica serovar Enteritidis str. 635290-58 Proteobacteria; Gammaproteobacteria; Enterobacterales; Enterobacteriaceae; Salmonella; Salmonella enterica; Salmonella enterica subsp. enterica; Salmonella enterica subsp. enterica serovar Enteritidis MTGDNTLITSHGINRRDFMKLCAALAATMGLSSKAAAEMAESVSNPQRPPVIWIGAQECT WP_000145429.1 hydrogenase 2 small subunit [Salmonella enterica] Length: 372\nNumber of predicted TMHs: 1\nExp number of AAs in TMHs: 24.51309\nExp number, first 60 AAs: 0.2682\nTotal prob of N-in: 0.14402\noutside 1 331\nTMhelix 332 354\ninside 355 372

23343 GCF_000329665.1_ASM32966v1 Salmonella enterica subsp. enterica serovar Enteritidis str. 635290-58 Proteobacteria; Gammaproteobacteria; Enterobacterales; Enterobacteriaceae; Salmonella; Salmonella enterica; Salmonella enterica subsp. enterica; Salmonella enterica subsp. enterica serovar Enteritidis MNNEETFYQAMRRKGVTRRSFLKFCSLAATSLGLGAGMTPKIAWALENKPRIPVVWIHGL WP_001058307.1 hydrogenase [Salmonella enterica] Length: 372\nNumber of predicted TMHs: 2\nExp number of AAs in TMHs: 36.51738\nExp number, first 60 AAs: 19.72849\nTotal prob of N-in: 0.96170\nPOSSIBLE N-term signal sequence\ninside 1 19\nTMhelix 20 39\noutside 40 325\nTMhelix 326 348\ninside 349 372

23344 GCF_000329965.1_ASM32996v1 Salmonella enterica subsp. enterica serovar Enteritidis str. 638970-15 Proteobacteria; Gammaproteobacteria; Enterobacterales; Enterobacteriaceae; Salmonella; Salmonella enterica; Salmonella enterica subsp. enterica; Salmonella enterica subsp. enterica serovar Enteritidis MTGDNTLITSHGINRRDFMKLCAALAATMGLSSKAAAEMAESVSNPQRPPVIWIGAQECT WP_000145429.1 hydrogenase 2 small subunit [Salmonella enterica] Length: 372\nNumber of predicted TMHs: 1\nExp number of AAs in TMHs: 24.51309\nExp number, first 60 AAs: 0.2682\nTotal prob of N-in: 0.14402\noutside 1 331\nTMhelix 332 354\ninside 355 372

23345 GCF_000329965.1_ASM32996v1 Salmonella enterica subsp. enterica serovar Enteritidis str. 638970-15 Proteobacteria; Gammaproteobacteria; Enterobacterales; Enterobacteriaceae; Salmonella; Salmonella enterica; Salmonella enterica subsp. enterica; Salmonella enterica subsp. enterica serovar Enteritidis MNNEETFYQAMRRKGVTRRSFLKFCSLAATSLGLGAGMTPKIAWALENKPRIPVVWIHGL WP_001058307.1 hydrogenase [Salmonella enterica] Length: 372\nNumber of predicted TMHs: 2\nExp number of AAs in TMHs: 36.51738\nExp number, first 60 AAs: 19.72849\nTotal prob of N-in: 0.96170\nPOSSIBLE N-term signal sequence\ninside 1 19\nTMhelix 20 39\noutside 40 325\nTMhelix 326 348\ninside 349 372

23346 GCF_000280095.1_ASM28009v1 Salmonella enterica subsp. enterica serovar Enteritidis str. 639016-6 Proteobacteria; Gammaproteobacteria; Enterobacterales; Enterobacteriaceae; Salmonella; Salmonella enterica; Salmonella enterica subsp. enterica; Salmonella enterica subsp. enterica serovar Enteritidis MTGDNTLITSHGINRRDFMKLCAALAATMGLSSKAAAEMAESVSNPQRPPVIWIGAQECT WP_000145429.1 hydrogenase 2 small subunit [Salmonella enterica] Length: 372\nNumber of predicted TMHs: 1\nExp

number of AAs in TMHs: 24.51309\nExp number, first 60 AAs: 0.2682\nTotal prob of N-in: 0.14402\noutside 1 331\nTMhelix 332 354\ninside 355 372

23347 GCF_000280095.1_ASM28009v1 Salmonella enterica subsp. enterica serovar Enteritidis str. 639016-6 Proteobacteria; Gammaproteobacteria; Enterobacterales; Enterobacteriaceae; Salmonella; Salmonella enterica; Salmonella enterica subsp. enterica; Salmonella enterica subsp. enterica serovar Enteritidis MNNEETFYQAMRRKGVTRRSFLKFCSLAATSLGLGAGMTPKIAWALENKPRIPVVWIHGL WP_001058307.1 hydrogenase [Salmonella enterica] Length: 372\nNumber of predicted TMHs: 2\nExp number of AAs in TMHs: 36.51738\nExp number, first 60 AAs: 19.72849\nTotal prob of N-in: 0.96170\nPOSSIBLE N-term signal sequence\ninside 1 19\nTMhelix 20 39\noutside 40 325\nTMhelix 326 348\ninside 349 372

23348 GCF_000280275.1_ASM28027v1 Salmonella enterica subsp. enterica serovar Enteritidis str. 639672-46 Proteobacteria; Gammaproteobacteria; Enterobacterales; Enterobacteriaceae; Salmonella; Salmonella enterica; Salmonella enterica subsp. enterica; Salmonella enterica subsp. enterica serovar Enteritidis MTGDNTLITSHGINRRDFMKLCAALAATMGLSSKAAAEMAESVSNPQRPPVIWIGAQECT WP_000145429.1 hydrogenase 2 small subunit [Salmonella enterica] Length: 372\nNumber of predicted TMHs: 1\nExp number of AAs in TMHs: 24.51309\nExp number, first 60 AAs: 0.2682\nTotal prob of N-in: 0.14402\noutside 1 331\nTMhelix 332 354\ninside 355 372

23349 GCF_000280275.1_ASM28027v1 Salmonella enterica subsp. enterica serovar Enteritidis str. 639672-46 Proteobacteria; Gammaproteobacteria; Enterobacterales; Enterobacteriaceae; Salmonella; Salmonella enterica; Salmonella enterica subsp. enterica; Salmonella enterica subsp. enterica serovar Enteritidis MNNEETFYQAMRRKGVTRRSFLKFCSLAATSLGLGAGMTPKIAWALENKPRIPVVWIHGL WP_001058307.1 hydrogenase [Salmonella enterica] Length: 372\nNumber of predicted TMHs: 2\nExp number of AAs in TMHs: 36.51738\nExp number, first 60 AAs: 19.72849\nTotal prob of N-in: 0.96170\nPOSSIBLE N-term signal sequence\ninside 1 19\nTMhelix 20 39\noutside 40 325\nTMhelix 326 348\ninside 349 372

23350 GCF_000280295.1_ASM28029v1 Salmonella enterica subsp. enterica serovar Enteritidis str. 639672-50 Proteobacteria; Gammaproteobacteria; Enterobacterales; Enterobacteriaceae; Salmonella; Salmonella enterica; Salmonella enterica subsp. enterica; Salmonella enterica subsp. enterica serovar Enteritidis MNNEETFYQAMRRKGVTRRSFLKFCSLAATSLGLGAGMTPKIAWALENKPRIPVVWIHGL WP_001058307.1 hydrogenase [Salmonella enterica] Length: 372\nNumber of predicted TMHs: 2\nExp number of AAs in TMHs: 36.51738\nExp number, first 60 AAs: 19.72849\nTotal prob of N-in: 0.96170\nPOSSIBLE N-term signal sequence\ninside 1 19\nTMhelix 20 39\noutside 40 325\nTMhelix 326 348\ninside 349 372

23351 GCF_000280295.1_ASM28029v1 Salmonella enterica subsp. enterica serovar Enteritidis str. 639672-50 Proteobacteria; Gammaproteobacteria; Enterobacterales; Enterobacteriaceae; Salmonella; Salmonella enterica; Salmonella enterica subsp. enterica; Salmonella enterica subsp. enterica serovar Enteritidis MTGDNTLITSHGINRRDFMKLCAALAATMGLSSKAAAEMAESVSNPQRPPVIWIGAQECT WP_000145429.1 hydrogenase 2 small subunit [Salmonella enterica] Length: 372\nNumber of predicted TMHs: 1\nExp number of AAs in TMHs: 24.51309\nExp number, first 60 AAs: 0.2682\nTotal prob of N-in: 0.14402\noutside 1 331\nTMhelix 332 354\ninside 355 372

23352 GCF_000280115.1_ASM28011v1 Salmonella enterica subsp. enterica serovar Enteritidis str. 640631 Proteobacteria; Gammaproteobacteria; Enterobacterales; Enterobacteriaceae; Salmonella; Salmonella enterica; Salmonella enterica subsp. enterica; Salmonella enterica subsp. enterica serovar Enteritidis MTGDNTLITSHGINRRDFMKLCAALAATMGLSSKAAAEMAESVSNPQRPPVIWIGAQECT WP_000145429.1 hydrogenase 2 small subunit [Salmonella enterica] Length: 372\nNumber of predicted TMHs: 1\nExp number of AAs in TMHs: 24.51309\nExp number, first 60 AAs: 0.2682\nTotal prob of N-in: 0.14402\noutside 1 331\nTMhelix 332 354\ninside 355 372

23353 GCF_000280115.1_ASM28011v1 Salmonella enterica subsp. enterica serovar Enteritidis str. 640631 Proteobacteria; Gammaproteobacteria; Enterobacterales; Enterobacteriaceae; Salmonella; Salmonella enterica; Salmonella enterica subsp. enterica; Salmonella enterica subsp. enterica serovar Enteritidis MNNEETFYQAMRRKGVTRRSFLKFCSLAATSLGLGAGMTPKIAWALENKPRIPVVWIHGL WP_001058307.1 hydrogenase [Salmonella enterica] Length: 372\nNumber of predicted TMHs: 2\nExp number of AAs in TMHs: 36.51738\nExp number, first 60 AAs: 19.72849\nTotal prob of N-in: 0.96170\nPOSSIBLE N-term signal sequence\ninside 1 19\nTMhelix 20 39\noutside 40 325\nTMhelix 326 348\ninside 349 372

23354 GCF_000330105.1_ASM33010v1 Salmonella enterica subsp. enterica serovar Enteritidis str. 642044 4-1 Proteobacteria; Gammaproteobacteria; Enterobacterales; Enterobacteriaceae; Salmonella; Salmonella enterica;

Salmonella enterica subsp. enterica; Salmonella enterica subsp. enterica serovar Enteritidis
MTGDNTLITSHGINRRDFMKLCAALAATMGLSSKAAAEMAESVSNPQRPPVIWIGAQECT WP_000145429.1
hydrogenase 2 small subunit [Salmonella enterica] Length: 372\nNumber of predicted TMHs: 1\nExp number of AAs in TMHs: 24.51309\nExp number, first 60 AAs: 0.2682\nTotal prob of N-in: 0.14402\noutside 1 331\nTMhelix 332 354\ninside 355 372

23355 GCF_000330105.1_ASM33010v1 Salmonella enterica subsp. enterica serovar Enteritidis str. 642044 4-1
Proteobacteria; Gammaproteobacteria; Enterobacterales; Enterobacteriaceae; Salmonella; Salmonella enterica; Salmonella enterica subsp. enterica; Salmonella enterica subsp. enterica serovar Enteritidis
MNNEETFYQAMRRKGVTRRSFLKFCSLAATSLGLGAGMTPKIAWALENKPRIPVVWIHGL WP_001058307.1
hydrogenase [Salmonella enterica] Length: 372\nNumber of predicted TMHs: 2\nExp number of AAs in TMHs: 36.51738\nExp number, first 60 AAs: 19.72849\nTotal prob of N-in: 0.96170\nPOSSIBLE N-term signal sequence\ninside 1 19\nTMhelix 20 39\noutside 40 325\nTMhelix 326 348\ninside 349 372

23356 GCF_000330125.1_ASM33012v1 Salmonella enterica subsp. enterica serovar Enteritidis str. 642046 4-7
Proteobacteria; Gammaproteobacteria; Enterobacterales; Enterobacteriaceae; Salmonella; Salmonella enterica; Salmonella enterica subsp. enterica; Salmonella enterica subsp. enterica serovar Enteritidis
MNNEETFYQAMRRKGVTRRSFLKFCSLAATSLGLGAGMTPKIAWALENKPRIPVVWIHGL WP_001058307.1
hydrogenase [Salmonella enterica] Length: 372\nNumber of predicted TMHs: 2\nExp number of AAs in TMHs: 36.51738\nExp number, first 60 AAs: 19.72849\nTotal prob of N-in: 0.96170\nPOSSIBLE N-term signal sequence\ninside 1 19\nTMhelix 20 39\noutside 40 325\nTMhelix 326 348\ninside 349 372

23357 GCF_000330125.1_ASM33012v1 Salmonella enterica subsp. enterica serovar Enteritidis str. 642046 4-7
Proteobacteria; Gammaproteobacteria; Enterobacterales; Enterobacteriaceae; Salmonella; Salmonella enterica; Salmonella enterica subsp. enterica; Salmonella enterica subsp. enterica serovar Enteritidis
MTGDNTLITSHGINRRDFMKLCAALAATMGLSSKAAAEMAESVSNPQRPPVIWIGAQECT WP_000145429.1
hydrogenase 2 small subunit [Salmonella enterica] Length: 372\nNumber of predicted TMHs: 1\nExp number of AAs in TMHs: 24.51309\nExp number, first 60 AAs: 0.2682\nTotal prob of N-in: 0.14402\noutside 1 331\nTMhelix 332 354\ninside 355 372

23358 GCF_000330145.1_ASM33014v1 Salmonella enterica subsp. enterica serovar Enteritidis str. 648898 4-5
Proteobacteria; Gammaproteobacteria; Enterobacterales; Enterobacteriaceae; Salmonella; Salmonella enterica; Salmonella enterica subsp. enterica; Salmonella enterica subsp. enterica serovar Enteritidis
MNNEETFYQAMRRKGVTRRSFLKFCSLAATSLGLGAGMTPKIAWALENKPRIPVVWIHGL WP_001058307.1
hydrogenase [Salmonella enterica] Length: 372\nNumber of predicted TMHs: 2\nExp number of AAs in TMHs: 36.51738\nExp number, first 60 AAs: 19.72849\nTotal prob of N-in: 0.96170\nPOSSIBLE N-term signal sequence\ninside 1 19\nTMhelix 20 39\noutside 40 325\nTMhelix 326 348\ninside 349 372

23359 GCF_000330145.1_ASM33014v1 Salmonella enterica subsp. enterica serovar Enteritidis str. 648898 4-5
Proteobacteria; Gammaproteobacteria; Enterobacterales; Enterobacteriaceae; Salmonella; Salmonella enterica; Salmonella enterica subsp. enterica; Salmonella enterica subsp. enterica serovar Enteritidis
MTGDNTLITSHGINRRDFMKLCAALAATMGLSSKAAAEMAESVSNPQRPPVIWIGAQECT WP_000145429.1
hydrogenase 2 small subunit [Salmonella enterica] Length: 372\nNumber of predicted TMHs: 1\nExp number of AAs in TMHs: 24.51309\nExp number, first 60 AAs: 0.2682\nTotal prob of N-in: 0.14402\noutside 1 331\nTMhelix 332 354\ninside 355 372

23360 GCF_000330165.1_ASM33016v1 Salmonella enterica subsp. enterica serovar Enteritidis str. 648899 3-17
Proteobacteria; Gammaproteobacteria; Enterobacterales; Enterobacteriaceae; Salmonella; Salmonella enterica; Salmonella enterica subsp. enterica; Salmonella enterica subsp. enterica serovar Enteritidis
MNNEETFYQAMRRKGVTRRSFLKFCSLAATSLGLGAGMTPKIAWALENKPRIPVVWIHGL WP_001058307.1
hydrogenase [Salmonella enterica] Length: 372\nNumber of predicted TMHs: 2\nExp number of AAs in TMHs: 36.51738\nExp number, first 60 AAs: 19.72849\nTotal prob of N-in: 0.96170\nPOSSIBLE N-term signal sequence\ninside 1 19\nTMhelix 20 39\noutside 40 325\nTMhelix 326 348\ninside 349 372

23361 GCF_000330165.1_ASM33016v1 Salmonella enterica subsp. enterica serovar Enteritidis str. 648899 3-17
Proteobacteria; Gammaproteobacteria; Enterobacterales; Enterobacteriaceae; Salmonella; Salmonella enterica; Salmonella enterica subsp. enterica; Salmonella enterica subsp. enterica serovar Enteritidis
MTGDNTLITSHGINRRDFMKLCAALAATMGLSSKAAAEMAESVSNPQRPPVIWIGAQECT WP_000145429.1
hydrogenase 2 small subunit [Salmonella enterica] Length: 372\nNumber of predicted TMHs: 1\nExp number of AAs in TMHs: 24.51309\nExp number, first 60 AAs: 0.2682\nTotal prob of N-in: 0.14402\noutside 1 331\nTMhelix 332 354\ninside 355 372

23362 GCF_000330185.1_ASM33018v1 Salmonella enterica subsp. enterica serovar Enteritidis str. 648900 1-16 Proteobacteria; Gammaproteobacteria; Enterobacterales; Enterobacteriaceae; Salmonella; Salmonella enterica; Salmonella enterica subsp. enterica; Salmonella enterica subsp. enterica serovar Enteritidis MNNEETFYQAMRRKGVTRRSFLKFCSLAATSLGLGAGMTPKIAWALENKPRIPVVWIHGL WP_001058307.1 hydrogenase [Salmonella enterica] Length: 372\nNumber of predicted TMHs: 2\nExp number of AAs in TMHs: 36.51738\nExp number, first 60 AAs: 19.72849\nTotal prob of N-in: 0.96170\nPOSSIBLE N-term signal sequence\ninside 1 19\nTMhelix 20 39\noutside 40 325\nTMhelix 326 348\ninside 349 372

23363 GCF_000330185.1_ASM33018v1 Salmonella enterica subsp. enterica serovar Enteritidis str. 648900 1-16 Proteobacteria; Gammaproteobacteria; Enterobacterales; Enterobacteriaceae; Salmonella; Salmonella enterica; Salmonella enterica subsp. enterica; Salmonella enterica subsp. enterica serovar Enteritidis MTGDNTLITSHGINRRDFMKLCAALAATMGLSSKAAAEMAESVSNPQRPPVIWIGAQECT WP_000145429.1 hydrogenase 2 small subunit [Salmonella enterica] Length: 372\nNumber of predicted TMHs: 1\nExp number of AAs in TMHs: 24.51309\nExp number, first 60 AAs: 0.2682\nTotal prob of N-in: 0.14402\noutside 1 331\nTMhelix 332 354\ninside 355 372

23364 GCF_000330205.1_ASM33020v1 Salmonella enterica subsp. enterica serovar Enteritidis str. 648901 1-17 Proteobacteria; Gammaproteobacteria; Enterobacterales; Enterobacteriaceae; Salmonella; Salmonella enterica; Salmonella enterica subsp. enterica; Salmonella enterica subsp. enterica serovar Enteritidis MNNEETFYQAMRRKGVTRRSFLKFCSLAATSLGLGAGMTPKIAWALENKPRIPVVWIHGL WP_001058307.1 hydrogenase [Salmonella enterica] Length: 372\nNumber of predicted TMHs: 2\nExp number of AAs in TMHs: 36.51738\nExp number, first 60 AAs: 19.72849\nTotal prob of N-in: 0.96170\nPOSSIBLE N-term signal sequence\ninside 1 19\nTMhelix 20 39\noutside 40 325\nTMhelix 326 348\ninside 349 372

23365 GCF_000330205.1_ASM33020v1 Salmonella enterica subsp. enterica serovar Enteritidis str. 648901 1-17 Proteobacteria; Gammaproteobacteria; Enterobacterales; Enterobacteriaceae; Salmonella; Salmonella enterica; Salmonella enterica subsp. enterica; Salmonella enterica subsp. enterica serovar Enteritidis MTGDNTLITSHGINRRDFMKLCAALAATMGLSSKAAAEMAESVSNPQRPPVIWIGAQECT WP_000145429.1 hydrogenase 2 small subunit [Salmonella enterica] Length: 372\nNumber of predicted TMHs: 1\nExp number of AAs in TMHs: 24.51309\nExp number, first 60 AAs: 0.2682\nTotal prob of N-in: 0.14402\noutside 1 331\nTMhelix 332 354\ninside 355 372

23366 GCF_000330385.1_ASM33038v1 Salmonella enterica subsp. enterica serovar Enteritidis str. 648901 16-16 Proteobacteria; Gammaproteobacteria; Enterobacterales; Enterobacteriaceae; Salmonella; Salmonella enterica; Salmonella enterica subsp. enterica; Salmonella enterica subsp. enterica serovar Enteritidis MTGDNTLITSHGINRRDFMKLCAALAATMGLSSKAAAEMAESVSNPQRPPVIWIGAQECT WP_000145429.1 hydrogenase 2 small subunit [Salmonella enterica] Length: 372\nNumber of predicted TMHs: 1\nExp number of AAs in TMHs: 24.51309\nExp number, first 60 AAs: 0.2682\nTotal prob of N-in: 0.14402\noutside 1 331\nTMhelix 332 354\ninside 355 372

23367 GCF_000330385.1_ASM33038v1 Salmonella enterica subsp. enterica serovar Enteritidis str. 648901 16-16 Proteobacteria; Gammaproteobacteria; Enterobacterales; Enterobacteriaceae; Salmonella; Salmonella enterica; Salmonella enterica subsp. enterica; Salmonella enterica subsp. enterica serovar Enteritidis MNNEETFYQAMRRKGVTRRSFLKFCSLAATSLGLGAGMTPKIAWALENKPRIPVVWIHGL WP_001058307.1 hydrogenase [Salmonella enterica] Length: 372\nNumber of predicted TMHs: 2\nExp number of AAs in TMHs: 36.51738\nExp number, first 60 AAs: 19.72849\nTotal prob of N-in: 0.96170\nPOSSIBLE N-term signal sequence\ninside 1 19\nTMhelix 20 39\noutside 40 325\nTMhelix 326 348\ninside 349 372

23368 GCF_000280435.1_ASM28043v1 Salmonella enterica subsp. enterica serovar Enteritidis str. 648901 6-18 Proteobacteria; Gammaproteobacteria; Enterobacterales; Enterobacteriaceae; Salmonella; Salmonella enterica; Salmonella enterica subsp. enterica; Salmonella enterica subsp. enterica serovar Enteritidis MTGDNTLITSHGINRRDFMKLCAALAATMGLSSKAAAEMAESVSNPQRPPVIWIGAQECT WP_000145429.1 hydrogenase 2 small subunit [Salmonella enterica] Length: 372\nNumber of predicted TMHs: 1\nExp number of AAs in TMHs: 24.51309\nExp number, first 60 AAs: 0.2682\nTotal prob of N-in: 0.14402\noutside 1 331\nTMhelix 332 354\ninside 355 372

23369 GCF_000280435.1_ASM28043v1 Salmonella enterica subsp. enterica serovar Enteritidis str. 648901 6-18 Proteobacteria; Gammaproteobacteria; Enterobacterales; Enterobacteriaceae; Salmonella; Salmonella enterica; Salmonella enterica subsp. enterica; Salmonella enterica subsp. enterica serovar Enteritidis MNNEETFYQAMRRKGVTRRSFLKFCSLAATSLGLGAGMTPKIAWALENKPRIPVVWIHGL WP_001058307.1 hydrogenase [Salmonella enterica] Length: 372\nNumber of predicted TMHs: 2\nExp number of AAs in

TMHs: 36.51738\nExp number, first 60 AAs: 19.72849\nTotal prob of N-in: 0.96170\nPOSSIBLE N-term signal sequence\ninside 1 19\nTMhelix 20 39\noutside 40 325\nTMhelix 326 348\ninside 349 372

23370 GCF_000330245.1_ASM33024v1 Salmonella enterica subsp. enterica serovar Enteritidis str. 648902 6-8 Proteobacteria; Gammaproteobacteria; Enterobacterales; Enterobacteriaceae; Salmonella; Salmonella enterica; Salmonella enterica subsp. enterica; Salmonella enterica subsp. enterica serovar Enteritidis MTGDNTLITSHGINRRDFMKLCAALAATMGLSSKAAAEMAESVSNPQRPPVIWIGAQECT WP_000145429.1 hydrogenase 2 small subunit [Salmonella enterica] Length: 372\nNumber of predicted TMHs: 1\nExp number of AAs in TMHs: 24.51309\nExp number, first 60 AAs: 0.2682\nTotal prob of N-in: 0.14402\noutside 1 331\nTMhelix 332 354\ninside 355 372

23371 GCF_000330245.1_ASM33024v1 Salmonella enterica subsp. enterica serovar Enteritidis str. 648902 6-8 Proteobacteria; Gammaproteobacteria; Enterobacterales; Enterobacteriaceae; Salmonella; Salmonella enterica; Salmonella enterica subsp. enterica; Salmonella enterica subsp. enterica serovar Enteritidis MNNEETFYQAMRRKGVTRRSFLKFCSLAATSLGLGAGMTPKIAWALENKPRIPVVWIHGL WP_001058307.1 hydrogenase [Salmonella enterica] Length: 372\nNumber of predicted TMHs: 2\nExp number of AAs in TMHs: 36.51738\nExp number, first 60 AAs: 19.72849\nTotal prob of N-in: 0.96170\nPOSSIBLE N-term signal sequence\ninside 1 19\nTMhelix 20 39\noutside 40 325\nTMhelix 326 348\ninside 349 372

23372 GCF_000330265.1_ASM33026v1 Salmonella enterica subsp. enterica serovar Enteritidis str. 648903 1-6 Proteobacteria; Gammaproteobacteria; Enterobacterales; Enterobacteriaceae; Salmonella; Salmonella enterica; Salmonella enterica subsp. enterica; Salmonella enterica subsp. enterica serovar Enteritidis MNNEETFYQAMRRKGVTRRSFLKFCSLAATSLGLGAGMTPKIAWALENKPRIPVVWIHGL WP_001058307.1 hydrogenase [Salmonella enterica] Length: 372\nNumber of predicted TMHs: 2\nExp number of AAs in TMHs: 36.51738\nExp number, first 60 AAs: 19.72849\nTotal prob of N-in: 0.96170\nPOSSIBLE N-term signal sequence\ninside 1 19\nTMhelix 20 39\noutside 40 325\nTMhelix 326 348\ninside 349 372

23373 GCF_000330265.1_ASM33026v1 Salmonella enterica subsp. enterica serovar Enteritidis str. 648903 1-6 Proteobacteria; Gammaproteobacteria; Enterobacterales; Enterobacteriaceae; Salmonella; Salmonella enterica; Salmonella enterica subsp. enterica; Salmonella enterica subsp. enterica serovar Enteritidis MTGDNTLITSHGINRRDFMKLCAALAATMGLSSKAAAEMAESVSNPQRPPVIWIGAQECT WP_000145429.1 hydrogenase 2 small subunit [Salmonella enterica] Length: 372\nNumber of predicted TMHs: 1\nExp number of AAs in TMHs: 24.51309\nExp number, first 60 AAs: 0.2682\nTotal prob of N-in: 0.14402\noutside 1 331\nTMhelix 332 354\ninside 355 372

23374 GCF_000330285.1_ASM33028v1 Salmonella enterica subsp. enterica serovar Enteritidis str. 648904 3-6 Proteobacteria; Gammaproteobacteria; Enterobacterales; Enterobacteriaceae; Salmonella; Salmonella enterica; Salmonella enterica subsp. enterica; Salmonella enterica subsp. enterica serovar Enteritidis MNNEETFYQAMRRKGVTRRSFLKFCSLAATSLGLGAGMTPKIAWALENKPRIPVVWIHGL WP_001058307.1 hydrogenase [Salmonella enterica] Length: 372\nNumber of predicted TMHs: 2\nExp number of AAs in TMHs: 36.51738\nExp number, first 60 AAs: 19.72849\nTotal prob of N-in: 0.96170\nPOSSIBLE N-term signal sequence\ninside 1 19\nTMhelix 20 39\noutside 40 325\nTMhelix 326 348\ninside 349 372

23375 GCF_000330285.1_ASM33028v1 Salmonella enterica subsp. enterica serovar Enteritidis str. 648904 3-6 Proteobacteria; Gammaproteobacteria; Enterobacterales; Enterobacteriaceae; Salmonella; Salmonella enterica; Salmonella enterica subsp. enterica; Salmonella enterica subsp. enterica serovar Enteritidis MTGDNTLITSHGINRRDFMKLCAALAATMGLSSKAAAEMAESVSNPQRPPVIWIGAQECT WP_000145429.1 hydrogenase 2 small subunit [Salmonella enterica] Length: 372\nNumber of predicted TMHs: 1\nExp number of AAs in TMHs: 24.51309\nExp number, first 60 AAs: 0.2682\nTotal prob of N-in: 0.14402\noutside 1 331\nTMhelix 332 354\ninside 355 372

23376 GCF_000280415.1_ASM28041v1 Salmonella enterica subsp. enterica serovar Enteritidis str. 648905 5-18 Proteobacteria; Gammaproteobacteria; Enterobacterales; Enterobacteriaceae; Salmonella; Salmonella enterica; Salmonella enterica subsp. enterica; Salmonella enterica subsp. enterica serovar Enteritidis MTGDNTLITSHGINRRDFMKLCAALAATMGLSSKAAAEMAESVSNPQRPPVIWIGAQECT WP_000145429.1 hydrogenase 2 small subunit [Salmonella enterica] Length: 372\nNumber of predicted TMHs: 1\nExp number of AAs in TMHs: 24.51309\nExp number, first 60 AAs: 0.2682\nTotal prob of N-in: 0.14402\noutside 1 331\nTMhelix 332 354\ninside 355 372

23377 GCF_000280415.1_ASM28041v1 Salmonella enterica subsp. enterica serovar Enteritidis str. 648905 5-18 Proteobacteria; Gammaproteobacteria; Enterobacterales; Enterobacteriaceae; Salmonella; Salmonella enterica;

Salmonella enterica subsp. enterica; Salmonella enterica subsp. enterica serovar Enteritidis
MNNEETFYQAMRRKGVTRRSFLKFCSLAATSLGLGAGMTPKIAWALENKPRIPVVWIHGL WP_001058307.1
hydrogenase [Salmonella enterica] Length: 372\nNumber of predicted TMHs: 2\nExp number of AAs in TMHs: 36.51738\nExp number, first 60 AAs: 19.72849\nTotal prob of N-in: 0.96170\nPOSSIBLE N-term signal sequence\ninside 1 19\nTMhelix 20 39\noutside 40 325\nTMhelix 326 348\ninside 349 372

23378 GCF_000330305.1_ASM33030v1 Salmonella enterica subsp. enterica serovar Enteritidis str. 653049 13-19
Proteobacteria; Gammaproteobacteria; Enterobacterales; Enterobacteriaceae; Salmonella; Salmonella enterica; Salmonella enterica subsp. enterica; Salmonella enterica subsp. enterica serovar Enteritidis
MTGDNTLITSHGINRRDFMKLCAALAATMGLSSKAAAEMAESVSNPQRPPVIWIGAQECT WP_000145429.1
hydrogenase 2 small subunit [Salmonella enterica] Length: 372\nNumber of predicted TMHs: 1\nExp number of AAs in TMHs: 24.51309\nExp number, first 60 AAs: 0.2682\nTotal prob of N-in: 0.14402\noutside 1 331\nTMhelix 332 354\ninside 355 372

23379 GCF_000330305.1_ASM33030v1 Salmonella enterica subsp. enterica serovar Enteritidis str. 653049 13-19
Proteobacteria; Gammaproteobacteria; Enterobacterales; Enterobacteriaceae; Salmonella; Salmonella enterica; Salmonella enterica subsp. enterica; Salmonella enterica subsp. enterica serovar Enteritidis
MNNEETFYQAMRRKGVTRRSFLKFCSLAATSLGLGAGMTPKIAWALENKPRIPVVWIHGL WP_001058307.1
hydrogenase [Salmonella enterica] Length: 372\nNumber of predicted TMHs: 2\nExp number of AAs in TMHs: 36.51738\nExp number, first 60 AAs: 19.72849\nTotal prob of N-in: 0.96170\nPOSSIBLE N-term signal sequence\ninside 1 19\nTMhelix 20 39\noutside 40 325\nTMhelix 326 348\ninside 349 372

23380 GCF_000280135.1_ASM28013v1 Salmonella enterica subsp. enterica serovar Enteritidis str. 77-0424
Proteobacteria; Gammaproteobacteria; Enterobacterales; Enterobacteriaceae; Salmonella; Salmonella enterica; Salmonella enterica subsp. enterica; Salmonella enterica subsp. enterica serovar Enteritidis
MTGDNTLITSHGINRRDFMKLCAALAATMGLSSKAAAEMAESVSNPQRPPVIWIGAQECT WP_000145429.1
hydrogenase 2 small subunit [Salmonella enterica] Length: 372\nNumber of predicted TMHs: 1\nExp number of AAs in TMHs: 24.51309\nExp number, first 60 AAs: 0.2682\nTotal prob of N-in: 0.14402\noutside 1 331\nTMhelix 332 354\ninside 355 372

23381 GCF_000280135.1_ASM28013v1 Salmonella enterica subsp. enterica serovar Enteritidis str. 77-0424
Proteobacteria; Gammaproteobacteria; Enterobacterales; Enterobacteriaceae; Salmonella; Salmonella enterica; Salmonella enterica subsp. enterica; Salmonella enterica subsp. enterica serovar Enteritidis
MNNEETFYQAMRRKGVTRRSFLKFCSLAATSLGLGAGMTPKIAWALENKPRIPVVWIHGL WP_001058307.1
hydrogenase [Salmonella enterica] Length: 372\nNumber of predicted TMHs: 2\nExp number of AAs in TMHs: 36.51738\nExp number, first 60 AAs: 19.72849\nTotal prob of N-in: 0.96170\nPOSSIBLE N-term signal sequence\ninside 1 19\nTMhelix 20 39\noutside 40 325\nTMhelix 326 348\ninside 349 372

23382 GCF_000280315.2_ASM28031v2 Salmonella enterica subsp. enterica serovar Enteritidis str. 77-1427
Proteobacteria; Gammaproteobacteria; Enterobacterales; Enterobacteriaceae; Salmonella; Salmonella enterica; Salmonella enterica subsp. enterica; Salmonella enterica subsp. enterica serovar Enteritidis
MTGDNTLITSHGINRRDFMKLCAALAATMGLSSKAAAEMAESVSNPQRPPVIWIGAQECT WP_000145429.1
hydrogenase 2 small subunit [Salmonella enterica] Length: 372\nNumber of predicted TMHs: 1\nExp number of AAs in TMHs: 24.51309\nExp number, first 60 AAs: 0.2682\nTotal prob of N-in: 0.14402\noutside 1 331\nTMhelix 332 354\ninside 355 372

23383 GCF_000280315.2_ASM28031v2 Salmonella enterica subsp. enterica serovar Enteritidis str. 77-1427
Proteobacteria; Gammaproteobacteria; Enterobacterales; Enterobacteriaceae; Salmonella; Salmonella enterica; Salmonella enterica subsp. enterica; Salmonella enterica subsp. enterica serovar Enteritidis
MNNEETFYQAMRRKGVTRRSFLKFCSLAATSLGLGAGMTPKIAWALENKPRIPVVWIHGL WP_001058307.1
hydrogenase [Salmonella enterica] Length: 372\nNumber of predicted TMHs: 2\nExp number of AAs in TMHs: 36.51738\nExp number, first 60 AAs: 19.72849\nTotal prob of N-in: 0.96170\nPOSSIBLE N-term signal sequence\ninside 1 19\nTMhelix 20 39\noutside 40 325\nTMhelix 326 348\ninside 349 372

23384 GCF_000280335.1_ASM28033v1 Salmonella enterica subsp. enterica serovar Enteritidis str. 77-2659
Proteobacteria; Gammaproteobacteria; Enterobacterales; Enterobacteriaceae; Salmonella; Salmonella enterica; Salmonella enterica subsp. enterica; Salmonella enterica subsp. enterica serovar Enteritidis
MNNEETFYQAMRRKGVTRRSFLKFCSLAATSLGLGAGMTPKIAWALENKPRIPVVWIHGL WP_001058307.1
hydrogenase [Salmonella enterica] Length: 372\nNumber of predicted TMHs: 2\nExp number of AAs in TMHs: 36.51738\nExp number, first 60 AAs: 19.72849\nTotal prob of N-in: 0.96170\nPOSSIBLE N-term signal sequence\ninside 1 19\nTMhelix 20 39\noutside 40 325\nTMhelix 326 348\ninside 349 372

23385 GCF_000280335.1_ASM28033v1 Salmonella enterica subsp. enterica serovar Enteritidis str. 77-2659 Proteobacteria; Gammaproteobacteria; Enterobacterales; Enterobacteriaceae; Salmonella; Salmonella enterica; Salmonella enterica subsp. enterica; Salmonella enterica subsp. enterica serovar Enteritidis MTGDNTLITSHGINRRDFMKLCAALAATMGLSSKAAAEMAESVSNPQRPPVIWIGAQECT WP_000145429.1 hydrogenase 2 small subunit [Salmonella enterica] Length: 372\nNumber of predicted TMHs: 1\nExp number of AAs in TMHs: 24.51309\nExp number, first 60 AAs: 0.2682\nTotal prob of N-in: 0.14402\noutside 1 331\nTMhelix 332 354\ninside 355 372

23386 GCF_000280355.1_ASM28035v1 Salmonella enterica subsp. enterica serovar Enteritidis str. 78-1757 Proteobacteria; Gammaproteobacteria; Enterobacterales; Enterobacteriaceae; Salmonella; Salmonella enterica; Salmonella enterica subsp. enterica; Salmonella enterica subsp. enterica serovar Enteritidis MTGDNTLITSHGINRRDFMKLCAALAATMGLSSKAAAEMAESVSNPQRPPVIWIGAQECT WP_000145429.1 hydrogenase 2 small subunit [Salmonella enterica] Length: 372\nNumber of predicted TMHs: 1\nExp number of AAs in TMHs: 24.51309\nExp number, first 60 AAs: 0.2682\nTotal prob of N-in: 0.14402\noutside 1 331\nTMhelix 332 354\ninside 355 372

23387 GCF_000280355.1_ASM28035v1 Salmonella enterica subsp. enterica serovar Enteritidis str. 78-1757 Proteobacteria; Gammaproteobacteria; Enterobacterales; Enterobacteriaceae; Salmonella; Salmonella enterica; Salmonella enterica subsp. enterica; Salmonella enterica subsp. enterica serovar Enteritidis MNNEETFYQAMRRKGVTRRSFLKFCSLAATSLGLGAGMTPKIAWALENKPRIPVVWIHGL WP_001058307.1 hydrogenase [Salmonella enterica] Length: 372\nNumber of predicted TMHs: 2\nExp number of AAs in TMHs: 36.51738\nExp number, first 60 AAs: 19.72849\nTotal prob of N-in: 0.96170\nPOSSIBLE N-term signal sequence\ninside 1 19\nTMhelix 20 39\noutside 40 325\nTMhelix 326 348\ninside 349 372

23388 GCF_000330465.1_ASM33046v1 Salmonella enterica subsp. enterica serovar Enteritidis str. 81-2625 Proteobacteria; Gammaproteobacteria; Enterobacterales; Enterobacteriaceae; Salmonella; Salmonella enterica; Salmonella enterica subsp. enterica; Salmonella enterica subsp. enterica serovar Enteritidis MNNEETFYQAMRRKGVTRRSFLKFCSLAATSLGLGAGMTPKIAWALENKPRIPVVWIHGL WP_001058307.1 hydrogenase [Salmonella enterica] Length: 372\nNumber of predicted TMHs: 2\nExp number of AAs in TMHs: 36.51738\nExp number, first 60 AAs: 19.72849\nTotal prob of N-in: 0.96170\nPOSSIBLE N-term signal sequence\ninside 1 19\nTMhelix 20 39\noutside 40 325\nTMhelix 326 348\ninside 349 372

23389 GCF_000330465.1_ASM33046v1 Salmonella enterica subsp. enterica serovar Enteritidis str. 81-2625 Proteobacteria; Gammaproteobacteria; Enterobacterales; Enterobacteriaceae; Salmonella; Salmonella enterica; Salmonella enterica subsp. enterica; Salmonella enterica subsp. enterica serovar Enteritidis MTGDNTLITSHGINRRDFMKLCAALAATMGLSSKAAAEMAESVSNPQRPPVIWIGAQECT WP_000145429.1 hydrogenase 2 small subunit [Salmonella enterica] Length: 372\nNumber of predicted TMHs: 1\nExp number of AAs in TMHs: 24.51309\nExp number, first 60 AAs: 0.2682\nTotal prob of N-in: 0.14402\noutside 1 331\nTMhelix 332 354\ninside 355 372

23390 GCF_000280395.1_ASM28039v1 Salmonella enterica subsp. enterica serovar Enteritidis str. 8b-1 Proteobacteria; Gammaproteobacteria; Enterobacterales; Enterobacteriaceae; Salmonella; Salmonella enterica; Salmonella enterica subsp. enterica; Salmonella enterica subsp. enterica serovar Enteritidis MNNEETFYQAMRRKGVTRRSFLKFCSLAATSLGLGAGMTPKIAWALENKPRIPVVWIHGL WP_001058307.1 hydrogenase [Salmonella enterica] Length: 372\nNumber of predicted TMHs: 2\nExp number of AAs in TMHs: 36.51738\nExp number, first 60 AAs: 19.72849\nTotal prob of N-in: 0.96170\nPOSSIBLE N-term signal sequence\ninside 1 19\nTMhelix 20 39\noutside 40 325\nTMhelix 326 348\ninside 349 372

23391 GCF_000280395.1_ASM28039v1 Salmonella enterica subsp. enterica serovar Enteritidis str. 8b-1 Proteobacteria; Gammaproteobacteria; Enterobacterales; Enterobacteriaceae; Salmonella; Salmonella enterica; Salmonella enterica subsp. enterica; Salmonella enterica subsp. enterica serovar Enteritidis MTGDNTLITSHGINRRDFMKLCAALAATMGLSSKAAAEMAESVSNPQRPPVIWIGAQECT WP_000145429.1 hydrogenase 2 small subunit [Salmonella enterica] Length: 372\nNumber of predicted TMHs: 1\nExp number of AAs in TMHs: 24.51309\nExp number, first 60 AAs: 0.2682\nTotal prob of N-in: 0.14402\noutside 1 331\nTMhelix 332 354\ninside 355 372

23392 GCF_000329285.1_ASM32928v1 Salmonella enterica subsp. enterica serovar Enteritidis str. CDC_2010K_0895 Proteobacteria; Gammaproteobacteria; Enterobacterales; Enterobacteriaceae; Salmonella; Salmonella enterica; Salmonella enterica subsp. enterica; Salmonella enterica subsp. enterica serovar Enteritidis MNNEETFYQAMRRKGVTRRSFLKFCSLAATSLGLGAGMTPKIAWALENKPRIPVVWIHGL WP_001058307.1 hydrogenase [Salmonella enterica] Length: 372\nNumber of predicted TMHs: 2\nExp number of AAs in

TMHs: 36.51738\nExp number, first 60 AAs: 19.72849\nTotal prob of N-in: 0.96170\nPOSSIBLE N-term signal sequence\ninside 1 19\nTMhelix 20 39\noutside 40 325\nTMhelix 326 348\ninside 349 372

23393 GCF_000329285.1_ASM32928v1 Salmonella enterica subsp. enterica serovar Enteritidis str. CDC_2010K_0895 Proteobacteria; Gammaproteobacteria; Enterobacterales; Enterobacteriaceae; Salmonella; Salmonella enterica; Salmonella enterica subsp. enterica; Salmonella enterica subsp. enterica serovar Enteritidis MTGDNTLITSHGINRRDFMKLCAALAATMGLSSKAAAEMAESVSNPQRPPVIWIGAQECT WP_000145429.1 hydrogenase 2 small subunit [Salmonella enterica] Length: 372\nNumber of predicted TMHs: 1\nExp number of AAs in TMHs: 24.51309\nExp number, first 60 AAs: 0.2682\nTotal prob of N-in: 0.14402\noutside 1 331\nTMhelix 332 354\ninside 355 372

23394 GCF_000329305.1_ASM32930v1 Salmonella enterica subsp. enterica serovar Enteritidis str. CDC_2010K_0899 Proteobacteria; Gammaproteobacteria; Enterobacterales; Enterobacteriaceae; Salmonella; Salmonella enterica; Salmonella enterica subsp. enterica; Salmonella enterica subsp. enterica serovar Enteritidis MNNEETFYQAMRRKGVTRRSFLKFCSLAATSLGLGAGMTPKIAWALENKPRIPVVWIHGL WP_001058307.1 hydrogenase [Salmonella enterica] Length: 372\nNumber of predicted TMHs: 2\nExp number of AAs in TMHs: 36.51738\nExp number, first 60 AAs: 19.72849\nTotal prob of N-in: 0.96170\nPOSSIBLE N-term signal sequence\ninside 1 19\nTMhelix 20 39\noutside 40 325\nTMhelix 326 348\ninside 349 372

23395 GCF_000329305.1_ASM32930v1 Salmonella enterica subsp. enterica serovar Enteritidis str. CDC_2010K_0899 Proteobacteria; Gammaproteobacteria; Enterobacterales; Enterobacteriaceae; Salmonella; Salmonella enterica; Salmonella enterica subsp. enterica; Salmonella enterica subsp. enterica serovar Enteritidis MTGDNTLITSHGINRRDFMKLCAALAATMGLSSKAAAEMAESVSNPQRPPVIWIGAQECT WP_000145429.1 hydrogenase 2 small subunit [Salmonella enterica] Length: 372\nNumber of predicted TMHs: 1\nExp number of AAs in TMHs: 24.51309\nExp number, first 60 AAs: 0.2682\nTotal prob of N-in: 0.14402\noutside 1 331\nTMhelix 332 354\ninside 355 372

23396 GCF_000329505.1_ASM32950v1 Salmonella enterica subsp. enterica serovar Enteritidis str. CDC_2010K_0956 Proteobacteria; Gammaproteobacteria; Enterobacterales; Enterobacteriaceae; Salmonella; Salmonella enterica; Salmonella enterica subsp. enterica; Salmonella enterica subsp. enterica serovar Enteritidis MTGDNTLITSHGINRRDFMKLCAALAATMGLSSKAAAEMAESVSNPQRPPVIWIGAQECT WP_000145429.1 hydrogenase 2 small subunit [Salmonella enterica] Length: 372\nNumber of predicted TMHs: 1\nExp number of AAs in TMHs: 24.51309\nExp number, first 60 AAs: 0.2682\nTotal prob of N-in: 0.14402\noutside 1 331\nTMhelix 332 354\ninside 355 372

23397 GCF_000329505.1_ASM32950v1 Salmonella enterica subsp. enterica serovar Enteritidis str. CDC_2010K_0956 Proteobacteria; Gammaproteobacteria; Enterobacterales; Enterobacteriaceae; Salmonella; Salmonella enterica; Salmonella enterica subsp. enterica; Salmonella enterica subsp. enterica serovar Enteritidis MNNEETFYQAMRRKGVTRRSFLKFCSLAATSLGLGAGMTPKIAWALENKPRIPVVWIHGL WP_001058307.1 hydrogenase [Salmonella enterica] Length: 372\nNumber of predicted TMHs: 2\nExp number of AAs in TMHs: 36.51738\nExp number, first 60 AAs: 19.72849\nTotal prob of N-in: 0.96170\nPOSSIBLE N-term signal sequence\ninside 1 19\nTMhelix 20 39\noutside 40 325\nTMhelix 326 348\ninside 349 372

23398 GCF_000329365.2_ASM32936v2 Salmonella enterica subsp. enterica serovar Enteritidis str. CDC_2010K_0968 Proteobacteria; Gammaproteobacteria; Enterobacterales; Enterobacteriaceae; Salmonella; Salmonella enterica; Salmonella enterica subsp. enterica; Salmonella enterica subsp. enterica serovar Enteritidis MNNEETFYQAMRRKGVTRRSFLKFCSLAATSLGLGAGMTPKIAWALENKPRIPVVWIHGL WP_001058307.1 hydrogenase [Salmonella enterica] Length: 372\nNumber of predicted TMHs: 2\nExp number of AAs in TMHs: 36.51738\nExp number, first 60 AAs: 19.72849\nTotal prob of N-in: 0.96170\nPOSSIBLE N-term signal sequence\ninside 1 19\nTMhelix 20 39\noutside 40 325\nTMhelix 326 348\ninside 349 372

23399 GCF_000329365.2_ASM32936v2 Salmonella enterica subsp. enterica serovar Enteritidis str. CDC_2010K_0968 Proteobacteria; Gammaproteobacteria; Enterobacterales; Enterobacteriaceae; Salmonella; Salmonella enterica; Salmonella enterica subsp. enterica; Salmonella enterica subsp. enterica serovar Enteritidis MTGDNTLITSHGINRRDFMKLCAALAATMGLSSKAAAEMAESVSNPQRPPVIWIGAQECT WP_000145429.1 hydrogenase 2 small subunit [Salmonella enterica] Length: 372\nNumber of predicted TMHs: 1\nExp number of AAs in TMHs: 24.51309\nExp number, first 60 AAs: 0.2682\nTotal prob of N-in: 0.14402\noutside 1 331\nTMhelix 332 354\ninside 355 372

23400 GCF_000329245.1_ASM32924v1 Salmonella enterica subsp. enterica serovar Enteritidis str. CDC_2010K_1010 Proteobacteria; Gammaproteobacteria; Enterobacterales; Enterobacteriaceae; Salmonella; Salmonella

enterica; Salmonella enterica subsp. enterica; Salmonella enterica subsp. enterica serovar Enteritidis
MNNEETFYQAMRRKGVTRRSFLKFCSLAATSLGLGAGMTPKIAWALENKPRIPVVWIHGL WP_001058307.1
hydrogenase [Salmonella enterica] Length: 372\nNumber of predicted TMHs: 2\nExp number of AAs in TMHs: 36.51738\nExp number, first 60 AAs: 19.72849\nTotal prob of N-in: 0.96170\nPOSSIBLE N-term signal sequence\ninside 1 19\nTMhelix 20 39\noutside 40 325\nTMhelix 326 348\ninside 349 372

23401 GCF_000329245.1_ASM32924v1 Salmonella enterica subsp. enterica serovar Enteritidis str. CDC_2010K_1010 Proteobacteria; Gammaproteobacteria; Enterobacterales; Enterobacteriaceae; Salmonella; Salmonella enterica; Salmonella enterica subsp. enterica; Salmonella enterica subsp. enterica serovar Enteritidis
MTGDNTLITSHGINRRDFMKLCAALAATMGLSSKAAAEMAESVSNPQRPPVIWIGAQECT WP_000145429.1
hydrogenase 2 small subunit [Salmonella enterica] Length: 372\nNumber of predicted TMHs: 1\nExp number of AAs in TMHs: 24.51309\nExp number, first 60 AAs: 0.2682\nTotal prob of N-in: 0.14402\noutside 1 331\nTMhelix 332 354\ninside 355 372

23402 GCF_000329225.1_ASM32922v1 Salmonella enterica subsp. enterica serovar Enteritidis str. CDC_2010K_1018 Proteobacteria; Gammaproteobacteria; Enterobacterales; Enterobacteriaceae; Salmonella; Salmonella enterica; Salmonella enterica subsp. enterica; Salmonella enterica subsp. enterica serovar Enteritidis
MNNEETFYQAMRRKGVTRRSFLKFCSLAATSLGLGAGMTPKIAWALENKPRIPVVWIHGL WP_001058307.1
hydrogenase [Salmonella enterica] Length: 372\nNumber of predicted TMHs: 2\nExp number of AAs in TMHs: 36.51738\nExp number, first 60 AAs: 19.72849\nTotal prob of N-in: 0.96170\nPOSSIBLE N-term signal sequence\ninside 1 19\nTMhelix 20 39\noutside 40 325\nTMhelix 326 348\ninside 349 372

23403 GCF_000329225.1_ASM32922v1 Salmonella enterica subsp. enterica serovar Enteritidis str. CDC_2010K_1018 Proteobacteria; Gammaproteobacteria; Enterobacterales; Enterobacteriaceae; Salmonella; Salmonella enterica; Salmonella enterica subsp. enterica; Salmonella enterica subsp. enterica serovar Enteritidis
MTGDNTLITSHGINRRDFMKLCAALAATMGLSSKAAAEMAESVSNPQRPPVIWIGAQECT WP_000145429.1
hydrogenase 2 small subunit [Salmonella enterica] Length: 372\nNumber of predicted TMHs: 1\nExp number of AAs in TMHs: 24.51309\nExp number, first 60 AAs: 0.2682\nTotal prob of N-in: 0.14402\noutside 1 331\nTMhelix 332 354\ninside 355 372

23404 GCF_000329165.1_ASM32916v1 Salmonella enterica subsp. enterica serovar Enteritidis str. CDC_2010K_1441 Proteobacteria; Gammaproteobacteria; Enterobacterales; Enterobacteriaceae; Salmonella; Salmonella enterica; Salmonella enterica subsp. enterica; Salmonella enterica subsp. enterica serovar Enteritidis
MTGDNTLITSHGINRRDFMKLCAALAATMGLSSKAAAEMAESVSNPQRPPVIWIGAQECT WP_000145429.1
hydrogenase 2 small subunit [Salmonella enterica] Length: 372\nNumber of predicted TMHs: 1\nExp number of AAs in TMHs: 24.51309\nExp number, first 60 AAs: 0.2682\nTotal prob of N-in: 0.14402\noutside 1 331\nTMhelix 332 354\ninside 355 372

23405 GCF_000329165.1_ASM32916v1 Salmonella enterica subsp. enterica serovar Enteritidis str. CDC_2010K_1441 Proteobacteria; Gammaproteobacteria; Enterobacterales; Enterobacteriaceae; Salmonella; Salmonella enterica; Salmonella enterica subsp. enterica; Salmonella enterica subsp. enterica serovar Enteritidis
MNNEETFYQAMRRKGVTRRSFLKFCSLAATSLGLGAGMTPKIAWALENKPRIPVVWIHGL WP_001058307.1
hydrogenase [Salmonella enterica] Length: 372\nNumber of predicted TMHs: 2\nExp number of AAs in TMHs: 36.51738\nExp number, first 60 AAs: 19.72849\nTotal prob of N-in: 0.96170\nPOSSIBLE N-term signal sequence\ninside 1 19\nTMhelix 20 39\noutside 40 325\nTMhelix 326 348\ninside 349 372

23406 GCF_000329385.1_ASM32938v1 Salmonella enterica subsp. enterica serovar Enteritidis str. CDC_2010K_1444 Proteobacteria; Gammaproteobacteria; Enterobacterales; Enterobacteriaceae; Salmonella; Salmonella enterica; Salmonella enterica subsp. enterica; Salmonella enterica subsp. enterica serovar Enteritidis
MTGDNTLITSHGINRRDFMKLCAALAATMGLSSKAAAEMAESVSNPQRPPVIWIGAQECT WP_000145429.1
hydrogenase 2 small subunit [Salmonella enterica] Length: 372\nNumber of predicted TMHs: 1\nExp number of AAs in TMHs: 24.51309\nExp number, first 60 AAs: 0.2682\nTotal prob of N-in: 0.14402\noutside 1 331\nTMhelix 332 354\ninside 355 372

23407 GCF_000329385.1_ASM32938v1 Salmonella enterica subsp. enterica serovar Enteritidis str. CDC_2010K_1444 Proteobacteria; Gammaproteobacteria; Enterobacterales; Enterobacteriaceae; Salmonella; Salmonella enterica; Salmonella enterica subsp. enterica; Salmonella enterica subsp. enterica serovar Enteritidis
MNNEETFYQAMRRKGVTRRSFLKFCSLAATSLGLGAGMTPKIAWALENKPRIPVVWIHGL WP_001058307.1
hydrogenase [Salmonella enterica] Length: 372\nNumber of predicted TMHs: 2\nExp number of AAs in TMHs: 36.51738\nExp number, first 60 AAs: 19.72849\nTotal prob of N-in: 0.96170\nPOSSIBLE N-term signal sequence\ninside 1 19\nTMhelix 20 39\noutside 40 325\nTMhelix 326 348\ninside 349 372

23408 GCF_000329405.1_ASM32940v1 Salmonella enterica subsp. enterica serovar Enteritidis str. CDC_2010K_1445 Proteobacteria; Gammaproteobacteria; Enterobacterales; Enterobacteriaceae; Salmonella; Salmonella enterica; Salmonella enterica subsp. enterica; Salmonella enterica subsp. enterica serovar Enteritidis MNNEETFYQAMRRKGVTRRSFLKFCSLAATSLGLGAGMTPKIAWALENKPRIPVVWIHGL WP_001058307.1 hydrogenase [Salmonella enterica] Length: 372\nNumber of predicted TMHs: 2\nExp number of AAs in TMHs: 36.51738\nExp number, first 60 AAs: 19.72849\nTotal prob of N-in: 0.96170\nPOSSIBLE N-term signal sequence\ninside 1 19\nTMhelix 20 39\noutside 40 325\nTMhelix 326 348\ninside 349 372

23409 GCF_000329405.1_ASM32940v1 Salmonella enterica subsp. enterica serovar Enteritidis str. CDC_2010K_1445 Proteobacteria; Gammaproteobacteria; Enterobacterales; Enterobacteriaceae; Salmonella; Salmonella enterica; Salmonella enterica subsp. enterica; Salmonella enterica subsp. enterica serovar Enteritidis MTGDNTLITSHGINRRDFMKLCAALAATMGLSSKAAAEMAESVSNPQRPPVIWIGAQECT WP_000145429.1 hydrogenase 2 small subunit [Salmonella enterica] Length: 372\nNumber of predicted TMHs: 1\nExp number of AAs in TMHs: 24.51309\nExp number, first 60 AAs: 0.2682\nTotal prob of N-in: 0.14402\noutside 1 331\nTMhelix 332 354\ninside 355 372

23410 GCF_000329525.1_ASM32952v1 Salmonella enterica subsp. enterica serovar Enteritidis str. CDC_2010K_1455 Proteobacteria; Gammaproteobacteria; Enterobacterales; Enterobacteriaceae; Salmonella; Salmonella enterica; Salmonella enterica subsp. enterica; Salmonella enterica subsp. enterica serovar Enteritidis MNNEETFYQAMRRKGVTRRSFLKFCSLAATSLGLGAGMTPKIAWALENKPRIPVVWIHGL WP_001058307.1 hydrogenase [Salmonella enterica] Length: 372\nNumber of predicted TMHs: 2\nExp number of AAs in TMHs: 36.51738\nExp number, first 60 AAs: 19.72849\nTotal prob of N-in: 0.96170\nPOSSIBLE N-term signal sequence\ninside 1 19\nTMhelix 20 39\noutside 40 325\nTMhelix 326 348\ninside 349 372

23411 GCF_000329525.1_ASM32952v1 Salmonella enterica subsp. enterica serovar Enteritidis str. CDC_2010K_1455 Proteobacteria; Gammaproteobacteria; Enterobacterales; Enterobacteriaceae; Salmonella; Salmonella enterica; Salmonella enterica subsp. enterica; Salmonella enterica subsp. enterica serovar Enteritidis MTGDNTLITSHGINRRDFMKLCAALAATMGLSSKAAAEMAESVSNPQRPPVIWIGAQECT WP_000145429.1 hydrogenase 2 small subunit [Salmonella enterica] Length: 372\nNumber of predicted TMHs: 1\nExp number of AAs in TMHs: 24.51309\nExp number, first 60 AAs: 0.2682\nTotal prob of N-in: 0.14402\noutside 1 331\nTMhelix 332 354\ninside 355 372

23412 GCF_000329325.1_ASM32932v1 Salmonella enterica subsp. enterica serovar Enteritidis str. CDC_2010K_1457 Proteobacteria; Gammaproteobacteria; Enterobacterales; Enterobacteriaceae; Salmonella; Salmonella enterica; Salmonella enterica subsp. enterica; Salmonella enterica subsp. enterica serovar Enteritidis MTGDNTLITSHGINRRDFMKLCAALAATMGLSSKAAAEMAESVSNPQRPPVIWIGAQECT WP_000145429.1 hydrogenase 2 small subunit [Salmonella enterica] Length: 372\nNumber of predicted TMHs: 1\nExp number of AAs in TMHs: 24.51309\nExp number, first 60 AAs: 0.2682\nTotal prob of N-in: 0.14402\noutside 1 331\nTMhelix 332 354\ninside 355 372

23413 GCF_000329325.1_ASM32932v1 Salmonella enterica subsp. enterica serovar Enteritidis str. CDC_2010K_1457 Proteobacteria; Gammaproteobacteria; Enterobacterales; Enterobacteriaceae; Salmonella; Salmonella enterica; Salmonella enterica subsp. enterica; Salmonella enterica subsp. enterica serovar Enteritidis MNNEETFYQAMRRKGVTRRSFLKFCSLAATSLGLGAGMTPKIAWALENKPRIPVVWIHGL WP_001058307.1 hydrogenase [Salmonella enterica] Length: 372\nNumber of predicted TMHs: 2\nExp number of AAs in TMHs: 36.51738\nExp number, first 60 AAs: 19.72849\nTotal prob of N-in: 0.96170\nPOSSIBLE N-term signal sequence\ninside 1 19\nTMhelix 20 39\noutside 40 325\nTMhelix 326 348\ninside 349 372

23414 GCF_000329145.1_ASM32914v1 Salmonella enterica subsp. enterica serovar Enteritidis str. CDC_2010K_1543 Proteobacteria; Gammaproteobacteria; Enterobacterales; Enterobacteriaceae; Salmonella; Salmonella enterica; Salmonella enterica subsp. enterica; Salmonella enterica subsp. enterica serovar Enteritidis MNNEETFYQAMRRKGVTRRSFLKFCSLAATSLGLGAGMTPKIAWALENKPRIPVVWIHGL WP_001058307.1 hydrogenase [Salmonella enterica] Length: 372\nNumber of predicted TMHs: 2\nExp number of AAs in TMHs: 36.51738\nExp number, first 60 AAs: 19.72849\nTotal prob of N-in: 0.96170\nPOSSIBLE N-term signal sequence\ninside 1 19\nTMhelix 20 39\noutside 40 325\nTMhelix 326 348\ninside 349 372

23415 GCF_000329145.1_ASM32914v1 Salmonella enterica subsp. enterica serovar Enteritidis str. CDC_2010K_1543 Proteobacteria; Gammaproteobacteria; Enterobacterales; Enterobacteriaceae; Salmonella; Salmonella enterica; Salmonella enterica subsp. enterica; Salmonella enterica subsp. enterica serovar Enteritidis MTGDNTLITSHGINRRDFMKLCAALAATMGLSSKAAAEMAESVSNPQRPPVIWIGAQECT WP_000145429.1 hydrogenase 2 small subunit [Salmonella enterica] Length: 372\nNumber of predicted TMHs: 1\nExp

number of AAs in TMHs: 24.51309\nExp number, first 60 AAs: 0.2682\nTotal prob of N-in: 0.14402\noutside 1 331\nTMhelix 332 354\ninside 355 372

23416 GCF_000329205.1_ASM32920v1 Salmonella enterica subsp. enterica serovar Enteritidis str. CDC_2010K_1558 Proteobacteria; Gammaproteobacteria; Enterobacterales; Enterobacteriaceae; Salmonella; Salmonella enterica; Salmonella enterica subsp. enterica; Salmonella enterica subsp. enterica serovar Enteritidis MNNEETFYQAMRRKGVTRRSFLKFCSLAATSLGLGAGMTPKIAWALENKPRIPVVWIHGL WP_001058307.1 hydrogenase [Salmonella enterica] Length: 372\nNumber of predicted TMHs: 2\nExp number of AAs in TMHs: 36.51738\nExp number, first 60 AAs: 19.72849\nTotal prob of N-in: 0.96170\nPOSSIBLE N-term signal sequence\ninside 1 19\nTMhelix 20 39\noutside 40 325\nTMhelix 326 348\ninside 349 372

23417 GCF_000329205.1_ASM32920v1 Salmonella enterica subsp. enterica serovar Enteritidis str. CDC_2010K_1558 Proteobacteria; Gammaproteobacteria; Enterobacterales; Enterobacteriaceae; Salmonella; Salmonella enterica; Salmonella enterica subsp. enterica; Salmonella enterica subsp. enterica serovar Enteritidis MTGDNTLITSHGINRRDFMKLCAALAATMGLSSKAAAEMAESVSNPQRPPVIWIGAQECT WP_000145429.1 hydrogenase 2 small subunit [Salmonella enterica] Length: 372\nNumber of predicted TMHs: 1\nExp number of AAs in TMHs: 24.51309\nExp number, first 60 AAs: 0.2682\nTotal prob of N-in: 0.14402\noutside 1 331\nTMhelix 332 354\ninside 355 372

23418 GCF_000329425.1_ASM32942v1 Salmonella enterica subsp. enterica serovar Enteritidis str. CDC_2010K_1559 Proteobacteria; Gammaproteobacteria; Enterobacterales; Enterobacteriaceae; Salmonella; Salmonella enterica; Salmonella enterica subsp. enterica; Salmonella enterica subsp. enterica serovar Enteritidis MTGDNTLITSHGINRRDFMKLCAALAATMGLSSKAAAEMAESVSNPQRPPVIWIGAQECT WP_000145429.1 hydrogenase 2 small subunit [Salmonella enterica] Length: 372\nNumber of predicted TMHs: 1\nExp number of AAs in TMHs: 24.51309\nExp number, first 60 AAs: 0.2682\nTotal prob of N-in: 0.14402\noutside 1 331\nTMhelix 332 354\ninside 355 372

23419 GCF_000329425.1_ASM32942v1 Salmonella enterica subsp. enterica serovar Enteritidis str. CDC_2010K_1559 Proteobacteria; Gammaproteobacteria; Enterobacterales; Enterobacteriaceae; Salmonella; Salmonella enterica; Salmonella enterica subsp. enterica; Salmonella enterica subsp. enterica serovar Enteritidis MNNEETFYQAMRRKGVTRRSFLKFCSLAATSLGLGAGMTPKIAWALENKPRIPVVWIHGL WP_001058307.1 hydrogenase [Salmonella enterica] Length: 372\nNumber of predicted TMHs: 2\nExp number of AAs in TMHs: 36.51738\nExp number, first 60 AAs: 19.72849\nTotal prob of N-in: 0.96170\nPOSSIBLE N-term signal sequence\ninside 1 19\nTMhelix 20 39\noutside 40 325\nTMhelix 326 348\ninside 349 372

23420 GCF_000329445.1_ASM32944v1 Salmonella enterica subsp. enterica serovar Enteritidis str. CDC_2010K_1565 Proteobacteria; Gammaproteobacteria; Enterobacterales; Enterobacteriaceae; Salmonella; Salmonella enterica; Salmonella enterica subsp. enterica; Salmonella enterica subsp. enterica serovar Enteritidis MTGDNTLITSHGINRRDFMKLCAALAATMGLSSKAAAEMAESVSNPQRPPVIWIGAQECT WP_000145429.1 hydrogenase 2 small subunit [Salmonella enterica] Length: 372\nNumber of predicted TMHs: 1\nExp number of AAs in TMHs: 24.51309\nExp number, first 60 AAs: 0.2682\nTotal prob of N-in: 0.14402\noutside 1 331\nTMhelix 332 354\ninside 355 372

23421 GCF_000329445.1_ASM32944v1 Salmonella enterica subsp. enterica serovar Enteritidis str. CDC_2010K_1565 Proteobacteria; Gammaproteobacteria; Enterobacterales; Enterobacteriaceae; Salmonella; Salmonella enterica; Salmonella enterica subsp. enterica; Salmonella enterica subsp. enterica serovar Enteritidis MNNEETFYQAMRRKGVTRRSFLKFCSLAATSLGLGAGMTPKIAWALENKPRIPVVWIHGL WP_001058307.1 hydrogenase [Salmonella enterica] Length: 372\nNumber of predicted TMHs: 2\nExp number of AAs in TMHs: 36.51738\nExp number, first 60 AAs: 19.72849\nTotal prob of N-in: 0.96170\nPOSSIBLE N-term signal sequence\ninside 1 19\nTMhelix 20 39\noutside 40 325\nTMhelix 326 348\ninside 349 372

23422 GCF_000329105.1_ASM32910v1 Salmonella enterica subsp. enterica serovar Enteritidis str. CDC_2010K_1566 Proteobacteria; Gammaproteobacteria; Enterobacterales; Enterobacteriaceae; Salmonella; Salmonella enterica; Salmonella enterica subsp. enterica; Salmonella enterica subsp. enterica serovar Enteritidis MTGDNTLITSHGINRRDFMKLCAALAATMGLSSKAAAEMAESVSNPQRPPVIWIGAQECT WP_000145429.1 hydrogenase 2 small subunit [Salmonella enterica] Length: 372\nNumber of predicted TMHs: 1\nExp number of AAs in TMHs: 24.51309\nExp number, first 60 AAs: 0.2682\nTotal prob of N-in: 0.14402\noutside 1 331\nTMhelix 332 354\ninside 355 372

23423 GCF_000329105.1_ASM32910v1 Salmonella enterica subsp. enterica serovar Enteritidis str. CDC_2010K_1566 Proteobacteria; Gammaproteobacteria; Enterobacterales; Enterobacteriaceae; Salmonella; Salmonella

enterica; Salmonella enterica subsp. enterica; Salmonella enterica subsp. enterica serovar Enteritidis
MNNEETFYQAMRRKGVTRRSFLKFCSLAATSLGLGAGMTPKIAWALENKPRIPVVWIHGL WP_001058307.1
hydrogenase [Salmonella enterica] Length: 372\nNumber of predicted TMHs: 2\nExp number of AAs in TMHs: 36.51738\nExp number, first 60 AAs: 19.72849\nTotal prob of N-in: 0.96170\nPOSSIBLE N-term signal sequence\ninside 1 19\nTMhelix 20 39\noutside 40 325\nTMhelix 326 348\ninside 349 372

23424 GCF_000329545.1_ASM32954v1 Salmonella enterica subsp. enterica serovar Enteritidis str. CDC_2010K_1575 Proteobacteria; Gammaproteobacteria; Enterobacterales; Enterobacteriaceae; Salmonella; Salmonella enterica; Salmonella enterica subsp. enterica; Salmonella enterica subsp. enterica serovar Enteritidis
MNNEETFYQAMRRKGVTRRSFLKFCSLAATSLGLGAGMTPKIAWALENKPRIPVVWIHGL WP_001058307.1
hydrogenase [Salmonella enterica] Length: 372\nNumber of predicted TMHs: 2\nExp number of AAs in TMHs: 36.51738\nExp number, first 60 AAs: 19.72849\nTotal prob of N-in: 0.96170\nPOSSIBLE N-term signal sequence\ninside 1 19\nTMhelix 20 39\noutside 40 325\nTMhelix 326 348\ninside 349 372

23425 GCF_000329545.1_ASM32954v1 Salmonella enterica subsp. enterica serovar Enteritidis str. CDC_2010K_1575 Proteobacteria; Gammaproteobacteria; Enterobacterales; Enterobacteriaceae; Salmonella; Salmonella enterica; Salmonella enterica subsp. enterica; Salmonella enterica subsp. enterica serovar Enteritidis
MTGDNTLITSHGINRRDFMKLCAALAATMGLSSKAAAEMAESVSNPQRPPVIWIGAQECT WP_000145429.1
hydrogenase 2 small subunit [Salmonella enterica] Length: 372\nNumber of predicted TMHs: 1\nExp number of AAs in TMHs: 24.51309\nExp number, first 60 AAs: 0.2682\nTotal prob of N-in: 0.14402\noutside 1 331\nTMhelix 332 354\ninside 355 372

23426 GCF_000329125.1_ASM32912v1 Salmonella enterica subsp. enterica serovar Enteritidis str. CDC_2010K_1580 Proteobacteria; Gammaproteobacteria; Enterobacterales; Enterobacteriaceae; Salmonella; Salmonella enterica; Salmonella enterica subsp. enterica; Salmonella enterica subsp. enterica serovar Enteritidis
MNNEETFYQAMRRKGVTRRSFLKFCSLAATSLGLGAGMTPKIAWALENKPRIPVVWIHGL WP_001058307.1
hydrogenase [Salmonella enterica] Length: 372\nNumber of predicted TMHs: 2\nExp number of AAs in TMHs: 36.51738\nExp number, first 60 AAs: 19.72849\nTotal prob of N-in: 0.96170\nPOSSIBLE N-term signal sequence\ninside 1 19\nTMhelix 20 39\noutside 40 325\nTMhelix 326 348\ninside 349 372

23427 GCF_000329125.1_ASM32912v1 Salmonella enterica subsp. enterica serovar Enteritidis str. CDC_2010K_1580 Proteobacteria; Gammaproteobacteria; Enterobacterales; Enterobacteriaceae; Salmonella; Salmonella enterica; Salmonella enterica subsp. enterica; Salmonella enterica subsp. enterica serovar Enteritidis
MTGDNTLITSHGINRRDFMKLCAALAATMGLSSKAAAEMAESVSNPQRPPVIWIGAQECT WP_000145429.1
hydrogenase 2 small subunit [Salmonella enterica] Length: 372\nNumber of predicted TMHs: 1\nExp number of AAs in TMHs: 24.51309\nExp number, first 60 AAs: 0.2682\nTotal prob of N-in: 0.14402\noutside 1 331\nTMhelix 332 354\ninside 355 372

23428 GCF_000329085.1_ASM32908v1 Salmonella enterica subsp. enterica serovar Enteritidis str. CDC_2010K_1594 Proteobacteria; Gammaproteobacteria; Enterobacterales; Enterobacteriaceae; Salmonella; Salmonella enterica; Salmonella enterica subsp. enterica; Salmonella enterica subsp. enterica serovar Enteritidis
MTGDNTLITSHGINRRDFMKLCAALAATMGLSSKAAAEMAESVSNPQRPPVIWIGAQECT WP_000145429.1
hydrogenase 2 small subunit [Salmonella enterica] Length: 372\nNumber of predicted TMHs: 1\nExp number of AAs in TMHs: 24.51309\nExp number, first 60 AAs: 0.2682\nTotal prob of N-in: 0.14402\noutside 1 331\nTMhelix 332 354\ninside 355 372

23429 GCF_000329085.1_ASM32908v1 Salmonella enterica subsp. enterica serovar Enteritidis str. CDC_2010K_1594 Proteobacteria; Gammaproteobacteria; Enterobacterales; Enterobacteriaceae; Salmonella; Salmonella enterica; Salmonella enterica subsp. enterica; Salmonella enterica subsp. enterica serovar Enteritidis
MNNEETFYQAMRRKGVTRRSFLKFCSLAATSLGLGAGMTPKIAWALENKPRIPVVWIHGL WP_001058307.1
hydrogenase [Salmonella enterica] Length: 372\nNumber of predicted TMHs: 2\nExp number of AAs in TMHs: 36.51738\nExp number, first 60 AAs: 19.72849\nTotal prob of N-in: 0.96170\nPOSSIBLE N-term signal sequence\ninside 1 19\nTMhelix 20 39\noutside 40 325\nTMhelix 326 348\ninside 349 372

23430 GCF_000329565.1_ASM32956v1 Salmonella enterica subsp. enterica serovar Enteritidis str. CDC_2010K_1725 Proteobacteria; Gammaproteobacteria; Enterobacterales; Enterobacteriaceae; Salmonella; Salmonella enterica; Salmonella enterica subsp. enterica; Salmonella enterica subsp. enterica serovar Enteritidis
MTGDNTLITSHGINRRDFMKLCAALAATMGLSSKAAAEMAESVSNPQRPPVIWIGAQECT WP_000145429.1
hydrogenase 2 small subunit [Salmonella enterica] Length: 372\nNumber of predicted TMHs: 1\nExp number of AAs in TMHs: 24.51309\nExp number, first 60 AAs: 0.2682\nTotal prob of N-in: 0.14402\noutside 1 331\nTMhelix 332 354\ninside 355 372

23431 GCF_000329565.1_ASM32956v1 Salmonella enterica subsp. enterica serovar Enteritidis str. CDC_2010K_1725 Proteobacteria; Gammaproteobacteria; Enterobacterales; Enterobacteriaceae; Salmonella; Salmonella enterica; Salmonella enterica subsp. enterica; Salmonella enterica subsp. enterica serovar Enteritidis
MNNEETFYQAMRRKGVTRRSFLKFCSLAATSLGLGAGMTPKIAWALENKPRIPVVWIHGL WP_001058307.1
hydrogenase [Salmonella enterica] Length: 372\nNumber of predicted TMHs: 2\nExp number of AAs in TMHs: 36.51738\nExp number, first 60 AAs: 19.72849\nTotal prob of N-in: 0.96170\nPOSSIBLE N-term signal sequence\ninside 1 19\nTMhelix 20 39\noutside 40 325\nTMhelix 326 348\ninside 349 372

23432 GCF_000329265.1_ASM32926v1 Salmonella enterica subsp. enterica serovar Enteritidis str. CDC_2010K_1729 Proteobacteria; Gammaproteobacteria; Enterobacterales; Enterobacteriaceae; Salmonella; Salmonella enterica; Salmonella enterica subsp. enterica; Salmonella enterica subsp. enterica serovar Enteritidis
MNNEETFYQAMRRKGVTRRSFLKFCSLAATSLGLGAGMTPKIAWALENKPRIPVVWIHGL WP_001058307.1
hydrogenase [Salmonella enterica] Length: 372\nNumber of predicted TMHs: 2\nExp number of AAs in TMHs: 36.51738\nExp number, first 60 AAs: 19.72849\nTotal prob of N-in: 0.96170\nPOSSIBLE N-term signal sequence\ninside 1 19\nTMhelix 20 39\noutside 40 325\nTMhelix 326 348\ninside 349 372

23433 GCF_000329265.1_ASM32926v1 Salmonella enterica subsp. enterica serovar Enteritidis str. CDC_2010K_1729 Proteobacteria; Gammaproteobacteria; Enterobacterales; Enterobacteriaceae; Salmonella; Salmonella enterica; Salmonella enterica subsp. enterica; Salmonella enterica subsp. enterica serovar Enteritidis
MTGDNTLITSHGINRRDFMKLCAALAATMGLSSKAAAEMAESVSNPQRPPVIWIGAQECT WP_000145429.1
hydrogenase 2 small subunit [Salmonella enterica] Length: 372\nNumber of predicted TMHs: 1\nExp number of AAs in TMHs: 24.51309\nExp number, first 60 AAs: 0.2682\nTotal prob of N-in: 0.14402\noutside 1 331\nTMhelix 332 354\ninside 355 372

23434 GCF_000329585.1_ASM32958v1 Salmonella enterica subsp. enterica serovar Enteritidis str. CDC_2010K_1745 Proteobacteria; Gammaproteobacteria; Enterobacterales; Enterobacteriaceae; Salmonella; Salmonella enterica; Salmonella enterica subsp. enterica; Salmonella enterica subsp. enterica serovar Enteritidis
MTGDNTLITSHGINRRDFMKLCAALAATMGLSSKAAAEMAESVSNPQRPPVIWIGAQECT WP_000145429.1
hydrogenase 2 small subunit [Salmonella enterica] Length: 372\nNumber of predicted TMHs: 1\nExp number of AAs in TMHs: 24.51309\nExp number, first 60 AAs: 0.2682\nTotal prob of N-in: 0.14402\noutside 1 331\nTMhelix 332 354\ninside 355 372

23435 GCF_000329585.1_ASM32958v1 Salmonella enterica subsp. enterica serovar Enteritidis str. CDC_2010K_1745 Proteobacteria; Gammaproteobacteria; Enterobacterales; Enterobacteriaceae; Salmonella; Salmonella enterica; Salmonella enterica subsp. enterica; Salmonella enterica subsp. enterica serovar Enteritidis
MNNEETFYQAMRRKGVTRRSFLKFCSLAATSLGLGAGMTPKIAWALENKPRIPVVWIHGL WP_001058307.1
hydrogenase [Salmonella enterica] Length: 372\nNumber of predicted TMHs: 2\nExp number of AAs in TMHs: 36.51738\nExp number, first 60 AAs: 19.72849\nTotal prob of N-in: 0.96170\nPOSSIBLE N-term signal sequence\ninside 1 19\nTMhelix 20 39\noutside 40 325\nTMhelix 326 348\ninside 349 372

23436 GCF_000329345.1_ASM32934v1 Salmonella enterica subsp. enterica serovar Enteritidis str. CDC_2010K_1747 Proteobacteria; Gammaproteobacteria; Enterobacterales; Enterobacteriaceae; Salmonella; Salmonella enterica; Salmonella enterica subsp. enterica; Salmonella enterica subsp. enterica serovar Enteritidis
MNNEETFYQAMRRKGVTRRSFLKFCSLAATSLGLGAGMTPKIAWALENKPRIPVVWIHGL WP_001058307.1
hydrogenase [Salmonella enterica] Length: 372\nNumber of predicted TMHs: 2\nExp number of AAs in TMHs: 36.51738\nExp number, first 60 AAs: 19.72849\nTotal prob of N-in: 0.96170\nPOSSIBLE N-term signal sequence\ninside 1 19\nTMhelix 20 39\noutside 40 325\nTMhelix 326 348\ninside 349 372

23437 GCF_000329345.1_ASM32934v1 Salmonella enterica subsp. enterica serovar Enteritidis str. CDC_2010K_1747 Proteobacteria; Gammaproteobacteria; Enterobacterales; Enterobacteriaceae; Salmonella; Salmonella enterica; Salmonella enterica subsp. enterica; Salmonella enterica subsp. enterica serovar Enteritidis
MTGDNTLITSHGINRRDFMKLCAALAATMGLSSKAAAEMAESVSNPQRPPVIWIGAQECT WP_000145429.1
hydrogenase 2 small subunit [Salmonella enterica] Length: 372\nNumber of predicted TMHs: 1\nExp number of AAs in TMHs: 24.51309\nExp number, first 60 AAs: 0.2682\nTotal prob of N-in: 0.14402\noutside 1 331\nTMhelix 332 354\ninside 355 372

23438 GCF_000329605.1_ASM32960v1 Salmonella enterica subsp. enterica serovar Enteritidis str. CDC_2010K_1791 Proteobacteria; Gammaproteobacteria; Enterobacterales; Enterobacteriaceae; Salmonella; Salmonella enterica; Salmonella enterica subsp. enterica; Salmonella enterica subsp. enterica serovar Enteritidis
MNNEETFYQAMRRKGVTRRSFLKFCSLAATSLGLGAGMTPKIAWALENKPRIPVVWIHGL WP_001058307.1
hydrogenase [Salmonella enterica] Length: 372\nNumber of predicted TMHs: 2\nExp number of AAs in

TMHs: 36.51738\nExp number, first 60 AAs: 19.72849\nTotal prob of N-in: 0.96170\nPOSSIBLE N-term signal sequence\ninside 1 19\nTMhelix 20 39\noutside 40 325\nTMhelix 326 348\ninside 349 372

23439 GCF_000329605.1_ASM32960v1 Salmonella enterica subsp. enterica serovar Enteritidis str. CDC_2010K_1791 Proteobacteria; Gammaproteobacteria; Enterobacterales; Enterobacteriaceae; Salmonella; Salmonella enterica; Salmonella enterica subsp. enterica; Salmonella enterica subsp. enterica serovar Enteritidis MTGDNTLITSHGINRRDFMKLCAALAATMGLSSKAAAEMAESVSNPQRPPVIWIGAQECT WP_000145429.1 hydrogenase 2 small subunit [Salmonella enterica] Length: 372\nNumber of predicted TMHs: 1\nExp number of AAs in TMHs: 24.51309\nExp number, first 60 AAs: 0.2682\nTotal prob of N-in: 0.14402\noutside 1 331\nTMhelix 332 354\ninside 355 372

23440 GCF_000329625.1_ASM32962v1 Salmonella enterica subsp. enterica serovar Enteritidis str. CDC_2010K_1795 Proteobacteria; Gammaproteobacteria; Enterobacterales; Enterobacteriaceae; Salmonella; Salmonella enterica; Salmonella enterica subsp. enterica; Salmonella enterica subsp. enterica serovar Enteritidis MTGDNTLITSHGINRRDFMKLCAALAATMGLSSKAAAEMAESVSNPQRPPVIWIGAQECT WP_000145429.1 hydrogenase 2 small subunit [Salmonella enterica] Length: 372\nNumber of predicted TMHs: 1\nExp number of AAs in TMHs: 24.51309\nExp number, first 60 AAs: 0.2682\nTotal prob of N-in: 0.14402\noutside 1 331\nTMhelix 332 354\ninside 355 372

23441 GCF_000329625.1_ASM32962v1 Salmonella enterica subsp. enterica serovar Enteritidis str. CDC_2010K_1795 Proteobacteria; Gammaproteobacteria; Enterobacterales; Enterobacteriaceae; Salmonella; Salmonella enterica; Salmonella enterica subsp. enterica; Salmonella enterica subsp. enterica serovar Enteritidis MNNEETFYQAMRRKGVTRRSFLKFCSLAATSLGLGAGMTPKIAWALENKPRIPVVWIHGL WP_001058307.1 hydrogenase [Salmonella enterica] Length: 372\nNumber of predicted TMHs: 2\nExp number of AAs in TMHs: 36.51738\nExp number, first 60 AAs: 19.72849\nTotal prob of N-in: 0.96170\nPOSSIBLE N-term signal sequence\ninside 1 19\nTMhelix 20 39\noutside 40 325\nTMhelix 326 348\ninside 349 372

23442 GCF_000329465.1_ASM32946v1 Salmonella enterica subsp. enterica serovar Enteritidis str. CDC_2010K_1808 Proteobacteria; Gammaproteobacteria; Enterobacterales; Enterobacteriaceae; Salmonella; Salmonella enterica; Salmonella enterica subsp. enterica; Salmonella enterica subsp. enterica serovar Enteritidis MTGDNTLITSHGINRRDFMKLCAALAATMGLSSKAAAEMAESVSNPQRPPVIWIGAQECT WP_000145429.1 hydrogenase 2 small subunit [Salmonella enterica] Length: 372\nNumber of predicted TMHs: 1\nExp number of AAs in TMHs: 24.51309\nExp number, first 60 AAs: 0.2682\nTotal prob of N-in: 0.14402\noutside 1 331\nTMhelix 332 354\ninside 355 372

23443 GCF_000329465.1_ASM32946v1 Salmonella enterica subsp. enterica serovar Enteritidis str. CDC_2010K_1808 Proteobacteria; Gammaproteobacteria; Enterobacterales; Enterobacteriaceae; Salmonella; Salmonella enterica; Salmonella enterica subsp. enterica; Salmonella enterica subsp. enterica serovar Enteritidis MNNEETFYQAMRRKGVTRRSFLKFCSLAATSLGLGAGMTPKIAWALENKPRIPVVWIHGL WP_001058307.1 hydrogenase [Salmonella enterica] Length: 372\nNumber of predicted TMHs: 2\nExp number of AAs in TMHs: 36.51738\nExp number, first 60 AAs: 19.72849\nTotal prob of N-in: 0.96170\nPOSSIBLE N-term signal sequence\ninside 1 19\nTMhelix 20 39\noutside 40 325\nTMhelix 326 348\ninside 349 372

23444 GCF_000329185.1_ASM32918v1 Salmonella enterica subsp. enterica serovar Enteritidis str. CDC_2010K_1810 Proteobacteria; Gammaproteobacteria; Enterobacterales; Enterobacteriaceae; Salmonella; Salmonella enterica; Salmonella enterica subsp. enterica; Salmonella enterica subsp. enterica serovar Enteritidis MTGDNTLITSHGINRRDFMKLCAALAATMGLSSKAAAEMAESVSNPQRPPVIWIGAQECT WP_000145429.1 hydrogenase 2 small subunit [Salmonella enterica] Length: 372\nNumber of predicted TMHs: 1\nExp number of AAs in TMHs: 24.51309\nExp number, first 60 AAs: 0.2682\nTotal prob of N-in: 0.14402\noutside 1 331\nTMhelix 332 354\ninside 355 372

23445 GCF_000329185.1_ASM32918v1 Salmonella enterica subsp. enterica serovar Enteritidis str. CDC_2010K_1810 Proteobacteria; Gammaproteobacteria; Enterobacterales; Enterobacteriaceae; Salmonella; Salmonella enterica; Salmonella enterica subsp. enterica; Salmonella enterica subsp. enterica serovar Enteritidis MNNEETFYQAMRRKGVTRRSFLKFCSLAATSLGLGAGMTPKIAWALENKPRIPVVWIHGL WP_001058307.1 hydrogenase [Salmonella enterica] Length: 372\nNumber of predicted TMHs: 2\nExp number of AAs in TMHs: 36.51738\nExp number, first 60 AAs: 19.72849\nTotal prob of N-in: 0.96170\nPOSSIBLE N-term signal sequence\ninside 1 19\nTMhelix 20 39\noutside 40 325\nTMhelix 326 348\ninside 349 372

23446 GCF_000329485.1_ASM32948v1 Salmonella enterica subsp. enterica serovar Enteritidis str. CDC_2010K_1811 Proteobacteria; Gammaproteobacteria; Enterobacterales; Enterobacteriaceae; Salmonella; Salmonella

enterica; Salmonella enterica subsp. enterica; Salmonella enterica subsp. enterica serovar Enteritidis
MTGDNTLITSHGINRRDFMKLCAALAATMGLSSKAAAEMAESVSNPQRPPVIWIGAQECT WP_000145429.1
hydrogenase 2 small subunit [Salmonella enterica] Length: 372\nNumber of predicted TMHs: 1\nExp number of AAs in TMHs: 24.51309\nExp number, first 60 AAs: 0.2682\nTotal prob of N-in: 0.14402\noutside 1 331\nTMhelix 332 354\ninside 355 372

23447 GCF_000329485.1_ASM32948v1 Salmonella enterica subsp. enterica serovar Enteritidis str. CDC_2010K_1811 Proteobacteria; Gammaproteobacteria; Enterobacterales; Enterobacteriaceae; Salmonella; Salmonella enterica; Salmonella enterica subsp. enterica; Salmonella enterica subsp. enterica serovar Enteritidis
MNNEETFYQAMRRKGVTRRSFLKFCSLAATSLGLGAGMTPKIAWALENKPRIPVVWIHGL WP_001058307.1
hydrogenase [Salmonella enterica] Length: 372\nNumber of predicted TMHs: 2\nExp number of AAs in TMHs: 36.51738\nExp number, first 60 AAs: 19.72849\nTotal prob of N-in: 0.96170\nPOSSIBLE N-term signal sequence\ninside 1 19\nTMhelix 20 39\noutside 40 325\nTMhelix 326 348\ninside 349 372

23448 GCF_000329045.1_ASM32904v1 Salmonella enterica subsp. enterica serovar Enteritidis str. CDC_2010K_1882 Proteobacteria; Gammaproteobacteria; Enterobacterales; Enterobacteriaceae; Salmonella; Salmonella enterica; Salmonella enterica subsp. enterica; Salmonella enterica subsp. enterica serovar Enteritidis
MNNEETFYQAMRRKGVTRRSFLKFCSLAATSLGLGAGMTPKIAWALENKPRIPVVWIHGL WP_001058307.1
hydrogenase [Salmonella enterica] Length: 372\nNumber of predicted TMHs: 2\nExp number of AAs in TMHs: 36.51738\nExp number, first 60 AAs: 19.72849\nTotal prob of N-in: 0.96170\nPOSSIBLE N-term signal sequence\ninside 1 19\nTMhelix 20 39\noutside 40 325\nTMhelix 326 348\ninside 349 372

23449 GCF_000329045.1_ASM32904v1 Salmonella enterica subsp. enterica serovar Enteritidis str. CDC_2010K_1882 Proteobacteria; Gammaproteobacteria; Enterobacterales; Enterobacteriaceae; Salmonella; Salmonella enterica; Salmonella enterica subsp. enterica; Salmonella enterica subsp. enterica serovar Enteritidis
MTGDNTLITSHGINRRDFMKLCAALAATMGLSSKAAAEMAESVSNPQRPPVIWIGAQECT WP_000145429.1
hydrogenase 2 small subunit [Salmonella enterica] Length: 372\nNumber of predicted TMHs: 1\nExp number of AAs in TMHs: 24.51309\nExp number, first 60 AAs: 0.2682\nTotal prob of N-in: 0.14402\noutside 1 331\nTMhelix 332 354\ninside 355 372

23450 GCF_000329065.1_ASM32906v1 Salmonella enterica subsp. enterica serovar Enteritidis str. CDC_2010K_1884 Proteobacteria; Gammaproteobacteria; Enterobacterales; Enterobacteriaceae; Salmonella; Salmonella enterica; Salmonella enterica subsp. enterica; Salmonella enterica subsp. enterica serovar Enteritidis
MTGDNTLITSHGINRRDFMKLCAALAATMGLSSKAAAEMAESVSNPQRPPVIWIGAQECT WP_000145429.1
hydrogenase 2 small subunit [Salmonella enterica] Length: 372\nNumber of predicted TMHs: 1\nExp number of AAs in TMHs: 24.51309\nExp number, first 60 AAs: 0.2682\nTotal prob of N-in: 0.14402\noutside 1 331\nTMhelix 332 354\ninside 355 372

23451 GCF_000329065.1_ASM32906v1 Salmonella enterica subsp. enterica serovar Enteritidis str. CDC_2010K_1884 Proteobacteria; Gammaproteobacteria; Enterobacterales; Enterobacteriaceae; Salmonella; Salmonella enterica; Salmonella enterica subsp. enterica; Salmonella enterica subsp. enterica serovar Enteritidis
MNNEETFYQAMRRKGVTRRSFLKFCSLAATSLGLGAGMTPKIAWALENKPRIPVVWIHGL WP_001058307.1
hydrogenase [Salmonella enterica] Length: 372\nNumber of predicted TMHs: 2\nExp number of AAs in TMHs: 36.51738\nExp number, first 60 AAs: 19.72849\nTotal prob of N-in: 0.96170\nPOSSIBLE N-term signal sequence\ninside 1 19\nTMhelix 20 39\noutside 40 325\nTMhelix 326 348\ninside 349 372

23452 GCF_000330005.1_ASM33000v1 Salmonella enterica subsp. enterica serovar Enteritidis str. CHS4
Proteobacteria; Gammaproteobacteria; Enterobacterales; Enterobacteriaceae; Salmonella; Salmonella enterica; Salmonella enterica subsp. enterica; Salmonella enterica subsp. enterica serovar Enteritidis
MNNEETFYQAMRRKGVTRRSFLKFCSLAATSLGLGAGMTPKIAWALENKPRIPVVWIHGL WP_001058307.1
hydrogenase [Salmonella enterica] Length: 372\nNumber of predicted TMHs: 2\nExp number of AAs in TMHs: 36.51738\nExp number, first 60 AAs: 19.72849\nTotal prob of N-in: 0.96170\nPOSSIBLE N-term signal sequence\ninside 1 19\nTMhelix 20 39\noutside 40 325\nTMhelix 326 348\ninside 349 372

23453 GCF_000330005.1_ASM33000v1 Salmonella enterica subsp. enterica serovar Enteritidis str. CHS4
Proteobacteria; Gammaproteobacteria; Enterobacterales; Enterobacteriaceae; Salmonella; Salmonella enterica; Salmonella enterica subsp. enterica; Salmonella enterica subsp. enterica serovar Enteritidis
MTGDNTLITSHGINRRDFMKLCAALAATMGLSSKAAAEMAESVSNPQRPPVIWIGAQECT WP_000145429.1
hydrogenase 2 small subunit [Salmonella enterica] Length: 372\nNumber of predicted TMHs: 1\nExp number of AAs in TMHs: 24.51309\nExp number, first 60 AAs: 0.2682\nTotal prob of N-in: 0.14402\noutside 1 331\nTMhelix 332 354\ninside 355 372

23454 GCF_000329025.1_ASM32902v1 Salmonella enterica subsp. enterica serovar Enteritidis str. CHS44 Proteobacteria; Gammaproteobacteria; Enterobacterales; Enterobacteriaceae; Salmonella; Salmonella enterica; Salmonella enterica subsp. enterica; Salmonella enterica subsp. enterica serovar Enteritidis MNNEETFYQAMRRKGVTRRSFLKFCSLAATSLGLGAGMTPKIAWALENKPRIPVVWIHGL WP_001058307.1 hydrogenase [Salmonella enterica] Length: 372\nNumber of predicted TMHs: 2\nExp number of AAs in TMHs: 36.51738\nExp number, first 60 AAs: 19.72849\nTotal prob of N-in: 0.96170\nPOSSIBLE N-term signal sequence\ninside 1 19\nTMhelix 20 39\noutside 40 325\nTMhelix 326 348\ninside 349 372

23455 GCF_000329025.1_ASM32902v1 Salmonella enterica subsp. enterica serovar Enteritidis str. CHS44 Proteobacteria; Gammaproteobacteria; Enterobacterales; Enterobacteriaceae; Salmonella; Salmonella enterica; Salmonella enterica subsp. enterica; Salmonella enterica subsp. enterica serovar Enteritidis MTGDNTLITSHGINRRDFMKLCAALAATMGLSSKAAAEMAESVSNPQRPPVIWIGAQECT WP_000145429.1 hydrogenase 2 small subunit [Salmonella enterica] Length: 372\nNumber of predicted TMHs: 1\nExp number of AAs in TMHs: 24.51309\nExp number, first 60 AAs: 0.2682\nTotal prob of N-in: 0.14402\noutside 1 331\nTMhelix 332 354\ninside 355 372

23456 GCF_000329825.1_ASM32982v1 Salmonella enterica subsp. enterica serovar Enteritidis str. CVM_56-3991 Proteobacteria; Gammaproteobacteria; Enterobacterales; Enterobacteriaceae; Salmonella; Salmonella enterica; Salmonella enterica subsp. enterica; Salmonella enterica subsp. enterica serovar Enteritidis MTGDNTLITSHGINRRDFMKLCAALAATMGLSSKAAAEMAESVSNPQRPPVIWIGAQECT WP_000145429.1 hydrogenase 2 small subunit [Salmonella enterica] Length: 372\nNumber of predicted TMHs: 1\nExp number of AAs in TMHs: 24.51309\nExp number, first 60 AAs: 0.2682\nTotal prob of N-in: 0.14402\noutside 1 331\nTMhelix 332 354\ninside 355 372

23457 GCF_000329825.1_ASM32982v1 Salmonella enterica subsp. enterica serovar Enteritidis str. CVM_56-3991 Proteobacteria; Gammaproteobacteria; Enterobacterales; Enterobacteriaceae; Salmonella; Salmonella enterica; Salmonella enterica subsp. enterica; Salmonella enterica subsp. enterica serovar Enteritidis MNNEETFYQAMRRKGVTRRSFLKFCSLAATSLGLGAGMTPKIAWALENKPRIPVVWIHGL WP_001058307.1 hydrogenase [Salmonella enterica] Length: 372\nNumber of predicted TMHs: 2\nExp number of AAs in TMHs: 36.51738\nExp number, first 60 AAs: 19.72849\nTotal prob of N-in: 0.96170\nPOSSIBLE N-term signal sequence\ninside 1 19\nTMhelix 20 39\noutside 40 325\nTMhelix 326 348\ninside 349 372

23458 GCF_000329945.1_ASM32994v1 Salmonella enterica subsp. enterica serovar Enteritidis str. CVM_69-4941 Proteobacteria; Gammaproteobacteria; Enterobacterales; Enterobacteriaceae; Salmonella; Salmonella enterica; Salmonella enterica subsp. enterica; Salmonella enterica subsp. enterica serovar Enteritidis MTGDNTLITSHGINRRDFMKLCAALAATMGLSSKAAAEMAESVSNPQRPPVIWIGAQECT WP_000145429.1 hydrogenase 2 small subunit [Salmonella enterica] Length: 372\nNumber of predicted TMHs: 1\nExp number of AAs in TMHs: 24.51309\nExp number, first 60 AAs: 0.2682\nTotal prob of N-in: 0.14402\noutside 1 331\nTMhelix 332 354\ninside 355 372

23459 GCF_000329945.1_ASM32994v1 Salmonella enterica subsp. enterica serovar Enteritidis str. CVM_69-4941 Proteobacteria; Gammaproteobacteria; Enterobacterales; Enterobacteriaceae; Salmonella; Salmonella enterica; Salmonella enterica subsp. enterica; Salmonella enterica subsp. enterica serovar Enteritidis MNNEETFYQAMRRKGVTRRSFLKFCSLAATSLGLGAGMTPKIAWALENKPRIPVVWIHGL WP_001058307.1 hydrogenase [Salmonella enterica] Length: 372\nNumber of predicted TMHs: 2\nExp number of AAs in TMHs: 36.51738\nExp number, first 60 AAs: 19.72849\nTotal prob of N-in: 0.96170\nPOSSIBLE N-term signal sequence\ninside 1 19\nTMhelix 20 39\noutside 40 325\nTMhelix 326 348\ninside 349 372

23460 GCF_000329845.1_ASM32984v1 Salmonella enterica subsp. enterica serovar Enteritidis str. CVM_76-3618 Proteobacteria; Gammaproteobacteria; Enterobacterales; Enterobacteriaceae; Salmonella; Salmonella enterica; Salmonella enterica subsp. enterica; Salmonella enterica subsp. enterica serovar Enteritidis MNNEETFYQAMRRKGVTRRSFLKFCSLAATSLGLGAGMTPKIAWALENKPRIPVVWIHGL WP_001058307.1 hydrogenase [Salmonella enterica] Length: 372\nNumber of predicted TMHs: 2\nExp number of AAs in TMHs: 36.51738\nExp number, first 60 AAs: 19.72849\nTotal prob of N-in: 0.96170\nPOSSIBLE N-term signal sequence\ninside 1 19\nTMhelix 20 39\noutside 40 325\nTMhelix 326 348\ninside 349 372

23461 GCF_000329845.1_ASM32984v1 Salmonella enterica subsp. enterica serovar Enteritidis str. CVM_76-3618 Proteobacteria; Gammaproteobacteria; Enterobacterales; Enterobacteriaceae; Salmonella; Salmonella enterica; Salmonella enterica subsp. enterica; Salmonella enterica subsp. enterica serovar Enteritidis MTGDNTLITSHGINRRDFMKLCAALAATMGLSSKAAAEMAESVSNPQRPPVIWIGAQECT WP_000145429.1 hydrogenase 2 small subunit [Salmonella enterica] Length: 372\nNumber of predicted TMHs: 1\nExp

number of AAs in TMHs: 24.51309\nExp number, first 60 AAs:  0.2682\nTotal prob of N-in:   0.14402\noutside   1   331\nTMhelix   332   354\ninside   355   372

23462 GCF_000329885.1_ASM32988v1 Salmonella enterica subsp. enterica serovar Enteritidis str. CVM_81-2490 Proteobacteria; Gammaproteobacteria; Enterobacterales; Enterobacteriaceae; Salmonella; Salmonella enterica; Salmonella enterica subsp. enterica; Salmonella enterica subsp. enterica serovar Enteritidis MNNEETFYQAMRRKGVTRRSFLKFCSLAATSLGLGAGMTPKIAWALENKPRIPVVWIHGL WP_001058307.1 hydrogenase [Salmonella enterica] Length: 372\nNumber of predicted TMHs:  2\nExp number of AAs in TMHs: 36.51738\nExp number, first 60 AAs:  19.72849\nTotal prob of N-in:   0.96170\nPOSSIBLE N-term signal sequence\ninside   1   19\nTMhelix   20   39\noutside   40   325\nTMhelix   326   348\ninside   349   372

23463 GCF_000329885.1_ASM32988v1 Salmonella enterica subsp. enterica serovar Enteritidis str. CVM_81-2490 Proteobacteria; Gammaproteobacteria; Enterobacterales; Enterobacteriaceae; Salmonella; Salmonella enterica; Salmonella enterica subsp. enterica; Salmonella enterica subsp. enterica serovar Enteritidis MTGDNTLITSHGINRRDFMKLCAALAATMGLSSKAAAEMAESVSNPQRPPVIWIGAQECT WP_000145429.1 hydrogenase 2 small subunit [Salmonella enterica] Length: 372\nNumber of predicted TMHs:  1\nExp number of AAs in TMHs: 24.51309\nExp number, first 60 AAs:  0.2682\nTotal prob of N-in:   0.14402\noutside   1   331\nTMhelix   332   354\ninside   355   372

23464 GCF_000329805.1_ASM32980v1 Salmonella enterica subsp. enterica serovar Enteritidis str. CVM_N202 Proteobacteria; Gammaproteobacteria; Enterobacterales; Enterobacteriaceae; Salmonella; Salmonella enterica; Salmonella enterica subsp. enterica; Salmonella enterica subsp. enterica serovar Enteritidis MNNEETFYQAMRRKGVTRRSFLKFCSLAATSLGLGAGMTPKIAWALENKPRIPVVWIHGL WP_001058307.1 hydrogenase [Salmonella enterica] Length: 372\nNumber of predicted TMHs:  2\nExp number of AAs in TMHs: 36.51738\nExp number, first 60 AAs:  19.72849\nTotal prob of N-in:   0.96170\nPOSSIBLE N-term signal sequence\ninside   1   19\nTMhelix   20   39\noutside   40   325\nTMhelix   326   348\ninside   349   372

23465 GCF_000329805.1_ASM32980v1 Salmonella enterica subsp. enterica serovar Enteritidis str. CVM_N202 Proteobacteria; Gammaproteobacteria; Enterobacterales; Enterobacteriaceae; Salmonella; Salmonella enterica; Salmonella enterica subsp. enterica; Salmonella enterica subsp. enterica serovar Enteritidis MTGDNTLITSHGINRRDFMKLCAALAATMGLSSKAAAEMAESVSNPQRPPVIWIGAQECT WP_000145429.1 hydrogenase 2 small subunit [Salmonella enterica] Length: 372\nNumber of predicted TMHs:  1\nExp number of AAs in TMHs: 24.51309\nExp number, first 60 AAs:  0.2682\nTotal prob of N-in:   0.14402\noutside   1   331\nTMhelix   332   354\ninside   355   372

23466 GCF_000623755.2_ASM62375v2 Salmonella enterica subsp. enterica serovar Enteritidis str. EC20090135 Proteobacteria; Gammaproteobacteria; Enterobacterales; Enterobacteriaceae; Salmonella; Salmonella enterica; Salmonella enterica subsp. enterica; Salmonella enterica subsp. enterica serovar Enteritidis MTGDNTLITSHGINRRDFMKLCAALAATMGLSSKAAAEMAESVSNPQRPPVIWIGAQECT WP_000145429.1 hydrogenase 2 small subunit [Salmonella enterica] Length: 372\nNumber of predicted TMHs:  1\nExp number of AAs in TMHs: 24.51309\nExp number, first 60 AAs:  0.2682\nTotal prob of N-in:   0.14402\noutside   1   331\nTMhelix   332   354\ninside   355   372

23467 GCF_000623755.2_ASM62375v2 Salmonella enterica subsp. enterica serovar Enteritidis str. EC20090135 Proteobacteria; Gammaproteobacteria; Enterobacterales; Enterobacteriaceae; Salmonella; Salmonella enterica; Salmonella enterica subsp. enterica; Salmonella enterica subsp. enterica serovar Enteritidis MNNEETFYQAMRRKGVTRRSFLKFCSLAATSLGLGAGMTPKIAWALENKPRIPVVWIHGL WP_001058307.1 hydrogenase [Salmonella enterica] Length: 372\nNumber of predicted TMHs:  2\nExp number of AAs in TMHs: 36.51738\nExp number, first 60 AAs:  19.72849\nTotal prob of N-in:   0.96170\nPOSSIBLE N-term signal sequence\ninside   1   19\nTMhelix   20   39\noutside   40   325\nTMhelix   326   348\ninside   349   372

23468 GCF_000623775.2_ASM62377v2 Salmonella enterica subsp. enterica serovar Enteritidis str. EC20090193 Proteobacteria; Gammaproteobacteria; Enterobacterales; Enterobacteriaceae; Salmonella; Salmonella enterica; Salmonella enterica subsp. enterica; Salmonella enterica subsp. enterica serovar Enteritidis MNNEETFYQAMRRKGVTRRSFLKFCSLAATSLGLGAGMTPKIAWALENKPRIPVVWIHGL WP_001058307.1 hydrogenase [Salmonella enterica] Length: 372\nNumber of predicted TMHs:  2\nExp number of AAs in TMHs: 36.51738\nExp number, first 60 AAs:  19.72849\nTotal prob of N-in:   0.96170\nPOSSIBLE N-term signal sequence\ninside   1   19\nTMhelix   20   39\noutside   40   325\nTMhelix   326   348\ninside   349   372

23469 GCF_000623775.2_ASM62377v2 Salmonella enterica subsp. enterica serovar Enteritidis str. EC20090193 Proteobacteria; Gammaproteobacteria; Enterobacterales; Enterobacteriaceae; Salmonella; Salmonella enterica;

Salmonella enterica subsp. enterica; Salmonella enterica subsp. enterica serovar Enteritidis
MTGDNTLITSHGINRRDFMKLCAALAATMGLSSKAAAEMAESVSNPQRPPVIWIGAQECT WP_000145429.1
hydrogenase 2 small subunit [Salmonella enterica] Length: 372\nNumber of predicted TMHs: 1\nExp number of AAs in TMHs: 24.51309\nExp number, first 60 AAs: 0.2682\nTotal prob of N-in: 0.14402\noutside 1 331\nTMhelix 332 354\ninside 355 372

23470 GCF_000625675.2_ASM62567v2 Salmonella enterica subsp. enterica serovar Enteritidis str. EC20090195
Proteobacteria; Gammaproteobacteria; Enterobacterales; Enterobacteriaceae; Salmonella; Salmonella enterica; Salmonella enterica subsp. enterica; Salmonella enterica subsp. enterica serovar Enteritidis
MNNEETFYQAMRRKGVTRRSFLKFCSLAATSLGLGAGMTPKIAWALENKPRIPVVWIHGL WP_001058307.1
hydrogenase [Salmonella enterica] Length: 372\nNumber of predicted TMHs: 2\nExp number of AAs in TMHs: 36.51738\nExp number, first 60 AAs: 19.72849\nTotal prob of N-in: 0.96170\nPOSSIBLE N-term signal sequence\ninside 1 19\nTMhelix 20 39\noutside 40 325\nTMhelix 326 348\ninside 349 372

23471 GCF_000625675.2_ASM62567v2 Salmonella enterica subsp. enterica serovar Enteritidis str. EC20090195
Proteobacteria; Gammaproteobacteria; Enterobacterales; Enterobacteriaceae; Salmonella; Salmonella enterica; Salmonella enterica subsp. enterica; Salmonella enterica subsp. enterica serovar Enteritidis
MTGDNTLITSHGINRRDFMKLCAALAATMGLSSKAAAEMAESVSNPQRPPVIWIGAQECT WP_000145429.1
hydrogenase 2 small subunit [Salmonella enterica] Length: 372\nNumber of predicted TMHs: 1\nExp number of AAs in TMHs: 24.51309\nExp number, first 60 AAs: 0.2682\nTotal prob of N-in: 0.14402\noutside 1 331\nTMhelix 332 354\ninside 355 372

23472 GCF_000623795.2_ASM62379v2 Salmonella enterica subsp. enterica serovar Enteritidis str. EC20090332
Proteobacteria; Gammaproteobacteria; Enterobacterales; Enterobacteriaceae; Salmonella; Salmonella enterica; Salmonella enterica subsp. enterica; Salmonella enterica subsp. enterica serovar Enteritidis
MTGDNTLITSHGINRRDFMKLCAALAATMGLSSKAAAEMAESVSNPQRPPVIWIGAQECT WP_000145429.1
hydrogenase 2 small subunit [Salmonella enterica] Length: 372\nNumber of predicted TMHs: 1\nExp number of AAs in TMHs: 24.51309\nExp number, first 60 AAs: 0.2682\nTotal prob of N-in: 0.14402\noutside 1 331\nTMhelix 332 354\ninside 355 372

23473 GCF_000623795.2_ASM62379v2 Salmonella enterica subsp. enterica serovar Enteritidis str. EC20090332
Proteobacteria; Gammaproteobacteria; Enterobacterales; Enterobacteriaceae; Salmonella; Salmonella enterica; Salmonella enterica subsp. enterica; Salmonella enterica subsp. enterica serovar Enteritidis
MNNEETFYQAMRRKGVTRRSFLKFCSLAATSLGLGAGMTPKIAWALENKPRIPVVWIHGL WP_001058307.1
hydrogenase [Salmonella enterica] Length: 372\nNumber of predicted TMHs: 2\nExp number of AAs in TMHs: 36.51738\nExp number, first 60 AAs: 19.72849\nTotal prob of N-in: 0.96170\nPOSSIBLE N-term signal sequence\ninside 1 19\nTMhelix 20 39\noutside 40 325\nTMhelix 326 348\ninside 349 372

23474 GCF_000625395.2_ASM62539v2 Salmonella enterica subsp. enterica serovar Enteritidis str. EC20090530
Proteobacteria; Gammaproteobacteria; Enterobacterales; Enterobacteriaceae; Salmonella; Salmonella enterica; Salmonella enterica subsp. enterica; Salmonella enterica subsp. enterica serovar Enteritidis
MTGDNTLITSHGINRRDFMKLCAALAATMGLSSKAAAEMAESVSNPQRPPVIWIGAQECT WP_000145429.1
hydrogenase 2 small subunit [Salmonella enterica] Length: 372\nNumber of predicted TMHs: 1\nExp number of AAs in TMHs: 24.51309\nExp number, first 60 AAs: 0.2682\nTotal prob of N-in: 0.14402\noutside 1 331\nTMhelix 332 354\ninside 355 372

23475 GCF_000625395.2_ASM62539v2 Salmonella enterica subsp. enterica serovar Enteritidis str. EC20090530
Proteobacteria; Gammaproteobacteria; Enterobacterales; Enterobacteriaceae; Salmonella; Salmonella enterica; Salmonella enterica subsp. enterica; Salmonella enterica subsp. enterica serovar Enteritidis
MNNEETFYQAMRRKGVTRRSFLKFCSLAATSLGLGAGMTPKIAWALENKPRIPVVWIHGL WP_001058307.1
hydrogenase [Salmonella enterica] Length: 372\nNumber of predicted TMHs: 2\nExp number of AAs in TMHs: 36.51738\nExp number, first 60 AAs: 19.72849\nTotal prob of N-in: 0.96170\nPOSSIBLE N-term signal sequence\ninside 1 19\nTMhelix 20 39\noutside 40 325\nTMhelix 326 348\ninside 349 372

23476 GCF_000626155.2_ASM62615v2 Salmonella enterica subsp. enterica serovar Enteritidis str. EC20090531
Proteobacteria; Gammaproteobacteria; Enterobacterales; Enterobacteriaceae; Salmonella; Salmonella enterica; Salmonella enterica subsp. enterica; Salmonella enterica subsp. enterica serovar Enteritidis
MTGDNTLITSHGINRRDFMKLCAALAATMGLSSKAAAEMAESVSNPQRPPVIWIGAQECT WP_000145429.1
hydrogenase 2 small subunit [Salmonella enterica] Length: 372\nNumber of predicted TMHs: 1\nExp number of AAs in TMHs: 24.51309\nExp number, first 60 AAs: 0.2682\nTotal prob of N-in: 0.14402\noutside 1 331\nTMhelix 332 354\ninside 355 372

23477 GCF_000626155.2_ASM62615v2 Salmonella enterica subsp. enterica serovar Enteritidis str. EC20090531 Proteobacteria; Gammaproteobacteria; Enterobacterales; Enterobacteriaceae; Salmonella; Salmonella enterica; Salmonella enterica subsp. enterica; Salmonella enterica subsp. enterica serovar Enteritidis MNNEETFYQAMRRKGVTRRSFLKFCSLAATSLGLGAGMTPKIAWALENKPRIPVVWIHGL WP_001058307.1 hydrogenase [Salmonella enterica] Length: 372\nNumber of predicted TMHs: 2\nExp number of AAs in TMHs: 36.51738\nExp number, first 60 AAs: 19.72849\nTotal prob of N-in: 0.96170\nPOSSIBLE N-term signal sequence\ninside 1 19\nTMhelix 20 39\noutside 40 325\nTMhelix 326 348\ninside 349 372

23478 GCF_000626275.2_ASM62627v2 Salmonella enterica subsp. enterica serovar Enteritidis str. EC20090641 Proteobacteria; Gammaproteobacteria; Enterobacterales; Enterobacteriaceae; Salmonella; Salmonella enterica; Salmonella enterica subsp. enterica; Salmonella enterica subsp. enterica serovar Enteritidis MTGDNTLITSHGINRRDFMKLCAALAATMGLSSKAAAEMAESVSNPQRPPVIWIGAQECT WP_000145429.1 hydrogenase 2 small subunit [Salmonella enterica] Length: 372\nNumber of predicted TMHs: 1\nExp number of AAs in TMHs: 24.51309\nExp number, first 60 AAs: 0.2682\nTotal prob of N-in: 0.14402\noutside 1 331\nTMhelix 332 354\ninside 355 372

23479 GCF_000626275.2_ASM62627v2 Salmonella enterica subsp. enterica serovar Enteritidis str. EC20090641 Proteobacteria; Gammaproteobacteria; Enterobacterales; Enterobacteriaceae; Salmonella; Salmonella enterica; Salmonella enterica subsp. enterica; Salmonella enterica subsp. enterica serovar Enteritidis MNNEETFYQAMRRKGVTRRSFLKFCSLAATSLGLGAGMTPKIAWALENKPRIPVVWIHGL WP_001058307.1 hydrogenase [Salmonella enterica] Length: 372\nNumber of predicted TMHs: 2\nExp number of AAs in TMHs: 36.51738\nExp number, first 60 AAs: 19.72849\nTotal prob of N-in: 0.96170\nPOSSIBLE N-term signal sequence\ninside 1 19\nTMhelix 20 39\noutside 40 325\nTMhelix 326 348\ninside 349 372

23480 GCF_000626255.1_ASM62625v1 Salmonella enterica subsp. enterica serovar Enteritidis str. EC20090698 Proteobacteria; Gammaproteobacteria; Enterobacterales; Enterobacteriaceae; Salmonella; Salmonella enterica; Salmonella enterica subsp. enterica; Salmonella enterica subsp. enterica serovar Enteritidis MNNEETFYQAMRRKGVTRRSFLKFCSLAATSLGLGAGMTPKIAWALENKPRIPVVWIHGL WP_001058307.1 hydrogenase [Salmonella enterica] Length: 372\nNumber of predicted TMHs: 2\nExp number of AAs in TMHs: 36.51738\nExp number, first 60 AAs: 19.72849\nTotal prob of N-in: 0.96170\nPOSSIBLE N-term signal sequence\ninside 1 19\nTMhelix 20 39\noutside 40 325\nTMhelix 326 348\ninside 349 372

23481 GCF_000626255.1_ASM62625v1 Salmonella enterica subsp. enterica serovar Enteritidis str. EC20090698 Proteobacteria; Gammaproteobacteria; Enterobacterales; Enterobacteriaceae; Salmonella; Salmonella enterica; Salmonella enterica subsp. enterica; Salmonella enterica subsp. enterica serovar Enteritidis MTGDNTLITSHGINRRDFMKLCAALAATMGLSSKAAAEMAESVSNPQRPPVIWIGAQECT WP_000145429.1 hydrogenase 2 small subunit [Salmonella enterica] Length: 372\nNumber of predicted TMHs: 1\nExp number of AAs in TMHs: 24.51309\nExp number, first 60 AAs: 0.2682\nTotal prob of N-in: 0.14402\noutside 1 331\nTMhelix 332 354\ninside 355 372

23482 GCF_000626135.2_ASM62613v2 Salmonella enterica subsp. enterica serovar Enteritidis str. EC20090884 Proteobacteria; Gammaproteobacteria; Enterobacterales; Enterobacteriaceae; Salmonella; Salmonella enterica; Salmonella enterica subsp. enterica; Salmonella enterica subsp. enterica serovar Enteritidis MTGDNTLITSHGINRRDFMKLCAALAATMGLSSKAAAEMAESVSNPQRPPVIWIGAQECT WP_000145429.1 hydrogenase 2 small subunit [Salmonella enterica] Length: 372\nNumber of predicted TMHs: 1\nExp number of AAs in TMHs: 24.51309\nExp number, first 60 AAs: 0.2682\nTotal prob of N-in: 0.14402\noutside 1 331\nTMhelix 332 354\ninside 355 372

23483 GCF_000626135.2_ASM62613v2 Salmonella enterica subsp. enterica serovar Enteritidis str. EC20090884 Proteobacteria; Gammaproteobacteria; Enterobacterales; Enterobacteriaceae; Salmonella; Salmonella enterica; Salmonella enterica subsp. enterica; Salmonella enterica subsp. enterica serovar Enteritidis MNNEETFYQAMRRKGVTRRSFLKFCSLAATSLGLGAGMTPKIAWALENKPRIPVVWIHGL WP_001058307.1 hydrogenase [Salmonella enterica] Length: 372\nNumber of predicted TMHs: 2\nExp number of AAs in TMHs: 36.51738\nExp number, first 60 AAs: 19.72849\nTotal prob of N-in: 0.96170\nPOSSIBLE N-term signal sequence\ninside 1 19\nTMhelix 20 39\noutside 40 325\nTMhelix 326 348\ninside 349 372

23484 GCF_000625415.2_ASM62541v2 Salmonella enterica subsp. enterica serovar Enteritidis str. EC20100088 Proteobacteria; Gammaproteobacteria; Enterobacterales; Enterobacteriaceae; Salmonella; Salmonella enterica; Salmonella enterica subsp. enterica; Salmonella enterica subsp. enterica serovar Enteritidis MNNEETFYQAMRRKGVTRRSFLKFCSLAATSLGLGAGMTPKIAWALENKPRIPVVWIHGL WP_001058307.1 hydrogenase [Salmonella enterica] Length: 372\nNumber of predicted TMHs: 2\nExp number of AAs in

TMHs: 36.51738\nExp number, first 60 AAs: 19.72849\nTotal prob of N-in: 0.96170\nPOSSIBLE N-term signal sequence\ninside 1 19\nTMhelix 20 39\noutside 40 325\nTMhelix 326 348\ninside 349 372

23485 GCF_000625415.2_ASM62541v2 Salmonella enterica subsp. enterica serovar Enteritidis str. EC20100088 Proteobacteria; Gammaproteobacteria; Enterobacterales; Enterobacteriaceae; Salmonella; Salmonella enterica; Salmonella enterica subsp. enterica; Salmonella enterica subsp. enterica serovar Enteritidis MTGDNTLITSHGINRRDFMKLCAALAATMGLSSKAAAEMAESVSNPQRPPVIWIGAQECT WP_000145429.1 hydrogenase 2 small subunit [Salmonella enterica] Length: 372\nNumber of predicted TMHs: 1\nExp number of AAs in TMHs: 24.51309\nExp number, first 60 AAs: 0.2682\nTotal prob of N-in: 0.14402\noutside 1 331\nTMhelix 332 354\ninside 355 372

23486 GCF_000625435.2_ASM62543v2 Salmonella enterica subsp. enterica serovar Enteritidis str. EC20100089 Proteobacteria; Gammaproteobacteria; Enterobacterales; Enterobacteriaceae; Salmonella; Salmonella enterica; Salmonella enterica subsp. enterica; Salmonella enterica subsp. enterica serovar Enteritidis MNNEETFYQAMRRKGVTRRSFLKFCSLAATSLGLGAGMTPKIAWALENKPRIPVVWIHGL WP_001058307.1 hydrogenase [Salmonella enterica] Length: 372\nNumber of predicted TMHs: 2\nExp number of AAs in TMHs: 36.51738\nExp number, first 60 AAs: 19.72849\nTotal prob of N-in: 0.96170\nPOSSIBLE N-term signal sequence\ninside 1 19\nTMhelix 20 39\noutside 40 325\nTMhelix 326 348\ninside 349 372

23487 GCF_000625435.2_ASM62543v2 Salmonella enterica subsp. enterica serovar Enteritidis str. EC20100089 Proteobacteria; Gammaproteobacteria; Enterobacterales; Enterobacteriaceae; Salmonella; Salmonella enterica; Salmonella enterica subsp. enterica; Salmonella enterica subsp. enterica serovar Enteritidis MTGDNTLITSHGINRRDFMKLCAALAATMGLSSKAAAEMAESVSNPQRPPVIWIGAQECT WP_000145429.1 hydrogenase 2 small subunit [Salmonella enterica] Length: 372\nNumber of predicted TMHs: 1\nExp number of AAs in TMHs: 24.51309\nExp number, first 60 AAs: 0.2682\nTotal prob of N-in: 0.14402\noutside 1 331\nTMhelix 332 354\ninside 355 372

23488 GCF_000968775.2_ASM96877v2 Salmonella enterica subsp. enterica serovar Enteritidis str. EC20100100 Proteobacteria; Gammaproteobacteria; Enterobacterales; Enterobacteriaceae; Salmonella; Salmonella enterica; Salmonella enterica subsp. enterica; Salmonella enterica subsp. enterica serovar Enteritidis MNNEETFYQAMRRKGVTRRSFLKFCSLAATSLGLGAGMTPKIAWALENKPRIPVVWIHGL WP_001058307.1 hydrogenase [Salmonella enterica] Length: 372\nNumber of predicted TMHs: 2\nExp number of AAs in TMHs: 36.51738\nExp number, first 60 AAs: 19.72849\nTotal prob of N-in: 0.96170\nPOSSIBLE N-term signal sequence\ninside 1 19\nTMhelix 20 39\noutside 40 325\nTMhelix 326 348\ninside 349 372

23489 GCF_000968775.2_ASM96877v2 Salmonella enterica subsp. enterica serovar Enteritidis str. EC20100100 Proteobacteria; Gammaproteobacteria; Enterobacterales; Enterobacteriaceae; Salmonella; Salmonella enterica; Salmonella enterica subsp. enterica; Salmonella enterica subsp. enterica serovar Enteritidis MTGDNTLITSHGINRRDFMKLCAALAATMGLSSKAAAEMAESVSNPQRPPVIWIGAQECT WP_000145429.1 hydrogenase 2 small subunit [Salmonella enterica] Length: 372\nNumber of predicted TMHs: 1\nExp number of AAs in TMHs: 24.51309\nExp number, first 60 AAs: 0.2682\nTotal prob of N-in: 0.14402\noutside 1 331\nTMhelix 332 354\ninside 355 372

23490 GCF_000626215.1_ASM62621v1 Salmonella enterica subsp. enterica serovar Enteritidis str. EC20100101 Proteobacteria; Gammaproteobacteria; Enterobacterales; Enterobacteriaceae; Salmonella; Salmonella enterica; Salmonella enterica subsp. enterica; Salmonella enterica subsp. enterica serovar Enteritidis MNNEETFYQAMRRKGVTRRSFLKFCSLAATSLGLGAGMTPKIAWALENKPRIPVVWIHGL WP_001058307.1 hydrogenase [Salmonella enterica] Length: 372\nNumber of predicted TMHs: 2\nExp number of AAs in TMHs: 36.51738\nExp number, first 60 AAs: 19.72849\nTotal prob of N-in: 0.96170\nPOSSIBLE N-term signal sequence\ninside 1 19\nTMhelix 20 39\noutside 40 325\nTMhelix 326 348\ninside 349 372

23491 GCF_000626215.1_ASM62621v1 Salmonella enterica subsp. enterica serovar Enteritidis str. EC20100101 Proteobacteria; Gammaproteobacteria; Enterobacterales; Enterobacteriaceae; Salmonella; Salmonella enterica; Salmonella enterica subsp. enterica; Salmonella enterica subsp. enterica serovar Enteritidis MTGDNTLITSHGINRRDFMKLCAALAATMGLSSKAAAEMAESVSNPQRPPVIWIGAQECT WP_000145429.1 hydrogenase 2 small subunit [Salmonella enterica] Length: 372\nNumber of predicted TMHs: 1\nExp number of AAs in TMHs: 24.51309\nExp number, first 60 AAs: 0.2682\nTotal prob of N-in: 0.14402\noutside 1 331\nTMhelix 332 354\ninside 355 372

23492 GCF_000626115.2_ASM62611v2 Salmonella enterica subsp. enterica serovar Enteritidis str. EC20100103 Proteobacteria; Gammaproteobacteria; Enterobacterales; Enterobacteriaceae; Salmonella; Salmonella enterica;

Salmonella enterica subsp. enterica; Salmonella enterica subsp. enterica serovar Enteritidis
MNNEETFYQAMRRKGVTRRSFLKFCSLAATSLGLGAGMTPKIAWALENKPRIPVVWIHGL WP_001058307.1
hydrogenase [Salmonella enterica] Length: 372\nNumber of predicted TMHs: 2\nExp number of AAs in TMHs: 36.51738\nExp number, first 60 AAs: 19.72849\nTotal prob of N-in: 0.96170\nPOSSIBLE N-term signal sequence\ninside 1 19\nTMhelix 20 39\noutside 40 325\nTMhelix 326 348\ninside 349 372

23493 GCF_000626115.2_ASM62611v2 Salmonella enterica subsp. enterica serovar Enteritidis str. EC20100103
Proteobacteria; Gammaproteobacteria; Enterobacterales; Enterobacteriaceae; Salmonella; Salmonella enterica; Salmonella enterica subsp. enterica; Salmonella enterica subsp. enterica serovar Enteritidis
MTGDNTLITSHGINRRDFMKLCAALAATMGLSSKAAAEMAESVSNPQRPPVIWIGAQECT WP_000145429.1
hydrogenase 2 small subunit [Salmonella enterica] Length: 372\nNumber of predicted TMHs: 1\nExp number of AAs in TMHs: 24.51309\nExp number, first 60 AAs: 0.2682\nTotal prob of N-in: 0.14402\noutside 1 331\nTMhelix 332 354\ninside 355 372

23494 GCF_000831045.2_ASM83104v2 Salmonella enterica subsp. enterica serovar Enteritidis str. EC20100130
Proteobacteria; Gammaproteobacteria; Enterobacterales; Enterobacteriaceae; Salmonella; Salmonella enterica; Salmonella enterica subsp. enterica; Salmonella enterica subsp. enterica serovar Enteritidis
MNNEETFYQAMRRKGVTRRSFLKFCSLAATSLGLGAGMTPKIAWALENKPRIPVVWIHGL WP_001058307.1
hydrogenase [Salmonella enterica] Length: 372\nNumber of predicted TMHs: 2\nExp number of AAs in TMHs: 36.51738\nExp number, first 60 AAs: 19.72849\nTotal prob of N-in: 0.96170\nPOSSIBLE N-term signal sequence\ninside 1 19\nTMhelix 20 39\noutside 40 325\nTMhelix 326 348\ninside 349 372

23495 GCF_000831045.2_ASM83104v2 Salmonella enterica subsp. enterica serovar Enteritidis str. EC20100130
Proteobacteria; Gammaproteobacteria; Enterobacterales; Enterobacteriaceae; Salmonella; Salmonella enterica; Salmonella enterica subsp. enterica; Salmonella enterica subsp. enterica serovar Enteritidis
MTGDNTLITSHGINRRDFMKLCAALAATMGLSSKAAAEMAESVSNPQRPPVIWIGAQECT WP_000145429.1
hydrogenase 2 small subunit [Salmonella enterica] Length: 372\nNumber of predicted TMHs: 1\nExp number of AAs in TMHs: 24.51309\nExp number, first 60 AAs: 0.2682\nTotal prob of N-in: 0.14402\noutside 1 331\nTMhelix 332 354\ninside 355 372

23496 GCF_000625715.2_ASM62571v2 Salmonella enterica subsp. enterica serovar Enteritidis str. EC20100131
Proteobacteria; Gammaproteobacteria; Enterobacterales; Enterobacteriaceae; Salmonella; Salmonella enterica; Salmonella enterica subsp. enterica; Salmonella enterica subsp. enterica serovar Enteritidis
MNNEETFYQAMRRKGVTRRSFLKFCSLAATSLGLGAGMTPKIAWALENKPRIPVVWIHGL WP_001058307.1
hydrogenase [Salmonella enterica] Length: 372\nNumber of predicted TMHs: 2\nExp number of AAs in TMHs: 36.51738\nExp number, first 60 AAs: 19.72849\nTotal prob of N-in: 0.96170\nPOSSIBLE N-term signal sequence\ninside 1 19\nTMhelix 20 39\noutside 40 325\nTMhelix 326 348\ninside 349 372

23497 GCF_000625715.2_ASM62571v2 Salmonella enterica subsp. enterica serovar Enteritidis str. EC20100131
Proteobacteria; Gammaproteobacteria; Enterobacterales; Enterobacteriaceae; Salmonella; Salmonella enterica; Salmonella enterica subsp. enterica; Salmonella enterica subsp. enterica serovar Enteritidis
MTGDNTLITSHGINRRDFMKLCAALAATMGLSSKAAAEMAESVSNPQRPPVIWIGAQECT WP_000145429.1
hydrogenase 2 small subunit [Salmonella enterica] Length: 372\nNumber of predicted TMHs: 1\nExp number of AAs in TMHs: 24.51309\nExp number, first 60 AAs: 0.2682\nTotal prob of N-in: 0.14402\noutside 1 331\nTMhelix 332 354\ninside 355 372

23498 GCF_000831025.2_ASM83102v2 Salmonella enterica subsp. enterica serovar Enteritidis str. EC20100134
Proteobacteria; Gammaproteobacteria; Enterobacterales; Enterobacteriaceae; Salmonella; Salmonella enterica; Salmonella enterica subsp. enterica; Salmonella enterica subsp. enterica serovar Enteritidis
MNNEETFYQAMRRKGVTRRSFLKFCSLAATSLGLGAGMTPKIAWALENKPRIPVVWIHGL WP_001058307.1
hydrogenase [Salmonella enterica] Length: 372\nNumber of predicted TMHs: 2\nExp number of AAs in TMHs: 36.51738\nExp number, first 60 AAs: 19.72849\nTotal prob of N-in: 0.96170\nPOSSIBLE N-term signal sequence\ninside 1 19\nTMhelix 20 39\noutside 40 325\nTMhelix 326 348\ninside 349 372

23499 GCF_000831025.2_ASM83102v2 Salmonella enterica subsp. enterica serovar Enteritidis str. EC20100134
Proteobacteria; Gammaproteobacteria; Enterobacterales; Enterobacteriaceae; Salmonella; Salmonella enterica; Salmonella enterica subsp. enterica; Salmonella enterica subsp. enterica serovar Enteritidis
MTGDNTLITSHGINRRDFMKLCAALAATMGLSSKAAAEMAESVSNPQRPPVIWIGAQECT WP_000145429.1
hydrogenase 2 small subunit [Salmonella enterica] Length: 372\nNumber of predicted TMHs: 1\nExp number of AAs in TMHs: 24.51309\nExp number, first 60 AAs: 0.2682\nTotal prob of N-in: 0.14402\noutside 1 331\nTMhelix 332 354\ninside 355 372

23500 GCF_000626695.2_ASM62669v2 Salmonella enterica subsp. enterica serovar Enteritidis str. EC20100325 Proteobacteria; Gammaproteobacteria; Enterobacterales; Enterobacteriaceae; Salmonella; Salmonella enterica; Salmonella enterica subsp. enterica; Salmonella enterica subsp. enterica serovar Enteritidis MTGDNTLITSHGINRRDFMKLCAALAATMGLSSKAAAEMAESVSNPQRPPVIWIGAQECT WP_000145429.1 hydrogenase 2 small subunit [Salmonella enterica] Length: 372\nNumber of predicted TMHs: 1\nExp number of AAs in TMHs: 24.51309\nExp number, first 60 AAs: 0.2682\nTotal prob of N-in: 0.14402\noutside 1 331\nTMhelix 332 354\ninside 355 372

23501 GCF_000626695.2_ASM62669v2 Salmonella enterica subsp. enterica serovar Enteritidis str. EC20100325 Proteobacteria; Gammaproteobacteria; Enterobacterales; Enterobacteriaceae; Salmonella; Salmonella enterica; Salmonella enterica subsp. enterica; Salmonella enterica subsp. enterica serovar Enteritidis MNNEETFYQAMRRKGVTRRSFLKFCSLAATSLGLGAGMTPKIAWALENKPRIPVVWIHGL WP_001058307.1 hydrogenase [Salmonella enterica] Length: 372\nNumber of predicted TMHs: 2\nExp number of AAs in TMHs: 36.51738\nExp number, first 60 AAs: 19.72849\nTotal prob of N-in: 0.96170\nPOSSIBLE N-term signal sequence\ninside 1 19\nTMhelix 20 39\noutside 40 325\nTMhelix 326 348\ninside 349 372

23502 GCF_000626235.1_ASM62623v1 Salmonella enterica subsp. enterica serovar Enteritidis str. EC20110221 Proteobacteria; Gammaproteobacteria; Enterobacterales; Enterobacteriaceae; Salmonella; Salmonella enterica; Salmonella enterica subsp. enterica; Salmonella enterica subsp. enterica serovar Enteritidis MNNEETFYQAMRRKGVTRRSFLKFCSLAATSLGLGAGMTPKIAWALENKPRIPVVWIHGL WP_001058307.1 hydrogenase [Salmonella enterica] Length: 372\nNumber of predicted TMHs: 2\nExp number of AAs in TMHs: 36.51738\nExp number, first 60 AAs: 19.72849\nTotal prob of N-in: 0.96170\nPOSSIBLE N-term signal sequence\ninside 1 19\nTMhelix 20 39\noutside 40 325\nTMhelix 326 348\ninside 349 372

23503 GCF_000626235.1_ASM62623v1 Salmonella enterica subsp. enterica serovar Enteritidis str. EC20110221 Proteobacteria; Gammaproteobacteria; Enterobacterales; Enterobacteriaceae; Salmonella; Salmonella enterica; Salmonella enterica subsp. enterica; Salmonella enterica subsp. enterica serovar Enteritidis MTGDNTLITSHGINRRDFMKLCAALAATMGLSSKAAAEMAESVSNPQRPPVIWIGAQECT WP_000145429.1 hydrogenase 2 small subunit [Salmonella enterica] Length: 372\nNumber of predicted TMHs: 1\nExp number of AAs in TMHs: 24.51309\nExp number, first 60 AAs: 0.2682\nTotal prob of N-in: 0.14402\noutside 1 331\nTMhelix 332 354\ninside 355 372

23504 GCF_000624275.2_ASM62427v2 Salmonella enterica subsp. enterica serovar Enteritidis str. EC20110222 Proteobacteria; Gammaproteobacteria; Enterobacterales; Enterobacteriaceae; Salmonella; Salmonella enterica; Salmonella enterica subsp. enterica; Salmonella enterica subsp. enterica serovar Enteritidis MTGDNTLITSHGINRRDFMKLCAALAATMGLSSKAAAEMAESVSNPQRPPVIWIGAQECT WP_000145429.1 hydrogenase 2 small subunit [Salmonella enterica] Length: 372\nNumber of predicted TMHs: 1\nExp number of AAs in TMHs: 24.51309\nExp number, first 60 AAs: 0.2682\nTotal prob of N-in: 0.14402\noutside 1 331\nTMhelix 332 354\ninside 355 372

23505 GCF_000624275.2_ASM62427v2 Salmonella enterica subsp. enterica serovar Enteritidis str. EC20110222 Proteobacteria; Gammaproteobacteria; Enterobacterales; Enterobacteriaceae; Salmonella; Salmonella enterica; Salmonella enterica subsp. enterica; Salmonella enterica subsp. enterica serovar Enteritidis MNNEETFYQAMRRKGVTRRSFLKFCSLAATSLGLGAGMTPKIAWALENKPRIPVVWIHGL WP_001058307.1 hydrogenase [Salmonella enterica] Length: 372\nNumber of predicted TMHs: 2\nExp number of AAs in TMHs: 36.51738\nExp number, first 60 AAs: 19.72849\nTotal prob of N-in: 0.96170\nPOSSIBLE N-term signal sequence\ninside 1 19\nTMhelix 20 39\noutside 40 325\nTMhelix 326 348\ninside 349 372

23506 GCF_000623175.2_ASM62317v2 Salmonella enterica subsp. enterica serovar Enteritidis str. EC20110223 Proteobacteria; Gammaproteobacteria; Enterobacterales; Enterobacteriaceae; Salmonella; Salmonella enterica; Salmonella enterica subsp. enterica; Salmonella enterica subsp. enterica serovar Enteritidis MTGDNTLITSHGINRRDFMKLCAALAATMGLSSKAAAEMAESVSNPQRPPVIWIGAQECT WP_000145429.1 hydrogenase 2 small subunit [Salmonella enterica] Length: 372\nNumber of predicted TMHs: 1\nExp number of AAs in TMHs: 24.51309\nExp number, first 60 AAs: 0.2682\nTotal prob of N-in: 0.14402\noutside 1 331\nTMhelix 332 354\ninside 355 372

23507 GCF_000623175.2_ASM62317v2 Salmonella enterica subsp. enterica serovar Enteritidis str. EC20110223 Proteobacteria; Gammaproteobacteria; Enterobacterales; Enterobacteriaceae; Salmonella; Salmonella enterica; Salmonella enterica subsp. enterica; Salmonella enterica subsp. enterica serovar Enteritidis MNNEETFYQAMRRKGVTRRSFLKFCSLAATSLGLGAGMTPKIAWALENKPRIPVVWIHGL WP_001058307.1 hydrogenase [Salmonella enterica] Length: 372\nNumber of predicted TMHs: 2\nExp number of AAs in

TMHs: 36.51738\nExp number, first 60 AAs: 19.72849\nTotal prob of N-in: 0.96170\nPOSSIBLE N-term signal sequence\ninside 1 19\nTMhelix 20 39\noutside 40 325\nTMhelix 326 348\ninside 349 372

23508 GCF_000626175.1_ASM62617v1 Salmonella enterica subsp. enterica serovar Enteritidis str. EC20110354 Proteobacteria; Gammaproteobacteria; Enterobacterales; Enterobacteriaceae; Salmonella; Salmonella enterica; Salmonella enterica subsp. enterica; Salmonella enterica subsp. enterica serovar Enteritidis MNNEETFYQAMRRKGVTRRSFLKFCSLAATSLGLGAGMTPKIAWALENKPRIPVVWIHGL WP_001058307.1 hydrogenase [Salmonella enterica] Length: 372\nNumber of predicted TMHs: 2\nExp number of AAs in TMHs: 36.51738\nExp number, first 60 AAs: 19.72849\nTotal prob of N-in: 0.96170\nPOSSIBLE N-term signal sequence\ninside 1 19\nTMhelix 20 39\noutside 40 325\nTMhelix 326 348\ninside 349 372

23509 GCF_000626175.1_ASM62617v1 Salmonella enterica subsp. enterica serovar Enteritidis str. EC20110354 Proteobacteria; Gammaproteobacteria; Enterobacterales; Enterobacteriaceae; Salmonella; Salmonella enterica; Salmonella enterica subsp. enterica; Salmonella enterica subsp. enterica serovar Enteritidis MTGDNTLITSHGINRRDFMKLCAALAATMGLSSKAAAEMAESVSNPQRPPVIWIGAQECT WP_000145429.1 hydrogenase 2 small subunit [Salmonella enterica] Length: 372\nNumber of predicted TMHs: 1\nExp number of AAs in TMHs: 24.51309\nExp number, first 60 AAs: 0.2682\nTotal prob of N-in: 0.14402\noutside 1 331\nTMhelix 332 354\ninside 355 372

23510 GCF_000626295.1_ASM62629v1 Salmonella enterica subsp. enterica serovar Enteritidis str. EC20110355 Proteobacteria; Gammaproteobacteria; Enterobacterales; Enterobacteriaceae; Salmonella; Salmonella enterica; Salmonella enterica subsp. enterica; Salmonella enterica subsp. enterica serovar Enteritidis MTGDNTLITSHGINRRDFMKLCAALAATMGLSSKAAAEMAESVSNPQRPPVIWIGAQECT WP_000145429.1 hydrogenase 2 small subunit [Salmonella enterica] Length: 372\nNumber of predicted TMHs: 1\nExp number of AAs in TMHs: 24.51309\nExp number, first 60 AAs: 0.2682\nTotal prob of N-in: 0.14402\noutside 1 331\nTMhelix 332 354\ninside 355 372

23511 GCF_000626295.1_ASM62629v1 Salmonella enterica subsp. enterica serovar Enteritidis str. EC20110355 Proteobacteria; Gammaproteobacteria; Enterobacterales; Enterobacteriaceae; Salmonella; Salmonella enterica; Salmonella enterica subsp. enterica; Salmonella enterica subsp. enterica serovar Enteritidis MNNEETFYQAMRRKGVTRRSFLKFCSLAATSLGLGAGMTPKIAWALENKPRIPVVWIHGL WP_001058307.1 hydrogenase [Salmonella enterica] Length: 372\nNumber of predicted TMHs: 2\nExp number of AAs in TMHs: 36.51738\nExp number, first 60 AAs: 19.72849\nTotal prob of N-in: 0.96170\nPOSSIBLE N-term signal sequence\ninside 1 19\nTMhelix 20 39\noutside 40 325\nTMhelix 326 348\ninside 349 372

23512 GCF_000623375.1_ASM62337v1 Salmonella enterica subsp. enterica serovar Enteritidis str. EC20110356 Proteobacteria; Gammaproteobacteria; Enterobacterales; Enterobacteriaceae; Salmonella; Salmonella enterica; Salmonella enterica subsp. enterica; Salmonella enterica subsp. enterica serovar Enteritidis MTGDNTLITSHGINRRDFMKLCAALAATMGLSSKAAAEMAESVSNPQRPPVIWIGAQECT WP_000145429.1 hydrogenase 2 small subunit [Salmonella enterica] Length: 372\nNumber of predicted TMHs: 1\nExp number of AAs in TMHs: 24.51309\nExp number, first 60 AAs: 0.2682\nTotal prob of N-in: 0.14402\noutside 1 331\nTMhelix 332 354\ninside 355 372

23513 GCF_000623375.1_ASM62337v1 Salmonella enterica subsp. enterica serovar Enteritidis str. EC20110356 Proteobacteria; Gammaproteobacteria; Enterobacterales; Enterobacteriaceae; Salmonella; Salmonella enterica; Salmonella enterica subsp. enterica; Salmonella enterica subsp. enterica serovar Enteritidis MNNEETFYQAMRRKGVTRRSFLKFCSLAATSLGLGAGMTPKIAWALENKPRIPVVWIHGL WP_001058307.1 hydrogenase [Salmonella enterica] Length: 372\nNumber of predicted TMHs: 2\nExp number of AAs in TMHs: 36.51738\nExp number, first 60 AAs: 19.72849\nTotal prob of N-in: 0.96170\nPOSSIBLE N-term signal sequence\ninside 1 19\nTMhelix 20 39\noutside 40 325\nTMhelix 326 348\ninside 349 372

23514 GCF_000623355.1_ASM62335v1 Salmonella enterica subsp. enterica serovar Enteritidis str. EC20110357 Proteobacteria; Gammaproteobacteria; Enterobacterales; Enterobacteriaceae; Salmonella; Salmonella enterica; Salmonella enterica subsp. enterica; Salmonella enterica subsp. enterica serovar Enteritidis MNNEETFYQAMRRKGVTRRSFLKFCSLAATSLGLGAGMTPKIAWALENKPRIPVVWIHGL WP_001058307.1 hydrogenase [Salmonella enterica] Length: 372\nNumber of predicted TMHs: 2\nExp number of AAs in TMHs: 36.51738\nExp number, first 60 AAs: 19.72849\nTotal prob of N-in: 0.96170\nPOSSIBLE N-term signal sequence\ninside 1 19\nTMhelix 20 39\noutside 40 325\nTMhelix 326 348\ninside 349 372

23515 GCF_000623355.1_ASM62335v1 Salmonella enterica subsp. enterica serovar Enteritidis str. EC20110357 Proteobacteria; Gammaproteobacteria; Enterobacterales; Enterobacteriaceae; Salmonella; Salmonella enterica;

Salmonella enterica subsp. enterica; Salmonella enterica subsp. enterica serovar Enteritidis
MTGDNTLITSHGINRRDFMKLCAALAATMGLSSKAAAEMAESVSNPQRPPVIWIGAQECT WP_000145429.1
hydrogenase 2 small subunit [Salmonella enterica] Length: 372\nNumber of predicted TMHs: 1\nExp number of AAs in TMHs: 24.51309\nExp number, first 60 AAs: 0.2682\nTotal prob of N-in: 0.14402\noutside 1 331\nTMhelix 332 354\ninside 355 372

23516 GCF_000623335.1_ASM62333v1 Salmonella enterica subsp. enterica serovar Enteritidis str. EC20110358
Proteobacteria; Gammaproteobacteria; Enterobacterales; Enterobacteriaceae; Salmonella; Salmonella enterica; Salmonella enterica subsp. enterica; Salmonella enterica subsp. enterica serovar Enteritidis
MTGDNTLITSHGINRRDFMKLCAALAATMGLSSKAAAEMAESVSNPQRPPVIWIGAQECT WP_000145429.1
hydrogenase 2 small subunit [Salmonella enterica] Length: 372\nNumber of predicted TMHs: 1\nExp number of AAs in TMHs: 24.51309\nExp number, first 60 AAs: 0.2682\nTotal prob of N-in: 0.14402\noutside 1 331\nTMhelix 332 354\ninside 355 372

23517 GCF_000623335.1_ASM62333v1 Salmonella enterica subsp. enterica serovar Enteritidis str. EC20110358
Proteobacteria; Gammaproteobacteria; Enterobacterales; Enterobacteriaceae; Salmonella; Salmonella enterica; Salmonella enterica subsp. enterica; Salmonella enterica subsp. enterica serovar Enteritidis
MNNEETFYQAMRRKGVTRRSFLKFCSLAATSLGLGAGMTPKIAWALENKPRIPVVWIHGL WP_001058307.1
hydrogenase [Salmonella enterica] Length: 372\nNumber of predicted TMHs: 2\nExp number of AAs in TMHs: 36.51738\nExp number, first 60 AAs: 19.72849\nTotal prob of N-in: 0.96170\nPOSSIBLE N-term signal sequence\ninside 1 19\nTMhelix 20 39\noutside 40 325\nTMhelix 326 348\ninside 349 372

23518 GCF_000623315.1_ASM62331v1 Salmonella enterica subsp. enterica serovar Enteritidis str. EC20110359
Proteobacteria; Gammaproteobacteria; Enterobacterales; Enterobacteriaceae; Salmonella; Salmonella enterica; Salmonella enterica subsp. enterica; Salmonella enterica subsp. enterica serovar Enteritidis
MNNEETFYQAMRRKGVTRRSFLKFCSLAATSLGLGAGMTPKIAWALENKPRIPVVWIHGL WP_001058307.1
hydrogenase [Salmonella enterica] Length: 372\nNumber of predicted TMHs: 2\nExp number of AAs in TMHs: 36.51738\nExp number, first 60 AAs: 19.72849\nTotal prob of N-in: 0.96170\nPOSSIBLE N-term signal sequence\ninside 1 19\nTMhelix 20 39\noutside 40 325\nTMhelix 326 348\ninside 349 372

23519 GCF_000623315.1_ASM62331v1 Salmonella enterica subsp. enterica serovar Enteritidis str. EC20110359
Proteobacteria; Gammaproteobacteria; Enterobacterales; Enterobacteriaceae; Salmonella; Salmonella enterica; Salmonella enterica subsp. enterica; Salmonella enterica subsp. enterica serovar Enteritidis
MTGDNTLITSHGINRRDFMKLCAALAATMGLSSKAAAEMAESVSNPQRPPVIWIGAQECT WP_000145429.1
hydrogenase 2 small subunit [Salmonella enterica] Length: 372\nNumber of predicted TMHs: 1\nExp number of AAs in TMHs: 24.51309\nExp number, first 60 AAs: 0.2682\nTotal prob of N-in: 0.14402\noutside 1 331\nTMhelix 332 354\ninside 355 372

23520 GCF_000623295.1_ASM62329v1 Salmonella enterica subsp. enterica serovar Enteritidis str. EC20110360
Proteobacteria; Gammaproteobacteria; Enterobacterales; Enterobacteriaceae; Salmonella; Salmonella enterica; Salmonella enterica subsp. enterica; Salmonella enterica subsp. enterica serovar Enteritidis
MTGDNTLITSHGINRRDFMKLCAALAATMGLSSKAAAEMAESVSNPQRPPVIWIGAQECT WP_000145429.1
hydrogenase 2 small subunit [Salmonella enterica] Length: 372\nNumber of predicted TMHs: 1\nExp number of AAs in TMHs: 24.51309\nExp number, first 60 AAs: 0.2682\nTotal prob of N-in: 0.14402\noutside 1 331\nTMhelix 332 354\ninside 355 372

23521 GCF_000623295.1_ASM62329v1 Salmonella enterica subsp. enterica serovar Enteritidis str. EC20110360
Proteobacteria; Gammaproteobacteria; Enterobacterales; Enterobacteriaceae; Salmonella; Salmonella enterica; Salmonella enterica subsp. enterica; Salmonella enterica subsp. enterica serovar Enteritidis
MNNEETFYQAMRRKGVTRRSFLKFCSLAATSLGLGAGMTPKIAWALENKPRIPVVWIHGL WP_001058307.1
hydrogenase [Salmonella enterica] Length: 372\nNumber of predicted TMHs: 2\nExp number of AAs in TMHs: 36.51738\nExp number, first 60 AAs: 19.72849\nTotal prob of N-in: 0.96170\nPOSSIBLE N-term signal sequence\ninside 1 19\nTMhelix 20 39\noutside 40 325\nTMhelix 326 348\ninside 349 372

23522 GCF_000623275.1_ASM62327v1 Salmonella enterica subsp. enterica serovar Enteritidis str. EC20110361
Proteobacteria; Gammaproteobacteria; Enterobacterales; Enterobacteriaceae; Salmonella; Salmonella enterica; Salmonella enterica subsp. enterica; Salmonella enterica subsp. enterica serovar Enteritidis
MNNEETFYQAMRRKGVTRRSFLKFCSLAATSLGLGAGMTPKIAWALENKPRIPVVWIHGL WP_001058307.1
hydrogenase [Salmonella enterica] Length: 372\nNumber of predicted TMHs: 2\nExp number of AAs in TMHs: 36.51738\nExp number, first 60 AAs: 19.72849\nTotal prob of N-in: 0.96170\nPOSSIBLE N-term signal sequence\ninside 1 19\nTMhelix 20 39\noutside 40 325\nTMhelix 326 348\ninside 349 372

23523 GCF_000623275.1_ASM62327v1 Salmonella enterica subsp. enterica serovar Enteritidis str. EC20110361 Proteobacteria; Gammaproteobacteria; Enterobacterales; Enterobacteriaceae; Salmonella; Salmonella enterica; Salmonella enterica subsp. enterica; Salmonella enterica subsp. enterica serovar Enteritidis MTGDNTLITSHGINRRDFMKLCAALAATMGLSSKAAAEMAESVSNPQRPPVIWIGAQECT WP_000145429.1 hydrogenase 2 small subunit [Salmonella enterica] Length: 372\nNumber of predicted TMHs: 1\nExp number of AAs in TMHs: 24.51309\nExp number, first 60 AAs: 0.2682\nTotal prob of N-in: 0.14402\noutside 1 331\nTMhelix 332 354\ninside 355 372

23524 GCF_000626375.1_ASM62637v1 Salmonella enterica subsp. enterica serovar Enteritidis str. EC20111095 Proteobacteria; Gammaproteobacteria; Enterobacterales; Enterobacteriaceae; Salmonella; Salmonella enterica; Salmonella enterica subsp. enterica; Salmonella enterica subsp. enterica serovar Enteritidis MTGDNTLITSHGINRRDFMKLCAALAATMGLSSKAAAEMAESVSNPQRPPVIWIGAQECT WP_000145429.1 hydrogenase 2 small subunit [Salmonella enterica] Length: 372\nNumber of predicted TMHs: 1\nExp number of AAs in TMHs: 24.51309\nExp number, first 60 AAs: 0.2682\nTotal prob of N-in: 0.14402\noutside 1 331\nTMhelix 332 354\ninside 355 372

23525 GCF_000626375.1_ASM62637v1 Salmonella enterica subsp. enterica serovar Enteritidis str. EC20111095 Proteobacteria; Gammaproteobacteria; Enterobacterales; Enterobacteriaceae; Salmonella; Salmonella enterica; Salmonella enterica subsp. enterica; Salmonella enterica subsp. enterica serovar Enteritidis MNNEETFYQAMRRKGVTRRSFLKFCSLAATSLGLGAGMTPKIAWALENKPRIPVVWIHGL WP_001058307.1 hydrogenase [Salmonella enterica] Length: 372\nNumber of predicted TMHs: 2\nExp number of AAs in TMHs: 36.51738\nExp number, first 60 AAs: 19.72849\nTotal prob of N-in: 0.96170\nPOSSIBLE N-term signal sequence\ninside 1 19\nTMhelix 20 39\noutside 40 325\nTMhelix 326 348\ninside 349 372

23526 GCF_000626355.1_ASM62635v1 Salmonella enterica subsp. enterica serovar Enteritidis str. EC20111174 Proteobacteria; Gammaproteobacteria; Enterobacterales; Enterobacteriaceae; Salmonella; Salmonella enterica; Salmonella enterica subsp. enterica; Salmonella enterica subsp. enterica serovar Enteritidis MTGDNTLITSHGINRRDFMKLCAALAATMGLSSKAAAEMAESVSNPQRPPVIWIGAQECT WP_000145429.1 hydrogenase 2 small subunit [Salmonella enterica] Length: 372\nNumber of predicted TMHs: 1\nExp number of AAs in TMHs: 24.51309\nExp number, first 60 AAs: 0.2682\nTotal prob of N-in: 0.14402\noutside 1 331\nTMhelix 332 354\ninside 355 372

23527 GCF_000626355.1_ASM62635v1 Salmonella enterica subsp. enterica serovar Enteritidis str. EC20111174 Proteobacteria; Gammaproteobacteria; Enterobacterales; Enterobacteriaceae; Salmonella; Salmonella enterica; Salmonella enterica subsp. enterica; Salmonella enterica subsp. enterica serovar Enteritidis MNNEETFYQAMRRKGVTRRSFLKFCSLAATSLGLGAGMTPKIAWALENKPRIPVVWIHGL WP_001058307.1 hydrogenase [Salmonella enterica] Length: 372\nNumber of predicted TMHs: 2\nExp number of AAs in TMHs: 36.51738\nExp number, first 60 AAs: 19.72849\nTotal prob of N-in: 0.96170\nPOSSIBLE N-term signal sequence\ninside 1 19\nTMhelix 20 39\noutside 40 325\nTMhelix 326 348\ninside 349 372

23528 GCF_000626335.1_ASM62633v1 Salmonella enterica subsp. enterica serovar Enteritidis str. EC20111175 Proteobacteria; Gammaproteobacteria; Enterobacterales; Enterobacteriaceae; Salmonella; Salmonella enterica; Salmonella enterica subsp. enterica; Salmonella enterica subsp. enterica serovar Enteritidis MNNEETFYQAMRRKGVTRRSFLKFCSLAATSLGLGAGMTPKIAWALENKPRIPVVWIHGL WP_001058307.1 hydrogenase [Salmonella enterica] Length: 372\nNumber of predicted TMHs: 2\nExp number of AAs in TMHs: 36.51738\nExp number, first 60 AAs: 19.72849\nTotal prob of N-in: 0.96170\nPOSSIBLE N-term signal sequence\ninside 1 19\nTMhelix 20 39\noutside 40 325\nTMhelix 326 348\ninside 349 372

23529 GCF_000626335.1_ASM62633v1 Salmonella enterica subsp. enterica serovar Enteritidis str. EC20111175 Proteobacteria; Gammaproteobacteria; Enterobacterales; Enterobacteriaceae; Salmonella; Salmonella enterica; Salmonella enterica subsp. enterica; Salmonella enterica subsp. enterica serovar Enteritidis MTGDNTLITSHGINRRDFMKLCAALAATMGLSSKAAAEMAESVSNPQRPPVIWIGAQECT WP_000145429.1 hydrogenase 2 small subunit [Salmonella enterica] Length: 372\nNumber of predicted TMHs: 1\nExp number of AAs in TMHs: 24.51309\nExp number, first 60 AAs: 0.2682\nTotal prob of N-in: 0.14402\noutside 1 331\nTMhelix 332 354\ninside 355 372

23530 GCF_000626555.2_ASM62655v2 Salmonella enterica subsp. enterica serovar Enteritidis str. EC20111510 Proteobacteria; Gammaproteobacteria; Enterobacterales; Enterobacteriaceae; Salmonella; Salmonella enterica; Salmonella enterica subsp. enterica; Salmonella enterica subsp. enterica serovar Enteritidis MTGDNTLITSHGINRRDFMKLCAALAATMGLSSKAAAEMAESVSNPQRPPVIWIGAQECT WP_000145429.1 hydrogenase 2 small subunit [Salmonella enterica] Length: 372\nNumber of predicted TMHs: 1\nExp

number of AAs in TMHs: 24.51309\nExp number, first 60 AAs: 0.2682\nTotal prob of N-in: 0.14402\noutside 1 331\nTMhelix 332 354\ninside 355 372

23531 GCF_000626555.2_ASM62655v2 Salmonella enterica subsp. enterica serovar Enteritidis str. EC20111510 Proteobacteria; Gammaproteobacteria; Enterobacterales; Enterobacteriaceae; Salmonella; Salmonella enterica; Salmonella enterica subsp. enterica; Salmonella enterica subsp. enterica serovar Enteritidis MNNEETFYQAMRRKGVTRRSFLKFCSLAATSLGLGAGMTPKIAWALENKPRIPVVWIHGL WP_001058307.1 hydrogenase [Salmonella enterica] Length: 372\nNumber of predicted TMHs: 2\nExp number of AAs in TMHs: 36.51738\nExp number, first 60 AAs: 19.72849\nTotal prob of N-in: 0.96170\nPOSSIBLE N-term signal sequence\ninside 1 19\nTMhelix 20 39\noutside 40 325\nTMhelix 326 348\ninside 349 372

23532 GCF_000624295.2_ASM62429v2 Salmonella enterica subsp. enterica serovar Enteritidis str. EC20111514 Proteobacteria; Gammaproteobacteria; Enterobacterales; Enterobacteriaceae; Salmonella; Salmonella enterica; Salmonella enterica subsp. enterica; Salmonella enterica subsp. enterica serovar Enteritidis MNNEETFYQAMRRKGVTRRSFLKFCSLAATSLGLGAGMTPKIAWALENKPRIPVVWIHGL WP_001058307.1 hydrogenase [Salmonella enterica] Length: 372\nNumber of predicted TMHs: 2\nExp number of AAs in TMHs: 36.51738\nExp number, first 60 AAs: 19.72849\nTotal prob of N-in: 0.96170\nPOSSIBLE N-term signal sequence\ninside 1 19\nTMhelix 20 39\noutside 40 325\nTMhelix 326 348\ninside 349 372

23533 GCF_000624295.2_ASM62429v2 Salmonella enterica subsp. enterica serovar Enteritidis str. EC20111514 Proteobacteria; Gammaproteobacteria; Enterobacterales; Enterobacteriaceae; Salmonella; Salmonella enterica; Salmonella enterica subsp. enterica; Salmonella enterica subsp. enterica serovar Enteritidis MTGDNTLITSHGINRRDFMKLCAALAATMGLSSKAAAEMAESVSNPQRPPVIWIGAQECT WP_000145429.1 hydrogenase 2 small subunit [Salmonella enterica] Length: 372\nNumber of predicted TMHs: 1\nExp number of AAs in TMHs: 24.51309\nExp number, first 60 AAs: 0.2682\nTotal prob of N-in: 0.14402\noutside 1 331\nTMhelix 332 354\ninside 355 372

23534 GCF_000624315.2_ASM62431v2 Salmonella enterica subsp. enterica serovar Enteritidis str. EC20111515 Proteobacteria; Gammaproteobacteria; Enterobacterales; Enterobacteriaceae; Salmonella; Salmonella enterica; Salmonella enterica subsp. enterica; Salmonella enterica subsp. enterica serovar Enteritidis MTGDNTLITSHGINRRDFMKLCAALAATMGLSSKAAAEMAESVSNPQRPPVIWIGAQECT WP_000145429.1 hydrogenase 2 small subunit [Salmonella enterica] Length: 372\nNumber of predicted TMHs: 1\nExp number of AAs in TMHs: 24.51309\nExp number, first 60 AAs: 0.2682\nTotal prob of N-in: 0.14402\noutside 1 331\nTMhelix 332 354\ninside 355 372

23535 GCF_000624315.2_ASM62431v2 Salmonella enterica subsp. enterica serovar Enteritidis str. EC20111515 Proteobacteria; Gammaproteobacteria; Enterobacterales; Enterobacteriaceae; Salmonella; Salmonella enterica; Salmonella enterica subsp. enterica; Salmonella enterica subsp. enterica serovar Enteritidis MNNEETFYQAMRRKGVTRRSFLKFCSLAATSLGLGAGMTPKIAWALENKPRIPVVWIHGL WP_001058307.1 hydrogenase [Salmonella enterica] Length: 372\nNumber of predicted TMHs: 2\nExp number of AAs in TMHs: 36.51738\nExp number, first 60 AAs: 19.72849\nTotal prob of N-in: 0.96170\nPOSSIBLE N-term signal sequence\ninside 1 19\nTMhelix 20 39\noutside 40 325\nTMhelix 326 348\ninside 349 372

23536 GCF_000624335.2_ASM62433v2 Salmonella enterica subsp. enterica serovar Enteritidis str. EC20111554 Proteobacteria; Gammaproteobacteria; Enterobacterales; Enterobacteriaceae; Salmonella; Salmonella enterica; Salmonella enterica subsp. enterica; Salmonella enterica subsp. enterica serovar Enteritidis MNNEETFYQAMRRKGVTRRSFLKFCSLAATSLGLGAGMTPKIAWALENKPRIPVVWIHGL WP_001058307.1 hydrogenase [Salmonella enterica] Length: 372\nNumber of predicted TMHs: 2\nExp number of AAs in TMHs: 36.51738\nExp number, first 60 AAs: 19.72849\nTotal prob of N-in: 0.96170\nPOSSIBLE N-term signal sequence\ninside 1 19\nTMhelix 20 39\noutside 40 325\nTMhelix 326 348\ninside 349 372

23537 GCF_000624335.2_ASM62433v2 Salmonella enterica subsp. enterica serovar Enteritidis str. EC20111554 Proteobacteria; Gammaproteobacteria; Enterobacterales; Enterobacteriaceae; Salmonella; Salmonella enterica; Salmonella enterica subsp. enterica; Salmonella enterica subsp. enterica serovar Enteritidis MTGDNTLITSHGINRRDFMKLCAALAATMGLSSKAAAEMAESVSNPQRPPVIWIGAQECT WP_000145429.1 hydrogenase 2 small subunit [Salmonella enterica] Length: 372\nNumber of predicted TMHs: 1\nExp number of AAs in TMHs: 24.51309\nExp number, first 60 AAs: 0.2682\nTotal prob of N-in: 0.14402\noutside 1 331\nTMhelix 332 354\ninside 355 372

23538 GCF_000624355.2_ASM62435v2 Salmonella enterica subsp. enterica serovar Enteritidis str. EC20111561 Proteobacteria; Gammaproteobacteria; Enterobacterales; Enterobacteriaceae; Salmonella; Salmonella enterica;

Salmonella enterica subsp. enterica; Salmonella enterica subsp. enterica serovar Enteritidis
MTGDNTLITSHGINRRDFMKLCAALAATMGLSSKAAAEMAESVSNPQRPPVIWIGAQECT WP_000145429.1
hydrogenase 2 small subunit [Salmonella enterica] Length: 372\nNumber of predicted TMHs: 1\nExp number of AAs in TMHs: 24.51309\nExp number, first 60 AAs: 0.2682\nTotal prob of N-in: 0.14402\noutside 1 331\nTMhelix 332 354\ninside 355 372

23539 GCF_000624355.2_ASM62435v2 Salmonella enterica subsp. enterica serovar Enteritidis str. EC20111561
Proteobacteria; Gammaproteobacteria; Enterobacterales; Enterobacteriaceae; Salmonella; Salmonella enterica; Salmonella enterica subsp. enterica; Salmonella enterica subsp. enterica serovar Enteritidis
MNNEETFYQAMRRKGVTRRSFLKFCSLAATSLGLGAGMTPKIAWALENKPRIPVVWIHGL WP_001058307.1
hydrogenase [Salmonella enterica] Length: 372\nNumber of predicted TMHs: 2\nExp number of AAs in TMHs: 36.51738\nExp number, first 60 AAs: 19.72849\nTotal prob of N-in: 0.96170\nPOSSIBLE N-term signal sequence\ninside 1 19\nTMhelix 20 39\noutside 40 325\nTMhelix 326 348\ninside 349 372

23540 GCF_000624375.2_ASM62437v2 Salmonella enterica subsp. enterica serovar Enteritidis str. EC20111576
Proteobacteria; Gammaproteobacteria; Enterobacterales; Enterobacteriaceae; Salmonella; Salmonella enterica; Salmonella enterica subsp. enterica; Salmonella enterica subsp. enterica serovar Enteritidis
MNNEETFYQAMRRKGVTRRSFLKFCSLAATSLGLGAGMTPKIAWALENKPRIPVVWIHGL WP_001058307.1
hydrogenase [Salmonella enterica] Length: 372\nNumber of predicted TMHs: 2\nExp number of AAs in TMHs: 36.51738\nExp number, first 60 AAs: 19.72849\nTotal prob of N-in: 0.96170\nPOSSIBLE N-term signal sequence\ninside 1 19\nTMhelix 20 39\noutside 40 325\nTMhelix 326 348\ninside 349 372

23541 GCF_000624375.2_ASM62437v2 Salmonella enterica subsp. enterica serovar Enteritidis str. EC20111576
Proteobacteria; Gammaproteobacteria; Enterobacterales; Enterobacteriaceae; Salmonella; Salmonella enterica; Salmonella enterica subsp. enterica; Salmonella enterica subsp. enterica serovar Enteritidis
MTGDNTLITSHGINRRDFMKLCAALAATMGLSSKAAAEMAESVSNPQRPPVIWIGAQECT WP_000145429.1
hydrogenase 2 small subunit [Salmonella enterica] Length: 372\nNumber of predicted TMHs: 1\nExp number of AAs in TMHs: 24.51309\nExp number, first 60 AAs: 0.2682\nTotal prob of N-in: 0.14402\noutside 1 331\nTMhelix 332 354\ninside 355 372

23542 GCF_000624395.2_ASM62439v2 Salmonella enterica subsp. enterica serovar Enteritidis str. EC20120002
Proteobacteria; Gammaproteobacteria; Enterobacterales; Enterobacteriaceae; Salmonella; Salmonella enterica; Salmonella enterica subsp. enterica; Salmonella enterica subsp. enterica serovar Enteritidis
MNNEETFYQAMRRKGVTRRSFLKFCSLAATSLGLGAGMTPKIAWALENKPRIPVVWIHGL WP_001058307.1
hydrogenase [Salmonella enterica] Length: 372\nNumber of predicted TMHs: 2\nExp number of AAs in TMHs: 36.51738\nExp number, first 60 AAs: 19.72849\nTotal prob of N-in: 0.96170\nPOSSIBLE N-term signal sequence\ninside 1 19\nTMhelix 20 39\noutside 40 325\nTMhelix 326 348\ninside 349 372

23543 GCF_000624395.2_ASM62439v2 Salmonella enterica subsp. enterica serovar Enteritidis str. EC20120002
Proteobacteria; Gammaproteobacteria; Enterobacterales; Enterobacteriaceae; Salmonella; Salmonella enterica; Salmonella enterica subsp. enterica; Salmonella enterica subsp. enterica serovar Enteritidis
MTGDNTLITSHGINRRDFMKLCAALAATMGLSSKAAAEMAESVSNPQRPPVIWIGAQECT WP_000145429.1
hydrogenase 2 small subunit [Salmonella enterica] Length: 372\nNumber of predicted TMHs: 1\nExp number of AAs in TMHs: 24.51309\nExp number, first 60 AAs: 0.2682\nTotal prob of N-in: 0.14402\noutside 1 331\nTMhelix 332 354\ninside 355 372

23544 GCF_000624415.2_ASM62441v2 Salmonella enterica subsp. enterica serovar Enteritidis str. EC20120003
Proteobacteria; Gammaproteobacteria; Enterobacterales; Enterobacteriaceae; Salmonella; Salmonella enterica; Salmonella enterica subsp. enterica; Salmonella enterica subsp. enterica serovar Enteritidis
MNNEETFYQAMRRKGVTRRSFLKFCSLAATSLGLGAGMTPKIAWALENKPRIPVVWIHGL WP_001058307.1
hydrogenase [Salmonella enterica] Length: 372\nNumber of predicted TMHs: 2\nExp number of AAs in TMHs: 36.51738\nExp number, first 60 AAs: 19.72849\nTotal prob of N-in: 0.96170\nPOSSIBLE N-term signal sequence\ninside 1 19\nTMhelix 20 39\noutside 40 325\nTMhelix 326 348\ninside 349 372

23545 GCF_000624415.2_ASM62441v2 Salmonella enterica subsp. enterica serovar Enteritidis str. EC20120003
Proteobacteria; Gammaproteobacteria; Enterobacterales; Enterobacteriaceae; Salmonella; Salmonella enterica; Salmonella enterica subsp. enterica; Salmonella enterica subsp. enterica serovar Enteritidis
MTGDNTLITSHGINRRDFMKLCAALAATMGLSSKAAAEMAESVSNPQRPPVIWIGAQECT WP_000145429.1
hydrogenase 2 small subunit [Salmonella enterica] Length: 372\nNumber of predicted TMHs: 1\nExp number of AAs in TMHs: 24.51309\nExp number, first 60 AAs: 0.2682\nTotal prob of N-in: 0.14402\noutside 1 331\nTMhelix 332 354\ninside 355 372

23546 GCF_000623195.2_ASM62319v2 Salmonella enterica subsp. enterica serovar Enteritidis str. EC20120005 Proteobacteria; Gammaproteobacteria; Enterobacterales; Enterobacteriaceae; Salmonella; Salmonella enterica; Salmonella enterica subsp. enterica; Salmonella enterica subsp. enterica serovar Enteritidis MTGDNTLITSHGINRRDFMKLCAALAATMGLSSKAAAEMAESVSNPQRPPVIWIGAQECT WP_000145429.1 hydrogenase 2 small subunit [Salmonella enterica] Length: 372\nNumber of predicted TMHs: 1\nExp number of AAs in TMHs: 24.51309\nExp number, first 60 AAs: 0.2682\nTotal prob of N-in: 0.14402\noutside 1 331\nTMhelix 332 354\ninside 355 372

23547 GCF_000623195.2_ASM62319v2 Salmonella enterica subsp. enterica serovar Enteritidis str. EC20120005 Proteobacteria; Gammaproteobacteria; Enterobacterales; Enterobacteriaceae; Salmonella; Salmonella enterica; Salmonella enterica subsp. enterica; Salmonella enterica subsp. enterica serovar Enteritidis MNNEETFYQAMRRKGVTRRSFLKFCSLAATSLGLGAGMTPKIAWALENKPRIPVVWIHGL WP_001058307.1 hydrogenase [Salmonella enterica] Length: 372\nNumber of predicted TMHs: 2\nExp number of AAs in TMHs: 36.51738\nExp number, first 60 AAs: 19.72849\nTotal prob of N-in: 0.96170\nPOSSIBLE N-term signal sequence\ninside 1 19\nTMhelix 20 39\noutside 40 325\nTMhelix 326 348\ninside 349 372

23548 GCF_000624435.2_ASM62443v2 Salmonella enterica subsp. enterica serovar Enteritidis str. EC20120007 Proteobacteria; Gammaproteobacteria; Enterobacterales; Enterobacteriaceae; Salmonella; Salmonella enterica; Salmonella enterica subsp. enterica; Salmonella enterica subsp. enterica serovar Enteritidis MNNEETFYQAMRRKGVTRRSFLKFCSLAATSLGLGAGMTPKIAWALENKPRIPVVWIHGL WP_001058307.1 hydrogenase [Salmonella enterica] Length: 372\nNumber of predicted TMHs: 2\nExp number of AAs in TMHs: 36.51738\nExp number, first 60 AAs: 19.72849\nTotal prob of N-in: 0.96170\nPOSSIBLE N-term signal sequence\ninside 1 19\nTMhelix 20 39\noutside 40 325\nTMhelix 326 348\ninside 349 372

23549 GCF_000624435.2_ASM62443v2 Salmonella enterica subsp. enterica serovar Enteritidis str. EC20120007 Proteobacteria; Gammaproteobacteria; Enterobacterales; Enterobacteriaceae; Salmonella; Salmonella enterica; Salmonella enterica subsp. enterica; Salmonella enterica subsp. enterica serovar Enteritidis MTGDNTLITSHGINRRDFMKLCAALAATMGLSSKAAAEMAESVSNPQRPPVIWIGAQECT WP_000145429.1 hydrogenase 2 small subunit [Salmonella enterica] Length: 372\nNumber of predicted TMHs: 1\nExp number of AAs in TMHs: 24.51309\nExp number, first 60 AAs: 0.2682\nTotal prob of N-in: 0.14402\noutside 1 331\nTMhelix 332 354\ninside 355 372

23550 GCF_000626195.1_ASM62619v1 Salmonella enterica subsp. enterica serovar Enteritidis str. EC20120008 Proteobacteria; Gammaproteobacteria; Enterobacterales; Enterobacteriaceae; Salmonella; Salmonella enterica; Salmonella enterica subsp. enterica; Salmonella enterica subsp. enterica serovar Enteritidis MNNEETFYQAMRRKGVTRRSFLKFCSLAATSLGLGAGMTPKIAWALENKPRIPVVWIHGL WP_001058307.1 hydrogenase [Salmonella enterica] Length: 372\nNumber of predicted TMHs: 2\nExp number of AAs in TMHs: 36.51738\nExp number, first 60 AAs: 19.72849\nTotal prob of N-in: 0.96170\nPOSSIBLE N-term signal sequence\ninside 1 19\nTMhelix 20 39\noutside 40 325\nTMhelix 326 348\ninside 349 372

23551 GCF_000626195.1_ASM62619v1 Salmonella enterica subsp. enterica serovar Enteritidis str. EC20120008 Proteobacteria; Gammaproteobacteria; Enterobacterales; Enterobacteriaceae; Salmonella; Salmonella enterica; Salmonella enterica subsp. enterica; Salmonella enterica subsp. enterica serovar Enteritidis MTGDNTLITSHGINRRDFMKLCAALAATMGLSSKAAAEMAESVSNPQRPPVIWIGAQECT WP_000145429.1 hydrogenase 2 small subunit [Salmonella enterica] Length: 372\nNumber of predicted TMHs: 1\nExp number of AAs in TMHs: 24.51309\nExp number, first 60 AAs: 0.2682\nTotal prob of N-in: 0.14402\noutside 1 331\nTMhelix 332 354\ninside 355 372

23552 GCF_000626395.1_ASM62639v2 Salmonella enterica subsp. enterica serovar Enteritidis str. EC20120009 Proteobacteria; Gammaproteobacteria; Enterobacterales; Enterobacteriaceae; Salmonella; Salmonella enterica; Salmonella enterica subsp. enterica; Salmonella enterica subsp. enterica serovar Enteritidis MNNEETFYQAMRRKGVTRRSFLKFCSLAATSLGLGAGMTPKIAWALENKPRIPVVWIHGL WP_001058307.1 hydrogenase [Salmonella enterica] Length: 372\nNumber of predicted TMHs: 2\nExp number of AAs in TMHs: 36.51738\nExp number, first 60 AAs: 19.72849\nTotal prob of N-in: 0.96170\nPOSSIBLE N-term signal sequence\ninside 1 19\nTMhelix 20 39\noutside 40 325\nTMhelix 326 348\ninside 349 372

23553 GCF_000626395.1_ASM62639v2 Salmonella enterica subsp. enterica serovar Enteritidis str. EC20120009 Proteobacteria; Gammaproteobacteria; Enterobacterales; Enterobacteriaceae; Salmonella; Salmonella enterica; Salmonella enterica subsp. enterica; Salmonella enterica subsp. enterica serovar Enteritidis MTGDNTLITSHGINRRDFMKLCAALAATMGLSSKAAAEMAESVSNPQRPPVIWIGAQECT WP_000145429.1 hydrogenase 2 small subunit [Salmonella enterica] Length: 372\nNumber of predicted TMHs: 1\nExp

number of AAs in TMHs: 24.51309\nExp number, first 60 AAs: 0.2682\nTotal prob of N-in: 0.14402\noutside 1 331\nTMhelix 332 354\ninside 355 372

23554 GCF_000625835.2_ASM62583v2 Salmonella enterica subsp. enterica serovar Enteritidis str. EC20120051 Proteobacteria; Gammaproteobacteria; Enterobacterales; Enterobacteriaceae; Salmonella; Salmonella enterica; Salmonella enterica subsp. enterica; Salmonella enterica subsp. enterica serovar Enteritidis MTGDNTLITSHGINRRDFMKLCAALAATMGLSSKAAAEMAESVSNPQRPPVIWIGAQECT WP_000145429.1 hydrogenase 2 small subunit [Salmonella enterica] Length: 372\nNumber of predicted TMHs: 1\nExp number of AAs in TMHs: 24.51309\nExp number, first 60 AAs: 0.2682\nTotal prob of N-in: 0.14402\noutside 1 331\nTMhelix 332 354\ninside 355 372

23555 GCF_000625835.2_ASM62583v2 Salmonella enterica subsp. enterica serovar Enteritidis str. EC20120051 Proteobacteria; Gammaproteobacteria; Enterobacterales; Enterobacteriaceae; Salmonella; Salmonella enterica; Salmonella enterica subsp. enterica; Salmonella enterica subsp. enterica serovar Enteritidis MNNEETFYQAMRRKGVTRRSFLKFCSLAATSLGLGAGMTPKIAWALENKPRIPVVWIHGL WP_001058307.1 hydrogenase [Salmonella enterica] Length: 372\nNumber of predicted TMHs: 2\nExp number of AAs in TMHs: 36.51738\nExp number, first 60 AAs: 19.72849\nTotal prob of N-in: 0.96170\nPOSSIBLE N-term signal sequence\ninside 1 19\nTMhelix 20 39\noutside 40 325\nTMhelix 326 348\ninside 349 372

23556 GCF_000626095.2_ASM62609v2 Salmonella enterica subsp. enterica serovar Enteritidis str. EC20120200 Proteobacteria; Gammaproteobacteria; Enterobacterales; Enterobacteriaceae; Salmonella; Salmonella enterica; Salmonella enterica subsp. enterica; Salmonella enterica subsp. enterica serovar Enteritidis MNNEETFYQAMRRKGVTRRSFLKFCSLAATSLGLGAGMTPKIAWALENKPRIPVVWIHGL WP_001058307.1 hydrogenase [Salmonella enterica] Length: 372\nNumber of predicted TMHs: 2\nExp number of AAs in TMHs: 36.51738\nExp number, first 60 AAs: 19.72849\nTotal prob of N-in: 0.96170\nPOSSIBLE N-term signal sequence\ninside 1 19\nTMhelix 20 39\noutside 40 325\nTMhelix 326 348\ninside 349 372

23557 GCF_000626095.2_ASM62609v2 Salmonella enterica subsp. enterica serovar Enteritidis str. EC20120200 Proteobacteria; Gammaproteobacteria; Enterobacterales; Enterobacteriaceae; Salmonella; Salmonella enterica; Salmonella enterica subsp. enterica; Salmonella enterica subsp. enterica serovar Enteritidis MTGDNTLITSHGINRRDFMKLCAALAATMGLSSKAAAEMAESVSNPQRPPVIWIGAQECT WP_000145429.1 hydrogenase 2 small subunit [Salmonella enterica] Length: 372\nNumber of predicted TMHs: 1\nExp number of AAs in TMHs: 24.51309\nExp number, first 60 AAs: 0.2682\nTotal prob of N-in: 0.14402\noutside 1 331\nTMhelix 332 354\ninside 355 372

23558 GCF_000624455.2_ASM62445v2 Salmonella enterica subsp. enterica serovar Enteritidis str. EC20120213 Proteobacteria; Gammaproteobacteria; Enterobacterales; Enterobacteriaceae; Salmonella; Salmonella enterica; Salmonella enterica subsp. enterica; Salmonella enterica subsp. enterica serovar Enteritidis MTGDNTLITSHGINRRDFMKLCAALAATMGLSSKAAAEMAESVSNPQRPPVIWIGAQECT WP_000145429.1 hydrogenase 2 small subunit [Salmonella enterica] Length: 372\nNumber of predicted TMHs: 1\nExp number of AAs in TMHs: 24.51309\nExp number, first 60 AAs: 0.2682\nTotal prob of N-in: 0.14402\noutside 1 331\nTMhelix 332 354\ninside 355 372

23559 GCF_000624455.2_ASM62445v2 Salmonella enterica subsp. enterica serovar Enteritidis str. EC20120213 Proteobacteria; Gammaproteobacteria; Enterobacterales; Enterobacteriaceae; Salmonella; Salmonella enterica; Salmonella enterica subsp. enterica; Salmonella enterica subsp. enterica serovar Enteritidis MNNEETFYQAMRRKGVTRRSFLKFCSLAATSLGLGAGMTPKIAWALENKPRIPVVWIHGL WP_001058307.1 hydrogenase [Salmonella enterica] Length: 372\nNumber of predicted TMHs: 2\nExp number of AAs in TMHs: 36.51738\nExp number, first 60 AAs: 19.72849\nTotal prob of N-in: 0.96170\nPOSSIBLE N-term signal sequence\ninside 1 19\nTMhelix 20 39\noutside 40 325\nTMhelix 326 348\ninside 349 372

23560 GCF_000624475.2_ASM62447v2 Salmonella enterica subsp. enterica serovar Enteritidis str. EC20120219 Proteobacteria; Gammaproteobacteria; Enterobacterales; Enterobacteriaceae; Salmonella; Salmonella enterica; Salmonella enterica subsp. enterica; Salmonella enterica subsp. enterica serovar Enteritidis MTGDNTLITSHGINRRDFMKLCAALAATMGLSSKAAAEMAESVSNPQRPPVIWIGAQECT WP_000145429.1 hydrogenase 2 small subunit [Salmonella enterica] Length: 372\nNumber of predicted TMHs: 1\nExp number of AAs in TMHs: 24.51309\nExp number, first 60 AAs: 0.2682\nTotal prob of N-in: 0.14402\noutside 1 331\nTMhelix 332 354\ninside 355 372

23561 GCF_000624475.2_ASM62447v2 Salmonella enterica subsp. enterica serovar Enteritidis str. EC20120219 Proteobacteria; Gammaproteobacteria; Enterobacterales; Enterobacteriaceae; Salmonella; Salmonella enterica;

Salmonella enterica subsp. enterica; Salmonella enterica subsp. enterica serovar Enteritidis
MNNEETFYQAMRRKGVTRRSFLKFCSLAATSLGLGAGMTPKIAWALENKPRIPVVWIHGL WP_001058307.1
hydrogenase [Salmonella enterica] Length: 372\nNumber of predicted TMHs: 2\nExp number of AAs in TMHs: 36.51738\nExp number, first 60 AAs: 19.72849\nTotal prob of N-in: 0.96170\nPOSSIBLE N-term signal sequence\ninside 1 19\nTMhelix 20 39\noutside 40 325\nTMhelix 326 348\ninside 349 372

23562 GCF_000624495.2_ASM62449v2 Salmonella enterica subsp. enterica serovar Enteritidis str. EC20120229
Proteobacteria; Gammaproteobacteria; Enterobacterales; Enterobacteriaceae; Salmonella; Salmonella enterica; Salmonella enterica subsp. enterica; Salmonella enterica subsp. enterica serovar Enteritidis
MNNEETFYQAMRRKGVTRRSFLKFCSLAATSLGLGAGMTPKIAWALENKPRIPVVWIHGL WP_001058307.1
hydrogenase [Salmonella enterica] Length: 372\nNumber of predicted TMHs: 2\nExp number of AAs in TMHs: 36.51738\nExp number, first 60 AAs: 19.72849\nTotal prob of N-in: 0.96170\nPOSSIBLE N-term signal sequence\ninside 1 19\nTMhelix 20 39\noutside 40 325\nTMhelix 326 348\ninside 349 372

23563 GCF_000624495.2_ASM62449v2 Salmonella enterica subsp. enterica serovar Enteritidis str. EC20120229
Proteobacteria; Gammaproteobacteria; Enterobacterales; Enterobacteriaceae; Salmonella; Salmonella enterica; Salmonella enterica subsp. enterica; Salmonella enterica subsp. enterica serovar Enteritidis
MTGDNTLITSHGINRRDFMKLCAALAATMGLSSKAAAEMAESVSNPQRPPVIWIGAQECT WP_000145429.1
hydrogenase 2 small subunit [Salmonella enterica] Length: 372\nNumber of predicted TMHs: 1\nExp number of AAs in TMHs: 24.51309\nExp number, first 60 AAs: 0.2682\nTotal prob of N-in: 0.14402\noutside 1 331\nTMhelix 332 354\ninside 355 372

23564 GCF_000624515.2_ASM62451v2 Salmonella enterica subsp. enterica serovar Enteritidis str. EC20120240
Proteobacteria; Gammaproteobacteria; Enterobacterales; Enterobacteriaceae; Salmonella; Salmonella enterica; Salmonella enterica subsp. enterica; Salmonella enterica subsp. enterica serovar Enteritidis
MTGDNTLITSHGINRRDFMKLCAALAATMGLSSKAAAEMAESVSNPQRPPVIWIGAQECT WP_000145429.1
hydrogenase 2 small subunit [Salmonella enterica] Length: 372\nNumber of predicted TMHs: 1\nExp number of AAs in TMHs: 24.51309\nExp number, first 60 AAs: 0.2682\nTotal prob of N-in: 0.14402\noutside 1 331\nTMhelix 332 354\ninside 355 372

23565 GCF_000624515.2_ASM62451v2 Salmonella enterica subsp. enterica serovar Enteritidis str. EC20120240
Proteobacteria; Gammaproteobacteria; Enterobacterales; Enterobacteriaceae; Salmonella; Salmonella enterica; Salmonella enterica subsp. enterica; Salmonella enterica subsp. enterica serovar Enteritidis
MNNEETFYQAMRRKGVTRRSFLKFCSLAATSLGLGAGMTPKIAWALENKPRIPVVWIHGL WP_001058307.1
hydrogenase [Salmonella enterica] Length: 372\nNumber of predicted TMHs: 2\nExp number of AAs in TMHs: 36.51738\nExp number, first 60 AAs: 19.72849\nTotal prob of N-in: 0.96170\nPOSSIBLE N-term signal sequence\ninside 1 19\nTMhelix 20 39\noutside 40 325\nTMhelix 326 348\ninside 349 372

23566 GCF_000624535.2_ASM62453v2 Salmonella enterica subsp. enterica serovar Enteritidis str. EC20120356
Proteobacteria; Gammaproteobacteria; Enterobacterales; Enterobacteriaceae; Salmonella; Salmonella enterica; Salmonella enterica subsp. enterica; Salmonella enterica subsp. enterica serovar Enteritidis
MTGDNTLITSHGINRRDFMKLCAALAATMGLSSKAAAEMAESVSNPQRPPVIWIGAQECT WP_000145429.1
hydrogenase 2 small subunit [Salmonella enterica] Length: 372\nNumber of predicted TMHs: 1\nExp number of AAs in TMHs: 24.51309\nExp number, first 60 AAs: 0.2682\nTotal prob of N-in: 0.14402\noutside 1 331\nTMhelix 332 354\ninside 355 372

23567 GCF_000624535.2_ASM62453v2 Salmonella enterica subsp. enterica serovar Enteritidis str. EC20120356
Proteobacteria; Gammaproteobacteria; Enterobacterales; Enterobacteriaceae; Salmonella; Salmonella enterica; Salmonella enterica subsp. enterica; Salmonella enterica subsp. enterica serovar Enteritidis
MNNEETFYQAMRRKGVTRRSFLKFCSLAATSLGLGAGMTPKIAWALENKPRIPVVWIHGL WP_001058307.1
hydrogenase [Salmonella enterica] Length: 372\nNumber of predicted TMHs: 2\nExp number of AAs in TMHs: 36.51738\nExp number, first 60 AAs: 19.72849\nTotal prob of N-in: 0.96170\nPOSSIBLE N-term signal sequence\ninside 1 19\nTMhelix 20 39\noutside 40 325\nTMhelix 326 348\ninside 349 372

23568 GCF_000624555.2_ASM62455v2 Salmonella enterica subsp. enterica serovar Enteritidis str. EC20120469
Proteobacteria; Gammaproteobacteria; Enterobacterales; Enterobacteriaceae; Salmonella; Salmonella enterica; Salmonella enterica subsp. enterica; Salmonella enterica subsp. enterica serovar Enteritidis
MTGDNTLITSHGINRRDFMKLCAALAATMGLSSKAAAEMAESVSNPQRPPVIWIGAQECT WP_000145429.1
hydrogenase 2 small subunit [Salmonella enterica] Length: 372\nNumber of predicted TMHs: 1\nExp number of AAs in TMHs: 24.51309\nExp number, first 60 AAs: 0.2682\nTotal prob of N-in: 0.14402\noutside 1 331\nTMhelix 332 354\ninside 355 372

23569 GCF_000624555.2_ASM62455v2 Salmonella enterica subsp. enterica serovar Enteritidis str. EC20120469 Proteobacteria; Gammaproteobacteria; Enterobacterales; Enterobacteriaceae; Salmonella; Salmonella enterica; Salmonella enterica subsp. enterica; Salmonella enterica subsp. enterica serovar Enteritidis MNNEETFYQAMRRKGVTRRSFLKFCSLAATSLGLGAGMTPKIAWALENKPRIPVVWIHGL WP_001058307.1 hydrogenase [Salmonella enterica] Length: 372\nNumber of predicted TMHs: 2\nExp number of AAs in TMHs: 36.51738\nExp number, first 60 AAs: 19.72849\nTotal prob of N-in: 0.96170\nPOSSIBLE N-term signal sequence\ninside 1 19\nTMhelix 20 39\noutside 40 325\nTMhelix 326 348\ninside 349 372

23570 GCF_000624575.2_ASM62457v2 Salmonella enterica subsp. enterica serovar Enteritidis str. EC20120496 Proteobacteria; Gammaproteobacteria; Enterobacterales; Enterobacteriaceae; Salmonella; Salmonella enterica; Salmonella enterica subsp. enterica; Salmonella enterica subsp. enterica serovar Enteritidis MTGDNTLITSHGINRRDFMKLCAALAATMGLSSKAAAEMAESVSNPQRPPVIWIGAQECT WP_000145429.1 hydrogenase 2 small subunit [Salmonella enterica] Length: 372\nNumber of predicted TMHs: 1\nExp number of AAs in TMHs: 24.51309\nExp number, first 60 AAs: 0.2682\nTotal prob of N-in: 0.14402\noutside 1 331\nTMhelix 332 354\ninside 355 372

23571 GCF_000624575.2_ASM62457v2 Salmonella enterica subsp. enterica serovar Enteritidis str. EC20120496 Proteobacteria; Gammaproteobacteria; Enterobacterales; Enterobacteriaceae; Salmonella; Salmonella enterica; Salmonella enterica subsp. enterica; Salmonella enterica subsp. enterica serovar Enteritidis MNNEETFYQAMRRKGVTRRSFLKFCSLAATSLGLGAGMTPKIAWALENKPRIPVVWIHGL WP_001058307.1 hydrogenase [Salmonella enterica] Length: 372\nNumber of predicted TMHs: 2\nExp number of AAs in TMHs: 36.51738\nExp number, first 60 AAs: 19.72849\nTotal prob of N-in: 0.96170\nPOSSIBLE N-term signal sequence\ninside 1 19\nTMhelix 20 39\noutside 40 325\nTMhelix 326 348\ninside 349 372

23572 GCF_000624595.2_ASM62459v2 Salmonella enterica subsp. enterica serovar Enteritidis str. EC20120497 Proteobacteria; Gammaproteobacteria; Enterobacterales; Enterobacteriaceae; Salmonella; Salmonella enterica; Salmonella enterica subsp. enterica; Salmonella enterica subsp. enterica serovar Enteritidis MTGDNTLITSHGINRRDFMKLCAALAATMGLSSKAAAEMAESVSNPQRPPVIWIGAQECT WP_000145429.1 hydrogenase 2 small subunit [Salmonella enterica] Length: 372\nNumber of predicted TMHs: 1\nExp number of AAs in TMHs: 24.51309\nExp number, first 60 AAs: 0.2682\nTotal prob of N-in: 0.14402\noutside 1 331\nTMhelix 332 354\ninside 355 372

23573 GCF_000624595.2_ASM62459v2 Salmonella enterica subsp. enterica serovar Enteritidis str. EC20120497 Proteobacteria; Gammaproteobacteria; Enterobacterales; Enterobacteriaceae; Salmonella; Salmonella enterica; Salmonella enterica subsp. enterica; Salmonella enterica subsp. enterica serovar Enteritidis MNNEETFYQAMRRKGVTRRSFLKFCSLAATSLGLGAGMTPKIAWALENKPRIPVVWIHGL WP_001058307.1 hydrogenase [Salmonella enterica] Length: 372\nNumber of predicted TMHs: 2\nExp number of AAs in TMHs: 36.51738\nExp number, first 60 AAs: 19.72849\nTotal prob of N-in: 0.96170\nPOSSIBLE N-term signal sequence\ninside 1 19\nTMhelix 20 39\noutside 40 325\nTMhelix 326 348\ninside 349 372

23574 GCF_000624615.2_ASM62461v2 Salmonella enterica subsp. enterica serovar Enteritidis str. EC20120498 Proteobacteria; Gammaproteobacteria; Enterobacterales; Enterobacteriaceae; Salmonella; Salmonella enterica; Salmonella enterica subsp. enterica; Salmonella enterica subsp. enterica serovar Enteritidis MNNEETFYQAMRRKGVTRRSFLKFCSLAATSLGLGAGMTPKIAWALENKPRIPVVWIHGL WP_001058307.1 hydrogenase [Salmonella enterica] Length: 372\nNumber of predicted TMHs: 2\nExp number of AAs in TMHs: 36.51738\nExp number, first 60 AAs: 19.72849\nTotal prob of N-in: 0.96170\nPOSSIBLE N-term signal sequence\ninside 1 19\nTMhelix 20 39\noutside 40 325\nTMhelix 326 348\ninside 349 372

23575 GCF_000624615.2_ASM62461v2 Salmonella enterica subsp. enterica serovar Enteritidis str. EC20120498 Proteobacteria; Gammaproteobacteria; Enterobacterales; Enterobacteriaceae; Salmonella; Salmonella enterica; Salmonella enterica subsp. enterica; Salmonella enterica subsp. enterica serovar Enteritidis MTGDNTLITSHGINRRDFMKLCAALAATMGLSSKAAAEMAESVSNPQRPPVIWIGAQECT WP_000145429.1 hydrogenase 2 small subunit [Salmonella enterica] Length: 372\nNumber of predicted TMHs: 1\nExp number of AAs in TMHs: 24.51309\nExp number, first 60 AAs: 0.2682\nTotal prob of N-in: 0.14402\noutside 1 331\nTMhelix 332 354\ninside 355 372

23576 GCF_000624635.2_ASM62463v2 Salmonella enterica subsp. enterica serovar Enteritidis str. EC20120505 Proteobacteria; Gammaproteobacteria; Enterobacterales; Enterobacteriaceae; Salmonella; Salmonella enterica; Salmonella enterica subsp. enterica; Salmonella enterica subsp. enterica serovar Enteritidis MNNEETFYQAMRRKGVTRRSFLKFCSLAATSLGLGAGMTPKIAWALENKPRIPVVWIHGL WP_001058307.1 hydrogenase [Salmonella enterica] Length: 372\nNumber of predicted TMHs: 2\nExp number of AAs in

TMHs: 36.51738\nExp number, first 60 AAs: 19.72849\nTotal prob of N-in: 0.96170\nPOSSIBLE N-term signal sequence\ninside 1 19\nTMhelix 20 39\noutside 40 325\nTMhelix 326 348\ninside 349 372

23577 GCF_000624635.2_ASM62463v2 Salmonella enterica subsp. enterica serovar Enteritidis str. EC20120505 Proteobacteria; Gammaproteobacteria; Enterobacterales; Enterobacteriaceae; Salmonella; Salmonella enterica; Salmonella enterica subsp. enterica; Salmonella enterica subsp. enterica serovar Enteritidis MTGDNTLITSHGINRRDFMKLCAALAATMGLSSKAAAEMAESVSNPQRPPVIWIGAQECT WP_000145429.1 hydrogenase 2 small subunit [Salmonella enterica] Length: 372\nNumber of predicted TMHs: 1\nExp number of AAs in TMHs: 24.51309\nExp number, first 60 AAs: 0.2682\nTotal prob of N-in: 0.14402\noutside 1 331\nTMhelix 332 354\ninside 355 372

23578 GCF_000624655.2_ASM62465v2 Salmonella enterica subsp. enterica serovar Enteritidis str. EC20120528 Proteobacteria; Gammaproteobacteria; Enterobacterales; Enterobacteriaceae; Salmonella; Salmonella enterica; Salmonella enterica subsp. enterica; Salmonella enterica subsp. enterica serovar Enteritidis MNNEETFYQAMRRKGVTRRSFLKFCSLAATSLGLGAGMTPKIAWALENKPRIPVVWIHGL WP_001058307.1 hydrogenase [Salmonella enterica] Length: 372\nNumber of predicted TMHs: 2\nExp number of AAs in TMHs: 36.51738\nExp number, first 60 AAs: 19.72849\nTotal prob of N-in: 0.96170\nPOSSIBLE N-term signal sequence\ninside 1 19\nTMhelix 20 39\noutside 40 325\nTMhelix 326 348\ninside 349 372

23579 GCF_000624655.2_ASM62465v2 Salmonella enterica subsp. enterica serovar Enteritidis str. EC20120528 Proteobacteria; Gammaproteobacteria; Enterobacterales; Enterobacteriaceae; Salmonella; Salmonella enterica; Salmonella enterica subsp. enterica; Salmonella enterica subsp. enterica serovar Enteritidis MTGDNTLITSHGINRRDFMKLCAALAATMGLSSKAAAEMAESVSNPQRPPVIWIGAQECT WP_000145429.1 hydrogenase 2 small subunit [Salmonella enterica] Length: 372\nNumber of predicted TMHs: 1\nExp number of AAs in TMHs: 24.51309\nExp number, first 60 AAs: 0.2682\nTotal prob of N-in: 0.14402\noutside 1 331\nTMhelix 332 354\ninside 355 372

23580 GCF_000623215.2_ASM62321v2 Salmonella enterica subsp. enterica serovar Enteritidis str. EC20120544 Proteobacteria; Gammaproteobacteria; Enterobacterales; Enterobacteriaceae; Salmonella; Salmonella enterica; Salmonella enterica subsp. enterica; Salmonella enterica subsp. enterica serovar Enteritidis MNNEETFYQAMRRKGVTRRSFLKFCSLAATSLGLGAGMTPKIAWALENKPRIPVVWIHGL WP_001058307.1 hydrogenase [Salmonella enterica] Length: 372\nNumber of predicted TMHs: 2\nExp number of AAs in TMHs: 36.51738\nExp number, first 60 AAs: 19.72849\nTotal prob of N-in: 0.96170\nPOSSIBLE N-term signal sequence\ninside 1 19\nTMhelix 20 39\noutside 40 325\nTMhelix 326 348\ninside 349 372

23581 GCF_000623215.2_ASM62321v2 Salmonella enterica subsp. enterica serovar Enteritidis str. EC20120544 Proteobacteria; Gammaproteobacteria; Enterobacterales; Enterobacteriaceae; Salmonella; Salmonella enterica; Salmonella enterica subsp. enterica; Salmonella enterica subsp. enterica serovar Enteritidis MTGDNTLITSHGINRRDFMKLCAALAATMGLSSKAAAEMAESVSNPQRPPVIWIGAQECT WP_000145429.1 hydrogenase 2 small subunit [Salmonella enterica] Length: 372\nNumber of predicted TMHs: 1\nExp number of AAs in TMHs: 24.51309\nExp number, first 60 AAs: 0.2682\nTotal prob of N-in: 0.14402\noutside 1 331\nTMhelix 332 354\ninside 355 372

23582 GCF_000623235.2_ASM62323v2 Salmonella enterica subsp. enterica serovar Enteritidis str. EC20120548 Proteobacteria; Gammaproteobacteria; Enterobacterales; Enterobacteriaceae; Salmonella; Salmonella enterica; Salmonella enterica subsp. enterica; Salmonella enterica subsp. enterica serovar Enteritidis MNNEETFYQAMRRKGVTRRSFLKFCSLAATSLGLGAGMTPKIAWALENKPRIPVVWIHGL WP_001058307.1 hydrogenase [Salmonella enterica] Length: 372\nNumber of predicted TMHs: 2\nExp number of AAs in TMHs: 36.51738\nExp number, first 60 AAs: 19.72849\nTotal prob of N-in: 0.96170\nPOSSIBLE N-term signal sequence\ninside 1 19\nTMhelix 20 39\noutside 40 325\nTMhelix 326 348\ninside 349 372

23583 GCF_000623235.2_ASM62323v2 Salmonella enterica subsp. enterica serovar Enteritidis str. EC20120548 Proteobacteria; Gammaproteobacteria; Enterobacterales; Enterobacteriaceae; Salmonella; Salmonella enterica; Salmonella enterica subsp. enterica; Salmonella enterica subsp. enterica serovar Enteritidis MTGDNTLITSHGINRRDFMKLCAALAATMGLSSKAAAEMAESVSNPQRPPVIWIGAQECT WP_000145429.1 hydrogenase 2 small subunit [Salmonella enterica] Length: 372\nNumber of predicted TMHs: 1\nExp number of AAs in TMHs: 24.51309\nExp number, first 60 AAs: 0.2682\nTotal prob of N-in: 0.14402\noutside 1 331\nTMhelix 332 354\ninside 355 372

23584 GCF_000623255.2_ASM62325v2 Salmonella enterica subsp. enterica serovar Enteritidis str. EC20120555 Proteobacteria; Gammaproteobacteria; Enterobacterales; Enterobacteriaceae; Salmonella; Salmonella enterica;

Salmonella enterica subsp. enterica; Salmonella enterica subsp. enterica serovar Enteritidis
MTGDNTLITSHGINRRDFMKLCAALAATMGLSSKAAAEMAESVSNPQRPPVIWIGAQECT WP_000145429.1
hydrogenase 2 small subunit [Salmonella enterica] Length: 372\nNumber of predicted TMHs: 1\nExp number of AAs in TMHs: 24.51309\nExp number, first 60 AAs: 0.2682\nTotal prob of N-in: 0.14402\noutside 1 331\nTMhelix 332 354\ninside 355 372

23585 GCF_000623255.2_ASM62325v2 Salmonella enterica subsp. enterica serovar Enteritidis str. EC20120555
Proteobacteria; Gammaproteobacteria; Enterobacterales; Enterobacteriaceae; Salmonella; Salmonella enterica; Salmonella enterica subsp. enterica; Salmonella enterica subsp. enterica serovar Enteritidis
MNNEETFYQAMRRKGVTRRSFLKFCSLAATSLGLGAGMTPKIAWALENKPRIPVVWIHGL WP_001058307.1
hydrogenase [Salmonella enterica] Length: 372\nNumber of predicted TMHs: 2\nExp number of AAs in TMHs: 36.51738\nExp number, first 60 AAs: 19.72849\nTotal prob of N-in: 0.96170\nPOSSIBLE N-term signal sequence\ninside 1 19\nTMhelix 20 39\noutside 40 325\nTMhelix 326 348\ninside 349 372

23586 GCF_000625455.2_ASM62545v2 Salmonella enterica subsp. enterica serovar Enteritidis str. EC20120580
Proteobacteria; Gammaproteobacteria; Enterobacterales; Enterobacteriaceae; Salmonella; Salmonella enterica; Salmonella enterica subsp. enterica; Salmonella enterica subsp. enterica serovar Enteritidis
MNNEETFYQAMRRKGVTRRSFLKFCSLAATSLGLGAGMTPKIAWALENKPRIPVVWIHGL WP_001058307.1
hydrogenase [Salmonella enterica] Length: 372\nNumber of predicted TMHs: 2\nExp number of AAs in TMHs: 36.51738\nExp number, first 60 AAs: 19.72849\nTotal prob of N-in: 0.96170\nPOSSIBLE N-term signal sequence\ninside 1 19\nTMhelix 20 39\noutside 40 325\nTMhelix 326 348\ninside 349 372

23587 GCF_000625455.2_ASM62545v2 Salmonella enterica subsp. enterica serovar Enteritidis str. EC20120580
Proteobacteria; Gammaproteobacteria; Enterobacterales; Enterobacteriaceae; Salmonella; Salmonella enterica; Salmonella enterica subsp. enterica; Salmonella enterica subsp. enterica serovar Enteritidis
MTGDNTLITSHGINRRDFMKLCAALAATMGLSSKAAAEMAESVSNPQRPPVIWIGAQECT WP_000145429.1
hydrogenase 2 small subunit [Salmonella enterica] Length: 372\nNumber of predicted TMHs: 1\nExp number of AAs in TMHs: 24.51309\nExp number, first 60 AAs: 0.2682\nTotal prob of N-in: 0.14402\noutside 1 331\nTMhelix 332 354\ninside 355 372

23588 GCF_000625475.2_ASM62547v2 Salmonella enterica subsp. enterica serovar Enteritidis str. EC20120581
Proteobacteria; Gammaproteobacteria; Enterobacterales; Enterobacteriaceae; Salmonella; Salmonella enterica; Salmonella enterica subsp. enterica; Salmonella enterica subsp. enterica serovar Enteritidis
MTGDNTLITSHGINRRDFMKLCAALAATMGLSSKAAAEMAESVSNPQRPPVIWIGAQECT WP_000145429.1
hydrogenase 2 small subunit [Salmonella enterica] Length: 372\nNumber of predicted TMHs: 1\nExp number of AAs in TMHs: 24.51309\nExp number, first 60 AAs: 0.2682\nTotal prob of N-in: 0.14402\noutside 1 331\nTMhelix 332 354\ninside 355 372

23589 GCF_000625475.2_ASM62547v2 Salmonella enterica subsp. enterica serovar Enteritidis str. EC20120581
Proteobacteria; Gammaproteobacteria; Enterobacterales; Enterobacteriaceae; Salmonella; Salmonella enterica; Salmonella enterica subsp. enterica; Salmonella enterica subsp. enterica serovar Enteritidis
MNNEETFYQAMRRKGVTRRSFLKFCSLAATSLGLGAGMTPKIAWALENKPRIPVVWIHGL WP_001058307.1
hydrogenase [Salmonella enterica] Length: 372\nNumber of predicted TMHs: 2\nExp number of AAs in TMHs: 36.51738\nExp number, first 60 AAs: 19.72849\nTotal prob of N-in: 0.96170\nPOSSIBLE N-term signal sequence\ninside 1 19\nTMhelix 20 39\noutside 40 325\nTMhelix 326 348\ninside 349 372

23590 GCF_000625495.2_ASM62549v2 Salmonella enterica subsp. enterica serovar Enteritidis str. EC20120590
Proteobacteria; Gammaproteobacteria; Enterobacterales; Enterobacteriaceae; Salmonella; Salmonella enterica; Salmonella enterica subsp. enterica; Salmonella enterica subsp. enterica serovar Enteritidis
MTGDNTLITSHGINRRDFMKLCAALAATMGLSSKAAAEMAESVSNPQRPPVIWIGAQECT WP_000145429.1
hydrogenase 2 small subunit [Salmonella enterica] Length: 372\nNumber of predicted TMHs: 1\nExp number of AAs in TMHs: 24.51309\nExp number, first 60 AAs: 0.2682\nTotal prob of N-in: 0.14402\noutside 1 331\nTMhelix 332 354\ninside 355 372

23591 GCF_000625495.2_ASM62549v2 Salmonella enterica subsp. enterica serovar Enteritidis str. EC20120590
Proteobacteria; Gammaproteobacteria; Enterobacterales; Enterobacteriaceae; Salmonella; Salmonella enterica; Salmonella enterica subsp. enterica; Salmonella enterica subsp. enterica serovar Enteritidis
MNNEETFYQAMRRKGVTRRSFLKFCSLAATSLGLGAGMTPKIAWALENKPRIPVVWIHGL WP_001058307.1
hydrogenase [Salmonella enterica] Length: 372\nNumber of predicted TMHs: 2\nExp number of AAs in TMHs: 36.51738\nExp number, first 60 AAs: 19.72849\nTotal prob of N-in: 0.96170\nPOSSIBLE N-term signal sequence\ninside 1 19\nTMhelix 20 39\noutside 40 325\nTMhelix 326 348\ninside 349 372

23592 GCF_000625515.2_ASM62551v2 Salmonella enterica subsp. enterica serovar Enteritidis str. EC20120597 Proteobacteria; Gammaproteobacteria; Enterobacterales; Enterobacteriaceae; Salmonella; Salmonella enterica; Salmonella enterica subsp. enterica; Salmonella enterica subsp. enterica serovar Enteritidis MTGDNTLITSHGINRRDFMKLCAALAATMGLSSKAAAEMAESVSNPQRPPVIWIGAQECT WP_000145429.1 hydrogenase 2 small subunit [Salmonella enterica] Length: 372\nNumber of predicted TMHs: 1\nExp number of AAs in TMHs: 24.51309\nExp number, first 60 AAs: 0.2682\nTotal prob of N-in: 0.14402\noutside 1 331\nTMhelix 332 354\ninside 355 372

23593 GCF_000625515.2_ASM62551v2 Salmonella enterica subsp. enterica serovar Enteritidis str. EC20120597 Proteobacteria; Gammaproteobacteria; Enterobacterales; Enterobacteriaceae; Salmonella; Salmonella enterica; Salmonella enterica subsp. enterica; Salmonella enterica subsp. enterica serovar Enteritidis MNNEETFYQAMRRKGVTRRSFLKFCSLAATSLGLGAGMTPKIAWALENKPRIPVVWIHGL WP_001058307.1 hydrogenase [Salmonella enterica] Length: 372\nNumber of predicted TMHs: 2\nExp number of AAs in TMHs: 36.51738\nExp number, first 60 AAs: 19.72849\nTotal prob of N-in: 0.96170\nPOSSIBLE N-term signal sequence\ninside 1 19\nTMhelix 20 39\noutside 40 325\nTMhelix 326 348\ninside 349 372

23594 GCF_000625755.1_ASM62575v2 Salmonella enterica subsp. enterica serovar Enteritidis str. EC20120677 Proteobacteria; Gammaproteobacteria; Enterobacterales; Enterobacteriaceae; Salmonella; Salmonella enterica; Salmonella enterica subsp. enterica; Salmonella enterica subsp. enterica serovar Enteritidis MTGDNTLITSHGINRRDFMKLCAALAATMGLSSKAAAEMAESVSNPQRPPVIWIGAQECT WP_000145429.1 hydrogenase 2 small subunit [Salmonella enterica] Length: 372\nNumber of predicted TMHs: 1\nExp number of AAs in TMHs: 24.51309\nExp number, first 60 AAs: 0.2682\nTotal prob of N-in: 0.14402\noutside 1 331\nTMhelix 332 354\ninside 355 372

23595 GCF_000625755.1_ASM62575v2 Salmonella enterica subsp. enterica serovar Enteritidis str. EC20120677 Proteobacteria; Gammaproteobacteria; Enterobacterales; Enterobacteriaceae; Salmonella; Salmonella enterica; Salmonella enterica subsp. enterica; Salmonella enterica subsp. enterica serovar Enteritidis MNNEETFYQAMRRKGVTRRSFLKFCSLAATSLGLGAGMTPKIAWALENKPRIPVVWIHGL WP_001058307.1 hydrogenase [Salmonella enterica] Length: 372\nNumber of predicted TMHs: 2\nExp number of AAs in TMHs: 36.51738\nExp number, first 60 AAs: 19.72849\nTotal prob of N-in: 0.96170\nPOSSIBLE N-term signal sequence\ninside 1 19\nTMhelix 20 39\noutside 40 325\nTMhelix 326 348\ninside 349 372

23596 GCF_000625535.1_ASM62553v2 Salmonella enterica subsp. enterica serovar Enteritidis str. EC20120685 Proteobacteria; Gammaproteobacteria; Enterobacterales; Enterobacteriaceae; Salmonella; Salmonella enterica; Salmonella enterica subsp. enterica; Salmonella enterica subsp. enterica serovar Enteritidis MNNEETFYQAMRRKGVTRRSFLKFCSLAATSLGLGAGMTPKIAWALENKPRIPVVWIHGL WP_001058307.1 hydrogenase [Salmonella enterica] Length: 372\nNumber of predicted TMHs: 2\nExp number of AAs in TMHs: 36.51738\nExp number, first 60 AAs: 19.72849\nTotal prob of N-in: 0.96170\nPOSSIBLE N-term signal sequence\ninside 1 19\nTMhelix 20 39\noutside 40 325\nTMhelix 326 348\ninside 349 372

23597 GCF_000625535.1_ASM62553v2 Salmonella enterica subsp. enterica serovar Enteritidis str. EC20120685 Proteobacteria; Gammaproteobacteria; Enterobacterales; Enterobacteriaceae; Salmonella; Salmonella enterica; Salmonella enterica subsp. enterica; Salmonella enterica subsp. enterica serovar Enteritidis MTGDNTLITSHGINRRDFMKLCAALAATMGLSSKAAAEMAESVSNPQRPPVIWIGAQECT WP_000145429.1 hydrogenase 2 small subunit [Salmonella enterica] Length: 372\nNumber of predicted TMHs: 1\nExp number of AAs in TMHs: 24.51309\nExp number, first 60 AAs: 0.2682\nTotal prob of N-in: 0.14402\noutside 1 331\nTMhelix 332 354\ninside 355 372

23598 GCF_000625555.2_ASM62555v2 Salmonella enterica subsp. enterica serovar Enteritidis str. EC20120686 Proteobacteria; Gammaproteobacteria; Enterobacterales; Enterobacteriaceae; Salmonella; Salmonella enterica; Salmonella enterica subsp. enterica; Salmonella enterica subsp. enterica serovar Enteritidis MNNEETFYQAMRRKGVTRRSFLKFCSLAATSLGLGAGMTPKIAWALENKPRIPVVWIHGL WP_001058307.1 hydrogenase [Salmonella enterica] Length: 372\nNumber of predicted TMHs: 2\nExp number of AAs in TMHs: 36.51738\nExp number, first 60 AAs: 19.72849\nTotal prob of N-in: 0.96170\nPOSSIBLE N-term signal sequence\ninside 1 19\nTMhelix 20 39\noutside 40 325\nTMhelix 326 348\ninside 349 372

23599 GCF_000625555.2_ASM62555v2 Salmonella enterica subsp. enterica serovar Enteritidis str. EC20120686 Proteobacteria; Gammaproteobacteria; Enterobacterales; Enterobacteriaceae; Salmonella; Salmonella enterica; Salmonella enterica subsp. enterica; Salmonella enterica subsp. enterica serovar Enteritidis MTGDNTLITSHGINRRDFMKLCAALAATMGLSSKAAAEMAESVSNPQRPPVIWIGAQECT WP_000145429.1 hydrogenase 2 small subunit [Salmonella enterica] Length: 372\nNumber of predicted TMHs: 1\nExp

number of AAs in TMHs: 24.51309\nExp number, first 60 AAs: 0.2682\nTotal prob of N-in: 0.14402\noutside 1 331\nTMhelix 332 354\ninside 355 372

23600 GCF_000625575.2_ASM62557v2 Salmonella enterica subsp. enterica serovar Enteritidis str. EC20120687 Proteobacteria; Gammaproteobacteria; Enterobacterales; Enterobacteriaceae; Salmonella; Salmonella enterica; Salmonella enterica subsp. enterica; Salmonella enterica subsp. enterica serovar Enteritidis MTGDNTLITSHGINRRDFMKLCAALAATMGLSSKAAAEMAESVSNPQRPPVIWIGAQECT WP_000145429.1 hydrogenase 2 small subunit [Salmonella enterica] Length: 372\nNumber of predicted TMHs: 1\nExp number of AAs in TMHs: 24.51309\nExp number, first 60 AAs: 0.2682\nTotal prob of N-in: 0.14402\noutside 1 331\nTMhelix 332 354\ninside 355 372

23601 GCF_000625575.2_ASM62557v2 Salmonella enterica subsp. enterica serovar Enteritidis str. EC20120687 Proteobacteria; Gammaproteobacteria; Enterobacterales; Enterobacteriaceae; Salmonella; Salmonella enterica; Salmonella enterica subsp. enterica; Salmonella enterica subsp. enterica serovar Enteritidis MNNEETFYQAMRRKGVTRRSFLKFCSLAATSLGLGAGMTPKIAWALENKPRIPVVWIHGL WP_001058307.1 hydrogenase [Salmonella enterica] Length: 372\nNumber of predicted TMHs: 2\nExp number of AAs in TMHs: 36.51738\nExp number, first 60 AAs: 19.72849\nTotal prob of N-in: 0.96170\nPOSSIBLE N-term signal sequence\ninside 1 19\nTMhelix 20 39\noutside 40 325\nTMhelix 326 348\ninside 349 372

23602 GCF_000625595.1_ASM62559v2 Salmonella enterica subsp. enterica serovar Enteritidis str. EC20120697 Proteobacteria; Gammaproteobacteria; Enterobacterales; Enterobacteriaceae; Salmonella; Salmonella enterica; Salmonella enterica subsp. enterica; Salmonella enterica subsp. enterica serovar Enteritidis MTGDNTLITSHGINRRDFMKLCAALAATMGLSSKAAAEMAESVSNPQRPPVIWIGAQECT WP_000145429.1 hydrogenase 2 small subunit [Salmonella enterica] Length: 372\nNumber of predicted TMHs: 1\nExp number of AAs in TMHs: 24.51309\nExp number, first 60 AAs: 0.2682\nTotal prob of N-in: 0.14402\noutside 1 331\nTMhelix 332 354\ninside 355 372

23603 GCF_000625595.1_ASM62559v2 Salmonella enterica subsp. enterica serovar Enteritidis str. EC20120697 Proteobacteria; Gammaproteobacteria; Enterobacterales; Enterobacteriaceae; Salmonella; Salmonella enterica; Salmonella enterica subsp. enterica; Salmonella enterica subsp. enterica serovar Enteritidis MNNEETFYQAMRRKGVTRRSFLKFCSLAATSLGLGAGMTPKIAWALENKPRIPVVWIHGL WP_001058307.1 hydrogenase [Salmonella enterica] Length: 372\nNumber of predicted TMHs: 2\nExp number of AAs in TMHs: 36.51738\nExp number, first 60 AAs: 19.72849\nTotal prob of N-in: 0.96170\nPOSSIBLE N-term signal sequence\ninside 1 19\nTMhelix 20 39\noutside 40 325\nTMhelix 326 348\ninside 349 372

23604 GCF_000625615.2_ASM62561v2 Salmonella enterica subsp. enterica serovar Enteritidis str. EC20120722 Proteobacteria; Gammaproteobacteria; Enterobacterales; Enterobacteriaceae; Salmonella; Salmonella enterica; Salmonella enterica subsp. enterica; Salmonella enterica subsp. enterica serovar Enteritidis MTGDNTLITSHGINRRDFMKLCAALAATMGLSSKAAAEMAESVSNPQRPPVIWIGAQECT WP_000145429.1 hydrogenase 2 small subunit [Salmonella enterica] Length: 372\nNumber of predicted TMHs: 1\nExp number of AAs in TMHs: 24.51309\nExp number, first 60 AAs: 0.2682\nTotal prob of N-in: 0.14402\noutside 1 331\nTMhelix 332 354\ninside 355 372

23605 GCF_000625615.2_ASM62561v2 Salmonella enterica subsp. enterica serovar Enteritidis str. EC20120722 Proteobacteria; Gammaproteobacteria; Enterobacterales; Enterobacteriaceae; Salmonella; Salmonella enterica; Salmonella enterica subsp. enterica; Salmonella enterica subsp. enterica serovar Enteritidis MNNEETFYQAMRRKGVTRRSFLKFCSLAATSLGLGAGMTPKIAWALENKPRIPVVWIHGL WP_001058307.1 hydrogenase [Salmonella enterica] Length: 372\nNumber of predicted TMHs: 2\nExp number of AAs in TMHs: 36.51738\nExp number, first 60 AAs: 19.72849\nTotal prob of N-in: 0.96170\nPOSSIBLE N-term signal sequence\ninside 1 19\nTMhelix 20 39\noutside 40 325\nTMhelix 326 348\ninside 349 372

23606 GCF_000624955.2_ASM62495v2 Salmonella enterica subsp. enterica serovar Enteritidis str. EC20120734 Proteobacteria; Gammaproteobacteria; Enterobacterales; Enterobacteriaceae; Salmonella; Salmonella enterica; Salmonella enterica subsp. enterica; Salmonella enterica subsp. enterica serovar Enteritidis MNNEETFYQAMRRKGVTRRSFLKFCSLAATSLGLGAGMTPKIAWALENKPRIPVVWIHGL WP_001058307.1 hydrogenase [Salmonella enterica] Length: 372\nNumber of predicted TMHs: 2\nExp number of AAs in TMHs: 36.51738\nExp number, first 60 AAs: 19.72849\nTotal prob of N-in: 0.96170\nPOSSIBLE N-term signal sequence\ninside 1 19\nTMhelix 20 39\noutside 40 325\nTMhelix 326 348\ninside 349 372

23607 GCF_000624955.2_ASM62495v2 Salmonella enterica subsp. enterica serovar Enteritidis str. EC20120734 Proteobacteria; Gammaproteobacteria; Enterobacterales; Enterobacteriaceae; Salmonella; Salmonella enterica;

Salmonella enterica subsp. enterica; Salmonella enterica subsp. enterica serovar Enteritidis
MTGDNTLITSHGINRRDFMKLCAALAATMGLSSKAAAEMAESVSNPQRPPVIWIGAQECT WP_000145429.1
hydrogenase 2 small subunit [Salmonella enterica] Length: 372\nNumber of predicted TMHs: 1\nExp number of AAs in TMHs: 24.51309\nExp number, first 60 AAs: 0.2682\nTotal prob of N-in: 0.14402\noutside 1 331\nTMhelix 332 354\ninside 355 372

23608 GCF_000624975.2_ASM62497v2 Salmonella enterica subsp. enterica serovar Enteritidis str. EC20120738
Proteobacteria; Gammaproteobacteria; Enterobacterales; Enterobacteriaceae; Salmonella; Salmonella enterica; Salmonella enterica subsp. enterica; Salmonella enterica subsp. enterica serovar Enteritidis
MNNEETFYQAMRRKGVTRRSFLKFCSLAATSLGLGAGMTPKIAWALENKPRIPVVWIHGL WP_001058307.1
hydrogenase [Salmonella enterica] Length: 372\nNumber of predicted TMHs: 2\nExp number of AAs in TMHs: 36.51738\nExp number, first 60 AAs: 19.72849\nTotal prob of N-in: 0.96170\nPOSSIBLE N-term signal sequence\ninside 1 19\nTMhelix 20 39\noutside 40 325\nTMhelix 326 348\ninside 349 372

23609 GCF_000624975.2_ASM62497v2 Salmonella enterica subsp. enterica serovar Enteritidis str. EC20120738
Proteobacteria; Gammaproteobacteria; Enterobacterales; Enterobacteriaceae; Salmonella; Salmonella enterica; Salmonella enterica subsp. enterica; Salmonella enterica subsp. enterica serovar Enteritidis
MTGDNTLITSHGINRRDFMKLCAALAATMGLSSKAAAEMAESVSNPQRPPVIWIGAQECT WP_000145429.1
hydrogenase 2 small subunit [Salmonella enterica] Length: 372\nNumber of predicted TMHs: 1\nExp number of AAs in TMHs: 24.51309\nExp number, first 60 AAs: 0.2682\nTotal prob of N-in: 0.14402\noutside 1 331\nTMhelix 332 354\ninside 355 372

23610 GCF_000624995.2_ASM62499v2 Salmonella enterica subsp. enterica serovar Enteritidis str. EC20120765
Proteobacteria; Gammaproteobacteria; Enterobacterales; Enterobacteriaceae; Salmonella; Salmonella enterica; Salmonella enterica subsp. enterica; Salmonella enterica subsp. enterica serovar Enteritidis
MNNEETFYQAMRRKGVTRRSFLKFCSLAATSLGLGAGMTPKIAWALENKPRIPVVWIHGL WP_001058307.1
hydrogenase [Salmonella enterica] Length: 372\nNumber of predicted TMHs: 2\nExp number of AAs in TMHs: 36.51738\nExp number, first 60 AAs: 19.72849\nTotal prob of N-in: 0.96170\nPOSSIBLE N-term signal sequence\ninside 1 19\nTMhelix 20 39\noutside 40 325\nTMhelix 326 348\ninside 349 372

23611 GCF_000624995.2_ASM62499v2 Salmonella enterica subsp. enterica serovar Enteritidis str. EC20120765
Proteobacteria; Gammaproteobacteria; Enterobacterales; Enterobacteriaceae; Salmonella; Salmonella enterica; Salmonella enterica subsp. enterica; Salmonella enterica subsp. enterica serovar Enteritidis
MTGDNTLITSHGINRRDFMKLCAALAATMGLSSKAAAEMAESVSNPQRPPVIWIGAQECT WP_000145429.1
hydrogenase 2 small subunit [Salmonella enterica] Length: 372\nNumber of predicted TMHs: 1\nExp number of AAs in TMHs: 24.51309\nExp number, first 60 AAs: 0.2682\nTotal prob of N-in: 0.14402\noutside 1 331\nTMhelix 332 354\ninside 355 372

23612 GCF_000625015.2_ASM62501v2 Salmonella enterica subsp. enterica serovar Enteritidis str. EC20120773
Proteobacteria; Gammaproteobacteria; Enterobacterales; Enterobacteriaceae; Salmonella; Salmonella enterica; Salmonella enterica subsp. enterica; Salmonella enterica subsp. enterica serovar Enteritidis
MTGDNTLITSHGINRRDFMKLCAALAATMGLSSKAAAEMAESVSNPQRPPVIWIGAQECT WP_000145429.1
hydrogenase 2 small subunit [Salmonella enterica] Length: 372\nNumber of predicted TMHs: 1\nExp number of AAs in TMHs: 24.51309\nExp number, first 60 AAs: 0.2682\nTotal prob of N-in: 0.14402\noutside 1 331\nTMhelix 332 354\ninside 355 372

23613 GCF_000625015.2_ASM62501v2 Salmonella enterica subsp. enterica serovar Enteritidis str. EC20120773
Proteobacteria; Gammaproteobacteria; Enterobacterales; Enterobacteriaceae; Salmonella; Salmonella enterica; Salmonella enterica subsp. enterica; Salmonella enterica subsp. enterica serovar Enteritidis
MNNEETFYQAMRRKGVTRRSFLKFCSLAATSLGLGAGMTPKIAWALENKPRIPVVWIHGL WP_001058307.1
hydrogenase [Salmonella enterica] Length: 372\nNumber of predicted TMHs: 2\nExp number of AAs in TMHs: 36.51738\nExp number, first 60 AAs: 19.72849\nTotal prob of N-in: 0.96170\nPOSSIBLE N-term signal sequence\ninside 1 19\nTMhelix 20 39\noutside 40 325\nTMhelix 326 348\ninside 349 372

23614 GCF_000625035.2_ASM62503v2 Salmonella enterica subsp. enterica serovar Enteritidis str. EC20120774
Proteobacteria; Gammaproteobacteria; Enterobacterales; Enterobacteriaceae; Salmonella; Salmonella enterica; Salmonella enterica subsp. enterica; Salmonella enterica subsp. enterica serovar Enteritidis
MNNEETFYQAMRRKGVTRRSFLKFCSLAATSLGLGAGMTPKIAWALENKPRIPVVWIHGL WP_001058307.1
hydrogenase [Salmonella enterica] Length: 372\nNumber of predicted TMHs: 2\nExp number of AAs in TMHs: 36.51738\nExp number, first 60 AAs: 19.72849\nTotal prob of N-in: 0.96170\nPOSSIBLE N-term signal sequence\ninside 1 19\nTMhelix 20 39\noutside 40 325\nTMhelix 326 348\ninside 349 372

23615 GCF_000625035.2_ASM62503v2 Salmonella enterica subsp. enterica serovar Enteritidis str. EC20120774 Proteobacteria; Gammaproteobacteria; Enterobacterales; Enterobacteriaceae; Salmonella; Salmonella enterica; Salmonella enterica subsp. enterica; Salmonella enterica subsp. enterica serovar Enteritidis MTGDNTLITSHGINRRDFMKLCAALAATMGLSSKAAAEMAESVSNPQRPPVIWIGAQECT WP_000145429.1 hydrogenase 2 small subunit [Salmonella enterica] Length: 372\nNumber of predicted TMHs: 1\nExp number of AAs in TMHs: 24.51309\nExp number, first 60 AAs: 0.2682\nTotal prob of N-in: 0.14402\noutside 1 331\nTMhelix 332 354\ninside 355 372

23616 GCF_000625055.2_ASM62505v2 Salmonella enterica subsp. enterica serovar Enteritidis str. EC20120775 Proteobacteria; Gammaproteobacteria; Enterobacterales; Enterobacteriaceae; Salmonella; Salmonella enterica; Salmonella enterica subsp. enterica; Salmonella enterica subsp. enterica serovar Enteritidis MNNEETFYQAMRRKGVTRRSFLKFCSLAATSLGLGAGMTPKIAWALENKPRIPVVWIHGL WP_001058307.1 hydrogenase [Salmonella enterica] Length: 372\nNumber of predicted TMHs: 2\nExp number of AAs in TMHs: 36.51738\nExp number, first 60 AAs: 19.72849\nTotal prob of N-in: 0.96170\nPOSSIBLE N-term signal sequence\ninside 1 19\nTMhelix 20 39\noutside 40 325\nTMhelix 326 348\ninside 349 372

23617 GCF_000625055.2_ASM62505v2 Salmonella enterica subsp. enterica serovar Enteritidis str. EC20120775 Proteobacteria; Gammaproteobacteria; Enterobacterales; Enterobacteriaceae; Salmonella; Salmonella enterica; Salmonella enterica subsp. enterica; Salmonella enterica subsp. enterica serovar Enteritidis MTGDNTLITSHGINRRDFMKLCAALAATMGLSSKAAAEMAESVSNPQRPPVIWIGAQECT WP_000145429.1 hydrogenase 2 small subunit [Salmonella enterica] Length: 372\nNumber of predicted TMHs: 1\nExp number of AAs in TMHs: 24.51309\nExp number, first 60 AAs: 0.2682\nTotal prob of N-in: 0.14402\noutside 1 331\nTMhelix 332 354\ninside 355 372

23618 GCF_000625075.2_ASM62507v2 Salmonella enterica subsp. enterica serovar Enteritidis str. EC20120776 Proteobacteria; Gammaproteobacteria; Enterobacterales; Enterobacteriaceae; Salmonella; Salmonella enterica; Salmonella enterica subsp. enterica; Salmonella enterica subsp. enterica serovar Enteritidis MNNEETFYQAMRRKGVTRRSFLKFCSLAATSLGLGAGMTPKIAWALENKPRIPVVWIHGL WP_001058307.1 hydrogenase [Salmonella enterica] Length: 372\nNumber of predicted TMHs: 2\nExp number of AAs in TMHs: 36.51738\nExp number, first 60 AAs: 19.72849\nTotal prob of N-in: 0.96170\nPOSSIBLE N-term signal sequence\ninside 1 19\nTMhelix 20 39\noutside 40 325\nTMhelix 326 348\ninside 349 372

23619 GCF_000625075.2_ASM62507v2 Salmonella enterica subsp. enterica serovar Enteritidis str. EC20120776 Proteobacteria; Gammaproteobacteria; Enterobacterales; Enterobacteriaceae; Salmonella; Salmonella enterica; Salmonella enterica subsp. enterica; Salmonella enterica subsp. enterica serovar Enteritidis MTGDNTLITSHGINRRDFMKLCAALAATMGLSSKAAAEMAESVSNPQRPPVIWIGAQECT WP_000145429.1 hydrogenase 2 small subunit [Salmonella enterica] Length: 372\nNumber of predicted TMHs: 1\nExp number of AAs in TMHs: 24.51309\nExp number, first 60 AAs: 0.2682\nTotal prob of N-in: 0.14402\noutside 1 331\nTMhelix 332 354\ninside 355 372

23620 GCF_000624155.1_ASM62415v1 Salmonella enterica subsp. enterica serovar Enteritidis str. EC20120916 Proteobacteria; Gammaproteobacteria; Enterobacterales; Enterobacteriaceae; Salmonella; Salmonella enterica; Salmonella enterica subsp. enterica; Salmonella enterica subsp. enterica serovar Enteritidis MTGDNTLITSHGINRRDFMKLCAALAATMGLSSKAAAEMAESVSNPQRPPVIWIGAQECT WP_000145429.1 hydrogenase 2 small subunit [Salmonella enterica] Length: 372\nNumber of predicted TMHs: 1\nExp number of AAs in TMHs: 24.51309\nExp number, first 60 AAs: 0.2682\nTotal prob of N-in: 0.14402\noutside 1 331\nTMhelix 332 354\ninside 355 372

23621 GCF_000624155.1_ASM62415v1 Salmonella enterica subsp. enterica serovar Enteritidis str. EC20120916 Proteobacteria; Gammaproteobacteria; Enterobacterales; Enterobacteriaceae; Salmonella; Salmonella enterica; Salmonella enterica subsp. enterica; Salmonella enterica subsp. enterica serovar Enteritidis MNNEETFYQAMRRKGVTRRSFLKFCSLAATSLGLGAGMTPKIAWALENKPRIPVVWIHGL WP_001058307.1 hydrogenase [Salmonella enterica] Length: 372\nNumber of predicted TMHs: 2\nExp number of AAs in TMHs: 36.51738\nExp number, first 60 AAs: 19.72849\nTotal prob of N-in: 0.96170\nPOSSIBLE N-term signal sequence\ninside 1 19\nTMhelix 20 39\noutside 40 325\nTMhelix 326 348\ninside 349 372

23622 GCF_000625095.2_ASM62509v2 Salmonella enterica subsp. enterica serovar Enteritidis str. EC20120917 Proteobacteria; Gammaproteobacteria; Enterobacterales; Enterobacteriaceae; Salmonella; Salmonella enterica; Salmonella enterica subsp. enterica; Salmonella enterica subsp. enterica serovar Enteritidis MTGDNTLITSHGINRRDFMKLCAALAATMGLSSKAAAEMAESVSNPQRPPVIWIGAQECT WP_000145429.1 hydrogenase 2 small subunit [Salmonella enterica] Length: 372\nNumber of predicted TMHs: 1\nExp

number of AAs in TMHs: 24.51309\nExp number, first 60 AAs: 0.2682\nTotal prob of N-in: 0.14402\noutside 1 331\nTMhelix 332 354\ninside 355 372

23623 GCF_000625095.2_ASM62509v2 Salmonella enterica subsp. enterica serovar Enteritidis str. EC20120917 Proteobacteria; Gammaproteobacteria; Enterobacterales; Enterobacteriaceae; Salmonella; Salmonella enterica; Salmonella enterica subsp. enterica; Salmonella enterica subsp. enterica serovar Enteritidis MNNEETFYQAMRRKGVTRRSFLKFCSLAATSLGLGAGMTPKIAWALENKPRIPVVWIHGL WP_001058307.1 hydrogenase [Salmonella enterica] Length: 372\nNumber of predicted TMHs: 2\nExp number of AAs in TMHs: 36.51738\nExp number, first 60 AAs: 19.72849\nTotal prob of N-in: 0.96170\nPOSSIBLE N-term signal sequence\ninside 1 19\nTMhelix 20 39\noutside 40 325\nTMhelix 326 348\ninside 349 372

23624 GCF_000625115.2_ASM62511v2 Salmonella enterica subsp. enterica serovar Enteritidis str. EC20120918 Proteobacteria; Gammaproteobacteria; Enterobacterales; Enterobacteriaceae; Salmonella; Salmonella enterica; Salmonella enterica subsp. enterica; Salmonella enterica subsp. enterica serovar Enteritidis MNNEETFYQAMRRKGVTRRSFLKFCSLAATSLGLGAGMTPKIAWALENKPRIPVVWIHGL WP_001058307.1 hydrogenase [Salmonella enterica] Length: 372\nNumber of predicted TMHs: 2\nExp number of AAs in TMHs: 36.51738\nExp number, first 60 AAs: 19.72849\nTotal prob of N-in: 0.96170\nPOSSIBLE N-term signal sequence\ninside 1 19\nTMhelix 20 39\noutside 40 325\nTMhelix 326 348\ninside 349 372

23625 GCF_000625115.2_ASM62511v2 Salmonella enterica subsp. enterica serovar Enteritidis str. EC20120918 Proteobacteria; Gammaproteobacteria; Enterobacterales; Enterobacteriaceae; Salmonella; Salmonella enterica; Salmonella enterica subsp. enterica; Salmonella enterica subsp. enterica serovar Enteritidis MTGDNTLITSHGINRRDFMKLCAALAATMGLSSKAAAEMAESVSNPQRPPVIWIGAQECT WP_000145429.1 hydrogenase 2 small subunit [Salmonella enterica] Length: 372\nNumber of predicted TMHs: 1\nExp number of AAs in TMHs: 24.51309\nExp number, first 60 AAs: 0.2682\nTotal prob of N-in: 0.14402\noutside 1 331\nTMhelix 332 354\ninside 355 372

23626 GCF_000625135.2_ASM62513v2 Salmonella enterica subsp. enterica serovar Enteritidis str. EC20120925 Proteobacteria; Gammaproteobacteria; Enterobacterales; Enterobacteriaceae; Salmonella; Salmonella enterica; Salmonella enterica subsp. enterica; Salmonella enterica subsp. enterica serovar Enteritidis MTGDNTLITSHGINRRDFMKLCAALAATMGLSSKAAAEMAESVSNPQRPPVIWIGAQECT WP_000145429.1 hydrogenase 2 small subunit [Salmonella enterica] Length: 372\nNumber of predicted TMHs: 1\nExp number of AAs in TMHs: 24.51309\nExp number, first 60 AAs: 0.2682\nTotal prob of N-in: 0.14402\noutside 1 331\nTMhelix 332 354\ninside 355 372

23627 GCF_000625135.2_ASM62513v2 Salmonella enterica subsp. enterica serovar Enteritidis str. EC20120925 Proteobacteria; Gammaproteobacteria; Enterobacterales; Enterobacteriaceae; Salmonella; Salmonella enterica; Salmonella enterica subsp. enterica; Salmonella enterica subsp. enterica serovar Enteritidis MNNEETFYQAMRRKGVTRRSFLKFCSLAATSLGLGAGMTPKIAWALENKPRIPVVWIHGL WP_001058307.1 hydrogenase [Salmonella enterica] Length: 372\nNumber of predicted TMHs: 2\nExp number of AAs in TMHs: 36.51738\nExp number, first 60 AAs: 19.72849\nTotal prob of N-in: 0.96170\nPOSSIBLE N-term signal sequence\ninside 1 19\nTMhelix 20 39\noutside 40 325\nTMhelix 326 348\ninside 349 372

23628 GCF_000625155.2_ASM62515v2 Salmonella enterica subsp. enterica serovar Enteritidis str. EC20120927 Proteobacteria; Gammaproteobacteria; Enterobacterales; Enterobacteriaceae; Salmonella; Salmonella enterica; Salmonella enterica subsp. enterica; Salmonella enterica subsp. enterica serovar Enteritidis MTGDNTLITSHGINRRDFMKLCAALAATMGLSSKAAAEMAESVSNPQRPPVIWIGAQECT WP_000145429.1 hydrogenase 2 small subunit [Salmonella enterica] Length: 372\nNumber of predicted TMHs: 1\nExp number of AAs in TMHs: 24.51309\nExp number, first 60 AAs: 0.2682\nTotal prob of N-in: 0.14402\noutside 1 331\nTMhelix 332 354\ninside 355 372

23629 GCF_000625155.2_ASM62515v2 Salmonella enterica subsp. enterica serovar Enteritidis str. EC20120927 Proteobacteria; Gammaproteobacteria; Enterobacterales; Enterobacteriaceae; Salmonella; Salmonella enterica; Salmonella enterica subsp. enterica; Salmonella enterica subsp. enterica serovar Enteritidis MNNEETFYQAMRRKGVTRRSFLKFCSLAATSLGLGAGMTPKIAWALENKPRIPVVWIHGL WP_001058307.1 hydrogenase [Salmonella enterica] Length: 372\nNumber of predicted TMHs: 2\nExp number of AAs in TMHs: 36.51738\nExp number, first 60 AAs: 19.72849\nTotal prob of N-in: 0.96170\nPOSSIBLE N-term signal sequence\ninside 1 19\nTMhelix 20 39\noutside 40 325\nTMhelix 326 348\ninside 349 372

23630 GCF_000626415.2_ASM62641v2 Salmonella enterica subsp. enterica serovar Enteritidis str. EC20120929 Proteobacteria; Gammaproteobacteria; Enterobacterales; Enterobacteriaceae; Salmonella; Salmonella enterica;

Salmonella enterica subsp. enterica; Salmonella enterica subsp. enterica serovar Enteritidis
MNNEETFYQAMRRKGVTRRSFLKFCSLAATSLGLGAGMTPKIAWALENKPRIPVVWIHGL WP_001058307.1
hydrogenase [Salmonella enterica] Length: 372\nNumber of predicted TMHs: 2\nExp number of AAs in TMHs: 36.51738\nExp number, first 60 AAs: 19.72849\nTotal prob of N-in: 0.96170\nPOSSIBLE N-term signal sequence\ninside 1 19\nTMhelix 20 39\noutside 40 325\nTMhelix 326 348\ninside 349 372

23631 GCF_000626415.2_ASM62641v2 Salmonella enterica subsp. enterica serovar Enteritidis str. EC20120929
Proteobacteria; Gammaproteobacteria; Enterobacterales; Enterobacteriaceae; Salmonella; Salmonella enterica; Salmonella enterica subsp. enterica; Salmonella enterica subsp. enterica serovar Enteritidis
MTGDNTLITSHGINRRDFMKLCAALAATMGLSSKAAAEMAESVSNPQRPPVIWIGAQECT WP_000145429.1
hydrogenase 2 small subunit [Salmonella enterica] Length: 372\nNumber of predicted TMHs: 1\nExp number of AAs in TMHs: 24.51309\nExp number, first 60 AAs: 0.2682\nTotal prob of N-in: 0.14402\noutside 1 331\nTMhelix 332 354\ninside 355 372

23632 GCF_000625175.2_ASM62517v2 Salmonella enterica subsp. enterica serovar Enteritidis str. EC20120963
Proteobacteria; Gammaproteobacteria; Enterobacterales; Enterobacteriaceae; Salmonella; Salmonella enterica; Salmonella enterica subsp. enterica; Salmonella enterica subsp. enterica serovar Enteritidis
MTGDNTLITSHGINRRDFMKLCAALAATMGLSSKAAAEMAESVSNPQRPPVIWIGAQECT WP_000145429.1
hydrogenase 2 small subunit [Salmonella enterica] Length: 372\nNumber of predicted TMHs: 1\nExp number of AAs in TMHs: 24.51309\nExp number, first 60 AAs: 0.2682\nTotal prob of N-in: 0.14402\noutside 1 331\nTMhelix 332 354\ninside 355 372

23633 GCF_000625175.2_ASM62517v2 Salmonella enterica subsp. enterica serovar Enteritidis str. EC20120963
Proteobacteria; Gammaproteobacteria; Enterobacterales; Enterobacteriaceae; Salmonella; Salmonella enterica; Salmonella enterica subsp. enterica; Salmonella enterica subsp. enterica serovar Enteritidis
MNNEETFYQAMRRKGVTRRSFLKFCSLAATSLGLGAGMTPKIAWALENKPRIPVVWIHGL WP_001058307.1
hydrogenase [Salmonella enterica] Length: 372\nNumber of predicted TMHs: 2\nExp number of AAs in TMHs: 36.51738\nExp number, first 60 AAs: 19.72849\nTotal prob of N-in: 0.96170\nPOSSIBLE N-term signal sequence\ninside 1 19\nTMhelix 20 39\noutside 40 325\nTMhelix 326 348\ninside 349 372

23634 GCF_000625195.2_ASM62519v2 Salmonella enterica subsp. enterica serovar Enteritidis str. EC20120968
Proteobacteria; Gammaproteobacteria; Enterobacterales; Enterobacteriaceae; Salmonella; Salmonella enterica; Salmonella enterica subsp. enterica; Salmonella enterica subsp. enterica serovar Enteritidis
MTGDNTLITSHGINRRDFMKLCAALAATMGLSSKAAAEMAESVSNPQRPPVIWIGAQECT WP_000145429.1
hydrogenase 2 small subunit [Salmonella enterica] Length: 372\nNumber of predicted TMHs: 1\nExp number of AAs in TMHs: 24.51309\nExp number, first 60 AAs: 0.2682\nTotal prob of N-in: 0.14402\noutside 1 331\nTMhelix 332 354\ninside 355 372

23635 GCF_000625195.2_ASM62519v2 Salmonella enterica subsp. enterica serovar Enteritidis str. EC20120968
Proteobacteria; Gammaproteobacteria; Enterobacterales; Enterobacteriaceae; Salmonella; Salmonella enterica; Salmonella enterica subsp. enterica; Salmonella enterica subsp. enterica serovar Enteritidis
MNNEETFYQAMRRKGVTRRSFLKFCSLAATSLGLGAGMTPKIAWALENKPRIPVVWIHGL WP_001058307.1
hydrogenase [Salmonella enterica] Length: 372\nNumber of predicted TMHs: 2\nExp number of AAs in TMHs: 36.51738\nExp number, first 60 AAs: 19.72849\nTotal prob of N-in: 0.96170\nPOSSIBLE N-term signal sequence\ninside 1 19\nTMhelix 20 39\noutside 40 325\nTMhelix 326 348\ninside 349 372

23636 GCF_000625215.2_ASM62521v2 Salmonella enterica subsp. enterica serovar Enteritidis str. EC20120969
Proteobacteria; Gammaproteobacteria; Enterobacterales; Enterobacteriaceae; Salmonella; Salmonella enterica; Salmonella enterica subsp. enterica; Salmonella enterica subsp. enterica serovar Enteritidis
MNNEETFYQAMRRKGVTRRSFLKFCSLAATSLGLGAGMTPKIAWALENKPRIPVVWIHGL WP_001058307.1
hydrogenase [Salmonella enterica] Length: 372\nNumber of predicted TMHs: 2\nExp number of AAs in TMHs: 36.51738\nExp number, first 60 AAs: 19.72849\nTotal prob of N-in: 0.96170\nPOSSIBLE N-term signal sequence\ninside 1 19\nTMhelix 20 39\noutside 40 325\nTMhelix 326 348\ninside 349 372

23637 GCF_000625215.2_ASM62521v2 Salmonella enterica subsp. enterica serovar Enteritidis str. EC20120969
Proteobacteria; Gammaproteobacteria; Enterobacterales; Enterobacteriaceae; Salmonella; Salmonella enterica; Salmonella enterica subsp. enterica; Salmonella enterica subsp. enterica serovar Enteritidis
MTGDNTLITSHGINRRDFMKLCAALAATMGLSSKAAAEMAESVSNPQRPPVIWIGAQECT WP_000145429.1
hydrogenase 2 small subunit [Salmonella enterica] Length: 372\nNumber of predicted TMHs: 1\nExp number of AAs in TMHs: 24.51309\nExp number, first 60 AAs: 0.2682\nTotal prob of N-in: 0.14402\noutside 1 331\nTMhelix 332 354\ninside 355 372

23638 GCF_000625235.2_ASM62523v2 Salmonella enterica subsp. enterica serovar Enteritidis str. EC20120970 Proteobacteria; Gammaproteobacteria; Enterobacterales; Enterobacteriaceae; Salmonella; Salmonella enterica; Salmonella enterica subsp. enterica; Salmonella enterica subsp. enterica serovar Enteritidis MNNEETFYQAMRRKGVTRRSFLKFCSLAATSLGLGAGMTPKIAWALENKPRIPVVWIHGL WP_001058307.1 hydrogenase [Salmonella enterica] Length: 372\nNumber of predicted TMHs: 2\nExp number of AAs in TMHs: 36.51738\nExp number, first 60 AAs: 19.72849\nTotal prob of N-in: 0.96170\nPOSSIBLE N-term signal sequence\ninside 1 19\nTMhelix 20 39\noutside 40 325\nTMhelix 326 348\ninside 349 372

23639 GCF_000625235.2_ASM62523v2 Salmonella enterica subsp. enterica serovar Enteritidis str. EC20120970 Proteobacteria; Gammaproteobacteria; Enterobacterales; Enterobacteriaceae; Salmonella; Salmonella enterica; Salmonella enterica subsp. enterica; Salmonella enterica subsp. enterica serovar Enteritidis MTGDNTLITSHGINRRDFMKLCAALAATMGLSSKAAAEMAESVSNPQRPPVIWIGAQECT WP_000145429.1 hydrogenase 2 small subunit [Salmonella enterica] Length: 372\nNumber of predicted TMHs: 1\nExp number of AAs in TMHs: 24.51309\nExp number, first 60 AAs: 0.2682\nTotal prob of N-in: 0.14402\noutside 1 331\nTMhelix 332 354\ninside 355 372

23640 GCF_000624675.1_ASM62467v2 Salmonella enterica subsp. enterica serovar Enteritidis str. EC20120994 Proteobacteria; Gammaproteobacteria; Enterobacterales; Enterobacteriaceae; Salmonella; Salmonella enterica; Salmonella enterica subsp. enterica; Salmonella enterica subsp. enterica serovar Enteritidis MNNEETFYQAMRRKGVTRRSFLKFCSLAATSLGLGAGMTPKIAWALENKPRIPVVWIHGL WP_001058307.1 hydrogenase [Salmonella enterica] Length: 372\nNumber of predicted TMHs: 2\nExp number of AAs in TMHs: 36.51738\nExp number, first 60 AAs: 19.72849\nTotal prob of N-in: 0.96170\nPOSSIBLE N-term signal sequence\ninside 1 19\nTMhelix 20 39\noutside 40 325\nTMhelix 326 348\ninside 349 372

23641 GCF_000624675.1_ASM62467v2 Salmonella enterica subsp. enterica serovar Enteritidis str. EC20120994 Proteobacteria; Gammaproteobacteria; Enterobacterales; Enterobacteriaceae; Salmonella; Salmonella enterica; Salmonella enterica subsp. enterica; Salmonella enterica subsp. enterica serovar Enteritidis MTGDNTLITSHGINRRDFMKLCAALAATMGLSSKAAAEMAESVSNPQRPPVIWIGAQECT WP_000145429.1 hydrogenase 2 small subunit [Salmonella enterica] Length: 372\nNumber of predicted TMHs: 1\nExp number of AAs in TMHs: 24.51309\nExp number, first 60 AAs: 0.2682\nTotal prob of N-in: 0.14402\noutside 1 331\nTMhelix 332 354\ninside 355 372

23642 GCF_000624695.1_ASM62469v2 Salmonella enterica subsp. enterica serovar Enteritidis str. EC20121004 Proteobacteria; Gammaproteobacteria; Enterobacterales; Enterobacteriaceae; Salmonella; Salmonella enterica; Salmonella enterica subsp. enterica; Salmonella enterica subsp. enterica serovar Enteritidis MNNEETFYQAMRRKGVTRRSFLKFCSLAATSLGLGAGMTPKIAWALENKPRIPVVWIHGL WP_001058307.1 hydrogenase [Salmonella enterica] Length: 372\nNumber of predicted TMHs: 2\nExp number of AAs in TMHs: 36.51738\nExp number, first 60 AAs: 19.72849\nTotal prob of N-in: 0.96170\nPOSSIBLE N-term signal sequence\ninside 1 19\nTMhelix 20 39\noutside 40 325\nTMhelix 326 348\ninside 349 372

23643 GCF_000624695.1_ASM62469v2 Salmonella enterica subsp. enterica serovar Enteritidis str. EC20121004 Proteobacteria; Gammaproteobacteria; Enterobacterales; Enterobacteriaceae; Salmonella; Salmonella enterica; Salmonella enterica subsp. enterica; Salmonella enterica subsp. enterica serovar Enteritidis MTGDNTLITSHGINRRDFMKLCAALAATMGLSSKAAAEMAESVSNPQRPPVIWIGAQECT WP_000145429.1 hydrogenase 2 small subunit [Salmonella enterica] Length: 372\nNumber of predicted TMHs: 1\nExp number of AAs in TMHs: 24.51309\nExp number, first 60 AAs: 0.2682\nTotal prob of N-in: 0.14402\noutside 1 331\nTMhelix 332 354\ninside 355 372

23644 GCF_000623055.1_ASM62305v2 Salmonella enterica subsp. enterica serovar Enteritidis str. EC20121175 Proteobacteria; Gammaproteobacteria; Enterobacterales; Enterobacteriaceae; Salmonella; Salmonella enterica; Salmonella enterica subsp. enterica; Salmonella enterica subsp. enterica serovar Enteritidis MTGDNTLITSHGINRRDFMKLCAALAATMGLSSKAAAEMAESVSNPQRPPVIWIGAQECT WP_000145429.1 hydrogenase 2 small subunit [Salmonella enterica] Length: 372\nNumber of predicted TMHs: 1\nExp number of AAs in TMHs: 24.51309\nExp number, first 60 AAs: 0.2682\nTotal prob of N-in: 0.14402\noutside 1 331\nTMhelix 332 354\ninside 355 372

23645 GCF_000623055.1_ASM62305v2 Salmonella enterica subsp. enterica serovar Enteritidis str. EC20121175 Proteobacteria; Gammaproteobacteria; Enterobacterales; Enterobacteriaceae; Salmonella; Salmonella enterica; Salmonella enterica subsp. enterica; Salmonella enterica subsp. enterica serovar Enteritidis MNNEETFYQAMRRKGVTRRSFLKFCSLAATSLGLGAGMTPKIAWALENKPRIPVVWIHGL WP_001058307.1 hydrogenase [Salmonella enterica] Length: 372\nNumber of predicted TMHs: 2\nExp number of AAs in

TMHs: 36.51738\nExp number, first 60 AAs: 19.72849\nTotal prob of N-in: 0.96170\nPOSSIBLE N-term signal sequence\ninside 1 19\nTMhelix 20 39\noutside 40 325\nTMhelix 326 348\ninside 349 372

23646 GCF_000623075.1_ASM62307v2 Salmonella enterica subsp. enterica serovar Enteritidis str. EC20121176 Proteobacteria; Gammaproteobacteria; Enterobacterales; Enterobacteriaceae; Salmonella; Salmonella enterica; Salmonella enterica subsp. enterica; Salmonella enterica subsp. enterica serovar Enteritidis MNNEETFYQAMRRKGVTRRSFLKFCSLAATSLGLGAGMTPKIAWALENKPRIPVVWIHGL WP_001058307.1 hydrogenase [Salmonella enterica] Length: 372\nNumber of predicted TMHs: 2\nExp number of AAs in TMHs: 36.51738\nExp number, first 60 AAs: 19.72849\nTotal prob of N-in: 0.96170\nPOSSIBLE N-term signal sequence\ninside 1 19\nTMhelix 20 39\noutside 40 325\nTMhelix 326 348\ninside 349 372

23647 GCF_000623075.1_ASM62307v2 Salmonella enterica subsp. enterica serovar Enteritidis str. EC20121176 Proteobacteria; Gammaproteobacteria; Enterobacterales; Enterobacteriaceae; Salmonella; Salmonella enterica; Salmonella enterica subsp. enterica; Salmonella enterica subsp. enterica serovar Enteritidis MTGDNTLITSHGINRRDFMKLCAALAATMGLSSKAAAEMAESVSNPQRPPVIWIGAQECT WP_000145429.1 hydrogenase 2 small subunit [Salmonella enterica] Length: 372\nNumber of predicted TMHs: 1\nExp number of AAs in TMHs: 24.51309\nExp number, first 60 AAs: 0.2682\nTotal prob of N-in: 0.14402\noutside 1 331\nTMhelix 332 354\ninside 355 372

23648 GCF_000624175.1_ASM62417v2 Salmonella enterica subsp. enterica serovar Enteritidis str. EC20121177 Proteobacteria; Gammaproteobacteria; Enterobacterales; Enterobacteriaceae; Salmonella; Salmonella enterica; Salmonella enterica subsp. enterica; Salmonella enterica subsp. enterica serovar Enteritidis MTGDNTLITSHGINRRDFMKLCAALAATMGLSSKAAAEMAESVSNPQRPPVIWIGAQECT WP_000145429.1 hydrogenase 2 small subunit [Salmonella enterica] Length: 372\nNumber of predicted TMHs: 1\nExp number of AAs in TMHs: 24.51309\nExp number, first 60 AAs: 0.2682\nTotal prob of N-in: 0.14402\noutside 1 331\nTMhelix 332 354\ninside 355 372

23649 GCF_000624175.1_ASM62417v2 Salmonella enterica subsp. enterica serovar Enteritidis str. EC20121177 Proteobacteria; Gammaproteobacteria; Enterobacterales; Enterobacteriaceae; Salmonella; Salmonella enterica; Salmonella enterica subsp. enterica; Salmonella enterica subsp. enterica serovar Enteritidis MNNEETFYQAMRRKGVTRRSFLKFCSLAATSLGLGAGMTPKIAWALENKPRIPVVWIHGL WP_001058307.1 hydrogenase [Salmonella enterica] Length: 372\nNumber of predicted TMHs: 2\nExp number of AAs in TMHs: 36.51738\nExp number, first 60 AAs: 19.72849\nTotal prob of N-in: 0.96170\nPOSSIBLE N-term signal sequence\ninside 1 19\nTMhelix 20 39\noutside 40 325\nTMhelix 326 348\ninside 349 372

23650 GCF_000623095.1_ASM62309v2 Salmonella enterica subsp. enterica serovar Enteritidis str. EC20121178 Proteobacteria; Gammaproteobacteria; Enterobacterales; Enterobacteriaceae; Salmonella; Salmonella enterica; Salmonella enterica subsp. enterica; Salmonella enterica subsp. enterica serovar Enteritidis MTGDNTLITSHGINRRDFMKLCAALAATMGLSSKAAAEMAESVSNPQRPPVIWIGAQECT WP_000145429.1 hydrogenase 2 small subunit [Salmonella enterica] Length: 372\nNumber of predicted TMHs: 1\nExp number of AAs in TMHs: 24.51309\nExp number, first 60 AAs: 0.2682\nTotal prob of N-in: 0.14402\noutside 1 331\nTMhelix 332 354\ninside 355 372

23651 GCF_000623095.1_ASM62309v2 Salmonella enterica subsp. enterica serovar Enteritidis str. EC20121178 Proteobacteria; Gammaproteobacteria; Enterobacterales; Enterobacteriaceae; Salmonella; Salmonella enterica; Salmonella enterica subsp. enterica; Salmonella enterica subsp. enterica serovar Enteritidis MNNEETFYQAMRRKGVTRRSFLKFCSLAATSLGLGAGMTPKIAWALENKPRIPVVWIHGL WP_001058307.1 hydrogenase [Salmonella enterica] Length: 372\nNumber of predicted TMHs: 2\nExp number of AAs in TMHs: 36.51738\nExp number, first 60 AAs: 19.72849\nTotal prob of N-in: 0.96170\nPOSSIBLE N-term signal sequence\ninside 1 19\nTMhelix 20 39\noutside 40 325\nTMhelix 326 348\ninside 349 372

23652 GCF_000623115.2_ASM62311v2 Salmonella enterica subsp. enterica serovar Enteritidis str. EC20121179 Proteobacteria; Gammaproteobacteria; Enterobacterales; Enterobacteriaceae; Salmonella; Salmonella enterica; Salmonella enterica subsp. enterica; Salmonella enterica subsp. enterica serovar Enteritidis MNNEETFYQAMRRKGVTRRSFLKFCSLAATSLGLGAGMTPKIAWALENKPRIPVVWIHGL WP_001058307.1 hydrogenase [Salmonella enterica] Length: 372\nNumber of predicted TMHs: 2\nExp number of AAs in TMHs: 36.51738\nExp number, first 60 AAs: 19.72849\nTotal prob of N-in: 0.96170\nPOSSIBLE N-term signal sequence\ninside 1 19\nTMhelix 20 39\noutside 40 325\nTMhelix 326 348\ninside 349 372

23653 GCF_000623115.2_ASM62311v2 Salmonella enterica subsp. enterica serovar Enteritidis str. EC20121179 Proteobacteria; Gammaproteobacteria; Enterobacterales; Enterobacteriaceae; Salmonella; Salmonella enterica;

Salmonella enterica subsp. enterica; Salmonella enterica subsp. enterica serovar Enteritidis
MTGDNTLITSHGINRRDFMKLCAALAATMGLSSKAAAEMAESVSNPQRPPVIWIGAQECT WP_000145429.1
hydrogenase 2 small subunit [Salmonella enterica] Length: 372\nNumber of predicted TMHs: 1\nExp number of AAs in TMHs: 24.51309\nExp number, first 60 AAs: 0.2682\nTotal prob of N-in: 0.14402\noutside 1 331\nTMhelix 332 354\ninside 355 372

23654 GCF_000623135.1_ASM62313v2 Salmonella enterica subsp. enterica serovar Enteritidis str. EC20121180
Proteobacteria; Gammaproteobacteria; Enterobacterales; Enterobacteriaceae; Salmonella; Salmonella enterica; Salmonella enterica subsp. enterica; Salmonella enterica subsp. enterica serovar Enteritidis
MNNEETFYQAMRRKGVTRRSFLKFCSLAATSLGLGAGMTPKIAWALENKPRIPVVWIHGL WP_001058307.1
hydrogenase [Salmonella enterica] Length: 372\nNumber of predicted TMHs: 2\nExp number of AAs in TMHs: 36.51738\nExp number, first 60 AAs: 19.72849\nTotal prob of N-in: 0.96170\nPOSSIBLE N-term signal sequence\ninside 1 19\nTMhelix 20 39\noutside 40 325\nTMhelix 326 348\ninside 349 372

23655 GCF_000623135.1_ASM62313v2 Salmonella enterica subsp. enterica serovar Enteritidis str. EC20121180
Proteobacteria; Gammaproteobacteria; Enterobacterales; Enterobacteriaceae; Salmonella; Salmonella enterica; Salmonella enterica subsp. enterica; Salmonella enterica subsp. enterica serovar Enteritidis
MTGDNTLITSHGINRRDFMKLCAALAATMGLSSKAAAEMAESVSNPQRPPVIWIGAQECT WP_000145429.1
hydrogenase 2 small subunit [Salmonella enterica] Length: 372\nNumber of predicted TMHs: 1\nExp number of AAs in TMHs: 24.51309\nExp number, first 60 AAs: 0.2682\nTotal prob of N-in: 0.14402\noutside 1 331\nTMhelix 332 354\ninside 355 372

23656 GCF_000624715.1_ASM62471v2 Salmonella enterica subsp. enterica serovar Enteritidis str. EC20121541
Proteobacteria; Gammaproteobacteria; Enterobacterales; Enterobacteriaceae; Salmonella; Salmonella enterica; Salmonella enterica subsp. enterica; Salmonella enterica subsp. enterica serovar Enteritidis
MTGDNTLITSHGINRRDFMKLCAALAATMGLSSKAAAEMAESVSNPQRPPVIWIGAQECT WP_000145429.1
hydrogenase 2 small subunit [Salmonella enterica] Length: 372\nNumber of predicted TMHs: 1\nExp number of AAs in TMHs: 24.51309\nExp number, first 60 AAs: 0.2682\nTotal prob of N-in: 0.14402\noutside 1 331\nTMhelix 332 354\ninside 355 372

23657 GCF_000624715.1_ASM62471v2 Salmonella enterica subsp. enterica serovar Enteritidis str. EC20121541
Proteobacteria; Gammaproteobacteria; Enterobacterales; Enterobacteriaceae; Salmonella; Salmonella enterica; Salmonella enterica subsp. enterica; Salmonella enterica subsp. enterica serovar Enteritidis
MNNEETFYQAMRRKGVTRRSFLKFCSLAATSLGLGAGMTPKIAWALENKPRIPVVWIHGL WP_001058307.1
hydrogenase [Salmonella enterica] Length: 372\nNumber of predicted TMHs: 2\nExp number of AAs in TMHs: 36.51738\nExp number, first 60 AAs: 19.72849\nTotal prob of N-in: 0.96170\nPOSSIBLE N-term signal sequence\ninside 1 19\nTMhelix 20 39\noutside 40 325\nTMhelix 326 348\ninside 349 372

23658 GCF_000624735.2_ASM62473v2 Salmonella enterica subsp. enterica serovar Enteritidis str. EC20121542
Proteobacteria; Gammaproteobacteria; Enterobacterales; Enterobacteriaceae; Salmonella; Salmonella enterica; Salmonella enterica subsp. enterica; Salmonella enterica subsp. enterica serovar Enteritidis
MNNEETFYQAMRRKGVTRRSFLKFCSLAATSLGLGAGMTPKIAWALENKPRIPVVWIHGL WP_001058307.1
hydrogenase [Salmonella enterica] Length: 372\nNumber of predicted TMHs: 2\nExp number of AAs in TMHs: 36.51738\nExp number, first 60 AAs: 19.72849\nTotal prob of N-in: 0.96170\nPOSSIBLE N-term signal sequence\ninside 1 19\nTMhelix 20 39\noutside 40 325\nTMhelix 326 348\ninside 349 372

23659 GCF_000624735.2_ASM62473v2 Salmonella enterica subsp. enterica serovar Enteritidis str. EC20121542
Proteobacteria; Gammaproteobacteria; Enterobacterales; Enterobacteriaceae; Salmonella; Salmonella enterica; Salmonella enterica subsp. enterica; Salmonella enterica subsp. enterica serovar Enteritidis
MTGDNTLITSHGINRRDFMKLCAALAATMGLSSKAAAEMAESVSNPQRPPVIWIGAQECT WP_000145429.1
hydrogenase 2 small subunit [Salmonella enterica] Length: 372\nNumber of predicted TMHs: 1\nExp number of AAs in TMHs: 24.51309\nExp number, first 60 AAs: 0.2682\nTotal prob of N-in: 0.14402\noutside 1 331\nTMhelix 332 354\ninside 355 372

23660 GCF_000624755.2_ASM62475v2 Salmonella enterica subsp. enterica serovar Enteritidis str. EC20121671
Proteobacteria; Gammaproteobacteria; Enterobacterales; Enterobacteriaceae; Salmonella; Salmonella enterica; Salmonella enterica subsp. enterica; Salmonella enterica subsp. enterica serovar Enteritidis
MNNEETFYQAMRRKGVTRRSFLKFCSLAATSLGLGAGMTPKIAWALENKPRIPVVWIHGL WP_001058307.1
hydrogenase [Salmonella enterica] Length: 372\nNumber of predicted TMHs: 2\nExp number of AAs in TMHs: 36.51738\nExp number, first 60 AAs: 19.72849\nTotal prob of N-in: 0.96170\nPOSSIBLE N-term signal sequence\ninside 1 19\nTMhelix 20 39\noutside 40 325\nTMhelix 326 348\ninside 349 372

23661 GCF_000624755.2_ASM62475v2 Salmonella enterica subsp. enterica serovar Enteritidis str. EC20121671 Proteobacteria; Gammaproteobacteria; Enterobacterales; Enterobacteriaceae; Salmonella; Salmonella enterica; Salmonella enterica subsp. enterica; Salmonella enterica subsp. enterica serovar Enteritidis MTGDNTLITSHGINRRDFMKLCAALAATMGLSSKAAAEMAESVSNPQRPPVIWIGAQECT WP_000145429.1 hydrogenase 2 small subunit [Salmonella enterica] Length: 372\nNumber of predicted TMHs: 1\nExp number of AAs in TMHs: 24.51309\nExp number, first 60 AAs: 0.2682\nTotal prob of N-in: 0.14402\noutside 1 331\nTMhelix 332 354\ninside 355 372

23662 GCF_000624775.2_ASM62477v2 Salmonella enterica subsp. enterica serovar Enteritidis str. EC20121672 Proteobacteria; Gammaproteobacteria; Enterobacterales; Enterobacteriaceae; Salmonella; Salmonella enterica; Salmonella enterica subsp. enterica; Salmonella enterica subsp. enterica serovar Enteritidis MTGDNTLITSHGINRRDFMKLCAALAATMGLSSKAAAEMAESVSNPQRPPVIWIGAQECT WP_000145429.1 hydrogenase 2 small subunit [Salmonella enterica] Length: 372\nNumber of predicted TMHs: 1\nExp number of AAs in TMHs: 24.51309\nExp number, first 60 AAs: 0.2682\nTotal prob of N-in: 0.14402\noutside 1 331\nTMhelix 332 354\ninside 355 372

23663 GCF_000624775.2_ASM62477v2 Salmonella enterica subsp. enterica serovar Enteritidis str. EC20121672 Proteobacteria; Gammaproteobacteria; Enterobacterales; Enterobacteriaceae; Salmonella; Salmonella enterica; Salmonella enterica subsp. enterica; Salmonella enterica subsp. enterica serovar Enteritidis MNNEETFYQAMRRKGVTRRSFLKFCSLAATSLGLGAGMTPKIAWALENKPRIPVVWIHGL WP_001058307.1 hydrogenase [Salmonella enterica] Length: 372\nNumber of predicted TMHs: 2\nExp number of AAs in TMHs: 36.51738\nExp number, first 60 AAs: 19.72849\nTotal prob of N-in: 0.96170\nPOSSIBLE N-term signal sequence\ninside 1 19\nTMhelix 20 39\noutside 40 325\nTMhelix 326 348\ninside 349 372

23664 GCF_000624795.2_ASM62479v2 Salmonella enterica subsp. enterica serovar Enteritidis str. EC20121689 Proteobacteria; Gammaproteobacteria; Enterobacterales; Enterobacteriaceae; Salmonella; Salmonella enterica; Salmonella enterica subsp. enterica; Salmonella enterica subsp. enterica serovar Enteritidis MTGDNTLITSHGINRRDFMKLCAALAATMGLSSKAAAEMAESVSNPQRPPVIWIGAQECT WP_000145429.1 hydrogenase 2 small subunit [Salmonella enterica] Length: 372\nNumber of predicted TMHs: 1\nExp number of AAs in TMHs: 24.51309\nExp number, first 60 AAs: 0.2682\nTotal prob of N-in: 0.14402\noutside 1 331\nTMhelix 332 354\ninside 355 372

23665 GCF_000624795.2_ASM62479v2 Salmonella enterica subsp. enterica serovar Enteritidis str. EC20121689 Proteobacteria; Gammaproteobacteria; Enterobacterales; Enterobacteriaceae; Salmonella; Salmonella enterica; Salmonella enterica subsp. enterica; Salmonella enterica subsp. enterica serovar Enteritidis MNNEETFYQAMRRKGVTRRSFLKFCSLAATSLGLGAGMTPKIAWALENKPRIPVVWIHGL WP_001058307.1 hydrogenase [Salmonella enterica] Length: 372\nNumber of predicted TMHs: 2\nExp number of AAs in TMHs: 36.51738\nExp number, first 60 AAs: 19.72849\nTotal prob of N-in: 0.96170\nPOSSIBLE N-term signal sequence\ninside 1 19\nTMhelix 20 39\noutside 40 325\nTMhelix 326 348\ninside 349 372

23666 GCF_000624835.2_ASM62483v2 Salmonella enterica subsp. enterica serovar Enteritidis str. EC20121744 Proteobacteria; Gammaproteobacteria; Enterobacterales; Enterobacteriaceae; Salmonella; Salmonella enterica; Salmonella enterica subsp. enterica; Salmonella enterica subsp. enterica serovar Enteritidis MNNEETFYQAMRRKGVTRRSFLKFCSLAATSLGLGAGMTPKIAWALENKPRIPVVWIHGL WP_001058307.1 hydrogenase [Salmonella enterica] Length: 372\nNumber of predicted TMHs: 2\nExp number of AAs in TMHs: 36.51738\nExp number, first 60 AAs: 19.72849\nTotal prob of N-in: 0.96170\nPOSSIBLE N-term signal sequence\ninside 1 19\nTMhelix 20 39\noutside 40 325\nTMhelix 326 348\ninside 349 372

23667 GCF_000624835.2_ASM62483v2 Salmonella enterica subsp. enterica serovar Enteritidis str. EC20121744 Proteobacteria; Gammaproteobacteria; Enterobacterales; Enterobacteriaceae; Salmonella; Salmonella enterica; Salmonella enterica subsp. enterica; Salmonella enterica subsp. enterica serovar Enteritidis MTGDNTLITSHGINRRDFMKLCAALAATMGLSSKAAAEMAESVSNPQRPPVIWIGAQECT WP_000145429.1 hydrogenase 2 small subunit [Salmonella enterica] Length: 372\nNumber of predicted TMHs: 1\nExp number of AAs in TMHs: 24.51309\nExp number, first 60 AAs: 0.2682\nTotal prob of N-in: 0.14402\noutside 1 331\nTMhelix 332 354\ninside 355 372

23668 GCF_000624855.2_ASM62485v2 Salmonella enterica subsp. enterica serovar Enteritidis str. EC20121746 Proteobacteria; Gammaproteobacteria; Enterobacterales; Enterobacteriaceae; Salmonella; Salmonella enterica; Salmonella enterica subsp. enterica; Salmonella enterica subsp. enterica serovar Enteritidis MTGDNTLITSHGINRRDFMKLCAALAATMGLSSKAAAEMAESVSNPQRPPVIWIGAQECT WP_000145429.1 hydrogenase 2 small subunit [Salmonella enterica] Length: 372\nNumber of predicted TMHs: 1\nExp

number of AAs in TMHs: 24.51309\nExp number, first 60 AAs: 0.2682\nTotal prob of N-in: 0.14402\noutside 1 331\nTMhelix 332 354\ninside 355 372

23669 GCF_000624855.2_ASM62485v2 Salmonella enterica subsp. enterica serovar Enteritidis str. EC20121746 Proteobacteria; Gammaproteobacteria; Enterobacterales; Enterobacteriaceae; Salmonella; Salmonella enterica; Salmonella enterica subsp. enterica; Salmonella enterica subsp. enterica serovar Enteritidis MNNEETFYQAMRRKGVTRRSFLKFCSLAATSLGLGAGMTPKIAWALENKPRIPVVWIHGL WP_001058307.1 hydrogenase [Salmonella enterica] Length: 372\nNumber of predicted TMHs: 2\nExp number of AAs in TMHs: 36.51738\nExp number, first 60 AAs: 19.72849\nTotal prob of N-in: 0.96170\nPOSSIBLE N-term signal sequence\ninside 1 19\nTMhelix 20 39\noutside 40 325\nTMhelix 326 348\ninside 349 372

23670 GCF_000626435.2_ASM62643v2 Salmonella enterica subsp. enterica serovar Enteritidis str. EC20121747 Proteobacteria; Gammaproteobacteria; Enterobacterales; Enterobacteriaceae; Salmonella; Salmonella enterica; Salmonella enterica subsp. enterica; Salmonella enterica subsp. enterica serovar Enteritidis MTGDNTLITSHGINRRDFMKLCAALAATMGLSSKAAAEMAESVSNPQRPPVIWIGAQECT WP_000145429.1 hydrogenase 2 small subunit [Salmonella enterica] Length: 372\nNumber of predicted TMHs: 1\nExp number of AAs in TMHs: 24.51309\nExp number, first 60 AAs: 0.2682\nTotal prob of N-in: 0.14402\noutside 1 331\nTMhelix 332 354\ninside 355 372

23671 GCF_000624875.2_ASM62487v2 Salmonella enterica subsp. enterica serovar Enteritidis str. EC20121748 Proteobacteria; Gammaproteobacteria; Enterobacterales; Enterobacteriaceae; Salmonella; Salmonella enterica; Salmonella enterica subsp. enterica; Salmonella enterica subsp. enterica serovar Enteritidis MTGDNTLITSHGINRRDFMKLCAALAATMGLSSKAAAEMAESVSNPQRPPVIWIGAQECT WP_000145429.1 hydrogenase 2 small subunit [Salmonella enterica] Length: 372\nNumber of predicted TMHs: 1\nExp number of AAs in TMHs: 24.51309\nExp number, first 60 AAs: 0.2682\nTotal prob of N-in: 0.14402\noutside 1 331\nTMhelix 332 354\ninside 355 372

23672 GCF_000624875.2_ASM62487v2 Salmonella enterica subsp. enterica serovar Enteritidis str. EC20121748 Proteobacteria; Gammaproteobacteria; Enterobacterales; Enterobacteriaceae; Salmonella; Salmonella enterica; Salmonella enterica subsp. enterica; Salmonella enterica subsp. enterica serovar Enteritidis MNNEETFYQAMRRKGVTRRSFLKFCSLAATSLGLGAGMTPKIAWALENKPRIPVVWIHGL WP_001058307.1 hydrogenase [Salmonella enterica] Length: 372\nNumber of predicted TMHs: 2\nExp number of AAs in TMHs: 36.51738\nExp number, first 60 AAs: 19.72849\nTotal prob of N-in: 0.96170\nPOSSIBLE N-term signal sequence\ninside 1 19\nTMhelix 20 39\noutside 40 325\nTMhelix 326 348\ninside 349 372

23673 GCF_000624895.2_ASM62489v2 Salmonella enterica subsp. enterica serovar Enteritidis str. EC20121750 Proteobacteria; Gammaproteobacteria; Enterobacterales; Enterobacteriaceae; Salmonella; Salmonella enterica; Salmonella enterica subsp. enterica; Salmonella enterica subsp. enterica serovar Enteritidis MTGDNTLITSHGINRRDFMKLCAALAATMGLSSKAAAEMAESVSNPQRPPVIWIGAQECT WP_000145429.1 hydrogenase 2 small subunit [Salmonella enterica] Length: 372\nNumber of predicted TMHs: 1\nExp number of AAs in TMHs: 24.51309\nExp number, first 60 AAs: 0.2682\nTotal prob of N-in: 0.14402\noutside 1 331\nTMhelix 332 354\ninside 355 372

23674 GCF_000624895.2_ASM62489v2 Salmonella enterica subsp. enterica serovar Enteritidis str. EC20121750 Proteobacteria; Gammaproteobacteria; Enterobacterales; Enterobacteriaceae; Salmonella; Salmonella enterica; Salmonella enterica subsp. enterica; Salmonella enterica subsp. enterica serovar Enteritidis MNNEETFYQAMRRKGVTRRSFLKFCSLAATSLGLGAGMTPKIAWALENKPRIPVVWIHGL WP_001058307.1 hydrogenase [Salmonella enterica] Length: 372\nNumber of predicted TMHs: 2\nExp number of AAs in TMHs: 36.51738\nExp number, first 60 AAs: 19.72849\nTotal prob of N-in: 0.96170\nPOSSIBLE N-term signal sequence\ninside 1 19\nTMhelix 20 39\noutside 40 325\nTMhelix 326 348\ninside 349 372

23675 GCF_000624915.2_ASM62491v2 Salmonella enterica subsp. enterica serovar Enteritidis str. EC20121751 Proteobacteria; Gammaproteobacteria; Enterobacterales; Enterobacteriaceae; Salmonella; Salmonella enterica; Salmonella enterica subsp. enterica; Salmonella enterica subsp. enterica serovar Enteritidis MTGDNTLITSHGINRRDFMKLCAALAATMGLSSKAAAEMAESVSNPQRPPVIWIGAQECT WP_000145429.1 hydrogenase 2 small subunit [Salmonella enterica] Length: 372\nNumber of predicted TMHs: 1\nExp number of AAs in TMHs: 24.51309\nExp number, first 60 AAs: 0.2682\nTotal prob of N-in: 0.14402\noutside 1 331\nTMhelix 332 354\ninside 355 372

23676 GCF_000624915.2_ASM62491v2 Salmonella enterica subsp. enterica serovar Enteritidis str. EC20121751 Proteobacteria; Gammaproteobacteria; Enterobacterales; Enterobacteriaceae; Salmonella; Salmonella enterica;

Salmonella enterica subsp. enterica; Salmonella enterica subsp. enterica serovar Enteritidis
MNNEETFYQAMRRKGVTRRSFLKFCSLAATSLGLGAGMTPKIAWALENKPRIPVVWIHGL WP_001058307.1
hydrogenase [Salmonella enterica] Length: 372\nNumber of predicted TMHs: 2\nExp number of AAs in TMHs: 36.51738\nExp number, first 60 AAs: 19.72849\nTotal prob of N-in: 0.96170\nPOSSIBLE N-term signal sequence\ninside 1 19\nTMhelix 20 39\noutside 40 325\nTMhelix 326 348\ninside 349 372

23677 GCF_000624935.2_ASM62493v2 Salmonella enterica subsp. enterica serovar Enteritidis str. EC20121753
Proteobacteria; Gammaproteobacteria; Enterobacterales; Enterobacteriaceae; Salmonella; Salmonella enterica; Salmonella enterica subsp. enterica; Salmonella enterica subsp. enterica serovar Enteritidis
MNNEETFYQAMRRKGVTRRSFLKFCSLAATSLGLGAGMTPKIAWALENKPRIPVVWIHGL WP_001058307.1
hydrogenase [Salmonella enterica] Length: 372\nNumber of predicted TMHs: 2\nExp number of AAs in TMHs: 36.51738\nExp number, first 60 AAs: 19.72849\nTotal prob of N-in: 0.96170\nPOSSIBLE N-term signal sequence\ninside 1 19\nTMhelix 20 39\noutside 40 325\nTMhelix 326 348\ninside 349 372

23678 GCF_000624935.2_ASM62493v2 Salmonella enterica subsp. enterica serovar Enteritidis str. EC20121753
Proteobacteria; Gammaproteobacteria; Enterobacterales; Enterobacteriaceae; Salmonella; Salmonella enterica; Salmonella enterica subsp. enterica; Salmonella enterica subsp. enterica serovar Enteritidis
MTGDNTLITSHGINRRDFMKLCAALAATMGLSSKAAAEMAESVSNPQRPPVIWIGAQECT WP_000145429.1
hydrogenase 2 small subunit [Salmonella enterica] Length: 372\nNumber of predicted TMHs: 1\nExp number of AAs in TMHs: 24.51309\nExp number, first 60 AAs: 0.2682\nTotal prob of N-in: 0.14402\noutside 1 331\nTMhelix 332 354\ninside 355 372

23679 GCF_000625775.2_ASM62577v2 Salmonella enterica subsp. enterica serovar Enteritidis str. EC20121765
Proteobacteria; Gammaproteobacteria; Enterobacterales; Enterobacteriaceae; Salmonella; Salmonella enterica; Salmonella enterica subsp. enterica; Salmonella enterica subsp. enterica serovar Enteritidis
MTGDNTLITSHGINRRDFMKLCAALAATMGLSSKAAAEMAESVSNPQRPPVIWIGAQECT WP_000145429.1
hydrogenase 2 small subunit [Salmonella enterica] Length: 372\nNumber of predicted TMHs: 1\nExp number of AAs in TMHs: 24.51309\nExp number, first 60 AAs: 0.2682\nTotal prob of N-in: 0.14402\noutside 1 331\nTMhelix 332 354\ninside 355 372

23680 GCF_000625775.2_ASM62577v2 Salmonella enterica subsp. enterica serovar Enteritidis str. EC20121765
Proteobacteria; Gammaproteobacteria; Enterobacterales; Enterobacteriaceae; Salmonella; Salmonella enterica; Salmonella enterica subsp. enterica; Salmonella enterica subsp. enterica serovar Enteritidis
MNNEETFYQAMRRKGVTRRSFLKFCSLAATSLGLGAGMTPKIAWALENKPRIPVVWIHGL WP_001058307.1
hydrogenase [Salmonella enterica] Length: 372\nNumber of predicted TMHs: 2\nExp number of AAs in TMHs: 36.51738\nExp number, first 60 AAs: 19.72849\nTotal prob of N-in: 0.96170\nPOSSIBLE N-term signal sequence\ninside 1 19\nTMhelix 20 39\noutside 40 325\nTMhelix 326 348\ninside 349 372

23681 GCF_000625255.2_ASM62525v2 Salmonella enterica subsp. enterica serovar Enteritidis str. EC20121812
Proteobacteria; Gammaproteobacteria; Enterobacterales; Enterobacteriaceae; Salmonella; Salmonella enterica; Salmonella enterica subsp. enterica; Salmonella enterica subsp. enterica serovar Enteritidis
MNNEETFYQAMRRKGVTRRSFLKFCSLAATSLGLGAGMTPKIAWALENKPRIPVVWIHGL WP_001058307.1
hydrogenase [Salmonella enterica] Length: 372\nNumber of predicted TMHs: 2\nExp number of AAs in TMHs: 36.51738\nExp number, first 60 AAs: 19.72849\nTotal prob of N-in: 0.96170\nPOSSIBLE N-term signal sequence\ninside 1 19\nTMhelix 20 39\noutside 40 325\nTMhelix 326 348\ninside 349 372

23682 GCF_000625255.2_ASM62525v2 Salmonella enterica subsp. enterica serovar Enteritidis str. EC20121812
Proteobacteria; Gammaproteobacteria; Enterobacterales; Enterobacteriaceae; Salmonella; Salmonella enterica; Salmonella enterica subsp. enterica; Salmonella enterica subsp. enterica serovar Enteritidis
MTGDNTLITSHGINRRDFMKLCAALAATMGLSSKAAAEMAESVSNPQRPPVIWIGAQECT WP_000145429.1
hydrogenase 2 small subunit [Salmonella enterica] Length: 372\nNumber of predicted TMHs: 1\nExp number of AAs in TMHs: 24.51309\nExp number, first 60 AAs: 0.2682\nTotal prob of N-in: 0.14402\noutside 1 331\nTMhelix 332 354\ninside 355 372

23683 GCF_000625275.2_ASM62527v2 Salmonella enterica subsp. enterica serovar Enteritidis str. EC20121825
Proteobacteria; Gammaproteobacteria; Enterobacterales; Enterobacteriaceae; Salmonella; Salmonella enterica; Salmonella enterica subsp. enterica; Salmonella enterica subsp. enterica serovar Enteritidis
MNNEETFYQAMRRKGVTRRSFLKFCSLAATSLGLGAGMTPKIAWALENKPRIPVVWIHGL WP_001058307.1
hydrogenase [Salmonella enterica] Length: 372\nNumber of predicted TMHs: 2\nExp number of AAs in TMHs: 36.51738\nExp number, first 60 AAs: 19.72849\nTotal prob of N-in: 0.96170\nPOSSIBLE N-term signal sequence\ninside 1 19\nTMhelix 20 39\noutside 40 325\nTMhelix 326 348\ninside 349 372

23684	GCF_000625275.2_ASM62527v2	Salmonella enterica subsp. enterica serovar Enteritidis str. EC20121825	Proteobacteria; Gammaproteobacteria; Enterobacterales; Enterobacteriaceae; Salmonella; Salmonella enterica; Salmonella enterica subsp. enterica; Salmonella enterica subsp. enterica serovar Enteritidis	MTGDNTLITSHGINRRDFMKLCAALAATMGLSSKAAAEMAESVSNPQRPPVIWIGAQECT	WP_000145429.1	hydrogenase 2 small subunit [Salmonella enterica]	Length: 372\nNumber of predicted TMHs: 1\nExp number of AAs in TMHs: 24.51309\nExp number, first 60 AAs: 0.2682\nTotal prob of N-in: 0.14402\noutside 1 331\nTMhelix 332 354\ninside 355 372

23685	GCF_000625295.2_ASM62529v2	Salmonella enterica subsp. enterica serovar Enteritidis str. EC20121826	Proteobacteria; Gammaproteobacteria; Enterobacterales; Enterobacteriaceae; Salmonella; Salmonella enterica; Salmonella enterica subsp. enterica; Salmonella enterica subsp. enterica serovar Enteritidis	MNNEETFYQAMRRKGVTRRSFLKFCSLAATSLGLGAGMTPKIAWALENKPRIPVVWIHGL	WP_001058307.1	hydrogenase [Salmonella enterica]	Length: 372\nNumber of predicted TMHs: 2\nExp number of AAs in TMHs: 36.51738\nExp number, first 60 AAs: 19.72849\nTotal prob of N-in: 0.96170\nPOSSIBLE N-term signal sequence\ninside 1 19\nTMhelix 20 39\noutside 40 325\nTMhelix 326 348\ninside 349 372

23686	GCF_000625295.2_ASM62529v2	Salmonella enterica subsp. enterica serovar Enteritidis str. EC20121826	Proteobacteria; Gammaproteobacteria; Enterobacterales; Enterobacteriaceae; Salmonella; Salmonella enterica; Salmonella enterica subsp. enterica; Salmonella enterica subsp. enterica serovar Enteritidis	MTGDNTLITSHGINRRDFMKLCAALAATMGLSSKAAAEMAESVSNPQRPPVIWIGAQECT	WP_000145429.1	hydrogenase 2 small subunit [Salmonella enterica]	Length: 372\nNumber of predicted TMHs: 1\nExp number of AAs in TMHs: 24.51309\nExp number, first 60 AAs: 0.2682\nTotal prob of N-in: 0.14402\noutside 1 331\nTMhelix 332 354\ninside 355 372

23687	GCF_000625315.2_ASM62531v2	Salmonella enterica subsp. enterica serovar Enteritidis str. EC20121969	Proteobacteria; Gammaproteobacteria; Enterobacterales; Enterobacteriaceae; Salmonella; Salmonella enterica; Salmonella enterica subsp. enterica; Salmonella enterica subsp. enterica serovar Enteritidis	MTGDNTLITSHGINRRDFMKLCAALAATMGLSSKAAAEMAESVSNPQRPPVIWIGAQECT	WP_000145429.1	hydrogenase 2 small subunit [Salmonella enterica]	Length: 372\nNumber of predicted TMHs: 1\nExp number of AAs in TMHs: 24.51309\nExp number, first 60 AAs: 0.2682\nTotal prob of N-in: 0.14402\noutside 1 331\nTMhelix 332 354\ninside 355 372

23688	GCF_000625315.2_ASM62531v2	Salmonella enterica subsp. enterica serovar Enteritidis str. EC20121969	Proteobacteria; Gammaproteobacteria; Enterobacterales; Enterobacteriaceae; Salmonella; Salmonella enterica; Salmonella enterica subsp. enterica; Salmonella enterica subsp. enterica serovar Enteritidis	MNNEETFYQAMRRKGVTRRSFLKFCSLAATSLGLGAGMTPKIAWALENKPRIPVVWIHGL	WP_001058307.1	hydrogenase [Salmonella enterica]	Length: 372\nNumber of predicted TMHs: 2\nExp number of AAs in TMHs: 36.51738\nExp number, first 60 AAs: 19.72849\nTotal prob of N-in: 0.96170\nPOSSIBLE N-term signal sequence\ninside 1 19\nTMhelix 20 39\noutside 40 325\nTMhelix 326 348\ninside 349 372

23689	GCF_000625335.2_ASM62533v2	Salmonella enterica subsp. enterica serovar Enteritidis str. EC20121970	Proteobacteria; Gammaproteobacteria; Enterobacterales; Enterobacteriaceae; Salmonella; Salmonella enterica; Salmonella enterica subsp. enterica; Salmonella enterica subsp. enterica serovar Enteritidis	MTGDNTLITSHGINRRDFMKLCAALAATMGLSSKAAAEMAESVSNPQRPPVIWIGAQECT	WP_000145429.1	hydrogenase 2 small subunit [Salmonella enterica]	Length: 372\nNumber of predicted TMHs: 1\nExp number of AAs in TMHs: 24.51309\nExp number, first 60 AAs: 0.2682\nTotal prob of N-in: 0.14402\noutside 1 331\nTMhelix 332 354\ninside 355 372

23690	GCF_000625335.2_ASM62533v2	Salmonella enterica subsp. enterica serovar Enteritidis str. EC20121970	Proteobacteria; Gammaproteobacteria; Enterobacterales; Enterobacteriaceae; Salmonella; Salmonella enterica; Salmonella enterica subsp. enterica; Salmonella enterica subsp. enterica serovar Enteritidis	MNNEETFYQAMRRKGVTRRSFLKFCSLAATSLGLGAGMTPKIAWALENKPRIPVVWIHGL	WP_001058307.1	hydrogenase [Salmonella enterica]	Length: 372\nNumber of predicted TMHs: 2\nExp number of AAs in TMHs: 36.51738\nExp number, first 60 AAs: 19.72849\nTotal prob of N-in: 0.96170\nPOSSIBLE N-term signal sequence\ninside 1 19\nTMhelix 20 39\noutside 40 325\nTMhelix 326 348\ninside 349 372

23691	GCF_000625635.2_ASM62563v2	Salmonella enterica subsp. enterica serovar Enteritidis str. EC20121976	Proteobacteria; Gammaproteobacteria; Enterobacterales; Enterobacteriaceae; Salmonella; Salmonella enterica; Salmonella enterica subsp. enterica; Salmonella enterica subsp. enterica serovar Enteritidis	MTGDNTLITSHGINRRDFMKLCAALAATMGLSSKAAAEMAESVSNPQRPPVIWIGAQECT	WP_000145429.1	hydrogenase 2 small subunit [Salmonella enterica]	Length: 372\nNumber of predicted TMHs: 1\nExp

number of AAs in TMHs: 24.51309\nExp number, first 60 AAs: 0.2682\nTotal prob of N-in: 0.14402\noutside 1 331\nTMhelix 332 354\ninside 355 372

23692 GCF_000625635.2_ASM62563v2 Salmonella enterica subsp. enterica serovar Enteritidis str. EC20121976 Proteobacteria; Gammaproteobacteria; Enterobacterales; Enterobacteriaceae; Salmonella; Salmonella enterica; Salmonella enterica subsp. enterica; Salmonella enterica subsp. enterica serovar Enteritidis MNNEETFYQAMRRKGVTRRSFLKFCSLAATSLGLGAGMTPKIAWALENKPRIPVVWIHGL WP_001058307.1 hydrogenase [Salmonella enterica] Length: 372\nNumber of predicted TMHs: 2\nExp number of AAs in TMHs: 36.51738\nExp number, first 60 AAs: 19.72849\nTotal prob of N-in: 0.96170\nPOSSIBLE N-term signal sequence\ninside 1 19\nTMhelix 20 39\noutside 40 325\nTMhelix 326 348\ninside 349 372

23693 GCF_000625795.2_ASM62579v2 Salmonella enterica subsp. enterica serovar Enteritidis str. EC20121986 Proteobacteria; Gammaproteobacteria; Enterobacterales; Enterobacteriaceae; Salmonella; Salmonella enterica; Salmonella enterica subsp. enterica; Salmonella enterica subsp. enterica serovar Enteritidis MNNEETFYQAMRRKGVTRRSFLKFCSLAATSLGLGAGMTPKIAWALENKPRIPVVWIHGL WP_001058307.1 hydrogenase [Salmonella enterica] Length: 372\nNumber of predicted TMHs: 2\nExp number of AAs in TMHs: 36.51738\nExp number, first 60 AAs: 19.72849\nTotal prob of N-in: 0.96170\nPOSSIBLE N-term signal sequence\ninside 1 19\nTMhelix 20 39\noutside 40 325\nTMhelix 326 348\ninside 349 372

23694 GCF_000625795.2_ASM62579v2 Salmonella enterica subsp. enterica serovar Enteritidis str. EC20121986 Proteobacteria; Gammaproteobacteria; Enterobacterales; Enterobacteriaceae; Salmonella; Salmonella enterica; Salmonella enterica subsp. enterica; Salmonella enterica subsp. enterica serovar Enteritidis MTGDNTLITSHGINRRDFMKLCAALAATMGLSSKAAAEMAESVSNPQRPPVIWIGAQECT WP_000145429.1 hydrogenase 2 small subunit [Salmonella enterica] Length: 372\nNumber of predicted TMHs: 1\nExp number of AAs in TMHs: 24.51309\nExp number, first 60 AAs: 0.2682\nTotal prob of N-in: 0.14402\noutside 1 331\nTMhelix 332 354\ninside 355 372

23695 GCF_000625815.2_ASM62581v2 Salmonella enterica subsp. enterica serovar Enteritidis str. EC20121989 Proteobacteria; Gammaproteobacteria; Enterobacterales; Enterobacteriaceae; Salmonella; Salmonella enterica; Salmonella enterica subsp. enterica; Salmonella enterica subsp. enterica serovar Enteritidis MNNEETFYQAMRRKGVTRRSFLKFCSLAATSLGLGAGMTPKIAWALENKPRIPVVWIHGL WP_001058307.1 hydrogenase [Salmonella enterica] Length: 372\nNumber of predicted TMHs: 2\nExp number of AAs in TMHs: 36.51738\nExp number, first 60 AAs: 19.72849\nTotal prob of N-in: 0.96170\nPOSSIBLE N-term signal sequence\ninside 1 19\nTMhelix 20 39\noutside 40 325\nTMhelix 326 348\ninside 349 372

23696 GCF_000625815.2_ASM62581v2 Salmonella enterica subsp. enterica serovar Enteritidis str. EC20121989 Proteobacteria; Gammaproteobacteria; Enterobacterales; Enterobacteriaceae; Salmonella; Salmonella enterica; Salmonella enterica subsp. enterica; Salmonella enterica subsp. enterica serovar Enteritidis MTGDNTLITSHGINRRDFMKLCAALAATMGLSSKAAAEMAESVSNPQRPPVIWIGAQECT WP_000145429.1 hydrogenase 2 small subunit [Salmonella enterica] Length: 372\nNumber of predicted TMHs: 1\nExp number of AAs in TMHs: 24.51309\nExp number, first 60 AAs: 0.2682\nTotal prob of N-in: 0.14402\noutside 1 331\nTMhelix 332 354\ninside 355 372

23697 GCF_000625875.2_ASM62587v2 Salmonella enterica subsp. enterica serovar Enteritidis str. EC20121990 Proteobacteria; Gammaproteobacteria; Enterobacterales; Enterobacteriaceae; Salmonella; Salmonella enterica; Salmonella enterica subsp. enterica; Salmonella enterica subsp. enterica serovar Enteritidis MTGDNTLITSHGINRRDFMKLCAALAATMGLSSKAAAEMAESVSNPQRPPVIWIGAQECT WP_000145429.1 hydrogenase 2 small subunit [Salmonella enterica] Length: 372\nNumber of predicted TMHs: 1\nExp number of AAs in TMHs: 24.51309\nExp number, first 60 AAs: 0.2682\nTotal prob of N-in: 0.14402\noutside 1 331\nTMhelix 332 354\ninside 355 372

23698 GCF_000625875.2_ASM62587v2 Salmonella enterica subsp. enterica serovar Enteritidis str. EC20121990 Proteobacteria; Gammaproteobacteria; Enterobacterales; Enterobacteriaceae; Salmonella; Salmonella enterica; Salmonella enterica subsp. enterica; Salmonella enterica subsp. enterica serovar Enteritidis MNNEETFYQAMRRKGVTRRSFLKFCSLAATSLGLGAGMTPKIAWALENKPRIPVVWIHGL WP_001058307.1 hydrogenase [Salmonella enterica] Length: 372\nNumber of predicted TMHs: 2\nExp number of AAs in TMHs: 36.51738\nExp number, first 60 AAs: 19.72849\nTotal prob of N-in: 0.96170\nPOSSIBLE N-term signal sequence\ninside 1 19\nTMhelix 20 39\noutside 40 325\nTMhelix 326 348\ninside 349 372

23699 GCF_000625895.2_ASM62589v2 Salmonella enterica subsp. enterica serovar Enteritidis str. EC20122022 Proteobacteria; Gammaproteobacteria; Enterobacterales; Enterobacteriaceae; Salmonella; Salmonella enterica;

Salmonella enterica subsp. enterica; Salmonella enterica subsp. enterica serovar Enteritidis
MTGDNTLITSHGINRRDFMKLCAALAATMGLSSKAAAEMAESVSNPQRPPVIWIGAQECT WP_000145429.1
hydrogenase 2 small subunit [Salmonella enterica] Length: 372\nNumber of predicted TMHs: 1\nExp number of AAs in TMHs: 24.51309\nExp number, first 60 AAs: 0.2682\nTotal prob of N-in: 0.14402\noutside 1 331\nTMhelix 332 354\ninside 355 372

23700 GCF_000625895.2_ASM62589v2 Salmonella enterica subsp. enterica serovar Enteritidis str. EC20122022
Proteobacteria; Gammaproteobacteria; Enterobacterales; Enterobacteriaceae; Salmonella; Salmonella enterica; Salmonella enterica subsp. enterica; Salmonella enterica subsp. enterica serovar Enteritidis
MNNEETFYQAMRRKGVTRRSFLKFCSLAATSLGLGAGMTPKIAWALENKPRIPVVWIHGL WP_001058307.1
hydrogenase [Salmonella enterica] Length: 372\nNumber of predicted TMHs: 2\nExp number of AAs in TMHs: 36.51738\nExp number, first 60 AAs: 19.72849\nTotal prob of N-in: 0.96170\nPOSSIBLE N-term signal sequence\ninside 1 19\nTMhelix 20 39\noutside 40 325\nTMhelix 326 348\ninside 349 372

23701 GCF_000625915.2_ASM62591v2 Salmonella enterica subsp. enterica serovar Enteritidis str. EC20122026
Proteobacteria; Gammaproteobacteria; Enterobacterales; Enterobacteriaceae; Salmonella; Salmonella enterica; Salmonella enterica subsp. enterica; Salmonella enterica subsp. enterica serovar Enteritidis
MTGDNTLITSHGINRRDFMKLCAALAATMGLSSKAAAEMAESVSNPQRPPVIWIGAQECT WP_000145429.1
hydrogenase 2 small subunit [Salmonella enterica] Length: 372\nNumber of predicted TMHs: 1\nExp number of AAs in TMHs: 24.51309\nExp number, first 60 AAs: 0.2682\nTotal prob of N-in: 0.14402\noutside 1 331\nTMhelix 332 354\ninside 355 372

23702 GCF_000625915.2_ASM62591v2 Salmonella enterica subsp. enterica serovar Enteritidis str. EC20122026
Proteobacteria; Gammaproteobacteria; Enterobacterales; Enterobacteriaceae; Salmonella; Salmonella enterica; Salmonella enterica subsp. enterica; Salmonella enterica subsp. enterica serovar Enteritidis
MNNEETFYQAMRRKGVTRRSFLKFCSLAATSLGLGAGMTPKIAWALENKPRIPVVWIHGL WP_001058307.1
hydrogenase [Salmonella enterica] Length: 372\nNumber of predicted TMHs: 2\nExp number of AAs in TMHs: 36.51738\nExp number, first 60 AAs: 19.72849\nTotal prob of N-in: 0.96170\nPOSSIBLE N-term signal sequence\ninside 1 19\nTMhelix 20 39\noutside 40 325\nTMhelix 326 348\ninside 349 372

23703 GCF_000625935.2_ASM62593v2 Salmonella enterica subsp. enterica serovar Enteritidis str. EC20122031
Proteobacteria; Gammaproteobacteria; Enterobacterales; Enterobacteriaceae; Salmonella; Salmonella enterica; Salmonella enterica subsp. enterica; Salmonella enterica subsp. enterica serovar Enteritidis
MNNEETFYQAMRRKGVTRRSFLKFCSLAATSLGLGAGMTPKIAWALENKPRIPVVWIHGL WP_001058307.1
hydrogenase [Salmonella enterica] Length: 372\nNumber of predicted TMHs: 2\nExp number of AAs in TMHs: 36.51738\nExp number, first 60 AAs: 19.72849\nTotal prob of N-in: 0.96170\nPOSSIBLE N-term signal sequence\ninside 1 19\nTMhelix 20 39\noutside 40 325\nTMhelix 326 348\ninside 349 372

23704 GCF_000625935.2_ASM62593v2 Salmonella enterica subsp. enterica serovar Enteritidis str. EC20122031
Proteobacteria; Gammaproteobacteria; Enterobacterales; Enterobacteriaceae; Salmonella; Salmonella enterica; Salmonella enterica subsp. enterica; Salmonella enterica subsp. enterica serovar Enteritidis
MTGDNTLITSHGINRRDFMKLCAALAATMGLSSKAAAEMAESVSNPQRPPVIWIGAQECT WP_000145429.1
hydrogenase 2 small subunit [Salmonella enterica] Length: 372\nNumber of predicted TMHs: 1\nExp number of AAs in TMHs: 24.51309\nExp number, first 60 AAs: 0.2682\nTotal prob of N-in: 0.14402\noutside 1 331\nTMhelix 332 354\ninside 355 372

23705 GCF_000625955.2_ASM62595v2 Salmonella enterica subsp. enterica serovar Enteritidis str. EC20122033
Proteobacteria; Gammaproteobacteria; Enterobacterales; Enterobacteriaceae; Salmonella; Salmonella enterica; Salmonella enterica subsp. enterica; Salmonella enterica subsp. enterica serovar Enteritidis
MTGDNTLITSHGINRRDFMKLCAALAATMGLSSKAAAEMAESVSNPQRPPVIWIGAQECT WP_000145429.1
hydrogenase 2 small subunit [Salmonella enterica] Length: 372\nNumber of predicted TMHs: 1\nExp number of AAs in TMHs: 24.51309\nExp number, first 60 AAs: 0.2682\nTotal prob of N-in: 0.14402\noutside 1 331\nTMhelix 332 354\ninside 355 372

23706 GCF_000625955.2_ASM62595v2 Salmonella enterica subsp. enterica serovar Enteritidis str. EC20122033
Proteobacteria; Gammaproteobacteria; Enterobacterales; Enterobacteriaceae; Salmonella; Salmonella enterica; Salmonella enterica subsp. enterica; Salmonella enterica subsp. enterica serovar Enteritidis
MNNEETFYQAMRRKGVTRRSFLKFCSLAATSLGLGAGMTPKIAWALENKPRIPVVWIHGL WP_001058307.1
hydrogenase [Salmonella enterica] Length: 372\nNumber of predicted TMHs: 2\nExp number of AAs in TMHs: 36.51738\nExp number, first 60 AAs: 19.72849\nTotal prob of N-in: 0.96170\nPOSSIBLE N-term signal sequence\ninside 1 19\nTMhelix 20 39\noutside 40 325\nTMhelix 326 348\ninside 349 372

23707	GCF_000625975.2_ASM62597v2	Salmonella enterica subsp. enterica serovar Enteritidis str. EC20122045	Proteobacteria; Gammaproteobacteria; Enterobacterales; Enterobacteriaceae; Salmonella; Salmonella enterica; Salmonella enterica subsp. enterica; Salmonella enterica subsp. enterica serovar Enteritidis	MTGDNTLITSHGINRRDFMKLCAALAATMGLSSKAAAEMAESVSNPQRPPVIWIGAQECT	WP_000145429.1	hydrogenase 2 small subunit [Salmonella enterica]	Length: 372\nNumber of predicted TMHs: 1\nExp number of AAs in TMHs: 24.51309\nExp number, first 60 AAs: 0.2682\nTotal prob of N-in: 0.14402\noutside 1 331\nTMhelix 332 354\ninside 355 372

23708	GCF_000625975.2_ASM62597v2	Salmonella enterica subsp. enterica serovar Enteritidis str. EC20122045	Proteobacteria; Gammaproteobacteria; Enterobacterales; Enterobacteriaceae; Salmonella; Salmonella enterica; Salmonella enterica subsp. enterica; Salmonella enterica subsp. enterica serovar Enteritidis	MNNEETFYQAMRRKGVTRRSFLKFCSLAATSLGLGAGMTPKIAWALENKPRIPVVWIHGL	WP_001058307.1	hydrogenase [Salmonella enterica]	Length: 372\nNumber of predicted TMHs: 2\nExp number of AAs in TMHs: 36.51738\nExp number, first 60 AAs: 19.72849\nTotal prob of N-in: 0.96170\nPOSSIBLE N-term signal sequence\ninside 1 19\nTMhelix 20 39\noutside 40 325\nTMhelix 326 348\ninside 349 372

23709	GCF_000626015.2_ASM62601v2	Salmonella enterica subsp. enterica serovar Enteritidis str. EC20130345	Proteobacteria; Gammaproteobacteria; Enterobacterales; Enterobacteriaceae; Salmonella; Salmonella enterica; Salmonella enterica subsp. enterica; Salmonella enterica subsp. enterica serovar Enteritidis	MTGDNTLITSHGINRRDFMKLCAALAATMGLSSKAAAEMAESVSNPQRPPVIWIGAQECT	WP_000145429.1	hydrogenase 2 small subunit [Salmonella enterica]	Length: 372\nNumber of predicted TMHs: 1\nExp number of AAs in TMHs: 24.51309\nExp number, first 60 AAs: 0.2682\nTotal prob of N-in: 0.14402\noutside 1 331\nTMhelix 332 354\ninside 355 372

23710	GCF_000626015.2_ASM62601v2	Salmonella enterica subsp. enterica serovar Enteritidis str. EC20130345	Proteobacteria; Gammaproteobacteria; Enterobacterales; Enterobacteriaceae; Salmonella; Salmonella enterica; Salmonella enterica subsp. enterica; Salmonella enterica subsp. enterica serovar Enteritidis	MNNEETFYQAMRRKGVTRRSFLKFCSLAATSLGLGAGMTPKIAWALENKPRIPVVWIHGL	WP_001058307.1	hydrogenase [Salmonella enterica]	Length: 372\nNumber of predicted TMHs: 2\nExp number of AAs in TMHs: 36.51738\nExp number, first 60 AAs: 19.72849\nTotal prob of N-in: 0.96170\nPOSSIBLE N-term signal sequence\ninside 1 19\nTMhelix 20 39\noutside 40 325\nTMhelix 326 348\ninside 349 372

23711	GCF_000626035.2_ASM62603v2	Salmonella enterica subsp. enterica serovar Enteritidis str. EC20130346	Proteobacteria; Gammaproteobacteria; Enterobacterales; Enterobacteriaceae; Salmonella; Salmonella enterica; Salmonella enterica subsp. enterica; Salmonella enterica subsp. enterica serovar Enteritidis	MTGDNTLITSHGINRRDFMKLCAALAATMGLSSKAAAEMAESVSNPQRPPVIWIGAQECT	WP_000145429.1	hydrogenase 2 small subunit [Salmonella enterica]	Length: 372\nNumber of predicted TMHs: 1\nExp number of AAs in TMHs: 24.51309\nExp number, first 60 AAs: 0.2682\nTotal prob of N-in: 0.14402\noutside 1 331\nTMhelix 332 354\ninside 355 372

23712	GCF_000626035.2_ASM62603v2	Salmonella enterica subsp. enterica serovar Enteritidis str. EC20130346	Proteobacteria; Gammaproteobacteria; Enterobacterales; Enterobacteriaceae; Salmonella; Salmonella enterica; Salmonella enterica subsp. enterica; Salmonella enterica subsp. enterica serovar Enteritidis	MNNEETFYQAMRRKGVTRRSFLKFCSLAATSLGLGAGMTPKIAWALENKPRIPVVWIHGL	WP_001058307.1	hydrogenase [Salmonella enterica]	Length: 372\nNumber of predicted TMHs: 2\nExp number of AAs in TMHs: 36.51738\nExp number, first 60 AAs: 19.72849\nTotal prob of N-in: 0.96170\nPOSSIBLE N-term signal sequence\ninside 1 19\nTMhelix 20 39\noutside 40 325\nTMhelix 326 348\ninside 349 372

23713	GCF_000626055.1_ASM62605v2	Salmonella enterica subsp. enterica serovar Enteritidis str. EC20130347	Proteobacteria; Gammaproteobacteria; Enterobacterales; Enterobacteriaceae; Salmonella; Salmonella enterica; Salmonella enterica subsp. enterica; Salmonella enterica subsp. enterica serovar Enteritidis	MNNEETFYQAMRRKGVTRRSFLKFCSLAATSLGLGAGMTPKIAWALENKPRIPVVWIHGL	WP_001058307.1	hydrogenase [Salmonella enterica]	Length: 372\nNumber of predicted TMHs: 2\nExp number of AAs in TMHs: 36.51738\nExp number, first 60 AAs: 19.72849\nTotal prob of N-in: 0.96170\nPOSSIBLE N-term signal sequence\ninside 1 19\nTMhelix 20 39\noutside 40 325\nTMhelix 326 348\ninside 349 372

23714	GCF_000626055.1_ASM62605v2	Salmonella enterica subsp. enterica serovar Enteritidis str. EC20130347	Proteobacteria; Gammaproteobacteria; Enterobacterales; Enterobacteriaceae; Salmonella; Salmonella enterica; Salmonella enterica subsp. enterica; Salmonella enterica subsp. enterica serovar Enteritidis	MTGDNTLITSHGINRRDFMKLCAALAATMGLSSKAAAEMAESVSNPQRPPVIWIGAQECT	WP_000145429.1	hydrogenase 2 small subunit [Salmonella enterica]	Length: 372\nNumber of predicted TMHs: 1\nExp

number of AAs in TMHs: 24.51309\nExp number, first 60 AAs: 0.2682\nTotal prob of N-in: 0.14402\noutside 1 331\nTMhelix 332 354\ninside 355 372

23715 GCF_000626075.2_ASM62607v2 Salmonella enterica subsp. enterica serovar Enteritidis str. EC20130348 Proteobacteria; Gammaproteobacteria; Enterobacterales; Enterobacteriaceae; Salmonella; Salmonella enterica; Salmonella enterica subsp. enterica; Salmonella enterica subsp. enterica serovar Enteritidis MNNEETFYQAMRRKGVTRRSFLKFCSLAATSLGLGAGMTPKIAWALENKPRIPVVWIHGL WP_001058307.1 hydrogenase [Salmonella enterica] Length: 372\nNumber of predicted TMHs: 2\nExp number of AAs in TMHs: 36.51738\nExp number, first 60 AAs: 19.72849\nTotal prob of N-in: 0.96170\nPOSSIBLE N-term signal sequence\ninside 1 19\nTMhelix 20 39\noutside 40 325\nTMhelix 326 348\ninside 349 372

23716 GCF_000626075.2_ASM62607v2 Salmonella enterica subsp. enterica serovar Enteritidis str. EC20130348 Proteobacteria; Gammaproteobacteria; Enterobacterales; Enterobacteriaceae; Salmonella; Salmonella enterica; Salmonella enterica subsp. enterica; Salmonella enterica subsp. enterica serovar Enteritidis MTGDNTLITSHGINRRDFMKLCAALAATMGLSSKAAAEMAESVSNPQRPPVIWIGAQECT WP_000145429.1 hydrogenase 2 small subunit [Salmonella enterica] Length: 372\nNumber of predicted TMHs: 1\nExp number of AAs in TMHs: 24.51309\nExp number, first 60 AAs: 0.2682\nTotal prob of N-in: 0.14402\noutside 1 331\nTMhelix 332 354\ninside 355 372

23717 GCF_000313715.1_ASM31371v2 Salmonella enterica subsp. enterica serovar Enteritidis str. LA5 Proteobacteria; Gammaproteobacteria; Enterobacterales; Enterobacteriaceae; Salmonella; Salmonella enterica; Salmonella enterica subsp. enterica; Salmonella enterica subsp. enterica serovar Enteritidis MNNEETFYQAMRRKGVTRRSFLKFCSLAATSLGLGAGMTPKIAWALENKPRIPVVWIHGL WP_001058307.1 hydrogenase [Salmonella enterica] Length: 372\nNumber of predicted TMHs: 2\nExp number of AAs in TMHs: 36.51738\nExp number, first 60 AAs: 19.72849\nTotal prob of N-in: 0.96170\nPOSSIBLE N-term signal sequence\ninside 1 19\nTMhelix 20 39\noutside 40 325\nTMhelix 326 348\ninside 349 372

23718 GCF_000313715.1_ASM31371v2 Salmonella enterica subsp. enterica serovar Enteritidis str. LA5 Proteobacteria; Gammaproteobacteria; Enterobacterales; Enterobacteriaceae; Salmonella; Salmonella enterica; Salmonella enterica subsp. enterica; Salmonella enterica subsp. enterica serovar Enteritidis MTGDNTLITSHGINRRDFMKLCAALAATMGLSSKAAAEMAESVSNPQRPPVIWIGAQECT WP_000145429.1 hydrogenase 2 small subunit [Salmonella enterica] Length: 372\nNumber of predicted TMHs: 1\nExp number of AAs in TMHs: 24.51309\nExp number, first 60 AAs: 0.2682\nTotal prob of N-in: 0.14402\noutside 1 331\nTMhelix 332 354\ninside 355 372

23719 GCF_000009505.1_ASM950v1 Salmonella enterica subsp. enterica serovar Enteritidis str. P125109 Proteobacteria; Gammaproteobacteria; Enterobacterales; Enterobacteriaceae; Salmonella; Salmonella enterica; Salmonella enterica subsp. enterica; Salmonella enterica subsp. enterica serovar Enteritidis MTGDNTLITSHGINRRDFMKLCAALAATMGLSSKAAAEMAESVSNPQRPPVIWIGAQECT WP_000145429.1 hydrogenase 2 small subunit [Salmonella enterica] Length: 372\nNumber of predicted TMHs: 1\nExp number of AAs in TMHs: 24.51309\nExp number, first 60 AAs: 0.2682\nTotal prob of N-in: 0.14402\noutside 1 331\nTMhelix 332 354\ninside 355 372

23720 GCF_000009505.1_ASM950v1 Salmonella enterica subsp. enterica serovar Enteritidis str. P125109 Proteobacteria; Gammaproteobacteria; Enterobacterales; Enterobacteriaceae; Salmonella; Salmonella enterica; Salmonella enterica subsp. enterica; Salmonella enterica subsp. enterica serovar Enteritidis MNNEETFYQAMRRKGVTRRSFLKFCSLAATSLGLGAGMTPKIAWALENKPRIPVVWIHGL WP_001058307.1 hydrogenase [Salmonella enterica] Length: 372\nNumber of predicted TMHs: 2\nExp number of AAs in TMHs: 36.51738\nExp number, first 60 AAs: 19.72849\nTotal prob of N-in: 0.96170\nPOSSIBLE N-term signal sequence\ninside 1 19\nTMhelix 20 39\noutside 40 325\nTMhelix 326 348\ninside 349 372

23721 GCF_000336115.1_ASM33611v1 Salmonella enterica subsp. enterica serovar Enteritidis str. PT23 Proteobacteria; Gammaproteobacteria; Enterobacterales; Enterobacteriaceae; Salmonella; Salmonella enterica; Salmonella enterica subsp. enterica; Salmonella enterica subsp. enterica serovar Enteritidis MNNEETFYQAMRRKGVTRRSFLKFCSLAATSLGLGAGMTPKIAWALENKPRIPVVWIHGL WP_001058307.1 hydrogenase [Salmonella enterica] Length: 372\nNumber of predicted TMHs: 2\nExp number of AAs in TMHs: 36.51738\nExp number, first 60 AAs: 19.72849\nTotal prob of N-in: 0.96170\nPOSSIBLE N-term signal sequence\ninside 1 19\nTMhelix 20 39\noutside 40 325\nTMhelix 326 348\ninside 349 372

23722 GCF_000336115.1_ASM33611v1 Salmonella enterica subsp. enterica serovar Enteritidis str. PT23 Proteobacteria; Gammaproteobacteria; Enterobacterales; Enterobacteriaceae; Salmonella; Salmonella enterica;

Salmonella enterica subsp. enterica; Salmonella enterica subsp. enterica serovar Enteritidis
MTGDNTLITSHGINRRDFMKLCAALAATMGLSSKAAAEMAESVSNPQRPPVIWIGAQECT WP_000145429.1
hydrogenase 2 small subunit [Salmonella enterica] Length: 372\nNumber of predicted TMHs: 1\nExp number of AAs in TMHs: 24.51309\nExp number, first 60 AAs: 0.2682\nTotal prob of N-in: 0.14402\noutside 1 331\nTMhelix 332 354\ninside 355 372

23723 GCF_000623555.2_ASM62355v2 Salmonella enterica subsp. enterica serovar Enteritidis str. SA19930684
Proteobacteria; Gammaproteobacteria; Enterobacterales; Enterobacteriaceae; Salmonella; Salmonella enterica; Salmonella enterica subsp. enterica; Salmonella enterica subsp. enterica serovar Enteritidis
MTGDNTLITSHGINRRDFMKLCAALAATMGLSSKAAAEMAESVSNPQRPPVIWIGAQECT WP_000145429.1
hydrogenase 2 small subunit [Salmonella enterica] Length: 372\nNumber of predicted TMHs: 1\nExp number of AAs in TMHs: 24.51309\nExp number, first 60 AAs: 0.2682\nTotal prob of N-in: 0.14402\noutside 1 331\nTMhelix 332 354\ninside 355 372

23724 GCF_000623555.2_ASM62355v2 Salmonella enterica subsp. enterica serovar Enteritidis str. SA19930684
Proteobacteria; Gammaproteobacteria; Enterobacterales; Enterobacteriaceae; Salmonella; Salmonella enterica; Salmonella enterica subsp. enterica; Salmonella enterica subsp. enterica serovar Enteritidis
MNNEETFYQAMRRKGVTRRSFLKFCSLAATSLGLGAGMTPKIAWALENKPRIPVVWIHGL WP_001058307.1
hydrogenase [Salmonella enterica] Length: 372\nNumber of predicted TMHs: 2\nExp number of AAs in TMHs: 36.51738\nExp number, first 60 AAs: 19.72849\nTotal prob of N-in: 0.96170\nPOSSIBLE N-term signal sequence\ninside 1 19\nTMhelix 20 39\noutside 40 325\nTMhelix 326 348\ninside 349 372

23725 GCF_000626455.1_ASM62645v1 Salmonella enterica subsp. enterica serovar Enteritidis str. SA19940857
Proteobacteria; Gammaproteobacteria; Enterobacterales; Enterobacteriaceae; Salmonella; Salmonella enterica; Salmonella enterica subsp. enterica; Salmonella enterica subsp. enterica serovar Enteritidis
MNNEETFYQAMRRKGVTRRSFLKFCSLAATSLGLGAGMTPKIAWALENKPRIPVVWIHGL WP_001058307.1
hydrogenase [Salmonella enterica] Length: 372\nNumber of predicted TMHs: 2\nExp number of AAs in TMHs: 36.51738\nExp number, first 60 AAs: 19.72849\nTotal prob of N-in: 0.96170\nPOSSIBLE N-term signal sequence\ninside 1 19\nTMhelix 20 39\noutside 40 325\nTMhelix 326 348\ninside 349 372

23726 GCF_000626455.1_ASM62645v1 Salmonella enterica subsp. enterica serovar Enteritidis str. SA19940857
Proteobacteria; Gammaproteobacteria; Enterobacterales; Enterobacteriaceae; Salmonella; Salmonella enterica; Salmonella enterica subsp. enterica; Salmonella enterica subsp. enterica serovar Enteritidis
MTGDNTLITSHGINRRDFMKLCAALAATMGLSSKAAAEMAESVSNPQRPPVIWIGAQECT WP_000145429.1
hydrogenase 2 small subunit [Salmonella enterica] Length: 372\nNumber of predicted TMHs: 1\nExp number of AAs in TMHs: 24.51309\nExp number, first 60 AAs: 0.2682\nTotal prob of N-in: 0.14402\noutside 1 331\nTMhelix 332 354\ninside 355 372

23727 GCF_000623575.2_ASM62357v2 Salmonella enterica subsp. enterica serovar Enteritidis str. SA19942384
Proteobacteria; Gammaproteobacteria; Enterobacterales; Enterobacteriaceae; Salmonella; Salmonella enterica; Salmonella enterica subsp. enterica; Salmonella enterica subsp. enterica serovar Enteritidis
MNNEETFYQAMRRKGVTRRSFLKFCSLAATSLGLGAGMTPKIAWALENKPRIPVVWIHGL WP_001058307.1
hydrogenase [Salmonella enterica] Length: 372\nNumber of predicted TMHs: 2\nExp number of AAs in TMHs: 36.51738\nExp number, first 60 AAs: 19.72849\nTotal prob of N-in: 0.96170\nPOSSIBLE N-term signal sequence\ninside 1 19\nTMhelix 20 39\noutside 40 325\nTMhelix 326 348\ninside 349 372

23728 GCF_000623575.2_ASM62357v2 Salmonella enterica subsp. enterica serovar Enteritidis str. SA19942384
Proteobacteria; Gammaproteobacteria; Enterobacterales; Enterobacteriaceae; Salmonella; Salmonella enterica; Salmonella enterica subsp. enterica; Salmonella enterica subsp. enterica serovar Enteritidis
MTGDNTLITSHGINRRDFMKLCAALAATMGLSSKAAAEMAESVSNPQRPPVIWIGAQECT WP_000145429.1
hydrogenase 2 small subunit [Salmonella enterica] Length: 372\nNumber of predicted TMHs: 1\nExp number of AAs in TMHs: 24.51309\nExp number, first 60 AAs: 0.2682\nTotal prob of N-in: 0.14402\noutside 1 331\nTMhelix 332 354\ninside 355 372

23729 GCF_000623595.2_ASM62359v2 Salmonella enterica subsp. enterica serovar Enteritidis str. SA19943269
Proteobacteria; Gammaproteobacteria; Enterobacterales; Enterobacteriaceae; Salmonella; Salmonella enterica; Salmonella enterica subsp. enterica; Salmonella enterica subsp. enterica serovar Enteritidis
MNNEETFYQAMRRKGVTRRSFLKFCSLAATSLGLGAGMTPKIAWALENKPRIPVVWIHGL WP_001058307.1
hydrogenase [Salmonella enterica] Length: 372\nNumber of predicted TMHs: 2\nExp number of AAs in TMHs: 36.51738\nExp number, first 60 AAs: 19.72849\nTotal prob of N-in: 0.96170\nPOSSIBLE N-term signal sequence\ninside 1 19\nTMhelix 20 39\noutside 40 325\nTMhelix 326 348\ninside 349 372

23730 GCF_000623595.2_ASM62359v2 Salmonella enterica subsp. enterica serovar Enteritidis str. SA19943269 Proteobacteria; Gammaproteobacteria; Enterobacterales; Enterobacteriaceae; Salmonella; Salmonella enterica; Salmonella enterica subsp. enterica; Salmonella enterica subsp. enterica serovar Enteritidis MTGDNTLITSHGINRRDFMKLCAALAATMGLSSKAAAEMAESVSNPQRPPVIWIGAQECT WP_000145429.1 hydrogenase 2 small subunit [Salmonella enterica] Length: 372\nNumber of predicted TMHs: 1\nExp number of AAs in TMHs: 24.51309\nExp number, first 60 AAs: 0.2682\nTotal prob of N-in: 0.14402\noutside 1 331\nTMhelix 332 354\ninside 355 372

23731 GCF_000623615.2_ASM62361v2 Salmonella enterica subsp. enterica serovar Enteritidis str. SA19960848 Proteobacteria; Gammaproteobacteria; Enterobacterales; Enterobacteriaceae; Salmonella; Salmonella enterica; Salmonella enterica subsp. enterica; Salmonella enterica subsp. enterica serovar Enteritidis MNNEETFYQAMRRKGVTRRSFLKFCSLAATSLGLGAGMTPKIAWALENKPRIPVVWIHGL WP_001058307.1 hydrogenase [Salmonella enterica] Length: 372\nNumber of predicted TMHs: 2\nExp number of AAs in TMHs: 36.51738\nExp number, first 60 AAs: 19.72849\nTotal prob of N-in: 0.96170\nPOSSIBLE N-term signal sequence\ninside 1 19\nTMhelix 20 39\noutside 40 325\nTMhelix 326 348\ninside 349 372

23732 GCF_000623615.2_ASM62361v2 Salmonella enterica subsp. enterica serovar Enteritidis str. SA19960848 Proteobacteria; Gammaproteobacteria; Enterobacterales; Enterobacteriaceae; Salmonella; Salmonella enterica; Salmonella enterica subsp. enterica; Salmonella enterica subsp. enterica serovar Enteritidis MTGDNTLITSHGINRRDFMKLCAALAATMGLSSKAAAEMAESVSNPQRPPVIWIGAQECT WP_000145429.1 hydrogenase 2 small subunit [Salmonella enterica] Length: 372\nNumber of predicted TMHs: 1\nExp number of AAs in TMHs: 24.51309\nExp number, first 60 AAs: 0.2682\nTotal prob of N-in: 0.14402\noutside 1 331\nTMhelix 332 354\ninside 355 372

23733 GCF_000623635.2_ASM62363v2 Salmonella enterica subsp. enterica serovar Enteritidis str. SA19961622 Proteobacteria; Gammaproteobacteria; Enterobacterales; Enterobacteriaceae; Salmonella; Salmonella enterica; Salmonella enterica subsp. enterica; Salmonella enterica subsp. enterica serovar Enteritidis MTGDNTLITSHGINRRDFMKLCAALAATMGLSSKAAAEMAESVSNPQRPPVIWIGAQECT WP_000145429.1 hydrogenase 2 small subunit [Salmonella enterica] Length: 372\nNumber of predicted TMHs: 1\nExp number of AAs in TMHs: 24.51309\nExp number, first 60 AAs: 0.2682\nTotal prob of N-in: 0.14402\noutside 1 331\nTMhelix 332 354\ninside 355 372

23734 GCF_000623635.2_ASM62363v2 Salmonella enterica subsp. enterica serovar Enteritidis str. SA19961622 Proteobacteria; Gammaproteobacteria; Enterobacterales; Enterobacteriaceae; Salmonella; Salmonella enterica; Salmonella enterica subsp. enterica; Salmonella enterica subsp. enterica serovar Enteritidis MNNEETFYQAMRRKGVTRRSFLKFCSLAATSLGLGAGMTPKIAWALENKPRIPVVWIHGL WP_001058307.1 hydrogenase [Salmonella enterica] Length: 372\nNumber of predicted TMHs: 2\nExp number of AAs in TMHs: 36.51738\nExp number, first 60 AAs: 19.72849\nTotal prob of N-in: 0.96170\nPOSSIBLE N-term signal sequence\ninside 1 19\nTMhelix 20 39\noutside 40 325\nTMhelix 326 348\ninside 349 372

23735 GCF_000623655.2_ASM62365v2 Salmonella enterica subsp. enterica serovar Enteritidis str. SA19970510 Proteobacteria; Gammaproteobacteria; Enterobacterales; Enterobacteriaceae; Salmonella; Salmonella enterica; Salmonella enterica subsp. enterica; Salmonella enterica subsp. enterica serovar Enteritidis MTGDNTLITSHGINRRDFMKLCAALAATMGLSSKAAAEMAESVSNPQRPPVIWIGAQECT WP_000145429.1 hydrogenase 2 small subunit [Salmonella enterica] Length: 372\nNumber of predicted TMHs: 1\nExp number of AAs in TMHs: 24.51309\nExp number, first 60 AAs: 0.2682\nTotal prob of N-in: 0.14402\noutside 1 331\nTMhelix 332 354\ninside 355 372

23736 GCF_000623655.2_ASM62365v2 Salmonella enterica subsp. enterica serovar Enteritidis str. SA19970510 Proteobacteria; Gammaproteobacteria; Enterobacterales; Enterobacteriaceae; Salmonella; Salmonella enterica; Salmonella enterica subsp. enterica; Salmonella enterica subsp. enterica serovar Enteritidis MNNEETFYQAMRRKGVTRRSFLKFCSLAATSLGLGAGMTPKIAWALENKPRIPVVWIHGL WP_001058307.1 hydrogenase [Salmonella enterica] Length: 372\nNumber of predicted TMHs: 2\nExp number of AAs in TMHs: 36.51738\nExp number, first 60 AAs: 19.72849\nTotal prob of N-in: 0.96170\nPOSSIBLE N-term signal sequence\ninside 1 19\nTMhelix 20 39\noutside 40 325\nTMhelix 326 348\ninside 349 372

23737 GCF_000623675.2_ASM62367v2 Salmonella enterica subsp. enterica serovar Enteritidis str. SA19970769 Proteobacteria; Gammaproteobacteria; Enterobacterales; Enterobacteriaceae; Salmonella; Salmonella enterica; Salmonella enterica subsp. enterica; Salmonella enterica subsp. enterica serovar Enteritidis MNNEETFYQAMRRKGVTRRSFLKFCSLAATSLGLGAGMTPKIAWALENKPRIPVVWIHGL WP_061202265.1 hydrogenase [Salmonella enterica] Length: 372\nNumber of predicted TMHs: 2\nExp number of AAs in

TMHs: 38.7219799999999\nExp number, first 60 AAs: 19.72878\nTotal prob of N-in: 0.96174\nPOSSIBLE N-term signal sequence\ninside 1 19\nTMhelix 20 39\noutside 40 325\nTMhelix 326 348\ninside 349 372

23738 GCF_000623675.2_ASM62367v2 Salmonella enterica subsp. enterica serovar Enteritidis str. SA19970769 Proteobacteria; Gammaproteobacteria; Enterobacterales; Enterobacteriaceae; Salmonella; Salmonella enterica; Salmonella enterica subsp. enterica; Salmonella enterica subsp. enterica serovar Enteritidis MTGDNTLITSHGINRRDFMKLCAALAATMGLSSKAAAEMAESVSNPQRPPVIWIGAQECT WP_000145429.1 hydrogenase 2 small subunit [Salmonella enterica] Length: 372\nNumber of predicted TMHs: 1\nExp number of AAs in TMHs: 24.51309\nExp number, first 60 AAs: 0.2682\nTotal prob of N-in: 0.14402\noutside 1 331\nTMhelix 332 354\ninside 355 372

23739 GCF_000623695.2_ASM62369v2 Salmonella enterica subsp. enterica serovar Enteritidis str. SA19971331 Proteobacteria; Gammaproteobacteria; Enterobacterales; Enterobacteriaceae; Salmonella; Salmonella enterica; Salmonella enterica subsp. enterica; Salmonella enterica subsp. enterica serovar Enteritidis MTGDNTLITSHGINRRDFMKLCAALAATMGLSSKAAAEMAESVSNPQRPPVIWIGAQECT WP_000145429.1 hydrogenase 2 small subunit [Salmonella enterica] Length: 372\nNumber of predicted TMHs: 1\nExp number of AAs in TMHs: 24.51309\nExp number, first 60 AAs: 0.2682\nTotal prob of N-in: 0.14402\noutside 1 331\nTMhelix 332 354\ninside 355 372

23740 GCF_000623695.2_ASM62369v2 Salmonella enterica subsp. enterica serovar Enteritidis str. SA19971331 Proteobacteria; Gammaproteobacteria; Enterobacterales; Enterobacteriaceae; Salmonella; Salmonella enterica; Salmonella enterica subsp. enterica; Salmonella enterica subsp. enterica serovar Enteritidis MNNEETFYQAMRRKGVTRRSFLKFCSLAATSLGLGAGMTPKIAWALENKPRIPVVWIHGL WP_001058307.1 hydrogenase [Salmonella enterica] Length: 372\nNumber of predicted TMHs: 2\nExp number of AAs in TMHs: 36.51738\nExp number, first 60 AAs: 19.72849\nTotal prob of N-in: 0.96170\nPOSSIBLE N-term signal sequence\ninside 1 19\nTMhelix 20 39\noutside 40 325\nTMhelix 326 348\ninside 349 372

23741 GCF_000623715.2_ASM62371v2 Salmonella enterica subsp. enterica serovar Enteritidis str. SA19980677 Proteobacteria; Gammaproteobacteria; Enterobacterales; Enterobacteriaceae; Salmonella; Salmonella enterica; Salmonella enterica subsp. enterica; Salmonella enterica subsp. enterica serovar Enteritidis MNNEETFYQAMRRKGVTRRSFLKFCSLAATSLGLGAGMTPKIAWALENKPRIPVVWIHGL WP_001058307.1 hydrogenase [Salmonella enterica] Length: 372\nNumber of predicted TMHs: 2\nExp number of AAs in TMHs: 36.51738\nExp number, first 60 AAs: 19.72849\nTotal prob of N-in: 0.96170\nPOSSIBLE N-term signal sequence\ninside 1 19\nTMhelix 20 39\noutside 40 325\nTMhelix 326 348\ninside 349 372

23742 GCF_000623715.2_ASM62371v2 Salmonella enterica subsp. enterica serovar Enteritidis str. SA19980677 Proteobacteria; Gammaproteobacteria; Enterobacterales; Enterobacteriaceae; Salmonella; Salmonella enterica; Salmonella enterica subsp. enterica; Salmonella enterica subsp. enterica serovar Enteritidis MTGDNTLITSHGINRRDFMKLCAALAATMGLSSKAAAEMAESVSNPQRPPVIWIGAQECT WP_000145429.1 hydrogenase 2 small subunit [Salmonella enterica] Length: 372\nNumber of predicted TMHs: 1\nExp number of AAs in TMHs: 24.51309\nExp number, first 60 AAs: 0.2682\nTotal prob of N-in: 0.14402\noutside 1 331\nTMhelix 332 354\ninside 355 372

23743 GCF_000623735.2_ASM62373v2 Salmonella enterica subsp. enterica serovar Enteritidis str. SA19981522 Proteobacteria; Gammaproteobacteria; Enterobacterales; Enterobacteriaceae; Salmonella; Salmonella enterica; Salmonella enterica subsp. enterica; Salmonella enterica subsp. enterica serovar Enteritidis MNNEETFYQAMRRKGVTRRSFLKFCSLAATSLGLGAGMTPKIAWALENKPRIPVVWIHGL WP_001058307.1 hydrogenase [Salmonella enterica] Length: 372\nNumber of predicted TMHs: 2\nExp number of AAs in TMHs: 36.51738\nExp number, first 60 AAs: 19.72849\nTotal prob of N-in: 0.96170\nPOSSIBLE N-term signal sequence\ninside 1 19\nTMhelix 20 39\noutside 40 325\nTMhelix 326 348\ninside 349 372

23744 GCF_000623735.2_ASM62373v2 Salmonella enterica subsp. enterica serovar Enteritidis str. SA19981522 Proteobacteria; Gammaproteobacteria; Enterobacterales; Enterobacteriaceae; Salmonella; Salmonella enterica; Salmonella enterica subsp. enterica; Salmonella enterica subsp. enterica serovar Enteritidis MTGDNTLITSHGINRRDFMKLCAALAATMGLSSKAAAEMAESVSNPQRPPVIWIGAQECT WP_000145429.1 hydrogenase 2 small subunit [Salmonella enterica] Length: 372\nNumber of predicted TMHs: 1\nExp number of AAs in TMHs: 24.51309\nExp number, first 60 AAs: 0.2682\nTotal prob of N-in: 0.14402\noutside 1 331\nTMhelix 332 354\ninside 355 372

23745 GCF_000623395.2_ASM62339v2 Salmonella enterica subsp. enterica serovar Enteritidis str. SA19981857 Proteobacteria; Gammaproteobacteria; Enterobacterales; Enterobacteriaceae; Salmonella; Salmonella enterica;

Salmonella enterica subsp. enterica; Salmonella enterica subsp. enterica serovar Enteritidis
MNNEETFYQAMRRKGVTRRSFLKFCSLAATSLGLGAGMTPKIAWALENKPRIPVVWIHGL WP_001058307.1
hydrogenase [Salmonella enterica] Length: 372\nNumber of predicted TMHs: 2\nExp number of AAs in TMHs: 36.51738\nExp number, first 60 AAs: 19.72849\nTotal prob of N-in: 0.96170\nPOSSIBLE N-term signal sequence\ninside 1 19\nTMhelix 20 39\noutside 40 325\nTMhelix 326 348\ninside 349 372

23746 GCF_000623395.2_ASM62339v2 Salmonella enterica subsp. enterica serovar Enteritidis str. SA19981857
Proteobacteria; Gammaproteobacteria; Enterobacterales; Enterobacteriaceae; Salmonella; Salmonella enterica; Salmonella enterica subsp. enterica; Salmonella enterica subsp. enterica serovar Enteritidis
MTGDNTLITSHGINRRDFMKLCAALAATMGLSSKAAAEMAESVSNPQRPPVIWIGAQECT WP_000145429.1
hydrogenase 2 small subunit [Salmonella enterica] Length: 372\nNumber of predicted TMHs: 1\nExp number of AAs in TMHs: 24.51309\nExp number, first 60 AAs: 0.2682\nTotal prob of N-in: 0.14402\noutside 1 331\nTMhelix 332 354\ninside 355 372

23747 GCF_000623415.2_ASM62341v2 Salmonella enterica subsp. enterica serovar Enteritidis str. SA19982831
Proteobacteria; Gammaproteobacteria; Enterobacterales; Enterobacteriaceae; Salmonella; Salmonella enterica; Salmonella enterica subsp. enterica; Salmonella enterica subsp. enterica serovar Enteritidis
MNNEETFYQAMRRKGVTRRSFLKFCSLAATSLGLGAGMTPKIAWALENKPRIPVVWIHGL WP_001058307.1
hydrogenase [Salmonella enterica] Length: 372\nNumber of predicted TMHs: 2\nExp number of AAs in TMHs: 36.51738\nExp number, first 60 AAs: 19.72849\nTotal prob of N-in: 0.96170\nPOSSIBLE N-term signal sequence\ninside 1 19\nTMhelix 20 39\noutside 40 325\nTMhelix 326 348\ninside 349 372

23748 GCF_000623415.2_ASM62341v2 Salmonella enterica subsp. enterica serovar Enteritidis str. SA19982831
Proteobacteria; Gammaproteobacteria; Enterobacterales; Enterobacteriaceae; Salmonella; Salmonella enterica; Salmonella enterica subsp. enterica; Salmonella enterica subsp. enterica serovar Enteritidis
MTGDNTLITSHGINRRDFMKLCAALAATMGLSSKAAAEMAESVSNPQRPPVIWIGAQECT WP_000145429.1
hydrogenase 2 small subunit [Salmonella enterica] Length: 372\nNumber of predicted TMHs: 1\nExp number of AAs in TMHs: 24.51309\nExp number, first 60 AAs: 0.2682\nTotal prob of N-in: 0.14402\noutside 1 331\nTMhelix 332 354\ninside 355 372

23749 GCF_000623435.2_ASM62343v2 Salmonella enterica subsp. enterica serovar Enteritidis str. SA19983126
Proteobacteria; Gammaproteobacteria; Enterobacterales; Enterobacteriaceae; Salmonella; Salmonella enterica; Salmonella enterica subsp. enterica; Salmonella enterica subsp. enterica serovar Enteritidis
MNNEETFYQAMRRKGVTRRSFLKFCSLAATSLGLGAGMTPKIAWALENKPRIPVVWIHGL WP_001058307.1
hydrogenase [Salmonella enterica] Length: 372\nNumber of predicted TMHs: 2\nExp number of AAs in TMHs: 36.51738\nExp number, first 60 AAs: 19.72849\nTotal prob of N-in: 0.96170\nPOSSIBLE N-term signal sequence\ninside 1 19\nTMhelix 20 39\noutside 40 325\nTMhelix 326 348\ninside 349 372

23750 GCF_000623435.2_ASM62343v2 Salmonella enterica subsp. enterica serovar Enteritidis str. SA19983126
Proteobacteria; Gammaproteobacteria; Enterobacterales; Enterobacteriaceae; Salmonella; Salmonella enterica; Salmonella enterica subsp. enterica; Salmonella enterica subsp. enterica serovar Enteritidis
MTGDNTLITSHGINRRDFMKLCAALAATMGLSSKAAAEMAESVSNPQRPPVIWIGAQECT WP_000145429.1
hydrogenase 2 small subunit [Salmonella enterica] Length: 372\nNumber of predicted TMHs: 1\nExp number of AAs in TMHs: 24.51309\nExp number, first 60 AAs: 0.2682\nTotal prob of N-in: 0.14402\noutside 1 331\nTMhelix 332 354\ninside 355 372

23751 GCF_000623455.2_ASM62345v2 Salmonella enterica subsp. enterica serovar Enteritidis str. SA19992322
Proteobacteria; Gammaproteobacteria; Enterobacterales; Enterobacteriaceae; Salmonella; Salmonella enterica; Salmonella enterica subsp. enterica; Salmonella enterica subsp. enterica serovar Enteritidis
MNNEETFYQAMRRKGVTRRSFLKFCSLAATSLGLGAGMTPKIAWALENKPRIPVVWIHGL WP_001058307.1
hydrogenase [Salmonella enterica] Length: 372\nNumber of predicted TMHs: 2\nExp number of AAs in TMHs: 36.51738\nExp number, first 60 AAs: 19.72849\nTotal prob of N-in: 0.96170\nPOSSIBLE N-term signal sequence\ninside 1 19\nTMhelix 20 39\noutside 40 325\nTMhelix 326 348\ninside 349 372

23752 GCF_000623455.2_ASM62345v2 Salmonella enterica subsp. enterica serovar Enteritidis str. SA19992322
Proteobacteria; Gammaproteobacteria; Enterobacterales; Enterobacteriaceae; Salmonella; Salmonella enterica; Salmonella enterica subsp. enterica; Salmonella enterica subsp. enterica serovar Enteritidis
MTGDNTLITSHGINRRDFMKLCAALAATMGLSSKAAAEMAESVSNPQRPPVIWIGAQECT WP_000145429.1
hydrogenase 2 small subunit [Salmonella enterica] Length: 372\nNumber of predicted TMHs: 1\nExp number of AAs in TMHs: 24.51309\nExp number, first 60 AAs: 0.2682\nTotal prob of N-in: 0.14402\noutside 1 331\nTMhelix 332 354\ninside 355 372

23753 GCF_000623475.1_ASM62347v2 Salmonella enterica subsp. enterica serovar Enteritidis str. SA19994216 Proteobacteria; Gammaproteobacteria; Enterobacterales; Enterobacteriaceae; Salmonella; Salmonella enterica; Salmonella enterica subsp. enterica; Salmonella enterica subsp. enterica serovar Enteritidis MNNEETFYQAMRRKGVTRRSFLKFCSLAATSLGLGAGMTPKIAWALENKPRIPVVWIHGL WP_001058307.1 hydrogenase [Salmonella enterica] Length: 372\nNumber of predicted TMHs: 2\nExp number of AAs in TMHs: 36.51738\nExp number, first 60 AAs: 19.72849\nTotal prob of N-in: 0.96170\nPOSSIBLE N-term signal sequence\ninside 1 19\nTMhelix 20 39\noutside 40 325\nTMhelix 326 348\ninside 349 372

23754 GCF_000623475.1_ASM62347v2 Salmonella enterica subsp. enterica serovar Enteritidis str. SA19994216 Proteobacteria; Gammaproteobacteria; Enterobacterales; Enterobacteriaceae; Salmonella; Salmonella enterica; Salmonella enterica subsp. enterica; Salmonella enterica subsp. enterica serovar Enteritidis MTGDNTLITSHGINRRDFMKLCAALAATMGLSSKAAAEMAESVSNPQRPPVIWIGAQECT WP_000145429.1 hydrogenase 2 small subunit [Salmonella enterica] Length: 372\nNumber of predicted TMHs: 1\nExp number of AAs in TMHs: 24.51309\nExp number, first 60 AAs: 0.2682\nTotal prob of N-in: 0.14402\noutside 1 331\nTMhelix 332 354\ninside 355 372

23755 GCF_000625855.1_ASM62585v2 Salmonella enterica subsp. enterica serovar Enteritidis str. SA20082034 Proteobacteria; Gammaproteobacteria; Enterobacterales; Enterobacteriaceae; Salmonella; Salmonella enterica; Salmonella enterica subsp. enterica; Salmonella enterica subsp. enterica serovar Enteritidis MTGDNTLITSHGINRRDFMKLCAALAATMGLSSKAAAEMAESVSNPQRPPVIWIGAQECT WP_000145429.1 hydrogenase 2 small subunit [Salmonella enterica] Length: 372\nNumber of predicted TMHs: 1\nExp number of AAs in TMHs: 24.51309\nExp number, first 60 AAs: 0.2682\nTotal prob of N-in: 0.14402\noutside 1 331\nTMhelix 332 354\ninside 355 372

23756 GCF_000625855.1_ASM62585v2 Salmonella enterica subsp. enterica serovar Enteritidis str. SA20082034 Proteobacteria; Gammaproteobacteria; Enterobacterales; Enterobacteriaceae; Salmonella; Salmonella enterica; Salmonella enterica subsp. enterica; Salmonella enterica subsp. enterica serovar Enteritidis MNNEETFYQAMRRKGVTRRSFLKFCSLAATSLGLGAGMTPKIAWALENKPRIPVVWIHGL WP_001058307.1 hydrogenase [Salmonella enterica] Length: 372\nNumber of predicted TMHs: 2\nExp number of AAs in TMHs: 36.51738\nExp number, first 60 AAs: 19.72849\nTotal prob of N-in: 0.96170\nPOSSIBLE N-term signal sequence\ninside 1 19\nTMhelix 20 39\noutside 40 325\nTMhelix 326 348\ninside 349 372

23757 GCF_000623495.2_ASM62349v2 Salmonella enterica subsp. enterica serovar Enteritidis str. SA20083456 Proteobacteria; Gammaproteobacteria; Enterobacterales; Enterobacteriaceae; Salmonella; Salmonella enterica; Salmonella enterica subsp. enterica; Salmonella enterica subsp. enterica serovar Enteritidis MNNEETFYQAMRRKGVTRRSFLKFCSLAATSLGLGAGMTPKIAWALENKPRIPVVWIHGL WP_001058307.1 hydrogenase [Salmonella enterica] Length: 372\nNumber of predicted TMHs: 2\nExp number of AAs in TMHs: 36.51738\nExp number, first 60 AAs: 19.72849\nTotal prob of N-in: 0.96170\nPOSSIBLE N-term signal sequence\ninside 1 19\nTMhelix 20 39\noutside 40 325\nTMhelix 326 348\ninside 349 372

23758 GCF_000623495.2_ASM62349v2 Salmonella enterica subsp. enterica serovar Enteritidis str. SA20083456 Proteobacteria; Gammaproteobacteria; Enterobacterales; Enterobacteriaceae; Salmonella; Salmonella enterica; Salmonella enterica subsp. enterica; Salmonella enterica subsp. enterica serovar Enteritidis MTGDNTLITSHGINRRDFMKLCAALAATMGLSSKAAAEMAESVSNPQRPPVIWIGAQECT WP_000145429.1 hydrogenase 2 small subunit [Salmonella enterica] Length: 372\nNumber of predicted TMHs: 1\nExp number of AAs in TMHs: 24.51309\nExp number, first 60 AAs: 0.2682\nTotal prob of N-in: 0.14402\noutside 1 331\nTMhelix 332 354\ninside 355 372

23759 GCF_000623515.2_ASM62351v2 Salmonella enterica subsp. enterica serovar Enteritidis str. SA20083636 Proteobacteria; Gammaproteobacteria; Enterobacterales; Enterobacteriaceae; Salmonella; Salmonella enterica; Salmonella enterica subsp. enterica; Salmonella enterica subsp. enterica serovar Enteritidis MNNEETFYQAMRRKGVTRRSFLKFCSLAATSLGLGAGMTPKIAWALENKPRIPVVWIHGL WP_001058307.1 hydrogenase [Salmonella enterica] Length: 372\nNumber of predicted TMHs: 2\nExp number of AAs in TMHs: 36.51738\nExp number, first 60 AAs: 19.72849\nTotal prob of N-in: 0.96170\nPOSSIBLE N-term signal sequence\ninside 1 19\nTMhelix 20 39\noutside 40 325\nTMhelix 326 348\ninside 349 372

23760 GCF_000623515.2_ASM62351v2 Salmonella enterica subsp. enterica serovar Enteritidis str. SA20083636 Proteobacteria; Gammaproteobacteria; Enterobacterales; Enterobacteriaceae; Salmonella; Salmonella enterica; Salmonella enterica subsp. enterica; Salmonella enterica subsp. enterica serovar Enteritidis MTGDNTLITSHGINRRDFMKLCAALAATMGLSSKAAAEMAESVSNPQRPPVIWIGAQECT WP_000145429.1 hydrogenase 2 small subunit [Salmonella enterica] Length: 372\nNumber of predicted TMHs: 1\nExp

number of AAs in TMHs: 24.51309\nExp number, first 60 AAs: 0.2682\nTotal prob of N-in: 0.14402\noutside 1 331\nTMhelix 332 354\ninside 355 372

23761 GCF_000623535.2_ASM62353v2 Salmonella enterica subsp. enterica serovar Enteritidis str. SA20084384 Proteobacteria; Gammaproteobacteria; Enterobacterales; Enterobacteriaceae; Salmonella; Salmonella enterica; Salmonella enterica subsp. enterica; Salmonella enterica subsp. enterica serovar Enteritidis MNNEETFYQAMRRKGVTRRSFLKFCSLAATSLGLGAGMTPKIAWALENKPRIPVVWIHGL WP_001058307.1 hydrogenase [Salmonella enterica] Length: 372\nNumber of predicted TMHs: 2\nExp number of AAs in TMHs: 36.51738\nExp number, first 60 AAs: 19.72849\nTotal prob of N-in: 0.96170\nPOSSIBLE N-term signal sequence\ninside 1 19\nTMhelix 20 39\noutside 40 325\nTMhelix 326 348\ninside 349 372

23762 GCF_000623535.2_ASM62353v2 Salmonella enterica subsp. enterica serovar Enteritidis str. SA20084384 Proteobacteria; Gammaproteobacteria; Enterobacterales; Enterobacteriaceae; Salmonella; Salmonella enterica; Salmonella enterica subsp. enterica; Salmonella enterica subsp. enterica serovar Enteritidis MTGDNTLITSHGINRRDFMKLCAALAATMGLSSKAAAEMAESVSNPQRPPVIWIGAQECT WP_000145429.1 hydrogenase 2 small subunit [Salmonella enterica] Length: 372\nNumber of predicted TMHs: 1\nExp number of AAs in TMHs: 24.51309\nExp number, first 60 AAs: 0.2682\nTotal prob of N-in: 0.14402\noutside 1 331\nTMhelix 332 354\ninside 355 372

23763 GCF_000626475.1_ASM62647v2 Salmonella enterica subsp. enterica serovar Enteritidis str. SA20084644 Proteobacteria; Gammaproteobacteria; Enterobacterales; Enterobacteriaceae; Salmonella; Salmonella enterica; Salmonella enterica subsp. enterica; Salmonella enterica subsp. enterica serovar Enteritidis MTGDNTLITSHGINRRDFMKLCAALAATMGLSSKAAAEMAESVSNPQRPPVIWIGAQECT WP_000145429.1 hydrogenase 2 small subunit [Salmonella enterica] Length: 372\nNumber of predicted TMHs: 1\nExp number of AAs in TMHs: 24.51309\nExp number, first 60 AAs: 0.2682\nTotal prob of N-in: 0.14402\noutside 1 331\nTMhelix 332 354\ninside 355 372

23764 GCF_000626475.1_ASM62647v2 Salmonella enterica subsp. enterica serovar Enteritidis str. SA20084644 Proteobacteria; Gammaproteobacteria; Enterobacterales; Enterobacteriaceae; Salmonella; Salmonella enterica; Salmonella enterica subsp. enterica; Salmonella enterica subsp. enterica serovar Enteritidis MNNEETFYQAMRRKGVTRRSFLKFCSLAATSLGLGAGMTPKIAWALENKPRIPVVWIHGL WP_001058307.1 hydrogenase [Salmonella enterica] Length: 372\nNumber of predicted TMHs: 2\nExp number of AAs in TMHs: 36.51738\nExp number, first 60 AAs: 19.72849\nTotal prob of N-in: 0.96170\nPOSSIBLE N-term signal sequence\ninside 1 19\nTMhelix 20 39\noutside 40 325\nTMhelix 326 348\ninside 349 372

23765 GCF_000626495.1_ASM62649v2 Salmonella enterica subsp. enterica serovar Enteritidis str. SA20084824 Proteobacteria; Gammaproteobacteria; Enterobacterales; Enterobacteriaceae; Salmonella; Salmonella enterica; Salmonella enterica subsp. enterica; Salmonella enterica subsp. enterica serovar Enteritidis MNNEETFYQAMRRKGVTRRSFLKFCSLAATSLGLGAGMTPKIAWALENKPRIPVVWIHGL WP_001058307.1 hydrogenase [Salmonella enterica] Length: 372\nNumber of predicted TMHs: 2\nExp number of AAs in TMHs: 36.51738\nExp number, first 60 AAs: 19.72849\nTotal prob of N-in: 0.96170\nPOSSIBLE N-term signal sequence\ninside 1 19\nTMhelix 20 39\noutside 40 325\nTMhelix 326 348\ninside 349 372

23766 GCF_000626495.1_ASM62649v2 Salmonella enterica subsp. enterica serovar Enteritidis str. SA20084824 Proteobacteria; Gammaproteobacteria; Enterobacterales; Enterobacteriaceae; Salmonella; Salmonella enterica; Salmonella enterica subsp. enterica; Salmonella enterica subsp. enterica serovar Enteritidis MTGDNTLITSHGINRRDFMKLCAALAATMGLSSKAAAEMAESVSNPQRPPVIWIGAQECT WP_000145429.1 hydrogenase 2 small subunit [Salmonella enterica] Length: 372\nNumber of predicted TMHs: 1\nExp number of AAs in TMHs: 24.51309\nExp number, first 60 AAs: 0.2682\nTotal prob of N-in: 0.14402\noutside 1 331\nTMhelix 332 354\ninside 355 372

23767 GCF_000625655.1_ASM62565v2 Salmonella enterica subsp. enterica serovar Enteritidis str. SA20085285 Proteobacteria; Gammaproteobacteria; Enterobacterales; Enterobacteriaceae; Salmonella; Salmonella enterica; Salmonella enterica subsp. enterica; Salmonella enterica subsp. enterica serovar Enteritidis MTGDNTLITSHGINRRDFMKLCAALAATMGLSSKAAAEMAESVSNPQRPPVIWIGAQECT WP_000145429.1 hydrogenase 2 small subunit [Salmonella enterica] Length: 372\nNumber of predicted TMHs: 1\nExp number of AAs in TMHs: 24.51309\nExp number, first 60 AAs: 0.2682\nTotal prob of N-in: 0.14402\noutside 1 331\nTMhelix 332 354\ninside 355 372

23768 GCF_000625655.1_ASM62565v2 Salmonella enterica subsp. enterica serovar Enteritidis str. SA20085285 Proteobacteria; Gammaproteobacteria; Enterobacterales; Enterobacteriaceae; Salmonella; Salmonella enterica;

Salmonella enterica subsp. enterica; Salmonella enterica subsp. enterica serovar Enteritidis
MNNEETFYQAMRRKGVTRRSFLKFCSLAATSLGLGAGMTPKIAWALENKPRIPVVWIHGL WP_001058307.1
hydrogenase [Salmonella enterica] Length: 372\nNumber of predicted TMHs: 2\nExp number of AAs in TMHs: 36.51738\nExp number, first 60 AAs: 19.72849\nTotal prob of N-in: 0.96170\nPOSSIBLE N-term signal sequence\ninside 1 19\nTMhelix 20 39\noutside 40 325\nTMhelix 326 348\ninside 349 372

23769 GCF_000625355.1_ASM62535v2 Salmonella enterica subsp. enterica serovar Enteritidis str. SA20090419
Proteobacteria; Gammaproteobacteria; Enterobacterales; Enterobacteriaceae; Salmonella; Salmonella enterica; Salmonella enterica subsp. enterica; Salmonella enterica subsp. enterica serovar Enteritidis
MNNEETFYQAMRRKGVTRRSFLKFCSLAATSLGLGAGMTPKIAWALENKPRIPVVWIHGL WP_001058307.1
hydrogenase [Salmonella enterica] Length: 372\nNumber of predicted TMHs: 2\nExp number of AAs in TMHs: 36.51738\nExp number, first 60 AAs: 19.72849\nTotal prob of N-in: 0.96170\nPOSSIBLE N-term signal sequence\ninside 1 19\nTMhelix 20 39\noutside 40 325\nTMhelix 326 348\ninside 349 372

23770 GCF_000625355.1_ASM62535v2 Salmonella enterica subsp. enterica serovar Enteritidis str. SA20090419
Proteobacteria; Gammaproteobacteria; Enterobacterales; Enterobacteriaceae; Salmonella; Salmonella enterica; Salmonella enterica subsp. enterica; Salmonella enterica subsp. enterica serovar Enteritidis
MTGDNTLITSHGINRRDFMKLCAALAATMGLSSKAAAEMAESVSNPQRPPVIWIGAQECT WP_000145429.1
hydrogenase 2 small subunit [Salmonella enterica] Length: 372\nNumber of predicted TMHs: 1\nExp number of AAs in TMHs: 24.51309\nExp number, first 60 AAs: 0.2682\nTotal prob of N-in: 0.14402\noutside 1 331\nTMhelix 332 354\ninside 355 372

23771 GCF_000625375.2_ASM62537v2 Salmonella enterica subsp. enterica serovar Enteritidis str. SA20090435
Proteobacteria; Gammaproteobacteria; Enterobacterales; Enterobacteriaceae; Salmonella; Salmonella enterica; Salmonella enterica subsp. enterica; Salmonella enterica subsp. enterica serovar Enteritidis
MTGDNTLITSHGINRRDFMKLCAALAATMGLSSKAAAEMAESVSNPQRPPVIWIGAQECT WP_000145429.1
hydrogenase 2 small subunit [Salmonella enterica] Length: 372\nNumber of predicted TMHs: 1\nExp number of AAs in TMHs: 24.51309\nExp number, first 60 AAs: 0.2682\nTotal prob of N-in: 0.14402\noutside 1 331\nTMhelix 332 354\ninside 355 372

23772 GCF_000625375.2_ASM62537v2 Salmonella enterica subsp. enterica serovar Enteritidis str. SA20090435
Proteobacteria; Gammaproteobacteria; Enterobacterales; Enterobacteriaceae; Salmonella; Salmonella enterica; Salmonella enterica subsp. enterica; Salmonella enterica subsp. enterica serovar Enteritidis
MNNEETFYQAMRRKGVTRRSFLKFCSLAATSLGLGAGMTPKIAWALENKPRIPVVWIHGL WP_001058307.1
hydrogenase [Salmonella enterica] Length: 372\nNumber of predicted TMHs: 2\nExp number of AAs in TMHs: 36.51738\nExp number, first 60 AAs: 19.72849\nTotal prob of N-in: 0.96170\nPOSSIBLE N-term signal sequence\ninside 1 19\nTMhelix 20 39\noutside 40 325\nTMhelix 326 348\ninside 349 372

23773 GCF_000623815.2_ASM62381v2 Salmonella enterica subsp. enterica serovar Enteritidis str. SA20090877
Proteobacteria; Gammaproteobacteria; Enterobacterales; Enterobacteriaceae; Salmonella; Salmonella enterica; Salmonella enterica subsp. enterica; Salmonella enterica subsp. enterica serovar Enteritidis
MNNEETFYQAMRRKGVTRRSFLKFCSLAATSLGLGAGMTPKIAWALENKPRIPVVWIHGL WP_001058307.1
hydrogenase [Salmonella enterica] Length: 372\nNumber of predicted TMHs: 2\nExp number of AAs in TMHs: 36.51738\nExp number, first 60 AAs: 19.72849\nTotal prob of N-in: 0.96170\nPOSSIBLE N-term signal sequence\ninside 1 19\nTMhelix 20 39\noutside 40 325\nTMhelix 326 348\ninside 349 372

23774 GCF_000623815.2_ASM62381v2 Salmonella enterica subsp. enterica serovar Enteritidis str. SA20090877
Proteobacteria; Gammaproteobacteria; Enterobacterales; Enterobacteriaceae; Salmonella; Salmonella enterica; Salmonella enterica subsp. enterica; Salmonella enterica subsp. enterica serovar Enteritidis
MTGDNTLITSHGINRRDFMKLCAALAATMGLSSKAAAEMAESVSNPQRPPVIWIGAQECT WP_000145429.1
hydrogenase 2 small subunit [Salmonella enterica] Length: 372\nNumber of predicted TMHs: 1\nExp number of AAs in TMHs: 24.51309\nExp number, first 60 AAs: 0.2682\nTotal prob of N-in: 0.14402\noutside 1 331\nTMhelix 332 354\ninside 355 372

23775 GCF_000623835.2_ASM62383v2 Salmonella enterica subsp. enterica serovar Enteritidis str. SA20091739
Proteobacteria; Gammaproteobacteria; Enterobacterales; Enterobacteriaceae; Salmonella; Salmonella enterica; Salmonella enterica subsp. enterica; Salmonella enterica subsp. enterica serovar Enteritidis
MNNEETFYQAMRRKGVTRRSFLKFCSLAATSLGLGAGMTPKIAWALENKPRIPVVWIHGL WP_001058307.1
hydrogenase [Salmonella enterica] Length: 372\nNumber of predicted TMHs: 2\nExp number of AAs in TMHs: 36.51738\nExp number, first 60 AAs: 19.72849\nTotal prob of N-in: 0.96170\nPOSSIBLE N-term signal sequence\ninside 1 19\nTMhelix 20 39\noutside 40 325\nTMhelix 326 348\ninside 349 372

23776 GCF_000623835.2_ASM62383v2 Salmonella enterica subsp. enterica serovar Enteritidis str. SA20091739 Proteobacteria; Gammaproteobacteria; Enterobacterales; Enterobacteriaceae; Salmonella; Salmonella enterica; Salmonella enterica subsp. enterica; Salmonella enterica subsp. enterica serovar Enteritidis MTGDNTLITSHGINRRDFMKLCAALAATMGLSSKAAAEMAESVSNPQRPPVIWIGAQECT WP_000145429.1 hydrogenase 2 small subunit [Salmonella enterica] Length: 372\nNumber of predicted TMHs: 1\nExp number of AAs in TMHs: 24.51309\nExp number, first 60 AAs: 0.2682\nTotal prob of N-in: 0.14402\noutside 1 331\nTMhelix 332 354\ninside 355 372

23777 GCF_000624195.2_ASM62419v2 Salmonella enterica subsp. enterica serovar Enteritidis str. SA20092320 Proteobacteria; Gammaproteobacteria; Enterobacterales; Enterobacteriaceae; Salmonella; Salmonella enterica; Salmonella enterica subsp. enterica; Salmonella enterica subsp. enterica serovar Enteritidis MNNEETFYQAMRRKGVTRRSFLKFCSLAATSLGLGAGMTPKIAWALENKPRIPVVWIHGL WP_001058307.1 hydrogenase [Salmonella enterica] Length: 372\nNumber of predicted TMHs: 2\nExp number of AAs in TMHs: 36.51738\nExp number, first 60 AAs: 19.72849\nTotal prob of N-in: 0.96170\nPOSSIBLE N-term signal sequence\ninside 1 19\nTMhelix 20 39\noutside 40 325\nTMhelix 326 348\ninside 349 372

23778 GCF_000624195.2_ASM62419v2 Salmonella enterica subsp. enterica serovar Enteritidis str. SA20092320 Proteobacteria; Gammaproteobacteria; Enterobacterales; Enterobacteriaceae; Salmonella; Salmonella enterica; Salmonella enterica subsp. enterica; Salmonella enterica subsp. enterica serovar Enteritidis MTGDNTLITSHGINRRDFMKLCAALAATMGLSSKAAAEMAESVSNPQRPPVIWIGAQECT WP_000145429.1 hydrogenase 2 small subunit [Salmonella enterica] Length: 372\nNumber of predicted TMHs: 1\nExp number of AAs in TMHs: 24.51309\nExp number, first 60 AAs: 0.2682\nTotal prob of N-in: 0.14402\noutside 1 331\nTMhelix 332 354\ninside 355 372

23779 GCF_000623155.2_ASM62315v2 Salmonella enterica subsp. enterica serovar Enteritidis str. SA20093266 Proteobacteria; Gammaproteobacteria; Enterobacterales; Enterobacteriaceae; Salmonella; Salmonella enterica; Salmonella enterica subsp. enterica; Salmonella enterica subsp. enterica serovar Enteritidis MTGDNTLITSHGINRRDFMKLCAALAATMGLSSKAAAEMAESVSNPQRPPVIWIGAQECT WP_000145429.1 hydrogenase 2 small subunit [Salmonella enterica] Length: 372\nNumber of predicted TMHs: 1\nExp number of AAs in TMHs: 24.51309\nExp number, first 60 AAs: 0.2682\nTotal prob of N-in: 0.14402\noutside 1 331\nTMhelix 332 354\ninside 355 372

23780 GCF_000623155.2_ASM62315v2 Salmonella enterica subsp. enterica serovar Enteritidis str. SA20093266 Proteobacteria; Gammaproteobacteria; Enterobacterales; Enterobacteriaceae; Salmonella; Salmonella enterica; Salmonella enterica subsp. enterica; Salmonella enterica subsp. enterica serovar Enteritidis MNNEETFYQAMRRKGVTRRSFLKFCSLAATSLGLGAGMTPKIAWALENKPRIPVVWIHGL WP_001058307.1 hydrogenase [Salmonella enterica] Length: 372\nNumber of predicted TMHs: 2\nExp number of AAs in TMHs: 36.51738\nExp number, first 60 AAs: 19.72849\nTotal prob of N-in: 0.96170\nPOSSIBLE N-term signal sequence\ninside 1 19\nTMhelix 20 39\noutside 40 325\nTMhelix 326 348\ninside 349 372

23781 GCF_000623855.2_ASM62385v2 Salmonella enterica subsp. enterica serovar Enteritidis str. SA20093421 Proteobacteria; Gammaproteobacteria; Enterobacterales; Enterobacteriaceae; Salmonella; Salmonella enterica; Salmonella enterica subsp. enterica; Salmonella enterica subsp. enterica serovar Enteritidis MNNEETFYQAMRRKGVTRRSFLKFCSLAATSLGLGAGMTPKIAWALENKPRIPVVWIHGL WP_001058307.1 hydrogenase [Salmonella enterica] Length: 372\nNumber of predicted TMHs: 2\nExp number of AAs in TMHs: 36.51738\nExp number, first 60 AAs: 19.72849\nTotal prob of N-in: 0.96170\nPOSSIBLE N-term signal sequence\ninside 1 19\nTMhelix 20 39\noutside 40 325\nTMhelix 326 348\ninside 349 372

23782 GCF_000623855.2_ASM62385v2 Salmonella enterica subsp. enterica serovar Enteritidis str. SA20093421 Proteobacteria; Gammaproteobacteria; Enterobacterales; Enterobacteriaceae; Salmonella; Salmonella enterica; Salmonella enterica subsp. enterica; Salmonella enterica subsp. enterica serovar Enteritidis MTGDNTLITSHGINRRDFMKLCAALAATMGLSSKAAAEMAESVSNPQRPPVIWIGAQECT WP_000145429.1 hydrogenase 2 small subunit [Salmonella enterica] Length: 372\nNumber of predicted TMHs: 1\nExp number of AAs in TMHs: 24.51309\nExp number, first 60 AAs: 0.2682\nTotal prob of N-in: 0.14402\noutside 1 331\nTMhelix 332 354\ninside 355 372

23783 GCF_000623875.2_ASM62387v2 Salmonella enterica subsp. enterica serovar Enteritidis str. SA20093430 Proteobacteria; Gammaproteobacteria; Enterobacterales; Enterobacteriaceae; Salmonella; Salmonella enterica; Salmonella enterica subsp. enterica; Salmonella enterica subsp. enterica serovar Enteritidis MNNEETFYQAMRRKGVTRRSFLKFCSLAATSLGLGAGMTPKIAWALENKPRIPVVWIHGL WP_001058307.1 hydrogenase [Salmonella enterica] Length: 372\nNumber of predicted TMHs: 2\nExp number of AAs in

TMHs: 36.51738\nExp number, first 60 AAs: 19.72849\nTotal prob of N-in: 0.96170\nPOSSIBLE N-term signal sequence\ninside 1 19\nTMhelix 20 39\noutside 40 325\nTMhelix 326 348\ninside 349 372

23784 GCF_000623875.2_ASM62387v2 Salmonella enterica subsp. enterica serovar Enteritidis str. SA20093430 Proteobacteria; Gammaproteobacteria; Enterobacterales; Enterobacteriaceae; Salmonella; Salmonella enterica; Salmonella enterica subsp. enterica; Salmonella enterica subsp. enterica serovar Enteritidis MTGDNTLITSHGINRRDFMKLCAALAATMGLSSKAAAEMAESVSNPQRPPVIWIGAQECT WP_000145429.1 hydrogenase 2 small subunit [Salmonella enterica] Length: 372\nNumber of predicted TMHs: 1\nExp number of AAs in TMHs: 24.51309\nExp number, first 60 AAs: 0.2682\nTotal prob of N-in: 0.14402\noutside 1 331\nTMhelix 332 354\ninside 355 372

23785 GCF_000623895.2_ASM62389v2 Salmonella enterica subsp. enterica serovar Enteritidis str. SA20093538 Proteobacteria; Gammaproteobacteria; Enterobacterales; Enterobacteriaceae; Salmonella; Salmonella enterica; Salmonella enterica subsp. enterica; Salmonella enterica subsp. enterica serovar Enteritidis MNNEETFYQAMRRKGVTRRSFLKFCSLAATSLGLGAGMTPKIAWALENKPRIPVVWIHGL WP_001058307.1 hydrogenase [Salmonella enterica] Length: 372\nNumber of predicted TMHs: 2\nExp number of AAs in TMHs: 36.51738\nExp number, first 60 AAs: 19.72849\nTotal prob of N-in: 0.96170\nPOSSIBLE N-term signal sequence\ninside 1 19\nTMhelix 20 39\noutside 40 325\nTMhelix 326 348\ninside 349 372

23786 GCF_000623895.2_ASM62389v2 Salmonella enterica subsp. enterica serovar Enteritidis str. SA20093538 Proteobacteria; Gammaproteobacteria; Enterobacterales; Enterobacteriaceae; Salmonella; Salmonella enterica; Salmonella enterica subsp. enterica; Salmonella enterica subsp. enterica serovar Enteritidis MTGDNTLITSHGINRRDFMKLCAALAATMGLSSKAAAEMAESVSNPQRPPVIWIGAQECT WP_000145429.1 hydrogenase 2 small subunit [Salmonella enterica] Length: 372\nNumber of predicted TMHs: 1\nExp number of AAs in TMHs: 24.51309\nExp number, first 60 AAs: 0.2682\nTotal prob of N-in: 0.14402\noutside 1 331\nTMhelix 332 354\ninside 355 372

23787 GCF_000623915.2_ASM62391v2 Salmonella enterica subsp. enterica serovar Enteritidis str. SA20093543 Proteobacteria; Gammaproteobacteria; Enterobacterales; Enterobacteriaceae; Salmonella; Salmonella enterica; Salmonella enterica subsp. enterica; Salmonella enterica subsp. enterica serovar Enteritidis MNNEETFYQAMRRKGVTRRSFLKFCSLAATSLGLGAGMTPKIAWALENKPRIPVVWIHGL WP_001058307.1 hydrogenase [Salmonella enterica] Length: 372\nNumber of predicted TMHs: 2\nExp number of AAs in TMHs: 36.51738\nExp number, first 60 AAs: 19.72849\nTotal prob of N-in: 0.96170\nPOSSIBLE N-term signal sequence\ninside 1 19\nTMhelix 20 39\noutside 40 325\nTMhelix 326 348\ninside 349 372

23788 GCF_000623915.2_ASM62391v2 Salmonella enterica subsp. enterica serovar Enteritidis str. SA20093543 Proteobacteria; Gammaproteobacteria; Enterobacterales; Enterobacteriaceae; Salmonella; Salmonella enterica; Salmonella enterica subsp. enterica; Salmonella enterica subsp. enterica serovar Enteritidis MTGDNTLITSHGINRRDFMKLCAALAATMGLSSKAAAEMAESVSNPQRPPVIWIGAQECT WP_000145429.1 hydrogenase 2 small subunit [Salmonella enterica] Length: 372\nNumber of predicted TMHs: 1\nExp number of AAs in TMHs: 24.51309\nExp number, first 60 AAs: 0.2682\nTotal prob of N-in: 0.14402\noutside 1 331\nTMhelix 332 354\ninside 355 372

23789 GCF_000623935.2_ASM62393v2 Salmonella enterica subsp. enterica serovar Enteritidis str. SA20093784 Proteobacteria; Gammaproteobacteria; Enterobacterales; Enterobacteriaceae; Salmonella; Salmonella enterica; Salmonella enterica subsp. enterica; Salmonella enterica subsp. enterica serovar Enteritidis MTGDNTLITSHGINRRDFMKLCAALAATMGLSSKAAAEMAESVSNPQRPPVIWIGAQECT WP_000145429.1 hydrogenase 2 small subunit [Salmonella enterica] Length: 372\nNumber of predicted TMHs: 1\nExp number of AAs in TMHs: 24.51309\nExp number, first 60 AAs: 0.2682\nTotal prob of N-in: 0.14402\noutside 1 331\nTMhelix 332 354\ninside 355 372

23790 GCF_000623935.2_ASM62393v2 Salmonella enterica subsp. enterica serovar Enteritidis str. SA20093784 Proteobacteria; Gammaproteobacteria; Enterobacterales; Enterobacteriaceae; Salmonella; Salmonella enterica; Salmonella enterica subsp. enterica; Salmonella enterica subsp. enterica serovar Enteritidis MNNEETFYQAMRRKGVTRRSFLKFCSLAATSLGLGAGMTPKIAWALENKPRIPVVWIHGL WP_001058307.1 hydrogenase [Salmonella enterica] Length: 372\nNumber of predicted TMHs: 2\nExp number of AAs in TMHs: 36.51738\nExp number, first 60 AAs: 19.72849\nTotal prob of N-in: 0.96170\nPOSSIBLE N-term signal sequence\ninside 1 19\nTMhelix 20 39\noutside 40 325\nTMhelix 326 348\ninside 349 372

23791 GCF_000623955.2_ASM62395v2 Salmonella enterica subsp. enterica serovar Enteritidis str. SA20093788 Proteobacteria; Gammaproteobacteria; Enterobacterales; Enterobacteriaceae; Salmonella; Salmonella enterica;

Salmonella enterica subsp. enterica; Salmonella enterica subsp. enterica serovar Enteritidis
MNNEETFYQAMRRKGVTRRSFLKFCSLAATSLGLGAGMTPKIAWALENKPRIPVVWIHGL WP_001058307.1
hydrogenase [Salmonella enterica] Length: 372\nNumber of predicted TMHs: 2\nExp number of AAs in TMHs: 36.51738\nExp number, first 60 AAs: 19.72849\nTotal prob of N-in: 0.96170\nPOSSIBLE N-term signal sequence\ninside 1 19\nTMhelix 20 39\noutside 40 325\nTMhelix 326 348\ninside 349 372

23792 GCF_000623955.2_ASM62395v2 Salmonella enterica subsp. enterica serovar Enteritidis str. SA20093788
Proteobacteria; Gammaproteobacteria; Enterobacterales; Enterobacteriaceae; Salmonella; Salmonella enterica; Salmonella enterica subsp. enterica; Salmonella enterica subsp. enterica serovar Enteritidis
MTGDNTLITSHGINRRDFMKLCAALAATMGLSSKAAAEMAESVSNPQRPPVIWIGAQECT WP_000145429.1
hydrogenase 2 small subunit [Salmonella enterica] Length: 372\nNumber of predicted TMHs: 1\nExp number of AAs in TMHs: 24.51309\nExp number, first 60 AAs: 0.2682\nTotal prob of N-in: 0.14402\noutside 1 331\nTMhelix 332 354\ninside 355 372

23793 GCF_000623975.2_ASM62397v2 Salmonella enterica subsp. enterica serovar Enteritidis str. SA20093950
Proteobacteria; Gammaproteobacteria; Enterobacterales; Enterobacteriaceae; Salmonella; Salmonella enterica; Salmonella enterica subsp. enterica; Salmonella enterica subsp. enterica serovar Enteritidis
MNNEETFYQAMRRKGVTRRSFLKFCSLAATSLGLGAGMTPKIAWALENKPRIPVVWIHGL WP_001058307.1
hydrogenase [Salmonella enterica] Length: 372\nNumber of predicted TMHs: 2\nExp number of AAs in TMHs: 36.51738\nExp number, first 60 AAs: 19.72849\nTotal prob of N-in: 0.96170\nPOSSIBLE N-term signal sequence\ninside 1 19\nTMhelix 20 39\noutside 40 325\nTMhelix 326 348\ninside 349 372

23794 GCF_000623975.2_ASM62397v2 Salmonella enterica subsp. enterica serovar Enteritidis str. SA20093950
Proteobacteria; Gammaproteobacteria; Enterobacterales; Enterobacteriaceae; Salmonella; Salmonella enterica; Salmonella enterica subsp. enterica; Salmonella enterica subsp. enterica serovar Enteritidis
MTGDNTLITSHGINRRDFMKLCAALAATMGLSSKAAAEMAESVSNPQRPPVIWIGAQECT WP_000145429.1
hydrogenase 2 small subunit [Salmonella enterica] Length: 372\nNumber of predicted TMHs: 1\nExp number of AAs in TMHs: 24.51309\nExp number, first 60 AAs: 0.2682\nTotal prob of N-in: 0.14402\noutside 1 331\nTMhelix 332 354\ninside 355 372

23795 GCF_000623995.2_ASM62399v2 Salmonella enterica subsp. enterica serovar Enteritidis str. SA20093977
Proteobacteria; Gammaproteobacteria; Enterobacterales; Enterobacteriaceae; Salmonella; Salmonella enterica; Salmonella enterica subsp. enterica; Salmonella enterica subsp. enterica serovar Enteritidis
MNNEETFYQAMRRKGVTRRSFLKFCSLAATSLGLGAGMTPKIAWALENKPRIPVVWIHGL WP_001058307.1
hydrogenase [Salmonella enterica] Length: 372\nNumber of predicted TMHs: 2\nExp number of AAs in TMHs: 36.51738\nExp number, first 60 AAs: 19.72849\nTotal prob of N-in: 0.96170\nPOSSIBLE N-term signal sequence\ninside 1 19\nTMhelix 20 39\noutside 40 325\nTMhelix 326 348\ninside 349 372

23796 GCF_000623995.2_ASM62399v2 Salmonella enterica subsp. enterica serovar Enteritidis str. SA20093977
Proteobacteria; Gammaproteobacteria; Enterobacterales; Enterobacteriaceae; Salmonella; Salmonella enterica; Salmonella enterica subsp. enterica; Salmonella enterica subsp. enterica serovar Enteritidis
MTGDNTLITSHGINRRDFMKLCAALAATMGLSSKAAAEMAESVSNPQRPPVIWIGAQECT WP_000145429.1
hydrogenase 2 small subunit [Salmonella enterica] Length: 372\nNumber of predicted TMHs: 1\nExp number of AAs in TMHs: 24.51309\nExp number, first 60 AAs: 0.2682\nTotal prob of N-in: 0.14402\noutside 1 331\nTMhelix 332 354\ninside 355 372

23797 GCF_000624015.2_ASM62401v2 Salmonella enterica subsp. enterica serovar Enteritidis str. SA20094079
Proteobacteria; Gammaproteobacteria; Enterobacterales; Enterobacteriaceae; Salmonella; Salmonella enterica; Salmonella enterica subsp. enterica; Salmonella enterica subsp. enterica serovar Enteritidis
MNNEETFYQAMRRKGVTRRSFLKFCSLAATSLGLGAGMTPKIAWALENKPRIPVVWIHGL WP_001058307.1
hydrogenase [Salmonella enterica] Length: 372\nNumber of predicted TMHs: 2\nExp number of AAs in TMHs: 36.51738\nExp number, first 60 AAs: 19.72849\nTotal prob of N-in: 0.96170\nPOSSIBLE N-term signal sequence\ninside 1 19\nTMhelix 20 39\noutside 40 325\nTMhelix 326 348\ninside 349 372

23798 GCF_000624015.2_ASM62401v2 Salmonella enterica subsp. enterica serovar Enteritidis str. SA20094079
Proteobacteria; Gammaproteobacteria; Enterobacterales; Enterobacteriaceae; Salmonella; Salmonella enterica; Salmonella enterica subsp. enterica; Salmonella enterica subsp. enterica serovar Enteritidis
MTGDNTLITSHGINRRDFMKLCAALAATMGLSSKAAAEMAESVSNPQRPPVIWIGAQECT WP_000145429.1
hydrogenase 2 small subunit [Salmonella enterica] Length: 372\nNumber of predicted TMHs: 1\nExp number of AAs in TMHs: 24.51309\nExp number, first 60 AAs: 0.2682\nTotal prob of N-in: 0.14402\noutside 1 331\nTMhelix 332 354\ninside 355 372

23799 GCF_000626515.1_ASM62651v2 Salmonella enterica subsp. enterica serovar Enteritidis str. SA20094177 Proteobacteria; Gammaproteobacteria; Enterobacterales; Enterobacteriaceae; Salmonella; Salmonella enterica; Salmonella enterica subsp. enterica; Salmonella enterica subsp. enterica serovar Enteritidis MTGDNTLITSHGINRRDFMKLCAALAATMGLSSKAAAEMAESVSNPQRPPVIWIGAQECT WP_000145429.1 hydrogenase 2 small subunit [Salmonella enterica] Length: 372\nNumber of predicted TMHs: 1\nExp number of AAs in TMHs: 24.51309\nExp number, first 60 AAs: 0.2682\nTotal prob of N-in: 0.14402\noutside 1 331\nTMhelix 332 354\ninside 355 372

23800 GCF_000626515.1_ASM62651v2 Salmonella enterica subsp. enterica serovar Enteritidis str. SA20094177 Proteobacteria; Gammaproteobacteria; Enterobacterales; Enterobacteriaceae; Salmonella; Salmonella enterica; Salmonella enterica subsp. enterica; Salmonella enterica subsp. enterica serovar Enteritidis MNNEETFYQAMRRKGVTRRSFLKFCSLAATSLGLGAGMTPKIAWALENKPRIPVVWIHGL WP_001058307.1 hydrogenase [Salmonella enterica] Length: 372\nNumber of predicted TMHs: 2\nExp number of AAs in TMHs: 36.51738\nExp number, first 60 AAs: 19.72849\nTotal prob of N-in: 0.96170\nPOSSIBLE N-term signal sequence\ninside 1 19\nTMhelix 20 39\noutside 40 325\nTMhelix 326 348\ninside 349 372

23801 GCF_000626535.1_ASM62653v2 Salmonella enterica subsp. enterica serovar Enteritidis str. SA20094301 Proteobacteria; Gammaproteobacteria; Enterobacterales; Enterobacteriaceae; Salmonella; Salmonella enterica; Salmonella enterica subsp. enterica; Salmonella enterica subsp. enterica serovar Enteritidis MTGDNTLITSHGINRRDFMKLCAALAATMGLSSKAAAEMAESVSNPQRPPVIWIGAQECT WP_000145429.1 hydrogenase 2 small subunit [Salmonella enterica] Length: 372\nNumber of predicted TMHs: 1\nExp number of AAs in TMHs: 24.51309\nExp number, first 60 AAs: 0.2682\nTotal prob of N-in: 0.14402\noutside 1 331\nTMhelix 332 354\ninside 355 372

23802 GCF_000626535.1_ASM62653v2 Salmonella enterica subsp. enterica serovar Enteritidis str. SA20094301 Proteobacteria; Gammaproteobacteria; Enterobacterales; Enterobacteriaceae; Salmonella; Salmonella enterica; Salmonella enterica subsp. enterica; Salmonella enterica subsp. enterica serovar Enteritidis MNNEETFYQAMRRKGVTRRSFLKFCSLAATSLGLGAGMTPKIAWALENKPRIPVVWIHGL WP_001058307.1 hydrogenase [Salmonella enterica] Length: 372\nNumber of predicted TMHs: 2\nExp number of AAs in TMHs: 36.51738\nExp number, first 60 AAs: 19.72849\nTotal prob of N-in: 0.96170\nPOSSIBLE N-term signal sequence\ninside 1 19\nTMhelix 20 39\noutside 40 325\nTMhelix 326 348\ninside 349 372

23803 GCF_000624035.2_ASM62403v2 Salmonella enterica subsp. enterica serovar Enteritidis str. SA20094350 Proteobacteria; Gammaproteobacteria; Enterobacterales; Enterobacteriaceae; Salmonella; Salmonella enterica; Salmonella enterica subsp. enterica; Salmonella enterica subsp. enterica serovar Enteritidis MNNEETFYQAMRRKGVTRRSFLKFCSLAATSLGLGAGMTPKIAWALENKPRIPVVWIHGL WP_001058307.1 hydrogenase [Salmonella enterica] Length: 372\nNumber of predicted TMHs: 2\nExp number of AAs in TMHs: 36.51738\nExp number, first 60 AAs: 19.72849\nTotal prob of N-in: 0.96170\nPOSSIBLE N-term signal sequence\ninside 1 19\nTMhelix 20 39\noutside 40 325\nTMhelix 326 348\ninside 349 372

23804 GCF_000624035.2_ASM62403v2 Salmonella enterica subsp. enterica serovar Enteritidis str. SA20094350 Proteobacteria; Gammaproteobacteria; Enterobacterales; Enterobacteriaceae; Salmonella; Salmonella enterica; Salmonella enterica subsp. enterica; Salmonella enterica subsp. enterica serovar Enteritidis MTGDNTLITSHGINRRDFMKLCAALAATMGLSSKAAAEMAESVSNPQRPPVIWIGAQECT WP_000145429.1 hydrogenase 2 small subunit [Salmonella enterica] Length: 372\nNumber of predicted TMHs: 1\nExp number of AAs in TMHs: 24.51309\nExp number, first 60 AAs: 0.2682\nTotal prob of N-in: 0.14402\noutside 1 331\nTMhelix 332 354\ninside 355 372

23805 GCF_000624055.2_ASM62405v2 Salmonella enterica subsp. enterica serovar Enteritidis str. SA20094352 Proteobacteria; Gammaproteobacteria; Enterobacterales; Enterobacteriaceae; Salmonella; Salmonella enterica; Salmonella enterica subsp. enterica; Salmonella enterica subsp. enterica serovar Enteritidis MTGDNTLITSHGINRRDFMKLCAALAATMGLSSKAAAEMAESVSNPQRPPVIWIGAQECT WP_000145429.1 hydrogenase 2 small subunit [Salmonella enterica] Length: 372\nNumber of predicted TMHs: 1\nExp number of AAs in TMHs: 24.51309\nExp number, first 60 AAs: 0.2682\nTotal prob of N-in: 0.14402\noutside 1 331\nTMhelix 332 354\ninside 355 372

23806 GCF_000624055.2_ASM62405v2 Salmonella enterica subsp. enterica serovar Enteritidis str. SA20094352 Proteobacteria; Gammaproteobacteria; Enterobacterales; Enterobacteriaceae; Salmonella; Salmonella enterica; Salmonella enterica subsp. enterica; Salmonella enterica subsp. enterica serovar Enteritidis MNNEETFYQAMRRKGVTRRSFLKFCSLAATSLGLGAGMTPKIAWALENKPRIPVVWIHGL WP_001058307.1 hydrogenase [Salmonella enterica] Length: 372\nNumber of predicted TMHs: 2\nExp number of AAs in

TMHs: 36.51738\nExp number, first 60 AAs: 19.72849\nTotal prob of N-in: 0.96170\nPOSSIBLE N-term signal sequence\ninside 1 19\nTMhelix 20 39\noutside 40 325\nTMhelix 326 348\ninside 349 372

23807 GCF_000624075.2_ASM62407v2 Salmonella enterica subsp. enterica serovar Enteritidis str. SA20094383 Proteobacteria; Gammaproteobacteria; Enterobacterales; Enterobacteriaceae; Salmonella; Salmonella enterica; Salmonella enterica subsp. enterica; Salmonella enterica subsp. enterica serovar Enteritidis MNNEETFYQAMRRKGVTRRSFLKFCSLAATSLGLGAGMTPKIAWALENKPRIPVVWIHGL WP_001058307.1 hydrogenase [Salmonella enterica] Length: 372\nNumber of predicted TMHs: 2\nExp number of AAs in TMHs: 36.51738\nExp number, first 60 AAs: 19.72849\nTotal prob of N-in: 0.96170\nPOSSIBLE N-term signal sequence\ninside 1 19\nTMhelix 20 39\noutside 40 325\nTMhelix 326 348\ninside 349 372

23808 GCF_000624075.2_ASM62407v2 Salmonella enterica subsp. enterica serovar Enteritidis str. SA20094383 Proteobacteria; Gammaproteobacteria; Enterobacterales; Enterobacteriaceae; Salmonella; Salmonella enterica; Salmonella enterica subsp. enterica; Salmonella enterica subsp. enterica serovar Enteritidis MTGDNTLITSHGINRRDFMKLCAALAATMGLSSKAAAEMAESVSNPQRPPVIWIGAQECT WP_000145429.1 hydrogenase 2 small subunit [Salmonella enterica] Length: 372\nNumber of predicted TMHs: 1\nExp number of AAs in TMHs: 24.51309\nExp number, first 60 AAs: 0.2682\nTotal prob of N-in: 0.14402\noutside 1 331\nTMhelix 332 354\ninside 355 372

23809 GCF_000624095.2_ASM62409v2 Salmonella enterica subsp. enterica serovar Enteritidis str. SA20094389 Proteobacteria; Gammaproteobacteria; Enterobacterales; Enterobacteriaceae; Salmonella; Salmonella enterica; Salmonella enterica subsp. enterica; Salmonella enterica subsp. enterica serovar Enteritidis MNNEETFYQAMRRKGVTRRSFLKFCSLAATSLGLGAGMTPKIAWALENKPRIPVVWIHGL WP_001058307.1 hydrogenase [Salmonella enterica] Length: 372\nNumber of predicted TMHs: 2\nExp number of AAs in TMHs: 36.51738\nExp number, first 60 AAs: 19.72849\nTotal prob of N-in: 0.96170\nPOSSIBLE N-term signal sequence\ninside 1 19\nTMhelix 20 39\noutside 40 325\nTMhelix 326 348\ninside 349 372

23810 GCF_000624095.2_ASM62409v2 Salmonella enterica subsp. enterica serovar Enteritidis str. SA20094389 Proteobacteria; Gammaproteobacteria; Enterobacterales; Enterobacteriaceae; Salmonella; Salmonella enterica; Salmonella enterica subsp. enterica; Salmonella enterica subsp. enterica serovar Enteritidis MTGDNTLITSHGINRRDFMKLCAALAATMGLSSKAAAEMAESVSNPQRPPVIWIGAQECT WP_000145429.1 hydrogenase 2 small subunit [Salmonella enterica] Length: 372\nNumber of predicted TMHs: 1\nExp number of AAs in TMHs: 24.51309\nExp number, first 60 AAs: 0.2682\nTotal prob of N-in: 0.14402\noutside 1 331\nTMhelix 332 354\ninside 355 372

23811 GCF_000624115.2_ASM62411v2 Salmonella enterica subsp. enterica serovar Enteritidis str. SA20094521 Proteobacteria; Gammaproteobacteria; Enterobacterales; Enterobacteriaceae; Salmonella; Salmonella enterica; Salmonella enterica subsp. enterica; Salmonella enterica subsp. enterica serovar Enteritidis MNNEETFYQAMRRKGVTRRSFLKFCSLAATSLGLGAGMTPKIAWALENKPRIPVVWIHGL WP_001058307.1 hydrogenase [Salmonella enterica] Length: 372\nNumber of predicted TMHs: 2\nExp number of AAs in TMHs: 36.51738\nExp number, first 60 AAs: 19.72849\nTotal prob of N-in: 0.96170\nPOSSIBLE N-term signal sequence\ninside 1 19\nTMhelix 20 39\noutside 40 325\nTMhelix 326 348\ninside 349 372

23812 GCF_000624115.2_ASM62411v2 Salmonella enterica subsp. enterica serovar Enteritidis str. SA20094521 Proteobacteria; Gammaproteobacteria; Enterobacterales; Enterobacteriaceae; Salmonella; Salmonella enterica; Salmonella enterica subsp. enterica; Salmonella enterica subsp. enterica serovar Enteritidis MTGDNTLITSHGINRRDFMKLCAALAATMGLSSKAAAEMAESVSNPQRPPVIWIGAQECT WP_000145429.1 hydrogenase 2 small subunit [Salmonella enterica] Length: 372\nNumber of predicted TMHs: 1\nExp number of AAs in TMHs: 24.51309\nExp number, first 60 AAs: 0.2682\nTotal prob of N-in: 0.14402\noutside 1 331\nTMhelix 332 354\ninside 355 372

23813 GCF_000624135.2_ASM62413v2 Salmonella enterica subsp. enterica serovar Enteritidis str. SA20094642 Proteobacteria; Gammaproteobacteria; Enterobacterales; Enterobacteriaceae; Salmonella; Salmonella enterica; Salmonella enterica subsp. enterica; Salmonella enterica subsp. enterica serovar Enteritidis MTGDNTLITSHGINRRDFMKLCAALAATMGLSSKAAAEMAESVSNPQRPPVIWIGAQECT WP_000145429.1 hydrogenase 2 small subunit [Salmonella enterica] Length: 372\nNumber of predicted TMHs: 1\nExp number of AAs in TMHs: 24.51309\nExp number, first 60 AAs: 0.2682\nTotal prob of N-in: 0.14402\noutside 1 331\nTMhelix 332 354\ninside 355 372

23814 GCF_000625695.2_ASM62569v2 Salmonella enterica subsp. enterica serovar Enteritidis str. SA20094682 Proteobacteria; Gammaproteobacteria; Enterobacterales; Enterobacteriaceae; Salmonella; Salmonella enterica;

Salmonella enterica subsp. enterica; Salmonella enterica subsp. enterica serovar Enteritidis
MTGDNTLITSHGINRRDFMKLCAALAATMGLSSKAAAEMAESVSNPQRPPVIWIGAQECT WP_000145429.1
hydrogenase 2 small subunit [Salmonella enterica] Length: 372\nNumber of predicted TMHs: 1\nExp number of AAs in TMHs: 24.51309\nExp number, first 60 AAs: 0.2682\nTotal prob of N-in: 0.14402\noutside 1 331\nTMhelix 332 354\ninside 355 372

23815 GCF_000624215.2_ASM62421v2 Salmonella enterica subsp. enterica serovar Enteritidis str. SA20094803
Proteobacteria; Gammaproteobacteria; Enterobacterales; Enterobacteriaceae; Salmonella; Salmonella enterica; Salmonella enterica subsp. enterica; Salmonella enterica subsp. enterica serovar Enteritidis
MNNEETFYQAMRRKGVTRRSFLKFCSLAATSLGLGAGMTPKIAWALENKPRIPVVWIHGL WP_001058307.1
hydrogenase [Salmonella enterica] Length: 372\nNumber of predicted TMHs: 2\nExp number of AAs in TMHs: 36.51738\nExp number, first 60 AAs: 19.72849\nTotal prob of N-in: 0.96170\nPOSSIBLE N-term signal sequence\ninside 1 19\nTMhelix 20 39\noutside 40 325\nTMhelix 326 348\ninside 349 372

23816 GCF_000624215.2_ASM62421v2 Salmonella enterica subsp. enterica serovar Enteritidis str. SA20094803
Proteobacteria; Gammaproteobacteria; Enterobacterales; Enterobacteriaceae; Salmonella; Salmonella enterica; Salmonella enterica subsp. enterica; Salmonella enterica subsp. enterica serovar Enteritidis
MTGDNTLITSHGINRRDFMKLCAALAATMGLSSKAAAEMAESVSNPQRPPVIWIGAQECT WP_000145429.1
hydrogenase 2 small subunit [Salmonella enterica] Length: 372\nNumber of predicted TMHs: 1\nExp number of AAs in TMHs: 24.51309\nExp number, first 60 AAs: 0.2682\nTotal prob of N-in: 0.14402\noutside 1 331\nTMhelix 332 354\ninside 355 372

23817 GCF_000624235.2_ASM62423v2 Salmonella enterica subsp. enterica serovar Enteritidis str. SA20095309
Proteobacteria; Gammaproteobacteria; Enterobacterales; Enterobacteriaceae; Salmonella; Salmonella enterica; Salmonella enterica subsp. enterica; Salmonella enterica subsp. enterica serovar Enteritidis
MNNEETFYQAMRRKGVTRRSFLKFCSLAATSLGLGAGMTPKIAWALENKPRIPVVWIHGL WP_001058307.1
hydrogenase [Salmonella enterica] Length: 372\nNumber of predicted TMHs: 2\nExp number of AAs in TMHs: 36.51738\nExp number, first 60 AAs: 19.72849\nTotal prob of N-in: 0.96170\nPOSSIBLE N-term signal sequence\ninside 1 19\nTMhelix 20 39\noutside 40 325\nTMhelix 326 348\ninside 349 372

23818 GCF_000624235.2_ASM62423v2 Salmonella enterica subsp. enterica serovar Enteritidis str. SA20095309
Proteobacteria; Gammaproteobacteria; Enterobacterales; Enterobacteriaceae; Salmonella; Salmonella enterica; Salmonella enterica subsp. enterica; Salmonella enterica subsp. enterica serovar Enteritidis
MTGDNTLITSHGINRRDFMKLCAALAATMGLSSKAAAEMAESVSNPQRPPVIWIGAQECT WP_000145429.1
hydrogenase 2 small subunit [Salmonella enterica] Length: 372\nNumber of predicted TMHs: 1\nExp number of AAs in TMHs: 24.51309\nExp number, first 60 AAs: 0.2682\nTotal prob of N-in: 0.14402\noutside 1 331\nTMhelix 332 354\ninside 355 372

23819 GCF_000624255.1_ASM62425v2 Salmonella enterica subsp. enterica serovar Enteritidis str. SA20095440
Proteobacteria; Gammaproteobacteria; Enterobacterales; Enterobacteriaceae; Salmonella; Salmonella enterica; Salmonella enterica subsp. enterica; Salmonella enterica subsp. enterica serovar Enteritidis
MNNEETFYQAMRRKGVTRRSFLKFCSLAATSLGLGAGMTPKIAWALENKPRIPVVWIHGL WP_001058307.1
hydrogenase [Salmonella enterica] Length: 372\nNumber of predicted TMHs: 2\nExp number of AAs in TMHs: 36.51738\nExp number, first 60 AAs: 19.72849\nTotal prob of N-in: 0.96170\nPOSSIBLE N-term signal sequence\ninside 1 19\nTMhelix 20 39\noutside 40 325\nTMhelix 326 348\ninside 349 372

23820 GCF_000624255.1_ASM62425v2 Salmonella enterica subsp. enterica serovar Enteritidis str. SA20095440
Proteobacteria; Gammaproteobacteria; Enterobacterales; Enterobacteriaceae; Salmonella; Salmonella enterica; Salmonella enterica subsp. enterica; Salmonella enterica subsp. enterica serovar Enteritidis
MTGDNTLITSHGINRRDFMKLCAALAATMGLSSKAAAEMAESVSNPQRPPVIWIGAQECT WP_000145429.1
hydrogenase 2 small subunit [Salmonella enterica] Length: 372\nNumber of predicted TMHs: 1\nExp number of AAs in TMHs: 24.51309\nExp number, first 60 AAs: 0.2682\nTotal prob of N-in: 0.14402\noutside 1 331\nTMhelix 332 354\ninside 355 372

23821 GCF_000625735.1_ASM62573v2 Salmonella enterica subsp. enterica serovar Enteritidis str. SA20100239
Proteobacteria; Gammaproteobacteria; Enterobacterales; Enterobacteriaceae; Salmonella; Salmonella enterica; Salmonella enterica subsp. enterica; Salmonella enterica subsp. enterica serovar Enteritidis
MTGDNTLITSHGINRRDFMKLCAALAATMGLSSKAAAEMAESVSNPQRPPVIWIGAQECT WP_000145429.1
hydrogenase 2 small subunit [Salmonella enterica] Length: 372\nNumber of predicted TMHs: 1\nExp number of AAs in TMHs: 24.51309\nExp number, first 60 AAs: 0.2682\nTotal prob of N-in: 0.14402\noutside 1 331\nTMhelix 332 354\ninside 355 372

23822 GCF_000625735.1_ASM62573v2 Salmonella enterica subsp. enterica serovar Enteritidis str. SA20100239 Proteobacteria; Gammaproteobacteria; Enterobacterales; Enterobacteriaceae; Salmonella; Salmonella enterica; Salmonella enterica subsp. enterica; Salmonella enterica subsp. enterica serovar Enteritidis MNNEETFYQAMRRKGVTRRSFLKFCSLAATSLGLGAGMTPKIAWALENKPRIPVVWIHGL WP_001058307.1 hydrogenase [Salmonella enterica] Length: 372\nNumber of predicted TMHs: 2\nExp number of AAs in TMHs: 36.51738\nExp number, first 60 AAs: 19.72849\nTotal prob of N-in: 0.96170\nPOSSIBLE N-term signal sequence\ninside 1 19\nTMhelix 20 39\noutside 40 325\nTMhelix 326 348\ninside 349 372

23823 GCF_000968795.1_ASM96879v2 Salmonella enterica subsp. enterica serovar Enteritidis str. SA20100349 Proteobacteria; Gammaproteobacteria; Enterobacterales; Enterobacteriaceae; Salmonella; Salmonella enterica; Salmonella enterica subsp. enterica; Salmonella enterica subsp. enterica serovar Enteritidis MNNEETFYQAMRRKGVTRRSFLKFCSLAATSLGLGAGMTPKIAWALENKPRIPVVWIHGL WP_001058307.1 hydrogenase [Salmonella enterica] Length: 372\nNumber of predicted TMHs: 2\nExp number of AAs in TMHs: 36.51738\nExp number, first 60 AAs: 19.72849\nTotal prob of N-in: 0.96170\nPOSSIBLE N-term signal sequence\ninside 1 19\nTMhelix 20 39\noutside 40 325\nTMhelix 326 348\ninside 349 372

23824 GCF_000968795.1_ASM96879v2 Salmonella enterica subsp. enterica serovar Enteritidis str. SA20100349 Proteobacteria; Gammaproteobacteria; Enterobacterales; Enterobacteriaceae; Salmonella; Salmonella enterica; Salmonella enterica subsp. enterica; Salmonella enterica subsp. enterica serovar Enteritidis MTGDNTLITSHGINRRDFMKLCAALAATMGLSSKAAAEMAESVSNPQRPPVIWIGAQECT WP_000145429.1 hydrogenase 2 small subunit [Salmonella enterica] Length: 372\nNumber of predicted TMHs: 1\nExp number of AAs in TMHs: 24.51309\nExp number, first 60 AAs: 0.2682\nTotal prob of N-in: 0.14402\noutside 1 331\nTMhelix 332 354\ninside 355 372

23825 GCF_000624815.2_ASM62481v2 Salmonella enterica subsp. enterica serovar Enteritidis str. SA20121703 Proteobacteria; Gammaproteobacteria; Enterobacterales; Enterobacteriaceae; Salmonella; Salmonella enterica; Salmonella enterica subsp. enterica; Salmonella enterica subsp. enterica serovar Enteritidis MNNEETFYQAMRRKGVTRRSFLKFCSLAATSLGLGAGMTPKIAWALENKPRIPVVWIHGL WP_001058307.1 hydrogenase [Salmonella enterica] Length: 372\nNumber of predicted TMHs: 2\nExp number of AAs in TMHs: 36.51738\nExp number, first 60 AAs: 19.72849\nTotal prob of N-in: 0.96170\nPOSSIBLE N-term signal sequence\ninside 1 19\nTMhelix 20 39\noutside 40 325\nTMhelix 326 348\ninside 349 372

23826 GCF_000624815.2_ASM62481v2 Salmonella enterica subsp. enterica serovar Enteritidis str. SA20121703 Proteobacteria; Gammaproteobacteria; Enterobacterales; Enterobacteriaceae; Salmonella; Salmonella enterica; Salmonella enterica subsp. enterica; Salmonella enterica subsp. enterica serovar Enteritidis MTGDNTLITSHGINRRDFMKLCAALAATMGLSSKAAAEMAESVSNPQRPPVIWIGAQECT WP_000145429.1 hydrogenase 2 small subunit [Salmonella enterica] Length: 372\nNumber of predicted TMHs: 1\nExp number of AAs in TMHs: 24.51309\nExp number, first 60 AAs: 0.2682\nTotal prob of N-in: 0.14402\noutside 1 331\nTMhelix 332 354\ninside 355 372

23827 GCF_000625995.2_ASM62599v2 Salmonella enterica subsp. enterica serovar Enteritidis str. SA20123395 Proteobacteria; Gammaproteobacteria; Enterobacterales; Enterobacteriaceae; Salmonella; Salmonella enterica; Salmonella enterica subsp. enterica; Salmonella enterica subsp. enterica serovar Enteritidis MTGDNTLITSHGINRRDFMKLCAALAATMGLSSKAAAEMAESVSNPQRPPVIWIGAQECT WP_000145429.1 hydrogenase 2 small subunit [Salmonella enterica] Length: 372\nNumber of predicted TMHs: 1\nExp number of AAs in TMHs: 24.51309\nExp number, first 60 AAs: 0.2682\nTotal prob of N-in: 0.14402\noutside 1 331\nTMhelix 332 354\ninside 355 372

23828 GCF_000625995.2_ASM62599v2 Salmonella enterica subsp. enterica serovar Enteritidis str. SA20123395 Proteobacteria; Gammaproteobacteria; Enterobacterales; Enterobacteriaceae; Salmonella; Salmonella enterica; Salmonella enterica subsp. enterica; Salmonella enterica subsp. enterica serovar Enteritidis MNNEETFYQAMRRKGVTRRSFLKFCSLAATSLGLGAGMTPKIAWALENKPRIPVVWIHGL WP_001058307.1 hydrogenase [Salmonella enterica] Length: 372\nNumber of predicted TMHs: 2\nExp number of AAs in TMHs: 36.51738\nExp number, first 60 AAs: 19.72849\nTotal prob of N-in: 0.96170\nPOSSIBLE N-term signal sequence\ninside 1 19\nTMhelix 20 39\noutside 40 325\nTMhelix 326 348\ninside 349 372

23829 GCF_000335855.1_ASM33585v1 Salmonella enterica subsp. enterica serovar Enteritidis str. SE10 Proteobacteria; Gammaproteobacteria; Enterobacterales; Enterobacteriaceae; Salmonella; Salmonella enterica; Salmonella enterica subsp. enterica; Salmonella enterica subsp. enterica serovar Enteritidis MTGDNTLITSHGINRRDFMKLCAALAATMGLSSKAAAEMAESVSNPQRPPVIWIGAQECT WP_000145429.1 hydrogenase 2 small subunit [Salmonella enterica] Length: 372\nNumber of predicted TMHs: 1\nExp

number of AAs in TMHs: 24.51309\nExp number, first 60 AAs: 0.2682\nTotal prob of N-in: 0.14402\noutside 1 331\nTMhelix 332 354\ninside 355 372

23830 GCF_000329785.1_ASM32978v1 Salmonella enterica subsp. enterica serovar Enteritidis str. SE15-1 Proteobacteria; Gammaproteobacteria; Enterobacterales; Enterobacteriaceae; Salmonella; Salmonella enterica; Salmonella enterica subsp. enterica; Salmonella enterica subsp. enterica serovar Enteritidis MNNEETFYQAMRRKGVTRRSFLKFCSLAATSLGLGAGMTPKIAWALENKPRIPVVWIHGL WP_001058307.1 hydrogenase [Salmonella enterica] Length: 372\nNumber of predicted TMHs: 2\nExp number of AAs in TMHs: 36.51738\nExp number, first 60 AAs: 19.72849\nTotal prob of N-in: 0.96170\nPOSSIBLE N-term signal sequence\ninside 1 19\nTMhelix 20 39\noutside 40 325\nTMhelix 326 348\ninside 349 372

23831 GCF_000329785.1_ASM32978v1 Salmonella enterica subsp. enterica serovar Enteritidis str. SE15-1 Proteobacteria; Gammaproteobacteria; Enterobacterales; Enterobacteriaceae; Salmonella; Salmonella enterica; Salmonella enterica subsp. enterica; Salmonella enterica subsp. enterica serovar Enteritidis MTGDNTLITSHGINRRDFMKLCAALAATMGLSSKAAAEMAESVSNPQRPPVIWIGAQECT WP_000145429.1 hydrogenase 2 small subunit [Salmonella enterica] Length: 372\nNumber of predicted TMHs: 1\nExp number of AAs in TMHs: 24.51309\nExp number, first 60 AAs: 0.2682\nTotal prob of N-in: 0.14402\noutside 1 331\nTMhelix 332 354\ninside 355 372

23832 GCF_000329005.1_ASM32900v1 Salmonella enterica subsp. enterica serovar Enteritidis str. SE30663 Proteobacteria; Gammaproteobacteria; Enterobacterales; Enterobacteriaceae; Salmonella; Salmonella enterica; Salmonella enterica subsp. enterica; Salmonella enterica subsp. enterica serovar Enteritidis MTGDNTLITSHGINRRDFMKLCAALAATMGLSSKAAAEMAESVSNPQRPPVIWIGAQECT WP_000145429.1 hydrogenase 2 small subunit [Salmonella enterica] Length: 372\nNumber of predicted TMHs: 1\nExp number of AAs in TMHs: 24.51309\nExp number, first 60 AAs: 0.2682\nTotal prob of N-in: 0.14402\noutside 1 331\nTMhelix 332 354\ninside 355 372

23833 GCF_000336055.1_ASM33605v1 Salmonella enterica subsp. enterica serovar Enteritidis str. SE8a Proteobacteria; Gammaproteobacteria; Enterobacterales; Enterobacteriaceae; Salmonella; Salmonella enterica; Salmonella enterica subsp. enterica; Salmonella enterica subsp. enterica serovar Enteritidis MNNEETFYQAMRRKGVTRRSFLKFCSLAATSLGLGAGMTPKIAWALENKPRIPVVWIHGL WP_001058307.1 hydrogenase [Salmonella enterica] Length: 372\nNumber of predicted TMHs: 2\nExp number of AAs in TMHs: 36.51738\nExp number, first 60 AAs: 19.72849\nTotal prob of N-in: 0.96170\nPOSSIBLE N-term signal sequence\ninside 1 19\nTMhelix 20 39\noutside 40 325\nTMhelix 326 348\ninside 349 372

23834 GCF_000336055.1_ASM33605v1 Salmonella enterica subsp. enterica serovar Enteritidis str. SE8a Proteobacteria; Gammaproteobacteria; Enterobacterales; Enterobacteriaceae; Salmonella; Salmonella enterica; Salmonella enterica subsp. enterica; Salmonella enterica subsp. enterica serovar Enteritidis MTGDNTLITSHGINRRDFMKLCAALAATMGLSSKAAAEMAESVSNPQRPPVIWIGAQECT WP_000145429.1 hydrogenase 2 small subunit [Salmonella enterica] Length: 372\nNumber of predicted TMHs: 1\nExp number of AAs in TMHs: 24.51309\nExp number, first 60 AAs: 0.2682\nTotal prob of N-in: 0.14402\noutside 1 331\nTMhelix 332 354\ninside 355 372

23835 GCF_000329905.1_ASM32990v1 Salmonella enterica subsp. enterica serovar Enteritidis str. SL909 Proteobacteria; Gammaproteobacteria; Enterobacterales; Enterobacteriaceae; Salmonella; Salmonella enterica; Salmonella enterica subsp. enterica; Salmonella enterica subsp. enterica serovar Enteritidis MNNEETFYQAMRRKGVTRRSFLKFCSLAATSLGLGAGMTPKIAWALENKPRIPVVWIHGL WP_001058307.1 hydrogenase [Salmonella enterica] Length: 372\nNumber of predicted TMHs: 2\nExp number of AAs in TMHs: 36.51738\nExp number, first 60 AAs: 19.72849\nTotal prob of N-in: 0.96170\nPOSSIBLE N-term signal sequence\ninside 1 19\nTMhelix 20 39\noutside 40 325\nTMhelix 326 348\ninside 349 372

23836 GCF_000329905.1_ASM32990v1 Salmonella enterica subsp. enterica serovar Enteritidis str. SL909 Proteobacteria; Gammaproteobacteria; Enterobacterales; Enterobacteriaceae; Salmonella; Salmonella enterica; Salmonella enterica subsp. enterica; Salmonella enterica subsp. enterica serovar Enteritidis MTGDNTLITSHGINRRDFMKLCAALAATMGLSSKAAAEMAESVSNPQRPPVIWIGAQECT WP_000145429.1 hydrogenase 2 small subunit [Salmonella enterica] Length: 372\nNumber of predicted TMHs: 1\nExp number of AAs in TMHs: 24.51309\nExp number, first 60 AAs: 0.2682\nTotal prob of N-in: 0.14402\noutside 1 331\nTMhelix 332 354\ninside 355 372

23837 GCF_000329925.1_ASM32992v1 Salmonella enterica subsp. enterica serovar Enteritidis str. SL913 Proteobacteria; Gammaproteobacteria; Enterobacterales; Enterobacteriaceae; Salmonella; Salmonella enterica;

Salmonella enterica subsp. enterica; Salmonella enterica subsp. enterica serovar Enteritidis
MTGDNTLITSHGINRRDFMKLCAALAATMGLSSKAAAEMAESVSNPQRPPVIWIGAQECT WP_000145429.1
hydrogenase 2 small subunit [Salmonella enterica] Length: 372\nNumber of predicted TMHs: 1\nExp number of AAs in TMHs: 24.51309\nExp number, first 60 AAs: 0.2682\nTotal prob of N-in: 0.14402\noutside 1 331\nTMhelix 332 354\ninside 355 372

23838 GCF_000329925.1_ASM32992v1 Salmonella enterica subsp. enterica serovar Enteritidis str. SL913
Proteobacteria; Gammaproteobacteria; Enterobacterales; Enterobacteriaceae; Salmonella; Salmonella enterica; Salmonella enterica subsp. enterica; Salmonella enterica subsp. enterica serovar Enteritidis
MNNEETFYQAMRRKGVTRRSFLKFCSLAATSLGLGAGMTPKIAWALENKPRIPVVWIHGL WP_001058307.1
hydrogenase [Salmonella enterica] Length: 372\nNumber of predicted TMHs: 2\nExp number of AAs in TMHs: 36.51738\nExp number, first 60 AAs: 19.72849\nTotal prob of N-in: 0.96170\nPOSSIBLE N-term signal sequence\ninside 1 19\nTMhelix 20 39\noutside 40 325\nTMhelix 326 348\ninside 349 372

23839 GCF_001448465.1_ASM144846v1 Salmonella enterica subsp. enterica serovar Fresno str. ST224
Proteobacteria; Gammaproteobacteria; Enterobacterales; Enterobacteriaceae; Salmonella; Salmonella enterica; Salmonella enterica subsp. enterica; Salmonella enterica subsp. enterica serovar Fresno
MNNEETFYQAMRRKGVTRRSFLKFCSLAATSLGLGAGMTPKIAWALENKPRIPVVWIHGL WP_001058311.1
hydrogenase [Salmonella enterica] Length: 372\nNumber of predicted TMHs: 2\nExp number of AAs in TMHs: 37.4699899999999\nExp number, first 60 AAs: 19.72699\nTotal prob of N-in: 0.96171\nPOSSIBLE N-term signal sequence\ninside 1 19\nTMhelix 20 39\noutside 40 325\nTMhelix 326 348\ninside 349 372

23840 GCF_001448465.1_ASM144846v1 Salmonella enterica subsp. enterica serovar Fresno str. ST224
Proteobacteria; Gammaproteobacteria; Enterobacterales; Enterobacteriaceae; Salmonella; Salmonella enterica; Salmonella enterica subsp. enterica; Salmonella enterica subsp. enterica serovar Fresno
MTGDNTLITSHGINRRDFMKLCAALAATMGLSSKAAAEMAESVSNPQRPPVIWIGAQECT WP_000145429.1
hydrogenase 2 small subunit [Salmonella enterica] Length: 372\nNumber of predicted TMHs: 1\nExp number of AAs in TMHs: 24.51309\nExp number, first 60 AAs: 0.2682\nTotal prob of N-in: 0.14402\noutside 1 331\nTMhelix 332 354\ninside 355 372

23841 GCF_000009525.1_ASM952v1 Salmonella enterica subsp. enterica serovar Gallinarum str. 287/91
Proteobacteria; Gammaproteobacteria; Enterobacterales; Enterobacteriaceae; Salmonella; Salmonella enterica; Salmonella enterica subsp. enterica; Salmonella enterica subsp. enterica serovar Gallinarum
MNNEETFYQAMRRKGVTRRSFLKFCSLAATSLGLGAGMTPKIAWALENKPRIPVVWIHGL WP_001058304.1
hydrogenase [Salmonella enterica] Length: 372\nNumber of predicted TMHs: 2\nExp number of AAs in TMHs: 36.51703\nExp number, first 60 AAs: 19.7285\nTotal prob of N-in: 0.96169\nPOSSIBLE N-term signal sequence\ninside 1 19\nTMhelix 20 39\noutside 40 325\nTMhelix 326 348\ninside 349 372

23842 GCF_000009525.1_ASM952v1 Salmonella enterica subsp. enterica serovar Gallinarum str. 287/91
Proteobacteria; Gammaproteobacteria; Enterobacterales; Enterobacteriaceae; Salmonella; Salmonella enterica; Salmonella enterica subsp. enterica; Salmonella enterica subsp. enterica serovar Gallinarum
MTGDNTLITSHGINRRDFMKLCAALAATMGLSSKAAAEMAESVSNPQRPPVIWIGAQECT WP_000145429.1
hydrogenase 2 small subunit [Salmonella enterica] Length: 372\nNumber of predicted TMHs: 1\nExp number of AAs in TMHs: 24.51309\nExp number, first 60 AAs: 0.2682\nTotal prob of N-in: 0.14402\noutside 1 331\nTMhelix 332 354\ninside 355 372

23843 GCF_000335995.2_ASM33599v2 Salmonella enterica subsp. enterica serovar Gallinarum str. 9184
Proteobacteria; Gammaproteobacteria; Enterobacterales; Enterobacteriaceae; Salmonella; Salmonella enterica; Salmonella enterica subsp. enterica; Salmonella enterica subsp. enterica serovar Gallinarum
MTGDNTLITSHGINRRDFMKLCAALAATMGLSSKAAAEMAESVSNPQRPPVIWIGAQECT WP_000145429.1
hydrogenase 2 small subunit [Salmonella enterica] Length: 372\nNumber of predicted TMHs: 1\nExp number of AAs in TMHs: 24.51309\nExp number, first 60 AAs: 0.2682\nTotal prob of N-in: 0.14402\noutside 1 331\nTMhelix 332 354\ninside 355 372

23844 GCF_000335995.2_ASM33599v2 Salmonella enterica subsp. enterica serovar Gallinarum str. 9184
Proteobacteria; Gammaproteobacteria; Enterobacterales; Enterobacteriaceae; Salmonella; Salmonella enterica; Salmonella enterica subsp. enterica; Salmonella enterica subsp. enterica serovar Gallinarum
MNNEETFYQAMRRKGVTRRSFLKFCSLAATSLGLGAGMTPKIAWALENKPRIPVVWIHGL WP_001058304.1
hydrogenase [Salmonella enterica] Length: 372\nNumber of predicted TMHs: 2\nExp number of AAs in TMHs: 36.51703\nExp number, first 60 AAs: 19.7285\nTotal prob of N-in: 0.96169\nPOSSIBLE N-term signal sequence\ninside 1 19\nTMhelix 20 39\noutside 40 325\nTMhelix 326 348\ninside 349 372

23845 GCF_000192125.1_ASM19212v1 Salmonella enterica subsp. enterica serovar Gallinarum str. SG9 Proteobacteria; Gammaproteobacteria; Enterobacterales; Enterobacteriaceae; Salmonella; Salmonella enterica; Salmonella enterica subsp. enterica; Salmonella enterica subsp. enterica serovar Gallinarum MNNEETFYQAMRRKGVTRRSFLKFCSLAATSLGLGAGMTPKIAWALENKPRIPVVWIHGL WP_001058304.1 hydrogenase [Salmonella enterica] Length: 372\nNumber of predicted TMHs: 2\nExp number of AAs in TMHs: 36.51703\nExp number, first 60 AAs: 19.7285\nTotal prob of N-in: 0.96169\nPOSSIBLE N-term signal sequence\ninside 1 19\nTMhelix 20 39\noutside 40 325\nTMhelix 326 348\ninside 349 372

23846 GCF_000192125.1_ASM19212v1 Salmonella enterica subsp. enterica serovar Gallinarum str. SG9 Proteobacteria; Gammaproteobacteria; Enterobacterales; Enterobacteriaceae; Salmonella; Salmonella enterica; Salmonella enterica subsp. enterica; Salmonella enterica subsp. enterica serovar Gallinarum MTGDNTLITSHGINRRDFMKLCAALAATMGLSSKAAAEMAESVSNPQRPPVIWIGAQECT WP_000145429.1 hydrogenase 2 small subunit [Salmonella enterica] Length: 372\nNumber of predicted TMHs: 1\nExp number of AAs in TMHs: 24.51309\nExp number, first 60 AAs: 0.2682\nTotal prob of N-in: 0.14402\noutside 1 331\nTMhelix 332 354\ninside 355 372

23847 GCF_000462995.1_ASM46299v1 Salmonella enterica subsp. enterica serovar Gallinarum/pullorum str. CDC1983-67 Proteobacteria; Gammaproteobacteria; Enterobacterales; Enterobacteriaceae; Salmonella; Salmonella enterica; Salmonella enterica subsp. enterica; Salmonella enterica subsp. enterica serovar Gallinarum/pullorum MNNEETFYQAMRRKGVTRRSFLKFCSLAATSLGLGAGMTPKIAWALENKPRIPVVWIHGL WP_001058309.1 hydrogenase [Salmonella enterica] Length: 372\nNumber of predicted TMHs: 2\nExp number of AAs in TMHs: 36.51725\nExp number, first 60 AAs: 19.7285\nTotal prob of N-in: 0.96170\nPOSSIBLE N-term signal sequence\ninside 1 19\nTMhelix 20 39\noutside 40 325\nTMhelix 326 348\ninside 349 372

23848 GCF_000462995.1_ASM46299v1 Salmonella enterica subsp. enterica serovar Gallinarum/pullorum str. CDC1983-67 Proteobacteria; Gammaproteobacteria; Enterobacterales; Enterobacteriaceae; Salmonella; Salmonella enterica; Salmonella enterica subsp. enterica; Salmonella enterica subsp. enterica serovar Gallinarum/pullorum MTGDNTLITSHGINRRDFMKLCAALAATMGLSSKAAAEMAESVSNPQRPPVIWIGAQECT WP_000145430.1 hydrogenase 2 small subunit [Salmonella enterica] Length: 372\nNumber of predicted TMHs: 1\nExp number of AAs in TMHs: 24.51306\nExp number, first 60 AAs: 0.2682\nTotal prob of N-in: 0.14402\noutside 1 331\nTMhelix 332 354\ninside 355 372

23849 GCF_000521885.1_SP-v1 Salmonella enterica subsp. enterica serovar Gallinarum/pullorum str. FCAV198 Proteobacteria; Gammaproteobacteria; Enterobacterales; Enterobacteriaceae; Salmonella; Salmonella enterica; Salmonella enterica subsp. enterica; Salmonella enterica subsp. enterica serovar Gallinarum/pullorum MNNEETFYQAMRRKGVTRRSFLKFCSLAATSLGLGAGMTPKIAWALENKPRIPVVWIHGL WP_001058309.1 hydrogenase [Salmonella enterica] Length: 372\nNumber of predicted TMHs: 2\nExp number of AAs in TMHs: 36.51725\nExp number, first 60 AAs: 19.7285\nTotal prob of N-in: 0.96170\nPOSSIBLE N-term signal sequence\ninside 1 19\nTMhelix 20 39\noutside 40 325\nTMhelix 326 348\ninside 349 372

23850 GCF_000521885.1_SP-v1 Salmonella enterica subsp. enterica serovar Gallinarum/pullorum str. FCAV198 Proteobacteria; Gammaproteobacteria; Enterobacterales; Enterobacteriaceae; Salmonella; Salmonella enterica; Salmonella enterica subsp. enterica; Salmonella enterica subsp. enterica serovar Gallinarum/pullorum MTGDNTLITSHGINRRDFMKLCAALAATMGLSSKAAAEMAESVSNPQRPPVIWIGAQECT WP_000145430.1 hydrogenase 2 small subunit [Salmonella enterica] Length: 372\nNumber of predicted TMHs: 1\nExp number of AAs in TMHs: 24.51306\nExp number, first 60 AAs: 0.2682\nTotal prob of N-in: 0.14402\noutside 1 331\nTMhelix 332 354\ninside 355 372

23851 GCF_000235545.1_ASM23554v1 Salmonella enterica subsp. enterica serovar Gallinarum/pullorum str. RKS5078 Proteobacteria; Gammaproteobacteria; Enterobacterales; Enterobacteriaceae; Salmonella; Salmonella enterica; Salmonella enterica subsp. enterica; Salmonella enterica subsp. enterica serovar Gallinarum/pullorum MNNEETFYQAMRRKGVTRRSFLKFCSLAATSLGLGAGMTPKIAWALENKPRIPVVWIHGL WP_001058309.1 hydrogenase [Salmonella enterica] Length: 372\nNumber of predicted TMHs: 2\nExp number of AAs in TMHs: 36.51725\nExp number, first 60 AAs: 19.7285\nTotal prob of N-in: 0.96170\nPOSSIBLE N-term signal sequence\ninside 1 19\nTMhelix 20 39\noutside 40 325\nTMhelix 326 348\ninside 349 372

23852 GCF_000235545.1_ASM23554v1 Salmonella enterica subsp. enterica serovar Gallinarum/pullorum str. RKS5078 Proteobacteria; Gammaproteobacteria; Enterobacterales; Enterobacteriaceae; Salmonella; Salmonella enterica; Salmonella enterica subsp. enterica; Salmonella enterica subsp. enterica serovar Gallinarum/pullorum MTGDNTLITSHGINRRDFMKLCAALAATMGLSSKAAAEMAESVSNPQRPPVIWIGAQECT WP_000145430.1 hydrogenase 2 small subunit [Salmonella enterica] Length: 372\nNumber of predicted TMHs: 1\nExp

number of AAs in TMHs: 24.51306\nExp number, first 60 AAs: 0.2682\nTotal prob of N-in: 0.14402\noutside 1 331\nTMhelix 332 354\ninside 355 372

23853 GCF_001448665.1_ASM144866v1 Salmonella enterica subsp. enterica serovar Gaminara str. SA20063285 Proteobacteria; Gammaproteobacteria; Enterobacterales; Enterobacteriaceae; Salmonella; Salmonella enterica; Salmonella enterica subsp. enterica; Salmonella enterica subsp. enterica serovar Gaminara MTGDNTLITSHGINRRDFMKLCAALAATMGLSSKAAAEMAESVSNPQRPPVIWIGAQECT WP_000145429.1 hydrogenase 2 small subunit [Salmonella enterica] Length: 372\nNumber of predicted TMHs: 1\nExp number of AAs in TMHs: 24.51309\nExp number, first 60 AAs: 0.2682\nTotal prob of N-in: 0.14402\noutside 1 331\nTMhelix 332 354\ninside 355 372

23854 GCF_001448665.1_ASM144866v1 Salmonella enterica subsp. enterica serovar Gaminara str. SA20063285 Proteobacteria; Gammaproteobacteria; Enterobacterales; Enterobacteriaceae; Salmonella; Salmonella enterica; Salmonella enterica subsp. enterica; Salmonella enterica subsp. enterica serovar Gaminara MNNEETFYQAMRRKGVTRRSFLKFCSLAATSLGLGAGMTPKIAWALENKPRIPVVWIHGL WP_001662439.1 hydrogenase [Salmonella enterica] Length: 372\nNumber of predicted TMHs: 2\nExp number of AAs in TMHs: 36.5172\nExp number, first 60 AAs: 19.72849\nTotal prob of N-in: 0.96169\nPOSSIBLE N-term signal sequence\ninside 1 19\nTMhelix 20 39\noutside 40 325\nTMhelix 326 348\ninside 349 372

23855 GCF_000487215.1_ASM48721v1 Salmonella enterica subsp. enterica serovar Give str. 564 Proteobacteria; Gammaproteobacteria; Enterobacterales; Enterobacteriaceae; Salmonella; Salmonella enterica; Salmonella enterica subsp. enterica; Salmonella enterica subsp. enterica serovar Give MNNEETFYQAMRRKGVTRRSFLKFCSLAATSLGLGAGMTPKIAWALENKPRIPVVWIHGL WP_001058312.1 hydrogenase [Salmonella enterica] Length: 372\nNumber of predicted TMHs: 2\nExp number of AAs in TMHs: 36.51738\nExp number, first 60 AAs: 19.72849\nTotal prob of N-in: 0.96170\nPOSSIBLE N-term signal sequence\ninside 1 19\nTMhelix 20 39\noutside 40 325\nTMhelix 326 348\ninside 349 372

23856 GCF_000487215.1_ASM48721v1 Salmonella enterica subsp. enterica serovar Give str. 564 Proteobacteria; Gammaproteobacteria; Enterobacterales; Enterobacteriaceae; Salmonella; Salmonella enterica; Salmonella enterica subsp. enterica; Salmonella enterica subsp. enterica serovar Give MTGDNTLITSHGINRRDFMKLCAALAATMGLSSKAAAEMAESVSNPQRPPVIWIGAQECT WP_000145429.1 hydrogenase 2 small subunit [Salmonella enterica] Length: 372\nNumber of predicted TMHs: 1\nExp number of AAs in TMHs: 24.51309\nExp number, first 60 AAs: 0.2682\nTotal prob of N-in: 0.14402\noutside 1 331\nTMhelix 332 354\ninside 355 372

23857 GCF_000505465.1_CFSAN004343_01.0Salmonella enterica subsp. enterica serovar Give var. 15 str. CFSAN004343 Proteobacteria; Gammaproteobacteria; Enterobacterales; Enterobacteriaceae; Salmonella; Salmonella enterica; Salmonella enterica subsp. enterica; Salmonella enterica subsp. enterica serovar Give MTGDNTLITSHGINRRDFMKLCAALAATMGLSSKAAAEMAESVSNPQRPPVIWIGAQECT WP_000145429.1 hydrogenase 2 small subunit [Salmonella enterica] Length: 372\nNumber of predicted TMHs: 1\nExp number of AAs in TMHs: 24.51309\nExp number, first 60 AAs: 0.2682\nTotal prob of N-in: 0.14402\noutside 1 331\nTMhelix 332 354\ninside 355 372

23858 GCF_000505465.1_CFSAN004343_01.0Salmonella enterica subsp. enterica serovar Give var. 15 str. CFSAN004343 Proteobacteria; Gammaproteobacteria; Enterobacterales; Enterobacteriaceae; Salmonella; Salmonella enterica; Salmonella enterica subsp. enterica; Salmonella enterica subsp. enterica serovar Give MNNEETFYQAMRRKGVTRRSFLKFCSLAATSLGLGAGMTPKIAWALENKPRIPVVWIHGL WP_001058312.1 hydrogenase [Salmonella enterica] Length: 372\nNumber of predicted TMHs: 2\nExp number of AAs in TMHs: 36.51738\nExp number, first 60 AAs: 19.72849\nTotal prob of N-in: 0.96170\nPOSSIBLE N-term signal sequence\ninside 1 19\nTMhelix 20 39\noutside 40 325\nTMhelix 326 348\ninside 349 372

23859 GCF_000293095.1_ASM29309v1 Salmonella enterica subsp. enterica serovar Hadar str. 2011K-0112 Proteobacteria; Gammaproteobacteria; Enterobacterales; Enterobacteriaceae; Salmonella; Salmonella enterica; Salmonella enterica subsp. enterica; Salmonella enterica subsp. enterica serovar Hadar MTGDNTLITSHGINRRDFMKLCAALAATMGLSSKAAAEMAESVSNPQRPPVIWIGAQECT WP_000145429.1 hydrogenase 2 small subunit [Salmonella enterica] Length: 372\nNumber of predicted TMHs: 1\nExp number of AAs in TMHs: 24.51309\nExp number, first 60 AAs: 0.2682\nTotal prob of N-in: 0.14402\noutside 1 331\nTMhelix 332 354\ninside 355 372

23860 GCF_000293095.1_ASM29309v1 Salmonella enterica subsp. enterica serovar Hadar str. 2011K-0112 Proteobacteria; Gammaproteobacteria; Enterobacterales; Enterobacteriaceae; Salmonella; Salmonella enterica;

Salmonella enterica subsp. enterica; Salmonella enterica subsp. enterica serovar Hadar
MNNEETFYQAMRRKGVTRRSFLKFCSLAATSLGLGAGMTPKIAWALENKPRIPVVWIHGL WP_001058308.1
hydrogenase [Salmonella enterica] Length: 372\nNumber of predicted TMHs: 2\nExp number of AAs in TMHs: 39.08204\nExp number, first 60 AAs: 19.72474\nTotal prob of N-in: 0.96174\nPOSSIBLE N-term signal sequence\ninside 1 19\nTMhelix 20 39\noutside 40 325\nTMhelix 326 348\ninside 349 372

23861 GCF_000412915.1_2012K-0266_Ver1 Salmonella enterica subsp. enterica serovar Hadar str. 2012K-0272
Proteobacteria; Gammaproteobacteria; Enterobacterales; Enterobacteriaceae; Salmonella; Salmonella enterica; Salmonella enterica subsp. enterica; Salmonella enterica subsp. enterica serovar Hadar
MTGDNTLITSHGINRRDFMKLCAALAATMGLSSKAAAEMAESVSNPQRPPVIWIGAQECT WP_000145429.1
hydrogenase 2 small subunit [Salmonella enterica] Length: 372\nNumber of predicted TMHs: 1\nExp number of AAs in TMHs: 24.51309\nExp number, first 60 AAs: 0.2682\nTotal prob of N-in: 0.14402\noutside 1 331\nTMhelix 332 354\ninside 355 372

23862 GCF_000487175.1_ASM48717v1 Salmonella enterica subsp. enterica serovar Hadar str. ATCC 51956
Proteobacteria; Gammaproteobacteria; Enterobacterales; Enterobacteriaceae; Salmonella; Salmonella enterica; Salmonella enterica subsp. enterica; Salmonella enterica subsp. enterica serovar Hadar
MTGDNTLITSHGINRRDFMKLCAALAATMGLSSKAAAEMAESVSNPQRPPVIWIGAQECT WP_000145429.1
hydrogenase 2 small subunit [Salmonella enterica] Length: 372\nNumber of predicted TMHs: 1\nExp number of AAs in TMHs: 24.51309\nExp number, first 60 AAs: 0.2682\nTotal prob of N-in: 0.14402\noutside 1 331\nTMhelix 332 354\ninside 355 372

23863 GCF_000487175.1_ASM48717v1 Salmonella enterica subsp. enterica serovar Hadar str. ATCC 51956
Proteobacteria; Gammaproteobacteria; Enterobacterales; Enterobacteriaceae; Salmonella; Salmonella enterica; Salmonella enterica subsp. enterica; Salmonella enterica subsp. enterica serovar Hadar
MNNEETFYQAMRRKGVTRRSFLKFCSLAATSLGLGAGMTPKIAWALENKPRIPVVWIHGL WP_001058308.1
hydrogenase [Salmonella enterica] Length: 372\nNumber of predicted TMHs: 2\nExp number of AAs in TMHs: 39.08204\nExp number, first 60 AAs: 19.72474\nTotal prob of N-in: 0.96174\nPOSSIBLE N-term signal sequence\ninside 1 19\nTMhelix 20 39\noutside 40 325\nTMhelix 326 348\ninside 349 372

23864 GCF_000171515.1_ASM17151v1 Salmonella enterica subsp. enterica serovar Hadar str. RI_05P066
Proteobacteria; Gammaproteobacteria; Enterobacterales; Enterobacteriaceae; Salmonella; Salmonella enterica; Salmonella enterica subsp. enterica; Salmonella enterica subsp. enterica serovar Hadar
MNNEETFYQAMRRKGVTRRSFLKFCSLAATSLGLGAGMTPKIAWALENKPRIPVVWIHGL WP_001058308.1
hydrogenase [Salmonella enterica] Length: 372\nNumber of predicted TMHs: 2\nExp number of AAs in TMHs: 39.08204\nExp number, first 60 AAs: 19.72474\nTotal prob of N-in: 0.96174\nPOSSIBLE N-term signal sequence\ninside 1 19\nTMhelix 20 39\noutside 40 325\nTMhelix 326 348\ninside 349 372

23865 GCF_000171515.1_ASM17151v1 Salmonella enterica subsp. enterica serovar Hadar str. RI_05P066
Proteobacteria; Gammaproteobacteria; Enterobacterales; Enterobacteriaceae; Salmonella; Salmonella enterica; Salmonella enterica subsp. enterica; Salmonella enterica subsp. enterica serovar Hadar
MTGDNTLITSHGINRRDFMKLCAALAATMGLSSKAAAEMAESVSNPQRPPVIWIGAQECT WP_000145429.1
hydrogenase 2 small subunit [Salmonella enterica] Length: 372\nNumber of predicted TMHs: 1\nExp number of AAs in TMHs: 24.51309\nExp number, first 60 AAs: 0.2682\nTotal prob of N-in: 0.14402\noutside 1 331\nTMhelix 332 354\ninside 355 372

23866 GCF_001448685.1_ASM144868v1 Salmonella enterica subsp. enterica serovar Hadar str. SA20026260
Proteobacteria; Gammaproteobacteria; Enterobacterales; Enterobacteriaceae; Salmonella; Salmonella enterica; Salmonella enterica subsp. enterica; Salmonella enterica subsp. enterica serovar Hadar
MNNEETFYQAMRRKGVTRRSFLKFCSLAATSLGLGAGMTPKIAWALENKPRIPVVWIHGL WP_001058308.1
hydrogenase [Salmonella enterica] Length: 372\nNumber of predicted TMHs: 2\nExp number of AAs in TMHs: 39.08204\nExp number, first 60 AAs: 19.72474\nTotal prob of N-in: 0.96174\nPOSSIBLE N-term signal sequence\ninside 1 19\nTMhelix 20 39\noutside 40 325\nTMhelix 326 348\ninside 349 372

23867 GCF_001448685.1_ASM144868v1 Salmonella enterica subsp. enterica serovar Hadar str. SA20026260
Proteobacteria; Gammaproteobacteria; Enterobacterales; Enterobacteriaceae; Salmonella; Salmonella enterica; Salmonella enterica subsp. enterica; Salmonella enterica subsp. enterica serovar Hadar
MTGDNTLITSHGINRRDFMKLCAALAATMGLSSKAAAEMAESVSNPQRPPVIWIGAQECT WP_000145429.1
hydrogenase 2 small subunit [Salmonella enterica] Length: 372\nNumber of predicted TMHs: 1\nExp number of AAs in TMHs: 24.51309\nExp number, first 60 AAs: 0.2682\nTotal prob of N-in: 0.14402\noutside 1 331\nTMhelix 332 354\ninside 355 372

23868 GCF_000293055.1_ASM29305v1 Salmonella enterica subsp. enterica serovar Hartford str. 06-0676 Proteobacteria; Gammaproteobacteria; Enterobacterales; Enterobacteriaceae; Salmonella; Salmonella enterica; Salmonella enterica subsp. enterica; Salmonella enterica subsp. enterica serovar Hartford MNNEETFYQAMRRKGVTRRSFLKFCSLAATSLGLGAGMTPKIAWALENKPRIPVVWIHGL WP_001058312.1 hydrogenase [Salmonella enterica] Length: 372\nNumber of predicted TMHs: 2\nExp number of AAs in TMHs: 36.51738\nExp number, first 60 AAs: 19.72849\nTotal prob of N-in: 0.96170\nPOSSIBLE N-term signal sequence\ninside 1 19\nTMhelix 20 39\noutside 40 325\nTMhelix 326 348\ninside 349 372

23869 GCF_000293055.1_ASM29305v1 Salmonella enterica subsp. enterica serovar Hartford str. 06-0676 Proteobacteria; Gammaproteobacteria; Enterobacterales; Enterobacteriaceae; Salmonella; Salmonella enterica; Salmonella enterica subsp. enterica; Salmonella enterica subsp. enterica serovar Hartford MTGDNTLITSHGINRRDFMKLCAALAATMGLSSKAAAEMAESVSNPQRPPVIWIGAQECT WP_000145429.1 hydrogenase 2 small subunit [Salmonella enterica] Length: 372\nNumber of predicted TMHs: 1\nExp number of AAs in TMHs: 24.51309\nExp number, first 60 AAs: 0.2682\nTotal prob of N-in: 0.14402\noutside 1 331\nTMhelix 332 354\ninside 355 372

23870 GCF_000412935.1_2012K-0273_Ver1 Salmonella enterica subsp. enterica serovar Hartford str. 2012K-0272 Proteobacteria; Gammaproteobacteria; Enterobacterales; Enterobacteriaceae; Salmonella; Salmonella enterica; Salmonella enterica subsp. enterica; Salmonella enterica subsp. enterica serovar Hartford MTGDNTLITSHGINRRDFMKLCAALAATMGLSSKAAAEMAESVSNPQRPPVIWIGAQECT WP_000145429.1 hydrogenase 2 small subunit [Salmonella enterica] Length: 372\nNumber of predicted TMHs: 1\nExp number of AAs in TMHs: 24.51309\nExp number, first 60 AAs: 0.2682\nTotal prob of N-in: 0.14402\noutside 1 331\nTMhelix 332 354\ninside 355 372

23871 GCF_000487875.1_ASM48787v1 Salmonella enterica subsp. enterica serovar Hartford str. CFSAN001075 Proteobacteria; Gammaproteobacteria; Enterobacterales; Enterobacteriaceae; Salmonella; Salmonella enterica; Salmonella enterica subsp. enterica; Salmonella enterica subsp. enterica serovar Hartford MNNEETFYQAMRRKGVTRRSFLKFCSLAATSLGLGAGMTPKIAWALENKPRIPVVWIHGL WP_001058312.1 hydrogenase [Salmonella enterica] Length: 372\nNumber of predicted TMHs: 2\nExp number of AAs in TMHs: 36.51738\nExp number, first 60 AAs: 19.72849\nTotal prob of N-in: 0.96170\nPOSSIBLE N-term signal sequence\ninside 1 19\nTMhelix 20 39\noutside 40 325\nTMhelix 326 348\ninside 349 372

23872 GCF_000487875.1_ASM48787v1 Salmonella enterica subsp. enterica serovar Hartford str. CFSAN001075 Proteobacteria; Gammaproteobacteria; Enterobacterales; Enterobacteriaceae; Salmonella; Salmonella enterica; Salmonella enterica subsp. enterica; Salmonella enterica subsp. enterica serovar Hartford MTGDNTLITSHGINRRDFMKLCAALAATMGLSSKAAAEMAESVSNPQRPPVIWIGAQECT WP_000145429.1 hydrogenase 2 small subunit [Salmonella enterica] Length: 372\nNumber of predicted TMHs: 1\nExp number of AAs in TMHs: 24.51309\nExp number, first 60 AAs: 0.2682\nTotal prob of N-in: 0.14402\noutside 1 331\nTMhelix 332 354\ninside 355 372

23873 GCF_000487735.1_ASM48773v1 Salmonella enterica subsp. enterica serovar Havana str. CFSAN001082 Proteobacteria; Gammaproteobacteria; Enterobacterales; Enterobacteriaceae; Salmonella; Salmonella enterica; Salmonella enterica subsp. enterica; Salmonella enterica subsp. enterica serovar Havana MTGDNTLITSHGINRRDFMKLCAALAATMGLSSKAAAEMAESVSNPQRPPVIWIGAQECT WP_000145429.1 hydrogenase 2 small subunit [Salmonella enterica] Length: 372\nNumber of predicted TMHs: 1\nExp number of AAs in TMHs: 24.51309\nExp number, first 60 AAs: 0.2682\nTotal prob of N-in: 0.14402\noutside 1 331\nTMhelix 332 354\ninside 355 372

23874 GCF_000487735.1_ASM48773v1 Salmonella enterica subsp. enterica serovar Havana str. CFSAN001082 Proteobacteria; Gammaproteobacteria; Enterobacterales; Enterobacteriaceae; Salmonella; Salmonella enterica; Salmonella enterica subsp. enterica; Salmonella enterica subsp. enterica serovar Havana MNNEETFYQAMRRKGVTRRSFLKFCSLAATSLGLGAGMTPKIAWALENKPRIPVVWIHGL WP_023254288.1 hydrogenase 1, small subunit [Salmonella enterica] Length: 372\nNumber of predicted TMHs: 2\nExp number of AAs in TMHs: 37.68168\nExp number, first 60 AAs: 19.73265\nTotal prob of N-in: 0.96171\nPOSSIBLE N-term signal sequence\ninside 1 19\nTMhelix 20 39\noutside 40 325\nTMhelix 326 348\ninside 349 372

23875 GCF_000603505.1_ASM60350v1 Salmonella enterica subsp. enterica serovar Heidelberg str. 21381 Proteobacteria; Gammaproteobacteria; Enterobacterales; Enterobacteriaceae; Salmonella; Salmonella enterica; Salmonella enterica subsp. enterica; Salmonella enterica subsp. enterica serovar Heidelberg MNNEETFYQAMRRKGVTRRSFLKFCSLAATSLGLGAGMTPKIAWALENKPRIPVVWIHGL WP_001058312.1 hydrogenase [Salmonella enterica] Length: 372\nNumber of predicted TMHs: 2\nExp number of AAs in

TMHs: 36.51738\nExp number, first 60 AAs: 19.72849\nTotal prob of N-in: 0.96170\nPOSSIBLE N-term signal sequence\ninside 1 19\nTMhelix 20 39\noutside 40 325\nTMhelix 326 348\ninside 349 372

23876 GCF_000603505.1_ASM60350v1 Salmonella enterica subsp. enterica serovar Heidelberg str. 21381 Proteobacteria; Gammaproteobacteria; Enterobacterales; Enterobacteriaceae; Salmonella; Salmonella enterica; Salmonella enterica subsp. enterica; Salmonella enterica subsp. enterica serovar Heidelberg MTGDNTLITSHGINRRDFMKLCAALAATMGLSSKAAAEMAESVSNPQRPPVIWIGAQECT WP_000145429.1 hydrogenase 2 small subunit [Salmonella enterica] Length: 372\nNumber of predicted TMHs: 1\nExp number of AAs in TMHs: 24.51309\nExp number, first 60 AAs: 0.2682\nTotal prob of N-in: 0.14402\noutside 1 331\nTMhelix 332 354\ninside 355 372

23877 GCF_000486085.1_ASM48608v1 Salmonella enterica subsp. enterica serovar Heidelberg str. 24390 Proteobacteria; Gammaproteobacteria; Enterobacterales; Enterobacteriaceae; Salmonella; Salmonella enterica; Salmonella enterica subsp. enterica; Salmonella enterica subsp. enterica serovar Heidelberg MNNEETFYQAMRRKGVTRRSFLKFCSLAATSLGLGAGMTPKIAWALENKPRIPVVWIHGL WP_001058312.1 hydrogenase [Salmonella enterica] Length: 372\nNumber of predicted TMHs: 2\nExp number of AAs in TMHs: 36.51738\nExp number, first 60 AAs: 19.72849\nTotal prob of N-in: 0.96170\nPOSSIBLE N-term signal sequence\ninside 1 19\nTMhelix 20 39\noutside 40 325\nTMhelix 326 348\ninside 349 372

23878 GCF_000486085.1_ASM48608v1 Salmonella enterica subsp. enterica serovar Heidelberg str. 24390 Proteobacteria; Gammaproteobacteria; Enterobacterales; Enterobacteriaceae; Salmonella; Salmonella enterica; Salmonella enterica subsp. enterica; Salmonella enterica subsp. enterica serovar Heidelberg MTGDNTLITSHGINRRDFMKLCAALAATMGLSSKAAAEMAESVSNPQRPPVIWIGAQECT WP_000145429.1 hydrogenase 2 small subunit [Salmonella enterica] Length: 372\nNumber of predicted TMHs: 1\nExp number of AAs in TMHs: 24.51309\nExp number, first 60 AAs: 0.2682\nTotal prob of N-in: 0.14402\noutside 1 331\nTMhelix 332 354\ninside 355 372

23879 GCF_000601415.1_CFSAN000309_01.0Salmonella enterica subsp. enterica serovar Heidelberg str. 29169 Proteobacteria; Gammaproteobacteria; Enterobacterales; Enterobacteriaceae; Salmonella; Salmonella enterica; Salmonella enterica subsp. enterica; Salmonella enterica subsp. enterica serovar Heidelberg MTGDNTLITSHGINRRDFMKLCAALAATMGLSSKAAAEMAESVSNPQRPPVIWIGAQECT WP_000145429.1 hydrogenase 2 small subunit [Salmonella enterica] Length: 372\nNumber of predicted TMHs: 1\nExp number of AAs in TMHs: 24.51309\nExp number, first 60 AAs: 0.2682\nTotal prob of N-in: 0.14402\noutside 1 331\nTMhelix 332 354\ninside 355 372

23880 GCF_000601415.1_CFSAN000309_01.0Salmonella enterica subsp. enterica serovar Heidelberg str. 29169 Proteobacteria; Gammaproteobacteria; Enterobacterales; Enterobacteriaceae; Salmonella; Salmonella enterica; Salmonella enterica subsp. enterica; Salmonella enterica subsp. enterica serovar Heidelberg MNNEETFYQAMRRKGVTRRSFLKFCSLAATSLGLGAGMTPKIAWALENKPRIPVVWIHGL WP_001058312.1 hydrogenase [Salmonella enterica] Length: 372\nNumber of predicted TMHs: 2\nExp number of AAs in TMHs: 36.51738\nExp number, first 60 AAs: 19.72849\nTotal prob of N-in: 0.96170\nPOSSIBLE N-term signal sequence\ninside 1 19\nTMhelix 20 39\noutside 40 325\nTMhelix 326 348\ninside 349 372

23881 GCF_000603725.1_ASM60372v1 Salmonella enterica subsp. enterica serovar Heidelberg str. 32507 Proteobacteria; Gammaproteobacteria; Enterobacterales; Enterobacteriaceae; Salmonella; Salmonella enterica; Salmonella enterica subsp. enterica; Salmonella enterica subsp. enterica serovar Heidelberg MNNEETFYQAMRRKGVTRRSFLKFCSLAATSLGLGAGMTPKIAWALENKPRIPVVWIHGL WP_001058312.1 hydrogenase [Salmonella enterica] Length: 372\nNumber of predicted TMHs: 2\nExp number of AAs in TMHs: 36.51738\nExp number, first 60 AAs: 19.72849\nTotal prob of N-in: 0.96170\nPOSSIBLE N-term signal sequence\ninside 1 19\nTMhelix 20 39\noutside 40 325\nTMhelix 326 348\ninside 349 372

23882 GCF_000603725.1_ASM60372v1 Salmonella enterica subsp. enterica serovar Heidelberg str. 32507 Proteobacteria; Gammaproteobacteria; Enterobacterales; Enterobacteriaceae; Salmonella; Salmonella enterica; Salmonella enterica subsp. enterica; Salmonella enterica subsp. enterica serovar Heidelberg MTGDNTLITSHGINRRDFMKLCAALAATMGLSSKAAAEMAESVSNPQRPPVIWIGAQECT WP_000145429.1 hydrogenase 2 small subunit [Salmonella enterica] Length: 372\nNumber of predicted TMHs: 1\nExp number of AAs in TMHs: 24.51309\nExp number, first 60 AAs: 0.2682\nTotal prob of N-in: 0.14402\noutside 1 331\nTMhelix 332 354\ninside 355 372

23883 GCF_000284755.1_ASM28475v1 Salmonella enterica subsp. enterica serovar Heidelberg str. 41563 Proteobacteria; Gammaproteobacteria; Enterobacterales; Enterobacteriaceae; Salmonella; Salmonella enterica;

Salmonella enterica subsp. enterica; Salmonella enterica subsp. enterica serovar Heidelberg
MNNEETFYQAMRRKGVTRRSFLKFCSLAATSLGLGAGMTPKIAWALENKPRIPVVWIHGL WP_001058312.1
hydrogenase [Salmonella enterica] Length: 372\nNumber of predicted TMHs: 2\nExp number of AAs in TMHs: 36.51738\nExp number, first 60 AAs: 19.72849\nTotal prob of N-in: 0.96170\nPOSSIBLE N-term signal sequence\ninside 1 19\nTMhelix 20 39\noutside 40 325\nTMhelix 326 348\ninside 349 372

23884 GCF_000284755.1_ASM28475v1 Salmonella enterica subsp. enterica serovar Heidelberg str. 41563
Proteobacteria; Gammaproteobacteria; Enterobacterales; Enterobacteriaceae; Salmonella; Salmonella enterica; Salmonella enterica subsp. enterica; Salmonella enterica subsp. enterica serovar Heidelberg
MTGDNTLITSHGINRRDFMKLCAALAATMGLSSKAAAEMAESVSNPQRPPVIWIGAQECT WP_000145429.1
hydrogenase 2 small subunit [Salmonella enterica] Length: 372\nNumber of predicted TMHs: 1\nExp number of AAs in TMHs: 24.51309\nExp number, first 60 AAs: 0.2682\nTotal prob of N-in: 0.14402\noutside 1 331\nTMhelix 332 354\ninside 355 372

23885 GCF_000284815.1_ASM28481v1 Salmonella enterica subsp. enterica serovar Heidelberg str. 41565
Proteobacteria; Gammaproteobacteria; Enterobacterales; Enterobacteriaceae; Salmonella; Salmonella enterica; Salmonella enterica subsp. enterica; Salmonella enterica subsp. enterica serovar Heidelberg
MNNEETFYQAMRRKGVTRRSFLKFCSLAATSLGLGAGMTPKIAWALENKPRIPVVWIHGL WP_001058312.1
hydrogenase [Salmonella enterica] Length: 372\nNumber of predicted TMHs: 2\nExp number of AAs in TMHs: 36.51738\nExp number, first 60 AAs: 19.72849\nTotal prob of N-in: 0.96170\nPOSSIBLE N-term signal sequence\ninside 1 19\nTMhelix 20 39\noutside 40 325\nTMhelix 326 348\ninside 349 372

23886 GCF_000284815.1_ASM28481v1 Salmonella enterica subsp. enterica serovar Heidelberg str. 41565
Proteobacteria; Gammaproteobacteria; Enterobacterales; Enterobacteriaceae; Salmonella; Salmonella enterica; Salmonella enterica subsp. enterica; Salmonella enterica subsp. enterica serovar Heidelberg
MTGDNTLITSHGINRRDFMKLCAALAATMGLSSKAAAEMAESVSNPQRPPVIWIGAQECT WP_000145429.1
hydrogenase 2 small subunit [Salmonella enterica] Length: 372\nNumber of predicted TMHs: 1\nExp number of AAs in TMHs: 24.51309\nExp number, first 60 AAs: 0.2682\nTotal prob of N-in: 0.14402\noutside 1 331\nTMhelix 332 354\ninside 355 372

23887 GCF_000284795.1_ASM28479v1 Salmonella enterica subsp. enterica serovar Heidelberg str. 41566
Proteobacteria; Gammaproteobacteria; Enterobacterales; Enterobacteriaceae; Salmonella; Salmonella enterica; Salmonella enterica subsp. enterica; Salmonella enterica subsp. enterica serovar Heidelberg
MTGDNTLITSHGINRRDFMKLCAALAATMGLSSKAAAEMAESVSNPQRPPVIWIGAQECT WP_000145429.1
hydrogenase 2 small subunit [Salmonella enterica] Length: 372\nNumber of predicted TMHs: 1\nExp number of AAs in TMHs: 24.51309\nExp number, first 60 AAs: 0.2682\nTotal prob of N-in: 0.14402\noutside 1 331\nTMhelix 332 354\ninside 355 372

23888 GCF_000284795.1_ASM28479v1 Salmonella enterica subsp. enterica serovar Heidelberg str. 41566
Proteobacteria; Gammaproteobacteria; Enterobacterales; Enterobacteriaceae; Salmonella; Salmonella enterica; Salmonella enterica subsp. enterica; Salmonella enterica subsp. enterica serovar Heidelberg
MNNEETFYQAMRRKGVTRRSFLKFCSLAATSLGLGAGMTPKIAWALENKPRIPVVWIHGL WP_001058312.1
hydrogenase [Salmonella enterica] Length: 372\nNumber of predicted TMHs: 2\nExp number of AAs in TMHs: 36.51738\nExp number, first 60 AAs: 19.72849\nTotal prob of N-in: 0.96170\nPOSSIBLE N-term signal sequence\ninside 1 19\nTMhelix 20 39\noutside 40 325\nTMhelix 326 348\ninside 349 372

23889 GCF_000603625.1_ASM60362v1 Salmonella enterica subsp. enterica serovar Heidelberg str. 41567
Proteobacteria; Gammaproteobacteria; Enterobacterales; Enterobacteriaceae; Salmonella; Salmonella enterica; Salmonella enterica subsp. enterica; Salmonella enterica subsp. enterica serovar Heidelberg
MNNEETFYQAMRRKGVTRRSFLKFCSLAATSLGLGAGMTPKIAWALENKPRIPVVWIHGL WP_001058312.1
hydrogenase [Salmonella enterica] Length: 372\nNumber of predicted TMHs: 2\nExp number of AAs in TMHs: 36.51738\nExp number, first 60 AAs: 19.72849\nTotal prob of N-in: 0.96170\nPOSSIBLE N-term signal sequence\ninside 1 19\nTMhelix 20 39\noutside 40 325\nTMhelix 326 348\ninside 349 372

23890 GCF_000603625.1_ASM60362v1 Salmonella enterica subsp. enterica serovar Heidelberg str. 41567
Proteobacteria; Gammaproteobacteria; Enterobacterales; Enterobacteriaceae; Salmonella; Salmonella enterica; Salmonella enterica subsp. enterica; Salmonella enterica subsp. enterica serovar Heidelberg
MTGDNTLITSHGINRRDFMKLCAALAATMGLSSKAAAEMAESVSNPQRPPVIWIGAQECT WP_000145429.1
hydrogenase 2 small subunit [Salmonella enterica] Length: 372\nNumber of predicted TMHs: 1\nExp number of AAs in TMHs: 24.51309\nExp number, first 60 AAs: 0.2682\nTotal prob of N-in: 0.14402\noutside 1 331\nTMhelix 332 354\ninside 355 372

23891 GCF_000284775.1_ASM28477v1 Salmonella enterica subsp. enterica serovar Heidelberg str. 41573 Proteobacteria; Gammaproteobacteria; Enterobacterales; Enterobacteriaceae; Salmonella; Salmonella enterica; Salmonella enterica subsp. enterica; Salmonella enterica subsp. enterica serovar Heidelberg MNNEETFYQAMRRKGVTRRSFLKFCSLAATSLGLGAGMTPKIAWALENKPRIPVVWIHGL WP_001058312.1 hydrogenase [Salmonella enterica] Length: 372\nNumber of predicted TMHs: 2\nExp number of AAs in TMHs: 36.51738\nExp number, first 60 AAs: 19.72849\nTotal prob of N-in: 0.96170\nPOSSIBLE N-term signal sequence\ninside 1 19\nTMhelix 20 39\noutside 40 325\nTMhelix 326 348\ninside 349 372

23892 GCF_000284775.1_ASM28477v1 Salmonella enterica subsp. enterica serovar Heidelberg str. 41573 Proteobacteria; Gammaproteobacteria; Enterobacterales; Enterobacteriaceae; Salmonella; Salmonella enterica; Salmonella enterica subsp. enterica; Salmonella enterica subsp. enterica serovar Heidelberg MTGDNTLITSHGINRRDFMKLCAALAATMGLSSKAAAEMAESVSNPQRPPVIWIGAQECT WP_000145429.1 hydrogenase 2 small subunit [Salmonella enterica] Length: 372\nNumber of predicted TMHs: 1\nExp number of AAs in TMHs: 24.51309\nExp number, first 60 AAs: 0.2682\nTotal prob of N-in: 0.14402\noutside 1 331\nTMhelix 332 354\ninside 355 372

23893 GCF_000430105.1_ASM43010v1 Salmonella enterica subsp. enterica serovar Heidelberg str. 41578 Proteobacteria; Gammaproteobacteria; Enterobacterales; Enterobacteriaceae; Salmonella; Salmonella enterica; Salmonella enterica subsp. enterica; Salmonella enterica subsp. enterica serovar Heidelberg MTGDNTLITSHGINRRDFMKLCAALAATMGLSSKAAAEMAESVSNPQRPPVIWIGAQECT WP_000145429.1 hydrogenase 2 small subunit [Salmonella enterica] Length: 372\nNumber of predicted TMHs: 1\nExp number of AAs in TMHs: 24.51309\nExp number, first 60 AAs: 0.2682\nTotal prob of N-in: 0.14402\noutside 1 331\nTMhelix 332 354\ninside 355 372

23894 GCF_000430105.1_ASM43010v1 Salmonella enterica subsp. enterica serovar Heidelberg str. 41578 Proteobacteria; Gammaproteobacteria; Enterobacterales; Enterobacteriaceae; Salmonella; Salmonella enterica; Salmonella enterica subsp. enterica; Salmonella enterica subsp. enterica serovar Heidelberg MNNEETFYQAMRRKGVTRRSFLKFCSLAATSLGLGAGMTPKIAWALENKPRIPVVWIHGL WP_001058312.1 hydrogenase [Salmonella enterica] Length: 372\nNumber of predicted TMHs: 2\nExp number of AAs in TMHs: 36.51738\nExp number, first 60 AAs: 19.72849\nTotal prob of N-in: 0.96170\nPOSSIBLE N-term signal sequence\ninside 1 19\nTMhelix 20 39\noutside 40 325\nTMhelix 326 348\ninside 349 372

23895 GCF_000256625.1_ASM25662v1 Salmonella enterica subsp. enterica serovar Heidelberg str. 41579 Proteobacteria; Gammaproteobacteria; Enterobacterales; Enterobacteriaceae; Salmonella; Salmonella enterica; Salmonella enterica subsp. enterica; Salmonella enterica subsp. enterica serovar Heidelberg MTGDNTLITSHGINRRDFMKLCAALAATMGLSSKAAAEMAESVSNPQRPPVIWIGAQECT WP_000145429.1 hydrogenase 2 small subunit [Salmonella enterica] Length: 372\nNumber of predicted TMHs: 1\nExp number of AAs in TMHs: 24.51309\nExp number, first 60 AAs: 0.2682\nTotal prob of N-in: 0.14402\noutside 1 331\nTMhelix 332 354\ninside 355 372

23896 GCF_000256625.1_ASM25662v1 Salmonella enterica subsp. enterica serovar Heidelberg str. 41579 Proteobacteria; Gammaproteobacteria; Enterobacterales; Enterobacteriaceae; Salmonella; Salmonella enterica; Salmonella enterica subsp. enterica; Salmonella enterica subsp. enterica serovar Heidelberg MNNEETFYQAMRRKGVTRRSFLKFCSLAATSLGLGAGMTPKIAWALENKPRIPVVWIHGL WP_001058312.1 hydrogenase [Salmonella enterica] Length: 372\nNumber of predicted TMHs: 2\nExp number of AAs in TMHs: 36.51738\nExp number, first 60 AAs: 19.72849\nTotal prob of N-in: 0.96170\nPOSSIBLE N-term signal sequence\ninside 1 19\nTMhelix 20 39\noutside 40 325\nTMhelix 326 348\ninside 349 372

23897 GCF_000603665.1_ASM60366v1 Salmonella enterica subsp. enterica serovar Heidelberg str. 41584 Proteobacteria; Gammaproteobacteria; Enterobacterales; Enterobacteriaceae; Salmonella; Salmonella enterica; Salmonella enterica subsp. enterica; Salmonella enterica subsp. enterica serovar Heidelberg MTGDNTLITSHGINRRDFMKLCAALAATMGLSSKAAAEMAESVSNPQRPPVIWIGAQECT WP_000145429.1 hydrogenase 2 small subunit [Salmonella enterica] Length: 372\nNumber of predicted TMHs: 1\nExp number of AAs in TMHs: 24.51309\nExp number, first 60 AAs: 0.2682\nTotal prob of N-in: 0.14402\noutside 1 331\nTMhelix 332 354\ninside 355 372

23898 GCF_000603665.1_ASM60366v1 Salmonella enterica subsp. enterica serovar Heidelberg str. 41584 Proteobacteria; Gammaproteobacteria; Enterobacterales; Enterobacteriaceae; Salmonella; Salmonella enterica; Salmonella enterica subsp. enterica; Salmonella enterica subsp. enterica serovar Heidelberg MNNEETFYQAMRRKGVTRRSFLKFCSLAATSLGLGAGMTPKIAWALENKPRIPVVWIHGL WP_001058312.1 hydrogenase [Salmonella enterica] Length: 372\nNumber of predicted TMHs: 2\nExp number of AAs in

TMHs: 36.51738\nExp number, first 60 AAs: 19.72849\nTotal prob of N-in: 0.96170\nPOSSIBLE N-term signal sequence\ninside 1 19\nTMhelix 20 39\noutside 40 325\nTMhelix 326 348\ninside 349 372

23899 GCF_000962315.1_ASM96231v1 Salmonella enterica subsp. enterica serovar Heidelberg str. 579083-10 Proteobacteria; Gammaproteobacteria; Enterobacterales; Enterobacteriaceae; Salmonella; Salmonella enterica; Salmonella enterica subsp. enterica; Salmonella enterica subsp. enterica serovar Heidelberg MTGDNTLITSHGINRRDFMKLCAALAATMGLSSKAAAEMAESVSNPQRPPVIWIGAQECT WP_000145429.1 hydrogenase 2 small subunit [Salmonella enterica] Length: 372\nNumber of predicted TMHs: 1\nExp number of AAs in TMHs: 24.51309\nExp number, first 60 AAs: 0.2682\nTotal prob of N-in: 0.14402\noutside 1 331\nTMhelix 332 354\ninside 355 372

23900 GCF_000962315.1_ASM96231v1 Salmonella enterica subsp. enterica serovar Heidelberg str. 579083-10 Proteobacteria; Gammaproteobacteria; Enterobacterales; Enterobacteriaceae; Salmonella; Salmonella enterica; Salmonella enterica subsp. enterica; Salmonella enterica subsp. enterica serovar Heidelberg MNNEETFYQAMRRKGVTRRSFLKFCSLAATSLGLGAGMTPKIAWALENKPRIPVVWIHGL WP_001058312.1 hydrogenase [Salmonella enterica] Length: 372\nNumber of predicted TMHs: 2\nExp number of AAs in TMHs: 36.51738\nExp number, first 60 AAs: 19.72849\nTotal prob of N-in: 0.96170\nPOSSIBLE N-term signal sequence\ninside 1 19\nTMhelix 20 39\noutside 40 325\nTMhelix 326 348\ninside 349 372

23901 GCF_000962075.1_ASM96207v1 Salmonella enterica subsp. enterica serovar Heidelberg str. 607309-34 Proteobacteria; Gammaproteobacteria; Enterobacterales; Enterobacteriaceae; Salmonella; Salmonella enterica; Salmonella enterica subsp. enterica; Salmonella enterica subsp. enterica serovar Heidelberg MNNEETFYQAMRRKGVTRRSFLKFCSLAATSLGLGAGMTPKIAWALENKPRIPVVWIHGL WP_001058312.1 hydrogenase [Salmonella enterica] Length: 372\nNumber of predicted TMHs: 2\nExp number of AAs in TMHs: 36.51738\nExp number, first 60 AAs: 19.72849\nTotal prob of N-in: 0.96170\nPOSSIBLE N-term signal sequence\ninside 1 19\nTMhelix 20 39\noutside 40 325\nTMhelix 326 348\ninside 349 372

23902 GCF_000962075.1_ASM96207v1 Salmonella enterica subsp. enterica serovar Heidelberg str. 607309-34 Proteobacteria; Gammaproteobacteria; Enterobacterales; Enterobacteriaceae; Salmonella; Salmonella enterica; Salmonella enterica subsp. enterica; Salmonella enterica subsp. enterica serovar Heidelberg MTGDNTLITSHGINRRDFMKLCAALAATMGLSSKAAAEMAESVSNPQRPPVIWIGAQECT WP_000145429.1 hydrogenase 2 small subunit [Salmonella enterica] Length: 372\nNumber of predicted TMHs: 1\nExp number of AAs in TMHs: 24.51309\nExp number, first 60 AAs: 0.2682\nTotal prob of N-in: 0.14402\noutside 1 331\nTMhelix 332 354\ninside 355 372

23903 GCF_000962025.1_ASM96202v1 Salmonella enterica subsp. enterica serovar Heidelberg str. 607309-5 Proteobacteria; Gammaproteobacteria; Enterobacterales; Enterobacteriaceae; Salmonella; Salmonella enterica; Salmonella enterica subsp. enterica; Salmonella enterica subsp. enterica serovar Heidelberg MTGDNTLITSHGINRRDFMKLCAALAATMGLSSKAAAEMAESVSNPQRPPVIWIGAQECT WP_000145429.1 hydrogenase 2 small subunit [Salmonella enterica] Length: 372\nNumber of predicted TMHs: 1\nExp number of AAs in TMHs: 24.51309\nExp number, first 60 AAs: 0.2682\nTotal prob of N-in: 0.14402\noutside 1 331\nTMhelix 332 354\ninside 355 372

23904 GCF_000962025.1_ASM96202v1 Salmonella enterica subsp. enterica serovar Heidelberg str. 607309-5 Proteobacteria; Gammaproteobacteria; Enterobacterales; Enterobacteriaceae; Salmonella; Salmonella enterica; Salmonella enterica subsp. enterica; Salmonella enterica subsp. enterica serovar Heidelberg MNNEETFYQAMRRKGVTRRSFLKFCSLAATSLGLGAGMTPKIAWALENKPRIPVVWIHGL WP_045720686.1 hydrogenase, partial [Salmonella enterica] Length: 370\nNumber of predicted TMHs: 2\nExp number of AAs in TMHs: 36.54535\nExp number, first 60 AAs: 19.72794\nTotal prob of N-in: 0.96170\nPOSSIBLE N-term signal sequence\ninside 1 19\nTMhelix 20 39\noutside 40 325\nTMhelix 326 348\ninside 349 370

23905 GCF_000962295.1_ASM96229v1 Salmonella enterica subsp. enterica serovar Heidelberg str. 622737-21 Proteobacteria; Gammaproteobacteria; Enterobacterales; Enterobacteriaceae; Salmonella; Salmonella enterica; Salmonella enterica subsp. enterica; Salmonella enterica subsp. enterica serovar Heidelberg MTGDNTLITSHGINRRDFMKLCAALAATMGLSSKAAAEMAESVSNPQRPPVIWIGAQECT WP_000145429.1 hydrogenase 2 small subunit [Salmonella enterica] Length: 372\nNumber of predicted TMHs: 1\nExp number of AAs in TMHs: 24.51309\nExp number, first 60 AAs: 0.2682\nTotal prob of N-in: 0.14402\noutside 1 331\nTMhelix 332 354\ninside 355 372

23906 GCF_000962085.1_ASM96208v1 Salmonella enterica subsp. enterica serovar Heidelberg str. 632675-24 Proteobacteria; Gammaproteobacteria; Enterobacterales; Enterobacteriaceae; Salmonella; Salmonella enterica;

Salmonella enterica subsp. enterica; Salmonella enterica subsp. enterica serovar Heidelberg
MNNEETFYQAMRRKGVTRRSFLKFCSLAATSLGLGAGMTPKIAWALENKPRIPVVWIHGL WP_001058312.1
hydrogenase [Salmonella enterica] Length: 372\nNumber of predicted TMHs: 2\nExp number of AAs in TMHs: 36.51738\nExp number, first 60 AAs: 19.72849\nTotal prob of N-in: 0.96170\nPOSSIBLE N-term signal sequence\ninside 1 19\nTMhelix 20 39\noutside 40 325\nTMhelix 326 348\ninside 349 372

23907 GCF_000962085.1_ASM96208v1 Salmonella enterica subsp. enterica serovar Heidelberg str. 632675-24
Proteobacteria; Gammaproteobacteria; Enterobacterales; Enterobacteriaceae; Salmonella; Salmonella enterica; Salmonella enterica subsp. enterica; Salmonella enterica subsp. enterica serovar Heidelberg
MTGDNTLITSHGINRRDFMKLCAALAATMGLSSKAAAEMAESVSNPQRPPVIWIGAQECT WP_000145429.1
hydrogenase 2 small subunit [Salmonella enterica] Length: 372\nNumber of predicted TMHs: 1\nExp number of AAs in TMHs: 24.51309\nExp number, first 60 AAs: 0.2682\nTotal prob of N-in: 0.14402\noutside 1 331\nTMhelix 332 354\ninside 355 372

23908 GCF_000700785.1_SEEH5111_1.0 Salmonella enterica subsp. enterica serovar Heidelberg str. 640151-11
Proteobacteria; Gammaproteobacteria; Enterobacterales; Enterobacteriaceae; Salmonella; Salmonella enterica; Salmonella enterica subsp. enterica; Salmonella enterica subsp. enterica serovar Heidelberg
MTGDNTLITSHGINRRDFMKLCAALAATMGLSSKAAAEMAESVSNPQRPPVIWIGAQECT WP_000145429.1
hydrogenase 2 small subunit [Salmonella enterica] Length: 372\nNumber of predicted TMHs: 1\nExp number of AAs in TMHs: 24.51309\nExp number, first 60 AAs: 0.2682\nTotal prob of N-in: 0.14402\noutside 1 331\nTMhelix 332 354\ninside 355 372

23909 GCF_000700785.1_SEEH5111_1.0 Salmonella enterica subsp. enterica serovar Heidelberg str. 640151-11
Proteobacteria; Gammaproteobacteria; Enterobacterales; Enterobacteriaceae; Salmonella; Salmonella enterica; Salmonella enterica subsp. enterica; Salmonella enterica subsp. enterica serovar Heidelberg
MNNEETFYQAMRRKGVTRRSFLKFCSLAATSLGLGAGMTPKIAWALENKPRIPVVWIHGL WP_001058312.1
hydrogenase [Salmonella enterica] Length: 372\nNumber of predicted TMHs: 2\nExp number of AAs in TMHs: 36.51738\nExp number, first 60 AAs: 19.72849\nTotal prob of N-in: 0.96170\nPOSSIBLE N-term signal sequence\ninside 1 19\nTMhelix 20 39\noutside 40 325\nTMhelix 326 348\ninside 349 372

23910 GCF_000962355.1_ASM96235v1 Salmonella enterica subsp. enterica serovar Heidelberg str. 670102-3
Proteobacteria; Gammaproteobacteria; Enterobacterales; Enterobacteriaceae; Salmonella; Salmonella enterica; Salmonella enterica subsp. enterica; Salmonella enterica subsp. enterica serovar Heidelberg
MTGDNTLITSHGINRRDFMKLCAALAATMGLSSKAAAEMAESVSNPQRPPVIWIGAQECT WP_000145429.1
hydrogenase 2 small subunit [Salmonella enterica] Length: 372\nNumber of predicted TMHs: 1\nExp number of AAs in TMHs: 24.51309\nExp number, first 60 AAs: 0.2682\nTotal prob of N-in: 0.14402\noutside 1 331\nTMhelix 332 354\ninside 355 372

23911 GCF_000962355.1_ASM96235v1 Salmonella enterica subsp. enterica serovar Heidelberg str. 670102-3
Proteobacteria; Gammaproteobacteria; Enterobacterales; Enterobacteriaceae; Salmonella; Salmonella enterica; Salmonella enterica subsp. enterica; Salmonella enterica subsp. enterica serovar Heidelberg
MNNEETFYQAMRRKGVTRRSFLKFCSLAATSLGLGAGMTPKIAWALENKPRIPVVWIHGL WP_001058312.1
hydrogenase [Salmonella enterica] Length: 372\nNumber of predicted TMHs: 2\nExp number of AAs in TMHs: 36.51738\nExp number, first 60 AAs: 19.72849\nTotal prob of N-in: 0.96170\nPOSSIBLE N-term signal sequence\ninside 1 19\nTMhelix 20 39\noutside 40 325\nTMhelix 326 348\ninside 349 372

23912 GCF_000962115.1_ASM96211v1 Salmonella enterica subsp. enterica serovar Heidelberg str. 670102-5
Proteobacteria; Gammaproteobacteria; Enterobacterales; Enterobacteriaceae; Salmonella; Salmonella enterica; Salmonella enterica subsp. enterica; Salmonella enterica subsp. enterica serovar Heidelberg
MNNEETFYQAMRRKGVTRRSFLKFCSLAATSLGLGAGMTPKIAWALENKPRIPVVWIHGL WP_001058312.1
hydrogenase [Salmonella enterica] Length: 372\nNumber of predicted TMHs: 2\nExp number of AAs in TMHs: 36.51738\nExp number, first 60 AAs: 19.72849\nTotal prob of N-in: 0.96170\nPOSSIBLE N-term signal sequence\ninside 1 19\nTMhelix 20 39\noutside 40 325\nTMhelix 326 348\ninside 349 372

23913 GCF_000962115.1_ASM96211v1 Salmonella enterica subsp. enterica serovar Heidelberg str. 670102-5
Proteobacteria; Gammaproteobacteria; Enterobacterales; Enterobacteriaceae; Salmonella; Salmonella enterica; Salmonella enterica subsp. enterica; Salmonella enterica subsp. enterica serovar Heidelberg
MTGDNTLITSHGINRRDFMKLCAALAATMGLSSKAAAEMAESVSNPQRPPVIWIGAQECT WP_000145429.1
hydrogenase 2 small subunit [Salmonella enterica] Length: 372\nNumber of predicted TMHs: 1\nExp number of AAs in TMHs: 24.51309\nExp number, first 60 AAs: 0.2682\nTotal prob of N-in: 0.14402\noutside 1 331\nTMhelix 332 354\ninside 355 372

23914 GCF_000962605.1_ASM96260v1 Salmonella enterica subsp. enterica serovar Heidelberg str. 82-1617 Proteobacteria; Gammaproteobacteria; Enterobacterales; Enterobacteriaceae; Salmonella; Salmonella enterica; Salmonella enterica subsp. enterica; Salmonella enterica subsp. enterica serovar Heidelberg MNNEETFYQAMRRKGVTRRSFLKFCSLAATSLGLGAGMTPKIAWALENKPRIPVVWIHGL WP_001058312.1 hydrogenase [Salmonella enterica] Length: 372\nNumber of predicted TMHs: 2\nExp number of AAs in TMHs: 36.51738\nExp number, first 60 AAs: 19.72849\nTotal prob of N-in: 0.96170\nPOSSIBLE N-term signal sequence\ninside 1 19\nTMhelix 20 39\noutside 40 325\nTMhelix 326 348\ninside 349 372

23915 GCF_000962605.1_ASM96260v1 Salmonella enterica subsp. enterica serovar Heidelberg str. 82-1617 Proteobacteria; Gammaproteobacteria; Enterobacterales; Enterobacteriaceae; Salmonella; Salmonella enterica; Salmonella enterica subsp. enterica; Salmonella enterica subsp. enterica serovar Heidelberg MTGDNTLITSHGINRRDFMKLCAALAATMGLSSKAAAEMAESVSNPQRPPVIWIGAQECT WP_000145429.1 hydrogenase 2 small subunit [Salmonella enterica] Length: 372\nNumber of predicted TMHs: 1\nExp number of AAs in TMHs: 24.51309\nExp number, first 60 AAs: 0.2682\nTotal prob of N-in: 0.14402\noutside 1 331\nTMhelix 332 354\ninside 355 372

23916 GCF_000486065.1_ASM48606v1 Salmonella enterica subsp. enterica serovar Heidelberg str. 82-2052 Proteobacteria; Gammaproteobacteria; Enterobacterales; Enterobacteriaceae; Salmonella; Salmonella enterica; Salmonella enterica subsp. enterica; Salmonella enterica subsp. enterica serovar Heidelberg MNNEETFYQAMRRKGVTRRSFLKFCSLAATSLGLGAGMTPKIAWALENKPRIPVVWIHGL WP_001058312.1 hydrogenase [Salmonella enterica] Length: 372\nNumber of predicted TMHs: 2\nExp number of AAs in TMHs: 36.51738\nExp number, first 60 AAs: 19.72849\nTotal prob of N-in: 0.96170\nPOSSIBLE N-term signal sequence\ninside 1 19\nTMhelix 20 39\noutside 40 325\nTMhelix 326 348\ninside 349 372

23917 GCF_000962445.1_ASM96244v1 Salmonella enterica subsp. enterica serovar Heidelberg str. 83-1068 Proteobacteria; Gammaproteobacteria; Enterobacterales; Enterobacteriaceae; Salmonella; Salmonella enterica; Salmonella enterica subsp. enterica; Salmonella enterica subsp. enterica serovar Heidelberg MTGDNTLITSHGINRRDFMKLCAALAATMGLSSKAAAEMAESVSNPQRPPVIWIGAQECT WP_000145429.1 hydrogenase 2 small subunit [Salmonella enterica] Length: 372\nNumber of predicted TMHs: 1\nExp number of AAs in TMHs: 24.51309\nExp number, first 60 AAs: 0.2682\nTotal prob of N-in: 0.14402\noutside 1 331\nTMhelix 332 354\ninside 355 372

23918 GCF_000962445.1_ASM96244v1 Salmonella enterica subsp. enterica serovar Heidelberg str. 83-1068 Proteobacteria; Gammaproteobacteria; Enterobacterales; Enterobacteriaceae; Salmonella; Salmonella enterica; Salmonella enterica subsp. enterica; Salmonella enterica subsp. enterica serovar Heidelberg MNNEETFYQAMRRKGVTRRSFLKFCSLAATSLGLGAGMTPKIAWALENKPRIPVVWIHGL WP_001058312.1 hydrogenase [Salmonella enterica] Length: 372\nNumber of predicted TMHs: 2\nExp number of AAs in TMHs: 36.51738\nExp number, first 60 AAs: 19.72849\nTotal prob of N-in: 0.96170\nPOSSIBLE N-term signal sequence\ninside 1 19\nTMhelix 20 39\noutside 40 325\nTMhelix 326 348\ninside 349 372

23919 GCF_000962625.1_ASM96262v1 Salmonella enterica subsp. enterica serovar Heidelberg str. 84-1004 Proteobacteria; Gammaproteobacteria; Enterobacterales; Enterobacteriaceae; Salmonella; Salmonella enterica; Salmonella enterica subsp. enterica; Salmonella enterica subsp. enterica serovar Heidelberg MTGDNTLITSHGINRRDFMKLCAALAATMGLSSKAAAEMAESVSNPQRPPVIWIGAQECT WP_000145429.1 hydrogenase 2 small subunit [Salmonella enterica] Length: 372\nNumber of predicted TMHs: 1\nExp number of AAs in TMHs: 24.51309\nExp number, first 60 AAs: 0.2682\nTotal prob of N-in: 0.14402\noutside 1 331\nTMhelix 332 354\ninside 355 372

23920 GCF_000962625.1_ASM96262v1 Salmonella enterica subsp. enterica serovar Heidelberg str. 84-1004 Proteobacteria; Gammaproteobacteria; Enterobacterales; Enterobacteriaceae; Salmonella; Salmonella enterica; Salmonella enterica subsp. enterica; Salmonella enterica subsp. enterica serovar Heidelberg MNNEETFYQAMRRKGVTRRSFLKFCSLAATSLGLGAGMTPKIAWALENKPRIPVVWIHGL WP_001058312.1 hydrogenase [Salmonella enterica] Length: 372\nNumber of predicted TMHs: 2\nExp number of AAs in TMHs: 36.51738\nExp number, first 60 AAs: 19.72849\nTotal prob of N-in: 0.96170\nPOSSIBLE N-term signal sequence\ninside 1 19\nTMhelix 20 39\noutside 40 325\nTMhelix 326 348\ninside 349 372

23921 GCF_000962655.1_ASM96265v1 Salmonella enterica subsp. enterica serovar Heidelberg str. 85-0486 Proteobacteria; Gammaproteobacteria; Enterobacterales; Enterobacteriaceae; Salmonella; Salmonella enterica; Salmonella enterica subsp. enterica; Salmonella enterica subsp. enterica serovar Heidelberg MNNEETFYQAMRRKGVTRRSFLKFCSLAATSLGLGAGMTPKIAWALENKPRIPVVWIHGL WP_001058312.1 hydrogenase [Salmonella enterica] Length: 372\nNumber of predicted TMHs: 2\nExp number of AAs in

TMHs: 36.51738\nExp number, first 60 AAs: 19.72849\nTotal prob of N-in: 0.96170\nPOSSIBLE N-term signal sequence\ninside 1 19\nTMhelix 20 39\noutside 40 325\nTMhelix 326 348\ninside 349 372

23922 GCF_000962655.1_ASM96265v1 Salmonella enterica subsp. enterica serovar Heidelberg str. 85-0486 Proteobacteria; Gammaproteobacteria; Enterobacterales; Enterobacteriaceae; Salmonella; Salmonella enterica; Salmonella enterica subsp. enterica; Salmonella enterica subsp. enterica serovar Heidelberg MTGDNTLITSHGINRRDFMKLCAALAATMGLSSKAAAEMAESVSNPQRPPVIWIGAQECT WP_000145429.1 hydrogenase 2 small subunit [Salmonella enterica] Length: 372\nNumber of predicted TMHs: 1\nExp number of AAs in TMHs: 24.51309\nExp number, first 60 AAs: 0.2682\nTotal prob of N-in: 0.14402\noutside 1 331\nTMhelix 332 354\ninside 355 372

23923 GCF_000962475.1_ASM96247v1 Salmonella enterica subsp. enterica serovar Heidelberg str. 86-0255 Proteobacteria; Gammaproteobacteria; Enterobacterales; Enterobacteriaceae; Salmonella; Salmonella enterica; Salmonella enterica subsp. enterica; Salmonella enterica subsp. enterica serovar Heidelberg MTGDNTLITSHGINRRDFMKLCAALAATMGLSSKAAAEMAESVSNPQRPPVIWIGAQECT WP_000145429.1 hydrogenase 2 small subunit [Salmonella enterica] Length: 372\nNumber of predicted TMHs: 1\nExp number of AAs in TMHs: 24.51309\nExp number, first 60 AAs: 0.2682\nTotal prob of N-in: 0.14402\noutside 1 331\nTMhelix 332 354\ninside 355 372

23924 GCF_000962475.1_ASM96247v1 Salmonella enterica subsp. enterica serovar Heidelberg str. 86-0255 Proteobacteria; Gammaproteobacteria; Enterobacterales; Enterobacteriaceae; Salmonella; Salmonella enterica; Salmonella enterica subsp. enterica; Salmonella enterica subsp. enterica serovar Heidelberg MNNEETFYQAMRRKGVTRRSFLKFCSLAATSLGLGAGMTPKIAWALENKPRIPVVWIHGL WP_001058312.1 hydrogenase [Salmonella enterica] Length: 372\nNumber of predicted TMHs: 2\nExp number of AAs in TMHs: 36.51738\nExp number, first 60 AAs: 19.72849\nTotal prob of N-in: 0.96170\nPOSSIBLE N-term signal sequence\ninside 1 19\nTMhelix 20 39\noutside 40 325\nTMhelix 326 348\ninside 349 372

23925 GCF_000962485.1_ASM96248v1 Salmonella enterica subsp. enterica serovar Heidelberg str. 87-0208 Proteobacteria; Gammaproteobacteria; Enterobacterales; Enterobacteriaceae; Salmonella; Salmonella enterica; Salmonella enterica subsp. enterica; Salmonella enterica subsp. enterica serovar Heidelberg MNNEETFYQAMRRKGVTRRSFLKFCSLAATSLGLGAGMTPKIAWALENKPRIPVVWIHGL WP_001058312.1 hydrogenase [Salmonella enterica] Length: 372\nNumber of predicted TMHs: 2\nExp number of AAs in TMHs: 36.51738\nExp number, first 60 AAs: 19.72849\nTotal prob of N-in: 0.96170\nPOSSIBLE N-term signal sequence\ninside 1 19\nTMhelix 20 39\noutside 40 325\nTMhelix 326 348\ninside 349 372

23926 GCF_000962485.1_ASM96248v1 Salmonella enterica subsp. enterica serovar Heidelberg str. 87-0208 Proteobacteria; Gammaproteobacteria; Enterobacterales; Enterobacteriaceae; Salmonella; Salmonella enterica; Salmonella enterica subsp. enterica; Salmonella enterica subsp. enterica serovar Heidelberg MTGDNTLITSHGINRRDFMKLCAALAATMGLSSKAAAEMAESVSNPQRPPVIWIGAQECT WP_000145429.1 hydrogenase 2 small subunit [Salmonella enterica] Length: 372\nNumber of predicted TMHs: 1\nExp number of AAs in TMHs: 24.51309\nExp number, first 60 AAs: 0.2682\nTotal prob of N-in: 0.14402\noutside 1 331\nTMhelix 332 354\ninside 355 372

23927 GCF_000962515.1_ASM96251v1 Salmonella enterica subsp. enterica serovar Heidelberg str. 88-0312 Proteobacteria; Gammaproteobacteria; Enterobacterales; Enterobacteriaceae; Salmonella; Salmonella enterica; Salmonella enterica subsp. enterica; Salmonella enterica subsp. enterica serovar Heidelberg MTGDNTLITSHGINRRDFMKLCAALAATMGLSSKAAAEMAESVSNPQRPPVIWIGAQECT WP_000145429.1 hydrogenase 2 small subunit [Salmonella enterica] Length: 372\nNumber of predicted TMHs: 1\nExp number of AAs in TMHs: 24.51309\nExp number, first 60 AAs: 0.2682\nTotal prob of N-in: 0.14402\noutside 1 331\nTMhelix 332 354\ninside 355 372

23928 GCF_000962515.1_ASM96251v1 Salmonella enterica subsp. enterica serovar Heidelberg str. 88-0312 Proteobacteria; Gammaproteobacteria; Enterobacterales; Enterobacteriaceae; Salmonella; Salmonella enterica; Salmonella enterica subsp. enterica; Salmonella enterica subsp. enterica serovar Heidelberg MNNEETFYQAMRRKGVTRRSFLKFCSLAATSLGLGAGMTPKIAWALENKPRIPVVWIHGL WP_001058312.1 hydrogenase [Salmonella enterica] Length: 372\nNumber of predicted TMHs: 2\nExp number of AAs in TMHs: 36.51738\nExp number, first 60 AAs: 19.72849\nTotal prob of N-in: 0.96170\nPOSSIBLE N-term signal sequence\ninside 1 19\nTMhelix 20 39\noutside 40 325\nTMhelix 326 348\ninside 349 372

23929 GCF_000962525.1_ASM96252v1 Salmonella enterica subsp. enterica serovar Heidelberg str. 89-0213 Proteobacteria; Gammaproteobacteria; Enterobacterales; Enterobacteriaceae; Salmonella; Salmonella enterica;

Salmonella enterica subsp. enterica; Salmonella enterica subsp. enterica serovar Heidelberg
MNNEETFYQAMRRKGVTRRSFLKFCSLAATSLGLGAGMTPKIAWALENKPRIPVVWIHGL WP_001058312.1
hydrogenase [Salmonella enterica] Length: 372\nNumber of predicted TMHs: 2\nExp number of AAs in TMHs: 36.51738\nExp number, first 60 AAs: 19.72849\nTotal prob of N-in: 0.96170\nPOSSIBLE N-term signal sequence\ninside 1 19\nTMhelix 20 39\noutside 40 325\nTMhelix 326 348\ninside 349 372

23930 GCF_000962525.1_ASM96252v1 Salmonella enterica subsp. enterica serovar Heidelberg str. 89-0213
Proteobacteria; Gammaproteobacteria; Enterobacterales; Enterobacteriaceae; Salmonella; Salmonella enterica; Salmonella enterica subsp. enterica; Salmonella enterica subsp. enterica serovar Heidelberg
MTGDNTLITSHGINRRDFMKLCAALAATMGLSSKAAAEMAESVSNPQRPPVIWIGAQECT WP_000145429.1
hydrogenase 2 small subunit [Salmonella enterica] Length: 372\nNumber of predicted TMHs: 1\nExp number of AAs in TMHs: 24.51309\nExp number, first 60 AAs: 0.2682\nTotal prob of N-in: 0.14402\noutside 1 331\nTMhelix 332 354\ninside 355 372

23931 GCF_000962675.1_ASM96267v1 Salmonella enterica subsp. enterica serovar Heidelberg str. 90-0318
Proteobacteria; Gammaproteobacteria; Enterobacterales; Enterobacteriaceae; Salmonella; Salmonella enterica; Salmonella enterica subsp. enterica; Salmonella enterica subsp. enterica serovar Heidelberg
MTGDNTLITSHGINRRDFMKLCAALAATMGLSSKAAAEMAESVSNPQRPPVIWIGAQECT WP_000145429.1
hydrogenase 2 small subunit [Salmonella enterica] Length: 372\nNumber of predicted TMHs: 1\nExp number of AAs in TMHs: 24.51309\nExp number, first 60 AAs: 0.2682\nTotal prob of N-in: 0.14402\noutside 1 331\nTMhelix 332 354\ninside 355 372

23932 GCF_000962675.1_ASM96267v1 Salmonella enterica subsp. enterica serovar Heidelberg str. 90-0318
Proteobacteria; Gammaproteobacteria; Enterobacterales; Enterobacteriaceae; Salmonella; Salmonella enterica; Salmonella enterica subsp. enterica; Salmonella enterica subsp. enterica serovar Heidelberg
MNNEETFYQAMRRKGVTRRSFLKFCSLAATSLGLGAGMTPKIAWALENKPRIPVVWIHGL WP_001058312.1
hydrogenase [Salmonella enterica] Length: 372\nNumber of predicted TMHs: 2\nExp number of AAs in TMHs: 36.51738\nExp number, first 60 AAs: 19.72849\nTotal prob of N-in: 0.96170\nPOSSIBLE N-term signal sequence\ninside 1 19\nTMhelix 20 39\noutside 40 325\nTMhelix 326 348\ninside 349 372

23933 GCF_000962565.1_ASM96256v1 Salmonella enterica subsp. enterica serovar Heidelberg str. 92-0138
Proteobacteria; Gammaproteobacteria; Enterobacterales; Enterobacteriaceae; Salmonella; Salmonella enterica; Salmonella enterica subsp. enterica; Salmonella enterica subsp. enterica serovar Heidelberg
MNNEETFYQAMRRKGVTRRSFLKFCSLAATSLGLGAGMTPKIAWALENKPRIPVVWIHGL WP_001058312.1
hydrogenase [Salmonella enterica] Length: 372\nNumber of predicted TMHs: 2\nExp number of AAs in TMHs: 36.51738\nExp number, first 60 AAs: 19.72849\nTotal prob of N-in: 0.96170\nPOSSIBLE N-term signal sequence\ninside 1 19\nTMhelix 20 39\noutside 40 325\nTMhelix 326 348\ninside 349 372

23934 GCF_000962565.1_ASM96256v1 Salmonella enterica subsp. enterica serovar Heidelberg str. 92-0138
Proteobacteria; Gammaproteobacteria; Enterobacterales; Enterobacteriaceae; Salmonella; Salmonella enterica; Salmonella enterica subsp. enterica; Salmonella enterica subsp. enterica serovar Heidelberg
MTGDNTLITSHGINRRDFMKLCAALAATMGLSSKAAAEMAESVSNPQRPPVIWIGAQECT WP_000145429.1
hydrogenase 2 small subunit [Salmonella enterica] Length: 372\nNumber of predicted TMHs: 1\nExp number of AAs in TMHs: 24.51309\nExp number, first 60 AAs: 0.2682\nTotal prob of N-in: 0.14402\noutside 1 331\nTMhelix 332 354\ninside 355 372

23935 GCF_000962685.1_ASM96268v1 Salmonella enterica subsp. enterica serovar Heidelberg str. 92-0144
Proteobacteria; Gammaproteobacteria; Enterobacterales; Enterobacteriaceae; Salmonella; Salmonella enterica; Salmonella enterica subsp. enterica; Salmonella enterica subsp. enterica serovar Heidelberg
MNNEETFYQAMRRKGVTRRSFLKFCSLAATSLGLGAGMTPKIAWALENKPRIPVVWIHGL WP_001058312.1
hydrogenase [Salmonella enterica] Length: 372\nNumber of predicted TMHs: 2\nExp number of AAs in TMHs: 36.51738\nExp number, first 60 AAs: 19.72849\nTotal prob of N-in: 0.96170\nPOSSIBLE N-term signal sequence\ninside 1 19\nTMhelix 20 39\noutside 40 325\nTMhelix 326 348\ninside 349 372

23936 GCF_000962685.1_ASM96268v1 Salmonella enterica subsp. enterica serovar Heidelberg str. 92-0144
Proteobacteria; Gammaproteobacteria; Enterobacterales; Enterobacteriaceae; Salmonella; Salmonella enterica; Salmonella enterica subsp. enterica; Salmonella enterica subsp. enterica serovar Heidelberg
MTGDNTLITSHGINRRDFMKLCAALAATMGLSSKAAAEMAESVSNPQRPPVIWIGAQECT WP_000145429.1
hydrogenase 2 small subunit [Salmonella enterica] Length: 372\nNumber of predicted TMHs: 1\nExp number of AAs in TMHs: 24.51309\nExp number, first 60 AAs: 0.2682\nTotal prob of N-in: 0.14402\noutside 1 331\nTMhelix 332 354\ninside 355 372

23937 GCF_000258365.1_ASM25836v1 Salmonella enterica subsp. enterica serovar Heidelberg str. B182 Proteobacteria; Gammaproteobacteria; Enterobacterales; Enterobacteriaceae; Salmonella; Salmonella enterica; Salmonella enterica subsp. enterica; Salmonella enterica subsp. enterica serovar Heidelberg MTGDNTLITSHGINRRDFMKLCAALAATMGLSSKAAAEMAESVSNPQRPPVIWIGAQECT WP_000145429.1 hydrogenase 2 small subunit [Salmonella enterica] Length: 372\nNumber of predicted TMHs: 1\nExp number of AAs in TMHs: 24.51309\nExp number, first 60 AAs: 0.2682\nTotal prob of N-in: 0.14402\noutside 1 331\nTMhelix 332 354\ninside 355 372

23938 GCF_000258365.1_ASM25836v1 Salmonella enterica subsp. enterica serovar Heidelberg str. B182 Proteobacteria; Gammaproteobacteria; Enterobacterales; Enterobacteriaceae; Salmonella; Salmonella enterica; Salmonella enterica subsp. enterica; Salmonella enterica subsp. enterica serovar Heidelberg MNNEETFYQAMRRKGVTRRSFLKFCSLAATSLGLGAGMTPKIAWALENKPRIPVVWIHGL WP_001058312.1 hydrogenase [Salmonella enterica] Length: 372\nNumber of predicted TMHs: 2\nExp number of AAs in TMHs: 36.51738\nExp number, first 60 AAs: 19.72849\nTotal prob of N-in: 0.96170\nPOSSIBLE N-term signal sequence\ninside 1 19\nTMhelix 20 39\noutside 40 325\nTMhelix 326 348\ninside 349 372

23939 GCF_001682115.1_ASM168211v1 Salmonella enterica subsp. enterica serovar Heidelberg str. CFSAN002063 Proteobacteria; Gammaproteobacteria; Enterobacterales; Enterobacteriaceae; Salmonella; Salmonella enterica; Salmonella enterica subsp. enterica; Salmonella enterica subsp. enterica serovar Heidelberg MNNEETFYQAMRRKGVTRRSFLKFCSLAATSLGLGAGMTPKIAWALENKPRIPVVWIHGL WP_001058312.1 hydrogenase [Salmonella enterica] Length: 372\nNumber of predicted TMHs: 2\nExp number of AAs in TMHs: 36.51738\nExp number, first 60 AAs: 19.72849\nTotal prob of N-in: 0.96170\nPOSSIBLE N-term signal sequence\ninside 1 19\nTMhelix 20 39\noutside 40 325\nTMhelix 326 348\ninside 349 372

23940 GCF_001682115.1_ASM168211v1 Salmonella enterica subsp. enterica serovar Heidelberg str. CFSAN002063 Proteobacteria; Gammaproteobacteria; Enterobacterales; Enterobacteriaceae; Salmonella; Salmonella enterica; Salmonella enterica subsp. enterica; Salmonella enterica subsp. enterica serovar Heidelberg MTGDNTLITSHGINRRDFMKLCAALAATMGLSSKAAAEMAESVSNPQRPPVIWIGAQECT WP_000145429.1 hydrogenase 2 small subunit [Salmonella enterica] Length: 372\nNumber of predicted TMHs: 1\nExp number of AAs in TMHs: 24.51309\nExp number, first 60 AAs: 0.2682\nTotal prob of N-in: 0.14402\noutside 1 331\nTMhelix 332 354\ninside 355 372

23941 GCF_000505705.1_ASM50570v1 Salmonella enterica subsp. enterica serovar Heidelberg str. CFSAN002064 Proteobacteria; Gammaproteobacteria; Enterobacterales; Enterobacteriaceae; Salmonella; Salmonella enterica; Salmonella enterica subsp. enterica; Salmonella enterica subsp. enterica serovar Heidelberg MTGDNTLITSHGINRRDFMKLCAALAATMGLSSKAAAEMAESVSNPQRPPVIWIGAQECT WP_000145429.1 hydrogenase 2 small subunit [Salmonella enterica] Length: 372\nNumber of predicted TMHs: 1\nExp number of AAs in TMHs: 24.51309\nExp number, first 60 AAs: 0.2682\nTotal prob of N-in: 0.14402\noutside 1 331\nTMhelix 332 354\ninside 355 372

23942 GCF_000505705.1_ASM50570v1 Salmonella enterica subsp. enterica serovar Heidelberg str. CFSAN002064 Proteobacteria; Gammaproteobacteria; Enterobacterales; Enterobacteriaceae; Salmonella; Salmonella enterica; Salmonella enterica subsp. enterica; Salmonella enterica subsp. enterica serovar Heidelberg MNNEETFYQAMRRKGVTRRSFLKFCSLAATSLGLGAGMTPKIAWALENKPRIPVVWIHGL WP_001058312.1 hydrogenase [Salmonella enterica] Length: 372\nNumber of predicted TMHs: 2\nExp number of AAs in TMHs: 36.51738\nExp number, first 60 AAs: 19.72849\nTotal prob of N-in: 0.96170\nPOSSIBLE N-term signal sequence\ninside 1 19\nTMhelix 20 39\noutside 40 325\nTMhelix 326 348\ninside 349 372

23943 GCF_000430085.2_ASM43008v2 Salmonella enterica subsp. enterica serovar Heidelberg str. CFSAN002069 Proteobacteria; Gammaproteobacteria; Enterobacterales; Enterobacteriaceae; Salmonella; Salmonella enterica; Salmonella enterica subsp. enterica; Salmonella enterica subsp. enterica serovar Heidelberg MNNEETFYQAMRRKGVTRRSFLKFCSLAATSLGLGAGMTPKIAWALENKPRIPVVWIHGL WP_001058312.1 hydrogenase [Salmonella enterica] Length: 372\nNumber of predicted TMHs: 2\nExp number of AAs in TMHs: 36.51738\nExp number, first 60 AAs: 19.72849\nTotal prob of N-in: 0.96170\nPOSSIBLE N-term signal sequence\ninside 1 19\nTMhelix 20 39\noutside 40 325\nTMhelix 326 348\ninside 349 372

23944 GCF_000430085.2_ASM43008v2 Salmonella enterica subsp. enterica serovar Heidelberg str. CFSAN002069 Proteobacteria; Gammaproteobacteria; Enterobacterales; Enterobacteriaceae; Salmonella; Salmonella enterica; Salmonella enterica subsp. enterica; Salmonella enterica subsp. enterica serovar Heidelberg MTGDNTLITSHGINRRDFMKLCAALAATMGLSSKAAAEMAESVSNPQRPPVIWIGAQECT WP_000145429.1 hydrogenase 2 small subunit [Salmonella enterica] Length: 372\nNumber of predicted TMHs: 1\nExp

number of AAs in TMHs: 24.51309\nExp number, first 60 AAs: 0.2682\nTotal prob of N-in: 0.14402\noutside 1 331\nTMhelix 332 354\ninside 355 372

23945 GCF_000293905.1_ASM29390v1 Salmonella enterica subsp. enterica serovar Heidelberg str. CFSAN00322 Proteobacteria; Gammaproteobacteria; Enterobacterales; Enterobacteriaceae; Salmonella; Salmonella enterica; Salmonella enterica subsp. enterica; Salmonella enterica subsp. enterica serovar Heidelberg MTGDNTLITSHGINRRDFMKLCAALAATMGLSSKAAAEMAESVSNPQRPPVIWIGAQECT WP_000145429.1 hydrogenase 2 small subunit [Salmonella enterica] Length: 372\nNumber of predicted TMHs: 1\nExp number of AAs in TMHs: 24.51309\nExp number, first 60 AAs: 0.2682\nTotal prob of N-in: 0.14402\noutside 1 331\nTMhelix 332 354\ninside 355 372

23946 GCF_000293905.1_ASM29390v1 Salmonella enterica subsp. enterica serovar Heidelberg str. CFSAN00322 Proteobacteria; Gammaproteobacteria; Enterobacterales; Enterobacteriaceae; Salmonella; Salmonella enterica; Salmonella enterica subsp. enterica; Salmonella enterica subsp. enterica serovar Heidelberg MNNEETFYQAMRRKGVTRRSFLKFCSLAATSLGLGAGMTPKIAWALENKPRIPVVWIHGL WP_001058312.1 hydrogenase [Salmonella enterica] Length: 372\nNumber of predicted TMHs: 2\nExp number of AAs in TMHs: 36.51738\nExp number, first 60 AAs: 19.72849\nTotal prob of N-in: 0.96170\nPOSSIBLE N-term signal sequence\ninside 1 19\nTMhelix 20 39\noutside 40 325\nTMhelix 326 348\ninside 349 372

23947 GCF_000962695.1_ASM96269v1 Salmonella enterica subsp. enterica serovar Heidelberg str. CFSAN00323 Proteobacteria; Gammaproteobacteria; Enterobacterales; Enterobacteriaceae; Salmonella; Salmonella enterica; Salmonella enterica subsp. enterica; Salmonella enterica subsp. enterica serovar Heidelberg MTGDNTLITSHGINRRDFMKLCAALAATMGLSSKAAAEMAESVSNPQRPPVIWIGAQECT WP_000145429.1 hydrogenase 2 small subunit [Salmonella enterica] Length: 372\nNumber of predicted TMHs: 1\nExp number of AAs in TMHs: 24.51309\nExp number, first 60 AAs: 0.2682\nTotal prob of N-in: 0.14402\noutside 1 331\nTMhelix 332 354\ninside 355 372

23948 GCF_000962695.1_ASM96269v1 Salmonella enterica subsp. enterica serovar Heidelberg str. CFSAN00323 Proteobacteria; Gammaproteobacteria; Enterobacterales; Enterobacteriaceae; Salmonella; Salmonella enterica; Salmonella enterica subsp. enterica; Salmonella enterica subsp. enterica serovar Heidelberg MNNEETFYQAMRRKGVTRRSFLKFCSLAATSLGLGAGMTPKIAWALENKPRIPVVWIHGL WP_001058312.1 hydrogenase [Salmonella enterica] Length: 372\nNumber of predicted TMHs: 2\nExp number of AAs in TMHs: 36.51738\nExp number, first 60 AAs: 19.72849\nTotal prob of N-in: 0.96170\nPOSSIBLE N-term signal sequence\ninside 1 19\nTMhelix 20 39\noutside 40 325\nTMhelix 326 348\ninside 349 372

23949 GCF_000962725.1_ASM96272v1 Salmonella enterica subsp. enterica serovar Heidelberg str. CFSAN00324 Proteobacteria; Gammaproteobacteria; Enterobacterales; Enterobacteriaceae; Salmonella; Salmonella enterica; Salmonella enterica subsp. enterica; Salmonella enterica subsp. enterica serovar Heidelberg MTGDNTLITSHGINRRDFMKLCAALAATMGLSSKAAAEMAESVSNPQRPPVIWIGAQECT WP_000145429.1 hydrogenase 2 small subunit [Salmonella enterica] Length: 372\nNumber of predicted TMHs: 1\nExp number of AAs in TMHs: 24.51309\nExp number, first 60 AAs: 0.2682\nTotal prob of N-in: 0.14402\noutside 1 331\nTMhelix 332 354\ninside 355 372

23950 GCF_000962725.1_ASM96272v1 Salmonella enterica subsp. enterica serovar Heidelberg str. CFSAN00324 Proteobacteria; Gammaproteobacteria; Enterobacterales; Enterobacteriaceae; Salmonella; Salmonella enterica; Salmonella enterica subsp. enterica; Salmonella enterica subsp. enterica serovar Heidelberg MNNEETFYQAMRRKGVTRRSFLKFCSLAATSLGLGAGMTPKIAWALENKPRIPVVWIHGL WP_001058312.1 hydrogenase [Salmonella enterica] Length: 372\nNumber of predicted TMHs: 2\nExp number of AAs in TMHs: 36.51738\nExp number, first 60 AAs: 19.72849\nTotal prob of N-in: 0.96170\nPOSSIBLE N-term signal sequence\ninside 1 19\nTMhelix 20 39\noutside 40 325\nTMhelix 326 348\ninside 349 372

23951 GCF_000293945.1_ASM29394v1 Salmonella enterica subsp. enterica serovar Heidelberg str. CFSAN00325 Proteobacteria; Gammaproteobacteria; Enterobacterales; Enterobacteriaceae; Salmonella; Salmonella enterica; Salmonella enterica subsp. enterica; Salmonella enterica subsp. enterica serovar Heidelberg MNNEETFYQAMRRKGVTRRSFLKFCSLAATSLGLGAGMTPKIAWALENKPRIPVVWIHGL WP_001058312.1 hydrogenase [Salmonella enterica] Length: 372\nNumber of predicted TMHs: 2\nExp number of AAs in TMHs: 36.51738\nExp number, first 60 AAs: 19.72849\nTotal prob of N-in: 0.96170\nPOSSIBLE N-term signal sequence\ninside 1 19\nTMhelix 20 39\noutside 40 325\nTMhelix 326 348\ninside 349 372

23952 GCF_000293945.1_ASM29394v1 Salmonella enterica subsp. enterica serovar Heidelberg str. CFSAN00325 Proteobacteria; Gammaproteobacteria; Enterobacterales; Enterobacteriaceae; Salmonella; Salmonella enterica;

Salmonella enterica subsp. enterica; Salmonella enterica subsp. enterica serovar Heidelberg
MTGDNTLITSHGINRRDFMKLCAALAATMGLSSKAAAEMAESVSNPQRPPVIWIGAQECT WP_000145429.1
hydrogenase 2 small subunit [Salmonella enterica] Length: 372\nNumber of predicted TMHs: 1\nExp number of AAs in TMHs: 24.51309\nExp number, first 60 AAs: 0.2682\nTotal prob of N-in: 0.14402\noutside 1 331\nTMhelix 332 354\ninside 355 372

23953 GCF_000293965.1_ASM29396v1 Salmonella enterica subsp. enterica serovar Heidelberg str. CFSAN00326
Proteobacteria; Gammaproteobacteria; Enterobacterales; Enterobacteriaceae; Salmonella; Salmonella enterica; Salmonella enterica subsp. enterica; Salmonella enterica subsp. enterica serovar Heidelberg
MTGDNTLITSHGINRRDFMKLCAALAATMGLSSKAAAEMAESVSNPQRPPVIWIGAQECT WP_000145429.1
hydrogenase 2 small subunit [Salmonella enterica] Length: 372\nNumber of predicted TMHs: 1\nExp number of AAs in TMHs: 24.51309\nExp number, first 60 AAs: 0.2682\nTotal prob of N-in: 0.14402\noutside 1 331\nTMhelix 332 354\ninside 355 372

23954 GCF_000293965.1_ASM29396v1 Salmonella enterica subsp. enterica serovar Heidelberg str. CFSAN00326
Proteobacteria; Gammaproteobacteria; Enterobacterales; Enterobacteriaceae; Salmonella; Salmonella enterica; Salmonella enterica subsp. enterica; Salmonella enterica subsp. enterica serovar Heidelberg
MNNEETFYQAMRRKGVTRRSFLKFCSLAATSLGLGAGMTPKIAWALENKPRIPVVWIHGL WP_001058312.1
hydrogenase [Salmonella enterica] Length: 372\nNumber of predicted TMHs: 2\nExp number of AAs in TMHs: 36.51738\nExp number, first 60 AAs: 19.72849\nTotal prob of N-in: 0.96170\nPOSSIBLE N-term signal sequence\ninside 1 19\nTMhelix 20 39\noutside 40 325\nTMhelix 326 348\ninside 349 372

23955 GCF_000962555.1_ASM96255v1 Salmonella enterica subsp. enterica serovar Heidelberg str. CFSAN00327
Proteobacteria; Gammaproteobacteria; Enterobacterales; Enterobacteriaceae; Salmonella; Salmonella enterica; Salmonella enterica subsp. enterica; Salmonella enterica subsp. enterica serovar Heidelberg
MNNEETFYQAMRRKGVTRRSFLKFCSLAATSLGLGAGMTPKIAWALENKPRIPVVWIHGL WP_001058312.1
hydrogenase [Salmonella enterica] Length: 372\nNumber of predicted TMHs: 2\nExp number of AAs in TMHs: 36.51738\nExp number, first 60 AAs: 19.72849\nTotal prob of N-in: 0.96170\nPOSSIBLE N-term signal sequence\ninside 1 19\nTMhelix 20 39\noutside 40 325\nTMhelix 326 348\ninside 349 372

23956 GCF_000962555.1_ASM96255v1 Salmonella enterica subsp. enterica serovar Heidelberg str. CFSAN00327
Proteobacteria; Gammaproteobacteria; Enterobacterales; Enterobacteriaceae; Salmonella; Salmonella enterica; Salmonella enterica subsp. enterica; Salmonella enterica subsp. enterica serovar Heidelberg
MTGDNTLITSHGINRRDFMKLCAALAATMGLSSKAAAEMAESVSNPQRPPVIWIGAQECT WP_000145429.1
hydrogenase 2 small subunit [Salmonella enterica] Length: 372\nNumber of predicted TMHs: 1\nExp number of AAs in TMHs: 24.51309\nExp number, first 60 AAs: 0.2682\nTotal prob of N-in: 0.14402\noutside 1 331\nTMhelix 332 354\ninside 355 372

23957 GCF_000293925.1_ASM29392v1 Salmonella enterica subsp. enterica serovar Heidelberg str. CFSAN00328
Proteobacteria; Gammaproteobacteria; Enterobacterales; Enterobacteriaceae; Salmonella; Salmonella enterica; Salmonella enterica subsp. enterica; Salmonella enterica subsp. enterica serovar Heidelberg
MNNEETFYQAMRRKGVTRRSFLKFCSLAATSLGLGAGMTPKIAWALENKPRIPVVWIHGL WP_001058312.1
hydrogenase [Salmonella enterica] Length: 372\nNumber of predicted TMHs: 2\nExp number of AAs in TMHs: 36.51738\nExp number, first 60 AAs: 19.72849\nTotal prob of N-in: 0.96170\nPOSSIBLE N-term signal sequence\ninside 1 19\nTMhelix 20 39\noutside 40 325\nTMhelix 326 348\ninside 349 372

23958 GCF_000293925.1_ASM29392v1 Salmonella enterica subsp. enterica serovar Heidelberg str. CFSAN00328
Proteobacteria; Gammaproteobacteria; Enterobacterales; Enterobacteriaceae; Salmonella; Salmonella enterica; Salmonella enterica subsp. enterica; Salmonella enterica subsp. enterica serovar Heidelberg
MTGDNTLITSHGINRRDFMKLCAALAATMGLSSKAAAEMAESVSNPQRPPVIWIGAQECT WP_000145429.1
hydrogenase 2 small subunit [Salmonella enterica] Length: 372\nNumber of predicted TMHs: 1\nExp number of AAs in TMHs: 24.51309\nExp number, first 60 AAs: 0.2682\nTotal prob of N-in: 0.14402\noutside 1 331\nTMhelix 332 354\ninside 355 372

23959 GCF_000603845.1_ASM60384v1 Salmonella enterica subsp. enterica serovar Heidelberg str. CVM20752
Proteobacteria; Gammaproteobacteria; Enterobacterales; Enterobacteriaceae; Salmonella; Salmonella enterica; Salmonella enterica subsp. enterica; Salmonella enterica subsp. enterica serovar Heidelberg
MTGDNTLITSHGINRRDFMKLCAALAATMGLSSKAAAEMAESVSNPQRPPVIWIGAQECT WP_000145429.1
hydrogenase 2 small subunit [Salmonella enterica] Length: 372\nNumber of predicted TMHs: 1\nExp number of AAs in TMHs: 24.51309\nExp number, first 60 AAs: 0.2682\nTotal prob of N-in: 0.14402\noutside 1 331\nTMhelix 332 354\ninside 355 372

23960 GCF_000603845.1_ASM60384v1 Salmonella enterica subsp. enterica serovar Heidelberg str. CVM20752 Proteobacteria; Gammaproteobacteria; Enterobacterales; Enterobacteriaceae; Salmonella; Salmonella enterica; Salmonella enterica subsp. enterica; Salmonella enterica subsp. enterica serovar Heidelberg MNNEETFYQAMRRKGVTRRSFLKFCSLAATSLGLGAGMTPKIAWALENKPRIPVVWIHGL WP_001058312.1 hydrogenase [Salmonella enterica] Length: 372\nNumber of predicted TMHs: 2\nExp number of AAs in TMHs: 36.51738\nExp number, first 60 AAs: 19.72849\nTotal prob of N-in: 0.96170\nPOSSIBLE N-term signal sequence\ninside 1 19\nTMhelix 20 39\noutside 40 325\nTMhelix 326 348\ninside 349 372

23961 GCF_000603805.1_ASM60380v1 Salmonella enterica subsp. enterica serovar Heidelberg str. CVM24359 Proteobacteria; Gammaproteobacteria; Enterobacterales; Enterobacteriaceae; Salmonella; Salmonella enterica; Salmonella enterica subsp. enterica; Salmonella enterica subsp. enterica serovar Heidelberg MNNEETFYQAMRRKGVTRRSFLKFCSLAATSLGLGAGMTPKIAWALENKPRIPVVWIHGL WP_001058312.1 hydrogenase [Salmonella enterica] Length: 372\nNumber of predicted TMHs: 2\nExp number of AAs in TMHs: 36.51738\nExp number, first 60 AAs: 19.72849\nTotal prob of N-in: 0.96170\nPOSSIBLE N-term signal sequence\ninside 1 19\nTMhelix 20 39\noutside 40 325\nTMhelix 326 348\ninside 349 372

23962 GCF_000603805.1_ASM60380v1 Salmonella enterica subsp. enterica serovar Heidelberg str. CVM24359 Proteobacteria; Gammaproteobacteria; Enterobacterales; Enterobacteriaceae; Salmonella; Salmonella enterica; Salmonella enterica subsp. enterica; Salmonella enterica subsp. enterica serovar Heidelberg MTGDNTLITSHGINRRDFMKLCAALAATMGLSSKAAAEMAESVSNPQRPPVIWIGAQECT WP_000145429.1 hydrogenase 2 small subunit [Salmonella enterica] Length: 372\nNumber of predicted TMHs: 1\nExp number of AAs in TMHs: 24.51309\nExp number, first 60 AAs: 0.2682\nTotal prob of N-in: 0.14402\noutside 1 331\nTMhelix 332 354\ninside 355 372

23963 GCF_000603825.1_ASM60382v1 Salmonella enterica subsp. enterica serovar Heidelberg str. CVM24388 Proteobacteria; Gammaproteobacteria; Enterobacterales; Enterobacteriaceae; Salmonella; Salmonella enterica; Salmonella enterica subsp. enterica; Salmonella enterica subsp. enterica serovar Heidelberg MTGDNTLITSHGINRRDFMKLCAALAATMGLSSKAAAEMAESVSNPQRPPVIWIGAQECT WP_000145429.1 hydrogenase 2 small subunit [Salmonella enterica] Length: 372\nNumber of predicted TMHs: 1\nExp number of AAs in TMHs: 24.51309\nExp number, first 60 AAs: 0.2682\nTotal prob of N-in: 0.14402\noutside 1 331\nTMhelix 332 354\ninside 355 372

23964 GCF_000603785.1_ASM60378v1 Salmonella enterica subsp. enterica serovar Heidelberg str. CVM24391 Proteobacteria; Gammaproteobacteria; Enterobacterales; Enterobacteriaceae; Salmonella; Salmonella enterica; Salmonella enterica subsp. enterica; Salmonella enterica subsp. enterica serovar Heidelberg MTGDNTLITSHGINRRDFMKLCAALAATMGLSSKAAAEMAESVSNPQRPPVIWIGAQECT WP_000145429.1 hydrogenase 2 small subunit [Salmonella enterica] Length: 372\nNumber of predicted TMHs: 1\nExp number of AAs in TMHs: 24.51309\nExp number, first 60 AAs: 0.2682\nTotal prob of N-in: 0.14402\noutside 1 331\nTMhelix 332 354\ninside 355 372

23965 GCF_000603785.1_ASM60378v1 Salmonella enterica subsp. enterica serovar Heidelberg str. CVM24391 Proteobacteria; Gammaproteobacteria; Enterobacterales; Enterobacteriaceae; Salmonella; Salmonella enterica; Salmonella enterica subsp. enterica; Salmonella enterica subsp. enterica serovar Heidelberg MNNEETFYQAMRRKGVTRRSFLKFCSLAATSLGLGAGMTPKIAWALENKPRIPVVWIHGL WP_001058312.1 hydrogenase [Salmonella enterica] Length: 372\nNumber of predicted TMHs: 2\nExp number of AAs in TMHs: 36.51738\nExp number, first 60 AAs: 19.72849\nTotal prob of N-in: 0.96170\nPOSSIBLE N-term signal sequence\ninside 1 19\nTMhelix 20 39\noutside 40 325\nTMhelix 326 348\ninside 349 372

23966 GCF_000601395.1_SEEH1536_1.0 Salmonella enterica subsp. enterica serovar Heidelberg str. N1536 Proteobacteria; Gammaproteobacteria; Enterobacterales; Enterobacteriaceae; Salmonella; Salmonella enterica; Salmonella enterica subsp. enterica; Salmonella enterica subsp. enterica serovar Heidelberg MNNEETFYQAMRRKGVTRRSFLKFCSLAATSLGLGAGMTPKIAWALENKPRIPVVWIHGL WP_001058312.1 hydrogenase [Salmonella enterica] Length: 372\nNumber of predicted TMHs: 2\nExp number of AAs in TMHs: 36.51738\nExp number, first 60 AAs: 19.72849\nTotal prob of N-in: 0.96170\nPOSSIBLE N-term signal sequence\ninside 1 19\nTMhelix 20 39\noutside 40 325\nTMhelix 326 348\ninside 349 372

23967 GCF_000601395.1_SEEH1536_1.0 Salmonella enterica subsp. enterica serovar Heidelberg str. N1536 Proteobacteria; Gammaproteobacteria; Enterobacterales; Enterobacteriaceae; Salmonella; Salmonella enterica; Salmonella enterica subsp. enterica; Salmonella enterica subsp. enterica serovar Heidelberg MTGDNTLITSHGINRRDFMKLCAALAATMGLSSKAAAEMAESVSNPQRPPVIWIGAQECT WP_000145429.1 hydrogenase 2 small subunit [Salmonella enterica] Length: 372\nNumber of predicted TMHs: 1\nExp

number of AAs in TMHs: 24.51309\nExp number, first 60 AAs: 0.2682\nTotal prob of N-in: 0.14402\noutside 1 331\nTMhelix 332 354\ninside 355 372

23968 GCF_000603565.1_ASM60356v1 Salmonella enterica subsp. enterica serovar Heidelberg str. N15757 Proteobacteria; Gammaproteobacteria; Enterobacterales; Enterobacteriaceae; Salmonella; Salmonella enterica; Salmonella enterica subsp. enterica; Salmonella enterica subsp. enterica serovar Heidelberg MNNEETFYQAMRRKGVTRRSFLKFCSLAATSLGLGAGMTPKIAWALENKPRIPVVWIHGL WP_001058312.1 hydrogenase [Salmonella enterica] Length: 372\nNumber of predicted TMHs: 2\nExp number of AAs in TMHs: 36.51738\nExp number, first 60 AAs: 19.72849\nTotal prob of N-in: 0.96170\nPOSSIBLE N-term signal sequence\ninside 1 19\nTMhelix 20 39\noutside 40 325\nTMhelix 326 348\ninside 349 372

23969 GCF_000603565.1_ASM60356v1 Salmonella enterica subsp. enterica serovar Heidelberg str. N15757 Proteobacteria; Gammaproteobacteria; Enterobacterales; Enterobacteriaceae; Salmonella; Salmonella enterica; Salmonella enterica subsp. enterica; Salmonella enterica subsp. enterica serovar Heidelberg MTGDNTLITSHGINRRDFMKLCAALAATMGLSSKAAAEMAESVSNPQRPPVIWIGAQECT WP_000145429.1 hydrogenase 2 small subunit [Salmonella enterica] Length: 372\nNumber of predicted TMHs: 1\nExp number of AAs in TMHs: 24.51309\nExp number, first 60 AAs: 0.2682\nTotal prob of N-in: 0.14402\noutside 1 331\nTMhelix 332 354\ninside 355 372

23970 GCF_000603605.1_ASM60360v1 Salmonella enterica subsp. enterica serovar Heidelberg str. N18393 Proteobacteria; Gammaproteobacteria; Enterobacterales; Enterobacteriaceae; Salmonella; Salmonella enterica; Salmonella enterica subsp. enterica; Salmonella enterica subsp. enterica serovar Heidelberg MTGDNTLITSHGINRRDFMKLCAALAATMGLSSKAAAEMAESVSNPQRPPVIWIGAQECT WP_000145429.1 hydrogenase 2 small subunit [Salmonella enterica] Length: 372\nNumber of predicted TMHs: 1\nExp number of AAs in TMHs: 24.51309\nExp number, first 60 AAs: 0.2682\nTotal prob of N-in: 0.14402\noutside 1 331\nTMhelix 332 354\ninside 355 372

23971 GCF_000603605.1_ASM60360v1 Salmonella enterica subsp. enterica serovar Heidelberg str. N18393 Proteobacteria; Gammaproteobacteria; Enterobacterales; Enterobacteriaceae; Salmonella; Salmonella enterica; Salmonella enterica subsp. enterica; Salmonella enterica subsp. enterica serovar Heidelberg MNNEETFYQAMRRKGVTRRSFLKFCSLAATSLGLGAGMTPKIAWALENKPRIPVVWIHGL WP_001058312.1 hydrogenase [Salmonella enterica] Length: 372\nNumber of predicted TMHs: 2\nExp number of AAs in TMHs: 36.51738\nExp number, first 60 AAs: 19.72849\nTotal prob of N-in: 0.96170\nPOSSIBLE N-term signal sequence\ninside 1 19\nTMhelix 20 39\noutside 40 325\nTMhelix 326 348\ninside 349 372

23972 GCF_000603865.1_ASM60386v1 Salmonella enterica subsp. enterica serovar Heidelberg str. N18413 Proteobacteria; Gammaproteobacteria; Enterobacterales; Enterobacteriaceae; Salmonella; Salmonella enterica; Salmonella enterica subsp. enterica; Salmonella enterica subsp. enterica serovar Heidelberg MNNEETFYQAMRRKGVTRRSFLKFCSLAATSLGLGAGMTPKIAWALENKPRIPVVWIHGL WP_001058312.1 hydrogenase [Salmonella enterica] Length: 372\nNumber of predicted TMHs: 2\nExp number of AAs in TMHs: 36.51738\nExp number, first 60 AAs: 19.72849\nTotal prob of N-in: 0.96170\nPOSSIBLE N-term signal sequence\ninside 1 19\nTMhelix 20 39\noutside 40 325\nTMhelix 326 348\ninside 349 372

23973 GCF_000603865.1_ASM60386v1 Salmonella enterica subsp. enterica serovar Heidelberg str. N18413 Proteobacteria; Gammaproteobacteria; Enterobacterales; Enterobacteriaceae; Salmonella; Salmonella enterica; Salmonella enterica subsp. enterica; Salmonella enterica subsp. enterica serovar Heidelberg MTGDNTLITSHGINRRDFMKLCAALAATMGLSSKAAAEMAESVSNPQRPPVIWIGAQECT WP_000145429.1 hydrogenase 2 small subunit [Salmonella enterica] Length: 372\nNumber of predicted TMHs: 1\nExp number of AAs in TMHs: 24.51309\nExp number, first 60 AAs: 0.2682\nTotal prob of N-in: 0.14402\noutside 1 331\nTMhelix 332 354\ninside 355 372

23974 GCF_000603885.1_ASM60388v1 Salmonella enterica subsp. enterica serovar Heidelberg str. N18440 Proteobacteria; Gammaproteobacteria; Enterobacterales; Enterobacteriaceae; Salmonella; Salmonella enterica; Salmonella enterica subsp. enterica; Salmonella enterica subsp. enterica serovar Heidelberg MTGDNTLITSHGINRRDFMKLCAALAATMGLSSKAAAEMAESVSNPQRPPVIWIGAQECT WP_000145429.1 hydrogenase 2 small subunit [Salmonella enterica] Length: 372\nNumber of predicted TMHs: 1\nExp number of AAs in TMHs: 24.51309\nExp number, first 60 AAs: 0.2682\nTotal prob of N-in: 0.14402\noutside 1 331\nTMhelix 332 354\ninside 355 372

23975 GCF_000603885.1_ASM60388v1 Salmonella enterica subsp. enterica serovar Heidelberg str. N18440 Proteobacteria; Gammaproteobacteria; Enterobacterales; Enterobacteriaceae; Salmonella; Salmonella enterica;

Salmonella enterica subsp. enterica; Salmonella enterica subsp. enterica serovar Heidelberg
MNNEETFYQAMRRKGVTRRSFLKFCSLAATSLGLGAGMTPKIAWALENKPRIPVVWIHGL WP_001058312.1
hydrogenase [Salmonella enterica] Length: 372\nNumber of predicted TMHs: 2\nExp number of AAs in TMHs: 36.51738\nExp number, first 60 AAs: 19.72849\nTotal prob of N-in: 0.96170\nPOSSIBLE N-term signal sequence\ninside 1 19\nTMhelix 20 39\noutside 40 325\nTMhelix 326 348\ninside 349 372

23976 GCF_000603905.1_ASM60390v1 Salmonella enterica subsp. enterica serovar Heidelberg str. N18453
Proteobacteria; Gammaproteobacteria; Enterobacterales; Enterobacteriaceae; Salmonella; Salmonella enterica; Salmonella enterica subsp. enterica; Salmonella enterica subsp. enterica serovar Heidelberg
MTGDNTLITSHGINRRDFMKLCAALAATMGLSSKAAAEMAESVSNPQRPPVIWIGAQECT WP_000145429.1
hydrogenase 2 small subunit [Salmonella enterica] Length: 372\nNumber of predicted TMHs: 1\nExp number of AAs in TMHs: 24.51309\nExp number, first 60 AAs: 0.2682\nTotal prob of N-in: 0.14402\noutside 1 331\nTMhelix 332 354\ninside 355 372

23977 GCF_000603905.1_ASM60390v1 Salmonella enterica subsp. enterica serovar Heidelberg str. N18453
Proteobacteria; Gammaproteobacteria; Enterobacterales; Enterobacteriaceae; Salmonella; Salmonella enterica; Salmonella enterica subsp. enterica; Salmonella enterica subsp. enterica serovar Heidelberg
MNNEETFYQAMRRKGVTRRSFLKFCSLAATSLGLGAGMTPKIAWALENKPRIPVVWIHGL WP_001058312.1
hydrogenase [Salmonella enterica] Length: 372\nNumber of predicted TMHs: 2\nExp number of AAs in TMHs: 36.51738\nExp number, first 60 AAs: 19.72849\nTotal prob of N-in: 0.96170\nPOSSIBLE N-term signal sequence\ninside 1 19\nTMhelix 20 39\noutside 40 325\nTMhelix 326 348\ninside 349 372

23978 GCF_000603545.1_ASM60354v1 Salmonella enterica subsp. enterica serovar Heidelberg str. N19871
Proteobacteria; Gammaproteobacteria; Enterobacterales; Enterobacteriaceae; Salmonella; Salmonella enterica; Salmonella enterica subsp. enterica; Salmonella enterica subsp. enterica serovar Heidelberg
MTGDNTLITSHGINRRDFMKLCAALAATMGLSSKAAAEMAESVSNPQRPPVIWIGAQECT WP_000145429.1
hydrogenase 2 small subunit [Salmonella enterica] Length: 372\nNumber of predicted TMHs: 1\nExp number of AAs in TMHs: 24.51309\nExp number, first 60 AAs: 0.2682\nTotal prob of N-in: 0.14402\noutside 1 331\nTMhelix 332 354\ninside 355 372

23979 GCF_000603545.1_ASM60354v1 Salmonella enterica subsp. enterica serovar Heidelberg str. N19871
Proteobacteria; Gammaproteobacteria; Enterobacterales; Enterobacteriaceae; Salmonella; Salmonella enterica; Salmonella enterica subsp. enterica; Salmonella enterica subsp. enterica serovar Heidelberg
MNNEETFYQAMRRKGVTRRSFLKFCSLAATSLGLGAGMTPKIAWALENKPRIPVVWIHGL WP_001058312.1
hydrogenase [Salmonella enterica] Length: 372\nNumber of predicted TMHs: 2\nExp number of AAs in TMHs: 36.51738\nExp number, first 60 AAs: 19.72849\nTotal prob of N-in: 0.96170\nPOSSIBLE N-term signal sequence\ninside 1 19\nTMhelix 20 39\noutside 40 325\nTMhelix 326 348\ninside 349 372

23980 GCF_000603365.1_ASM60336v1 Salmonella enterica subsp. enterica serovar Heidelberg str. N19992
Proteobacteria; Gammaproteobacteria; Enterobacterales; Enterobacteriaceae; Salmonella; Salmonella enterica; Salmonella enterica subsp. enterica; Salmonella enterica subsp. enterica serovar Heidelberg
MNNEETFYQAMRRKGVTRRSFLKFCSLAATSLGLGAGMTPKIAWALENKPRIPVVWIHGL WP_001058312.1
hydrogenase [Salmonella enterica] Length: 372\nNumber of predicted TMHs: 2\nExp number of AAs in TMHs: 36.51738\nExp number, first 60 AAs: 19.72849\nTotal prob of N-in: 0.96170\nPOSSIBLE N-term signal sequence\ninside 1 19\nTMhelix 20 39\noutside 40 325\nTMhelix 326 348\ninside 349 372

23981 GCF_000603365.1_ASM60336v1 Salmonella enterica subsp. enterica serovar Heidelberg str. N19992
Proteobacteria; Gammaproteobacteria; Enterobacterales; Enterobacteriaceae; Salmonella; Salmonella enterica; Salmonella enterica subsp. enterica; Salmonella enterica subsp. enterica serovar Heidelberg
MTGDNTLITSHGINRRDFMKLCAALAATMGLSSKAAAEMAESVSNPQRPPVIWIGAQECT WP_000145429.1
hydrogenase 2 small subunit [Salmonella enterica] Length: 372\nNumber of predicted TMHs: 1\nExp number of AAs in TMHs: 24.51309\nExp number, first 60 AAs: 0.2682\nTotal prob of N-in: 0.14402\noutside 1 331\nTMhelix 332 354\ninside 355 372

23982 GCF_000603765.1_ASM60376v1 Salmonella enterica subsp. enterica serovar Heidelberg str. N20134
Proteobacteria; Gammaproteobacteria; Enterobacterales; Enterobacteriaceae; Salmonella; Salmonella enterica; Salmonella enterica subsp. enterica; Salmonella enterica subsp. enterica serovar Heidelberg
MNNEETFYQAMRRKGVTRRSFLKFCSLAATSLGLGAGMTPKIAWALENKPRIPVVWIHGL WP_001058312.1
hydrogenase [Salmonella enterica] Length: 372\nNumber of predicted TMHs: 2\nExp number of AAs in TMHs: 36.51738\nExp number, first 60 AAs: 19.72849\nTotal prob of N-in: 0.96170\nPOSSIBLE N-term signal sequence\ninside 1 19\nTMhelix 20 39\noutside 40 325\nTMhelix 326 348\ninside 349 372

23983 GCF_000603765.1_ASM60376v1 Salmonella enterica subsp. enterica serovar Heidelberg str. N20134 Proteobacteria; Gammaproteobacteria; Enterobacterales; Enterobacteriaceae; Salmonella; Salmonella enterica; Salmonella enterica subsp. enterica; Salmonella enterica subsp. enterica serovar Heidelberg MTGDNTLITSHGINRRDFMKLCAALAATMGLSSKAAAEMAESVSNPQRPPVIWIGAQECT WP_000145429.1 hydrogenase 2 small subunit [Salmonella enterica] Length: 372\nNumber of predicted TMHs: 1\nExp number of AAs in TMHs: 24.51309\nExp number, first 60 AAs: 0.2682\nTotal prob of N-in: 0.14402\noutside 1 331\nTMhelix 332 354\ninside 355 372

23984 GCF_000603585.1_ASM60358v1 Salmonella enterica subsp. enterica serovar Heidelberg str. N26457 Proteobacteria; Gammaproteobacteria; Enterobacterales; Enterobacteriaceae; Salmonella; Salmonella enterica; Salmonella enterica subsp. enterica; Salmonella enterica subsp. enterica serovar Heidelberg MNNEETFYQAMRRKGVTRRSFLKFCSLAATSLGLGAGMTPKIAWALENKPRIPVVWIHGL WP_001058312.1 hydrogenase [Salmonella enterica] Length: 372\nNumber of predicted TMHs: 2\nExp number of AAs in TMHs: 36.51738\nExp number, first 60 AAs: 19.72849\nTotal prob of N-in: 0.96170\nPOSSIBLE N-term signal sequence\ninside 1 19\nTMhelix 20 39\noutside 40 325\nTMhelix 326 348\ninside 349 372

23985 GCF_000603585.1_ASM60358v1 Salmonella enterica subsp. enterica serovar Heidelberg str. N26457 Proteobacteria; Gammaproteobacteria; Enterobacterales; Enterobacteriaceae; Salmonella; Salmonella enterica; Salmonella enterica subsp. enterica; Salmonella enterica subsp. enterica serovar Heidelberg MTGDNTLITSHGINRRDFMKLCAALAATMGLSSKAAAEMAESVSNPQRPPVIWIGAQECT WP_000145429.1 hydrogenase 2 small subunit [Salmonella enterica] Length: 372\nNumber of predicted TMHs: 1\nExp number of AAs in TMHs: 24.51309\nExp number, first 60 AAs: 0.2682\nTotal prob of N-in: 0.14402\noutside 1 331\nTMhelix 332 354\ninside 355 372

23986 GCF_000603685.1_ASM60368v1 Salmonella enterica subsp. enterica serovar Heidelberg str. N29341 Proteobacteria; Gammaproteobacteria; Enterobacterales; Enterobacteriaceae; Salmonella; Salmonella enterica; Salmonella enterica subsp. enterica; Salmonella enterica subsp. enterica serovar Heidelberg MNNEETFYQAMRRKGVTRRSFLKFCSLAATSLGLGAGMTPKIAWALENKPRIPVVWIHGL WP_001058312.1 hydrogenase [Salmonella enterica] Length: 372\nNumber of predicted TMHs: 2\nExp number of AAs in TMHs: 36.51738\nExp number, first 60 AAs: 19.72849\nTotal prob of N-in: 0.96170\nPOSSIBLE N-term signal sequence\ninside 1 19\nTMhelix 20 39\noutside 40 325\nTMhelix 326 348\ninside 349 372

23987 GCF_000603685.1_ASM60368v1 Salmonella enterica subsp. enterica serovar Heidelberg str. N29341 Proteobacteria; Gammaproteobacteria; Enterobacterales; Enterobacteriaceae; Salmonella; Salmonella enterica; Salmonella enterica subsp. enterica; Salmonella enterica subsp. enterica serovar Heidelberg MTGDNTLITSHGINRRDFMKLCAALAATMGLSSKAAAEMAESVSNPQRPPVIWIGAQECT WP_000145429.1 hydrogenase 2 small subunit [Salmonella enterica] Length: 372\nNumber of predicted TMHs: 1\nExp number of AAs in TMHs: 24.51309\nExp number, first 60 AAs: 0.2682\nTotal prob of N-in: 0.14402\noutside 1 331\nTMhelix 332 354\ninside 355 372

23988 GCF_000603705.1_ASM60370v1 Salmonella enterica subsp. enterica serovar Heidelberg str. N30678 Proteobacteria; Gammaproteobacteria; Enterobacterales; Enterobacteriaceae; Salmonella; Salmonella enterica; Salmonella enterica subsp. enterica; Salmonella enterica subsp. enterica serovar Heidelberg MTGDNTLITSHGINRRDFMKLCAALAATMGLSSKAAAEMAESVSNPQRPPVIWIGAQECT WP_000145429.1 hydrogenase 2 small subunit [Salmonella enterica] Length: 372\nNumber of predicted TMHs: 1\nExp number of AAs in TMHs: 24.51309\nExp number, first 60 AAs: 0.2682\nTotal prob of N-in: 0.14402\noutside 1 331\nTMhelix 332 354\ninside 355 372

23989 GCF_000603705.1_ASM60370v1 Salmonella enterica subsp. enterica serovar Heidelberg str. N30678 Proteobacteria; Gammaproteobacteria; Enterobacterales; Enterobacteriaceae; Salmonella; Salmonella enterica; Salmonella enterica subsp. enterica; Salmonella enterica subsp. enterica serovar Heidelberg MNNEETFYQAMRRKGVTRRSFLKFCSLAATSLGLGAGMTPKIAWALENKPRIPVVWIHGL WP_001058312.1 hydrogenase [Salmonella enterica] Length: 372\nNumber of predicted TMHs: 2\nExp number of AAs in TMHs: 36.51738\nExp number, first 60 AAs: 19.72849\nTotal prob of N-in: 0.96170\nPOSSIBLE N-term signal sequence\ninside 1 19\nTMhelix 20 39\noutside 40 325\nTMhelix 326 348\ninside 349 372

23990 GCF_000603345.3_ASM60334v3 Salmonella enterica subsp. enterica serovar Heidelberg str. N418 Proteobacteria; Gammaproteobacteria; Enterobacterales; Enterobacteriaceae; Salmonella; Salmonella enterica; Salmonella enterica subsp. enterica; Salmonella enterica subsp. enterica serovar Heidelberg MTGDNTLITSHGINRRDFMKLCAALAATMGLSSKAAAEMAESVSNPQRPPVIWIGAQECT WP_000145429.1 hydrogenase 2 small subunit [Salmonella enterica] Length: 372\nNumber of predicted TMHs: 1\nExp

number of AAs in TMHs: 24.51309\nExp number, first 60 AAs: 0.2682\nTotal prob of N-in: 0.14402\noutside 1 331\nTMhelix 332 354\ninside 355 372

23991 GCF_000603345.3_ASM60334v3 Salmonella enterica subsp. enterica serovar Heidelberg str. N418 Proteobacteria; Gammaproteobacteria; Enterobacterales; Enterobacteriaceae; Salmonella; Salmonella enterica; Salmonella enterica subsp. enterica; Salmonella enterica subsp. enterica serovar Heidelberg MNNEETFYQAMRRKGVTRRSFLKFCSLAATSLGLGAGMTPKIAWALENKPRIPVVWIHGL WP_001058312.1 hydrogenase [Salmonella enterica] Length: 372\nNumber of predicted TMHs: 2\nExp number of AAs in TMHs: 36.51738\nExp number, first 60 AAs: 19.72849\nTotal prob of N-in: 0.96170\nPOSSIBLE N-term signal sequence\ninside 1 19\nTMhelix 20 39\noutside 40 325\nTMhelix 326 348\ninside 349 372

23992 GCF_000603525.1_ASM60352v1 Salmonella enterica subsp. enterica serovar Heidelberg str. N4630 Proteobacteria; Gammaproteobacteria; Enterobacterales; Enterobacteriaceae; Salmonella; Salmonella enterica; Salmonella enterica subsp. enterica; Salmonella enterica subsp. enterica serovar Heidelberg MTGDNTLITSHGINRRDFMKLCAALAATMGLSSKAAAEMAESVSNPQRPPVIWIGAQECT WP_000145429.1 hydrogenase 2 small subunit [Salmonella enterica] Length: 372\nNumber of predicted TMHs: 1\nExp number of AAs in TMHs: 24.51309\nExp number, first 60 AAs: 0.2682\nTotal prob of N-in: 0.14402\noutside 1 331\nTMhelix 332 354\ninside 355 372

23993 GCF_000603525.1_ASM60352v1 Salmonella enterica subsp. enterica serovar Heidelberg str. N4630 Proteobacteria; Gammaproteobacteria; Enterobacterales; Enterobacteriaceae; Salmonella; Salmonella enterica; Salmonella enterica subsp. enterica; Salmonella enterica subsp. enterica serovar Heidelberg MNNEETFYQAMRRKGVTRRSFLKFCSLAATSLGLGAGMTPKIAWALENKPRIPVVWIHGL WP_001058312.1 hydrogenase [Salmonella enterica] Length: 372\nNumber of predicted TMHs: 2\nExp number of AAs in TMHs: 36.51738\nExp number, first 60 AAs: 19.72849\nTotal prob of N-in: 0.96170\nPOSSIBLE N-term signal sequence\ninside 1 19\nTMhelix 20 39\noutside 40 325\nTMhelix 326 348\ninside 349 372

23994 GCF_000603485.1_ASM60348v1 Salmonella enterica subsp. enterica serovar Heidelberg str. N653 Proteobacteria; Gammaproteobacteria; Enterobacterales; Enterobacteriaceae; Salmonella; Salmonella enterica; Salmonella enterica subsp. enterica; Salmonella enterica subsp. enterica serovar Heidelberg MNNEETFYQAMRRKGVTRRSFLKFCSLAATSLGLGAGMTPKIAWALENKPRIPVVWIHGL WP_001058312.1 hydrogenase [Salmonella enterica] Length: 372\nNumber of predicted TMHs: 2\nExp number of AAs in TMHs: 36.51738\nExp number, first 60 AAs: 19.72849\nTotal prob of N-in: 0.96170\nPOSSIBLE N-term signal sequence\ninside 1 19\nTMhelix 20 39\noutside 40 325\nTMhelix 326 348\ninside 349 372

23995 GCF_000603485.1_ASM60348v1 Salmonella enterica subsp. enterica serovar Heidelberg str. N653 Proteobacteria; Gammaproteobacteria; Enterobacterales; Enterobacteriaceae; Salmonella; Salmonella enterica; Salmonella enterica subsp. enterica; Salmonella enterica subsp. enterica serovar Heidelberg MTGDNTLITSHGINRRDFMKLCAALAATMGLSSKAAAEMAESVSNPQRPPVIWIGAQECT WP_000145429.1 hydrogenase 2 small subunit [Salmonella enterica] Length: 372\nNumber of predicted TMHs: 1\nExp number of AAs in TMHs: 24.51309\nExp number, first 60 AAs: 0.2682\nTotal prob of N-in: 0.14402\noutside 1 331\nTMhelix 332 354\ninside 355 372

23996 GCF_000962015.1_ASM96201v1 Salmonella enterica subsp. enterica serovar Heidelberg str. RI-11-013193 Proteobacteria; Gammaproteobacteria; Enterobacterales; Enterobacteriaceae; Salmonella; Salmonella enterica; Salmonella enterica subsp. enterica; Salmonella enterica subsp. enterica serovar Heidelberg MNNEETFYQAMRRKGVTRRSFLKFCSLAATSLGLGAGMTPKIAWALENKPRIPVVWIHGL WP_001058312.1 hydrogenase [Salmonella enterica] Length: 372\nNumber of predicted TMHs: 2\nExp number of AAs in TMHs: 36.51738\nExp number, first 60 AAs: 19.72849\nTotal prob of N-in: 0.96170\nPOSSIBLE N-term signal sequence\ninside 1 19\nTMhelix 20 39\noutside 40 325\nTMhelix 326 348\ninside 349 372

23997 GCF_000962015.1_ASM96201v1 Salmonella enterica subsp. enterica serovar Heidelberg str. RI-11-013193 Proteobacteria; Gammaproteobacteria; Enterobacterales; Enterobacteriaceae; Salmonella; Salmonella enterica; Salmonella enterica subsp. enterica; Salmonella enterica subsp. enterica serovar Heidelberg MTGDNTLITSHGINRRDFMKLCAALAATMGLSSKAAAEMAESVSNPQRPPVIWIGAQECT WP_000145429.1 hydrogenase 2 small subunit [Salmonella enterica] Length: 372\nNumber of predicted TMHs: 1\nExp number of AAs in TMHs: 24.51309\nExp number, first 60 AAs: 0.2682\nTotal prob of N-in: 0.14402\noutside 1 331\nTMhelix 332 354\ninside 355 372

23998 GCF_000700765.1_SEEH3312_1.0 Salmonella enterica subsp. enterica serovar Heidelberg str. RI-11-013312 Proteobacteria; Gammaproteobacteria; Enterobacterales; Enterobacteriaceae; Salmonella; Salmonella enterica;

Salmonella enterica subsp. enterica; Salmonella enterica subsp. enterica serovar Heidelberg
MNNEETFYQAMRRKGVTRRSFLKFCSLAATSLGLGAGMTPKIAWALENKPRIPVVWIHGL WP_001058312.1
hydrogenase [Salmonella enterica] Length: 372\nNumber of predicted TMHs: 2\nExp number of AAs in TMHs: 36.51738\nExp number, first 60 AAs: 19.72849\nTotal prob of N-in: 0.96170\nPOSSIBLE N-term signal sequence\ninside 1 19\nTMhelix 20 39\noutside 40 325\nTMhelix 326 348\ninside 349 372

23999 GCF_000700765.1_SEEH3312_1.0 Salmonella enterica subsp. enterica serovar Heidelberg str. RI-11-013312
Proteobacteria; Gammaproteobacteria; Enterobacterales; Enterobacteriaceae; Salmonella; Salmonella enterica; Salmonella enterica subsp. enterica; Salmonella enterica subsp. enterica serovar Heidelberg
MTGDNTLITSHGINRRDFMKLCAALAATMGLSSKAAAEMAESVSNPQRPPVIWIGAQECT WP_000145429.1
hydrogenase 2 small subunit [Salmonella enterica] Length: 372\nNumber of predicted TMHs: 1\nExp number of AAs in TMHs: 24.51309\nExp number, first 60 AAs: 0.2682\nTotal prob of N-in: 0.14402\noutside 1 331\nTMhelix 332 354\ninside 355 372

24000 GCF_000699605.1_SEEH3988_1.0 Salmonella enterica subsp. enterica serovar Heidelberg str. RI-11-013988
Proteobacteria; Gammaproteobacteria; Enterobacterales; Enterobacteriaceae; Salmonella; Salmonella enterica; Salmonella enterica subsp. enterica; Salmonella enterica subsp. enterica serovar Heidelberg
MTGDNTLITSHGINRRDFMKLCAALAATMGLSSKAAAEMAESVSNPQRPPVIWIGAQECT WP_000145429.1
hydrogenase 2 small subunit [Salmonella enterica] Length: 372\nNumber of predicted TMHs: 1\nExp number of AAs in TMHs: 24.51309\nExp number, first 60 AAs: 0.2682\nTotal prob of N-in: 0.14402\noutside 1 331\nTMhelix 332 354\ninside 355 372

24001 GCF_000699605.1_SEEH3988_1.0 Salmonella enterica subsp. enterica serovar Heidelberg str. RI-11-013988
Proteobacteria; Gammaproteobacteria; Enterobacterales; Enterobacteriaceae; Salmonella; Salmonella enterica; Salmonella enterica subsp. enterica; Salmonella enterica subsp. enterica serovar Heidelberg
MNNEETFYQAMRRKGVTRRSFLKFCSLAATSLGLGAGMTPKIAWALENKPRIPVVWIHGL WP_001058312.1
hydrogenase [Salmonella enterica] Length: 372\nNumber of predicted TMHs: 2\nExp number of AAs in TMHs: 36.51738\nExp number, first 60 AAs: 19.72849\nTotal prob of N-in: 0.96170\nPOSSIBLE N-term signal sequence\ninside 1 19\nTMhelix 20 39\noutside 40 325\nTMhelix 326 348\ninside 349 372

24002 GCF_000699625.1_ASM69962v1 Salmonella enterica subsp. enterica serovar Heidelberg str. RI-11-014319
Proteobacteria; Gammaproteobacteria; Enterobacterales; Enterobacteriaceae; Salmonella; Salmonella enterica; Salmonella enterica subsp. enterica; Salmonella enterica subsp. enterica serovar Heidelberg
MTGDNTLITSHGINRRDFMKLCAALAATMGLSSKAAAEMAESVSNPQRPPVIWIGAQECT WP_000145429.1
hydrogenase 2 small subunit [Salmonella enterica] Length: 372\nNumber of predicted TMHs: 1\nExp number of AAs in TMHs: 24.51309\nExp number, first 60 AAs: 0.2682\nTotal prob of N-in: 0.14402\noutside 1 331\nTMhelix 332 354\ninside 355 372

24003 GCF_000699625.1_ASM69962v1 Salmonella enterica subsp. enterica serovar Heidelberg str. RI-11-014319
Proteobacteria; Gammaproteobacteria; Enterobacterales; Enterobacteriaceae; Salmonella; Salmonella enterica; Salmonella enterica subsp. enterica; Salmonella enterica subsp. enterica serovar Heidelberg
MNNEETFYQAMRRKGVTRRSFLKFCSLAATSLGLGAGMTPKIAWALENKPRIPVVWIHGL WP_001058312.1
hydrogenase [Salmonella enterica] Length: 372\nNumber of predicted TMHs: 2\nExp number of AAs in TMHs: 36.51738\nExp number, first 60 AAs: 19.72849\nTotal prob of N-in: 0.96170\nPOSSIBLE N-term signal sequence\ninside 1 19\nTMhelix 20 39\noutside 40 325\nTMhelix 326 348\ninside 349 372

24004 GCF_000603325.1_ASM60332v1 Salmonella enterica subsp. enterica serovar Heidelberg str. SARA31
Proteobacteria; Gammaproteobacteria; Enterobacterales; Enterobacteriaceae; Salmonella; Salmonella enterica; Salmonella enterica subsp. enterica; Salmonella enterica subsp. enterica serovar Heidelberg
MTGDNTLITSHGINRRDFMKLCAALAATMGLSSKAAAEMAESVSNPQRPPVIWIGAQECT WP_000145429.1
hydrogenase 2 small subunit [Salmonella enterica] Length: 372\nNumber of predicted TMHs: 1\nExp number of AAs in TMHs: 24.51309\nExp number, first 60 AAs: 0.2682\nTotal prob of N-in: 0.14402\noutside 1 331\nTMhelix 332 354\ninside 355 372

24005 GCF_000603325.1_ASM60332v1 Salmonella enterica subsp. enterica serovar Heidelberg str. SARA31
Proteobacteria; Gammaproteobacteria; Enterobacterales; Enterobacteriaceae; Salmonella; Salmonella enterica; Salmonella enterica subsp. enterica; Salmonella enterica subsp. enterica serovar Heidelberg
MNNEETFYQAMRRKGVTRRSFLKFCSLAATSLGLGAGMTPKIAWALENKPRIPVVWIHGL WP_001058312.1
hydrogenase [Salmonella enterica] Length: 372\nNumber of predicted TMHs: 2\nExp number of AAs in TMHs: 36.51738\nExp number, first 60 AAs: 19.72849\nTotal prob of N-in: 0.96170\nPOSSIBLE N-term signal sequence\ninside 1 19\nTMhelix 20 39\noutside 40 325\nTMhelix 326 348\ninside 349 372

24006 GCF_000603305.1_ASM60330v1 Salmonella enterica subsp. enterica serovar Heidelberg str. SARA32 Proteobacteria; Gammaproteobacteria; Enterobacterales; Enterobacteriaceae; Salmonella; Salmonella enterica; Salmonella enterica subsp. enterica; Salmonella enterica subsp. enterica serovar Heidelberg MNNEETFYQAMRRKGVTRRSFLKFCSLAATSLGLGAGMTPKIAWALENKPRIPVVWIHGL WP_001058312.1 hydrogenase [Salmonella enterica] Length: 372\nNumber of predicted TMHs: 2\nExp number of AAs in TMHs: 36.51738\nExp number, first 60 AAs: 19.72849\nTotal prob of N-in: 0.96170\nPOSSIBLE N-term signal sequence\ninside 1 19\nTMhelix 20 39\noutside 40 325\nTMhelix 326 348\ninside 349 372

24007 GCF_000603305.1_ASM60330v1 Salmonella enterica subsp. enterica serovar Heidelberg str. SARA32 Proteobacteria; Gammaproteobacteria; Enterobacterales; Enterobacteriaceae; Salmonella; Salmonella enterica; Salmonella enterica subsp. enterica; Salmonella enterica subsp. enterica serovar Heidelberg MTGDNTLITSHGINRRDFMKLCAALAATMGLSSKAAAEMAESVSNPQRPPVIWIGAQECT WP_000145429.1 hydrogenase 2 small subunit [Salmonella enterica] Length: 372\nNumber of predicted TMHs: 1\nExp number of AAs in TMHs: 24.51309\nExp number, first 60 AAs: 0.2682\nTotal prob of N-in: 0.14402\noutside 1 331\nTMhelix 332 354\ninside 355 372

24008 GCF_000486045.2_ASM48604v2 Salmonella enterica subsp. enterica serovar Heidelberg str. SARA35 Proteobacteria; Gammaproteobacteria; Enterobacterales; Enterobacteriaceae; Salmonella; Salmonella enterica; Salmonella enterica subsp. enterica; Salmonella enterica subsp. enterica serovar Heidelberg MNNEETFYQAMRRKGVTRRSFLKFCSLAATSLGLGAGMTPKIAWALENKPRIPVVWIHGL WP_001058312.1 hydrogenase [Salmonella enterica] Length: 372\nNumber of predicted TMHs: 2\nExp number of AAs in TMHs: 36.51738\nExp number, first 60 AAs: 19.72849\nTotal prob of N-in: 0.96170\nPOSSIBLE N-term signal sequence\ninside 1 19\nTMhelix 20 39\noutside 40 325\nTMhelix 326 348\ninside 349 372

24009 GCF_000486045.2_ASM48604v2 Salmonella enterica subsp. enterica serovar Heidelberg str. SARA35 Proteobacteria; Gammaproteobacteria; Enterobacterales; Enterobacteriaceae; Salmonella; Salmonella enterica; Salmonella enterica subsp. enterica; Salmonella enterica subsp. enterica serovar Heidelberg MTGDNTLITSHGINRRDFMKLCAALAATMGLSSKAAAEMAESVSNPQRPPVIWIGAQECT WP_000145429.1 hydrogenase 2 small subunit [Salmonella enterica] Length: 372\nNumber of predicted TMHs: 1\nExp number of AAs in TMHs: 24.51309\nExp number, first 60 AAs: 0.2682\nTotal prob of N-in: 0.14402\noutside 1 331\nTMhelix 332 354\ninside 355 372

24010 GCF_000962005.1_ASM96200v1 Salmonella enterica subsp. enterica serovar Heidelberg str. SARA36 Proteobacteria; Gammaproteobacteria; Enterobacterales; Enterobacteriaceae; Salmonella; Salmonella enterica; Salmonella enterica subsp. enterica; Salmonella enterica subsp. enterica serovar Heidelberg MTGDNTLITSHGINRRDFMKLCAALAATMGLSSKAAAEMAESVSNPQRPPVIWIGAQECT WP_000145429.1 hydrogenase 2 small subunit [Salmonella enterica] Length: 372\nNumber of predicted TMHs: 1\nExp number of AAs in TMHs: 24.51309\nExp number, first 60 AAs: 0.2682\nTotal prob of N-in: 0.14402\noutside 1 331\nTMhelix 332 354\ninside 355 372

24011 GCF_000962005.1_ASM96200v1 Salmonella enterica subsp. enterica serovar Heidelberg str. SARA36 Proteobacteria; Gammaproteobacteria; Enterobacterales; Enterobacteriaceae; Salmonella; Salmonella enterica; Salmonella enterica subsp. enterica; Salmonella enterica subsp. enterica serovar Heidelberg MNNEETFYQAMRRKGVTRRSFLKFCSLAATSLGLGAGMTPKIAWALENKPRIPVVWIHGL WP_001058312.1 hydrogenase [Salmonella enterica] Length: 372\nNumber of predicted TMHs: 2\nExp number of AAs in TMHs: 36.51738\nExp number, first 60 AAs: 19.72849\nTotal prob of N-in: 0.96170\nPOSSIBLE N-term signal sequence\ninside 1 19\nTMhelix 20 39\noutside 40 325\nTMhelix 326 348\ninside 349 372

24012 GCF_000603285.1_ASM60328v1 Salmonella enterica subsp. enterica serovar Heidelberg str. SARA37 Proteobacteria; Gammaproteobacteria; Enterobacterales; Enterobacteriaceae; Salmonella; Salmonella enterica; Salmonella enterica subsp. enterica; Salmonella enterica subsp. enterica serovar Heidelberg MNNEETFYQAMRRKGVTRRSFLKFCSLAATSLGLGAGMTPKIAWALENKPRIPVVWIHGL WP_001058312.1 hydrogenase [Salmonella enterica] Length: 372\nNumber of predicted TMHs: 2\nExp number of AAs in TMHs: 36.51738\nExp number, first 60 AAs: 19.72849\nTotal prob of N-in: 0.96170\nPOSSIBLE N-term signal sequence\ninside 1 19\nTMhelix 20 39\noutside 40 325\nTMhelix 326 348\ninside 349 372

24013 GCF_000603285.1_ASM60328v1 Salmonella enterica subsp. enterica serovar Heidelberg str. SARA37 Proteobacteria; Gammaproteobacteria; Enterobacterales; Enterobacteriaceae; Salmonella; Salmonella enterica; Salmonella enterica subsp. enterica; Salmonella enterica subsp. enterica serovar Heidelberg MTGDNTLITSHGINRRDFMKLCAALAATMGLSSKAAAEMAESVSNPQRPPVIWIGAQECT WP_000145429.1 hydrogenase 2 small subunit [Salmonella enterica] Length: 372\nNumber of predicted TMHs: 1\nExp

number of AAs in TMHs: 24.51309\nExp number, first 60 AAs: 0.2682\nTotal prob of N-in: 0.14402\noutside 1 331\nTMhelix 332 354\ninside 355 372

24014 GCF_000961995.1_ASM96199v1 Salmonella enterica subsp. enterica serovar Heidelberg str. SARA40 Proteobacteria; Gammaproteobacteria; Enterobacterales; Enterobacteriaceae; Salmonella; Salmonella enterica; Salmonella enterica subsp. enterica; Salmonella enterica subsp. enterica serovar Heidelberg MTGDNTLITSHGINRRDFMKLCAALAATMGLSSKAAAEMAESVSNPQRPPVIWIGAQECT WP_000145429.1 hydrogenase 2 small subunit [Salmonella enterica] Length: 372\nNumber of predicted TMHs: 1\nExp number of AAs in TMHs: 24.51309\nExp number, first 60 AAs: 0.2682\nTotal prob of N-in: 0.14402\noutside 1 331\nTMhelix 332 354\ninside 355 372

24015 GCF_000961995.1_ASM96199v1 Salmonella enterica subsp. enterica serovar Heidelberg str. SARA40 Proteobacteria; Gammaproteobacteria; Enterobacterales; Enterobacteriaceae; Salmonella; Salmonella enterica; Salmonella enterica subsp. enterica; Salmonella enterica subsp. enterica serovar Heidelberg MNNEETFYQAMRRKGVTRRSFLKFCSLAATSLGLGAGMTPKIAWALENKPRIPVVWIHGL WP_001058312.1 hydrogenase [Salmonella enterica] Length: 372\nNumber of predicted TMHs: 2\nExp number of AAs in TMHs: 36.51738\nExp number, first 60 AAs: 19.72849\nTotal prob of N-in: 0.96170\nPOSSIBLE N-term signal sequence\ninside 1 19\nTMhelix 20 39\noutside 40 325\nTMhelix 326 348\ninside 349 372

24016 GCF_000020705.1_ASM2070v1 Salmonella enterica subsp. enterica serovar Heidelberg str. SL476 Proteobacteria; Gammaproteobacteria; Enterobacterales; Enterobacteriaceae; Salmonella; Salmonella enterica; Salmonella enterica subsp. enterica; Salmonella enterica subsp. enterica serovar Heidelberg MNNEETFYQAMRRKGVTRRSFLKFCSLAATSLGLGAGMTPKIAWALENKPRIPVVWIHGL WP_001058312.1 hydrogenase [Salmonella enterica] Length: 372\nNumber of predicted TMHs: 2\nExp number of AAs in TMHs: 36.51738\nExp number, first 60 AAs: 19.72849\nTotal prob of N-in: 0.96170\nPOSSIBLE N-term signal sequence\ninside 1 19\nTMhelix 20 39\noutside 40 325\nTMhelix 326 348\ninside 349 372

24017 GCF_000020705.1_ASM2070v1 Salmonella enterica subsp. enterica serovar Heidelberg str. SL476 Proteobacteria; Gammaproteobacteria; Enterobacterales; Enterobacteriaceae; Salmonella; Salmonella enterica; Salmonella enterica subsp. enterica; Salmonella enterica subsp. enterica serovar Heidelberg MTGDNTLITSHGINRRDFMKLCAALAATMGLSSKAAAEMAESVSNPQRPPVIWIGAQECT WP_000145429.1 hydrogenase 2 small subunit [Salmonella enterica] Length: 372\nNumber of predicted TMHs: 1\nExp number of AAs in TMHs: 24.51309\nExp number, first 60 AAs: 0.2682\nTotal prob of N-in: 0.14402\noutside 1 331\nTMhelix 332 354\ninside 355 372

24018 GCF_000171315.1_ASM17131v1 Salmonella enterica subsp. enterica serovar Heidelberg str. SL486 Proteobacteria; Gammaproteobacteria; Enterobacterales; Enterobacteriaceae; Salmonella; Salmonella enterica; Salmonella enterica subsp. enterica; Salmonella enterica subsp. enterica serovar Heidelberg MTGDNTLITSHGINRRDFMKLCAALAATMGLSSKAAAEMAESVSNPQRPPVIWIGAQECT WP_000145429.1 hydrogenase 2 small subunit [Salmonella enterica] Length: 372\nNumber of predicted TMHs: 1\nExp number of AAs in TMHs: 24.51309\nExp number, first 60 AAs: 0.2682\nTotal prob of N-in: 0.14402\noutside 1 331\nTMhelix 332 354\ninside 355 372

24019 GCF_000171315.1_ASM17131v1 Salmonella enterica subsp. enterica serovar Heidelberg str. SL486 Proteobacteria; Gammaproteobacteria; Enterobacterales; Enterobacteriaceae; Salmonella; Salmonella enterica; Salmonella enterica subsp. enterica; Salmonella enterica subsp. enterica serovar Heidelberg MNNEETFYQAMRRKGVTRRSFLKFCSLAATSLGLGAGMTPKIAWALENKPRIPVVWIHGL WP_001058312.1 hydrogenase [Salmonella enterica] Length: 372\nNumber of predicted TMHs: 2\nExp number of AAs in TMHs: 36.51738\nExp number, first 60 AAs: 19.72849\nTotal prob of N-in: 0.96170\nPOSSIBLE N-term signal sequence\ninside 1 19\nTMhelix 20 39\noutside 40 325\nTMhelix 326 348\ninside 349 372

24020 GCF_001975385.1_ASM197538v1 Salmonella enterica subsp. enterica serovar Hillingdon str. N1529-D3 Proteobacteria; Gammaproteobacteria; Enterobacterales; Enterobacteriaceae; Salmonella; Salmonella enterica; Salmonella enterica subsp. enterica; Salmonella enterica subsp. enterica serovar Hillingdon MTGDNTLITSHGINRRDFMKLCAALAATMGLSSKAAAEMAESVSNPQRPPVIWIGAQECT WP_000145429.1 hydrogenase 2 small subunit [Salmonella enterica] Length: 372\nNumber of predicted TMHs: 1\nExp number of AAs in TMHs: 24.51309\nExp number, first 60 AAs: 0.2682\nTotal prob of N-in: 0.14402\noutside 1 331\nTMhelix 332 354\ninside 355 372

24021 GCF_001975385.1_ASM197538v1 Salmonella enterica subsp. enterica serovar Hillingdon str. N1529-D3 Proteobacteria; Gammaproteobacteria; Enterobacterales; Enterobacteriaceae; Salmonella; Salmonella enterica;

Salmonella enterica subsp. enterica; Salmonella enterica subsp. enterica serovar Hillingdon
MNNEETFYQAMRRKGVTRRSFLKFCSLAATSLGLGAGMTPKIAWALENKPRIPVVWIHGL WP_001058311.1 hydrogenase [Salmonella enterica] Length: 372\nNumber of predicted TMHs: 2\nExp number of AAs in TMHs: 37.4699899999999\nExp number, first 60 AAs: 19.72699\nTotal prob of N-in: 0.96171\nPOSSIBLE N-term signal sequence\ninside 1 19\nTMhelix 20 39\noutside 40 325\nTMhelix 326 348\ninside 349 372

24022 GCF_001448705.1_ASM144870v1 Salmonella enterica subsp. enterica serovar Hvittingfoss str. SA20014981 Proteobacteria; Gammaproteobacteria; Enterobacterales; Enterobacteriaceae; Salmonella; Salmonella enterica; Salmonella enterica subsp. enterica; Salmonella enterica subsp. enterica serovar Hvittingfoss
MNNEETFYQAMRRKGVTRRSFLKFCSLAATSLGLGAGMTPKIAWALENKPRIPVVWIHGL WP_001058307.1 hydrogenase [Salmonella enterica] Length: 372\nNumber of predicted TMHs: 2\nExp number of AAs in TMHs: 36.51738\nExp number, first 60 AAs: 19.72849\nTotal prob of N-in: 0.96170\nPOSSIBLE N-term signal sequence\ninside 1 19\nTMhelix 20 39\noutside 40 325\nTMhelix 326 348\ninside 349 372

24023 GCF_001448705.1_ASM144870v1 Salmonella enterica subsp. enterica serovar Hvittingfoss str. SA20014981 Proteobacteria; Gammaproteobacteria; Enterobacterales; Enterobacteriaceae; Salmonella; Salmonella enterica; Salmonella enterica subsp. enterica; Salmonella enterica subsp. enterica serovar Hvittingfoss
MTGDNTLITSHGINRRDFMKLCAALAATMGLSSKAAAEMAESVSNPQRPPVIWIGAQECT WP_000145429.1 hydrogenase 2 small subunit [Salmonella enterica] Length: 372\nNumber of predicted TMHs: 1\nExp number of AAs in TMHs: 24.51309\nExp number, first 60 AAs: 0.2682\nTotal prob of N-in: 0.14402\noutside 1 331\nTMhelix 332 354\ninside 355 372

24024 GCF_000487135.1_ASM48713v1 Salmonella enterica subsp. enterica serovar Indiana str. ATCC 51959 Proteobacteria; Gammaproteobacteria; Enterobacterales; Enterobacteriaceae; Salmonella; Salmonella enterica; Salmonella enterica subsp. enterica; Salmonella enterica subsp. enterica serovar Indiana
MTGDNTLITSHGINRRDFMKLCAALAATMGLSSKAAAEMAESVSNPQRPPVIWIGAQECT WP_000145429.1 hydrogenase 2 small subunit [Salmonella enterica] Length: 372\nNumber of predicted TMHs: 1\nExp number of AAs in TMHs: 24.51309\nExp number, first 60 AAs: 0.2682\nTotal prob of N-in: 0.14402\noutside 1 331\nTMhelix 332 354\ninside 355 372

24025 GCF_000487135.1_ASM48713v1 Salmonella enterica subsp. enterica serovar Indiana str. ATCC 51959 Proteobacteria; Gammaproteobacteria; Enterobacterales; Enterobacteriaceae; Salmonella; Salmonella enterica; Salmonella enterica subsp. enterica; Salmonella enterica subsp. enterica serovar Indiana
MNNEETFYQAMRRKGVTRRSFLKFCSLAATSLGLGAGMTPQIAWALENKPRIPVVWIHGL WP_023227615.1 hydrogenase 1, small subunit [Salmonella enterica] Length: 372\nNumber of predicted TMHs: 2\nExp number of AAs in TMHs: 39.8591699999999\nExp number, first 60 AAs: 22.13093\nTotal prob of N-in: 0.98639\nPOSSIBLE N-term signal sequence\ninside 1 20\nTMhelix 21 43\noutside 44 325\nTMhelix 326 348\ninside 349 372

24026 GCF_000487135.1_ASM48713v1 Salmonella enterica subsp. enterica serovar Indiana str. ATCC 51959 Proteobacteria; Gammaproteobacteria; Enterobacterales; Enterobacteriaceae; Salmonella; Salmonella enterica; Salmonella enterica subsp. enterica; Salmonella enterica subsp. enterica serovar Indiana
MLKFRTIFLKKIFTLKNSILARRGAVIVIVSAVFTSIMFFAHSWASDKEVAMTLSVLNSN WP_023226644.1 hypothetical protein [Salmonella enterica] Length: 204\nNumber of predicted TMHs: 2\nExp number of AAs in TMHs: 44.99526\nExp number, first 60 AAs: 22.33786\nTotal prob of N-in: 0.87083\nPOSSIBLE N-term signal sequence\ninside 1 19\nTMhelix 20 42\noutside 43 171\nTMhelix 172 194\ninside 195 204

24027 GCF_000506925.1_SI119944 Salmonella enterica subsp. enterica serovar Infantis str. 119944 Proteobacteria; Gammaproteobacteria; Enterobacterales; Enterobacteriaceae; Salmonella; Salmonella enterica; Salmonella enterica subsp. enterica; Salmonella enterica subsp. enterica serovar Infantis
MNNEETFYQAMRRKGVTRRSFLKFCSLAATSLGLGAGMTPKIAWALENKPRIPVVWIHGL WP_001058311.1 hydrogenase [Salmonella enterica] Length: 372\nNumber of predicted TMHs: 2\nExp number of AAs in TMHs: 37.4699899999999\nExp number, first 60 AAs: 19.72699\nTotal prob of N-in: 0.96171\nPOSSIBLE N-term signal sequence\ninside 1 19\nTMhelix 20 39\noutside 40 325\nTMhelix 326 348\ninside 349 372

24028 GCF_000506925.1_SI119944 Salmonella enterica subsp. enterica serovar Infantis str. 119944 Proteobacteria; Gammaproteobacteria; Enterobacterales; Enterobacteriaceae; Salmonella; Salmonella enterica; Salmonella enterica subsp. enterica; Salmonella enterica subsp. enterica serovar Infantis
MTGDNTLITSHGINRRDFMKLCAALAATMGLSSKAAAEMAESVSNPQRPPVIWIGAQECT WP_000145429.1 hydrogenase 2 small subunit [Salmonella enterica] Length: 372\nNumber of predicted TMHs: 1\nExp

number of AAs in TMHs: 24.51309\nExp number, first 60 AAs: 0.2682\nTotal prob of N-in: 0.14402\noutside 1 331\nTMhelix 332 354\ninside 355 372

24029 GCF_000506945.1_S._Infantis_strain_335-3 Salmonella enterica subsp. enterica serovar Infantis str. 335-3 Proteobacteria; Gammaproteobacteria; Enterobacterales; Enterobacteriaceae; Salmonella; Salmonella enterica; Salmonella enterica subsp. enterica; Salmonella enterica subsp. enterica serovar Infantis MNNEETFYQAMRRKGVTRRSFLKFCSLAATSLGLGAGMTPKIAWALENKPRIPVVWIHGL WP_001058311.1 hydrogenase [Salmonella enterica] Length: 372\nNumber of predicted TMHs: 2\nExp number of AAs in TMHs: 37.4699899999999\nExp number, first 60 AAs: 19.72699\nTotal prob of N-in: 0.96171\nPOSSIBLE N-term signal sequence\ninside 1 19\nTMhelix 20 39\noutside 40 325\nTMhelix 326 348\ninside 349 372

24030 GCF_000506945.1_S._Infantis_strain_335-3 Salmonella enterica subsp. enterica serovar Infantis str. 335-3 Proteobacteria; Gammaproteobacteria; Enterobacterales; Enterobacteriaceae; Salmonella; Salmonella enterica; Salmonella enterica subsp. enterica; Salmonella enterica subsp. enterica serovar Infantis MTGDNTLITSHGINRRDFMKLCAALAATMGLSSKAAAEMAESVSNPQRPPVIWIGAQECT WP_000145429.1 hydrogenase 2 small subunit [Salmonella enterica] Length: 372\nNumber of predicted TMHs: 1\nExp number of AAs in TMHs: 24.51309\nExp number, first 60 AAs: 0.2682\nTotal prob of N-in: 0.14402\noutside 1 331\nTMhelix 332 354\ninside 355 372

24031 GCF_001970895.1_ASM197089v1 Salmonella enterica subsp. enterica serovar Infantis str. CVM N13139 Proteobacteria; Gammaproteobacteria; Enterobacterales; Enterobacteriaceae; Salmonella; Salmonella enterica; Salmonella enterica subsp. enterica; Salmonella enterica subsp. enterica serovar Infantis MTGDNTLITSHGINRRDFMKLCAALAATMGLSSKAAAEMAESVSNPQRPPVIWIGAQECT WP_000145429.1 hydrogenase 2 small subunit [Salmonella enterica] Length: 372\nNumber of predicted TMHs: 1\nExp number of AAs in TMHs: 24.51309\nExp number, first 60 AAs: 0.2682\nTotal prob of N-in: 0.14402\noutside 1 331\nTMhelix 332 354\ninside 355 372

24032 GCF_001970895.1_ASM197089v1 Salmonella enterica subsp. enterica serovar Infantis str. CVM N13139 Proteobacteria; Gammaproteobacteria; Enterobacterales; Enterobacteriaceae; Salmonella; Salmonella enterica; Salmonella enterica subsp. enterica; Salmonella enterica subsp. enterica serovar Infantis MNNEETFYQAMRRKGVTRRSFLKFCSLAATSLGLGAGMTPKIAWALENKPRIPVVWIHGL WP_001058311.1 hydrogenase [Salmonella enterica] Length: 372\nNumber of predicted TMHs: 2\nExp number of AAs in TMHs: 37.4699899999999\nExp number, first 60 AAs: 19.72699\nTotal prob of N-in: 0.96171\nPOSSIBLE N-term signal sequence\ninside 1 19\nTMhelix 20 39\noutside 40 325\nTMhelix 326 348\ninside 349 372

24033 GCF_001970965.1_ASM197096v1 Salmonella enterica subsp. enterica serovar Infantis str. CVM N15228 Proteobacteria; Gammaproteobacteria; Enterobacterales; Enterobacteriaceae; Salmonella; Salmonella enterica; Salmonella enterica subsp. enterica; Salmonella enterica subsp. enterica serovar Infantis MNNEETFYQAMRRKGVTRRSFLKFCSLAATSLGLGAGMTPKIAWALENKPRIPVVWIHGL WP_001058311.1 hydrogenase [Salmonella enterica] Length: 372\nNumber of predicted TMHs: 2\nExp number of AAs in TMHs: 37.4699899999999\nExp number, first 60 AAs: 19.72699\nTotal prob of N-in: 0.96171\nPOSSIBLE N-term signal sequence\ninside 1 19\nTMhelix 20 39\noutside 40 325\nTMhelix 326 348\ninside 349 372

24034 GCF_001970965.1_ASM197096v1 Salmonella enterica subsp. enterica serovar Infantis str. CVM N15228 Proteobacteria; Gammaproteobacteria; Enterobacterales; Enterobacteriaceae; Salmonella; Salmonella enterica; Salmonella enterica subsp. enterica; Salmonella enterica subsp. enterica serovar Infantis MTGDNTLITSHGINRRDFMKLCAALAATMGLSSKAAAEMAESVSNPQRPPVIWIGAQECT WP_000145429.1 hydrogenase 2 small subunit [Salmonella enterica] Length: 372\nNumber of predicted TMHs: 1\nExp number of AAs in TMHs: 24.51309\nExp number, first 60 AAs: 0.2682\nTotal prob of N-in: 0.14402\noutside 1 331\nTMhelix 332 354\ninside 355 372

24035 GCF_001951705.1_ASM195170v1 Salmonella enterica subsp. enterica serovar Infantis str. CVM N15773 Proteobacteria; Gammaproteobacteria; Enterobacterales; Enterobacteriaceae; Salmonella; Salmonella enterica; Salmonella enterica subsp. enterica; Salmonella enterica subsp. enterica serovar Infantis MNNEETFYQAMRRKGVTRRSFLKFCSLAATSLGLGAGMTPKIAWALENKPRIPVVWIHGL WP_001058311.1 hydrogenase [Salmonella enterica] Length: 372\nNumber of predicted TMHs: 2\nExp number of AAs in TMHs: 37.4699899999999\nExp number, first 60 AAs: 19.72699\nTotal prob of N-in: 0.96171\nPOSSIBLE N-term signal sequence\ninside 1 19\nTMhelix 20 39\noutside 40 325\nTMhelix 326 348\ninside 349 372

24036 GCF_001951705.1_ASM195170v1 Salmonella enterica subsp. enterica serovar Infantis str. CVM N15773 Proteobacteria; Gammaproteobacteria; Enterobacterales; Enterobacteriaceae; Salmonella; Salmonella enterica;

Salmonella enterica subsp. enterica; Salmonella enterica subsp. enterica serovar Infantis
MTGDNTLITSHGINRRDFMKLCAALAATMGLSSKAAAEMAESVSNPQRPPVIWIGAQECT WP_000145429.1
hydrogenase 2 small subunit [Salmonella enterica] Length: 372\nNumber of predicted TMHs: 1\nExp number of AAs in TMHs: 24.51309\nExp number, first 60 AAs: 0.2682\nTotal prob of N-in: 0.14402\noutside 1 331\nTMhelix 332 354\ninside 355 372

24037 GCF_001951675.1_ASM195167v1 Salmonella enterica subsp. enterica serovar Infantis str. CVM N19983
Proteobacteria; Gammaproteobacteria; Enterobacterales; Enterobacteriaceae; Salmonella; Salmonella enterica; Salmonella enterica subsp. enterica; Salmonella enterica subsp. enterica serovar Infantis
MTGDNTLITSHGINRRDFMKLCAALAATMGLSSKAAAEMAESVSNPQRPPVIWIGAQECT WP_000145429.1
hydrogenase 2 small subunit [Salmonella enterica] Length: 372\nNumber of predicted TMHs: 1\nExp number of AAs in TMHs: 24.51309\nExp number, first 60 AAs: 0.2682\nTotal prob of N-in: 0.14402\noutside 1 331\nTMhelix 332 354\ninside 355 372

24038 GCF_001951675.1_ASM195167v1 Salmonella enterica subsp. enterica serovar Infantis str. CVM N19983
Proteobacteria; Gammaproteobacteria; Enterobacterales; Enterobacteriaceae; Salmonella; Salmonella enterica; Salmonella enterica subsp. enterica; Salmonella enterica subsp. enterica serovar Infantis
MNNEETFYQAMRRKGVTRRSFLKFCSLAATSLGLGAGMTPKIAWALENKPRIPVVWIHGL WP_001058311.1
hydrogenase [Salmonella enterica] Length: 372\nNumber of predicted TMHs: 2\nExp number of AAs in TMHs: 37.4699899999999\nExp number, first 60 AAs: 19.72699\nTotal prob of N-in: 0.96171\nPOSSIBLE N-term signal sequence\ninside 1 19\nTMhelix 20 39\noutside 40 325\nTMhelix 326 348\ninside 349 372

24039 GCF_001951685.1_ASM195168v1 Salmonella enterica subsp. enterica serovar Infantis str. CVM N20078
Proteobacteria; Gammaproteobacteria; Enterobacterales; Enterobacteriaceae; Salmonella; Salmonella enterica; Salmonella enterica subsp. enterica; Salmonella enterica subsp. enterica serovar Infantis
MTGDNTLITSHGINRRDFMKLCAALAATMGLSSKAAAEMAESVSNPQRPPVIWIGAQECT WP_000145429.1
hydrogenase 2 small subunit [Salmonella enterica] Length: 372\nNumber of predicted TMHs: 1\nExp number of AAs in TMHs: 24.51309\nExp number, first 60 AAs: 0.2682\nTotal prob of N-in: 0.14402\noutside 1 331\nTMhelix 332 354\ninside 355 372

24040 GCF_001951685.1_ASM195168v1 Salmonella enterica subsp. enterica serovar Infantis str. CVM N20078
Proteobacteria; Gammaproteobacteria; Enterobacterales; Enterobacteriaceae; Salmonella; Salmonella enterica; Salmonella enterica subsp. enterica; Salmonella enterica subsp. enterica serovar Infantis
MNNEETFYQAMRRKGVTRRSFLKFCSLAATSLGLGAGMTPKIAWALENKPRIPVVWIHGL WP_001058311.1
hydrogenase [Salmonella enterica] Length: 372\nNumber of predicted TMHs: 2\nExp number of AAs in TMHs: 37.4699899999999\nExp number, first 60 AAs: 19.72699\nTotal prob of N-in: 0.96171\nPOSSIBLE N-term signal sequence\ninside 1 19\nTMhelix 20 39\noutside 40 325\nTMhelix 326 348\ninside 349 372

24041 GCF_001951755.1_ASM195175v1 Salmonella enterica subsp. enterica serovar Infantis str. CVM N20272
Proteobacteria; Gammaproteobacteria; Enterobacterales; Enterobacteriaceae; Salmonella; Salmonella enterica; Salmonella enterica subsp. enterica; Salmonella enterica subsp. enterica serovar Infantis
MTGDNTLITSHGINRRDFMKLCAALAATMGLSSKAAAEMAESVSNPQRPPVIWIGAQECT WP_000145429.1
hydrogenase 2 small subunit [Salmonella enterica] Length: 372\nNumber of predicted TMHs: 1\nExp number of AAs in TMHs: 24.51309\nExp number, first 60 AAs: 0.2682\nTotal prob of N-in: 0.14402\noutside 1 331\nTMhelix 332 354\ninside 355 372

24042 GCF_001951755.1_ASM195175v1 Salmonella enterica subsp. enterica serovar Infantis str. CVM N20272
Proteobacteria; Gammaproteobacteria; Enterobacterales; Enterobacteriaceae; Salmonella; Salmonella enterica; Salmonella enterica subsp. enterica; Salmonella enterica subsp. enterica serovar Infantis
MNNEETFYQAMRRKGVTRRSFLKFCSLAATSLGLGAGMTPKIAWALENKPRIPVVWIHGL WP_052901769.1
hydrogenase [Salmonella enterica] Length: 372\nNumber of predicted TMHs: 2\nExp number of AAs in TMHs: 37.42632\nExp number, first 60 AAs: 19.72508\nTotal prob of N-in: 0.96168\nPOSSIBLE N-term signal sequence\ninside 1 19\nTMhelix 20 39\noutside 40 325\nTMhelix 326 348\ninside 349 372

24043 GCF_001951765.1_ASM195176v1 Salmonella enterica subsp. enterica serovar Infantis str. CVM N23532
Proteobacteria; Gammaproteobacteria; Enterobacterales; Enterobacteriaceae; Salmonella; Salmonella enterica; Salmonella enterica subsp. enterica; Salmonella enterica subsp. enterica serovar Infantis
MNNEETFYQAMRRKGVTRRSFLKFCSLAATSLGLGAGMTPKIAWALENKPRIPVVWIHGL WP_001058311.1
hydrogenase [Salmonella enterica] Length: 372\nNumber of predicted TMHs: 2\nExp number of AAs in TMHs: 37.4699899999999\nExp number, first 60 AAs: 19.72699\nTotal prob of N-in: 0.96171\nPOSSIBLE N-term signal sequence\ninside 1 19\nTMhelix 20 39\noutside 40 325\nTMhelix 326 348\ninside 349 372

24044 GCF_001951765.1_ASM195176v1 Salmonella enterica subsp. enterica serovar Infantis str. CVM N23532 Proteobacteria; Gammaproteobacteria; Enterobacterales; Enterobacteriaceae; Salmonella; Salmonella enterica; Salmonella enterica subsp. enterica; Salmonella enterica subsp. enterica serovar Infantis MTGDNTLITSHGINRRDFMKLCAALAATMGLSSKAAAEMAESVSNPQRPPVIWIGAQECT WP_000145429.1 hydrogenase 2 small subunit [Salmonella enterica] Length: 372\nNumber of predicted TMHs: 1\nExp number of AAs in TMHs: 24.51309\nExp number, first 60 AAs: 0.2682\nTotal prob of N-in: 0.14402\noutside 1 331\nTMhelix 332 354\ninside 355 372

24045 GCF_001951695.1_ASM195169v1 Salmonella enterica subsp. enterica serovar Infantis str. CVM N23771 Proteobacteria; Gammaproteobacteria; Enterobacterales; Enterobacteriaceae; Salmonella; Salmonella enterica; Salmonella enterica subsp. enterica; Salmonella enterica subsp. enterica serovar Infantis MTGDNTLITSHGINRRDFMKLCAALAATMGLSSKAAAEMAESVSNPQRPPVIWIGAQECT WP_000145429.1 hydrogenase 2 small subunit [Salmonella enterica] Length: 372\nNumber of predicted TMHs: 1\nExp number of AAs in TMHs: 24.51309\nExp number, first 60 AAs: 0.2682\nTotal prob of N-in: 0.14402\noutside 1 331\nTMhelix 332 354\ninside 355 372

24046 GCF_001951695.1_ASM195169v1 Salmonella enterica subsp. enterica serovar Infantis str. CVM N23771 Proteobacteria; Gammaproteobacteria; Enterobacterales; Enterobacteriaceae; Salmonella; Salmonella enterica; Salmonella enterica subsp. enterica; Salmonella enterica subsp. enterica serovar Infantis MNNEETFYQAMRRKGVTRRSFLKFCSLAATSLGLGAGMTPKIAWALENKPRIPVVWIHGL WP_052901769.1 hydrogenase [Salmonella enterica] Length: 372\nNumber of predicted TMHs: 2\nExp number of AAs in TMHs: 37.42632\nExp number, first 60 AAs: 19.72508\nTotal prob of N-in: 0.96168\nPOSSIBLE N-term signal sequence\ninside 1 19\nTMhelix 20 39\noutside 40 325\nTMhelix 326 348\ninside 349 372

24047 GCF_001951775.1_ASM195177v1 Salmonella enterica subsp. enterica serovar Infantis str. CVM N23791 Proteobacteria; Gammaproteobacteria; Enterobacterales; Enterobacteriaceae; Salmonella; Salmonella enterica; Salmonella enterica subsp. enterica; Salmonella enterica subsp. enterica serovar Infantis MTGDNTLITSHGINRRDFMKLCAALAATMGLSSKAAAEMAESVSNPQRPPVIWIGAQECT WP_000145429.1 hydrogenase 2 small subunit [Salmonella enterica] Length: 372\nNumber of predicted TMHs: 1\nExp number of AAs in TMHs: 24.51309\nExp number, first 60 AAs: 0.2682\nTotal prob of N-in: 0.14402\noutside 1 331\nTMhelix 332 354\ninside 355 372

24048 GCF_001951775.1_ASM195177v1 Salmonella enterica subsp. enterica serovar Infantis str. CVM N23791 Proteobacteria; Gammaproteobacteria; Enterobacterales; Enterobacteriaceae; Salmonella; Salmonella enterica; Salmonella enterica subsp. enterica; Salmonella enterica subsp. enterica serovar Infantis MNNEETFYQAMRRKGVTRRSFLKFCSLAATSLGLGAGMTPKIAWALENKPRIPVVWIHGL WP_001058311.1 hydrogenase [Salmonella enterica] Length: 372\nNumber of predicted TMHs: 2\nExp number of AAs in TMHs: 37.4699899999999\nExp number, first 60 AAs: 19.72699\nTotal prob of N-in: 0.96171\nPOSSIBLE N-term signal sequence\ninside 1 19\nTMhelix 20 39\noutside 40 325\nTMhelix 326 348\ninside 349 372

24049 GCF_001951835.1_ASM195183v1 Salmonella enterica subsp. enterica serovar Infantis str. CVM N26093 Proteobacteria; Gammaproteobacteria; Enterobacterales; Enterobacteriaceae; Salmonella; Salmonella enterica; Salmonella enterica subsp. enterica; Salmonella enterica subsp. enterica serovar Infantis MTGDNTLITSHGINRRDFMKLCAALAATMGLSSKAAAEMAESVSNPQRPPVIWIGAQECT WP_000145429.1 hydrogenase 2 small subunit [Salmonella enterica] Length: 372\nNumber of predicted TMHs: 1\nExp number of AAs in TMHs: 24.51309\nExp number, first 60 AAs: 0.2682\nTotal prob of N-in: 0.14402\noutside 1 331\nTMhelix 332 354\ninside 355 372

24050 GCF_001951835.1_ASM195183v1 Salmonella enterica subsp. enterica serovar Infantis str. CVM N26093 Proteobacteria; Gammaproteobacteria; Enterobacterales; Enterobacteriaceae; Salmonella; Salmonella enterica; Salmonella enterica subsp. enterica; Salmonella enterica subsp. enterica serovar Infantis MNNEETFYQAMRRKGVTRRSFLKFCSLAATSLGLGAGMTPKIAWALENKPRIPVVWIHGL WP_001058311.1 hydrogenase [Salmonella enterica] Length: 372\nNumber of predicted TMHs: 2\nExp number of AAs in TMHs: 37.4699899999999\nExp number, first 60 AAs: 19.72699\nTotal prob of N-in: 0.96171\nPOSSIBLE N-term signal sequence\ninside 1 19\nTMhelix 20 39\noutside 40 325\nTMhelix 326 348\ninside 349 372

24051 GCF_001951785.1_ASM195178v1 Salmonella enterica subsp. enterica serovar Infantis str. CVM N29304 Proteobacteria; Gammaproteobacteria; Enterobacterales; Enterobacteriaceae; Salmonella; Salmonella enterica; Salmonella enterica subsp. enterica; Salmonella enterica subsp. enterica serovar Infantis MNNEETFYQAMRRKGVTRRSFLKFCSLAATSLGLGAGMTPKIAWALENKPRIPVVWIHGL WP_001058311.1 hydrogenase [Salmonella enterica] Length: 372\nNumber of predicted TMHs: 2\nExp number of AAs in

TMHs: 37.4699899999999\nExp number, first 60 AAs:  19.72699\nTotal prob of N-in:   0.96171\nPOSSIBLE N-term signal sequence\ninside   1   19\nTMhelix   20   39\noutside   40   325\nTMhelix   326   348\ninside   349   372

24052 GCF_001951785.1_ASM195178v1 Salmonella enterica subsp. enterica serovar Infantis str. CVM N29304 Proteobacteria; Gammaproteobacteria; Enterobacterales; Enterobacteriaceae; Salmonella; Salmonella enterica; Salmonella enterica subsp. enterica; Salmonella enterica subsp. enterica serovar Infantis MTGDNTLITSHGINRRDFMKLCAALAATMGLSSKAAAEMAESVSNPQRPPVIWIGAQECT WP_000145429.1 hydrogenase 2 small subunit [Salmonella enterica] Length: 372\nNumber of predicted TMHs:  1\nExp number of AAs in TMHs: 24.51309\nExp number, first 60 AAs:  0.2682\nTotal prob of N-in:   0.14402\noutside   1   331\nTMhelix   332   354\ninside   355   372

24053 GCF_001970975.1_CLC_Genomic_Workbench Salmonella enterica subsp. enterica serovar Infantis str. CVM N29379 Proteobacteria; Gammaproteobacteria; Enterobacterales; Enterobacteriaceae; Salmonella; Salmonella enterica; Salmonella enterica subsp. enterica; Salmonella enterica subsp. enterica serovar Infantis MTGDNTLITSHGINRRDFMKLCAALAATMGLSSKAAAEMAESVSNPQRPPVIWIGAQECT WP_000145429.1 hydrogenase 2 small subunit [Salmonella enterica] Length: 372\nNumber of predicted TMHs:  1\nExp number of AAs in TMHs: 24.51309\nExp number, first 60 AAs:  0.2682\nTotal prob of N-in:   0.14402\noutside   1   331\nTMhelix   332   354\ninside   355   372

24054 GCF_001970975.1_CLC_Genomic_Workbench Salmonella enterica subsp. enterica serovar Infantis str. CVM N29379 Proteobacteria; Gammaproteobacteria; Enterobacterales; Enterobacteriaceae; Salmonella; Salmonella enterica; Salmonella enterica subsp. enterica; Salmonella enterica subsp. enterica serovar Infantis MNNEETFYQAMRRKGVTRRSFLKFCSLAATSLGLGAGMTPKIAWALENKPRIPVVWIHGL WP_052901769.1 hydrogenase [Salmonella enterica] Length: 372\nNumber of predicted TMHs:  2\nExp number of AAs in TMHs: 37.42632\nExp number, first 60 AAs:  19.72508\nTotal prob of N-in:   0.96168\nPOSSIBLE N-term signal sequence\ninside   1   19\nTMhelix   20   39\noutside   40   325\nTMhelix   326   348\ninside   349   372

24055 GCF_001970985.1_CLC_Genomic_Workbench Salmonella enterica subsp. enterica serovar Infantis str. CVM N31594 Proteobacteria; Gammaproteobacteria; Enterobacterales; Enterobacteriaceae; Salmonella; Salmonella enterica; Salmonella enterica subsp. enterica; Salmonella enterica subsp. enterica serovar Infantis MNNEETFYQAMRRKGVTRRSFLKFCSLAATSLGLGAGMTPKIAWALENKPRIPVVWIHGL WP_001058311.1 hydrogenase [Salmonella enterica] Length: 372\nNumber of predicted TMHs:  2\nExp number of AAs in TMHs: 37.4699899999999\nExp number, first 60 AAs:  19.72699\nTotal prob of N-in:   0.96171\nPOSSIBLE N-term signal sequence\ninside   1   19\nTMhelix   20   39\noutside   40   325\nTMhelix   326   348\ninside   349   372

24056 GCF_001970985.1_CLC_Genomic_Workbench Salmonella enterica subsp. enterica serovar Infantis str. CVM N31594 Proteobacteria; Gammaproteobacteria; Enterobacterales; Enterobacteriaceae; Salmonella; Salmonella enterica; Salmonella enterica subsp. enterica; Salmonella enterica subsp. enterica serovar Infantis MTGDNTLITSHGINRRDFMKLCAALAATMGLSSKAAAEMAESVSNPQRPPVIWIGAQECT WP_000145429.1 hydrogenase 2 small subunit [Salmonella enterica] Length: 372\nNumber of predicted TMHs:  1\nExp number of AAs in TMHs: 24.51309\nExp number, first 60 AAs:  0.2682\nTotal prob of N-in:   0.14402\noutside   1   331\nTMhelix   332   354\ninside   355   372

24057 GCF_001971045.1_CLC_Genomic_Workbench Salmonella enterica subsp. enterica serovar Infantis str. CVM N32760 Proteobacteria; Gammaproteobacteria; Enterobacterales; Enterobacteriaceae; Salmonella; Salmonella enterica; Salmonella enterica subsp. enterica; Salmonella enterica subsp. enterica serovar Infantis MNNEETFYQAMRRKGVTRRSFLKFCSLAATSLGLGAGMTPKIAWALENKPRIPVVWIHGL WP_001058311.1 hydrogenase [Salmonella enterica] Length: 372\nNumber of predicted TMHs:  2\nExp number of AAs in TMHs: 37.4699899999999\nExp number, first 60 AAs:  19.72699\nTotal prob of N-in:   0.96171\nPOSSIBLE N-term signal sequence\ninside   1   19\nTMhelix   20   39\noutside   40   325\nTMhelix   326   348\ninside   349   372

24058 GCF_001971045.1_CLC_Genomic_Workbench Salmonella enterica subsp. enterica serovar Infantis str. CVM N32760 Proteobacteria; Gammaproteobacteria; Enterobacterales; Enterobacteriaceae; Salmonella; Salmonella enterica; Salmonella enterica subsp. enterica; Salmonella enterica subsp. enterica serovar Infantis MTGDNTLITSHGINRRDFMKLCAALAATMGLSSKAAAEMAESVSNPQRPPVIWIGAQECT WP_000145429.1 hydrogenase 2 small subunit [Salmonella enterica] Length: 372\nNumber of predicted TMHs:  1\nExp number of AAs in TMHs: 24.51309\nExp number, first 60 AAs:  0.2682\nTotal prob of N-in:   0.14402\noutside   1   331\nTMhelix   332   354\ninside   355   372

24059 GCF_001971055.1_CLC_Genomic_Workbench Salmonella enterica subsp. enterica serovar Infantis str. CVM N35518 Proteobacteria; Gammaproteobacteria; Enterobacterales; Enterobacteriaceae; Salmonella; Salmonella enterica;

Salmonella enterica subsp. enterica; Salmonella enterica subsp. enterica serovar Infantis
MNNEETFYQAMRRKGVTRRSFLKFCSLAATSLGLGAGMTPKIAWALENKPRIPVVWIHGL WP_001058311.1
hydrogenase [Salmonella enterica] Length: 372\nNumber of predicted TMHs: 2\nExp number of AAs in TMHs: 37.4699899999999\nExp number, first 60 AAs: 19.72699\nTotal prob of N-in: 0.96171\nPOSSIBLE N-term signal sequence\ninside 1 19\nTMhelix 20 39\noutside 40 325\nTMhelix 326 348\ninside 349 372

24060 GCF_001971055.1_CLC_Genomic_Workbench Salmonella enterica subsp. enterica serovar Infantis str. CVM N35518 Proteobacteria; Gammaproteobacteria; Enterobacterales; Enterobacteriaceae; Salmonella; Salmonella enterica; Salmonella enterica subsp. enterica; Salmonella enterica subsp. enterica serovar Infantis
MTGDNTLITSHGINRRDFMKLCAALAATMGLSSKAAAEMAESVSNPQRPPVIWIGAQECT WP_000145429.1
hydrogenase 2 small subunit [Salmonella enterica] Length: 372\nNumber of predicted TMHs: 1\nExp number of AAs in TMHs: 24.51309\nExp number, first 60 AAs: 0.2682\nTotal prob of N-in: 0.14402\noutside 1 331\nTMhelix 332 354\ninside 355 372

24061 GCF_001952165.1_ASM195216v1 Salmonella enterica subsp. enterica serovar Infantis str. CVM N38848 Proteobacteria; Gammaproteobacteria; Enterobacterales; Enterobacteriaceae; Salmonella; Salmonella enterica; Salmonella enterica subsp. enterica; Salmonella enterica subsp. enterica serovar Infantis
MNNEETFYQAMRRKGVTRRSFLKFCSLAATSLGLGAGMTPKIAWALENKPRIPVVWIHGL WP_001058311.1
hydrogenase [Salmonella enterica] Length: 372\nNumber of predicted TMHs: 2\nExp number of AAs in TMHs: 37.4699899999999\nExp number, first 60 AAs: 19.72699\nTotal prob of N-in: 0.96171\nPOSSIBLE N-term signal sequence\ninside 1 19\nTMhelix 20 39\noutside 40 325\nTMhelix 326 348\ninside 349 372

24062 GCF_001952165.1_ASM195216v1 Salmonella enterica subsp. enterica serovar Infantis str. CVM N38848 Proteobacteria; Gammaproteobacteria; Enterobacterales; Enterobacteriaceae; Salmonella; Salmonella enterica; Salmonella enterica subsp. enterica; Salmonella enterica subsp. enterica serovar Infantis
MTGDNTLITSHGINRRDFMKLCAALAATMGLSSKAAAEMAESVSNPQRPPVIWIGAQECT WP_000145429.1
hydrogenase 2 small subunit [Salmonella enterica] Length: 372\nNumber of predicted TMHs: 1\nExp number of AAs in TMHs: 24.51309\nExp number, first 60 AAs: 0.2682\nTotal prob of N-in: 0.14402\noutside 1 331\nTMhelix 332 354\ninside 355 372

24063 GCF_001952215.1_ASM195221v1 Salmonella enterica subsp. enterica serovar Infantis str. CVM N41903 Proteobacteria; Gammaproteobacteria; Enterobacterales; Enterobacteriaceae; Salmonella; Salmonella enterica; Salmonella enterica subsp. enterica; Salmonella enterica subsp. enterica serovar Infantis
MTGDNTLITSHGINRRDFMKLCAALAATMGLSSKAAAEMAESVSNPQRPPVIWIGAQECT WP_000145429.1
hydrogenase 2 small subunit [Salmonella enterica] Length: 372\nNumber of predicted TMHs: 1\nExp number of AAs in TMHs: 24.51309\nExp number, first 60 AAs: 0.2682\nTotal prob of N-in: 0.14402\noutside 1 331\nTMhelix 332 354\ninside 355 372

24064 GCF_001952215.1_ASM195221v1 Salmonella enterica subsp. enterica serovar Infantis str. CVM N41903 Proteobacteria; Gammaproteobacteria; Enterobacterales; Enterobacteriaceae; Salmonella; Salmonella enterica; Salmonella enterica subsp. enterica; Salmonella enterica subsp. enterica serovar Infantis
MNNEETFYQAMRRKGVTRRSFLKFCSLAATSLGLGAGMTPKIAWALENKPRIPVVWIHGL WP_001058311.1
hydrogenase [Salmonella enterica] Length: 372\nNumber of predicted TMHs: 2\nExp number of AAs in TMHs: 37.4699899999999\nExp number, first 60 AAs: 19.72699\nTotal prob of N-in: 0.96171\nPOSSIBLE N-term signal sequence\ninside 1 19\nTMhelix 20 39\noutside 40 325\nTMhelix 326 348\ninside 349 372

24065 GCF_001952225.1_ASM195222v1 Salmonella enterica subsp. enterica serovar Infantis str. CVM N42236 Proteobacteria; Gammaproteobacteria; Enterobacterales; Enterobacteriaceae; Salmonella; Salmonella enterica; Salmonella enterica subsp. enterica; Salmonella enterica subsp. enterica serovar Infantis
MNNEETFYQAMRRKGVTRRSFLKFCSLAATSLGLGAGMTPKIAWALENKPRIPVVWIHGL WP_001058311.1
hydrogenase [Salmonella enterica] Length: 372\nNumber of predicted TMHs: 2\nExp number of AAs in TMHs: 37.4699899999999\nExp number, first 60 AAs: 19.72699\nTotal prob of N-in: 0.96171\nPOSSIBLE N-term signal sequence\ninside 1 19\nTMhelix 20 39\noutside 40 325\nTMhelix 326 348\ninside 349 372

24066 GCF_001952225.1_ASM195222v1 Salmonella enterica subsp. enterica serovar Infantis str. CVM N42236 Proteobacteria; Gammaproteobacteria; Enterobacterales; Enterobacteriaceae; Salmonella; Salmonella enterica; Salmonella enterica subsp. enterica; Salmonella enterica subsp. enterica serovar Infantis
MTGDNTLITSHGINRRDFMKLCAALAATMGLSSKAAAEMAESVSNPQRPPVIWIGAQECT WP_000145429.1
hydrogenase 2 small subunit [Salmonella enterica] Length: 372\nNumber of predicted TMHs: 1\nExp number of AAs in TMHs: 24.51309\nExp number, first 60 AAs: 0.2682\nTotal prob of N-in: 0.14402\noutside 1 331\nTMhelix 332 354\ninside 355 372

24067 GCF_001952205.1_ASM195220v1 Salmonella enterica subsp. enterica serovar Infantis str. CVM N42459 Proteobacteria; Gammaproteobacteria; Enterobacterales; Enterobacteriaceae; Salmonella; Salmonella enterica; Salmonella enterica subsp. enterica; Salmonella enterica subsp. enterica serovar Infantis MTGDNTLITSHGINRRDFMKLCAALAATMGLSSKAAAEMAESVSNPQRPPVIWIGAQECT WP_000145429.1 hydrogenase 2 small subunit [Salmonella enterica] Length: 372\nNumber of predicted TMHs: 1\nExp number of AAs in TMHs: 24.51309\nExp number, first 60 AAs: 0.2682\nTotal prob of N-in: 0.14402\noutside 1 331\nTMhelix 332 354\ninside 355 372

24068 GCF_001952205.1_ASM195220v1 Salmonella enterica subsp. enterica serovar Infantis str. CVM N42459 Proteobacteria; Gammaproteobacteria; Enterobacterales; Enterobacteriaceae; Salmonella; Salmonella enterica; Salmonella enterica subsp. enterica; Salmonella enterica subsp. enterica serovar Infantis MNNEETFYQAMRRKGVTRRSFLKFCSLAATSLGLGAGMTPKIAWALENKPRIPVVWIHGL WP_052901769.1 hydrogenase [Salmonella enterica] Length: 372\nNumber of predicted TMHs: 2\nExp number of AAs in TMHs: 37.42632\nExp number, first 60 AAs: 19.72508\nTotal prob of N-in: 0.96168\nPOSSIBLE N-term signal sequence\ninside 1 19\nTMhelix 20 39\noutside 40 325\nTMhelix 326 348\ninside 349 372

24069 GCF_000230875.1_Salmonella_enterica_SARB27-2.0 Salmonella enterica subsp. enterica serovar Infantis str. SARB27 Proteobacteria; Gammaproteobacteria; Enterobacterales; Enterobacteriaceae; Salmonella; Salmonella enterica; Salmonella enterica subsp. enterica; Salmonella enterica subsp. enterica serovar Infantis MTGDNTLITSHGINRRDFMKLCAALAATMGLSSKAAAEMAESVSNPQRPPVIWIGAQECT WP_000145431.1 hydrogenase 2 small subunit [Salmonella enterica] Length: 372\nNumber of predicted TMHs: 1\nExp number of AAs in TMHs: 24.51505\nExp number, first 60 AAs: 0.2682\nTotal prob of N-in: 0.14406\noutside 1 331\nTMhelix 332 354\ninside 355 372

24070 GCF_000230875.1_Salmonella_enterica_SARB27-2.0 Salmonella enterica subsp. enterica serovar Infantis str. SARB27 Proteobacteria; Gammaproteobacteria; Enterobacterales; Enterobacteriaceae; Salmonella; Salmonella enterica; Salmonella enterica subsp. enterica; Salmonella enterica subsp. enterica serovar Infantis MNNEETFYQAMRRKGVTRRSFLKFCSLAATSLGLGAGMTPKIAWALENKPRIPVVWIHGL WP_001662439.1 hydrogenase [Salmonella enterica] Length: 372\nNumber of predicted TMHs: 2\nExp number of AAs in TMHs: 36.5172\nExp number, first 60 AAs: 19.72849\nTotal prob of N-in: 0.96169\nPOSSIBLE N-term signal sequence\ninside 1 19\nTMhelix 20 39\noutside 40 325\nTMhelix 326 348\ninside 349 372

24071 GCF_000487155.2_ASM48715v2 Salmonella enterica subsp. enterica serovar Inverness str. ATCC 10720 Proteobacteria; Gammaproteobacteria; Enterobacterales; Enterobacteriaceae; Salmonella; Salmonella enterica; Salmonella enterica subsp. enterica; Salmonella enterica subsp. enterica serovar Inverness MNNEETFYQAMRRKGVTRRSFLKFCSLAATSLGLGAGMTPKIAWALENKPRIPVVWIHGL WP_001058312.1 hydrogenase [Salmonella enterica] Length: 372\nNumber of predicted TMHs: 2\nExp number of AAs in TMHs: 36.51738\nExp number, first 60 AAs: 19.72849\nTotal prob of N-in: 0.96170\nPOSSIBLE N-term signal sequence\ninside 1 19\nTMhelix 20 39\noutside 40 325\nTMhelix 326 348\ninside 349 372

24072 GCF_000487155.2_ASM48715v2 Salmonella enterica subsp. enterica serovar Inverness str. ATCC 10720 Proteobacteria; Gammaproteobacteria; Enterobacterales; Enterobacteriaceae; Salmonella; Salmonella enterica; Salmonella enterica subsp. enterica; Salmonella enterica subsp. enterica serovar Inverness MTGDNTLITSHGINRRDFMKLCAALAATMGLSSKAAAEMAESVSNPQRPPVIWIGAQECT WP_000145429.1 hydrogenase 2 small subunit [Salmonella enterica] Length: 372\nNumber of predicted TMHs: 1\nExp number of AAs in TMHs: 24.51309\nExp number, first 60 AAs: 0.2682\nTotal prob of N-in: 0.14402\noutside 1 331\nTMhelix 332 354\ninside 355 372

24073 GCF_001448495.1_ASM144849v1 Salmonella enterica subsp. enterica serovar Itami str. SA20014991 Proteobacteria; Gammaproteobacteria; Enterobacterales; Enterobacteriaceae; Salmonella; Salmonella enterica; Salmonella enterica subsp. enterica; Salmonella enterica subsp. enterica serovar Itami MTGDNTLITSHGINRRDFMKLCAALAATMGLSSKAAAEMAESVSNPQRPPVIWIGAQECT WP_000145429.1 hydrogenase 2 small subunit [Salmonella enterica] Length: 372\nNumber of predicted TMHs: 1\nExp number of AAs in TMHs: 24.51309\nExp number, first 60 AAs: 0.2682\nTotal prob of N-in: 0.14402\noutside 1 331\nTMhelix 332 354\ninside 355 372

24074 GCF_001448495.1_ASM144849v1 Salmonella enterica subsp. enterica serovar Itami str. SA20014991 Proteobacteria; Gammaproteobacteria; Enterobacterales; Enterobacteriaceae; Salmonella; Salmonella enterica; Salmonella enterica subsp. enterica; Salmonella enterica subsp. enterica serovar Itami MNNEETFYQAMRRKGVTRRSFLKFCSLAATSLGLGAGMTPKIAWALENKPRIPVVWIHGL WP_001058312.1 hydrogenase [Salmonella enterica] Length: 372\nNumber of predicted TMHs: 2\nExp number of AAs in

TMHs: 36.51738\nExp number, first 60 AAs: 19.72849\nTotal prob of N-in: 0.96170\nPOSSIBLE N-term signal sequence\ninside 1 19\nTMhelix 20 39\noutside 40 325\nTMhelix 326 348\ninside 349 372

24075 GCF_000487095.1_ASM48709v1 Salmonella enterica subsp. enterica serovar Javiana str. 10721 Proteobacteria; Gammaproteobacteria; Enterobacterales; Enterobacteriaceae; Salmonella; Salmonella enterica; Salmonella enterica subsp. enterica; Salmonella enterica subsp. enterica serovar Javiana MNNEETFYQAMRRKGVTRRSFLKFCSLAATSLGLGAGMTPKIAWALENKPRIPVVWIHGL WP_001058312.1 hydrogenase [Salmonella enterica] Length: 372\nNumber of predicted TMHs: 2\nExp number of AAs in TMHs: 36.51738\nExp number, first 60 AAs: 19.72849\nTotal prob of N-in: 0.96170\nPOSSIBLE N-term signal sequence\ninside 1 19\nTMhelix 20 39\noutside 40 325\nTMhelix 326 348\ninside 349 372

24076 GCF_000487095.1_ASM48709v1 Salmonella enterica subsp. enterica serovar Javiana str. 10721 Proteobacteria; Gammaproteobacteria; Enterobacterales; Enterobacteriaceae; Salmonella; Salmonella enterica; Salmonella enterica subsp. enterica; Salmonella enterica subsp. enterica serovar Javiana MTGDNTLITSHGINRRDFMKLCAALAATMGLSSKAAAEMAESVSNPQRPPVIWIGAQECT WP_000145431.1 hydrogenase 2 small subunit [Salmonella enterica] Length: 372\nNumber of predicted TMHs: 1\nExp number of AAs in TMHs: 24.51505\nExp number, first 60 AAs: 0.2682\nTotal prob of N-in: 0.14406\noutside 1 331\nTMhelix 332 354\ninside 355 372

24077 GCF_000335955.1_ASM33595v1 Salmonella enterica subsp. enterica serovar Javiana str. ATCC BAA-1593 Proteobacteria; Gammaproteobacteria; Enterobacterales; Enterobacteriaceae; Salmonella; Salmonella enterica; Salmonella enterica subsp. enterica; Salmonella enterica subsp. enterica serovar Javiana MNNEETFYQAMRRKGVTRRSFLKFCSLAATSLGLGAGMTPKIAWALENKPRIPVVWIHGL WP_001058312.1 hydrogenase [Salmonella enterica] Length: 372\nNumber of predicted TMHs: 2\nExp number of AAs in TMHs: 36.51738\nExp number, first 60 AAs: 19.72849\nTotal prob of N-in: 0.96170\nPOSSIBLE N-term signal sequence\ninside 1 19\nTMhelix 20 39\noutside 40 325\nTMhelix 326 348\ninside 349 372

24078 GCF_000335955.1_ASM33595v1 Salmonella enterica subsp. enterica serovar Javiana str. ATCC BAA-1593 Proteobacteria; Gammaproteobacteria; Enterobacterales; Enterobacteriaceae; Salmonella; Salmonella enterica; Salmonella enterica subsp. enterica; Salmonella enterica subsp. enterica serovar Javiana MTGDNTLITSHGINRRDFMKLCAALAATMGLSSKAAAEMAESVSNPQRPPVIWIGAQECT WP_000145429.1 hydrogenase 2 small subunit [Salmonella enterica] Length: 372\nNumber of predicted TMHs: 1\nExp number of AAs in TMHs: 24.51309\nExp number, first 60 AAs: 0.2682\nTotal prob of N-in: 0.14402\noutside 1 331\nTMhelix 332 354\ninside 355 372

24079 GCF_001707555.1_ASM170755v1 Salmonella enterica subsp. enterica serovar Javiana str. CFSAN000905 Proteobacteria; Gammaproteobacteria; Enterobacterales; Enterobacteriaceae; Salmonella; Salmonella enterica; Salmonella enterica subsp. enterica; Salmonella enterica subsp. enterica serovar Javiana MNNEETFYQAMRRKGVTRRSFLKFCSLAATSLGLGAGMTPKIAWALENKPRIPVVWIHGL WP_001058312.1 hydrogenase [Salmonella enterica] Length: 372\nNumber of predicted TMHs: 2\nExp number of AAs in TMHs: 36.51738\nExp number, first 60 AAs: 19.72849\nTotal prob of N-in: 0.96170\nPOSSIBLE N-term signal sequence\ninside 1 19\nTMhelix 20 39\noutside 40 325\nTMhelix 326 348\ninside 349 372

24080 GCF_001707555.1_ASM170755v1 Salmonella enterica subsp. enterica serovar Javiana str. CFSAN000905 Proteobacteria; Gammaproteobacteria; Enterobacterales; Enterobacteriaceae; Salmonella; Salmonella enterica; Salmonella enterica subsp. enterica; Salmonella enterica subsp. enterica serovar Javiana MTGDNTLITSHGINRRDFMKLCAALAATMGLSSKAAAEMAESVSNPQRPPVIWIGAQECT WP_000145429.1 hydrogenase 2 small subunit [Salmonella enterica] Length: 372\nNumber of predicted TMHs: 1\nExp number of AAs in TMHs: 24.51309\nExp number, first 60 AAs: 0.2682\nTotal prob of N-in: 0.14402\noutside 1 331\nTMhelix 332 354\ninside 355 372

24081 GCF_000341425.1_ASM34142v1 Salmonella enterica subsp. enterica serovar Javiana str. CFSAN001992 Proteobacteria; Gammaproteobacteria; Enterobacterales; Enterobacteriaceae; Salmonella; Salmonella enterica; Salmonella enterica subsp. enterica; Salmonella enterica subsp. enterica serovar Javiana MNNEETFYQAMRRKGVTRRSFLKFCSLAATSLGLGAGMTPKIAWALENKPRIPVVWIHGL WP_001058312.1 hydrogenase [Salmonella enterica] Length: 372\nNumber of predicted TMHs: 2\nExp number of AAs in TMHs: 36.51738\nExp number, first 60 AAs: 19.72849\nTotal prob of N-in: 0.96170\nPOSSIBLE N-term signal sequence\ninside 1 19\nTMhelix 20 39\noutside 40 325\nTMhelix 326 348\ninside 349 372

24082 GCF_000341425.1_ASM34142v1 Salmonella enterica subsp. enterica serovar Javiana str. CFSAN001992 Proteobacteria; Gammaproteobacteria; Enterobacterales; Enterobacteriaceae; Salmonella; Salmonella enterica;

Salmonella enterica subsp. enterica; Salmonella enterica subsp. enterica serovar Javiana
MTGDNTLITSHGINRRDFMKLCAALAATMGLSSKAAAEMAESVSNPQRPPVIWIGAQECT WP_000145429.1
hydrogenase 2 small subunit [Salmonella enterica] Length: 372\nNumber of predicted TMHs: 1\nExp number of AAs in TMHs: 24.51309\nExp number, first 60 AAs: 0.2682\nTotal prob of N-in: 0.14402\noutside 1 331\nTMhelix 332 354\ninside 355 372

24083 GCF_000171255.1_ASM17125v1 Salmonella enterica subsp. enterica serovar Javiana str. GA_MM04042433
Proteobacteria; Gammaproteobacteria; Enterobacterales; Enterobacteriaceae; Salmonella; Salmonella enterica; Salmonella enterica subsp. enterica; Salmonella enterica subsp. enterica serovar Javiana
MNNEETFYQAMRRKGVTRRSFLKFCSLAATSLGLGAGMTPKIAWALENKPRIPVVWIHGL WP_001058312.1
hydrogenase [Salmonella enterica] Length: 372\nNumber of predicted TMHs: 2\nExp number of AAs in TMHs: 36.51738\nExp number, first 60 AAs: 19.72849\nTotal prob of N-in: 0.96170\nPOSSIBLE N-term signal sequence\ninside 1 19\nTMhelix 20 39\noutside 40 325\nTMhelix 326 348\ninside 349 372

24084 GCF_000171255.1_ASM17125v1 Salmonella enterica subsp. enterica serovar Javiana str. GA_MM04042433
Proteobacteria; Gammaproteobacteria; Enterobacterales; Enterobacteriaceae; Salmonella; Salmonella enterica; Salmonella enterica subsp. enterica; Salmonella enterica subsp. enterica serovar Javiana
MTGDNTLITSHGINRRDFMKLCAALAATMGLSSKAAAEMAESVSNPQRPPVIWIGAQECT WP_000145429.1
hydrogenase 2 small subunit [Salmonella enterica] Length: 372\nNumber of predicted TMHs: 1\nExp number of AAs in TMHs: 24.51309\nExp number, first 60 AAs: 0.2682\nTotal prob of N-in: 0.14402\noutside 1 331\nTMhelix 332 354\ninside 355 372

24085 GCF_000487115.1_ASM48711v1 Salmonella enterica subsp. enterica serovar Javiana str. PRS_2010_0720
Proteobacteria; Gammaproteobacteria; Enterobacterales; Enterobacteriaceae; Salmonella; Salmonella enterica; Salmonella enterica subsp. enterica; Salmonella enterica subsp. enterica serovar Javiana
MTGDNTLITSHGINRRDFMKLCAALAATMGLSSKAAAEMAESVSNPQRPPVIWIGAQECT WP_000145429.1
hydrogenase 2 small subunit [Salmonella enterica] Length: 372\nNumber of predicted TMHs: 1\nExp number of AAs in TMHs: 24.51309\nExp number, first 60 AAs: 0.2682\nTotal prob of N-in: 0.14402\noutside 1 331\nTMhelix 332 354\ninside 355 372

24086 GCF_000487115.1_ASM48711v1 Salmonella enterica subsp. enterica serovar Javiana str. PRS_2010_0720
Proteobacteria; Gammaproteobacteria; Enterobacterales; Enterobacteriaceae; Salmonella; Salmonella enterica; Salmonella enterica subsp. enterica; Salmonella enterica subsp. enterica serovar Javiana
MNNEETFYQAMRRKGVTRRSFLKFCSLAATSLGLGAGMTPKIAWALENKPRIPVVWIHGL WP_001058312.1
hydrogenase [Salmonella enterica] Length: 372\nNumber of predicted TMHs: 2\nExp number of AAs in TMHs: 36.51738\nExp number, first 60 AAs: 19.72849\nTotal prob of N-in: 0.96170\nPOSSIBLE N-term signal sequence\ninside 1 19\nTMhelix 20 39\noutside 40 325\nTMhelix 326 348\ninside 349 372

24087 GCF_001975405.1_ASM197540v1 Salmonella enterica subsp. enterica serovar Johannesburg str. ST203
Proteobacteria; Gammaproteobacteria; Enterobacterales; Enterobacteriaceae; Salmonella; Salmonella enterica; Salmonella enterica subsp. enterica; Salmonella enterica subsp. enterica serovar Johannesburg
MNNEETFYQAMRRKGVTRRSFLKFCSLAATSLGLGAGMTPKIAWALENKPRIPVVWIHGL WP_023198237.1
hydrogenase 1, small subunit [Salmonella enterica] Length: 372\nNumber of predicted TMHs: 2\nExp number of AAs in TMHs: 36.39423\nExp number, first 60 AAs: 19.72877\nTotal prob of N-in: 0.96162\nPOSSIBLE N-term signal sequence\ninside 1 19\nTMhelix 20 39\noutside 40 325\nTMhelix 326 348\ninside 349 372

24088 GCF_001975405.1_ASM197540v1 Salmonella enterica subsp. enterica serovar Johannesburg str. ST203
Proteobacteria; Gammaproteobacteria; Enterobacterales; Enterobacteriaceae; Salmonella; Salmonella enterica; Salmonella enterica subsp. enterica; Salmonella enterica subsp. enterica serovar Johannesburg
MTGDNTLITSHGINRRDFMKLCAALAATMGLSSKAAAEMAESVSNPQRPPVIWIGAQECT WP_000145429.1
hydrogenase 2 small subunit [Salmonella enterica] Length: 372\nNumber of predicted TMHs: 1\nExp number of AAs in TMHs: 24.51309\nExp number, first 60 AAs: 0.2682\nTotal prob of N-in: 0.14402\noutside 1 331\nTMhelix 332 354\ninside 355 372

24089 GCF_000484035.1_ASM48403v1 Salmonella enterica subsp. enterica serovar Kentucky str. 0253
Proteobacteria; Gammaproteobacteria; Enterobacterales; Enterobacteriaceae; Salmonella; Salmonella enterica; Salmonella enterica subsp. enterica; Salmonella enterica subsp. enterica serovar Kentucky
MNNEETFYQAMRRKGVTRRSFLKFCSLAATSLGLGAGMTPKIAWALENKPRIPVVWIHGL WP_001058311.1
hydrogenase [Salmonella enterica] Length: 372\nNumber of predicted TMHs: 2\nExp number of AAs in TMHs: 37.4699899999999\nExp number, first 60 AAs: 19.72699\nTotal prob of N-in: 0.96171\nPOSSIBLE N-term signal sequence\ninside 1 19\nTMhelix 20 39\noutside 40 325\nTMhelix 326 348\ninside 349 372

24090 GCF_000484035.1_ASM48403v1 Salmonella enterica subsp. enterica serovar Kentucky str. 0253 Proteobacteria; Gammaproteobacteria; Enterobacterales; Enterobacteriaceae; Salmonella; Salmonella enterica; Salmonella enterica subsp. enterica; Salmonella enterica subsp. enterica serovar Kentucky MTGDNTLITSHGINRRDFMKLCAALAATMGLSSKAAAEMAESVSNPQRPPVIWIGAQECT WP_000145429.1 hydrogenase 2 small subunit [Salmonella enterica] Length: 372\nNumber of predicted TMHs: 1\nExp number of AAs in TMHs: 24.51309\nExp number, first 60 AAs: 0.2682\nTotal prob of N-in: 0.14402\noutside 1 331\nTMhelix 332 354\ninside 355 372

24091 GCF_000474415.1_CFSAN001162_01.0Salmonella enterica subsp. enterica serovar Kentucky str. 13562 Proteobacteria; Gammaproteobacteria; Enterobacterales; Enterobacteriaceae; Salmonella; Salmonella enterica; Salmonella enterica subsp. enterica; Salmonella enterica subsp. enterica serovar Kentucky MTGDNTLITSHGINRRDFMKLCAALAATMGLSSKAAAEMAESVSNPQRPPVIWIGAQECT WP_000145429.1 hydrogenase 2 small subunit [Salmonella enterica] Length: 372\nNumber of predicted TMHs: 1\nExp number of AAs in TMHs: 24.51309\nExp number, first 60 AAs: 0.2682\nTotal prob of N-in: 0.14402\noutside 1 331\nTMhelix 332 354\ninside 355 372

24092 GCF_000474415.1_CFSAN001162_01.0Salmonella enterica subsp. enterica serovar Kentucky str. 13562 Proteobacteria; Gammaproteobacteria; Enterobacterales; Enterobacteriaceae; Salmonella; Salmonella enterica; Salmonella enterica subsp. enterica; Salmonella enterica subsp. enterica serovar Kentucky MNNEETFYQAMRRKGVTRRSFLKFCSLAATSLGLGAGMTPKIAWALENKPRIPVVWIHGL WP_001058311.1 hydrogenase [Salmonella enterica] Length: 372\nNumber of predicted TMHs: 2\nExp number of AAs in TMHs: 37.4699899999999\nExp number, first 60 AAs: 19.72699\nTotal prob of N-in: 0.96171\nPOSSIBLE N-term signal sequence\ninside 1 19\nTMhelix 20 39\noutside 40 325\nTMhelix 326 348\ninside 349 372

24093 GCF_000474455.1_CFSAN001166_01.0Salmonella enterica subsp. enterica serovar Kentucky str. 20793 Proteobacteria; Gammaproteobacteria; Enterobacterales; Enterobacteriaceae; Salmonella; Salmonella enterica; Salmonella enterica subsp. enterica; Salmonella enterica subsp. enterica serovar Kentucky MTGDNTLITSHGINRRDFMKLCAALAATMGLSSKAAAEMAESVSNPQRPPVIWIGAQECT WP_000145429.1 hydrogenase 2 small subunit [Salmonella enterica] Length: 372\nNumber of predicted TMHs: 1\nExp number of AAs in TMHs: 24.51309\nExp number, first 60 AAs: 0.2682\nTotal prob of N-in: 0.14402\noutside 1 331\nTMhelix 332 354\ninside 355 372

24094 GCF_000474455.1_CFSAN001166_01.0Salmonella enterica subsp. enterica serovar Kentucky str. 20793 Proteobacteria; Gammaproteobacteria; Enterobacterales; Enterobacteriaceae; Salmonella; Salmonella enterica; Salmonella enterica subsp. enterica; Salmonella enterica subsp. enterica serovar Kentucky MNNEETFYQAMRRKGVTRRSFLKFCSLAATSLGLGAGMTPKIAWALENKPRIPVVWIHGL WP_001058311.1 hydrogenase [Salmonella enterica] Length: 372\nNumber of predicted TMHs: 2\nExp number of AAs in TMHs: 37.4699899999999\nExp number, first 60 AAs: 19.72699\nTotal prob of N-in: 0.96171\nPOSSIBLE N-term signal sequence\ninside 1 19\nTMhelix 20 39\noutside 40 325\nTMhelix 326 348\ninside 349 372

24095 GCF_000474295.1_CFSAN001164_01.0Salmonella enterica subsp. enterica serovar Kentucky str. 22694 Proteobacteria; Gammaproteobacteria; Enterobacterales; Enterobacteriaceae; Salmonella; Salmonella enterica; Salmonella enterica subsp. enterica; Salmonella enterica subsp. enterica serovar Kentucky MTGDNTLITSHGINRRDFMKLCAALAATMGLSSKAAAEMAESVSNPQRPPVIWIGAQECT WP_000145429.1 hydrogenase 2 small subunit [Salmonella enterica] Length: 372\nNumber of predicted TMHs: 1\nExp number of AAs in TMHs: 24.51309\nExp number, first 60 AAs: 0.2682\nTotal prob of N-in: 0.14402\noutside 1 331\nTMhelix 332 354\ninside 355 372

24096 GCF_000474295.1_CFSAN001164_01.0Salmonella enterica subsp. enterica serovar Kentucky str. 22694 Proteobacteria; Gammaproteobacteria; Enterobacterales; Enterobacteriaceae; Salmonella; Salmonella enterica; Salmonella enterica subsp. enterica; Salmonella enterica subsp. enterica serovar Kentucky MNNEETFYQAMRRKGVTRRSFLKFCSLAATSLGLGAGMTPKIAWALENKPRIPVVWIHGL WP_001058311.1 hydrogenase [Salmonella enterica] Length: 372\nNumber of predicted TMHs: 2\nExp number of AAs in TMHs: 37.4699899999999\nExp number, first 60 AAs: 19.72699\nTotal prob of N-in: 0.96171\nPOSSIBLE N-term signal sequence\ninside 1 19\nTMhelix 20 39\noutside 40 325\nTMhelix 326 348\ninside 349 372

24097 GCF_000474435.1_CFSAN001163_01.0Salmonella enterica subsp. enterica serovar Kentucky str. 29166 Proteobacteria; Gammaproteobacteria; Enterobacterales; Enterobacteriaceae; Salmonella; Salmonella enterica; Salmonella enterica subsp. enterica; Salmonella enterica subsp. enterica serovar Kentucky MTGDNTLITSHGINRRDFMKLCAALAATMGLSSKAAAEMAESVSNPQRPPVIWIGAQECT WP_000145429.1 hydrogenase 2 small subunit [Salmonella enterica] Length: 372\nNumber of predicted TMHs: 1\nExp

number of AAs in TMHs: 24.51309\nExp number, first 60 AAs: 0.2682\nTotal prob of N-in: 0.14402\noutside 1 331\nTMhelix 332 354\ninside 355 372

24098 GCF_000474435.1_CFSAN001163_01.0Salmonella enterica subsp. enterica serovar Kentucky str. 29166 Proteobacteria; Gammaproteobacteria; Enterobacterales; Enterobacteriaceae; Salmonella; Salmonella enterica; Salmonella enterica subsp. enterica; Salmonella enterica subsp. enterica serovar Kentucky MNNEETFYQAMRRKGVTRRSFLKFCSLAATSLGLGAGMTPKIAWALENKPRIPVVWIHGL WP_001058311.1 hydrogenase [Salmonella enterica] Length: 372\nNumber of predicted TMHs: 2\nExp number of AAs in TMHs: 37.4699899999999\nExp number, first 60 AAs: 19.72699\nTotal prob of N-in: 0.96171\nPOSSIBLE N-term signal sequence\ninside 1 19\nTMhelix 20 39\noutside 40 325\nTMhelix 326 348\ninside 349 372

24099 GCF_000336135.1_ASM33613v1 Salmonella enterica subsp. enterica serovar Kentucky str. 29439 Proteobacteria; Gammaproteobacteria; Enterobacterales; Enterobacteriaceae; Salmonella; Salmonella enterica; Salmonella enterica subsp. enterica; Salmonella enterica subsp. enterica serovar Kentucky MNNEETFYQAMRRKGVTRRSFLKFCSLAATSLGLGAGMTPKIAWALENKPRIPVVWIHGL WP_001058311.1 hydrogenase [Salmonella enterica] Length: 372\nNumber of predicted TMHs: 2\nExp number of AAs in TMHs: 37.4699899999999\nExp number, first 60 AAs: 19.72699\nTotal prob of N-in: 0.96171\nPOSSIBLE N-term signal sequence\ninside 1 19\nTMhelix 20 39\noutside 40 325\nTMhelix 326 348\ninside 349 372

24100 GCF_000336135.1_ASM33613v1 Salmonella enterica subsp. enterica serovar Kentucky str. 29439 Proteobacteria; Gammaproteobacteria; Enterobacterales; Enterobacteriaceae; Salmonella; Salmonella enterica; Salmonella enterica subsp. enterica; Salmonella enterica subsp. enterica serovar Kentucky MTGDNTLITSHGINRRDFMKLCAALAATMGLSSKAAAEMAESVSNPQRPPVIWIGAQECT WP_000145429.1 hydrogenase 2 small subunit [Salmonella enterica] Length: 372\nNumber of predicted TMHs: 1\nExp number of AAs in TMHs: 24.51309\nExp number, first 60 AAs: 0.2682\nTotal prob of N-in: 0.14402\noutside 1 331\nTMhelix 332 354\ninside 355 372

24101 GCF_000487075.1_ASM48707v1 Salmonella enterica subsp. enterica serovar Kentucky str. 5349 Proteobacteria; Gammaproteobacteria; Enterobacterales; Enterobacteriaceae; Salmonella; Salmonella enterica; Salmonella enterica subsp. enterica; Salmonella enterica subsp. enterica serovar Kentucky MTGDNTLITSHGINRRDFMKLCAALAATMGLSSKAAAEMAESVSNPQRPPVIWIGAQECT WP_000145429.1 hydrogenase 2 small subunit [Salmonella enterica] Length: 372\nNumber of predicted TMHs: 1\nExp number of AAs in TMHs: 24.51309\nExp number, first 60 AAs: 0.2682\nTotal prob of N-in: 0.14402\noutside 1 331\nTMhelix 332 354\ninside 355 372

24102 GCF_000487075.1_ASM48707v1 Salmonella enterica subsp. enterica serovar Kentucky str. 5349 Proteobacteria; Gammaproteobacteria; Enterobacterales; Enterobacteriaceae; Salmonella; Salmonella enterica; Salmonella enterica subsp. enterica; Salmonella enterica subsp. enterica serovar Kentucky MNNEETFYQAMRRKGVTRRSFLKFCSLAATSLGLGAGMTPKIAWALENKPRIPVVWIHGL WP_001058311.1 hydrogenase [Salmonella enterica] Length: 372\nNumber of predicted TMHs: 2\nExp number of AAs in TMHs: 37.4699899999999\nExp number, first 60 AAs: 19.72699\nTotal prob of N-in: 0.96171\nPOSSIBLE N-term signal sequence\ninside 1 19\nTMhelix 20 39\noutside 40 325\nTMhelix 326 348\ninside 349 372

24103 GCF_000171275.1_ASM17127v1 Salmonella enterica subsp. enterica serovar Kentucky str. CDC 191 Proteobacteria; Gammaproteobacteria; Enterobacterales; Enterobacteriaceae; Salmonella; Salmonella enterica; Salmonella enterica subsp. enterica; Salmonella enterica subsp. enterica serovar Kentucky MTGDNTLITSHGINRRDFMKLCAALAATMGLSSKAAAEMAESVSNPQRPPVIWIGAQECT WP_000145429.1 hydrogenase 2 small subunit [Salmonella enterica] Length: 372\nNumber of predicted TMHs: 1\nExp number of AAs in TMHs: 24.51309\nExp number, first 60 AAs: 0.2682\nTotal prob of N-in: 0.14402\noutside 1 331\nTMhelix 332 354\ninside 355 372

24104 GCF_000171275.1_ASM17127v1 Salmonella enterica subsp. enterica serovar Kentucky str. CDC 191 Proteobacteria; Gammaproteobacteria; Enterobacterales; Enterobacteriaceae; Salmonella; Salmonella enterica; Salmonella enterica subsp. enterica; Salmonella enterica subsp. enterica serovar Kentucky MNNEETFYQAMRRKGVTRRSFLKFCSLAATSLGLGAGMTPKIAWALENKPRIPVVWIHGL WP_001058311.1 hydrogenase [Salmonella enterica] Length: 372\nNumber of predicted TMHs: 2\nExp number of AAs in TMHs: 37.4699899999999\nExp number, first 60 AAs: 19.72699\nTotal prob of N-in: 0.96171\nPOSSIBLE N-term signal sequence\ninside 1 19\nTMhelix 20 39\noutside 40 325\nTMhelix 326 348\ninside 349 372

24105 GCF_000170195.2_ASM17019v2 Salmonella enterica subsp. enterica serovar Kentucky str. CVM29188 Proteobacteria; Gammaproteobacteria; Enterobacterales; Enterobacteriaceae; Salmonella; Salmonella enterica;

Salmonella enterica subsp. enterica; Salmonella enterica subsp. enterica serovar Kentucky
MTGDNTLITSHGINRRDFMKLCAALAATMGLSSKAAAEMAESVSNPQRPPVIWIGAQECT WP_000145429.1
hydrogenase 2 small subunit [Salmonella enterica] Length: 372\nNumber of predicted TMHs: 1\nExp number of AAs in TMHs: 24.51309\nExp number, first 60 AAs: 0.2682\nTotal prob of N-in: 0.14402\noutside 1 331\nTMhelix 332 354\ninside 355 372

24106 GCF_000170195.2_ASM17019v2 Salmonella enterica subsp. enterica serovar Kentucky str. CVM29188
Proteobacteria; Gammaproteobacteria; Enterobacterales; Enterobacteriaceae; Salmonella; Salmonella enterica; Salmonella enterica subsp. enterica; Salmonella enterica subsp. enterica serovar Kentucky
MNNEETFYQAMRRKGVTRRSFLKFCSLAATSLGLGAGMTPKIAWALENKPRIPVVWIHGL WP_001058311.1
hydrogenase [Salmonella enterica] Length: 372\nNumber of predicted TMHs: 2\nExp number of AAs in TMHs: 37.4699899999999\nExp number, first 60 AAs: 19.72699\nTotal prob of N-in: 0.96171\nPOSSIBLE N-term signal sequence\ninside 1 19\nTMhelix 20 39\noutside 40 325\nTMhelix 326 348\ninside 349 372

24107 GCF_000474315.1_CFSAN001165_01.0Salmonella enterica subsp. enterica serovar Kentucky str. N312
Proteobacteria; Gammaproteobacteria; Enterobacterales; Enterobacteriaceae; Salmonella; Salmonella enterica; Salmonella enterica subsp. enterica; Salmonella enterica subsp. enterica serovar Kentucky
MNNEETFYQAMRRKGVTRRSFLKFCSLAATSLGLGAGMTPKIAWALENKPRIPVVWIHGL WP_001058311.1
hydrogenase [Salmonella enterica] Length: 372\nNumber of predicted TMHs: 2\nExp number of AAs in TMHs: 37.4699899999999\nExp number, first 60 AAs: 19.72699\nTotal prob of N-in: 0.96171\nPOSSIBLE N-term signal sequence\ninside 1 19\nTMhelix 20 39\noutside 40 325\nTMhelix 326 348\ninside 349 372

24108 GCF_000474315.1_CFSAN001165_01.0Salmonella enterica subsp. enterica serovar Kentucky str. N312
Proteobacteria; Gammaproteobacteria; Enterobacterales; Enterobacteriaceae; Salmonella; Salmonella enterica; Salmonella enterica subsp. enterica; Salmonella enterica subsp. enterica serovar Kentucky
MTGDNTLITSHGINRRDFMKLCAALAATMGLSSKAAAEMAESVSNPQRPPVIWIGAQECT WP_000145429.1
hydrogenase 2 small subunit [Salmonella enterica] Length: 372\nNumber of predicted TMHs: 1\nExp number of AAs in TMHs: 24.51309\nExp number, first 60 AAs: 0.2682\nTotal prob of N-in: 0.14402\noutside 1 331\nTMhelix 332 354\ninside 355 372

24109 GCF_001448485.1_ASM144848v1 Salmonella enterica subsp. enterica serovar Kentucky str. SA20030505
Proteobacteria; Gammaproteobacteria; Enterobacterales; Enterobacteriaceae; Salmonella; Salmonella enterica; Salmonella enterica subsp. enterica; Salmonella enterica subsp. enterica serovar Kentucky
MNNEETFYQAMRRKGVTRRSFLKFCSLAATSLGLGAGMTPKIAWALENKPRIPVVWIHGL WP_001058311.1
hydrogenase [Salmonella enterica] Length: 372\nNumber of predicted TMHs: 2\nExp number of AAs in TMHs: 37.4699899999999\nExp number, first 60 AAs: 19.72699\nTotal prob of N-in: 0.96171\nPOSSIBLE N-term signal sequence\ninside 1 19\nTMhelix 20 39\noutside 40 325\nTMhelix 326 348\ninside 349 372

24110 GCF_001448485.1_ASM144848v1 Salmonella enterica subsp. enterica serovar Kentucky str. SA20030505
Proteobacteria; Gammaproteobacteria; Enterobacterales; Enterobacteriaceae; Salmonella; Salmonella enterica; Salmonella enterica subsp. enterica; Salmonella enterica subsp. enterica serovar Kentucky
MTGDNTLITSHGINRRDFMKLCAALAATMGLSSKAAAEMAESVSNPQRPPVIWIGAQECT WP_000145429.1
hydrogenase 2 small subunit [Salmonella enterica] Length: 372\nNumber of predicted TMHs: 1\nExp number of AAs in TMHs: 24.51309\nExp number, first 60 AAs: 0.2682\nTotal prob of N-in: 0.14402\noutside 1 331\nTMhelix 332 354\ninside 355 372

24111 GCF_001975425.1_ASM197542v1 Salmonella enterica subsp. enterica serovar Koessen str. S-1501
Proteobacteria; Gammaproteobacteria; Enterobacterales; Enterobacteriaceae; Salmonella; Salmonella enterica; Salmonella enterica subsp. enterica; Salmonella enterica subsp. enterica serovar Koessen
MTGDNTLITSHGINRRDFMKLCAALAATMGLSSKAAAEMAESVSNPQRPPVIWIGAQECT WP_000145429.1
hydrogenase 2 small subunit [Salmonella enterica] Length: 372\nNumber of predicted TMHs: 1\nExp number of AAs in TMHs: 24.51309\nExp number, first 60 AAs: 0.2682\nTotal prob of N-in: 0.14402\noutside 1 331\nTMhelix 332 354\ninside 355 372

24112 GCF_001975425.1_ASM197542v1 Salmonella enterica subsp. enterica serovar Koessen str. S-1501
Proteobacteria; Gammaproteobacteria; Enterobacterales; Enterobacteriaceae; Salmonella; Salmonella enterica; Salmonella enterica subsp. enterica; Salmonella enterica subsp. enterica serovar Koessen
MNNEETFYQAMRRKGVTRRSFLKFCSLAATSLGLGAGMTPKIAWALENKPRIPVVWIHGL WP_001058312.1
hydrogenase [Salmonella enterica] Length: 372\nNumber of predicted TMHs: 2\nExp number of AAs in TMHs: 36.51738\nExp number, first 60 AAs: 19.72849\nTotal prob of N-in: 0.96170\nPOSSIBLE N-term signal sequence\ninside 1 19\nTMhelix 20 39\noutside 40 325\nTMhelix 326 348\ninside 349 372

24113 GCF_001975445.1_ASM197544v1 Salmonella enterica subsp. enterica serovar Krefeld str. SA20030536 Proteobacteria; Gammaproteobacteria; Enterobacterales; Enterobacteriaceae; Salmonella; Salmonella enterica; Salmonella enterica subsp. enterica; Salmonella enterica subsp. enterica serovar Krefeld MTGDNTLITSHGINRRDFMKLCAALAATMGLSSKAAAEMAESVSNPQRPPVIWIGAQECT WP_000145429.1 hydrogenase 2 small subunit [Salmonella enterica] Length: 372\nNumber of predicted TMHs: 1\nExp number of AAs in TMHs: 24.51309\nExp number, first 60 AAs: 0.2682\nTotal prob of N-in: 0.14402\noutside 1 331\nTMhelix 332 354\ninside 355 372

24114 GCF_001975445.1_ASM197544v1 Salmonella enterica subsp. enterica serovar Krefeld str. SA20030536 Proteobacteria; Gammaproteobacteria; Enterobacterales; Enterobacteriaceae; Salmonella; Salmonella enterica; Salmonella enterica subsp. enterica; Salmonella enterica subsp. enterica serovar Krefeld MNNEETFYQAMRRKGVTRRSFLKFCSLAATSLGLGAGMTPKIAWALENKPRIPVVWIHGL WP_001058312.1 hydrogenase [Salmonella enterica] Length: 372\nNumber of predicted TMHs: 2\nExp number of AAs in TMHs: 36.51738\nExp number, first 60 AAs: 19.72849\nTotal prob of N-in: 0.96170\nPOSSIBLE N-term signal sequence\ninside 1 19\nTMhelix 20 39\noutside 40 325\nTMhelix 326 348\ninside 349 372

24115 GCF_000487855.1_ASM48785v1 Salmonella enterica subsp. enterica serovar Litchfield str. CFSAN001076 Proteobacteria; Gammaproteobacteria; Enterobacterales; Enterobacteriaceae; Salmonella; Salmonella enterica; Salmonella enterica subsp. enterica; Salmonella enterica subsp. enterica serovar Litchfield MNNEETFYQAMRRKGVTRRSFLKFCSLAATSLGLGAGMTPKIAWALENKPRIPVVWIHGL WP_001058312.1 hydrogenase [Salmonella enterica] Length: 372\nNumber of predicted TMHs: 2\nExp number of AAs in TMHs: 36.51738\nExp number, first 60 AAs: 19.72849\nTotal prob of N-in: 0.96170\nPOSSIBLE N-term signal sequence\ninside 1 19\nTMhelix 20 39\noutside 40 325\nTMhelix 326 348\ninside 349 372

24116 GCF_000487855.1_ASM48785v1 Salmonella enterica subsp. enterica serovar Litchfield str. CFSAN001076 Proteobacteria; Gammaproteobacteria; Enterobacterales; Enterobacteriaceae; Salmonella; Salmonella enterica; Salmonella enterica subsp. enterica; Salmonella enterica subsp. enterica serovar Litchfield MTGDNTLITSHGINRRDFMKLCAALAATMGLSSKAAAEMAESVSNPQRPPVIWIGAQECT WP_000145429.1 hydrogenase 2 small subunit [Salmonella enterica] Length: 372\nNumber of predicted TMHs: 1\nExp number of AAs in TMHs: 24.51309\nExp number, first 60 AAs: 0.2682\nTotal prob of N-in: 0.14402\noutside 1 331\nTMhelix 332 354\ninside 355 372

24117 GCF_001975515.1_ASM197551v1 Salmonella enterica subsp. enterica serovar Manchester str. ST278 Proteobacteria; Gammaproteobacteria; Enterobacterales; Enterobacteriaceae; Salmonella; Salmonella enterica; Salmonella enterica subsp. enterica; Salmonella enterica subsp. enterica serovar Manchester MNNEETFYQAMRRKGVTRRSFLKFCSLAATSLGLGAGMTPKIAWALENKPRIPVVWIHGL WP_001058311.1 hydrogenase [Salmonella enterica] Length: 372\nNumber of predicted TMHs: 2\nExp number of AAs in TMHs: 37.4699899999999\nExp number, first 60 AAs: 19.72699\nTotal prob of N-in: 0.96171\nPOSSIBLE N-term signal sequence\ninside 1 19\nTMhelix 20 39\noutside 40 325\nTMhelix 326 348\ninside 349 372

24118 GCF_001975515.1_ASM197551v1 Salmonella enterica subsp. enterica serovar Manchester str. ST278 Proteobacteria; Gammaproteobacteria; Enterobacterales; Enterobacteriaceae; Salmonella; Salmonella enterica; Salmonella enterica subsp. enterica; Salmonella enterica subsp. enterica serovar Manchester MTGDNTLITSHGINRRDFMKLCAALAATMGLSSKAAAEMAESVSNPQRPPVIWIGAQECT WP_000145429.1 hydrogenase 2 small subunit [Salmonella enterica] Length: 372\nNumber of predicted TMHs: 1\nExp number of AAs in TMHs: 24.51309\nExp number, first 60 AAs: 0.2682\nTotal prob of N-in: 0.14402\noutside 1 331\nTMhelix 332 354\ninside 355 372

24119 GCF_000430975.2_ASM43097v2 Salmonella enterica subsp. enterica serovar Manhattan str. 111113 Proteobacteria; Gammaproteobacteria; Enterobacterales; Enterobacteriaceae; Salmonella; Salmonella enterica; Salmonella enterica subsp. enterica; Salmonella enterica subsp. enterica serovar Manhattan MNNEETFYQAMRRKGVTRRSFLKFCSLAATSLGLGAGMTPKIAWALENKPRIPVVWIHGL WP_001058311.1 hydrogenase [Salmonella enterica] Length: 372\nNumber of predicted TMHs: 2\nExp number of AAs in TMHs: 37.4699899999999\nExp number, first 60 AAs: 19.72699\nTotal prob of N-in: 0.96171\nPOSSIBLE N-term signal sequence\ninside 1 19\nTMhelix 20 39\noutside 40 325\nTMhelix 326 348\ninside 349 372

24120 GCF_000430975.2_ASM43097v2 Salmonella enterica subsp. enterica serovar Manhattan str. 111113 Proteobacteria; Gammaproteobacteria; Enterobacterales; Enterobacteriaceae; Salmonella; Salmonella enterica; Salmonella enterica subsp. enterica; Salmonella enterica subsp. enterica serovar Manhattan MTGDNTLITSHGINRRDFMKLCAALAATMGLSSKAAAEMAESVSNPQRPPVIWIGAQECT WP_000145429.1 hydrogenase 2 small subunit [Salmonella enterica] Length: 372\nNumber of predicted TMHs: 1\nExp

number of AAs in TMHs: 24.51309\nExp number, first 60 AAs: 0.2682\nTotal prob of N-in: 0.14402\noutside 1 331\nTMhelix 332 354\ninside 355 372

24121 GCF_000487815.1_ASM48781v1 Salmonella enterica subsp. enterica serovar Manhattan str. CFSAN001078 Proteobacteria; Gammaproteobacteria; Enterobacterales; Enterobacteriaceae; Salmonella; Salmonella enterica; Salmonella enterica subsp. enterica; Salmonella enterica subsp. enterica serovar Manhattan MTGDNTLITSHGINRRDFMKLCAALAATMGLSSKAAAEMAESVSNPQRPPVIWIGAQECT WP_000145429.1 hydrogenase 2 small subunit [Salmonella enterica] Length: 372\nNumber of predicted TMHs: 1\nExp number of AAs in TMHs: 24.51309\nExp number, first 60 AAs: 0.2682\nTotal prob of N-in: 0.14402\noutside 1 331\nTMhelix 332 354\ninside 355 372

24122 GCF_000487815.1_ASM48781v1 Salmonella enterica subsp. enterica serovar Manhattan str. CFSAN001078 Proteobacteria; Gammaproteobacteria; Enterobacterales; Enterobacteriaceae; Salmonella; Salmonella enterica; Salmonella enterica subsp. enterica; Salmonella enterica subsp. enterica serovar Manhattan MNNEETFYQAMRRKGVTRRSFLKFCSLAATSLGLGAGMTPKIAWALENKPRIPVVWIHGL WP_001058311.1 hydrogenase [Salmonella enterica] Length: 372\nNumber of predicted TMHs: 2\nExp number of AAs in TMHs: 37.4699899999999\nExp number, first 60 AAs: 19.72699\nTotal prob of N-in: 0.96171\nPOSSIBLE N-term signal sequence\ninside 1 19\nTMhelix 20 39\noutside 40 325\nTMhelix 326 348\ninside 349 372

24123 GCF_001448765.1_ASM144876v1 Salmonella enterica subsp. enterica serovar Manhattan str. SA20034532 Proteobacteria; Gammaproteobacteria; Enterobacterales; Enterobacteriaceae; Salmonella; Salmonella enterica; Salmonella enterica subsp. enterica; Salmonella enterica subsp. enterica serovar Manhattan MTGDNTLITSHGINRRDFMKLCAALAATMGLSSKAAAEMAESVSNPQRPPVIWIGAQECT WP_000145429.1 hydrogenase 2 small subunit [Salmonella enterica] Length: 372\nNumber of predicted TMHs: 1\nExp number of AAs in TMHs: 24.51309\nExp number, first 60 AAs: 0.2682\nTotal prob of N-in: 0.14402\noutside 1 331\nTMhelix 332 354\ninside 355 372

24124 GCF_001448765.1_ASM144876v1 Salmonella enterica subsp. enterica serovar Manhattan str. SA20034532 Proteobacteria; Gammaproteobacteria; Enterobacterales; Enterobacteriaceae; Salmonella; Salmonella enterica; Salmonella enterica subsp. enterica; Salmonella enterica subsp. enterica serovar Manhattan MNNEETFYQAMRRKGVTRRSFLKFCSLAATSLGLGAGMTPKIAWALENKPRIPVVWIHGL WP_001058311.1 hydrogenase [Salmonella enterica] Length: 372\nNumber of predicted TMHs: 2\nExp number of AAs in TMHs: 37.4699899999999\nExp number, first 60 AAs: 19.72699\nTotal prob of N-in: 0.96171\nPOSSIBLE N-term signal sequence\ninside 1 19\nTMhelix 20 39\noutside 40 325\nTMhelix 326 348\ninside 349 372

24125 GCF_000316015.1_2009K-0807v1 Salmonella enterica subsp. enterica serovar Mbandaka str. 2009K-0807 Proteobacteria; Gammaproteobacteria; Enterobacterales; Enterobacteriaceae; Salmonella; Salmonella enterica; Salmonella enterica subsp. enterica; Salmonella enterica subsp. enterica serovar Mbandaka MTGDNTLITSHGINRRDFMKLCAALAATMGLSSKAAAEMAESVSNPQRPPVIWIGAQECT WP_000145429.1 hydrogenase 2 small subunit [Salmonella enterica] Length: 372\nNumber of predicted TMHs: 1\nExp number of AAs in TMHs: 24.51309\nExp number, first 60 AAs: 0.2682\nTotal prob of N-in: 0.14402\noutside 1 331\nTMhelix 332 354\ninside 355 372

24126 GCF_000316015.1_2009K-0807v1 Salmonella enterica subsp. enterica serovar Mbandaka str. 2009K-0807 Proteobacteria; Gammaproteobacteria; Enterobacterales; Enterobacteriaceae; Salmonella; Salmonella enterica; Salmonella enterica subsp. enterica; Salmonella enterica subsp. enterica serovar Mbandaka MNNEETFYQAMRRKGVTRRSFLKFCSLAATSLGLGAGMTPKIAWALENKPRIPVVWIHGL WP_001058311.1 hydrogenase [Salmonella enterica] Length: 372\nNumber of predicted TMHs: 2\nExp number of AAs in TMHs: 37.4699899999999\nExp number, first 60 AAs: 19.72699\nTotal prob of N-in: 0.96171\nPOSSIBLE N-term signal sequence\ninside 1 19\nTMhelix 20 39\noutside 40 325\nTMhelix 326 348\ninside 349 372

24127 GCF_000416295.1_2012K-0273_Ver1 Salmonella enterica subsp. enterica serovar Mbandaka str. 2012K-0273 Proteobacteria; Gammaproteobacteria; Enterobacterales; Enterobacteriaceae; Salmonella; Salmonella enterica; Salmonella enterica subsp. enterica; Salmonella enterica subsp. enterica serovar Mbandaka MTGDNTLITSHGINRRDFMKLCAALAATMGLSSKAAAEMAESVSNPQRPPVIWIGAQECT WP_000145429.1 hydrogenase 2 small subunit [Salmonella enterica] Length: 372\nNumber of predicted TMHs: 1\nExp number of AAs in TMHs: 24.51309\nExp number, first 60 AAs: 0.2682\nTotal prob of N-in: 0.14402\noutside 1 331\nTMhelix 332 354\ninside 355 372

24128 GCF_000486915.2_ASM48691v2 Salmonella enterica subsp. enterica serovar Mbandaka str. ATCC 51958 Proteobacteria; Gammaproteobacteria; Enterobacterales; Enterobacteriaceae; Salmonella; Salmonella enterica;

Salmonella enterica subsp. enterica; Salmonella enterica subsp. enterica serovar Mbandaka
MTGDNTLITSHGINRRDFMKLCAALAATMGLSSKAAAEMAESVSNPQRPPVIWIGAQECT WP_000145429.1
hydrogenase 2 small subunit [Salmonella enterica] Length: 372\nNumber of predicted TMHs: 1\nExp number of AAs in TMHs: 24.51309\nExp number, first 60 AAs: 0.2682\nTotal prob of N-in: 0.14402\noutside 1 331\nTMhelix 332 354\ninside 355 372

24129 GCF_000486915.2_ASM48691v2 Salmonella enterica subsp. enterica serovar Mbandaka str. ATCC 51958
Proteobacteria; Gammaproteobacteria; Enterobacterales; Enterobacteriaceae; Salmonella; Salmonella enterica; Salmonella enterica subsp. enterica; Salmonella enterica subsp. enterica serovar Mbandaka
MNNEETFYQAMRRKGVTRRSFLKFCSLAATSLGLGAGMTPKIAWALENKPRIPVVWIHGL WP_001058310.1
hydrogenase [Salmonella enterica] Length: 372\nNumber of predicted TMHs: 2\nExp number of AAs in TMHs: 37.4567299999999\nExp number, first 60 AAs: 19.72694\nTotal prob of N-in: 0.96171\nPOSSIBLE N-term signal sequence\ninside 1 19\nTMhelix 20 39\noutside 40 325\nTMhelix 326 348\ninside 349 372

24130 GCF_000486935.1_ASM48693v1 Salmonella enterica subsp. enterica serovar Miami str. 1923
Proteobacteria; Gammaproteobacteria; Enterobacterales; Enterobacteriaceae; Salmonella; Salmonella enterica; Salmonella enterica subsp. enterica; Salmonella enterica subsp. enterica serovar Miami
MTGDNTLITSHGINRRDFMKLCAALAATMGLSSKAAAEMAESVSNPQRPPVIWIGAQECT WP_000145429.1
hydrogenase 2 small subunit [Salmonella enterica] Length: 372\nNumber of predicted TMHs: 1\nExp number of AAs in TMHs: 24.51309\nExp number, first 60 AAs: 0.2682\nTotal prob of N-in: 0.14402\noutside 1 331\nTMhelix 332 354\ninside 355 372

24131 GCF_000486935.1_ASM48693v1 Salmonella enterica subsp. enterica serovar Miami str. 1923
Proteobacteria; Gammaproteobacteria; Enterobacterales; Enterobacteriaceae; Salmonella; Salmonella enterica; Salmonella enterica subsp. enterica; Salmonella enterica subsp. enterica serovar Miami
MNNEETFYQAMRRKGVTRRSFLKFCSLAATSLGLGAGMTPKIAWALENKPRIPVVWIHGL WP_001662439.1
hydrogenase [Salmonella enterica] Length: 372\nNumber of predicted TMHs: 2\nExp number of AAs in TMHs: 36.5172\nExp number, first 60 AAs: 19.72849\nTotal prob of N-in: 0.96169\nPOSSIBLE N-term signal sequence\ninside 1 19\nTMhelix 20 39\noutside 40 325\nTMhelix 326 348\ninside 349 372

24132 GCF_001448785.1_ASM144878v1 Salmonella enterica subsp. enterica serovar Milwaukee str. SA19950795
Proteobacteria; Gammaproteobacteria; Enterobacterales; Enterobacteriaceae; Salmonella; Salmonella enterica; Salmonella enterica subsp. enterica; Salmonella enterica subsp. enterica serovar Milwaukee
MNNEETFYQAMRRKGVTRRSFLKFCSLAATSLGLGAGMTPKIAWALENKPRIPVVWIHGL WP_001058311.1
hydrogenase [Salmonella enterica] Length: 372\nNumber of predicted TMHs: 2\nExp number of AAs in TMHs: 37.4699899999999\nExp number, first 60 AAs: 19.72699\nTotal prob of N-in: 0.96171\nPOSSIBLE N-term signal sequence\ninside 1 19\nTMhelix 20 39\noutside 40 325\nTMhelix 326 348\ninside 349 372

24133 GCF_001448785.1_ASM144878v1 Salmonella enterica subsp. enterica serovar Milwaukee str. SA19950795
Proteobacteria; Gammaproteobacteria; Enterobacterales; Enterobacteriaceae; Salmonella; Salmonella enterica; Salmonella enterica subsp. enterica; Salmonella enterica subsp. enterica serovar Milwaukee
MTGDNTLITSHGINRRDFMKLCAALAATMGLSSKAAAEMAESVSNPQRPPVIWIGAQECT WP_000145429.1
hydrogenase 2 small subunit [Salmonella enterica] Length: 372\nNumber of predicted TMHs: 1\nExp number of AAs in TMHs: 24.51309\nExp number, first 60 AAs: 0.2682\nTotal prob of N-in: 0.14402\noutside 1 331\nTMhelix 332 354\ninside 355 372

24134 GCF_000486855.2_ASM48685v2 Salmonella enterica subsp. enterica serovar Minnesota str. ATCC 49284
Proteobacteria; Gammaproteobacteria; Enterobacterales; Enterobacteriaceae; Salmonella; Salmonella enterica; Salmonella enterica subsp. enterica; Salmonella enterica subsp. enterica serovar Minnesota
MTGDNTLITSHGINRRDFMKLCAALAATMGLSSKAAAEMAESVSNPQRPPVIWIGAQECT WP_000145429.1
hydrogenase 2 small subunit [Salmonella enterica] Length: 372\nNumber of predicted TMHs: 1\nExp number of AAs in TMHs: 24.51309\nExp number, first 60 AAs: 0.2682\nTotal prob of N-in: 0.14402\noutside 1 331\nTMhelix 332 354\ninside 355 372

24135 GCF_000293115.1_ASM29311v1 Salmonella enterica subsp. enterica serovar Mississippi str. 2010K-1406
Proteobacteria; Gammaproteobacteria; Enterobacterales; Enterobacteriaceae; Salmonella; Salmonella enterica; Salmonella enterica subsp. enterica; Salmonella enterica subsp. enterica serovar Mississippi
MNNEETFYQAMRRKGVTRRSFLKFCSLAATSLGLGAGMTPKIAWALENKPRIPVVWIHGL WP_001058312.1
hydrogenase [Salmonella enterica] Length: 372\nNumber of predicted TMHs: 2\nExp number of AAs in TMHs: 36.51738\nExp number, first 60 AAs: 19.72849\nTotal prob of N-in: 0.96170\nPOSSIBLE N-term signal sequence\ninside 1 19\nTMhelix 20 39\noutside 40 325\nTMhelix 326 348\ninside 349 372

24136 GCF_000293115.1_ASM29311v1 Salmonella enterica subsp. enterica serovar Mississippi str. 2010K-1406 Proteobacteria; Gammaproteobacteria; Enterobacterales; Enterobacteriaceae; Salmonella; Salmonella enterica; Salmonella enterica subsp. enterica; Salmonella enterica subsp. enterica serovar Mississippi MTGDNTLITSHGINRRDFMKLCAALAATMGLSSKAAAEMAESVSNPQRPPVIWIGAQECT WP_000145429.1 hydrogenase 2 small subunit [Salmonella enterica] Length: 372\nNumber of predicted TMHs: 1\nExp number of AAs in TMHs: 24.51309\nExp number, first 60 AAs: 0.2682\nTotal prob of N-in: 0.14402\noutside 1 331\nTMhelix 332 354\ninside 355 372

24137 GCF_000188535.1_ASM18853v2 Salmonella enterica subsp. enterica serovar Montevideo str. 19N Proteobacteria; Gammaproteobacteria; Enterobacterales; Enterobacteriaceae; Salmonella; Salmonella enterica; Salmonella enterica subsp. enterica; Salmonella enterica subsp. enterica serovar Montevideo MNNEETFYQAMRRKGVTRRSFLKFCSLAATSLGLGAGMTPKIAWALENKPRIPVVWIHGL WP_001058312.1 hydrogenase [Salmonella enterica] Length: 372\nNumber of predicted TMHs: 2\nExp number of AAs in TMHs: 36.51738\nExp number, first 60 AAs: 19.72849\nTotal prob of N-in: 0.96170\nPOSSIBLE N-term signal sequence\ninside 1 19\nTMhelix 20 39\noutside 40 325\nTMhelix 326 348\ninside 349 372

24138 GCF_000188535.1_ASM18853v2 Salmonella enterica subsp. enterica serovar Montevideo str. 19N Proteobacteria; Gammaproteobacteria; Enterobacterales; Enterobacteriaceae; Salmonella; Salmonella enterica; Salmonella enterica subsp. enterica; Salmonella enterica subsp. enterica serovar Montevideo MTGDNTLITSHGINRRDFMKLCAALAATMGLSSKAAAEMAESVSNPQRPPVIWIGAQECT WP_000145429.1 hydrogenase 2 small subunit [Salmonella enterica] Length: 372\nNumber of predicted TMHs: 1\nExp number of AAs in TMHs: 24.51309\nExp number, first 60 AAs: 0.2682\nTotal prob of N-in: 0.14402\noutside 1 331\nTMhelix 332 354\ninside 355 372

24139 GCF_000189075.1_ASM18907v2 Salmonella enterica subsp. enterica serovar Montevideo str. 2009083312 Proteobacteria; Gammaproteobacteria; Enterobacterales; Enterobacteriaceae; Salmonella; Salmonella enterica; Salmonella enterica subsp. enterica; Salmonella enterica subsp. enterica serovar Montevideo MTGDNTLITSHGINRRDFMKLCAALAATMGLSSKAAAEMAESVSNPQRPPVIWIGAQECT WP_000145429.1 hydrogenase 2 small subunit [Salmonella enterica] Length: 372\nNumber of predicted TMHs: 1\nExp number of AAs in TMHs: 24.51309\nExp number, first 60 AAs: 0.2682\nTotal prob of N-in: 0.14402\noutside 1 331\nTMhelix 332 354\ninside 355 372

24140 GCF_000189075.1_ASM18907v2 Salmonella enterica subsp. enterica serovar Montevideo str. 2009083312 Proteobacteria; Gammaproteobacteria; Enterobacterales; Enterobacteriaceae; Salmonella; Salmonella enterica; Salmonella enterica subsp. enterica; Salmonella enterica subsp. enterica serovar Montevideo MNNEETFYQAMRRKGVTRRSFLKFCSLAATSLGLGAGMTPKIAWALENKPRIPVVWIHGL WP_001058312.1 hydrogenase [Salmonella enterica] Length: 372\nNumber of predicted TMHs: 2\nExp number of AAs in TMHs: 36.51738\nExp number, first 60 AAs: 19.72849\nTotal prob of N-in: 0.96170\nPOSSIBLE N-term signal sequence\ninside 1 19\nTMhelix 20 39\noutside 40 325\nTMhelix 326 348\ninside 349 372

24141 GCF_000189095.1_ASM18909v2 Salmonella enterica subsp. enterica serovar Montevideo str. 2009085258 Proteobacteria; Gammaproteobacteria; Enterobacterales; Enterobacteriaceae; Salmonella; Salmonella enterica; Salmonella enterica subsp. enterica; Salmonella enterica subsp. enterica serovar Montevideo MNNEETFYQAMRRKGVTRRSFLKFCSLAATSLGLGAGMTPKIAWALENKPRIPVVWIHGL WP_001058312.1 hydrogenase [Salmonella enterica] Length: 372\nNumber of predicted TMHs: 2\nExp number of AAs in TMHs: 36.51738\nExp number, first 60 AAs: 19.72849\nTotal prob of N-in: 0.96170\nPOSSIBLE N-term signal sequence\ninside 1 19\nTMhelix 20 39\noutside 40 325\nTMhelix 326 348\ninside 349 372

24142 GCF_000189095.1_ASM18909v2 Salmonella enterica subsp. enterica serovar Montevideo str. 2009085258 Proteobacteria; Gammaproteobacteria; Enterobacterales; Enterobacteriaceae; Salmonella; Salmonella enterica; Salmonella enterica subsp. enterica; Salmonella enterica subsp. enterica serovar Montevideo MTGDNTLITSHGINRRDFMKLCAALAATMGLSSKAAAEMAESVSNPQRPPVIWIGAQECT WP_000145429.1 hydrogenase 2 small subunit [Salmonella enterica] Length: 372\nNumber of predicted TMHs: 1\nExp number of AAs in TMHs: 24.51309\nExp number, first 60 AAs: 0.2682\nTotal prob of N-in: 0.14402\noutside 1 331\nTMhelix 332 354\ninside 355 372

24143 GCF_000238475.1_ASM23847v2 Salmonella enterica subsp. enterica serovar Montevideo str. 29N Proteobacteria; Gammaproteobacteria; Enterobacterales; Enterobacteriaceae; Salmonella; Salmonella enterica; Salmonella enterica subsp. enterica; Salmonella enterica subsp. enterica serovar Montevideo MTGDNTLITSHGINRRDFMKLCAALAATMGLSSKAAAEMAESVSNPQRPPVIWIGAQECT WP_000145429.1 hydrogenase 2 small subunit [Salmonella enterica] Length: 372\nNumber of predicted TMHs: 1\nExp

number of AAs in TMHs: 24.51309\nExp number, first 60 AAs: 0.2682\nTotal prob of N-in: 0.14402\noutside 1 331\nTMhelix 332 354\ninside 355 372

24144 GCF_000238475.1_ASM23847v2 Salmonella enterica subsp. enterica serovar Montevideo str. 29N Proteobacteria; Gammaproteobacteria; Enterobacterales; Enterobacteriaceae; Salmonella; Salmonella enterica; Salmonella enterica subsp. enterica; Salmonella enterica subsp. enterica serovar Montevideo MNNEETFYQAMRRKGVTRRSFLKFCSLAATSLGLGAGMTPKIAWALENKPRIPVVWIHGL WP_001058312.1 hydrogenase [Salmonella enterica] Length: 372\nNumber of predicted TMHs: 2\nExp number of AAs in TMHs: 36.51738\nExp number, first 60 AAs: 19.72849\nTotal prob of N-in: 0.96170\nPOSSIBLE N-term signal sequence\ninside 1 19\nTMhelix 20 39\noutside 40 325\nTMhelix 326 348\ninside 349 372

24145 GCF_000189115.1_ASM18911v2 Salmonella enterica subsp. enterica serovar Montevideo str. 315731156 Proteobacteria; Gammaproteobacteria; Enterobacterales; Enterobacteriaceae; Salmonella; Salmonella enterica; Salmonella enterica subsp. enterica; Salmonella enterica subsp. enterica serovar Montevideo MNNEETFYQAMRRKGVTRRSFLKFCSLAATSLGLGAGMTPKIAWALENKPRIPVVWIHGL WP_001058312.1 hydrogenase [Salmonella enterica] Length: 372\nNumber of predicted TMHs: 2\nExp number of AAs in TMHs: 36.51738\nExp number, first 60 AAs: 19.72849\nTotal prob of N-in: 0.96170\nPOSSIBLE N-term signal sequence\ninside 1 19\nTMhelix 20 39\noutside 40 325\nTMhelix 326 348\ninside 349 372

24146 GCF_000189115.1_ASM18911v2 Salmonella enterica subsp. enterica serovar Montevideo str. 315731156 Proteobacteria; Gammaproteobacteria; Enterobacterales; Enterobacteriaceae; Salmonella; Salmonella enterica; Salmonella enterica subsp. enterica; Salmonella enterica subsp. enterica serovar Montevideo MTGDNTLITSHGINRRDFMKLCAALAATMGLSSKAAAEMAESVSNPQRPPVIWIGAQECT WP_000145429.1 hydrogenase 2 small subunit [Salmonella enterica] Length: 372\nNumber of predicted TMHs: 1\nExp number of AAs in TMHs: 24.51309\nExp number, first 60 AAs: 0.2682\nTotal prob of N-in: 0.14402\noutside 1 331\nTMhelix 332 354\ninside 355 372

24147 GCF_000188335.1_ASM18833v2 Salmonella enterica subsp. enterica serovar Montevideo str. 315996572 Proteobacteria; Gammaproteobacteria; Enterobacterales; Enterobacteriaceae; Salmonella; Salmonella enterica; Salmonella enterica subsp. enterica; Salmonella enterica subsp. enterica serovar Montevideo MTGDNTLITSHGINRRDFMKLCAALAATMGLSSKAAAEMAESVSNPQRPPVIWIGAQECT WP_000145429.1 hydrogenase 2 small subunit [Salmonella enterica] Length: 372\nNumber of predicted TMHs: 1\nExp number of AAs in TMHs: 24.51309\nExp number, first 60 AAs: 0.2682\nTotal prob of N-in: 0.14402\noutside 1 331\nTMhelix 332 354\ninside 355 372

24148 GCF_000188335.1_ASM18833v2 Salmonella enterica subsp. enterica serovar Montevideo str. 315996572 Proteobacteria; Gammaproteobacteria; Enterobacterales; Enterobacteriaceae; Salmonella; Salmonella enterica; Salmonella enterica subsp. enterica; Salmonella enterica subsp. enterica serovar Montevideo MNNEETFYQAMRRKGVTRRSFLKFCSLAATSLGLGAGMTPKIAWALENKPRIPVVWIHGL WP_001058312.1 hydrogenase [Salmonella enterica] Length: 372\nNumber of predicted TMHs: 2\nExp number of AAs in TMHs: 36.51738\nExp number, first 60 AAs: 19.72849\nTotal prob of N-in: 0.96170\nPOSSIBLE N-term signal sequence\ninside 1 19\nTMhelix 20 39\noutside 40 325\nTMhelix 326 348\ninside 349 372

24149 GCF_000335975.1_ASM33597v1 Salmonella enterica subsp. enterica serovar Montevideo str. 316111868 Proteobacteria; Gammaproteobacteria; Enterobacterales; Enterobacteriaceae; Salmonella; Salmonella enterica; Salmonella enterica subsp. enterica; Salmonella enterica subsp. enterica serovar Montevideo MNNEETFYQAMRRKGVTRRSFLKFCSLAATSLGLGAGMTPKIAWALENKPRIPVVWIHGL WP_001058312.1 hydrogenase [Salmonella enterica] Length: 372\nNumber of predicted TMHs: 2\nExp number of AAs in TMHs: 36.51738\nExp number, first 60 AAs: 19.72849\nTotal prob of N-in: 0.96170\nPOSSIBLE N-term signal sequence\ninside 1 19\nTMhelix 20 39\noutside 40 325\nTMhelix 326 348\ninside 349 372

24150 GCF_000335975.1_ASM33597v1 Salmonella enterica subsp. enterica serovar Montevideo str. 316111868 Proteobacteria; Gammaproteobacteria; Enterobacterales; Enterobacteriaceae; Salmonella; Salmonella enterica; Salmonella enterica subsp. enterica; Salmonella enterica subsp. enterica serovar Montevideo MTGDNTLITSHGINRRDFMKLCAALAATMGLSSKAAAEMAESVSNPQRPPVIWIGAQECT WP_000145429.1 hydrogenase 2 small subunit [Salmonella enterica] Length: 372\nNumber of predicted TMHs: 1\nExp number of AAs in TMHs: 24.51309\nExp number, first 60 AAs: 0.2682\nTotal prob of N-in: 0.14402\noutside 1 331\nTMhelix 332 354\ninside 355 372

24151 GCF_000188615.1_ASM18861v2 Salmonella enterica subsp. enterica serovar Montevideo str. 366867 Proteobacteria; Gammaproteobacteria; Enterobacterales; Enterobacteriaceae; Salmonella; Salmonella enterica;

Salmonella enterica subsp. enterica; Salmonella enterica subsp. enterica serovar Montevideo
MTGDNTLITSHGINRRDFMKLCAALAATMGLSSKAAAEMAESVSNPQRPPVIWIGAQECT WP_000145429.1
hydrogenase 2 small subunit [Salmonella enterica] Length: 372\nNumber of predicted TMHs: 1\nExp number of AAs in TMHs: 24.51309\nExp number, first 60 AAs: 0.2682\nTotal prob of N-in: 0.14402\noutside 1 331\nTMhelix 332 354\ninside 355 372

24152 GCF_000188615.1_ASM18861v2 Salmonella enterica subsp. enterica serovar Montevideo str. 366867
Proteobacteria; Gammaproteobacteria; Enterobacterales; Enterobacteriaceae; Salmonella; Salmonella enterica; Salmonella enterica subsp. enterica; Salmonella enterica subsp. enterica serovar Montevideo
MNNEETFYQAMRRKGVTRRSFLKFCSLAATSLGLGAGMTPKIAWALENKPRIPVVWIHGL WP_001058312.1
hydrogenase [Salmonella enterica] Length: 372\nNumber of predicted TMHs: 2\nExp number of AAs in TMHs: 36.51738\nExp number, first 60 AAs: 19.72849\nTotal prob of N-in: 0.96170\nPOSSIBLE N-term signal sequence\ninside 1 19\nTMhelix 20 39\noutside 40 325\nTMhelix 326 348\ninside 349 372

24153 GCF_000188635.1_ASM18863v2 Salmonella enterica subsp. enterica serovar Montevideo str. 413180
Proteobacteria; Gammaproteobacteria; Enterobacterales; Enterobacteriaceae; Salmonella; Salmonella enterica; Salmonella enterica subsp. enterica; Salmonella enterica subsp. enterica serovar Montevideo
MTGDNTLITSHGINRRDFMKLCAALAATMGLSSKAAAEMAESVSNPQRPPVIWIGAQECT WP_000145429.1
hydrogenase 2 small subunit [Salmonella enterica] Length: 372\nNumber of predicted TMHs: 1\nExp number of AAs in TMHs: 24.51309\nExp number, first 60 AAs: 0.2682\nTotal prob of N-in: 0.14402\noutside 1 331\nTMhelix 332 354\ninside 355 372

24154 GCF_000188635.1_ASM18863v2 Salmonella enterica subsp. enterica serovar Montevideo str. 413180
Proteobacteria; Gammaproteobacteria; Enterobacterales; Enterobacteriaceae; Salmonella; Salmonella enterica; Salmonella enterica subsp. enterica; Salmonella enterica subsp. enterica serovar Montevideo
MNNEETFYQAMRRKGVTRRSFLKFCSLAATSLGLGAGMTPKIAWALENKPRIPVVWIHGL WP_001058312.1
hydrogenase [Salmonella enterica] Length: 372\nNumber of predicted TMHs: 2\nExp number of AAs in TMHs: 36.51738\nExp number, first 60 AAs: 19.72849\nTotal prob of N-in: 0.96170\nPOSSIBLE N-term signal sequence\ninside 1 19\nTMhelix 20 39\noutside 40 325\nTMhelix 326 348\ninside 349 372

24155 GCF_000188595.1_ASM18859v2 Salmonella enterica subsp. enterica serovar Montevideo str. 414877
Proteobacteria; Gammaproteobacteria; Enterobacterales; Enterobacteriaceae; Salmonella; Salmonella enterica; Salmonella enterica subsp. enterica; Salmonella enterica subsp. enterica serovar Montevideo
MTGDNTLITSHGINRRDFMKLCAALAATMGLSSKAAAEMAESVSNPQRPPVIWIGAQECT WP_000145429.1
hydrogenase 2 small subunit [Salmonella enterica] Length: 372\nNumber of predicted TMHs: 1\nExp number of AAs in TMHs: 24.51309\nExp number, first 60 AAs: 0.2682\nTotal prob of N-in: 0.14402\noutside 1 331\nTMhelix 332 354\ninside 355 372

24156 GCF_000188595.1_ASM18859v2 Salmonella enterica subsp. enterica serovar Montevideo str. 414877
Proteobacteria; Gammaproteobacteria; Enterobacterales; Enterobacteriaceae; Salmonella; Salmonella enterica; Salmonella enterica subsp. enterica; Salmonella enterica subsp. enterica serovar Montevideo
MNNEETFYQAMRRKGVTRRSFLKFCSLAATSLGLGAGMTPKIAWALENKPRIPVVWIHGL WP_001058312.1
hydrogenase [Salmonella enterica] Length: 372\nNumber of predicted TMHs: 2\nExp number of AAs in TMHs: 36.51738\nExp number, first 60 AAs: 19.72849\nTotal prob of N-in: 0.96170\nPOSSIBLE N-term signal sequence\ninside 1 19\nTMhelix 20 39\noutside 40 325\nTMhelix 326 348\ninside 349 372

24157 GCF_000238455.1_ASM23845v2 Salmonella enterica subsp. enterica serovar Montevideo str. 42N
Proteobacteria; Gammaproteobacteria; Enterobacterales; Enterobacteriaceae; Salmonella; Salmonella enterica; Salmonella enterica subsp. enterica; Salmonella enterica subsp. enterica serovar Montevideo
MNNEETFYQAMRRKGVTRRSFLKFCSLAATSLGLGAGMTPKIAWALENKPRIPVVWIHGL WP_001058312.1
hydrogenase [Salmonella enterica] Length: 372\nNumber of predicted TMHs: 2\nExp number of AAs in TMHs: 36.51738\nExp number, first 60 AAs: 19.72849\nTotal prob of N-in: 0.96170\nPOSSIBLE N-term signal sequence\ninside 1 19\nTMhelix 20 39\noutside 40 325\nTMhelix 326 348\ninside 349 372

24158 GCF_000238455.1_ASM23845v2 Salmonella enterica subsp. enterica serovar Montevideo str. 42N
Proteobacteria; Gammaproteobacteria; Enterobacterales; Enterobacteriaceae; Salmonella; Salmonella enterica; Salmonella enterica subsp. enterica; Salmonella enterica subsp. enterica serovar Montevideo
MTGDNTLITSHGINRRDFMKLCAALAATMGLSSKAAAEMAESVSNPQRPPVIWIGAQECT WP_000145429.1
hydrogenase 2 small subunit [Salmonella enterica] Length: 372\nNumber of predicted TMHs: 1\nExp number of AAs in TMHs: 24.51309\nExp number, first 60 AAs: 0.2682\nTotal prob of N-in: 0.14402\noutside 1 331\nTMhelix 332 354\ninside 355 372

24159 GCF_000238435.1_ASM23843v2 Salmonella enterica subsp. enterica serovar Montevideo str. 4441 H Proteobacteria; Gammaproteobacteria; Enterobacterales; Enterobacteriaceae; Salmonella; Salmonella enterica; Salmonella enterica subsp. enterica; Salmonella enterica subsp. enterica serovar Montevideo MTGDNTLITSHGINRRDFMKLCAALAATMGLSSKAAAEMAESVSNPQRPPVIWIGAQECT WP_000145429.1 hydrogenase 2 small subunit [Salmonella enterica] Length: 372\nNumber of predicted TMHs: 1\nExp number of AAs in TMHs: 24.51309\nExp number, first 60 AAs: 0.2682\nTotal prob of N-in: 0.14402\noutside 1 331\nTMhelix 332 354\ninside 355 372

24160 GCF_000238435.1_ASM23843v2 Salmonella enterica subsp. enterica serovar Montevideo str. 4441 H Proteobacteria; Gammaproteobacteria; Enterobacterales; Enterobacteriaceae; Salmonella; Salmonella enterica; Salmonella enterica subsp. enterica; Salmonella enterica subsp. enterica serovar Montevideo MNNEETFYQAMRRKGVTRRSFLKFCSLAATSLGLGAGMTPKIAWALENKPRIPVVWIHGL WP_001058312.1 hydrogenase [Salmonella enterica] Length: 372\nNumber of predicted TMHs: 2\nExp number of AAs in TMHs: 36.51738\nExp number, first 60 AAs: 19.72849\nTotal prob of N-in: 0.96170\nPOSSIBLE N-term signal sequence\ninside 1 19\nTMhelix 20 39\noutside 40 325\nTMhelix 326 348\ninside 349 372

24161 GCF_000188655.1_ASM18865v2 Salmonella enterica subsp. enterica serovar Montevideo str. 446600 Proteobacteria; Gammaproteobacteria; Enterobacterales; Enterobacteriaceae; Salmonella; Salmonella enterica; Salmonella enterica subsp. enterica; Salmonella enterica subsp. enterica serovar Montevideo MNNEETFYQAMRRKGVTRRSFLKFCSLAATSLGLGAGMTPKIAWALENKPRIPVVWIHGL WP_001058312.1 hydrogenase [Salmonella enterica] Length: 372\nNumber of predicted TMHs: 2\nExp number of AAs in TMHs: 36.51738\nExp number, first 60 AAs: 19.72849\nTotal prob of N-in: 0.96170\nPOSSIBLE N-term signal sequence\ninside 1 19\nTMhelix 20 39\noutside 40 325\nTMhelix 326 348\ninside 349 372

24162 GCF_000188655.1_ASM18865v2 Salmonella enterica subsp. enterica serovar Montevideo str. 446600 Proteobacteria; Gammaproteobacteria; Enterobacterales; Enterobacteriaceae; Salmonella; Salmonella enterica; Salmonella enterica subsp. enterica; Salmonella enterica subsp. enterica serovar Montevideo MTGDNTLITSHGINRRDFMKLCAALAATMGLSSKAAAEMAESVSNPQRPPVIWIGAQECT WP_000145429.1 hydrogenase 2 small subunit [Salmonella enterica] Length: 372\nNumber of predicted TMHs: 1\nExp number of AAs in TMHs: 24.51309\nExp number, first 60 AAs: 0.2682\nTotal prob of N-in: 0.14402\noutside 1 331\nTMhelix 332 354\ninside 355 372

24163 GCF_000188355.1_ASM18835v2 Salmonella enterica subsp. enterica serovar Montevideo str. 495297-1 Proteobacteria; Gammaproteobacteria; Enterobacterales; Enterobacteriaceae; Salmonella; Salmonella enterica; Salmonella enterica subsp. enterica; Salmonella enterica subsp. enterica serovar Montevideo MTGDNTLITSHGINRRDFMKLCAALAATMGLSSKAAAEMAESVSNPQRPPVIWIGAQECT WP_000145429.1 hydrogenase 2 small subunit [Salmonella enterica] Length: 372\nNumber of predicted TMHs: 1\nExp number of AAs in TMHs: 24.51309\nExp number, first 60 AAs: 0.2682\nTotal prob of N-in: 0.14402\noutside 1 331\nTMhelix 332 354\ninside 355 372

24164 GCF_000188355.1_ASM18835v2 Salmonella enterica subsp. enterica serovar Montevideo str. 495297-1 Proteobacteria; Gammaproteobacteria; Enterobacterales; Enterobacteriaceae; Salmonella; Salmonella enterica; Salmonella enterica subsp. enterica; Salmonella enterica subsp. enterica serovar Montevideo MNNEETFYQAMRRKGVTRRSFLKFCSLAATSLGLGAGMTPKIAWALENKPRIPVVWIHGL WP_001058312.1 hydrogenase [Salmonella enterica] Length: 372\nNumber of predicted TMHs: 2\nExp number of AAs in TMHs: 36.51738\nExp number, first 60 AAs: 19.72849\nTotal prob of N-in: 0.96170\nPOSSIBLE N-term signal sequence\ninside 1 19\nTMhelix 20 39\noutside 40 325\nTMhelix 326 348\ninside 349 372

24165 GCF_000188375.1_ASM18837v2 Salmonella enterica subsp. enterica serovar Montevideo str. 495297-3 Proteobacteria; Gammaproteobacteria; Enterobacterales; Enterobacteriaceae; Salmonella; Salmonella enterica; Salmonella enterica subsp. enterica; Salmonella enterica subsp. enterica serovar Montevideo MNNEETFYQAMRRKGVTRRSFLKFCSLAATSLGLGAGMTPKIAWALENKPRIPVVWIHGL WP_001058312.1 hydrogenase [Salmonella enterica] Length: 372\nNumber of predicted TMHs: 2\nExp number of AAs in TMHs: 36.51738\nExp number, first 60 AAs: 19.72849\nTotal prob of N-in: 0.96170\nPOSSIBLE N-term signal sequence\ninside 1 19\nTMhelix 20 39\noutside 40 325\nTMhelix 326 348\ninside 349 372

24166 GCF_000188375.1_ASM18837v2 Salmonella enterica subsp. enterica serovar Montevideo str. 495297-3 Proteobacteria; Gammaproteobacteria; Enterobacterales; Enterobacteriaceae; Salmonella; Salmonella enterica; Salmonella enterica subsp. enterica; Salmonella enterica subsp. enterica serovar Montevideo MTGDNTLITSHGINRRDFMKLCAALAATMGLSSKAAAEMAESVSNPQRPPVIWIGAQECT WP_000145429.1 hydrogenase 2 small subunit [Salmonella enterica] Length: 372\nNumber of predicted TMHs: 1\nExp

number of AAs in TMHs: 24.51309\nExp number, first 60 AAs: 0.2682\nTotal prob of N-in: 0.14402\noutside 1 331\nTMhelix 332 354\ninside 355 372

24167 GCF_000188395.1_ASM18839v2 Salmonella enterica subsp. enterica serovar Montevideo str. 495297-4 Proteobacteria; Gammaproteobacteria; Enterobacterales; Enterobacteriaceae; Salmonella; Salmonella enterica; Salmonella enterica subsp. enterica; Salmonella enterica subsp. enterica serovar Montevideo MTGDNTLITSHGINRRDFMKLCAALAATMGLSSKAAAEMAESVSNPQRPPVIWIGAQECT WP_000145429.1 hydrogenase 2 small subunit [Salmonella enterica] Length: 372\nNumber of predicted TMHs: 1\nExp number of AAs in TMHs: 24.51309\nExp number, first 60 AAs: 0.2682\nTotal prob of N-in: 0.14402\noutside 1 331\nTMhelix 332 354\ninside 355 372

24168 GCF_000188395.1_ASM18839v2 Salmonella enterica subsp. enterica serovar Montevideo str. 495297-4 Proteobacteria; Gammaproteobacteria; Enterobacterales; Enterobacteriaceae; Salmonella; Salmonella enterica; Salmonella enterica subsp. enterica; Salmonella enterica subsp. enterica serovar Montevideo MNNEETFYQAMRRKGVTRRSFLKFCSLAATSLGLGAGMTPKIAWALENKPRIPVVWIHGL WP_001058312.1 hydrogenase [Salmonella enterica] Length: 372\nNumber of predicted TMHs: 2\nExp number of AAs in TMHs: 36.51738\nExp number, first 60 AAs: 19.72849\nTotal prob of N-in: 0.96170\nPOSSIBLE N-term signal sequence\ninside 1 19\nTMhelix 20 39\noutside 40 325\nTMhelix 326 348\ninside 349 372

24169 GCF_000188955.2_ASM18895v5 Salmonella enterica subsp. enterica serovar Montevideo str. 507440-20 Proteobacteria; Gammaproteobacteria; Enterobacterales; Enterobacteriaceae; Salmonella; Salmonella enterica; Salmonella enterica subsp. enterica; Salmonella enterica subsp. enterica serovar Montevideo MNNEETFYQAMRRKGVTRRSFLKFCSLAATSLGLGAGMTPKIAWALENKPRIPVVWIHGL WP_001058312.1 hydrogenase [Salmonella enterica] Length: 372\nNumber of predicted TMHs: 2\nExp number of AAs in TMHs: 36.51738\nExp number, first 60 AAs: 19.72849\nTotal prob of N-in: 0.96170\nPOSSIBLE N-term signal sequence\ninside 1 19\nTMhelix 20 39\noutside 40 325\nTMhelix 326 348\ninside 349 372

24170 GCF_000188955.2_ASM18895v5 Salmonella enterica subsp. enterica serovar Montevideo str. 507440-20 Proteobacteria; Gammaproteobacteria; Enterobacterales; Enterobacteriaceae; Salmonella; Salmonella enterica; Salmonella enterica subsp. enterica; Salmonella enterica subsp. enterica serovar Montevideo MTGDNTLITSHGINRRDFMKLCAALAATMGLSSKAAAEMAESVSNPQRPPVIWIGAQECT WP_000145429.1 hydrogenase 2 small subunit [Salmonella enterica] Length: 372\nNumber of predicted TMHs: 1\nExp number of AAs in TMHs: 24.51309\nExp number, first 60 AAs: 0.2682\nTotal prob of N-in: 0.14402\noutside 1 331\nTMhelix 332 354\ninside 355 372

24171 GCF_000188415.1_ASM18841v2 Salmonella enterica subsp. enterica serovar Montevideo str. 515920-1 Proteobacteria; Gammaproteobacteria; Enterobacterales; Enterobacteriaceae; Salmonella; Salmonella enterica; Salmonella enterica subsp. enterica; Salmonella enterica subsp. enterica serovar Montevideo MNNEETFYQAMRRKGVTRRSFLKFCSLAATSLGLGAGMTPKIAWALENKPRIPVVWIHGL WP_001058312.1 hydrogenase [Salmonella enterica] Length: 372\nNumber of predicted TMHs: 2\nExp number of AAs in TMHs: 36.51738\nExp number, first 60 AAs: 19.72849\nTotal prob of N-in: 0.96170\nPOSSIBLE N-term signal sequence\ninside 1 19\nTMhelix 20 39\noutside 40 325\nTMhelix 326 348\ninside 349 372

24172 GCF_000188415.1_ASM18841v2 Salmonella enterica subsp. enterica serovar Montevideo str. 515920-1 Proteobacteria; Gammaproteobacteria; Enterobacterales; Enterobacteriaceae; Salmonella; Salmonella enterica; Salmonella enterica subsp. enterica; Salmonella enterica subsp. enterica serovar Montevideo MTGDNTLITSHGINRRDFMKLCAALAATMGLSSKAAAEMAESVSNPQRPPVIWIGAQECT WP_000145429.1 hydrogenase 2 small subunit [Salmonella enterica] Length: 372\nNumber of predicted TMHs: 1\nExp number of AAs in TMHs: 24.51309\nExp number, first 60 AAs: 0.2682\nTotal prob of N-in: 0.14402\noutside 1 331\nTMhelix 332 354\ninside 355 372

24173 GCF_000188435.1_ASM18843v2 Salmonella enterica subsp. enterica serovar Montevideo str. 515920-2 Proteobacteria; Gammaproteobacteria; Enterobacterales; Enterobacteriaceae; Salmonella; Salmonella enterica; Salmonella enterica subsp. enterica; Salmonella enterica subsp. enterica serovar Montevideo MNNEETFYQAMRRKGVTRRSFLKFCSLAATSLGLGAGMTPKIAWALENKPRIPVVWIHGL WP_001058312.1 hydrogenase [Salmonella enterica] Length: 372\nNumber of predicted TMHs: 2\nExp number of AAs in TMHs: 36.51738\nExp number, first 60 AAs: 19.72849\nTotal prob of N-in: 0.96170\nPOSSIBLE N-term signal sequence\ninside 1 19\nTMhelix 20 39\noutside 40 325\nTMhelix 326 348\ninside 349 372

24174 GCF_000188435.1_ASM18843v2 Salmonella enterica subsp. enterica serovar Montevideo str. 515920-2 Proteobacteria; Gammaproteobacteria; Enterobacterales; Enterobacteriaceae; Salmonella; Salmonella enterica;

Salmonella enterica subsp. enterica; Salmonella enterica subsp. enterica serovar Montevideo
MTGDNTLITSHGINRRDFMKLCAALAATMGLSSKAAAEMAESVSNPQRPPVIWIGAQECT WP_000145429.1
hydrogenase 2 small subunit [Salmonella enterica] Length: 372\nNumber of predicted TMHs: 1\nExp number of AAs in TMHs: 24.51309\nExp number, first 60 AAs: 0.2682\nTotal prob of N-in: 0.14402\noutside 1 331\nTMhelix 332 354\ninside 355 372

24175 GCF_000188455.1_ASM18845v2 Salmonella enterica subsp. enterica serovar Montevideo str. 531954
Proteobacteria; Gammaproteobacteria; Enterobacterales; Enterobacteriaceae; Salmonella; Salmonella enterica; Salmonella enterica subsp. enterica; Salmonella enterica subsp. enterica serovar Montevideo
MNNEETFYQAMRRKGVTRRSFLKFCSLAATSLGLGAGMTPKIAWALENKPRIPVVWIHGL WP_001058312.1
hydrogenase [Salmonella enterica] Length: 372\nNumber of predicted TMHs: 2\nExp number of AAs in TMHs: 36.51738\nExp number, first 60 AAs: 19.72849\nTotal prob of N-in: 0.96170\nPOSSIBLE N-term signal sequence\ninside 1 19\nTMhelix 20 39\noutside 40 325\nTMhelix 326 348\ninside 349 372

24176 GCF_000188455.1_ASM18845v2 Salmonella enterica subsp. enterica serovar Montevideo str. 531954
Proteobacteria; Gammaproteobacteria; Enterobacterales; Enterobacteriaceae; Salmonella; Salmonella enterica; Salmonella enterica subsp. enterica; Salmonella enterica subsp. enterica serovar Montevideo
MTGDNTLITSHGINRRDFMKLCAALAATMGLSSKAAAEMAESVSNPQRPPVIWIGAQECT WP_000145429.1
hydrogenase 2 small subunit [Salmonella enterica] Length: 372\nNumber of predicted TMHs: 1\nExp number of AAs in TMHs: 24.51309\nExp number, first 60 AAs: 0.2682\nTotal prob of N-in: 0.14402\noutside 1 331\nTMhelix 332 354\ninside 355 372

24177 GCF_000188915.1_ASM18891v2 Salmonella enterica subsp. enterica serovar Montevideo str. 556150-1
Proteobacteria; Gammaproteobacteria; Enterobacterales; Enterobacteriaceae; Salmonella; Salmonella enterica; Salmonella enterica subsp. enterica; Salmonella enterica subsp. enterica serovar Montevideo
MNNEETFYQAMRRKGVTRRSFLKFCSLAATSLGLGAGMTPKIAWALENKPRIPVVWIHGL WP_001058312.1
hydrogenase [Salmonella enterica] Length: 372\nNumber of predicted TMHs: 2\nExp number of AAs in TMHs: 36.51738\nExp number, first 60 AAs: 19.72849\nTotal prob of N-in: 0.96170\nPOSSIBLE N-term signal sequence\ninside 1 19\nTMhelix 20 39\noutside 40 325\nTMhelix 326 348\ninside 349 372

24178 GCF_000188915.1_ASM18891v2 Salmonella enterica subsp. enterica serovar Montevideo str. 556150-1
Proteobacteria; Gammaproteobacteria; Enterobacterales; Enterobacteriaceae; Salmonella; Salmonella enterica; Salmonella enterica subsp. enterica; Salmonella enterica subsp. enterica serovar Montevideo
MTGDNTLITSHGINRRDFMKLCAALAATMGLSSKAAAEMAESVSNPQRPPVIWIGAQECT WP_000145429.1
hydrogenase 2 small subunit [Salmonella enterica] Length: 372\nNumber of predicted TMHs: 1\nExp number of AAs in TMHs: 24.51309\nExp number, first 60 AAs: 0.2682\nTotal prob of N-in: 0.14402\noutside 1 331\nTMhelix 332 354\ninside 355 372

24179 GCF_000188975.1_ASM18897v2 Salmonella enterica subsp. enterica serovar Montevideo str. 556152
Proteobacteria; Gammaproteobacteria; Enterobacterales; Enterobacteriaceae; Salmonella; Salmonella enterica; Salmonella enterica subsp. enterica; Salmonella enterica subsp. enterica serovar Montevideo
MTGDNTLITSHGINRRDFMKLCAALAATMGLSSKAAAEMAESVSNPQRPPVIWIGAQECT WP_000145429.1
hydrogenase 2 small subunit [Salmonella enterica] Length: 372\nNumber of predicted TMHs: 1\nExp number of AAs in TMHs: 24.51309\nExp number, first 60 AAs: 0.2682\nTotal prob of N-in: 0.14402\noutside 1 331\nTMhelix 332 354\ninside 355 372

24180 GCF_000188975.1_ASM18897v2 Salmonella enterica subsp. enterica serovar Montevideo str. 556152
Proteobacteria; Gammaproteobacteria; Enterobacterales; Enterobacteriaceae; Salmonella; Salmonella enterica; Salmonella enterica subsp. enterica; Salmonella enterica subsp. enterica serovar Montevideo
MNNEETFYQAMRRKGVTRRSFLKFCSLAATSLGLGAGMTPKIAWALENKPRIPVVWIHGL WP_001058312.1
hydrogenase [Salmonella enterica] Length: 372\nNumber of predicted TMHs: 2\nExp number of AAs in TMHs: 36.51738\nExp number, first 60 AAs: 19.72849\nTotal prob of N-in: 0.96170\nPOSSIBLE N-term signal sequence\ninside 1 19\nTMhelix 20 39\noutside 40 325\nTMhelix 326 348\ninside 349 372

24181 GCF_000188895.1_ASM18889v2 Salmonella enterica subsp. enterica serovar Montevideo str. 609458-1
Proteobacteria; Gammaproteobacteria; Enterobacterales; Enterobacteriaceae; Salmonella; Salmonella enterica; Salmonella enterica subsp. enterica; Salmonella enterica subsp. enterica serovar Montevideo
MNNEETFYQAMRRKGVTRRSFLKFCSLAATSLGLGAGMTPKIAWALENKPRIPVVWIHGL WP_001058312.1
hydrogenase [Salmonella enterica] Length: 372\nNumber of predicted TMHs: 2\nExp number of AAs in TMHs: 36.51738\nExp number, first 60 AAs: 19.72849\nTotal prob of N-in: 0.96170\nPOSSIBLE N-term signal sequence\ninside 1 19\nTMhelix 20 39\noutside 40 325\nTMhelix 326 348\ninside 349 372

24182 GCF_000188895.1_ASM18889v2 Salmonella enterica subsp. enterica serovar Montevideo str. 609458-1 Proteobacteria; Gammaproteobacteria; Enterobacterales; Enterobacteriaceae; Salmonella; Salmonella enterica; Salmonella enterica subsp. enterica; Salmonella enterica subsp. enterica serovar Montevideo MTGDNTLITSHGINRRDFMKLCAALAATMGLSSKAAAEMAESVSNPQRPPVIWIGAQECT WP_000145429.1 hydrogenase 2 small subunit [Salmonella enterica] Length: 372\nNumber of predicted TMHs: 1\nExp number of AAs in TMHs: 24.51309\nExp number, first 60 AAs: 0.2682\nTotal prob of N-in: 0.14402\noutside 1 331\nTMhelix 332 354\ninside 355 372

24183 GCF_001707585.1_ASM170758v1 Salmonella enterica subsp. enterica serovar Montevideo str. 609458-2 Proteobacteria; Gammaproteobacteria; Enterobacterales; Enterobacteriaceae; Salmonella; Salmonella enterica; Salmonella enterica subsp. enterica; Salmonella enterica subsp. enterica serovar Montevideo MTGDNTLITSHGINRRDFMKLCAALAATMGLSSKAAAEMAESVSNPQRPPVIWIGAQECT WP_000145429.1 hydrogenase 2 small subunit [Salmonella enterica] Length: 372\nNumber of predicted TMHs: 1\nExp number of AAs in TMHs: 24.51309\nExp number, first 60 AAs: 0.2682\nTotal prob of N-in: 0.14402\noutside 1 331\nTMhelix 332 354\ninside 355 372

24184 GCF_001707585.1_ASM170758v1 Salmonella enterica subsp. enterica serovar Montevideo str. 609458-2 Proteobacteria; Gammaproteobacteria; Enterobacterales; Enterobacteriaceae; Salmonella; Salmonella enterica; Salmonella enterica subsp. enterica; Salmonella enterica subsp. enterica serovar Montevideo MNNEETFYQAMRRKGVTRRSFLKFCSLAATSLGLGAGMTPKIAWALENKPRIPVVWIHGL WP_001058312.1 hydrogenase [Salmonella enterica] Length: 372\nNumber of predicted TMHs: 2\nExp number of AAs in TMHs: 36.51738\nExp number, first 60 AAs: 19.72849\nTotal prob of N-in: 0.96170\nPOSSIBLE N-term signal sequence\ninside 1 19\nTMhelix 20 39\noutside 40 325\nTMhelix 326 348\ninside 349 372

24185 GCF_000188935.1_ASM18893v2 Salmonella enterica subsp. enterica serovar Montevideo str. 609460 Proteobacteria; Gammaproteobacteria; Enterobacterales; Enterobacteriaceae; Salmonella; Salmonella enterica; Salmonella enterica subsp. enterica; Salmonella enterica subsp. enterica serovar Montevideo MTGDNTLITSHGINRRDFMKLCAALAATMGLSSKAAAEMAESVSNPQRPPVIWIGAQECT WP_000145429.1 hydrogenase 2 small subunit [Salmonella enterica] Length: 372\nNumber of predicted TMHs: 1\nExp number of AAs in TMHs: 24.51309\nExp number, first 60 AAs: 0.2682\nTotal prob of N-in: 0.14402\noutside 1 331\nTMhelix 332 354\ninside 355 372

24186 GCF_000188935.1_ASM18893v2 Salmonella enterica subsp. enterica serovar Montevideo str. 609460 Proteobacteria; Gammaproteobacteria; Enterobacterales; Enterobacteriaceae; Salmonella; Salmonella enterica; Salmonella enterica subsp. enterica; Salmonella enterica subsp. enterica serovar Montevideo MNNEETFYQAMRRKGVTRRSFLKFCSLAATSLGLGAGMTPKIAWALENKPRIPVVWIHGL WP_001058312.1 hydrogenase [Salmonella enterica] Length: 372\nNumber of predicted TMHs: 2\nExp number of AAs in TMHs: 36.51738\nExp number, first 60 AAs: 19.72849\nTotal prob of N-in: 0.96170\nPOSSIBLE N-term signal sequence\ninside 1 19\nTMhelix 20 39\noutside 40 325\nTMhelix 326 348\ninside 349 372

24187 GCF_000240785.1_ASM24078v2 Salmonella enterica subsp. enterica serovar Montevideo str. 80959-06 Proteobacteria; Gammaproteobacteria; Enterobacterales; Enterobacteriaceae; Salmonella; Salmonella enterica; Salmonella enterica subsp. enterica; Salmonella enterica subsp. enterica serovar Montevideo MNNEETFYQAMRRKGVTRRSFLKFCSLAATSLGLGAGMTPKIAWALENKPRIPVVWIHGL WP_001058312.1 hydrogenase [Salmonella enterica] Length: 372\nNumber of predicted TMHs: 2\nExp number of AAs in TMHs: 36.51738\nExp number, first 60 AAs: 19.72849\nTotal prob of N-in: 0.96170\nPOSSIBLE N-term signal sequence\ninside 1 19\nTMhelix 20 39\noutside 40 325\nTMhelix 326 348\ninside 349 372

24188 GCF_000240785.1_ASM24078v2 Salmonella enterica subsp. enterica serovar Montevideo str. 80959-06 Proteobacteria; Gammaproteobacteria; Enterobacterales; Enterobacteriaceae; Salmonella; Salmonella enterica; Salmonella enterica subsp. enterica; Salmonella enterica subsp. enterica serovar Montevideo MTGDNTLITSHGINRRDFMKLCAALAATMGLSSKAAAEMAESVSNPQRPPVIWIGAQECT WP_000145429.1 hydrogenase 2 small subunit [Salmonella enterica] Length: 372\nNumber of predicted TMHs: 1\nExp number of AAs in TMHs: 24.51309\nExp number, first 60 AAs: 0.2682\nTotal prob of N-in: 0.14402\noutside 1 331\nTMhelix 332 354\ninside 355 372

24189 GCF_000188555.1_ASM18855v2 Salmonella enterica subsp. enterica serovar Montevideo str. 81038-01 Proteobacteria; Gammaproteobacteria; Enterobacterales; Enterobacteriaceae; Salmonella; Salmonella enterica; Salmonella enterica subsp. enterica; Salmonella enterica subsp. enterica serovar Montevideo MNNEETFYQAMRRKGVTRRSFLKFCSLAATSLGLGAGMTPKIAWALENKPRIPVVWIHGL WP_001058312.1 hydrogenase [Salmonella enterica] Length: 372\nNumber of predicted TMHs: 2\nExp number of AAs in

TMHs: 36.51738\nExp number, first 60 AAs: 19.72849\nTotal prob of N-in: 0.96170\nPOSSIBLE N-term signal sequence\ninside 1 19\nTMhelix 20 39\noutside 40 325\nTMhelix 326 348\ninside 349 372

24190 GCF_000188555.1_ASM18855v2 Salmonella enterica subsp. enterica serovar Montevideo str. 81038-01 Proteobacteria; Gammaproteobacteria; Enterobacterales; Enterobacteriaceae; Salmonella; Salmonella enterica; Salmonella enterica subsp. enterica; Salmonella enterica subsp. enterica serovar Montevideo MTGDNTLITSHGINRRDFMKLCAALAATMGLSSKAAAEMAESVSNPQRPPVIWIGAQECT WP_000145429.1 hydrogenase 2 small subunit [Salmonella enterica] Length: 372\nNumber of predicted TMHs: 1\nExp number of AAs in TMHs: 24.51309\nExp number, first 60 AAs: 0.2682\nTotal prob of N-in: 0.14402\noutside 1 331\nTMhelix 332 354\ninside 355 372

24191 GCF_000486895.1_ASM48689v1 Salmonella enterica subsp. enterica serovar Montevideo str. 8387 Proteobacteria; Gammaproteobacteria; Enterobacterales; Enterobacteriaceae; Salmonella; Salmonella enterica; Salmonella enterica subsp. enterica; Salmonella enterica subsp. enterica serovar Montevideo MNNEETFYQAMRRKGVTRRSFLKFCSLAATSLGLGAGMTPKIAWALENKPRIPVVWIHGL WP_001058312.1 hydrogenase [Salmonella enterica] Length: 372\nNumber of predicted TMHs: 2\nExp number of AAs in TMHs: 36.51738\nExp number, first 60 AAs: 19.72849\nTotal prob of N-in: 0.96170\nPOSSIBLE N-term signal sequence\ninside 1 19\nTMhelix 20 39\noutside 40 325\nTMhelix 326 348\ninside 349 372

24192 GCF_000486895.1_ASM48689v1 Salmonella enterica subsp. enterica serovar Montevideo str. 8387 Proteobacteria; Gammaproteobacteria; Enterobacterales; Enterobacteriaceae; Salmonella; Salmonella enterica; Salmonella enterica subsp. enterica; Salmonella enterica subsp. enterica serovar Montevideo MTGDNTLITSHGINRRDFMKLCAALAATMGLSSKAAAEMAESVSNPQRPPVIWIGAQECT WP_000145429.1 hydrogenase 2 small subunit [Salmonella enterica] Length: 372\nNumber of predicted TMHs: 1\nExp number of AAs in TMHs: 24.51309\nExp number, first 60 AAs: 0.2682\nTotal prob of N-in: 0.14402\noutside 1 331\nTMhelix 332 354\ninside 355 372

24193 GCF_000238535.1_ASM23853v2 Salmonella enterica subsp. enterica serovar Montevideo str. ATCC BAA710 Proteobacteria; Gammaproteobacteria; Enterobacterales; Enterobacteriaceae; Salmonella; Salmonella enterica; Salmonella enterica subsp. enterica; Salmonella enterica subsp. enterica serovar Montevideo MTGDNTLITSHGINRRDFMKLCAALAATMGLSSKAAAEMAESVSNPQRPPVIWIGAQECT WP_000145429.1 hydrogenase 2 small subunit [Salmonella enterica] Length: 372\nNumber of predicted TMHs: 1\nExp number of AAs in TMHs: 24.51309\nExp number, first 60 AAs: 0.2682\nTotal prob of N-in: 0.14402\noutside 1 331\nTMhelix 332 354\ninside 355 372

24194 GCF_000238535.1_ASM23853v2 Salmonella enterica subsp. enterica serovar Montevideo str. ATCC BAA710 Proteobacteria; Gammaproteobacteria; Enterobacterales; Enterobacteriaceae; Salmonella; Salmonella enterica; Salmonella enterica subsp. enterica; Salmonella enterica subsp. enterica serovar Montevideo MNNEETFYQAMRRKGVTRRSFLKFCSLAATSLGLGAGMTPKIAWALENKPRIPVVWIHGL WP_001058312.1 hydrogenase [Salmonella enterica] Length: 372\nNumber of predicted TMHs: 2\nExp number of AAs in TMHs: 36.51738\nExp number, first 60 AAs: 19.72849\nTotal prob of N-in: 0.96170\nPOSSIBLE N-term signal sequence\ninside 1 19\nTMhelix 20 39\noutside 40 325\nTMhelix 326 348\ninside 349 372

24195 GCF_000188515.1_ASM18851v2 Salmonella enterica subsp. enterica serovar Montevideo str. CASC_09SCPH15965 Proteobacteria; Gammaproteobacteria; Enterobacterales; Enterobacteriaceae; Salmonella; Salmonella enterica; Salmonella enterica subsp. enterica; Salmonella enterica subsp. enterica serovar Montevideo MTGDNTLITSHGINRRDFMKLCAALAATMGLSSKAAAEMAESVSNPQRPPVIWIGAQECT WP_000145429.1 hydrogenase 2 small subunit [Salmonella enterica] Length: 372\nNumber of predicted TMHs: 1\nExp number of AAs in TMHs: 24.51309\nExp number, first 60 AAs: 0.2682\nTotal prob of N-in: 0.14402\noutside 1 331\nTMhelix 332 354\ninside 355 372

24196 GCF_000188515.1_ASM18851v2 Salmonella enterica subsp. enterica serovar Montevideo str. CASC_09SCPH15965 Proteobacteria; Gammaproteobacteria; Enterobacterales; Enterobacteriaceae; Salmonella; Salmonella enterica; Salmonella enterica subsp. enterica; Salmonella enterica subsp. enterica serovar Montevideo MNNEETFYQAMRRKGVTRRSFLKFCSLAATSLGLGAGMTPKIAWALENKPRIPVVWIHGL WP_001058312.1 hydrogenase [Salmonella enterica] Length: 372\nNumber of predicted TMHs: 2\nExp number of AAs in TMHs: 36.51738\nExp number, first 60 AAs: 19.72849\nTotal prob of N-in: 0.96170\nPOSSIBLE N-term signal sequence\ninside 1 19\nTMhelix 20 39\noutside 40 325\nTMhelix 326 348\ninside 349 372

24197 GCF_000505525.1_CFSAN004346_01.0Salmonella enterica subsp. enterica serovar Montevideo str. CFSAN004346 Proteobacteria; Gammaproteobacteria; Enterobacterales; Enterobacteriaceae; Salmonella; Salmonella

enterica; Salmonella enterica subsp. enterica; Salmonella enterica subsp. enterica serovar Montevideo
MNNEETFYQAMRRKGVTRRSFLKFCSLAATSLGLGAGMTPKIAWALENKPRIPVVWIHGL WP_001058307.1
hydrogenase [Salmonella enterica] Length: 372\nNumber of predicted TMHs: 2\nExp number of AAs in TMHs: 36.51738\nExp number, first 60 AAs: 19.72849\nTotal prob of N-in: 0.96170\nPOSSIBLE N-term signal sequence\ninside 1 19\nTMhelix 20 39\noutside 40 325\nTMhelix 326 348\ninside 349 372

24198 GCF_000505525.1_CFSAN004346_01.0 Salmonella enterica subsp. enterica serovar Montevideo str. CFSAN004346 Proteobacteria; Gammaproteobacteria; Enterobacterales; Enterobacteriaceae; Salmonella; Salmonella enterica; Salmonella enterica subsp. enterica; Salmonella enterica subsp. enterica serovar Montevideo
MTGDNTLITSHGINRRDFMKLCAALAATMGLSSKAAAEMAESVSNPQRPPVIWIGAQECT WP_000145429.1
hydrogenase 2 small subunit [Salmonella enterica] Length: 372\nNumber of predicted TMHs: 1\nExp number of AAs in TMHs: 24.51309\nExp number, first 60 AAs: 0.2682\nTotal prob of N-in: 0.14402\noutside 1 331\nTMhelix 332 354\ninside 355 372

24199 GCF_000240825.1_ASM24082v2 Salmonella enterica subsp. enterica serovar Montevideo str. CT_02035318 Proteobacteria; Gammaproteobacteria; Enterobacterales; Enterobacteriaceae; Salmonella; Salmonella enterica; Salmonella enterica subsp. enterica; Salmonella enterica subsp. enterica serovar Montevideo
MTGDNTLITSHGINRRDFMKLCAALAATMGLSSKAAAEMAESVSNPQRPPVIWIGAQECT WP_000145429.1
hydrogenase 2 small subunit [Salmonella enterica] Length: 372\nNumber of predicted TMHs: 1\nExp number of AAs in TMHs: 24.51309\nExp number, first 60 AAs: 0.2682\nTotal prob of N-in: 0.14402\noutside 1 331\nTMhelix 332 354\ninside 355 372

24200 GCF_000240825.1_ASM24082v2 Salmonella enterica subsp. enterica serovar Montevideo str. CT_02035318 Proteobacteria; Gammaproteobacteria; Enterobacterales; Enterobacteriaceae; Salmonella; Salmonella enterica; Salmonella enterica subsp. enterica; Salmonella enterica subsp. enterica serovar Montevideo
MNNEETFYQAMRRKGVTRRSFLKFCSLAATSLGLGAGMTPKIAWALENKPRIPVVWIHGL WP_001058312.1
hydrogenase [Salmonella enterica] Length: 372\nNumber of predicted TMHs: 2\nExp number of AAs in TMHs: 36.51738\nExp number, first 60 AAs: 19.72849\nTotal prob of N-in: 0.96170\nPOSSIBLE N-term signal sequence\ninside 1 19\nTMhelix 20 39\noutside 40 325\nTMhelix 326 348\ninside 349 372

24201 GCF_000240845.1_ASM24084v2 Salmonella enterica subsp. enterica serovar Montevideo str. CT_02035320 Proteobacteria; Gammaproteobacteria; Enterobacterales; Enterobacteriaceae; Salmonella; Salmonella enterica; Salmonella enterica subsp. enterica; Salmonella enterica subsp. enterica serovar Montevideo
MTGDNTLITSHGINRRDFMKLCAALAATMGLSSKAAAEMAESVSNPQRPPVIWIGAQECT WP_000145429.1
hydrogenase 2 small subunit [Salmonella enterica] Length: 372\nNumber of predicted TMHs: 1\nExp number of AAs in TMHs: 24.51309\nExp number, first 60 AAs: 0.2682\nTotal prob of N-in: 0.14402\noutside 1 331\nTMhelix 332 354\ninside 355 372

24202 GCF_000240845.1_ASM24084v2 Salmonella enterica subsp. enterica serovar Montevideo str. CT_02035320 Proteobacteria; Gammaproteobacteria; Enterobacterales; Enterobacteriaceae; Salmonella; Salmonella enterica; Salmonella enterica subsp. enterica; Salmonella enterica subsp. enterica serovar Montevideo
MNNEETFYQAMRRKGVTRRSFLKFCSLAATSLGLGAGMTPKIAWALENKPRIPVVWIHGL WP_001058312.1
hydrogenase [Salmonella enterica] Length: 372\nNumber of predicted TMHs: 2\nExp number of AAs in TMHs: 36.51738\nExp number, first 60 AAs: 19.72849\nTotal prob of N-in: 0.96170\nPOSSIBLE N-term signal sequence\ninside 1 19\nTMhelix 20 39\noutside 40 325\nTMhelix 326 348\ninside 349 372

24203 GCF_000240865.1_ASM24086v2 Salmonella enterica subsp. enterica serovar Montevideo str. CT_02035321 Proteobacteria; Gammaproteobacteria; Enterobacterales; Enterobacteriaceae; Salmonella; Salmonella enterica; Salmonella enterica subsp. enterica; Salmonella enterica subsp. enterica serovar Montevideo
MNNEETFYQAMRRKGVTRRSFLKFCSLAATSLGLGAGMTPKIAWALENKPRIPVVWIHGL WP_001058312.1
hydrogenase [Salmonella enterica] Length: 372\nNumber of predicted TMHs: 2\nExp number of AAs in TMHs: 36.51738\nExp number, first 60 AAs: 19.72849\nTotal prob of N-in: 0.96170\nPOSSIBLE N-term signal sequence\ninside 1 19\nTMhelix 20 39\noutside 40 325\nTMhelix 326 348\ninside 349 372

24204 GCF_000240865.1_ASM24086v2 Salmonella enterica subsp. enterica serovar Montevideo str. CT_02035321 Proteobacteria; Gammaproteobacteria; Enterobacterales; Enterobacteriaceae; Salmonella; Salmonella enterica; Salmonella enterica subsp. enterica; Salmonella enterica subsp. enterica serovar Montevideo
MTGDNTLITSHGINRRDFMKLCAALAATMGLSSKAAAEMAESVSNPQRPPVIWIGAQECT WP_000145429.1
hydrogenase 2 small subunit [Salmonella enterica] Length: 372\nNumber of predicted TMHs: 1\nExp number of AAs in TMHs: 24.51309\nExp number, first 60 AAs: 0.2682\nTotal prob of N-in: 0.14402\noutside 1 331\nTMhelix 332 354\ninside 355 372

24205 GCF_000240885.1_ASM24088v2 Salmonella enterica subsp. enterica serovar Montevideo str. CT_02035327 Proteobacteria; Gammaproteobacteria; Enterobacterales; Enterobacteriaceae; Salmonella; Salmonella enterica; Salmonella enterica subsp. enterica; Salmonella enterica subsp. enterica serovar Montevideo MTGDNTLITSHGINRRDFMKLCAALAATMGLSSKAAAEMAESVSNPQRPPVIWIGAQECT WP_000145429.1 hydrogenase 2 small subunit [Salmonella enterica] Length: 372\nNumber of predicted TMHs: 1\nExp number of AAs in TMHs: 24.51309\nExp number, first 60 AAs: 0.2682\nTotal prob of N-in: 0.14402\noutside 1 331\nTMhelix 332 354\ninside 355 372

24206 GCF_000240885.1_ASM24088v2 Salmonella enterica subsp. enterica serovar Montevideo str. CT_02035327 Proteobacteria; Gammaproteobacteria; Enterobacterales; Enterobacteriaceae; Salmonella; Salmonella enterica; Salmonella enterica subsp. enterica; Salmonella enterica subsp. enterica serovar Montevideo MNNEETFYQAMRRKGVTRRSFLKFCSLAATSLGLGAGMTPKIAWALENKPRIPVVWIHGL WP_001058312.1 hydrogenase [Salmonella enterica] Length: 372\nNumber of predicted TMHs: 2\nExp number of AAs in TMHs: 36.51738\nExp number, first 60 AAs: 19.72849\nTotal prob of N-in: 0.96170\nPOSSIBLE N-term signal sequence\ninside 1 19\nTMhelix 20 39\noutside 40 325\nTMhelix 326 348\ninside 349 372

24207 GCF_000189135.1_ASM18913v2 Salmonella enterica subsp. enterica serovar Montevideo str. IA_2009159199 Proteobacteria; Gammaproteobacteria; Enterobacterales; Enterobacteriaceae; Salmonella; Salmonella enterica; Salmonella enterica subsp. enterica; Salmonella enterica subsp. enterica serovar Montevideo MNNEETFYQAMRRKGVTRRSFLKFCSLAATSLGLGAGMTPKIAWALENKPRIPVVWIHGL WP_001058312.1 hydrogenase [Salmonella enterica] Length: 372\nNumber of predicted TMHs: 2\nExp number of AAs in TMHs: 36.51738\nExp number, first 60 AAs: 19.72849\nTotal prob of N-in: 0.96170\nPOSSIBLE N-term signal sequence\ninside 1 19\nTMhelix 20 39\noutside 40 325\nTMhelix 326 348\ninside 349 372

24208 GCF_000189135.1_ASM18913v2 Salmonella enterica subsp. enterica serovar Montevideo str. IA_2009159199 Proteobacteria; Gammaproteobacteria; Enterobacterales; Enterobacteriaceae; Salmonella; Salmonella enterica; Salmonella enterica subsp. enterica; Salmonella enterica subsp. enterica serovar Montevideo MTGDNTLITSHGINRRDFMKLCAALAATMGLSSKAAAEMAESVSNPQRPPVIWIGAQECT WP_000145429.1 hydrogenase 2 small subunit [Salmonella enterica] Length: 372\nNumber of predicted TMHs: 1\nExp number of AAs in TMHs: 24.51309\nExp number, first 60 AAs: 0.2682\nTotal prob of N-in: 0.14402\noutside 1 331\nTMhelix 332 354\ninside 355 372

24209 GCF_000189155.1_ASM18915v2 Salmonella enterica subsp. enterica serovar Montevideo str. IA_2010008282 Proteobacteria; Gammaproteobacteria; Enterobacterales; Enterobacteriaceae; Salmonella; Salmonella enterica; Salmonella enterica subsp. enterica; Salmonella enterica subsp. enterica serovar Montevideo MTGDNTLITSHGINRRDFMKLCAALAATMGLSSKAAAEMAESVSNPQRPPVIWIGAQECT WP_000145429.1 hydrogenase 2 small subunit [Salmonella enterica] Length: 372\nNumber of predicted TMHs: 1\nExp number of AAs in TMHs: 24.51309\nExp number, first 60 AAs: 0.2682\nTotal prob of N-in: 0.14402\noutside 1 331\nTMhelix 332 354\ninside 355 372

24210 GCF_000189155.1_ASM18915v2 Salmonella enterica subsp. enterica serovar Montevideo str. IA_2010008282 Proteobacteria; Gammaproteobacteria; Enterobacterales; Enterobacteriaceae; Salmonella; Salmonella enterica; Salmonella enterica subsp. enterica; Salmonella enterica subsp. enterica serovar Montevideo MNNEETFYQAMRRKGVTRRSFLKFCSLAATSLGLGAGMTPKIAWALENKPRIPVVWIHGL WP_001058312.1 hydrogenase [Salmonella enterica] Length: 372\nNumber of predicted TMHs: 2\nExp number of AAs in TMHs: 36.51738\nExp number, first 60 AAs: 19.72849\nTotal prob of N-in: 0.96170\nPOSSIBLE N-term signal sequence\ninside 1 19\nTMhelix 20 39\noutside 40 325\nTMhelix 326 348\ninside 349 372

24211 GCF_000189175.1_ASM18917v2 Salmonella enterica subsp. enterica serovar Montevideo str. IA_2010008283 Proteobacteria; Gammaproteobacteria; Enterobacterales; Enterobacteriaceae; Salmonella; Salmonella enterica; Salmonella enterica subsp. enterica; Salmonella enterica subsp. enterica serovar Montevideo MTGDNTLITSHGINRRDFMKLCAALAATMGLSSKAAAEMAESVSNPQRPPVIWIGAQECT WP_000145429.1 hydrogenase 2 small subunit [Salmonella enterica] Length: 372\nNumber of predicted TMHs: 1\nExp number of AAs in TMHs: 24.51309\nExp number, first 60 AAs: 0.2682\nTotal prob of N-in: 0.14402\noutside 1 331\nTMhelix 332 354\ninside 355 372

24212 GCF_000189175.1_ASM18917v2 Salmonella enterica subsp. enterica serovar Montevideo str. IA_2010008283 Proteobacteria; Gammaproteobacteria; Enterobacterales; Enterobacteriaceae; Salmonella; Salmonella enterica; Salmonella enterica subsp. enterica; Salmonella enterica subsp. enterica serovar Montevideo MNNEETFYQAMRRKGVTRRSFLKFCSLAATSLGLGAGMTPKIAWALENKPRIPVVWIHGL WP_001058312.1 hydrogenase [Salmonella enterica] Length: 372\nNumber of predicted TMHs: 2\nExp number of AAs in

TMHs: 36.51738\nExp number, first 60 AAs: 19.72849\nTotal prob of N-in: 0.96170\nPOSSIBLE N-term signal sequence\ninside 1 19\nTMhelix 20 39\noutside 40 325\nTMhelix 326 348\ninside 349 372

24213 GCF_000189195.1_ASM18919v2 Salmonella enterica subsp. enterica serovar Montevideo str. IA_2010008284 Proteobacteria; Gammaproteobacteria; Enterobacterales; Enterobacteriaceae; Salmonella; Salmonella enterica; Salmonella enterica subsp. enterica; Salmonella enterica subsp. enterica serovar Montevideo MNNEETFYQAMRRKGVTRRSFLKFCSLAATSLGLGAGMTPKIAWALENKPRIPVVWIHGL WP_001058312.1 hydrogenase [Salmonella enterica] Length: 372\nNumber of predicted TMHs: 2\nExp number of AAs in TMHs: 36.51738\nExp number, first 60 AAs: 19.72849\nTotal prob of N-in: 0.96170\nPOSSIBLE N-term signal sequence\ninside 1 19\nTMhelix 20 39\noutside 40 325\nTMhelix 326 348\ninside 349 372

24214 GCF_000189195.1_ASM18919v2 Salmonella enterica subsp. enterica serovar Montevideo str. IA_2010008284 Proteobacteria; Gammaproteobacteria; Enterobacterales; Enterobacteriaceae; Salmonella; Salmonella enterica; Salmonella enterica subsp. enterica; Salmonella enterica subsp. enterica serovar Montevideo MTGDNTLITSHGINRRDFMKLCAALAATMGLSSKAAAEMAESVSNPQRPPVIWIGAQECT WP_000145429.1 hydrogenase 2 small subunit [Salmonella enterica] Length: 372\nNumber of predicted TMHs: 1\nExp number of AAs in TMHs: 24.51309\nExp number, first 60 AAs: 0.2682\nTotal prob of N-in: 0.14402\noutside 1 331\nTMhelix 332 354\ninside 355 372

24215 GCF_000189215.1_ASM18921v2 Salmonella enterica subsp. enterica serovar Montevideo str. IA_2010008285 Proteobacteria; Gammaproteobacteria; Enterobacterales; Enterobacteriaceae; Salmonella; Salmonella enterica; Salmonella enterica subsp. enterica; Salmonella enterica subsp. enterica serovar Montevideo MNNEETFYQAMRRKGVTRRSFLKFCSLAATSLGLGAGMTPKIAWALENKPRIPVVWIHGL WP_001058312.1 hydrogenase [Salmonella enterica] Length: 372\nNumber of predicted TMHs: 2\nExp number of AAs in TMHs: 36.51738\nExp number, first 60 AAs: 19.72849\nTotal prob of N-in: 0.96170\nPOSSIBLE N-term signal sequence\ninside 1 19\nTMhelix 20 39\noutside 40 325\nTMhelix 326 348\ninside 349 372

24216 GCF_000189215.1_ASM18921v2 Salmonella enterica subsp. enterica serovar Montevideo str. IA_2010008285 Proteobacteria; Gammaproteobacteria; Enterobacterales; Enterobacteriaceae; Salmonella; Salmonella enterica; Salmonella enterica subsp. enterica; Salmonella enterica subsp. enterica serovar Montevideo MTGDNTLITSHGINRRDFMKLCAALAATMGLSSKAAAEMAESVSNPQRPPVIWIGAQECT WP_000145429.1 hydrogenase 2 small subunit [Salmonella enterica] Length: 372\nNumber of predicted TMHs: 1\nExp number of AAs in TMHs: 24.51309\nExp number, first 60 AAs: 0.2682\nTotal prob of N-in: 0.14402\noutside 1 331\nTMhelix 332 354\ninside 355 372

24217 GCF_000242795.1_ASM24279v2 Salmonella enterica subsp. enterica serovar Montevideo str. IA_2010008286 Proteobacteria; Gammaproteobacteria; Enterobacterales; Enterobacteriaceae; Salmonella; Salmonella enterica; Salmonella enterica subsp. enterica; Salmonella enterica subsp. enterica serovar Montevideo MTGDNTLITSHGINRRDFMKLCAALAATMGLSSKAAAEMAESVSNPQRPPVIWIGAQECT WP_000145429.1 hydrogenase 2 small subunit [Salmonella enterica] Length: 372\nNumber of predicted TMHs: 1\nExp number of AAs in TMHs: 24.51309\nExp number, first 60 AAs: 0.2682\nTotal prob of N-in: 0.14402\noutside 1 331\nTMhelix 332 354\ninside 355 372

24218 GCF_000242795.1_ASM24279v2 Salmonella enterica subsp. enterica serovar Montevideo str. IA_2010008286 Proteobacteria; Gammaproteobacteria; Enterobacterales; Enterobacteriaceae; Salmonella; Salmonella enterica; Salmonella enterica subsp. enterica; Salmonella enterica subsp. enterica serovar Montevideo MNNEETFYQAMRRKGVTRRSFLKFCSLAATSLGLGAGMTPKIAWALENKPRIPVVWIHGL WP_001058312.1 hydrogenase [Salmonella enterica] Length: 372\nNumber of predicted TMHs: 2\nExp number of AAs in TMHs: 36.51738\nExp number, first 60 AAs: 19.72849\nTotal prob of N-in: 0.96170\nPOSSIBLE N-term signal sequence\ninside 1 19\nTMhelix 20 39\noutside 40 325\nTMhelix 326 348\ninside 349 372

24219 GCF_000189235.1_ASM18923v2 Salmonella enterica subsp. enterica serovar Montevideo str. IA_2010008287 Proteobacteria; Gammaproteobacteria; Enterobacterales; Enterobacteriaceae; Salmonella; Salmonella enterica; Salmonella enterica subsp. enterica; Salmonella enterica subsp. enterica serovar Montevideo MTGDNTLITSHGINRRDFMKLCAALAATMGLSSKAAAEMAESVSNPQRPPVIWIGAQECT WP_000145429.1 hydrogenase 2 small subunit [Salmonella enterica] Length: 372\nNumber of predicted TMHs: 1\nExp number of AAs in TMHs: 24.51309\nExp number, first 60 AAs: 0.2682\nTotal prob of N-in: 0.14402\noutside 1 331\nTMhelix 332 354\ninside 355 372

24220 GCF_000189235.1_ASM18923v2 Salmonella enterica subsp. enterica serovar Montevideo str. IA_2010008287 Proteobacteria; Gammaproteobacteria; Enterobacterales; Enterobacteriaceae; Salmonella; Salmonella

enterica; Salmonella enterica subsp. enterica; Salmonella enterica subsp. enterica serovar Montevideo
MNNEETFYQAMRRKGVTRRSFLKFCSLAATSLGLGAGMTPKIAWALENKPRIPVVWIHGL WP_001058312.1
hydrogenase [Salmonella enterica] Length: 372\nNumber of predicted TMHs: 2\nExp number of AAs in TMHs: 36.51738\nExp number, first 60 AAs: 19.72849\nTotal prob of N-in: 0.96170\nPOSSIBLE N-term signal sequence\ninside 1 19\nTMhelix 20 39\noutside 40 325\nTMhelix 326 348\ninside 349 372

24221 GCF_000238515.1_ASM23851v2 Salmonella enterica subsp. enterica serovar Montevideo str. LQC 10
Proteobacteria; Gammaproteobacteria; Enterobacterales; Enterobacteriaceae; Salmonella; Salmonella enterica; Salmonella enterica subsp. enterica; Salmonella enterica subsp. enterica serovar Montevideo
MTGDNTLITSHGINRRDFMKLCAALAATMGLSSKAAAEMAESVSNPQRPPVIWIGAQECT WP_000145429.1
hydrogenase 2 small subunit [Salmonella enterica] Length: 372\nNumber of predicted TMHs: 1\nExp number of AAs in TMHs: 24.51309\nExp number, first 60 AAs: 0.2682\nTotal prob of N-in: 0.14402\noutside 1 331\nTMhelix 332 354\ninside 355 372

24222 GCF_000238515.1_ASM23851v2 Salmonella enterica subsp. enterica serovar Montevideo str. LQC 10
Proteobacteria; Gammaproteobacteria; Enterobacterales; Enterobacteriaceae; Salmonella; Salmonella enterica; Salmonella enterica subsp. enterica; Salmonella enterica subsp. enterica serovar Montevideo
MNNEETFYQAMRRKGVTRRSFLKFCSLAATSLGLGAGMTPKIAWALENKPRIPVVWIHGL WP_001058312.1
hydrogenase [Salmonella enterica] Length: 372\nNumber of predicted TMHs: 2\nExp number of AAs in TMHs: 36.51738\nExp number, first 60 AAs: 19.72849\nTotal prob of N-in: 0.96170\nPOSSIBLE N-term signal sequence\ninside 1 19\nTMhelix 20 39\noutside 40 325\nTMhelix 326 348\ninside 349 372

24223 GCF_000188995.1_ASM18899v2 Salmonella enterica subsp. enterica serovar Montevideo str. MB101509-0077 Proteobacteria; Gammaproteobacteria; Enterobacterales; Enterobacteriaceae; Salmonella; Salmonella enterica; Salmonella enterica subsp. enterica; Salmonella enterica subsp. enterica serovar Montevideo
MNNEETFYQAMRRKGVTRRSFLKFCSLAATSLGLGAGMTPKIAWALENKPRIPVVWIHGL WP_001058312.1
hydrogenase [Salmonella enterica] Length: 372\nNumber of predicted TMHs: 2\nExp number of AAs in TMHs: 36.51738\nExp number, first 60 AAs: 19.72849\nTotal prob of N-in: 0.96170\nPOSSIBLE N-term signal sequence\ninside 1 19\nTMhelix 20 39\noutside 40 325\nTMhelix 326 348\ninside 349 372

24224 GCF_000188995.1_ASM18899v2 Salmonella enterica subsp. enterica serovar Montevideo str. MB101509-0077 Proteobacteria; Gammaproteobacteria; Enterobacterales; Enterobacteriaceae; Salmonella; Salmonella enterica; Salmonella enterica subsp. enterica; Salmonella enterica subsp. enterica serovar Montevideo
MTGDNTLITSHGINRRDFMKLCAALAATMGLSSKAAAEMAESVSNPQRPPVIWIGAQECT WP_000145429.1
hydrogenase 2 small subunit [Salmonella enterica] Length: 372\nNumber of predicted TMHs: 1\nExp number of AAs in TMHs: 24.51309\nExp number, first 60 AAs: 0.2682\nTotal prob of N-in: 0.14402\noutside 1 331\nTMhelix 332 354\ninside 355 372

24225 GCF_000189015.1_ASM18901v2 Salmonella enterica subsp. enterica serovar Montevideo str. MB102109-0047 Proteobacteria; Gammaproteobacteria; Enterobacterales; Enterobacteriaceae; Salmonella; Salmonella enterica; Salmonella enterica subsp. enterica; Salmonella enterica subsp. enterica serovar Montevideo
MTGDNTLITSHGINRRDFMKLCAALAATMGLSSKAAAEMAESVSNPQRPPVIWIGAQECT WP_000145429.1
hydrogenase 2 small subunit [Salmonella enterica] Length: 372\nNumber of predicted TMHs: 1\nExp number of AAs in TMHs: 24.51309\nExp number, first 60 AAs: 0.2682\nTotal prob of N-in: 0.14402\noutside 1 331\nTMhelix 332 354\ninside 355 372

24226 GCF_000189015.1_ASM18901v2 Salmonella enterica subsp. enterica serovar Montevideo str. MB102109-0047 Proteobacteria; Gammaproteobacteria; Enterobacterales; Enterobacteriaceae; Salmonella; Salmonella enterica; Salmonella enterica subsp. enterica; Salmonella enterica subsp. enterica serovar Montevideo
MNNEETFYQAMRRKGVTRRSFLKFCSLAATSLGLGAGMTPKIAWALENKPRIPVVWIHGL WP_001058312.1
hydrogenase [Salmonella enterica] Length: 372\nNumber of predicted TMHs: 2\nExp number of AAs in TMHs: 36.51738\nExp number, first 60 AAs: 19.72849\nTotal prob of N-in: 0.96170\nPOSSIBLE N-term signal sequence\ninside 1 19\nTMhelix 20 39\noutside 40 325\nTMhelix 326 348\ninside 349 372

24227 GCF_000189035.1_ASM18903v2 Salmonella enterica subsp. enterica serovar Montevideo str. MB110209-0055 Proteobacteria; Gammaproteobacteria; Enterobacterales; Enterobacteriaceae; Salmonella; Salmonella enterica; Salmonella enterica subsp. enterica; Salmonella enterica subsp. enterica serovar Montevideo
MTGDNTLITSHGINRRDFMKLCAALAATMGLSSKAAAEMAESVSNPQRPPVIWIGAQECT WP_000145429.1
hydrogenase 2 small subunit [Salmonella enterica] Length: 372\nNumber of predicted TMHs: 1\nExp number of AAs in TMHs: 24.51309\nExp number, first 60 AAs: 0.2682\nTotal prob of N-in: 0.14402\noutside 1 331\nTMhelix 332 354\ninside 355 372

24228	GCF_000189035.1_ASM18903v2	Salmonella enterica subsp. enterica serovar Montevideo str. MB110209-0055	Proteobacteria; Gammaproteobacteria; Enterobacterales; Enterobacteriaceae; Salmonella; Salmonella enterica; Salmonella enterica subsp. enterica; Salmonella enterica subsp. enterica serovar Montevideo
MNNEETFYQAMRRKGVTRRSFLKFCSLAATSLGLGAGMTPKIAWALENKPRIPVVWIHGL	WP_001058312.1
hydrogenase [Salmonella enterica]	Length: 372\nNumber of predicted TMHs:  2\nExp number of AAs in TMHs: 36.51738\nExp number, first 60 AAs:  19.72849\nTotal prob of N-in:   0.96170\nPOSSIBLE N-term signal sequence\ninside   1   19\nTMhelix   20   39\noutside   40   325\nTMhelix   326   348\ninside   349   372

24229	GCF_000189055.1_ASM18905v2	Salmonella enterica subsp. enterica serovar Montevideo str. MB111609-0052	Proteobacteria; Gammaproteobacteria; Enterobacterales; Enterobacteriaceae; Salmonella; Salmonella enterica; Salmonella enterica subsp. enterica; Salmonella enterica subsp. enterica serovar Montevideo
MTGDNTLITSHGINRRDFMKLCAALAATMGLSSKAAAEMAESVSNPQRPPVIWIGAQECT	WP_000145429.1
hydrogenase 2 small subunit [Salmonella enterica]	Length: 372\nNumber of predicted TMHs:  1\nExp number of AAs in TMHs: 24.51309\nExp number, first 60 AAs:  0.2682\nTotal prob of N-in:   0.14402\noutside   1   331\nTMhelix   332   354\ninside   355   372

24230	GCF_000189055.1_ASM18905v2	Salmonella enterica subsp. enterica serovar Montevideo str. MB111609-0052	Proteobacteria; Gammaproteobacteria; Enterobacterales; Enterobacteriaceae; Salmonella; Salmonella enterica; Salmonella enterica subsp. enterica; Salmonella enterica subsp. enterica serovar Montevideo
MNNEETFYQAMRRKGVTRRSFLKFCSLAATSLGLGAGMTPKIAWALENKPRIPVVWIHGL	WP_001058312.1
hydrogenase [Salmonella enterica]	Length: 372\nNumber of predicted TMHs:  2\nExp number of AAs in TMHs: 36.51738\nExp number, first 60 AAs:  19.72849\nTotal prob of N-in:   0.96170\nPOSSIBLE N-term signal sequence\ninside   1   19\nTMhelix   20   39\noutside   40   325\nTMhelix   326   348\ninside   349   372

24231	GCF_000188575.1_ASM18857v2	Salmonella enterica subsp. enterica serovar Montevideo str. MD_MDA09249507	Proteobacteria; Gammaproteobacteria; Enterobacterales; Enterobacteriaceae; Salmonella; Salmonella enterica; Salmonella enterica subsp. enterica; Salmonella enterica subsp. enterica serovar Montevideo
MNNEETFYQAMRRKGVTRRSFLKFCSLAATSLGLGAGMTPKIAWALENKPRIPVVWIHGL	WP_001058312.1
hydrogenase [Salmonella enterica]	Length: 372\nNumber of predicted TMHs:  2\nExp number of AAs in TMHs: 36.51738\nExp number, first 60 AAs:  19.72849\nTotal prob of N-in:   0.96170\nPOSSIBLE N-term signal sequence\ninside   1   19\nTMhelix   20   39\noutside   40   325\nTMhelix   326   348\ninside   349   372

24232	GCF_000188575.1_ASM18857v2	Salmonella enterica subsp. enterica serovar Montevideo str. MD_MDA09249507	Proteobacteria; Gammaproteobacteria; Enterobacterales; Enterobacteriaceae; Salmonella; Salmonella enterica; Salmonella enterica subsp. enterica; Salmonella enterica subsp. enterica serovar Montevideo
MTGDNTLITSHGINRRDFMKLCAALAATMGLSSKAAAEMAESVSNPQRPPVIWIGAQECT	WP_000145429.1
hydrogenase 2 small subunit [Salmonella enterica]	Length: 372\nNumber of predicted TMHs:  1\nExp number of AAs in TMHs: 24.51309\nExp number, first 60 AAs:  0.2682\nTotal prob of N-in:   0.14402\noutside   1   331\nTMhelix   332   354\ninside   355   372

24233	GCF_000188475.1_ASM18847v2	Salmonella enterica subsp. enterica serovar Montevideo str. NC_MB110209-0054	Proteobacteria; Gammaproteobacteria; Enterobacterales; Enterobacteriaceae; Salmonella; Salmonella enterica; Salmonella enterica subsp. enterica; Salmonella enterica subsp. enterica serovar Montevideo
MTGDNTLITSHGINRRDFMKLCAALAATMGLSSKAAAEMAESVSNPQRPPVIWIGAQECT	WP_000145429.1
hydrogenase 2 small subunit [Salmonella enterica]	Length: 372\nNumber of predicted TMHs:  1\nExp number of AAs in TMHs: 24.51309\nExp number, first 60 AAs:  0.2682\nTotal prob of N-in:   0.14402\noutside   1   331\nTMhelix   332   354\ninside   355   372

24234	GCF_000188475.1_ASM18847v2	Salmonella enterica subsp. enterica serovar Montevideo str. NC_MB110209-0054	Proteobacteria; Gammaproteobacteria; Enterobacterales; Enterobacteriaceae; Salmonella; Salmonella enterica; Salmonella enterica subsp. enterica; Salmonella enterica subsp. enterica serovar Montevideo
MNNEETFYQAMRRKGVTRRSFLKFCSLAATSLGLGAGMTPKIAWALENKPRIPVVWIHGL	WP_001058312.1
hydrogenase [Salmonella enterica]	Length: 372\nNumber of predicted TMHs:  2\nExp number of AAs in TMHs: 36.51738\nExp number, first 60 AAs:  19.72849\nTotal prob of N-in:   0.96170\nPOSSIBLE N-term signal sequence\ninside   1   19\nTMhelix   20   39\noutside   40   325\nTMhelix   326   348\ninside   349   372

24235	GCF_000188495.1_ASM18849v2	Salmonella enterica subsp. enterica serovar Montevideo str. OH_2009072675	Proteobacteria; Gammaproteobacteria; Enterobacterales; Enterobacteriaceae; Salmonella; Salmonella enterica; Salmonella enterica subsp. enterica; Salmonella enterica subsp. enterica serovar Montevideo
MNNEETFYQAMRRKGVTRRSFLKFCSLAATSLGLGAGMTPKIAWALENKPRIPVVWIHGL	WP_001058312.1
hydrogenase [Salmonella enterica]	Length: 372\nNumber of predicted TMHs:  2\nExp number of AAs in

TMHs: 36.51738\nExp number, first 60 AAs: 19.72849\nTotal prob of N-in: 0.96170\nPOSSIBLE N-term signal sequence\ninside 1 19\nTMhelix 20 39\noutside 40 325\nTMhelix 326 348\ninside 349 372

24236 GCF_000188495.1_ASM18849v2 Salmonella enterica subsp. enterica serovar Montevideo str. OH_2009072675 Proteobacteria; Gammaproteobacteria; Enterobacterales; Enterobacteriaceae; Salmonella; Salmonella enterica; Salmonella enterica subsp. enterica; Salmonella enterica subsp. enterica serovar Montevideo MTGDNTLITSHGINRRDFMKLCAALAATMGLSSKAAAEMAESVSNPQRPPVIWIGAQECT WP_000145429.1 hydrogenase 2 small subunit [Salmonella enterica] Length: 372\nNumber of predicted TMHs: 1\nExp number of AAs in TMHs: 24.51309\nExp number, first 60 AAs: 0.2682\nTotal prob of N-in: 0.14402\noutside 1 331\nTMhelix 332 354\ninside 355 372

24237 GCF_000238495.1_ASM23849v2 Salmonella enterica subsp. enterica serovar Montevideo str. SARB30 Proteobacteria; Gammaproteobacteria; Enterobacterales; Enterobacteriaceae; Salmonella; Salmonella enterica; Salmonella enterica subsp. enterica; Salmonella enterica subsp. enterica serovar Montevideo MNNEETFYQAMRRKGVTRRSFLKFCSLAATSLGLGAGMTPKIAWALENKPRIPVVWIHGL WP_001058312.1 hydrogenase [Salmonella enterica] Length: 372\nNumber of predicted TMHs: 2\nExp number of AAs in TMHs: 36.51738\nExp number, first 60 AAs: 19.72849\nTotal prob of N-in: 0.96170\nPOSSIBLE N-term signal sequence\ninside 1 19\nTMhelix 20 39\noutside 40 325\nTMhelix 326 348\ninside 349 372

24238 GCF_000238495.1_ASM23849v2 Salmonella enterica subsp. enterica serovar Montevideo str. SARB30 Proteobacteria; Gammaproteobacteria; Enterobacterales; Enterobacteriaceae; Salmonella; Salmonella enterica; Salmonella enterica subsp. enterica; Salmonella enterica subsp. enterica serovar Montevideo MTGDNTLITSHGINRRDFMKLCAALAATMGLSSKAAAEMAESVSNPQRPPVIWIGAQECT WP_000145429.1 hydrogenase 2 small subunit [Salmonella enterica] Length: 372\nNumber of predicted TMHs: 1\nExp number of AAs in TMHs: 24.51309\nExp number, first 60 AAs: 0.2682\nTotal prob of N-in: 0.14402\noutside 1 331\nTMhelix 332 354\ninside 355 372

24239 GCF_000238555.1_ASM23855v2 Salmonella enterica subsp. enterica serovar Montevideo str. SARB31 Proteobacteria; Gammaproteobacteria; Enterobacterales; Enterobacteriaceae; Salmonella; Salmonella enterica; Salmonella enterica subsp. enterica; Salmonella enterica subsp. enterica serovar Montevideo MTGDNTLITSHGINRRDFMKLCAALAATMGLSSKAAAEMAESVSNPQRPPVIWIGAQECT WP_000145429.1 hydrogenase 2 small subunit [Salmonella enterica] Length: 372\nNumber of predicted TMHs: 1\nExp number of AAs in TMHs: 24.51309\nExp number, first 60 AAs: 0.2682\nTotal prob of N-in: 0.14402\noutside 1 331\nTMhelix 332 354\ninside 355 372

24240 GCF_000238555.1_ASM23855v2 Salmonella enterica subsp. enterica serovar Montevideo str. SARB31 Proteobacteria; Gammaproteobacteria; Enterobacterales; Enterobacteriaceae; Salmonella; Salmonella enterica; Salmonella enterica subsp. enterica; Salmonella enterica subsp. enterica serovar Montevideo MNNEETFYQAMRRKGVTRRSFLKFCSLAATSLGLGAGMTPKIAWALENKPRIPVVWIHGL WP_001058312.1 hydrogenase [Salmonella enterica] Length: 372\nNumber of predicted TMHs: 2\nExp number of AAs in TMHs: 36.51738\nExp number, first 60 AAs: 19.72849\nTotal prob of N-in: 0.96170\nPOSSIBLE N-term signal sequence\ninside 1 19\nTMhelix 20 39\noutside 40 325\nTMhelix 326 348\ninside 349 372

24241 GCF_000940975.1_ASM94097v1 Salmonella enterica subsp. enterica serovar Montevideo str. USDA-ARS-USMARC-1903 Proteobacteria; Gammaproteobacteria; Enterobacterales; Enterobacteriaceae; Salmonella; Salmonella enterica; Salmonella enterica subsp. enterica; Salmonella enterica subsp. enterica serovar Montevideo MTGDNTLITSHGINRRDFMKLCAALAATMGLSSKAAAEMAESVSNPQRPPVIWIGAQECT WP_000145429.1 hydrogenase 2 small subunit [Salmonella enterica] Length: 372\nNumber of predicted TMHs: 1\nExp number of AAs in TMHs: 24.51309\nExp number, first 60 AAs: 0.2682\nTotal prob of N-in: 0.14402\noutside 1 331\nTMhelix 332 354\ninside 355 372

24242 GCF_000940975.1_ASM94097v1 Salmonella enterica subsp. enterica serovar Montevideo str. USDA-ARS-USMARC-1903 Proteobacteria; Gammaproteobacteria; Enterobacterales; Enterobacteriaceae; Salmonella; Salmonella enterica; Salmonella enterica subsp. enterica; Salmonella enterica subsp. enterica serovar Montevideo MNNEETFYQAMRRKGVTRRSFLKFCSLAATSLGLGAGMTPKIAWALENKPRIPVVWIHGL WP_044782462.1 hydrogenase [Salmonella enterica] Length: 372\nNumber of predicted TMHs: 2\nExp number of AAs in TMHs: 36.51702\nExp number, first 60 AAs: 19.72849\nTotal prob of N-in: 0.96170\nPOSSIBLE N-term signal sequence\ninside 1 19\nTMhelix 20 39\noutside 40 325\nTMhelix 326 348\ninside 349 372

24243 GCF_000973665.2_ASM97366v2 Salmonella enterica subsp. enterica serovar Montevideo str. USDA-ARS-USMARC-1921 Proteobacteria; Gammaproteobacteria; Enterobacterales; Enterobacteriaceae; Salmonella; Salmonella

enterica; Salmonella enterica subsp. enterica; Salmonella enterica subsp. enterica serovar Montevideo
MTGDNTLITSHGINRRDFMKLCAALAATMGLSSKAAAEMAESVSNPQRPPVIWIGAQECT WP_000145429.1
hydrogenase 2 small subunit [Salmonella enterica] Length: 372\nNumber of predicted TMHs: 1\nExp number of AAs in TMHs: 24.51309\nExp number, first 60 AAs: 0.2682\nTotal prob of N-in: 0.14402\noutside 1 331\nTMhelix 332 354\ninside 355 372

24244 GCF_000973665.2_ASM97366v2 Salmonella enterica subsp. enterica serovar Montevideo str. USDA-ARS-USMARC-1921 Proteobacteria; Gammaproteobacteria; Enterobacterales; Enterobacteriaceae; Salmonella; Salmonella enterica; Salmonella enterica subsp. enterica; Salmonella enterica subsp. enterica serovar Montevideo
MNNEETFYQAMRRKGVTRRSFLKFCSLAATSLGLGAGMTPKIAWALENKPRIPVVWIHGL WP_044782462.1
hydrogenase [Salmonella enterica] Length: 372\nNumber of predicted TMHs: 2\nExp number of AAs in TMHs: 36.51702\nExp number, first 60 AAs: 19.72849\nTotal prob of N-in: 0.96170\nPOSSIBLE N-term signal sequence\ninside 1 19\nTMhelix 20 39\noutside 40 325\nTMhelix 326 348\ninside 349 372

24245 GCF_001975555.1_ASM197555v1 Salmonella enterica subsp. enterica serovar Moscow str. S-1843
Proteobacteria; Gammaproteobacteria; Enterobacterales; Enterobacteriaceae; Salmonella; Salmonella enterica; Salmonella enterica subsp. enterica; Salmonella enterica subsp. enterica serovar Moscow
MNNEETFYQAMRRKGVTRRSFLKFCSLAATSLGLGAGMTPKIAWALENKPRIPVVWIHGL WP_001058307.1
hydrogenase [Salmonella enterica] Length: 372\nNumber of predicted TMHs: 2\nExp number of AAs in TMHs: 36.51738\nExp number, first 60 AAs: 19.72849\nTotal prob of N-in: 0.96170\nPOSSIBLE N-term signal sequence\ninside 1 19\nTMhelix 20 39\noutside 40 325\nTMhelix 326 348\ninside 349 372

24246 GCF_001975555.1_ASM197555v1 Salmonella enterica subsp. enterica serovar Moscow str. S-1843
Proteobacteria; Gammaproteobacteria; Enterobacterales; Enterobacteriaceae; Salmonella; Salmonella enterica; Salmonella enterica subsp. enterica; Salmonella enterica subsp. enterica serovar Moscow
MTGDNTLITSHGINRRDFMKLCAALAATMGLSSKAAAEMAESVSNPQRPPVIWIGAQECT WP_000145429.1
hydrogenase 2 small subunit [Salmonella enterica] Length: 372\nNumber of predicted TMHs: 1\nExp number of AAs in TMHs: 24.51309\nExp number, first 60 AAs: 0.2682\nTotal prob of N-in: 0.14402\noutside 1 331\nTMhelix 332 354\ninside 355 372

24247 GCF_000412795.1_08-1125_Ver1 Salmonella enterica subsp. enterica serovar Muenchen str. 08-1125
Proteobacteria; Gammaproteobacteria; Enterobacterales; Enterobacteriaceae; Salmonella; Salmonella enterica; Salmonella enterica subsp. enterica; Salmonella enterica subsp. enterica serovar Muenchen
MTGDNTLITSHGINRRDFMKLCAALAATMGLSSKAAAEMAESVSNPQRPPVIWIGAQECT WP_000145429.1
hydrogenase 2 small subunit [Salmonella enterica] Length: 372\nNumber of predicted TMHs: 1\nExp number of AAs in TMHs: 24.51309\nExp number, first 60 AAs: 0.2682\nTotal prob of N-in: 0.14402\noutside 1 331\nTMhelix 332 354\ninside 355 372

24248 GCF_000412815.1_2009K-0951_Ver1 Salmonella enterica subsp. enterica serovar Muenchen str. 2009K-0951
Proteobacteria; Gammaproteobacteria; Enterobacterales; Enterobacteriaceae; Salmonella; Salmonella enterica; Salmonella enterica subsp. enterica; Salmonella enterica subsp. enterica serovar Muenchen
MTGDNTLITSHGINRRDFMKLCAALAATMGLSSKAAAEMAESVSNPQRPPVIWIGAQECT WP_000145429.1
hydrogenase 2 small subunit [Salmonella enterica] Length: 372\nNumber of predicted TMHs: 1\nExp number of AAs in TMHs: 24.51309\nExp number, first 60 AAs: 0.2682\nTotal prob of N-in: 0.14402\noutside 1 331\nTMhelix 332 354\ninside 355 372

24249 GCF_000486325.1_ASM48632v1 Salmonella enterica subsp. enterica serovar Muenchen str. ATCC 8388
Proteobacteria; Gammaproteobacteria; Enterobacterales; Enterobacteriaceae; Salmonella; Salmonella enterica; Salmonella enterica subsp. enterica; Salmonella enterica subsp. enterica serovar Muenchen
MTGDNTLITSHGINRRDFMKLCAALAATMGLSSKAAAEMAESVSNPQRPPVIWIGAQECT WP_000145429.1
hydrogenase 2 small subunit [Salmonella enterica] Length: 372\nNumber of predicted TMHs: 1\nExp number of AAs in TMHs: 24.51309\nExp number, first 60 AAs: 0.2682\nTotal prob of N-in: 0.14402\noutside 1 331\nTMhelix 332 354\ninside 355 372

24250 GCF_000486325.1_ASM48632v1 Salmonella enterica subsp. enterica serovar Muenchen str. ATCC 8388
Proteobacteria; Gammaproteobacteria; Enterobacterales; Enterobacteriaceae; Salmonella; Salmonella enterica; Salmonella enterica subsp. enterica; Salmonella enterica subsp. enterica serovar Muenchen
MNNEETFYQAMRRKGVTRRSFLKFCSLAATSLGLGAGMTPKIAWALENKPRIPVVWIHGL WP_001058311.1
hydrogenase [Salmonella enterica] Length: 372\nNumber of predicted TMHs: 2\nExp number of AAs in TMHs: 37.4699899999999\nExp number, first 60 AAs: 19.72699\nTotal prob of N-in: 0.96171\nPOSSIBLE N-term signal sequence\ninside 1 19\nTMhelix 20 39\noutside 40 325\nTMhelix 326 348\ninside 349 372

24251	GCF_000486955.1_ASM48695v1	Salmonella enterica subsp. enterica serovar Muenchen str. baa1674	Proteobacteria; Gammaproteobacteria; Enterobacterales; Enterobacteriaceae; Salmonella; Salmonella enterica; Salmonella enterica subsp. enterica; Salmonella enterica subsp. enterica serovar Muenchen	MNNEETFYQAMRRKGVTRRSFLKFCSLAATSLGLGAGMTPKIAWALENKPRIPVVWIHGL	WP_001058312.1	hydrogenase [Salmonella enterica]	Length: 372\nNumber of predicted TMHs:  2\nExp number of AAs in TMHs: 36.51738\nExp number, first 60 AAs:  19.72849\nTotal prob of N-in:   0.96170\nPOSSIBLE N-term signal sequence\ninside   1   19\nTMhelix   20   39\noutside   40   325\nTMhelix   326   348\ninside   349   372

24252	GCF_000486955.1_ASM48695v1	Salmonella enterica subsp. enterica serovar Muenchen str. baa1674	Proteobacteria; Gammaproteobacteria; Enterobacterales; Enterobacteriaceae; Salmonella; Salmonella enterica; Salmonella enterica subsp. enterica; Salmonella enterica subsp. enterica serovar Muenchen	MTGDNTLITSHGINRRDFMKLCAALAATMGLSSKAAAEMAESVSNPQRPPVIWIGAQECT	WP_023221305.1	hydrogenase 2 small subunit [Salmonella enterica]	Length: 372\nNumber of predicted TMHs:  1\nExp number of AAs in TMHs: 24.51314\nExp number, first 60 AAs:  0.26822\nTotal prob of N-in:   0.14403\noutside   1   331\nTMhelix   332   354\ninside   355   372

24253	GCF_000487035.2_ASM48703v2	Salmonella enterica subsp. enterica serovar Muenster str. 0315	Proteobacteria; Gammaproteobacteria; Enterobacterales; Enterobacteriaceae; Salmonella; Salmonella enterica; Salmonella enterica subsp. enterica; Salmonella enterica subsp. enterica serovar Muenster	MTGDNTLITSHGINRRDFMKLCAALAATMGLSSKAAAEMAESVSNPQRPPVIWIGAQECT	WP_000145431.1	hydrogenase 2 small subunit [Salmonella enterica]	Length: 372\nNumber of predicted TMHs:  1\nExp number of AAs in TMHs: 24.51505\nExp number, first 60 AAs:  0.2682\nTotal prob of N-in:   0.14406\noutside   1   331\nTMhelix   332   354\ninside   355   372

24254	GCF_000487035.2_ASM48703v2	Salmonella enterica subsp. enterica serovar Muenster str. 0315	Proteobacteria; Gammaproteobacteria; Enterobacterales; Enterobacteriaceae; Salmonella; Salmonella enterica; Salmonella enterica subsp. enterica; Salmonella enterica subsp. enterica serovar Muenster	MNNEETFYQAMRRKGVTRRSFLKFCSLAATSLGLGAGMTPKIAWALENKPRIPVVWIHGL	WP_001058312.1	hydrogenase [Salmonella enterica]	Length: 372\nNumber of predicted TMHs:  2\nExp number of AAs in TMHs: 36.51738\nExp number, first 60 AAs:  19.72849\nTotal prob of N-in:   0.96170\nPOSSIBLE N-term signal sequence\ninside   1   19\nTMhelix   20   39\noutside   40   325\nTMhelix   326   348\ninside   349   372

24255	GCF_000487015.1_ASM48701v1	Salmonella enterica subsp. enterica serovar Muenster str. 420	Proteobacteria; Gammaproteobacteria; Enterobacterales; Enterobacteriaceae; Salmonella; Salmonella enterica; Salmonella enterica subsp. enterica; Salmonella enterica subsp. enterica serovar Muenster	MTGDNTLITSHGINRRDFMKLCAALAATMGLSSKAAAEMAESVSNPQRPPVIWIGAQECT	WP_000145431.1	hydrogenase 2 small subunit [Salmonella enterica]	Length: 372\nNumber of predicted TMHs:  1\nExp number of AAs in TMHs: 24.51505\nExp number, first 60 AAs:  0.2682\nTotal prob of N-in:   0.14406\noutside   1   331\nTMhelix   332   354\ninside   355   372

24256	GCF_000487015.1_ASM48701v1	Salmonella enterica subsp. enterica serovar Muenster str. 420	Proteobacteria; Gammaproteobacteria; Enterobacterales; Enterobacteriaceae; Salmonella; Salmonella enterica; Salmonella enterica subsp. enterica; Salmonella enterica subsp. enterica serovar Muenster	MNNEETFYQAMRRKGVTRRSFLKFCSLAATSLGLGAGMTPKIAWALENKPRIPVVWIHGL	WP_001058312.1	hydrogenase [Salmonella enterica]	Length: 372\nNumber of predicted TMHs:  2\nExp number of AAs in TMHs: 36.51738\nExp number, first 60 AAs:  19.72849\nTotal prob of N-in:   0.96170\nPOSSIBLE N-term signal sequence\ninside   1   19\nTMhelix   20   39\noutside   40   325\nTMhelix   326   348\ninside   349   372

24257	GCF_000483855.1_ASM48385v1	Salmonella enterica subsp. enterica serovar Muenster str. 660	Proteobacteria; Gammaproteobacteria; Enterobacterales; Enterobacteriaceae; Salmonella; Salmonella enterica; Salmonella enterica subsp. enterica; Salmonella enterica subsp. enterica serovar Muenster	MTGDNTLITSHGINRRDFMKLCAALAATMGLSSKAAAEMAESVSNPQRPPVIWIGAQECT	WP_000145431.1	hydrogenase 2 small subunit [Salmonella enterica]	Length: 372\nNumber of predicted TMHs:  1\nExp number of AAs in TMHs: 24.51505\nExp number, first 60 AAs:  0.2682\nTotal prob of N-in:   0.14406\noutside   1   331\nTMhelix   332   354\ninside   355   372

24258	GCF_000483855.1_ASM48385v1	Salmonella enterica subsp. enterica serovar Muenster str. 660	Proteobacteria; Gammaproteobacteria; Enterobacterales; Enterobacteriaceae; Salmonella; Salmonella enterica; Salmonella enterica subsp. enterica; Salmonella enterica subsp. enterica serovar Muenster	MNNEETFYQAMRRKGVTRRSFLKFCSLAATSLGLGAGMTPKIAWALENKPRIPVVWIHGL	WP_001058312.1	hydrogenase [Salmonella enterica]	Length: 372\nNumber of predicted TMHs:  2\nExp number of AAs in

TMHs: 36.51738\nExp number, first 60 AAs: 19.72849\nTotal prob of N-in: 0.96170\nPOSSIBLE N-term signal sequence\ninside 1 19\nTMhelix 20 39\noutside 40 325\nTMhelix 326 348\ninside 349 372

24259 GCF_000505485.1_CFSAN004344_01.0Salmonella enterica subsp. enterica serovar Muenster str. CFSAN004344 Proteobacteria; Gammaproteobacteria; Enterobacterales; Enterobacteriaceae; Salmonella; Salmonella enterica; Salmonella enterica subsp. enterica; Salmonella enterica subsp. enterica serovar Muenster MTGDNTLITSHGINRRDFMKLCAALAATMGLSSKAAAEMAESVSNPQRPPVIWIGAQECT WP_000145429.1 hydrogenase 2 small subunit [Salmonella enterica] Length: 372\nNumber of predicted TMHs: 1\nExp number of AAs in TMHs: 24.51309\nExp number, first 60 AAs: 0.2682\nTotal prob of N-in: 0.14402\noutside 1 331\nTMhelix 332 354\ninside 355 372

24260 GCF_000599425.1_Salmonella_enterica_enterica_Namur Salmonella enterica subsp. enterica serovar Namur str. 05-2929 Proteobacteria; Gammaproteobacteria; Enterobacterales; Enterobacteriaceae; Salmonella; Salmonella enterica; Salmonella enterica subsp. enterica; Salmonella enterica subsp. enterica serovar Namur MTGDNTLITSHGINRRDFMKLCAALAATMGLSSKAAAEMAESVSNPQRPPVIWIGAQECT WP_000145429.1 hydrogenase 2 small subunit [Salmonella enterica] Length: 372\nNumber of predicted TMHs: 1\nExp number of AAs in TMHs: 24.51309\nExp number, first 60 AAs: 0.2682\nTotal prob of N-in: 0.14402\noutside 1 331\nTMhelix 332 354\ninside 355 372

24261 GCF_000599425.1_Salmonella_enterica_enterica_Namur Salmonella enterica subsp. enterica serovar Namur str. 05-2929 Proteobacteria; Gammaproteobacteria; Enterobacterales; Enterobacteriaceae; Salmonella; Salmonella enterica; Salmonella enterica subsp. enterica; Salmonella enterica subsp. enterica serovar Namur MNNEETFYQAMRRKGVTRRSFLKFCSLAATSLGLGAGMTPKIAWALENKPRIPVVWIHGL WP_001058311.1 hydrogenase [Salmonella enterica] Length: 372\nNumber of predicted TMHs: 2\nExp number of AAs in TMHs: 37.4699899999999\nExp number, first 60 AAs: 19.72699\nTotal prob of N-in: 0.96171\nPOSSIBLE N-term signal sequence\ninside 1 19\nTMhelix 20 39\noutside 40 325\nTMhelix 326 348\ninside 349 372

24262 GCF_000487675.1_ASM48767v1 Salmonella enterica subsp. enterica serovar Nchanga str. CFSAN001091 Proteobacteria; Gammaproteobacteria; Enterobacterales; Enterobacteriaceae; Salmonella; Salmonella enterica; Salmonella enterica subsp. enterica; Salmonella enterica subsp. enterica serovar Nchanga MTGDNTLITSHGINRRDFMKLCAALAATMGLSSKAAAEMAESVSNPQRPPVIWIGAQECT WP_000145429.1 hydrogenase 2 small subunit [Salmonella enterica] Length: 372\nNumber of predicted TMHs: 1\nExp number of AAs in TMHs: 24.51309\nExp number, first 60 AAs: 0.2682\nTotal prob of N-in: 0.14402\noutside 1 331\nTMhelix 332 354\ninside 355 372

24263 GCF_000487655.1_ASM48765v1 Salmonella enterica subsp. enterica serovar Nchanga str. CFSAN001092 Proteobacteria; Gammaproteobacteria; Enterobacterales; Enterobacteriaceae; Salmonella; Salmonella enterica; Salmonella enterica subsp. enterica; Salmonella enterica subsp. enterica serovar Nchanga MTGDNTLITSHGINRRDFMKLCAALAATMGLSSKAAAEMAESVSNPQRPPVIWIGAQECT WP_000145429.1 hydrogenase 2 small subunit [Salmonella enterica] Length: 372\nNumber of predicted TMHs: 1\nExp number of AAs in TMHs: 24.51309\nExp number, first 60 AAs: 0.2682\nTotal prob of N-in: 0.14402\noutside 1 331\nTMhelix 332 354\ninside 355 372

24264 GCF_000487655.1_ASM48765v1 Salmonella enterica subsp. enterica serovar Nchanga str. CFSAN001092 Proteobacteria; Gammaproteobacteria; Enterobacterales; Enterobacteriaceae; Salmonella; Salmonella enterica; Salmonella enterica subsp. enterica; Salmonella enterica subsp. enterica serovar Nchanga MNNEETFYQAMRRKGVTRRSFLKFCSLAATSLGLGAGMTPKIAWALENKPRIPVVWIHGL WP_001058312.1 hydrogenase [Salmonella enterica] Length: 372\nNumber of predicted TMHs: 2\nExp number of AAs in TMHs: 36.51738\nExp number, first 60 AAs: 19.72849\nTotal prob of N-in: 0.96170\nPOSSIBLE N-term signal sequence\ninside 1 19\nTMhelix 20 39\noutside 40 325\nTMhelix 326 348\ninside 349 372

24265 GCF_000474375.1_CFSAN001193_01.0Salmonella enterica subsp. enterica serovar Newport str. #11-2 Proteobacteria; Gammaproteobacteria; Enterobacterales; Enterobacteriaceae; Salmonella; Salmonella enterica; Salmonella enterica subsp. enterica; Salmonella enterica subsp. enterica serovar Newport MNNEETFYQAMRRKGVTRRSFLKFCSLAATSLGLGAGMTPKIAWALENKPRIPVVWIHGL WP_001058312.1 hydrogenase [Salmonella enterica] Length: 372\nNumber of predicted TMHs: 2\nExp number of AAs in TMHs: 36.51738\nExp number, first 60 AAs: 19.72849\nTotal prob of N-in: 0.96170\nPOSSIBLE N-term signal sequence\ninside 1 19\nTMhelix 20 39\noutside 40 325\nTMhelix 326 348\ninside 349 372

24266 GCF_000474375.1_CFSAN001193_01.0Salmonella enterica subsp. enterica serovar Newport str. #11-2 Proteobacteria; Gammaproteobacteria; Enterobacterales; Enterobacteriaceae; Salmonella; Salmonella enterica;

Salmonella enterica subsp. enterica; Salmonella enterica subsp. enterica serovar Newport
MTGDNTLITSHGINRRDFMKLCAALAATMGLSSKAAAEMAESVSNPQRPPVIWIGAQECT WP_000145429.1
hydrogenase 2 small subunit [Salmonella enterica] Length: 372\nNumber of predicted TMHs: 1\nExp number of AAs in TMHs: 24.51309\nExp number, first 60 AAs: 0.2682\nTotal prob of N-in: 0.14402\noutside 1 331\nTMhelix 332 354\ninside 355 372

24267 GCF_000474535.1_CFSAN001192_01.0Salmonella enterica subsp. enterica serovar Newport str. #11-3
Proteobacteria; Gammaproteobacteria; Enterobacterales; Enterobacteriaceae; Salmonella; Salmonella enterica; Salmonella enterica subsp. enterica; Salmonella enterica subsp. enterica serovar Newport
MTGDNTLITSHGINRRDFMKLCAALAATMGLSSKAAAEMAESVSNPQRPPVIWIGAQECT WP_000145429.1
hydrogenase 2 small subunit [Salmonella enterica] Length: 372\nNumber of predicted TMHs: 1\nExp number of AAs in TMHs: 24.51309\nExp number, first 60 AAs: 0.2682\nTotal prob of N-in: 0.14402\noutside 1 331\nTMhelix 332 354\ninside 355 372

24268 GCF_000474535.1_CFSAN001192_01.0Salmonella enterica subsp. enterica serovar Newport str. #11-3
Proteobacteria; Gammaproteobacteria; Enterobacterales; Enterobacteriaceae; Salmonella; Salmonella enterica; Salmonella enterica subsp. enterica; Salmonella enterica subsp. enterica serovar Newport
MNNEETFYQAMRRKGVTRRSFLKFCSLAATSLGLGAGMTPKIAWALENKPRIPVVWIHGL WP_001058312.1
hydrogenase [Salmonella enterica] Length: 372\nNumber of predicted TMHs: 2\nExp number of AAs in TMHs: 36.51738\nExp number, first 60 AAs: 19.72849\nTotal prob of N-in: 0.96170\nPOSSIBLE N-term signal sequence\ninside 1 19\nTMhelix 20 39\noutside 40 325\nTMhelix 326 348\ninside 349 372

24269 GCF_000474515.1_CFSAN001191_01.0Salmonella enterica subsp. enterica serovar Newport str. #11-4
Proteobacteria; Gammaproteobacteria; Enterobacterales; Enterobacteriaceae; Salmonella; Salmonella enterica; Salmonella enterica subsp. enterica; Salmonella enterica subsp. enterica serovar Newport
MTGDNTLITSHGINRRDFMKLCAALAATMGLSSKAAAEMAESVSNPQRPPVIWIGAQECT WP_000145429.1
hydrogenase 2 small subunit [Salmonella enterica] Length: 372\nNumber of predicted TMHs: 1\nExp number of AAs in TMHs: 24.51309\nExp number, first 60 AAs: 0.2682\nTotal prob of N-in: 0.14402\noutside 1 331\nTMhelix 332 354\ninside 355 372

24270 GCF_000474515.1_CFSAN001191_01.0Salmonella enterica subsp. enterica serovar Newport str. #11-4
Proteobacteria; Gammaproteobacteria; Enterobacterales; Enterobacteriaceae; Salmonella; Salmonella enterica; Salmonella enterica subsp. enterica; Salmonella enterica subsp. enterica serovar Newport
MNNEETFYQAMRRKGVTRRSFLKFCSLAATSLGLGAGMTPKIAWALENKPRIPVVWIHGL WP_001058312.1
hydrogenase [Salmonella enterica] Length: 372\nNumber of predicted TMHs: 2\nExp number of AAs in TMHs: 36.51738\nExp number, first 60 AAs: 19.72849\nTotal prob of N-in: 0.96170\nPOSSIBLE N-term signal sequence\ninside 1 19\nTMhelix 20 39\noutside 40 325\nTMhelix 326 348\ninside 349 372

24271 GCF_001051615.1_ASM105161v1 Salmonella enterica subsp. enterica serovar Newport str. 36799
Proteobacteria; Gammaproteobacteria; Enterobacterales; Enterobacteriaceae; Salmonella; Salmonella enterica; Salmonella enterica subsp. enterica; Salmonella enterica subsp. enterica serovar Newport
MTGDNTLITSHGINRRDFMKLCAALAATMGLSSKAAAEMAESVSNPQRPPVIWIGAQECT WP_000145429.1
hydrogenase 2 small subunit [Salmonella enterica] Length: 372\nNumber of predicted TMHs: 1\nExp number of AAs in TMHs: 24.51309\nExp number, first 60 AAs: 0.2682\nTotal prob of N-in: 0.14402\noutside 1 331\nTMhelix 332 354\ninside 355 372

24272 GCF_001051615.1_ASM105161v1 Salmonella enterica subsp. enterica serovar Newport str. 36799
Proteobacteria; Gammaproteobacteria; Enterobacterales; Enterobacteriaceae; Salmonella; Salmonella enterica; Salmonella enterica subsp. enterica; Salmonella enterica subsp. enterica serovar Newport
MNNEETFYQAMRRKGVTRRSFLKFCSLAATSLGLGAGMTPKIAWALENKPRIPVVWIHGL WP_001058312.1
hydrogenase [Salmonella enterica] Length: 372\nNumber of predicted TMHs: 2\nExp number of AAs in TMHs: 36.51738\nExp number, first 60 AAs: 19.72849\nTotal prob of N-in: 0.96170\nPOSSIBLE N-term signal sequence\ninside 1 19\nTMhelix 20 39\noutside 40 325\nTMhelix 326 348\ninside 349 372

24273 GCF_001051555.1_ASM105155v1 Salmonella enterica subsp. enterica serovar Newport str. 36801
Proteobacteria; Gammaproteobacteria; Enterobacterales; Enterobacteriaceae; Salmonella; Salmonella enterica; Salmonella enterica subsp. enterica; Salmonella enterica subsp. enterica serovar Newport
MTGDNTLITSHGINRRDFMKLCAALAATMGLSSKAAAEMAESVSNPQRPPVIWIGAQECT WP_000145429.1
hydrogenase 2 small subunit [Salmonella enterica] Length: 372\nNumber of predicted TMHs: 1\nExp number of AAs in TMHs: 24.51309\nExp number, first 60 AAs: 0.2682\nTotal prob of N-in: 0.14402\noutside 1 331\nTMhelix 332 354\ninside 355 372

24274	GCF_001051555.1_ASM105155v1	Salmonella enterica subsp. enterica serovar Newport str. 36801	Proteobacteria; Gammaproteobacteria; Enterobacterales; Enterobacteriaceae; Salmonella; Salmonella enterica; Salmonella enterica subsp. enterica; Salmonella enterica subsp. enterica serovar Newport	MNNEETFYQAMRRKGVTRRSFLKFCSLAATSLGLGAGMTPKIAWALENKPRIPVVWIHGL	WP_001058312.1	hydrogenase [Salmonella enterica]	Length: 372\nNumber of predicted TMHs:  2\nExp number of AAs in TMHs: 36.51738\nExp number, first 60 AAs:  19.72849\nTotal prob of N-in:   0.96170\nPOSSIBLE N-term signal sequence\ninside   1   19\nTMhelix   20   39\noutside   40   325\nTMhelix   326   348\ninside   349   372

24275	GCF_001051605.1_ASM105160v1	Salmonella enterica subsp. enterica serovar Newport str. 36802	Proteobacteria; Gammaproteobacteria; Enterobacterales; Enterobacteriaceae; Salmonella; Salmonella enterica; Salmonella enterica subsp. enterica; Salmonella enterica subsp. enterica serovar Newport	MTGDNTLITSHGINRRDFMKLCAALAATMGLSSKAAAEMAESVSNPQRPPVIWIGAQECT	WP_000145429.1	hydrogenase 2 small subunit [Salmonella enterica]	Length: 372\nNumber of predicted TMHs:  1\nExp number of AAs in TMHs: 24.51309\nExp number, first 60 AAs:  0.2682\nTotal prob of N-in:   0.14402\noutside   1   331\nTMhelix   332   354\ninside   355   372

24276	GCF_001051605.1_ASM105160v1	Salmonella enterica subsp. enterica serovar Newport str. 36802	Proteobacteria; Gammaproteobacteria; Enterobacterales; Enterobacteriaceae; Salmonella; Salmonella enterica; Salmonella enterica subsp. enterica; Salmonella enterica subsp. enterica serovar Newport	MNNEETFYQAMRRKGVTRRSFLKFCSLAATSLGLGAGMTPKIAWALENKPRIPVVWIHGL	WP_001058312.1	hydrogenase [Salmonella enterica]	Length: 372\nNumber of predicted TMHs:  2\nExp number of AAs in TMHs: 36.51738\nExp number, first 60 AAs:  19.72849\nTotal prob of N-in:   0.96170\nPOSSIBLE N-term signal sequence\ninside   1   19\nTMhelix   20   39\noutside   40   325\nTMhelix   326   348\ninside   349   372

24277	GCF_001051795.1_ASM105179v1	Salmonella enterica subsp. enterica serovar Newport str. 36803	Proteobacteria; Gammaproteobacteria; Enterobacterales; Enterobacteriaceae; Salmonella; Salmonella enterica; Salmonella enterica subsp. enterica; Salmonella enterica subsp. enterica serovar Newport	MTGDNTLITSHGINRRDFMKLCAALAATMGLSSKAAAEMAESVSNPQRPPVIWIGAQECT	WP_000145429.1	hydrogenase 2 small subunit [Salmonella enterica]	Length: 372\nNumber of predicted TMHs:  1\nExp number of AAs in TMHs: 24.51309\nExp number, first 60 AAs:  0.2682\nTotal prob of N-in:   0.14402\noutside   1   331\nTMhelix   332   354\ninside   355   372

24278	GCF_001051795.1_ASM105179v1	Salmonella enterica subsp. enterica serovar Newport str. 36803	Proteobacteria; Gammaproteobacteria; Enterobacterales; Enterobacteriaceae; Salmonella; Salmonella enterica; Salmonella enterica subsp. enterica; Salmonella enterica subsp. enterica serovar Newport	MNNEETFYQAMRRKGVTRRSFLKFCSLAATSLGLGAGMTPKIAWALENKPRIPVVWIHGL	WP_001058312.1	hydrogenase [Salmonella enterica]	Length: 372\nNumber of predicted TMHs:  2\nExp number of AAs in TMHs: 36.51738\nExp number, first 60 AAs:  19.72849\nTotal prob of N-in:   0.96170\nPOSSIBLE N-term signal sequence\ninside   1   19\nTMhelix   20   39\noutside   40   325\nTMhelix   326   348\ninside   349   372

24279	GCF_001051545.1_ASM105154v1	Salmonella enterica subsp. enterica serovar Newport str. 36804	Proteobacteria; Gammaproteobacteria; Enterobacterales; Enterobacteriaceae; Salmonella; Salmonella enterica; Salmonella enterica subsp. enterica; Salmonella enterica subsp. enterica serovar Newport	MNNEETFYQAMRRKGVTRRSFLKFCSLAATSLGLGAGMTPKIAWALENKPRIPVVWIHGL	WP_001058312.1	hydrogenase [Salmonella enterica]	Length: 372\nNumber of predicted TMHs:  2\nExp number of AAs in TMHs: 36.51738\nExp number, first 60 AAs:  19.72849\nTotal prob of N-in:   0.96170\nPOSSIBLE N-term signal sequence\ninside   1   19\nTMhelix   20   39\noutside   40   325\nTMhelix   326   348\ninside   349   372

24280	GCF_001051545.1_ASM105154v1	Salmonella enterica subsp. enterica serovar Newport str. 36804	Proteobacteria; Gammaproteobacteria; Enterobacterales; Enterobacteriaceae; Salmonella; Salmonella enterica; Salmonella enterica subsp. enterica; Salmonella enterica subsp. enterica serovar Newport	MTGDNTLITSHGINRRDFMKLCAALAATMGLSSKAAAEMAESVSNPQRPPVIWIGAQECT	WP_000145429.1	hydrogenase 2 small subunit [Salmonella enterica]	Length: 372\nNumber of predicted TMHs:  1\nExp number of AAs in TMHs: 24.51309\nExp number, first 60 AAs:  0.2682\nTotal prob of N-in:   0.14402\noutside   1   331\nTMhelix   332   354\ninside   355   372

24281	GCF_001051535.1_ASM105153v1	Salmonella enterica subsp. enterica serovar Newport str. 36805	Proteobacteria; Gammaproteobacteria; Enterobacterales; Enterobacteriaceae; Salmonella; Salmonella enterica; Salmonella enterica subsp. enterica; Salmonella enterica subsp. enterica serovar Newport	MNNEETFYQAMRRKGVTRRSFLKFCSLAATSLGLGAGMTPKIAWALENKPRIPVVWIHGL	WP_001058312.1	hydrogenase [Salmonella enterica]	Length: 372\nNumber of predicted TMHs:  2\nExp number of AAs in

TMHs: 36.51738\nExp number, first 60 AAs: 19.72849\nTotal prob of N-in: 0.96170\nPOSSIBLE N-term signal sequence\ninside 1 19\nTMhelix 20 39\noutside 40 325\nTMhelix 326 348\ninside 349 372

24282 GCF_001051535.1_ASM105153v1 Salmonella enterica subsp. enterica serovar Newport str. 36805 Proteobacteria; Gammaproteobacteria; Enterobacterales; Enterobacteriaceae; Salmonella; Salmonella enterica; Salmonella enterica subsp. enterica; Salmonella enterica subsp. enterica serovar Newport MTGDNTLITSHGINRRDFMKLCAALAATMGLSSKAAAEMAESVSNPQRPPVIWIGAQECT WP_000145429.1 hydrogenase 2 small subunit [Salmonella enterica] Length: 372\nNumber of predicted TMHs: 1\nExp number of AAs in TMHs: 24.51309\nExp number, first 60 AAs: 0.2682\nTotal prob of N-in: 0.14402\noutside 1 331\nTMhelix 332 354\ninside 355 372

24283 GCF_001051525.1_ASM105152v1 Salmonella enterica subsp. enterica serovar Newport str. 36807 Proteobacteria; Gammaproteobacteria; Enterobacterales; Enterobacteriaceae; Salmonella; Salmonella enterica; Salmonella enterica subsp. enterica; Salmonella enterica subsp. enterica serovar Newport MTGDNTLITSHGINRRDFMKLCAALAATMGLSSKAAAEMAESVSNPQRPPVIWIGAQECT WP_000145429.1 hydrogenase 2 small subunit [Salmonella enterica] Length: 372\nNumber of predicted TMHs: 1\nExp number of AAs in TMHs: 24.51309\nExp number, first 60 AAs: 0.2682\nTotal prob of N-in: 0.14402\noutside 1 331\nTMhelix 332 354\ninside 355 372

24284 GCF_001051525.1_ASM105152v1 Salmonella enterica subsp. enterica serovar Newport str. 36807 Proteobacteria; Gammaproteobacteria; Enterobacterales; Enterobacteriaceae; Salmonella; Salmonella enterica; Salmonella enterica subsp. enterica; Salmonella enterica subsp. enterica serovar Newport MNNEETFYQAMRRKGVTRRSFLKFCSLAATSLGLGAGMTPKIAWALENKPRIPVVWIHGL WP_001058312.1 hydrogenase [Salmonella enterica] Length: 372\nNumber of predicted TMHs: 2\nExp number of AAs in TMHs: 36.51738\nExp number, first 60 AAs: 19.72849\nTotal prob of N-in: 0.96170\nPOSSIBLE N-term signal sequence\ninside 1 19\nTMhelix 20 39\noutside 40 325\nTMhelix 326 348\ninside 349 372

24285 GCF_000484215.1_ASM48421v1 Salmonella enterica subsp. enterica serovar Newport str. 637564_17 Proteobacteria; Gammaproteobacteria; Enterobacterales; Enterobacteriaceae; Salmonella; Salmonella enterica; Salmonella enterica subsp. enterica; Salmonella enterica subsp. enterica serovar Newport MTGDNTLITSHGINRRDFMKLCAALAATMGLSSKAAAEMAESVSNPQRPPVIWIGAQECT WP_000145429.1 hydrogenase 2 small subunit [Salmonella enterica] Length: 372\nNumber of predicted TMHs: 1\nExp number of AAs in TMHs: 24.51309\nExp number, first 60 AAs: 0.2682\nTotal prob of N-in: 0.14402\noutside 1 331\nTMhelix 332 354\ninside 355 372

24286 GCF_000484215.1_ASM48421v1 Salmonella enterica subsp. enterica serovar Newport str. 637564_17 Proteobacteria; Gammaproteobacteria; Enterobacterales; Enterobacteriaceae; Salmonella; Salmonella enterica; Salmonella enterica subsp. enterica; Salmonella enterica subsp. enterica serovar Newport MNNEETFYQAMRRKGVTRRSFLKFCSLAATSLGLGAGMTPKIAWALENKPRIPVVWIHGL WP_001058312.1 hydrogenase [Salmonella enterica] Length: 372\nNumber of predicted TMHs: 2\nExp number of AAs in TMHs: 36.51738\nExp number, first 60 AAs: 19.72849\nTotal prob of N-in: 0.96170\nPOSSIBLE N-term signal sequence\ninside 1 19\nTMhelix 20 39\noutside 40 325\nTMhelix 326 348\ninside 349 372

24287 GCF_001051765.1_CFSAN000927_1.0 Salmonella enterica subsp. enterica serovar Newport str. A182RVB Proteobacteria; Gammaproteobacteria; Enterobacterales; Enterobacteriaceae; Salmonella; Salmonella enterica; Salmonella enterica subsp. enterica; Salmonella enterica subsp. enterica serovar Newport MTGDNTLITSHGINRRDFMKLCAALAATMGLSSKAAAEMAESVSNPQRPPVIWIGAQECT WP_000145429.1 hydrogenase 2 small subunit [Salmonella enterica] Length: 372\nNumber of predicted TMHs: 1\nExp number of AAs in TMHs: 24.51309\nExp number, first 60 AAs: 0.2682\nTotal prob of N-in: 0.14402\noutside 1 331\nTMhelix 332 354\ninside 355 372

24288 GCF_001051765.1_CFSAN000927_1.0 Salmonella enterica subsp. enterica serovar Newport str. A182RVB Proteobacteria; Gammaproteobacteria; Enterobacterales; Enterobacteriaceae; Salmonella; Salmonella enterica; Salmonella enterica subsp. enterica; Salmonella enterica subsp. enterica serovar Newport MNNEETFYQAMRRKGVTRRSFLKFCSLAATSLGLGAGMTPKIAWALENKPRIPVVWIHGL WP_001058312.1 hydrogenase [Salmonella enterica] Length: 372\nNumber of predicted TMHs: 2\nExp number of AAs in TMHs: 36.51738\nExp number, first 60 AAs: 19.72849\nTotal prob of N-in: 0.96170\nPOSSIBLE N-term signal sequence\ninside 1 19\nTMhelix 20 39\noutside 40 325\nTMhelix 326 348\ninside 349 372

24289 GCF_000973685.1_ASM97368v1 Salmonella enterica subsp. enterica serovar Newport str. CDC 2010K-2159 Proteobacteria; Gammaproteobacteria; Enterobacterales; Enterobacteriaceae; Salmonella; Salmonella enterica;

Salmonella enterica subsp. enterica; Salmonella enterica subsp. enterica serovar Newport
MNNEETFYQAMRRKGVTRRSFLKFCSLAATSLGLGAGMTPKIAWALENKPRIPVVWIHGL WP_001058312.1
hydrogenase [Salmonella enterica] Length: 372\nNumber of predicted TMHs: 2\nExp number of AAs in TMHs: 36.51738\nExp number, first 60 AAs: 19.72849\nTotal prob of N-in: 0.96170\nPOSSIBLE N-term signal sequence\ninside 1 19\nTMhelix 20 39\noutside 40 325\nTMhelix 326 348\ninside 349 372

24290 GCF_000973685.1_ASM97368v1 Salmonella enterica subsp. enterica serovar Newport str. CDC 2010K-2159
Proteobacteria; Gammaproteobacteria; Enterobacterales; Enterobacteriaceae; Salmonella; Salmonella enterica; Salmonella enterica subsp. enterica; Salmonella enterica subsp. enterica serovar Newport
MTGDNTLITSHGINRRDFMKLCAALAATMGLSSKAAAEMAESVSNPQRPPVIWIGAQECT WP_000145429.1
hydrogenase 2 small subunit [Salmonella enterica] Length: 372\nNumber of predicted TMHs: 1\nExp number of AAs in TMHs: 24.51309\nExp number, first 60 AAs: 0.2682\nTotal prob of N-in: 0.14402\noutside 1 331\nTMhelix 332 354\ninside 355 372

24291 GCF_000272915.1_ASM27291v1 Salmonella enterica subsp. enterica serovar Newport str. CVM 19443
Proteobacteria; Gammaproteobacteria; Enterobacterales; Enterobacteriaceae; Salmonella; Salmonella enterica; Salmonella enterica subsp. enterica; Salmonella enterica subsp. enterica serovar Newport
MTGDNTLITSHGINRRDFMKLCAALAATMGLSSKAAAEMAESVSNPQRPPVIWIGAQECT WP_000145429.1
hydrogenase 2 small subunit [Salmonella enterica] Length: 372\nNumber of predicted TMHs: 1\nExp number of AAs in TMHs: 24.51309\nExp number, first 60 AAs: 0.2682\nTotal prob of N-in: 0.14402\noutside 1 331\nTMhelix 332 354\ninside 355 372

24292 GCF_000272915.1_ASM27291v1 Salmonella enterica subsp. enterica serovar Newport str. CVM 19443
Proteobacteria; Gammaproteobacteria; Enterobacterales; Enterobacteriaceae; Salmonella; Salmonella enterica; Salmonella enterica subsp. enterica; Salmonella enterica subsp. enterica serovar Newport
MNNEETFYQAMRRKGVTRRSFLKFCSLAATSLGLGAGMTPKIAWALENKPRIPVVWIHGL WP_001058312.1
hydrogenase [Salmonella enterica] Length: 372\nNumber of predicted TMHs: 2\nExp number of AAs in TMHs: 36.51738\nExp number, first 60 AAs: 19.72849\nTotal prob of N-in: 0.96170\nPOSSIBLE N-term signal sequence\ninside 1 19\nTMhelix 20 39\noutside 40 325\nTMhelix 326 348\ninside 349 372

24293 GCF_000272575.1_ASM27257v1 Salmonella enterica subsp. enterica serovar Newport str. CVM 19447
Proteobacteria; Gammaproteobacteria; Enterobacterales; Enterobacteriaceae; Salmonella; Salmonella enterica; Salmonella enterica subsp. enterica; Salmonella enterica subsp. enterica serovar Newport
MTGDNTLITSHGINRRDFMKLCAALAATMGLSSKAAAEMAESVSNPQRPPVIWIGAQECT WP_000145429.1
hydrogenase 2 small subunit [Salmonella enterica] Length: 372\nNumber of predicted TMHs: 1\nExp number of AAs in TMHs: 24.51309\nExp number, first 60 AAs: 0.2682\nTotal prob of N-in: 0.14402\noutside 1 331\nTMhelix 332 354\ninside 355 372

24294 GCF_000272575.1_ASM27257v1 Salmonella enterica subsp. enterica serovar Newport str. CVM 19447
Proteobacteria; Gammaproteobacteria; Enterobacterales; Enterobacteriaceae; Salmonella; Salmonella enterica; Salmonella enterica subsp. enterica; Salmonella enterica subsp. enterica serovar Newport
MNNEETFYQAMRRKGVTRRSFLKFCSLAATSLGLGAGMTPKIAWALENKPRIPVVWIHGL WP_001058312.1
hydrogenase [Salmonella enterica] Length: 372\nNumber of predicted TMHs: 2\nExp number of AAs in TMHs: 36.51738\nExp number, first 60 AAs: 19.72849\nTotal prob of N-in: 0.96170\nPOSSIBLE N-term signal sequence\ninside 1 19\nTMhelix 20 39\noutside 40 325\nTMhelix 326 348\ninside 349 372

24295 GCF_000272695.1_ASM27269v1 Salmonella enterica subsp. enterica serovar Newport str. CVM 19449
Proteobacteria; Gammaproteobacteria; Enterobacterales; Enterobacteriaceae; Salmonella; Salmonella enterica; Salmonella enterica subsp. enterica; Salmonella enterica subsp. enterica serovar Newport
MTGDNTLITSHGINRRDFMKLCAALAATMGLSSKAAAEMAESVSNPQRPPVIWIGAQECT WP_000145429.1
hydrogenase 2 small subunit [Salmonella enterica] Length: 372\nNumber of predicted TMHs: 1\nExp number of AAs in TMHs: 24.51309\nExp number, first 60 AAs: 0.2682\nTotal prob of N-in: 0.14402\noutside 1 331\nTMhelix 332 354\ninside 355 372

24296 GCF_000272695.1_ASM27269v1 Salmonella enterica subsp. enterica serovar Newport str. CVM 19449
Proteobacteria; Gammaproteobacteria; Enterobacterales; Enterobacteriaceae; Salmonella; Salmonella enterica; Salmonella enterica subsp. enterica; Salmonella enterica subsp. enterica serovar Newport
MNNEETFYQAMRRKGVTRRSFLKFCSLAATSLGLGAGMTPKIAWALENKPRIPVVWIHGL WP_001058312.1
hydrogenase [Salmonella enterica] Length: 372\nNumber of predicted TMHs: 2\nExp number of AAs in TMHs: 36.51738\nExp number, first 60 AAs: 19.72849\nTotal prob of N-in: 0.96170\nPOSSIBLE N-term signal sequence\ninside 1 19\nTMhelix 20 39\noutside 40 325\nTMhelix 326 348\ninside 349 372

24297 GCF_000272975.1_ASM27297v1 Salmonella enterica subsp. enterica serovar Newport str. CVM 19470 Proteobacteria; Gammaproteobacteria; Enterobacterales; Enterobacteriaceae; Salmonella; Salmonella enterica; Salmonella enterica subsp. enterica; Salmonella enterica subsp. enterica serovar Newport MNNEETFYQAMRRKGVTRRSFLKFCSLAATSLGLGAGMTPKIAWALENKPRIPVVWIHGL WP_001058312.1 hydrogenase [Salmonella enterica] Length: 372\nNumber of predicted TMHs: 2\nExp number of AAs in TMHs: 36.51738\nExp number, first 60 AAs: 19.72849\nTotal prob of N-in: 0.96170\nPOSSIBLE N-term signal sequence\ninside 1 19\nTMhelix 20 39\noutside 40 325\nTMhelix 326 348\ninside 349 372

24298 GCF_000272975.1_ASM27297v1 Salmonella enterica subsp. enterica serovar Newport str. CVM 19470 Proteobacteria; Gammaproteobacteria; Enterobacterales; Enterobacteriaceae; Salmonella; Salmonella enterica; Salmonella enterica subsp. enterica; Salmonella enterica subsp. enterica serovar Newport MTGDNTLITSHGINRRDFMKLCAALAATMGLSSKAAAEMAESVSNPQRPPVIWIGAQECT WP_000145429.1 hydrogenase 2 small subunit [Salmonella enterica] Length: 372\nNumber of predicted TMHs: 1\nExp number of AAs in TMHs: 24.51309\nExp number, first 60 AAs: 0.2682\nTotal prob of N-in: 0.14402\noutside 1 331\nTMhelix 332 354\ninside 355 372

24299 GCF_000272855.1_ASM27285v1 Salmonella enterica subsp. enterica serovar Newport str. CVM 19536 Proteobacteria; Gammaproteobacteria; Enterobacterales; Enterobacteriaceae; Salmonella; Salmonella enterica; Salmonella enterica subsp. enterica; Salmonella enterica subsp. enterica serovar Newport MNNEETFYQAMRRKGVTRRSFLKFCSLAATSLGLGAGMTPKIAWALENKPRIPVVWIHGL WP_001058312.1 hydrogenase [Salmonella enterica] Length: 372\nNumber of predicted TMHs: 2\nExp number of AAs in TMHs: 36.51738\nExp number, first 60 AAs: 19.72849\nTotal prob of N-in: 0.96170\nPOSSIBLE N-term signal sequence\ninside 1 19\nTMhelix 20 39\noutside 40 325\nTMhelix 326 348\ninside 349 372

24300 GCF_000272855.1_ASM27285v1 Salmonella enterica subsp. enterica serovar Newport str. CVM 19536 Proteobacteria; Gammaproteobacteria; Enterobacterales; Enterobacteriaceae; Salmonella; Salmonella enterica; Salmonella enterica subsp. enterica; Salmonella enterica subsp. enterica serovar Newport MTGDNTLITSHGINRRDFMKLCAALAATMGLSSKAAAEMAESVSNPQRPPVIWIGAQECT WP_000145429.1 hydrogenase 2 small subunit [Salmonella enterica] Length: 372\nNumber of predicted TMHs: 1\nExp number of AAs in TMHs: 24.51309\nExp number, first 60 AAs: 0.2682\nTotal prob of N-in: 0.14402\noutside 1 331\nTMhelix 332 354\ninside 355 372

24301 GCF_000272595.1_ASM27259v1 Salmonella enterica subsp. enterica serovar Newport str. CVM 19567 Proteobacteria; Gammaproteobacteria; Enterobacterales; Enterobacteriaceae; Salmonella; Salmonella enterica; Salmonella enterica subsp. enterica; Salmonella enterica subsp. enterica serovar Newport MTGDNTLITSHGINRRDFMKLCAALAATMGLSSKAAAEMAESVSNPQRPPVIWIGAQECT WP_000145429.1 hydrogenase 2 small subunit [Salmonella enterica] Length: 372\nNumber of predicted TMHs: 1\nExp number of AAs in TMHs: 24.51309\nExp number, first 60 AAs: 0.2682\nTotal prob of N-in: 0.14402\noutside 1 331\nTMhelix 332 354\ninside 355 372

24302 GCF_000272595.1_ASM27259v1 Salmonella enterica subsp. enterica serovar Newport str. CVM 19567 Proteobacteria; Gammaproteobacteria; Enterobacterales; Enterobacteriaceae; Salmonella; Salmonella enterica; Salmonella enterica subsp. enterica; Salmonella enterica subsp. enterica serovar Newport MNNEETFYQAMRRKGVTRRSFLKFCSLAATSLGLGAGMTPKIAWALENKPRIPVVWIHGL WP_001058312.1 hydrogenase [Salmonella enterica] Length: 372\nNumber of predicted TMHs: 2\nExp number of AAs in TMHs: 36.51738\nExp number, first 60 AAs: 19.72849\nTotal prob of N-in: 0.96170\nPOSSIBLE N-term signal sequence\ninside 1 19\nTMhelix 20 39\noutside 40 325\nTMhelix 326 348\ninside 349 372

24303 GCF_000272955.1_ASM27295v1 Salmonella enterica subsp. enterica serovar Newport str. CVM 19593 Proteobacteria; Gammaproteobacteria; Enterobacterales; Enterobacteriaceae; Salmonella; Salmonella enterica; Salmonella enterica subsp. enterica; Salmonella enterica subsp. enterica serovar Newport MNNEETFYQAMRRKGVTRRSFLKFCSLAATSLGLGAGMTPKIAWALENKPRIPVVWIHGL WP_001058312.1 hydrogenase [Salmonella enterica] Length: 372\nNumber of predicted TMHs: 2\nExp number of AAs in TMHs: 36.51738\nExp number, first 60 AAs: 19.72849\nTotal prob of N-in: 0.96170\nPOSSIBLE N-term signal sequence\ninside 1 19\nTMhelix 20 39\noutside 40 325\nTMhelix 326 348\ninside 349 372

24304 GCF_000272955.1_ASM27295v1 Salmonella enterica subsp. enterica serovar Newport str. CVM 19593 Proteobacteria; Gammaproteobacteria; Enterobacterales; Enterobacteriaceae; Salmonella; Salmonella enterica; Salmonella enterica subsp. enterica; Salmonella enterica subsp. enterica serovar Newport MTGDNTLITSHGINRRDFMKLCAALAATMGLSSKAAAEMAESVSNPQRPPVIWIGAQECT WP_000145429.1 hydrogenase 2 small subunit [Salmonella enterica] Length: 372\nNumber of predicted TMHs: 1\nExp

number of AAs in TMHs: 24.51309\nExp number, first 60 AAs: 0.2682\nTotal prob of N-in: 0.14402\noutside 1 331\nTMhelix 332 354\ninside 355 372

24305 GCF_000272755.3_ASM27275v3 Salmonella enterica subsp. enterica serovar Newport str. CVM 21538 Proteobacteria; Gammaproteobacteria; Enterobacterales; Enterobacteriaceae; Salmonella; Salmonella enterica; Salmonella enterica subsp. enterica; Salmonella enterica subsp. enterica serovar Newport MTGDNTLITSHGINRRDFMKLCAALAATMGLSSKAAAEMAESVSNPQRPPVIWIGAQECT WP_000145429.1 hydrogenase 2 small subunit [Salmonella enterica] Length: 372\nNumber of predicted TMHs: 1\nExp number of AAs in TMHs: 24.51309\nExp number, first 60 AAs: 0.2682\nTotal prob of N-in: 0.14402\noutside 1 331\nTMhelix 332 354\ninside 355 372

24306 GCF_000272755.3_ASM27275v3 Salmonella enterica subsp. enterica serovar Newport str. CVM 21538 Proteobacteria; Gammaproteobacteria; Enterobacterales; Enterobacteriaceae; Salmonella; Salmonella enterica; Salmonella enterica subsp. enterica; Salmonella enterica subsp. enterica serovar Newport MNNEETFYQAMRRKGVTRRSFLKFCSLAATSLGLGAGMTPKIAWALENKPRIPVVWIHGL WP_001058312.1 hydrogenase [Salmonella enterica] Length: 372\nNumber of predicted TMHs: 2\nExp number of AAs in TMHs: 36.51738\nExp number, first 60 AAs: 19.72849\nTotal prob of N-in: 0.96170\nPOSSIBLE N-term signal sequence\ninside 1 19\nTMhelix 20 39\noutside 40 325\nTMhelix 326 348\ninside 349 372

24307 GCF_000272655.1_ASM27265v1 Salmonella enterica subsp. enterica serovar Newport str. CVM 21539 Proteobacteria; Gammaproteobacteria; Enterobacterales; Enterobacteriaceae; Salmonella; Salmonella enterica; Salmonella enterica subsp. enterica; Salmonella enterica subsp. enterica serovar Newport MNNEETFYQAMRRKGVTRRSFLKFCSLAATSLGLGAGMTPKIAWALENKPRIPVVWIHGL WP_001058312.1 hydrogenase [Salmonella enterica] Length: 372\nNumber of predicted TMHs: 2\nExp number of AAs in TMHs: 36.51738\nExp number, first 60 AAs: 19.72849\nTotal prob of N-in: 0.96170\nPOSSIBLE N-term signal sequence\ninside 1 19\nTMhelix 20 39\noutside 40 325\nTMhelix 326 348\ninside 349 372

24308 GCF_000272655.1_ASM27265v1 Salmonella enterica subsp. enterica serovar Newport str. CVM 21539 Proteobacteria; Gammaproteobacteria; Enterobacterales; Enterobacteriaceae; Salmonella; Salmonella enterica; Salmonella enterica subsp. enterica; Salmonella enterica subsp. enterica serovar Newport MTGDNTLITSHGINRRDFMKLCAALAATMGLSSKAAAEMAESVSNPQRPPVIWIGAQECT WP_000145429.1 hydrogenase 2 small subunit [Salmonella enterica] Length: 372\nNumber of predicted TMHs: 1\nExp number of AAs in TMHs: 24.51309\nExp number, first 60 AAs: 0.2682\nTotal prob of N-in: 0.14402\noutside 1 331\nTMhelix 332 354\ninside 355 372

24309 GCF_000272715.3_ASM27271v3 Salmonella enterica subsp. enterica serovar Newport str. CVM 21550 Proteobacteria; Gammaproteobacteria; Enterobacterales; Enterobacteriaceae; Salmonella; Salmonella enterica; Salmonella enterica subsp. enterica; Salmonella enterica subsp. enterica serovar Newport MTGDNTLITSHGINRRDFMKLCAALAATMGLSSKAAAEMAESVSNPQRPPVIWIGAQECT WP_000145429.1 hydrogenase 2 small subunit [Salmonella enterica] Length: 372\nNumber of predicted TMHs: 1\nExp number of AAs in TMHs: 24.51309\nExp number, first 60 AAs: 0.2682\nTotal prob of N-in: 0.14402\noutside 1 331\nTMhelix 332 354\ninside 355 372

24310 GCF_000272715.3_ASM27271v3 Salmonella enterica subsp. enterica serovar Newport str. CVM 21550 Proteobacteria; Gammaproteobacteria; Enterobacterales; Enterobacteriaceae; Salmonella; Salmonella enterica; Salmonella enterica subsp. enterica; Salmonella enterica subsp. enterica serovar Newport MNNEETFYQAMRRKGVTRRSFLKFCSLAATSLGLGAGMTPKIAWALENKPRIPVVWIHGL WP_001058312.1 hydrogenase [Salmonella enterica] Length: 372\nNumber of predicted TMHs: 2\nExp number of AAs in TMHs: 36.51738\nExp number, first 60 AAs: 19.72849\nTotal prob of N-in: 0.96170\nPOSSIBLE N-term signal sequence\ninside 1 19\nTMhelix 20 39\noutside 40 325\nTMhelix 326 348\ninside 349 372

24311 GCF_000272555.1_ASM27255v1 Salmonella enterica subsp. enterica serovar Newport str. CVM 21559 Proteobacteria; Gammaproteobacteria; Enterobacterales; Enterobacteriaceae; Salmonella; Salmonella enterica; Salmonella enterica subsp. enterica; Salmonella enterica subsp. enterica serovar Newport MNNEETFYQAMRRKGVTRRSFLKFCSLAATSLGLGAGMTPKIAWALENKPRIPVVWIHGL WP_001058312.1 hydrogenase [Salmonella enterica] Length: 372\nNumber of predicted TMHs: 2\nExp number of AAs in TMHs: 36.51738\nExp number, first 60 AAs: 19.72849\nTotal prob of N-in: 0.96170\nPOSSIBLE N-term signal sequence\ninside 1 19\nTMhelix 20 39\noutside 40 325\nTMhelix 326 348\ninside 349 372

24312 GCF_000272555.1_ASM27255v1 Salmonella enterica subsp. enterica serovar Newport str. CVM 21559 Proteobacteria; Gammaproteobacteria; Enterobacterales; Enterobacteriaceae; Salmonella; Salmonella enterica;

Salmonella enterica subsp. enterica; Salmonella enterica subsp. enterica serovar Newport
MTGDNTLITSHGINRRDFMKLCAALAATMGLSSKAAAEMAESVSNPQRPPVIWIGAQECT WP_000145429.1
hydrogenase 2 small subunit [Salmonella enterica] Length: 372\nNumber of predicted TMHs: 1\nExp number of AAs in TMHs: 24.51309\nExp number, first 60 AAs: 0.2682\nTotal prob of N-in: 0.14402\noutside 1 331\nTMhelix 332 354\ninside 355 372

24313 GCF_000272775.3_ASM27277v3 Salmonella enterica subsp. enterica serovar Newport str. CVM 22425
Proteobacteria; Gammaproteobacteria; Enterobacterales; Enterobacteriaceae; Salmonella; Salmonella enterica; Salmonella enterica subsp. enterica; Salmonella enterica subsp. enterica serovar Newport
MNNEETFYQAMRRKGVTRRSFLKFCSLAATSLGLGAGMTPKIAWALENKPRIPVVWIHGL WP_001058312.1
hydrogenase [Salmonella enterica] Length: 372\nNumber of predicted TMHs: 2\nExp number of AAs in TMHs: 36.51738\nExp number, first 60 AAs: 19.72849\nTotal prob of N-in: 0.96170\nPOSSIBLE N-term signal sequence\ninside 1 19\nTMhelix 20 39\noutside 40 325\nTMhelix 326 348\ninside 349 372

24314 GCF_000272775.3_ASM27277v3 Salmonella enterica subsp. enterica serovar Newport str. CVM 22425
Proteobacteria; Gammaproteobacteria; Enterobacterales; Enterobacteriaceae; Salmonella; Salmonella enterica; Salmonella enterica subsp. enterica; Salmonella enterica subsp. enterica serovar Newport
MTGDNTLITSHGINRRDFMKLCAALAATMGLSSKAAAEMAESVSNPQRPPVIWIGAQECT WP_000145429.1
hydrogenase 2 small subunit [Salmonella enterica] Length: 372\nNumber of predicted TMHs: 1\nExp number of AAs in TMHs: 24.51309\nExp number, first 60 AAs: 0.2682\nTotal prob of N-in: 0.14402\noutside 1 331\nTMhelix 332 354\ninside 355 372

24315 GCF_000272895.2_ASM27289v3 Salmonella enterica subsp. enterica serovar Newport str. CVM 22462
Proteobacteria; Gammaproteobacteria; Enterobacterales; Enterobacteriaceae; Salmonella; Salmonella enterica; Salmonella enterica subsp. enterica; Salmonella enterica subsp. enterica serovar Newport
MTGDNTLITSHGINRRDFMKLCAALAATMGLSSKAAAEMAESVSNPQRPPVIWIGAQECT WP_000145429.1
hydrogenase 2 small subunit [Salmonella enterica] Length: 372\nNumber of predicted TMHs: 1\nExp number of AAs in TMHs: 24.51309\nExp number, first 60 AAs: 0.2682\nTotal prob of N-in: 0.14402\noutside 1 331\nTMhelix 332 354\ninside 355 372

24316 GCF_000272895.2_ASM27289v3 Salmonella enterica subsp. enterica serovar Newport str. CVM 22462
Proteobacteria; Gammaproteobacteria; Enterobacterales; Enterobacteriaceae; Salmonella; Salmonella enterica; Salmonella enterica subsp. enterica; Salmonella enterica subsp. enterica serovar Newport
MNNEETFYQAMRRKGVTRRSFLKFCSLAATSLGLGAGMTPKIAWALENKPRIPVVWIHGL WP_001058312.1
hydrogenase [Salmonella enterica] Length: 372\nNumber of predicted TMHs: 2\nExp number of AAs in TMHs: 36.51738\nExp number, first 60 AAs: 19.72849\nTotal prob of N-in: 0.96170\nPOSSIBLE N-term signal sequence\ninside 1 19\nTMhelix 20 39\noutside 40 325\nTMhelix 326 348\ninside 349 372

24317 GCF_000272735.3_ASM27273v3 Salmonella enterica subsp. enterica serovar Newport str. CVM 22513
Proteobacteria; Gammaproteobacteria; Enterobacterales; Enterobacteriaceae; Salmonella; Salmonella enterica; Salmonella enterica subsp. enterica; Salmonella enterica subsp. enterica serovar Newport
MNNEETFYQAMRRKGVTRRSFLKFCSLAATSLGLGAGMTPKIAWALENKPRIPVVWIHGL WP_001058312.1
hydrogenase [Salmonella enterica] Length: 372\nNumber of predicted TMHs: 2\nExp number of AAs in TMHs: 36.51738\nExp number, first 60 AAs: 19.72849\nTotal prob of N-in: 0.96170\nPOSSIBLE N-term signal sequence\ninside 1 19\nTMhelix 20 39\noutside 40 325\nTMhelix 326 348\ninside 349 372

24318 GCF_000272735.3_ASM27273v3 Salmonella enterica subsp. enterica serovar Newport str. CVM 22513
Proteobacteria; Gammaproteobacteria; Enterobacterales; Enterobacteriaceae; Salmonella; Salmonella enterica; Salmonella enterica subsp. enterica; Salmonella enterica subsp. enterica serovar Newport
MTGDNTLITSHGINRRDFMKLCAALAATMGLSSKAAAEMAESVSNPQRPPVIWIGAQECT WP_000145429.1
hydrogenase 2 small subunit [Salmonella enterica] Length: 372\nNumber of predicted TMHs: 1\nExp number of AAs in TMHs: 24.51309\nExp number, first 60 AAs: 0.2682\nTotal prob of N-in: 0.14402\noutside 1 331\nTMhelix 332 354\ninside 355 372

24319 GCF_000272675.1_ASM27267v1 Salmonella enterica subsp. enterica serovar Newport str. CVM 33953
Proteobacteria; Gammaproteobacteria; Enterobacterales; Enterobacteriaceae; Salmonella; Salmonella enterica; Salmonella enterica subsp. enterica; Salmonella enterica subsp. enterica serovar Newport
MNNEETFYQAMRRKGVTRRSFLKFCSLAATSLGLGAGMTPKIAWALENKPRIPVVWIHGL WP_001058312.1
hydrogenase [Salmonella enterica] Length: 372\nNumber of predicted TMHs: 2\nExp number of AAs in TMHs: 36.51738\nExp number, first 60 AAs: 19.72849\nTotal prob of N-in: 0.96170\nPOSSIBLE N-term signal sequence\ninside 1 19\nTMhelix 20 39\noutside 40 325\nTMhelix 326 348\ninside 349 372

24320 GCF_000272675.1_ASM27267v1 Salmonella enterica subsp. enterica serovar Newport str. CVM 33953 Proteobacteria; Gammaproteobacteria; Enterobacterales; Enterobacteriaceae; Salmonella; Salmonella enterica; Salmonella enterica subsp. enterica; Salmonella enterica subsp. enterica serovar Newport MTGDNTLITSHGINRRDFMKLCAALAATMGLSSKAAAEMAESVSNPQRPPVIWIGAQECT WP_000145429.1 hydrogenase 2 small subunit [Salmonella enterica] Length: 372\nNumber of predicted TMHs: 1\nExp number of AAs in TMHs: 24.51309\nExp number, first 60 AAs: 0.2682\nTotal prob of N-in: 0.14402\noutside 1 331\nTMhelix 332 354\ninside 355 372

24321 GCF_000272515.1_ASM27251v1 Salmonella enterica subsp. enterica serovar Newport str. CVM 35185 Proteobacteria; Gammaproteobacteria; Enterobacterales; Enterobacteriaceae; Salmonella; Salmonella enterica; Salmonella enterica subsp. enterica; Salmonella enterica subsp. enterica serovar Newport MNNEETFYQAMRRKGVTRRSFLKFCSLAATSLGLGAGMTPKIAWALENKPRIPVVWIHGL WP_001058312.1 hydrogenase [Salmonella enterica] Length: 372\nNumber of predicted TMHs: 2\nExp number of AAs in TMHs: 36.51738\nExp number, first 60 AAs: 19.72849\nTotal prob of N-in: 0.96170\nPOSSIBLE N-term signal sequence\ninside 1 19\nTMhelix 20 39\noutside 40 325\nTMhelix 326 348\ninside 349 372

24322 GCF_000272515.1_ASM27251v1 Salmonella enterica subsp. enterica serovar Newport str. CVM 35185 Proteobacteria; Gammaproteobacteria; Enterobacterales; Enterobacteriaceae; Salmonella; Salmonella enterica; Salmonella enterica subsp. enterica; Salmonella enterica subsp. enterica serovar Newport MTGDNTLITSHGINRRDFMKLCAALAATMGLSSKAAAEMAESVSNPQRPPVIWIGAQECT WP_000145429.1 hydrogenase 2 small subunit [Salmonella enterica] Length: 372\nNumber of predicted TMHs: 1\nExp number of AAs in TMHs: 24.51309\nExp number, first 60 AAs: 0.2682\nTotal prob of N-in: 0.14402\noutside 1 331\nTMhelix 332 354\ninside 355 372

24323 GCF_000272535.1_ASM27253v1 Salmonella enterica subsp. enterica serovar Newport str. CVM 35188 Proteobacteria; Gammaproteobacteria; Enterobacterales; Enterobacteriaceae; Salmonella; Salmonella enterica; Salmonella enterica subsp. enterica; Salmonella enterica subsp. enterica serovar Newport MTGDNTLITSHGINRRDFMKLCAALAATMGLSSKAAAEMAESVSNPQRPPVIWIGAQECT WP_000145429.1 hydrogenase 2 small subunit [Salmonella enterica] Length: 372\nNumber of predicted TMHs: 1\nExp number of AAs in TMHs: 24.51309\nExp number, first 60 AAs: 0.2682\nTotal prob of N-in: 0.14402\noutside 1 331\nTMhelix 332 354\ninside 355 372

24324 GCF_000272535.1_ASM27253v1 Salmonella enterica subsp. enterica serovar Newport str. CVM 35188 Proteobacteria; Gammaproteobacteria; Enterobacterales; Enterobacteriaceae; Salmonella; Salmonella enterica; Salmonella enterica subsp. enterica; Salmonella enterica subsp. enterica serovar Newport MNNEETFYQAMRRKGVTRRSFLKFCSLAATSLGLGAGMTPKIAWALENKPRIPVVWIHGL WP_001058312.1 hydrogenase [Salmonella enterica] Length: 372\nNumber of predicted TMHs: 2\nExp number of AAs in TMHs: 36.51738\nExp number, first 60 AAs: 19.72849\nTotal prob of N-in: 0.96170\nPOSSIBLE N-term signal sequence\ninside 1 19\nTMhelix 20 39\noutside 40 325\nTMhelix 326 348\ninside 349 372

24325 GCF_000272635.1_ASM27263v1 Salmonella enterica subsp. enterica serovar Newport str. CVM 35199 Proteobacteria; Gammaproteobacteria; Enterobacterales; Enterobacteriaceae; Salmonella; Salmonella enterica; Salmonella enterica subsp. enterica; Salmonella enterica subsp. enterica serovar Newport MNNEETFYQAMRRKGVTRRSFLKFCSLAATSLGLGAGMTPKIAWALENKPRIPVVWIHGL WP_001058312.1 hydrogenase [Salmonella enterica] Length: 372\nNumber of predicted TMHs: 2\nExp number of AAs in TMHs: 36.51738\nExp number, first 60 AAs: 19.72849\nTotal prob of N-in: 0.96170\nPOSSIBLE N-term signal sequence\ninside 1 19\nTMhelix 20 39\noutside 40 325\nTMhelix 326 348\ninside 349 372

24326 GCF_000272635.1_ASM27263v1 Salmonella enterica subsp. enterica serovar Newport str. CVM 35199 Proteobacteria; Gammaproteobacteria; Enterobacterales; Enterobacteriaceae; Salmonella; Salmonella enterica; Salmonella enterica subsp. enterica; Salmonella enterica subsp. enterica serovar Newport MTGDNTLITSHGINRRDFMKLCAALAATMGLSSKAAAEMAESVSNPQRPPVIWIGAQECT WP_000145429.1 hydrogenase 2 small subunit [Salmonella enterica] Length: 372\nNumber of predicted TMHs: 1\nExp number of AAs in TMHs: 24.51309\nExp number, first 60 AAs: 0.2682\nTotal prob of N-in: 0.14402\noutside 1 331\nTMhelix 332 354\ninside 355 372

24327 GCF_000272615.1_ASM27261v1 Salmonella enterica subsp. enterica serovar Newport str. CVM 35202 Proteobacteria; Gammaproteobacteria; Enterobacterales; Enterobacteriaceae; Salmonella; Salmonella enterica; Salmonella enterica subsp. enterica; Salmonella enterica subsp. enterica serovar Newport MNNEETFYQAMRRKGVTRRSFLKFCSLAATSLGLGAGMTPKIAWALENKPRIPVVWIHGL WP_001058312.1 hydrogenase [Salmonella enterica] Length: 372\nNumber of predicted TMHs: 2\nExp number of AAs in

TMHs: 36.51738\nExp number, first 60 AAs: 19.72849\nTotal prob of N-in: 0.96170\nPOSSIBLE N-term signal sequence\ninside 1 19\nTMhelix 20 39\noutside 40 325\nTMhelix 326 348\ninside 349 372

24328 GCF_000272615.1_ASM27261v1 Salmonella enterica subsp. enterica serovar Newport str. CVM 35202 Proteobacteria; Gammaproteobacteria; Enterobacterales; Enterobacteriaceae; Salmonella; Salmonella enterica; Salmonella enterica subsp. enterica; Salmonella enterica subsp. enterica serovar Newport MTGDNTLITSHGINRRDFMKLCAALAATMGLSSKAAAEMAESVSNPQRPPVIWIGAQECT WP_000145429.1 hydrogenase 2 small subunit [Salmonella enterica] Length: 372\nNumber of predicted TMHs: 1\nExp number of AAs in TMHs: 24.51309\nExp number, first 60 AAs: 0.2682\nTotal prob of N-in: 0.14402\noutside 1 331\nTMhelix 332 354\ninside 355 372

24329 GCF_000272935.1_ASM27293v1 Salmonella enterica subsp. enterica serovar Newport str. CVM 37978 Proteobacteria; Gammaproteobacteria; Enterobacterales; Enterobacteriaceae; Salmonella; Salmonella enterica; Salmonella enterica subsp. enterica; Salmonella enterica subsp. enterica serovar Newport MNNEETFYQAMRRKGVTRRSFLKFCSLAATSLGLGAGMTPKIAWALENKPRIPVVWIHGL WP_001058312.1 hydrogenase [Salmonella enterica] Length: 372\nNumber of predicted TMHs: 2\nExp number of AAs in TMHs: 36.51738\nExp number, first 60 AAs: 19.72849\nTotal prob of N-in: 0.96170\nPOSSIBLE N-term signal sequence\ninside 1 19\nTMhelix 20 39\noutside 40 325\nTMhelix 326 348\ninside 349 372

24330 GCF_000272935.1_ASM27293v1 Salmonella enterica subsp. enterica serovar Newport str. CVM 37978 Proteobacteria; Gammaproteobacteria; Enterobacterales; Enterobacteriaceae; Salmonella; Salmonella enterica; Salmonella enterica subsp. enterica; Salmonella enterica subsp. enterica serovar Newport MTGDNTLITSHGINRRDFMKLCAALAATMGLSSKAAAEMAESVSNPQRPPVIWIGAQECT WP_000145429.1 hydrogenase 2 small subunit [Salmonella enterica] Length: 372\nNumber of predicted TMHs: 1\nExp number of AAs in TMHs: 24.51309\nExp number, first 60 AAs: 0.2682\nTotal prob of N-in: 0.14402\noutside 1 331\nTMhelix 332 354\ninside 355 372

24331 GCF_000272875.1_ASM27287v1 Salmonella enterica subsp. enterica serovar Newport str. CVM 4176 Proteobacteria; Gammaproteobacteria; Enterobacterales; Enterobacteriaceae; Salmonella; Salmonella enterica; Salmonella enterica subsp. enterica; Salmonella enterica subsp. enterica serovar Newport MTGDNTLITSHGINRRDFMKLCAALAATMGLSSKAAAEMAESVSNPQRPPVIWIGAQECT WP_000145429.1 hydrogenase 2 small subunit [Salmonella enterica] Length: 372\nNumber of predicted TMHs: 1\nExp number of AAs in TMHs: 24.51309\nExp number, first 60 AAs: 0.2682\nTotal prob of N-in: 0.14402\noutside 1 331\nTMhelix 332 354\ninside 355 372

24332 GCF_000272875.1_ASM27287v1 Salmonella enterica subsp. enterica serovar Newport str. CVM 4176 Proteobacteria; Gammaproteobacteria; Enterobacterales; Enterobacteriaceae; Salmonella; Salmonella enterica; Salmonella enterica subsp. enterica; Salmonella enterica subsp. enterica serovar Newport MNNEETFYQAMRRKGVTRRSFLKFCSLAATSLGLGAGMTPKIAWALENKPRIPVVWIHGL WP_001058312.1 hydrogenase [Salmonella enterica] Length: 372\nNumber of predicted TMHs: 2\nExp number of AAs in TMHs: 36.51738\nExp number, first 60 AAs: 19.72849\nTotal prob of N-in: 0.96170\nPOSSIBLE N-term signal sequence\ninside 1 19\nTMhelix 20 39\noutside 40 325\nTMhelix 326 348\ninside 349 372

24333 GCF_000272835.3_ASM27283v3 Salmonella enterica subsp. enterica serovar Newport str. CVM N1543 Proteobacteria; Gammaproteobacteria; Enterobacterales; Enterobacteriaceae; Salmonella; Salmonella enterica; Salmonella enterica subsp. enterica; Salmonella enterica subsp. enterica serovar Newport MTGDNTLITSHGINRRDFMKLCAALAATMGLSSKAAAEMAESVSNPQRPPVIWIGAQECT WP_000145429.1 hydrogenase 2 small subunit [Salmonella enterica] Length: 372\nNumber of predicted TMHs: 1\nExp number of AAs in TMHs: 24.51309\nExp number, first 60 AAs: 0.2682\nTotal prob of N-in: 0.14402\noutside 1 331\nTMhelix 332 354\ninside 355 372

24334 GCF_000272835.3_ASM27283v3 Salmonella enterica subsp. enterica serovar Newport str. CVM N1543 Proteobacteria; Gammaproteobacteria; Enterobacterales; Enterobacteriaceae; Salmonella; Salmonella enterica; Salmonella enterica subsp. enterica; Salmonella enterica subsp. enterica serovar Newport MNNEETFYQAMRRKGVTRRSFLKFCSLAATSLGLGAGMTPKIAWALENKPRIPVVWIHGL WP_001058312.1 hydrogenase [Salmonella enterica] Length: 372\nNumber of predicted TMHs: 2\nExp number of AAs in TMHs: 36.51738\nExp number, first 60 AAs: 19.72849\nTotal prob of N-in: 0.96170\nPOSSIBLE N-term signal sequence\ninside 1 19\nTMhelix 20 39\noutside 40 325\nTMhelix 326 348\ninside 349 372

24335 GCF_000272815.2_ASM27281v2 Salmonella enterica subsp. enterica serovar Newport str. CVM N18486 Proteobacteria; Gammaproteobacteria; Enterobacterales; Enterobacteriaceae; Salmonella; Salmonella enterica;

Salmonella enterica subsp. enterica; Salmonella enterica subsp. enterica serovar Newport
MNNEETFYQAMRRKGVTRRSFLKFCSLAATSLGLGAGMTPKIAWALENKPRIPVVWIHGL WP_001058312.1
hydrogenase [Salmonella enterica] Length: 372\nNumber of predicted TMHs: 2\nExp number of AAs in TMHs: 36.51738\nExp number, first 60 AAs: 19.72849\nTotal prob of N-in: 0.96170\nPOSSIBLE N-term signal sequence\ninside 1 19\nTMhelix 20 39\noutside 40 325\nTMhelix 326 348\ninside 349 372

24336 GCF_000272815.2_ASM27281v2 Salmonella enterica subsp. enterica serovar Newport str. CVM N18486
Proteobacteria; Gammaproteobacteria; Enterobacterales; Enterobacteriaceae; Salmonella; Salmonella enterica; Salmonella enterica subsp. enterica; Salmonella enterica subsp. enterica serovar Newport
MTGDNTLITSHGINRRDFMKLCAALAATMGLSSKAAAEMAESVSNPQRPPVIWIGAQECT WP_000145429.1
hydrogenase 2 small subunit [Salmonella enterica] Length: 372\nNumber of predicted TMHs: 1\nExp number of AAs in TMHs: 24.51309\nExp number, first 60 AAs: 0.2682\nTotal prob of N-in: 0.14402\noutside 1 331\nTMhelix 332 354\ninside 355 372

24337 GCF_001051695.1_CFSAN000862_1.0 Salmonella enterica subsp. enterica serovar Newport str. DC_10-446
Proteobacteria; Gammaproteobacteria; Enterobacterales; Enterobacteriaceae; Salmonella; Salmonella enterica; Salmonella enterica subsp. enterica; Salmonella enterica subsp. enterica serovar Newport
MNNEETFYQAMRRKGVTRRSFLKFCSLAATSLGLGAGMTPKIAWALENKPRIPVVWIHGL WP_001058312.1
hydrogenase [Salmonella enterica] Length: 372\nNumber of predicted TMHs: 2\nExp number of AAs in TMHs: 36.51738\nExp number, first 60 AAs: 19.72849\nTotal prob of N-in: 0.96170\nPOSSIBLE N-term signal sequence\ninside 1 19\nTMhelix 20 39\noutside 40 325\nTMhelix 326 348\ninside 349 372

24338 GCF_001051695.1_CFSAN000862_1.0 Salmonella enterica subsp. enterica serovar Newport str. DC_10-446
Proteobacteria; Gammaproteobacteria; Enterobacterales; Enterobacteriaceae; Salmonella; Salmonella enterica; Salmonella enterica subsp. enterica; Salmonella enterica subsp. enterica serovar Newport
MTGDNTLITSHGINRRDFMKLCAALAATMGLSSKAAAEMAESVSNPQRPPVIWIGAQECT WP_000145429.1
hydrogenase 2 small subunit [Salmonella enterica] Length: 372\nNumber of predicted TMHs: 1\nExp number of AAs in TMHs: 24.51309\nExp number, first 60 AAs: 0.2682\nTotal prob of N-in: 0.14402\noutside 1 331\nTMhelix 332 354\ninside 355 372

24339 GCF_001051685.1_CFSAN000863_1.0 Salmonella enterica subsp. enterica serovar Newport str. DC_10-449
Proteobacteria; Gammaproteobacteria; Enterobacterales; Enterobacteriaceae; Salmonella; Salmonella enterica; Salmonella enterica subsp. enterica; Salmonella enterica subsp. enterica serovar Newport
MTGDNTLITSHGINRRDFMKLCAALAATMGLSSKAAAEMAESVSNPQRPPVIWIGAQECT WP_000145429.1
hydrogenase 2 small subunit [Salmonella enterica] Length: 372\nNumber of predicted TMHs: 1\nExp number of AAs in TMHs: 24.51309\nExp number, first 60 AAs: 0.2682\nTotal prob of N-in: 0.14402\noutside 1 331\nTMhelix 332 354\ninside 355 372

24340 GCF_001051685.1_CFSAN000863_1.0 Salmonella enterica subsp. enterica serovar Newport str. DC_10-449
Proteobacteria; Gammaproteobacteria; Enterobacterales; Enterobacteriaceae; Salmonella; Salmonella enterica; Salmonella enterica subsp. enterica; Salmonella enterica subsp. enterica serovar Newport
MNNEETFYQAMRRKGVTRRSFLKFCSLAATSLGLGAGMTPKIAWALENKPRIPVVWIHGL WP_001058312.1
hydrogenase [Salmonella enterica] Length: 372\nNumber of predicted TMHs: 2\nExp number of AAs in TMHs: 36.51738\nExp number, first 60 AAs: 19.72849\nTotal prob of N-in: 0.96170\nPOSSIBLE N-term signal sequence\ninside 1 19\nTMhelix 20 39\noutside 40 325\nTMhelix 326 348\ninside 349 372

24341 GCF_001051715.1_CFSAN000864_1.0 Salmonella enterica subsp. enterica serovar Newport str. DC_10-450
Proteobacteria; Gammaproteobacteria; Enterobacterales; Enterobacteriaceae; Salmonella; Salmonella enterica; Salmonella enterica subsp. enterica; Salmonella enterica subsp. enterica serovar Newport
MNNEETFYQAMRRKGVTRRSFLKFCSLAATSLGLGAGMTPKIAWALENKPRIPVVWIHGL WP_001058312.1
hydrogenase [Salmonella enterica] Length: 372\nNumber of predicted TMHs: 2\nExp number of AAs in TMHs: 36.51738\nExp number, first 60 AAs: 19.72849\nTotal prob of N-in: 0.96170\nPOSSIBLE N-term signal sequence\ninside 1 19\nTMhelix 20 39\noutside 40 325\nTMhelix 326 348\ninside 349 372

24342 GCF_001051715.1_CFSAN000864_1.0 Salmonella enterica subsp. enterica serovar Newport str. DC_10-450
Proteobacteria; Gammaproteobacteria; Enterobacterales; Enterobacteriaceae; Salmonella; Salmonella enterica; Salmonella enterica subsp. enterica; Salmonella enterica subsp. enterica serovar Newport
MTGDNTLITSHGINRRDFMKLCAALAATMGLSSKAAAEMAESVSNPQRPPVIWIGAQECT WP_000145429.1
hydrogenase 2 small subunit [Salmonella enterica] Length: 372\nNumber of predicted TMHs: 1\nExp number of AAs in TMHs: 24.51309\nExp number, first 60 AAs: 0.2682\nTotal prob of N-in: 0.14402\noutside 1 331\nTMhelix 332 354\ninside 355 372

24343 GCF_000344885.1_SOAPdenovo1.05 Salmonella enterica subsp. enterica serovar Newport str. Henan_3 Proteobacteria; Gammaproteobacteria; Enterobacterales; Enterobacteriaceae; Salmonella; Salmonella enterica; Salmonella enterica subsp. enterica; Salmonella enterica subsp. enterica serovar Newport MTGDNTLITSHGINRRDFMKLCAALAATMGLSSKAAAEMAESVSNPQRPPVIWIGAQECT WP_000145429.1 hydrogenase 2 small subunit [Salmonella enterica] Length: 372\nNumber of predicted TMHs: 1\nExp number of AAs in TMHs: 24.51309\nExp number, first 60 AAs: 0.2682\nTotal prob of N-in: 0.14402\noutside 1 331\nTMhelix 332 354\ninside 355 372

24344 GCF_000344885.1_SOAPdenovo1.05 Salmonella enterica subsp. enterica serovar Newport str. Henan_3 Proteobacteria; Gammaproteobacteria; Enterobacterales; Enterobacteriaceae; Salmonella; Salmonella enterica; Salmonella enterica subsp. enterica; Salmonella enterica subsp. enterica serovar Newport MNNEETFYQAMRRKGVTRRSFLKFCSLAATSLGLGAGMTPKIAWALENKPRIPVVWIHGL WP_001058312.1 hydrogenase [Salmonella enterica] Length: 372\nNumber of predicted TMHs: 2\nExp number of AAs in TMHs: 36.51738\nExp number, first 60 AAs: 19.72849\nTotal prob of N-in: 0.96170\nPOSSIBLE N-term signal sequence\ninside 1 19\nTMhelix 20 39\noutside 40 325\nTMhelix 326 348\ninside 349 372

24345 GCF_000344865.1_SOAPdenovo1.05 Salmonella enterica subsp. enterica serovar Newport str. JS09102 Proteobacteria; Gammaproteobacteria; Enterobacterales; Enterobacteriaceae; Salmonella; Salmonella enterica; Salmonella enterica subsp. enterica; Salmonella enterica subsp. enterica serovar Newport MNNEETFYQAMRRKGVTRRSFLKFCSLAATSLGLGAGMTPKIAWALENKPRIPVVWIHGL WP_001058312.1 hydrogenase [Salmonella enterica] Length: 372\nNumber of predicted TMHs: 2\nExp number of AAs in TMHs: 36.51738\nExp number, first 60 AAs: 19.72849\nTotal prob of N-in: 0.96170\nPOSSIBLE N-term signal sequence\ninside 1 19\nTMhelix 20 39\noutside 40 325\nTMhelix 326 348\ninside 349 372

24346 GCF_000344865.1_SOAPdenovo1.05 Salmonella enterica subsp. enterica serovar Newport str. JS09102 Proteobacteria; Gammaproteobacteria; Enterobacterales; Enterobacteriaceae; Salmonella; Salmonella enterica; Salmonella enterica subsp. enterica; Salmonella enterica subsp. enterica serovar Newport MTGDNTLITSHGINRRDFMKLCAALAATMGLSSKAAAEMAESVSNPQRPPVIWIGAQECT WP_000145429.1 hydrogenase 2 small subunit [Salmonella enterica] Length: 372\nNumber of predicted TMHs: 1\nExp number of AAs in TMHs: 24.51309\nExp number, first 60 AAs: 0.2682\nTotal prob of N-in: 0.14402\noutside 1 331\nTMhelix 332 354\ninside 355 372

24347 GCF_000271885.2_ASM27188v2 Salmonella enterica subsp. enterica serovar Newport str. Levine 1 Proteobacteria; Gammaproteobacteria; Enterobacterales; Enterobacteriaceae; Salmonella; Salmonella enterica; Salmonella enterica subsp. enterica; Salmonella enterica subsp. enterica serovar Newport MNNEETFYQAMRRKGVTRRSFLKFCSLAATSLGLGAGMTPKIAWALENKPRIPVVWIHGL WP_001058312.1 hydrogenase [Salmonella enterica] Length: 372\nNumber of predicted TMHs: 2\nExp number of AAs in TMHs: 36.51738\nExp number, first 60 AAs: 19.72849\nTotal prob of N-in: 0.96170\nPOSSIBLE N-term signal sequence\ninside 1 19\nTMhelix 20 39\noutside 40 325\nTMhelix 326 348\ninside 349 372

24348 GCF_000271885.2_ASM27188v2 Salmonella enterica subsp. enterica serovar Newport str. Levine 1 Proteobacteria; Gammaproteobacteria; Enterobacterales; Enterobacteriaceae; Salmonella; Salmonella enterica; Salmonella enterica subsp. enterica; Salmonella enterica subsp. enterica serovar Newport MTGDNTLITSHGINRRDFMKLCAALAATMGLSSKAAAEMAESVSNPQRPPVIWIGAQECT WP_000145429.1 hydrogenase 2 small subunit [Salmonella enterica] Length: 372\nNumber of predicted TMHs: 1\nExp number of AAs in TMHs: 24.51309\nExp number, first 60 AAs: 0.2682\nTotal prob of N-in: 0.14402\noutside 1 331\nTMhelix 332 354\ninside 355 372

24349 GCF_000271905.2_ASM27190v2 Salmonella enterica subsp. enterica serovar Newport str. Levine 15 Proteobacteria; Gammaproteobacteria; Enterobacterales; Enterobacteriaceae; Salmonella; Salmonella enterica; Salmonella enterica subsp. enterica; Salmonella enterica subsp. enterica serovar Newport MNNEETFYQAMRRKGVTRRSFLKFCSLAATSLGLGAGMTPKIAWALENKPRIPVVWIHGL WP_001058312.1 hydrogenase [Salmonella enterica] Length: 372\nNumber of predicted TMHs: 2\nExp number of AAs in TMHs: 36.51738\nExp number, first 60 AAs: 19.72849\nTotal prob of N-in: 0.96170\nPOSSIBLE N-term signal sequence\ninside 1 19\nTMhelix 20 39\noutside 40 325\nTMhelix 326 348\ninside 349 372

24350 GCF_000271905.2_ASM27190v2 Salmonella enterica subsp. enterica serovar Newport str. Levine 15 Proteobacteria; Gammaproteobacteria; Enterobacterales; Enterobacteriaceae; Salmonella; Salmonella enterica; Salmonella enterica subsp. enterica; Salmonella enterica subsp. enterica serovar Newport MTGDNTLITSHGINRRDFMKLCAALAATMGLSSKAAAEMAESVSNPQRPPVIWIGAQECT WP_000145429.1 hydrogenase 2 small subunit [Salmonella enterica] Length: 372\nNumber of predicted TMHs: 1\nExp

number of AAs in TMHs: 24.51309\nExp number, first 60 AAs: 0.2682\nTotal prob of N-in: 0.14402\noutside 1 331\nTMhelix 332 354\ninside 355 372

24351 GCF_001051705.1_CFSAN000861_1.0 Salmonella enterica subsp. enterica serovar Newport str. MA_10EN1469 Proteobacteria; Gammaproteobacteria; Enterobacterales; Enterobacteriaceae; Salmonella; Salmonella enterica; Salmonella enterica subsp. enterica; Salmonella enterica subsp. enterica serovar Newport MTGDNTLITSHGINRRDFMKLCAALAATMGLSSKAAAEMAESVSNPQRPPVIWIGAQECT WP_000145429.1 hydrogenase 2 small subunit [Salmonella enterica] Length: 372\nNumber of predicted TMHs: 1\nExp number of AAs in TMHs: 24.51309\nExp number, first 60 AAs: 0.2682\nTotal prob of N-in: 0.14402\noutside 1 331\nTMhelix 332 354\ninside 355 372

24352 GCF_001051705.1_CFSAN000861_1.0 Salmonella enterica subsp. enterica serovar Newport str. MA_10EN1469 Proteobacteria; Gammaproteobacteria; Enterobacterales; Enterobacteriaceae; Salmonella; Salmonella enterica; Salmonella enterica subsp. enterica; Salmonella enterica subsp. enterica serovar Newport MNNEETFYQAMRRKGVTRRSFLKFCSLAATSLGLGAGMTPKIAWALENKPRIPVVWIHGL WP_001058312.1 hydrogenase [Salmonella enterica] Length: 372\nNumber of predicted TMHs: 2\nExp number of AAs in TMHs: 36.51738\nExp number, first 60 AAs: 19.72849\nTotal prob of N-in: 0.96170\nPOSSIBLE N-term signal sequence\ninside 1 19\nTMhelix 20 39\noutside 40 325\nTMhelix 326 348\ninside 349 372

24353 GCF_001051775.1_CFSAN000928_1.0 Salmonella enterica subsp. enterica serovar Newport str. Pond080-2TTA Proteobacteria; Gammaproteobacteria; Enterobacterales; Enterobacteriaceae; Salmonella; Salmonella enterica; Salmonella enterica subsp. enterica; Salmonella enterica subsp. enterica serovar Newport MTGDNTLITSHGINRRDFMKLCAALAATMGLSSKAAAEMAESVSNPQRPPVIWIGAQECT WP_000145429.1 hydrogenase 2 small subunit [Salmonella enterica] Length: 372\nNumber of predicted TMHs: 1\nExp number of AAs in TMHs: 24.51309\nExp number, first 60 AAs: 0.2682\nTotal prob of N-in: 0.14402\noutside 1 331\nTMhelix 332 354\ninside 355 372

24354 GCF_001051775.1_CFSAN000928_1.0 Salmonella enterica subsp. enterica serovar Newport str. Pond080-2TTA Proteobacteria; Gammaproteobacteria; Enterobacterales; Enterobacteriaceae; Salmonella; Salmonella enterica; Salmonella enterica subsp. enterica; Salmonella enterica subsp. enterica serovar Newport MNNEETFYQAMRRKGVTRRSFLKFCSLAATSLGLGAGMTPKIAWALENKPRIPVVWIHGL WP_001058312.1 hydrogenase [Salmonella enterica] Length: 372\nNumber of predicted TMHs: 2\nExp number of AAs in TMHs: 36.51738\nExp number, first 60 AAs: 19.72849\nTotal prob of N-in: 0.96170\nPOSSIBLE N-term signal sequence\ninside 1 19\nTMhelix 20 39\noutside 40 325\nTMhelix 326 348\ninside 349 372

24355 GCF_001051845.1_CFSAN001243_1.0 Salmonella enterica subsp. enterica serovar Newport str. PRS_2010_0624 Proteobacteria; Gammaproteobacteria; Enterobacterales; Enterobacteriaceae; Salmonella; Salmonella enterica; Salmonella enterica subsp. enterica; Salmonella enterica subsp. enterica serovar Newport MTGDNTLITSHGINRRDFMKLCAALAATMGLSSKAAAEMAESVSNPQRPPVIWIGAQECT WP_000145429.1 hydrogenase 2 small subunit [Salmonella enterica] Length: 372\nNumber of predicted TMHs: 1\nExp number of AAs in TMHs: 24.51309\nExp number, first 60 AAs: 0.2682\nTotal prob of N-in: 0.14402\noutside 1 331\nTMhelix 332 354\ninside 355 372

24356 GCF_001051845.1_CFSAN001243_1.0 Salmonella enterica subsp. enterica serovar Newport str. PRS_2010_0624 Proteobacteria; Gammaproteobacteria; Enterobacterales; Enterobacteriaceae; Salmonella; Salmonella enterica; Salmonella enterica subsp. enterica; Salmonella enterica subsp. enterica serovar Newport MNNEETFYQAMRRKGVTRRSFLKFCSLAATSLGLGAGMTPKIAWALENKPRIPVVWIHGL WP_001058312.1 hydrogenase [Salmonella enterica] Length: 372\nNumber of predicted TMHs: 2\nExp number of AAs in TMHs: 36.51738\nExp number, first 60 AAs: 19.72849\nTotal prob of N-in: 0.96170\nPOSSIBLE N-term signal sequence\ninside 1 19\nTMhelix 20 39\noutside 40 325\nTMhelix 326 348\ninside 349 372

24357 GCF_000483955.1_ASM48395v1 Salmonella enterica subsp. enterica serovar Newport str. RI_10P068 Proteobacteria; Gammaproteobacteria; Enterobacterales; Enterobacteriaceae; Salmonella; Salmonella enterica; Salmonella enterica subsp. enterica; Salmonella enterica subsp. enterica serovar Newport MTGDNTLITSHGINRRDFMKLCAALAATMGLSSKAAAEMAESVSNPQRPPVIWIGAQECT WP_000145429.1 hydrogenase 2 small subunit [Salmonella enterica] Length: 372\nNumber of predicted TMHs: 1\nExp number of AAs in TMHs: 24.51309\nExp number, first 60 AAs: 0.2682\nTotal prob of N-in: 0.14402\noutside 1 331\nTMhelix 332 354\ninside 355 372

24358 GCF_000483955.1_ASM48395v1 Salmonella enterica subsp. enterica serovar Newport str. RI_10P068 Proteobacteria; Gammaproteobacteria; Enterobacterales; Enterobacteriaceae; Salmonella; Salmonella enterica;

Salmonella enterica subsp. enterica; Salmonella enterica subsp. enterica serovar Newport
MNNEETFYQAMRRKGVTRRSFLKFCSLAATSLGLGAGMTPKIAWALENKPRIPVVWIHGL WP_001058312.1
hydrogenase [Salmonella enterica] Length: 372\nNumber of predicted TMHs: 2\nExp number of AAs in TMHs: 36.51738\nExp number, first 60 AAs: 19.72849\nTotal prob of N-in: 0.96170\nPOSSIBLE N-term signal sequence\ninside 1 19\nTMhelix 20 39\noutside 40 325\nTMhelix 326 348\ninside 349 372

24359 GCF_000483775.1_ASM48377v1 Salmonella enterica subsp. enterica serovar Newport str. RI_10P069
Proteobacteria; Gammaproteobacteria; Enterobacterales; Enterobacteriaceae; Salmonella; Salmonella enterica; Salmonella enterica subsp. enterica; Salmonella enterica subsp. enterica serovar Newport
MNNEETFYQAMRRKGVTRRSFLKFCSLAATSLGLGAGMTPKIAWALENKPRIPVVWIHGL WP_001058312.1
hydrogenase [Salmonella enterica] Length: 372\nNumber of predicted TMHs: 2\nExp number of AAs in TMHs: 36.51738\nExp number, first 60 AAs: 19.72849\nTotal prob of N-in: 0.96170\nPOSSIBLE N-term signal sequence\ninside 1 19\nTMhelix 20 39\noutside 40 325\nTMhelix 326 348\ninside 349 372

24360 GCF_000483775.1_ASM48377v1 Salmonella enterica subsp. enterica serovar Newport str. RI_10P069
Proteobacteria; Gammaproteobacteria; Enterobacterales; Enterobacteriaceae; Salmonella; Salmonella enterica; Salmonella enterica subsp. enterica; Salmonella enterica subsp. enterica serovar Newport
MTGDNTLITSHGINRRDFMKLCAALAATMGLSSKAAAEMAESVSNPQRPPVIWIGAQECT WP_000145429.1
hydrogenase 2 small subunit [Salmonella enterica] Length: 372\nNumber of predicted TMHs: 1\nExp number of AAs in TMHs: 24.51309\nExp number, first 60 AAs: 0.2682\nTotal prob of N-in: 0.14402\noutside 1 331\nTMhelix 332 354\ninside 355 372

24361 GCF_001051815.1_CFSAN000929_1.0 Salmonella enterica subsp. enterica serovar Newport str. S103RVX
Proteobacteria; Gammaproteobacteria; Enterobacterales; Enterobacteriaceae; Salmonella; Salmonella enterica; Salmonella enterica subsp. enterica; Salmonella enterica subsp. enterica serovar Newport
MTGDNTLITSHGINRRDFMKLCAALAATMGLSSKAAAEMAESVSNPQRPPVIWIGAQECT WP_000145429.1
hydrogenase 2 small subunit [Salmonella enterica] Length: 372\nNumber of predicted TMHs: 1\nExp number of AAs in TMHs: 24.51309\nExp number, first 60 AAs: 0.2682\nTotal prob of N-in: 0.14402\noutside 1 331\nTMhelix 332 354\ninside 355 372

24362 GCF_001051815.1_CFSAN000929_1.0 Salmonella enterica subsp. enterica serovar Newport str. S103RVX
Proteobacteria; Gammaproteobacteria; Enterobacterales; Enterobacteriaceae; Salmonella; Salmonella enterica; Salmonella enterica subsp. enterica; Salmonella enterica subsp. enterica serovar Newport
MNNEETFYQAMRRKGVTRRSFLKFCSLAATSLGLGAGMTPKIAWALENKPRIPVVWIHGL WP_001058312.1
hydrogenase [Salmonella enterica] Length: 372\nNumber of predicted TMHs: 2\nExp number of AAs in TMHs: 36.51738\nExp number, first 60 AAs: 19.72849\nTotal prob of N-in: 0.96170\nPOSSIBLE N-term signal sequence\ninside 1 19\nTMhelix 20 39\noutside 40 325\nTMhelix 326 348\ninside 349 372

24363 GCF_000344825.1_SOAPdenovo1.05 Salmonella enterica subsp. enterica serovar Newport str. SH111077
Proteobacteria; Gammaproteobacteria; Enterobacterales; Enterobacteriaceae; Salmonella; Salmonella enterica; Salmonella enterica subsp. enterica; Salmonella enterica subsp. enterica serovar Newport
MNNEETFYQAMRRKGVTRRSFLKFCSLAATSLGLGAGMTPKIAWALENKPRIPVVWIHGL WP_001058312.1
hydrogenase [Salmonella enterica] Length: 372\nNumber of predicted TMHs: 2\nExp number of AAs in TMHs: 36.51738\nExp number, first 60 AAs: 19.72849\nTotal prob of N-in: 0.96170\nPOSSIBLE N-term signal sequence\ninside 1 19\nTMhelix 20 39\noutside 40 325\nTMhelix 326 348\ninside 349 372

24364 GCF_000344825.1_SOAPdenovo1.05 Salmonella enterica subsp. enterica serovar Newport str. SH111077
Proteobacteria; Gammaproteobacteria; Enterobacterales; Enterobacteriaceae; Salmonella; Salmonella enterica; Salmonella enterica subsp. enterica; Salmonella enterica subsp. enterica serovar Newport
MTGDNTLITSHGINRRDFMKLCAALAATMGLSSKAAAEMAESVSNPQRPPVIWIGAQECT WP_000145429.1
hydrogenase 2 small subunit [Salmonella enterica] Length: 372\nNumber of predicted TMHs: 1\nExp number of AAs in TMHs: 24.51309\nExp number, first 60 AAs: 0.2682\nTotal prob of N-in: 0.14402\noutside 1 331\nTMhelix 332 354\ninside 355 372

24365 GCF_000344845.1_SOAPdenovo1.05 Salmonella enterica subsp. enterica serovar Newport str. Shandong_3
Proteobacteria; Gammaproteobacteria; Enterobacterales; Enterobacteriaceae; Salmonella; Salmonella enterica; Salmonella enterica subsp. enterica; Salmonella enterica subsp. enterica serovar Newport
MTGDNTLITSHGINRRDFMKLCAALAATMGLSSKAAAEMAESVSNPQRPPVIWIGAQECT WP_000145429.1
hydrogenase 2 small subunit [Salmonella enterica] Length: 372\nNumber of predicted TMHs: 1\nExp number of AAs in TMHs: 24.51309\nExp number, first 60 AAs: 0.2682\nTotal prob of N-in: 0.14402\noutside 1 331\nTMhelix 332 354\ninside 355 372

24366 GCF_000344845.1_SOAPdenovo1.05 Salmonella enterica subsp. enterica serovar Newport str. Shandong_3 Proteobacteria; Gammaproteobacteria; Enterobacterales; Enterobacteriaceae; Salmonella; Salmonella enterica; Salmonella enterica subsp. enterica; Salmonella enterica subsp. enterica serovar Newport MNNEETFYQAMRRKGVTRRSFLKFCSLAATSLGLGAGMTPKIAWALENKPRIPVVWIHGL WP_001058312.1 hydrogenase [Salmonella enterica] Length: 372\nNumber of predicted TMHs: 2\nExp number of AAs in TMHs: 36.51738\nExp number, first 60 AAs: 19.72849\nTotal prob of N-in: 0.96170\nPOSSIBLE N-term signal sequence\ninside 1 19\nTMhelix 20 39\noutside 40 325\nTMhelix 326 348\ninside 349 372

24367 GCF_002109145.1_S_Newport_SHSN001_v01 Salmonella enterica subsp. enterica serovar Newport str. SHSN001 Proteobacteria; Gammaproteobacteria; Enterobacterales; Enterobacteriaceae; Salmonella; Salmonella enterica; Salmonella enterica subsp. enterica; Salmonella enterica subsp. enterica serovar Newport MTGDNTLITSHGINRRDFMKLCAALAATMGLSSKAAAEMAESVSNPQRPPVIWIGAQECT WP_000145429.1 hydrogenase 2 small subunit [Salmonella enterica] Length: 372\nNumber of predicted TMHs: 1\nExp number of AAs in TMHs: 24.51309\nExp number, first 60 AAs: 0.2682\nTotal prob of N-in: 0.14402\noutside 1 331\nTMhelix 332 354\ninside 355 372

24368 GCF_002109145.1_S_Newport_SHSN001_v01 Salmonella enterica subsp. enterica serovar Newport str. SHSN001 Proteobacteria; Gammaproteobacteria; Enterobacterales; Enterobacteriaceae; Salmonella; Salmonella enterica; Salmonella enterica subsp. enterica; Salmonella enterica subsp. enterica serovar Newport MNNEETFYQAMRRKGVTRRSFLKFCSLAATSLGLGAGMTPKIAWALENKPRIPVVWIHGL WP_001058312.1 hydrogenase [Salmonella enterica] Length: 372\nNumber of predicted TMHs: 2\nExp number of AAs in TMHs: 36.51738\nExp number, first 60 AAs: 19.72849\nTotal prob of N-in: 0.96170\nPOSSIBLE N-term signal sequence\ninside 1 19\nTMhelix 20 39\noutside 40 325\nTMhelix 326 348\ninside 349 372

24369 GCF_002109185.1_S_Newport_SHSN002_v01 Salmonella enterica subsp. enterica serovar Newport str. SHSN002 Proteobacteria; Gammaproteobacteria; Enterobacterales; Enterobacteriaceae; Salmonella; Salmonella enterica; Salmonella enterica subsp. enterica; Salmonella enterica subsp. enterica serovar Newport MNNEETFYQAMRRKGVTRRSFLKFCSLAATSLGLGAGMTPKIAWALENKPRIPVVWIHGL WP_001058312.1 hydrogenase [Salmonella enterica] Length: 372\nNumber of predicted TMHs: 2\nExp number of AAs in TMHs: 36.51738\nExp number, first 60 AAs: 19.72849\nTotal prob of N-in: 0.96170\nPOSSIBLE N-term signal sequence\ninside 1 19\nTMhelix 20 39\noutside 40 325\nTMhelix 326 348\ninside 349 372

24370 GCF_002109185.1_S_Newport_SHSN002_v01 Salmonella enterica subsp. enterica serovar Newport str. SHSN002 Proteobacteria; Gammaproteobacteria; Enterobacterales; Enterobacteriaceae; Salmonella; Salmonella enterica; Salmonella enterica subsp. enterica; Salmonella enterica subsp. enterica serovar Newport MTGDNTLITSHGINRRDFMKLCAALAATMGLSSKAAAEMAESVSNPQRPPVIWIGAQECT WP_000145429.1 hydrogenase 2 small subunit [Salmonella enterica] Length: 372\nNumber of predicted TMHs: 1\nExp number of AAs in TMHs: 24.51309\nExp number, first 60 AAs: 0.2682\nTotal prob of N-in: 0.14402\noutside 1 331\nTMhelix 332 354\ninside 355 372

24371 GCF_002109075.1_S_Newport_SHSN003_v01 Salmonella enterica subsp. enterica serovar Newport str. SHSN003 Proteobacteria; Gammaproteobacteria; Enterobacterales; Enterobacteriaceae; Salmonella; Salmonella enterica; Salmonella enterica subsp. enterica; Salmonella enterica subsp. enterica serovar Newport MNNEETFYQAMRRKGVTRRSFLKFCSLAATSLGLGAGMTPKIAWALENKPRIPVVWIHGL WP_001058312.1 hydrogenase [Salmonella enterica] Length: 372\nNumber of predicted TMHs: 2\nExp number of AAs in TMHs: 36.51738\nExp number, first 60 AAs: 19.72849\nTotal prob of N-in: 0.96170\nPOSSIBLE N-term signal sequence\ninside 1 19\nTMhelix 20 39\noutside 40 325\nTMhelix 326 348\ninside 349 372

24372 GCF_002109075.1_S_Newport_SHSN003_v01 Salmonella enterica subsp. enterica serovar Newport str. SHSN003 Proteobacteria; Gammaproteobacteria; Enterobacterales; Enterobacteriaceae; Salmonella; Salmonella enterica; Salmonella enterica subsp. enterica; Salmonella enterica subsp. enterica serovar Newport MTGDNTLITSHGINRRDFMKLCAALAATMGLSSKAAAEMAESVSNPQRPPVIWIGAQECT WP_000145429.1 hydrogenase 2 small subunit [Salmonella enterica] Length: 372\nNumber of predicted TMHs: 1\nExp number of AAs in TMHs: 24.51309\nExp number, first 60 AAs: 0.2682\nTotal prob of N-in: 0.14402\noutside 1 331\nTMhelix 332 354\ninside 355 372

24373 GCF_002109045.1_S_Newport_SHSN004_v01 Salmonella enterica subsp. enterica serovar Newport str. SHSN004 Proteobacteria; Gammaproteobacteria; Enterobacterales; Enterobacteriaceae; Salmonella; Salmonella enterica; Salmonella enterica subsp. enterica; Salmonella enterica subsp. enterica serovar Newport MTGDNTLITSHGINRRDFMKLCAALAATMGLSSKAAAEMAESVSNPQRPPVIWIGAQECT WP_000145429.1 hydrogenase 2 small subunit [Salmonella enterica] Length: 372\nNumber of predicted TMHs: 1\nExp

number of AAs in TMHs: 24.51309\nExp number, first 60 AAs: 0.2682\nTotal prob of N-in: 0.14402\noutside 1 331\nTMhelix 332 354\ninside 355 372

24374 GCF_002109045.1_S_Newport_SHSN004_v01 Salmonella enterica subsp. enterica serovar Newport str. SHSN004 Proteobacteria; Gammaproteobacteria; Enterobacterales; Enterobacteriaceae; Salmonella; Salmonella enterica; Salmonella enterica subsp. enterica; Salmonella enterica subsp. enterica serovar Newport MNNEETFYQAMRRKGVTRRSFLKFCSLAATSLGLGAGMTPKIAWALENKPRIPVVWIHGL WP_001058312.1 hydrogenase [Salmonella enterica] Length: 372\nNumber of predicted TMHs: 2\nExp number of AAs in TMHs: 36.51738\nExp number, first 60 AAs: 19.72849\nTotal prob of N-in: 0.96170\nPOSSIBLE N-term signal sequence\ninside 1 19\nTMhelix 20 39\noutside 40 325\nTMhelix 326 348\ninside 349 372

24375 GCF_002109205.1_S_Newport_SHSN005_v01 Salmonella enterica subsp. enterica serovar Newport str. SHSN005 Proteobacteria; Gammaproteobacteria; Enterobacterales; Enterobacteriaceae; Salmonella; Salmonella enterica; Salmonella enterica subsp. enterica; Salmonella enterica subsp. enterica serovar Newport MNNEETFYQAMRRKGVTRRSFLKFCSLAATSLGLGAGMTPKIAWALENKPRIPVVWIHGL WP_001058312.1 hydrogenase [Salmonella enterica] Length: 372\nNumber of predicted TMHs: 2\nExp number of AAs in TMHs: 36.51738\nExp number, first 60 AAs: 19.72849\nTotal prob of N-in: 0.96170\nPOSSIBLE N-term signal sequence\ninside 1 19\nTMhelix 20 39\noutside 40 325\nTMhelix 326 348\ninside 349 372

24376 GCF_002109205.1_S_Newport_SHSN005_v01 Salmonella enterica subsp. enterica serovar Newport str. SHSN005 Proteobacteria; Gammaproteobacteria; Enterobacterales; Enterobacteriaceae; Salmonella; Salmonella enterica; Salmonella enterica subsp. enterica; Salmonella enterica subsp. enterica serovar Newport MTGDNTLITSHGINRRDFMKLCAALAATMGLSSKAAAEMAESVSNPQRPPVIWIGAQECT WP_000145429.1 hydrogenase 2 small subunit [Salmonella enterica] Length: 372\nNumber of predicted TMHs: 1\nExp number of AAs in TMHs: 24.51309\nExp number, first 60 AAs: 0.2682\nTotal prob of N-in: 0.14402\noutside 1 331\nTMhelix 332 354\ninside 355 372

24377 GCF_002109215.1_S_Newport_SHSN006_v01 Salmonella enterica subsp. enterica serovar Newport str. SHSN006 Proteobacteria; Gammaproteobacteria; Enterobacterales; Enterobacteriaceae; Salmonella; Salmonella enterica; Salmonella enterica subsp. enterica; Salmonella enterica subsp. enterica serovar Newport MNNEETFYQAMRRKGVTRRSFLKFCSLAATSLGLGAGMTPKIAWALENKPRIPVVWIHGL WP_001058312.1 hydrogenase [Salmonella enterica] Length: 372\nNumber of predicted TMHs: 2\nExp number of AAs in TMHs: 36.51738\nExp number, first 60 AAs: 19.72849\nTotal prob of N-in: 0.96170\nPOSSIBLE N-term signal sequence\ninside 1 19\nTMhelix 20 39\noutside 40 325\nTMhelix 326 348\ninside 349 372

24378 GCF_002109215.1_S_Newport_SHSN006_v01 Salmonella enterica subsp. enterica serovar Newport str. SHSN006 Proteobacteria; Gammaproteobacteria; Enterobacterales; Enterobacteriaceae; Salmonella; Salmonella enterica; Salmonella enterica subsp. enterica; Salmonella enterica subsp. enterica serovar Newport MTGDNTLITSHGINRRDFMKLCAALAATMGLSSKAAAEMAESVSNPQRPPVIWIGAQECT WP_000145429.1 hydrogenase 2 small subunit [Salmonella enterica] Length: 372\nNumber of predicted TMHs: 1\nExp number of AAs in TMHs: 24.51309\nExp number, first 60 AAs: 0.2682\nTotal prob of N-in: 0.14402\noutside 1 331\nTMhelix 332 354\ninside 355 372

24379 GCF_002109245.1_S_Newport_SHSN007_v01 Salmonella enterica subsp. enterica serovar Newport str. SHSN007 Proteobacteria; Gammaproteobacteria; Enterobacterales; Enterobacteriaceae; Salmonella; Salmonella enterica; Salmonella enterica subsp. enterica; Salmonella enterica subsp. enterica serovar Newport MNNEETFYQAMRRKGVTRRSFLKFCSLAATSLGLGAGMTPKIAWALENKPRIPVVWIHGL WP_001058312.1 hydrogenase [Salmonella enterica] Length: 372\nNumber of predicted TMHs: 2\nExp number of AAs in TMHs: 36.51738\nExp number, first 60 AAs: 19.72849\nTotal prob of N-in: 0.96170\nPOSSIBLE N-term signal sequence\ninside 1 19\nTMhelix 20 39\noutside 40 325\nTMhelix 326 348\ninside 349 372

24380 GCF_002109245.1_S_Newport_SHSN007_v01 Salmonella enterica subsp. enterica serovar Newport str. SHSN007 Proteobacteria; Gammaproteobacteria; Enterobacterales; Enterobacteriaceae; Salmonella; Salmonella enterica; Salmonella enterica subsp. enterica; Salmonella enterica subsp. enterica serovar Newport MTGDNTLITSHGINRRDFMKLCAALAATMGLSSKAAAEMAESVSNPQRPPVIWIGAQECT WP_000145429.1 hydrogenase 2 small subunit [Salmonella enterica] Length: 372\nNumber of predicted TMHs: 1\nExp number of AAs in TMHs: 24.51309\nExp number, first 60 AAs: 0.2682\nTotal prob of N-in: 0.14402\noutside 1 331\nTMhelix 332 354\ninside 355 372

24381 GCF_002109105.1_S_Newport_SHSN008_v01 Salmonella enterica subsp. enterica serovar Newport str. SHSN008 Proteobacteria; Gammaproteobacteria; Enterobacterales; Enterobacteriaceae; Salmonella; Salmonella enterica;

Salmonella enterica subsp. enterica; Salmonella enterica subsp. enterica serovar Newport
MNNEETFYQAMRRKGVTRRSFLKFCSLAATSLGLGAGMTPKIAWALENKPRIPVVWIHGL WP_001058311.1
hydrogenase [Salmonella enterica] Length: 372\nNumber of predicted TMHs: 2\nExp number of AAs in TMHs: 37.4699899999999\nExp number, first 60 AAs: 19.72699\nTotal prob of N-in: 0.96171\nPOSSIBLE N-term signal sequence\ninside 1 19\nTMhelix 20 39\noutside 40 325\nTMhelix 326 348\ninside 349 372

24382 GCF_002109105.1_S_Newport_SHSN008_v01 Salmonella enterica subsp. enterica serovar Newport str. SHSN008 Proteobacteria; Gammaproteobacteria; Enterobacterales; Enterobacteriaceae; Salmonella; Salmonella enterica; Salmonella enterica subsp. enterica; Salmonella enterica subsp. enterica serovar Newport
MTGDNTLITSHGINRRDFMKLCAALAATMGLSSKAAAEMAESVSNPQRPPVIWIGAQECT WP_000145429.1
hydrogenase 2 small subunit [Salmonella enterica] Length: 372\nNumber of predicted TMHs: 1\nExp number of AAs in TMHs: 24.51309\nExp number, first 60 AAs: 0.2682\nTotal prob of N-in: 0.14402\noutside 1 331\nTMhelix 332 354\ninside 355 372

24383 GCF_002109255.1_S_Newport_SHSN009_v01 Salmonella enterica subsp. enterica serovar Newport str. SHSN009 Proteobacteria; Gammaproteobacteria; Enterobacterales; Enterobacteriaceae; Salmonella; Salmonella enterica; Salmonella enterica subsp. enterica; Salmonella enterica subsp. enterica serovar Newport
MNNEETFYQAMRRKGVTRRSFLKFCSLAATSLGLGAGMTPKIAWALENKPRIPVVWIHGL WP_001058312.1
hydrogenase [Salmonella enterica] Length: 372\nNumber of predicted TMHs: 2\nExp number of AAs in TMHs: 36.51738\nExp number, first 60 AAs: 19.72849\nTotal prob of N-in: 0.96170\nPOSSIBLE N-term signal sequence\ninside 1 19\nTMhelix 20 39\noutside 40 325\nTMhelix 326 348\ninside 349 372

24384 GCF_002109255.1_S_Newport_SHSN009_v01 Salmonella enterica subsp. enterica serovar Newport str. SHSN009 Proteobacteria; Gammaproteobacteria; Enterobacterales; Enterobacteriaceae; Salmonella; Salmonella enterica; Salmonella enterica subsp. enterica; Salmonella enterica subsp. enterica serovar Newport
MTGDNTLITSHGINRRDFMKLCAALAATMGLSSKAAAEMAESVSNPQRPPVIWIGAQECT WP_000145429.1
hydrogenase 2 small subunit [Salmonella enterica] Length: 372\nNumber of predicted TMHs: 1\nExp number of AAs in TMHs: 24.51309\nExp number, first 60 AAs: 0.2682\nTotal prob of N-in: 0.14402\noutside 1 331\nTMhelix 332 354\ninside 355 372

24385 GCF_002109055.1_S_Newport_SHSN010_v01 Salmonella enterica subsp. enterica serovar Newport str. SHSN010 Proteobacteria; Gammaproteobacteria; Enterobacterales; Enterobacteriaceae; Salmonella; Salmonella enterica; Salmonella enterica subsp. enterica; Salmonella enterica subsp. enterica serovar Newport
MTGDNTLITSHGINRRDFMKLCAALAATMGLSSKAAAEMAESVSNPQRPPVIWIGAQECT WP_000145429.1
hydrogenase 2 small subunit [Salmonella enterica] Length: 372\nNumber of predicted TMHs: 1\nExp number of AAs in TMHs: 24.51309\nExp number, first 60 AAs: 0.2682\nTotal prob of N-in: 0.14402\noutside 1 331\nTMhelix 332 354\ninside 355 372

24386 GCF_002109055.1_S_Newport_SHSN010_v01 Salmonella enterica subsp. enterica serovar Newport str. SHSN010 Proteobacteria; Gammaproteobacteria; Enterobacterales; Enterobacteriaceae; Salmonella; Salmonella enterica; Salmonella enterica subsp. enterica; Salmonella enterica subsp. enterica serovar Newport
MNNEETFYQAMRRKGVTRRSFLKFCSLAATSLGLGAGMTPKIAWALENKPRIPVVWIHGL WP_001058312.1
hydrogenase [Salmonella enterica] Length: 372\nNumber of predicted TMHs: 2\nExp number of AAs in TMHs: 36.51738\nExp number, first 60 AAs: 19.72849\nTotal prob of N-in: 0.96170\nPOSSIBLE N-term signal sequence\ninside 1 19\nTMhelix 20 39\noutside 40 325\nTMhelix 326 348\ninside 349 372

24387 GCF_002109285.1_S_Newport_SHSN011_v01 Salmonella enterica subsp. enterica serovar Newport str. SHSN011 Proteobacteria; Gammaproteobacteria; Enterobacterales; Enterobacteriaceae; Salmonella; Salmonella enterica; Salmonella enterica subsp. enterica; Salmonella enterica subsp. enterica serovar Newport
MTGDNTLITSHGINRRDFMKLCAALAATMGLSSKAAAEMAESVSNPQRPPVIWIGAQECT WP_000145429.1
hydrogenase 2 small subunit [Salmonella enterica] Length: 372\nNumber of predicted TMHs: 1\nExp number of AAs in TMHs: 24.51309\nExp number, first 60 AAs: 0.2682\nTotal prob of N-in: 0.14402\noutside 1 331\nTMhelix 332 354\ninside 355 372

24388 GCF_002109285.1_S_Newport_SHSN011_v01 Salmonella enterica subsp. enterica serovar Newport str. SHSN011 Proteobacteria; Gammaproteobacteria; Enterobacterales; Enterobacteriaceae; Salmonella; Salmonella enterica; Salmonella enterica subsp. enterica; Salmonella enterica subsp. enterica serovar Newport
MNNEETFYQAMRRKGVTRRSFLKFCSLAATSLGLGAGMTPKIAWALENKPRIPVVWIHGL WP_001058312.1
hydrogenase [Salmonella enterica] Length: 372\nNumber of predicted TMHs: 2\nExp number of AAs in TMHs: 36.51738\nExp number, first 60 AAs: 19.72849\nTotal prob of N-in: 0.96170\nPOSSIBLE N-term signal sequence\ninside 1 19\nTMhelix 20 39\noutside 40 325\nTMhelix 326 348\ninside 349 372

24389 GCF_002109125.1_S_Newport_SHSN012_v01 Salmonella enterica subsp. enterica serovar Newport str. SHSN012 Proteobacteria; Gammaproteobacteria; Enterobacterales; Enterobacteriaceae; Salmonella; Salmonella enterica; Salmonella enterica subsp. enterica; Salmonella enterica subsp. enterica serovar Newport
MTGDNTLITSHGINRRDFMKLCAALAATMGLSSKAAAEMAESVSNPQRPPVIWIGAQECT WP_000145429.1
hydrogenase 2 small subunit [Salmonella enterica] Length: 372\nNumber of predicted TMHs: 1\nExp number of AAs in TMHs: 24.51309\nExp number, first 60 AAs: 0.2682\nTotal prob of N-in: 0.14402\noutside 1 331\nTMhelix 332 354\ninside 355 372

24390 GCF_002109135.1_S_Newport_SHSN013_v01 Salmonella enterica subsp. enterica serovar Newport str. SHSN013 Proteobacteria; Gammaproteobacteria; Enterobacterales; Enterobacteriaceae; Salmonella; Salmonella enterica; Salmonella enterica subsp. enterica; Salmonella enterica subsp. enterica serovar Newport
MNNEETFYQAMRRKGVTRRSFLKFCSLAATSLGLGAGMTPKIAWALENKPRIPVVWIHGL WP_001058312.1
hydrogenase [Salmonella enterica] Length: 372\nNumber of predicted TMHs: 2\nExp number of AAs in TMHs: 36.51738\nExp number, first 60 AAs: 19.72849\nTotal prob of N-in: 0.96170\nPOSSIBLE N-term signal sequence\ninside 1 19\nTMhelix 20 39\noutside 40 325\nTMhelix 326 348\ninside 349 372

24391 GCF_002109135.1_S_Newport_SHSN013_v01 Salmonella enterica subsp. enterica serovar Newport str. SHSN013 Proteobacteria; Gammaproteobacteria; Enterobacterales; Enterobacteriaceae; Salmonella; Salmonella enterica; Salmonella enterica subsp. enterica; Salmonella enterica subsp. enterica serovar Newport
MTGDNTLITSHGINRRDFMKLCAALAATMGLSSKAAAEMAESVSNPQRPPVIWIGAQECT WP_000145429.1
hydrogenase 2 small subunit [Salmonella enterica] Length: 372\nNumber of predicted TMHs: 1\nExp number of AAs in TMHs: 24.51309\nExp number, first 60 AAs: 0.2682\nTotal prob of N-in: 0.14402\noutside 1 331\nTMhelix 332 354\ninside 355 372

24392 GCF_002108925.1_S_Newport_SHSN014_v01 Salmonella enterica subsp. enterica serovar Newport str. SHSN014 Proteobacteria; Gammaproteobacteria; Enterobacterales; Enterobacteriaceae; Salmonella; Salmonella enterica; Salmonella enterica subsp. enterica; Salmonella enterica subsp. enterica serovar Newport
MNNEETFYQAMRRKGVTRRSFLKFCSLAATSLGLGAGMTPKIAWALENKPRIPVVWIHGL WP_001058312.1
hydrogenase [Salmonella enterica] Length: 372\nNumber of predicted TMHs: 2\nExp number of AAs in TMHs: 36.51738\nExp number, first 60 AAs: 19.72849\nTotal prob of N-in: 0.96170\nPOSSIBLE N-term signal sequence\ninside 1 19\nTMhelix 20 39\noutside 40 325\nTMhelix 326 348\ninside 349 372

24393 GCF_002108925.1_S_Newport_SHSN014_v01 Salmonella enterica subsp. enterica serovar Newport str. SHSN014 Proteobacteria; Gammaproteobacteria; Enterobacterales; Enterobacteriaceae; Salmonella; Salmonella enterica; Salmonella enterica subsp. enterica; Salmonella enterica subsp. enterica serovar Newport
MTGDNTLITSHGINRRDFMKLCAALAATMGLSSKAAAEMAESVSNPQRPPVIWIGAQECT WP_000145429.1
hydrogenase 2 small subunit [Salmonella enterica] Length: 372\nNumber of predicted TMHs: 1\nExp number of AAs in TMHs: 24.51309\nExp number, first 60 AAs: 0.2682\nTotal prob of N-in: 0.14402\noutside 1 331\nTMhelix 332 354\ninside 355 372

24394 GCF_000751335.1_Salmonella_enterica_subsp._enterica_strain_100TC3 Salmonella enterica subsp. enterica serovar Newport str. SL254 Proteobacteria; Gammaproteobacteria; Enterobacterales; Enterobacteriaceae; Salmonella; Salmonella enterica; Salmonella enterica subsp. enterica; Salmonella enterica subsp. enterica serovar Newport
MTGDNTLITSHGINRRDFMKLCAALAATMGLSSKAAAEMAESVSNPQRPPVIWIGAQECT WP_000145429.1
hydrogenase 2 small subunit [Salmonella enterica] Length: 372\nNumber of predicted TMHs: 1\nExp number of AAs in TMHs: 24.51309\nExp number, first 60 AAs: 0.2682\nTotal prob of N-in: 0.14402\noutside 1 331\nTMhelix 332 354\ninside 355 372

24395 GCF_000751335.1_Salmonella_enterica_subsp._enterica_strain_100TC3 Salmonella enterica subsp. enterica serovar Newport str. SL254 Proteobacteria; Gammaproteobacteria; Enterobacterales; Enterobacteriaceae; Salmonella; Salmonella enterica; Salmonella enterica subsp. enterica; Salmonella enterica subsp. enterica serovar Newport
MNNEETFYQAMRRKGVTRRSFLKFCSLAATSLGLGAGMTPKIAWALENKPRIPVVWIHGL WP_001058312.1
hydrogenase [Salmonella enterica] Length: 372\nNumber of predicted TMHs: 2\nExp number of AAs in TMHs: 36.51738\nExp number, first 60 AAs: 19.72849\nTotal prob of N-in: 0.96170\nPOSSIBLE N-term signal sequence\ninside 1 19\nTMhelix 20 39\noutside 40 325\nTMhelix 326 348\ninside 349 372

24396 GCF_000016045.1_ASM1604v1 Salmonella enterica subsp. enterica serovar Newport str. SL254
Proteobacteria; Gammaproteobacteria; Enterobacterales; Enterobacteriaceae; Salmonella; Salmonella enterica; Salmonella enterica subsp. enterica; Salmonella enterica subsp. enterica serovar Newport
MNNEETFYQAMRRKGVTRRSFLKFCSLAATSLGLGAGMTPKIAWALENKPRIPVVWIHGL WP_001058312.1
hydrogenase [Salmonella enterica] Length: 372\nNumber of predicted TMHs: 2\nExp number of AAs in

TMHs: 36.51738\nExp number, first 60 AAs: 19.72849\nTotal prob of N-in: 0.96170\nPOSSIBLE N-term signal sequence\ninside 1 19\nTMhelix 20 39\noutside 40 325\nTMhelix 326 348\ninside 349 372

24397 GCF_000016045.1_ASM1604v1 Salmonella enterica subsp. enterica serovar Newport str. SL254 Proteobacteria; Gammaproteobacteria; Enterobacterales; Enterobacteriaceae; Salmonella; Salmonella enterica; Salmonella enterica subsp. enterica; Salmonella enterica subsp. enterica serovar Newport MTGDNTLITSHGINRRDFMKLCAALAATMGLSSKAAAEMAESVSNPQRPPVIWIGAQECT WP_000145429.1 hydrogenase 2 small subunit [Salmonella enterica] Length: 372\nNumber of predicted TMHs: 1\nExp number of AAs in TMHs: 24.51309\nExp number, first 60 AAs: 0.2682\nTotal prob of N-in: 0.14402\noutside 1 331\nTMhelix 332 354\ninside 355 372

24398 GCF_000171415.1_ASM17141v1 Salmonella enterica subsp. enterica serovar Newport str. SL317 Proteobacteria; Gammaproteobacteria; Enterobacterales; Enterobacteriaceae; Salmonella; Salmonella enterica; Salmonella enterica subsp. enterica; Salmonella enterica subsp. enterica serovar Newport MNNEETFYQAMRRKGVTRRSFLKFCSLAATSLGLGAGMTPKIAWALENKPRIPVVWIHGL WP_001058312.1 hydrogenase [Salmonella enterica] Length: 372\nNumber of predicted TMHs: 2\nExp number of AAs in TMHs: 36.51738\nExp number, first 60 AAs: 19.72849\nTotal prob of N-in: 0.96170\nPOSSIBLE N-term signal sequence\ninside 1 19\nTMhelix 20 39\noutside 40 325\nTMhelix 326 348\ninside 349 372

24399 GCF_000171415.1_ASM17141v1 Salmonella enterica subsp. enterica serovar Newport str. SL317 Proteobacteria; Gammaproteobacteria; Enterobacterales; Enterobacteriaceae; Salmonella; Salmonella enterica; Salmonella enterica subsp. enterica; Salmonella enterica subsp. enterica serovar Newport MTGDNTLITSHGINRRDFMKLCAALAATMGLSSKAAAEMAESVSNPQRPPVIWIGAQECT WP_000145429.1 hydrogenase 2 small subunit [Salmonella enterica] Length: 372\nNumber of predicted TMHs: 1\nExp number of AAs in TMHs: 24.51309\nExp number, first 60 AAs: 0.2682\nTotal prob of N-in: 0.14402\noutside 1 331\nTMhelix 332 354\ninside 355 372

24400 GCF_000940935.1_ASM94093v1 Salmonella enterica subsp. enterica serovar Newport str. USDA-ARS-USMARC-1927 Proteobacteria; Gammaproteobacteria; Enterobacterales; Enterobacteriaceae; Salmonella; Salmonella enterica; Salmonella enterica subsp. enterica; Salmonella enterica subsp. enterica serovar Newport MNNEETFYQAMRRKGVTRRSFLKFCSLAATSLGLGAGMTPKIAWALENKPRIPVVWIHGL WP_001058312.1 hydrogenase [Salmonella enterica] Length: 372\nNumber of predicted TMHs: 2\nExp number of AAs in TMHs: 36.51738\nExp number, first 60 AAs: 19.72849\nTotal prob of N-in: 0.96170\nPOSSIBLE N-term signal sequence\ninside 1 19\nTMhelix 20 39\noutside 40 325\nTMhelix 326 348\ninside 349 372

24401 GCF_000940935.1_ASM94093v1 Salmonella enterica subsp. enterica serovar Newport str. USDA-ARS-USMARC-1927 Proteobacteria; Gammaproteobacteria; Enterobacterales; Enterobacteriaceae; Salmonella; Salmonella enterica; Salmonella enterica subsp. enterica; Salmonella enterica subsp. enterica serovar Newport MTGDNTLITSHGINRRDFMKLCAALAATMGLSSKAAAEMAESVSNPQRPPVIWIGAQECT WP_000145429.1 hydrogenase 2 small subunit [Salmonella enterica] Length: 372\nNumber of predicted TMHs: 1\nExp number of AAs in TMHs: 24.51309\nExp number, first 60 AAs: 0.2682\nTotal prob of N-in: 0.14402\noutside 1 331\nTMhelix 332 354\ninside 355 372

24402 GCF_000442415.1_ASM44241v1 Salmonella enterica subsp. enterica serovar Newport str. USMARC-S3124.1 Proteobacteria; Gammaproteobacteria; Enterobacterales; Enterobacteriaceae; Salmonella; Salmonella enterica; Salmonella enterica subsp. enterica; Salmonella enterica subsp. enterica serovar Newport MTGDNTLITSHGINRRDFMKLCAALAATMGLSSKAAAEMAESVSNPQRPPVIWIGAQECT WP_000145429.1 hydrogenase 2 small subunit [Salmonella enterica] Length: 372\nNumber of predicted TMHs: 1\nExp number of AAs in TMHs: 24.51309\nExp number, first 60 AAs: 0.2682\nTotal prob of N-in: 0.14402\noutside 1 331\nTMhelix 332 354\ninside 355 372

24403 GCF_000442415.1_ASM44241v1 Salmonella enterica subsp. enterica serovar Newport str. USMARC-S3124.1 Proteobacteria; Gammaproteobacteria; Enterobacterales; Enterobacteriaceae; Salmonella; Salmonella enterica; Salmonella enterica subsp. enterica; Salmonella enterica subsp. enterica serovar Newport MNNEETFYQAMRRKGVTRRSFLKFCSLAATSLGLGAGMTPKIAWALENKPRIPVVWIHGL WP_001058312.1 hydrogenase [Salmonella enterica] Length: 372\nNumber of predicted TMHs: 2\nExp number of AAs in TMHs: 36.51738\nExp number, first 60 AAs: 19.72849\nTotal prob of N-in: 0.96170\nPOSSIBLE N-term signal sequence\ninside 1 19\nTMhelix 20 39\noutside 40 325\nTMhelix 326 348\ninside 349 372

24404 GCF_001051625.1_CFSAN000859_1.0 Salmonella enterica subsp. enterica serovar Newport str. VA_R100804798 Proteobacteria; Gammaproteobacteria; Enterobacterales; Enterobacteriaceae; Salmonella; Salmonella enterica;

Salmonella enterica subsp. enterica; Salmonella enterica subsp. enterica serovar Newport
MTGDNTLITSHGINRRDFMKLCAALAATMGLSSKAAAEMAESVSNPQRPPVIWIGAQECT WP_000145429.1
hydrogenase 2 small subunit [Salmonella enterica] Length: 372\nNumber of predicted TMHs: 1\nExp number of AAs in TMHs: 24.51309\nExp number, first 60 AAs: 0.2682\nTotal prob of N-in: 0.14402\noutside 1 331\nTMhelix 332 354\ninside 355 372

24405 GCF_001051625.1_CFSAN000859_1.0 Salmonella enterica subsp. enterica serovar Newport str. VA_R100804798
Proteobacteria; Gammaproteobacteria; Enterobacterales; Enterobacteriaceae; Salmonella; Salmonella enterica; Salmonella enterica subsp. enterica; Salmonella enterica subsp. enterica serovar Newport
MNNEETFYQAMRRKGVTRRSFLKFCSLAATSLGLGAGMTPKIAWALENKPRIPVVWIHGL WP_001058312.1
hydrogenase [Salmonella enterica] Length: 372\nNumber of predicted TMHs: 2\nExp number of AAs in TMHs: 36.51738\nExp number, first 60 AAs: 19.72849\nTotal prob of N-in: 0.96170\nPOSSIBLE N-term signal sequence\ninside 1 19\nTMhelix 20 39\noutside 40 325\nTMhelix 326 348\ninside 349 372

24406 GCF_001051635.1_CFSAN000860_1.0 Salmonella enterica subsp. enterica serovar Newport str. VA_R100808502
Proteobacteria; Gammaproteobacteria; Enterobacterales; Enterobacteriaceae; Salmonella; Salmonella enterica; Salmonella enterica subsp. enterica; Salmonella enterica subsp. enterica serovar Newport
MNNEETFYQAMRRKGVTRRSFLKFCSLAATSLGLGAGMTPKIAWALENKPRIPVVWIHGL WP_001058312.1
hydrogenase [Salmonella enterica] Length: 372\nNumber of predicted TMHs: 2\nExp number of AAs in TMHs: 36.51738\nExp number, first 60 AAs: 19.72849\nTotal prob of N-in: 0.96170\nPOSSIBLE N-term signal sequence\ninside 1 19\nTMhelix 20 39\noutside 40 325\nTMhelix 326 348\ninside 349 372

24407 GCF_001051635.1_CFSAN000860_1.0 Salmonella enterica subsp. enterica serovar Newport str. VA_R100808502
Proteobacteria; Gammaproteobacteria; Enterobacterales; Enterobacteriaceae; Salmonella; Salmonella enterica; Salmonella enterica subsp. enterica; Salmonella enterica subsp. enterica serovar Newport
MTGDNTLITSHGINRRDFMKLCAALAATMGLSSKAAAEMAESVSNPQRPPVIWIGAQECT WP_000145429.1
hydrogenase 2 small subunit [Salmonella enterica] Length: 372\nNumber of predicted TMHs: 1\nExp number of AAs in TMHs: 24.51309\nExp number, first 60 AAs: 0.2682\nTotal prob of N-in: 0.14402\noutside 1 331\nTMhelix 332 354\ninside 355 372

24408 GCF_000483715.2_ASM48371v2 Salmonella enterica subsp. enterica serovar Newport str. WA_14882
Proteobacteria; Gammaproteobacteria; Enterobacterales; Enterobacteriaceae; Salmonella; Salmonella enterica; Salmonella enterica subsp. enterica; Salmonella enterica subsp. enterica serovar Newport
MTGDNTLITSHGINRRDFMKLCAALAATMGLSSKAAAEMAESVSNPQRPPVIWIGAQECT WP_000145429.1
hydrogenase 2 small subunit [Salmonella enterica] Length: 372\nNumber of predicted TMHs: 1\nExp number of AAs in TMHs: 24.51309\nExp number, first 60 AAs: 0.2682\nTotal prob of N-in: 0.14402\noutside 1 331\nTMhelix 332 354\ninside 355 372

24409 GCF_000483715.2_ASM48371v2 Salmonella enterica subsp. enterica serovar Newport str. WA_14882
Proteobacteria; Gammaproteobacteria; Enterobacterales; Enterobacteriaceae; Salmonella; Salmonella enterica; Salmonella enterica subsp. enterica; Salmonella enterica subsp. enterica serovar Newport
MNNEETFYQAMRRKGVTRRSFLKFCSLAATSLGLGAGMTPKIAWALENKPRIPVVWIHGL WP_001058312.1
hydrogenase [Salmonella enterica] Length: 372\nNumber of predicted TMHs: 2\nExp number of AAs in TMHs: 36.51738\nExp number, first 60 AAs: 19.72849\nTotal prob of N-in: 0.96170\nPOSSIBLE N-term signal sequence\ninside 1 19\nTMhelix 20 39\noutside 40 325\nTMhelix 326 348\ninside 349 372

24410 GCF_000483675.1_ASM48367v1 Salmonella enterica subsp. enterica serovar Newport str. WA_14885
Proteobacteria; Gammaproteobacteria; Enterobacterales; Enterobacteriaceae; Salmonella; Salmonella enterica; Salmonella enterica subsp. enterica; Salmonella enterica subsp. enterica serovar Newport
MTGDNTLITSHGINRRDFMKLCAALAATMGLSSKAAAEMAESVSNPQRPPVIWIGAQECT WP_000145429.1
hydrogenase 2 small subunit [Salmonella enterica] Length: 372\nNumber of predicted TMHs: 1\nExp number of AAs in TMHs: 24.51309\nExp number, first 60 AAs: 0.2682\nTotal prob of N-in: 0.14402\noutside 1 331\nTMhelix 332 354\ninside 355 372

24411 GCF_000483675.1_ASM48367v1 Salmonella enterica subsp. enterica serovar Newport str. WA_14885
Proteobacteria; Gammaproteobacteria; Enterobacterales; Enterobacteriaceae; Salmonella; Salmonella enterica; Salmonella enterica subsp. enterica; Salmonella enterica subsp. enterica serovar Newport
MNNEETFYQAMRRKGVTRRSFLKFCSLAATSLGLGAGMTPKIAWALENKPRIPVVWIHGL WP_001058312.1
hydrogenase [Salmonella enterica] Length: 372\nNumber of predicted TMHs: 2\nExp number of AAs in TMHs: 36.51738\nExp number, first 60 AAs: 19.72849\nTotal prob of N-in: 0.96170\nPOSSIBLE N-term signal sequence\ninside 1 19\nTMhelix 20 39\noutside 40 325\nTMhelix 326 348\ninside 349 372

24412 GCF_000487835.1_ASM48783v1 Salmonella enterica subsp. enterica serovar Norwich str. CFSAN001077 Proteobacteria; Gammaproteobacteria; Enterobacterales; Enterobacteriaceae; Salmonella; Salmonella enterica; Salmonella enterica subsp. enterica; Salmonella enterica subsp. enterica serovar Norwich MTGDNTLITSHGINRRDFMKLCAALAATMGLSSKAAAEMAESVSNPQRPPVIWIGAQECT WP_000145429.1 hydrogenase 2 small subunit [Salmonella enterica] Length: 372\nNumber of predicted TMHs: 1\nExp number of AAs in TMHs: 24.51309\nExp number, first 60 AAs: 0.2682\nTotal prob of N-in: 0.14402\noutside 1 331\nTMhelix 332 354\ninside 355 372

24413 GCF_000487835.1_ASM48783v1 Salmonella enterica subsp. enterica serovar Norwich str. CFSAN001077 Proteobacteria; Gammaproteobacteria; Enterobacterales; Enterobacteriaceae; Salmonella; Salmonella enterica; Salmonella enterica subsp. enterica; Salmonella enterica subsp. enterica serovar Norwich MNNEETFYQAMRRKGVTRRSFLKFCSLAATSLGLGAGMTPKIAWALENKPRIPVVWIHGL WP_023260323.1 hydrogenase 1, small subunit [Salmonella enterica] Length: 372\nNumber of predicted TMHs: 2\nExp number of AAs in TMHs: 37.4697199999999\nExp number, first 60 AAs: 19.72699\nTotal prob of N-in: 0.96171\nPOSSIBLE N-term signal sequence\ninside 1 19\nTMhelix 20 39\noutside 40 325\nTMhelix 326 348\ninside 349 372

24414 GCF_000487795.1_ASM48779v1 Salmonella enterica subsp. enterica serovar Ohio str. CFSAN001079 Proteobacteria; Gammaproteobacteria; Enterobacterales; Enterobacteriaceae; Salmonella; Salmonella enterica; Salmonella enterica subsp. enterica; Salmonella enterica subsp. enterica serovar Ohio MTGDNTLITSHGINRRDFMKLCAALAATMGLSSKAAAEMAESVSNPQRPPVIWIGAQECT WP_000145429.1 hydrogenase 2 small subunit [Salmonella enterica] Length: 372\nNumber of predicted TMHs: 1\nExp number of AAs in TMHs: 24.51309\nExp number, first 60 AAs: 0.2682\nTotal prob of N-in: 0.14402\noutside 1 331\nTMhelix 332 354\ninside 355 372

24415 GCF_000487795.1_ASM48779v1 Salmonella enterica subsp. enterica serovar Ohio str. CFSAN001079 Proteobacteria; Gammaproteobacteria; Enterobacterales; Enterobacteriaceae; Salmonella; Salmonella enterica; Salmonella enterica subsp. enterica; Salmonella enterica subsp. enterica serovar Ohio MNNEETFYQAMRRKGVTRRSFLKFCSLATTSLGLGAGMTPKIAWALENKPRIPVVWIHGL WP_023259200.1 hydrogenase 1, small subunit [Salmonella enterica] Length: 372\nNumber of predicted TMHs: 2\nExp number of AAs in TMHs: 35.98973\nExp number, first 60 AAs: 19.19893\nTotal prob of N-in: 0.93662\nPOSSIBLE N-term signal sequence\ninside 1 19\nTMhelix 20 39\noutside 40 325\nTMhelix 326 348\ninside 349 372

24416 GCF_002208255.1_ASM220825v1 Salmonella enterica subsp. enterica serovar Onderstepoort str. SA20060086 Proteobacteria; Gammaproteobacteria; Enterobacterales; Enterobacteriaceae; Salmonella; Salmonella enterica; Salmonella enterica subsp. enterica; Salmonella enterica subsp. Enterica serovar Onderstepoort MNNEETFYQAMRRKGVTRRSFLKFCSLAATSLGLGAGMTPKIAWALENKPRIPVVWIHGL WP_001058312.1 hydrogenase [Salmonella enterica] Length: 372\nNumber of predicted TMHs: 2\nExp number of AAs in TMHs: 36.51738\nExp number, first 60 AAs: 19.72849\nTotal prob of N-in: 0.96170\nPOSSIBLE N-term signal sequence\ninside 1 19\nTMhelix 20 39\noutside 40 325\nTMhelix 326 348\ninside 349 372

24417 GCF_002208255.1_ASM220825v1 Salmonella enterica subsp. enterica serovar Onderstepoort str. SA20060086 Proteobacteria; Gammaproteobacteria; Enterobacterales; Enterobacteriaceae; Salmonella; Salmonella enterica; Salmonella enterica subsp. enterica; Salmonella enterica subsp. Enterica serovar Onderstepoort MTGDNTLITSHGINRRDFMKLCAALAATMGLSSKAAAEMAESVSNPQRPPVIWIGAQECT WP_000145429.1 hydrogenase 2 small subunit [Salmonella enterica] Length: 372\nNumber of predicted TMHs: 1\nExp number of AAs in TMHs: 24.51309\nExp number, first 60 AAs: 0.2682\nTotal prob of N-in: 0.14402\noutside 1 331\nTMhelix 332 354\ninside 355 372

24418 GCF_000486795.1_ASM48679v1 Salmonella enterica subsp. enterica serovar Oranienburg str. 701 Proteobacteria; Gammaproteobacteria; Enterobacterales; Enterobacteriaceae; Salmonella; Salmonella enterica; Salmonella enterica subsp. enterica; Salmonella enterica subsp. enterica serovar Oranienburg MTGDNTLITSHGINRRDFMKLCAALAATMGLSSKAAAEMAESVSNPQRPPVIWIGAQECT WP_000145431.1 hydrogenase 2 small subunit [Salmonella enterica] Length: 372\nNumber of predicted TMHs: 1\nExp number of AAs in TMHs: 24.51505\nExp number, first 60 AAs: 0.2682\nTotal prob of N-in: 0.14406\noutside 1 331\nTMhelix 332 354\ninside 355 372

24419 GCF_000486795.1_ASM48679v1 Salmonella enterica subsp. enterica serovar Oranienburg str. 701 Proteobacteria; Gammaproteobacteria; Enterobacterales; Enterobacteriaceae; Salmonella; Salmonella enterica; Salmonella enterica subsp. enterica; Salmonella enterica subsp. enterica serovar Oranienburg MNNEETFYQAMRRKGVTRRSFLKFCSLAATSLGLGAGMTPKIAWALENKPRIPVVWIHGL WP_001058312.1

hydrogenase [Salmonella enterica] Length: 372\nNumber of predicted TMHs: 2\nExp number of AAs in TMHs: 36.51738\nExp number, first 60 AAs: 19.72849\nTotal prob of N-in: 0.96170\nPOSSIBLE N-term signal sequence\ninside 1 19\nTMhelix 20 39\noutside 40 325\nTMhelix 326 348\ninside 349 372

24420 GCF_000486765.2_ASM48676v2 Salmonella enterica subsp. enterica serovar Panama str. ATCC 7378 Proteobacteria; Gammaproteobacteria; Enterobacterales; Enterobacteriaceae; Salmonella; Salmonella enterica; Salmonella enterica subsp. enterica; Salmonella enterica subsp. enterica serovar Panama MTGDNTLITSHGINRRDFMKLCAALAATMGLSSKAAAEMAESVSNPQRPPVIWIGAQECT WP_000145429.1 hydrogenase 2 small subunit [Salmonella enterica] Length: 372\nNumber of predicted TMHs: 1\nExp number of AAs in TMHs: 24.51309\nExp number, first 60 AAs: 0.2682\nTotal prob of N-in: 0.14402\noutside 1 331\nTMhelix 332 354\ninside 355 372

24421 GCF_000486765.2_ASM48676v2 Salmonella enterica subsp. enterica serovar Panama str. ATCC 7378 Proteobacteria; Gammaproteobacteria; Enterobacterales; Enterobacteriaceae; Salmonella; Salmonella enterica; Salmonella enterica subsp. enterica; Salmonella enterica subsp. enterica serovar Panama MNNEETFYQAMRRKGVTRRSFLKFCSLAATSLGLGAGMTPKIAWALENKPRIPVVWIHGL WP_001058312.1 hydrogenase [Salmonella enterica] Length: 372\nNumber of predicted TMHs: 2\nExp number of AAs in TMHs: 36.51738\nExp number, first 60 AAs: 19.72849\nTotal prob of N-in: 0.96170\nPOSSIBLE N-term signal sequence\ninside 1 19\nTMhelix 20 39\noutside 40 325\nTMhelix 326 348\ninside 349 372

24422 GCF_001448795.1_ASM144879v1 Salmonella enterica subsp. enterica serovar Panama str. SA20030878 Proteobacteria; Gammaproteobacteria; Enterobacterales; Enterobacteriaceae; Salmonella; Salmonella enterica; Salmonella enterica subsp. enterica; Salmonella enterica subsp. enterica serovar Panama MTGDNTLITSHGINRRDFMKLCAALAATMGLSSKAAAEMAESVSNPQRPPVIWIGAQECT WP_000145429.1 hydrogenase 2 small subunit [Salmonella enterica] Length: 372\nNumber of predicted TMHs: 1\nExp number of AAs in TMHs: 24.51309\nExp number, first 60 AAs: 0.2682\nTotal prob of N-in: 0.14402\noutside 1 331\nTMhelix 332 354\ninside 355 372

24423 GCF_001448795.1_ASM144879v1 Salmonella enterica subsp. enterica serovar Panama str. SA20030878 Proteobacteria; Gammaproteobacteria; Enterobacterales; Enterobacteriaceae; Salmonella; Salmonella enterica; Salmonella enterica subsp. enterica; Salmonella enterica subsp. enterica serovar Panama MNNEETFYQAMRRKGVTRRSFLKFCSLAATSLGLGAGMTPKIAWALENKPRIPVVWIHGL WP_001058312.1 hydrogenase [Salmonella enterica] Length: 372\nNumber of predicted TMHs: 2\nExp number of AAs in TMHs: 36.51738\nExp number, first 60 AAs: 19.72849\nTotal prob of N-in: 0.96170\nPOSSIBLE N-term signal sequence\ninside 1 19\nTMhelix 20 39\noutside 40 325\nTMhelix 326 348\ninside 349 372

24424 GCF_000026565.1_ASM2656v1 Salmonella enterica subsp. enterica serovar Paratyphi A str. AKU_12601 Proteobacteria; Gammaproteobacteria; Enterobacterales; Enterobacteriaceae; Salmonella; Salmonella enterica; Salmonella enterica subsp. enterica; Salmonella enterica subsp. enterica serovar Paratyphi A MTGDNTLITSHGINRRDFMKLCAALAATMGLSSKAAAEMAESVSNPQRPPVIWIGAQECT WP_000145427.1 hydrogenase 2 small subunit [Salmonella enterica] Length: 372\nNumber of predicted TMHs: 1\nExp number of AAs in TMHs: 24.48576\nExp number, first 60 AAs: 0.2692\nTotal prob of N-in: 0.14251\noutside 1 331\nTMhelix 332 354\ninside 355 372

24425 GCF_000486725.2_ASM48672v2 Salmonella enterica subsp. enterica serovar Paratyphi A str. ATCC 11511 Proteobacteria; Gammaproteobacteria; Enterobacterales; Enterobacteriaceae; Salmonella; Salmonella enterica; Salmonella enterica subsp. enterica; Salmonella enterica subsp. enterica serovar Paratyphi A MTGDNTLITSHGINRRDFMKLCAALAATMGLSSKAAAEMAESVSNPQRPPVIWIGAQECT WP_000145427.1 hydrogenase 2 small subunit [Salmonella enterica] Length: 372\nNumber of predicted TMHs: 1\nExp number of AAs in TMHs: 24.48576\nExp number, first 60 AAs: 0.2692\nTotal prob of N-in: 0.14251\noutside 1 331\nTMhelix 332 354\ninside 355 372

24426 GCF_000011885.1_ASM1188v1 Salmonella enterica subsp. enterica serovar Paratyphi A str. ATCC 9150 Proteobacteria; Gammaproteobacteria; Enterobacterales; Enterobacteriaceae; Salmonella; Salmonella enterica; Salmonella enterica subsp. enterica; Salmonella enterica subsp. enterica serovar Paratyphi A MTGDNTLITSHGINRRDFMKLCAALAATMGLSSKAAAEMAESVSNPQRPPVIWIGAQECT WP_000145427.1 hydrogenase 2 small subunit [Salmonella enterica] Length: 372\nNumber of predicted TMHs: 1\nExp number of AAs in TMHs: 24.48576\nExp number, first 60 AAs: 0.2692\nTotal prob of N-in: 0.14251\noutside 1 331\nTMhelix 332 354\ninside 355 372

24427 GCF_000411815.1_ASM41181v1 Salmonella enterica subsp. enterica serovar Paratyphi A str. GXS2268 Proteobacteria; Gammaproteobacteria; Enterobacterales; Enterobacteriaceae; Salmonella; Salmonella enterica; Salmonella enterica subsp. enterica; Salmonella enterica subsp. enterica serovar Paratyphi A MTGDNTLITSHGINRRDFMKLCAALAATMGLSSKAAAEMAESVSNPQRPPVIWIGAQECT WP_000145427.1 hydrogenase 2 small subunit [Salmonella enterica] Length: 372\nNumber of predicted TMHs: 1\nExp number of AAs in TMHs: 24.48576\nExp number, first 60 AAs: 0.2692\nTotal prob of N-in: 0.14251\noutside 1 331\nTMhelix 332 354\ninside 355 372

24428 GCF_000411795.1_ASM41179v1 Salmonella enterica subsp. enterica serovar Paratyphi A str. GZ9A00052 Proteobacteria; Gammaproteobacteria; Enterobacterales; Enterobacteriaceae; Salmonella; Salmonella enterica; Salmonella enterica subsp. enterica; Salmonella enterica subsp. enterica serovar Paratyphi A MTGDNTLITSHGINRRDFMKLCAALAATMGLSSKAAAEMAESVSNPQRPPVIWIGAQECT WP_000145427.1 hydrogenase 2 small subunit [Salmonella enterica] Length: 372\nNumber of predicted TMHs: 1\nExp number of AAs in TMHs: 24.48576\nExp number, first 60 AAs: 0.2692\nTotal prob of N-in: 0.14251\noutside 1 331\nTMhelix 332 354\ninside 355 372

24429 GCF_000411835.1_ASM41183v1 Salmonella enterica subsp. enterica serovar Paratyphi A str. JX05-19 Proteobacteria; Gammaproteobacteria; Enterobacterales; Enterobacteriaceae; Salmonella; Salmonella enterica; Salmonella enterica subsp. enterica; Salmonella enterica subsp. enterica serovar Paratyphi A MTGDNTLITSHGINRRDFMKLCAALAATMGLSSKAAAEMAESVSNPQRPPVIWIGAQECT WP_000145427.1 hydrogenase 2 small subunit [Salmonella enterica] Length: 372\nNumber of predicted TMHs: 1\nExp number of AAs in TMHs: 24.48576\nExp number, first 60 AAs: 0.2692\nTotal prob of N-in: 0.14251\noutside 1 331\nTMhelix 332 354\ninside 355 372

24430 GCF_001448825.1_ASM144882v1 Salmonella enterica subsp. enterica serovar Paratyphi A str. SA19950809 Proteobacteria; Gammaproteobacteria; Enterobacterales; Enterobacteriaceae; Salmonella; Salmonella enterica; Salmonella enterica subsp. enterica; Salmonella enterica subsp. enterica serovar Paratyphi A MTGDNTLITSHGINRRDFMKLCAALAATMGLSSKAAAEMAESVSNPQRPPVIWIGAQECT WP_000145427.1 hydrogenase 2 small subunit [Salmonella enterica] Length: 372\nNumber of predicted TMHs: 1\nExp number of AAs in TMHs: 24.48576\nExp number, first 60 AAs: 0.2692\nTotal prob of N-in: 0.14251\noutside 1 331\nTMhelix 332 354\ninside 355 372

24431 GCF_000411855.1_ASM41185v1 Salmonella enterica subsp. enterica serovar Paratyphi A str. YN09620 Proteobacteria; Gammaproteobacteria; Enterobacterales; Enterobacteriaceae; Salmonella; Salmonella enterica; Salmonella enterica subsp. enterica; Salmonella enterica subsp. enterica serovar Paratyphi A MTGDNTLITSHGINRRDFMKLCAALAATMGLSSKAAAEMAESVSNPQRPPVIWIGAQECT WP_000145427.1 hydrogenase 2 small subunit [Salmonella enterica] Length: 372\nNumber of predicted TMHs: 1\nExp number of AAs in TMHs: 24.48576\nExp number, first 60 AAs: 0.2692\nTotal prob of N-in: 0.14251\noutside 1 331\nTMhelix 332 354\ninside 355 372

24432 GCF_000411875.1_ASM41187v1 Salmonella enterica subsp. enterica serovar Paratyphi A str. ZJ98-53 Proteobacteria; Gammaproteobacteria; Enterobacterales; Enterobacteriaceae; Salmonella; Salmonella enterica; Salmonella enterica subsp. enterica; Salmonella enterica subsp. enterica serovar Paratyphi A MTGDNTLITSHGINRRDFMKLCAALAATMGLSSKAAAEMAESVSNPQRPPVIWIGAQECT WP_000145427.1 hydrogenase 2 small subunit [Salmonella enterica] Length: 372\nNumber of predicted TMHs: 1\nExp number of AAs in TMHs: 24.48576\nExp number, first 60 AAs: 0.2692\nTotal prob of N-in: 0.14251\noutside 1 331\nTMhelix 332 354\ninside 355 372

24433 GCF_900086565.1_39641_H02 Salmonella enterica subsp. enterica serovar Java Proteobacteria; Gammaproteobacteria; Enterobacterales; Enterobacteriaceae; Salmonella; Salmonella enterica; Salmonella enterica subsp. enterica; Salmonella enterica subsp. enterica serovar Paratyphi B MTGDNTLITSHGINRRDFMKLCAALAATMGLSSKAAAEMAESVSNPQRPPVIWIGAQECT WP_000145429.1 hydrogenase 2 small subunit [Salmonella enterica] Length: 372\nNumber of predicted TMHs: 1\nExp number of AAs in TMHs: 24.51309\nExp number, first 60 AAs: 0.2682\nTotal prob of N-in: 0.14402\noutside 1 331\nTMhelix 332 354\ninside 355 372

24434 GCF_002063335.1_ASM206333v1 Salmonella enterica subsp. enterica serovar Java Proteobacteria; Gammaproteobacteria; Enterobacterales; Enterobacteriaceae; Salmonella; Salmonella enterica; Salmonella enterica subsp. enterica; Salmonella enterica subsp. enterica serovar Paratyphi B MTGDNTLITSHGINRRDFMKLCAALAATMGLSSKAAAEMAESVSNPQRPPVIWIGAQECT WP_000145429.1 hydrogenase 2 small subunit [Salmonella enterica] Length: 372\nNumber of predicted TMHs: 1\nExp

number of AAs in TMHs: 24.51309\nExp number, first 60 AAs: 0.2682\nTotal prob of N-in: 0.14402\noutside 1 331\nTMhelix 332 354\ninside 355 372

24435 GCF_002064125.1_ASM206412v1 Salmonella enterica subsp. enterica serovar Java Proteobacteria; Gammaproteobacteria; Enterobacterales; Enterobacteriaceae; Salmonella; Salmonella enterica; Salmonella enterica subsp. enterica; Salmonella enterica subsp. enterica serovar Paratyphi B
MNNEETFYQAMRRKGVTRRSFLKFCSLAATSLGLGAGMTPKIAWALENKPRIPVVWIHGL WP_001058311.1 hydrogenase [Salmonella enterica] Length: 372\nNumber of predicted TMHs: 2\nExp number of AAs in TMHs: 37.4699899999999\nExp number, first 60 AAs: 19.72699\nTotal prob of N-in: 0.96171\nPOSSIBLE N-term signal sequence\ninside 1 19\nTMhelix 20 39\noutside 40 325\nTMhelix 326 348\ninside 349 372

24436 GCF_002064125.1_ASM206412v1 Salmonella enterica subsp. enterica serovar Java Proteobacteria; Gammaproteobacteria; Enterobacterales; Enterobacteriaceae; Salmonella; Salmonella enterica; Salmonella enterica subsp. enterica; Salmonella enterica subsp. enterica serovar Paratyphi B
MTGDNTLITSHGINRRDFMKLCAALAATMGLSSKAAAEMAESVSNPQRPPVIWIGAQECT WP_000145429.1 hydrogenase 2 small subunit [Salmonella enterica] Length: 372\nNumber of predicted TMHs: 1\nExp number of AAs in TMHs: 24.51309\nExp number, first 60 AAs: 0.2682\nTotal prob of N-in: 0.14402\noutside 1 331\nTMhelix 332 354\ninside 355 372

24437 GCF_002063335.1_ASM206333v1 Salmonella enterica subsp. enterica serovar Java Proteobacteria; Gammaproteobacteria; Enterobacterales; Enterobacteriaceae; Salmonella; Salmonella enterica; Salmonella enterica subsp. enterica; Salmonella enterica subsp. enterica serovar Paratyphi B
MNNEETFYQAMRRKGVTRRSFLKFCSLAATSLGLGAGMTPKIAWALENKPRIPVVWIHGL WP_001058311.1 hydrogenase [Salmonella enterica] Length: 372\nNumber of predicted TMHs: 2\nExp number of AAs in TMHs: 37.4699899999999\nExp number, first 60 AAs: 19.72699\nTotal prob of N-in: 0.96171\nPOSSIBLE N-term signal sequence\ninside 1 19\nTMhelix 20 39\noutside 40 325\nTMhelix 326 348\ninside 349 372

24438 GCF_002064135.1_ASM206413v1 Salmonella enterica subsp. enterica serovar Java Proteobacteria; Gammaproteobacteria; Enterobacterales; Enterobacteriaceae; Salmonella; Salmonella enterica; Salmonella enterica subsp. enterica; Salmonella enterica subsp. enterica serovar Paratyphi B
MTGDNTLITSHGINRRDFMKLCAALAATMGLSSKAAAEMAESVSNPQRPPVIWIGAQECT WP_000145429.1 hydrogenase 2 small subunit [Salmonella enterica] Length: 372\nNumber of predicted TMHs: 1\nExp number of AAs in TMHs: 24.51309\nExp number, first 60 AAs: 0.2682\nTotal prob of N-in: 0.14402\noutside 1 331\nTMhelix 332 354\ninside 355 372

24439 GCF_002064135.1_ASM206413v1 Salmonella enterica subsp. enterica serovar Java Proteobacteria; Gammaproteobacteria; Enterobacterales; Enterobacteriaceae; Salmonella; Salmonella enterica; Salmonella enterica subsp. enterica; Salmonella enterica subsp. enterica serovar Paratyphi B
MNNEETFYQAMRRKGVTRRSFLKFCSLAATSLGLGAGMTPKIAWALENKPRIPVVWIHGL WP_001058311.1 hydrogenase [Salmonella enterica] Length: 372\nNumber of predicted TMHs: 2\nExp number of AAs in TMHs: 37.4699899999999\nExp number, first 60 AAs: 19.72699\nTotal prob of N-in: 0.96171\nPOSSIBLE N-term signal sequence\ninside 1 19\nTMhelix 20 39\noutside 40 325\nTMhelix 326 348\ninside 349 372

24440 GCF_900086565.1_39641_H02 Salmonella enterica subsp. enterica serovar Java Proteobacteria; Gammaproteobacteria; Enterobacterales; Enterobacteriaceae; Salmonella; Salmonella enterica; Salmonella enterica subsp. enterica; Salmonella enterica subsp. enterica serovar Paratyphi B
MNNEETFYQAMRRKGVTRRSFLKFCSLAATSLGLGAGMTPKIAWALENKPRIPVVWIHGL WP_001058307.1 hydrogenase [Salmonella enterica] Length: 372\nNumber of predicted TMHs: 2\nExp number of AAs in TMHs: 36.51738\nExp number, first 60 AAs: 19.72849\nTotal prob of N-in: 0.96170\nPOSSIBLE N-term signal sequence\ninside 1 19\nTMhelix 20 39\noutside 40 325\nTMhelix 326 348\ninside 349 372

24441 GCF_000486685.1_ASM48668v1 Salmonella enterica subsp. enterica serovar Paratyphi B str. ATCC 10719 Proteobacteria; Gammaproteobacteria; Enterobacterales; Enterobacteriaceae; Salmonella; Salmonella enterica; Salmonella enterica subsp. enterica; Salmonella enterica subsp. enterica serovar Paratyphi B
MTGDNTLITSHGINRRDFMKLCAALAATMGLSSKAAAEMAESVSNPQRPPVIWIGAQECT WP_000145429.1 hydrogenase 2 small subunit [Salmonella enterica] Length: 372\nNumber of predicted TMHs: 1\nExp number of AAs in TMHs: 24.51309\nExp number, first 60 AAs: 0.2682\nTotal prob of N-in: 0.14402\noutside 1 331\nTMhelix 332 354\ninside 355 372

24442 GCF_000486685.1_ASM48668v1 Salmonella enterica subsp. enterica serovar Paratyphi B str. ATCC 10719 Proteobacteria; Gammaproteobacteria; Enterobacterales; Enterobacteriaceae; Salmonella; Salmonella enterica;

Salmonella enterica subsp. enterica; Salmonella enterica subsp. enterica serovar Paratyphi B
MNNEETFYQAMRRKGVTRRSFLKFCSLAATSLGLGAGMTPKIAWALENKPRIPVVWIHGL WP_001058312.1
hydrogenase [Salmonella enterica] Length: 372\nNumber of predicted TMHs: 2\nExp number of AAs in TMHs: 36.51738\nExp number, first 60 AAs: 19.72849\nTotal prob of N-in: 0.96170\nPOSSIBLE N-term signal sequence\ninside 1 19\nTMhelix 20 39\noutside 40 325\nTMhelix 326 348\ninside 349 372

24443 GCF_000486645.1_ASM48664v1 Salmonella enterica subsp. enterica serovar Paratyphi B str. ATCC 19940
Proteobacteria; Gammaproteobacteria; Enterobacterales; Enterobacteriaceae; Salmonella; Salmonella enterica; Salmonella enterica subsp. enterica; Salmonella enterica subsp. enterica serovar Paratyphi B
MNNEETFYQAMRRKGVTRRSFLKFCSLAATSLGLGAGMTPKIAWVLENKPRIPVVWIHGL WP_023210509.1
hydrogenase 1, small subunit [Salmonella enterica] Length: 372\nNumber of predicted TMHs: 2\nExp number of AAs in TMHs: 36.5381799999999\nExp number, first 60 AAs: 19.73808\nTotal prob of N-in: 0.95980\nPOSSIBLE N-term signal sequence\ninside 1 19\nTMhelix 20 39\noutside 40 325\nTMhelix 326 348\ninside 349 372

24444 GCF_000486645.1_ASM48664v1 Salmonella enterica subsp. enterica serovar Paratyphi B str. ATCC 19940
Proteobacteria; Gammaproteobacteria; Enterobacterales; Enterobacteriaceae; Salmonella; Salmonella enterica; Salmonella enterica subsp. enterica; Salmonella enterica subsp. enterica serovar Paratyphi B
MTGDNTLITSHGINRRDFMKLCAALAATMGLSSKAAAEMAESVSNPQRPPVIWIGAQECT WP_000145429.1
hydrogenase 2 small subunit [Salmonella enterica] Length: 372\nNumber of predicted TMHs: 1\nExp number of AAs in TMHs: 24.51309\nExp number, first 60 AAs: 0.2682\nTotal prob of N-in: 0.14402\noutside 1 331\nTMhelix 332 354\ninside 355 372

24445 GCF_000486625.1_ASM48662v1 Salmonella enterica subsp. enterica serovar Paratyphi B str. ATCC 51962
Proteobacteria; Gammaproteobacteria; Enterobacterales; Enterobacteriaceae; Salmonella; Salmonella enterica; Salmonella enterica subsp. enterica; Salmonella enterica subsp. enterica serovar Paratyphi B
MNNEETFYQAMRRKGVTRRSFLKFCSLAATSLGLGAGMTPKIAWALENKPRIPVVWIHGL WP_001058307.1
hydrogenase [Salmonella enterica] Length: 372\nNumber of predicted TMHs: 2\nExp number of AAs in TMHs: 36.51738\nExp number, first 60 AAs: 19.72849\nTotal prob of N-in: 0.96170\nPOSSIBLE N-term signal sequence\ninside 1 19\nTMhelix 20 39\noutside 40 325\nTMhelix 326 348\ninside 349 372

24446 GCF_000486625.1_ASM48662v1 Salmonella enterica subsp. enterica serovar Paratyphi B str. ATCC 51962
Proteobacteria; Gammaproteobacteria; Enterobacterales; Enterobacteriaceae; Salmonella; Salmonella enterica; Salmonella enterica subsp. enterica; Salmonella enterica subsp. enterica serovar Paratyphi B
MTGDNTLITSHGINRRDFMKLCAALAATMGLSSKAAAEMAESVSNPQRPPVIWIGAQECT WP_000145429.1
hydrogenase 2 small subunit [Salmonella enterica] Length: 372\nNumber of predicted TMHs: 1\nExp number of AAs in TMHs: 24.51309\nExp number, first 60 AAs: 0.2682\nTotal prob of N-in: 0.14402\noutside 1 331\nTMhelix 332 354\ninside 355 372

24447 GCF_000486665.1_ASM48666v1 Salmonella enterica subsp. enterica serovar Paratyphi B str. ATCC 8759
Proteobacteria; Gammaproteobacteria; Enterobacterales; Enterobacteriaceae; Salmonella; Salmonella enterica; Salmonella enterica subsp. enterica; Salmonella enterica subsp. enterica serovar Paratyphi B
MNNEETFYQAMRRKGVTRRSFLKFCSLAATSLGLGAGMTPKIAWALENKPRIPVVWIHGL WP_001058307.1
hydrogenase [Salmonella enterica] Length: 372\nNumber of predicted TMHs: 2\nExp number of AAs in TMHs: 36.51738\nExp number, first 60 AAs: 19.72849\nTotal prob of N-in: 0.96170\nPOSSIBLE N-term signal sequence\ninside 1 19\nTMhelix 20 39\noutside 40 325\nTMhelix 326 348\ninside 349 372

24448 GCF_000486665.1_ASM48666v1 Salmonella enterica subsp. enterica serovar Paratyphi B str. ATCC 8759
Proteobacteria; Gammaproteobacteria; Enterobacterales; Enterobacteriaceae; Salmonella; Salmonella enterica; Salmonella enterica subsp. enterica; Salmonella enterica subsp. enterica serovar Paratyphi B
MTGDNTLITSHGINRRDFMKLCAALAATMGLSSKAAAEMAESVSNPQRPPVIWIGAQECT WP_000145429.1
hydrogenase 2 small subunit [Salmonella enterica] Length: 372\nNumber of predicted TMHs: 1\nExp number of AAs in TMHs: 24.51309\nExp number, first 60 AAs: 0.2682\nTotal prob of N-in: 0.14402\noutside 1 331\nTMhelix 332 354\ninside 355 372

24449 GCF_000486705.1_ASM48670v1 Salmonella enterica subsp. enterica serovar Paratyphi B str. ATCC BAA-1585 Proteobacteria; Gammaproteobacteria; Enterobacterales; Enterobacteriaceae; Salmonella; Salmonella enterica; Salmonella enterica subsp. enterica; Salmonella enterica subsp. enterica serovar Paratyphi B
MTGDNTLITSHGINRRDFMKLCAALAATMGLSSKAAAEMAESVSNPQRPPVIWIGAQECT WP_000145429.1
hydrogenase 2 small subunit [Salmonella enterica] Length: 372\nNumber of predicted TMHs: 1\nExp

number of AAs in TMHs: 24.51309\nExp number, first 60 AAs: 0.2682\nTotal prob of N-in: 0.14402\noutside 1 331\nTMhelix 332 354\ninside 355 372

24450 GCF_000486705.1_ASM48670v1 Salmonella enterica subsp. enterica serovar Paratyphi B str. ATCC BAA-1585 Proteobacteria; Gammaproteobacteria; Enterobacterales; Enterobacteriaceae; Salmonella; Salmonella enterica; Salmonella enterica subsp. enterica; Salmonella enterica subsp. enterica serovar Paratyphi B MNNEETFYQAMRRKGVTRRSFLKFCSLAATSLGLGAGMTPKIAWALENKPRIPVVWIHGL WP_001058312.1 hydrogenase [Salmonella enterica] Length: 372\nNumber of predicted TMHs: 2\nExp number of AAs in TMHs: 36.51738\nExp number, first 60 AAs: 19.72849\nTotal prob of N-in: 0.96170\nPOSSIBLE N-term signal sequence\ninside 1 19\nTMhelix 20 39\noutside 40 325\nTMhelix 326 348\ninside 349 372

24451 GCF_000484015.1_ASM48401v1 Salmonella enterica subsp. enterica serovar Paratyphi B str. SARA61 Proteobacteria; Gammaproteobacteria; Enterobacterales; Enterobacteriaceae; Salmonella; Salmonella enterica; Salmonella enterica subsp. enterica; Salmonella enterica subsp. enterica serovar Paratyphi B MTGDNTLITSHGINRRDFMKLCAALAATMGLSSKAAAEMAESVSNPQRPPVIWIGAQECT WP_000145429.1 hydrogenase 2 small subunit [Salmonella enterica] Length: 372\nNumber of predicted TMHs: 1\nExp number of AAs in TMHs: 24.51309\nExp number, first 60 AAs: 0.2682\nTotal prob of N-in: 0.14402\noutside 1 331\nTMhelix 332 354\ninside 355 372

24452 GCF_000484015.1_ASM48401v1 Salmonella enterica subsp. enterica serovar Paratyphi B str. SARA61 Proteobacteria; Gammaproteobacteria; Enterobacterales; Enterobacteriaceae; Salmonella; Salmonella enterica; Salmonella enterica subsp. enterica; Salmonella enterica subsp. enterica serovar Paratyphi B MNNEETFYQAMRRKGVTRRSFLKFCSLAATSLGLSAGMTPKIAWALENKPRIPVVWIHGL WP_001058316.1 hydrogenase [Salmonella enterica] Length: 372\nNumber of predicted TMHs: 2\nExp number of AAs in TMHs: 35.76397\nExp number, first 60 AAs: 18.97201\nTotal prob of N-in: 0.92785\nPOSSIBLE N-term signal sequence\ninside 1 19\nTMhelix 20 39\noutside 40 325\nTMhelix 326 348\ninside 349 372

24453 GCF_000486185.1_ASM48618v1 Salmonella enterica subsp. enterica serovar Paratyphi B str. SARA62 Proteobacteria; Gammaproteobacteria; Enterobacterales; Enterobacteriaceae; Salmonella; Salmonella enterica; Salmonella enterica subsp. enterica; Salmonella enterica subsp. enterica serovar Paratyphi B MTGDNTLITSHGINRRDFMKLCAALAATMGLSSKAAAEMAESVSNPQRPPVIWIGAQECT WP_000145429.1 hydrogenase 2 small subunit [Salmonella enterica] Length: 372\nNumber of predicted TMHs: 1\nExp number of AAs in TMHs: 24.51309\nExp number, first 60 AAs: 0.2682\nTotal prob of N-in: 0.14402\noutside 1 331\nTMhelix 332 354\ninside 355 372

24454 GCF_000486185.1_ASM48618v1 Salmonella enterica subsp. enterica serovar Paratyphi B str. SARA62 Proteobacteria; Gammaproteobacteria; Enterobacterales; Enterobacteriaceae; Salmonella; Salmonella enterica; Salmonella enterica subsp. enterica; Salmonella enterica subsp. enterica serovar Paratyphi B MNNEETFYQAMRRKGVTRRSFLKFCSLAATSLGLGAGMTPKIAWALENKPRIPVVWIHGL WP_001058312.1 hydrogenase [Salmonella enterica] Length: 372\nNumber of predicted TMHs: 2\nExp number of AAs in TMHs: 36.51738\nExp number, first 60 AAs: 19.72849\nTotal prob of N-in: 0.96170\nPOSSIBLE N-term signal sequence\ninside 1 19\nTMhelix 20 39\noutside 40 325\nTMhelix 326 348\ninside 349 372

24455 GCF_000018705.1_ASM1870v1 Salmonella enterica subsp. enterica serovar Paratyphi B str. SPB7 Proteobacteria; Gammaproteobacteria; Enterobacterales; Enterobacteriaceae; Salmonella; Salmonella enterica; Salmonella enterica subsp. enterica; Salmonella enterica subsp. enterica serovar Paratyphi B MTGDNTLITSHGINRRDFMKLCAALAATMGLSSKAAAEMAESVSNPQRPPVIWIGAQECT WP_000145429.1 hydrogenase 2 small subunit [Salmonella enterica] Length: 372\nNumber of predicted TMHs: 1\nExp number of AAs in TMHs: 24.51309\nExp number, first 60 AAs: 0.2682\nTotal prob of N-in: 0.14402\noutside 1 331\nTMhelix 332 354\ninside 355 372

24456 GCF_000018705.1_ASM1870v1 Salmonella enterica subsp. enterica serovar Paratyphi B str. SPB7 Proteobacteria; Gammaproteobacteria; Enterobacterales; Enterobacteriaceae; Salmonella; Salmonella enterica; Salmonella enterica subsp. enterica; Salmonella enterica subsp. enterica serovar Paratyphi B MNNEETFYQAMRRKGVTRRSFLKFCSLAATSLGLGAGMTPKIAWALENKPRIPVVWIHGL WP_001058312.1 hydrogenase [Salmonella enterica] Length: 372\nNumber of predicted TMHs: 2\nExp number of AAs in TMHs: 36.51738\nExp number, first 60 AAs: 19.72849\nTotal prob of N-in: 0.96170\nPOSSIBLE N-term signal sequence\ninside 1 19\nTMhelix 20 39\noutside 40 325\nTMhelix 326 348\ninside 349 372

24457 GCF_000018385.1_ASM1838v1 Salmonella enterica subsp. enterica serovar Paratyphi C str. RKS4594 Proteobacteria; Gammaproteobacteria; Enterobacterales; Enterobacteriaceae; Salmonella; Salmonella enterica;

Salmonella enterica subsp. enterica; Salmonella enterica subsp. enterica serovar Paratyphi C
MNNEETFYQAMRRKGVTRRSFLKFCSLAATSLGLGAGMTPKIAWALENKPRIPVVWIHGL WP_001058311.1
hydrogenase [Salmonella enterica] Length: 372\nNumber of predicted TMHs: 2\nExp number of AAs in TMHs: 37.4699899999999\nExp number, first 60 AAs: 19.72699\nTotal prob of N-in: 0.96171\nPOSSIBLE N-term signal sequence\ninside 1 19\nTMhelix 20 39\noutside 40 325\nTMhelix 326 348\ninside 349 372

24458 GCF_000018385.1_ASM1838v1 Salmonella enterica subsp. enterica serovar Paratyphi C str. RKS4594
Proteobacteria; Gammaproteobacteria; Enterobacterales; Enterobacteriaceae; Salmonella; Salmonella enterica; Salmonella enterica subsp. enterica; Salmonella enterica subsp. enterica serovar Paratyphi C
MTGDNTLITSHGINRRDFMKLCAALAATMGLSSKAAAEMAESVSNPQRPPVIWIGAQECT WP_000145429.1
hydrogenase 2 small subunit [Salmonella enterica] Length: 372\nNumber of predicted TMHs: 1\nExp number of AAs in TMHs: 24.51309\nExp number, first 60 AAs: 0.2682\nTotal prob of N-in: 0.14402\noutside 1 331\nTMhelix 332 354\ninside 355 372

24459 GCF_000240905.1_ASM24090v2 Salmonella enterica subsp. enterica serovar Pomona str. ATCC 10729
Proteobacteria; Gammaproteobacteria; Enterobacterales; Enterobacteriaceae; Salmonella; Salmonella enterica; Salmonella enterica subsp. enterica; Salmonella enterica subsp. enterica serovar Pomona
MTGDNTLITSHGINRRDFMKLCAALAATMGLSSKAAAEMAESVSNPQRPPVIWIGAQECT WP_000145431.1
hydrogenase 2 small subunit [Salmonella enterica] Length: 372\nNumber of predicted TMHs: 1\nExp number of AAs in TMHs: 24.51505\nExp number, first 60 AAs: 0.2682\nTotal prob of N-in: 0.14406\noutside 1 331\nTMhelix 332 354\ninside 355 372

24460 GCF_000412855.1_2010K-2244_Ver1 Salmonella enterica subsp. enterica serovar Poona str. 2010K-2244
Proteobacteria; Gammaproteobacteria; Enterobacterales; Enterobacteriaceae; Salmonella; Salmonella enterica; Salmonella enterica subsp. enterica; Salmonella enterica subsp. enterica serovar Poona
MTGDNTLITSHGINRRDFMKLCAALAATMGLSSKAAAEMAESVSNPQRPPVIWIGAQECT WP_000145429.1
hydrogenase 2 small subunit [Salmonella enterica] Length: 372\nNumber of predicted TMHs: 1\nExp number of AAs in TMHs: 24.51309\nExp number, first 60 AAs: 0.2682\nTotal prob of N-in: 0.14402\noutside 1 331\nTMhelix 332 354\ninside 355 372

24461 GCF_000493295.1_ASM49329v1 Salmonella enterica subsp. enterica serovar Poona str. ATCC BAA-1673
Proteobacteria; Gammaproteobacteria; Enterobacterales; Enterobacteriaceae; Salmonella; Salmonella enterica; Salmonella enterica subsp. enterica; Salmonella enterica subsp. enterica serovar Poona
MNNEETFYQAMRRKGVTRRSFLKFCSLAATSLGLGAGMTPKIAWALENKPRIPVVWIHGL WP_023213673.1
hydrogenase 1, small subunit [Salmonella enterica] Length: 372\nNumber of predicted TMHs: 2\nExp number of AAs in TMHs: 36.51753\nExp number, first 60 AAs: 19.72849\nTotal prob of N-in: 0.96170\nPOSSIBLE N-term signal sequence\ninside 1 19\nTMhelix 20 39\noutside 40 325\nTMhelix 326 348\ninside 349 372

24462 GCF_000493295.1_ASM49329v1 Salmonella enterica subsp. enterica serovar Poona str. ATCC BAA-1673
Proteobacteria; Gammaproteobacteria; Enterobacterales; Enterobacteriaceae; Salmonella; Salmonella enterica; Salmonella enterica subsp. enterica; Salmonella enterica subsp. enterica serovar Poona
MTGDNTLITSHGINRRDFMKLCAALAATMGLSSKAAAEMAESVSNPQRPPVIWIGAQECT WP_000145429.1
hydrogenase 2 small subunit [Salmonella enterica] Length: 372\nNumber of predicted TMHs: 1\nExp number of AAs in TMHs: 24.51309\nExp number, first 60 AAs: 0.2682\nTotal prob of N-in: 0.14402\noutside 1 331\nTMhelix 332 354\ninside 355 372

24463 GCF_000484295.1_ASM48429v1 Salmonella enterica subsp. enterica serovar Pullorum str. 13036
Proteobacteria; Gammaproteobacteria; Enterobacterales; Enterobacteriaceae; Salmonella; Salmonella enterica; Salmonella enterica subsp. enterica; Salmonella enterica subsp. enterica serovar Pullorum
MNNEETFYQAMRRKGVTRRSFLKFCSLAATSLGLGAGMTPKIAWALENKPRIPVVWIHGL WP_001058309.1
hydrogenase [Salmonella enterica] Length: 372\nNumber of predicted TMHs: 2\nExp number of AAs in TMHs: 36.51725\nExp number, first 60 AAs: 19.7285\nTotal prob of N-in: 0.96170\nPOSSIBLE N-term signal sequence\ninside 1 19\nTMhelix 20 39\noutside 40 325\nTMhelix 326 348\ninside 349 372

24464 GCF_000484295.1_ASM48429v1 Salmonella enterica subsp. enterica serovar Pullorum str. 13036
Proteobacteria; Gammaproteobacteria; Enterobacterales; Enterobacteriaceae; Salmonella; Salmonella enterica; Salmonella enterica subsp. enterica; Salmonella enterica subsp. enterica serovar Pullorum
MTGDNTLITSHGINRRDFMKLCAALAATMGLSSKAAAEMAESVSNPQRPPVIWIGAQECT WP_000145430.1
hydrogenase 2 small subunit [Salmonella enterica] Length: 372\nNumber of predicted TMHs: 1\nExp number of AAs in TMHs: 24.51306\nExp number, first 60 AAs: 0.2682\nTotal prob of N-in: 0.14402\noutside 1 331\nTMhelix 332 354\ninside 355 372

24465 GCF_000486745.1_ASM48674v1 Salmonella enterica subsp. enterica serovar Pullorum str. 19945 Proteobacteria; Gammaproteobacteria; Enterobacterales; Enterobacteriaceae; Salmonella; Salmonella enterica; Salmonella enterica subsp. enterica; Salmonella enterica subsp. enterica serovar Pullorum MNNEETFYQAMRRKGVTRRSFLKFCSLAATSLGLGAGMTPKIAWALENKPRIPVVWIHGL WP_001058309.1 hydrogenase [Salmonella enterica] Length: 372\nNumber of predicted TMHs: 2\nExp number of AAs in TMHs: 36.51725\nExp number, first 60 AAs: 19.7285\nTotal prob of N-in: 0.96170\nPOSSIBLE N-term signal sequence\ninside 1 19\nTMhelix 20 39\noutside 40 325\nTMhelix 326 348\ninside 349 372

24466 GCF_000486745.1_ASM48674v1 Salmonella enterica subsp. enterica serovar Pullorum str. 19945 Proteobacteria; Gammaproteobacteria; Enterobacterales; Enterobacteriaceae; Salmonella; Salmonella enterica; Salmonella enterica subsp. enterica; Salmonella enterica subsp. enterica serovar Pullorum MTGDNTLITSHGINRRDFMKLCAALAATMGLSSKAAAEMAESVSNPQRPPVIWIGAQECT WP_000145430.1 hydrogenase 2 small subunit [Salmonella enterica] Length: 372\nNumber of predicted TMHs: 1\nExp number of AAs in TMHs: 24.51306\nExp number, first 60 AAs: 0.2682\nTotal prob of N-in: 0.14402\noutside 1 331\nTMhelix 332 354\ninside 355 372

24467 GCF_000330485.2_ASM33048v2 Salmonella enterica subsp. enterica serovar Pullorum str. ATCC 9120 Proteobacteria; Gammaproteobacteria; Enterobacterales; Enterobacteriaceae; Salmonella; Salmonella enterica; Salmonella enterica subsp. enterica; Salmonella enterica subsp. enterica serovar Pullorum MNNEETFYQAMRRKGVTRRSFLKFCSLAATSLGLGAGMTPKIAWALENKPRIPVVWIHGL WP_001058309.1 hydrogenase [Salmonella enterica] Length: 372\nNumber of predicted TMHs: 2\nExp number of AAs in TMHs: 36.51725\nExp number, first 60 AAs: 19.7285\nTotal prob of N-in: 0.96170\nPOSSIBLE N-term signal sequence\ninside 1 19\nTMhelix 20 39\noutside 40 325\nTMhelix 326 348\ninside 349 372

24468 GCF_000330485.2_ASM33048v2 Salmonella enterica subsp. enterica serovar Pullorum str. ATCC 9120 Proteobacteria; Gammaproteobacteria; Enterobacterales; Enterobacteriaceae; Salmonella; Salmonella enterica; Salmonella enterica subsp. enterica; Salmonella enterica subsp. enterica serovar Pullorum MTGDNTLITSHGINRRDFMKLCAALAATMGLSSKAAAEMAESVSNPQRPPVIWIGAQECT WP_000145430.1 hydrogenase 2 small subunit [Salmonella enterica] Length: 372\nNumber of predicted TMHs: 1\nExp number of AAs in TMHs: 24.51306\nExp number, first 60 AAs: 0.2682\nTotal prob of N-in: 0.14402\noutside 1 331\nTMhelix 332 354\ninside 355 372

24469 GCF_002208315.1_ASM220831v1 Salmonella enterica subsp. enterica serovar Quebec str. S-1267 Proteobacteria; Gammaproteobacteria; Enterobacterales; Enterobacteriaceae; Salmonella; Salmonella enterica; Salmonella enterica subsp. enterica; Salmonella enterica subsp. enterica serovar Quebec MNNEETFYQAMRRKGVTRRSFLKFCSLAATSLGLGAGMTPKIAWALENKPRIPVVWIHGL WP_001058312.1 hydrogenase [Salmonella enterica] Length: 372\nNumber of predicted TMHs: 2\nExp number of AAs in TMHs: 36.51738\nExp number, first 60 AAs: 19.72849\nTotal prob of N-in: 0.96170\nPOSSIBLE N-term signal sequence\ninside 1 19\nTMhelix 20 39\noutside 40 325\nTMhelix 326 348\ninside 349 372

24470 GCF_002208315.1_ASM220831v1 Salmonella enterica subsp. enterica serovar Quebec str. S-1267 Proteobacteria; Gammaproteobacteria; Enterobacterales; Enterobacteriaceae; Salmonella; Salmonella enterica; Salmonella enterica subsp. enterica; Salmonella enterica subsp. enterica serovar Quebec MTGDNTLITSHGINRRDFMKLCAALAATMGLSSKAAAEMAESVSNPQRPPVIWIGAQECT WP_000145429.1 hydrogenase 2 small subunit [Salmonella enterica] Length: 372\nNumber of predicted TMHs: 1\nExp number of AAs in TMHs: 24.51309\nExp number, first 60 AAs: 0.2682\nTotal prob of N-in: 0.14402\noutside 1 331\nTMhelix 332 354\ninside 355 372

24471 GCF_000335835.1_ASM33583v1 Salmonella enterica subsp. enterica serovar Rissen str. 150 Proteobacteria; Gammaproteobacteria; Enterobacterales; Enterobacteriaceae; Salmonella; Salmonella enterica; Salmonella enterica subsp. enterica; Salmonella enterica subsp. enterica serovar Rissen MNNEETFYQAMRRKGVTRRSFLKFCSLAATSLGLSAGMTPKIAWALENKPRIPVVWIHGL WP_001058316.1 hydrogenase [Salmonella enterica] Length: 372\nNumber of predicted TMHs: 2\nExp number of AAs in TMHs: 35.76397\nExp number, first 60 AAs: 18.97201\nTotal prob of N-in: 0.92785\nPOSSIBLE N-term signal sequence\ninside 1 19\nTMhelix 20 39\noutside 40 325\nTMhelix 326 348\ninside 349 372

24472 GCF_000335835.1_ASM33583v1 Salmonella enterica subsp. enterica serovar Rissen str. 150 Proteobacteria; Gammaproteobacteria; Enterobacterales; Enterobacteriaceae; Salmonella; Salmonella enterica; Salmonella enterica subsp. enterica; Salmonella enterica subsp. enterica serovar Rissen MTGDNTLITSHGINRRDFMKLCAALAATMGLSSKAAAEMAESVSNPQRPPVIWIGAQECT WP_000145429.1 hydrogenase 2 small subunit [Salmonella enterica] Length: 372\nNumber of predicted TMHs: 1\nExp

number of AAs in TMHs: 24.51309\nExp number, first 60 AAs: 0.2682\nTotal prob of N-in: 0.14402\noutside 1 331\nTMhelix 332 354\ninside 355 372

24473 GCF_000486585.2_ASM48658v2 Salmonella enterica subsp. enterica serovar Rubislaw str. ATCC 10717 Proteobacteria; Gammaproteobacteria; Enterobacterales; Enterobacteriaceae; Salmonella; Salmonella enterica; Salmonella enterica subsp. enterica; Salmonella enterica subsp. enterica serovar Rubislaw MTGDNTLITSHGINRRDFMKLCAALAATMGLSSKAAAEMAESVSNPQRPPVIWIGAQECT WP_000145429.1 hydrogenase 2 small subunit [Salmonella enterica] Length: 372\nNumber of predicted TMHs: 1\nExp number of AAs in TMHs: 24.51309\nExp number, first 60 AAs: 0.2682\nTotal prob of N-in: 0.14402\noutside 1 331\nTMhelix 332 354\ninside 355 372

24474 GCF_000486585.2_ASM48658v2 Salmonella enterica subsp. enterica serovar Rubislaw str. ATCC 10717 Proteobacteria; Gammaproteobacteria; Enterobacterales; Enterobacteriaceae; Salmonella; Salmonella enterica; Salmonella enterica subsp. enterica; Salmonella enterica subsp. enterica serovar Rubislaw MNNEETFYQAMRRKGVTRRSFLKFCSLAATSLGLGAGMTPKIAWALENKPRIPVVWIHGL WP_001058312.1 hydrogenase [Salmonella enterica] Length: 372\nNumber of predicted TMHs: 2\nExp number of AAs in TMHs: 36.51738\nExp number, first 60 AAs: 19.72849\nTotal prob of N-in: 0.96170\nPOSSIBLE N-term signal sequence\ninside 1 19\nTMhelix 20 39\noutside 40 325\nTMhelix 326 348\ninside 349 372

24475 GCF_001448845.1_ASM144884v1 Salmonella enterica subsp. enterica serovar Rubislaw str. SA20030553 Proteobacteria; Gammaproteobacteria; Enterobacterales; Enterobacteriaceae; Salmonella; Salmonella enterica; Salmonella enterica subsp. enterica; Salmonella enterica subsp. enterica serovar Rubislaw MTGDNTLITSHGINRRDFMKLCAALAATMGLSSKAAAEMAESVSNPQRPPVIWIGAQECT WP_000145429.1 hydrogenase 2 small subunit [Salmonella enterica] Length: 372\nNumber of predicted TMHs: 1\nExp number of AAs in TMHs: 24.51309\nExp number, first 60 AAs: 0.2682\nTotal prob of N-in: 0.14402\noutside 1 331\nTMhelix 332 354\ninside 355 372

24476 GCF_000483975.1_ASM48397v1 Salmonella enterica subsp. enterica serovar Saintpaul str. 9712 Proteobacteria; Gammaproteobacteria; Enterobacterales; Enterobacteriaceae; Salmonella; Salmonella enterica; Salmonella enterica subsp. enterica; Salmonella enterica subsp. enterica serovar Saintpaul MTGDNTLITSHGINRRDFMKLCAALAATMGLSSKAAAEMAESVSNPQRPPVIWIGAQECT WP_000145429.1 hydrogenase 2 small subunit [Salmonella enterica] Length: 372\nNumber of predicted TMHs: 1\nExp number of AAs in TMHs: 24.51309\nExp number, first 60 AAs: 0.2682\nTotal prob of N-in: 0.14402\noutside 1 331\nTMhelix 332 354\ninside 355 372

24477 GCF_000483975.1_ASM48397v1 Salmonella enterica subsp. enterica serovar Saintpaul str. 9712 Proteobacteria; Gammaproteobacteria; Enterobacterales; Enterobacteriaceae; Salmonella; Salmonella enterica; Salmonella enterica subsp. enterica; Salmonella enterica subsp. enterica serovar Saintpaul MNNEETFYQAMRRKGVTRRSFLKFCSLAATSLGLGAGMTPKIAWALENKPRIPVVWIHGL WP_001058312.1 hydrogenase [Salmonella enterica] Length: 372\nNumber of predicted TMHs: 2\nExp number of AAs in TMHs: 36.51738\nExp number, first 60 AAs: 19.72849\nTotal prob of N-in: 0.96170\nPOSSIBLE N-term signal sequence\ninside 1 19\nTMhelix 20 39\noutside 40 325\nTMhelix 326 348\ninside 349 372

24478 GCF_000486545.1_ASM48654v1 Salmonella enterica subsp. enterica serovar Saintpaul str. JO2008 Proteobacteria; Gammaproteobacteria; Enterobacterales; Enterobacteriaceae; Salmonella; Salmonella enterica; Salmonella enterica subsp. enterica; Salmonella enterica subsp. enterica serovar Saintpaul MTGDNTLITSHGINRRDFMKLCAALAATMGLSSKAAAEMAESVSNPQRPPVIWIGAQECT WP_000145429.1 hydrogenase 2 small subunit [Salmonella enterica] Length: 372\nNumber of predicted TMHs: 1\nExp number of AAs in TMHs: 24.51309\nExp number, first 60 AAs: 0.2682\nTotal prob of N-in: 0.14402\noutside 1 331\nTMhelix 332 354\ninside 355 372

24479 GCF_000486545.1_ASM48654v1 Salmonella enterica subsp. enterica serovar Saintpaul str. JO2008 Proteobacteria; Gammaproteobacteria; Enterobacterales; Enterobacteriaceae; Salmonella; Salmonella enterica; Salmonella enterica subsp. enterica; Salmonella enterica subsp. enterica serovar Saintpaul MNNEETFYQAMRRKGVTRRSFLKFCSLAATSLGLGAGMTPKIAWALENKPRIPVVWIHGL WP_001058312.1 hydrogenase [Salmonella enterica] Length: 372\nNumber of predicted TMHs: 2\nExp number of AAs in TMHs: 36.51738\nExp number, first 60 AAs: 19.72849\nTotal prob of N-in: 0.96170\nPOSSIBLE N-term signal sequence\ninside 1 19\nTMhelix 20 39\noutside 40 325\nTMhelix 326 348\ninside 349 372

24480 GCF_000170215.1_ASM17021v1 Salmonella enterica subsp. enterica serovar Saintpaul str. SARA23 Proteobacteria; Gammaproteobacteria; Enterobacterales; Enterobacteriaceae; Salmonella; Salmonella enterica;

Salmonella enterica subsp. enterica; Salmonella enterica subsp. enterica serovar Saintpaul
MNNEETFYQAMRRKGVTRRSFLKFCSLAATSLGLGAGMTPKIAWALENKPRIPVVWIHGL WP_001058312.1
hydrogenase [Salmonella enterica] Length: 372\nNumber of predicted TMHs: 2\nExp number of AAs in TMHs: 36.51738\nExp number, first 60 AAs: 19.72849\nTotal prob of N-in: 0.96170\nPOSSIBLE N-term signal sequence\ninside 1 19\nTMhelix 20 39\noutside 40 325\nTMhelix 326 348\ninside 349 372

24481 GCF_000170215.1_ASM17021v1 Salmonella enterica subsp. enterica serovar Saintpaul str. SARA23
Proteobacteria; Gammaproteobacteria; Enterobacterales; Enterobacteriaceae; Salmonella; Salmonella enterica; Salmonella enterica subsp. enterica; Salmonella enterica subsp. enterica serovar Saintpaul
MTGDNTLITSHGINRRDFMKLCAALAATMGLSSKAAAEMAESVSNPQRPPVIWIGAQECT WP_000145429.1
hydrogenase 2 small subunit [Salmonella enterica] Length: 372\nNumber of predicted TMHs: 1\nExp number of AAs in TMHs: 24.51309\nExp number, first 60 AAs: 0.2682\nTotal prob of N-in: 0.14402\noutside 1 331\nTMhelix 332 354\ninside 355 372

24482 GCF_000486165.2_ASM48616v2 Salmonella enterica subsp. enterica serovar Saintpaul str. SARA26
Proteobacteria; Gammaproteobacteria; Enterobacterales; Enterobacteriaceae; Salmonella; Salmonella enterica; Salmonella enterica subsp. enterica; Salmonella enterica subsp. enterica serovar Saintpaul
MNNEETFYQAMRRKGVTRRSFLKFCSLAATSLGLGAGMTPKIAWALENKPRIPVVWIHGL WP_001058312.1
hydrogenase [Salmonella enterica] Length: 372\nNumber of predicted TMHs: 2\nExp number of AAs in TMHs: 36.51738\nExp number, first 60 AAs: 19.72849\nTotal prob of N-in: 0.96170\nPOSSIBLE N-term signal sequence\ninside 1 19\nTMhelix 20 39\noutside 40 325\nTMhelix 326 348\ninside 349 372

24483 GCF_000486165.2_ASM48616v2 Salmonella enterica subsp. enterica serovar Saintpaul str. SARA26
Proteobacteria; Gammaproteobacteria; Enterobacterales; Enterobacteriaceae; Salmonella; Salmonella enterica; Salmonella enterica subsp. enterica; Salmonella enterica subsp. enterica serovar Saintpaul
MTGDNTLITSHGINRRDFMKLCAALAATMGLSSKAAAEMAESVSNPQRPPVIWIGAQECT WP_000145429.1
hydrogenase 2 small subunit [Salmonella enterica] Length: 372\nNumber of predicted TMHs: 1\nExp number of AAs in TMHs: 24.51309\nExp number, first 60 AAs: 0.2682\nTotal prob of N-in: 0.14402\noutside 1 331\nTMhelix 332 354\ninside 355 372

24484 GCF_000170235.1_ASM17023v1 Salmonella enterica subsp. enterica serovar Saintpaul str. SARA29
Proteobacteria; Gammaproteobacteria; Enterobacterales; Enterobacteriaceae; Salmonella; Salmonella enterica; Salmonella enterica subsp. enterica; Salmonella enterica subsp. enterica serovar Saintpaul
MNNEETFYQAMRRKGVTRRSFLKFCSLAATSLGLGAGMTPKIAWALENKPRIPVVWIHGL WP_001058311.1
hydrogenase [Salmonella enterica] Length: 372\nNumber of predicted TMHs: 2\nExp number of AAs in TMHs: 37.4699899999999\nExp number, first 60 AAs: 19.72699\nTotal prob of N-in: 0.96171\nPOSSIBLE N-term signal sequence\ninside 1 19\nTMhelix 20 39\noutside 40 325\nTMhelix 326 348\ninside 349 372

24485 GCF_000170235.1_ASM17023v1 Salmonella enterica subsp. enterica serovar Saintpaul str. SARA29
Proteobacteria; Gammaproteobacteria; Enterobacterales; Enterobacteriaceae; Salmonella; Salmonella enterica; Salmonella enterica subsp. enterica; Salmonella enterica subsp. enterica serovar Saintpaul
MTGDNTLITSHGINRRDFMKLCAALAATMGLSSKAAAEMAESVSNPQRPPVIWIGAQECT WP_000145429.1
hydrogenase 2 small subunit [Salmonella enterica] Length: 372\nNumber of predicted TMHs: 1\nExp number of AAs in TMHs: 24.51309\nExp number, first 60 AAs: 0.2682\nTotal prob of N-in: 0.14402\noutside 1 331\nTMhelix 332 354\ninside 355 372

24486 GCF_000020745.1_ASM2074v1 Salmonella enterica subsp. enterica serovar Schwarzengrund str. CVM19633 Proteobacteria; Gammaproteobacteria; Enterobacterales; Enterobacteriaceae; Salmonella; Salmonella enterica; Salmonella enterica subsp. enterica; Salmonella enterica subsp. enterica serovar Schwarzengrund
MTGDNTLITSHGINRRDFMKLCAALAATMGLSSKAAAEMAESVSNPQRPPVIWIGAQECT WP_000145431.1
hydrogenase 2 small subunit [Salmonella enterica] Length: 372\nNumber of predicted TMHs: 1\nExp number of AAs in TMHs: 24.51505\nExp number, first 60 AAs: 0.2682\nTotal prob of N-in: 0.14406\noutside 1 331\nTMhelix 332 354\ninside 355 372

24487 GCF_000020745.1_ASM2074v1 Salmonella enterica subsp. enterica serovar Schwarzengrund str. CVM19633 Proteobacteria; Gammaproteobacteria; Enterobacterales; Enterobacteriaceae; Salmonella; Salmonella enterica; Salmonella enterica subsp. enterica; Salmonella enterica subsp. enterica serovar Schwarzengrund
MNNEETFYQAMRRKGVTRRSFLKFCSLAATSLGLGAGMTPKIAWALENKPRIPVVWIHGL WP_001058312.1
hydrogenase [Salmonella enterica] Length: 372\nNumber of predicted TMHs: 2\nExp number of AAs in TMHs: 36.51738\nExp number, first 60 AAs: 19.72849\nTotal prob of N-in: 0.96170\nPOSSIBLE N-term signal sequence\ninside 1 19\nTMhelix 20 39\noutside 40 325\nTMhelix 326 348\ninside 349 372

24488 GCF_000171295.1_ASM17129v1 Salmonella enterica subsp. enterica serovar Schwarzengrund str. SL480 Proteobacteria; Gammaproteobacteria; Enterobacterales; Enterobacteriaceae; Salmonella; Salmonella enterica; Salmonella enterica subsp. enterica; Salmonella enterica subsp. enterica serovar Schwarzengrund MTGDNTLITSHGINRRDFMKLCAALAATMGLSSKAAAEMAESVSNPQRPPVIWIGAQECT WP_000145431.1 hydrogenase 2 small subunit [Salmonella enterica] Length: 372\nNumber of predicted TMHs: 1\nExp number of AAs in TMHs: 24.51505\nExp number, first 60 AAs: 0.2682\nTotal prob of N-in: 0.14406\noutside 1 331\nTMhelix 332 354\ninside 355 372

24489 GCF_000171295.1_ASM17129v1 Salmonella enterica subsp. enterica serovar Schwarzengrund str. SL480 Proteobacteria; Gammaproteobacteria; Enterobacterales; Enterobacteriaceae; Salmonella; Salmonella enterica; Salmonella enterica subsp. enterica; Salmonella enterica subsp. enterica serovar Schwarzengrund MNNEETFYQAMRRKGVTRRSFLKFCSLAATSLGLGAGMTPKIAWALENKPRIPVVWIHGL WP_001058312.1 hydrogenase [Salmonella enterica] Length: 372\nNumber of predicted TMHs: 2\nExp number of AAs in TMHs: 36.51738\nExp number, first 60 AAs: 19.72849\nTotal prob of N-in: 0.96170\nPOSSIBLE N-term signal sequence\ninside 1 19\nTMhelix 20 39\noutside 40 325\nTMhelix 326 348\ninside 349 372

24490 GCF_000486975.1_ASM48697v1 Salmonella enterica subsp. enterica serovar Senftenberg str. 316235162 Proteobacteria; Gammaproteobacteria; Enterobacterales; Enterobacteriaceae; Salmonella; Salmonella enterica; Salmonella enterica subsp. enterica; Salmonella enterica subsp. enterica serovar Senftenberg MNNEETFYQAMRRKGVTRRSFLKFCSLAATSLGLGAGMTPKIAWALENKPRIPVVWIHGL WP_001058312.1 hydrogenase [Salmonella enterica] Length: 372\nNumber of predicted TMHs: 2\nExp number of AAs in TMHs: 36.51738\nExp number, first 60 AAs: 19.72849\nTotal prob of N-in: 0.96170\nPOSSIBLE N-term signal sequence\ninside 1 19\nTMhelix 20 39\noutside 40 325\nTMhelix 326 348\ninside 349 372

24491 GCF_000486975.1_ASM48697v1 Salmonella enterica subsp. enterica serovar Senftenberg str. 316235162 Proteobacteria; Gammaproteobacteria; Enterobacterales; Enterobacteriaceae; Salmonella; Salmonella enterica; Salmonella enterica subsp. enterica; Salmonella enterica subsp. enterica serovar Senftenberg MTGDNTLITSHGINRRDFMKLCAALAATMGLSSKAAAEMAESVSNPQRPPVIWIGAQECT WP_000145429.1 hydrogenase 2 small subunit [Salmonella enterica] Length: 372\nNumber of predicted TMHs: 1\nExp number of AAs in TMHs: 24.51309\nExp number, first 60 AAs: 0.2682\nTotal prob of N-in: 0.14402\noutside 1 331\nTMhelix 332 354\ninside 355 372

24492 GCF_000483995.1_ASM48399v1 Salmonella enterica subsp. enterica serovar Senftenberg str. 361154004 Proteobacteria; Gammaproteobacteria; Enterobacterales; Enterobacteriaceae; Salmonella; Salmonella enterica; Salmonella enterica subsp. enterica; Salmonella enterica subsp. enterica serovar Senftenberg MNNEETFYQAMRRKGVTRRSFLKFCSLAATSLGLGAGMTPKIAWALENKPRIPVVWIHGL WP_001058312.1 hydrogenase [Salmonella enterica] Length: 372\nNumber of predicted TMHs: 2\nExp number of AAs in TMHs: 36.51738\nExp number, first 60 AAs: 19.72849\nTotal prob of N-in: 0.96170\nPOSSIBLE N-term signal sequence\ninside 1 19\nTMhelix 20 39\noutside 40 325\nTMhelix 326 348\ninside 349 372

24493 GCF_000483995.1_ASM48399v1 Salmonella enterica subsp. enterica serovar Senftenberg str. 361154004 Proteobacteria; Gammaproteobacteria; Enterobacterales; Enterobacteriaceae; Salmonella; Salmonella enterica; Salmonella enterica subsp. enterica; Salmonella enterica subsp. enterica serovar Senftenberg MTGDNTLITSHGINRRDFMKLCAALAATMGLSSKAAAEMAESVSNPQRPPVIWIGAQECT WP_000145429.1 hydrogenase 2 small subunit [Salmonella enterica] Length: 372\nNumber of predicted TMHs: 1\nExp number of AAs in TMHs: 24.51309\nExp number, first 60 AAs: 0.2682\nTotal prob of N-in: 0.14402\noutside 1 331\nTMhelix 332 354\ninside 355 372

24494 GCF_000483915.1_ASM48391v1 Salmonella enterica subsp. enterica serovar Senftenberg str. 423984-1 Proteobacteria; Gammaproteobacteria; Enterobacterales; Enterobacteriaceae; Salmonella; Salmonella enterica; Salmonella enterica subsp. enterica; Salmonella enterica subsp. enterica serovar Senftenberg MTGDNTLITSHGINRRDFMKLCAALAATMGLSSKAAAEMAESVSNPQRPPVIWIGAQECT WP_000145429.1 hydrogenase 2 small subunit [Salmonella enterica] Length: 372\nNumber of predicted TMHs: 1\nExp number of AAs in TMHs: 24.51309\nExp number, first 60 AAs: 0.2682\nTotal prob of N-in: 0.14402\noutside 1 331\nTMhelix 332 354\ninside 355 372

24495 GCF_000483915.1_ASM48391v1 Salmonella enterica subsp. enterica serovar Senftenberg str. 423984-1 Proteobacteria; Gammaproteobacteria; Enterobacterales; Enterobacteriaceae; Salmonella; Salmonella enterica; Salmonella enterica subsp. enterica; Salmonella enterica subsp. enterica serovar Senftenberg MNNEETFYQAMRRKGVTRRSFLKFCSLAATSLGLGAGMTPKIAWALENKPRIPVVWIHGL WP_001058312.1 hydrogenase [Salmonella enterica] Length: 372\nNumber of predicted TMHs: 2\nExp number of AAs in

TMHs: 36.51738\nExp number, first 60 AAs: 19.72849\nTotal prob of N-in: 0.96170\nPOSSIBLE N-term signal sequence\ninside 1 19\nTMhelix 20 39\noutside 40 325\nTMhelix 326 348\ninside 349 372

24496 GCF_000486875.1_ASM48687v1 Salmonella enterica subsp. enterica serovar Senftenberg str. 423984-2 Proteobacteria; Gammaproteobacteria; Enterobacterales; Enterobacteriaceae; Salmonella; Salmonella enterica; Salmonella enterica subsp. enterica; Salmonella enterica subsp. enterica serovar Senftenberg MTGDNTLITSHGINRRDFMKLCAALAATMGLSSKAAAEMAESVSNPQRPPVIWIGAQECT WP_000145429.1 hydrogenase 2 small subunit [Salmonella enterica] Length: 372\nNumber of predicted TMHs: 1\nExp number of AAs in TMHs: 24.51309\nExp number, first 60 AAs: 0.2682\nTotal prob of N-in: 0.14402\noutside 1 331\nTMhelix 332 354\ninside 355 372

24497 GCF_000486875.1_ASM48687v1 Salmonella enterica subsp. enterica serovar Senftenberg str. 423984-2 Proteobacteria; Gammaproteobacteria; Enterobacterales; Enterobacteriaceae; Salmonella; Salmonella enterica; Salmonella enterica subsp. enterica; Salmonella enterica subsp. enterica serovar Senftenberg MNNEETFYQAMRRKGVTRRSFLKFCSLAATSLGLGAGMTPKIAWALENKPRIPVVWIHGL WP_001058312.1 hydrogenase [Salmonella enterica] Length: 372\nNumber of predicted TMHs: 2\nExp number of AAs in TMHs: 36.51738\nExp number, first 60 AAs: 19.72849\nTotal prob of N-in: 0.96170\nPOSSIBLE N-term signal sequence\ninside 1 19\nTMhelix 20 39\noutside 40 325\nTMhelix 326 348\ninside 349 372

24498 GCF_000486505.1_ASM48650v1 Salmonella enterica subsp. enterica serovar Senftenberg str. 604314 Proteobacteria; Gammaproteobacteria; Enterobacterales; Enterobacteriaceae; Salmonella; Salmonella enterica; Salmonella enterica subsp. enterica; Salmonella enterica subsp. enterica serovar Senftenberg MNNEETFYQAMRRKGVTRRSFLKFCSLAATSLGLGAGMTPKIAWALENKPRIPVVWIHGL WP_001058312.1 hydrogenase [Salmonella enterica] Length: 372\nNumber of predicted TMHs: 2\nExp number of AAs in TMHs: 36.51738\nExp number, first 60 AAs: 19.72849\nTotal prob of N-in: 0.96170\nPOSSIBLE N-term signal sequence\ninside 1 19\nTMhelix 20 39\noutside 40 325\nTMhelix 326 348\ninside 349 372

24499 GCF_000486505.1_ASM48650v1 Salmonella enterica subsp. enterica serovar Senftenberg str. 604314 Proteobacteria; Gammaproteobacteria; Enterobacterales; Enterobacteriaceae; Salmonella; Salmonella enterica; Salmonella enterica subsp. enterica; Salmonella enterica subsp. enterica serovar Senftenberg MTGDNTLITSHGINRRDFMKLCAALAATMGLSSKAAAEMAESVSNPQRPPVIWIGAQECT WP_000145429.1 hydrogenase 2 small subunit [Salmonella enterica] Length: 372\nNumber of predicted TMHs: 1\nExp number of AAs in TMHs: 24.51309\nExp number, first 60 AAs: 0.2682\nTotal prob of N-in: 0.14402\noutside 1 331\nTMhelix 332 354\ninside 355 372

24500 GCF_000486465.1_ASM48646v1 Salmonella enterica subsp. enterica serovar Senftenberg str. ATCC 8400 Proteobacteria; Gammaproteobacteria; Enterobacterales; Enterobacteriaceae; Salmonella; Salmonella enterica; Salmonella enterica subsp. enterica; Salmonella enterica subsp. enterica serovar Senftenberg MTGDNTLITSHGINRRDFMKLCAALAATMGLSSKAAAEMAESVSNPQRPPVIWIGAQECT WP_000145429.1 hydrogenase 2 small subunit [Salmonella enterica] Length: 372\nNumber of predicted TMHs: 1\nExp number of AAs in TMHs: 24.51309\nExp number, first 60 AAs: 0.2682\nTotal prob of N-in: 0.14402\noutside 1 331\nTMhelix 332 354\ninside 355 372

24501 GCF_000486465.1_ASM48646v1 Salmonella enterica subsp. enterica serovar Senftenberg str. ATCC 8400 Proteobacteria; Gammaproteobacteria; Enterobacterales; Enterobacteriaceae; Salmonella; Salmonella enterica; Salmonella enterica subsp. enterica; Salmonella enterica subsp. enterica serovar Senftenberg MNNEETFYQAMRRKGVTRRSFLKFCSLAATSLGLGAGMTPKIAWALENKPRIPVVWIHGL WP_001058311.1 hydrogenase [Salmonella enterica] Length: 372\nNumber of predicted TMHs: 2\nExp number of AAs in TMHs: 37.4699899999999\nExp number, first 60 AAs: 19.72699\nTotal prob of N-in: 0.96171\nPOSSIBLE N-term signal sequence\ninside 1 19\nTMhelix 20 39\noutside 40 325\nTMhelix 326 348\ninside 349 372

24502 GCF_000484435.1_ASM48443v1 Salmonella enterica subsp. enterica serovar Senftenberg str. NC_MB012510-0038 Proteobacteria; Gammaproteobacteria; Enterobacterales; Enterobacteriaceae; Salmonella; Salmonella enterica; Salmonella enterica subsp. enterica; Salmonella enterica subsp. enterica serovar Senftenberg MTGDNTLITSHGINRRDFMKLCAALAATMGLSSKAAAEMAESVSNPQRPPVIWIGAQECT WP_000145429.1 hydrogenase 2 small subunit [Salmonella enterica] Length: 372\nNumber of predicted TMHs: 1\nExp number of AAs in TMHs: 24.51309\nExp number, first 60 AAs: 0.2682\nTotal prob of N-in: 0.14402\noutside 1 331\nTMhelix 332 354\ninside 355 372

24503 GCF_000484435.1_ASM48443v1 Salmonella enterica subsp. enterica serovar Senftenberg str. NC_MB012510-0038 Proteobacteria; Gammaproteobacteria; Enterobacterales; Enterobacteriaceae; Salmonella;

Salmonella enterica; Salmonella enterica subsp. enterica; Salmonella enterica subsp. enterica serovar Senftenberg
MNNEETFYQAMRRKGVTRRSFLKFCSLAATSLGLGAGMTPKIAWALENKPRIPVVWIHGL WP_001058312.1
hydrogenase [Salmonella enterica] Length: 372\nNumber of predicted TMHs: 2\nExp number of AAs in TMHs: 36.51738\nExp number, first 60 AAs: 19.72849\nTotal prob of N-in: 0.96170\nPOSSIBLE N-term signal sequence\ninside 1 19\nTMhelix 20 39\noutside 40 325\nTMhelix 326 348\ninside 349 372

24504 GCF_000263295.1_SE-SS209 Salmonella enterica subsp. enterica serovar Senftenberg str. SS209
Proteobacteria; Gammaproteobacteria; Enterobacterales; Enterobacteriaceae; Salmonella; Salmonella enterica; Salmonella enterica subsp. enterica; Salmonella enterica subsp. enterica serovar Senftenberg
MNNEETFYQAMRRKGVTRRSFLKFCSLAATSLGLGAGMTPKIAWALENKPRIPVVWIHGL WP_001058312.1
hydrogenase [Salmonella enterica] Length: 372\nNumber of predicted TMHs: 2\nExp number of AAs in TMHs: 36.51738\nExp number, first 60 AAs: 19.72849\nTotal prob of N-in: 0.96170\nPOSSIBLE N-term signal sequence\ninside 1 19\nTMhelix 20 39\noutside 40 325\nTMhelix 326 348\ninside 349 372

24505 GCF_000263295.1_SE-SS209 Salmonella enterica subsp. enterica serovar Senftenberg str. SS209
Proteobacteria; Gammaproteobacteria; Enterobacterales; Enterobacteriaceae; Salmonella; Salmonella enterica; Salmonella enterica subsp. enterica; Salmonella enterica subsp. enterica serovar Senftenberg
MTGDNTLITSHGINRRDFMKLCAALAATMGLSSKAAAEMAESVSNPQRPPVIWIGAQECT WP_000145429.1
hydrogenase 2 small subunit [Salmonella enterica] Length: 372\nNumber of predicted TMHs: 1\nExp number of AAs in TMHs: 24.51309\nExp number, first 60 AAs: 0.2682\nTotal prob of N-in: 0.14402\noutside 1 331\nTMhelix 332 354\ninside 355 372

24506 GCF_000486445.2_ASM48644v2 Salmonella enterica subsp. enterica serovar Sloterdijk str. ATCC 15791
Proteobacteria; Gammaproteobacteria; Enterobacterales; Enterobacteriaceae; Salmonella; Salmonella enterica; Salmonella enterica subsp. enterica; Salmonella enterica subsp. enterica serovar Sloterdijk
MTGDNTLITSHGINRRDFMKLCAALAATMGLSSKAAAEMAESVSNPQRPPVIWIGAQECT WP_000145429.1
hydrogenase 2 small subunit [Salmonella enterica] Length: 372\nNumber of predicted TMHs: 1\nExp number of AAs in TMHs: 24.51309\nExp number, first 60 AAs: 0.2682\nTotal prob of N-in: 0.14402\noutside 1 331\nTMhelix 332 354\ninside 355 372

24507 GCF_000486445.2_ASM48644v2 Salmonella enterica subsp. enterica serovar Sloterdijk str. ATCC 15791
Proteobacteria; Gammaproteobacteria; Enterobacterales; Enterobacteriaceae; Salmonella; Salmonella enterica; Salmonella enterica subsp. enterica; Salmonella enterica subsp. enterica serovar Sloterdijk
MNNEETFYQAMRRKGVTRRSFLKFCSLAATSLGLGAGMTPKIAWALENKPRIPVVWIHGL WP_001058311.1
hydrogenase [Salmonella enterica] Length: 372\nNumber of predicted TMHs: 2\nExp number of AAs in TMHs: 37.4699899999999\nExp number, first 60 AAs: 19.72699\nTotal prob of N-in: 0.96171\nPOSSIBLE N-term signal sequence\ninside 1 19\nTMhelix 20 39\noutside 40 325\nTMhelix 326 348\ninside 349 372

24508 GCF_000486565.1_ASM48656v1 Salmonella enterica subsp. enterica serovar Soerenga str. 695
Proteobacteria; Gammaproteobacteria; Enterobacterales; Enterobacteriaceae; Salmonella; Salmonella enterica; Salmonella enterica subsp. enterica; Salmonella enterica subsp. enterica serovar Soerenga
MTGDNTLITSHGINRRDFMKLCAALAATMGLSSKAAAEMAESVSNPQRPPVIWIGAQECT WP_000145429.1
hydrogenase 2 small subunit [Salmonella enterica] Length: 372\nNumber of predicted TMHs: 1\nExp number of AAs in TMHs: 24.51309\nExp number, first 60 AAs: 0.2682\nTotal prob of N-in: 0.14402\noutside 1 331\nTMhelix 332 354\ninside 355 372

24509 GCF_000486565.1_ASM48656v1 Salmonella enterica subsp. enterica serovar Soerenga str. 695
Proteobacteria; Gammaproteobacteria; Enterobacterales; Enterobacteriaceae; Salmonella; Salmonella enterica; Salmonella enterica subsp. enterica; Salmonella enterica subsp. enterica serovar Soerenga
MNNRRGLIAVLATLITFSLSATPVLANPGNGNGGGHGNNAANQGNNGNGHKGNAGQKTEH WP_023206755.1
hypothetical protein [Salmonella enterica] Length: 159\nNumber of predicted TMHs: 2\nExp number of AAs in TMHs: 42.06129\nExp number, first 60 AAs: 20.61241\nTotal prob of N-in: 0.99707\nPOSSIBLE N-term signal sequence\ninside 1 6\nTMhelix 7 26\noutside 27 134\nTMhelix 135 157\ninside 158 159

24510 GCF_000486565.1_ASM48656v1 Salmonella enterica subsp. enterica serovar Soerenga str. 695
Proteobacteria; Gammaproteobacteria; Enterobacterales; Enterobacteriaceae; Salmonella; Salmonella enterica; Salmonella enterica subsp. enterica; Salmonella enterica subsp. enterica serovar Soerenga
MNNEETFYQAMRRKGVTRRSFLKFCSLAATSLGLSAGMTPKIAWALENKPRIPVVWIHGL WP_001058316.1
hydrogenase [Salmonella enterica] Length: 372\nNumber of predicted TMHs: 2\nExp number of AAs in TMHs: 35.76397\nExp number, first 60 AAs: 18.97201\nTotal prob of N-in: 0.92785\nPOSSIBLE N-term signal sequence\ninside 1 19\nTMhelix 20 39\noutside 40 325\nTMhelix 326 348\ninside 349 372

24511 GCF_000412835.1_06-0538_Ver1 Salmonella enterica subsp. enterica serovar Stanley str. 06-0538 Proteobacteria; Gammaproteobacteria; Enterobacterales; Enterobacteriaceae; Salmonella; Salmonella enterica; Salmonella enterica subsp. enterica; Salmonella enterica subsp. enterica serovar Stanley MTGDNTLITSHGINRRDFMKLCAALAATMGLSSKAAAEMAESVSNPQRPPVIWIGAQECT WP_000145429.1 hydrogenase 2 small subunit [Salmonella enterica] Length: 372\nNumber of predicted TMHs: 1\nExp number of AAs in TMHs: 24.51309\nExp number, first 60 AAs: 0.2682\nTotal prob of N-in: 0.14402\noutside 1 331\nTMhelix 332 354\ninside 355 372

24512 GCF_000412835.1_06-0538_Ver1 Salmonella enterica subsp. enterica serovar Stanley str. 06-0538 Proteobacteria; Gammaproteobacteria; Enterobacterales; Enterobacteriaceae; Salmonella; Salmonella enterica; Salmonella enterica subsp. enterica; Salmonella enterica subsp. enterica serovar Stanley MNNEETFYQAMRRKGVTRRSFLKFCSLAATSLGLGAGMTPKIAWALENKPRIPVVWIHGL WP_001058310.1 hydrogenase [Salmonella enterica] Length: 372\nNumber of predicted TMHs: 2\nExp number of AAs in TMHs: 37.4567299999999\nExp number, first 60 AAs: 19.72694\nTotal prob of N-in: 0.96171\nPOSSIBLE N-term signal sequence\ninside 1 19\nTMhelix 20 39\noutside 40 325\nTMhelix 326 348\ninside 349 372

24513 GCF_000486485.1_ASM48648v1 Salmonella enterica subsp. enterica serovar Stanley str. ATCC 7308 Proteobacteria; Gammaproteobacteria; Enterobacterales; Enterobacteriaceae; Salmonella; Salmonella enterica; Salmonella enterica subsp. enterica; Salmonella enterica subsp. enterica serovar Stanley MNNEETFYQAMRRKGVTRRSFLKFCSLAATSLGLGAGMTPKIAWALENKPRIPVVWIHGL WP_001058310.1 hydrogenase [Salmonella enterica] Length: 372\nNumber of predicted TMHs: 2\nExp number of AAs in TMHs: 37.4567299999999\nExp number, first 60 AAs: 19.72694\nTotal prob of N-in: 0.96171\nPOSSIBLE N-term signal sequence\ninside 1 19\nTMhelix 20 39\noutside 40 325\nTMhelix 326 348\ninside 349 372

24514 GCF_000486485.1_ASM48648v1 Salmonella enterica subsp. enterica serovar Stanley str. ATCC 7308 Proteobacteria; Gammaproteobacteria; Enterobacterales; Enterobacteriaceae; Salmonella; Salmonella enterica; Salmonella enterica subsp. enterica; Salmonella enterica subsp. enterica serovar Stanley MTGDNTLITSHGINRRDFMKLCAALAATMGLSSKAAAEMAESVSNPQRPPVIWIGAQECT WP_000145429.1 hydrogenase 2 small subunit [Salmonella enterica] Length: 372\nNumber of predicted TMHs: 1\nExp number of AAs in TMHs: 24.51309\nExp number, first 60 AAs: 0.2682\nTotal prob of N-in: 0.14402\noutside 1 331\nTMhelix 332 354\ninside 355 372

24515 GCF_000487935.2_ASM48793v2 Salmonella enterica subsp. enterica serovar Stanleyville str. CFSAN000624 Proteobacteria; Gammaproteobacteria; Enterobacterales; Enterobacteriaceae; Salmonella; Salmonella enterica; Salmonella enterica subsp. enterica; Salmonella enterica subsp. enterica serovar Stanleyville MNNEETFYQAMRRKGVTRRSFLKFCSLAATSLGLGAGMTPKIAWALENKPRIPVVWIHGL WP_001058312.1 hydrogenase [Salmonella enterica] Length: 372\nNumber of predicted TMHs: 2\nExp number of AAs in TMHs: 36.51738\nExp number, first 60 AAs: 19.72849\nTotal prob of N-in: 0.96170\nPOSSIBLE N-term signal sequence\ninside 1 19\nTMhelix 20 39\noutside 40 325\nTMhelix 326 348\ninside 349 372

24516 GCF_000487935.2_ASM48793v2 Salmonella enterica subsp. enterica serovar Stanleyville str. CFSAN000624 Proteobacteria; Gammaproteobacteria; Enterobacterales; Enterobacteriaceae; Salmonella; Salmonella enterica; Salmonella enterica subsp. enterica; Salmonella enterica subsp. enterica serovar Stanleyville MTGDNTLITSHGINRRDFMKLCAALAATMGLSSKAAAEMAESVSNPQRPPVIWIGAQECT WP_000145429.1 hydrogenase 2 small subunit [Salmonella enterica] Length: 372\nNumber of predicted TMHs: 1\nExp number of AAs in TMHs: 24.51309\nExp number, first 60 AAs: 0.2682\nTotal prob of N-in: 0.14402\noutside 1 331\nTMhelix 332 354\ninside 355 372

24517 GCF_001535365.1_ASM153536v1 Salmonella enterica subsp. enterica serovar Tennessee str. 2691 Proteobacteria; Gammaproteobacteria; Enterobacterales; Enterobacteriaceae; Salmonella; Salmonella enterica; Salmonella enterica subsp. enterica; Salmonella enterica subsp. enterica serovar Tennessee MNNEETFYQAMRRKGVTRRSFLKFCSLAATSLGLGAGMTPKIAWALENKPRIPVVWIHGL WP_001058312.1 hydrogenase [Salmonella enterica] Length: 372\nNumber of predicted TMHs: 2\nExp number of AAs in TMHs: 36.51738\nExp number, first 60 AAs: 19.72849\nTotal prob of N-in: 0.96170\nPOSSIBLE N-term signal sequence\ninside 1 19\nTMhelix 20 39\noutside 40 325\nTMhelix 326 348\ninside 349 372

24518 GCF_001535365.1_ASM153536v1 Salmonella enterica subsp. enterica serovar Tennessee str. 2691 Proteobacteria; Gammaproteobacteria; Enterobacterales; Enterobacteriaceae; Salmonella; Salmonella enterica; Salmonella enterica subsp. enterica; Salmonella enterica subsp. enterica serovar Tennessee MTGDNTLITSHGINRRDFMKLCAALAATMGLSSKAAAEMAESVSNPQRPPVIWIGAQECT WP_000145429.1 hydrogenase 2 small subunit [Salmonella enterica] Length: 372\nNumber of predicted TMHs: 1\nExp

number of AAs in TMHs: 24.51309\nExp number, first 60 AAs: 0.2682\nTotal prob of N-in: 0.14402\noutside 1 331\nTMhelix 332 354\ninside 355 372

24519 GCF_000336175.1_ASM33617v1 Salmonella enterica subsp. enterica serovar Tennessee str. 4535 Proteobacteria; Gammaproteobacteria; Enterobacterales; Enterobacteriaceae; Salmonella; Salmonella enterica; Salmonella enterica subsp. enterica; Salmonella enterica subsp. enterica serovar Tennessee MTGDNTLITSHGINRRDFMKLCAALAATMGLSSKAAAEMAESVSNPQRPPVIWIGAQECT WP_000145429.1 hydrogenase 2 small subunit [Salmonella enterica] Length: 372\nNumber of predicted TMHs: 1\nExp number of AAs in TMHs: 24.51309\nExp number, first 60 AAs: 0.2682\nTotal prob of N-in: 0.14402\noutside 1 331\nTMhelix 332 354\ninside 355 372

24520 GCF_000336175.1_ASM33617v1 Salmonella enterica subsp. enterica serovar Tennessee str. 4535 Proteobacteria; Gammaproteobacteria; Enterobacterales; Enterobacteriaceae; Salmonella; Salmonella enterica; Salmonella enterica subsp. enterica; Salmonella enterica subsp. enterica serovar Tennessee MNNEETFYQAMRRKGVTRRSFLKFCSLAATSLGLGAGMTPKIAWALENKPRIPVVWIHGL WP_001058312.1 hydrogenase [Salmonella enterica] Length: 372\nNumber of predicted TMHs: 2\nExp number of AAs in TMHs: 36.51738\nExp number, first 60 AAs: 19.72849\nTotal prob of N-in: 0.96170\nPOSSIBLE N-term signal sequence\ninside 1 19\nTMhelix 20 39\noutside 40 325\nTMhelix 326 348\ninside 349 372

24521 GCF_000353585.1_S._enterica_Tennessee_CDC07-0191 Salmonella enterica subsp. enterica serovar Tennessee str. CDC07-0191 Proteobacteria; Gammaproteobacteria; Enterobacterales; Enterobacteriaceae; Salmonella; Salmonella enterica; Salmonella enterica subsp. enterica; Salmonella enterica subsp. enterica serovar Tennessee MNNEETFYQAMRRKGVTRRSFLKFCSLAATSLGLGAGMTPKIAWALENKPRIPVVWIHGL WP_001058312.1 hydrogenase [Salmonella enterica] Length: 372\nNumber of predicted TMHs: 2\nExp number of AAs in TMHs: 36.51738\nExp number, first 60 AAs: 19.72849\nTotal prob of N-in: 0.96170\nPOSSIBLE N-term signal sequence\ninside 1 19\nTMhelix 20 39\noutside 40 325\nTMhelix 326 348\ninside 349 372

24522 GCF_000173695.1_ASM17369v1 Salmonella enterica subsp. enterica serovar Tennessee str. CDC07-0191 Proteobacteria; Gammaproteobacteria; Enterobacterales; Enterobacteriaceae; Salmonella; Salmonella enterica; Salmonella enterica subsp. enterica; Salmonella enterica subsp. enterica serovar Tennessee MTGDNTLITSHGINRRDFMKLCAALAATMGLSSKAAAEMAESVSNPQRPPVIWIGAQECT WP_000145429.1 hydrogenase 2 small subunit [Salmonella enterica] Length: 372\nNumber of predicted TMHs: 1\nExp number of AAs in TMHs: 24.51309\nExp number, first 60 AAs: 0.2682\nTotal prob of N-in: 0.14402\noutside 1 331\nTMhelix 332 354\ninside 355 372

24523 GCF_000173695.1_ASM17369v1 Salmonella enterica subsp. enterica serovar Tennessee str. CDC07-0191 Proteobacteria; Gammaproteobacteria; Enterobacterales; Enterobacteriaceae; Salmonella; Salmonella enterica; Salmonella enterica subsp. enterica; Salmonella enterica subsp. enterica serovar Tennessee MNNEETFYQAMRRKGVTRRSFLKFCSLAATSLGLGAGMTPKIAWALENKPRIPVVWIHGL WP_001058312.1 hydrogenase [Salmonella enterica] Length: 372\nNumber of predicted TMHs: 2\nExp number of AAs in TMHs: 36.51738\nExp number, first 60 AAs: 19.72849\nTotal prob of N-in: 0.96170\nPOSSIBLE N-term signal sequence\ninside 1 19\nTMhelix 20 39\noutside 40 325\nTMhelix 326 348\ninside 349 372

24524 GCF_000353585.1_S._enterica_Tennessee_CDC07-0191 Salmonella enterica subsp. enterica serovar Tennessee str. CDC07-0191 Proteobacteria; Gammaproteobacteria; Enterobacterales; Enterobacteriaceae; Salmonella; Salmonella enterica; Salmonella enterica subsp. enterica; Salmonella enterica subsp. enterica serovar Tennessee MTGDNTLITSHGINRRDFMKLCAALAATMGLSSKAAAEMAESVSNPQRPPVIWIGAQECT WP_000145429.1 hydrogenase 2 small subunit [Salmonella enterica] Length: 372\nNumber of predicted TMHs: 1\nExp number of AAs in TMHs: 24.51309\nExp number, first 60 AAs: 0.2682\nTotal prob of N-in: 0.14402\noutside 1 331\nTMhelix 332 354\ninside 355 372

24525 GCF_000486405.2_ASM48640v2 Salmonella enterica subsp. enterica serovar Tennessee str. TXSC_TXSC08-19 Proteobacteria; Gammaproteobacteria; Enterobacterales; Enterobacteriaceae; Salmonella; Salmonella enterica; Salmonella enterica subsp. enterica; Salmonella enterica subsp. enterica serovar Tennessee MNNEETFYQAMRRKGVTRRSFLKFCSLAATSLGLGAGMTPKIAWALENKPRIPVVWIHGL WP_001058312.1 hydrogenase [Salmonella enterica] Length: 372\nNumber of predicted TMHs: 2\nExp number of AAs in TMHs: 36.51738\nExp number, first 60 AAs: 19.72849\nTotal prob of N-in: 0.96170\nPOSSIBLE N-term signal sequence\ninside 1 19\nTMhelix 20 39\noutside 40 325\nTMhelix 326 348\ninside 349 372

24526 GCF_000486405.2_ASM48640v2 Salmonella enterica subsp. enterica serovar Tennessee str. TXSC_TXSC08-19 Proteobacteria; Gammaproteobacteria; Enterobacterales; Enterobacteriaceae; Salmonella; Salmonella enterica;

Salmonella enterica subsp. enterica; Salmonella enterica subsp. enterica serovar Tennessee
MTGDNTLITSHGINRRDFMKLCAALAATMGLSSKAAAEMAESVSNPQRPPVIWIGAQECT WP_000145429.1
hydrogenase 2 small subunit [Salmonella enterica] Length: 372\nNumber of predicted TMHs: 1\nExp number of AAs in TMHs: 24.51309\nExp number, first 60 AAs: 0.2682\nTotal prob of N-in: 0.14402\noutside 1 331\nTMhelix 332 354\ninside 355 372

24527 GCF_000315995.1_2009K-1863v1 Salmonella enterica subsp. enterica serovar Thompson str. 2010K-1863
Proteobacteria; Gammaproteobacteria; Enterobacterales; Enterobacteriaceae; Salmonella; Salmonella enterica; Salmonella enterica subsp. enterica; Salmonella enterica subsp. enterica serovar Thompson
MTGDNTLITSHGINRRDFMKLCAALAATMGLSSKAAAEMAESVSNPQRPPVIWIGAQECT WP_000145429.1
hydrogenase 2 small subunit [Salmonella enterica] Length: 372\nNumber of predicted TMHs: 1\nExp number of AAs in TMHs: 24.51309\nExp number, first 60 AAs: 0.2682\nTotal prob of N-in: 0.14402\noutside 1 331\nTMhelix 332 354\ninside 355 372

24528 GCF_000315995.1_2009K-1863v1 Salmonella enterica subsp. enterica serovar Thompson str. 2010K-1863
Proteobacteria; Gammaproteobacteria; Enterobacterales; Enterobacteriaceae; Salmonella; Salmonella enterica; Salmonella enterica subsp. enterica; Salmonella enterica subsp. enterica serovar Thompson
MNNEETFYQAMRRKGVTRRSFLKFCSLAATSLGLGAGMTPKIAWALENKPRIPVVWIHGL WP_001058310.1
hydrogenase [Salmonella enterica] Length: 372\nNumber of predicted TMHs: 2\nExp number of AAs in TMHs: 37.4567299999999\nExp number, first 60 AAs: 19.72694\nTotal prob of N-in: 0.96171\nPOSSIBLE N-term signal sequence\ninside 1 19\nTMhelix 20 39\noutside 40 325\nTMhelix 326 348\ninside 349 372

24529 GCF_000486365.2_ASM48636v2 Salmonella enterica subsp. enterica serovar Thompson str. ATCC 8391
Proteobacteria; Gammaproteobacteria; Enterobacterales; Enterobacteriaceae; Salmonella; Salmonella enterica; Salmonella enterica subsp. enterica; Salmonella enterica subsp. enterica serovar Thompson
MTGDNTLITSHGINRRDFMKLCAALAATMGLSSKAAAEMAESVSNPQRPPVIWIGAQECT WP_000145429.1
hydrogenase 2 small subunit [Salmonella enterica] Length: 372\nNumber of predicted TMHs: 1\nExp number of AAs in TMHs: 24.51309\nExp number, first 60 AAs: 0.2682\nTotal prob of N-in: 0.14402\noutside 1 331\nTMhelix 332 354\ninside 355 372

24530 GCF_000486365.2_ASM48636v2 Salmonella enterica subsp. enterica serovar Thompson str. ATCC 8391
Proteobacteria; Gammaproteobacteria; Enterobacterales; Enterobacteriaceae; Salmonella; Salmonella enterica; Salmonella enterica subsp. enterica; Salmonella enterica subsp. enterica serovar Thompson
MNNEETFYQAMRRKGVTRRSFLKFCSLAATSLGLGAGMTPKIAWALENKPRIPVVWIHGL WP_001058310.1
hydrogenase [Salmonella enterica] Length: 372\nNumber of predicted TMHs: 2\nExp number of AAs in TMHs: 37.4567299999999\nExp number, first 60 AAs: 19.72694\nTotal prob of N-in: 0.96171\nPOSSIBLE N-term signal sequence\ninside 1 19\nTMhelix 20 39\noutside 40 325\nTMhelix 326 348\ninside 349 372

24531 GCF_000473275.1_ASM47327v1 Salmonella enterica subsp. enterica serovar Thompson str. RM6836
Proteobacteria; Gammaproteobacteria; Enterobacterales; Enterobacteriaceae; Salmonella; Salmonella enterica; Salmonella enterica subsp. enterica; Salmonella enterica subsp. enterica serovar Thompson
MNNEETFYQAMRRKGVTRRSFLKFCSLAATSLGLGAGMTPKIAWALENKPRIPVVWIHGL WP_001058310.1
hydrogenase [Salmonella enterica] Length: 372\nNumber of predicted TMHs: 2\nExp number of AAs in TMHs: 37.4567299999999\nExp number, first 60 AAs: 19.72694\nTotal prob of N-in: 0.96171\nPOSSIBLE N-term signal sequence\ninside 1 19\nTMhelix 20 39\noutside 40 325\nTMhelix 326 348\ninside 349 372

24532 GCF_000473275.1_ASM47327v1 Salmonella enterica subsp. enterica serovar Thompson str. RM6836
Proteobacteria; Gammaproteobacteria; Enterobacterales; Enterobacteriaceae; Salmonella; Salmonella enterica; Salmonella enterica subsp. enterica; Salmonella enterica subsp. enterica serovar Thompson
MTGDNTLITSHGINRRDFMKLCAALAATMGLSSKAAAEMAESVSNPQRPPVIWIGAQECT WP_000145429.1
hydrogenase 2 small subunit [Salmonella enterica] Length: 372\nNumber of predicted TMHs: 1\nExp number of AAs in TMHs: 24.51309\nExp number, first 60 AAs: 0.2682\nTotal prob of N-in: 0.14402\noutside 1 331\nTMhelix 332 354\ninside 355 372

24533 GCF_000256015.1_SalTypBL196_1.0 Salmonella enterica subsp. enterica serovar Typhi str. BL196
Proteobacteria; Gammaproteobacteria; Enterobacterales; Enterobacteriaceae; Salmonella; Salmonella enterica; Salmonella enterica subsp. enterica; Salmonella enterica subsp. enterica serovar Typhi
MTGDNTLITSHGINRRDFMKLCAALAATMGLSSKAAAEMAESVSNPQRPPVIWIGAQECT WP_000145429.1
hydrogenase 2 small subunit [Salmonella enterica] Length: 372\nNumber of predicted TMHs: 1\nExp number of AAs in TMHs: 24.51309\nExp number, first 60 AAs: 0.2682\nTotal prob of N-in: 0.14402\noutside 1 331\nTMhelix 332 354\ninside 355 372

24534 GCF_000282875.1_SalTypSTCR0044_1.0 Salmonella enterica subsp. enterica serovar Typhi str. CR0044 Proteobacteria; Gammaproteobacteria; Enterobacterales; Enterobacteriaceae; Salmonella; Salmonella enterica; Salmonella enterica subsp. enterica; Salmonella enterica subsp. enterica serovar Typhi MTGDNTLITSHGINRRDFMKLCAALAATMGLSSKAAAEMAESVSNPQRPPVIWIGAQECT WP_000145429.1 hydrogenase 2 small subunit [Salmonella enterica] Length: 372\nNumber of predicted TMHs: 1\nExp number of AAs in TMHs: 24.51309\nExp number, first 60 AAs: 0.2682\nTotal prob of N-in: 0.14402\noutside 1 331\nTMhelix 332 354\ninside 355 372

24535 GCF_000282855.1_SalTypCR0063_1.0 Salmonella enterica subsp. enterica serovar Typhi str. CR0063 Proteobacteria; Gammaproteobacteria; Enterobacterales; Enterobacteriaceae; Salmonella; Salmonella enterica; Salmonella enterica subsp. enterica; Salmonella enterica subsp. enterica serovar Typhi MTGDNTLITSHGINRRDFMKLCAALAATMGLSSKAAAEMAESVSNPQRPPVIWIGAQECT WP_000145429.1 hydrogenase 2 small subunit [Salmonella enterica] Length: 372\nNumber of predicted TMHs: 1\nExp number of AAs in TMHs: 24.51309\nExp number, first 60 AAs: 0.2682\nTotal prob of N-in: 0.14402\noutside 1 331\nTMhelix 332 354\ninside 355 372

24536 GCF_000195995.1_ASM19599v1 Salmonella enterica subsp. enterica serovar Typhi str. CT18 Proteobacteria; Gammaproteobacteria; Enterobacterales; Enterobacteriaceae; Salmonella; Salmonella enterica; Salmonella enterica subsp. enterica; Salmonella enterica subsp. enterica serovar Typhi MTGDNTLITSHGINRRDFMKLCAALAATMGLSSKAAAEMAESVSNPQRPPVIWIGAQECT NP_457545.1 hydrogenase-2 small subunit [Salmonella enterica subsp. enterica serovar Typhi str. CT18] Length: 372\nNumber of predicted TMHs: 1\nExp number of AAs in TMHs: 24.51309\nExp number, first 60 AAs: 0.2682\nTotal prob of N-in: 0.14402\noutside 1 331\nTMhelix 332 354\ninside 355 372

24537 GCF_000245535.1_ASM24553v1 Salmonella enterica subsp. enterica serovar Typhi str. P-stx-12 Proteobacteria; Gammaproteobacteria; Enterobacterales; Enterobacteriaceae; Salmonella; Salmonella enterica; Salmonella enterica subsp. enterica; Salmonella enterica subsp. enterica serovar Typhi MTGDNTLITSHGINRRDFMKLCAALAATMGLSSKAAAEMAESVSNPQRPPVIWIGAQECT WP_000145429.1 hydrogenase 2 small subunit [Salmonella enterica] Length: 372\nNumber of predicted TMHs: 1\nExp number of AAs in TMHs: 24.51309\nExp number, first 60 AAs: 0.2682\nTotal prob of N-in: 0.14402\noutside 1 331\nTMhelix 332 354\ninside 355 372

24538 GCF_000267565.1_YKP860805.1 Salmonella enterica subsp. enterica serovar Typhi str. ST0208 Proteobacteria; Gammaproteobacteria; Enterobacterales; Enterobacteriaceae; Salmonella; Salmonella enterica; Salmonella enterica subsp. enterica; Salmonella enterica subsp. enterica serovar Typhi MTGDNTLITSHGINRRDFMKLCAALAATMGLSSKAAAEMAESVSNPQRPPVIWIGAQECT WP_000145429.1 hydrogenase 2 small subunit [Salmonella enterica] Length: 372\nNumber of predicted TMHs: 1\nExp number of AAs in TMHs: 24.51309\nExp number, first 60 AAs: 0.2682\nTotal prob of N-in: 0.14402\noutside 1 331\nTMhelix 332 354\ninside 355 372

24539 GCF_000523415.2_STH2370v1 Salmonella enterica subsp. enterica serovar Typhi str. STH2370 Proteobacteria; Gammaproteobacteria; Enterobacterales; Enterobacteriaceae; Salmonella; Salmonella enterica; Salmonella enterica subsp. enterica; Salmonella enterica subsp. enterica serovar Typhi MTGDNTLITSHGINRRDFMKLCAALAATMGLSSKAAAEMAESVSNPQRPPVIWIGAQECT WP_000145429.1 hydrogenase 2 small subunit [Salmonella enterica] Length: 372\nNumber of predicted TMHs: 1\nExp number of AAs in TMHs: 24.51309\nExp number, first 60 AAs: 0.2682\nTotal prob of N-in: 0.14402\noutside 1 331\nTMhelix 332 354\ninside 355 372

24540 GCF_000007545.1_ASM754v1 Salmonella enterica subsp. enterica serovar Typhi str. Ty2 Proteobacteria; Gammaproteobacteria; Enterobacterales; Enterobacteriaceae; Salmonella; Salmonella enterica; Salmonella enterica subsp. enterica; Salmonella enterica subsp. enterica serovar Typhi MTGDNTLITSHGINRRDFMKLCAALAATMGLSSKAAAEMAESVSNPQRPPVIWIGAQECT WP_000145429.1 hydrogenase 2 small subunit [Salmonella enterica] Length: 372\nNumber of predicted TMHs: 1\nExp number of AAs in TMHs: 24.51309\nExp number, first 60 AAs: 0.2682\nTotal prob of N-in: 0.14402\noutside 1 331\nTMhelix 332 354\ninside 355 372

24541 GCF_000385905.1_ASM38590v1 Salmonella enterica subsp. enterica serovar Typhi str. Ty21a Proteobacteria; Gammaproteobacteria; Enterobacterales; Enterobacteriaceae; Salmonella; Salmonella enterica; Salmonella enterica subsp. enterica; Salmonella enterica subsp. enterica serovar Typhi MTGDNTLITSHGINRRDFMKLCAALAATMGLSSKAAAEMAESVSNPQRPPVIWIGAQECT WP_000145429.1 hydrogenase 2 small subunit [Salmonella enterica] Length: 372\nNumber of predicted TMHs: 1\nExp

number of AAs in TMHs: 24.51309\nExp number, first 60 AAs: 0.2682\nTotal prob of N-in: 0.14402\noutside 1 331\nTMhelix 332 354\ninside 355 372

24542 GCF_000262635.1_SalTypUJ308A_1.0 Salmonella enterica subsp. enterica serovar Typhi str. UJ308A Proteobacteria; Gammaproteobacteria; Enterobacterales; Enterobacteriaceae; Salmonella; Salmonella enterica; Salmonella enterica subsp. enterica; Salmonella enterica subsp. enterica serovar Typhi MTGDNTLITSHGINRRDFMKLCAALAATMGLSSKAAAEMAESVSNPQRPPVIWIGAQECT WP_000145429.1 hydrogenase 2 small subunit [Salmonella enterica] Length: 372\nNumber of predicted TMHs: 1\nExp number of AAs in TMHs: 24.51309\nExp number, first 60 AAs: 0.2682\nTotal prob of N-in: 0.14402\noutside 1 331\nTMhelix 332 354\ninside 355 372

24543 GCF_000262615.1_SalTypUJ816A_1.0 Salmonella enterica subsp. enterica serovar Typhi str. UJ816A Proteobacteria; Gammaproteobacteria; Enterobacterales; Enterobacteriaceae; Salmonella; Salmonella enterica; Salmonella enterica subsp. enterica; Salmonella enterica subsp. enterica serovar Typhi MTGDNTLITSHGINRRDFMKLCAALAATMGLSSKAAAEMAESVSNPQRPPVIWIGAQECT WP_000145429.1 hydrogenase 2 small subunit [Salmonella enterica] Length: 372\nNumber of predicted TMHs: 1\nExp number of AAs in TMHs: 24.51309\nExp number, first 60 AAs: 0.2682\nTotal prob of N-in: 0.14402\noutside 1 331\nTMhelix 332 354\ninside 355 372

24544 GCF_000648475.2_104772_ver_1 Salmonella enterica subsp. enterica serovar Typhimurium str. 104772 Proteobacteria; Gammaproteobacteria; Enterobacterales; Enterobacteriaceae; Salmonella; Salmonella enterica; Salmonella enterica subsp. enterica; Salmonella enterica subsp. enterica serovar Typhimurium MTGDNTLITSHGINRRDFMKLCAALAATMGLSSKAAAEMAESVSNPQRPPVIWIGAQECT WP_000145429.1 hydrogenase 2 small subunit [Salmonella enterica] Length: 372\nNumber of predicted TMHs: 1\nExp number of AAs in TMHs: 24.51309\nExp number, first 60 AAs: 0.2682\nTotal prob of N-in: 0.14402\noutside 1 331\nTMhelix 332 354\ninside 355 372

24545 GCF_000648475.2_104772_ver_1 Salmonella enterica subsp. enterica serovar Typhimurium str. 104772 Proteobacteria; Gammaproteobacteria; Enterobacterales; Enterobacteriaceae; Salmonella; Salmonella enterica; Salmonella enterica subsp. enterica; Salmonella enterica subsp. enterica serovar Typhimurium MNNEETFYQAMRRKGVTRRSFLKFCSLAATSLGLGAGMTPKIAWALENKPRIPVVWIHGL WP_001058312.1 hydrogenase [Salmonella enterica] Length: 372\nNumber of predicted TMHs: 2\nExp number of AAs in TMHs: 36.51738\nExp number, first 60 AAs: 19.72849\nTotal prob of N-in: 0.96170\nPOSSIBLE N-term signal sequence\ninside 1 19\nTMhelix 20 39\noutside 40 325\nTMhelix 326 348\ninside 349 372

24546 GCF_000648495.2_108402_Ver_1 Salmonella enterica subsp. enterica serovar Typhimurium str. 108402 Proteobacteria; Gammaproteobacteria; Enterobacterales; Enterobacteriaceae; Salmonella; Salmonella enterica; Salmonella enterica subsp. enterica; Salmonella enterica subsp. enterica serovar Typhimurium MNNEETFYQAMRRKGVTRRSFLKFCSLAATSLGLGAGMTPKIAWALENKPRIPVVWIHGL WP_001058312.1 hydrogenase [Salmonella enterica] Length: 372\nNumber of predicted TMHs: 2\nExp number of AAs in TMHs: 36.51738\nExp number, first 60 AAs: 19.72849\nTotal prob of N-in: 0.96170\nPOSSIBLE N-term signal sequence\ninside 1 19\nTMhelix 20 39\noutside 40 325\nTMhelix 326 348\ninside 349 372

24547 GCF_000648495.2_108402_Ver_1 Salmonella enterica subsp. enterica serovar Typhimurium str. 108402 Proteobacteria; Gammaproteobacteria; Enterobacterales; Enterobacteriaceae; Salmonella; Salmonella enterica; Salmonella enterica subsp. enterica; Salmonella enterica subsp. enterica serovar Typhimurium MTGDNTLITSHGINRRDFMKLCAALAATMGLSSKAAAEMAESVSNPQRPPVIWIGAQECT WP_000145429.1 hydrogenase 2 small subunit [Salmonella enterica] Length: 372\nNumber of predicted TMHs: 1\nExp number of AAs in TMHs: 24.51309\nExp number, first 60 AAs: 0.2682\nTotal prob of N-in: 0.14402\noutside 1 331\nTMhelix 332 354\ninside 355 372

24548 GCF_000648455.2_116045_Ver_1 Salmonella enterica subsp. enterica serovar Typhimurium str. 116045 Proteobacteria; Gammaproteobacteria; Enterobacterales; Enterobacteriaceae; Salmonella; Salmonella enterica; Salmonella enterica subsp. enterica; Salmonella enterica subsp. enterica serovar Typhimurium MTGDNTLITSHGINRRDFMKLCAALAATMGLSSKAAAEMAESVSNPQRPPVIWIGAQECT WP_000145429.1 hydrogenase 2 small subunit [Salmonella enterica] Length: 372\nNumber of predicted TMHs: 1\nExp number of AAs in TMHs: 24.51309\nExp number, first 60 AAs: 0.2682\nTotal prob of N-in: 0.14402\noutside 1 331\nTMhelix 332 354\ninside 355 372

24549 GCF_000648455.2_116045_Ver_1 Salmonella enterica subsp. enterica serovar Typhimurium str. 116045 Proteobacteria; Gammaproteobacteria; Enterobacterales; Enterobacteriaceae; Salmonella; Salmonella enterica;

Salmonella enterica subsp. enterica; Salmonella enterica subsp. enterica serovar Typhimurium
MNNEETFYQAMRRKGVTRRSFLKFCSLAATSLGLGAGMTPKIAWALENKPRIPVVWIHGL WP_001058312.1
hydrogenase [Salmonella enterica] Length: 372\nNumber of predicted TMHs: 2\nExp number of AAs in TMHs: 36.51738\nExp number, first 60 AAs: 19.72849\nTotal prob of N-in: 0.96170\nPOSSIBLE N-term signal sequence\ninside 1 19\nTMhelix 20 39\noutside 40 325\nTMhelix 326 348\ninside 349 372

24550 GCF_000474555.1_CFSAN001198_01.0Salmonella enterica subsp. enterica serovar Typhimurium str. 34502
Proteobacteria; Gammaproteobacteria; Enterobacterales; Enterobacteriaceae; Salmonella; Salmonella enterica; Salmonella enterica subsp. enterica; Salmonella enterica subsp. enterica serovar Typhimurium
MTGDNTLITSHGINRRDFMKLCAALAATMGLSSKAAAEMAESVSNPQRPPVIWIGAQECT WP_000145429.1
hydrogenase 2 small subunit [Salmonella enterica] Length: 372\nNumber of predicted TMHs: 1\nExp number of AAs in TMHs: 24.51309\nExp number, first 60 AAs: 0.2682\nTotal prob of N-in: 0.14402\noutside 1 331\nTMhelix 332 354\ninside 355 372

24551 GCF_000474555.1_CFSAN001198_01.0Salmonella enterica subsp. enterica serovar Typhimurium str. 34502
Proteobacteria; Gammaproteobacteria; Enterobacterales; Enterobacteriaceae; Salmonella; Salmonella enterica; Salmonella enterica subsp. enterica; Salmonella enterica subsp. enterica serovar Typhimurium
MNNEETFYQAMRRKGVTRRSFLKFCSLAATSLGLGAGMTPKIAWALENKPRIPVVWIHGL WP_001058312.1
hydrogenase [Salmonella enterica] Length: 372\nNumber of predicted TMHs: 2\nExp number of AAs in TMHs: 36.51738\nExp number, first 60 AAs: 19.72849\nTotal prob of N-in: 0.96170\nPOSSIBLE N-term signal sequence\ninside 1 19\nTMhelix 20 39\noutside 40 325\nTMhelix 326 348\ninside 349 372

24552 GCF_000474395.1_CFSAN001199_01.0Salmonella enterica subsp. enterica serovar Typhimurium str. 35423
Proteobacteria; Gammaproteobacteria; Enterobacterales; Enterobacteriaceae; Salmonella; Salmonella enterica; Salmonella enterica subsp. enterica; Salmonella enterica subsp. enterica serovar Typhimurium
MTGDNTLITSHGINRRDFMKLCAALAATMGLSSKAAAEMAESVSNPQRPPVIWIGAQECT WP_000145429.1
hydrogenase 2 small subunit [Salmonella enterica] Length: 372\nNumber of predicted TMHs: 1\nExp number of AAs in TMHs: 24.51309\nExp number, first 60 AAs: 0.2682\nTotal prob of N-in: 0.14402\noutside 1 331\nTMhelix 332 354\ninside 355 372

24553 GCF_000474395.1_CFSAN001199_01.0Salmonella enterica subsp. enterica serovar Typhimurium str. 35423
Proteobacteria; Gammaproteobacteria; Enterobacterales; Enterobacteriaceae; Salmonella; Salmonella enterica; Salmonella enterica subsp. enterica; Salmonella enterica subsp. enterica serovar Typhimurium
MNNEETFYQAMRRKGVTRRSFLKFCSLAATSLGLGAGMTPKIAWALENKPRIPVVWIHGL WP_001058312.1
hydrogenase [Salmonella enterica] Length: 372\nNumber of predicted TMHs: 2\nExp number of AAs in TMHs: 36.51738\nExp number, first 60 AAs: 19.72849\nTotal prob of N-in: 0.96170\nPOSSIBLE N-term signal sequence\ninside 1 19\nTMhelix 20 39\noutside 40 325\nTMhelix 326 348\ninside 349 372

24554 GCF_000474575.1_CFSAN001200_01.0Salmonella enterica subsp. enterica serovar Typhimurium str. 36618
Proteobacteria; Gammaproteobacteria; Enterobacterales; Enterobacteriaceae; Salmonella; Salmonella enterica; Salmonella enterica subsp. enterica; Salmonella enterica subsp. enterica serovar Typhimurium
MNNEETFYQAMRRKGVTRRSFLKFCSLAATSLGLGAGMTPKIAWALENKPRIPVVWIHGL WP_001058312.1
hydrogenase [Salmonella enterica] Length: 372\nNumber of predicted TMHs: 2\nExp number of AAs in TMHs: 36.51738\nExp number, first 60 AAs: 19.72849\nTotal prob of N-in: 0.96170\nPOSSIBLE N-term signal sequence\ninside 1 19\nTMhelix 20 39\noutside 40 325\nTMhelix 326 348\ninside 349 372

24555 GCF_000474575.1_CFSAN001200_01.0Salmonella enterica subsp. enterica serovar Typhimurium str. 36618
Proteobacteria; Gammaproteobacteria; Enterobacterales; Enterobacteriaceae; Salmonella; Salmonella enterica; Salmonella enterica subsp. enterica; Salmonella enterica subsp. enterica serovar Typhimurium
MTGDNTLITSHGINRRDFMKLCAALAATMGLSSKAAAEMAESVSNPQRPPVIWIGAQECT WP_000145429.1
hydrogenase 2 small subunit [Salmonella enterica] Length: 372\nNumber of predicted TMHs: 1\nExp number of AAs in TMHs: 24.51309\nExp number, first 60 AAs: 0.2682\nTotal prob of N-in: 0.14402\noutside 1 331\nTMhelix 332 354\ninside 355 372

24556 GCF_000252875.1_ASM25287v1 Salmonella enterica subsp. enterica serovar Typhimurium str. 798
Proteobacteria; Gammaproteobacteria; Enterobacterales; Enterobacteriaceae; Salmonella; Salmonella enterica; Salmonella enterica subsp. enterica; Salmonella enterica subsp. enterica serovar Typhimurium
MNNEETFYQAMRRKGVTRRSFLKFCSLAATSLGLGAGMTPKIAWALENKPRIPVVWIHGL WP_001058312.1
hydrogenase [Salmonella enterica] Length: 372\nNumber of predicted TMHs: 2\nExp number of AAs in TMHs: 36.51738\nExp number, first 60 AAs: 19.72849\nTotal prob of N-in: 0.96170\nPOSSIBLE N-term signal sequence\ninside 1 19\nTMhelix 20 39\noutside 40 325\nTMhelix 326 348\ninside 349 372

24557 GCF_000252875.1_ASM25287v1 Salmonella enterica subsp. enterica serovar Typhimurium str. 798 Proteobacteria; Gammaproteobacteria; Enterobacterales; Enterobacteriaceae; Salmonella; Salmonella enterica; Salmonella enterica subsp. enterica; Salmonella enterica subsp. enterica serovar Typhimurium MTGDNTLITSHGINRRDFMKLCAALAATMGLSSKAAAEMAESVSNPQRPPVIWIGAQECT WP_000145429.1 hydrogenase 2 small subunit [Salmonella enterica] Length: 372\nNumber of predicted TMHs: 1\nExp number of AAs in TMHs: 24.51309\nExp number, first 60 AAs: 0.2682\nTotal prob of N-in: 0.14402\noutside 1 331\nTMhelix 332 354\ninside 355 372

24558 GCF_000786025.1_CFSAN001197_01.0Salmonella enterica subsp. enterica serovar Typhimurium str. 86-0368 Proteobacteria; Gammaproteobacteria; Enterobacterales; Enterobacteriaceae; Salmonella; Salmonella enterica; Salmonella enterica subsp. enterica; Salmonella enterica subsp. enterica serovar Typhimurium MNNEETFYQAMRRKGVTRRSFLKFCSLAATSLGLGAGMTPKIAWALENKPRIPVVWIHGL WP_001058312.1 hydrogenase [Salmonella enterica] Length: 372\nNumber of predicted TMHs: 2\nExp number of AAs in TMHs: 36.51738\nExp number, first 60 AAs: 19.72849\nTotal prob of N-in: 0.96170\nPOSSIBLE N-term signal sequence\ninside 1 19\nTMhelix 20 39\noutside 40 325\nTMhelix 326 348\ninside 349 372

24559 GCF_000786025.1_CFSAN001197_01.0Salmonella enterica subsp. enterica serovar Typhimurium str. 86-0368 Proteobacteria; Gammaproteobacteria; Enterobacterales; Enterobacteriaceae; Salmonella; Salmonella enterica; Salmonella enterica subsp. enterica; Salmonella enterica subsp. enterica serovar Typhimurium MTGDNTLITSHGINRRDFMKLCAALAATMGLSSKAAAEMAESVSNPQRPPVIWIGAQECT WP_000145429.1 hydrogenase 2 small subunit [Salmonella enterica] Length: 372\nNumber of predicted TMHs: 1\nExp number of AAs in TMHs: 24.51309\nExp number, first 60 AAs: 0.2682\nTotal prob of N-in: 0.14402\noutside 1 331\nTMhelix 332 354\ninside 355 372

24560 GCF_000648415.2_95799_Ver_1 Salmonella enterica subsp. enterica serovar Typhimurium str. 95799 Proteobacteria; Gammaproteobacteria; Enterobacterales; Enterobacteriaceae; Salmonella; Salmonella enterica; Salmonella enterica subsp. enterica; Salmonella enterica subsp. enterica serovar Typhimurium MTGDNTLITSHGINRRDFMKLCAALAATMGLSSKAAAEMAESVSNPQRPPVIWIGAQECT WP_000145429.1 hydrogenase 2 small subunit [Salmonella enterica] Length: 372\nNumber of predicted TMHs: 1\nExp number of AAs in TMHs: 24.51309\nExp number, first 60 AAs: 0.2682\nTotal prob of N-in: 0.14402\noutside 1 331\nTMhelix 332 354\ninside 355 372

24561 GCF_000648415.2_95799_Ver_1 Salmonella enterica subsp. enterica serovar Typhimurium str. 95799 Proteobacteria; Gammaproteobacteria; Enterobacterales; Enterobacteriaceae; Salmonella; Salmonella enterica; Salmonella enterica subsp. enterica; Salmonella enterica subsp. enterica serovar Typhimurium MNNEETFYQAMRRKGVTRRSFLKFCSLAATSLGLGAGMTPKIAWALENKPRIPVVWIHGL WP_001058312.1 hydrogenase [Salmonella enterica] Length: 372\nNumber of predicted TMHs: 2\nExp number of AAs in TMHs: 36.51738\nExp number, first 60 AAs: 19.72849\nTotal prob of N-in: 0.96170\nPOSSIBLE N-term signal sequence\ninside 1 19\nTMhelix 20 39\noutside 40 325\nTMhelix 326 348\ninside 349 372

24562 GCF_000648435.2_98346_ver1 Salmonella enterica subsp. enterica serovar Typhimurium str. 98346 Proteobacteria; Gammaproteobacteria; Enterobacterales; Enterobacteriaceae; Salmonella; Salmonella enterica; Salmonella enterica subsp. enterica; Salmonella enterica subsp. enterica serovar Typhimurium MTGDNTLITSHGINRRDFMKLCAALAATMGLSSKAAAEMAESVSNPQRPPVIWIGAQECT WP_000145429.1 hydrogenase 2 small subunit [Salmonella enterica] Length: 372\nNumber of predicted TMHs: 1\nExp number of AAs in TMHs: 24.51309\nExp number, first 60 AAs: 0.2682\nTotal prob of N-in: 0.14402\noutside 1 331\nTMhelix 332 354\ninside 355 372

24563 GCF_000648435.2_98346_ver1 Salmonella enterica subsp. enterica serovar Typhimurium str. 98346 Proteobacteria; Gammaproteobacteria; Enterobacterales; Enterobacteriaceae; Salmonella; Salmonella enterica; Salmonella enterica subsp. enterica; Salmonella enterica subsp. enterica serovar Typhimurium MNNEETFYQAMRRKGVTRRSFLKFCSLAATSLGLGAGMTPKIAWALENKPRIPVVWIHGL WP_001058312.1 hydrogenase [Salmonella enterica] Length: 372\nNumber of predicted TMHs: 2\nExp number of AAs in TMHs: 36.51738\nExp number, first 60 AAs: 19.72849\nTotal prob of N-in: 0.96170\nPOSSIBLE N-term signal sequence\ninside 1 19\nTMhelix 20 39\noutside 40 325\nTMhelix 326 348\ninside 349 372

24564 GCF_000486105.1_ASM48610v1 Salmonella enterica subsp. enterica serovar Typhimurium str. AZ 057 Proteobacteria; Gammaproteobacteria; Enterobacterales; Enterobacteriaceae; Salmonella; Salmonella enterica; Salmonella enterica subsp. enterica; Salmonella enterica subsp. enterica serovar Typhimurium MNNEETFYQAMRRKGVTRRSFLKFCSLAATSLGLGAGMTPKIAWALENKPRIPVVWIHGL WP_001058312.1 hydrogenase [Salmonella enterica] Length: 372\nNumber of predicted TMHs: 2\nExp number of AAs in

TMHs: 36.51738\nExp number, first 60 AAs: 19.72849\nTotal prob of N-in: 0.96170\nPOSSIBLE N-term signal sequence\ninside 1 19\nTMhelix 20 39\noutside 40 325\nTMhelix 326 348\ninside 349 372

24565 GCF_000486105.1_ASM48610v1 Salmonella enterica subsp. enterica serovar Typhimurium str. AZ 057 Proteobacteria; Gammaproteobacteria; Enterobacterales; Enterobacteriaceae; Salmonella; Salmonella enterica; Salmonella enterica subsp. enterica; Salmonella enterica subsp. enterica serovar Typhimurium MTGDNTLITSHGINRRDFMKLCAALAATMGLSSKAAAEMAESVSNPQRPPVIWIGAQECT WP_000145429.1 hydrogenase 2 small subunit [Salmonella enterica] Length: 372\nNumber of predicted TMHs: 1\nExp number of AAs in TMHs: 24.51309\nExp number, first 60 AAs: 0.2682\nTotal prob of N-in: 0.14402\noutside 1 331\nTMhelix 332 354\ninside 355 372

24566 GCF_000336155.1_ASM33615v1 Salmonella enterica subsp. enterica serovar Typhimurium str. CDC_2009K1153 Proteobacteria; Gammaproteobacteria; Enterobacterales; Enterobacteriaceae; Salmonella; Salmonella enterica; Salmonella enterica subsp. enterica; Salmonella enterica subsp. enterica serovar Typhimurium MTGDNTLITSHGINRRDFMKLCAALAATMGLSSKAAAEMAESVSNPQRPPVIWIGAQECT WP_000145429.1 hydrogenase 2 small subunit [Salmonella enterica] Length: 372\nNumber of predicted TMHs: 1\nExp number of AAs in TMHs: 24.51309\nExp number, first 60 AAs: 0.2682\nTotal prob of N-in: 0.14402\noutside 1 331\nTMhelix 332 354\ninside 355 372

24567 GCF_000336155.1_ASM33615v1 Salmonella enterica subsp. enterica serovar Typhimurium str. CDC_2009K1153 Proteobacteria; Gammaproteobacteria; Enterobacterales; Enterobacteriaceae; Salmonella; Salmonella enterica; Salmonella enterica subsp. enterica; Salmonella enterica subsp. enterica serovar Typhimurium MNNEETFYQAMRRKGVTRRSFLKFCSLAATSLGLGAGMTPKIAWALENKPRIPVVWIHGL WP_001058312.1 hydrogenase [Salmonella enterica] Length: 372\nNumber of predicted TMHs: 2\nExp number of AAs in TMHs: 36.51738\nExp number, first 60 AAs: 19.72849\nTotal prob of N-in: 0.96170\nPOSSIBLE N-term signal sequence\ninside 1 19\nTMhelix 20 39\noutside 40 325\nTMhelix 326 348\ninside 349 372

24568 GCF_000474475.1_CFSAN001184_01.0Salmonella enterica subsp. enterica serovar Typhimurium str. CDC_2009K1158 Proteobacteria; Gammaproteobacteria; Enterobacterales; Enterobacteriaceae; Salmonella; Salmonella enterica; Salmonella enterica subsp. enterica; Salmonella enterica subsp. enterica serovar Typhimurium MTGDNTLITSHGINRRDFMKLCAALAATMGLSSKAAAEMAESVSNPQRPPVIWIGAQECT WP_000145429.1 hydrogenase 2 small subunit [Salmonella enterica] Length: 372\nNumber of predicted TMHs: 1\nExp number of AAs in TMHs: 24.51309\nExp number, first 60 AAs: 0.2682\nTotal prob of N-in: 0.14402\noutside 1 331\nTMhelix 332 354\ninside 355 372

24569 GCF_000474475.1_CFSAN001184_01.0Salmonella enterica subsp. enterica serovar Typhimurium str. CDC_2009K1158 Proteobacteria; Gammaproteobacteria; Enterobacterales; Enterobacteriaceae; Salmonella; Salmonella enterica; Salmonella enterica subsp. enterica; Salmonella enterica subsp. enterica serovar Typhimurium MNNEETFYQAMRRKGVTRRSFLKFCSLAATSLGLGAGMTPKIAWALENKPRIPVVWIHGL WP_001058312.1 hydrogenase [Salmonella enterica] Length: 372\nNumber of predicted TMHs: 2\nExp number of AAs in TMHs: 36.51738\nExp number, first 60 AAs: 19.72849\nTotal prob of N-in: 0.96170\nPOSSIBLE N-term signal sequence\ninside 1 19\nTMhelix 20 39\noutside 40 325\nTMhelix 326 348\ninside 349 372

24570 GCF_000474495.1_CFSAN001187_01.0Salmonella enterica subsp. enterica serovar Typhimurium str. CDC_2009K1277 Proteobacteria; Gammaproteobacteria; Enterobacterales; Enterobacteriaceae; Salmonella; Salmonella enterica; Salmonella enterica subsp. enterica; Salmonella enterica subsp. enterica serovar Typhimurium MNNEETFYQAMRRKGVTRRSFLKFCSLAATSLGLGAGMTPKIAWALENKPRIPVVWIHGL WP_001058312.1 hydrogenase [Salmonella enterica] Length: 372\nNumber of predicted TMHs: 2\nExp number of AAs in TMHs: 36.51738\nExp number, first 60 AAs: 19.72849\nTotal prob of N-in: 0.96170\nPOSSIBLE N-term signal sequence\ninside 1 19\nTMhelix 20 39\noutside 40 325\nTMhelix 326 348\ninside 349 372

24571 GCF_000474495.1_CFSAN001187_01.0Salmonella enterica subsp. enterica serovar Typhimurium str. CDC_2009K1277 Proteobacteria; Gammaproteobacteria; Enterobacterales; Enterobacteriaceae; Salmonella; Salmonella enterica; Salmonella enterica subsp. enterica; Salmonella enterica subsp. enterica serovar Typhimurium MTGDNTLITSHGINRRDFMKLCAALAATMGLSSKAAAEMAESVSNPQRPPVIWIGAQECT WP_000145429.1 hydrogenase 2 small subunit [Salmonella enterica] Length: 372\nNumber of predicted TMHs: 1\nExp number of AAs in TMHs: 24.51309\nExp number, first 60 AAs: 0.2682\nTotal prob of N-in: 0.14402\noutside 1 331\nTMhelix 332 354\ninside 355 372

24572 GCF_000474355.1_CFSAN001186_01.0Salmonella enterica subsp. enterica serovar Typhimurium str. CDC_2009K1283 Proteobacteria; Gammaproteobacteria; Enterobacterales; Enterobacteriaceae; Salmonella; Salmonella

enterica; Salmonella enterica subsp. enterica; Salmonella enterica subsp. enterica serovar Typhimurium
MTGDNTLITSHGINRRDFMKLCAALAATMGLSSKAAAEMAESVSNPQRPPVIWIGAQECT WP_000145429.1
hydrogenase 2 small subunit [Salmonella enterica] Length: 372\nNumber of predicted TMHs: 1\nExp number of AAs in TMHs: 24.51309\nExp number, first 60 AAs: 0.2682\nTotal prob of N-in: 0.14402\noutside 1 331\nTMhelix 332 354\ninside 355 372

24573 GCF_000474355.1_CFSAN001186_01.0Salmonella enterica subsp. enterica serovar Typhimurium str. CDC_2009K1283 Proteobacteria; Gammaproteobacteria; Enterobacterales; Enterobacteriaceae; Salmonella; Salmonella enterica; Salmonella enterica subsp. enterica; Salmonella enterica subsp. enterica serovar Typhimurium
MNNEETFYQAMRRKGVTRRSFLKFCSLAATSLGLGAGMTPKIAWALENKPRIPVVWIHGL WP_001058312.1
hydrogenase [Salmonella enterica] Length: 372\nNumber of predicted TMHs: 2\nExp number of AAs in TMHs: 36.51738\nExp number, first 60 AAs: 19.72849\nTotal prob of N-in: 0.96170\nPOSSIBLE N-term signal sequence\ninside 1 19\nTMhelix 20 39\noutside 40 325\nTMhelix 326 348\ninside 349 372

24574 GCF_000474335.1_CFSAN001183_01.0Salmonella enterica subsp. enterica serovar Typhimurium str. CDC_2009K1288 Proteobacteria; Gammaproteobacteria; Enterobacterales; Enterobacteriaceae; Salmonella; Salmonella enterica; Salmonella enterica subsp. enterica; Salmonella enterica subsp. enterica serovar Typhimurium
MTGDNTLITSHGINRRDFMKLCAALAATMGLSSKAAAEMAESVSNPQRPPVIWIGAQECT WP_000145429.1
hydrogenase 2 small subunit [Salmonella enterica] Length: 372\nNumber of predicted TMHs: 1\nExp number of AAs in TMHs: 24.51309\nExp number, first 60 AAs: 0.2682\nTotal prob of N-in: 0.14402\noutside 1 331\nTMhelix 332 354\ninside 355 372

24575 GCF_000474335.1_CFSAN001183_01.0Salmonella enterica subsp. enterica serovar Typhimurium str. CDC_2009K1288 Proteobacteria; Gammaproteobacteria; Enterobacterales; Enterobacteriaceae; Salmonella; Salmonella enterica; Salmonella enterica subsp. enterica; Salmonella enterica subsp. enterica serovar Typhimurium
MNNEETFYQAMRRKGVTRRSFLKFCSLAATSLGLGAGMTPKIAWALENKPRIPVVWIHGL WP_001058312.1
hydrogenase [Salmonella enterica] Length: 372\nNumber of predicted TMHs: 2\nExp number of AAs in TMHs: 36.51738\nExp number, first 60 AAs: 19.72849\nTotal prob of N-in: 0.96170\nPOSSIBLE N-term signal sequence\ninside 1 19\nTMhelix 20 39\noutside 40 325\nTMhelix 326 348\ninside 349 372

24576 GCF_001623805.1_ASM162380v1 Salmonella enterica subsp. enterica serovar Typhimurium str. CDC 2009K-1640 Proteobacteria; Gammaproteobacteria; Enterobacterales; Enterobacteriaceae; Salmonella; Salmonella enterica; Salmonella enterica subsp. enterica; Salmonella enterica subsp. enterica serovar Typhimurium
MNNEETFYQAMRRKGVTRRSFLKFCSLAATSLGLGAGMTPKIAWALENKPRIPVVWIHGL WP_001058312.1
hydrogenase [Salmonella enterica] Length: 372\nNumber of predicted TMHs: 2\nExp number of AAs in TMHs: 36.51738\nExp number, first 60 AAs: 19.72849\nTotal prob of N-in: 0.96170\nPOSSIBLE N-term signal sequence\ninside 1 19\nTMhelix 20 39\noutside 40 325\nTMhelix 326 348\ninside 349 372

24577 GCF_001623805.1_ASM162380v1 Salmonella enterica subsp. enterica serovar Typhimurium str. CDC 2009K-1640 Proteobacteria; Gammaproteobacteria; Enterobacterales; Enterobacteriaceae; Salmonella; Salmonella enterica; Salmonella enterica subsp. enterica; Salmonella enterica subsp. enterica serovar Typhimurium
MTGDNTLITSHGINRRDFMKLCAALAATMGLSSKAAAEMAESVSNPQRPPVIWIGAQECT WP_000145429.1
hydrogenase 2 small subunit [Salmonella enterica] Length: 372\nNumber of predicted TMHs: 1\nExp number of AAs in TMHs: 24.51309\nExp number, first 60 AAs: 0.2682\nTotal prob of N-in: 0.14402\noutside 1 331\nTMhelix 332 354\ninside 355 372

24578 GCF_001623775.1_ASM162377v1 Salmonella enterica subsp. enterica serovar Typhimurium str. CDC 2009K-2059 Proteobacteria; Gammaproteobacteria; Enterobacterales; Enterobacteriaceae; Salmonella; Salmonella enterica; Salmonella enterica subsp. enterica; Salmonella enterica subsp. enterica serovar Typhimurium
MNNEETFYQAMRRKGVTRRSFLKFCSLAATSLGLGAGMTPKIAWALENKPRIPVVWIHGL WP_001058312.1
hydrogenase [Salmonella enterica] Length: 372\nNumber of predicted TMHs: 2\nExp number of AAs in TMHs: 36.51738\nExp number, first 60 AAs: 19.72849\nTotal prob of N-in: 0.96170\nPOSSIBLE N-term signal sequence\ninside 1 19\nTMhelix 20 39\noutside 40 325\nTMhelix 326 348\ninside 349 372

24579 GCF_001623775.1_ASM162377v1 Salmonella enterica subsp. enterica serovar Typhimurium str. CDC 2009K-2059 Proteobacteria; Gammaproteobacteria; Enterobacterales; Enterobacteriaceae; Salmonella; Salmonella enterica; Salmonella enterica subsp. enterica; Salmonella enterica subsp. enterica serovar Typhimurium
MTGDNTLITSHGINRRDFMKLCAALAATMGLSSKAAAEMAESVSNPQRPPVIWIGAQECT WP_000145429.1
hydrogenase 2 small subunit [Salmonella enterica] Length: 372\nNumber of predicted TMHs: 1\nExp number of AAs in TMHs: 24.51309\nExp number, first 60 AAs: 0.2682\nTotal prob of N-in: 0.14402\noutside 1 331\nTMhelix 332 354\ninside 355 372

24580	GCF_001623645.1_ASM162364v1	Salmonella enterica subsp. enterica serovar Typhimurium str. CDC 2010K-1587	Proteobacteria; Gammaproteobacteria; Enterobacterales; Enterobacteriaceae; Salmonella; Salmonella enterica; Salmonella enterica subsp. enterica; Salmonella enterica subsp. enterica serovar Typhimurium	MNNEETFYQAMRRKGVTRRSFLKFCSLAATSLGLGAGMTPKIAWALENKPRIPVVWIHGL	WP_001058312.1	hydrogenase [Salmonella enterica]	Length: 372\nNumber of predicted TMHs: 2\nExp number of AAs in TMHs: 36.51738\nExp number, first 60 AAs: 19.72849\nTotal prob of N-in: 0.96170\nPOSSIBLE N-term signal sequence\ninside 1 19\nTMhelix 20 39\noutside 40 325\nTMhelix 326 348\ninside 349 372

24581	GCF_001623645.1_ASM162364v1	Salmonella enterica subsp. enterica serovar Typhimurium str. CDC 2010K-1587	Proteobacteria; Gammaproteobacteria; Enterobacterales; Enterobacteriaceae; Salmonella; Salmonella enterica; Salmonella enterica subsp. enterica; Salmonella enterica subsp. enterica serovar Typhimurium	MTGDNTLITSHGINRRDFMKLCAALAATMGLSSKAAAEMAESVSNPQRPPVIWIGAQECT	WP_000145429.1	hydrogenase 2 small subunit [Salmonella enterica]	Length: 372\nNumber of predicted TMHs: 1\nExp number of AAs in TMHs: 24.51309\nExp number, first 60 AAs: 0.2682\nTotal prob of N-in: 0.14402\noutside 1 331\nTMhelix 332 354\ninside 355 372

24582	GCF_000973645.1_ASM97364v1	Salmonella enterica subsp. enterica serovar Typhimurium str. CDC 2011K-0870	Proteobacteria; Gammaproteobacteria; Enterobacterales; Enterobacteriaceae; Salmonella; Salmonella enterica; Salmonella enterica subsp. enterica; Salmonella enterica subsp. enterica serovar Typhimurium	MTGDNTLITSHGINRRDFMKLCAALAATMGLSSKAAAEMAESVSNPQRPPVIWIGAQECT	WP_000145429.1	hydrogenase 2 small subunit [Salmonella enterica]	Length: 372\nNumber of predicted TMHs: 1\nExp number of AAs in TMHs: 24.51309\nExp number, first 60 AAs: 0.2682\nTotal prob of N-in: 0.14402\noutside 1 331\nTMhelix 332 354\ninside 355 372

24583	GCF_000973645.1_ASM97364v1	Salmonella enterica subsp. enterica serovar Typhimurium str. CDC 2011K-0870	Proteobacteria; Gammaproteobacteria; Enterobacterales; Enterobacteriaceae; Salmonella; Salmonella enterica; Salmonella enterica subsp. enterica; Salmonella enterica subsp. enterica serovar Typhimurium	MNNEETFYQAMRRKGVTRRSFLKFCSLAATSLGLGAGMTPKIAWALENKPRIPVVWIHGL	WP_001058312.1	hydrogenase [Salmonella enterica]	Length: 372\nNumber of predicted TMHs: 2\nExp number of AAs in TMHs: 36.51738\nExp number, first 60 AAs: 19.72849\nTotal prob of N-in: 0.96170\nPOSSIBLE N-term signal sequence\ninside 1 19\nTMhelix 20 39\noutside 40 325\nTMhelix 326 348\ninside 349 372

24584	GCF_001623685.1_ASM162368v1	Salmonella enterica subsp. enterica serovar Typhimurium str. CDC 2011K-1702	Proteobacteria; Gammaproteobacteria; Enterobacterales; Enterobacteriaceae; Salmonella; Salmonella enterica; Salmonella enterica subsp. enterica; Salmonella enterica subsp. enterica serovar Typhimurium	MTGDNTLITSHGINRRDFMKLCAALAATMGLSSKAAAEMAESVSNPQRPPVIWIGAQECT	WP_000145429.1	hydrogenase 2 small subunit [Salmonella enterica]	Length: 372\nNumber of predicted TMHs: 1\nExp number of AAs in TMHs: 24.51309\nExp number, first 60 AAs: 0.2682\nTotal prob of N-in: 0.14402\noutside 1 331\nTMhelix 332 354\ninside 355 372

24585	GCF_001623685.1_ASM162368v1	Salmonella enterica subsp. enterica serovar Typhimurium str. CDC 2011K-1702	Proteobacteria; Gammaproteobacteria; Enterobacterales; Enterobacteriaceae; Salmonella; Salmonella enterica; Salmonella enterica subsp. enterica; Salmonella enterica subsp. enterica serovar Typhimurium	MNNEETFYQAMRRKGVTRRSFLKFCSLAATSLGLGAGMTPKIAWALENKPRIPVVWIHGL	WP_001058312.1	hydrogenase [Salmonella enterica]	Length: 372\nNumber of predicted TMHs: 2\nExp number of AAs in TMHs: 36.51738\nExp number, first 60 AAs: 19.72849\nTotal prob of N-in: 0.96170\nPOSSIBLE N-term signal sequence\ninside 1 19\nTMhelix 20 39\noutside 40 325\nTMhelix 326 348\ninside 349 372

24586	GCF_001623825.2_ASM162382v2	Salmonella enterica subsp. enterica serovar Typhimurium str. CDC H2662	Proteobacteria; Gammaproteobacteria; Enterobacterales; Enterobacteriaceae; Salmonella; Salmonella enterica; Salmonella enterica subsp. enterica; Salmonella enterica subsp. enterica serovar Typhimurium	MNNEETFYQAMRRKGVTRRSFLKFCSLAATSLGLGAGMTPKIAWALENKPRIPVVWIHGL	WP_001058312.1	hydrogenase [Salmonella enterica]	Length: 372\nNumber of predicted TMHs: 2\nExp number of AAs in TMHs: 36.51738\nExp number, first 60 AAs: 19.72849\nTotal prob of N-in: 0.96170\nPOSSIBLE N-term signal sequence\ninside 1 19\nTMhelix 20 39\noutside 40 325\nTMhelix 326 348\ninside 349 372

24587	GCF_001623825.2_ASM162382v2	Salmonella enterica subsp. enterica serovar Typhimurium str. CDC H2662	Proteobacteria; Gammaproteobacteria; Enterobacterales; Enterobacteriaceae; Salmonella; Salmonella enterica; Salmonella enterica subsp. enterica; Salmonella enterica subsp. enterica serovar Typhimurium	MTGDNTLITSHGINRRDFMKLCAALAATMGLSSKAAAEMAESVSNPQRPPVIWIGAQECT	WP_000145429.1	hydrogenase 2 small subunit [Salmonella enterica]	Length: 372\nNumber of predicted TMHs: 1\nExp

number of AAs in TMHs: 24.51309\nExp number, first 60 AAs: 0.2682\nTotal prob of N-in: 0.14402\noutside 1 331\nTMhelix 332 354\ninside 355 372

24588 GCF_000027025.1_ASM2702v1 Salmonella enterica subsp. enterica serovar Typhimurium str. D23580 Proteobacteria; Gammaproteobacteria; Enterobacterales; Enterobacteriaceae; Salmonella; Salmonella enterica; Salmonella enterica subsp. enterica; Salmonella enterica subsp. enterica serovar Typhimurium MTGDNTLITSHGINRRDFMKLCAALAATMGLSSKAAAEMAESVSNPQRPPVIWIGAQECT WP_000145429.1 hydrogenase 2 small subunit [Salmonella enterica] Length: 372\nNumber of predicted TMHs: 1\nExp number of AAs in TMHs: 24.51309\nExp number, first 60 AAs: 0.2682\nTotal prob of N-in: 0.14402\noutside 1 331\nTMhelix 332 354\ninside 355 372

24589 GCF_000027025.1_ASM2702v1 Salmonella enterica subsp. enterica serovar Typhimurium str. D23580 Proteobacteria; Gammaproteobacteria; Enterobacterales; Enterobacteriaceae; Salmonella; Salmonella enterica; Salmonella enterica subsp. enterica; Salmonella enterica subsp. enterica serovar Typhimurium MNNEETFYQAMRRKGVTRRSFLKFCSLAATSLGLGAGMTPKIAWALENKPRIPVVWIHGL WP_001058312.1 hydrogenase [Salmonella enterica] Length: 372\nNumber of predicted TMHs: 2\nExp number of AAs in TMHs: 36.51738\nExp number, first 60 AAs: 19.72849\nTotal prob of N-in: 0.96170\nPOSSIBLE N-term signal sequence\ninside 1 19\nTMhelix 20 39\noutside 40 325\nTMhelix 326 348\ninside 349 372

24590 GCF_001218505.1_5285_7_7Salmonella enterica subsp. enterica serovar Typhimurium str. DT104 Proteobacteria; Gammaproteobacteria; Enterobacterales; Enterobacteriaceae; Salmonella; Salmonella enterica; Salmonella enterica subsp. enterica; Salmonella enterica subsp. enterica serovar Typhimurium MTGDNTLITSHGINRRDFMKLCAALAATMGLSSKAAAEMAESVSNPQRPPVIWIGAQECT WP_000145429.1 hydrogenase 2 small subunit [Salmonella enterica] Length: 372\nNumber of predicted TMHs: 1\nExp number of AAs in TMHs: 24.51309\nExp number, first 60 AAs: 0.2682\nTotal prob of N-in: 0.14402\noutside 1 331\nTMhelix 332 354\ninside 355 372

24591 GCF_001218305.1_5299_6_3Salmonella enterica subsp. enterica serovar Typhimurium str. DT104 Proteobacteria; Gammaproteobacteria; Enterobacterales; Enterobacteriaceae; Salmonella; Salmonella enterica; Salmonella enterica subsp. enterica; Salmonella enterica subsp. enterica serovar Typhimurium MNNEETFYQAMRRKGVTRRSFLKFCSLAATSLGLGAGMTPKIAWALENKPRIPVVWIHGL WP_001058312.1 hydrogenase [Salmonella enterica] Length: 372\nNumber of predicted TMHs: 2\nExp number of AAs in TMHs: 36.51738\nExp number, first 60 AAs: 19.72849\nTotal prob of N-in: 0.96170\nPOSSIBLE N-term signal sequence\ninside 1 19\nTMhelix 20 39\noutside 40 325\nTMhelix 326 348\ninside 349 372

24592 GCF_001222225.1_5275_5_11 Salmonella enterica subsp. enterica serovar Typhimurium str. DT104 Proteobacteria; Gammaproteobacteria; Enterobacterales; Enterobacteriaceae; Salmonella; Salmonella enterica; Salmonella enterica subsp. enterica; Salmonella enterica subsp. enterica serovar Typhimurium MNNEETFYQAMRRKGVTRRSFLKFCSLAATSLGLGAGMTPKIAWALENKPRIPVVWIHGL WP_001058312.1 hydrogenase [Salmonella enterica] Length: 372\nNumber of predicted TMHs: 2\nExp number of AAs in TMHs: 36.51738\nExp number, first 60 AAs: 19.72849\nTotal prob of N-in: 0.96170\nPOSSIBLE N-term signal sequence\ninside 1 19\nTMhelix 20 39\noutside 40 325\nTMhelix 326 348\ninside 349 372

24593 GCF_001218265.1_8036_3_79 Salmonella enterica subsp. enterica serovar Typhimurium str. DT104 Proteobacteria; Gammaproteobacteria; Enterobacterales; Enterobacteriaceae; Salmonella; Salmonella enterica; Salmonella enterica subsp. enterica; Salmonella enterica subsp. enterica serovar Typhimurium MNNEETFYQAMRRKGVTRRSFLKFCSLAATSLGLGAGMTPKIAWALENKPRIPVVWIHGL WP_001058312.1 hydrogenase [Salmonella enterica] Length: 372\nNumber of predicted TMHs: 2\nExp number of AAs in TMHs: 36.51738\nExp number, first 60 AAs: 19.72849\nTotal prob of N-in: 0.96170\nPOSSIBLE N-term signal sequence\ninside 1 19\nTMhelix 20 39\noutside 40 325\nTMhelix 326 348\ninside 349 372

24594 GCF_001218105.1_5285_6_3Salmonella enterica subsp. enterica serovar Typhimurium str. DT104 Proteobacteria; Gammaproteobacteria; Enterobacterales; Enterobacteriaceae; Salmonella; Salmonella enterica; Salmonella enterica subsp. enterica; Salmonella enterica subsp. enterica serovar Typhimurium MNNEETFYQAMRRKGVTRRSFLKFCSLAATSLGLGAGMTPKIAWALENKPRIPVVWIHGL WP_001058312.1 hydrogenase [Salmonella enterica] Length: 372\nNumber of predicted TMHs: 2\nExp number of AAs in TMHs: 36.51738\nExp number, first 60 AAs: 19.72849\nTotal prob of N-in: 0.96170\nPOSSIBLE N-term signal sequence\ninside 1 19\nTMhelix 20 39\noutside 40 325\nTMhelix 326 348\ninside 349 372

24595 GCF_001218145.1_5295_2_12 Salmonella enterica subsp. enterica serovar Typhimurium str. DT104 Proteobacteria; Gammaproteobacteria; Enterobacterales; Enterobacteriaceae; Salmonella; Salmonella enterica;

Salmonella enterica subsp. enterica; Salmonella enterica subsp. enterica serovar Typhimurium
MTGDNTLITSHGINRRDFMKLCAALAATMGLSSKAAAEMAESVSNPQRPPVIWIGAQECT WP_000145429.1
hydrogenase 2 small subunit [Salmonella enterica] Length: 372\nNumber of predicted TMHs: 1\nExp number of AAs in TMHs: 24.51309\nExp number, first 60 AAs: 0.2682\nTotal prob of N-in: 0.14402\noutside 1 331\nTMhelix 332 354\ninside 355 372

24596 GCF_001218065.1_8080_3_2Salmonella enterica subsp. enterica serovar Typhimurium str. DT104
Proteobacteria; Gammaproteobacteria; Enterobacterales; Enterobacteriaceae; Salmonella; Salmonella enterica; Salmonella enterica subsp. enterica; Salmonella enterica subsp. enterica serovar Typhimurium
MTGDNTLITSHGINRRDFMKLCAALAATMGLSSKAAAEMAESVSNPQRPPVIWIGAQECT WP_000145429.1
hydrogenase 2 small subunit [Salmonella enterica] Length: 372\nNumber of predicted TMHs: 1\nExp number of AAs in TMHs: 24.51309\nExp number, first 60 AAs: 0.2682\nTotal prob of N-in: 0.14402\noutside 1 331\nTMhelix 332 354\ninside 355 372

24597 GCF_001217945.1_8036_3_37 Salmonella enterica subsp. enterica serovar Typhimurium str. DT104
Proteobacteria; Gammaproteobacteria; Enterobacterales; Enterobacteriaceae; Salmonella; Salmonella enterica; Salmonella enterica subsp. enterica; Salmonella enterica subsp. enterica serovar Typhimurium
MNNEETFYQAMRRKGVTRRSFLKFCSLAATSLGLGAGMTPKIAWALENKPRIPVVWIHGL WP_001058312.1
hydrogenase [Salmonella enterica] Length: 372\nNumber of predicted TMHs: 2\nExp number of AAs in TMHs: 36.51738\nExp number, first 60 AAs: 19.72849\nTotal prob of N-in: 0.96170\nPOSSIBLE N-term signal sequence\ninside 1 19\nTMhelix 20 39\noutside 40 325\nTMhelix 326 348\ninside 349 372

24598 GCF_001217905.1_5275_7_3Salmonella enterica subsp. enterica serovar Typhimurium str. DT104
Proteobacteria; Gammaproteobacteria; Enterobacterales; Enterobacteriaceae; Salmonella; Salmonella enterica; Salmonella enterica subsp. enterica; Salmonella enterica subsp. enterica serovar Typhimurium
MNNEETFYQAMRRKGVTRRSFLKFCSLAATSLGLGAGMTPKIAWALENKPRIPVVWIHGL WP_001058312.1
hydrogenase [Salmonella enterica] Length: 372\nNumber of predicted TMHs: 2\nExp number of AAs in TMHs: 36.51738\nExp number, first 60 AAs: 19.72849\nTotal prob of N-in: 0.96170\nPOSSIBLE N-term signal sequence\ninside 1 19\nTMhelix 20 39\noutside 40 325\nTMhelix 326 348\ninside 349 372

24599 GCF_001217865.1_8080_3_17 Salmonella enterica subsp. enterica serovar Typhimurium str. DT104
Proteobacteria; Gammaproteobacteria; Enterobacterales; Enterobacteriaceae; Salmonella; Salmonella enterica; Salmonella enterica subsp. enterica; Salmonella enterica subsp. enterica serovar Typhimurium
MTGDNTLITSHGINRRDFMKLCAALAATMGLSSKAAAEMAESVSNPQRPPVIWIGAQECT WP_000145429.1
hydrogenase 2 small subunit [Salmonella enterica] Length: 372\nNumber of predicted TMHs: 1\nExp number of AAs in TMHs: 24.51309\nExp number, first 60 AAs: 0.2682\nTotal prob of N-in: 0.14402\noutside 1 331\nTMhelix 332 354\ninside 355 372

24600 GCF_001217385.1_5285_7_11 Salmonella enterica subsp. enterica serovar Typhimurium str. DT104
Proteobacteria; Gammaproteobacteria; Enterobacterales; Enterobacteriaceae; Salmonella; Salmonella enterica; Salmonella enterica subsp. enterica; Salmonella enterica subsp. enterica serovar Typhimurium
MNNEETFYQAMRRKGVTRRSFLKFCSLAATSLGLGAGMTPKIAWALENKPRIPVVWIHGL WP_001058312.1
hydrogenase [Salmonella enterica] Length: 372\nNumber of predicted TMHs: 2\nExp number of AAs in TMHs: 36.51738\nExp number, first 60 AAs: 19.72849\nTotal prob of N-in: 0.96170\nPOSSIBLE N-term signal sequence\ninside 1 19\nTMhelix 20 39\noutside 40 325\nTMhelix 326 348\ninside 349 372

24601 GCF_001222185.1_5285_7_1Salmonella enterica subsp. enterica serovar Typhimurium str. DT104
Proteobacteria; Gammaproteobacteria; Enterobacterales; Enterobacteriaceae; Salmonella; Salmonella enterica; Salmonella enterica subsp. enterica; Salmonella enterica subsp. enterica serovar Typhimurium
MNNEETFYQAMRRKGVTRRSFLKFCSLAATSLGLGAGMTPKIAWALENKPRIPVVWIHGL WP_001058312.1
hydrogenase [Salmonella enterica] Length: 372\nNumber of predicted TMHs: 2\nExp number of AAs in TMHs: 36.51738\nExp number, first 60 AAs: 19.72849\nTotal prob of N-in: 0.96170\nPOSSIBLE N-term signal sequence\ninside 1 19\nTMhelix 20 39\noutside 40 325\nTMhelix 326 348\ninside 349 372

24602 GCF_001222345.1_5285_6_5Salmonella enterica subsp. enterica serovar Typhimurium str. DT104
Proteobacteria; Gammaproteobacteria; Enterobacterales; Enterobacteriaceae; Salmonella; Salmonella enterica; Salmonella enterica subsp. enterica; Salmonella enterica subsp. enterica serovar Typhimurium
MNNEETFYQAMRRKGVTRRSFLKFCSLAATSLGLGAGMTPKIAWALENKPRIPVVWIHGL WP_001058312.1
hydrogenase [Salmonella enterica] Length: 372\nNumber of predicted TMHs: 2\nExp number of AAs in TMHs: 36.51738\nExp number, first 60 AAs: 19.72849\nTotal prob of N-in: 0.96170\nPOSSIBLE N-term signal sequence\ninside 1 19\nTMhelix 20 39\noutside 40 325\nTMhelix 326 348\ninside 349 372

24603 GCF_001222005.1_8036_3_63 Salmonella enterica subsp. enterica serovar Typhimurium str. DT104 Proteobacteria; Gammaproteobacteria; Enterobacterales; Enterobacteriaceae; Salmonella; Salmonella enterica; Salmonella enterica subsp. enterica; Salmonella enterica subsp. enterica serovar Typhimurium MNNEETFYQAMRRKGVTRRSFLKFCSLAATSLGLGAGMTPKIAWALENKPRIPVVWIHGL WP_001058312.1 hydrogenase [Salmonella enterica] Length: 372\nNumber of predicted TMHs: 2\nExp number of AAs in TMHs: 36.51738\nExp number, first 60 AAs: 19.72849\nTotal prob of N-in: 0.96170\nPOSSIBLE N-term signal sequence\ninside 1 19\nTMhelix 20 39\noutside 40 325\nTMhelix 326 348\ninside 349 372

24604 GCF_001222385.1_5295_2_6Salmonella enterica subsp. enterica serovar Typhimurium str. DT104 Proteobacteria; Gammaproteobacteria; Enterobacterales; Enterobacteriaceae; Salmonella; Salmonella enterica; Salmonella enterica subsp. enterica; Salmonella enterica subsp. enterica serovar Typhimurium MTGDNTLITSHGINRRDFMKLCAALAATMGLSSKAAAEMAESVSNPQRPPVIWIGAQECT WP_000145429.1 hydrogenase 2 small subunit [Salmonella enterica] Length: 372\nNumber of predicted TMHs: 1\nExp number of AAs in TMHs: 24.51309\nExp number, first 60 AAs: 0.2682\nTotal prob of N-in: 0.14402\noutside 1 331\nTMhelix 332 354\ninside 355 372

24605 GCF_001217825.1_5275_2_4Salmonella enterica subsp. enterica serovar Typhimurium str. DT104 Proteobacteria; Gammaproteobacteria; Enterobacterales; Enterobacteriaceae; Salmonella; Salmonella enterica; Salmonella enterica subsp. enterica; Salmonella enterica subsp. enterica serovar Typhimurium MTGDNTLITSHGINRRDFMKLCAALAATMGLSSKAAAEMAESVSNPQRPPVIWIGAQECT WP_000145429.1 hydrogenase 2 small subunit [Salmonella enterica] Length: 372\nNumber of predicted TMHs: 1\nExp number of AAs in TMHs: 24.51309\nExp number, first 60 AAs: 0.2682\nTotal prob of N-in: 0.14402\noutside 1 331\nTMhelix 332 354\ninside 355 372

24606 GCF_001095425.1_8490_2_87 Salmonella enterica subsp. enterica serovar Typhimurium str. DT104 Proteobacteria; Gammaproteobacteria; Enterobacterales; Enterobacteriaceae; Salmonella; Salmonella enterica; Salmonella enterica subsp. enterica; Salmonella enterica subsp. enterica serovar Typhimurium MTGDNTLITSHGINRRDFMKLCAALAATMGLSSKAAAEMAESVSNPQRPPVIWIGAQECT WP_000145429.1 hydrogenase 2 small subunit [Salmonella enterica] Length: 372\nNumber of predicted TMHs: 1\nExp number of AAs in TMHs: 24.51309\nExp number, first 60 AAs: 0.2682\nTotal prob of N-in: 0.14402\noutside 1 331\nTMhelix 332 354\ninside 355 372

24607 GCF_001137885.1_8490_2_64 Salmonella enterica subsp. enterica serovar Typhimurium str. DT104 Proteobacteria; Gammaproteobacteria; Enterobacterales; Enterobacteriaceae; Salmonella; Salmonella enterica; Salmonella enterica subsp. enterica; Salmonella enterica subsp. enterica serovar Typhimurium MNNEETFYQAMRRKGVTRRSFLKFCSLAATSLGLGAGMTPKIAWALENKPRIPVVWIHGL WP_001058312.1 hydrogenase [Salmonella enterica] Length: 372\nNumber of predicted TMHs: 2\nExp number of AAs in TMHs: 36.51738\nExp number, first 60 AAs: 19.72849\nTotal prob of N-in: 0.96170\nPOSSIBLE N-term signal sequence\ninside 1 19\nTMhelix 20 39\noutside 40 325\nTMhelix 326 348\ninside 349 372

24608 GCF_001222505.1_5388_1_8Salmonella enterica subsp. enterica serovar Typhimurium str. DT104 Proteobacteria; Gammaproteobacteria; Enterobacterales; Enterobacteriaceae; Salmonella; Salmonella enterica; Salmonella enterica subsp. enterica; Salmonella enterica subsp. enterica serovar Typhimurium MNNEETFYQAMRRKGVTRRSFLKFCSLAATSLGLGAGMTPKIAWALENKPRIPVVWIHGL WP_001058312.1 hydrogenase [Salmonella enterica] Length: 372\nNumber of predicted TMHs: 2\nExp number of AAs in TMHs: 36.51738\nExp number, first 60 AAs: 19.72849\nTotal prob of N-in: 0.96170\nPOSSIBLE N-term signal sequence\ninside 1 19\nTMhelix 20 39\noutside 40 325\nTMhelix 326 348\ninside 349 372

24609 GCF_001140725.1_8490_2_51 Salmonella enterica subsp. enterica serovar Typhimurium str. DT104 Proteobacteria; Gammaproteobacteria; Enterobacterales; Enterobacteriaceae; Salmonella; Salmonella enterica; Salmonella enterica subsp. enterica; Salmonella enterica subsp. enterica serovar Typhimurium MTGDNTLITSHGINRRDFMKLCAALAATMGLSSKAAAEMAESVSNPQRPPVIWIGAQECT WP_000145429.1 hydrogenase 2 small subunit [Salmonella enterica] Length: 372\nNumber of predicted TMHs: 1\nExp number of AAs in TMHs: 24.51309\nExp number, first 60 AAs: 0.2682\nTotal prob of N-in: 0.14402\noutside 1 331\nTMhelix 332 354\ninside 355 372

24610 GCF_001222585.1_8080_3_18 Salmonella enterica subsp. enterica serovar Typhimurium str. DT104 Proteobacteria; Gammaproteobacteria; Enterobacterales; Enterobacteriaceae; Salmonella; Salmonella enterica; Salmonella enterica subsp. enterica; Salmonella enterica subsp. enterica serovar Typhimurium MNNEETFYQAMRRKGVTRRSFLKFCSLAATSLGLGAGMTPKIAWALENKPRIPVVWIHGL WP_001058312.1 hydrogenase [Salmonella enterica] Length: 372\nNumber of predicted TMHs: 2\nExp number of AAs in

TMHs: 36.51738\nExp number, first 60 AAs: 19.72849\nTotal prob of N-in: 0.96170\nPOSSIBLE N-term signal sequence\ninside 1 19\nTMhelix 20 39\noutside 40 325\nTMhelix 326 348\ninside 349 372

24611 GCF_001222625.1_5285_7_2Salmonella enterica subsp. enterica serovar Typhimurium str. DT104 Proteobacteria; Gammaproteobacteria; Enterobacterales; Enterobacteriaceae; Salmonella; Salmonella enterica; Salmonella enterica subsp. enterica; Salmonella enterica subsp. enterica serovar Typhimurium MTGDNTLITSHGINRRDFMKLCAALAATMGLSSKAAAEMAESVSNPQRPPVIWIGAQECT WP_000145429.1 hydrogenase 2 small subunit [Salmonella enterica] Length: 372\nNumber of predicted TMHs: 1\nExp number of AAs in TMHs: 24.51309\nExp number, first 60 AAs: 0.2682\nTotal prob of N-in: 0.14402\noutside 1 331\nTMhelix 332 354\ninside 355 372

24612 GCF_001222665.1_8080_3_72 Salmonella enterica subsp. enterica serovar Typhimurium str. DT104 Proteobacteria; Gammaproteobacteria; Enterobacterales; Enterobacteriaceae; Salmonella; Salmonella enterica; Salmonella enterica subsp. enterica; Salmonella enterica subsp. enterica serovar Typhimurium MTGDNTLITSHGINRRDFMKLCAALAATMGLSSKAAAEMAESVSNPQRPPVIWIGAQECT WP_000145429.1 hydrogenase 2 small subunit [Salmonella enterica] Length: 372\nNumber of predicted TMHs: 1\nExp number of AAs in TMHs: 24.51309\nExp number, first 60 AAs: 0.2682\nTotal prob of N-in: 0.14402\noutside 1 331\nTMhelix 332 354\ninside 355 372

24613 GCF_001172545.1_8490_2_78 Salmonella enterica subsp. enterica serovar Typhimurium str. DT104 Proteobacteria; Gammaproteobacteria; Enterobacterales; Enterobacteriaceae; Salmonella; Salmonella enterica; Salmonella enterica subsp. enterica; Salmonella enterica subsp. enterica serovar Typhimurium MTGDNTLITSHGINRRDFMKLCAALAATMGLSSKAAAEMAESVSNPQRPPVIWIGAQECT WP_000145429.1 hydrogenase 2 small subunit [Salmonella enterica] Length: 372\nNumber of predicted TMHs: 1\nExp number of AAs in TMHs: 24.51309\nExp number, first 60 AAs: 0.2682\nTotal prob of N-in: 0.14402\noutside 1 331\nTMhelix 332 354\ninside 355 372

24614 GCF_001222725.1_8080_3_79 Salmonella enterica subsp. enterica serovar Typhimurium str. DT104 Proteobacteria; Gammaproteobacteria; Enterobacterales; Enterobacteriaceae; Salmonella; Salmonella enterica; Salmonella enterica subsp. enterica; Salmonella enterica subsp. enterica serovar Typhimurium MTGDNTLITSHGINRRDFMKLCAALAATMGLSSKAAAEMAESVSNPQRPPVIWIGAQECT WP_000145429.1 hydrogenase 2 small subunit [Salmonella enterica] Length: 372\nNumber of predicted TMHs: 1\nExp number of AAs in TMHs: 24.51309\nExp number, first 60 AAs: 0.2682\nTotal prob of N-in: 0.14402\noutside 1 331\nTMhelix 332 354\ninside 355 372

24615 GCF_001222825.1_8036_3_39 Salmonella enterica subsp. enterica serovar Typhimurium str. DT104 Proteobacteria; Gammaproteobacteria; Enterobacterales; Enterobacteriaceae; Salmonella; Salmonella enterica; Salmonella enterica subsp. enterica; Salmonella enterica subsp. enterica serovar Typhimurium MNNEETFYQAMRRKGVTRRSFLKFCSLAATSLGLGAGMTPKIAWALENKPRIPVVWIHGL WP_001058312.1 hydrogenase [Salmonella enterica] Length: 372\nNumber of predicted TMHs: 2\nExp number of AAs in TMHs: 36.51738\nExp number, first 60 AAs: 19.72849\nTotal prob of N-in: 0.96170\nPOSSIBLE N-term signal sequence\ninside 1 19\nTMhelix 20 39\noutside 40 325\nTMhelix 326 348\ninside 349 372

24616 GCF_001223185.1_5275_2_9Salmonella enterica subsp. enterica serovar Typhimurium str. DT104 Proteobacteria; Gammaproteobacteria; Enterobacterales; Enterobacteriaceae; Salmonella; Salmonella enterica; Salmonella enterica subsp. enterica; Salmonella enterica subsp. enterica serovar Typhimurium MTGDNTLITSHGINRRDFMKLCAALAATMGLSSKAAAEMAESVSNPQRPPVIWIGAQECT WP_000145429.1 hydrogenase 2 small subunit [Salmonella enterica] Length: 372\nNumber of predicted TMHs: 1\nExp number of AAs in TMHs: 24.51309\nExp number, first 60 AAs: 0.2682\nTotal prob of N-in: 0.14402\noutside 1 331\nTMhelix 332 354\ninside 355 372

24617 GCF_001223185.1_5275_2_9Salmonella enterica subsp. enterica serovar Typhimurium str. DT104 Proteobacteria; Gammaproteobacteria; Enterobacterales; Enterobacteriaceae; Salmonella; Salmonella enterica; Salmonella enterica subsp. enterica; Salmonella enterica subsp. enterica serovar Typhimurium MNNEETFYQAMRRKGVTRRSFLKFCSLAATSLGLGAGMTPKIAWALENKPRIPVVWIHGL WP_001058312.1 hydrogenase [Salmonella enterica] Length: 372\nNumber of predicted TMHs: 2\nExp number of AAs in TMHs: 36.51738\nExp number, first 60 AAs: 19.72849\nTotal prob of N-in: 0.96170\nPOSSIBLE N-term signal sequence\ninside 1 19\nTMhelix 20 39\noutside 40 325\nTMhelix 326 348\ninside 349 372

24618 GCF_001223025.1_5275_3_6Salmonella enterica subsp. enterica serovar Typhimurium str. DT104 Proteobacteria; Gammaproteobacteria; Enterobacterales; Enterobacteriaceae; Salmonella; Salmonella enterica;

Salmonella enterica subsp. enterica; Salmonella enterica subsp. enterica serovar Typhimurium
MNNEETFYQAMRRKGVTRRSFLKFCSLAATSLGLGAGMTPKIAWALENKPRIPVVWIHGL WP_001058312.1
hydrogenase [Salmonella enterica] Length: 372\nNumber of predicted TMHs: 2\nExp number of AAs in TMHs: 36.51738\nExp number, first 60 AAs: 19.72849\nTotal prob of N-in: 0.96170\nPOSSIBLE N-term signal sequence\ninside 1 19\nTMhelix 20 39\noutside 40 325\nTMhelix 326 348\ninside 349 372

24619 GCF_001223225.1_5275_5_5Salmonella enterica subsp. enterica serovar Typhimurium str. DT104
Proteobacteria; Gammaproteobacteria; Enterobacterales; Enterobacteriaceae; Salmonella; Salmonella enterica; Salmonella enterica subsp. enterica; Salmonella enterica subsp. enterica serovar Typhimurium
MTGDNTLITSHGINRRDFMKLCAALAATMGLSSKAAAEMAESVSNPQRPPVIWIGAQECT WP_000145429.1
hydrogenase 2 small subunit [Salmonella enterica] Length: 372\nNumber of predicted TMHs: 1\nExp number of AAs in TMHs: 24.51309\nExp number, first 60 AAs: 0.2682\nTotal prob of N-in: 0.14402\noutside 1 331\nTMhelix 332 354\ninside 355 372

24620 GCF_001223285.1_8080_3_90 Salmonella enterica subsp. enterica serovar Typhimurium str. DT104
Proteobacteria; Gammaproteobacteria; Enterobacterales; Enterobacteriaceae; Salmonella; Salmonella enterica; Salmonella enterica subsp. enterica; Salmonella enterica subsp. enterica serovar Typhimurium
MTGDNTLITSHGINRRDFMKLCAALAATMGLSSKAAAEMAESVSNPQRPPVIWIGAQECT WP_000145429.1
hydrogenase 2 small subunit [Salmonella enterica] Length: 372\nNumber of predicted TMHs: 1\nExp number of AAs in TMHs: 24.51309\nExp number, first 60 AAs: 0.2682\nTotal prob of N-in: 0.14402\noutside 1 331\nTMhelix 332 354\ninside 355 372

24621 GCF_001326255.1_5388_1_10 Salmonella enterica subsp. enterica serovar Typhimurium str. DT104
Proteobacteria; Gammaproteobacteria; Enterobacterales; Enterobacteriaceae; Salmonella; Salmonella enterica; Salmonella enterica subsp. enterica; Salmonella enterica subsp. enterica serovar Typhimurium
MTGDNTLITSHGINRRDFMKLCAALAATMGLSSKAAAEMAESVSNPQRPPVIWIGAQECT WP_000145429.1
hydrogenase 2 small subunit [Salmonella enterica] Length: 372\nNumber of predicted TMHs: 1\nExp number of AAs in TMHs: 24.51309\nExp number, first 60 AAs: 0.2682\nTotal prob of N-in: 0.14402\noutside 1 331\nTMhelix 332 354\ninside 355 372

24622 GCF_001214485.1_8036_3_68 Salmonella enterica subsp. enterica serovar Typhimurium str. DT104
Proteobacteria; Gammaproteobacteria; Enterobacterales; Enterobacteriaceae; Salmonella; Salmonella enterica; Salmonella enterica subsp. enterica; Salmonella enterica subsp. enterica serovar Typhimurium
MTGDNTLITSHGINRRDFMKLCAALAATMGLSSKAAAEMAESVSNPQRPPVIWIGAQECT WP_000145429.1
hydrogenase 2 small subunit [Salmonella enterica] Length: 372\nNumber of predicted TMHs: 1\nExp number of AAs in TMHs: 24.51309\nExp number, first 60 AAs: 0.2682\nTotal prob of N-in: 0.14402\noutside 1 331\nTMhelix 332 354\ninside 355 372

24623 GCF_001214745.1_5275_2_6Salmonella enterica subsp. enterica serovar Typhimurium str. DT104
Proteobacteria; Gammaproteobacteria; Enterobacterales; Enterobacteriaceae; Salmonella; Salmonella enterica; Salmonella enterica subsp. enterica; Salmonella enterica subsp. enterica serovar Typhimurium
MTGDNTLITSHGINRRDFMKLCAALAATMGLSSKAAAEMAESVSNPQRPPVIWIGAQECT WP_000145429.1
hydrogenase 2 small subunit [Salmonella enterica] Length: 372\nNumber of predicted TMHs: 1\nExp number of AAs in TMHs: 24.51309\nExp number, first 60 AAs: 0.2682\nTotal prob of N-in: 0.14402\noutside 1 331\nTMhelix 332 354\ninside 355 372

24624 GCF_001214865.1_9065_1_3Salmonella enterica subsp. enterica serovar Typhimurium str. DT104
Proteobacteria; Gammaproteobacteria; Enterobacterales; Enterobacteriaceae; Salmonella; Salmonella enterica; Salmonella enterica subsp. enterica; Salmonella enterica subsp. enterica serovar Typhimurium
MTGDNTLITSHGINRRDFMKLCAALAATMGLSSKAAAEMAESVSNPQRPPVIWIGAQECT WP_000145429.1
hydrogenase 2 small subunit [Salmonella enterica] Length: 372\nNumber of predicted TMHs: 1\nExp number of AAs in TMHs: 24.51309\nExp number, first 60 AAs: 0.2682\nTotal prob of N-in: 0.14402\noutside 1 331\nTMhelix 332 354\ninside 355 372

24625 GCF_001214485.1_8036_3_68 Salmonella enterica subsp. enterica serovar Typhimurium str. DT104
Proteobacteria; Gammaproteobacteria; Enterobacterales; Enterobacteriaceae; Salmonella; Salmonella enterica; Salmonella enterica subsp. enterica; Salmonella enterica subsp. enterica serovar Typhimurium
MNNEETFYQAMRRKGVTRRSFLKFCSLAATSLGLGAGMTPKIAWALENKPRIPVVWIHGL WP_001058312.1
hydrogenase [Salmonella enterica] Length: 372\nNumber of predicted TMHs: 2\nExp number of AAs in TMHs: 36.51738\nExp number, first 60 AAs: 19.72849\nTotal prob of N-in: 0.96170\nPOSSIBLE N-term signal sequence\ninside 1 19\nTMhelix 20 39\noutside 40 325\nTMhelix 326 348\ninside 349 372

24626 GCF_001215665.1_8080_3_65 Salmonella enterica subsp. enterica serovar Typhimurium str. DT104 Proteobacteria; Gammaproteobacteria; Enterobacterales; Enterobacteriaceae; Salmonella; Salmonella enterica; Salmonella enterica subsp. enterica; Salmonella enterica subsp. enterica serovar Typhimurium MNNEETFYQAMRRKGVTRRSFLKFCSLAATSLGLGAGMTPKIAWALENKPRIPVVWIHGL WP_001058312.1 hydrogenase [Salmonella enterica] Length: 372\nNumber of predicted TMHs: 2\nExp number of AAs in TMHs: 36.51738\nExp number, first 60 AAs: 19.72849\nTotal prob of N-in: 0.96170\nPOSSIBLE N-term signal sequence\ninside 1 19\nTMhelix 20 39\noutside 40 325\nTMhelix 326 348\ninside 349 372

24627 GCF_001215725.1_5275_3_8Salmonella enterica subsp. enterica serovar Typhimurium str. DT104 Proteobacteria; Gammaproteobacteria; Enterobacterales; Enterobacteriaceae; Salmonella; Salmonella enterica; Salmonella enterica subsp. enterica; Salmonella enterica subsp. enterica serovar Typhimurium MNNEETFYQAMRRKGVTRRSFLKFCSLAATSLGLGAGMTPKIAWALENKPRIPVVWIHGL WP_001058312.1 hydrogenase [Salmonella enterica] Length: 372\nNumber of predicted TMHs: 2\nExp number of AAs in TMHs: 36.51738\nExp number, first 60 AAs: 19.72849\nTotal prob of N-in: 0.96170\nPOSSIBLE N-term signal sequence\ninside 1 19\nTMhelix 20 39\noutside 40 325\nTMhelix 326 348\ninside 349 372

24628 GCF_001215925.1_8036_3_66 Salmonella enterica subsp. enterica serovar Typhimurium str. DT104 Proteobacteria; Gammaproteobacteria; Enterobacterales; Enterobacteriaceae; Salmonella; Salmonella enterica; Salmonella enterica subsp. enterica; Salmonella enterica subsp. enterica serovar Typhimurium MTGDNTLITSHGINRRDFMKLCAALAATMGLSSKAAAEMAESVSNPQRPPVIWIGAQECT WP_000145429.1 hydrogenase 2 small subunit [Salmonella enterica] Length: 372\nNumber of predicted TMHs: 1\nExp number of AAs in TMHs: 24.51309\nExp number, first 60 AAs: 0.2682\nTotal prob of N-in: 0.14402\noutside 1 331\nTMhelix 332 354\ninside 355 372

24629 GCF_001215925.1_8036_3_66 Salmonella enterica subsp. enterica serovar Typhimurium str. DT104 Proteobacteria; Gammaproteobacteria; Enterobacterales; Enterobacteriaceae; Salmonella; Salmonella enterica; Salmonella enterica subsp. enterica; Salmonella enterica subsp. enterica serovar Typhimurium MNNEETFYQAMRRKGVTRRSFLKFCSLAATSLGLGAGMTPKIAWALENKPRIPVVWIHGL WP_001058312.1 hydrogenase [Salmonella enterica] Length: 372\nNumber of predicted TMHs: 2\nExp number of AAs in TMHs: 36.51738\nExp number, first 60 AAs: 19.72849\nTotal prob of N-in: 0.96170\nPOSSIBLE N-term signal sequence\ninside 1 19\nTMhelix 20 39\noutside 40 325\nTMhelix 326 348\ninside 349 372

24630 GCF_001566035.1_8080_3_37 Salmonella enterica subsp. enterica serovar Typhimurium str. DT104 Proteobacteria; Gammaproteobacteria; Enterobacterales; Enterobacteriaceae; Salmonella; Salmonella enterica; Salmonella enterica subsp. enterica; Salmonella enterica subsp. enterica serovar Typhimurium MNNEETFYQAMRRKGVTRRSFLKFCSLAATSLGLGAGMTPKIAWALENKPRIPVVWIHGL WP_001058312.1 hydrogenase [Salmonella enterica] Length: 372\nNumber of predicted TMHs: 2\nExp number of AAs in TMHs: 36.51738\nExp number, first 60 AAs: 19.72849\nTotal prob of N-in: 0.96170\nPOSSIBLE N-term signal sequence\ninside 1 19\nTMhelix 20 39\noutside 40 325\nTMhelix 326 348\ninside 349 372

24631 GCF_001326255.1_5388_1_10 Salmonella enterica subsp. enterica serovar Typhimurium str. DT104 Proteobacteria; Gammaproteobacteria; Enterobacterales; Enterobacteriaceae; Salmonella; Salmonella enterica; Salmonella enterica subsp. enterica; Salmonella enterica subsp. enterica serovar Typhimurium MNNEETFYQAMRRKGVTRRSFLKFCSLAATSLGLGAGMTPKIAWALENKPRIPVVWIHGL WP_001058312.1 hydrogenase [Salmonella enterica] Length: 372\nNumber of predicted TMHs: 2\nExp number of AAs in TMHs: 36.51738\nExp number, first 60 AAs: 19.72849\nTotal prob of N-in: 0.96170\nPOSSIBLE N-term signal sequence\ninside 1 19\nTMhelix 20 39\noutside 40 325\nTMhelix 326 348\ninside 349 372

24632 GCF_001217545.1_8036_3_43 Salmonella enterica subsp. enterica serovar Typhimurium str. DT104 Proteobacteria; Gammaproteobacteria; Enterobacterales; Enterobacteriaceae; Salmonella; Salmonella enterica; Salmonella enterica subsp. enterica; Salmonella enterica subsp. enterica serovar Typhimurium MTGDNTLITSHGINRRDFMKLCAALAATMGLSSKAAAEMAESVSNPQRPPVIWIGAQECT WP_000145429.1 hydrogenase 2 small subunit [Salmonella enterica] Length: 372\nNumber of predicted TMHs: 1\nExp number of AAs in TMHs: 24.51309\nExp number, first 60 AAs: 0.2682\nTotal prob of N-in: 0.14402\noutside 1 331\nTMhelix 332 354\ninside 355 372

24633 GCF_001216385.1_8080_3_44 Salmonella enterica subsp. enterica serovar Typhimurium str. DT104 Proteobacteria; Gammaproteobacteria; Enterobacterales; Enterobacteriaceae; Salmonella; Salmonella enterica; Salmonella enterica subsp. enterica; Salmonella enterica subsp. enterica serovar Typhimurium MNNEETFYQAMRRKGVTRRSFLKFCSLAATSLGLGAGMTPKIAWALENKPRIPVVWIHGL WP_001058312.1 hydrogenase [Salmonella enterica] Length: 372\nNumber of predicted TMHs: 2\nExp number of AAs in

TMHs: 36.51738\nExp number, first 60 AAs:  19.72849\nTotal prob of N-in:   0.96170\nPOSSIBLE N-term signal sequence\ninside   1   19\nTMhelix   20   39\noutside   40   325\nTMhelix   326   348\ninside   349   372

24634	GCF_001216785.1_8080_3_76	Salmonella enterica subsp. enterica serovar Typhimurium str. DT104	Proteobacteria; Gammaproteobacteria; Enterobacterales; Enterobacteriaceae; Salmonella; Salmonella enterica; Salmonella enterica subsp. enterica; Salmonella enterica subsp. enterica serovar Typhimurium	MTGDNTLITSHGINRRDFMKLCAALAATMGLSSKAAAEMAESVSNPQRPPVIWIGAQECT	WP_000145429.1	hydrogenase 2 small subunit [Salmonella enterica]	Length: 372\nNumber of predicted TMHs:  1\nExp number of AAs in TMHs: 24.51309\nExp number, first 60 AAs:  0.2682\nTotal prob of N-in:   0.14402\noutside   1   331\nTMhelix   332   354\ninside   355   372

24635	GCF_001216785.1_8080_3_76	Salmonella enterica subsp. enterica serovar Typhimurium str. DT104	Proteobacteria; Gammaproteobacteria; Enterobacterales; Enterobacteriaceae; Salmonella; Salmonella enterica; Salmonella enterica subsp. enterica; Salmonella enterica subsp. enterica serovar Typhimurium	MNNEETFYQAMRRKGVTRRSFLKFCSLAATSLGLGAGMTPKIAWALENKPRIPVVWIHGL	WP_001058312.1	hydrogenase [Salmonella enterica]	Length: 372\nNumber of predicted TMHs:  2\nExp number of AAs in TMHs: 36.51738\nExp number, first 60 AAs:  19.72849\nTotal prob of N-in:   0.96170\nPOSSIBLE N-term signal sequence\ninside   1   19\nTMhelix   20   39\noutside   40   325\nTMhelix   326   348\ninside   349   372

24636	GCF_001217305.1_8080_3_24	Salmonella enterica subsp. enterica serovar Typhimurium str. DT104	Proteobacteria; Gammaproteobacteria; Enterobacterales; Enterobacteriaceae; Salmonella; Salmonella enterica; Salmonella enterica subsp. enterica; Salmonella enterica subsp. enterica serovar Typhimurium	MNNEETFYQAMRRKGVTRRSFLKFCSLAATSLGLGAGMTPKIAWALENKPRIPVVWIHGL	WP_001058312.1	hydrogenase [Salmonella enterica]	Length: 372\nNumber of predicted TMHs:  2\nExp number of AAs in TMHs: 36.51738\nExp number, first 60 AAs:  19.72849\nTotal prob of N-in:   0.96170\nPOSSIBLE N-term signal sequence\ninside   1   19\nTMhelix   20   39\noutside   40   325\nTMhelix   326   348\ninside   349   372

24637	GCF_001217825.1_5275_2_4Salmonella enterica subsp. enterica serovar Typhimurium str. DT104	Proteobacteria; Gammaproteobacteria; Enterobacterales; Enterobacteriaceae; Salmonella; Salmonella enterica; Salmonella enterica subsp. enterica; Salmonella enterica subsp. enterica serovar Typhimurium	MNNEETFYQAMRRKGVTRRSFLKFCSLAATSLGLGAGMTPKIAWALENKPRIPVVWIHGL	WP_001058312.1	hydrogenase [Salmonella enterica]	Length: 372\nNumber of predicted TMHs:  2\nExp number of AAs in TMHs: 36.51738\nExp number, first 60 AAs:  19.72849\nTotal prob of N-in:   0.96170\nPOSSIBLE N-term signal sequence\ninside   1   19\nTMhelix   20   39\noutside   40   325\nTMhelix   326   348\ninside   349   372

24638	GCF_001216465.1_8036_3_69	Salmonella enterica subsp. enterica serovar Typhimurium str. DT104	Proteobacteria; Gammaproteobacteria; Enterobacterales; Enterobacteriaceae; Salmonella; Salmonella enterica; Salmonella enterica subsp. enterica; Salmonella enterica subsp. enterica serovar Typhimurium	MNNEETFYQAMRRKGVTRRSFLKFCSLAATSLGLGAGMTPKIAWALENKPRIPVVWIHGL	WP_001058312.1	hydrogenase [Salmonella enterica]	Length: 372\nNumber of predicted TMHs:  2\nExp number of AAs in TMHs: 36.51738\nExp number, first 60 AAs:  19.72849\nTotal prob of N-in:   0.96170\nPOSSIBLE N-term signal sequence\ninside   1   19\nTMhelix   20   39\noutside   40   325\nTMhelix   326   348\ninside   349   372

24639	GCF_001216985.1_5275_8_3Salmonella enterica subsp. enterica serovar Typhimurium str. DT104	Proteobacteria; Gammaproteobacteria; Enterobacterales; Enterobacteriaceae; Salmonella; Salmonella enterica; Salmonella enterica subsp. enterica; Salmonella enterica subsp. enterica serovar Typhimurium	MTGDNTLITSHGINRRDFMKLCAALAATMGLSSKAAAEMAESVSNPQRPPVIWIGAQECT	WP_000145429.1	hydrogenase 2 small subunit [Salmonella enterica]	Length: 372\nNumber of predicted TMHs:  1\nExp number of AAs in TMHs: 24.51309\nExp number, first 60 AAs:  0.2682\nTotal prob of N-in:   0.14402\noutside   1   331\nTMhelix   332   354\ninside   355   372

24640	GCF_001217705.1_8036_3_62	Salmonella enterica subsp. enterica serovar Typhimurium str. DT104	Proteobacteria; Gammaproteobacteria; Enterobacterales; Enterobacteriaceae; Salmonella; Salmonella enterica; Salmonella enterica subsp. enterica; Salmonella enterica subsp. enterica serovar Typhimurium	MNNEETFYQAMRRKGVTRRSFLKFCSLAATSLGLGAGMTPKIAWALENKPRIPVVWIHGL	WP_001058312.1	hydrogenase [Salmonella enterica]	Length: 372\nNumber of predicted TMHs:  2\nExp number of AAs in TMHs: 36.51738\nExp number, first 60 AAs:  19.72849\nTotal prob of N-in:   0.96170\nPOSSIBLE N-term signal sequence\ninside   1   19\nTMhelix   20   39\noutside   40   325\nTMhelix   326   348\ninside   349   372

24641	GCF_001217085.1_5275_3_3Salmonella enterica subsp. enterica serovar Typhimurium str. DT104	Proteobacteria; Gammaproteobacteria; Enterobacterales; Enterobacteriaceae; Salmonella; Salmonella enterica;

Salmonella enterica subsp. enterica; Salmonella enterica subsp. enterica serovar Typhimurium
MTGDNTLITSHGINRRDFMKLCAALAATMGLSSKAAAEMAESVSNPQRPPVIWIGAQECT WP_000145429.1
hydrogenase 2 small subunit [Salmonella enterica] Length: 372\nNumber of predicted TMHs: 1\nExp number of AAs in TMHs: 24.51309\nExp number, first 60 AAs: 0.2682\nTotal prob of N-in: 0.14402\noutside 1 331\nTMhelix 332 354\ninside 355 372

24642 GCF_001220105.1_8080_3_7Salmonella enterica subsp. enterica serovar Typhimurium str. DT104
Proteobacteria; Gammaproteobacteria; Enterobacterales; Enterobacteriaceae; Salmonella; Salmonella enterica; Salmonella enterica subsp. enterica; Salmonella enterica subsp. enterica serovar Typhimurium
MTGDNTLITSHGINRRDFMKLCAALAATMGLSSKAAAEMAESVSNPQRPPVIWIGAQECT WP_000145429.1
hydrogenase 2 small subunit [Salmonella enterica] Length: 372\nNumber of predicted TMHs: 1\nExp number of AAs in TMHs: 24.51309\nExp number, first 60 AAs: 0.2682\nTotal prob of N-in: 0.14402\noutside 1 331\nTMhelix 332 354\ninside 355 372

24643 GCF_001219945.1_5285_6_2Salmonella enterica subsp. enterica serovar Typhimurium str. DT104
Proteobacteria; Gammaproteobacteria; Enterobacterales; Enterobacteriaceae; Salmonella; Salmonella enterica; Salmonella enterica subsp. enterica; Salmonella enterica subsp. enterica serovar Typhimurium
MNNEETFYQAMRRKGVTRRSFLKFCSLAATSLGLGAGMTPKIAWALENKPRIPVVWIHGL WP_001058312.1
hydrogenase [Salmonella enterica] Length: 372\nNumber of predicted TMHs: 2\nExp number of AAs in TMHs: 36.51738\nExp number, first 60 AAs: 19.72849\nTotal prob of N-in: 0.96170\nPOSSIBLE N-term signal sequence\ninside 1 19\nTMhelix 20 39\noutside 40 325\nTMhelix 326 348\ninside 349 372

24644 GCF_001219905.1_8036_3_54 Salmonella enterica subsp. enterica serovar Typhimurium str. DT104
Proteobacteria; Gammaproteobacteria; Enterobacterales; Enterobacteriaceae; Salmonella; Salmonella enterica; Salmonella enterica subsp. enterica; Salmonella enterica subsp. enterica serovar Typhimurium
MNNEETFYQAMRRKGVTRRSFLKFCSLAATSLGLGAGMTPKIAWALENKPRIPVVWIHGL WP_001058312.1
hydrogenase [Salmonella enterica] Length: 372\nNumber of predicted TMHs: 2\nExp number of AAs in TMHs: 36.51738\nExp number, first 60 AAs: 19.72849\nTotal prob of N-in: 0.96170\nPOSSIBLE N-term signal sequence\ninside 1 19\nTMhelix 20 39\noutside 40 325\nTMhelix 326 348\ninside 349 372

24645 GCF_001219725.1_8080_3_35 Salmonella enterica subsp. enterica serovar Typhimurium str. DT104
Proteobacteria; Gammaproteobacteria; Enterobacterales; Enterobacteriaceae; Salmonella; Salmonella enterica; Salmonella enterica subsp. enterica; Salmonella enterica subsp. enterica serovar Typhimurium
MNNEETFYQAMRRKGVTRRSFLKFCSLAATSLGLGAGMTPKIAWALENKPRIPVVWIHGL WP_001058312.1
hydrogenase [Salmonella enterica] Length: 372\nNumber of predicted TMHs: 2\nExp number of AAs in TMHs: 36.51738\nExp number, first 60 AAs: 19.72849\nTotal prob of N-in: 0.96170\nPOSSIBLE N-term signal sequence\ninside 1 19\nTMhelix 20 39\noutside 40 325\nTMhelix 326 348\ninside 349 372

24646 GCF_001219725.1_8080_3_35 Salmonella enterica subsp. enterica serovar Typhimurium str. DT104
Proteobacteria; Gammaproteobacteria; Enterobacterales; Enterobacteriaceae; Salmonella; Salmonella enterica; Salmonella enterica subsp. enterica; Salmonella enterica subsp. enterica serovar Typhimurium
MTGDNTLITSHGINRRDFMKLCAALAATMGLSSKAAAEMAESVSNPQRPPVIWIGAQECT WP_000145429.1
hydrogenase 2 small subunit [Salmonella enterica] Length: 372\nNumber of predicted TMHs: 1\nExp number of AAs in TMHs: 24.51309\nExp number, first 60 AAs: 0.2682\nTotal prob of N-in: 0.14402\noutside 1 331\nTMhelix 332 354\ninside 355 372

24647 GCF_001219685.1_5285_6_1Salmonella enterica subsp. enterica serovar Typhimurium str. DT104
Proteobacteria; Gammaproteobacteria; Enterobacterales; Enterobacteriaceae; Salmonella; Salmonella enterica; Salmonella enterica subsp. enterica; Salmonella enterica subsp. enterica serovar Typhimurium
MTGDNTLITSHGINRRDFMKLCAALAATMGLSSKAAAEMAESVSNPQRPPVIWIGAQECT WP_000145429.1
hydrogenase 2 small subunit [Salmonella enterica] Length: 372\nNumber of predicted TMHs: 1\nExp number of AAs in TMHs: 24.51309\nExp number, first 60 AAs: 0.2682\nTotal prob of N-in: 0.14402\noutside 1 331\nTMhelix 332 354\ninside 355 372

24648 GCF_001219565.1_8080_3_74 Salmonella enterica subsp. enterica serovar Typhimurium str. DT104
Proteobacteria; Gammaproteobacteria; Enterobacterales; Enterobacteriaceae; Salmonella; Salmonella enterica; Salmonella enterica subsp. enterica; Salmonella enterica subsp. enterica serovar Typhimurium
MTGDNTLITSHGINRRDFMKLCAALAATMGLSSKAAAEMAESVSNPQRPPVIWIGAQECT WP_000145429.1
hydrogenase 2 small subunit [Salmonella enterica] Length: 372\nNumber of predicted TMHs: 1\nExp number of AAs in TMHs: 24.51309\nExp number, first 60 AAs: 0.2682\nTotal prob of N-in: 0.14402\noutside 1 331\nTMhelix 332 354\ninside 355 372

24649 GCF_001219525.1_9065_1_4Salmonella enterica subsp. enterica serovar Typhimurium str. DT104 Proteobacteria; Gammaproteobacteria; Enterobacterales; Enterobacteriaceae; Salmonella; Salmonella enterica; Salmonella enterica subsp. enterica; Salmonella enterica subsp. enterica serovar Typhimurium MTGDNTLITSHGINRRDFMKLCAALAATMGLSSKAAAEMAESVSNPQRPPVIWIGAQECT WP_000145429.1 hydrogenase 2 small subunit [Salmonella enterica] Length: 372\nNumber of predicted TMHs: 1\nExp number of AAs in TMHs: 24.51309\nExp number, first 60 AAs: 0.2682\nTotal prob of N-in: 0.14402\noutside 1 331\nTMhelix 332 354\ninside 355 372

24650 GCF_001219485.1_5275_6_4Salmonella enterica subsp. enterica serovar Typhimurium str. DT104 Proteobacteria; Gammaproteobacteria; Enterobacterales; Enterobacteriaceae; Salmonella; Salmonella enterica; Salmonella enterica subsp. enterica; Salmonella enterica subsp. enterica serovar Typhimurium MTGDNTLITSHGINRRDFMKLCAALAATMGLSSKAAAEMAESVSNPQRPPVIWIGAQECT WP_000145429.1 hydrogenase 2 small subunit [Salmonella enterica] Length: 372\nNumber of predicted TMHs: 1\nExp number of AAs in TMHs: 24.51309\nExp number, first 60 AAs: 0.2682\nTotal prob of N-in: 0.14402\noutside 1 331\nTMhelix 332 354\ninside 355 372

24651 GCF_001219005.1_8036_3_47 Salmonella enterica subsp. enterica serovar Typhimurium str. DT104 Proteobacteria; Gammaproteobacteria; Enterobacterales; Enterobacteriaceae; Salmonella; Salmonella enterica; Salmonella enterica subsp. enterica; Salmonella enterica subsp. enterica serovar Typhimurium MNNEETFYQAMRRKGVTRRSFLKFCSLAATSLGLGAGMTPKIAWALENKPRIPVVWIHGL WP_001058312.1 hydrogenase [Salmonella enterica] Length: 372\nNumber of predicted TMHs: 2\nExp number of AAs in TMHs: 36.51738\nExp number, first 60 AAs: 19.72849\nTotal prob of N-in: 0.96170\nPOSSIBLE N-term signal sequence\ninside 1 19\nTMhelix 20 39\noutside 40 325\nTMhelix 326 348\ninside 349 372

24652 GCF_001220165.1_5275_2_7Salmonella enterica subsp. enterica serovar Typhimurium str. DT104 Proteobacteria; Gammaproteobacteria; Enterobacterales; Enterobacteriaceae; Salmonella; Salmonella enterica; Salmonella enterica subsp. enterica; Salmonella enterica subsp. enterica serovar Typhimurium MTGDNTLITSHGINRRDFMKLCAALAATMGLSSKAAAEMAESVSNPQRPPVIWIGAQECT WP_000145429.1 hydrogenase 2 small subunit [Salmonella enterica] Length: 372\nNumber of predicted TMHs: 1\nExp number of AAs in TMHs: 24.51309\nExp number, first 60 AAs: 0.2682\nTotal prob of N-in: 0.14402\noutside 1 331\nTMhelix 332 354\ninside 355 372

24653 GCF_001220225.1_9066_1_3Salmonella enterica subsp. enterica serovar Typhimurium str. DT104 Proteobacteria; Gammaproteobacteria; Enterobacterales; Enterobacteriaceae; Salmonella; Salmonella enterica; Salmonella enterica subsp. enterica; Salmonella enterica subsp. enterica serovar Typhimurium MTGDNTLITSHGINRRDFMKLCAALAATMGLSSKAAAEMAESVSNPQRPPVIWIGAQECT WP_000145429.1 hydrogenase 2 small subunit [Salmonella enterica] Length: 372\nNumber of predicted TMHs: 1\nExp number of AAs in TMHs: 24.51309\nExp number, first 60 AAs: 0.2682\nTotal prob of N-in: 0.14402\noutside 1 331\nTMhelix 332 354\ninside 355 372

24654 GCF_001220505.1_8036_3_64 Salmonella enterica subsp. enterica serovar Typhimurium str. DT104 Proteobacteria; Gammaproteobacteria; Enterobacterales; Enterobacteriaceae; Salmonella; Salmonella enterica; Salmonella enterica subsp. enterica; Salmonella enterica subsp. enterica serovar Typhimurium MTGDNTLITSHGINRRDFMKLCAALAATMGLSSKAAAEMAESVSNPQRPPVIWIGAQECT WP_000145429.1 hydrogenase 2 small subunit [Salmonella enterica] Length: 372\nNumber of predicted TMHs: 1\nExp number of AAs in TMHs: 24.51309\nExp number, first 60 AAs: 0.2682\nTotal prob of N-in: 0.14402\noutside 1 331\nTMhelix 332 354\ninside 355 372

24655 GCF_001220545.1_5275_2_1Salmonella enterica subsp. enterica serovar Typhimurium str. DT104 Proteobacteria; Gammaproteobacteria; Enterobacterales; Enterobacteriaceae; Salmonella; Salmonella enterica; Salmonella enterica subsp. enterica; Salmonella enterica subsp. enterica serovar Typhimurium MTGDNTLITSHGINRRDFMKLCAALAATMGLSSKAAAEMAESVSNPQRPPVIWIGAQECT WP_000145429.1 hydrogenase 2 small subunit [Salmonella enterica] Length: 372\nNumber of predicted TMHs: 1\nExp number of AAs in TMHs: 24.51309\nExp number, first 60 AAs: 0.2682\nTotal prob of N-in: 0.14402\noutside 1 331\nTMhelix 332 354\ninside 355 372

24656 GCF_001220505.1_8036_3_64 Salmonella enterica subsp. enterica serovar Typhimurium str. DT104 Proteobacteria; Gammaproteobacteria; Enterobacterales; Enterobacteriaceae; Salmonella; Salmonella enterica; Salmonella enterica subsp. enterica; Salmonella enterica subsp. enterica serovar Typhimurium MNNEETFYQAMRRKGVTRRSFLKFCSLAATSLGLGAGMTPKIAWALENKPRIPVVWIHGL WP_001058312.1 hydrogenase [Salmonella enterica] Length: 372\nNumber of predicted TMHs: 2\nExp number of AAs in

TMHs: 36.51738\nExp number, first 60 AAs: 19.72849\nTotal prob of N-in: 0.96170\nPOSSIBLE N-term signal sequence\ninside 1 19\nTMhelix 20 39\noutside 40 325\nTMhelix 326 348\ninside 349 372

24657 GCF_001218865.1_5285_7_12 Salmonella enterica subsp. enterica serovar Typhimurium str. DT104 Proteobacteria; Gammaproteobacteria; Enterobacterales; Enterobacteriaceae; Salmonella; Salmonella enterica; Salmonella enterica subsp. enterica; Salmonella enterica subsp. enterica serovar Typhimurium MNNEETFYQAMRRKGVTRRSFLKFCSLAATSLGLGAGMTPKIAWALENKPRIPVVWIHGL WP_001058312.1 hydrogenase [Salmonella enterica] Length: 372\nNumber of predicted TMHs: 2\nExp number of AAs in TMHs: 36.51738\nExp number, first 60 AAs: 19.72849\nTotal prob of N-in: 0.96170\nPOSSIBLE N-term signal sequence\ninside 1 19\nTMhelix 20 39\noutside 40 325\nTMhelix 326 348\ninside 349 372

24658 GCF_001218785.1_8080_3_45 Salmonella enterica subsp. enterica serovar Typhimurium str. DT104 Proteobacteria; Gammaproteobacteria; Enterobacterales; Enterobacteriaceae; Salmonella; Salmonella enterica; Salmonella enterica subsp. enterica; Salmonella enterica subsp. enterica serovar Typhimurium MNNEETFYQAMRRKGVTRRSFLKFCSLAATSLGLGAGMTPKIAWALENKPRIPVVWIHGL WP_001058312.1 hydrogenase [Salmonella enterica] Length: 372\nNumber of predicted TMHs: 2\nExp number of AAs in TMHs: 36.51738\nExp number, first 60 AAs: 19.72849\nTotal prob of N-in: 0.96170\nPOSSIBLE N-term signal sequence\ninside 1 19\nTMhelix 20 39\noutside 40 325\nTMhelix 326 348\ninside 349 372

24659 GCF_001220545.1_5275_2_1Salmonella enterica subsp. enterica serovar Typhimurium str. DT104 Proteobacteria; Gammaproteobacteria; Enterobacterales; Enterobacteriaceae; Salmonella; Salmonella enterica; Salmonella enterica subsp. enterica; Salmonella enterica subsp. enterica serovar Typhimurium MNNEETFYQAMRRKGVTRRSFLKFCSLAATSLGLGAGMTPKIAWALENKPRIPVVWIHGL WP_001058312.1 hydrogenase [Salmonella enterica] Length: 372\nNumber of predicted TMHs: 2\nExp number of AAs in TMHs: 36.51738\nExp number, first 60 AAs: 19.72849\nTotal prob of N-in: 0.96170\nPOSSIBLE N-term signal sequence\ninside 1 19\nTMhelix 20 39\noutside 40 325\nTMhelix 326 348\ninside 349 372

24660 GCF_001220365.1_9066_1_1Salmonella enterica subsp. enterica serovar Typhimurium str. DT104 Proteobacteria; Gammaproteobacteria; Enterobacterales; Enterobacteriaceae; Salmonella; Salmonella enterica; Salmonella enterica subsp. enterica; Salmonella enterica subsp. enterica serovar Typhimurium MNNEETFYQAMRRKGVTRRSFLKFCSLAATSLGLGAGMTPKIAWALENKPRIPVVWIHGL WP_001058312.1 hydrogenase [Salmonella enterica] Length: 372\nNumber of predicted TMHs: 2\nExp number of AAs in TMHs: 36.51738\nExp number, first 60 AAs: 19.72849\nTotal prob of N-in: 0.96170\nPOSSIBLE N-term signal sequence\ninside 1 19\nTMhelix 20 39\noutside 40 325\nTMhelix 326 348\ninside 349 372

24661 GCF_001219085.1_8036_3_58 Salmonella enterica subsp. enterica serovar Typhimurium str. DT104 Proteobacteria; Gammaproteobacteria; Enterobacterales; Enterobacteriaceae; Salmonella; Salmonella enterica; Salmonella enterica subsp. enterica; Salmonella enterica subsp. enterica serovar Typhimurium MTGDNTLITSHGINRRDFMKLCAALAATMGLSSKAAAEMAESVSNPQRPPVIWIGAQECT WP_000145429.1 hydrogenase 2 small subunit [Salmonella enterica] Length: 372\nNumber of predicted TMHs: 1\nExp number of AAs in TMHs: 24.51309\nExp number, first 60 AAs: 0.2682\nTotal prob of N-in: 0.14402\noutside 1 331\nTMhelix 332 354\ninside 355 372

24662 GCF_001218745.1_5275_6_10 Salmonella enterica subsp. enterica serovar Typhimurium str. DT104 Proteobacteria; Gammaproteobacteria; Enterobacterales; Enterobacteriaceae; Salmonella; Salmonella enterica; Salmonella enterica subsp. enterica; Salmonella enterica subsp. enterica serovar Typhimurium MNNEETFYQAMRRKGVTRRSFLKFCSLAATSLGLGAGMTPKIAWALENKPRIPVVWIHGL WP_001058312.1 hydrogenase [Salmonella enterica] Length: 372\nNumber of predicted TMHs: 2\nExp number of AAs in TMHs: 36.51738\nExp number, first 60 AAs: 19.72849\nTotal prob of N-in: 0.96170\nPOSSIBLE N-term signal sequence\ninside 1 19\nTMhelix 20 39\noutside 40 325\nTMhelix 326 348\ninside 349 372

24663 GCF_001220405.1_9065_1_12 Salmonella enterica subsp. enterica serovar Typhimurium str. DT104 Proteobacteria; Gammaproteobacteria; Enterobacterales; Enterobacteriaceae; Salmonella; Salmonella enterica; Salmonella enterica subsp. enterica; Salmonella enterica subsp. enterica serovar Typhimurium MNNEETFYQAMRRKGVTRRSFLKFCSLAATSLGLGAGMTPKIAWALENKPRIPVVWIHGL WP_001058312.1 hydrogenase [Salmonella enterica] Length: 372\nNumber of predicted TMHs: 2\nExp number of AAs in TMHs: 36.51738\nExp number, first 60 AAs: 19.72849\nTotal prob of N-in: 0.96170\nPOSSIBLE N-term signal sequence\ninside 1 19\nTMhelix 20 39\noutside 40 325\nTMhelix 326 348\ninside 349 372

24664 GCF_001221125.1_5275_8_5Salmonella enterica subsp. enterica serovar Typhimurium str. DT104 Proteobacteria; Gammaproteobacteria; Enterobacterales; Enterobacteriaceae; Salmonella; Salmonella enterica;

Salmonella enterica subsp. enterica; Salmonella enterica subsp. enterica serovar Typhimurium
MTGDNTLITSHGINRRDFMKLCAALAATMGLSSKAAAEMAESVSNPQRPPVIWIGAQECT WP_000145429.1
hydrogenase 2 small subunit [Salmonella enterica] Length: 372\nNumber of predicted TMHs: 1\nExp number of AAs in TMHs: 24.51309\nExp number, first 60 AAs: 0.2682\nTotal prob of N-in: 0.14402\noutside 1 331\nTMhelix 332 354\ninside 355 372

24665 GCF_001221125.1_5275_8_5Salmonella enterica subsp. enterica serovar Typhimurium str. DT104
Proteobacteria; Gammaproteobacteria; Enterobacterales; Enterobacteriaceae; Salmonella; Salmonella enterica; Salmonella enterica subsp. enterica; Salmonella enterica subsp. enterica serovar Typhimurium
MNNEETFYQAMRRKGVTRRSFLKFCSLAATSLGLGAGMTPKIAWALENKPRIPVVWIHGL WP_001058312.1
hydrogenase [Salmonella enterica] Length: 372\nNumber of predicted TMHs: 2\nExp number of AAs in TMHs: 36.51738\nExp number, first 60 AAs: 19.72849\nTotal prob of N-in: 0.96170\nPOSSIBLE N-term signal sequence\ninside 1 19\nTMhelix 20 39\noutside 40 325\nTMhelix 326 348\ninside 349 372

24666 GCF_001220865.1_5299_6_11 Salmonella enterica subsp. enterica serovar Typhimurium str. DT104
Proteobacteria; Gammaproteobacteria; Enterobacterales; Enterobacteriaceae; Salmonella; Salmonella enterica; Salmonella enterica subsp. enterica; Salmonella enterica subsp. enterica serovar Typhimurium
MNNEETFYQAMRRKGVTRRSFLKFCSLAATSLGLGAGMTPKIAWALENKPRIPVVWIHGL WP_001058312.1
hydrogenase [Salmonella enterica] Length: 372\nNumber of predicted TMHs: 2\nExp number of AAs in TMHs: 36.51738\nExp number, first 60 AAs: 19.72849\nTotal prob of N-in: 0.96170\nPOSSIBLE N-term signal sequence\ninside 1 19\nTMhelix 20 39\noutside 40 325\nTMhelix 326 348\ninside 349 372

24667 GCF_001326195.1_9066_1_8Salmonella enterica subsp. enterica serovar Typhimurium str. DT104
Proteobacteria; Gammaproteobacteria; Enterobacterales; Enterobacteriaceae; Salmonella; Salmonella enterica; Salmonella enterica subsp. enterica; Salmonella enterica subsp. enterica serovar Typhimurium
MNNEETFYQAMRRKGVTRRSFLKFCSLAATSLGLGAGMTPKIAWALENKPRIPVVWIHGL WP_001058312.1
hydrogenase [Salmonella enterica] Length: 372\nNumber of predicted TMHs: 2\nExp number of AAs in TMHs: 36.51738\nExp number, first 60 AAs: 19.72849\nTotal prob of N-in: 0.96170\nPOSSIBLE N-term signal sequence\ninside 1 19\nTMhelix 20 39\noutside 40 325\nTMhelix 326 348\ninside 349 372

24668 GCF_001220725.1_8080_3_94 Salmonella enterica subsp. enterica serovar Typhimurium str. DT104
Proteobacteria; Gammaproteobacteria; Enterobacterales; Enterobacteriaceae; Salmonella; Salmonella enterica; Salmonella enterica subsp. enterica; Salmonella enterica subsp. enterica serovar Typhimurium
MNNEETFYQAMRRKGVTRRSFLKFCSLAATSLGLGAGMTPKIAWALENKPRIPVVWIHGL WP_001058312.1
hydrogenase [Salmonella enterica] Length: 372\nNumber of predicted TMHs: 2\nExp number of AAs in TMHs: 36.51738\nExp number, first 60 AAs: 19.72849\nTotal prob of N-in: 0.96170\nPOSSIBLE N-term signal sequence\ninside 1 19\nTMhelix 20 39\noutside 40 325\nTMhelix 326 348\ninside 349 372

24669 GCF_001221205.1_8080_3_67 Salmonella enterica subsp. enterica serovar Typhimurium str. DT104
Proteobacteria; Gammaproteobacteria; Enterobacterales; Enterobacteriaceae; Salmonella; Salmonella enterica; Salmonella enterica subsp. enterica; Salmonella enterica subsp. enterica serovar Typhimurium
MTGDNTLITSHGINRRDFMKLCAALAATMGLSSKAAAEMAESVSNPQRPPVIWIGAQECT WP_000145429.1
hydrogenase 2 small subunit [Salmonella enterica] Length: 372\nNumber of predicted TMHs: 1\nExp number of AAs in TMHs: 24.51309\nExp number, first 60 AAs: 0.2682\nTotal prob of N-in: 0.14402\noutside 1 331\nTMhelix 332 354\ninside 355 372

24670 GCF_001221205.1_8080_3_67 Salmonella enterica subsp. enterica serovar Typhimurium str. DT104
Proteobacteria; Gammaproteobacteria; Enterobacterales; Enterobacteriaceae; Salmonella; Salmonella enterica; Salmonella enterica subsp. enterica; Salmonella enterica subsp. enterica serovar Typhimurium
MNNEETFYQAMRRKGVTRRSFLKFCSLAATSLGLGAGMTPKIAWALENKPRIPVVWIHGL WP_001058312.1
hydrogenase [Salmonella enterica] Length: 372\nNumber of predicted TMHs: 2\nExp number of AAs in TMHs: 36.51738\nExp number, first 60 AAs: 19.72849\nTotal prob of N-in: 0.96170\nPOSSIBLE N-term signal sequence\ninside 1 19\nTMhelix 20 39\noutside 40 325\nTMhelix 326 348\ninside 349 372

24671 GCF_001221585.1_5275_7_9Salmonella enterica subsp. enterica serovar Typhimurium str. DT104
Proteobacteria; Gammaproteobacteria; Enterobacterales; Enterobacteriaceae; Salmonella; Salmonella enterica; Salmonella enterica subsp. enterica; Salmonella enterica subsp. enterica serovar Typhimurium
MTGDNTLITSHGINRRDFMKLCAALAATMGLSSKAAAEMAESVSNPQRPPVIWIGAQECT WP_000145429.1
hydrogenase 2 small subunit [Salmonella enterica] Length: 372\nNumber of predicted TMHs: 1\nExp number of AAs in TMHs: 24.51309\nExp number, first 60 AAs: 0.2682\nTotal prob of N-in: 0.14402\noutside 1 331\nTMhelix 332 354\ninside 355 372

24672 GCF_001218685.1_5299_3_6Salmonella enterica subsp. enterica serovar Typhimurium str. DT104 Proteobacteria; Gammaproteobacteria; Enterobacterales; Enterobacteriaceae; Salmonella; Salmonella enterica; Salmonella enterica subsp. enterica; Salmonella enterica subsp. enterica serovar Typhimurium MTGDNTLITSHGINRRDFMKLCAALAATMGLSSKAAAEMAESVSNPQRPPVIWIGAQECT WP_000145429.1 hydrogenase 2 small subunit [Salmonella enterica] Length: 372\nNumber of predicted TMHs: 1\nExp number of AAs in TMHs: 24.51309\nExp number, first 60 AAs: 0.2682\nTotal prob of N-in: 0.14402\noutside 1 331\nTMhelix 332 354\ninside 355 372

24673 GCF_001221625.1_5275_6_11 Salmonella enterica subsp. enterica serovar Typhimurium str. DT104 Proteobacteria; Gammaproteobacteria; Enterobacterales; Enterobacteriaceae; Salmonella; Salmonella enterica; Salmonella enterica subsp. enterica; Salmonella enterica subsp. enterica serovar Typhimurium MTGDNTLITSHGINRRDFMKLCAALAATMGLSSKAAAEMAESVSNPQRPPVIWIGAQECT WP_000145429.1 hydrogenase 2 small subunit [Salmonella enterica] Length: 372\nNumber of predicted TMHs: 1\nExp number of AAs in TMHs: 24.51309\nExp number, first 60 AAs: 0.2682\nTotal prob of N-in: 0.14402\noutside 1 331\nTMhelix 332 354\ninside 355 372

24674 GCF_001218565.1_8080_3_66 Salmonella enterica subsp. enterica serovar Typhimurium str. DT104 Proteobacteria; Gammaproteobacteria; Enterobacterales; Enterobacteriaceae; Salmonella; Salmonella enterica; Salmonella enterica subsp. enterica; Salmonella enterica subsp. enterica serovar Typhimurium MTGDNTLITSHGINRRDFMKLCAALAATMGLSSKAAAEMAESVSNPQRPPVIWIGAQECT WP_000145429.1 hydrogenase 2 small subunit [Salmonella enterica] Length: 372\nNumber of predicted TMHs: 1\nExp number of AAs in TMHs: 24.51309\nExp number, first 60 AAs: 0.2682\nTotal prob of N-in: 0.14402\noutside 1 331\nTMhelix 332 354\ninside 355 372

24675 GCF_001221665.1_9065_1_2Salmonella enterica subsp. enterica serovar Typhimurium str. DT104 Proteobacteria; Gammaproteobacteria; Enterobacterales; Enterobacteriaceae; Salmonella; Salmonella enterica; Salmonella enterica subsp. enterica; Salmonella enterica subsp. enterica serovar Typhimurium MTGDNTLITSHGINRRDFMKLCAALAATMGLSSKAAAEMAESVSNPQRPPVIWIGAQECT WP_000145429.1 hydrogenase 2 small subunit [Salmonella enterica] Length: 372\nNumber of predicted TMHs: 1\nExp number of AAs in TMHs: 24.51309\nExp number, first 60 AAs: 0.2682\nTotal prob of N-in: 0.14402\noutside 1 331\nTMhelix 332 354\ninside 355 372

24676 GCF_001221725.1_5275_1_6Salmonella enterica subsp. enterica serovar Typhimurium str. DT104 Proteobacteria; Gammaproteobacteria; Enterobacterales; Enterobacteriaceae; Salmonella; Salmonella enterica; Salmonella enterica subsp. enterica; Salmonella enterica subsp. enterica serovar Typhimurium MTGDNTLITSHGINRRDFMKLCAALAATMGLSSKAAAEMAESVSNPQRPPVIWIGAQECT WP_000145429.1 hydrogenase 2 small subunit [Salmonella enterica] Length: 372\nNumber of predicted TMHs: 1\nExp number of AAs in TMHs: 24.51309\nExp number, first 60 AAs: 0.2682\nTotal prob of N-in: 0.14402\noutside 1 331\nTMhelix 332 354\ninside 355 372

24677 GCF_001221585.1_5275_7_9Salmonella enterica subsp. enterica serovar Typhimurium str. DT104 Proteobacteria; Gammaproteobacteria; Enterobacterales; Enterobacteriaceae; Salmonella; Salmonella enterica; Salmonella enterica subsp. enterica; Salmonella enterica subsp. enterica serovar Typhimurium MNNEETFYQAMRRKGVTRRSFLKFCSLAATSLGLGAGMTPKIAWALENKPRIPVVWIHGL WP_001058312.1 hydrogenase [Salmonella enterica] Length: 372\nNumber of predicted TMHs: 2\nExp number of AAs in TMHs: 36.51738\nExp number, first 60 AAs: 19.72849\nTotal prob of N-in: 0.96170\nPOSSIBLE N-term signal sequence\ninside 1 19\nTMhelix 20 39\noutside 40 325\nTMhelix 326 348\ninside 349 372

24678 GCF_001222345.1_5285_6_5Salmonella enterica subsp. enterica serovar Typhimurium str. DT104 Proteobacteria; Gammaproteobacteria; Enterobacterales; Enterobacteriaceae; Salmonella; Salmonella enterica; Salmonella enterica subsp. enterica; Salmonella enterica subsp. enterica serovar Typhimurium MTGDNTLITSHGINRRDFMKLCAALAATMGLSSKAAAEMAESVSNPQRPPVIWIGAQECT WP_000145429.1 hydrogenase 2 small subunit [Salmonella enterica] Length: 372\nNumber of predicted TMHs: 1\nExp number of AAs in TMHs: 24.51309\nExp number, first 60 AAs: 0.2682\nTotal prob of N-in: 0.14402\noutside 1 331\nTMhelix 332 354\ninside 355 372

24679 GCF_001220025.1_5275_8_8Salmonella enterica subsp. enterica serovar Typhimurium str. DT104 Proteobacteria; Gammaproteobacteria; Enterobacterales; Enterobacteriaceae; Salmonella; Salmonella enterica; Salmonella enterica subsp. enterica; Salmonella enterica subsp. enterica serovar Typhimurium MTGDNTLITSHGINRRDFMKLCAALAATMGLSSKAAAEMAESVSNPQRPPVIWIGAQECT WP_000145429.1 hydrogenase 2 small subunit [Salmonella enterica] Length: 372\nNumber of predicted TMHs: 1\nExp

number of AAs in TMHs: 24.51309\nExp number, first 60 AAs: 0.2682\nTotal prob of N-in: 0.14402\noutside 1 331\nTMhelix 332 354\ninside 355 372

24680 GCF_001221265.1_8080_3_39 Salmonella enterica subsp. enterica serovar Typhimurium str. DT104 Proteobacteria; Gammaproteobacteria; Enterobacterales; Enterobacteriaceae; Salmonella; Salmonella enterica; Salmonella enterica subsp. enterica; Salmonella enterica subsp. enterica serovar Typhimurium MNNEETFYQAMRRKGVTRRSFLKFCSLAATSLGLGAGMTPKIAWALENKPRIPVVWIHGL WP_001058312.1 hydrogenase [Salmonella enterica] Length: 372\nNumber of predicted TMHs: 2\nExp number of AAs in TMHs: 36.51738\nExp number, first 60 AAs: 19.72849\nTotal prob of N-in: 0.96170\nPOSSIBLE N-term signal sequence\ninside 1 19\nTMhelix 20 39\noutside 40 325\nTMhelix 326 348\ninside 349 372

24681 GCF_001221445.1_5299_6_6Salmonella enterica subsp. enterica serovar Typhimurium str. DT104 Proteobacteria; Gammaproteobacteria; Enterobacterales; Enterobacteriaceae; Salmonella; Salmonella enterica; Salmonella enterica subsp. enterica; Salmonella enterica subsp. enterica serovar Typhimurium MTGDNTLITSHGINRRDFMKLCAALAATMGLSSKAAAEMAESVSNPQRPPVIWIGAQECT WP_000145429.1 hydrogenase 2 small subunit [Salmonella enterica] Length: 372\nNumber of predicted TMHs: 1\nExp number of AAs in TMHs: 24.51309\nExp number, first 60 AAs: 0.2682\nTotal prob of N-in: 0.14402\noutside 1 331\nTMhelix 332 354\ninside 355 372

24682 GCF_001221505.1_8036_3_56 Salmonella enterica subsp. enterica serovar Typhimurium str. DT104 Proteobacteria; Gammaproteobacteria; Enterobacterales; Enterobacteriaceae; Salmonella; Salmonella enterica; Salmonella enterica subsp. enterica; Salmonella enterica subsp. enterica serovar Typhimurium MNNEETFYQAMRRKGVTRRSFLKFCSLAATSLGLGAGMTPKIAWALENKPRIPVVWIHGL WP_001058312.1 hydrogenase [Salmonella enterica] Length: 372\nNumber of predicted TMHs: 2\nExp number of AAs in TMHs: 36.51738\nExp number, first 60 AAs: 19.72849\nTotal prob of N-in: 0.96170\nPOSSIBLE N-term signal sequence\ninside 1 19\nTMhelix 20 39\noutside 40 325\nTMhelix 326 348\ninside 349 372

24683 GCF_001221305.1_5275_8_2Salmonella enterica subsp. enterica serovar Typhimurium str. DT104 Proteobacteria; Gammaproteobacteria; Enterobacterales; Enterobacteriaceae; Salmonella; Salmonella enterica; Salmonella enterica subsp. enterica; Salmonella enterica subsp. enterica serovar Typhimurium MNNEETFYQAMRRKGVTRRSFLKFCSLAATSLGLGAGMTPKIAWALENKPRIPVVWIHGL WP_001058312.1 hydrogenase [Salmonella enterica] Length: 372\nNumber of predicted TMHs: 2\nExp number of AAs in TMHs: 36.51738\nExp number, first 60 AAs: 19.72849\nTotal prob of N-in: 0.96170\nPOSSIBLE N-term signal sequence\ninside 1 19\nTMhelix 20 39\noutside 40 325\nTMhelix 326 348\ninside 349 372

24684 GCF_001221565.1_8036_3_85 Salmonella enterica subsp. enterica serovar Typhimurium str. DT104 Proteobacteria; Gammaproteobacteria; Enterobacterales; Enterobacteriaceae; Salmonella; Salmonella enterica; Salmonella enterica subsp. enterica; Salmonella enterica subsp. enterica serovar Typhimurium MTGDNTLITSHGINRRDFMKLCAALAATMGLSSKAAAEMAESVSNPQRPPVIWIGAQECT WP_000145429.1 hydrogenase 2 small subunit [Salmonella enterica] Length: 372\nNumber of predicted TMHs: 1\nExp number of AAs in TMHs: 24.51309\nExp number, first 60 AAs: 0.2682\nTotal prob of N-in: 0.14402\noutside 1 331\nTMhelix 332 354\ninside 355 372

24685 GCF_001221645.1_8080_3_22 Salmonella enterica subsp. enterica serovar Typhimurium str. DT104 Proteobacteria; Gammaproteobacteria; Enterobacterales; Enterobacteriaceae; Salmonella; Salmonella enterica; Salmonella enterica subsp. enterica; Salmonella enterica subsp. enterica serovar Typhimurium MTGDNTLITSHGINRRDFMKLCAALAATMGLSSKAAAEMAESVSNPQRPPVIWIGAQECT WP_000145429.1 hydrogenase 2 small subunit [Salmonella enterica] Length: 372\nNumber of predicted TMHs: 1\nExp number of AAs in TMHs: 24.51309\nExp number, first 60 AAs: 0.2682\nTotal prob of N-in: 0.14402\noutside 1 331\nTMhelix 332 354\ninside 355 372

24686 GCF_001221605.1_5299_6_10 Salmonella enterica subsp. enterica serovar Typhimurium str. DT104 Proteobacteria; Gammaproteobacteria; Enterobacterales; Enterobacteriaceae; Salmonella; Salmonella enterica; Salmonella enterica subsp. enterica; Salmonella enterica subsp. enterica serovar Typhimurium MTGDNTLITSHGINRRDFMKLCAALAATMGLSSKAAAEMAESVSNPQRPPVIWIGAQECT WP_000145429.1 hydrogenase 2 small subunit [Salmonella enterica] Length: 372\nNumber of predicted TMHs: 1\nExp number of AAs in TMHs: 24.51309\nExp number, first 60 AAs: 0.2682\nTotal prob of N-in: 0.14402\noutside 1 331\nTMhelix 332 354\ninside 355 372

24687 GCF_001221685.1_8080_3_64 Salmonella enterica subsp. enterica serovar Typhimurium str. DT104 Proteobacteria; Gammaproteobacteria; Enterobacterales; Enterobacteriaceae; Salmonella; Salmonella enterica;

Salmonella enterica subsp. enterica; Salmonella enterica subsp. enterica serovar Typhimurium
MTGDNTLITSHGINRRDFMKLCAALAATMGLSSKAAAEMAESVSNPQRPPVIWIGAQECT WP_000145429.1
hydrogenase 2 small subunit [Salmonella enterica] Length: 372\nNumber of predicted TMHs: 1\nExp number of AAs in TMHs: 24.51309\nExp number, first 60 AAs: 0.2682\nTotal prob of N-in: 0.14402\noutside 1 331\nTMhelix 332 354\ninside 355 372

24688 GCF_001221605.1_5299_6_10 Salmonella enterica subsp. enterica serovar Typhimurium str. DT104
Proteobacteria; Gammaproteobacteria; Enterobacterales; Enterobacteriaceae; Salmonella; Salmonella enterica; Salmonella enterica subsp. enterica; Salmonella enterica subsp. enterica serovar Typhimurium
MNNEETFYQAMRRKGVTRRSFLKFCSLAATSLGLGAGMTPKIAWALENKPRIPVVWIHGL WP_001058312.1
hydrogenase [Salmonella enterica] Length: 372\nNumber of predicted TMHs: 2\nExp number of AAs in TMHs: 36.51738\nExp number, first 60 AAs: 19.72849\nTotal prob of N-in: 0.96170\nPOSSIBLE N-term signal sequence\ninside 1 19\nTMhelix 20 39\noutside 40 325\nTMhelix 326 348\ninside 349 372

24689 GCF_001221325.1_8080_3_16 Salmonella enterica subsp. enterica serovar Typhimurium str. DT104
Proteobacteria; Gammaproteobacteria; Enterobacterales; Enterobacteriaceae; Salmonella; Salmonella enterica; Salmonella enterica subsp. enterica; Salmonella enterica subsp. enterica serovar Typhimurium
MNNEETFYQAMRRKGVTRRSFLKFCSLAATSLGLGAGMTPKIAWALENKPRIPVVWIHGL WP_001058312.1
hydrogenase [Salmonella enterica] Length: 372\nNumber of predicted TMHs: 2\nExp number of AAs in TMHs: 36.51738\nExp number, first 60 AAs: 19.72849\nTotal prob of N-in: 0.96170\nPOSSIBLE N-term signal sequence\ninside 1 19\nTMhelix 20 39\noutside 40 325\nTMhelix 326 348\ninside 349 372

24690 GCF_001221345.1_8080_3_12 Salmonella enterica subsp. enterica serovar Typhimurium str. DT104
Proteobacteria; Gammaproteobacteria; Enterobacterales; Enterobacteriaceae; Salmonella; Salmonella enterica; Salmonella enterica subsp. enterica; Salmonella enterica subsp. enterica serovar Typhimurium
MNNEETFYQAMRRKGVTRRSFLKFCSLAATSLGLGAGMTPKIAWALENKPRIPVVWIHGL WP_001058312.1
hydrogenase [Salmonella enterica] Length: 372\nNumber of predicted TMHs: 2\nExp number of AAs in TMHs: 36.51738\nExp number, first 60 AAs: 19.72849\nTotal prob of N-in: 0.96170\nPOSSIBLE N-term signal sequence\ninside 1 19\nTMhelix 20 39\noutside 40 325\nTMhelix 326 348\ninside 349 372

24691 GCF_001221385.1_9066_1_7Salmonella enterica subsp. enterica serovar Typhimurium str. DT104
Proteobacteria; Gammaproteobacteria; Enterobacterales; Enterobacteriaceae; Salmonella; Salmonella enterica; Salmonella enterica subsp. enterica; Salmonella enterica subsp. enterica serovar Typhimurium
MNNEETFYQAMRRKGVTRRSFLKFCSLAATSLGLGAGMTPKIAWALENKPRIPVVWIHGL WP_001058312.1
hydrogenase [Salmonella enterica] Length: 372\nNumber of predicted TMHs: 2\nExp number of AAs in TMHs: 36.51738\nExp number, first 60 AAs: 19.72849\nTotal prob of N-in: 0.96170\nPOSSIBLE N-term signal sequence\ninside 1 19\nTMhelix 20 39\noutside 40 325\nTMhelix 326 348\ninside 349 372

24692 GCF_001216845.1_5299_3_8Salmonella enterica subsp. enterica serovar Typhimurium str. DT104
Proteobacteria; Gammaproteobacteria; Enterobacterales; Enterobacteriaceae; Salmonella; Salmonella enterica; Salmonella enterica subsp. enterica; Salmonella enterica subsp. enterica serovar Typhimurium
MNNEETFYQAMRRKGVTRRSFLKFCSLAATSLGLGAGMTPKIAWALENKPRIPVVWIHGL WP_001058312.1
hydrogenase [Salmonella enterica] Length: 372\nNumber of predicted TMHs: 2\nExp number of AAs in TMHs: 36.51738\nExp number, first 60 AAs: 19.72849\nTotal prob of N-in: 0.96170\nPOSSIBLE N-term signal sequence\ninside 1 19\nTMhelix 20 39\noutside 40 325\nTMhelix 326 348\ninside 349 372

24693 GCF_001221565.1_8036_3_85 Salmonella enterica subsp. enterica serovar Typhimurium str. DT104
Proteobacteria; Gammaproteobacteria; Enterobacterales; Enterobacteriaceae; Salmonella; Salmonella enterica; Salmonella enterica subsp. enterica; Salmonella enterica subsp. enterica serovar Typhimurium
MNNEETFYQAMRRKGVTRRSFLKFCSLAATSLGLGAGMTPKIAWALENKPRIPVVWIHGL WP_001058312.1
hydrogenase [Salmonella enterica] Length: 372\nNumber of predicted TMHs: 2\nExp number of AAs in TMHs: 36.51738\nExp number, first 60 AAs: 19.72849\nTotal prob of N-in: 0.96170\nPOSSIBLE N-term signal sequence\ninside 1 19\nTMhelix 20 39\noutside 40 325\nTMhelix 326 348\ninside 349 372

24694 GCF_001221645.1_8080_3_22 Salmonella enterica subsp. enterica serovar Typhimurium str. DT104
Proteobacteria; Gammaproteobacteria; Enterobacterales; Enterobacteriaceae; Salmonella; Salmonella enterica; Salmonella enterica subsp. enterica; Salmonella enterica subsp. enterica serovar Typhimurium
MNNEETFYQAMRRKGVTRRSFLKFCSLAATSLGLGAGMTPKIAWALENKPRIPVVWIHGL WP_001058312.1
hydrogenase [Salmonella enterica] Length: 372\nNumber of predicted TMHs: 2\nExp number of AAs in TMHs: 36.51738\nExp number, first 60 AAs: 19.72849\nTotal prob of N-in: 0.96170\nPOSSIBLE N-term signal sequence\ninside 1 19\nTMhelix 20 39\noutside 40 325\nTMhelix 326 348\ninside 349 372

24695 GCF_001221685.1_8080_3_64 Salmonella enterica subsp. enterica serovar Typhimurium str. DT104 Proteobacteria; Gammaproteobacteria; Enterobacterales; Enterobacteriaceae; Salmonella; Salmonella enterica; Salmonella enterica subsp. enterica; Salmonella enterica subsp. enterica serovar Typhimurium MNNEETFYQAMRRKGVTRRSFLKFCSLAATSLGLGAGMTPKIAWALENKPRIPVVWIHGL WP_001058312.1 hydrogenase [Salmonella enterica] Length: 372\nNumber of predicted TMHs: 2\nExp number of AAs in TMHs: 36.51738\nExp number, first 60 AAs: 19.72849\nTotal prob of N-in: 0.96170\nPOSSIBLE N-term signal sequence\ninside 1 19\nTMhelix 20 39\noutside 40 325\nTMhelix 326 348\ninside 349 372

24696 GCF_001221705.1_8080_3_93 Salmonella enterica subsp. enterica serovar Typhimurium str. DT104 Proteobacteria; Gammaproteobacteria; Enterobacterales; Enterobacteriaceae; Salmonella; Salmonella enterica; Salmonella enterica subsp. enterica; Salmonella enterica subsp. enterica serovar Typhimurium MTGDNTLITSHGINRRDFMKLCAALAATMGLSSKAAAEMAESVSNPQRPPVIWIGAQECT WP_000145429.1 hydrogenase 2 small subunit [Salmonella enterica] Length: 372\nNumber of predicted TMHs: 1\nExp number of AAs in TMHs: 24.51309\nExp number, first 60 AAs: 0.2682\nTotal prob of N-in: 0.14402\noutside 1 331\nTMhelix 332 354\ninside 355 372

24697 GCF_001221745.1_8080_3_75 Salmonella enterica subsp. enterica serovar Typhimurium str. DT104 Proteobacteria; Gammaproteobacteria; Enterobacterales; Enterobacteriaceae; Salmonella; Salmonella enterica; Salmonella enterica subsp. enterica; Salmonella enterica subsp. enterica serovar Typhimurium MTGDNTLITSHGINRRDFMKLCAALAATMGLSSKAAAEMAESVSNPQRPPVIWIGAQECT WP_000145429.1 hydrogenase 2 small subunit [Salmonella enterica] Length: 372\nNumber of predicted TMHs: 1\nExp number of AAs in TMHs: 24.51309\nExp number, first 60 AAs: 0.2682\nTotal prob of N-in: 0.14402\noutside 1 331\nTMhelix 332 354\ninside 355 372

24698 GCF_001221785.1_8080_3_60 Salmonella enterica subsp. enterica serovar Typhimurium str. DT104 Proteobacteria; Gammaproteobacteria; Enterobacterales; Enterobacteriaceae; Salmonella; Salmonella enterica; Salmonella enterica subsp. enterica; Salmonella enterica subsp. enterica serovar Typhimurium MTGDNTLITSHGINRRDFMKLCAALAATMGLSSKAAAEMAESVSNPQRPPVIWIGAQECT WP_000145429.1 hydrogenase 2 small subunit [Salmonella enterica] Length: 372\nNumber of predicted TMHs: 1\nExp number of AAs in TMHs: 24.51309\nExp number, first 60 AAs: 0.2682\nTotal prob of N-in: 0.14402\noutside 1 331\nTMhelix 332 354\ninside 355 372

24699 GCF_001221845.1_8080_3_55 Salmonella enterica subsp. enterica serovar Typhimurium str. DT104 Proteobacteria; Gammaproteobacteria; Enterobacterales; Enterobacteriaceae; Salmonella; Salmonella enterica; Salmonella enterica subsp. enterica; Salmonella enterica subsp. enterica serovar Typhimurium MTGDNTLITSHGINRRDFMKLCAALAATMGLSSKAAAEMAESVSNPQRPPVIWIGAQECT WP_000145429.1 hydrogenase 2 small subunit [Salmonella enterica] Length: 372\nNumber of predicted TMHs: 1\nExp number of AAs in TMHs: 24.51309\nExp number, first 60 AAs: 0.2682\nTotal prob of N-in: 0.14402\noutside 1 331\nTMhelix 332 354\ninside 355 372

24700 GCF_001221925.1_8080_3_9Salmonella enterica subsp. enterica serovar Typhimurium str. DT104 Proteobacteria; Gammaproteobacteria; Enterobacterales; Enterobacteriaceae; Salmonella; Salmonella enterica; Salmonella enterica subsp. enterica; Salmonella enterica subsp. enterica serovar Typhimurium MTGDNTLITSHGINRRDFMKLCAALAATMGLSSKAAAEMAESVSNPQRPPVIWIGAQECT WP_000145429.1 hydrogenase 2 small subunit [Salmonella enterica] Length: 372\nNumber of predicted TMHs: 1\nExp number of AAs in TMHs: 24.51309\nExp number, first 60 AAs: 0.2682\nTotal prob of N-in: 0.14402\noutside 1 331\nTMhelix 332 354\ninside 355 372

24701 GCF_001326275.1_5275_1_5Salmonella enterica subsp. enterica serovar Typhimurium str. DT104 Proteobacteria; Gammaproteobacteria; Enterobacterales; Enterobacteriaceae; Salmonella; Salmonella enterica; Salmonella enterica subsp. enterica; Salmonella enterica subsp. enterica serovar Typhimurium MNNEETFYQAMRRKGVTRRSFLKFCSLAATSLGLGAGMTPKIAWALENKPRIPVVWIHGL WP_001058312.1 hydrogenase [Salmonella enterica] Length: 372\nNumber of predicted TMHs: 2\nExp number of AAs in TMHs: 36.51738\nExp number, first 60 AAs: 19.72849\nTotal prob of N-in: 0.96170\nPOSSIBLE N-term signal sequence\ninside 1 19\nTMhelix 20 39\noutside 40 325\nTMhelix 326 348\ninside 349 372

24702 GCF_001221885.1_9065_1_1Salmonella enterica subsp. enterica serovar Typhimurium str. DT104 Proteobacteria; Gammaproteobacteria; Enterobacterales; Enterobacteriaceae; Salmonella; Salmonella enterica; Salmonella enterica subsp. enterica; Salmonella enterica subsp. enterica serovar Typhimurium MTGDNTLITSHGINRRDFMKLCAALAATMGLSSKAAAEMAESVSNPQRPPVIWIGAQECT WP_000145429.1 hydrogenase 2 small subunit [Salmonella enterica] Length: 372\nNumber of predicted TMHs: 1\nExp

number of AAs in TMHs: 24.51309\nExp number, first 60 AAs: 0.2682\nTotal prob of N-in: 0.14402\noutside 1 331\nTMhelix 332 354\ninside 355 372

24703 GCF_001221885.1_9065_1_1Salmonella enterica subsp. enterica serovar Typhimurium str. DT104 Proteobacteria; Gammaproteobacteria; Enterobacterales; Enterobacteriaceae; Salmonella; Salmonella enterica; Salmonella enterica subsp. enterica; Salmonella enterica subsp. enterica serovar Typhimurium MNNEETFYQAMRRKGVTRRSFLKFCSLAATSLGLGAGMTPKIAWALENKPRIPVVWIHGL WP_001058312.1 hydrogenase [Salmonella enterica] Length: 372\nNumber of predicted TMHs: 2\nExp number of AAs in TMHs: 36.51738\nExp number, first 60 AAs: 19.72849\nTotal prob of N-in: 0.96170\nPOSSIBLE N-term signal sequence\ninside 1 19\nTMhelix 20 39\noutside 40 325\nTMhelix 326 348\ninside 349 372

24704 GCF_001221705.1_8080_3_93 Salmonella enterica subsp. enterica serovar Typhimurium str. DT104 Proteobacteria; Gammaproteobacteria; Enterobacterales; Enterobacteriaceae; Salmonella; Salmonella enterica; Salmonella enterica subsp. enterica; Salmonella enterica subsp. enterica serovar Typhimurium MNNEETFYQAMRRKGVTRRSFLKFCSLAATSLGLGAGMTPKIAWALENKPRIPVVWIHGL WP_001058312.1 hydrogenase [Salmonella enterica] Length: 372\nNumber of predicted TMHs: 2\nExp number of AAs in TMHs: 36.51738\nExp number, first 60 AAs: 19.72849\nTotal prob of N-in: 0.96170\nPOSSIBLE N-term signal sequence\ninside 1 19\nTMhelix 20 39\noutside 40 325\nTMhelix 326 348\ninside 349 372

24705 GCF_001216585.1_5388_1_12 Salmonella enterica subsp. enterica serovar Typhimurium str. DT104 Proteobacteria; Gammaproteobacteria; Enterobacterales; Enterobacteriaceae; Salmonella; Salmonella enterica; Salmonella enterica subsp. enterica; Salmonella enterica subsp. enterica serovar Typhimurium MNNEETFYQAMRRKGVTRRSFLKFCSLAATSLGLGAGMTPKIAWALENKPRIPVVWIHGL WP_001058312.1 hydrogenase [Salmonella enterica] Length: 372\nNumber of predicted TMHs: 2\nExp number of AAs in TMHs: 36.51738\nExp number, first 60 AAs: 19.72849\nTotal prob of N-in: 0.96170\nPOSSIBLE N-term signal sequence\ninside 1 19\nTMhelix 20 39\noutside 40 325\nTMhelix 326 348\ninside 349 372

24706 GCF_001221745.1_8080_3_75 Salmonella enterica subsp. enterica serovar Typhimurium str. DT104 Proteobacteria; Gammaproteobacteria; Enterobacterales; Enterobacteriaceae; Salmonella; Salmonella enterica; Salmonella enterica subsp. enterica; Salmonella enterica subsp. enterica serovar Typhimurium MNNEETFYQAMRRKGVTRRSFLKFCSLAATSLGLGAGMTPKIAWALENKPRIPVVWIHGL WP_001058312.1 hydrogenase [Salmonella enterica] Length: 372\nNumber of predicted TMHs: 2\nExp number of AAs in TMHs: 36.51738\nExp number, first 60 AAs: 19.72849\nTotal prob of N-in: 0.96170\nPOSSIBLE N-term signal sequence\ninside 1 19\nTMhelix 20 39\noutside 40 325\nTMhelix 326 348\ninside 349 372

24707 GCF_001221985.1_8036_3_70 Salmonella enterica subsp. enterica serovar Typhimurium str. DT104 Proteobacteria; Gammaproteobacteria; Enterobacterales; Enterobacteriaceae; Salmonella; Salmonella enterica; Salmonella enterica subsp. enterica; Salmonella enterica subsp. enterica serovar Typhimurium MTGDNTLITSHGINRRDFMKLCAALAATMGLSSKAAAEMAESVSNPQRPPVIWIGAQECT WP_000145429.1 hydrogenase 2 small subunit [Salmonella enterica] Length: 372\nNumber of predicted TMHs: 1\nExp number of AAs in TMHs: 24.51309\nExp number, first 60 AAs: 0.2682\nTotal prob of N-in: 0.14402\noutside 1 331\nTMhelix 332 354\ninside 355 372

24708 GCF_001222025.1_8080_3_42 Salmonella enterica subsp. enterica serovar Typhimurium str. DT104 Proteobacteria; Gammaproteobacteria; Enterobacterales; Enterobacteriaceae; Salmonella; Salmonella enterica; Salmonella enterica subsp. enterica; Salmonella enterica subsp. enterica serovar Typhimurium MTGDNTLITSHGINRRDFMKLCAALAATMGLSSKAAAEMAESVSNPQRPPVIWIGAQECT WP_000145429.1 hydrogenase 2 small subunit [Salmonella enterica] Length: 372\nNumber of predicted TMHs: 1\nExp number of AAs in TMHs: 24.51309\nExp number, first 60 AAs: 0.2682\nTotal prob of N-in: 0.14402\noutside 1 331\nTMhelix 332 354\ninside 355 372

24709 GCF_001222085.1_8080_3_3Salmonella enterica subsp. enterica serovar Typhimurium str. DT104 Proteobacteria; Gammaproteobacteria; Enterobacterales; Enterobacteriaceae; Salmonella; Salmonella enterica; Salmonella enterica subsp. enterica; Salmonella enterica subsp. enterica serovar Typhimurium MTGDNTLITSHGINRRDFMKLCAALAATMGLSSKAAAEMAESVSNPQRPPVIWIGAQECT WP_000145429.1 hydrogenase 2 small subunit [Salmonella enterica] Length: 372\nNumber of predicted TMHs: 1\nExp number of AAs in TMHs: 24.51309\nExp number, first 60 AAs: 0.2682\nTotal prob of N-in: 0.14402\noutside 1 331\nTMhelix 332 354\ninside 355 372

24710 GCF_001222165.1_8036_3_78 Salmonella enterica subsp. enterica serovar Typhimurium str. DT104 Proteobacteria; Gammaproteobacteria; Enterobacterales; Enterobacteriaceae; Salmonella; Salmonella enterica;

Salmonella enterica subsp. enterica; Salmonella enterica subsp. enterica serovar Typhimurium
MTGDNTLITSHGINRRDFMKLCAALAATMGLSSKAAAEMAESVSNPQRPPVIWIGAQECT WP_000145429.1
hydrogenase 2 small subunit [Salmonella enterica] Length: 372\nNumber of predicted TMHs: 1\nExp number of AAs in TMHs: 24.51309\nExp number, first 60 AAs: 0.2682\nTotal prob of N-in: 0.14402\noutside 1 331\nTMhelix 332 354\ninside 355 372

24711 GCF_001222145.1_5299_3_9Salmonella enterica subsp. enterica serovar Typhimurium str. DT104
Proteobacteria; Gammaproteobacteria; Enterobacterales; Enterobacteriaceae; Salmonella; Salmonella enterica; Salmonella enterica subsp. enterica; Salmonella enterica subsp. enterica serovar Typhimurium
MTGDNTLITSHGINRRDFMKLCAALAATMGLSSKAAAEMAESVSNPQRPPVIWIGAQECT WP_000145429.1
hydrogenase 2 small subunit [Salmonella enterica] Length: 372\nNumber of predicted TMHs: 1\nExp number of AAs in TMHs: 24.51309\nExp number, first 60 AAs: 0.2682\nTotal prob of N-in: 0.14402\noutside 1 331\nTMhelix 332 354\ninside 355 372

24712 GCF_001326135.1_8080_3_62 Salmonella enterica subsp. enterica serovar Typhimurium str. DT104
Proteobacteria; Gammaproteobacteria; Enterobacterales; Enterobacteriaceae; Salmonella; Salmonella enterica; Salmonella enterica subsp. enterica; Salmonella enterica subsp. enterica serovar Typhimurium
MNNEETFYQAMRRKGVTRRSFLKFCSLAATSLGLGAGMTPKIAWALENKPRIPVVWIHGL WP_001058312.1
hydrogenase [Salmonella enterica] Length: 372\nNumber of predicted TMHs: 2\nExp number of AAs in TMHs: 36.51738\nExp number, first 60 AAs: 19.72849\nTotal prob of N-in: 0.96170\nPOSSIBLE N-term signal sequence\ninside 1 19\nTMhelix 20 39\noutside 40 325\nTMhelix 326 348\ninside 349 372

24713 GCF_001222205.1_5275_1_1Salmonella enterica subsp. enterica serovar Typhimurium str. DT104
Proteobacteria; Gammaproteobacteria; Enterobacterales; Enterobacteriaceae; Salmonella; Salmonella enterica; Salmonella enterica subsp. enterica; Salmonella enterica subsp. enterica serovar Typhimurium
MTGDNTLITSHGINRRDFMKLCAALAATMGLSSKAAAEMAESVSNPQRPPVIWIGAQECT WP_000145429.1
hydrogenase 2 small subunit [Salmonella enterica] Length: 372\nNumber of predicted TMHs: 1\nExp number of AAs in TMHs: 24.51309\nExp number, first 60 AAs: 0.2682\nTotal prob of N-in: 0.14402\noutside 1 331\nTMhelix 332 354\ninside 355 372

24714 GCF_001222205.1_5275_1_1Salmonella enterica subsp. enterica serovar Typhimurium str. DT104
Proteobacteria; Gammaproteobacteria; Enterobacterales; Enterobacteriaceae; Salmonella; Salmonella enterica; Salmonella enterica subsp. enterica; Salmonella enterica subsp. enterica serovar Typhimurium
MNNEETFYQAMRRKGVTRRSFLKFCSLAATSLGLGAGMTPKIAWALENKPRIPVVWIHGL WP_001058312.1
hydrogenase [Salmonella enterica] Length: 372\nNumber of predicted TMHs: 2\nExp number of AAs in TMHs: 36.51738\nExp number, first 60 AAs: 19.72849\nTotal prob of N-in: 0.96170\nPOSSIBLE N-term signal sequence\ninside 1 19\nTMhelix 20 39\noutside 40 325\nTMhelix 326 348\ninside 349 372

24715 GCF_001221845.1_8080_3_55 Salmonella enterica subsp. enterica serovar Typhimurium str. DT104
Proteobacteria; Gammaproteobacteria; Enterobacterales; Enterobacteriaceae; Salmonella; Salmonella enterica; Salmonella enterica subsp. enterica; Salmonella enterica subsp. enterica serovar Typhimurium
MNNEETFYQAMRRKGVTRRSFLKFCSLAATSLGLGAGMTPKIAWALENKPRIPVVWIHGL WP_001058312.1
hydrogenase [Salmonella enterica] Length: 372\nNumber of predicted TMHs: 2\nExp number of AAs in TMHs: 36.51738\nExp number, first 60 AAs: 19.72849\nTotal prob of N-in: 0.96170\nPOSSIBLE N-term signal sequence\ninside 1 19\nTMhelix 20 39\noutside 40 325\nTMhelix 326 348\ninside 349 372

24716 GCF_001221925.1_8080_3_9Salmonella enterica subsp. enterica serovar Typhimurium str. DT104
Proteobacteria; Gammaproteobacteria; Enterobacterales; Enterobacteriaceae; Salmonella; Salmonella enterica; Salmonella enterica subsp. enterica; Salmonella enterica subsp. enterica serovar Typhimurium
MNNEETFYQAMRRKGVTRRSFLKFCSLAATSLGLGAGMTPKIAWALENKPRIPVVWIHGL WP_001058312.1
hydrogenase [Salmonella enterica] Length: 372\nNumber of predicted TMHs: 2\nExp number of AAs in TMHs: 36.51738\nExp number, first 60 AAs: 19.72849\nTotal prob of N-in: 0.96170\nPOSSIBLE N-term signal sequence\ninside 1 19\nTMhelix 20 39\noutside 40 325\nTMhelix 326 348\ninside 349 372

24717 GCF_001221985.1_8036_3_70 Salmonella enterica subsp. enterica serovar Typhimurium str. DT104
Proteobacteria; Gammaproteobacteria; Enterobacterales; Enterobacteriaceae; Salmonella; Salmonella enterica; Salmonella enterica subsp. enterica; Salmonella enterica subsp. enterica serovar Typhimurium
MNNEETFYQAMRRKGVTRRSFLKFCSLAATSLGLGAGMTPKIAWALENKPRIPVVWIHGL WP_001058312.1
hydrogenase [Salmonella enterica] Length: 372\nNumber of predicted TMHs: 2\nExp number of AAs in TMHs: 36.51738\nExp number, first 60 AAs: 19.72849\nTotal prob of N-in: 0.96170\nPOSSIBLE N-term signal sequence\ninside 1 19\nTMhelix 20 39\noutside 40 325\nTMhelix 326 348\ninside 349 372

24718 GCF_001222145.1_5299_3_9Salmonella enterica subsp. enterica serovar Typhimurium str. DT104 Proteobacteria; Gammaproteobacteria; Enterobacterales; Enterobacteriaceae; Salmonella; Salmonella enterica; Salmonella enterica subsp. enterica; Salmonella enterica subsp. enterica serovar Typhimurium MNNEETFYQAMRRKGVTRRSFLKFCSLAATSLGLGAGMTPKIAWALENKPRIPVVWIHGL WP_001058312.1 hydrogenase [Salmonella enterica] Length: 372\nNumber of predicted TMHs: 2\nExp number of AAs in TMHs: 36.51738\nExp number, first 60 AAs: 19.72849\nTotal prob of N-in: 0.96170\nPOSSIBLE N-term signal sequence\ninside 1 19\nTMhelix 20 39\noutside 40 325\nTMhelix 326 348\ninside 349 372

24719 GCF_001222085.1_8080_3_3Salmonella enterica subsp. enterica serovar Typhimurium str. DT104 Proteobacteria; Gammaproteobacteria; Enterobacterales; Enterobacteriaceae; Salmonella; Salmonella enterica; Salmonella enterica subsp. enterica; Salmonella enterica subsp. enterica serovar Typhimurium MNNEETFYQAMRRKGVTRRSFLKFCSLAATSLGLGAGMTPKIAWALENKPRIPVVWIHGL WP_001058312.1 hydrogenase [Salmonella enterica] Length: 372\nNumber of predicted TMHs: 2\nExp number of AAs in TMHs: 36.51738\nExp number, first 60 AAs: 19.72849\nTotal prob of N-in: 0.96170\nPOSSIBLE N-term signal sequence\ninside 1 19\nTMhelix 20 39\noutside 40 325\nTMhelix 326 348\ninside 349 372

24720 GCF_001222165.1_8036_3_78 Salmonella enterica subsp. enterica serovar Typhimurium str. DT104 Proteobacteria; Gammaproteobacteria; Enterobacterales; Enterobacteriaceae; Salmonella; Salmonella enterica; Salmonella enterica subsp. enterica; Salmonella enterica subsp. enterica serovar Typhimurium MNNEETFYQAMRRKGVTRRSFLKFCSLAATSLGLGAGMTPKIAWALENKPRIPVVWIHGL WP_001058312.1 hydrogenase [Salmonella enterica] Length: 372\nNumber of predicted TMHs: 2\nExp number of AAs in TMHs: 36.51738\nExp number, first 60 AAs: 19.72849\nTotal prob of N-in: 0.96170\nPOSSIBLE N-term signal sequence\ninside 1 19\nTMhelix 20 39\noutside 40 325\nTMhelix 326 348\ninside 349 372

24721 GCF_001222025.1_8080_3_42 Salmonella enterica subsp. enterica serovar Typhimurium str. DT104 Proteobacteria; Gammaproteobacteria; Enterobacterales; Enterobacteriaceae; Salmonella; Salmonella enterica; Salmonella enterica subsp. enterica; Salmonella enterica subsp. enterica serovar Typhimurium MNNEETFYQAMRRKGVTRRSFLKFCSLAATSLGLGAGMTPKIAWALENKPRIPVVWIHGL WP_001058312.1 hydrogenase [Salmonella enterica] Length: 372\nNumber of predicted TMHs: 2\nExp number of AAs in TMHs: 36.51738\nExp number, first 60 AAs: 19.72849\nTotal prob of N-in: 0.96170\nPOSSIBLE N-term signal sequence\ninside 1 19\nTMhelix 20 39\noutside 40 325\nTMhelix 326 348\ninside 349 372

24722 GCF_001221785.1_8080_3_60 Salmonella enterica subsp. enterica serovar Typhimurium str. DT104 Proteobacteria; Gammaproteobacteria; Enterobacterales; Enterobacteriaceae; Salmonella; Salmonella enterica; Salmonella enterica subsp. enterica; Salmonella enterica subsp. enterica serovar Typhimurium MNNEETFYQAMRRKGVTRRSFLKFCSLAATSLGLGAGMTPKIAWALENKPRIPVVWIHGL WP_050196264.1 hydrogenase [Salmonella enterica] Length: 372\nNumber of predicted TMHs: 2\nExp number of AAs in TMHs: 36.76337\nExp number, first 60 AAs: 19.73298\nTotal prob of N-in: 0.96170\nPOSSIBLE N-term signal sequence\ninside 1 19\nTMhelix 20 39\noutside 40 325\nTMhelix 326 348\ninside 349 372

24723 GCF_001222245.1_8080_3_52 Salmonella enterica subsp. enterica serovar Typhimurium str. DT104 Proteobacteria; Gammaproteobacteria; Enterobacterales; Enterobacteriaceae; Salmonella; Salmonella enterica; Salmonella enterica subsp. enterica; Salmonella enterica subsp. enterica serovar Typhimurium MTGDNTLITSHGINRRDFMKLCAALAATMGLSSKAAAEMAESVSNPQRPPVIWIGAQECT WP_000145429.1 hydrogenase 2 small subunit [Salmonella enterica] Length: 372\nNumber of predicted TMHs: 1\nExp number of AAs in TMHs: 24.51309\nExp number, first 60 AAs: 0.2682\nTotal prob of N-in: 0.14402\noutside 1 331\nTMhelix 332 354\ninside 355 372

24724 GCF_001222325.1_8080_3_73 Salmonella enterica subsp. enterica serovar Typhimurium str. DT104 Proteobacteria; Gammaproteobacteria; Enterobacterales; Enterobacteriaceae; Salmonella; Salmonella enterica; Salmonella enterica subsp. enterica; Salmonella enterica subsp. enterica serovar Typhimurium MTGDNTLITSHGINRRDFMKLCAALAATMGLSSKAAAEMAESVSNPQRPPVIWIGAQECT WP_000145429.1 hydrogenase 2 small subunit [Salmonella enterica] Length: 372\nNumber of predicted TMHs: 1\nExp number of AAs in TMHs: 24.51309\nExp number, first 60 AAs: 0.2682\nTotal prob of N-in: 0.14402\noutside 1 331\nTMhelix 332 354\ninside 355 372

24725 GCF_001222365.1_8080_3_85 Salmonella enterica subsp. enterica serovar Typhimurium str. DT104 Proteobacteria; Gammaproteobacteria; Enterobacterales; Enterobacteriaceae; Salmonella; Salmonella enterica; Salmonella enterica subsp. enterica; Salmonella enterica subsp. enterica serovar Typhimurium MTGDNTLITSHGINRRDFMKLCAALAATMGLSSKAAAEMAESVSNPQRPPVIWIGAQECT WP_000145429.1 hydrogenase 2 small subunit [Salmonella enterica] Length: 372\nNumber of predicted TMHs: 1\nExp

number of AAs in TMHs: 24.51309\nExp number, first 60 AAs:  0.2682\nTotal prob of N-in:   0.14402\noutside   1   331\nTMhelix   332   354\ninside   355   372

24726 GCF_001222405.1_8036_3_84 Salmonella enterica subsp. enterica serovar Typhimurium str. DT104 Proteobacteria; Gammaproteobacteria; Enterobacterales; Enterobacteriaceae; Salmonella; Salmonella enterica; Salmonella enterica subsp. enterica; Salmonella enterica subsp. enterica serovar Typhimurium MTGDNTLITSHGINRRDFMKLCAALAATMGLSSKAAAEMAESVSNPQRPPVIWIGAQECT WP_000145429.1 hydrogenase 2 small subunit [Salmonella enterica] Length: 372\nNumber of predicted TMHs:  1\nExp number of AAs in TMHs: 24.51309\nExp number, first 60 AAs:  0.2682\nTotal prob of N-in:   0.14402\noutside   1   331\nTMhelix   332   354\ninside   355   372

24727 GCF_001222525.1_8080_3_27 Salmonella enterica subsp. enterica serovar Typhimurium str. DT104 Proteobacteria; Gammaproteobacteria; Enterobacterales; Enterobacteriaceae; Salmonella; Salmonella enterica; Salmonella enterica subsp. enterica; Salmonella enterica subsp. enterica serovar Typhimurium MTGDNTLITSHGINRRDFMKLCAALAATMGLSSKAAAEMAESVSNPQRPPVIWIGAQECT WP_000145429.1 hydrogenase 2 small subunit [Salmonella enterica] Length: 372\nNumber of predicted TMHs:  1\nExp number of AAs in TMHs: 24.51309\nExp number, first 60 AAs:  0.2682\nTotal prob of N-in:   0.14402\noutside   1   331\nTMhelix   332   354\ninside   355   372

24728 GCF_001326175.1_9066_1_9Salmonella enterica subsp. enterica serovar Typhimurium str. DT104 Proteobacteria; Gammaproteobacteria; Enterobacterales; Enterobacteriaceae; Salmonella; Salmonella enterica; Salmonella enterica subsp. enterica; Salmonella enterica subsp. enterica serovar Typhimurium MNNEETFYQAMRRKGVTRRSFLKFCSLAATSLGLGAGMTPKIAWALENKPRIPVVWIHGL WP_001058312.1 hydrogenase [Salmonella enterica] Length: 372\nNumber of predicted TMHs:  2\nExp number of AAs in TMHs: 36.51738\nExp number, first 60 AAs:  19.72849\nTotal prob of N-in:   0.96170\nPOSSIBLE N-term signal sequence\ninside   1   19\nTMhelix   20   39\noutside   40   325\nTMhelix   326   348\ninside   349   372

24729 GCF_001222485.1_5295_2_11 Salmonella enterica subsp. enterica serovar Typhimurium str. DT104 Proteobacteria; Gammaproteobacteria; Enterobacterales; Enterobacteriaceae; Salmonella; Salmonella enterica; Salmonella enterica subsp. enterica; Salmonella enterica subsp. enterica serovar Typhimurium MTGDNTLITSHGINRRDFMKLCAALAATMGLSSKAAAEMAESVSNPQRPPVIWIGAQECT WP_000145429.1 hydrogenase 2 small subunit [Salmonella enterica] Length: 372\nNumber of predicted TMHs:  1\nExp number of AAs in TMHs: 24.51309\nExp number, first 60 AAs:  0.2682\nTotal prob of N-in:   0.14402\noutside   1   331\nTMhelix   332   354\ninside   355   372

24730 GCF_001222485.1_5295_2_11 Salmonella enterica subsp. enterica serovar Typhimurium str. DT104 Proteobacteria; Gammaproteobacteria; Enterobacterales; Enterobacteriaceae; Salmonella; Salmonella enterica; Salmonella enterica subsp. enterica; Salmonella enterica subsp. enterica serovar Typhimurium MNNEETFYQAMRRKGVTRRSFLKFCSLAATSLGLGAGMTPKIAWALENKPRIPVVWIHGL WP_001058312.1 hydrogenase [Salmonella enterica] Length: 372\nNumber of predicted TMHs:  2\nExp number of AAs in TMHs: 36.51738\nExp number, first 60 AAs:  19.72849\nTotal prob of N-in:   0.96170\nPOSSIBLE N-term signal sequence\ninside   1   19\nTMhelix   20   39\noutside   40   325\nTMhelix   326   348\ninside   349   372

24731 GCF_001222965.1_8036_3_72 Salmonella enterica subsp. enterica serovar Typhimurium str. DT104 Proteobacteria; Gammaproteobacteria; Enterobacterales; Enterobacteriaceae; Salmonella; Salmonella enterica; Salmonella enterica subsp. enterica; Salmonella enterica subsp. enterica serovar Typhimurium MTGDNTLITSHGINRRDFMKLCAALAATMGLSSKAAAEMAESVSNPQRPPVIWIGAQECT WP_000145429.1 hydrogenase 2 small subunit [Salmonella enterica] Length: 372\nNumber of predicted TMHs:  1\nExp number of AAs in TMHs: 24.51309\nExp number, first 60 AAs:  0.2682\nTotal prob of N-in:   0.14402\noutside   1   331\nTMhelix   332   354\ninside   355   372

24732 GCF_001222245.1_8080_3_52 Salmonella enterica subsp. enterica serovar Typhimurium str. DT104 Proteobacteria; Gammaproteobacteria; Enterobacterales; Enterobacteriaceae; Salmonella; Salmonella enterica; Salmonella enterica subsp. enterica; Salmonella enterica subsp. enterica serovar Typhimurium MNNEETFYQAMRRKGVTRRSFLKFCSLAATSLGLGAGMTPKIAWALENKPRIPVVWIHGL WP_001058312.1 hydrogenase [Salmonella enterica] Length: 372\nNumber of predicted TMHs:  2\nExp number of AAs in TMHs: 36.51738\nExp number, first 60 AAs:  19.72849\nTotal prob of N-in:   0.96170\nPOSSIBLE N-term signal sequence\ninside   1   19\nTMhelix   20   39\noutside   40   325\nTMhelix   326   348\ninside   349   372

24733 GCF_001222565.1_8036_3_40 Salmonella enterica subsp. enterica serovar Typhimurium str. DT104 Proteobacteria; Gammaproteobacteria; Enterobacterales; Enterobacteriaceae; Salmonella; Salmonella enterica;

Salmonella enterica subsp. enterica; Salmonella enterica subsp. enterica serovar Typhimurium
MTGDNTLITSHGINRRDFMKLCAALAATMGLSSKAAAEMAESVSNPQRPPVIWIGAQECT WP_000145429.1
hydrogenase 2 small subunit [Salmonella enterica] Length: 372\nNumber of predicted TMHs: 1\nExp number of AAs in TMHs: 24.51309\nExp number, first 60 AAs: 0.2682\nTotal prob of N-in: 0.14402\noutside 1 331\nTMhelix 332 354\ninside 355 372

24734 GCF_001222705.1_5295_2_9Salmonella enterica subsp. enterica serovar Typhimurium str. DT104
Proteobacteria; Gammaproteobacteria; Enterobacterales; Enterobacteriaceae; Salmonella; Salmonella enterica; Salmonella enterica subsp. enterica; Salmonella enterica subsp. enterica serovar Typhimurium
MTGDNTLITSHGINRRDFMKLCAALAATMGLSSKAAAEMAESVSNPQRPPVIWIGAQECT WP_000145429.1
hydrogenase 2 small subunit [Salmonella enterica] Length: 372\nNumber of predicted TMHs: 1\nExp number of AAs in TMHs: 24.51309\nExp number, first 60 AAs: 0.2682\nTotal prob of N-in: 0.14402\noutside 1 331\nTMhelix 332 354\ninside 355 372

24735 GCF_001222605.1_8080_3_63 Salmonella enterica subsp. enterica serovar Typhimurium str. DT104
Proteobacteria; Gammaproteobacteria; Enterobacterales; Enterobacteriaceae; Salmonella; Salmonella enterica; Salmonella enterica subsp. enterica; Salmonella enterica subsp. enterica serovar Typhimurium
MTGDNTLITSHGINRRDFMKLCAALAATMGLSSKAAAEMAESVSNPQRPPVIWIGAQECT WP_000145429.1
hydrogenase 2 small subunit [Salmonella enterica] Length: 372\nNumber of predicted TMHs: 1\nExp number of AAs in TMHs: 24.51309\nExp number, first 60 AAs: 0.2682\nTotal prob of N-in: 0.14402\noutside 1 331\nTMhelix 332 354\ninside 355 372

24736 GCF_001222605.1_8080_3_63 Salmonella enterica subsp. enterica serovar Typhimurium str. DT104
Proteobacteria; Gammaproteobacteria; Enterobacterales; Enterobacteriaceae; Salmonella; Salmonella enterica; Salmonella enterica subsp. enterica; Salmonella enterica subsp. enterica serovar Typhimurium
MNNEETFYQAMRRKGVTRRSFLKFCSLAATSLGLGAGMTPKIAWALENKPRIPVVWIHGL WP_001058312.1
hydrogenase [Salmonella enterica] Length: 372\nNumber of predicted TMHs: 2\nExp number of AAs in TMHs: 36.51738\nExp number, first 60 AAs: 19.72849\nTotal prob of N-in: 0.96170\nPOSSIBLE N-term signal sequence\ninside 1 19\nTMhelix 20 39\noutside 40 325\nTMhelix 326 348\ninside 349 372

24737 GCF_001222705.1_5295_2_9Salmonella enterica subsp. enterica serovar Typhimurium str. DT104
Proteobacteria; Gammaproteobacteria; Enterobacterales; Enterobacteriaceae; Salmonella; Salmonella enterica; Salmonella enterica subsp. enterica; Salmonella enterica subsp. enterica serovar Typhimurium
MNNEETFYQAMRRKGVTRRSFLKFCSLAATSLGLGAGMTPKIAWALENKPRIPVVWIHGL WP_001058312.1
hydrogenase [Salmonella enterica] Length: 372\nNumber of predicted TMHs: 2\nExp number of AAs in TMHs: 36.51738\nExp number, first 60 AAs: 19.72849\nTotal prob of N-in: 0.96170\nPOSSIBLE N-term signal sequence\ninside 1 19\nTMhelix 20 39\noutside 40 325\nTMhelix 326 348\ninside 349 372

24738 GCF_001222325.1_8080_3_73 Salmonella enterica subsp. enterica serovar Typhimurium str. DT104
Proteobacteria; Gammaproteobacteria; Enterobacterales; Enterobacteriaceae; Salmonella; Salmonella enterica; Salmonella enterica subsp. enterica; Salmonella enterica subsp. enterica serovar Typhimurium
MNNEETFYQAMRRKGVTRRSFLKFCSLAATSLGLGAGMTPKIAWALENKPRIPVVWIHGL WP_001058312.1
hydrogenase [Salmonella enterica] Length: 372\nNumber of predicted TMHs: 2\nExp number of AAs in TMHs: 36.51738\nExp number, first 60 AAs: 19.72849\nTotal prob of N-in: 0.96170\nPOSSIBLE N-term signal sequence\ninside 1 19\nTMhelix 20 39\noutside 40 325\nTMhelix 326 348\ninside 349 372

24739 GCF_001222765.1_5275_2_3Salmonella enterica subsp. enterica serovar Typhimurium str. DT104
Proteobacteria; Gammaproteobacteria; Enterobacterales; Enterobacteriaceae; Salmonella; Salmonella enterica; Salmonella enterica subsp. enterica; Salmonella enterica subsp. enterica serovar Typhimurium
MTGDNTLITSHGINRRDFMKLCAALAATMGLSSKAAAEMAESVSNPQRPPVIWIGAQECT WP_000145429.1
hydrogenase 2 small subunit [Salmonella enterica] Length: 372\nNumber of predicted TMHs: 1\nExp number of AAs in TMHs: 24.51309\nExp number, first 60 AAs: 0.2682\nTotal prob of N-in: 0.14402\noutside 1 331\nTMhelix 332 354\ninside 355 372

24740 GCF_001222645.1_5299_3_7Salmonella enterica subsp. enterica serovar Typhimurium str. DT104
Proteobacteria; Gammaproteobacteria; Enterobacterales; Enterobacteriaceae; Salmonella; Salmonella enterica; Salmonella enterica subsp. enterica; Salmonella enterica subsp. enterica serovar Typhimurium
MTGDNTLITSHGINRRDFMKLCAALAATMGLSSKAAAEMAESVSNPQRPPVIWIGAQECT WP_000145429.1
hydrogenase 2 small subunit [Salmonella enterica] Length: 372\nNumber of predicted TMHs: 1\nExp number of AAs in TMHs: 24.51309\nExp number, first 60 AAs: 0.2682\nTotal prob of N-in: 0.14402\noutside 1 331\nTMhelix 332 354\ninside 355 372

24741 GCF_001222785.1_8080_3_84 Salmonella enterica subsp. enterica serovar Typhimurium str. DT104 Proteobacteria; Gammaproteobacteria; Enterobacterales; Enterobacteriaceae; Salmonella; Salmonella enterica; Salmonella enterica subsp. enterica; Salmonella enterica subsp. enterica serovar Typhimurium MTGDNTLITSHGINRRDFMKLCAALAATMGLSSKAAAEMAESVSNPQRPPVIWIGAQECT WP_000145429.1 hydrogenase 2 small subunit [Salmonella enterica] Length: 372\nNumber of predicted TMHs: 1\nExp number of AAs in TMHs: 24.51309\nExp number, first 60 AAs: 0.2682\nTotal prob of N-in: 0.14402\noutside 1 331\nTMhelix 332 354\ninside 355 372

24742 GCF_001222645.1_5299_3_7Salmonella enterica subsp. enterica serovar Typhimurium str. DT104 Proteobacteria; Gammaproteobacteria; Enterobacterales; Enterobacteriaceae; Salmonella; Salmonella enterica; Salmonella enterica subsp. enterica; Salmonella enterica subsp. enterica serovar Typhimurium MNNEETFYQAMRRKGVTRRSFLKFCSLAATSLGLGAGMTPKIAWALENKPRIPVVWIHGL WP_001058312.1 hydrogenase [Salmonella enterica] Length: 372\nNumber of predicted TMHs: 2\nExp number of AAs in TMHs: 36.51738\nExp number, first 60 AAs: 19.72849\nTotal prob of N-in: 0.96170\nPOSSIBLE N-term signal sequence\ninside 1 19\nTMhelix 20 39\noutside 40 325\nTMhelix 326 348\ninside 349 372

24743 GCF_001222765.1_5275_2_3Salmonella enterica subsp. enterica serovar Typhimurium str. DT104 Proteobacteria; Gammaproteobacteria; Enterobacterales; Enterobacteriaceae; Salmonella; Salmonella enterica; Salmonella enterica subsp. enterica; Salmonella enterica subsp. enterica serovar Typhimurium MNNEETFYQAMRRKGVTRRSFLKFCSLAATSLGLGAGMTPKIAWALENKPRIPVVWIHGL WP_001058312.1 hydrogenase [Salmonella enterica] Length: 372\nNumber of predicted TMHs: 2\nExp number of AAs in TMHs: 36.51738\nExp number, first 60 AAs: 19.72849\nTotal prob of N-in: 0.96170\nPOSSIBLE N-term signal sequence\ninside 1 19\nTMhelix 20 39\noutside 40 325\nTMhelix 326 348\ninside 349 372

24744 GCF_001222365.1_8080_3_85 Salmonella enterica subsp. enterica serovar Typhimurium str. DT104 Proteobacteria; Gammaproteobacteria; Enterobacterales; Enterobacteriaceae; Salmonella; Salmonella enterica; Salmonella enterica subsp. enterica; Salmonella enterica subsp. enterica serovar Typhimurium MNNEETFYQAMRRKGVTRRSFLKFCSLAATSLGLGAGMTPKIAWALENKPRIPVVWIHGL WP_001058312.1 hydrogenase [Salmonella enterica] Length: 372\nNumber of predicted TMHs: 2\nExp number of AAs in TMHs: 36.51738\nExp number, first 60 AAs: 19.72849\nTotal prob of N-in: 0.96170\nPOSSIBLE N-term signal sequence\ninside 1 19\nTMhelix 20 39\noutside 40 325\nTMhelix 326 348\ninside 349 372

24745 GCF_001222405.1_8036_3_84 Salmonella enterica subsp. enterica serovar Typhimurium str. DT104 Proteobacteria; Gammaproteobacteria; Enterobacterales; Enterobacteriaceae; Salmonella; Salmonella enterica; Salmonella enterica subsp. enterica; Salmonella enterica subsp. enterica serovar Typhimurium MNNEETFYQAMRRKGVTRRSFLKFCSLAATSLGLGAGMTPKIAWALENKPRIPVVWIHGL WP_001058312.1 hydrogenase [Salmonella enterica] Length: 372\nNumber of predicted TMHs: 2\nExp number of AAs in TMHs: 36.51738\nExp number, first 60 AAs: 19.72849\nTotal prob of N-in: 0.96170\nPOSSIBLE N-term signal sequence\ninside 1 19\nTMhelix 20 39\noutside 40 325\nTMhelix 326 348\ninside 349 372

24746 GCF_001222525.1_8080_3_27 Salmonella enterica subsp. enterica serovar Typhimurium str. DT104 Proteobacteria; Gammaproteobacteria; Enterobacterales; Enterobacteriaceae; Salmonella; Salmonella enterica; Salmonella enterica subsp. enterica; Salmonella enterica subsp. enterica serovar Typhimurium MNNEETFYQAMRRKGVTRRSFLKFCSLAATSLGLGAGMTPKIAWALENKPRIPVVWIHGL WP_001058312.1 hydrogenase [Salmonella enterica] Length: 372\nNumber of predicted TMHs: 2\nExp number of AAs in TMHs: 36.51738\nExp number, first 60 AAs: 19.72849\nTotal prob of N-in: 0.96170\nPOSSIBLE N-term signal sequence\ninside 1 19\nTMhelix 20 39\noutside 40 325\nTMhelix 326 348\ninside 349 372

24747 GCF_001222785.1_8080_3_84 Salmonella enterica subsp. enterica serovar Typhimurium str. DT104 Proteobacteria; Gammaproteobacteria; Enterobacterales; Enterobacteriaceae; Salmonella; Salmonella enterica; Salmonella enterica subsp. enterica; Salmonella enterica subsp. enterica serovar Typhimurium MNNEETFYQAMRRKGVTRRSFLKFCSLAATSLGLGAGMTPKIAWALENKPRIPVVWIHGL WP_001058312.1 hydrogenase [Salmonella enterica] Length: 372\nNumber of predicted TMHs: 2\nExp number of AAs in TMHs: 36.51738\nExp number, first 60 AAs: 19.72849\nTotal prob of N-in: 0.96170\nPOSSIBLE N-term signal sequence\ninside 1 19\nTMhelix 20 39\noutside 40 325\nTMhelix 326 348\ninside 349 372

24748 GCF_001222865.1_5275_6_12 Salmonella enterica subsp. enterica serovar Typhimurium str. DT104 Proteobacteria; Gammaproteobacteria; Enterobacterales; Enterobacteriaceae; Salmonella; Salmonella enterica; Salmonella enterica subsp. enterica; Salmonella enterica subsp. enterica serovar Typhimurium MTGDNTLITSHGINRRDFMKLCAALAATMGLSSKAAAEMAESVSNPQRPPVIWIGAQECT WP_000145429.1 hydrogenase 2 small subunit [Salmonella enterica] Length: 372\nNumber of predicted TMHs: 1\nExp

number of AAs in TMHs: 24.51309\nExp number, first 60 AAs: 0.2682\nTotal prob of N-in: 0.14402\noutside 1 331\nTMhelix 332 354\ninside 355 372

24749 GCF_001222905.1_5275_8_1Salmonella enterica subsp. enterica serovar Typhimurium str. DT104 Proteobacteria; Gammaproteobacteria; Enterobacterales; Enterobacteriaceae; Salmonella; Salmonella enterica; Salmonella enterica subsp. enterica; Salmonella enterica subsp. enterica serovar Typhimurium MTGDNTLITSHGINRRDFMKLCAALAATMGLSSKAAAEMAESVSNPQRPPVIWIGAQECT WP_000145429.1 hydrogenase 2 small subunit [Salmonella enterica] Length: 372\nNumber of predicted TMHs: 1\nExp number of AAs in TMHs: 24.51309\nExp number, first 60 AAs: 0.2682\nTotal prob of N-in: 0.14402\noutside 1 331\nTMhelix 332 354\ninside 355 372

24750 GCF_001223045.1_8080_3_1Salmonella enterica subsp. enterica serovar Typhimurium str. DT104 Proteobacteria; Gammaproteobacteria; Enterobacterales; Enterobacteriaceae; Salmonella; Salmonella enterica; Salmonella enterica subsp. enterica; Salmonella enterica subsp. enterica serovar Typhimurium MTGDNTLITSHGINRRDFMKLCAALAATMGLSSKAAAEMAESVSNPQRPPVIWIGAQECT WP_000145429.1 hydrogenase 2 small subunit [Salmonella enterica] Length: 372\nNumber of predicted TMHs: 1\nExp number of AAs in TMHs: 24.51309\nExp number, first 60 AAs: 0.2682\nTotal prob of N-in: 0.14402\noutside 1 331\nTMhelix 332 354\ninside 355 372

24751 GCF_001223085.1_8080_3_58 Salmonella enterica subsp. enterica serovar Typhimurium str. DT104 Proteobacteria; Gammaproteobacteria; Enterobacterales; Enterobacteriaceae; Salmonella; Salmonella enterica; Salmonella enterica subsp. enterica; Salmonella enterica subsp. enterica serovar Typhimurium MTGDNTLITSHGINRRDFMKLCAALAATMGLSSKAAAEMAESVSNPQRPPVIWIGAQECT WP_000145429.1 hydrogenase 2 small subunit [Salmonella enterica] Length: 372\nNumber of predicted TMHs: 1\nExp number of AAs in TMHs: 24.51309\nExp number, first 60 AAs: 0.2682\nTotal prob of N-in: 0.14402\noutside 1 331\nTMhelix 332 354\ninside 355 372

24752 GCF_001350135.1_8490_2_85 Salmonella enterica subsp. enterica serovar Typhimurium str. DT104 Proteobacteria; Gammaproteobacteria; Enterobacterales; Enterobacteriaceae; Salmonella; Salmonella enterica; Salmonella enterica subsp. enterica; Salmonella enterica subsp. enterica serovar Typhimurium MNNEETFYQAMRRKGVTRRSFLKFCSLAATSLGLGAGMTPKIAWALENKPRIPVVWIHGL WP_001058312.1 hydrogenase [Salmonella enterica] Length: 372\nNumber of predicted TMHs: 2\nExp number of AAs in TMHs: 36.51738\nExp number, first 60 AAs: 19.72849\nTotal prob of N-in: 0.96170\nPOSSIBLE N-term signal sequence\ninside 1 19\nTMhelix 20 39\noutside 40 325\nTMhelix 326 348\ninside 349 372

24753 GCF_001223105.1_5285_7_6Salmonella enterica subsp. enterica serovar Typhimurium str. DT104 Proteobacteria; Gammaproteobacteria; Enterobacterales; Enterobacteriaceae; Salmonella; Salmonella enterica; Salmonella enterica subsp. enterica; Salmonella enterica subsp. enterica serovar Typhimurium MTGDNTLITSHGINRRDFMKLCAALAATMGLSSKAAAEMAESVSNPQRPPVIWIGAQECT WP_000145429.1 hydrogenase 2 small subunit [Salmonella enterica] Length: 372\nNumber of predicted TMHs: 1\nExp number of AAs in TMHs: 24.51309\nExp number, first 60 AAs: 0.2682\nTotal prob of N-in: 0.14402\noutside 1 331\nTMhelix 332 354\ninside 355 372

24754 GCF_001223105.1_5285_7_6Salmonella enterica subsp. enterica serovar Typhimurium str. DT104 Proteobacteria; Gammaproteobacteria; Enterobacterales; Enterobacteriaceae; Salmonella; Salmonella enterica; Salmonella enterica subsp. enterica; Salmonella enterica subsp. enterica serovar Typhimurium MNNEETFYQAMRRKGVTRRSFLKFCSLAATSLGLGAGMTPKIAWALENKPRIPVVWIHGL WP_001058312.1 hydrogenase [Salmonella enterica] Length: 372\nNumber of predicted TMHs: 2\nExp number of AAs in TMHs: 36.51738\nExp number, first 60 AAs: 19.72849\nTotal prob of N-in: 0.96170\nPOSSIBLE N-term signal sequence\ninside 1 19\nTMhelix 20 39\noutside 40 325\nTMhelix 326 348\ninside 349 372

24755 GCF_001222865.1_5275_6_12 Salmonella enterica subsp. enterica serovar Typhimurium str. DT104 Proteobacteria; Gammaproteobacteria; Enterobacterales; Enterobacteriaceae; Salmonella; Salmonella enterica; Salmonella enterica subsp. enterica; Salmonella enterica subsp. enterica serovar Typhimurium MNNEETFYQAMRRKGVTRRSFLKFCSLAATSLGLGAGMTPKIAWALENKPRIPVVWIHGL WP_001058312.1 hydrogenase [Salmonella enterica] Length: 372\nNumber of predicted TMHs: 2\nExp number of AAs in TMHs: 36.51738\nExp number, first 60 AAs: 19.72849\nTotal prob of N-in: 0.96170\nPOSSIBLE N-term signal sequence\ninside 1 19\nTMhelix 20 39\noutside 40 325\nTMhelix 326 348\ninside 349 372

24756 GCF_001217425.1_5285_6_7Salmonella enterica subsp. enterica serovar Typhimurium str. DT104 Proteobacteria; Gammaproteobacteria; Enterobacterales; Enterobacteriaceae; Salmonella; Salmonella enterica;

Salmonella enterica subsp. enterica; Salmonella enterica subsp. enterica serovar Typhimurium
MTGDNTLITSHGINRRDFMKLCAALAATMGLSSKAAAEMAESVSNPQRPPVIWIGAQECT WP_000145429.1
hydrogenase 2 small subunit [Salmonella enterica] Length: 372\nNumber of predicted TMHs: 1\nExp number of AAs in TMHs: 24.51309\nExp number, first 60 AAs: 0.2682\nTotal prob of N-in: 0.14402\noutside 1 331\nTMhelix 332 354\ninside 355 372

24757 GCF_001223165.1_8080_3_23 Salmonella enterica subsp. enterica serovar Typhimurium str. DT104
Proteobacteria; Gammaproteobacteria; Enterobacterales; Enterobacteriaceae; Salmonella; Salmonella enterica; Salmonella enterica subsp. enterica; Salmonella enterica subsp. enterica serovar Typhimurium
MTGDNTLITSHGINRRDFMKLCAALAATMGLSSKAAAEMAESVSNPQRPPVIWIGAQECT WP_000145429.1
hydrogenase 2 small subunit [Salmonella enterica] Length: 372\nNumber of predicted TMHs: 1\nExp number of AAs in TMHs: 24.51309\nExp number, first 60 AAs: 0.2682\nTotal prob of N-in: 0.14402\noutside 1 331\nTMhelix 332 354\ninside 355 372

24758 GCF_001223205.1_8080_3_68 Salmonella enterica subsp. enterica serovar Typhimurium str. DT104
Proteobacteria; Gammaproteobacteria; Enterobacterales; Enterobacteriaceae; Salmonella; Salmonella enterica; Salmonella enterica subsp. enterica; Salmonella enterica subsp. enterica serovar Typhimurium
MTGDNTLITSHGINRRDFMKLCAALAATMGLSSKAAAEMAESVSNPQRPPVIWIGAQECT WP_000145429.1
hydrogenase 2 small subunit [Salmonella enterica] Length: 372\nNumber of predicted TMHs: 1\nExp number of AAs in TMHs: 24.51309\nExp number, first 60 AAs: 0.2682\nTotal prob of N-in: 0.14402\noutside 1 331\nTMhelix 332 354\ninside 355 372

24759 GCF_001223245.1_5285_6_11 Salmonella enterica subsp. enterica serovar Typhimurium str. DT104
Proteobacteria; Gammaproteobacteria; Enterobacterales; Enterobacteriaceae; Salmonella; Salmonella enterica; Salmonella enterica subsp. enterica; Salmonella enterica subsp. enterica serovar Typhimurium
MTGDNTLITSHGINRRDFMKLCAALAATMGLSSKAAAEMAESVSNPQRPPVIWIGAQECT WP_000145429.1
hydrogenase 2 small subunit [Salmonella enterica] Length: 372\nNumber of predicted TMHs: 1\nExp number of AAs in TMHs: 24.51309\nExp number, first 60 AAs: 0.2682\nTotal prob of N-in: 0.14402\noutside 1 331\nTMhelix 332 354\ninside 355 372

24760 GCF_001223265.1_8036_3_77 Salmonella enterica subsp. enterica serovar Typhimurium str. DT104
Proteobacteria; Gammaproteobacteria; Enterobacterales; Enterobacteriaceae; Salmonella; Salmonella enterica; Salmonella enterica subsp. enterica; Salmonella enterica subsp. enterica serovar Typhimurium
MTGDNTLITSHGINRRDFMKLCAALAATMGLSSKAAAEMAESVSNPQRPPVIWIGAQECT WP_000145429.1
hydrogenase 2 small subunit [Salmonella enterica] Length: 372\nNumber of predicted TMHs: 1\nExp number of AAs in TMHs: 24.51309\nExp number, first 60 AAs: 0.2682\nTotal prob of N-in: 0.14402\noutside 1 331\nTMhelix 332 354\ninside 355 372

24761 GCF_001223165.1_8080_3_23 Salmonella enterica subsp. enterica serovar Typhimurium str. DT104
Proteobacteria; Gammaproteobacteria; Enterobacterales; Enterobacteriaceae; Salmonella; Salmonella enterica; Salmonella enterica subsp. enterica; Salmonella enterica subsp. enterica serovar Typhimurium
MNNEETFYQAMRRKGVTRRSFLKFCSLAATSLGLGAGMTPKIAWALENKPRIPVVWIHGL WP_001058312.1
hydrogenase [Salmonella enterica] Length: 372\nNumber of predicted TMHs: 2\nExp number of AAs in TMHs: 36.51738\nExp number, first 60 AAs: 19.72849\nTotal prob of N-in: 0.96170\nPOSSIBLE N-term signal sequence\ninside 1 19\nTMhelix 20 39\noutside 40 325\nTMhelix 326 348\ninside 349 372

24762 GCF_001222905.1_5275_8_1Salmonella enterica subsp. enterica serovar Typhimurium str. DT104
Proteobacteria; Gammaproteobacteria; Enterobacterales; Enterobacteriaceae; Salmonella; Salmonella enterica; Salmonella enterica subsp. enterica; Salmonella enterica subsp. enterica serovar Typhimurium
MNNEETFYQAMRRKGVTRRSFLKFCSLAATSLGLGAGMTPKIAWALENKPRIPVVWIHGL WP_001058312.1
hydrogenase [Salmonella enterica] Length: 372\nNumber of predicted TMHs: 2\nExp number of AAs in TMHs: 36.51738\nExp number, first 60 AAs: 19.72849\nTotal prob of N-in: 0.96170\nPOSSIBLE N-term signal sequence\ninside 1 19\nTMhelix 20 39\noutside 40 325\nTMhelix 326 348\ninside 349 372

24763 GCF_001223305.1_5275_7_1Salmonella enterica subsp. enterica serovar Typhimurium str. DT104
Proteobacteria; Gammaproteobacteria; Enterobacterales; Enterobacteriaceae; Salmonella; Salmonella enterica; Salmonella enterica subsp. enterica; Salmonella enterica subsp. enterica serovar Typhimurium
MTGDNTLITSHGINRRDFMKLCAALAATMGLSSKAAAEMAESVSNPQRPPVIWIGAQECT WP_000145429.1
hydrogenase 2 small subunit [Salmonella enterica] Length: 372\nNumber of predicted TMHs: 1\nExp number of AAs in TMHs: 24.51309\nExp number, first 60 AAs: 0.2682\nTotal prob of N-in: 0.14402\noutside 1 331\nTMhelix 332 354\ninside 355 372

24764 GCF_001223305.1_5275_7_1Salmonella enterica subsp. enterica serovar Typhimurium str. DT104 Proteobacteria; Gammaproteobacteria; Enterobacterales; Enterobacteriaceae; Salmonella; Salmonella enterica; Salmonella enterica subsp. enterica; Salmonella enterica subsp. enterica serovar Typhimurium MNNEETFYQAMRRKGVTRRSFLKFCSLAATSLGLGAGMTPKIAWALENKPRIPVVWIHGL WP_001058312.1 hydrogenase [Salmonella enterica] Length: 372\nNumber of predicted TMHs: 2\nExp number of AAs in TMHs: 36.51738\nExp number, first 60 AAs: 19.72849\nTotal prob of N-in: 0.96170\nPOSSIBLE N-term signal sequence\ninside 1 19\nTMhelix 20 39\noutside 40 325\nTMhelix 326 348\ninside 349 372

24765 GCF_001222965.1_8036_3_72 Salmonella enterica subsp. enterica serovar Typhimurium str. DT104 Proteobacteria; Gammaproteobacteria; Enterobacterales; Enterobacteriaceae; Salmonella; Salmonella enterica; Salmonella enterica subsp. enterica; Salmonella enterica subsp. enterica serovar Typhimurium MNNEETFYQAMRRKGVTRRSFLKFCSLAATSLGLGAGMTPKIAWALENKPRIPVVWIHGL WP_001058312.1 hydrogenase [Salmonella enterica] Length: 372\nNumber of predicted TMHs: 2\nExp number of AAs in TMHs: 36.51738\nExp number, first 60 AAs: 19.72849\nTotal prob of N-in: 0.96170\nPOSSIBLE N-term signal sequence\ninside 1 19\nTMhelix 20 39\noutside 40 325\nTMhelix 326 348\ninside 349 372

24766 GCF_001223345.1_8036_3_73 Salmonella enterica subsp. enterica serovar Typhimurium str. DT104 Proteobacteria; Gammaproteobacteria; Enterobacterales; Enterobacteriaceae; Salmonella; Salmonella enterica; Salmonella enterica subsp. enterica; Salmonella enterica subsp. enterica serovar Typhimurium MTGDNTLITSHGINRRDFMKLCAALAATMGLSSKAAAEMAESVSNPQRPPVIWIGAQECT WP_000145429.1 hydrogenase 2 small subunit [Salmonella enterica] Length: 372\nNumber of predicted TMHs: 1\nExp number of AAs in TMHs: 24.51309\nExp number, first 60 AAs: 0.2682\nTotal prob of N-in: 0.14402\noutside 1 331\nTMhelix 332 354\ninside 355 372

24767 GCF_001223345.1_8036_3_73 Salmonella enterica subsp. enterica serovar Typhimurium str. DT104 Proteobacteria; Gammaproteobacteria; Enterobacterales; Enterobacteriaceae; Salmonella; Salmonella enterica; Salmonella enterica subsp. enterica; Salmonella enterica subsp. enterica serovar Typhimurium MNNEETFYQAMRRKGVTRRSFLKFCSLAATSLGLGAGMTPKIAWALENKPRIPVVWIHGL WP_001058312.1 hydrogenase [Salmonella enterica] Length: 372\nNumber of predicted TMHs: 2\nExp number of AAs in TMHs: 36.51738\nExp number, first 60 AAs: 19.72849\nTotal prob of N-in: 0.96170\nPOSSIBLE N-term signal sequence\ninside 1 19\nTMhelix 20 39\noutside 40 325\nTMhelix 326 348\ninside 349 372

24768 GCF_001223045.1_8080_3_1Salmonella enterica subsp. enterica serovar Typhimurium str. DT104 Proteobacteria; Gammaproteobacteria; Enterobacterales; Enterobacteriaceae; Salmonella; Salmonella enterica; Salmonella enterica subsp. enterica; Salmonella enterica subsp. enterica serovar Typhimurium MNNEETFYQAMRRKGVTRRSFLKFCSLAATSLGLGAGMTPKIAWALENKPRIPVVWIHGL WP_001058312.1 hydrogenase [Salmonella enterica] Length: 372\nNumber of predicted TMHs: 2\nExp number of AAs in TMHs: 36.51738\nExp number, first 60 AAs: 19.72849\nTotal prob of N-in: 0.96170\nPOSSIBLE N-term signal sequence\ninside 1 19\nTMhelix 20 39\noutside 40 325\nTMhelix 326 348\ninside 349 372

24769 GCF_001223085.1_8080_3_58 Salmonella enterica subsp. enterica serovar Typhimurium str. DT104 Proteobacteria; Gammaproteobacteria; Enterobacterales; Enterobacteriaceae; Salmonella; Salmonella enterica; Salmonella enterica subsp. enterica; Salmonella enterica subsp. enterica serovar Typhimurium MNNEETFYQAMRRKGVTRRSFLKFCSLAATSLGLGAGMTPKIAWALENKPRIPVVWIHGL WP_001058312.1 hydrogenase [Salmonella enterica] Length: 372\nNumber of predicted TMHs: 2\nExp number of AAs in TMHs: 36.51738\nExp number, first 60 AAs: 19.72849\nTotal prob of N-in: 0.96170\nPOSSIBLE N-term signal sequence\ninside 1 19\nTMhelix 20 39\noutside 40 325\nTMhelix 326 348\ninside 349 372

24770 GCF_001223205.1_8080_3_68 Salmonella enterica subsp. enterica serovar Typhimurium str. DT104 Proteobacteria; Gammaproteobacteria; Enterobacterales; Enterobacteriaceae; Salmonella; Salmonella enterica; Salmonella enterica subsp. enterica; Salmonella enterica subsp. enterica serovar Typhimurium MNNEETFYQAMRRKGVTRRSFLKFCSLAATSLGLGAGMTPKIAWALENKPRIPVVWIHGL WP_001058312.1 hydrogenase [Salmonella enterica] Length: 372\nNumber of predicted TMHs: 2\nExp number of AAs in TMHs: 36.51738\nExp number, first 60 AAs: 19.72849\nTotal prob of N-in: 0.96170\nPOSSIBLE N-term signal sequence\ninside 1 19\nTMhelix 20 39\noutside 40 325\nTMhelix 326 348\ninside 349 372

24771 GCF_001223245.1_5285_6_11 Salmonella enterica subsp. enterica serovar Typhimurium str. DT104 Proteobacteria; Gammaproteobacteria; Enterobacterales; Enterobacteriaceae; Salmonella; Salmonella enterica; Salmonella enterica subsp. enterica; Salmonella enterica subsp. enterica serovar Typhimurium MNNEETFYQAMRRKGVTRRSFLKFCSLAATSLGLGAGMTPKIAWALENKPRIPVVWIHGL WP_001058312.1 hydrogenase [Salmonella enterica] Length: 372\nNumber of predicted TMHs: 2\nExp number of AAs in

TMHs: 36.51738\nExp number, first 60 AAs: 19.72849\nTotal prob of N-in: 0.96170\nPOSSIBLE N-term signal sequence\ninside 1 19\nTMhelix 20 39\noutside 40 325\nTMhelix 326 348\ninside 349 372

24772 GCF_001223265.1_8036_3_77 Salmonella enterica subsp. enterica serovar Typhimurium str. DT104 Proteobacteria; Gammaproteobacteria; Enterobacterales; Enterobacteriaceae; Salmonella; Salmonella enterica; Salmonella enterica subsp. enterica; Salmonella enterica subsp. enterica serovar Typhimurium MNNEETFYQAMRRKGVTRRSFLKFCSLAATSLGLGAGMTPKIAWALENKPRIPVVWIHGL WP_001058312.1 hydrogenase [Salmonella enterica] Length: 372\nNumber of predicted TMHs: 2\nExp number of AAs in TMHs: 36.51738\nExp number, first 60 AAs: 19.72849\nTotal prob of N-in: 0.96170\nPOSSIBLE N-term signal sequence\ninside 1 19\nTMhelix 20 39\noutside 40 325\nTMhelix 326 348\ninside 349 372

24773 GCF_001223405.1_5275_2_10 Salmonella enterica subsp. enterica serovar Typhimurium str. DT104 Proteobacteria; Gammaproteobacteria; Enterobacterales; Enterobacteriaceae; Salmonella; Salmonella enterica; Salmonella enterica subsp. enterica; Salmonella enterica subsp. enterica serovar Typhimurium MTGDNTLITSHGINRRDFMKLCAALAATMGLSSKAAAEMAESVSNPQRPPVIWIGAQECT WP_000145429.1 hydrogenase 2 small subunit [Salmonella enterica] Length: 372\nNumber of predicted TMHs: 1\nExp number of AAs in TMHs: 24.51309\nExp number, first 60 AAs: 0.2682\nTotal prob of N-in: 0.14402\noutside 1 331\nTMhelix 332 354\ninside 355 372

24774 GCF_001223445.1_5275_6_9Salmonella enterica subsp. enterica serovar Typhimurium str. DT104 Proteobacteria; Gammaproteobacteria; Enterobacterales; Enterobacteriaceae; Salmonella; Salmonella enterica; Salmonella enterica subsp. enterica; Salmonella enterica subsp. enterica serovar Typhimurium MTGDNTLITSHGINRRDFMKLCAALAATMGLSSKAAAEMAESVSNPQRPPVIWIGAQECT WP_000145429.1 hydrogenase 2 small subunit [Salmonella enterica] Length: 372\nNumber of predicted TMHs: 1\nExp number of AAs in TMHs: 24.51309\nExp number, first 60 AAs: 0.2682\nTotal prob of N-in: 0.14402\noutside 1 331\nTMhelix 332 354\ninside 355 372

24775 GCF_001223505.1_8036_3_81 Salmonella enterica subsp. enterica serovar Typhimurium str. DT104 Proteobacteria; Gammaproteobacteria; Enterobacterales; Enterobacteriaceae; Salmonella; Salmonella enterica; Salmonella enterica subsp. enterica; Salmonella enterica subsp. enterica serovar Typhimurium MTGDNTLITSHGINRRDFMKLCAALAATMGLSSKAAAEMAESVSNPQRPPVIWIGAQECT WP_000145429.1 hydrogenase 2 small subunit [Salmonella enterica] Length: 372\nNumber of predicted TMHs: 1\nExp number of AAs in TMHs: 24.51309\nExp number, first 60 AAs: 0.2682\nTotal prob of N-in: 0.14402\noutside 1 331\nTMhelix 332 354\ninside 355 372

24776 GCF_001223545.1_8036_3_49 Salmonella enterica subsp. enterica serovar Typhimurium str. DT104 Proteobacteria; Gammaproteobacteria; Enterobacterales; Enterobacteriaceae; Salmonella; Salmonella enterica; Salmonella enterica subsp. enterica; Salmonella enterica subsp. enterica serovar Typhimurium MTGDNTLITSHGINRRDFMKLCAALAATMGLSSKAAAEMAESVSNPQRPPVIWIGAQECT WP_000145429.1 hydrogenase 2 small subunit [Salmonella enterica] Length: 372\nNumber of predicted TMHs: 1\nExp number of AAs in TMHs: 24.51309\nExp number, first 60 AAs: 0.2682\nTotal prob of N-in: 0.14402\noutside 1 331\nTMhelix 332 354\ninside 355 372

24777 GCF_001223445.1_5275_6_9Salmonella enterica subsp. enterica serovar Typhimurium str. DT104 Proteobacteria; Gammaproteobacteria; Enterobacterales; Enterobacteriaceae; Salmonella; Salmonella enterica; Salmonella enterica subsp. enterica; Salmonella enterica subsp. enterica serovar Typhimurium MNNEETFYQAMRRKGVTRRSFLKFCSLAATSLGLGAGMTPKIAWALENKPRIPVVWIHGL WP_001058312.1 hydrogenase [Salmonella enterica] Length: 372\nNumber of predicted TMHs: 2\nExp number of AAs in TMHs: 36.51738\nExp number, first 60 AAs: 19.72849\nTotal prob of N-in: 0.96170\nPOSSIBLE N-term signal sequence\ninside 1 19\nTMhelix 20 39\noutside 40 325\nTMhelix 326 348\ninside 349 372

24778 GCF_001223405.1_5275_2_10 Salmonella enterica subsp. enterica serovar Typhimurium str. DT104 Proteobacteria; Gammaproteobacteria; Enterobacterales; Enterobacteriaceae; Salmonella; Salmonella enterica; Salmonella enterica subsp. enterica; Salmonella enterica subsp. enterica serovar Typhimurium MNNEETFYQAMRRKGVTRRSFLKFCSLAATSLGLGAGMTPKIAWALENKPRIPVVWIHGL WP_001058312.1 hydrogenase [Salmonella enterica] Length: 372\nNumber of predicted TMHs: 2\nExp number of AAs in TMHs: 36.51738\nExp number, first 60 AAs: 19.72849\nTotal prob of N-in: 0.96170\nPOSSIBLE N-term signal sequence\ninside 1 19\nTMhelix 20 39\noutside 40 325\nTMhelix 326 348\ninside 349 372

24779 GCF_001326155.1_9066_1_10 Salmonella enterica subsp. enterica serovar Typhimurium str. DT104 Proteobacteria; Gammaproteobacteria; Enterobacterales; Enterobacteriaceae; Salmonella; Salmonella enterica;

Salmonella enterica subsp. enterica; Salmonella enterica subsp. enterica serovar Typhimurium
MNNEETFYQAMRRKGVTRRSFLKFCSLAATSLGLGAGMTPKIAWALENKPRIPVVWIHGL WP_001058312.1
hydrogenase [Salmonella enterica] Length: 372\nNumber of predicted TMHs: 2\nExp number of AAs in TMHs: 36.51738\nExp number, first 60 AAs: 19.72849\nTotal prob of N-in: 0.96170\nPOSSIBLE N-term signal sequence\ninside 1 19\nTMhelix 20 39\noutside 40 325\nTMhelix 326 348\ninside 349 372

24780 GCF_001217345.1_5275_6_8Salmonella enterica subsp. enterica serovar Typhimurium str. DT104
Proteobacteria; Gammaproteobacteria; Enterobacterales; Enterobacteriaceae; Salmonella; Salmonella enterica; Salmonella enterica subsp. enterica; Salmonella enterica subsp. enterica serovar Typhimurium
MTGDNTLITSHGINRRDFMKLCAALAATMGLSSKAAAEMAESVSNPQRPPVIWIGAQECT WP_000145429.1
hydrogenase 2 small subunit [Salmonella enterica] Length: 372\nNumber of predicted TMHs: 1\nExp number of AAs in TMHs: 24.51309\nExp number, first 60 AAs: 0.2682\nTotal prob of N-in: 0.14402\noutside 1 331\nTMhelix 332 354\ninside 355 372

24781 GCF_001217205.1_8080_3_61 Salmonella enterica subsp. enterica serovar Typhimurium str. DT104
Proteobacteria; Gammaproteobacteria; Enterobacterales; Enterobacteriaceae; Salmonella; Salmonella enterica; Salmonella enterica subsp. enterica; Salmonella enterica subsp. enterica serovar Typhimurium
MTGDNTLITSHGINRRDFMKLCAALAATMGLSSKAAAEMAESVSNPQRPPVIWIGAQECT WP_000145429.1
hydrogenase 2 small subunit [Salmonella enterica] Length: 372\nNumber of predicted TMHs: 1\nExp number of AAs in TMHs: 24.51309\nExp number, first 60 AAs: 0.2682\nTotal prob of N-in: 0.14402\noutside 1 331\nTMhelix 332 354\ninside 355 372

24782 GCF_001326215.1_9065_1_5Salmonella enterica subsp. enterica serovar Typhimurium str. DT104
Proteobacteria; Gammaproteobacteria; Enterobacterales; Enterobacteriaceae; Salmonella; Salmonella enterica; Salmonella enterica subsp. enterica; Salmonella enterica subsp. enterica serovar Typhimurium
MNNEETFYQAMRRKGVTRRSFLKFCSLAATSLGLGAGMTPKIAWALENKPRIPVVWIHGL WP_001058312.1
hydrogenase [Salmonella enterica] Length: 372\nNumber of predicted TMHs: 2\nExp number of AAs in TMHs: 36.51738\nExp number, first 60 AAs: 19.72849\nTotal prob of N-in: 0.96170\nPOSSIBLE N-term signal sequence\ninside 1 19\nTMhelix 20 39\noutside 40 325\nTMhelix 326 348\ninside 349 372

24783 GCF_001350175.1_8490_2_80 Salmonella enterica subsp. enterica serovar Typhimurium str. DT104
Proteobacteria; Gammaproteobacteria; Enterobacterales; Enterobacteriaceae; Salmonella; Salmonella enterica; Salmonella enterica subsp. enterica; Salmonella enterica subsp. enterica serovar Typhimurium
MNNEETFYQAMRRKGVTRRSFLKFCSLAATSLGLGAGMTPKIAWALENKPRIPVVWIHGL WP_001058312.1
hydrogenase [Salmonella enterica] Length: 372\nNumber of predicted TMHs: 2\nExp number of AAs in TMHs: 36.51738\nExp number, first 60 AAs: 19.72849\nTotal prob of N-in: 0.96170\nPOSSIBLE N-term signal sequence\ninside 1 19\nTMhelix 20 39\noutside 40 325\nTMhelix 326 348\ninside 349 372

24784 GCF_001223505.1_8036_3_81 Salmonella enterica subsp. enterica serovar Typhimurium str. DT104
Proteobacteria; Gammaproteobacteria; Enterobacterales; Enterobacteriaceae; Salmonella; Salmonella enterica; Salmonella enterica subsp. enterica; Salmonella enterica subsp. enterica serovar Typhimurium
MNNEETFYQAMRRKGVTRRSFLKFCSLAATSLGLGAGMTPKIAWALENKPRIPVVWIHGL WP_001058312.1
hydrogenase [Salmonella enterica] Length: 372\nNumber of predicted TMHs: 2\nExp number of AAs in TMHs: 36.51738\nExp number, first 60 AAs: 19.72849\nTotal prob of N-in: 0.96170\nPOSSIBLE N-term signal sequence\ninside 1 19\nTMhelix 20 39\noutside 40 325\nTMhelix 326 348\ninside 349 372

24785 GCF_001326235.1_9066_1_2Salmonella enterica subsp. enterica serovar Typhimurium str. DT104
Proteobacteria; Gammaproteobacteria; Enterobacterales; Enterobacteriaceae; Salmonella; Salmonella enterica; Salmonella enterica subsp. enterica; Salmonella enterica subsp. enterica serovar Typhimurium
MNNEETFYQAMRRKGVTRRSFLKFCSLAATSLGLGAGMTPKIAWALENKPRIPVVWIHGL WP_001058312.1
hydrogenase [Salmonella enterica] Length: 372\nNumber of predicted TMHs: 2\nExp number of AAs in TMHs: 36.51738\nExp number, first 60 AAs: 19.72849\nTotal prob of N-in: 0.96170\nPOSSIBLE N-term signal sequence\ninside 1 19\nTMhelix 20 39\noutside 40 325\nTMhelix 326 348\ninside 349 372

24786 GCF_001223545.1_8036_3_49 Salmonella enterica subsp. enterica serovar Typhimurium str. DT104
Proteobacteria; Gammaproteobacteria; Enterobacterales; Enterobacteriaceae; Salmonella; Salmonella enterica; Salmonella enterica subsp. enterica; Salmonella enterica subsp. enterica serovar Typhimurium
MNNEETFYQAMRRKGVTRRSFLKFCSLAATSLGLGAGMTPKIAWALENKPRIPVVWIHGL WP_001058312.1
hydrogenase [Salmonella enterica] Length: 372\nNumber of predicted TMHs: 2\nExp number of AAs in TMHs: 36.51738\nExp number, first 60 AAs: 19.72849\nTotal prob of N-in: 0.96170\nPOSSIBLE N-term signal sequence\ninside 1 19\nTMhelix 20 39\noutside 40 325\nTMhelix 326 348\ninside 349 372

24787 GCF_001217265.1_5299_3_3Salmonella enterica subsp. enterica serovar Typhimurium str. DT104 Proteobacteria; Gammaproteobacteria; Enterobacterales; Enterobacteriaceae; Salmonella; Salmonella enterica; Salmonella enterica subsp. enterica; Salmonella enterica subsp. enterica serovar Typhimurium MTGDNTLITSHGINRRDFMKLCAALAATMGLSSKAAAEMAESVSNPQRPPVIWIGAQECT WP_000145429.1 hydrogenase 2 small subunit [Salmonella enterica] Length: 372\nNumber of predicted TMHs: 1\nExp number of AAs in TMHs: 24.51309\nExp number, first 60 AAs: 0.2682\nTotal prob of N-in: 0.14402\noutside 1 331\nTMhelix 332 354\ninside 355 372

24788 GCF_001217045.1_5275_3_5Salmonella enterica subsp. enterica serovar Typhimurium str. DT104 Proteobacteria; Gammaproteobacteria; Enterobacterales; Enterobacteriaceae; Salmonella; Salmonella enterica; Salmonella enterica subsp. enterica; Salmonella enterica subsp. enterica serovar Typhimurium MTGDNTLITSHGINRRDFMKLCAALAATMGLSSKAAAEMAESVSNPQRPPVIWIGAQECT WP_000145429.1 hydrogenase 2 small subunit [Salmonella enterica] Length: 372\nNumber of predicted TMHs: 1\nExp number of AAs in TMHs: 24.51309\nExp number, first 60 AAs: 0.2682\nTotal prob of N-in: 0.14402\noutside 1 331\nTMhelix 332 354\ninside 355 372

24789 GCF_001350235.1_8490_2_81 Salmonella enterica subsp. enterica serovar Typhimurium str. DT104 Proteobacteria; Gammaproteobacteria; Enterobacterales; Enterobacteriaceae; Salmonella; Salmonella enterica; Salmonella enterica subsp. enterica; Salmonella enterica subsp. enterica serovar Typhimurium MNNEETFYQAMRRKGVTRRSFLKFCSLAATSLGLGAGMTPKIAWALENKPRIPVVWIHGL WP_050196264.1 hydrogenase [Salmonella enterica] Length: 372\nNumber of predicted TMHs: 2\nExp number of AAs in TMHs: 36.76337\nExp number, first 60 AAs: 19.73298\nTotal prob of N-in: 0.96170\nPOSSIBLE N-term signal sequence\ninside 1 19\nTMhelix 20 39\noutside 40 325\nTMhelix 326 348\ninside 349 372

24790 GCF_001350215.1_8490_2_66 Salmonella enterica subsp. enterica serovar Typhimurium str. DT104 Proteobacteria; Gammaproteobacteria; Enterobacterales; Enterobacteriaceae; Salmonella; Salmonella enterica; Salmonella enterica subsp. enterica; Salmonella enterica subsp. enterica serovar Typhimurium MNNEETFYQAMRRKGVTRRSFLKFCSLAATSLGLGAGMTPKIAWALENKPRIPVVWIHGL WP_001058312.1 hydrogenase [Salmonella enterica] Length: 372\nNumber of predicted TMHs: 2\nExp number of AAs in TMHs: 36.51738\nExp number, first 60 AAs: 19.72849\nTotal prob of N-in: 0.96170\nPOSSIBLE N-term signal sequence\ninside 1 19\nTMhelix 20 39\noutside 40 325\nTMhelix 326 348\ninside 349 372

24791 GCF_001217025.1_5275_8_11 Salmonella enterica subsp. enterica serovar Typhimurium str. DT104 Proteobacteria; Gammaproteobacteria; Enterobacterales; Enterobacteriaceae; Salmonella; Salmonella enterica; Salmonella enterica subsp. enterica; Salmonella enterica subsp. enterica serovar Typhimurium MTGDNTLITSHGINRRDFMKLCAALAATMGLSSKAAAEMAESVSNPQRPPVIWIGAQECT WP_000145429.1 hydrogenase 2 small subunit [Salmonella enterica] Length: 372\nNumber of predicted TMHs: 1\nExp number of AAs in TMHs: 24.51309\nExp number, first 60 AAs: 0.2682\nTotal prob of N-in: 0.14402\noutside 1 331\nTMhelix 332 354\ninside 355 372

24792 GCF_001216765.1_8036_3_82 Salmonella enterica subsp. enterica serovar Typhimurium str. DT104 Proteobacteria; Gammaproteobacteria; Enterobacterales; Enterobacteriaceae; Salmonella; Salmonella enterica; Salmonella enterica subsp. enterica; Salmonella enterica subsp. enterica serovar Typhimurium MTGDNTLITSHGINRRDFMKLCAALAATMGLSSKAAAEMAESVSNPQRPPVIWIGAQECT WP_000145429.1 hydrogenase 2 small subunit [Salmonella enterica] Length: 372\nNumber of predicted TMHs: 1\nExp number of AAs in TMHs: 24.51309\nExp number, first 60 AAs: 0.2682\nTotal prob of N-in: 0.14402\noutside 1 331\nTMhelix 332 354\ninside 355 372

24793 GCF_001216645.1_5275_3_11 Salmonella enterica subsp. enterica serovar Typhimurium str. DT104 Proteobacteria; Gammaproteobacteria; Enterobacterales; Enterobacteriaceae; Salmonella; Salmonella enterica; Salmonella enterica subsp. enterica; Salmonella enterica subsp. enterica serovar Typhimurium MTGDNTLITSHGINRRDFMKLCAALAATMGLSSKAAAEMAESVSNPQRPPVIWIGAQECT WP_000145429.1 hydrogenase 2 small subunit [Salmonella enterica] Length: 372\nNumber of predicted TMHs: 1\nExp number of AAs in TMHs: 24.51309\nExp number, first 60 AAs: 0.2682\nTotal prob of N-in: 0.14402\noutside 1 331\nTMhelix 332 354\ninside 355 372

24794 GCF_001216845.1_5299_3_8Salmonella enterica subsp. enterica serovar Typhimurium str. DT104 Proteobacteria; Gammaproteobacteria; Enterobacterales; Enterobacteriaceae; Salmonella; Salmonella enterica; Salmonella enterica subsp. enterica; Salmonella enterica subsp. enterica serovar Typhimurium MTGDNTLITSHGINRRDFMKLCAALAATMGLSSKAAAEMAESVSNPQRPPVIWIGAQECT WP_000145429.1 hydrogenase 2 small subunit [Salmonella enterica] Length: 372\nNumber of predicted TMHs: 1\nExp

number of AAs in TMHs: 24.51309\nExp number, first 60 AAs: 0.2682\nTotal prob of N-in: 0.14402\noutside 1 331\nTMhelix 332 354\ninside 355 372

24795 GCF_001216705.1_5275_6_7Salmonella enterica subsp. enterica serovar Typhimurium str. DT104 Proteobacteria; Gammaproteobacteria; Enterobacterales; Enterobacteriaceae; Salmonella; Salmonella enterica; Salmonella enterica subsp. enterica; Salmonella enterica subsp. enterica serovar Typhimurium MTGDNTLITSHGINRRDFMKLCAALAATMGLSSKAAAEMAESVSNPQRPPVIWIGAQECT WP_000145429.1 hydrogenase 2 small subunit [Salmonella enterica] Length: 372\nNumber of predicted TMHs: 1\nExp number of AAs in TMHs: 24.51309\nExp number, first 60 AAs: 0.2682\nTotal prob of N-in: 0.14402\noutside 1 331\nTMhelix 332 354\ninside 355 372

24796 GCF_001216585.1_5388_1_12 Salmonella enterica subsp. enterica serovar Typhimurium str. DT104 Proteobacteria; Gammaproteobacteria; Enterobacterales; Enterobacteriaceae; Salmonella; Salmonella enterica; Salmonella enterica subsp. enterica; Salmonella enterica subsp. enterica serovar Typhimurium MTGDNTLITSHGINRRDFMKLCAALAATMGLSSKAAAEMAESVSNPQRPPVIWIGAQECT WP_000145429.1 hydrogenase 2 small subunit [Salmonella enterica] Length: 372\nNumber of predicted TMHs: 1\nExp number of AAs in TMHs: 24.51309\nExp number, first 60 AAs: 0.2682\nTotal prob of N-in: 0.14402\noutside 1 331\nTMhelix 332 354\ninside 355 372

24797 GCF_001216545.1_5285_6_8Salmonella enterica subsp. enterica serovar Typhimurium str. DT104 Proteobacteria; Gammaproteobacteria; Enterobacterales; Enterobacteriaceae; Salmonella; Salmonella enterica; Salmonella enterica subsp. enterica; Salmonella enterica subsp. enterica serovar Typhimurium MTGDNTLITSHGINRRDFMKLCAALAATMGLSSKAAAEMAESVSNPQRPPVIWIGAQECT WP_000145429.1 hydrogenase 2 small subunit [Salmonella enterica] Length: 372\nNumber of predicted TMHs: 1\nExp number of AAs in TMHs: 24.51309\nExp number, first 60 AAs: 0.2682\nTotal prob of N-in: 0.14402\noutside 1 331\nTMhelix 332 354\ninside 355 372

24798 GCF_001215685.1_5275_3_12 Salmonella enterica subsp. enterica serovar Typhimurium str. DT104 Proteobacteria; Gammaproteobacteria; Enterobacterales; Enterobacteriaceae; Salmonella; Salmonella enterica; Salmonella enterica subsp. enterica; Salmonella enterica subsp. enterica serovar Typhimurium MNNEETFYQAMRRKGVTRRSFLKFCSLAATSLGLGAGMTPKIAWALENKPRIPVVWIHGL WP_001058312.1 hydrogenase [Salmonella enterica] Length: 372\nNumber of predicted TMHs: 2\nExp number of AAs in TMHs: 36.51738\nExp number, first 60 AAs: 19.72849\nTotal prob of N-in: 0.96170\nPOSSIBLE N-term signal sequence\ninside 1 19\nTMhelix 20 39\noutside 40 325\nTMhelix 326 348\ninside 349 372

24799 GCF_001216445.1_8036_3_52 Salmonella enterica subsp. enterica serovar Typhimurium str. DT104 Proteobacteria; Gammaproteobacteria; Enterobacterales; Enterobacteriaceae; Salmonella; Salmonella enterica; Salmonella enterica subsp. enterica; Salmonella enterica subsp. enterica serovar Typhimurium MNNEETFYQAMRRKGVTRRSFLKFCSLAATSLGLGAGMTPKIAWALENKPRIPVVWIHGL WP_001058312.1 hydrogenase [Salmonella enterica] Length: 372\nNumber of predicted TMHs: 2\nExp number of AAs in TMHs: 36.51738\nExp number, first 60 AAs: 19.72849\nTotal prob of N-in: 0.96170\nPOSSIBLE N-term signal sequence\ninside 1 19\nTMhelix 20 39\noutside 40 325\nTMhelix 326 348\ninside 349 372

24800 GCF_001215765.1_8080_3_20 Salmonella enterica subsp. enterica serovar Typhimurium str. DT104 Proteobacteria; Gammaproteobacteria; Enterobacterales; Enterobacteriaceae; Salmonella; Salmonella enterica; Salmonella enterica subsp. enterica; Salmonella enterica subsp. enterica serovar Typhimurium MNNEETFYQAMRRKGVTRRSFLKFCSLAATSLGLGAGMTPKIAWALENKPRIPVVWIHGL WP_001058312.1 hydrogenase [Salmonella enterica] Length: 372\nNumber of predicted TMHs: 2\nExp number of AAs in TMHs: 36.51738\nExp number, first 60 AAs: 19.72849\nTotal prob of N-in: 0.96170\nPOSSIBLE N-term signal sequence\ninside 1 19\nTMhelix 20 39\noutside 40 325\nTMhelix 326 348\ninside 349 372

24801 GCF_001216425.1_5275_7_7Salmonella enterica subsp. enterica serovar Typhimurium str. DT104 Proteobacteria; Gammaproteobacteria; Enterobacterales; Enterobacteriaceae; Salmonella; Salmonella enterica; Salmonella enterica subsp. enterica; Salmonella enterica subsp. enterica serovar Typhimurium MNNEETFYQAMRRKGVTRRSFLKFCSLAATSLGLGAGMTPKIAWALENKPRIPVVWIHGL WP_001058312.1 hydrogenase [Salmonella enterica] Length: 372\nNumber of predicted TMHs: 2\nExp number of AAs in TMHs: 36.51738\nExp number, first 60 AAs: 19.72849\nTotal prob of N-in: 0.96170\nPOSSIBLE N-term signal sequence\ninside 1 19\nTMhelix 20 39\noutside 40 325\nTMhelix 326 348\ninside 349 372

24802 GCF_001216345.1_8080_3_30 Salmonella enterica subsp. enterica serovar Typhimurium str. DT104 Proteobacteria; Gammaproteobacteria; Enterobacterales; Enterobacteriaceae; Salmonella; Salmonella enterica;

Salmonella enterica subsp. enterica; Salmonella enterica subsp. enterica serovar Typhimurium
MNNEETFYQAMRRKGVTRRSFLKFCSLAATSLGLGAGMTPKIAWALENKPRIPVVWIHGL WP_001058312.1 hydrogenase [Salmonella enterica] Length: 372\nNumber of predicted TMHs: 2\nExp number of AAs in TMHs: 36.51738\nExp number, first 60 AAs: 19.72849\nTotal prob of N-in: 0.96170\nPOSSIBLE N-term signal sequence\ninside 1 19\nTMhelix 20 39\noutside 40 325\nTMhelix 326 348\ninside 349 372

24803 GCF_001215865.1_8080_3_26 Salmonella enterica subsp. enterica serovar Typhimurium str. DT104 Proteobacteria; Gammaproteobacteria; Enterobacterales; Enterobacteriaceae; Salmonella; Salmonella enterica; Salmonella enterica subsp. enterica; Salmonella enterica subsp. enterica serovar Typhimurium
MNNEETFYQAMRRKGVTRRSFLKFCSLAATSLGLGAGMTPKIAWALENKPRIPVVWIHGL WP_001058312.1 hydrogenase [Salmonella enterica] Length: 372\nNumber of predicted TMHs: 2\nExp number of AAs in TMHs: 36.51738\nExp number, first 60 AAs: 19.72849\nTotal prob of N-in: 0.96170\nPOSSIBLE N-term signal sequence\ninside 1 19\nTMhelix 20 39\noutside 40 325\nTMhelix 326 348\ninside 349 372

24804 GCF_001215525.1_5275_1_9Salmonella enterica subsp. enterica serovar Typhimurium str. DT104 Proteobacteria; Gammaproteobacteria; Enterobacterales; Enterobacteriaceae; Salmonella; Salmonella enterica; Salmonella enterica subsp. enterica; Salmonella enterica subsp. enterica serovar Typhimurium
MNNEETFYQAMRRKGVTRRSFLKFCSLAATSLGLGAGMTPKIAWALENKPRIPVVWIHGL WP_001058312.1 hydrogenase [Salmonella enterica] Length: 372\nNumber of predicted TMHs: 2\nExp number of AAs in TMHs: 36.51738\nExp number, first 60 AAs: 19.72849\nTotal prob of N-in: 0.96170\nPOSSIBLE N-term signal sequence\ninside 1 19\nTMhelix 20 39\noutside 40 325\nTMhelix 326 348\ninside 349 372

24805 GCF_001215885.1_5275_1_4Salmonella enterica subsp. enterica serovar Typhimurium str. DT104 Proteobacteria; Gammaproteobacteria; Enterobacterales; Enterobacteriaceae; Salmonella; Salmonella enterica; Salmonella enterica subsp. enterica; Salmonella enterica subsp. enterica serovar Typhimurium
MNNEETFYQAMRRKGVTRRSFLKFCSLAATSLGLGAGMTPKIAWALENKPRIPVVWIHGL WP_001058312.1 hydrogenase [Salmonella enterica] Length: 372\nNumber of predicted TMHs: 2\nExp number of AAs in TMHs: 36.51738\nExp number, first 60 AAs: 19.72849\nTotal prob of N-in: 0.96170\nPOSSIBLE N-term signal sequence\ninside 1 19\nTMhelix 20 39\noutside 40 325\nTMhelix 326 348\ninside 349 372

24806 GCF_001216125.1_8036_3_61 Salmonella enterica subsp. enterica serovar Typhimurium str. DT104 Proteobacteria; Gammaproteobacteria; Enterobacterales; Enterobacteriaceae; Salmonella; Salmonella enterica; Salmonella enterica subsp. enterica; Salmonella enterica subsp. enterica serovar Typhimurium
MNNEETFYQAMRRKGVTRRSFLKFCSLAATSLGLGAGMTPKIAWALENKPRIPVVWIHGL WP_001058312.1 hydrogenase [Salmonella enterica] Length: 372\nNumber of predicted TMHs: 2\nExp number of AAs in TMHs: 36.51738\nExp number, first 60 AAs: 19.72849\nTotal prob of N-in: 0.96170\nPOSSIBLE N-term signal sequence\ninside 1 19\nTMhelix 20 39\noutside 40 325\nTMhelix 326 348\ninside 349 372

24807 GCF_001215985.1_5275_5_6Salmonella enterica subsp. enterica serovar Typhimurium str. DT104 Proteobacteria; Gammaproteobacteria; Enterobacterales; Enterobacteriaceae; Salmonella; Salmonella enterica; Salmonella enterica subsp. enterica; Salmonella enterica subsp. enterica serovar Typhimurium
MNNEETFYQAMRRKGVTRRSFLKFCSLAATSLGLGAGMTPKIAWALENKPRIPVVWIHGL WP_001058312.1 hydrogenase [Salmonella enterica] Length: 372\nNumber of predicted TMHs: 2\nExp number of AAs in TMHs: 36.51738\nExp number, first 60 AAs: 19.72849\nTotal prob of N-in: 0.96170\nPOSSIBLE N-term signal sequence\ninside 1 19\nTMhelix 20 39\noutside 40 325\nTMhelix 326 348\ninside 349 372

24808 GCF_001215585.1_5295_2_8Salmonella enterica subsp. enterica serovar Typhimurium str. DT104 Proteobacteria; Gammaproteobacteria; Enterobacterales; Enterobacteriaceae; Salmonella; Salmonella enterica; Salmonella enterica subsp. enterica; Salmonella enterica subsp. enterica serovar Typhimurium
MNNEETFYQAMRRKGVTRRSFLKFCSLAATSLGLGAGMTPKIAWALENKPRIPVVWIHGL WP_001058312.1 hydrogenase [Salmonella enterica] Length: 372\nNumber of predicted TMHs: 2\nExp number of AAs in TMHs: 36.51738\nExp number, first 60 AAs: 19.72849\nTotal prob of N-in: 0.96170\nPOSSIBLE N-term signal sequence\ninside 1 19\nTMhelix 20 39\noutside 40 325\nTMhelix 326 348\ninside 349 372

24809 GCF_001215685.1_5275_3_12 Salmonella enterica subsp. enterica serovar Typhimurium str. DT104 Proteobacteria; Gammaproteobacteria; Enterobacterales; Enterobacteriaceae; Salmonella; Salmonella enterica; Salmonella enterica subsp. enterica; Salmonella enterica subsp. enterica serovar Typhimurium
MTGDNTLITSHGINRRDFMKLCAALAATMGLSSKAAAEMAESVSNPQRPPVIWIGAQECT WP_000145429.1 hydrogenase 2 small subunit [Salmonella enterica] Length: 372\nNumber of predicted TMHs: 1\nExp number of AAs in TMHs: 24.51309\nExp number, first 60 AAs: 0.2682\nTotal prob of N-in: 0.14402\noutside 1 331\nTMhelix 332 354\ninside 355 372

24810 GCF_001216425.1_5275_7_7Salmonella enterica subsp. enterica serovar Typhimurium str. DT104 Proteobacteria; Gammaproteobacteria; Enterobacterales; Enterobacteriaceae; Salmonella; Salmonella enterica; Salmonella enterica subsp. enterica; Salmonella enterica subsp. enterica serovar Typhimurium MTGDNTLITSHGINRRDFMKLCAALAATMGLSSKAAAEMAESVSNPQRPPVIWIGAQECT WP_000145429.1 hydrogenase 2 small subunit [Salmonella enterica] Length: 372\nNumber of predicted TMHs: 1\nExp number of AAs in TMHs: 24.51309\nExp number, first 60 AAs: 0.2682\nTotal prob of N-in: 0.14402\noutside 1 331\nTMhelix 332 354\ninside 355 372

24811 GCF_001350275.1_8490_2_82 Salmonella enterica subsp. enterica serovar Typhimurium str. DT104 Proteobacteria; Gammaproteobacteria; Enterobacterales; Enterobacteriaceae; Salmonella; Salmonella enterica; Salmonella enterica subsp. enterica; Salmonella enterica subsp. enterica serovar Typhimurium MNNEETFYQAMRRKGVTRRSFLKFCSLAATSLGLGAGMTPKIAWALENKPRIPVVWIHGL WP_001058312.1 hydrogenase [Salmonella enterica] Length: 372\nNumber of predicted TMHs: 2\nExp number of AAs in TMHs: 36.51738\nExp number, first 60 AAs: 19.72849\nTotal prob of N-in: 0.96170\nPOSSIBLE N-term signal sequence\ninside 1 19\nTMhelix 20 39\noutside 40 325\nTMhelix 326 348\ninside 349 372

24812 GCF_001216445.1_8036_3_52 Salmonella enterica subsp. enterica serovar Typhimurium str. DT104 Proteobacteria; Gammaproteobacteria; Enterobacterales; Enterobacteriaceae; Salmonella; Salmonella enterica; Salmonella enterica subsp. enterica; Salmonella enterica subsp. enterica serovar Typhimurium MTGDNTLITSHGINRRDFMKLCAALAATMGLSSKAAAEMAESVSNPQRPPVIWIGAQECT WP_000145429.1 hydrogenase 2 small subunit [Salmonella enterica] Length: 372\nNumber of predicted TMHs: 1\nExp number of AAs in TMHs: 24.51309\nExp number, first 60 AAs: 0.2682\nTotal prob of N-in: 0.14402\noutside 1 331\nTMhelix 332 354\ninside 355 372

24813 GCF_001216345.1_8080_3_30 Salmonella enterica subsp. enterica serovar Typhimurium str. DT104 Proteobacteria; Gammaproteobacteria; Enterobacterales; Enterobacteriaceae; Salmonella; Salmonella enterica; Salmonella enterica subsp. enterica; Salmonella enterica subsp. enterica serovar Typhimurium MTGDNTLITSHGINRRDFMKLCAALAATMGLSSKAAAEMAESVSNPQRPPVIWIGAQECT WP_000145429.1 hydrogenase 2 small subunit [Salmonella enterica] Length: 372\nNumber of predicted TMHs: 1\nExp number of AAs in TMHs: 24.51309\nExp number, first 60 AAs: 0.2682\nTotal prob of N-in: 0.14402\noutside 1 331\nTMhelix 332 354\ninside 355 372

24814 GCF_001216125.1_8036_3_61 Salmonella enterica subsp. enterica serovar Typhimurium str. DT104 Proteobacteria; Gammaproteobacteria; Enterobacterales; Enterobacteriaceae; Salmonella; Salmonella enterica; Salmonella enterica subsp. enterica; Salmonella enterica subsp. enterica serovar Typhimurium MTGDNTLITSHGINRRDFMKLCAALAATMGLSSKAAAEMAESVSNPQRPPVIWIGAQECT WP_000145429.1 hydrogenase 2 small subunit [Salmonella enterica] Length: 372\nNumber of predicted TMHs: 1\nExp number of AAs in TMHs: 24.51309\nExp number, first 60 AAs: 0.2682\nTotal prob of N-in: 0.14402\noutside 1 331\nTMhelix 332 354\ninside 355 372

24815 GCF_001215885.1_5275_1_4Salmonella enterica subsp. enterica serovar Typhimurium str. DT104 Proteobacteria; Gammaproteobacteria; Enterobacterales; Enterobacteriaceae; Salmonella; Salmonella enterica; Salmonella enterica subsp. enterica; Salmonella enterica subsp. enterica serovar Typhimurium MTGDNTLITSHGINRRDFMKLCAALAATMGLSSKAAAEMAESVSNPQRPPVIWIGAQECT WP_000145429.1 hydrogenase 2 small subunit [Salmonella enterica] Length: 372\nNumber of predicted TMHs: 1\nExp number of AAs in TMHs: 24.51309\nExp number, first 60 AAs: 0.2682\nTotal prob of N-in: 0.14402\noutside 1 331\nTMhelix 332 354\ninside 355 372

24816 GCF_001215765.1_8080_3_20 Salmonella enterica subsp. enterica serovar Typhimurium str. DT104 Proteobacteria; Gammaproteobacteria; Enterobacterales; Enterobacteriaceae; Salmonella; Salmonella enterica; Salmonella enterica subsp. enterica; Salmonella enterica subsp. enterica serovar Typhimurium MTGDNTLITSHGINRRDFMKLCAALAATMGLSSKAAAEMAESVSNPQRPPVIWIGAQECT WP_000145429.1 hydrogenase 2 small subunit [Salmonella enterica] Length: 372\nNumber of predicted TMHs: 1\nExp number of AAs in TMHs: 24.51309\nExp number, first 60 AAs: 0.2682\nTotal prob of N-in: 0.14402\noutside 1 331\nTMhelix 332 354\ninside 355 372

24817 GCF_001215585.1_5295_2_8Salmonella enterica subsp. enterica serovar Typhimurium str. DT104 Proteobacteria; Gammaproteobacteria; Enterobacterales; Enterobacteriaceae; Salmonella; Salmonella enterica; Salmonella enterica subsp. enterica; Salmonella enterica subsp. enterica serovar Typhimurium MTGDNTLITSHGINRRDFMKLCAALAATMGLSSKAAAEMAESVSNPQRPPVIWIGAQECT WP_000145429.1 hydrogenase 2 small subunit [Salmonella enterica] Length: 372\nNumber of predicted TMHs: 1\nExp

number of AAs in TMHs: 24.51309\nExp number, first 60 AAs: 0.2682\nTotal prob of N-in: 0.14402\noutside 1 331\nTMhelix 332 354\ninside 355 372

24818 GCF_001216485.1_5275_2_12 Salmonella enterica subsp. enterica serovar Typhimurium str. DT104 Proteobacteria; Gammaproteobacteria; Enterobacterales; Enterobacteriaceae; Salmonella; Salmonella enterica; Salmonella enterica subsp. enterica; Salmonella enterica subsp. enterica serovar Typhimurium MTGDNTLITSHGINRRDFMKLCAALAATMGLSSKAAAEMAESVSNPQRPPVIWIGAQECT WP_000145429.1 hydrogenase 2 small subunit [Salmonella enterica] Length: 372\nNumber of predicted TMHs: 1\nExp number of AAs in TMHs: 24.51309\nExp number, first 60 AAs: 0.2682\nTotal prob of N-in: 0.14402\noutside 1 331\nTMhelix 332 354\ninside 355 372

24819 GCF_001215865.1_8080_3_26 Salmonella enterica subsp. enterica serovar Typhimurium str. DT104 Proteobacteria; Gammaproteobacteria; Enterobacterales; Enterobacteriaceae; Salmonella; Salmonella enterica; Salmonella enterica subsp. enterica; Salmonella enterica subsp. enterica serovar Typhimurium MTGDNTLITSHGINRRDFMKLCAALAATMGLSSKAAAEMAESVSNPQRPPVIWIGAQECT WP_000145429.1 hydrogenase 2 small subunit [Salmonella enterica] Length: 372\nNumber of predicted TMHs: 1\nExp number of AAs in TMHs: 24.51309\nExp number, first 60 AAs: 0.2682\nTotal prob of N-in: 0.14402\noutside 1 331\nTMhelix 332 354\ninside 355 372

24820 GCF_001215985.1_5275_5_6Salmonella enterica subsp. enterica serovar Typhimurium str. DT104 Proteobacteria; Gammaproteobacteria; Enterobacterales; Enterobacteriaceae; Salmonella; Salmonella enterica; Salmonella enterica subsp. enterica; Salmonella enterica subsp. enterica serovar Typhimurium MTGDNTLITSHGINRRDFMKLCAALAATMGLSSKAAAEMAESVSNPQRPPVIWIGAQECT WP_000145429.1 hydrogenase 2 small subunit [Salmonella enterica] Length: 372\nNumber of predicted TMHs: 1\nExp number of AAs in TMHs: 24.51309\nExp number, first 60 AAs: 0.2682\nTotal prob of N-in: 0.14402\noutside 1 331\nTMhelix 332 354\ninside 355 372

24821 GCF_001215525.1_5275_1_9Salmonella enterica subsp. enterica serovar Typhimurium str. DT104 Proteobacteria; Gammaproteobacteria; Enterobacterales; Enterobacteriaceae; Salmonella; Salmonella enterica; Salmonella enterica subsp. enterica; Salmonella enterica subsp. enterica serovar Typhimurium MTGDNTLITSHGINRRDFMKLCAALAATMGLSSKAAAEMAESVSNPQRPPVIWIGAQECT WP_000145429.1 hydrogenase 2 small subunit [Salmonella enterica] Length: 372\nNumber of predicted TMHs: 1\nExp number of AAs in TMHs: 24.51309\nExp number, first 60 AAs: 0.2682\nTotal prob of N-in: 0.14402\noutside 1 331\nTMhelix 332 354\ninside 355 372

24822 GCF_001350035.1_8490_2_58 Salmonella enterica subsp. enterica serovar Typhimurium str. DT104 Proteobacteria; Gammaproteobacteria; Enterobacterales; Enterobacteriaceae; Salmonella; Salmonella enterica; Salmonella enterica subsp. enterica; Salmonella enterica subsp. enterica serovar Typhimurium MTGDNTLITSHGINRRDFMKLCAALAATMGLSSKAAAEMAESVSNPQRPPVIWIGAQECT WP_000145429.1 hydrogenase 2 small subunit [Salmonella enterica] Length: 372\nNumber of predicted TMHs: 1\nExp number of AAs in TMHs: 24.51309\nExp number, first 60 AAs: 0.2682\nTotal prob of N-in: 0.14402\noutside 1 331\nTMhelix 332 354\ninside 355 372

24823 GCF_001214765.1_8080_3_54 Salmonella enterica subsp. enterica serovar Typhimurium str. DT104 Proteobacteria; Gammaproteobacteria; Enterobacterales; Enterobacteriaceae; Salmonella; Salmonella enterica; Salmonella enterica subsp. enterica; Salmonella enterica subsp. enterica serovar Typhimurium MNNEETFYQAMRRKGVTRRSFLKFCSLAATSLGLGAGMTPKIAWALENKPRIPVVWIHGL WP_001058312.1 hydrogenase [Salmonella enterica] Length: 372\nNumber of predicted TMHs: 2\nExp number of AAs in TMHs: 36.51738\nExp number, first 60 AAs: 19.72849\nTotal prob of N-in: 0.96170\nPOSSIBLE N-term signal sequence\ninside 1 19\nTMhelix 20 39\noutside 40 325\nTMhelix 326 348\ninside 349 372

24824 GCF_001214965.1_8080_3_86 Salmonella enterica subsp. enterica serovar Typhimurium str. DT104 Proteobacteria; Gammaproteobacteria; Enterobacterales; Enterobacteriaceae; Salmonella; Salmonella enterica; Salmonella enterica subsp. enterica; Salmonella enterica subsp. enterica serovar Typhimurium MNNEETFYQAMRRKGVTRRSFLKFCSLAATSLGLGAGMTPKIAWALENKPRIPVVWIHGL WP_001058312.1 hydrogenase [Salmonella enterica] Length: 372\nNumber of predicted TMHs: 2\nExp number of AAs in TMHs: 36.51738\nExp number, first 60 AAs: 19.72849\nTotal prob of N-in: 0.96170\nPOSSIBLE N-term signal sequence\ninside 1 19\nTMhelix 20 39\noutside 40 325\nTMhelix 326 348\ninside 349 372

24825 GCF_001214545.1_5299_6_9Salmonella enterica subsp. enterica serovar Typhimurium str. DT104 Proteobacteria; Gammaproteobacteria; Enterobacterales; Enterobacteriaceae; Salmonella; Salmonella enterica;

Salmonella enterica subsp. enterica; Salmonella enterica subsp. enterica serovar Typhimurium
MNNEETFYQAMRRKGVTRRSFLKFCSLAATSLGLGAGMTPKIAWALENKPRIPVVWIHGL WP_001058312.1
hydrogenase [Salmonella enterica] Length: 372\nNumber of predicted TMHs: 2\nExp number of AAs in TMHs: 36.51738\nExp number, first 60 AAs: 19.72849\nTotal prob of N-in: 0.96170\nPOSSIBLE N-term signal sequence\ninside 1 19\nTMhelix 20 39\noutside 40 325\nTMhelix 326 348\ninside 349 372

24826 GCF_001214705.1_8080_3_36 Salmonella enterica subsp. enterica serovar Typhimurium str. DT104
Proteobacteria; Gammaproteobacteria; Enterobacterales; Enterobacteriaceae; Salmonella; Salmonella enterica; Salmonella enterica subsp. enterica; Salmonella enterica subsp. enterica serovar Typhimurium
MNNEETFYQAMRRKGVTRRSFLKFCSLAATSLGLGAGMTPKIAWALENKPRIPVVWIHGL WP_001058312.1
hydrogenase [Salmonella enterica] Length: 372\nNumber of predicted TMHs: 2\nExp number of AAs in TMHs: 36.51738\nExp number, first 60 AAs: 19.72849\nTotal prob of N-in: 0.96170\nPOSSIBLE N-term signal sequence\ninside 1 19\nTMhelix 20 39\noutside 40 325\nTMhelix 326 348\ninside 349 372

24827 GCF_001214425.1_8036_3_65 Salmonella enterica subsp. enterica serovar Typhimurium str. DT104
Proteobacteria; Gammaproteobacteria; Enterobacterales; Enterobacteriaceae; Salmonella; Salmonella enterica; Salmonella enterica subsp. enterica; Salmonella enterica subsp. enterica serovar Typhimurium
MNNEETFYQAMRRKGVTRRSFLKFCSLAATSLGLGAGMTPKIAWALENKPRIPVVWIHGL WP_001058312.1
hydrogenase [Salmonella enterica] Length: 372\nNumber of predicted TMHs: 2\nExp number of AAs in TMHs: 36.51738\nExp number, first 60 AAs: 19.72849\nTotal prob of N-in: 0.96170\nPOSSIBLE N-term signal sequence\ninside 1 19\nTMhelix 20 39\noutside 40 325\nTMhelix 326 348\ninside 349 372

24828 GCF_001215105.1_5275_8_9Salmonella enterica subsp. enterica serovar Typhimurium str. DT104
Proteobacteria; Gammaproteobacteria; Enterobacterales; Enterobacteriaceae; Salmonella; Salmonella enterica; Salmonella enterica subsp. enterica; Salmonella enterica subsp. enterica serovar Typhimurium
MNNEETFYQAMRRKGVTRRSFLKFCSLAATSLGLGAGMTPKIAWALENKPRIPVVWIHGL WP_001058312.1
hydrogenase [Salmonella enterica] Length: 372\nNumber of predicted TMHs: 2\nExp number of AAs in TMHs: 36.51738\nExp number, first 60 AAs: 19.72849\nTotal prob of N-in: 0.96170\nPOSSIBLE N-term signal sequence\ninside 1 19\nTMhelix 20 39\noutside 40 325\nTMhelix 326 348\ninside 349 372

24829 GCF_001215505.1_8080_3_53 Salmonella enterica subsp. enterica serovar Typhimurium str. DT104
Proteobacteria; Gammaproteobacteria; Enterobacterales; Enterobacteriaceae; Salmonella; Salmonella enterica; Salmonella enterica subsp. enterica; Salmonella enterica subsp. enterica serovar Typhimurium
MNNEETFYQAMRRKGVTRRSFLKFCSLAATSLGLGAGMTPKIAWALENKPRIPVVWIHGL WP_001058312.1
hydrogenase [Salmonella enterica] Length: 372\nNumber of predicted TMHs: 2\nExp number of AAs in TMHs: 36.51738\nExp number, first 60 AAs: 19.72849\nTotal prob of N-in: 0.96170\nPOSSIBLE N-term signal sequence\ninside 1 19\nTMhelix 20 39\noutside 40 325\nTMhelix 326 348\ninside 349 372

24830 GCF_001215365.1_8080_3_46 Salmonella enterica subsp. enterica serovar Typhimurium str. DT104
Proteobacteria; Gammaproteobacteria; Enterobacterales; Enterobacteriaceae; Salmonella; Salmonella enterica; Salmonella enterica subsp. enterica; Salmonella enterica subsp. enterica serovar Typhimurium
MNNEETFYQAMRRKGVTRRSFLKFCSLAATSLGLGAGMTPKIAWALENKPRIPVVWIHGL WP_001058312.1
hydrogenase [Salmonella enterica] Length: 372\nNumber of predicted TMHs: 2\nExp number of AAs in TMHs: 36.51738\nExp number, first 60 AAs: 19.72849\nTotal prob of N-in: 0.96170\nPOSSIBLE N-term signal sequence\ninside 1 19\nTMhelix 20 39\noutside 40 325\nTMhelix 326 348\ninside 349 372

24831 GCF_001215105.1_5275_8_9Salmonella enterica subsp. enterica serovar Typhimurium str. DT104
Proteobacteria; Gammaproteobacteria; Enterobacterales; Enterobacteriaceae; Salmonella; Salmonella enterica; Salmonella enterica subsp. enterica; Salmonella enterica subsp. enterica serovar Typhimurium
MTGDNTLITSHGINRRDFMKLCAALAATMGLSSKAAAEMAESVSNPQRPPVIWIGAQECT WP_000145429.1
hydrogenase 2 small subunit [Salmonella enterica] Length: 372\nNumber of predicted TMHs: 1\nExp number of AAs in TMHs: 24.51309\nExp number, first 60 AAs: 0.2682\nTotal prob of N-in: 0.14402\noutside 1 331\nTMhelix 332 354\ninside 355 372

24832 GCF_001215505.1_8080_3_53 Salmonella enterica subsp. enterica serovar Typhimurium str. DT104
Proteobacteria; Gammaproteobacteria; Enterobacterales; Enterobacteriaceae; Salmonella; Salmonella enterica; Salmonella enterica subsp. enterica; Salmonella enterica subsp. enterica serovar Typhimurium
MTGDNTLITSHGINRRDFMKLCAALAATMGLSSKAAAEMAESVSNPQRPPVIWIGAQECT WP_000145429.1
hydrogenase 2 small subunit [Salmonella enterica] Length: 372\nNumber of predicted TMHs: 1\nExp number of AAs in TMHs: 24.51309\nExp number, first 60 AAs: 0.2682\nTotal prob of N-in: 0.14402\noutside 1 331\nTMhelix 332 354\ninside 355 372

24833 GCF_001215365.1_8080_3_46 Salmonella enterica subsp. enterica serovar Typhimurium str. DT104 Proteobacteria; Gammaproteobacteria; Enterobacterales; Enterobacteriaceae; Salmonella; Salmonella enterica; Salmonella enterica subsp. enterica; Salmonella enterica subsp. enterica serovar Typhimurium MTGDNTLITSHGINRRDFMKLCAALAATMGLSSKAAAEMAESVSNPQRPPVIWIGAQECT WP_000145429.1 hydrogenase 2 small subunit [Salmonella enterica] Length: 372\nNumber of predicted TMHs: 1\nExp number of AAs in TMHs: 24.51309\nExp number, first 60 AAs: 0.2682\nTotal prob of N-in: 0.14402\noutside 1 331\nTMhelix 332 354\ninside 355 372

24834 GCF_001214965.1_8080_3_86 Salmonella enterica subsp. enterica serovar Typhimurium str. DT104 Proteobacteria; Gammaproteobacteria; Enterobacterales; Enterobacteriaceae; Salmonella; Salmonella enterica; Salmonella enterica subsp. enterica; Salmonella enterica subsp. enterica serovar Typhimurium MTGDNTLITSHGINRRDFMKLCAALAATMGLSSKAAAEMAESVSNPQRPPVIWIGAQECT WP_000145429.1 hydrogenase 2 small subunit [Salmonella enterica] Length: 372\nNumber of predicted TMHs: 1\nExp number of AAs in TMHs: 24.51309\nExp number, first 60 AAs: 0.2682\nTotal prob of N-in: 0.14402\noutside 1 331\nTMhelix 332 354\ninside 355 372

24835 GCF_001214765.1_8080_3_54 Salmonella enterica subsp. enterica serovar Typhimurium str. DT104 Proteobacteria; Gammaproteobacteria; Enterobacterales; Enterobacteriaceae; Salmonella; Salmonella enterica; Salmonella enterica subsp. enterica; Salmonella enterica subsp. enterica serovar Typhimurium MTGDNTLITSHGINRRDFMKLCAALAATMGLSSKAAAEMAESVSNPQRPPVIWIGAQECT WP_000145429.1 hydrogenase 2 small subunit [Salmonella enterica] Length: 372\nNumber of predicted TMHs: 1\nExp number of AAs in TMHs: 24.51309\nExp number, first 60 AAs: 0.2682\nTotal prob of N-in: 0.14402\noutside 1 331\nTMhelix 332 354\ninside 355 372

24836 GCF_001214705.1_8080_3_36 Salmonella enterica subsp. enterica serovar Typhimurium str. DT104 Proteobacteria; Gammaproteobacteria; Enterobacterales; Enterobacteriaceae; Salmonella; Salmonella enterica; Salmonella enterica subsp. enterica; Salmonella enterica subsp. enterica serovar Typhimurium MTGDNTLITSHGINRRDFMKLCAALAATMGLSSKAAAEMAESVSNPQRPPVIWIGAQECT WP_000145429.1 hydrogenase 2 small subunit [Salmonella enterica] Length: 372\nNumber of predicted TMHs: 1\nExp number of AAs in TMHs: 24.51309\nExp number, first 60 AAs: 0.2682\nTotal prob of N-in: 0.14402\noutside 1 331\nTMhelix 332 354\ninside 355 372

24837 GCF_001214545.1_5299_6_9Salmonella enterica subsp. enterica serovar Typhimurium str. DT104 Proteobacteria; Gammaproteobacteria; Enterobacterales; Enterobacteriaceae; Salmonella; Salmonella enterica; Salmonella enterica subsp. enterica; Salmonella enterica subsp. enterica serovar Typhimurium MTGDNTLITSHGINRRDFMKLCAALAATMGLSSKAAAEMAESVSNPQRPPVIWIGAQECT WP_000145429.1 hydrogenase 2 small subunit [Salmonella enterica] Length: 372\nNumber of predicted TMHs: 1\nExp number of AAs in TMHs: 24.51309\nExp number, first 60 AAs: 0.2682\nTotal prob of N-in: 0.14402\noutside 1 331\nTMhelix 332 354\ninside 355 372

24838 GCF_001214425.1_8036_3_65 Salmonella enterica subsp. enterica serovar Typhimurium str. DT104 Proteobacteria; Gammaproteobacteria; Enterobacterales; Enterobacteriaceae; Salmonella; Salmonella enterica; Salmonella enterica subsp. enterica; Salmonella enterica subsp. enterica serovar Typhimurium MTGDNTLITSHGINRRDFMKLCAALAATMGLSSKAAAEMAESVSNPQRPPVIWIGAQECT WP_000145429.1 hydrogenase 2 small subunit [Salmonella enterica] Length: 372\nNumber of predicted TMHs: 1\nExp number of AAs in TMHs: 24.51309\nExp number, first 60 AAs: 0.2682\nTotal prob of N-in: 0.14402\noutside 1 331\nTMhelix 332 354\ninside 355 372

24839 GCF_001214325.1_5388_1_7Salmonella enterica subsp. enterica serovar Typhimurium str. DT104 Proteobacteria; Gammaproteobacteria; Enterobacterales; Enterobacteriaceae; Salmonella; Salmonella enterica; Salmonella enterica subsp. enterica; Salmonella enterica subsp. enterica serovar Typhimurium MTGDNTLITSHGINRRDFMKLCAALAATMGLSSKAAAEMAESVSNPQRPPVIWIGAQECT WP_000145429.1 hydrogenase 2 small subunit [Salmonella enterica] Length: 372\nNumber of predicted TMHs: 1\nExp number of AAs in TMHs: 24.51309\nExp number, first 60 AAs: 0.2682\nTotal prob of N-in: 0.14402\noutside 1 331\nTMhelix 332 354\ninside 355 372

24840 GCF_001172145.1_8490_2_61 Salmonella enterica subsp. enterica serovar Typhimurium str. DT104 Proteobacteria; Gammaproteobacteria; Enterobacterales; Enterobacteriaceae; Salmonella; Salmonella enterica; Salmonella enterica subsp. enterica; Salmonella enterica subsp. enterica serovar Typhimurium MNNEETFYQAMRRKGVTRRSFLKFCSLAATSLGLGAGMTPKIAWALENKPRIPVVWIHGL WP_001058312.1 hydrogenase [Salmonella enterica] Length: 372\nNumber of predicted TMHs: 2\nExp number of AAs in

TMHs: 36.51738\nExp number, first 60 AAs: 19.72849\nTotal prob of N-in: 0.96170\nPOSSIBLE N-term signal sequence\ninside 1 19\nTMhelix 20 39\noutside 40 325\nTMhelix 326 348\ninside 349 372

24841 GCF_001172145.1_8490_2_61 Salmonella enterica subsp. enterica serovar Typhimurium str. DT104 Proteobacteria; Gammaproteobacteria; Enterobacterales; Enterobacteriaceae; Salmonella; Salmonella enterica; Salmonella enterica subsp. enterica; Salmonella enterica subsp. enterica serovar Typhimurium MTGDNTLITSHGINRRDFMKLCAALAATMGLSSKAAAEMAESVSNPQRPPVIWIGAQECT WP_000145429.1 hydrogenase 2 small subunit [Salmonella enterica] Length: 372\nNumber of predicted TMHs: 1\nExp number of AAs in TMHs: 24.51309\nExp number, first 60 AAs: 0.2682\nTotal prob of N-in: 0.14402\noutside 1 331\nTMhelix 332 354\ninside 355 372

24842 GCF_001164765.1_8490_2_92 Salmonella enterica subsp. enterica serovar Typhimurium str. DT104 Proteobacteria; Gammaproteobacteria; Enterobacterales; Enterobacteriaceae; Salmonella; Salmonella enterica; Salmonella enterica subsp. enterica; Salmonella enterica subsp. enterica serovar Typhimurium MNNEETFYQAMRRKGVTRRSFLKFCSLAATSLGLGAGMTPKIAWALENKPRIPVVWIHGL WP_001058312.1 hydrogenase [Salmonella enterica] Length: 372\nNumber of predicted TMHs: 2\nExp number of AAs in TMHs: 36.51738\nExp number, first 60 AAs: 19.72849\nTotal prob of N-in: 0.96170\nPOSSIBLE N-term signal sequence\ninside 1 19\nTMhelix 20 39\noutside 40 325\nTMhelix 326 348\ninside 349 372

24843 GCF_001164765.1_8490_2_92 Salmonella enterica subsp. enterica serovar Typhimurium str. DT104 Proteobacteria; Gammaproteobacteria; Enterobacterales; Enterobacteriaceae; Salmonella; Salmonella enterica; Salmonella enterica subsp. enterica; Salmonella enterica subsp. enterica serovar Typhimurium MTGDNTLITSHGINRRDFMKLCAALAATMGLSSKAAAEMAESVSNPQRPPVIWIGAQECT WP_000145429.1 hydrogenase 2 small subunit [Salmonella enterica] Length: 372\nNumber of predicted TMHs: 1\nExp number of AAs in TMHs: 24.51309\nExp number, first 60 AAs: 0.2682\nTotal prob of N-in: 0.14402\noutside 1 331\nTMhelix 332 354\ninside 355 372

24844 GCF_001162405.1_8490_2_53 Salmonella enterica subsp. enterica serovar Typhimurium str. DT104 Proteobacteria; Gammaproteobacteria; Enterobacterales; Enterobacteriaceae; Salmonella; Salmonella enterica; Salmonella enterica subsp. enterica; Salmonella enterica subsp. enterica serovar Typhimurium MNNEETFYQAMRRKGVTRRSFLKFCSLAATSLGLGAGMTPKIAWALENKPRIPVVWIHGL WP_001058312.1 hydrogenase [Salmonella enterica] Length: 372\nNumber of predicted TMHs: 2\nExp number of AAs in TMHs: 36.51738\nExp number, first 60 AAs: 19.72849\nTotal prob of N-in: 0.96170\nPOSSIBLE N-term signal sequence\ninside 1 19\nTMhelix 20 39\noutside 40 325\nTMhelix 326 348\ninside 349 372

24845 GCF_001161925.1_8490_2_65 Salmonella enterica subsp. enterica serovar Typhimurium str. DT104 Proteobacteria; Gammaproteobacteria; Enterobacterales; Enterobacteriaceae; Salmonella; Salmonella enterica; Salmonella enterica subsp. enterica; Salmonella enterica subsp. enterica serovar Typhimurium MNNEETFYQAMRRKGVTRRSFLKFCSLAATSLGLGAGMTPKIAWALENKPRIPVVWIHGL WP_001058312.1 hydrogenase [Salmonella enterica] Length: 372\nNumber of predicted TMHs: 2\nExp number of AAs in TMHs: 36.51738\nExp number, first 60 AAs: 19.72849\nTotal prob of N-in: 0.96170\nPOSSIBLE N-term signal sequence\ninside 1 19\nTMhelix 20 39\noutside 40 325\nTMhelix 326 348\ninside 349 372

24846 GCF_001350055.1_8490_2_67 Salmonella enterica subsp. enterica serovar Typhimurium str. DT104 Proteobacteria; Gammaproteobacteria; Enterobacterales; Enterobacteriaceae; Salmonella; Salmonella enterica; Salmonella enterica subsp. enterica; Salmonella enterica subsp. enterica serovar Typhimurium MTGDNTLITSHGINRRDFMKLCAALAATMGLSSKAAAEMAESVSNPQRPPVIWIGAQECT WP_000145429.1 hydrogenase 2 small subunit [Salmonella enterica] Length: 372\nNumber of predicted TMHs: 1\nExp number of AAs in TMHs: 24.51309\nExp number, first 60 AAs: 0.2682\nTotal prob of N-in: 0.14402\noutside 1 331\nTMhelix 332 354\ninside 355 372

24847 GCF_001162405.1_8490_2_53 Salmonella enterica subsp. enterica serovar Typhimurium str. DT104 Proteobacteria; Gammaproteobacteria; Enterobacterales; Enterobacteriaceae; Salmonella; Salmonella enterica; Salmonella enterica subsp. enterica; Salmonella enterica subsp. enterica serovar Typhimurium MTGDNTLITSHGINRRDFMKLCAALAATMGLSSKAAAEMAESVSNPQRPPVIWIGAQECT WP_000145429.1 hydrogenase 2 small subunit [Salmonella enterica] Length: 372\nNumber of predicted TMHs: 1\nExp number of AAs in TMHs: 24.51309\nExp number, first 60 AAs: 0.2682\nTotal prob of N-in: 0.14402\noutside 1 331\nTMhelix 332 354\ninside 355 372

24848 GCF_001161925.1_8490_2_65 Salmonella enterica subsp. enterica serovar Typhimurium str. DT104 Proteobacteria; Gammaproteobacteria; Enterobacterales; Enterobacteriaceae; Salmonella; Salmonella enterica;

Salmonella enterica subsp. enterica; Salmonella enterica subsp. enterica serovar Typhimurium
MTGDNTLITSHGINRRDFMKLCAALAATMGLSSKAAAEMAESVSNPQRPPVIWIGAQECT WP_000145429.1
hydrogenase 2 small subunit [Salmonella enterica] Length: 372\nNumber of predicted TMHs: 1\nExp number of AAs in TMHs: 24.51309\nExp number, first 60 AAs: 0.2682\nTotal prob of N-in: 0.14402\noutside 1 331\nTMhelix 332 354\ninside 355 372

24849 GCF_001154585.1_8490_2_47 Salmonella enterica subsp. enterica serovar Typhimurium str. DT104
Proteobacteria; Gammaproteobacteria; Enterobacterales; Enterobacteriaceae; Salmonella; Salmonella enterica; Salmonella enterica subsp. enterica; Salmonella enterica subsp. enterica serovar Typhimurium
MNNEETFYQAMRRKGVTRRSFLKFCSLAATSLGLGAGMTPKIAWALENKPRIPVVWIHGL WP_001058312.1
hydrogenase [Salmonella enterica] Length: 372\nNumber of predicted TMHs: 2\nExp number of AAs in TMHs: 36.51738\nExp number, first 60 AAs: 19.72849\nTotal prob of N-in: 0.96170\nPOSSIBLE N-term signal sequence\ninside 1 19\nTMhelix 20 39\noutside 40 325\nTMhelix 326 348\ninside 349 372

24850 GCF_001155485.1_8490_2_73 Salmonella enterica subsp. enterica serovar Typhimurium str. DT104
Proteobacteria; Gammaproteobacteria; Enterobacterales; Enterobacteriaceae; Salmonella; Salmonella enterica; Salmonella enterica subsp. enterica; Salmonella enterica subsp. enterica serovar Typhimurium
MNNEETFYQAMRRKGVTRRSFLKFCSLAATSLGLGAGMTPKIAWALENKPRIPVVWIHGL WP_001058312.1
hydrogenase [Salmonella enterica] Length: 372\nNumber of predicted TMHs: 2\nExp number of AAs in TMHs: 36.51738\nExp number, first 60 AAs: 19.72849\nTotal prob of N-in: 0.96170\nPOSSIBLE N-term signal sequence\ninside 1 19\nTMhelix 20 39\noutside 40 325\nTMhelix 326 348\ninside 349 372

24851 GCF_001155485.1_8490_2_73 Salmonella enterica subsp. enterica serovar Typhimurium str. DT104
Proteobacteria; Gammaproteobacteria; Enterobacterales; Enterobacteriaceae; Salmonella; Salmonella enterica; Salmonella enterica subsp. enterica; Salmonella enterica subsp. enterica serovar Typhimurium
MTGDNTLITSHGINRRDFMKLCAALAATMGLSSKAAAEMAESVSNPQRPPVIWIGAQECT WP_000145429.1
hydrogenase 2 small subunit [Salmonella enterica] Length: 372\nNumber of predicted TMHs: 1\nExp number of AAs in TMHs: 24.51309\nExp number, first 60 AAs: 0.2682\nTotal prob of N-in: 0.14402\noutside 1 331\nTMhelix 332 354\ninside 355 372

24852 GCF_001154585.1_8490_2_47 Salmonella enterica subsp. enterica serovar Typhimurium str. DT104
Proteobacteria; Gammaproteobacteria; Enterobacterales; Enterobacteriaceae; Salmonella; Salmonella enterica; Salmonella enterica subsp. enterica; Salmonella enterica subsp. enterica serovar Typhimurium
MTGDNTLITSHGINRRDFMKLCAALAATMGLSSKAAAEMAESVSNPQRPPVIWIGAQECT WP_000145429.1
hydrogenase 2 small subunit [Salmonella enterica] Length: 372\nNumber of predicted TMHs: 1\nExp number of AAs in TMHs: 24.51309\nExp number, first 60 AAs: 0.2682\nTotal prob of N-in: 0.14402\noutside 1 331\nTMhelix 332 354\ninside 355 372

24853 GCF_001153005.1_8490_2_84 Salmonella enterica subsp. enterica serovar Typhimurium str. DT104
Proteobacteria; Gammaproteobacteria; Enterobacterales; Enterobacteriaceae; Salmonella; Salmonella enterica; Salmonella enterica subsp. enterica; Salmonella enterica subsp. enterica serovar Typhimurium
MNNEETFYQAMRRKGVTRRSFLKFCSLAATSLGLGAGMTPKIAWALENKPRIPVVWIHGL WP_001058312.1
hydrogenase [Salmonella enterica] Length: 372\nNumber of predicted TMHs: 2\nExp number of AAs in TMHs: 36.51738\nExp number, first 60 AAs: 19.72849\nTotal prob of N-in: 0.96170\nPOSSIBLE N-term signal sequence\ninside 1 19\nTMhelix 20 39\noutside 40 325\nTMhelix 326 348\ninside 349 372

24854 GCF_001153005.1_8490_2_84 Salmonella enterica subsp. enterica serovar Typhimurium str. DT104
Proteobacteria; Gammaproteobacteria; Enterobacterales; Enterobacteriaceae; Salmonella; Salmonella enterica; Salmonella enterica subsp. enterica; Salmonella enterica subsp. enterica serovar Typhimurium
MTGDNTLITSHGINRRDFMKLCAALAATMGLSSKAAAEMAESVSNPQRPPVIWIGAQECT WP_000145429.1
hydrogenase 2 small subunit [Salmonella enterica] Length: 372\nNumber of predicted TMHs: 1\nExp number of AAs in TMHs: 24.51309\nExp number, first 60 AAs: 0.2682\nTotal prob of N-in: 0.14402\noutside 1 331\nTMhelix 332 354\ninside 355 372

24855 GCF_001146465.1_8490_2_74 Salmonella enterica subsp. enterica serovar Typhimurium str. DT104
Proteobacteria; Gammaproteobacteria; Enterobacterales; Enterobacteriaceae; Salmonella; Salmonella enterica; Salmonella enterica subsp. enterica; Salmonella enterica subsp. enterica serovar Typhimurium
MNNEETFYQAMRRKGVTRRSFLKFCSLAATSLGLGAGMTPKIAWALENKPRIPVVWIHGL WP_001058312.1
hydrogenase [Salmonella enterica] Length: 372\nNumber of predicted TMHs: 2\nExp number of AAs in TMHs: 36.51738\nExp number, first 60 AAs: 19.72849\nTotal prob of N-in: 0.96170\nPOSSIBLE N-term signal sequence\ninside 1 19\nTMhelix 20 39\noutside 40 325\nTMhelix 326 348\ninside 349 372

24856 GCF_001146465.1_8490_2_74 Salmonella enterica subsp. enterica serovar Typhimurium str. DT104 Proteobacteria; Gammaproteobacteria; Enterobacterales; Enterobacteriaceae; Salmonella; Salmonella enterica; Salmonella enterica subsp. enterica; Salmonella enterica subsp. enterica serovar Typhimurium MTGDNTLITSHGINRRDFMKLCAALAATMGLSSKAAAEMAESVSNPQRPPVIWIGAQECT WP_000145429.1 hydrogenase 2 small subunit [Salmonella enterica] Length: 372\nNumber of predicted TMHs: 1\nExp number of AAs in TMHs: 24.51309\nExp number, first 60 AAs: 0.2682\nTotal prob of N-in: 0.14402\noutside 1 331\nTMhelix 332 354\ninside 355 372

24857 GCF_001139325.1_8490_2_90 Salmonella enterica subsp. enterica serovar Typhimurium str. DT104 Proteobacteria; Gammaproteobacteria; Enterobacterales; Enterobacteriaceae; Salmonella; Salmonella enterica; Salmonella enterica subsp. enterica; Salmonella enterica subsp. enterica serovar Typhimurium MNNEETFYQAMRRKGVTRRSFLKFCSLAATSLGLGAGMTPKIAWALENKPRIPVVWIHGL WP_001058312.1 hydrogenase [Salmonella enterica] Length: 372\nNumber of predicted TMHs: 2\nExp number of AAs in TMHs: 36.51738\nExp number, first 60 AAs: 19.72849\nTotal prob of N-in: 0.96170\nPOSSIBLE N-term signal sequence\ninside 1 19\nTMhelix 20 39\noutside 40 325\nTMhelix 326 348\ninside 349 372

24858 GCF_001139325.1_8490_2_90 Salmonella enterica subsp. enterica serovar Typhimurium str. DT104 Proteobacteria; Gammaproteobacteria; Enterobacterales; Enterobacteriaceae; Salmonella; Salmonella enterica; Salmonella enterica subsp. enterica; Salmonella enterica subsp. enterica serovar Typhimurium MTGDNTLITSHGINRRDFMKLCAALAATMGLSSKAAAEMAESVSNPQRPPVIWIGAQECT WP_000145429.1 hydrogenase 2 small subunit [Salmonella enterica] Length: 372\nNumber of predicted TMHs: 1\nExp number of AAs in TMHs: 24.51309\nExp number, first 60 AAs: 0.2682\nTotal prob of N-in: 0.14402\noutside 1 331\nTMhelix 332 354\ninside 355 372

24859 GCF_001350095.1_8490_2_79 Salmonella enterica subsp. enterica serovar Typhimurium str. DT104 Proteobacteria; Gammaproteobacteria; Enterobacterales; Enterobacteriaceae; Salmonella; Salmonella enterica; Salmonella enterica subsp. enterica; Salmonella enterica subsp. enterica serovar Typhimurium MTGDNTLITSHGINRRDFMKLCAALAATMGLSSKAAAEMAESVSNPQRPPVIWIGAQECT WP_000145429.1 hydrogenase 2 small subunit [Salmonella enterica] Length: 372\nNumber of predicted TMHs: 1\nExp number of AAs in TMHs: 24.51309\nExp number, first 60 AAs: 0.2682\nTotal prob of N-in: 0.14402\noutside 1 331\nTMhelix 332 354\ninside 355 372

24860 GCF_001135785.1_8490_2_63 Salmonella enterica subsp. enterica serovar Typhimurium str. DT104 Proteobacteria; Gammaproteobacteria; Enterobacterales; Enterobacteriaceae; Salmonella; Salmonella enterica; Salmonella enterica subsp. enterica; Salmonella enterica subsp. enterica serovar Typhimurium MNNEETFYQAMRRKGVTRRSFLKFCSLAATSLGLGAGMTPKIAWALENKPRIPVVWIHGL WP_001058312.1 hydrogenase [Salmonella enterica] Length: 372\nNumber of predicted TMHs: 2\nExp number of AAs in TMHs: 36.51738\nExp number, first 60 AAs: 19.72849\nTotal prob of N-in: 0.96170\nPOSSIBLE N-term signal sequence\ninside 1 19\nTMhelix 20 39\noutside 40 325\nTMhelix 326 348\ninside 349 372

24861 GCF_001135785.1_8490_2_63 Salmonella enterica subsp. enterica serovar Typhimurium str. DT104 Proteobacteria; Gammaproteobacteria; Enterobacterales; Enterobacteriaceae; Salmonella; Salmonella enterica; Salmonella enterica subsp. enterica; Salmonella enterica subsp. enterica serovar Typhimurium MTGDNTLITSHGINRRDFMKLCAALAATMGLSSKAAAEMAESVSNPQRPPVIWIGAQECT WP_000145429.1 hydrogenase 2 small subunit [Salmonella enterica] Length: 372\nNumber of predicted TMHs: 1\nExp number of AAs in TMHs: 24.51309\nExp number, first 60 AAs: 0.2682\nTotal prob of N-in: 0.14402\noutside 1 331\nTMhelix 332 354\ninside 355 372

24862 GCF_001350135.1_8490_2_85 Salmonella enterica subsp. enterica serovar Typhimurium str. DT104 Proteobacteria; Gammaproteobacteria; Enterobacterales; Enterobacteriaceae; Salmonella; Salmonella enterica; Salmonella enterica subsp. enterica; Salmonella enterica subsp. enterica serovar Typhimurium MTGDNTLITSHGINRRDFMKLCAALAATMGLSSKAAAEMAESVSNPQRPPVIWIGAQECT WP_000145429.1 hydrogenase 2 small subunit [Salmonella enterica] Length: 372\nNumber of predicted TMHs: 1\nExp number of AAs in TMHs: 24.51309\nExp number, first 60 AAs: 0.2682\nTotal prob of N-in: 0.14402\noutside 1 331\nTMhelix 332 354\ninside 355 372

24863 GCF_001131525.1_8490_2_83 Salmonella enterica subsp. enterica serovar Typhimurium str. DT104 Proteobacteria; Gammaproteobacteria; Enterobacterales; Enterobacteriaceae; Salmonella; Salmonella enterica; Salmonella enterica subsp. enterica; Salmonella enterica subsp. enterica serovar Typhimurium MNNEETFYQAMRRKGVTRRSFLKFCSLAATSLGLGAGMTPKIAWALENKPRIPVVWIHGL WP_001058312.1 hydrogenase [Salmonella enterica] Length: 372\nNumber of predicted TMHs: 2\nExp number of AAs in

TMHs: 36.51738\nExp number, first 60 AAs: 19.72849\nTotal prob of N-in: 0.96170\nPOSSIBLE N-term signal sequence\ninside 1 19\nTMhelix 20 39\noutside 40 325\nTMhelix 326 348\ninside 349 372

24864 GCF_001130925.1_8490_2_77 Salmonella enterica subsp. enterica serovar Typhimurium str. DT104 Proteobacteria; Gammaproteobacteria; Enterobacterales; Enterobacteriaceae; Salmonella; Salmonella enterica; Salmonella enterica subsp. enterica; Salmonella enterica subsp. enterica serovar Typhimurium MNNEETFYQAMRRKGVTRRSFLKFCSLAATSLGLGAGMTPKIAWALENKPRIPVVWIHGL WP_001058312.1 hydrogenase [Salmonella enterica] Length: 372\nNumber of predicted TMHs: 2\nExp number of AAs in TMHs: 36.51738\nExp number, first 60 AAs: 19.72849\nTotal prob of N-in: 0.96170\nPOSSIBLE N-term signal sequence\ninside 1 19\nTMhelix 20 39\noutside 40 325\nTMhelix 326 348\ninside 349 372

24865 GCF_001131525.1_8490_2_83 Salmonella enterica subsp. enterica serovar Typhimurium str. DT104 Proteobacteria; Gammaproteobacteria; Enterobacterales; Enterobacteriaceae; Salmonella; Salmonella enterica; Salmonella enterica subsp. enterica; Salmonella enterica subsp. enterica serovar Typhimurium MTGDNTLITSHGINRRDFMKLCAALAATMGLSSKAAAEMAESVSNPQRPPVIWIGAQECT WP_000145429.1 hydrogenase 2 small subunit [Salmonella enterica] Length: 372\nNumber of predicted TMHs: 1\nExp number of AAs in TMHs: 24.51309\nExp number, first 60 AAs: 0.2682\nTotal prob of N-in: 0.14402\noutside 1 331\nTMhelix 332 354\ninside 355 372

24866 GCF_001130925.1_8490_2_77 Salmonella enterica subsp. enterica serovar Typhimurium str. DT104 Proteobacteria; Gammaproteobacteria; Enterobacterales; Enterobacteriaceae; Salmonella; Salmonella enterica; Salmonella enterica subsp. enterica; Salmonella enterica subsp. enterica serovar Typhimurium MTGDNTLITSHGINRRDFMKLCAALAATMGLSSKAAAEMAESVSNPQRPPVIWIGAQECT WP_000145429.1 hydrogenase 2 small subunit [Salmonella enterica] Length: 372\nNumber of predicted TMHs: 1\nExp number of AAs in TMHs: 24.51309\nExp number, first 60 AAs: 0.2682\nTotal prob of N-in: 0.14402\noutside 1 331\nTMhelix 332 354\ninside 355 372

24867 GCF_001129385.1_8490_2_56 Salmonella enterica subsp. enterica serovar Typhimurium str. DT104 Proteobacteria; Gammaproteobacteria; Enterobacterales; Enterobacteriaceae; Salmonella; Salmonella enterica; Salmonella enterica subsp. enterica; Salmonella enterica subsp. enterica serovar Typhimurium MNNEETFYQAMRRKGVTRRSFLKFCSLAATSLGLGAGMTPKIAWALENKPRIPVVWIHGL WP_001058312.1 hydrogenase [Salmonella enterica] Length: 372\nNumber of predicted TMHs: 2\nExp number of AAs in TMHs: 36.51738\nExp number, first 60 AAs: 19.72849\nTotal prob of N-in: 0.96170\nPOSSIBLE N-term signal sequence\ninside 1 19\nTMhelix 20 39\noutside 40 325\nTMhelix 326 348\ninside 349 372

24868 GCF_001130025.1_8490_2_93 Salmonella enterica subsp. enterica serovar Typhimurium str. DT104 Proteobacteria; Gammaproteobacteria; Enterobacterales; Enterobacteriaceae; Salmonella; Salmonella enterica; Salmonella enterica subsp. enterica; Salmonella enterica subsp. enterica serovar Typhimurium MNNEETFYQAMRRKGVTRRSFLKFCSLAATSLGLGAGMTPKIAWALENKPRIPVVWIHGL WP_001058312.1 hydrogenase [Salmonella enterica] Length: 372\nNumber of predicted TMHs: 2\nExp number of AAs in TMHs: 36.51738\nExp number, first 60 AAs: 19.72849\nTotal prob of N-in: 0.96170\nPOSSIBLE N-term signal sequence\ninside 1 19\nTMhelix 20 39\noutside 40 325\nTMhelix 326 348\ninside 349 372

24869 GCF_001130025.1_8490_2_93 Salmonella enterica subsp. enterica serovar Typhimurium str. DT104 Proteobacteria; Gammaproteobacteria; Enterobacterales; Enterobacteriaceae; Salmonella; Salmonella enterica; Salmonella enterica subsp. enterica; Salmonella enterica subsp. enterica serovar Typhimurium MTGDNTLITSHGINRRDFMKLCAALAATMGLSSKAAAEMAESVSNPQRPPVIWIGAQECT WP_000145429.1 hydrogenase 2 small subunit [Salmonella enterica] Length: 372\nNumber of predicted TMHs: 1\nExp number of AAs in TMHs: 24.51309\nExp number, first 60 AAs: 0.2682\nTotal prob of N-in: 0.14402\noutside 1 331\nTMhelix 332 354\ninside 355 372

24870 GCF_001129385.1_8490_2_56 Salmonella enterica subsp. enterica serovar Typhimurium str. DT104 Proteobacteria; Gammaproteobacteria; Enterobacterales; Enterobacteriaceae; Salmonella; Salmonella enterica; Salmonella enterica subsp. enterica; Salmonella enterica subsp. enterica serovar Typhimurium MTGDNTLITSHGINRRDFMKLCAALAATMGLSSKAAAEMAESVSNPQRPPVIWIGAQECT WP_000145429.1 hydrogenase 2 small subunit [Salmonella enterica] Length: 372\nNumber of predicted TMHs: 1\nExp number of AAs in TMHs: 24.51309\nExp number, first 60 AAs: 0.2682\nTotal prob of N-in: 0.14402\noutside 1 331\nTMhelix 332 354\ninside 355 372

24871 GCF_001127885.1_8490_2_71 Salmonella enterica subsp. enterica serovar Typhimurium str. DT104 Proteobacteria; Gammaproteobacteria; Enterobacterales; Enterobacteriaceae; Salmonella; Salmonella enterica;

Salmonella enterica subsp. enterica; Salmonella enterica subsp. enterica serovar Typhimurium
MTGDNTLITSHGINRRDFMKLCAALAATMGLSSKAAAEMAESVSNPQRPPVIWIGAQECT WP_000145429.1
hydrogenase 2 small subunit [Salmonella enterica] Length: 372\nNumber of predicted TMHs: 1\nExp number of AAs in TMHs: 24.51309\nExp number, first 60 AAs: 0.2682\nTotal prob of N-in: 0.14402\noutside 1 331\nTMhelix 332 354\ninside 355 372

24872 GCF_001125725.1_8490_2_57 Salmonella enterica subsp. enterica serovar Typhimurium str. DT104
Proteobacteria; Gammaproteobacteria; Enterobacterales; Enterobacteriaceae; Salmonella; Salmonella enterica; Salmonella enterica subsp. enterica; Salmonella enterica subsp. enterica serovar Typhimurium
MNNEETFYQAMRRKGVTRRSFLKFCSLAATSLGLGAGMTPKIAWALENKPRIPVVWIHGL WP_001058312.1
hydrogenase [Salmonella enterica] Length: 372\nNumber of predicted TMHs: 2\nExp number of AAs in TMHs: 36.51738\nExp number, first 60 AAs: 19.72849\nTotal prob of N-in: 0.96170\nPOSSIBLE N-term signal sequence\ninside 1 19\nTMhelix 20 39\noutside 40 325\nTMhelix 326 348\ninside 349 372

24873 GCF_001125725.1_8490_2_57 Salmonella enterica subsp. enterica serovar Typhimurium str. DT104
Proteobacteria; Gammaproteobacteria; Enterobacterales; Enterobacteriaceae; Salmonella; Salmonella enterica; Salmonella enterica subsp. enterica; Salmonella enterica subsp. enterica serovar Typhimurium
MTGDNTLITSHGINRRDFMKLCAALAATMGLSSKAAAEMAESVSNPQRPPVIWIGAQECT WP_000145429.1
hydrogenase 2 small subunit [Salmonella enterica] Length: 372\nNumber of predicted TMHs: 1\nExp number of AAs in TMHs: 24.51309\nExp number, first 60 AAs: 0.2682\nTotal prob of N-in: 0.14402\noutside 1 331\nTMhelix 332 354\ninside 355 372

24874 GCF_001120065.1_8490_2_59 Salmonella enterica subsp. enterica serovar Typhimurium str. DT104
Proteobacteria; Gammaproteobacteria; Enterobacterales; Enterobacteriaceae; Salmonella; Salmonella enterica; Salmonella enterica subsp. enterica; Salmonella enterica subsp. enterica serovar Typhimurium
MNNEETFYQAMRRKGVTRRSFLKFCSLAATSLGLGAGMTPKIAWALENKPRIPVVWIHGL WP_001058312.1
hydrogenase [Salmonella enterica] Length: 372\nNumber of predicted TMHs: 2\nExp number of AAs in TMHs: 36.51738\nExp number, first 60 AAs: 19.72849\nTotal prob of N-in: 0.96170\nPOSSIBLE N-term signal sequence\ninside 1 19\nTMhelix 20 39\noutside 40 325\nTMhelix 326 348\ninside 349 372

24875 GCF_001119985.1_8490_2_89 Salmonella enterica subsp. enterica serovar Typhimurium str. DT104
Proteobacteria; Gammaproteobacteria; Enterobacterales; Enterobacteriaceae; Salmonella; Salmonella enterica; Salmonella enterica subsp. enterica; Salmonella enterica subsp. enterica serovar Typhimurium
MNNEETFYQAMRRKGVTRRSFLKFCSLAATSLGLGAGMTPKIAWALENKPRIPVVWIHGL WP_001058312.1
hydrogenase [Salmonella enterica] Length: 372\nNumber of predicted TMHs: 2\nExp number of AAs in TMHs: 36.51738\nExp number, first 60 AAs: 19.72849\nTotal prob of N-in: 0.96170\nPOSSIBLE N-term signal sequence\ninside 1 19\nTMhelix 20 39\noutside 40 325\nTMhelix 326 348\ninside 349 372

24876 GCF_001350175.1_8490_2_80 Salmonella enterica subsp. enterica serovar Typhimurium str. DT104
Proteobacteria; Gammaproteobacteria; Enterobacterales; Enterobacteriaceae; Salmonella; Salmonella enterica; Salmonella enterica subsp. enterica; Salmonella enterica subsp. enterica serovar Typhimurium
MTGDNTLITSHGINRRDFMKLCAALAATMGLSSKAAAEMAESVSNPQRPPVIWIGAQECT WP_000145429.1
hydrogenase 2 small subunit [Salmonella enterica] Length: 372\nNumber of predicted TMHs: 1\nExp number of AAs in TMHs: 24.51309\nExp number, first 60 AAs: 0.2682\nTotal prob of N-in: 0.14402\noutside 1 331\nTMhelix 332 354\ninside 355 372

24877 GCF_001120665.1_8490_2_91 Salmonella enterica subsp. enterica serovar Typhimurium str. DT104
Proteobacteria; Gammaproteobacteria; Enterobacterales; Enterobacteriaceae; Salmonella; Salmonella enterica; Salmonella enterica subsp. enterica; Salmonella enterica subsp. enterica serovar Typhimurium
MNNEETFYQAMRRKGVTRRSFLKFCSLAATSLGLGAGMTPKIAWALENKPRIPVVWIHGL WP_001058312.1
hydrogenase [Salmonella enterica] Length: 372\nNumber of predicted TMHs: 2\nExp number of AAs in TMHs: 36.51738\nExp number, first 60 AAs: 19.72849\nTotal prob of N-in: 0.96170\nPOSSIBLE N-term signal sequence\ninside 1 19\nTMhelix 20 39\noutside 40 325\nTMhelix 326 348\ninside 349 372

24878 GCF_001120665.1_8490_2_91 Salmonella enterica subsp. enterica serovar Typhimurium str. DT104
Proteobacteria; Gammaproteobacteria; Enterobacterales; Enterobacteriaceae; Salmonella; Salmonella enterica; Salmonella enterica subsp. enterica; Salmonella enterica subsp. enterica serovar Typhimurium
MTGDNTLITSHGINRRDFMKLCAALAATMGLSSKAAAEMAESVSNPQRPPVIWIGAQECT WP_000145429.1
hydrogenase 2 small subunit [Salmonella enterica] Length: 372\nNumber of predicted TMHs: 1\nExp number of AAs in TMHs: 24.51309\nExp number, first 60 AAs: 0.2682\nTotal prob of N-in: 0.14402\noutside 1 331\nTMhelix 332 354\ninside 355 372

24879 GCF_001120065.1_8490_2_59 Salmonella enterica subsp. enterica serovar Typhimurium str. DT104 Proteobacteria; Gammaproteobacteria; Enterobacterales; Enterobacteriaceae; Salmonella; Salmonella enterica; Salmonella enterica subsp. enterica; Salmonella enterica subsp. enterica serovar Typhimurium MTGDNTLITSHGINRRDFMKLCAALAATMGLSSKAAAEMAESVSNPQRPPVIWIGAQECT WP_000145429.1 hydrogenase 2 small subunit [Salmonella enterica] Length: 372\nNumber of predicted TMHs: 1\nExp number of AAs in TMHs: 24.51309\nExp number, first 60 AAs: 0.2682\nTotal prob of N-in: 0.14402\noutside 1 331\nTMhelix 332 354\ninside 355 372

24880 GCF_001119985.1_8490_2_89 Salmonella enterica subsp. enterica serovar Typhimurium str. DT104 Proteobacteria; Gammaproteobacteria; Enterobacterales; Enterobacteriaceae; Salmonella; Salmonella enterica; Salmonella enterica subsp. enterica; Salmonella enterica subsp. enterica serovar Typhimurium MTGDNTLITSHGINRRDFMKLCAALAATMGLSSKAAAEMAESVSNPQRPPVIWIGAQECT WP_000145429.1 hydrogenase 2 small subunit [Salmonella enterica] Length: 372\nNumber of predicted TMHs: 1\nExp number of AAs in TMHs: 24.51309\nExp number, first 60 AAs: 0.2682\nTotal prob of N-in: 0.14402\noutside 1 331\nTMhelix 332 354\ninside 355 372

24881 GCF_001116925.1_8490_2_62 Salmonella enterica subsp. enterica serovar Typhimurium str. DT104 Proteobacteria; Gammaproteobacteria; Enterobacterales; Enterobacteriaceae; Salmonella; Salmonella enterica; Salmonella enterica subsp. enterica; Salmonella enterica subsp. enterica serovar Typhimurium MNNEETFYQAMRRKGVTRRSFLKFCSLAATSLGLGAGMTPKIAWALENKPRIPVVWIHGL WP_050196264.1 hydrogenase [Salmonella enterica] Length: 372\nNumber of predicted TMHs: 2\nExp number of AAs in TMHs: 36.76337\nExp number, first 60 AAs: 19.73298\nTotal prob of N-in: 0.96170\nPOSSIBLE N-term signal sequence\ninside 1 19\nTMhelix 20 39\noutside 40 325\nTMhelix 326 348\ninside 349 372

24882 GCF_001116925.1_8490_2_62 Salmonella enterica subsp. enterica serovar Typhimurium str. DT104 Proteobacteria; Gammaproteobacteria; Enterobacterales; Enterobacteriaceae; Salmonella; Salmonella enterica; Salmonella enterica subsp. enterica; Salmonella enterica subsp. enterica serovar Typhimurium MTGDNTLITSHGINRRDFMKLCAALAATMGLSSKAAAEMAESVSNPQRPPVIWIGAQECT WP_000145429.1 hydrogenase 2 small subunit [Salmonella enterica] Length: 372\nNumber of predicted TMHs: 1\nExp number of AAs in TMHs: 24.51309\nExp number, first 60 AAs: 0.2682\nTotal prob of N-in: 0.14402\noutside 1 331\nTMhelix 332 354\ninside 355 372

24883 GCF_001111905.1_8490_2_68 Salmonella enterica subsp. enterica serovar Typhimurium str. DT104 Proteobacteria; Gammaproteobacteria; Enterobacterales; Enterobacteriaceae; Salmonella; Salmonella enterica; Salmonella enterica subsp. enterica; Salmonella enterica subsp. enterica serovar Typhimurium MNNEETFYQAMRRKGVTRRSFLKFCSLAATSLGLGAGMTPKIAWALENKPRIPVVWIHGL WP_001058312.1 hydrogenase [Salmonella enterica] Length: 372\nNumber of predicted TMHs: 2\nExp number of AAs in TMHs: 36.51738\nExp number, first 60 AAs: 19.72849\nTotal prob of N-in: 0.96170\nPOSSIBLE N-term signal sequence\ninside 1 19\nTMhelix 20 39\noutside 40 325\nTMhelix 326 348\ninside 349 372

24884 GCF_001111905.1_8490_2_68 Salmonella enterica subsp. enterica serovar Typhimurium str. DT104 Proteobacteria; Gammaproteobacteria; Enterobacterales; Enterobacteriaceae; Salmonella; Salmonella enterica; Salmonella enterica subsp. enterica; Salmonella enterica subsp. enterica serovar Typhimurium MTGDNTLITSHGINRRDFMKLCAALAATMGLSSKAAAEMAESVSNPQRPPVIWIGAQECT WP_000145429.1 hydrogenase 2 small subunit [Salmonella enterica] Length: 372\nNumber of predicted TMHs: 1\nExp number of AAs in TMHs: 24.51309\nExp number, first 60 AAs: 0.2682\nTotal prob of N-in: 0.14402\noutside 1 331\nTMhelix 332 354\ninside 355 372

24885 GCF_001110745.1_8490_2_69 Salmonella enterica subsp. enterica serovar Typhimurium str. DT104 Proteobacteria; Gammaproteobacteria; Enterobacterales; Enterobacteriaceae; Salmonella; Salmonella enterica; Salmonella enterica subsp. enterica; Salmonella enterica subsp. enterica serovar Typhimurium MNNEETFYQAMRRKGVTRRSFLKFCSLAATSLGLGAGMTPKIAWALENKPRIPVVWIHGL WP_001058312.1 hydrogenase [Salmonella enterica] Length: 372\nNumber of predicted TMHs: 2\nExp number of AAs in TMHs: 36.51738\nExp number, first 60 AAs: 19.72849\nTotal prob of N-in: 0.96170\nPOSSIBLE N-term signal sequence\ninside 1 19\nTMhelix 20 39\noutside 40 325\nTMhelix 326 348\ninside 349 372

24886 GCF_001110745.1_8490_2_69 Salmonella enterica subsp. enterica serovar Typhimurium str. DT104 Proteobacteria; Gammaproteobacteria; Enterobacterales; Enterobacteriaceae; Salmonella; Salmonella enterica; Salmonella enterica subsp. enterica; Salmonella enterica subsp. enterica serovar Typhimurium MTGDNTLITSHGINRRDFMKLCAALAATMGLSSKAAAEMAESVSNPQRPPVIWIGAQECT WP_000145429.1 hydrogenase 2 small subunit [Salmonella enterica] Length: 372\nNumber of predicted TMHs: 1\nExp

number of AAs in TMHs: 24.51309\nExp number, first 60 AAs: 0.2682\nTotal prob of N-in: 0.14402\noutside 1 331\nTMhelix 332 354\ninside 355 372

24887 GCF_001105145.1_8490_2_50 Salmonella enterica subsp. enterica serovar Typhimurium str. DT104 Proteobacteria; Gammaproteobacteria; Enterobacterales; Enterobacteriaceae; Salmonella; Salmonella enterica; Salmonella enterica subsp. enterica; Salmonella enterica subsp. enterica serovar Typhimurium MNNEETFYQAMRRKGVTRRSFLKFCSLAATSLGLGAGMTPKIAWALENKPRIPVVWIHGL WP_001058312.1 hydrogenase [Salmonella enterica] Length: 372\nNumber of predicted TMHs: 2\nExp number of AAs in TMHs: 36.51738\nExp number, first 60 AAs: 19.72849\nTotal prob of N-in: 0.96170\nPOSSIBLE N-term signal sequence\ninside 1 19\nTMhelix 20 39\noutside 40 325\nTMhelix 326 348\ninside 349 372

24888 GCF_001105145.1_8490_2_50 Salmonella enterica subsp. enterica serovar Typhimurium str. DT104 Proteobacteria; Gammaproteobacteria; Enterobacterales; Enterobacteriaceae; Salmonella; Salmonella enterica; Salmonella enterica subsp. enterica; Salmonella enterica subsp. enterica serovar Typhimurium MTGDNTLITSHGINRRDFMKLCAALAATMGLSSKAAAEMAESVSNPQRPPVIWIGAQECT WP_000145429.1 hydrogenase 2 small subunit [Salmonella enterica] Length: 372\nNumber of predicted TMHs: 1\nExp number of AAs in TMHs: 24.51309\nExp number, first 60 AAs: 0.2682\nTotal prob of N-in: 0.14402\noutside 1 331\nTMhelix 332 354\ninside 355 372

24889 GCF_001102665.1_8490_2_49 Salmonella enterica subsp. enterica serovar Typhimurium str. DT104 Proteobacteria; Gammaproteobacteria; Enterobacterales; Enterobacteriaceae; Salmonella; Salmonella enterica; Salmonella enterica subsp. enterica; Salmonella enterica subsp. enterica serovar Typhimurium MNNEETFYQAMRRKGVTRRSFLKFCSLAATSLGLGAGMTPKIAWALENKPRIPVVWIHGL WP_001058312.1 hydrogenase [Salmonella enterica] Length: 372\nNumber of predicted TMHs: 2\nExp number of AAs in TMHs: 36.51738\nExp number, first 60 AAs: 19.72849\nTotal prob of N-in: 0.96170\nPOSSIBLE N-term signal sequence\ninside 1 19\nTMhelix 20 39\noutside 40 325\nTMhelix 326 348\ninside 349 372

24890 GCF_001350215.1_8490_2_66 Salmonella enterica subsp. enterica serovar Typhimurium str. DT104 Proteobacteria; Gammaproteobacteria; Enterobacterales; Enterobacteriaceae; Salmonella; Salmonella enterica; Salmonella enterica subsp. enterica; Salmonella enterica subsp. enterica serovar Typhimurium MTGDNTLITSHGINRRDFMKLCAALAATMGLSSKAAAEMAESVSNPQRPPVIWIGAQECT WP_000145429.1 hydrogenase 2 small subunit [Salmonella enterica] Length: 372\nNumber of predicted TMHs: 1\nExp number of AAs in TMHs: 24.51309\nExp number, first 60 AAs: 0.2682\nTotal prob of N-in: 0.14402\noutside 1 331\nTMhelix 332 354\ninside 355 372

24891 GCF_001102665.1_8490_2_49 Salmonella enterica subsp. enterica serovar Typhimurium str. DT104 Proteobacteria; Gammaproteobacteria; Enterobacterales; Enterobacteriaceae; Salmonella; Salmonella enterica; Salmonella enterica subsp. enterica; Salmonella enterica subsp. enterica serovar Typhimurium MTGDNTLITSHGINRRDFMKLCAALAATMGLSSKAAAEMAESVSNPQRPPVIWIGAQECT WP_000145429.1 hydrogenase 2 small subunit [Salmonella enterica] Length: 372\nNumber of predicted TMHs: 1\nExp number of AAs in TMHs: 24.51309\nExp number, first 60 AAs: 0.2682\nTotal prob of N-in: 0.14402\noutside 1 331\nTMhelix 332 354\ninside 355 372

24892 GCF_001097045.1_8490_2_94 Salmonella enterica subsp. enterica serovar Typhimurium str. DT104 Proteobacteria; Gammaproteobacteria; Enterobacterales; Enterobacteriaceae; Salmonella; Salmonella enterica; Salmonella enterica subsp. enterica; Salmonella enterica subsp. enterica serovar Typhimurium MNNEETFYQAMRRKGVTRRSFLKFCSLAATSLGLGAGMTPKIAWALENKPRIPVVWIHGL WP_001058312.1 hydrogenase [Salmonella enterica] Length: 372\nNumber of predicted TMHs: 2\nExp number of AAs in TMHs: 36.51738\nExp number, first 60 AAs: 19.72849\nTotal prob of N-in: 0.96170\nPOSSIBLE N-term signal sequence\ninside 1 19\nTMhelix 20 39\noutside 40 325\nTMhelix 326 348\ninside 349 372

24893 GCF_001096565.1_8490_2_48 Salmonella enterica subsp. enterica serovar Typhimurium str. DT104 Proteobacteria; Gammaproteobacteria; Enterobacterales; Enterobacteriaceae; Salmonella; Salmonella enterica; Salmonella enterica subsp. enterica; Salmonella enterica subsp. enterica serovar Typhimurium MNNEETFYQAMRRKGVTRRSFLKFCSLAATSLGLGAGMTPKIAWALENKPRIPVVWIHGL WP_001058312.1 hydrogenase [Salmonella enterica] Length: 372\nNumber of predicted TMHs: 2\nExp number of AAs in TMHs: 36.51738\nExp number, first 60 AAs: 19.72849\nTotal prob of N-in: 0.96170\nPOSSIBLE N-term signal sequence\ninside 1 19\nTMhelix 20 39\noutside 40 325\nTMhelix 326 348\ninside 349 372

24894 GCF_001097045.1_8490_2_94 Salmonella enterica subsp. enterica serovar Typhimurium str. DT104 Proteobacteria; Gammaproteobacteria; Enterobacterales; Enterobacteriaceae; Salmonella; Salmonella enterica;

Salmonella enterica subsp. enterica; Salmonella enterica subsp. enterica serovar Typhimurium
MTGDNTLITSHGINRRDFMKLCAALAATMGLSSKAAAEMAESVSNPQRPPVIWIGAQECT WP_000145429.1
hydrogenase 2 small subunit [Salmonella enterica] Length: 372\nNumber of predicted TMHs: 1\nExp number of AAs in TMHs: 24.51309\nExp number, first 60 AAs: 0.2682\nTotal prob of N-in: 0.14402\noutside 1 331\nTMhelix 332 354\ninside 355 372

24895 GCF_001096565.1_8490_2_48 Salmonella enterica subsp. enterica serovar Typhimurium str. DT104
Proteobacteria; Gammaproteobacteria; Enterobacterales; Enterobacteriaceae; Salmonella; Salmonella enterica; Salmonella enterica subsp. enterica; Salmonella enterica subsp. enterica serovar Typhimurium
MTGDNTLITSHGINRRDFMKLCAALAATMGLSSKAAAEMAESVSNPQRPPVIWIGAQECT WP_000145429.1
hydrogenase 2 small subunit [Salmonella enterica] Length: 372\nNumber of predicted TMHs: 1\nExp number of AAs in TMHs: 24.51309\nExp number, first 60 AAs: 0.2682\nTotal prob of N-in: 0.14402\noutside 1 331\nTMhelix 332 354\ninside 355 372

24896 GCF_001093545.1_8490_2_60 Salmonella enterica subsp. enterica serovar Typhimurium str. DT104
Proteobacteria; Gammaproteobacteria; Enterobacterales; Enterobacteriaceae; Salmonella; Salmonella enterica; Salmonella enterica subsp. enterica; Salmonella enterica subsp. enterica serovar Typhimurium
MNNEETFYQAMRRKGVTRRSFLKFCSLAATSLGLGAGMTPKIAWALENKPRIPVVWIHGL WP_001058312.1
hydrogenase [Salmonella enterica] Length: 372\nNumber of predicted TMHs: 2\nExp number of AAs in TMHs: 36.51738\nExp number, first 60 AAs: 19.72849\nTotal prob of N-in: 0.96170\nPOSSIBLE N-term signal sequence\ninside 1 19\nTMhelix 20 39\noutside 40 325\nTMhelix 326 348\ninside 349 372

24897 GCF_001093545.1_8490_2_60 Salmonella enterica subsp. enterica serovar Typhimurium str. DT104
Proteobacteria; Gammaproteobacteria; Enterobacterales; Enterobacteriaceae; Salmonella; Salmonella enterica; Salmonella enterica subsp. enterica; Salmonella enterica subsp. enterica serovar Typhimurium
MTGDNTLITSHGINRRDFMKLCAALAATMGLSSKAAAEMAESVSNPQRPPVIWIGAQECT WP_000145429.1
hydrogenase 2 small subunit [Salmonella enterica] Length: 372\nNumber of predicted TMHs: 1\nExp number of AAs in TMHs: 24.51309\nExp number, first 60 AAs: 0.2682\nTotal prob of N-in: 0.14402\noutside 1 331\nTMhelix 332 354\ninside 355 372

24898 GCF_001091445.1_8490_2_44 Salmonella enterica subsp. enterica serovar Typhimurium str. DT104
Proteobacteria; Gammaproteobacteria; Enterobacterales; Enterobacteriaceae; Salmonella; Salmonella enterica; Salmonella enterica subsp. enterica; Salmonella enterica subsp. enterica serovar Typhimurium
MNNEETFYQAMRRKGVTRRSFLKFCSLAATSLGLGAGMTPKIAWALENKPRIPVVWIHGL WP_001058312.1
hydrogenase [Salmonella enterica] Length: 372\nNumber of predicted TMHs: 2\nExp number of AAs in TMHs: 36.51738\nExp number, first 60 AAs: 19.72849\nTotal prob of N-in: 0.96170\nPOSSIBLE N-term signal sequence\ninside 1 19\nTMhelix 20 39\noutside 40 325\nTMhelix 326 348\ninside 349 372

24899 GCF_001091445.1_8490_2_44 Salmonella enterica subsp. enterica serovar Typhimurium str. DT104
Proteobacteria; Gammaproteobacteria; Enterobacterales; Enterobacteriaceae; Salmonella; Salmonella enterica; Salmonella enterica subsp. enterica; Salmonella enterica subsp. enterica serovar Typhimurium
MTGDNTLITSHGINRRDFMKLCAALAATMGLSSKAAAEMAESVSNPQRPPVIWIGAQECT WP_000145429.1
hydrogenase 2 small subunit [Salmonella enterica] Length: 372\nNumber of predicted TMHs: 1\nExp number of AAs in TMHs: 24.51309\nExp number, first 60 AAs: 0.2682\nTotal prob of N-in: 0.14402\noutside 1 331\nTMhelix 332 354\ninside 355 372

24900 GCF_001085685.1_8490_2_75 Salmonella enterica subsp. enterica serovar Typhimurium str. DT104
Proteobacteria; Gammaproteobacteria; Enterobacterales; Enterobacteriaceae; Salmonella; Salmonella enterica; Salmonella enterica subsp. enterica; Salmonella enterica subsp. enterica serovar Typhimurium
MNNEETFYQAMRRKGVTRRSFLKFCSLAATSLGLGAGMTPKIAWALENKPRIPVVWIHGL WP_050196264.1
hydrogenase [Salmonella enterica] Length: 372\nNumber of predicted TMHs: 2\nExp number of AAs in TMHs: 36.76337\nExp number, first 60 AAs: 19.73298\nTotal prob of N-in: 0.96170\nPOSSIBLE N-term signal sequence\ninside 1 19\nTMhelix 20 39\noutside 40 325\nTMhelix 326 348\ninside 349 372

24901 GCF_001085685.1_8490_2_75 Salmonella enterica subsp. enterica serovar Typhimurium str. DT104
Proteobacteria; Gammaproteobacteria; Enterobacterales; Enterobacteriaceae; Salmonella; Salmonella enterica; Salmonella enterica subsp. enterica; Salmonella enterica subsp. enterica serovar Typhimurium
MTGDNTLITSHGINRRDFMKLCAALAATMGLSSKAAAEMAESVSNPQRPPVIWIGAQECT WP_000145429.1
hydrogenase 2 small subunit [Salmonella enterica] Length: 372\nNumber of predicted TMHs: 1\nExp number of AAs in TMHs: 24.51309\nExp number, first 60 AAs: 0.2682\nTotal prob of N-in: 0.14402\noutside 1 331\nTMhelix 332 354\ninside 355 372

24902 GCF_001085145.1_8490_2_76 Salmonella enterica subsp. enterica serovar Typhimurium str. DT104 Proteobacteria; Gammaproteobacteria; Enterobacterales; Enterobacteriaceae; Salmonella; Salmonella enterica; Salmonella enterica subsp. enterica; Salmonella enterica subsp. enterica serovar Typhimurium MNNEETFYQAMRRKGVTRRSFLKFCSLAATSLGLGAGMTPKIAWALENKPRIPVVWIHGL WP_001058312.1 hydrogenase [Salmonella enterica] Length: 372\nNumber of predicted TMHs: 2\nExp number of AAs in TMHs: 36.51738\nExp number, first 60 AAs: 19.72849\nTotal prob of N-in: 0.96170\nPOSSIBLE N-term signal sequence\ninside 1 19\nTMhelix 20 39\noutside 40 325\nTMhelix 326 348\ninside 349 372

24903 GCF_001085145.1_8490_2_76 Salmonella enterica subsp. enterica serovar Typhimurium str. DT104 Proteobacteria; Gammaproteobacteria; Enterobacterales; Enterobacteriaceae; Salmonella; Salmonella enterica; Salmonella enterica subsp. enterica; Salmonella enterica subsp. enterica serovar Typhimurium MTGDNTLITSHGINRRDFMKLCAALAATMGLSSKAAAEMAESVSNPQRPPVIWIGAQECT WP_000145429.1 hydrogenase 2 small subunit [Salmonella enterica] Length: 372\nNumber of predicted TMHs: 1\nExp number of AAs in TMHs: 24.51309\nExp number, first 60 AAs: 0.2682\nTotal prob of N-in: 0.14402\noutside 1 331\nTMhelix 332 354\ninside 355 372

24904 GCF_001350235.1_8490_2_81 Salmonella enterica subsp. enterica serovar Typhimurium str. DT104 Proteobacteria; Gammaproteobacteria; Enterobacterales; Enterobacteriaceae; Salmonella; Salmonella enterica; Salmonella enterica subsp. enterica; Salmonella enterica subsp. enterica serovar Typhimurium MTGDNTLITSHGINRRDFMKLCAALAATMGLSSKAAAEMAESVSNPQRPPVIWIGAQECT WP_000145429.1 hydrogenase 2 small subunit [Salmonella enterica] Length: 372\nNumber of predicted TMHs: 1\nExp number of AAs in TMHs: 24.51309\nExp number, first 60 AAs: 0.2682\nTotal prob of N-in: 0.14402\noutside 1 331\nTMhelix 332 354\ninside 355 372

24905 GCF_001219645.1_5295_2_7Salmonella enterica subsp. enterica serovar Typhimurium str. DT104 Proteobacteria; Gammaproteobacteria; Enterobacterales; Enterobacteriaceae; Salmonella; Salmonella enterica; Salmonella enterica subsp. enterica; Salmonella enterica subsp. enterica serovar Typhimurium MNNEETFYQAMRRKGVTRRSFLKFCSLAATSLGLGAGMTPKIAWALENKPRIPVVWIHGL WP_001058312.1 hydrogenase [Salmonella enterica] Length: 372\nNumber of predicted TMHs: 2\nExp number of AAs in TMHs: 36.51738\nExp number, first 60 AAs: 19.72849\nTotal prob of N-in: 0.96170\nPOSSIBLE N-term signal sequence\ninside 1 19\nTMhelix 20 39\noutside 40 325\nTMhelix 326 348\ninside 349 372

24906 GCF_001219965.1_5299_3_5Salmonella enterica subsp. enterica serovar Typhimurium str. DT104 Proteobacteria; Gammaproteobacteria; Enterobacterales; Enterobacteriaceae; Salmonella; Salmonella enterica; Salmonella enterica subsp. enterica; Salmonella enterica subsp. enterica serovar Typhimurium MTGDNTLITSHGINRRDFMKLCAALAATMGLSSKAAAEMAESVSNPQRPPVIWIGAQECT WP_000145429.1 hydrogenase 2 small subunit [Salmonella enterica] Length: 372\nNumber of predicted TMHs: 1\nExp number of AAs in TMHs: 24.51309\nExp number, first 60 AAs: 0.2682\nTotal prob of N-in: 0.14402\noutside 1 331\nTMhelix 332 354\ninside 355 372

24907 GCF_001219965.1_5299_3_5Salmonella enterica subsp. enterica serovar Typhimurium str. DT104 Proteobacteria; Gammaproteobacteria; Enterobacterales; Enterobacteriaceae; Salmonella; Salmonella enterica; Salmonella enterica subsp. enterica; Salmonella enterica subsp. enterica serovar Typhimurium MNNEETFYQAMRRKGVTRRSFLKFCSLAATSLGLGAGMTPKIAWALENKPRIPVVWIHGL WP_001058312.1 hydrogenase [Salmonella enterica] Length: 372\nNumber of predicted TMHs: 2\nExp number of AAs in TMHs: 36.51738\nExp number, first 60 AAs: 19.72849\nTotal prob of N-in: 0.96170\nPOSSIBLE N-term signal sequence\ninside 1 19\nTMhelix 20 39\noutside 40 325\nTMhelix 326 348\ninside 349 372

24908 GCF_001219665.1_5388_1_6Salmonella enterica subsp. enterica serovar Typhimurium str. DT104 Proteobacteria; Gammaproteobacteria; Enterobacterales; Enterobacteriaceae; Salmonella; Salmonella enterica; Salmonella enterica subsp. enterica; Salmonella enterica subsp. enterica serovar Typhimurium MNNEETFYQAMRRKGVTRRSFLKFCSLAATSLGLGAGMTPKIAWALENKPRIPVVWIHGL WP_001058312.1 hydrogenase [Salmonella enterica] Length: 372\nNumber of predicted TMHs: 2\nExp number of AAs in TMHs: 36.51738\nExp number, first 60 AAs: 19.72849\nTotal prob of N-in: 0.96170\nPOSSIBLE N-term signal sequence\ninside 1 19\nTMhelix 20 39\noutside 40 325\nTMhelix 326 348\ninside 349 372

24909 GCF_001219705.1_8080_3_83 Salmonella enterica subsp. enterica serovar Typhimurium str. DT104 Proteobacteria; Gammaproteobacteria; Enterobacterales; Enterobacteriaceae; Salmonella; Salmonella enterica; Salmonella enterica subsp. enterica; Salmonella enterica subsp. enterica serovar Typhimurium MNNEETFYQAMRRKGVTRRSFLKFCSLAATSLGLGAGMTPKIAWALENKPRIPVVWIHGL WP_001058312.1 hydrogenase [Salmonella enterica] Length: 372\nNumber of predicted TMHs: 2\nExp number of AAs in

TMHs: 36.51738\nExp number, first 60 AAs:  19.72849\nTotal prob of N-in:   0.96170\nPOSSIBLE N-term signal sequence\ninside   1   19\nTMhelix   20   39\noutside   40   325\nTMhelix   326   348\ninside   349   372

24910 GCF_001219845.1_5299_3_11 Salmonella enterica subsp. enterica serovar Typhimurium str. DT104 Proteobacteria; Gammaproteobacteria; Enterobacterales; Enterobacteriaceae; Salmonella; Salmonella enterica; Salmonella enterica subsp. enterica; Salmonella enterica subsp. enterica serovar Typhimurium MNNEETFYQAMRRKGVTRRSFLKFCSLAATSLGLGAGMTPKIAWALENKPRIPVVWIHGL WP_001058312.1 hydrogenase [Salmonella enterica] Length: 372\nNumber of predicted TMHs:  2\nExp number of AAs in TMHs: 36.51738\nExp number, first 60 AAs:  19.72849\nTotal prob of N-in:   0.96170\nPOSSIBLE N-term signal sequence\ninside   1   19\nTMhelix   20   39\noutside   40   325\nTMhelix   326   348\ninside   349   372

24911 GCF_001219765.1_8080_3_57 Salmonella enterica subsp. enterica serovar Typhimurium str. DT104 Proteobacteria; Gammaproteobacteria; Enterobacterales; Enterobacteriaceae; Salmonella; Salmonella enterica; Salmonella enterica subsp. enterica; Salmonella enterica subsp. enterica serovar Typhimurium MNNEETFYQAMRRKGVTRRSFLKFCSLAATSLGLGAGMTPKIAWALENKPRIPVVWIHGL WP_001058312.1 hydrogenase [Salmonella enterica] Length: 372\nNumber of predicted TMHs:  2\nExp number of AAs in TMHs: 36.51738\nExp number, first 60 AAs:  19.72849\nTotal prob of N-in:   0.96170\nPOSSIBLE N-term signal sequence\ninside   1   19\nTMhelix   20   39\noutside   40   325\nTMhelix   326   348\ninside   349   372

24912 GCF_001220065.1_5299_3_10 Salmonella enterica subsp. enterica serovar Typhimurium str. DT104 Proteobacteria; Gammaproteobacteria; Enterobacterales; Enterobacteriaceae; Salmonella; Salmonella enterica; Salmonella enterica subsp. enterica; Salmonella enterica subsp. enterica serovar Typhimurium MNNEETFYQAMRRKGVTRRSFLKFCSLAATSLGLGAGMTPKIAWALENKPRIPVVWIHGL WP_001058312.1 hydrogenase [Salmonella enterica] Length: 372\nNumber of predicted TMHs:  2\nExp number of AAs in TMHs: 36.51738\nExp number, first 60 AAs:  19.72849\nTotal prob of N-in:   0.96170\nPOSSIBLE N-term signal sequence\ninside   1   19\nTMhelix   20   39\noutside   40   325\nTMhelix   326   348\ninside   349   372

24913 GCF_001219805.1_8036_3_75 Salmonella enterica subsp. enterica serovar Typhimurium str. DT104 Proteobacteria; Gammaproteobacteria; Enterobacterales; Enterobacteriaceae; Salmonella; Salmonella enterica; Salmonella enterica subsp. enterica; Salmonella enterica subsp. enterica serovar Typhimurium MNNEETFYQAMRRKGVTRRSFLKFCSLAATSLGLGAGMTPKIAWALENKPRIPVVWIHGL WP_001058312.1 hydrogenase [Salmonella enterica] Length: 372\nNumber of predicted TMHs:  2\nExp number of AAs in TMHs: 36.51738\nExp number, first 60 AAs:  19.72849\nTotal prob of N-in:   0.96170\nPOSSIBLE N-term signal sequence\ninside   1   19\nTMhelix   20   39\noutside   40   325\nTMhelix   326   348\ninside   349   372

24914 GCF_001219885.1_8080_3_38 Salmonella enterica subsp. enterica serovar Typhimurium str. DT104 Proteobacteria; Gammaproteobacteria; Enterobacterales; Enterobacteriaceae; Salmonella; Salmonella enterica; Salmonella enterica subsp. enterica; Salmonella enterica subsp. enterica serovar Typhimurium MNNEETFYQAMRRKGVTRRSFLKFCSLAATSLGLGAGMTPKIAWALENKPRIPVVWIHGL WP_001058312.1 hydrogenase [Salmonella enterica] Length: 372\nNumber of predicted TMHs:  2\nExp number of AAs in TMHs: 36.51738\nExp number, first 60 AAs:  19.72849\nTotal prob of N-in:   0.96170\nPOSSIBLE N-term signal sequence\ninside   1   19\nTMhelix   20   39\noutside   40   325\nTMhelix   326   348\ninside   349   372

24915 GCF_001219925.1_5275_5_1Salmonella enterica subsp. enterica serovar Typhimurium str. DT104 Proteobacteria; Gammaproteobacteria; Enterobacterales; Enterobacteriaceae; Salmonella; Salmonella enterica; Salmonella enterica subsp. enterica; Salmonella enterica subsp. enterica serovar Typhimurium MNNEETFYQAMRRKGVTRRSFLKFCSLAATSLGLGAGMTPKIAWALENKPRIPVVWIHGL WP_001058312.1 hydrogenase [Salmonella enterica] Length: 372\nNumber of predicted TMHs:  2\nExp number of AAs in TMHs: 36.51738\nExp number, first 60 AAs:  19.72849\nTotal prob of N-in:   0.96170\nPOSSIBLE N-term signal sequence\ninside   1   19\nTMhelix   20   39\noutside   40   325\nTMhelix   326   348\ninside   349   372

24916 GCF_001219985.1_8080_3_6Salmonella enterica subsp. enterica serovar Typhimurium str. DT104 Proteobacteria; Gammaproteobacteria; Enterobacterales; Enterobacteriaceae; Salmonella; Salmonella enterica; Salmonella enterica subsp. enterica; Salmonella enterica subsp. enterica serovar Typhimurium MNNEETFYQAMRRKGVTRRSFLKFCSLAATSLGLGAGMTPKIAWALENKPRIPVVWIHGL WP_001058312.1 hydrogenase [Salmonella enterica] Length: 372\nNumber of predicted TMHs:  2\nExp number of AAs in TMHs: 36.51738\nExp number, first 60 AAs:  19.72849\nTotal prob of N-in:   0.96170\nPOSSIBLE N-term signal sequence\ninside   1   19\nTMhelix   20   39\noutside   40   325\nTMhelix   326   348\ninside   349   372

24917 GCF_001220125.1_8080_3_91 Salmonella enterica subsp. enterica serovar Typhimurium str. DT104 Proteobacteria; Gammaproteobacteria; Enterobacterales; Enterobacteriaceae; Salmonella; Salmonella enterica;

Salmonella enterica subsp. enterica; Salmonella enterica subsp. enterica serovar Typhimurium
MNNEETFYQAMRRKGVTRRSFLKFCSLAATSLGLGAGMTPKIAWALENKPRIPVVWIHGL WP_001058312.1
hydrogenase [Salmonella enterica] Length: 372\nNumber of predicted TMHs: 2\nExp number of AAs in TMHs: 36.51738\nExp number, first 60 AAs: 19.72849\nTotal prob of N-in: 0.96170\nPOSSIBLE N-term signal sequence\ninside 1 19\nTMhelix 20 39\noutside 40 325\nTMhelix 326 348\ninside 349 372

24918 GCF_001220205.1_8080_3_77 Salmonella enterica subsp. enterica serovar Typhimurium str. DT104
Proteobacteria; Gammaproteobacteria; Enterobacterales; Enterobacteriaceae; Salmonella; Salmonella enterica; Salmonella enterica subsp. enterica; Salmonella enterica subsp. enterica serovar Typhimurium
MTGDNTLITSHGINRRDFMKLCAALAATMGLSSKAAAEMAESVSNPQRPPVIWIGAQECT WP_000145429.1
hydrogenase 2 small subunit [Salmonella enterica] Length: 372\nNumber of predicted TMHs: 1\nExp number of AAs in TMHs: 24.51309\nExp number, first 60 AAs: 0.2682\nTotal prob of N-in: 0.14402\noutside 1 331\nTMhelix 332 354\ninside 355 372

24919 GCF_001220245.1_8080_3_96 Salmonella enterica subsp. enterica serovar Typhimurium str. DT104
Proteobacteria; Gammaproteobacteria; Enterobacterales; Enterobacteriaceae; Salmonella; Salmonella enterica; Salmonella enterica subsp. enterica; Salmonella enterica subsp. enterica serovar Typhimurium
MTGDNTLITSHGINRRDFMKLCAALAATMGLSSKAAAEMAESVSNPQRPPVIWIGAQECT WP_000145429.1
hydrogenase 2 small subunit [Salmonella enterica] Length: 372\nNumber of predicted TMHs: 1\nExp number of AAs in TMHs: 24.51309\nExp number, first 60 AAs: 0.2682\nTotal prob of N-in: 0.14402\noutside 1 331\nTMhelix 332 354\ninside 355 372

24920 GCF_001220265.1_5275_2_11 Salmonella enterica subsp. enterica serovar Typhimurium str. DT104
Proteobacteria; Gammaproteobacteria; Enterobacterales; Enterobacteriaceae; Salmonella; Salmonella enterica; Salmonella enterica subsp. enterica; Salmonella enterica subsp. enterica serovar Typhimurium
MTGDNTLITSHGINRRDFMKLCAALAATMGLSSKAAAEMAESVSNPQRPPVIWIGAQECT WP_000145429.1
hydrogenase 2 small subunit [Salmonella enterica] Length: 372\nNumber of predicted TMHs: 1\nExp number of AAs in TMHs: 24.51309\nExp number, first 60 AAs: 0.2682\nTotal prob of N-in: 0.14402\noutside 1 331\nTMhelix 332 354\ninside 355 372

24921 GCF_001220205.1_8080_3_77 Salmonella enterica subsp. enterica serovar Typhimurium str. DT104
Proteobacteria; Gammaproteobacteria; Enterobacterales; Enterobacteriaceae; Salmonella; Salmonella enterica; Salmonella enterica subsp. enterica; Salmonella enterica subsp. enterica serovar Typhimurium
MNNEETFYQAMRRKGVTRRSFLKFCSLAATSLGLGAGMTPKIAWALENKPRIPVVWIHGL WP_001058312.1
hydrogenase [Salmonella enterica] Length: 372\nNumber of predicted TMHs: 2\nExp number of AAs in TMHs: 36.51738\nExp number, first 60 AAs: 19.72849\nTotal prob of N-in: 0.96170\nPOSSIBLE N-term signal sequence\ninside 1 19\nTMhelix 20 39\noutside 40 325\nTMhelix 326 348\ninside 349 372

24922 GCF_001220445.1_8036_3_50 Salmonella enterica subsp. enterica serovar Typhimurium str. DT104
Proteobacteria; Gammaproteobacteria; Enterobacterales; Enterobacteriaceae; Salmonella; Salmonella enterica; Salmonella enterica subsp. enterica; Salmonella enterica subsp. enterica serovar Typhimurium
MTGDNTLITSHGINRRDFMKLCAALAATMGLSSKAAAEMAESVSNPQRPPVIWIGAQECT WP_000145429.1
hydrogenase 2 small subunit [Salmonella enterica] Length: 372\nNumber of predicted TMHs: 1\nExp number of AAs in TMHs: 24.51309\nExp number, first 60 AAs: 0.2682\nTotal prob of N-in: 0.14402\noutside 1 331\nTMhelix 332 354\ninside 355 372

24923 GCF_001220305.1_8080_3_40 Salmonella enterica subsp. enterica serovar Typhimurium str. DT104
Proteobacteria; Gammaproteobacteria; Enterobacterales; Enterobacteriaceae; Salmonella; Salmonella enterica; Salmonella enterica subsp. enterica; Salmonella enterica subsp. enterica serovar Typhimurium
MTGDNTLITSHGINRRDFMKLCAALAATMGLSSKAAAEMAESVSNPQRPPVIWIGAQECT WP_000145429.1
hydrogenase 2 small subunit [Salmonella enterica] Length: 372\nNumber of predicted TMHs: 1\nExp number of AAs in TMHs: 24.51309\nExp number, first 60 AAs: 0.2682\nTotal prob of N-in: 0.14402\noutside 1 331\nTMhelix 332 354\ninside 355 372

24924 GCF_001220585.1_5299_6_7Salmonella enterica subsp. enterica serovar Typhimurium str. DT104
Proteobacteria; Gammaproteobacteria; Enterobacterales; Enterobacteriaceae; Salmonella; Salmonella enterica; Salmonella enterica subsp. enterica; Salmonella enterica subsp. enterica serovar Typhimurium
MTGDNTLITSHGINRRDFMKLCAALAATMGLSSKAAAEMAESVSNPQRPPVIWIGAQECT WP_000145429.1
hydrogenase 2 small subunit [Salmonella enterica] Length: 372\nNumber of predicted TMHs: 1\nExp number of AAs in TMHs: 24.51309\nExp number, first 60 AAs: 0.2682\nTotal prob of N-in: 0.14402\noutside 1 331\nTMhelix 332 354\ninside 355 372

24925 GCF_001220525.1_5295_2_1Salmonella enterica subsp. enterica serovar Typhimurium str. DT104 Proteobacteria; Gammaproteobacteria; Enterobacterales; Enterobacteriaceae; Salmonella; Salmonella enterica; Salmonella enterica subsp. enterica; Salmonella enterica subsp. enterica serovar Typhimurium MTGDNTLITSHGINRRDFMKLCAALAATMGLSSKAAAEMAESVSNPQRPPVIWIGAQECT WP_000145429.1 hydrogenase 2 small subunit [Salmonella enterica] Length: 372\nNumber of predicted TMHs: 1\nExp number of AAs in TMHs: 24.51309\nExp number, first 60 AAs: 0.2682\nTotal prob of N-in: 0.14402\noutside 1 331\nTMhelix 332 354\ninside 355 372

24926 GCF_001220385.1_5275_7_6Salmonella enterica subsp. enterica serovar Typhimurium str. DT104 Proteobacteria; Gammaproteobacteria; Enterobacterales; Enterobacteriaceae; Salmonella; Salmonella enterica; Salmonella enterica subsp. enterica; Salmonella enterica subsp. enterica serovar Typhimurium MTGDNTLITSHGINRRDFMKLCAALAATMGLSSKAAAEMAESVSNPQRPPVIWIGAQECT WP_000145429.1 hydrogenase 2 small subunit [Salmonella enterica] Length: 372\nNumber of predicted TMHs: 1\nExp number of AAs in TMHs: 24.51309\nExp number, first 60 AAs: 0.2682\nTotal prob of N-in: 0.14402\noutside 1 331\nTMhelix 332 354\ninside 355 372

24927 GCF_001220465.1_8036_3_80 Salmonella enterica subsp. enterica serovar Typhimurium str. DT104 Proteobacteria; Gammaproteobacteria; Enterobacterales; Enterobacteriaceae; Salmonella; Salmonella enterica; Salmonella enterica subsp. enterica; Salmonella enterica subsp. enterica serovar Typhimurium MTGDNTLITSHGINRRDFMKLCAALAATMGLSSKAAAEMAESVSNPQRPPVIWIGAQECT WP_000145429.1 hydrogenase 2 small subunit [Salmonella enterica] Length: 372\nNumber of predicted TMHs: 1\nExp number of AAs in TMHs: 24.51309\nExp number, first 60 AAs: 0.2682\nTotal prob of N-in: 0.14402\noutside 1 331\nTMhelix 332 354\ninside 355 372

24928 GCF_001220625.1_5295_2_10 Salmonella enterica subsp. enterica serovar Typhimurium str. DT104 Proteobacteria; Gammaproteobacteria; Enterobacterales; Enterobacteriaceae; Salmonella; Salmonella enterica; Salmonella enterica subsp. enterica; Salmonella enterica subsp. enterica serovar Typhimurium MTGDNTLITSHGINRRDFMKLCAALAATMGLSSKAAAEMAESVSNPQRPPVIWIGAQECT WP_000145429.1 hydrogenase 2 small subunit [Salmonella enterica] Length: 372\nNumber of predicted TMHs: 1\nExp number of AAs in TMHs: 24.51309\nExp number, first 60 AAs: 0.2682\nTotal prob of N-in: 0.14402\noutside 1 331\nTMhelix 332 354\ninside 355 372

24929 GCF_001220705.1_5275_5_10 Salmonella enterica subsp. enterica serovar Typhimurium str. DT104 Proteobacteria; Gammaproteobacteria; Enterobacterales; Enterobacteriaceae; Salmonella; Salmonella enterica; Salmonella enterica subsp. enterica; Salmonella enterica subsp. enterica serovar Typhimurium MTGDNTLITSHGINRRDFMKLCAALAATMGLSSKAAAEMAESVSNPQRPPVIWIGAQECT WP_000145429.1 hydrogenase 2 small subunit [Salmonella enterica] Length: 372\nNumber of predicted TMHs: 1\nExp number of AAs in TMHs: 24.51309\nExp number, first 60 AAs: 0.2682\nTotal prob of N-in: 0.14402\noutside 1 331\nTMhelix 332 354\ninside 355 372

24930 GCF_001220245.1_8080_3_96 Salmonella enterica subsp. enterica serovar Typhimurium str. DT104 Proteobacteria; Gammaproteobacteria; Enterobacterales; Enterobacteriaceae; Salmonella; Salmonella enterica; Salmonella enterica subsp. enterica; Salmonella enterica subsp. enterica serovar Typhimurium MNNEETFYQAMRRKGVTRRSFLKFCSLAATSLGLGAGMTPKIAWALENKPRIPVVWIHGL WP_001058312.1 hydrogenase [Salmonella enterica] Length: 372\nNumber of predicted TMHs: 2\nExp number of AAs in TMHs: 36.51738\nExp number, first 60 AAs: 19.72849\nTotal prob of N-in: 0.96170\nPOSSIBLE N-term signal sequence\ninside 1 19\nTMhelix 20 39\noutside 40 325\nTMhelix 326 348\ninside 349 372

24931 GCF_001220265.1_5275_2_11 Salmonella enterica subsp. enterica serovar Typhimurium str. DT104 Proteobacteria; Gammaproteobacteria; Enterobacterales; Enterobacteriaceae; Salmonella; Salmonella enterica; Salmonella enterica subsp. enterica; Salmonella enterica subsp. enterica serovar Typhimurium MNNEETFYQAMRRKGVTRRSFLKFCSLAATSLGLGAGMTPKIAWALENKPRIPVVWIHGL WP_001058312.1 hydrogenase [Salmonella enterica] Length: 372\nNumber of predicted TMHs: 2\nExp number of AAs in TMHs: 36.51738\nExp number, first 60 AAs: 19.72849\nTotal prob of N-in: 0.96170\nPOSSIBLE N-term signal sequence\ninside 1 19\nTMhelix 20 39\noutside 40 325\nTMhelix 326 348\ninside 349 372

24932 GCF_001220585.1_5299_6_7Salmonella enterica subsp. enterica serovar Typhimurium str. DT104 Proteobacteria; Gammaproteobacteria; Enterobacterales; Enterobacteriaceae; Salmonella; Salmonella enterica; Salmonella enterica subsp. enterica; Salmonella enterica subsp. enterica serovar Typhimurium MNNEETFYQAMRRKGVTRRSFLKFCSLAATSLGLGAGMTPKIAWALENKPRIPVVWIHGL WP_001058312.1 hydrogenase [Salmonella enterica] Length: 372\nNumber of predicted TMHs: 2\nExp number of AAs in

TMHs: 36.51738\nExp number, first 60 AAs: 19.72849\nTotal prob of N-in: 0.96170\nPOSSIBLE N-term signal sequence\ninside 1 19\nTMhelix 20 39\noutside 40 325\nTMhelix 326 348\ninside 349 372

24933 GCF_001220445.1_8036_3_50 Salmonella enterica subsp. enterica serovar Typhimurium str. DT104 Proteobacteria; Gammaproteobacteria; Enterobacterales; Enterobacteriaceae; Salmonella; Salmonella enterica; Salmonella enterica subsp. enterica; Salmonella enterica subsp. enterica serovar Typhimurium MNNEETFYQAMRRKGVTRRSFLKFCSLAATSLGLGAGMTPKIAWALENKPRIPVVWIHGL WP_001058312.1 hydrogenase [Salmonella enterica] Length: 372\nNumber of predicted TMHs: 2\nExp number of AAs in TMHs: 36.51738\nExp number, first 60 AAs: 19.72849\nTotal prob of N-in: 0.96170\nPOSSIBLE N-term signal sequence\ninside 1 19\nTMhelix 20 39\noutside 40 325\nTMhelix 326 348\ninside 349 372

24934 GCF_001220345.1_8080_3_50 Salmonella enterica subsp. enterica serovar Typhimurium str. DT104 Proteobacteria; Gammaproteobacteria; Enterobacterales; Enterobacteriaceae; Salmonella; Salmonella enterica; Salmonella enterica subsp. enterica; Salmonella enterica subsp. enterica serovar Typhimurium MNNEETFYQAMRRKGVTRRSFLKFCSLAATSLGLGAGMTPKIAWALENKPRIPVVWIHGL WP_001058312.1 hydrogenase [Salmonella enterica] Length: 372\nNumber of predicted TMHs: 2\nExp number of AAs in TMHs: 36.51738\nExp number, first 60 AAs: 19.72849\nTotal prob of N-in: 0.96170\nPOSSIBLE N-term signal sequence\ninside 1 19\nTMhelix 20 39\noutside 40 325\nTMhelix 326 348\ninside 349 372

24935 GCF_001220625.1_5295_2_10 Salmonella enterica subsp. enterica serovar Typhimurium str. DT104 Proteobacteria; Gammaproteobacteria; Enterobacterales; Enterobacteriaceae; Salmonella; Salmonella enterica; Salmonella enterica subsp. enterica; Salmonella enterica subsp. enterica serovar Typhimurium MNNEETFYQAMRRKGVTRRSFLKFCSLAATSLGLGAGMTPKIAWALENKPRIPVVWIHGL WP_001058312.1 hydrogenase [Salmonella enterica] Length: 372\nNumber of predicted TMHs: 2\nExp number of AAs in TMHs: 36.51738\nExp number, first 60 AAs: 19.72849\nTotal prob of N-in: 0.96170\nPOSSIBLE N-term signal sequence\ninside 1 19\nTMhelix 20 39\noutside 40 325\nTMhelix 326 348\ninside 349 372

24936 GCF_001220425.1_8036_3_83 Salmonella enterica subsp. enterica serovar Typhimurium str. DT104 Proteobacteria; Gammaproteobacteria; Enterobacterales; Enterobacteriaceae; Salmonella; Salmonella enterica; Salmonella enterica subsp. enterica; Salmonella enterica subsp. enterica serovar Typhimurium MNNEETFYQAMRRKGVTRRSFLKFCSLAATSLGLGAGMTPKIAWALENKPRIPVVWIHGL WP_001058312.1 hydrogenase [Salmonella enterica] Length: 372\nNumber of predicted TMHs: 2\nExp number of AAs in TMHs: 36.51738\nExp number, first 60 AAs: 19.72849\nTotal prob of N-in: 0.96170\nPOSSIBLE N-term signal sequence\ninside 1 19\nTMhelix 20 39\noutside 40 325\nTMhelix 326 348\ninside 349 372

24937 GCF_001220525.1_5295_2_1Salmonella enterica subsp. enterica serovar Typhimurium str. DT104 Proteobacteria; Gammaproteobacteria; Enterobacterales; Enterobacteriaceae; Salmonella; Salmonella enterica; Salmonella enterica subsp. enterica; Salmonella enterica subsp. enterica serovar Typhimurium MNNEETFYQAMRRKGVTRRSFLKFCSLAATSLGLGAGMTPKIAWALENKPRIPVVWIHGL WP_001058312.1 hydrogenase [Salmonella enterica] Length: 372\nNumber of predicted TMHs: 2\nExp number of AAs in TMHs: 36.51738\nExp number, first 60 AAs: 19.72849\nTotal prob of N-in: 0.96170\nPOSSIBLE N-term signal sequence\ninside 1 19\nTMhelix 20 39\noutside 40 325\nTMhelix 326 348\ninside 349 372

24938 GCF_001220385.1_5275_7_6Salmonella enterica subsp. enterica serovar Typhimurium str. DT104 Proteobacteria; Gammaproteobacteria; Enterobacterales; Enterobacteriaceae; Salmonella; Salmonella enterica; Salmonella enterica subsp. enterica; Salmonella enterica subsp. enterica serovar Typhimurium MNNEETFYQAMRRKGVTRRSFLKFCSLAATSLGLGAGMTPKIAWALENKPRIPVVWIHGL WP_001058312.1 hydrogenase [Salmonella enterica] Length: 372\nNumber of predicted TMHs: 2\nExp number of AAs in TMHs: 36.51738\nExp number, first 60 AAs: 19.72849\nTotal prob of N-in: 0.96170\nPOSSIBLE N-term signal sequence\ninside 1 19\nTMhelix 20 39\noutside 40 325\nTMhelix 326 348\ninside 349 372

24939 GCF_001220305.1_8080_3_40 Salmonella enterica subsp. enterica serovar Typhimurium str. DT104 Proteobacteria; Gammaproteobacteria; Enterobacterales; Enterobacteriaceae; Salmonella; Salmonella enterica; Salmonella enterica subsp. enterica; Salmonella enterica subsp. enterica serovar Typhimurium MNNEETFYQAMRRKGVTRRSFLKFCSLAATSLGLGAGMTPKIAWALENKPRIPVVWIHGL WP_001058312.1 hydrogenase [Salmonella enterica] Length: 372\nNumber of predicted TMHs: 2\nExp number of AAs in TMHs: 36.51738\nExp number, first 60 AAs: 19.72849\nTotal prob of N-in: 0.96170\nPOSSIBLE N-term signal sequence\ninside 1 19\nTMhelix 20 39\noutside 40 325\nTMhelix 326 348\ninside 349 372

24940 GCF_001220465.1_8036_3_80 Salmonella enterica subsp. enterica serovar Typhimurium str. DT104 Proteobacteria; Gammaproteobacteria; Enterobacterales; Enterobacteriaceae; Salmonella; Salmonella enterica;

Salmonella enterica subsp. enterica; Salmonella enterica subsp. enterica serovar Typhimurium
MNNEETFYQAMRRKGVTRRSFLKFCSLAATSLGLGAGMTPKIAWALENKPRIPVVWIHGL WP_001058312.1
hydrogenase [Salmonella enterica] Length: 372\nNumber of predicted TMHs: 2\nExp number of AAs in TMHs: 36.51738\nExp number, first 60 AAs: 19.72849\nTotal prob of N-in: 0.96170\nPOSSIBLE N-term signal sequence\ninside 1 19\nTMhelix 20 39\noutside 40 325\nTMhelix 326 348\ninside 349 372

24941 GCF_001220665.1_5299_3_2Salmonella enterica subsp. enterica serovar Typhimurium str. DT104
Proteobacteria; Gammaproteobacteria; Enterobacterales; Enterobacteriaceae; Salmonella; Salmonella enterica; Salmonella enterica subsp. enterica; Salmonella enterica subsp. enterica serovar Typhimurium
MTGDNTLITSHGINRRDFMKLCAALAATMGLSSKAAAEMAESVSNPQRPPVIWIGAQECT WP_000145429.1
hydrogenase 2 small subunit [Salmonella enterica] Length: 372\nNumber of predicted TMHs: 1\nExp number of AAs in TMHs: 24.51309\nExp number, first 60 AAs: 0.2682\nTotal prob of N-in: 0.14402\noutside 1 331\nTMhelix 332 354\ninside 355 372

24942 GCF_001220785.1_8036_3_42 Salmonella enterica subsp. enterica serovar Typhimurium str. DT104
Proteobacteria; Gammaproteobacteria; Enterobacterales; Enterobacteriaceae; Salmonella; Salmonella enterica; Salmonella enterica subsp. enterica; Salmonella enterica subsp. enterica serovar Typhimurium
MTGDNTLITSHGINRRDFMKLCAALAATMGLSSKAAAEMAESVSNPQRPPVIWIGAQECT WP_000145429.1
hydrogenase 2 small subunit [Salmonella enterica] Length: 372\nNumber of predicted TMHs: 1\nExp number of AAs in TMHs: 24.51309\nExp number, first 60 AAs: 0.2682\nTotal prob of N-in: 0.14402\noutside 1 331\nTMhelix 332 354\ninside 355 372

24943 GCF_001220665.1_5299_3_2Salmonella enterica subsp. enterica serovar Typhimurium str. DT104
Proteobacteria; Gammaproteobacteria; Enterobacterales; Enterobacteriaceae; Salmonella; Salmonella enterica; Salmonella enterica subsp. enterica; Salmonella enterica subsp. enterica serovar Typhimurium
MNNEETFYQAMRRKGVTRRSFLKFCSLAATSLGLGAGMTPKIAWALENKPRIPVVWIHGL WP_001058312.1
hydrogenase [Salmonella enterica] Length: 372\nNumber of predicted TMHs: 2\nExp number of AAs in TMHs: 36.51738\nExp number, first 60 AAs: 19.72849\nTotal prob of N-in: 0.96170\nPOSSIBLE N-term signal sequence\ninside 1 19\nTMhelix 20 39\noutside 40 325\nTMhelix 326 348\ninside 349 372

24944 GCF_001220885.1_8036_3_86 Salmonella enterica subsp. enterica serovar Typhimurium str. DT104
Proteobacteria; Gammaproteobacteria; Enterobacterales; Enterobacteriaceae; Salmonella; Salmonella enterica; Salmonella enterica subsp. enterica; Salmonella enterica subsp. enterica serovar Typhimurium
MTGDNTLITSHGINRRDFMKLCAALAATMGLSSKAAAEMAESVSNPQRPPVIWIGAQECT WP_000145429.1
hydrogenase 2 small subunit [Salmonella enterica] Length: 372\nNumber of predicted TMHs: 1\nExp number of AAs in TMHs: 24.51309\nExp number, first 60 AAs: 0.2682\nTotal prob of N-in: 0.14402\noutside 1 331\nTMhelix 332 354\ninside 355 372

24945 GCF_001220945.1_8036_3_51 Salmonella enterica subsp. enterica serovar Typhimurium str. DT104
Proteobacteria; Gammaproteobacteria; Enterobacterales; Enterobacteriaceae; Salmonella; Salmonella enterica; Salmonella enterica subsp. enterica; Salmonella enterica subsp. enterica serovar Typhimurium
MTGDNTLITSHGINRRDFMKLCAALAATMGLSSKAAAEMAESVSNPQRPPVIWIGAQECT WP_000145429.1
hydrogenase 2 small subunit [Salmonella enterica] Length: 372\nNumber of predicted TMHs: 1\nExp number of AAs in TMHs: 24.51309\nExp number, first 60 AAs: 0.2682\nTotal prob of N-in: 0.14402\noutside 1 331\nTMhelix 332 354\ninside 355 372

24946 GCF_001221105.1_8036_3_53 Salmonella enterica subsp. enterica serovar Typhimurium str. DT104
Proteobacteria; Gammaproteobacteria; Enterobacterales; Enterobacteriaceae; Salmonella; Salmonella enterica; Salmonella enterica subsp. enterica; Salmonella enterica subsp. enterica serovar Typhimurium
MTGDNTLITSHGINRRDFMKLCAALAATMGLSSKAAAEMAESVSNPQRPPVIWIGAQECT WP_000145429.1
hydrogenase 2 small subunit [Salmonella enterica] Length: 372\nNumber of predicted TMHs: 1\nExp number of AAs in TMHs: 24.51309\nExp number, first 60 AAs: 0.2682\nTotal prob of N-in: 0.14402\noutside 1 331\nTMhelix 332 354\ninside 355 372

24947 GCF_001220905.1_8080_3_43 Salmonella enterica subsp. enterica serovar Typhimurium str. DT104
Proteobacteria; Gammaproteobacteria; Enterobacterales; Enterobacteriaceae; Salmonella; Salmonella enterica; Salmonella enterica subsp. enterica; Salmonella enterica subsp. enterica serovar Typhimurium
MTGDNTLITSHGINRRDFMKLCAALAATMGLSSKAAAEMAESVSNPQRPPVIWIGAQECT WP_000145429.1
hydrogenase 2 small subunit [Salmonella enterica] Length: 372\nNumber of predicted TMHs: 1\nExp number of AAs in TMHs: 24.51309\nExp number, first 60 AAs: 0.2682\nTotal prob of N-in: 0.14402\noutside 1 331\nTMhelix 332 354\ninside 355 372

24948 GCF_001221065.1_5299_6_4Salmonella enterica subsp. enterica serovar Typhimurium str. DT104 Proteobacteria; Gammaproteobacteria; Enterobacterales; Enterobacteriaceae; Salmonella; Salmonella enterica; Salmonella enterica subsp. enterica; Salmonella enterica subsp. enterica serovar Typhimurium MTGDNTLITSHGINRRDFMKLCAALAATMGLSSKAAAEMAESVSNPQRPPVIWIGAQECT WP_000145429.1 hydrogenase 2 small subunit [Salmonella enterica] Length: 372\nNumber of predicted TMHs: 1\nExp number of AAs in TMHs: 24.51309\nExp number, first 60 AAs: 0.2682\nTotal prob of N-in: 0.14402\noutside 1 331\nTMhelix 332 354\ninside 355 372

24949 GCF_001221185.1_8080_3_31 Salmonella enterica subsp. enterica serovar Typhimurium str. DT104 Proteobacteria; Gammaproteobacteria; Enterobacterales; Enterobacteriaceae; Salmonella; Salmonella enterica; Salmonella enterica subsp. enterica; Salmonella enterica subsp. enterica serovar Typhimurium MTGDNTLITSHGINRRDFMKLCAALAATMGLSSKAAAEMAESVSNPQRPPVIWIGAQECT WP_000145429.1 hydrogenase 2 small subunit [Salmonella enterica] Length: 372\nNumber of predicted TMHs: 1\nExp number of AAs in TMHs: 24.51309\nExp number, first 60 AAs: 0.2682\nTotal prob of N-in: 0.14402\noutside 1 331\nTMhelix 332 354\ninside 355 372

24950 GCF_001221225.1_8036_3_71 Salmonella enterica subsp. enterica serovar Typhimurium str. DT104 Proteobacteria; Gammaproteobacteria; Enterobacterales; Enterobacteriaceae; Salmonella; Salmonella enterica; Salmonella enterica subsp. enterica; Salmonella enterica subsp. enterica serovar Typhimurium MTGDNTLITSHGINRRDFMKLCAALAATMGLSSKAAAEMAESVSNPQRPPVIWIGAQECT WP_000145429.1 hydrogenase 2 small subunit [Salmonella enterica] Length: 372\nNumber of predicted TMHs: 1\nExp number of AAs in TMHs: 24.51309\nExp number, first 60 AAs: 0.2682\nTotal prob of N-in: 0.14402\noutside 1 331\nTMhelix 332 354\ninside 355 372

24951 GCF_001221025.1_5275_8_10 Salmonella enterica subsp. enterica serovar Typhimurium str. DT104 Proteobacteria; Gammaproteobacteria; Enterobacterales; Enterobacteriaceae; Salmonella; Salmonella enterica; Salmonella enterica subsp. enterica; Salmonella enterica subsp. enterica serovar Typhimurium MTGDNTLITSHGINRRDFMKLCAALAATMGLSSKAAAEMAESVSNPQRPPVIWIGAQECT WP_000145429.1 hydrogenase 2 small subunit [Salmonella enterica] Length: 372\nNumber of predicted TMHs: 1\nExp number of AAs in TMHs: 24.51309\nExp number, first 60 AAs: 0.2682\nTotal prob of N-in: 0.14402\noutside 1 331\nTMhelix 332 354\ninside 355 372

24952 GCF_001220705.1_5275_5_10 Salmonella enterica subsp. enterica serovar Typhimurium str. DT104 Proteobacteria; Gammaproteobacteria; Enterobacterales; Enterobacteriaceae; Salmonella; Salmonella enterica; Salmonella enterica subsp. enterica; Salmonella enterica subsp. enterica serovar Typhimurium MNNEETFYQAMRRKGVTRRSFLKFCSLAATSLGLGAGMTPKIAWALENKPRIPVVWIHGL WP_001058312.1 hydrogenase [Salmonella enterica] Length: 372\nNumber of predicted TMHs: 2\nExp number of AAs in TMHs: 36.51738\nExp number, first 60 AAs: 19.72849\nTotal prob of N-in: 0.96170\nPOSSIBLE N-term signal sequence\ninside 1 19\nTMhelix 20 39\noutside 40 325\nTMhelix 326 348\ninside 349 372

24953 GCF_001220985.1_5275_5_2Salmonella enterica subsp. enterica serovar Typhimurium str. DT104 Proteobacteria; Gammaproteobacteria; Enterobacterales; Enterobacteriaceae; Salmonella; Salmonella enterica; Salmonella enterica subsp. enterica; Salmonella enterica subsp. enterica serovar Typhimurium MTGDNTLITSHGINRRDFMKLCAALAATMGLSSKAAAEMAESVSNPQRPPVIWIGAQECT WP_000145429.1 hydrogenase 2 small subunit [Salmonella enterica] Length: 372\nNumber of predicted TMHs: 1\nExp number of AAs in TMHs: 24.51309\nExp number, first 60 AAs: 0.2682\nTotal prob of N-in: 0.14402\noutside 1 331\nTMhelix 332 354\ninside 355 372

24954 GCF_001326235.1_9066_1_2Salmonella enterica subsp. enterica serovar Typhimurium str. DT104 Proteobacteria; Gammaproteobacteria; Enterobacterales; Enterobacteriaceae; Salmonella; Salmonella enterica; Salmonella enterica subsp. enterica; Salmonella enterica subsp. enterica serovar Typhimurium MTGDNTLITSHGINRRDFMKLCAALAATMGLSSKAAAEMAESVSNPQRPPVIWIGAQECT WP_000145429.1 hydrogenase 2 small subunit [Salmonella enterica] Length: 372\nNumber of predicted TMHs: 1\nExp number of AAs in TMHs: 24.51309\nExp number, first 60 AAs: 0.2682\nTotal prob of N-in: 0.14402\noutside 1 331\nTMhelix 332 354\ninside 355 372

24955 GCF_001220985.1_5275_5_2Salmonella enterica subsp. enterica serovar Typhimurium str. DT104 Proteobacteria; Gammaproteobacteria; Enterobacterales; Enterobacteriaceae; Salmonella; Salmonella enterica; Salmonella enterica subsp. enterica; Salmonella enterica subsp. enterica serovar Typhimurium MNNEETFYQAMRRKGVTRRSFLKFCSLAATSLGLGAGMTPKIAWALENKPRIPVVWIHGL WP_001058312.1 hydrogenase [Salmonella enterica] Length: 372\nNumber of predicted TMHs: 2\nExp number of AAs in

TMHs: 36.51738\nExp number, first 60 AAs: 19.72849\nTotal prob of N-in: 0.96170\nPOSSIBLE N-term signal sequence\ninside 1 19\nTMhelix 20 39\noutside 40 325\nTMhelix 326 348\ninside 349 372

24956 GCF_001220785.1_8036_3_42 Salmonella enterica subsp. enterica serovar Typhimurium str. DT104 Proteobacteria; Gammaproteobacteria; Enterobacterales; Enterobacteriaceae; Salmonella; Salmonella enterica; Salmonella enterica subsp. enterica; Salmonella enterica subsp. enterica serovar Typhimurium MNNEETFYQAMRRKGVTRRSFLKFCSLAATSLGLGAGMTPKIAWALENKPRIPVVWIHGL WP_001058312.1 hydrogenase [Salmonella enterica] Length: 372\nNumber of predicted TMHs: 2\nExp number of AAs in TMHs: 36.51738\nExp number, first 60 AAs: 19.72849\nTotal prob of N-in: 0.96170\nPOSSIBLE N-term signal sequence\ninside 1 19\nTMhelix 20 39\noutside 40 325\nTMhelix 326 348\ninside 349 372

24957 GCF_001220945.1_8036_3_51 Salmonella enterica subsp. enterica serovar Typhimurium str. DT104 Proteobacteria; Gammaproteobacteria; Enterobacterales; Enterobacteriaceae; Salmonella; Salmonella enterica; Salmonella enterica subsp. enterica; Salmonella enterica subsp. enterica serovar Typhimurium MNNEETFYQAMRRKGVTRRSFLKFCSLAATSLGLGAGMTPKIAWALENKPRIPVVWIHGL WP_001058312.1 hydrogenase [Salmonella enterica] Length: 372\nNumber of predicted TMHs: 2\nExp number of AAs in TMHs: 36.51738\nExp number, first 60 AAs: 19.72849\nTotal prob of N-in: 0.96170\nPOSSIBLE N-term signal sequence\ninside 1 19\nTMhelix 20 39\noutside 40 325\nTMhelix 326 348\ninside 349 372

24958 GCF_001220885.1_8036_3_86 Salmonella enterica subsp. enterica serovar Typhimurium str. DT104 Proteobacteria; Gammaproteobacteria; Enterobacterales; Enterobacteriaceae; Salmonella; Salmonella enterica; Salmonella enterica subsp. enterica; Salmonella enterica subsp. enterica serovar Typhimurium MNNEETFYQAMRRKGVTRRSFLKFCSLAATSLGLGAGMTPKIAWALENKPRIPVVWIHGL WP_001058312.1 hydrogenase [Salmonella enterica] Length: 372\nNumber of predicted TMHs: 2\nExp number of AAs in TMHs: 36.51738\nExp number, first 60 AAs: 19.72849\nTotal prob of N-in: 0.96170\nPOSSIBLE N-term signal sequence\ninside 1 19\nTMhelix 20 39\noutside 40 325\nTMhelix 326 348\ninside 349 372

24959 GCF_001221105.1_8036_3_53 Salmonella enterica subsp. enterica serovar Typhimurium str. DT104 Proteobacteria; Gammaproteobacteria; Enterobacterales; Enterobacteriaceae; Salmonella; Salmonella enterica; Salmonella enterica subsp. enterica; Salmonella enterica subsp. enterica serovar Typhimurium MNNEETFYQAMRRKGVTRRSFLKFCSLAATSLGLGAGMTPKIAWALENKPRIPVVWIHGL WP_001058312.1 hydrogenase [Salmonella enterica] Length: 372\nNumber of predicted TMHs: 2\nExp number of AAs in TMHs: 36.51738\nExp number, first 60 AAs: 19.72849\nTotal prob of N-in: 0.96170\nPOSSIBLE N-term signal sequence\ninside 1 19\nTMhelix 20 39\noutside 40 325\nTMhelix 326 348\ninside 349 372

24960 GCF_001220905.1_8080_3_43 Salmonella enterica subsp. enterica serovar Typhimurium str. DT104 Proteobacteria; Gammaproteobacteria; Enterobacterales; Enterobacteriaceae; Salmonella; Salmonella enterica; Salmonella enterica subsp. enterica; Salmonella enterica subsp. enterica serovar Typhimurium MNNEETFYQAMRRKGVTRRSFLKFCSLAATSLGLGAGMTPKIAWALENKPRIPVVWIHGL WP_001058312.1 hydrogenase [Salmonella enterica] Length: 372\nNumber of predicted TMHs: 2\nExp number of AAs in TMHs: 36.51738\nExp number, first 60 AAs: 19.72849\nTotal prob of N-in: 0.96170\nPOSSIBLE N-term signal sequence\ninside 1 19\nTMhelix 20 39\noutside 40 325\nTMhelix 326 348\ninside 349 372

24961 GCF_001221025.1_5275_8_10 Salmonella enterica subsp. enterica serovar Typhimurium str. DT104 Proteobacteria; Gammaproteobacteria; Enterobacterales; Enterobacteriaceae; Salmonella; Salmonella enterica; Salmonella enterica subsp. enterica; Salmonella enterica subsp. enterica serovar Typhimurium MNNEETFYQAMRRKGVTRRSFLKFCSLAATSLGLGAGMTPKIAWALENKPRIPVVWIHGL WP_001058312.1 hydrogenase [Salmonella enterica] Length: 372\nNumber of predicted TMHs: 2\nExp number of AAs in TMHs: 36.51738\nExp number, first 60 AAs: 19.72849\nTotal prob of N-in: 0.96170\nPOSSIBLE N-term signal sequence\ninside 1 19\nTMhelix 20 39\noutside 40 325\nTMhelix 326 348\ninside 349 372

24962 GCF_001221185.1_8080_3_31 Salmonella enterica subsp. enterica serovar Typhimurium str. DT104 Proteobacteria; Gammaproteobacteria; Enterobacterales; Enterobacteriaceae; Salmonella; Salmonella enterica; Salmonella enterica subsp. enterica; Salmonella enterica subsp. enterica serovar Typhimurium MNNEETFYQAMRRKGVTRRSFLKFCSLAATSLGLGAGMTPKIAWALENKPRIPVVWIHGL WP_001058312.1 hydrogenase [Salmonella enterica] Length: 372\nNumber of predicted TMHs: 2\nExp number of AAs in TMHs: 36.51738\nExp number, first 60 AAs: 19.72849\nTotal prob of N-in: 0.96170\nPOSSIBLE N-term signal sequence\ninside 1 19\nTMhelix 20 39\noutside 40 325\nTMhelix 326 348\ninside 349 372

24963 GCF_001221225.1_8036_3_71 Salmonella enterica subsp. enterica serovar Typhimurium str. DT104 Proteobacteria; Gammaproteobacteria; Enterobacterales; Enterobacteriaceae; Salmonella; Salmonella enterica;

Salmonella enterica subsp. enterica; Salmonella enterica subsp. enterica serovar Typhimurium
MNNEETFYQAMRRKGVTRRSFLKFCSLAATSLGLGAGMTPKIAWALENKPRIPVVWIHGL WP_001058312.1
hydrogenase [Salmonella enterica] Length: 372\nNumber of predicted TMHs: 2\nExp number of AAs in TMHs: 36.51738\nExp number, first 60 AAs: 19.72849\nTotal prob of N-in: 0.96170\nPOSSIBLE N-term signal sequence\ninside 1 19\nTMhelix 20 39\noutside 40 325\nTMhelix 326 348\ninside 349 372

24964 GCF_001221065.1_5299_6_4Salmonella enterica subsp. enterica serovar Typhimurium str. DT104
Proteobacteria; Gammaproteobacteria; Enterobacterales; Enterobacteriaceae; Salmonella; Salmonella enterica; Salmonella enterica subsp. enterica; Salmonella enterica subsp. enterica serovar Typhimurium
MNNEETFYQAMRRKGVTRRSFLKFCSLAATSLGLGAGMTPKIAWALENKPRIPVVWIHGL WP_001058312.1
hydrogenase [Salmonella enterica] Length: 372\nNumber of predicted TMHs: 2\nExp number of AAs in TMHs: 36.51738\nExp number, first 60 AAs: 19.72849\nTotal prob of N-in: 0.96170\nPOSSIBLE N-term signal sequence\ninside 1 19\nTMhelix 20 39\noutside 40 325\nTMhelix 326 348\ninside 349 372

24965 GCF_001221265.1_8080_3_39 Salmonella enterica subsp. enterica serovar Typhimurium str. DT104
Proteobacteria; Gammaproteobacteria; Enterobacterales; Enterobacteriaceae; Salmonella; Salmonella enterica; Salmonella enterica subsp. enterica; Salmonella enterica subsp. enterica serovar Typhimurium
MTGDNTLITSHGINRRDFMKLCAALAATMGLSSKAAAEMAESVSNPQRPPVIWIGAQECT WP_000145429.1
hydrogenase 2 small subunit [Salmonella enterica] Length: 372\nNumber of predicted TMHs: 1\nExp number of AAs in TMHs: 24.51309\nExp number, first 60 AAs: 0.2682\nTotal prob of N-in: 0.14402\noutside 1 331\nTMhelix 332 354\ninside 355 372

24966 GCF_001221305.1_5275_8_2Salmonella enterica subsp. enterica serovar Typhimurium str. DT104
Proteobacteria; Gammaproteobacteria; Enterobacterales; Enterobacteriaceae; Salmonella; Salmonella enterica; Salmonella enterica subsp. enterica; Salmonella enterica subsp. enterica serovar Typhimurium
MTGDNTLITSHGINRRDFMKLCAALAATMGLSSKAAAEMAESVSNPQRPPVIWIGAQECT WP_000145429.1
hydrogenase 2 small subunit [Salmonella enterica] Length: 372\nNumber of predicted TMHs: 1\nExp number of AAs in TMHs: 24.51309\nExp number, first 60 AAs: 0.2682\nTotal prob of N-in: 0.14402\noutside 1 331\nTMhelix 332 354\ninside 355 372

24967 GCF_001221325.1_8080_3_16 Salmonella enterica subsp. enterica serovar Typhimurium str. DT104
Proteobacteria; Gammaproteobacteria; Enterobacterales; Enterobacteriaceae; Salmonella; Salmonella enterica; Salmonella enterica subsp. enterica; Salmonella enterica subsp. enterica serovar Typhimurium
MTGDNTLITSHGINRRDFMKLCAALAATMGLSSKAAAEMAESVSNPQRPPVIWIGAQECT WP_000145429.1
hydrogenase 2 small subunit [Salmonella enterica] Length: 372\nNumber of predicted TMHs: 1\nExp number of AAs in TMHs: 24.51309\nExp number, first 60 AAs: 0.2682\nTotal prob of N-in: 0.14402\noutside 1 331\nTMhelix 332 354\ninside 355 372

24968 GCF_001221345.1_8080_3_12 Salmonella enterica subsp. enterica serovar Typhimurium str. DT104
Proteobacteria; Gammaproteobacteria; Enterobacterales; Enterobacteriaceae; Salmonella; Salmonella enterica; Salmonella enterica subsp. enterica; Salmonella enterica subsp. enterica serovar Typhimurium
MTGDNTLITSHGINRRDFMKLCAALAATMGLSSKAAAEMAESVSNPQRPPVIWIGAQECT WP_000145429.1
hydrogenase 2 small subunit [Salmonella enterica] Length: 372\nNumber of predicted TMHs: 1\nExp number of AAs in TMHs: 24.51309\nExp number, first 60 AAs: 0.2682\nTotal prob of N-in: 0.14402\noutside 1 331\nTMhelix 332 354\ninside 355 372

24969 GCF_001221385.1_9066_1_7Salmonella enterica subsp. enterica serovar Typhimurium str. DT104
Proteobacteria; Gammaproteobacteria; Enterobacterales; Enterobacteriaceae; Salmonella; Salmonella enterica; Salmonella enterica subsp. enterica; Salmonella enterica subsp. enterica serovar Typhimurium
MTGDNTLITSHGINRRDFMKLCAALAATMGLSSKAAAEMAESVSNPQRPPVIWIGAQECT WP_000145429.1
hydrogenase 2 small subunit [Salmonella enterica] Length: 372\nNumber of predicted TMHs: 1\nExp number of AAs in TMHs: 24.51309\nExp number, first 60 AAs: 0.2682\nTotal prob of N-in: 0.14402\noutside 1 331\nTMhelix 332 354\ninside 355 372

24970 GCF_001326275.1_5275_1_5Salmonella enterica subsp. enterica serovar Typhimurium str. DT104
Proteobacteria; Gammaproteobacteria; Enterobacterales; Enterobacteriaceae; Salmonella; Salmonella enterica; Salmonella enterica subsp. enterica; Salmonella enterica subsp. enterica serovar Typhimurium
MTGDNTLITSHGINRRDFMKLCAALAATMGLSSKAAAEMAESVSNPQRPPVIWIGAQECT WP_000145429.1
hydrogenase 2 small subunit [Salmonella enterica] Length: 372\nNumber of predicted TMHs: 1\nExp number of AAs in TMHs: 24.51309\nExp number, first 60 AAs: 0.2682\nTotal prob of N-in: 0.14402\noutside 1 331\nTMhelix 332 354\ninside 355 372

24971 GCF_001221425.1_5275_5_9Salmonella enterica subsp. enterica serovar Typhimurium str. DT104 Proteobacteria; Gammaproteobacteria; Enterobacterales; Enterobacteriaceae; Salmonella; Salmonella enterica; Salmonella enterica subsp. enterica; Salmonella enterica subsp. enterica serovar Typhimurium MTGDNTLITSHGINRRDFMKLCAALAATMGLSSKAAAEMAESVSNPQRPPVIWIGAQECT WP_000145429.1 hydrogenase 2 small subunit [Salmonella enterica] Length: 372\nNumber of predicted TMHs: 1\nExp number of AAs in TMHs: 24.51309\nExp number, first 60 AAs: 0.2682\nTotal prob of N-in: 0.14402\noutside 1 331\nTMhelix 332 354\ninside 355 372

24972 GCF_001221505.1_8036_3_56 Salmonella enterica subsp. enterica serovar Typhimurium str. DT104 Proteobacteria; Gammaproteobacteria; Enterobacterales; Enterobacteriaceae; Salmonella; Salmonella enterica; Salmonella enterica subsp. enterica; Salmonella enterica subsp. enterica serovar Typhimurium MTGDNTLITSHGINRRDFMKLCAALAATMGLSSKAAAEMAESVSNPQRPPVIWIGAQECT WP_000145429.1 hydrogenase 2 small subunit [Salmonella enterica] Length: 372\nNumber of predicted TMHs: 1\nExp number of AAs in TMHs: 24.51309\nExp number, first 60 AAs: 0.2682\nTotal prob of N-in: 0.14402\noutside 1 331\nTMhelix 332 354\ninside 355 372

24973 GCF_001350275.1_8490_2_82 Salmonella enterica subsp. enterica serovar Typhimurium str. DT104 Proteobacteria; Gammaproteobacteria; Enterobacterales; Enterobacteriaceae; Salmonella; Salmonella enterica; Salmonella enterica subsp. enterica; Salmonella enterica subsp. enterica serovar Typhimurium MTGDNTLITSHGINRRDFMKLCAALAATMGLSSKAAAEMAESVSNPQRPPVIWIGAQECT WP_000145429.1 hydrogenase 2 small subunit [Salmonella enterica] Length: 372\nNumber of predicted TMHs: 1\nExp number of AAs in TMHs: 24.51309\nExp number, first 60 AAs: 0.2682\nTotal prob of N-in: 0.14402\noutside 1 331\nTMhelix 332 354\ninside 355 372

24974 GCF_001350255.1_8490_2_72 Salmonella enterica subsp. enterica serovar Typhimurium str. DT104 Proteobacteria; Gammaproteobacteria; Enterobacterales; Enterobacteriaceae; Salmonella; Salmonella enterica; Salmonella enterica subsp. enterica; Salmonella enterica subsp. enterica serovar Typhimurium MTGDNTLITSHGINRRDFMKLCAALAATMGLSSKAAAEMAESVSNPQRPPVIWIGAQECT WP_000145429.1 hydrogenase 2 small subunit [Salmonella enterica] Length: 372\nNumber of predicted TMHs: 1\nExp number of AAs in TMHs: 24.51309\nExp number, first 60 AAs: 0.2682\nTotal prob of N-in: 0.14402\noutside 1 331\nTMhelix 332 354\ninside 355 372

24975 GCF_001350255.1_8490_2_72 Salmonella enterica subsp. enterica serovar Typhimurium str. DT104 Proteobacteria; Gammaproteobacteria; Enterobacterales; Enterobacteriaceae; Salmonella; Salmonella enterica; Salmonella enterica subsp. enterica; Salmonella enterica subsp. enterica serovar Typhimurium MNNEETFYQAMRRKGVTRRSFLKFCSLAATSLGLGAGMTPKIAWALENKPRIPVVWIHGL WP_001058312.1 hydrogenase [Salmonella enterica] Length: 372\nNumber of predicted TMHs: 2\nExp number of AAs in TMHs: 36.51738\nExp number, first 60 AAs: 19.72849\nTotal prob of N-in: 0.96170\nPOSSIBLE N-term signal sequence\ninside 1 19\nTMhelix 20 39\noutside 40 325\nTMhelix 326 348\ninside 349 372

24976 GCF_001350035.1_8490_2_58 Salmonella enterica subsp. enterica serovar Typhimurium str. DT104 Proteobacteria; Gammaproteobacteria; Enterobacterales; Enterobacteriaceae; Salmonella; Salmonella enterica; Salmonella enterica subsp. enterica; Salmonella enterica subsp. enterica serovar Typhimurium MNNEETFYQAMRRKGVTRRSFLKFCSLAATSLGLGAGMTPKIAWALENKPRIPVVWIHGL WP_001058312.1 hydrogenase [Salmonella enterica] Length: 372\nNumber of predicted TMHs: 2\nExp number of AAs in TMHs: 36.51738\nExp number, first 60 AAs: 19.72849\nTotal prob of N-in: 0.96170\nPOSSIBLE N-term signal sequence\ninside 1 19\nTMhelix 20 39\noutside 40 325\nTMhelix 326 348\ninside 349 372

24977 GCF_001216645.1_5275_3_11 Salmonella enterica subsp. enterica serovar Typhimurium str. DT104 Proteobacteria; Gammaproteobacteria; Enterobacterales; Enterobacteriaceae; Salmonella; Salmonella enterica; Salmonella enterica subsp. enterica; Salmonella enterica subsp. enterica serovar Typhimurium MNNEETFYQAMRRKGVTRRSFLKFCSLAATSLGLGAGMTPKIAWALENKPRIPVVWIHGL WP_001058312.1 hydrogenase [Salmonella enterica] Length: 372\nNumber of predicted TMHs: 2\nExp number of AAs in TMHs: 36.51738\nExp number, first 60 AAs: 19.72849\nTotal prob of N-in: 0.96170\nPOSSIBLE N-term signal sequence\ninside 1 19\nTMhelix 20 39\noutside 40 325\nTMhelix 326 348\ninside 349 372

24978 GCF_001217265.1_5299_3_3Salmonella enterica subsp. enterica serovar Typhimurium str. DT104 Proteobacteria; Gammaproteobacteria; Enterobacterales; Enterobacteriaceae; Salmonella; Salmonella enterica; Salmonella enterica subsp. enterica; Salmonella enterica subsp. enterica serovar Typhimurium MNNEETFYQAMRRKGVTRRSFLKFCSLAATSLGLGAGMTPKIAWALENKPRIPVVWIHGL WP_001058312.1 hydrogenase [Salmonella enterica] Length: 372\nNumber of predicted TMHs: 2\nExp number of AAs in

TMHs: 36.51738\nExp number, first 60 AAs: 19.72849\nTotal prob of N-in: 0.96170\nPOSSIBLE N-term signal sequence\ninside 1 19\nTMhelix 20 39\noutside 40 325\nTMhelix 326 348\ninside 349 372

24979 GCF_001217025.1_5275_8_11 Salmonella enterica subsp. enterica serovar Typhimurium str. DT104 Proteobacteria; Gammaproteobacteria; Enterobacterales; Enterobacteriaceae; Salmonella; Salmonella enterica; Salmonella enterica subsp. enterica; Salmonella enterica subsp. enterica serovar Typhimurium MNNEETFYQAMRRKGVTRRSFLKFCSLAATSLGLGAGMTPKIAWALENKPRIPVVWIHGL WP_001058312.1 hydrogenase [Salmonella enterica] Length: 372\nNumber of predicted TMHs: 2\nExp number of AAs in TMHs: 36.51738\nExp number, first 60 AAs: 19.72849\nTotal prob of N-in: 0.96170\nPOSSIBLE N-term signal sequence\ninside 1 19\nTMhelix 20 39\noutside 40 325\nTMhelix 326 348\ninside 349 372

24980 GCF_001217045.1_5275_3_5Salmonella enterica subsp. enterica serovar Typhimurium str. DT104 Proteobacteria; Gammaproteobacteria; Enterobacterales; Enterobacteriaceae; Salmonella; Salmonella enterica; Salmonella enterica subsp. enterica; Salmonella enterica subsp. enterica serovar Typhimurium MNNEETFYQAMRRKGVTRRSFLKFCSLAATSLGLGAGMTPKIAWALENKPRIPVVWIHGL WP_001058312.1 hydrogenase [Salmonella enterica] Length: 372\nNumber of predicted TMHs: 2\nExp number of AAs in TMHs: 36.51738\nExp number, first 60 AAs: 19.72849\nTotal prob of N-in: 0.96170\nPOSSIBLE N-term signal sequence\ninside 1 19\nTMhelix 20 39\noutside 40 325\nTMhelix 326 348\ninside 349 372

24981 GCF_001216765.1_8036_3_82 Salmonella enterica subsp. enterica serovar Typhimurium str. DT104 Proteobacteria; Gammaproteobacteria; Enterobacterales; Enterobacteriaceae; Salmonella; Salmonella enterica; Salmonella enterica subsp. enterica; Salmonella enterica subsp. enterica serovar Typhimurium MNNEETFYQAMRRKGVTRRSFLKFCSLAATSLGLGAGMTPKIAWALENKPRIPVVWIHGL WP_001058312.1 hydrogenase [Salmonella enterica] Length: 372\nNumber of predicted TMHs: 2\nExp number of AAs in TMHs: 36.51738\nExp number, first 60 AAs: 19.72849\nTotal prob of N-in: 0.96170\nPOSSIBLE N-term signal sequence\ninside 1 19\nTMhelix 20 39\noutside 40 325\nTMhelix 326 348\ninside 349 372

24982 GCF_001217205.1_8080_3_61 Salmonella enterica subsp. enterica serovar Typhimurium str. DT104 Proteobacteria; Gammaproteobacteria; Enterobacterales; Enterobacteriaceae; Salmonella; Salmonella enterica; Salmonella enterica subsp. enterica; Salmonella enterica subsp. enterica serovar Typhimurium MNNEETFYQAMRRKGVTRRSFLKFCSLAATSLGLGAGMTPKIAWALENKPRIPVVWIHGL WP_001058312.1 hydrogenase [Salmonella enterica] Length: 372\nNumber of predicted TMHs: 2\nExp number of AAs in TMHs: 36.51738\nExp number, first 60 AAs: 19.72849\nTotal prob of N-in: 0.96170\nPOSSIBLE N-term signal sequence\ninside 1 19\nTMhelix 20 39\noutside 40 325\nTMhelix 326 348\ninside 349 372

24983 GCF_001217425.1_5285_6_7Salmonella enterica subsp. enterica serovar Typhimurium str. DT104 Proteobacteria; Gammaproteobacteria; Enterobacterales; Enterobacteriaceae; Salmonella; Salmonella enterica; Salmonella enterica subsp. enterica; Salmonella enterica subsp. enterica serovar Typhimurium MNNEETFYQAMRRKGVTRRSFLKFCSLAATSLGLGAGMTPKIAWALENKPRIPVVWIHGL WP_001058312.1 hydrogenase [Salmonella enterica] Length: 372\nNumber of predicted TMHs: 2\nExp number of AAs in TMHs: 36.51738\nExp number, first 60 AAs: 19.72849\nTotal prob of N-in: 0.96170\nPOSSIBLE N-term signal sequence\ninside 1 19\nTMhelix 20 39\noutside 40 325\nTMhelix 326 348\ninside 349 372

24984 GCF_001217445.1_5275_5_3Salmonella enterica subsp. enterica serovar Typhimurium str. DT104 Proteobacteria; Gammaproteobacteria; Enterobacterales; Enterobacteriaceae; Salmonella; Salmonella enterica; Salmonella enterica subsp. enterica; Salmonella enterica subsp. enterica serovar Typhimurium MTGDNTLITSHGINRRDFMKLCAALAATMGLSSKAAAEMAESVSNPQRPPVIWIGAQECT WP_000145429.1 hydrogenase 2 small subunit [Salmonella enterica] Length: 372\nNumber of predicted TMHs: 1\nExp number of AAs in TMHs: 24.51309\nExp number, first 60 AAs: 0.2682\nTotal prob of N-in: 0.14402\noutside 1 331\nTMhelix 332 354\ninside 355 372

24985 GCF_001217625.1_9065_1_11 Salmonella enterica subsp. enterica serovar Typhimurium str. DT104 Proteobacteria; Gammaproteobacteria; Enterobacterales; Enterobacteriaceae; Salmonella; Salmonella enterica; Salmonella enterica subsp. enterica; Salmonella enterica subsp. enterica serovar Typhimurium MTGDNTLITSHGINRRDFMKLCAALAATMGLSSKAAAEMAESVSNPQRPPVIWIGAQECT WP_000145429.1 hydrogenase 2 small subunit [Salmonella enterica] Length: 372\nNumber of predicted TMHs: 1\nExp number of AAs in TMHs: 24.51309\nExp number, first 60 AAs: 0.2682\nTotal prob of N-in: 0.14402\noutside 1 331\nTMhelix 332 354\ninside 355 372

24986 GCF_001217585.1_8036_3_46 Salmonella enterica subsp. enterica serovar Typhimurium str. DT104 Proteobacteria; Gammaproteobacteria; Enterobacterales; Enterobacteriaceae; Salmonella; Salmonella enterica;

Salmonella enterica subsp. enterica; Salmonella enterica subsp. enterica serovar Typhimurium
MTGDNTLITSHGINRRDFMKLCAALAATMGLSSKAAAEMAESVSNPQRPPVIWIGAQECT WP_000145429.1
hydrogenase 2 small subunit [Salmonella enterica] Length: 372\nNumber of predicted TMHs: 1\nExp number of AAs in TMHs: 24.51309\nExp number, first 60 AAs: 0.2682\nTotal prob of N-in: 0.14402\noutside 1 331\nTMhelix 332 354\ninside 355 372

24987 GCF_001217665.1_9065_1_8Salmonella enterica subsp. enterica serovar Typhimurium str. DT104
Proteobacteria; Gammaproteobacteria; Enterobacterales; Enterobacteriaceae; Salmonella; Salmonella enterica; Salmonella enterica subsp. enterica; Salmonella enterica subsp. enterica serovar Typhimurium
MTGDNTLITSHGINRRDFMKLCAALAATMGLSSKAAAEMAESVSNPQRPPVIWIGAQECT WP_000145429.1
hydrogenase 2 small subunit [Salmonella enterica] Length: 372\nNumber of predicted TMHs: 1\nExp number of AAs in TMHs: 24.51309\nExp number, first 60 AAs: 0.2682\nTotal prob of N-in: 0.14402\noutside 1 331\nTMhelix 332 354\ninside 355 372

24988 GCF_001217765.1_5295_2_2Salmonella enterica subsp. enterica serovar Typhimurium str. DT104
Proteobacteria; Gammaproteobacteria; Enterobacterales; Enterobacteriaceae; Salmonella; Salmonella enterica; Salmonella enterica subsp. enterica; Salmonella enterica subsp. enterica serovar Typhimurium
MTGDNTLITSHGINRRDFMKLCAALAATMGLSSKAAAEMAESVSNPQRPPVIWIGAQECT WP_000145429.1
hydrogenase 2 small subunit [Salmonella enterica] Length: 372\nNumber of predicted TMHs: 1\nExp number of AAs in TMHs: 24.51309\nExp number, first 60 AAs: 0.2682\nTotal prob of N-in: 0.14402\noutside 1 331\nTMhelix 332 354\ninside 355 372

24989 GCF_001350055.1_8490_2_67 Salmonella enterica subsp. enterica serovar Typhimurium str. DT104
Proteobacteria; Gammaproteobacteria; Enterobacterales; Enterobacteriaceae; Salmonella; Salmonella enterica; Salmonella enterica subsp. enterica; Salmonella enterica subsp. enterica serovar Typhimurium
MNNEETFYQAMRRKGVTRRSFLKFCSLAATSLGLGAGMTPKIAWALENKPRIPVVWIHGL WP_001058312.1
hydrogenase [Salmonella enterica] Length: 372\nNumber of predicted TMHs: 2\nExp number of AAs in TMHs: 36.51738\nExp number, first 60 AAs: 19.72849\nTotal prob of N-in: 0.96170\nPOSSIBLE N-term signal sequence\ninside 1 19\nTMhelix 20 39\noutside 40 325\nTMhelix 326 348\ninside 349 372

24990 GCF_001217785.1_8080_3_47 Salmonella enterica subsp. enterica serovar Typhimurium str. DT104
Proteobacteria; Gammaproteobacteria; Enterobacterales; Enterobacteriaceae; Salmonella; Salmonella enterica; Salmonella enterica subsp. enterica; Salmonella enterica subsp. enterica serovar Typhimurium
MTGDNTLITSHGINRRDFMKLCAALAATMGLSSKAAAEMAESVSNPQRPPVIWIGAQECT WP_000145429.1
hydrogenase 2 small subunit [Salmonella enterica] Length: 372\nNumber of predicted TMHs: 1\nExp number of AAs in TMHs: 24.51309\nExp number, first 60 AAs: 0.2682\nTotal prob of N-in: 0.14402\noutside 1 331\nTMhelix 332 354\ninside 355 372

24991 GCF_001217885.1_9066_1_11 Salmonella enterica subsp. enterica serovar Typhimurium str. DT104
Proteobacteria; Gammaproteobacteria; Enterobacterales; Enterobacteriaceae; Salmonella; Salmonella enterica; Salmonella enterica subsp. enterica; Salmonella enterica subsp. enterica serovar Typhimurium
MTGDNTLITSHGINRRDFMKLCAALAATMGLSSKAAAEMAESVSNPQRPPVIWIGAQECT WP_000145429.1
hydrogenase 2 small subunit [Salmonella enterica] Length: 372\nNumber of predicted TMHs: 1\nExp number of AAs in TMHs: 24.51309\nExp number, first 60 AAs: 0.2682\nTotal prob of N-in: 0.14402\noutside 1 331\nTMhelix 332 354\ninside 355 372

24992 GCF_001217965.1_9066_1_4Salmonella enterica subsp. enterica serovar Typhimurium str. DT104
Proteobacteria; Gammaproteobacteria; Enterobacterales; Enterobacteriaceae; Salmonella; Salmonella enterica; Salmonella enterica subsp. enterica; Salmonella enterica subsp. enterica serovar Typhimurium
MTGDNTLITSHGINRRDFMKLCAALAATMGLSSKAAAEMAESVSNPQRPPVIWIGAQECT WP_000145429.1
hydrogenase 2 small subunit [Salmonella enterica] Length: 372\nNumber of predicted TMHs: 1\nExp number of AAs in TMHs: 24.51309\nExp number, first 60 AAs: 0.2682\nTotal prob of N-in: 0.14402\noutside 1 331\nTMhelix 332 354\ninside 355 372

24993 GCF_001217925.1_9065_1_6Salmonella enterica subsp. enterica serovar Typhimurium str. DT104
Proteobacteria; Gammaproteobacteria; Enterobacterales; Enterobacteriaceae; Salmonella; Salmonella enterica; Salmonella enterica subsp. enterica; Salmonella enterica subsp. enterica serovar Typhimurium
MTGDNTLITSHGINRRDFMKLCAALAATMGLSSKAAAEMAESVSNPQRPPVIWIGAQECT WP_000145429.1
hydrogenase 2 small subunit [Salmonella enterica] Length: 372\nNumber of predicted TMHs: 1\nExp number of AAs in TMHs: 24.51309\nExp number, first 60 AAs: 0.2682\nTotal prob of N-in: 0.14402\noutside 1 331\nTMhelix 332 354\ninside 355 372

24994 GCF_001217845.1_8080_3_71 Salmonella enterica subsp. enterica serovar Typhimurium str. DT104
Proteobacteria; Gammaproteobacteria; Enterobacterales; Enterobacteriaceae; Salmonella; Salmonella enterica; Salmonella enterica subsp. enterica; Salmonella enterica subsp. enterica serovar Typhimurium
MTGDNTLITSHGINRRDFMKLCAALAATMGLSSKAAAEMAESVSNPQRPPVIWIGAQECT WP_000145429.1
hydrogenase 2 small subunit [Salmonella enterica] Length: 372\nNumber of predicted TMHs: 1\nExp number of AAs in TMHs: 24.51309\nExp number, first 60 AAs: 0.2682\nTotal prob of N-in: 0.14402\noutside 1 331\nTMhelix 332 354\ninside 355 372

24995 GCF_001217985.1_8080_3_89 Salmonella enterica subsp. enterica serovar Typhimurium str. DT104
Proteobacteria; Gammaproteobacteria; Enterobacterales; Enterobacteriaceae; Salmonella; Salmonella enterica; Salmonella enterica subsp. enterica; Salmonella enterica subsp. enterica serovar Typhimurium
MTGDNTLITSHGINRRDFMKLCAALAATMGLSSKAAAEMAESVSNPQRPPVIWIGAQECT WP_000145429.1
hydrogenase 2 small subunit [Salmonella enterica] Length: 372\nNumber of predicted TMHs: 1\nExp number of AAs in TMHs: 24.51309\nExp number, first 60 AAs: 0.2682\nTotal prob of N-in: 0.14402\noutside 1 331\nTMhelix 332 354\ninside 355 372

24996 GCF_001218025.1_8080_3_49 Salmonella enterica subsp. enterica serovar Typhimurium str. DT104
Proteobacteria; Gammaproteobacteria; Enterobacterales; Enterobacteriaceae; Salmonella; Salmonella enterica; Salmonella enterica subsp. enterica; Salmonella enterica subsp. enterica serovar Typhimurium
MTGDNTLITSHGINRRDFMKLCAALAATMGLSSKAAAEMAESVSNPQRPPVIWIGAQECT WP_000145429.1
hydrogenase 2 small subunit [Salmonella enterica] Length: 372\nNumber of predicted TMHs: 1\nExp number of AAs in TMHs: 24.51309\nExp number, first 60 AAs: 0.2682\nTotal prob of N-in: 0.14402\noutside 1 331\nTMhelix 332 354\ninside 355 372

24997 GCF_001217445.1_5275_5_3Salmonella enterica subsp. enterica serovar Typhimurium str. DT104
Proteobacteria; Gammaproteobacteria; Enterobacterales; Enterobacteriaceae; Salmonella; Salmonella enterica; Salmonella enterica subsp. enterica; Salmonella enterica subsp. enterica serovar Typhimurium
MNNEETFYQAMRRKGVTRRSFLKFCSLAATSLGLGAGMTPKIAWALENKPRIPVVWIHGL WP_001058312.1
hydrogenase [Salmonella enterica] Length: 372\nNumber of predicted TMHs: 2\nExp number of AAs in TMHs: 36.51738\nExp number, first 60 AAs: 19.72849\nTotal prob of N-in: 0.96170\nPOSSIBLE N-term signal sequence\ninside 1 19\nTMhelix 20 39\noutside 40 325\nTMhelix 326 348\ninside 349 372

24998 GCF_001217625.1_9065_1_11 Salmonella enterica subsp. enterica serovar Typhimurium str. DT104
Proteobacteria; Gammaproteobacteria; Enterobacterales; Enterobacteriaceae; Salmonella; Salmonella enterica; Salmonella enterica subsp. enterica; Salmonella enterica subsp. enterica serovar Typhimurium
MNNEETFYQAMRRKGVTRRSFLKFCSLAATSLGLGAGMTPKIAWALENKPRIPVVWIHGL WP_001058312.1
hydrogenase [Salmonella enterica] Length: 372\nNumber of predicted TMHs: 2\nExp number of AAs in TMHs: 36.51738\nExp number, first 60 AAs: 19.72849\nTotal prob of N-in: 0.96170\nPOSSIBLE N-term signal sequence\ninside 1 19\nTMhelix 20 39\noutside 40 325\nTMhelix 326 348\ninside 349 372

24999 GCF_001217665.1_9065_1_8Salmonella enterica subsp. enterica serovar Typhimurium str. DT104
Proteobacteria; Gammaproteobacteria; Enterobacterales; Enterobacteriaceae; Salmonella; Salmonella enterica; Salmonella enterica subsp. enterica; Salmonella enterica subsp. enterica serovar Typhimurium
MNNEETFYQAMRRKGVTRRSFLKFCSLAATSLGLGAGMTPKIAWALENKPRIPVVWIHGL WP_001058312.1
hydrogenase [Salmonella enterica] Length: 372\nNumber of predicted TMHs: 2\nExp number of AAs in TMHs: 36.51738\nExp number, first 60 AAs: 19.72849\nTotal prob of N-in: 0.96170\nPOSSIBLE N-term signal sequence\ninside 1 19\nTMhelix 20 39\noutside 40 325\nTMhelix 326 348\ninside 349 372

25000 GCF_001217585.1_8036_3_46 Salmonella enterica subsp. enterica serovar Typhimurium str. DT104
Proteobacteria; Gammaproteobacteria; Enterobacterales; Enterobacteriaceae; Salmonella; Salmonella enterica; Salmonella enterica subsp. enterica; Salmonella enterica subsp. enterica serovar Typhimurium
MNNEETFYQAMRRKGVTRRSFLKFCSLAATSLGLGAGMTPKIAWALENKPRIPVVWIHGL WP_001058312.1
hydrogenase [Salmonella enterica] Length: 372\nNumber of predicted TMHs: 2\nExp number of AAs in TMHs: 36.51738\nExp number, first 60 AAs: 19.72849\nTotal prob of N-in: 0.96170\nPOSSIBLE N-term signal sequence\ninside 1 19\nTMhelix 20 39\noutside 40 325\nTMhelix 326 348\ninside 349 372

25001 GCF_001217965.1_9066_1_4Salmonella enterica subsp. enterica serovar Typhimurium str. DT104
Proteobacteria; Gammaproteobacteria; Enterobacterales; Enterobacteriaceae; Salmonella; Salmonella enterica; Salmonella enterica subsp. enterica; Salmonella enterica subsp. enterica serovar Typhimurium
MNNEETFYQAMRRKGVTRRSFLKFCSLAATSLGLGAGMTPKIAWALENKPRIPVVWIHGL WP_001058312.1
hydrogenase [Salmonella enterica] Length: 372\nNumber of predicted TMHs: 2\nExp number of AAs in

TMHs: 36.51738\nExp number, first 60 AAs: 19.72849\nTotal prob of N-in: 0.96170\nPOSSIBLE N-term signal sequence\ninside 1 19\nTMhelix 20 39\noutside 40 325\nTMhelix 326 348\ninside 349 372

25002 GCF_001217885.1_9066_1_11 Salmonella enterica subsp. enterica serovar Typhimurium str. DT104 Proteobacteria; Gammaproteobacteria; Enterobacterales; Enterobacteriaceae; Salmonella; Salmonella enterica; Salmonella enterica subsp. enterica; Salmonella enterica subsp. enterica serovar Typhimurium MNNEETFYQAMRRKGVTRRSFLKFCSLAATSLGLGAGMTPKIAWALENKPRIPVVWIHGL WP_001058312.1 hydrogenase [Salmonella enterica] Length: 372\nNumber of predicted TMHs: 2\nExp number of AAs in TMHs: 36.51738\nExp number, first 60 AAs: 19.72849\nTotal prob of N-in: 0.96170\nPOSSIBLE N-term signal sequence\ninside 1 19\nTMhelix 20 39\noutside 40 325\nTMhelix 326 348\ninside 349 372

25003 GCF_001217765.1_5295_2_2Salmonella enterica subsp. enterica serovar Typhimurium str. DT104 Proteobacteria; Gammaproteobacteria; Enterobacterales; Enterobacteriaceae; Salmonella; Salmonella enterica; Salmonella enterica subsp. enterica; Salmonella enterica subsp. enterica serovar Typhimurium MNNEETFYQAMRRKGVTRRSFLKFCSLAATSLGLGAGMTPKIAWALENKPRIPVVWIHGL WP_001058312.1 hydrogenase [Salmonella enterica] Length: 372\nNumber of predicted TMHs: 2\nExp number of AAs in TMHs: 36.51738\nExp number, first 60 AAs: 19.72849\nTotal prob of N-in: 0.96170\nPOSSIBLE N-term signal sequence\ninside 1 19\nTMhelix 20 39\noutside 40 325\nTMhelix 326 348\ninside 349 372

25004 GCF_001217785.1_8080_3_47 Salmonella enterica subsp. enterica serovar Typhimurium str. DT104 Proteobacteria; Gammaproteobacteria; Enterobacterales; Enterobacteriaceae; Salmonella; Salmonella enterica; Salmonella enterica subsp. enterica; Salmonella enterica subsp. enterica serovar Typhimurium MNNEETFYQAMRRKGVTRRSFLKFCSLAATSLGLGAGMTPKIAWALENKPRIPVVWIHGL WP_001058312.1 hydrogenase [Salmonella enterica] Length: 372\nNumber of predicted TMHs: 2\nExp number of AAs in TMHs: 36.51738\nExp number, first 60 AAs: 19.72849\nTotal prob of N-in: 0.96170\nPOSSIBLE N-term signal sequence\ninside 1 19\nTMhelix 20 39\noutside 40 325\nTMhelix 326 348\ninside 349 372

25005 GCF_001217845.1_8080_3_71 Salmonella enterica subsp. enterica serovar Typhimurium str. DT104 Proteobacteria; Gammaproteobacteria; Enterobacterales; Enterobacteriaceae; Salmonella; Salmonella enterica; Salmonella enterica subsp. enterica; Salmonella enterica subsp. enterica serovar Typhimurium MNNEETFYQAMRRKGVTRRSFLKFCSLAATSLGLGAGMTPKIAWALENKPRIPVVWIHGL WP_001058312.1 hydrogenase [Salmonella enterica] Length: 372\nNumber of predicted TMHs: 2\nExp number of AAs in TMHs: 36.51738\nExp number, first 60 AAs: 19.72849\nTotal prob of N-in: 0.96170\nPOSSIBLE N-term signal sequence\ninside 1 19\nTMhelix 20 39\noutside 40 325\nTMhelix 326 348\ninside 349 372

25006 GCF_001217985.1_8080_3_89 Salmonella enterica subsp. enterica serovar Typhimurium str. DT104 Proteobacteria; Gammaproteobacteria; Enterobacterales; Enterobacteriaceae; Salmonella; Salmonella enterica; Salmonella enterica subsp. enterica; Salmonella enterica subsp. enterica serovar Typhimurium MNNEETFYQAMRRKGVTRRSFLKFCSLAATSLGLGAGMTPKIAWALENKPRIPVVWIHGL WP_001058312.1 hydrogenase [Salmonella enterica] Length: 372\nNumber of predicted TMHs: 2\nExp number of AAs in TMHs: 36.51738\nExp number, first 60 AAs: 19.72849\nTotal prob of N-in: 0.96170\nPOSSIBLE N-term signal sequence\ninside 1 19\nTMhelix 20 39\noutside 40 325\nTMhelix 326 348\ninside 349 372

25007 GCF_001217925.1_9065_1_6Salmonella enterica subsp. enterica serovar Typhimurium str. DT104 Proteobacteria; Gammaproteobacteria; Enterobacterales; Enterobacteriaceae; Salmonella; Salmonella enterica; Salmonella enterica subsp. enterica; Salmonella enterica subsp. enterica serovar Typhimurium MNNEETFYQAMRRKGVTRRSFLKFCSLAATSLGLGAGMTPKIAWALENKPRIPVVWIHGL WP_001058312.1 hydrogenase [Salmonella enterica] Length: 372\nNumber of predicted TMHs: 2\nExp number of AAs in TMHs: 36.51738\nExp number, first 60 AAs: 19.72849\nTotal prob of N-in: 0.96170\nPOSSIBLE N-term signal sequence\ninside 1 19\nTMhelix 20 39\noutside 40 325\nTMhelix 326 348\ninside 349 372

25008 GCF_001218025.1_8080_3_49 Salmonella enterica subsp. enterica serovar Typhimurium str. DT104 Proteobacteria; Gammaproteobacteria; Enterobacterales; Enterobacteriaceae; Salmonella; Salmonella enterica; Salmonella enterica subsp. enterica; Salmonella enterica subsp. enterica serovar Typhimurium MNNEETFYQAMRRKGVTRRSFLKFCSLAATSLGLGAGMTPKIAWALENKPRIPVVWIHGL WP_001058312.1 hydrogenase [Salmonella enterica] Length: 372\nNumber of predicted TMHs: 2\nExp number of AAs in TMHs: 36.51738\nExp number, first 60 AAs: 19.72849\nTotal prob of N-in: 0.96170\nPOSSIBLE N-term signal sequence\ninside 1 19\nTMhelix 20 39\noutside 40 325\nTMhelix 326 348\ninside 349 372

25009 GCF_001218125.1_5295_2_3Salmonella enterica subsp. enterica serovar Typhimurium str. DT104 Proteobacteria; Gammaproteobacteria; Enterobacterales; Enterobacteriaceae; Salmonella; Salmonella enterica;

Salmonella enterica subsp. enterica; Salmonella enterica subsp. enterica serovar Typhimurium MTGDNTLITSHGINRRDFMKLCAALAATMGLSSKAAAEMAESVSNPQRPPVIWIGAQECT WP_000145429.1 hydrogenase 2 small subunit [Salmonella enterica] Length: 372\nNumber of predicted TMHs: 1\nExp number of AAs in TMHs: 24.51309\nExp number, first 60 AAs: 0.2682\nTotal prob of N-in: 0.14402\noutside 1 331\nTMhelix 332 354\ninside 355 372

25010 GCF_001218085.1_5275_7_8Salmonella enterica subsp. enterica serovar Typhimurium str. DT104 Proteobacteria; Gammaproteobacteria; Enterobacterales; Enterobacteriaceae; Salmonella; Salmonella enterica; Salmonella enterica subsp. enterica; Salmonella enterica subsp. enterica serovar Typhimurium MTGDNTLITSHGINRRDFMKLCAALAATMGLSSKAAAEMAESVSNPQRPPVIWIGAQECT WP_000145429.1 hydrogenase 2 small subunit [Salmonella enterica] Length: 372\nNumber of predicted TMHs: 1\nExp number of AAs in TMHs: 24.51309\nExp number, first 60 AAs: 0.2682\nTotal prob of N-in: 0.14402\noutside 1 331\nTMhelix 332 354\ninside 355 372

25011 GCF_001218225.1_8080_3_95 Salmonella enterica subsp. enterica serovar Typhimurium str. DT104 Proteobacteria; Gammaproteobacteria; Enterobacterales; Enterobacteriaceae; Salmonella; Salmonella enterica; Salmonella enterica subsp. enterica; Salmonella enterica subsp. enterica serovar Typhimurium MTGDNTLITSHGINRRDFMKLCAALAATMGLSSKAAAEMAESVSNPQRPPVIWIGAQECT WP_000145429.1 hydrogenase 2 small subunit [Salmonella enterica] Length: 372\nNumber of predicted TMHs: 1\nExp number of AAs in TMHs: 24.51309\nExp number, first 60 AAs: 0.2682\nTotal prob of N-in: 0.14402\noutside 1 331\nTMhelix 332 354\ninside 355 372

25012 GCF_001218245.1_9065_1_7Salmonella enterica subsp. enterica serovar Typhimurium str. DT104 Proteobacteria; Gammaproteobacteria; Enterobacterales; Enterobacteriaceae; Salmonella; Salmonella enterica; Salmonella enterica subsp. enterica; Salmonella enterica subsp. enterica serovar Typhimurium MTGDNTLITSHGINRRDFMKLCAALAATMGLSSKAAAEMAESVSNPQRPPVIWIGAQECT WP_000145429.1 hydrogenase 2 small subunit [Salmonella enterica] Length: 372\nNumber of predicted TMHs: 1\nExp number of AAs in TMHs: 24.51309\nExp number, first 60 AAs: 0.2682\nTotal prob of N-in: 0.14402\noutside 1 331\nTMhelix 332 354\ninside 355 372

25013 GCF_001218405.1_5299_6_12 Salmonella enterica subsp. enterica serovar Typhimurium str. DT104 Proteobacteria; Gammaproteobacteria; Enterobacterales; Enterobacteriaceae; Salmonella; Salmonella enterica; Salmonella enterica subsp. enterica; Salmonella enterica subsp. enterica serovar Typhimurium MTGDNTLITSHGINRRDFMKLCAALAATMGLSSKAAAEMAESVSNPQRPPVIWIGAQECT WP_000145429.1 hydrogenase 2 small subunit [Salmonella enterica] Length: 372\nNumber of predicted TMHs: 1\nExp number of AAs in TMHs: 24.51309\nExp number, first 60 AAs: 0.2682\nTotal prob of N-in: 0.14402\noutside 1 331\nTMhelix 332 354\ninside 355 372

25014 GCF_001218365.1_8080_3_34 Salmonella enterica subsp. enterica serovar Typhimurium str. DT104 Proteobacteria; Gammaproteobacteria; Enterobacterales; Enterobacteriaceae; Salmonella; Salmonella enterica; Salmonella enterica subsp. enterica; Salmonella enterica subsp. enterica serovar Typhimurium MTGDNTLITSHGINRRDFMKLCAALAATMGLSSKAAAEMAESVSNPQRPPVIWIGAQECT WP_000145429.1 hydrogenase 2 small subunit [Salmonella enterica] Length: 372\nNumber of predicted TMHs: 1\nExp number of AAs in TMHs: 24.51309\nExp number, first 60 AAs: 0.2682\nTotal prob of N-in: 0.14402\noutside 1 331\nTMhelix 332 354\ninside 355 372

25015 GCF_001218285.1_5285_6_12 Salmonella enterica subsp. enterica serovar Typhimurium str. DT104 Proteobacteria; Gammaproteobacteria; Enterobacterales; Enterobacteriaceae; Salmonella; Salmonella enterica; Salmonella enterica subsp. enterica; Salmonella enterica subsp. enterica serovar Typhimurium MTGDNTLITSHGINRRDFMKLCAALAATMGLSSKAAAEMAESVSNPQRPPVIWIGAQECT WP_000145429.1 hydrogenase 2 small subunit [Salmonella enterica] Length: 372\nNumber of predicted TMHs: 1\nExp number of AAs in TMHs: 24.51309\nExp number, first 60 AAs: 0.2682\nTotal prob of N-in: 0.14402\noutside 1 331\nTMhelix 332 354\ninside 355 372

25016 GCF_001218325.1_5388_1_5Salmonella enterica subsp. enterica serovar Typhimurium str. DT104 Proteobacteria; Gammaproteobacteria; Enterobacterales; Enterobacteriaceae; Salmonella; Salmonella enterica; Salmonella enterica subsp. enterica; Salmonella enterica subsp. enterica serovar Typhimurium MTGDNTLITSHGINRRDFMKLCAALAATMGLSSKAAAEMAESVSNPQRPPVIWIGAQECT WP_000145429.1 hydrogenase 2 small subunit [Salmonella enterica] Length: 372\nNumber of predicted TMHs: 1\nExp number of AAs in TMHs: 24.51309\nExp number, first 60 AAs: 0.2682\nTotal prob of N-in: 0.14402\noutside 1 331\nTMhelix 332 354\ninside 355 372

25017 GCF_001218425.1_8080_3_41 Salmonella enterica subsp. enterica serovar Typhimurium str. DT104 Proteobacteria; Gammaproteobacteria; Enterobacterales; Enterobacteriaceae; Salmonella; Salmonella enterica; Salmonella enterica subsp. enterica; Salmonella enterica subsp. enterica serovar Typhimurium MTGDNTLITSHGINRRDFMKLCAALAATMGLSSKAAAEMAESVSNPQRPPVIWIGAQECT WP_000145429.1 hydrogenase 2 small subunit [Salmonella enterica] Length: 372\nNumber of predicted TMHs: 1\nExp number of AAs in TMHs: 24.51309\nExp number, first 60 AAs: 0.2682\nTotal prob of N-in: 0.14402\noutside 1 331\nTMhelix 332 354\ninside 355 372

25018 GCF_001218545.1_5388_1_3Salmonella enterica subsp. enterica serovar Typhimurium str. DT104 Proteobacteria; Gammaproteobacteria; Enterobacterales; Enterobacteriaceae; Salmonella; Salmonella enterica; Salmonella enterica subsp. enterica; Salmonella enterica subsp. enterica serovar Typhimurium MTGDNTLITSHGINRRDFMKLCAALAATMGLSSKAAAEMAESVSNPQRPPVIWIGAQECT WP_000145429.1 hydrogenase 2 small subunit [Salmonella enterica] Length: 372\nNumber of predicted TMHs: 1\nExp number of AAs in TMHs: 24.51309\nExp number, first 60 AAs: 0.2682\nTotal prob of N-in: 0.14402\noutside 1 331\nTMhelix 332 354\ninside 355 372

25019 GCF_001218465.1_8080_3_8Salmonella enterica subsp. enterica serovar Typhimurium str. DT104 Proteobacteria; Gammaproteobacteria; Enterobacterales; Enterobacteriaceae; Salmonella; Salmonella enterica; Salmonella enterica subsp. enterica; Salmonella enterica subsp. enterica serovar Typhimurium MTGDNTLITSHGINRRDFMKLCAALAATMGLSSKAAAEMAESVSNPQRPPVIWIGAQECT WP_000145429.1 hydrogenase 2 small subunit [Salmonella enterica] Length: 372\nNumber of predicted TMHs: 1\nExp number of AAs in TMHs: 24.51309\nExp number, first 60 AAs: 0.2682\nTotal prob of N-in: 0.14402\noutside 1 331\nTMhelix 332 354\ninside 355 372

25020 GCF_001218585.1_5388_1_11 Salmonella enterica subsp. enterica serovar Typhimurium str. DT104 Proteobacteria; Gammaproteobacteria; Enterobacterales; Enterobacteriaceae; Salmonella; Salmonella enterica; Salmonella enterica subsp. enterica; Salmonella enterica subsp. enterica serovar Typhimurium MTGDNTLITSHGINRRDFMKLCAALAATMGLSSKAAAEMAESVSNPQRPPVIWIGAQECT WP_000145429.1 hydrogenase 2 small subunit [Salmonella enterica] Length: 372\nNumber of predicted TMHs: 1\nExp number of AAs in TMHs: 24.51309\nExp number, first 60 AAs: 0.2682\nTotal prob of N-in: 0.14402\noutside 1 331\nTMhelix 332 354\ninside 355 372

25021 GCF_001218625.1_5299_6_2Salmonella enterica subsp. enterica serovar Typhimurium str. DT104 Proteobacteria; Gammaproteobacteria; Enterobacterales; Enterobacteriaceae; Salmonella; Salmonella enterica; Salmonella enterica subsp. enterica; Salmonella enterica subsp. enterica serovar Typhimurium MTGDNTLITSHGINRRDFMKLCAALAATMGLSSKAAAEMAESVSNPQRPPVIWIGAQECT WP_000145429.1 hydrogenase 2 small subunit [Salmonella enterica] Length: 372\nNumber of predicted TMHs: 1\nExp number of AAs in TMHs: 24.51309\nExp number, first 60 AAs: 0.2682\nTotal prob of N-in: 0.14402\noutside 1 331\nTMhelix 332 354\ninside 355 372

25022 GCF_001218085.1_5275_7_8Salmonella enterica subsp. enterica serovar Typhimurium str. DT104 Proteobacteria; Gammaproteobacteria; Enterobacterales; Enterobacteriaceae; Salmonella; Salmonella enterica; Salmonella enterica subsp. enterica; Salmonella enterica subsp. enterica serovar Typhimurium MNNEETFYQAMRRKGVTRRSFLKFCSLAATSLGLGAGMTPKIAWALENKPRIPVVWIHGL WP_001058312.1 hydrogenase [Salmonella enterica] Length: 372\nNumber of predicted TMHs: 2\nExp number of AAs in TMHs: 36.51738\nExp number, first 60 AAs: 19.72849\nTotal prob of N-in: 0.96170\nPOSSIBLE N-term signal sequence\ninside 1 19\nTMhelix 20 39\noutside 40 325\nTMhelix 326 348\ninside 349 372

25023 GCF_001218125.1_5295_2_3Salmonella enterica subsp. enterica serovar Typhimurium str. DT104 Proteobacteria; Gammaproteobacteria; Enterobacterales; Enterobacteriaceae; Salmonella; Salmonella enterica; Salmonella enterica subsp. enterica; Salmonella enterica subsp. enterica serovar Typhimurium MNNEETFYQAMRRKGVTRRSFLKFCSLAATSLGLGAGMTPKIAWALENKPRIPVVWIHGL WP_001058312.1 hydrogenase [Salmonella enterica] Length: 372\nNumber of predicted TMHs: 2\nExp number of AAs in TMHs: 36.51738\nExp number, first 60 AAs: 19.72849\nTotal prob of N-in: 0.96170\nPOSSIBLE N-term signal sequence\ninside 1 19\nTMhelix 20 39\noutside 40 325\nTMhelix 326 348\ninside 349 372

25024 GCF_001218225.1_8080_3_95 Salmonella enterica subsp. enterica serovar Typhimurium str. DT104 Proteobacteria; Gammaproteobacteria; Enterobacterales; Enterobacteriaceae; Salmonella; Salmonella enterica; Salmonella enterica subsp. enterica; Salmonella enterica subsp. enterica serovar Typhimurium MNNEETFYQAMRRKGVTRRSFLKFCSLAATSLGLGAGMTPKIAWALENKPRIPVVWIHGL WP_001058312.1 hydrogenase [Salmonella enterica] Length: 372\nNumber of predicted TMHs: 2\nExp number of AAs in

TMHs: 36.51738\nExp number, first 60 AAs:  19.72849\nTotal prob of N-in:   0.96170\nPOSSIBLE N-term signal sequence\ninside   1   19\nTMhelix   20   39\noutside   40   325\nTMhelix   326   348\ninside   349   372

25025 GCF_001218245.1_9065_1_7Salmonella enterica subsp. enterica serovar Typhimurium str. DT104 Proteobacteria; Gammaproteobacteria; Enterobacterales; Enterobacteriaceae; Salmonella; Salmonella enterica; Salmonella enterica subsp. enterica; Salmonella enterica subsp. enterica serovar Typhimurium MNNEETFYQAMRRKGVTRRSFLKFCSLAATSLGLGAGMTPKIAWALENKPRIPVVWIHGL WP_001058312.1 hydrogenase [Salmonella enterica] Length: 372\nNumber of predicted TMHs:  2\nExp number of AAs in TMHs: 36.51738\nExp number, first 60 AAs:  19.72849\nTotal prob of N-in:   0.96170\nPOSSIBLE N-term signal sequence\ninside   1   19\nTMhelix   20   39\noutside   40   325\nTMhelix   326   348\ninside   349   372

25026 GCF_001218325.1_5388_1_5Salmonella enterica subsp. enterica serovar Typhimurium str. DT104 Proteobacteria; Gammaproteobacteria; Enterobacterales; Enterobacteriaceae; Salmonella; Salmonella enterica; Salmonella enterica subsp. enterica; Salmonella enterica subsp. enterica serovar Typhimurium MNNEETFYQAMRRKGVTRRSFLKFCSLAATSLGLGAGMTPKIAWALENKPRIPVVWIHGL WP_001058312.1 hydrogenase [Salmonella enterica] Length: 372\nNumber of predicted TMHs:  2\nExp number of AAs in TMHs: 36.51738\nExp number, first 60 AAs:  19.72849\nTotal prob of N-in:   0.96170\nPOSSIBLE N-term signal sequence\ninside   1   19\nTMhelix   20   39\noutside   40   325\nTMhelix   326   348\ninside   349   372

25027 GCF_001218405.1_5299_6_12 Salmonella enterica subsp. enterica serovar Typhimurium str. DT104 Proteobacteria; Gammaproteobacteria; Enterobacterales; Enterobacteriaceae; Salmonella; Salmonella enterica; Salmonella enterica subsp. enterica; Salmonella enterica subsp. enterica serovar Typhimurium MNNEETFYQAMRRKGVTRRSFLKFCSLAATSLGLGAGMTPKIAWALENKPRIPVVWIHGL WP_001058312.1 hydrogenase [Salmonella enterica] Length: 372\nNumber of predicted TMHs:  2\nExp number of AAs in TMHs: 36.51738\nExp number, first 60 AAs:  19.72849\nTotal prob of N-in:   0.96170\nPOSSIBLE N-term signal sequence\ninside   1   19\nTMhelix   20   39\noutside   40   325\nTMhelix   326   348\ninside   349   372

25028 GCF_001218365.1_8080_3_34 Salmonella enterica subsp. enterica serovar Typhimurium str. DT104 Proteobacteria; Gammaproteobacteria; Enterobacterales; Enterobacteriaceae; Salmonella; Salmonella enterica; Salmonella enterica subsp. enterica; Salmonella enterica subsp. enterica serovar Typhimurium MNNEETFYQAMRRKGVTRRSFLKFCSLAATSLGLGAGMTPKIAWALENKPRIPVVWIHGL WP_001058312.1 hydrogenase [Salmonella enterica] Length: 372\nNumber of predicted TMHs:  2\nExp number of AAs in TMHs: 36.51738\nExp number, first 60 AAs:  19.72849\nTotal prob of N-in:   0.96170\nPOSSIBLE N-term signal sequence\ninside   1   19\nTMhelix   20   39\noutside   40   325\nTMhelix   326   348\ninside   349   372

25029 GCF_001218425.1_8080_3_41 Salmonella enterica subsp. enterica serovar Typhimurium str. DT104 Proteobacteria; Gammaproteobacteria; Enterobacterales; Enterobacteriaceae; Salmonella; Salmonella enterica; Salmonella enterica subsp. enterica; Salmonella enterica subsp. enterica serovar Typhimurium MNNEETFYQAMRRKGVTRRSFLKFCSLAATSLGLGAGMTPKIAWALENKPRIPVVWIHGL WP_001058312.1 hydrogenase [Salmonella enterica] Length: 372\nNumber of predicted TMHs:  2\nExp number of AAs in TMHs: 36.51738\nExp number, first 60 AAs:  19.72849\nTotal prob of N-in:   0.96170\nPOSSIBLE N-term signal sequence\ninside   1   19\nTMhelix   20   39\noutside   40   325\nTMhelix   326   348\ninside   349   372

25030 GCF_001218285.1_5285_6_12 Salmonella enterica subsp. enterica serovar Typhimurium str. DT104 Proteobacteria; Gammaproteobacteria; Enterobacterales; Enterobacteriaceae; Salmonella; Salmonella enterica; Salmonella enterica subsp. enterica; Salmonella enterica subsp. enterica serovar Typhimurium MNNEETFYQAMRRKGVTRRSFLKFCSLAATSLGLGAGMTPKIAWALENKPRIPVVWIHGL WP_001058312.1 hydrogenase [Salmonella enterica] Length: 372\nNumber of predicted TMHs:  2\nExp number of AAs in TMHs: 36.51738\nExp number, first 60 AAs:  19.72849\nTotal prob of N-in:   0.96170\nPOSSIBLE N-term signal sequence\ninside   1   19\nTMhelix   20   39\noutside   40   325\nTMhelix   326   348\ninside   349   372

25031 GCF_001218545.1_5388_1_3Salmonella enterica subsp. enterica serovar Typhimurium str. DT104 Proteobacteria; Gammaproteobacteria; Enterobacterales; Enterobacteriaceae; Salmonella; Salmonella enterica; Salmonella enterica subsp. enterica; Salmonella enterica subsp. enterica serovar Typhimurium MNNEETFYQAMRRKGVTRRSFLKFCSLAATSLGLGAGMTPKIAWALENKPRIPVVWIHGL WP_001058312.1 hydrogenase [Salmonella enterica] Length: 372\nNumber of predicted TMHs:  2\nExp number of AAs in TMHs: 36.51738\nExp number, first 60 AAs:  19.72849\nTotal prob of N-in:   0.96170\nPOSSIBLE N-term signal sequence\ninside   1   19\nTMhelix   20   39\noutside   40   325\nTMhelix   326   348\ninside   349   372

25032 GCF_001218465.1_8080_3_8Salmonella enterica subsp. enterica serovar Typhimurium str. DT104 Proteobacteria; Gammaproteobacteria; Enterobacterales; Enterobacteriaceae; Salmonella; Salmonella enterica;

Salmonella enterica subsp. enterica; Salmonella enterica subsp. enterica serovar Typhimurium
MNNEETFYQAMRRKGVTRRSFLKFCSLAATSLGLGAGMTPKIAWALENKPRIPVVWIHGL WP_001058312.1
hydrogenase [Salmonella enterica] Length: 372\nNumber of predicted TMHs: 2\nExp number of AAs in TMHs: 36.51738\nExp number, first 60 AAs: 19.72849\nTotal prob of N-in: 0.96170\nPOSSIBLE N-term signal sequence\ninside 1 19\nTMhelix 20 39\noutside 40 325\nTMhelix 326 348\ninside 349 372

25033 GCF_001218585.1_5388_1_11 Salmonella enterica subsp. enterica serovar Typhimurium str. DT104
Proteobacteria; Gammaproteobacteria; Enterobacterales; Enterobacteriaceae; Salmonella; Salmonella enterica; Salmonella enterica subsp. enterica; Salmonella enterica subsp. enterica serovar Typhimurium
MNNEETFYQAMRRKGVTRRSFLKFCSLAATSLGLGAGMTPKIAWALENKPRIPVVWIHGL WP_001058312.1
hydrogenase [Salmonella enterica] Length: 372\nNumber of predicted TMHs: 2\nExp number of AAs in TMHs: 36.51738\nExp number, first 60 AAs: 19.72849\nTotal prob of N-in: 0.96170\nPOSSIBLE N-term signal sequence\ninside 1 19\nTMhelix 20 39\noutside 40 325\nTMhelix 326 348\ninside 349 372

25034 GCF_001218665.1_8080_3_69 Salmonella enterica subsp. enterica serovar Typhimurium str. DT104
Proteobacteria; Gammaproteobacteria; Enterobacterales; Enterobacteriaceae; Salmonella; Salmonella enterica; Salmonella enterica subsp. enterica; Salmonella enterica subsp. enterica serovar Typhimurium
MTGDNTLITSHGINRRDFMKLCAALAATMGLSSKAAAEMAESVSNPQRPPVIWIGAQECT WP_000145429.1
hydrogenase 2 small subunit [Salmonella enterica] Length: 372\nNumber of predicted TMHs: 1\nExp number of AAs in TMHs: 24.51309\nExp number, first 60 AAs: 0.2682\nTotal prob of N-in: 0.14402\noutside 1 331\nTMhelix 332 354\ninside 355 372

25035 GCF_001218725.1_5275_3_4Salmonella enterica subsp. enterica serovar Typhimurium str. DT104
Proteobacteria; Gammaproteobacteria; Enterobacterales; Enterobacteriaceae; Salmonella; Salmonella enterica; Salmonella enterica subsp. enterica; Salmonella enterica subsp. enterica serovar Typhimurium
MTGDNTLITSHGINRRDFMKLCAALAATMGLSSKAAAEMAESVSNPQRPPVIWIGAQECT WP_000145429.1
hydrogenase 2 small subunit [Salmonella enterica] Length: 372\nNumber of predicted TMHs: 1\nExp number of AAs in TMHs: 24.51309\nExp number, first 60 AAs: 0.2682\nTotal prob of N-in: 0.14402\noutside 1 331\nTMhelix 332 354\ninside 355 372

25036 GCF_001218805.1_5285_7_3Salmonella enterica subsp. enterica serovar Typhimurium str. DT104
Proteobacteria; Gammaproteobacteria; Enterobacterales; Enterobacteriaceae; Salmonella; Salmonella enterica; Salmonella enterica subsp. enterica; Salmonella enterica subsp. enterica serovar Typhimurium
MTGDNTLITSHGINRRDFMKLCAALAATMGLSSKAAAEMAESVSNPQRPPVIWIGAQECT WP_000145429.1
hydrogenase 2 small subunit [Salmonella enterica] Length: 372\nNumber of predicted TMHs: 1\nExp number of AAs in TMHs: 24.51309\nExp number, first 60 AAs: 0.2682\nTotal prob of N-in: 0.14402\noutside 1 331\nTMhelix 332 354\ninside 355 372

25037 GCF_001218765.1_5299_3_12 Salmonella enterica subsp. enterica serovar Typhimurium str. DT104
Proteobacteria; Gammaproteobacteria; Enterobacterales; Enterobacteriaceae; Salmonella; Salmonella enterica; Salmonella enterica subsp. enterica; Salmonella enterica subsp. enterica serovar Typhimurium
MTGDNTLITSHGINRRDFMKLCAALAATMGLSSKAAAEMAESVSNPQRPPVIWIGAQECT WP_000145429.1
hydrogenase 2 small subunit [Salmonella enterica] Length: 372\nNumber of predicted TMHs: 1\nExp number of AAs in TMHs: 24.51309\nExp number, first 60 AAs: 0.2682\nTotal prob of N-in: 0.14402\noutside 1 331\nTMhelix 332 354\ninside 355 372

25038 GCF_001350095.1_8490_2_79 Salmonella enterica subsp. enterica serovar Typhimurium str. DT104
Proteobacteria; Gammaproteobacteria; Enterobacterales; Enterobacteriaceae; Salmonella; Salmonella enterica; Salmonella enterica subsp. enterica; Salmonella enterica subsp. enterica serovar Typhimurium
MNNEETFYQAMRRKGVTRRSFLKFCSLAATSLGLGAGMTPKIAWALENKPRIPVVWIHGL WP_001058312.1
hydrogenase [Salmonella enterica] Length: 372\nNumber of predicted TMHs: 2\nExp number of AAs in TMHs: 36.51738\nExp number, first 60 AAs: 19.72849\nTotal prob of N-in: 0.96170\nPOSSIBLE N-term signal sequence\ninside 1 19\nTMhelix 20 39\noutside 40 325\nTMhelix 326 348\ninside 349 372

25039 GCF_001218825.1_8080_3_14 Salmonella enterica subsp. enterica serovar Typhimurium str. DT104
Proteobacteria; Gammaproteobacteria; Enterobacterales; Enterobacteriaceae; Salmonella; Salmonella enterica; Salmonella enterica subsp. enterica; Salmonella enterica subsp. enterica serovar Typhimurium
MTGDNTLITSHGINRRDFMKLCAALAATMGLSSKAAAEMAESVSNPQRPPVIWIGAQECT WP_000145429.1
hydrogenase 2 small subunit [Salmonella enterica] Length: 372\nNumber of predicted TMHs: 1\nExp number of AAs in TMHs: 24.51309\nExp number, first 60 AAs: 0.2682\nTotal prob of N-in: 0.14402\noutside 1 331\nTMhelix 332 354\ninside 355 372

25040 GCF_001218885.1_8080_3_82 Salmonella enterica subsp. enterica serovar Typhimurium str. DT104 Proteobacteria; Gammaproteobacteria; Enterobacterales; Enterobacteriaceae; Salmonella; Salmonella enterica; Salmonella enterica subsp. enterica; Salmonella enterica subsp. enterica serovar Typhimurium MTGDNTLITSHGINRRDFMKLCAALAATMGLSSKAAAEMAESVSNPQRPPVIWIGAQECT WP_000145429.1 hydrogenase 2 small subunit [Salmonella enterica] Length: 372\nNumber of predicted TMHs: 1\nExp number of AAs in TMHs: 24.51309\nExp number, first 60 AAs: 0.2682\nTotal prob of N-in: 0.14402\noutside 1 331\nTMhelix 332 354\ninside 355 372

25041 GCF_001218845.1_5275_5_7Salmonella enterica subsp. enterica serovar Typhimurium str. DT104 Proteobacteria; Gammaproteobacteria; Enterobacterales; Enterobacteriaceae; Salmonella; Salmonella enterica; Salmonella enterica subsp. enterica; Salmonella enterica subsp. enterica serovar Typhimurium MTGDNTLITSHGINRRDFMKLCAALAATMGLSSKAAAEMAESVSNPQRPPVIWIGAQECT WP_000145429.1 hydrogenase 2 small subunit [Salmonella enterica] Length: 372\nNumber of predicted TMHs: 1\nExp number of AAs in TMHs: 24.51309\nExp number, first 60 AAs: 0.2682\nTotal prob of N-in: 0.14402\noutside 1 331\nTMhelix 332 354\ninside 355 372

25042 GCF_001218965.1_5275_5_12 Salmonella enterica subsp. enterica serovar Typhimurium str. DT104 Proteobacteria; Gammaproteobacteria; Enterobacterales; Enterobacteriaceae; Salmonella; Salmonella enterica; Salmonella enterica subsp. enterica; Salmonella enterica subsp. enterica serovar Typhimurium MTGDNTLITSHGINRRDFMKLCAALAATMGLSSKAAAEMAESVSNPQRPPVIWIGAQECT WP_000145429.1 hydrogenase 2 small subunit [Salmonella enterica] Length: 372\nNumber of predicted TMHs: 1\nExp number of AAs in TMHs: 24.51309\nExp number, first 60 AAs: 0.2682\nTotal prob of N-in: 0.14402\noutside 1 331\nTMhelix 332 354\ninside 355 372

25043 GCF_001218985.1_8036_3_57 Salmonella enterica subsp. enterica serovar Typhimurium str. DT104 Proteobacteria; Gammaproteobacteria; Enterobacterales; Enterobacteriaceae; Salmonella; Salmonella enterica; Salmonella enterica subsp. enterica; Salmonella enterica subsp. enterica serovar Typhimurium MTGDNTLITSHGINRRDFMKLCAALAATMGLSSKAAAEMAESVSNPQRPPVIWIGAQECT WP_000145429.1 hydrogenase 2 small subunit [Salmonella enterica] Length: 372\nNumber of predicted TMHs: 1\nExp number of AAs in TMHs: 24.51309\nExp number, first 60 AAs: 0.2682\nTotal prob of N-in: 0.14402\noutside 1 331\nTMhelix 332 354\ninside 355 372

25044 GCF_001219025.1_8080_3_29 Salmonella enterica subsp. enterica serovar Typhimurium str. DT104 Proteobacteria; Gammaproteobacteria; Enterobacterales; Enterobacteriaceae; Salmonella; Salmonella enterica; Salmonella enterica subsp. enterica; Salmonella enterica subsp. enterica serovar Typhimurium MTGDNTLITSHGINRRDFMKLCAALAATMGLSSKAAAEMAESVSNPQRPPVIWIGAQECT WP_000145429.1 hydrogenase 2 small subunit [Salmonella enterica] Length: 372\nNumber of predicted TMHs: 1\nExp number of AAs in TMHs: 24.51309\nExp number, first 60 AAs: 0.2682\nTotal prob of N-in: 0.14402\noutside 1 331\nTMhelix 332 354\ninside 355 372

25045 GCF_001219065.1_5285_7_9Salmonella enterica subsp. enterica serovar Typhimurium str. DT104 Proteobacteria; Gammaproteobacteria; Enterobacterales; Enterobacteriaceae; Salmonella; Salmonella enterica; Salmonella enterica subsp. enterica; Salmonella enterica subsp. enterica serovar Typhimurium MTGDNTLITSHGINRRDFMKLCAALAATMGLSSKAAAEMAESVSNPQRPPVIWIGAQECT WP_000145429.1 hydrogenase 2 small subunit [Salmonella enterica] Length: 372\nNumber of predicted TMHs: 1\nExp number of AAs in TMHs: 24.51309\nExp number, first 60 AAs: 0.2682\nTotal prob of N-in: 0.14402\noutside 1 331\nTMhelix 332 354\ninside 355 372

25046 GCF_001218625.1_5299_6_2Salmonella enterica subsp. enterica serovar Typhimurium str. DT104 Proteobacteria; Gammaproteobacteria; Enterobacterales; Enterobacteriaceae; Salmonella; Salmonella enterica; Salmonella enterica subsp. enterica; Salmonella enterica subsp. enterica serovar Typhimurium MNNEETFYQAMRRKGVTRRSFLKFCSLAATSLGLGAGMTPKIAWALENKPRIPVVWIHGL WP_001058312.1 hydrogenase [Salmonella enterica] Length: 372\nNumber of predicted TMHs: 2\nExp number of AAs in TMHs: 36.51738\nExp number, first 60 AAs: 19.72849\nTotal prob of N-in: 0.96170\nPOSSIBLE N-term signal sequence\ninside 1 19\nTMhelix 20 39\noutside 40 325\nTMhelix 326 348\ninside 349 372

25047 GCF_001326135.1_8080_3_62 Salmonella enterica subsp. enterica serovar Typhimurium str. DT104 Proteobacteria; Gammaproteobacteria; Enterobacterales; Enterobacteriaceae; Salmonella; Salmonella enterica; Salmonella enterica subsp. enterica; Salmonella enterica subsp. enterica serovar Typhimurium MTGDNTLITSHGINRRDFMKLCAALAATMGLSSKAAAEMAESVSNPQRPPVIWIGAQECT WP_000145429.1 hydrogenase 2 small subunit [Salmonella enterica] Length: 372\nNumber of predicted TMHs: 1\nExp

number of AAs in TMHs: 24.51309\nExp number, first 60 AAs: 0.2682\nTotal prob of N-in: 0.14402\noutside 1 331\nTMhelix 332 354\ninside 355 372

25048 GCF_001218665.1_8080_3_69 Salmonella enterica subsp. enterica serovar Typhimurium str. DT104 Proteobacteria; Gammaproteobacteria; Enterobacterales; Enterobacteriaceae; Salmonella; Salmonella enterica; Salmonella enterica subsp. enterica; Salmonella enterica subsp. enterica serovar Typhimurium MNNEETFYQAMRRKGVTRRSFLKFCSLAATSLGLGAGMTPKIAWALENKPRIPVVWIHGL WP_001058312.1 hydrogenase [Salmonella enterica] Length: 372\nNumber of predicted TMHs: 2\nExp number of AAs in TMHs: 36.51738\nExp number, first 60 AAs: 19.72849\nTotal prob of N-in: 0.96170\nPOSSIBLE N-term signal sequence\ninside 1 19\nTMhelix 20 39\noutside 40 325\nTMhelix 326 348\ninside 349 372

25049 GCF_001218725.1_5275_3_4Salmonella enterica subsp. enterica serovar Typhimurium str. DT104 Proteobacteria; Gammaproteobacteria; Enterobacterales; Enterobacteriaceae; Salmonella; Salmonella enterica; Salmonella enterica subsp. enterica; Salmonella enterica subsp. enterica serovar Typhimurium MNNEETFYQAMRRKGVTRRSFLKFCSLAATSLGLGAGMTPKIAWALENKPRIPVVWIHGL WP_001058312.1 hydrogenase [Salmonella enterica] Length: 372\nNumber of predicted TMHs: 2\nExp number of AAs in TMHs: 36.51738\nExp number, first 60 AAs: 19.72849\nTotal prob of N-in: 0.96170\nPOSSIBLE N-term signal sequence\ninside 1 19\nTMhelix 20 39\noutside 40 325\nTMhelix 326 348\ninside 349 372

25050 GCF_001218805.1_5285_7_3Salmonella enterica subsp. enterica serovar Typhimurium str. DT104 Proteobacteria; Gammaproteobacteria; Enterobacterales; Enterobacteriaceae; Salmonella; Salmonella enterica; Salmonella enterica subsp. enterica; Salmonella enterica subsp. enterica serovar Typhimurium MNNEETFYQAMRRKGVTRRSFLKFCSLAATSLGLGAGMTPKIAWALENKPRIPVVWIHGL WP_001058312.1 hydrogenase [Salmonella enterica] Length: 372\nNumber of predicted TMHs: 2\nExp number of AAs in TMHs: 36.51738\nExp number, first 60 AAs: 19.72849\nTotal prob of N-in: 0.96170\nPOSSIBLE N-term signal sequence\ninside 1 19\nTMhelix 20 39\noutside 40 325\nTMhelix 326 348\ninside 349 372

25051 GCF_001218825.1_8080_3_14 Salmonella enterica subsp. enterica serovar Typhimurium str. DT104 Proteobacteria; Gammaproteobacteria; Enterobacterales; Enterobacteriaceae; Salmonella; Salmonella enterica; Salmonella enterica subsp. enterica; Salmonella enterica subsp. enterica serovar Typhimurium MNNEETFYQAMRRKGVTRRSFLKFCSLAATSLGLGAGMTPKIAWALENKPRIPVVWIHGL WP_001058312.1 hydrogenase [Salmonella enterica] Length: 372\nNumber of predicted TMHs: 2\nExp number of AAs in TMHs: 36.51738\nExp number, first 60 AAs: 19.72849\nTotal prob of N-in: 0.96170\nPOSSIBLE N-term signal sequence\ninside 1 19\nTMhelix 20 39\noutside 40 325\nTMhelix 326 348\ninside 349 372

25052 GCF_001218765.1_5299_3_12 Salmonella enterica subsp. enterica serovar Typhimurium str. DT104 Proteobacteria; Gammaproteobacteria; Enterobacterales; Enterobacteriaceae; Salmonella; Salmonella enterica; Salmonella enterica subsp. enterica; Salmonella enterica subsp. enterica serovar Typhimurium MNNEETFYQAMRRKGVTRRSFLKFCSLAATSLGLGAGMTPKIAWALENKPRIPVVWIHGL WP_001058312.1 hydrogenase [Salmonella enterica] Length: 372\nNumber of predicted TMHs: 2\nExp number of AAs in TMHs: 36.51738\nExp number, first 60 AAs: 19.72849\nTotal prob of N-in: 0.96170\nPOSSIBLE N-term signal sequence\ninside 1 19\nTMhelix 20 39\noutside 40 325\nTMhelix 326 348\ninside 349 372

25053 GCF_001218885.1_8080_3_82 Salmonella enterica subsp. enterica serovar Typhimurium str. DT104 Proteobacteria; Gammaproteobacteria; Enterobacterales; Enterobacteriaceae; Salmonella; Salmonella enterica; Salmonella enterica subsp. enterica; Salmonella enterica subsp. enterica serovar Typhimurium MNNEETFYQAMRRKGVTRRSFLKFCSLAATSLGLGAGMTPKIAWALENKPRIPVVWIHGL WP_001058312.1 hydrogenase [Salmonella enterica] Length: 372\nNumber of predicted TMHs: 2\nExp number of AAs in TMHs: 36.51738\nExp number, first 60 AAs: 19.72849\nTotal prob of N-in: 0.96170\nPOSSIBLE N-term signal sequence\ninside 1 19\nTMhelix 20 39\noutside 40 325\nTMhelix 326 348\ninside 349 372

25054 GCF_001218845.1_5275_5_7Salmonella enterica subsp. enterica serovar Typhimurium str. DT104 Proteobacteria; Gammaproteobacteria; Enterobacterales; Enterobacteriaceae; Salmonella; Salmonella enterica; Salmonella enterica subsp. enterica; Salmonella enterica subsp. enterica serovar Typhimurium MNNEETFYQAMRRKGVTRRSFLKFCSLAATSLGLGAGMTPKIAWALENKPRIPVVWIHGL WP_001058312.1 hydrogenase [Salmonella enterica] Length: 372\nNumber of predicted TMHs: 2\nExp number of AAs in TMHs: 36.51738\nExp number, first 60 AAs: 19.72849\nTotal prob of N-in: 0.96170\nPOSSIBLE N-term signal sequence\ninside 1 19\nTMhelix 20 39\noutside 40 325\nTMhelix 326 348\ninside 349 372

25055 GCF_001218965.1_5275_5_12 Salmonella enterica subsp. enterica serovar Typhimurium str. DT104 Proteobacteria; Gammaproteobacteria; Enterobacterales; Enterobacteriaceae; Salmonella; Salmonella enterica;

Salmonella enterica subsp. enterica; Salmonella enterica subsp. enterica serovar Typhimurium
MNNEETFYQAMRRKGVTRRSFLKFCSLAATSLGLGAGMTPKIAWALENKPRIPVVWIHGL WP_001058312.1
hydrogenase [Salmonella enterica] Length: 372\nNumber of predicted TMHs: 2\nExp number of AAs in TMHs: 36.51738\nExp number, first 60 AAs: 19.72849\nTotal prob of N-in: 0.96170\nPOSSIBLE N-term signal sequence\ninside 1 19\nTMhelix 20 39\noutside 40 325\nTMhelix 326 348\ninside 349 372

25056 GCF_001218985.1_8036_3_57 Salmonella enterica subsp. enterica serovar Typhimurium str. DT104
Proteobacteria; Gammaproteobacteria; Enterobacterales; Enterobacteriaceae; Salmonella; Salmonella enterica; Salmonella enterica subsp. enterica; Salmonella enterica subsp. enterica serovar Typhimurium
MNNEETFYQAMRRKGVTRRSFLKFCSLAATSLGLGAGMTPKIAWALENKPRIPVVWIHGL WP_001058312.1
hydrogenase [Salmonella enterica] Length: 372\nNumber of predicted TMHs: 2\nExp number of AAs in TMHs: 36.51738\nExp number, first 60 AAs: 19.72849\nTotal prob of N-in: 0.96170\nPOSSIBLE N-term signal sequence\ninside 1 19\nTMhelix 20 39\noutside 40 325\nTMhelix 326 348\ninside 349 372

25057 GCF_001219025.1_8080_3_29 Salmonella enterica subsp. enterica serovar Typhimurium str. DT104
Proteobacteria; Gammaproteobacteria; Enterobacterales; Enterobacteriaceae; Salmonella; Salmonella enterica; Salmonella enterica subsp. enterica; Salmonella enterica subsp. enterica serovar Typhimurium
MNNEETFYQAMRRKGVTRRSFLKFCSLAATSLGLGAGMTPKIAWALENKPRIPVVWIHGL WP_001058312.1
hydrogenase [Salmonella enterica] Length: 372\nNumber of predicted TMHs: 2\nExp number of AAs in TMHs: 36.51738\nExp number, first 60 AAs: 19.72849\nTotal prob of N-in: 0.96170\nPOSSIBLE N-term signal sequence\ninside 1 19\nTMhelix 20 39\noutside 40 325\nTMhelix 326 348\ninside 349 372

25058 GCF_001219065.1_5285_7_9Salmonella enterica subsp. enterica serovar Typhimurium str. DT104
Proteobacteria; Gammaproteobacteria; Enterobacterales; Enterobacteriaceae; Salmonella; Salmonella enterica; Salmonella enterica subsp. enterica; Salmonella enterica subsp. enterica serovar Typhimurium
MNNEETFYQAMRRKGVTRRSFLKFCSLAATSLGLGAGMTPKIAWALENKPRIPVVWIHGL WP_001058312.1
hydrogenase [Salmonella enterica] Length: 372\nNumber of predicted TMHs: 2\nExp number of AAs in TMHs: 36.51738\nExp number, first 60 AAs: 19.72849\nTotal prob of N-in: 0.96170\nPOSSIBLE N-term signal sequence\ninside 1 19\nTMhelix 20 39\noutside 40 325\nTMhelix 326 348\ninside 349 372

25059 GCF_001219105.1_5285_7_10 Salmonella enterica subsp. enterica serovar Typhimurium str. DT104
Proteobacteria; Gammaproteobacteria; Enterobacterales; Enterobacteriaceae; Salmonella; Salmonella enterica; Salmonella enterica subsp. enterica; Salmonella enterica subsp. enterica serovar Typhimurium
MTGDNTLITSHGINRRDFMKLCAALAATMGLSSKAAAEMAESVSNPQRPPVIWIGAQECT WP_000145429.1
hydrogenase 2 small subunit [Salmonella enterica] Length: 372\nNumber of predicted TMHs: 1\nExp number of AAs in TMHs: 24.51309\nExp number, first 60 AAs: 0.2682\nTotal prob of N-in: 0.14402\noutside 1 331\nTMhelix 332 354\ninside 355 372

25060 GCF_001219145.1_5295_2_4Salmonella enterica subsp. enterica serovar Typhimurium str. DT104
Proteobacteria; Gammaproteobacteria; Enterobacterales; Enterobacteriaceae; Salmonella; Salmonella enterica; Salmonella enterica subsp. enterica; Salmonella enterica subsp. enterica serovar Typhimurium
MTGDNTLITSHGINRRDFMKLCAALAATMGLSSKAAAEMAESVSNPQRPPVIWIGAQECT WP_000145429.1
hydrogenase 2 small subunit [Salmonella enterica] Length: 372\nNumber of predicted TMHs: 1\nExp number of AAs in TMHs: 24.51309\nExp number, first 60 AAs: 0.2682\nTotal prob of N-in: 0.14402\noutside 1 331\nTMhelix 332 354\ninside 355 372

25061 GCF_001219165.1_5285_7_8Salmonella enterica subsp. enterica serovar Typhimurium str. DT104
Proteobacteria; Gammaproteobacteria; Enterobacterales; Enterobacteriaceae; Salmonella; Salmonella enterica; Salmonella enterica subsp. enterica; Salmonella enterica subsp. enterica serovar Typhimurium
MTGDNTLITSHGINRRDFMKLCAALAATMGLSSKAAAEMAESVSNPQRPPVIWIGAQECT WP_000145429.1
hydrogenase 2 small subunit [Salmonella enterica] Length: 372\nNumber of predicted TMHs: 1\nExp number of AAs in TMHs: 24.51309\nExp number, first 60 AAs: 0.2682\nTotal prob of N-in: 0.14402\noutside 1 331\nTMhelix 332 354\ninside 355 372

25062 GCF_001219205.1_8036_3_48 Salmonella enterica subsp. enterica serovar Typhimurium str. DT104
Proteobacteria; Gammaproteobacteria; Enterobacterales; Enterobacteriaceae; Salmonella; Salmonella enterica; Salmonella enterica subsp. enterica; Salmonella enterica subsp. enterica serovar Typhimurium
MTGDNTLITSHGINRRDFMKLCAALAATMGLSSKAAAEMAESVSNPQRPPVIWIGAQECT WP_000145429.1
hydrogenase 2 small subunit [Salmonella enterica] Length: 372\nNumber of predicted TMHs: 1\nExp number of AAs in TMHs: 24.51309\nExp number, first 60 AAs: 0.2682\nTotal prob of N-in: 0.14402\noutside 1 331\nTMhelix 332 354\ninside 355 372

25063 GCF_001219265.1_5275_2_5Salmonella enterica subsp. enterica serovar Typhimurium str. DT104 Proteobacteria; Gammaproteobacteria; Enterobacterales; Enterobacteriaceae; Salmonella; Salmonella enterica; Salmonella enterica subsp. enterica; Salmonella enterica subsp. enterica serovar Typhimurium MTGDNTLITSHGINRRDFMKLCAALAATMGLSSKAAAEMAESVSNPQRPPVIWIGAQECT WP_000145429.1 hydrogenase 2 small subunit [Salmonella enterica] Length: 372\nNumber of predicted TMHs: 1\nExp number of AAs in TMHs: 24.51309\nExp number, first 60 AAs: 0.2682\nTotal prob of N-in: 0.14402\noutside 1 331\nTMhelix 332 354\ninside 355 372

25064 GCF_001326175.1_9066_1_9Salmonella enterica subsp. enterica serovar Typhimurium str. DT104 Proteobacteria; Gammaproteobacteria; Enterobacterales; Enterobacteriaceae; Salmonella; Salmonella enterica; Salmonella enterica subsp. enterica; Salmonella enterica subsp. enterica serovar Typhimurium MTGDNTLITSHGINRRDFMKLCAALAATMGLSSKAAAEMAESVSNPQRPPVIWIGAQECT WP_000145429.1 hydrogenase 2 small subunit [Salmonella enterica] Length: 372\nNumber of predicted TMHs: 1\nExp number of AAs in TMHs: 24.51309\nExp number, first 60 AAs: 0.2682\nTotal prob of N-in: 0.14402\noutside 1 331\nTMhelix 332 354\ninside 355 372

25065 GCF_001219305.1_5275_7_4Salmonella enterica subsp. enterica serovar Typhimurium str. DT104 Proteobacteria; Gammaproteobacteria; Enterobacterales; Enterobacteriaceae; Salmonella; Salmonella enterica; Salmonella enterica subsp. enterica; Salmonella enterica subsp. enterica serovar Typhimurium MTGDNTLITSHGINRRDFMKLCAALAATMGLSSKAAAEMAESVSNPQRPPVIWIGAQECT WP_000145429.1 hydrogenase 2 small subunit [Salmonella enterica] Length: 372\nNumber of predicted TMHs: 1\nExp number of AAs in TMHs: 24.51309\nExp number, first 60 AAs: 0.2682\nTotal prob of N-in: 0.14402\noutside 1 331\nTMhelix 332 354\ninside 355 372

25066 GCF_001219405.1_5275_8_7Salmonella enterica subsp. enterica serovar Typhimurium str. DT104 Proteobacteria; Gammaproteobacteria; Enterobacterales; Enterobacteriaceae; Salmonella; Salmonella enterica; Salmonella enterica subsp. enterica; Salmonella enterica subsp. enterica serovar Typhimurium MTGDNTLITSHGINRRDFMKLCAALAATMGLSSKAAAEMAESVSNPQRPPVIWIGAQECT WP_000145429.1 hydrogenase 2 small subunit [Salmonella enterica] Length: 372\nNumber of predicted TMHs: 1\nExp number of AAs in TMHs: 24.51309\nExp number, first 60 AAs: 0.2682\nTotal prob of N-in: 0.14402\noutside 1 331\nTMhelix 332 354\ninside 355 372

25067 GCF_001219425.1_5275_6_5Salmonella enterica subsp. enterica serovar Typhimurium str. DT104 Proteobacteria; Gammaproteobacteria; Enterobacterales; Enterobacteriaceae; Salmonella; Salmonella enterica; Salmonella enterica subsp. enterica; Salmonella enterica subsp. enterica serovar Typhimurium MTGDNTLITSHGINRRDFMKLCAALAATMGLSSKAAAEMAESVSNPQRPPVIWIGAQECT WP_000145429.1 hydrogenase 2 small subunit [Salmonella enterica] Length: 372\nNumber of predicted TMHs: 1\nExp number of AAs in TMHs: 24.51309\nExp number, first 60 AAs: 0.2682\nTotal prob of N-in: 0.14402\noutside 1 331\nTMhelix 332 354\ninside 355 372

25068 GCF_001219505.1_5299_3_1Salmonella enterica subsp. enterica serovar Typhimurium str. DT104 Proteobacteria; Gammaproteobacteria; Enterobacterales; Enterobacteriaceae; Salmonella; Salmonella enterica; Salmonella enterica subsp. enterica; Salmonella enterica subsp. enterica serovar Typhimurium MTGDNTLITSHGINRRDFMKLCAALAATMGLSSKAAAEMAESVSNPQRPPVIWIGAQECT WP_000145429.1 hydrogenase 2 small subunit [Salmonella enterica] Length: 372\nNumber of predicted TMHs: 1\nExp number of AAs in TMHs: 24.51309\nExp number, first 60 AAs: 0.2682\nTotal prob of N-in: 0.14402\noutside 1 331\nTMhelix 332 354\ninside 355 372

25069 GCF_001219465.1_8080_3_92 Salmonella enterica subsp. enterica serovar Typhimurium str. DT104 Proteobacteria; Gammaproteobacteria; Enterobacterales; Enterobacteriaceae; Salmonella; Salmonella enterica; Salmonella enterica subsp. enterica; Salmonella enterica subsp. enterica serovar Typhimurium MTGDNTLITSHGINRRDFMKLCAALAATMGLSSKAAAEMAESVSNPQRPPVIWIGAQECT WP_000145429.1 hydrogenase 2 small subunit [Salmonella enterica] Length: 372\nNumber of predicted TMHs: 1\nExp number of AAs in TMHs: 24.51309\nExp number, first 60 AAs: 0.2682\nTotal prob of N-in: 0.14402\noutside 1 331\nTMhelix 332 354\ninside 355 372

25070 GCF_001219405.1_5275_8_7Salmonella enterica subsp. enterica serovar Typhimurium str. DT104 Proteobacteria; Gammaproteobacteria; Enterobacterales; Enterobacteriaceae; Salmonella; Salmonella enterica; Salmonella enterica subsp. enterica; Salmonella enterica subsp. enterica serovar Typhimurium MNNEETFYQAMRRKGVTRRSFLKFCSLAATSLGLGAGMTPKIAWALENKPRIPVVWIHGL WP_001058312.1 hydrogenase [Salmonella enterica] Length: 372\nNumber of predicted TMHs: 2\nExp number of AAs in

TMHs: 36.51738\nExp number, first 60 AAs: 19.72849\nTotal prob of N-in: 0.96170\nPOSSIBLE N-term signal sequence\ninside 1 19\nTMhelix 20 39\noutside 40 325\nTMhelix 326 348\ninside 349 372

25071 GCF_001326155.1_9066_1_10 Salmonella enterica subsp. enterica serovar Typhimurium str. DT104 Proteobacteria; Gammaproteobacteria; Enterobacterales; Enterobacteriaceae; Salmonella; Salmonella enterica; Salmonella enterica subsp. enterica; Salmonella enterica subsp. enterica serovar Typhimurium MTGDNTLITSHGINRRDFMKLCAALAATMGLSSKAAAEMAESVSNPQRPPVIWIGAQECT WP_000145429.1 hydrogenase 2 small subunit [Salmonella enterica] Length: 372\nNumber of predicted TMHs: 1\nExp number of AAs in TMHs: 24.51309\nExp number, first 60 AAs: 0.2682\nTotal prob of N-in: 0.14402\noutside 1 331\nTMhelix 332 354\ninside 355 372

25072 GCF_001219145.1_5295_2_4Salmonella enterica subsp. enterica serovar Typhimurium str. DT104 Proteobacteria; Gammaproteobacteria; Enterobacterales; Enterobacteriaceae; Salmonella; Salmonella enterica; Salmonella enterica subsp. enterica; Salmonella enterica subsp. enterica serovar Typhimurium MNNEETFYQAMRRKGVTRRSFLKFCSLAATSLGLGAGMTPKIAWALENKPRIPVVWIHGL WP_001058312.1 hydrogenase [Salmonella enterica] Length: 372\nNumber of predicted TMHs: 2\nExp number of AAs in TMHs: 36.51738\nExp number, first 60 AAs: 19.72849\nTotal prob of N-in: 0.96170\nPOSSIBLE N-term signal sequence\ninside 1 19\nTMhelix 20 39\noutside 40 325\nTMhelix 326 348\ninside 349 372

25073 GCF_001216545.1_5285_6_8Salmonella enterica subsp. enterica serovar Typhimurium str. DT104 Proteobacteria; Gammaproteobacteria; Enterobacterales; Enterobacteriaceae; Salmonella; Salmonella enterica; Salmonella enterica subsp. enterica; Salmonella enterica subsp. enterica serovar Typhimurium MNNEETFYQAMRRKGVTRRSFLKFCSLAATSLGLGAGMTPKIAWALENKPRIPVVWIHGL WP_001058312.1 hydrogenase [Salmonella enterica] Length: 372\nNumber of predicted TMHs: 2\nExp number of AAs in TMHs: 36.51738\nExp number, first 60 AAs: 19.72849\nTotal prob of N-in: 0.96170\nPOSSIBLE N-term signal sequence\ninside 1 19\nTMhelix 20 39\noutside 40 325\nTMhelix 326 348\ninside 349 372

25074 GCF_001219545.1_5275_3_7Salmonella enterica subsp. enterica serovar Typhimurium str. DT104 Proteobacteria; Gammaproteobacteria; Enterobacterales; Enterobacteriaceae; Salmonella; Salmonella enterica; Salmonella enterica subsp. enterica; Salmonella enterica subsp. enterica serovar Typhimurium MTGDNTLITSHGINRRDFMKLCAALAATMGLSSKAAAEMAESVSNPQRPPVIWIGAQECT WP_000145429.1 hydrogenase 2 small subunit [Salmonella enterica] Length: 372\nNumber of predicted TMHs: 1\nExp number of AAs in TMHs: 24.51309\nExp number, first 60 AAs: 0.2682\nTotal prob of N-in: 0.14402\noutside 1 331\nTMhelix 332 354\ninside 355 372

25075 GCF_001219585.1_8080_3_5Salmonella enterica subsp. enterica serovar Typhimurium str. DT104 Proteobacteria; Gammaproteobacteria; Enterobacterales; Enterobacteriaceae; Salmonella; Salmonella enterica; Salmonella enterica subsp. enterica; Salmonella enterica subsp. enterica serovar Typhimurium MTGDNTLITSHGINRRDFMKLCAALAATMGLSSKAAAEMAESVSNPQRPPVIWIGAQECT WP_000145429.1 hydrogenase 2 small subunit [Salmonella enterica] Length: 372\nNumber of predicted TMHs: 1\nExp number of AAs in TMHs: 24.51309\nExp number, first 60 AAs: 0.2682\nTotal prob of N-in: 0.14402\noutside 1 331\nTMhelix 332 354\ninside 355 372

25076 GCF_001219585.1_8080_3_5Salmonella enterica subsp. enterica serovar Typhimurium str. DT104 Proteobacteria; Gammaproteobacteria; Enterobacterales; Enterobacteriaceae; Salmonella; Salmonella enterica; Salmonella enterica subsp. enterica; Salmonella enterica subsp. enterica serovar Typhimurium MNNEETFYQAMRRKGVTRRSFLKFCSLAATSLGLGAGMTPKIAWALENKPRIPVVWIHGL WP_001058312.1 hydrogenase [Salmonella enterica] Length: 372\nNumber of predicted TMHs: 2\nExp number of AAs in TMHs: 36.51738\nExp number, first 60 AAs: 19.72849\nTotal prob of N-in: 0.96170\nPOSSIBLE N-term signal sequence\ninside 1 19\nTMhelix 20 39\noutside 40 325\nTMhelix 326 348\ninside 349 372

25077 GCF_001219545.1_5275_3_7Salmonella enterica subsp. enterica serovar Typhimurium str. DT104 Proteobacteria; Gammaproteobacteria; Enterobacterales; Enterobacteriaceae; Salmonella; Salmonella enterica; Salmonella enterica subsp. enterica; Salmonella enterica subsp. enterica serovar Typhimurium MNNEETFYQAMRRKGVTRRSFLKFCSLAATSLGLGAGMTPKIAWALENKPRIPVVWIHGL WP_001058312.1 hydrogenase [Salmonella enterica] Length: 372\nNumber of predicted TMHs: 2\nExp number of AAs in TMHs: 36.51738\nExp number, first 60 AAs: 19.72849\nTotal prob of N-in: 0.96170\nPOSSIBLE N-term signal sequence\ninside 1 19\nTMhelix 20 39\noutside 40 325\nTMhelix 326 348\ninside 349 372

25078 GCF_001219105.1_5285_7_10 Salmonella enterica subsp. enterica serovar Typhimurium str. DT104 Proteobacteria; Gammaproteobacteria; Enterobacterales; Enterobacteriaceae; Salmonella; Salmonella enterica;

Salmonella enterica subsp. enterica; Salmonella enterica subsp. enterica serovar Typhimurium
MNNEETFYQAMRRKGVTRRSFLKFCSLAATSLGLGAGMTPKIAWALENKPRIPVVWIHGL WP_001058312.1 hydrogenase [Salmonella enterica] Length: 372\nNumber of predicted TMHs: 2\nExp number of AAs in TMHs: 36.51738\nExp number, first 60 AAs: 19.72849\nTotal prob of N-in: 0.96170\nPOSSIBLE N-term signal sequence\ninside 1 19\nTMhelix 20 39\noutside 40 325\nTMhelix 326 348\ninside 349 372

25079 GCF_001219165.1_5285_7_8Salmonella enterica subsp. enterica serovar Typhimurium str. DT104 Proteobacteria; Gammaproteobacteria; Enterobacterales; Enterobacteriaceae; Salmonella; Salmonella enterica; Salmonella enterica subsp. enterica; Salmonella enterica subsp. enterica serovar Typhimurium
MNNEETFYQAMRRKGVTRRSFLKFCSLAATSLGLGAGMTPKIAWALENKPRIPVVWIHGL WP_001058312.1 hydrogenase [Salmonella enterica] Length: 372\nNumber of predicted TMHs: 2\nExp number of AAs in TMHs: 36.51738\nExp number, first 60 AAs: 19.72849\nTotal prob of N-in: 0.96170\nPOSSIBLE N-term signal sequence\ninside 1 19\nTMhelix 20 39\noutside 40 325\nTMhelix 326 348\ninside 349 372

25080 GCF_001219205.1_8036_3_48 Salmonella enterica subsp. enterica serovar Typhimurium str. DT104 Proteobacteria; Gammaproteobacteria; Enterobacterales; Enterobacteriaceae; Salmonella; Salmonella enterica; Salmonella enterica subsp. enterica; Salmonella enterica subsp. enterica serovar Typhimurium
MNNEETFYQAMRRKGVTRRSFLKFCSLAATSLGLGAGMTPKIAWALENKPRIPVVWIHGL WP_001058312.1 hydrogenase [Salmonella enterica] Length: 372\nNumber of predicted TMHs: 2\nExp number of AAs in TMHs: 36.51738\nExp number, first 60 AAs: 19.72849\nTotal prob of N-in: 0.96170\nPOSSIBLE N-term signal sequence\ninside 1 19\nTMhelix 20 39\noutside 40 325\nTMhelix 326 348\ninside 349 372

25081 GCF_001219265.1_5275_2_5Salmonella enterica subsp. enterica serovar Typhimurium str. DT104 Proteobacteria; Gammaproteobacteria; Enterobacterales; Enterobacteriaceae; Salmonella; Salmonella enterica; Salmonella enterica subsp. enterica; Salmonella enterica subsp. enterica serovar Typhimurium
MNNEETFYQAMRRKGVTRRSFLKFCSLAATSLGLGAGMTPKIAWALENKPRIPVVWIHGL WP_001058312.1 hydrogenase [Salmonella enterica] Length: 372\nNumber of predicted TMHs: 2\nExp number of AAs in TMHs: 36.51738\nExp number, first 60 AAs: 19.72849\nTotal prob of N-in: 0.96170\nPOSSIBLE N-term signal sequence\ninside 1 19\nTMhelix 20 39\noutside 40 325\nTMhelix 326 348\ninside 349 372

25082 GCF_001219305.1_5275_7_4Salmonella enterica subsp. enterica serovar Typhimurium str. DT104 Proteobacteria; Gammaproteobacteria; Enterobacterales; Enterobacteriaceae; Salmonella; Salmonella enterica; Salmonella enterica subsp. enterica; Salmonella enterica subsp. enterica serovar Typhimurium
MNNEETFYQAMRRKGVTRRSFLKFCSLAATSLGLGAGMTPKIAWALENKPRIPVVWIHGL WP_001058312.1 hydrogenase [Salmonella enterica] Length: 372\nNumber of predicted TMHs: 2\nExp number of AAs in TMHs: 36.51738\nExp number, first 60 AAs: 19.72849\nTotal prob of N-in: 0.96170\nPOSSIBLE N-term signal sequence\ninside 1 19\nTMhelix 20 39\noutside 40 325\nTMhelix 326 348\ninside 349 372

25083 GCF_001219425.1_5275_6_5Salmonella enterica subsp. enterica serovar Typhimurium str. DT104 Proteobacteria; Gammaproteobacteria; Enterobacterales; Enterobacteriaceae; Salmonella; Salmonella enterica; Salmonella enterica subsp. enterica; Salmonella enterica subsp. enterica serovar Typhimurium
MNNEETFYQAMRRKGVTRRSFLKFCSLAATSLGLGAGMTPKIAWALENKPRIPVVWIHGL WP_001058312.1 hydrogenase [Salmonella enterica] Length: 372\nNumber of predicted TMHs: 2\nExp number of AAs in TMHs: 36.51738\nExp number, first 60 AAs: 19.72849\nTotal prob of N-in: 0.96170\nPOSSIBLE N-term signal sequence\ninside 1 19\nTMhelix 20 39\noutside 40 325\nTMhelix 326 348\ninside 349 372

25084 GCF_001219465.1_8080_3_92 Salmonella enterica subsp. enterica serovar Typhimurium str. DT104 Proteobacteria; Gammaproteobacteria; Enterobacterales; Enterobacteriaceae; Salmonella; Salmonella enterica; Salmonella enterica subsp. enterica; Salmonella enterica subsp. enterica serovar Typhimurium
MNNEETFYQAMRRKGVTRRSFLKFCSLAATSLGLGAGMTPKIAWALENKPRIPVVWIHGL WP_001058312.1 hydrogenase [Salmonella enterica] Length: 372\nNumber of predicted TMHs: 2\nExp number of AAs in TMHs: 36.51738\nExp number, first 60 AAs: 19.72849\nTotal prob of N-in: 0.96170\nPOSSIBLE N-term signal sequence\ninside 1 19\nTMhelix 20 39\noutside 40 325\nTMhelix 326 348\ninside 349 372

25085 GCF_001219505.1_5299_3_1Salmonella enterica subsp. enterica serovar Typhimurium str. DT104 Proteobacteria; Gammaproteobacteria; Enterobacterales; Enterobacteriaceae; Salmonella; Salmonella enterica; Salmonella enterica subsp. enterica; Salmonella enterica subsp. enterica serovar Typhimurium
MNNEETFYQAMRRKGVTRRSFLKFCSLAATSLGLGAGMTPKIAWALENKPRIPVVWIHGL WP_050954650.1 hydrogenase [Salmonella enterica] Length: 372\nNumber of predicted TMHs: 2\nExp number of AAs in TMHs: 36.4666\nExp number, first 60 AAs: 19.72983\nTotal prob of N-in: 0.96168\nPOSSIBLE N-term signal sequence\ninside 1 19\nTMhelix 20 39\noutside 40 325\nTMhelix 326 348\ninside 349 372

25086 GCF_001219645.1_5295_2_7Salmonella enterica subsp. enterica serovar Typhimurium str. DT104 Proteobacteria; Gammaproteobacteria; Enterobacterales; Enterobacteriaceae; Salmonella; Salmonella enterica; Salmonella enterica subsp. enterica; Salmonella enterica subsp. enterica serovar Typhimurium MTGDNTLITSHGINRRDFMKLCAALAATMGLSSKAAAEMAESVSNPQRPPVIWIGAQECT WP_000145429.1 hydrogenase 2 small subunit [Salmonella enterica] Length: 372\nNumber of predicted TMHs: 1\nExp number of AAs in TMHs: 24.51309\nExp number, first 60 AAs: 0.2682\nTotal prob of N-in: 0.14402\noutside 1 331\nTMhelix 332 354\ninside 355 372

25087 GCF_001219665.1_5388_1_6Salmonella enterica subsp. enterica serovar Typhimurium str. DT104 Proteobacteria; Gammaproteobacteria; Enterobacterales; Enterobacteriaceae; Salmonella; Salmonella enterica; Salmonella enterica subsp. enterica; Salmonella enterica subsp. enterica serovar Typhimurium MTGDNTLITSHGINRRDFMKLCAALAATMGLSSKAAAEMAESVSNPQRPPVIWIGAQECT WP_000145429.1 hydrogenase 2 small subunit [Salmonella enterica] Length: 372\nNumber of predicted TMHs: 1\nExp number of AAs in TMHs: 24.51309\nExp number, first 60 AAs: 0.2682\nTotal prob of N-in: 0.14402\noutside 1 331\nTMhelix 332 354\ninside 355 372

25088 GCF_001219705.1_8080_3_83 Salmonella enterica subsp. enterica serovar Typhimurium str. DT104 Proteobacteria; Gammaproteobacteria; Enterobacterales; Enterobacteriaceae; Salmonella; Salmonella enterica; Salmonella enterica subsp. enterica; Salmonella enterica subsp. enterica serovar Typhimurium MTGDNTLITSHGINRRDFMKLCAALAATMGLSSKAAAEMAESVSNPQRPPVIWIGAQECT WP_000145429.1 hydrogenase 2 small subunit [Salmonella enterica] Length: 372\nNumber of predicted TMHs: 1\nExp number of AAs in TMHs: 24.51309\nExp number, first 60 AAs: 0.2682\nTotal prob of N-in: 0.14402\noutside 1 331\nTMhelix 332 354\ninside 355 372

25089 GCF_001219765.1_8080_3_57 Salmonella enterica subsp. enterica serovar Typhimurium str. DT104 Proteobacteria; Gammaproteobacteria; Enterobacterales; Enterobacteriaceae; Salmonella; Salmonella enterica; Salmonella enterica subsp. enterica; Salmonella enterica subsp. enterica serovar Typhimurium MTGDNTLITSHGINRRDFMKLCAALAATMGLSSKAAAEMAESVSNPQRPPVIWIGAQECT WP_000145429.1 hydrogenase 2 small subunit [Salmonella enterica] Length: 372\nNumber of predicted TMHs: 1\nExp number of AAs in TMHs: 24.51309\nExp number, first 60 AAs: 0.2682\nTotal prob of N-in: 0.14402\noutside 1 331\nTMhelix 332 354\ninside 355 372

25090 GCF_001219805.1_8036_3_75 Salmonella enterica subsp. enterica serovar Typhimurium str. DT104 Proteobacteria; Gammaproteobacteria; Enterobacterales; Enterobacteriaceae; Salmonella; Salmonella enterica; Salmonella enterica subsp. enterica; Salmonella enterica subsp. enterica serovar Typhimurium MTGDNTLITSHGINRRDFMKLCAALAATMGLSSKAAAEMAESVSNPQRPPVIWIGAQECT WP_000145429.1 hydrogenase 2 small subunit [Salmonella enterica] Length: 372\nNumber of predicted TMHs: 1\nExp number of AAs in TMHs: 24.51309\nExp number, first 60 AAs: 0.2682\nTotal prob of N-in: 0.14402\noutside 1 331\nTMhelix 332 354\ninside 355 372

25091 GCF_001326215.1_9065_1_5Salmonella enterica subsp. enterica serovar Typhimurium str. DT104 Proteobacteria; Gammaproteobacteria; Enterobacterales; Enterobacteriaceae; Salmonella; Salmonella enterica; Salmonella enterica subsp. enterica; Salmonella enterica subsp. enterica serovar Typhimurium MTGDNTLITSHGINRRDFMKLCAALAATMGLSSKAAAEMAESVSNPQRPPVIWIGAQECT WP_000145429.1 hydrogenase 2 small subunit [Salmonella enterica] Length: 372\nNumber of predicted TMHs: 1\nExp number of AAs in TMHs: 24.51309\nExp number, first 60 AAs: 0.2682\nTotal prob of N-in: 0.14402\noutside 1 331\nTMhelix 332 354\ninside 355 372

25092 GCF_001219845.1_5299_3_11 Salmonella enterica subsp. enterica serovar Typhimurium str. DT104 Proteobacteria; Gammaproteobacteria; Enterobacterales; Enterobacteriaceae; Salmonella; Salmonella enterica; Salmonella enterica subsp. enterica; Salmonella enterica subsp. enterica serovar Typhimurium MTGDNTLITSHGINRRDFMKLCAALAATMGLSSKAAAEMAESVSNPQRPPVIWIGAQECT WP_000145429.1 hydrogenase 2 small subunit [Salmonella enterica] Length: 372\nNumber of predicted TMHs: 1\nExp number of AAs in TMHs: 24.51309\nExp number, first 60 AAs: 0.2682\nTotal prob of N-in: 0.14402\noutside 1 331\nTMhelix 332 354\ninside 355 372

25093 GCF_001219885.1_8080_3_38 Salmonella enterica subsp. enterica serovar Typhimurium str. DT104 Proteobacteria; Gammaproteobacteria; Enterobacterales; Enterobacteriaceae; Salmonella; Salmonella enterica; Salmonella enterica subsp. enterica; Salmonella enterica subsp. enterica serovar Typhimurium MTGDNTLITSHGINRRDFMKLCAALAATMGLSSKAAAEMAESVSNPQRPPVIWIGAQECT WP_000145429.1 hydrogenase 2 small subunit [Salmonella enterica] Length: 372\nNumber of predicted TMHs: 1\nExp

number of AAs in TMHs: 24.51309\nExp number, first 60 AAs: 0.2682\nTotal prob of N-in: 0.14402\noutside 1 331\nTMhelix 332 354\ninside 355 372

25094 GCF_001220065.1_5299_3_10 Salmonella enterica subsp. enterica serovar Typhimurium str. DT104 Proteobacteria; Gammaproteobacteria; Enterobacterales; Enterobacteriaceae; Salmonella; Salmonella enterica; Salmonella enterica subsp. enterica; Salmonella enterica subsp. enterica serovar Typhimurium MTGDNTLITSHGINRRDFMKLCAALAATMGLSSKAAAEMAESVSNPQRPPVIWIGAQECT WP_000145429.1 hydrogenase 2 small subunit [Salmonella enterica] Length: 372\nNumber of predicted TMHs: 1\nExp number of AAs in TMHs: 24.51309\nExp number, first 60 AAs: 0.2682\nTotal prob of N-in: 0.14402\noutside 1 331\nTMhelix 332 354\ninside 355 372

25095 GCF_001219985.1_8080_3_6Salmonella enterica subsp. enterica serovar Typhimurium str. DT104 Proteobacteria; Gammaproteobacteria; Enterobacterales; Enterobacteriaceae; Salmonella; Salmonella enterica; Salmonella enterica subsp. enterica; Salmonella enterica subsp. enterica serovar Typhimurium MTGDNTLITSHGINRRDFMKLCAALAATMGLSSKAAAEMAESVSNPQRPPVIWIGAQECT WP_000145429.1 hydrogenase 2 small subunit [Salmonella enterica] Length: 372\nNumber of predicted TMHs: 1\nExp number of AAs in TMHs: 24.51309\nExp number, first 60 AAs: 0.2682\nTotal prob of N-in: 0.14402\noutside 1 331\nTMhelix 332 354\ninside 355 372

25096 GCF_001219925.1_5275_5_1Salmonella enterica subsp. enterica serovar Typhimurium str. DT104 Proteobacteria; Gammaproteobacteria; Enterobacterales; Enterobacteriaceae; Salmonella; Salmonella enterica; Salmonella enterica subsp. enterica; Salmonella enterica subsp. enterica serovar Typhimurium MTGDNTLITSHGINRRDFMKLCAALAATMGLSSKAAAEMAESVSNPQRPPVIWIGAQECT WP_000145429.1 hydrogenase 2 small subunit [Salmonella enterica] Length: 372\nNumber of predicted TMHs: 1\nExp number of AAs in TMHs: 24.51309\nExp number, first 60 AAs: 0.2682\nTotal prob of N-in: 0.14402\noutside 1 331\nTMhelix 332 354\ninside 355 372

25097 GCF_001220125.1_8080_3_91 Salmonella enterica subsp. enterica serovar Typhimurium str. DT104 Proteobacteria; Gammaproteobacteria; Enterobacterales; Enterobacteriaceae; Salmonella; Salmonella enterica; Salmonella enterica subsp. enterica; Salmonella enterica subsp. enterica serovar Typhimurium MTGDNTLITSHGINRRDFMKLCAALAATMGLSSKAAAEMAESVSNPQRPPVIWIGAQECT WP_000145429.1 hydrogenase 2 small subunit [Salmonella enterica] Length: 372\nNumber of predicted TMHs: 1\nExp number of AAs in TMHs: 24.51309\nExp number, first 60 AAs: 0.2682\nTotal prob of N-in: 0.14402\noutside 1 331\nTMhelix 332 354\ninside 355 372

25098 GCF_001220425.1_8036_3_83 Salmonella enterica subsp. enterica serovar Typhimurium str. DT104 Proteobacteria; Gammaproteobacteria; Enterobacterales; Enterobacteriaceae; Salmonella; Salmonella enterica; Salmonella enterica subsp. enterica; Salmonella enterica subsp. enterica serovar Typhimurium MTGDNTLITSHGINRRDFMKLCAALAATMGLSSKAAAEMAESVSNPQRPPVIWIGAQECT WP_000145429.1 hydrogenase 2 small subunit [Salmonella enterica] Length: 372\nNumber of predicted TMHs: 1\nExp number of AAs in TMHs: 24.51309\nExp number, first 60 AAs: 0.2682\nTotal prob of N-in: 0.14402\noutside 1 331\nTMhelix 332 354\ninside 355 372

25099 GCF_001220345.1_8080_3_50 Salmonella enterica subsp. enterica serovar Typhimurium str. DT104 Proteobacteria; Gammaproteobacteria; Enterobacterales; Enterobacteriaceae; Salmonella; Salmonella enterica; Salmonella enterica subsp. enterica; Salmonella enterica subsp. enterica serovar Typhimurium MTGDNTLITSHGINRRDFMKLCAALAATMGLSSKAAAEMAESVSNPQRPPVIWIGAQECT WP_000145429.1 hydrogenase 2 small subunit [Salmonella enterica] Length: 372\nNumber of predicted TMHs: 1\nExp number of AAs in TMHs: 24.51309\nExp number, first 60 AAs: 0.2682\nTotal prob of N-in: 0.14402\noutside 1 331\nTMhelix 332 354\ninside 355 372

25100 GCF_001095425.1_8490_2_87 Salmonella enterica subsp. enterica serovar Typhimurium str. DT104 Proteobacteria; Gammaproteobacteria; Enterobacterales; Enterobacteriaceae; Salmonella; Salmonella enterica; Salmonella enterica subsp. enterica; Salmonella enterica subsp. enterica serovar Typhimurium MNNEETFYQAMRRKGVTRRSFLKFCSLAATSLGLGAGMTPKIAWALENKPRIPVVWIHGL WP_001058312.1 hydrogenase [Salmonella enterica] Length: 372\nNumber of predicted TMHs: 2\nExp number of AAs in TMHs: 36.51738\nExp number, first 60 AAs: 19.72849\nTotal prob of N-in: 0.96170\nPOSSIBLE N-term signal sequence\ninside 1 19\nTMhelix 20 39\noutside 40 325\nTMhelix 326 348\ninside 349 372

25101 GCF_001350115.1_8490_2_45 Salmonella enterica subsp. enterica serovar Typhimurium str. DT104 Proteobacteria; Gammaproteobacteria; Enterobacterales; Enterobacteriaceae; Salmonella; Salmonella enterica;

Salmonella enterica subsp. enterica; Salmonella enterica subsp. enterica serovar Typhimurium
MNNEETFYQAMRRKGVTRRSFLKFCSLAATSLGLGAGMTPKIAWALENKPRIPVVWIHGL WP_001058312.1
hydrogenase [Salmonella enterica] Length: 372\nNumber of predicted TMHs: 2\nExp number of AAs in TMHs: 36.51738\nExp number, first 60 AAs: 19.72849\nTotal prob of N-in: 0.96170\nPOSSIBLE N-term signal sequence\ninside 1 19\nTMhelix 20 39\noutside 40 325\nTMhelix 326 348\ninside 349 372

25102 GCF_001350155.1_8490_2_54 Salmonella enterica subsp. enterica serovar Typhimurium str. DT104
Proteobacteria; Gammaproteobacteria; Enterobacterales; Enterobacteriaceae; Salmonella; Salmonella enterica; Salmonella enterica subsp. enterica; Salmonella enterica subsp. enterica serovar Typhimurium
MTGDNTLITSHGINRRDFMKLCAALAATMGLSSKAAAEMAESVSNPQRPPVIWIGAQECT WP_000145429.1
hydrogenase 2 small subunit [Salmonella enterica] Length: 372\nNumber of predicted TMHs: 1\nExp number of AAs in TMHs: 24.51309\nExp number, first 60 AAs: 0.2682\nTotal prob of N-in: 0.14402\noutside 1 331\nTMhelix 332 354\ninside 355 372

25103 GCF_001350115.1_8490_2_45 Salmonella enterica subsp. enterica serovar Typhimurium str. DT104
Proteobacteria; Gammaproteobacteria; Enterobacterales; Enterobacteriaceae; Salmonella; Salmonella enterica; Salmonella enterica subsp. enterica; Salmonella enterica subsp. enterica serovar Typhimurium
MTGDNTLITSHGINRRDFMKLCAALAATMGLSSKAAAEMAESVSNPQRPPVIWIGAQECT WP_000145429.1
hydrogenase 2 small subunit [Salmonella enterica] Length: 372\nNumber of predicted TMHs: 1\nExp number of AAs in TMHs: 24.51309\nExp number, first 60 AAs: 0.2682\nTotal prob of N-in: 0.14402\noutside 1 331\nTMhelix 332 354\ninside 355 372

25104 GCF_001350075.1_8490_2_55 Salmonella enterica subsp. enterica serovar Typhimurium str. DT104
Proteobacteria; Gammaproteobacteria; Enterobacterales; Enterobacteriaceae; Salmonella; Salmonella enterica; Salmonella enterica subsp. enterica; Salmonella enterica subsp. enterica serovar Typhimurium
MTGDNTLITSHGINRRDFMKLCAALAATMGLSSKAAAEMAESVSNPQRPPVIWIGAQECT WP_000145429.1
hydrogenase 2 small subunit [Salmonella enterica] Length: 372\nNumber of predicted TMHs: 1\nExp number of AAs in TMHs: 24.51309\nExp number, first 60 AAs: 0.2682\nTotal prob of N-in: 0.14402\noutside 1 331\nTMhelix 332 354\ninside 355 372

25105 GCF_001223465.1_5275_8_12 Salmonella enterica subsp. enterica serovar Typhimurium str. DT104
Proteobacteria; Gammaproteobacteria; Enterobacterales; Enterobacteriaceae; Salmonella; Salmonella enterica; Salmonella enterica subsp. enterica; Salmonella enterica subsp. enterica serovar Typhimurium
MTGDNTLITSHGINRRDFMKLCAALAATMGLSSKAAAEMAESVSNPQRPPVIWIGAQECT WP_000145429.1
hydrogenase 2 small subunit [Salmonella enterica] Length: 372\nNumber of predicted TMHs: 1\nExp number of AAs in TMHs: 24.51309\nExp number, first 60 AAs: 0.2682\nTotal prob of N-in: 0.14402\noutside 1 331\nTMhelix 332 354\ninside 355 372

25106 GCF_001223525.1_8080_3_81 Salmonella enterica subsp. enterica serovar Typhimurium str. DT104
Proteobacteria; Gammaproteobacteria; Enterobacterales; Enterobacteriaceae; Salmonella; Salmonella enterica; Salmonella enterica subsp. enterica; Salmonella enterica subsp. enterica serovar Typhimurium
MTGDNTLITSHGINRRDFMKLCAALAATMGLSSKAAAEMAESVSNPQRPPVIWIGAQECT WP_000145429.1
hydrogenase 2 small subunit [Salmonella enterica] Length: 372\nNumber of predicted TMHs: 1\nExp number of AAs in TMHs: 24.51309\nExp number, first 60 AAs: 0.2682\nTotal prob of N-in: 0.14402\noutside 1 331\nTMhelix 332 354\ninside 355 372

25107 GCF_001223225.1_5275_5_5Salmonella enterica subsp. enterica serovar Typhimurium str. DT104
Proteobacteria; Gammaproteobacteria; Enterobacterales; Enterobacteriaceae; Salmonella; Salmonella enterica; Salmonella enterica subsp. enterica; Salmonella enterica subsp. enterica serovar Typhimurium
MNNEETFYQAMRRKGVTRRSFLKFCSLAATSLGLGAGMTPKIAWALENKPRIPVVWIHGL WP_001058312.1
hydrogenase [Salmonella enterica] Length: 372\nNumber of predicted TMHs: 2\nExp number of AAs in TMHs: 36.51738\nExp number, first 60 AAs: 19.72849\nTotal prob of N-in: 0.96170\nPOSSIBLE N-term signal sequence\ninside 1 19\nTMhelix 20 39\noutside 40 325\nTMhelix 326 348\ninside 349 372

25108 GCF_001223365.1_8080_3_70 Salmonella enterica subsp. enterica serovar Typhimurium str. DT104
Proteobacteria; Gammaproteobacteria; Enterobacterales; Enterobacteriaceae; Salmonella; Salmonella enterica; Salmonella enterica subsp. enterica; Salmonella enterica subsp. enterica serovar Typhimurium
MTGDNTLITSHGINRRDFMKLCAALAATMGLSSKAAAEMAESVSNPQRPPVIWIGAQECT WP_000145429.1
hydrogenase 2 small subunit [Salmonella enterica] Length: 372\nNumber of predicted TMHs: 1\nExp number of AAs in TMHs: 24.51309\nExp number, first 60 AAs: 0.2682\nTotal prob of N-in: 0.14402\noutside 1 331\nTMhelix 332 354\ninside 355 372

25109 GCF_001223325.1_5299_6_1Salmonella enterica subsp. enterica serovar Typhimurium str. DT104 Proteobacteria; Gammaproteobacteria; Enterobacterales; Enterobacteriaceae; Salmonella; Salmonella enterica; Salmonella enterica subsp. enterica; Salmonella enterica subsp. enterica serovar Typhimurium MTGDNTLITSHGINRRDFMKLCAALAATMGLSSKAAAEMAESVSNPQRPPVIWIGAQECT WP_000145429.1 hydrogenase 2 small subunit [Salmonella enterica] Length: 372\nNumber of predicted TMHs: 1\nExp number of AAs in TMHs: 24.51309\nExp number, first 60 AAs: 0.2682\nTotal prob of N-in: 0.14402\noutside 1 331\nTMhelix 332 354\ninside 355 372

25110 GCF_001222945.1_5275_1_7Salmonella enterica subsp. enterica serovar Typhimurium str. DT104 Proteobacteria; Gammaproteobacteria; Enterobacterales; Enterobacteriaceae; Salmonella; Salmonella enterica; Salmonella enterica subsp. enterica; Salmonella enterica subsp. enterica serovar Typhimurium MTGDNTLITSHGINRRDFMKLCAALAATMGLSSKAAAEMAESVSNPQRPPVIWIGAQECT WP_000145429.1 hydrogenase 2 small subunit [Salmonella enterica] Length: 372\nNumber of predicted TMHs: 1\nExp number of AAs in TMHs: 24.51309\nExp number, first 60 AAs: 0.2682\nTotal prob of N-in: 0.14402\noutside 1 331\nTMhelix 332 354\ninside 355 372

25111 GCF_001222885.1_5275_2_2Salmonella enterica subsp. enterica serovar Typhimurium str. DT104 Proteobacteria; Gammaproteobacteria; Enterobacterales; Enterobacteriaceae; Salmonella; Salmonella enterica; Salmonella enterica subsp. enterica; Salmonella enterica subsp. enterica serovar Typhimurium MTGDNTLITSHGINRRDFMKLCAALAATMGLSSKAAAEMAESVSNPQRPPVIWIGAQECT WP_000145429.1 hydrogenase 2 small subunit [Salmonella enterica] Length: 372\nNumber of predicted TMHs: 1\nExp number of AAs in TMHs: 24.51309\nExp number, first 60 AAs: 0.2682\nTotal prob of N-in: 0.14402\noutside 1 331\nTMhelix 332 354\ninside 355 372

25112 GCF_001222665.1_8080_3_72 Salmonella enterica subsp. enterica serovar Typhimurium str. DT104 Proteobacteria; Gammaproteobacteria; Enterobacterales; Enterobacteriaceae; Salmonella; Salmonella enterica; Salmonella enterica subsp. enterica; Salmonella enterica subsp. enterica serovar Typhimurium MNNEETFYQAMRRKGVTRRSFLKFCSLAATSLGLGAGMTPKIAWALENKPRIPVVWIHGL WP_001058312.1 hydrogenase [Salmonella enterica] Length: 372\nNumber of predicted TMHs: 2\nExp number of AAs in TMHs: 36.51738\nExp number, first 60 AAs: 19.72849\nTotal prob of N-in: 0.96170\nPOSSIBLE N-term signal sequence\ninside 1 19\nTMhelix 20 39\noutside 40 325\nTMhelix 326 348\ninside 349 372

25113 GCF_001222825.1_8036_3_39 Salmonella enterica subsp. enterica serovar Typhimurium str. DT104 Proteobacteria; Gammaproteobacteria; Enterobacterales; Enterobacteriaceae; Salmonella; Salmonella enterica; Salmonella enterica subsp. enterica; Salmonella enterica subsp. enterica serovar Typhimurium MTGDNTLITSHGINRRDFMKLCAALAATMGLSSKAAAEMAESVSNPQRPPVIWIGAQECT WP_000145429.1 hydrogenase 2 small subunit [Salmonella enterica] Length: 372\nNumber of predicted TMHs: 1\nExp number of AAs in TMHs: 24.51309\nExp number, first 60 AAs: 0.2682\nTotal prob of N-in: 0.14402\noutside 1 331\nTMhelix 332 354\ninside 355 372

25114 GCF_001222625.1_5285_7_2Salmonella enterica subsp. enterica serovar Typhimurium str. DT104 Proteobacteria; Gammaproteobacteria; Enterobacterales; Enterobacteriaceae; Salmonella; Salmonella enterica; Salmonella enterica subsp. enterica; Salmonella enterica subsp. enterica serovar Typhimurium MNNEETFYQAMRRKGVTRRSFLKFCSLAATSLGLGAGMTPKIAWALENKPRIPVVWIHGL WP_001058312.1 hydrogenase [Salmonella enterica] Length: 372\nNumber of predicted TMHs: 2\nExp number of AAs in TMHs: 36.51738\nExp number, first 60 AAs: 19.72849\nTotal prob of N-in: 0.96170\nPOSSIBLE N-term signal sequence\ninside 1 19\nTMhelix 20 39\noutside 40 325\nTMhelix 326 348\ninside 349 372

25115 GCF_001222225.1_5275_5_11 Salmonella enterica subsp. enterica serovar Typhimurium str. DT104 Proteobacteria; Gammaproteobacteria; Enterobacterales; Enterobacteriaceae; Salmonella; Salmonella enterica; Salmonella enterica subsp. enterica; Salmonella enterica subsp. enterica serovar Typhimurium MTGDNTLITSHGINRRDFMKLCAALAATMGLSSKAAAEMAESVSNPQRPPVIWIGAQECT WP_000145429.1 hydrogenase 2 small subunit [Salmonella enterica] Length: 372\nNumber of predicted TMHs: 1\nExp number of AAs in TMHs: 24.51309\nExp number, first 60 AAs: 0.2682\nTotal prob of N-in: 0.14402\noutside 1 331\nTMhelix 332 354\ninside 355 372

25116 GCF_001222185.1_5285_7_1Salmonella enterica subsp. enterica serovar Typhimurium str. DT104 Proteobacteria; Gammaproteobacteria; Enterobacterales; Enterobacteriaceae; Salmonella; Salmonella enterica; Salmonella enterica subsp. enterica; Salmonella enterica subsp. enterica serovar Typhimurium MTGDNTLITSHGINRRDFMKLCAALAATMGLSSKAAAEMAESVSNPQRPPVIWIGAQECT WP_000145429.1 hydrogenase 2 small subunit [Salmonella enterica] Length: 372\nNumber of predicted TMHs: 1\nExp

number of AAs in TMHs: 24.51309\nExp number, first 60 AAs: 0.2682\nTotal prob of N-in: 0.14402\noutside 1 331\nTMhelix 332 354\ninside 355 372

25117 GCF_001222065.1_5388_1_4Salmonella enterica subsp. enterica serovar Typhimurium str. DT104 Proteobacteria; Gammaproteobacteria; Enterobacterales; Enterobacteriaceae; Salmonella; Salmonella enterica; Salmonella enterica subsp. enterica; Salmonella enterica subsp. enterica serovar Typhimurium MTGDNTLITSHGINRRDFMKLCAALAATMGLSSKAAAEMAESVSNPQRPPVIWIGAQECT WP_000145429.1 hydrogenase 2 small subunit [Salmonella enterica] Length: 372\nNumber of predicted TMHs: 1\nExp number of AAs in TMHs: 24.51309\nExp number, first 60 AAs: 0.2682\nTotal prob of N-in: 0.14402\noutside 1 331\nTMhelix 332 354\ninside 355 372

25118 GCF_001221945.1_8080_3_21 Salmonella enterica subsp. enterica serovar Typhimurium str. DT104 Proteobacteria; Gammaproteobacteria; Enterobacterales; Enterobacteriaceae; Salmonella; Salmonella enterica; Salmonella enterica subsp. enterica; Salmonella enterica subsp. enterica serovar Typhimurium MNNEETFYQAMRRKGVTRRSFLKFCSLAATSLGLGAGMTPKIAWALENKPRIPVVWIHGL WP_001058312.1 hydrogenase [Salmonella enterica] Length: 372\nNumber of predicted TMHs: 2\nExp number of AAs in TMHs: 36.51738\nExp number, first 60 AAs: 19.72849\nTotal prob of N-in: 0.96170\nPOSSIBLE N-term signal sequence\ninside 1 19\nTMhelix 20 39\noutside 40 325\nTMhelix 326 348\ninside 349 372

25119 GCF_001222005.1_8036_3_63 Salmonella enterica subsp. enterica serovar Typhimurium str. DT104 Proteobacteria; Gammaproteobacteria; Enterobacterales; Enterobacteriaceae; Salmonella; Salmonella enterica; Salmonella enterica subsp. enterica; Salmonella enterica subsp. enterica serovar Typhimurium MTGDNTLITSHGINRRDFMKLCAALAATMGLSSKAAAEMAESVSNPQRPPVIWIGAQECT WP_000145429.1 hydrogenase 2 small subunit [Salmonella enterica] Length: 372\nNumber of predicted TMHs: 1\nExp number of AAs in TMHs: 24.51309\nExp number, first 60 AAs: 0.2682\nTotal prob of N-in: 0.14402\noutside 1 331\nTMhelix 332 354\ninside 355 372

25120 GCF_001221805.1_5285_6_10 Salmonella enterica subsp. enterica serovar Typhimurium str. DT104 Proteobacteria; Gammaproteobacteria; Enterobacterales; Enterobacteriaceae; Salmonella; Salmonella enterica; Salmonella enterica subsp. enterica; Salmonella enterica subsp. enterica serovar Typhimurium MTGDNTLITSHGINRRDFMKLCAALAATMGLSSKAAAEMAESVSNPQRPPVIWIGAQECT WP_000145429.1 hydrogenase 2 small subunit [Salmonella enterica] Length: 372\nNumber of predicted TMHs: 1\nExp number of AAs in TMHs: 24.51309\nExp number, first 60 AAs: 0.2682\nTotal prob of N-in: 0.14402\noutside 1 331\nTMhelix 332 354\ninside 355 372

25121 GCF_001221545.1_8080_3_80 Salmonella enterica subsp. enterica serovar Typhimurium str. DT104 Proteobacteria; Gammaproteobacteria; Enterobacterales; Enterobacteriaceae; Salmonella; Salmonella enterica; Salmonella enterica subsp. enterica; Salmonella enterica subsp. enterica serovar Typhimurium MNNEETFYQAMRRKGVTRRSFLKFCSLAATSLGLGAGMTPKIAWALENKPRIPVVWIHGL WP_001058312.1 hydrogenase [Salmonella enterica] Length: 372\nNumber of predicted TMHs: 2\nExp number of AAs in TMHs: 36.51738\nExp number, first 60 AAs: 19.72849\nTotal prob of N-in: 0.96170\nPOSSIBLE N-term signal sequence\ninside 1 19\nTMhelix 20 39\noutside 40 325\nTMhelix 326 348\ninside 349 372

25122 GCF_001221765.1_5295_2_5Salmonella enterica subsp. enterica serovar Typhimurium str. DT104 Proteobacteria; Gammaproteobacteria; Enterobacterales; Enterobacteriaceae; Salmonella; Salmonella enterica; Salmonella enterica subsp. enterica; Salmonella enterica subsp. enterica serovar Typhimurium MNNEETFYQAMRRKGVTRRSFLKFCSLAATSLGLGAGMTPKIAWALENKPRIPVVWIHGL WP_001058312.1 hydrogenase [Salmonella enterica] Length: 372\nNumber of predicted TMHs: 2\nExp number of AAs in TMHs: 36.51738\nExp number, first 60 AAs: 19.72849\nTotal prob of N-in: 0.96170\nPOSSIBLE N-term signal sequence\ninside 1 19\nTMhelix 20 39\noutside 40 325\nTMhelix 326 348\ninside 349 372

25123 GCF_001221765.1_5295_2_5Salmonella enterica subsp. enterica serovar Typhimurium str. DT104 Proteobacteria; Gammaproteobacteria; Enterobacterales; Enterobacteriaceae; Salmonella; Salmonella enterica; Salmonella enterica subsp. enterica; Salmonella enterica subsp. enterica serovar Typhimurium MTGDNTLITSHGINRRDFMKLCAALAATMGLSSKAAAEMAESVSNPQRPPVIWIGAQECT WP_000145429.1 hydrogenase 2 small subunit [Salmonella enterica] Length: 372\nNumber of predicted TMHs: 1\nExp number of AAs in TMHs: 24.51309\nExp number, first 60 AAs: 0.2682\nTotal prob of N-in: 0.14402\noutside 1 331\nTMhelix 332 354\ninside 355 372

25124 GCF_001221405.1_8036_3_41 Salmonella enterica subsp. enterica serovar Typhimurium str. DT104 Proteobacteria; Gammaproteobacteria; Enterobacterales; Enterobacteriaceae; Salmonella; Salmonella enterica;

Salmonella enterica subsp. enterica; Salmonella enterica subsp. enterica serovar Typhimurium
MNNEETFYQAMRRKGVTRRSFLKFCSLAATSLGLGAGMTPKIAWALENKPRIPVVWIHGL WP_001058312.1
hydrogenase [Salmonella enterica] Length: 372\nNumber of predicted TMHs: 2\nExp number of AAs in TMHs: 36.51738\nExp number, first 60 AAs: 19.72849\nTotal prob of N-in: 0.96170\nPOSSIBLE N-term signal sequence\ninside 1 19\nTMhelix 20 39\noutside 40 325\nTMhelix 326 348\ninside 349 372

25125 GCF_001221485.1_5275_6_1Salmonella enterica subsp. enterica serovar Typhimurium str. DT104
Proteobacteria; Gammaproteobacteria; Enterobacterales; Enterobacteriaceae; Salmonella; Salmonella enterica; Salmonella enterica subsp. enterica; Salmonella enterica subsp. enterica serovar Typhimurium
MNNEETFYQAMRRKGVTRRSFLKFCSLAATSLGLGAGMTPKIAWALENKPRIPVVWIHGL WP_001058312.1
hydrogenase [Salmonella enterica] Length: 372\nNumber of predicted TMHs: 2\nExp number of AAs in TMHs: 36.51738\nExp number, first 60 AAs: 19.72849\nTotal prob of N-in: 0.96170\nPOSSIBLE N-term signal sequence\ninside 1 19\nTMhelix 20 39\noutside 40 325\nTMhelix 326 348\ninside 349 372

25126 GCF_001221365.1_8036_3_45 Salmonella enterica subsp. enterica serovar Typhimurium str. DT104
Proteobacteria; Gammaproteobacteria; Enterobacterales; Enterobacteriaceae; Salmonella; Salmonella enterica; Salmonella enterica subsp. enterica; Salmonella enterica subsp. enterica serovar Typhimurium
MNNEETFYQAMRRKGVTRRSFLKFCSLAATSLGLGAGMTPKIAWALENKPRIPVVWIHGL WP_001058312.1
hydrogenase [Salmonella enterica] Length: 372\nNumber of predicted TMHs: 2\nExp number of AAs in TMHs: 36.51738\nExp number, first 60 AAs: 19.72849\nTotal prob of N-in: 0.96170\nPOSSIBLE N-term signal sequence\ninside 1 19\nTMhelix 20 39\noutside 40 325\nTMhelix 326 348\ninside 349 372

25127 GCF_001221545.1_8080_3_80 Salmonella enterica subsp. enterica serovar Typhimurium str. DT104
Proteobacteria; Gammaproteobacteria; Enterobacterales; Enterobacteriaceae; Salmonella; Salmonella enterica; Salmonella enterica subsp. enterica; Salmonella enterica subsp. enterica serovar Typhimurium
MTGDNTLITSHGINRRDFMKLCAALAATMGLSSKAAAEMAESVSNPQRPPVIWIGAQECT WP_000145429.1
hydrogenase 2 small subunit [Salmonella enterica] Length: 372\nNumber of predicted TMHs: 1\nExp number of AAs in TMHs: 24.51309\nExp number, first 60 AAs: 0.2682\nTotal prob of N-in: 0.14402\noutside 1 331\nTMhelix 332 354\ninside 355 372

25128 GCF_001221405.1_8036_3_41 Salmonella enterica subsp. enterica serovar Typhimurium str. DT104
Proteobacteria; Gammaproteobacteria; Enterobacterales; Enterobacteriaceae; Salmonella; Salmonella enterica; Salmonella enterica subsp. enterica; Salmonella enterica subsp. enterica serovar Typhimurium
MTGDNTLITSHGINRRDFMKLCAALAATMGLSSKAAAEMAESVSNPQRPPVIWIGAQECT WP_000145429.1
hydrogenase 2 small subunit [Salmonella enterica] Length: 372\nNumber of predicted TMHs: 1\nExp number of AAs in TMHs: 24.51309\nExp number, first 60 AAs: 0.2682\nTotal prob of N-in: 0.14402\noutside 1 331\nTMhelix 332 354\ninside 355 372

25129 GCF_001221485.1_5275_6_1Salmonella enterica subsp. enterica serovar Typhimurium str. DT104
Proteobacteria; Gammaproteobacteria; Enterobacterales; Enterobacteriaceae; Salmonella; Salmonella enterica; Salmonella enterica subsp. enterica; Salmonella enterica subsp. enterica serovar Typhimurium
MTGDNTLITSHGINRRDFMKLCAALAATMGLSSKAAAEMAESVSNPQRPPVIWIGAQECT WP_000145429.1
hydrogenase 2 small subunit [Salmonella enterica] Length: 372\nNumber of predicted TMHs: 1\nExp number of AAs in TMHs: 24.51309\nExp number, first 60 AAs: 0.2682\nTotal prob of N-in: 0.14402\noutside 1 331\nTMhelix 332 354\ninside 355 372

25130 GCF_001221365.1_8036_3_45 Salmonella enterica subsp. enterica serovar Typhimurium str. DT104
Proteobacteria; Gammaproteobacteria; Enterobacterales; Enterobacteriaceae; Salmonella; Salmonella enterica; Salmonella enterica subsp. enterica; Salmonella enterica subsp. enterica serovar Typhimurium
MTGDNTLITSHGINRRDFMKLCAALAATMGLSSKAAAEMAESVSNPQRPPVIWIGAQECT WP_000145429.1
hydrogenase 2 small subunit [Salmonella enterica] Length: 372\nNumber of predicted TMHs: 1\nExp number of AAs in TMHs: 24.51309\nExp number, first 60 AAs: 0.2682\nTotal prob of N-in: 0.14402\noutside 1 331\nTMhelix 332 354\ninside 355 372

25131 GCF_001221045.1_5285_6_4Salmonella enterica subsp. enterica serovar Typhimurium str. DT104
Proteobacteria; Gammaproteobacteria; Enterobacterales; Enterobacteriaceae; Salmonella; Salmonella enterica; Salmonella enterica subsp. enterica; Salmonella enterica subsp. enterica serovar Typhimurium
MNNEETFYQAMRRKGVTRRSFLKFCSLAATSLGLGAGMTPKIAWALENKPRIPVVWIHGL WP_001058312.1
hydrogenase [Salmonella enterica] Length: 372\nNumber of predicted TMHs: 2\nExp number of AAs in TMHs: 36.51738\nExp number, first 60 AAs: 19.72849\nTotal prob of N-in: 0.96170\nPOSSIBLE N-term signal sequence\ninside 1 19\nTMhelix 20 39\noutside 40 325\nTMhelix 326 348\ninside 349 372

25132 GCF_001220685.1_8080_3_33 Salmonella enterica subsp. enterica serovar Typhimurium str. DT104
Proteobacteria; Gammaproteobacteria; Enterobacterales; Enterobacteriaceae; Salmonella; Salmonella enterica; Salmonella enterica subsp. enterica; Salmonella enterica subsp. enterica serovar Typhimurium
MNNEETFYQAMRRKGVTRRSFLKFCSLAATSLGLGAGMTPKIAWALENKPRIPVVWIHGL WP_001058312.1
hydrogenase [Salmonella enterica] Length: 372\nNumber of predicted TMHs: 2\nExp number of AAs in TMHs: 36.51738\nExp number, first 60 AAs: 19.72849\nTotal prob of N-in: 0.96170\nPOSSIBLE N-term signal sequence\ninside 1 19\nTMhelix 20 39\noutside 40 325\nTMhelix 326 348\ninside 349 372

25133 GCF_001220965.1_5275_3_9Salmonella enterica subsp. enterica serovar Typhimurium str. DT104
Proteobacteria; Gammaproteobacteria; Enterobacterales; Enterobacteriaceae; Salmonella; Salmonella enterica; Salmonella enterica subsp. enterica; Salmonella enterica subsp. enterica serovar Typhimurium
MNNEETFYQAMRRKGVTRRSFLKFCSLAATSLGLGAGMTPKIAWALENKPRIPVVWIHGL WP_001058312.1
hydrogenase [Salmonella enterica] Length: 372\nNumber of predicted TMHs: 2\nExp number of AAs in TMHs: 36.51738\nExp number, first 60 AAs: 19.72849\nTotal prob of N-in: 0.96170\nPOSSIBLE N-term signal sequence\ninside 1 19\nTMhelix 20 39\noutside 40 325\nTMhelix 326 348\ninside 349 372

25134 GCF_001220925.1_8080_3_15 Salmonella enterica subsp. enterica serovar Typhimurium str. DT104
Proteobacteria; Gammaproteobacteria; Enterobacterales; Enterobacteriaceae; Salmonella; Salmonella enterica; Salmonella enterica subsp. enterica; Salmonella enterica subsp. enterica serovar Typhimurium
MNNEETFYQAMRRKGVTRRSFLKFCSLAATSLGLGAGMTPKIAWALENKPRIPVVWIHGL WP_001058312.1
hydrogenase [Salmonella enterica] Length: 372\nNumber of predicted TMHs: 2\nExp number of AAs in TMHs: 36.51738\nExp number, first 60 AAs: 19.72849\nTotal prob of N-in: 0.96170\nPOSSIBLE N-term signal sequence\ninside 1 19\nTMhelix 20 39\noutside 40 325\nTMhelix 326 348\ninside 349 372

25135 GCF_001221045.1_5285_6_4Salmonella enterica subsp. enterica serovar Typhimurium str. DT104
Proteobacteria; Gammaproteobacteria; Enterobacterales; Enterobacteriaceae; Salmonella; Salmonella enterica; Salmonella enterica subsp. enterica; Salmonella enterica subsp. enterica serovar Typhimurium
MTGDNTLITSHGINRRDFMKLCAALAATMGLSSKAAAEMAESVSNPQRPPVIWIGAQECT WP_000145429.1
hydrogenase 2 small subunit [Salmonella enterica] Length: 372\nNumber of predicted TMHs: 1\nExp number of AAs in TMHs: 24.51309\nExp number, first 60 AAs: 0.2682\nTotal prob of N-in: 0.14402\noutside 1 331\nTMhelix 332 354\ninside 355 372

25136 GCF_001221005.1_8080_3_48 Salmonella enterica subsp. enterica serovar Typhimurium str. DT104
Proteobacteria; Gammaproteobacteria; Enterobacterales; Enterobacteriaceae; Salmonella; Salmonella enterica; Salmonella enterica subsp. enterica; Salmonella enterica subsp. enterica serovar Typhimurium
MTGDNTLITSHGINRRDFMKLCAALAATMGLSSKAAAEMAESVSNPQRPPVIWIGAQECT WP_000145429.1
hydrogenase 2 small subunit [Salmonella enterica] Length: 372\nNumber of predicted TMHs: 1\nExp number of AAs in TMHs: 24.51309\nExp number, first 60 AAs: 0.2682\nTotal prob of N-in: 0.14402\noutside 1 331\nTMhelix 332 354\ninside 355 372

25137 GCF_001220965.1_5275_3_9Salmonella enterica subsp. enterica serovar Typhimurium str. DT104
Proteobacteria; Gammaproteobacteria; Enterobacterales; Enterobacteriaceae; Salmonella; Salmonella enterica; Salmonella enterica subsp. enterica; Salmonella enterica subsp. enterica serovar Typhimurium
MTGDNTLITSHGINRRDFMKLCAALAATMGLSSKAAAEMAESVSNPQRPPVIWIGAQECT WP_000145429.1
hydrogenase 2 small subunit [Salmonella enterica] Length: 372\nNumber of predicted TMHs: 1\nExp number of AAs in TMHs: 24.51309\nExp number, first 60 AAs: 0.2682\nTotal prob of N-in: 0.14402\noutside 1 331\nTMhelix 332 354\ninside 355 372

25138 GCF_001220325.1_8080_3_78 Salmonella enterica subsp. enterica serovar Typhimurium str. DT104
Proteobacteria; Gammaproteobacteria; Enterobacterales; Enterobacteriaceae; Salmonella; Salmonella enterica; Salmonella enterica subsp. enterica; Salmonella enterica subsp. enterica serovar Typhimurium
MNNEETFYQAMRRKGVTRRSFLKFCSLAATSLGLGAGMTPKIAWALENKPRIPVVWIHGL WP_001058312.1
hydrogenase [Salmonella enterica] Length: 372\nNumber of predicted TMHs: 2\nExp number of AAs in TMHs: 36.51738\nExp number, first 60 AAs: 19.72849\nTotal prob of N-in: 0.96170\nPOSSIBLE N-term signal sequence\ninside 1 19\nTMhelix 20 39\noutside 40 325\nTMhelix 326 348\ninside 349 372

25139 GCF_001220285.1_8080_3_88 Salmonella enterica subsp. enterica serovar Typhimurium str. DT104
Proteobacteria; Gammaproteobacteria; Enterobacterales; Enterobacteriaceae; Salmonella; Salmonella enterica; Salmonella enterica subsp. enterica; Salmonella enterica subsp. enterica serovar Typhimurium
MNNEETFYQAMRRKGVTRRSFLKFCSLAATSLGLGAGMTPKIAWALENKPRIPVVWIHGL WP_001058312.1
hydrogenase [Salmonella enterica] Length: 372\nNumber of predicted TMHs: 2\nExp number of AAs in

TMHs: 36.51738\nExp number, first 60 AAs: 19.72849\nTotal prob of N-in: 0.96170\nPOSSIBLE N-term signal sequence\ninside 1 19\nTMhelix 20 39\noutside 40 325\nTMhelix 326 348\ninside 349 372

25140 GCF_001220165.1_5275_2_7Salmonella enterica subsp. enterica serovar Typhimurium str. DT104 Proteobacteria; Gammaproteobacteria; Enterobacterales; Enterobacteriaceae; Salmonella; Salmonella enterica; Salmonella enterica subsp. enterica; Salmonella enterica subsp. enterica serovar Typhimurium MNNEETFYQAMRRKGVTRRSFLKFCSLAATSLGLGAGMTPKIAWALENKPRIPVVWIHGL WP_001058312.1 hydrogenase [Salmonella enterica] Length: 372\nNumber of predicted TMHs: 2\nExp number of AAs in TMHs: 36.51738\nExp number, first 60 AAs: 19.72849\nTotal prob of N-in: 0.96170\nPOSSIBLE N-term signal sequence\ninside 1 19\nTMhelix 20 39\noutside 40 325\nTMhelix 326 348\ninside 349 372

25141 GCF_001220105.1_8080_3_7Salmonella enterica subsp. enterica serovar Typhimurium str. DT104 Proteobacteria; Gammaproteobacteria; Enterobacterales; Enterobacteriaceae; Salmonella; Salmonella enterica; Salmonella enterica subsp. enterica; Salmonella enterica subsp. enterica serovar Typhimurium MNNEETFYQAMRRKGVTRRSFLKFCSLAATSLGLGAGMTPKIAWALENKPRIPVVWIHGL WP_001058312.1 hydrogenase [Salmonella enterica] Length: 372\nNumber of predicted TMHs: 2\nExp number of AAs in TMHs: 36.51738\nExp number, first 60 AAs: 19.72849\nTotal prob of N-in: 0.96170\nPOSSIBLE N-term signal sequence\ninside 1 19\nTMhelix 20 39\noutside 40 325\nTMhelix 326 348\ninside 349 372

25142 GCF_001219825.1_8036_3_60 Salmonella enterica subsp. enterica serovar Typhimurium str. DT104 Proteobacteria; Gammaproteobacteria; Enterobacterales; Enterobacteriaceae; Salmonella; Salmonella enterica; Salmonella enterica subsp. enterica; Salmonella enterica subsp. enterica serovar Typhimurium MTGDNTLITSHGINRRDFMKLCAALAATMGLSSKAAAEMAESVSNPQRPPVIWIGAQECT WP_000145429.1 hydrogenase 2 small subunit [Salmonella enterica] Length: 372\nNumber of predicted TMHs: 1\nExp number of AAs in TMHs: 24.51309\nExp number, first 60 AAs: 0.2682\nTotal prob of N-in: 0.14402\noutside 1 331\nTMhelix 332 354\ninside 355 372

25143 GCF_001219565.1_8080_3_74 Salmonella enterica subsp. enterica serovar Typhimurium str. DT104 Proteobacteria; Gammaproteobacteria; Enterobacterales; Enterobacteriaceae; Salmonella; Salmonella enterica; Salmonella enterica subsp. enterica; Salmonella enterica subsp. enterica serovar Typhimurium MNNEETFYQAMRRKGVTRRSFLKFCSLAATSLGLGAGMTPKIAWALENKPRIPVVWIHGL WP_001058312.1 hydrogenase [Salmonella enterica] Length: 372\nNumber of predicted TMHs: 2\nExp number of AAs in TMHs: 36.51738\nExp number, first 60 AAs: 19.72849\nTotal prob of N-in: 0.96170\nPOSSIBLE N-term signal sequence\ninside 1 19\nTMhelix 20 39\noutside 40 325\nTMhelix 326 348\ninside 349 372

25144 GCF_001219525.1_9065_1_4Salmonella enterica subsp. enterica serovar Typhimurium str. DT104 Proteobacteria; Gammaproteobacteria; Enterobacterales; Enterobacteriaceae; Salmonella; Salmonella enterica; Salmonella enterica subsp. enterica; Salmonella enterica subsp. enterica serovar Typhimurium MNNEETFYQAMRRKGVTRRSFLKFCSLAATSLGLGAGMTPKIAWALENKPRIPVVWIHGL WP_001058312.1 hydrogenase [Salmonella enterica] Length: 372\nNumber of predicted TMHs: 2\nExp number of AAs in TMHs: 36.51738\nExp number, first 60 AAs: 19.72849\nTotal prob of N-in: 0.96170\nPOSSIBLE N-term signal sequence\ninside 1 19\nTMhelix 20 39\noutside 40 325\nTMhelix 326 348\ninside 349 372

25145 GCF_001219485.1_5275_6_4Salmonella enterica subsp. enterica serovar Typhimurium str. DT104 Proteobacteria; Gammaproteobacteria; Enterobacterales; Enterobacteriaceae; Salmonella; Salmonella enterica; Salmonella enterica subsp. enterica; Salmonella enterica subsp. enterica serovar Typhimurium MNNEETFYQAMRRKGVTRRSFLKFCSLAATSLGLGAGMTPKIAWALENKPRIPVVWIHGL WP_001058312.1 hydrogenase [Salmonella enterica] Length: 372\nNumber of predicted TMHs: 2\nExp number of AAs in TMHs: 36.51738\nExp number, first 60 AAs: 19.72849\nTotal prob of N-in: 0.96170\nPOSSIBLE N-term signal sequence\ninside 1 19\nTMhelix 20 39\noutside 40 325\nTMhelix 326 348\ninside 349 372

25146 GCF_001219445.1_8036_3_74 Salmonella enterica subsp. enterica serovar Typhimurium str. DT104 Proteobacteria; Gammaproteobacteria; Enterobacterales; Enterobacteriaceae; Salmonella; Salmonella enterica; Salmonella enterica subsp. enterica; Salmonella enterica subsp. enterica serovar Typhimurium MNNEETFYQAMRRKGVTRRSFLKFCSLAATSLGLGAGMTPKIAWALENKPRIPVVWIHGL WP_001058312.1 hydrogenase [Salmonella enterica] Length: 372\nNumber of predicted TMHs: 2\nExp number of AAs in TMHs: 36.51738\nExp number, first 60 AAs: 19.72849\nTotal prob of N-in: 0.96170\nPOSSIBLE N-term signal sequence\ninside 1 19\nTMhelix 20 39\noutside 40 325\nTMhelix 326 348\ninside 349 372

25147 GCF_001219125.1_5275_8_4Salmonella enterica subsp. enterica serovar Typhimurium str. DT104 Proteobacteria; Gammaproteobacteria; Enterobacterales; Enterobacteriaceae; Salmonella; Salmonella enterica;

Salmonella enterica subsp. enterica; Salmonella enterica subsp. enterica serovar Typhimurium
MNNEETFYQAMRRKGVTRRSFLKFCSLAATSLGLGAGMTPKIAWALENKPRIPVVWIHGL WP_001058312.1
hydrogenase [Salmonella enterica] Length: 372\nNumber of predicted TMHs: 2\nExp number of AAs in TMHs: 36.51738\nExp number, first 60 AAs: 19.72849\nTotal prob of N-in: 0.96170\nPOSSIBLE N-term signal sequence\ninside 1 19\nTMhelix 20 39\noutside 40 325\nTMhelix 326 348\ninside 349 372

25148 GCF_001219285.1_5388_1_2Salmonella enterica subsp. enterica serovar Typhimurium str. DT104
Proteobacteria; Gammaproteobacteria; Enterobacterales; Enterobacteriaceae; Salmonella; Salmonella enterica; Salmonella enterica subsp. enterica; Salmonella enterica subsp. enterica serovar Typhimurium
MTGDNTLITSHGINRRDFMKLCAALAATMGLSSKAAAEMAESVSNPQRPPVIWIGAQECT WP_000145429.1
hydrogenase 2 small subunit [Salmonella enterica] Length: 372\nNumber of predicted TMHs: 1\nExp number of AAs in TMHs: 24.51309\nExp number, first 60 AAs: 0.2682\nTotal prob of N-in: 0.14402\noutside 1 331\nTMhelix 332 354\ninside 355 372

25149 GCF_001219225.1_5275_3_1Salmonella enterica subsp. enterica serovar Typhimurium str. DT104
Proteobacteria; Gammaproteobacteria; Enterobacterales; Enterobacteriaceae; Salmonella; Salmonella enterica; Salmonella enterica subsp. enterica; Salmonella enterica subsp. enterica serovar Typhimurium
MTGDNTLITSHGINRRDFMKLCAALAATMGLSSKAAAEMAESVSNPQRPPVIWIGAQECT WP_000145429.1
hydrogenase 2 small subunit [Salmonella enterica] Length: 372\nNumber of predicted TMHs: 1\nExp number of AAs in TMHs: 24.51309\nExp number, first 60 AAs: 0.2682\nTotal prob of N-in: 0.14402\noutside 1 331\nTMhelix 332 354\ninside 355 372

25150 GCF_001219185.1_8036_3_55 Salmonella enterica subsp. enterica serovar Typhimurium str. DT104
Proteobacteria; Gammaproteobacteria; Enterobacterales; Enterobacteriaceae; Salmonella; Salmonella enterica; Salmonella enterica subsp. enterica; Salmonella enterica subsp. enterica serovar Typhimurium
MTGDNTLITSHGINRRDFMKLCAALAATMGLSSKAAAEMAESVSNPQRPPVIWIGAQECT WP_000145429.1
hydrogenase 2 small subunit [Salmonella enterica] Length: 372\nNumber of predicted TMHs: 1\nExp number of AAs in TMHs: 24.51309\nExp number, first 60 AAs: 0.2682\nTotal prob of N-in: 0.14402\noutside 1 331\nTMhelix 332 354\ninside 355 372

25151 GCF_001219125.1_5275_8_4Salmonella enterica subsp. enterica serovar Typhimurium str. DT104
Proteobacteria; Gammaproteobacteria; Enterobacterales; Enterobacteriaceae; Salmonella; Salmonella enterica; Salmonella enterica subsp. enterica; Salmonella enterica subsp. enterica serovar Typhimurium
MTGDNTLITSHGINRRDFMKLCAALAATMGLSSKAAAEMAESVSNPQRPPVIWIGAQECT WP_000145429.1
hydrogenase 2 small subunit [Salmonella enterica] Length: 372\nNumber of predicted TMHs: 1\nExp number of AAs in TMHs: 24.51309\nExp number, first 60 AAs: 0.2682\nTotal prob of N-in: 0.14402\noutside 1 331\nTMhelix 332 354\ninside 355 372

25152 GCF_001218925.1_5275_7_5Salmonella enterica subsp. enterica serovar Typhimurium str. DT104
Proteobacteria; Gammaproteobacteria; Enterobacterales; Enterobacteriaceae; Salmonella; Salmonella enterica; Salmonella enterica subsp. enterica; Salmonella enterica subsp. enterica serovar Typhimurium
MNNEETFYQAMRRKGVTRRSFLKFCSLAATSLGLGAGMTPKIAWALENKPRIPVVWIHGL WP_001058312.1
hydrogenase [Salmonella enterica] Length: 372\nNumber of predicted TMHs: 2\nExp number of AAs in TMHs: 36.51738\nExp number, first 60 AAs: 19.72849\nTotal prob of N-in: 0.96170\nPOSSIBLE N-term signal sequence\ninside 1 19\nTMhelix 20 39\noutside 40 325\nTMhelix 326 348\ninside 349 372

25153 GCF_001218745.1_5275_6_10 Salmonella enterica subsp. enterica serovar Typhimurium str. DT104
Proteobacteria; Gammaproteobacteria; Enterobacterales; Enterobacteriaceae; Salmonella; Salmonella enterica; Salmonella enterica subsp. enterica; Salmonella enterica subsp. enterica serovar Typhimurium
MTGDNTLITSHGINRRDFMKLCAALAATMGLSSKAAAEMAESVSNPQRPPVIWIGAQECT WP_000145429.1
hydrogenase 2 small subunit [Salmonella enterica] Length: 372\nNumber of predicted TMHs: 1\nExp number of AAs in TMHs: 24.51309\nExp number, first 60 AAs: 0.2682\nTotal prob of N-in: 0.14402\noutside 1 331\nTMhelix 332 354\ninside 355 372

25154 GCF_001218605.1_5275_1_11 Salmonella enterica subsp. enterica serovar Typhimurium str. DT104
Proteobacteria; Gammaproteobacteria; Enterobacterales; Enterobacteriaceae; Salmonella; Salmonella enterica; Salmonella enterica subsp. enterica; Salmonella enterica subsp. enterica serovar Typhimurium
MNNEETFYQAMRRKGVTRRSFLKFCSLAATSLGLGAGMTPKIAWALENKPRIPVVWIHGL WP_001058312.1
hydrogenase [Salmonella enterica] Length: 372\nNumber of predicted TMHs: 2\nExp number of AAs in TMHs: 36.51738\nExp number, first 60 AAs: 19.72849\nTotal prob of N-in: 0.96170\nPOSSIBLE N-term signal sequence\ninside 1 19\nTMhelix 20 39\noutside 40 325\nTMhelix 326 348\ninside 349 372

25155 GCF_001218685.1_5299_3_6Salmonella enterica subsp. enterica serovar Typhimurium str. DT104 Proteobacteria; Gammaproteobacteria; Enterobacterales; Enterobacteriaceae; Salmonella; Salmonella enterica; Salmonella enterica subsp. enterica; Salmonella enterica subsp. enterica serovar Typhimurium MNNEETFYQAMRRKGVTRRSFLKFCSLAATSLGLGAGMTPKIAWALENKPRIPVVWIHGL WP_001058312.1 hydrogenase [Salmonella enterica] Length: 372\nNumber of predicted TMHs: 2\nExp number of AAs in TMHs: 36.51738\nExp number, first 60 AAs: 19.72849\nTotal prob of N-in: 0.96170\nPOSSIBLE N-term signal sequence\ninside 1 19\nTMhelix 20 39\noutside 40 325\nTMhelix 326 348\ninside 349 372

25156 GCF_001218565.1_8080_3_66 Salmonella enterica subsp. enterica serovar Typhimurium str. DT104 Proteobacteria; Gammaproteobacteria; Enterobacterales; Enterobacteriaceae; Salmonella; Salmonella enterica; Salmonella enterica subsp. enterica; Salmonella enterica subsp. enterica serovar Typhimurium MNNEETFYQAMRRKGVTRRSFLKFCSLAATSLGLGAGMTPKIAWALENKPRIPVVWIHGL WP_001058312.1 hydrogenase [Salmonella enterica] Length: 372\nNumber of predicted TMHs: 2\nExp number of AAs in TMHs: 36.51738\nExp number, first 60 AAs: 19.72849\nTotal prob of N-in: 0.96170\nPOSSIBLE N-term signal sequence\ninside 1 19\nTMhelix 20 39\noutside 40 325\nTMhelix 326 348\ninside 349 372

25157 GCF_001218505.1_5285_7_7Salmonella enterica subsp. enterica serovar Typhimurium str. DT104 Proteobacteria; Gammaproteobacteria; Enterobacterales; Enterobacteriaceae; Salmonella; Salmonella enterica; Salmonella enterica subsp. enterica; Salmonella enterica subsp. enterica serovar Typhimurium MNNEETFYQAMRRKGVTRRSFLKFCSLAATSLGLGAGMTPKIAWALENKPRIPVVWIHGL WP_001058312.1 hydrogenase [Salmonella enterica] Length: 372\nNumber of predicted TMHs: 2\nExp number of AAs in TMHs: 36.51738\nExp number, first 60 AAs: 19.72849\nTotal prob of N-in: 0.96170\nPOSSIBLE N-term signal sequence\ninside 1 19\nTMhelix 20 39\noutside 40 325\nTMhelix 326 348\ninside 349 372

25158 GCF_001218605.1_5275_1_11 Salmonella enterica subsp. enterica serovar Typhimurium str. DT104 Proteobacteria; Gammaproteobacteria; Enterobacterales; Enterobacteriaceae; Salmonella; Salmonella enterica; Salmonella enterica subsp. enterica; Salmonella enterica subsp. enterica serovar Typhimurium MTGDNTLITSHGINRRDFMKLCAALAATMGLSSKAAAEMAESVSNPQRPPVIWIGAQECT WP_000145429.1 hydrogenase 2 small subunit [Salmonella enterica] Length: 372\nNumber of predicted TMHs: 1\nExp number of AAs in TMHs: 24.51309\nExp number, first 60 AAs: 0.2682\nTotal prob of N-in: 0.14402\noutside 1 331\nTMhelix 332 354\ninside 355 372

25159 GCF_001218345.1_8080_3_25 Salmonella enterica subsp. enterica serovar Typhimurium str. DT104 Proteobacteria; Gammaproteobacteria; Enterobacterales; Enterobacteriaceae; Salmonella; Salmonella enterica; Salmonella enterica subsp. enterica; Salmonella enterica subsp. enterica serovar Typhimurium MNNEETFYQAMRRKGVTRRSFLKFCSLAATSLGLGAGMTPKIAWALENKPRIPVVWIHGL WP_001058312.1 hydrogenase [Salmonella enterica] Length: 372\nNumber of predicted TMHs: 2\nExp number of AAs in TMHs: 36.51738\nExp number, first 60 AAs: 19.72849\nTotal prob of N-in: 0.96170\nPOSSIBLE N-term signal sequence\ninside 1 19\nTMhelix 20 39\noutside 40 325\nTMhelix 326 348\ninside 349 372

25160 GCF_001218345.1_8080_3_25 Salmonella enterica subsp. enterica serovar Typhimurium str. DT104 Proteobacteria; Gammaproteobacteria; Enterobacterales; Enterobacteriaceae; Salmonella; Salmonella enterica; Salmonella enterica subsp. enterica; Salmonella enterica subsp. enterica serovar Typhimurium MTGDNTLITSHGINRRDFMKLCAALAATMGLSSKAAAEMAESVSNPQRPPVIWIGAQECT WP_000145429.1 hydrogenase 2 small subunit [Salmonella enterica] Length: 372\nNumber of predicted TMHs: 1\nExp number of AAs in TMHs: 24.51309\nExp number, first 60 AAs: 0.2682\nTotal prob of N-in: 0.14402\noutside 1 331\nTMhelix 332 354\ninside 355 372

25161 GCF_001218305.1_5299_6_3Salmonella enterica subsp. enterica serovar Typhimurium str. DT104 Proteobacteria; Gammaproteobacteria; Enterobacterales; Enterobacteriaceae; Salmonella; Salmonella enterica; Salmonella enterica subsp. enterica; Salmonella enterica subsp. enterica serovar Typhimurium MTGDNTLITSHGINRRDFMKLCAALAATMGLSSKAAAEMAESVSNPQRPPVIWIGAQECT WP_000145429.1 hydrogenase 2 small subunit [Salmonella enterica] Length: 372\nNumber of predicted TMHs: 1\nExp number of AAs in TMHs: 24.51309\nExp number, first 60 AAs: 0.2682\nTotal prob of N-in: 0.14402\noutside 1 331\nTMhelix 332 354\ninside 355 372

25162 GCF_001218265.1_8036_3_79 Salmonella enterica subsp. enterica serovar Typhimurium str. DT104 Proteobacteria; Gammaproteobacteria; Enterobacterales; Enterobacteriaceae; Salmonella; Salmonella enterica; Salmonella enterica subsp. enterica; Salmonella enterica subsp. enterica serovar Typhimurium MTGDNTLITSHGINRRDFMKLCAALAATMGLSSKAAAEMAESVSNPQRPPVIWIGAQECT WP_000145429.1 hydrogenase 2 small subunit [Salmonella enterica] Length: 372\nNumber of predicted TMHs: 1\nExp

number of AAs in TMHs: 24.51309\nExp number, first 60 AAs: 0.2682\nTotal prob of N-in: 0.14402\noutside 1 331\nTMhelix 332 354\ninside 355 372

25163 GCF_001218105.1_5285_6_3Salmonella enterica subsp. enterica serovar Typhimurium str. DT104 Proteobacteria; Gammaproteobacteria; Enterobacterales; Enterobacteriaceae; Salmonella; Salmonella enterica; Salmonella enterica subsp. enterica; Salmonella enterica subsp. enterica serovar Typhimurium MTGDNTLITSHGINRRDFMKLCAALAATMGLSSKAAAEMAESVSNPQRPPVIWIGAQECT WP_000145429.1 hydrogenase 2 small subunit [Salmonella enterica] Length: 372\nNumber of predicted TMHs: 1\nExp number of AAs in TMHs: 24.51309\nExp number, first 60 AAs: 0.2682\nTotal prob of N-in: 0.14402\noutside 1 331\nTMhelix 332 354\ninside 355 372

25164 GCF_001217705.1_8036_3_62 Salmonella enterica subsp. enterica serovar Typhimurium str. DT104 Proteobacteria; Gammaproteobacteria; Enterobacterales; Enterobacteriaceae; Salmonella; Salmonella enterica; Salmonella enterica subsp. enterica; Salmonella enterica subsp. enterica serovar Typhimurium MTGDNTLITSHGINRRDFMKLCAALAATMGLSSKAAAEMAESVSNPQRPPVIWIGAQECT WP_000145429.1 hydrogenase 2 small subunit [Salmonella enterica] Length: 372\nNumber of predicted TMHs: 1\nExp number of AAs in TMHs: 24.51309\nExp number, first 60 AAs: 0.2682\nTotal prob of N-in: 0.14402\noutside 1 331\nTMhelix 332 354\ninside 355 372

25165 GCF_001217245.1_8080_3_4Salmonella enterica subsp. enterica serovar Typhimurium str. DT104 Proteobacteria; Gammaproteobacteria; Enterobacterales; Enterobacteriaceae; Salmonella; Salmonella enterica; Salmonella enterica subsp. enterica; Salmonella enterica subsp. enterica serovar Typhimurium MNNEETFYQAMRRKGVTRRSFLKFCSLAATSLGLGAGMTPKIAWALENKPRIPVVWIHGL WP_001058312.1 hydrogenase [Salmonella enterica] Length: 372\nNumber of predicted TMHs: 2\nExp number of AAs in TMHs: 36.51738\nExp number, first 60 AAs: 19.72849\nTotal prob of N-in: 0.96170\nPOSSIBLE N-term signal sequence\ninside 1 19\nTMhelix 20 39\noutside 40 325\nTMhelix 326 348\ninside 349 372

25166 GCF_001217645.1_5275_3_10 Salmonella enterica subsp. enterica serovar Typhimurium str. DT104 Proteobacteria; Gammaproteobacteria; Enterobacterales; Enterobacteriaceae; Salmonella; Salmonella enterica; Salmonella enterica subsp. enterica; Salmonella enterica subsp. enterica serovar Typhimurium MNNEETFYQAMRRKGVTRRSFLKFCSLAATSLGLGAGMTPKIAWALENKPRIPVVWIHGL WP_001058312.1 hydrogenase [Salmonella enterica] Length: 372\nNumber of predicted TMHs: 2\nExp number of AAs in TMHs: 36.51738\nExp number, first 60 AAs: 19.72849\nTotal prob of N-in: 0.96170\nPOSSIBLE N-term signal sequence\ninside 1 19\nTMhelix 20 39\noutside 40 325\nTMhelix 326 348\ninside 349 372

25167 GCF_001217645.1_5275_3_10 Salmonella enterica subsp. enterica serovar Typhimurium str. DT104 Proteobacteria; Gammaproteobacteria; Enterobacterales; Enterobacteriaceae; Salmonella; Salmonella enterica; Salmonella enterica subsp. enterica; Salmonella enterica subsp. enterica serovar Typhimurium MTGDNTLITSHGINRRDFMKLCAALAATMGLSSKAAAEMAESVSNPQRPPVIWIGAQECT WP_000145429.1 hydrogenase 2 small subunit [Salmonella enterica] Length: 372\nNumber of predicted TMHs: 1\nExp number of AAs in TMHs: 24.51309\nExp number, first 60 AAs: 0.2682\nTotal prob of N-in: 0.14402\noutside 1 331\nTMhelix 332 354\ninside 355 372

25168 GCF_001217605.1_5275_1_12 Salmonella enterica subsp. enterica serovar Typhimurium str. DT104 Proteobacteria; Gammaproteobacteria; Enterobacterales; Enterobacteriaceae; Salmonella; Salmonella enterica; Salmonella enterica subsp. enterica; Salmonella enterica subsp. enterica serovar Typhimurium MTGDNTLITSHGINRRDFMKLCAALAATMGLSSKAAAEMAESVSNPQRPPVIWIGAQECT WP_000145429.1 hydrogenase 2 small subunit [Salmonella enterica] Length: 372\nNumber of predicted TMHs: 1\nExp number of AAs in TMHs: 24.51309\nExp number, first 60 AAs: 0.2682\nTotal prob of N-in: 0.14402\noutside 1 331\nTMhelix 332 354\ninside 355 372

25169 GCF_001216205.1_8036_3_76 Salmonella enterica subsp. enterica serovar Typhimurium str. DT104 Proteobacteria; Gammaproteobacteria; Enterobacterales; Enterobacteriaceae; Salmonella; Salmonella enterica; Salmonella enterica subsp. enterica; Salmonella enterica subsp. enterica serovar Typhimurium MNNEETFYQAMRRKGVTRRSFLKFCSLAATSLGLGAGMTPKIAWALENKPRIPVVWIHGL WP_001058312.1 hydrogenase [Salmonella enterica] Length: 372\nNumber of predicted TMHs: 2\nExp number of AAs in TMHs: 36.51738\nExp number, first 60 AAs: 19.72849\nTotal prob of N-in: 0.96170\nPOSSIBLE N-term signal sequence\ninside 1 19\nTMhelix 20 39\noutside 40 325\nTMhelix 326 348\ninside 349 372

25170 GCF_001216605.1_5285_6_9Salmonella enterica subsp. enterica serovar Typhimurium str. DT104 Proteobacteria; Gammaproteobacteria; Enterobacterales; Enterobacteriaceae; Salmonella; Salmonella enterica;

Salmonella enterica subsp. enterica; Salmonella enterica subsp. enterica serovar Typhimurium
MNNEETFYQAMRRKGVTRRSFLKFCSLAATSLGLGAGMTPKIAWALENKPRIPVVWIHGL WP_001058312.1
hydrogenase [Salmonella enterica] Length: 372\nNumber of predicted TMHs: 2\nExp number of AAs in TMHs: 36.51738\nExp number, first 60 AAs: 19.72849\nTotal prob of N-in: 0.96170\nPOSSIBLE N-term signal sequence\ninside 1 19\nTMhelix 20 39\noutside 40 325\nTMhelix 326 348\ninside 349 372

25171 GCF_001216605.1_5285_6_9Salmonella enterica subsp. enterica serovar Typhimurium str. DT104
Proteobacteria; Gammaproteobacteria; Enterobacterales; Enterobacteriaceae; Salmonella; Salmonella enterica; Salmonella enterica subsp. enterica; Salmonella enterica subsp. enterica serovar Typhimurium
MTGDNTLITSHGINRRDFMKLCAALAATMGLSSKAAAEMAESVSNPQRPPVIWIGAQECT WP_000145429.1
hydrogenase 2 small subunit [Salmonella enterica] Length: 372\nNumber of predicted TMHs: 1\nExp number of AAs in TMHs: 24.51309\nExp number, first 60 AAs: 0.2682\nTotal prob of N-in: 0.14402\noutside 1 331\nTMhelix 332 354\ninside 355 372

25172 GCF_001216085.1_8080_3_87 Salmonella enterica subsp. enterica serovar Typhimurium str. DT104
Proteobacteria; Gammaproteobacteria; Enterobacterales; Enterobacteriaceae; Salmonella; Salmonella enterica; Salmonella enterica subsp. enterica; Salmonella enterica subsp. enterica serovar Typhimurium
MNNEETFYQAMRRKGVTRRSFLKFCSLAATSLGLGAGMTPKIAWALENKPRIPVVWIHGL WP_001058312.1
hydrogenase [Salmonella enterica] Length: 372\nNumber of predicted TMHs: 2\nExp number of AAs in TMHs: 36.51738\nExp number, first 60 AAs: 19.72849\nTotal prob of N-in: 0.96170\nPOSSIBLE N-term signal sequence\ninside 1 19\nTMhelix 20 39\noutside 40 325\nTMhelix 326 348\ninside 349 372

25173 GCF_001215785.1_5275_1_8Salmonella enterica subsp. enterica serovar Typhimurium str. DT104
Proteobacteria; Gammaproteobacteria; Enterobacterales; Enterobacteriaceae; Salmonella; Salmonella enterica; Salmonella enterica subsp. enterica; Salmonella enterica subsp. enterica serovar Typhimurium
MTGDNTLITSHGINRRDFMKLCAALAATMGLSSKAAAEMAESVSNPQRPPVIWIGAQECT WP_000145429.1
hydrogenase 2 small subunit [Salmonella enterica] Length: 372\nNumber of predicted TMHs: 1\nExp number of AAs in TMHs: 24.51309\nExp number, first 60 AAs: 0.2682\nTotal prob of N-in: 0.14402\noutside 1 331\nTMhelix 332 354\ninside 355 372

25174 GCF_001215725.1_5275_3_8Salmonella enterica subsp. enterica serovar Typhimurium str. DT104
Proteobacteria; Gammaproteobacteria; Enterobacterales; Enterobacteriaceae; Salmonella; Salmonella enterica; Salmonella enterica subsp. enterica; Salmonella enterica subsp. enterica serovar Typhimurium
MTGDNTLITSHGINRRDFMKLCAALAATMGLSSKAAAEMAESVSNPQRPPVIWIGAQECT WP_000145429.1
hydrogenase 2 small subunit [Salmonella enterica] Length: 372\nNumber of predicted TMHs: 1\nExp number of AAs in TMHs: 24.51309\nExp number, first 60 AAs: 0.2682\nTotal prob of N-in: 0.14402\noutside 1 331\nTMhelix 332 354\ninside 355 372

25175 GCF_001215665.1_8080_3_65 Salmonella enterica subsp. enterica serovar Typhimurium str. DT104
Proteobacteria; Gammaproteobacteria; Enterobacterales; Enterobacteriaceae; Salmonella; Salmonella enterica; Salmonella enterica subsp. enterica; Salmonella enterica subsp. enterica serovar Typhimurium
MTGDNTLITSHGINRRDFMKLCAALAATMGLSSKAAAEMAESVSNPQRPPVIWIGAQECT WP_000145429.1
hydrogenase 2 small subunit [Salmonella enterica] Length: 372\nNumber of predicted TMHs: 1\nExp number of AAs in TMHs: 24.51309\nExp number, first 60 AAs: 0.2682\nTotal prob of N-in: 0.14402\noutside 1 331\nTMhelix 332 354\ninside 355 372

25176 GCF_001214865.1_9065_1_3Salmonella enterica subsp. enterica serovar Typhimurium str. DT104
Proteobacteria; Gammaproteobacteria; Enterobacterales; Enterobacteriaceae; Salmonella; Salmonella enterica; Salmonella enterica subsp. enterica; Salmonella enterica subsp. enterica serovar Typhimurium
MNNEETFYQAMRRKGVTRRSFLKFCSLAATSLGLGAGMTPKIAWALENKPRIPVVWIHGL WP_001058312.1
hydrogenase [Salmonella enterica] Length: 372\nNumber of predicted TMHs: 2\nExp number of AAs in TMHs: 36.51738\nExp number, first 60 AAs: 19.72849\nTotal prob of N-in: 0.96170\nPOSSIBLE N-term signal sequence\ninside 1 19\nTMhelix 20 39\noutside 40 325\nTMhelix 326 348\ninside 349 372

25177 GCF_001215545.1_5275_6_2Salmonella enterica subsp. enterica serovar Typhimurium str. DT104
Proteobacteria; Gammaproteobacteria; Enterobacterales; Enterobacteriaceae; Salmonella; Salmonella enterica; Salmonella enterica subsp. enterica; Salmonella enterica subsp. enterica serovar Typhimurium
MNNEETFYQAMRRKGVTRRSFLKFCSLAATSLGLGAGMTPKIAWALENKPRIPVVWIHGL WP_001058312.1
hydrogenase [Salmonella enterica] Length: 372\nNumber of predicted TMHs: 2\nExp number of AAs in TMHs: 36.51738\nExp number, first 60 AAs: 19.72849\nTotal prob of N-in: 0.96170\nPOSSIBLE N-term signal sequence\ninside 1 19\nTMhelix 20 39\noutside 40 325\nTMhelix 326 348\ninside 349 372

25178 GCF_001215545.1_5275_6_2Salmonella enterica subsp. enterica serovar Typhimurium str. DT104 Proteobacteria; Gammaproteobacteria; Enterobacterales; Enterobacteriaceae; Salmonella; Salmonella enterica; Salmonella enterica subsp. enterica; Salmonella enterica subsp. enterica serovar Typhimurium MTGDNTLITSHGINRRDFMKLCAALAATMGLSSKAAAEMAESVSNPQRPPVIWIGAQECT WP_000145429.1 hydrogenase 2 small subunit [Salmonella enterica] Length: 372\nNumber of predicted TMHs: 1\nExp number of AAs in TMHs: 24.51309\nExp number, first 60 AAs: 0.2682\nTotal prob of N-in: 0.14402\noutside 1 331\nTMhelix 332 354\ninside 355 372

25179 GCF_001172545.1_8490_2_78 Salmonella enterica subsp. enterica serovar Typhimurium str. DT104 Proteobacteria; Gammaproteobacteria; Enterobacterales; Enterobacteriaceae; Salmonella; Salmonella enterica; Salmonella enterica subsp. enterica; Salmonella enterica subsp. enterica serovar Typhimurium MNNEETFYQAMRRKGVTRRSFLKFCSLAATSLGLGAGMTPKIAWALENKPRIPVVWIHGL WP_001058312.1 hydrogenase [Salmonella enterica] Length: 372\nNumber of predicted TMHs: 2\nExp number of AAs in TMHs: 36.51738\nExp number, first 60 AAs: 19.72849\nTotal prob of N-in: 0.96170\nPOSSIBLE N-term signal sequence\ninside 1 19\nTMhelix 20 39\noutside 40 325\nTMhelix 326 348\ninside 349 372

25180 GCF_001164125.1_8490_2_52 Salmonella enterica subsp. enterica serovar Typhimurium str. DT104 Proteobacteria; Gammaproteobacteria; Enterobacterales; Enterobacteriaceae; Salmonella; Salmonella enterica; Salmonella enterica subsp. enterica; Salmonella enterica subsp. enterica serovar Typhimurium MTGDNTLITSHGINRRDFMKLCAALAATMGLSSKAAAEMAESVSNPQRPPVIWIGAQECT WP_000145429.1 hydrogenase 2 small subunit [Salmonella enterica] Length: 372\nNumber of predicted TMHs: 1\nExp number of AAs in TMHs: 24.51309\nExp number, first 60 AAs: 0.2682\nTotal prob of N-in: 0.14402\noutside 1 331\nTMhelix 332 354\ninside 355 372

25181 GCF_001159405.1_8490_2_88 Salmonella enterica subsp. enterica serovar Typhimurium str. DT104 Proteobacteria; Gammaproteobacteria; Enterobacterales; Enterobacteriaceae; Salmonella; Salmonella enterica; Salmonella enterica subsp. enterica; Salmonella enterica subsp. enterica serovar Typhimurium MNNEETFYQAMRRKGVTRRSFLKFCSLAATSLGLGAGMTPKIAWALENKPRIPVVWIHGL WP_001058312.1 hydrogenase [Salmonella enterica] Length: 372\nNumber of predicted TMHs: 2\nExp number of AAs in TMHs: 36.51738\nExp number, first 60 AAs: 19.72849\nTotal prob of N-in: 0.96170\nPOSSIBLE N-term signal sequence\ninside 1 19\nTMhelix 20 39\noutside 40 325\nTMhelix 326 348\ninside 349 372

25182 GCF_001159405.1_8490_2_88 Salmonella enterica subsp. enterica serovar Typhimurium str. DT104 Proteobacteria; Gammaproteobacteria; Enterobacterales; Enterobacteriaceae; Salmonella; Salmonella enterica; Salmonella enterica subsp. enterica; Salmonella enterica subsp. enterica serovar Typhimurium MTGDNTLITSHGINRRDFMKLCAALAATMGLSSKAAAEMAESVSNPQRPPVIWIGAQECT WP_000145429.1 hydrogenase 2 small subunit [Salmonella enterica] Length: 372\nNumber of predicted TMHs: 1\nExp number of AAs in TMHs: 24.51309\nExp number, first 60 AAs: 0.2682\nTotal prob of N-in: 0.14402\noutside 1 331\nTMhelix 332 354\ninside 355 372

25183 GCF_001154285.1_8490_2_70 Salmonella enterica subsp. enterica serovar Typhimurium str. DT104 Proteobacteria; Gammaproteobacteria; Enterobacterales; Enterobacteriaceae; Salmonella; Salmonella enterica; Salmonella enterica subsp. enterica; Salmonella enterica subsp. enterica serovar Typhimurium MNNEETFYQAMRRKGVTRRSFLKFCSLAATSLGLGAGMTPKIAWALENKPRIPVVWIHGL WP_001058312.1 hydrogenase [Salmonella enterica] Length: 372\nNumber of predicted TMHs: 2\nExp number of AAs in TMHs: 36.51738\nExp number, first 60 AAs: 19.72849\nTotal prob of N-in: 0.96170\nPOSSIBLE N-term signal sequence\ninside 1 19\nTMhelix 20 39\noutside 40 325\nTMhelix 326 348\ninside 349 372

25184 GCF_001154285.1_8490_2_70 Salmonella enterica subsp. enterica serovar Typhimurium str. DT104 Proteobacteria; Gammaproteobacteria; Enterobacterales; Enterobacteriaceae; Salmonella; Salmonella enterica; Salmonella enterica subsp. enterica; Salmonella enterica subsp. enterica serovar Typhimurium MTGDNTLITSHGINRRDFMKLCAALAATMGLSSKAAAEMAESVSNPQRPPVIWIGAQECT WP_000145429.1 hydrogenase 2 small subunit [Salmonella enterica] Length: 372\nNumber of predicted TMHs: 1\nExp number of AAs in TMHs: 24.51309\nExp number, first 60 AAs: 0.2682\nTotal prob of N-in: 0.14402\noutside 1 331\nTMhelix 332 354\ninside 355 372

25185 GCF_001140725.1_8490_2_51 Salmonella enterica subsp. enterica serovar Typhimurium str. DT104 Proteobacteria; Gammaproteobacteria; Enterobacterales; Enterobacteriaceae; Salmonella; Salmonella enterica; Salmonella enterica subsp. enterica; Salmonella enterica subsp. enterica serovar Typhimurium MNNEETFYQAMRRKGVTRRSFLKFCSLAATSLGLGAGMTPKIAWALENKPRIPVVWIHGL WP_001058312.1 hydrogenase [Salmonella enterica] Length: 372\nNumber of predicted TMHs: 2\nExp number of AAs in

TMHs: 36.51738\nExp number, first 60 AAs: 19.72849\nTotal prob of N-in: 0.96170\nPOSSIBLE N-term signal sequence\ninside 1 19\nTMhelix 20 39\noutside 40 325\nTMhelix 326 348\ninside 349 372

25186 GCF_001137885.1_8490_2_64 Salmonella enterica subsp. enterica serovar Typhimurium str. DT104 Proteobacteria; Gammaproteobacteria; Enterobacterales; Enterobacteriaceae; Salmonella; Salmonella enterica; Salmonella enterica subsp. enterica; Salmonella enterica subsp. enterica serovar Typhimurium MTGDNTLITSHGINRRDFMKLCAALAATMGLSSKAAAEMAESVSNPQRPPVIWIGAQECT WP_000145429.1 hydrogenase 2 small subunit [Salmonella enterica] Length: 372\nNumber of predicted TMHs: 1\nExp number of AAs in TMHs: 24.51309\nExp number, first 60 AAs: 0.2682\nTotal prob of N-in: 0.14402\noutside 1 331\nTMhelix 332 354\ninside 355 372

25187 GCF_001566155.1_8080_3_32 Salmonella enterica subsp. enterica serovar Typhimurium str. DT104 Proteobacteria; Gammaproteobacteria; Enterobacterales; Enterobacteriaceae; Salmonella; Salmonella enterica; Salmonella enterica subsp. enterica; Salmonella enterica subsp. enterica serovar Typhimurium MNNEETFYQAMRRKGVTRRSFLKFCSLAATSLGLGAGMTPKIAWALENKPRIPVVWIHGL WP_001058312.1 hydrogenase [Salmonella enterica] Length: 372\nNumber of predicted TMHs: 2\nExp number of AAs in TMHs: 36.51738\nExp number, first 60 AAs: 19.72849\nTotal prob of N-in: 0.96170\nPOSSIBLE N-term signal sequence\ninside 1 19\nTMhelix 20 39\noutside 40 325\nTMhelix 326 348\ninside 349 372

25188 GCF_001565975.1_8080_3_10 Salmonella enterica subsp. enterica serovar Typhimurium str. DT104 Proteobacteria; Gammaproteobacteria; Enterobacterales; Enterobacteriaceae; Salmonella; Salmonella enterica; Salmonella enterica subsp. enterica; Salmonella enterica subsp. enterica serovar Typhimurium MTGDNTLITSHGINRRDFMKLCAALAATMGLSSKAAAEMAESVSNPQRPPVIWIGAQECT WP_000145429.1 hydrogenase 2 small subunit [Salmonella enterica] Length: 372\nNumber of predicted TMHs: 1\nExp number of AAs in TMHs: 24.51309\nExp number, first 60 AAs: 0.2682\nTotal prob of N-in: 0.14402\noutside 1 331\nTMhelix 332 354\ninside 355 372

25189 GCF_001566115.1_5275_1_10 Salmonella enterica subsp. enterica serovar Typhimurium str. DT104 Proteobacteria; Gammaproteobacteria; Enterobacterales; Enterobacteriaceae; Salmonella; Salmonella enterica; Salmonella enterica subsp. enterica; Salmonella enterica subsp. enterica serovar Typhimurium MTGDNTLITSHGINRRDFMKLCAALAATMGLSSKAAAEMAESVSNPQRPPVIWIGAQECT WP_000145429.1 hydrogenase 2 small subunit [Salmonella enterica] Length: 372\nNumber of predicted TMHs: 1\nExp number of AAs in TMHs: 24.51309\nExp number, first 60 AAs: 0.2682\nTotal prob of N-in: 0.14402\noutside 1 331\nTMhelix 332 354\ninside 355 372

25190 GCF_001566095.1_8080_3_59 Salmonella enterica subsp. enterica serovar Typhimurium str. DT104 Proteobacteria; Gammaproteobacteria; Enterobacterales; Enterobacteriaceae; Salmonella; Salmonella enterica; Salmonella enterica subsp. enterica; Salmonella enterica subsp. enterica serovar Typhimurium MNNEETFYQAMRRKGVTRRSFLKFCSLAATSLGLGAGMTPKIAWALENKPRIPVVWIHGL WP_001058312.1 hydrogenase [Salmonella enterica] Length: 372\nNumber of predicted TMHs: 2\nExp number of AAs in TMHs: 36.51738\nExp number, first 60 AAs: 19.72849\nTotal prob of N-in: 0.96170\nPOSSIBLE N-term signal sequence\ninside 1 19\nTMhelix 20 39\noutside 40 325\nTMhelix 326 348\ninside 349 372

25191 GCF_001566175.1_5275_7_2Salmonella enterica subsp. enterica serovar Typhimurium str. DT104 Proteobacteria; Gammaproteobacteria; Enterobacterales; Enterobacteriaceae; Salmonella; Salmonella enterica; Salmonella enterica subsp. enterica; Salmonella enterica subsp. enterica serovar Typhimurium MNNEETFYQAMRRKGVTRRSFLKFCSLAATSLGLGAGMTPKIAWALENKPRIPVVWIHGL WP_001058312.1 hydrogenase [Salmonella enterica] Length: 372\nNumber of predicted TMHs: 2\nExp number of AAs in TMHs: 36.51738\nExp number, first 60 AAs: 19.72849\nTotal prob of N-in: 0.96170\nPOSSIBLE N-term signal sequence\ninside 1 19\nTMhelix 20 39\noutside 40 325\nTMhelix 326 348\ninside 349 372

25192 GCF_001566175.1_5275_7_2Salmonella enterica subsp. enterica serovar Typhimurium str. DT104 Proteobacteria; Gammaproteobacteria; Enterobacterales; Enterobacteriaceae; Salmonella; Salmonella enterica; Salmonella enterica subsp. enterica; Salmonella enterica subsp. enterica serovar Typhimurium MTGDNTLITSHGINRRDFMKLCAALAATMGLSSKAAAEMAESVSNPQRPPVIWIGAQECT WP_000145429.1 hydrogenase 2 small subunit [Salmonella enterica] Length: 372\nNumber of predicted TMHs: 1\nExp number of AAs in TMHs: 24.51309\nExp number, first 60 AAs: 0.2682\nTotal prob of N-in: 0.14402\noutside 1 331\nTMhelix 332 354\ninside 355 372

25193 GCF_001566055.1_5275_1_2Salmonella enterica subsp. enterica serovar Typhimurium str. DT104 Proteobacteria; Gammaproteobacteria; Enterobacterales; Enterobacteriaceae; Salmonella; Salmonella enterica;

Salmonella enterica subsp. enterica; Salmonella enterica subsp. enterica serovar Typhimurium
MNNEETFYQAMRRKGVTRRSFLKFCSLAATSLGLGAGMTPKIAWALENKPRIPVVWIHGL WP_001058312.1
hydrogenase [Salmonella enterica] Length: 372\nNumber of predicted TMHs: 2\nExp number of AAs in TMHs: 36.51738\nExp number, first 60 AAs: 19.72849\nTotal prob of N-in: 0.96170\nPOSSIBLE N-term signal sequence\ninside 1 19\nTMhelix 20 39\noutside 40 325\nTMhelix 326 348\ninside 349 372

25194 GCF_001566115.1_5275_1_10 Salmonella enterica subsp. enterica serovar Typhimurium str. DT104
Proteobacteria; Gammaproteobacteria; Enterobacterales; Enterobacteriaceae; Salmonella; Salmonella enterica; Salmonella enterica subsp. enterica; Salmonella enterica subsp. enterica serovar Typhimurium
MNNEETFYQAMRRKGVTRRSFLKFCSLAATSLGLGAGMTPKIAWALENKPRIPVVWIHGL WP_001058312.1
hydrogenase [Salmonella enterica] Length: 372\nNumber of predicted TMHs: 2\nExp number of AAs in TMHs: 36.51738\nExp number, first 60 AAs: 19.72849\nTotal prob of N-in: 0.96170\nPOSSIBLE N-term signal sequence\ninside 1 19\nTMhelix 20 39\noutside 40 325\nTMhelix 326 348\ninside 349 372

25195 GCF_001565995.1_9065_1_9Salmonella enterica subsp. enterica serovar Typhimurium str. DT104
Proteobacteria; Gammaproteobacteria; Enterobacterales; Enterobacteriaceae; Salmonella; Salmonella enterica; Salmonella enterica subsp. enterica; Salmonella enterica subsp. enterica serovar Typhimurium
MNNEETFYQAMRRKGVTRRSFLKFCSLAATSLGLGAGMTPKIAWALENKPRIPVVWIHGL WP_001058312.1
hydrogenase [Salmonella enterica] Length: 372\nNumber of predicted TMHs: 2\nExp number of AAs in TMHs: 36.51738\nExp number, first 60 AAs: 19.72849\nTotal prob of N-in: 0.96170\nPOSSIBLE N-term signal sequence\ninside 1 19\nTMhelix 20 39\noutside 40 325\nTMhelix 326 348\ninside 349 372

25196 GCF_001565935.1_9066_1_12 Salmonella enterica subsp. enterica serovar Typhimurium str. DT104
Proteobacteria; Gammaproteobacteria; Enterobacterales; Enterobacteriaceae; Salmonella; Salmonella enterica; Salmonella enterica subsp. enterica; Salmonella enterica subsp. enterica serovar Typhimurium
MNNEETFYQAMRRKGVTRRSFLKFCSLAATSLGLGAGMTPKIAWALENKPRIPVVWIHGL WP_001058312.1
hydrogenase [Salmonella enterica] Length: 372\nNumber of predicted TMHs: 2\nExp number of AAs in TMHs: 36.51738\nExp number, first 60 AAs: 19.72849\nTotal prob of N-in: 0.96170\nPOSSIBLE N-term signal sequence\ninside 1 19\nTMhelix 20 39\noutside 40 325\nTMhelix 326 348\ninside 349 372

25197 GCF_001565955.1_5299_6_5Salmonella enterica subsp. enterica serovar Typhimurium str. DT104
Proteobacteria; Gammaproteobacteria; Enterobacterales; Enterobacteriaceae; Salmonella; Salmonella enterica; Salmonella enterica subsp. enterica; Salmonella enterica subsp. enterica serovar Typhimurium
MNNEETFYQAMRRKGVTRRSFLKFCSLAATSLGLGAGMTPKIAWALENKPRIPVVWIHGL WP_001058312.1
hydrogenase [Salmonella enterica] Length: 372\nNumber of predicted TMHs: 2\nExp number of AAs in TMHs: 36.51738\nExp number, first 60 AAs: 19.72849\nTotal prob of N-in: 0.96170\nPOSSIBLE N-term signal sequence\ninside 1 19\nTMhelix 20 39\noutside 40 325\nTMhelix 326 348\ninside 349 372

25198 GCF_001566095.1_8080_3_59 Salmonella enterica subsp. enterica serovar Typhimurium str. DT104
Proteobacteria; Gammaproteobacteria; Enterobacterales; Enterobacteriaceae; Salmonella; Salmonella enterica; Salmonella enterica subsp. enterica; Salmonella enterica subsp. enterica serovar Typhimurium
MTGDNTLITSHGINRRDFMKLCAALAATMGLSSKAAAEMAESVSNPQRPPVIWIGAQECT WP_000145429.1
hydrogenase 2 small subunit [Salmonella enterica] Length: 372\nNumber of predicted TMHs: 1\nExp number of AAs in TMHs: 24.51309\nExp number, first 60 AAs: 0.2682\nTotal prob of N-in: 0.14402\noutside 1 331\nTMhelix 332 354\ninside 355 372

25199 GCF_001566055.1_5275_1_2Salmonella enterica subsp. enterica serovar Typhimurium str. DT104
Proteobacteria; Gammaproteobacteria; Enterobacterales; Enterobacteriaceae; Salmonella; Salmonella enterica; Salmonella enterica subsp. enterica; Salmonella enterica subsp. enterica serovar Typhimurium
MTGDNTLITSHGINRRDFMKLCAALAATMGLSSKAAAEMAESVSNPQRPPVIWIGAQECT WP_000145429.1
hydrogenase 2 small subunit [Salmonella enterica] Length: 372\nNumber of predicted TMHs: 1\nExp number of AAs in TMHs: 24.51309\nExp number, first 60 AAs: 0.2682\nTotal prob of N-in: 0.14402\noutside 1 331\nTMhelix 332 354\ninside 355 372

25200 GCF_001565995.1_9065_1_9Salmonella enterica subsp. enterica serovar Typhimurium str. DT104
Proteobacteria; Gammaproteobacteria; Enterobacterales; Enterobacteriaceae; Salmonella; Salmonella enterica; Salmonella enterica subsp. enterica; Salmonella enterica subsp. enterica serovar Typhimurium
MTGDNTLITSHGINRRDFMKLCAALAATMGLSSKAAAEMAESVSNPQRPPVIWIGAQECT WP_000145429.1
hydrogenase 2 small subunit [Salmonella enterica] Length: 372\nNumber of predicted TMHs: 1\nExp number of AAs in TMHs: 24.51309\nExp number, first 60 AAs: 0.2682\nTotal prob of N-in: 0.14402\noutside 1 331\nTMhelix 332 354\ninside 355 372

25201 GCF_001565955.1_5299_6_5Salmonella enterica subsp. enterica serovar Typhimurium str. DT104 Proteobacteria; Gammaproteobacteria; Enterobacterales; Enterobacteriaceae; Salmonella; Salmonella enterica; Salmonella enterica subsp. enterica; Salmonella enterica subsp. enterica serovar Typhimurium MTGDNTLITSHGINRRDFMKLCAALAATMGLSSKAAAEMAESVSNPQRPPVIWIGAQECT WP_000145429.1 hydrogenase 2 small subunit [Salmonella enterica] Length: 372\nNumber of predicted TMHs: 1\nExp number of AAs in TMHs: 24.51309\nExp number, first 60 AAs: 0.2682\nTotal prob of N-in: 0.14402\noutside 1 331\nTMhelix 332 354\ninside 355 372

25202 GCF_001565935.1_9066_1_12 Salmonella enterica subsp. enterica serovar Typhimurium str. DT104 Proteobacteria; Gammaproteobacteria; Enterobacterales; Enterobacteriaceae; Salmonella; Salmonella enterica; Salmonella enterica subsp. enterica; Salmonella enterica subsp. enterica serovar Typhimurium MTGDNTLITSHGINRRDFMKLCAALAATMGLSSKAAAEMAESVSNPQRPPVIWIGAQECT WP_000145429.1 hydrogenase 2 small subunit [Salmonella enterica] Length: 372\nNumber of predicted TMHs: 1\nExp number of AAs in TMHs: 24.51309\nExp number, first 60 AAs: 0.2682\nTotal prob of N-in: 0.14402\noutside 1 331\nTMhelix 332 354\ninside 355 372

25203 GCF_000493675.1_DT104 Salmonella enterica subsp. enterica serovar Typhimurium str. DT104 Proteobacteria; Gammaproteobacteria; Enterobacterales; Enterobacteriaceae; Salmonella; Salmonella enterica; Salmonella enterica subsp. enterica; Salmonella enterica subsp. enterica serovar Typhimurium MTGDNTLITSHGINRRDFMKLCAALAATMGLSSKAAAEMAESVSNPQRPPVIWIGAQECT WP_000145429.1 hydrogenase 2 small subunit [Salmonella enterica] Length: 372\nNumber of predicted TMHs: 1\nExp number of AAs in TMHs: 24.51309\nExp number, first 60 AAs: 0.2682\nTotal prob of N-in: 0.14402\noutside 1 331\nTMhelix 332 354\ninside 355 372

25204 GCF_000493675.1_DT104 Salmonella enterica subsp. enterica serovar Typhimurium str. DT104 Proteobacteria; Gammaproteobacteria; Enterobacterales; Enterobacteriaceae; Salmonella; Salmonella enterica; Salmonella enterica subsp. enterica; Salmonella enterica subsp. enterica serovar Typhimurium MNNEETFYQAMRRKGVTRRSFLKFCSLAATSLGLGAGMTPKIAWALENKPRIPVVWIHGL WP_001058312.1 hydrogenase [Salmonella enterica] Length: 372\nNumber of predicted TMHs: 2\nExp number of AAs in TMHs: 36.51738\nExp number, first 60 AAs: 19.72849\nTotal prob of N-in: 0.96170\nPOSSIBLE N-term signal sequence\ninside 1 19\nTMhelix 20 39\noutside 40 325\nTMhelix 326 348\ninside 349 372

25205 GCF_001221285.1_8080_3_19 Salmonella enterica subsp. enterica serovar Typhimurium str. DT104 Proteobacteria; Gammaproteobacteria; Enterobacterales; Enterobacteriaceae; Salmonella; Salmonella enterica; Salmonella enterica subsp. enterica; Salmonella enterica subsp. enterica serovar Typhimurium MNNEETFYQAMRRKGVTRRSFLKFCSLAATSLGLGAGMTPKIAWALENKPRIPVVWIHGL WP_001058312.1 hydrogenase [Salmonella enterica] Length: 372\nNumber of predicted TMHs: 2\nExp number of AAs in TMHs: 36.51738\nExp number, first 60 AAs: 19.72849\nTotal prob of N-in: 0.96170\nPOSSIBLE N-term signal sequence\ninside 1 19\nTMhelix 20 39\noutside 40 325\nTMhelix 326 348\ninside 349 372

25206 GCF_001221805.1_5285_6_10 Salmonella enterica subsp. enterica serovar Typhimurium str. DT104 Proteobacteria; Gammaproteobacteria; Enterobacterales; Enterobacteriaceae; Salmonella; Salmonella enterica; Salmonella enterica subsp. enterica; Salmonella enterica subsp. enterica serovar Typhimurium MNNEETFYQAMRRKGVTRRSFLKFCSLAATSLGLGAGMTPKIAWALENKPRIPVVWIHGL WP_001058312.1 hydrogenase [Salmonella enterica] Length: 372\nNumber of predicted TMHs: 2\nExp number of AAs in TMHs: 36.51738\nExp number, first 60 AAs: 19.72849\nTotal prob of N-in: 0.96170\nPOSSIBLE N-term signal sequence\ninside 1 19\nTMhelix 20 39\noutside 40 325\nTMhelix 326 348\ninside 349 372

25207 GCF_001221245.1_5285_7_4Salmonella enterica subsp. enterica serovar Typhimurium str. DT104 Proteobacteria; Gammaproteobacteria; Enterobacterales; Enterobacteriaceae; Salmonella; Salmonella enterica; Salmonella enterica subsp. enterica; Salmonella enterica subsp. enterica serovar Typhimurium MNNEETFYQAMRRKGVTRRSFLKFCSLAATSLGLGAGMTPKIAWALENKPRIPVVWIHGL WP_001058312.1 hydrogenase [Salmonella enterica] Length: 372\nNumber of predicted TMHs: 2\nExp number of AAs in TMHs: 36.51738\nExp number, first 60 AAs: 19.72849\nTotal prob of N-in: 0.96170\nPOSSIBLE N-term signal sequence\ninside 1 19\nTMhelix 20 39\noutside 40 325\nTMhelix 326 348\ninside 349 372

25208 GCF_001219445.1_8036_3_74 Salmonella enterica subsp. enterica serovar Typhimurium str. DT104 Proteobacteria; Gammaproteobacteria; Enterobacterales; Enterobacteriaceae; Salmonella; Salmonella enterica; Salmonella enterica subsp. enterica; Salmonella enterica subsp. enterica serovar Typhimurium MTGDNTLITSHGINRRDFMKLCAALAATMGLSSKAAAEMAESVSNPQRPPVIWIGAQECT WP_000145429.1 hydrogenase 2 small subunit [Salmonella enterica] Length: 372\nNumber of predicted TMHs: 1\nExp

number of AAs in TMHs: 24.51309\nExp number, first 60 AAs: 0.2682\nTotal prob of N-in: 0.14402\noutside 1 331\nTMhelix 332 354\ninside 355 372

25209 GCF_001221905.1_5275_2_8Salmonella enterica subsp. enterica serovar Typhimurium str. DT104 Proteobacteria; Gammaproteobacteria; Enterobacterales; Enterobacteriaceae; Salmonella; Salmonella enterica; Salmonella enterica subsp. enterica; Salmonella enterica subsp. enterica serovar Typhimurium MNNEETFYQAMRRKGVTRRSFLKFCSLAATSLGLGAGMTPKIAWALENKPRIPVVWIHGL WP_001058312.1 hydrogenase [Salmonella enterica] Length: 372\nNumber of predicted TMHs: 2\nExp number of AAs in TMHs: 36.51738\nExp number, first 60 AAs: 19.72849\nTotal prob of N-in: 0.96170\nPOSSIBLE N-term signal sequence\ninside 1 19\nTMhelix 20 39\noutside 40 325\nTMhelix 326 348\ninside 349 372

25210 GCF_001219285.1_5388_1_2Salmonella enterica subsp. enterica serovar Typhimurium str. DT104 Proteobacteria; Gammaproteobacteria; Enterobacterales; Enterobacteriaceae; Salmonella; Salmonella enterica; Salmonella enterica subsp. enterica; Salmonella enterica subsp. enterica serovar Typhimurium MNNEETFYQAMRRKGVTRRSFLKFCSLAATSLGLGAGMTPKIAWALENKPRIPVVWIHGL WP_001058312.1 hydrogenase [Salmonella enterica] Length: 372\nNumber of predicted TMHs: 2\nExp number of AAs in TMHs: 36.51738\nExp number, first 60 AAs: 19.72849\nTotal prob of N-in: 0.96170\nPOSSIBLE N-term signal sequence\ninside 1 19\nTMhelix 20 39\noutside 40 325\nTMhelix 326 348\ninside 349 372

25211 GCF_001219365.1_8080_3_56 Salmonella enterica subsp. enterica serovar Typhimurium str. DT104 Proteobacteria; Gammaproteobacteria; Enterobacterales; Enterobacteriaceae; Salmonella; Salmonella enterica; Salmonella enterica subsp. enterica; Salmonella enterica subsp. enterica serovar Typhimurium MNNEETFYQAMRRKGVTRRSFLKFCSLAATSLGLGAGMTPKIAWALENKPRIPVVWIHGL WP_001058312.1 hydrogenase [Salmonella enterica] Length: 372\nNumber of predicted TMHs: 2\nExp number of AAs in TMHs: 36.51738\nExp number, first 60 AAs: 19.72849\nTotal prob of N-in: 0.96170\nPOSSIBLE N-term signal sequence\ninside 1 19\nTMhelix 20 39\noutside 40 325\nTMhelix 326 348\ninside 349 372

25212 GCF_001221665.1_9065_1_2Salmonella enterica subsp. enterica serovar Typhimurium str. DT104 Proteobacteria; Gammaproteobacteria; Enterobacterales; Enterobacteriaceae; Salmonella; Salmonella enterica; Salmonella enterica subsp. enterica; Salmonella enterica subsp. enterica serovar Typhimurium MNNEETFYQAMRRKGVTRRSFLKFCSLAATSLGLGAGMTPKIAWALENKPRIPVVWIHGL WP_001058312.1 hydrogenase [Salmonella enterica] Length: 372\nNumber of predicted TMHs: 2\nExp number of AAs in TMHs: 36.51738\nExp number, first 60 AAs: 19.72849\nTotal prob of N-in: 0.96170\nPOSSIBLE N-term signal sequence\ninside 1 19\nTMhelix 20 39\noutside 40 325\nTMhelix 326 348\ninside 349 372

25213 GCF_001219365.1_8080_3_56 Salmonella enterica subsp. enterica serovar Typhimurium str. DT104 Proteobacteria; Gammaproteobacteria; Enterobacterales; Enterobacteriaceae; Salmonella; Salmonella enterica; Salmonella enterica subsp. enterica; Salmonella enterica subsp. enterica serovar Typhimurium MTGDNTLITSHGINRRDFMKLCAALAATMGLSSKAAAEMAESVSNPQRPPVIWIGAQECT WP_000145429.1 hydrogenase 2 small subunit [Salmonella enterica] Length: 372\nNumber of predicted TMHs: 1\nExp number of AAs in TMHs: 24.51309\nExp number, first 60 AAs: 0.2682\nTotal prob of N-in: 0.14402\noutside 1 331\nTMhelix 332 354\ninside 355 372

25214 GCF_001219225.1_5275_3_1Salmonella enterica subsp. enterica serovar Typhimurium str. DT104 Proteobacteria; Gammaproteobacteria; Enterobacterales; Enterobacteriaceae; Salmonella; Salmonella enterica; Salmonella enterica subsp. enterica; Salmonella enterica subsp. enterica serovar Typhimurium MNNEETFYQAMRRKGVTRRSFLKFCSLAATSLGLGAGMTPKIAWALENKPRIPVVWIHGL WP_001058312.1 hydrogenase [Salmonella enterica] Length: 372\nNumber of predicted TMHs: 2\nExp number of AAs in TMHs: 36.51738\nExp number, first 60 AAs: 19.72849\nTotal prob of N-in: 0.96170\nPOSSIBLE N-term signal sequence\ninside 1 19\nTMhelix 20 39\noutside 40 325\nTMhelix 326 348\ninside 349 372

25215 GCF_001219185.1_8036_3_55 Salmonella enterica subsp. enterica serovar Typhimurium str. DT104 Proteobacteria; Gammaproteobacteria; Enterobacterales; Enterobacteriaceae; Salmonella; Salmonella enterica; Salmonella enterica subsp. enterica; Salmonella enterica subsp. enterica serovar Typhimurium MNNEETFYQAMRRKGVTRRSFLKFCSLAATSLGLGAGMTPKIAWALENKPRIPVVWIHGL WP_001058312.1 hydrogenase [Salmonella enterica] Length: 372\nNumber of predicted TMHs: 2\nExp number of AAs in TMHs: 36.51738\nExp number, first 60 AAs: 19.72849\nTotal prob of N-in: 0.96170\nPOSSIBLE N-term signal sequence\ninside 1 19\nTMhelix 20 39\noutside 40 325\nTMhelix 326 348\ninside 349 372

25216 GCF_001219045.1_5275_6_6Salmonella enterica subsp. enterica serovar Typhimurium str. DT104 Proteobacteria; Gammaproteobacteria; Enterobacterales; Enterobacteriaceae; Salmonella; Salmonella enterica;

Salmonella enterica subsp. enterica; Salmonella enterica subsp. enterica serovar Typhimurium
MNNEETFYQAMRRKGVTRRSFLKFCSLAATSLGLGAGMTPKIAWALENKPRIPVVWIHGL WP_001058312.1
hydrogenase [Salmonella enterica] Length: 372\nNumber of predicted TMHs: 2\nExp number of AAs in TMHs: 36.51738\nExp number, first 60 AAs: 19.72849\nTotal prob of N-in: 0.96170\nPOSSIBLE N-term signal sequence\ninside 1 19\nTMhelix 20 39\noutside 40 325\nTMhelix 326 348\ninside 349 372

25217 GCF_001219045.1_5275_6_6Salmonella enterica subsp. enterica serovar Typhimurium str. DT104
Proteobacteria; Gammaproteobacteria; Enterobacterales; Enterobacteriaceae; Salmonella; Salmonella enterica; Salmonella enterica subsp. enterica; Salmonella enterica subsp. enterica serovar Typhimurium
MTGDNTLITSHGINRRDFMKLCAALAATMGLSSKAAAEMAESVSNPQRPPVIWIGAQECT WP_000145429.1
hydrogenase 2 small subunit [Salmonella enterica] Length: 372\nNumber of predicted TMHs: 1\nExp number of AAs in TMHs: 24.51309\nExp number, first 60 AAs: 0.2682\nTotal prob of N-in: 0.14402\noutside 1 331\nTMhelix 332 354\ninside 355 372

25218 GCF_001218925.1_5275_7_5Salmonella enterica subsp. enterica serovar Typhimurium str. DT104
Proteobacteria; Gammaproteobacteria; Enterobacterales; Enterobacteriaceae; Salmonella; Salmonella enterica; Salmonella enterica subsp. enterica; Salmonella enterica subsp. enterica serovar Typhimurium
MTGDNTLITSHGINRRDFMKLCAALAATMGLSSKAAAEMAESVSNPQRPPVIWIGAQECT WP_000145429.1
hydrogenase 2 small subunit [Salmonella enterica] Length: 372\nNumber of predicted TMHs: 1\nExp number of AAs in TMHs: 24.51309\nExp number, first 60 AAs: 0.2682\nTotal prob of N-in: 0.14402\noutside 1 331\nTMhelix 332 354\ninside 355 372

25219 GCF_001219005.1_8036_3_47 Salmonella enterica subsp. enterica serovar Typhimurium str. DT104
Proteobacteria; Gammaproteobacteria; Enterobacterales; Enterobacteriaceae; Salmonella; Salmonella enterica; Salmonella enterica subsp. enterica; Salmonella enterica subsp. enterica serovar Typhimurium
MTGDNTLITSHGINRRDFMKLCAALAATMGLSSKAAAEMAESVSNPQRPPVIWIGAQECT WP_000145429.1
hydrogenase 2 small subunit [Salmonella enterica] Length: 372\nNumber of predicted TMHs: 1\nExp number of AAs in TMHs: 24.51309\nExp number, first 60 AAs: 0.2682\nTotal prob of N-in: 0.14402\noutside 1 331\nTMhelix 332 354\ninside 355 372

25220 GCF_001218865.1_5285_7_12 Salmonella enterica subsp. enterica serovar Typhimurium str. DT104
Proteobacteria; Gammaproteobacteria; Enterobacterales; Enterobacteriaceae; Salmonella; Salmonella enterica; Salmonella enterica subsp. enterica; Salmonella enterica subsp. enterica serovar Typhimurium
MTGDNTLITSHGINRRDFMKLCAALAATMGLSSKAAAEMAESVSNPQRPPVIWIGAQECT WP_000145429.1
hydrogenase 2 small subunit [Salmonella enterica] Length: 372\nNumber of predicted TMHs: 1\nExp number of AAs in TMHs: 24.51309\nExp number, first 60 AAs: 0.2682\nTotal prob of N-in: 0.14402\noutside 1 331\nTMhelix 332 354\ninside 355 372

25221 GCF_001218785.1_8080_3_45 Salmonella enterica subsp. enterica serovar Typhimurium str. DT104
Proteobacteria; Gammaproteobacteria; Enterobacterales; Enterobacteriaceae; Salmonella; Salmonella enterica; Salmonella enterica subsp. enterica; Salmonella enterica subsp. enterica serovar Typhimurium
MTGDNTLITSHGINRRDFMKLCAALAATMGLSSKAAAEMAESVSNPQRPPVIWIGAQECT WP_000145429.1
hydrogenase 2 small subunit [Salmonella enterica] Length: 372\nNumber of predicted TMHs: 1\nExp number of AAs in TMHs: 24.51309\nExp number, first 60 AAs: 0.2682\nTotal prob of N-in: 0.14402\noutside 1 331\nTMhelix 332 354\ninside 355 372

25222 GCF_001218145.1_5295_2_12 Salmonella enterica subsp. enterica serovar Typhimurium str. DT104
Proteobacteria; Gammaproteobacteria; Enterobacterales; Enterobacteriaceae; Salmonella; Salmonella enterica; Salmonella enterica subsp. enterica; Salmonella enterica subsp. enterica serovar Typhimurium
MNNEETFYQAMRRKGVTRRSFLKFCSLAATSLGLGAGMTPKIAWALENKPRIPVVWIHGL WP_001058312.1
hydrogenase [Salmonella enterica] Length: 372\nNumber of predicted TMHs: 2\nExp number of AAs in TMHs: 36.51738\nExp number, first 60 AAs: 19.72849\nTotal prob of N-in: 0.96170\nPOSSIBLE N-term signal sequence\ninside 1 19\nTMhelix 20 39\noutside 40 325\nTMhelix 326 348\ninside 349 372

25223 GCF_001221245.1_5285_7_4Salmonella enterica subsp. enterica serovar Typhimurium str. DT104
Proteobacteria; Gammaproteobacteria; Enterobacterales; Enterobacteriaceae; Salmonella; Salmonella enterica; Salmonella enterica subsp. enterica; Salmonella enterica subsp. enterica serovar Typhimurium
MTGDNTLITSHGINRRDFMKLCAALAATMGLSSKAAAEMAESVSNPQRPPVIWIGAQECT WP_000145429.1
hydrogenase 2 small subunit [Salmonella enterica] Length: 372\nNumber of predicted TMHs: 1\nExp number of AAs in TMHs: 24.51309\nExp number, first 60 AAs: 0.2682\nTotal prob of N-in: 0.14402\noutside 1 331\nTMhelix 332 354\ninside 355 372

25224	GCF_001218445.1_9066_1_5Salmonella enterica subsp. enterica serovar Typhimurium str. DT104	Proteobacteria; Gammaproteobacteria; Enterobacterales; Enterobacteriaceae; Salmonella; Salmonella enterica; Salmonella enterica subsp. enterica; Salmonella enterica subsp. enterica serovar Typhimurium	MNNEETFYQAMRRKGVTRRSFLKFCSLAATSLGLGAGMTPKIAWALENKPRIPVVWIHGL	WP_001058312.1	hydrogenase [Salmonella enterica]	Length: 372\nNumber of predicted TMHs:  2\nExp number of AAs in TMHs: 36.51738\nExp number, first 60 AAs:  19.72849\nTotal prob of N-in:   0.96170\nPOSSIBLE N-term signal sequence\ninside   1   19\nTMhelix   20   39\noutside   40   325\nTMhelix   326   348\ninside   349   372

25225	GCF_001218385.1_5299_3_4Salmonella enterica subsp. enterica serovar Typhimurium str. DT104	Proteobacteria; Gammaproteobacteria; Enterobacterales; Enterobacteriaceae; Salmonella; Salmonella enterica; Salmonella enterica subsp. enterica; Salmonella enterica subsp. enterica serovar Typhimurium	MNNEETFYQAMRRKGVTRRSFLKFCSLAATSLGLGAGMTPKIAWALENKPRIPVVWIHGL	WP_001058312.1	hydrogenase [Salmonella enterica]	Length: 372\nNumber of predicted TMHs:  2\nExp number of AAs in TMHs: 36.51738\nExp number, first 60 AAs:  19.72849\nTotal prob of N-in:   0.96170\nPOSSIBLE N-term signal sequence\ninside   1   19\nTMhelix   20   39\noutside   40   325\nTMhelix   326   348\ninside   349   372

25226	GCF_001218445.1_9066_1_5Salmonella enterica subsp. enterica serovar Typhimurium str. DT104	Proteobacteria; Gammaproteobacteria; Enterobacterales; Enterobacteriaceae; Salmonella; Salmonella enterica; Salmonella enterica subsp. enterica; Salmonella enterica subsp. enterica serovar Typhimurium	MTGDNTLITSHGINRRDFMKLCAALAATMGLSSKAAAEMAESVSNPQRPPVIWIGAQECT	WP_000145429.1	hydrogenase 2 small subunit [Salmonella enterica]	Length: 372\nNumber of predicted TMHs:  1\nExp number of AAs in TMHs: 24.51309\nExp number, first 60 AAs:  0.2682\nTotal prob of N-in:   0.14402\noutside   1   331\nTMhelix   332   354\ninside   355   372

25227	GCF_001220285.1_8080_3_88	Salmonella enterica subsp. enterica serovar Typhimurium str. DT104	Proteobacteria; Gammaproteobacteria; Enterobacterales; Enterobacteriaceae; Salmonella; Salmonella enterica; Salmonella enterica subsp. enterica; Salmonella enterica subsp. enterica serovar Typhimurium	MTGDNTLITSHGINRRDFMKLCAALAATMGLSSKAAAEMAESVSNPQRPPVIWIGAQECT	WP_000145429.1	hydrogenase 2 small subunit [Salmonella enterica]	Length: 372\nNumber of predicted TMHs:  1\nExp number of AAs in TMHs: 24.51309\nExp number, first 60 AAs:  0.2682\nTotal prob of N-in:   0.14402\noutside   1   331\nTMhelix   332   354\ninside   355   372

25228	GCF_001565975.1_8080_3_10	Salmonella enterica subsp. enterica serovar Typhimurium str. DT104	Proteobacteria; Gammaproteobacteria; Enterobacterales; Enterobacteriaceae; Salmonella; Salmonella enterica; Salmonella enterica subsp. enterica; Salmonella enterica subsp. enterica serovar Typhimurium	MNNEETFYQAMRRKGVTRRSFLKFCSLAATSLGLGAGMTPKIAWALENKPRIPVVWIHGL	WP_001058312.1	hydrogenase [Salmonella enterica]	Length: 372\nNumber of predicted TMHs:  2\nExp number of AAs in TMHs: 36.51738\nExp number, first 60 AAs:  19.72849\nTotal prob of N-in:   0.96170\nPOSSIBLE N-term signal sequence\ninside   1   19\nTMhelix   20   39\noutside   40   325\nTMhelix   326   348\ninside   349   372

25229	GCF_001220025.1_5275_8_8Salmonella enterica subsp. enterica serovar Typhimurium str. DT104	Proteobacteria; Gammaproteobacteria; Enterobacterales; Enterobacteriaceae; Salmonella; Salmonella enterica; Salmonella enterica subsp. enterica; Salmonella enterica subsp. enterica serovar Typhimurium	MNNEETFYQAMRRKGVTRRSFLKFCSLAATSLGLGAGMTPKIAWALENKPRIPVVWIHGL	WP_001058312.1	hydrogenase [Salmonella enterica]	Length: 372\nNumber of predicted TMHs:  2\nExp number of AAs in TMHs: 36.51738\nExp number, first 60 AAs:  19.72849\nTotal prob of N-in:   0.96170\nPOSSIBLE N-term signal sequence\ninside   1   19\nTMhelix   20   39\noutside   40   325\nTMhelix   326   348\ninside   349   372

25230	GCF_001220645.1_8036_3_67	Salmonella enterica subsp. enterica serovar Typhimurium str. DT104	Proteobacteria; Gammaproteobacteria; Enterobacterales; Enterobacteriaceae; Salmonella; Salmonella enterica; Salmonella enterica subsp. enterica; Salmonella enterica subsp. enterica serovar Typhimurium	MTGDNTLITSHGINRRDFMKLCAALAATMGLSSKAAAEMAESVSNPQRPPVIWIGAQECT	WP_000145429.1	hydrogenase 2 small subunit [Salmonella enterica]	Length: 372\nNumber of predicted TMHs:  1\nExp number of AAs in TMHs: 24.51309\nExp number, first 60 AAs:  0.2682\nTotal prob of N-in:   0.14402\noutside   1   331\nTMhelix   332   354\ninside   355   372

25231	GCF_001220685.1_8080_3_33	Salmonella enterica subsp. enterica serovar Typhimurium str. DT104	Proteobacteria; Gammaproteobacteria; Enterobacterales; Enterobacteriaceae; Salmonella; Salmonella enterica; Salmonella enterica subsp. enterica; Salmonella enterica subsp. enterica serovar Typhimurium	MTGDNTLITSHGINRRDFMKLCAALAATMGLSSKAAAEMAESVSNPQRPPVIWIGAQECT	WP_000145429.1	hydrogenase 2 small subunit [Salmonella enterica]	Length: 372\nNumber of predicted TMHs:  1\nExp

number of AAs in TMHs: 24.51309\nExp number, first 60 AAs: 0.2682\nTotal prob of N-in: 0.14402\noutside 1 331\nTMhelix 332 354\ninside 355 372

25232 GCF_001220645.1_8036_3_67 Salmonella enterica subsp. enterica serovar Typhimurium str. DT104 Proteobacteria; Gammaproteobacteria; Enterobacterales; Enterobacteriaceae; Salmonella; Salmonella enterica; Salmonella enterica subsp. enterica; Salmonella enterica subsp. enterica serovar Typhimurium MNNEETFYQAMRRKGVTRRSFLKFCSLAATSLGLGAGMTPKIAWALENKPRIPVVWIHGL WP_001058312.1 hydrogenase [Salmonella enterica] Length: 372\nNumber of predicted TMHs: 2\nExp number of AAs in TMHs: 36.51738\nExp number, first 60 AAs: 19.72849\nTotal prob of N-in: 0.96170\nPOSSIBLE N-term signal sequence\ninside 1 19\nTMhelix 20 39\noutside 40 325\nTMhelix 326 348\ninside 349 372

25233 GCF_001220725.1_8080_3_94 Salmonella enterica subsp. enterica serovar Typhimurium str. DT104 Proteobacteria; Gammaproteobacteria; Enterobacterales; Enterobacteriaceae; Salmonella; Salmonella enterica; Salmonella enterica subsp. enterica; Salmonella enterica subsp. enterica serovar Typhimurium MTGDNTLITSHGINRRDFMKLCAALAATMGLSSKAAAEMAESVSNPQRPPVIWIGAQECT WP_000145429.1 hydrogenase 2 small subunit [Salmonella enterica] Length: 372\nNumber of predicted TMHs: 1\nExp number of AAs in TMHs: 24.51309\nExp number, first 60 AAs: 0.2682\nTotal prob of N-in: 0.14402\noutside 1 331\nTMhelix 332 354\ninside 355 372

25234 GCF_001220865.1_5299_6_11 Salmonella enterica subsp. enterica serovar Typhimurium str. DT104 Proteobacteria; Gammaproteobacteria; Enterobacterales; Enterobacteriaceae; Salmonella; Salmonella enterica; Salmonella enterica subsp. enterica; Salmonella enterica subsp. enterica serovar Typhimurium MTGDNTLITSHGINRRDFMKLCAALAATMGLSSKAAAEMAESVSNPQRPPVIWIGAQECT WP_000145429.1 hydrogenase 2 small subunit [Salmonella enterica] Length: 372\nNumber of predicted TMHs: 1\nExp number of AAs in TMHs: 24.51309\nExp number, first 60 AAs: 0.2682\nTotal prob of N-in: 0.14402\noutside 1 331\nTMhelix 332 354\ninside 355 372

25235 GCF_001220925.1_8080_3_15 Salmonella enterica subsp. enterica serovar Typhimurium str. DT104 Proteobacteria; Gammaproteobacteria; Enterobacterales; Enterobacteriaceae; Salmonella; Salmonella enterica; Salmonella enterica subsp. enterica; Salmonella enterica subsp. enterica serovar Typhimurium MTGDNTLITSHGINRRDFMKLCAALAATMGLSSKAAAEMAESVSNPQRPPVIWIGAQECT WP_000145429.1 hydrogenase 2 small subunit [Salmonella enterica] Length: 372\nNumber of predicted TMHs: 1\nExp number of AAs in TMHs: 24.51309\nExp number, first 60 AAs: 0.2682\nTotal prob of N-in: 0.14402\noutside 1 331\nTMhelix 332 354\ninside 355 372

25236 GCF_001350075.1_8490_2_55 Salmonella enterica subsp. enterica serovar Typhimurium str. DT104 Proteobacteria; Gammaproteobacteria; Enterobacterales; Enterobacteriaceae; Salmonella; Salmonella enterica; Salmonella enterica subsp. enterica; Salmonella enterica subsp. enterica serovar Typhimurium MNNEETFYQAMRRKGVTRRSFLKFCSLAATSLGLGAGMTPKIAWALENKPRIPVVWIHGL WP_001058312.1 hydrogenase [Salmonella enterica] Length: 372\nNumber of predicted TMHs: 2\nExp number of AAs in TMHs: 36.51738\nExp number, first 60 AAs: 19.72849\nTotal prob of N-in: 0.96170\nPOSSIBLE N-term signal sequence\ninside 1 19\nTMhelix 20 39\noutside 40 325\nTMhelix 326 348\ninside 349 372

25237 GCF_001350155.1_8490_2_54 Salmonella enterica subsp. enterica serovar Typhimurium str. DT104 Proteobacteria; Gammaproteobacteria; Enterobacterales; Enterobacteriaceae; Salmonella; Salmonella enterica; Salmonella enterica subsp. enterica; Salmonella enterica subsp. enterica serovar Typhimurium MNNEETFYQAMRRKGVTRRSFLKFCSLAATSLGLGAGMTPKIAWALENKPRIPVVWIHGL WP_050196264.1 hydrogenase [Salmonella enterica] Length: 372\nNumber of predicted TMHs: 2\nExp number of AAs in TMHs: 36.76337\nExp number, first 60 AAs: 19.73298\nTotal prob of N-in: 0.96170\nPOSSIBLE N-term signal sequence\ninside 1 19\nTMhelix 20 39\noutside 40 325\nTMhelix 326 348\ninside 349 372

25238 GCF_001350195.1_8490_2_46 Salmonella enterica subsp. enterica serovar Typhimurium str. DT104 Proteobacteria; Gammaproteobacteria; Enterobacterales; Enterobacteriaceae; Salmonella; Salmonella enterica; Salmonella enterica subsp. enterica; Salmonella enterica subsp. enterica serovar Typhimurium MTGDNTLITSHGINRRDFMKLCAALAATMGLSSKAAAEMAESVSNPQRPPVIWIGAQECT WP_000145429.1 hydrogenase 2 small subunit [Salmonella enterica] Length: 372\nNumber of predicted TMHs: 1\nExp number of AAs in TMHs: 24.51309\nExp number, first 60 AAs: 0.2682\nTotal prob of N-in: 0.14402\noutside 1 331\nTMhelix 332 354\ninside 355 372

25239 GCF_001326195.1_9066_1_8Salmonella enterica subsp. enterica serovar Typhimurium str. DT104 Proteobacteria; Gammaproteobacteria; Enterobacterales; Enterobacteriaceae; Salmonella; Salmonella enterica;

Salmonella enterica subsp. enterica; Salmonella enterica subsp. enterica serovar Typhimurium
MTGDNTLITSHGINRRDFMKLCAALAATMGLSSKAAAEMAESVSNPQRPPVIWIGAQECT WP_000145429.1
hydrogenase 2 small subunit [Salmonella enterica] Length: 372\nNumber of predicted TMHs: 1\nExp number of AAs in TMHs: 24.51309\nExp number, first 60 AAs: 0.2682\nTotal prob of N-in: 0.14402\noutside 1 331\nTMhelix 332 354\ninside 355 372

25240 GCF_001221005.1_8080_3_48 Salmonella enterica subsp. enterica serovar Typhimurium str. DT104
Proteobacteria; Gammaproteobacteria; Enterobacterales; Enterobacteriaceae; Salmonella; Salmonella enterica; Salmonella enterica subsp. enterica; Salmonella enterica subsp. enterica serovar Typhimurium
MNNEETFYQAMRRKGVTRRSFLKFCSLAATSLGLGAGMTPKIAWALENKPRIPVVWIHGL WP_001058312.1
hydrogenase [Salmonella enterica] Length: 372\nNumber of predicted TMHs: 2\nExp number of AAs in TMHs: 36.51738\nExp number, first 60 AAs: 19.72849\nTotal prob of N-in: 0.96170\nPOSSIBLE N-term signal sequence\ninside 1 19\nTMhelix 20 39\noutside 40 325\nTMhelix 326 348\ninside 349 372

25241 GCF_001218385.1_5299_3_4Salmonella enterica subsp. enterica serovar Typhimurium str. DT104
Proteobacteria; Gammaproteobacteria; Enterobacterales; Enterobacteriaceae; Salmonella; Salmonella enterica; Salmonella enterica subsp. enterica; Salmonella enterica subsp. enterica serovar Typhimurium
MTGDNTLITSHGINRRDFMKLCAALAATMGLSSKAAAEMAESVSNPQRPPVIWIGAQECT WP_000145429.1
hydrogenase 2 small subunit [Salmonella enterica] Length: 372\nNumber of predicted TMHs: 1\nExp number of AAs in TMHs: 24.51309\nExp number, first 60 AAs: 0.2682\nTotal prob of N-in: 0.14402\noutside 1 331\nTMhelix 332 354\ninside 355 372

25242 GCF_001218065.1_8080_3_2Salmonella enterica subsp. enterica serovar Typhimurium str. DT104
Proteobacteria; Gammaproteobacteria; Enterobacterales; Enterobacteriaceae; Salmonella; Salmonella enterica; Salmonella enterica subsp. enterica; Salmonella enterica subsp. enterica serovar Typhimurium
MNNEETFYQAMRRKGVTRRSFLKFCSLAATSLGLGAGMTPKIAWALENKPRIPVVWIHGL WP_001058312.1
hydrogenase [Salmonella enterica] Length: 372\nNumber of predicted TMHs: 2\nExp number of AAs in TMHs: 36.51738\nExp number, first 60 AAs: 19.72849\nTotal prob of N-in: 0.96170\nPOSSIBLE N-term signal sequence\ninside 1 19\nTMhelix 20 39\noutside 40 325\nTMhelix 326 348\ninside 349 372

25243 GCF_001221285.1_8080_3_19 Salmonella enterica subsp. enterica serovar Typhimurium str. DT104
Proteobacteria; Gammaproteobacteria; Enterobacterales; Enterobacteriaceae; Salmonella; Salmonella enterica; Salmonella enterica subsp. enterica; Salmonella enterica subsp. enterica serovar Typhimurium
MTGDNTLITSHGINRRDFMKLCAALAATMGLSSKAAAEMAESVSNPQRPPVIWIGAQECT WP_000145429.1
hydrogenase 2 small subunit [Salmonella enterica] Length: 372\nNumber of predicted TMHs: 1\nExp number of AAs in TMHs: 24.51309\nExp number, first 60 AAs: 0.2682\nTotal prob of N-in: 0.14402\noutside 1 331\nTMhelix 332 354\ninside 355 372

25244 GCF_001221725.1_5275_1_6Salmonella enterica subsp. enterica serovar Typhimurium str. DT104
Proteobacteria; Gammaproteobacteria; Enterobacterales; Enterobacteriaceae; Salmonella; Salmonella enterica; Salmonella enterica subsp. enterica; Salmonella enterica subsp. enterica serovar Typhimurium
MNNEETFYQAMRRKGVTRRSFLKFCSLAATSLGLGAGMTPKIAWALENKPRIPVVWIHGL WP_001058312.1
hydrogenase [Salmonella enterica] Length: 372\nNumber of predicted TMHs: 2\nExp number of AAs in TMHs: 36.51738\nExp number, first 60 AAs: 19.72849\nTotal prob of N-in: 0.96170\nPOSSIBLE N-term signal sequence\ninside 1 19\nTMhelix 20 39\noutside 40 325\nTMhelix 326 348\ninside 349 372

25245 GCF_001217945.1_8036_3_37 Salmonella enterica subsp. enterica serovar Typhimurium str. DT104
Proteobacteria; Gammaproteobacteria; Enterobacterales; Enterobacteriaceae; Salmonella; Salmonella enterica; Salmonella enterica subsp. enterica; Salmonella enterica subsp. enterica serovar Typhimurium
MTGDNTLITSHGINRRDFMKLCAALAATMGLSSKAAAEMAESVSNPQRPPVIWIGAQECT WP_000145429.1
hydrogenase 2 small subunit [Salmonella enterica] Length: 372\nNumber of predicted TMHs: 1\nExp number of AAs in TMHs: 24.51309\nExp number, first 60 AAs: 0.2682\nTotal prob of N-in: 0.14402\noutside 1 331\nTMhelix 332 354\ninside 355 372

25246 GCF_001217905.1_5275_7_3Salmonella enterica subsp. enterica serovar Typhimurium str. DT104
Proteobacteria; Gammaproteobacteria; Enterobacterales; Enterobacteriaceae; Salmonella; Salmonella enterica; Salmonella enterica subsp. enterica; Salmonella enterica subsp. enterica serovar Typhimurium
MTGDNTLITSHGINRRDFMKLCAALAATMGLSSKAAAEMAESVSNPQRPPVIWIGAQECT WP_000145429.1
hydrogenase 2 small subunit [Salmonella enterica] Length: 372\nNumber of predicted TMHs: 1\nExp number of AAs in TMHs: 24.51309\nExp number, first 60 AAs: 0.2682\nTotal prob of N-in: 0.14402\noutside 1 331\nTMhelix 332 354\ninside 355 372

25247	GCF_001217605.1_5275_1_12	Salmonella enterica subsp. enterica serovar Typhimurium str. DT104	Proteobacteria; Gammaproteobacteria; Enterobacterales; Enterobacteriaceae; Salmonella; Salmonella enterica; Salmonella enterica subsp. enterica; Salmonella enterica subsp. enterica serovar Typhimurium	MNNEETFYQAMRRKGVTRRSFLKFCSLAATSLGLGAGMTPKIAWALENKPRIPVVWIHGL	WP_001058312.1	hydrogenase [Salmonella enterica]	Length: 372\nNumber of predicted TMHs:  2\nExp number of AAs in TMHs: 36.51738\nExp number, first 60 AAs:  19.72849\nTotal prob of N-in:   0.96170\nPOSSIBLE N-term signal sequence\ninside   1   19\nTMhelix   20   39\noutside   40   325\nTMhelix   326   348\ninside   349   372

25248	GCF_001217545.1_8036_3_43	Salmonella enterica subsp. enterica serovar Typhimurium str. DT104	Proteobacteria; Gammaproteobacteria; Enterobacterales; Enterobacteriaceae; Salmonella; Salmonella enterica; Salmonella enterica subsp. enterica; Salmonella enterica subsp. enterica serovar Typhimurium	MNNEETFYQAMRRKGVTRRSFLKFCSLAATSLGLGAGMTPKIAWALENKPRIPVVWIHGL	WP_001058312.1	hydrogenase [Salmonella enterica]	Length: 372\nNumber of predicted TMHs:  2\nExp number of AAs in TMHs: 36.51738\nExp number, first 60 AAs:  19.72849\nTotal prob of N-in:   0.96170\nPOSSIBLE N-term signal sequence\ninside   1   19\nTMhelix   20   39\noutside   40   325\nTMhelix   326   348\ninside   349   372

25249	GCF_001217865.1_8080_3_17	Salmonella enterica subsp. enterica serovar Typhimurium str. DT104	Proteobacteria; Gammaproteobacteria; Enterobacterales; Enterobacteriaceae; Salmonella; Salmonella enterica; Salmonella enterica subsp. enterica; Salmonella enterica subsp. enterica serovar Typhimurium	MNNEETFYQAMRRKGVTRRSFLKFCSLAATSLGLGAGMTPKIAWALENKPRIPVVWIHGL	WP_001058312.1	hydrogenase [Salmonella enterica]	Length: 372\nNumber of predicted TMHs:  2\nExp number of AAs in TMHs: 36.51738\nExp number, first 60 AAs:  19.72849\nTotal prob of N-in:   0.96170\nPOSSIBLE N-term signal sequence\ninside   1   19\nTMhelix   20   39\noutside   40   325\nTMhelix   326   348\ninside   349   372

25250	GCF_001217385.1_5285_7_11	Salmonella enterica subsp. enterica serovar Typhimurium str. DT104	Proteobacteria; Gammaproteobacteria; Enterobacterales; Enterobacteriaceae; Salmonella; Salmonella enterica; Salmonella enterica subsp. enterica; Salmonella enterica subsp. enterica serovar Typhimurium	MTGDNTLITSHGINRRDFMKLCAALAATMGLSSKAAAEMAESVSNPQRPPVIWIGAQECT	WP_000145429.1	hydrogenase 2 small subunit [Salmonella enterica]	Length: 372\nNumber of predicted TMHs:  1\nExp number of AAs in TMHs: 24.51309\nExp number, first 60 AAs:  0.2682\nTotal prob of N-in:   0.14402\noutside   1   331\nTMhelix   332   354\ninside   355   372

25251	GCF_001217305.1_8080_3_24	Salmonella enterica subsp. enterica serovar Typhimurium str. DT104	Proteobacteria; Gammaproteobacteria; Enterobacterales; Enterobacteriaceae; Salmonella; Salmonella enterica; Salmonella enterica subsp. enterica; Salmonella enterica subsp. enterica serovar Typhimurium	MTGDNTLITSHGINRRDFMKLCAALAATMGLSSKAAAEMAESVSNPQRPPVIWIGAQECT	WP_000145429.1	hydrogenase 2 small subunit [Salmonella enterica]	Length: 372\nNumber of predicted TMHs:  1\nExp number of AAs in TMHs: 24.51309\nExp number, first 60 AAs:  0.2682\nTotal prob of N-in:   0.14402\noutside   1   331\nTMhelix   332   354\ninside   355   372

25252	GCF_001217245.1_8080_3_4Salmonella enterica subsp. enterica serovar Typhimurium str. DT104	Proteobacteria; Gammaproteobacteria; Enterobacterales; Enterobacteriaceae; Salmonella; Salmonella enterica; Salmonella enterica subsp. enterica; Salmonella enterica subsp. enterica serovar Typhimurium	MTGDNTLITSHGINRRDFMKLCAALAATMGLSSKAAAEMAESVSNPQRPPVIWIGAQECT	WP_000145429.1	hydrogenase 2 small subunit [Salmonella enterica]	Length: 372\nNumber of predicted TMHs:  1\nExp number of AAs in TMHs: 24.51309\nExp number, first 60 AAs:  0.2682\nTotal prob of N-in:   0.14402\noutside   1   331\nTMhelix   332   354\ninside   355   372

25253	GCF_001216565.1_9066_1_6Salmonella enterica subsp. enterica serovar Typhimurium str. DT104	Proteobacteria; Gammaproteobacteria; Enterobacterales; Enterobacteriaceae; Salmonella; Salmonella enterica; Salmonella enterica subsp. enterica; Salmonella enterica subsp. enterica serovar Typhimurium	MNNEETFYQAMRRKGVTRRSFLKFCSLAATSLGLGAGMTPKIAWALENKPRIPVVWIHGL	WP_001058312.1	hydrogenase [Salmonella enterica]	Length: 372\nNumber of predicted TMHs:  2\nExp number of AAs in TMHs: 36.51738\nExp number, first 60 AAs:  19.72849\nTotal prob of N-in:   0.96170\nPOSSIBLE N-term signal sequence\ninside   1   19\nTMhelix   20   39\noutside   40   325\nTMhelix   326   348\ninside   349   372

25254	GCF_001217085.1_5275_3_3Salmonella enterica subsp. enterica serovar Typhimurium str. DT104	Proteobacteria; Gammaproteobacteria; Enterobacterales; Enterobacteriaceae; Salmonella; Salmonella enterica; Salmonella enterica subsp. enterica; Salmonella enterica subsp. enterica serovar Typhimurium	MNNEETFYQAMRRKGVTRRSFLKFCSLAATSLGLGAGMTPKIAWALENKPRIPVVWIHGL	WP_001058312.1	hydrogenase [Salmonella enterica]	Length: 372\nNumber of predicted TMHs:  2\nExp number of AAs in

TMHs: 36.51738\nExp number, first 60 AAs: 19.72849\nTotal prob of N-in: 0.96170\nPOSSIBLE N-term signal sequence\ninside 1 19\nTMhelix 20 39\noutside 40 325\nTMhelix 326 348\ninside 349 372

25255 GCF_001216985.1_5275_8_3Salmonella enterica subsp. enterica serovar Typhimurium str. DT104 Proteobacteria; Gammaproteobacteria; Enterobacterales; Enterobacteriaceae; Salmonella; Salmonella enterica; Salmonella enterica subsp. enterica; Salmonella enterica subsp. enterica serovar Typhimurium MNNEETFYQAMRRKGVTRRSFLKFCSLAATSLGLGAGMTPKIAWALENKPRIPVVWIHGL WP_001058312.1 hydrogenase [Salmonella enterica] Length: 372\nNumber of predicted TMHs: 2\nExp number of AAs in TMHs: 36.51738\nExp number, first 60 AAs: 19.72849\nTotal prob of N-in: 0.96170\nPOSSIBLE N-term signal sequence\ninside 1 19\nTMhelix 20 39\noutside 40 325\nTMhelix 326 348\ninside 349 372

25256 GCF_001221945.1_8080_3_21 Salmonella enterica subsp. enterica serovar Typhimurium str. DT104 Proteobacteria; Gammaproteobacteria; Enterobacterales; Enterobacteriaceae; Salmonella; Salmonella enterica; Salmonella enterica subsp. enterica; Salmonella enterica subsp. enterica serovar Typhimurium MTGDNTLITSHGINRRDFMKLCAALAATMGLSSKAAAEMAESVSNPQRPPVIWIGAQECT WP_000145429.1 hydrogenase 2 small subunit [Salmonella enterica] Length: 372\nNumber of predicted TMHs: 1\nExp number of AAs in TMHs: 24.51309\nExp number, first 60 AAs: 0.2682\nTotal prob of N-in: 0.14402\noutside 1 331\nTMhelix 332 354\ninside 355 372

25257 GCF_001216565.1_9066_1_6Salmonella enterica subsp. enterica serovar Typhimurium str. DT104 Proteobacteria; Gammaproteobacteria; Enterobacterales; Enterobacteriaceae; Salmonella; Salmonella enterica; Salmonella enterica subsp. enterica; Salmonella enterica subsp. enterica serovar Typhimurium MTGDNTLITSHGINRRDFMKLCAALAATMGLSSKAAAEMAESVSNPQRPPVIWIGAQECT WP_000145429.1 hydrogenase 2 small subunit [Salmonella enterica] Length: 372\nNumber of predicted TMHs: 1\nExp number of AAs in TMHs: 24.51309\nExp number, first 60 AAs: 0.2682\nTotal prob of N-in: 0.14402\noutside 1 331\nTMhelix 332 354\ninside 355 372

25258 GCF_001216465.1_8036_3_69 Salmonella enterica subsp. enterica serovar Typhimurium str. DT104 Proteobacteria; Gammaproteobacteria; Enterobacterales; Enterobacteriaceae; Salmonella; Salmonella enterica; Salmonella enterica subsp. enterica; Salmonella enterica subsp. enterica serovar Typhimurium MTGDNTLITSHGINRRDFMKLCAALAATMGLSSKAAAEMAESVSNPQRPPVIWIGAQECT WP_000145429.1 hydrogenase 2 small subunit [Salmonella enterica] Length: 372\nNumber of predicted TMHs: 1\nExp number of AAs in TMHs: 24.51309\nExp number, first 60 AAs: 0.2682\nTotal prob of N-in: 0.14402\noutside 1 331\nTMhelix 332 354\ninside 355 372

25259 GCF_001216385.1_8080_3_44 Salmonella enterica subsp. enterica serovar Typhimurium str. DT104 Proteobacteria; Gammaproteobacteria; Enterobacterales; Enterobacteriaceae; Salmonella; Salmonella enterica; Salmonella enterica subsp. enterica; Salmonella enterica subsp. enterica serovar Typhimurium MTGDNTLITSHGINRRDFMKLCAALAATMGLSSKAAAEMAESVSNPQRPPVIWIGAQECT WP_000145429.1 hydrogenase 2 small subunit [Salmonella enterica] Length: 372\nNumber of predicted TMHs: 1\nExp number of AAs in TMHs: 24.51309\nExp number, first 60 AAs: 0.2682\nTotal prob of N-in: 0.14402\noutside 1 331\nTMhelix 332 354\ninside 355 372

25260 GCF_001216205.1_8036_3_76 Salmonella enterica subsp. enterica serovar Typhimurium str. DT104 Proteobacteria; Gammaproteobacteria; Enterobacterales; Enterobacteriaceae; Salmonella; Salmonella enterica; Salmonella enterica subsp. enterica; Salmonella enterica subsp. enterica serovar Typhimurium MTGDNTLITSHGINRRDFMKLCAALAATMGLSSKAAAEMAESVSNPQRPPVIWIGAQECT WP_000145429.1 hydrogenase 2 small subunit [Salmonella enterica] Length: 372\nNumber of predicted TMHs: 1\nExp number of AAs in TMHs: 24.51309\nExp number, first 60 AAs: 0.2682\nTotal prob of N-in: 0.14402\noutside 1 331\nTMhelix 332 354\ninside 355 372

25261 GCF_001216085.1_8080_3_87 Salmonella enterica subsp. enterica serovar Typhimurium str. DT104 Proteobacteria; Gammaproteobacteria; Enterobacterales; Enterobacteriaceae; Salmonella; Salmonella enterica; Salmonella enterica subsp. enterica; Salmonella enterica subsp. enterica serovar Typhimurium MTGDNTLITSHGINRRDFMKLCAALAATMGLSSKAAAEMAESVSNPQRPPVIWIGAQECT WP_000145429.1 hydrogenase 2 small subunit [Salmonella enterica] Length: 372\nNumber of predicted TMHs: 1\nExp number of AAs in TMHs: 24.51309\nExp number, first 60 AAs: 0.2682\nTotal prob of N-in: 0.14402\noutside 1 331\nTMhelix 332 354\ninside 355 372

25262 GCF_001215785.1_5275_1_8Salmonella enterica subsp. enterica serovar Typhimurium str. DT104 Proteobacteria; Gammaproteobacteria; Enterobacterales; Enterobacteriaceae; Salmonella; Salmonella enterica;

Salmonella enterica subsp. enterica; Salmonella enterica subsp. enterica serovar Typhimurium
MNNEETFYQAMRRKGVTRRSFLKFCSLAATSLGLGAGMTPKIAWALENKPRIPVVWIHGL WP_001058312.1
hydrogenase [Salmonella enterica] Length: 372\nNumber of predicted TMHs: 2\nExp number of AAs in TMHs: 36.51738\nExp number, first 60 AAs: 19.72849\nTotal prob of N-in: 0.96170\nPOSSIBLE N-term signal sequence\ninside 1 19\nTMhelix 20 39\noutside 40 325\nTMhelix 326 348\ninside 349 372

25263 GCF_001214745.1_5275_2_6Salmonella enterica subsp. enterica serovar Typhimurium str. DT104
Proteobacteria; Gammaproteobacteria; Enterobacterales; Enterobacteriaceae; Salmonella; Salmonella enterica; Salmonella enterica subsp. enterica; Salmonella enterica subsp. enterica serovar Typhimurium
MNNEETFYQAMRRKGVTRRSFLKFCSLAATSLGLGAGMTPKIAWALENKPRIPVVWIHGL WP_001058312.1
hydrogenase [Salmonella enterica] Length: 372\nNumber of predicted TMHs: 2\nExp number of AAs in TMHs: 36.51738\nExp number, first 60 AAs: 19.72849\nTotal prob of N-in: 0.96170\nPOSSIBLE N-term signal sequence\ninside 1 19\nTMhelix 20 39\noutside 40 325\nTMhelix 326 348\ninside 349 372

25264 GCF_001215405.1_5275_7_10 Salmonella enterica subsp. enterica serovar Typhimurium str. DT104
Proteobacteria; Gammaproteobacteria; Enterobacterales; Enterobacteriaceae; Salmonella; Salmonella enterica; Salmonella enterica subsp. enterica; Salmonella enterica subsp. enterica serovar Typhimurium
MNNEETFYQAMRRKGVTRRSFLKFCSLAATSLGLGAGMTPKIAWALENKPRIPVVWIHGL WP_001058312.1
hydrogenase [Salmonella enterica] Length: 372\nNumber of predicted TMHs: 2\nExp number of AAs in TMHs: 36.51738\nExp number, first 60 AAs: 19.72849\nTotal prob of N-in: 0.96170\nPOSSIBLE N-term signal sequence\ninside 1 19\nTMhelix 20 39\noutside 40 325\nTMhelix 326 348\ninside 349 372

25265 GCF_001215145.1_8080_3_51 Salmonella enterica subsp. enterica serovar Typhimurium str. DT104
Proteobacteria; Gammaproteobacteria; Enterobacterales; Enterobacteriaceae; Salmonella; Salmonella enterica; Salmonella enterica subsp. enterica; Salmonella enterica subsp. enterica serovar Typhimurium
MNNEETFYQAMRRKGVTRRSFLKFCSLAATSLGLGAGMTPKIAWALENKPRIPVVWIHGL WP_001058312.1
hydrogenase [Salmonella enterica] Length: 372\nNumber of predicted TMHs: 2\nExp number of AAs in TMHs: 36.51738\nExp number, first 60 AAs: 19.72849\nTotal prob of N-in: 0.96170\nPOSSIBLE N-term signal sequence\ninside 1 19\nTMhelix 20 39\noutside 40 325\nTMhelix 326 348\ninside 349 372

25266 GCF_001215405.1_5275_7_10 Salmonella enterica subsp. enterica serovar Typhimurium str. DT104
Proteobacteria; Gammaproteobacteria; Enterobacterales; Enterobacteriaceae; Salmonella; Salmonella enterica; Salmonella enterica subsp. enterica; Salmonella enterica subsp. enterica serovar Typhimurium
MTGDNTLITSHGINRRDFMKLCAALAATMGLSSKAAAEMAESVSNPQRPPVIWIGAQECT WP_000145429.1
hydrogenase 2 small subunit [Salmonella enterica] Length: 372\nNumber of predicted TMHs: 1\nExp number of AAs in TMHs: 24.51309\nExp number, first 60 AAs: 0.2682\nTotal prob of N-in: 0.14402\noutside 1 331\nTMhelix 332 354\ninside 355 372

25267 GCF_001215145.1_8080_3_51 Salmonella enterica subsp. enterica serovar Typhimurium str. DT104
Proteobacteria; Gammaproteobacteria; Enterobacterales; Enterobacteriaceae; Salmonella; Salmonella enterica; Salmonella enterica subsp. enterica; Salmonella enterica subsp. enterica serovar Typhimurium
MTGDNTLITSHGINRRDFMKLCAALAATMGLSSKAAAEMAESVSNPQRPPVIWIGAQECT WP_000145429.1
hydrogenase 2 small subunit [Salmonella enterica] Length: 372\nNumber of predicted TMHs: 1\nExp number of AAs in TMHs: 24.51309\nExp number, first 60 AAs: 0.2682\nTotal prob of N-in: 0.14402\noutside 1 331\nTMhelix 332 354\ninside 355 372

25268 GCF_001214365.1_8080_3_11 Salmonella enterica subsp. enterica serovar Typhimurium str. DT104
Proteobacteria; Gammaproteobacteria; Enterobacterales; Enterobacteriaceae; Salmonella; Salmonella enterica; Salmonella enterica subsp. enterica; Salmonella enterica subsp. enterica serovar Typhimurium
MNNEETFYQAMRRKGVTRRSFLKFCSLAATSLGLGAGMTPKIAWALENKPRIPVVWIHGL WP_001058312.1
hydrogenase [Salmonella enterica] Length: 372\nNumber of predicted TMHs: 2\nExp number of AAs in TMHs: 36.51738\nExp number, first 60 AAs: 19.72849\nTotal prob of N-in: 0.96170\nPOSSIBLE N-term signal sequence\ninside 1 19\nTMhelix 20 39\noutside 40 325\nTMhelix 326 348\ninside 349 372

25269 GCF_001214365.1_8080_3_11 Salmonella enterica subsp. enterica serovar Typhimurium str. DT104
Proteobacteria; Gammaproteobacteria; Enterobacterales; Enterobacteriaceae; Salmonella; Salmonella enterica; Salmonella enterica subsp. enterica; Salmonella enterica subsp. enterica serovar Typhimurium
MTGDNTLITSHGINRRDFMKLCAALAATMGLSSKAAAEMAESVSNPQRPPVIWIGAQECT WP_000145429.1
hydrogenase 2 small subunit [Salmonella enterica] Length: 372\nNumber of predicted TMHs: 1\nExp number of AAs in TMHs: 24.51309\nExp number, first 60 AAs: 0.2682\nTotal prob of N-in: 0.14402\noutside 1 331\nTMhelix 332 354\ninside 355 372

25270 GCF_001164125.1_8490_2_52 Salmonella enterica subsp. enterica serovar Typhimurium str. DT104 Proteobacteria; Gammaproteobacteria; Enterobacterales; Enterobacteriaceae; Salmonella; Salmonella enterica; Salmonella enterica subsp. enterica; Salmonella enterica subsp. enterica serovar Typhimurium MNNEETFYQAMRRKGVTRRSFLKFCSLAATSLGLGAGMTPKIAWALENKPRIPVVWIHGL WP_001058312.1 hydrogenase [Salmonella enterica] Length: 372\nNumber of predicted TMHs: 2\nExp number of AAs in TMHs: 36.51738\nExp number, first 60 AAs: 19.72849\nTotal prob of N-in: 0.96170\nPOSSIBLE N-term signal sequence\ninside 1 19\nTMhelix 20 39\noutside 40 325\nTMhelix 326 348\ninside 349 372

25271 GCF_001222465.1_5275_6_3Salmonella enterica subsp. enterica serovar Typhimurium str. DT104 Proteobacteria; Gammaproteobacteria; Enterobacterales; Enterobacteriaceae; Salmonella; Salmonella enterica; Salmonella enterica subsp. enterica; Salmonella enterica subsp. enterica serovar Typhimurium MTGDNTLITSHGINRRDFMKLCAALAATMGLSSKAAAEMAESVSNPQRPPVIWIGAQECT WP_000145429.1 hydrogenase 2 small subunit [Salmonella enterica] Length: 372\nNumber of predicted TMHs: 1\nExp number of AAs in TMHs: 24.51309\nExp number, first 60 AAs: 0.2682\nTotal prob of N-in: 0.14402\noutside 1 331\nTMhelix 332 354\ninside 355 372

25272 GCF_001219865.1_5388_1_1Salmonella enterica subsp. enterica serovar Typhimurium str. DT104 Proteobacteria; Gammaproteobacteria; Enterobacterales; Enterobacteriaceae; Salmonella; Salmonella enterica; Salmonella enterica subsp. enterica; Salmonella enterica subsp. enterica serovar Typhimurium MTGDNTLITSHGINRRDFMKLCAALAATMGLSSKAAAEMAESVSNPQRPPVIWIGAQECT WP_000145429.1 hydrogenase 2 small subunit [Salmonella enterica] Length: 372\nNumber of predicted TMHs: 1\nExp number of AAs in TMHs: 24.51309\nExp number, first 60 AAs: 0.2682\nTotal prob of N-in: 0.14402\noutside 1 331\nTMhelix 332 354\ninside 355 372

25273 GCF_001222385.1_5295_2_6Salmonella enterica subsp. enterica serovar Typhimurium str. DT104 Proteobacteria; Gammaproteobacteria; Enterobacterales; Enterobacteriaceae; Salmonella; Salmonella enterica; Salmonella enterica subsp. enterica; Salmonella enterica subsp. enterica serovar Typhimurium MNNEETFYQAMRRKGVTRRSFLKFCSLAATSLGLGAGMTPKIAWALENKPRIPVVWIHGL WP_001058312.1 hydrogenase [Salmonella enterica] Length: 372\nNumber of predicted TMHs: 2\nExp number of AAs in TMHs: 36.51738\nExp number, first 60 AAs: 19.72849\nTotal prob of N-in: 0.96170\nPOSSIBLE N-term signal sequence\ninside 1 19\nTMhelix 20 39\noutside 40 325\nTMhelix 326 348\ninside 349 372

25274 GCF_001222465.1_5275_6_3Salmonella enterica subsp. enterica serovar Typhimurium str. DT104 Proteobacteria; Gammaproteobacteria; Enterobacterales; Enterobacteriaceae; Salmonella; Salmonella enterica; Salmonella enterica subsp. enterica; Salmonella enterica subsp. enterica serovar Typhimurium MNNEETFYQAMRRKGVTRRSFLKFCSLAATSLGLGAGMTPKIAWALENKPRIPVVWIHGL WP_001058312.1 hydrogenase [Salmonella enterica] Length: 372\nNumber of predicted TMHs: 2\nExp number of AAs in TMHs: 36.51738\nExp number, first 60 AAs: 19.72849\nTotal prob of N-in: 0.96170\nPOSSIBLE N-term signal sequence\ninside 1 19\nTMhelix 20 39\noutside 40 325\nTMhelix 326 348\ninside 349 372

25275 GCF_001222065.1_5388_1_4Salmonella enterica subsp. enterica serovar Typhimurium str. DT104 Proteobacteria; Gammaproteobacteria; Enterobacterales; Enterobacteriaceae; Salmonella; Salmonella enterica; Salmonella enterica subsp. enterica; Salmonella enterica subsp. enterica serovar Typhimurium MNNEETFYQAMRRKGVTRRSFLKFCSLAATSLGLGAGMTPKIAWALENKPRIPVVWIHGL WP_001058312.1 hydrogenase [Salmonella enterica] Length: 372\nNumber of predicted TMHs: 2\nExp number of AAs in TMHs: 36.51738\nExp number, first 60 AAs: 19.72849\nTotal prob of N-in: 0.96170\nPOSSIBLE N-term signal sequence\ninside 1 19\nTMhelix 20 39\noutside 40 325\nTMhelix 326 348\ninside 349 372

25276 GCF_001222505.1_5388_1_8Salmonella enterica subsp. enterica serovar Typhimurium str. DT104 Proteobacteria; Gammaproteobacteria; Enterobacterales; Enterobacteriaceae; Salmonella; Salmonella enterica; Salmonella enterica subsp. enterica; Salmonella enterica subsp. enterica serovar Typhimurium MTGDNTLITSHGINRRDFMKLCAALAATMGLSSKAAAEMAESVSNPQRPPVIWIGAQECT WP_000145429.1 hydrogenase 2 small subunit [Salmonella enterica] Length: 372\nNumber of predicted TMHs: 1\nExp number of AAs in TMHs: 24.51309\nExp number, first 60 AAs: 0.2682\nTotal prob of N-in: 0.14402\noutside 1 331\nTMhelix 332 354\ninside 355 372

25277 GCF_001219825.1_8036_3_60 Salmonella enterica subsp. enterica serovar Typhimurium str. DT104 Proteobacteria; Gammaproteobacteria; Enterobacterales; Enterobacteriaceae; Salmonella; Salmonella enterica; Salmonella enterica subsp. enterica; Salmonella enterica subsp. enterica serovar Typhimurium MNNEETFYQAMRRKGVTRRSFLKFCSLAATSLGLGAGMTPKIAWALENKPRIPVVWIHGL WP_001058312.1 hydrogenase [Salmonella enterica] Length: 372\nNumber of predicted TMHs: 2\nExp number of AAs in

TMHs: 36.51738\nExp number, first 60 AAs: 19.72849\nTotal prob of N-in: 0.96170\nPOSSIBLE N-term signal sequence\ninside 1 19\nTMhelix 20 39\noutside 40 325\nTMhelix 326 348\ninside 349 372

25278 GCF_001222585.1_8080_3_18 Salmonella enterica subsp. enterica serovar Typhimurium str. DT104 Proteobacteria; Gammaproteobacteria; Enterobacterales; Enterobacteriaceae; Salmonella; Salmonella enterica; Salmonella enterica subsp. enterica; Salmonella enterica subsp. enterica serovar Typhimurium MTGDNTLITSHGINRRDFMKLCAALAATMGLSSKAAAEMAESVSNPQRPPVIWIGAQECT WP_000145429.1 hydrogenase 2 small subunit [Salmonella enterica] Length: 372\nNumber of predicted TMHs: 1\nExp number of AAs in TMHs: 24.51309\nExp number, first 60 AAs: 0.2682\nTotal prob of N-in: 0.14402\noutside 1 331\nTMhelix 332 354\ninside 355 372

25279 GCF_001219865.1_5388_1_1Salmonella enterica subsp. enterica serovar Typhimurium str. DT104 Proteobacteria; Gammaproteobacteria; Enterobacterales; Enterobacteriaceae; Salmonella; Salmonella enterica; Salmonella enterica subsp. enterica; Salmonella enterica subsp. enterica serovar Typhimurium MNNEETFYQAMRRKGVTRRSFLKFCSLAATSLGLGAGMTPKIAWALENKPRIPVVWIHGL WP_001058312.1 hydrogenase [Salmonella enterica] Length: 372\nNumber of predicted TMHs: 2\nExp number of AAs in TMHs: 36.51738\nExp number, first 60 AAs: 19.72849\nTotal prob of N-in: 0.96170\nPOSSIBLE N-term signal sequence\ninside 1 19\nTMhelix 20 39\noutside 40 325\nTMhelix 326 348\ninside 349 372

25280 GCF_001223025.1_5275_3_6Salmonella enterica subsp. enterica serovar Typhimurium str. DT104 Proteobacteria; Gammaproteobacteria; Enterobacterales; Enterobacteriaceae; Salmonella; Salmonella enterica; Salmonella enterica subsp. enterica; Salmonella enterica subsp. enterica serovar Typhimurium MTGDNTLITSHGINRRDFMKLCAALAATMGLSSKAAAEMAESVSNPQRPPVIWIGAQECT WP_000145429.1 hydrogenase 2 small subunit [Salmonella enterica] Length: 372\nNumber of predicted TMHs: 1\nExp number of AAs in TMHs: 24.51309\nExp number, first 60 AAs: 0.2682\nTotal prob of N-in: 0.14402\noutside 1 331\nTMhelix 332 354\ninside 355 372

25281 GCF_001222885.1_5275_2_2Salmonella enterica subsp. enterica serovar Typhimurium str. DT104 Proteobacteria; Gammaproteobacteria; Enterobacterales; Enterobacteriaceae; Salmonella; Salmonella enterica; Salmonella enterica subsp. enterica; Salmonella enterica subsp. enterica serovar Typhimurium MNNEETFYQAMRRKGVTRRSFLKFCSLAATSLGLGAGMTPKIAWALENKPRIPVVWIHGL WP_001058312.1 hydrogenase [Salmonella enterica] Length: 372\nNumber of predicted TMHs: 2\nExp number of AAs in TMHs: 36.51738\nExp number, first 60 AAs: 19.72849\nTotal prob of N-in: 0.96170\nPOSSIBLE N-term signal sequence\ninside 1 19\nTMhelix 20 39\noutside 40 325\nTMhelix 326 348\ninside 349 372

25282 GCF_001219685.1_5285_6_1Salmonella enterica subsp. enterica serovar Typhimurium str. DT104 Proteobacteria; Gammaproteobacteria; Enterobacterales; Enterobacteriaceae; Salmonella; Salmonella enterica; Salmonella enterica subsp. enterica; Salmonella enterica subsp. enterica serovar Typhimurium MNNEETFYQAMRRKGVTRRSFLKFCSLAATSLGLGAGMTPKIAWALENKPRIPVVWIHGL WP_001058312.1 hydrogenase [Salmonella enterica] Length: 372\nNumber of predicted TMHs: 2\nExp number of AAs in TMHs: 36.51738\nExp number, first 60 AAs: 19.72849\nTotal prob of N-in: 0.96170\nPOSSIBLE N-term signal sequence\ninside 1 19\nTMhelix 20 39\noutside 40 325\nTMhelix 326 348\ninside 349 372

25283 GCF_001222725.1_8080_3_79 Salmonella enterica subsp. enterica serovar Typhimurium str. DT104 Proteobacteria; Gammaproteobacteria; Enterobacterales; Enterobacteriaceae; Salmonella; Salmonella enterica; Salmonella enterica subsp. enterica; Salmonella enterica subsp. enterica serovar Typhimurium MNNEETFYQAMRRKGVTRRSFLKFCSLAATSLGLGAGMTPKIAWALENKPRIPVVWIHGL WP_001058312.1 hydrogenase [Salmonella enterica] Length: 372\nNumber of predicted TMHs: 2\nExp number of AAs in TMHs: 36.51738\nExp number, first 60 AAs: 19.72849\nTotal prob of N-in: 0.96170\nPOSSIBLE N-term signal sequence\ninside 1 19\nTMhelix 20 39\noutside 40 325\nTMhelix 326 348\ninside 349 372

25284 GCF_001223125.1_8080_3_28 Salmonella enterica subsp. enterica serovar Typhimurium str. DT104 Proteobacteria; Gammaproteobacteria; Enterobacterales; Enterobacteriaceae; Salmonella; Salmonella enterica; Salmonella enterica subsp. enterica; Salmonella enterica subsp. enterica serovar Typhimurium MTGDNTLITSHGINRRDFMKLCAALAATMGLSSKAAAEMAESVSNPQRPPVIWIGAQECT WP_000145429.1 hydrogenase 2 small subunit [Salmonella enterica] Length: 372\nNumber of predicted TMHs: 1\nExp number of AAs in TMHs: 24.51309\nExp number, first 60 AAs: 0.2682\nTotal prob of N-in: 0.14402\noutside 1 331\nTMhelix 332 354\ninside 355 372

25285 GCF_001223125.1_8080_3_28 Salmonella enterica subsp. enterica serovar Typhimurium str. DT104 Proteobacteria; Gammaproteobacteria; Enterobacterales; Enterobacteriaceae; Salmonella; Salmonella enterica;

Salmonella enterica subsp. enterica; Salmonella enterica subsp. enterica serovar Typhimurium
MNNEETFYQAMRRKGVTRRSFLKFCSLAATSLGLGAGMTPKIAWALENKPRIPVVWIHGL WP_001058312.1
hydrogenase [Salmonella enterica] Length: 372\nNumber of predicted TMHs: 2\nExp number of AAs in TMHs: 36.51738\nExp number, first 60 AAs: 19.72849\nTotal prob of N-in: 0.96170\nPOSSIBLE N-term signal sequence\ninside 1 19\nTMhelix 20 39\noutside 40 325\nTMhelix 326 348\ninside 349 372

25286 GCF_001219905.1_8036_3_54 Salmonella enterica subsp. enterica serovar Typhimurium str. DT104
Proteobacteria; Gammaproteobacteria; Enterobacterales; Enterobacteriaceae; Salmonella; Salmonella enterica; Salmonella enterica subsp. enterica; Salmonella enterica subsp. enterica serovar Typhimurium
MTGDNTLITSHGINRRDFMKLCAALAATMGLSSKAAAEMAESVSNPQRPPVIWIGAQECT WP_000145429.1
hydrogenase 2 small subunit [Salmonella enterica] Length: 372\nNumber of predicted TMHs: 1\nExp number of AAs in TMHs: 24.51309\nExp number, first 60 AAs: 0.2682\nTotal prob of N-in: 0.14402\noutside 1 331\nTMhelix 332 354\ninside 355 372

25287 GCF_001223525.1_8080_3_81 Salmonella enterica subsp. enterica serovar Typhimurium str. DT104
Proteobacteria; Gammaproteobacteria; Enterobacterales; Enterobacteriaceae; Salmonella; Salmonella enterica; Salmonella enterica subsp. enterica; Salmonella enterica subsp. enterica serovar Typhimurium
MNNEETFYQAMRRKGVTRRSFLKFCSLAATSLGLGAGMTPKIAWALENKPRIPVVWIHGL WP_001058312.1
hydrogenase [Salmonella enterica] Length: 372\nNumber of predicted TMHs: 2\nExp number of AAs in TMHs: 36.51738\nExp number, first 60 AAs: 19.72849\nTotal prob of N-in: 0.96170\nPOSSIBLE N-term signal sequence\ninside 1 19\nTMhelix 20 39\noutside 40 325\nTMhelix 326 348\ninside 349 372

25288 GCF_001219945.1_5285_6_2Salmonella enterica subsp. enterica serovar Typhimurium str. DT104
Proteobacteria; Gammaproteobacteria; Enterobacterales; Enterobacteriaceae; Salmonella; Salmonella enterica; Salmonella enterica subsp. enterica; Salmonella enterica subsp. enterica serovar Typhimurium
MTGDNTLITSHGINRRDFMKLCAALAATMGLSSKAAAEMAESVSNPQRPPVIWIGAQECT WP_000145429.1
hydrogenase 2 small subunit [Salmonella enterica] Length: 372\nNumber of predicted TMHs: 1\nExp number of AAs in TMHs: 24.51309\nExp number, first 60 AAs: 0.2682\nTotal prob of N-in: 0.14402\noutside 1 331\nTMhelix 332 354\ninside 355 372

25289 GCF_001223465.1_5275_8_12 Salmonella enterica subsp. enterica serovar Typhimurium str. DT104
Proteobacteria; Gammaproteobacteria; Enterobacterales; Enterobacteriaceae; Salmonella; Salmonella enterica; Salmonella enterica subsp. enterica; Salmonella enterica subsp. enterica serovar Typhimurium
MNNEETFYQAMRRKGVTRRSFLKFCSLAATSLGLGAGMTPKIAWALENKPRIPVVWIHGL WP_001058312.1
hydrogenase [Salmonella enterica] Length: 372\nNumber of predicted TMHs: 2\nExp number of AAs in TMHs: 36.51738\nExp number, first 60 AAs: 19.72849\nTotal prob of N-in: 0.96170\nPOSSIBLE N-term signal sequence\ninside 1 19\nTMhelix 20 39\noutside 40 325\nTMhelix 326 348\ninside 349 372

25290 GCF_001223285.1_8080_3_90 Salmonella enterica subsp. enterica serovar Typhimurium str. DT104
Proteobacteria; Gammaproteobacteria; Enterobacterales; Enterobacteriaceae; Salmonella; Salmonella enterica; Salmonella enterica subsp. enterica; Salmonella enterica subsp. enterica serovar Typhimurium
MNNEETFYQAMRRKGVTRRSFLKFCSLAATSLGLGAGMTPKIAWALENKPRIPVVWIHGL WP_001058312.1
hydrogenase [Salmonella enterica] Length: 372\nNumber of predicted TMHs: 2\nExp number of AAs in TMHs: 36.51738\nExp number, first 60 AAs: 19.72849\nTotal prob of N-in: 0.96170\nPOSSIBLE N-term signal sequence\ninside 1 19\nTMhelix 20 39\noutside 40 325\nTMhelix 326 348\ninside 349 372

25291 GCF_001223325.1_5299_6_1Salmonella enterica subsp. enterica serovar Typhimurium str. DT104
Proteobacteria; Gammaproteobacteria; Enterobacterales; Enterobacteriaceae; Salmonella; Salmonella enterica; Salmonella enterica subsp. enterica; Salmonella enterica subsp. enterica serovar Typhimurium
MNNEETFYQAMRRKGVTRRSFLKFCSLAATSLGLGAGMTPKIAWALENKPRIPVVWIHGL WP_001058312.1
hydrogenase [Salmonella enterica] Length: 372\nNumber of predicted TMHs: 2\nExp number of AAs in TMHs: 36.51738\nExp number, first 60 AAs: 19.72849\nTotal prob of N-in: 0.96170\nPOSSIBLE N-term signal sequence\ninside 1 19\nTMhelix 20 39\noutside 40 325\nTMhelix 326 348\ninside 349 372

25292 GCF_001221905.1_5275_2_8Salmonella enterica subsp. enterica serovar Typhimurium str. DT104
Proteobacteria; Gammaproteobacteria; Enterobacterales; Enterobacteriaceae; Salmonella; Salmonella enterica; Salmonella enterica subsp. enterica; Salmonella enterica subsp. enterica serovar Typhimurium
MTGDNTLITSHGINRRDFMKLCAALAATMGLSSKAAAEMAESVSNPQRPPVIWIGAQECT WP_000145429.1
hydrogenase 2 small subunit [Salmonella enterica] Length: 372\nNumber of predicted TMHs: 1\nExp number of AAs in TMHs: 24.51309\nExp number, first 60 AAs: 0.2682\nTotal prob of N-in: 0.14402\noutside 1 331\nTMhelix 332 354\ninside 355 372

25293	GCF_001223365.1_8080_3_70	Salmonella enterica subsp. enterica serovar Typhimurium str. DT104	Proteobacteria; Gammaproteobacteria; Enterobacterales; Enterobacteriaceae; Salmonella; Salmonella enterica; Salmonella enterica subsp. enterica; Salmonella enterica subsp. enterica serovar Typhimurium	MNNEETFYQAMRRKGVTRRSFLKFCSLAATSLGLGAGMTPKIAWALENKPRIPVVWIHGL	WP_001058312.1	hydrogenase [Salmonella enterica]	Length: 372\nNumber of predicted TMHs:  2\nExp number of AAs in TMHs: 36.51738\nExp number, first 60 AAs:  19.72849\nTotal prob of N-in:   0.96170\nPOSSIBLE N-term signal sequence\ninside   1   19\nTMhelix   20   39\noutside   40   325\nTMhelix   326   348\ninside   349   372

25294	GCF_001566155.1_8080_3_32	Salmonella enterica subsp. enterica serovar Typhimurium str. DT104	Proteobacteria; Gammaproteobacteria; Enterobacterales; Enterobacteriaceae; Salmonella; Salmonella enterica; Salmonella enterica subsp. enterica; Salmonella enterica subsp. enterica serovar Typhimurium	MTGDNTLITSHGINRRDFMKLCAALAATMGLSSKAAAEMAESVSNPQRPPVIWIGAQECT	WP_000145429.1	hydrogenase 2 small subunit [Salmonella enterica]	Length: 372\nNumber of predicted TMHs:  1\nExp number of AAs in TMHs: 24.51309\nExp number, first 60 AAs:  0.2682\nTotal prob of N-in:   0.14402\noutside   1   331\nTMhelix   332   354\ninside   355   372

25295	GCF_001220325.1_8080_3_78	Salmonella enterica subsp. enterica serovar Typhimurium str. DT104	Proteobacteria; Gammaproteobacteria; Enterobacterales; Enterobacteriaceae; Salmonella; Salmonella enterica; Salmonella enterica subsp. enterica; Salmonella enterica subsp. enterica serovar Typhimurium	MTGDNTLITSHGINRRDFMKLCAALAATMGLSSKAAAEMAESVSNPQRPPVIWIGAQECT	WP_000145429.1	hydrogenase 2 small subunit [Salmonella enterica]	Length: 372\nNumber of predicted TMHs:  1\nExp number of AAs in TMHs: 24.51309\nExp number, first 60 AAs:  0.2682\nTotal prob of N-in:   0.14402\noutside   1   331\nTMhelix   332   354\ninside   355   372

25296	GCF_001566035.1_8080_3_37	Salmonella enterica subsp. enterica serovar Typhimurium str. DT104	Proteobacteria; Gammaproteobacteria; Enterobacterales; Enterobacteriaceae; Salmonella; Salmonella enterica; Salmonella enterica subsp. enterica; Salmonella enterica subsp. enterica serovar Typhimurium	MTGDNTLITSHGINRRDFMKLCAALAATMGLSSKAAAEMAESVSNPQRPPVIWIGAQECT	WP_000145429.1	hydrogenase 2 small subunit [Salmonella enterica]	Length: 372\nNumber of predicted TMHs:  1\nExp number of AAs in TMHs: 24.51309\nExp number, first 60 AAs:  0.2682\nTotal prob of N-in:   0.14402\noutside   1   331\nTMhelix   332   354\ninside   355   372

25297	GCF_001220365.1_9066_1_1Salmonella enterica subsp. enterica serovar Typhimurium str. DT104	Proteobacteria; Gammaproteobacteria; Enterobacterales; Enterobacteriaceae; Salmonella; Salmonella enterica; Salmonella enterica subsp. enterica; Salmonella enterica subsp. enterica serovar Typhimurium	MTGDNTLITSHGINRRDFMKLCAALAATMGLSSKAAAEMAESVSNPQRPPVIWIGAQECT	WP_000145429.1	hydrogenase 2 small subunit [Salmonella enterica]	Length: 372\nNumber of predicted TMHs:  1\nExp number of AAs in TMHs: 24.51309\nExp number, first 60 AAs:  0.2682\nTotal prob of N-in:   0.14402\noutside   1   331\nTMhelix   332   354\ninside   355   372

25298	GCF_001220405.1_9065_1_12	Salmonella enterica subsp. enterica serovar Typhimurium str. DT104	Proteobacteria; Gammaproteobacteria; Enterobacterales; Enterobacteriaceae; Salmonella; Salmonella enterica; Salmonella enterica subsp. enterica; Salmonella enterica subsp. enterica serovar Typhimurium	MTGDNTLITSHGINRRDFMKLCAALAATMGLSSKAAAEMAESVSNPQRPPVIWIGAQECT	WP_000145429.1	hydrogenase 2 small subunit [Salmonella enterica]	Length: 372\nNumber of predicted TMHs:  1\nExp number of AAs in TMHs: 24.51309\nExp number, first 60 AAs:  0.2682\nTotal prob of N-in:   0.14402\noutside   1   331\nTMhelix   332   354\ninside   355   372

25299	GCF_001220225.1_9066_1_3Salmonella enterica subsp. enterica serovar Typhimurium str. DT104	Proteobacteria; Gammaproteobacteria; Enterobacterales; Enterobacteriaceae; Salmonella; Salmonella enterica; Salmonella enterica subsp. enterica; Salmonella enterica subsp. enterica serovar Typhimurium	MNNEETFYQAMRRKGVTRRSFLKFCSLAATSLGLGAGMTPKIAWALENKPRIPVVWIHGL	WP_001058312.1	hydrogenase [Salmonella enterica]	Length: 372\nNumber of predicted TMHs:  2\nExp number of AAs in TMHs: 36.51738\nExp number, first 60 AAs:  19.72849\nTotal prob of N-in:   0.96170\nPOSSIBLE N-term signal sequence\ninside   1   19\nTMhelix   20   39\noutside   40   325\nTMhelix   326   348\ninside   349   372

25300	GCF_000493535.2_DT2	Salmonella enterica subsp. enterica serovar Typhimurium str. DT2	Proteobacteria; Gammaproteobacteria; Enterobacterales; Enterobacteriaceae; Salmonella; Salmonella enterica; Salmonella enterica subsp. enterica; Salmonella enterica subsp. enterica serovar Typhimurium	MNNEETFYQAMRRKGVTRRSFLKFCSLAATSLGLGAGMTPKIAWALENKPRIPVVWIHGL	WP_001058312.1	hydrogenase [Salmonella enterica]	Length: 372\nNumber of predicted TMHs:  2\nExp number of AAs in

TMHs: 36.51738\nExp number, first 60 AAs: 19.72849\nTotal prob of N-in: 0.96170\nPOSSIBLE N-term signal sequence\ninside 1 19\nTMhelix 20 39\noutside 40 325\nTMhelix 326 348\ninside 349 372

25301 GCF_000493535.2_DT2 Salmonella enterica subsp. enterica serovar Typhimurium str. DT2 Proteobacteria; Gammaproteobacteria; Enterobacterales; Enterobacteriaceae; Salmonella; Salmonella enterica; Salmonella enterica subsp. enterica; Salmonella enterica subsp. enterica serovar Typhimurium MTGDNTLITSHGINRRDFMKLCAALAATMGLSSKAAAEMAESVSNPQRPPVIWIGAQECT WP_022562441.1 hydrogenase-2 small chain protein [Salmonella enterica] Length: 372\nNumber of predicted TMHs: 1\nExp number of AAs in TMHs: 24.51309\nExp number, first 60 AAs: 0.2682\nTotal prob of N-in: 0.14402\noutside 1 331\nTMhelix 332 354\ninside 355 372

25302 GCF_000828595.1_ASM82859v1 Salmonella enterica subsp. enterica serovar Typhimurium str. L-3553 Proteobacteria; Gammaproteobacteria; Enterobacterales; Enterobacteriaceae; Salmonella; Salmonella enterica; Salmonella enterica subsp. enterica; Salmonella enterica subsp. enterica serovar Typhimurium MNNEETFYQAMRRKGVTRRSFLKFCSLAATSLGLGAGMTPKIAWALENKPRIPVVWIHGL WP_001058312.1 hydrogenase [Salmonella enterica] Length: 372\nNumber of predicted TMHs: 2\nExp number of AAs in TMHs: 36.51738\nExp number, first 60 AAs: 19.72849\nTotal prob of N-in: 0.96170\nPOSSIBLE N-term signal sequence\ninside 1 19\nTMhelix 20 39\noutside 40 325\nTMhelix 326 348\ninside 349 372

25303 GCF_000828595.1_ASM82859v1 Salmonella enterica subsp. enterica serovar Typhimurium str. L-3553 Proteobacteria; Gammaproteobacteria; Enterobacterales; Enterobacteriaceae; Salmonella; Salmonella enterica; Salmonella enterica subsp. enterica; Salmonella enterica subsp. enterica serovar Typhimurium MTGDNTLITSHGINRRDFMKLCAALAATMGLSSKAAAEMAESVSNPQRPPVIWIGAQECT WP_000145429.1 hydrogenase 2 small subunit [Salmonella enterica] Length: 372\nNumber of predicted TMHs: 1\nExp number of AAs in TMHs: 24.51309\nExp number, first 60 AAs: 0.2682\nTotal prob of N-in: 0.14402\noutside 1 331\nTMhelix 332 354\ninside 355 372

25304 GCF_000006945.2_ASM694v2 Salmonella enterica subsp. enterica serovar Typhimurium str. LT2 Proteobacteria; Gammaproteobacteria; Enterobacterales; Enterobacteriaceae; Salmonella; Salmonella enterica; Salmonella enterica subsp. enterica; Salmonella enterica subsp. enterica serovar Typhimurium MTGDNTLITSHGINRRDFMKLCAALAATMGLSSKAAAEMAESVSNPQRPPVIWIGAQECT NP_462065.1 hydrogenase 2 small subunit [Salmonella enterica subsp. enterica serovar Typhimurium str. LT2] Length: 372\nNumber of predicted TMHs: 1\nExp number of AAs in TMHs: 24.51309\nExp number, first 60 AAs: 0.2682\nTotal prob of N-in: 0.14402\noutside 1 331\nTMhelix 332 354\ninside 355 372

25305 GCF_000006945.2_ASM694v2 Salmonella enterica subsp. enterica serovar Typhimurium str. LT2 Proteobacteria; Gammaproteobacteria; Enterobacterales; Enterobacteriaceae; Salmonella; Salmonella enterica; Salmonella enterica subsp. enterica; Salmonella enterica subsp. enterica serovar Typhimurium MNNEETFYQAMRRKGVTRRSFLKFCSLAATSLGLGAGMTPKIAWALENKPRIPVVWIHGL NP_460742.1 [Ni/Fe] hydrogenase small subunit [Salmonella enterica subsp. enterica serovar Typhimurium str. LT2] Length: 372\nNumber of predicted TMHs: 2\nExp number of AAs in TMHs: 36.51738\nExp number, first 60 AAs: 19.72849\nTotal prob of N-in: 0.96170\nPOSSIBLE N-term signal sequence\ninside 1 19\nTMhelix 20 39\noutside 40 325\nTMhelix 326 348\ninside 349 372

25306 GCF_000486345.2_ASM48634v2 Salmonella enterica subsp. enterica serovar Typhimurium str. SARA13 Proteobacteria; Gammaproteobacteria; Enterobacterales; Enterobacteriaceae; Salmonella; Salmonella enterica; Salmonella enterica subsp. enterica; Salmonella enterica subsp. enterica serovar Typhimurium MNNEETFYQAMRRKGVTRRSFLKFCSLAATSLGLGAGMTPKIAWALENKPRIPVVWIHGL WP_023198237.1 hydrogenase 1, small subunit [Salmonella enterica] Length: 372\nNumber of predicted TMHs: 2\nExp number of AAs in TMHs: 36.39423\nExp number, first 60 AAs: 19.72877\nTotal prob of N-in: 0.96162\nPOSSIBLE N-term signal sequence\ninside 1 19\nTMhelix 20 39\noutside 40 325\nTMhelix 326 348\ninside 349 372

25307 GCF_000486345.2_ASM48634v2 Salmonella enterica subsp. enterica serovar Typhimurium str. SARA13 Proteobacteria; Gammaproteobacteria; Enterobacterales; Enterobacteriaceae; Salmonella; Salmonella enterica; Salmonella enterica subsp. enterica; Salmonella enterica subsp. enterica serovar Typhimurium MTGDNTLITSHGINRRDFMKLCAALAATMGLSSKAAAEMAESVSNPQRPPVIWIGAQECT WP_000145429.1 hydrogenase 2 small subunit [Salmonella enterica] Length: 372\nNumber of predicted TMHs: 1\nExp number of AAs in TMHs: 24.51309\nExp number, first 60 AAs: 0.2682\nTotal prob of N-in: 0.14402\noutside 1 331\nTMhelix 332 354\ninside 355 372

25308 GCF_000210855.2_ASM21085v2 Salmonella enterica subsp. enterica serovar Typhimurium str. SL1344 Proteobacteria; Gammaproteobacteria; Enterobacterales; Enterobacteriaceae; Salmonella; Salmonella enterica; Salmonella enterica subsp. enterica; Salmonella enterica subsp. enterica serovar Typhimurium MNNEETFYQAMRRKGVTRRSFLKFCSLAATSLGLGAGMTPKIAWALENKPRIPVVWIHGL WP_001058312.1 hydrogenase [Salmonella enterica] Length: 372\nNumber of predicted TMHs: 2\nExp number of AAs in TMHs: 36.51738\nExp number, first 60 AAs: 19.72849\nTotal prob of N-in: 0.96170\nPOSSIBLE N-term signal sequence\ninside 1 19\nTMhelix 20 39\noutside 40 325\nTMhelix 326 348\ninside 349 372

25309 GCF_000210855.2_ASM21085v2 Salmonella enterica subsp. enterica serovar Typhimurium str. SL1344 Proteobacteria; Gammaproteobacteria; Enterobacterales; Enterobacteriaceae; Salmonella; Salmonella enterica; Salmonella enterica subsp. enterica; Salmonella enterica subsp. enterica serovar Typhimurium MTGDNTLITSHGINRRDFMKLCAALAATMGLSSKAAAEMAESVSNPQRPPVIWIGAQECT WP_000145429.1 hydrogenase 2 small subunit [Salmonella enterica] Length: 372\nNumber of predicted TMHs: 1\nExp number of AAs in TMHs: 24.51309\nExp number, first 60 AAs: 0.2682\nTotal prob of N-in: 0.14402\noutside 1 331\nTMhelix 332 354\ninside 355 372

25310 GCF_001295495.1_ASM129549v1 Salmonella enterica subsp. enterica serovar Typhimurium str. ST1489 Proteobacteria; Gammaproteobacteria; Enterobacterales; Enterobacteriaceae; Salmonella; Salmonella enterica; Salmonella enterica subsp. enterica; Salmonella enterica subsp. enterica serovar Typhimurium MNNEETFYQAMRRKGVTRRSFLKFCSLAATSLGLGAGMTPKIAWALENKPRIPVVWIHGL WP_001058312.1 hydrogenase [Salmonella enterica] Length: 372\nNumber of predicted TMHs: 2\nExp number of AAs in TMHs: 36.51738\nExp number, first 60 AAs: 19.72849\nTotal prob of N-in: 0.96170\nPOSSIBLE N-term signal sequence\ninside 1 19\nTMhelix 20 39\noutside 40 325\nTMhelix 326 348\ninside 349 372

25311 GCF_001295495.1_ASM129549v1 Salmonella enterica subsp. enterica serovar Typhimurium str. ST1489 Proteobacteria; Gammaproteobacteria; Enterobacterales; Enterobacteriaceae; Salmonella; Salmonella enterica; Salmonella enterica subsp. enterica; Salmonella enterica subsp. enterica serovar Typhimurium MTGDNTLITSHGINRRDFMKLCAALAATMGLSSKAAAEMAESVSNPQRPPVIWIGAQECT WP_000145429.1 hydrogenase 2 small subunit [Salmonella enterica] Length: 372\nNumber of predicted TMHs: 1\nExp number of AAs in TMHs: 24.51309\nExp number, first 60 AAs: 0.2682\nTotal prob of N-in: 0.14402\noutside 1 331\nTMhelix 332 354\ninside 355 372

25312 GCF_000465195.1_ST1489_for_the_version_1_of_Salmonella_enterica_subsp._enterica_serovar_Typhimurium_ST1489 Salmonella enterica subsp. enterica serovar Typhimurium str. ST1489 Proteobacteria; Gammaproteobacteria; Enterobacterales; Enterobacteriaceae; Salmonella; Salmonella enterica; Salmonella enterica subsp. enterica; Salmonella enterica subsp. enterica serovar Typhimurium MTGDNTLITSHGINRRDFMKLCAALAATMGLSSKAAAEMAESVSNPQRPPVIWIGAQECT WP_000145429.1 hydrogenase 2 small subunit [Salmonella enterica] Length: 372\nNumber of predicted TMHs: 1\nExp number of AAs in TMHs: 24.51309\nExp number, first 60 AAs: 0.2682\nTotal prob of N-in: 0.14402\noutside 1 331\nTMhelix 332 354\ninside 355 372

25313 GCF_000465195.1_ST1489_for_the_version_1_of_Salmonella_enterica_subsp._enterica_serovar_Typhimurium_ST1489 Salmonella enterica subsp. enterica serovar Typhimurium str. ST1489 Proteobacteria; Gammaproteobacteria; Enterobacterales; Enterobacteriaceae; Salmonella; Salmonella enterica; Salmonella enterica subsp. enterica; Salmonella enterica subsp. enterica serovar Typhimurium MNNEETFYQAMRRKGVTRRSFLKFCSLAATSLGLGAGMTPKIAWALENKPRIPVVWIHGL WP_001058312.1 hydrogenase [Salmonella enterica] Length: 372\nNumber of predicted TMHs: 2\nExp number of AAs in TMHs: 36.51738\nExp number, first 60 AAs: 19.72849\nTotal prob of N-in: 0.96170\nPOSSIBLE N-term signal sequence\ninside 1 19\nTMhelix 20 39\noutside 40 325\nTMhelix 326 348\ninside 349 372

25314 GCF_000292755.1_ASM29275v1 Salmonella enterica subsp. enterica serovar Typhimurium str. ST1660/06 Proteobacteria; Gammaproteobacteria; Enterobacterales; Enterobacteriaceae; Salmonella; Salmonella enterica; Salmonella enterica subsp. enterica; Salmonella enterica subsp. enterica serovar Typhimurium MNNEETFYQAMRRKGVTRRSFLKFCSLAATSLGLGAGMTPKIAWALENKPRIPVVWIHGL WP_001058312.1 hydrogenase [Salmonella enterica] Length: 372\nNumber of predicted TMHs: 2\nExp number of AAs in TMHs: 36.51738\nExp number, first 60 AAs: 19.72849\nTotal prob of N-in: 0.96170\nPOSSIBLE N-term signal sequence\ninside 1 19\nTMhelix 20 39\noutside 40 325\nTMhelix 326 348\ninside 349 372

25315 GCF_000292755.1_ASM29275v1 Salmonella enterica subsp. enterica serovar Typhimurium str. ST1660/06 Proteobacteria; Gammaproteobacteria; Enterobacterales; Enterobacteriaceae; Salmonella; Salmonella enterica; Salmonella enterica subsp. enterica; Salmonella enterica subsp. enterica serovar Typhimurium MTGDNTLITSHGINRRDFMKLCAALAATMGLSSKAAAEMAESVSNPQRPPVIWIGAQECT WP_000145429.1 hydrogenase 2 small subunit [Salmonella enterica] Length: 372\nNumber of predicted TMHs: 1\nExp number of AAs in TMHs: 24.51309\nExp number, first 60 AAs: 0.2682\nTotal prob of N-in: 0.14402\noutside 1 331\nTMhelix 332 354\ninside 355 372

25316 GCF_000188735.1_ASM18873v1 Salmonella enterica subsp. enterica serovar Typhimurium str. ST4/74 Proteobacteria; Gammaproteobacteria; Enterobacterales; Enterobacteriaceae; Salmonella; Salmonella enterica; Salmonella enterica subsp. enterica; Salmonella enterica subsp. enterica serovar Typhimurium MTGDNTLITSHGINRRDFMKLCAALAATMGLSSKAAAEMAESVSNPQRPPVIWIGAQECT WP_000145429.1 hydrogenase 2 small subunit [Salmonella enterica] Length: 372\nNumber of predicted TMHs: 1\nExp number of AAs in TMHs: 24.51309\nExp number, first 60 AAs: 0.2682\nTotal prob of N-in: 0.14402\noutside 1 331\nTMhelix 332 354\ninside 355 372

25317 GCF_000188735.1_ASM18873v1 Salmonella enterica subsp. enterica serovar Typhimurium str. ST4/74 Proteobacteria; Gammaproteobacteria; Enterobacterales; Enterobacteriaceae; Salmonella; Salmonella enterica; Salmonella enterica subsp. enterica; Salmonella enterica subsp. enterica serovar Typhimurium MNNEETFYQAMRRKGVTRRSFLKFCSLAATSLGLGAGMTPKIAWALENKPRIPVVWIHGL WP_001058312.1 hydrogenase [Salmonella enterica] Length: 372\nNumber of predicted TMHs: 2\nExp number of AAs in TMHs: 36.51738\nExp number, first 60 AAs: 19.72849\nTotal prob of N-in: 0.96170\nPOSSIBLE N-term signal sequence\ninside 1 19\nTMhelix 20 39\noutside 40 325\nTMhelix 326 348\ninside 349 372

25318 GCF_000468255.1_ST4848_v1 Salmonella enterica subsp. enterica serovar Typhimurium str. ST4848 Proteobacteria; Gammaproteobacteria; Enterobacterales; Enterobacteriaceae; Salmonella; Salmonella enterica; Salmonella enterica subsp. enterica; Salmonella enterica subsp. enterica serovar Typhimurium MNNEETFYQAMRRKGVTRRSFLKFCSLAATSLGLGAGMTPKIAWALENKPRIPVVWIHGL WP_001058312.1 hydrogenase [Salmonella enterica] Length: 372\nNumber of predicted TMHs: 2\nExp number of AAs in TMHs: 36.51738\nExp number, first 60 AAs: 19.72849\nTotal prob of N-in: 0.96170\nPOSSIBLE N-term signal sequence\ninside 1 19\nTMhelix 20 39\noutside 40 325\nTMhelix 326 348\ninside 349 372

25319 GCF_000468255.1_ST4848_v1 Salmonella enterica subsp. enterica serovar Typhimurium str. ST4848 Proteobacteria; Gammaproteobacteria; Enterobacterales; Enterobacteriaceae; Salmonella; Salmonella enterica; Salmonella enterica subsp. enterica; Salmonella enterica subsp. enterica serovar Typhimurium MTGDNTLITSHGINRRDFMKLCAALAATMGLSSKAAAEMAESVSNPQRPPVIWIGAQECT WP_000145429.1 hydrogenase 2 small subunit [Salmonella enterica] Length: 372\nNumber of predicted TMHs: 1\nExp number of AAs in TMHs: 24.51309\nExp number, first 60 AAs: 0.2682\nTotal prob of N-in: 0.14402\noutside 1 331\nTMhelix 332 354\ninside 355 372

25320 GCF_000465175.1_ST78896_for_the_version_1_of_Salmonella_enterica_subsp._enterica_serovar_Typhimurium_ST78896Salmonella enterica subsp. enterica serovar Typhimurium str. ST78896 Proteobacteria; Gammaproteobacteria; Enterobacterales; Enterobacteriaceae; Salmonella; Salmonella enterica; Salmonella enterica subsp. enterica; Salmonella enterica subsp. enterica serovar Typhimurium MTGDNTLITSHGINRRDFMKLCAALAATMGLSSKAAAEMAESVSNPQRPPVIWIGAQECT WP_000145429.1 hydrogenase 2 small subunit [Salmonella enterica] Length: 372\nNumber of predicted TMHs: 1\nExp number of AAs in TMHs: 24.51309\nExp number, first 60 AAs: 0.2682\nTotal prob of N-in: 0.14402\noutside 1 331\nTMhelix 332 354\ninside 355 372

25321 GCF_000465175.1_ST78896_for_the_version_1_of_Salmonella_enterica_subsp._enterica_serovar_Typhimurium_ST78896Salmonella enterica subsp. enterica serovar Typhimurium str. ST78896 Proteobacteria; Gammaproteobacteria; Enterobacterales; Enterobacteriaceae; Salmonella; Salmonella enterica; Salmonella enterica subsp. enterica; Salmonella enterica subsp. enterica serovar Typhimurium MNNEETFYQAMRRKGVTRRSFLKFCSLAATSLGLGAGMTPKIAWALENKPRIPVVWIHGL WP_001058312.1 hydrogenase [Salmonella enterica] Length: 372\nNumber of predicted TMHs: 2\nExp number of AAs in TMHs: 36.51738\nExp number, first 60 AAs: 19.72849\nTotal prob of N-in: 0.96170\nPOSSIBLE N-term signal sequence\ninside 1 19\nTMhelix 20 39\noutside 40 325\nTMhelix 326 348\ninside 349 372

25322 GCF_000312825.2_STm1.2 Salmonella enterica subsp. enterica serovar Typhimurium str. STm1 Proteobacteria; Gammaproteobacteria; Enterobacterales; Enterobacteriaceae; Salmonella; Salmonella enterica; Salmonella enterica subsp. enterica; Salmonella enterica subsp. enterica serovar Typhimurium MTGDNTLITSHGINRRDFMKLCAALAATMGLSSKAAAEMAESVSNPQRPPVIWIGAQECT WP_000145429.1 hydrogenase 2 small subunit [Salmonella enterica] Length: 372\nNumber of predicted TMHs: 1\nExp number of AAs in TMHs: 24.51309\nExp number, first 60 AAs: 0.2682\nTotal prob of N-in: 0.14402\noutside 1 331\nTMhelix 332 354\ninside 355 372

25323 GCF_000312825.2_STm1.2 Salmonella enterica subsp. enterica serovar Typhimurium str. STm1 Proteobacteria; Gammaproteobacteria; Enterobacterales; Enterobacteriaceae; Salmonella; Salmonella enterica; Salmonella enterica subsp. enterica; Salmonella enterica subsp. enterica serovar Typhimurium MNNEETFYQAMRRKGVTRRSFLKFCSLAATSLGLGAGMTPKIAWALENKPRIPVVWIHGL WP_001058312.1 hydrogenase [Salmonella enterica] Length: 372\nNumber of predicted TMHs: 2\nExp number of AAs in TMHs: 36.51738\nExp number, first 60 AAs: 19.72849\nTotal prob of N-in: 0.96170\nPOSSIBLE N-term signal sequence\ninside 1 19\nTMhelix 20 39\noutside 40 325\nTMhelix 326 348\ninside 349 372

25324 GCF_000312905.2_STm10.2 Salmonella enterica subsp. enterica serovar Typhimurium str. STm10 Proteobacteria; Gammaproteobacteria; Enterobacterales; Enterobacteriaceae; Salmonella; Salmonella enterica; Salmonella enterica subsp. enterica; Salmonella enterica subsp. enterica serovar Typhimurium MTGDNTLITSHGINRRDFMKLCAALAATMGLSSKAAAEMAESVSNPQRPPVIWIGAQECT WP_000145429.1 hydrogenase 2 small subunit [Salmonella enterica] Length: 372\nNumber of predicted TMHs: 1\nExp number of AAs in TMHs: 24.51309\nExp number, first 60 AAs: 0.2682\nTotal prob of N-in: 0.14402\noutside 1 331\nTMhelix 332 354\ninside 355 372

25325 GCF_000312905.2_STm10.2 Salmonella enterica subsp. enterica serovar Typhimurium str. STm10 Proteobacteria; Gammaproteobacteria; Enterobacterales; Enterobacteriaceae; Salmonella; Salmonella enterica; Salmonella enterica subsp. enterica; Salmonella enterica subsp. enterica serovar Typhimurium MNNEETFYQAMRRKGVTRRSFLKFCSLAATSLGLGAGMTPKIAWALENKPRIPVVWIHGL WP_001058312.1 hydrogenase [Salmonella enterica] Length: 372\nNumber of predicted TMHs: 2\nExp number of AAs in TMHs: 36.51738\nExp number, first 60 AAs: 19.72849\nTotal prob of N-in: 0.96170\nPOSSIBLE N-term signal sequence\ninside 1 19\nTMhelix 20 39\noutside 40 325\nTMhelix 326 348\ninside 349 372

25326 GCF_000312765.2_STm11.2 Salmonella enterica subsp. enterica serovar Typhimurium str. STm11 Proteobacteria; Gammaproteobacteria; Enterobacterales; Enterobacteriaceae; Salmonella; Salmonella enterica; Salmonella enterica subsp. enterica; Salmonella enterica subsp. enterica serovar Typhimurium MNNEETFYQAMRRKGVTRRSFLKFCSLAATSLGLGAGMTPKIAWALENKPRIPVVWIHGL WP_001058312.1 hydrogenase [Salmonella enterica] Length: 372\nNumber of predicted TMHs: 2\nExp number of AAs in TMHs: 36.51738\nExp number, first 60 AAs: 19.72849\nTotal prob of N-in: 0.96170\nPOSSIBLE N-term signal sequence\ninside 1 19\nTMhelix 20 39\noutside 40 325\nTMhelix 326 348\ninside 349 372

25327 GCF_000312765.2_STm11.2 Salmonella enterica subsp. enterica serovar Typhimurium str. STm11 Proteobacteria; Gammaproteobacteria; Enterobacterales; Enterobacteriaceae; Salmonella; Salmonella enterica; Salmonella enterica subsp. enterica; Salmonella enterica subsp. enterica serovar Typhimurium MTGDNTLITSHGINRRDFMKLCAALAATMGLSSKAAAEMAESVSNPQRPPVIWIGAQECT WP_000145429.1 hydrogenase 2 small subunit [Salmonella enterica] Length: 372\nNumber of predicted TMHs: 1\nExp number of AAs in TMHs: 24.51309\nExp number, first 60 AAs: 0.2682\nTotal prob of N-in: 0.14402\noutside 1 331\nTMhelix 332 354\ninside 355 372

25328 GCF_000312785.2_STm12.2 Salmonella enterica subsp. enterica serovar Typhimurium str. STm12 Proteobacteria; Gammaproteobacteria; Enterobacterales; Enterobacteriaceae; Salmonella; Salmonella enterica; Salmonella enterica subsp. enterica; Salmonella enterica subsp. enterica serovar Typhimurium MTGDNTLITSHGINRRDFMKLCAALAATMGLSSKAAAEMAESVSNPQRPPVIWIGAQECT WP_000145429.1 hydrogenase 2 small subunit [Salmonella enterica] Length: 372\nNumber of predicted TMHs: 1\nExp number of AAs in TMHs: 24.51309\nExp number, first 60 AAs: 0.2682\nTotal prob of N-in: 0.14402\noutside 1 331\nTMhelix 332 354\ninside 355 372

25329 GCF_000312785.2_STm12.2 Salmonella enterica subsp. enterica serovar Typhimurium str. STm12 Proteobacteria; Gammaproteobacteria; Enterobacterales; Enterobacteriaceae; Salmonella; Salmonella enterica; Salmonella enterica subsp. enterica; Salmonella enterica subsp. enterica serovar Typhimurium MNNEETFYQAMRRKGVTRRSFLKFCSLAATSLGLGAGMTPKIAWALENKPRIPVVWIHGL WP_001058312.1 hydrogenase [Salmonella enterica] Length: 372\nNumber of predicted TMHs: 2\nExp number of AAs in

TMHs: 36.51738\nExp number, first 60 AAs: 19.72849\nTotal prob of N-in: 0.96170\nPOSSIBLE N-term signal sequence\ninside 1 19\nTMhelix 20 39\noutside 40 325\nTMhelix 326 348\ninside 349 372

25330 GCF_000314915.1_ASM31491v1 Salmonella enterica subsp. enterica serovar Typhimurium str. STm2 Proteobacteria; Gammaproteobacteria; Enterobacterales; Enterobacteriaceae; Salmonella; Salmonella enterica; Salmonella enterica subsp. enterica; Salmonella enterica subsp. enterica serovar Typhimurium MTGDNTLITSHGINRRDFMKLCAALAATMGLSSKAAAEMAESVSNPQRPPVIWIGAQECT WP_000145429.1 hydrogenase 2 small subunit [Salmonella enterica] Length: 372\nNumber of predicted TMHs: 1\nExp number of AAs in TMHs: 24.51309\nExp number, first 60 AAs: 0.2682\nTotal prob of N-in: 0.14402\noutside 1 331\nTMhelix 332 354\ninside 355 372

25331 GCF_000312865.2_STm3.2 Salmonella enterica subsp. enterica serovar Typhimurium str. STm3 Proteobacteria; Gammaproteobacteria; Enterobacterales; Enterobacteriaceae; Salmonella; Salmonella enterica; Salmonella enterica subsp. enterica; Salmonella enterica subsp. enterica serovar Typhimurium MTGDNTLITSHGINRRDFMKLCAALAATMGLSSKAAAEMAESVSNPQRPPVIWIGAQECT WP_000145429.1 hydrogenase 2 small subunit [Salmonella enterica] Length: 372\nNumber of predicted TMHs: 1\nExp number of AAs in TMHs: 24.51309\nExp number, first 60 AAs: 0.2682\nTotal prob of N-in: 0.14402\noutside 1 331\nTMhelix 332 354\ninside 355 372

25332 GCF_000312865.2_STm3.2 Salmonella enterica subsp. enterica serovar Typhimurium str. STm3 Proteobacteria; Gammaproteobacteria; Enterobacterales; Enterobacteriaceae; Salmonella; Salmonella enterica; Salmonella enterica subsp. enterica; Salmonella enterica subsp. enterica serovar Typhimurium MNNEETFYQAMRRKGVTRRSFLKFCSLAATSLGLGAGMTPKIAWALENKPRIPVVWIHGL WP_001058312.1 hydrogenase [Salmonella enterica] Length: 372\nNumber of predicted TMHs: 2\nExp number of AAs in TMHs: 36.51738\nExp number, first 60 AAs: 19.72849\nTotal prob of N-in: 0.96170\nPOSSIBLE N-term signal sequence\ninside 1 19\nTMhelix 20 39\noutside 40 325\nTMhelix 326 348\ninside 349 372

25333 GCF_000312885.2_STm4.2 Salmonella enterica subsp. enterica serovar Typhimurium str. STm4 Proteobacteria; Gammaproteobacteria; Enterobacterales; Enterobacteriaceae; Salmonella; Salmonella enterica; Salmonella enterica subsp. enterica; Salmonella enterica subsp. enterica serovar Typhimurium MNNEETFYQAMRRKGVTRRSFLKFCSLAATSLGLGAGMTPKIAWALENKPRIPVVWIHGL WP_001058312.1 hydrogenase [Salmonella enterica] Length: 372\nNumber of predicted TMHs: 2\nExp number of AAs in TMHs: 36.51738\nExp number, first 60 AAs: 19.72849\nTotal prob of N-in: 0.96170\nPOSSIBLE N-term signal sequence\ninside 1 19\nTMhelix 20 39\noutside 40 325\nTMhelix 326 348\ninside 349 372

25334 GCF_000312885.2_STm4.2 Salmonella enterica subsp. enterica serovar Typhimurium str. STm4 Proteobacteria; Gammaproteobacteria; Enterobacterales; Enterobacteriaceae; Salmonella; Salmonella enterica; Salmonella enterica subsp. enterica; Salmonella enterica subsp. enterica serovar Typhimurium MTGDNTLITSHGINRRDFMKLCAALAATMGLSSKAAAEMAESVSNPQRPPVIWIGAQECT WP_000145429.1 hydrogenase 2 small subunit [Salmonella enterica] Length: 372\nNumber of predicted TMHs: 1\nExp number of AAs in TMHs: 24.51309\nExp number, first 60 AAs: 0.2682\nTotal prob of N-in: 0.14402\noutside 1 331\nTMhelix 332 354\ninside 355 372

25335 GCF_000312805.2_STm5.2 Salmonella enterica subsp. enterica serovar Typhimurium str. STm5 Proteobacteria; Gammaproteobacteria; Enterobacterales; Enterobacteriaceae; Salmonella; Salmonella enterica; Salmonella enterica subsp. enterica; Salmonella enterica subsp. enterica serovar Typhimurium MTGDNTLITSHGINRRDFMKLCAALAATMGLSSKAAAEMAESVSNPQRPPVIWIGAQECT WP_000145429.1 hydrogenase 2 small subunit [Salmonella enterica] Length: 372\nNumber of predicted TMHs: 1\nExp number of AAs in TMHs: 24.51309\nExp number, first 60 AAs: 0.2682\nTotal prob of N-in: 0.14402\noutside 1 331\nTMhelix 332 354\ninside 355 372

25336 GCF_000312745.2_STm6.2 Salmonella enterica subsp. enterica serovar Typhimurium str. STm6 Proteobacteria; Gammaproteobacteria; Enterobacterales; Enterobacteriaceae; Salmonella; Salmonella enterica; Salmonella enterica subsp. enterica; Salmonella enterica subsp. enterica serovar Typhimurium MNNEETFYQAMRRKGVTRRSFLKFCSLAATSLGLGAGMTPKIAWALENKPRIPVVWIHGL WP_001058312.1 hydrogenase [Salmonella enterica] Length: 372\nNumber of predicted TMHs: 2\nExp number of AAs in TMHs: 36.51738\nExp number, first 60 AAs: 19.72849\nTotal prob of N-in: 0.96170\nPOSSIBLE N-term signal sequence\ninside 1 19\nTMhelix 20 39\noutside 40 325\nTMhelix 326 348\ninside 349 372

25337 GCF_000312745.2_STm6.2 Salmonella enterica subsp. enterica serovar Typhimurium str. STm6 Proteobacteria; Gammaproteobacteria; Enterobacterales; Enterobacteriaceae; Salmonella; Salmonella enterica;

Salmonella enterica subsp. enterica; Salmonella enterica subsp. enterica serovar Typhimurium
MTGDNTLITSHGINRRDFMKLCAALAATMGLSSKAAAEMAESVSNPQRPPVIWIGAQECT WP_000145429.1
hydrogenase 2 small subunit [Salmonella enterica] Length: 372\nNumber of predicted TMHs: 1\nExp number of AAs in TMHs: 24.51309\nExp number, first 60 AAs: 0.2682\nTotal prob of N-in: 0.14402\noutside 1 331\nTMhelix 332 354\ninside 355 372

25338 GCF_000444835.1_STm7.1 Salmonella enterica subsp. enterica serovar Typhimurium str. STm7
Proteobacteria; Gammaproteobacteria; Enterobacterales; Enterobacteriaceae; Salmonella; Salmonella enterica; Salmonella enterica subsp. enterica; Salmonella enterica subsp. enterica serovar Typhimurium
MTGDNTLITSHGINRRDFMKLCAALAATMGLSSKAAAEMAESVSNPQRPPVIWIGAQECT WP_000145429.1
hydrogenase 2 small subunit [Salmonella enterica] Length: 372\nNumber of predicted TMHs: 1\nExp number of AAs in TMHs: 24.51309\nExp number, first 60 AAs: 0.2682\nTotal prob of N-in: 0.14402\noutside 1 331\nTMhelix 332 354\ninside 355 372

25339 GCF_000444835.1_STm7.1 Salmonella enterica subsp. enterica serovar Typhimurium str. STm7
Proteobacteria; Gammaproteobacteria; Enterobacterales; Enterobacteriaceae; Salmonella; Salmonella enterica; Salmonella enterica subsp. enterica; Salmonella enterica subsp. enterica serovar Typhimurium
MNNEETFYQAMRRKGVTRRSFLKFCSLAATSLGLGAGMTPKIAWALENKPRIPVVWIHGL WP_001058312.1
hydrogenase [Salmonella enterica] Length: 372\nNumber of predicted TMHs: 2\nExp number of AAs in TMHs: 36.51738\nExp number, first 60 AAs: 19.72849\nTotal prob of N-in: 0.96170\nPOSSIBLE N-term signal sequence\ninside 1 19\nTMhelix 20 39\noutside 40 325\nTMhelix 326 348\ninside 349 372

25340 GCF_000312845.2_STm8.2 Salmonella enterica subsp. enterica serovar Typhimurium str. STm8
Proteobacteria; Gammaproteobacteria; Enterobacterales; Enterobacteriaceae; Salmonella; Salmonella enterica; Salmonella enterica subsp. enterica; Salmonella enterica subsp. enterica serovar Typhimurium
MNNEETFYQAMRRKGVTRRSFLKFCSLAATSLGLGAGMTPKIAWALENKPRIPVVWIHGL WP_001058312.1
hydrogenase [Salmonella enterica] Length: 372\nNumber of predicted TMHs: 2\nExp number of AAs in TMHs: 36.51738\nExp number, first 60 AAs: 19.72849\nTotal prob of N-in: 0.96170\nPOSSIBLE N-term signal sequence\ninside 1 19\nTMhelix 20 39\noutside 40 325\nTMhelix 326 348\ninside 349 372

25341 GCF_000312845.2_STm8.2 Salmonella enterica subsp. enterica serovar Typhimurium str. STm8
Proteobacteria; Gammaproteobacteria; Enterobacterales; Enterobacteriaceae; Salmonella; Salmonella enterica; Salmonella enterica subsp. enterica; Salmonella enterica subsp. enterica serovar Typhimurium
MTGDNTLITSHGINRRDFMKLCAALAATMGLSSKAAAEMAESVSNPQRPPVIWIGAQECT WP_000145429.1
hydrogenase 2 small subunit [Salmonella enterica] Length: 372\nNumber of predicted TMHs: 1\nExp number of AAs in TMHs: 24.51309\nExp number, first 60 AAs: 0.2682\nTotal prob of N-in: 0.14402\noutside 1 331\nTMhelix 332 354\ninside 355 372

25342 GCF_000319795.2_STm9.2 Salmonella enterica subsp. enterica serovar Typhimurium str. STm9
Proteobacteria; Gammaproteobacteria; Enterobacterales; Enterobacteriaceae; Salmonella; Salmonella enterica; Salmonella enterica subsp. enterica; Salmonella enterica subsp. enterica serovar Typhimurium
MNNEETFYQAMRRKGVTRRSFLKFCSLAATSLGLGAGMTPKIAWALENKPRIPVVWIHGL WP_001058312.1
hydrogenase [Salmonella enterica] Length: 372\nNumber of predicted TMHs: 2\nExp number of AAs in TMHs: 36.51738\nExp number, first 60 AAs: 19.72849\nTotal prob of N-in: 0.96170\nPOSSIBLE N-term signal sequence\ninside 1 19\nTMhelix 20 39\noutside 40 325\nTMhelix 326 348\ninside 349 372

25343 GCF_000319795.2_STm9.2 Salmonella enterica subsp. enterica serovar Typhimurium str. STm9
Proteobacteria; Gammaproteobacteria; Enterobacterales; Enterobacteriaceae; Salmonella; Salmonella enterica; Salmonella enterica subsp. enterica; Salmonella enterica subsp. enterica serovar Typhimurium
MTGDNTLITSHGINRRDFMKLCAALAATMGLSSKAAAEMAESVSNPQRPPVIWIGAQECT WP_000145429.1
hydrogenase 2 small subunit [Salmonella enterica] Length: 372\nNumber of predicted TMHs: 1\nExp number of AAs in TMHs: 24.51309\nExp number, first 60 AAs: 0.2682\nTotal prob of N-in: 0.14402\noutside 1 331\nTMhelix 332 354\ninside 355 372

25344 GCF_000283735.1_ASM28373v1 Salmonella enterica subsp. enterica serovar Typhimurium str. T000240
Proteobacteria; Gammaproteobacteria; Enterobacterales; Enterobacteriaceae; Salmonella; Salmonella enterica; Salmonella enterica subsp. enterica; Salmonella enterica subsp. enterica serovar Typhimurium
MNNEETFYQAMRRKGVTRRSFLKFCSLAATSLGLGAGMTPKIAWALENKPRIPVVWIHGL WP_001058312.1
hydrogenase [Salmonella enterica] Length: 372\nNumber of predicted TMHs: 2\nExp number of AAs in TMHs: 36.51738\nExp number, first 60 AAs: 19.72849\nTotal prob of N-in: 0.96170\nPOSSIBLE N-term signal sequence\ninside 1 19\nTMhelix 20 39\noutside 40 325\nTMhelix 326 348\ninside 349 372

25345 GCF_000283735.1_ASM28373v1 Salmonella enterica subsp. enterica serovar Typhimurium str. T000240 Proteobacteria; Gammaproteobacteria; Enterobacterales; Enterobacteriaceae; Salmonella; Salmonella enterica; Salmonella enterica subsp. enterica; Salmonella enterica subsp. enterica serovar Typhimurium MTGDNTLITSHGINRRDFMKLCAALAATMGLSSKAAAEMAESVSNPQRPPVIWIGAQECT WP_000145429.1 hydrogenase 2 small subunit [Salmonella enterica] Length: 372\nNumber of predicted TMHs: 1\nExp number of AAs in TMHs: 24.51309\nExp number, first 60 AAs: 0.2682\nTotal prob of N-in: 0.14402\noutside 1 331\nTMhelix 332 354\ninside 355 372

25346 GCF_000187785.1_ASM18778v2 Salmonella enterica subsp. enterica serovar Typhimurium str. TN061786 Proteobacteria; Gammaproteobacteria; Enterobacterales; Enterobacteriaceae; Salmonella; Salmonella enterica; Salmonella enterica subsp. enterica; Salmonella enterica subsp. enterica serovar Typhimurium MTGDNTLITSHGINRRDFMKLCAALAATMGLSSKAAAEMAESVSNPQRPPVIWIGAQECT WP_000145429.1 hydrogenase 2 small subunit [Salmonella enterica] Length: 372\nNumber of predicted TMHs: 1\nExp number of AAs in TMHs: 24.51309\nExp number, first 60 AAs: 0.2682\nTotal prob of N-in: 0.14402\noutside 1 331\nTMhelix 332 354\ninside 355 372

25347 GCF_000187785.1_ASM18778v2 Salmonella enterica subsp. enterica serovar Typhimurium str. TN061786 Proteobacteria; Gammaproteobacteria; Enterobacterales; Enterobacteriaceae; Salmonella; Salmonella enterica; Salmonella enterica subsp. enterica; Salmonella enterica subsp. enterica serovar Typhimurium MNNEETFYQAMRRKGVTRRSFLKFCSLAATSLGLGAGMTPKIAWALENKPRIPVVWIHGL WP_001058312.1 hydrogenase [Salmonella enterica] Length: 372\nNumber of predicted TMHs: 2\nExp number of AAs in TMHs: 36.51738\nExp number, first 60 AAs: 19.72849\nTotal prob of N-in: 0.96170\nPOSSIBLE N-term signal sequence\ninside 1 19\nTMhelix 20 39\noutside 40 325\nTMhelix 326 348\ninside 349 372

25348 GCF_000380325.1_ASM38032v1 Salmonella enterica subsp. enterica serovar Typhimurium str. U288 Proteobacteria; Gammaproteobacteria; Enterobacterales; Enterobacteriaceae; Salmonella; Salmonella enterica; Salmonella enterica subsp. enterica; Salmonella enterica subsp. enterica serovar Typhimurium MNNEETFYQAMRRKGVTRRSFLKFCSLAATSLGLGAGMTPKIAWALENKPRIPVVWIHGL WP_001058312.1 hydrogenase [Salmonella enterica] Length: 372\nNumber of predicted TMHs: 2\nExp number of AAs in TMHs: 36.51738\nExp number, first 60 AAs: 19.72849\nTotal prob of N-in: 0.96170\nPOSSIBLE N-term signal sequence\ninside 1 19\nTMhelix 20 39\noutside 40 325\nTMhelix 326 348\ninside 349 372

25349 GCF_000380325.1_ASM38032v1 Salmonella enterica subsp. enterica serovar Typhimurium str. U288 Proteobacteria; Gammaproteobacteria; Enterobacterales; Enterobacteriaceae; Salmonella; Salmonella enterica; Salmonella enterica subsp. enterica; Salmonella enterica subsp. enterica serovar Typhimurium MTGDNTLITSHGINRRDFMKLCAALAATMGLSSKAAAEMAESVSNPQRPPVIWIGAQECT WP_015589585.1 hydrogenase 2 small subunit [Salmonella enterica] Length: 372\nNumber of predicted TMHs: 1\nExp number of AAs in TMHs: 24.51304\nExp number, first 60 AAs: 0.2682\nTotal prob of N-in: 0.14402\noutside 1 331\nTMhelix 332 354\ninside 355 372

25350 GCF_000213635.1_ASM21363v1 Salmonella enterica subsp. enterica serovar Typhimurium str. UK-1 Proteobacteria; Gammaproteobacteria; Enterobacterales; Enterobacteriaceae; Salmonella; Salmonella enterica; Salmonella enterica subsp. enterica; Salmonella enterica subsp. enterica serovar Typhimurium MTGDNTLITSHGINRRDFMKLCAALAATMGLSSKAAAEMAESVSNPQRPPVIWIGAQECT WP_000145429.1 hydrogenase 2 small subunit [Salmonella enterica] Length: 372\nNumber of predicted TMHs: 1\nExp number of AAs in TMHs: 24.51309\nExp number, first 60 AAs: 0.2682\nTotal prob of N-in: 0.14402\noutside 1 331\nTMhelix 332 354\ninside 355 372

25351 GCF_000213635.1_ASM21363v1 Salmonella enterica subsp. enterica serovar Typhimurium str. UK-1 Proteobacteria; Gammaproteobacteria; Enterobacterales; Enterobacteriaceae; Salmonella; Salmonella enterica; Salmonella enterica subsp. enterica; Salmonella enterica subsp. enterica serovar Typhimurium MNNEETFYQAMRRKGVTRRSFLKFCSLAATSLGLGAGMTPKIAWALENKPRIPVVWIHGL WP_001058312.1 hydrogenase [Salmonella enterica] Length: 372\nNumber of predicted TMHs: 2\nExp number of AAs in TMHs: 36.51738\nExp number, first 60 AAs: 19.72849\nTotal prob of N-in: 0.96170\nPOSSIBLE N-term signal sequence\ninside 1 19\nTMhelix 20 39\noutside 40 325\nTMhelix 326 348\ninside 349 372

25352 GCF_001623725.1_ASM162372v1 Salmonella enterica subsp. enterica serovar Typhimurium str. USDA-ARS-USMARC-1808 Proteobacteria; Gammaproteobacteria; Enterobacterales; Enterobacteriaceae; Salmonella; Salmonella enterica; Salmonella enterica subsp. enterica; Salmonella enterica subsp. enterica serovar Typhimurium MTGDNTLITSHGINRRDFMKLCAALAATMGLSSKAAAEMAESVSNPQRPPVIWIGAQECT WP_000145429.1 hydrogenase 2 small subunit [Salmonella enterica] Length: 372\nNumber of predicted TMHs: 1\nExp

number of AAs in TMHs: 24.51309\nExp number, first 60 AAs: 0.2682\nTotal prob of N-in: 0.14402\noutside 1 331\nTMhelix 332 354\ninside 355 372

25353 GCF_001623725.1_ASM162372v1 Salmonella enterica subsp. enterica serovar Typhimurium str. USDA-ARS-USMARC-1808 Proteobacteria; Gammaproteobacteria; Enterobacterales; Enterobacteriaceae; Salmonella; Salmonella enterica; Salmonella enterica subsp. enterica; Salmonella enterica subsp. enterica serovar Typhimurium MNNEETFYQAMRRKGVTRRSFLKFCSLAATSLGLGAGMTPKIAWALENKPRIPVVWIHGL WP_001058312.1 hydrogenase [Salmonella enterica] Length: 372\nNumber of predicted TMHs: 2\nExp number of AAs in TMHs: 36.51738\nExp number, first 60 AAs: 19.72849\nTotal prob of N-in: 0.96170\nPOSSIBLE N-term signal sequence\ninside 1 19\nTMhelix 20 39\noutside 40 325\nTMhelix 326 348\ninside 349 372

25354 GCF_001623745.2_ASM162374v2 Salmonella enterica subsp. enterica serovar Typhimurium str. USDA-ARS-USMARC-1810 Proteobacteria; Gammaproteobacteria; Enterobacterales; Enterobacteriaceae; Salmonella; Salmonella enterica; Salmonella enterica subsp. enterica; Salmonella enterica subsp. enterica serovar Typhimurium MNNEETFYQAMRRKGVTRRSFLKFCSLAATSLGLGAGMTPKIAWALENKPRIPVVWIHGL WP_001058312.1 hydrogenase [Salmonella enterica] Length: 372\nNumber of predicted TMHs: 2\nExp number of AAs in TMHs: 36.51738\nExp number, first 60 AAs: 19.72849\nTotal prob of N-in: 0.96170\nPOSSIBLE N-term signal sequence\ninside 1 19\nTMhelix 20 39\noutside 40 325\nTMhelix 326 348\ninside 349 372

25355 GCF_001623745.2_ASM162374v2 Salmonella enterica subsp. enterica serovar Typhimurium str. USDA-ARS-USMARC-1810 Proteobacteria; Gammaproteobacteria; Enterobacterales; Enterobacteriaceae; Salmonella; Salmonella enterica; Salmonella enterica subsp. enterica; Salmonella enterica subsp. enterica serovar Typhimurium MTGDNTLITSHGINRRDFMKLCAALAATMGLSSKAAAEMAESVSNPQRPPVIWIGAQECT WP_000145429.1 hydrogenase 2 small subunit [Salmonella enterica] Length: 372\nNumber of predicted TMHs: 1\nExp number of AAs in TMHs: 24.51309\nExp number, first 60 AAs: 0.2682\nTotal prob of N-in: 0.14402\noutside 1 331\nTMhelix 332 354\ninside 355 372

25356 GCF_001623705.1_ASM162370v1 Salmonella enterica subsp. enterica serovar Typhimurium str. USDA-ARS-USMARC-1880 Proteobacteria; Gammaproteobacteria; Enterobacterales; Enterobacteriaceae; Salmonella; Salmonella enterica; Salmonella enterica subsp. enterica; Salmonella enterica subsp. enterica serovar Typhimurium MNNEETFYQAMRRKGVTRRSFLKFCSLAATSLGLGAGMTPKIAWALENKPRIPVVWIHGL WP_001058312.1 hydrogenase [Salmonella enterica] Length: 372\nNumber of predicted TMHs: 2\nExp number of AAs in TMHs: 36.51738\nExp number, first 60 AAs: 19.72849\nTotal prob of N-in: 0.96170\nPOSSIBLE N-term signal sequence\ninside 1 19\nTMhelix 20 39\noutside 40 325\nTMhelix 326 348\ninside 349 372

25357 GCF_001623705.1_ASM162370v1 Salmonella enterica subsp. enterica serovar Typhimurium str. USDA-ARS-USMARC-1880 Proteobacteria; Gammaproteobacteria; Enterobacterales; Enterobacteriaceae; Salmonella; Salmonella enterica; Salmonella enterica subsp. enterica; Salmonella enterica subsp. enterica serovar Typhimurium MTGDNTLITSHGINRRDFMKLCAALAATMGLSSKAAAEMAESVSNPQRPPVIWIGAQECT WP_000145429.1 hydrogenase 2 small subunit [Salmonella enterica] Length: 372\nNumber of predicted TMHs: 1\nExp number of AAs in TMHs: 24.51309\nExp number, first 60 AAs: 0.2682\nTotal prob of N-in: 0.14402\noutside 1 331\nTMhelix 332 354\ninside 355 372

25358 GCF_001623845.1_ASM162384v1 Salmonella enterica subsp. enterica serovar Typhimurium str. USDA-ARS-USMARC-1896 Proteobacteria; Gammaproteobacteria; Enterobacterales; Enterobacteriaceae; Salmonella; Salmonella enterica; Salmonella enterica subsp. enterica; Salmonella enterica subsp. enterica serovar Typhimurium MTGDNTLITSHGINRRDFMKLCAALAATMGLSSKAAAEMAESVSNPQRPPVIWIGAQECT WP_000145429.1 hydrogenase 2 small subunit [Salmonella enterica] Length: 372\nNumber of predicted TMHs: 1\nExp number of AAs in TMHs: 24.51309\nExp number, first 60 AAs: 0.2682\nTotal prob of N-in: 0.14402\noutside 1 331\nTMhelix 332 354\ninside 355 372

25359 GCF_001623845.1_ASM162384v1 Salmonella enterica subsp. enterica serovar Typhimurium str. USDA-ARS-USMARC-1896 Proteobacteria; Gammaproteobacteria; Enterobacterales; Enterobacteriaceae; Salmonella; Salmonella enterica; Salmonella enterica subsp. enterica; Salmonella enterica subsp. enterica serovar Typhimurium MNNEETFYQAMRRKGVTRRSFLKFCSLAATSLGLGAGMTPKIAWALENKPRIPVVWIHGL WP_001058312.1 hydrogenase [Salmonella enterica] Length: 372\nNumber of predicted TMHs: 2\nExp number of AAs in TMHs: 36.51738\nExp number, first 60 AAs: 19.72849\nTotal prob of N-in: 0.96170\nPOSSIBLE N-term signal sequence\ninside 1 19\nTMhelix 20 39\noutside 40 325\nTMhelix 326 348\ninside 349 372

25360 GCF_001623765.2_ASM162376v2 Salmonella enterica subsp. enterica serovar Typhimurium str. USDA-ARS-USMARC-1898 Proteobacteria; Gammaproteobacteria; Enterobacterales; Enterobacteriaceae; Salmonella; Salmonella

enterica; Salmonella enterica subsp. enterica; Salmonella enterica subsp. enterica serovar Typhimurium
MNNEETFYQAMRRKGVTRRSFLKFCSLAATSLGLGAGMTPKIAWALENKPRIPVVWIHGL WP_001058312.1
hydrogenase [Salmonella enterica] Length: 372\nNumber of predicted TMHs: 2\nExp number of AAs in TMHs: 36.51738\nExp number, first 60 AAs: 19.72849\nTotal prob of N-in: 0.96170\nPOSSIBLE N-term signal sequence\ninside 1 19\nTMhelix 20 39\noutside 40 325\nTMhelix 326 348\ninside 349 372

25361 GCF_001623765.2_ASM162376v2 Salmonella enterica subsp. enterica serovar Typhimurium str. USDA-ARS-USMARC-1898 Proteobacteria; Gammaproteobacteria; Enterobacterales; Enterobacteriaceae; Salmonella; Salmonella enterica; Salmonella enterica subsp. enterica; Salmonella enterica subsp. enterica serovar Typhimurium
MTGDNTLITSHGINRRDFMKLCAALAATMGLSSKAAAEMAESVSNPQRPPVIWIGAQECT WP_000145429.1
hydrogenase 2 small subunit [Salmonella enterica] Length: 372\nNumber of predicted TMHs: 1\nExp number of AAs in TMHs: 24.51309\nExp number, first 60 AAs: 0.2682\nTotal prob of N-in: 0.14402\noutside 1 331\nTMhelix 332 354\ninside 355 372

25362 GCF_000941015.2_ASM94101v2 Salmonella enterica subsp. enterica serovar Typhimurium str. USDA-ARS-USMARC-1899 Proteobacteria; Gammaproteobacteria; Enterobacterales; Enterobacteriaceae; Salmonella; Salmonella enterica; Salmonella enterica subsp. enterica; Salmonella enterica subsp. enterica serovar Typhimurium
MNNEETFYQAMRRKGVTRRSFLKFCSLAATSLGLGAGMTPKIAWALENKPRIPVVWIHGL WP_001058312.1
hydrogenase [Salmonella enterica] Length: 372\nNumber of predicted TMHs: 2\nExp number of AAs in TMHs: 36.51738\nExp number, first 60 AAs: 19.72849\nTotal prob of N-in: 0.96170\nPOSSIBLE N-term signal sequence\ninside 1 19\nTMhelix 20 39\noutside 40 325\nTMhelix 326 348\ninside 349 372

25363 GCF_000941015.2_ASM94101v2 Salmonella enterica subsp. enterica serovar Typhimurium str. USDA-ARS-USMARC-1899 Proteobacteria; Gammaproteobacteria; Enterobacterales; Enterobacteriaceae; Salmonella; Salmonella enterica; Salmonella enterica subsp. enterica; Salmonella enterica subsp. enterica serovar Typhimurium
MTGDNTLITSHGINRRDFMKLCAALAATMGLSSKAAAEMAESVSNPQRPPVIWIGAQECT WP_000145429.1
hydrogenase 2 small subunit [Salmonella enterica] Length: 372\nNumber of predicted TMHs: 1\nExp number of AAs in TMHs: 24.51309\nExp number, first 60 AAs: 0.2682\nTotal prob of N-in: 0.14402\noutside 1 331\nTMhelix 332 354\ninside 355 372

25364 GCF_001271565.1_Salmonella_enterica_CVM_N45391_v1.0 Salmonella enterica subsp. enterica serovar Typhimurium var. 5- Proteobacteria; Gammaproteobacteria; Enterobacterales; Enterobacteriaceae; Salmonella; Salmonella enterica; Salmonella enterica subsp. enterica; Salmonella enterica subsp. enterica serovar Typhimurium
MNNEETFYQAMRRKGVTRRSFLKFCSLAATSLGLGAGMTPKIAWALENKPRIPVVWIHGL WP_001058312.1
hydrogenase [Salmonella enterica] Length: 372\nNumber of predicted TMHs: 2\nExp number of AAs in TMHs: 36.51738\nExp number, first 60 AAs: 19.72849\nTotal prob of N-in: 0.96170\nPOSSIBLE N-term signal sequence\ninside 1 19\nTMhelix 20 39\noutside 40 325\nTMhelix 326 348\ninside 349 372

25365 GCF_001272045.1_Salmonella_enterica_CVM_N44717_v1.0 Salmonella enterica subsp. enterica serovar Typhimurium var. 5- Proteobacteria; Gammaproteobacteria; Enterobacterales; Enterobacteriaceae; Salmonella; Salmonella enterica; Salmonella enterica subsp. enterica; Salmonella enterica subsp. enterica serovar Typhimurium
MTGDNTLITSHGINRRDFMKLCAALAATMGLSSKAAAEMAESVSNPQRPPVIWIGAQECT WP_000145429.1
hydrogenase 2 small subunit [Salmonella enterica] Length: 372\nNumber of predicted TMHs: 1\nExp number of AAs in TMHs: 24.51309\nExp number, first 60 AAs: 0.2682\nTotal prob of N-in: 0.14402\noutside 1 331\nTMhelix 332 354\ninside 355 372

25366 GCF_001272095.1_Salmonella_enterica_CVM_N46826_v1.0 Salmonella enterica subsp. enterica serovar Typhimurium var. 5- Proteobacteria; Gammaproteobacteria; Enterobacterales; Enterobacteriaceae; Salmonella; Salmonella enterica; Salmonella enterica subsp. enterica; Salmonella enterica subsp. enterica serovar Typhimurium
MTGDNTLITSHGINRRDFMKLCAALAATMGLSSKAAAEMAESVSNPQRPPVIWIGAQECT WP_000145429.1
hydrogenase 2 small subunit [Salmonella enterica] Length: 372\nNumber of predicted TMHs: 1\nExp number of AAs in TMHs: 24.51309\nExp number, first 60 AAs: 0.2682\nTotal prob of N-in: 0.14402\noutside 1 331\nTMhelix 332 354\ninside 355 372

25367 GCF_001271505.1_Salmonella_enterica_CVM_N45390_v1.0 Salmonella enterica subsp. enterica serovar Typhimurium var. 5- Proteobacteria; Gammaproteobacteria; Enterobacterales; Enterobacteriaceae; Salmonella; Salmonella enterica; Salmonella enterica subsp. enterica; Salmonella enterica subsp. enterica serovar Typhimurium
MNNEETFYQAMRRKGVTRRSFLKFCSLAATSLGLGAGMTPKIAWALENKPRIPVVWIHGL WP_001058312.1
hydrogenase [Salmonella enterica] Length: 372\nNumber of predicted TMHs: 2\nExp number of AAs in TMHs: 36.51738\nExp number, first 60 AAs: 19.72849\nTotal prob of N-in: 0.96170\nPOSSIBLE N-term signal sequence\ninside 1 19\nTMhelix 20 39\noutside 40 325\nTMhelix 326 348\ninside 349 372

25368 GCF_001271565.1_Salmonella_enterica_CVM_N45391_v1.0 Salmonella enterica subsp. enterica serovar Typhimurium var. 5- Proteobacteria; Gammaproteobacteria; Enterobacterales; Enterobacteriaceae; Salmonella; Salmonella enterica; Salmonella enterica subsp. enterica; Salmonella enterica subsp. enterica serovar Typhimurium MTGDNTLITSHGINRRDFMKLCAALAATMGLSSKAAAEMAESVSNPQRPPVIWIGAQECT WP_000145429.1 hydrogenase 2 small subunit [Salmonella enterica] Length: 372\nNumber of predicted TMHs: 1\nExp number of AAs in TMHs: 24.51309\nExp number, first 60 AAs: 0.2682\nTotal prob of N-in: 0.14402\noutside 1 331\nTMhelix 332 354\ninside 355 372

25369 GCF_001271505.1_Salmonella_enterica_CVM_N45390_v1.0 Salmonella enterica subsp. enterica serovar Typhimurium var. 5- Proteobacteria; Gammaproteobacteria; Enterobacterales; Enterobacteriaceae; Salmonella; Salmonella enterica; Salmonella enterica subsp. enterica; Salmonella enterica subsp. enterica serovar Typhimurium MTGDNTLITSHGINRRDFMKLCAALAATMGLSSKAAAEMAESVSNPQRPPVIWIGAQECT WP_000145429.1 hydrogenase 2 small subunit [Salmonella enterica] Length: 372\nNumber of predicted TMHs: 1\nExp number of AAs in TMHs: 24.51309\nExp number, first 60 AAs: 0.2682\nTotal prob of N-in: 0.14402\noutside 1 331\nTMhelix 332 354\ninside 355 372

25370 GCF_001272045.1_Salmonella_enterica_CVM_N44717_v1.0 Salmonella enterica subsp. enterica serovar Typhimurium var. 5- Proteobacteria; Gammaproteobacteria; Enterobacterales; Enterobacteriaceae; Salmonella; Salmonella enterica; Salmonella enterica subsp. enterica; Salmonella enterica subsp. enterica serovar Typhimurium MNNEETFYQAMRRKGVTRRSFLKFCSLAATSLGLGAGMTPKIAWALENKPRIPVVWIHGL WP_001058312.1 hydrogenase [Salmonella enterica] Length: 372\nNumber of predicted TMHs: 2\nExp number of AAs in TMHs: 36.51738\nExp number, first 60 AAs: 19.72849\nTotal prob of N-in: 0.96170\nPOSSIBLE N-term signal sequence\ninside 1 19\nTMhelix 20 39\noutside 40 325\nTMhelix 326 348\ninside 349 372

25371 GCF_001247275.1_Salmonella_enterica_CVM_N51303_v1.0 Salmonella enterica subsp. enterica serovar Typhimurium var. 5- Proteobacteria; Gammaproteobacteria; Enterobacterales; Enterobacteriaceae; Salmonella; Salmonella enterica; Salmonella enterica subsp. enterica; Salmonella enterica subsp. enterica serovar Typhimurium MNNEETFYQAMRRKGVTRRSFLKFCSLAATSLGLGAGMTPKIAWALENKPRIPVVWIHGL WP_001058312.1 hydrogenase [Salmonella enterica] Length: 372\nNumber of predicted TMHs: 2\nExp number of AAs in TMHs: 36.51738\nExp number, first 60 AAs: 19.72849\nTotal prob of N-in: 0.96170\nPOSSIBLE N-term signal sequence\ninside 1 19\nTMhelix 20 39\noutside 40 325\nTMhelix 326 348\ninside 349 372

25372 GCF_001247275.1_Salmonella_enterica_CVM_N51303_v1.0 Salmonella enterica subsp. enterica serovar Typhimurium var. 5- Proteobacteria; Gammaproteobacteria; Enterobacterales; Enterobacteriaceae; Salmonella; Salmonella enterica; Salmonella enterica subsp. enterica; Salmonella enterica subsp. enterica serovar Typhimurium MTGDNTLITSHGINRRDFMKLCAALAATMGLSSKAAAEMAESVSNPQRPPVIWIGAQECT WP_000145429.1 hydrogenase 2 small subunit [Salmonella enterica] Length: 372\nNumber of predicted TMHs: 1\nExp number of AAs in TMHs: 24.51309\nExp number, first 60 AAs: 0.2682\nTotal prob of N-in: 0.14402\noutside 1 331\nTMhelix 332 354\ninside 355 372

25373 GCF_001246625.1_Salmonella_enterica_CVM_N51272_v1.0 Salmonella enterica subsp. enterica serovar Typhimurium var. 5- Proteobacteria; Gammaproteobacteria; Enterobacterales; Enterobacteriaceae; Salmonella; Salmonella enterica; Salmonella enterica subsp. enterica; Salmonella enterica subsp. enterica serovar Typhimurium MNNEETFYQAMRRKGVTRRSFLKFCSLAATSLGLGAGMTPKIAWALENKPRIPVVWIHGL WP_001058312.1 hydrogenase [Salmonella enterica] Length: 372\nNumber of predicted TMHs: 2\nExp number of AAs in TMHs: 36.51738\nExp number, first 60 AAs: 19.72849\nTotal prob of N-in: 0.96170\nPOSSIBLE N-term signal sequence\ninside 1 19\nTMhelix 20 39\noutside 40 325\nTMhelix 326 348\ninside 349 372

25374 GCF_001246625.1_Salmonella_enterica_CVM_N51272_v1.0 Salmonella enterica subsp. enterica serovar Typhimurium var. 5- Proteobacteria; Gammaproteobacteria; Enterobacterales; Enterobacteriaceae; Salmonella; Salmonella enterica; Salmonella enterica subsp. enterica; Salmonella enterica subsp. enterica serovar Typhimurium MTGDNTLITSHGINRRDFMKLCAALAATMGLSSKAAAEMAESVSNPQRPPVIWIGAQECT WP_000145429.1 hydrogenase 2 small subunit [Salmonella enterica] Length: 372\nNumber of predicted TMHs: 1\nExp number of AAs in TMHs: 24.51309\nExp number, first 60 AAs: 0.2682\nTotal prob of N-in: 0.14402\noutside 1 331\nTMhelix 332 354\ninside 355 372

25375 GCF_001246565.1_Salmonella_enterica_CVM_N51268_v1.0 Salmonella enterica subsp. enterica serovar Typhimurium var. 5- Proteobacteria; Gammaproteobacteria; Enterobacterales; Enterobacteriaceae; Salmonella; Salmonella enterica; Salmonella enterica subsp. enterica; Salmonella enterica subsp. enterica serovar Typhimurium MNNEETFYQAMRRKGVTRRSFLKFCSLAATSLGLGAGMTPKIAWALENKPRIPVVWIHGL WP_001058312.1 hydrogenase [Salmonella enterica] Length: 372\nNumber of predicted TMHs: 2\nExp number of AAs in TMHs: 36.51738\nExp number, first 60 AAs: 19.72849\nTotal prob of N-in: 0.96170\nPOSSIBLE N-term signal sequence\ninside 1 19\nTMhelix 20 39\noutside 40 325\nTMhelix 326 348\ninside 349 372

TMHs: 36.51738\nExp number, first 60 AAs: 19.72849\nTotal prob of N-in: 0.96170\nPOSSIBLE N-term signal sequence\ninside 1 19\nTMhelix 20 39\noutside 40 325\nTMhelix 326 348\ninside 349 372

25376 GCF_001246565.1_Salmonella_enterica_CVM_N51268_v1.0 Salmonella enterica subsp. enterica serovar Typhimurium var. 5- Proteobacteria; Gammaproteobacteria; Enterobacterales; Enterobacteriaceae; Salmonella; Salmonella enterica; Salmonella enterica subsp. enterica; Salmonella enterica subsp. enterica serovar Typhimurium MTGDNTLITSHGINRRDFMKLCAALAATMGLSSKAAAEMAESVSNPQRPPVIWIGAQECT WP_000145429.1 hydrogenase 2 small subunit [Salmonella enterica] Length: 372\nNumber of predicted TMHs: 1\nExp number of AAs in TMHs: 24.51309\nExp number, first 60 AAs: 0.2682\nTotal prob of N-in: 0.14402\noutside 1 331\nTMhelix 332 354\ninside 355 372

25377 GCF_001245535.1_Salmonella_enterica_CVM_N50433_v1.0 Salmonella enterica subsp. enterica serovar Typhimurium var. 5- Proteobacteria; Gammaproteobacteria; Enterobacterales; Enterobacteriaceae; Salmonella; Salmonella enterica; Salmonella enterica subsp. enterica; Salmonella enterica subsp. enterica serovar Typhimurium MNNEETFYQAMRRKGVTRRSFLKFCSLAATSLGLGAGMTPKIAWALENKPRIPVVWIHGL WP_001058312.1 hydrogenase [Salmonella enterica] Length: 372\nNumber of predicted TMHs: 2\nExp number of AAs in TMHs: 36.51738\nExp number, first 60 AAs: 19.72849\nTotal prob of N-in: 0.96170\nPOSSIBLE N-term signal sequence\ninside 1 19\nTMhelix 20 39\noutside 40 325\nTMhelix 326 348\ninside 349 372

25378 GCF_001272095.1_Salmonella_enterica_CVM_N46826_v1.0 Salmonella enterica subsp. enterica serovar Typhimurium var. 5- Proteobacteria; Gammaproteobacteria; Enterobacterales; Enterobacteriaceae; Salmonella; Salmonella enterica; Salmonella enterica subsp. enterica; Salmonella enterica subsp. enterica serovar Typhimurium MNNEETFYQAMRRKGVTRRSFLKFCSLAATSLGLGAGMTPKIAWALENKPRIPVVWIHGL WP_001058312.1 hydrogenase [Salmonella enterica] Length: 372\nNumber of predicted TMHs: 2\nExp number of AAs in TMHs: 36.51738\nExp number, first 60 AAs: 19.72849\nTotal prob of N-in: 0.96170\nPOSSIBLE N-term signal sequence\ninside 1 19\nTMhelix 20 39\noutside 40 325\nTMhelix 326 348\ninside 349 372

25379 GCF_001245535.1_Salmonella_enterica_CVM_N50433_v1.0 Salmonella enterica subsp. enterica serovar Typhimurium var. 5- Proteobacteria; Gammaproteobacteria; Enterobacterales; Enterobacteriaceae; Salmonella; Salmonella enterica; Salmonella enterica subsp. enterica; Salmonella enterica subsp. enterica serovar Typhimurium MTGDNTLITSHGINRRDFMKLCAALAATMGLSSKAAAEMAESVSNPQRPPVIWIGAQECT WP_000145429.1 hydrogenase 2 small subunit [Salmonella enterica] Length: 372\nNumber of predicted TMHs: 1\nExp number of AAs in TMHs: 24.51309\nExp number, first 60 AAs: 0.2682\nTotal prob of N-in: 0.14402\noutside 1 331\nTMhelix 332 354\ninside 355 372

25380 GCF_001240475.1_Salmonella_enterica_CVM_N43469_v1.0 Salmonella enterica subsp. enterica serovar Typhimurium var. 5- Proteobacteria; Gammaproteobacteria; Enterobacterales; Enterobacteriaceae; Salmonella; Salmonella enterica; Salmonella enterica subsp. enterica; Salmonella enterica subsp. enterica serovar Typhimurium MNNEETFYQAMRRKGVTRRSFLKFCSLAATSLGLGAGMTPKIAWALENKPRIPVVWIHGL WP_001058312.1 hydrogenase [Salmonella enterica] Length: 372\nNumber of predicted TMHs: 2\nExp number of AAs in TMHs: 36.51738\nExp number, first 60 AAs: 19.72849\nTotal prob of N-in: 0.96170\nPOSSIBLE N-term signal sequence\ninside 1 19\nTMhelix 20 39\noutside 40 325\nTMhelix 326 348\ninside 349 372

25381 GCF_001271835.1_Salmonella_enterica_CVM_N46848-R_v1.0 Salmonella enterica subsp. enterica serovar Typhimurium var. 5- Proteobacteria; Gammaproteobacteria; Enterobacterales; Enterobacteriaceae; Salmonella; Salmonella enterica; Salmonella enterica subsp. enterica; Salmonella enterica subsp. enterica serovar Typhimurium MTGDNTLITSHGINRRDFMKLCAALAATMGLSSKAAAEMAESVSNPQRPPVIWIGAQECT WP_000145429.1 hydrogenase 2 small subunit [Salmonella enterica] Length: 372\nNumber of predicted TMHs: 1\nExp number of AAs in TMHs: 24.51309\nExp number, first 60 AAs: 0.2682\nTotal prob of N-in: 0.14402\noutside 1 331\nTMhelix 332 354\ninside 355 372

25382 GCF_001240785.1_Salmonella_enterica_CVM_N43822_v1.0 Salmonella enterica subsp. enterica serovar Typhimurium var. 5- Proteobacteria; Gammaproteobacteria; Enterobacterales; Enterobacteriaceae; Salmonella; Salmonella enterica; Salmonella enterica subsp. enterica; Salmonella enterica subsp. enterica serovar Typhimurium MNNEETFYQAMRRKGVTRRSFLKFCSLAATSLGLGAGMTPKIAWALENKPRIPVVWIHGL WP_001058312.1 hydrogenase [Salmonella enterica] Length: 372\nNumber of predicted TMHs: 2\nExp number of AAs in TMHs: 36.51738\nExp number, first 60 AAs: 19.72849\nTotal prob of N-in: 0.96170\nPOSSIBLE N-term signal sequence\ninside 1 19\nTMhelix 20 39\noutside 40 325\nTMhelix 326 348\ninside 349 372

25383 GCF_001240935.1_Salmonella_enterica_CVM_N43830-R_v1.0 Salmonella enterica subsp. enterica serovar Typhimurium var. 5- Proteobacteria; Gammaproteobacteria; Enterobacterales; Enterobacteriaceae; Salmonella;

Salmonella enterica; Salmonella enterica subsp. enterica; Salmonella enterica subsp. enterica serovar Typhimurium MTGDNTLITSHGINRRDFMKLCAALAATMGLSSKAAAEMAESVSNPQRPPVIWIGAQECT WP_000145429.1 hydrogenase 2 small subunit [Salmonella enterica] Length: 372\nNumber of predicted TMHs: 1\nExp number of AAs in TMHs: 24.51309\nExp number, first 60 AAs: 0.2682\nTotal prob of N-in: 0.14402\noutside 1 331\nTMhelix 332 354\ninside 355 372

25384 GCF_001240825.1_Salmonella_enterica_CVM_N43823_v1.0 Salmonella enterica subsp. enterica serovar Typhimurium var. 5- Proteobacteria; Gammaproteobacteria; Enterobacterales; Enterobacteriaceae; Salmonella; Salmonella enterica; Salmonella enterica subsp. enterica; Salmonella enterica subsp. enterica serovar Typhimurium MTGDNTLITSHGINRRDFMKLCAALAATMGLSSKAAAEMAESVSNPQRPPVIWIGAQECT WP_000145429.1 hydrogenase 2 small subunit [Salmonella enterica] Length: 372\nNumber of predicted TMHs: 1\nExp number of AAs in TMHs: 24.51309\nExp number, first 60 AAs: 0.2682\nTotal prob of N-in: 0.14402\noutside 1 331\nTMhelix 332 354\ninside 355 372

25385 GCF_001240785.1_Salmonella_enterica_CVM_N43822_v1.0 Salmonella enterica subsp. enterica serovar Typhimurium var. 5- Proteobacteria; Gammaproteobacteria; Enterobacterales; Enterobacteriaceae; Salmonella; Salmonella enterica; Salmonella enterica subsp. enterica; Salmonella enterica subsp. enterica serovar Typhimurium MTGDNTLITSHGINRRDFMKLCAALAATMGLSSKAAAEMAESVSNPQRPPVIWIGAQECT WP_000145429.1 hydrogenase 2 small subunit [Salmonella enterica] Length: 372\nNumber of predicted TMHs: 1\nExp number of AAs in TMHs: 24.51309\nExp number, first 60 AAs: 0.2682\nTotal prob of N-in: 0.14402\noutside 1 331\nTMhelix 332 354\ninside 355 372

25386 GCF_001271835.1_Salmonella_enterica_CVM_N46848-R_v1.0 Salmonella enterica subsp. enterica serovar Typhimurium var. 5- Proteobacteria; Gammaproteobacteria; Enterobacterales; Enterobacteriaceae; Salmonella; Salmonella enterica; Salmonella enterica subsp. enterica; Salmonella enterica subsp. enterica serovar Typhimurium MNNEETFYQAMRRKGVTRRSFLKFCSLAATSLGLGAGMTPKIAWALENKPRIPVVWIHGL WP_001058312.1 hydrogenase [Salmonella enterica] Length: 372\nNumber of predicted TMHs: 2\nExp number of AAs in TMHs: 36.51738\nExp number, first 60 AAs: 19.72849\nTotal prob of N-in: 0.96170\nPOSSIBLE N-term signal sequence\ninside 1 19\nTMhelix 20 39\noutside 40 325\nTMhelix 326 348\ninside 349 372

25387 GCF_001244705.1_Salmonella_enterica_CVM_N48689_v1.0 Salmonella enterica subsp. enterica serovar Typhimurium var. 5- Proteobacteria; Gammaproteobacteria; Enterobacterales; Enterobacteriaceae; Salmonella; Salmonella enterica; Salmonella enterica subsp. enterica; Salmonella enterica subsp. enterica serovar Typhimurium MNNEETFYQAMRRKGVTRRSFLKFCSLAATSLGLGAGMTPKIAWALENKPRIPVVWIHGL WP_001058312.1 hydrogenase [Salmonella enterica] Length: 372\nNumber of predicted TMHs: 2\nExp number of AAs in TMHs: 36.51738\nExp number, first 60 AAs: 19.72849\nTotal prob of N-in: 0.96170\nPOSSIBLE N-term signal sequence\ninside 1 19\nTMhelix 20 39\noutside 40 325\nTMhelix 326 348\ninside 349 372

25388 GCF_001244385.1_Salmonella_enterica_CVM_N48680_v1.0 Salmonella enterica subsp. enterica serovar Typhimurium var. 5- Proteobacteria; Gammaproteobacteria; Enterobacterales; Enterobacteriaceae; Salmonella; Salmonella enterica; Salmonella enterica subsp. enterica; Salmonella enterica subsp. enterica serovar Typhimurium MNNEETFYQAMRRKGVTRRSFLKFCSLAATSLGLGAGMTPKIAWALENKPRIPVVWIHGL WP_001058312.1 hydrogenase [Salmonella enterica] Length: 372\nNumber of predicted TMHs: 2\nExp number of AAs in TMHs: 36.51738\nExp number, first 60 AAs: 19.72849\nTotal prob of N-in: 0.96170\nPOSSIBLE N-term signal sequence\ninside 1 19\nTMhelix 20 39\noutside 40 325\nTMhelix 326 348\ninside 349 372

25389 GCF_001244255.1_Salmonella_enterica_CVM_N48676_v1.0 Salmonella enterica subsp. enterica serovar Typhimurium var. 5- Proteobacteria; Gammaproteobacteria; Enterobacterales; Enterobacteriaceae; Salmonella; Salmonella enterica; Salmonella enterica subsp. enterica; Salmonella enterica subsp. enterica serovar Typhimurium MNNEETFYQAMRRKGVTRRSFLKFCSLAATSLGLGAGMTPKIAWALENKPRIPVVWIHGL WP_001058312.1 hydrogenase [Salmonella enterica] Length: 372\nNumber of predicted TMHs: 2\nExp number of AAs in TMHs: 36.51738\nExp number, first 60 AAs: 19.72849\nTotal prob of N-in: 0.96170\nPOSSIBLE N-term signal sequence\ninside 1 19\nTMhelix 20 39\noutside 40 325\nTMhelix 326 348\ninside 349 372

25390 GCF_001244705.1_Salmonella_enterica_CVM_N48689_v1.0 Salmonella enterica subsp. enterica serovar Typhimurium var. 5- Proteobacteria; Gammaproteobacteria; Enterobacterales; Enterobacteriaceae; Salmonella; Salmonella enterica; Salmonella enterica subsp. enterica; Salmonella enterica subsp. enterica serovar Typhimurium MTGDNTLITSHGINRRDFMKLCAALAATMGLSSKAAAEMAESVSNPQRPPVIWIGAQECT WP_000145429.1 hydrogenase 2 small subunit [Salmonella enterica] Length: 372\nNumber of predicted TMHs: 1\nExp number of AAs in TMHs: 24.51309\nExp number, first 60 AAs: 0.2682\nTotal prob of N-in: 0.14402\noutside 1 331\nTMhelix 332 354\ninside 355 372

25391 GCF_001272465.1_Salmonella_enterica_CVM_N51269_v1.0 Salmonella enterica subsp. enterica serovar Typhimurium var. 5- Proteobacteria; Gammaproteobacteria; Enterobacterales; Enterobacteriaceae; Salmonella; Salmonella enterica; Salmonella enterica subsp. enterica; Salmonella enterica subsp. enterica serovar Typhimurium MTGDNTLITSHGINRRDFMKLCAALAATMGLSSKAAAEMAESVSNPQRPPVIWIGAQECT WP_000145429.1 hydrogenase 2 small subunit [Salmonella enterica] Length: 372\nNumber of predicted TMHs: 1\nExp number of AAs in TMHs: 24.51309\nExp number, first 60 AAs: 0.2682\nTotal prob of N-in: 0.14402\noutside 1 331\nTMhelix 332 354\ninside 355 372

25392 GCF_001244385.1_Salmonella_enterica_CVM_N48680_v1.0 Salmonella enterica subsp. enterica serovar Typhimurium var. 5- Proteobacteria; Gammaproteobacteria; Enterobacterales; Enterobacteriaceae; Salmonella; Salmonella enterica; Salmonella enterica subsp. enterica; Salmonella enterica subsp. enterica serovar Typhimurium MTGDNTLITSHGINRRDFMKLCAALAATMGLSSKAAAEMAESVSNPQRPPVIWIGAQECT WP_000145429.1 hydrogenase 2 small subunit [Salmonella enterica] Length: 372\nNumber of predicted TMHs: 1\nExp number of AAs in TMHs: 24.51309\nExp number, first 60 AAs: 0.2682\nTotal prob of N-in: 0.14402\noutside 1 331\nTMhelix 332 354\ninside 355 372

25393 GCF_001272465.1_Salmonella_enterica_CVM_N51269_v1.0 Salmonella enterica subsp. enterica serovar Typhimurium var. 5- Proteobacteria; Gammaproteobacteria; Enterobacterales; Enterobacteriaceae; Salmonella; Salmonella enterica; Salmonella enterica subsp. enterica; Salmonella enterica subsp. enterica serovar Typhimurium MNNEETFYQAMRRKGVTRRSFLKFCSLAATSLGLGAGMTPKIAWALENKPRIPVVWIHGL WP_001058312.1 hydrogenase [Salmonella enterica] Length: 372\nNumber of predicted TMHs: 2\nExp number of AAs in TMHs: 36.51738\nExp number, first 60 AAs: 19.72849\nTotal prob of N-in: 0.96170\nPOSSIBLE N-term signal sequence\ninside 1 19\nTMhelix 20 39\noutside 40 325\nTMhelix 326 348\ninside 349 372

25394 GCF_001244255.1_Salmonella_enterica_CVM_N48676_v1.0 Salmonella enterica subsp. enterica serovar Typhimurium var. 5- Proteobacteria; Gammaproteobacteria; Enterobacterales; Enterobacteriaceae; Salmonella; Salmonella enterica; Salmonella enterica subsp. enterica; Salmonella enterica subsp. enterica serovar Typhimurium MTGDNTLITSHGINRRDFMKLCAALAATMGLSSKAAAEMAESVSNPQRPPVIWIGAQECT WP_000145429.1 hydrogenase 2 small subunit [Salmonella enterica] Length: 372\nNumber of predicted TMHs: 1\nExp number of AAs in TMHs: 24.51309\nExp number, first 60 AAs: 0.2682\nTotal prob of N-in: 0.14402\noutside 1 331\nTMhelix 332 354\ninside 355 372

25395 GCF_001243815.1_Salmonella_enterica_CVM_N47715_v1.0 Salmonella enterica subsp. enterica serovar Typhimurium var. 5- Proteobacteria; Gammaproteobacteria; Enterobacterales; Enterobacteriaceae; Salmonella; Salmonella enterica; Salmonella enterica subsp. enterica; Salmonella enterica subsp. enterica serovar Typhimurium MNNEETFYQAMRRKGVTRRSFLKFCSLAATSLGLGAGMTPKIAWALENKPRIPVVWIHGL WP_001058312.1 hydrogenase [Salmonella enterica] Length: 372\nNumber of predicted TMHs: 2\nExp number of AAs in TMHs: 36.51738\nExp number, first 60 AAs: 19.72849\nTotal prob of N-in: 0.96170\nPOSSIBLE N-term signal sequence\ninside 1 19\nTMhelix 20 39\noutside 40 325\nTMhelix 326 348\ninside 349 372

25396 GCF_001243815.1_Salmonella_enterica_CVM_N47715_v1.0 Salmonella enterica subsp. enterica serovar Typhimurium var. 5- Proteobacteria; Gammaproteobacteria; Enterobacterales; Enterobacteriaceae; Salmonella; Salmonella enterica; Salmonella enterica subsp. enterica; Salmonella enterica subsp. enterica serovar Typhimurium MTGDNTLITSHGINRRDFMKLCAALAATMGLSSKAAAEMAESVSNPQRPPVIWIGAQECT WP_000145429.1 hydrogenase 2 small subunit [Salmonella enterica] Length: 372\nNumber of predicted TMHs: 1\nExp number of AAs in TMHs: 24.51309\nExp number, first 60 AAs: 0.2682\nTotal prob of N-in: 0.14402\noutside 1 331\nTMhelix 332 354\ninside 355 372

25397 GCF_001243695.1_Salmonella_enterica_CVM_N46860_v1.0 Salmonella enterica subsp. enterica serovar Typhimurium var. 5- Proteobacteria; Gammaproteobacteria; Enterobacterales; Enterobacteriaceae; Salmonella; Salmonella enterica; Salmonella enterica subsp. enterica; Salmonella enterica subsp. enterica serovar Typhimurium MNNEETFYQAMRRKGVTRRSFLKFCSLAATSLGLGAGMTPKIAWALENKPRIPVVWIHGL WP_001058312.1 hydrogenase [Salmonella enterica] Length: 372\nNumber of predicted TMHs: 2\nExp number of AAs in TMHs: 36.51738\nExp number, first 60 AAs: 19.72849\nTotal prob of N-in: 0.96170\nPOSSIBLE N-term signal sequence\ninside 1 19\nTMhelix 20 39\noutside 40 325\nTMhelix 326 348\ninside 349 372

25398 GCF_001243695.1_Salmonella_enterica_CVM_N46860_v1.0 Salmonella enterica subsp. enterica serovar Typhimurium var. 5- Proteobacteria; Gammaproteobacteria; Enterobacterales; Enterobacteriaceae; Salmonella; Salmonella enterica; Salmonella enterica subsp. enterica; Salmonella enterica subsp. enterica serovar Typhimurium MTGDNTLITSHGINRRDFMKLCAALAATMGLSSKAAAEMAESVSNPQRPPVIWIGAQECT WP_000145429.1 hydrogenase 2 small subunit [Salmonella enterica] Length: 372\nNumber of predicted TMHs: 1\nExp

number of AAs in TMHs: 24.51309\nExp number, first 60 AAs: 0.2682\nTotal prob of N-in: 0.14402\noutside 1 331\nTMhelix 332 354\ninside 355 372

25399 GCF_001243505.1_Salmonella_enterica_CVM_N46851-R_v1.0 Salmonella enterica subsp. enterica serovar Typhimurium var. 5- Proteobacteria; Gammaproteobacteria; Enterobacterales; Enterobacteriaceae; Salmonella; Salmonella enterica; Salmonella enterica subsp. enterica; Salmonella enterica subsp. enterica serovar Typhimurium MNNEETFYQAMRRKGVTRRSFLKFCSLAATSLGLGAGMTPKIAWALENKPRIPVVWIHGL WP_001058312.1 hydrogenase [Salmonella enterica] Length: 372\nNumber of predicted TMHs: 2\nExp number of AAs in TMHs: 36.51738\nExp number, first 60 AAs: 19.72849\nTotal prob of N-in: 0.96170\nPOSSIBLE N-term signal sequence\ninside 1 19\nTMhelix 20 39\noutside 40 325\nTMhelix 326 348\ninside 349 372

25400 GCF_001243555.1_Salmonella_enterica_CVM_N46852_v1.0 Salmonella enterica subsp. enterica serovar Typhimurium var. 5- Proteobacteria; Gammaproteobacteria; Enterobacterales; Enterobacteriaceae; Salmonella; Salmonella enterica; Salmonella enterica subsp. enterica; Salmonella enterica subsp. enterica serovar Typhimurium MNNEETFYQAMRRKGVTRRSFLKFCSLAATSLGLGAGMTPKIAWALENKPRIPVVWIHGL WP_001058312.1 hydrogenase [Salmonella enterica] Length: 372\nNumber of predicted TMHs: 2\nExp number of AAs in TMHs: 36.51738\nExp number, first 60 AAs: 19.72849\nTotal prob of N-in: 0.96170\nPOSSIBLE N-term signal sequence\ninside 1 19\nTMhelix 20 39\noutside 40 325\nTMhelix 326 348\ninside 349 372

25401 GCF_001243555.1_Salmonella_enterica_CVM_N46852_v1.0 Salmonella enterica subsp. enterica serovar Typhimurium var. 5- Proteobacteria; Gammaproteobacteria; Enterobacterales; Enterobacteriaceae; Salmonella; Salmonella enterica; Salmonella enterica subsp. enterica; Salmonella enterica subsp. enterica serovar Typhimurium MTGDNTLITSHGINRRDFMKLCAALAATMGLSSKAAAEMAESVSNPQRPPVIWIGAQECT WP_000145429.1 hydrogenase 2 small subunit [Salmonella enterica] Length: 372\nNumber of predicted TMHs: 1\nExp number of AAs in TMHs: 24.51309\nExp number, first 60 AAs: 0.2682\nTotal prob of N-in: 0.14402\noutside 1 331\nTMhelix 332 354\ninside 355 372

25402 GCF_001243425.1_Salmonella_enterica_CVM_N46845-R_v1.0 Salmonella enterica subsp. enterica serovar Typhimurium var. 5- Proteobacteria; Gammaproteobacteria; Enterobacterales; Enterobacteriaceae; Salmonella; Salmonella enterica; Salmonella enterica subsp. enterica; Salmonella enterica subsp. enterica serovar Typhimurium MNNEETFYQAMRRKGVTRRSFLKFCSLAATSLGLGAGMTPKIAWALENKPRIPVVWIHGL WP_001058312.1 hydrogenase [Salmonella enterica] Length: 372\nNumber of predicted TMHs: 2\nExp number of AAs in TMHs: 36.51738\nExp number, first 60 AAs: 19.72849\nTotal prob of N-in: 0.96170\nPOSSIBLE N-term signal sequence\ninside 1 19\nTMhelix 20 39\noutside 40 325\nTMhelix 326 348\ninside 349 372

25403 GCF_001243505.1_Salmonella_enterica_CVM_N46851-R_v1.0 Salmonella enterica subsp. enterica serovar Typhimurium var. 5- Proteobacteria; Gammaproteobacteria; Enterobacterales; Enterobacteriaceae; Salmonella; Salmonella enterica; Salmonella enterica subsp. enterica; Salmonella enterica subsp. enterica serovar Typhimurium MTGDNTLITSHGINRRDFMKLCAALAATMGLSSKAAAEMAESVSNPQRPPVIWIGAQECT WP_000145429.1 hydrogenase 2 small subunit [Salmonella enterica] Length: 372\nNumber of predicted TMHs: 1\nExp number of AAs in TMHs: 24.51309\nExp number, first 60 AAs: 0.2682\nTotal prob of N-in: 0.14402\noutside 1 331\nTMhelix 332 354\ninside 355 372

25404 GCF_001243425.1_Salmonella_enterica_CVM_N46845-R_v1.0 Salmonella enterica subsp. enterica serovar Typhimurium var. 5- Proteobacteria; Gammaproteobacteria; Enterobacterales; Enterobacteriaceae; Salmonella; Salmonella enterica; Salmonella enterica subsp. enterica; Salmonella enterica subsp. enterica serovar Typhimurium MTGDNTLITSHGINRRDFMKLCAALAATMGLSSKAAAEMAESVSNPQRPPVIWIGAQECT WP_000145429.1 hydrogenase 2 small subunit [Salmonella enterica] Length: 372\nNumber of predicted TMHs: 1\nExp number of AAs in TMHs: 24.51309\nExp number, first 60 AAs: 0.2682\nTotal prob of N-in: 0.14402\noutside 1 331\nTMhelix 332 354\ninside 355 372

25405 GCF_001243095.1_Salmonella_enterica_CVM_N46824_v1.0 Salmonella enterica subsp. enterica serovar Typhimurium var. 5- Proteobacteria; Gammaproteobacteria; Enterobacterales; Enterobacteriaceae; Salmonella; Salmonella enterica; Salmonella enterica subsp. enterica; Salmonella enterica subsp. enterica serovar Typhimurium MNNEETFYQAMRRKGVTRRSFLKFCSLAATSLGLGAGMTPKIAWALENKPRIPVVWIHGL WP_001058312.1 hydrogenase [Salmonella enterica] Length: 372\nNumber of predicted TMHs: 2\nExp number of AAs in TMHs: 36.51738\nExp number, first 60 AAs: 19.72849\nTotal prob of N-in: 0.96170\nPOSSIBLE N-term signal sequence\ninside 1 19\nTMhelix 20 39\noutside 40 325\nTMhelix 326 348\ninside 349 372

25406 GCF_001243095.1_Salmonella_enterica_CVM_N46824_v1.0 Salmonella enterica subsp. enterica serovar Typhimurium var. 5- Proteobacteria; Gammaproteobacteria; Enterobacterales; Enterobacteriaceae; Salmonella;

Salmonella enterica; Salmonella enterica subsp. enterica; Salmonella enterica subsp. enterica serovar Typhimurium MTGDNTLITSHGINRRDFMKLCAALAATMGLSSKAAAEMAESVSNPQRPPVIWIGAQECT WP_000145429.1 hydrogenase 2 small subunit [Salmonella enterica] Length: 372\nNumber of predicted TMHs: 1\nExp number of AAs in TMHs: 24.51309\nExp number, first 60 AAs: 0.2682\nTotal prob of N-in: 0.14402\noutside 1 331\nTMhelix 332 354\ninside 355 372

25407 GCF_001242345.1_Salmonella_enterica_CVM_N45936_v1.0 Salmonella enterica subsp. enterica serovar Typhimurium var. 5- Proteobacteria; Gammaproteobacteria; Enterobacterales; Enterobacteriaceae; Salmonella; Salmonella enterica; Salmonella enterica subsp. enterica; Salmonella enterica subsp. enterica serovar Typhimurium MNNEETFYQAMRRKGVTRRSFLKFCSLAATSLGLGAGMTPKIAWALENKPRIPVVWIHGL WP_001058312.1 hydrogenase [Salmonella enterica] Length: 372\nNumber of predicted TMHs: 2\nExp number of AAs in TMHs: 36.51738\nExp number, first 60 AAs: 19.72849\nTotal prob of N-in: 0.96170\nPOSSIBLE N-term signal sequence\ninside 1 19\nTMhelix 20 39\noutside 40 325\nTMhelix 326 348\ninside 349 372

25408 GCF_001242275.1_Salmonella_enterica_CVM_N45935_v1.0 Salmonella enterica subsp. enterica serovar Typhimurium var. 5- Proteobacteria; Gammaproteobacteria; Enterobacterales; Enterobacteriaceae; Salmonella; Salmonella enterica; Salmonella enterica subsp. enterica; Salmonella enterica subsp. enterica serovar Typhimurium MNNEETFYQAMRRKGVTRRSFLKFCSLAATSLGLGAGMTPKIAWALENKPRIPVVWIHGL WP_001058312.1 hydrogenase [Salmonella enterica] Length: 372\nNumber of predicted TMHs: 2\nExp number of AAs in TMHs: 36.51738\nExp number, first 60 AAs: 19.72849\nTotal prob of N-in: 0.96170\nPOSSIBLE N-term signal sequence\ninside 1 19\nTMhelix 20 39\noutside 40 325\nTMhelix 326 348\ninside 349 372

25409 GCF_001242345.1_Salmonella_enterica_CVM_N45936_v1.0 Salmonella enterica subsp. enterica serovar Typhimurium var. 5- Proteobacteria; Gammaproteobacteria; Enterobacterales; Enterobacteriaceae; Salmonella; Salmonella enterica; Salmonella enterica subsp. enterica; Salmonella enterica subsp. enterica serovar Typhimurium MTGDNTLITSHGINRRDFMKLCAALAATMGLSSKAAAEMAESVSNPQRPPVIWIGAQECT WP_000145429.1 hydrogenase 2 small subunit [Salmonella enterica] Length: 372\nNumber of predicted TMHs: 1\nExp number of AAs in TMHs: 24.51309\nExp number, first 60 AAs: 0.2682\nTotal prob of N-in: 0.14402\noutside 1 331\nTMhelix 332 354\ninside 355 372

25410 GCF_001242275.1_Salmonella_enterica_CVM_N45935_v1.0 Salmonella enterica subsp. enterica serovar Typhimurium var. 5- Proteobacteria; Gammaproteobacteria; Enterobacterales; Enterobacteriaceae; Salmonella; Salmonella enterica; Salmonella enterica subsp. enterica; Salmonella enterica subsp. enterica serovar Typhimurium MTGDNTLITSHGINRRDFMKLCAALAATMGLSSKAAAEMAESVSNPQRPPVIWIGAQECT WP_000145429.1 hydrogenase 2 small subunit [Salmonella enterica] Length: 372\nNumber of predicted TMHs: 1\nExp number of AAs in TMHs: 24.51309\nExp number, first 60 AAs: 0.2682\nTotal prob of N-in: 0.14402\noutside 1 331\nTMhelix 332 354\ninside 355 372

25411 GCF_001241905.1_Salmonella_enterica_CVM_N45404_v1.0 Salmonella enterica subsp. enterica serovar Typhimurium var. 5- Proteobacteria; Gammaproteobacteria; Enterobacterales; Enterobacteriaceae; Salmonella; Salmonella enterica; Salmonella enterica subsp. enterica; Salmonella enterica subsp. enterica serovar Typhimurium MNNEETFYQAMRRKGVTRRSFLKFCSLAATSLGLGAGMTPKIAWALENKPRIPVVWIHGL WP_001058312.1 hydrogenase [Salmonella enterica] Length: 372\nNumber of predicted TMHs: 2\nExp number of AAs in TMHs: 36.51738\nExp number, first 60 AAs: 19.72849\nTotal prob of N-in: 0.96170\nPOSSIBLE N-term signal sequence\ninside 1 19\nTMhelix 20 39\noutside 40 325\nTMhelix 326 348\ninside 349 372

25412 GCF_001241905.1_Salmonella_enterica_CVM_N45404_v1.0 Salmonella enterica subsp. enterica serovar Typhimurium var. 5- Proteobacteria; Gammaproteobacteria; Enterobacterales; Enterobacteriaceae; Salmonella; Salmonella enterica; Salmonella enterica subsp. enterica; Salmonella enterica subsp. enterica serovar Typhimurium MTGDNTLITSHGINRRDFMKLCAALAATMGLSSKAAAEMAESVSNPQRPPVIWIGAQECT WP_000145429.1 hydrogenase 2 small subunit [Salmonella enterica] Length: 372\nNumber of predicted TMHs: 1\nExp number of AAs in TMHs: 24.51309\nExp number, first 60 AAs: 0.2682\nTotal prob of N-in: 0.14402\noutside 1 331\nTMhelix 332 354\ninside 355 372

25413 GCF_001241605.1_Salmonella_enterica_CVM_N44716_v1.0 Salmonella enterica subsp. enterica serovar Typhimurium var. 5- Proteobacteria; Gammaproteobacteria; Enterobacterales; Enterobacteriaceae; Salmonella; Salmonella enterica; Salmonella enterica subsp. enterica; Salmonella enterica subsp. enterica serovar Typhimurium MNNEETFYQAMRRKGVTRRSFLKFCSLAATSLGLGAGMTPKIAWALENKPRIPVVWIHGL WP_001058312.1 hydrogenase [Salmonella enterica] Length: 372\nNumber of predicted TMHs: 2\nExp number of AAs in TMHs: 36.51738\nExp number, first 60 AAs: 19.72849\nTotal prob of N-in: 0.96170\nPOSSIBLE N-term signal sequence\ninside 1 19\nTMhelix 20 39\noutside 40 325\nTMhelix 326 348\ninside 349 372

25414 GCF_001241555.1_Salmonella_enterica_CVM_N44714_v1.0 Salmonella enterica subsp. enterica serovar Typhimurium var. 5- Proteobacteria; Gammaproteobacteria; Enterobacterales; Enterobacteriaceae; Salmonella; Salmonella enterica; Salmonella enterica subsp. enterica; Salmonella enterica subsp. enterica serovar Typhimurium MNNEETFYQAMRRKGVTRRSFLKFCSLAATSLGLGAGMTPKIAWALENKPRIPVVWIHGL WP_001058312.1 hydrogenase [Salmonella enterica] Length: 372\nNumber of predicted TMHs: 2\nExp number of AAs in TMHs: 36.51738\nExp number, first 60 AAs: 19.72849\nTotal prob of N-in: 0.96170\nPOSSIBLE N-term signal sequence\ninside 1 19\nTMhelix 20 39\noutside 40 325\nTMhelix 326 348\ninside 349 372

25415 GCF_001241605.1_Salmonella_enterica_CVM_N44716_v1.0 Salmonella enterica subsp. enterica serovar Typhimurium var. 5- Proteobacteria; Gammaproteobacteria; Enterobacterales; Enterobacteriaceae; Salmonella; Salmonella enterica; Salmonella enterica subsp. enterica; Salmonella enterica subsp. enterica serovar Typhimurium MTGDNTLITSHGINRRDFMKLCAALAATMGLSSKAAAEMAESVSNPQRPPVIWIGAQECT WP_000145429.1 hydrogenase 2 small subunit [Salmonella enterica] Length: 372\nNumber of predicted TMHs: 1\nExp number of AAs in TMHs: 24.51309\nExp number, first 60 AAs: 0.2682\nTotal prob of N-in: 0.14402\noutside 1 331\nTMhelix 332 354\ninside 355 372

25416 GCF_001241555.1_Salmonella_enterica_CVM_N44714_v1.0 Salmonella enterica subsp. enterica serovar Typhimurium var. 5- Proteobacteria; Gammaproteobacteria; Enterobacterales; Enterobacteriaceae; Salmonella; Salmonella enterica; Salmonella enterica subsp. enterica; Salmonella enterica subsp. enterica serovar Typhimurium MTGDNTLITSHGINRRDFMKLCAALAATMGLSSKAAAEMAESVSNPQRPPVIWIGAQECT WP_000145429.1 hydrogenase 2 small subunit [Salmonella enterica] Length: 372\nNumber of predicted TMHs: 1\nExp number of AAs in TMHs: 24.51309\nExp number, first 60 AAs: 0.2682\nTotal prob of N-in: 0.14402\noutside 1 331\nTMhelix 332 354\ninside 355 372

25417 GCF_001241515.1_Salmonella_enterica_CVM_N44713_v1.0 Salmonella enterica subsp. enterica serovar Typhimurium var. 5- Proteobacteria; Gammaproteobacteria; Enterobacterales; Enterobacteriaceae; Salmonella; Salmonella enterica; Salmonella enterica subsp. enterica; Salmonella enterica subsp. enterica serovar Typhimurium MTGDNTLITSHGINRRDFMKLCAALAATMGLSSKAAAEMAESVSNPQRPPVIWIGAQECT WP_000145429.1 hydrogenase 2 small subunit [Salmonella enterica] Length: 372\nNumber of predicted TMHs: 1\nExp number of AAs in TMHs: 24.51309\nExp number, first 60 AAs: 0.2682\nTotal prob of N-in: 0.14402\noutside 1 331\nTMhelix 332 354\ninside 355 372

25418 GCF_001241165.1_Salmonella_enterica_CVM_N44697-R_v1.0 Salmonella enterica subsp. enterica serovar Typhimurium var. 5- Proteobacteria; Gammaproteobacteria; Enterobacterales; Enterobacteriaceae; Salmonella; Salmonella enterica; Salmonella enterica subsp. enterica; Salmonella enterica subsp. enterica serovar Typhimurium MNNEETFYQAMRRKGVTRRSFLKFCSLAATSLGLGAGMTPKIAWALENKPRIPVVWIHGL WP_001058312.1 hydrogenase [Salmonella enterica] Length: 372\nNumber of predicted TMHs: 2\nExp number of AAs in TMHs: 36.51738\nExp number, first 60 AAs: 19.72849\nTotal prob of N-in: 0.96170\nPOSSIBLE N-term signal sequence\ninside 1 19\nTMhelix 20 39\noutside 40 325\nTMhelix 326 348\ninside 349 372

25419 GCF_001241165.1_Salmonella_enterica_CVM_N44697-R_v1.0 Salmonella enterica subsp. enterica serovar Typhimurium var. 5- Proteobacteria; Gammaproteobacteria; Enterobacterales; Enterobacteriaceae; Salmonella; Salmonella enterica; Salmonella enterica subsp. enterica; Salmonella enterica subsp. enterica serovar Typhimurium MTGDNTLITSHGINRRDFMKLCAALAATMGLSSKAAAEMAESVSNPQRPPVIWIGAQECT WP_000145429.1 hydrogenase 2 small subunit [Salmonella enterica] Length: 372\nNumber of predicted TMHs: 1\nExp number of AAs in TMHs: 24.51309\nExp number, first 60 AAs: 0.2682\nTotal prob of N-in: 0.14402\noutside 1 331\nTMhelix 332 354\ninside 355 372

25420 GCF_001240975.1_Salmonella_enterica_CVM_N43832_v1.0 Salmonella enterica subsp. enterica serovar Typhimurium var. 5- Proteobacteria; Gammaproteobacteria; Enterobacterales; Enterobacteriaceae; Salmonella; Salmonella enterica; Salmonella enterica subsp. enterica; Salmonella enterica subsp. enterica serovar Typhimurium MNNEETFYQAMRRKGVTRRSFLKFCSLAATSLGLGAGMTPKIAWALENKPRIPVVWIHGL WP_001058312.1 hydrogenase [Salmonella enterica] Length: 372\nNumber of predicted TMHs: 2\nExp number of AAs in TMHs: 36.51738\nExp number, first 60 AAs: 19.72849\nTotal prob of N-in: 0.96170\nPOSSIBLE N-term signal sequence\ninside 1 19\nTMhelix 20 39\noutside 40 325\nTMhelix 326 348\ninside 349 372

25421 GCF_001240475.1_Salmonella_enterica_CVM_N43469_v1.0 Salmonella enterica subsp. enterica serovar Typhimurium var. 5- Proteobacteria; Gammaproteobacteria; Enterobacterales; Enterobacteriaceae; Salmonella; Salmonella enterica; Salmonella enterica subsp. enterica; Salmonella enterica subsp. enterica serovar Typhimurium MTGDNTLITSHGINRRDFMKLCAALAATMGLSSKAAAEMAESVSNPQRPPVIWIGAQECT WP_000145429.1 hydrogenase 2 small subunit [Salmonella enterica] Length: 372\nNumber of predicted TMHs: 1\nExp

number of AAs in TMHs: 24.51309\nExp number, first 60 AAs: 0.2682\nTotal prob of N-in: 0.14402\noutside 1 331\nTMhelix 332 354\ninside 355 372

25422 GCF_001240975.1_Salmonella_enterica_CVM_N43832_v1.0 Salmonella enterica subsp. enterica serovar Typhimurium var. 5- Proteobacteria; Gammaproteobacteria; Enterobacterales; Enterobacteriaceae; Salmonella; Salmonella enterica; Salmonella enterica subsp. enterica; Salmonella enterica subsp. enterica serovar Typhimurium MTGDNTLITSHGINRRDFMKLCAALAATMGLSSKAAAEMAESVSNPQRPPVIWIGAQECT WP_000145429.1 hydrogenase 2 small subunit [Salmonella enterica] Length: 372\nNumber of predicted TMHs: 1\nExp number of AAs in TMHs: 24.51309\nExp number, first 60 AAs: 0.2682\nTotal prob of N-in: 0.14402\noutside 1 331\nTMhelix 332 354\ninside 355 372

25423 GCF_001240935.1_Salmonella_enterica_CVM_N43830-R_v1.0 Salmonella enterica subsp. enterica serovar Typhimurium var. 5- Proteobacteria; Gammaproteobacteria; Enterobacterales; Enterobacteriaceae; Salmonella; Salmonella enterica; Salmonella enterica subsp. enterica; Salmonella enterica subsp. enterica serovar Typhimurium MNNEETFYQAMRRKGVTRRSFLKFCSLAATSLGLGAGMTPKIAWALENKPRIPVVWIHGL WP_001058312.1 hydrogenase [Salmonella enterica] Length: 372\nNumber of predicted TMHs: 2\nExp number of AAs in TMHs: 36.51738\nExp number, first 60 AAs: 19.72849\nTotal prob of N-in: 0.96170\nPOSSIBLE N-term signal sequence\ninside 1 19\nTMhelix 20 39\noutside 40 325\nTMhelix 326 348\ninside 349 372

25424 GCF_001240825.1_Salmonella_enterica_CVM_N43823_v1.0 Salmonella enterica subsp. enterica serovar Typhimurium var. 5- Proteobacteria; Gammaproteobacteria; Enterobacterales; Enterobacteriaceae; Salmonella; Salmonella enterica; Salmonella enterica subsp. enterica; Salmonella enterica subsp. enterica serovar Typhimurium MNNEETFYQAMRRKGVTRRSFLKFCSLAATSLGLGAGMTPKIAWALENKPRIPVVWIHGL WP_001058312.1 hydrogenase [Salmonella enterica] Length: 372\nNumber of predicted TMHs: 2\nExp number of AAs in TMHs: 36.51738\nExp number, first 60 AAs: 19.72849\nTotal prob of N-in: 0.96170\nPOSSIBLE N-term signal sequence\ninside 1 19\nTMhelix 20 39\noutside 40 325\nTMhelix 326 348\ninside 349 372

25425 GCF_001244105.1_Salmonella_enterica_CVM_N47728_v1.0 Salmonella enterica subsp. enterica serovar Typhimurium var. 5- Proteobacteria; Gammaproteobacteria; Enterobacterales; Enterobacteriaceae; Salmonella; Salmonella enterica; Salmonella enterica subsp. enterica; Salmonella enterica subsp. enterica serovar Typhimurium MNNEETFYQAMRRKGVTRRSFLKFCSLAATSLGLGAGMTPKIAWALENKPRIPVVWIHGL WP_001058312.1 hydrogenase [Salmonella enterica] Length: 372\nNumber of predicted TMHs: 2\nExp number of AAs in TMHs: 36.51738\nExp number, first 60 AAs: 19.72849\nTotal prob of N-in: 0.96170\nPOSSIBLE N-term signal sequence\ninside 1 19\nTMhelix 20 39\noutside 40 325\nTMhelix 326 348\ninside 349 372

25426 GCF_001272475.1_Salmonella_enterica_CVM_N51296_v1.0 Salmonella enterica subsp. enterica serovar Typhimurium var. 5- Proteobacteria; Gammaproteobacteria; Enterobacterales; Enterobacteriaceae; Salmonella; Salmonella enterica; Salmonella enterica subsp. enterica; Salmonella enterica subsp. enterica serovar Typhimurium MTGDNTLITSHGINRRDFMKLCAALAATMGLSSKAAAEMAESVSNPQRPPVIWIGAQECT WP_000145429.1 hydrogenase 2 small subunit [Salmonella enterica] Length: 372\nNumber of predicted TMHs: 1\nExp number of AAs in TMHs: 24.51309\nExp number, first 60 AAs: 0.2682\nTotal prob of N-in: 0.14402\noutside 1 331\nTMhelix 332 354\ninside 355 372

25427 GCF_001272115.1_Salmonella_enterica_CVM_N46840_v1.0 Salmonella enterica subsp. enterica serovar Typhimurium var. 5- Proteobacteria; Gammaproteobacteria; Enterobacterales; Enterobacteriaceae; Salmonella; Salmonella enterica; Salmonella enterica subsp. enterica; Salmonella enterica subsp. enterica serovar Typhimurium MTGDNTLITSHGINRRDFMKLCAALAATMGLSSKAAAEMAESVSNPQRPPVIWIGAQECT WP_000145429.1 hydrogenase 2 small subunit [Salmonella enterica] Length: 372\nNumber of predicted TMHs: 1\nExp number of AAs in TMHs: 24.51309\nExp number, first 60 AAs: 0.2682\nTotal prob of N-in: 0.14402\noutside 1 331\nTMhelix 332 354\ninside 355 372

25428 GCF_001272035.1_Salmonella_enterica_CVM_N45410_v1.0 Salmonella enterica subsp. enterica serovar Typhimurium var. 5- Proteobacteria; Gammaproteobacteria; Enterobacterales; Enterobacteriaceae; Salmonella; Salmonella enterica; Salmonella enterica subsp. enterica; Salmonella enterica subsp. enterica serovar Typhimurium MTGDNTLITSHGINRRDFMKLCAALAATMGLSSKAAAEMAESVSNPQRPPVIWIGAQECT WP_000145429.1 hydrogenase 2 small subunit [Salmonella enterica] Length: 372\nNumber of predicted TMHs: 1\nExp number of AAs in TMHs: 24.51309\nExp number, first 60 AAs: 0.2682\nTotal prob of N-in: 0.14402\noutside 1 331\nTMhelix 332 354\ninside 355 372

25429 GCF_001480225.1_Salmonella_enterica_CVM_N40384-R_v1.0 Salmonella enterica subsp. enterica serovar Typhimurium var. 5- Proteobacteria; Gammaproteobacteria; Enterobacterales; Enterobacteriaceae; Salmonella;

Salmonella enterica; Salmonella enterica subsp. enterica; Salmonella enterica subsp. enterica serovar Typhimurium
MTGDNTLITSHGINRRDFMKLCAALAATMGLSSKAAAEMAESVSNPQRPPVIWIGAQECT WP_000145429.1
hydrogenase 2 small subunit [Salmonella enterica] Length: 372\nNumber of predicted TMHs: 1\nExp number of AAs in TMHs: 24.51309\nExp number, first 60 AAs: 0.2682\nTotal prob of N-in: 0.14402\noutside 1 331\nTMhelix 332 354\ninside 355 372

25430 GCF_001271745.1_Salmonella_enterica_CVM_N46841_v1.0 Salmonella enterica subsp. enterica serovar Typhimurium var. 5- Proteobacteria; Gammaproteobacteria; Enterobacterales; Enterobacteriaceae; Salmonella; Salmonella enterica; Salmonella enterica subsp. enterica; Salmonella enterica subsp. enterica serovar Typhimurium
MNNEETFYQAMRRKGVTRRSFLKFCSLAATSLGLGAGMTPKIAWALENKPRIPVVWIHGL WP_001058312.1
hydrogenase [Salmonella enterica] Length: 372\nNumber of predicted TMHs: 2\nExp number of AAs in TMHs: 36.51738\nExp number, first 60 AAs: 19.72849\nTotal prob of N-in: 0.96170\nPOSSIBLE N-term signal sequence\ninside 1 19\nTMhelix 20 39\noutside 40 325\nTMhelix 326 348\ninside 349 372

25431 GCF_001271715.1_Salmonella_enterica_CVM_N46828_v1.0 Salmonella enterica subsp. enterica serovar Typhimurium var. 5- Proteobacteria; Gammaproteobacteria; Enterobacterales; Enterobacteriaceae; Salmonella; Salmonella enterica; Salmonella enterica subsp. enterica; Salmonella enterica subsp. enterica serovar Typhimurium
MTGDNTLITSHGINRRDFMKLCAALAATMGLSSKAAAEMAESVSNPQRPPVIWIGAQECT WP_000145429.1
hydrogenase 2 small subunit [Salmonella enterica] Length: 372\nNumber of predicted TMHs: 1\nExp number of AAs in TMHs: 24.51309\nExp number, first 60 AAs: 0.2682\nTotal prob of N-in: 0.14402\noutside 1 331\nTMhelix 332 354\ninside 355 372

25432 GCF_001247265.1_Salmonella_enterica_CVM_N51301_v1.0 Salmonella enterica subsp. enterica serovar Typhimurium var. 5- Proteobacteria; Gammaproteobacteria; Enterobacterales; Enterobacteriaceae; Salmonella; Salmonella enterica; Salmonella enterica subsp. enterica; Salmonella enterica subsp. enterica serovar Typhimurium
MNNEETFYQAMRRKGVTRRSFLKFCSLAATSLGLGAGMTPKIAWALENKPRIPVVWIHGL WP_001058312.1
hydrogenase [Salmonella enterica] Length: 372\nNumber of predicted TMHs: 2\nExp number of AAs in TMHs: 36.51738\nExp number, first 60 AAs: 19.72849\nTotal prob of N-in: 0.96170\nPOSSIBLE N-term signal sequence\ninside 1 19\nTMhelix 20 39\noutside 40 325\nTMhelix 326 348\ninside 349 372

25433 GCF_001243105.1_Salmonella_enterica_CVM_N46825_v1.0 Salmonella enterica subsp. enterica serovar Typhimurium var. 5- Proteobacteria; Gammaproteobacteria; Enterobacterales; Enterobacteriaceae; Salmonella; Salmonella enterica; Salmonella enterica subsp. enterica; Salmonella enterica subsp. enterica serovar Typhimurium
MTGDNTLITSHGINRRDFMKLCAALAATMGLSSKAAAEMAESVSNPQRPPVIWIGAQECT WP_000145429.1
hydrogenase 2 small subunit [Salmonella enterica] Length: 372\nNumber of predicted TMHs: 1\nExp number of AAs in TMHs: 24.51309\nExp number, first 60 AAs: 0.2682\nTotal prob of N-in: 0.14402\noutside 1 331\nTMhelix 332 354\ninside 355 372

25434 GCF_001477745.1_Salmonella_enterica_CVM_N29315-SQ_v1.0 Salmonella enterica subsp. enterica serovar Typhimurium var. 5- Proteobacteria; Gammaproteobacteria; Enterobacterales; Enterobacteriaceae; Salmonella; Salmonella enterica; Salmonella enterica subsp. enterica; Salmonella enterica subsp. enterica serovar Typhimurium
MTGDNTLITSHGINRRDFMKLCAALAATMGLSSKAAAEMAESVSNPQRPPVIWIGAQECT WP_000145429.1
hydrogenase 2 small subunit [Salmonella enterica] Length: 372\nNumber of predicted TMHs: 1\nExp number of AAs in TMHs: 24.51309\nExp number, first 60 AAs: 0.2682\nTotal prob of N-in: 0.14402\noutside 1 331\nTMhelix 332 354\ninside 355 372

25435 GCF_001478445.1_Salmonella_enterica_CVM_N38944-SQ_v1.0 Salmonella enterica subsp. enterica serovar Typhimurium var. 5- Proteobacteria; Gammaproteobacteria; Enterobacterales; Enterobacteriaceae; Salmonella; Salmonella enterica; Salmonella enterica subsp. enterica; Salmonella enterica subsp. enterica serovar Typhimurium
MTGDNTLITSHGINRRDFMKLCAALAATMGLSSKAAAEMAESVSNPQRPPVIWIGAQECT WP_000145429.1
hydrogenase 2 small subunit [Salmonella enterica] Length: 372\nNumber of predicted TMHs: 1\nExp number of AAs in TMHs: 24.51309\nExp number, first 60 AAs: 0.2682\nTotal prob of N-in: 0.14402\noutside 1 331\nTMhelix 332 354\ninside 355 372

25436 GCF_001478325.1_Salmonella_enterica_CVM_N38858-SQ_v1.0 Salmonella enterica subsp. enterica serovar Typhimurium var. 5- Proteobacteria; Gammaproteobacteria; Enterobacterales; Enterobacteriaceae; Salmonella; Salmonella enterica; Salmonella enterica subsp. enterica; Salmonella enterica subsp. enterica serovar Typhimurium
MNNEETFYQAMRRKGVTRRSFLKFCSLAATSLGLGAGMTPKIAWALENKPRIPVVWIHGL WP_001058312.1
hydrogenase [Salmonella enterica] Length: 372\nNumber of predicted TMHs: 2\nExp number of AAs in TMHs: 36.51738\nExp number, first 60 AAs: 19.72849\nTotal prob of N-in: 0.96170\nPOSSIBLE N-term signal sequence\ninside 1 19\nTMhelix 20 39\noutside 40 325\nTMhelix 326 348\ninside 349 372

25437 GCF_001243255.1_Salmonella_enterica_CVM_N46834_v1.0 Salmonella enterica subsp. enterica serovar Typhimurium var. 5- Proteobacteria; Gammaproteobacteria; Enterobacterales; Enterobacteriaceae; Salmonella; Salmonella enterica; Salmonella enterica subsp. enterica; Salmonella enterica subsp. enterica serovar Typhimurium MTGDNTLITSHGINRRDFMKLCAALAATMGLSSKAAAEMAESVSNPQRPPVIWIGAQECT WP_000145429.1 hydrogenase 2 small subunit [Salmonella enterica] Length: 372\nNumber of predicted TMHs: 1\nExp number of AAs in TMHs: 24.51309\nExp number, first 60 AAs: 0.2682\nTotal prob of N-in: 0.14402\noutside 1 331\nTMhelix 332 354\ninside 355 372

25438 GCF_001477745.1_Salmonella_enterica_CVM_N29315-SQ_v1.0 Salmonella enterica subsp. enterica serovar Typhimurium var. 5- Proteobacteria; Gammaproteobacteria; Enterobacterales; Enterobacteriaceae; Salmonella; Salmonella enterica; Salmonella enterica subsp. enterica; Salmonella enterica subsp. enterica serovar Typhimurium MNNEETFYQAMRRKGVTRRSFLKFCSLAATSLGLGAGMTPKIAWALENKPRIPVVWIHGL WP_001058312.1 hydrogenase [Salmonella enterica] Length: 372\nNumber of predicted TMHs: 2\nExp number of AAs in TMHs: 36.51738\nExp number, first 60 AAs: 19.72849\nTotal prob of N-in: 0.96170\nPOSSIBLE N-term signal sequence\ninside 1 19\nTMhelix 20 39\noutside 40 325\nTMhelix 326 348\ninside 349 372

25439 GCF_001243255.1_Salmonella_enterica_CVM_N46834_v1.0 Salmonella enterica subsp. enterica serovar Typhimurium var. 5- Proteobacteria; Gammaproteobacteria; Enterobacterales; Enterobacteriaceae; Salmonella; Salmonella enterica; Salmonella enterica subsp. enterica; Salmonella enterica subsp. enterica serovar Typhimurium MNNEETFYQAMRRKGVTRRSFLKFCSLAATSLGLGAGMTPKIAWALENKPRIPVVWIHGL WP_001058312.1 hydrogenase [Salmonella enterica] Length: 372\nNumber of predicted TMHs: 2\nExp number of AAs in TMHs: 36.51738\nExp number, first 60 AAs: 19.72849\nTotal prob of N-in: 0.96170\nPOSSIBLE N-term signal sequence\ninside 1 19\nTMhelix 20 39\noutside 40 325\nTMhelix 326 348\ninside 349 372

25440 GCF_002032065.1_ASM203206v1 Salmonella enterica subsp. enterica serovar Typhimurium var. 5- Proteobacteria; Gammaproteobacteria; Enterobacterales; Enterobacteriaceae; Salmonella; Salmonella enterica; Salmonella enterica subsp. enterica; Salmonella enterica subsp. enterica serovar Typhimurium MNNEETFYQAMRRKGVTRRSFLKFCSLAATSLGLGAGMTPKIAWALENKPRIPVVWIHGL WP_079839574.1 [Ni/Fe] hydrogenase small subunit [Salmonella enterica] Length: 372\nNumber of predicted TMHs: 2\nExp number of AAs in TMHs: 36.47233\nExp number, first 60 AAs: 19.72898\nTotal prob of N-in: 0.96168\nPOSSIBLE N-term signal sequence\ninside 1 19\nTMhelix 20 39\noutside 40 325\nTMhelix 326 348\ninside 349 372

25441 GCF_002032065.1_ASM203206v1 Salmonella enterica subsp. enterica serovar Typhimurium var. 5- Proteobacteria; Gammaproteobacteria; Enterobacterales; Enterobacteriaceae; Salmonella; Salmonella enterica; Salmonella enterica subsp. enterica; Salmonella enterica subsp. enterica serovar Typhimurium MTGDNTLITSHGINRRDFMKLCAALAATMGLSSKAAAEMAESVSNPQRPPVIWIGAQECT WP_000145429.1 hydrogenase 2 small subunit [Salmonella enterica] Length: 372\nNumber of predicted TMHs: 1\nExp number of AAs in TMHs: 24.51309\nExp number, first 60 AAs: 0.2682\nTotal prob of N-in: 0.14402\noutside 1 331\nTMhelix 332 354\ninside 355 372

25442 GCF_002031265.1_ASM203126v1 Salmonella enterica subsp. enterica serovar Typhimurium var. 5- Proteobacteria; Gammaproteobacteria; Enterobacterales; Enterobacteriaceae; Salmonella; Salmonella enterica; Salmonella enterica subsp. enterica; Salmonella enterica subsp. enterica serovar Typhimurium MNNEETFYQAMRRKGVTRRSFLKFCSLAATSLGLGAGMTPKIAWALENKPRIPVVWIHGL WP_001058312.1 hydrogenase [Salmonella enterica] Length: 372\nNumber of predicted TMHs: 2\nExp number of AAs in TMHs: 36.51738\nExp number, first 60 AAs: 19.72849\nTotal prob of N-in: 0.96170\nPOSSIBLE N-term signal sequence\ninside 1 19\nTMhelix 20 39\noutside 40 325\nTMhelix 326 348\ninside 349 372

25443 GCF_001240965.1_Salmonella_enterica_CVM_N43831_v1.0 Salmonella enterica subsp. enterica serovar Typhimurium var. 5- Proteobacteria; Gammaproteobacteria; Enterobacterales; Enterobacteriaceae; Salmonella; Salmonella enterica; Salmonella enterica subsp. enterica; Salmonella enterica subsp. enterica serovar Typhimurium MTGDNTLITSHGINRRDFMKLCAALAATMGLSSKAAAEMAESVSNPQRPPVIWIGAQECT WP_000145429.1 hydrogenase 2 small subunit [Salmonella enterica] Length: 372\nNumber of predicted TMHs: 1\nExp number of AAs in TMHs: 24.51309\nExp number, first 60 AAs: 0.2682\nTotal prob of N-in: 0.14402\noutside 1 331\nTMhelix 332 354\ninside 355 372

25444 GCF_001241075.1_Salmonella_enterica_CVM_N44692_v1.0 Salmonella enterica subsp. enterica serovar Typhimurium var. 5- Proteobacteria; Gammaproteobacteria; Enterobacterales; Enterobacteriaceae; Salmonella; Salmonella enterica; Salmonella enterica subsp. enterica; Salmonella enterica subsp. enterica serovar Typhimurium MTGDNTLITSHGINRRDFMKLCAALAATMGLSSKAAAEMAESVSNPQRPPVIWIGAQECT WP_000145429.1 hydrogenase 2 small subunit [Salmonella enterica] Length: 372\nNumber of predicted TMHs: 1\nExp

number of AAs in TMHs: 24.51309\nExp number, first 60 AAs: 0.2682\nTotal prob of N-in: 0.14402\noutside 1 331\nTMhelix 332 354\ninside 355 372

25445 GCF_001240965.1_Salmonella_enterica_CVM_N43831_v1.0 Salmonella enterica subsp. enterica serovar Typhimurium var. 5- Proteobacteria; Gammaproteobacteria; Enterobacterales; Enterobacteriaceae; Salmonella; Salmonella enterica; Salmonella enterica subsp. enterica; Salmonella enterica subsp. enterica serovar Typhimurium MNNEETFYQAMRRKGVTRRSFLKFCSLAATSLGLGAGMTPKIAWALENKPRIPVVWIHGL WP_001058312.1 hydrogenase [Salmonella enterica] Length: 372\nNumber of predicted TMHs: 2\nExp number of AAs in TMHs: 36.51738\nExp number, first 60 AAs: 19.72849\nTotal prob of N-in: 0.96170\nPOSSIBLE N-term signal sequence\ninside 1 19\nTMhelix 20 39\noutside 40 325\nTMhelix 326 348\ninside 349 372

25446 GCF_001241585.1_Salmonella_enterica_CVM_N44715_v1.0 Salmonella enterica subsp. enterica serovar Typhimurium var. 5- Proteobacteria; Gammaproteobacteria; Enterobacterales; Enterobacteriaceae; Salmonella; Salmonella enterica; Salmonella enterica subsp. enterica; Salmonella enterica subsp. enterica serovar Typhimurium MTGDNTLITSHGINRRDFMKLCAALAATMGLSSKAAAEMAESVSNPQRPPVIWIGAQECT WP_000145429.1 hydrogenase 2 small subunit [Salmonella enterica] Length: 372\nNumber of predicted TMHs: 1\nExp number of AAs in TMHs: 24.51309\nExp number, first 60 AAs: 0.2682\nTotal prob of N-in: 0.14402\noutside 1 331\nTMhelix 332 354\ninside 355 372

25447 GCF_001241505.1_Salmonella_enterica_CVM_N44712_v1.0 Salmonella enterica subsp. enterica serovar Typhimurium var. 5- Proteobacteria; Gammaproteobacteria; Enterobacterales; Enterobacteriaceae; Salmonella; Salmonella enterica; Salmonella enterica subsp. enterica; Salmonella enterica subsp. enterica serovar Typhimurium MNNEETFYQAMRRKGVTRRSFLKFCSLAATSLGLGAGMTPKIAWALENKPRIPVVWIHGL WP_001058312.1 hydrogenase [Salmonella enterica] Length: 372\nNumber of predicted TMHs: 2\nExp number of AAs in TMHs: 36.51738\nExp number, first 60 AAs: 19.72849\nTotal prob of N-in: 0.96170\nPOSSIBLE N-term signal sequence\ninside 1 19\nTMhelix 20 39\noutside 40 325\nTMhelix 326 348\ninside 349 372

25448 GCF_001241585.1_Salmonella_enterica_CVM_N44715_v1.0 Salmonella enterica subsp. enterica serovar Typhimurium var. 5- Proteobacteria; Gammaproteobacteria; Enterobacterales; Enterobacteriaceae; Salmonella; Salmonella enterica; Salmonella enterica subsp. enterica; Salmonella enterica subsp. enterica serovar Typhimurium MNNEETFYQAMRRKGVTRRSFLKFCSLAATSLGLGAGMTPKIAWALENKPRIPVVWIHGL WP_001058312.1 hydrogenase [Salmonella enterica] Length: 372\nNumber of predicted TMHs: 2\nExp number of AAs in TMHs: 36.51738\nExp number, first 60 AAs: 19.72849\nTotal prob of N-in: 0.96170\nPOSSIBLE N-term signal sequence\ninside 1 19\nTMhelix 20 39\noutside 40 325\nTMhelix 326 348\ninside 349 372

25449 GCF_001479395.1_Salmonella_enterica_CVM_N32056-R_v1.0 Salmonella enterica subsp. enterica serovar Typhimurium var. 5- Proteobacteria; Gammaproteobacteria; Enterobacterales; Enterobacteriaceae; Salmonella; Salmonella enterica; Salmonella enterica subsp. enterica; Salmonella enterica subsp. enterica serovar Typhimurium MTGDNTLITSHGINRRDFMKLCAALAATMGLSSKAAAEMAESVSNPQRPPVIWIGAQECT WP_000145429.1 hydrogenase 2 small subunit [Salmonella enterica] Length: 372\nNumber of predicted TMHs: 1\nExp number of AAs in TMHs: 24.51309\nExp number, first 60 AAs: 0.2682\nTotal prob of N-in: 0.14402\noutside 1 331\nTMhelix 332 354\ninside 355 372

25450 GCF_001245495.1_Salmonella_enterica_CVM_N50430_v1.0 Salmonella enterica subsp. enterica serovar Typhimurium var. 5- Proteobacteria; Gammaproteobacteria; Enterobacterales; Enterobacteriaceae; Salmonella; Salmonella enterica; Salmonella enterica subsp. enterica; Salmonella enterica subsp. enterica serovar Typhimurium MTGDNTLITSHGINRRDFMKLCAALAATMGLSSKAAAEMAESVSNPQRPPVIWIGAQECT WP_000145429.1 hydrogenase 2 small subunit [Salmonella enterica] Length: 372\nNumber of predicted TMHs: 1\nExp number of AAs in TMHs: 24.51309\nExp number, first 60 AAs: 0.2682\nTotal prob of N-in: 0.14402\noutside 1 331\nTMhelix 332 354\ninside 355 372

25451 GCF_001241975.1_Salmonella_enterica_CVM_N45408_v1.0 Salmonella enterica subsp. enterica serovar Typhimurium var. 5- Proteobacteria; Gammaproteobacteria; Enterobacterales; Enterobacteriaceae; Salmonella; Salmonella enterica; Salmonella enterica subsp. enterica; Salmonella enterica subsp. enterica serovar Typhimurium MTGDNTLITSHGINRRDFMKLCAALAATMGLSSKAAAEMAESVSNPQRPPVIWIGAQECT WP_000145429.1 hydrogenase 2 small subunit [Salmonella enterica] Length: 372\nNumber of predicted TMHs: 1\nExp number of AAs in TMHs: 24.51309\nExp number, first 60 AAs: 0.2682\nTotal prob of N-in: 0.14402\noutside 1 331\nTMhelix 332 354\ninside 355 372

25452 GCF_001242395.1_Salmonella_enterica_CVM_N45938_v1.0 Salmonella enterica subsp. enterica serovar Typhimurium var. 5- Proteobacteria; Gammaproteobacteria; Enterobacterales; Enterobacteriaceae; Salmonella;

Salmonella enterica; Salmonella enterica subsp. enterica; Salmonella enterica subsp. enterica serovar Typhimurium
MTGDNTLITSHGINRRDFMKLCAALAATMGLSSKAAAEMAESVSNPQRPPVIWIGAQECT WP_000145429.1
hydrogenase 2 small subunit [Salmonella enterica] Length: 372\nNumber of predicted TMHs: 1\nExp number of AAs in TMHs: 24.51309\nExp number, first 60 AAs: 0.2682\nTotal prob of N-in: 0.14402\noutside 1 331\nTMhelix 332 354\ninside 355 372

25453 GCF_001242565.1_Salmonella_enterica_CVM_N45950_v1.0 Salmonella enterica subsp. enterica serovar Typhimurium var. 5- Proteobacteria; Gammaproteobacteria; Enterobacterales; Enterobacteriaceae; Salmonella; Salmonella enterica; Salmonella enterica subsp. enterica; Salmonella enterica subsp. enterica serovar Typhimurium
MTGDNTLITSHGINRRDFMKLCAALAATMGLSSKAAAEMAESVSNPQRPPVIWIGAQECT WP_000145429.1
hydrogenase 2 small subunit [Salmonella enterica] Length: 372\nNumber of predicted TMHs: 1\nExp number of AAs in TMHs: 24.51309\nExp number, first 60 AAs: 0.2682\nTotal prob of N-in: 0.14402\noutside 1 331\nTMhelix 332 354\ninside 355 372

25454 GCF_002030225.1_ASM203022v1 Salmonella enterica subsp. enterica serovar Typhimurium var. 5-
Proteobacteria; Gammaproteobacteria; Enterobacterales; Enterobacteriaceae; Salmonella; Salmonella enterica; Salmonella enterica subsp. enterica; Salmonella enterica subsp. enterica serovar Typhimurium
MTGDNTLITSHGINRRDFMKLCAALAATMGLSSKAAAEMAESVSNPQRPPVIWIGAQECT WP_000145429.1
hydrogenase 2 small subunit [Salmonella enterica] Length: 372\nNumber of predicted TMHs: 1\nExp number of AAs in TMHs: 24.51309\nExp number, first 60 AAs: 0.2682\nTotal prob of N-in: 0.14402\noutside 1 331\nTMhelix 332 354\ninside 355 372

25455 GCF_002058005.1_ASM205800v1 Salmonella enterica subsp. enterica serovar Typhimurium var. 5-
Proteobacteria; Gammaproteobacteria; Enterobacterales; Enterobacteriaceae; Salmonella; Salmonella enterica; Salmonella enterica subsp. enterica; Salmonella enterica subsp. enterica serovar Typhimurium
MTGDNTLITSHGINRRDFMKLCAALAATMGLSSKAAAEMAESVSNPQRPPVIWIGAQECT WP_000145429.1
hydrogenase 2 small subunit [Salmonella enterica] Length: 372\nNumber of predicted TMHs: 1\nExp number of AAs in TMHs: 24.51309\nExp number, first 60 AAs: 0.2682\nTotal prob of N-in: 0.14402\noutside 1 331\nTMhelix 332 354\ninside 355 372

25456 GCF_002058595.1_ASM205859v1 Salmonella enterica subsp. enterica serovar Typhimurium var. 5-
Proteobacteria; Gammaproteobacteria; Enterobacterales; Enterobacteriaceae; Salmonella; Salmonella enterica; Salmonella enterica subsp. enterica; Salmonella enterica subsp. enterica serovar Typhimurium
MNNEETFYQAMRRKGVTRRSFLKFCSLAATSLGLGAGMTPKIAWALENKPRIPVVWIHGL WP_001058312.1
hydrogenase [Salmonella enterica] Length: 372\nNumber of predicted TMHs: 2\nExp number of AAs in TMHs: 36.51738\nExp number, first 60 AAs: 19.72849\nTotal prob of N-in: 0.96170\nPOSSIBLE N-term signal sequence\ninside 1 19\nTMhelix 20 39\noutside 40 325\nTMhelix 326 348\ninside 349 372

25457 GCF_002058595.1_ASM205859v1 Salmonella enterica subsp. enterica serovar Typhimurium var. 5-
Proteobacteria; Gammaproteobacteria; Enterobacterales; Enterobacteriaceae; Salmonella; Salmonella enterica; Salmonella enterica subsp. enterica; Salmonella enterica subsp. enterica serovar Typhimurium
MTGDNTLITSHGINRRDFMKLCAALAATMGLSSKAAAEMAESVSNPQRPPVIWIGAQECT WP_000145429.1
hydrogenase 2 small subunit [Salmonella enterica] Length: 372\nNumber of predicted TMHs: 1\nExp number of AAs in TMHs: 24.51309\nExp number, first 60 AAs: 0.2682\nTotal prob of N-in: 0.14402\noutside 1 331\nTMhelix 332 354\ninside 355 372

25458 GCF_002058005.1_ASM205800v1 Salmonella enterica subsp. enterica serovar Typhimurium var. 5-
Proteobacteria; Gammaproteobacteria; Enterobacterales; Enterobacteriaceae; Salmonella; Salmonella enterica; Salmonella enterica subsp. enterica; Salmonella enterica subsp. enterica serovar Typhimurium
MNNEETFYQAMRRKGVTRRSFLKFCSLAATSLGLGAGMTPKIAWALENKPRIPVVWIHGL WP_001058312.1
hydrogenase [Salmonella enterica] Length: 372\nNumber of predicted TMHs: 2\nExp number of AAs in TMHs: 36.51738\nExp number, first 60 AAs: 19.72849\nTotal prob of N-in: 0.96170\nPOSSIBLE N-term signal sequence\ninside 1 19\nTMhelix 20 39\noutside 40 325\nTMhelix 326 348\ninside 349 372

25459 GCF_002035635.1_ASM203563v1 Salmonella enterica subsp. enterica serovar Typhimurium var. 5-
Proteobacteria; Gammaproteobacteria; Enterobacterales; Enterobacteriaceae; Salmonella; Salmonella enterica; Salmonella enterica subsp. enterica; Salmonella enterica subsp. enterica serovar Typhimurium
MNNEETFYQAMRRKGVTRRSFLKFCSLAATSLGLGAGMTPKIAWALENKPRIPVVWIHGL WP_001058312.1
hydrogenase [Salmonella enterica] Length: 372\nNumber of predicted TMHs: 2\nExp number of AAs in TMHs: 36.51738\nExp number, first 60 AAs: 19.72849\nTotal prob of N-in: 0.96170\nPOSSIBLE N-term signal sequence\ninside 1 19\nTMhelix 20 39\noutside 40 325\nTMhelix 326 348\ninside 349 372

25460 GCF_002035635.1_ASM203563v1 Salmonella enterica subsp. enterica serovar Typhimurium var. 5- Proteobacteria; Gammaproteobacteria; Enterobacterales; Enterobacteriaceae; Salmonella; Salmonella enterica; Salmonella enterica subsp. enterica; Salmonella enterica subsp. enterica serovar Typhimurium MTGDNTLITSHGINRRDFMKLCAALAATMGLSSKAAAEMAESVSNPQRPPVIWIGAQECT WP_000145429.1 hydrogenase 2 small subunit [Salmonella enterica] Length: 372\nNumber of predicted TMHs: 1\nExp number of AAs in TMHs: 24.51309\nExp number, first 60 AAs: 0.2682\nTotal prob of N-in: 0.14402\noutside 1 331\nTMhelix 332 354\ninside 355 372

25461 GCF_002033135.1_ASM203313v1 Salmonella enterica subsp. enterica serovar Typhimurium var. 5- Proteobacteria; Gammaproteobacteria; Enterobacterales; Enterobacteriaceae; Salmonella; Salmonella enterica; Salmonella enterica subsp. enterica; Salmonella enterica subsp. enterica serovar Typhimurium MNNEETFYQAMRRKGVTRRSFLKFCSLAATSLGLGAGMTPKIAWALENKPRIPVVWIHGL WP_001058312.1 hydrogenase [Salmonella enterica] Length: 372\nNumber of predicted TMHs: 2\nExp number of AAs in TMHs: 36.51738\nExp number, first 60 AAs: 19.72849\nTotal prob of N-in: 0.96170\nPOSSIBLE N-term signal sequence\ninside 1 19\nTMhelix 20 39\noutside 40 325\nTMhelix 326 348\ninside 349 372

25462 GCF_002033065.1_ASM203306v1 Salmonella enterica subsp. enterica serovar Typhimurium var. 5- Proteobacteria; Gammaproteobacteria; Enterobacterales; Enterobacteriaceae; Salmonella; Salmonella enterica; Salmonella enterica subsp. enterica; Salmonella enterica subsp. enterica serovar Typhimurium MNNEETFYQAMRRKGVTRRSFLKFCSLAATSLGLGAGMTPKIAWALENKPRIPVVWIHGL WP_001058312.1 hydrogenase [Salmonella enterica] Length: 372\nNumber of predicted TMHs: 2\nExp number of AAs in TMHs: 36.51738\nExp number, first 60 AAs: 19.72849\nTotal prob of N-in: 0.96170\nPOSSIBLE N-term signal sequence\ninside 1 19\nTMhelix 20 39\noutside 40 325\nTMhelix 326 348\ninside 349 372

25463 GCF_002033135.1_ASM203313v1 Salmonella enterica subsp. enterica serovar Typhimurium var. 5- Proteobacteria; Gammaproteobacteria; Enterobacterales; Enterobacteriaceae; Salmonella; Salmonella enterica; Salmonella enterica subsp. enterica; Salmonella enterica subsp. enterica serovar Typhimurium MTGDNTLITSHGINRRDFMKLCAALAATMGLSSKAAAEMAESVSNPQRPPVIWIGAQECT WP_000145429.1 hydrogenase 2 small subunit [Salmonella enterica] Length: 372\nNumber of predicted TMHs: 1\nExp number of AAs in TMHs: 24.51309\nExp number, first 60 AAs: 0.2682\nTotal prob of N-in: 0.14402\noutside 1 331\nTMhelix 332 354\ninside 355 372

25464 GCF_002032875.1_ASM203287v1 Salmonella enterica subsp. enterica serovar Typhimurium var. 5- Proteobacteria; Gammaproteobacteria; Enterobacterales; Enterobacteriaceae; Salmonella; Salmonella enterica; Salmonella enterica subsp. enterica; Salmonella enterica subsp. enterica serovar Typhimurium MTGDNTLITSHGINRRDFMKLCAALAATMGLSSKAAAEMAESVSNPQRPPVIWIGAQECT WP_000145429.1 hydrogenase 2 small subunit [Salmonella enterica] Length: 372\nNumber of predicted TMHs: 1\nExp number of AAs in TMHs: 24.51309\nExp number, first 60 AAs: 0.2682\nTotal prob of N-in: 0.14402\noutside 1 331\nTMhelix 332 354\ninside 355 372

25465 GCF_002032705.1_ASM203270v1 Salmonella enterica subsp. enterica serovar Typhimurium var. 5- Proteobacteria; Gammaproteobacteria; Enterobacterales; Enterobacteriaceae; Salmonella; Salmonella enterica; Salmonella enterica subsp. enterica; Salmonella enterica subsp. enterica serovar Typhimurium MTGDNTLITSHGINRRDFMKLCAALAATMGLSSKAAAEMAESVSNPQRPPVIWIGAQECT WP_000145429.1 hydrogenase 2 small subunit [Salmonella enterica] Length: 372\nNumber of predicted TMHs: 1\nExp number of AAs in TMHs: 24.51309\nExp number, first 60 AAs: 0.2682\nTotal prob of N-in: 0.14402\noutside 1 331\nTMhelix 332 354\ninside 355 372

25466 GCF_002032795.1_ASM203279v1 Salmonella enterica subsp. enterica serovar Typhimurium var. 5- Proteobacteria; Gammaproteobacteria; Enterobacterales; Enterobacteriaceae; Salmonella; Salmonella enterica; Salmonella enterica subsp. enterica; Salmonella enterica subsp. enterica serovar Typhimurium MTGDNTLITSHGINRRDFMKLCAALAATMGLSSKAAAEMAESVSNPQRPPVIWIGAQECT WP_000145429.1 hydrogenase 2 small subunit [Salmonella enterica] Length: 372\nNumber of predicted TMHs: 1\nExp number of AAs in TMHs: 24.51309\nExp number, first 60 AAs: 0.2682\nTotal prob of N-in: 0.14402\noutside 1 331\nTMhelix 332 354\ninside 355 372

25467 GCF_002031565.1_ASM203156v1 Salmonella enterica subsp. enterica serovar Typhimurium var. 5- Proteobacteria; Gammaproteobacteria; Enterobacterales; Enterobacteriaceae; Salmonella; Salmonella enterica; Salmonella enterica subsp. enterica; Salmonella enterica subsp. enterica serovar Typhimurium MNNEETFYQAMRRKGVTRRSFLKFCSLAATSLGLGAGMTPKIAWALENKPRIPVVWIHGL WP_001058312.1 hydrogenase [Salmonella enterica] Length: 372\nNumber of predicted TMHs: 2\nExp number of AAs in

TMHs: 36.51738\nExp number, first 60 AAs: 19.72849\nTotal prob of N-in: 0.96170\nPOSSIBLE N-term signal sequence\ninside 1 19\nTMhelix 20 39\noutside 40 325\nTMhelix 326 348\ninside 349 372

25468 GCF_002031405.1_ASM203140v1 Salmonella enterica subsp. enterica serovar Typhimurium var. 5- Proteobacteria; Gammaproteobacteria; Enterobacterales; Enterobacteriaceae; Salmonella; Salmonella enterica; Salmonella enterica subsp. enterica; Salmonella enterica subsp. enterica serovar Typhimurium MNNEETFYQAMRRKGVTRRSFLKFCSLAATSLGLGAGMTPKIAWALENKPRIPVVWIHGL WP_001058312.1 hydrogenase [Salmonella enterica] Length: 372\nNumber of predicted TMHs: 2\nExp number of AAs in TMHs: 36.51738\nExp number, first 60 AAs: 19.72849\nTotal prob of N-in: 0.96170\nPOSSIBLE N-term signal sequence\ninside 1 19\nTMhelix 20 39\noutside 40 325\nTMhelix 326 348\ninside 349 372

25469 GCF_002031525.1_ASM203152v1 Salmonella enterica subsp. enterica serovar Typhimurium var. 5- Proteobacteria; Gammaproteobacteria; Enterobacterales; Enterobacteriaceae; Salmonella; Salmonella enterica; Salmonella enterica subsp. enterica; Salmonella enterica subsp. enterica serovar Typhimurium MNNEETFYQAMRRKGVTRRSFLKFCSLAATSLGLGAGMTPKIAWALENKPRIPVVWIHGL WP_001058312.1 hydrogenase [Salmonella enterica] Length: 372\nNumber of predicted TMHs: 2\nExp number of AAs in TMHs: 36.51738\nExp number, first 60 AAs: 19.72849\nTotal prob of N-in: 0.96170\nPOSSIBLE N-term signal sequence\ninside 1 19\nTMhelix 20 39\noutside 40 325\nTMhelix 326 348\ninside 349 372

25470 GCF_002031405.1_ASM203140v1 Salmonella enterica subsp. enterica serovar Typhimurium var. 5- Proteobacteria; Gammaproteobacteria; Enterobacterales; Enterobacteriaceae; Salmonella; Salmonella enterica; Salmonella enterica subsp. enterica; Salmonella enterica subsp. enterica serovar Typhimurium MTGDNTLITSHGINRRDFMKLCAALAATMGLSSKAAAEMAESVSNPQRPPVIWIGAQECT WP_000145429.1 hydrogenase 2 small subunit [Salmonella enterica] Length: 372\nNumber of predicted TMHs: 1\nExp number of AAs in TMHs: 24.51309\nExp number, first 60 AAs: 0.2682\nTotal prob of N-in: 0.14402\noutside 1 331\nTMhelix 332 354\ninside 355 372

25471 GCF_002030995.1_ASM203099v1 Salmonella enterica subsp. enterica serovar Typhimurium var. 5- Proteobacteria; Gammaproteobacteria; Enterobacterales; Enterobacteriaceae; Salmonella; Salmonella enterica; Salmonella enterica subsp. enterica; Salmonella enterica subsp. enterica serovar Typhimurium MTGDNTLITSHGINRRDFMKLCAALAATMGLSSKAAAEMAESVSNPQRPPVIWIGAQECT WP_000145429.1 hydrogenase 2 small subunit [Salmonella enterica] Length: 372\nNumber of predicted TMHs: 1\nExp number of AAs in TMHs: 24.51309\nExp number, first 60 AAs: 0.2682\nTotal prob of N-in: 0.14402\noutside 1 331\nTMhelix 332 354\ninside 355 372

25472 GCF_002030775.1_ASM203077v1 Salmonella enterica subsp. enterica serovar Typhimurium var. 5- Proteobacteria; Gammaproteobacteria; Enterobacterales; Enterobacteriaceae; Salmonella; Salmonella enterica; Salmonella enterica subsp. enterica; Salmonella enterica subsp. enterica serovar Typhimurium MNNEETFYQAMRRKGVTRRSFLKFCSLAATSLGLGAGMTPKIAWALENKPRIPVVWIHGL WP_001058312.1 hydrogenase [Salmonella enterica] Length: 372\nNumber of predicted TMHs: 2\nExp number of AAs in TMHs: 36.51738\nExp number, first 60 AAs: 19.72849\nTotal prob of N-in: 0.96170\nPOSSIBLE N-term signal sequence\ninside 1 19\nTMhelix 20 39\noutside 40 325\nTMhelix 326 348\ninside 349 372

25473 GCF_002030605.1_ASM203060v1 Salmonella enterica subsp. enterica serovar Typhimurium var. 5- Proteobacteria; Gammaproteobacteria; Enterobacterales; Enterobacteriaceae; Salmonella; Salmonella enterica; Salmonella enterica subsp. enterica; Salmonella enterica subsp. enterica serovar Typhimurium MNNEETFYQAMRRKGVTRRSFLKFCSLAATSLGLGAGMTPKIAWALENKPRIPVVWIHGL WP_001058312.1 hydrogenase [Salmonella enterica] Length: 372\nNumber of predicted TMHs: 2\nExp number of AAs in TMHs: 36.51738\nExp number, first 60 AAs: 19.72849\nTotal prob of N-in: 0.96170\nPOSSIBLE N-term signal sequence\ninside 1 19\nTMhelix 20 39\noutside 40 325\nTMhelix 326 348\ninside 349 372

25474 GCF_001478445.1_Salmonella_enterica_CVM_N38944-SQ_v1.0 Salmonella enterica subsp. enterica serovar Typhimurium var. 5- Proteobacteria; Gammaproteobacteria; Enterobacterales; Enterobacteriaceae; Salmonella; Salmonella enterica; Salmonella enterica subsp. enterica; Salmonella enterica subsp. enterica serovar Typhimurium MNNEETFYQAMRRKGVTRRSFLKFCSLAATSLGLGAGMTPKIAWALENKPRIPVVWIHGL WP_001058312.1 hydrogenase [Salmonella enterica] Length: 372\nNumber of predicted TMHs: 2\nExp number of AAs in TMHs: 36.51738\nExp number, first 60 AAs: 19.72849\nTotal prob of N-in: 0.96170\nPOSSIBLE N-term signal sequence\ninside 1 19\nTMhelix 20 39\noutside 40 325\nTMhelix 326 348\ninside 349 372

25475 GCF_001478045.1_Salmonella_enterica_CVM_N30693-SQ_v1.0 Salmonella enterica subsp. enterica serovar Typhimurium var. 5- Proteobacteria; Gammaproteobacteria; Enterobacterales; Enterobacteriaceae; Salmonella;

Salmonella enterica; Salmonella enterica subsp. enterica; Salmonella enterica subsp. enterica serovar Typhimurium MTGDNTLITSHGINRRDFMKLCAALAATMGLSSKAAAEMAESVSNPQRPPVIWIGAQECT WP_000145429.1 hydrogenase 2 small subunit [Salmonella enterica] Length: 372\nNumber of predicted TMHs: 1\nExp number of AAs in TMHs: 24.51309\nExp number, first 60 AAs: 0.2682\nTotal prob of N-in: 0.14402\noutside 1 331\nTMhelix 332 354\ninside 355 372

25476 GCF_001272475.1_Salmonella_enterica_CVM_N51296_v1.0 Salmonella enterica subsp. enterica serovar Typhimurium var. 5- Proteobacteria; Gammaproteobacteria; Enterobacterales; Enterobacteriaceae; Salmonella; Salmonella enterica; Salmonella enterica subsp. enterica; Salmonella enterica subsp. enterica serovar Typhimurium MNNEETFYQAMRRKGVTRRSFLKFCSLAATSLGLGAGMTPKIAWALENKPRIPVVWIHGL WP_001058312.1 hydrogenase [Salmonella enterica] Length: 372\nNumber of predicted TMHs: 2\nExp number of AAs in TMHs: 36.51738\nExp number, first 60 AAs: 19.72849\nTotal prob of N-in: 0.96170\nPOSSIBLE N-term signal sequence\ninside 1 19\nTMhelix 20 39\noutside 40 325\nTMhelix 326 348\ninside 349 372

25477 GCF_001272115.1_Salmonella_enterica_CVM_N46840_v1.0 Salmonella enterica subsp. enterica serovar Typhimurium var. 5- Proteobacteria; Gammaproteobacteria; Enterobacterales; Enterobacteriaceae; Salmonella; Salmonella enterica; Salmonella enterica subsp. enterica; Salmonella enterica subsp. enterica serovar Typhimurium MNNEETFYQAMRRKGVTRRSFLKFCSLAATSLGLGAGMTPKIAWALENKPRIPVVWIHGL WP_001058312.1 hydrogenase [Salmonella enterica] Length: 372\nNumber of predicted TMHs: 2\nExp number of AAs in TMHs: 36.51738\nExp number, first 60 AAs: 19.72849\nTotal prob of N-in: 0.96170\nPOSSIBLE N-term signal sequence\ninside 1 19\nTMhelix 20 39\noutside 40 325\nTMhelix 326 348\ninside 349 372

25478 GCF_001272035.1_Salmonella_enterica_CVM_N45410_v1.0 Salmonella enterica subsp. enterica serovar Typhimurium var. 5- Proteobacteria; Gammaproteobacteria; Enterobacterales; Enterobacteriaceae; Salmonella; Salmonella enterica; Salmonella enterica subsp. enterica; Salmonella enterica subsp. enterica serovar Typhimurium MNNEETFYQAMRRKGVTRRSFLKFCSLAATSLGLGAGMTPKIAWALENKPRIPVVWIHGL WP_001058312.1 hydrogenase [Salmonella enterica] Length: 372\nNumber of predicted TMHs: 2\nExp number of AAs in TMHs: 36.51738\nExp number, first 60 AAs: 19.72849\nTotal prob of N-in: 0.96170\nPOSSIBLE N-term signal sequence\ninside 1 19\nTMhelix 20 39\noutside 40 325\nTMhelix 326 348\ninside 349 372

25479 GCF_001271715.1_Salmonella_enterica_CVM_N46828_v1.0 Salmonella enterica subsp. enterica serovar Typhimurium var. 5- Proteobacteria; Gammaproteobacteria; Enterobacterales; Enterobacteriaceae; Salmonella; Salmonella enterica; Salmonella enterica subsp. enterica; Salmonella enterica subsp. enterica serovar Typhimurium MNNEETFYQAMRRKGVTRRSFLKFCSLAATSLGLGAGMTPKIAWALENKPRIPVVWIHGL WP_001058312.1 hydrogenase [Salmonella enterica] Length: 372\nNumber of predicted TMHs: 2\nExp number of AAs in TMHs: 36.51738\nExp number, first 60 AAs: 19.72849\nTotal prob of N-in: 0.96170\nPOSSIBLE N-term signal sequence\ninside 1 19\nTMhelix 20 39\noutside 40 325\nTMhelix 326 348\ninside 349 372

25480 GCF_001271795.1_Salmonella_enterica_CVM_N46843_v1.0 Salmonella enterica subsp. enterica serovar Typhimurium var. 5- Proteobacteria; Gammaproteobacteria; Enterobacterales; Enterobacteriaceae; Salmonella; Salmonella enterica; Salmonella enterica subsp. enterica; Salmonella enterica subsp. enterica serovar Typhimurium MNNEETFYQAMRRKGVTRRSFLKFCSLAATSLGLGAGMTPKIAWALENKPRIPVVWIHGL WP_001058312.1 hydrogenase [Salmonella enterica] Length: 372\nNumber of predicted TMHs: 2\nExp number of AAs in TMHs: 36.51738\nExp number, first 60 AAs: 19.72849\nTotal prob of N-in: 0.96170\nPOSSIBLE N-term signal sequence\ninside 1 19\nTMhelix 20 39\noutside 40 325\nTMhelix 326 348\ninside 349 372

25481 GCF_001271795.1_Salmonella_enterica_CVM_N46843_v1.0 Salmonella enterica subsp. enterica serovar Typhimurium var. 5- Proteobacteria; Gammaproteobacteria; Enterobacterales; Enterobacteriaceae; Salmonella; Salmonella enterica; Salmonella enterica subsp. enterica; Salmonella enterica subsp. enterica serovar Typhimurium MTGDNTLITSHGINRRDFMKLCAALAATMGLSSKAAAEMAESVSNPQRPPVIWIGAQECT WP_000145429.1 hydrogenase 2 small subunit [Salmonella enterica] Length: 372\nNumber of predicted TMHs: 1\nExp number of AAs in TMHs: 24.51309\nExp number, first 60 AAs: 0.2682\nTotal prob of N-in: 0.14402\noutside 1 331\nTMhelix 332 354\ninside 355 372

25482 GCF_001271745.1_Salmonella_enterica_CVM_N46841_v1.0 Salmonella enterica subsp. enterica serovar Typhimurium var. 5- Proteobacteria; Gammaproteobacteria; Enterobacterales; Enterobacteriaceae; Salmonella; Salmonella enterica; Salmonella enterica subsp. enterica; Salmonella enterica subsp. enterica serovar Typhimurium MTGDNTLITSHGINRRDFMKLCAALAATMGLSSKAAAEMAESVSNPQRPPVIWIGAQECT WP_000145429.1 hydrogenase 2 small subunit [Salmonella enterica] Length: 372\nNumber of predicted TMHs: 1\nExp number of AAs in TMHs: 24.51309\nExp number, first 60 AAs: 0.2682\nTotal prob of N-in: 0.14402\noutside 1 331\nTMhelix 332 354\ninside 355 372

25483 GCF_001247205.1_Salmonella_enterica_CVM_N51300_v1.0 Salmonella enterica subsp. enterica serovar Typhimurium var. 5- Proteobacteria; Gammaproteobacteria; Enterobacterales; Enterobacteriaceae; Salmonella; Salmonella enterica; Salmonella enterica subsp. enterica; Salmonella enterica subsp. enterica serovar Typhimurium MTGDNTLITSHGINRRDFMKLCAALAATMGLSSKAAAEMAESVSNPQRPPVIWIGAQECT WP_000145429.1 hydrogenase 2 small subunit [Salmonella enterica] Length: 372\nNumber of predicted TMHs: 1\nExp number of AAs in TMHs: 24.51309\nExp number, first 60 AAs: 0.2682\nTotal prob of N-in: 0.14402\noutside 1 331\nTMhelix 332 354\ninside 355 372

25484 GCF_001246695.1_Salmonella_enterica_CVM_N51275_v1.0 Salmonella enterica subsp. enterica serovar Typhimurium var. 5- Proteobacteria; Gammaproteobacteria; Enterobacterales; Enterobacteriaceae; Salmonella; Salmonella enterica; Salmonella enterica subsp. enterica; Salmonella enterica subsp. enterica serovar Typhimurium MNNEETFYQAMRRKGVTRRSFLKFCSLAATSLGLGAGMTPKIAWALENKPRIPVVWIHGL WP_001058312.1 hydrogenase [Salmonella enterica] Length: 372\nNumber of predicted TMHs: 2\nExp number of AAs in TMHs: 36.51738\nExp number, first 60 AAs: 19.72849\nTotal prob of N-in: 0.96170\nPOSSIBLE N-term signal sequence\ninside 1 19\nTMhelix 20 39\noutside 40 325\nTMhelix 326 348\ninside 349 372

25485 GCF_001246695.1_Salmonella_enterica_CVM_N51275_v1.0 Salmonella enterica subsp. enterica serovar Typhimurium var. 5- Proteobacteria; Gammaproteobacteria; Enterobacterales; Enterobacteriaceae; Salmonella; Salmonella enterica; Salmonella enterica subsp. enterica; Salmonella enterica subsp. enterica serovar Typhimurium MTGDNTLITSHGINRRDFMKLCAALAATMGLSSKAAAEMAESVSNPQRPPVIWIGAQECT WP_000145429.1 hydrogenase 2 small subunit [Salmonella enterica] Length: 372\nNumber of predicted TMHs: 1\nExp number of AAs in TMHs: 24.51309\nExp number, first 60 AAs: 0.2682\nTotal prob of N-in: 0.14402\noutside 1 331\nTMhelix 332 354\ninside 355 372

25486 GCF_001246265.1_Salmonella_enterica_CVM_N51255_v1.0 Salmonella enterica subsp. enterica serovar Typhimurium var. 5- Proteobacteria; Gammaproteobacteria; Enterobacterales; Enterobacteriaceae; Salmonella; Salmonella enterica; Salmonella enterica subsp. enterica; Salmonella enterica subsp. enterica serovar Typhimurium MTGDNTLITSHGINRRDFMKLCAALAATMGLSSKAAAEMAESVSNPQRPPVIWIGAQECT WP_000145429.1 hydrogenase 2 small subunit [Salmonella enterica] Length: 372\nNumber of predicted TMHs: 1\nExp number of AAs in TMHs: 24.51309\nExp number, first 60 AAs: 0.2682\nTotal prob of N-in: 0.14402\noutside 1 331\nTMhelix 332 354\ninside 355 372

25487 GCF_001245415.1_Salmonella_enterica_CVM_N50425_v1.0 Salmonella enterica subsp. enterica serovar Typhimurium var. 5- Proteobacteria; Gammaproteobacteria; Enterobacterales; Enterobacteriaceae; Salmonella; Salmonella enterica; Salmonella enterica subsp. enterica; Salmonella enterica subsp. enterica serovar Typhimurium MTGDNTLITSHGINRRDFMKLCAALAATMGLSSKAAAEMAESVSNPQRPPVIWIGAQECT WP_052935412.1 hydrogenase 2 small subunit [Salmonella enterica] Length: 372\nNumber of predicted TMHs: 1\nExp number of AAs in TMHs: 24.51307\nExp number, first 60 AAs: 0.2682\nTotal prob of N-in: 0.14402\noutside 1 331\nTMhelix 332 354\ninside 355 372

25488 GCF_001245125.1_Salmonella_enterica_CVM_N48708_v1.0 Salmonella enterica subsp. enterica serovar Typhimurium var. 5- Proteobacteria; Gammaproteobacteria; Enterobacterales; Enterobacteriaceae; Salmonella; Salmonella enterica; Salmonella enterica subsp. enterica; Salmonella enterica subsp. enterica serovar Typhimurium MNNEETFYQAMRRKGVTRRSFLKFCSLAATSLGLGAGMTPKIAWALENKPRIPVVWIHGL WP_001058312.1 hydrogenase [Salmonella enterica] Length: 372\nNumber of predicted TMHs: 2\nExp number of AAs in TMHs: 36.51738\nExp number, first 60 AAs: 19.72849\nTotal prob of N-in: 0.96170\nPOSSIBLE N-term signal sequence\ninside 1 19\nTMhelix 20 39\noutside 40 325\nTMhelix 326 348\ninside 349 372

25489 GCF_001244335.1_Salmonella_enterica_CVM_N48679_v1.0 Salmonella enterica subsp. enterica serovar Typhimurium var. 5- Proteobacteria; Gammaproteobacteria; Enterobacterales; Enterobacteriaceae; Salmonella; Salmonella enterica; Salmonella enterica subsp. enterica; Salmonella enterica subsp. enterica serovar Typhimurium MTGDNTLITSHGINRRDFMKLCAALAATMGLSSKAAAEMAESVSNPQRPPVIWIGAQECT WP_000145429.1 hydrogenase 2 small subunit [Salmonella enterica] Length: 372\nNumber of predicted TMHs: 1\nExp number of AAs in TMHs: 24.51309\nExp number, first 60 AAs: 0.2682\nTotal prob of N-in: 0.14402\noutside 1 331\nTMhelix 332 354\ninside 355 372

25490 GCF_001244285.1_Salmonella_enterica_CVM_N48677_v1.0 Salmonella enterica subsp. enterica serovar Typhimurium var. 5- Proteobacteria; Gammaproteobacteria; Enterobacterales; Enterobacteriaceae; Salmonella; Salmonella enterica; Salmonella enterica subsp. enterica; Salmonella enterica subsp. enterica serovar Typhimurium MTGDNTLITSHGINRRDFMKLCAALAATMGLSSKAAAEMAESVSNPQRPPVIWIGAQECT WP_000145429.1 hydrogenase 2 small subunit [Salmonella enterica] Length: 372\nNumber of predicted TMHs: 1\nExp

number of AAs in TMHs: 24.51309\nExp number, first 60 AAs: 0.2682\nTotal prob of N-in: 0.14402\noutside 1 331\nTMhelix 332 354\ninside 355 372

25491 GCF_001244105.1_Salmonella_enterica_CVM_N47728_v1.0 Salmonella enterica subsp. enterica serovar Typhimurium var. 5- Proteobacteria; Gammaproteobacteria; Enterobacterales; Enterobacteriaceae; Salmonella; Salmonella enterica; Salmonella enterica subsp. enterica; Salmonella enterica subsp. enterica serovar Typhimurium MTGDNTLITSHGINRRDFMKLCAALAATMGLSSKAAAEMAESVSNPQRPPVIWIGAQECT WP_000145429.1 hydrogenase 2 small subunit [Salmonella enterica] Length: 372\nNumber of predicted TMHs: 1\nExp number of AAs in TMHs: 24.51309\nExp number, first 60 AAs: 0.2682\nTotal prob of N-in: 0.14402\noutside 1 331\nTMhelix 332 354\ninside 355 372

25492 GCF_001243395.1_Salmonella_enterica_CVM_N46844_v1.0 Salmonella enterica subsp. enterica serovar Typhimurium var. 5- Proteobacteria; Gammaproteobacteria; Enterobacterales; Enterobacteriaceae; Salmonella; Salmonella enterica; Salmonella enterica subsp. enterica; Salmonella enterica subsp. enterica serovar Typhimurium MNNEETFYQAMRRKGVTRRSFLKFCSLAATSLGLGAGMTPKIAWALENKPRIPVVWIHGL WP_001058312.1 hydrogenase [Salmonella enterica] Length: 372\nNumber of predicted TMHs: 2\nExp number of AAs in TMHs: 36.51738\nExp number, first 60 AAs: 19.72849\nTotal prob of N-in: 0.96170\nPOSSIBLE N-term signal sequence\ninside 1 19\nTMhelix 20 39\noutside 40 325\nTMhelix 326 348\ninside 349 372

25493 GCF_001242565.1_Salmonella_enterica_CVM_N45950_v1.0 Salmonella enterica subsp. enterica serovar Typhimurium var. 5- Proteobacteria; Gammaproteobacteria; Enterobacterales; Enterobacteriaceae; Salmonella; Salmonella enterica; Salmonella enterica subsp. enterica; Salmonella enterica subsp. enterica serovar Typhimurium MNNEETFYQAMRRKGVTRRSFLKFCSLAATSLGLGAGMTPKIAWALENKPRIPVVWIHGL WP_001058312.1 hydrogenase [Salmonella enterica] Length: 372\nNumber of predicted TMHs: 2\nExp number of AAs in TMHs: 36.51738\nExp number, first 60 AAs: 19.72849\nTotal prob of N-in: 0.96170\nPOSSIBLE N-term signal sequence\ninside 1 19\nTMhelix 20 39\noutside 40 325\nTMhelix 326 348\ninside 349 372

25494 GCF_001242395.1_Salmonella_enterica_CVM_N45938_v1.0 Salmonella enterica subsp. enterica serovar Typhimurium var. 5- Proteobacteria; Gammaproteobacteria; Enterobacterales; Enterobacteriaceae; Salmonella; Salmonella enterica; Salmonella enterica subsp. enterica; Salmonella enterica subsp. enterica serovar Typhimurium MNNEETFYQAMRRKGVTRRSFLKFCSLAATSLGLGAGMTPKIAWALENKPRIPVVWIHGL WP_001058312.1 hydrogenase [Salmonella enterica] Length: 372\nNumber of predicted TMHs: 2\nExp number of AAs in TMHs: 36.51738\nExp number, first 60 AAs: 19.72849\nTotal prob of N-in: 0.96170\nPOSSIBLE N-term signal sequence\ninside 1 19\nTMhelix 20 39\noutside 40 325\nTMhelix 326 348\ninside 349 372

25495 GCF_001241945.1_Salmonella_enterica_CVM_N45405_v1.0 Salmonella enterica subsp. enterica serovar Typhimurium var. 5- Proteobacteria; Gammaproteobacteria; Enterobacterales; Enterobacteriaceae; Salmonella; Salmonella enterica; Salmonella enterica subsp. enterica; Salmonella enterica subsp. enterica serovar Typhimurium MTGDNTLITSHGINRRDFMKLCAALAATMGLSSKAAAEMAESVSNPQRPPVIWIGAQECT WP_000145429.1 hydrogenase 2 small subunit [Salmonella enterica] Length: 372\nNumber of predicted TMHs: 1\nExp number of AAs in TMHs: 24.51309\nExp number, first 60 AAs: 0.2682\nTotal prob of N-in: 0.14402\noutside 1 331\nTMhelix 332 354\ninside 355 372

25496 GCF_001241895.1_Salmonella_enterica_CVM_N45403_v1.0 Salmonella enterica subsp. enterica serovar Typhimurium var. 5- Proteobacteria; Gammaproteobacteria; Enterobacterales; Enterobacteriaceae; Salmonella; Salmonella enterica; Salmonella enterica subsp. enterica; Salmonella enterica subsp. enterica serovar Typhimurium MTGDNTLITSHGINRRDFMKLCAALAATMGLSSKAAAEMAESVSNPQRPPVIWIGAQECT WP_000145429.1 hydrogenase 2 small subunit [Salmonella enterica] Length: 372\nNumber of predicted TMHs: 1\nExp number of AAs in TMHs: 24.51309\nExp number, first 60 AAs: 0.2682\nTotal prob of N-in: 0.14402\noutside 1 331\nTMhelix 332 354\ninside 355 372

25497 GCF_001241505.1_Salmonella_enterica_CVM_N44712_v1.0 Salmonella enterica subsp. enterica serovar Typhimurium var. 5- Proteobacteria; Gammaproteobacteria; Enterobacterales; Enterobacteriaceae; Salmonella; Salmonella enterica; Salmonella enterica subsp. enterica; Salmonella enterica subsp. enterica serovar Typhimurium MTGDNTLITSHGINRRDFMKLCAALAATMGLSSKAAAEMAESVSNPQRPPVIWIGAQECT WP_000145429.1 hydrogenase 2 small subunit [Salmonella enterica] Length: 372\nNumber of predicted TMHs: 1\nExp number of AAs in TMHs: 24.51309\nExp number, first 60 AAs: 0.2682\nTotal prob of N-in: 0.14402\noutside 1 331\nTMhelix 332 354\ninside 355 372

25498 GCF_001241425.1_Salmonella_enterica_CVM_N44709_v1.0 Salmonella enterica subsp. enterica serovar Typhimurium var. 5- Proteobacteria; Gammaproteobacteria; Enterobacterales; Enterobacteriaceae; Salmonella;

Salmonella enterica; Salmonella enterica subsp. enterica; Salmonella enterica subsp. enterica serovar Typhimurium
MTGDNTLITSHGINRRDFMKLCAALAATMGLSSKAAAEMAESVSNPQRPPVIWIGAQECT WP_000145429.1
hydrogenase 2 small subunit [Salmonella enterica] Length: 372\nNumber of predicted TMHs: 1\nExp number of AAs in TMHs: 24.51309\nExp number, first 60 AAs: 0.2682\nTotal prob of N-in: 0.14402\noutside 1 331\nTMhelix 332 354\ninside 355 372

25499 GCF_001240805.1_Salmonella_enterica_CVM_N43824_v1.0 Salmonella enterica subsp. enterica serovar Typhimurium var. 5- Proteobacteria; Gammaproteobacteria; Enterobacterales; Enterobacteriaceae; Salmonella; Salmonella enterica; Salmonella enterica subsp. enterica; Salmonella enterica subsp. enterica serovar Typhimurium
MNNEETFYQAMRRKGVTRRSFLKFCSLAATSLGLGAGMTPKIAWALENKPRIPVVWIHGL WP_001058312.1
hydrogenase [Salmonella enterica] Length: 372\nNumber of predicted TMHs: 2\nExp number of AAs in TMHs: 36.51738\nExp number, first 60 AAs: 19.72849\nTotal prob of N-in: 0.96170\nPOSSIBLE N-term signal sequence\ninside 1 19\nTMhelix 20 39\noutside 40 325\nTMhelix 326 348\ninside 349 372

25500 GCF_002031265.1_ASM203126v1 Salmonella enterica subsp. enterica serovar Typhimurium var. 5- Proteobacteria; Gammaproteobacteria; Enterobacterales; Enterobacteriaceae; Salmonella; Salmonella enterica; Salmonella enterica subsp. enterica; Salmonella enterica subsp. enterica serovar Typhimurium
MTGDNTLITSHGINRRDFMKLCAALAATMGLSSKAAAEMAESVSNPQRPPVIWIGAQECT WP_000145429.1
hydrogenase 2 small subunit [Salmonella enterica] Length: 372\nNumber of predicted TMHs: 1\nExp number of AAs in TMHs: 24.51309\nExp number, first 60 AAs: 0.2682\nTotal prob of N-in: 0.14402\noutside 1 331\nTMhelix 332 354\ninside 355 372

25501 GCF_002032165.1_ASM203216v1 Salmonella enterica subsp. enterica serovar Typhimurium var. 5- Proteobacteria; Gammaproteobacteria; Enterobacterales; Enterobacteriaceae; Salmonella; Salmonella enterica; Salmonella enterica subsp. enterica; Salmonella enterica subsp. enterica serovar Typhimurium
MTGDNTLITSHGINRRDFMKLCAALAATMGLSSKAAAEMAESVSNPQRPPVIWIGAQECT WP_000145429.1
hydrogenase 2 small subunit [Salmonella enterica] Length: 372\nNumber of predicted TMHs: 1\nExp number of AAs in TMHs: 24.51309\nExp number, first 60 AAs: 0.2682\nTotal prob of N-in: 0.14402\noutside 1 331\nTMhelix 332 354\ninside 355 372

25502 GCF_002032165.1_ASM203216v1 Salmonella enterica subsp. enterica serovar Typhimurium var. 5- Proteobacteria; Gammaproteobacteria; Enterobacterales; Enterobacteriaceae; Salmonella; Salmonella enterica; Salmonella enterica subsp. enterica; Salmonella enterica subsp. enterica serovar Typhimurium
MNNEETFYQAMRRKGVTRRSFLKFCSLAATSLGLGAGMTPKIAWALENKPRIPVVWIHGL WP_001058312.1
hydrogenase [Salmonella enterica] Length: 372\nNumber of predicted TMHs: 2\nExp number of AAs in TMHs: 36.51738\nExp number, first 60 AAs: 19.72849\nTotal prob of N-in: 0.96170\nPOSSIBLE N-term signal sequence\ninside 1 19\nTMhelix 20 39\noutside 40 325\nTMhelix 326 348\ninside 349 372

25503 GCF_002033165.1_ASM203316v1 Salmonella enterica subsp. enterica serovar Typhimurium var. 5- Proteobacteria; Gammaproteobacteria; Enterobacterales; Enterobacteriaceae; Salmonella; Salmonella enterica; Salmonella enterica subsp. enterica; Salmonella enterica subsp. enterica serovar Typhimurium
MTGDNTLITSHGINRRDFMKLCAALAATMGLSSKAAAEMAESVSNPQRPPVIWIGAQECT WP_000145429.1
hydrogenase 2 small subunit [Salmonella enterica] Length: 372\nNumber of predicted TMHs: 1\nExp number of AAs in TMHs: 24.51309\nExp number, first 60 AAs: 0.2682\nTotal prob of N-in: 0.14402\noutside 1 331\nTMhelix 332 354\ninside 355 372

25504 GCF_001481235.1_Salmonella_enterica_CVM_N42518-SQ_v1.0 Salmonella enterica subsp. enterica serovar Typhimurium var. 5- Proteobacteria; Gammaproteobacteria; Enterobacterales; Enterobacteriaceae; Salmonella; Salmonella enterica; Salmonella enterica subsp. enterica; Salmonella enterica subsp. enterica serovar Typhimurium
MNNEETFYQAMRRKGVTRRSFLKFCSLAATSLGLGAGMTPKIAWALENKPRIPVVWIHGL WP_001058312.1
hydrogenase [Salmonella enterica] Length: 372\nNumber of predicted TMHs: 2\nExp number of AAs in TMHs: 36.51738\nExp number, first 60 AAs: 19.72849\nTotal prob of N-in: 0.96170\nPOSSIBLE N-term signal sequence\ninside 1 19\nTMhelix 20 39\noutside 40 325\nTMhelix 326 348\ninside 349 372

25505 GCF_001481235.1_Salmonella_enterica_CVM_N42518-SQ_v1.0 Salmonella enterica subsp. enterica serovar Typhimurium var. 5- Proteobacteria; Gammaproteobacteria; Enterobacterales; Enterobacteriaceae; Salmonella; Salmonella enterica; Salmonella enterica subsp. enterica; Salmonella enterica subsp. enterica serovar Typhimurium
MTGDNTLITSHGINRRDFMKLCAALAATMGLSSKAAAEMAESVSNPQRPPVIWIGAQECT WP_000145429.1
hydrogenase 2 small subunit [Salmonella enterica] Length: 372\nNumber of predicted TMHs: 1\nExp number of AAs in TMHs: 24.51309\nExp number, first 60 AAs: 0.2682\nTotal prob of N-in: 0.14402\noutside 1 331\nTMhelix 332 354\ninside 355 372

25506 GCF_001479395.1_Salmonella_enterica_CVM_N32056-R_v1.0 Salmonella enterica subsp. enterica serovar Typhimurium var. 5- Proteobacteria; Gammaproteobacteria; Enterobacterales; Enterobacteriaceae; Salmonella; Salmonella enterica; Salmonella enterica subsp. enterica; Salmonella enterica subsp. enterica serovar Typhimurium MNNEETFYQAMRRKGVTRRSFLKFCSLAATSLGLGAGMTPKIAWALENKPRIPVVWIHGL WP_001058312.1 hydrogenase [Salmonella enterica] Length: 372\nNumber of predicted TMHs: 2\nExp number of AAs in TMHs: 36.51738\nExp number, first 60 AAs: 19.72849\nTotal prob of N-in: 0.96170\nPOSSIBLE N-term signal sequence\ninside 1 19\nTMhelix 20 39\noutside 40 325\nTMhelix 326 348\ninside 349 372

25507 GCF_001241975.1_Salmonella_enterica_CVM_N45408_v1.0 Salmonella enterica subsp. enterica serovar Typhimurium var. 5- Proteobacteria; Gammaproteobacteria; Enterobacterales; Enterobacteriaceae; Salmonella; Salmonella enterica; Salmonella enterica subsp. enterica; Salmonella enterica subsp. enterica serovar Typhimurium MNNEETFYQAMRRKGVTRRSFLKFCSLAATSLGLGAGMTPKIAWALENKPRIPVVWIHGL WP_001058312.1 hydrogenase [Salmonella enterica] Length: 372\nNumber of predicted TMHs: 2\nExp number of AAs in TMHs: 36.51738\nExp number, first 60 AAs: 19.72849\nTotal prob of N-in: 0.96170\nPOSSIBLE N-term signal sequence\ninside 1 19\nTMhelix 20 39\noutside 40 325\nTMhelix 326 348\ninside 349 372

25508 GCF_001478285.1_Salmonella_enterica_CVM_N37950-SQ_v1.0 Salmonella enterica subsp. enterica serovar Typhimurium var. 5- Proteobacteria; Gammaproteobacteria; Enterobacterales; Enterobacteriaceae; Salmonella; Salmonella enterica; Salmonella enterica subsp. enterica; Salmonella enterica subsp. enterica serovar Typhimurium MNNEETFYQAMRRKGVTRRSFLKFCSLAATSLGLGAGMTPKIAWALENKPRIPVVWIHGL WP_001058312.1 hydrogenase [Salmonella enterica] Length: 372\nNumber of predicted TMHs: 2\nExp number of AAs in TMHs: 36.51738\nExp number, first 60 AAs: 19.72849\nTotal prob of N-in: 0.96170\nPOSSIBLE N-term signal sequence\ninside 1 19\nTMhelix 20 39\noutside 40 325\nTMhelix 326 348\ninside 349 372

25509 GCF_001478285.1_Salmonella_enterica_CVM_N37950-SQ_v1.0 Salmonella enterica subsp. enterica serovar Typhimurium var. 5- Proteobacteria; Gammaproteobacteria; Enterobacterales; Enterobacteriaceae; Salmonella; Salmonella enterica; Salmonella enterica subsp. enterica; Salmonella enterica subsp. enterica serovar Typhimurium MTGDNTLITSHGINRRDFMKLCAALAATMGLSSKAAAEMAESVSNPQRPPVIWIGAQECT WP_000145429.1 hydrogenase 2 small subunit [Salmonella enterica] Length: 372\nNumber of predicted TMHs: 1\nExp number of AAs in TMHs: 24.51309\nExp number, first 60 AAs: 0.2682\nTotal prob of N-in: 0.14402\noutside 1 331\nTMhelix 332 354\ninside 355 372

25510 GCF_001479365.1_Salmonella_enterica_CVM_N32051-SQ_v1.0 Salmonella enterica subsp. enterica serovar Typhimurium var. 5- Proteobacteria; Gammaproteobacteria; Enterobacterales; Enterobacteriaceae; Salmonella; Salmonella enterica; Salmonella enterica subsp. enterica; Salmonella enterica subsp. enterica serovar Typhimurium MTGDNTLITSHGINRRDFMKLCAALAATMGLSSKAAAEMAESVSNPQRPPVIWIGAQECT WP_000145429.1 hydrogenase 2 small subunit [Salmonella enterica] Length: 372\nNumber of predicted TMHs: 1\nExp number of AAs in TMHs: 24.51309\nExp number, first 60 AAs: 0.2682\nTotal prob of N-in: 0.14402\noutside 1 331\nTMhelix 332 354\ninside 355 372

25511 GCF_001479365.1_Salmonella_enterica_CVM_N32051-SQ_v1.0 Salmonella enterica subsp. enterica serovar Typhimurium var. 5- Proteobacteria; Gammaproteobacteria; Enterobacterales; Enterobacteriaceae; Salmonella; Salmonella enterica; Salmonella enterica subsp. enterica; Salmonella enterica subsp. enterica serovar Typhimurium MNNEETFYQAMRRKGVTRRSFLKFCSLAATSLGLGAGMTPKIAWALENKPRIPVVWIHGL WP_001058312.1 hydrogenase [Salmonella enterica] Length: 372\nNumber of predicted TMHs: 2\nExp number of AAs in TMHs: 36.51738\nExp number, first 60 AAs: 19.72849\nTotal prob of N-in: 0.96170\nPOSSIBLE N-term signal sequence\ninside 1 19\nTMhelix 20 39\noutside 40 325\nTMhelix 326 348\ninside 349 372

25512 GCF_001479695.1_Salmonella_enterica_CVM_N38234-SQ_v1.0 Salmonella enterica subsp. enterica serovar Typhimurium var. 5- Proteobacteria; Gammaproteobacteria; Enterobacterales; Enterobacteriaceae; Salmonella; Salmonella enterica; Salmonella enterica subsp. enterica; Salmonella enterica subsp. enterica serovar Typhimurium MTGDNTLITSHGINRRDFMKLCAALAATMGLSSKAAAEMAESVSNPQRPPVIWIGAQECT WP_000145429.1 hydrogenase 2 small subunit [Salmonella enterica] Length: 372\nNumber of predicted TMHs: 1\nExp number of AAs in TMHs: 24.51309\nExp number, first 60 AAs: 0.2682\nTotal prob of N-in: 0.14402\noutside 1 331\nTMhelix 332 354\ninside 355 372

25513 GCF_001479695.1_Salmonella_enterica_CVM_N38234-SQ_v1.0 Salmonella enterica subsp. enterica serovar Typhimurium var. 5- Proteobacteria; Gammaproteobacteria; Enterobacterales; Enterobacteriaceae; Salmonella; Salmonella enterica; Salmonella enterica subsp. enterica; Salmonella enterica subsp. enterica serovar Typhimurium MNNEETFYQAMRRKGVTRRSFLKFCSLAATSLGLGAGMTPKIAWALENKPRIPVVWIHGL WP_001058312.1 hydrogenase [Salmonella enterica] Length: 372\nNumber of predicted TMHs: 2\nExp number of AAs in

TMHs: 36.51738\nExp number, first 60 AAs: 19.72849\nTotal prob of N-in: 0.96170\nPOSSIBLE N-term signal sequence\ninside 1 19\nTMhelix 20 39\noutside 40 325\nTMhelix 326 348\ninside 349 372

25514 GCF_001480225.1_Salmonella_enterica_CVM_N40384-R_v1.0 Salmonella enterica subsp. enterica serovar Typhimurium var. 5- Proteobacteria; Gammaproteobacteria; Enterobacterales; Enterobacteriaceae; Salmonella; Salmonella enterica; Salmonella enterica subsp. enterica; Salmonella enterica subsp. enterica serovar Typhimurium MNNEETFYQAMRRKGVTRRSFLKFCSLAATSLGLGAGMTPKIAWALENKPRIPVVWIHGL WP_058802265.1 hydrogenase [Salmonella enterica] Length: 372\nNumber of predicted TMHs: 2\nExp number of AAs in TMHs: 36.51188\nExp number, first 60 AAs: 19.73066\nTotal prob of N-in: 0.96172\nPOSSIBLE N-term signal sequence\ninside 1 19\nTMhelix 20 39\noutside 40 325\nTMhelix 326 348\ninside 349 372

25515 GCF_001480795.1_Salmonella_enterica_CVM_N38229-SQ_v1.0 Salmonella enterica subsp. enterica serovar Typhimurium var. 5- Proteobacteria; Gammaproteobacteria; Enterobacterales; Enterobacteriaceae; Salmonella; Salmonella enterica; Salmonella enterica subsp. enterica; Salmonella enterica subsp. enterica serovar Typhimurium MTGDNTLITSHGINRRDFMKLCAALAATMGLSSKAAAEMAESVSNPQRPPVIWIGAQECT WP_000145429.1 hydrogenase 2 small subunit [Salmonella enterica] Length: 372\nNumber of predicted TMHs: 1\nExp number of AAs in TMHs: 24.51309\nExp number, first 60 AAs: 0.2682\nTotal prob of N-in: 0.14402\noutside 1 331\nTMhelix 332 354\ninside 355 372

25516 GCF_001480795.1_Salmonella_enterica_CVM_N38229-SQ_v1.0 Salmonella enterica subsp. enterica serovar Typhimurium var. 5- Proteobacteria; Gammaproteobacteria; Enterobacterales; Enterobacteriaceae; Salmonella; Salmonella enterica; Salmonella enterica subsp. enterica; Salmonella enterica subsp. enterica serovar Typhimurium MNNEETFYQAMRRKGVTRRSFLKFCSLAATSLGLGAGMTPKIAWALENKPRIPVVWIHGL WP_001058312.1 hydrogenase [Salmonella enterica] Length: 372\nNumber of predicted TMHs: 2\nExp number of AAs in TMHs: 36.51738\nExp number, first 60 AAs: 19.72849\nTotal prob of N-in: 0.96170\nPOSSIBLE N-term signal sequence\ninside 1 19\nTMhelix 20 39\noutside 40 325\nTMhelix 326 348\ninside 349 372

25517 GCF_001271825.1_Salmonella_enterica_CVM_N46861_v1.0 Salmonella enterica subsp. enterica serovar Typhimurium var. 5- Proteobacteria; Gammaproteobacteria; Enterobacterales; Enterobacteriaceae; Salmonella; Salmonella enterica; Salmonella enterica subsp. enterica; Salmonella enterica subsp. enterica serovar Typhimurium MNNEETFYQAMRRKGVTRRSFLKFCSLAATSLGLGAGMTPKIAWALENKPRIPVVWIHGL WP_001058312.1 hydrogenase [Salmonella enterica] Length: 372\nNumber of predicted TMHs: 2\nExp number of AAs in TMHs: 36.51738\nExp number, first 60 AAs: 19.72849\nTotal prob of N-in: 0.96170\nPOSSIBLE N-term signal sequence\ninside 1 19\nTMhelix 20 39\noutside 40 325\nTMhelix 326 348\ninside 349 372

25518 GCF_001271825.1_Salmonella_enterica_CVM_N46861_v1.0 Salmonella enterica subsp. enterica serovar Typhimurium var. 5- Proteobacteria; Gammaproteobacteria; Enterobacterales; Enterobacteriaceae; Salmonella; Salmonella enterica; Salmonella enterica subsp. enterica; Salmonella enterica subsp. enterica serovar Typhimurium MTGDNTLITSHGINRRDFMKLCAALAATMGLSSKAAAEMAESVSNPQRPPVIWIGAQECT WP_000145429.1 hydrogenase 2 small subunit [Salmonella enterica] Length: 372\nNumber of predicted TMHs: 1\nExp number of AAs in TMHs: 24.51309\nExp number, first 60 AAs: 0.2682\nTotal prob of N-in: 0.14402\noutside 1 331\nTMhelix 332 354\ninside 355 372

25519 GCF_001247205.1_Salmonella_enterica_CVM_N51300_v1.0 Salmonella enterica subsp. enterica serovar Typhimurium var. 5- Proteobacteria; Gammaproteobacteria; Enterobacterales; Enterobacteriaceae; Salmonella; Salmonella enterica; Salmonella enterica subsp. enterica; Salmonella enterica subsp. enterica serovar Typhimurium MNNEETFYQAMRRKGVTRRSFLKFCSLAATSLGLGAGMTPKIAWALENKPRIPVVWIHGL WP_001058312.1 hydrogenase [Salmonella enterica] Length: 372\nNumber of predicted TMHs: 2\nExp number of AAs in TMHs: 36.51738\nExp number, first 60 AAs: 19.72849\nTotal prob of N-in: 0.96170\nPOSSIBLE N-term signal sequence\ninside 1 19\nTMhelix 20 39\noutside 40 325\nTMhelix 326 348\ninside 349 372

25520 GCF_001247265.1_Salmonella_enterica_CVM_N51301_v1.0 Salmonella enterica subsp. enterica serovar Typhimurium var. 5- Proteobacteria; Gammaproteobacteria; Enterobacterales; Enterobacteriaceae; Salmonella; Salmonella enterica; Salmonella enterica subsp. enterica; Salmonella enterica subsp. enterica serovar Typhimurium MTGDNTLITSHGINRRDFMKLCAALAATMGLSSKAAAEMAESVSNPQRPPVIWIGAQECT WP_000145429.1 hydrogenase 2 small subunit [Salmonella enterica] Length: 372\nNumber of predicted TMHs: 1\nExp number of AAs in TMHs: 24.51309\nExp number, first 60 AAs: 0.2682\nTotal prob of N-in: 0.14402\noutside 1 331\nTMhelix 332 354\ninside 355 372

25521 GCF_001246265.1_Salmonella_enterica_CVM_N51255_v1.0 Salmonella enterica subsp. enterica serovar Typhimurium var. 5- Proteobacteria; Gammaproteobacteria; Enterobacterales; Enterobacteriaceae; Salmonella;

Salmonella enterica; Salmonella enterica subsp. enterica; Salmonella enterica subsp. enterica serovar Typhimurium
MNNEETFYQAMRRKGVTRRSFLKFCSLAATSLGLGAGMTPKIAWALENKPRIPVVWIHGL WP_001058312.1
hydrogenase [Salmonella enterica] Length: 372\nNumber of predicted TMHs: 2\nExp number of AAs in TMHs: 36.51738\nExp number, first 60 AAs: 19.72849\nTotal prob of N-in: 0.96170\nPOSSIBLE N-term signal sequence\ninside 1 19\nTMhelix 20 39\noutside 40 325\nTMhelix 326 348\ninside 349 372

25522 GCF_001245495.1_Salmonella_enterica_CVM_N50430_v1.0 Salmonella enterica subsp. enterica serovar Typhimurium var. 5- Proteobacteria; Gammaproteobacteria; Enterobacterales; Enterobacteriaceae; Salmonella; Salmonella enterica; Salmonella enterica subsp. enterica; Salmonella enterica subsp. enterica serovar Typhimurium
MNNEETFYQAMRRKGVTRRSFLKFCSLAATSLGLGAGMTPKIAWALENKPRIPVVWIHGL WP_001058312.1
hydrogenase [Salmonella enterica] Length: 372\nNumber of predicted TMHs: 2\nExp number of AAs in TMHs: 36.51738\nExp number, first 60 AAs: 19.72849\nTotal prob of N-in: 0.96170\nPOSSIBLE N-term signal sequence\ninside 1 19\nTMhelix 20 39\noutside 40 325\nTMhelix 326 348\ninside 349 372

25523 GCF_001245415.1_Salmonella_enterica_CVM_N50425_v1.0 Salmonella enterica subsp. enterica serovar Typhimurium var. 5- Proteobacteria; Gammaproteobacteria; Enterobacterales; Enterobacteriaceae; Salmonella; Salmonella enterica; Salmonella enterica subsp. enterica; Salmonella enterica subsp. enterica serovar Typhimurium
MNNEETFYQAMRRKGVTRRSFLKFCSLAATSLGLGAGMTPKIAWALENKPRIPVVWIHGL WP_001058312.1
hydrogenase [Salmonella enterica] Length: 372\nNumber of predicted TMHs: 2\nExp number of AAs in TMHs: 36.51738\nExp number, first 60 AAs: 19.72849\nTotal prob of N-in: 0.96170\nPOSSIBLE N-term signal sequence\ninside 1 19\nTMhelix 20 39\noutside 40 325\nTMhelix 326 348\ninside 349 372

25524 GCF_001245125.1_Salmonella_enterica_CVM_N48708_v1.0 Salmonella enterica subsp. enterica serovar Typhimurium var. 5- Proteobacteria; Gammaproteobacteria; Enterobacterales; Enterobacteriaceae; Salmonella; Salmonella enterica; Salmonella enterica subsp. enterica; Salmonella enterica subsp. enterica serovar Typhimurium
MTGDNTLITSHGINRRDFMKLCAALAATMGLSSKAAAEMAESVSNPQRPPVIWIGAQECT WP_000145429.1
hydrogenase 2 small subunit [Salmonella enterica] Length: 372\nNumber of predicted TMHs: 1\nExp number of AAs in TMHs: 24.51309\nExp number, first 60 AAs: 0.2682\nTotal prob of N-in: 0.14402\noutside 1 331\nTMhelix 332 354\ninside 355 372

25525 GCF_001244335.1_Salmonella_enterica_CVM_N48679_v1.0 Salmonella enterica subsp. enterica serovar Typhimurium var. 5- Proteobacteria; Gammaproteobacteria; Enterobacterales; Enterobacteriaceae; Salmonella; Salmonella enterica; Salmonella enterica subsp. enterica; Salmonella enterica subsp. enterica serovar Typhimurium
MNNEETFYQAMRRKGVTRRSFLKFCSLAATSLGLGAGMTPKIAWALENKPRIPVVWIHGL WP_001058312.1
hydrogenase [Salmonella enterica] Length: 372\nNumber of predicted TMHs: 2\nExp number of AAs in TMHs: 36.51738\nExp number, first 60 AAs: 19.72849\nTotal prob of N-in: 0.96170\nPOSSIBLE N-term signal sequence\ninside 1 19\nTMhelix 20 39\noutside 40 325\nTMhelix 326 348\ninside 349 372

25526 GCF_001244695.1_Salmonella_enterica_CVM_N48690_v1.0 Salmonella enterica subsp. enterica serovar Typhimurium var. 5- Proteobacteria; Gammaproteobacteria; Enterobacterales; Enterobacteriaceae; Salmonella; Salmonella enterica; Salmonella enterica subsp. enterica; Salmonella enterica subsp. enterica serovar Typhimurium
MNNEETFYQAMRRKGVTRRSFLKFCSLAATSLGLGAGMTPKIAWALENKPRIPVVWIHGL WP_001058312.1
hydrogenase [Salmonella enterica] Length: 372\nNumber of predicted TMHs: 2\nExp number of AAs in TMHs: 36.51738\nExp number, first 60 AAs: 19.72849\nTotal prob of N-in: 0.96170\nPOSSIBLE N-term signal sequence\ninside 1 19\nTMhelix 20 39\noutside 40 325\nTMhelix 326 348\ninside 349 372

25527 GCF_001244695.1_Salmonella_enterica_CVM_N48690_v1.0 Salmonella enterica subsp. enterica serovar Typhimurium var. 5- Proteobacteria; Gammaproteobacteria; Enterobacterales; Enterobacteriaceae; Salmonella; Salmonella enterica; Salmonella enterica subsp. enterica; Salmonella enterica subsp. enterica serovar Typhimurium
MTGDNTLITSHGINRRDFMKLCAALAATMGLSSKAAAEMAESVSNPQRPPVIWIGAQECT WP_000145429.1
hydrogenase 2 small subunit [Salmonella enterica] Length: 372\nNumber of predicted TMHs: 1\nExp number of AAs in TMHs: 24.51309\nExp number, first 60 AAs: 0.2682\nTotal prob of N-in: 0.14402\noutside 1 331\nTMhelix 332 354\ninside 355 372

25528 GCF_001244285.1_Salmonella_enterica_CVM_N48677_v1.0 Salmonella enterica subsp. enterica serovar Typhimurium var. 5- Proteobacteria; Gammaproteobacteria; Enterobacterales; Enterobacteriaceae; Salmonella; Salmonella enterica; Salmonella enterica subsp. enterica; Salmonella enterica subsp. enterica serovar Typhimurium
MNNEETFYQAMRRKGVTRRSFLKFCSLAATSLGLGAGMTPKIAWALENKPRIPVVWIHGL WP_001058312.1
hydrogenase [Salmonella enterica] Length: 372\nNumber of predicted TMHs: 2\nExp number of AAs in TMHs: 36.51738\nExp number, first 60 AAs: 19.72849\nTotal prob of N-in: 0.96170\nPOSSIBLE N-term signal sequence\ninside 1 19\nTMhelix 20 39\noutside 40 325\nTMhelix 326 348\ninside 349 372

25529 GCF_001243395.1_Salmonella_enterica_CVM_N46844_v1.0 Salmonella enterica subsp. enterica serovar Typhimurium var. 5- Proteobacteria; Gammaproteobacteria; Enterobacterales; Enterobacteriaceae; Salmonella; Salmonella enterica; Salmonella enterica subsp. enterica; Salmonella enterica subsp. enterica serovar Typhimurium MTGDNTLITSHGINRRDFMKLCAALAATMGLSSKAAAEMAESVSNPQRPPVIWIGAQECT WP_000145429.1 hydrogenase 2 small subunit [Salmonella enterica] Length: 372\nNumber of predicted TMHs: 1\nExp number of AAs in TMHs: 24.51309\nExp number, first 60 AAs: 0.2682\nTotal prob of N-in: 0.14402\noutside 1 331\nTMhelix 332 354\ninside 355 372

25530 GCF_001243105.1_Salmonella_enterica_CVM_N46825_v1.0 Salmonella enterica subsp. enterica serovar Typhimurium var. 5- Proteobacteria; Gammaproteobacteria; Enterobacterales; Enterobacteriaceae; Salmonella; Salmonella enterica; Salmonella enterica subsp. enterica; Salmonella enterica subsp. enterica serovar Typhimurium MNNEETFYQAMRRKGVTRRSFLKFCSLAATSLGLGAGMTPKIAWALENKPRIPVVWIHGL WP_001058312.1 hydrogenase [Salmonella enterica] Length: 372\nNumber of predicted TMHs: 2\nExp number of AAs in TMHs: 36.51738\nExp number, first 60 AAs: 19.72849\nTotal prob of N-in: 0.96170\nPOSSIBLE N-term signal sequence\ninside 1 19\nTMhelix 20 39\noutside 40 325\nTMhelix 326 348\ninside 349 372

25531 GCF_001478045.1_Salmonella_enterica_CVM_N30693-SQ_v1.0 Salmonella enterica subsp. enterica serovar Typhimurium var. 5- Proteobacteria; Gammaproteobacteria; Enterobacterales; Enterobacteriaceae; Salmonella; Salmonella enterica; Salmonella enterica subsp. enterica; Salmonella enterica subsp. enterica serovar Typhimurium MNNEETFYQAMRRKGVTRRSFLKFCSLAATSLGLGAGMTPKIAWALENKPRIPVVWIHGL WP_001058312.1 hydrogenase [Salmonella enterica] Length: 372\nNumber of predicted TMHs: 2\nExp number of AAs in TMHs: 36.51738\nExp number, first 60 AAs: 19.72849\nTotal prob of N-in: 0.96170\nPOSSIBLE N-term signal sequence\ninside 1 19\nTMhelix 20 39\noutside 40 325\nTMhelix 326 348\ninside 349 372

25532 GCF_001241895.1_Salmonella_enterica_CVM_N45403_v1.0 Salmonella enterica subsp. enterica serovar Typhimurium var. 5- Proteobacteria; Gammaproteobacteria; Enterobacterales; Enterobacteriaceae; Salmonella; Salmonella enterica; Salmonella enterica subsp. enterica; Salmonella enterica subsp. enterica serovar Typhimurium MNNEETFYQAMRRKGVTRRSFLKFCSLAATSLGLGAGMTPKIAWALENKPRIPVVWIHGL WP_001058312.1 hydrogenase [Salmonella enterica] Length: 372\nNumber of predicted TMHs: 2\nExp number of AAs in TMHs: 36.51738\nExp number, first 60 AAs: 19.72849\nTotal prob of N-in: 0.96170\nPOSSIBLE N-term signal sequence\ninside 1 19\nTMhelix 20 39\noutside 40 325\nTMhelix 326 348\ninside 349 372

25533 GCF_001241945.1_Salmonella_enterica_CVM_N45405_v1.0 Salmonella enterica subsp. enterica serovar Typhimurium var. 5- Proteobacteria; Gammaproteobacteria; Enterobacterales; Enterobacteriaceae; Salmonella; Salmonella enterica; Salmonella enterica subsp. enterica; Salmonella enterica subsp. enterica serovar Typhimurium MNNEETFYQAMRRKGVTRRSFLKFCSLAATSLGLGAGMTPKIAWALENKPRIPVVWIHGL WP_001058312.1 hydrogenase [Salmonella enterica] Length: 372\nNumber of predicted TMHs: 2\nExp number of AAs in TMHs: 36.51738\nExp number, first 60 AAs: 19.72849\nTotal prob of N-in: 0.96170\nPOSSIBLE N-term signal sequence\ninside 1 19\nTMhelix 20 39\noutside 40 325\nTMhelix 326 348\ninside 349 372

25534 GCF_001241425.1_Salmonella_enterica_CVM_N44709_v1.0 Salmonella enterica subsp. enterica serovar Typhimurium var. 5- Proteobacteria; Gammaproteobacteria; Enterobacterales; Enterobacteriaceae; Salmonella; Salmonella enterica; Salmonella enterica subsp. enterica; Salmonella enterica subsp. enterica serovar Typhimurium MNNEETFYQAMRRKGVTRRSFLKFCSLAATSLGLGAGMTPKIAWALENKPRIPVVWIHGL WP_001058312.1 hydrogenase [Salmonella enterica] Length: 372\nNumber of predicted TMHs: 2\nExp number of AAs in TMHs: 36.51738\nExp number, first 60 AAs: 19.72849\nTotal prob of N-in: 0.96170\nPOSSIBLE N-term signal sequence\ninside 1 19\nTMhelix 20 39\noutside 40 325\nTMhelix 326 348\ninside 349 372

25535 GCF_001241075.1_Salmonella_enterica_CVM_N44692_v1.0 Salmonella enterica subsp. enterica serovar Typhimurium var. 5- Proteobacteria; Gammaproteobacteria; Enterobacterales; Enterobacteriaceae; Salmonella; Salmonella enterica; Salmonella enterica subsp. enterica; Salmonella enterica subsp. enterica serovar Typhimurium MNNEETFYQAMRRKGVTRRSFLKFCSLAATSLGLGAGMTPKIAWALENKPRIPVVWIHGL WP_001058312.1 hydrogenase [Salmonella enterica] Length: 372\nNumber of predicted TMHs: 2\nExp number of AAs in TMHs: 36.51738\nExp number, first 60 AAs: 19.72849\nTotal prob of N-in: 0.96170\nPOSSIBLE N-term signal sequence\ninside 1 19\nTMhelix 20 39\noutside 40 325\nTMhelix 326 348\ninside 349 372

25536 GCF_001240805.1_Salmonella_enterica_CVM_N43824_v1.0 Salmonella enterica subsp. enterica serovar Typhimurium var. 5- Proteobacteria; Gammaproteobacteria; Enterobacterales; Enterobacteriaceae; Salmonella; Salmonella enterica; Salmonella enterica subsp. enterica; Salmonella enterica subsp. enterica serovar Typhimurium MTGDNTLITSHGINRRDFMKLCAALAATMGLSSKAAAEMAESVSNPQRPPVIWIGAQECT WP_000145429.1 hydrogenase 2 small subunit [Salmonella enterica] Length: 372\nNumber of predicted TMHs: 1\nExp

number of AAs in TMHs: 24.51309\nExp number, first 60 AAs: 0.2682\nTotal prob of N-in: 0.14402\noutside 1 331\nTMhelix 332 354\ninside 355 372

25537 GCF_002030225.1_ASM203022v1 Salmonella enterica subsp. enterica serovar Typhimurium var. 5- Proteobacteria; Gammaproteobacteria; Enterobacterales; Enterobacteriaceae; Salmonella; Salmonella enterica; Salmonella enterica subsp. enterica; Salmonella enterica subsp. enterica serovar Typhimurium MNNEETFYQAMRRKGVTRRSFLKFCSLAATSLGLGAGMTPKIAWALENKPRIPVVWIHGL WP_001058312.1 hydrogenase [Salmonella enterica] Length: 372\nNumber of predicted TMHs: 2\nExp number of AAs in TMHs: 36.51738\nExp number, first 60 AAs: 19.72849\nTotal prob of N-in: 0.96170\nPOSSIBLE N-term signal sequence\ninside 1 19\nTMhelix 20 39\noutside 40 325\nTMhelix 326 348\ninside 349 372

25538 GCF_002033165.1_ASM203316v1 Salmonella enterica subsp. enterica serovar Typhimurium var. 5- Proteobacteria; Gammaproteobacteria; Enterobacterales; Enterobacteriaceae; Salmonella; Salmonella enterica; Salmonella enterica subsp. enterica; Salmonella enterica subsp. enterica serovar Typhimurium MNNEETFYQAMRRKGVTRRSFLKFCSLAATSLGLGAGMTPKIAWALENKPRIPVVWIHGL WP_001058312.1 hydrogenase [Salmonella enterica] Length: 372\nNumber of predicted TMHs: 2\nExp number of AAs in TMHs: 36.51738\nExp number, first 60 AAs: 19.72849\nTotal prob of N-in: 0.96170\nPOSSIBLE N-term signal sequence\ninside 1 19\nTMhelix 20 39\noutside 40 325\nTMhelix 326 348\ninside 349 372

25539 GCF_001478325.1_Salmonella_enterica_CVM_N38858-SQ_v1.0 Salmonella enterica subsp. enterica serovar Typhimurium var. 5- Proteobacteria; Gammaproteobacteria; Enterobacterales; Enterobacteriaceae; Salmonella; Salmonella enterica; Salmonella enterica subsp. enterica; Salmonella enterica subsp. enterica serovar Typhimurium MTGDNTLITSHGINRRDFMKLCAALAATMGLSSKAAAEMAESVSNPQRPPVIWIGAQECT WP_000145429.1 hydrogenase 2 small subunit [Salmonella enterica] Length: 372\nNumber of predicted TMHs: 1\nExp number of AAs in TMHs: 24.51309\nExp number, first 60 AAs: 0.2682\nTotal prob of N-in: 0.14402\noutside 1 331\nTMhelix 332 354\ninside 355 372

25540 GCF_001480105.1_Salmonella_enterica_CVM_N38943-SQ_v1.0 Salmonella enterica subsp. enterica serovar Typhimurium var. 5- Proteobacteria; Gammaproteobacteria; Enterobacterales; Enterobacteriaceae; Salmonella; Salmonella enterica; Salmonella enterica subsp. enterica; Salmonella enterica subsp. enterica serovar Typhimurium MNNEETFYQAMRRKGVTRRSFLKFCSLAATSLGLGAGMTPKIAWALENKPRIPVVWIHGL WP_001058312.1 hydrogenase [Salmonella enterica] Length: 372\nNumber of predicted TMHs: 2\nExp number of AAs in TMHs: 36.51738\nExp number, first 60 AAs: 19.72849\nTotal prob of N-in: 0.96170\nPOSSIBLE N-term signal sequence\ninside 1 19\nTMhelix 20 39\noutside 40 325\nTMhelix 326 348\ninside 349 372

25541 GCF_001480025.1_Salmonella_enterica_CVM_N38903-SQ_v1.0 Salmonella enterica subsp. enterica serovar Typhimurium var. 5- Proteobacteria; Gammaproteobacteria; Enterobacterales; Enterobacteriaceae; Salmonella; Salmonella enterica; Salmonella enterica subsp. enterica; Salmonella enterica subsp. enterica serovar Typhimurium MNNEETFYQAMRRKGVTRRSFLKFCSLAATSLGLGAGMTPKIAWALENKPRIPVVWIHGL WP_001058312.1 hydrogenase [Salmonella enterica] Length: 372\nNumber of predicted TMHs: 2\nExp number of AAs in TMHs: 36.51738\nExp number, first 60 AAs: 19.72849\nTotal prob of N-in: 0.96170\nPOSSIBLE N-term signal sequence\ninside 1 19\nTMhelix 20 39\noutside 40 325\nTMhelix 326 348\ninside 349 372

25542 GCF_001480265.1_Salmonella_enterica_CVM_N40416-SQ_v1.0 Salmonella enterica subsp. enterica serovar Typhimurium var. 5- Proteobacteria; Gammaproteobacteria; Enterobacterales; Enterobacteriaceae; Salmonella; Salmonella enterica; Salmonella enterica subsp. enterica; Salmonella enterica subsp. enterica serovar Typhimurium MNNEETFYQAMRRKGVTRRSFLKFCSLAATSLGLGAGMTPKIAWALENKPRIPVVWIHGL WP_001058312.1 hydrogenase [Salmonella enterica] Length: 372\nNumber of predicted TMHs: 2\nExp number of AAs in TMHs: 36.51738\nExp number, first 60 AAs: 19.72849\nTotal prob of N-in: 0.96170\nPOSSIBLE N-term signal sequence\ninside 1 19\nTMhelix 20 39\noutside 40 325\nTMhelix 326 348\ninside 349 372

25543 GCF_001480205.1_Salmonella_enterica_CVM_N40379-SQ_v1.0 Salmonella enterica subsp. enterica serovar Typhimurium var. 5- Proteobacteria; Gammaproteobacteria; Enterobacterales; Enterobacteriaceae; Salmonella; Salmonella enterica; Salmonella enterica subsp. enterica; Salmonella enterica subsp. enterica serovar Typhimurium MNNEETFYQAMRRKGVTRRSFLKFCSLAATSLGLGAGMTPKIAWALENKPRIPVVWIHGL WP_001058312.1 hydrogenase [Salmonella enterica] Length: 372\nNumber of predicted TMHs: 2\nExp number of AAs in TMHs: 36.51738\nExp number, first 60 AAs: 19.72849\nTotal prob of N-in: 0.96170\nPOSSIBLE N-term signal sequence\ninside 1 19\nTMhelix 20 39\noutside 40 325\nTMhelix 326 348\ninside 349 372

25544 GCF_001480265.1_Salmonella_enterica_CVM_N40416-SQ_v1.0 Salmonella enterica subsp. enterica serovar Typhimurium var. 5- Proteobacteria; Gammaproteobacteria; Enterobacterales; Enterobacteriaceae; Salmonella;

Salmonella enterica; Salmonella enterica subsp. enterica; Salmonella enterica subsp. enterica serovar Typhimurium MTGDNTLITSHGINRRDFMKLCAALAATMGLSSKAAAEMAESVSNPQRPPVIWIGAQECT WP_000145429.1 hydrogenase 2 small subunit [Salmonella enterica] Length: 372\nNumber of predicted TMHs: 1\nExp number of AAs in TMHs: 24.51309\nExp number, first 60 AAs: 0.2682\nTotal prob of N-in: 0.14402\noutside 1 331\nTMhelix 332 354\ninside 355 372

25545 GCF_001480145.1_Salmonella_enterica_CVM_N40368-SQ_v1.0 Salmonella enterica subsp. enterica serovar Typhimurium var. 5- Proteobacteria; Gammaproteobacteria; Enterobacterales; Enterobacteriaceae; Salmonella; Salmonella enterica; Salmonella enterica subsp. enterica; Salmonella enterica subsp. enterica serovar Typhimurium MTGDNTLITSHGINRRDFMKLCAALAATMGLSSKAAAEMAESVSNPQRPPVIWIGAQECT WP_000145429.1 hydrogenase 2 small subunit [Salmonella enterica] Length: 372\nNumber of predicted TMHs: 1\nExp number of AAs in TMHs: 24.51309\nExp number, first 60 AAs: 0.2682\nTotal prob of N-in: 0.14402\noutside 1 331\nTMhelix 332 354\ninside 355 372

25546 GCF_001480105.1_Salmonella_enterica_CVM_N38943-SQ_v1.0 Salmonella enterica subsp. enterica serovar Typhimurium var. 5- Proteobacteria; Gammaproteobacteria; Enterobacterales; Enterobacteriaceae; Salmonella; Salmonella enterica; Salmonella enterica subsp. enterica; Salmonella enterica subsp. enterica serovar Typhimurium MTGDNTLITSHGINRRDFMKLCAALAATMGLSSKAAAEMAESVSNPQRPPVIWIGAQECT WP_000145429.1 hydrogenase 2 small subunit [Salmonella enterica] Length: 372\nNumber of predicted TMHs: 1\nExp number of AAs in TMHs: 24.51309\nExp number, first 60 AAs: 0.2682\nTotal prob of N-in: 0.14402\noutside 1 331\nTMhelix 332 354\ninside 355 372

25547 GCF_001480285.1_Salmonella_enterica_CVM_N40935-SQ_v1.0 Salmonella enterica subsp. enterica serovar Typhimurium var. 5- Proteobacteria; Gammaproteobacteria; Enterobacterales; Enterobacteriaceae; Salmonella; Salmonella enterica; Salmonella enterica subsp. enterica; Salmonella enterica subsp. enterica serovar Typhimurium MNNEETFYQAMRRKGVTRRSFLKFCSLAATSLGLGAGMTPKIAWALENKPRIPVVWIHGL WP_001058312.1 hydrogenase [Salmonella enterica] Length: 372\nNumber of predicted TMHs: 2\nExp number of AAs in TMHs: 36.51738\nExp number, first 60 AAs: 19.72849\nTotal prob of N-in: 0.96170\nPOSSIBLE N-term signal sequence\ninside 1 19\nTMhelix 20 39\noutside 40 325\nTMhelix 326 348\ninside 349 372

25548 GCF_001480285.1_Salmonella_enterica_CVM_N40935-SQ_v1.0 Salmonella enterica subsp. enterica serovar Typhimurium var. 5- Proteobacteria; Gammaproteobacteria; Enterobacterales; Enterobacteriaceae; Salmonella; Salmonella enterica; Salmonella enterica subsp. enterica; Salmonella enterica subsp. enterica serovar Typhimurium MTGDNTLITSHGINRRDFMKLCAALAATMGLSSKAAAEMAESVSNPQRPPVIWIGAQECT WP_000145429.1 hydrogenase 2 small subunit [Salmonella enterica] Length: 372\nNumber of predicted TMHs: 1\nExp number of AAs in TMHs: 24.51309\nExp number, first 60 AAs: 0.2682\nTotal prob of N-in: 0.14402\noutside 1 331\nTMhelix 332 354\ninside 355 372

25549 GCF_001480145.1_Salmonella_enterica_CVM_N40368-SQ_v1.0 Salmonella enterica subsp. enterica serovar Typhimurium var. 5- Proteobacteria; Gammaproteobacteria; Enterobacterales; Enterobacteriaceae; Salmonella; Salmonella enterica; Salmonella enterica subsp. enterica; Salmonella enterica subsp. enterica serovar Typhimurium MNNEETFYQAMRRKGVTRRSFLKFCSLAATSLGLGAGMTPKIAWALENKPRIPVVWIHGL WP_001058312.1 hydrogenase [Salmonella enterica] Length: 372\nNumber of predicted TMHs: 2\nExp number of AAs in TMHs: 36.51738\nExp number, first 60 AAs: 19.72849\nTotal prob of N-in: 0.96170\nPOSSIBLE N-term signal sequence\ninside 1 19\nTMhelix 20 39\noutside 40 325\nTMhelix 326 348\ninside 349 372

25550 GCF_001480385.1_Salmonella_enterica_CVM_N41904-SQ_v1.0 Salmonella enterica subsp. enterica serovar Typhimurium var. 5- Proteobacteria; Gammaproteobacteria; Enterobacterales; Enterobacteriaceae; Salmonella; Salmonella enterica; Salmonella enterica subsp. enterica; Salmonella enterica subsp. enterica serovar Typhimurium MTGDNTLITSHGINRRDFMKLCAALAATMGLSSKAAAEMAESVSNPQRPPVIWIGAQECT WP_000145429.1 hydrogenase 2 small subunit [Salmonella enterica] Length: 372\nNumber of predicted TMHs: 1\nExp number of AAs in TMHs: 24.51309\nExp number, first 60 AAs: 0.2682\nTotal prob of N-in: 0.14402\noutside 1 331\nTMhelix 332 354\ninside 355 372

25551 GCF_001479155.1_Salmonella_enterica_CVM_N31411-SQ_v1.0 Salmonella enterica subsp. enterica serovar Typhimurium var. 5- Proteobacteria; Gammaproteobacteria; Enterobacterales; Enterobacteriaceae; Salmonella; Salmonella enterica; Salmonella enterica subsp. enterica; Salmonella enterica subsp. enterica serovar Typhimurium MNNEETFYQAMRRKGVTRRSFLKFCSLAATSLGLGAGMTPKIAWALENKPRIPVVWIHGL WP_001058312.1 hydrogenase [Salmonella enterica] Length: 372\nNumber of predicted TMHs: 2\nExp number of AAs in TMHs: 36.51738\nExp number, first 60 AAs: 19.72849\nTotal prob of N-in: 0.96170\nPOSSIBLE N-term signal sequence\ninside 1 19\nTMhelix 20 39\noutside 40 325\nTMhelix 326 348\ninside 349 372

25552 GCF_001479155.1_Salmonella_enterica_CVM_N31411-SQ_v1.0 Salmonella enterica subsp. enterica serovar Typhimurium var. 5- Proteobacteria; Gammaproteobacteria; Enterobacterales; Enterobacteriaceae; Salmonella; Salmonella enterica; Salmonella enterica subsp. enterica; Salmonella enterica subsp. enterica serovar Typhimurium MTGDNTLITSHGINRRDFMKLCAALAATMGLSSKAAAEMAESVSNPQRPPVIWIGAQECT WP_000145429.1 hydrogenase 2 small subunit [Salmonella enterica] Length: 372\nNumber of predicted TMHs: 1\nExp number of AAs in TMHs: 24.51309\nExp number, first 60 AAs: 0.2682\nTotal prob of N-in: 0.14402\noutside 1 331\nTMhelix 332 354\ninside 355 372

25553 GCF_001479065.1_Salmonella_enterica_CVM_N31392-SQ_v1.0 Salmonella enterica subsp. enterica serovar Typhimurium var. 5- Proteobacteria; Gammaproteobacteria; Enterobacterales; Enterobacteriaceae; Salmonella; Salmonella enterica; Salmonella enterica subsp. enterica; Salmonella enterica subsp. enterica serovar Typhimurium MNNEETFYQAMRRKGVTRRSFLKFCSLAATSLGLGAGMTPKIAWALENKPRIPVVWIHGL WP_001058312.1 hydrogenase [Salmonella enterica] Length: 372\nNumber of predicted TMHs: 2\nExp number of AAs in TMHs: 36.51738\nExp number, first 60 AAs: 19.72849\nTotal prob of N-in: 0.96170\nPOSSIBLE N-term signal sequence\ninside 1 19\nTMhelix 20 39\noutside 40 325\nTMhelix 326 348\ninside 349 372

25554 GCF_001479065.1_Salmonella_enterica_CVM_N31392-SQ_v1.0 Salmonella enterica subsp. enterica serovar Typhimurium var. 5- Proteobacteria; Gammaproteobacteria; Enterobacterales; Enterobacteriaceae; Salmonella; Salmonella enterica; Salmonella enterica subsp. enterica; Salmonella enterica subsp. enterica serovar Typhimurium MTGDNTLITSHGINRRDFMKLCAALAATMGLSSKAAAEMAESVSNPQRPPVIWIGAQECT WP_000145429.1 hydrogenase 2 small subunit [Salmonella enterica] Length: 372\nNumber of predicted TMHs: 1\nExp number of AAs in TMHs: 24.51309\nExp number, first 60 AAs: 0.2682\nTotal prob of N-in: 0.14402\noutside 1 331\nTMhelix 332 354\ninside 355 372

25555 GCF_001478885.1_Salmonella_enterica_CVM_N30655-SQ_v1.0 Salmonella enterica subsp. enterica serovar Typhimurium var. 5- Proteobacteria; Gammaproteobacteria; Enterobacterales; Enterobacteriaceae; Salmonella; Salmonella enterica; Salmonella enterica subsp. enterica; Salmonella enterica subsp. enterica serovar Typhimurium MNNEETFYQAMRRKGVTRRSFLKFCSLAATSLGLGAGMTPKIAWALENKPRIPVVWIHGL WP_001058312.1 hydrogenase [Salmonella enterica] Length: 372\nNumber of predicted TMHs: 2\nExp number of AAs in TMHs: 36.51738\nExp number, first 60 AAs: 19.72849\nTotal prob of N-in: 0.96170\nPOSSIBLE N-term signal sequence\ninside 1 19\nTMhelix 20 39\noutside 40 325\nTMhelix 326 348\ninside 349 372

25556 GCF_001478885.1_Salmonella_enterica_CVM_N30655-SQ_v1.0 Salmonella enterica subsp. enterica serovar Typhimurium var. 5- Proteobacteria; Gammaproteobacteria; Enterobacterales; Enterobacteriaceae; Salmonella; Salmonella enterica; Salmonella enterica subsp. enterica; Salmonella enterica subsp. enterica serovar Typhimurium MTGDNTLITSHGINRRDFMKLCAALAATMGLSSKAAAEMAESVSNPQRPPVIWIGAQECT WP_000145429.1 hydrogenase 2 small subunit [Salmonella enterica] Length: 372\nNumber of predicted TMHs: 1\nExp number of AAs in TMHs: 24.51309\nExp number, first 60 AAs: 0.2682\nTotal prob of N-in: 0.14402\noutside 1 331\nTMhelix 332 354\ninside 355 372

25557 GCF_001478625.1_Salmonella_enterica_CVM_N29343-SQ_v1.0 Salmonella enterica subsp. enterica serovar Typhimurium var. 5- Proteobacteria; Gammaproteobacteria; Enterobacterales; Enterobacteriaceae; Salmonella; Salmonella enterica; Salmonella enterica subsp. enterica; Salmonella enterica subsp. enterica serovar Typhimurium MNNEETFYQAMRRKGVTRRSFLKFCSLAATSLGLGAGMTPKIAWALENKPRIPVVWIHGL WP_001058312.1 hydrogenase [Salmonella enterica] Length: 372\nNumber of predicted TMHs: 2\nExp number of AAs in TMHs: 36.51738\nExp number, first 60 AAs: 19.72849\nTotal prob of N-in: 0.96170\nPOSSIBLE N-term signal sequence\ninside 1 19\nTMhelix 20 39\noutside 40 325\nTMhelix 326 348\ninside 349 372

25558 GCF_001478625.1_Salmonella_enterica_CVM_N29343-SQ_v1.0 Salmonella enterica subsp. enterica serovar Typhimurium var. 5- Proteobacteria; Gammaproteobacteria; Enterobacterales; Enterobacteriaceae; Salmonella; Salmonella enterica; Salmonella enterica subsp. enterica; Salmonella enterica subsp. enterica serovar Typhimurium MTGDNTLITSHGINRRDFMKLCAALAATMGLSSKAAAEMAESVSNPQRPPVIWIGAQECT WP_000145429.1 hydrogenase 2 small subunit [Salmonella enterica] Length: 372\nNumber of predicted TMHs: 1\nExp number of AAs in TMHs: 24.51309\nExp number, first 60 AAs: 0.2682\nTotal prob of N-in: 0.14402\noutside 1 331\nTMhelix 332 354\ninside 355 372

25559 GCF_001477855.1_Salmonella_enterica_CVM_N29350-SQ_v1.0 Salmonella enterica subsp. enterica serovar Typhimurium var. 5- Proteobacteria; Gammaproteobacteria; Enterobacterales; Enterobacteriaceae; Salmonella; Salmonella enterica; Salmonella enterica subsp. enterica; Salmonella enterica subsp. enterica serovar Typhimurium MNNEETFYQAMRRKGVTRRSFLKFCSLAATSLGLGAGMTPKIAWALENKPRIPVVWIHGL WP_001058312.1 hydrogenase [Salmonella enterica] Length: 372\nNumber of predicted TMHs: 2\nExp number of AAs in

TMHs: 36.51738\nExp number, first 60 AAs: 19.72849\nTotal prob of N-in: 0.96170\nPOSSIBLE N-term signal sequence\ninside 1 19\nTMhelix 20 39\noutside 40 325\nTMhelix 326 348\ninside 349 372

25560 GCF_001480385.1_Salmonella_enterica_CVM_N41904-SQ_v1.0 Salmonella enterica subsp. enterica serovar Typhimurium var. 5- Proteobacteria; Gammaproteobacteria; Enterobacterales; Enterobacteriaceae; Salmonella; Salmonella enterica; Salmonella enterica subsp. enterica; Salmonella enterica subsp. enterica serovar Typhimurium MNNEETFYQAMRRKGVTRRSFLKFCSLAATSLGLGAGMTPKIAWALENKPRIPVVWIHGL WP_001058312.1 hydrogenase [Salmonella enterica] Length: 372\nNumber of predicted TMHs: 2\nExp number of AAs in TMHs: 36.51738\nExp number, first 60 AAs: 19.72849\nTotal prob of N-in: 0.96170\nPOSSIBLE N-term signal sequence\ninside 1 19\nTMhelix 20 39\noutside 40 325\nTMhelix 326 348\ninside 349 372

25561 GCF_001477855.1_Salmonella_enterica_CVM_N29350-SQ_v1.0 Salmonella enterica subsp. enterica serovar Typhimurium var. 5- Proteobacteria; Gammaproteobacteria; Enterobacterales; Enterobacteriaceae; Salmonella; Salmonella enterica; Salmonella enterica subsp. enterica; Salmonella enterica subsp. enterica serovar Typhimurium MTGDNTLITSHGINRRDFMKLCAALAATMGLSSKAAAEMAESVSNPQRPPVIWIGAQECT WP_000145429.1 hydrogenase 2 small subunit [Salmonella enterica] Length: 372\nNumber of predicted TMHs: 1\nExp number of AAs in TMHs: 24.51309\nExp number, first 60 AAs: 0.2682\nTotal prob of N-in: 0.14402\noutside 1 331\nTMhelix 332 354\ninside 355 372

25562 GCF_001480775.1_Salmonella_enterica_CVM_N37941-SQ_v1.0 Salmonella enterica subsp. enterica serovar Typhimurium var. 5- Proteobacteria; Gammaproteobacteria; Enterobacterales; Enterobacteriaceae; Salmonella; Salmonella enterica; Salmonella enterica subsp. enterica; Salmonella enterica subsp. enterica serovar Typhimurium MTGDNTLITSHGINRRDFMKLCAALAATMGLSSKAAAEMAESVSNPQRPPVIWIGAQECT WP_000145429.1 hydrogenase 2 small subunit [Salmonella enterica] Length: 372\nNumber of predicted TMHs: 1\nExp number of AAs in TMHs: 24.51309\nExp number, first 60 AAs: 0.2682\nTotal prob of N-in: 0.14402\noutside 1 331\nTMhelix 332 354\ninside 355 372

25563 GCF_001480775.1_Salmonella_enterica_CVM_N37941-SQ_v1.0 Salmonella enterica subsp. enterica serovar Typhimurium var. 5- Proteobacteria; Gammaproteobacteria; Enterobacterales; Enterobacteriaceae; Salmonella; Salmonella enterica; Salmonella enterica subsp. enterica; Salmonella enterica subsp. enterica serovar Typhimurium MNNEETFYQAMRRKGVTRRSFLKFCSLAATSLGLGAGMTPKIAWALENKPRIPVVWIHGL WP_001058312.1 hydrogenase [Salmonella enterica] Length: 372\nNumber of predicted TMHs: 2\nExp number of AAs in TMHs: 36.51738\nExp number, first 60 AAs: 19.72849\nTotal prob of N-in: 0.96170\nPOSSIBLE N-term signal sequence\ninside 1 19\nTMhelix 20 39\noutside 40 325\nTMhelix 326 348\ninside 349 372

25564 GCF_001480955.1_Salmonella_enterica_CVM_N40413-SQ_v1.0 Salmonella enterica subsp. enterica serovar Typhimurium var. 5- Proteobacteria; Gammaproteobacteria; Enterobacterales; Enterobacteriaceae; Salmonella; Salmonella enterica; Salmonella enterica subsp. enterica; Salmonella enterica subsp. enterica serovar Typhimurium MTGDNTLITSHGINRRDFMKLCAALAATMGLSSKAAAEMAESVSNPQRPPVIWIGAQECT WP_000145429.1 hydrogenase 2 small subunit [Salmonella enterica] Length: 372\nNumber of predicted TMHs: 1\nExp number of AAs in TMHs: 24.51309\nExp number, first 60 AAs: 0.2682\nTotal prob of N-in: 0.14402\noutside 1 331\nTMhelix 332 354\ninside 355 372

25565 GCF_001480985.1_Salmonella_enterica_CVM_N41906-R_v1.0 Salmonella enterica subsp. enterica serovar Typhimurium var. 5- Proteobacteria; Gammaproteobacteria; Enterobacterales; Enterobacteriaceae; Salmonella; Salmonella enterica; Salmonella enterica subsp. enterica; Salmonella enterica subsp. enterica serovar Typhimurium MTGDNTLITSHGINRRDFMKLCAALAATMGLSSKAAAEMAESVSNPQRPPVIWIGAQECT WP_000145429.1 hydrogenase 2 small subunit [Salmonella enterica] Length: 372\nNumber of predicted TMHs: 1\nExp number of AAs in TMHs: 24.51309\nExp number, first 60 AAs: 0.2682\nTotal prob of N-in: 0.14402\noutside 1 331\nTMhelix 332 354\ninside 355 372

25566 GCF_001481175.1_Salmonella_enterica_CVM_N42492_v1.0 Salmonella enterica subsp. enterica serovar Typhimurium var. 5- Proteobacteria; Gammaproteobacteria; Enterobacterales; Enterobacteriaceae; Salmonella; Salmonella enterica; Salmonella enterica subsp. enterica; Salmonella enterica subsp. enterica serovar Typhimurium MTGDNTLITSHGINRRDFMKLCAALAATMGLSSKAAAEMAESVSNPQRPPVIWIGAQECT WP_000145429.1 hydrogenase 2 small subunit [Salmonella enterica] Length: 372\nNumber of predicted TMHs: 1\nExp number of AAs in TMHs: 24.51309\nExp number, first 60 AAs: 0.2682\nTotal prob of N-in: 0.14402\noutside 1 331\nTMhelix 332 354\ninside 355 372

25567 GCF_001481175.1_Salmonella_enterica_CVM_N42492_v1.0 Salmonella enterica subsp. enterica serovar Typhimurium var. 5- Proteobacteria; Gammaproteobacteria; Enterobacterales; Enterobacteriaceae; Salmonella;

Salmonella enterica; Salmonella enterica subsp. enterica; Salmonella enterica subsp. enterica serovar Typhimurium
MNNEETFYQAMRRKGVTRRSFLKFCSLAATSLGLGAGMTPKIAWALENKPRIPVVWIHGL WP_001058312.1
hydrogenase [Salmonella enterica] Length: 372\nNumber of predicted TMHs: 2\nExp number of AAs in TMHs: 36.51738\nExp number, first 60 AAs: 19.72849\nTotal prob of N-in: 0.96170\nPOSSIBLE N-term signal sequence\ninside 1 19\nTMhelix 20 39\noutside 40 325\nTMhelix 326 348\ninside 349 372

25568 GCF_001480955.1_Salmonella_enterica_CVM_N40413-SQ_v1.0 Salmonella enterica subsp. enterica serovar Typhimurium var. 5- Proteobacteria; Gammaproteobacteria; Enterobacterales; Enterobacteriaceae; Salmonella; Salmonella enterica; Salmonella enterica subsp. enterica; Salmonella enterica subsp. enterica serovar Typhimurium
MNNEETFYQAMRRKGVTRRSFLKFCSLAATSLGLGAGMTPKIAWALENKPRIPVVWIHGL WP_001058312.1
hydrogenase [Salmonella enterica] Length: 372\nNumber of predicted TMHs: 2\nExp number of AAs in TMHs: 36.51738\nExp number, first 60 AAs: 19.72849\nTotal prob of N-in: 0.96170\nPOSSIBLE N-term signal sequence\ninside 1 19\nTMhelix 20 39\noutside 40 325\nTMhelix 326 348\ninside 349 372

25569 GCF_001480985.1_Salmonella_enterica_CVM_N41906-R_v1.0 Salmonella enterica subsp. enterica serovar Typhimurium var. 5- Proteobacteria; Gammaproteobacteria; Enterobacterales; Enterobacteriaceae; Salmonella; Salmonella enterica; Salmonella enterica subsp. enterica; Salmonella enterica subsp. enterica serovar Typhimurium
MNNEETFYQAMRRKGVTRRSFLKFCSLAATSLGLGAGMTPKIAWALENKPRIPVVWIHGL WP_001058312.1
hydrogenase [Salmonella enterica] Length: 372\nNumber of predicted TMHs: 2\nExp number of AAs in TMHs: 36.51738\nExp number, first 60 AAs: 19.72849\nTotal prob of N-in: 0.96170\nPOSSIBLE N-term signal sequence\ninside 1 19\nTMhelix 20 39\noutside 40 325\nTMhelix 326 348\ninside 349 372

25570 GCF_002030605.1_ASM203060v1 Salmonella enterica subsp. enterica serovar Typhimurium var. 5- Proteobacteria; Gammaproteobacteria; Enterobacterales; Enterobacteriaceae; Salmonella; Salmonella enterica; Salmonella enterica subsp. enterica; Salmonella enterica subsp. enterica serovar Typhimurium
MTGDNTLITSHGINRRDFMKLCAALAATMGLSSKAAAEMAESVSNPQRPPVIWIGAQECT WP_000145429.1
hydrogenase 2 small subunit [Salmonella enterica] Length: 372\nNumber of predicted TMHs: 1\nExp number of AAs in TMHs: 24.51309\nExp number, first 60 AAs: 0.2682\nTotal prob of N-in: 0.14402\noutside 1 331\nTMhelix 332 354\ninside 355 372

25571 GCF_002030775.1_ASM203077v1 Salmonella enterica subsp. enterica serovar Typhimurium var. 5- Proteobacteria; Gammaproteobacteria; Enterobacterales; Enterobacteriaceae; Salmonella; Salmonella enterica; Salmonella enterica subsp. enterica; Salmonella enterica subsp. enterica serovar Typhimurium
MTGDNTLITSHGINRRDFMKLCAALAATMGLSSKAAAEMAESVSNPQRPPVIWIGAQECT WP_000145429.1
hydrogenase 2 small subunit [Salmonella enterica] Length: 372\nNumber of predicted TMHs: 1\nExp number of AAs in TMHs: 24.51309\nExp number, first 60 AAs: 0.2682\nTotal prob of N-in: 0.14402\noutside 1 331\nTMhelix 332 354\ninside 355 372

25572 GCF_002030995.1_ASM203099v1 Salmonella enterica subsp. enterica serovar Typhimurium var. 5- Proteobacteria; Gammaproteobacteria; Enterobacterales; Enterobacteriaceae; Salmonella; Salmonella enterica; Salmonella enterica subsp. enterica; Salmonella enterica subsp. enterica serovar Typhimurium
MNNEETFYQAMRRKGVTRRSFLKFCSLAATSLGLGAGMTPKIAWALENKPRIPVVWIHGL WP_001058312.1
hydrogenase [Salmonella enterica] Length: 372\nNumber of predicted TMHs: 2\nExp number of AAs in TMHs: 36.51738\nExp number, first 60 AAs: 19.72849\nTotal prob of N-in: 0.96170\nPOSSIBLE N-term signal sequence\ninside 1 19\nTMhelix 20 39\noutside 40 325\nTMhelix 326 348\ninside 349 372

25573 GCF_002031525.1_ASM203152v1 Salmonella enterica subsp. enterica serovar Typhimurium var. 5- Proteobacteria; Gammaproteobacteria; Enterobacterales; Enterobacteriaceae; Salmonella; Salmonella enterica; Salmonella enterica subsp. enterica; Salmonella enterica subsp. enterica serovar Typhimurium
MTGDNTLITSHGINRRDFMKLCAALAATMGLSSKAAAEMAESVSNPQRPPVIWIGAQECT WP_000145429.1
hydrogenase 2 small subunit [Salmonella enterica] Length: 372\nNumber of predicted TMHs: 1\nExp number of AAs in TMHs: 24.51309\nExp number, first 60 AAs: 0.2682\nTotal prob of N-in: 0.14402\noutside 1 331\nTMhelix 332 354\ninside 355 372

25574 GCF_002031565.1_ASM203156v1 Salmonella enterica subsp. enterica serovar Typhimurium var. 5- Proteobacteria; Gammaproteobacteria; Enterobacterales; Enterobacteriaceae; Salmonella; Salmonella enterica; Salmonella enterica subsp. enterica; Salmonella enterica subsp. enterica serovar Typhimurium
MTGDNTLITSHGINRRDFMKLCAALAATMGLSSKAAAEMAESVSNPQRPPVIWIGAQECT WP_000145429.1
hydrogenase 2 small subunit [Salmonella enterica] Length: 372\nNumber of predicted TMHs: 1\nExp number of AAs in TMHs: 24.51309\nExp number, first 60 AAs: 0.2682\nTotal prob of N-in: 0.14402\noutside 1 331\nTMhelix 332 354\ninside 355 372

25575 GCF_002031925.1_ASM203192v1 Salmonella enterica subsp. enterica serovar Typhimurium var. 5- Proteobacteria; Gammaproteobacteria; Enterobacterales; Enterobacteriaceae; Salmonella; Salmonella enterica; Salmonella enterica subsp. enterica; Salmonella enterica subsp. enterica serovar Typhimurium MTGDNTLITSHGINRRDFMKLCAALAATMGLSSKAAAEMAESVSNPQRPPVIWIGAQECT WP_000145429.1 hydrogenase 2 small subunit [Salmonella enterica] Length: 372\nNumber of predicted TMHs: 1\nExp number of AAs in TMHs: 24.51309\nExp number, first 60 AAs: 0.2682\nTotal prob of N-in: 0.14402\noutside 1 331\nTMhelix 332 354\ninside 355 372

25576 GCF_002031955.1_ASM203195v1 Salmonella enterica subsp. enterica serovar Typhimurium var. 5- Proteobacteria; Gammaproteobacteria; Enterobacterales; Enterobacteriaceae; Salmonella; Salmonella enterica; Salmonella enterica subsp. enterica; Salmonella enterica subsp. enterica serovar Typhimurium MTGDNTLITSHGINRRDFMKLCAALAATMGLSSKAAAEMAESVSNPQRPPVIWIGAQECT WP_000145429.1 hydrogenase 2 small subunit [Salmonella enterica] Length: 372\nNumber of predicted TMHs: 1\nExp number of AAs in TMHs: 24.51309\nExp number, first 60 AAs: 0.2682\nTotal prob of N-in: 0.14402\noutside 1 331\nTMhelix 332 354\ninside 355 372

25577 GCF_002031925.1_ASM203192v1 Salmonella enterica subsp. enterica serovar Typhimurium var. 5- Proteobacteria; Gammaproteobacteria; Enterobacterales; Enterobacteriaceae; Salmonella; Salmonella enterica; Salmonella enterica subsp. enterica; Salmonella enterica subsp. enterica serovar Typhimurium MNNEETFYQAMRRKGVTRRSFLKFCSLAATSLGLGAGMTPKIAWALENKPRIPVVWIHGL WP_001058312.1 hydrogenase [Salmonella enterica] Length: 372\nNumber of predicted TMHs: 2\nExp number of AAs in TMHs: 36.51738\nExp number, first 60 AAs: 19.72849\nTotal prob of N-in: 0.96170\nPOSSIBLE N-term signal sequence\ninside 1 19\nTMhelix 20 39\noutside 40 325\nTMhelix 326 348\ninside 349 372

25578 GCF_002031955.1_ASM203195v1 Salmonella enterica subsp. enterica serovar Typhimurium var. 5- Proteobacteria; Gammaproteobacteria; Enterobacterales; Enterobacteriaceae; Salmonella; Salmonella enterica; Salmonella enterica subsp. enterica; Salmonella enterica subsp. enterica serovar Typhimurium MNNEETFYQAMRRKGVTRRSFLKFCSLAATSLGLGAGMTPKIAWALENKPRIPVVWIHGL WP_001058312.1 hydrogenase [Salmonella enterica] Length: 372\nNumber of predicted TMHs: 2\nExp number of AAs in TMHs: 36.51738\nExp number, first 60 AAs: 19.72849\nTotal prob of N-in: 0.96170\nPOSSIBLE N-term signal sequence\ninside 1 19\nTMhelix 20 39\noutside 40 325\nTMhelix 326 348\ninside 349 372

25579 GCF_002032435.1_ASM203243v1 Salmonella enterica subsp. enterica serovar Typhimurium var. 5- Proteobacteria; Gammaproteobacteria; Enterobacterales; Enterobacteriaceae; Salmonella; Salmonella enterica; Salmonella enterica subsp. enterica; Salmonella enterica subsp. enterica serovar Typhimurium MTGDNTLITSHGINRRDFMKLCAALAATMGLSSKAAAEMAESVSNPQRPPVIWIGAQECT WP_000145429.1 hydrogenase 2 small subunit [Salmonella enterica] Length: 372\nNumber of predicted TMHs: 1\nExp number of AAs in TMHs: 24.51309\nExp number, first 60 AAs: 0.2682\nTotal prob of N-in: 0.14402\noutside 1 331\nTMhelix 332 354\ninside 355 372

25580 GCF_002032435.1_ASM203243v1 Salmonella enterica subsp. enterica serovar Typhimurium var. 5- Proteobacteria; Gammaproteobacteria; Enterobacterales; Enterobacteriaceae; Salmonella; Salmonella enterica; Salmonella enterica subsp. enterica; Salmonella enterica subsp. enterica serovar Typhimurium MNNEETFYQAMRRKGVTRRSFLKFCSLAATSLGLGAGMTPKIAWALENKPRIPVVWIHGL WP_001058312.1 hydrogenase [Salmonella enterica] Length: 372\nNumber of predicted TMHs: 2\nExp number of AAs in TMHs: 36.51738\nExp number, first 60 AAs: 19.72849\nTotal prob of N-in: 0.96170\nPOSSIBLE N-term signal sequence\ninside 1 19\nTMhelix 20 39\noutside 40 325\nTMhelix 326 348\ninside 349 372

25581 GCF_002032875.1_ASM203287v1 Salmonella enterica subsp. enterica serovar Typhimurium var. 5- Proteobacteria; Gammaproteobacteria; Enterobacterales; Enterobacteriaceae; Salmonella; Salmonella enterica; Salmonella enterica subsp. enterica; Salmonella enterica subsp. enterica serovar Typhimurium MNNEETFYQAMRRKGVTRRSFLKFCSLAATSLGLGAGMTPKIAWALENKPRIPVVWIHGL WP_001058312.1 hydrogenase [Salmonella enterica] Length: 372\nNumber of predicted TMHs: 2\nExp number of AAs in TMHs: 36.51738\nExp number, first 60 AAs: 19.72849\nTotal prob of N-in: 0.96170\nPOSSIBLE N-term signal sequence\ninside 1 19\nTMhelix 20 39\noutside 40 325\nTMhelix 326 348\ninside 349 372

25582 GCF_002032705.1_ASM203270v1 Salmonella enterica subsp. enterica serovar Typhimurium var. 5- Proteobacteria; Gammaproteobacteria; Enterobacterales; Enterobacteriaceae; Salmonella; Salmonella enterica; Salmonella enterica subsp. enterica; Salmonella enterica subsp. enterica serovar Typhimurium MNNEETFYQAMRRKGVTRRSFLKFCSLAATSLGLGAGMTPKIAWALENKPRIPVVWIHGL WP_001058312.1 hydrogenase [Salmonella enterica] Length: 372\nNumber of predicted TMHs: 2\nExp number of AAs in

TMHs: 36.51738\nExp number, first 60 AAs: 19.72849\nTotal prob of N-in: 0.96170\nPOSSIBLE N-term signal sequence\ninside 1 19\nTMhelix 20 39\noutside 40 325\nTMhelix 326 348\ninside 349 372

25583 GCF_002032795.1_ASM203279v1 Salmonella enterica subsp. enterica serovar Typhimurium var. 5- Proteobacteria; Gammaproteobacteria; Enterobacterales; Enterobacteriaceae; Salmonella; Salmonella enterica; Salmonella enterica subsp. enterica; Salmonella enterica subsp. enterica serovar Typhimurium MNNEETFYQAMRRKGVTRRSFLKFCSLAATSLGLGAGMTPKIAWALENKPRIPVVWIHGL WP_079805161.1 [Ni/Fe] hydrogenase small subunit [Salmonella enterica] Length: 372\nNumber of predicted TMHs: 2\nExp number of AAs in TMHs: 36.9928999999999\nExp number, first 60 AAs: 19.72755\nTotal prob of N-in: 0.96172\nPOSSIBLE N-term signal sequence\ninside 1 19\nTMhelix 20 39\noutside 40 325\nTMhelix 326 348\ninside 349 372

25584 GCF_002033065.1_ASM203306v1 Salmonella enterica subsp. enterica serovar Typhimurium var. 5- Proteobacteria; Gammaproteobacteria; Enterobacterales; Enterobacteriaceae; Salmonella; Salmonella enterica; Salmonella enterica subsp. enterica; Salmonella enterica subsp. enterica serovar Typhimurium MTGDNTLITSHGINRRDFMKLCAALAATMGLSSKAAAEMAESVSNPQRPPVIWIGAQECT WP_000145429.1 hydrogenase 2 small subunit [Salmonella enterica] Length: 372\nNumber of predicted TMHs: 1\nExp number of AAs in TMHs: 24.51309\nExp number, first 60 AAs: 0.2682\nTotal prob of N-in: 0.14402\noutside 1 331\nTMhelix 332 354\ninside 355 372

25585 GCF_002058075.1_ASM205807v1 Salmonella enterica subsp. enterica serovar Typhimurium var. 5- Proteobacteria; Gammaproteobacteria; Enterobacterales; Enterobacteriaceae; Salmonella; Salmonella enterica; Salmonella enterica subsp. enterica; Salmonella enterica subsp. enterica serovar Typhimurium MTGDNTLITSHGINRRDFMKLCAALAATMGLSSKAAAEMAESVSNPQRPPVIWIGAQECT WP_000145429.1 hydrogenase 2 small subunit [Salmonella enterica] Length: 372\nNumber of predicted TMHs: 1\nExp number of AAs in TMHs: 24.51309\nExp number, first 60 AAs: 0.2682\nTotal prob of N-in: 0.14402\noutside 1 331\nTMhelix 332 354\ninside 355 372

25586 GCF_001480025.1_Salmonella_enterica_CVM_N38903-SQ_v1.0 Salmonella enterica subsp. enterica serovar Typhimurium var. 5- Proteobacteria; Gammaproteobacteria; Enterobacterales; Enterobacteriaceae; Salmonella; Salmonella enterica; Salmonella enterica subsp. enterica; Salmonella enterica subsp. enterica serovar Typhimurium MTGDNTLITSHGINRRDFMKLCAALAATMGLSSKAAAEMAESVSNPQRPPVIWIGAQECT WP_000145429.1 hydrogenase 2 small subunit [Salmonella enterica] Length: 372\nNumber of predicted TMHs: 1\nExp number of AAs in TMHs: 24.51309\nExp number, first 60 AAs: 0.2682\nTotal prob of N-in: 0.14402\noutside 1 331\nTMhelix 332 354\ninside 355 372

25587 GCF_001479925.1_Salmonella_enterica_CVM_N38873-R_v1.0 Salmonella enterica subsp. enterica serovar Typhimurium var. 5- Proteobacteria; Gammaproteobacteria; Enterobacterales; Enterobacteriaceae; Salmonella; Salmonella enterica; Salmonella enterica subsp. enterica; Salmonella enterica subsp. enterica serovar Typhimurium MNNEETFYQAMRRKGVTRRSFLKFCSLAATSLGLGAGMTPKIAWALENKPRIPVVWIHGL WP_001058312.1 hydrogenase [Salmonella enterica] Length: 372\nNumber of predicted TMHs: 2\nExp number of AAs in TMHs: 36.51738\nExp number, first 60 AAs: 19.72849\nTotal prob of N-in: 0.96170\nPOSSIBLE N-term signal sequence\ninside 1 19\nTMhelix 20 39\noutside 40 325\nTMhelix 326 348\ninside 349 372

25588 GCF_001479925.1_Salmonella_enterica_CVM_N38873-R_v1.0 Salmonella enterica subsp. enterica serovar Typhimurium var. 5- Proteobacteria; Gammaproteobacteria; Enterobacterales; Enterobacteriaceae; Salmonella; Salmonella enterica; Salmonella enterica subsp. enterica; Salmonella enterica subsp. enterica serovar Typhimurium MTGDNTLITSHGINRRDFMKLCAALAATMGLSSKAAAEMAESVSNPQRPPVIWIGAQECT WP_000145429.1 hydrogenase 2 small subunit [Salmonella enterica] Length: 372\nNumber of predicted TMHs: 1\nExp number of AAs in TMHs: 24.51309\nExp number, first 60 AAs: 0.2682\nTotal prob of N-in: 0.14402\noutside 1 331\nTMhelix 332 354\ninside 355 372

25589 GCF_001480205.1_Salmonella_enterica_CVM_N40379-SQ_v1.0 Salmonella enterica subsp. enterica serovar Typhimurium var. 5- Proteobacteria; Gammaproteobacteria; Enterobacterales; Enterobacteriaceae; Salmonella; Salmonella enterica; Salmonella enterica subsp. enterica; Salmonella enterica subsp. enterica serovar Typhimurium MTGDNTLITSHGINRRDFMKLCAALAATMGLSSKAAAEMAESVSNPQRPPVIWIGAQECT WP_000145429.1 hydrogenase 2 small subunit [Salmonella enterica] Length: 372\nNumber of predicted TMHs: 1\nExp number of AAs in TMHs: 24.51309\nExp number, first 60 AAs: 0.2682\nTotal prob of N-in: 0.14402\noutside 1 331\nTMhelix 332 354\ninside 355 372

25590 GCF_000505085.1_CFSAN001284_01.0Salmonella enterica subsp. enterica serovar Typhimurium var. Copenhagen str. 0084 Proteobacteria; Gammaproteobacteria; Enterobacterales; Enterobacteriaceae; Salmonella; Salmonella enterica; Salmonella enterica subsp. enterica; Salmonella enterica subsp. enterica serovar Typhimurium MTGDNTLITSHGINRRDFMKLCAALAATMGLSSKAAAEMAESVSNPQRPPVIWIGAQECT WP_000145429.1 hydrogenase 2 small subunit [Salmonella enterica] Length: 372\nNumber of predicted TMHs: 1\nExp number of AAs in TMHs: 24.51309\nExp number, first 60 AAs: 0.2682\nTotal prob of N-in: 0.14402\noutside 1 331\nTMhelix 332 354\ninside 355 372

25591 GCF_000505085.1_CFSAN001284_01.0Salmonella enterica subsp. enterica serovar Typhimurium var. Copenhagen str. 0084 Proteobacteria; Gammaproteobacteria; Enterobacterales; Enterobacteriaceae; Salmonella; Salmonella enterica; Salmonella enterica subsp. enterica; Salmonella enterica subsp. enterica serovar Typhimurium MNNEETFYQAMRRKGVTRRSFLKFCSLAATSLGLGAGMTPKIAWALENKPRIPVVWIHGL WP_001058312.1 hydrogenase [Salmonella enterica] Length: 372\nNumber of predicted TMHs: 2\nExp number of AAs in TMHs: 36.51738\nExp number, first 60 AAs: 19.72849\nTotal prob of N-in: 0.96170\nPOSSIBLE N-term signal sequence\ninside 1 19\nTMhelix 20 39\noutside 40 325\nTMhelix 326 348\ninside 349 372

25592 GCF_000022165.1_ASM2216v1 Salmonella enterica subsp. enterica serovar Typhimurium str. 14028S Proteobacteria; Gammaproteobacteria; Enterobacterales; Enterobacteriaceae; Salmonella; Salmonella enterica; Salmonella enterica subsp. enterica; Salmonella enterica subsp. enterica serovar Typhimurium; Salmonella enterica subsp. enterica serovar Typhimurium str. LT2 MTGDNTLITSHGINRRDFMKLCAALAATMGLSSKAAAEMAESVSNPQRPPVIWIGAQECT WP_000145429.1 hydrogenase 2 small subunit [Salmonella enterica] Length: 372\nNumber of predicted TMHs: 1\nExp number of AAs in TMHs: 24.51309\nExp number, first 60 AAs: 0.2682\nTotal prob of N-in: 0.14402\noutside 1 331\nTMhelix 332 354\ninside 355 372

25593 GCF_000022165.1_ASM2216v1 Salmonella enterica subsp. enterica serovar Typhimurium str. 14028S Proteobacteria; Gammaproteobacteria; Enterobacterales; Enterobacteriaceae; Salmonella; Salmonella enterica; Salmonella enterica subsp. enterica; Salmonella enterica subsp. enterica serovar Typhimurium; Salmonella enterica subsp. enterica serovar Typhimurium str. LT2 MNNEETFYQAMRRKGVTRRSFLKFCSLAATSLGLGAGMTPKIAWALENKPRIPVVWIHGL WP_001058312.1 hydrogenase [Salmonella enterica] Length: 372\nNumber of predicted TMHs: 2\nExp number of AAs in TMHs: 36.51738\nExp number, first 60 AAs: 19.72849\nTotal prob of N-in: 0.96170\nPOSSIBLE N-term signal sequence\ninside 1 19\nTMhelix 20 39\noutside 40 325\nTMhelix 326 348\ninside 349 372

25594 GCF_000336215.1_ASM33621v1 Salmonella enterica subsp. enterica serovar Typhimurium str. LT2-4 Proteobacteria; Gammaproteobacteria; Enterobacterales; Enterobacteriaceae; Salmonella; Salmonella enterica; Salmonella enterica subsp. enterica; Salmonella enterica subsp. enterica serovar Typhimurium; Salmonella enterica subsp. enterica serovar Typhimurium str. LT2 MTGDNTLITSHGINRRDFMKLCAALAATMGLSSKAAAEMAESVSNPQRPPVIWIGAQECT WP_000145429.1 hydrogenase 2 small subunit [Salmonella enterica] Length: 372\nNumber of predicted TMHs: 1\nExp number of AAs in TMHs: 24.51309\nExp number, first 60 AAs: 0.2682\nTotal prob of N-in: 0.14402\noutside 1 331\nTMhelix 332 354\ninside 355 372

25595 GCF_000336215.1_ASM33621v1 Salmonella enterica subsp. enterica serovar Typhimurium str. LT2-4 Proteobacteria; Gammaproteobacteria; Enterobacterales; Enterobacteriaceae; Salmonella; Salmonella enterica; Salmonella enterica subsp. enterica; Salmonella enterica subsp. enterica serovar Typhimurium; Salmonella enterica subsp. enterica serovar Typhimurium str. LT2 MNNEETFYQAMRRKGVTRRSFLKFCSLAATSLGLGAGMTPKIAWALENKPRIPVVWIHGL WP_001058312.1 hydrogenase [Salmonella enterica] Length: 372\nNumber of predicted TMHs: 2\nExp number of AAs in TMHs: 36.51738\nExp number, first 60 AAs: 19.72849\nTotal prob of N-in: 0.96170\nPOSSIBLE N-term signal sequence\ninside 1 19\nTMhelix 20 39\noutside 40 325\nTMhelix 326 348\ninside 349 372

25596 GCF_000336195.1_ASM33619v1 Salmonella enterica subsp. enterica serovar Typhimurium str. LT2-4_delta.ramA::kan Proteobacteria; Gammaproteobacteria; Enterobacterales; Enterobacteriaceae; Salmonella; Salmonella enterica; Salmonella enterica subsp. enterica; Salmonella enterica subsp. enterica serovar Typhimurium; Salmonella enterica subsp. enterica serovar Typhimurium str. LT2 MNNEETFYQAMRRKGVTRRSFLKFCSLAATSLGLGAGMTPKIAWALENKPRIPVVWIHGL WP_001058312.1 hydrogenase [Salmonella enterica] Length: 372\nNumber of predicted TMHs: 2\nExp number of AAs in TMHs: 36.51738\nExp number, first 60 AAs: 19.72849\nTotal prob of N-in: 0.96170\nPOSSIBLE N-term signal sequence\ninside 1 19\nTMhelix 20 39\noutside 40 325\nTMhelix 326 348\ninside 349 372

25597 GCF_000336195.1_ASM33619v1 Salmonella enterica subsp. enterica serovar Typhimurium str. LT2-4_delta.ramA::kan Proteobacteria; Gammaproteobacteria; Enterobacterales; Enterobacteriaceae; Salmonella; Salmonella enterica; Salmonella enterica subsp. enterica; Salmonella enterica subsp. enterica serovar Typhimurium; Salmonella enterica subsp. enterica serovar Typhimurium str. LT2

MTGDNTLITSHGINRRDFMKLCAALAATMGLSSKAAAEMAESVSNPQRPPVIWIGAQECT WP_000145429.1 hydrogenase 2 small subunit [Salmonella enterica] Length: 372\nNumber of predicted TMHs: 1\nExp number of AAs in TMHs: 24.51309\nExp number, first 60 AAs: 0.2682\nTotal prob of N-in: 0.14402\noutside 1 331\nTMhelix 332 354\ninside 355 372

25598 GCF_000430145.2_ASM43014v3 Salmonella enterica subsp. enterica serovar Typhimurium var. 5- str. CFSAN001921 Proteobacteria; Gammaproteobacteria; Enterobacterales; Enterobacteriaceae; Salmonella; Salmonella enterica; Salmonella enterica subsp. enterica; Salmonella enterica subsp. enterica serovar Typhimurium; Salmonella enterica subsp. enterica serovar Typhimurium var. 5-
MTGDNTLITSHGINRRDFMKLCAALAATMGLSSKAAAEMAESVSNPQRPPVIWIGAQECT WP_000145429.1 hydrogenase 2 small subunit [Salmonella enterica] Length: 372\nNumber of predicted TMHs: 1\nExp number of AAs in TMHs: 24.51309\nExp number, first 60 AAs: 0.2682\nTotal prob of N-in: 0.14402\noutside 1 331\nTMhelix 332 354\ninside 355 372

25599 GCF_000430145.2_ASM43014v3 Salmonella enterica subsp. enterica serovar Typhimurium var. 5- str. CFSAN001921 Proteobacteria; Gammaproteobacteria; Enterobacterales; Enterobacteriaceae; Salmonella; Salmonella enterica; Salmonella enterica subsp. enterica; Salmonella enterica subsp. enterica serovar Typhimurium; Salmonella enterica subsp. enterica serovar Typhimurium var. 5-
MNNEETFYQAMRRKGVTRRSFLKFCSLAATSLGLGAGMTPKIAWALENKPRIPVVWIHGL WP_001058312.1 hydrogenase [Salmonella enterica] Length: 372\nNumber of predicted TMHs: 2\nExp number of AAs in TMHs: 36.51738\nExp number, first 60 AAs: 19.72849\nTotal prob of N-in: 0.96170\nPOSSIBLE N-term signal sequence\ninside 1 19\nTMhelix 20 39\noutside 40 325\nTMhelix 326 348\ninside 349 372

25600 GCF_000505505.1_CFSAN004345_01.0Salmonella enterica subsp. enterica serovar Typhimurium var. 5- str. CFSAN004345 Proteobacteria; Gammaproteobacteria; Enterobacterales; Enterobacteriaceae; Salmonella; Salmonella enterica; Salmonella enterica subsp. enterica; Salmonella enterica subsp. enterica serovar Typhimurium; Salmonella enterica subsp. enterica serovar Typhimurium var. 5-
MTGDNTLITSHGINRRDFMKLCAALAATMGLSSKAAAEMAESVSNPQRPPVIWIGAQECT WP_000145431.1 hydrogenase 2 small subunit [Salmonella enterica] Length: 372\nNumber of predicted TMHs: 1\nExp number of AAs in TMHs: 24.51505\nExp number, first 60 AAs: 0.2682\nTotal prob of N-in: 0.14406\noutside 1 331\nTMhelix 332 354\ninside 355 372

25601 GCF_000505505.1_CFSAN004345_01.0Salmonella enterica subsp. enterica serovar Typhimurium var. 5- str. CFSAN004345 Proteobacteria; Gammaproteobacteria; Enterobacterales; Enterobacteriaceae; Salmonella; Salmonella enterica; Salmonella enterica subsp. enterica; Salmonella enterica subsp. enterica serovar Typhimurium; Salmonella enterica subsp. enterica serovar Typhimurium var. 5-
MNNEETFYQAMRRKGVTRRSFLKFCSLAATSLGLGAGMTPKIAWALENKPRIPVVWIHGL WP_001058312.1 hydrogenase [Salmonella enterica] Length: 372\nNumber of predicted TMHs: 2\nExp number of AAs in TMHs: 36.51738\nExp number, first 60 AAs: 19.72849\nTotal prob of N-in: 0.96170\nPOSSIBLE N-term signal sequence\ninside 1 19\nTMhelix 20 39\noutside 40 325\nTMhelix 326 348\ninside 349 372

25602 GCF_000486305.1_ASM48630v1 Salmonella enterica subsp. enterica serovar Urbana str. ATCC 9261 Proteobacteria; Gammaproteobacteria; Enterobacterales; Enterobacteriaceae; Salmonella; Salmonella enterica; Salmonella enterica subsp. enterica; Salmonella enterica subsp. enterica serovar Urbana
MTRRSFLKFCSLAATSLGLGAGMTPKIAWALENKPRIPVVWIHGLECTCCTESFIRSSHP WP_065304937.1 hydrogenase [Salmonella enterica] Length: 357\nNumber of predicted TMHs: 1\nExp number of AAs in TMHs: 19.92219\nExp number, first 60 AAs: 3.08454\nTotal prob of N-in: 0.16445\noutside 1 310\nTMhelix 311 333\ninside 334 357

25603 GCF_000486305.1_ASM48630v1 Salmonella enterica subsp. enterica serovar Urbana str. ATCC 9261 Proteobacteria; Gammaproteobacteria; Enterobacterales; Enterobacteriaceae; Salmonella; Salmonella enterica; Salmonella enterica subsp. enterica; Salmonella enterica subsp. enterica serovar Urbana
MTGDNTLITSHGINRRDFMKLCAALAATMGLSSKAAAEMAESVSNPQRPPVIWIGAQECT WP_000145429.1 hydrogenase 2 small subunit [Salmonella enterica] Length: 372\nNumber of predicted TMHs: 1\nExp number of AAs in TMHs: 24.51309\nExp number, first 60 AAs: 0.2682\nTotal prob of N-in: 0.14402\noutside 1 331\nTMhelix 332 354\ninside 355 372

25604 GCF_000171535.2_ASM17153v2 Salmonella enterica subsp. enterica serovar Virchow str. SL491 Proteobacteria; Gammaproteobacteria; Enterobacterales; Enterobacteriaceae; Salmonella; Salmonella enterica; Salmonella enterica subsp. enterica; Salmonella enterica subsp. enterica serovar Virchow
MNNEETFYQAMRRKGVTRRSFLKFCSLAATSLGLGAGMTPKIAWALENKPRIPVVWIHGL WP_001058310.1 hydrogenase [Salmonella enterica] Length: 372\nNumber of predicted TMHs: 2\nExp number of AAs in

TMHs: 37.4567299999999\nExp number, first 60 AAs: 19.72694\nTotal prob of N-in: 0.96171\nPOSSIBLE N-term signal sequence\ninside 1 19\nTMhelix 20 39\noutside 40 325\nTMhelix 326 348\ninside 349 372

25605 GCF_000171535.2_ASM17153v2 Salmonella enterica subsp. enterica serovar Virchow str. SL491 Proteobacteria; Gammaproteobacteria; Enterobacterales; Enterobacteriaceae; Salmonella; Salmonella enterica; Salmonella enterica subsp. enterica; Salmonella enterica subsp. enterica serovar Virchow MTGDNTLITSHGINRRDFMKLCAALAATMGLSSKAAAEMAESVSNPQRPPVIWIGAQECT WP_000145429.1 hydrogenase 2 small subunit [Salmonella enterica] Length: 372\nNumber of predicted TMHs: 1\nExp number of AAs in TMHs: 24.51309\nExp number, first 60 AAs: 0.2682\nTotal prob of N-in: 0.14402\noutside 1 331\nTMhelix 332 354\ninside 355 372

25606 GCF_000516855.1_SvirSVQ1_v1.0 Salmonella enterica subsp. enterica serovar Virchow str. SVQ1 Proteobacteria; Gammaproteobacteria; Enterobacterales; Enterobacteriaceae; Salmonella; Salmonella enterica; Salmonella enterica subsp. enterica; Salmonella enterica subsp. enterica serovar Virchow MTGDNTLITSHGINRRDFMKLCAALAATMGLSSKAAAEMAESVSNPQRPPVIWIGAQECT WP_000145429.1 hydrogenase 2 small subunit [Salmonella enterica] Length: 372\nNumber of predicted TMHs: 1\nExp number of AAs in TMHs: 24.51309\nExp number, first 60 AAs: 0.2682\nTotal prob of N-in: 0.14402\noutside 1 331\nTMhelix 332 354\ninside 355 372

25607 GCF_000516855.1_SvirSVQ1_v1.0 Salmonella enterica subsp. enterica serovar Virchow str. SVQ1 Proteobacteria; Gammaproteobacteria; Enterobacterales; Enterobacteriaceae; Salmonella; Salmonella enterica; Salmonella enterica subsp. enterica; Salmonella enterica subsp. enterica serovar Virchow MNNEETFYQAMRRKGVTRRSFLKFCSLAATSLGLGAGMTPKIAWALENKPRIPVVWIHGL WP_001058310.1 hydrogenase [Salmonella enterica] Length: 372\nNumber of predicted TMHs: 2\nExp number of AAs in TMHs: 37.4567299999999\nExp number, first 60 AAs: 19.72694\nTotal prob of N-in: 0.96171\nPOSSIBLE N-term signal sequence\ninside 1 19\nTMhelix 20 39\noutside 40 325\nTMhelix 326 348\ninside 349 372

25608 GCF_001448885.1_ASM144888v1 Salmonella enterica subsp. enterica serovar Virginia str. SA19971529 Proteobacteria; Gammaproteobacteria; Enterobacterales; Enterobacteriaceae; Salmonella; Salmonella enterica; Salmonella enterica subsp. enterica; Salmonella enterica subsp. enterica serovar Virginia MTGDNTLITSHGINRRDFMKLCAALAATMGLSSKAAAEMAESVSNPQRPPVIWIGAQECT WP_000145429.1 hydrogenase 2 small subunit [Salmonella enterica] Length: 372\nNumber of predicted TMHs: 1\nExp number of AAs in TMHs: 24.51309\nExp number, first 60 AAs: 0.2682\nTotal prob of N-in: 0.14402\noutside 1 331\nTMhelix 332 354\ninside 355 372

25609 GCF_001448885.1_ASM144888v1 Salmonella enterica subsp. enterica serovar Virginia str. SA19971529 Proteobacteria; Gammaproteobacteria; Enterobacterales; Enterobacteriaceae; Salmonella; Salmonella enterica; Salmonella enterica subsp. enterica; Salmonella enterica subsp. enterica serovar Virginia MNNEETFYQAMRRKGVTRRSFLKFCSLAATSLGLGAGMTPKIAWALENKPRIPVVWIHGL WP_001058312.1 hydrogenase [Salmonella enterica] Length: 372\nNumber of predicted TMHs: 2\nExp number of AAs in TMHs: 36.51738\nExp number, first 60 AAs: 19.72849\nTotal prob of N-in: 0.96170\nPOSSIBLE N-term signal sequence\ninside 1 19\nTMhelix 20 39\noutside 40 325\nTMhelix 326 348\ninside 349 372

25610 GCF_001975625.1_ASM197562v1 Salmonella enterica subsp. enterica serovar Wandsworth str. SA20092095 Proteobacteria; Gammaproteobacteria; Enterobacterales; Enterobacteriaceae; Salmonella; Salmonella enterica; Salmonella enterica subsp. enterica; Salmonella enterica subsp. enterica serovar Wandsworth MNNEETFYQAMRRKGVTRRSFLKFCSLAATSLGLGAGMTPKIAWALENKPRIPVVWIHGL WP_001058312.1 hydrogenase [Salmonella enterica] Length: 372\nNumber of predicted TMHs: 2\nExp number of AAs in TMHs: 36.51738\nExp number, first 60 AAs: 19.72849\nTotal prob of N-in: 0.96170\nPOSSIBLE N-term signal sequence\ninside 1 19\nTMhelix 20 39\noutside 40 325\nTMhelix 326 348\ninside 349 372

25611 GCF_001975625.1_ASM197562v1 Salmonella enterica subsp. enterica serovar Wandsworth str. SA20092095 Proteobacteria; Gammaproteobacteria; Enterobacterales; Enterobacteriaceae; Salmonella; Salmonella enterica; Salmonella enterica subsp. enterica; Salmonella enterica subsp. enterica serovar Wandsworth MTGDNTLITSHGINRRDFMKLCAALAATMGLSSKAAAEMAESVSNPQRPPVIWIGAQECT WP_000145429.1 hydrogenase 2 small subunit [Salmonella enterica] Length: 372\nNumber of predicted TMHs: 1\nExp number of AAs in TMHs: 24.51309\nExp number, first 60 AAs: 0.2682\nTotal prob of N-in: 0.14402\noutside 1 331\nTMhelix 332 354\ninside 355 372

25612 GCF_001612515.1_ASM161251v1 Salmonella enterica subsp. enterica serovar Weltevreden str. 1655 Proteobacteria; Gammaproteobacteria; Enterobacterales; Enterobacteriaceae; Salmonella; Salmonella enterica;

Salmonella enterica subsp. enterica; Salmonella enterica subsp. enterica serovar Weltevreden
MTGDNTLITSHGINRRDFMKLCAALAATMGLSSKAAAEMAESVSNPQRPPVIWIGAQECT WP_000145429.1
hydrogenase 2 small subunit [Salmonella enterica] Length: 372\nNumber of predicted TMHs: 1\nExp number of AAs in TMHs: 24.51309\nExp number, first 60 AAs: 0.2682\nTotal prob of N-in: 0.14402\noutside 1 331\nTMhelix 332 354\ninside 355 372

25613 GCF_001612515.1_ASM161251v1 Salmonella enterica subsp. enterica serovar Weltevreden str. 1655
Proteobacteria; Gammaproteobacteria; Enterobacterales; Enterobacteriaceae; Salmonella; Salmonella enterica; Salmonella enterica subsp. enterica; Salmonella enterica subsp. enterica serovar Weltevreden
MNNEETFYQAMRRKGVTRRSFLKFCSLAATSLGLGAGMTPKIAWALENKPRIPVVWIHGL WP_001058307.1
hydrogenase [Salmonella enterica] Length: 372\nNumber of predicted TMHs: 2\nExp number of AAs in TMHs: 36.51738\nExp number, first 60 AAs: 19.72849\nTotal prob of N-in: 0.96170\nPOSSIBLE N-term signal sequence\ninside 1 19\nTMhelix 20 39\noutside 40 325\nTMhelix 326 348\ninside 349 372

25614 GCF_000171495.1_ASM17149v1 Salmonella enterica subsp. enterica serovar Weltevreden str. HI_N05-537
Proteobacteria; Gammaproteobacteria; Enterobacterales; Enterobacteriaceae; Salmonella; Salmonella enterica; Salmonella enterica subsp. enterica; Salmonella enterica subsp. enterica serovar Weltevreden
MNNEETFYQAMRRKGVTRRSFLKFCSLAATSLGLGAGMTPKIAWALENKPRIPVVWIHGL WP_001058307.1
hydrogenase [Salmonella enterica] Length: 372\nNumber of predicted TMHs: 2\nExp number of AAs in TMHs: 36.51738\nExp number, first 60 AAs: 19.72849\nTotal prob of N-in: 0.96170\nPOSSIBLE N-term signal sequence\ninside 1 19\nTMhelix 20 39\noutside 40 325\nTMhelix 326 348\ninside 349 372

25615 GCF_000171495.1_ASM17149v1 Salmonella enterica subsp. enterica serovar Weltevreden str. HI_N05-537
Proteobacteria; Gammaproteobacteria; Enterobacterales; Enterobacteriaceae; Salmonella; Salmonella enterica; Salmonella enterica subsp. enterica; Salmonella enterica subsp. enterica serovar Weltevreden
MTGDNTLITSHGINRRDFMKLCAALAATMGLSSKAAAEMAESVSNPQRPPVIWIGAQECT WP_000145429.1
hydrogenase 2 small subunit [Salmonella enterica] Length: 372\nNumber of predicted TMHs: 1\nExp number of AAs in TMHs: 24.51309\nExp number, first 60 AAs: 0.2682\nTotal prob of N-in: 0.14402\noutside 1 331\nTMhelix 332 354\ninside 355 372

25616 GCF_001448905.1_ASM144890v1 Salmonella enterica subsp. enterica serovar Weslaco str. 247K
Proteobacteria; Gammaproteobacteria; Enterobacterales; Enterobacteriaceae; Salmonella; Salmonella enterica; Salmonella enterica subsp. enterica; Salmonella enterica subsp. enterica serovar Weslaco
MNNEETFYQAIRRKGVTRRSFLKFCSLAATSLGLGAGMTPKIAWALENKPRIPVVWIHGL WP_058110568.1
hydrogenase [Salmonella enterica] Length: 372\nNumber of predicted TMHs: 2\nExp number of AAs in TMHs: 35.65465\nExp number, first 60 AAs: 18.86174\nTotal prob of N-in: 0.91931\nPOSSIBLE N-term signal sequence\ninside 1 19\nTMhelix 20 39\noutside 40 325\nTMhelix 326 348\ninside 349 372

25617 GCF_001448905.1_ASM144890v1 Salmonella enterica subsp. enterica serovar Weslaco str. 247K
Proteobacteria; Gammaproteobacteria; Enterobacterales; Enterobacteriaceae; Salmonella; Salmonella enterica; Salmonella enterica subsp. enterica; Salmonella enterica subsp. enterica serovar Weslaco
MTGDNTLITSHGINRRDFMKLCAALAATMGLSSKAAAEMAESVSNPQRPPVIWIGAQECT WP_000145429.1
hydrogenase 2 small subunit [Salmonella enterica] Length: 372\nNumber of predicted TMHs: 1\nExp number of AAs in TMHs: 24.51309\nExp number, first 60 AAs: 0.2682\nTotal prob of N-in: 0.14402\noutside 1 331\nTMhelix 332 354\ninside 355 372

25618 GCF_000486265.1_ASM48626v1 Salmonella enterica subsp. enterica serovar Worthington str. ATCC 9607
Proteobacteria; Gammaproteobacteria; Enterobacterales; Enterobacteriaceae; Salmonella; Salmonella enterica; Salmonella enterica subsp. enterica; Salmonella enterica subsp. enterica serovar Worthington
MTGDNTLITSHGINRRDFMKLCAALAATMGLSSKAAAEMAESVSNPQRPPVIWIGAQECT WP_000145429.1
hydrogenase 2 small subunit [Salmonella enterica] Length: 372\nNumber of predicted TMHs: 1\nExp number of AAs in TMHs: 24.51309\nExp number, first 60 AAs: 0.2682\nTotal prob of N-in: 0.14402\noutside 1 331\nTMhelix 332 354\ninside 355 372

25619 GCF_001975645.1_ASM197564v1 Salmonella enterica subsp. enterica serovar Yovokome str. S-1850
Proteobacteria; Gammaproteobacteria; Enterobacterales; Enterobacteriaceae; Salmonella; Salmonella enterica; Salmonella enterica subsp. enterica; Salmonella enterica subsp. enterica serovar Yovokome
MNNEETFYQAMRRKGVTRRSFLKFCSLAATSLGLGAGMTPKIAWALENKPRIPVVWIHGL WP_001058311.1
hydrogenase [Salmonella enterica] Length: 372\nNumber of predicted TMHs: 2\nExp number of AAs in TMHs: 37.4699899999999\nExp number, first 60 AAs: 19.72699\nTotal prob of N-in: 0.96171\nPOSSIBLE N-term signal sequence\ninside 1 19\nTMhelix 20 39\noutside 40 325\nTMhelix 326 348\ninside 349 372

25620 GCF_001975645.1_ASM197564v1 Salmonella enterica subsp. enterica serovar Yovokome str. S-1850 Proteobacteria; Gammaproteobacteria; Enterobacterales; Enterobacteriaceae; Salmonella; Salmonella enterica; Salmonella enterica subsp. enterica; Salmonella enterica subsp. enterica serovar Yovokome MTGDNTLITSHGINRRDFMKLCAALAATMGLSSKAAAEMAESVSNPQRPPVIWIGAQECT WP_000145429.1 hydrogenase 2 small subunit [Salmonella enterica] Length: 372\nNumber of predicted TMHs: 1\nExp number of AAs in TMHs: 24.51309\nExp number, first 60 AAs: 0.2682\nTotal prob of N-in: 0.14402\noutside 1 331\nTMhelix 332 354\ninside 355 372

25621 GCF_002037585.1_ASM203758v1 Salmonella enterica subsp. houtenae serovar 11:z4,z23:- Proteobacteria; Gammaproteobacteria; Enterobacterales; Enterobacteriaceae; Salmonella; Salmonella enterica; Salmonella enterica subsp. houtenae MTGDNTLITSHGINRRDFMKLCAALAATMGLSSKAAAEMAESVSNPQRPPVIWIGAQECT WP_000145429.1 hydrogenase 2 small subunit [Salmonella enterica] Length: 372\nNumber of predicted TMHs: 1\nExp number of AAs in TMHs: 24.51309\nExp number, first 60 AAs: 0.2682\nTotal prob of N-in: 0.14402\noutside 1 331\nTMhelix 332 354\ninside 355 372

25622 GCF_002037585.1_ASM203758v1 Salmonella enterica subsp. houtenae serovar 11:z4,z23:- Proteobacteria; Gammaproteobacteria; Enterobacterales; Enterobacteriaceae; Salmonella; Salmonella enterica; Salmonella enterica subsp. houtenae MNNEETFYQAMRRKGVTRRSFLKFCSLAATSLGLGAGMTPKIAWALENKPRIPVVWIHGL WP_001058310.1 hydrogenase [Salmonella enterica] Length: 372\nNumber of predicted TMHs: 2\nExp number of AAs in TMHs: 37.4567299999999\nExp number, first 60 AAs: 19.72694\nTotal prob of N-in: 0.96171\nPOSSIBLE N-term signal sequence\ninside 1 19\nTMhelix 20 39\noutside 40 325\nTMhelix 326 348\ninside 349 372

25623 GCF_002035325.1_ASM203532v1 Salmonella enterica subsp. houtenae serovar 16:z4,z32:- Proteobacteria; Gammaproteobacteria; Enterobacterales; Enterobacteriaceae; Salmonella; Salmonella enterica; Salmonella enterica subsp. houtenae MNNEETFYQAMRRKGVTRRSFLKFCSLATTSLGLGAGMTPKIAWALENKPRIPVVWIHGL WP_001058317.1 hydrogenase [Salmonella enterica] Length: 372\nNumber of predicted TMHs: 2\nExp number of AAs in TMHs: 36.4727299999999\nExp number, first 60 AAs: 19.19744\nTotal prob of N-in: 0.93666\nPOSSIBLE N-term signal sequence\ninside 1 19\nTMhelix 20 39\noutside 40 325\nTMhelix 326 348\ninside 349 372

25624 GCF_002035325.1_ASM203532v1 Salmonella enterica subsp. houtenae serovar 16:z4,z32:- Proteobacteria; Gammaproteobacteria; Enterobacterales; Enterobacteriaceae; Salmonella; Salmonella enterica; Salmonella enterica subsp. houtenae MTGDNTLITSHGINRRDFMKLCAALAATMGLSSKAAAEMAESVSNPQRPPVIWIGAQECT WP_000145429.1 hydrogenase 2 small subunit [Salmonella enterica] Length: 372\nNumber of predicted TMHs: 1\nExp number of AAs in TMHs: 24.51309\nExp number, first 60 AAs: 0.2682\nTotal prob of N-in: 0.14402\noutside 1 331\nTMhelix 332 354\ninside 355 372

25625 GCF_002127815.1_ASM212781v1 Salmonella enterica subsp. houtenae serovar 40gt:- Proteobacteria; Gammaproteobacteria; Enterobacterales; Enterobacteriaceae; Salmonella; Salmonella enterica; Salmonella enterica subsp. houtenae MTGDNTLITSHGINRRDFMKLCAALAATMGLSSKAAAEMAESVSNPQRPPVIWIGAQECT WP_000145429.1 hydrogenase 2 small subunit [Salmonella enterica] Length: 372\nNumber of predicted TMHs: 1\nExp number of AAs in TMHs: 24.51309\nExp number, first 60 AAs: 0.2682\nTotal prob of N-in: 0.14402\noutside 1 331\nTMhelix 332 354\ninside 355 372

25626 GCF_001953555.1_ASM195355v1 Salmonella enterica subsp. houtenae serovar 40gt:- Proteobacteria; Gammaproteobacteria; Enterobacterales; Enterobacteriaceae; Salmonella; Salmonella enterica; Salmonella enterica subsp. houtenae MNNEETFYQAMRRKGVTRRSFLKFCSLAATSLGLGAGMTPKIAWALENKPRIPVVWIHGL WP_023187532.1 hydrogenase 1, small subunit [Salmonella enterica] Length: 372\nNumber of predicted TMHs: 2\nExp number of AAs in TMHs: 36.9997\nExp number, first 60 AAs: 19.72759\nTotal prob of N-in: 0.96171\nPOSSIBLE N-term signal sequence\ninside 1 19\nTMhelix 20 39\noutside 40 325\nTMhelix 326 348\ninside 349 372

25627 GCF_001953555.1_ASM195355v1 Salmonella enterica subsp. houtenae serovar 40gt:- Proteobacteria; Gammaproteobacteria; Enterobacterales; Enterobacteriaceae; Salmonella; Salmonella enterica; Salmonella enterica subsp. houtenae MTGDNTLITSHGINRRDFMKLCAALAATMGLSSKAAAEMAESVSNPQRPPVIWIGAQECT WP_000145429.1 hydrogenase 2 small subunit [Salmonella enterica] Length: 372\nNumber of predicted TMHs: 1\nExp number of AAs in TMHs: 24.51309\nExp number, first 60 AAs: 0.2682\nTotal prob of N-in: 0.14402\noutside 1 331\nTMhelix 332 354\ninside 355 372

25628 GCF_002127815.1_ASM212781v1 Salmonella enterica subsp. houtenae serovar 40gt:- Proteobacteria; Gammaproteobacteria; Enterobacterales; Enterobacteriaceae; Salmonella; Salmonella enterica; Salmonella enterica subsp. houtenae MNNEETFYQAMRRKGVTRRSFLKFCSLAATSLGLGAGMTPKIAWALENKPRIPVVWIHGL WP_023187532.1 hydrogenase 1, small subunit [Salmonella enterica] Length: 372\nNumber of predicted TMHs: 2\nExp number of AAs in TMHs: 36.9997\nExp number, first 60 AAs: 19.72759\nTotal prob of N-in: 0.96171\nPOSSIBLE N-term signal sequence\ninside 1 19\nTMhelix 20 39\noutside 40 325\nTMhelix 326 348\ninside 349 372

25629 GCF_001953515.1_ASM195351v1 Salmonella enterica subsp. houtenae serovar 40gt:- Proteobacteria; Gammaproteobacteria; Enterobacterales; Enterobacteriaceae; Salmonella; Salmonella enterica; Salmonella enterica subsp. houtenae MTGDNTLITSHGINRRDFMKLCAALAATMGLSSKAAAEMAESVSNPQRPPVIWIGAQECT WP_000145429.1 hydrogenase 2 small subunit [Salmonella enterica] Length: 372\nNumber of predicted TMHs: 1\nExp number of AAs in TMHs: 24.51309\nExp number, first 60 AAs: 0.2682\nTotal prob of N-in: 0.14402\noutside 1 331\nTMhelix 332 354\ninside 355 372

25630 GCF_001953525.1_ASM195352v1 Salmonella enterica subsp. houtenae serovar 40gt:- Proteobacteria; Gammaproteobacteria; Enterobacterales; Enterobacteriaceae; Salmonella; Salmonella enterica; Salmonella enterica subsp. houtenae MTGDNTLITSHGINRRDFMKLCAALAATMGLSSKAAAEMAESVSNPQRPPVIWIGAQECT WP_000145429.1 hydrogenase 2 small subunit [Salmonella enterica] Length: 372\nNumber of predicted TMHs: 1\nExp number of AAs in TMHs: 24.51309\nExp number, first 60 AAs: 0.2682\nTotal prob of N-in: 0.14402\noutside 1 331\nTMhelix 332 354\ninside 355 372

25631 GCF_001953515.1_ASM195351v1 Salmonella enterica subsp. houtenae serovar 40gt:- Proteobacteria; Gammaproteobacteria; Enterobacterales; Enterobacteriaceae; Salmonella; Salmonella enterica; Salmonella enterica subsp. houtenae MNNEETFYQAMRRKGVTRRSFLKFCSLAATSLGLGAGMTPKIAWALENKPRIPVVWIHGL WP_023187532.1 hydrogenase 1, small subunit [Salmonella enterica] Length: 372\nNumber of predicted TMHs: 2\nExp number of AAs in TMHs: 36.9997\nExp number, first 60 AAs: 19.72759\nTotal prob of N-in: 0.96171\nPOSSIBLE N-term signal sequence\ninside 1 19\nTMhelix 20 39\noutside 40 325\nTMhelix 326 348\ninside 349 372

25632 GCF_001953525.1_ASM195352v1 Salmonella enterica subsp. houtenae serovar 40gt:- Proteobacteria; Gammaproteobacteria; Enterobacterales; Enterobacteriaceae; Salmonella; Salmonella enterica; Salmonella enterica subsp. houtenae MNNEETFYQAMRRKGVTRRSFLKFCSLAATSLGLGAGMTPKIAWALENKPRIPVVWIHGL WP_023187532.1 hydrogenase 1, small subunit [Salmonella enterica] Length: 372\nNumber of predicted TMHs: 2\nExp number of AAs in TMHs: 36.9997\nExp number, first 60 AAs: 19.72759\nTotal prob of N-in: 0.96171\nPOSSIBLE N-term signal sequence\ninside 1 19\nTMhelix 20 39\noutside 40 325\nTMhelix 326 348\ninside 349 372

25633 GCF_002106115.1_ASM210611v1 Salmonella enterica subsp. houtenae serovar 40:z4,z32:- Proteobacteria; Gammaproteobacteria; Enterobacterales; Enterobacteriaceae; Salmonella; Salmonella enterica; Salmonella enterica subsp. houtenae MTGDNTLITSHGINRRDFMKLCAALAATMGLSSKAAAEMAESVSNPQRPPVIWIGAQECT WP_000145429.1 hydrogenase 2 small subunit [Salmonella enterica] Length: 372\nNumber of predicted TMHs: 1\nExp number of AAs in TMHs: 24.51309\nExp number, first 60 AAs: 0.2682\nTotal prob of N-in: 0.14402\noutside 1 331\nTMhelix 332 354\ninside 355 372

25634 GCF_002106585.1_ASM210658v1 Salmonella enterica subsp. houtenae serovar 40:z4,z32:- Proteobacteria; Gammaproteobacteria; Enterobacterales; Enterobacteriaceae; Salmonella; Salmonella enterica; Salmonella enterica subsp. houtenae MNNEETFYQAMRRKGVTRRSFLKFCSLATTSLGLGAGMTPKIAWALENKPRIPVVWIHGL WP_085335578.1 [Ni/Fe] hydrogenase small subunit [Salmonella enterica] Length: 372\nNumber of predicted TMHs: 2\nExp number of AAs in TMHs: 36.4727299999999\nExp number, first 60 AAs: 19.19744\nTotal prob of N-in: 0.93666\nPOSSIBLE N-term signal sequence\ninside 1 19\nTMhelix 20 39\noutside 40 325\nTMhelix 326 348\ninside 349 372

25635 GCF_002106415.1_ASM210641v1 Salmonella enterica subsp. houtenae serovar 40:z4,z32:- Proteobacteria; Gammaproteobacteria; Enterobacterales; Enterobacteriaceae; Salmonella; Salmonella enterica; Salmonella enterica subsp. houtenae MTGDNTLITSHGINRRDFMKLCAALAATMGLSSKAAAEMAESVSNPQRPPVIWIGAQECT WP_000145429.1 hydrogenase 2 small subunit [Salmonella enterica] Length: 372\nNumber of predicted TMHs: 1\nExp number of AAs in TMHs: 24.51309\nExp number, first 60 AAs: 0.2682\nTotal prob of N-in: 0.14402\noutside 1 331\nTMhelix 332 354\ninside 355 372

25636	GCF_002106115.1_ASM210611v1	Salmonella enterica subsp. houtenae serovar 40:z4,z32:-	Proteobacteria; Gammaproteobacteria; Enterobacterales; Enterobacteriaceae; Salmonella; Salmonella enterica; Salmonella enterica subsp. houtenae	MNNEETFYQAMRRKGVTRRSFLKFCSLATTSLGLGAGMTPKIAWALENKPRIPVVWIHGL	WP_085335578.1	[Ni/Fe] hydrogenase small subunit [Salmonella enterica]	Length: 372\nNumber of predicted TMHs: 2\nExp number of AAs in TMHs: 36.4727299999999\nExp number, first 60 AAs: 19.19744\nTotal prob of N-in: 0.93666\nPOSSIBLE N-term signal sequence\ninside 1 19\nTMhelix 20 39\noutside 40 325\nTMhelix 326 348\ninside 349 372

25637	GCF_002105995.1_ASM210599v1	Salmonella enterica subsp. houtenae serovar 40:z4,z32:-	Proteobacteria; Gammaproteobacteria; Enterobacterales; Enterobacteriaceae; Salmonella; Salmonella enterica; Salmonella enterica subsp. houtenae	MTGDNTLITSHGINRRDFMKLCAALAATMGLSSKAAAEMAESVSNPQRPPVIWIGAQECT	WP_000145429.1	hydrogenase 2 small subunit [Salmonella enterica]	Length: 372\nNumber of predicted TMHs: 1\nExp number of AAs in TMHs: 24.51309\nExp number, first 60 AAs: 0.2682\nTotal prob of N-in: 0.14402\noutside 1 331\nTMhelix 332 354\ninside 355 372

25638	GCF_002106445.1_ASM210644v1	Salmonella enterica subsp. houtenae serovar 40:z4,z32:-	Proteobacteria; Gammaproteobacteria; Enterobacterales; Enterobacteriaceae; Salmonella; Salmonella enterica; Salmonella enterica subsp. houtenae	MNNEETFYQAMRRKGVTRRSFLKFCSLATTSLGLGAGMTPKIAWALENKPRIPVVWIHGL	WP_085335578.1	[Ni/Fe] hydrogenase small subunit [Salmonella enterica]	Length: 372\nNumber of predicted TMHs: 2\nExp number of AAs in TMHs: 36.4727299999999\nExp number, first 60 AAs: 19.19744\nTotal prob of N-in: 0.93666\nPOSSIBLE N-term signal sequence\ninside 1 19\nTMhelix 20 39\noutside 40 325\nTMhelix 326 348\ninside 349 372

25639	GCF_002105995.1_ASM210599v1	Salmonella enterica subsp. houtenae serovar 40:z4,z32:-	Proteobacteria; Gammaproteobacteria; Enterobacterales; Enterobacteriaceae; Salmonella; Salmonella enterica; Salmonella enterica subsp. houtenae	MNNEETFYQAMRRKGVTRRSFLKFCSLATTSLGLGAGMTPKIAWALENKPRIPVVWIHGL	WP_085335578.1	[Ni/Fe] hydrogenase small subunit [Salmonella enterica]	Length: 372\nNumber of predicted TMHs: 2\nExp number of AAs in TMHs: 36.4727299999999\nExp number, first 60 AAs: 19.19744\nTotal prob of N-in: 0.93666\nPOSSIBLE N-term signal sequence\ninside 1 19\nTMhelix 20 39\noutside 40 325\nTMhelix 326 348\ninside 349 372

25640	GCF_002106445.1_ASM210644v1	Salmonella enterica subsp. houtenae serovar 40:z4,z32:-	Proteobacteria; Gammaproteobacteria; Enterobacterales; Enterobacteriaceae; Salmonella; Salmonella enterica; Salmonella enterica subsp. houtenae	MTGDNTLITSHGINRRDFMKLCAALAATMGLSSKAAAEMAESVSNPQRPPVIWIGAQECT	WP_000145429.1	hydrogenase 2 small subunit [Salmonella enterica]	Length: 372\nNumber of predicted TMHs: 1\nExp number of AAs in TMHs: 24.51309\nExp number, first 60 AAs: 0.2682\nTotal prob of N-in: 0.14402\noutside 1 331\nTMhelix 332 354\ninside 355 372

25641	GCF_002106585.1_ASM210658v1	Salmonella enterica subsp. houtenae serovar 40:z4,z32:-	Proteobacteria; Gammaproteobacteria; Enterobacterales; Enterobacteriaceae; Salmonella; Salmonella enterica; Salmonella enterica subsp. houtenae	MTGDNTLITSHGINRRDFMKLCAALAATMGLSSKAAAEMAESVSNPQRPPVIWIGAQECT	WP_000145429.1	hydrogenase 2 small subunit [Salmonella enterica]	Length: 372\nNumber of predicted TMHs: 1\nExp number of AAs in TMHs: 24.51309\nExp number, first 60 AAs: 0.2682\nTotal prob of N-in: 0.14402\noutside 1 331\nTMhelix 332 354\ninside 355 372

25642	GCF_002106415.1_ASM210641v1	Salmonella enterica subsp. houtenae serovar 40:z4,z32:-	Proteobacteria; Gammaproteobacteria; Enterobacterales; Enterobacteriaceae; Salmonella; Salmonella enterica; Salmonella enterica subsp. houtenae	MNNEETFYQAMRRKGVTRRSFLKFCSLATTSLGLGAGMTPKIAWALENKPRIPVVWIHGL	WP_085335578.1	[Ni/Fe] hydrogenase small subunit [Salmonella enterica]	Length: 372\nNumber of predicted TMHs: 2\nExp number of AAs in TMHs: 36.4727299999999\nExp number, first 60 AAs: 19.19744\nTotal prob of N-in: 0.93666\nPOSSIBLE N-term signal sequence\ninside 1 19\nTMhelix 20 39\noutside 40 325\nTMhelix 326 348\ninside 349 372

25643	GCF_002044835.1_ASM204483v1	Salmonella enterica subsp. houtenae serovar 44:z4,z23:-	Proteobacteria; Gammaproteobacteria; Enterobacterales; Enterobacteriaceae; Salmonella; Salmonella enterica; Salmonella enterica subsp. houtenae	MNNEETFYQAMRRKGVTRRSFLKFCSLAATSLGLGAGMTPKIAWALENKPRIPVVWIHGL	WP_023187532.1	hydrogenase 1, small subunit [Salmonella enterica]	Length: 372\nNumber of predicted TMHs: 2\nExp number of AAs in TMHs: 36.9997\nExp number, first 60 AAs: 19.72759\nTotal prob of N-in: 0.96171\nPOSSIBLE N-term signal sequence\ninside 1 19\nTMhelix 20 39\noutside 40 325\nTMhelix 326 348\ninside 349 372

25644 GCF_002044835.1_ASM204483v1 Salmonella enterica subsp. houtenae serovar 44:z4,z23:- Proteobacteria; Gammaproteobacteria; Enterobacterales; Enterobacteriaceae; Salmonella; Salmonella enterica; Salmonella enterica subsp. houtenae MTGDNTLITSHGINRRDFMKLCAALAATMGLSSKAAAEMAESVSNPQRPPVIWIGAQECT WP_000145429.1 hydrogenase 2 small subunit [Salmonella enterica] Length: 372\nNumber of predicted TMHs: 1\nExp number of AAs in TMHs: 24.51309\nExp number, first 60 AAs: 0.2682\nTotal prob of N-in: 0.14402\noutside 1 331\nTMhelix 332 354\ninside 355 372

25645 GCF_002045125.1_ASM204512v1 Salmonella enterica subsp. houtenae serovar 44:z4,z23:- Proteobacteria; Gammaproteobacteria; Enterobacterales; Enterobacteriaceae; Salmonella; Salmonella enterica; Salmonella enterica subsp. houtenae MNNEETFYQAMRRKGVTRRSFLKFCSLAATSLGLGAGMTPKIAWALENKPRIPVVWIHGL WP_023187532.1 hydrogenase 1, small subunit [Salmonella enterica] Length: 372\nNumber of predicted TMHs: 2\nExp number of AAs in TMHs: 36.9997\nExp number, first 60 AAs: 19.72759\nTotal prob of N-in: 0.96171\nPOSSIBLE N-term signal sequence\ninside 1 19\nTMhelix 20 39\noutside 40 325\nTMhelix 326 348\ninside 349 372

25646 GCF_002044905.1_ASM204490v1 Salmonella enterica subsp. houtenae serovar 44:z4,z23:- Proteobacteria; Gammaproteobacteria; Enterobacterales; Enterobacteriaceae; Salmonella; Salmonella enterica; Salmonella enterica subsp. houtenae MNNEETFYQAMRRKGVTRRSFLKFCSLAATSLGLGAGMTPKIAWALENKPRIPVVWIHGL WP_023187532.1 hydrogenase 1, small subunit [Salmonella enterica] Length: 372\nNumber of predicted TMHs: 2\nExp number of AAs in TMHs: 36.9997\nExp number, first 60 AAs: 19.72759\nTotal prob of N-in: 0.96171\nPOSSIBLE N-term signal sequence\ninside 1 19\nTMhelix 20 39\noutside 40 325\nTMhelix 326 348\ninside 349 372

25647 GCF_002045125.1_ASM204512v1 Salmonella enterica subsp. houtenae serovar 44:z4,z23:- Proteobacteria; Gammaproteobacteria; Enterobacterales; Enterobacteriaceae; Salmonella; Salmonella enterica; Salmonella enterica subsp. houtenae MTGDNTLITSHGINRRDFMKLCAALAATMGLSSKAAAEMAESVSNPQRPPVIWIGAQECT WP_000145429.1 hydrogenase 2 small subunit [Salmonella enterica] Length: 372\nNumber of predicted TMHs: 1\nExp number of AAs in TMHs: 24.51309\nExp number, first 60 AAs: 0.2682\nTotal prob of N-in: 0.14402\noutside 1 331\nTMhelix 332 354\ninside 355 372

25648 GCF_002044905.1_ASM204490v1 Salmonella enterica subsp. houtenae serovar 44:z4,z23:- Proteobacteria; Gammaproteobacteria; Enterobacterales; Enterobacteriaceae; Salmonella; Salmonella enterica; Salmonella enterica subsp. houtenae MTGDNTLITSHGINRRDFMKLCAALAATMGLSSKAAAEMAESVSNPQRPPVIWIGAQECT WP_000145429.1 hydrogenase 2 small subunit [Salmonella enterica] Length: 372\nNumber of predicted TMHs: 1\nExp number of AAs in TMHs: 24.51309\nExp number, first 60 AAs: 0.2682\nTotal prob of N-in: 0.14402\noutside 1 331\nTMhelix 332 354\ninside 355 372

25649 GCF_002047425.1_ASM204742v1 Salmonella enterica subsp. houtenae serovar 44:z4,z23:- Proteobacteria; Gammaproteobacteria; Enterobacterales; Enterobacteriaceae; Salmonella; Salmonella enterica; Salmonella enterica subsp. houtenae MNNEETFYQAMRRKGVTRRSFLKFCSLAATSLGLGAGMTPKIAWALENKPRIPVVWIHGL WP_023187532.1 hydrogenase 1, small subunit [Salmonella enterica] Length: 372\nNumber of predicted TMHs: 2\nExp number of AAs in TMHs: 36.9997\nExp number, first 60 AAs: 19.72759\nTotal prob of N-in: 0.96171\nPOSSIBLE N-term signal sequence\ninside 1 19\nTMhelix 20 39\noutside 40 325\nTMhelix 326 348\ninside 349 372

25650 GCF_002047425.1_ASM204742v1 Salmonella enterica subsp. houtenae serovar 44:z4,z23:- Proteobacteria; Gammaproteobacteria; Enterobacterales; Enterobacteriaceae; Salmonella; Salmonella enterica; Salmonella enterica subsp. houtenae MTGDNTLITSHGINRRDFMKLCAALAATMGLSSKAAAEMAESVSNPQRPPVIWIGAQECT WP_000145429.1 hydrogenase 2 small subunit [Salmonella enterica] Length: 372\nNumber of predicted TMHs: 1\nExp number of AAs in TMHs: 24.51309\nExp number, first 60 AAs: 0.2682\nTotal prob of N-in: 0.14402\noutside 1 331\nTMhelix 332 354\ninside 355 372

25651 GCF_002045945.1_ASM204594v1 Salmonella enterica subsp. houtenae serovar 44:z4,z23:- Proteobacteria; Gammaproteobacteria; Enterobacterales; Enterobacteriaceae; Salmonella; Salmonella enterica; Salmonella enterica subsp. houtenae MTGDNTLITSHGINRRDFMKLCAALAATMGLSSKAAAEMAESVSNPQRPPVIWIGAQECT WP_000145429.1 hydrogenase 2 small subunit [Salmonella enterica] Length: 372\nNumber of predicted TMHs: 1\nExp number of AAs in TMHs: 24.51309\nExp number, first 60 AAs: 0.2682\nTotal prob of N-in: 0.14402\noutside 1 331\nTMhelix 332 354\ninside 355 372

25652 GCF_002045945.1_ASM204594v1 Salmonella enterica subsp. houtenae serovar 44:z4,z23:- Proteobacteria; Gammaproteobacteria; Enterobacterales; Enterobacteriaceae; Salmonella; Salmonella enterica;

Salmonella enterica subsp. houtenae MNNEETFYQAMRRKGVTRRSFLKFCSLAATSLGLGAGMTPKIAWALENKPRIPVVWIHGL WP_023187532.1 hydrogenase 1, small subunit [Salmonella enterica] Length: 372\nNumber of predicted TMHs: 2\nExp number of AAs in TMHs: 36.9997\nExp number, first 60 AAs: 19.72759\nTotal prob of N-in: 0.96171\nPOSSIBLE N-term signal sequence\ninside 1 19\nTMhelix 20 39\noutside 40 325\nTMhelix 326 348\ninside 349 372

25653 GCF_002035365.1_ASM203536v1 Salmonella enterica subsp. houtenae serovar 45:g,z51:- Proteobacteria; Gammaproteobacteria; Enterobacterales; Enterobacteriaceae; Salmonella; Salmonella enterica; Salmonella enterica subsp. houtenae MNNEETFYQAMRRKGVTRRSFLKFCSLAATSLGLGAGMTPKIAWALENKPRIPVVWIHGL WP_023187532.1 hydrogenase 1, small subunit [Salmonella enterica] Length: 372\nNumber of predicted TMHs: 2\nExp number of AAs in TMHs: 36.9997\nExp number, first 60 AAs: 19.72759\nTotal prob of N-in: 0.96171\nPOSSIBLE N-term signal sequence\ninside 1 19\nTMhelix 20 39\noutside 40 325\nTMhelix 326 348\ninside 349 372

25654 GCF_002035365.1_ASM203536v1 Salmonella enterica subsp. houtenae serovar 45:g,z51:- Proteobacteria; Gammaproteobacteria; Enterobacterales; Enterobacteriaceae; Salmonella; Salmonella enterica; Salmonella enterica subsp. houtenae MTGDNTLITSHGINRRDFMKLCAALAATMGLSSKAAAEMAESVSNPQRPPVIWIGAQECT WP_000145429.1 hydrogenase 2 small subunit [Salmonella enterica] Length: 372\nNumber of predicted TMHs: 1\nExp number of AAs in TMHs: 24.51309\nExp number, first 60 AAs: 0.2682\nTotal prob of N-in: 0.14402\noutside 1 331\nTMhelix 332 354\ninside 355 372

25655 GCF_002046205.1_ASM204620v1 Salmonella enterica subsp. houtenae serovar 48:g,z51:- Proteobacteria; Gammaproteobacteria; Enterobacterales; Enterobacteriaceae; Salmonella; Salmonella enterica; Salmonella enterica subsp. houtenae MNNEETFYQAMRRKGVTRRSFLKFCSLATTSLGLGAGMTPKIAWALENKPRIPVVWIHGL WP_001058317.1 hydrogenase [Salmonella enterica] Length: 372\nNumber of predicted TMHs: 2\nExp number of AAs in TMHs: 36.4727299999999\nExp number, first 60 AAs: 19.19744\nTotal prob of N-in: 0.93666\nPOSSIBLE N-term signal sequence\ninside 1 19\nTMhelix 20 39\noutside 40 325\nTMhelix 326 348\ninside 349 372

25656 GCF_002033735.1_ASM203373v1 Salmonella enterica subsp. houtenae serovar 48:g,z51:- Proteobacteria; Gammaproteobacteria; Enterobacterales; Enterobacteriaceae; Salmonella; Salmonella enterica; Salmonella enterica subsp. houtenae MNNEETFYQAMRRKGVTRRSFLKFCSLAATSLGLGAGMTPKIAWALENKPRIPVVWIHGL WP_001058306.1 hydrogenase [Salmonella enterica] Length: 372\nNumber of predicted TMHs: 2\nExp number of AAs in TMHs: 36.5540099999999\nExp number, first 60 AAs: 19.72732\nTotal prob of N-in: 0.96171\nPOSSIBLE N-term signal sequence\ninside 1 19\nTMhelix 20 39\noutside 40 325\nTMhelix 326 348\ninside 349 372

25657 GCF_002046345.1_ASM204634v1 Salmonella enterica subsp. houtenae serovar 48:g,z51:- Proteobacteria; Gammaproteobacteria; Enterobacterales; Enterobacteriaceae; Salmonella; Salmonella enterica; Salmonella enterica subsp. houtenae MTGDNTLITSHGINRRDFMKLCAALAATMGLSSKAAAEMAESVSNPQRPPVIWIGAQECT WP_000145429.1 hydrogenase 2 small subunit [Salmonella enterica] Length: 372\nNumber of predicted TMHs: 1\nExp number of AAs in TMHs: 24.51309\nExp number, first 60 AAs: 0.2682\nTotal prob of N-in: 0.14402\noutside 1 331\nTMhelix 332 354\ninside 355 372

25658 GCF_002033735.1_ASM203373v1 Salmonella enterica subsp. houtenae serovar 48:g,z51:- Proteobacteria; Gammaproteobacteria; Enterobacterales; Enterobacteriaceae; Salmonella; Salmonella enterica; Salmonella enterica subsp. houtenae MTGDNTLITSHGINRRDFMKLCAALAATMGLSSKAAAEMAESVSNPQRPPVIWIGAQECT WP_023248548.1 hydrogenase 2 small subunit [Salmonella enterica] Length: 372\nNumber of predicted TMHs: 1\nExp number of AAs in TMHs: 24.51281\nExp number, first 60 AAs: 0.26822\nTotal prob of N-in: 0.14402\noutside 1 331\nTMhelix 332 354\ninside 355 372

25659 GCF_002035575.1_ASM203557v1 Salmonella enterica subsp. houtenae serovar 48:g,z51:- Proteobacteria; Gammaproteobacteria; Enterobacterales; Enterobacteriaceae; Salmonella; Salmonella enterica; Salmonella enterica subsp. houtenae MTGDNTLITSHGINRRDFMKLCAALAATMGLSSKAAAEMAESVSNPQRPPVIWIGAQECT WP_000145429.1 hydrogenase 2 small subunit [Salmonella enterica] Length: 372\nNumber of predicted TMHs: 1\nExp number of AAs in TMHs: 24.51309\nExp number, first 60 AAs: 0.2682\nTotal prob of N-in: 0.14402\noutside 1 331\nTMhelix 332 354\ninside 355 372

25660 GCF_002046565.1_ASM204656v1 Salmonella enterica subsp. houtenae serovar 48:g,z51:- Proteobacteria; Gammaproteobacteria; Enterobacterales; Enterobacteriaceae; Salmonella; Salmonella enterica; Salmonella enterica subsp. houtenae MTGDNTLITSHGINRRDFMKLCAALAATMGLSSKAAAEMAESVSNPQRPPVIWIGAQECT

WP_000145429.1 hydrogenase 2 small subunit [Salmonella enterica] Length: 372\nNumber of predicted TMHs: 1\nExp number of AAs in TMHs: 24.51309\nExp number, first 60 AAs: 0.2682\nTotal prob of N-in: 0.14402\noutside 1 331\nTMhelix 332 354\ninside 355 372

25661 GCF_002033835.1_ASM203383v1 Salmonella enterica subsp. houtenae serovar 48:g,z51:- Proteobacteria; Gammaproteobacteria; Enterobacterales; Enterobacteriaceae; Salmonella; Salmonella enterica; Salmonella enterica subsp. houtenae MTGDNTLITSHGINRRDFMKLCAALAATMGLSSKAAAEMAESVSNPQRPPVIWIGAQECT WP_000145429.1 hydrogenase 2 small subunit [Salmonella enterica] Length: 372\nNumber of predicted TMHs: 1\nExp number of AAs in TMHs: 24.51309\nExp number, first 60 AAs: 0.2682\nTotal prob of N-in: 0.14402\noutside 1 331\nTMhelix 332 354\ninside 355 372

25662 GCF_002033805.1_ASM203380v1 Salmonella enterica subsp. houtenae serovar 48:g,z51:- Proteobacteria; Gammaproteobacteria; Enterobacterales; Enterobacteriaceae; Salmonella; Salmonella enterica; Salmonella enterica subsp. houtenae MNNEETFYQAMRRKGVTRRSFLKFCSLATTSLGLGAGMTPKIAWALENKPRIPVVWIHGL WP_001058317.1 hydrogenase [Salmonella enterica] Length: 372\nNumber of predicted TMHs: 2\nExp number of AAs in TMHs: 36.4727299999999\nExp number, first 60 AAs: 19.19744\nTotal prob of N-in: 0.93666\nPOSSIBLE N-term signal sequence\ninside 1 19\nTMhelix 20 39\noutside 40 325\nTMhelix 326 348\ninside 349 372

25663 GCF_002033805.1_ASM203380v1 Salmonella enterica subsp. houtenae serovar 48:g,z51:- Proteobacteria; Gammaproteobacteria; Enterobacterales; Enterobacteriaceae; Salmonella; Salmonella enterica; Salmonella enterica subsp. houtenae MTGDNTLITSHGINRRDFMKLCAALAATMGLSSKAAAEMAESVSNPQRPPVIWIGAQECT WP_000145429.1 hydrogenase 2 small subunit [Salmonella enterica] Length: 372\nNumber of predicted TMHs: 1\nExp number of AAs in TMHs: 24.51309\nExp number, first 60 AAs: 0.2682\nTotal prob of N-in: 0.14402\noutside 1 331\nTMhelix 332 354\ninside 355 372

25664 GCF_002035575.1_ASM203557v1 Salmonella enterica subsp. houtenae serovar 48:g,z51:- Proteobacteria; Gammaproteobacteria; Enterobacterales; Enterobacteriaceae; Salmonella; Salmonella enterica; Salmonella enterica subsp. houtenae MNNEETFYQAMRRKGVTRRSFLKFCSLATTSLGLGAGMTPKIAWALENKPRIPVVWIHGL WP_001058317.1 hydrogenase [Salmonella enterica] Length: 372\nNumber of predicted TMHs: 2\nExp number of AAs in TMHs: 36.4727299999999\nExp number, first 60 AAs: 19.19744\nTotal prob of N-in: 0.93666\nPOSSIBLE N-term signal sequence\ninside 1 19\nTMhelix 20 39\noutside 40 325\nTMhelix 326 348\ninside 349 372

25665 GCF_002046345.1_ASM204634v1 Salmonella enterica subsp. houtenae serovar 48:g,z51:- Proteobacteria; Gammaproteobacteria; Enterobacterales; Enterobacteriaceae; Salmonella; Salmonella enterica; Salmonella enterica subsp. houtenae MNNEETFYQAMRRKGVTRRSFLKFCSLATTSLGLGAGMTPKIAWALENKPRIPVVWIHGL WP_001058317.1 hydrogenase [Salmonella enterica] Length: 372\nNumber of predicted TMHs: 2\nExp number of AAs in TMHs: 36.4727299999999\nExp number, first 60 AAs: 19.19744\nTotal prob of N-in: 0.93666\nPOSSIBLE N-term signal sequence\ninside 1 19\nTMhelix 20 39\noutside 40 325\nTMhelix 326 348\ninside 349 372

25666 GCF_002046205.1_ASM204620v1 Salmonella enterica subsp. houtenae serovar 48:g,z51:- Proteobacteria; Gammaproteobacteria; Enterobacterales; Enterobacteriaceae; Salmonella; Salmonella enterica; Salmonella enterica subsp. houtenae MTGDNTLITSHGINRRDFMKLCAALAATMGLSSKAAAEMAESVSNPQRPPVIWIGAQECT WP_000145429.1 hydrogenase 2 small subunit [Salmonella enterica] Length: 372\nNumber of predicted TMHs: 1\nExp number of AAs in TMHs: 24.51309\nExp number, first 60 AAs: 0.2682\nTotal prob of N-in: 0.14402\noutside 1 331\nTMhelix 332 354\ninside 355 372

25667 GCF_002046565.1_ASM204656v1 Salmonella enterica subsp. houtenae serovar 48:g,z51:- Proteobacteria; Gammaproteobacteria; Enterobacterales; Enterobacteriaceae; Salmonella; Salmonella enterica; Salmonella enterica subsp. houtenae MNNEETFYQAMRRKGVTRRSFLKFCSLATTSLGLGAGMTPKIAWALENKPRIPVVWIHGL WP_001058317.1 hydrogenase [Salmonella enterica] Length: 372\nNumber of predicted TMHs: 2\nExp number of AAs in TMHs: 36.4727299999999\nExp number, first 60 AAs: 19.19744\nTotal prob of N-in: 0.93666\nPOSSIBLE N-term signal sequence\ninside 1 19\nTMhelix 20 39\noutside 40 325\nTMhelix 326 348\ninside 349 372

25668 GCF_002045345.1_ASM204534v1 Salmonella enterica subsp. houtenae serovar 48:g,z51:- Proteobacteria; Gammaproteobacteria; Enterobacterales; Enterobacteriaceae; Salmonella; Salmonella enterica; Salmonella enterica subsp. houtenae MNNEETFYQAMRRKGVTRRSFLKFCSLATTSLGLGAGMTPKIAWALENKPRIPVVWIHGL WP_001058317.1 hydrogenase [Salmonella enterica] Length: 372\nNumber of predicted TMHs: 2\nExp

number of AAs in TMHs: 36.4727299999999\nExp number, first 60 AAs: 19.19744\nTotal prob of N-in: 0.93666\nPOSSIBLE N-term signal sequence\ninside 1 19\nTMhelix 20 39\noutside 40 325\nTMhelix 326 348\ninside 349 372

25669 GCF_002046405.1_ASM204640v1 Salmonella enterica subsp. houtenae serovar 48:g,z51:- Proteobacteria; Gammaproteobacteria; Enterobacterales; Enterobacteriaceae; Salmonella; Salmonella enterica; Salmonella enterica subsp. houtenae MTGDNTLITSHGINRRDFMKLCAALAATMGLSSKAAAEMAESVSNPQRPPVIWIGAQECT WP_000145429.1 hydrogenase 2 small subunit [Salmonella enterica] Length: 372\nNumber of predicted TMHs: 1\nExp number of AAs in TMHs: 24.51309\nExp number, first 60 AAs: 0.2682\nTotal prob of N-in: 0.14402\noutside 1 331\nTMhelix 332 354\ninside 355 372

25670 GCF_002045345.1_ASM204534v1 Salmonella enterica subsp. houtenae serovar 48:g,z51:- Proteobacteria; Gammaproteobacteria; Enterobacterales; Enterobacteriaceae; Salmonella; Salmonella enterica; Salmonella enterica subsp. houtenae MTGDNTLITSHGINRRDFMKLCAALAATMGLSSKAAAEMAESVSNPQRPPVIWIGAQECT WP_000145429.1 hydrogenase 2 small subunit [Salmonella enterica] Length: 372\nNumber of predicted TMHs: 1\nExp number of AAs in TMHs: 24.51309\nExp number, first 60 AAs: 0.2682\nTotal prob of N-in: 0.14402\noutside 1 331\nTMhelix 332 354\ninside 355 372

25671 GCF_002033835.1_ASM203383v1 Salmonella enterica subsp. houtenae serovar 48:g,z51:- Proteobacteria; Gammaproteobacteria; Enterobacterales; Enterobacteriaceae; Salmonella; Salmonella enterica; Salmonella enterica subsp. houtenae MNNEETFYQAMRRKGVTRRSFLKFCSLATTSLGLGAGMTPKIAWALENKPRIPVVWIHGL WP_001058317.1 hydrogenase [Salmonella enterica] Length: 372\nNumber of predicted TMHs: 2\nExp number of AAs in TMHs: 36.4727299999999\nExp number, first 60 AAs: 19.19744\nTotal prob of N-in: 0.93666\nPOSSIBLE N-term signal sequence\ninside 1 19\nTMhelix 20 39\noutside 40 325\nTMhelix 326 348\ninside 349 372

25672 GCF_002046405.1_ASM204640v1 Salmonella enterica subsp. houtenae serovar 48:g,z51:- Proteobacteria; Gammaproteobacteria; Enterobacterales; Enterobacteriaceae; Salmonella; Salmonella enterica; Salmonella enterica subsp. houtenae MNNEETFYQAMRRKGVTRRSFLKFCSLATTSLGLGAGMTPKIAWALENKPRIPVVWIHGL WP_001058317.1 hydrogenase [Salmonella enterica] Length: 372\nNumber of predicted TMHs: 2\nExp number of AAs in TMHs: 36.4727299999999\nExp number, first 60 AAs: 19.19744\nTotal prob of N-in: 0.93666\nPOSSIBLE N-term signal sequence\ninside 1 19\nTMhelix 20 39\noutside 40 325\nTMhelix 326 348\ninside 349 372

25673 GCF_002045205.1_ASM204520v1 Salmonella enterica subsp. houtenae serovar 50:g,z51:- Proteobacteria; Gammaproteobacteria; Enterobacterales; Enterobacteriaceae; Salmonella; Salmonella enterica; Salmonella enterica subsp. houtenae MTGDNTLITSHGINRRDFMKLCAALAATMGLSSKAAAEMAESVSNPQRPPVIWIGAQECT WP_000145429.1 hydrogenase 2 small subunit [Salmonella enterica] Length: 372\nNumber of predicted TMHs: 1\nExp number of AAs in TMHs: 24.51309\nExp number, first 60 AAs: 0.2682\nTotal prob of N-in: 0.14402\noutside 1 331\nTMhelix 332 354\ninside 355 372

25674 GCF_002045905.1_ASM204590v1 Salmonella enterica subsp. houtenae serovar 50:g,z51:- Proteobacteria; Gammaproteobacteria; Enterobacterales; Enterobacteriaceae; Salmonella; Salmonella enterica; Salmonella enterica subsp. houtenae MNNEETFYQAMRRKGVTRRSFLKFCSLAATSLGLGAGMTPKIAWALENKPRIPVVWIHGL WP_023187532.1 hydrogenase 1, small subunit [Salmonella enterica] Length: 372\nNumber of predicted TMHs: 2\nExp number of AAs in TMHs: 36.9997\nExp number, first 60 AAs: 19.72759\nTotal prob of N-in: 0.96171\nPOSSIBLE N-term signal sequence\ninside 1 19\nTMhelix 20 39\noutside 40 325\nTMhelix 326 348\ninside 349 372

25675 GCF_002044985.1_ASM204498v1 Salmonella enterica subsp. houtenae serovar 50:g,z51:- Proteobacteria; Gammaproteobacteria; Enterobacterales; Enterobacteriaceae; Salmonella; Salmonella enterica; Salmonella enterica subsp. houtenae MTGDNTLITSHGINRRDFMKLCAALAATMGLSSKAAAEMAESVSNPQRPPVIWIGAQECT WP_000145429.1 hydrogenase 2 small subunit [Salmonella enterica] Length: 372\nNumber of predicted TMHs: 1\nExp number of AAs in TMHs: 24.51309\nExp number, first 60 AAs: 0.2682\nTotal prob of N-in: 0.14402\noutside 1 331\nTMhelix 332 354\ninside 355 372

25676 GCF_002044985.1_ASM204498v1 Salmonella enterica subsp. houtenae serovar 50:g,z51:- Proteobacteria; Gammaproteobacteria; Enterobacterales; Enterobacteriaceae; Salmonella; Salmonella enterica; Salmonella enterica subsp. houtenae MNNEETFYQAMRRKGVTRRSFLKFCSLAATSLGLGAGMTPKIAWALENKPRIPVVWIHGL WP_023187532.1 hydrogenase 1, small subunit [Salmonella enterica] Length: 372\nNumber of predicted TMHs: 2\nExp number of AAs in TMHs: 36.9997\nExp number, first 60 AAs: 19.72759\nTotal prob of N-in:

0.96171\nPOSSIBLE N-term signal sequence\ninside 1 19\nTMhelix 20 39\noutside 40 325\nTMhelix 326 348\ninside 349 372

25677 GCF_002045205.1_ASM204520v1 Salmonella enterica subsp. houtenae serovar 50:g,z51:- Proteobacteria; Gammaproteobacteria; Enterobacterales; Enterobacteriaceae; Salmonella; Salmonella enterica; Salmonella enterica subsp. houtenae MNNEETFYQAMRRKGVTRRSFLKFCSLAATSLGLGAGMTPKIAWALENKPRIPVVWIHGL WP_023187532.1 hydrogenase 1, small subunit [Salmonella enterica] Length: 372\nNumber of predicted TMHs: 2\nExp number of AAs in TMHs: 36.9997\nExp number, first 60 AAs: 19.72759\nTotal prob of N-in: 0.96171\nPOSSIBLE N-term signal sequence\ninside 1 19\nTMhelix 20 39\noutside 40 325\nTMhelix 326 348\ninside 349 372

25678 GCF_002045755.1_ASM204575v1 Salmonella enterica subsp. houtenae serovar 50:g,z51:- Proteobacteria; Gammaproteobacteria; Enterobacterales; Enterobacteriaceae; Salmonella; Salmonella enterica; Salmonella enterica subsp. houtenae MTGDNTLITSHGINRRDFMKLCAALAATMGLSSKAAAEMAESVSNPQRPPVIWIGAQECT WP_000145429.1 hydrogenase 2 small subunit [Salmonella enterica] Length: 372\nNumber of predicted TMHs: 1\nExp number of AAs in TMHs: 24.51309\nExp number, first 60 AAs: 0.2682\nTotal prob of N-in: 0.14402\noutside 1 331\nTMhelix 332 354\ninside 355 372

25679 GCF_002045905.1_ASM204590v1 Salmonella enterica subsp. houtenae serovar 50:g,z51:- Proteobacteria; Gammaproteobacteria; Enterobacterales; Enterobacteriaceae; Salmonella; Salmonella enterica; Salmonella enterica subsp. houtenae MTGDNTLITSHGINRRDFMKLCAALAATMGLSSKAAAEMAESVSNPQRPPVIWIGAQECT WP_000145429.1 hydrogenase 2 small subunit [Salmonella enterica] Length: 372\nNumber of predicted TMHs: 1\nExp number of AAs in TMHs: 24.51309\nExp number, first 60 AAs: 0.2682\nTotal prob of N-in: 0.14402\noutside 1 331\nTMhelix 332 354\ninside 355 372

25680 GCF_002045755.1_ASM204575v1 Salmonella enterica subsp. houtenae serovar 50:g,z51:- Proteobacteria; Gammaproteobacteria; Enterobacterales; Enterobacteriaceae; Salmonella; Salmonella enterica; Salmonella enterica subsp. houtenae MNNEETFYQAMRRKGVTRRSFLKFCSLAATSLGLGAGMTPKIAWALENKPRIPVVWIHGL WP_023187532.1 hydrogenase 1, small subunit [Salmonella enterica] Length: 372\nNumber of predicted TMHs: 2\nExp number of AAs in TMHs: 36.9997\nExp number, first 60 AAs: 19.72759\nTotal prob of N-in: 0.96171\nPOSSIBLE N-term signal sequence\ninside 1 19\nTMhelix 20 39\noutside 40 325\nTMhelix 326 348\ninside 349 372

25681 GCF_002046485.1_ASM204648v1 Salmonella enterica subsp. houtenae serovar 50:z4,z23:- Proteobacteria; Gammaproteobacteria; Enterobacterales; Enterobacteriaceae; Salmonella; Salmonella enterica; Salmonella enterica subsp. houtenae MNNEETFYQAMRRKGVTRRSFLKFCSLATTSLGLGAGMTPKIAWALENKPRIPVVWIHGL WP_001058317.1 hydrogenase [Salmonella enterica] Length: 372\nNumber of predicted TMHs: 2\nExp number of AAs in TMHs: 36.4727299999999\nExp number, first 60 AAs: 19.19744\nTotal prob of N-in: 0.93666\nPOSSIBLE N-term signal sequence\ninside 1 19\nTMhelix 20 39\noutside 40 325\nTMhelix 326 348\ninside 349 372

25682 GCF_002046355.1_ASM204635v1 Salmonella enterica subsp. houtenae serovar 50:z4,z23:- Proteobacteria; Gammaproteobacteria; Enterobacterales; Enterobacteriaceae; Salmonella; Salmonella enterica; Salmonella enterica subsp. houtenae MNNEETFYQAMRRKGVTRRSFLKFCSLATTSLGLGAGMTPKIAWALENKPRIPVVWIHGL WP_001058317.1 hydrogenase [Salmonella enterica] Length: 372\nNumber of predicted TMHs: 2\nExp number of AAs in TMHs: 36.4727299999999\nExp number, first 60 AAs: 19.19744\nTotal prob of N-in: 0.93666\nPOSSIBLE N-term signal sequence\ninside 1 19\nTMhelix 20 39\noutside 40 325\nTMhelix 326 348\ninside 349 372

25683 GCF_002046465.1_ASM204646v1 Salmonella enterica subsp. houtenae serovar 50:z4,z23:- Proteobacteria; Gammaproteobacteria; Enterobacterales; Enterobacteriaceae; Salmonella; Salmonella enterica; Salmonella enterica subsp. houtenae MTGDNTLITSHGINRRDFMKLCAALAATMGLSSKAAAEMAESVSNPQRPPVIWIGAQECT WP_000145429.1 hydrogenase 2 small subunit [Salmonella enterica] Length: 372\nNumber of predicted TMHs: 1\nExp number of AAs in TMHs: 24.51309\nExp number, first 60 AAs: 0.2682\nTotal prob of N-in: 0.14402\noutside 1 331\nTMhelix 332 354\ninside 355 372

25684 GCF_002046045.1_ASM204604v1 Salmonella enterica subsp. houtenae serovar 50:z4,z23:- Proteobacteria; Gammaproteobacteria; Enterobacterales; Enterobacteriaceae; Salmonella; Salmonella enterica; Salmonella enterica subsp. houtenae MNNEETFYQAMRRKGVTRRSFLKFCSLAATSLGLGAGMTPKIAWALENKPRIPVVWIHGL WP_023187532.1 hydrogenase 1, small subunit [Salmonella enterica] Length: 372\nNumber of predicted TMHs: 2\nExp number of AAs in TMHs: 36.9997\nExp number, first 60 AAs: 19.72759\nTotal prob of N-in:

0.96171\nPOSSIBLE N-term signal sequence\ninside 1 19\nTMhelix 20 39\noutside 40 325\nTMhelix 326 348\ninside 349 372

25685 GCF_002046465.1_ASM204646v1 Salmonella enterica subsp. houtenae serovar 50:z4,z23:- Proteobacteria; Gammaproteobacteria; Enterobacterales; Enterobacteriaceae; Salmonella; Salmonella enterica; Salmonella enterica subsp. houtenae MNNEETFYQAMRRKGVTRRSFLKFCSLATTSLGLGAGMTPKIAWALENKPRIPVVWIHGL WP_001058317.1 hydrogenase [Salmonella enterica] Length: 372\nNumber of predicted TMHs: 2\nExp number of AAs in TMHs: 36.4727299999999\nExp number, first 60 AAs: 19.19744\nTotal prob of N-in: 0.93666\nPOSSIBLE N-term signal sequence\ninside 1 19\nTMhelix 20 39\noutside 40 325\nTMhelix 326 348\ninside 349 372

25686 GCF_002046045.1_ASM204604v1 Salmonella enterica subsp. houtenae serovar 50:z4,z23:- Proteobacteria; Gammaproteobacteria; Enterobacterales; Enterobacteriaceae; Salmonella; Salmonella enterica; Salmonella enterica subsp. houtenae MTGDNTLITSHGINRRDFMKLCAALAATMGLSSKAAAEMAESVSNPQRPPVIWIGAQECT WP_000145429.1 hydrogenase 2 small subunit [Salmonella enterica] Length: 372\nNumber of predicted TMHs: 1\nExp number of AAs in TMHs: 24.51309\nExp number, first 60 AAs: 0.2682\nTotal prob of N-in: 0.14402\noutside 1 331\nTMhelix 332 354\ninside 355 372

25687 GCF_002046095.1_ASM204609v1 Salmonella enterica subsp. houtenae serovar 50:z4,z23:- Proteobacteria; Gammaproteobacteria; Enterobacterales; Enterobacteriaceae; Salmonella; Salmonella enterica; Salmonella enterica subsp. houtenae MTGDNTLITSHGINRRDFMKLCAALAATMGLSSKAAAEMAESVSNPQRPPVIWIGAQECT WP_000145429.1 hydrogenase 2 small subunit [Salmonella enterica] Length: 372\nNumber of predicted TMHs: 1\nExp number of AAs in TMHs: 24.51309\nExp number, first 60 AAs: 0.2682\nTotal prob of N-in: 0.14402\noutside 1 331\nTMhelix 332 354\ninside 355 372

25688 GCF_002046095.1_ASM204609v1 Salmonella enterica subsp. houtenae serovar 50:z4,z23:- Proteobacteria; Gammaproteobacteria; Enterobacterales; Enterobacteriaceae; Salmonella; Salmonella enterica; Salmonella enterica subsp. houtenae MNNEETFYQAMRRKGVTRRSFLKFCSLAATSLGLGAGMTPKIAWALENKPRIPVVWIHGL WP_023187532.1 hydrogenase 1, small subunit [Salmonella enterica] Length: 372\nNumber of predicted TMHs: 2\nExp number of AAs in TMHs: 36.9997\nExp number, first 60 AAs: 19.72759\nTotal prob of N-in: 0.96171\nPOSSIBLE N-term signal sequence\ninside 1 19\nTMhelix 20 39\noutside 40 325\nTMhelix 326 348\ninside 349 372

25689 GCF_002046485.1_ASM204648v1 Salmonella enterica subsp. houtenae serovar 50:z4,z23:- Proteobacteria; Gammaproteobacteria; Enterobacterales; Enterobacteriaceae; Salmonella; Salmonella enterica; Salmonella enterica subsp. houtenae MTGDNTLITSHGINRRDFMKLCAALAATMGLSSKAAAEMAESVSNPQRPPVIWIGAQECT WP_000145429.1 hydrogenase 2 small subunit [Salmonella enterica] Length: 372\nNumber of predicted TMHs: 1\nExp number of AAs in TMHs: 24.51309\nExp number, first 60 AAs: 0.2682\nTotal prob of N-in: 0.14402\noutside 1 331\nTMhelix 332 354\ninside 355 372

25690 GCF_002046355.1_ASM204635v1 Salmonella enterica subsp. houtenae serovar 50:z4,z23:- Proteobacteria; Gammaproteobacteria; Enterobacterales; Enterobacteriaceae; Salmonella; Salmonella enterica; Salmonella enterica subsp. houtenae MTGDNTLITSHGINRRDFMKLCAALAATMGLSSKAAAEMAESVSNPQRPPVIWIGAQECT WP_079896656.1 hydrogenase 2 small subunit [Salmonella enterica] Length: 372\nNumber of predicted TMHs: 1\nExp number of AAs in TMHs: 24.52378\nExp number, first 60 AAs: 0.2682\nTotal prob of N-in: 0.14424\noutside 1 331\nTMhelix 332 354\ninside 355 372

25691 GCF_000251025.1_Salmonella_enterica_IV_05-0642-2.0.1 Salmonella enterica subsp. houtenae str. ATCC BAA-1581 Proteobacteria; Gammaproteobacteria; Enterobacterales; Enterobacteriaceae; Salmonella; Salmonella enterica; Salmonella enterica subsp. houtenae MTGDNTLITSHGINRRDFMKLCAALAATMGLSSKAAAEMAESVSNPQRPPVIWIGAQECT WP_000145429.1 hydrogenase 2 small subunit [Salmonella enterica] Length: 372\nNumber of predicted TMHs: 1\nExp number of AAs in TMHs: 24.51309\nExp number, first 60 AAs: 0.2682\nTotal prob of N-in: 0.14402\noutside 1 331\nTMhelix 332 354\ninside 355 372

25692 GCF_000251025.1_Salmonella_enterica_IV_05-0642-2.0.1 Salmonella enterica subsp. houtenae str. ATCC BAA-1581 Proteobacteria; Gammaproteobacteria; Enterobacterales; Enterobacteriaceae; Salmonella; Salmonella enterica; Salmonella enterica subsp. houtenae MNNEETFYQAMRRKGVTRRSFLKFCSLATTSLGLGAGMTPKIAWALENKPRIPVVWIHGL WP_001058317.1 hydrogenase [Salmonella enterica] Length: 372\nNumber of predicted TMHs: 2\nExp number of AAs in TMHs: 36.4727299999999\nExp number, first 60 AAs: 19.19744\nTotal prob of N-in: 0.93666\nPOSSIBLE N-term signal sequence\ninside 1 19\nTMhelix 20 39\noutside 40 325\nTMhelix 326 348\ninside 349 372

25693 GCF_000376525.1_W41 Salmonella enterica subsp. houtenae serovar 16:z4,z32:-- str. RKS3027 Proteobacteria; Gammaproteobacteria; Enterobacterales; Enterobacteriaceae; Salmonella; Salmonella enterica; Salmonella enterica subsp. houtenae; Salmonella enterica subsp. houtenae serovar 16 MTGDNTLITSHGINRRDFMKLCAALAATMGLSSKAAAEMAESVSNPQRPPVIWIGAQECT WP_000145429.1 hydrogenase 2 small subunit [Salmonella enterica] Length: 372\nNumber of predicted TMHs: 1\nExp number of AAs in TMHs: 24.51309\nExp number, first 60 AAs: 0.2682\nTotal prob of N-in: 0.14402\noutside 1 331\nTMhelix 332 354\ninside 355 372

25694 GCF_000376525.1_W41 Salmonella enterica subsp. houtenae serovar 16:z4,z32:-- str. RKS3027 Proteobacteria; Gammaproteobacteria; Enterobacterales; Enterobacteriaceae; Salmonella; Salmonella enterica; Salmonella enterica subsp. houtenae; Salmonella enterica subsp. houtenae serovar 16 MNNEETFYQAMRRKGVTRRSFLKFCSLATTSLGLGAGMTPKIAWALENKPRIPVVWIHGL WP_001058317.1 hydrogenase [Salmonella enterica] Length: 372\nNumber of predicted TMHs: 2\nExp number of AAs in TMHs: 36.4727299999999\nExp number, first 60 AAs: 19.19744\nTotal prob of N-in: 0.93666\nPOSSIBLE N-term signal sequence\ninside 1 19\nTMhelix 20 39\noutside 40 325\nTMhelix 326 348\ninside 349 372

25695 GCF_000486245.1_ASM48624v1 Salmonella enterica subsp. houtenae serovar 50:g,z51:- str. 01-0133 Proteobacteria; Gammaproteobacteria; Enterobacterales; Enterobacteriaceae; Salmonella; Salmonella enterica; Salmonella enterica subsp. houtenae; Salmonella enterica subsp. houtenae serovar 50 MTGDNTLITSHGINRRDFMKLCAALAATMGLSSKAAAEMAESVSNPQRPPVIWIGAQECT WP_000145429.1 hydrogenase 2 small subunit [Salmonella enterica] Length: 372\nNumber of predicted TMHs: 1\nExp number of AAs in TMHs: 24.51309\nExp number, first 60 AAs: 0.2682\nTotal prob of N-in: 0.14402\noutside 1 331\nTMhelix 332 354\ninside 355 372

25696 GCF_000486245.1_ASM48624v1 Salmonella enterica subsp. houtenae serovar 50:g,z51:- str. 01-0133 Proteobacteria; Gammaproteobacteria; Enterobacterales; Enterobacteriaceae; Salmonella; Salmonella enterica; Salmonella enterica subsp. houtenae; Salmonella enterica subsp. houtenae serovar 50 MNNEETFYQAMRRKGVTRRSFLKFCSLAATSLGLGAGMTPKIAWALENKPRIPVVWIHGL WP_023187532.1 hydrogenase 1, small subunit [Salmonella enterica] Length: 372\nNumber of predicted TMHs: 2\nExp number of AAs in TMHs: 36.9997\nExp number, first 60 AAs: 19.72759\nTotal prob of N-in: 0.96171\nPOSSIBLE N-term signal sequence\ninside 1 19\nTMhelix 20 39\noutside 40 325\nTMhelix 326 348\ninside 349 372

25697 GCF_002035225.1_ASM203522v1 Salmonella enterica subsp. indica serovar 11:b:1,7 Proteobacteria; Gammaproteobacteria; Enterobacterales; Enterobacteriaceae; Salmonella; Salmonella enterica; Salmonella enterica subsp. indica MTGDNTLITSHGINRRDFMKLCAALAATMGLSSKAAAEMAESVSNPQRPPVIWIGAQECT WP_079776618.1 MULTISPECIES: hydrogenase 2 small subunit [Salmonella] Length: 372\nNumber of predicted TMHs: 1\nExp number of AAs in TMHs: 24.65705\nExp number, first 60 AAs: 0.26805\nTotal prob of N-in: 0.13761\noutside 1 331\nTMhelix 332 354\ninside 355 372

25698 GCF_002066875.1_ASM206687v1 Salmonella enterica subsp. indica serovar 11:b:e,n,x Proteobacteria; Gammaproteobacteria; Enterobacterales; Enterobacteriaceae; Salmonella; Salmonella enterica; Salmonella enterica subsp. indica MNNEETFYQAMRRKGVTRRSFLKFCSLAATSLGLGAGMTPKIAWALENKPRIPVVWIHGL WP_080249058.1 [Ni/Fe] hydrogenase small subunit [Salmonella enterica] Length: 372\nNumber of predicted TMHs: 2\nExp number of AAs in TMHs: 36.54989\nExp number, first 60 AAs: 19.72756\nTotal prob of N-in: 0.96171\nPOSSIBLE N-term signal sequence\ninside 1 19\nTMhelix 20 39\noutside 40 325\nTMhelix 326 348\ninside 349 372

25699 GCF_002066875.1_ASM206687v1 Salmonella enterica subsp. indica serovar 11:b:e,n,x Proteobacteria; Gammaproteobacteria; Enterobacterales; Enterobacteriaceae; Salmonella; Salmonella enterica; Salmonella enterica subsp. indica MTGDNTLITSHGINRRDFMKLCAALAATMGLSSKAAAEMAESVSNPQRPPVIWIGAQECT WP_079776618.1 MULTISPECIES: hydrogenase 2 small subunit [Salmonella] Length: 372\nNumber of predicted TMHs: 1\nExp number of AAs in TMHs: 24.65705\nExp number, first 60 AAs: 0.26805\nTotal prob of N-in: 0.13761\noutside 1 331\nTMhelix 332 354\ninside 355 372

25700 GCF_002066855.1_ASM206685v1 Salmonella enterica subsp. indica serovar 1,6,14:a:e,n,x Proteobacteria; Gammaproteobacteria; Enterobacterales; Enterobacteriaceae; Salmonella; Salmonella enterica; Salmonella enterica subsp. indica MTGDNTLITSHGINRRDFMKLCAALAATMGLSSKAAAEMAESVSNPQRPPVIWIGAQECT WP_079776618.1 MULTISPECIES: hydrogenase 2 small subunit [Salmonella] Length: 372\nNumber of predicted TMHs: 1\nExp number of AAs in TMHs: 24.65705\nExp number, first 60 AAs: 0.26805\nTotal prob of N-in: 0.13761\noutside 1 331\nTMhelix 332 354\ninside 355 372

25701	GCF_002035485.1_ASM203548v1	Salmonella enterica subsp. indica serovar 6,14,25:z10:1,(2),7	Proteobacteria; Gammaproteobacteria; Enterobacterales; Enterobacteriaceae; Salmonella; Salmonella enterica; Salmonella enterica subsp. indica	MTGDNTLITSHDINRRDFMKLCAALAATMGLSSKAAAEMAESVSNPQRPPVIWIGAQECT	WP_023185463.1	hydrogenase 2 small subunit [Salmonella enterica]	Length: 372\nNumber of predicted TMHs: 1\nExp number of AAs in TMHs: 24.68266\nExp number, first 60 AAs: 0.29511\nTotal prob of N-in: 0.13876\noutside 1 331\nTMhelix 332 354\ninside 355 372

25702	GCF_002035485.1_ASM203548v1	Salmonella enterica subsp. indica serovar 6,14,25:z10:1,(2),7	Proteobacteria; Gammaproteobacteria; Enterobacterales; Enterobacteriaceae; Salmonella; Salmonella enterica; Salmonella enterica subsp. indica	MNNEETFYQAMRRKGVTRRSFLKFCSLAATSLGLGAGMTPKIAWALENKPRIPVVWIHGL	WP_023185544.1	hydrogenase 1, small subunit [Salmonella enterica]	Length: 372\nNumber of predicted TMHs: 2\nExp number of AAs in TMHs: 36.55018\nExp number, first 60 AAs: 19.72756\nTotal prob of N-in: 0.96171\nPOSSIBLE N-term signal sequence\ninside 1 19\nTMhelix 20 39\noutside 40 325\nTMhelix 326 348\ninside 349 372

25703	GCF_000486225.1_ASM48622v1	Salmonella enterica subsp. indica serovar 6,14,25:z10:1,(2),7 str. 1121	Proteobacteria; Gammaproteobacteria; Enterobacterales; Enterobacteriaceae; Salmonella; Salmonella enterica; Salmonella enterica subsp. indica; Salmonella enterica subsp. indica serovar 6,14,25	MNNEETFYQAMRRKGVTRRSFLKFCSLAATSLGLGAGMTPKIAWALENKPRIPVVWIHGL	WP_023185544.1	hydrogenase 1, small subunit [Salmonella enterica]	Length: 372\nNumber of predicted TMHs: 2\nExp number of AAs in TMHs: 36.55018\nExp number, first 60 AAs: 19.72756\nTotal prob of N-in: 0.96171\nPOSSIBLE N-term signal sequence\ninside 1 19\nTMhelix 20 39\noutside 40 325\nTMhelix 326 348\ninside 349 372

25704	GCF_000486225.1_ASM48622v1	Salmonella enterica subsp. indica serovar 6,14,25:z10:1,(2),7 str. 1121	Proteobacteria; Gammaproteobacteria; Enterobacterales; Enterobacteriaceae; Salmonella; Salmonella enterica; Salmonella enterica subsp. indica; Salmonella enterica subsp. indica serovar 6,14,25	MTGDNTLITSHDINRRDFMKLCAALAATMGLSSKAAAEMAESVSNPQRPPVIWIGAQECT	WP_023185463.1	hydrogenase 2 small subunit [Salmonella enterica]	Length: 372\nNumber of predicted TMHs: 1\nExp number of AAs in TMHs: 24.68266\nExp number, first 60 AAs: 0.29511\nTotal prob of N-in: 0.13876\noutside 1 331\nTMhelix 332 354\ninside 355 372

25705	GCF_002035875.1_ASM203587v1	Salmonella enterica subsp. salamae serovar 13,22:z29:enx	Proteobacteria; Gammaproteobacteria; Enterobacterales; Enterobacteriaceae; Salmonella; Salmonella enterica; Salmonella enterica subsp. salamae	MTGDNTLITSHGINRRDFMKLCAALAATMGLSSKAAAEMAESVSNPQRPPVIWIGAQECT	WP_000145432.1	hydrogenase 2 small subunit [Salmonella enterica]	Length: 372\nNumber of predicted TMHs: 1\nExp number of AAs in TMHs: 24.5101\nExp number, first 60 AAs: 0.2682\nTotal prob of N-in: 0.14397\noutside 1 331\nTMhelix 332 354\ninside 355 372

25706	GCF_002035875.1_ASM203587v1	Salmonella enterica subsp. salamae serovar 13,22:z29:enx	Proteobacteria; Gammaproteobacteria; Enterobacterales; Enterobacteriaceae; Salmonella; Salmonella enterica; Salmonella enterica subsp. salamae	MNNEETFYQAMRRKGVTRRSFLKFCSLAATSLGLGAGMTPKIAWALENKPRIPVVWIHGL	WP_001058314.1	hydrogenase [Salmonella enterica]	Length: 372\nNumber of predicted TMHs: 2\nExp number of AAs in TMHs: 36.3110299999999\nExp number, first 60 AAs: 19.72774\nTotal prob of N-in: 0.96169\nPOSSIBLE N-term signal sequence\ninside 1 19\nTMhelix 20 39\noutside 40 325\nTMhelix 326 348\ninside 349 372

25707	GCF_002036105.1_ASM203610v1	Salmonella enterica subsp. salamae serovar 47:d:z39	Proteobacteria; Gammaproteobacteria; Enterobacterales; Enterobacteriaceae; Salmonella; Salmonella enterica; Salmonella enterica subsp. salamae	MTGDNTLITSHGINRRDFMKLCAALAATMGLSSKAAAEMAESVSNPQRPPVIWIGAQECT	WP_000145432.1	hydrogenase 2 small subunit [Salmonella enterica]	Length: 372\nNumber of predicted TMHs: 1\nExp number of AAs in TMHs: 24.5101\nExp number, first 60 AAs: 0.2682\nTotal prob of N-in: 0.14397\noutside 1 331\nTMhelix 332 354\ninside 355 372

25708	GCF_002036105.1_ASM203610v1	Salmonella enterica subsp. salamae serovar 47:d:z39	Proteobacteria; Gammaproteobacteria; Enterobacterales; Enterobacteriaceae; Salmonella; Salmonella enterica; Salmonella enterica subsp. salamae	MNNEETFYQAMRRKGVTRRSFLKFCSLAATSLGLGAGMTPKIAWALENKPRIPVVWIHGL	WP_001058314.1	hydrogenase [Salmonella enterica]	Length: 372\nNumber of predicted TMHs: 2\nExp number of AAs in TMHs: 36.3110299999999\nExp number, first 60 AAs: 19.72774\nTotal prob of N-in: 0.96169\nPOSSIBLE N-term signal sequence\ninside 1 19\nTMhelix 20 39\noutside 40 325\nTMhelix 326 348\ninside 349 372

25709	GCF_002036095.1_ASM203609v1	Salmonella enterica subsp. salamae serovar 48:d:z6	Proteobacteria; Gammaproteobacteria; Enterobacterales; Enterobacteriaceae; Salmonella; Salmonella enterica; Salmonella enterica subsp. salamae	MNNEETFYQAMRRKGVTRRSFLKFCSLAATSLGLGAGMTPKIAWALENKPRIPVVWIHGL	WP_001058314.1	hydrogenase [Salmonella enterica]	Length: 372\nNumber of predicted TMHs:  2\nExp number of AAs in TMHs: 36.3110299999999\nExp number, first 60 AAs:  19.72774\nTotal prob of N-in: 0.96169\nPOSSIBLE N-term signal sequence\ninside   1   19\nTMhelix   20   39\noutside   40   325\nTMhelix   326   348\ninside   349   372

25710	GCF_002036095.1_ASM203609v1	Salmonella enterica subsp. salamae serovar 48:d:z6	Proteobacteria; Gammaproteobacteria; Enterobacterales; Enterobacteriaceae; Salmonella; Salmonella enterica; Salmonella enterica subsp. salamae	MTGDNTLITSHGINRRDFMKLCAALAVTMGLSSKAAAEMAESVSNPQRPPVIWIGAQECT	WP_079815268.1	hydrogenase 2 small subunit [Salmonella enterica]	Length: 372\nNumber of predicted TMHs:  1\nExp number of AAs in TMHs: 24.69423\nExp number, first 60 AAs:  0.46183\nTotal prob of N-in: 0.15218\noutside   1   331\nTMhelix   332   354\ninside   355   372

25711	GCF_002036085.1_ASM203608v1	Salmonella enterica subsp. salamae serovar 50:b:z6	Proteobacteria; Gammaproteobacteria; Enterobacterales; Enterobacteriaceae; Salmonella; Salmonella enterica; Salmonella enterica subsp. salamae	MTGDNTLITSHGINRRDFMKLCAALAATMGLSSKAAAEMAESVSNPQRPPVIWIGAQECT	WP_000145432.1	hydrogenase 2 small subunit [Salmonella enterica]	Length: 372\nNumber of predicted TMHs:  1\nExp number of AAs in TMHs: 24.5101\nExp number, first 60 AAs:  0.2682\nTotal prob of N-in: 0.14397\noutside   1   331\nTMhelix   332   354\ninside   355   372

25712	GCF_002036085.1_ASM203608v1	Salmonella enterica subsp. salamae serovar 50:b:z6	Proteobacteria; Gammaproteobacteria; Enterobacterales; Enterobacteriaceae; Salmonella; Salmonella enterica; Salmonella enterica subsp. salamae	MNNEETFYQAMRRKGVTRRSFLKFCSLAATSLGLGAGMTPKIAWALENKPRIPVVWIHGL	WP_001058314.1	hydrogenase [Salmonella enterica]	Length: 372\nNumber of predicted TMHs:  2\nExp number of AAs in TMHs: 36.3110299999999\nExp number, first 60 AAs:  19.72774\nTotal prob of N-in: 0.96169\nPOSSIBLE N-term signal sequence\ninside   1   19\nTMhelix   20   39\noutside   40   325\nTMhelix   326   348\ninside   349   372

25713	GCF_002035885.1_ASM203588v1	Salmonella enterica subsp. salamae serovar 53:lz28:z39	Proteobacteria; Gammaproteobacteria; Enterobacterales; Enterobacteriaceae; Salmonella; Salmonella enterica; Salmonella enterica subsp. salamae	MTGDNTLITSHGINRRDFMKLCAALAATMGLSSKAAAEMAESVSNPQRPPVIWIGAQECT	WP_000145432.1	hydrogenase 2 small subunit [Salmonella enterica]	Length: 372\nNumber of predicted TMHs:  1\nExp number of AAs in TMHs: 24.5101\nExp number, first 60 AAs:  0.2682\nTotal prob of N-in: 0.14397\noutside   1   331\nTMhelix   332   354\ninside   355   372

25714	GCF_002035885.1_ASM203588v1	Salmonella enterica subsp. salamae serovar 53:lz28:z39	Proteobacteria; Gammaproteobacteria; Enterobacterales; Enterobacteriaceae; Salmonella; Salmonella enterica; Salmonella enterica subsp. salamae	MNNEETFYQAMRRKGVTRRSFLKFCSLAATSLGLGAGMTPKIAWALENKPRIPVVWIHGL	WP_001058314.1	hydrogenase [Salmonella enterica]	Length: 372\nNumber of predicted TMHs:  2\nExp number of AAs in TMHs: 36.3110299999999\nExp number, first 60 AAs:  19.72774\nTotal prob of N-in: 0.96169\nPOSSIBLE N-term signal sequence\ninside   1   19\nTMhelix   20   39\noutside   40   325\nTMhelix   326   348\ninside   349   372

25715	GCF_002065095.1_ASM206509v1	Salmonella enterica subsp. salamae serovar 55:k:z39	Proteobacteria; Gammaproteobacteria; Enterobacterales; Enterobacteriaceae; Salmonella; Salmonella enterica; Salmonella enterica subsp. salamae	MNNEETFYQAMRRKGVTRRSFLKFCSLAATSLGLGAGMTPKIAWALENKPRIPVVWIHGL	WP_080227380.1	[Ni/Fe] hydrogenase small subunit [Salmonella enterica]	Length: 372\nNumber of predicted TMHs:  2\nExp number of AAs in TMHs: 36.98541\nExp number, first 60 AAs:  19.72756\nTotal prob of N-in: 0.96171\nPOSSIBLE N-term signal sequence\ninside   1   19\nTMhelix   20   39\noutside   40   325\nTMhelix   326   348\ninside   349   372

25716	GCF_002065095.1_ASM206509v1	Salmonella enterica subsp. salamae serovar 55:k:z39	Proteobacteria; Gammaproteobacteria; Enterobacterales; Enterobacteriaceae; Salmonella; Salmonella enterica; Salmonella enterica subsp. salamae	MTGDNTLITSHGINRRDFMKLCAALAATMGLSSKAAAEMAESVSNPQRPPVIWIGAQECT	WP_023248548.1	hydrogenase 2 small subunit [Salmonella enterica]	Length: 372\nNumber of predicted TMHs:  1\nExp number of AAs in TMHs: 24.51281\nExp number, first 60 AAs:  0.26822\nTotal prob of N-in: 0.14402\noutside   1   331\nTMhelix   332   354\ninside   355   372

25717 GCF_002034565.1_ASM203456v1 Salmonella enterica subsp. salamae serovar 58:d:z6 Proteobacteria; Gammaproteobacteria; Enterobacterales; Enterobacteriaceae; Salmonella; Salmonella enterica; Salmonella enterica subsp. salamae MNNEETFYQAMRRKGVTRRSFLKFCSLAATSLGLGAGMTPKIAWALENKPRIPVVWIHGL WP_001058314.1 hydrogenase [Salmonella enterica] Length: 372\nNumber of predicted TMHs: 2\nExp number of AAs in TMHs: 36.3110299999999\nExp number, first 60 AAs: 19.72774\nTotal prob of N-in: 0.96169\nPOSSIBLE N-term signal sequence\ninside 1 19\nTMhelix 20 39\noutside 40 325\nTMhelix 326 348\ninside 349 372

25718 GCF_002034565.1_ASM203456v1 Salmonella enterica subsp. salamae serovar 58:d:z6 Proteobacteria; Gammaproteobacteria; Enterobacterales; Enterobacteriaceae; Salmonella; Salmonella enterica; Salmonella enterica subsp. salamae MTGDNTLITSHGINRRDFMKLCAALAATMGLSSKAAAEMAESVSNPQRPPVIWIGAQECT WP_000145432.1 hydrogenase 2 small subunit [Salmonella enterica] Length: 372\nNumber of predicted TMHs: 1\nExp number of AAs in TMHs: 24.5101\nExp number, first 60 AAs: 0.2682\nTotal prob of N-in: 0.14397\noutside 1 331\nTMhelix 332 354\ninside 355 372

25719 GCF_002036155.1_ASM203615v1 Salmonella enterica subsp. salamae serovar 58:l,z13,z28:z6 Proteobacteria; Gammaproteobacteria; Enterobacterales; Enterobacteriaceae; Salmonella; Salmonella enterica; Salmonella enterica subsp. salamae MTGDNTLITSHGINRRDFMKLCAALAATMGLSSKAAAEMAESVSNPQRPPVIWIGAQECT WP_000145432.1 hydrogenase 2 small subunit [Salmonella enterica] Length: 372\nNumber of predicted TMHs: 1\nExp number of AAs in TMHs: 24.5101\nExp number, first 60 AAs: 0.2682\nTotal prob of N-in: 0.14397\noutside 1 331\nTMhelix 332 354\ninside 355 372

25720 GCF_002036155.1_ASM203615v1 Salmonella enterica subsp. salamae serovar 58:l,z13,z28:z6 Proteobacteria; Gammaproteobacteria; Enterobacterales; Enterobacteriaceae; Salmonella; Salmonella enterica; Salmonella enterica subsp. salamae MNNEETFYQAMRRKGVTRRSFLKFCSLAATSLGLGAGMTPKIAWALENKPRIPVVWIHGL WP_001058314.1 hydrogenase [Salmonella enterica] Length: 372\nNumber of predicted TMHs: 2\nExp number of AAs in TMHs: 36.3110299999999\nExp number, first 60 AAs: 19.72774\nTotal prob of N-in: 0.96169\nPOSSIBLE N-term signal sequence\ninside 1 19\nTMhelix 20 39\noutside 40 325\nTMhelix 326 348\ninside 349 372

25721 GCF_000308035.1_ASM30803v1 Salmonella enterica subsp. salamae str. 3588/07 Proteobacteria; Gammaproteobacteria; Enterobacterales; Enterobacteriaceae; Salmonella; Salmonella enterica; Salmonella enterica subsp. salamae MNNEETFYQAMRRKGVTRRSFLKFCSLAATSLGLGAGMTPKIAWALENKPRIPVVWIHGL WP_001058314.1 hydrogenase [Salmonella enterica] Length: 372\nNumber of predicted TMHs: 2\nExp number of AAs in TMHs: 36.3110299999999\nExp number, first 60 AAs: 19.72774\nTotal prob of N-in: 0.96169\nPOSSIBLE N-term signal sequence\ninside 1 19\nTMhelix 20 39\noutside 40 325\nTMhelix 326 348\ninside 349 372

25722 GCF_000308035.1_ASM30803v1 Salmonella enterica subsp. salamae str. 3588/07 Proteobacteria; Gammaproteobacteria; Enterobacterales; Enterobacteriaceae; Salmonella; Salmonella enterica; Salmonella enterica subsp. salamae MTGDNTLITSHGINRRDFMKLCAALAATMGLSSKAAAEMAESVSNPQRPPVIWIGAQECT WP_000145432.1 hydrogenase 2 small subunit [Salmonella enterica] Length: 372\nNumber of predicted TMHs: 1\nExp number of AAs in TMHs: 24.5101\nExp number, first 60 AAs: 0.2682\nTotal prob of N-in: 0.14397\noutside 1 331\nTMhelix 332 354\ninside 355 372

25723 GCF_000416565.1_Salmonella_enterica_subsp._salamae_strain_DMA-1 Salmonella enterica subsp. salamae str. DMA-1Proteobacteria; Gammaproteobacteria; Enterobacterales; Enterobacteriaceae; Salmonella; Salmonella enterica; Salmonella enterica subsp. salamae MNNEETFYQAMRRKGVTRRSFLKFCSLAATSLGLGAGMTPKIAWALENKPRIPVVWIHGL WP_001058312.1 hydrogenase [Salmonella enterica] Length: 372\nNumber of predicted TMHs: 2\nExp number of AAs in TMHs: 36.51738\nExp number, first 60 AAs: 19.72849\nTotal prob of N-in: 0.96170\nPOSSIBLE N-term signal sequence\ninside 1 19\nTMhelix 20 39\noutside 40 325\nTMhelix 326 348\ninside 349 372

25724 GCF_000416565.1_Salmonella_enterica_subsp._salamae_strain_DMA-1 Salmonella enterica subsp. salamae str. DMA-1Proteobacteria; Gammaproteobacteria; Enterobacterales; Enterobacteriaceae; Salmonella; Salmonella enterica; Salmonella enterica subsp. salamae MTGDNTLITSHGINRRDFMKLCAALAATMGLSSKAAAEMAESVSNPQRPPVIWIGAQECT WP_000145429.1 hydrogenase 2 small subunit [Salmonella enterica] Length: 372\nNumber of predicted TMHs: 1\nExp number of AAs in TMHs: 24.51309\nExp number, first 60 AAs: 0.2682\nTotal prob of N-in: 0.14402\noutside 1 331\nTMhelix 332 354\ninside 355 372

25725 GCF_001448865.1_ASM144886v1 Salmonella enterica subsp. salamae serovar 56:z10:e,n,x str. 1369-73 Proteobacteria; Gammaproteobacteria; Enterobacterales; Enterobacteriaceae; Salmonella; Salmonella enterica; Salmonella enterica subsp. salamae; Salmonella enterica subsp. salamae serovar 56 MNNEETFYQAMRRKGVTRRSFLKFCSLAATSLGLGAGMTPKIAWALENKPRIPVVWIHGL WP_001058314.1 hydrogenase [Salmonella enterica] Length: 372\nNumber of predicted TMHs: 2\nExp number of AAs in TMHs: 36.3110299999999\nExp number, first 60 AAs: 19.72774\nTotal prob of N-in: 0.96169\nPOSSIBLE N-term signal sequence\ninside 1 19\nTMhelix 20 39\noutside 40 325\nTMhelix 326 348\ninside 349 372

25726 GCF_001448865.1_ASM144886v1 Salmonella enterica subsp. salamae serovar 56:z10:e,n,x str. 1369-73 Proteobacteria; Gammaproteobacteria; Enterobacterales; Enterobacteriaceae; Salmonella; Salmonella enterica; Salmonella enterica subsp. salamae; Salmonella enterica subsp. salamae serovar 56 MTGDNTLITSHGINRRDFMKLCAALAATMGLSSKAAAEMAESVSNPQRPPVIWIGAQECT WP_000145432.1 hydrogenase 2 small subunit [Salmonella enterica] Length: 372\nNumber of predicted TMHs: 1\nExp number of AAs in TMHs: 24.5101\nExp number, first 60 AAs: 0.2682\nTotal prob of N-in: 0.14397\noutside 1 331\nTMhelix 332 354\ninside 355 372

25727 GCF_000486145.1_ASM48614v1 Salmonella enterica subsp. salamae serovar 58:l,z13,z28:z6 str. 00-0163 Proteobacteria; Gammaproteobacteria; Enterobacterales; Enterobacteriaceae; Salmonella; Salmonella enterica; Salmonella enterica subsp. salamae; Salmonella enterica subsp. salamae serovar 58 MTGDNTLITSHGINRRDFMKLCAALAATMGLSSKAAAEMAESVSNPQRPPVIWIGAQECT WP_000145432.1 hydrogenase 2 small subunit [Salmonella enterica] Length: 372\nNumber of predicted TMHs: 1\nExp number of AAs in TMHs: 24.5101\nExp number, first 60 AAs: 0.2682\nTotal prob of N-in: 0.14397\noutside 1 331\nTMhelix 332 354\ninside 355 372

25728 GCF_000486145.1_ASM48614v1 Salmonella enterica subsp. salamae serovar 58:l,z13,z28:z6 str. 00-0163 Proteobacteria; Gammaproteobacteria; Enterobacterales; Enterobacteriaceae; Salmonella; Salmonella enterica; Salmonella enterica subsp. salamae; Salmonella enterica subsp. salamae serovar 58 MNNEETFYQAMRRKGVTRRSFLKFCSLAATSLGLGAGMTPKIAWALENKPRIPVVWIHGL WP_001058314.1 hydrogenase [Salmonella enterica] Length: 372\nNumber of predicted TMHs: 2\nExp number of AAs in TMHs: 36.3110299999999\nExp number, first 60 AAs: 19.72774\nTotal prob of N-in: 0.96169\nPOSSIBLE N-term signal sequence\ninside 1 19\nTMhelix 20 39\noutside 40 325\nTMhelix 326 348\ninside 349 372

25729 GCF_002034545.1_ASM203454v1 Salmonella enterica subsp. VII serovar 1,40:g,z51:-- Proteobacteria; Gammaproteobacteria; Enterobacterales; Enterobacteriaceae; Salmonella; Salmonella enterica; Salmonella enterica subsp. VII MTGDNTLITSHGINRRDFMKLCAALAATMGLSSKAAAEMAESVSNPQRPPVIWIGAQECT WP_000145429.1 hydrogenase 2 small subunit [Salmonella enterica] Length: 372\nNumber of predicted TMHs: 1\nExp number of AAs in TMHs: 24.51309\nExp number, first 60 AAs: 0.2682\nTotal prob of N-in: 0.14402\noutside 1 331\nTMhelix 332 354\ninside 355 372

25730 GCF_002034545.1_ASM203454v1 Salmonella enterica subsp. VII serovar 1,40:g,z51:-- Proteobacteria; Gammaproteobacteria; Enterobacterales; Enterobacteriaceae; Salmonella; Salmonella enterica; Salmonella enterica subsp. VII MNNEETFYQAMRRKGVTRRSFLKFCSLAATSLGLGAGMTPKIAWALENKPRIPVVWIHGL WP_079805161.1 [Ni/Fe] hydrogenase small subunit [Salmonella enterica] Length: 372\nNumber of predicted TMHs: 2\nExp number of AAs in TMHs: 36.9928999999999\nExp number, first 60 AAs: 19.72755\nTotal prob of N-in: 0.96172\nPOSSIBLE N-term signal sequence\ninside 1 19\nTMhelix 20 39\noutside 40 325\nTMhelix 326 348\ninside 349 372

25731 GCF_001906355.1_ASM190635v1 Shigella boydii Proteobacteria; Gammaproteobacteria; Enterobacterales; Enterobacteriaceae; Shigella MNNEETFYQAMRRQGVTRRSFLKYCSLAATSLGLGAGMAPKIAWALENKPRIPVVWIHGL WP_001058323.1 MULTISPECIES: hydrogenase-1 small chain [Proteobacteria] Length: 372\nNumber of predicted TMHs: 2\nExp number of AAs in TMHs: 37.2608499999999\nExp number, first 60 AAs: 19.52923\nTotal prob of N-in: 0.94351\nPOSSIBLE N-term signal sequence\ninside 1 19\nTMhelix 20 39\noutside 40 325\nTMhelix 326 348\ninside 349 372

25732 GCF_001905915.1_ASM190591v1 Shigella boydii Proteobacteria; Gammaproteobacteria; Enterobacterales; Enterobacteriaceae; Shigella MTGDNTLIHSHGINRRDFMKLCAALAATMGLSSKAAAEMAESVTNPQRPPVIWIGAQECT WP_000145413.1 MULTISPECIES: hydrogenase 2 small subunit [Shigella] Length: 372\nNumber of predicted TMHs: 1\nExp number of AAs in TMHs: 23.72614\nExp number, first 60 AAs: 0.22038\nTotal prob of N-in: 0.13004\noutside 1 331\nTMhelix 332 354\ninside 355 372

25733 GCF_001905975.1_ASM190597v1 Shigella boydii Proteobacteria; Gammaproteobacteria; Enterobacterales; Enterobacteriaceae; Shigella MTGDNTLIHSHGINRRDFMKLCAALAATMGLSSKAAAEMAESVTNPQRPPVIWIGAQECT WP_000145413.1 MULTISPECIES: hydrogenase 2 small subunit [Shigella] Length: 372\nNumber of predicted TMHs: 1\nExp number of AAs in TMHs: 23.72614\nExp number, first 60 AAs: 0.22038\nTotal prob of N-in: 0.13004\noutside 1 331\nTMhelix 332 354\ninside 355 372

25734 GCF_001905975.1_ASM190597v1 Shigella boydii Proteobacteria; Gammaproteobacteria; Enterobacterales; Enterobacteriaceae; Shigella MNNEETFYQAMRRQGVTRRSFLKYCSLAATSLGLGAGMAPKIAWALENKPRIPVVWIHGL WP_001058323.1 MULTISPECIES: hydrogenase-1 small chain [Proteobacteria] Length: 372\nNumber of predicted TMHs: 2\nExp number of AAs in TMHs: 37.2608499999999\nExp number, first 60 AAs: 19.52923\nTotal prob of N-in: 0.94351\nPOSSIBLE N-term signal sequence\ninside 1 19\nTMhelix 20 39\noutside 40 325\nTMhelix 326 348\ninside 349 372

25735 GCF_001905915.1_ASM190591v1 Shigella boydii Proteobacteria; Gammaproteobacteria; Enterobacterales; Enterobacteriaceae; Shigella MNNEETFYQAMRRQGVTRRSFLKYCSLAATSLGLGAGMAPKIAWALENKPRIPVVWIHGL WP_001058323.1 MULTISPECIES: hydrogenase-1 small chain [Proteobacteria] Length: 372\nNumber of predicted TMHs: 2\nExp number of AAs in TMHs: 37.2608499999999\nExp number, first 60 AAs: 19.52923\nTotal prob of N-in: 0.94351\nPOSSIBLE N-term signal sequence\ninside 1 19\nTMhelix 20 39\noutside 40 325\nTMhelix 326 348\ninside 349 372

25736 GCF_001061295.1_ASM106129v1 Shigella boydii Proteobacteria; Gammaproteobacteria; Enterobacterales; Enterobacteriaceae; Shigella MTGDNTLIHSHGINRRDFMKLCAALAATMGLSSKAAAEMAESVTNPQRPPVIWIGAQECT WP_000145410.1 MULTISPECIES: hydrogenase-2 small chain [Proteobacteria] Length: 372\nNumber of predicted TMHs: 1\nExp number of AAs in TMHs: 23.72226\nExp number, first 60 AAs: 0.22038\nTotal prob of N-in: 0.12994\noutside 1 331\nTMhelix 332 354\ninside 355 372

25737 GCF_001906205.1_ASM190620v1 Shigella boydii Proteobacteria; Gammaproteobacteria; Enterobacterales; Enterobacteriaceae; Shigella MNNEETFYQAMRRQGVTRRSFLKYCSLAATSLGLGAGMAPKIAWALENKPRIPVVWIHGL WP_001058323.1 MULTISPECIES: hydrogenase-1 small chain [Proteobacteria] Length: 372\nNumber of predicted TMHs: 2\nExp number of AAs in TMHs: 37.2608499999999\nExp number, first 60 AAs: 19.52923\nTotal prob of N-in: 0.94351\nPOSSIBLE N-term signal sequence\ninside 1 19\nTMhelix 20 39\noutside 40 325\nTMhelix 326 348\ninside 349 372

25738 GCF_001906055.1_ASM190605v1 Shigella boydii Proteobacteria; Gammaproteobacteria; Enterobacterales; Enterobacteriaceae; Shigella MNNEETFYQAMRRQGVTRRSFLKYCSLAATSLGLGAGMAPKIAWALENKPRIPVVWIHGL WP_001058323.1 MULTISPECIES: hydrogenase-1 small chain [Proteobacteria] Length: 372\nNumber of predicted TMHs: 2\nExp number of AAs in TMHs: 37.2608499999999\nExp number, first 60 AAs: 19.52923\nTotal prob of N-in: 0.94351\nPOSSIBLE N-term signal sequence\ninside 1 19\nTMhelix 20 39\noutside 40 325\nTMhelix 326 348\ninside 349 372

25739 GCF_001906265.1_ASM190626v1 Shigella boydii Proteobacteria; Gammaproteobacteria; Enterobacterales; Enterobacteriaceae; Shigella MTGDNTLIHSHGINRRDFMKLCAALAATMGLSSKAAAEMAESVTNPQRPPVIWIGAQECT WP_000145410.1 MULTISPECIES: hydrogenase-2 small chain [Proteobacteria] Length: 372\nNumber of predicted TMHs: 1\nExp number of AAs in TMHs: 23.72226\nExp number, first 60 AAs: 0.22038\nTotal prob of N-in: 0.12994\noutside 1 331\nTMhelix 332 354\ninside 355 372

25740 GCF_001906265.1_ASM190626v1 Shigella boydii Proteobacteria; Gammaproteobacteria; Enterobacterales; Enterobacteriaceae; Shigella MNNEETFYQAMRRQGVTRRSFLKYCSLAATSLGLGAGMAPKIAWALENKPRIPVVWIHGL WP_001058323.1 MULTISPECIES: hydrogenase-1 small chain [Proteobacteria] Length: 372\nNumber of predicted TMHs: 2\nExp number of AAs in TMHs: 37.2608499999999\nExp number, first 60 AAs: 19.52923\nTotal prob of N-in: 0.94351\nPOSSIBLE N-term signal sequence\ninside 1 19\nTMhelix 20 39\noutside 40 325\nTMhelix 326 348\ninside 349 372

25741 GCF_001906115.1_ASM190611v1 Shigella boydii Proteobacteria; Gammaproteobacteria; Enterobacterales; Enterobacteriaceae; Shigella
MNNEETFYQAMRRQGVTRRSFLKYCSLAATSLGLGAGMAPKIAWALENKPRIPVVWIHGL WP_001058323.1
MULTISPECIES: hydrogenase-1 small chain [Proteobacteria] Length: 372\nNumber of predicted TMHs: 2\nExp number of AAs in TMHs: 37.2608499999999\nExp number, first 60 AAs: 19.52923\nTotal prob of N-in: 0.94351\nPOSSIBLE N-term signal sequence\ninside 1 19\nTMhelix 20 39\noutside 40 325\nTMhelix 326 348\ninside 349 372

25742 GCF_001906145.1_ASM190614v1 Shigella boydii Proteobacteria; Gammaproteobacteria; Enterobacterales; Enterobacteriaceae; Shigella
MNNEETFYQAMRRQGVTRRSFLKYCSLAATSLGLGAGMAPKIAWALENKPRIPVVWIHGL WP_001058323.1
MULTISPECIES: hydrogenase-1 small chain [Proteobacteria] Length: 372\nNumber of predicted TMHs: 2\nExp number of AAs in TMHs: 37.2608499999999\nExp number, first 60 AAs: 19.52923\nTotal prob of N-in: 0.94351\nPOSSIBLE N-term signal sequence\ninside 1 19\nTMhelix 20 39\noutside 40 325\nTMhelix 326 348\ninside 349 372

25743 GCF_001906295.1_ASM190629v1 Shigella boydii Proteobacteria; Gammaproteobacteria; Enterobacterales; Enterobacteriaceae; Shigella
MNNEETFYQAMRRQGVTRRSFLKYCSLAATSLGLGAGMAPKIAWALENKPRIPVVWIHGL WP_001058323.1
MULTISPECIES: hydrogenase-1 small chain [Proteobacteria] Length: 372\nNumber of predicted TMHs: 2\nExp number of AAs in TMHs: 37.2608499999999\nExp number, first 60 AAs: 19.52923\nTotal prob of N-in: 0.94351\nPOSSIBLE N-term signal sequence\ninside 1 19\nTMhelix 20 39\noutside 40 325\nTMhelix 326 348\ninside 349 372

25744 GCF_001933075.2_ASM193307v2 Shigella boydii Proteobacteria; Gammaproteobacteria; Enterobacterales; Enterobacteriaceae; Shigella
MNNEETFYQAMRRQGVTRRSFLKYCSLAATSLGLGAGMAPKIAWALENKPRIPVVWIHGL WP_001058323.1
MULTISPECIES: hydrogenase-1 small chain [Proteobacteria] Length: 372\nNumber of predicted TMHs: 2\nExp number of AAs in TMHs: 37.2608499999999\nExp number, first 60 AAs: 19.52923\nTotal prob of N-in: 0.94351\nPOSSIBLE N-term signal sequence\ninside 1 19\nTMhelix 20 39\noutside 40 325\nTMhelix 326 348\ninside 349 372

25745 GCF_001906445.1_ASM190644v1 Shigella boydii Proteobacteria; Gammaproteobacteria; Enterobacterales; Enterobacteriaceae; Shigella
MNNEETFYQAMRRQGVTRRSFLKYCSLAATSLGLGAGMAPKIAWALENKPRIPVVWIHGL WP_001058323.1
MULTISPECIES: hydrogenase-1 small chain [Proteobacteria] Length: 372\nNumber of predicted TMHs: 2\nExp number of AAs in TMHs: 37.2608499999999\nExp number, first 60 AAs: 19.52923\nTotal prob of N-in: 0.94351\nPOSSIBLE N-term signal sequence\ninside 1 19\nTMhelix 20 39\noutside 40 325\nTMhelix 326 348\ninside 349 372

25746 GCF_001906135.1_ASM190613v1 Shigella boydii Proteobacteria; Gammaproteobacteria; Enterobacterales; Enterobacteriaceae; Shigella
MTGDNTLIHSHGINRRDFMKLCAALAATMGLSSKAAAEMAESVTNPQRPPVIWIGAQECT WP_000145410.1
MULTISPECIES: hydrogenase-2 small chain [Proteobacteria] Length: 372\nNumber of predicted TMHs: 1\nExp number of AAs in TMHs: 23.72226\nExp number, first 60 AAs: 0.22038\nTotal prob of N-in: 0.12994\noutside 1 331\nTMhelix 332 354\ninside 355 372

25747 GCF_001906185.1_ASM190618v1 Shigella boydii Proteobacteria; Gammaproteobacteria; Enterobacterales; Enterobacteriaceae; Shigella
MTGDNTLIHSHGINRRDFMKLCAALAATMGLSSKAAAEMAESVTNPQRPPVIWIGAQECT WP_000145410.1
MULTISPECIES: hydrogenase-2 small chain [Proteobacteria] Length: 372\nNumber of predicted TMHs: 1\nExp number of AAs in TMHs: 23.72226\nExp number, first 60 AAs: 0.22038\nTotal prob of N-in: 0.12994\noutside 1 331\nTMhelix 332 354\ninside 355 372

25748 GCF_001906135.1_ASM190613v1 Shigella boydii Proteobacteria; Gammaproteobacteria; Enterobacterales; Enterobacteriaceae; Shigella
MNNEETFYQAMRRQGVTRRSFLKYCSLAATSLGLGAGMAPKIAWALENKPRIPVVWIHGL WP_078166434.1
[Ni/Fe] hydrogenase small subunit [Shigella boydii] Length: 372\nNumber of predicted TMHs: 2\nExp number of AAs in TMHs: 37.25829\nExp number, first 60 AAs: 19.52923\nTotal prob of N-in: 0.94351\nPOSSIBLE N-term signal sequence\ninside 1 19\nTMhelix 20 39\noutside 40 325\nTMhelix 326 348\ninside 349 372

25749	GCF_001906185.1_ASM190618v1	Shigella boydii	Proteobacteria; Gammaproteobacteria; Enterobacterales; Enterobacteriaceae; Shigella
MNNEETFYQAMRRQGVTRRSFLKYCSLAATSLGLGAGMAPKIAWALENKPRIPVVWIHGL	WP_001058323.1
MULTISPECIES: hydrogenase-1 small chain [Proteobacteria]	Length: 372\nNumber of predicted TMHs: 2\nExp number of AAs in TMHs: 37.2608499999999\nExp number, first 60 AAs: 19.52923\nTotal prob of N-in: 0.94351\nPOSSIBLE N-term signal sequence\ninside 1 19\nTMhelix 20 39\noutside 40 325\nTMhelix 326 348\ninside 349 372

25750	GCF_001906105.1_ASM190610v1	Shigella boydii	Proteobacteria; Gammaproteobacteria; Enterobacterales; Enterobacteriaceae; Shigella
MNNEETFYQAMRRQGVTRRSFLKYCSLAATSLGLGVGMAPKIAWALENKPRIPVVWIHGL	WP_073714848.1
hydrogenase [Shigella boydii]	Length: 372\nNumber of predicted TMHs: 2\nExp number of AAs in TMHs: 37.83336\nExp number, first 60 AAs: 20.10829\nTotal prob of N-in: 0.96228\nPOSSIBLE N-term signal sequence\ninside 1 20\nTMhelix 21 43\noutside 44 325\nTMhelix 326 348\ninside 349 372

25751	GCF_001906275.1_ASM190627v1	Shigella boydii	Proteobacteria; Gammaproteobacteria; Enterobacterales; Enterobacteriaceae; Shigella
MNNEETFYQAMRRQGVTRRSFLKYCSLAATSLGLGAGMAPKIAWALENKPRIPVVWIHGL	WP_001058323.1
MULTISPECIES: hydrogenase-1 small chain [Proteobacteria]	Length: 372\nNumber of predicted TMHs: 2\nExp number of AAs in TMHs: 37.2608499999999\nExp number, first 60 AAs: 19.52923\nTotal prob of N-in: 0.94351\nPOSSIBLE N-term signal sequence\ninside 1 19\nTMhelix 20 39\noutside 40 325\nTMhelix 326 348\ninside 349 372

25752	GCF_001906365.1_100706	Shigella boydii	Proteobacteria; Gammaproteobacteria; Enterobacterales; Enterobacteriaceae; Shigella	MNNEETFYQAMRRQGVTRRSFLKYCSLAATSLGLGAGMAPKIAWALENKPRIPVVWIHGL
WP_001058323.1	MULTISPECIES: hydrogenase-1 small chain [Proteobacteria]	Length: 372\nNumber of predicted TMHs: 2\nExp number of AAs in TMHs: 37.2608499999999\nExp number, first 60 AAs: 19.52923\nTotal prob of N-in: 0.94351\nPOSSIBLE N-term signal sequence\ninside 1 19\nTMhelix 20 39\noutside 40 325\nTMhelix 326 348\ninside 349 372

25753	GCF_001061275.1_ASM106127v1	Shigella boydii	Proteobacteria; Gammaproteobacteria; Enterobacterales; Enterobacteriaceae; Shigella
MTGDNTLIHSHGINRRDFMKLCAALAATMGLSSKAAAEMAESVTNPQRPPVIWIGAQECT	WP_000145410.1
MULTISPECIES: hydrogenase-2 small chain [Proteobacteria]	Length: 372\nNumber of predicted TMHs: 1\nExp number of AAs in TMHs: 23.72226\nExp number, first 60 AAs: 0.22038\nTotal prob of N-in: 0.12994\noutside 1 331\nTMhelix 332 354\ninside 355 372

25754	GCF_001906285.1_ASM190628v1	Shigella boydii	Proteobacteria; Gammaproteobacteria; Enterobacterales; Enterobacteriaceae; Shigella
MNNEETFYQAMRRQGVTRRSFLKYCSLAATSLGLGAGMAPKIAWALENKPRIPVVWIHGL	WP_001058323.1
MULTISPECIES: hydrogenase-1 small chain [Proteobacteria]	Length: 372\nNumber of predicted TMHs: 2\nExp number of AAs in TMHs: 37.2608499999999\nExp number, first 60 AAs: 19.52923\nTotal prob of N-in: 0.94351\nPOSSIBLE N-term signal sequence\ninside 1 19\nTMhelix 20 39\noutside 40 325\nTMhelix 326 348\ninside 349 372

25755	GCF_001906385.1_ASM190638v1	Shigella boydii	Proteobacteria; Gammaproteobacteria; Enterobacterales; Enterobacteriaceae; Shigella
MNNEETFYQAMRRQGVTRRSFLKYCSLAATSLGLGAGMAPKIAWALENKPRIPVVWIHGL	WP_073692392.1
hydrogenase [Shigella boydii]	Length: 372\nNumber of predicted TMHs: 2\nExp number of AAs in TMHs: 37.2732\nExp number, first 60 AAs: 19.52926\nTotal prob of N-in: 0.94351\nPOSSIBLE N-term signal sequence\ninside 1 19\nTMhelix 20 39\noutside 40 325\nTMhelix 326 348\ninside 349 372

25756	GCF_001076355.1_ASM107635v1	Shigella boydii	Proteobacteria; Gammaproteobacteria; Enterobacterales; Enterobacteriaceae; Shigella
MNNEETFYQAMRRQGVTRRSFLKYCSLAATSLGLGAGMAPKIAWALENKPRIPVVWIHGL	WP_001058323.1
MULTISPECIES: hydrogenase-1 small chain [Proteobacteria]	Length: 372\nNumber of predicted TMHs: 2\nExp number of AAs in TMHs: 37.2608499999999\nExp number, first 60 AAs: 19.52923\nTotal prob of N-in: 0.94351\nPOSSIBLE N-term signal sequence\ninside 1 19\nTMhelix 20 39\noutside 40 325\nTMhelix 326 348\ninside 349 372

25757 GCF_001062045.1_ASM106204v1 Shigella boydii Proteobacteria; Gammaproteobacteria; Enterobacterales; Enterobacteriaceae; Shigella
MNKRRTTLTVLALIASLGLSSAPALADKGGNGNGNGNGHGNSGNNGNHGNNGNHGNKGNK WP_049200367.1
MULTISPECIES: membrane protein [Enterobacterales] Length: 164\nNumber of predicted TMHs: 2\nExp number of AAs in TMHs: 30.51526\nExp number, first 60 AAs: 14.42673\nTotal prob of N-in: 0.94579\nPOSSIBLE N-term signal sequence\ninside 1 6\nTMhelix 7 26\noutside 27 144\nTMhelix 145 163\ninside 164 164

25758 GCF_001906045.1_ASM190604v1 Shigella boydii Proteobacteria; Gammaproteobacteria; Enterobacterales; Enterobacteriaceae; Shigella
MNNEETFYQAMRRQGVTRRSFLKYCSLAATSLGLGAGMAPKIAWALENKPRIPVVWIHGL WP_001058323.1
MULTISPECIES: hydrogenase-1 small chain [Proteobacteria] Length: 372\nNumber of predicted TMHs: 2\nExp number of AAs in TMHs: 37.2608499999999\nExp number, first 60 AAs: 19.52923\nTotal prob of N-in: 0.94351\nPOSSIBLE N-term signal sequence\ninside 1 19\nTMhelix 20 39\noutside 40 325\nTMhelix 326 348\ninside 349 372

25759 GCF_001061115.1_ASM106111v1 Shigella boydii Proteobacteria; Gammaproteobacteria; Enterobacterales; Enterobacteriaceae; Shigella
MTGDNTLIHSHGINRRDFMKLCAALAATMGLSSKAAAEMAESVTNPQRPPVIWIGAQECT WP_000145410.1
MULTISPECIES: hydrogenase-2 small chain [Proteobacteria] Length: 372\nNumber of predicted TMHs: 1\nExp number of AAs in TMHs: 23.72226\nExp number, first 60 AAs: 0.22038\nTotal prob of N-in: 0.12994\noutside 1 331\nTMhelix 332 354\ninside 355 372

25760 GCF_001060475.1_ASM106047v1 Shigella boydii Proteobacteria; Gammaproteobacteria; Enterobacterales; Enterobacteriaceae; Shigella
MNKRRTTLTVLALIASLGLSSAPALADKGGNGNGNGNGHGNSGNNGNHGNNGNHGNKGNK WP_049200367.1
MULTISPECIES: membrane protein [Enterobacterales] Length: 164\nNumber of predicted TMHs: 2\nExp number of AAs in TMHs: 30.51526\nExp number, first 60 AAs: 14.42673\nTotal prob of N-in: 0.94579\nPOSSIBLE N-term signal sequence\ninside 1 6\nTMhelix 7 26\noutside 27 144\nTMhelix 145 163\ninside 164 164

25761 GCF_001905965.1_ASM190596v1 Shigella boydii Proteobacteria; Gammaproteobacteria; Enterobacterales; Enterobacteriaceae; Shigella
MNNEETFYQAMRRQGVTRRSFLKYCSLAATSLGLGAGMAPKIAWALENKPRIPVVWIHGL WP_001058323.1
MULTISPECIES: hydrogenase-1 small chain [Proteobacteria] Length: 372\nNumber of predicted TMHs: 2\nExp number of AAs in TMHs: 37.2608499999999\nExp number, first 60 AAs: 19.52923\nTotal prob of N-in: 0.94351\nPOSSIBLE N-term signal sequence\ninside 1 19\nTMhelix 20 39\noutside 40 325\nTMhelix 326 348\ninside 349 372

25762 GCF_001064675.1_ASM106467v1 Shigella boydii Proteobacteria; Gammaproteobacteria; Enterobacterales; Enterobacteriaceae; Shigella
MTGDNTLIHSHGINRRDFMKLCAALAATMGLSSKAAAEMAESVTNPQRPPVIWIGAQECT WP_000145410.1
MULTISPECIES: hydrogenase-2 small chain [Proteobacteria] Length: 372\nNumber of predicted TMHs: 1\nExp number of AAs in TMHs: 23.72226\nExp number, first 60 AAs: 0.22038\nTotal prob of N-in: 0.12994\noutside 1 331\nTMhelix 332 354\ninside 355 372

25763 GCF_001076355.1_ASM107635v1 Shigella boydii Proteobacteria; Gammaproteobacteria; Enterobacterales; Enterobacteriaceae; Shigella
MTGDNTLIHSHGINRRDFMKLCAALAATMGLSSKAAAEMAESVTNPQRPPVIWIGAQECT WP_000145410.1
MULTISPECIES: hydrogenase-2 small chain [Proteobacteria] Length: 372\nNumber of predicted TMHs: 1\nExp number of AAs in TMHs: 23.72226\nExp number, first 60 AAs: 0.22038\nTotal prob of N-in: 0.12994\noutside 1 331\nTMhelix 332 354\ninside 355 372

25764 GCF_001905965.1_ASM190596v1 Shigella boydii Proteobacteria; Gammaproteobacteria; Enterobacterales; Enterobacteriaceae; Shigella
MTGDNTLIHSHGINRRDFMKLCAALAATMGLSSKAAAEMAESVTNPQRPPVIWIGAQECT WP_000145413.1
MULTISPECIES: hydrogenase 2 small subunit [Shigella] Length: 372\nNumber of predicted TMHs: 1\nExp number of AAs in TMHs: 23.72614\nExp number, first 60 AAs: 0.22038\nTotal prob of N-in: 0.13004\noutside 1 331\nTMhelix 332 354\ninside 355 372

25765 GCF_001905945.1_ASM190594v1 Shigella boydii Proteobacteria; Gammaproteobacteria; Enterobacterales; Enterobacteriaceae; Shigella
MTGDNTLIHSHGINRRDFMKLCAALAATMGLSSKAAAEMAESVTNPQRPPVIWIGAQECT WP_000145410.1

MULTISPECIES: hydrogenase-2 small chain [Proteobacteria] Length: 372\nNumber of predicted TMHs: 1\nExp number of AAs in TMHs: 23.72226\nExp number, first 60 AAs: 0.22038\nTotal prob of N-in: 0.12994\noutside 1 331\nTMhelix 332 354\ninside 355 372

25766 GCF_001905945.1_ASM190594v1 Shigella boydii Proteobacteria; Gammaproteobacteria; Enterobacterales; Enterobacteriaceae; Shigella
MNNEETFYQAMRRQGVTRRSFLKYCSLAATSLGLGAGMAPKIAWALENKPRIPVVWIHGL WP_001058323.1
MULTISPECIES: hydrogenase-1 small chain [Proteobacteria] Length: 372\nNumber of predicted TMHs: 2\nExp number of AAs in TMHs: 37.2608499999999\nExp number, first 60 AAs: 19.52923\nTotal prob of N-in: 0.94351\nPOSSIBLE N-term signal sequence\ninside 1 19\nTMhelix 20 39\noutside 40 325\nTMhelix 326 348\ninside 349 372

25767 GCF_001061115.1_ASM106111v1 Shigella boydii Proteobacteria; Gammaproteobacteria; Enterobacterales; Enterobacteriaceae; Shigella
MNNEETFYQAMRRQGVTRRSFLKYCSLAATSLGLGAGMAPKIAWALENKPRIPVVWIHGL WP_001295940.1
MULTISPECIES: hydrogenase [Enterobacteriaceae] Length: 372\nNumber of predicted TMHs: 2\nExp number of AAs in TMHs: 37.2608899999999\nExp number, first 60 AAs: 19.52923\nTotal prob of N-in: 0.94351\nPOSSIBLE N-term signal sequence\ninside 1 19\nTMhelix 20 39\noutside 40 325\nTMhelix 326 348\ninside 349 372

25768 GCF_001905995.1_ASM190599v1 Shigella boydii Proteobacteria; Gammaproteobacteria; Enterobacterales; Enterobacteriaceae; Shigella
MTGDNTLIHSHGINRRDFMKLCAALAATMGLSSKAAAEMAESVTNPQRPPVIWIGAQECT WP_000145410.1
MULTISPECIES: hydrogenase-2 small chain [Proteobacteria] Length: 372\nNumber of predicted TMHs: 1\nExp number of AAs in TMHs: 23.72226\nExp number, first 60 AAs: 0.22038\nTotal prob of N-in: 0.12994\noutside 1 331\nTMhelix 332 354\ninside 355 372

25769 GCF_001906045.1_ASM190604v1 Shigella boydii Proteobacteria; Gammaproteobacteria; Enterobacterales; Enterobacteriaceae; Shigella
MTGDNTLIHSHGINRRDFMKLCAALAATMGLSSKAAAEMAESVTNPQRPPVIWIGAQECT WP_000145410.1
MULTISPECIES: hydrogenase-2 small chain [Proteobacteria] Length: 372\nNumber of predicted TMHs: 1\nExp number of AAs in TMHs: 23.72226\nExp number, first 60 AAs: 0.22038\nTotal prob of N-in: 0.12994\noutside 1 331\nTMhelix 332 354\ninside 355 372

25770 GCF_001906065.1_ASM190606v1 Shigella boydii Proteobacteria; Gammaproteobacteria; Enterobacterales; Enterobacteriaceae; Shigella
MTGDNTLIHSHGINRRDFMKLCAALAATMGLSSKAAAEMAESVTNPQRPPVIWIGAQECT WP_000145410.1
MULTISPECIES: hydrogenase-2 small chain [Proteobacteria] Length: 372\nNumber of predicted TMHs: 1\nExp number of AAs in TMHs: 23.72226\nExp number, first 60 AAs: 0.22038\nTotal prob of N-in: 0.12994\noutside 1 331\nTMhelix 332 354\ninside 355 372

25771 GCF_001906105.1_ASM190610v1 Shigella boydii Proteobacteria; Gammaproteobacteria; Enterobacterales; Enterobacteriaceae; Shigella
MTGDNTLIHSHGINRRDFMKLCAALAATMGLSSKAAAEMAESVTNPQRPPVIWIGAQECT WP_000145410.1
MULTISPECIES: hydrogenase-2 small chain [Proteobacteria] Length: 372\nNumber of predicted TMHs: 1\nExp number of AAs in TMHs: 23.72226\nExp number, first 60 AAs: 0.22038\nTotal prob of N-in: 0.12994\noutside 1 331\nTMhelix 332 354\ninside 355 372

25772 GCF_001905995.1_ASM190599v1 Shigella boydii Proteobacteria; Gammaproteobacteria; Enterobacterales; Enterobacteriaceae; Shigella
MNNEETFYQAMRRQGVTRRSFLKYCSLAATSLGLGAGMAPKIAWALENKPRIPVVWIHGL WP_001058323.1
MULTISPECIES: hydrogenase-1 small chain [Proteobacteria] Length: 372\nNumber of predicted TMHs: 2\nExp number of AAs in TMHs: 37.2608499999999\nExp number, first 60 AAs: 19.52923\nTotal prob of N-in: 0.94351\nPOSSIBLE N-term signal sequence\ninside 1 19\nTMhelix 20 39\noutside 40 325\nTMhelix 326 348\ninside 349 372

25773 GCF_001906065.1_ASM190606v1 Shigella boydii Proteobacteria; Gammaproteobacteria; Enterobacterales; Enterobacteriaceae; Shigella
MNNEETFYQAMRRQGVTRRSFLKYCSLAATSLGLGAGMAPKIAWALENKPRIPVVWIHGL WP_001058323.1
MULTISPECIES: hydrogenase-1 small chain [Proteobacteria] Length: 372\nNumber of predicted TMHs: 2\nExp number of AAs in TMHs: 37.2608499999999\nExp number, first 60 AAs: 19.52923\nTotal prob of N-in:

0.94351\nPOSSIBLE N-term signal sequence\ninside 1 19\nTMhelix 20 39\noutside 40 325\nTMhelix 326 348\ninside 349 372

25774 GCF_001906195.1_ASM190619v1 Shigella boydii Proteobacteria; Gammaproteobacteria; Enterobacterales; Enterobacteriaceae; Shigella MTGDNTLIHSHGINRRDFMKLCAALAATMGLSSKAAAEMAESVTNPQRPPVIWIGAQECT WP_000145410.1 MULTISPECIES: hydrogenase-2 small chain [Proteobacteria] Length: 372\nNumber of predicted TMHs: 1\nExp number of AAs in TMHs: 23.72226\nExp number, first 60 AAs: 0.22038\nTotal prob of N-in: 0.12994\noutside 1 331\nTMhelix 332 354\ninside 355 372

25775 GCF_001906225.1_ASM190622v1 Shigella boydii Proteobacteria; Gammaproteobacteria; Enterobacterales; Enterobacteriaceae; Shigella MTGDNTLIHSHGINRRDFMKLCAALAATMGLSSKAAAEMAESVTNPQRPPVIWIGAQECT WP_000145410.1 MULTISPECIES: hydrogenase-2 small chain [Proteobacteria] Length: 372\nNumber of predicted TMHs: 1\nExp number of AAs in TMHs: 23.72226\nExp number, first 60 AAs: 0.22038\nTotal prob of N-in: 0.12994\noutside 1 331\nTMhelix 332 354\ninside 355 372

25776 GCF_001906345.1_ASM190634v1 Shigella boydii Proteobacteria; Gammaproteobacteria; Enterobacterales; Enterobacteriaceae; Shigella MTGDNTLIHSHGINRRDFMKLCAALAATMGLSSKAAAEMAESVTNPQRPPVIWIGAQECT WP_000145413.1 MULTISPECIES: hydrogenase 2 small subunit [Shigella] Length: 372\nNumber of predicted TMHs: 1\nExp number of AAs in TMHs: 23.72614\nExp number, first 60 AAs: 0.22038\nTotal prob of N-in: 0.13004\noutside 1 331\nTMhelix 332 354\ninside 355 372

25777 GCF_001906275.1_ASM190627v1 Shigella boydii Proteobacteria; Gammaproteobacteria; Enterobacterales; Enterobacteriaceae; Shigella MTGDNTLIHSHGINRRDFMKLCAALAATMGLSSKAAAEMAESVTNPQRPPVIWIGAQECT WP_000145410.1 MULTISPECIES: hydrogenase-2 small chain [Proteobacteria] Length: 372\nNumber of predicted TMHs: 1\nExp number of AAs in TMHs: 23.72226\nExp number, first 60 AAs: 0.22038\nTotal prob of N-in: 0.12994\noutside 1 331\nTMhelix 332 354\ninside 355 372

25778 GCF_001906285.1_ASM190628v1 Shigella boydii Proteobacteria; Gammaproteobacteria; Enterobacterales; Enterobacteriaceae; Shigella MTGDNTLIHSHGINRRDFMKLCAALAATMGLSSKAAAEMAESVTNPQRPPVIWIGAQECT WP_000145410.1 MULTISPECIES: hydrogenase-2 small chain [Proteobacteria] Length: 372\nNumber of predicted TMHs: 1\nExp number of AAs in TMHs: 23.72226\nExp number, first 60 AAs: 0.22038\nTotal prob of N-in: 0.12994\noutside 1 331\nTMhelix 332 354\ninside 355 372

25779 GCF_001906365.1_100706 Shigella boydii Proteobacteria; Gammaproteobacteria; Enterobacterales; Enterobacteriaceae; Shigella MTGDNTLIHSHGINRRDFMKLCAALAATMGLSSKAAAEMAESVTNPQRPPVIWIGAQECT WP_000145421.1 MULTISPECIES: hydrogenase 2 small subunit [Shigella] Length: 372\nNumber of predicted TMHs: 1\nExp number of AAs in TMHs: 22.90256\nExp number, first 60 AAs: 0.2236\nTotal prob of N-in: 0.09847\noutside 1 331\nTMhelix 332 354\ninside 355 372

25780 GCF_001906385.1_ASM190638v1 Shigella boydii Proteobacteria; Gammaproteobacteria; Enterobacterales; Enterobacteriaceae; Shigella MTGDNTLIHSHGINRRDFMKLCAALAATMGLSSKAAAEMAESVTNPQRPPVIWIGAQECT WP_000145410.1 MULTISPECIES: hydrogenase-2 small chain [Proteobacteria] Length: 372\nNumber of predicted TMHs: 1\nExp number of AAs in TMHs: 23.72226\nExp number, first 60 AAs: 0.22038\nTotal prob of N-in: 0.12994\noutside 1 331\nTMhelix 332 354\ninside 355 372

25781 GCF_001906195.1_ASM190619v1 Shigella boydii Proteobacteria; Gammaproteobacteria; Enterobacterales; Enterobacteriaceae; Shigella MNNEETFYQAMRRQGVTRRSFLKYCSLAATSLGLGAGMAPKIAWALENKPRIPVVWIHGL WP_001058323.1 MULTISPECIES: hydrogenase-1 small chain [Proteobacteria] Length: 372\nNumber of predicted TMHs: 2\nExp number of AAs in TMHs: 37.2608499999999\nExp number, first 60 AAs: 19.52923\nTotal prob of N-in: 0.94351\nPOSSIBLE N-term signal sequence\ninside 1 19\nTMhelix 20 39\noutside 40 325\nTMhelix 326 348\ninside 349 372

25782 GCF_001906435.1_ASM190643v1 Shigella boydii Proteobacteria; Gammaproteobacteria; Enterobacterales; Enterobacteriaceae; Shigella

MTGDNTLIHSHGINRRDFMKLCAALAATMGLSSKAAAEMAESVTNPQRPPVIWIGAQECT WP_001685818.1 MULTISPECIES: hydrogenase-2 small chain [Enterobacteriaceae] Length: 372\nNumber of predicted TMHs: 1\nExp number of AAs in TMHs: 23.82505\nExp number, first 60 AAs: 0.21974\nTotal prob of N-in: 0.13379\noutside 1 331\nTMhelix 332 354\ninside 355 372

25783 GCF_001933015.2_ASM193301v2 Shigella boydii Proteobacteria; Gammaproteobacteria; Enterobacterales; Enterobacteriaceae; Shigella MTGDNTLIHSHGINRRDFMKLCAALAATMGLSSKAAAEMAESVTNPQRPPVIWIGAQECT WP_000145410.1 MULTISPECIES: hydrogenase-2 small chain [Proteobacteria] Length: 372\nNumber of predicted TMHs: 1\nExp number of AAs in TMHs: 23.72226\nExp number, first 60 AAs: 0.22038\nTotal prob of N-in: 0.12994\noutside 1 331\nTMhelix 332 354\ninside 355 372

25784 GCF_002205865.1_ASM220586v1 Shigella boydii Proteobacteria; Gammaproteobacteria; Enterobacterales; Enterobacteriaceae; Shigella MTGDNTLIHSHGINRRDFMKLCAALAATMGLSSKAAAEMAESVTNPQRPPVIWIGAQECT WP_000145410.1 MULTISPECIES: hydrogenase-2 small chain [Proteobacteria] Length: 372\nNumber of predicted TMHs: 1\nExp number of AAs in TMHs: 23.72226\nExp number, first 60 AAs: 0.22038\nTotal prob of N-in: 0.12994\noutside 1 331\nTMhelix 332 354\ninside 355 372

25785 GCF_002205865.1_ASM220586v1 Shigella boydii Proteobacteria; Gammaproteobacteria; Enterobacterales; Enterobacteriaceae; Shigella MNNEETFYQAMRRQGVTRRSFLKYCSLAATSLGLGAGMAPKIAWALENKPRIPVVWIHGL WP_001058323.1 MULTISPECIES: hydrogenase-1 small chain [Proteobacteria] Length: 372\nNumber of predicted TMHs: 2\nExp number of AAs in TMHs: 37.2608499999999\nExp number, first 60 AAs: 19.52923\nTotal prob of N-in: 0.94351\nPOSSIBLE N-term signal sequence\ninside 1 19\nTMhelix 20 39\noutside 40 325\nTMhelix 326 348\ninside 349 372

25786 GCF_001933015.2_ASM193301v2 Shigella boydii Proteobacteria; Gammaproteobacteria; Enterobacterales; Enterobacteriaceae; Shigella MNNEETFYQAMRRQGVTRRSFLKYCSLAATSLGLGAGMAPKIAWALENKPRIPVVWIHGL WP_001058323.1 MULTISPECIES: hydrogenase-1 small chain [Proteobacteria] Length: 372\nNumber of predicted TMHs: 2\nExp number of AAs in TMHs: 37.2608499999999\nExp number, first 60 AAs: 19.52923\nTotal prob of N-in: 0.94351\nPOSSIBLE N-term signal sequence\ninside 1 19\nTMhelix 20 39\noutside 40 325\nTMhelix 326 348\ninside 349 372

25787 GCF_001906345.1_ASM190634v1 Shigella boydii Proteobacteria; Gammaproteobacteria; Enterobacterales; Enterobacteriaceae; Shigella MNNEETFYQAMRRQGVTRRSFLKYCSLAATSLGLGAGMAPKIAWALENKPRIPVVWIHGL WP_001058323.1 MULTISPECIES: hydrogenase-1 small chain [Proteobacteria] Length: 372\nNumber of predicted TMHs: 2\nExp number of AAs in TMHs: 37.2608499999999\nExp number, first 60 AAs: 19.52923\nTotal prob of N-in: 0.94351\nPOSSIBLE N-term signal sequence\ninside 1 19\nTMhelix 20 39\noutside 40 325\nTMhelix 326 348\ninside 349 372

25788 GCF_001906225.1_ASM190622v1 Shigella boydii Proteobacteria; Gammaproteobacteria; Enterobacterales; Enterobacteriaceae; Shigella MNNEETFYQAMRRQGVTRRSFLKYCSLAATSLGLGAGMAPKIAWALENKPRIPVVWIHGL WP_001058323.1 MULTISPECIES: hydrogenase-1 small chain [Proteobacteria] Length: 372\nNumber of predicted TMHs: 2\nExp number of AAs in TMHs: 37.2608499999999\nExp number, first 60 AAs: 19.52923\nTotal prob of N-in: 0.94351\nPOSSIBLE N-term signal sequence\ninside 1 19\nTMhelix 20 39\noutside 40 325\nTMhelix 326 348\ninside 349 372

25789 GCF_001906435.1_ASM190643v1 Shigella boydii Proteobacteria; Gammaproteobacteria; Enterobacterales; Enterobacteriaceae; Shigella MNNEETFYQAMRRQGVTRRSFLKYCSLAATSLGLGAGMAPKIAWALENKPRIPVVWIHGL WP_001058323.1 MULTISPECIES: hydrogenase-1 small chain [Proteobacteria] Length: 372\nNumber of predicted TMHs: 2\nExp number of AAs in TMHs: 37.2608499999999\nExp number, first 60 AAs: 19.52923\nTotal prob of N-in: 0.94351\nPOSSIBLE N-term signal sequence\ninside 1 19\nTMhelix 20 39\noutside 40 325\nTMhelix 326 348\ninside 349 372

25790 GCF_001933075.2_ASM193307v2 Shigella boydii Proteobacteria; Gammaproteobacteria; Enterobacterales; Enterobacteriaceae; Shigella

MTGDNTLIHSHGINRRDFMKLCAALAATMGLSSKAAAEMAESVTNPQRPPVIWIGAQECT WP_000145410.1 MULTISPECIES: hydrogenase-2 small chain [Proteobacteria] Length: 372\nNumber of predicted TMHs: 1\nExp number of AAs in TMHs: 23.72226\nExp number, first 60 AAs: 0.22038\nTotal prob of N-in: 0.12994\noutside 1 331\nTMhelix 332 354\ninside 355 372

25791 GCF_001906115.1_ASM190611v1 Shigella boydii Proteobacteria; Gammaproteobacteria; Enterobacterales; Enterobacteriaceae; Shigella MTGDNTLIHSHGINRRDFMKLCAALAATMGLSSKAAAEMAESVTNPQRPPVIWIGAQECT WP_000145410.1 MULTISPECIES: hydrogenase-2 small chain [Proteobacteria] Length: 372\nNumber of predicted TMHs: 1\nExp number of AAs in TMHs: 23.72226\nExp number, first 60 AAs: 0.22038\nTotal prob of N-in: 0.12994\noutside 1 331\nTMhelix 332 354\ninside 355 372

25792 GCF_001063095.1_ASM106309v1 Shigella boydii Proteobacteria; Gammaproteobacteria; Enterobacterales; Enterobacteriaceae; Shigella MNNEETFYQAMRRQGVTRRSFLKYCSLAATSLGLGAGMAPKIAWALENKPRIPVVWIHGL WP_001058323.1 MULTISPECIES: hydrogenase-1 small chain [Proteobacteria] Length: 372\nNumber of predicted TMHs: 2\nExp number of AAs in TMHs: 37.2608499999999\nExp number, first 60 AAs: 19.52923\nTotal prob of N-in: 0.94351\nPOSSIBLE N-term signal sequence\ninside 1 19\nTMhelix 20 39\noutside 40 325\nTMhelix 326 348\ninside 349 372

25793 GCF_001906145.1_ASM190614v1 Shigella boydii Proteobacteria; Gammaproteobacteria; Enterobacterales; Enterobacteriaceae; Shigella MTGDNTLIHSHGINRRDFMKLCAALAATMGLSSKAAAEMAESVTNPQRPPVIWIGAQECT WP_000145410.1 MULTISPECIES: hydrogenase-2 small chain [Proteobacteria] Length: 372\nNumber of predicted TMHs: 1\nExp number of AAs in TMHs: 23.72226\nExp number, first 60 AAs: 0.22038\nTotal prob of N-in: 0.12994\noutside 1 331\nTMhelix 332 354\ninside 355 372

25794 GCF_001906025.1_ASM190602v1 Shigella boydii Proteobacteria; Gammaproteobacteria; Enterobacterales; Enterobacteriaceae; Shigella MNNEETFYQAMRRQGVTRRSFLKYCSLAATSLGLGAGMAPKIAWALENKPRIPVVWIHGL WP_001058323.1 MULTISPECIES: hydrogenase-1 small chain [Proteobacteria] Length: 372\nNumber of predicted TMHs: 2\nExp number of AAs in TMHs: 37.2608499999999\nExp number, first 60 AAs: 19.52923\nTotal prob of N-in: 0.94351\nPOSSIBLE N-term signal sequence\ninside 1 19\nTMhelix 20 39\noutside 40 325\nTMhelix 326 348\ninside 349 372

25795 GCF_001906205.1_ASM190620v1 Shigella boydii Proteobacteria; Gammaproteobacteria; Enterobacterales; Enterobacteriaceae; Shigella MTGDNTLIHSHGINRRDFMKLCAALAATMGLSSKAAAEMAESVTNPQRPPVIWIGAQECT WP_000145410.1 MULTISPECIES: hydrogenase-2 small chain [Proteobacteria] Length: 372\nNumber of predicted TMHs: 1\nExp number of AAs in TMHs: 23.72226\nExp number, first 60 AAs: 0.22038\nTotal prob of N-in: 0.12994\noutside 1 331\nTMhelix 332 354\ninside 355 372

25796 GCF_001906295.1_ASM190629v1 Shigella boydii Proteobacteria; Gammaproteobacteria; Enterobacterales; Enterobacteriaceae; Shigella MTGDNTLIHSHGINRRDFMKLCAALAATMGLSSKAAAEMAESVTNPQRPPVIWIGAQECT WP_001685818.1 MULTISPECIES: hydrogenase-2 small chain [Enterobacteriaceae] Length: 372\nNumber of predicted TMHs: 1\nExp number of AAs in TMHs: 23.82505\nExp number, first 60 AAs: 0.21974\nTotal prob of N-in: 0.13379\noutside 1 331\nTMhelix 332 354\ninside 355 372

25797 GCF_001063095.1_ASM106309v1 Shigella boydii Proteobacteria; Gammaproteobacteria; Enterobacterales; Enterobacteriaceae; Shigella MTGDNTLIHSHGINRRDFMKLCAALATTMGLSSKAAAEMAESVTNPQRPPVIWIGAQECT WP_023149629.1 MULTISPECIES: hydrogenase-2 small chain [Enterobacteriaceae] Length: 372\nNumber of predicted TMHs: 1\nExp number of AAs in TMHs: 23.63603\nExp number, first 60 AAs: 0.12971\nTotal prob of N-in: 0.12608\noutside 1 331\nTMhelix 332 354\ninside 355 372

25798 GCF_001906355.1_ASM190635v1 Shigella boydii Proteobacteria; Gammaproteobacteria; Enterobacterales; Enterobacteriaceae; Shigella MTGDNTLIHSHGINRRDFMKLCAALAATMGLSSKAAAEMAESVTNPQRPPVIWIGAQECT WP_000145410.1 MULTISPECIES: hydrogenase-2 small chain [Proteobacteria] Length: 372\nNumber of predicted TMHs:

1\nExp number of AAs in TMHs: 23.72226\nExp number, first 60 AAs: 0.22038\nTotal prob of N-in: 0.12994\noutside 1 331\nTMhelix 332 354\ninside 355 372

25799 GCF_001906425.1_ASM190642v1 Shigella boydii Proteobacteria; Gammaproteobacteria; Enterobacterales; Enterobacteriaceae; Shigella
MTGDNTLIHSHGINRRDFMKLCAALAATMGLSSKAAAEMAESVTNPQRPPVIWIGAQECT WP_000145410.1
MULTISPECIES: hydrogenase-2 small chain [Proteobacteria] Length: 372\nNumber of predicted TMHs: 1\nExp number of AAs in TMHs: 23.72226\nExp number, first 60 AAs: 0.22038\nTotal prob of N-in: 0.12994\noutside 1 331\nTMhelix 332 354\ninside 355 372

25800 GCF_001906445.1_ASM190644v1 Shigella boydii Proteobacteria; Gammaproteobacteria; Enterobacterales; Enterobacteriaceae; Shigella
MTGDNTLIHSHGINRRDFMKLCAALAATMGLSSKAAAEMAESVTNPQRPPVIWIGAQECT WP_000145410.1
MULTISPECIES: hydrogenase-2 small chain [Proteobacteria] Length: 372\nNumber of predicted TMHs: 1\nExp number of AAs in TMHs: 23.72226\nExp number, first 60 AAs: 0.22038\nTotal prob of N-in: 0.12994\noutside 1 331\nTMhelix 332 354\ninside 355 372

25801 GCF_001906025.1_ASM190602v1 Shigella boydii Proteobacteria; Gammaproteobacteria; Enterobacterales; Enterobacteriaceae; Shigella
MTGDNTLIHSHGINRRDFMKLCAALAATMGLSSKAAAEMAESVTNPQRPPVIWIGAQECT WP_000145410.1
MULTISPECIES: hydrogenase-2 small chain [Proteobacteria] Length: 372\nNumber of predicted TMHs: 1\nExp number of AAs in TMHs: 23.72226\nExp number, first 60 AAs: 0.22038\nTotal prob of N-in: 0.12994\noutside 1 331\nTMhelix 332 354\ninside 355 372

25802 GCF_001906055.1_ASM190605v1 Shigella boydii Proteobacteria; Gammaproteobacteria; Enterobacterales; Enterobacteriaceae; Shigella
MTGDNTLIHSHGINRRDFMKLCAALAATMGLSSKAAAEMAESVTNPQRPPVIWIGAQECT WP_000145410.1
MULTISPECIES: hydrogenase-2 small chain [Proteobacteria] Length: 372\nNumber of predicted TMHs: 1\nExp number of AAs in TMHs: 23.72226\nExp number, first 60 AAs: 0.22038\nTotal prob of N-in: 0.12994\noutside 1 331\nTMhelix 332 354\ninside 355 372

25803 GCF_001060695.1_ASM106069v1 Shigella boydii Proteobacteria; Gammaproteobacteria; Enterobacterales; Enterobacteriaceae; Shigella
MNKRRTTLTVLALIASLGLSSAPALADKGGNGNGNGNGHGNSGNNGNHGNNGNHGNKGNK WP_049200367.1
MULTISPECIES: membrane protein [Enterobacterales] Length: 164\nNumber of predicted TMHs: 2\nExp number of AAs in TMHs: 30.51526\nExp number, first 60 AAs: 14.42673\nTotal prob of N-in: 0.94579\nPOSSIBLE N-term signal sequence\ninside 1 6\nTMhelix 7 26\noutside 27 144\nTMhelix 145 163\ninside 164 164

25804 GCF_001933095.2_ASM193309v2 Shigella dysenteriae Proteobacteria; Gammaproteobacteria; Enterobacterales; Enterobacteriaceae; Shigella
MNNEETFYQAMRRQGVTRRSFLKYCSLAATSLGLGAGMAPKIAWALENKPRIPVVWIHGL WP_001058323.1
MULTISPECIES: hydrogenase-1 small chain [Proteobacteria] Length: 372\nNumber of predicted TMHs: 2\nExp number of AAs in TMHs: 37.2608499999999\nExp number, first 60 AAs: 19.52923\nTotal prob of N-in: 0.94351\nPOSSIBLE N-term signal sequence\ninside 1 19\nTMhelix 20 39\noutside 40 325\nTMhelix 326 348\ninside 349 372

25805 GCF_001933095.2_ASM193309v2 Shigella dysenteriae Proteobacteria; Gammaproteobacteria; Enterobacterales; Enterobacteriaceae; Shigella
MTGDNTLIHSHGINRRDFMKLCAALAATMGLSSKAAAEMAESVTNPQRPPVIWIGAQECT WP_000145410.1
MULTISPECIES: hydrogenase-2 small chain [Proteobacteria] Length: 372\nNumber of predicted TMHs: 1\nExp number of AAs in TMHs: 23.72226\nExp number, first 60 AAs: 0.22038\nTotal prob of N-in: 0.12994\noutside 1 331\nTMhelix 332 354\ninside 355 372

25806 GCF_001932995.2_ASM193299v2 Shigella dysenteriae Proteobacteria; Gammaproteobacteria; Enterobacterales; Enterobacteriaceae; Shigella
MTGDNTLIHSHGINRRDFMKLCAALAATMGLSSKAAAEMAESVTNPQRPPVIWIGAQECT WP_000145410.1
MULTISPECIES: hydrogenase-2 small chain [Proteobacteria] Length: 372\nNumber of predicted TMHs: 1\nExp number of AAs in TMHs: 23.72226\nExp number, first 60 AAs: 0.22038\nTotal prob of N-in: 0.12994\noutside 1 331\nTMhelix 332 354\ninside 355 372

25807 GCF_001932975.2_ASM193297v2 Shigella dysenteriae Proteobacteria; Gammaproteobacteria; Enterobacterales; Enterobacteriaceae; Shigella
MTGDNTLIHSHGINRRDFMKLCAALAATMGLSSKAAAEMAESVTNPQRPPVIWIGAQECT WP_000145410.1
MULTISPECIES: hydrogenase-2 small chain [Proteobacteria] Length: 372\nNumber of predicted TMHs: 1\nExp number of AAs in TMHs: 23.72226\nExp number, first 60 AAs: 0.22038\nTotal prob of N-in: 0.12994\noutside 1 331\nTMhelix 332 354\ninside 355 372

25808 GCF_001932975.2_ASM193297v2 Shigella dysenteriae Proteobacteria; Gammaproteobacteria; Enterobacterales; Enterobacteriaceae; Shigella
MNNEETFYQAMRRQGVTRRSFLKYCSLAATSLGLGAGMAPKIAWALENKPRIPVVWIHGL WP_001058323.1
MULTISPECIES: hydrogenase-1 small chain [Proteobacteria] Length: 372\nNumber of predicted TMHs: 2\nExp number of AAs in TMHs: 37.2608499999999\nExp number, first 60 AAs: 19.52923\nTotal prob of N-in: 0.94351\nPOSSIBLE N-term signal sequence\ninside 1 19\nTMhelix 20 39\noutside 40 325\nTMhelix 326 348\ninside 349 372

25809 GCF_001932995.2_ASM193299v2 Shigella dysenteriae Proteobacteria; Gammaproteobacteria; Enterobacterales; Enterobacteriaceae; Shigella
MNNEETFYQAMRRQGVTRRSFLKYCSLAATSLGLGAGMAPKIAWALENKPRIPVVWIHGL WP_004999783.1
MULTISPECIES: hydrogenase [Enterobacteriaceae] Length: 372\nNumber of predicted TMHs: 2\nExp number of AAs in TMHs: 37.2630799999999\nExp number, first 60 AAs: 19.52922\nTotal prob of N-in: 0.94351\nPOSSIBLE N-term signal sequence\ninside 1 19\nTMhelix 20 39\noutside 40 325\nTMhelix 326 348\ninside 349 372

25810 GCF_002174695.1_ASM217469v1 Shigella flexneri Proteobacteria; Gammaproteobacteria; Enterobacterales; Enterobacteriaceae; Shigella
MTGDNTLIHSHGINRRDFMKLCAALAATMGLSSKAAAEMAESVTNPQRPPVIWIGAQECT WP_000145421.1
MULTISPECIES: hydrogenase 2 small subunit [Shigella] Length: 372\nNumber of predicted TMHs: 1\nExp number of AAs in TMHs: 22.90256\nExp number, first 60 AAs: 0.2236\nTotal prob of N-in: 0.09847\noutside 1 331\nTMhelix 332 354\ninside 355 372

25811 GCF_002174725.1_ASM217472v1 Shigella flexneri Proteobacteria; Gammaproteobacteria; Enterobacterales; Enterobacteriaceae; Shigella
MTGDNTLIHSHGINRRDFMKLCAALAATMGLSSKAAAEMAESVTNPQRPPVIWIGAQECT WP_000145410.1
MULTISPECIES: hydrogenase-2 small chain [Proteobacteria] Length: 372\nNumber of predicted TMHs: 1\nExp number of AAs in TMHs: 23.72226\nExp number, first 60 AAs: 0.22038\nTotal prob of N-in: 0.12994\noutside 1 331\nTMhelix 332 354\ninside 355 372

25812 GCF_001933235.2_ASM193323v2 Shigella flexneri Proteobacteria; Gammaproteobacteria; Enterobacterales; Enterobacteriaceae; Shigella
MTGDNTLIHSHGINRRDFMKLCAALAATMGLSSKAAAEMAESVTNPQRPPVIWIGAQECT WP_000145421.1
MULTISPECIES: hydrogenase 2 small subunit [Shigella] Length: 372\nNumber of predicted TMHs: 1\nExp number of AAs in TMHs: 22.90256\nExp number, first 60 AAs: 0.2236\nTotal prob of N-in: 0.09847\noutside 1 331\nTMhelix 332 354\ninside 355 372

25813 GCF_000754175.1_SFC Shigella flexneri Proteobacteria; Gammaproteobacteria; Enterobacterales; Enterobacteriaceae; Shigella MTGDNTLIHSHGINRRDFMKLCAALAATMGLSSKAAAEMAESVTNPQRPPVIWIGAQECT
WP_000145421.1 MULTISPECIES: hydrogenase 2 small subunit [Shigella] Length: 372\nNumber of predicted TMHs: 1\nExp number of AAs in TMHs: 22.90256\nExp number, first 60 AAs: 0.2236\nTotal prob of N-in: 0.09847\noutside 1 331\nTMhelix 332 354\ninside 355 372

25814 GCF_001933135.2_ASM193313v2 Shigella flexneri Proteobacteria; Gammaproteobacteria; Enterobacterales; Enterobacteriaceae; Shigella
MTGDNTLIHSHGINRRDFMKLCAALAATMGLSSKAAAEMAESVTNPQRPPVIWIGAQECT WP_000145421.1
MULTISPECIES: hydrogenase 2 small subunit [Shigella] Length: 372\nNumber of predicted TMHs: 1\nExp number of AAs in TMHs: 22.90256\nExp number, first 60 AAs: 0.2236\nTotal prob of N-in: 0.09847\noutside 1 331\nTMhelix 332 354\ninside 355 372

25815 GCF_001933165.2_ASM193316v2 Shigella flexneri Proteobacteria; Gammaproteobacteria; Enterobacterales; Enterobacteriaceae; Shigella
MTGDNTLIHSHGINRRDFMKLCAALAATMGLSSKAAAEMAESVTNPQRPPVIWIGAQECT WP_000145421.1
MULTISPECIES: hydrogenase 2 small subunit [Shigella] Length: 372\nNumber of predicted TMHs: 1\nExp number

number of AAs in TMHs: 22.90256\nExp number, first 60 AAs: 0.2236\nTotal prob of N-in: 0.09847\noutside 1 331\nTMhelix 332 354\ninside 355 372

25816 GCF_002154165.1_ASM215416v1 Shigella flexneri Proteobacteria; Gammaproteobacteria; Enterobacterales; Enterobacteriaceae; Shigella
MTGDNTLIHSHGINRRDFMKLCAALAATMGLSSKAAAEMAESVTNPQRPPVIWIGAQECT WP_000145421.1
MULTISPECIES: hydrogenase 2 small subunit [Shigella] Length: 372\nNumber of predicted TMHs: 1\nExp number of AAs in TMHs: 22.90256\nExp number, first 60 AAs: 0.2236\nTotal prob of N-in: 0.09847\noutside 1 331\nTMhelix 332 354\ninside 355 372

25817 GCF_001933215.2_ASM193321v2 Shigella flexneri Proteobacteria; Gammaproteobacteria; Enterobacterales; Enterobacteriaceae; Shigella
MTGDNTLIHSHGINRRDFMKLCAALAATMGLSSKAAAEMAESVTNPQRPPVIWIGAQECT WP_000145421.1
MULTISPECIES: hydrogenase 2 small subunit [Shigella] Length: 372\nNumber of predicted TMHs: 1\nExp number of AAs in TMHs: 22.90256\nExp number, first 60 AAs: 0.2236\nTotal prob of N-in: 0.09847\noutside 1 331\nTMhelix 332 354\ninside 355 372

25818 GCF_001649215.1_ASM164921v1 Shigella flexneri Proteobacteria; Gammaproteobacteria; Enterobacterales; Enterobacteriaceae; Shigella
MNNEETFYQAMRRQGVTRRSFLKYCSLAATSLGLGAGMAPKIAWALENKPRIPVVWIHGL WP_001058323.1
MULTISPECIES: hydrogenase-1 small chain [Proteobacteria] Length: 372\nNumber of predicted TMHs: 2\nExp number of AAs in TMHs: 37.2608499999999\nExp number, first 60 AAs: 19.52923\nTotal prob of N-in: 0.94351\nPOSSIBLE N-term signal sequence\ninside 1 19\nTMhelix 20 39\noutside 40 325\nTMhelix 326 348\ninside 349 372

25819 GCF_000710235.1_ASM71023v1 Shigella flexneri Proteobacteria; Gammaproteobacteria; Enterobacterales; Enterobacteriaceae; Shigella
MTGDNTLIHSHGINRRDFMKLCAALAATMGLSSKAAAEMAESVTNPQRPPVIWIGAQECT WP_000145421.1
MULTISPECIES: hydrogenase 2 small subunit [Shigella] Length: 372\nNumber of predicted TMHs: 1\nExp number of AAs in TMHs: 22.90256\nExp number, first 60 AAs: 0.2236\nTotal prob of N-in: 0.09847\noutside 1 331\nTMhelix 332 354\ninside 355 372

25820 GCF_001649215.1_ASM164921v1 Shigella flexneri Proteobacteria; Gammaproteobacteria; Enterobacterales; Enterobacteriaceae; Shigella
MTGDNTLIHSHGINRRDFMKLCAALAATMGLSSKAAAEMAESVTNPQRPPVIWIGAQECT WP_000145421.1
MULTISPECIES: hydrogenase 2 small subunit [Shigella] Length: 372\nNumber of predicted TMHs: 1\nExp number of AAs in TMHs: 22.90256\nExp number, first 60 AAs: 0.2236\nTotal prob of N-in: 0.09847\noutside 1 331\nTMhelix 332 354\ninside 355 372

25821 GCF_001649345.1_ASM164934v1 Shigella flexneri Proteobacteria; Gammaproteobacteria; Enterobacterales; Enterobacteriaceae; Shigella
MTGDNTLIHSHGINRRDFMKLCAALAATMGLSSKAAAEMAESVTNPQRPPVIWIGAQECT WP_000145421.1
MULTISPECIES: hydrogenase 2 small subunit [Shigella] Length: 372\nNumber of predicted TMHs: 1\nExp number of AAs in TMHs: 22.90256\nExp number, first 60 AAs: 0.2236\nTotal prob of N-in: 0.09847\noutside 1 331\nTMhelix 332 354\ninside 355 372

25822 GCF_001649295.1_ASM164929v1 Shigella flexneri Proteobacteria; Gammaproteobacteria; Enterobacterales; Enterobacteriaceae; Shigella
MTGDNTLIHSHGINRRDFMKLCAALAATMGLSSKAAAEMAESVTNPQRPPVIWIGAQECT WP_000145421.1
MULTISPECIES: hydrogenase 2 small subunit [Shigella] Length: 372\nNumber of predicted TMHs: 1\nExp number of AAs in TMHs: 22.90256\nExp number, first 60 AAs: 0.2236\nTotal prob of N-in: 0.09847\noutside 1 331\nTMhelix 332 354\ninside 355 372

25823 GCF_001933055.2_ASM193305v2 Shigella flexneri Proteobacteria; Gammaproteobacteria; Enterobacterales; Enterobacteriaceae; Shigella
MTGDNTLIHSHGINRRDFMKLCAALAATMGLSSKAAAEMAESVTNPQRPPVIWIGAQECT WP_000145421.1
MULTISPECIES: hydrogenase 2 small subunit [Shigella] Length: 372\nNumber of predicted TMHs: 1\nExp number of AAs in TMHs: 22.90256\nExp number, first 60 AAs: 0.2236\nTotal prob of N-in: 0.09847\noutside 1 331\nTMhelix 332 354\ninside 355 372

25824 GCF_001933155.2_ASM193315v2 Shigella flexneri Proteobacteria; Gammaproteobacteria; Enterobacterales; Enterobacteriaceae; Shigella
MTGDNTLIHSHGINRRDFMKLCAALAATMGLSSKAAAEMAESVTNPQRPPVIWIGAQECT WP_000145421.1
MULTISPECIES: hydrogenase 2 small subunit [Shigella] Length: 372\nNumber of predicted TMHs: 1\nExp number of AAs in TMHs: 22.90256\nExp number, first 60 AAs: 0.2236\nTotal prob of N-in: 0.09847\noutside 1 331\nTMhelix 332 354\ninside 355 372

25825 GCF_001933055.2_ASM193305v2 Shigella flexneri Proteobacteria; Gammaproteobacteria; Enterobacterales; Enterobacteriaceae; Shigella
MNNEETFYQAMRRQGVTRRSFLKYCSLAATSLGLGAGMAPKIAWALENKPRIPVVWIHGL WP_001058323.1
MULTISPECIES: hydrogenase-1 small chain [Proteobacteria] Length: 372\nNumber of predicted TMHs: 2\nExp number of AAs in TMHs: 37.2608499999999\nExp number, first 60 AAs: 19.52923\nTotal prob of N-in: 0.94351\nPOSSIBLE N-term signal sequence\ninside 1 19\nTMhelix 20 39\noutside 40 325\nTMhelix 326 348\ninside 349 372

25826 GCF_001933155.2_ASM193315v2 Shigella flexneri Proteobacteria; Gammaproteobacteria; Enterobacterales; Enterobacteriaceae; Shigella
MNNEETFYQAMRRQGVTRRSFLKYCSLAATSLGLGAGMAPKIAWALENKPRIPVVWIHGL WP_001058323.1
MULTISPECIES: hydrogenase-1 small chain [Proteobacteria] Length: 372\nNumber of predicted TMHs: 2\nExp number of AAs in TMHs: 37.2608499999999\nExp number, first 60 AAs: 19.52923\nTotal prob of N-in: 0.94351\nPOSSIBLE N-term signal sequence\ninside 1 19\nTMhelix 20 39\noutside 40 325\nTMhelix 326 348\ninside 349 372

25827 GCF_001649345.1_ASM164934v1 Shigella flexneri Proteobacteria; Gammaproteobacteria; Enterobacterales; Enterobacteriaceae; Shigella
MNNEETFYQAMRRQGVTRRSFLKYCSLAATSLGLGAGMAPKIAWALENKPRIPVVWIHGL WP_001058323.1
MULTISPECIES: hydrogenase-1 small chain [Proteobacteria] Length: 372\nNumber of predicted TMHs: 2\nExp number of AAs in TMHs: 37.2608499999999\nExp number, first 60 AAs: 19.52923\nTotal prob of N-in: 0.94351\nPOSSIBLE N-term signal sequence\ninside 1 19\nTMhelix 20 39\noutside 40 325\nTMhelix 326 348\ninside 349 372

25828 GCF_001649295.1_ASM164929v1 Shigella flexneri Proteobacteria; Gammaproteobacteria; Enterobacterales; Enterobacteriaceae; Shigella
MNNEETFYQAMRRQGVTRRSFLKYCSLAATSLGLGAGMAPKIAWALENKPRIPVVWIHGL WP_001058323.1
MULTISPECIES: hydrogenase-1 small chain [Proteobacteria] Length: 372\nNumber of predicted TMHs: 2\nExp number of AAs in TMHs: 37.2608499999999\nExp number, first 60 AAs: 19.52923\nTotal prob of N-in: 0.94351\nPOSSIBLE N-term signal sequence\ninside 1 19\nTMhelix 20 39\noutside 40 325\nTMhelix 326 348\ninside 349 372

25829 GCF_001933175.2_ASM193317v2 Shigella flexneri Proteobacteria; Gammaproteobacteria; Enterobacterales; Enterobacteriaceae; Shigella
MTGDNTLIHSHGINRRDFMKLCAALAATMGLSSKAAAEMAESVTNPQRPPVIWIGAQECT WP_000145410.1
MULTISPECIES: hydrogenase-2 small chain [Proteobacteria] Length: 372\nNumber of predicted TMHs: 1\nExp number of AAs in TMHs: 23.72226\nExp number, first 60 AAs: 0.22038\nTotal prob of N-in: 0.12994\noutside 1 331\nTMhelix 332 354\ninside 355 372

25830 GCF_001933255.2_ASM193325v2 Shigella flexneri Proteobacteria; Gammaproteobacteria; Enterobacterales; Enterobacteriaceae; Shigella
MTGDNTLIHSHGINRRDFMKLCAALAATMGLSSKAAAEMAESVTNPQRPPVIWIGAQECT WP_000145410.1
MULTISPECIES: hydrogenase-2 small chain [Proteobacteria] Length: 372\nNumber of predicted TMHs: 1\nExp number of AAs in TMHs: 23.72226\nExp number, first 60 AAs: 0.22038\nTotal prob of N-in: 0.12994\noutside 1 331\nTMhelix 332 354\ninside 355 372

25831 GCF_002174615.1_ASM217461v1 Shigella flexneri Proteobacteria; Gammaproteobacteria; Enterobacterales; Enterobacteriaceae; Shigella
MTGDNTLIHSHGINRRDFMKLCAALAATMGLSSKAAAEMAESVTNPQRPPVIWIGAQECT WP_000145410.1
MULTISPECIES: hydrogenase-2 small chain [Proteobacteria] Length: 372\nNumber of predicted TMHs: 1\nExp number of AAs in TMHs: 23.72226\nExp number, first 60 AAs: 0.22038\nTotal prob of N-in: 0.12994\noutside 1 331\nTMhelix 332 354\ninside 355 372

25832 GCF_001933175.2_ASM193317v2 Shigella flexneri Proteobacteria; Gammaproteobacteria; Enterobacterales; Enterobacteriaceae; Shigella
MNNEETFYQAMRRQGVTRRSFLKYCSLAATSLGLGAGMAPKIAWALENKPRIPVVWIHGL WP_001058323.1
MULTISPECIES: hydrogenase-1 small chain [Proteobacteria] Length: 372\nNumber of predicted TMHs: 2\nExp number of AAs in TMHs: 37.2608499999999\nExp number, first 60 AAs: 19.52923\nTotal prob of N-in: 0.94351\nPOSSIBLE N-term signal sequence\ninside 1 19\nTMhelix 20 39\noutside 40 325\nTMhelix 326 348\ninside 349 372

25833 GCF_002174615.1_ASM217461v1 Shigella flexneri Proteobacteria; Gammaproteobacteria; Enterobacterales; Enterobacteriaceae; Shigella
MNNEETFYQAMRRQGVTRRSFLKYCSLAATSLGLGAGMAPKIAWALENKPRIPVVWIHGL WP_001058323.1
MULTISPECIES: hydrogenase-1 small chain [Proteobacteria] Length: 372\nNumber of predicted TMHs: 2\nExp number of AAs in TMHs: 37.2608499999999\nExp number, first 60 AAs: 19.52923\nTotal prob of N-in: 0.94351\nPOSSIBLE N-term signal sequence\ninside 1 19\nTMhelix 20 39\noutside 40 325\nTMhelix 326 348\ninside 349 372

25834 GCF_001933255.2_ASM193325v2 Shigella flexneri Proteobacteria; Gammaproteobacteria; Enterobacterales; Enterobacteriaceae; Shigella
MNNEETFYQAMRRQGVTRRSFLKYCSLAATSLGLGAGMAPKIAWALENKPRIPVVWIHGL WP_001058323.1
MULTISPECIES: hydrogenase-1 small chain [Proteobacteria] Length: 372\nNumber of predicted TMHs: 2\nExp number of AAs in TMHs: 37.2608499999999\nExp number, first 60 AAs: 19.52923\nTotal prob of N-in: 0.94351\nPOSSIBLE N-term signal sequence\ninside 1 19\nTMhelix 20 39\noutside 40 325\nTMhelix 326 348\ninside 349 372

25835 GCF_002154165.1_ASM215416v1 Shigella flexneri Proteobacteria; Gammaproteobacteria; Enterobacterales; Enterobacteriaceae; Shigella
MNNEETFYQAMRRQGVTRRSFLKYCSLAATSLGLGAGMAPKIAWALENKPRIPVVWIHGL WP_001058323.1
MULTISPECIES: hydrogenase-1 small chain [Proteobacteria] Length: 372\nNumber of predicted TMHs: 2\nExp number of AAs in TMHs: 37.2608499999999\nExp number, first 60 AAs: 19.52923\nTotal prob of N-in: 0.94351\nPOSSIBLE N-term signal sequence\ninside 1 19\nTMhelix 20 39\noutside 40 325\nTMhelix 326 348\ninside 349 372

25836 GCF_000783735.1_ASM78373v1 Shigella flexneri Proteobacteria; Gammaproteobacteria; Enterobacterales; Enterobacteriaceae; Shigella
MNNEETFYQAMRRQGVTRRSFLKYCSLAATSLGLGAGMAPKIAWALENKPRIPVVWIHGL WP_025758242.1
hydrogenase [Shigella flexneri] Length: 372\nNumber of predicted TMHs: 2\nExp number of AAs in TMHs: 37.0188199999999\nExp number, first 60 AAs: 19.52908\nTotal prob of N-in: 0.94338\nPOSSIBLE N-term signal sequence\ninside 1 19\nTMhelix 20 39\noutside 40 325\nTMhelix 326 348\ninside 349 372

25837 GCF_000754175.1_SFC Shigella flexneri Proteobacteria; Gammaproteobacteria; Enterobacterales; Enterobacteriaceae; Shigella MNNEETFYQAMRRQGVTRRSFLKYCSLAATSLGLGAGMAPKIAWALENKPRIPVVWIHGL
WP_001058323.1 MULTISPECIES: hydrogenase-1 small chain [Proteobacteria] Length: 372\nNumber of predicted TMHs: 2\nExp number of AAs in TMHs: 37.2608499999999\nExp number, first 60 AAs: 19.52923\nTotal prob of N-in: 0.94351\nPOSSIBLE N-term signal sequence\ninside 1 19\nTMhelix 20 39\noutside 40 325\nTMhelix 326 348\ninside 349 372

25838 GCF_001933085.2_ASM193308v2 Shigella flexneri Proteobacteria; Gammaproteobacteria; Enterobacterales; Enterobacteriaceae; Shigella
MNNEETFYQAMRRQGVTRRSFLKYCSLAATSLGLGAGMAPKIAWALENKPRIPVVWIHGL WP_075328580.1
[Ni/Fe] hydrogenase small subunit [Shigella flexneri] Length: 372\nNumber of predicted TMHs: 2\nExp number of AAs in TMHs: 37.0193199999999\nExp number, first 60 AAs: 19.52908\nTotal prob of N-in: 0.94338\nPOSSIBLE N-term signal sequence\ninside 1 19\nTMhelix 20 39\noutside 40 325\nTMhelix 326 348\ninside 349 372

25839 GCF_001648955.1_ASM164895v1 Shigella flexneri Proteobacteria; Gammaproteobacteria; Enterobacterales; Enterobacteriaceae; Shigella
MTGDNTLIHSHGINRRDFMKLCAALAATMGLSSKAAAEMAESVTNPQRPPVIWIGAQECT WP_000145421.1
MULTISPECIES: hydrogenase 2 small subunit [Shigella] Length: 372\nNumber of predicted TMHs: 1\nExp number of AAs in TMHs: 22.90256\nExp number, first 60 AAs: 0.2236\nTotal prob of N-in: 0.09847\noutside 1 331\nTMhelix 332 354\ninside 355 372

25840	GCF_001648955.1_ASM164895v1	Shigella flexneri	Proteobacteria; Gammaproteobacteria; Enterobacterales; Enterobacteriaceae; Shigella
MNNEETFYQAMRRQGVTRRSFLKYCSLAATSLGLGAGMAPKIAWALENKPRIPVVWIHGL	WP_001058323.1
MULTISPECIES: hydrogenase-1 small chain [Proteobacteria]	Length: 372\nNumber of predicted TMHs: 2\nExp number of AAs in TMHs: 37.2608499999999\nExp number, first 60 AAs: 19.52923\nTotal prob of N-in: 0.94351\nPOSSIBLE N-term signal sequence\ninside 1 19\nTMhelix 20 39\noutside 40 325\nTMhelix 326 348\ninside 349 372

25841	GCF_001649175.1_ASM164917v1	Shigella flexneri	Proteobacteria; Gammaproteobacteria; Enterobacterales; Enterobacteriaceae; Shigella
MTGDNTLIHSHGINRRDFMKLCAALAATMGLSSKAAAEMAESVTNPQRPPVIWIGAQECT	WP_000145421.1
MULTISPECIES: hydrogenase 2 small subunit [Shigella]	Length: 372\nNumber of predicted TMHs: 1\nExp number of AAs in TMHs: 22.90256\nExp number, first 60 AAs: 0.2236\nTotal prob of N-in: 0.09847\noutside 1 331\nTMhelix 332 354\ninside 355 372

25842	GCF_002174695.1_ASM217469v1	Shigella flexneri	Proteobacteria; Gammaproteobacteria; Enterobacterales; Enterobacteriaceae; Shigella
MNNEETFYQAMRRQGVTRRSFLKYCSLAATSLGLGAGMAPKIAWALENKPRIPVVWIHGL	WP_001058323.1
MULTISPECIES: hydrogenase-1 small chain [Proteobacteria]	Length: 372\nNumber of predicted TMHs: 2\nExp number of AAs in TMHs: 37.2608499999999\nExp number, first 60 AAs: 19.52923\nTotal prob of N-in: 0.94351\nPOSSIBLE N-term signal sequence\ninside 1 19\nTMhelix 20 39\noutside 40 325\nTMhelix 326 348\ninside 349 372

25843	GCF_001649175.1_ASM164917v1	Shigella flexneri	Proteobacteria; Gammaproteobacteria; Enterobacterales; Enterobacteriaceae; Shigella
MNNEETFYQAMRRQGVTRRSFLKYCSLAATSLGLGAGMAPKIAWALENKPRIPVVWIHGL	WP_001058323.1
MULTISPECIES: hydrogenase-1 small chain [Proteobacteria]	Length: 372\nNumber of predicted TMHs: 2\nExp number of AAs in TMHs: 37.2608499999999\nExp number, first 60 AAs: 19.52923\nTotal prob of N-in: 0.94351\nPOSSIBLE N-term signal sequence\ninside 1 19\nTMhelix 20 39\noutside 40 325\nTMhelix 326 348\ninside 349 372

25844	GCF_001649275.1_ASM164927v1	Shigella flexneri	Proteobacteria; Gammaproteobacteria; Enterobacterales; Enterobacteriaceae; Shigella
MTGDNTLIHSHGINRRDFMKLCAALAATMGLSSKAAAEMAESVTNPQRPPVIWIGAQECT	WP_000145421.1
MULTISPECIES: hydrogenase 2 small subunit [Shigella]	Length: 372\nNumber of predicted TMHs: 1\nExp number of AAs in TMHs: 22.90256\nExp number, first 60 AAs: 0.2236\nTotal prob of N-in: 0.09847\noutside 1 331\nTMhelix 332 354\ninside 355 372

25845	GCF_001649335.1_ASM164933v1	Shigella flexneri	Proteobacteria; Gammaproteobacteria; Enterobacterales; Enterobacteriaceae; Shigella
MNNEETFYQAMRRQGVTRRSFLKYCSLAATSLGLGAGMAPKIAWALENKPRIPVVWIHGL	WP_001058323.1
MULTISPECIES: hydrogenase-1 small chain [Proteobacteria]	Length: 372\nNumber of predicted TMHs: 2\nExp number of AAs in TMHs: 37.2608499999999\nExp number, first 60 AAs: 19.52923\nTotal prob of N-in: 0.94351\nPOSSIBLE N-term signal sequence\ninside 1 19\nTMhelix 20 39\noutside 40 325\nTMhelix 326 348\ninside 349 372

25846	GCF_001933235.2_ASM193323v2	Shigella flexneri	Proteobacteria; Gammaproteobacteria; Enterobacterales; Enterobacteriaceae; Shigella
MNNEETFYQAMRRQGVTRRSFLKYCSLAATSLGLGAGMAPKIAWALENKPRIPVVWIHGL	WP_001058323.1
MULTISPECIES: hydrogenase-1 small chain [Proteobacteria]	Length: 372\nNumber of predicted TMHs: 2\nExp number of AAs in TMHs: 37.2608499999999\nExp number, first 60 AAs: 19.52923\nTotal prob of N-in: 0.94351\nPOSSIBLE N-term signal sequence\ninside 1 19\nTMhelix 20 39\noutside 40 325\nTMhelix 326 348\ninside 349 372

25847	GCF_001933085.2_ASM193308v2	Shigella flexneri	Proteobacteria; Gammaproteobacteria; Enterobacterales; Enterobacteriaceae; Shigella
MTGDNTLIHSHGINRRDFMKLCAALAATMGLSSKAAAEMAESVTNPQRPPVIWIGAQECT	WP_000145413.1
MULTISPECIES: hydrogenase 2 small subunit [Shigella]	Length: 372\nNumber of predicted TMHs: 1\nExp number of AAs in TMHs: 23.72614\nExp number, first 60 AAs: 0.22038\nTotal prob of N-in: 0.13004\noutside 1 331\nTMhelix 332 354\ninside 355 372

25848 GCF_001649275.1_ASM164927v1 Shigella flexneri Proteobacteria; Gammaproteobacteria; Enterobacterales; Enterobacteriaceae; Shigella
MNNEETFYQAMRRQGVTRRSFLKYCSLAATSLGLGAGMAPKIAWALENKPRIPVVWIHGL WP_001058323.1
MULTISPECIES: hydrogenase-1 small chain [Proteobacteria] Length: 372\nNumber of predicted TMHs: 2\nExp number of AAs in TMHs: 37.2608499999999\nExp number, first 60 AAs: 19.52923\nTotal prob of N-in: 0.94351\nPOSSIBLE N-term signal sequence\ninside 1 19\nTMhelix 20 39\noutside 40 325\nTMhelix 326 348\ninside 349 372

25849 GCF_002174725.1_ASM217472v1 Shigella flexneri Proteobacteria; Gammaproteobacteria; Enterobacterales; Enterobacteriaceae; Shigella
MNNEETFYQAMRRQGVTRRSFLKYCSLAATSLGLGAGMAPKIAWALENKPRIPVVWIHGL WP_001058323.1
MULTISPECIES: hydrogenase-1 small chain [Proteobacteria] Length: 372\nNumber of predicted TMHs: 2\nExp number of AAs in TMHs: 37.2608499999999\nExp number, first 60 AAs: 19.52923\nTotal prob of N-in: 0.94351\nPOSSIBLE N-term signal sequence\ninside 1 19\nTMhelix 20 39\noutside 40 325\nTMhelix 326 348\ninside 349 372

25850 GCF_001649335.1_ASM164933v1 Shigella flexneri Proteobacteria; Gammaproteobacteria; Enterobacterales; Enterobacteriaceae; Shigella
MTGDNTLIHSHGINRRDFMKLCAALAATMGLSSKAAAEMAESVTNPQRPPVIWIGAQECT WP_000145421.1
MULTISPECIES: hydrogenase 2 small subunit [Shigella] Length: 372\nNumber of predicted TMHs: 1\nExp number of AAs in TMHs: 22.90256\nExp number, first 60 AAs: 0.2236\nTotal prob of N-in: 0.09847\noutside 1 331\nTMhelix 332 354\ninside 355 372

25851 GCF_001933215.2_ASM193321v2 Shigella flexneri Proteobacteria; Gammaproteobacteria; Enterobacterales; Enterobacteriaceae; Shigella
MNNEETFYQAMRRQGVTRRSFLKYCSLAATSLGLGAGMAPKIAWALENKPRIPVVWIHGL WP_001058323.1
MULTISPECIES: hydrogenase-1 small chain [Proteobacteria] Length: 372\nNumber of predicted TMHs: 2\nExp number of AAs in TMHs: 37.2608499999999\nExp number, first 60 AAs: 19.52923\nTotal prob of N-in: 0.94351\nPOSSIBLE N-term signal sequence\ninside 1 19\nTMhelix 20 39\noutside 40 325\nTMhelix 326 348\ninside 349 372

25852 GCF_001933165.2_ASM193316v2 Shigella flexneri Proteobacteria; Gammaproteobacteria; Enterobacterales; Enterobacteriaceae; Shigella
MNNEETFYQAMRRQGVTRRSFLKYCSLAATSLGLGAGMAPKIAWALENKPRIPVVWIHGL WP_001058323.1
MULTISPECIES: hydrogenase-1 small chain [Proteobacteria] Length: 372\nNumber of predicted TMHs: 2\nExp number of AAs in TMHs: 37.2608499999999\nExp number, first 60 AAs: 19.52923\nTotal prob of N-in: 0.94351\nPOSSIBLE N-term signal sequence\ninside 1 19\nTMhelix 20 39\noutside 40 325\nTMhelix 326 348\ninside 349 372

25853 GCF_001933135.2_ASM193313v2 Shigella flexneri Proteobacteria; Gammaproteobacteria; Enterobacterales; Enterobacteriaceae; Shigella
MNNEETFYQAMRRQGVTRRSFLKYCSLAATSLGLGAGMAPKIAWALENKPRIPVVWIHGL WP_025746233.1
hydrogenase [Shigella flexneri] Length: 372\nNumber of predicted TMHs: 2\nExp number of AAs in TMHs: 38.0705399999999\nExp number, first 60 AAs: 19.53132\nTotal prob of N-in: 0.94350\nPOSSIBLE N-term signal sequence\ninside 1 19\nTMhelix 20 39\noutside 40 325\nTMhelix 326 348\ninside 349 372

25854 GCF_000783735.1_ASM78373v1 Shigella flexneri Proteobacteria; Gammaproteobacteria; Enterobacterales; Enterobacteriaceae; Shigella
MTGDNTLIHSHGINRRDFMKLCAALAATMGLSSKAAAEMAESVTNPQRPPVIWIGAQECT WP_000145413.1
MULTISPECIES: hydrogenase 2 small subunit [Shigella] Length: 372\nNumber of predicted TMHs: 1\nExp number of AAs in TMHs: 23.72614\nExp number, first 60 AAs: 0.22038\nTotal prob of N-in: 0.13004\noutside 1 331\nTMhelix 332 354\ninside 355 372

25855 GCF_000710235.1_ASM71023v1 Shigella flexneri Proteobacteria; Gammaproteobacteria; Enterobacterales; Enterobacteriaceae; Shigella
MNNEETFYQAMRRQGVTRRSFLKYCSLAATSLGLGAGMAPKIAWALENKPRIPVVWIHGL WP_001058323.1
MULTISPECIES: hydrogenase-1 small chain [Proteobacteria] Length: 372\nNumber of predicted TMHs: 2\nExp number of AAs in TMHs: 37.2608499999999\nExp number, first 60 AAs: 19.52923\nTotal prob of N-in: 0.94351\nPOSSIBLE N-term signal sequence\ninside 1 19\nTMhelix 20 39\noutside 40 325\nTMhelix 326 348\ninside 349 372

25856 GCF_001249635.1_5236_2_9Shigella sonnei Proteobacteria; Gammaproteobacteria; Enterobacterales; Enterobacteriaceae; Shigella MTGDNTLIHSHGINRRDFMKLCAALAATMGLSSKAAAEMAESVTNPQRPPVIWIGAQECT WP_000145410.1 MULTISPECIES: hydrogenase-2 small chain [Proteobacteria] Length: 372\nNumber of predicted TMHs: 1\nExp number of AAs in TMHs: 23.72226\nExp number, first 60 AAs: 0.22038\nTotal prob of N-in: 0.12994\noutside 1 331\nTMhelix 332 354\ninside 355 372

25857 GCF_000783795.1_ASM78379v1 Shigella sonnei Proteobacteria; Gammaproteobacteria; Enterobacterales; Enterobacteriaceae; Shigella MTGDNTLIHSHGINRRDFMKLCAALAATMGLSSKAAAEMAESVTNPQRPPVIWIGAQECT WP_000145410.1 MULTISPECIES: hydrogenase-2 small chain [Proteobacteria] Length: 372\nNumber of predicted TMHs: 1\nExp number of AAs in TMHs: 23.72226\nExp number, first 60 AAs: 0.22038\nTotal prob of N-in: 0.12994\noutside 1 331\nTMhelix 332 354\ninside 355 372

25858 GCF_000783795.1_ASM78379v1 Shigella sonnei Proteobacteria; Gammaproteobacteria; Enterobacterales; Enterobacteriaceae; Shigella MNNEETFYQAMRRQGVTRRSFLKYCSLAATSLGLGAGMAPKIAWALENKPRIPVVWIHGL WP_005139779.1 hydrogenase [Shigella sonnei] Length: 372\nNumber of predicted TMHs: 2\nExp number of AAs in TMHs: 37.02628\nExp number, first 60 AAs: 19.39684\nTotal prob of N-in: 0.94313\nPOSSIBLE N-term signal sequence\ninside 1 19\nTMhelix 20 39\noutside 40 325\nTMhelix 326 348\ninside 349 372

25859 GCF_001063115.1_ASM106311v1 Shigella sonnei Proteobacteria; Gammaproteobacteria; Enterobacterales; Enterobacteriaceae; Shigella MTGDNTLIHSHGINRRDFMKLCAALAATMGLSSKAAAEMAESVTNPQRPPVIWIGAQECT WP_000145410.1 MULTISPECIES: hydrogenase-2 small chain [Proteobacteria] Length: 372\nNumber of predicted TMHs: 1\nExp number of AAs in TMHs: 23.72226\nExp number, first 60 AAs: 0.22038\nTotal prob of N-in: 0.12994\noutside 1 331\nTMhelix 332 354\ninside 355 372

25860 GCF_001063115.1_ASM106311v1 Shigella sonnei Proteobacteria; Gammaproteobacteria; Enterobacterales; Enterobacteriaceae; Shigella MNNEETFYQAMRRQGVTRRSFLKYCSLAATSLGLGAGMAPKIAWALENKPRIPVVWIHGL WP_001058323.1 MULTISPECIES: hydrogenase-1 small chain [Proteobacteria] Length: 372\nNumber of predicted TMHs: 2\nExp number of AAs in TMHs: 37.2608499999999\nExp number, first 60 AAs: 19.52923\nTotal prob of N-in: 0.94351\nPOSSIBLE N-term signal sequence\ninside 1 19\nTMhelix 20 39\noutside 40 325\nTMhelix 326 348\ninside 349 372

25861 GCF_001245685.1_12971_8_30 Shigella sonnei Proteobacteria; Gammaproteobacteria; Enterobacterales; Enterobacteriaceae; Shigella MTGDNTLIHSHGINRRDFMKLCAALAATMGLSSKAAAEMAESVTNPQRPPVIWIGAQECT WP_000145410.1 MULTISPECIES: hydrogenase-2 small chain [Proteobacteria] Length: 372\nNumber of predicted TMHs: 1\nExp number of AAs in TMHs: 23.72226\nExp number, first 60 AAs: 0.22038\nTotal prob of N-in: 0.12994\noutside 1 331\nTMhelix 332 354\ninside 355 372

25862 GCF_001245685.1_12971_8_30 Shigella sonnei Proteobacteria; Gammaproteobacteria; Enterobacterales; Enterobacteriaceae; Shigella MNNEETFYQAMRRQGVTRRSFLKYCSLAATSLGLGAGMAPKIAWALENKPRIPVVWIHGL WP_005139779.1 hydrogenase [Shigella sonnei] Length: 372\nNumber of predicted TMHs: 2\nExp number of AAs in TMHs: 37.02628\nExp number, first 60 AAs: 19.39684\nTotal prob of N-in: 0.94313\nPOSSIBLE N-term signal sequence\ninside 1 19\nTMhelix 20 39\noutside 40 325\nTMhelix 326 348\ninside 349 372

25863 GCF_001246015.1_5417_1_1Shigella sonnei Proteobacteria; Gammaproteobacteria; Enterobacterales; Enterobacteriaceae; Shigella MNNEETFYQAMRRQGVTRRSFLKYCSLAATSLGLGAGMAPKIAWALENKPRIPVVWIHGL WP_005139779.1 hydrogenase [Shigella sonnei] Length: 372\nNumber of predicted TMHs: 2\nExp number of AAs in TMHs: 37.02628\nExp number, first 60 AAs: 19.39684\nTotal prob of N-in: 0.94313\nPOSSIBLE N-term signal sequence\ninside 1 19\nTMhelix 20 39\noutside 40 325\nTMhelix 326 348\ninside 349 372

25864 GCF_001246385.1_5417_3_1Shigella sonnei Proteobacteria; Gammaproteobacteria; Enterobacterales; Enterobacteriaceae; Shigella MNNEETFYQAMRRQGVTRRSFLKYCSLAATSLGLGAGMAPKIAWALENKPRIPVVWIHGL WP_005139779.1 hydrogenase [Shigella sonnei] Length: 372\nNumber of predicted TMHs: 2\nExp number of AAs in TMHs: 37.02628\nExp number, first 60 AAs: 19.39684\nTotal prob of N-in: 0.94313\nPOSSIBLE N-term signal sequence\ninside 1 19\nTMhelix 20 39\noutside 40 325\nTMhelix 326 348\ninside 349 372

25865 GCF_001246585.1_5236_7_10 Shigella sonnei Proteobacteria; Gammaproteobacteria; Enterobacterales; Enterobacteriaceae; Shigella MNNEETFYQAMRRQGVTRRSFLKYCSLAATSLGLGAGMAPKIAWALENKPRIPVVWIHGL WP_005139779.1 hydrogenase [Shigella sonnei] Length: 372\nNumber of predicted TMHs: 2\nExp number of AAs in TMHs: 37.02628\nExp number, first 60 AAs: 19.39684\nTotal prob of N-in: 0.94313\nPOSSIBLE N-term signal sequence\ninside 1 19\nTMhelix 20 39\noutside 40 325\nTMhelix 326 348\ninside 349 372

25866 GCF_001247385.1_8489_2_91 Shigella sonnei Proteobacteria; Gammaproteobacteria; Enterobacterales; Enterobacteriaceae; Shigella MTGDNTLIHSHGINRRDFMKLCAALAATMGLSSKAAAEMAESVTNPQRPPVIWIGAQECT WP_000145410.1 MULTISPECIES: hydrogenase-2 small chain [Proteobacteria] Length: 372\nNumber of predicted TMHs: 1\nExp number of AAs in TMHs: 23.72226\nExp number, first 60 AAs: 0.22038\nTotal prob of N-in: 0.12994\noutside 1 331\nTMhelix 332 354\ninside 355 372

25867 GCF_001246675.1_6551_1_10 Shigella sonnei Proteobacteria; Gammaproteobacteria; Enterobacterales; Enterobacteriaceae; Shigella MNNEETFYQAMRRQGVTRRSFLKYCSLAATSLGLGAGMAPKIAWALENKPRIPVVWIHGL WP_005139779.1 hydrogenase [Shigella sonnei] Length: 372\nNumber of predicted TMHs: 2\nExp number of AAs in TMHs: 37.02628\nExp number, first 60 AAs: 19.39684\nTotal prob of N-in: 0.94313\nPOSSIBLE N-term signal sequence\ninside 1 19\nTMhelix 20 39\noutside 40 325\nTMhelix 326 348\ninside 349 372

25868 GCF_001246975.1_9870_6_25 Shigella sonnei Proteobacteria; Gammaproteobacteria; Enterobacterales; Enterobacteriaceae; Shigella MNNEETFYQAMRRQGVTRRSFLKYCSLAATSLGLGAGMAPKIAWALENKPRIPVVWIHGL WP_005139779.1 hydrogenase [Shigella sonnei] Length: 372\nNumber of predicted TMHs: 2\nExp number of AAs in TMHs: 37.02628\nExp number, first 60 AAs: 19.39684\nTotal prob of N-in: 0.94313\nPOSSIBLE N-term signal sequence\ninside 1 19\nTMhelix 20 39\noutside 40 325\nTMhelix 326 348\ninside 349 372

25869 GCF_001247925.1_6350_1_5Shigella sonnei Proteobacteria; Gammaproteobacteria; Enterobacterales; Enterobacteriaceae; Shigella MTGDNTLIHSHGINRRDFMKLCAALAATMGLSSKAAAEMAESVTNPQRPPVIWIGAQECT WP_000145410.1 MULTISPECIES: hydrogenase-2 small chain [Proteobacteria] Length: 372\nNumber of predicted TMHs: 1\nExp number of AAs in TMHs: 23.72226\nExp number, first 60 AAs: 0.22038\nTotal prob of N-in: 0.12994\noutside 1 331\nTMhelix 332 354\ninside 355 372

25870 GCF_001247975.1_12971_8_32 Shigella sonnei Proteobacteria; Gammaproteobacteria; Enterobacterales; Enterobacteriaceae; Shigella MTGDNTLIHSHGINRRDFMKLCAALAATMGLSSKAAAEMAESVTNPQRPPVIWIGAQECT WP_000145410.1 MULTISPECIES: hydrogenase-2 small chain [Proteobacteria] Length: 372\nNumber of predicted TMHs: 1\nExp number of AAs in TMHs: 23.72226\nExp number, first 60 AAs: 0.22038\nTotal prob of N-in: 0.12994\noutside 1 331\nTMhelix 332 354\ninside 355 372

25871 GCF_001247805.1_8290_4_90 Shigella sonnei Proteobacteria; Gammaproteobacteria; Enterobacterales; Enterobacteriaceae; Shigella MTGDNTLIHSHGINRRDFMKLCAALAATMGLSSKAAAEMAESVTNPQRPPVIWIGAQECT WP_000145410.1 MULTISPECIES: hydrogenase-2 small chain [Proteobacteria] Length: 372\nNumber of predicted TMHs: 1\nExp number of AAs in TMHs: 23.72226\nExp number, first 60 AAs: 0.22038\nTotal prob of N-in: 0.12994\noutside 1 331\nTMhelix 332 354\ninside 355 372

25872 GCF_001247995.1_8290_4_42 Shigella sonnei Proteobacteria; Gammaproteobacteria; Enterobacterales; Enterobacteriaceae; Shigella MTGDNTLIHSHGINRRDFMKLCAALAATMGLSSKAAAEMAESVTNPQRPPVIWIGAQECT WP_000145410.1 MULTISPECIES: hydrogenase-2 small chain [Proteobacteria] Length: 372\nNumber of predicted TMHs: 1\nExp number of AAs in TMHs: 23.72226\nExp number, first 60 AAs: 0.22038\nTotal prob of N-in: 0.12994\noutside 1 331\nTMhelix 332 354\ninside 355 372

25873 GCF_001247905.1_12971_8_78 Shigella sonnei Proteobacteria; Gammaproteobacteria; Enterobacterales; Enterobacteriaceae; Shigella MTGDNTLIHSHGINRRDFMKLCAALAATMGLSSKAAAEMAESVTNPQRPPVIWIGAQECT WP_000145410.1 MULTISPECIES: hydrogenase-2 small chain [Proteobacteria] Length: 372\nNumber of predicted TMHs: 1\nExp number of AAs in TMHs: 23.72226\nExp number, first 60 AAs: 0.22038\nTotal prob of N-in: 0.12994\noutside 1 331\nTMhelix 332 354\ninside 355 372

25874 GCF_001247155.1_6350_1_16 Shigella sonnei Proteobacteria; Gammaproteobacteria; Enterobacterales; Enterobacteriaceae; Shigella MNNEETFYQAMRRQGVTRRSFLKYCSLAATSLGLGAGMAPKIAWALENKPRIPVVWIHGL WP_005139779.1 hydrogenase [Shigella sonnei] Length: 372\nNumber of predicted TMHs: 2\nExp number of AAs in TMHs: 37.02628\nExp number, first 60 AAs: 19.39684\nTotal prob of N-in: 0.94313\nPOSSIBLE N-term signal sequence\ninside 1 19\nTMhelix 20 39\noutside 40 325\nTMhelix 326 348\ninside 349 372

25875 GCF_001248155.1_12971_8_75 Shigella sonnei Proteobacteria; Gammaproteobacteria; Enterobacterales; Enterobacteriaceae; Shigella MTGDNTLIHSHGINRRDFMKLCAALAATMGLSSKAAAEMAESVTNPQRPPVIWIGAQECT WP_000145410.1 MULTISPECIES: hydrogenase-2 small chain [Proteobacteria] Length: 372\nNumber of predicted TMHs: 1\nExp number of AAs in TMHs: 23.72226\nExp number, first 60 AAs: 0.22038\nTotal prob of N-in: 0.12994\noutside 1 331\nTMhelix 332 354\ninside 355 372

25876 GCF_001248215.1_Sg_201401278_GTAGAG_L001 Shigella sonnei Proteobacteria; Gammaproteobacteria; Enterobacterales; Enterobacteriaceae; Shigella MTGDNTLIHSHGINRRDFMKLCAALAATMGLSSKAAAEMAESVTNPQRPPVIWIGAQECT WP_000145410.1 MULTISPECIES: hydrogenase-2 small chain [Proteobacteria] Length: 372\nNumber of predicted TMHs: 1\nExp number of AAs in TMHs: 23.72226\nExp number, first 60 AAs: 0.22038\nTotal prob of N-in: 0.12994\noutside 1 331\nTMhelix 332 354\ninside 355 372

25877 GCF_001247925.1_6350_1_5Shigella sonnei Proteobacteria; Gammaproteobacteria; Enterobacterales; Enterobacteriaceae; Shigella MNNEETFYQAMRRQGVTRRSFLKYCSLAATSLGLGAGMAPKIAWALENKPRIPVVWIHGL WP_005139779.1 hydrogenase [Shigella sonnei] Length: 372\nNumber of predicted TMHs: 2\nExp number of AAs in TMHs: 37.02628\nExp number, first 60 AAs: 19.39684\nTotal prob of N-in: 0.94313\nPOSSIBLE N-term signal sequence\ninside 1 19\nTMhelix 20 39\noutside 40 325\nTMhelix 326 348\ninside 349 372

25878 GCF_001247805.1_8290_4_90 Shigella sonnei Proteobacteria; Gammaproteobacteria; Enterobacterales; Enterobacteriaceae; Shigella MNNEETFYQAMRRQGVTRRSFLKYCSLAATSLGLGAGMAPKIAWALENKPRIPVVWIHGL WP_005139779.1 hydrogenase [Shigella sonnei] Length: 372\nNumber of predicted TMHs: 2\nExp number of AAs in TMHs: 37.02628\nExp number, first 60 AAs: 19.39684\nTotal prob of N-in: 0.94313\nPOSSIBLE N-term signal sequence\ninside 1 19\nTMhelix 20 39\noutside 40 325\nTMhelix 326 348\ninside 349 372

25879 GCF_001247995.1_8290_4_42 Shigella sonnei Proteobacteria; Gammaproteobacteria; Enterobacterales; Enterobacteriaceae; Shigella MNNEETFYQAMRRQGVTRRSFLKYCSLAATSLGLGAGMAPKIAWALENKPRIPVVWIHGL WP_005139779.1 hydrogenase [Shigella sonnei] Length: 372\nNumber of predicted TMHs: 2\nExp number of AAs in TMHs: 37.02628\nExp number, first 60 AAs: 19.39684\nTotal prob of N-in: 0.94313\nPOSSIBLE N-term signal sequence\ninside 1 19\nTMhelix 20 39\noutside 40 325\nTMhelix 326 348\ninside 349 372

25880 GCF_001247975.1_12971_8_32 Shigella sonnei Proteobacteria; Gammaproteobacteria; Enterobacterales; Enterobacteriaceae; Shigella MNNEETFYQAMRRQGVTRRSFLKYCSLAATSLGLGAGMAPKIAWALENKPRIPVVWIHGL WP_005139779.1 hydrogenase [Shigella sonnei] Length: 372\nNumber of predicted TMHs: 2\nExp number of AAs in TMHs: 37.02628\nExp number, first 60 AAs: 19.39684\nTotal prob of N-in: 0.94313\nPOSSIBLE N-term signal sequence\ninside 1 19\nTMhelix 20 39\noutside 40 325\nTMhelix 326 348\ninside 349 372

25881 GCF_001247905.1_12971_8_78 Shigella sonnei Proteobacteria; Gammaproteobacteria; Enterobacterales; Enterobacteriaceae; Shigella MNNEETFYQAMRRQGVTRRSFLKYCSLAATSLGLGAGMAPKIAWALENKPRIPVVWIHGL WP_005139779.1 hydrogenase [Shigella sonnei] Length: 372\nNumber of predicted TMHs: 2\nExp number of AAs in TMHs: 37.02628\nExp number, first 60 AAs: 19.39684\nTotal prob of N-in: 0.94313\nPOSSIBLE N-term signal sequence\ninside 1 19\nTMhelix 20 39\noutside 40 325\nTMhelix 326 348\ninside 349 372

25882 GCF_001248345.1_8290_4_87 Shigella sonnei Proteobacteria; Gammaproteobacteria; Enterobacterales; Enterobacteriaceae; Shigella MTGDNTLIHSHGINRRDFMKLCAALAATMGLSSKAAAEMAESVTNPQRPPVIWIGAQECT WP_000145410.1 MULTISPECIES: hydrogenase-2 small chain [Proteobacteria] Length: 372\nNumber of predicted TMHs: 1\nExp number of AAs in TMHs: 23.72226\nExp number, first 60 AAs: 0.22038\nTotal prob of N-in: 0.12994\noutside 1 331\nTMhelix 332 354\ninside 355 372

25883 GCF_001248385.1_6551_1_3Shigella sonnei Proteobacteria; Gammaproteobacteria; Enterobacterales; Enterobacteriaceae; Shigella MTGDNTLIHSHGINRRDFMKLCAALAATMGLSSKAAAEMAESVTNPQRPPVIWIGAQECT WP_000145410.1 MULTISPECIES: hydrogenase-2 small chain [Proteobacteria] Length: 372\nNumber of predicted TMHs: 1\nExp number of AAs in TMHs: 23.72226\nExp number, first 60 AAs: 0.22038\nTotal prob of N-in: 0.12994\noutside 1 331\nTMhelix 332 354\ninside 355 372

25884 GCF_001248425.1_H14134026501 Shigella sonnei Proteobacteria; Gammaproteobacteria; Enterobacterales; Enterobacteriaceae; Shigella MTGDNTLIHSHGINRRDFMKLCAALAATMGLSSKAAAEMAESVTNPQRPPVIWIGAQECT WP_000145410.1 MULTISPECIES: hydrogenase-2 small chain [Proteobacteria] Length: 372\nNumber of predicted TMHs: 1\nExp number of AAs in TMHs: 23.72226\nExp number, first 60 AAs: 0.22038\nTotal prob of N-in: 0.12994\noutside 1 331\nTMhelix 332 354\ninside 355 372

25885 GCF_001248445.1_Sg_201401573_GTGGCC_L001 Shigella sonnei Proteobacteria; Gammaproteobacteria; Enterobacterales; Enterobacteriaceae; Shigella MTGDNTLIHSHGINRRDFMKLCAALAATMGLSSKAAAEMAESVTNPQRPPVIWIGAQECT WP_000145410.1 MULTISPECIES: hydrogenase-2 small chain [Proteobacteria] Length: 372\nNumber of predicted TMHs: 1\nExp number of AAs in TMHs: 23.72226\nExp number, first 60 AAs: 0.22038\nTotal prob of N-in: 0.12994\noutside 1 331\nTMhelix 332 354\ninside 355 372

25886 GCF_001248525.1_Sg_201400893_GGCTAC_L001 Shigella sonnei Proteobacteria; Gammaproteobacteria; Enterobacterales; Enterobacteriaceae; Shigella MTGDNTLIHSHGINRRDFMKLCAALAATMGLSSKAAAEMAESVTNPQRPPVIWIGAQECT WP_000145410.1 MULTISPECIES: hydrogenase-2 small chain [Proteobacteria] Length: 372\nNumber of predicted TMHs: 1\nExp number of AAs in TMHs: 23.72226\nExp number, first 60 AAs: 0.22038\nTotal prob of N-in: 0.12994\noutside 1 331\nTMhelix 332 354\ninside 355 372

25887 GCF_001248565.1_6246_5_1Shigella sonnei Proteobacteria; Gammaproteobacteria; Enterobacterales; Enterobacteriaceae; Shigella MTGDNTLIHSHGINRRDFMKLCAALAATMGLSSKAAAEMAESVTNPQRPPVIWIGAQECT WP_000145410.1 MULTISPECIES: hydrogenase-2 small chain [Proteobacteria] Length: 372\nNumber of predicted TMHs: 1\nExp number of AAs in TMHs: 23.72226\nExp number, first 60 AAs: 0.22038\nTotal prob of N-in: 0.12994\noutside 1 331\nTMhelix 332 354\ninside 355 372

25888 GCF_001248585.1_H14086038101 Shigella sonnei Proteobacteria; Gammaproteobacteria; Enterobacterales; Enterobacteriaceae; Shigella MTGDNTLIHSHGINRRDFMKLCAALAATMGLSSKAAAEMAESVTNPQRPPVIWIGAQECT WP_000145410.1 MULTISPECIES: hydrogenase-2 small chain [Proteobacteria] Length: 372\nNumber of predicted TMHs: 1\nExp number of AAs in TMHs: 23.72226\nExp number, first 60 AAs: 0.22038\nTotal prob of N-in: 0.12994\noutside 1 331\nTMhelix 332 354\ninside 355 372

25889 GCF_001248625.1_5236_8_8Shigella sonnei Proteobacteria; Gammaproteobacteria; Enterobacterales; Enterobacteriaceae; Shigella MTGDNTLIHSHGINRRDFMKLCAALAATMGLSSKAAAEMAESVTNPQRPPVIWIGAQECT WP_000145410.1 MULTISPECIES: hydrogenase-2 small chain [Proteobacteria] Length: 372\nNumber of predicted TMHs: 1\nExp number of AAs in TMHs: 23.72226\nExp number, first 60 AAs: 0.22038\nTotal prob of N-in: 0.12994\noutside 1 331\nTMhelix 332 354\ninside 355 372

25890 GCF_001248685.1_8290_4_4Shigella sonnei Proteobacteria; Gammaproteobacteria; Enterobacterales; Enterobacteriaceae; Shigella MTGDNTLIHSHGINRRDFMKLCAALAATMGLSSKAAAEMAESVTNPQRPPVIWIGAQECT WP_000145410.1 MULTISPECIES: hydrogenase-2 small chain [Proteobacteria] Length: 372\nNumber of predicted TMHs: 1\nExp number of AAs in TMHs: 23.72226\nExp number, first 60 AAs: 0.22038\nTotal prob of N-in: 0.12994\noutside 1 331\nTMhelix 332 354\ninside 355 372

25891 GCF_001248625.1_5236_8_8Shigella sonnei Proteobacteria; Gammaproteobacteria; Enterobacterales; Enterobacteriaceae; Shigella MNNEETFYQAMRRQGVTRRSFLKYCSLAATSLGLGAGMAPKIAWALENKPRIPVVWIHGL WP_005139779.1 hydrogenase [Shigella sonnei] Length: 372\nNumber of predicted TMHs: 2\nExp number of AAs in TMHs: 37.02628\nExp number, first 60 AAs: 19.39684\nTotal prob of N-in: 0.94313\nPOSSIBLE N-term signal sequence\ninside 1 19\nTMhelix 20 39\noutside 40 325\nTMhelix 326 348\ninside 349 372

25892 GCF_001248785.1_5417_3_8Shigella sonnei Proteobacteria; Gammaproteobacteria; Enterobacterales; Enterobacteriaceae; Shigella MNNEETFYQAMRRQGVTRRSFLKYCSLAATSLGLGAGMAPKIAWALENKPRIPVVWIHGL WP_005139779.1 hydrogenase [Shigella sonnei] Length: 372\nNumber of predicted TMHs: 2\nExp

number of AAs in TMHs: 37.02628\nExp number, first 60 AAs: 19.39684\nTotal prob of N-in: 0.94313\nPOSSIBLE N-term signal sequence\ninside 1 19\nTMhelix 20 39\noutside 40 325\nTMhelix 326 348\ninside 349 372

25893 GCF_001248745.1_H14146070501 Shigella sonnei Proteobacteria; Gammaproteobacteria; Enterobacterales; Enterobacteriaceae; Shigella MNNEETFYQAMRRQGVTRRSFLKYCSLAATSLGLGAGMAPKIAWALENKPRIPVVWIHGL WP_052962775.1 hydrogenase [Shigella sonnei] Length: 372\nNumber of predicted TMHs: 2\nExp number of AAs in TMHs: 37.09248\nExp number, first 60 AAs: 19.39296\nTotal prob of N-in: 0.94316\nPOSSIBLE N-term signal sequence\ninside 1 19\nTMhelix 20 39\noutside 40 325\nTMhelix 326 348\ninside 349 372

25894 GCF_001248425.1_H14134026501 Shigella sonnei Proteobacteria; Gammaproteobacteria; Enterobacterales; Enterobacteriaceae; Shigella MNNEETFYQAMRRQGVTRRSFLKYCSLAATSLGLGAGMAPKIAWALENKPRIPVVWIHGL WP_052962775.1 hydrogenase [Shigella sonnei] Length: 372\nNumber of predicted TMHs: 2\nExp number of AAs in TMHs: 37.09248\nExp number, first 60 AAs: 19.39296\nTotal prob of N-in: 0.94316\nPOSSIBLE N-term signal sequence\ninside 1 19\nTMhelix 20 39\noutside 40 325\nTMhelix 326 348\ninside 349 372

25895 GCF_001248445.1_Sg_201401573_GTGGCC_L001 Shigella sonnei Proteobacteria; Gammaproteobacteria; Enterobacterales; Enterobacteriaceae; Shigella MNNEETFYQAMRRQGVTRRSFLKYCSLAATSLGLGAGMAPKIAWALENKPRIPVVWIHGL WP_005139779.1 hydrogenase [Shigella sonnei] Length: 372\nNumber of predicted TMHs: 2\nExp number of AAs in TMHs: 37.02628\nExp number, first 60 AAs: 19.39684\nTotal prob of N-in: 0.94313\nPOSSIBLE N-term signal sequence\ninside 1 19\nTMhelix 20 39\noutside 40 325\nTMhelix 326 348\ninside 349 372

25896 GCF_001248565.1_6246_5_1Shigella sonnei Proteobacteria; Gammaproteobacteria; Enterobacterales; Enterobacteriaceae; Shigella MNNEETFYQAMRRQGVTRRSFLKYCSLAATSLGLGAGMAPKIAWALENKPRIPVVWIHGL WP_005139779.1 hydrogenase [Shigella sonnei] Length: 372\nNumber of predicted TMHs: 2\nExp number of AAs in TMHs: 37.02628\nExp number, first 60 AAs: 19.39684\nTotal prob of N-in: 0.94313\nPOSSIBLE N-term signal sequence\ninside 1 19\nTMhelix 20 39\noutside 40 325\nTMhelix 326 348\ninside 349 372

25897 GCF_001248525.1_Sg_201400893_GGCTAC_L001 Shigella sonnei Proteobacteria; Gammaproteobacteria; Enterobacterales; Enterobacteriaceae; Shigella MNNEETFYQAMRRQGVTRRSFLKYCSLAATSLGLGAGMAPKIAWALENKPRIPVVWIHGL WP_005139779.1 hydrogenase [Shigella sonnei] Length: 372\nNumber of predicted TMHs: 2\nExp number of AAs in TMHs: 37.02628\nExp number, first 60 AAs: 19.39684\nTotal prob of N-in: 0.94313\nPOSSIBLE N-term signal sequence\ninside 1 19\nTMhelix 20 39\noutside 40 325\nTMhelix 326 348\ninside 349 372

25898 GCF_001248585.1_H14086038101 Shigella sonnei Proteobacteria; Gammaproteobacteria; Enterobacterales; Enterobacteriaceae; Shigella MNNEETFYQAMRRQGVTRRSFLKYCSLAATSLGLGAGMAPKIAWALENKPRIPVVWIHGL WP_005139779.1 hydrogenase [Shigella sonnei] Length: 372\nNumber of predicted TMHs: 2\nExp number of AAs in TMHs: 37.02628\nExp number, first 60 AAs: 19.39684\nTotal prob of N-in: 0.94313\nPOSSIBLE N-term signal sequence\ninside 1 19\nTMhelix 20 39\noutside 40 325\nTMhelix 326 348\ninside 349 372

25899 GCF_001248805.1_5008_8_1Shigella sonnei Proteobacteria; Gammaproteobacteria; Enterobacterales; Enterobacteriaceae; Shigella MTGDNTLIHSHGINRRDFMKLCAALAATMGLSSKAAAEMAESVTNPQRPPVIWIGAQECT WP_000145410.1 MULTISPECIES: hydrogenase-2 small chain [Proteobacteria] Length: 372\nNumber of predicted TMHs: 1\nExp number of AAs in TMHs: 23.72226\nExp number, first 60 AAs: 0.22038\nTotal prob of N-in: 0.12994\noutside 1 331\nTMhelix 332 354\ninside 355 372

25900 GCF_001248885.1_12971_8_64 Shigella sonnei Proteobacteria; Gammaproteobacteria; Enterobacterales; Enterobacteriaceae; Shigella MTGDNTLIHSHGINRRDFMKLCAALAATMGLSSKAAAEMAESVTNPQRPPVIWIGAQECT WP_000145410.1 MULTISPECIES: hydrogenase-2 small chain [Proteobacteria] Length: 372\nNumber of predicted TMHs: 1\nExp number of AAs in TMHs: 23.72226\nExp number, first 60 AAs: 0.22038\nTotal prob of N-in: 0.12994\noutside 1 331\nTMhelix 332 354\ninside 355 372

25901 GCF_001248985.1_H14090035601 Shigella sonnei Proteobacteria; Gammaproteobacteria; Enterobacterales; Enterobacteriaceae; Shigella MTGDNTLIHSHGINRRDFMKLCAALAATMGLSSKAAAEMAESVTNPQRPPVIWIGAQECT WP_000145410.1 MULTISPECIES: hydrogenase-2 small chain [Proteobacteria] Length: 372\nNumber of predicted TMHs:

1\nExp number of AAs in TMHs: 23.72226\nExp number, first 60 AAs: 0.22038\nTotal prob of N-in: 0.12994\noutside 1 331\nTMhelix 332 354\ninside 355 372

25902 GCF_001249025.1_H14090035401 Shigella sonnei Proteobacteria; Gammaproteobacteria; Enterobacterales; Enterobacteriaceae; Shigella MTGDNTLIHSHGINRRDFMKLCAALAATMGLSSKAAAEMAESVTNPQRPPVIWIGAQECT WP_000145410.1 MULTISPECIES: hydrogenase-2 small chain [Proteobacteria] Length: 372\nNumber of predicted TMHs: 1\nExp number of AAs in TMHs: 23.72226\nExp number, first 60 AAs: 0.22038\nTotal prob of N-in: 0.12994\noutside 1 331\nTMhelix 332 354\ninside 355 372

25903 GCF_001249065.1_5236_8_9Shigella sonnei Proteobacteria; Gammaproteobacteria; Enterobacterales; Enterobacteriaceae; Shigella MTGDNTLIHSHGINRRDFMKLCAALAATMGLSSKAAAEMAESVTNPQRPPVIWIGAQECT WP_000145410.1 MULTISPECIES: hydrogenase-2 small chain [Proteobacteria] Length: 372\nNumber of predicted TMHs: 1\nExp number of AAs in TMHs: 23.72226\nExp number, first 60 AAs: 0.22038\nTotal prob of N-in: 0.12994\noutside 1 331\nTMhelix 332 354\ninside 355 372

25904 GCF_001249115.1_12971_8_80 Shigella sonnei Proteobacteria; Gammaproteobacteria; Enterobacterales; Enterobacteriaceae; Shigella MTGDNTLIHSHGINRRDFMKLCAALAATMGLSSKAAAEMAESVTNPQRPPVIWIGAQECT WP_000145410.1 MULTISPECIES: hydrogenase-2 small chain [Proteobacteria] Length: 372\nNumber of predicted TMHs: 1\nExp number of AAs in TMHs: 23.72226\nExp number, first 60 AAs: 0.22038\nTotal prob of N-in: 0.12994\noutside 1 331\nTMhelix 332 354\ninside 355 372

25905 GCF_001249155.1_8489_2_85 Shigella sonnei Proteobacteria; Gammaproteobacteria; Enterobacterales; Enterobacteriaceae; Shigella MTGDNTLIHSHGINRRDFMKLCAALAATMGLSSKAAAEMAESVTNPQRPPVIWIGAQECT WP_000145410.1 MULTISPECIES: hydrogenase-2 small chain [Proteobacteria] Length: 372\nNumber of predicted TMHs: 1\nExp number of AAs in TMHs: 23.72226\nExp number, first 60 AAs: 0.22038\nTotal prob of N-in: 0.12994\noutside 1 331\nTMhelix 332 354\ninside 355 372

25906 GCF_001249215.1_8290_4_53 Shigella sonnei Proteobacteria; Gammaproteobacteria; Enterobacterales; Enterobacteriaceae; Shigella MTGDNTLIHSHGINRRDFMKLCAALAATMGLSSKAAAEMAESVTNPQRPPVIWIGAQECT WP_000145410.1 MULTISPECIES: hydrogenase-2 small chain [Proteobacteria] Length: 372\nNumber of predicted TMHs: 1\nExp number of AAs in TMHs: 23.72226\nExp number, first 60 AAs: 0.22038\nTotal prob of N-in: 0.12994\noutside 1 331\nTMhelix 332 354\ninside 355 372

25907 GCF_001249275.1_Sg_201312374_TCATTC_L001Shigella sonnei Proteobacteria; Gammaproteobacteria; Enterobacterales; Enterobacteriaceae; Shigella MTGDNTLIHSHGINRRDFMKLCAALAATMGLSSKAAAEMAESVTNPQRPPVIWIGAQECT WP_000145410.1 MULTISPECIES: hydrogenase-2 small chain [Proteobacteria] Length: 372\nNumber of predicted TMHs: 1\nExp number of AAs in TMHs: 23.72226\nExp number, first 60 AAs: 0.22038\nTotal prob of N-in: 0.12994\noutside 1 331\nTMhelix 332 354\ninside 355 372

25908 GCF_001249235.1_5417_3_3Shigella sonnei Proteobacteria; Gammaproteobacteria; Enterobacterales; Enterobacteriaceae; Shigella MTGDNTLIHSHGINRRDFMKLCAALAATMGLSSKAAAEMAESVTNPQRPPVIWIGAQECT WP_000145410.1 MULTISPECIES: hydrogenase-2 small chain [Proteobacteria] Length: 372\nNumber of predicted TMHs: 1\nExp number of AAs in TMHs: 23.72226\nExp number, first 60 AAs: 0.22038\nTotal prob of N-in: 0.12994\noutside 1 331\nTMhelix 332 354\ninside 355 372

25909 GCF_001249215.1_8290_4_53 Shigella sonnei Proteobacteria; Gammaproteobacteria; Enterobacterales; Enterobacteriaceae; Shigella MNNEETFYQAMRRQGVTRRSFLKYCSLAATSLGLGAGMAPKIAWALENKPRIPVVWIHGL WP_005139779.1 hydrogenase [Shigella sonnei] Length: 372\nNumber of predicted TMHs: 2\nExp number of AAs in TMHs: 37.02628\nExp number, first 60 AAs: 19.39684\nTotal prob of N-in: 0.94313\nPOSSIBLE N-term signal sequence\ninside 1 19\nTMhelix 20 39\noutside 40 325\nTMhelix 326 348\ninside 349 372

25910 GCF_001249275.1_Sg_201312374_TCATTC_L001Shigella sonnei Proteobacteria; Gammaproteobacteria; Enterobacterales; Enterobacteriaceae; Shigella MNNEETFYQAMRRQGVTRRSFLKYCSLAATSLGLGAGMAPKIAWALENKPRIPVVWIHGL WP_005139779.1 hydrogenase [Shigella sonnei] Length: 372\nNumber of predicted TMHs: 2\nExp number of AAs in

TMHs: 37.02628\nExp number, first 60 AAs: 19.39684\nTotal prob of N-in: 0.94313\nPOSSIBLE N-term signal sequence\ninside 1 19\nTMhelix 20 39\noutside 40 325\nTMhelix 326 348\ninside 349 372

25911 GCF_001249235.1_5417_3_3Shigella sonnei Proteobacteria; Gammaproteobacteria; Enterobacterales; Enterobacteriaceae; Shigella MNNEETFYQAMRRQGVTRRSFLKYCSLAATSLGLGAGMAPKIAWALENKPRIPVVWIHGL WP_005139779.1 hydrogenase [Shigella sonnei] Length: 372\nNumber of predicted TMHs: 2\nExp number of AAs in TMHs: 37.02628\nExp number, first 60 AAs: 19.39684\nTotal prob of N-in: 0.94313\nPOSSIBLE N-term signal sequence\ninside 1 19\nTMhelix 20 39\noutside 40 325\nTMhelix 326 348\ninside 349 372

25912 GCF_001248885.1_12971_8_64 Shigella sonnei Proteobacteria; Gammaproteobacteria; Enterobacterales; Enterobacteriaceae; Shigella MNNEETFYQAMRRQGVTRRSFLKYCSLAATSLGLGAGMAPKIAWALENKPRIPVVWIHGL WP_005139779.1 hydrogenase [Shigella sonnei] Length: 372\nNumber of predicted TMHs: 2\nExp number of AAs in TMHs: 37.02628\nExp number, first 60 AAs: 19.39684\nTotal prob of N-in: 0.94313\nPOSSIBLE N-term signal sequence\ninside 1 19\nTMhelix 20 39\noutside 40 325\nTMhelix 326 348\ninside 349 372

25913 GCF_001248805.1_5008_8_1Shigella sonnei Proteobacteria; Gammaproteobacteria; Enterobacterales; Enterobacteriaceae; Shigella MNNEETFYQAMRRQGVTRRSFLKYCSLAATSLGLGAGMAPKIAWALENKPRIPVVWIHGL WP_005139779.1 hydrogenase [Shigella sonnei] Length: 372\nNumber of predicted TMHs: 2\nExp number of AAs in TMHs: 37.02628\nExp number, first 60 AAs: 19.39684\nTotal prob of N-in: 0.94313\nPOSSIBLE N-term signal sequence\ninside 1 19\nTMhelix 20 39\noutside 40 325\nTMhelix 326 348\ninside 349 372

25914 GCF_001249395.1_9870_6_6Shigella sonnei Proteobacteria; Gammaproteobacteria; Enterobacterales; Enterobacteriaceae; Shigella MNNEETFYQAMRRQGVTRRSFLKYCSLAATSLGLGAGMAPKIAWALENKPRIPVVWIHGL WP_005139779.1 hydrogenase [Shigella sonnei] Length: 372\nNumber of predicted TMHs: 2\nExp number of AAs in TMHs: 37.02628\nExp number, first 60 AAs: 19.39684\nTotal prob of N-in: 0.94313\nPOSSIBLE N-term signal sequence\ninside 1 19\nTMhelix 20 39\noutside 40 325\nTMhelix 326 348\ninside 349 372

25915 GCF_001249115.1_12971_8_80 Shigella sonnei Proteobacteria; Gammaproteobacteria; Enterobacterales; Enterobacteriaceae; Shigella MNNEETFYQAMRRQGVTRRSFLKYCSLAATSLGLGAGMAPKIAWALENKPRIPVVWIHGL WP_005139779.1 hydrogenase [Shigella sonnei] Length: 372\nNumber of predicted TMHs: 2\nExp number of AAs in TMHs: 37.02628\nExp number, first 60 AAs: 19.39684\nTotal prob of N-in: 0.94313\nPOSSIBLE N-term signal sequence\ninside 1 19\nTMhelix 20 39\noutside 40 325\nTMhelix 326 348\ninside 349 372

25916 GCF_001249155.1_8489_2_85 Shigella sonnei Proteobacteria; Gammaproteobacteria; Enterobacterales; Enterobacteriaceae; Shigella MNNEETFYQAMRRQGVTRRSFLKYCSLAATSLGLGAGMAPKIAWALENKPRIPVVWIHGL WP_005139779.1 hydrogenase [Shigella sonnei] Length: 372\nNumber of predicted TMHs: 2\nExp number of AAs in TMHs: 37.02628\nExp number, first 60 AAs: 19.39684\nTotal prob of N-in: 0.94313\nPOSSIBLE N-term signal sequence\ninside 1 19\nTMhelix 20 39\noutside 40 325\nTMhelix 326 348\ninside 349 372

25917 GCF_001249415.1_9870_6_40 Shigella sonnei Proteobacteria; Gammaproteobacteria; Enterobacterales; Enterobacteriaceae; Shigella MTGDNTLIHSHGINRRDFMKLCAALAATMGLSSKAAAEMAESVTNPQRPPVIWIGAQECT WP_000145410.1 MULTISPECIES: hydrogenase-2 small chain [Proteobacteria] Length: 372\nNumber of predicted TMHs: 1\nExp number of AAs in TMHs: 23.72226\nExp number, first 60 AAs: 0.22038\nTotal prob of N-in: 0.12994\noutside 1 331\nTMhelix 332 354\ninside 355 372

25918 GCF_001249455.1_12844_1_58 Shigella sonnei Proteobacteria; Gammaproteobacteria; Enterobacterales; Enterobacteriaceae; Shigella MTGDNTLIHSHGINRRDFMKLCAALAATMGLSSKAAAEMAESVTNPQRPPVIWIGAQECT WP_000145410.1 MULTISPECIES: hydrogenase-2 small chain [Proteobacteria] Length: 372\nNumber of predicted TMHs: 1\nExp number of AAs in TMHs: 23.72226\nExp number, first 60 AAs: 0.22038\nTotal prob of N-in: 0.12994\noutside 1 331\nTMhelix 332 354\ninside 355 372

25919 GCF_001249495.1_9870_6_30 Shigella sonnei Proteobacteria; Gammaproteobacteria; Enterobacterales; Enterobacteriaceae; Shigella MTGDNTLIHSHGINRRDFMKLCAALAATMGLSSKAAAEMAESVTNPQRPPVIWIGAQECT WP_000145410.1 MULTISPECIES: hydrogenase-2 small chain [Proteobacteria] Length: 372\nNumber of predicted TMHs:

1\nExp number of AAs in TMHs: 23.72226\nExp number, first 60 AAs: 0.22038\nTotal prob of N-in: 0.12994\noutside 1 331\nTMhelix 332 354\ninside 355 372

25920 GCF_001249535.1_8489_2_82 Shigella sonnei Proteobacteria; Gammaproteobacteria; Enterobacterales; Enterobacteriaceae; Shigella
MTGDNTLIHSHGINRRDFMKLCAALAATMGLSSKAAAEMAESVTNPQRPPVIWIGAQECT WP_000145410.1
MULTISPECIES: hydrogenase-2 small chain [Proteobacteria] Length: 372\nNumber of predicted TMHs: 1\nExp number of AAs in TMHs: 23.72226\nExp number, first 60 AAs: 0.22038\nTotal prob of N-in: 0.12994\noutside 1 331\nTMhelix 332 354\ninside 355 372

25921 GCF_001249575.1_8289_7_94 Shigella sonnei Proteobacteria; Gammaproteobacteria; Enterobacterales; Enterobacteriaceae; Shigella
MTGDNTLIHSHGINRRDFMKLCAALAATMGLSSKAAAEMAESVTNPQRPPVIWIGAQECT WP_000145410.1
MULTISPECIES: hydrogenase-2 small chain [Proteobacteria] Length: 372\nNumber of predicted TMHs: 1\nExp number of AAs in TMHs: 23.72226\nExp number, first 60 AAs: 0.22038\nTotal prob of N-in: 0.12994\noutside 1 331\nTMhelix 332 354\ninside 355 372

25922 GCF_001249615.1_Sg_201400696_ACTTGA_L001 Shigella sonnei Proteobacteria; Gammaproteobacteria; Enterobacterales; Enterobacteriaceae; Shigella
MTGDNTLIHSHGINRRDFMKLCAALAATMGLSSKAAAEMAESVTNPQRPPVIWIGAQECT WP_000145410.1
MULTISPECIES: hydrogenase-2 small chain [Proteobacteria] Length: 372\nNumber of predicted TMHs: 1\nExp number of AAs in TMHs: 23.72226\nExp number, first 60 AAs: 0.22038\nTotal prob of N-in: 0.12994\noutside 1 331\nTMhelix 332 354\ninside 355 372

25923 GCF_001249675.1_7655_8_15 Shigella sonnei Proteobacteria; Gammaproteobacteria; Enterobacterales; Enterobacteriaceae; Shigella
MTGDNTLIHSHGINRRDFMKLCAALAATMGLSSKAAAEMAESVTNPQRPPVIWIGAQECT WP_000145410.1
MULTISPECIES: hydrogenase-2 small chain [Proteobacteria] Length: 372\nNumber of predicted TMHs: 1\nExp number of AAs in TMHs: 23.72226\nExp number, first 60 AAs: 0.22038\nTotal prob of N-in: 0.12994\noutside 1 331\nTMhelix 332 354\ninside 355 372

25924 GCF_001249735.1_8489_2_86 Shigella sonnei Proteobacteria; Gammaproteobacteria; Enterobacterales; Enterobacteriaceae; Shigella
MTGDNTLIHSHGINRRDFMKLCAALAATMGLSSKAAAEMAESVTNPQRPPVIWIGAQECT WP_000145410.1
MULTISPECIES: hydrogenase-2 small chain [Proteobacteria] Length: 372\nNumber of predicted TMHs: 1\nExp number of AAs in TMHs: 23.72226\nExp number, first 60 AAs: 0.22038\nTotal prob of N-in: 0.12994\noutside 1 331\nTMhelix 332 354\ninside 355 372

25925 GCF_001249675.1_7655_8_15 Shigella sonnei Proteobacteria; Gammaproteobacteria; Enterobacterales; Enterobacteriaceae; Shigella
MNNEETFYQAMRRQGVTRRSFLKYCSLAATSLGLGAGMAPKIAWALENKPRIPVVWIHGL WP_005139779.1
hydrogenase [Shigella sonnei] Length: 372\nNumber of predicted TMHs: 2\nExp number of AAs in TMHs: 37.02628\nExp number, first 60 AAs: 19.39684\nTotal prob of N-in: 0.94313\nPOSSIBLE N-term signal sequence\ninside 1 19\nTMhelix 20 39\noutside 40 325\nTMhelix 326 348\ninside 349 372

25926 GCF_001249735.1_8489_2_86 Shigella sonnei Proteobacteria; Gammaproteobacteria; Enterobacterales; Enterobacteriaceae; Shigella
MNNEETFYQAMRRQGVTRRSFLKYCSLAATSLGLGAGMAPKIAWALENKPRIPVVWIHGL WP_005139779.1
hydrogenase [Shigella sonnei] Length: 372\nNumber of predicted TMHs: 2\nExp number of AAs in TMHs: 37.02628\nExp number, first 60 AAs: 19.39684\nTotal prob of N-in: 0.94313\nPOSSIBLE N-term signal sequence\ninside 1 19\nTMhelix 20 39\noutside 40 325\nTMhelix 326 348\ninside 349 372

25927 GCF_001249875.1_5236_5_2Shigella sonnei Proteobacteria; Gammaproteobacteria; Enterobacterales; Enterobacteriaceae; Shigella MNNEETFYQAMRRQGVTRRSFLKYCSLAATSLGLGAGMAPKIAWALENKPRIPVVWIHGL
WP_005139779.1 hydrogenase [Shigella sonnei] Length: 372\nNumber of predicted TMHs: 2\nExp number of AAs in TMHs: 37.02628\nExp number, first 60 AAs: 19.39684\nTotal prob of N-in: 0.94313\nPOSSIBLE N-term signal sequence\ninside 1 19\nTMhelix 20 39\noutside 40 325\nTMhelix 326 348\ninside 349 372

25928 GCF_001249575.1_8289_7_94 Shigella sonnei Proteobacteria; Gammaproteobacteria; Enterobacterales; Enterobacteriaceae; Shigella
MNNEETFYQAMRRQGVTRRSFLKYCSLAATSLGLGAGMAPKIAWALENKPRIPVVWIHGL WP_005139779.1

hydrogenase [Shigella sonnei] Length: 372\nNumber of predicted TMHs: 2\nExp number of AAs in TMHs: 37.02628\nExp number, first 60 AAs: 19.39684\nTotal prob of N-in: 0.94313\nPOSSIBLE N-term signal sequence\ninside 1 19\nTMhelix 20 39\noutside 40 325\nTMhelix 326 348\ninside 349 372

25929 GCF_001249895.1_12971_8_33 Shigella sonnei Proteobacteria; Gammaproteobacteria; Enterobacterales; Enterobacteriaceae; Shigella MTGDNTLIHSHGINRRDFMKLCAALAATMGLSSKAAAEMAESVTNPQRPPVIWIGAQECT WP_000145410.1 MULTISPECIES: hydrogenase-2 small chain [Proteobacteria] Length: 372\nNumber of predicted TMHs: 1\nExp number of AAs in TMHs: 23.72226\nExp number, first 60 AAs: 0.22038\nTotal prob of N-in: 0.12994\noutside 1 331\nTMhelix 332 354\ninside 355 372

25930 GCF_001249895.1_12971_8_33 Shigella sonnei Proteobacteria; Gammaproteobacteria; Enterobacterales; Enterobacteriaceae; Shigella MNNEETFYQAMRRQGVTRRSFLKYCSLAATSLGLGAGMAPKIAWALENKPRIPVVWIHGL WP_005139779.1 hydrogenase [Shigella sonnei] Length: 372\nNumber of predicted TMHs: 2\nExp number of AAs in TMHs: 37.02628\nExp number, first 60 AAs: 19.39684\nTotal prob of N-in: 0.94313\nPOSSIBLE N-term signal sequence\ninside 1 19\nTMhelix 20 39\noutside 40 325\nTMhelix 326 348\ninside 349 372

25931 GCF_001249615.1_Sg_201400696_ACTTGA_L001 Shigella sonnei Proteobacteria; Gammaproteobacteria; Enterobacterales; Enterobacteriaceae; Shigella MNNEETFYQAMRRQGVTRRSFLKYCSLAATSLGLGAGMAPKIAWALENKPRIPVVWIHGL WP_005139779.1 hydrogenase [Shigella sonnei] Length: 372\nNumber of predicted TMHs: 2\nExp number of AAs in TMHs: 37.02628\nExp number, first 60 AAs: 19.39684\nTotal prob of N-in: 0.94313\nPOSSIBLE N-term signal sequence\ninside 1 19\nTMhelix 20 39\noutside 40 325\nTMhelix 326 348\ninside 349 372

25932 GCF_001249635.1_5236_2_9Shigella sonnei Proteobacteria; Gammaproteobacteria; Enterobacterales; Enterobacteriaceae; Shigella MNNEETFYQAMRRQGVTRRSFLKYCSLAATSLGLGAGMAPKIAWALENKPRIPVVWIHGL WP_005139779.1 hydrogenase [Shigella sonnei] Length: 372\nNumber of predicted TMHs: 2\nExp number of AAs in TMHs: 37.02628\nExp number, first 60 AAs: 19.39684\nTotal prob of N-in: 0.94313\nPOSSIBLE N-term signal sequence\ninside 1 19\nTMhelix 20 39\noutside 40 325\nTMhelix 326 348\ninside 349 372

25933 GCF_001249935.1_H14140035801 Shigella sonnei Proteobacteria; Gammaproteobacteria; Enterobacterales; Enterobacteriaceae; Shigella MTGDNTLIHSHGINRRDFMKLCAALAATMGLSSKAAAEMAESVTNPQRPPVIWIGAQECT WP_000145410.1 MULTISPECIES: hydrogenase-2 small chain [Proteobacteria] Length: 372\nNumber of predicted TMHs: 1\nExp number of AAs in TMHs: 23.72226\nExp number, first 60 AAs: 0.22038\nTotal prob of N-in: 0.12994\noutside 1 331\nTMhelix 332 354\ninside 355 372

25934 GCF_001249755.1_H14088031401 Shigella sonnei Proteobacteria; Gammaproteobacteria; Enterobacterales; Enterobacteriaceae; Shigella MNNEETFYQAMRRQGVTRRSFLKYCSLAATSLGLGAGMAPKIAWALENKPRIPVVWIHGL WP_005139779.1 hydrogenase [Shigella sonnei] Length: 372\nNumber of predicted TMHs: 2\nExp number of AAs in TMHs: 37.02628\nExp number, first 60 AAs: 19.39684\nTotal prob of N-in: 0.94313\nPOSSIBLE N-term signal sequence\ninside 1 19\nTMhelix 20 39\noutside 40 325\nTMhelix 326 348\ninside 349 372

25935 GCF_001249975.1_5236_2_7Shigella sonnei Proteobacteria; Gammaproteobacteria; Enterobacterales; Enterobacteriaceae; Shigella MTGDNTLIHSHGINRRDFMKLCAALAATMGLSSKAAAEMAESVTNPQRPPVIWIGAQECT WP_000145410.1 MULTISPECIES: hydrogenase-2 small chain [Proteobacteria] Length: 372\nNumber of predicted TMHs: 1\nExp number of AAs in TMHs: 23.72226\nExp number, first 60 AAs: 0.22038\nTotal prob of N-in: 0.12994\noutside 1 331\nTMhelix 332 354\ninside 355 372

25936 GCF_001250055.1_7655_8_12 Shigella sonnei Proteobacteria; Gammaproteobacteria; Enterobacterales; Enterobacteriaceae; Shigella MTGDNTLIHSHGINRRDFMKLCAALAATMGLSSKAAAEMAESVTNPQRPPVIWIGAQECT WP_000145410.1 MULTISPECIES: hydrogenase-2 small chain [Proteobacteria] Length: 372\nNumber of predicted TMHs: 1\nExp number of AAs in TMHs: 23.72226\nExp number, first 60 AAs: 0.22038\nTotal prob of N-in: 0.12994\noutside 1 331\nTMhelix 332 354\ninside 355 372

25937 GCF_001250075.1_8489_2_92 Shigella sonnei Proteobacteria; Gammaproteobacteria; Enterobacterales; Enterobacteriaceae; Shigella MTGDNTLIHSHGINRRDFMKLCAALAATMGLSSKAAAEMAESVTNPQRPPVIWIGAQECT WP_000145410.1

MULTISPECIES: hydrogenase-2 small chain [Proteobacteria] Length: 372\nNumber of predicted TMHs: 1\nExp number of AAs in TMHs: 23.72226\nExp number, first 60 AAs: 0.22038\nTotal prob of N-in: 0.12994\noutside 1 331\nTMhelix 332 354\ninside 355 372

25938 GCF_001250115.1_12971_8_18 Shigella sonnei Proteobacteria; Gammaproteobacteria; Enterobacterales; Enterobacteriaceae; Shigella MTGDNTLIHSHGINRRDFMKLCAALAATMGLSSKAAAEMAESVTNPQRPPVIWIGAQECT WP_000145410.1 MULTISPECIES: hydrogenase-2 small chain [Proteobacteria] Length: 372\nNumber of predicted TMHs: 1\nExp number of AAs in TMHs: 23.72226\nExp number, first 60 AAs: 0.22038\nTotal prob of N-in: 0.12994\noutside 1 331\nTMhelix 332 354\ninside 355 372

25939 GCF_001250155.1_5236_2_2Shigella sonnei Proteobacteria; Gammaproteobacteria; Enterobacterales; Enterobacteriaceae; Shigella MTGDNTLIHSHGINRRDFMKLCAALAATMGLSSKAAAEMAESVTNPQRPPVIWIGAQECT WP_000145410.1 MULTISPECIES: hydrogenase-2 small chain [Proteobacteria] Length: 372\nNumber of predicted TMHs: 1\nExp number of AAs in TMHs: 23.72226\nExp number, first 60 AAs: 0.22038\nTotal prob of N-in: 0.12994\noutside 1 331\nTMhelix 332 354\ninside 355 372

25940 GCF_001250175.1_8289_7_61 Shigella sonnei Proteobacteria; Gammaproteobacteria; Enterobacterales; Enterobacteriaceae; Shigella MTGDNTLIHSHGINRRDFMKLCAALAATMGLSSKAAAEMAESVTNPQRPPVIWIGAQECT WP_000145410.1 MULTISPECIES: hydrogenase-2 small chain [Proteobacteria] Length: 372\nNumber of predicted TMHs: 1\nExp number of AAs in TMHs: 23.72226\nExp number, first 60 AAs: 0.22038\nTotal prob of N-in: 0.12994\noutside 1 331\nTMhelix 332 354\ninside 355 372

25941 GCF_001250275.1_8290_4_55 Shigella sonnei Proteobacteria; Gammaproteobacteria; Enterobacterales; Enterobacteriaceae; Shigella MTGDNTLIHSHGINRRDFMKLCAALAATMGLSSKAAAEMAESVTNPQRPPVIWIGAQECT WP_000145410.1 MULTISPECIES: hydrogenase-2 small chain [Proteobacteria] Length: 372\nNumber of predicted TMHs: 1\nExp number of AAs in TMHs: 23.72226\nExp number, first 60 AAs: 0.22038\nTotal prob of N-in: 0.12994\noutside 1 331\nTMhelix 332 354\ninside 355 372

25942 GCF_001250075.1_8489_2_92 Shigella sonnei Proteobacteria; Gammaproteobacteria; Enterobacterales; Enterobacteriaceae; Shigella MNNEETFYQAMRRQGVTRRSFLKYCSLAATSLGLGAGMAPKIAWALENKPRIPVVWIHGL WP_005139779.1 hydrogenase [Shigella sonnei] Length: 372\nNumber of predicted TMHs: 2\nExp number of AAs in TMHs: 37.02628\nExp number, first 60 AAs: 19.39684\nTotal prob of N-in: 0.94313\nPOSSIBLE N-term signal sequence\ninside 1 19\nTMhelix 20 39\noutside 40 325\nTMhelix 326 348\ninside 349 372

25943 GCF_001250335.1_8290_4_14 Shigella sonnei Proteobacteria; Gammaproteobacteria; Enterobacterales; Enterobacteriaceae; Shigella MTGDNTLIHSHGINRRDFMKLCAALAATMGLSSKAAAEMAESVTNPQRPPVIWIGAQECT WP_000145410.1 MULTISPECIES: hydrogenase-2 small chain [Proteobacteria] Length: 372\nNumber of predicted TMHs: 1\nExp number of AAs in TMHs: 23.72226\nExp number, first 60 AAs: 0.22038\nTotal prob of N-in: 0.12994\noutside 1 331\nTMhelix 332 354\ninside 355 372

25944 GCF_001250375.1_5417_2_3Shigella sonnei Proteobacteria; Gammaproteobacteria; Enterobacterales; Enterobacteriaceae; Shigella MTGDNTLIHSHGINRRDFMKLCAALAATMGLSSKAAAEMAESVTNPQRPPVIWIGAQECT WP_000145410.1 MULTISPECIES: hydrogenase-2 small chain [Proteobacteria] Length: 372\nNumber of predicted TMHs: 1\nExp number of AAs in TMHs: 23.72226\nExp number, first 60 AAs: 0.22038\nTotal prob of N-in: 0.12994\noutside 1 331\nTMhelix 332 354\ninside 355 372

25945 GCF_001250375.1_5417_2_3Shigella sonnei Proteobacteria; Gammaproteobacteria; Enterobacterales; Enterobacteriaceae; Shigella MNNEETFYQAMRRQGVTRRSFLKYCSLAATSLGLGAGMAPKIAWALENKPRIPVVWIHGL WP_005139779.1 hydrogenase [Shigella sonnei] Length: 372\nNumber of predicted TMHs: 2\nExp number of AAs in TMHs: 37.02628\nExp number, first 60 AAs: 19.39684\nTotal prob of N-in: 0.94313\nPOSSIBLE N-term signal sequence\ninside 1 19\nTMhelix 20 39\noutside 40 325\nTMhelix 326 348\ninside 349 372

25946 GCF_001250335.1_8290_4_14 Shigella sonnei Proteobacteria; Gammaproteobacteria; Enterobacterales; Enterobacteriaceae; Shigella MNNEETFYQAMRRQGVTRRSFLKYCSLAATSLGLGAGMAPKIAWALENKPRIPVVWIHGL WP_005139779.1 hydrogenase [Shigella sonnei] Length: 372\nNumber of predicted TMHs: 2\nExp number of AAs in

TMHs: 37.02628\nExp number, first 60 AAs: 19.39684\nTotal prob of N-in: 0.94313\nPOSSIBLE N-term signal sequence\ninside 1 19\nTMhelix 20 39\noutside 40 325\nTMhelix 326 348\ninside 349 372

25947 GCF_001250115.1_12971_8_18 Shigella sonnei Proteobacteria; Gammaproteobacteria; Enterobacterales; Enterobacteriaceae; Shigella MNNEETFYQAMRRQGVTRRSFLKYCSLAATSLGLGAGMAPKIAWALENKPRIPVVWIHGL WP_005139779.1 hydrogenase [Shigella sonnei] Length: 372\nNumber of predicted TMHs: 2\nExp number of AAs in TMHs: 37.02628\nExp number, first 60 AAs: 19.39684\nTotal prob of N-in: 0.94313\nPOSSIBLE N-term signal sequence\ninside 1 19\nTMhelix 20 39\noutside 40 325\nTMhelix 326 348\ninside 349 372

25948 GCF_001250415.1_6350_1_3Shigella sonnei Proteobacteria; Gammaproteobacteria; Enterobacterales; Enterobacteriaceae; Shigella MTGDNTLIHSHGINRRDFMKLCAALAATMGLSSKAAAEMAESVTNPQRPPVIWIGAQECT WP_000145410.1 MULTISPECIES: hydrogenase-2 small chain [Proteobacteria] Length: 372\nNumber of predicted TMHs: 1\nExp number of AAs in TMHs: 23.72226\nExp number, first 60 AAs: 0.22038\nTotal prob of N-in: 0.12994\noutside 1 331\nTMhelix 332 354\ninside 355 372

25949 GCF_001250415.1_6350_1_3Shigella sonnei Proteobacteria; Gammaproteobacteria; Enterobacterales; Enterobacteriaceae; Shigella MNNEETFYQAMRRQGVTRRSFLKYCSLAATSLGLGAGMAPKIAWALENKPRIPVVWIHGL WP_005139779.1 hydrogenase [Shigella sonnei] Length: 372\nNumber of predicted TMHs: 2\nExp number of AAs in TMHs: 37.02628\nExp number, first 60 AAs: 19.39684\nTotal prob of N-in: 0.94313\nPOSSIBLE N-term signal sequence\ninside 1 19\nTMhelix 20 39\noutside 40 325\nTMhelix 326 348\ninside 349 372

25950 GCF_001250155.1_5236_2_2Shigella sonnei Proteobacteria; Gammaproteobacteria; Enterobacterales; Enterobacteriaceae; Shigella MNNEETFYQAMRRQGVTRRSFLKYCSLAATSLGLGAGMAPKIAWALENKPRIPVVWIHGL WP_005139779.1 hydrogenase [Shigella sonnei] Length: 372\nNumber of predicted TMHs: 2\nExp number of AAs in TMHs: 37.02628\nExp number, first 60 AAs: 19.39684\nTotal prob of N-in: 0.94313\nPOSSIBLE N-term signal sequence\ninside 1 19\nTMhelix 20 39\noutside 40 325\nTMhelix 326 348\ninside 349 372

25951 GCF_001250475.1_H14058015001 Shigella sonnei Proteobacteria; Gammaproteobacteria; Enterobacterales; Enterobacteriaceae; Shigella MTGDNTLIHSHGINRRDFMKLCAALAATMGLSSKAAAEMAESVTNPQRPPVIWIGAQECT WP_000145410.1 MULTISPECIES: hydrogenase-2 small chain [Proteobacteria] Length: 372\nNumber of predicted TMHs: 1\nExp number of AAs in TMHs: 23.72226\nExp number, first 60 AAs: 0.22038\nTotal prob of N-in: 0.12994\noutside 1 331\nTMhelix 332 354\ninside 355 372

25952 GCF_001250515.1_6350_1_2Shigella sonnei Proteobacteria; Gammaproteobacteria; Enterobacterales; Enterobacteriaceae; Shigella MTGDNTLIHSHGINRRDFMKLCAALAATMGLSSKAAAEMAESVTNPQRPPVIWIGAQECT WP_000145410.1 MULTISPECIES: hydrogenase-2 small chain [Proteobacteria] Length: 372\nNumber of predicted TMHs: 1\nExp number of AAs in TMHs: 23.72226\nExp number, first 60 AAs: 0.22038\nTotal prob of N-in: 0.12994\noutside 1 331\nTMhelix 332 354\ninside 355 372

25953 GCF_001250555.1_6350_1_10 Shigella sonnei Proteobacteria; Gammaproteobacteria; Enterobacterales; Enterobacteriaceae; Shigella MTGDNTLIHSHGINRRDFMKLCAALAATMGLSSKAAAEMAESVTNPQRPPVIWIGAQECT WP_000145410.1 MULTISPECIES: hydrogenase-2 small chain [Proteobacteria] Length: 372\nNumber of predicted TMHs: 1\nExp number of AAs in TMHs: 23.72226\nExp number, first 60 AAs: 0.22038\nTotal prob of N-in: 0.12994\noutside 1 331\nTMhelix 332 354\ninside 355 372

25954 GCF_001250575.1_12971_8_23 Shigella sonnei Proteobacteria; Gammaproteobacteria; Enterobacterales; Enterobacteriaceae; Shigella MTGDNTLIHSHGINRRDFMKLCAALAATMGLSSKAAAEMAESVTNPQRPPVIWIGAQECT WP_000145410.1 MULTISPECIES: hydrogenase-2 small chain [Proteobacteria] Length: 372\nNumber of predicted TMHs: 1\nExp number of AAs in TMHs: 23.72226\nExp number, first 60 AAs: 0.22038\nTotal prob of N-in: 0.12994\noutside 1 331\nTMhelix 332 354\ninside 355 372

25955 GCF_001250635.1_8290_5_12 Shigella sonnei Proteobacteria; Gammaproteobacteria; Enterobacterales; Enterobacteriaceae; Shigella MTGDNTLIHSHGINRRDFMKLCAALAATMGLSSKAAAEMAESVTNPQRPPVIWIGAQECT WP_000145410.1 MULTISPECIES: hydrogenase-2 small chain [Proteobacteria] Length: 372\nNumber of predicted TMHs: 1\nExp number of AAs in TMHs: 23.72226\nExp number, first 60 AAs: 0.22038\nTotal prob of N-in: 0.12994\noutside 1 331\nTMhelix 332 354\ninside 355 372

25956 GCF_001250655.1_5008_8_8Shigella sonnei Proteobacteria; Gammaproteobacteria; Enterobacterales; Enterobacteriaceae; Shigella MTGDNTLIHSHGINRRDFMKLCAALAATMGLSSKAAAEMAESVTNPQRPPVIWIGAQECT WP_000145410.1 MULTISPECIES: hydrogenase-2 small chain [Proteobacteria] Length: 372\nNumber of predicted TMHs: 1\nExp number of AAs in TMHs: 23.72226\nExp number, first 60 AAs: 0.22038\nTotal prob of N-in: 0.12994\noutside 1 331\nTMhelix 332 354\ninside 355 372

25957 GCF_001250695.1_H14154048001 Shigella sonnei Proteobacteria; Gammaproteobacteria; Enterobacterales; Enterobacteriaceae; Shigella MTGDNTLIHSHGINRRDFMKLCAALAATMGLSSKAAAEMAESVTNPQRPPVIWIGAQECT WP_000145410.1 MULTISPECIES: hydrogenase-2 small chain [Proteobacteria] Length: 372\nNumber of predicted TMHs: 1\nExp number of AAs in TMHs: 23.72226\nExp number, first 60 AAs: 0.22038\nTotal prob of N-in: 0.12994\noutside 1 331\nTMhelix 332 354\ninside 355 372

25958 GCF_001250475.1_H14058015001 Shigella sonnei Proteobacteria; Gammaproteobacteria; Enterobacterales; Enterobacteriaceae; Shigella MNNEETFYQAMRRQGVTRRSFLKYCSLAATSLGLGAGMAPKIAWALENKPRIPVVWIHGL WP_005139779.1 hydrogenase [Shigella sonnei] Length: 372\nNumber of predicted TMHs: 2\nExp number of AAs in TMHs: 37.02628\nExp number, first 60 AAs: 19.39684\nTotal prob of N-in: 0.94313\nPOSSIBLE N-term signal sequence\ninside 1 19\nTMhelix 20 39\noutside 40 325\nTMhelix 326 348\ninside 349 372

25959 GCF_001250735.1_5417_1_5Shigella sonnei Proteobacteria; Gammaproteobacteria; Enterobacterales; Enterobacteriaceae; Shigella MTGDNTLIHSHGINRRDFMKLCAALAATMGLSSKAAAEMAESVTNPQRPPVIWIGAQECT WP_000145410.1 MULTISPECIES: hydrogenase-2 small chain [Proteobacteria] Length: 372\nNumber of predicted TMHs: 1\nExp number of AAs in TMHs: 23.72226\nExp number, first 60 AAs: 0.22038\nTotal prob of N-in: 0.12994\noutside 1 331\nTMhelix 332 354\ninside 355 372

25960 GCF_001250755.1_7655_8_9Shigella sonnei Proteobacteria; Gammaproteobacteria; Enterobacterales; Enterobacteriaceae; Shigella MTGDNTLIHSHGINRRDFMKLCAALAATMGLSSKAAAEMAESVTNPQRPPVIWIGAQECT WP_000145410.1 MULTISPECIES: hydrogenase-2 small chain [Proteobacteria] Length: 372\nNumber of predicted TMHs: 1\nExp number of AAs in TMHs: 23.72226\nExp number, first 60 AAs: 0.22038\nTotal prob of N-in: 0.12994\noutside 1 331\nTMhelix 332 354\ninside 355 372

25961 GCF_001250735.1_5417_1_5Shigella sonnei Proteobacteria; Gammaproteobacteria; Enterobacterales; Enterobacteriaceae; Shigella MNNEETFYQAMRRQGVTRRSFLKYCSLAATSLGLGAGMAPKIAWALENKPRIPVVWIHGL WP_005139779.1 hydrogenase [Shigella sonnei] Length: 372\nNumber of predicted TMHs: 2\nExp number of AAs in TMHs: 37.02628\nExp number, first 60 AAs: 19.39684\nTotal prob of N-in: 0.94313\nPOSSIBLE N-term signal sequence\ninside 1 19\nTMhelix 20 39\noutside 40 325\nTMhelix 326 348\ninside 349 372

25962 GCF_001250755.1_7655_8_9Shigella sonnei Proteobacteria; Gammaproteobacteria; Enterobacterales; Enterobacteriaceae; Shigella MNNEETFYQAMRRQGVTRRSFLKYCSLAATSLGLGAGMAPKIAWALENKPRIPVVWIHGL WP_005139779.1 hydrogenase [Shigella sonnei] Length: 372\nNumber of predicted TMHs: 2\nExp number of AAs in TMHs: 37.02628\nExp number, first 60 AAs: 19.39684\nTotal prob of N-in: 0.94313\nPOSSIBLE N-term signal sequence\ninside 1 19\nTMhelix 20 39\noutside 40 325\nTMhelix 326 348\ninside 349 372

25963 GCF_001250515.1_6350_1_2Shigella sonnei Proteobacteria; Gammaproteobacteria; Enterobacterales; Enterobacteriaceae; Shigella MNNEETFYQAMRRQGVTRRSFLKYCSLAATSLGLGAGMAPKIAWALENKPRIPVVWIHGL WP_005139779.1 hydrogenase [Shigella sonnei] Length: 372\nNumber of predicted TMHs: 2\nExp number of AAs in TMHs: 37.02628\nExp number, first 60 AAs: 19.39684\nTotal prob of N-in: 0.94313\nPOSSIBLE N-term signal sequence\ninside 1 19\nTMhelix 20 39\noutside 40 325\nTMhelix 326 348\ninside 349 372

25964 GCF_001250875.1_Sg_201401463_GTCCGC_L001 Shigella sonnei Proteobacteria; Gammaproteobacteria; Enterobacterales; Enterobacteriaceae; Shigella MNNEETFYQAMRRQGVTRRSFLKYCSLAATSLGLGAGMAPKIAWALENKPRIPVVWIHGL WP_005139779.1 hydrogenase [Shigella sonnei] Length: 372\nNumber of predicted TMHs: 2\nExp number of AAs in TMHs: 37.02628\nExp number, first 60 AAs: 19.39684\nTotal prob of N-in: 0.94313\nPOSSIBLE N-term signal sequence\ninside 1 19\nTMhelix 20 39\noutside 40 325\nTMhelix 326 348\ninside 349 372

25965 GCF_001250655.1_5008_8_8Shigella sonnei Proteobacteria; Gammaproteobacteria; Enterobacterales; Enterobacteriaceae; Shigella MNNEETFYQAMRRQGVTRRSFLKYCSLAATSLGLGAGMAPKIAWALENKPRIPVVWIHGL WP_005139779.1 hydrogenase [Shigella sonnei] Length: 372\nNumber of predicted TMHs: 2\nExp

number of AAs in TMHs: 37.02628\nExp number, first 60 AAs: 19.39684\nTotal prob of N-in: 0.94313\nPOSSIBLE N-term signal sequence\ninside 1 19\nTMhelix 20 39\noutside 40 325\nTMhelix 326 348\ninside 349 372

25966 GCF_001250635.1_8290_5_12 Shigella sonnei Proteobacteria; Gammaproteobacteria; Enterobacterales; Enterobacteriaceae; Shigella
MNNEETFYQAMRRQGVTRRSFLKYCSLAATSLGLGAGMAPKIAWALENKPRIPVVWIHGL WP_005139779.1
hydrogenase [Shigella sonnei] Length: 372\nNumber of predicted TMHs: 2\nExp number of AAs in TMHs: 37.02628\nExp number, first 60 AAs: 19.39684\nTotal prob of N-in: 0.94313\nPOSSIBLE N-term signal sequence\ninside 1 19\nTMhelix 20 39\noutside 40 325\nTMhelix 326 348\ninside 349 372

25967 GCF_001250955.1_12971_8_39 Shigella sonnei Proteobacteria; Gammaproteobacteria; Enterobacterales; Enterobacteriaceae; Shigella
MTGDNTLIHSHGINRRDFMKLCAALAATMGLSSKAAAEMAESVTNPQRPPVIWIGAQECT WP_000145410.1
MULTISPECIES: hydrogenase-2 small chain [Proteobacteria] Length: 372\nNumber of predicted TMHs: 1\nExp number of AAs in TMHs: 23.72226\nExp number, first 60 AAs: 0.22038\nTotal prob of N-in: 0.12994\noutside 1 331\nTMhelix 332 354\ninside 355 372

25968 GCF_001250995.1_8290_4_31 Shigella sonnei Proteobacteria; Gammaproteobacteria; Enterobacterales; Enterobacteriaceae; Shigella
MTGDNTLIHSHGINRRDFMKLCAALAATMGLSSKAAAEMAESVTNPQRPPVIWIGAQECT WP_000145410.1
MULTISPECIES: hydrogenase-2 small chain [Proteobacteria] Length: 372\nNumber of predicted TMHs: 1\nExp number of AAs in TMHs: 23.72226\nExp number, first 60 AAs: 0.22038\nTotal prob of N-in: 0.12994\noutside 1 331\nTMhelix 332 354\ninside 355 372

25969 GCF_001251015.1_12971_8_46 Shigella sonnei Proteobacteria; Gammaproteobacteria; Enterobacterales; Enterobacteriaceae; Shigella
MTGDNTLIHSHGINRRDFMKLCAALAATMGLSSKAAAEMAESVTNPQRPPVIWIGAQECT WP_000145410.1
MULTISPECIES: hydrogenase-2 small chain [Proteobacteria] Length: 372\nNumber of predicted TMHs: 1\nExp number of AAs in TMHs: 23.72226\nExp number, first 60 AAs: 0.22038\nTotal prob of N-in: 0.12994\noutside 1 331\nTMhelix 332 354\ninside 355 372

25970 GCF_001251055.1_8489_2_84 Shigella sonnei Proteobacteria; Gammaproteobacteria; Enterobacterales; Enterobacteriaceae; Shigella
MTGDNTLIHSHGINRRDFMKLCAALAATMGLSSKAAAEMAESVTNPQRPPVIWIGAQECT WP_000145410.1
MULTISPECIES: hydrogenase-2 small chain [Proteobacteria] Length: 372\nNumber of predicted TMHs: 1\nExp number of AAs in TMHs: 23.72226\nExp number, first 60 AAs: 0.22038\nTotal prob of N-in: 0.12994\noutside 1 331\nTMhelix 332 354\ninside 355 372

25971 GCF_001251135.1_H14136026101 Shigella sonnei Proteobacteria; Gammaproteobacteria; Enterobacterales; Enterobacteriaceae; Shigella
MTGDNTLIHSHGINRRDFMKLCAALAATMGLSSKAAAEMAESVTNPQRPPVIWIGAQECT WP_000145410.1
MULTISPECIES: hydrogenase-2 small chain [Proteobacteria] Length: 372\nNumber of predicted TMHs: 1\nExp number of AAs in TMHs: 23.72226\nExp number, first 60 AAs: 0.22038\nTotal prob of N-in: 0.12994\noutside 1 331\nTMhelix 332 354\ninside 355 372

25972 GCF_001251075.1_8289_7_58 Shigella sonnei Proteobacteria; Gammaproteobacteria; Enterobacterales; Enterobacteriaceae; Shigella
MTGDNTLIHSHGINRRDFMKLCAALAATMGLSSKAAAEMAESVTNPQRPPVIWIGAQECT WP_000145410.1
MULTISPECIES: hydrogenase-2 small chain [Proteobacteria] Length: 372\nNumber of predicted TMHs: 1\nExp number of AAs in TMHs: 23.72226\nExp number, first 60 AAs: 0.22038\nTotal prob of N-in: 0.12994\noutside 1 331\nTMhelix 332 354\ninside 355 372

25973 GCF_001251195.1_12971_8_50 Shigella sonnei Proteobacteria; Gammaproteobacteria; Enterobacterales; Enterobacteriaceae; Shigella
MTGDNTLIHSHGINRRDFMKLCAALAATMGLSSKAAAEMAESVTNPQRPPVIWIGAQECT WP_000145410.1
MULTISPECIES: hydrogenase-2 small chain [Proteobacteria] Length: 372\nNumber of predicted TMHs: 1\nExp number of AAs in TMHs: 23.72226\nExp number, first 60 AAs: 0.22038\nTotal prob of N-in: 0.12994\noutside 1 331\nTMhelix 332 354\ninside 355 372

25974 GCF_001250955.1_12971_8_39 Shigella sonnei Proteobacteria; Gammaproteobacteria; Enterobacterales; Enterobacteriaceae; Shigella

MNNEETFYQAMRRQGVTRRSFLKYCSLAATSLGLGAGMAPKIAWALENKPRIPVVWIHGL WP_005139779.1 hydrogenase [Shigella sonnei] Length: 372\nNumber of predicted TMHs: 2\nExp number of AAs in TMHs: 37.02628\nExp number, first 60 AAs: 19.39684\nTotal prob of N-in: 0.94313\nPOSSIBLE N-term signal sequence\ninside 1 19\nTMhelix 20 39\noutside 40 325\nTMhelix 326 348\ninside 349 372

25975 GCF_001251235.1_5236_2_12 Shigella sonnei Proteobacteria; Gammaproteobacteria; Enterobacterales; Enterobacteriaceae; Shigella MTGDNTLIHSHGINRRDFMKLCAALAATMGLSSKAAAEMAESVTNPQRPPVIWIGAQECT WP_000145410.1 MULTISPECIES: hydrogenase-2 small chain [Proteobacteria] Length: 372\nNumber of predicted TMHs: 1\nExp number of AAs in TMHs: 23.72226\nExp number, first 60 AAs: 0.22038\nTotal prob of N-in: 0.12994\noutside 1 331\nTMhelix 332 354\ninside 355 372

25976 GCF_001251255.1_5236_5_12 Shigella sonnei Proteobacteria; Gammaproteobacteria; Enterobacterales; Enterobacteriaceae; Shigella MTGDNTLIHSHGINRRDFMKLCAALAATMGLSSKAAAEMAESVTNPQRPPVIWIGAQECT WP_000145410.1 MULTISPECIES: hydrogenase-2 small chain [Proteobacteria] Length: 372\nNumber of predicted TMHs: 1\nExp number of AAs in TMHs: 23.72226\nExp number, first 60 AAs: 0.22038\nTotal prob of N-in: 0.12994\noutside 1 331\nTMhelix 332 354\ninside 355 372

25977 GCF_001251235.1_5236_2_12 Shigella sonnei Proteobacteria; Gammaproteobacteria; Enterobacterales; Enterobacteriaceae; Shigella MNNEETFYQAMRRQGVTRRSFLKYCSLAATSLGLGAGMAPKIAWALENKPRIPVVWIHGL WP_005139779.1 hydrogenase [Shigella sonnei] Length: 372\nNumber of predicted TMHs: 2\nExp number of AAs in TMHs: 37.02628\nExp number, first 60 AAs: 19.39684\nTotal prob of N-in: 0.94313\nPOSSIBLE N-term signal sequence\ninside 1 19\nTMhelix 20 39\noutside 40 325\nTMhelix 326 348\ninside 349 372

25978 GCF_001251355.1_8290_4_44 Shigella sonnei Proteobacteria; Gammaproteobacteria; Enterobacterales; Enterobacteriaceae; Shigella MTGDNTLIHSHGINRRDFMKLCAALAATMGLSSKAAAEMAESVTNPQRPPVIWIGAQECT WP_000145410.1 MULTISPECIES: hydrogenase-2 small chain [Proteobacteria] Length: 372\nNumber of predicted TMHs: 1\nExp number of AAs in TMHs: 23.72226\nExp number, first 60 AAs: 0.22038\nTotal prob of N-in: 0.12994\noutside 1 331\nTMhelix 332 354\ninside 355 372

25979 GCF_001251295.1_6350_1_9Shigella sonnei Proteobacteria; Gammaproteobacteria; Enterobacterales; Enterobacteriaceae; Shigella MNNEETFYQAMRRQGVTRRSFLKYCSLAATSLGLGAGMAPKIAWALENKPRIPVVWIHGL WP_005139779.1 hydrogenase [Shigella sonnei] Length: 372\nNumber of predicted TMHs: 2\nExp number of AAs in TMHs: 37.02628\nExp number, first 60 AAs: 19.39684\nTotal prob of N-in: 0.94313\nPOSSIBLE N-term signal sequence\ninside 1 19\nTMhelix 20 39\noutside 40 325\nTMhelix 326 348\ninside 349 372

25980 GCF_001251335.1_7353_1_22 Shigella sonnei Proteobacteria; Gammaproteobacteria; Enterobacterales; Enterobacteriaceae; Shigella MNNEETFYQAMRRQGVTRRSFLKYCSLAATSLGLGAGMAPKIAWALENKPRIPVVWIHGL WP_005139779.1 hydrogenase [Shigella sonnei] Length: 372\nNumber of predicted TMHs: 2\nExp number of AAs in TMHs: 37.02628\nExp number, first 60 AAs: 19.39684\nTotal prob of N-in: 0.94313\nPOSSIBLE N-term signal sequence\ninside 1 19\nTMhelix 20 39\noutside 40 325\nTMhelix 326 348\ninside 349 372

25981 GCF_001251355.1_8290_4_44 Shigella sonnei Proteobacteria; Gammaproteobacteria; Enterobacterales; Enterobacteriaceae; Shigella MNNEETFYQAMRRQGVTRRSFLKYCSLAATSLGLGAGMAPKIAWALENKPRIPVVWIHGL WP_005139779.1 hydrogenase [Shigella sonnei] Length: 372\nNumber of predicted TMHs: 2\nExp number of AAs in TMHs: 37.02628\nExp number, first 60 AAs: 19.39684\nTotal prob of N-in: 0.94313\nPOSSIBLE N-term signal sequence\ninside 1 19\nTMhelix 20 39\noutside 40 325\nTMhelix 326 348\ninside 349 372

25982 GCF_001251075.1_8289_7_58 Shigella sonnei Proteobacteria; Gammaproteobacteria; Enterobacterales; Enterobacteriaceae; Shigella MNNEETFYQAMRRQGVTRRSFLKYCSLAATSLGLGAGMAPKIAWALENKPRIPVVWIHGL WP_005139779.1 hydrogenase [Shigella sonnei] Length: 372\nNumber of predicted TMHs: 2\nExp number of AAs in TMHs: 37.02628\nExp number, first 60 AAs: 19.39684\nTotal prob of N-in: 0.94313\nPOSSIBLE N-term signal sequence\ninside 1 19\nTMhelix 20 39\noutside 40 325\nTMhelix 326 348\ninside 349 372

25983 GCF_001251395.1_5008_7_4Shigella sonnei Proteobacteria; Gammaproteobacteria; Enterobacterales; Enterobacteriaceae; Shigella MTGDNTLIHSHGINRRDFMKLCAALAATMGLSSKAAAEMAESVTNPQRPPVIWIGAQECT WP_000145410.1 MULTISPECIES: hydrogenase-2 small chain [Proteobacteria] Length: 372\nNumber of predicted TMHs: 1\nExp number of AAs in TMHs: 23.72226\nExp number, first 60 AAs: 0.22038\nTotal prob of N-in: 0.12994\noutside 1 331\nTMhelix 332 354\ninside 355 372

25984 GCF_001251455.1_8289_7_38 Shigella sonnei Proteobacteria; Gammaproteobacteria; Enterobacterales; Enterobacteriaceae; Shigella MTGDNTLIHSHGINRRDFMKLCAALAATMGLSSKAAAEMAESVTNPQRPPVIWIGAQECT WP_000145410.1 MULTISPECIES: hydrogenase-2 small chain [Proteobacteria] Length: 372\nNumber of predicted TMHs: 1\nExp number of AAs in TMHs: 23.72226\nExp number, first 60 AAs: 0.22038\nTotal prob of N-in: 0.12994\noutside 1 331\nTMhelix 332 354\ninside 355 372

25985 GCF_001251535.1_8290_4_38 Shigella sonnei Proteobacteria; Gammaproteobacteria; Enterobacterales; Enterobacteriaceae; Shigella MTGDNTLIHSHGINRRDFMKLCAALAATMGLSSKAAAEMAESVTNPQRPPVIWIGAQECT WP_000145410.1 MULTISPECIES: hydrogenase-2 small chain [Proteobacteria] Length: 372\nNumber of predicted TMHs: 1\nExp number of AAs in TMHs: 23.72226\nExp number, first 60 AAs: 0.22038\nTotal prob of N-in: 0.12994\noutside 1 331\nTMhelix 332 354\ninside 355 372

25986 GCF_001251395.1_5008_7_4Shigella sonnei Proteobacteria; Gammaproteobacteria; Enterobacterales; Enterobacteriaceae; Shigella MNNEETFYQAMRRQGVTRRSFLKYCSLAATSLGLGAGMAPKIAWALENKPRIPVVWIHGL WP_005139779.1 hydrogenase [Shigella sonnei] Length: 372\nNumber of predicted TMHs: 2\nExp number of AAs in TMHs: 37.02628\nExp number, first 60 AAs: 19.39684\nTotal prob of N-in: 0.94313\nPOSSIBLE N-term signal sequence\ninside 1 19\nTMhelix 20 39\noutside 40 325\nTMhelix 326 348\ninside 349 372

25987 GCF_001251455.1_8289_7_38 Shigella sonnei Proteobacteria; Gammaproteobacteria; Enterobacterales; Enterobacteriaceae; Shigella MNNEETFYQAMRRQGVTRRSFLKYCSLAATSLGLGAGMAPKIAWALENKPRIPVVWIHGL WP_005139779.1 hydrogenase [Shigella sonnei] Length: 372\nNumber of predicted TMHs: 2\nExp number of AAs in TMHs: 37.02628\nExp number, first 60 AAs: 19.39684\nTotal prob of N-in: 0.94313\nPOSSIBLE N-term signal sequence\ninside 1 19\nTMhelix 20 39\noutside 40 325\nTMhelix 326 348\ninside 349 372

25988 GCF_001251535.1_8290_4_38 Shigella sonnei Proteobacteria; Gammaproteobacteria; Enterobacterales; Enterobacteriaceae; Shigella MNNEETFYQAMRRQGVTRRSFLKYCSLAATSLGLGAGMAPKIAWALENKPRIPVVWIHGL WP_005139779.1 hydrogenase [Shigella sonnei] Length: 372\nNumber of predicted TMHs: 2\nExp number of AAs in TMHs: 37.02628\nExp number, first 60 AAs: 19.39684\nTotal prob of N-in: 0.94313\nPOSSIBLE N-term signal sequence\ninside 1 19\nTMhelix 20 39\noutside 40 325\nTMhelix 326 348\ninside 349 372

25989 GCF_001251195.1_12971_8_50 Shigella sonnei Proteobacteria; Gammaproteobacteria; Enterobacterales; Enterobacteriaceae; Shigella MNNEETFYQAMRRQGVTRRSFLKYCSLAATSLGLGAGMAPKIAWALENKPRIPVVWIHGL WP_052962775.1 hydrogenase [Shigella sonnei] Length: 372\nNumber of predicted TMHs: 2\nExp number of AAs in TMHs: 37.09248\nExp number, first 60 AAs: 19.39296\nTotal prob of N-in: 0.94316\nPOSSIBLE N-term signal sequence\ninside 1 19\nTMhelix 20 39\noutside 40 325\nTMhelix 326 348\ninside 349 372

25990 GCF_001251575.1_8289_7_84 Shigella sonnei Proteobacteria; Gammaproteobacteria; Enterobacterales; Enterobacteriaceae; Shigella MTGDNTLIHSHGINRRDFMKLCAALAATMGLSSKAAAEMAESVTNPQRPPVIWIGAQECT WP_000145410.1 MULTISPECIES: hydrogenase-2 small chain [Proteobacteria] Length: 372\nNumber of predicted TMHs: 1\nExp number of AAs in TMHs: 23.72226\nExp number, first 60 AAs: 0.22038\nTotal prob of N-in: 0.12994\noutside 1 331\nTMhelix 332 354\ninside 355 372

25991 GCF_001251615.1_H14128025901 Shigella sonnei Proteobacteria; Gammaproteobacteria; Enterobacterales; Enterobacteriaceae; Shigella MTGDNTLIHSHGINRRDFMKLCAALAATMGLSSKAAAEMAESVTNPQRPPVIWIGAQECT WP_000145410.1 MULTISPECIES: hydrogenase-2 small chain [Proteobacteria] Length: 372\nNumber of predicted TMHs: 1\nExp number of AAs in TMHs: 23.72226\nExp number, first 60 AAs: 0.22038\nTotal prob of N-in: 0.12994\noutside 1 331\nTMhelix 332 354\ninside 355 372

25992 GCF_001251655.1_8289_7_50 Shigella sonnei Proteobacteria; Gammaproteobacteria; Enterobacterales; Enterobacteriaceae; Shigella
MTGDNTLIHSHGINRRDFMKLCAALAATMGLSSKAAAEMAESVTNPQRPPVIWIGAQECT WP_000145410.1
MULTISPECIES: hydrogenase-2 small chain [Proteobacteria] Length: 372\nNumber of predicted TMHs: 1\nExp number of AAs in TMHs: 23.72226\nExp number, first 60 AAs: 0.22038\nTotal prob of N-in: 0.12994\noutside 1 331\nTMhelix 332 354\ninside 355 372

25993 GCF_001251695.1_8289_7_52 Shigella sonnei Proteobacteria; Gammaproteobacteria; Enterobacterales; Enterobacteriaceae; Shigella
MTGDNTLIHSHGINRRDFMKLCAALAATMGLSSKAAAEMAESVTNPQRPPVIWIGAQECT WP_000145410.1
MULTISPECIES: hydrogenase-2 small chain [Proteobacteria] Length: 372\nNumber of predicted TMHs: 1\nExp number of AAs in TMHs: 23.72226\nExp number, first 60 AAs: 0.22038\nTotal prob of N-in: 0.12994\noutside 1 331\nTMhelix 332 354\ninside 355 372

25994 GCF_001251715.1_9870_6_41 Shigella sonnei Proteobacteria; Gammaproteobacteria; Enterobacterales; Enterobacteriaceae; Shigella
MTGDNTLIHSHGINRRDFMKLCAALAATMGLSSKAAAEMAESVTNPQRPPVIWIGAQECT WP_000145410.1
MULTISPECIES: hydrogenase-2 small chain [Proteobacteria] Length: 372\nNumber of predicted TMHs: 1\nExp number of AAs in TMHs: 23.72226\nExp number, first 60 AAs: 0.22038\nTotal prob of N-in: 0.12994\noutside 1 331\nTMhelix 332 354\ninside 355 372

25995 GCF_001251755.1_12971_8_9 Shigella sonnei Proteobacteria; Gammaproteobacteria; Enterobacterales; Enterobacteriaceae; Shigella
MTGDNTLIHSHGINRRDFMKLCAALAATMGLSSKAAAEMAESVTNPQRPPVIWIGAQECT WP_000145410.1
MULTISPECIES: hydrogenase-2 small chain [Proteobacteria] Length: 372\nNumber of predicted TMHs: 1\nExp number of AAs in TMHs: 23.72226\nExp number, first 60 AAs: 0.22038\nTotal prob of N-in: 0.12994\noutside 1 331\nTMhelix 332 354\ninside 355 372

25996 GCF_001251815.1_9870_6_15 Shigella sonnei Proteobacteria; Gammaproteobacteria; Enterobacterales; Enterobacteriaceae; Shigella
MTGDNTLIHSHGINRRDFMKLCAALAATMGLSSKAAAEMAESVTNPQRPPVIWIGAQECT WP_000145410.1
MULTISPECIES: hydrogenase-2 small chain [Proteobacteria] Length: 372\nNumber of predicted TMHs: 1\nExp number of AAs in TMHs: 23.72226\nExp number, first 60 AAs: 0.22038\nTotal prob of N-in: 0.12994\noutside 1 331\nTMhelix 332 354\ninside 355 372

25997 GCF_001251835.1_9870_7_11 Shigella sonnei Proteobacteria; Gammaproteobacteria; Enterobacterales; Enterobacteriaceae; Shigella
MTGDNTLIHSHGINRRDFMKLCAALAATMGLSSKAAAEMAESVTNPQRPPVIWIGAQECT WP_000145410.1
MULTISPECIES: hydrogenase-2 small chain [Proteobacteria] Length: 372\nNumber of predicted TMHs: 1\nExp number of AAs in TMHs: 23.72226\nExp number, first 60 AAs: 0.22038\nTotal prob of N-in: 0.12994\noutside 1 331\nTMhelix 332 354\ninside 355 372

25998 GCF_001251835.1_9870_7_11 Shigella sonnei Proteobacteria; Gammaproteobacteria; Enterobacterales; Enterobacteriaceae; Shigella
MNNEETFYQAMRRQGVTRRSFLKYCSLAATSLGLGAGMAPKIAWALENKPRIPVVWIHGL WP_005139779.1
hydrogenase [Shigella sonnei] Length: 372\nNumber of predicted TMHs: 2\nExp number of AAs in TMHs: 37.02628\nExp number, first 60 AAs: 19.39684\nTotal prob of N-in: 0.94313\nPOSSIBLE N-term signal sequence\ninside 1 19\nTMhelix 20 39\noutside 40 325\nTMhelix 326 348\ninside 349 372

25999 GCF_001251695.1_8289_7_52 Shigella sonnei Proteobacteria; Gammaproteobacteria; Enterobacterales; Enterobacteriaceae; Shigella
MNNEETFYQAMRRQGVTRRSFLKYCSLAATSLGLGAGMAPKIAWALENKPRIPVVWIHGL WP_005139779.1
hydrogenase [Shigella sonnei] Length: 372\nNumber of predicted TMHs: 2\nExp number of AAs in TMHs: 37.02628\nExp number, first 60 AAs: 19.39684\nTotal prob of N-in: 0.94313\nPOSSIBLE N-term signal sequence\ninside 1 19\nTMhelix 20 39\noutside 40 325\nTMhelix 326 348\ninside 349 372

26000 GCF_001251715.1_9870_6_41 Shigella sonnei Proteobacteria; Gammaproteobacteria; Enterobacterales; Enterobacteriaceae; Shigella
MNNEETFYQAMRRQGVTRRSFLKYCSLAATSLGLGAGMAPKIAWALENKPRIPVVWIHGL WP_005139779.1
hydrogenase [Shigella sonnei] Length: 372\nNumber of predicted TMHs: 2\nExp number of AAs in

TMHs: 37.02628\nExp number, first 60 AAs: 19.39684\nTotal prob of N-in: 0.94313\nPOSSIBLE N-term signal sequence\ninside 1 19\nTMhelix 20 39\noutside 40 325\nTMhelix 326 348\ninside 349 372

26001 GCF_001251995.1_7655_8_17 Shigella sonnei Proteobacteria; Gammaproteobacteria; Enterobacterales; Enterobacteriaceae; Shigella MTGDNTLIHSHGINRRDFMKLCAALAATMGLSSKAAAEMAESVTNPQRPPVIWIGAQECT WP_000145410.1 MULTISPECIES: hydrogenase-2 small chain [Proteobacteria] Length: 372\nNumber of predicted TMHs: 1\nExp number of AAs in TMHs: 23.72226\nExp number, first 60 AAs: 0.22038\nTotal prob of N-in: 0.12994\noutside 1 331\nTMhelix 332 354\ninside 355 372

26002 GCF_001251995.1_7655_8_17 Shigella sonnei Proteobacteria; Gammaproteobacteria; Enterobacterales; Enterobacteriaceae; Shigella MNNEETFYQAMRRQGVTRRSFLKYCSLAATSLGLGAGMAPKIAWALENKPRIPVVWIHGL WP_005139779.1 hydrogenase [Shigella sonnei] Length: 372\nNumber of predicted TMHs: 2\nExp number of AAs in TMHs: 37.02628\nExp number, first 60 AAs: 19.39684\nTotal prob of N-in: 0.94313\nPOSSIBLE N-term signal sequence\ninside 1 19\nTMhelix 20 39\noutside 40 325\nTMhelix 326 348\ninside 349 372

26003 GCF_001251755.1_12971_8_9 Shigella sonnei Proteobacteria; Gammaproteobacteria; Enterobacterales; Enterobacteriaceae; Shigella MNNEETFYQAMRRQGVTRRSFLKYCSLAATSLGLGAGMAPKIAWALENKPRIPVVWIHGL WP_005139779.1 hydrogenase [Shigella sonnei] Length: 372\nNumber of predicted TMHs: 2\nExp number of AAs in TMHs: 37.02628\nExp number, first 60 AAs: 19.39684\nTotal prob of N-in: 0.94313\nPOSSIBLE N-term signal sequence\ninside 1 19\nTMhelix 20 39\noutside 40 325\nTMhelix 326 348\ninside 349 372

26004 GCF_001251815.1_9870_6_15 Shigella sonnei Proteobacteria; Gammaproteobacteria; Enterobacterales; Enterobacteriaceae; Shigella MNNEETFYQAMRRQGVTRRSFLKYCSLAATSLGLGAGMAPKIAWALENKPRIPVVWIHGL WP_005139779.1 hydrogenase [Shigella sonnei] Length: 372\nNumber of predicted TMHs: 2\nExp number of AAs in TMHs: 37.02628\nExp number, first 60 AAs: 19.39684\nTotal prob of N-in: 0.94313\nPOSSIBLE N-term signal sequence\ninside 1 19\nTMhelix 20 39\noutside 40 325\nTMhelix 326 348\ninside 349 372

26005 GCF_001252035.1_5417_2_2Shigella sonnei Proteobacteria; Gammaproteobacteria; Enterobacterales; Enterobacteriaceae; Shigella MTGDNTLIHSHGINRRDFMKLCAALAATMGLSSKAAAEMAESVTNPQRPPVIWIGAQECT WP_000145410.1 MULTISPECIES: hydrogenase-2 small chain [Proteobacteria] Length: 372\nNumber of predicted TMHs: 1\nExp number of AAs in TMHs: 23.72226\nExp number, first 60 AAs: 0.22038\nTotal prob of N-in: 0.12994\noutside 1 331\nTMhelix 332 354\ninside 355 372

26006 GCF_001252095.1_Sg_201401096_AGTTCC_L001 Shigella sonnei Proteobacteria; Gammaproteobacteria; Enterobacterales; Enterobacteriaceae; Shigella MTGDNTLIHSHGINRRDFMKLCAALAATMGLSSKAAAEMAESVTNPQRPPVIWIGAQECT WP_000145410.1 MULTISPECIES: hydrogenase-2 small chain [Proteobacteria] Length: 372\nNumber of predicted TMHs: 1\nExp number of AAs in TMHs: 23.72226\nExp number, first 60 AAs: 0.22038\nTotal prob of N-in: 0.12994\noutside 1 331\nTMhelix 332 354\ninside 355 372

26007 GCF_001252135.1_5236_6_9Shigella sonnei Proteobacteria; Gammaproteobacteria; Enterobacterales; Enterobacteriaceae; Shigella MTGDNTLIHSHGINRRDFMKLCAALAATMGLSSKAAAEMAESVTNPQRPPVIWIGAQECT WP_000145410.1 MULTISPECIES: hydrogenase-2 small chain [Proteobacteria] Length: 372\nNumber of predicted TMHs: 1\nExp number of AAs in TMHs: 23.72226\nExp number, first 60 AAs: 0.22038\nTotal prob of N-in: 0.12994\noutside 1 331\nTMhelix 332 354\ninside 355 372

26008 GCF_001252175.1_5236_7_8Shigella sonnei Proteobacteria; Gammaproteobacteria; Enterobacterales; Enterobacteriaceae; Shigella MTGDNTLIHSHGINRRDFMKLCAALAATMGLSSKAAAEMAESVTNPQRPPVIWIGAQECT WP_000145410.1 MULTISPECIES: hydrogenase-2 small chain [Proteobacteria] Length: 372\nNumber of predicted TMHs: 1\nExp number of AAs in TMHs: 23.72226\nExp number, first 60 AAs: 0.22038\nTotal prob of N-in: 0.12994\noutside 1 331\nTMhelix 332 354\ninside 355 372

26009 GCF_001252255.1_Sg_201400855_TAGCTT_L001 Shigella sonnei Proteobacteria; Gammaproteobacteria; Enterobacterales; Enterobacteriaceae; Shigella MTGDNTLIHSHGINRRDFMKLCAALAATMGLSSKAAAEMAESVTNPQRPPVIWIGAQECT WP_000145410.1 MULTISPECIES: hydrogenase-2 small chain [Proteobacteria] Length: 372\nNumber of predicted TMHs:

1\nExp number of AAs in TMHs: 23.72226\nExp number, first 60 AAs: 0.22038\nTotal prob of N-in: 0.12994\noutside 1 331\nTMhelix 332 354\ninside 355 372

26010 GCF_001252295.1_5236_8_10 Shigella sonnei Proteobacteria; Gammaproteobacteria; Enterobacterales; Enterobacteriaceae; Shigella MTGDNTLIHSHGINRRDFMKLCAALAATMGLSSKAAAEMAESVTNPQRPPVIWIGAQECT WP_000145410.1 MULTISPECIES: hydrogenase-2 small chain [Proteobacteria] Length: 372\nNumber of predicted TMHs: 1\nExp number of AAs in TMHs: 23.72226\nExp number, first 60 AAs: 0.22038\nTotal prob of N-in: 0.12994\noutside 1 331\nTMhelix 332 354\ninside 355 372

26011 GCF_001252315.1_9870_7_8Shigella sonnei Proteobacteria; Gammaproteobacteria; Enterobacterales; Enterobacteriaceae; Shigella MTGDNTLIHSHGINRRDFMKLCAALAATMGLSSKAAAEMAESVTNPQRPPVIWIGAQECT WP_000145410.1 MULTISPECIES: hydrogenase-2 small chain [Proteobacteria] Length: 372\nNumber of predicted TMHs: 1\nExp number of AAs in TMHs: 23.72226\nExp number, first 60 AAs: 0.22038\nTotal prob of N-in: 0.12994\noutside 1 331\nTMhelix 332 354\ninside 355 372

26012 GCF_001252135.1_5236_6_9Shigella sonnei Proteobacteria; Gammaproteobacteria; Enterobacterales; Enterobacteriaceae; Shigella MNNEETFYQAMRRQGVTRRSFLKYCSLAATSLGLGAGMAPKIAWALENKPRIPVVWIHGL WP_005139779.1 hydrogenase [Shigella sonnei] Length: 372\nNumber of predicted TMHs: 2\nExp number of AAs in TMHs: 37.02628\nExp number, first 60 AAs: 19.39684\nTotal prob of N-in: 0.94313\nPOSSIBLE N-term signal sequence\ninside 1 19\nTMhelix 20 39\noutside 40 325\nTMhelix 326 348\ninside 349 372

26013 GCF_001252415.1_12971_8_47 Shigella sonnei Proteobacteria; Gammaproteobacteria; Enterobacterales; Enterobacteriaceae; Shigella MTGDNTLIHSHGINRRDFMKLCAALAATMGLSSKAAAEMAESVTNPQRPPVIWIGAQECT WP_000145410.1 MULTISPECIES: hydrogenase-2 small chain [Proteobacteria] Length: 372\nNumber of predicted TMHs: 1\nExp number of AAs in TMHs: 23.72226\nExp number, first 60 AAs: 0.22038\nTotal prob of N-in: 0.12994\noutside 1 331\nTMhelix 332 354\ninside 355 372

26014 GCF_001252455.1_12971_8_54 Shigella sonnei Proteobacteria; Gammaproteobacteria; Enterobacterales; Enterobacteriaceae; Shigella MTGDNTLIHSHGINRRDFMKLCAALAATMGLSSKAAAEMAESVTNPQRPPVIWIGAQECT WP_000145410.1 MULTISPECIES: hydrogenase-2 small chain [Proteobacteria] Length: 372\nNumber of predicted TMHs: 1\nExp number of AAs in TMHs: 23.72226\nExp number, first 60 AAs: 0.22038\nTotal prob of N-in: 0.12994\noutside 1 331\nTMhelix 332 354\ninside 355 372

26015 GCF_001252415.1_12971_8_47 Shigella sonnei Proteobacteria; Gammaproteobacteria; Enterobacterales; Enterobacteriaceae; Shigella MNNEETFYQAMRRQGVTRRSFLKYCSLAATSLGLGAGMAPKIAWALENKPRIPVVWIHGL WP_005139779.1 hydrogenase [Shigella sonnei] Length: 372\nNumber of predicted TMHs: 2\nExp number of AAs in TMHs: 37.02628\nExp number, first 60 AAs: 19.39684\nTotal prob of N-in: 0.94313\nPOSSIBLE N-term signal sequence\ninside 1 19\nTMhelix 20 39\noutside 40 325\nTMhelix 326 348\ninside 349 372

26016 GCF_001252395.1_12971_8_61 Shigella sonnei Proteobacteria; Gammaproteobacteria; Enterobacterales; Enterobacteriaceae; Shigella MNNEETFYQAMRRQGVTRRSFLKYCSLAATSLGLGAGMAPKIAWALENKPRIPVVWIHGL WP_005139779.1 hydrogenase [Shigella sonnei] Length: 372\nNumber of predicted TMHs: 2\nExp number of AAs in TMHs: 37.02628\nExp number, first 60 AAs: 19.39684\nTotal prob of N-in: 0.94313\nPOSSIBLE N-term signal sequence\ninside 1 19\nTMhelix 20 39\noutside 40 325\nTMhelix 326 348\ninside 349 372

26017 GCF_001252455.1_12971_8_54 Shigella sonnei Proteobacteria; Gammaproteobacteria; Enterobacterales; Enterobacteriaceae; Shigella MNNEETFYQAMRRQGVTRRSFLKYCSLAATSLGLGAGMAPKIAWALENKPRIPVVWIHGL WP_052962775.1 hydrogenase [Shigella sonnei] Length: 372\nNumber of predicted TMHs: 2\nExp number of AAs in TMHs: 37.09248\nExp number, first 60 AAs: 19.39296\nTotal prob of N-in: 0.94316\nPOSSIBLE N-term signal sequence\ninside 1 19\nTMhelix 20 39\noutside 40 325\nTMhelix 326 348\ninside 349 372

26018 GCF_001252255.1_Sg_201400855_TAGCTT_L001 Shigella sonnei Proteobacteria; Gammaproteobacteria; Enterobacterales; Enterobacteriaceae; Shigella MNNEETFYQAMRRQGVTRRSFLKYCSLAATSLGLGAGMAPKIAWALENKPRIPVVWIHGL WP_005139779.1 hydrogenase [Shigella sonnei] Length: 372\nNumber of predicted TMHs: 2\nExp number of AAs in

TMHs: 37.02628\nExp number, first 60 AAs: 19.39684\nTotal prob of N-in: 0.94313\nPOSSIBLE N-term signal sequence\ninside 1 19\nTMhelix 20 39\noutside 40 325\nTMhelix 326 348\ninside 349 372

26019 GCF_001252295.1_5236_8_10 Shigella sonnei Proteobacteria; Gammaproteobacteria; Enterobacterales; Enterobacteriaceae; Shigella MNNEETFYQAMRRQGVTRRSFLKYCSLAATSLGLGAGMAPKIAWALENKPRIPVVWIHGL WP_005139779.1 hydrogenase [Shigella sonnei] Length: 372\nNumber of predicted TMHs: 2\nExp number of AAs in TMHs: 37.02628\nExp number, first 60 AAs: 19.39684\nTotal prob of N-in: 0.94313\nPOSSIBLE N-term signal sequence\ninside 1 19\nTMhelix 20 39\noutside 40 325\nTMhelix 326 348\ninside 349 372

26020 GCF_001252515.1_6350_1_18 Shigella sonnei Proteobacteria; Gammaproteobacteria; Enterobacterales; Enterobacteriaceae; Shigella MTGDNTLIHSHGINRRDFMKLCAALAATMGLSSKAAAEMAESVTNPQRPPVIWIGAQECT WP_000145410.1 MULTISPECIES: hydrogenase-2 small chain [Proteobacteria] Length: 372\nNumber of predicted TMHs: 1\nExp number of AAs in TMHs: 23.72226\nExp number, first 60 AAs: 0.22038\nTotal prob of N-in: 0.12994\noutside 1 331\nTMhelix 332 354\ninside 355 372

26021 GCF_001252555.1_H14132053801 Shigella sonnei Proteobacteria; Gammaproteobacteria; Enterobacterales; Enterobacteriaceae; Shigella MTGDNTLIHSHGINRRDFMKLCAALAATMGLSSKAAAEMAESVTNPQRPPVIWIGAQECT WP_000145410.1 MULTISPECIES: hydrogenase-2 small chain [Proteobacteria] Length: 372\nNumber of predicted TMHs: 1\nExp number of AAs in TMHs: 23.72226\nExp number, first 60 AAs: 0.22038\nTotal prob of N-in: 0.12994\noutside 1 331\nTMhelix 332 354\ninside 355 372

26022 GCF_001252515.1_6350_1_18 Shigella sonnei Proteobacteria; Gammaproteobacteria; Enterobacterales; Enterobacteriaceae; Shigella MNNEETFYQAMRRQGVTRRSFLKYCSLAATSLGLGAGMAPKIAWALENKPRIPVVWIHGL WP_005139779.1 hydrogenase [Shigella sonnei] Length: 372\nNumber of predicted TMHs: 2\nExp number of AAs in TMHs: 37.02628\nExp number, first 60 AAs: 19.39684\nTotal prob of N-in: 0.94313\nPOSSIBLE N-term signal sequence\ninside 1 19\nTMhelix 20 39\noutside 40 325\nTMhelix 326 348\ninside 349 372

26023 GCF_001252555.1_H14132053801 Shigella sonnei Proteobacteria; Gammaproteobacteria; Enterobacterales; Enterobacteriaceae; Shigella MNNEETFYQAMRRQGVTRRSFLKYCSLAATSLGLGAGMAPKIAWALENKPRIPVVWIHGL WP_052962775.1 hydrogenase [Shigella sonnei] Length: 372\nNumber of predicted TMHs: 2\nExp number of AAs in TMHs: 37.09248\nExp number, first 60 AAs: 19.39296\nTotal prob of N-in: 0.94316\nPOSSIBLE N-term signal sequence\ninside 1 19\nTMhelix 20 39\noutside 40 325\nTMhelix 326 348\ninside 349 372

26024 GCF_001252315.1_9870_7_8Shigella sonnei Proteobacteria; Gammaproteobacteria; Enterobacterales; Enterobacteriaceae; Shigella MNNEETFYQAMRRQGVTRRSFLKYCSLAATSLGLGAGMAPKIAWALENKPRIPVVWIHGL WP_005139779.1 hydrogenase [Shigella sonnei] Length: 372\nNumber of predicted TMHs: 2\nExp number of AAs in TMHs: 37.02628\nExp number, first 60 AAs: 19.39684\nTotal prob of N-in: 0.94313\nPOSSIBLE N-term signal sequence\ninside 1 19\nTMhelix 20 39\noutside 40 325\nTMhelix 326 348\ninside 349 372

26025 GCF_001252595.1_6246_5_6Shigella sonnei Proteobacteria; Gammaproteobacteria; Enterobacterales; Enterobacteriaceae; Shigella MTGDNTLIHSHGINRRDFMKLCAALAATMGLSSKAAAEMAESVTNPQRPPVIWIGAQECT WP_000145410.1 MULTISPECIES: hydrogenase-2 small chain [Proteobacteria] Length: 372\nNumber of predicted TMHs: 1\nExp number of AAs in TMHs: 23.72226\nExp number, first 60 AAs: 0.22038\nTotal prob of N-in: 0.12994\noutside 1 331\nTMhelix 332 354\ninside 355 372

26026 GCF_001252635.1_8290_4_39 Shigella sonnei Proteobacteria; Gammaproteobacteria; Enterobacterales; Enterobacteriaceae; Shigella MTGDNTLIHSHGINRRDFMKLCAALAATMGLSSKAAAEMAESVTNPQRPPVIWIGAQECT WP_000145410.1 MULTISPECIES: hydrogenase-2 small chain [Proteobacteria] Length: 372\nNumber of predicted TMHs: 1\nExp number of AAs in TMHs: 23.72226\nExp number, first 60 AAs: 0.22038\nTotal prob of N-in: 0.12994\noutside 1 331\nTMhelix 332 354\ninside 355 372

26027 GCF_001252715.1_12971_8_37 Shigella sonnei Proteobacteria; Gammaproteobacteria; Enterobacterales; Enterobacteriaceae; Shigella MTGDNTLIHSHGINRRDFMKLCAALAATMGLSSKAAAEMAESVTNPQRPPVIWIGAQECT WP_000145410.1 MULTISPECIES: hydrogenase-2 small chain [Proteobacteria] Length: 372\nNumber of predicted TMHs:

1\nExp number of AAs in TMHs: 23.72226\nExp number, first 60 AAs: 0.22038\nTotal prob of N-in: 0.12994\noutside 1 331\nTMhelix 332 354\ninside 355 372

26028 GCF_001252955.1_8289_7_72 Shigella sonnei Proteobacteria; Gammaproteobacteria; Enterobacterales; Enterobacteriaceae; Shigella MNNEETFYQAMRRQGVTRRSFLKYCSLAATSLGLGAGMAPKIAWALENKPRIPVVWIHGL WP_005139779.1 hydrogenase [Shigella sonnei] Length: 372\nNumber of predicted TMHs: 2\nExp number of AAs in TMHs: 37.02628\nExp number, first 60 AAs: 19.39684\nTotal prob of N-in: 0.94313\nPOSSIBLE N-term signal sequence\ninside 1 19\nTMhelix 20 39\noutside 40 325\nTMhelix 326 348\ninside 349 372

26029 GCF_001252995.1_5236_3_6Shigella sonnei Proteobacteria; Gammaproteobacteria; Enterobacterales; Enterobacteriaceae; Shigella MNNEETFYQAMRRQGVTRRSFLKYCSLAATSLGLGAGMAPKIAWALENKPRIPVVWIHGL WP_005139779.1 hydrogenase [Shigella sonnei] Length: 372\nNumber of predicted TMHs: 2\nExp number of AAs in TMHs: 37.02628\nExp number, first 60 AAs: 19.39684\nTotal prob of N-in: 0.94313\nPOSSIBLE N-term signal sequence\ninside 1 19\nTMhelix 20 39\noutside 40 325\nTMhelix 326 348\ninside 349 372

26030 GCF_001252635.1_8290_4_39 Shigella sonnei Proteobacteria; Gammaproteobacteria; Enterobacterales; Enterobacteriaceae; Shigella MNNEETFYQAMRRQGVTRRSFLKYCSLAATSLGLGAGMAPKIAWALENKPRIPVVWIHGL WP_005139779.1 hydrogenase [Shigella sonnei] Length: 372\nNumber of predicted TMHs: 2\nExp number of AAs in TMHs: 37.02628\nExp number, first 60 AAs: 19.39684\nTotal prob of N-in: 0.94313\nPOSSIBLE N-term signal sequence\ninside 1 19\nTMhelix 20 39\noutside 40 325\nTMhelix 326 348\ninside 349 372

26031 GCF_001252715.1_12971_8_37 Shigella sonnei Proteobacteria; Gammaproteobacteria; Enterobacterales; Enterobacteriaceae; Shigella MNNEETFYQAMRRQGVTRRSFLKYCSLAATSLGLGAGMAPKIAWALENKPRIPVVWIHGL WP_005139779.1 hydrogenase [Shigella sonnei] Length: 372\nNumber of predicted TMHs: 2\nExp number of AAs in TMHs: 37.02628\nExp number, first 60 AAs: 19.39684\nTotal prob of N-in: 0.94313\nPOSSIBLE N-term signal sequence\ninside 1 19\nTMhelix 20 39\noutside 40 325\nTMhelix 326 348\ninside 349 372

26032 GCF_001253075.1_8290_4_96 Shigella sonnei Proteobacteria; Gammaproteobacteria; Enterobacterales; Enterobacteriaceae; Shigella MTGDNTLIHSHGINRRDFMKLCAALAATMGLSSKAAAEMAESVTNPQRPPVIWIGAQECT WP_000145410.1 MULTISPECIES: hydrogenase-2 small chain [Proteobacteria] Length: 372\nNumber of predicted TMHs: 1\nExp number of AAs in TMHs: 23.72226\nExp number, first 60 AAs: 0.22038\nTotal prob of N-in: 0.12994\noutside 1 331\nTMhelix 332 354\ninside 355 372

26033 GCF_001253095.1_5236_6_6Shigella sonnei Proteobacteria; Gammaproteobacteria; Enterobacterales; Enterobacteriaceae; Shigella MTGDNTLIHSHGINRRDFMKLCAALAATMGLSSKAAAEMAESVTNPQRPPVIWIGAQECT WP_000145410.1 MULTISPECIES: hydrogenase-2 small chain [Proteobacteria] Length: 372\nNumber of predicted TMHs: 1\nExp number of AAs in TMHs: 23.72226\nExp number, first 60 AAs: 0.22038\nTotal prob of N-in: 0.12994\noutside 1 331\nTMhelix 332 354\ninside 355 372

26034 GCF_001253095.1_5236_6_6Shigella sonnei Proteobacteria; Gammaproteobacteria; Enterobacterales; Enterobacteriaceae; Shigella MNNEETFYQAMRRQGVTRRSFLKYCSLAATSLGLGAGMAPKIAWALENKPRIPVVWIHGL WP_005139779.1 hydrogenase [Shigella sonnei] Length: 372\nNumber of predicted TMHs: 2\nExp number of AAs in TMHs: 37.02628\nExp number, first 60 AAs: 19.39684\nTotal prob of N-in: 0.94313\nPOSSIBLE N-term signal sequence\ninside 1 19\nTMhelix 20 39\noutside 40 325\nTMhelix 326 348\ninside 349 372

26035 GCF_001253075.1_8290_4_96 Shigella sonnei Proteobacteria; Gammaproteobacteria; Enterobacterales; Enterobacteriaceae; Shigella MNNEETFYQAMRRQGVTRRSFLKYCSLAATSLGLGAGMAPKIAWALENKPRIPVVWIHGL WP_005139779.1 hydrogenase [Shigella sonnei] Length: 372\nNumber of predicted TMHs: 2\nExp number of AAs in TMHs: 37.02628\nExp number, first 60 AAs: 19.39684\nTotal prob of N-in: 0.94313\nPOSSIBLE N-term signal sequence\ninside 1 19\nTMhelix 20 39\noutside 40 325\nTMhelix 326 348\ninside 349 372

26036 GCF_001252755.1_12830_1_41 Shigella sonnei Proteobacteria; Gammaproteobacteria; Enterobacterales; Enterobacteriaceae; Shigella MNNEETFYQAMRRQGVTRRSFLKYCSLAATSLGLGAGMAPKIAWALENKPRIPVVWIHGL WP_005139779.1 hydrogenase [Shigella sonnei] Length: 372\nNumber of predicted TMHs: 2\nExp number of AAs in

TMHs: 37.02628\nExp number, first 60 AAs: 19.39684\nTotal prob of N-in: 0.94313\nPOSSIBLE N-term signal sequence\ninside 1 19\nTMhelix 20 39\noutside 40 325\nTMhelix 326 348\ninside 349 372

26037 GCF_001252655.1_5417_2_12 Shigella sonnei Proteobacteria; Gammaproteobacteria; Enterobacterales; Enterobacteriaceae; Shigella MNNEETFYQAMRRQGVTRRSFLKYCSLAATSLGLGAGMAPKIAWALENKPRIPVVWIHGL WP_005139779.1 hydrogenase [Shigella sonnei] Length: 372\nNumber of predicted TMHs: 2\nExp number of AAs in TMHs: 37.02628\nExp number, first 60 AAs: 19.39684\nTotal prob of N-in: 0.94313\nPOSSIBLE N-term signal sequence\ninside 1 19\nTMhelix 20 39\noutside 40 325\nTMhelix 326 348\ninside 349 372

26038 GCF_001253135.1_9870_6_23 Shigella sonnei Proteobacteria; Gammaproteobacteria; Enterobacterales; Enterobacteriaceae; Shigella MTGDNTLIHSHGINRRDFMKLCAALAATMGLSSKAAAEMAESVTNPQRPPVIWIGAQECT WP_000145410.1 MULTISPECIES: hydrogenase-2 small chain [Proteobacteria] Length: 372\nNumber of predicted TMHs: 1\nExp number of AAs in TMHs: 23.72226\nExp number, first 60 AAs: 0.22038\nTotal prob of N-in: 0.12994\noutside 1 331\nTMhelix 332 354\ninside 355 372

26039 GCF_001253255.1_H14126044301 Shigella sonnei Proteobacteria; Gammaproteobacteria; Enterobacterales; Enterobacteriaceae; Shigella MTGDNTLIHSHGINRRDFMKLCAALAATMGLSSKAAAEMAESVTNPQRPPVIWIGAQECT WP_000145410.1 MULTISPECIES: hydrogenase-2 small chain [Proteobacteria] Length: 372\nNumber of predicted TMHs: 1\nExp number of AAs in TMHs: 23.72226\nExp number, first 60 AAs: 0.22038\nTotal prob of N-in: 0.12994\noutside 1 331\nTMhelix 332 354\ninside 355 372

26040 GCF_001253255.1_H14126044301 Shigella sonnei Proteobacteria; Gammaproteobacteria; Enterobacterales; Enterobacteriaceae; Shigella MNNEETFYQAMRRQGVTRRSFLKYCSLAATSLGLGAGMAPKIAWALENKPRIPVVWIHGL WP_005139779.1 hydrogenase [Shigella sonnei] Length: 372\nNumber of predicted TMHs: 2\nExp number of AAs in TMHs: 37.02628\nExp number, first 60 AAs: 19.39684\nTotal prob of N-in: 0.94313\nPOSSIBLE N-term signal sequence\ninside 1 19\nTMhelix 20 39\noutside 40 325\nTMhelix 326 348\ninside 349 372

26041 GCF_001253135.1_9870_6_23 Shigella sonnei Proteobacteria; Gammaproteobacteria; Enterobacterales; Enterobacteriaceae; Shigella MNNEETFYQAMRRQGVTRRSFLKYCSLAATSLGLGAGMAPKIAWALENKPRIPVVWIHGL WP_005139779.1 hydrogenase [Shigella sonnei] Length: 372\nNumber of predicted TMHs: 2\nExp number of AAs in TMHs: 37.02628\nExp number, first 60 AAs: 19.39684\nTotal prob of N-in: 0.94313\nPOSSIBLE N-term signal sequence\ninside 1 19\nTMhelix 20 39\noutside 40 325\nTMhelix 326 348\ninside 349 372

26042 GCF_001253675.1_5417_3_2Shigella sonnei Proteobacteria; Gammaproteobacteria; Enterobacterales; Enterobacteriaceae; Shigella MTGDNTLIHSHGINRRDFMKLCAALAATMGLSSKAAAEMAESVTNPQRPPVIWIGAQECT WP_000145410.1 MULTISPECIES: hydrogenase-2 small chain [Proteobacteria] Length: 372\nNumber of predicted TMHs: 1\nExp number of AAs in TMHs: 23.72226\nExp number, first 60 AAs: 0.22038\nTotal prob of N-in: 0.12994\noutside 1 331\nTMhelix 332 354\ninside 355 372

26043 GCF_001253675.1_5417_3_2Shigella sonnei Proteobacteria; Gammaproteobacteria; Enterobacterales; Enterobacteriaceae; Shigella MNNEETFYQAMRRQGVTRRSFLKYCSLAATSLGLGAGMAPKIAWALENKPRIPVVWIHGL WP_005139779.1 hydrogenase [Shigella sonnei] Length: 372\nNumber of predicted TMHs: 2\nExp number of AAs in TMHs: 37.02628\nExp number, first 60 AAs: 19.39684\nTotal prob of N-in: 0.94313\nPOSSIBLE N-term signal sequence\ninside 1 19\nTMhelix 20 39\noutside 40 325\nTMhelix 326 348\ninside 349 372

26044 GCF_001253275.1_8290_4_17 Shigella sonnei Proteobacteria; Gammaproteobacteria; Enterobacterales; Enterobacteriaceae; Shigella MNNEETFYQAMRRQGVTRRSFLKYCSLAATSLGLGAGMAPKIAWALENKPRIPVVWIHGL WP_005139779.1 hydrogenase [Shigella sonnei] Length: 372\nNumber of predicted TMHs: 2\nExp number of AAs in TMHs: 37.02628\nExp number, first 60 AAs: 19.39684\nTotal prob of N-in: 0.94313\nPOSSIBLE N-term signal sequence\ninside 1 19\nTMhelix 20 39\noutside 40 325\nTMhelix 326 348\ninside 349 372

26045 GCF_001253355.1_6246_5_4Shigella sonnei Proteobacteria; Gammaproteobacteria; Enterobacterales; Enterobacteriaceae; Shigella MNNEETFYQAMRRQGVTRRSFLKYCSLAATSLGLGAGMAPKIAWALENKPRIPVVWIHGL WP_005139779.1 hydrogenase [Shigella sonnei] Length: 372\nNumber of predicted TMHs: 2\nExp

number of AAs in TMHs: 37.02628\nExp number, first 60 AAs: 19.39684\nTotal prob of N-in: 0.94313\nPOSSIBLE N-term signal sequence\ninside 1 19\nTMhelix 20 39\noutside 40 325\nTMhelix 326 348\ninside 349 372

26046 GCF_001253395.1_Sg_201403963_CACTCA_L001 Shigella sonnei Proteobacteria; Gammaproteobacteria; Enterobacterales; Enterobacteriaceae; Shigella
MNNEETFYQAMRRQGVTRRSFLKYCSLAATSLGLGAGMAPKIAWALENKPRIPVVWIHGL WP_005139779.1
hydrogenase [Shigella sonnei] Length: 372\nNumber of predicted TMHs: 2\nExp number of AAs in TMHs: 37.02628\nExp number, first 60 AAs: 19.39684\nTotal prob of N-in: 0.94313\nPOSSIBLE N-term signal sequence\ninside 1 19\nTMhelix 20 39\noutside 40 325\nTMhelix 326 348\ninside 349 372

26047 GCF_001253415.1_6350_1_11 Shigella sonnei Proteobacteria; Gammaproteobacteria; Enterobacterales; Enterobacteriaceae; Shigella
MNNEETFYQAMRRQGVTRRSFLKYCSLAATSLGLGAGMAPKIAWALENKPRIPVVWIHGL WP_005139779.1
hydrogenase [Shigella sonnei] Length: 372\nNumber of predicted TMHs: 2\nExp number of AAs in TMHs: 37.02628\nExp number, first 60 AAs: 19.39684\nTotal prob of N-in: 0.94313\nPOSSIBLE N-term signal sequence\ninside 1 19\nTMhelix 20 39\noutside 40 325\nTMhelix 326 348\ninside 349 372

26048 GCF_001253715.1_8289_7_92 Shigella sonnei Proteobacteria; Gammaproteobacteria; Enterobacterales; Enterobacteriaceae; Shigella
MTGDNTLIHSHGINRRDFMKLCAALAATMGLSSKAAAEMAESVTNPQRPPVIWIGAQECT WP_000145410.1
MULTISPECIES: hydrogenase-2 small chain [Proteobacteria] Length: 372\nNumber of predicted TMHs: 1\nExp number of AAs in TMHs: 23.72226\nExp number, first 60 AAs: 0.22038\nTotal prob of N-in: 0.12994\noutside 1 331\nTMhelix 332 354\ninside 355 372

26049 GCF_001253775.1_8289_7_95 Shigella sonnei Proteobacteria; Gammaproteobacteria; Enterobacterales; Enterobacteriaceae; Shigella
MTGDNTLIHSHGINRRDFMKLCAALAATMGLSSKAAAEMAESVTNPQRPPVIWIGAQECT WP_000145410.1
MULTISPECIES: hydrogenase-2 small chain [Proteobacteria] Length: 372\nNumber of predicted TMHs: 1\nExp number of AAs in TMHs: 23.72226\nExp number, first 60 AAs: 0.22038\nTotal prob of N-in: 0.12994\noutside 1 331\nTMhelix 332 354\ninside 355 372

26050 GCF_001253815.1_8289_7_42 Shigella sonnei Proteobacteria; Gammaproteobacteria; Enterobacterales; Enterobacteriaceae; Shigella
MTGDNTLIHSHGINRRDFMKLCAALAATMGLSSKAAAEMAESVTNPQRPPVIWIGAQECT WP_000145410.1
MULTISPECIES: hydrogenase-2 small chain [Proteobacteria] Length: 372\nNumber of predicted TMHs: 1\nExp number of AAs in TMHs: 23.72226\nExp number, first 60 AAs: 0.22038\nTotal prob of N-in: 0.12994\noutside 1 331\nTMhelix 332 354\ninside 355 372

26051 GCF_001253855.1_7353_1_24 Shigella sonnei Proteobacteria; Gammaproteobacteria; Enterobacterales; Enterobacteriaceae; Shigella
MTGDNTLIHSHGINRRDFMKLCAALAATMGLSSKAAAEMAESVTNPQRPPVIWIGAQECT WP_000145410.1
MULTISPECIES: hydrogenase-2 small chain [Proteobacteria] Length: 372\nNumber of predicted TMHs: 1\nExp number of AAs in TMHs: 23.72226\nExp number, first 60 AAs: 0.22038\nTotal prob of N-in: 0.12994\noutside 1 331\nTMhelix 332 354\ninside 355 372

26052 GCF_001253715.1_8289_7_92 Shigella sonnei Proteobacteria; Gammaproteobacteria; Enterobacterales; Enterobacteriaceae; Shigella
MNNEETFYQAMRRQGVTRRSFLKYCSLAATSLGLGAGMAPKIAWALENKPRIPVVWIHGL WP_005139779.1
hydrogenase [Shigella sonnei] Length: 372\nNumber of predicted TMHs: 2\nExp number of AAs in TMHs: 37.02628\nExp number, first 60 AAs: 19.39684\nTotal prob of N-in: 0.94313\nPOSSIBLE N-term signal sequence\ninside 1 19\nTMhelix 20 39\noutside 40 325\nTMhelix 326 348\ninside 349 372

26053 GCF_001253815.1_8289_7_42 Shigella sonnei Proteobacteria; Gammaproteobacteria; Enterobacterales; Enterobacteriaceae; Shigella
MNNEETFYQAMRRQGVTRRSFLKYCSLAATSLGLGAGMAPKIAWALENKPRIPVVWIHGL WP_005139779.1
hydrogenase [Shigella sonnei] Length: 372\nNumber of predicted TMHs: 2\nExp number of AAs in TMHs: 37.02628\nExp number, first 60 AAs: 19.39684\nTotal prob of N-in: 0.94313\nPOSSIBLE N-term signal sequence\ninside 1 19\nTMhelix 20 39\noutside 40 325\nTMhelix 326 348\ninside 349 372

26054 GCF_001253515.1_12971_8_22 Shigella sonnei Proteobacteria; Gammaproteobacteria; Enterobacterales; Enterobacteriaceae; Shigella

MNNEETFYQAMRRQGVTRRSFLKYCSLAATSLGLGAGMAPKIAWALENKPRIPVVWIHGL WP_005139779.1 hydrogenase [Shigella sonnei] Length: 372\nNumber of predicted TMHs: 2\nExp number of AAs in TMHs: 37.02628\nExp number, first 60 AAs: 19.39684\nTotal prob of N-in: 0.94313\nPOSSIBLE N-term signal sequence\ninside 1 19\nTMhelix 20 39\noutside 40 325\nTMhelix 326 348\ninside 349 372

26055 GCF_001253895.1_5236_7_9Shigella sonnei Proteobacteria; Gammaproteobacteria; Enterobacterales; Enterobacteriaceae; Shigella MTGDNTLIHSHGINRRDFMKLCAALAATMGLSSKAAAEMAESVTNPQRPPVIWIGAQECT WP_000145410.1 MULTISPECIES: hydrogenase-2 small chain [Proteobacteria] Length: 372\nNumber of predicted TMHs: 1\nExp number of AAs in TMHs: 23.72226\nExp number, first 60 AAs: 0.22038\nTotal prob of N-in: 0.12994\noutside 1 331\nTMhelix 332 354\ninside 355 372

26056 GCF_001253995.1_8289_7_75 Shigella sonnei Proteobacteria; Gammaproteobacteria; Enterobacterales; Enterobacteriaceae; Shigella MNNEETFYQAMRRQGVTRRSFLKYCSLAATSLGLGAGMAPKIAWALENKPRIPVVWIHGL WP_005139779.1 hydrogenase [Shigella sonnei] Length: 372\nNumber of predicted TMHs: 2\nExp number of AAs in TMHs: 37.02628\nExp number, first 60 AAs: 19.39684\nTotal prob of N-in: 0.94313\nPOSSIBLE N-term signal sequence\ninside 1 19\nTMhelix 20 39\noutside 40 325\nTMhelix 326 348\ninside 349 372

26057 GCF_001253635.1_5008_7_2Shigella sonnei Proteobacteria; Gammaproteobacteria; Enterobacterales; Enterobacteriaceae; Shigella MNNEETFYQAMRRQGVTRRSFLKYCSLAATSLGLGAGMAPKIAWALENKPRIPVVWIHGL WP_005139779.1 hydrogenase [Shigella sonnei] Length: 372\nNumber of predicted TMHs: 2\nExp number of AAs in TMHs: 37.02628\nExp number, first 60 AAs: 19.39684\nTotal prob of N-in: 0.94313\nPOSSIBLE N-term signal sequence\ninside 1 19\nTMhelix 20 39\noutside 40 325\nTMhelix 326 348\ninside 349 372

26058 GCF_001254035.1_5236_2_11 Shigella sonnei Proteobacteria; Gammaproteobacteria; Enterobacterales; Enterobacteriaceae; Shigella MTGDNTLIHSHGINRRDFMKLCAALAATMGLSSKAAAEMAESVTNPQRPPVIWIGAQECT WP_000145410.1 MULTISPECIES: hydrogenase-2 small chain [Proteobacteria] Length: 372\nNumber of predicted TMHs: 1\nExp number of AAs in TMHs: 23.72226\nExp number, first 60 AAs: 0.22038\nTotal prob of N-in: 0.12994\noutside 1 331\nTMhelix 332 354\ninside 355 372

26059 GCF_001253775.1_8289_7_95 Shigella sonnei Proteobacteria; Gammaproteobacteria; Enterobacterales; Enterobacteriaceae; Shigella MNNEETFYQAMRRQGVTRRSFLKYCSLAATSLGLGAGMAPKIAWALENKPRIPVVWIHGL WP_005139779.1 hydrogenase [Shigella sonnei] Length: 372\nNumber of predicted TMHs: 2\nExp number of AAs in TMHs: 37.02628\nExp number, first 60 AAs: 19.39684\nTotal prob of N-in: 0.94313\nPOSSIBLE N-term signal sequence\ninside 1 19\nTMhelix 20 39\noutside 40 325\nTMhelix 326 348\ninside 349 372

26060 GCF_001253855.1_7353_1_24 Shigella sonnei Proteobacteria; Gammaproteobacteria; Enterobacterales; Enterobacteriaceae; Shigella MNNEETFYQAMRRQGVTRRSFLKYCSLAATSLGLGAGMAPKIAWALENKPRIPVVWIHGL WP_005139779.1 hydrogenase [Shigella sonnei] Length: 372\nNumber of predicted TMHs: 2\nExp number of AAs in TMHs: 37.02628\nExp number, first 60 AAs: 19.39684\nTotal prob of N-in: 0.94313\nPOSSIBLE N-term signal sequence\ninside 1 19\nTMhelix 20 39\noutside 40 325\nTMhelix 326 348\ninside 349 372

26061 GCF_001254075.1_12971_8_57 Shigella sonnei Proteobacteria; Gammaproteobacteria; Enterobacterales; Enterobacteriaceae; Shigella MTGDNTLIHSHGINRRDFMKLCAALAATMGLSSKAAAEMAESVTNPQRPPVIWIGAQECT WP_000145410.1 MULTISPECIES: hydrogenase-2 small chain [Proteobacteria] Length: 372\nNumber of predicted TMHs: 1\nExp number of AAs in TMHs: 23.72226\nExp number, first 60 AAs: 0.22038\nTotal prob of N-in: 0.12994\noutside 1 331\nTMhelix 332 354\ninside 355 372

26062 GCF_001254155.1_5008_7_7Shigella sonnei Proteobacteria; Gammaproteobacteria; Enterobacterales; Enterobacteriaceae; Shigella MTGDNTLIHSHGINRRDFMKLCAALAATMGLSSKAAAEMAESVTNPQRPPVIWIGAQECT WP_000145410.1 MULTISPECIES: hydrogenase-2 small chain [Proteobacteria] Length: 372\nNumber of predicted TMHs: 1\nExp number of AAs in TMHs: 23.72226\nExp number, first 60 AAs: 0.22038\nTotal prob of N-in: 0.12994\noutside 1 331\nTMhelix 332 354\ninside 355 372

26063 GCF_001254195.1_6551_1_8Shigella sonnei Proteobacteria; Gammaproteobacteria; Enterobacterales; Enterobacteriaceae; Shigella MTGDNTLIHSHGINRRDFMKLCAALAATMGLSSKAAAEMAESVTNPQRPPVIWIGAQECT WP_000145410.1 MULTISPECIES: hydrogenase-2 small chain [Proteobacteria] Length: 372\nNumber of

predicted TMHs: 1\nExp number of AAs in TMHs: 23.72226\nExp number, first 60 AAs: 0.22038\nTotal prob of N-in: 0.12994\noutside 1 331\nTMhelix 332 354\ninside 355 372

26064 GCF_001254215.1_8290_4_33 Shigella sonnei Proteobacteria; Gammaproteobacteria; Enterobacterales; Enterobacteriaceae; Shigella MTGDNTLIHSHGINRRDFMKLCAALAATMGLSSKAAAEMAESVTNPQRPPVIWIGAQECT WP_000145410.1 MULTISPECIES: hydrogenase-2 small chain [Proteobacteria] Length: 372\nNumber of predicted TMHs: 1\nExp number of AAs in TMHs: 23.72226\nExp number, first 60 AAs: 0.22038\nTotal prob of N-in: 0.12994\noutside 1 331\nTMhelix 332 354\ninside 355 372

26065 GCF_001254315.1_H14128036101 Shigella sonnei Proteobacteria; Gammaproteobacteria; Enterobacterales; Enterobacteriaceae; Shigella MTGDNTLIHSHGINRRDFMKLCAALAATMGLSSKAAAEMAESVTNPQRPPVIWIGAQECT WP_000145410.1 MULTISPECIES: hydrogenase-2 small chain [Proteobacteria] Length: 372\nNumber of predicted TMHs: 1\nExp number of AAs in TMHs: 23.72226\nExp number, first 60 AAs: 0.22038\nTotal prob of N-in: 0.12994\noutside 1 331\nTMhelix 332 354\ninside 355 372

26066 GCF_001254275.1_Sg_201312542_TCGAAG_L001 Shigella sonnei Proteobacteria; Gammaproteobacteria; Enterobacterales; Enterobacteriaceae; Shigella MTGDNTLIHSHGINRRDFMKLCAALAATMGLSSKAAAEMAESVTNPQRPPVIWIGAQECT WP_000145410.1 MULTISPECIES: hydrogenase-2 small chain [Proteobacteria] Length: 372\nNumber of predicted TMHs: 1\nExp number of AAs in TMHs: 23.72226\nExp number, first 60 AAs: 0.22038\nTotal prob of N-in: 0.12994\noutside 1 331\nTMhelix 332 354\ninside 355 372

26067 GCF_001254375.1_8290_4_37 Shigella sonnei Proteobacteria; Gammaproteobacteria; Enterobacterales; Enterobacteriaceae; Shigella MTGDNTLIHSHGINRRDFMKLCAALAATMGLSSKAAAEMAESVTNPQRPPVIWIGAQECT WP_000145410.1 MULTISPECIES: hydrogenase-2 small chain [Proteobacteria] Length: 372\nNumber of predicted TMHs: 1\nExp number of AAs in TMHs: 23.72226\nExp number, first 60 AAs: 0.22038\nTotal prob of N-in: 0.12994\noutside 1 331\nTMhelix 332 354\ninside 355 372

26068 GCF_001254495.1_5008_7_6Shigella sonnei Proteobacteria; Gammaproteobacteria; Enterobacterales; Enterobacteriaceae; Shigella MTGDNTLIHSHGINRRDFMKLCAALAATMGLSSKAAAEMAESVTNPQRPPVIWIGAQECT WP_000145410.1 MULTISPECIES: hydrogenase-2 small chain [Proteobacteria] Length: 372\nNumber of predicted TMHs: 1\nExp number of AAs in TMHs: 23.72226\nExp number, first 60 AAs: 0.22038\nTotal prob of N-in: 0.12994\noutside 1 331\nTMhelix 332 354\ninside 355 372

26069 GCF_001254455.1_8290_4_92 Shigella sonnei Proteobacteria; Gammaproteobacteria; Enterobacterales; Enterobacteriaceae; Shigella MTGDNTLIHSHGINRRDFMKLCAALAATMGLSSKAAAEMAESVTNPQRPPVIWIGAQECT WP_000145410.1 MULTISPECIES: hydrogenase-2 small chain [Proteobacteria] Length: 372\nNumber of predicted TMHs: 1\nExp number of AAs in TMHs: 23.72226\nExp number, first 60 AAs: 0.22038\nTotal prob of N-in: 0.12994\noutside 1 331\nTMhelix 332 354\ninside 355 372

26070 GCF_001254495.1_5008_7_6Shigella sonnei Proteobacteria; Gammaproteobacteria; Enterobacterales; Enterobacteriaceae; Shigella MNNEETFYQAMRRQGVTRRSFLKYCSLAATSLGLGAGMAPKIAWALENKPRIPVVWIHGL WP_005139779.1 hydrogenase [Shigella sonnei] Length: 372\nNumber of predicted TMHs: 2\nExp number of AAs in TMHs: 37.02628\nExp number, first 60 AAs: 19.39684\nTotal prob of N-in: 0.94313\nPOSSIBLE N-term signal sequence\ninside 1 19\nTMhelix 20 39\noutside 40 325\nTMhelix 326 348\ninside 349 372

26071 GCF_001254695.1_H14142025101 Shigella sonnei Proteobacteria; Gammaproteobacteria; Enterobacterales; Enterobacteriaceae; Shigella MNNEETFYQAMRRQGVTRRSFLKYCSLAATSLGLGAGMAPKIAWALENKPRIPVVWIHGL WP_052962775.1 hydrogenase [Shigella sonnei] Length: 372\nNumber of predicted TMHs: 2\nExp number of AAs in TMHs: 37.09248\nExp number, first 60 AAs: 19.39296\nTotal prob of N-in: 0.94316\nPOSSIBLE N-term signal sequence\ninside 1 19\nTMhelix 20 39\noutside 40 325\nTMhelix 326 348\ninside 349 372

26072 GCF_001254595.1_8289_7_37 Shigella sonnei Proteobacteria; Gammaproteobacteria; Enterobacterales; Enterobacteriaceae; Shigella MNNEETFYQAMRRQGVTRRSFLKYCSLAATSLGLGAGMAPKIAWALENKPRIPVVWIHGL WP_005139779.1 hydrogenase [Shigella sonnei] Length: 372\nNumber of predicted TMHs: 2\nExp number of AAs in

TMHs: 37.02628\nExp number, first 60 AAs: 19.39684\nTotal prob of N-in: 0.94313\nPOSSIBLE N-term signal sequence\ninside 1 19\nTMhelix 20 39\noutside 40 325\nTMhelix 326 348\ninside 349 372

26073 GCF_001254075.1_12971_8_57 Shigella sonnei Proteobacteria; Gammaproteobacteria; Enterobacterales; Enterobacteriaceae; Shigella MNNEETFYQAMRRQGVTRRSFLKYCSLAATSLGLGAGMAPKIAWALENKPRIPVVWIHGL WP_052962775.1 hydrogenase [Shigella sonnei] Length: 372\nNumber of predicted TMHs: 2\nExp number of AAs in TMHs: 37.09248\nExp number, first 60 AAs: 19.39296\nTotal prob of N-in: 0.94316\nPOSSIBLE N-term signal sequence\ninside 1 19\nTMhelix 20 39\noutside 40 325\nTMhelix 326 348\ninside 349 372

26074 GCF_001254315.1_H14128036101 Shigella sonnei Proteobacteria; Gammaproteobacteria; Enterobacterales; Enterobacteriaceae; Shigella MNNEETFYQAMRRQGVTRRSFLKYCSLAATSLGLGAGMAPKIAWALENKPRIPVVWIHGL WP_005139779.1 hydrogenase [Shigella sonnei] Length: 372\nNumber of predicted TMHs: 2\nExp number of AAs in TMHs: 37.02628\nExp number, first 60 AAs: 19.39684\nTotal prob of N-in: 0.94313\nPOSSIBLE N-term signal sequence\ninside 1 19\nTMhelix 20 39\noutside 40 325\nTMhelix 326 348\ninside 349 372

26075 GCF_001254215.1_8290_4_33 Shigella sonnei Proteobacteria; Gammaproteobacteria; Enterobacterales; Enterobacteriaceae; Shigella MNNEETFYQAMRRQGVTRRSFLKYCSLAATSLGLGAGMAPKIAWALENKPRIPVVWIHGL WP_005139779.1 hydrogenase [Shigella sonnei] Length: 372\nNumber of predicted TMHs: 2\nExp number of AAs in TMHs: 37.02628\nExp number, first 60 AAs: 19.39684\nTotal prob of N-in: 0.94313\nPOSSIBLE N-term signal sequence\ninside 1 19\nTMhelix 20 39\noutside 40 325\nTMhelix 326 348\ninside 349 372

26076 GCF_001254275.1_Sg_201312542_TCGAAG_L001 Shigella sonnei Proteobacteria; Gammaproteobacteria; Enterobacterales; Enterobacteriaceae; Shigella MNNEETFYQAMRRQGVTRRSFLKYCSLAATSLGLGAGMAPKIAWALENKPRIPVVWIHGL WP_005139779.1 hydrogenase [Shigella sonnei] Length: 372\nNumber of predicted TMHs: 2\nExp number of AAs in TMHs: 37.02628\nExp number, first 60 AAs: 19.39684\nTotal prob of N-in: 0.94313\nPOSSIBLE N-term signal sequence\ninside 1 19\nTMhelix 20 39\noutside 40 325\nTMhelix 326 348\ninside 349 372

26077 GCF_001254375.1_8290_4_37 Shigella sonnei Proteobacteria; Gammaproteobacteria; Enterobacterales; Enterobacteriaceae; Shigella MNNEETFYQAMRRQGVTRRSFLKYCSLAATSLGLGAGMAPKIAWALENKPRIPVVWIHGL WP_005139779.1 hydrogenase [Shigella sonnei] Length: 372\nNumber of predicted TMHs: 2\nExp number of AAs in TMHs: 37.02628\nExp number, first 60 AAs: 19.39684\nTotal prob of N-in: 0.94313\nPOSSIBLE N-term signal sequence\ninside 1 19\nTMhelix 20 39\noutside 40 325\nTMhelix 326 348\ninside 349 372

26078 GCF_001254775.1_8289_7_47 Shigella sonnei Proteobacteria; Gammaproteobacteria; Enterobacterales; Enterobacteriaceae; Shigella MTGDNTLIHSHGINRRDFMKLCAALAATMGLSSKAAAEMAESVTNPQRPPVIWIGAQECT WP_000145410.1 MULTISPECIES: hydrogenase-2 small chain [Proteobacteria] Length: 372\nNumber of predicted TMHs: 1\nExp number of AAs in TMHs: 23.72226\nExp number, first 60 AAs: 0.22038\nTotal prob of N-in: 0.12994\noutside 1 331\nTMhelix 332 354\ninside 355 372

26079 GCF_001254835.1_6350_1_15 Shigella sonnei Proteobacteria; Gammaproteobacteria; Enterobacterales; Enterobacteriaceae; Shigella MTGDNTLIHSHGINRRDFMKLCAALAATMGLSSKAAAEMAESVTNPQRPPVIWIGAQECT WP_000145410.1 MULTISPECIES: hydrogenase-2 small chain [Proteobacteria] Length: 372\nNumber of predicted TMHs: 1\nExp number of AAs in TMHs: 23.72226\nExp number, first 60 AAs: 0.22038\nTotal prob of N-in: 0.12994\noutside 1 331\nTMhelix 332 354\ninside 355 372

26080 GCF_001254875.1_12971_8_56 Shigella sonnei Proteobacteria; Gammaproteobacteria; Enterobacterales; Enterobacteriaceae; Shigella MTGDNTLIHSHGINRRDFMKLCAALAATMGLSSKAAAEMAESVTNPQRPPVIWIGAQECT WP_000145410.1 MULTISPECIES: hydrogenase-2 small chain [Proteobacteria] Length: 372\nNumber of predicted TMHs: 1\nExp number of AAs in TMHs: 23.72226\nExp number, first 60 AAs: 0.22038\nTotal prob of N-in: 0.12994\noutside 1 331\nTMhelix 332 354\ninside 355 372

26081 GCF_001254975.1_5417_1_4Shigella sonnei Proteobacteria; Gammaproteobacteria; Enterobacterales; Enterobacteriaceae; Shigella MTGDNTLIHSHGINRRDFMKLCAALAATMGLSSKAAAEMAESVTNPQRPPVIWIGAQECT

WP_000145410.1 MULTISPECIES: hydrogenase-2 small chain [Proteobacteria] Length: 372\nNumber of predicted TMHs: 1\nExp number of AAs in TMHs: 23.72226\nExp number, first 60 AAs: 0.22038\nTotal prob of N-in: 0.12994\noutside 1 331\nTMhelix 332 354\ninside 355 372

26082 GCF_001255015.1_8290_4_6Shigella sonnei Proteobacteria; Gammaproteobacteria; Enterobacterales; Enterobacteriaceae; Shigella MTGDNTLIHSHGINRRDFMKLCAALAATMGLSSKAAAEMAESVTNPQRPPVIWIGAQECT WP_000145410.1 MULTISPECIES: hydrogenase-2 small chain [Proteobacteria] Length: 372\nNumber of predicted TMHs: 1\nExp number of AAs in TMHs: 23.72226\nExp number, first 60 AAs: 0.22038\nTotal prob of N-in: 0.12994\noutside 1 331\nTMhelix 332 354\ninside 355 372

26083 GCF_001255035.1_12971_8_26 Shigella sonnei Proteobacteria; Gammaproteobacteria; Enterobacterales; Enterobacteriaceae; Shigella MTGDNTLIHSHGINRRDFMKLCAALAATMGLSSKAAAEMAESVTNPQRPPVIWIGAQECT WP_000145410.1 MULTISPECIES: hydrogenase-2 small chain [Proteobacteria] Length: 372\nNumber of predicted TMHs: 1\nExp number of AAs in TMHs: 23.72226\nExp number, first 60 AAs: 0.22038\nTotal prob of N-in: 0.12994\noutside 1 331\nTMhelix 332 354\ninside 355 372

26084 GCF_001255075.1_12971_8_63 Shigella sonnei Proteobacteria; Gammaproteobacteria; Enterobacterales; Enterobacteriaceae; Shigella MTGDNTLIHSHGINRRDFMKLCAALAATMGLSSKAAAEMAESVTNPQRPPVIWIGAQECT WP_000145410.1 MULTISPECIES: hydrogenase-2 small chain [Proteobacteria] Length: 372\nNumber of predicted TMHs: 1\nExp number of AAs in TMHs: 23.72226\nExp number, first 60 AAs: 0.22038\nTotal prob of N-in: 0.12994\noutside 1 331\nTMhelix 332 354\ninside 355 372

26085 GCF_001066285.1_ASM106628v1 Shigella sonnei Proteobacteria; Gammaproteobacteria; Enterobacterales; Enterobacteriaceae; Shigella MNKRRTTLTVLALIASLGLSSAPALADKGGNGNGNGNGHGNSGNHGNSGNHGNNGNKGNK WP_033646351.1 MULTISPECIES: membrane protein [Enterobacterales] Length: 161\nNumber of predicted TMHs: 2\nExp number of AAs in TMHs: 31.11104\nExp number, first 60 AAs: 14.67675\nTotal prob of N-in: 0.95023\nPOSSIBLE N-term signal sequence\ninside 1 6\nTMhelix 7 26\noutside 27 141\nTMhelix 142 160\ninside 161 161

26086 GCF_001246975.1_9870_6_25 Shigella sonnei Proteobacteria; Gammaproteobacteria; Enterobacterales; Enterobacteriaceae; Shigella MTGDNTLIHSHGINRRDFMKLCAALAATMGLSSKAAAEMAESVTNPQRPPVIWIGAQECT WP_000145410.1 MULTISPECIES: hydrogenase-2 small chain [Proteobacteria] Length: 372\nNumber of predicted TMHs: 1\nExp number of AAs in TMHs: 23.72226\nExp number, first 60 AAs: 0.22038\nTotal prob of N-in: 0.12994\noutside 1 331\nTMhelix 332 354\ninside 355 372

26087 GCF_001255115.1_9870_6_64 Shigella sonnei Proteobacteria; Gammaproteobacteria; Enterobacterales; Enterobacteriaceae; Shigella MTGDNTLIHSHGINRRDFMKLCAALAATMGLSSKAAAEMAESVTNPQRPPVIWIGAQECT WP_000145410.1 MULTISPECIES: hydrogenase-2 small chain [Proteobacteria] Length: 372\nNumber of predicted TMHs: 1\nExp number of AAs in TMHs: 23.72226\nExp number, first 60 AAs: 0.22038\nTotal prob of N-in: 0.12994\noutside 1 331\nTMhelix 332 354\ninside 355 372

26088 GCF_900159035.1_16703_8_9 Shigella sonnei Proteobacteria; Gammaproteobacteria; Enterobacterales; Enterobacteriaceae; Shigella MTGDNTLIHSHGINRRDFMKLCAALAATMGLSSKAAAEMAESVTNPQRPPVIWIGAQECT WP_000145410.1 MULTISPECIES: hydrogenase-2 small chain [Proteobacteria] Length: 372\nNumber of predicted TMHs: 1\nExp number of AAs in TMHs: 23.72226\nExp number, first 60 AAs: 0.22038\nTotal prob of N-in: 0.12994\noutside 1 331\nTMhelix 332 354\ninside 355 372

26089 GCF_900159055.1_16703_8_11 Shigella sonnei Proteobacteria; Gammaproteobacteria; Enterobacterales; Enterobacteriaceae; Shigella MTGDNTLIHSHGINRRDFMKLCAALAATMGLSSKAAAEMAESVTNPQRPPVIWIGAQECT WP_000145410.1 MULTISPECIES: hydrogenase-2 small chain [Proteobacteria] Length: 372\nNumber of predicted TMHs: 1\nExp number of AAs in TMHs: 23.72226\nExp number, first 60 AAs: 0.22038\nTotal prob of N-in: 0.12994\noutside 1 331\nTMhelix 332 354\ninside 355 372

26090 GCF_900159055.1_16703_8_11 Shigella sonnei Proteobacteria; Gammaproteobacteria; Enterobacterales; Enterobacteriaceae; Shigella

MNNEETFYQAMRRQGVTRRSFLKYCSLAATSLGLGAGMAPKIAWALENKPRIPVVWIHGL WP_005139779.1 hydrogenase [Shigella sonnei] Length: 372\nNumber of predicted TMHs: 2\nExp number of AAs in TMHs: 37.02628\nExp number, first 60 AAs: 19.39684\nTotal prob of N-in: 0.94313\nPOSSIBLE N-term signal sequence\ninside 1 19\nTMhelix 20 39\noutside 40 325\nTMhelix 326 348\ninside 349 372

26091 GCF_900158945.1_9803_4_95 Shigella sonnei Proteobacteria; Gammaproteobacteria; Enterobacterales; Enterobacteriaceae; Shigella MNNEETFYQAMRRQGVTRRSFLKYCSLAATSLGLGAGMAPKIAWALENKPRIPVVWIHGL WP_005139779.1 hydrogenase [Shigella sonnei] Length: 372\nNumber of predicted TMHs: 2\nExp number of AAs in TMHs: 37.02628\nExp number, first 60 AAs: 19.39684\nTotal prob of N-in: 0.94313\nPOSSIBLE N-term signal sequence\ninside 1 19\nTMhelix 20 39\noutside 40 325\nTMhelix 326 348\ninside 349 372

26092 GCF_900158955.1_9803_4_96 Shigella sonnei Proteobacteria; Gammaproteobacteria; Enterobacterales; Enterobacteriaceae; Shigella MNNEETFYQAMRRQGVTRRSFLKYCSLAATSLGLGAGMAPKIAWALENKPRIPVVWIHGL WP_005139779.1 hydrogenase [Shigella sonnei] Length: 372\nNumber of predicted TMHs: 2\nExp number of AAs in TMHs: 37.02628\nExp number, first 60 AAs: 19.39684\nTotal prob of N-in: 0.94313\nPOSSIBLE N-term signal sequence\ninside 1 19\nTMhelix 20 39\noutside 40 325\nTMhelix 326 348\ninside 349 372

26093 GCF_900159115.1_16703_8_18 Shigella sonnei Proteobacteria; Gammaproteobacteria; Enterobacterales; Enterobacteriaceae; Shigella MTGDNTLIHSHGINRRDFMKLCAALAATMGLSSKAAAEMAESVTNPQRPPVIWIGAQECT WP_000145410.1 MULTISPECIES: hydrogenase-2 small chain [Proteobacteria] Length: 372\nNumber of predicted TMHs: 1\nExp number of AAs in TMHs: 23.72226\nExp number, first 60 AAs: 0.22038\nTotal prob of N-in: 0.12994\noutside 1 331\nTMhelix 332 354\ninside 355 372

26094 GCF_900159095.1_16703_8_15 Shigella sonnei Proteobacteria; Gammaproteobacteria; Enterobacterales; Enterobacteriaceae; Shigella MTGDNTLIHSHGINRRDFMKLCAALAATMGLSSKAAAEMAESVTNPQRPPVIWIGAQECT WP_000145410.1 MULTISPECIES: hydrogenase-2 small chain [Proteobacteria] Length: 372\nNumber of predicted TMHs: 1\nExp number of AAs in TMHs: 23.72226\nExp number, first 60 AAs: 0.22038\nTotal prob of N-in: 0.12994\noutside 1 331\nTMhelix 332 354\ninside 355 372

26095 GCF_900159125.1_16703_8_19 Shigella sonnei Proteobacteria; Gammaproteobacteria; Enterobacterales; Enterobacteriaceae; Shigella MTGDNTLIHSHGINRRDFMKLCAALAATMGLSSKAAAEMAESVTNPQRPPVIWIGAQECT WP_000145410.1 MULTISPECIES: hydrogenase-2 small chain [Proteobacteria] Length: 372\nNumber of predicted TMHs: 1\nExp number of AAs in TMHs: 23.72226\nExp number, first 60 AAs: 0.22038\nTotal prob of N-in: 0.12994\noutside 1 331\nTMhelix 332 354\ninside 355 372

26096 GCF_900159145.1_16703_8_21 Shigella sonnei Proteobacteria; Gammaproteobacteria; Enterobacterales; Enterobacteriaceae; Shigella MTGDNTLIHSHGINRRDFMKLCAALAATMGLSSKAAAEMAESVTNPQRPPVIWIGAQECT WP_000145410.1 MULTISPECIES: hydrogenase-2 small chain [Proteobacteria] Length: 372\nNumber of predicted TMHs: 1\nExp number of AAs in TMHs: 23.72226\nExp number, first 60 AAs: 0.22038\nTotal prob of N-in: 0.12994\noutside 1 331\nTMhelix 332 354\ninside 355 372

26097 GCF_900159115.1_16703_8_18 Shigella sonnei Proteobacteria; Gammaproteobacteria; Enterobacterales; Enterobacteriaceae; Shigella MNNEETFYQAMRRQGVTRRSFLKYCSLAATSLGLGAGMAPKIAWALENKPRIPVVWIHGL WP_005139779.1 hydrogenase [Shigella sonnei] Length: 372\nNumber of predicted TMHs: 2\nExp number of AAs in TMHs: 37.02628\nExp number, first 60 AAs: 19.39684\nTotal prob of N-in: 0.94313\nPOSSIBLE N-term signal sequence\ninside 1 19\nTMhelix 20 39\noutside 40 325\nTMhelix 326 348\ninside 349 372

26098 GCF_900159095.1_16703_8_15 Shigella sonnei Proteobacteria; Gammaproteobacteria; Enterobacterales; Enterobacteriaceae; Shigella MNNEETFYQAMRRQGVTRRSFLKYCSLAATSLGLGAGMAPKIAWALENKPRIPVVWIHGL WP_005139779.1 hydrogenase [Shigella sonnei] Length: 372\nNumber of predicted TMHs: 2\nExp number of AAs in TMHs: 37.02628\nExp number, first 60 AAs: 19.39684\nTotal prob of N-in: 0.94313\nPOSSIBLE N-term signal sequence\ninside 1 19\nTMhelix 20 39\noutside 40 325\nTMhelix 326 348\ninside 349 372

26099 GCF_900158995.1_16703_8_5 Shigella sonnei Proteobacteria; Gammaproteobacteria; Enterobacterales; Enterobacteriaceae; Shigella
MNNEETFYQAMRRQGVTRRSFLKYCSLAATSLGLGAGMAPKIAWALENKPRIPVVWIHGL WP_005139779.1
hydrogenase [Shigella sonnei] Length: 372\nNumber of predicted TMHs: 2\nExp number of AAs in TMHs: 37.02628\nExp number, first 60 AAs: 19.39684\nTotal prob of N-in: 0.94313\nPOSSIBLE N-term signal sequence\ninside 1 19\nTMhelix 20 39\noutside 40 325\nTMhelix 326 348\ninside 349 372

26100 GCF_900158975.1_16703_8_2 Shigella sonnei Proteobacteria; Gammaproteobacteria; Enterobacterales; Enterobacteriaceae; Shigella
MNNEETFYQAMRRQGVTRRSFLKYCSLAATSLGLGAGMAPKIAWALENKPRIPVVWIHGL WP_005139779.1
hydrogenase [Shigella sonnei] Length: 372\nNumber of predicted TMHs: 2\nExp number of AAs in TMHs: 37.02628\nExp number, first 60 AAs: 19.39684\nTotal prob of N-in: 0.94313\nPOSSIBLE N-term signal sequence\ninside 1 19\nTMhelix 20 39\noutside 40 325\nTMhelix 326 348\ninside 349 372

26101 GCF_900159005.1_16703_8_6 Shigella sonnei Proteobacteria; Gammaproteobacteria; Enterobacterales; Enterobacteriaceae; Shigella
MNNEETFYQAMRRQGVTRRSFLKYCSLAATSLGLGAGMAPKIAWALENKPRIPVVWIHGL WP_005139779.1
hydrogenase [Shigella sonnei] Length: 372\nNumber of predicted TMHs: 2\nExp number of AAs in TMHs: 37.02628\nExp number, first 60 AAs: 19.39684\nTotal prob of N-in: 0.94313\nPOSSIBLE N-term signal sequence\ninside 1 19\nTMhelix 20 39\noutside 40 325\nTMhelix 326 348\ninside 349 372

26102 GCF_900159155.1_16703_8_22 Shigella sonnei Proteobacteria; Gammaproteobacteria; Enterobacterales; Enterobacteriaceae; Shigella
MTGDNTLIHSHGINRRDFMKLCAALAATMGLSSKAAAEMAESVTNPQRPPVIWIGAQECT WP_000145410.1
MULTISPECIES: hydrogenase-2 small chain [Proteobacteria] Length: 372\nNumber of predicted TMHs: 1\nExp number of AAs in TMHs: 23.72226\nExp number, first 60 AAs: 0.22038\nTotal prob of N-in: 0.12994\noutside 1 331\nTMhelix 332 354\ninside 355 372

26103 GCF_900159155.1_16703_8_22 Shigella sonnei Proteobacteria; Gammaproteobacteria; Enterobacterales; Enterobacteriaceae; Shigella
MNNEETFYQAMRRQGVTRRSFLKYCSLAATSLGLGAGMAPKIAWALENKPRIPVVWIHGL WP_005139779.1
hydrogenase [Shigella sonnei] Length: 372\nNumber of predicted TMHs: 2\nExp number of AAs in TMHs: 37.02628\nExp number, first 60 AAs: 19.39684\nTotal prob of N-in: 0.94313\nPOSSIBLE N-term signal sequence\ninside 1 19\nTMhelix 20 39\noutside 40 325\nTMhelix 326 348\ninside 349 372

26104 GCF_900159025.1_16703_8_8 Shigella sonnei Proteobacteria; Gammaproteobacteria; Enterobacterales; Enterobacteriaceae; Shigella
MNNEETFYQAMRRQGVTRRSFLKYCSLAATSLGLGAGMAPKIAWALENKPRIPVVWIHGL WP_005139779.1
hydrogenase [Shigella sonnei] Length: 372\nNumber of predicted TMHs: 2\nExp number of AAs in TMHs: 37.02628\nExp number, first 60 AAs: 19.39684\nTotal prob of N-in: 0.94313\nPOSSIBLE N-term signal sequence\ninside 1 19\nTMhelix 20 39\noutside 40 325\nTMhelix 326 348\ninside 349 372

26105 GCF_900159035.1_16703_8_9 Shigella sonnei Proteobacteria; Gammaproteobacteria; Enterobacterales; Enterobacteriaceae; Shigella
MNNEETFYQAMRRQGVTRRSFLKYCSLAATSLGLGAGMAPKIAWALENKPRIPVVWIHGL WP_005139779.1
hydrogenase [Shigella sonnei] Length: 372\nNumber of predicted TMHs: 2\nExp number of AAs in TMHs: 37.02628\nExp number, first 60 AAs: 19.39684\nTotal prob of N-in: 0.94313\nPOSSIBLE N-term signal sequence\ninside 1 19\nTMhelix 20 39\noutside 40 325\nTMhelix 326 348\ninside 349 372

26106 GCF_900159175.1_16703_8_24 Shigella sonnei Proteobacteria; Gammaproteobacteria; Enterobacterales; Enterobacteriaceae; Shigella
MTGDNTLIHSHGINRRDFMKLCAALAATMGLSSKAAAEMAESVTNPQRPPVIWIGAQECT WP_000145410.1
MULTISPECIES: hydrogenase-2 small chain [Proteobacteria] Length: 372\nNumber of predicted TMHs: 1\nExp number of AAs in TMHs: 23.72226\nExp number, first 60 AAs: 0.22038\nTotal prob of N-in: 0.12994\noutside 1 331\nTMhelix 332 354\ninside 355 372

26107 GCF_900159205.1_16703_8_27 Shigella sonnei Proteobacteria; Gammaproteobacteria; Enterobacterales; Enterobacteriaceae; Shigella
MTGDNTLIHSHGINRRDFMKLCAALAATMGLSSKAAAEMAESVTNPQRPPVIWIGAQECT WP_000145410.1
MULTISPECIES: hydrogenase-2 small chain [Proteobacteria] Length: 372\nNumber of predicted TMHs:

1\nExp number of AAs in TMHs: 23.72226\nExp number, first 60 AAs: 0.22038\nTotal prob of N-in: 0.12994\noutside 1 331\nTMhelix 332 354\ninside 355 372

26108 GCF_900159225.1_16703_8_29 Shigella sonnei Proteobacteria; Gammaproteobacteria; Enterobacterales; Enterobacteriaceae; Shigella
MTGDNTLIHSHGINRRDFMKLCAALAATMGLSSKAAAEMAESVTNPQRPPVIWIGAQECT WP_000145410.1
MULTISPECIES: hydrogenase-2 small chain [Proteobacteria] Length: 372\nNumber of predicted TMHs: 1\nExp number of AAs in TMHs: 23.72226\nExp number, first 60 AAs: 0.22038\nTotal prob of N-in: 0.12994\noutside 1 331\nTMhelix 332 354\ninside 355 372

26109 GCF_900159195.1_16703_8_26 Shigella sonnei Proteobacteria; Gammaproteobacteria; Enterobacterales; Enterobacteriaceae; Shigella
MTGDNTLIHSHGINRRDFMKLCAALAATMGLSSKAAAEMAESVTNPQRPPVIWIGAQECT WP_000145410.1
MULTISPECIES: hydrogenase-2 small chain [Proteobacteria] Length: 372\nNumber of predicted TMHs: 1\nExp number of AAs in TMHs: 23.72226\nExp number, first 60 AAs: 0.22038\nTotal prob of N-in: 0.12994\noutside 1 331\nTMhelix 332 354\ninside 355 372

26110 GCF_900159235.1_16703_8_30 Shigella sonnei Proteobacteria; Gammaproteobacteria; Enterobacterales; Enterobacteriaceae; Shigella
MTGDNTLIHSHGINRRDFMKLCAALAATMGLSSKAAAEMAESVTNPQRPPVIWIGAQECT WP_000145410.1
MULTISPECIES: hydrogenase-2 small chain [Proteobacteria] Length: 372\nNumber of predicted TMHs: 1\nExp number of AAs in TMHs: 23.72226\nExp number, first 60 AAs: 0.22038\nTotal prob of N-in: 0.12994\noutside 1 331\nTMhelix 332 354\ninside 355 372

26111 GCF_900159175.1_16703_8_24 Shigella sonnei Proteobacteria; Gammaproteobacteria; Enterobacterales; Enterobacteriaceae; Shigella
MNNEETFYQAMRRQGVTRRSFLKYCSLAATSLGLGAGMAPKIAWALENKPRIPVVWIHGL WP_005139779.1
hydrogenase [Shigella sonnei] Length: 372\nNumber of predicted TMHs: 2\nExp number of AAs in TMHs: 37.02628\nExp number, first 60 AAs: 19.39684\nTotal prob of N-in: 0.94313\nPOSSIBLE N-term signal sequence\ninside 1 19\nTMhelix 20 39\noutside 40 325\nTMhelix 326 348\ninside 349 372

26112 GCF_900159125.1_16703_8_19 Shigella sonnei Proteobacteria; Gammaproteobacteria; Enterobacterales; Enterobacteriaceae; Shigella
MNNEETFYQAMRRQGVTRRSFLKYCSLAATSLGLGAGMAPKIAWALENKPRIPVVWIHGL WP_005139779.1
hydrogenase [Shigella sonnei] Length: 372\nNumber of predicted TMHs: 2\nExp number of AAs in TMHs: 37.02628\nExp number, first 60 AAs: 19.39684\nTotal prob of N-in: 0.94313\nPOSSIBLE N-term signal sequence\ninside 1 19\nTMhelix 20 39\noutside 40 325\nTMhelix 326 348\ninside 349 372

26113 GCF_900159255.1_16703_8_32 Shigella sonnei Proteobacteria; Gammaproteobacteria; Enterobacterales; Enterobacteriaceae; Shigella
MTGDNTLIHSHGINRRDFMKLCAALAATMGLSSKAAAEMAESVTNPQRPPVIWIGAQECT WP_000145410.1
MULTISPECIES: hydrogenase-2 small chain [Proteobacteria] Length: 372\nNumber of predicted TMHs: 1\nExp number of AAs in TMHs: 23.72226\nExp number, first 60 AAs: 0.22038\nTotal prob of N-in: 0.12994\noutside 1 331\nTMhelix 332 354\ninside 355 372

26114 GCF_900159255.1_16703_8_32 Shigella sonnei Proteobacteria; Gammaproteobacteria; Enterobacterales; Enterobacteriaceae; Shigella
MNNEETFYQAMRRQGVTRRSFLKYCSLAATSLGLGAGMAPKIAWALENKPRIPVVWIHGL WP_005139779.1
hydrogenase [Shigella sonnei] Length: 372\nNumber of predicted TMHs: 2\nExp number of AAs in TMHs: 37.02628\nExp number, first 60 AAs: 19.39684\nTotal prob of N-in: 0.94313\nPOSSIBLE N-term signal sequence\ninside 1 19\nTMhelix 20 39\noutside 40 325\nTMhelix 326 348\ninside 349 372

26115 GCF_900159145.1_16703_8_21 Shigella sonnei Proteobacteria; Gammaproteobacteria; Enterobacterales; Enterobacteriaceae; Shigella
MNNEETFYQAMRRQGVTRRSFLKYCSLAATSLGLGAGMAPKIAWALENKPRIPVVWIHGL WP_005139779.1
hydrogenase [Shigella sonnei] Length: 372\nNumber of predicted TMHs: 2\nExp number of AAs in TMHs: 37.02628\nExp number, first 60 AAs: 19.39684\nTotal prob of N-in: 0.94313\nPOSSIBLE N-term signal sequence\ninside 1 19\nTMhelix 20 39\noutside 40 325\nTMhelix 326 348\ninside 349 372

26116 GCF_900159275.1_16703_8_34 Shigella sonnei Proteobacteria; Gammaproteobacteria; Enterobacterales; Enterobacteriaceae; Shigella

MTGDNTLIHSHGINRRDFMKLCAALAATMGLSSKAAAEMAESVTNPQRPPVIWIGAQECT WP_000145410.1 MULTISPECIES: hydrogenase-2 small chain [Proteobacteria] Length: 372\nNumber of predicted TMHs: 1\nExp number of AAs in TMHs: 23.72226\nExp number, first 60 AAs: 0.22038\nTotal prob of N-in: 0.12994\noutside 1 331\nTMhelix 332 354\ninside 355 372

26117 GCF_900159285.1_16703_8_35 Shigella sonnei Proteobacteria; Gammaproteobacteria; Enterobacterales; Enterobacteriaceae; Shigella MTGDNTLIHSHGINRRDFMKLCAALAATMGLSSKAAAEMAESVTNPQRPPVIWIGAQECT WP_000145410.1 MULTISPECIES: hydrogenase-2 small chain [Proteobacteria] Length: 372\nNumber of predicted TMHs: 1\nExp number of AAs in TMHs: 23.72226\nExp number, first 60 AAs: 0.22038\nTotal prob of N-in: 0.12994\noutside 1 331\nTMhelix 332 354\ninside 355 372

26118 GCF_900159315.1_16703_8_39 Shigella sonnei Proteobacteria; Gammaproteobacteria; Enterobacterales; Enterobacteriaceae; Shigella MTGDNTLIHSHGINRRDFMKLCAALAATMGLSSKAAAEMAESVTNPQRPPVIWIGAQECT WP_000145410.1 MULTISPECIES: hydrogenase-2 small chain [Proteobacteria] Length: 372\nNumber of predicted TMHs: 1\nExp number of AAs in TMHs: 23.72226\nExp number, first 60 AAs: 0.22038\nTotal prob of N-in: 0.12994\noutside 1 331\nTMhelix 332 354\ninside 355 372

26119 GCF_900159305.1_16703_8_38 Shigella sonnei Proteobacteria; Gammaproteobacteria; Enterobacterales; Enterobacteriaceae; Shigella MTGDNTLIHSHGINRRDFMKLCAALAATMGLSSKAAAEMAESVTNPQRPPVIWIGAQECT WP_000145410.1 MULTISPECIES: hydrogenase-2 small chain [Proteobacteria] Length: 372\nNumber of predicted TMHs: 1\nExp number of AAs in TMHs: 23.72226\nExp number, first 60 AAs: 0.22038\nTotal prob of N-in: 0.12994\noutside 1 331\nTMhelix 332 354\ninside 355 372

26120 GCF_900159275.1_16703_8_34 Shigella sonnei Proteobacteria; Gammaproteobacteria; Enterobacterales; Enterobacteriaceae; Shigella MNNEETFYQAMRRQGVTRRSFLKYCSLAATSLGLGAGMAPKIAWALENKPRIPVVWIHGL WP_005139779.1 hydrogenase [Shigella sonnei] Length: 372\nNumber of predicted TMHs: 2\nExp number of AAs in TMHs: 37.02628\nExp number, first 60 AAs: 19.39684\nTotal prob of N-in: 0.94313\nPOSSIBLE N-term signal sequence\ninside 1 19\nTMhelix 20 39\noutside 40 325\nTMhelix 326 348\ninside 349 372

26121 GCF_900159305.1_16703_8_38 Shigella sonnei Proteobacteria; Gammaproteobacteria; Enterobacterales; Enterobacteriaceae; Shigella MNNEETFYQAMRRQGVTRRSFLKYCSLAATSLGLGAGMAPKIAWALENKPRIPVVWIHGL WP_005139779.1 hydrogenase [Shigella sonnei] Length: 372\nNumber of predicted TMHs: 2\nExp number of AAs in TMHs: 37.02628\nExp number, first 60 AAs: 19.39684\nTotal prob of N-in: 0.94313\nPOSSIBLE N-term signal sequence\ninside 1 19\nTMhelix 20 39\noutside 40 325\nTMhelix 326 348\ninside 349 372

26122 GCF_900159195.1_16703_8_26 Shigella sonnei Proteobacteria; Gammaproteobacteria; Enterobacterales; Enterobacteriaceae; Shigella MNNEETFYQAMRRQGVTRRSFLKYCSLAATSLGLGAGMAPKIAWALENKPRIPVVWIHGL WP_005139779.1 hydrogenase [Shigella sonnei] Length: 372\nNumber of predicted TMHs: 2\nExp number of AAs in TMHs: 37.02628\nExp number, first 60 AAs: 19.39684\nTotal prob of N-in: 0.94313\nPOSSIBLE N-term signal sequence\ninside 1 19\nTMhelix 20 39\noutside 40 325\nTMhelix 326 348\ninside 349 372

26123 GCF_900159365.1_8489_1_59 Shigella sonnei Proteobacteria; Gammaproteobacteria; Enterobacterales; Enterobacteriaceae; Shigella MTGDNTLIHSHGINRRDFMKLCAALAATMGLSSKAAAEMAESVTNPQRPPVIWIGAQECT WP_000145410.1 MULTISPECIES: hydrogenase-2 small chain [Proteobacteria] Length: 372\nNumber of predicted TMHs: 1\nExp number of AAs in TMHs: 23.72226\nExp number, first 60 AAs: 0.22038\nTotal prob of N-in: 0.12994\noutside 1 331\nTMhelix 332 354\ninside 355 372

26124 GCF_900159205.1_16703_8_27 Shigella sonnei Proteobacteria; Gammaproteobacteria; Enterobacterales; Enterobacteriaceae; Shigella MNNEETFYQAMRRQGVTRRSFLKYCSLAATSLGLGAGMAPKIAWALENKPRIPVVWIHGL WP_005139779.1 hydrogenase [Shigella sonnei] Length: 372\nNumber of predicted TMHs: 2\nExp number of AAs in TMHs: 37.02628\nExp number, first 60 AAs: 19.39684\nTotal prob of N-in: 0.94313\nPOSSIBLE N-term signal sequence\ninside 1 19\nTMhelix 20 39\noutside 40 325\nTMhelix 326 348\ninside 349 372

26125 GCF_900159225.1_16703_8_29 Shigella sonnei Proteobacteria; Gammaproteobacteria; Enterobacterales; Enterobacteriaceae; Shigella
MNNEETFYQAMRRQGVTRRSFLKYCSLAATSLGLGAGMAPKIAWALENKPRIPVVWIHGL WP_005139779.1
hydrogenase [Shigella sonnei] Length: 372\nNumber of predicted TMHs: 2\nExp number of AAs in TMHs: 37.02628\nExp number, first 60 AAs: 19.39684\nTotal prob of N-in: 0.94313\nPOSSIBLE N-term signal sequence\ninside 1 19\nTMhelix 20 39\noutside 40 325\nTMhelix 326 348\ninside 349 372

26126 GCF_900159335.1_16703_8_45 Shigella sonnei Proteobacteria; Gammaproteobacteria; Enterobacterales; Enterobacteriaceae; Shigella
MTGDNTLIHSHGINRRDFMKLCAALAATMGLSSKAAAEMAESVTNPQRPPVIWIGAQECT WP_000145410.1
MULTISPECIES: hydrogenase-2 small chain [Proteobacteria] Length: 372\nNumber of predicted TMHs: 1\nExp number of AAs in TMHs: 23.72226\nExp number, first 60 AAs: 0.22038\nTotal prob of N-in: 0.12994\noutside 1 331\nTMhelix 332 354\ninside 355 372

26127 GCF_900159355.1_16703_8_51 Shigella sonnei Proteobacteria; Gammaproteobacteria; Enterobacterales; Enterobacteriaceae; Shigella
MTGDNTLIHSHGINRRDFMKLCAALAATMGLSSKAAAEMAESVTNPQRPPVIWIGAQECT WP_000145410.1
MULTISPECIES: hydrogenase-2 small chain [Proteobacteria] Length: 372\nNumber of predicted TMHs: 1\nExp number of AAs in TMHs: 23.72226\nExp number, first 60 AAs: 0.22038\nTotal prob of N-in: 0.12994\noutside 1 331\nTMhelix 332 354\ninside 355 372

26128 GCF_900159335.1_16703_8_45 Shigella sonnei Proteobacteria; Gammaproteobacteria; Enterobacterales; Enterobacteriaceae; Shigella
MNNEETFYQAMRRQGVTRRSFLKYCSLAATSLGLGAGMAPKIAWALENKPRIPVVWIHGL WP_005139779.1
hydrogenase [Shigella sonnei] Length: 372\nNumber of predicted TMHs: 2\nExp number of AAs in TMHs: 37.02628\nExp number, first 60 AAs: 19.39684\nTotal prob of N-in: 0.94313\nPOSSIBLE N-term signal sequence\ninside 1 19\nTMhelix 20 39\noutside 40 325\nTMhelix 326 348\ninside 349 372

26129 GCF_900159355.1_16703_8_51 Shigella sonnei Proteobacteria; Gammaproteobacteria; Enterobacterales; Enterobacteriaceae; Shigella
MNNEETFYQAMRRQGVTRRSFLKYCSLAATSLGLGAGMAPKIAWALENKPRIPVVWIHGL WP_005139779.1
hydrogenase [Shigella sonnei] Length: 372\nNumber of predicted TMHs: 2\nExp number of AAs in TMHs: 37.02628\nExp number, first 60 AAs: 19.39684\nTotal prob of N-in: 0.94313\nPOSSIBLE N-term signal sequence\ninside 1 19\nTMhelix 20 39\noutside 40 325\nTMhelix 326 348\ninside 349 372

26130 GCF_900159235.1_16703_8_30 Shigella sonnei Proteobacteria; Gammaproteobacteria; Enterobacterales; Enterobacteriaceae; Shigella
MNNEETFYQAMRRQGVTRRSFLKYCSLAATSLGLGAGMAPKIAWALENKPRIPVVWIHGL WP_005139779.1
hydrogenase [Shigella sonnei] Length: 372\nNumber of predicted TMHs: 2\nExp number of AAs in TMHs: 37.02628\nExp number, first 60 AAs: 19.39684\nTotal prob of N-in: 0.94313\nPOSSIBLE N-term signal sequence\ninside 1 19\nTMhelix 20 39\noutside 40 325\nTMhelix 326 348\ninside 349 372

26131 GCF_900159385.1_8489_1_61 Shigella sonnei Proteobacteria; Gammaproteobacteria; Enterobacterales; Enterobacteriaceae; Shigella
MTGDNTLIHSHGINRRDFMKLCAALAATMGLSSKAAAEMAESVTNPQRPPVIWIGAQECT WP_000145410.1
MULTISPECIES: hydrogenase-2 small chain [Proteobacteria] Length: 372\nNumber of predicted TMHs: 1\nExp number of AAs in TMHs: 23.72226\nExp number, first 60 AAs: 0.22038\nTotal prob of N-in: 0.12994\noutside 1 331\nTMhelix 332 354\ninside 355 372

26132 GCF_900159395.1_8489_1_62 Shigella sonnei Proteobacteria; Gammaproteobacteria; Enterobacterales; Enterobacteriaceae; Shigella
MTGDNTLIHSHGINRRDFMKLCAALAATMGLSSKAAAEMAESVTNPQRPPVIWIGAQECT WP_000145410.1
MULTISPECIES: hydrogenase-2 small chain [Proteobacteria] Length: 372\nNumber of predicted TMHs: 1\nExp number of AAs in TMHs: 23.72226\nExp number, first 60 AAs: 0.22038\nTotal prob of N-in: 0.12994\noutside 1 331\nTMhelix 332 354\ninside 355 372

26133 GCF_900159415.1_8489_1_64 Shigella sonnei Proteobacteria; Gammaproteobacteria; Enterobacterales; Enterobacteriaceae; Shigella
MTGDNTLIHSHGINRRDFMKLCAALAATMGLSSKAAAEMAESVTNPQRPPVIWIGAQECT WP_000145410.1
MULTISPECIES: hydrogenase-2 small chain [Proteobacteria] Length: 372\nNumber of predicted TMHs:

1\nExp number of AAs in TMHs: 23.72226\nExp number, first 60 AAs: 0.22038\nTotal prob of N-in: 0.12994\noutside 1 331\nTMhelix 332 354\ninside 355 372

26134 GCF_900159365.1_8489_1_59 Shigella sonnei Proteobacteria; Gammaproteobacteria; Enterobacterales; Enterobacteriaceae; Shigella
MNNEETFYQAMRRQGVTRRSFLKYCSLAATSLGLGAGMAPKIAWALENKPRIPVVWIHGL WP_005139779.1
hydrogenase [Shigella sonnei] Length: 372\nNumber of predicted TMHs: 2\nExp number of AAs in TMHs: 37.02628\nExp number, first 60 AAs: 19.39684\nTotal prob of N-in: 0.94313\nPOSSIBLE N-term signal sequence\ninside 1 19\nTMhelix 20 39\noutside 40 325\nTMhelix 326 348\ninside 349 372

26135 GCF_900159385.1_8489_1_61 Shigella sonnei Proteobacteria; Gammaproteobacteria; Enterobacterales; Enterobacteriaceae; Shigella
MNNEETFYQAMRRQGVTRRSFLKYCSLAATSLGLGAGMAPKIAWALENKPRIPVVWIHGL WP_005139779.1
hydrogenase [Shigella sonnei] Length: 372\nNumber of predicted TMHs: 2\nExp number of AAs in TMHs: 37.02628\nExp number, first 60 AAs: 19.39684\nTotal prob of N-in: 0.94313\nPOSSIBLE N-term signal sequence\ninside 1 19\nTMhelix 20 39\noutside 40 325\nTMhelix 326 348\ninside 349 372

26136 GCF_900159285.1_16703_8_35 Shigella sonnei Proteobacteria; Gammaproteobacteria; Enterobacterales; Enterobacteriaceae; Shigella
MNNEETFYQAMRRQGVTRRSFLKYCSLAATSLGLGAGMAPKIAWALENKPRIPVVWIHGL WP_005139779.1
hydrogenase [Shigella sonnei] Length: 372\nNumber of predicted TMHs: 2\nExp number of AAs in TMHs: 37.02628\nExp number, first 60 AAs: 19.39684\nTotal prob of N-in: 0.94313\nPOSSIBLE N-term signal sequence\ninside 1 19\nTMhelix 20 39\noutside 40 325\nTMhelix 326 348\ninside 349 372

26137 GCF_900159575.1_8489_1_81 Shigella sonnei Proteobacteria; Gammaproteobacteria; Enterobacterales; Enterobacteriaceae; Shigella
MTGDNTLIHSHGINRRDFMKLCAALAATMGLSSKAAAEMAESVTNPQRPPVIWIGAQECT WP_000145410.1
MULTISPECIES: hydrogenase-2 small chain [Proteobacteria] Length: 372\nNumber of predicted TMHs: 1\nExp number of AAs in TMHs: 23.72226\nExp number, first 60 AAs: 0.22038\nTotal prob of N-in: 0.12994\noutside 1 331\nTMhelix 332 354\ninside 355 372

26138 GCF_900159315.1_16703_8_39 Shigella sonnei Proteobacteria; Gammaproteobacteria; Enterobacterales; Enterobacteriaceae; Shigella
MNNEETFYQAMRRQGVTRRSFLKYCSLAATSLGLGAGMAPKIAWALENKPRIPVVWIHGL WP_005139779.1
hydrogenase [Shigella sonnei] Length: 372\nNumber of predicted TMHs: 2\nExp number of AAs in TMHs: 37.02628\nExp number, first 60 AAs: 19.39684\nTotal prob of N-in: 0.94313\nPOSSIBLE N-term signal sequence\ninside 1 19\nTMhelix 20 39\noutside 40 325\nTMhelix 326 348\ninside 349 372

26139 GCF_900159435.1_8489_1_66 Shigella sonnei Proteobacteria; Gammaproteobacteria; Enterobacterales; Enterobacteriaceae; Shigella
MTGDNTLIHSHGINRRDFMKLCAALAATMGLSSKAAAEMAESVTNPQRPPVIWIGAQECT WP_000145410.1
MULTISPECIES: hydrogenase-2 small chain [Proteobacteria] Length: 372\nNumber of predicted TMHs: 1\nExp number of AAs in TMHs: 23.72226\nExp number, first 60 AAs: 0.22038\nTotal prob of N-in: 0.12994\noutside 1 331\nTMhelix 332 354\ninside 355 372

26140 GCF_900159455.1_8489_1_68 Shigella sonnei Proteobacteria; Gammaproteobacteria; Enterobacterales; Enterobacteriaceae; Shigella
MTGDNTLIHSHGINRRDFMKLCAALAATMGLSSKAAAEMAESVTNPQRPPVIWIGAQECT WP_000145410.1
MULTISPECIES: hydrogenase-2 small chain [Proteobacteria] Length: 372\nNumber of predicted TMHs: 1\nExp number of AAs in TMHs: 23.72226\nExp number, first 60 AAs: 0.22038\nTotal prob of N-in: 0.12994\noutside 1 331\nTMhelix 332 354\ninside 355 372

26141 GCF_900159465.1_8489_1_69 Shigella sonnei Proteobacteria; Gammaproteobacteria; Enterobacterales; Enterobacteriaceae; Shigella
MTGDNTLIHSHGINRRDFMKLCAALAATMGLSSKAAAEMAESVTNPQRPPVIWIGAQECT WP_000145410.1
MULTISPECIES: hydrogenase-2 small chain [Proteobacteria] Length: 372\nNumber of predicted TMHs: 1\nExp number of AAs in TMHs: 23.72226\nExp number, first 60 AAs: 0.22038\nTotal prob of N-in: 0.12994\noutside 1 331\nTMhelix 332 354\ninside 355 372

26142 GCF_900159495.1_8489_1_74 Shigella sonnei Proteobacteria; Gammaproteobacteria; Enterobacterales; Enterobacteriaceae; Shigella

MTGDNTLIHSHGINRRDFMKLCAALAATMGLSSKAAAEMAESVTNPQRPPVIWIGAQECT WP_000145410.1 MULTISPECIES: hydrogenase-2 small chain [Proteobacteria] Length: 372\nNumber of predicted TMHs: 1\nExp number of AAs in TMHs: 23.72226\nExp number, first 60 AAs: 0.22038\nTotal prob of N-in: 0.12994\noutside 1 331\nTMhelix 332 354\ninside 355 372

26143 GCF_900159505.1_8489_1_73 Shigella sonnei Proteobacteria; Gammaproteobacteria; Enterobacterales; Enterobacteriaceae; Shigella MTGDNTLIHSHGINRRDFMKLCAALAATMGLSSKAAAEMAESVTNPQRPPVIWIGAQECT WP_000145410.1 MULTISPECIES: hydrogenase-2 small chain [Proteobacteria] Length: 372\nNumber of predicted TMHs: 1\nExp number of AAs in TMHs: 23.72226\nExp number, first 60 AAs: 0.22038\nTotal prob of N-in: 0.12994\noutside 1 331\nTMhelix 332 354\ninside 355 372

26144 GCF_900159525.1_8489_1_76 Shigella sonnei Proteobacteria; Gammaproteobacteria; Enterobacterales; Enterobacteriaceae; Shigella MTGDNTLIHSHGINRRDFMKLCAALAATMGLSSKAAAEMAESVTNPQRPPVIWIGAQECT WP_000145410.1 MULTISPECIES: hydrogenase-2 small chain [Proteobacteria] Length: 372\nNumber of predicted TMHs: 1\nExp number of AAs in TMHs: 23.72226\nExp number, first 60 AAs: 0.22038\nTotal prob of N-in: 0.12994\noutside 1 331\nTMhelix 332 354\ninside 355 372

26145 GCF_900159395.1_8489_1_62 Shigella sonnei Proteobacteria; Gammaproteobacteria; Enterobacterales; Enterobacteriaceae; Shigella MNNEETFYQAMRRQGVTRRSFLKYCSLAATSLGLGAGMAPKIAWALENKPRIPVVWIHGL WP_005139779.1 hydrogenase [Shigella sonnei] Length: 372\nNumber of predicted TMHs: 2\nExp number of AAs in TMHs: 37.02628\nExp number, first 60 AAs: 19.39684\nTotal prob of N-in: 0.94313\nPOSSIBLE N-term signal sequence\ninside 1 19\nTMhelix 20 39\noutside 40 325\nTMhelix 326 348\ninside 349 372

26146 GCF_900159415.1_8489_1_64 Shigella sonnei Proteobacteria; Gammaproteobacteria; Enterobacterales; Enterobacteriaceae; Shigella MNNEETFYQAMRRQGVTRRSFLKYCSLAATSLGLGAGMAPKIAWALENKPRIPVVWIHGL WP_005139779.1 hydrogenase [Shigella sonnei] Length: 372\nNumber of predicted TMHs: 2\nExp number of AAs in TMHs: 37.02628\nExp number, first 60 AAs: 19.39684\nTotal prob of N-in: 0.94313\nPOSSIBLE N-term signal sequence\ninside 1 19\nTMhelix 20 39\noutside 40 325\nTMhelix 326 348\ninside 349 372

26147 GCF_900159545.1_8489_1_78 Shigella sonnei Proteobacteria; Gammaproteobacteria; Enterobacterales; Enterobacteriaceae; Shigella MTGDNTLIHSHGINRRDFMKLCAALAATMGLSSKAAAEMAESVTNPQRPPVIWIGAQECT WP_000145410.1 MULTISPECIES: hydrogenase-2 small chain [Proteobacteria] Length: 372\nNumber of predicted TMHs: 1\nExp number of AAs in TMHs: 23.72226\nExp number, first 60 AAs: 0.22038\nTotal prob of N-in: 0.12994\noutside 1 331\nTMhelix 332 354\ninside 355 372

26148 GCF_900159555.1_8489_1_79 Shigella sonnei Proteobacteria; Gammaproteobacteria; Enterobacterales; Enterobacteriaceae; Shigella MTGDNTLIHSHGINRRDFMKLCAALAATMGLSSKAAAEMAESVTNPQRPPVIWIGAQECT WP_000145410.1 MULTISPECIES: hydrogenase-2 small chain [Proteobacteria] Length: 372\nNumber of predicted TMHs: 1\nExp number of AAs in TMHs: 23.72226\nExp number, first 60 AAs: 0.22038\nTotal prob of N-in: 0.12994\noutside 1 331\nTMhelix 332 354\ninside 355 372

26149 GCF_900159545.1_8489_1_78 Shigella sonnei Proteobacteria; Gammaproteobacteria; Enterobacterales; Enterobacteriaceae; Shigella MNNEETFYQAMRRQGVTRRSFLKYCSLAATSLGLGAGMAPKIAWALENKPRIPVVWIHGL WP_005139779.1 hydrogenase [Shigella sonnei] Length: 372\nNumber of predicted TMHs: 2\nExp number of AAs in TMHs: 37.02628\nExp number, first 60 AAs: 19.39684\nTotal prob of N-in: 0.94313\nPOSSIBLE N-term signal sequence\ninside 1 19\nTMhelix 20 39\noutside 40 325\nTMhelix 326 348\ninside 349 372

26150 GCF_900159555.1_8489_1_79 Shigella sonnei Proteobacteria; Gammaproteobacteria; Enterobacterales; Enterobacteriaceae; Shigella MNNEETFYQAMRRQGVTRRSFLKYCSLAATSLGLGAGMAPKIAWALENKPRIPVVWIHGL WP_005139779.1 hydrogenase [Shigella sonnei] Length: 372\nNumber of predicted TMHs: 2\nExp number of AAs in TMHs: 37.02628\nExp number, first 60 AAs: 19.39684\nTotal prob of N-in: 0.94313\nPOSSIBLE N-term signal sequence\ninside 1 19\nTMhelix 20 39\noutside 40 325\nTMhelix 326 348\ninside 349 372

26151 GCF_900159455.1_8489_1_68 Shigella sonnei Proteobacteria; Gammaproteobacteria; Enterobacterales; Enterobacteriaceae; Shigella
MNNEETFYQAMRRQGVTRRSFLKYCSLAATSLGLGAGMAPKIAWALENKPRIPVVWIHGL WP_005139779.1
hydrogenase [Shigella sonnei] Length: 372\nNumber of predicted TMHs: 2\nExp number of AAs in TMHs: 37.02628\nExp number, first 60 AAs: 19.39684\nTotal prob of N-in: 0.94313\nPOSSIBLE N-term signal sequence\ninside 1 19\nTMhelix 20 39\noutside 40 325\nTMhelix 326 348\ninside 349 372

26152 GCF_900159575.1_8489_1_81 Shigella sonnei Proteobacteria; Gammaproteobacteria; Enterobacterales; Enterobacteriaceae; Shigella
MNNEETFYQAMRRQGVTRRSFLKYCSLAATSLGLGAGMAPKIAWALENKPRIPVVWIHGL WP_005139779.1
hydrogenase [Shigella sonnei] Length: 372\nNumber of predicted TMHs: 2\nExp number of AAs in TMHs: 37.02628\nExp number, first 60 AAs: 19.39684\nTotal prob of N-in: 0.94313\nPOSSIBLE N-term signal sequence\ninside 1 19\nTMhelix 20 39\noutside 40 325\nTMhelix 326 348\ninside 349 372

26153 GCF_900159435.1_8489_1_66 Shigella sonnei Proteobacteria; Gammaproteobacteria; Enterobacterales; Enterobacteriaceae; Shigella
MNNEETFYQAMRRQGVTRRSFLKYCSLAATSLGLGAGMAPKIAWALENKPRIPVVWIHGL WP_005139779.1
hydrogenase [Shigella sonnei] Length: 372\nNumber of predicted TMHs: 2\nExp number of AAs in TMHs: 37.02628\nExp number, first 60 AAs: 19.39684\nTotal prob of N-in: 0.94313\nPOSSIBLE N-term signal sequence\ninside 1 19\nTMhelix 20 39\noutside 40 325\nTMhelix 326 348\ninside 349 372

26154 GCF_900159465.1_8489_1_69 Shigella sonnei Proteobacteria; Gammaproteobacteria; Enterobacterales; Enterobacteriaceae; Shigella
MNNEETFYQAMRRQGVTRRSFLKYCSLAATSLGLGAGMAPKIAWALENKPRIPVVWIHGL WP_005139779.1
hydrogenase [Shigella sonnei] Length: 372\nNumber of predicted TMHs: 2\nExp number of AAs in TMHs: 37.02628\nExp number, first 60 AAs: 19.39684\nTotal prob of N-in: 0.94313\nPOSSIBLE N-term signal sequence\ninside 1 19\nTMhelix 20 39\noutside 40 325\nTMhelix 326 348\ninside 349 372

26155 GCF_900159595.1_8489_1_82 Shigella sonnei Proteobacteria; Gammaproteobacteria; Enterobacterales; Enterobacteriaceae; Shigella
MTGDNTLIHSHGINRRDFMKLCAALAATMGLSSKAAAEMAESVTNPQRPPVIWIGAQECT WP_000145410.1
MULTISPECIES: hydrogenase-2 small chain [Proteobacteria] Length: 372\nNumber of predicted TMHs: 1\nExp number of AAs in TMHs: 23.72226\nExp number, first 60 AAs: 0.22038\nTotal prob of N-in: 0.12994\noutside 1 331\nTMhelix 332 354\ninside 355 372

26156 GCF_900159605.1_8489_1_84 Shigella sonnei Proteobacteria; Gammaproteobacteria; Enterobacterales; Enterobacteriaceae; Shigella
MTGDNTLIHSHGINRRDFMKLCAALAATMGLSSKAAAEMAESVTNPQRPPVIWIGAQECT WP_000145410.1
MULTISPECIES: hydrogenase-2 small chain [Proteobacteria] Length: 372\nNumber of predicted TMHs: 1\nExp number of AAs in TMHs: 23.72226\nExp number, first 60 AAs: 0.22038\nTotal prob of N-in: 0.12994\noutside 1 331\nTMhelix 332 354\ninside 355 372

26157 GCF_900159625.1_8489_1_87 Shigella sonnei Proteobacteria; Gammaproteobacteria; Enterobacterales; Enterobacteriaceae; Shigella
MTGDNTLIHSHGINRRDFMKLCAALAATMGLSSKAAAEMAESVTNPQRPPVIWIGAQECT WP_000145410.1
MULTISPECIES: hydrogenase-2 small chain [Proteobacteria] Length: 372\nNumber of predicted TMHs: 1\nExp number of AAs in TMHs: 23.72226\nExp number, first 60 AAs: 0.22038\nTotal prob of N-in: 0.12994\noutside 1 331\nTMhelix 332 354\ninside 355 372

26158 GCF_900159595.1_8489_1_82 Shigella sonnei Proteobacteria; Gammaproteobacteria; Enterobacterales; Enterobacteriaceae; Shigella
MNNEETFYQAMRRQGVTRRSFLKYCSLAATSLGLGAGMAPKIAWALENKPRIPVVWIHGL WP_005139779.1
hydrogenase [Shigella sonnei] Length: 372\nNumber of predicted TMHs: 2\nExp number of AAs in TMHs: 37.02628\nExp number, first 60 AAs: 19.39684\nTotal prob of N-in: 0.94313\nPOSSIBLE N-term signal sequence\ninside 1 19\nTMhelix 20 39\noutside 40 325\nTMhelix 326 348\ninside 349 372

26159 GCF_900159605.1_8489_1_84 Shigella sonnei Proteobacteria; Gammaproteobacteria; Enterobacterales; Enterobacteriaceae; Shigella
MNNEETFYQAMRRQGVTRRSFLKYCSLAATSLGLGAGMAPKIAWALENKPRIPVVWIHGL WP_005139779.1
hydrogenase [Shigella sonnei] Length: 372\nNumber of predicted TMHs: 2\nExp number of AAs in TMHs: 37.02628\nExp number, first 60 AAs: 19.39684\nTotal prob of N-in: 0.94313\nPOSSIBLE N-term signal sequence\ninside 1 19\nTMhelix 20 39\noutside 40 325\nTMhelix 326 348\ninside 349 372

TMHs: 37.02628\nExp number, first 60 AAs:  19.39684\nTotal prob of N-in:   0.94313\nPOSSIBLE N-term signal sequence\ninside   1   19\nTMhelix   20   39\noutside   40   325\nTMhelix   326   348\ninside   349   372

26160 GCF_900159625.1_8489_1_87 Shigella sonnei Proteobacteria; Gammaproteobacteria; Enterobacterales; Enterobacteriaceae; Shigella
MNNEETFYQAMRRQGVTRRSFLKYCSLAATSLGLGAGMAPKIAWALENKPRIPVVWIHGL WP_005139779.1
hydrogenase [Shigella sonnei] Length: 372\nNumber of predicted TMHs:  2\nExp number of AAs in TMHs: 37.02628\nExp number, first 60 AAs:  19.39684\nTotal prob of N-in:   0.94313\nPOSSIBLE N-term signal sequence\ninside   1   19\nTMhelix   20   39\noutside   40   325\nTMhelix   326   348\ninside   349   372

26161 GCF_900159525.1_8489_1_76 Shigella sonnei Proteobacteria; Gammaproteobacteria; Enterobacterales; Enterobacteriaceae; Shigella
MNNEETFYQAMRRQGVTRRSFLKYCSLAATSLGLGAGMAPKIAWALENKPRIPVVWIHGL WP_005139779.1
hydrogenase [Shigella sonnei] Length: 372\nNumber of predicted TMHs:  2\nExp number of AAs in TMHs: 37.02628\nExp number, first 60 AAs:  19.39684\nTotal prob of N-in:   0.94313\nPOSSIBLE N-term signal sequence\ninside   1   19\nTMhelix   20   39\noutside   40   325\nTMhelix   326   348\ninside   349   372

26162 GCF_900159495.1_8489_1_74 Shigella sonnei Proteobacteria; Gammaproteobacteria; Enterobacterales; Enterobacteriaceae; Shigella
MNNEETFYQAMRRQGVTRRSFLKYCSLAATSLGLGAGMAPKIAWALENKPRIPVVWIHGL WP_005139779.1
hydrogenase [Shigella sonnei] Length: 372\nNumber of predicted TMHs:  2\nExp number of AAs in TMHs: 37.02628\nExp number, first 60 AAs:  19.39684\nTotal prob of N-in:   0.94313\nPOSSIBLE N-term signal sequence\ninside   1   19\nTMhelix   20   39\noutside   40   325\nTMhelix   326   348\ninside   349   372

26163 GCF_900159645.1_8489_1_89 Shigella sonnei Proteobacteria; Gammaproteobacteria; Enterobacterales; Enterobacteriaceae; Shigella
MTGDNTLIHSHGINRRDFMKLCAALAATMGLSSKAAAEMAESVTNPQRPPVIWIGAQECT WP_000145410.1
MULTISPECIES: hydrogenase-2 small chain [Proteobacteria] Length: 372\nNumber of predicted TMHs: 1\nExp number of AAs in TMHs: 23.72226\nExp number, first 60 AAs:  0.22038\nTotal prob of N-in:   0.12994\noutside   1   331\nTMhelix   332   354\ninside   355   372

26164 GCF_900159665.1_8489_1_91 Shigella sonnei Proteobacteria; Gammaproteobacteria; Enterobacterales; Enterobacteriaceae; Shigella
MTGDNTLIHSHGINRRDFMKLCAALAATMGLSSKAAAEMAESVTNPQRPPVIWIGAQECT WP_000145410.1
MULTISPECIES: hydrogenase-2 small chain [Proteobacteria] Length: 372\nNumber of predicted TMHs: 1\nExp number of AAs in TMHs: 23.72226\nExp number, first 60 AAs:  0.22038\nTotal prob of N-in:   0.12994\noutside   1   331\nTMhelix   332   354\ninside   355   372

26165 GCF_900159665.1_8489_1_91 Shigella sonnei Proteobacteria; Gammaproteobacteria; Enterobacterales; Enterobacteriaceae; Shigella
MNNEETFYQAMRRQGVTRRSFLKYCSLAATSLGLGAGMAPKIAWALENKPRIPVVWIHGL WP_005139779.1
hydrogenase [Shigella sonnei] Length: 372\nNumber of predicted TMHs:  2\nExp number of AAs in TMHs: 37.02628\nExp number, first 60 AAs:  19.39684\nTotal prob of N-in:   0.94313\nPOSSIBLE N-term signal sequence\ninside   1   19\nTMhelix   20   39\noutside   40   325\nTMhelix   326   348\ninside   349   372

26166 GCF_900159645.1_8489_1_89 Shigella sonnei Proteobacteria; Gammaproteobacteria; Enterobacterales; Enterobacteriaceae; Shigella
MNNEETFYQAMRRQGVTRRSFLKYCSLAATSLGLGAGMAPKIAWALENKPRIPVVWIHGL WP_005139779.1
hydrogenase [Shigella sonnei] Length: 372\nNumber of predicted TMHs:  2\nExp number of AAs in TMHs: 37.02628\nExp number, first 60 AAs:  19.39684\nTotal prob of N-in:   0.94313\nPOSSIBLE N-term signal sequence\ninside   1   19\nTMhelix   20   39\noutside   40   325\nTMhelix   326   348\ninside   349   372

26167 GCF_900159505.1_8489_1_73 Shigella sonnei Proteobacteria; Gammaproteobacteria; Enterobacterales; Enterobacteriaceae; Shigella
MNNEETFYQAMRRQGVTRRSFLKYCSLAATSLGLGAGMAPKIAWALENKPRIPVVWIHGL WP_005139779.1
hydrogenase [Shigella sonnei] Length: 372\nNumber of predicted TMHs:  2\nExp number of AAs in TMHs: 37.02628\nExp number, first 60 AAs:  19.39684\nTotal prob of N-in:   0.94313\nPOSSIBLE N-term signal sequence\ninside   1   19\nTMhelix   20   39\noutside   40   325\nTMhelix   326   348\ninside   349   372

26168 GCF_900159675.1_8489_1_92 Shigella sonnei Proteobacteria; Gammaproteobacteria; Enterobacterales; Enterobacteriaceae; Shigella

MTGDNTLIHSHGINRRDFMKLCAALAATMGLSSKAAAEMAESVTNPQRPPVIWIGAQECT WP_000145410.1 MULTISPECIES: hydrogenase-2 small chain [Proteobacteria] Length: 372\nNumber of predicted TMHs: 1\nExp number of AAs in TMHs: 23.72226\nExp number, first 60 AAs: 0.22038\nTotal prob of N-in: 0.12994\noutside 1 331\nTMhelix 332 354\ninside 355 372

26169 GCF_900159695.1_8489_1_94 Shigella sonnei Proteobacteria; Gammaproteobacteria; Enterobacterales; Enterobacteriaceae; Shigella MTGDNTLIHSHGINRRDFMKLCAALAATMGLSSKAAAEMAESVTNPQRPPVIWIGAQECT WP_000145410.1 MULTISPECIES: hydrogenase-2 small chain [Proteobacteria] Length: 372\nNumber of predicted TMHs: 1\nExp number of AAs in TMHs: 23.72226\nExp number, first 60 AAs: 0.22038\nTotal prob of N-in: 0.12994\noutside 1 331\nTMhelix 332 354\ninside 355 372

26170 GCF_900159715.1_8403_8_1Shigella sonnei Proteobacteria; Gammaproteobacteria; Enterobacterales; Enterobacteriaceae; Shigella MTGDNTLIHSHGINRRDFMKLCAALAATMGLSSKAAAEMAESVTNPQRPPVIWIGAQECT WP_000145410.1 MULTISPECIES: hydrogenase-2 small chain [Proteobacteria] Length: 372\nNumber of predicted TMHs: 1\nExp number of AAs in TMHs: 23.72226\nExp number, first 60 AAs: 0.22038\nTotal prob of N-in: 0.12994\noutside 1 331\nTMhelix 332 354\ninside 355 372

26171 GCF_900159725.1_8403_8_2Shigella sonnei Proteobacteria; Gammaproteobacteria; Enterobacterales; Enterobacteriaceae; Shigella MTGDNTLIHSHGINRRDFMKLCAALAATMGLSSKAAAEMAESVTNPQRPPVIWIGAQECT WP_000145410.1 MULTISPECIES: hydrogenase-2 small chain [Proteobacteria] Length: 372\nNumber of predicted TMHs: 1\nExp number of AAs in TMHs: 23.72226\nExp number, first 60 AAs: 0.22038\nTotal prob of N-in: 0.12994\noutside 1 331\nTMhelix 332 354\ninside 355 372

26172 GCF_900159745.1_8403_8_4Shigella sonnei Proteobacteria; Gammaproteobacteria; Enterobacterales; Enterobacteriaceae; Shigella MTGDNTLIHSHGINRRDFMKLCAALAATMGLSSKAAAEMAESVTNPQRPPVIWIGAQECT WP_000145410.1 MULTISPECIES: hydrogenase-2 small chain [Proteobacteria] Length: 372\nNumber of predicted TMHs: 1\nExp number of AAs in TMHs: 23.72226\nExp number, first 60 AAs: 0.22038\nTotal prob of N-in: 0.12994\noutside 1 331\nTMhelix 332 354\ninside 355 372

26173 GCF_900159765.1_8403_8_6Shigella sonnei Proteobacteria; Gammaproteobacteria; Enterobacterales; Enterobacteriaceae; Shigella MTGDNTLIHSHGINRRDFMKLCAALAATMGLSSKAAAEMAESVTNPQRPPVIWIGAQECT WP_000145410.1 MULTISPECIES: hydrogenase-2 small chain [Proteobacteria] Length: 372\nNumber of predicted TMHs: 1\nExp number of AAs in TMHs: 23.72226\nExp number, first 60 AAs: 0.22038\nTotal prob of N-in: 0.12994\noutside 1 331\nTMhelix 332 354\ninside 355 372

26174 GCF_900159775.1_8403_8_7Shigella sonnei Proteobacteria; Gammaproteobacteria; Enterobacterales; Enterobacteriaceae; Shigella MTGDNTLIHSHGINRRDFMKLCAALAATMGLSSKAAAEMAESVTNPQRPPVIWIGAQECT WP_000145410.1 MULTISPECIES: hydrogenase-2 small chain [Proteobacteria] Length: 372\nNumber of predicted TMHs: 1\nExp number of AAs in TMHs: 23.72226\nExp number, first 60 AAs: 0.22038\nTotal prob of N-in: 0.12994\noutside 1 331\nTMhelix 332 354\ninside 355 372

26175 GCF_900159675.1_8489_1_92 Shigella sonnei Proteobacteria; Gammaproteobacteria; Enterobacterales; Enterobacteriaceae; Shigella MNNEETFYQAMRRQGVTRRSFLKYCSLAATSLGLGAGMAPKIAWALENKPRIPVVWIHGL WP_005139779.1 hydrogenase [Shigella sonnei] Length: 372\nNumber of predicted TMHs: 2\nExp number of AAs in TMHs: 37.02628\nExp number, first 60 AAs: 19.39684\nTotal prob of N-in: 0.94313\nPOSSIBLE N-term signal sequence\ninside 1 19\nTMhelix 20 39\noutside 40 325\nTMhelix 326 348\ninside 349 372

26176 GCF_900159795.1_8403_8_9Shigella sonnei Proteobacteria; Gammaproteobacteria; Enterobacterales; Enterobacteriaceae; Shigella MTGDNTLIHSHGINRRDFMKLCAALAATMGLSSKAAAEMAESVTNPQRPPVIWIGAQECT WP_000145410.1 MULTISPECIES: hydrogenase-2 small chain [Proteobacteria] Length: 372\nNumber of predicted TMHs: 1\nExp number of AAs in TMHs: 23.72226\nExp number, first 60 AAs: 0.22038\nTotal prob of N-in: 0.12994\noutside 1 331\nTMhelix 332 354\ninside 355 372

26177 GCF_900159815.1_8403_8_11 Shigella sonnei Proteobacteria; Gammaproteobacteria; Enterobacterales; Enterobacteriaceae; Shigella MTGDNTLIHSHGINRRDFMKLCAALAATMGLSSKAAAEMAESVTNPQRPPVIWIGAQECT WP_000145410.1 MULTISPECIES: hydrogenase-2 small chain [Proteobacteria] Length: 372\nNumber of predicted TMHs: 1\nExp number of AAs in TMHs: 23.72226\nExp number, first 60 AAs: 0.22038\nTotal prob of N-in: 0.12994\noutside 1 331\nTMhelix 332 354\ninside 355 372

26178 GCF_900159835.1_8403_8_13 Shigella sonnei Proteobacteria; Gammaproteobacteria; Enterobacterales; Enterobacteriaceae; Shigella MTGDNTLIHSHGINRRDFMKLCAALAATMGLSSKAAAEMAESVTNPQRPPVIWIGAQECT WP_000145410.1 MULTISPECIES: hydrogenase-2 small chain [Proteobacteria] Length: 372\nNumber of predicted TMHs: 1\nExp number of AAs in TMHs: 23.72226\nExp number, first 60 AAs: 0.22038\nTotal prob of N-in: 0.12994\noutside 1 331\nTMhelix 332 354\ninside 355 372

26179 GCF_900159795.1_8403_8_9Shigella sonnei Proteobacteria; Gammaproteobacteria; Enterobacterales; Enterobacteriaceae; Shigella MNNEETFYQAMRRQGVTRRSFLKYCSLAATSLGLGAGMAPKIAWALENKPRIPVVWIHGL WP_005139779.1 hydrogenase [Shigella sonnei] Length: 372\nNumber of predicted TMHs: 2\nExp number of AAs in TMHs: 37.02628\nExp number, first 60 AAs: 19.39684\nTotal prob of N-in: 0.94313\nPOSSIBLE N-term signal sequence\ninside 1 19\nTMhelix 20 39\noutside 40 325\nTMhelix 326 348\ninside 349 372

26180 GCF_900159835.1_8403_8_13 Shigella sonnei Proteobacteria; Gammaproteobacteria; Enterobacterales; Enterobacteriaceae; Shigella MNNEETFYQAMRRQGVTRRSFLKYCSLAATSLGLGAGMAPKIAWALENKPRIPVVWIHGL WP_005139779.1 hydrogenase [Shigella sonnei] Length: 372\nNumber of predicted TMHs: 2\nExp number of AAs in TMHs: 37.02628\nExp number, first 60 AAs: 19.39684\nTotal prob of N-in: 0.94313\nPOSSIBLE N-term signal sequence\ninside 1 19\nTMhelix 20 39\noutside 40 325\nTMhelix 326 348\ninside 349 372

26181 GCF_900159815.1_8403_8_11 Shigella sonnei Proteobacteria; Gammaproteobacteria; Enterobacterales; Enterobacteriaceae; Shigella MNNEETFYQAMRRQGVTRRSFLKYCSLAATSLGLGAGMAPKIAWALENKPRIPVVWIHGL WP_005139779.1 hydrogenase [Shigella sonnei] Length: 372\nNumber of predicted TMHs: 2\nExp number of AAs in TMHs: 37.02628\nExp number, first 60 AAs: 19.39684\nTotal prob of N-in: 0.94313\nPOSSIBLE N-term signal sequence\ninside 1 19\nTMhelix 20 39\noutside 40 325\nTMhelix 326 348\ninside 349 372

26182 GCF_900159695.1_8489_1_94 Shigella sonnei Proteobacteria; Gammaproteobacteria; Enterobacterales; Enterobacteriaceae; Shigella MNNEETFYQAMRRQGVTRRSFLKYCSLAATSLGLGAGMAPKIAWALENKPRIPVVWIHGL WP_005139779.1 hydrogenase [Shigella sonnei] Length: 372\nNumber of predicted TMHs: 2\nExp number of AAs in TMHs: 37.02628\nExp number, first 60 AAs: 19.39684\nTotal prob of N-in: 0.94313\nPOSSIBLE N-term signal sequence\ninside 1 19\nTMhelix 20 39\noutside 40 325\nTMhelix 326 348\ninside 349 372

26183 GCF_900159715.1_8403_8_1Shigella sonnei Proteobacteria; Gammaproteobacteria; Enterobacterales; Enterobacteriaceae; Shigella MNNEETFYQAMRRQGVTRRSFLKYCSLAATSLGLGAGMAPKIAWALENKPRIPVVWIHGL WP_005139779.1 hydrogenase [Shigella sonnei] Length: 372\nNumber of predicted TMHs: 2\nExp number of AAs in TMHs: 37.02628\nExp number, first 60 AAs: 19.39684\nTotal prob of N-in: 0.94313\nPOSSIBLE N-term signal sequence\ninside 1 19\nTMhelix 20 39\noutside 40 325\nTMhelix 326 348\ninside 349 372

26184 GCF_900159845.1_8403_8_15 Shigella sonnei Proteobacteria; Gammaproteobacteria; Enterobacterales; Enterobacteriaceae; Shigella MTGDNTLIHSHGINRRDFMKLCAALAATMGLSSKAAAEMAESVTNPQRPPVIWIGAQECT WP_000145410.1 MULTISPECIES: hydrogenase-2 small chain [Proteobacteria] Length: 372\nNumber of predicted TMHs: 1\nExp number of AAs in TMHs: 23.72226\nExp number, first 60 AAs: 0.22038\nTotal prob of N-in: 0.12994\noutside 1 331\nTMhelix 332 354\ninside 355 372

26185 GCF_900159865.1_8403_8_16 Shigella sonnei Proteobacteria; Gammaproteobacteria; Enterobacterales; Enterobacteriaceae; Shigella MTGDNTLIHSHGINRRDFMKLCAALAATMGLSSKAAAEMAESVTNPQRPPVIWIGAQECT WP_000145410.1 MULTISPECIES: hydrogenase-2 small chain [Proteobacteria] Length: 372\nNumber of predicted TMHs: 1\nExp number of AAs in TMHs: 23.72226\nExp number, first 60 AAs: 0.22038\nTotal prob of N-in: 0.12994\noutside 1 331\nTMhelix 332 354\ninside 355 372

26186 GCF_900159845.1_8403_8_15 Shigella sonnei Proteobacteria; Gammaproteobacteria; Enterobacterales; Enterobacteriaceae; Shigella MNNEETFYQAMRRQGVTRRSFLKYCSLAATSLGLGAGMAPKIAWALENKPRIPVVWIHGL WP_005139779.1 hydrogenase [Shigella sonnei] Length: 372\nNumber of predicted TMHs: 2\nExp number of AAs in TMHs: 37.02628\nExp number, first 60 AAs: 19.39684\nTotal prob of N-in: 0.94313\nPOSSIBLE N-term signal sequence\ninside 1 19\nTMhelix 20 39\noutside 40 325\nTMhelix 326 348\ninside 349 372

26187 GCF_900159865.1_8403_8_16 Shigella sonnei Proteobacteria; Gammaproteobacteria; Enterobacterales; Enterobacteriaceae; Shigella MNNEETFYQAMRRQGVTRRSFLKYCSLAATSLGLGAGMAPKIAWALENKPRIPVVWIHGL WP_005139779.1 hydrogenase [Shigella sonnei] Length: 372\nNumber of predicted TMHs: 2\nExp number of AAs in TMHs: 37.02628\nExp number, first 60 AAs: 19.39684\nTotal prob of N-in: 0.94313\nPOSSIBLE N-term signal sequence\ninside 1 19\nTMhelix 20 39\noutside 40 325\nTMhelix 326 348\ninside 349 372

26188 GCF_900159725.1_8403_8_2Shigella sonnei Proteobacteria; Gammaproteobacteria; Enterobacterales; Enterobacteriaceae; Shigella MNNEETFYQAMRRQGVTRRSFLKYCSLAATSLGLGAGMAPKIAWALENKPRIPVVWIHGL WP_005139779.1 hydrogenase [Shigella sonnei] Length: 372\nNumber of predicted TMHs: 2\nExp number of AAs in TMHs: 37.02628\nExp number, first 60 AAs: 19.39684\nTotal prob of N-in: 0.94313\nPOSSIBLE N-term signal sequence\ninside 1 19\nTMhelix 20 39\noutside 40 325\nTMhelix 326 348\ninside 349 372

26189 GCF_900159885.1_8403_8_18 Shigella sonnei Proteobacteria; Gammaproteobacteria; Enterobacterales; Enterobacteriaceae; Shigella MTGDNTLIHSHGINRRDFMKLCAALAATMGLSSKAAAEMAESVTNPQRPPVIWIGAQECT WP_000145410.1 MULTISPECIES: hydrogenase-2 small chain [Proteobacteria] Length: 372\nNumber of predicted TMHs: 1\nExp number of AAs in TMHs: 23.72226\nExp number, first 60 AAs: 0.22038\nTotal prob of N-in: 0.12994\noutside 1 331\nTMhelix 332 354\ninside 355 372

26190 GCF_900159885.1_8403_8_18 Shigella sonnei Proteobacteria; Gammaproteobacteria; Enterobacterales; Enterobacteriaceae; Shigella MNNEETFYQAMRRQGVTRRSFLKYCSLAATSLGLGAGMAPKIAWALENKPRIPVVWIHGL WP_005139779.1 hydrogenase [Shigella sonnei] Length: 372\nNumber of predicted TMHs: 2\nExp number of AAs in TMHs: 37.02628\nExp number, first 60 AAs: 19.39684\nTotal prob of N-in: 0.94313\nPOSSIBLE N-term signal sequence\ninside 1 19\nTMhelix 20 39\noutside 40 325\nTMhelix 326 348\ninside 349 372

26191 GCF_900159765.1_8403_8_6Shigella sonnei Proteobacteria; Gammaproteobacteria; Enterobacterales; Enterobacteriaceae; Shigella MNNEETFYQAMRRQGVTRRSFLKYCSLAATSLGLGAGMAPKIAWALENKPRIPVVWIHGL WP_005139779.1 hydrogenase [Shigella sonnei] Length: 372\nNumber of predicted TMHs: 2\nExp number of AAs in TMHs: 37.02628\nExp number, first 60 AAs: 19.39684\nTotal prob of N-in: 0.94313\nPOSSIBLE N-term signal sequence\ninside 1 19\nTMhelix 20 39\noutside 40 325\nTMhelix 326 348\ninside 349 372

26192 GCF_900159745.1_8403_8_4Shigella sonnei Proteobacteria; Gammaproteobacteria; Enterobacterales; Enterobacteriaceae; Shigella MNNEETFYQAMRRQGVTRRSFLKYCSLAATSLGLGAGMAPKIAWALENKPRIPVVWIHGL WP_005139779.1 hydrogenase [Shigella sonnei] Length: 372\nNumber of predicted TMHs: 2\nExp number of AAs in TMHs: 37.02628\nExp number, first 60 AAs: 19.39684\nTotal prob of N-in: 0.94313\nPOSSIBLE N-term signal sequence\ninside 1 19\nTMhelix 20 39\noutside 40 325\nTMhelix 326 348\ninside 349 372

26193 GCF_900159775.1_8403_8_7Shigella sonnei Proteobacteria; Gammaproteobacteria; Enterobacterales; Enterobacteriaceae; Shigella MNNEETFYQAMRRQGVTRRSFLKYCSLAATSLGLGAGMAPKIAWALENKPRIPVVWIHGL WP_005139779.1 hydrogenase [Shigella sonnei] Length: 372\nNumber of predicted TMHs: 2\nExp number of AAs in TMHs: 37.02628\nExp number, first 60 AAs: 19.39684\nTotal prob of N-in: 0.94313\nPOSSIBLE N-term signal sequence\ninside 1 19\nTMhelix 20 39\noutside 40 325\nTMhelix 326 348\ninside 349 372

26194 GCF_900159915.1_8403_8_21 Shigella sonnei Proteobacteria; Gammaproteobacteria; Enterobacterales; Enterobacteriaceae; Shigella MTGDNTLIHSHGINRRDFMKLCAALAATMGLSSKAAAEMAESVTNPQRPPVIWIGAQECT WP_000145410.1 MULTISPECIES: hydrogenase-2 small chain [Proteobacteria] Length: 372\nNumber of predicted TMHs: 1\nExp number of AAs in TMHs: 23.72226\nExp number, first 60 AAs: 0.22038\nTotal prob of N-in: 0.12994\noutside 1 331\nTMhelix 332 354\ninside 355 372

26195 GCF_900159925.1_8403_8_22 Shigella sonnei Proteobacteria; Gammaproteobacteria; Enterobacterales; Enterobacteriaceae; Shigella MTGDNTLIHSHGINRRDFMKLCAALAATMGLSSKAAAEMAESVTNPQRPPVIWIGAQECT WP_000145410.1 MULTISPECIES: hydrogenase-2 small chain [Proteobacteria] Length: 372\nNumber of predicted TMHs: 1\nExp number of AAs in TMHs: 23.72226\nExp number, first 60 AAs: 0.22038\nTotal prob of N-in: 0.12994\noutside 1 331\nTMhelix 332 354\ninside 355 372

26196 GCF_900159945.1_8403_8_24 Shigella sonnei Proteobacteria; Gammaproteobacteria; Enterobacterales; Enterobacteriaceae; Shigella

MTGDNTLIHSHGINRRDFMKLCAALAATMGLSSKAAAEMAESVTNPQRPPVIWIGAQECT WP_000145410.1 MULTISPECIES: hydrogenase-2 small chain [Proteobacteria] Length: 372\nNumber of predicted TMHs: 1\nExp number of AAs in TMHs: 23.72226\nExp number, first 60 AAs: 0.22038\nTotal prob of N-in: 0.12994\noutside 1 331\nTMhelix 332 354\ninside 355 372

26197 GCF_900159965.1_8403_8_26 Shigella sonnei Proteobacteria; Gammaproteobacteria; Enterobacterales; Enterobacteriaceae; Shigella MTGDNTLIHSHGINRRDFMKLCAALAATMGLSSKAAAEMAESVTNPQRPPVIWIGAQECT WP_000145410.1 MULTISPECIES: hydrogenase-2 small chain [Proteobacteria] Length: 372\nNumber of predicted TMHs: 1\nExp number of AAs in TMHs: 23.72226\nExp number, first 60 AAs: 0.22038\nTotal prob of N-in: 0.12994\noutside 1 331\nTMhelix 332 354\ninside 355 372

26198 GCF_900159975.1_8403_8_27 Shigella sonnei Proteobacteria; Gammaproteobacteria; Enterobacterales; Enterobacteriaceae; Shigella MTGDNTLIHSHGINRRDFMKLCAALAATMGLSSKAAAEMAESVTNPQRPPVIWIGAQECT WP_000145410.1 MULTISPECIES: hydrogenase-2 small chain [Proteobacteria] Length: 372\nNumber of predicted TMHs: 1\nExp number of AAs in TMHs: 23.72226\nExp number, first 60 AAs: 0.22038\nTotal prob of N-in: 0.12994\noutside 1 331\nTMhelix 332 354\ninside 355 372

26199 GCF_900159995.1_8403_8_34 Shigella sonnei Proteobacteria; Gammaproteobacteria; Enterobacterales; Enterobacteriaceae; Shigella MTGDNTLIHSHGINRRDFMKLCAALAATMGLSSKAAAEMAESVTNPQRPPVIWIGAQECT WP_000145410.1 MULTISPECIES: hydrogenase-2 small chain [Proteobacteria] Length: 372\nNumber of predicted TMHs: 1\nExp number of AAs in TMHs: 23.72226\nExp number, first 60 AAs: 0.22038\nTotal prob of N-in: 0.12994\noutside 1 331\nTMhelix 332 354\ninside 355 372

26200 GCF_900160015.1_8403_8_30 Shigella sonnei Proteobacteria; Gammaproteobacteria; Enterobacterales; Enterobacteriaceae; Shigella MTGDNTLIHSHGINRRDFMKLCAALAATMGLSSKAAAEMAESVTNPQRPPVIWIGAQECT WP_000145410.1 MULTISPECIES: hydrogenase-2 small chain [Proteobacteria] Length: 372\nNumber of predicted TMHs: 1\nExp number of AAs in TMHs: 23.72226\nExp number, first 60 AAs: 0.22038\nTotal prob of N-in: 0.12994\noutside 1 331\nTMhelix 332 354\ninside 355 372

26201 GCF_900159915.1_8403_8_21 Shigella sonnei Proteobacteria; Gammaproteobacteria; Enterobacterales; Enterobacteriaceae; Shigella MNNEETFYQAMRRQGVTRRSFLKYCSLAATSLGLGAGMAPKIAWALENKPRIPVVWIHGL WP_005139779.1 hydrogenase [Shigella sonnei] Length: 372\nNumber of predicted TMHs: 2\nExp number of AAs in TMHs: 37.02628\nExp number, first 60 AAs: 19.39684\nTotal prob of N-in: 0.94313\nPOSSIBLE N-term signal sequence\ninside 1 19\nTMhelix 20 39\noutside 40 325\nTMhelix 326 348\ninside 349 372

26202 GCF_900160085.1_8403_8_40 Shigella sonnei Proteobacteria; Gammaproteobacteria; Enterobacterales; Enterobacteriaceae; Shigella MTGDNTLIHSHGINRRDFMKLCAALAATMGLSSKAAAEMAESVTNPQRPPVIWIGAQECT WP_000145410.1 MULTISPECIES: hydrogenase-2 small chain [Proteobacteria] Length: 372\nNumber of predicted TMHs: 1\nExp number of AAs in TMHs: 23.72226\nExp number, first 60 AAs: 0.22038\nTotal prob of N-in: 0.12994\noutside 1 331\nTMhelix 332 354\ninside 355 372

26203 GCF_900159925.1_8403_8_22 Shigella sonnei Proteobacteria; Gammaproteobacteria; Enterobacterales; Enterobacteriaceae; Shigella MNNEETFYQAMRRQGVTRRSFLKYCSLAATSLGLGAGMAPKIAWALENKPRIPVVWIHGL WP_005139779.1 hydrogenase [Shigella sonnei] Length: 372\nNumber of predicted TMHs: 2\nExp number of AAs in TMHs: 37.02628\nExp number, first 60 AAs: 19.39684\nTotal prob of N-in: 0.94313\nPOSSIBLE N-term signal sequence\ninside 1 19\nTMhelix 20 39\noutside 40 325\nTMhelix 326 348\ninside 349 372

26204 GCF_900160025.1_8403_8_33 Shigella sonnei Proteobacteria; Gammaproteobacteria; Enterobacterales; Enterobacteriaceae; Shigella MTGDNTLIHSHGINRRDFMKLCAALAATMGLSSKAAAEMAESVTNPQRPPVIWIGAQECT WP_000145410.1 MULTISPECIES: hydrogenase-2 small chain [Proteobacteria] Length: 372\nNumber of predicted TMHs: 1\nExp number of AAs in TMHs: 23.72226\nExp number, first 60 AAs: 0.22038\nTotal prob of N-in: 0.12994\noutside 1 331\nTMhelix 332 354\ninside 355 372

26205 GCF_900160045.1_8403_8_32 Shigella sonnei Proteobacteria; Gammaproteobacteria; Enterobacterales; Enterobacteriaceae; Shigella
MTGDNTLIHSHGINRRDFMKLCAALAATMGLSSKAAAEMAESVTNPQRPPVIWIGAQECT WP_000145410.1
MULTISPECIES: hydrogenase-2 small chain [Proteobacteria] Length: 372\nNumber of predicted TMHs: 1\nExp number of AAs in TMHs: 23.72226\nExp number, first 60 AAs: 0.22038\nTotal prob of N-in: 0.12994\noutside 1 331\nTMhelix 332 354\ninside 355 372

26206 GCF_900160045.1_8403_8_32 Shigella sonnei Proteobacteria; Gammaproteobacteria; Enterobacterales; Enterobacteriaceae; Shigella
MNNEETFYQAMRRQGVTRRSFLKYCSLAATSLGLGAGMAPKIAWALENKPRIPVVWIHGL WP_005139779.1
hydrogenase [Shigella sonnei] Length: 372\nNumber of predicted TMHs: 2\nExp number of AAs in TMHs: 37.02628\nExp number, first 60 AAs: 19.39684\nTotal prob of N-in: 0.94313\nPOSSIBLE N-term signal sequence\ninside 1 19\nTMhelix 20 39\noutside 40 325\nTMhelix 326 348\ninside 349 372

26207 GCF_900160025.1_8403_8_33 Shigella sonnei Proteobacteria; Gammaproteobacteria; Enterobacterales; Enterobacteriaceae; Shigella
MNNEETFYQAMRRQGVTRRSFLKYCSLAATSLGLGAGMAPKIAWALENKPRIPVVWIHGL WP_005139779.1
hydrogenase [Shigella sonnei] Length: 372\nNumber of predicted TMHs: 2\nExp number of AAs in TMHs: 37.02628\nExp number, first 60 AAs: 19.39684\nTotal prob of N-in: 0.94313\nPOSSIBLE N-term signal sequence\ninside 1 19\nTMhelix 20 39\noutside 40 325\nTMhelix 326 348\ninside 349 372

26208 GCF_900159945.1_8403_8_24 Shigella sonnei Proteobacteria; Gammaproteobacteria; Enterobacterales; Enterobacteriaceae; Shigella
MNNEETFYQAMRRQGVTRRSFLKYCSLAATSLGLGAGMAPKIAWALENKPRIPVVWIHGL WP_005139779.1
hydrogenase [Shigella sonnei] Length: 372\nNumber of predicted TMHs: 2\nExp number of AAs in TMHs: 37.02628\nExp number, first 60 AAs: 19.39684\nTotal prob of N-in: 0.94313\nPOSSIBLE N-term signal sequence\ninside 1 19\nTMhelix 20 39\noutside 40 325\nTMhelix 326 348\ninside 349 372

26209 GCF_900160065.1_8403_8_37 Shigella sonnei Proteobacteria; Gammaproteobacteria; Enterobacterales; Enterobacteriaceae; Shigella
MTGDNTLIHSHGINRRDFMKLCAALAATMGLSSKAAAEMAESVTNPQRPPVIWIGAQECT WP_000145410.1
MULTISPECIES: hydrogenase-2 small chain [Proteobacteria] Length: 372\nNumber of predicted TMHs: 1\nExp number of AAs in TMHs: 23.72226\nExp number, first 60 AAs: 0.22038\nTotal prob of N-in: 0.12994\noutside 1 331\nTMhelix 332 354\ninside 355 372

26210 GCF_900160085.1_8403_8_40 Shigella sonnei Proteobacteria; Gammaproteobacteria; Enterobacterales; Enterobacteriaceae; Shigella
MNNEETFYQAMRRQGVTRRSFLKYCSLAATSLGLGAGMAPKIAWALENKPRIPVVWIHGL WP_005139779.1
hydrogenase [Shigella sonnei] Length: 372\nNumber of predicted TMHs: 2\nExp number of AAs in TMHs: 37.02628\nExp number, first 60 AAs: 19.39684\nTotal prob of N-in: 0.94313\nPOSSIBLE N-term signal sequence\ninside 1 19\nTMhelix 20 39\noutside 40 325\nTMhelix 326 348\ninside 349 372

26211 GCF_900160065.1_8403_8_37 Shigella sonnei Proteobacteria; Gammaproteobacteria; Enterobacterales; Enterobacteriaceae; Shigella
MNNEETFYQAMRRQGVTRRSFLKYCSLAATSLGLGAGMAPKIAWALENKPRIPVVWIHGL WP_005139779.1
hydrogenase [Shigella sonnei] Length: 372\nNumber of predicted TMHs: 2\nExp number of AAs in TMHs: 37.02628\nExp number, first 60 AAs: 19.39684\nTotal prob of N-in: 0.94313\nPOSSIBLE N-term signal sequence\ninside 1 19\nTMhelix 20 39\noutside 40 325\nTMhelix 326 348\ninside 349 372

26212 GCF_900159965.1_8403_8_26 Shigella sonnei Proteobacteria; Gammaproteobacteria; Enterobacterales; Enterobacteriaceae; Shigella
MNNEETFYQAMRRQGVTRRSFLKYCSLAATSLGLGAGMAPKIAWALENKPRIPVVWIHGL WP_005139779.1
hydrogenase [Shigella sonnei] Length: 372\nNumber of predicted TMHs: 2\nExp number of AAs in TMHs: 37.02628\nExp number, first 60 AAs: 19.39684\nTotal prob of N-in: 0.94313\nPOSSIBLE N-term signal sequence\ninside 1 19\nTMhelix 20 39\noutside 40 325\nTMhelix 326 348\ninside 349 372

26213 GCF_900159975.1_8403_8_27 Shigella sonnei Proteobacteria; Gammaproteobacteria; Enterobacterales; Enterobacteriaceae; Shigella
MNNEETFYQAMRRQGVTRRSFLKYCSLAATSLGLGAGMAPKIAWALENKPRIPVVWIHGL WP_005139779.1
hydrogenase [Shigella sonnei] Length: 372\nNumber of predicted TMHs: 2\nExp number of AAs in

TMHs: 37.02628\nExp number, first 60 AAs: 19.39684\nTotal prob of N-in: 0.94313\nPOSSIBLE N-term signal sequence\ninside 1 19\nTMhelix 20 39\noutside 40 325\nTMhelix 326 348\ninside 349 372

26214 GCF_900159995.1_8403_8_34 Shigella sonnei Proteobacteria; Gammaproteobacteria; Enterobacterales; Enterobacteriaceae; Shigella
MNNEETFYQAMRRQGVTRRSFLKYCSLAATSLGLGAGMAPKIAWALENKPRIPVVWIHGL WP_005139779.1
hydrogenase [Shigella sonnei] Length: 372\nNumber of predicted TMHs: 2\nExp number of AAs in TMHs: 37.02628\nExp number, first 60 AAs: 19.39684\nTotal prob of N-in: 0.94313\nPOSSIBLE N-term signal sequence\ninside 1 19\nTMhelix 20 39\noutside 40 325\nTMhelix 326 348\ninside 349 372

26215 GCF_900160095.1_8403_8_39 Shigella sonnei Proteobacteria; Gammaproteobacteria; Enterobacterales; Enterobacteriaceae; Shigella
MTGDNTLIHSHGINRRDFMKLCAALAATMGLSSKAAAEMAESVTNPQRPPVIWIGAQECT WP_000145410.1
MULTISPECIES: hydrogenase-2 small chain [Proteobacteria] Length: 372\nNumber of predicted TMHs: 1\nExp number of AAs in TMHs: 23.72226\nExp number, first 60 AAs: 0.22038\nTotal prob of N-in: 0.12994\noutside 1 331\nTMhelix 332 354\ninside 355 372

26216 GCF_900160115.1_8403_8_42 Shigella sonnei Proteobacteria; Gammaproteobacteria; Enterobacterales; Enterobacteriaceae; Shigella
MTGDNTLIHSHGINRRDFMKLCAALAATMGLSSKAAAEMAESVTNPQRPPVIWIGAQECT WP_000145410.1
MULTISPECIES: hydrogenase-2 small chain [Proteobacteria] Length: 372\nNumber of predicted TMHs: 1\nExp number of AAs in TMHs: 23.72226\nExp number, first 60 AAs: 0.22038\nTotal prob of N-in: 0.12994\noutside 1 331\nTMhelix 332 354\ninside 355 372

26217 GCF_900160095.1_8403_8_39 Shigella sonnei Proteobacteria; Gammaproteobacteria; Enterobacterales; Enterobacteriaceae; Shigella
MNNEETFYQAMRRQGVTRRSFLKYCSLAATSLGLGAGMAPKIAWALENKPRIPVVWIHGL WP_005139779.1
hydrogenase [Shigella sonnei] Length: 372\nNumber of predicted TMHs: 2\nExp number of AAs in TMHs: 37.02628\nExp number, first 60 AAs: 19.39684\nTotal prob of N-in: 0.94313\nPOSSIBLE N-term signal sequence\ninside 1 19\nTMhelix 20 39\noutside 40 325\nTMhelix 326 348\ninside 349 372

26218 GCF_900160115.1_8403_8_42 Shigella sonnei Proteobacteria; Gammaproteobacteria; Enterobacterales; Enterobacteriaceae; Shigella
MNNEETFYQAMRRQGVTRRSFLKYCSLAATSLGLGAGMAPKIAWALENKPRIPVVWIHGL WP_005139779.1
hydrogenase [Shigella sonnei] Length: 372\nNumber of predicted TMHs: 2\nExp number of AAs in TMHs: 37.02628\nExp number, first 60 AAs: 19.39684\nTotal prob of N-in: 0.94313\nPOSSIBLE N-term signal sequence\ninside 1 19\nTMhelix 20 39\noutside 40 325\nTMhelix 326 348\ninside 349 372

26219 GCF_900160015.1_8403_8_30 Shigella sonnei Proteobacteria; Gammaproteobacteria; Enterobacterales; Enterobacteriaceae; Shigella
MNNEETFYQAMRRQGVTRRSFLKYCSLAATSLGLGAGMAPKIAWALENKPRIPVVWIHGL WP_005139779.1
hydrogenase [Shigella sonnei] Length: 372\nNumber of predicted TMHs: 2\nExp number of AAs in TMHs: 37.02628\nExp number, first 60 AAs: 19.39684\nTotal prob of N-in: 0.94313\nPOSSIBLE N-term signal sequence\ninside 1 19\nTMhelix 20 39\noutside 40 325\nTMhelix 326 348\ninside 349 372

26220 GCF_900160135.1_8403_8_44 Shigella sonnei Proteobacteria; Gammaproteobacteria; Enterobacterales; Enterobacteriaceae; Shigella
MTGDNTLIHSHGINRRDFMKLCAALAATMGLSSKAAAEMAESVTNPQRPPVIWIGAQECT WP_000145410.1
MULTISPECIES: hydrogenase-2 small chain [Proteobacteria] Length: 372\nNumber of predicted TMHs: 1\nExp number of AAs in TMHs: 23.72226\nExp number, first 60 AAs: 0.22038\nTotal prob of N-in: 0.12994\noutside 1 331\nTMhelix 332 354\ninside 355 372

26221 GCF_900160145.1_8403_8_45 Shigella sonnei Proteobacteria; Gammaproteobacteria; Enterobacterales; Enterobacteriaceae; Shigella
MTGDNTLIHSHGINRRDFMKLCAALAATMGLSSKAAAEMAESVTNPQRPPVIWIGAQECT WP_000145410.1
MULTISPECIES: hydrogenase-2 small chain [Proteobacteria] Length: 372\nNumber of predicted TMHs: 1\nExp number of AAs in TMHs: 23.72226\nExp number, first 60 AAs: 0.22038\nTotal prob of N-in: 0.12994\noutside 1 331\nTMhelix 332 354\ninside 355 372

26222 GCF_900160165.1_8403_8_48 Shigella sonnei Proteobacteria; Gammaproteobacteria; Enterobacterales; Enterobacteriaceae; Shigella

MTGDNTLIHSHGINRRDFMKLCAALAATMGLSSKAAAEMAESVTNPQRPPVIWIGAQECT WP_000145410.1 MULTISPECIES: hydrogenase-2 small chain [Proteobacteria] Length: 372\nNumber of predicted TMHs: 1\nExp number of AAs in TMHs: 23.72226\nExp number, first 60 AAs: 0.22038\nTotal prob of N-in: 0.12994\noutside 1 331\nTMhelix 332 354\ninside 355 372

26223 GCF_900160185.1_8403_8_49 Shigella sonnei Proteobacteria; Gammaproteobacteria; Enterobacterales; Enterobacteriaceae; Shigella MTGDNTLIHSHGINRRDFMKLCAALAATMGLSSKAAAEMAESVTNPQRPPVIWIGAQECT WP_000145410.1 MULTISPECIES: hydrogenase-2 small chain [Proteobacteria] Length: 372\nNumber of predicted TMHs: 1\nExp number of AAs in TMHs: 23.72226\nExp number, first 60 AAs: 0.22038\nTotal prob of N-in: 0.12994\noutside 1 331\nTMhelix 332 354\ninside 355 372

26224 GCF_900160195.1_8403_8_50 Shigella sonnei Proteobacteria; Gammaproteobacteria; Enterobacterales; Enterobacteriaceae; Shigella MTGDNTLIHSHGINRRDFMKLCAALAATMGLSSKAAAEMAESVTNPQRPPVIWIGAQECT WP_000145410.1 MULTISPECIES: hydrogenase-2 small chain [Proteobacteria] Length: 372\nNumber of predicted TMHs: 1\nExp number of AAs in TMHs: 23.72226\nExp number, first 60 AAs: 0.22038\nTotal prob of N-in: 0.12994\noutside 1 331\nTMhelix 332 354\ninside 355 372

26225 GCF_900160215.1_8403_8_52 Shigella sonnei Proteobacteria; Gammaproteobacteria; Enterobacterales; Enterobacteriaceae; Shigella MTGDNTLIHSHGINRRDFMKLCAALAATMGLSSKAAAEMAESVTNPQRPPVIWIGAQECT WP_000145410.1 MULTISPECIES: hydrogenase-2 small chain [Proteobacteria] Length: 372\nNumber of predicted TMHs: 1\nExp number of AAs in TMHs: 23.72226\nExp number, first 60 AAs: 0.22038\nTotal prob of N-in: 0.12994\noutside 1 331\nTMhelix 332 354\ninside 355 372

26226 GCF_900160235.1_8403_8_54 Shigella sonnei Proteobacteria; Gammaproteobacteria; Enterobacterales; Enterobacteriaceae; Shigella MTGDNTLIHSHGINRRDFMKLCAALAATMGLSSKAAAEMAESVTNPQRPPVIWIGAQECT WP_000145410.1 MULTISPECIES: hydrogenase-2 small chain [Proteobacteria] Length: 372\nNumber of predicted TMHs: 1\nExp number of AAs in TMHs: 23.72226\nExp number, first 60 AAs: 0.22038\nTotal prob of N-in: 0.12994\noutside 1 331\nTMhelix 332 354\ninside 355 372

26227 GCF_900160135.1_8403_8_44 Shigella sonnei Proteobacteria; Gammaproteobacteria; Enterobacterales; Enterobacteriaceae; Shigella MNNEETFYQAMRRQGVTRRSFLKYCSLAATSLGLGAGMAPKIAWALENKPRIPVVWIHGL WP_005139779.1 hydrogenase [Shigella sonnei] Length: 372\nNumber of predicted TMHs: 2\nExp number of AAs in TMHs: 37.02628\nExp number, first 60 AAs: 19.39684\nTotal prob of N-in: 0.94313\nPOSSIBLE N-term signal sequence\ninside 1 19\nTMhelix 20 39\noutside 40 325\nTMhelix 326 348\ninside 349 372

26228 GCF_900160245.1_8403_8_55 Shigella sonnei Proteobacteria; Gammaproteobacteria; Enterobacterales; Enterobacteriaceae; Shigella MTGDNTLIHSHGINRRDFMKLCAALAATMGLSSKAAAEMAESVTNPQRPPVIWIGAQECT WP_000145410.1 MULTISPECIES: hydrogenase-2 small chain [Proteobacteria] Length: 372\nNumber of predicted TMHs: 1\nExp number of AAs in TMHs: 23.72226\nExp number, first 60 AAs: 0.22038\nTotal prob of N-in: 0.12994\noutside 1 331\nTMhelix 332 354\ninside 355 372

26229 GCF_900160265.1_8403_8_57 Shigella sonnei Proteobacteria; Gammaproteobacteria; Enterobacterales; Enterobacteriaceae; Shigella MTGDNTLIHSHGINRRDFMKLCAALAATMGLSSKAAAEMAESVTNPQRPPVIWIGAQECT WP_000145410.1 MULTISPECIES: hydrogenase-2 small chain [Proteobacteria] Length: 372\nNumber of predicted TMHs: 1\nExp number of AAs in TMHs: 23.72226\nExp number, first 60 AAs: 0.22038\nTotal prob of N-in: 0.12994\noutside 1 331\nTMhelix 332 354\ninside 355 372

26230 GCF_900160285.1_8403_8_60 Shigella sonnei Proteobacteria; Gammaproteobacteria; Enterobacterales; Enterobacteriaceae; Shigella MTGDNTLIHSHGINRRDFMKLCAALAATMGLSSKAAAEMAESVTNPQRPPVIWIGAQECT WP_000145410.1 MULTISPECIES: hydrogenase-2 small chain [Proteobacteria] Length: 372\nNumber of predicted TMHs: 1\nExp number of AAs in TMHs: 23.72226\nExp number, first 60 AAs: 0.22038\nTotal prob of N-in: 0.12994\noutside 1 331\nTMhelix 332 354\ninside 355 372

26231	GCF_900160305.1_8403_8_61	Shigella sonnei	Proteobacteria; Gammaproteobacteria; Enterobacterales; Enterobacteriaceae; Shigella
MTGDNTLIHSHGINRRDFMKLCAALAATMGLSSKAAAEMAESVTNPQRPPVIWIGAQECT	WP_000145410.1
MULTISPECIES: hydrogenase-2 small chain [Proteobacteria]	Length: 372\nNumber of predicted TMHs: 1\nExp number of AAs in TMHs: 23.72226\nExp number, first 60 AAs: 0.22038\nTotal prob of N-in: 0.12994\noutside 1 331\nTMhelix 332 354\ninside 355 372

26232	GCF_900160245.1_8403_8_55	Shigella sonnei	Proteobacteria; Gammaproteobacteria; Enterobacterales; Enterobacteriaceae; Shigella
MNNEETFYQAMRRQGVTRRSFLKYCSLAATSLGLGAGMAPKIAWALENKPRIPVVWIHGL	WP_005139779.1
hydrogenase [Shigella sonnei]	Length: 372\nNumber of predicted TMHs: 2\nExp number of AAs in TMHs: 37.02628\nExp number, first 60 AAs: 19.39684\nTotal prob of N-in: 0.94313\nPOSSIBLE N-term signal sequence\ninside 1 19\nTMhelix 20 39\noutside 40 325\nTMhelix 326 348\ninside 349 372

26233	GCF_900160285.1_8403_8_60	Shigella sonnei	Proteobacteria; Gammaproteobacteria; Enterobacterales; Enterobacteriaceae; Shigella
MNNEETFYQAMRRQGVTRRSFLKYCSLAATSLGLGAGMAPKIAWALENKPRIPVVWIHGL	WP_005139779.1
hydrogenase [Shigella sonnei]	Length: 372\nNumber of predicted TMHs: 2\nExp number of AAs in TMHs: 37.02628\nExp number, first 60 AAs: 19.39684\nTotal prob of N-in: 0.94313\nPOSSIBLE N-term signal sequence\ninside 1 19\nTMhelix 20 39\noutside 40 325\nTMhelix 326 348\ninside 349 372

26234	GCF_900160265.1_8403_8_57	Shigella sonnei	Proteobacteria; Gammaproteobacteria; Enterobacterales; Enterobacteriaceae; Shigella
MNNEETFYQAMRRQGVTRRSFLKYCSLAATSLGLGAGMAPKIAWALENKPRIPVVWIHGL	WP_005139779.1
hydrogenase [Shigella sonnei]	Length: 372\nNumber of predicted TMHs: 2\nExp number of AAs in TMHs: 37.02628\nExp number, first 60 AAs: 19.39684\nTotal prob of N-in: 0.94313\nPOSSIBLE N-term signal sequence\ninside 1 19\nTMhelix 20 39\noutside 40 325\nTMhelix 326 348\ninside 349 372

26235	GCF_900160145.1_8403_8_45	Shigella sonnei	Proteobacteria; Gammaproteobacteria; Enterobacterales; Enterobacteriaceae; Shigella
MNNEETFYQAMRRQGVTRRSFLKYCSLAATSLGLGAGMAPKIAWALENKPRIPVVWIHGL	WP_005139779.1
hydrogenase [Shigella sonnei]	Length: 372\nNumber of predicted TMHs: 2\nExp number of AAs in TMHs: 37.02628\nExp number, first 60 AAs: 19.39684\nTotal prob of N-in: 0.94313\nPOSSIBLE N-term signal sequence\ninside 1 19\nTMhelix 20 39\noutside 40 325\nTMhelix 326 348\ninside 349 372

26236	GCF_900160185.1_8403_8_49	Shigella sonnei	Proteobacteria; Gammaproteobacteria; Enterobacterales; Enterobacteriaceae; Shigella
MNNEETFYQAMRRQGVTRRSFLKYCSLAATSLGLGAGMAPKIAWALENKPRIPVVWIHGL	WP_005139779.1
hydrogenase [Shigella sonnei]	Length: 372\nNumber of predicted TMHs: 2\nExp number of AAs in TMHs: 37.02628\nExp number, first 60 AAs: 19.39684\nTotal prob of N-in: 0.94313\nPOSSIBLE N-term signal sequence\ninside 1 19\nTMhelix 20 39\noutside 40 325\nTMhelix 326 348\ninside 349 372

26237	GCF_900160195.1_8403_8_50	Shigella sonnei	Proteobacteria; Gammaproteobacteria; Enterobacterales; Enterobacteriaceae; Shigella
MNNEETFYQAMRRQGVTRRSFLKYCSLAATSLGLGAGMAPKIAWALENKPRIPVVWIHGL	WP_005139779.1
hydrogenase [Shigella sonnei]	Length: 372\nNumber of predicted TMHs: 2\nExp number of AAs in TMHs: 37.02628\nExp number, first 60 AAs: 19.39684\nTotal prob of N-in: 0.94313\nPOSSIBLE N-term signal sequence\ninside 1 19\nTMhelix 20 39\noutside 40 325\nTMhelix 326 348\ninside 349 372

26238	GCF_900160165.1_8403_8_48	Shigella sonnei	Proteobacteria; Gammaproteobacteria; Enterobacterales; Enterobacteriaceae; Shigella
MNNEETFYQAMRRQGVTRRSFLKYCSLAATSLGLGAGMAPKIAWALENKPRIPVVWIHGL	WP_005139779.1
hydrogenase [Shigella sonnei]	Length: 372\nNumber of predicted TMHs: 2\nExp number of AAs in TMHs: 37.02628\nExp number, first 60 AAs: 19.39684\nTotal prob of N-in: 0.94313\nPOSSIBLE N-term signal sequence\ninside 1 19\nTMhelix 20 39\noutside 40 325\nTMhelix 326 348\ninside 349 372

26239	GCF_900160335.1_8403_8_64	Shigella sonnei	Proteobacteria; Gammaproteobacteria; Enterobacterales; Enterobacteriaceae; Shigella
MTGDNTLIHSHGINRRDFMKLCAALAATMGLSSKAAAEMAESVTNPQRPPVIWIGAQECT	WP_000145410.1
MULTISPECIES: hydrogenase-2 small chain [Proteobacteria]	Length: 372\nNumber of predicted TMHs:

1\nExp number of AAs in TMHs: 23.72226\nExp number, first 60 AAs: 0.22038\nTotal prob of N-in: 0.12994\noutside 1 331\nTMhelix 332 354\ninside 355 372

26240 GCF_900160355.1_8403_8_66 Shigella sonnei Proteobacteria; Gammaproteobacteria; Enterobacterales; Enterobacteriaceae; Shigella
MTGDNTLIHSHGINRRDFMKLCAALAATMGLSSKAAAEMAESVTNPQRPPVIWIGAQECT WP_000145410.1
MULTISPECIES: hydrogenase-2 small chain [Proteobacteria] Length: 372\nNumber of predicted TMHs: 1\nExp number of AAs in TMHs: 23.72226\nExp number, first 60 AAs: 0.22038\nTotal prob of N-in: 0.12994\noutside 1 331\nTMhelix 332 354\ninside 355 372

26241 GCF_900160375.1_8403_8_68 Shigella sonnei Proteobacteria; Gammaproteobacteria; Enterobacterales; Enterobacteriaceae; Shigella
MTGDNTLIHSHGINRRDFMKLCAALAATMGLSSKAAAEMAESVTNPQRPPVIWIGAQECT WP_000145410.1
MULTISPECIES: hydrogenase-2 small chain [Proteobacteria] Length: 372\nNumber of predicted TMHs: 1\nExp number of AAs in TMHs: 23.72226\nExp number, first 60 AAs: 0.22038\nTotal prob of N-in: 0.12994\noutside 1 331\nTMhelix 332 354\ninside 355 372

26242 GCF_900160305.1_8403_8_61 Shigella sonnei Proteobacteria; Gammaproteobacteria; Enterobacterales; Enterobacteriaceae; Shigella
MNNEETFYQAMRRQGVTRRSFLKYCSLAATSLGLGAGMAPKIAWALENKPRIPVVWIHGL WP_005139779.1
hydrogenase [Shigella sonnei] Length: 372\nNumber of predicted TMHs: 2\nExp number of AAs in TMHs: 37.02628\nExp number, first 60 AAs: 19.39684\nTotal prob of N-in: 0.94313\nPOSSIBLE N-term signal sequence\ninside 1 19\nTMhelix 20 39\noutside 40 325\nTMhelix 326 348\ninside 349 372

26243 GCF_900160335.1_8403_8_64 Shigella sonnei Proteobacteria; Gammaproteobacteria; Enterobacterales; Enterobacteriaceae; Shigella
MNNEETFYQAMRRQGVTRRSFLKYCSLAATSLGLGAGMAPKIAWALENKPRIPVVWIHGL WP_005139779.1
hydrogenase [Shigella sonnei] Length: 372\nNumber of predicted TMHs: 2\nExp number of AAs in TMHs: 37.02628\nExp number, first 60 AAs: 19.39684\nTotal prob of N-in: 0.94313\nPOSSIBLE N-term signal sequence\ninside 1 19\nTMhelix 20 39\noutside 40 325\nTMhelix 326 348\ninside 349 372

26244 GCF_900160215.1_8403_8_52 Shigella sonnei Proteobacteria; Gammaproteobacteria; Enterobacterales; Enterobacteriaceae; Shigella
MNNEETFYQAMRRQGVTRRSFLKYCSLAATSLGLGAGMAPKIAWALENKPRIPVVWIHGL WP_005139779.1
hydrogenase [Shigella sonnei] Length: 372\nNumber of predicted TMHs: 2\nExp number of AAs in TMHs: 37.02628\nExp number, first 60 AAs: 19.39684\nTotal prob of N-in: 0.94313\nPOSSIBLE N-term signal sequence\ninside 1 19\nTMhelix 20 39\noutside 40 325\nTMhelix 326 348\ninside 349 372

26245 GCF_900162795.1_9789_6_88 Shigella sonnei Proteobacteria; Gammaproteobacteria; Enterobacterales; Enterobacteriaceae; Shigella
MTGDNTLIHSHGINRRDFMKLCAALAATMGLSSKAAAEMAESVTNPQRPPVIWIGAQECT WP_000145410.1
MULTISPECIES: hydrogenase-2 small chain [Proteobacteria] Length: 372\nNumber of predicted TMHs: 1\nExp number of AAs in TMHs: 23.72226\nExp number, first 60 AAs: 0.22038\nTotal prob of N-in: 0.12994\noutside 1 331\nTMhelix 332 354\ninside 355 372

26246 GCF_900160235.1_8403_8_54 Shigella sonnei Proteobacteria; Gammaproteobacteria; Enterobacterales; Enterobacteriaceae; Shigella
MNNEETFYQAMRRQGVTRRSFLKYCSLAATSLGLGAGMAPKIAWALENKPRIPVVWIHGL WP_005139779.1
hydrogenase [Shigella sonnei] Length: 372\nNumber of predicted TMHs: 2\nExp number of AAs in TMHs: 37.02628\nExp number, first 60 AAs: 19.39684\nTotal prob of N-in: 0.94313\nPOSSIBLE N-term signal sequence\ninside 1 19\nTMhelix 20 39\noutside 40 325\nTMhelix 326 348\ninside 349 372

26247 GCF_900160385.1_8403_8_69 Shigella sonnei Proteobacteria; Gammaproteobacteria; Enterobacterales; Enterobacteriaceae; Shigella
MTGDNTLIHSHGINRRDFMKLCAALAATMGLSSKAAAEMAESVTNPQRPPVIWIGAQECT WP_000145410.1
MULTISPECIES: hydrogenase-2 small chain [Proteobacteria] Length: 372\nNumber of predicted TMHs: 1\nExp number of AAs in TMHs: 23.72226\nExp number, first 60 AAs: 0.22038\nTotal prob of N-in: 0.12994\noutside 1 331\nTMhelix 332 354\ninside 355 372

26248 GCF_900160405.1_8403_8_71 Shigella sonnei Proteobacteria; Gammaproteobacteria; Enterobacterales; Enterobacteriaceae; Shigella

MTGDNTLIHSHGINRRDFMKLCAALAATMGLSSKAAAEMAESVTNPQRPPVIWIGAQECT WP_000145410.1 MULTISPECIES: hydrogenase-2 small chain [Proteobacteria] Length: 372\nNumber of predicted TMHs: 1\nExp number of AAs in TMHs: 23.72226\nExp number, first 60 AAs: 0.22038\nTotal prob of N-in: 0.12994\noutside 1 331\nTMhelix 332 354\ninside 355 372

26249 GCF_900160425.1_8403_8_73 Shigella sonnei Proteobacteria; Gammaproteobacteria; Enterobacterales; Enterobacteriaceae; Shigella MTGDNTLIHSHGINRRDFMKLCAALAATMGLSSKAAAEMAESVTNPQRPPVIWIGAQECT WP_000145410.1 MULTISPECIES: hydrogenase-2 small chain [Proteobacteria] Length: 372\nNumber of predicted TMHs: 1\nExp number of AAs in TMHs: 23.72226\nExp number, first 60 AAs: 0.22038\nTotal prob of N-in: 0.12994\noutside 1 331\nTMhelix 332 354\ninside 355 372

26250 GCF_900160385.1_8403_8_69 Shigella sonnei Proteobacteria; Gammaproteobacteria; Enterobacterales; Enterobacteriaceae; Shigella MNNEETFYQAMRRQGVTRRSFLKYCSLAATSLGLGAGMAPKIAWALENKPRIPVVWIHGL WP_005139779.1 hydrogenase [Shigella sonnei] Length: 372\nNumber of predicted TMHs: 2\nExp number of AAs in TMHs: 37.02628\nExp number, first 60 AAs: 19.39684\nTotal prob of N-in: 0.94313\nPOSSIBLE N-term signal sequence\ninside 1 19\nTMhelix 20 39\noutside 40 325\nTMhelix 326 348\ninside 349 372

26251 GCF_900160405.1_8403_8_71 Shigella sonnei Proteobacteria; Gammaproteobacteria; Enterobacterales; Enterobacteriaceae; Shigella MNNEETFYQAMRRQGVTRRSFLKYCSLAATSLGLGAGMAPKIAWALENKPRIPVVWIHGL WP_005139779.1 hydrogenase [Shigella sonnei] Length: 372\nNumber of predicted TMHs: 2\nExp number of AAs in TMHs: 37.02628\nExp number, first 60 AAs: 19.39684\nTotal prob of N-in: 0.94313\nPOSSIBLE N-term signal sequence\ninside 1 19\nTMhelix 20 39\noutside 40 325\nTMhelix 326 348\ninside 349 372

26252 GCF_900160425.1_8403_8_73 Shigella sonnei Proteobacteria; Gammaproteobacteria; Enterobacterales; Enterobacteriaceae; Shigella MNNEETFYQAMRRQGVTRRSFLKYCSLAATSLGLGAGMAPKIAWALENKPRIPVVWIHGL WP_005139779.1 hydrogenase [Shigella sonnei] Length: 372\nNumber of predicted TMHs: 2\nExp number of AAs in TMHs: 37.02628\nExp number, first 60 AAs: 19.39684\nTotal prob of N-in: 0.94313\nPOSSIBLE N-term signal sequence\ninside 1 19\nTMhelix 20 39\noutside 40 325\nTMhelix 326 348\ninside 349 372

26253 GCF_900160355.1_8403_8_66 Shigella sonnei Proteobacteria; Gammaproteobacteria; Enterobacterales; Enterobacteriaceae; Shigella MNNEETFYQAMRRQGVTRRSFLKYCSLAATSLGLGAGMAPKIAWALENKPRIPVVWIHGL WP_005139779.1 hydrogenase [Shigella sonnei] Length: 372\nNumber of predicted TMHs: 2\nExp number of AAs in TMHs: 37.02628\nExp number, first 60 AAs: 19.39684\nTotal prob of N-in: 0.94313\nPOSSIBLE N-term signal sequence\ninside 1 19\nTMhelix 20 39\noutside 40 325\nTMhelix 326 348\ninside 349 372

26254 GCF_900160435.1_8403_8_74 Shigella sonnei Proteobacteria; Gammaproteobacteria; Enterobacterales; Enterobacteriaceae; Shigella MTGDNTLIHSHGINRRDFMKLCAALAATMGLSSKAAAEMAESVTNPQRPPVIWIGAQECT WP_000145410.1 MULTISPECIES: hydrogenase-2 small chain [Proteobacteria] Length: 372\nNumber of predicted TMHs: 1\nExp number of AAs in TMHs: 23.72226\nExp number, first 60 AAs: 0.22038\nTotal prob of N-in: 0.12994\noutside 1 331\nTMhelix 332 354\ninside 355 372

26255 GCF_900160455.1_8403_8_77 Shigella sonnei Proteobacteria; Gammaproteobacteria; Enterobacterales; Enterobacteriaceae; Shigella MTGDNTLIHSHGINRRDFMKLCAALAATMGLSSKAAAEMAESVTNPQRPPVIWIGAQECT WP_000145410.1 MULTISPECIES: hydrogenase-2 small chain [Proteobacteria] Length: 372\nNumber of predicted TMHs: 1\nExp number of AAs in TMHs: 23.72226\nExp number, first 60 AAs: 0.22038\nTotal prob of N-in: 0.12994\noutside 1 331\nTMhelix 332 354\ninside 355 372

26256 GCF_900160475.1_8403_8_79 Shigella sonnei Proteobacteria; Gammaproteobacteria; Enterobacterales; Enterobacteriaceae; Shigella MTGDNTLIHSHGINRRDFMKLCAALAATMGLSSKAAAEMAESVTNPQRPPVIWIGAQECT WP_000145410.1 MULTISPECIES: hydrogenase-2 small chain [Proteobacteria] Length: 372\nNumber of predicted TMHs: 1\nExp number of AAs in TMHs: 23.72226\nExp number, first 60 AAs: 0.22038\nTotal prob of N-in: 0.12994\noutside 1 331\nTMhelix 332 354\ninside 355 372

26257 GCF_900160435.1_8403_8_74 Shigella sonnei Proteobacteria; Gammaproteobacteria; Enterobacterales; Enterobacteriaceae; Shigella
MNNEETFYQAMRRQGVTRRSFLKYCSLAATSLGLGAGMAPKIAWALENKPRIPVVWIHGL WP_005139779.1
hydrogenase [Shigella sonnei] Length: 372\nNumber of predicted TMHs: 2\nExp number of AAs in TMHs: 37.02628\nExp number, first 60 AAs: 19.39684\nTotal prob of N-in: 0.94313\nPOSSIBLE N-term signal sequence\ninside 1 19\nTMhelix 20 39\noutside 40 325\nTMhelix 326 348\ninside 349 372

26258 GCF_900160455.1_8403_8_77 Shigella sonnei Proteobacteria; Gammaproteobacteria; Enterobacterales; Enterobacteriaceae; Shigella
MNNEETFYQAMRRQGVTRRSFLKYCSLAATSLGLGAGMAPKIAWALENKPRIPVVWIHGL WP_005139779.1
hydrogenase [Shigella sonnei] Length: 372\nNumber of predicted TMHs: 2\nExp number of AAs in TMHs: 37.02628\nExp number, first 60 AAs: 19.39684\nTotal prob of N-in: 0.94313\nPOSSIBLE N-term signal sequence\ninside 1 19\nTMhelix 20 39\noutside 40 325\nTMhelix 326 348\ninside 349 372

26259 GCF_900160475.1_8403_8_79 Shigella sonnei Proteobacteria; Gammaproteobacteria; Enterobacterales; Enterobacteriaceae; Shigella
MNNEETFYQAMRRQGVTRRSFLKYCSLAATSLGLGAGMAPKIAWALENKPRIPVVWIHGL WP_005139779.1
hydrogenase [Shigella sonnei] Length: 372\nNumber of predicted TMHs: 2\nExp number of AAs in TMHs: 37.02628\nExp number, first 60 AAs: 19.39684\nTotal prob of N-in: 0.94313\nPOSSIBLE N-term signal sequence\ninside 1 19\nTMhelix 20 39\noutside 40 325\nTMhelix 326 348\ninside 349 372

26260 GCF_900160375.1_8403_8_68 Shigella sonnei Proteobacteria; Gammaproteobacteria; Enterobacterales; Enterobacteriaceae; Shigella
MNNEETFYQAMRRQGVTRRSFLKYCSLAATSLGLGAGMAPKIAWALENKPRIPVVWIHGL WP_005139779.1
hydrogenase [Shigella sonnei] Length: 372\nNumber of predicted TMHs: 2\nExp number of AAs in TMHs: 37.02628\nExp number, first 60 AAs: 19.39684\nTotal prob of N-in: 0.94313\nPOSSIBLE N-term signal sequence\ninside 1 19\nTMhelix 20 39\noutside 40 325\nTMhelix 326 348\ninside 349 372

26261 GCF_900160485.1_8403_8_84 Shigella sonnei Proteobacteria; Gammaproteobacteria; Enterobacterales; Enterobacteriaceae; Shigella
MTGDNTLIHSHGINRRDFMKLCAALAATMGLSSKAAAEMAESVTNPQRPPVIWIGAQECT WP_000145410.1
MULTISPECIES: hydrogenase-2 small chain [Proteobacteria] Length: 372\nNumber of predicted TMHs: 1\nExp number of AAs in TMHs: 23.72226\nExp number, first 60 AAs: 0.22038\nTotal prob of N-in: 0.12994\noutside 1 331\nTMhelix 332 354\ninside 355 372

26262 GCF_900160505.1_8403_8_85 Shigella sonnei Proteobacteria; Gammaproteobacteria; Enterobacterales; Enterobacteriaceae; Shigella
MTGDNTLIHSHGINRRDFMKLCAALAATMGLSSKAAAEMAESVTNPQRPPVIWIGAQECT WP_000145410.1
MULTISPECIES: hydrogenase-2 small chain [Proteobacteria] Length: 372\nNumber of predicted TMHs: 1\nExp number of AAs in TMHs: 23.72226\nExp number, first 60 AAs: 0.22038\nTotal prob of N-in: 0.12994\noutside 1 331\nTMhelix 332 354\ninside 355 372

26263 GCF_900160525.1_8403_8_86 Shigella sonnei Proteobacteria; Gammaproteobacteria; Enterobacterales; Enterobacteriaceae; Shigella
MTGDNTLIHSHGINRRDFMKLCAALAATMGLSSKAAAEMAESVTNPQRPPVIWIGAQECT WP_000145410.1
MULTISPECIES: hydrogenase-2 small chain [Proteobacteria] Length: 372\nNumber of predicted TMHs: 1\nExp number of AAs in TMHs: 23.72226\nExp number, first 60 AAs: 0.22038\nTotal prob of N-in: 0.12994\noutside 1 331\nTMhelix 332 354\ninside 355 372

26264 GCF_900160555.1_8403_8_88 Shigella sonnei Proteobacteria; Gammaproteobacteria; Enterobacterales; Enterobacteriaceae; Shigella
MTGDNTLIHSHGINRRDFMKLCAALAATMGLSSKAAAEMAESVTNPQRPPVIWIGAQECT WP_000145410.1
MULTISPECIES: hydrogenase-2 small chain [Proteobacteria] Length: 372\nNumber of predicted TMHs: 1\nExp number of AAs in TMHs: 23.72226\nExp number, first 60 AAs: 0.22038\nTotal prob of N-in: 0.12994\noutside 1 331\nTMhelix 332 354\ninside 355 372

26265 GCF_900160545.1_8403_8_87 Shigella sonnei Proteobacteria; Gammaproteobacteria; Enterobacterales; Enterobacteriaceae; Shigella
MTGDNTLIHSHGINRRDFMKLCAALAATMGLSSKAAAEMAESVTNPQRPPVIWIGAQECT WP_000145410.1
MULTISPECIES: hydrogenase-2 small chain [Proteobacteria] Length: 372\nNumber of predicted TMHs:

1\nExp number of AAs in TMHs: 23.72226\nExp number, first 60 AAs: 0.22038\nTotal prob of N-in: 0.12994\noutside 1 331\nTMhelix 332 354\ninside 355 372

26266 GCF_900160575.1_8403_8_90 Shigella sonnei Proteobacteria; Gammaproteobacteria; Enterobacterales; Enterobacteriaceae; Shigella
MTGDNTLIHSHGINRRDFMKLCAALAATMGLSSKAAAEMAESVTNPQRPPVIWIGAQECT WP_000145410.1
MULTISPECIES: hydrogenase-2 small chain [Proteobacteria] Length: 372\nNumber of predicted TMHs: 1\nExp number of AAs in TMHs: 23.72226\nExp number, first 60 AAs: 0.22038\nTotal prob of N-in: 0.12994\noutside 1 331\nTMhelix 332 354\ninside 355 372

26267 GCF_900160595.1_8403_8_93 Shigella sonnei Proteobacteria; Gammaproteobacteria; Enterobacterales; Enterobacteriaceae; Shigella
MTGDNTLIHSHGINRRDFMKLCAALAATMGLSSKAAAEMAESVTNPQRPPVIWIGAQECT WP_000145410.1
MULTISPECIES: hydrogenase-2 small chain [Proteobacteria] Length: 372\nNumber of predicted TMHs: 1\nExp number of AAs in TMHs: 23.72226\nExp number, first 60 AAs: 0.22038\nTotal prob of N-in: 0.12994\noutside 1 331\nTMhelix 332 354\ninside 355 372

26268 GCF_900160505.1_8403_8_85 Shigella sonnei Proteobacteria; Gammaproteobacteria; Enterobacterales; Enterobacteriaceae; Shigella
MNNEETFYQAMRRQGVTRRSFLKYCSLAATSLGLGAGMAPKIAWALENKPRIPVVWIHGL WP_005139779.1
hydrogenase [Shigella sonnei] Length: 372\nNumber of predicted TMHs: 2\nExp number of AAs in TMHs: 37.02628\nExp number, first 60 AAs: 19.39684\nTotal prob of N-in: 0.94313\nPOSSIBLE N-term signal sequence\ninside 1 19\nTMhelix 20 39\noutside 40 325\nTMhelix 326 348\ninside 349 372

26269 GCF_900160485.1_8403_8_84 Shigella sonnei Proteobacteria; Gammaproteobacteria; Enterobacterales; Enterobacteriaceae; Shigella
MNNEETFYQAMRRQGVTRRSFLKYCSLAATSLGLGAGMAPKIAWALENKPRIPVVWIHGL WP_005139779.1
hydrogenase [Shigella sonnei] Length: 372\nNumber of predicted TMHs: 2\nExp number of AAs in TMHs: 37.02628\nExp number, first 60 AAs: 19.39684\nTotal prob of N-in: 0.94313\nPOSSIBLE N-term signal sequence\ninside 1 19\nTMhelix 20 39\noutside 40 325\nTMhelix 326 348\ninside 349 372

26270 GCF_900160625.1_8403_8_95 Shigella sonnei Proteobacteria; Gammaproteobacteria; Enterobacterales; Enterobacteriaceae; Shigella
MTGDNTLIHSHGINRRDFMKLCAALAATMGLSSKAAAEMAESVTNPQRPPVIWIGAQECT WP_000145410.1
MULTISPECIES: hydrogenase-2 small chain [Proteobacteria] Length: 372\nNumber of predicted TMHs: 1\nExp number of AAs in TMHs: 23.72226\nExp number, first 60 AAs: 0.22038\nTotal prob of N-in: 0.12994\noutside 1 331\nTMhelix 332 354\ninside 355 372

26271 GCF_900160605.1_8403_8_92 Shigella sonnei Proteobacteria; Gammaproteobacteria; Enterobacterales; Enterobacteriaceae; Shigella
MTGDNTLIHSHGINRRDFMKLCAALAATMGLSSKAAAEMAESVTNPQRPPVIWIGAQECT WP_000145410.1
MULTISPECIES: hydrogenase-2 small chain [Proteobacteria] Length: 372\nNumber of predicted TMHs: 1\nExp number of AAs in TMHs: 23.72226\nExp number, first 60 AAs: 0.22038\nTotal prob of N-in: 0.12994\noutside 1 331\nTMhelix 332 354\ninside 355 372

26272 GCF_900160605.1_8403_8_92 Shigella sonnei Proteobacteria; Gammaproteobacteria; Enterobacterales; Enterobacteriaceae; Shigella
MNNEETFYQAMRRQGVTRRSFLKYCSLAATSLGLGAGMAPKIAWALENKPRIPVVWIHGL WP_005139779.1
hydrogenase [Shigella sonnei] Length: 372\nNumber of predicted TMHs: 2\nExp number of AAs in TMHs: 37.02628\nExp number, first 60 AAs: 19.39684\nTotal prob of N-in: 0.94313\nPOSSIBLE N-term signal sequence\ninside 1 19\nTMhelix 20 39\noutside 40 325\nTMhelix 326 348\ninside 349 372

26273 GCF_900160625.1_8403_8_95 Shigella sonnei Proteobacteria; Gammaproteobacteria; Enterobacterales; Enterobacteriaceae; Shigella
MNNEETFYQAMRRQGVTRRSFLKYCSLAATSLGLGAGMAPKIAWALENKPRIPVVWIHGL WP_005139779.1
hydrogenase [Shigella sonnei] Length: 372\nNumber of predicted TMHs: 2\nExp number of AAs in TMHs: 37.02628\nExp number, first 60 AAs: 19.39684\nTotal prob of N-in: 0.94313\nPOSSIBLE N-term signal sequence\ninside 1 19\nTMhelix 20 39\noutside 40 325\nTMhelix 326 348\ninside 349 372

26274 GCF_900160525.1_8403_8_86 Shigella sonnei Proteobacteria; Gammaproteobacteria; Enterobacterales; Enterobacteriaceae; Shigella

MNNEETFYQAMRRQGVTRRSFLKYCSLAATSLGLGAGMAPKIAWALENKPRIPVVWIHGL WP_005139779.1 hydrogenase [Shigella sonnei] Length: 372\nNumber of predicted TMHs: 2\nExp number of AAs in TMHs: 37.02628\nExp number, first 60 AAs: 19.39684\nTotal prob of N-in: 0.94313\nPOSSIBLE N-term signal sequence\ninside 1 19\nTMhelix 20 39\noutside 40 325\nTMhelix 326 348\ninside 349 372

26275 GCF_900160555.1_8403_8_88 Shigella sonnei Proteobacteria; Gammaproteobacteria; Enterobacterales; Enterobacteriaceae; Shigella MNNEETFYQAMRRQGVTRRSFLKYCSLAATSLGLGAGMAPKIAWALENKPRIPVVWIHGL WP_005139779.1 hydrogenase [Shigella sonnei] Length: 372\nNumber of predicted TMHs: 2\nExp number of AAs in TMHs: 37.02628\nExp number, first 60 AAs: 19.39684\nTotal prob of N-in: 0.94313\nPOSSIBLE N-term signal sequence\ninside 1 19\nTMhelix 20 39\noutside 40 325\nTMhelix 326 348\ninside 349 372

26276 GCF_900160545.1_8403_8_87 Shigella sonnei Proteobacteria; Gammaproteobacteria; Enterobacterales; Enterobacteriaceae; Shigella MNNEETFYQAMRRQGVTRRSFLKYCSLAATSLGLGAGMAPKIAWALENKPRIPVVWIHGL WP_005139779.1 hydrogenase [Shigella sonnei] Length: 372\nNumber of predicted TMHs: 2\nExp number of AAs in TMHs: 37.02628\nExp number, first 60 AAs: 19.39684\nTotal prob of N-in: 0.94313\nPOSSIBLE N-term signal sequence\ninside 1 19\nTMhelix 20 39\noutside 40 325\nTMhelix 326 348\ninside 349 372

26277 GCF_900162735.1_8489_1_72 Shigella sonnei Proteobacteria; Gammaproteobacteria; Enterobacterales; Enterobacteriaceae; Shigella MTGDNTLIHSHGINRRDFMKLCAALAATMGLSSKAAAEMAESVTNPQRPPVIWIGAQECT WP_000145410.1 MULTISPECIES: hydrogenase-2 small chain [Proteobacteria] Length: 372\nNumber of predicted TMHs: 1\nExp number of AAs in TMHs: 23.72226\nExp number, first 60 AAs: 0.22038\nTotal prob of N-in: 0.12994\noutside 1 331\nTMhelix 332 354\ninside 355 372

26278 GCF_900162735.1_8489_1_72 Shigella sonnei Proteobacteria; Gammaproteobacteria; Enterobacterales; Enterobacteriaceae; Shigella MNNEETFYQAMRRQGVTRRSFLKYCSLAATSLGLGAGMAPKIAWALENKPRIPVVWIHGL WP_005139779.1 hydrogenase [Shigella sonnei] Length: 372\nNumber of predicted TMHs: 2\nExp number of AAs in TMHs: 37.02628\nExp number, first 60 AAs: 19.39684\nTotal prob of N-in: 0.94313\nPOSSIBLE N-term signal sequence\ninside 1 19\nTMhelix 20 39\noutside 40 325\nTMhelix 326 348\ninside 349 372

26279 GCF_900160575.1_8403_8_90 Shigella sonnei Proteobacteria; Gammaproteobacteria; Enterobacterales; Enterobacteriaceae; Shigella MNNEETFYQAMRRQGVTRRSFLKYCSLAATSLGLGAGMAPKIAWALENKPRIPVVWIHGL WP_005139779.1 hydrogenase [Shigella sonnei] Length: 372\nNumber of predicted TMHs: 2\nExp number of AAs in TMHs: 37.02628\nExp number, first 60 AAs: 19.39684\nTotal prob of N-in: 0.94313\nPOSSIBLE N-term signal sequence\ninside 1 19\nTMhelix 20 39\noutside 40 325\nTMhelix 326 348\ninside 349 372

26280 GCF_900162755.1_8403_8_80 Shigella sonnei Proteobacteria; Gammaproteobacteria; Enterobacterales; Enterobacteriaceae; Shigella MTGDNTLIHSHGINRRDFMKLCAALAATMGLSSKAAAEMAESVTNPQRPPVIWIGAQECT WP_000145410.1 MULTISPECIES: hydrogenase-2 small chain [Proteobacteria] Length: 372\nNumber of predicted TMHs: 1\nExp number of AAs in TMHs: 23.72226\nExp number, first 60 AAs: 0.22038\nTotal prob of N-in: 0.12994\noutside 1 331\nTMhelix 332 354\ninside 355 372

26281 GCF_900162775.1_8403_8_29 Shigella sonnei Proteobacteria; Gammaproteobacteria; Enterobacterales; Enterobacteriaceae; Shigella MTGDNTLIHSHGINRRDFMKLCAALAATMGLSSKAAAEMAESVTNPQRPPVIWIGAQECT WP_000145410.1 MULTISPECIES: hydrogenase-2 small chain [Proteobacteria] Length: 372\nNumber of predicted TMHs: 1\nExp number of AAs in TMHs: 23.72226\nExp number, first 60 AAs: 0.22038\nTotal prob of N-in: 0.12994\noutside 1 331\nTMhelix 332 354\ninside 355 372

26282 GCF_900163485.1_8403_8_76 Shigella sonnei Proteobacteria; Gammaproteobacteria; Enterobacterales; Enterobacteriaceae; Shigella MTGDNTLIHSHGINRRDFMKLCAALAATMGLSSKAAAEMAESVTNPQRPPVIWIGAQECT WP_000145410.1 MULTISPECIES: hydrogenase-2 small chain [Proteobacteria] Length: 372\nNumber of predicted TMHs: 1\nExp number of AAs in TMHs: 23.72226\nExp number, first 60 AAs: 0.22038\nTotal prob of N-in: 0.12994\noutside 1 331\nTMhelix 332 354\ninside 355 372

26283 GCF_900162775.1_8403_8_29 Shigella sonnei Proteobacteria; Gammaproteobacteria; Enterobacterales; Enterobacteriaceae; Shigella
MNNEETFYQAMRRQGVTRRSFLKYCSLAATSLGLGAGMAPKIAWALENKPRIPVVWIHGL WP_005139779.1
hydrogenase [Shigella sonnei] Length: 372\nNumber of predicted TMHs: 2\nExp number of AAs in TMHs: 37.02628\nExp number, first 60 AAs: 19.39684\nTotal prob of N-in: 0.94313\nPOSSIBLE N-term signal sequence\ninside 1 19\nTMhelix 20 39\noutside 40 325\nTMhelix 326 348\ninside 349 372

26284 GCF_900162755.1_8403_8_80 Shigella sonnei Proteobacteria; Gammaproteobacteria; Enterobacterales; Enterobacteriaceae; Shigella
MNNEETFYQAMRRQGVTRRSFLKYCSLAATSLGLGAGMAPKIAWALENKPRIPVVWIHGL WP_005139779.1
hydrogenase [Shigella sonnei] Length: 372\nNumber of predicted TMHs: 2\nExp number of AAs in TMHs: 37.02628\nExp number, first 60 AAs: 19.39684\nTotal prob of N-in: 0.94313\nPOSSIBLE N-term signal sequence\ninside 1 19\nTMhelix 20 39\noutside 40 325\nTMhelix 326 348\ninside 349 372

26285 GCF_900160595.1_8403_8_93 Shigella sonnei Proteobacteria; Gammaproteobacteria; Enterobacterales; Enterobacteriaceae; Shigella
MNNEETFYQAMRRQGVTRRSFLKYCSLAATSLGLGAGMAPKIAWALENKPRIPVVWIHGL WP_005139779.1
hydrogenase [Shigella sonnei] Length: 372\nNumber of predicted TMHs: 2\nExp number of AAs in TMHs: 37.02628\nExp number, first 60 AAs: 19.39684\nTotal prob of N-in: 0.94313\nPOSSIBLE N-term signal sequence\ninside 1 19\nTMhelix 20 39\noutside 40 325\nTMhelix 326 348\ninside 349 372

26286 GCF_900162795.1_9789_6_88 Shigella sonnei Proteobacteria; Gammaproteobacteria; Enterobacterales; Enterobacteriaceae; Shigella
MNNEETFYQAMRRQGVTRRSFLKYCSLAATSLGLGAGMAPKIAWALENKPRIPVVWIHGL WP_005139779.1
hydrogenase [Shigella sonnei] Length: 372\nNumber of predicted TMHs: 2\nExp number of AAs in TMHs: 37.02628\nExp number, first 60 AAs: 19.39684\nTotal prob of N-in: 0.94313\nPOSSIBLE N-term signal sequence\ninside 1 19\nTMhelix 20 39\noutside 40 325\nTMhelix 326 348\ninside 349 372

26287 GCF_900163485.1_8403_8_76 Shigella sonnei Proteobacteria; Gammaproteobacteria; Enterobacterales; Enterobacteriaceae; Shigella
MNNEETFYQAMRRQGVTRRSFLKYCSLAATSLGLGAGMAPKIAWALENKPRIPVVWIHGL WP_005139779.1
hydrogenase [Shigella sonnei] Length: 372\nNumber of predicted TMHs: 2\nExp number of AAs in TMHs: 37.02628\nExp number, first 60 AAs: 19.39684\nTotal prob of N-in: 0.94313\nPOSSIBLE N-term signal sequence\ninside 1 19\nTMhelix 20 39\noutside 40 325\nTMhelix 326 348\ninside 349 372

26288 GCF_002174605.1_ASM217460v1 Shigella sonnei Proteobacteria; Gammaproteobacteria; Enterobacterales; Enterobacteriaceae; Shigella
MTGDNTLIHSHGINRRDFMKLCAALAATMGLSSKAAAEMAESVTNPQRPPVIWIGAQECT WP_000145410.1
MULTISPECIES: hydrogenase-2 small chain [Proteobacteria] Length: 372\nNumber of predicted TMHs: 1\nExp number of AAs in TMHs: 23.72226\nExp number, first 60 AAs: 0.22038\nTotal prob of N-in: 0.12994\noutside 1 331\nTMhelix 332 354\ninside 355 372

26289 GCF_900157475.1_9789_6_40 Shigella sonnei Proteobacteria; Gammaproteobacteria; Enterobacterales; Enterobacteriaceae; Shigella
MTGDNTLIHSHGINRRDFMKLCAALAATMGLSSKAAAEMAESVTNPQRPPVIWIGAQECT WP_000145410.1
MULTISPECIES: hydrogenase-2 small chain [Proteobacteria] Length: 372\nNumber of predicted TMHs: 1\nExp number of AAs in TMHs: 23.72226\nExp number, first 60 AAs: 0.22038\nTotal prob of N-in: 0.12994\noutside 1 331\nTMhelix 332 354\ninside 355 372

26290 GCF_900157485.1_9789_6_35 Shigella sonnei Proteobacteria; Gammaproteobacteria; Enterobacterales; Enterobacteriaceae; Shigella
MTGDNTLIHSHGINRRDFMKLCAALAATMGLSSKAAAEMAESVTNPQRPPVIWIGAQECT WP_000145410.1
MULTISPECIES: hydrogenase-2 small chain [Proteobacteria] Length: 372\nNumber of predicted TMHs: 1\nExp number of AAs in TMHs: 23.72226\nExp number, first 60 AAs: 0.22038\nTotal prob of N-in: 0.12994\noutside 1 331\nTMhelix 332 354\ninside 355 372

26291 GCF_900157485.1_9789_6_35 Shigella sonnei Proteobacteria; Gammaproteobacteria; Enterobacterales; Enterobacteriaceae; Shigella
MNNEETFYQAMRRQGVTRRSFLKYCSLAATSLGLGAGMAPKIAWALENKPRIPVVWIHGL WP_005139779.1
hydrogenase [Shigella sonnei] Length: 372\nNumber of predicted TMHs: 2\nExp number of AAs in

TMHs: 37.02628\nExp number, first 60 AAs: 19.39684\nTotal prob of N-in: 0.94313\nPOSSIBLE N-term signal sequence\ninside 1 19\nTMhelix 20 39\noutside 40 325\nTMhelix 326 348\ninside 349 372

26292 GCF_001689285.1_ASM168928v1 Shigella sonnei Proteobacteria; Gammaproteobacteria; Enterobacterales; Enterobacteriaceae; Shigella
MNNEETFYQAMRRQGVTRRSFLKYCSLAATSLGLGAGMAPKIAWALENKPRIPVVWIHGL WP_005139779.1
hydrogenase [Shigella sonnei] Length: 372\nNumber of predicted TMHs: 2\nExp number of AAs in TMHs: 37.02628\nExp number, first 60 AAs: 19.39684\nTotal prob of N-in: 0.94313\nPOSSIBLE N-term signal sequence\ninside 1 19\nTMhelix 20 39\noutside 40 325\nTMhelix 326 348\ninside 349 372

26293 GCF_001689345.1_ASM168934v1 Shigella sonnei Proteobacteria; Gammaproteobacteria; Enterobacterales; Enterobacteriaceae; Shigella
MNNEETFYQAMRRQGVTRRSFLKYCSLAATSLGLGAGMAPKIAWALENKPRIPVVWIHGL WP_005139779.1
hydrogenase [Shigella sonnei] Length: 372\nNumber of predicted TMHs: 2\nExp number of AAs in TMHs: 37.02628\nExp number, first 60 AAs: 19.39684\nTotal prob of N-in: 0.94313\nPOSSIBLE N-term signal sequence\ninside 1 19\nTMhelix 20 39\noutside 40 325\nTMhelix 326 348\ninside 349 372

26294 GCF_001689295.1_ASM168929v1 Shigella sonnei Proteobacteria; Gammaproteobacteria; Enterobacterales; Enterobacteriaceae; Shigella
MNNEETFYQAMRRQGVTRRSFLKYCSLAATSLGLGAGMAPKIAWALENKPRIPVVWIHGL WP_005139779.1
hydrogenase [Shigella sonnei] Length: 372\nNumber of predicted TMHs: 2\nExp number of AAs in TMHs: 37.02628\nExp number, first 60 AAs: 19.39684\nTotal prob of N-in: 0.94313\nPOSSIBLE N-term signal sequence\ninside 1 19\nTMhelix 20 39\noutside 40 325\nTMhelix 326 348\ninside 349 372

26295 GCF_001689365.1_ASM168936v1 Shigella sonnei Proteobacteria; Gammaproteobacteria; Enterobacterales; Enterobacteriaceae; Shigella
MNNEETFYQAMRRQGVTRRSFLKYCSLAATSLGLGAGMAPKIAWALENKPRIPVVWIHGL WP_005139779.1
hydrogenase [Shigella sonnei] Length: 372\nNumber of predicted TMHs: 2\nExp number of AAs in TMHs: 37.02628\nExp number, first 60 AAs: 19.39684\nTotal prob of N-in: 0.94313\nPOSSIBLE N-term signal sequence\ninside 1 19\nTMhelix 20 39\noutside 40 325\nTMhelix 326 348\ninside 349 372

26296 GCF_002174605.1_ASM217460v1 Shigella sonnei Proteobacteria; Gammaproteobacteria; Enterobacterales; Enterobacteriaceae; Shigella
MNNEETFYQAMRRQGVTRRSFLKYCSLAATSLGLGAGMAPKIAWALENKPRIPVVWIHGL WP_005139779.1
hydrogenase [Shigella sonnei] Length: 372\nNumber of predicted TMHs: 2\nExp number of AAs in TMHs: 37.02628\nExp number, first 60 AAs: 19.39684\nTotal prob of N-in: 0.94313\nPOSSIBLE N-term signal sequence\ninside 1 19\nTMhelix 20 39\noutside 40 325\nTMhelix 326 348\ninside 349 372

26297 GCF_001933245.2_ASM193324v2 Shigella sonnei Proteobacteria; Gammaproteobacteria; Enterobacterales; Enterobacteriaceae; Shigella
MNNEETFYQAMRRQGVTRRSFLKYCSLAATSLGLGAGMAPKIAWALENKPRIPVVWIHGL WP_005139779.1
hydrogenase [Shigella sonnei] Length: 372\nNumber of predicted TMHs: 2\nExp number of AAs in TMHs: 37.02628\nExp number, first 60 AAs: 19.39684\nTotal prob of N-in: 0.94313\nPOSSIBLE N-term signal sequence\ninside 1 19\nTMhelix 20 39\noutside 40 325\nTMhelix 326 348\ninside 349 372

26298 GCF_900157515.1_9789_6_37 Shigella sonnei Proteobacteria; Gammaproteobacteria; Enterobacterales; Enterobacteriaceae; Shigella
MTGDNTLIHSHGINRRDFMKLCAALAATMGLSSKAAAEMAESVTNPQRPPVIWIGAQECT WP_000145410.1
MULTISPECIES: hydrogenase-2 small chain [Proteobacteria] Length: 372\nNumber of predicted TMHs: 1\nExp number of AAs in TMHs: 23.72226\nExp number, first 60 AAs: 0.22038\nTotal prob of N-in: 0.12994\noutside 1 331\nTMhelix 332 354\ninside 355 372

26299 GCF_900157525.1_9789_6_18 Shigella sonnei Proteobacteria; Gammaproteobacteria; Enterobacterales; Enterobacteriaceae; Shigella
MTGDNTLIHSHGINRRDFMKLCAALAATMGLSSKAAAEMAESVTNPQRPPVIWIGAQECT WP_000145410.1
MULTISPECIES: hydrogenase-2 small chain [Proteobacteria] Length: 372\nNumber of predicted TMHs: 1\nExp number of AAs in TMHs: 23.72226\nExp number, first 60 AAs: 0.22038\nTotal prob of N-in: 0.12994\noutside 1 331\nTMhelix 332 354\ninside 355 372

26300 GCF_900157545.1_9789_6_39 Shigella sonnei Proteobacteria; Gammaproteobacteria; Enterobacterales; Enterobacteriaceae; Shigella

MTGDNTLIHSHGINRRDFMKLCAALAATMGLSSKAAAEMAESVTNPQRPPVIWIGAQECT WP_000145410.1 MULTISPECIES: hydrogenase-2 small chain [Proteobacteria] Length: 372\nNumber of predicted TMHs: 1\nExp number of AAs in TMHs: 23.72226\nExp number, first 60 AAs: 0.22038\nTotal prob of N-in: 0.12994\noutside 1 331\nTMhelix 332 354\ninside 355 372

26301 GCF_900157525.1_9789_6_18 Shigella sonnei Proteobacteria; Gammaproteobacteria; Enterobacterales; Enterobacteriaceae; Shigella MNNEETFYQAMRRQGVTRRSFLKYCSLAATSLGLGAGMAPKIAWALENKPRIPVVWIHGL WP_005139779.1 hydrogenase [Shigella sonnei] Length: 372\nNumber of predicted TMHs: 2\nExp number of AAs in TMHs: 37.02628\nExp number, first 60 AAs: 19.39684\nTotal prob of N-in: 0.94313\nPOSSIBLE N-term signal sequence\ninside 1 19\nTMhelix 20 39\noutside 40 325\nTMhelix 326 348\ninside 349 372

26302 GCF_900157515.1_9789_6_37 Shigella sonnei Proteobacteria; Gammaproteobacteria; Enterobacterales; Enterobacteriaceae; Shigella MNNEETFYQAMRRQGVTRRSFLKYCSLAATSLGLGAGMAPKIAWALENKPRIPVVWIHGL WP_005139779.1 hydrogenase [Shigella sonnei] Length: 372\nNumber of predicted TMHs: 2\nExp number of AAs in TMHs: 37.02628\nExp number, first 60 AAs: 19.39684\nTotal prob of N-in: 0.94313\nPOSSIBLE N-term signal sequence\ninside 1 19\nTMhelix 20 39\noutside 40 325\nTMhelix 326 348\ninside 349 372

26303 GCF_900157545.1_9789_6_39 Shigella sonnei Proteobacteria; Gammaproteobacteria; Enterobacterales; Enterobacteriaceae; Shigella MNNEETFYQAMRRQGVTRRSFLKYCSLAATSLGLGAGMAPKIAWALENKPRIPVVWIHGL WP_005139779.1 hydrogenase [Shigella sonnei] Length: 372\nNumber of predicted TMHs: 2\nExp number of AAs in TMHs: 37.02628\nExp number, first 60 AAs: 19.39684\nTotal prob of N-in: 0.94313\nPOSSIBLE N-term signal sequence\ninside 1 19\nTMhelix 20 39\noutside 40 325\nTMhelix 326 348\ninside 349 372

26304 GCF_900157475.1_9789_6_40 Shigella sonnei Proteobacteria; Gammaproteobacteria; Enterobacterales; Enterobacteriaceae; Shigella MNNEETFYQAMRRQGVTRRSFLKYCSLAATSLGLGAGMAPKIAWALENKPRIPVVWIHGL WP_005139779.1 hydrogenase [Shigella sonnei] Length: 372\nNumber of predicted TMHs: 2\nExp number of AAs in TMHs: 37.02628\nExp number, first 60 AAs: 19.39684\nTotal prob of N-in: 0.94313\nPOSSIBLE N-term signal sequence\ninside 1 19\nTMhelix 20 39\noutside 40 325\nTMhelix 326 348\ninside 349 372

26305 GCF_900157575.1_9789_6_71 Shigella sonnei Proteobacteria; Gammaproteobacteria; Enterobacterales; Enterobacteriaceae; Shigella MTGDNTLIHSHGINRRDFMKLCAALAATMGLSSKAAAEMAESVTNPQRPPVIWIGAQECT WP_000145410.1 MULTISPECIES: hydrogenase-2 small chain [Proteobacteria] Length: 372\nNumber of predicted TMHs: 1\nExp number of AAs in TMHs: 23.72226\nExp number, first 60 AAs: 0.22038\nTotal prob of N-in: 0.12994\noutside 1 331\nTMhelix 332 354\ninside 355 372

26306 GCF_900157565.1_9789_6_53 Shigella sonnei Proteobacteria; Gammaproteobacteria; Enterobacterales; Enterobacteriaceae; Shigella MTGDNTLIHSHGINRRDFMKLCAALAATMGLSSKAAAEMAESVTNPQRPPVIWIGAQECT WP_000145410.1 MULTISPECIES: hydrogenase-2 small chain [Proteobacteria] Length: 372\nNumber of predicted TMHs: 1\nExp number of AAs in TMHs: 23.72226\nExp number, first 60 AAs: 0.22038\nTotal prob of N-in: 0.12994\noutside 1 331\nTMhelix 332 354\ninside 355 372

26307 GCF_900157595.1_9789_6_59 Shigella sonnei Proteobacteria; Gammaproteobacteria; Enterobacterales; Enterobacteriaceae; Shigella MTGDNTLIHSHGINRRDFMKLCAALAATMGLSSKAAAEMAESVTNPQRPPVIWIGAQECT WP_000145410.1 MULTISPECIES: hydrogenase-2 small chain [Proteobacteria] Length: 372\nNumber of predicted TMHs: 1\nExp number of AAs in TMHs: 23.72226\nExp number, first 60 AAs: 0.22038\nTotal prob of N-in: 0.12994\noutside 1 331\nTMhelix 332 354\ninside 355 372

26308 GCF_900157625.1_9789_6_77 Shigella sonnei Proteobacteria; Gammaproteobacteria; Enterobacterales; Enterobacteriaceae; Shigella MTGDNTLIHSHGINRRDFMKLCAALAATMGLSSKAAAEMAESVTNPQRPPVIWIGAQECT WP_000145410.1 MULTISPECIES: hydrogenase-2 small chain [Proteobacteria] Length: 372\nNumber of predicted TMHs: 1\nExp number of AAs in TMHs: 23.72226\nExp number, first 60 AAs: 0.22038\nTotal prob of N-in: 0.12994\noutside 1 331\nTMhelix 332 354\ninside 355 372

26309 GCF_900157605.1_9789_6_22 Shigella sonnei Proteobacteria; Gammaproteobacteria; Enterobacterales; Enterobacteriaceae; Shigella
MTGDNTLIHSHGINRRDFMKLCAALAATMGLSSKAAAEMAESVTNPQRPPVIWIGAQECT WP_000145410.1
MULTISPECIES: hydrogenase-2 small chain [Proteobacteria] Length: 372\nNumber of predicted TMHs: 1\nExp number of AAs in TMHs: 23.72226\nExp number, first 60 AAs: 0.22038\nTotal prob of N-in: 0.12994\noutside 1 331\nTMhelix 332 354\ninside 355 372

26310 GCF_900157645.1_9789_6_32 Shigella sonnei Proteobacteria; Gammaproteobacteria; Enterobacterales; Enterobacteriaceae; Shigella
MTGDNTLIHSHGINRRDFMKLCAALAATMGLSSKAAAEMAESVTNPQRPPVIWIGAQECT WP_000145410.1
MULTISPECIES: hydrogenase-2 small chain [Proteobacteria] Length: 372\nNumber of predicted TMHs: 1\nExp number of AAs in TMHs: 23.72226\nExp number, first 60 AAs: 0.22038\nTotal prob of N-in: 0.12994\noutside 1 331\nTMhelix 332 354\ninside 355 372

26311 GCF_900157655.1_9789_6_66 Shigella sonnei Proteobacteria; Gammaproteobacteria; Enterobacterales; Enterobacteriaceae; Shigella
MTGDNTLIHSHGINRRDFMKLCAALAATMGLSSKAAAEMAESVTNPQRPPVIWIGAQECT WP_000145410.1
MULTISPECIES: hydrogenase-2 small chain [Proteobacteria] Length: 372\nNumber of predicted TMHs: 1\nExp number of AAs in TMHs: 23.72226\nExp number, first 60 AAs: 0.22038\nTotal prob of N-in: 0.12994\noutside 1 331\nTMhelix 332 354\ninside 355 372

26312 GCF_900157675.1_9789_6_31 Shigella sonnei Proteobacteria; Gammaproteobacteria; Enterobacterales; Enterobacteriaceae; Shigella
MTGDNTLIHSHGINRRDFMKLCAALAATMGLSSKAAAEMAESVTNPQRPPVIWIGAQECT WP_000145410.1
MULTISPECIES: hydrogenase-2 small chain [Proteobacteria] Length: 372\nNumber of predicted TMHs: 1\nExp number of AAs in TMHs: 23.72226\nExp number, first 60 AAs: 0.22038\nTotal prob of N-in: 0.12994\noutside 1 331\nTMhelix 332 354\ninside 355 372

26313 GCF_900157675.1_9789_6_31 Shigella sonnei Proteobacteria; Gammaproteobacteria; Enterobacterales; Enterobacteriaceae; Shigella
MNNEETFYQAMRRQGVTRRSFLKYCSLAATSLGLGAGMAPKIAWALENKPRIPVVWIHGL WP_005139779.1
hydrogenase [Shigella sonnei] Length: 372\nNumber of predicted TMHs: 2\nExp number of AAs in TMHs: 37.02628\nExp number, first 60 AAs: 19.39684\nTotal prob of N-in: 0.94313\nPOSSIBLE N-term signal sequence\ninside 1 19\nTMhelix 20 39\noutside 40 325\nTMhelix 326 348\ninside 349 372

26314 GCF_900157565.1_9789_6_53 Shigella sonnei Proteobacteria; Gammaproteobacteria; Enterobacterales; Enterobacteriaceae; Shigella
MNNEETFYQAMRRQGVTRRSFLKYCSLAATSLGLGAGMAPKIAWALENKPRIPVVWIHGL WP_005139779.1
hydrogenase [Shigella sonnei] Length: 372\nNumber of predicted TMHs: 2\nExp number of AAs in TMHs: 37.02628\nExp number, first 60 AAs: 19.39684\nTotal prob of N-in: 0.94313\nPOSSIBLE N-term signal sequence\ninside 1 19\nTMhelix 20 39\noutside 40 325\nTMhelix 326 348\ninside 349 372

26315 GCF_900157575.1_9789_6_71 Shigella sonnei Proteobacteria; Gammaproteobacteria; Enterobacterales; Enterobacteriaceae; Shigella
MNNEETFYQAMRRQGVTRRSFLKYCSLAATSLGLGAGMAPKIAWALENKPRIPVVWIHGL WP_005139779.1
hydrogenase [Shigella sonnei] Length: 372\nNumber of predicted TMHs: 2\nExp number of AAs in TMHs: 37.02628\nExp number, first 60 AAs: 19.39684\nTotal prob of N-in: 0.94313\nPOSSIBLE N-term signal sequence\ninside 1 19\nTMhelix 20 39\noutside 40 325\nTMhelix 326 348\ninside 349 372

26316 GCF_900157645.1_9789_6_32 Shigella sonnei Proteobacteria; Gammaproteobacteria; Enterobacterales; Enterobacteriaceae; Shigella
MNNEETFYQAMRRQGVTRRSFLKYCSLAATSLGLGAGMAPKIAWALENKPRIPVVWIHGL WP_005139779.1
hydrogenase [Shigella sonnei] Length: 372\nNumber of predicted TMHs: 2\nExp number of AAs in TMHs: 37.02628\nExp number, first 60 AAs: 19.39684\nTotal prob of N-in: 0.94313\nPOSSIBLE N-term signal sequence\ninside 1 19\nTMhelix 20 39\noutside 40 325\nTMhelix 326 348\ninside 349 372

26317 GCF_900157605.1_9789_6_22 Shigella sonnei Proteobacteria; Gammaproteobacteria; Enterobacterales; Enterobacteriaceae; Shigella
MNNEETFYQAMRRQGVTRRSFLKYCSLAATSLGLGAGMAPKIAWALENKPRIPVVWIHGL WP_005139779.1
hydrogenase [Shigella sonnei] Length: 372\nNumber of predicted TMHs: 2\nExp number of AAs in

TMHs: 37.02628\nExp number, first 60 AAs: 19.39684\nTotal prob of N-in: 0.94313\nPOSSIBLE N-term signal sequence\ninside 1 19\nTMhelix 20 39\noutside 40 325\nTMhelix 326 348\ninside 349 372

26318 GCF_900157595.1_9789_6_59 Shigella sonnei Proteobacteria; Gammaproteobacteria; Enterobacterales; Enterobacteriaceae; Shigella
MNNEETFYQAMRRQGVTRRSFLKYCSLAATSLGLGAGMAPKIAWALENKPRIPVVWIHGL WP_005139779.1
hydrogenase [Shigella sonnei] Length: 372\nNumber of predicted TMHs: 2\nExp number of AAs in TMHs: 37.02628\nExp number, first 60 AAs: 19.39684\nTotal prob of N-in: 0.94313\nPOSSIBLE N-term signal sequence\ninside 1 19\nTMhelix 20 39\noutside 40 325\nTMhelix 326 348\ninside 349 372

26319 GCF_900157625.1_9789_6_77 Shigella sonnei Proteobacteria; Gammaproteobacteria; Enterobacterales; Enterobacteriaceae; Shigella
MNNEETFYQAMRRQGVTRRSFLKYCSLAATSLGLGAGMAPKIAWALENKPRIPVVWIHGL WP_005139779.1
hydrogenase [Shigella sonnei] Length: 372\nNumber of predicted TMHs: 2\nExp number of AAs in TMHs: 37.02628\nExp number, first 60 AAs: 19.39684\nTotal prob of N-in: 0.94313\nPOSSIBLE N-term signal sequence\ninside 1 19\nTMhelix 20 39\noutside 40 325\nTMhelix 326 348\ninside 349 372

26320 GCF_900157685.1_9789_6_72 Shigella sonnei Proteobacteria; Gammaproteobacteria; Enterobacterales; Enterobacteriaceae; Shigella
MTGDNTLIHSHGINRRDFMKLCAALAATMGLSSKAAAEMAESVTNPQRPPVIWIGAQECT WP_000145410.1
MULTISPECIES: hydrogenase-2 small chain [Proteobacteria] Length: 372\nNumber of predicted TMHs: 1\nExp number of AAs in TMHs: 23.72226\nExp number, first 60 AAs: 0.22038\nTotal prob of N-in: 0.12994\noutside 1 331\nTMhelix 332 354\ninside 355 372

26321 GCF_900157705.1_9789_6_70 Shigella sonnei Proteobacteria; Gammaproteobacteria; Enterobacterales; Enterobacteriaceae; Shigella
MTGDNTLIHSHGINRRDFMKLCAALAATMGLSSKAAAEMAESVTNPQRPPVIWIGAQECT WP_000145410.1
MULTISPECIES: hydrogenase-2 small chain [Proteobacteria] Length: 372\nNumber of predicted TMHs: 1\nExp number of AAs in TMHs: 23.72226\nExp number, first 60 AAs: 0.22038\nTotal prob of N-in: 0.12994\noutside 1 331\nTMhelix 332 354\ninside 355 372

26322 GCF_900157725.1_9789_6_61 Shigella sonnei Proteobacteria; Gammaproteobacteria; Enterobacterales; Enterobacteriaceae; Shigella
MTGDNTLIHSHGINRRDFMKLCAALAATMGLSSKAAAEMAESVTNPQRPPVIWIGAQECT WP_000145410.1
MULTISPECIES: hydrogenase-2 small chain [Proteobacteria] Length: 372\nNumber of predicted TMHs: 1\nExp number of AAs in TMHs: 23.72226\nExp number, first 60 AAs: 0.22038\nTotal prob of N-in: 0.12994\noutside 1 331\nTMhelix 332 354\ninside 355 372

26323 GCF_900157735.1_9789_6_69 Shigella sonnei Proteobacteria; Gammaproteobacteria; Enterobacterales; Enterobacteriaceae; Shigella
MTGDNTLIHSHGINRRDFMKLCAALAATMGLSSKAAAEMAESVTNPQRPPVIWIGAQECT WP_000145410.1
MULTISPECIES: hydrogenase-2 small chain [Proteobacteria] Length: 372\nNumber of predicted TMHs: 1\nExp number of AAs in TMHs: 23.72226\nExp number, first 60 AAs: 0.22038\nTotal prob of N-in: 0.12994\noutside 1 331\nTMhelix 332 354\ninside 355 372

26324 GCF_900157685.1_9789_6_72 Shigella sonnei Proteobacteria; Gammaproteobacteria; Enterobacterales; Enterobacteriaceae; Shigella
MNNEETFYQAMRRQGVTRRSFLKYCSLAATSLGLGAGMAPKIAWALENKPRIPVVWIHGL WP_005139779.1
hydrogenase [Shigella sonnei] Length: 372\nNumber of predicted TMHs: 2\nExp number of AAs in TMHs: 37.02628\nExp number, first 60 AAs: 19.39684\nTotal prob of N-in: 0.94313\nPOSSIBLE N-term signal sequence\ninside 1 19\nTMhelix 20 39\noutside 40 325\nTMhelix 326 348\ninside 349 372

26325 GCF_900157725.1_9789_6_61 Shigella sonnei Proteobacteria; Gammaproteobacteria; Enterobacterales; Enterobacteriaceae; Shigella
MNNEETFYQAMRRQGVTRRSFLKYCSLAATSLGLGAGMAPKIAWALENKPRIPVVWIHGL WP_005139779.1
hydrogenase [Shigella sonnei] Length: 372\nNumber of predicted TMHs: 2\nExp number of AAs in TMHs: 37.02628\nExp number, first 60 AAs: 19.39684\nTotal prob of N-in: 0.94313\nPOSSIBLE N-term signal sequence\ninside 1 19\nTMhelix 20 39\noutside 40 325\nTMhelix 326 348\ninside 349 372

26326 GCF_900157655.1_9789_6_66 Shigella sonnei Proteobacteria; Gammaproteobacteria; Enterobacterales; Enterobacteriaceae; Shigella

MNNEETFYQAMRRQGVTRRSFLKYCSLAATSLGLGAGMAPKIAWALENKPRIPVVWIHGL WP_005139779.1 hydrogenase [Shigella sonnei] Length: 372\nNumber of predicted TMHs: 2\nExp number of AAs in TMHs: 37.02628\nExp number, first 60 AAs: 19.39684\nTotal prob of N-in: 0.94313\nPOSSIBLE N-term signal sequence\ninside 1 19\nTMhelix 20 39\noutside 40 325\nTMhelix 326 348\ninside 349 372

26327 GCF_900157765.1_9789_6_48 Shigella sonnei Proteobacteria; Gammaproteobacteria; Enterobacterales; Enterobacteriaceae; Shigella MTGDNTLIHSHGINRRDFMKLCAALAATMGLSSKAAAEMAESVTNPQRPPVIWIGAQECT WP_000145410.1 MULTISPECIES: hydrogenase-2 small chain [Proteobacteria] Length: 372\nNumber of predicted TMHs: 1\nExp number of AAs in TMHs: 23.72226\nExp number, first 60 AAs: 0.22038\nTotal prob of N-in: 0.12994\noutside 1 331\nTMhelix 332 354\ninside 355 372

26328 GCF_900157755.1_9789_6_11 Shigella sonnei Proteobacteria; Gammaproteobacteria; Enterobacterales; Enterobacteriaceae; Shigella MTGDNTLIHSHGINRRDFMKLCAALAATMGLSSKAAAEMAESVTNPQRPPVIWIGAQECT WP_000145410.1 MULTISPECIES: hydrogenase-2 small chain [Proteobacteria] Length: 372\nNumber of predicted TMHs: 1\nExp number of AAs in TMHs: 23.72226\nExp number, first 60 AAs: 0.22038\nTotal prob of N-in: 0.12994\noutside 1 331\nTMhelix 332 354\ninside 355 372

26329 GCF_900157785.1_9789_6_56 Shigella sonnei Proteobacteria; Gammaproteobacteria; Enterobacterales; Enterobacteriaceae; Shigella MTGDNTLIHSHGINRRDFMKLCAALAATMGLSSKAAAEMAESVTNPQRPPVIWIGAQECT WP_000145410.1 MULTISPECIES: hydrogenase-2 small chain [Proteobacteria] Length: 372\nNumber of predicted TMHs: 1\nExp number of AAs in TMHs: 23.72226\nExp number, first 60 AAs: 0.22038\nTotal prob of N-in: 0.12994\noutside 1 331\nTMhelix 332 354\ninside 355 372

26330 GCF_900157805.1_9789_6_23 Shigella sonnei Proteobacteria; Gammaproteobacteria; Enterobacterales; Enterobacteriaceae; Shigella MTGDNTLIHSHGINRRDFMKLCAALAATMGLSSKAAAEMAESVTNPQRPPVIWIGAQECT WP_000145410.1 MULTISPECIES: hydrogenase-2 small chain [Proteobacteria] Length: 372\nNumber of predicted TMHs: 1\nExp number of AAs in TMHs: 23.72226\nExp number, first 60 AAs: 0.22038\nTotal prob of N-in: 0.12994\noutside 1 331\nTMhelix 332 354\ninside 355 372

26331 GCF_900157835.1_9789_6_38 Shigella sonnei Proteobacteria; Gammaproteobacteria; Enterobacterales; Enterobacteriaceae; Shigella MTGDNTLIHSHGINRRDFMKLCAALAATMGLSSKAAAEMAESVTNPQRPPVIWIGAQECT WP_000145410.1 MULTISPECIES: hydrogenase-2 small chain [Proteobacteria] Length: 372\nNumber of predicted TMHs: 1\nExp number of AAs in TMHs: 23.72226\nExp number, first 60 AAs: 0.22038\nTotal prob of N-in: 0.12994\noutside 1 331\nTMhelix 332 354\ninside 355 372

26332 GCF_900157815.1_9789_6_17 Shigella sonnei Proteobacteria; Gammaproteobacteria; Enterobacterales; Enterobacteriaceae; Shigella MTGDNTLIHSHGINRRDFMKLCAALAATMGLSSKAAAEMAESVTNPQRPPVIWIGAQECT WP_000145410.1 MULTISPECIES: hydrogenase-2 small chain [Proteobacteria] Length: 372\nNumber of predicted TMHs: 1\nExp number of AAs in TMHs: 23.72226\nExp number, first 60 AAs: 0.22038\nTotal prob of N-in: 0.12994\noutside 1 331\nTMhelix 332 354\ninside 355 372

26333 GCF_900157855.1_9789_6_83 Shigella sonnei Proteobacteria; Gammaproteobacteria; Enterobacterales; Enterobacteriaceae; Shigella MTGDNTLIHSHGINRRDFMKLCAALAATMGLSSKAAAEMAESVTNPQRPPVIWIGAQECT WP_000145410.1 MULTISPECIES: hydrogenase-2 small chain [Proteobacteria] Length: 372\nNumber of predicted TMHs: 1\nExp number of AAs in TMHs: 23.72226\nExp number, first 60 AAs: 0.22038\nTotal prob of N-in: 0.12994\noutside 1 331\nTMhelix 332 354\ninside 355 372

26334 GCF_900157865.1_9789_6_21 Shigella sonnei Proteobacteria; Gammaproteobacteria; Enterobacterales; Enterobacteriaceae; Shigella MTGDNTLIHSHGINRRDFMKLCAALAATMGLSSKAAAEMAESVTNPQRPPVIWIGAQECT WP_000145410.1 MULTISPECIES: hydrogenase-2 small chain [Proteobacteria] Length: 372\nNumber of predicted TMHs: 1\nExp number of AAs in TMHs: 23.72226\nExp number, first 60 AAs: 0.22038\nTotal prob of N-in: 0.12994\noutside 1 331\nTMhelix 332 354\ninside 355 372

26335 GCF_900157865.1_9789_6_21 Shigella sonnei Proteobacteria; Gammaproteobacteria; Enterobacterales; Enterobacteriaceae; Shigella
MNNEETFYQAMRRQGVTRRSFLKYCSLAATSLGLGAGMAPKIAWALENKPRIPVVWIHGL WP_005139779.1
hydrogenase [Shigella sonnei] Length: 372\nNumber of predicted TMHs: 2\nExp number of AAs in TMHs: 37.02628\nExp number, first 60 AAs: 19.39684\nTotal prob of N-in: 0.94313\nPOSSIBLE N-term signal sequence\ninside 1 19\nTMhelix 20 39\noutside 40 325\nTMhelix 326 348\ninside 349 372

26336 GCF_900157705.1_9789_6_70 Shigella sonnei Proteobacteria; Gammaproteobacteria; Enterobacterales; Enterobacteriaceae; Shigella
MNNEETFYQAMRRQGVTRRSFLKYCSLAATSLGLGAGMAPKIAWALENKPRIPVVWIHGL WP_005139779.1
hydrogenase [Shigella sonnei] Length: 372\nNumber of predicted TMHs: 2\nExp number of AAs in TMHs: 37.02628\nExp number, first 60 AAs: 19.39684\nTotal prob of N-in: 0.94313\nPOSSIBLE N-term signal sequence\ninside 1 19\nTMhelix 20 39\noutside 40 325\nTMhelix 326 348\ninside 349 372

26337 GCF_900157735.1_9789_6_69 Shigella sonnei Proteobacteria; Gammaproteobacteria; Enterobacterales; Enterobacteriaceae; Shigella
MNNEETFYQAMRRQGVTRRSFLKYCSLAATSLGLGAGMAPKIAWALENKPRIPVVWIHGL WP_005139779.1
hydrogenase [Shigella sonnei] Length: 372\nNumber of predicted TMHs: 2\nExp number of AAs in TMHs: 37.02628\nExp number, first 60 AAs: 19.39684\nTotal prob of N-in: 0.94313\nPOSSIBLE N-term signal sequence\ninside 1 19\nTMhelix 20 39\noutside 40 325\nTMhelix 326 348\ninside 349 372

26338 GCF_900157755.1_9789_6_11 Shigella sonnei Proteobacteria; Gammaproteobacteria; Enterobacterales; Enterobacteriaceae; Shigella
MNNEETFYQAMRRQGVTRRSFLKYCSLAATSLGLGAGMAPKIAWALENKPRIPVVWIHGL WP_005139779.1
hydrogenase [Shigella sonnei] Length: 372\nNumber of predicted TMHs: 2\nExp number of AAs in TMHs: 37.02628\nExp number, first 60 AAs: 19.39684\nTotal prob of N-in: 0.94313\nPOSSIBLE N-term signal sequence\ninside 1 19\nTMhelix 20 39\noutside 40 325\nTMhelix 326 348\ninside 349 372

26339 GCF_900157765.1_9789_6_48 Shigella sonnei Proteobacteria; Gammaproteobacteria; Enterobacterales; Enterobacteriaceae; Shigella
MNNEETFYQAMRRQGVTRRSFLKYCSLAATSLGLGAGMAPKIAWALENKPRIPVVWIHGL WP_005139779.1
hydrogenase [Shigella sonnei] Length: 372\nNumber of predicted TMHs: 2\nExp number of AAs in TMHs: 37.02628\nExp number, first 60 AAs: 19.39684\nTotal prob of N-in: 0.94313\nPOSSIBLE N-term signal sequence\ninside 1 19\nTMhelix 20 39\noutside 40 325\nTMhelix 326 348\ninside 349 372

26340 GCF_900157785.1_9789_6_56 Shigella sonnei Proteobacteria; Gammaproteobacteria; Enterobacterales; Enterobacteriaceae; Shigella
MNNEETFYQAMRRQGVTRRSFLKYCSLAATSLGLGAGMAPKIAWALENKPRIPVVWIHGL WP_005139779.1
hydrogenase [Shigella sonnei] Length: 372\nNumber of predicted TMHs: 2\nExp number of AAs in TMHs: 37.02628\nExp number, first 60 AAs: 19.39684\nTotal prob of N-in: 0.94313\nPOSSIBLE N-term signal sequence\ninside 1 19\nTMhelix 20 39\noutside 40 325\nTMhelix 326 348\ninside 349 372

26341 GCF_900157815.1_9789_6_17 Shigella sonnei Proteobacteria; Gammaproteobacteria; Enterobacterales; Enterobacteriaceae; Shigella
MNNEETFYQAMRRQGVTRRSFLKYCSLAATSLGLGAGMAPKIAWALENKPRIPVVWIHGL WP_005139779.1
hydrogenase [Shigella sonnei] Length: 372\nNumber of predicted TMHs: 2\nExp number of AAs in TMHs: 37.02628\nExp number, first 60 AAs: 19.39684\nTotal prob of N-in: 0.94313\nPOSSIBLE N-term signal sequence\ninside 1 19\nTMhelix 20 39\noutside 40 325\nTMhelix 326 348\ninside 349 372

26342 GCF_900157805.1_9789_6_23 Shigella sonnei Proteobacteria; Gammaproteobacteria; Enterobacterales; Enterobacteriaceae; Shigella
MNNEETFYQAMRRQGVTRRSFLKYCSLAATSLGLGAGMAPKIAWALENKPRIPVVWIHGL WP_005139779.1
hydrogenase [Shigella sonnei] Length: 372\nNumber of predicted TMHs: 2\nExp number of AAs in TMHs: 37.02628\nExp number, first 60 AAs: 19.39684\nTotal prob of N-in: 0.94313\nPOSSIBLE N-term signal sequence\ninside 1 19\nTMhelix 20 39\noutside 40 325\nTMhelix 326 348\ninside 349 372

26343 GCF_900157855.1_9789_6_83 Shigella sonnei Proteobacteria; Gammaproteobacteria; Enterobacterales; Enterobacteriaceae; Shigella
MNNEETFYQAMRRQGVTRRSFLKYCSLAATSLGLGAGMAPKIAWALENKPRIPVVWIHGL WP_005139779.1
hydrogenase [Shigella sonnei] Length: 372\nNumber of predicted TMHs: 2\nExp number of AAs in

TMHs: 37.02628\nExp number, first 60 AAs: 19.39684\nTotal prob of N-in: 0.94313\nPOSSIBLE N-term signal sequence\ninside 1 19\nTMhelix 20 39\noutside 40 325\nTMhelix 326 348\ninside 349 372

26344 GCF_900157905.1_9789_6_36 Shigella sonnei Proteobacteria; Gammaproteobacteria; Enterobacterales; Enterobacteriaceae; Shigella
MTGDNTLIHSHGINRRDFMKLCAALAATMGLSSKAAAEMAESVTNPQRPPVIWIGAQECT WP_000145410.1
MULTISPECIES: hydrogenase-2 small chain [Proteobacteria] Length: 372\nNumber of predicted TMHs: 1\nExp number of AAs in TMHs: 23.72226\nExp number, first 60 AAs: 0.22038\nTotal prob of N-in: 0.12994\noutside 1 331\nTMhelix 332 354\ninside 355 372

26345 GCF_900157905.1_9789_6_36 Shigella sonnei Proteobacteria; Gammaproteobacteria; Enterobacterales; Enterobacteriaceae; Shigella
MNNEETFYQAMRRQGVTRRSFLKYCSLAATSLGLGAGMAPKIAWALENKPRIPVVWIHGL WP_005139779.1
hydrogenase [Shigella sonnei] Length: 372\nNumber of predicted TMHs: 2\nExp number of AAs in TMHs: 37.02628\nExp number, first 60 AAs: 19.39684\nTotal prob of N-in: 0.94313\nPOSSIBLE N-term signal sequence\ninside 1 19\nTMhelix 20 39\noutside 40 325\nTMhelix 326 348\ninside 349 372

26346 GCF_900157835.1_9789_6_38 Shigella sonnei Proteobacteria; Gammaproteobacteria; Enterobacterales; Enterobacteriaceae; Shigella
MNNEETFYQAMRRQGVTRRSFLKYCSLAATSLGLGAGMAPKIAWALENKPRIPVVWIHGL WP_005139779.1
hydrogenase [Shigella sonnei] Length: 372\nNumber of predicted TMHs: 2\nExp number of AAs in TMHs: 37.02628\nExp number, first 60 AAs: 19.39684\nTotal prob of N-in: 0.94313\nPOSSIBLE N-term signal sequence\ninside 1 19\nTMhelix 20 39\noutside 40 325\nTMhelix 326 348\ninside 349 372

26347 GCF_900157955.1_9789_6_42 Shigella sonnei Proteobacteria; Gammaproteobacteria; Enterobacterales; Enterobacteriaceae; Shigella
MTGDNTLIHSHGINRRDFMKLCAALAATMGLSSKAAAEMAESVTNPQRPPVIWIGAQECT WP_000145410.1
MULTISPECIES: hydrogenase-2 small chain [Proteobacteria] Length: 372\nNumber of predicted TMHs: 1\nExp number of AAs in TMHs: 23.72226\nExp number, first 60 AAs: 0.22038\nTotal prob of N-in: 0.12994\noutside 1 331\nTMhelix 332 354\ninside 355 372

26348 GCF_900157925.1_9789_6_44 Shigella sonnei Proteobacteria; Gammaproteobacteria; Enterobacterales; Enterobacteriaceae; Shigella
MTGDNTLIHSHGINRRDFMKLCAALAATMGLSSKAAAEMAESVTNPQRPPVIWIGAQECT WP_000145410.1
MULTISPECIES: hydrogenase-2 small chain [Proteobacteria] Length: 372\nNumber of predicted TMHs: 1\nExp number of AAs in TMHs: 23.72226\nExp number, first 60 AAs: 0.22038\nTotal prob of N-in: 0.12994\noutside 1 331\nTMhelix 332 354\ninside 355 372

26349 GCF_900157935.1_9789_6_16 Shigella sonnei Proteobacteria; Gammaproteobacteria; Enterobacterales; Enterobacteriaceae; Shigella
MTGDNTLIHSHGINRRDFMKLCAALAATMGLSSKAAAEMAESVTNPQRPPVIWIGAQECT WP_000145410.1
MULTISPECIES: hydrogenase-2 small chain [Proteobacteria] Length: 372\nNumber of predicted TMHs: 1\nExp number of AAs in TMHs: 23.72226\nExp number, first 60 AAs: 0.22038\nTotal prob of N-in: 0.12994\noutside 1 331\nTMhelix 332 354\ninside 355 372

26350 GCF_900157975.1_9789_6_96 Shigella sonnei Proteobacteria; Gammaproteobacteria; Enterobacterales; Enterobacteriaceae; Shigella
MTGDNTLIHSHGINRRDFMKLCAALAATMGLSSKAAAEMAESVTNPQRPPVIWIGAQECT WP_000145410.1
MULTISPECIES: hydrogenase-2 small chain [Proteobacteria] Length: 372\nNumber of predicted TMHs: 1\nExp number of AAs in TMHs: 23.72226\nExp number, first 60 AAs: 0.22038\nTotal prob of N-in: 0.12994\noutside 1 331\nTMhelix 332 354\ninside 355 372

26351 GCF_900157985.1_9789_6_95 Shigella sonnei Proteobacteria; Gammaproteobacteria; Enterobacterales; Enterobacteriaceae; Shigella
MTGDNTLIHSHGINRRDFMKLCAALAATMGLSSKAAAEMAESVTNPQRPPVIWIGAQECT WP_000145410.1
MULTISPECIES: hydrogenase-2 small chain [Proteobacteria] Length: 372\nNumber of predicted TMHs: 1\nExp number of AAs in TMHs: 23.72226\nExp number, first 60 AAs: 0.22038\nTotal prob of N-in: 0.12994\noutside 1 331\nTMhelix 332 354\ninside 355 372

26352 GCF_900158005.1_9789_6_93 Shigella sonnei Proteobacteria; Gammaproteobacteria; Enterobacterales; Enterobacteriaceae; Shigella

MTGDNTLIHSHGINRRDFMKLCAALAATMGLSSKAAAEMAESVTNPQRPPVIWIGAQECT WP_000145410.1 MULTISPECIES: hydrogenase-2 small chain [Proteobacteria] Length: 372\nNumber of predicted TMHs: 1\nExp number of AAs in TMHs: 23.72226\nExp number, first 60 AAs: 0.22038\nTotal prob of N-in: 0.12994\noutside 1 331\nTMhelix 332 354\ninside 355 372

26353 GCF_900158055.1_9803_4_1Shigella sonnei Proteobacteria; Gammaproteobacteria; Enterobacterales; Enterobacteriaceae; Shigella MTGDNTLIHSHGINRRDFMKLCAALAATMGLSSKAAAEMAESVTNPQRPPVIWIGAQECT WP_000145410.1 MULTISPECIES: hydrogenase-2 small chain [Proteobacteria] Length: 372\nNumber of predicted TMHs: 1\nExp number of AAs in TMHs: 23.72226\nExp number, first 60 AAs: 0.22038\nTotal prob of N-in: 0.12994\noutside 1 331\nTMhelix 332 354\ninside 355 372

26354 GCF_900158015.1_9789_6_92 Shigella sonnei Proteobacteria; Gammaproteobacteria; Enterobacterales; Enterobacteriaceae; Shigella MTGDNTLIHSHGINRRDFMKLCAALAATMGLSSKAAAEMAESVTNPQRPPVIWIGAQECT WP_000145410.1 MULTISPECIES: hydrogenase-2 small chain [Proteobacteria] Length: 372\nNumber of predicted TMHs: 1\nExp number of AAs in TMHs: 23.72226\nExp number, first 60 AAs: 0.22038\nTotal prob of N-in: 0.12994\noutside 1 331\nTMhelix 332 354\ninside 355 372

26355 GCF_900158035.1_9803_4_2Shigella sonnei Proteobacteria; Gammaproteobacteria; Enterobacterales; Enterobacteriaceae; Shigella MTGDNTLIHSHGINRRDFMKLCAALAATMGLSSKAAAEMAESVTNPQRPPVIWIGAQECT WP_000145410.1 MULTISPECIES: hydrogenase-2 small chain [Proteobacteria] Length: 372\nNumber of predicted TMHs: 1\nExp number of AAs in TMHs: 23.72226\nExp number, first 60 AAs: 0.22038\nTotal prob of N-in: 0.12994\noutside 1 331\nTMhelix 332 354\ninside 355 372

26356 GCF_900158065.1_9789_6_89 Shigella sonnei Proteobacteria; Gammaproteobacteria; Enterobacterales; Enterobacteriaceae; Shigella MTGDNTLIHSHGINRRDFMKLCAALAATMGLSSKAAAEMAESVTNPQRPPVIWIGAQECT WP_000145410.1 MULTISPECIES: hydrogenase-2 small chain [Proteobacteria] Length: 372\nNumber of predicted TMHs: 1\nExp number of AAs in TMHs: 23.72226\nExp number, first 60 AAs: 0.22038\nTotal prob of N-in: 0.12994\noutside 1 331\nTMhelix 332 354\ninside 355 372

26357 GCF_900158015.1_9789_6_92 Shigella sonnei Proteobacteria; Gammaproteobacteria; Enterobacterales; Enterobacteriaceae; Shigella MNNEETFYQAMRRQGVTRRSFLKYCSLAATSLGLGAGMAPKIAWALENKPRIPVVWIHGL WP_005139779.1 hydrogenase [Shigella sonnei] Length: 372\nNumber of predicted TMHs: 2\nExp number of AAs in TMHs: 37.02628\nExp number, first 60 AAs: 19.39684\nTotal prob of N-in: 0.94313\nPOSSIBLE N-term signal sequence\ninside 1 19\nTMhelix 20 39\noutside 40 325\nTMhelix 326 348\ninside 349 372

26358 GCF_900158035.1_9803_4_2Shigella sonnei Proteobacteria; Gammaproteobacteria; Enterobacterales; Enterobacteriaceae; Shigella MNNEETFYQAMRRQGVTRRSFLKYCSLAATSLGLGAGMAPKIAWALENKPRIPVVWIHGL WP_005139779.1 hydrogenase [Shigella sonnei] Length: 372\nNumber of predicted TMHs: 2\nExp number of AAs in TMHs: 37.02628\nExp number, first 60 AAs: 19.39684\nTotal prob of N-in: 0.94313\nPOSSIBLE N-term signal sequence\ninside 1 19\nTMhelix 20 39\noutside 40 325\nTMhelix 326 348\ninside 349 372

26359 GCF_900158065.1_9789_6_89 Shigella sonnei Proteobacteria; Gammaproteobacteria; Enterobacterales; Enterobacteriaceae; Shigella MNNEETFYQAMRRQGVTRRSFLKYCSLAATSLGLGAGMAPKIAWALENKPRIPVVWIHGL WP_005139779.1 hydrogenase [Shigella sonnei] Length: 372\nNumber of predicted TMHs: 2\nExp number of AAs in TMHs: 37.02628\nExp number, first 60 AAs: 19.39684\nTotal prob of N-in: 0.94313\nPOSSIBLE N-term signal sequence\ninside 1 19\nTMhelix 20 39\noutside 40 325\nTMhelix 326 348\ninside 349 372

26360 GCF_900157935.1_9789_6_16 Shigella sonnei Proteobacteria; Gammaproteobacteria; Enterobacterales; Enterobacteriaceae; Shigella MNNEETFYQAMRRQGVTRRSFLKYCSLAATSLGLGAGMAPKIAWALENKPRIPVVWIHGL WP_005139779.1 hydrogenase [Shigella sonnei] Length: 372\nNumber of predicted TMHs: 2\nExp number of AAs in TMHs: 37.02628\nExp number, first 60 AAs: 19.39684\nTotal prob of N-in: 0.94313\nPOSSIBLE N-term signal sequence\ninside 1 19\nTMhelix 20 39\noutside 40 325\nTMhelix 326 348\ninside 349 372

26361 GCF_900157925.1_9789_6_44 Shigella sonnei Proteobacteria; Gammaproteobacteria; Enterobacterales; Enterobacteriaceae; Shigella MNNEETFYQAMRRQGVTRRSFLKYCSLAATSLGLGAGMAPKIAWALENKPRIPVVWIHGL WP_005139779.1

hydrogenase [Shigella sonnei] Length: 372\nNumber of predicted TMHs: 2\nExp number of AAs in TMHs: 37.02628\nExp number, first 60 AAs: 19.39684\nTotal prob of N-in: 0.94313\nPOSSIBLE N-term signal sequence\ninside 1 19\nTMhelix 20 39\noutside 40 325\nTMhelix 326 348\ninside 349 372

26362 GCF_900157975.1_9789_6_96 Shigella sonnei Proteobacteria; Gammaproteobacteria; Enterobacterales; Enterobacteriaceae; Shigella MNNEETFYQAMRRQGVTRRSFLKYCSLAATSLGLGAGMAPKIAWALENKPRIPVVWIHGL WP_005139779.1 hydrogenase [Shigella sonnei] Length: 372\nNumber of predicted TMHs: 2\nExp number of AAs in TMHs: 37.02628\nExp number, first 60 AAs: 19.39684\nTotal prob of N-in: 0.94313\nPOSSIBLE N-term signal sequence\ninside 1 19\nTMhelix 20 39\noutside 40 325\nTMhelix 326 348\ninside 349 372

26363 GCF_900157985.1_9789_6_95 Shigella sonnei Proteobacteria; Gammaproteobacteria; Enterobacterales; Enterobacteriaceae; Shigella MNNEETFYQAMRRQGVTRRSFLKYCSLAATSLGLGAGMAPKIAWALENKPRIPVVWIHGL WP_005139779.1 hydrogenase [Shigella sonnei] Length: 372\nNumber of predicted TMHs: 2\nExp number of AAs in TMHs: 37.02628\nExp number, first 60 AAs: 19.39684\nTotal prob of N-in: 0.94313\nPOSSIBLE N-term signal sequence\ninside 1 19\nTMhelix 20 39\noutside 40 325\nTMhelix 326 348\ninside 349 372

26364 GCF_900158085.1_9803_4_4Shigella sonnei Proteobacteria; Gammaproteobacteria; Enterobacterales; Enterobacteriaceae; Shigella MTGDNTLIHSHGINRRDFMKLCAALAATMGLSSKAAAEMAESVTNPQRPPVIWIGAQECT WP_000145410.1 MULTISPECIES: hydrogenase-2 small chain [Proteobacteria] Length: 372\nNumber of predicted TMHs: 1\nExp number of AAs in TMHs: 23.72226\nExp number, first 60 AAs: 0.22038\nTotal prob of N-in: 0.12994\noutside 1 331\nTMhelix 332 354\ninside 355 372

26365 GCF_900158085.1_9803_4_4Shigella sonnei Proteobacteria; Gammaproteobacteria; Enterobacterales; Enterobacteriaceae; Shigella MNNEETFYQAMRRQGVTRRSFLKYCSLAATSLGLGAGMAPKIAWALENKPRIPVVWIHGL WP_005139779.1 hydrogenase [Shigella sonnei] Length: 372\nNumber of predicted TMHs: 2\nExp number of AAs in TMHs: 37.02628\nExp number, first 60 AAs: 19.39684\nTotal prob of N-in: 0.94313\nPOSSIBLE N-term signal sequence\ninside 1 19\nTMhelix 20 39\noutside 40 325\nTMhelix 326 348\ninside 349 372

26366 GCF_900158005.1_9789_6_93 Shigella sonnei Proteobacteria; Gammaproteobacteria; Enterobacterales; Enterobacteriaceae; Shigella MNNEETFYQAMRRQGVTRRSFLKYCSLAATSLGLGAGMAPKIAWALENKPRIPVVWIHGL WP_005139779.1 hydrogenase [Shigella sonnei] Length: 372\nNumber of predicted TMHs: 2\nExp number of AAs in TMHs: 37.02628\nExp number, first 60 AAs: 19.39684\nTotal prob of N-in: 0.94313\nPOSSIBLE N-term signal sequence\ninside 1 19\nTMhelix 20 39\noutside 40 325\nTMhelix 326 348\ninside 349 372

26367 GCF_900158105.1_9803_4_6Shigella sonnei Proteobacteria; Gammaproteobacteria; Enterobacterales; Enterobacteriaceae; Shigella MTGDNTLIHSHGINRRDFMKLCAALAATMGLSSKAAAEMAESVTNPQRPPVIWIGAQECT WP_000145410.1 MULTISPECIES: hydrogenase-2 small chain [Proteobacteria] Length: 372\nNumber of predicted TMHs: 1\nExp number of AAs in TMHs: 23.72226\nExp number, first 60 AAs: 0.22038\nTotal prob of N-in: 0.12994\noutside 1 331\nTMhelix 332 354\ninside 355 372

26368 GCF_900158115.1_9803_4_7Shigella sonnei Proteobacteria; Gammaproteobacteria; Enterobacterales; Enterobacteriaceae; Shigella MTGDNTLIHSHGINRRDFMKLCAALAATMGLSSKAAAEMAESVTNPQRPPVIWIGAQECT WP_000145410.1 MULTISPECIES: hydrogenase-2 small chain [Proteobacteria] Length: 372\nNumber of predicted TMHs: 1\nExp number of AAs in TMHs: 23.72226\nExp number, first 60 AAs: 0.22038\nTotal prob of N-in: 0.12994\noutside 1 331\nTMhelix 332 354\ninside 355 372

26369 GCF_900158105.1_9803_4_6Shigella sonnei Proteobacteria; Gammaproteobacteria; Enterobacterales; Enterobacteriaceae; Shigella MNNEETFYQAMRRQGVTRRSFLKYCSLAATSLGLGAGMAPKIAWALENKPRIPVVWIHGL WP_005139779.1 hydrogenase [Shigella sonnei] Length: 372\nNumber of predicted TMHs: 2\nExp number of AAs in TMHs: 37.02628\nExp number, first 60 AAs: 19.39684\nTotal prob of N-in: 0.94313\nPOSSIBLE N-term signal sequence\ninside 1 19\nTMhelix 20 39\noutside 40 325\nTMhelix 326 348\ninside 349 372

26370 GCF_900158115.1_9803_4_7Shigella sonnei Proteobacteria; Gammaproteobacteria; Enterobacterales; Enterobacteriaceae; Shigella MNNEETFYQAMRRQGVTRRSFLKYCSLAATSLGLGAGMAPKIAWALENKPRIPVVWIHGL WP_005139779.1 hydrogenase [Shigella sonnei] Length: 372\nNumber of predicted TMHs: 2\nExp number of AAs in TMHs: 37.02628\nExp number, first 60 AAs: 19.39684\nTotal prob of N-in: 0.94313\nPOSSIBLE N-term signal sequence\ninside 1 19\nTMhelix 20 39\noutside 40 325\nTMhelix 326 348\ninside 349 372

26371 GCF_900158055.1_9803_4_1Shigella sonnei Proteobacteria; Gammaproteobacteria; Enterobacterales; Enterobacteriaceae; Shigella MNNEETFYQAMRRQGVTRRSFLKYCSLAATSLGLGAGMAPKIAWALENKPRIPVVWIHGL WP_005139779.1 hydrogenase [Shigella sonnei] Length: 372\nNumber of predicted TMHs: 2\nExp number of AAs in TMHs: 37.02628\nExp number, first 60 AAs: 19.39684\nTotal prob of N-in: 0.94313\nPOSSIBLE N-term signal sequence\ninside 1 19\nTMhelix 20 39\noutside 40 325\nTMhelix 326 348\ninside 349 372

26372 GCF_900158135.1_9803_4_10 Shigella sonnei Proteobacteria; Gammaproteobacteria; Enterobacterales; Enterobacteriaceae; Shigella MTGDNTLIHSHGINRRDFMKLCAALAATMGLSSKAAAEMAESVTNPQRPPVIWIGAQECT WP_000145410.1 MULTISPECIES: hydrogenase-2 small chain [Proteobacteria] Length: 372\nNumber of predicted TMHs: 1\nExp number of AAs in TMHs: 23.72226\nExp number, first 60 AAs: 0.22038\nTotal prob of N-in: 0.12994\noutside 1 331\nTMhelix 332 354\ninside 355 372

26373 GCF_900158155.1_9803_4_9Shigella sonnei Proteobacteria; Gammaproteobacteria; Enterobacterales; Enterobacteriaceae; Shigella MTGDNTLIHSHGINRRDFMKLCAALAATMGLSSKAAAEMAESVTNPQRPPVIWIGAQECT WP_000145410.1 MULTISPECIES: hydrogenase-2 small chain [Proteobacteria] Length: 372\nNumber of predicted TMHs: 1\nExp number of AAs in TMHs: 23.72226\nExp number, first 60 AAs: 0.22038\nTotal prob of N-in: 0.12994\noutside 1 331\nTMhelix 332 354\ninside 355 372

26374 GCF_900158165.1_9803_4_13 Shigella sonnei Proteobacteria; Gammaproteobacteria; Enterobacterales; Enterobacteriaceae; Shigella MTGDNTLIHSHGINRRDFMKLCAALAATMGLSSKAAAEMAESVTNPQRPPVIWIGAQECT WP_000145410.1 MULTISPECIES: hydrogenase-2 small chain [Proteobacteria] Length: 372\nNumber of predicted TMHs: 1\nExp number of AAs in TMHs: 23.72226\nExp number, first 60 AAs: 0.22038\nTotal prob of N-in: 0.12994\noutside 1 331\nTMhelix 332 354\ninside 355 372

26375 GCF_900158185.1_9803_4_15 Shigella sonnei Proteobacteria; Gammaproteobacteria; Enterobacterales; Enterobacteriaceae; Shigella MTGDNTLIHSHGINRRDFMKLCAALAATMGLSSKAAAEMAESVTNPQRPPVIWIGAQECT WP_000145410.1 MULTISPECIES: hydrogenase-2 small chain [Proteobacteria] Length: 372\nNumber of predicted TMHs: 1\nExp number of AAs in TMHs: 23.72226\nExp number, first 60 AAs: 0.22038\nTotal prob of N-in: 0.12994\noutside 1 331\nTMhelix 332 354\ninside 355 372

26376 GCF_900158195.1_9803_4_16 Shigella sonnei Proteobacteria; Gammaproteobacteria; Enterobacterales; Enterobacteriaceae; Shigella MTGDNTLIHSHGINRRDFMKLCAALAATMGLSSKAAAEMAESVTNPQRPPVIWIGAQECT WP_000145410.1 MULTISPECIES: hydrogenase-2 small chain [Proteobacteria] Length: 372\nNumber of predicted TMHs: 1\nExp number of AAs in TMHs: 23.72226\nExp number, first 60 AAs: 0.22038\nTotal prob of N-in: 0.12994\noutside 1 331\nTMhelix 332 354\ninside 355 372

26377 GCF_900158215.1_9803_4_19 Shigella sonnei Proteobacteria; Gammaproteobacteria; Enterobacterales; Enterobacteriaceae; Shigella MTGDNTLIHSHGINRRDFMKLCAALAATMGLSSKAAAEMAESVTNPQRPPVIWIGAQECT WP_000145410.1 MULTISPECIES: hydrogenase-2 small chain [Proteobacteria] Length: 372\nNumber of predicted TMHs: 1\nExp number of AAs in TMHs: 23.72226\nExp number, first 60 AAs: 0.22038\nTotal prob of N-in: 0.12994\noutside 1 331\nTMhelix 332 354\ninside 355 372

26378 GCF_900158235.1_9803_4_20 Shigella sonnei Proteobacteria; Gammaproteobacteria; Enterobacterales; Enterobacteriaceae; Shigella MTGDNTLIHSHGINRRDFMKLCAALAATMGLSSKAAAEMAESVTNPQRPPVIWIGAQECT WP_000145410.1 MULTISPECIES: hydrogenase-2 small chain [Proteobacteria] Length: 372\nNumber of predicted TMHs: 1\nExp number of AAs in TMHs: 23.72226\nExp number, first 60 AAs: 0.22038\nTotal prob of N-in: 0.12994\noutside 1 331\nTMhelix 332 354\ninside 355 372

26379 GCF_900158245.1_9803_4_21 Shigella sonnei Proteobacteria; Gammaproteobacteria; Enterobacterales; Enterobacteriaceae; Shigella MTGDNTLIHSHGINRRDFMKLCAALAATMGLSSKAAAEMAESVTNPQRPPVIWIGAQECT WP_000145410.1 MULTISPECIES: hydrogenase-2 small chain [Proteobacteria] Length: 372\nNumber of predicted TMHs: 1\nExp number of AAs in TMHs: 23.72226\nExp number, first 60 AAs: 0.22038\nTotal prob of N-in: 0.12994\noutside 1 331\nTMhelix 332 354\ninside 355 372

26380 GCF_900158265.1_9803_4_23 Shigella sonnei Proteobacteria; Gammaproteobacteria; Enterobacterales; Enterobacteriaceae; Shigella MTGDNTLIHSHGINRRDFMKLCAALAATMGLSSKAAAEMAESVTNPQRPPVIWIGAQECT WP_000145410.1 MULTISPECIES: hydrogenase-2 small chain [Proteobacteria] Length: 372\nNumber of predicted TMHs: 1\nExp number of AAs in TMHs: 23.72226\nExp number, first 60 AAs: 0.22038\nTotal prob of N-in: 0.12994\noutside 1 331\nTMhelix 332 354\ninside 355 372

26381 GCF_900158245.1_9803_4_21 Shigella sonnei Proteobacteria; Gammaproteobacteria; Enterobacterales; Enterobacteriaceae; Shigella MNNEETFYQAMRRQGVTRRSFLKYCSLAATSLGLGAGMAPKIAWALENKPRIPVVWIHGL WP_005139779.1 hydrogenase [Shigella sonnei] Length: 372\nNumber of predicted TMHs: 2\nExp number of AAs in TMHs: 37.02628\nExp number, first 60 AAs: 19.39684\nTotal prob of N-in: 0.94313\nPOSSIBLE N-term signal sequence\ninside 1 19\nTMhelix 20 39\noutside 40 325\nTMhelix 326 348\ninside 349 372

26382 GCF_900158265.1_9803_4_23 Shigella sonnei Proteobacteria; Gammaproteobacteria; Enterobacterales; Enterobacteriaceae; Shigella MNNEETFYQAMRRQGVTRRSFLKYCSLAATSLGLGAGMAPKIAWALENKPRIPVVWIHGL WP_005139779.1 hydrogenase [Shigella sonnei] Length: 372\nNumber of predicted TMHs: 2\nExp number of AAs in TMHs: 37.02628\nExp number, first 60 AAs: 19.39684\nTotal prob of N-in: 0.94313\nPOSSIBLE N-term signal sequence\ninside 1 19\nTMhelix 20 39\noutside 40 325\nTMhelix 326 348\ninside 349 372

26383 GCF_900158135.1_9803_4_10 Shigella sonnei Proteobacteria; Gammaproteobacteria; Enterobacterales; Enterobacteriaceae; Shigella MNNEETFYQAMRRQGVTRRSFLKYCSLAATSLGLGAGMAPKIAWALENKPRIPVVWIHGL WP_005139779.1 hydrogenase [Shigella sonnei] Length: 372\nNumber of predicted TMHs: 2\nExp number of AAs in TMHs: 37.02628\nExp number, first 60 AAs: 19.39684\nTotal prob of N-in: 0.94313\nPOSSIBLE N-term signal sequence\ninside 1 19\nTMhelix 20 39\noutside 40 325\nTMhelix 326 348\ninside 349 372

26384 GCF_900158155.1_9803_4_9Shigella sonnei Proteobacteria; Gammaproteobacteria; Enterobacterales; Enterobacteriaceae; Shigella MNNEETFYQAMRRQGVTRRSFLKYCSLAATSLGLGAGMAPKIAWALENKPRIPVVWIHGL WP_005139779.1 hydrogenase [Shigella sonnei] Length: 372\nNumber of predicted TMHs: 2\nExp number of AAs in TMHs: 37.02628\nExp number, first 60 AAs: 19.39684\nTotal prob of N-in: 0.94313\nPOSSIBLE N-term signal sequence\ninside 1 19\nTMhelix 20 39\noutside 40 325\nTMhelix 326 348\ninside 349 372

26385 GCF_900158185.1_9803_4_15 Shigella sonnei Proteobacteria; Gammaproteobacteria; Enterobacterales; Enterobacteriaceae; Shigella MNNEETFYQAMRRQGVTRRSFLKYCSLAATSLGLGAGMAPKIAWALENKPRIPVVWIHGL WP_005139779.1 hydrogenase [Shigella sonnei] Length: 372\nNumber of predicted TMHs: 2\nExp number of AAs in TMHs: 37.02628\nExp number, first 60 AAs: 19.39684\nTotal prob of N-in: 0.94313\nPOSSIBLE N-term signal sequence\ninside 1 19\nTMhelix 20 39\noutside 40 325\nTMhelix 326 348\ninside 349 372

26386 GCF_900158165.1_9803_4_13 Shigella sonnei Proteobacteria; Gammaproteobacteria; Enterobacterales; Enterobacteriaceae; Shigella MNNEETFYQAMRRQGVTRRSFLKYCSLAATSLGLGAGMAPKIAWALENKPRIPVVWIHGL WP_005139779.1 hydrogenase [Shigella sonnei] Length: 372\nNumber of predicted TMHs: 2\nExp number of AAs in TMHs: 37.02628\nExp number, first 60 AAs: 19.39684\nTotal prob of N-in: 0.94313\nPOSSIBLE N-term signal sequence\ninside 1 19\nTMhelix 20 39\noutside 40 325\nTMhelix 326 348\ninside 349 372

26387 GCF_900158195.1_9803_4_16 Shigella sonnei Proteobacteria; Gammaproteobacteria; Enterobacterales; Enterobacteriaceae; Shigella MNNEETFYQAMRRQGVTRRSFLKYCSLAATSLGLGAGMAPKIAWALENKPRIPVVWIHGL WP_005139779.1 hydrogenase [Shigella sonnei] Length: 372\nNumber of predicted TMHs: 2\nExp number of AAs in TMHs: 37.02628\nExp number, first 60 AAs: 19.39684\nTotal prob of N-in: 0.94313\nPOSSIBLE N-term signal sequence\ninside 1 19\nTMhelix 20 39\noutside 40 325\nTMhelix 326 348\ninside 349 372

26388 GCF_900158305.1_9803_4_28 Shigella sonnei Proteobacteria; Gammaproteobacteria; Enterobacterales; Enterobacteriaceae; Shigella MTGDNTLIHSHGINRRDFMKLCAALAATMGLSSKAAAEMAESVTNPQRPPVIWIGAQECT WP_000145410.1 MULTISPECIES: hydrogenase-2 small chain [Proteobacteria] Length: 372\nNumber of predicted TMHs: 1\nExp number of AAs in TMHs: 23.72226\nExp number, first 60 AAs: 0.22038\nTotal prob of N-in: 0.12994\noutside 1 331\nTMhelix 332 354\ninside 355 372

26389 GCF_900158305.1_9803_4_28 Shigella sonnei Proteobacteria; Gammaproteobacteria; Enterobacterales; Enterobacteriaceae; Shigella
MNNEETFYQAMRRQGVTRRSFLKYCSLAATSLGLGAGMAPKIAWALENKPRIPVVWIHGL WP_005139779.1
hydrogenase [Shigella sonnei] Length: 372\nNumber of predicted TMHs: 2\nExp number of AAs in TMHs: 37.02628\nExp number, first 60 AAs: 19.39684\nTotal prob of N-in: 0.94313\nPOSSIBLE N-term signal sequence\ninside 1 19\nTMhelix 20 39\noutside 40 325\nTMhelix 326 348\ninside 349 372

26390 GCF_900158385.1_9803_4_36 Shigella sonnei Proteobacteria; Gammaproteobacteria; Enterobacterales; Enterobacteriaceae; Shigella
MTGDNTLIHSHGINRRDFMKLCAALAATMGLSSKAAAEMAESVTNPQRPPVIWIGAQECT WP_000145410.1
MULTISPECIES: hydrogenase-2 small chain [Proteobacteria] Length: 372\nNumber of predicted TMHs: 1\nExp number of AAs in TMHs: 23.72226\nExp number, first 60 AAs: 0.22038\nTotal prob of N-in: 0.12994\noutside 1 331\nTMhelix 332 354\ninside 355 372

26391 GCF_900158215.1_9803_4_19 Shigella sonnei Proteobacteria; Gammaproteobacteria; Enterobacterales; Enterobacteriaceae; Shigella
MNNEETFYQAMRRQGVTRRSFLKYCSLAATSLGLGAGMAPKIAWALENKPRIPVVWIHGL WP_005139779.1
hydrogenase [Shigella sonnei] Length: 372\nNumber of predicted TMHs: 2\nExp number of AAs in TMHs: 37.02628\nExp number, first 60 AAs: 19.39684\nTotal prob of N-in: 0.94313\nPOSSIBLE N-term signal sequence\ninside 1 19\nTMhelix 20 39\noutside 40 325\nTMhelix 326 348\ninside 349 372

26392 GCF_900158325.1_9803_4_30 Shigella sonnei Proteobacteria; Gammaproteobacteria; Enterobacterales; Enterobacteriaceae; Shigella
MTGDNTLIHSHGINRRDFMKLCAALAATMGLSSKAAAEMAESVTNPQRPPVIWIGAQECT WP_000145410.1
MULTISPECIES: hydrogenase-2 small chain [Proteobacteria] Length: 372\nNumber of predicted TMHs: 1\nExp number of AAs in TMHs: 23.72226\nExp number, first 60 AAs: 0.22038\nTotal prob of N-in: 0.12994\noutside 1 331\nTMhelix 332 354\ninside 355 372

26393 GCF_900158325.1_9803_4_30 Shigella sonnei Proteobacteria; Gammaproteobacteria; Enterobacterales; Enterobacteriaceae; Shigella
MNNEETFYQAMRRQGVTRRSFLKYCSLAATSLGLGAGMAPKIAWALENKPRIPVVWIHGL WP_005139779.1
hydrogenase [Shigella sonnei] Length: 372\nNumber of predicted TMHs: 2\nExp number of AAs in TMHs: 37.02628\nExp number, first 60 AAs: 19.39684\nTotal prob of N-in: 0.94313\nPOSSIBLE N-term signal sequence\ninside 1 19\nTMhelix 20 39\noutside 40 325\nTMhelix 326 348\ninside 349 372

26394 GCF_900158235.1_9803_4_20 Shigella sonnei Proteobacteria; Gammaproteobacteria; Enterobacterales; Enterobacteriaceae; Shigella
MNNEETFYQAMRRQGVTRRSFLKYCSLAATSLGLGAGMAPKIAWALENKPRIPVVWIHGL WP_005139779.1
hydrogenase [Shigella sonnei] Length: 372\nNumber of predicted TMHs: 2\nExp number of AAs in TMHs: 37.02628\nExp number, first 60 AAs: 19.39684\nTotal prob of N-in: 0.94313\nPOSSIBLE N-term signal sequence\ninside 1 19\nTMhelix 20 39\noutside 40 325\nTMhelix 326 348\ninside 349 372

26395 GCF_900158335.1_9803_4_31 Shigella sonnei Proteobacteria; Gammaproteobacteria; Enterobacterales; Enterobacteriaceae; Shigella
MTGDNTLIHSHGINRRDFMKLCAALAATMGLSSKAAAEMAESVTNPQRPPVIWIGAQECT WP_000145410.1
MULTISPECIES: hydrogenase-2 small chain [Proteobacteria] Length: 372\nNumber of predicted TMHs: 1\nExp number of AAs in TMHs: 23.72226\nExp number, first 60 AAs: 0.22038\nTotal prob of N-in: 0.12994\noutside 1 331\nTMhelix 332 354\ninside 355 372

26396 GCF_900158355.1_9803_4_32 Shigella sonnei Proteobacteria; Gammaproteobacteria; Enterobacterales; Enterobacteriaceae; Shigella
MTGDNTLIHSHGINRRDFMKLCAALAATMGLSSKAAAEMAESVTNPQRPPVIWIGAQECT WP_000145410.1
MULTISPECIES: hydrogenase-2 small chain [Proteobacteria] Length: 372\nNumber of predicted TMHs: 1\nExp number of AAs in TMHs: 23.72226\nExp number, first 60 AAs: 0.22038\nTotal prob of N-in: 0.12994\noutside 1 331\nTMhelix 332 354\ninside 355 372

26397 GCF_900158365.1_9803_4_34 Shigella sonnei Proteobacteria; Gammaproteobacteria; Enterobacterales; Enterobacteriaceae; Shigella
MTGDNTLIHSHGINRRDFMKLCAALAATMGLSSKAAAEMAESVTNPQRPPVIWIGAQECT WP_000145410.1
MULTISPECIES: hydrogenase-2 small chain [Proteobacteria] Length: 372\nNumber of predicted TMHs:

1\nExp number of AAs in TMHs: 23.72226\nExp number, first 60 AAs: 0.22038\nTotal prob of N-in: 0.12994\noutside 1 331\nTMhelix 332 354\ninside 355 372

26398 GCF_900158405.1_9803_4_38 Shigella sonnei Proteobacteria; Gammaproteobacteria; Enterobacterales; Enterobacteriaceae; Shigella
MTGDNTLIHSHGINRRDFMKLCAALAATMGLSSKAAAEMAESVTNPQRPPVIWIGAQECT WP_000145410.1
MULTISPECIES: hydrogenase-2 small chain [Proteobacteria] Length: 372\nNumber of predicted TMHs: 1\nExp number of AAs in TMHs: 23.72226\nExp number, first 60 AAs: 0.22038\nTotal prob of N-in: 0.12994\noutside 1 331\nTMhelix 332 354\ninside 355 372

26399 GCF_900158415.1_9803_4_39 Shigella sonnei Proteobacteria; Gammaproteobacteria; Enterobacterales; Enterobacteriaceae; Shigella
MTGDNTLIHSHGINRRDFMKLCAALAATMGLSSKAAAEMAESVTNPQRPPVIWIGAQECT WP_000145410.1
MULTISPECIES: hydrogenase-2 small chain [Proteobacteria] Length: 372\nNumber of predicted TMHs: 1\nExp number of AAs in TMHs: 23.72226\nExp number, first 60 AAs: 0.22038\nTotal prob of N-in: 0.12994\noutside 1 331\nTMhelix 332 354\ninside 355 372

26400 GCF_900158435.1_9803_4_41 Shigella sonnei Proteobacteria; Gammaproteobacteria; Enterobacterales; Enterobacteriaceae; Shigella
MTGDNTLIHSHGINRRDFMKLCAALAATMGLSSKAAAEMAESVTNPQRPPVIWIGAQECT WP_000145410.1
MULTISPECIES: hydrogenase-2 small chain [Proteobacteria] Length: 372\nNumber of predicted TMHs: 1\nExp number of AAs in TMHs: 23.72226\nExp number, first 60 AAs: 0.22038\nTotal prob of N-in: 0.12994\noutside 1 331\nTMhelix 332 354\ninside 355 372

26401 GCF_900158455.1_9803_4_43 Shigella sonnei Proteobacteria; Gammaproteobacteria; Enterobacterales; Enterobacteriaceae; Shigella
MTGDNTLIHSHGINRRDFMKLCAALAATMGLSSKAAAEMAESVTNPQRPPVIWIGAQECT WP_000145410.1
MULTISPECIES: hydrogenase-2 small chain [Proteobacteria] Length: 372\nNumber of predicted TMHs: 1\nExp number of AAs in TMHs: 23.72226\nExp number, first 60 AAs: 0.22038\nTotal prob of N-in: 0.12994\noutside 1 331\nTMhelix 332 354\ninside 355 372

26402 GCF_900158455.1_9803_4_43 Shigella sonnei Proteobacteria; Gammaproteobacteria; Enterobacterales; Enterobacteriaceae; Shigella
MNNEETFYQAMRRQGVTRRSFLKYCSLAATSLGLGAGMAPKIAWALENKPRIPVVWIHGL WP_005139779.1
hydrogenase [Shigella sonnei] Length: 372\nNumber of predicted TMHs: 2\nExp number of AAs in TMHs: 37.02628\nExp number, first 60 AAs: 19.39684\nTotal prob of N-in: 0.94313\nPOSSIBLE N-term signal sequence\ninside 1 19\nTMhelix 20 39\noutside 40 325\nTMhelix 326 348\ninside 349 372

26403 GCF_900158335.1_9803_4_31 Shigella sonnei Proteobacteria; Gammaproteobacteria; Enterobacterales; Enterobacteriaceae; Shigella
MNNEETFYQAMRRQGVTRRSFLKYCSLAATSLGLGAGMAPKIAWALENKPRIPVVWIHGL WP_005139779.1
hydrogenase [Shigella sonnei] Length: 372\nNumber of predicted TMHs: 2\nExp number of AAs in TMHs: 37.02628\nExp number, first 60 AAs: 19.39684\nTotal prob of N-in: 0.94313\nPOSSIBLE N-term signal sequence\ninside 1 19\nTMhelix 20 39\noutside 40 325\nTMhelix 326 348\ninside 349 372

26404 GCF_900158355.1_9803_4_32 Shigella sonnei Proteobacteria; Gammaproteobacteria; Enterobacterales; Enterobacteriaceae; Shigella
MNNEETFYQAMRRQGVTRRSFLKYCSLAATSLGLGAGMAPKIAWALENKPRIPVVWIHGL WP_005139779.1
hydrogenase [Shigella sonnei] Length: 372\nNumber of predicted TMHs: 2\nExp number of AAs in TMHs: 37.02628\nExp number, first 60 AAs: 19.39684\nTotal prob of N-in: 0.94313\nPOSSIBLE N-term signal sequence\ninside 1 19\nTMhelix 20 39\noutside 40 325\nTMhelix 326 348\ninside 349 372

26405 GCF_900158465.1_9803_4_44 Shigella sonnei Proteobacteria; Gammaproteobacteria; Enterobacterales; Enterobacteriaceae; Shigella
MTGDNTLIHSHGINRRDFMKLCAALAATMGLSSKAAAEMAESVTNPQRPPVIWIGAQECT WP_000145410.1
MULTISPECIES: hydrogenase-2 small chain [Proteobacteria] Length: 372\nNumber of predicted TMHs: 1\nExp number of AAs in TMHs: 23.72226\nExp number, first 60 AAs: 0.22038\nTotal prob of N-in: 0.12994\noutside 1 331\nTMhelix 332 354\ninside 355 372

26406 GCF_900158465.1_9803_4_44 Shigella sonnei Proteobacteria; Gammaproteobacteria; Enterobacterales; Enterobacteriaceae; Shigella

MNNEETFYQAMRRQGVTRRSFLKYCSLAATSLGLGAGMAPKIAWALENKPRIPVVWIHGL WP_005139779.1 hydrogenase [Shigella sonnei] Length: 372\nNumber of predicted TMHs: 2\nExp number of AAs in TMHs: 37.02628\nExp number, first 60 AAs: 19.39684\nTotal prob of N-in: 0.94313\nPOSSIBLE N-term signal sequence\ninside 1 19\nTMhelix 20 39\noutside 40 325\nTMhelix 326 348\ninside 349 372

26407 GCF_900158365.1_9803_4_34 Shigella sonnei Proteobacteria; Gammaproteobacteria; Enterobacterales; Enterobacteriaceae; Shigella MNNEETFYQAMRRQGVTRRSFLKYCSLAATSLGLGAGMAPKIAWALENKPRIPVVWIHGL WP_005139779.1 hydrogenase [Shigella sonnei] Length: 372\nNumber of predicted TMHs: 2\nExp number of AAs in TMHs: 37.02628\nExp number, first 60 AAs: 19.39684\nTotal prob of N-in: 0.94313\nPOSSIBLE N-term signal sequence\ninside 1 19\nTMhelix 20 39\noutside 40 325\nTMhelix 326 348\ninside 349 372

26408 GCF_900158385.1_9803_4_36 Shigella sonnei Proteobacteria; Gammaproteobacteria; Enterobacterales; Enterobacteriaceae; Shigella MNNEETFYQAMRRQGVTRRSFLKYCSLAATSLGLGAGMAPKIAWALENKPRIPVVWIHGL WP_005139779.1 hydrogenase [Shigella sonnei] Length: 372\nNumber of predicted TMHs: 2\nExp number of AAs in TMHs: 37.02628\nExp number, first 60 AAs: 19.39684\nTotal prob of N-in: 0.94313\nPOSSIBLE N-term signal sequence\ninside 1 19\nTMhelix 20 39\noutside 40 325\nTMhelix 326 348\ninside 349 372

26409 GCF_900158415.1_9803_4_39 Shigella sonnei Proteobacteria; Gammaproteobacteria; Enterobacterales; Enterobacteriaceae; Shigella MNNEETFYQAMRRQGVTRRSFLKYCSLAATSLGLGAGMAPKIAWALENKPRIPVVWIHGL WP_005139779.1 hydrogenase [Shigella sonnei] Length: 372\nNumber of predicted TMHs: 2\nExp number of AAs in TMHs: 37.02628\nExp number, first 60 AAs: 19.39684\nTotal prob of N-in: 0.94313\nPOSSIBLE N-term signal sequence\ninside 1 19\nTMhelix 20 39\noutside 40 325\nTMhelix 326 348\ninside 349 372

26410 GCF_900158405.1_9803_4_38 Shigella sonnei Proteobacteria; Gammaproteobacteria; Enterobacterales; Enterobacteriaceae; Shigella MNNEETFYQAMRRQGVTRRSFLKYCSLAATSLGLGAGMAPKIAWALENKPRIPVVWIHGL WP_005139779.1 hydrogenase [Shigella sonnei] Length: 372\nNumber of predicted TMHs: 2\nExp number of AAs in TMHs: 37.02628\nExp number, first 60 AAs: 19.39684\nTotal prob of N-in: 0.94313\nPOSSIBLE N-term signal sequence\ninside 1 19\nTMhelix 20 39\noutside 40 325\nTMhelix 326 348\ninside 349 372

26411 GCF_900158485.1_9803_4_47 Shigella sonnei Proteobacteria; Gammaproteobacteria; Enterobacterales; Enterobacteriaceae; Shigella MTGDNTLIHSHGINRRDFMKLCAALAATMGLSSKAAAEMAESVTNPQRPPVIWIGAQECT WP_000145410.1 MULTISPECIES: hydrogenase-2 small chain [Proteobacteria] Length: 372\nNumber of predicted TMHs: 1\nExp number of AAs in TMHs: 23.72226\nExp number, first 60 AAs: 0.22038\nTotal prob of N-in: 0.12994\noutside 1 331\nTMhelix 332 354\ninside 355 372

26412 GCF_900158505.1_9803_4_49 Shigella sonnei Proteobacteria; Gammaproteobacteria; Enterobacterales; Enterobacteriaceae; Shigella MTGDNTLIHSHGINRRDFMKLCAALAATMGLSSKAAAEMAESVTNPQRPPVIWIGAQECT WP_000145410.1 MULTISPECIES: hydrogenase-2 small chain [Proteobacteria] Length: 372\nNumber of predicted TMHs: 1\nExp number of AAs in TMHs: 23.72226\nExp number, first 60 AAs: 0.22038\nTotal prob of N-in: 0.12994\noutside 1 331\nTMhelix 332 354\ninside 355 372

26413 GCF_900158515.1_9803_4_50 Shigella sonnei Proteobacteria; Gammaproteobacteria; Enterobacterales; Enterobacteriaceae; Shigella MTGDNTLIHSHGINRRDFMKLCAALAATMGLSSKAAAEMAESVTNPQRPPVIWIGAQECT WP_000145410.1 MULTISPECIES: hydrogenase-2 small chain [Proteobacteria] Length: 372\nNumber of predicted TMHs: 1\nExp number of AAs in TMHs: 23.72226\nExp number, first 60 AAs: 0.22038\nTotal prob of N-in: 0.12994\noutside 1 331\nTMhelix 332 354\ninside 355 372

26414 GCF_900158535.1_9803_4_52 Shigella sonnei Proteobacteria; Gammaproteobacteria; Enterobacterales; Enterobacteriaceae; Shigella MTGDNTLIHSHGINRRDFMKLCAALAATMGLSSKAAAEMAESVTNPQRPPVIWIGAQECT WP_000145410.1 MULTISPECIES: hydrogenase-2 small chain [Proteobacteria] Length: 372\nNumber of predicted TMHs: 1\nExp number of AAs in TMHs: 23.72226\nExp number, first 60 AAs: 0.22038\nTotal prob of N-in: 0.12994\noutside 1 331\nTMhelix 332 354\ninside 355 372

26415 GCF_900158485.1_9803_4_47 Shigella sonnei Proteobacteria; Gammaproteobacteria; Enterobacterales; Enterobacteriaceae; Shigella
MNNEETFYQAMRRQGVTRRSFLKYCSLAATSLGLGAGMAPKIAWALENKPRIPVVWIHGL WP_005139779.1
hydrogenase [Shigella sonnei] Length: 372\nNumber of predicted TMHs: 2\nExp number of AAs in TMHs: 37.02628\nExp number, first 60 AAs: 19.39684\nTotal prob of N-in: 0.94313\nPOSSIBLE N-term signal sequence\ninside 1 19\nTMhelix 20 39\noutside 40 325\nTMhelix 326 348\ninside 349 372

26416 GCF_900158505.1_9803_4_49 Shigella sonnei Proteobacteria; Gammaproteobacteria; Enterobacterales; Enterobacteriaceae; Shigella
MNNEETFYQAMRRQGVTRRSFLKYCSLAATSLGLGAGMAPKIAWALENKPRIPVVWIHGL WP_005139779.1
hydrogenase [Shigella sonnei] Length: 372\nNumber of predicted TMHs: 2\nExp number of AAs in TMHs: 37.02628\nExp number, first 60 AAs: 19.39684\nTotal prob of N-in: 0.94313\nPOSSIBLE N-term signal sequence\ninside 1 19\nTMhelix 20 39\noutside 40 325\nTMhelix 326 348\ninside 349 372

26417 GCF_900158435.1_9803_4_41 Shigella sonnei Proteobacteria; Gammaproteobacteria; Enterobacterales; Enterobacteriaceae; Shigella
MNNEETFYQAMRRQGVTRRSFLKYCSLAATSLGLGAGMAPKIAWALENKPRIPVVWIHGL WP_005139779.1
hydrogenase [Shigella sonnei] Length: 372\nNumber of predicted TMHs: 2\nExp number of AAs in TMHs: 37.02628\nExp number, first 60 AAs: 19.39684\nTotal prob of N-in: 0.94313\nPOSSIBLE N-term signal sequence\ninside 1 19\nTMhelix 20 39\noutside 40 325\nTMhelix 326 348\ninside 349 372

26418 GCF_900158545.1_9803_4_53 Shigella sonnei Proteobacteria; Gammaproteobacteria; Enterobacterales; Enterobacteriaceae; Shigella
MTGDNTLIHSHGINRRDFMKLCAALAATMGLSSKAAAEMAESVTNPQRPPVIWIGAQECT WP_000145410.1
MULTISPECIES: hydrogenase-2 small chain [Proteobacteria] Length: 372\nNumber of predicted TMHs: 1\nExp number of AAs in TMHs: 23.72226\nExp number, first 60 AAs: 0.22038\nTotal prob of N-in: 0.12994\noutside 1 331\nTMhelix 332 354\ninside 355 372

26419 GCF_900158565.1_9803_4_55 Shigella sonnei Proteobacteria; Gammaproteobacteria; Enterobacterales; Enterobacteriaceae; Shigella
MTGDNTLIHSHGINRRDFMKLCAALAATMGLSSKAAAEMAESVTNPQRPPVIWIGAQECT WP_000145410.1
MULTISPECIES: hydrogenase-2 small chain [Proteobacteria] Length: 372\nNumber of predicted TMHs: 1\nExp number of AAs in TMHs: 23.72226\nExp number, first 60 AAs: 0.22038\nTotal prob of N-in: 0.12994\noutside 1 331\nTMhelix 332 354\ninside 355 372

26420 GCF_900158585.1_9803_4_58 Shigella sonnei Proteobacteria; Gammaproteobacteria; Enterobacterales; Enterobacteriaceae; Shigella
MTGDNTLIHSHGINRRDFMKLCAALAATMGLSSKAAAEMAESVTNPQRPPVIWIGAQECT WP_000145410.1
MULTISPECIES: hydrogenase-2 small chain [Proteobacteria] Length: 372\nNumber of predicted TMHs: 1\nExp number of AAs in TMHs: 23.72226\nExp number, first 60 AAs: 0.22038\nTotal prob of N-in: 0.12994\noutside 1 331\nTMhelix 332 354\ninside 355 372

26421 GCF_900158615.1_9803_4_61 Shigella sonnei Proteobacteria; Gammaproteobacteria; Enterobacterales; Enterobacteriaceae; Shigella
MTGDNTLIHSHGINRRDFMKLCAALAATMGLSSKAAAEMAESVTNPQRPPVIWIGAQECT WP_000145410.1
MULTISPECIES: hydrogenase-2 small chain [Proteobacteria] Length: 372\nNumber of predicted TMHs: 1\nExp number of AAs in TMHs: 23.72226\nExp number, first 60 AAs: 0.22038\nTotal prob of N-in: 0.12994\noutside 1 331\nTMhelix 332 354\ninside 355 372

26422 GCF_900158595.1_9803_4_59 Shigella sonnei Proteobacteria; Gammaproteobacteria; Enterobacterales; Enterobacteriaceae; Shigella
MTGDNTLIHSHGINRRDFMKLCAALAATMGLSSKAAAEMAESVTNPQRPPVIWIGAQECT WP_000145410.1
MULTISPECIES: hydrogenase-2 small chain [Proteobacteria] Length: 372\nNumber of predicted TMHs: 1\nExp number of AAs in TMHs: 23.72226\nExp number, first 60 AAs: 0.22038\nTotal prob of N-in: 0.12994\noutside 1 331\nTMhelix 332 354\ninside 355 372

26423 GCF_900158635.1_9803_4_63 Shigella sonnei Proteobacteria; Gammaproteobacteria; Enterobacterales; Enterobacteriaceae; Shigella
MTGDNTLIHSHGINRRDFMKLCAALAATMGLSSKAAAEMAESVTNPQRPPVIWIGAQECT WP_000145410.1
MULTISPECIES: hydrogenase-2 small chain [Proteobacteria] Length: 372\nNumber of predicted TMHs:

1\nExp number of AAs in TMHs: 23.72226\nExp number, first 60 AAs: 0.22038\nTotal prob of N-in: 0.12994\noutside 1 331\nTMhelix 332 354\ninside 355 372

26424 GCF_900158645.1_9803_4_64 Shigella sonnei Proteobacteria; Gammaproteobacteria; Enterobacterales; Enterobacteriaceae; Shigella
MTGDNTLIHSHGINRRDFMKLCAALAATMGLSSKAAAEMAESVTNPQRPPVIWIGAQECT WP_000145410.1
MULTISPECIES: hydrogenase-2 small chain [Proteobacteria] Length: 372\nNumber of predicted TMHs: 1\nExp number of AAs in TMHs: 23.72226\nExp number, first 60 AAs: 0.22038\nTotal prob of N-in: 0.12994\noutside 1 331\nTMhelix 332 354\ninside 355 372

26425 GCF_900158515.1_9803_4_50 Shigella sonnei Proteobacteria; Gammaproteobacteria; Enterobacterales; Enterobacteriaceae; Shigella
MNNEETFYQAMRRQGVTRRSFLKYCSLAATSLGLGAGMAPKIAWALENKPRIPVVWIHGL WP_005139779.1
hydrogenase [Shigella sonnei] Length: 372\nNumber of predicted TMHs: 2\nExp number of AAs in TMHs: 37.02628\nExp number, first 60 AAs: 19.39684\nTotal prob of N-in: 0.94313\nPOSSIBLE N-term signal sequence\ninside 1 19\nTMhelix 20 39\noutside 40 325\nTMhelix 326 348\ninside 349 372

26426 GCF_900158735.1_9803_4_73 Shigella sonnei Proteobacteria; Gammaproteobacteria; Enterobacterales; Enterobacteriaceae; Shigella
MTGDNTLIHSHGINRRDFMKLCAALAATMGLSSKAAAEMAESVTNPQRPPVIWIGAQECT WP_000145410.1
MULTISPECIES: hydrogenase-2 small chain [Proteobacteria] Length: 372\nNumber of predicted TMHs: 1\nExp number of AAs in TMHs: 23.72226\nExp number, first 60 AAs: 0.22038\nTotal prob of N-in: 0.12994\noutside 1 331\nTMhelix 332 354\ninside 355 372

26427 GCF_900158535.1_9803_4_52 Shigella sonnei Proteobacteria; Gammaproteobacteria; Enterobacterales; Enterobacteriaceae; Shigella
MNNEETFYQAMRRQGVTRRSFLKYCSLAATSLGLGAGMAPKIAWALENKPRIPVVWIHGL WP_005139779.1
hydrogenase [Shigella sonnei] Length: 372\nNumber of predicted TMHs: 2\nExp number of AAs in TMHs: 37.02628\nExp number, first 60 AAs: 19.39684\nTotal prob of N-in: 0.94313\nPOSSIBLE N-term signal sequence\ninside 1 19\nTMhelix 20 39\noutside 40 325\nTMhelix 326 348\ninside 349 372

26428 GCF_900158665.1_9803_4_66 Shigella sonnei Proteobacteria; Gammaproteobacteria; Enterobacterales; Enterobacteriaceae; Shigella
MTGDNTLIHSHGINRRDFMKLCAALAATMGLSSKAAAEMAESVTNPQRPPVIWIGAQECT WP_000145410.1
MULTISPECIES: hydrogenase-2 small chain [Proteobacteria] Length: 372\nNumber of predicted TMHs: 1\nExp number of AAs in TMHs: 23.72226\nExp number, first 60 AAs: 0.22038\nTotal prob of N-in: 0.12994\noutside 1 331\nTMhelix 332 354\ninside 355 372

26429 GCF_900158665.1_9803_4_66 Shigella sonnei Proteobacteria; Gammaproteobacteria; Enterobacterales; Enterobacteriaceae; Shigella
MNNEETFYQAMRRQGVTRRSFLKYCSLAATSLGLGAGMAPKIAWALENKPRIPVVWIHGL WP_005139779.1
hydrogenase [Shigella sonnei] Length: 372\nNumber of predicted TMHs: 2\nExp number of AAs in TMHs: 37.02628\nExp number, first 60 AAs: 19.39684\nTotal prob of N-in: 0.94313\nPOSSIBLE N-term signal sequence\ninside 1 19\nTMhelix 20 39\noutside 40 325\nTMhelix 326 348\ninside 349 372

26430 GCF_900158545.1_9803_4_53 Shigella sonnei Proteobacteria; Gammaproteobacteria; Enterobacterales; Enterobacteriaceae; Shigella
MNNEETFYQAMRRQGVTRRSFLKYCSLAATSLGLGAGMAPKIAWALENKPRIPVVWIHGL WP_005139779.1
hydrogenase [Shigella sonnei] Length: 372\nNumber of predicted TMHs: 2\nExp number of AAs in TMHs: 37.02628\nExp number, first 60 AAs: 19.39684\nTotal prob of N-in: 0.94313\nPOSSIBLE N-term signal sequence\ninside 1 19\nTMhelix 20 39\noutside 40 325\nTMhelix 326 348\ninside 349 372

26431 GCF_900158565.1_9803_4_55 Shigella sonnei Proteobacteria; Gammaproteobacteria; Enterobacterales; Enterobacteriaceae; Shigella
MNNEETFYQAMRRQGVTRRSFLKYCSLAATSLGLGAGMAPKIAWALENKPRIPVVWIHGL WP_005139779.1
hydrogenase [Shigella sonnei] Length: 372\nNumber of predicted TMHs: 2\nExp number of AAs in TMHs: 37.02628\nExp number, first 60 AAs: 19.39684\nTotal prob of N-in: 0.94313\nPOSSIBLE N-term signal sequence\ninside 1 19\nTMhelix 20 39\noutside 40 325\nTMhelix 326 348\ninside 349 372

26432 GCF_900158615.1_9803_4_61 Shigella sonnei Proteobacteria; Gammaproteobacteria; Enterobacterales; Enterobacteriaceae; Shigella

MNNEETFYQAMRRQGVTRRSFLKYCSLAATSLGLGAGMAPKIAWALENKPRIPVVWIHGL WP_005139779.1 hydrogenase [Shigella sonnei] Length: 372\nNumber of predicted TMHs: 2\nExp number of AAs in TMHs: 37.02628\nExp number, first 60 AAs: 19.39684\nTotal prob of N-in: 0.94313\nPOSSIBLE N-term signal sequence\ninside 1 19\nTMhelix 20 39\noutside 40 325\nTMhelix 326 348\ninside 349 372

26433 GCF_900158695.1_9803_4_69 Shigella sonnei Proteobacteria; Gammaproteobacteria; Enterobacterales; Enterobacteriaceae; Shigella MTGDNTLIHSHGINRRDFMKLCAALAATMGLSSKAAAEMAESVTNPQRPPVIWIGAQECT WP_000145410.1 MULTISPECIES: hydrogenase-2 small chain [Proteobacteria] Length: 372\nNumber of predicted TMHs: 1\nExp number of AAs in TMHs: 23.72226\nExp number, first 60 AAs: 0.22038\nTotal prob of N-in: 0.12994\noutside 1 331\nTMhelix 332 354\ninside 355 372

26434 GCF_900158695.1_9803_4_69 Shigella sonnei Proteobacteria; Gammaproteobacteria; Enterobacterales; Enterobacteriaceae; Shigella MNNEETFYQAMRRQGVTRRSFLKYCSLAATSLGLGAGMAPKIAWALENKPRIPVVWIHGL WP_005139779.1 hydrogenase [Shigella sonnei] Length: 372\nNumber of predicted TMHs: 2\nExp number of AAs in TMHs: 37.02628\nExp number, first 60 AAs: 19.39684\nTotal prob of N-in: 0.94313\nPOSSIBLE N-term signal sequence\ninside 1 19\nTMhelix 20 39\noutside 40 325\nTMhelix 326 348\ninside 349 372

26435 GCF_900158595.1_9803_4_59 Shigella sonnei Proteobacteria; Gammaproteobacteria; Enterobacterales; Enterobacteriaceae; Shigella MNNEETFYQAMRRQGVTRRSFLKYCSLAATSLGLGAGMAPKIAWALENKPRIPVVWIHGL WP_005139779.1 hydrogenase [Shigella sonnei] Length: 372\nNumber of predicted TMHs: 2\nExp number of AAs in TMHs: 37.02628\nExp number, first 60 AAs: 19.39684\nTotal prob of N-in: 0.94313\nPOSSIBLE N-term signal sequence\ninside 1 19\nTMhelix 20 39\noutside 40 325\nTMhelix 326 348\ninside 349 372

26436 GCF_900158715.1_9803_4_71 Shigella sonnei Proteobacteria; Gammaproteobacteria; Enterobacterales; Enterobacteriaceae; Shigella MTGDNTLIHSHGINRRDFMKLCAALAATMGLSSKAAAEMAESVTNPQRPPVIWIGAQECT WP_000145410.1 MULTISPECIES: hydrogenase-2 small chain [Proteobacteria] Length: 372\nNumber of predicted TMHs: 1\nExp number of AAs in TMHs: 23.72226\nExp number, first 60 AAs: 0.22038\nTotal prob of N-in: 0.12994\noutside 1 331\nTMhelix 332 354\ninside 355 372

26437 GCF_900158715.1_9803_4_71 Shigella sonnei Proteobacteria; Gammaproteobacteria; Enterobacterales; Enterobacteriaceae; Shigella MNNEETFYQAMRRQGVTRRSFLKYCSLAATSLGLGAGMAPKIAWALENKPRIPVVWIHGL WP_005139779.1 hydrogenase [Shigella sonnei] Length: 372\nNumber of predicted TMHs: 2\nExp number of AAs in TMHs: 37.02628\nExp number, first 60 AAs: 19.39684\nTotal prob of N-in: 0.94313\nPOSSIBLE N-term signal sequence\ninside 1 19\nTMhelix 20 39\noutside 40 325\nTMhelix 326 348\ninside 349 372

26438 GCF_900158635.1_9803_4_63 Shigella sonnei Proteobacteria; Gammaproteobacteria; Enterobacterales; Enterobacteriaceae; Shigella MNNEETFYQAMRRQGVTRRSFLKYCSLAATSLGLGAGMAPKIAWALENKPRIPVVWIHGL WP_005139779.1 hydrogenase [Shigella sonnei] Length: 372\nNumber of predicted TMHs: 2\nExp number of AAs in TMHs: 37.02628\nExp number, first 60 AAs: 19.39684\nTotal prob of N-in: 0.94313\nPOSSIBLE N-term signal sequence\ninside 1 19\nTMhelix 20 39\noutside 40 325\nTMhelix 326 348\ninside 349 372

26439 GCF_900158735.1_9803_4_73 Shigella sonnei Proteobacteria; Gammaproteobacteria; Enterobacterales; Enterobacteriaceae; Shigella MNNEETFYQAMRRQGVTRRSFLKYCSLAATSLGLGAGMAPKIAWALENKPRIPVVWIHGL WP_005139779.1 hydrogenase [Shigella sonnei] Length: 372\nNumber of predicted TMHs: 2\nExp number of AAs in TMHs: 37.02628\nExp number, first 60 AAs: 19.39684\nTotal prob of N-in: 0.94313\nPOSSIBLE N-term signal sequence\ninside 1 19\nTMhelix 20 39\noutside 40 325\nTMhelix 326 348\ninside 349 372

26440 GCF_900158585.1_9803_4_58 Shigella sonnei Proteobacteria; Gammaproteobacteria; Enterobacterales; Enterobacteriaceae; Shigella MNNEETFYQAMRRQGVTRRSFLKYCSLAATSLGLGAGMAPKIAWALENKPRIPVVWIHGL WP_005139779.1 hydrogenase [Shigella sonnei] Length: 372\nNumber of predicted TMHs: 2\nExp number of AAs in TMHs: 37.02628\nExp number, first 60 AAs: 19.39684\nTotal prob of N-in: 0.94313\nPOSSIBLE N-term signal sequence\ninside 1 19\nTMhelix 20 39\noutside 40 325\nTMhelix 326 348\ninside 349 372

26441 GCF_900158645.1_9803_4_64 Shigella sonnei Proteobacteria; Gammaproteobacteria; Enterobacterales; Enterobacteriaceae; Shigella
MNNEETFYQAMRRQGVTRRSFLKYCSLAATSLGLGAGMAPKIAWALENKPRIPVVWIHGL WP_005139779.1
hydrogenase [Shigella sonnei] Length: 372\nNumber of predicted TMHs: 2\nExp number of AAs in TMHs: 37.02628\nExp number, first 60 AAs: 19.39684\nTotal prob of N-in: 0.94313\nPOSSIBLE N-term signal sequence\ninside 1 19\nTMhelix 20 39\noutside 40 325\nTMhelix 326 348\ninside 349 372

26442 GCF_900158745.1_9803_4_74 Shigella sonnei Proteobacteria; Gammaproteobacteria; Enterobacterales; Enterobacteriaceae; Shigella
MTGDNTLIHSHGINRRDFMKLCAALAATMGLSSKAAAEMAESVTNPQRPPVIWIGAQECT WP_000145410.1
MULTISPECIES: hydrogenase-2 small chain [Proteobacteria] Length: 372\nNumber of predicted TMHs: 1\nExp number of AAs in TMHs: 23.72226\nExp number, first 60 AAs: 0.22038\nTotal prob of N-in: 0.12994\noutside 1 331\nTMhelix 332 354\ninside 355 372

26443 GCF_900158765.1_9803_4_76 Shigella sonnei Proteobacteria; Gammaproteobacteria; Enterobacterales; Enterobacteriaceae; Shigella
MTGDNTLIHSHGINRRDFMKLCAALAATMGLSSKAAAEMAESVTNPQRPPVIWIGAQECT WP_000145410.1
MULTISPECIES: hydrogenase-2 small chain [Proteobacteria] Length: 372\nNumber of predicted TMHs: 1\nExp number of AAs in TMHs: 23.72226\nExp number, first 60 AAs: 0.22038\nTotal prob of N-in: 0.12994\noutside 1 331\nTMhelix 332 354\ninside 355 372

26444 GCF_900158775.1_9803_4_77 Shigella sonnei Proteobacteria; Gammaproteobacteria; Enterobacterales; Enterobacteriaceae; Shigella
MTGDNTLIHSHGINRRDFMKLCAALAATMGLSSKAAAEMAESVTNPQRPPVIWIGAQECT WP_000145410.1
MULTISPECIES: hydrogenase-2 small chain [Proteobacteria] Length: 372\nNumber of predicted TMHs: 1\nExp number of AAs in TMHs: 23.72226\nExp number, first 60 AAs: 0.22038\nTotal prob of N-in: 0.12994\noutside 1 331\nTMhelix 332 354\ninside 355 372

26445 GCF_900158795.1_9803_4_79 Shigella sonnei Proteobacteria; Gammaproteobacteria; Enterobacterales; Enterobacteriaceae; Shigella
MTGDNTLIHSHGINRRDFMKLCAALAATMGLSSKAAAEMAESVTNPQRPPVIWIGAQECT WP_000145410.1
MULTISPECIES: hydrogenase-2 small chain [Proteobacteria] Length: 372\nNumber of predicted TMHs: 1\nExp number of AAs in TMHs: 23.72226\nExp number, first 60 AAs: 0.22038\nTotal prob of N-in: 0.12994\noutside 1 331\nTMhelix 332 354\ninside 355 372

26446 GCF_900158815.1_9803_4_81 Shigella sonnei Proteobacteria; Gammaproteobacteria; Enterobacterales; Enterobacteriaceae; Shigella
MTGDNTLIHSHGINRRDFMKLCAALAATMGLSSKAAAEMAESVTNPQRPPVIWIGAQECT WP_000145410.1
MULTISPECIES: hydrogenase-2 small chain [Proteobacteria] Length: 372\nNumber of predicted TMHs: 1\nExp number of AAs in TMHs: 23.72226\nExp number, first 60 AAs: 0.22038\nTotal prob of N-in: 0.12994\noutside 1 331\nTMhelix 332 354\ninside 355 372

26447 GCF_900158825.1_9803_4_82 Shigella sonnei Proteobacteria; Gammaproteobacteria; Enterobacterales; Enterobacteriaceae; Shigella
MTGDNTLIHSHGINRRDFMKLCAALAATMGLSSKAAAEMAESVTNPQRPPVIWIGAQECT WP_000145410.1
MULTISPECIES: hydrogenase-2 small chain [Proteobacteria] Length: 372\nNumber of predicted TMHs: 1\nExp number of AAs in TMHs: 23.72226\nExp number, first 60 AAs: 0.22038\nTotal prob of N-in: 0.12994\noutside 1 331\nTMhelix 332 354\ninside 355 372

26448 GCF_900158845.1_9803_4_84 Shigella sonnei Proteobacteria; Gammaproteobacteria; Enterobacterales; Enterobacteriaceae; Shigella
MTGDNTLIHSHGINRRDFMKLCAALAATMGLSSKAAAEMAESVTNPQRPPVIWIGAQECT WP_000145410.1
MULTISPECIES: hydrogenase-2 small chain [Proteobacteria] Length: 372\nNumber of predicted TMHs: 1\nExp number of AAs in TMHs: 23.72226\nExp number, first 60 AAs: 0.22038\nTotal prob of N-in: 0.12994\noutside 1 331\nTMhelix 332 354\ninside 355 372

26449 GCF_900158865.1_9803_4_86 Shigella sonnei Proteobacteria; Gammaproteobacteria; Enterobacterales; Enterobacteriaceae; Shigella
MTGDNTLIHSHGINRRDFMKLCAALAATMGLSSKAAAEMAESVTNPQRPPVIWIGAQECT WP_000145410.1
MULTISPECIES: hydrogenase-2 small chain [Proteobacteria] Length: 372\nNumber of predicted TMHs:

1\nExp number of AAs in TMHs: 23.72226\nExp number, first 60 AAs: 0.22038\nTotal prob of N-in: 0.12994\noutside 1 331\nTMhelix 332 354\ninside 355 372

26450 GCF_900158865.1_9803_4_86 Shigella sonnei Proteobacteria; Gammaproteobacteria; Enterobacterales; Enterobacteriaceae; Shigella MNNEETFYQAMRRQGVTRRSFLKYCSLAATSLGLGAGMAPKIAWALENKPRIPVVWIHGL WP_005139779.1 hydrogenase [Shigella sonnei] Length: 372\nNumber of predicted TMHs: 2\nExp number of AAs in TMHs: 37.02628\nExp number, first 60 AAs: 19.39684\nTotal prob of N-in: 0.94313\nPOSSIBLE N-term signal sequence\ninside 1 19\nTMhelix 20 39\noutside 40 325\nTMhelix 326 348\ninside 349 372

26451 GCF_900158745.1_9803_4_74 Shigella sonnei Proteobacteria; Gammaproteobacteria; Enterobacterales; Enterobacteriaceae; Shigella MNNEETFYQAMRRQGVTRRSFLKYCSLAATSLGLGAGMAPKIAWALENKPRIPVVWIHGL WP_005139779.1 hydrogenase [Shigella sonnei] Length: 372\nNumber of predicted TMHs: 2\nExp number of AAs in TMHs: 37.02628\nExp number, first 60 AAs: 19.39684\nTotal prob of N-in: 0.94313\nPOSSIBLE N-term signal sequence\ninside 1 19\nTMhelix 20 39\noutside 40 325\nTMhelix 326 348\ninside 349 372

26452 GCF_900158765.1_9803_4_76 Shigella sonnei Proteobacteria; Gammaproteobacteria; Enterobacterales; Enterobacteriaceae; Shigella MNNEETFYQAMRRQGVTRRSFLKYCSLAATSLGLGAGMAPKIAWALENKPRIPVVWIHGL WP_005139779.1 hydrogenase [Shigella sonnei] Length: 372\nNumber of predicted TMHs: 2\nExp number of AAs in TMHs: 37.02628\nExp number, first 60 AAs: 19.39684\nTotal prob of N-in: 0.94313\nPOSSIBLE N-term signal sequence\ninside 1 19\nTMhelix 20 39\noutside 40 325\nTMhelix 326 348\ninside 349 372

26453 GCF_900158775.1_9803_4_77 Shigella sonnei Proteobacteria; Gammaproteobacteria; Enterobacterales; Enterobacteriaceae; Shigella MNNEETFYQAMRRQGVTRRSFLKYCSLAATSLGLGAGMAPKIAWALENKPRIPVVWIHGL WP_005139779.1 hydrogenase [Shigella sonnei] Length: 372\nNumber of predicted TMHs: 2\nExp number of AAs in TMHs: 37.02628\nExp number, first 60 AAs: 19.39684\nTotal prob of N-in: 0.94313\nPOSSIBLE N-term signal sequence\ninside 1 19\nTMhelix 20 39\noutside 40 325\nTMhelix 326 348\ninside 349 372

26454 GCF_900158875.1_9803_4_87 Shigella sonnei Proteobacteria; Gammaproteobacteria; Enterobacterales; Enterobacteriaceae; Shigella MTGDNTLIHSHGINRRDFMKLCAALAATMGLSSKAAAEMAESVTNPQRPPVIWIGAQECT WP_000145410.1 MULTISPECIES: hydrogenase-2 small chain [Proteobacteria] Length: 372\nNumber of predicted TMHs: 1\nExp number of AAs in TMHs: 23.72226\nExp number, first 60 AAs: 0.22038\nTotal prob of N-in: 0.12994\noutside 1 331\nTMhelix 332 354\ninside 355 372

26455 GCF_900158875.1_9803_4_87 Shigella sonnei Proteobacteria; Gammaproteobacteria; Enterobacterales; Enterobacteriaceae; Shigella MNNEETFYQAMRRQGVTRRSFLKYCSLAATSLGLGAGMAPKIAWALENKPRIPVVWIHGL WP_005139779.1 hydrogenase [Shigella sonnei] Length: 372\nNumber of predicted TMHs: 2\nExp number of AAs in TMHs: 37.02628\nExp number, first 60 AAs: 19.39684\nTotal prob of N-in: 0.94313\nPOSSIBLE N-term signal sequence\ninside 1 19\nTMhelix 20 39\noutside 40 325\nTMhelix 326 348\ninside 349 372

26456 GCF_900158815.1_9803_4_81 Shigella sonnei Proteobacteria; Gammaproteobacteria; Enterobacterales; Enterobacteriaceae; Shigella MNNEETFYQAMRRQGVTRRSFLKYCSLAATSLGLGAGMAPKIAWALENKPRIPVVWIHGL WP_005139779.1 hydrogenase [Shigella sonnei] Length: 372\nNumber of predicted TMHs: 2\nExp number of AAs in TMHs: 37.02628\nExp number, first 60 AAs: 19.39684\nTotal prob of N-in: 0.94313\nPOSSIBLE N-term signal sequence\ninside 1 19\nTMhelix 20 39\noutside 40 325\nTMhelix 326 348\ninside 349 372

26457 GCF_900158895.1_9803_4_90 Shigella sonnei Proteobacteria; Gammaproteobacteria; Enterobacterales; Enterobacteriaceae; Shigella MTGDNTLIHSHGINRRDFMKLCAALAATMGLSSKAAAEMAESVTNPQRPPVIWIGAQECT WP_000145410.1 MULTISPECIES: hydrogenase-2 small chain [Proteobacteria] Length: 372\nNumber of predicted TMHs: 1\nExp number of AAs in TMHs: 23.72226\nExp number, first 60 AAs: 0.22038\nTotal prob of N-in: 0.12994\noutside 1 331\nTMhelix 332 354\ninside 355 372

26458 GCF_900158905.1_9803_4_91 Shigella sonnei Proteobacteria; Gammaproteobacteria; Enterobacterales; Enterobacteriaceae; Shigella

MTGDNTLIHSHGINRRDFMKLCAALAATMGLSSKAAAEMAESVTNPQRPPVIWIGAQECT WP_000145410.1 MULTISPECIES: hydrogenase-2 small chain [Proteobacteria] Length: 372\nNumber of predicted TMHs: 1\nExp number of AAs in TMHs: 23.72226\nExp number, first 60 AAs: 0.22038\nTotal prob of N-in: 0.12994\noutside 1 331\nTMhelix 332 354\ninside 355 372

26459 GCF_900158925.1_9803_4_93 Shigella sonnei Proteobacteria; Gammaproteobacteria; Enterobacterales; Enterobacteriaceae; Shigella MTGDNTLIHSHGINRRDFMKLCAALAATMGLSSKAAAEMAESVTNPQRPPVIWIGAQECT WP_000145410.1 MULTISPECIES: hydrogenase-2 small chain [Proteobacteria] Length: 372\nNumber of predicted TMHs: 1\nExp number of AAs in TMHs: 23.72226\nExp number, first 60 AAs: 0.22038\nTotal prob of N-in: 0.12994\noutside 1 331\nTMhelix 332 354\ninside 355 372

26460 GCF_900158895.1_9803_4_90 Shigella sonnei Proteobacteria; Gammaproteobacteria; Enterobacterales; Enterobacteriaceae; Shigella MNNEETFYQAMRRQGVTRRSFLKYCSLAATSLGLGAGMAPKIAWALENKPRIPVVWIHGL WP_005139779.1 hydrogenase [Shigella sonnei] Length: 372\nNumber of predicted TMHs: 2\nExp number of AAs in TMHs: 37.02628\nExp number, first 60 AAs: 19.39684\nTotal prob of N-in: 0.94313\nPOSSIBLE N-term signal sequence\ninside 1 19\nTMhelix 20 39\noutside 40 325\nTMhelix 326 348\ninside 349 372

26461 GCF_900158925.1_9803_4_93 Shigella sonnei Proteobacteria; Gammaproteobacteria; Enterobacterales; Enterobacteriaceae; Shigella MNNEETFYQAMRRQGVTRRSFLKYCSLAATSLGLGAGMAPKIAWALENKPRIPVVWIHGL WP_005139779.1 hydrogenase [Shigella sonnei] Length: 372\nNumber of predicted TMHs: 2\nExp number of AAs in TMHs: 37.02628\nExp number, first 60 AAs: 19.39684\nTotal prob of N-in: 0.94313\nPOSSIBLE N-term signal sequence\ninside 1 19\nTMhelix 20 39\noutside 40 325\nTMhelix 326 348\ninside 349 372

26462 GCF_900158905.1_9803_4_91 Shigella sonnei Proteobacteria; Gammaproteobacteria; Enterobacterales; Enterobacteriaceae; Shigella MNNEETFYQAMRRQGVTRRSFLKYCSLAATSLGLGAGMAPKIAWALENKPRIPVVWIHGL WP_005139779.1 hydrogenase [Shigella sonnei] Length: 372\nNumber of predicted TMHs: 2\nExp number of AAs in TMHs: 37.02628\nExp number, first 60 AAs: 19.39684\nTotal prob of N-in: 0.94313\nPOSSIBLE N-term signal sequence\ninside 1 19\nTMhelix 20 39\noutside 40 325\nTMhelix 326 348\ninside 349 372

26463 GCF_900158795.1_9803_4_79 Shigella sonnei Proteobacteria; Gammaproteobacteria; Enterobacterales; Enterobacteriaceae; Shigella MNNEETFYQAMRRQGVTRRSFLKYCSLAATSLGLGAGMAPKIAWALENKPRIPVVWIHGL WP_005139779.1 hydrogenase [Shigella sonnei] Length: 372\nNumber of predicted TMHs: 2\nExp number of AAs in TMHs: 37.02628\nExp number, first 60 AAs: 19.39684\nTotal prob of N-in: 0.94313\nPOSSIBLE N-term signal sequence\ninside 1 19\nTMhelix 20 39\noutside 40 325\nTMhelix 326 348\ninside 349 372

26464 GCF_900158845.1_9803_4_84 Shigella sonnei Proteobacteria; Gammaproteobacteria; Enterobacterales; Enterobacteriaceae; Shigella MNNEETFYQAMRRQGVTRRSFLKYCSLAATSLGLGAGMAPKIAWALENKPRIPVVWIHGL WP_005139779.1 hydrogenase [Shigella sonnei] Length: 372\nNumber of predicted TMHs: 2\nExp number of AAs in TMHs: 37.02628\nExp number, first 60 AAs: 19.39684\nTotal prob of N-in: 0.94313\nPOSSIBLE N-term signal sequence\ninside 1 19\nTMhelix 20 39\noutside 40 325\nTMhelix 326 348\ninside 349 372

26465 GCF_900158825.1_9803_4_82 Shigella sonnei Proteobacteria; Gammaproteobacteria; Enterobacterales; Enterobacteriaceae; Shigella MNNEETFYQAMRRQGVTRRSFLKYCSLAATSLGLGAGMAPKIAWALENKPRIPVVWIHGL WP_005139779.1 hydrogenase [Shigella sonnei] Length: 372\nNumber of predicted TMHs: 2\nExp number of AAs in TMHs: 37.02628\nExp number, first 60 AAs: 19.39684\nTotal prob of N-in: 0.94313\nPOSSIBLE N-term signal sequence\ninside 1 19\nTMhelix 20 39\noutside 40 325\nTMhelix 326 348\ninside 349 372

26466 GCF_900158945.1_9803_4_95 Shigella sonnei Proteobacteria; Gammaproteobacteria; Enterobacterales; Enterobacteriaceae; Shigella MTGDNTLIHSHGINRRDFMKLCAALAATMGLSSKAAAEMAESVTNPQRPPVIWIGAQECT WP_000145410.1 MULTISPECIES: hydrogenase-2 small chain [Proteobacteria] Length: 372\nNumber of predicted TMHs: 1\nExp number of AAs in TMHs: 23.72226\nExp number, first 60 AAs: 0.22038\nTotal prob of N-in: 0.12994\noutside 1 331\nTMhelix 332 354\ninside 355 372

26467 GCF_900158955.1_9803_4_96 Shigella sonnei Proteobacteria; Gammaproteobacteria; Enterobacterales; Enterobacteriaceae; Shigella MTGDNTLIHSHGINRRDFMKLCAALAATMGLSSKAAAEMAESVTNPQRPPVIWIGAQECT WP_000145410.1 MULTISPECIES: hydrogenase-2 small chain [Proteobacteria] Length: 372\nNumber of predicted TMHs: 1\nExp number of AAs in TMHs: 23.72226\nExp number, first 60 AAs: 0.22038\nTotal prob of N-in: 0.12994\noutside 1 331\nTMhelix 332 354\ninside 355 372

26468 GCF_900158975.1_16703_8_2 Shigella sonnei Proteobacteria; Gammaproteobacteria; Enterobacterales; Enterobacteriaceae; Shigella MTGDNTLIHSHGINRRDFMKLCAALAATMGLSSKAAAEMAESVTNPQRPPVIWIGAQECT WP_000145410.1 MULTISPECIES: hydrogenase-2 small chain [Proteobacteria] Length: 372\nNumber of predicted TMHs: 1\nExp number of AAs in TMHs: 23.72226\nExp number, first 60 AAs: 0.22038\nTotal prob of N-in: 0.12994\noutside 1 331\nTMhelix 332 354\ninside 355 372

26469 GCF_900158995.1_16703_8_5 Shigella sonnei Proteobacteria; Gammaproteobacteria; Enterobacterales; Enterobacteriaceae; Shigella MTGDNTLIHSHGINRRDFMKLCAALAATMGLSSKAAAEMAESVTNPQRPPVIWIGAQECT WP_000145410.1 MULTISPECIES: hydrogenase-2 small chain [Proteobacteria] Length: 372\nNumber of predicted TMHs: 1\nExp number of AAs in TMHs: 23.72226\nExp number, first 60 AAs: 0.22038\nTotal prob of N-in: 0.12994\noutside 1 331\nTMhelix 332 354\ninside 355 372

26470 GCF_900159005.1_16703_8_6 Shigella sonnei Proteobacteria; Gammaproteobacteria; Enterobacterales; Enterobacteriaceae; Shigella MTGDNTLIHSHGINRRDFMKLCAALAATMGLSSKAAAEMAESVTNPQRPPVIWIGAQECT WP_000145410.1 MULTISPECIES: hydrogenase-2 small chain [Proteobacteria] Length: 372\nNumber of predicted TMHs: 1\nExp number of AAs in TMHs: 23.72226\nExp number, first 60 AAs: 0.22038\nTotal prob of N-in: 0.12994\noutside 1 331\nTMhelix 332 354\ninside 355 372

26471 GCF_900159025.1_16703_8_8 Shigella sonnei Proteobacteria; Gammaproteobacteria; Enterobacterales; Enterobacteriaceae; Shigella MTGDNTLIHSHGINRRDFMKLCAALAATMGLSSKAAAEMAESVTNPQRPPVIWIGAQECT WP_000145410.1 MULTISPECIES: hydrogenase-2 small chain [Proteobacteria] Length: 372\nNumber of predicted TMHs: 1\nExp number of AAs in TMHs: 23.72226\nExp number, first 60 AAs: 0.22038\nTotal prob of N-in: 0.12994\noutside 1 331\nTMhelix 332 354\ninside 355 372

26472 GCF_001246015.1_5417_1_1Shigella sonnei Proteobacteria; Gammaproteobacteria; Enterobacterales; Enterobacteriaceae; Shigella MTGDNTLIHSHGINRRDFMKLCAALAATMGLSSKAAAEMAESVTNPQRPPVIWIGAQECT WP_000145410.1 MULTISPECIES: hydrogenase-2 small chain [Proteobacteria] Length: 372\nNumber of predicted TMHs: 1\nExp number of AAs in TMHs: 23.72226\nExp number, first 60 AAs: 0.22038\nTotal prob of N-in: 0.12994\noutside 1 331\nTMhelix 332 354\ninside 355 372

26473 GCF_001245885.1_8290_4_91 Shigella sonnei Proteobacteria; Gammaproteobacteria; Enterobacterales; Enterobacteriaceae; Shigella MTGDNTLIHSHGINRRDFMKLCAALAATMGLSSKAAAEMAESVTNPQRPPVIWIGAQECT WP_000145410.1 MULTISPECIES: hydrogenase-2 small chain [Proteobacteria] Length: 372\nNumber of predicted TMHs: 1\nExp number of AAs in TMHs: 23.72226\nExp number, first 60 AAs: 0.22038\nTotal prob of N-in: 0.12994\noutside 1 331\nTMhelix 332 354\ninside 355 372

26474 GCF_001246385.1_5417_3_1Shigella sonnei Proteobacteria; Gammaproteobacteria; Enterobacterales; Enterobacteriaceae; Shigella MTGDNTLIHSHGINRRDFMKLCAALAATMGLSSKAAAEMAESVTNPQRPPVIWIGAQECT WP_000145410.1 MULTISPECIES: hydrogenase-2 small chain [Proteobacteria] Length: 372\nNumber of predicted TMHs: 1\nExp number of AAs in TMHs: 23.72226\nExp number, first 60 AAs: 0.22038\nTotal prob of N-in: 0.12994\noutside 1 331\nTMhelix 332 354\ninside 355 372

26475 GCF_001246585.1_5236_7_10 Shigella sonnei Proteobacteria; Gammaproteobacteria; Enterobacterales; Enterobacteriaceae; Shigella MTGDNTLIHSHGINRRDFMKLCAALAATMGLSSKAAAEMAESVTNPQRPPVIWIGAQECT WP_000145410.1 MULTISPECIES: hydrogenase-2 small chain [Proteobacteria] Length: 372\nNumber of predicted TMHs: 1\nExp number of AAs in TMHs: 23.72226\nExp number, first 60 AAs: 0.22038\nTotal prob of N-in: 0.12994\noutside 1 331\nTMhelix 332 354\ninside 355 372

26476 GCF_001246675.1_6551_1_10 Shigella sonnei Proteobacteria; Gammaproteobacteria; Enterobacterales; Enterobacteriaceae; Shigella
MTGDNTLIHSHGINRRDFMKLCAALAATMGLSSKAAAEMAESVTNPQRPPVIWIGAQECT WP_000145410.1
MULTISPECIES: hydrogenase-2 small chain [Proteobacteria] Length: 372\nNumber of predicted TMHs: 1\nExp number of AAs in TMHs: 23.72226\nExp number, first 60 AAs: 0.22038\nTotal prob of N-in: 0.12994\noutside 1 331\nTMhelix 332 354\ninside 355 372

26477 GCF_001247155.1_6350_1_16 Shigella sonnei Proteobacteria; Gammaproteobacteria; Enterobacterales; Enterobacteriaceae; Shigella
MTGDNTLIHSHGINRRDFMKLCAALAATMGLSSKAAAEMAESVTNPQRPPVIWIGAQECT WP_000145410.1
MULTISPECIES: hydrogenase-2 small chain [Proteobacteria] Length: 372\nNumber of predicted TMHs: 1\nExp number of AAs in TMHs: 23.72226\nExp number, first 60 AAs: 0.22038\nTotal prob of N-in: 0.12994\noutside 1 331\nTMhelix 332 354\ninside 355 372

26478 GCF_001245885.1_8290_4_91 Shigella sonnei Proteobacteria; Gammaproteobacteria; Enterobacterales; Enterobacteriaceae; Shigella
MNNEETFYQAMRRQGVTRRSFLKYCSLAATSLGLGAGMAPKIAWALENKPRIPVVWIHGL WP_005139779.1
hydrogenase [Shigella sonnei] Length: 372\nNumber of predicted TMHs: 2\nExp number of AAs in TMHs: 37.02628\nExp number, first 60 AAs: 19.39684\nTotal prob of N-in: 0.94313\nPOSSIBLE N-term signal sequence\ninside 1 19\nTMhelix 20 39\noutside 40 325\nTMhelix 326 348\ninside 349 372

26479 GCF_001248155.1_12971_8_75 Shigella sonnei Proteobacteria; Gammaproteobacteria; Enterobacterales; Enterobacteriaceae; Shigella
MNNEETFYQAMRRQGVTRRSFLKYCSLAATSLGLGAGMAPKIAWALENKPRIPVVWIHGL WP_052962775.1
hydrogenase [Shigella sonnei] Length: 372\nNumber of predicted TMHs: 2\nExp number of AAs in TMHs: 37.09248\nExp number, first 60 AAs: 19.39296\nTotal prob of N-in: 0.94316\nPOSSIBLE N-term signal sequence\ninside 1 19\nTMhelix 20 39\noutside 40 325\nTMhelix 326 348\ninside 349 372

26480 GCF_001248215.1_Sg_201401278_GTAGAG_L001 Shigella sonnei Proteobacteria; Gammaproteobacteria; Enterobacterales; Enterobacteriaceae; Shigella
MNNEETFYQAMRRQGVTRRSFLKYCSLAATSLGLGAGMAPKIAWALENKPRIPVVWIHGL WP_005139779.1
hydrogenase [Shigella sonnei] Length: 372\nNumber of predicted TMHs: 2\nExp number of AAs in TMHs: 37.02628\nExp number, first 60 AAs: 19.39684\nTotal prob of N-in: 0.94313\nPOSSIBLE N-term signal sequence\ninside 1 19\nTMhelix 20 39\noutside 40 325\nTMhelix 326 348\ninside 349 372

26481 GCF_001247385.1_8489_2_91 Shigella sonnei Proteobacteria; Gammaproteobacteria; Enterobacterales; Enterobacteriaceae; Shigella
MNNEETFYQAMRRQGVTRRSFLKYCSLAATSLGLGAGMAPKIAWALENKPRIPVVWIHGL WP_005139779.1
hydrogenase [Shigella sonnei] Length: 372\nNumber of predicted TMHs: 2\nExp number of AAs in TMHs: 37.02628\nExp number, first 60 AAs: 19.39684\nTotal prob of N-in: 0.94313\nPOSSIBLE N-term signal sequence\ninside 1 19\nTMhelix 20 39\noutside 40 325\nTMhelix 326 348\ninside 349 372

26482 GCF_001248255.1_5236_8_2Shigella sonnei Proteobacteria; Gammaproteobacteria; Enterobacterales; Enterobacteriaceae; Shigella MTGDNTLIHSHGINRRDFMKLCAALAATMGLSSKAAAEMAESVTNPQRPPVIWIGAQECT
WP_000145410.1 MULTISPECIES: hydrogenase-2 small chain [Proteobacteria] Length: 372\nNumber of predicted TMHs: 1\nExp number of AAs in TMHs: 23.72226\nExp number, first 60 AAs: 0.22038\nTotal prob of N-in: 0.12994\noutside 1 331\nTMhelix 332 354\ninside 355 372

26483 GCF_001248295.1_H14056026101 Shigella sonnei Proteobacteria; Gammaproteobacteria; Enterobacterales; Enterobacteriaceae; Shigella
MTGDNTLIHSHGINRRDFMKLCAALAATMGLSSKAAAEMAESVTNPQRPPVIWIGAQECT WP_000145410.1
MULTISPECIES: hydrogenase-2 small chain [Proteobacteria] Length: 372\nNumber of predicted TMHs: 1\nExp number of AAs in TMHs: 23.72226\nExp number, first 60 AAs: 0.22038\nTotal prob of N-in: 0.12994\noutside 1 331\nTMhelix 332 354\ninside 355 372

26484 GCF_001248295.1_H14056026101 Shigella sonnei Proteobacteria; Gammaproteobacteria; Enterobacterales; Enterobacteriaceae; Shigella
MNNEETFYQAMRRQGVTRRSFLKYCSLAATSLGLGAGMAPKIAWALENKPRIPVVWIHGL WP_005139779.1
hydrogenase [Shigella sonnei] Length: 372\nNumber of predicted TMHs: 2\nExp number of AAs in TMHs: 37.02628\nExp number, first 60 AAs: 19.39684\nTotal prob of N-in: 0.94313\nPOSSIBLE N-term signal sequence\ninside 1 19\nTMhelix 20 39\noutside 40 325\nTMhelix 326 348\ninside 349 372

26485 GCF_001248255.1_5236_8_2Shigella sonnei Proteobacteria; Gammaproteobacteria; Enterobacterales; Enterobacteriaceae; Shigella MNNEETFYQAMRRQGVTRRSFLKYCSLAATSLGLGAGMAPKIAWALENKPRIPVVWIHGL WP_005139779.1 hydrogenase [Shigella sonnei] Length: 372\nNumber of predicted TMHs: 2\nExp number of AAs in TMHs: 37.02628\nExp number, first 60 AAs: 19.39684\nTotal prob of N-in: 0.94313\nPOSSIBLE N-term signal sequence\ninside 1 19\nTMhelix 20 39\noutside 40 325\nTMhelix 326 348\ninside 349 372

26486 GCF_001248685.1_8290_4_4Shigella sonnei Proteobacteria; Gammaproteobacteria; Enterobacterales; Enterobacteriaceae; Shigella MNNEETFYQAMRRQGVTRRSFLKYCSLAATSLGLGAGMAPKIAWALENKPRIPVVWIHGL WP_005139779.1 hydrogenase [Shigella sonnei] Length: 372\nNumber of predicted TMHs: 2\nExp number of AAs in TMHs: 37.02628\nExp number, first 60 AAs: 19.39684\nTotal prob of N-in: 0.94313\nPOSSIBLE N-term signal sequence\ninside 1 19\nTMhelix 20 39\noutside 40 325\nTMhelix 326 348\ninside 349 372

26487 GCF_001248385.1_6551_1_3Shigella sonnei Proteobacteria; Gammaproteobacteria; Enterobacterales; Enterobacteriaceae; Shigella MNNEETFYQAMRRQGVTRRSFLKYCSLAATSLGLGAGMAPKIAWALENKPRIPVVWIHGL WP_005139779.1 hydrogenase [Shigella sonnei] Length: 372\nNumber of predicted TMHs: 2\nExp number of AAs in TMHs: 37.02628\nExp number, first 60 AAs: 19.39684\nTotal prob of N-in: 0.94313\nPOSSIBLE N-term signal sequence\ninside 1 19\nTMhelix 20 39\noutside 40 325\nTMhelix 326 348\ninside 349 372

26488 GCF_001248705.1_8290_4_40 Shigella sonnei Proteobacteria; Gammaproteobacteria; Enterobacterales; Enterobacteriaceae; Shigella MTGDNTLIHSHGINRRDFMKLCAALAATMGLSSKAAAEMAESVTNPQRPPVIWIGAQECT WP_000145410.1 MULTISPECIES: hydrogenase-2 small chain [Proteobacteria] Length: 372\nNumber of predicted TMHs: 1\nExp number of AAs in TMHs: 23.72226\nExp number, first 60 AAs: 0.22038\nTotal prob of N-in: 0.12994\noutside 1 331\nTMhelix 332 354\ninside 355 372

26489 GCF_001248705.1_8290_4_40 Shigella sonnei Proteobacteria; Gammaproteobacteria; Enterobacterales; Enterobacteriaceae; Shigella MNNEETFYQAMRRQGVTRRSFLKYCSLAATSLGLGAGMAPKIAWALENKPRIPVVWIHGL WP_005139779.1 hydrogenase [Shigella sonnei] Length: 372\nNumber of predicted TMHs: 2\nExp number of AAs in TMHs: 37.02628\nExp number, first 60 AAs: 19.39684\nTotal prob of N-in: 0.94313\nPOSSIBLE N-term signal sequence\ninside 1 19\nTMhelix 20 39\noutside 40 325\nTMhelix 326 348\ninside 349 372

26490 GCF_001248345.1_8290_4_87 Shigella sonnei Proteobacteria; Gammaproteobacteria; Enterobacterales; Enterobacteriaceae; Shigella MNNEETFYQAMRRQGVTRRSFLKYCSLAATSLGLGAGMAPKIAWALENKPRIPVVWIHGL WP_005139779.1 hydrogenase [Shigella sonnei] Length: 372\nNumber of predicted TMHs: 2\nExp number of AAs in TMHs: 37.02628\nExp number, first 60 AAs: 19.39684\nTotal prob of N-in: 0.94313\nPOSSIBLE N-term signal sequence\ninside 1 19\nTMhelix 20 39\noutside 40 325\nTMhelix 326 348\ninside 349 372

26491 GCF_001248785.1_5417_3_8Shigella sonnei Proteobacteria; Gammaproteobacteria; Enterobacterales; Enterobacteriaceae; Shigella MTGDNTLIHSHGINRRDFMKLCAALAATMGLSSKAAAEMAESVTNPQRPPVIWIGAQECT WP_000145410.1 MULTISPECIES: hydrogenase-2 small chain [Proteobacteria] Length: 372\nNumber of predicted TMHs: 1\nExp number of AAs in TMHs: 23.72226\nExp number, first 60 AAs: 0.22038\nTotal prob of N-in: 0.12994\noutside 1 331\nTMhelix 332 354\ninside 355 372

26492 GCF_001248745.1_H14146070501 Shigella sonnei Proteobacteria; Gammaproteobacteria; Enterobacterales; Enterobacteriaceae; Shigella MTGDNTLIHSHGINRRDFMKLCAALAATMGLSSKAAAEMAESVTNPQRPPVIWIGAQECT WP_000145410.1 MULTISPECIES: hydrogenase-2 small chain [Proteobacteria] Length: 372\nNumber of predicted TMHs: 1\nExp number of AAs in TMHs: 23.72226\nExp number, first 60 AAs: 0.22038\nTotal prob of N-in: 0.12994\noutside 1 331\nTMhelix 332 354\ninside 355 372

26493 GCF_001249335.1_12830_1_37 Shigella sonnei Proteobacteria; Gammaproteobacteria; Enterobacterales; Enterobacteriaceae; Shigella MTGDNTLIHSHGINRRDFMKLCAALAATMGLSSKAAAEMAESVTNPQRPPVIWIGAQECT WP_000145410.1 MULTISPECIES: hydrogenase-2 small chain [Proteobacteria] Length: 372\nNumber of predicted TMHs: 1\nExp number of AAs in TMHs: 23.72226\nExp number, first 60 AAs: 0.22038\nTotal prob of N-in: 0.12994\noutside 1 331\nTMhelix 332 354\ninside 355 372

26494 GCF_001249335.1_12830_1_37 Shigella sonnei Proteobacteria; Gammaproteobacteria; Enterobacterales; Enterobacteriaceae; Shigella

MNNEETFYQAMRRQGVTRRSFLKYCSLAATSLGLGAGMAPKIAWALENKPRIPVVWIHGL WP_005139779.1 hydrogenase [Shigella sonnei] Length: 372\nNumber of predicted TMHs: 2\nExp number of AAs in TMHs: 37.02628\nExp number, first 60 AAs: 19.39684\nTotal prob of N-in: 0.94313\nPOSSIBLE N-term signal sequence\ninside 1 19\nTMhelix 20 39\noutside 40 325\nTMhelix 326 348\ninside 349 372

26495 GCF_001249025.1_H14090035401 Shigella sonnei Proteobacteria; Gammaproteobacteria; Enterobacterales; Enterobacteriaceae; Shigella MNNEETFYQAMRRQGVTRRSFLKYCSLAATSLGLGAGMAPKIAWALENKPRIPVVWIHGL WP_005139779.1 hydrogenase [Shigella sonnei] Length: 372\nNumber of predicted TMHs: 2\nExp number of AAs in TMHs: 37.02628\nExp number, first 60 AAs: 19.39684\nTotal prob of N-in: 0.94313\nPOSSIBLE N-term signal sequence\ninside 1 19\nTMhelix 20 39\noutside 40 325\nTMhelix 326 348\ninside 349 372

26496 GCF_001249355.1_12971_8_69 Shigella sonnei Proteobacteria; Gammaproteobacteria; Enterobacterales; Enterobacteriaceae; Shigella MTGDNTLIHSHGINRRDFMKLCAALAATMGLSSKAAAEMAESVTNPQRPPVIWIGAQECT WP_000145410.1 MULTISPECIES: hydrogenase-2 small chain [Proteobacteria] Length: 372\nNumber of predicted TMHs: 1\nExp number of AAs in TMHs: 23.72226\nExp number, first 60 AAs: 0.22038\nTotal prob of N-in: 0.12994\noutside 1 331\nTMhelix 332 354\ninside 355 372

26497 GCF_001249355.1_12971_8_69 Shigella sonnei Proteobacteria; Gammaproteobacteria; Enterobacterales; Enterobacteriaceae; Shigella MNNEETFYQAMRRQGVTRRSFLKYCSLAATSLGLGAGMAPKIAWALENKPRIPVVWIHGL WP_005139779.1 hydrogenase [Shigella sonnei] Length: 372\nNumber of predicted TMHs: 2\nExp number of AAs in TMHs: 37.02628\nExp number, first 60 AAs: 19.39684\nTotal prob of N-in: 0.94313\nPOSSIBLE N-term signal sequence\ninside 1 19\nTMhelix 20 39\noutside 40 325\nTMhelix 326 348\ninside 349 372

26498 GCF_001249065.1_5236_8_9Shigella sonnei Proteobacteria; Gammaproteobacteria; Enterobacterales; Enterobacteriaceae; Shigella MNNEETFYQAMRRQGVTRRSFLKYCSLAATSLGLGAGMAPKIAWALENKPRIPVVWIHGL WP_005139779.1 hydrogenase [Shigella sonnei] Length: 372\nNumber of predicted TMHs: 2\nExp number of AAs in TMHs: 37.02628\nExp number, first 60 AAs: 19.39684\nTotal prob of N-in: 0.94313\nPOSSIBLE N-term signal sequence\ninside 1 19\nTMhelix 20 39\noutside 40 325\nTMhelix 326 348\ninside 349 372

26499 GCF_001249395.1_9870_6_6Shigella sonnei Proteobacteria; Gammaproteobacteria; Enterobacterales; Enterobacteriaceae; Shigella MTGDNTLIHSHGINRRDFMKLCAALAATMGLSSKAAAEMAESVTNPQRPPVIWIGAQECT WP_000145410.1 MULTISPECIES: hydrogenase-2 small chain [Proteobacteria] Length: 372\nNumber of predicted TMHs: 1\nExp number of AAs in TMHs: 23.72226\nExp number, first 60 AAs: 0.22038\nTotal prob of N-in: 0.12994\noutside 1 331\nTMhelix 332 354\ninside 355 372

26500 GCF_001249415.1_9870_6_40 Shigella sonnei Proteobacteria; Gammaproteobacteria; Enterobacterales; Enterobacteriaceae; Shigella MNNEETFYQAMRRQGVTRRSFLKYCSLAATSLGLGAGMAPKIAWALENKPRIPVVWIHGL WP_005139779.1 hydrogenase [Shigella sonnei] Length: 372\nNumber of predicted TMHs: 2\nExp number of AAs in TMHs: 37.02628\nExp number, first 60 AAs: 19.39684\nTotal prob of N-in: 0.94313\nPOSSIBLE N-term signal sequence\ninside 1 19\nTMhelix 20 39\noutside 40 325\nTMhelix 326 348\ninside 349 372

26501 GCF_001249455.1_12844_1_58 Shigella sonnei Proteobacteria; Gammaproteobacteria; Enterobacterales; Enterobacteriaceae; Shigella MNNEETFYQAMRRQGVTRRSFLKYCSLAATSLGLGAGMAPKIAWALENKPRIPVVWIHGL WP_005139779.1 hydrogenase [Shigella sonnei] Length: 372\nNumber of predicted TMHs: 2\nExp number of AAs in TMHs: 37.02628\nExp number, first 60 AAs: 19.39684\nTotal prob of N-in: 0.94313\nPOSSIBLE N-term signal sequence\ninside 1 19\nTMhelix 20 39\noutside 40 325\nTMhelix 326 348\ninside 349 372

26502 GCF_001249495.1_9870_6_30 Shigella sonnei Proteobacteria; Gammaproteobacteria; Enterobacterales; Enterobacteriaceae; Shigella MNNEETFYQAMRRQGVTRRSFLKYCSLAATSLGLGAGMAPKIAWALENKPRIPVVWIHGL WP_005139779.1 hydrogenase [Shigella sonnei] Length: 372\nNumber of predicted TMHs: 2\nExp number of AAs in TMHs: 37.02628\nExp number, first 60 AAs: 19.39684\nTotal prob of N-in: 0.94313\nPOSSIBLE N-term signal sequence\ninside 1 19\nTMhelix 20 39\noutside 40 325\nTMhelix 326 348\ninside 349 372

26503 GCF_001249535.1_8489_2_82 Shigella sonnei Proteobacteria; Gammaproteobacteria; Enterobacterales; Enterobacteriaceae; Shigella

MNNEETFYQAMRRQGVTRRSFLKYCSLAATSLGLGAGMAPKIAWALENKPRIPVVWIHGL WP_005139779.1 hydrogenase [Shigella sonnei] Length: 372\nNumber of predicted TMHs: 2\nExp number of AAs in TMHs: 37.02628\nExp number, first 60 AAs: 19.39684\nTotal prob of N-in: 0.94313\nPOSSIBLE N-term signal sequence\ninside 1 19\nTMhelix 20 39\noutside 40 325\nTMhelix 326 348\ninside 349 372

26504 GCF_001249835.1_12971_8_44 Shigella sonnei Proteobacteria; Gammaproteobacteria; Enterobacterales; Enterobacteriaceae; Shigella MTGDNTLIHSHGINRRDFMKLCAALAATMGLSSKAAAEMAESVTNPQRPPVIWIGAQECT WP_000145410.1 MULTISPECIES: hydrogenase-2 small chain [Proteobacteria] Length: 372\nNumber of predicted TMHs: 1\nExp number of AAs in TMHs: 23.72226\nExp number, first 60 AAs: 0.22038\nTotal prob of N-in: 0.12994\noutside 1 331\nTMhelix 332 354\ninside 355 372

26505 GCF_001249875.1_5236_5_2Shigella sonnei Proteobacteria; Gammaproteobacteria; Enterobacterales; Enterobacteriaceae; Shigella MTGDNTLIHSHGINRRDFMKLCAALAATMGLSSKAAAEMAESVTNPQRPPVIWIGAQECT WP_000145410.1 MULTISPECIES: hydrogenase-2 small chain [Proteobacteria] Length: 372\nNumber of predicted TMHs: 1\nExp number of AAs in TMHs: 23.72226\nExp number, first 60 AAs: 0.22038\nTotal prob of N-in: 0.12994\noutside 1 331\nTMhelix 332 354\ninside 355 372

26506 GCF_001249835.1_12971_8_44 Shigella sonnei Proteobacteria; Gammaproteobacteria; Enterobacterales; Enterobacteriaceae; Shigella MNNEETFYQAMRRQGVTRRSFLKYCSLAATSLGLGAGMAPKIAWALENKPRIPVVWIHGL WP_005139779.1 hydrogenase [Shigella sonnei] Length: 372\nNumber of predicted TMHs: 2\nExp number of AAs in TMHs: 37.02628\nExp number, first 60 AAs: 19.39684\nTotal prob of N-in: 0.94313\nPOSSIBLE N-term signal sequence\ninside 1 19\nTMhelix 20 39\noutside 40 325\nTMhelix 326 348\ninside 349 372

26507 GCF_001250175.1_8289_7_61 Shigella sonnei Proteobacteria; Gammaproteobacteria; Enterobacterales; Enterobacteriaceae; Shigella MNNEETFYQAMRRQGVTRRSFLKYCSLAATSLGLGAGMAPKIAWALENKPRIPVVWIHGL WP_005139779.1 hydrogenase [Shigella sonnei] Length: 372\nNumber of predicted TMHs: 2\nExp number of AAs in TMHs: 37.02628\nExp number, first 60 AAs: 19.39684\nTotal prob of N-in: 0.94313\nPOSSIBLE N-term signal sequence\ninside 1 19\nTMhelix 20 39\noutside 40 325\nTMhelix 326 348\ninside 349 372

26508 GCF_001250275.1_8290_4_55 Shigella sonnei Proteobacteria; Gammaproteobacteria; Enterobacterales; Enterobacteriaceae; Shigella MNNEETFYQAMRRQGVTRRSFLKYCSLAATSLGLGAGMAPKIAWALENKPRIPVVWIHGL WP_005139779.1 hydrogenase [Shigella sonnei] Length: 372\nNumber of predicted TMHs: 2\nExp number of AAs in TMHs: 37.02628\nExp number, first 60 AAs: 19.39684\nTotal prob of N-in: 0.94313\nPOSSIBLE N-term signal sequence\ninside 1 19\nTMhelix 20 39\noutside 40 325\nTMhelix 326 348\ninside 349 372

26509 GCF_001249935.1_H14140035801 Shigella sonnei Proteobacteria; Gammaproteobacteria; Enterobacterales; Enterobacteriaceae; Shigella MNNEETFYQAMRRQGVTRRSFLKYCSLAATSLGLGAGMAPKIAWALENKPRIPVVWIHGL WP_052962775.1 hydrogenase [Shigella sonnei] Length: 372\nNumber of predicted TMHs: 2\nExp number of AAs in TMHs: 37.09248\nExp number, first 60 AAs: 19.39296\nTotal prob of N-in: 0.94316\nPOSSIBLE N-term signal sequence\ninside 1 19\nTMhelix 20 39\noutside 40 325\nTMhelix 326 348\ninside 349 372

26510 GCF_001250315.1_8289_7_89 Shigella sonnei Proteobacteria; Gammaproteobacteria; Enterobacterales; Enterobacteriaceae; Shigella MTGDNTLIHSHGINRRDFMKLCAALAATMGLSSKAAAEMAESVTNPQRPPVIWIGAQECT WP_000145410.1 MULTISPECIES: hydrogenase-2 small chain [Proteobacteria] Length: 372\nNumber of predicted TMHs: 1\nExp number of AAs in TMHs: 23.72226\nExp number, first 60 AAs: 0.22038\nTotal prob of N-in: 0.12994\noutside 1 331\nTMhelix 332 354\ninside 355 372

26511 GCF_001250315.1_8289_7_89 Shigella sonnei Proteobacteria; Gammaproteobacteria; Enterobacterales; Enterobacteriaceae; Shigella MNNEETFYQAMRRQGVTRRSFLKYCSLAATSLGLGAGMAPKIAWALENKPRIPVVWIHGL WP_005139779.1 hydrogenase [Shigella sonnei] Length: 372\nNumber of predicted TMHs: 2\nExp number of AAs in TMHs: 37.02628\nExp number, first 60 AAs: 19.39684\nTotal prob of N-in: 0.94313\nPOSSIBLE N-term signal sequence\ninside 1 19\nTMhelix 20 39\noutside 40 325\nTMhelix 326 348\ninside 349 372

26512 GCF_001249975.1_5236_2_7Shigella sonnei Proteobacteria; Gammaproteobacteria; Enterobacterales; Enterobacteriaceae; Shigella MNNEETFYQAMRRQGVTRRSFLKYCSLAATSLGLGAGMAPKIAWALENKPRIPVVWIHGL WP_005139779.1 hydrogenase [Shigella sonnei] Length: 372\nNumber of predicted TMHs: 2\nExp number of AAs in TMHs: 37.02628\nExp number, first 60 AAs: 19.39684\nTotal prob of N-in: 0.94313\nPOSSIBLE N-term signal sequence\ninside 1 19\nTMhelix 20 39\noutside 40 325\nTMhelix 326 348\ninside 349 372

26513 GCF_001250055.1_7655_8_12 Shigella sonnei Proteobacteria; Gammaproteobacteria; Enterobacterales; Enterobacteriaceae; Shigella MNNEETFYQAMRRQGVTRRSFLKYCSLAATSLGLGAGMAPKIAWALENKPRIPVVWIHGL WP_005139779.1 hydrogenase [Shigella sonnei] Length: 372\nNumber of predicted TMHs: 2\nExp number of AAs in TMHs: 37.02628\nExp number, first 60 AAs: 19.39684\nTotal prob of N-in: 0.94313\nPOSSIBLE N-term signal sequence\ninside 1 19\nTMhelix 20 39\noutside 40 325\nTMhelix 326 348\ninside 349 372

26514 GCF_001250555.1_6350_1_10 Shigella sonnei Proteobacteria; Gammaproteobacteria; Enterobacterales; Enterobacteriaceae; Shigella MNNEETFYQAMRRQGVTRRSFLKYCSLAATSLGLGAGMAPKIAWALENKPRIPVVWIHGL WP_005139779.1 hydrogenase [Shigella sonnei] Length: 372\nNumber of predicted TMHs: 2\nExp number of AAs in TMHs: 37.02628\nExp number, first 60 AAs: 19.39684\nTotal prob of N-in: 0.94313\nPOSSIBLE N-term signal sequence\ninside 1 19\nTMhelix 20 39\noutside 40 325\nTMhelix 326 348\ninside 349 372

26515 GCF_001250815.1_Sg_201402408_ACTGAT_L001 Shigella sonnei Proteobacteria; Gammaproteobacteria; Enterobacterales; Enterobacteriaceae; Shigella MTGDNTLIHSHGINRRDFMKLCAALAATMGLSSKAAAEMAESVTNPQRPPVIWIGAQECT WP_000145410.1 MULTISPECIES: hydrogenase-2 small chain [Proteobacteria] Length: 372\nNumber of predicted TMHs: 1\nExp number of AAs in TMHs: 23.72226\nExp number, first 60 AAs: 0.22038\nTotal prob of N-in: 0.12994\noutside 1 331\nTMhelix 332 354\ninside 355 372

26516 GCF_001250815.1_Sg_201402408_ACTGAT_L001 Shigella sonnei Proteobacteria; Gammaproteobacteria; Enterobacterales; Enterobacteriaceae; Shigella MNNEETFYQAMRRQGVTRRSFLKYCSLAATSLGLGAGMAPKIAWALENKPRIPVVWIHGL WP_005139779.1 hydrogenase [Shigella sonnei] Length: 372\nNumber of predicted TMHs: 2\nExp number of AAs in TMHs: 37.02628\nExp number, first 60 AAs: 19.39684\nTotal prob of N-in: 0.94313\nPOSSIBLE N-term signal sequence\ninside 1 19\nTMhelix 20 39\noutside 40 325\nTMhelix 326 348\ninside 349 372

26517 GCF_001250575.1_12971_8_23 Shigella sonnei Proteobacteria; Gammaproteobacteria; Enterobacterales; Enterobacteriaceae; Shigella MNNEETFYQAMRRQGVTRRSFLKYCSLAATSLGLGAGMAPKIAWALENKPRIPVVWIHGL WP_005139779.1 hydrogenase [Shigella sonnei] Length: 372\nNumber of predicted TMHs: 2\nExp number of AAs in TMHs: 37.02628\nExp number, first 60 AAs: 19.39684\nTotal prob of N-in: 0.94313\nPOSSIBLE N-term signal sequence\ninside 1 19\nTMhelix 20 39\noutside 40 325\nTMhelix 326 348\ninside 349 372

26518 GCF_001250855.1_7655_8_13 Shigella sonnei Proteobacteria; Gammaproteobacteria; Enterobacterales; Enterobacteriaceae; Shigella MTGDNTLIHSHGINRRDFMKLCAALAATMGLSSKAAAEMAESVTNPQRPPVIWIGAQECT WP_000145410.1 MULTISPECIES: hydrogenase-2 small chain [Proteobacteria] Length: 372\nNumber of predicted TMHs: 1\nExp number of AAs in TMHs: 23.72226\nExp number, first 60 AAs: 0.22038\nTotal prob of N-in: 0.12994\noutside 1 331\nTMhelix 332 354\ninside 355 372

26519 GCF_001250875.1_Sg_201401463_GTCCGC_L001 Shigella sonnei Proteobacteria; Gammaproteobacteria; Enterobacterales; Enterobacteriaceae; Shigella MTGDNTLIHSHGINRRDFMKLCAALAATMGLSSKAAAEMAESVTNPQRPPVIWIGAQECT WP_000145410.1 MULTISPECIES: hydrogenase-2 small chain [Proteobacteria] Length: 372\nNumber of predicted TMHs: 1\nExp number of AAs in TMHs: 23.72226\nExp number, first 60 AAs: 0.22038\nTotal prob of N-in: 0.12994\noutside 1 331\nTMhelix 332 354\ninside 355 372

26520 GCF_001250855.1_7655_8_13 Shigella sonnei Proteobacteria; Gammaproteobacteria; Enterobacterales; Enterobacteriaceae; Shigella MNNEETFYQAMRRQGVTRRSFLKYCSLAATSLGLGAGMAPKIAWALENKPRIPVVWIHGL WP_005139779.1 hydrogenase [Shigella sonnei] Length: 372\nNumber of predicted TMHs: 2\nExp number of AAs in TMHs: 37.02628\nExp number, first 60 AAs: 19.39684\nTotal prob of N-in: 0.94313\nPOSSIBLE N-term signal sequence\ninside 1 19\nTMhelix 20 39\noutside 40 325\nTMhelix 326 348\ninside 349 372

26521 GCF_001251255.1_5236_5_12 Shigella sonnei Proteobacteria; Gammaproteobacteria; Enterobacterales; Enterobacteriaceae; Shigella MNNEETFYQAMRRQGVTRRSFLKYCSLAATSLGLGAGMAPKIAWALENKPRIPVVWIHGL WP_005139779.1 hydrogenase [Shigella sonnei] Length: 372\nNumber of predicted TMHs: 2\nExp number of AAs in TMHs: 37.02628\nExp number, first 60 AAs: 19.39684\nTotal prob of N-in: 0.94313\nPOSSIBLE N-term signal sequence\ninside 1 19\nTMhelix 20 39\noutside 40 325\nTMhelix 326 348\ninside 349 372

26522 GCF_001250995.1_8290_4_31 Shigella sonnei Proteobacteria; Gammaproteobacteria; Enterobacterales; Enterobacteriaceae; Shigella MNNEETFYQAMRRQGVTRRSFLKYCSLAATSLGLGAGMAPKIAWALENKPRIPVVWIHGL WP_005139779.1 hydrogenase [Shigella sonnei] Length: 372\nNumber of predicted TMHs: 2\nExp number of AAs in TMHs: 37.02628\nExp number, first 60 AAs: 19.39684\nTotal prob of N-in: 0.94313\nPOSSIBLE N-term signal sequence\ninside 1 19\nTMhelix 20 39\noutside 40 325\nTMhelix 326 348\ninside 349 372

26523 GCF_001251015.1_12971_8_46 Shigella sonnei Proteobacteria; Gammaproteobacteria; Enterobacterales; Enterobacteriaceae; Shigella MNNEETFYQAMRRQGVTRRSFLKYCSLAATSLGLGAGMAPKIAWALENKPRIPVVWIHGL WP_005139779.1 hydrogenase [Shigella sonnei] Length: 372\nNumber of predicted TMHs: 2\nExp number of AAs in TMHs: 37.02628\nExp number, first 60 AAs: 19.39684\nTotal prob of N-in: 0.94313\nPOSSIBLE N-term signal sequence\ninside 1 19\nTMhelix 20 39\noutside 40 325\nTMhelix 326 348\ninside 349 372

26524 GCF_001251055.1_8489_2_84 Shigella sonnei Proteobacteria; Gammaproteobacteria; Enterobacterales; Enterobacteriaceae; Shigella MNNEETFYQAMRRQGVTRRSFLKYCSLAATSLGLGAGMAPKIAWALENKPRIPVVWIHGL WP_005139779.1 hydrogenase [Shigella sonnei] Length: 372\nNumber of predicted TMHs: 2\nExp number of AAs in TMHs: 37.02628\nExp number, first 60 AAs: 19.39684\nTotal prob of N-in: 0.94313\nPOSSIBLE N-term signal sequence\ninside 1 19\nTMhelix 20 39\noutside 40 325\nTMhelix 326 348\ninside 349 372

26525 GCF_001251135.1_H14136026101 Shigella sonnei Proteobacteria; Gammaproteobacteria; Enterobacterales; Enterobacteriaceae; Shigella MNNEETFYQAMRRQGVTRRSFLKYCSLAATSLGLGAGMAPKIAWALENKPRIPVVWIHGL WP_005139779.1 hydrogenase [Shigella sonnei] Length: 372\nNumber of predicted TMHs: 2\nExp number of AAs in TMHs: 37.02628\nExp number, first 60 AAs: 19.39684\nTotal prob of N-in: 0.94313\nPOSSIBLE N-term signal sequence\ninside 1 19\nTMhelix 20 39\noutside 40 325\nTMhelix 326 348\ninside 349 372

26526 GCF_001251295.1_6350_1_9Shigella sonnei Proteobacteria; Gammaproteobacteria; Enterobacterales; Enterobacteriaceae; Shigella MTGDNTLIHSHGINRRDFMKLCAALAATMGLSSKAAAEMAESVTNPQRPPVIWIGAQECT WP_000145410.1 MULTISPECIES: hydrogenase-2 small chain [Proteobacteria] Length: 372\nNumber of predicted TMHs: 1\nExp number of AAs in TMHs: 23.72226\nExp number, first 60 AAs: 0.22038\nTotal prob of N-in: 0.12994\noutside 1 331\nTMhelix 332 354\ninside 355 372

26527 GCF_001251335.1_7353_1_22 Shigella sonnei Proteobacteria; Gammaproteobacteria; Enterobacterales; Enterobacteriaceae; Shigella MTGDNTLIHSHGINRRDFMKLCAALAATMGLSSKAAAEMAESVTNPQRPPVIWIGAQECT WP_000145410.1 MULTISPECIES: hydrogenase-2 small chain [Proteobacteria] Length: 372\nNumber of predicted TMHs: 1\nExp number of AAs in TMHs: 23.72226\nExp number, first 60 AAs: 0.22038\nTotal prob of N-in: 0.12994\noutside 1 331\nTMhelix 332 354\ninside 355 372

26528 GCF_001251575.1_8289_7_84 Shigella sonnei Proteobacteria; Gammaproteobacteria; Enterobacterales; Enterobacteriaceae; Shigella MNNEETFYQAMRRQGVTRRSFLKYCSLAATSLGLGAGMAPKIAWALENKPRIPVVWIHGL WP_005139779.1 hydrogenase [Shigella sonnei] Length: 372\nNumber of predicted TMHs: 2\nExp number of AAs in TMHs: 37.02628\nExp number, first 60 AAs: 19.39684\nTotal prob of N-in: 0.94313\nPOSSIBLE N-term signal sequence\ninside 1 19\nTMhelix 20 39\noutside 40 325\nTMhelix 326 348\ninside 349 372

26529 GCF_001251875.1_8289_7_51 Shigella sonnei Proteobacteria; Gammaproteobacteria; Enterobacterales; Enterobacteriaceae; Shigella MTGDNTLIHSHGINRRDFMKLCAALAATMGLSSKAAAEMAESVTNPQRPPVIWIGAQECT WP_000145410.1 MULTISPECIES: hydrogenase-2 small chain [Proteobacteria] Length: 372\nNumber of predicted TMHs: 1\nExp number of AAs in TMHs: 23.72226\nExp number, first 60 AAs: 0.22038\nTotal prob of N-in: 0.12994\noutside 1 331\nTMhelix 332 354\ninside 355 372

26530 GCF_001251875.1_8289_7_51 Shigella sonnei Proteobacteria; Gammaproteobacteria; Enterobacterales; Enterobacteriaceae; Shigella MNNEETFYQAMRRQGVTRRSFLKYCSLAATSLGLGAGMAPKIAWALENKPRIPVVWIHGL WP_005139779.1 hydrogenase [Shigella sonnei] Length: 372\nNumber of predicted TMHs: 2\nExp number of AAs in TMHs: 37.02628\nExp number, first 60 AAs: 19.39684\nTotal prob of N-in: 0.94313\nPOSSIBLE N-term signal sequence\ninside 1 19\nTMhelix 20 39\noutside 40 325\nTMhelix 326 348\ninside 349 372

26531 GCF_001251615.1_H14128025901 Shigella sonnei Proteobacteria; Gammaproteobacteria; Enterobacterales; Enterobacteriaceae; Shigella MNNEETFYQAMRRQGVTRRSFLKYCSLAATSLGLGAGMAPKIAWALENKPRIPVVWIHGL WP_005139779.1 hydrogenase [Shigella sonnei] Length: 372\nNumber of predicted TMHs: 2\nExp number of AAs in TMHs: 37.02628\nExp number, first 60 AAs: 19.39684\nTotal prob of N-in: 0.94313\nPOSSIBLE N-term signal sequence\ninside 1 19\nTMhelix 20 39\noutside 40 325\nTMhelix 326 348\ninside 349 372

26532 GCF_001251655.1_8289_7_50 Shigella sonnei Proteobacteria; Gammaproteobacteria; Enterobacterales; Enterobacteriaceae; Shigella MNNEETFYQAMRRQGVTRRSFLKYCSLAATSLGLGAGMAPKIAWALENKPRIPVVWIHGL WP_005139779.1 hydrogenase [Shigella sonnei] Length: 372\nNumber of predicted TMHs: 2\nExp number of AAs in TMHs: 37.02628\nExp number, first 60 AAs: 19.39684\nTotal prob of N-in: 0.94313\nPOSSIBLE N-term signal sequence\ninside 1 19\nTMhelix 20 39\noutside 40 325\nTMhelix 326 348\ninside 349 372

26533 GCF_001251955.1_5008_8_11 Shigella sonnei Proteobacteria; Gammaproteobacteria; Enterobacterales; Enterobacteriaceae; Shigella MTGDNTLIHSHGINRRDFMKLCAALAATMGLSSKAAAEMAESVTNPQRPPVIWIGAQECT WP_000145410.1 MULTISPECIES: hydrogenase-2 small chain [Proteobacteria] Length: 372\nNumber of predicted TMHs: 1\nExp number of AAs in TMHs: 23.72226\nExp number, first 60 AAs: 0.22038\nTotal prob of N-in: 0.12994\noutside 1 331\nTMhelix 332 354\ninside 355 372

26534 GCF_001251955.1_5008_8_11 Shigella sonnei Proteobacteria; Gammaproteobacteria; Enterobacterales; Enterobacteriaceae; Shigella MNNEETFYQAMRRQGVTRRSFLKYCSLAATSLGLGAGMAPKIAWALENKPRIPVVWIHGL WP_005139779.1 hydrogenase [Shigella sonnei] Length: 372\nNumber of predicted TMHs: 2\nExp number of AAs in TMHs: 37.02628\nExp number, first 60 AAs: 19.39684\nTotal prob of N-in: 0.94313\nPOSSIBLE N-term signal sequence\ninside 1 19\nTMhelix 20 39\noutside 40 325\nTMhelix 326 348\ninside 349 372

26535 GCF_001252215.1_H14124064101 Shigella sonnei Proteobacteria; Gammaproteobacteria; Enterobacterales; Enterobacteriaceae; Shigella MNNEETFYQAMRRQGVTRRSFLKYCSLAATSLGLGAGMAPKIAWALENKPRIPVVWIHGL WP_005139779.1 hydrogenase [Shigella sonnei] Length: 372\nNumber of predicted TMHs: 2\nExp number of AAs in TMHs: 37.02628\nExp number, first 60 AAs: 19.39684\nTotal prob of N-in: 0.94313\nPOSSIBLE N-term signal sequence\ninside 1 19\nTMhelix 20 39\noutside 40 325\nTMhelix 326 348\ninside 349 372

26536 GCF_001252035.1_5417_2_2Shigella sonnei Proteobacteria; Gammaproteobacteria; Enterobacterales; Enterobacteriaceae; Shigella MNNEETFYQAMRRQGVTRRSFLKYCSLAATSLGLGAGMAPKIAWALENKPRIPVVWIHGL WP_005139779.1 hydrogenase [Shigella sonnei] Length: 372\nNumber of predicted TMHs: 2\nExp number of AAs in TMHs: 37.02628\nExp number, first 60 AAs: 19.39684\nTotal prob of N-in: 0.94313\nPOSSIBLE N-term signal sequence\ninside 1 19\nTMhelix 20 39\noutside 40 325\nTMhelix 326 348\ninside 349 372

26537 GCF_001252355.1_5236_6_7Shigella sonnei Proteobacteria; Gammaproteobacteria; Enterobacterales; Enterobacteriaceae; Shigella MTGDNTLIHSHGINRRDFMKLCAALAATMGLSSKAAAEMAESVTNPQRPPVIWIGAQECT WP_000145410.1 MULTISPECIES: hydrogenase-2 small chain [Proteobacteria] Length: 372\nNumber of predicted TMHs: 1\nExp number of AAs in TMHs: 23.72226\nExp number, first 60 AAs: 0.22038\nTotal prob of N-in: 0.12994\noutside 1 331\nTMhelix 332 354\ninside 355 372

26538 GCF_001252355.1_5236_6_7Shigella sonnei Proteobacteria; Gammaproteobacteria; Enterobacterales; Enterobacteriaceae; Shigella MNNEETFYQAMRRQGVTRRSFLKYCSLAATSLGLGAGMAPKIAWALENKPRIPVVWIHGL WP_005139779.1 hydrogenase [Shigella sonnei] Length: 372\nNumber of predicted TMHs: 2\nExp number of AAs in TMHs: 37.02628\nExp number, first 60 AAs: 19.39684\nTotal prob of N-in: 0.94313\nPOSSIBLE N-term signal sequence\ninside 1 19\nTMhelix 20 39\noutside 40 325\nTMhelix 326 348\ninside 349 372

26539 GCF_001252175.1_5236_7_8Shigella sonnei Proteobacteria; Gammaproteobacteria; Enterobacterales; Enterobacteriaceae; Shigella MNNEETFYQAMRRQGVTRRSFLKYCSLAATSLGLGAGMAPKIAWALENKPRIPVVWIHGL WP_005139779.1 hydrogenase [Shigella sonnei] Length: 372\nNumber of predicted TMHs: 2\nExp number of AAs in TMHs: 37.02628\nExp number, first 60 AAs: 19.39684\nTotal prob of N-in: 0.94313\nPOSSIBLE N-term signal sequence\ninside 1 19\nTMhelix 20 39\noutside 40 325\nTMhelix 326 348\ninside 349 372

26540 GCF_001252095.1_Sg_201401096_AGTTCC_L001 Shigella sonnei Proteobacteria; Gammaproteobacteria; Enterobacterales; Enterobacteriaceae; Shigella MNNEETFYQAMRRQGVTRRSFLKYCSLAATSLGLGAGMAPKIAWALENKPRIPVVWIHGL WP_005139779.1 hydrogenase [Shigella sonnei] Length: 372\nNumber of predicted TMHs: 2\nExp number of AAs in TMHs: 37.02628\nExp number, first 60 AAs: 19.39684\nTotal prob of N-in: 0.94313\nPOSSIBLE N-term signal sequence\ninside 1 19\nTMhelix 20 39\noutside 40 325\nTMhelix 326 348\ninside 349 372

26541 GCF_001252395.1_12971_8_61 Shigella sonnei Proteobacteria; Gammaproteobacteria; Enterobacterales; Enterobacteriaceae; Shigella MTGDNTLIHSHGINRRDFMKLCAALAATMGLSSKAAAEMAESVTNPQRPPVIWIGAQECT WP_000145410.1 MULTISPECIES: hydrogenase-2 small chain [Proteobacteria] Length: 372\nNumber of predicted TMHs: 1\nExp number of AAs in TMHs: 23.72226\nExp number, first 60 AAs: 0.22038\nTotal prob of N-in: 0.12994\noutside 1 331\nTMhelix 332 354\ninside 355 372

26542 GCF_001252655.1_5417_2_12 Shigella sonnei Proteobacteria; Gammaproteobacteria; Enterobacterales; Enterobacteriaceae; Shigella MTGDNTLIHSHGINRRDFMKLCAALAATMGLSSKAAAEMAESVTNPQRPPVIWIGAQECT WP_000145410.1 MULTISPECIES: hydrogenase-2 small chain [Proteobacteria] Length: 372\nNumber of predicted TMHs: 1\nExp number of AAs in TMHs: 23.72226\nExp number, first 60 AAs: 0.22038\nTotal prob of N-in: 0.12994\noutside 1 331\nTMhelix 332 354\ninside 355 372

26543 GCF_001252755.1_12830_1_41 Shigella sonnei Proteobacteria; Gammaproteobacteria; Enterobacterales; Enterobacteriaceae; Shigella MTGDNTLIHSHGINRRDFMKLCAALAATMGLSSKAAAEMAESVTNPQRPPVIWIGAQECT WP_000145410.1 MULTISPECIES: hydrogenase-2 small chain [Proteobacteria] Length: 372\nNumber of predicted TMHs: 1\nExp number of AAs in TMHs: 23.72226\nExp number, first 60 AAs: 0.22038\nTotal prob of N-in: 0.12994\noutside 1 331\nTMhelix 332 354\ninside 355 372

26544 GCF_001252795.1_6551_1_9Shigella sonnei Proteobacteria; Gammaproteobacteria; Enterobacterales; Enterobacteriaceae; Shigella MTGDNTLIHSHGINRRDFMKLCAALAATMGLSSKAAAEMAESVTNPQRPPVIWIGAQECT WP_000145410.1 MULTISPECIES: hydrogenase-2 small chain [Proteobacteria] Length: 372\nNumber of predicted TMHs: 1\nExp number of AAs in TMHs: 23.72226\nExp number, first 60 AAs: 0.22038\nTotal prob of N-in: 0.12994\noutside 1 331\nTMhelix 332 354\ninside 355 372

26545 GCF_001252595.1_6246_5_6Shigella sonnei Proteobacteria; Gammaproteobacteria; Enterobacterales; Enterobacteriaceae; Shigella MNNEETFYQAMRRQGVTRRSFLKYCSLAATSLGLGAGMAPKIAWALENKPRIPVVWIHGL WP_005139779.1 hydrogenase [Shigella sonnei] Length: 372\nNumber of predicted TMHs: 2\nExp number of AAs in TMHs: 37.02628\nExp number, first 60 AAs: 19.39684\nTotal prob of N-in: 0.94313\nPOSSIBLE N-term signal sequence\ninside 1 19\nTMhelix 20 39\noutside 40 325\nTMhelix 326 348\ninside 349 372

26546 GCF_001252835.1_6246_5_2Shigella sonnei Proteobacteria; Gammaproteobacteria; Enterobacterales; Enterobacteriaceae; Shigella MTGDNTLIHSHGINRRDFMKLCAALAATMGLSSKAAAEMAESVTNPQRPPVIWIGAQECT WP_000145410.1 MULTISPECIES: hydrogenase-2 small chain [Proteobacteria] Length: 372\nNumber of predicted TMHs: 1\nExp number of AAs in TMHs: 23.72226\nExp number, first 60 AAs: 0.22038\nTotal prob of N-in: 0.12994\noutside 1 331\nTMhelix 332 354\ninside 355 372

26547 GCF_001252955.1_8289_7_72 Shigella sonnei Proteobacteria; Gammaproteobacteria; Enterobacterales; Enterobacteriaceae; Shigella MTGDNTLIHSHGINRRDFMKLCAALAATMGLSSKAAAEMAESVTNPQRPPVIWIGAQECT WP_000145410.1 MULTISPECIES: hydrogenase-2 small chain [Proteobacteria] Length: 372\nNumber of predicted TMHs: 1\nExp number of AAs in TMHs: 23.72226\nExp number, first 60 AAs: 0.22038\nTotal prob of N-in: 0.12994\noutside 1 331\nTMhelix 332 354\ninside 355 372

26548 GCF_001252995.1_5236_3_6Shigella sonnei Proteobacteria; Gammaproteobacteria; Enterobacterales; Enterobacteriaceae; Shigella MTGDNTLIHSHGINRRDFMKLCAALAATMGLSSKAAAEMAESVTNPQRPPVIWIGAQECT

WP_000145410.1 MULTISPECIES: hydrogenase-2 small chain [Proteobacteria] Length: 372\nNumber of predicted TMHs: 1\nExp number of AAs in TMHs: 23.72226\nExp number, first 60 AAs: 0.22038\nTotal prob of N-in: 0.12994\noutside 1 331\nTMhelix 332 354\ninside 355 372

26549 GCF_001252835.1_6246_5_2Shigella sonnei Proteobacteria; Gammaproteobacteria; Enterobacterales; Enterobacteriaceae; Shigella MNNEETFYQAMRRQGVTRRSFLKYCSLAATSLGLGAGMAPKIAWALENKPRIPVVWIHGL WP_005139779.1 hydrogenase [Shigella sonnei] Length: 372\nNumber of predicted TMHs: 2\nExp number of AAs in TMHs: 37.02628\nExp number, first 60 AAs: 19.39684\nTotal prob of N-in: 0.94313\nPOSSIBLE N-term signal sequence\ninside 1 19\nTMhelix 20 39\noutside 40 325\nTMhelix 326 348\ninside 349 372

26550 GCF_001252795.1_6551_1_9Shigella sonnei Proteobacteria; Gammaproteobacteria; Enterobacterales; Enterobacteriaceae; Shigella MNNEETFYQAMRRQGVTRRSFLKYCSLAATSLGLGAGMAPKIAWALENKPRIPVVWIHGL WP_005139779.1 hydrogenase [Shigella sonnei] Length: 372\nNumber of predicted TMHs: 2\nExp number of AAs in TMHs: 37.02628\nExp number, first 60 AAs: 19.39684\nTotal prob of N-in: 0.94313\nPOSSIBLE N-term signal sequence\ninside 1 19\nTMhelix 20 39\noutside 40 325\nTMhelix 326 348\ninside 349 372

26551 GCF_001253275.1_8290_4_17 Shigella sonnei Proteobacteria; Gammaproteobacteria; Enterobacterales; Enterobacteriaceae; Shigella MTGDNTLIHSHGINRRDFMKLCAALAATMGLSSKAAAEMAESVTNPQRPPVIWIGAQECT WP_000145410.1 MULTISPECIES: hydrogenase-2 small chain [Proteobacteria] Length: 372\nNumber of predicted TMHs: 1\nExp number of AAs in TMHs: 23.72226\nExp number, first 60 AAs: 0.22038\nTotal prob of N-in: 0.12994\noutside 1 331\nTMhelix 332 354\ninside 355 372

26552 GCF_001253355.1_6246_5_4Shigella sonnei Proteobacteria; Gammaproteobacteria; Enterobacterales; Enterobacteriaceae; Shigella MTGDNTLIHSHGINRRDFMKLCAALAATMGLSSKAAAEMAESVTNPQRPPVIWIGAQECT WP_000145410.1 MULTISPECIES: hydrogenase-2 small chain [Proteobacteria] Length: 372\nNumber of predicted TMHs: 1\nExp number of AAs in TMHs: 23.72226\nExp number, first 60 AAs: 0.22038\nTotal prob of N-in: 0.12994\noutside 1 331\nTMhelix 332 354\ninside 355 372

26553 GCF_001253395.1_Sg_201403963_CACTCA_L001 Shigella sonnei Proteobacteria; Gammaproteobacteria; Enterobacterales; Enterobacteriaceae; Shigella MTGDNTLIHSHGINRRDFMKLCAALAATMGLSSKAAAEMAESVTNPQRPPVIWIGAQECT WP_000145410.1 MULTISPECIES: hydrogenase-2 small chain [Proteobacteria] Length: 372\nNumber of predicted TMHs: 1\nExp number of AAs in TMHs: 23.72226\nExp number, first 60 AAs: 0.22038\nTotal prob of N-in: 0.12994\noutside 1 331\nTMhelix 332 354\ninside 355 372

26554 GCF_001253415.1_6350_1_11 Shigella sonnei Proteobacteria; Gammaproteobacteria; Enterobacterales; Enterobacteriaceae; Shigella MTGDNTLIHSHGINRRDFMKLCAALAATMGLSSKAAAEMAESVTNPQRPPVIWIGAQECT WP_000145410.1 MULTISPECIES: hydrogenase-2 small chain [Proteobacteria] Length: 372\nNumber of predicted TMHs: 1\nExp number of AAs in TMHs: 23.72226\nExp number, first 60 AAs: 0.22038\nTotal prob of N-in: 0.12994\noutside 1 331\nTMhelix 332 354\ninside 355 372

26555 GCF_001253515.1_12971_8_22 Shigella sonnei Proteobacteria; Gammaproteobacteria; Enterobacterales; Enterobacteriaceae; Shigella MTGDNTLIHSHGINRRDFMKLCAALAATMGLSSKAAAEMAESVTNPQRPPVIWIGAQECT WP_000145410.1 MULTISPECIES: hydrogenase-2 small chain [Proteobacteria] Length: 372\nNumber of predicted TMHs: 1\nExp number of AAs in TMHs: 23.72226\nExp number, first 60 AAs: 0.22038\nTotal prob of N-in: 0.12994\noutside 1 331\nTMhelix 332 354\ninside 355 372

26556 GCF_001253595.1_12971_8_66 Shigella sonnei Proteobacteria; Gammaproteobacteria; Enterobacterales; Enterobacteriaceae; Shigella MTGDNTLIHSHGINRRDFMKLCAALAATMGLSSKAAAEMAESVTNPQRPPVIWIGAQECT WP_000145410.1 MULTISPECIES: hydrogenase-2 small chain [Proteobacteria] Length: 372\nNumber of predicted TMHs: 1\nExp number of AAs in TMHs: 23.72226\nExp number, first 60 AAs: 0.22038\nTotal prob of N-in: 0.12994\noutside 1 331\nTMhelix 332 354\ninside 355 372

26557 GCF_001253635.1_5008_7_2Shigella sonnei Proteobacteria; Gammaproteobacteria; Enterobacterales; Enterobacteriaceae; Shigella MTGDNTLIHSHGINRRDFMKLCAALAATMGLSSKAAAEMAESVTNPQRPPVIWIGAQECT WP_000145410.1 MULTISPECIES: hydrogenase-2 small chain [Proteobacteria] Length: 372\nNumber of

predicted TMHs: 1\nExp number of AAs in TMHs: 23.72226\nExp number, first 60 AAs: 0.22038\nTotal prob of N-in: 0.12994\noutside 1 331\nTMhelix 332 354\ninside 355 372

26558 GCF_001253895.1_5236_7_9Shigella sonnei Proteobacteria; Gammaproteobacteria; Enterobacterales; Enterobacteriaceae; Shigella MNNEETFYQAMRRQGVTRRSFLKYCSLAATSLGLGAGMAPKIAWALENKPRIPVVWIHGL WP_005139779.1 hydrogenase [Shigella sonnei] Length: 372\nNumber of predicted TMHs: 2\nExp number of AAs in TMHs: 37.02628\nExp number, first 60 AAs: 19.39684\nTotal prob of N-in: 0.94313\nPOSSIBLE N-term signal sequence\ninside 1 19\nTMhelix 20 39\noutside 40 325\nTMhelix 326 348\ninside 349 372

26559 GCF_001253595.1_12971_8_66 Shigella sonnei Proteobacteria; Gammaproteobacteria; Enterobacterales; Enterobacteriaceae; Shigella MNNEETFYQAMRRQGVTRRSFLKYCSLAATSLGLGAGMAPKIAWALENKPRIPVVWIHGL WP_005139779.1 hydrogenase [Shigella sonnei] Length: 372\nNumber of predicted TMHs: 2\nExp number of AAs in TMHs: 37.02628\nExp number, first 60 AAs: 19.39684\nTotal prob of N-in: 0.94313\nPOSSIBLE N-term signal sequence\ninside 1 19\nTMhelix 20 39\noutside 40 325\nTMhelix 326 348\ninside 349 372

26560 GCF_001253935.1_5236_3_4Shigella sonnei Proteobacteria; Gammaproteobacteria; Enterobacterales; Enterobacteriaceae; Shigella MTGDNTLIHSHGINRRDFMKLCAALAATMGLSSKAAAEMAESVTNPQRPPVIWIGAQECT WP_000145410.1 MULTISPECIES: hydrogenase-2 small chain [Proteobacteria] Length: 372\nNumber of predicted TMHs: 1\nExp number of AAs in TMHs: 23.72226\nExp number, first 60 AAs: 0.22038\nTotal prob of N-in: 0.12994\noutside 1 331\nTMhelix 332 354\ninside 355 372

26561 GCF_001253955.1_8290_4_18 Shigella sonnei Proteobacteria; Gammaproteobacteria; Enterobacterales; Enterobacteriaceae; Shigella MTGDNTLIHSHGINRRDFMKLCAALAATMGLSSKAAAEMAESVTNPQRPPVIWIGAQECT WP_000145410.1 MULTISPECIES: hydrogenase-2 small chain [Proteobacteria] Length: 372\nNumber of predicted TMHs: 1\nExp number of AAs in TMHs: 23.72226\nExp number, first 60 AAs: 0.22038\nTotal prob of N-in: 0.12994\noutside 1 331\nTMhelix 332 354\ninside 355 372

26562 GCF_001253995.1_8289_7_75 Shigella sonnei Proteobacteria; Gammaproteobacteria; Enterobacterales; Enterobacteriaceae; Shigella MTGDNTLIHSHGINRRDFMKLCAALAATMGLSSKAAAEMAESVTNPQRPPVIWIGAQECT WP_000145410.1 MULTISPECIES: hydrogenase-2 small chain [Proteobacteria] Length: 372\nNumber of predicted TMHs: 1\nExp number of AAs in TMHs: 23.72226\nExp number, first 60 AAs: 0.22038\nTotal prob of N-in: 0.12994\noutside 1 331\nTMhelix 332 354\ninside 355 372

26563 GCF_001253935.1_5236_3_4Shigella sonnei Proteobacteria; Gammaproteobacteria; Enterobacterales; Enterobacteriaceae; Shigella MNNEETFYQAMRRQGVTRRSFLKYCSLAATSLGLGAGMAPKIAWALENKPRIPVVWIHGL WP_005139779.1 hydrogenase [Shigella sonnei] Length: 372\nNumber of predicted TMHs: 2\nExp number of AAs in TMHs: 37.02628\nExp number, first 60 AAs: 19.39684\nTotal prob of N-in: 0.94313\nPOSSIBLE N-term signal sequence\ninside 1 19\nTMhelix 20 39\noutside 40 325\nTMhelix 326 348\ninside 349 372

26564 GCF_001253955.1_8290_4_18 Shigella sonnei Proteobacteria; Gammaproteobacteria; Enterobacterales; Enterobacteriaceae; Shigella MNNEETFYQAMRRQGVTRRSFLKYCSLAATSLGLGAGMAPKIAWALENKPRIPVVWIHGL WP_005139779.1 hydrogenase [Shigella sonnei] Length: 372\nNumber of predicted TMHs: 2\nExp number of AAs in TMHs: 37.02628\nExp number, first 60 AAs: 19.39684\nTotal prob of N-in: 0.94313\nPOSSIBLE N-term signal sequence\ninside 1 19\nTMhelix 20 39\noutside 40 325\nTMhelix 326 348\ninside 349 372

26565 GCF_001254455.1_8290_4_92 Shigella sonnei Proteobacteria; Gammaproteobacteria; Enterobacterales; Enterobacteriaceae; Shigella MNNEETFYQAMRRQGVTRRSFLKYCSLAATSLGLGAGMAPKIAWALENKPRIPVVWIHGL WP_005139779.1 hydrogenase [Shigella sonnei] Length: 372\nNumber of predicted TMHs: 2\nExp number of AAs in TMHs: 37.02628\nExp number, first 60 AAs: 19.39684\nTotal prob of N-in: 0.94313\nPOSSIBLE N-term signal sequence\ninside 1 19\nTMhelix 20 39\noutside 40 325\nTMhelix 326 348\ninside 349 372

26566 GCF_001254155.1_5008_7_7Shigella sonnei Proteobacteria; Gammaproteobacteria; Enterobacterales; Enterobacteriaceae; Shigella MNNEETFYQAMRRQGVTRRSFLKYCSLAATSLGLGAGMAPKIAWALENKPRIPVVWIHGL WP_005139779.1 hydrogenase [Shigella sonnei] Length: 372\nNumber of predicted TMHs: 2\nExp number of AAs in TMHs: 37.02628\nExp number, first 60 AAs: 19.39684\nTotal prob of N-in: 0.94313\nPOSSIBLE N-term signal sequence\ninside 1 19\nTMhelix 20 39\noutside 40 325\nTMhelix 326 348\ninside 349 372

26567 GCF_001254515.1_H14050007401 Shigella sonnei Proteobacteria; Gammaproteobacteria; Enterobacterales; Enterobacteriaceae; Shigella MTGDNTLIHSHGINRRDFMKLCAALAATMGLSSKAAAEMAESVTNPQRPPVIWIGAQECT WP_000145410.1 MULTISPECIES: hydrogenase-2 small chain [Proteobacteria] Length: 372\nNumber of predicted TMHs: 1\nExp number of AAs in TMHs: 23.72226\nExp number, first 60 AAs: 0.22038\nTotal prob of N-in: 0.12994\noutside 1 331\nTMhelix 332 354\ninside 355 372

26568 GCF_001254515.1_H14050007401 Shigella sonnei Proteobacteria; Gammaproteobacteria; Enterobacterales; Enterobacteriaceae; Shigella MNNEETFYQAMRRQGVTRRSFLKYCSLAATSLGLGAGMAPKIAWALENKPRIPVVWIHGL WP_005139779.1 hydrogenase [Shigella sonnei] Length: 372\nNumber of predicted TMHs: 2\nExp number of AAs in TMHs: 37.02628\nExp number, first 60 AAs: 19.39684\nTotal prob of N-in: 0.94313\nPOSSIBLE N-term signal sequence\ninside 1 19\nTMhelix 20 39\noutside 40 325\nTMhelix 326 348\ninside 349 372

26569 GCF_001254195.1_6551_1_8Shigella sonnei Proteobacteria; Gammaproteobacteria; Enterobacterales; Enterobacteriaceae; Shigella MNNEETFYQAMRRQGVTRRSFLKYCSLAATSLGLGAGMAPKIAWALENKPRIPVVWIHGL WP_005139779.1 hydrogenase [Shigella sonnei] Length: 372\nNumber of predicted TMHs: 2\nExp number of AAs in TMHs: 37.02628\nExp number, first 60 AAs: 19.39684\nTotal prob of N-in: 0.94313\nPOSSIBLE N-term signal sequence\ninside 1 19\nTMhelix 20 39\noutside 40 325\nTMhelix 326 348\ninside 349 372

26570 GCF_001254595.1_8289_7_37 Shigella sonnei Proteobacteria; Gammaproteobacteria; Enterobacterales; Enterobacteriaceae; Shigella MTGDNTLIHSHGINRRDFMKLCAALAATMGLSSKAAAEMAESVTNPQRPPVIWIGAQECT WP_000145410.1 MULTISPECIES: hydrogenase-2 small chain [Proteobacteria] Length: 372\nNumber of predicted TMHs: 1\nExp number of AAs in TMHs: 23.72226\nExp number, first 60 AAs: 0.22038\nTotal prob of N-in: 0.12994\noutside 1 331\nTMhelix 332 354\ninside 355 372

26571 GCF_001254695.1_H14142025101 Shigella sonnei Proteobacteria; Gammaproteobacteria; Enterobacterales; Enterobacteriaceae; Shigella MTGDNTLIHSHGINRRDFMKLCAALAATMGLSSKAAAEMAESVTNPQRPPVIWIGAQECT WP_000145410.1 MULTISPECIES: hydrogenase-2 small chain [Proteobacteria] Length: 372\nNumber of predicted TMHs: 1\nExp number of AAs in TMHs: 23.72226\nExp number, first 60 AAs: 0.22038\nTotal prob of N-in: 0.12994\noutside 1 331\nTMhelix 332 354\ninside 355 372

26572 GCF_001255135.1_5236_5_9Shigella sonnei Proteobacteria; Gammaproteobacteria; Enterobacterales; Enterobacteriaceae; Shigella MTGDNTLIHSHGINRRDFMKLCAALAATMGLSSKAAAEMAESVTNPQRPPVIWIGAQECT WP_000145410.1 MULTISPECIES: hydrogenase-2 small chain [Proteobacteria] Length: 372\nNumber of predicted TMHs: 1\nExp number of AAs in TMHs: 23.72226\nExp number, first 60 AAs: 0.22038\nTotal prob of N-in: 0.12994\noutside 1 331\nTMhelix 332 354\ninside 355 372

26573 GCF_001255115.1_9870_6_64 Shigella sonnei Proteobacteria; Gammaproteobacteria; Enterobacterales; Enterobacteriaceae; Shigella MNNEETFYQAMRRQGVTRRSFLKYCSLAATSLGLGAGMAPKIAWALENKPRIPVVWIHGL WP_005139779.1 hydrogenase [Shigella sonnei] Length: 372\nNumber of predicted TMHs: 2\nExp number of AAs in TMHs: 37.02628\nExp number, first 60 AAs: 19.39684\nTotal prob of N-in: 0.94313\nPOSSIBLE N-term signal sequence\ninside 1 19\nTMhelix 20 39\noutside 40 325\nTMhelix 326 348\ninside 349 372

26574 GCF_001255135.1_5236_5_9Shigella sonnei Proteobacteria; Gammaproteobacteria; Enterobacterales; Enterobacteriaceae; Shigella MNNEETFYQAMRRQGVTRRSFLKYCSLAATSLGLGAGMAPKIAWALENKPRIPVVWIHGL WP_005139779.1 hydrogenase [Shigella sonnei] Length: 372\nNumber of predicted TMHs: 2\nExp number of AAs in TMHs: 37.02628\nExp number, first 60 AAs: 19.39684\nTotal prob of N-in: 0.94313\nPOSSIBLE N-term signal sequence\ninside 1 19\nTMhelix 20 39\noutside 40 325\nTMhelix 326 348\ninside 349 372

26575 GCF_001254835.1_6350_1_15 Shigella sonnei Proteobacteria; Gammaproteobacteria; Enterobacterales; Enterobacteriaceae; Shigella MNNEETFYQAMRRQGVTRRSFLKYCSLAATSLGLGAGMAPKIAWALENKPRIPVVWIHGL WP_005139779.1 hydrogenase [Shigella sonnei] Length: 372\nNumber of predicted TMHs: 2\nExp number of AAs in TMHs: 37.02628\nExp number, first 60 AAs: 19.39684\nTotal prob of N-in: 0.94313\nPOSSIBLE N-term signal sequence\ninside 1 19\nTMhelix 20 39\noutside 40 325\nTMhelix 326 348\ninside 349 372

26576 GCF_001255195.1_5236_2_6Shigella sonnei Proteobacteria; Gammaproteobacteria; Enterobacterales; Enterobacteriaceae; Shigella MTGDNTLIHSHGINRRDFMKLCAALAATMGLSSKAAAEMAESVTNPQRPPVIWIGAQECT WP_000145410.1 MULTISPECIES: hydrogenase-2 small chain [Proteobacteria] Length: 372\nNumber of predicted TMHs: 1\nExp number of AAs in TMHs: 23.72226\nExp number, first 60 AAs: 0.22038\nTotal prob of N-in: 0.12994\noutside 1 331\nTMhelix 332 354\ninside 355 372

26577 GCF_001255255.1_12971_8_65 Shigella sonnei Proteobacteria; Gammaproteobacteria; Enterobacterales; Enterobacteriaceae; Shigella MTGDNTLIHSHGINRRDFMKLCAALAATMGLSSKAAAEMAESVTNPQRPPVIWIGAQECT WP_000145410.1 MULTISPECIES: hydrogenase-2 small chain [Proteobacteria] Length: 372\nNumber of predicted TMHs: 1\nExp number of AAs in TMHs: 23.72226\nExp number, first 60 AAs: 0.22038\nTotal prob of N-in: 0.12994\noutside 1 331\nTMhelix 332 354\ninside 355 372

26578 GCF_001254715.1_H14126044401 Shigella sonnei Proteobacteria; Gammaproteobacteria; Enterobacterales; Enterobacteriaceae; Shigella TGDNTLIHSHGINRRDFMKLCAALAATMGLSSKAAAEMAESVTNPQRPPVIWIGAQECTG WP_065406877.1 hydrogenase 2 small subunit, partial [Shigella sonnei] Length: 371\nNumber of predicted TMHs: 1\nExp number of AAs in TMHs: 23.87942\nExp number, first 60 AAs: 0.21256\nTotal prob of N-in: 0.13793\noutside 1 330\nTMhelix 331 353\ninside 354 371

26579 GCF_001255195.1_5236_2_6Shigella sonnei Proteobacteria; Gammaproteobacteria; Enterobacterales; Enterobacteriaceae; Shigella MNNEETFYQAMRRQGVTRRSFLKYCSLAATSLGLGAGMAPKIAWALENKPRIPVVWIHGL WP_005139779.1 hydrogenase [Shigella sonnei] Length: 372\nNumber of predicted TMHs: 2\nExp number of AAs in TMHs: 37.02628\nExp number, first 60 AAs: 19.39684\nTotal prob of N-in: 0.94313\nPOSSIBLE N-term signal sequence\ninside 1 19\nTMhelix 20 39\noutside 40 325\nTMhelix 326 348\ninside 349 372

26580 GCF_001255255.1_12971_8_65 Shigella sonnei Proteobacteria; Gammaproteobacteria; Enterobacterales; Enterobacteriaceae; Shigella MNNEETFYQAMRRQGVTRRSFLKYCSLAATSLGLGAGMAPKIAWALENKPRIPVVWIHGL WP_005139779.1 hydrogenase [Shigella sonnei] Length: 372\nNumber of predicted TMHs: 2\nExp number of AAs in TMHs: 37.02628\nExp number, first 60 AAs: 19.39684\nTotal prob of N-in: 0.94313\nPOSSIBLE N-term signal sequence\ninside 1 19\nTMhelix 20 39\noutside 40 325\nTMhelix 326 348\ninside 349 372

26581 GCF_001254775.1_8289_7_47 Shigella sonnei Proteobacteria; Gammaproteobacteria; Enterobacterales; Enterobacteriaceae; Shigella MNNEETFYQAMRRQGVTRRSFLKYCSLAATSLGLGAGMAPKIAWALENKPRIPVVWIHGL WP_005139779.1 hydrogenase [Shigella sonnei] Length: 372\nNumber of predicted TMHs: 2\nExp number of AAs in TMHs: 37.02628\nExp number, first 60 AAs: 19.39684\nTotal prob of N-in: 0.94313\nPOSSIBLE N-term signal sequence\ninside 1 19\nTMhelix 20 39\noutside 40 325\nTMhelix 326 348\ninside 349 372

26582 GCF_001254875.1_12971_8_56 Shigella sonnei Proteobacteria; Gammaproteobacteria; Enterobacterales; Enterobacteriaceae; Shigella MNNEETFYQAMRRQGVTRRSFLKYCSLAATSLGLGAGMAPKIAWALENKPRIPVVWIHGL WP_005139779.1 hydrogenase [Shigella sonnei] Length: 372\nNumber of predicted TMHs: 2\nExp number of AAs in TMHs: 37.02628\nExp number, first 60 AAs: 19.39684\nTotal prob of N-in: 0.94313\nPOSSIBLE N-term signal sequence\ninside 1 19\nTMhelix 20 39\noutside 40 325\nTMhelix 326 348\ninside 349 372

26583 GCF_001255275.1_5417_1_2Shigella sonnei Proteobacteria; Gammaproteobacteria; Enterobacterales; Enterobacteriaceae; Shigella MTGDNTLIHSHGINRRDFMKLCAALAATMGLSSKAAAEMAESVTNPQRPPVIWIGAQECT WP_000145410.1 MULTISPECIES: hydrogenase-2 small chain [Proteobacteria] Length: 372\nNumber of predicted TMHs: 1\nExp number of AAs in TMHs: 23.72226\nExp number, first 60 AAs: 0.22038\nTotal prob of N-in: 0.12994\noutside 1 331\nTMhelix 332 354\ninside 355 372

26584 GCF_001255275.1_5417_1_2Shigella sonnei Proteobacteria; Gammaproteobacteria; Enterobacterales; Enterobacteriaceae; Shigella MNNEETFYQAMRRQGVTRRSFLKYCSLAATSLGLGAGMAPKIAWALENKPRIPVVWIHGL WP_005139779.1 hydrogenase [Shigella sonnei] Length: 372\nNumber of predicted TMHs: 2\nExp number of AAs in TMHs: 37.02628\nExp number, first 60 AAs: 19.39684\nTotal prob of N-in: 0.94313\nPOSSIBLE N-term signal sequence\ninside 1 19\nTMhelix 20 39\noutside 40 325\nTMhelix 326 348\ninside 349 372

26585 GCF_001255015.1_8290_4_6Shigella sonnei Proteobacteria; Gammaproteobacteria; Enterobacterales; Enterobacteriaceae; Shigella MNNEETFYQAMRRQGVTRRSFLKYCSLAATSLGLGAGMAPKIAWALENKPRIPVVWIHGL

WP_005139779.1 hydrogenase [Shigella sonnei] Length: 372\nNumber of predicted TMHs: 2\nExp number of AAs in TMHs: 37.02628\nExp number, first 60 AAs: 19.39684\nTotal prob of N-in: 0.94313\nPOSSIBLE N-term signal sequence\ninside 1 19\nTMhelix 20 39\noutside 40 325\nTMhelix 326 348\ninside 349 372

26586 GCF_001255035.1_12971_8_26 Shigella sonnei Proteobacteria; Gammaproteobacteria; Enterobacterales; Enterobacteriaceae; Shigella MNNEETFYQAMRRQGVTRRSFLKYCSLAATSLGLGAGMAPKIAWALENKPRIPVVWIHGL WP_005139779.1 hydrogenase [Shigella sonnei] Length: 372\nNumber of predicted TMHs: 2\nExp number of AAs in TMHs: 37.02628\nExp number, first 60 AAs: 19.39684\nTotal prob of N-in: 0.94313\nPOSSIBLE N-term signal sequence\ninside 1 19\nTMhelix 20 39\noutside 40 325\nTMhelix 326 348\ninside 349 372

26587 GCF_001255335.1_8489_2_94 Shigella sonnei Proteobacteria; Gammaproteobacteria; Enterobacterales; Enterobacteriaceae; Shigella MTGDNTLIHSHGINRRDFMKLCAALAATMGLSSKAAAEMAESVTNPQRPPVIWIGAQECT WP_000145410.1 MULTISPECIES: hydrogenase-2 small chain [Proteobacteria] Length: 372\nNumber of predicted TMHs: 1\nExp number of AAs in TMHs: 23.72226\nExp number, first 60 AAs: 0.22038\nTotal prob of N-in: 0.12994\noutside 1 331\nTMhelix 332 354\ninside 355 372

26588 GCF_001255335.1_8489_2_94 Shigella sonnei Proteobacteria; Gammaproteobacteria; Enterobacterales; Enterobacteriaceae; Shigella MNNEETFYQAMRRQGVTRRSFLKYCSLAATSLGLGAGMAPKIAWALENKPRIPVVWIHGL WP_005139779.1 hydrogenase [Shigella sonnei] Length: 372\nNumber of predicted TMHs: 2\nExp number of AAs in TMHs: 37.02628\nExp number, first 60 AAs: 19.39684\nTotal prob of N-in: 0.94313\nPOSSIBLE N-term signal sequence\ninside 1 19\nTMhelix 20 39\noutside 40 325\nTMhelix 326 348\ninside 349 372

26589 GCF_001255075.1_12971_8_63 Shigella sonnei Proteobacteria; Gammaproteobacteria; Enterobacterales; Enterobacteriaceae; Shigella MNNEETFYQAMRRQGVTRRSFLKYCSLAATSLGLGAGMAPKIAWALENKPRIPVVWIHGL WP_005139779.1 hydrogenase [Shigella sonnei] Length: 372\nNumber of predicted TMHs: 2\nExp number of AAs in TMHs: 37.02628\nExp number, first 60 AAs: 19.39684\nTotal prob of N-in: 0.94313\nPOSSIBLE N-term signal sequence\ninside 1 19\nTMhelix 20 39\noutside 40 325\nTMhelix 326 348\ninside 349 372

26590 GCF_001255375.1_H14090035501 Shigella sonnei Proteobacteria; Gammaproteobacteria; Enterobacterales; Enterobacteriaceae; Shigella MTGDNTLIHSHGINRRDFMKLCAALAATMGLSSKAAAEMAESVTNPQRPPVIWIGAQECT WP_000145410.1 MULTISPECIES: hydrogenase-2 small chain [Proteobacteria] Length: 372\nNumber of predicted TMHs: 1\nExp number of AAs in TMHs: 23.72226\nExp number, first 60 AAs: 0.22038\nTotal prob of N-in: 0.12994\noutside 1 331\nTMhelix 332 354\ninside 355 372

26591 GCF_001255455.1_Sg_201401879_GGTAGC_L001 Shigella sonnei Proteobacteria; Gammaproteobacteria; Enterobacterales; Enterobacteriaceae; Shigella MTGDNTLIHSHGINRRDFMKLCAALAATMGLSSKAAAEMAESVTNPQRPPVIWIGAQECT WP_000145410.1 MULTISPECIES: hydrogenase-2 small chain [Proteobacteria] Length: 372\nNumber of predicted TMHs: 1\nExp number of AAs in TMHs: 23.72226\nExp number, first 60 AAs: 0.22038\nTotal prob of N-in: 0.12994\noutside 1 331\nTMhelix 332 354\ninside 355 372

26592 GCF_001255435.1_5008_8_7Shigella sonnei Proteobacteria; Gammaproteobacteria; Enterobacterales; Enterobacteriaceae; Shigella MTGDNTLIHSHGINRRDFMKLCAALAATMGLSSKAAAEMAESVTNPQRPPVIWIGAQECT WP_000145410.1 MULTISPECIES: hydrogenase-2 small chain [Proteobacteria] Length: 372\nNumber of predicted TMHs: 1\nExp number of AAs in TMHs: 23.72226\nExp number, first 60 AAs: 0.22038\nTotal prob of N-in: 0.12994\noutside 1 331\nTMhelix 332 354\ninside 355 372

26593 GCF_001255555.1_H14052014601 Shigella sonnei Proteobacteria; Gammaproteobacteria; Enterobacterales; Enterobacteriaceae; Shigella MTGDNTLIHSHGINRRDFMKLCAALAATMGLSSKAAAEMAESVTNPQRPPVIWIGAQECT WP_000145410.1 MULTISPECIES: hydrogenase-2 small chain [Proteobacteria] Length: 372\nNumber of predicted TMHs: 1\nExp number of AAs in TMHs: 23.72226\nExp number, first 60 AAs: 0.22038\nTotal prob of N-in: 0.12994\noutside 1 331\nTMhelix 332 354\ninside 355 372

26594 GCF_001255515.1_12850_1_42 Shigella sonnei Proteobacteria; Gammaproteobacteria; Enterobacterales; Enterobacteriaceae; Shigella

MTGDNTLIHSHGINRRDFMKLCAALAATMGLSSKAAAEMAESVTNPQRPPVIWIGAQECT WP_000145410.1 MULTISPECIES: hydrogenase-2 small chain [Proteobacteria] Length: 372\nNumber of predicted TMHs: 1\nExp number of AAs in TMHs: 23.72226\nExp number, first 60 AAs: 0.22038\nTotal prob of N-in: 0.12994\noutside 1 331\nTMhelix 332 354\ninside 355 372

26595 GCF_001255595.1_H14060002901 Shigella sonnei Proteobacteria; Gammaproteobacteria; Enterobacterales; Enterobacteriaceae; Shigella MTGDNTLIHSHGINRRDFMKLCAALAATMGLSSKAAAEMAESVTNPQRPPVIWIGAQECT WP_000145410.1 MULTISPECIES: hydrogenase-2 small chain [Proteobacteria] Length: 372\nNumber of predicted TMHs: 1\nExp number of AAs in TMHs: 23.72226\nExp number, first 60 AAs: 0.22038\nTotal prob of N-in: 0.12994\noutside 1 331\nTMhelix 332 354\ninside 355 372

26596 GCF_001255635.1_8290_4_29 Shigella sonnei Proteobacteria; Gammaproteobacteria; Enterobacterales; Enterobacteriaceae; Shigella MTGDNTLIHSHGINRRDFMKLCAALAATMGLSSKAAAEMAESVTNPQRPPVIWIGAQECT WP_000145410.1 MULTISPECIES: hydrogenase-2 small chain [Proteobacteria] Length: 372\nNumber of predicted TMHs: 1\nExp number of AAs in TMHs: 23.72226\nExp number, first 60 AAs: 0.22038\nTotal prob of N-in: 0.12994\noutside 1 331\nTMhelix 332 354\ninside 355 372

26597 GCF_001255375.1_H14090035501 Shigella sonnei Proteobacteria; Gammaproteobacteria; Enterobacterales; Enterobacteriaceae; Shigella MNNEETFYQAMRRQGVTRRSFLKYCSLAATSLGLGAGMAPKIAWALENKPRIPVVWIHGL WP_005139779.1 hydrogenase [Shigella sonnei] Length: 372\nNumber of predicted TMHs: 2\nExp number of AAs in TMHs: 37.02628\nExp number, first 60 AAs: 19.39684\nTotal prob of N-in: 0.94313\nPOSSIBLE N-term signal sequence\ninside 1 19\nTMhelix 20 39\noutside 40 325\nTMhelix 326 348\ninside 349 372

26598 GCF_001255675.1_12850_1_50 Shigella sonnei Proteobacteria; Gammaproteobacteria; Enterobacterales; Enterobacteriaceae; Shigella MTGDNTLIHSHGINRRDFMKLCAALAATMGLSSKAAAEMAESVTNPQRPPVIWIGAQECT WP_000145410.1 MULTISPECIES: hydrogenase-2 small chain [Proteobacteria] Length: 372\nNumber of predicted TMHs: 1\nExp number of AAs in TMHs: 23.72226\nExp number, first 60 AAs: 0.22038\nTotal prob of N-in: 0.12994\noutside 1 331\nTMhelix 332 354\ninside 355 372

26599 GCF_001255695.1_12830_1_33 Shigella sonnei Proteobacteria; Gammaproteobacteria; Enterobacterales; Enterobacteriaceae; Shigella MTGDNTLIHSHGINRRDFMKLCAALAATMGLSSKAAAEMAESVTNPQRPPVIWIGAQECT WP_000145410.1 MULTISPECIES: hydrogenase-2 small chain [Proteobacteria] Length: 372\nNumber of predicted TMHs: 1\nExp number of AAs in TMHs: 23.72226\nExp number, first 60 AAs: 0.22038\nTotal prob of N-in: 0.12994\noutside 1 331\nTMhelix 332 354\ninside 355 372

26600 GCF_001255735.1_12971_8_12 Shigella sonnei Proteobacteria; Gammaproteobacteria; Enterobacterales; Enterobacteriaceae; Shigella MTGDNTLIHSHGINRRDFMKLCAALAATMGLSSKAAAEMAESVTNPQRPPVIWIGAQECT WP_000145410.1 MULTISPECIES: hydrogenase-2 small chain [Proteobacteria] Length: 372\nNumber of predicted TMHs: 1\nExp number of AAs in TMHs: 23.72226\nExp number, first 60 AAs: 0.22038\nTotal prob of N-in: 0.12994\noutside 1 331\nTMhelix 332 354\ninside 355 372

26601 GCF_001255675.1_12850_1_50 Shigella sonnei Proteobacteria; Gammaproteobacteria; Enterobacterales; Enterobacteriaceae; Shigella MNNEETFYQAMRRQGVTRRSFLKYCSLAATSLGLGAGMAPKIAWALENKPRIPVVWIHGL WP_005139779.1 hydrogenase [Shigella sonnei] Length: 372\nNumber of predicted TMHs: 2\nExp number of AAs in TMHs: 37.02628\nExp number, first 60 AAs: 19.39684\nTotal prob of N-in: 0.94313\nPOSSIBLE N-term signal sequence\ninside 1 19\nTMhelix 20 39\noutside 40 325\nTMhelix 326 348\ninside 349 372

26602 GCF_001255695.1_12830_1_33 Shigella sonnei Proteobacteria; Gammaproteobacteria; Enterobacterales; Enterobacteriaceae; Shigella MNNEETFYQAMRRQGVTRRSFLKYCSLAATSLGLGAGMAPKIAWALENKPRIPVVWIHGL WP_005139779.1 hydrogenase [Shigella sonnei] Length: 372\nNumber of predicted TMHs: 2\nExp number of AAs in TMHs: 37.02628\nExp number, first 60 AAs: 19.39684\nTotal prob of N-in: 0.94313\nPOSSIBLE N-term signal sequence\ninside 1 19\nTMhelix 20 39\noutside 40 325\nTMhelix 326 348\ninside 349 372

26603	GCF_001255735.1_12971_8_12	Shigella sonnei	Proteobacteria; Gammaproteobacteria; Enterobacterales; Enterobacteriaceae; Shigella	MNNEETFYQAMRRQGVTRRSFLKYCSLAATSLGLGAGMAPKIAWALENKPRIPVVWIHGL	WP_005139779.1	hydrogenase [Shigella sonnei]	Length: 372\nNumber of predicted TMHs:  2\nExp number of AAs in TMHs: 37.02628\nExp number, first 60 AAs:  19.39684\nTotal prob of N-in:   0.94313\nPOSSIBLE N-term signal sequence\ninside   1   19\nTMhelix   20   39\noutside   40   325\nTMhelix   326   348\ninside   349   372

26604	GCF_001255435.1_5008_8_7Shigella sonnei	Proteobacteria; Gammaproteobacteria; Enterobacterales; Enterobacteriaceae; Shigella	MNNEETFYQAMRRQGVTRRSFLKYCSLAATSLGLGAGMAPKIAWALENKPRIPVVWIHGL	WP_005139779.1	hydrogenase [Shigella sonnei]	Length: 372\nNumber of predicted TMHs:  2\nExp number of AAs in TMHs: 37.02628\nExp number, first 60 AAs:  19.39684\nTotal prob of N-in:   0.94313\nPOSSIBLE N-term signal sequence\ninside   1   19\nTMhelix   20   39\noutside   40   325\nTMhelix   326   348\ninside   349   372

26605	GCF_001255515.1_12850_1_42	Shigella sonnei	Proteobacteria; Gammaproteobacteria; Enterobacterales; Enterobacteriaceae; Shigella	MNNEETFYQAMRRQGVTRRSFLKYCSLAATSLGLGAGMAPKIAWALENKPRIPVVWIHGL	WP_005139779.1	hydrogenase [Shigella sonnei]	Length: 372\nNumber of predicted TMHs:  2\nExp number of AAs in TMHs: 37.02628\nExp number, first 60 AAs:  19.39684\nTotal prob of N-in:   0.94313\nPOSSIBLE N-term signal sequence\ninside   1   19\nTMhelix   20   39\noutside   40   325\nTMhelix   326   348\ninside   349   372

26606	GCF_001255455.1_Sg_201401879_GGTAGC_L001	Shigella sonnei	Proteobacteria; Gammaproteobacteria; Enterobacterales; Enterobacteriaceae; Shigella	MNNEETFYQAMRRQGVTRRSFLKYCSLAATSLGLGAGMAPKIAWALENKPRIPVVWIHGL	WP_005139779.1	hydrogenase [Shigella sonnei]	Length: 372\nNumber of predicted TMHs:  2\nExp number of AAs in TMHs: 37.02628\nExp number, first 60 AAs:  19.39684\nTotal prob of N-in:   0.94313\nPOSSIBLE N-term signal sequence\ninside   1   19\nTMhelix   20   39\noutside   40   325\nTMhelix   326   348\ninside   349   372

26607	GCF_001255755.1_12844_1_54	Shigella sonnei	Proteobacteria; Gammaproteobacteria; Enterobacterales; Enterobacteriaceae; Shigella	MTGDNTLIHSHGINRRDFMKLCAALAATMGLSSKAAAEMAESVTNPQRPPVIWIGAQECT	WP_000145410.1	MULTISPECIES: hydrogenase-2 small chain [Proteobacteria]	Length: 372\nNumber of predicted TMHs:  1\nExp number of AAs in TMHs: 23.72226\nExp number, first 60 AAs:  0.22038\nTotal prob of N-in:   0.12994\noutside   1   331\nTMhelix   332   354\ninside   355   372

26608	GCF_001255755.1_12844_1_54	Shigella sonnei	Proteobacteria; Gammaproteobacteria; Enterobacterales; Enterobacteriaceae; Shigella	MNNEETFYQAMRRQGVTRRSFLKYCSLAATSLGLGAGMAPKIAWALENKPRIPVVWIHGL	WP_005139779.1	hydrogenase [Shigella sonnei]	Length: 372\nNumber of predicted TMHs:  2\nExp number of AAs in TMHs: 37.02628\nExp number, first 60 AAs:  19.39684\nTotal prob of N-in:   0.94313\nPOSSIBLE N-term signal sequence\ninside   1   19\nTMhelix   20   39\noutside   40   325\nTMhelix   326   348\ninside   349   372

26609	GCF_001255595.1_H14060002901	Shigella sonnei	Proteobacteria; Gammaproteobacteria; Enterobacterales; Enterobacteriaceae; Shigella	MNNEETFYQAMRRQGVTRRSFLKYCSLAATSLGLGAGMAPKIAWALENKPRIPVVWIHGL	WP_005139779.1	hydrogenase [Shigella sonnei]	Length: 372\nNumber of predicted TMHs:  2\nExp number of AAs in TMHs: 37.02628\nExp number, first 60 AAs:  19.39684\nTotal prob of N-in:   0.94313\nPOSSIBLE N-term signal sequence\ninside   1   19\nTMhelix   20   39\noutside   40   325\nTMhelix   326   348\ninside   349   372

26610	GCF_001255815.1_8289_7_64	Shigella sonnei	Proteobacteria; Gammaproteobacteria; Enterobacterales; Enterobacteriaceae; Shigella	MTGDNTLIHSHGINRRDFMKLCAALAATMGLSSKAAAEMAESVTNPQRPPVIWIGAQECT	WP_000145410.1	MULTISPECIES: hydrogenase-2 small chain [Proteobacteria]	Length: 372\nNumber of predicted TMHs:  1\nExp number of AAs in TMHs: 23.72226\nExp number, first 60 AAs:  0.22038\nTotal prob of N-in:   0.12994\noutside   1   331\nTMhelix   332   354\ninside   355   372

26611	GCF_001255875.1_5236_1_3Shigella sonnei	Proteobacteria; Gammaproteobacteria; Enterobacterales; Enterobacteriaceae; Shigella	MTGDNTLIHSHGINRRDFMKLCAALAATMGLSSKAAAEMAESVTNPQRPPVIWIGAQECT	WP_000145410.1	MULTISPECIES: hydrogenase-2 small chain [Proteobacteria]	Length: 372\nNumber of predicted TMHs:  1\nExp number of AAs in TMHs: 23.72226\nExp number, first 60 AAs:  0.22038\nTotal prob of N-in:   0.12994\noutside   1   331\nTMhelix   332   354\ninside   355   372

26612 GCF_001255815.1_8289_7_64 Shigella sonnei Proteobacteria; Gammaproteobacteria; Enterobacterales; Enterobacteriaceae; Shigella MNNEETFYQAMRRQGVTRRSFLKYCSLAATSLGLGAGMAPKIAWALENKPRIPVVWIHGL WP_005139779.1 hydrogenase [Shigella sonnei] Length: 372\nNumber of predicted TMHs: 2\nExp number of AAs in TMHs: 37.02628\nExp number, first 60 AAs: 19.39684\nTotal prob of N-in: 0.94313\nPOSSIBLE N-term signal sequence\ninside 1 19\nTMhelix 20 39\noutside 40 325\nTMhelix 326 348\ninside 349 372

26613 GCF_001255875.1_5236_1_3Shigella sonnei Proteobacteria; Gammaproteobacteria; Enterobacterales; Enterobacteriaceae; Shigella MNNEETFYQAMRRQGVTRRSFLKYCSLAATSLGLGAGMAPKIAWALENKPRIPVVWIHGL WP_005139779.1 hydrogenase [Shigella sonnei] Length: 372\nNumber of predicted TMHs: 2\nExp number of AAs in TMHs: 37.02628\nExp number, first 60 AAs: 19.39684\nTotal prob of N-in: 0.94313\nPOSSIBLE N-term signal sequence\ninside 1 19\nTMhelix 20 39\noutside 40 325\nTMhelix 326 348\ninside 349 372

26614 GCF_001255635.1_8290_4_29 Shigella sonnei Proteobacteria; Gammaproteobacteria; Enterobacterales; Enterobacteriaceae; Shigella MNNEETFYQAMRRQGVTRRSFLKYCSLAATSLGLGAGMAPKIAWALENKPRIPVVWIHGL WP_005139779.1 hydrogenase [Shigella sonnei] Length: 372\nNumber of predicted TMHs: 2\nExp number of AAs in TMHs: 37.02628\nExp number, first 60 AAs: 19.39684\nTotal prob of N-in: 0.94313\nPOSSIBLE N-term signal sequence\ninside 1 19\nTMhelix 20 39\noutside 40 325\nTMhelix 326 348\ninside 349 372

26615 GCF_001255895.1_Sg_201401842_GAGTGG_L001 Shigella sonnei Proteobacteria; Gammaproteobacteria; Enterobacterales; Enterobacteriaceae; Shigella MTGDNTLIHSHGINRRDFMKLCAALAATMGLSSKAAAEMAESVTNPQRPPVIWIGAQECT WP_000145410.1 MULTISPECIES: hydrogenase-2 small chain [Proteobacteria] Length: 372\nNumber of predicted TMHs: 1\nExp number of AAs in TMHs: 23.72226\nExp number, first 60 AAs: 0.22038\nTotal prob of N-in: 0.12994\noutside 1 331\nTMhelix 332 354\ninside 355 372

26616 GCF_001255955.1_5417_2_9Shigella sonnei Proteobacteria; Gammaproteobacteria; Enterobacterales; Enterobacteriaceae; Shigella MTGDNTLIHSHGINRRDFMKLCAALAATMGLSSKAAAEMAESVTNPQRPPVIWIGAQECT WP_000145410.1 MULTISPECIES: hydrogenase-2 small chain [Proteobacteria] Length: 372\nNumber of predicted TMHs: 1\nExp number of AAs in TMHs: 23.72226\nExp number, first 60 AAs: 0.22038\nTotal prob of N-in: 0.12994\noutside 1 331\nTMhelix 332 354\ninside 355 372

26617 GCF_001255995.1_12850_1_48 Shigella sonnei Proteobacteria; Gammaproteobacteria; Enterobacterales; Enterobacteriaceae; Shigella MTGDNTLIHSHGINRRDFMKLCAALAATMGLSSKAAAEMAESVTNPQRPPVIWIGAQECT WP_000145410.1 MULTISPECIES: hydrogenase-2 small chain [Proteobacteria] Length: 372\nNumber of predicted TMHs: 1\nExp number of AAs in TMHs: 23.72226\nExp number, first 60 AAs: 0.22038\nTotal prob of N-in: 0.12994\noutside 1 331\nTMhelix 332 354\ninside 355 372

26618 GCF_001256035.1_H14138024701 Shigella sonnei Proteobacteria; Gammaproteobacteria; Enterobacterales; Enterobacteriaceae; Shigella MTGDNTLIHSHGINRRDFMKLCAALAATMGLSSKAAAEMAESVTNPQRPPVIWIGAQECT WP_000145410.1 MULTISPECIES: hydrogenase-2 small chain [Proteobacteria] Length: 372\nNumber of predicted TMHs: 1\nExp number of AAs in TMHs: 23.72226\nExp number, first 60 AAs: 0.22038\nTotal prob of N-in: 0.12994\noutside 1 331\nTMhelix 332 354\ninside 355 372

26619 GCF_001256075.1_12971_8_53 Shigella sonnei Proteobacteria; Gammaproteobacteria; Enterobacterales; Enterobacteriaceae; Shigella MTGDNTLIHSHGINRRDFMKLCAALAATMGLSSKAAAEMAESVTNPQRPPVIWIGAQECT WP_000145410.1 MULTISPECIES: hydrogenase-2 small chain [Proteobacteria] Length: 372\nNumber of predicted TMHs: 1\nExp number of AAs in TMHs: 23.72226\nExp number, first 60 AAs: 0.22038\nTotal prob of N-in: 0.12994\noutside 1 331\nTMhelix 332 354\ninside 355 372

26620 GCF_001256135.1_9870_6_62 Shigella sonnei Proteobacteria; Gammaproteobacteria; Enterobacterales; Enterobacteriaceae; Shigella MTGDNTLIHSHGINRRDFMKLCAALAATMGLSSKAAAEMAESVTNPQRPPVIWIGAQECT WP_000145410.1 MULTISPECIES: hydrogenase-2 small chain [Proteobacteria] Length: 372\nNumber of predicted TMHs: 1\nExp number of AAs in TMHs: 23.72226\nExp number, first 60 AAs: 0.22038\nTotal prob of N-in: 0.12994\noutside 1 331\nTMhelix 332 354\ninside 355 372

26621 GCF_001256115.1_8289_7_87 Shigella sonnei Proteobacteria; Gammaproteobacteria; Enterobacterales; Enterobacteriaceae; Shigella MTGDNTLIHSHGINRRDFMKLCAALAATMGLSSKAAAEMAESVTNPQRPPVIWIGAQECT WP_000145410.1 MULTISPECIES: hydrogenase-2 small chain [Proteobacteria] Length: 372\nNumber of predicted TMHs: 1\nExp number of AAs in TMHs: 23.72226\nExp number, first 60 AAs: 0.22038\nTotal prob of N-in: 0.12994\noutside 1 331\nTMhelix 332 354\ninside 355 372

26622 GCF_001256175.1_6496_8_12 Shigella sonnei Proteobacteria; Gammaproteobacteria; Enterobacterales; Enterobacteriaceae; Shigella MTGDNTLIHSHGINRRDFMKLCAALAATMGLSSKAAAEMAESVTNPQRPPVIWIGAQECT WP_000145410.1 MULTISPECIES: hydrogenase-2 small chain [Proteobacteria] Length: 372\nNumber of predicted TMHs: 1\nExp number of AAs in TMHs: 23.72226\nExp number, first 60 AAs: 0.22038\nTotal prob of N-in: 0.12994\noutside 1 331\nTMhelix 332 354\ninside 355 372

26623 GCF_001256215.1_H14086038001 Shigella sonnei Proteobacteria; Gammaproteobacteria; Enterobacterales; Enterobacteriaceae; Shigella MTGDNTLIHSHGINRRDFMKLCAALAATMGLSSKAAAEMAESVTNPQRPPVIWIGAQECT WP_000145410.1 MULTISPECIES: hydrogenase-2 small chain [Proteobacteria] Length: 372\nNumber of predicted TMHs: 1\nExp number of AAs in TMHs: 23.72226\nExp number, first 60 AAs: 0.22038\nTotal prob of N-in: 0.12994\noutside 1 331\nTMhelix 332 354\ninside 355 372

26624 GCF_001256175.1_6496_8_12 Shigella sonnei Proteobacteria; Gammaproteobacteria; Enterobacterales; Enterobacteriaceae; Shigella MNNEETFYQAMRRQGVTRRSFLKYCSLAATSLGLGAGMAPKIAWALENKPRIPVVWIHGL WP_005139779.1 hydrogenase [Shigella sonnei] Length: 372\nNumber of predicted TMHs: 2\nExp number of AAs in TMHs: 37.02628\nExp number, first 60 AAs: 19.39684\nTotal prob of N-in: 0.94313\nPOSSIBLE N-term signal sequence\ninside 1 19\nTMhelix 20 39\noutside 40 325\nTMhelix 326 348\ninside 349 372

26625 GCF_001256215.1_H14086038001 Shigella sonnei Proteobacteria; Gammaproteobacteria; Enterobacterales; Enterobacteriaceae; Shigella MNNEETFYQAMRRQGVTRRSFLKYCSLAATSLGLGAGMAPKIAWALENKPRIPVVWIHGL WP_005139779.1 hydrogenase [Shigella sonnei] Length: 372\nNumber of predicted TMHs: 2\nExp number of AAs in TMHs: 37.02628\nExp number, first 60 AAs: 19.39684\nTotal prob of N-in: 0.94313\nPOSSIBLE N-term signal sequence\ninside 1 19\nTMhelix 20 39\noutside 40 325\nTMhelix 326 348\ninside 349 372

26626 GCF_001255895.1_Sg_201401842_GAGTGG_L001 Shigella sonnei Proteobacteria; Gammaproteobacteria; Enterobacterales; Enterobacteriaceae; Shigella MNNEETFYQAMRRQGVTRRSFLKYCSLAATSLGLGAGMAPKIAWALENKPRIPVVWIHGL WP_005139779.1 hydrogenase [Shigella sonnei] Length: 372\nNumber of predicted TMHs: 2\nExp number of AAs in TMHs: 37.02628\nExp number, first 60 AAs: 19.39684\nTotal prob of N-in: 0.94313\nPOSSIBLE N-term signal sequence\ninside 1 19\nTMhelix 20 39\noutside 40 325\nTMhelix 326 348\ninside 349 372

26627 GCF_001255955.1_5417_2_9Shigella sonnei Proteobacteria; Gammaproteobacteria; Enterobacterales; Enterobacteriaceae; Shigella MNNEETFYQAMRRQGVTRRSFLKYCSLAATSLGLGAGMAPKIAWALENKPRIPVVWIHGL WP_005139779.1 hydrogenase [Shigella sonnei] Length: 372\nNumber of predicted TMHs: 2\nExp number of AAs in TMHs: 37.02628\nExp number, first 60 AAs: 19.39684\nTotal prob of N-in: 0.94313\nPOSSIBLE N-term signal sequence\ninside 1 19\nTMhelix 20 39\noutside 40 325\nTMhelix 326 348\ninside 349 372

26628 GCF_001255995.1_12850_1_48 Shigella sonnei Proteobacteria; Gammaproteobacteria; Enterobacterales; Enterobacteriaceae; Shigella MNNEETFYQAMRRQGVTRRSFLKYCSLAATSLGLGAGMAPKIAWALENKPRIPVVWIHGL WP_005139779.1 hydrogenase [Shigella sonnei] Length: 372\nNumber of predicted TMHs: 2\nExp number of AAs in TMHs: 37.02628\nExp number, first 60 AAs: 19.39684\nTotal prob of N-in: 0.94313\nPOSSIBLE N-term signal sequence\ninside 1 19\nTMhelix 20 39\noutside 40 325\nTMhelix 326 348\ninside 349 372

26629 GCF_001256035.1_H14138024701 Shigella sonnei Proteobacteria; Gammaproteobacteria; Enterobacterales; Enterobacteriaceae; Shigella MNNEETFYQAMRRQGVTRRSFLKYCSLAATSLGLGAGMAPKIAWALENKPRIPVVWIHGL WP_052962775.1 hydrogenase [Shigella sonnei] Length: 372\nNumber of predicted TMHs: 2\nExp number of AAs in TMHs: 37.09248\nExp number, first 60 AAs: 19.39296\nTotal prob of N-in: 0.94316\nPOSSIBLE N-term signal sequence\ninside 1 19\nTMhelix 20 39\noutside 40 325\nTMhelix 326 348\ninside 349 372

26630 GCF_001256235.1_12971_8_79 Shigella sonnei Proteobacteria; Gammaproteobacteria; Enterobacterales; Enterobacteriaceae; Shigella
MTGDNTLIHSHGINRRDFMKLCAALAATMGLSSKAAAEMAESVTNPQRPPVIWIGAQECT WP_000145410.1
MULTISPECIES: hydrogenase-2 small chain [Proteobacteria] Length: 372\nNumber of predicted TMHs: 1\nExp number of AAs in TMHs: 23.72226\nExp number, first 60 AAs: 0.22038\nTotal prob of N-in: 0.12994\noutside 1 331\nTMhelix 332 354\ninside 355 372

26631 GCF_001256295.1_9870_6_14 Shigella sonnei Proteobacteria; Gammaproteobacteria; Enterobacterales; Enterobacteriaceae; Shigella
MTGDNTLIHSHGINRRDFMKLCAALAATMGLSSKAAAEMAESVTNPQRPPVIWIGAQECT WP_000145410.1
MULTISPECIES: hydrogenase-2 small chain [Proteobacteria] Length: 372\nNumber of predicted TMHs: 1\nExp number of AAs in TMHs: 23.72226\nExp number, first 60 AAs: 0.22038\nTotal prob of N-in: 0.12994\noutside 1 331\nTMhelix 332 354\ninside 355 372

26632 GCF_001256235.1_12971_8_79 Shigella sonnei Proteobacteria; Gammaproteobacteria; Enterobacterales; Enterobacteriaceae; Shigella
MNNEETFYQAMRRQGVTRRSFLKYCSLAATSLGLGAGMAPKIAWALENKPRIPVVWIHGL WP_005139779.1
hydrogenase [Shigella sonnei] Length: 372\nNumber of predicted TMHs: 2\nExp number of AAs in TMHs: 37.02628\nExp number, first 60 AAs: 19.39684\nTotal prob of N-in: 0.94313\nPOSSIBLE N-term signal sequence\ninside 1 19\nTMhelix 20 39\noutside 40 325\nTMhelix 326 348\ninside 349 372

26633 GCF_001256295.1_9870_6_14 Shigella sonnei Proteobacteria; Gammaproteobacteria; Enterobacterales; Enterobacteriaceae; Shigella
MNNEETFYQAMRRQGVTRRSFLKYCSLAATSLGLGAGMAPKIAWALENKPRIPVVWIHGL WP_005139779.1
hydrogenase [Shigella sonnei] Length: 372\nNumber of predicted TMHs: 2\nExp number of AAs in TMHs: 37.02628\nExp number, first 60 AAs: 19.39684\nTotal prob of N-in: 0.94313\nPOSSIBLE N-term signal sequence\ninside 1 19\nTMhelix 20 39\noutside 40 325\nTMhelix 326 348\ninside 349 372

26634 GCF_001256135.1_9870_6_62 Shigella sonnei Proteobacteria; Gammaproteobacteria; Enterobacterales; Enterobacteriaceae; Shigella
MNNEETFYQAMRRQGVTRRSFLKYCSLAATSLGLGAGMAPKIAWALENKPRIPVVWIHGL WP_005139779.1
hydrogenase [Shigella sonnei] Length: 372\nNumber of predicted TMHs: 2\nExp number of AAs in TMHs: 37.02628\nExp number, first 60 AAs: 19.39684\nTotal prob of N-in: 0.94313\nPOSSIBLE N-term signal sequence\ninside 1 19\nTMhelix 20 39\noutside 40 325\nTMhelix 326 348\ninside 349 372

26635 GCF_001256115.1_8289_7_87 Shigella sonnei Proteobacteria; Gammaproteobacteria; Enterobacterales; Enterobacteriaceae; Shigella
MNNEETFYQAMRRQGVTRRSFLKYCSLAATSLGLGAGMAPKIAWALENKPRIPVVWIHGL WP_005139779.1
hydrogenase [Shigella sonnei] Length: 372\nNumber of predicted TMHs: 2\nExp number of AAs in TMHs: 37.02628\nExp number, first 60 AAs: 19.39684\nTotal prob of N-in: 0.94313\nPOSSIBLE N-term signal sequence\ninside 1 19\nTMhelix 20 39\noutside 40 325\nTMhelix 326 348\ninside 349 372

26636 GCF_001256315.1_7655_8_8Shigella sonnei Proteobacteria; Gammaproteobacteria; Enterobacterales; Enterobacteriaceae; Shigella MTGDNTLIHSHGINRRDFMKLCAALAATMGLSSKAAAEMAESVTNPQRPPVIWIGAQECT
WP_000145410.1 MULTISPECIES: hydrogenase-2 small chain [Proteobacteria] Length: 372\nNumber of predicted TMHs: 1\nExp number of AAs in TMHs: 23.72226\nExp number, first 60 AAs: 0.22038\nTotal prob of N-in: 0.12994\noutside 1 331\nTMhelix 332 354\ninside 355 372

26637 GCF_001256375.1_5236_2_1Shigella sonnei Proteobacteria; Gammaproteobacteria; Enterobacterales; Enterobacteriaceae; Shigella MTGDNTLIHSHGINRRDFMKLCAALAATMGLSSKAAAEMAESVTNPQRPPVIWIGAQECT
WP_000145410.1 MULTISPECIES: hydrogenase-2 small chain [Proteobacteria] Length: 372\nNumber of predicted TMHs: 1\nExp number of AAs in TMHs: 23.72226\nExp number, first 60 AAs: 0.22038\nTotal prob of N-in: 0.12994\noutside 1 331\nTMhelix 332 354\ninside 355 372

26638 GCF_001256375.1_5236_2_1Shigella sonnei Proteobacteria; Gammaproteobacteria; Enterobacterales; Enterobacteriaceae; Shigella MNNEETFYQAMRRQGVTRRSFLKYCSLAATSLGLGAGMAPKIAWALENKPRIPVVWIHGL
WP_005139779.1 hydrogenase [Shigella sonnei] Length: 372\nNumber of predicted TMHs: 2\nExp number of AAs in TMHs: 37.02628\nExp number, first 60 AAs: 19.39684\nTotal prob of N-in: 0.94313\nPOSSIBLE N-term signal sequence\ninside 1 19\nTMhelix 20 39\noutside 40 325\nTMhelix 326 348\ninside 349 372

26639 GCF_001256315.1_7655_8_8Shigella sonnei Proteobacteria; Gammaproteobacteria; Enterobacterales; Enterobacteriaceae; Shigella MNNEETFYQAMRRQGVTRRSFLKYCSLAATSLGLGAGMAPKIAWALENKPRIPVVWIHGL WP_005139779.1 hydrogenase [Shigella sonnei] Length: 372\nNumber of predicted TMHs: 2\nExp number of AAs in TMHs: 37.02628\nExp number, first 60 AAs: 19.39684\nTotal prob of N-in: 0.94313\nPOSSIBLE N-term signal sequence\ninside 1 19\nTMhelix 20 39\noutside 40 325\nTMhelix 326 348\ninside 349 372

26640 GCF_001256075.1_12971_8_53 Shigella sonnei Proteobacteria; Gammaproteobacteria; Enterobacterales; Enterobacteriaceae; Shigella MNNEETFYQAMRRQGVTRRSFLKYCSLAATSLGLGAGMAPKIAWALENKPRIPVVWIHGL WP_052962775.1 hydrogenase [Shigella sonnei] Length: 372\nNumber of predicted TMHs: 2\nExp number of AAs in TMHs: 37.09248\nExp number, first 60 AAs: 19.39296\nTotal prob of N-in: 0.94316\nPOSSIBLE N-term signal sequence\ninside 1 19\nTMhelix 20 39\noutside 40 325\nTMhelix 326 348\ninside 349 372

26641 GCF_001256415.1_12971_8_62 Shigella sonnei Proteobacteria; Gammaproteobacteria; Enterobacterales; Enterobacteriaceae; Shigella MTGDNTLIHSHGINRRDFMKLCAALAATMGLSSKAAAEMAESVTNPQRPPVIWIGAQECT WP_000145410.1 MULTISPECIES: hydrogenase-2 small chain [Proteobacteria] Length: 372\nNumber of predicted TMHs: 1\nExp number of AAs in TMHs: 23.72226\nExp number, first 60 AAs: 0.22038\nTotal prob of N-in: 0.12994\noutside 1 331\nTMhelix 332 354\ninside 355 372

26642 GCF_001256435.1_Sg_201405327_CTAGCT_L001 Shigella sonnei Proteobacteria; Gammaproteobacteria; Enterobacterales; Enterobacteriaceae; Shigella MTGDNTLIHSHGINRRDFMKLCAALAATMGLSSKAAAEMAESVTNPQRPPVIWIGAQECT WP_000145410.1 MULTISPECIES: hydrogenase-2 small chain [Proteobacteria] Length: 372\nNumber of predicted TMHs: 1\nExp number of AAs in TMHs: 23.72226\nExp number, first 60 AAs: 0.22038\nTotal prob of N-in: 0.12994\noutside 1 331\nTMhelix 332 354\ninside 355 372

26643 GCF_001256475.1_8290_4_48 Shigella sonnei Proteobacteria; Gammaproteobacteria; Enterobacterales; Enterobacteriaceae; Shigella MTGDNTLIHSHGINRRDFMKLCAALAATMGLSSKAAAEMAESVTNPQRPPVIWIGAQECT WP_000145410.1 MULTISPECIES: hydrogenase-2 small chain [Proteobacteria] Length: 372\nNumber of predicted TMHs: 1\nExp number of AAs in TMHs: 23.72226\nExp number, first 60 AAs: 0.22038\nTotal prob of N-in: 0.12994\noutside 1 331\nTMhelix 332 354\ninside 355 372

26644 GCF_001256495.1_12844_1_56 Shigella sonnei Proteobacteria; Gammaproteobacteria; Enterobacterales; Enterobacteriaceae; Shigella MTGDNTLIHSHGINRRDFMKLCAALAATMGLSSKAAAEMAESVTNPQRPPVIWIGAQECT WP_000145410.1 MULTISPECIES: hydrogenase-2 small chain [Proteobacteria] Length: 372\nNumber of predicted TMHs: 1\nExp number of AAs in TMHs: 23.72226\nExp number, first 60 AAs: 0.22038\nTotal prob of N-in: 0.12994\noutside 1 331\nTMhelix 332 354\ninside 355 372

26645 GCF_001256555.1_5236_1_6Shigella sonnei Proteobacteria; Gammaproteobacteria; Enterobacterales; Enterobacteriaceae; Shigella MTGDNTLIHSHGINRRDFMKLCAALAATMGLSSKAAAEMAESVTNPQRPPVIWIGAQECT WP_000145410.1 MULTISPECIES: hydrogenase-2 small chain [Proteobacteria] Length: 372\nNumber of predicted TMHs: 1\nExp number of AAs in TMHs: 23.72226\nExp number, first 60 AAs: 0.22038\nTotal prob of N-in: 0.12994\noutside 1 331\nTMhelix 332 354\ninside 355 372

26646 GCF_001256635.1_8289_7_59 Shigella sonnei Proteobacteria; Gammaproteobacteria; Enterobacterales; Enterobacteriaceae; Shigella MTGDNTLIHSHGINRRDFMKLCAALAATMGLSSKAAAEMAESVTNPQRPPVIWIGAQECT WP_000145410.1 MULTISPECIES: hydrogenase-2 small chain [Proteobacteria] Length: 372\nNumber of predicted TMHs: 1\nExp number of AAs in TMHs: 23.72226\nExp number, first 60 AAs: 0.22038\nTotal prob of N-in: 0.12994\noutside 1 331\nTMhelix 332 354\ninside 355 372

26647 GCF_001256615.1_12971_8_13 Shigella sonnei Proteobacteria; Gammaproteobacteria; Enterobacterales; Enterobacteriaceae; Shigella MTGDNTLIHSHGINRRDFMKLCAALAATMGLSSKAAAEMAESVTNPQRPPVIWIGAQECT WP_000145410.1 MULTISPECIES: hydrogenase-2 small chain [Proteobacteria] Length: 372\nNumber of predicted TMHs: 1\nExp number of AAs in TMHs: 23.72226\nExp number, first 60 AAs: 0.22038\nTotal prob of N-in: 0.12994\noutside 1 331\nTMhelix 332 354\ninside 355 372

26648 GCF_001256435.1_Sg_201405327_CTAGCT_L001 Shigella sonnei Proteobacteria; Gammaproteobacteria; Enterobacterales; Enterobacteriaceae; Shigella MNNEETFYQAMRRQGVTRRSFLKYCSLAATSLGLGAGMAPKIAWALENKPRIPVVWIHGL WP_005139779.1 hydrogenase [Shigella sonnei] Length: 372\nNumber of predicted TMHs: 2\nExp number of AAs in TMHs: 37.02628\nExp number, first 60 AAs: 19.39684\nTotal prob of N-in: 0.94313\nPOSSIBLE N-term signal sequence\ninside 1 19\nTMhelix 20 39\noutside 40 325\nTMhelix 326 348\ninside 349 372

26649 GCF_001256415.1_12971_8_62 Shigella sonnei Proteobacteria; Gammaproteobacteria; Enterobacterales; Enterobacteriaceae; Shigella MNNEETFYQAMRRQGVTRRSFLKYCSLAATSLGLGAGMAPKIAWALENKPRIPVVWIHGL WP_005139779.1 hydrogenase [Shigella sonnei] Length: 372\nNumber of predicted TMHs: 2\nExp number of AAs in TMHs: 37.02628\nExp number, first 60 AAs: 19.39684\nTotal prob of N-in: 0.94313\nPOSSIBLE N-term signal sequence\ninside 1 19\nTMhelix 20 39\noutside 40 325\nTMhelix 326 348\ninside 349 372

26650 GCF_001256735.1_H14132054001 Shigella sonnei Proteobacteria; Gammaproteobacteria; Enterobacterales; Enterobacteriaceae; Shigella MTGDNTLIHSHGINRRDFMKLCAALAATMGLSSKAAAEMAESVTNPQRPPVIWIGAQECT WP_000145410.1 MULTISPECIES: hydrogenase-2 small chain [Proteobacteria] Length: 372\nNumber of predicted TMHs: 1\nExp number of AAs in TMHs: 23.72226\nExp number, first 60 AAs: 0.22038\nTotal prob of N-in: 0.12994\noutside 1 331\nTMhelix 332 354\ninside 355 372

26651 GCF_001256775.1_7655_8_6Shigella sonnei Proteobacteria; Gammaproteobacteria; Enterobacterales; Enterobacteriaceae; Shigella MTGDNTLIHSHGINRRDFMKLCAALAATMGLSSKAAAEMAESVTNPQRPPVIWIGAQECT WP_000145410.1 MULTISPECIES: hydrogenase-2 small chain [Proteobacteria] Length: 372\nNumber of predicted TMHs: 1\nExp number of AAs in TMHs: 23.72226\nExp number, first 60 AAs: 0.22038\nTotal prob of N-in: 0.12994\noutside 1 331\nTMhelix 332 354\ninside 355 372

26652 GCF_001256815.1_Sg_201401237_CCGTCC_L001 Shigella sonnei Proteobacteria; Gammaproteobacteria; Enterobacterales; Enterobacteriaceae; Shigella MTGDNTLIHSHGINRRDFMKLCAALAATMGLSSKAAAEMAESVTNPQRPPVIWIGAQECT WP_000145410.1 MULTISPECIES: hydrogenase-2 small chain [Proteobacteria] Length: 372\nNumber of predicted TMHs: 1\nExp number of AAs in TMHs: 23.72226\nExp number, first 60 AAs: 0.22038\nTotal prob of N-in: 0.12994\noutside 1 331\nTMhelix 332 354\ninside 355 372

26653 GCF_001256875.1_Sg_201400003_CGATGT_L001 Shigella sonnei Proteobacteria; Gammaproteobacteria; Enterobacterales; Enterobacteriaceae; Shigella MTGDNTLIHSHGINRRDFMKLCAALAATMGLSSKAAAEMAESVTNPQRPPVIWIGAQECT WP_000145410.1 MULTISPECIES: hydrogenase-2 small chain [Proteobacteria] Length: 372\nNumber of predicted TMHs: 1\nExp number of AAs in TMHs: 23.72226\nExp number, first 60 AAs: 0.22038\nTotal prob of N-in: 0.12994\noutside 1 331\nTMhelix 332 354\ninside 355 372

26654 GCF_001256735.1_H14132054001 Shigella sonnei Proteobacteria; Gammaproteobacteria; Enterobacterales; Enterobacteriaceae; Shigella MNNEETFYQAMRRQGVTRRSFLKYCSLAATSLGLGAGMAPKIAWALENKPRIPVVWIHGL WP_005139779.1 hydrogenase [Shigella sonnei] Length: 372\nNumber of predicted TMHs: 2\nExp number of AAs in TMHs: 37.02628\nExp number, first 60 AAs: 19.39684\nTotal prob of N-in: 0.94313\nPOSSIBLE N-term signal sequence\ninside 1 19\nTMhelix 20 39\noutside 40 325\nTMhelix 326 348\ninside 349 372

26655 GCF_001256815.1_Sg_201401237_CCGTCC_L001 Shigella sonnei Proteobacteria; Gammaproteobacteria; Enterobacterales; Enterobacteriaceae; Shigella MNNEETFYQAMRRQGVTRRSFLKYCSLAATSLGLGAGMAPKIAWALENKPRIPVVWIHGL WP_005139779.1 hydrogenase [Shigella sonnei] Length: 372\nNumber of predicted TMHs: 2\nExp number of AAs in TMHs: 37.02628\nExp number, first 60 AAs: 19.39684\nTotal prob of N-in: 0.94313\nPOSSIBLE N-term signal sequence\ninside 1 19\nTMhelix 20 39\noutside 40 325\nTMhelix 326 348\ninside 349 372

26656 GCF_001256475.1_8290_4_48 Shigella sonnei Proteobacteria; Gammaproteobacteria; Enterobacterales; Enterobacteriaceae; Shigella MNNEETFYQAMRRQGVTRRSFLKYCSLAATSLGLGAGMAPKIAWALENKPRIPVVWIHGL WP_005139779.1 hydrogenase [Shigella sonnei] Length: 372\nNumber of predicted TMHs: 2\nExp number of AAs in TMHs: 37.02628\nExp number, first 60 AAs: 19.39684\nTotal prob of N-in: 0.94313\nPOSSIBLE N-term signal sequence\ninside 1 19\nTMhelix 20 39\noutside 40 325\nTMhelix 326 348\ninside 349 372

26657 GCF_001256495.1_12844_1_56 Shigella sonnei Proteobacteria; Gammaproteobacteria; Enterobacterales; Enterobacteriaceae; Shigella MNNEETFYQAMRRQGVTRRSFLKYCSLAATSLGLGAGMAPKIAWALENKPRIPVVWIHGL WP_005139779.1 hydrogenase [Shigella sonnei] Length: 372\nNumber of predicted TMHs: 2\nExp number of AAs in TMHs: 37.02628\nExp number, first 60 AAs: 19.39684\nTotal prob of N-in: 0.94313\nPOSSIBLE N-term signal sequence\ninside 1 19\nTMhelix 20 39\noutside 40 325\nTMhelix 326 348\ninside 349 372

26658 GCF_001256915.1_12850_1_44 Shigella sonnei Proteobacteria; Gammaproteobacteria; Enterobacterales; Enterobacteriaceae; Shigella MTGDNTLIHSHGINRRDFMKLCAALAATMGLSSKAAAEMAESVTNPQRPPVIWIGAQECT WP_000145410.1 MULTISPECIES: hydrogenase-2 small chain [Proteobacteria] Length: 372\nNumber of predicted TMHs: 1\nExp number of AAs in TMHs: 23.72226\nExp number, first 60 AAs: 0.22038\nTotal prob of N-in: 0.12994\noutside 1 331\nTMhelix 332 354\ninside 355 372

26659 GCF_001256915.1_12850_1_44 Shigella sonnei Proteobacteria; Gammaproteobacteria; Enterobacterales; Enterobacteriaceae; Shigella MNNEETFYQAMRRQGVTRRSFLKYCSLAATSLGLGAGMAPKIAWALENKPRIPVVWIHGL WP_005139779.1 hydrogenase [Shigella sonnei] Length: 372\nNumber of predicted TMHs: 2\nExp number of AAs in TMHs: 37.02628\nExp number, first 60 AAs: 19.39684\nTotal prob of N-in: 0.94313\nPOSSIBLE N-term signal sequence\ninside 1 19\nTMhelix 20 39\noutside 40 325\nTMhelix 326 348\ninside 349 372

26660 GCF_001256635.1_8289_7_59 Shigella sonnei Proteobacteria; Gammaproteobacteria; Enterobacterales; Enterobacteriaceae; Shigella MNNEETFYQAMRRQGVTRRSFLKYCSLAATSLGLGAGMAPKIAWALENKPRIPVVWIHGL WP_005139779.1 hydrogenase [Shigella sonnei] Length: 372\nNumber of predicted TMHs: 2\nExp number of AAs in TMHs: 37.02628\nExp number, first 60 AAs: 19.39684\nTotal prob of N-in: 0.94313\nPOSSIBLE N-term signal sequence\ninside 1 19\nTMhelix 20 39\noutside 40 325\nTMhelix 326 348\ninside 349 372

26661 GCF_001256555.1_5236_1_6Shigella sonnei Proteobacteria; Gammaproteobacteria; Enterobacterales; Enterobacteriaceae; Shigella MNNEETFYQAMRRQGVTRRSFLKYCSLAATSLGLGAGMAPKIAWALENKPRIPVVWIHGL WP_005139779.1 hydrogenase [Shigella sonnei] Length: 372\nNumber of predicted TMHs: 2\nExp number of AAs in TMHs: 37.02628\nExp number, first 60 AAs: 19.39684\nTotal prob of N-in: 0.94313\nPOSSIBLE N-term signal sequence\ninside 1 19\nTMhelix 20 39\noutside 40 325\nTMhelix 326 348\ninside 349 372

26662 GCF_001256615.1_12971_8_13 Shigella sonnei Proteobacteria; Gammaproteobacteria; Enterobacterales; Enterobacteriaceae; Shigella MNNEETFYQAMRRQGVTRRSFLKYCSLAATSLGLGAGMAPKIAWALENKPRIPVVWIHGL WP_005139779.1 hydrogenase [Shigella sonnei] Length: 372\nNumber of predicted TMHs: 2\nExp number of AAs in TMHs: 37.02628\nExp number, first 60 AAs: 19.39684\nTotal prob of N-in: 0.94313\nPOSSIBLE N-term signal sequence\ninside 1 19\nTMhelix 20 39\noutside 40 325\nTMhelix 326 348\ninside 349 372

26663 GCF_001256935.1_Sg_201401464_GTGAAA_L001 Shigella sonnei Proteobacteria; Gammaproteobacteria; Enterobacterales; Enterobacteriaceae; Shigella MTGDNTLIHSHGINRRDFMKLCAALAATMGLSSKAAAEMAESVTNPQRPPVIWIGAQECT WP_000145410.1 MULTISPECIES: hydrogenase-2 small chain [Proteobacteria] Length: 372\nNumber of predicted TMHs: 1\nExp number of AAs in TMHs: 23.72226\nExp number, first 60 AAs: 0.22038\nTotal prob of N-in: 0.12994\noutside 1 331\nTMhelix 332 354\ninside 355 372

26664 GCF_001256995.1_8289_7_96 Shigella sonnei Proteobacteria; Gammaproteobacteria; Enterobacterales; Enterobacteriaceae; Shigella MTGDNTLIHSHGINRRDFMKLCAALAATMGLSSKAAAEMAESVTNPQRPPVIWIGAQECT WP_000145410.1 MULTISPECIES: hydrogenase-2 small chain [Proteobacteria] Length: 372\nNumber of predicted TMHs: 1\nExp number of AAs in TMHs: 23.72226\nExp number, first 60 AAs: 0.22038\nTotal prob of N-in: 0.12994\noutside 1 331\nTMhelix 332 354\ninside 355 372

26665 GCF_001257055.1_8289_7_40 Shigella sonnei Proteobacteria; Gammaproteobacteria; Enterobacterales; Enterobacteriaceae; Shigella MTGDNTLIHSHGINRRDFMKLCAALAATMGLSSKAAAEMAESVTNPQRPPVIWIGAQECT WP_000145410.1 MULTISPECIES: hydrogenase-2 small chain [Proteobacteria] Length: 372\nNumber of predicted TMHs: 1\nExp number of AAs in TMHs: 23.72226\nExp number, first 60 AAs: 0.22038\nTotal prob of N-in: 0.12994\noutside 1 331\nTMhelix 332 354\ninside 355 372

26666 GCF_001257095.1_8289_7_41 Shigella sonnei Proteobacteria; Gammaproteobacteria; Enterobacterales; Enterobacteriaceae; Shigella MTGDNTLIHSHGINRRDFMKLCAALAATMGLSSKAAAEMAESVTNPQRPPVIWIGAQECT WP_000145410.1 MULTISPECIES: hydrogenase-2 small chain [Proteobacteria] Length: 372\nNumber of predicted TMHs: 1\nExp number of AAs in TMHs: 23.72226\nExp number, first 60 AAs: 0.22038\nTotal prob of N-in: 0.12994\noutside 1 331\nTMhelix 332 354\ninside 355 372

26667 GCF_001257135.1_Sg_201311838_TAATCG_L001 Shigella sonnei Proteobacteria; Gammaproteobacteria; Enterobacterales; Enterobacteriaceae; Shigella MTGDNTLIHSHGINRRDFMKLCAALAATMGLSSKAAAEMAESVTNPQRPPVIWIGAQECT WP_000145410.1 MULTISPECIES: hydrogenase-2 small chain [Proteobacteria] Length: 372\nNumber of predicted TMHs: 1\nExp number of AAs in TMHs: 23.72226\nExp number, first 60 AAs: 0.22038\nTotal prob of N-in: 0.12994\noutside 1 331\nTMhelix 332 354\ninside 355 372

26668 GCF_001257175.1_8290_4_11 Shigella sonnei Proteobacteria; Gammaproteobacteria; Enterobacterales; Enterobacteriaceae; Shigella MTGDNTLIHSHGINRRDFMKLCAALAATMGLSSKAAAEMAESVTNPQRPPVIWIGAQECT WP_000145410.1 MULTISPECIES: hydrogenase-2 small chain [Proteobacteria] Length: 372\nNumber of predicted TMHs: 1\nExp number of AAs in TMHs: 23.72226\nExp number, first 60 AAs: 0.22038\nTotal prob of N-in: 0.12994\noutside 1 331\nTMhelix 332 354\ninside 355 372

26669 GCF_001258195.1_9870_6_4Shigella sonnei Proteobacteria; Gammaproteobacteria; Enterobacterales; Enterobacteriaceae; Shigella MTGDNTLIHSHGINRRDFMKLCAALAATMGLSSKAAAEMAESVTNPQRPPVIWIGAQECT WP_000145410.1 MULTISPECIES: hydrogenase-2 small chain [Proteobacteria] Length: 372\nNumber of predicted TMHs: 1\nExp number of AAs in TMHs: 23.72226\nExp number, first 60 AAs: 0.22038\nTotal prob of N-in: 0.12994\noutside 1 331\nTMhelix 332 354\ninside 355 372

26670 GCF_001256775.1_7655_8_6Shigella sonnei Proteobacteria; Gammaproteobacteria; Enterobacterales; Enterobacteriaceae; Shigella MNNEETFYQAMRRQGVTRRSFLKYCSLAATSLGLGAGMAPKIAWALENKPRIPVVWIHGL WP_005139779.1 hydrogenase [Shigella sonnei] Length: 372\nNumber of predicted TMHs: 2\nExp number of AAs in TMHs: 37.02628\nExp number, first 60 AAs: 19.39684\nTotal prob of N-in: 0.94313\nPOSSIBLE N-term signal sequence\ninside 1 19\nTMhelix 20 39\noutside 40 325\nTMhelix 326 348\ninside 349 372

26671 GCF_001257195.1_12971_8_7 Shigella sonnei Proteobacteria; Gammaproteobacteria; Enterobacterales; Enterobacteriaceae; Shigella MTGDNTLIHSHGINRRDFMKLCAALAATMGLSSKAAAEMAESVTNPQRPPVIWIGAQECT WP_000145410.1 MULTISPECIES: hydrogenase-2 small chain [Proteobacteria] Length: 372\nNumber of predicted TMHs: 1\nExp number of AAs in TMHs: 23.72226\nExp number, first 60 AAs: 0.22038\nTotal prob of N-in: 0.12994\noutside 1 331\nTMhelix 332 354\ninside 355 372

26672 GCF_001257195.1_12971_8_7 Shigella sonnei Proteobacteria; Gammaproteobacteria; Enterobacterales; Enterobacteriaceae; Shigella MNNEETFYQAMRRQGVTRRSFLKYCSLAATSLGLGAGMAPKIAWALENKPRIPVVWIHGL WP_005139779.1 hydrogenase [Shigella sonnei] Length: 372\nNumber of predicted TMHs: 2\nExp number of AAs in TMHs: 37.02628\nExp number, first 60 AAs: 19.39684\nTotal prob of N-in: 0.94313\nPOSSIBLE N-term signal sequence\ninside 1 19\nTMhelix 20 39\noutside 40 325\nTMhelix 326 348\ninside 349 372

26673 GCF_001256875.1_Sg_201400003_CGATGT_L001 Shigella sonnei Proteobacteria; Gammaproteobacteria; Enterobacterales; Enterobacteriaceae; Shigella MNNEETFYQAMRRQGVTRRSFLKYCSLAATSLGLGAGMAPKIAWALENKPRIPVVWIHGL WP_005139779.1 hydrogenase [Shigella sonnei] Length: 372\nNumber of predicted TMHs: 2\nExp number of AAs in TMHs: 37.02628\nExp number, first 60 AAs: 19.39684\nTotal prob of N-in: 0.94313\nPOSSIBLE N-term signal sequence\ninside 1 19\nTMhelix 20 39\noutside 40 325\nTMhelix 326 348\ninside 349 372

26674 GCF_001256935.1_Sg_201401464_GTGAAA_L001 Shigella sonnei Proteobacteria; Gammaproteobacteria; Enterobacterales; Enterobacteriaceae; Shigella MNNEETFYQAMRRQGVTRRSFLKYCSLAATSLGLGAGMAPKIAWALENKPRIPVVWIHGL WP_005139779.1 hydrogenase [Shigella sonnei] Length: 372\nNumber of predicted TMHs: 2\nExp number of AAs in TMHs: 37.02628\nExp number, first 60 AAs: 19.39684\nTotal prob of N-in: 0.94313\nPOSSIBLE N-term signal sequence\ninside 1 19\nTMhelix 20 39\noutside 40 325\nTMhelix 326 348\ninside 349 372

26675 GCF_001256995.1_8289_7_96 Shigella sonnei Proteobacteria; Gammaproteobacteria; Enterobacterales; Enterobacteriaceae; Shigella MNNEETFYQAMRRQGVTRRSFLKYCSLAATSLGLGAGMAPKIAWALENKPRIPVVWIHGL WP_005139779.1 hydrogenase [Shigella sonnei] Length: 372\nNumber of predicted TMHs: 2\nExp number of AAs in TMHs: 37.02628\nExp number, first 60 AAs: 19.39684\nTotal prob of N-in: 0.94313\nPOSSIBLE N-term signal sequence\ninside 1 19\nTMhelix 20 39\noutside 40 325\nTMhelix 326 348\ninside 349 372

26676 GCF_001257275.1_12971_8_5 Shigella sonnei Proteobacteria; Gammaproteobacteria; Enterobacterales; Enterobacteriaceae; Shigella MTGDNTLIHSHGINRRDFMKLCAALAATMGLSSKAAAEMAESVTNPQRPPVIWIGAQECT WP_000145410.1 MULTISPECIES: hydrogenase-2 small chain [Proteobacteria] Length: 372\nNumber of predicted TMHs: 1\nExp number of AAs in TMHs: 23.72226\nExp number, first 60 AAs: 0.22038\nTotal prob of N-in: 0.12994\noutside 1 331\nTMhelix 332 354\ninside 355 372

26677 GCF_001257335.1_9870_7_2Shigella sonnei Proteobacteria; Gammaproteobacteria; Enterobacterales; Enterobacteriaceae; Shigella MTGDNTLIHSHGINRRDFMKLCAALAATMGLSSKAAAEMAESVTNPQRPPVIWIGAQECT WP_000145410.1 MULTISPECIES: hydrogenase-2 small chain [Proteobacteria] Length: 372\nNumber of predicted TMHs: 1\nExp number of AAs in TMHs: 23.72226\nExp number, first 60 AAs: 0.22038\nTotal prob of N-in: 0.12994\noutside 1 331\nTMhelix 332 354\ninside 355 372

26678 GCF_001257415.1_5236_5_1Shigella sonnei Proteobacteria; Gammaproteobacteria; Enterobacterales; Enterobacteriaceae; Shigella MTGDNTLIHSHGINRRDFMKLCAALAATMGLSSKAAAEMAESVTNPQRPPVIWIGAQECT WP_000145410.1 MULTISPECIES: hydrogenase-2 small chain [Proteobacteria] Length: 372\nNumber of predicted TMHs: 1\nExp number of AAs in TMHs: 23.72226\nExp number, first 60 AAs: 0.22038\nTotal prob of N-in: 0.12994\noutside 1 331\nTMhelix 332 354\ninside 355 372

26679 GCF_001257455.1_8290_5_5Shigella sonnei Proteobacteria; Gammaproteobacteria; Enterobacterales; Enterobacteriaceae; Shigella MTGDNTLIHSHGINRRDFMKLCAALAATMGLSSKAAAEMAESVTNPQRPPVIWIGAQECT WP_000145410.1 MULTISPECIES: hydrogenase-2 small chain [Proteobacteria] Length: 372\nNumber of predicted TMHs: 1\nExp number of AAs in TMHs: 23.72226\nExp number, first 60 AAs: 0.22038\nTotal prob of N-in: 0.12994\noutside 1 331\nTMhelix 332 354\ninside 355 372

26680 GCF_001257375.1_12971_8_14 Shigella sonnei Proteobacteria; Gammaproteobacteria; Enterobacterales; Enterobacteriaceae; Shigella MTGDNTLIHSHGINRRDFMKLCAALAATMGLSSKAAAEMAESVTNPQRPPVIWIGAQECT WP_000145410.1 MULTISPECIES: hydrogenase-2 small chain [Proteobacteria] Length: 372\nNumber of predicted TMHs: 1\nExp number of AAs in TMHs: 23.72226\nExp number, first 60 AAs: 0.22038\nTotal prob of N-in: 0.12994\noutside 1 331\nTMhelix 332 354\ninside 355 372

26681 GCF_001257275.1_12971_8_5 Shigella sonnei Proteobacteria; Gammaproteobacteria; Enterobacterales; Enterobacteriaceae; Shigella MNNEETFYQAMRRQGVTRRSFLKYCSLAATSLGLGAGMAPKIAWALENKPRIPVVWIHGL WP_005139779.1 hydrogenase [Shigella sonnei] Length: 372\nNumber of predicted TMHs: 2\nExp number of AAs in TMHs: 37.02628\nExp number, first 60 AAs: 19.39684\nTotal prob of N-in: 0.94313\nPOSSIBLE N-term signal sequence\ninside 1 19\nTMhelix 20 39\noutside 40 325\nTMhelix 326 348\ninside 349 372

26682 GCF_001258735.1_9870_6_20 Shigella sonnei Proteobacteria; Gammaproteobacteria; Enterobacterales; Enterobacteriaceae; Shigella MTGDNTLIHSHGINRRDFMKLCAALAATMGLSSKAAAEMAESVTNPQRPPVIWIGAQECT WP_000145410.1 MULTISPECIES: hydrogenase-2 small chain [Proteobacteria] Length: 372\nNumber of predicted TMHs: 1\nExp number of AAs in TMHs: 23.72226\nExp number, first 60 AAs: 0.22038\nTotal prob of N-in: 0.12994\noutside 1 331\nTMhelix 332 354\ninside 355 372

26683 GCF_001257055.1_8289_7_40 Shigella sonnei Proteobacteria; Gammaproteobacteria; Enterobacterales; Enterobacteriaceae; Shigella MNNEETFYQAMRRQGVTRRSFLKYCSLAATSLGLGAGMAPKIAWALENKPRIPVVWIHGL WP_005139779.1 hydrogenase [Shigella sonnei] Length: 372\nNumber of predicted TMHs: 2\nExp number of AAs in TMHs: 37.02628\nExp number, first 60 AAs: 19.39684\nTotal prob of N-in: 0.94313\nPOSSIBLE N-term signal sequence\ninside 1 19\nTMhelix 20 39\noutside 40 325\nTMhelix 326 348\ninside 349 372

26684 GCF_001257095.1_8289_7_41 Shigella sonnei Proteobacteria; Gammaproteobacteria; Enterobacterales; Enterobacteriaceae; Shigella
MNNEETFYQAMRRQGVTRRSFLKYCSLAATSLGLGAGMAPKIAWALENKPRIPVVWIHGL WP_005139779.1
hydrogenase [Shigella sonnei] Length: 372\nNumber of predicted TMHs: 2\nExp number of AAs in TMHs: 37.02628\nExp number, first 60 AAs: 19.39684\nTotal prob of N-in: 0.94313\nPOSSIBLE N-term signal sequence\ninside 1 19\nTMhelix 20 39\noutside 40 325\nTMhelix 326 348\ninside 349 372

26685 GCF_001257135.1_Sg_201311838_TAATCG_L001 Shigella sonnei Proteobacteria; Gammaproteobacteria; Enterobacterales; Enterobacteriaceae; Shigella
MNNEETFYQAMRRQGVTRRSFLKYCSLAATSLGLGAGMAPKIAWALENKPRIPVVWIHGL WP_005139779.1
hydrogenase [Shigella sonnei] Length: 372\nNumber of predicted TMHs: 2\nExp number of AAs in TMHs: 37.02628\nExp number, first 60 AAs: 19.39684\nTotal prob of N-in: 0.94313\nPOSSIBLE N-term signal sequence\ninside 1 19\nTMhelix 20 39\noutside 40 325\nTMhelix 326 348\ninside 349 372

26686 GCF_001257175.1_8290_4_11 Shigella sonnei Proteobacteria; Gammaproteobacteria; Enterobacterales; Enterobacteriaceae; Shigella
MNNEETFYQAMRRQGVTRRSFLKYCSLAATSLGLGAGMAPKIAWALENKPRIPVVWIHGL WP_005139779.1
hydrogenase [Shigella sonnei] Length: 372\nNumber of predicted TMHs: 2\nExp number of AAs in TMHs: 37.02628\nExp number, first 60 AAs: 19.39684\nTotal prob of N-in: 0.94313\nPOSSIBLE N-term signal sequence\ninside 1 19\nTMhelix 20 39\noutside 40 325\nTMhelix 326 348\ninside 349 372

26687 GCF_001257475.1_Sg_201400029_TGACCA_L001 Shigella sonnei Proteobacteria; Gammaproteobacteria; Enterobacterales; Enterobacteriaceae; Shigella
MTGDNTLIHSHGINRRDFMKLCAALAATMGLSSKAAAEMAESVTNPQRPPVIWIGAQECT WP_000145410.1
MULTISPECIES: hydrogenase-2 small chain [Proteobacteria] Length: 372\nNumber of predicted TMHs: 1\nExp number of AAs in TMHs: 23.72226\nExp number, first 60 AAs: 0.22038\nTotal prob of N-in: 0.12994\noutside 1 331\nTMhelix 332 354\ninside 355 372

26688 GCF_001257695.1_5417_3_9Shigella sonnei Proteobacteria; Gammaproteobacteria; Enterobacterales; Enterobacteriaceae; Shigella MTGDNTLIHSHGINRRDFMKLCAALAATMGLSSKAAAEMAESVTNPQRPPVIWIGAQECT
WP_000145410.1 MULTISPECIES: hydrogenase-2 small chain [Proteobacteria] Length: 372\nNumber of predicted TMHs: 1\nExp number of AAs in TMHs: 23.72226\nExp number, first 60 AAs: 0.22038\nTotal prob of N-in: 0.12994\noutside 1 331\nTMhelix 332 354\ninside 355 372

26689 GCF_001257475.1_Sg_201400029_TGACCA_L001 Shigella sonnei Proteobacteria; Gammaproteobacteria; Enterobacterales; Enterobacteriaceae; Shigella
MNNEETFYQAMRRQGVTRRSFLKYCSLAATSLGLGAGMAPKIAWALENKPRIPVVWIHGL WP_005139779.1
hydrogenase [Shigella sonnei] Length: 372\nNumber of predicted TMHs: 2\nExp number of AAs in TMHs: 37.02628\nExp number, first 60 AAs: 19.39684\nTotal prob of N-in: 0.94313\nPOSSIBLE N-term signal sequence\ninside 1 19\nTMhelix 20 39\noutside 40 325\nTMhelix 326 348\ninside 349 372

26690 GCF_001257695.1_5417_3_9Shigella sonnei Proteobacteria; Gammaproteobacteria; Enterobacterales; Enterobacteriaceae; Shigella MNNEETFYQAMRRQGVTRRSFLKYCSLAATSLGLGAGMAPKIAWALENKPRIPVVWIHGL
WP_005139779.1 hydrogenase [Shigella sonnei] Length: 372\nNumber of predicted TMHs: 2\nExp number of AAs in TMHs: 37.02628\nExp number, first 60 AAs: 19.39684\nTotal prob of N-in: 0.94313\nPOSSIBLE N-term signal sequence\ninside 1 19\nTMhelix 20 39\noutside 40 325\nTMhelix 326 348\ninside 349 372

26691 GCF_001257335.1_9870_7_2Shigella sonnei Proteobacteria; Gammaproteobacteria; Enterobacterales; Enterobacteriaceae; Shigella MNNEETFYQAMRRQGVTRRSFLKYCSLAATSLGLGAGMAPKIAWALENKPRIPVVWIHGL
WP_005139779.1 hydrogenase [Shigella sonnei] Length: 372\nNumber of predicted TMHs: 2\nExp number of AAs in TMHs: 37.02628\nExp number, first 60 AAs: 19.39684\nTotal prob of N-in: 0.94313\nPOSSIBLE N-term signal sequence\ninside 1 19\nTMhelix 20 39\noutside 40 325\nTMhelix 326 348\ninside 349 372

26692 GCF_001257715.1_6350_1_19 Shigella sonnei Proteobacteria; Gammaproteobacteria; Enterobacterales; Enterobacteriaceae; Shigella
MTGDNTLIHSHGINRRDFMKLCAALAATMGLSSKAAAEMAESVTNPQRPPVIWIGAQECT WP_000145410.1
MULTISPECIES: hydrogenase-2 small chain [Proteobacteria] Length: 372\nNumber of predicted TMHs: 1\nExp number of AAs in TMHs: 23.72226\nExp number, first 60 AAs: 0.22038\nTotal prob of N-in: 0.12994\noutside 1 331\nTMhelix 332 354\ninside 355 372

26693 GCF_001257755.1_Sg_201312273_TATAAT_L001 Shigella sonnei Proteobacteria; Gammaproteobacteria; Enterobacterales; Enterobacteriaceae; Shigella MTGDNTLIHSHGINRRDFMKLCAALAATMGLSSKAAAEMAESVTNPQRPPVIWIGAQECT WP_000145410.1 MULTISPECIES: hydrogenase-2 small chain [Proteobacteria] Length: 372\nNumber of predicted TMHs: 1\nExp number of AAs in TMHs: 23.72226\nExp number, first 60 AAs: 0.22038\nTotal prob of N-in: 0.12994\noutside 1 331\nTMhelix 332 354\ninside 355 372

26694 GCF_001257715.1_6350_1_19 Shigella sonnei Proteobacteria; Gammaproteobacteria; Enterobacterales; Enterobacteriaceae; Shigella MNNEETFYQAMRRQGVTRRSFLKYCSLAATSLGLGAGMAPKIAWALENKPRIPVVWIHGL WP_005139779.1 hydrogenase [Shigella sonnei] Length: 372\nNumber of predicted TMHs: 2\nExp number of AAs in TMHs: 37.02628\nExp number, first 60 AAs: 19.39684\nTotal prob of N-in: 0.94313\nPOSSIBLE N-term signal sequence\ninside 1 19\nTMhelix 20 39\noutside 40 325\nTMhelix 326 348\ninside 349 372

26695 GCF_001257755.1_Sg_201312273_TATAAT_L001 Shigella sonnei Proteobacteria; Gammaproteobacteria; Enterobacterales; Enterobacteriaceae; Shigella MNNEETFYQAMRRQGVTRRSFLKYCSLAATSLGLGAGMAPKIAWALENKPRIPVVWIHGL WP_005139779.1 hydrogenase [Shigella sonnei] Length: 372\nNumber of predicted TMHs: 2\nExp number of AAs in TMHs: 37.02628\nExp number, first 60 AAs: 19.39684\nTotal prob of N-in: 0.94313\nPOSSIBLE N-term signal sequence\ninside 1 19\nTMhelix 20 39\noutside 40 325\nTMhelix 326 348\ninside 349 372

26696 GCF_001257415.1_5236_5_1Shigella sonnei Proteobacteria; Gammaproteobacteria; Enterobacterales; Enterobacteriaceae; Shigella MNNEETFYQAMRRQGVTRRSFLKYCSLAATSLGLGAGMAPKIAWALENKPRIPVVWIHGL WP_005139779.1 hydrogenase [Shigella sonnei] Length: 372\nNumber of predicted TMHs: 2\nExp number of AAs in TMHs: 37.02628\nExp number, first 60 AAs: 19.39684\nTotal prob of N-in: 0.94313\nPOSSIBLE N-term signal sequence\ninside 1 19\nTMhelix 20 39\noutside 40 325\nTMhelix 326 348\ninside 349 372

26697 GCF_001257795.1_7655_8_16 Shigella sonnei Proteobacteria; Gammaproteobacteria; Enterobacterales; Enterobacteriaceae; Shigella MTGDNTLIHSHGINRRDFMKLCAALAATMGLSSKAAAEMAESVTNPQRPPVIWIGAQECT WP_000145410.1 MULTISPECIES: hydrogenase-2 small chain [Proteobacteria] Length: 372\nNumber of predicted TMHs: 1\nExp number of AAs in TMHs: 23.72226\nExp number, first 60 AAs: 0.22038\nTotal prob of N-in: 0.12994\noutside 1 331\nTMhelix 332 354\ninside 355 372

26698 GCF_001257815.1_5417_2_4Shigella sonnei Proteobacteria; Gammaproteobacteria; Enterobacterales; Enterobacteriaceae; Shigella MTGDNTLIHSHGINRRDFMKLCAALAATMGLSSKAAAEMAESVTNPQRPPVIWIGAQECT WP_000145410.1 MULTISPECIES: hydrogenase-2 small chain [Proteobacteria] Length: 372\nNumber of predicted TMHs: 1\nExp number of AAs in TMHs: 23.72226\nExp number, first 60 AAs: 0.22038\nTotal prob of N-in: 0.12994\noutside 1 331\nTMhelix 332 354\ninside 355 372

26699 GCF_001257795.1_7655_8_16 Shigella sonnei Proteobacteria; Gammaproteobacteria; Enterobacterales; Enterobacteriaceae; Shigella MNNEETFYQAMRRQGVTRRSFLKYCSLAATSLGLGAGMAPKIAWALENKPRIPVVWIHGL WP_005139779.1 hydrogenase [Shigella sonnei] Length: 372\nNumber of predicted TMHs: 2\nExp number of AAs in TMHs: 37.02628\nExp number, first 60 AAs: 19.39684\nTotal prob of N-in: 0.94313\nPOSSIBLE N-term signal sequence\ninside 1 19\nTMhelix 20 39\noutside 40 325\nTMhelix 326 348\ninside 349 372

26700 GCF_001257815.1_5417_2_4Shigella sonnei Proteobacteria; Gammaproteobacteria; Enterobacterales; Enterobacteriaceae; Shigella MNNEETFYQAMRRQGVTRRSFLKYCSLAATSLGLGAGMAPKIAWALENKPRIPVVWIHGL WP_005139779.1 hydrogenase [Shigella sonnei] Length: 372\nNumber of predicted TMHs: 2\nExp number of AAs in TMHs: 37.02628\nExp number, first 60 AAs: 19.39684\nTotal prob of N-in: 0.94313\nPOSSIBLE N-term signal sequence\ninside 1 19\nTMhelix 20 39\noutside 40 325\nTMhelix 326 348\ninside 349 372

26701 GCF_001257455.1_8290_5_5Shigella sonnei Proteobacteria; Gammaproteobacteria; Enterobacterales; Enterobacteriaceae; Shigella MNNEETFYQAMRRQGVTRRSFLKYCSLAATSLGLGAGMAPKIAWALENKPRIPVVWIHGL WP_005139779.1 hydrogenase [Shigella sonnei] Length: 372\nNumber of predicted TMHs: 2\nExp number of AAs in TMHs: 37.02628\nExp number, first 60 AAs: 19.39684\nTotal prob of N-in: 0.94313\nPOSSIBLE N-term signal sequence\ninside 1 19\nTMhelix 20 39\noutside 40 325\nTMhelix 326 348\ninside 349 372

26702 GCF_001257875.1_5236_3_2Shigella sonnei Proteobacteria; Gammaproteobacteria; Enterobacterales; Enterobacteriaceae; Shigella MTGDNTLIHSHGINRRDFMKLCAALAATMGLSSKAAAEMAESVTNPQRPPVIWIGAQECT

WP_000145410.1 MULTISPECIES: hydrogenase-2 small chain [Proteobacteria] Length: 372\nNumber of predicted TMHs: 1\nExp number of AAs in TMHs: 23.72226\nExp number, first 60 AAs: 0.22038\nTotal prob of N-in: 0.12994\noutside 1 331\nTMhelix 332 354\ninside 355 372

26703 GCF_001257875.1_5236_3_2Shigella sonnei Proteobacteria; Gammaproteobacteria; Enterobacterales; Enterobacteriaceae; Shigella MNNEETFYQAMRRQGVTRRSFLKYCSLAATSLGLGAGMAPKIAWALENKPRIPVVWIHGL WP_005139779.1 hydrogenase [Shigella sonnei] Length: 372\nNumber of predicted TMHs: 2\nExp number of AAs in TMHs: 37.02628\nExp number, first 60 AAs: 19.39684\nTotal prob of N-in: 0.94313\nPOSSIBLE N-term signal sequence\ninside 1 19\nTMhelix 20 39\noutside 40 325\nTMhelix 326 348\ninside 349 372

26704 GCF_001257375.1_12971_8_14 Shigella sonnei Proteobacteria; Gammaproteobacteria; Enterobacterales; Enterobacteriaceae; Shigella MNNEETFYQAMRRQGVTRRSFLKYCSLAATSLGLGAGMAPKIAWALENKPRIPVVWIHGL WP_005139779.1 hydrogenase [Shigella sonnei] Length: 372\nNumber of predicted TMHs: 2\nExp number of AAs in TMHs: 37.02628\nExp number, first 60 AAs: 19.39684\nTotal prob of N-in: 0.94313\nPOSSIBLE N-term signal sequence\ninside 1 19\nTMhelix 20 39\noutside 40 325\nTMhelix 326 348\ninside 349 372

26705 GCF_001257935.1_5236_1_1Shigella sonnei Proteobacteria; Gammaproteobacteria; Enterobacterales; Enterobacteriaceae; Shigella MTGDNTLIHSHGINRRDFMKLCAALAATMGLSSKAAAEMAESVTNPQRPPVIWIGAQECT WP_000145410.1 MULTISPECIES: hydrogenase-2 small chain [Proteobacteria] Length: 372\nNumber of predicted TMHs: 1\nExp number of AAs in TMHs: 23.72226\nExp number, first 60 AAs: 0.22038\nTotal prob of N-in: 0.12994\noutside 1 331\nTMhelix 332 354\ninside 355 372

26706 GCF_001257955.1_9870_6_7Shigella sonnei Proteobacteria; Gammaproteobacteria; Enterobacterales; Enterobacteriaceae; Shigella MTGDNTLIHSHGINRRDFMKLCAALAATMGLSSKAAAEMAESVTNPQRPPVIWIGAQECT WP_000145410.1 MULTISPECIES: hydrogenase-2 small chain [Proteobacteria] Length: 372\nNumber of predicted TMHs: 1\nExp number of AAs in TMHs: 23.72226\nExp number, first 60 AAs: 0.22038\nTotal prob of N-in: 0.12994\noutside 1 331\nTMhelix 332 354\ninside 355 372

26707 GCF_001258015.1_8289_7_80 Shigella sonnei Proteobacteria; Gammaproteobacteria; Enterobacterales; Enterobacteriaceae; Shigella MTGDNTLIHSHGINRRDFMKLCAALAATMGLSSKAAAEMAESVTNPQRPPVIWIGAQECT WP_000145410.1 MULTISPECIES: hydrogenase-2 small chain [Proteobacteria] Length: 372\nNumber of predicted TMHs: 1\nExp number of AAs in TMHs: 23.72226\nExp number, first 60 AAs: 0.22038\nTotal prob of N-in: 0.12994\noutside 1 331\nTMhelix 332 354\ninside 355 372

26708 GCF_001258075.1_8290_4_94 Shigella sonnei Proteobacteria; Gammaproteobacteria; Enterobacterales; Enterobacteriaceae; Shigella MTGDNTLIHSHGINRRDFMKLCAALAATMGLSSKAAAEMAESVTNPQRPPVIWIGAQECT WP_000145410.1 MULTISPECIES: hydrogenase-2 small chain [Proteobacteria] Length: 372\nNumber of predicted TMHs: 1\nExp number of AAs in TMHs: 23.72226\nExp number, first 60 AAs: 0.22038\nTotal prob of N-in: 0.12994\noutside 1 331\nTMhelix 332 354\ninside 355 372

26709 GCF_001258095.1_Sg_201405358_CTCAGA_L001 Shigella sonnei Proteobacteria; Gammaproteobacteria; Enterobacterales; Enterobacteriaceae; Shigella MTGDNTLIHSHGINRRDFMKLCAALAATMGLSSKAAAEMAESVTNPQRPPVIWIGAQECT WP_000145410.1 MULTISPECIES: hydrogenase-2 small chain [Proteobacteria] Length: 372\nNumber of predicted TMHs: 1\nExp number of AAs in TMHs: 23.72226\nExp number, first 60 AAs: 0.22038\nTotal prob of N-in: 0.12994\noutside 1 331\nTMhelix 332 354\ninside 355 372

26710 GCF_001258135.1_8290_4_41 Shigella sonnei Proteobacteria; Gammaproteobacteria; Enterobacterales; Enterobacteriaceae; Shigella MTGDNTLIHSHGINRRDFMKLCAALAATMGLSSKAAAEMAESVTNPQRPPVIWIGAQECT WP_000145410.1 MULTISPECIES: hydrogenase-2 small chain [Proteobacteria] Length: 372\nNumber of predicted TMHs: 1\nExp number of AAs in TMHs: 23.72226\nExp number, first 60 AAs: 0.22038\nTotal prob of N-in: 0.12994\noutside 1 331\nTMhelix 332 354\ninside 355 372

26711 GCF_001258175.1_5417_1_10 Shigella sonnei Proteobacteria; Gammaproteobacteria; Enterobacterales; Enterobacteriaceae; Shigella MTGDNTLIHSHGINRRDFMKLCAALAATMGLSSKAAAEMAESVTNPQRPPVIWIGAQECT WP_000145410.1 MULTISPECIES: hydrogenase-2 small chain [Proteobacteria] Length: 372\nNumber of predicted TMHs:

1\nExp number of AAs in TMHs: 23.72226\nExp number, first 60 AAs: 0.22038\nTotal prob of N-in: 0.12994\noutside 1 331\nTMhelix 332 354\ninside 355 372

26712 GCF_001257935.1_5236_1_1Shigella sonnei Proteobacteria; Gammaproteobacteria; Enterobacterales; Enterobacteriaceae; Shigella MNNEETFYQAMRRQGVTRRSFLKYCSLAATSLGLGAGMAPKIAWALENKPRIPVVWIHGL WP_005139779.1 hydrogenase [Shigella sonnei] Length: 372\nNumber of predicted TMHs: 2\nExp number of AAs in TMHs: 37.02628\nExp number, first 60 AAs: 19.39684\nTotal prob of N-in: 0.94313\nPOSSIBLE N-term signal sequence\ninside 1 19\nTMhelix 20 39\noutside 40 325\nTMhelix 326 348\ninside 349 372

26713 GCF_001257955.1_9870_6_7Shigella sonnei Proteobacteria; Gammaproteobacteria; Enterobacterales; Enterobacteriaceae; Shigella MNNEETFYQAMRRQGVTRRSFLKYCSLAATSLGLGAGMAPKIAWALENKPRIPVVWIHGL WP_005139779.1 hydrogenase [Shigella sonnei] Length: 372\nNumber of predicted TMHs: 2\nExp number of AAs in TMHs: 37.02628\nExp number, first 60 AAs: 19.39684\nTotal prob of N-in: 0.94313\nPOSSIBLE N-term signal sequence\ninside 1 19\nTMhelix 20 39\noutside 40 325\nTMhelix 326 348\ninside 349 372

26714 GCF_001258015.1_8289_7_80 Shigella sonnei Proteobacteria; Gammaproteobacteria; Enterobacterales; Enterobacteriaceae; Shigella MNNEETFYQAMRRQGVTRRSFLKYCSLAATSLGLGAGMAPKIAWALENKPRIPVVWIHGL WP_005139779.1 hydrogenase [Shigella sonnei] Length: 372\nNumber of predicted TMHs: 2\nExp number of AAs in TMHs: 37.02628\nExp number, first 60 AAs: 19.39684\nTotal prob of N-in: 0.94313\nPOSSIBLE N-term signal sequence\ninside 1 19\nTMhelix 20 39\noutside 40 325\nTMhelix 326 348\ninside 349 372

26715 GCF_001258235.1_H14072036401 Shigella sonnei Proteobacteria; Gammaproteobacteria; Enterobacterales; Enterobacteriaceae; Shigella MTGDNTLIHSHGINRRDFMKLCAALAATMGLSSKAAAEMAESVTNPQRPPVIWIGAQECT WP_000145410.1 MULTISPECIES: hydrogenase-2 small chain [Proteobacteria] Length: 372\nNumber of predicted TMHs: 1\nExp number of AAs in TMHs: 23.72226\nExp number, first 60 AAs: 0.22038\nTotal prob of N-in: 0.12994\noutside 1 331\nTMhelix 332 354\ninside 355 372

26716 GCF_001258255.1_6350_1_14 Shigella sonnei Proteobacteria; Gammaproteobacteria; Enterobacterales; Enterobacteriaceae; Shigella MTGDNTLIHSHGINRRDFMKLCAALAATMGLSSKAAAEMAESVTNPQRPPVIWIGAQECT WP_000145410.1 MULTISPECIES: hydrogenase-2 small chain [Proteobacteria] Length: 372\nNumber of predicted TMHs: 1\nExp number of AAs in TMHs: 23.72226\nExp number, first 60 AAs: 0.22038\nTotal prob of N-in: 0.12994\noutside 1 331\nTMhelix 332 354\ninside 355 372

26717 GCF_001258255.1_6350_1_14 Shigella sonnei Proteobacteria; Gammaproteobacteria; Enterobacterales; Enterobacteriaceae; Shigella MNNEETFYQAMRRQGVTRRSFLKYCSLAATSLGLGAGMAPKIAWALENKPRIPVVWIHGL WP_005139779.1 hydrogenase [Shigella sonnei] Length: 372\nNumber of predicted TMHs: 2\nExp number of AAs in TMHs: 37.02628\nExp number, first 60 AAs: 19.39684\nTotal prob of N-in: 0.94313\nPOSSIBLE N-term signal sequence\ninside 1 19\nTMhelix 20 39\noutside 40 325\nTMhelix 326 348\ninside 349 372

26718 GCF_001258075.1_8290_4_94 Shigella sonnei Proteobacteria; Gammaproteobacteria; Enterobacterales; Enterobacteriaceae; Shigella MNNEETFYQAMRRQGVTRRSFLKYCSLAATSLGLGAGMAPKIAWALENKPRIPVVWIHGL WP_005139779.1 hydrogenase [Shigella sonnei] Length: 372\nNumber of predicted TMHs: 2\nExp number of AAs in TMHs: 37.02628\nExp number, first 60 AAs: 19.39684\nTotal prob of N-in: 0.94313\nPOSSIBLE N-term signal sequence\ninside 1 19\nTMhelix 20 39\noutside 40 325\nTMhelix 326 348\ninside 349 372

26719 GCF_001258295.1_8289_7_70 Shigella sonnei Proteobacteria; Gammaproteobacteria; Enterobacterales; Enterobacteriaceae; Shigella MTGDNTLIHSHGINRRDFMKLCAALAATMGLSSKAAAEMAESVTNPQRPPVIWIGAQECT WP_000145410.1 MULTISPECIES: hydrogenase-2 small chain [Proteobacteria] Length: 372\nNumber of predicted TMHs: 1\nExp number of AAs in TMHs: 23.72226\nExp number, first 60 AAs: 0.22038\nTotal prob of N-in: 0.12994\noutside 1 331\nTMhelix 332 354\ninside 355 372

26720 GCF_001258295.1_8289_7_70 Shigella sonnei Proteobacteria; Gammaproteobacteria; Enterobacterales; Enterobacteriaceae; Shigella MNNEETFYQAMRRQGVTRRSFLKYCSLAATSLGLGAGMAPKIAWALENKPRIPVVWIHGL WP_005139779.1 hydrogenase [Shigella sonnei] Length: 372\nNumber of predicted TMHs: 2\nExp number of AAs in

TMHs: 37.02628\nExp number, first 60 AAs: 19.39684\nTotal prob of N-in: 0.94313\nPOSSIBLE N-term signal sequence\ninside 1 19\nTMhelix 20 39\noutside 40 325\nTMhelix 326 348\ninside 349 372

26721 GCF_001258095.1_Sg_201405358_CTCAGA_L001 Shigella sonnei Proteobacteria; Gammaproteobacteria; Enterobacterales; Enterobacteriaceae; Shigella
MNNEETFYQAMRRQGVTRRSFLKYCSLAATSLGLGAGMAPKIAWALENKPRIPVVWIHGL WP_005139779.1
hydrogenase [Shigella sonnei] Length: 372\nNumber of predicted TMHs: 2\nExp number of AAs in TMHs: 37.02628\nExp number, first 60 AAs: 19.39684\nTotal prob of N-in: 0.94313\nPOSSIBLE N-term signal sequence\ninside 1 19\nTMhelix 20 39\noutside 40 325\nTMhelix 326 348\ninside 349 372

26722 GCF_001258335.1_Sg_201402425_ATGAGC_L001 Shigella sonnei Proteobacteria; Gammaproteobacteria; Enterobacterales; Enterobacteriaceae; Shigella
MTGDNTLIHSHGINRRDFMKLCAALAATMGLSSKAAAEMAESVTNPQRPPVIWIGAQECT WP_000145410.1
MULTISPECIES: hydrogenase-2 small chain [Proteobacteria] Length: 372\nNumber of predicted TMHs: 1\nExp number of AAs in TMHs: 23.72226\nExp number, first 60 AAs: 0.22038\nTotal prob of N-in: 0.12994\noutside 1 331\nTMhelix 332 354\ninside 355 372

26723 GCF_001258335.1_Sg_201402425_ATGAGC_L001 Shigella sonnei Proteobacteria; Gammaproteobacteria; Enterobacterales; Enterobacteriaceae; Shigella
MNNEETFYQAMRRQGVTRRSFLKYCSLAATSLGLGAGMAPKIAWALENKPRIPVVWIHGL WP_005139779.1
hydrogenase [Shigella sonnei] Length: 372\nNumber of predicted TMHs: 2\nExp number of AAs in TMHs: 37.02628\nExp number, first 60 AAs: 19.39684\nTotal prob of N-in: 0.94313\nPOSSIBLE N-term signal sequence\ninside 1 19\nTMhelix 20 39\noutside 40 325\nTMhelix 326 348\ninside 349 372

26724 GCF_001258135.1_8290_4_41 Shigella sonnei Proteobacteria; Gammaproteobacteria; Enterobacterales; Enterobacteriaceae; Shigella
MNNEETFYQAMRRQGVTRRSFLKYCSLAATSLGLGAGMAPKIAWALENKPRIPVVWIHGL WP_005139779.1
hydrogenase [Shigella sonnei] Length: 372\nNumber of predicted TMHs: 2\nExp number of AAs in TMHs: 37.02628\nExp number, first 60 AAs: 19.39684\nTotal prob of N-in: 0.94313\nPOSSIBLE N-term signal sequence\ninside 1 19\nTMhelix 20 39\noutside 40 325\nTMhelix 326 348\ninside 349 372

26725 GCF_001258355.1_H14152025501 Shigella sonnei Proteobacteria; Gammaproteobacteria; Enterobacterales; Enterobacteriaceae; Shigella
MTGDNTLIHSHGINRRDFMKLCAALAATMGLSSKAAAEMAESVTNPQRPPVIWIGAQECT WP_000145410.1
MULTISPECIES: hydrogenase-2 small chain [Proteobacteria] Length: 372\nNumber of predicted TMHs: 1\nExp number of AAs in TMHs: 23.72226\nExp number, first 60 AAs: 0.22038\nTotal prob of N-in: 0.12994\noutside 1 331\nTMhelix 332 354\ninside 355 372

26726 GCF_001258395.1_12850_1_46 Shigella sonnei Proteobacteria; Gammaproteobacteria; Enterobacterales; Enterobacteriaceae; Shigella
MTGDNTLIHSHGINRRDFMKLCAALAATMGLSSKAAAEMAESVTNPQRPPVIWIGAQECT WP_000145410.1
MULTISPECIES: hydrogenase-2 small chain [Proteobacteria] Length: 372\nNumber of predicted TMHs: 1\nExp number of AAs in TMHs: 23.72226\nExp number, first 60 AAs: 0.22038\nTotal prob of N-in: 0.12994\noutside 1 331\nTMhelix 332 354\ninside 355 372

26727 GCF_001258355.1_H14152025501 Shigella sonnei Proteobacteria; Gammaproteobacteria; Enterobacterales; Enterobacteriaceae; Shigella
MNNEETFYQAMRRQGVTRRSFLKYCSLAATSLGLGAGMAPKIAWALENKPRIPVVWIHGL WP_005139779.1
hydrogenase [Shigella sonnei] Length: 372\nNumber of predicted TMHs: 2\nExp number of AAs in TMHs: 37.02628\nExp number, first 60 AAs: 19.39684\nTotal prob of N-in: 0.94313\nPOSSIBLE N-term signal sequence\ninside 1 19\nTMhelix 20 39\noutside 40 325\nTMhelix 326 348\ninside 349 372

26728 GCF_001258395.1_12850_1_46 Shigella sonnei Proteobacteria; Gammaproteobacteria; Enterobacterales; Enterobacteriaceae; Shigella
MNNEETFYQAMRRQGVTRRSFLKYCSLAATSLGLGAGMAPKIAWALENKPRIPVVWIHGL WP_005139779.1
hydrogenase [Shigella sonnei] Length: 372\nNumber of predicted TMHs: 2\nExp number of AAs in TMHs: 37.02628\nExp number, first 60 AAs: 19.39684\nTotal prob of N-in: 0.94313\nPOSSIBLE N-term signal sequence\ninside 1 19\nTMhelix 20 39\noutside 40 325\nTMhelix 326 348\ninside 349 372

26729 GCF_001258175.1_5417_1_10 Shigella sonnei Proteobacteria; Gammaproteobacteria; Enterobacterales; Enterobacteriaceae; Shigella

MNNEETFYQAMRRQGVTRRSFLKYCSLAATSLGLGAGMAPKIAWALENKPRIPVVWIHGL WP_005139779.1 hydrogenase [Shigella sonnei] Length: 372\nNumber of predicted TMHs: 2\nExp number of AAs in TMHs: 37.02628\nExp number, first 60 AAs: 19.39684\nTotal prob of N-in: 0.94313\nPOSSIBLE N-term signal sequence\ninside 1 19\nTMhelix 20 39\noutside 40 325\nTMhelix 326 348\ninside 349 372

26730 GCF_001258195.1_9870_6_4Shigella sonnei Proteobacteria; Gammaproteobacteria; Enterobacterales; Enterobacteriaceae; Shigella MNNEETFYQAMRRQGVTRRSFLKYCSLAATSLGLGAGMAPKIAWALENKPRIPVVWIHGL WP_005139779.1 hydrogenase [Shigella sonnei] Length: 372\nNumber of predicted TMHs: 2\nExp number of AAs in TMHs: 37.02628\nExp number, first 60 AAs: 19.39684\nTotal prob of N-in: 0.94313\nPOSSIBLE N-term signal sequence\ninside 1 19\nTMhelix 20 39\noutside 40 325\nTMhelix 326 348\ninside 349 372

26731 GCF_001258435.1_5236_6_1Shigella sonnei Proteobacteria; Gammaproteobacteria; Enterobacterales; Enterobacteriaceae; Shigella MTGDNTLIHSHGINRRDFMKLCAALAATMGLSSKAAAEMAESVTNPQRPPVIWIGAQECT WP_000145410.1 MULTISPECIES: hydrogenase-2 small chain [Proteobacteria] Length: 372\nNumber of predicted TMHs: 1\nExp number of AAs in TMHs: 23.72226\nExp number, first 60 AAs: 0.22038\nTotal prob of N-in: 0.12994\noutside 1 331\nTMhelix 332 354\ninside 355 372

26732 GCF_001258475.1_12971_8_41 Shigella sonnei Proteobacteria; Gammaproteobacteria; Enterobacterales; Enterobacteriaceae; Shigella MTGDNTLIHSHGINRRDFMKLCAALAATMGLSSKAAAEMAESVTNPQRPPVIWIGAQECT WP_000145410.1 MULTISPECIES: hydrogenase-2 small chain [Proteobacteria] Length: 372\nNumber of predicted TMHs: 1\nExp number of AAs in TMHs: 23.72226\nExp number, first 60 AAs: 0.22038\nTotal prob of N-in: 0.12994\noutside 1 331\nTMhelix 332 354\ninside 355 372

26733 GCF_001258515.1_H14120049201 Shigella sonnei Proteobacteria; Gammaproteobacteria; Enterobacterales; Enterobacteriaceae; Shigella MTGDNTLIHSHGINRRDFMKLCAALAATMGLSSKAAAEMAESVTNPQRPPVIWIGAQECT WP_000145410.1 MULTISPECIES: hydrogenase-2 small chain [Proteobacteria] Length: 372\nNumber of predicted TMHs: 1\nExp number of AAs in TMHs: 23.72226\nExp number, first 60 AAs: 0.22038\nTotal prob of N-in: 0.12994\noutside 1 331\nTMhelix 332 354\ninside 355 372

26734 GCF_001258595.1_9870_6_21 Shigella sonnei Proteobacteria; Gammaproteobacteria; Enterobacterales; Enterobacteriaceae; Shigella MTGDNTLIHSHGINRRDFMKLCAALAATMGLSSKAAAEMAESVTNPQRPPVIWIGAQECT WP_000145410.1 MULTISPECIES: hydrogenase-2 small chain [Proteobacteria] Length: 372\nNumber of predicted TMHs: 1\nExp number of AAs in TMHs: 23.72226\nExp number, first 60 AAs: 0.22038\nTotal prob of N-in: 0.12994\noutside 1 331\nTMhelix 332 354\ninside 355 372

26735 GCF_001258635.1_Sg_201405357_CTATAC_L001 Shigella sonnei Proteobacteria; Gammaproteobacteria; Enterobacterales; Enterobacteriaceae; Shigella MTGDNTLIHSHGINRRDFMKLCAALAATMGLSSKAAAEMAESVTNPQRPPVIWIGAQECT WP_000145410.1 MULTISPECIES: hydrogenase-2 small chain [Proteobacteria] Length: 372\nNumber of predicted TMHs: 1\nExp number of AAs in TMHs: 23.72226\nExp number, first 60 AAs: 0.22038\nTotal prob of N-in: 0.12994\noutside 1 331\nTMhelix 332 354\ninside 355 372

26736 GCF_001258655.1_9870_6_37 Shigella sonnei Proteobacteria; Gammaproteobacteria; Enterobacterales; Enterobacteriaceae; Shigella MTGDNTLIHSHGINRRDFMKLCAALAATMGLSSKAAAEMAESVTNPQRPPVIWIGAQECT WP_000145410.1 MULTISPECIES: hydrogenase-2 small chain [Proteobacteria] Length: 372\nNumber of predicted TMHs: 1\nExp number of AAs in TMHs: 23.72226\nExp number, first 60 AAs: 0.22038\nTotal prob of N-in: 0.12994\noutside 1 331\nTMhelix 332 354\ninside 355 372

26737 GCF_001258695.1_8289_7_86 Shigella sonnei Proteobacteria; Gammaproteobacteria; Enterobacterales; Enterobacteriaceae; Shigella MTGDNTLIHSHGINRRDFMKLCAALAATMGLSSKAAAEMAESVTNPQRPPVIWIGAQECT WP_000145410.1 MULTISPECIES: hydrogenase-2 small chain [Proteobacteria] Length: 372\nNumber of predicted TMHs: 1\nExp number of AAs in TMHs: 23.72226\nExp number, first 60 AAs: 0.22038\nTotal prob of N-in: 0.12994\noutside 1 331\nTMhelix 332 354\ninside 355 372

26738 GCF_001258755.1_12971_8_27 Shigella sonnei Proteobacteria; Gammaproteobacteria; Enterobacterales; Enterobacteriaceae; Shigella

MTGDNTLIHSHGINRRDFMKLCAALAATMGLSSKAAAEMAESVTNPQRPPVIWIGAQECT WP_000145410.1 MULTISPECIES: hydrogenase-2 small chain [Proteobacteria] Length: 372\nNumber of predicted TMHs: 1\nExp number of AAs in TMHs: 23.72226\nExp number, first 60 AAs: 0.22038\nTotal prob of N-in: 0.12994\noutside 1 331\nTMhelix 332 354\ninside 355 372

26739 GCF_001258755.1_12971_8_27 Shigella sonnei Proteobacteria; Gammaproteobacteria; Enterobacterales; Enterobacteriaceae; Shigella MNNEETFYQAMRRQGVTRRSFLKYCSLAATSLGLGAGMAPKIAWALENKPRIPVVWIHGL WP_005139779.1 hydrogenase [Shigella sonnei] Length: 372\nNumber of predicted TMHs: 2\nExp number of AAs in TMHs: 37.02628\nExp number, first 60 AAs: 19.39684\nTotal prob of N-in: 0.94313\nPOSSIBLE N-term signal sequence\ninside 1 19\nTMhelix 20 39\noutside 40 325\nTMhelix 326 348\ninside 349 372

26740 GCF_001258475.1_12971_8_41 Shigella sonnei Proteobacteria; Gammaproteobacteria; Enterobacterales; Enterobacteriaceae; Shigella MNNEETFYQAMRRQGVTRRSFLKYCSLAATSLGLGAGMAPKIAWALENKPRIPVVWIHGL WP_005139779.1 hydrogenase [Shigella sonnei] Length: 372\nNumber of predicted TMHs: 2\nExp number of AAs in TMHs: 37.02628\nExp number, first 60 AAs: 19.39684\nTotal prob of N-in: 0.94313\nPOSSIBLE N-term signal sequence\ninside 1 19\nTMhelix 20 39\noutside 40 325\nTMhelix 326 348\ninside 349 372

26741 GCF_001258435.1_5236_6_1Shigella sonnei Proteobacteria; Gammaproteobacteria; Enterobacterales; Enterobacteriaceae; Shigella MNNEETFYQAMRRQGVTRRSFLKYCSLAATSLGLGAGMAPKIAWALENKPRIPVVWIHGL WP_005139779.1 hydrogenase [Shigella sonnei] Length: 372\nNumber of predicted TMHs: 2\nExp number of AAs in TMHs: 37.02628\nExp number, first 60 AAs: 19.39684\nTotal prob of N-in: 0.94313\nPOSSIBLE N-term signal sequence\ninside 1 19\nTMhelix 20 39\noutside 40 325\nTMhelix 326 348\ninside 349 372

26742 GCF_001258815.1_5417_2_7Shigella sonnei Proteobacteria; Gammaproteobacteria; Enterobacterales; Enterobacteriaceae; Shigella MTGDNTLIHSHGINRRDFMKLCAALAATMGLSSKAAAEMAESVTNPQRPPVIWIGAQECT WP_000145410.1 MULTISPECIES: hydrogenase-2 small chain [Proteobacteria] Length: 372\nNumber of predicted TMHs: 1\nExp number of AAs in TMHs: 23.72226\nExp number, first 60 AAs: 0.22038\nTotal prob of N-in: 0.12994\noutside 1 331\nTMhelix 332 354\ninside 355 372

26743 GCF_001258815.1_5417_2_7Shigella sonnei Proteobacteria; Gammaproteobacteria; Enterobacterales; Enterobacteriaceae; Shigella MNNEETFYQAMRRQGVTRRSFLKYCSLAATSLGLGAGMAPKIAWALENKPRIPVVWIHGL WP_005139779.1 hydrogenase [Shigella sonnei] Length: 372\nNumber of predicted TMHs: 2\nExp number of AAs in TMHs: 37.02628\nExp number, first 60 AAs: 19.39684\nTotal prob of N-in: 0.94313\nPOSSIBLE N-term signal sequence\ninside 1 19\nTMhelix 20 39\noutside 40 325\nTMhelix 326 348\ninside 349 372

26744 GCF_001258515.1_H14120049201 Shigella sonnei Proteobacteria; Gammaproteobacteria; Enterobacterales; Enterobacteriaceae; Shigella MNNEETFYQAMRRQGVTRRSFLKYCSLAATSLGLGAGMAPKIAWALENKPRIPVVWIHGL WP_005139779.1 hydrogenase [Shigella sonnei] Length: 372\nNumber of predicted TMHs: 2\nExp number of AAs in TMHs: 37.02628\nExp number, first 60 AAs: 19.39684\nTotal prob of N-in: 0.94313\nPOSSIBLE N-term signal sequence\ninside 1 19\nTMhelix 20 39\noutside 40 325\nTMhelix 326 348\ninside 349 372

26745 GCF_001258835.1_8289_7_78 Shigella sonnei Proteobacteria; Gammaproteobacteria; Enterobacterales; Enterobacteriaceae; Shigella MTGDNTLIHSHGINRRDFMKLCAALAATMGLSSKAAAEMAESVTNPQRPPVIWIGAQECT WP_000145410.1 MULTISPECIES: hydrogenase-2 small chain [Proteobacteria] Length: 372\nNumber of predicted TMHs: 1\nExp number of AAs in TMHs: 23.72226\nExp number, first 60 AAs: 0.22038\nTotal prob of N-in: 0.12994\noutside 1 331\nTMhelix 332 354\ninside 355 372

26746 GCF_001258835.1_8289_7_78 Shigella sonnei Proteobacteria; Gammaproteobacteria; Enterobacterales; Enterobacteriaceae; Shigella MNNEETFYQAMRRQGVTRRSFLKYCSLAATSLGLGAGMAPKIAWALENKPRIPVVWIHGL WP_005139779.1 hydrogenase [Shigella sonnei] Length: 372\nNumber of predicted TMHs: 2\nExp number of AAs in TMHs: 37.02628\nExp number, first 60 AAs: 19.39684\nTotal prob of N-in: 0.94313\nPOSSIBLE N-term signal sequence\ninside 1 19\nTMhelix 20 39\noutside 40 325\nTMhelix 326 348\ninside 349 372

26747 GCF_001258595.1_9870_6_21 Shigella sonnei Proteobacteria; Gammaproteobacteria; Enterobacterales; Enterobacteriaceae; Shigella MNNEETFYQAMRRQGVTRRSFLKYCSLAATSLGLGAGMAPKIAWALENKPRIPVVWIHGL WP_005139779.1

hydrogenase [Shigella sonnei] Length: 372\nNumber of predicted TMHs: 2\nExp number of AAs in TMHs: 37.02628\nExp number, first 60 AAs: 19.39684\nTotal prob of N-in: 0.94313\nPOSSIBLE N-term signal sequence\ninside 1 19\nTMhelix 20 39\noutside 40 325\nTMhelix 326 348\ninside 349 372

26748 GCF_001258895.1_9870_7_9Shigella sonnei Proteobacteria; Gammaproteobacteria; Enterobacterales; Enterobacteriaceae; Shigella MTGDNTLIHSHGINRRDFMKLCAALAATMGLSSKAAAEMAESVTNPQRPPVIWIGAQECT WP_000145410.1 MULTISPECIES: hydrogenase-2 small chain [Proteobacteria] Length: 372\nNumber of predicted TMHs: 1\nExp number of AAs in TMHs: 23.72226\nExp number, first 60 AAs: 0.22038\nTotal prob of N-in: 0.12994\noutside 1 331\nTMhelix 332 354\ninside 355 372

26749 GCF_001258895.1_9870_7_9Shigella sonnei Proteobacteria; Gammaproteobacteria; Enterobacterales; Enterobacteriaceae; Shigella MNNEETFYQAMRRQGVTRRSFLKYCSLAATSLGLGAGMAPKIAWALENKPRIPVVWIHGL WP_005139779.1 hydrogenase [Shigella sonnei] Length: 372\nNumber of predicted TMHs: 2\nExp number of AAs in TMHs: 37.02628\nExp number, first 60 AAs: 19.39684\nTotal prob of N-in: 0.94313\nPOSSIBLE N-term signal sequence\ninside 1 19\nTMhelix 20 39\noutside 40 325\nTMhelix 326 348\ninside 349 372

26750 GCF_001258635.1_Sg_201405357_CTATAC_L001 Shigella sonnei Proteobacteria; Gammaproteobacteria; Enterobacterales; Enterobacteriaceae; Shigella MNNEETFYQAMRRQGVTRRSFLKYCSLAATSLGLGAGMAPKIAWALENKPRIPVVWIHGL WP_005139779.1 hydrogenase [Shigella sonnei] Length: 372\nNumber of predicted TMHs: 2\nExp number of AAs in TMHs: 37.02628\nExp number, first 60 AAs: 19.39684\nTotal prob of N-in: 0.94313\nPOSSIBLE N-term signal sequence\ninside 1 19\nTMhelix 20 39\noutside 40 325\nTMhelix 326 348\ninside 349 372

26751 GCF_001258735.1_9870_6_20 Shigella sonnei Proteobacteria; Gammaproteobacteria; Enterobacterales; Enterobacteriaceae; Shigella MNNEETFYQAMRRQGVTRRSFLKYCSLAATSLGLGAGMAPKIAWALENKPRIPVVWIHGL WP_005139779.1 hydrogenase [Shigella sonnei] Length: 372\nNumber of predicted TMHs: 2\nExp number of AAs in TMHs: 37.02628\nExp number, first 60 AAs: 19.39684\nTotal prob of N-in: 0.94313\nPOSSIBLE N-term signal sequence\ninside 1 19\nTMhelix 20 39\noutside 40 325\nTMhelix 326 348\ninside 349 372

26752 GCF_001258695.1_8289_7_86 Shigella sonnei Proteobacteria; Gammaproteobacteria; Enterobacterales; Enterobacteriaceae; Shigella MNNEETFYQAMRRQGVTRRSFLKYCSLAATSLGLGAGMAPKIAWALENKPRIPVVWIHGL WP_005139779.1 hydrogenase [Shigella sonnei] Length: 372\nNumber of predicted TMHs: 2\nExp number of AAs in TMHs: 37.02628\nExp number, first 60 AAs: 19.39684\nTotal prob of N-in: 0.94313\nPOSSIBLE N-term signal sequence\ninside 1 19\nTMhelix 20 39\noutside 40 325\nTMhelix 326 348\ninside 349 372

26753 GCF_001258655.1_9870_6_37 Shigella sonnei Proteobacteria; Gammaproteobacteria; Enterobacterales; Enterobacteriaceae; Shigella MNNEETFYQAMRRQGVTRRSFLKYCSLAATSLGLGAGMAPKIAWALENKPRIPVVWIHGL WP_005139779.1 hydrogenase [Shigella sonnei] Length: 372\nNumber of predicted TMHs: 2\nExp number of AAs in TMHs: 37.02628\nExp number, first 60 AAs: 19.39684\nTotal prob of N-in: 0.94313\nPOSSIBLE N-term signal sequence\ninside 1 19\nTMhelix 20 39\noutside 40 325\nTMhelix 326 348\ninside 349 372

26754 GCF_001258935.1_H14128036001 Shigella sonnei Proteobacteria; Gammaproteobacteria; Enterobacterales; Enterobacteriaceae; Shigella MTGDNTLIHSHGINRRDFMKLCAALAATMGLSSKAAAEMAESVTNPQRPPVIWIGAQECT WP_000145410.1 MULTISPECIES: hydrogenase-2 small chain [Proteobacteria] Length: 372\nNumber of predicted TMHs: 1\nExp number of AAs in TMHs: 23.72226\nExp number, first 60 AAs: 0.22038\nTotal prob of N-in: 0.12994\noutside 1 331\nTMhelix 332 354\ninside 355 372

26755 GCF_001259015.1_6551_1_11 Shigella sonnei Proteobacteria; Gammaproteobacteria; Enterobacterales; Enterobacteriaceae; Shigella MTGDNTLIHSHGINRRDFMKLCAALAATMGLSSKAAAEMAESVTNPQRPPVIWIGAQECT WP_000145410.1 MULTISPECIES: hydrogenase-2 small chain [Proteobacteria] Length: 372\nNumber of predicted TMHs: 1\nExp number of AAs in TMHs: 23.72226\nExp number, first 60 AAs: 0.22038\nTotal prob of N-in: 0.12994\noutside 1 331\nTMhelix 332 354\ninside 355 372

26756 GCF_001259035.1_12971_8_52 Shigella sonnei Proteobacteria; Gammaproteobacteria; Enterobacterales; Enterobacteriaceae; Shigella MTGDNTLIHSHGINRRDFMKLCAALAATMGLSSKAAAEMAESVTNPQRPPVIWIGAQECT WP_000145410.1

MULTISPECIES: hydrogenase-2 small chain [Proteobacteria] Length: 372\nNumber of predicted TMHs: 1\nExp number of AAs in TMHs: 23.72226\nExp number, first 60 AAs: 0.22038\nTotal prob of N-in: 0.12994\noutside 1 331\nTMhelix 332 354\ninside 355 372

26757 GCF_001259095.1_8290_4_49 Shigella sonnei Proteobacteria; Gammaproteobacteria; Enterobacterales; Enterobacteriaceae; Shigella MTGDNTLIHSHGINRRDFMKLCAALAATMGLSSKAAAEMAESVTNPQRPPVIWIGAQECT WP_000145410.1 MULTISPECIES: hydrogenase-2 small chain [Proteobacteria] Length: 372\nNumber of predicted TMHs: 1\nExp number of AAs in TMHs: 23.72226\nExp number, first 60 AAs: 0.22038\nTotal prob of N-in: 0.12994\noutside 1 331\nTMhelix 332 354\ninside 355 372

26758 GCF_001259115.1_8290_4_88 Shigella sonnei Proteobacteria; Gammaproteobacteria; Enterobacterales; Enterobacteriaceae; Shigella MTGDNTLIHSHGINRRDFMKLCAALAATMGLSSKAAAEMAESVTNPQRPPVIWIGAQECT WP_000145410.1 MULTISPECIES: hydrogenase-2 small chain [Proteobacteria] Length: 372\nNumber of predicted TMHs: 1\nExp number of AAs in TMHs: 23.72226\nExp number, first 60 AAs: 0.22038\nTotal prob of N-in: 0.12994\noutside 1 331\nTMhelix 332 354\ninside 355 372

26759 GCF_001259175.1_5008_8_9Shigella sonnei Proteobacteria; Gammaproteobacteria; Enterobacterales; Enterobacteriaceae; Shigella MTGDNTLIHSHGINRRDFMKLCAALAATMGLSSKAAAEMAESVTNPQRPPVIWIGAQECT WP_000145410.1 MULTISPECIES: hydrogenase-2 small chain [Proteobacteria] Length: 372\nNumber of predicted TMHs: 1\nExp number of AAs in TMHs: 23.72226\nExp number, first 60 AAs: 0.22038\nTotal prob of N-in: 0.12994\noutside 1 331\nTMhelix 332 354\ninside 355 372

26760 GCF_001259255.1_12830_1_40 Shigella sonnei Proteobacteria; Gammaproteobacteria; Enterobacterales; Enterobacteriaceae; Shigella MTGDNTLIHSHGINRRDFMKLCAALAATMGLSSKAAAEMAESVTNPQRPPVIWIGAQECT WP_000145410.1 MULTISPECIES: hydrogenase-2 small chain [Proteobacteria] Length: 372\nNumber of predicted TMHs: 1\nExp number of AAs in TMHs: 23.72226\nExp number, first 60 AAs: 0.22038\nTotal prob of N-in: 0.12994\noutside 1 331\nTMhelix 332 354\ninside 355 372

26761 GCF_001259215.1_12850_1_51 Shigella sonnei Proteobacteria; Gammaproteobacteria; Enterobacterales; Enterobacteriaceae; Shigella MTGDNTLIHSHGINRRDFMKLCAALAATMGLSSKAAAEMAESVTNPQRPPVIWIGAQECT WP_000145410.1 MULTISPECIES: hydrogenase-2 small chain [Proteobacteria] Length: 372\nNumber of predicted TMHs: 1\nExp number of AAs in TMHs: 23.72226\nExp number, first 60 AAs: 0.22038\nTotal prob of N-in: 0.12994\noutside 1 331\nTMhelix 332 354\ninside 355 372

26762 GCF_001259295.1_12830_1_34 Shigella sonnei Proteobacteria; Gammaproteobacteria; Enterobacterales; Enterobacteriaceae; Shigella MTGDNTLIHSHGINRRDFMKLCAALAATMGLSSKAAAEMAESVTNPQRPPVIWIGAQECT WP_000145410.1 MULTISPECIES: hydrogenase-2 small chain [Proteobacteria] Length: 372\nNumber of predicted TMHs: 1\nExp number of AAs in TMHs: 23.72226\nExp number, first 60 AAs: 0.22038\nTotal prob of N-in: 0.12994\noutside 1 331\nTMhelix 332 354\ninside 355 372

26763 GCF_001259355.1_6350_1_12 Shigella sonnei Proteobacteria; Gammaproteobacteria; Enterobacterales; Enterobacteriaceae; Shigella MTGDNTLIHSHGINRRDFMKLCAALAATMGLSSKAAAEMAESVTNPQRPPVIWIGAQECT WP_000145410.1 MULTISPECIES: hydrogenase-2 small chain [Proteobacteria] Length: 372\nNumber of predicted TMHs: 1\nExp number of AAs in TMHs: 23.72226\nExp number, first 60 AAs: 0.22038\nTotal prob of N-in: 0.12994\noutside 1 331\nTMhelix 332 354\ninside 355 372

26764 GCF_001259295.1_12830_1_34 Shigella sonnei Proteobacteria; Gammaproteobacteria; Enterobacterales; Enterobacteriaceae; Shigella MNNEETFYQAMRRQGVTRRSFLKYCSLAATSLGLGAGMAPKIAWALENKPRIPVVWIHGL WP_005139779.1 hydrogenase [Shigella sonnei] Length: 372\nNumber of predicted TMHs: 2\nExp number of AAs in TMHs: 37.02628\nExp number, first 60 AAs: 19.39684\nTotal prob of N-in: 0.94313\nPOSSIBLE N-term signal sequence\ninside 1 19\nTMhelix 20 39\noutside 40 325\nTMhelix 326 348\ninside 349 372

26765 GCF_001259215.1_12850_1_51 Shigella sonnei Proteobacteria; Gammaproteobacteria; Enterobacterales; Enterobacteriaceae; Shigella

MNNEETFYQAMRRQGVTRRSFLKYCSLAATSLGLGAGMAPKIAWALENKPRIPVVWIHGL WP_005139779.1 hydrogenase [Shigella sonnei] Length: 372\nNumber of predicted TMHs: 2\nExp number of AAs in TMHs: 37.02628\nExp number, first 60 AAs: 19.39684\nTotal prob of N-in: 0.94313\nPOSSIBLE N-term signal sequence\ninside 1 19\nTMhelix 20 39\noutside 40 325\nTMhelix 326 348\ninside 349 372

26766 GCF_001259355.1_6350_1_12 Shigella sonnei Proteobacteria; Gammaproteobacteria; Enterobacterales; Enterobacteriaceae; Shigella MNNEETFYQAMRRQGVTRRSFLKYCSLAATSLGLGAGMAPKIAWALENKPRIPVVWIHGL WP_005139779.1 hydrogenase [Shigella sonnei] Length: 372\nNumber of predicted TMHs: 2\nExp number of AAs in TMHs: 37.02628\nExp number, first 60 AAs: 19.39684\nTotal prob of N-in: 0.94313\nPOSSIBLE N-term signal sequence\ninside 1 19\nTMhelix 20 39\noutside 40 325\nTMhelix 326 348\ninside 349 372

26767 GCF_001259015.1_6551_1_11 Shigella sonnei Proteobacteria; Gammaproteobacteria; Enterobacterales; Enterobacteriaceae; Shigella MNNEETFYQAMRRQGVTRRSFLKYCSLAATSLGLGAGMAPKIAWALENKPRIPVVWIHGL WP_005139779.1 hydrogenase [Shigella sonnei] Length: 372\nNumber of predicted TMHs: 2\nExp number of AAs in TMHs: 37.02628\nExp number, first 60 AAs: 19.39684\nTotal prob of N-in: 0.94313\nPOSSIBLE N-term signal sequence\ninside 1 19\nTMhelix 20 39\noutside 40 325\nTMhelix 326 348\ninside 349 372

26768 GCF_001259395.1_8290_5_10 Shigella sonnei Proteobacteria; Gammaproteobacteria; Enterobacterales; Enterobacteriaceae; Shigella MTGDNTLIHSHGINRRDFMKLCAALAATMGLSSKAAAEMAESVTNPQRPPVIWIGAQECT WP_000145410.1 MULTISPECIES: hydrogenase-2 small chain [Proteobacteria] Length: 372\nNumber of predicted TMHs: 1\nExp number of AAs in TMHs: 23.72226\nExp number, first 60 AAs: 0.22038\nTotal prob of N-in: 0.12994\noutside 1 331\nTMhelix 332 354\ninside 355 372

26769 GCF_001259395.1_8290_5_10 Shigella sonnei Proteobacteria; Gammaproteobacteria; Enterobacterales; Enterobacteriaceae; Shigella MNNEETFYQAMRRQGVTRRSFLKYCSLAATSLGLGAGMAPKIAWALENKPRIPVVWIHGL WP_005139779.1 hydrogenase [Shigella sonnei] Length: 372\nNumber of predicted TMHs: 2\nExp number of AAs in TMHs: 37.02628\nExp number, first 60 AAs: 19.39684\nTotal prob of N-in: 0.94313\nPOSSIBLE N-term signal sequence\ninside 1 19\nTMhelix 20 39\noutside 40 325\nTMhelix 326 348\ninside 349 372

26770 GCF_001259095.1_8290_4_49 Shigella sonnei Proteobacteria; Gammaproteobacteria; Enterobacterales; Enterobacteriaceae; Shigella MNNEETFYQAMRRQGVTRRSFLKYCSLAATSLGLGAGMAPKIAWALENKPRIPVVWIHGL WP_005139779.1 hydrogenase [Shigella sonnei] Length: 372\nNumber of predicted TMHs: 2\nExp number of AAs in TMHs: 37.02628\nExp number, first 60 AAs: 19.39684\nTotal prob of N-in: 0.94313\nPOSSIBLE N-term signal sequence\ninside 1 19\nTMhelix 20 39\noutside 40 325\nTMhelix 326 348\ninside 349 372

26771 GCF_001259175.1_5008_8_9Shigella sonnei Proteobacteria; Gammaproteobacteria; Enterobacterales; Enterobacteriaceae; Shigella MNNEETFYQAMRRQGVTRRSFLKYCSLAATSLGLGAGMAPKIAWALENKPRIPVVWIHGL WP_005139779.1 hydrogenase [Shigella sonnei] Length: 372\nNumber of predicted TMHs: 2\nExp number of AAs in TMHs: 37.02628\nExp number, first 60 AAs: 19.39684\nTotal prob of N-in: 0.94313\nPOSSIBLE N-term signal sequence\ninside 1 19\nTMhelix 20 39\noutside 40 325\nTMhelix 326 348\ninside 349 372

26772 GCF_001259035.1_12971_8_52 Shigella sonnei Proteobacteria; Gammaproteobacteria; Enterobacterales; Enterobacteriaceae; Shigella MNNEETFYQAMRRQGVTRRSFLKYCSLAATSLGLGAGMAPKIAWALENKPRIPVVWIHGL WP_052962775.1 hydrogenase [Shigella sonnei] Length: 372\nNumber of predicted TMHs: 2\nExp number of AAs in TMHs: 37.09248\nExp number, first 60 AAs: 19.39296\nTotal prob of N-in: 0.94316\nPOSSIBLE N-term signal sequence\ninside 1 19\nTMhelix 20 39\noutside 40 325\nTMhelix 326 348\ninside 349 372

26773 GCF_001259515.1_5417_2_6Shigella sonnei Proteobacteria; Gammaproteobacteria; Enterobacterales; Enterobacteriaceae; Shigella MTGDNTLIHSHGINRRDFMKLCAALAATMGLSSKAAAEMAESVTNPQRPPVIWIGAQECT WP_000145410.1 MULTISPECIES: hydrogenase-2 small chain [Proteobacteria] Length: 372\nNumber of predicted TMHs: 1\nExp number of AAs in TMHs: 23.72226\nExp number, first 60 AAs: 0.22038\nTotal prob of N-in: 0.12994\noutside 1 331\nTMhelix 332 354\ninside 355 372

26774 GCF_001259435.1_5008_8_3Shigella sonnei Proteobacteria; Gammaproteobacteria; Enterobacterales; Enterobacteriaceae; Shigella MTGDNTLIHSHGINRRDFMKLCAALAATMGLSSKAAAEMAESVTNPQRPPVIWIGAQECT

WP_000145410.1 MULTISPECIES: hydrogenase-2 small chain [Proteobacteria] Length: 372\nNumber of predicted TMHs: 1\nExp number of AAs in TMHs: 23.72226\nExp number, first 60 AAs: 0.22038\nTotal prob of N-in: 0.12994\noutside 1 331\nTMhelix 332 354\ninside 355 372

26775 GCF_001259515.1_5417_2_6Shigella sonnei Proteobacteria; Gammaproteobacteria; Enterobacterales; Enterobacteriaceae; Shigella MNNEETFYQAMRRQGVTRRSFLKYCSLAATSLGLGAGMAPKIAWALENKPRIPVVWIHGL WP_005139779.1 hydrogenase [Shigella sonnei] Length: 372\nNumber of predicted TMHs: 2\nExp number of AAs in TMHs: 37.02628\nExp number, first 60 AAs: 19.39684\nTotal prob of N-in: 0.94313\nPOSSIBLE N-term signal sequence\ninside 1 19\nTMhelix 20 39\noutside 40 325\nTMhelix 326 348\ninside 349 372

26776 GCF_001259435.1_5008_8_3Shigella sonnei Proteobacteria; Gammaproteobacteria; Enterobacterales; Enterobacteriaceae; Shigella MNNEETFYQAMRRQGVTRRSFLKYCSLAATSLGLGAGMAPKIAWALENKPRIPVVWIHGL WP_005139779.1 hydrogenase [Shigella sonnei] Length: 372\nNumber of predicted TMHs: 2\nExp number of AAs in TMHs: 37.02628\nExp number, first 60 AAs: 19.39684\nTotal prob of N-in: 0.94313\nPOSSIBLE N-term signal sequence\ninside 1 19\nTMhelix 20 39\noutside 40 325\nTMhelix 326 348\ninside 349 372

26777 GCF_001259115.1_8290_4_88 Shigella sonnei Proteobacteria; Gammaproteobacteria; Enterobacterales; Enterobacteriaceae; Shigella MNNEETFYQAMRRQGVTRRSFLKYCSLAATSLGLGAGMAPKIAWALENKPRIPVVWIHGL WP_005139779.1 hydrogenase [Shigella sonnei] Length: 372\nNumber of predicted TMHs: 2\nExp number of AAs in TMHs: 37.02628\nExp number, first 60 AAs: 19.39684\nTotal prob of N-in: 0.94313\nPOSSIBLE N-term signal sequence\ninside 1 19\nTMhelix 20 39\noutside 40 325\nTMhelix 326 348\ninside 349 372

26778 GCF_001259615.1_5236_8_6Shigella sonnei Proteobacteria; Gammaproteobacteria; Enterobacterales; Enterobacteriaceae; Shigella MTGDNTLIHSHGINRRDFMKLCAALAATMGLSSKAAAEMAESVTNPQRPPVIWIGAQECT WP_000145410.1 MULTISPECIES: hydrogenase-2 small chain [Proteobacteria] Length: 372\nNumber of predicted TMHs: 1\nExp number of AAs in TMHs: 23.72226\nExp number, first 60 AAs: 0.22038\nTotal prob of N-in: 0.12994\noutside 1 331\nTMhelix 332 354\ninside 355 372

26779 GCF_001259615.1_5236_8_6Shigella sonnei Proteobacteria; Gammaproteobacteria; Enterobacterales; Enterobacteriaceae; Shigella MNNEETFYQAMRRQGVTRRSFLKYCSLAATSLGLGAGMAPKIAWALENKPRIPVVWIHGL WP_005139779.1 hydrogenase [Shigella sonnei] Length: 372\nNumber of predicted TMHs: 2\nExp number of AAs in TMHs: 37.02628\nExp number, first 60 AAs: 19.39684\nTotal prob of N-in: 0.94313\nPOSSIBLE N-term signal sequence\ninside 1 19\nTMhelix 20 39\noutside 40 325\nTMhelix 326 348\ninside 349 372

26780 GCF_001259255.1_12830_1_40 Shigella sonnei Proteobacteria; Gammaproteobacteria; Enterobacterales; Enterobacteriaceae; Shigella MNNEETFYQAMRRQGVTRRSFLKYCSLAATSLGLGAGMAPKIAWALENKPRIPVVWIHGL WP_005139779.1 hydrogenase [Shigella sonnei] Length: 372\nNumber of predicted TMHs: 2\nExp number of AAs in TMHs: 37.02628\nExp number, first 60 AAs: 19.39684\nTotal prob of N-in: 0.94313\nPOSSIBLE N-term signal sequence\ninside 1 19\nTMhelix 20 39\noutside 40 325\nTMhelix 326 348\ninside 349 372

26781 GCF_001259655.1_5417_2_11 Shigella sonnei Proteobacteria; Gammaproteobacteria; Enterobacterales; Enterobacteriaceae; Shigella MTGDNTLIHSHGINRRDFMKLCAALAATMGLSSKAAAEMAESVTNPQRPPVIWIGAQECT WP_000145410.1 MULTISPECIES: hydrogenase-2 small chain [Proteobacteria] Length: 372\nNumber of predicted TMHs: 1\nExp number of AAs in TMHs: 23.72226\nExp number, first 60 AAs: 0.22038\nTotal prob of N-in: 0.12994\noutside 1 331\nTMhelix 332 354\ninside 355 372

26782 GCF_001259695.1_12971_8_28 Shigella sonnei Proteobacteria; Gammaproteobacteria; Enterobacterales; Enterobacteriaceae; Shigella MTGDNTLIHSHGINRRDFMKLCAALAATMGLSSKAAAEMAESVTNPQRPPVIWIGAQECT WP_000145410.1 MULTISPECIES: hydrogenase-2 small chain [Proteobacteria] Length: 372\nNumber of predicted TMHs: 1\nExp number of AAs in TMHs: 23.72226\nExp number, first 60 AAs: 0.22038\nTotal prob of N-in: 0.12994\noutside 1 331\nTMhelix 332 354\ninside 355 372

26783 GCF_001259755.1_8289_7_83 Shigella sonnei Proteobacteria; Gammaproteobacteria; Enterobacterales; Enterobacteriaceae; Shigella MTGDNTLIHSHGINRRDFMKLCAALAATMGLSSKAAAEMAESVTNPQRPPVIWIGAQECT WP_000145410.1 MULTISPECIES: hydrogenase-2 small chain [Proteobacteria] Length: 372\nNumber of predicted TMHs:

1\nExp number of AAs in TMHs: 23.72226\nExp number, first 60 AAs: 0.22038\nTotal prob of N-in: 0.12994\noutside 1 331\nTMhelix 332 354\ninside 355 372

26784 GCF_001259775.1_12971_8_42 Shigella sonnei Proteobacteria; Gammaproteobacteria; Enterobacterales; Enterobacteriaceae; Shigella
MTGDNTLIHSHGINRRDFMKLCAALAATMGLSSKAAAEMAESVTNPQRPPVIWIGAQECT WP_000145410.1
MULTISPECIES: hydrogenase-2 small chain [Proteobacteria] Length: 372\nNumber of predicted TMHs: 1\nExp number of AAs in TMHs: 23.72226\nExp number, first 60 AAs: 0.22038\nTotal prob of N-in: 0.12994\noutside 1 331\nTMhelix 332 354\ninside 355 372

26785 GCF_001259835.1_8290_4_89 Shigella sonnei Proteobacteria; Gammaproteobacteria; Enterobacterales; Enterobacteriaceae; Shigella
MTGDNTLIHSHGINRRDFMKLCAALAATMGLSSKAAAEMAESVTNPQRPPVIWIGAQECT WP_000145410.1
MULTISPECIES: hydrogenase-2 small chain [Proteobacteria] Length: 372\nNumber of predicted TMHs: 1\nExp number of AAs in TMHs: 23.72226\nExp number, first 60 AAs: 0.22038\nTotal prob of N-in: 0.12994\noutside 1 331\nTMhelix 332 354\ninside 355 372

26786 GCF_001259895.1_8290_4_34 Shigella sonnei Proteobacteria; Gammaproteobacteria; Enterobacterales; Enterobacteriaceae; Shigella
MTGDNTLIHSHGINRRDFMKLCAALAATMGLSSKAAAEMAESVTNPQRPPVIWIGAQECT WP_000145410.1
MULTISPECIES: hydrogenase-2 small chain [Proteobacteria] Length: 372\nNumber of predicted TMHs: 1\nExp number of AAs in TMHs: 23.72226\nExp number, first 60 AAs: 0.22038\nTotal prob of N-in: 0.12994\noutside 1 331\nTMhelix 332 354\ninside 355 372

26787 GCF_001259955.1_12830_1_32 Shigella sonnei Proteobacteria; Gammaproteobacteria; Enterobacterales; Enterobacteriaceae; Shigella
MTGDNTLIHSHGINRRDFMKLCAALAATMGLSSKAAAEMAESVTNPQRPPVIWIGAQECT WP_000145410.1
MULTISPECIES: hydrogenase-2 small chain [Proteobacteria] Length: 372\nNumber of predicted TMHs: 1\nExp number of AAs in TMHs: 23.72226\nExp number, first 60 AAs: 0.22038\nTotal prob of N-in: 0.12994\noutside 1 331\nTMhelix 332 354\ninside 355 372

26788 GCF_001260015.1_12830_1_39 Shigella sonnei Proteobacteria; Gammaproteobacteria; Enterobacterales; Enterobacteriaceae; Shigella
MTGDNTLIHSHGINRRDFMKLCAALAATMGLSSKAAAEMAESVTNPQRPPVIWIGAQECT WP_000145410.1
MULTISPECIES: hydrogenase-2 small chain [Proteobacteria] Length: 372\nNumber of predicted TMHs: 1\nExp number of AAs in TMHs: 23.72226\nExp number, first 60 AAs: 0.22038\nTotal prob of N-in: 0.12994\noutside 1 331\nTMhelix 332 354\ninside 355 372

26789 GCF_001260095.1_Sg_201401743_CGTACG_L001 Shigella sonnei Proteobacteria; Gammaproteobacteria; Enterobacterales; Enterobacteriaceae; Shigella
MTGDNTLIHSHGINRRDFMKLCAALAATMGLSSKAAAEMAESVTNPQRPPVIWIGAQECT WP_000145410.1
MULTISPECIES: hydrogenase-2 small chain [Proteobacteria] Length: 372\nNumber of predicted TMHs: 1\nExp number of AAs in TMHs: 23.72226\nExp number, first 60 AAs: 0.22038\nTotal prob of N-in: 0.12994\noutside 1 331\nTMhelix 332 354\ninside 355 372

26790 GCF_001260015.1_12830_1_39 Shigella sonnei Proteobacteria; Gammaproteobacteria; Enterobacterales; Enterobacteriaceae; Shigella
MNNEETFYQAMRRQGVTRRSFLKYCSLAATSLGLGAGMAPKIAWALENKPRIPVVWIHGL WP_052985843.1
hydrogenase [Shigella sonnei] Length: 372\nNumber of predicted TMHs: 2\nExp number of AAs in TMHs: 37.38943\nExp number, first 60 AAs: 19.40855\nTotal prob of N-in: 0.94303\nPOSSIBLE N-term signal sequence\ninside 1 19\nTMhelix 20 39\noutside 40 325\nTMhelix 326 348\ninside 349 372

26791 GCF_001260095.1_Sg_201401743_CGTACG_L001 Shigella sonnei Proteobacteria; Gammaproteobacteria; Enterobacterales; Enterobacteriaceae; Shigella
MNNEETFYQAMRRQGVTRRSFLKYCSLAATSLGLGAGMAPKIAWALENKPRIPVVWIHGL WP_005139779.1
hydrogenase [Shigella sonnei] Length: 372\nNumber of predicted TMHs: 2\nExp number of AAs in TMHs: 37.02628\nExp number, first 60 AAs: 19.39684\nTotal prob of N-in: 0.94313\nPOSSIBLE N-term signal sequence\ninside 1 19\nTMhelix 20 39\noutside 40 325\nTMhelix 326 348\ninside 349 372

26792 GCF_001259655.1_5417_2_11 Shigella sonnei Proteobacteria; Gammaproteobacteria; Enterobacterales; Enterobacteriaceae; Shigella

MNNEETFYQAMRRQGVTRRSFLKYCSLAATSLGLGAGMAPKIAWALENKPRIPVVWIHGL WP_005139779.1 hydrogenase [Shigella sonnei] Length: 372\nNumber of predicted TMHs: 2\nExp number of AAs in TMHs: 37.02628\nExp number, first 60 AAs: 19.39684\nTotal prob of N-in: 0.94313\nPOSSIBLE N-term signal sequence\ninside 1 19\nTMhelix 20 39\noutside 40 325\nTMhelix 326 348\ninside 349 372

26793 GCF_001260115.1_8289_7_66 Shigella sonnei Proteobacteria; Gammaproteobacteria; Enterobacterales; Enterobacteriaceae; Shigella MTGDNTLIHSHGINRRDFMKLCAALAATMGLSSKAAAEMAESVTNPQRPPVIWIGAQECT WP_000145410.1 MULTISPECIES: hydrogenase-2 small chain [Proteobacteria] Length: 372\nNumber of predicted TMHs: 1\nExp number of AAs in TMHs: 23.72226\nExp number, first 60 AAs: 0.22038\nTotal prob of N-in: 0.12994\noutside 1 331\nTMhelix 332 354\ninside 355 372

26794 GCF_001260115.1_8289_7_66 Shigella sonnei Proteobacteria; Gammaproteobacteria; Enterobacterales; Enterobacteriaceae; Shigella MNNEETFYQAMRRQGVTRRSFLKYCSLAATSLGLGAGMAPKIAWALENKPRIPVVWIHGL WP_005139779.1 hydrogenase [Shigella sonnei] Length: 372\nNumber of predicted TMHs: 2\nExp number of AAs in TMHs: 37.02628\nExp number, first 60 AAs: 19.39684\nTotal prob of N-in: 0.94313\nPOSSIBLE N-term signal sequence\ninside 1 19\nTMhelix 20 39\noutside 40 325\nTMhelix 326 348\ninside 349 372

26795 GCF_001259695.1_12971_8_28 Shigella sonnei Proteobacteria; Gammaproteobacteria; Enterobacterales; Enterobacteriaceae; Shigella MNNEETFYQAMRRQGVTRRSFLKYCSLAATSLGLGAGMAPKIAWALENKPRIPVVWIHGL WP_005139779.1 hydrogenase [Shigella sonnei] Length: 372\nNumber of predicted TMHs: 2\nExp number of AAs in TMHs: 37.02628\nExp number, first 60 AAs: 19.39684\nTotal prob of N-in: 0.94313\nPOSSIBLE N-term signal sequence\ninside 1 19\nTMhelix 20 39\noutside 40 325\nTMhelix 326 348\ninside 349 372

26796 GCF_001260155.1_9870_6_33 Shigella sonnei Proteobacteria; Gammaproteobacteria; Enterobacterales; Enterobacteriaceae; Shigella MTGDNTLIHSHGINRRDFMKLCAALAATMGLSSKAAAEMAESVTNPQRPPVIWIGAQECT WP_000145410.1 MULTISPECIES: hydrogenase-2 small chain [Proteobacteria] Length: 372\nNumber of predicted TMHs: 1\nExp number of AAs in TMHs: 23.72226\nExp number, first 60 AAs: 0.22038\nTotal prob of N-in: 0.12994\noutside 1 331\nTMhelix 332 354\ninside 355 372

26797 GCF_001260155.1_9870_6_33 Shigella sonnei Proteobacteria; Gammaproteobacteria; Enterobacterales; Enterobacteriaceae; Shigella MNNEETFYQAMRRQGVTRRSFLKYCSLAATSLGLGAGMAPKIAWALENKPRIPVVWIHGL WP_005139779.1 hydrogenase [Shigella sonnei] Length: 372\nNumber of predicted TMHs: 2\nExp number of AAs in TMHs: 37.02628\nExp number, first 60 AAs: 19.39684\nTotal prob of N-in: 0.94313\nPOSSIBLE N-term signal sequence\ninside 1 19\nTMhelix 20 39\noutside 40 325\nTMhelix 326 348\ninside 349 372

26798 GCF_001259755.1_8289_7_83 Shigella sonnei Proteobacteria; Gammaproteobacteria; Enterobacterales; Enterobacteriaceae; Shigella MNNEETFYQAMRRQGVTRRSFLKYCSLAATSLGLGAGMAPKIAWALENKPRIPVVWIHGL WP_005139779.1 hydrogenase [Shigella sonnei] Length: 372\nNumber of predicted TMHs: 2\nExp number of AAs in TMHs: 37.02628\nExp number, first 60 AAs: 19.39684\nTotal prob of N-in: 0.94313\nPOSSIBLE N-term signal sequence\ninside 1 19\nTMhelix 20 39\noutside 40 325\nTMhelix 326 348\ninside 349 372

26799 GCF_001259775.1_12971_8_42 Shigella sonnei Proteobacteria; Gammaproteobacteria; Enterobacterales; Enterobacteriaceae; Shigella MNNEETFYQAMRRQGVTRRSFLKYCSLAATSLGLGAGMAPKIAWALENKPRIPVVWIHGL WP_005139779.1 hydrogenase [Shigella sonnei] Length: 372\nNumber of predicted TMHs: 2\nExp number of AAs in TMHs: 37.02628\nExp number, first 60 AAs: 19.39684\nTotal prob of N-in: 0.94313\nPOSSIBLE N-term signal sequence\ninside 1 19\nTMhelix 20 39\noutside 40 325\nTMhelix 326 348\ninside 349 372

26800 GCF_001259955.1_12830_1_32 Shigella sonnei Proteobacteria; Gammaproteobacteria; Enterobacterales; Enterobacteriaceae; Shigella MNNEETFYQAMRRQGVTRRSFLKYCSLAATSLGLGAGMAPKIAWALENKPRIPVVWIHGL WP_005139779.1 hydrogenase [Shigella sonnei] Length: 372\nNumber of predicted TMHs: 2\nExp number of AAs in TMHs: 37.02628\nExp number, first 60 AAs: 19.39684\nTotal prob of N-in: 0.94313\nPOSSIBLE N-term signal sequence\ninside 1 19\nTMhelix 20 39\noutside 40 325\nTMhelix 326 348\ninside 349 372

26801 GCF_001259835.1_8290_4_89 Shigella sonnei Proteobacteria; Gammaproteobacteria; Enterobacterales; Enterobacteriaceae; Shigella
MNNEETFYQAMRRQGVTRRSFLKYCSLAATSLGLGAGMAPKIAWALENKPRIPVVWIHGL WP_005139779.1
hydrogenase [Shigella sonnei] Length: 372\nNumber of predicted TMHs: 2\nExp number of AAs in TMHs: 37.02628\nExp number, first 60 AAs: 19.39684\nTotal prob of N-in: 0.94313\nPOSSIBLE N-term signal sequence\ninside 1 19\nTMhelix 20 39\noutside 40 325\nTMhelix 326 348\ninside 349 372

26802 GCF_001259895.1_8290_4_34 Shigella sonnei Proteobacteria; Gammaproteobacteria; Enterobacterales; Enterobacteriaceae; Shigella
MNNEETFYQAMRRQGVTRRSFLKYCSLAATSLGLGAGMAPKIAWALENKPRIPVVWIHGL WP_005139779.1
hydrogenase [Shigella sonnei] Length: 372\nNumber of predicted TMHs: 2\nExp number of AAs in TMHs: 37.02628\nExp number, first 60 AAs: 19.39684\nTotal prob of N-in: 0.94313\nPOSSIBLE N-term signal sequence\ninside 1 19\nTMhelix 20 39\noutside 40 325\nTMhelix 326 348\ninside 349 372

26803 GCF_001260235.1_5236_1_7Shigella sonnei Proteobacteria; Gammaproteobacteria; Enterobacterales; Enterobacteriaceae; Shigella MTGDNTLIHSHGINRRDFMKLCAALAATMGLSSKAAAEMAESVTNPQRPPVIWIGAQECT
WP_000145410.1 MULTISPECIES: hydrogenase-2 small chain [Proteobacteria] Length: 372\nNumber of predicted TMHs: 1\nExp number of AAs in TMHs: 23.72226\nExp number, first 60 AAs: 0.22038\nTotal prob of N-in: 0.12994\noutside 1 331\nTMhelix 332 354\ninside 355 372

26804 GCF_001260195.1_6551_1_5Shigella sonnei Proteobacteria; Gammaproteobacteria; Enterobacterales; Enterobacteriaceae; Shigella MTGDNTLIHSHGINRRDFMKLCAALAATMGLSSKAAAEMAESVTNPQRPPVIWIGAQECT
WP_000145410.1 MULTISPECIES: hydrogenase-2 small chain [Proteobacteria] Length: 372\nNumber of predicted TMHs: 1\nExp number of AAs in TMHs: 23.72226\nExp number, first 60 AAs: 0.22038\nTotal prob of N-in: 0.12994\noutside 1 331\nTMhelix 332 354\ninside 355 372

26805 GCF_001260275.1_7655_8_11 Shigella sonnei Proteobacteria; Gammaproteobacteria; Enterobacterales; Enterobacteriaceae; Shigella
MTGDNTLIHSHGINRRDFMKLCAALAATMGLSSKAAAEMAESVTNPQRPPVIWIGAQECT WP_000145410.1
MULTISPECIES: hydrogenase-2 small chain [Proteobacteria] Length: 372\nNumber of predicted TMHs: 1\nExp number of AAs in TMHs: 23.72226\nExp number, first 60 AAs: 0.22038\nTotal prob of N-in: 0.12994\noutside 1 331\nTMhelix 332 354\ninside 355 372

26806 GCF_001260315.1_5008_8_2Shigella sonnei Proteobacteria; Gammaproteobacteria; Enterobacterales; Enterobacteriaceae; Shigella MTGDNTLIHSHGINRRDFMKLCAALAATMGLSSKAAAEMAESVTNPQRPPVIWIGAQECT
WP_000145410.1 MULTISPECIES: hydrogenase-2 small chain [Proteobacteria] Length: 372\nNumber of predicted TMHs: 1\nExp number of AAs in TMHs: 23.72226\nExp number, first 60 AAs: 0.22038\nTotal prob of N-in: 0.12994\noutside 1 331\nTMhelix 332 354\ninside 355 372

26807 GCF_001260375.1_H14112034001 Shigella sonnei Proteobacteria; Gammaproteobacteria; Enterobacterales; Enterobacteriaceae; Shigella
MTGDNTLIHSHGINRRDFMKLCAALAATMGLSSKAAAEMAESVTNPQRPPVIWIGAQECT WP_000145410.1
MULTISPECIES: hydrogenase-2 small chain [Proteobacteria] Length: 372\nNumber of predicted TMHs: 1\nExp number of AAs in TMHs: 23.72226\nExp number, first 60 AAs: 0.22038\nTotal prob of N-in: 0.12994\noutside 1 331\nTMhelix 332 354\ninside 355 372

26808 GCF_001260415.1_5236_7_7Shigella sonnei Proteobacteria; Gammaproteobacteria; Enterobacterales; Enterobacteriaceae; Shigella MTGDNTLIHSHGINRRDFMKLCAALAATMGLSSKAAAEMAESVTNPQRPPVIWIGAQECT
WP_000145410.1 MULTISPECIES: hydrogenase-2 small chain [Proteobacteria] Length: 372\nNumber of predicted TMHs: 1\nExp number of AAs in TMHs: 23.72226\nExp number, first 60 AAs: 0.22038\nTotal prob of N-in: 0.12994\noutside 1 331\nTMhelix 332 354\ninside 355 372

26809 GCF_001260455.1_5417_2_10 Shigella sonnei Proteobacteria; Gammaproteobacteria; Enterobacterales; Enterobacteriaceae; Shigella
MTGDNTLIHSHGINRRDFMKLCAALAATMGLSSKAAAEMAESVTNPQRPPVIWIGAQECT WP_000145410.1
MULTISPECIES: hydrogenase-2 small chain [Proteobacteria] Length: 372\nNumber of predicted TMHs: 1\nExp number of AAs in TMHs: 23.72226\nExp number, first 60 AAs: 0.22038\nTotal prob of N-in: 0.12994\noutside 1 331\nTMhelix 332 354\ninside 355 372

26810 GCF_001260495.1_8289_7_81 Shigella sonnei Proteobacteria; Gammaproteobacteria; Enterobacterales; Enterobacteriaceae; Shigella

MTGDNTLIHSHGINRRDFMKLCAALAATMGLSSKAAAEMAESVTNPQRPPVIWIGAQECT WP_000145410.1 MULTISPECIES: hydrogenase-2 small chain [Proteobacteria] Length: 372\nNumber of predicted TMHs: 1\nExp number of AAs in TMHs: 23.72226\nExp number, first 60 AAs: 0.22038\nTotal prob of N-in: 0.12994\noutside 1 331\nTMhelix 332 354\ninside 355 372

26811 GCF_001260535.1_5417_1_12 Shigella sonnei Proteobacteria; Gammaproteobacteria; Enterobacterales; Enterobacteriaceae; Shigella MTGDNTLIHSHGINRRDFMKLCAALAATMGLSSKAAAEMAESVTNPQRPPVIWIGAQECT WP_000145410.1 MULTISPECIES: hydrogenase-2 small chain [Proteobacteria] Length: 372\nNumber of predicted TMHs: 1\nExp number of AAs in TMHs: 23.72226\nExp number, first 60 AAs: 0.22038\nTotal prob of N-in: 0.12994\noutside 1 331\nTMhelix 332 354\ninside 355 372

26812 GCF_001260555.1_8290_4_45 Shigella sonnei Proteobacteria; Gammaproteobacteria; Enterobacterales; Enterobacteriaceae; Shigella MTGDNTLIHSHGINRRDFMKLCAALAATMGLSSKAAAEMAESVTNPQRPPVIWIGAQECT WP_000145410.1 MULTISPECIES: hydrogenase-2 small chain [Proteobacteria] Length: 372\nNumber of predicted TMHs: 1\nExp number of AAs in TMHs: 23.72226\nExp number, first 60 AAs: 0.22038\nTotal prob of N-in: 0.12994\noutside 1 331\nTMhelix 332 354\ninside 355 372

26813 GCF_001260495.1_8289_7_81 Shigella sonnei Proteobacteria; Gammaproteobacteria; Enterobacterales; Enterobacteriaceae; Shigella MNNEETFYQAMRRQGVTRRSFLKYCSLAATSLGLGAGMAPKIAWALENKPRIPVVWIHGL WP_005139779.1 hydrogenase [Shigella sonnei] Length: 372\nNumber of predicted TMHs: 2\nExp number of AAs in TMHs: 37.02628\nExp number, first 60 AAs: 19.39684\nTotal prob of N-in: 0.94313\nPOSSIBLE N-term signal sequence\ninside 1 19\nTMhelix 20 39\noutside 40 325\nTMhelix 326 348\ninside 349 372

26814 GCF_001260535.1_5417_1_12 Shigella sonnei Proteobacteria; Gammaproteobacteria; Enterobacterales; Enterobacteriaceae; Shigella MNNEETFYQAMRRQGVTRRSFLKYCSLAATSLGLGAGMAPKIAWALENKPRIPVVWIHGL WP_005139779.1 hydrogenase [Shigella sonnei] Length: 372\nNumber of predicted TMHs: 2\nExp number of AAs in TMHs: 37.02628\nExp number, first 60 AAs: 19.39684\nTotal prob of N-in: 0.94313\nPOSSIBLE N-term signal sequence\ninside 1 19\nTMhelix 20 39\noutside 40 325\nTMhelix 326 348\ninside 349 372

26815 GCF_001260555.1_8290_4_45 Shigella sonnei Proteobacteria; Gammaproteobacteria; Enterobacterales; Enterobacteriaceae; Shigella MNNEETFYQAMRRQGVTRRSFLKYCSLAATSLGLGAGMAPKIAWALENKPRIPVVWIHGL WP_005139779.1 hydrogenase [Shigella sonnei] Length: 372\nNumber of predicted TMHs: 2\nExp number of AAs in TMHs: 37.02628\nExp number, first 60 AAs: 19.39684\nTotal prob of N-in: 0.94313\nPOSSIBLE N-term signal sequence\ninside 1 19\nTMhelix 20 39\noutside 40 325\nTMhelix 326 348\ninside 349 372

26816 GCF_001260195.1_6551_1_5Shigella sonnei Proteobacteria; Gammaproteobacteria; Enterobacterales; Enterobacteriaceae; Shigella MNNEETFYQAMRRQGVTRRSFLKYCSLAATSLGLGAGMAPKIAWALENKPRIPVVWIHGL WP_005139779.1 hydrogenase [Shigella sonnei] Length: 372\nNumber of predicted TMHs: 2\nExp number of AAs in TMHs: 37.02628\nExp number, first 60 AAs: 19.39684\nTotal prob of N-in: 0.94313\nPOSSIBLE N-term signal sequence\ninside 1 19\nTMhelix 20 39\noutside 40 325\nTMhelix 326 348\ninside 349 372

26817 GCF_001260235.1_5236_1_7Shigella sonnei Proteobacteria; Gammaproteobacteria; Enterobacterales; Enterobacteriaceae; Shigella MNNEETFYQAMRRQGVTRRSFLKYCSLAATSLGLGAGMAPKIAWALENKPRIPVVWIHGL WP_005139779.1 hydrogenase [Shigella sonnei] Length: 372\nNumber of predicted TMHs: 2\nExp number of AAs in TMHs: 37.02628\nExp number, first 60 AAs: 19.39684\nTotal prob of N-in: 0.94313\nPOSSIBLE N-term signal sequence\ninside 1 19\nTMhelix 20 39\noutside 40 325\nTMhelix 326 348\ninside 349 372

26818 GCF_001260275.1_7655_8_11 Shigella sonnei Proteobacteria; Gammaproteobacteria; Enterobacterales; Enterobacteriaceae; Shigella MNNEETFYQAMRRQGVTRRSFLKYCSLAATSLGLGAGMAPKIAWALENKPRIPVVWIHGL WP_005139779.1 hydrogenase [Shigella sonnei] Length: 372\nNumber of predicted TMHs: 2\nExp number of AAs in TMHs: 37.02628\nExp number, first 60 AAs: 19.39684\nTotal prob of N-in: 0.94313\nPOSSIBLE N-term signal sequence\ninside 1 19\nTMhelix 20 39\noutside 40 325\nTMhelix 326 348\ninside 349 372

26819 GCF_001260315.1_5008_8_2Shigella sonnei Proteobacteria; Gammaproteobacteria; Enterobacterales; Enterobacteriaceae; Shigella MNNEETFYQAMRRQGVTRRSFLKYCSLAATSLGLGAGMAPKIAWALENKPRIPVVWIHGL

WP_005139779.1 hydrogenase [Shigella sonnei] Length: 372\nNumber of predicted TMHs: 2\nExp number of AAs in TMHs: 37.02628\nExp number, first 60 AAs: 19.39684\nTotal prob of N-in: 0.94313\nPOSSIBLE N-term signal sequence\ninside 1 19\nTMhelix 20 39\noutside 40 325\nTMhelix 326 348\ninside 349 372

26820 GCF_001260635.1_8289_7_76 Shigella sonnei Proteobacteria; Gammaproteobacteria; Enterobacterales; Enterobacteriaceae; Shigella MTGDNTLIHSHGINRRDFMKLCAALAATMGLSSKAAAEMAESVTNPQRPPVIWIGAQECT WP_000145410.1 MULTISPECIES: hydrogenase-2 small chain [Proteobacteria] Length: 372\nNumber of predicted TMHs: 1\nExp number of AAs in TMHs: 23.72226\nExp number, first 60 AAs: 0.22038\nTotal prob of N-in: 0.12994\noutside 1 331\nTMhelix 332 354\ninside 355 372

26821 GCF_001260635.1_8289_7_76 Shigella sonnei Proteobacteria; Gammaproteobacteria; Enterobacterales; Enterobacteriaceae; Shigella MNNEETFYQAMRRQGVTRRSFLKYCSLAATSLGLGAGMAPKIAWALENKPRIPVVWIHGL WP_005139779.1 hydrogenase [Shigella sonnei] Length: 372\nNumber of predicted TMHs: 2\nExp number of AAs in TMHs: 37.02628\nExp number, first 60 AAs: 19.39684\nTotal prob of N-in: 0.94313\nPOSSIBLE N-term signal sequence\ninside 1 19\nTMhelix 20 39\noutside 40 325\nTMhelix 326 348\ninside 349 372

26822 GCF_001260415.1_5236_7_7Shigella sonnei Proteobacteria; Gammaproteobacteria; Enterobacterales; Enterobacteriaceae; Shigella MNNEETFYQAMRRQGVTRRSFLKYCSLAATSLGLGAGMAPKIAWALENKPRIPVVWIHGL WP_005139779.1 hydrogenase [Shigella sonnei] Length: 372\nNumber of predicted TMHs: 2\nExp number of AAs in TMHs: 37.02628\nExp number, first 60 AAs: 19.39684\nTotal prob of N-in: 0.94313\nPOSSIBLE N-term signal sequence\ninside 1 19\nTMhelix 20 39\noutside 40 325\nTMhelix 326 348\ninside 349 372

26823 GCF_001260375.1_H14112034001 Shigella sonnei Proteobacteria; Gammaproteobacteria; Enterobacterales; Enterobacteriaceae; Shigella MNNEETFYQAMRRQGVTRRSFLKYCSLAATSLGLGAGMAPKIAWALENKPRIPVVWIHGL WP_005139779.1 hydrogenase [Shigella sonnei] Length: 372\nNumber of predicted TMHs: 2\nExp number of AAs in TMHs: 37.02628\nExp number, first 60 AAs: 19.39684\nTotal prob of N-in: 0.94313\nPOSSIBLE N-term signal sequence\ninside 1 19\nTMhelix 20 39\noutside 40 325\nTMhelix 326 348\ninside 349 372

26824 GCF_001260675.1_5236_5_4Shigella sonnei Proteobacteria; Gammaproteobacteria; Enterobacterales; Enterobacteriaceae; Shigella MTGDNTLIHSHGINRRDFMKLCAALAATMGLSSKAAAEMAESVTNPQRPPVIWIGAQECT WP_000145410.1 MULTISPECIES: hydrogenase-2 small chain [Proteobacteria] Length: 372\nNumber of predicted TMHs: 1\nExp number of AAs in TMHs: 23.72226\nExp number, first 60 AAs: 0.22038\nTotal prob of N-in: 0.12994\noutside 1 331\nTMhelix 332 354\ninside 355 372

26825 GCF_001260715.1_7353_1_20 Shigella sonnei Proteobacteria; Gammaproteobacteria; Enterobacterales; Enterobacteriaceae; Shigella MTGDNTLIHSHGINRRDFMKLCAALAATMGLSSKAAAEMAESVTNPQRPPVIWIGAQECT WP_000145410.1 MULTISPECIES: hydrogenase-2 small chain [Proteobacteria] Length: 372\nNumber of predicted TMHs: 1\nExp number of AAs in TMHs: 23.72226\nExp number, first 60 AAs: 0.22038\nTotal prob of N-in: 0.12994\noutside 1 331\nTMhelix 332 354\ninside 355 372

26826 GCF_001260675.1_5236_5_4Shigella sonnei Proteobacteria; Gammaproteobacteria; Enterobacterales; Enterobacteriaceae; Shigella MNNEETFYQAMRRQGVTRRSFLKYCSLAATSLGLGAGMAPKIAWALENKPRIPVVWIHGL WP_005139779.1 hydrogenase [Shigella sonnei] Length: 372\nNumber of predicted TMHs: 2\nExp number of AAs in TMHs: 37.02628\nExp number, first 60 AAs: 19.39684\nTotal prob of N-in: 0.94313\nPOSSIBLE N-term signal sequence\ninside 1 19\nTMhelix 20 39\noutside 40 325\nTMhelix 326 348\ninside 349 372

26827 GCF_001260715.1_7353_1_20 Shigella sonnei Proteobacteria; Gammaproteobacteria; Enterobacterales; Enterobacteriaceae; Shigella MNNEETFYQAMRRQGVTRRSFLKYCSLAATSLGLGAGMAPKIAWALENKPRIPVVWIHGL WP_005139779.1 hydrogenase [Shigella sonnei] Length: 372\nNumber of predicted TMHs: 2\nExp number of AAs in TMHs: 37.02628\nExp number, first 60 AAs: 19.39684\nTotal prob of N-in: 0.94313\nPOSSIBLE N-term signal sequence\ninside 1 19\nTMhelix 20 39\noutside 40 325\nTMhelix 326 348\ninside 349 372

26828 GCF_001260455.1_5417_2_10 Shigella sonnei Proteobacteria; Gammaproteobacteria; Enterobacterales; Enterobacteriaceae; Shigella MNNEETFYQAMRRQGVTRRSFLKYCSLAATSLGLGAGMAPKIAWALENKPRIPVVWIHGL WP_005139779.1 hydrogenase [Shigella sonnei] Length: 372\nNumber of predicted TMHs: 2\nExp number of AAs in

TMHs: 37.02628\nExp number, first 60 AAs: 19.39684\nTotal prob of N-in: 0.94313\nPOSSIBLE N-term signal sequence\ninside 1 19\nTMhelix 20 39\noutside 40 325\nTMhelix 326 348\ninside 349 372

26829 GCF_001260755.1_5417_3_6Shigella sonnei Proteobacteria; Gammaproteobacteria; Enterobacterales; Enterobacteriaceae; Shigella MTGDNTLIHSHGINRRDFMKLCAALAATMGLSSKAAAEMAESVTNPQRPPVIWIGAQECT WP_000145410.1 MULTISPECIES: hydrogenase-2 small chain [Proteobacteria] Length: 372\nNumber of predicted TMHs: 1\nExp number of AAs in TMHs: 23.72226\nExp number, first 60 AAs: 0.22038\nTotal prob of N-in: 0.12994\noutside 1 331\nTMhelix 332 354\ninside 355 372

26830 GCF_001260795.1_5008_7_10 Shigella sonnei Proteobacteria; Gammaproteobacteria; Enterobacterales; Enterobacteriaceae; Shigella MTGDNTLIHSHGINRRDFMKLCAALAATMGLSSKAAAEMAESVTNPQRPPVIWIGAQECT WP_000145410.1 MULTISPECIES: hydrogenase-2 small chain [Proteobacteria] Length: 372\nNumber of predicted TMHs: 1\nExp number of AAs in TMHs: 23.72226\nExp number, first 60 AAs: 0.22038\nTotal prob of N-in: 0.12994\noutside 1 331\nTMhelix 332 354\ninside 355 372

26831 GCF_001260815.1_7353_1_17 Shigella sonnei Proteobacteria; Gammaproteobacteria; Enterobacterales; Enterobacteriaceae; Shigella MTGDNTLIHSHGINRRDFMKLCAALAATMGLSSKAAAEMAESVTNPQRPPVIWIGAQECT WP_000145410.1 MULTISPECIES: hydrogenase-2 small chain [Proteobacteria] Length: 372\nNumber of predicted TMHs: 1\nExp number of AAs in TMHs: 23.72226\nExp number, first 60 AAs: 0.22038\nTotal prob of N-in: 0.12994\noutside 1 331\nTMhelix 332 354\ninside 355 372

26832 GCF_001260855.1_8289_7_71 Shigella sonnei Proteobacteria; Gammaproteobacteria; Enterobacterales; Enterobacteriaceae; Shigella MTGDNTLIHSHGINRRDFMKLCAALAATMGLSSKAAAEMAESVTNPQRPPVIWIGAQECT WP_000145410.1 MULTISPECIES: hydrogenase-2 small chain [Proteobacteria] Length: 372\nNumber of predicted TMHs: 1\nExp number of AAs in TMHs: 23.72226\nExp number, first 60 AAs: 0.22038\nTotal prob of N-in: 0.12994\noutside 1 331\nTMhelix 332 354\ninside 355 372

26833 GCF_001260895.1_12971_8_36 Shigella sonnei Proteobacteria; Gammaproteobacteria; Enterobacterales; Enterobacteriaceae; Shigella MTGDNTLIHSHGINRRDFMKLCAALAATMGLSSKAAAEMAESVTNPQRPPVIWIGAQECT WP_000145410.1 MULTISPECIES: hydrogenase-2 small chain [Proteobacteria] Length: 372\nNumber of predicted TMHs: 1\nExp number of AAs in TMHs: 23.72226\nExp number, first 60 AAs: 0.22038\nTotal prob of N-in: 0.12994\noutside 1 331\nTMhelix 332 354\ninside 355 372

26834 GCF_001260935.1_8289_7_57 Shigella sonnei Proteobacteria; Gammaproteobacteria; Enterobacterales; Enterobacteriaceae; Shigella MTGDNTLIHSHGINRRDFMKLCAALAATMGLSSKAAAEMAESVTNPQRPPVIWIGAQECT WP_000145410.1 MULTISPECIES: hydrogenase-2 small chain [Proteobacteria] Length: 372\nNumber of predicted TMHs: 1\nExp number of AAs in TMHs: 23.72226\nExp number, first 60 AAs: 0.22038\nTotal prob of N-in: 0.12994\noutside 1 331\nTMhelix 332 354\ninside 355 372

26835 GCF_001260975.1_9870_6_3Shigella sonnei Proteobacteria; Gammaproteobacteria; Enterobacterales; Enterobacteriaceae; Shigella MTGDNTLIHSHGINRRDFMKLCAALAATMGLSSKAAAEMAESVTNPQRPPVIWIGAQECT WP_000145410.1 MULTISPECIES: hydrogenase-2 small chain [Proteobacteria] Length: 372\nNumber of predicted TMHs: 1\nExp number of AAs in TMHs: 23.72226\nExp number, first 60 AAs: 0.22038\nTotal prob of N-in: 0.12994\noutside 1 331\nTMhelix 332 354\ninside 355 372

26836 GCF_001261035.1_8289_7_90 Shigella sonnei Proteobacteria; Gammaproteobacteria; Enterobacterales; Enterobacteriaceae; Shigella MTGDNTLIHSHGINRRDFMKLCAALAATMGLSSKAAAEMAESVTNPQRPPVIWIGAQECT WP_000145410.1 MULTISPECIES: hydrogenase-2 small chain [Proteobacteria] Length: 372\nNumber of predicted TMHs: 1\nExp number of AAs in TMHs: 23.72226\nExp number, first 60 AAs: 0.22038\nTotal prob of N-in: 0.12994\noutside 1 331\nTMhelix 332 354\ninside 355 372

26837 GCF_001261055.1_9870_7_10 Shigella sonnei Proteobacteria; Gammaproteobacteria; Enterobacterales; Enterobacteriaceae; Shigella MTGDNTLIHSHGINRRDFMKLCAALAATMGLSSKAAAEMAESVTNPQRPPVIWIGAQECT WP_000145410.1 MULTISPECIES: hydrogenase-2 small chain [Proteobacteria] Length: 372\nNumber of predicted TMHs:

1\nExp number of AAs in TMHs: 23.72226\nExp number, first 60 AAs: 0.22038\nTotal prob of N-in: 0.12994\noutside 1 331\nTMhelix 332 354\ninside 355 372

26838 GCF_001261155.1_12971_8_51 Shigella sonnei Proteobacteria; Gammaproteobacteria; Enterobacterales; Enterobacteriaceae; Shigella MTGDNTLIHSHGINRRDFMKLCAALAATMGLSSKAAAEMAESVTNPQRPPVIWIGAQECT WP_000145410.1 MULTISPECIES: hydrogenase-2 small chain [Proteobacteria] Length: 372\nNumber of predicted TMHs: 1\nExp number of AAs in TMHs: 23.72226\nExp number, first 60 AAs: 0.22038\nTotal prob of N-in: 0.12994\noutside 1 331\nTMhelix 332 354\ninside 355 372

26839 GCF_001261035.1_8289_7_90 Shigella sonnei Proteobacteria; Gammaproteobacteria; Enterobacterales; Enterobacteriaceae; Shigella MNNEETFYQAMRRQGVTRRSFLKYCSLAATSLGLGAGMAPKIAWALENKPRIPVVWIHGL WP_005139779.1 hydrogenase [Shigella sonnei] Length: 372\nNumber of predicted TMHs: 2\nExp number of AAs in TMHs: 37.02628\nExp number, first 60 AAs: 19.39684\nTotal prob of N-in: 0.94313\nPOSSIBLE N-term signal sequence\ninside 1 19\nTMhelix 20 39\noutside 40 325\nTMhelix 326 348\ninside 349 372

26840 GCF_001261055.1_9870_7_10 Shigella sonnei Proteobacteria; Gammaproteobacteria; Enterobacterales; Enterobacteriaceae; Shigella MNNEETFYQAMRRQGVTRRSFLKYCSLAATSLGLGAGMAPKIAWALENKPRIPVVWIHGL WP_005139779.1 hydrogenase [Shigella sonnei] Length: 372\nNumber of predicted TMHs: 2\nExp number of AAs in TMHs: 37.02628\nExp number, first 60 AAs: 19.39684\nTotal prob of N-in: 0.94313\nPOSSIBLE N-term signal sequence\ninside 1 19\nTMhelix 20 39\noutside 40 325\nTMhelix 326 348\ninside 349 372

26841 GCF_001261155.1_12971_8_51 Shigella sonnei Proteobacteria; Gammaproteobacteria; Enterobacterales; Enterobacteriaceae; Shigella MNNEETFYQAMRRQGVTRRSFLKYCSLAATSLGLGAGMAPKIAWALENKPRIPVVWIHGL WP_052962775.1 hydrogenase [Shigella sonnei] Length: 372\nNumber of predicted TMHs: 2\nExp number of AAs in TMHs: 37.09248\nExp number, first 60 AAs: 19.39296\nTotal prob of N-in: 0.94316\nPOSSIBLE N-term signal sequence\ninside 1 19\nTMhelix 20 39\noutside 40 325\nTMhelix 326 348\ninside 349 372

26842 GCF_001260815.1_7353_1_17 Shigella sonnei Proteobacteria; Gammaproteobacteria; Enterobacterales; Enterobacteriaceae; Shigella MNNEETFYQAMRRQGVTRRSFLKYCSLAATSLGLGAGMAPKIAWALENKPRIPVVWIHGL WP_005139779.1 hydrogenase [Shigella sonnei] Length: 372\nNumber of predicted TMHs: 2\nExp number of AAs in TMHs: 37.02628\nExp number, first 60 AAs: 19.39684\nTotal prob of N-in: 0.94313\nPOSSIBLE N-term signal sequence\ninside 1 19\nTMhelix 20 39\noutside 40 325\nTMhelix 326 348\ninside 349 372

26843 GCF_001260755.1_5417_3_6Shigella sonnei Proteobacteria; Gammaproteobacteria; Enterobacterales; Enterobacteriaceae; Shigella MNNEETFYQAMRRQGVTRRSFLKYCSLAATSLGLGAGMAPKIAWALENKPRIPVVWIHGL WP_005139779.1 hydrogenase [Shigella sonnei] Length: 372\nNumber of predicted TMHs: 2\nExp number of AAs in TMHs: 37.02628\nExp number, first 60 AAs: 19.39684\nTotal prob of N-in: 0.94313\nPOSSIBLE N-term signal sequence\ninside 1 19\nTMhelix 20 39\noutside 40 325\nTMhelix 326 348\ninside 349 372

26844 GCF_001260795.1_5008_7_10 Shigella sonnei Proteobacteria; Gammaproteobacteria; Enterobacterales; Enterobacteriaceae; Shigella MNNEETFYQAMRRQGVTRRSFLKYCSLAATSLGLGAGMAPKIAWALENKPRIPVVWIHGL WP_005139779.1 hydrogenase [Shigella sonnei] Length: 372\nNumber of predicted TMHs: 2\nExp number of AAs in TMHs: 37.02628\nExp number, first 60 AAs: 19.39684\nTotal prob of N-in: 0.94313\nPOSSIBLE N-term signal sequence\ninside 1 19\nTMhelix 20 39\noutside 40 325\nTMhelix 326 348\ninside 349 372

26845 GCF_001260855.1_8289_7_71 Shigella sonnei Proteobacteria; Gammaproteobacteria; Enterobacterales; Enterobacteriaceae; Shigella MNNEETFYQAMRRQGVTRRSFLKYCSLAATSLGLGAGMAPKIAWALENKPRIPVVWIHGL WP_005139779.1 hydrogenase [Shigella sonnei] Length: 372\nNumber of predicted TMHs: 2\nExp number of AAs in TMHs: 37.02628\nExp number, first 60 AAs: 19.39684\nTotal prob of N-in: 0.94313\nPOSSIBLE N-term signal sequence\ninside 1 19\nTMhelix 20 39\noutside 40 325\nTMhelix 326 348\ninside 349 372

26846 GCF_001261195.1_H14140035901 Shigella sonnei Proteobacteria; Gammaproteobacteria; Enterobacterales; Enterobacteriaceae; Shigella MTGDNTLIHSHGINRRDFMKLCAALAATMGLSSKAAAEMAESVTNPQRPPVIWIGAQECT WP_000145410.1

MULTISPECIES: hydrogenase-2 small chain [Proteobacteria] Length: 372\nNumber of predicted TMHs: 1\nExp number of AAs in TMHs: 23.72226\nExp number, first 60 AAs: 0.22038\nTotal prob of N-in: 0.12994\noutside 1 331\nTMhelix 332 354\ninside 355 372

26847 GCF_001261215.1_5008_7_11 Shigella sonnei Proteobacteria; Gammaproteobacteria; Enterobacterales; Enterobacteriaceae; Shigella MTGDNTLIHSHGINRRDFMKLCAALAATMGLSSKAAAEMAESVTNPQRPPVIWIGAQECT WP_000145410.1 MULTISPECIES: hydrogenase-2 small chain [Proteobacteria] Length: 372\nNumber of predicted TMHs: 1\nExp number of AAs in TMHs: 23.72226\nExp number, first 60 AAs: 0.22038\nTotal prob of N-in: 0.12994\noutside 1 331\nTMhelix 332 354\ninside 355 372

26848 GCF_001261215.1_5008_7_11 Shigella sonnei Proteobacteria; Gammaproteobacteria; Enterobacterales; Enterobacteriaceae; Shigella MNNEETFYQAMRRQGVTRRSFLKYCSLAATSLGLGAGMAPKIAWALENKPRIPVVWIHGL WP_005139779.1 hydrogenase [Shigella sonnei] Length: 372\nNumber of predicted TMHs: 2\nExp number of AAs in TMHs: 37.02628\nExp number, first 60 AAs: 19.39684\nTotal prob of N-in: 0.94313\nPOSSIBLE N-term signal sequence\ninside 1 19\nTMhelix 20 39\noutside 40 325\nTMhelix 326 348\ninside 349 372

26849 GCF_001261195.1_H14140035901 Shigella sonnei Proteobacteria; Gammaproteobacteria; Enterobacterales; Enterobacteriaceae; Shigella MNNEETFYQAMRRQGVTRRSFLKYCSLAATSLGLGAGMAPKIAWALENKPRIPVVWIHGL WP_052962775.1 hydrogenase [Shigella sonnei] Length: 372\nNumber of predicted TMHs: 2\nExp number of AAs in TMHs: 37.09248\nExp number, first 60 AAs: 19.39296\nTotal prob of N-in: 0.94316\nPOSSIBLE N-term signal sequence\ninside 1 19\nTMhelix 20 39\noutside 40 325\nTMhelix 326 348\ninside 349 372

26850 GCF_001260895.1_12971_8_36 Shigella sonnei Proteobacteria; Gammaproteobacteria; Enterobacterales; Enterobacteriaceae; Shigella MNNEETFYQAMRRQGVTRRSFLKYCSLAATSLGLGAGMAPKIAWALENKPRIPVVWIHGL WP_005139779.1 hydrogenase [Shigella sonnei] Length: 372\nNumber of predicted TMHs: 2\nExp number of AAs in TMHs: 37.02628\nExp number, first 60 AAs: 19.39684\nTotal prob of N-in: 0.94313\nPOSSIBLE N-term signal sequence\ninside 1 19\nTMhelix 20 39\noutside 40 325\nTMhelix 326 348\ninside 349 372

26851 GCF_001260935.1_8289_7_57 Shigella sonnei Proteobacteria; Gammaproteobacteria; Enterobacterales; Enterobacteriaceae; Shigella MNNEETFYQAMRRQGVTRRSFLKYCSLAATSLGLGAGMAPKIAWALENKPRIPVVWIHGL WP_005139779.1 hydrogenase [Shigella sonnei] Length: 372\nNumber of predicted TMHs: 2\nExp number of AAs in TMHs: 37.02628\nExp number, first 60 AAs: 19.39684\nTotal prob of N-in: 0.94313\nPOSSIBLE N-term signal sequence\ninside 1 19\nTMhelix 20 39\noutside 40 325\nTMhelix 326 348\ninside 349 372

26852 GCF_001260975.1_9870_6_3Shigella sonnei Proteobacteria; Gammaproteobacteria; Enterobacterales; Enterobacteriaceae; Shigella MNNEETFYQAMRRQGVTRRSFLKYCSLAATSLGLGAGMAPKIAWALENKPRIPVVWIHGL WP_005139779.1 hydrogenase [Shigella sonnei] Length: 372\nNumber of predicted TMHs: 2\nExp number of AAs in TMHs: 37.02628\nExp number, first 60 AAs: 19.39684\nTotal prob of N-in: 0.94313\nPOSSIBLE N-term signal sequence\ninside 1 19\nTMhelix 20 39\noutside 40 325\nTMhelix 326 348\ninside 349 372

26853 GCF_001261315.1_6350_1_8Shigella sonnei Proteobacteria; Gammaproteobacteria; Enterobacterales; Enterobacteriaceae; Shigella MTGDNTLIHSHGINRRDFMKLCAALAATMGLSSKAAAEMAESVTNPQRPPVIWIGAQECT WP_000145410.1 MULTISPECIES: hydrogenase-2 small chain [Proteobacteria] Length: 372\nNumber of predicted TMHs: 1\nExp number of AAs in TMHs: 23.72226\nExp number, first 60 AAs: 0.22038\nTotal prob of N-in: 0.12994\noutside 1 331\nTMhelix 332 354\ninside 355 372

26854 GCF_001261255.1_8289_7_85 Shigella sonnei Proteobacteria; Gammaproteobacteria; Enterobacterales; Enterobacteriaceae; Shigella MTGDNTLIHSHGINRRDFMKLCAALAATMGLSSKAAAEMAESVTNPQRPPVIWIGAQECT WP_000145410.1 MULTISPECIES: hydrogenase-2 small chain [Proteobacteria] Length: 372\nNumber of predicted TMHs: 1\nExp number of AAs in TMHs: 23.72226\nExp number, first 60 AAs: 0.22038\nTotal prob of N-in: 0.12994\noutside 1 331\nTMhelix 332 354\ninside 355 372

26855 GCF_001261355.1_9870_6_22 Shigella sonnei Proteobacteria; Gammaproteobacteria; Enterobacterales; Enterobacteriaceae; Shigella MTGDNTLIHSHGINRRDFMKLCAALAATMGLSSKAAAEMAESVTNPQRPPVIWIGAQECT WP_000145410.1

MULTISPECIES: hydrogenase-2 small chain [Proteobacteria]	Length: 372\nNumber of predicted TMHs: 1\nExp number of AAs in TMHs: 23.72226\nExp number, first 60 AAs: 0.22038\nTotal prob of N-in: 0.12994\noutside 1 331\nTMhelix 332 354\ninside 355 372

26856 GCF_001261395.1_H14146070601 Shigella sonnei Proteobacteria; Gammaproteobacteria; Enterobacterales; Enterobacteriaceae; Shigella MTGDNTLIHSHGINRRDFMKLCAALAATMGLSSKAAAEMAESVTNPQRPPVIWIGAQECT WP_000145410.1 MULTISPECIES: hydrogenase-2 small chain [Proteobacteria] Length: 372\nNumber of predicted TMHs: 1\nExp number of AAs in TMHs: 23.72226\nExp number, first 60 AAs: 0.22038\nTotal prob of N-in: 0.12994\noutside 1 331\nTMhelix 332 354\ninside 355 372

26857 GCF_001261435.1_Sg_201402995_ATTCCT_L001Shigella sonnei Proteobacteria; Gammaproteobacteria; Enterobacterales; Enterobacteriaceae; Shigella MTGDNTLIHSHGINRRDFMKLCAALAATMGLSSKAAAEMAESVTNPQRPPVIWIGAQECT WP_000145410.1 MULTISPECIES: hydrogenase-2 small chain [Proteobacteria] Length: 372\nNumber of predicted TMHs: 1\nExp number of AAs in TMHs: 23.72226\nExp number, first 60 AAs: 0.22038\nTotal prob of N-in: 0.12994\noutside 1 331\nTMhelix 332 354\ninside 355 372

26858 GCF_001261455.1_Sg_201405193_CATTTT_L001Shigella sonnei Proteobacteria; Gammaproteobacteria; Enterobacterales; Enterobacteriaceae; Shigella MTGDNTLIHSHGINRRDFMKLCAALAATMGLSSKAAAEMAESVTNPQRPPVIWIGAQECT WP_000145410.1 MULTISPECIES: hydrogenase-2 small chain [Proteobacteria] Length: 372\nNumber of predicted TMHs: 1\nExp number of AAs in TMHs: 23.72226\nExp number, first 60 AAs: 0.22038\nTotal prob of N-in: 0.12994\noutside 1 331\nTMhelix 332 354\ninside 355 372

26859 GCF_001261595.1_6350_1_20 Shigella sonnei Proteobacteria; Gammaproteobacteria; Enterobacterales; Enterobacteriaceae; Shigella MTGDNTLIHSHGINRRDFMKLCAALAATMGLSSKAAAEMAESVTNPQRPPVIWIGAQECT WP_000145410.1 MULTISPECIES: hydrogenase-2 small chain [Proteobacteria] Length: 372\nNumber of predicted TMHs: 1\nExp number of AAs in TMHs: 23.72226\nExp number, first 60 AAs: 0.22038\nTotal prob of N-in: 0.12994\noutside 1 331\nTMhelix 332 354\ninside 355 372

26860 GCF_001261495.1_6350_1_4Shigella sonnei Proteobacteria; Gammaproteobacteria; Enterobacterales; Enterobacteriaceae; Shigella MTGDNTLIHSHGINRRDFMKLCAALAATMGLSSKAAAEMAESVTNPQRPPVIWIGAQECT WP_000145410.1 MULTISPECIES: hydrogenase-2 small chain [Proteobacteria] Length: 372\nNumber of predicted TMHs: 1\nExp number of AAs in TMHs: 23.72226\nExp number, first 60 AAs: 0.22038\nTotal prob of N-in: 0.12994\noutside 1 331\nTMhelix 332 354\ninside 355 372

26861 GCF_001261495.1_6350_1_4Shigella sonnei Proteobacteria; Gammaproteobacteria; Enterobacterales; Enterobacteriaceae; Shigella MNNEETFYQAMRRQGVTRRSFLKYCSLAATSLGLGAGMAPKIAWALENKPRIPVVWIHGL WP_005139779.1 hydrogenase [Shigella sonnei] Length: 372\nNumber of predicted TMHs: 2\nExp number of AAs in TMHs: 37.02628\nExp number, first 60 AAs: 19.39684\nTotal prob of N-in: 0.94313\nPOSSIBLE N-term signal sequence\ninside 1 19\nTMhelix 20 39\noutside 40 325\nTMhelix 326 348\ninside 349 372

26862 GCF_001261315.1_6350_1_8Shigella sonnei Proteobacteria; Gammaproteobacteria; Enterobacterales; Enterobacteriaceae; Shigella MNNEETFYQAMRRQGVTRRSFLKYCSLAATSLGLGAGMAPKIAWALENKPRIPVVWIHGL WP_005139779.1 hydrogenase [Shigella sonnei] Length: 372\nNumber of predicted TMHs: 2\nExp number of AAs in TMHs: 37.02628\nExp number, first 60 AAs: 19.39684\nTotal prob of N-in: 0.94313\nPOSSIBLE N-term signal sequence\ninside 1 19\nTMhelix 20 39\noutside 40 325\nTMhelix 326 348\ninside 349 372

26863 GCF_001261615.1_12971_8_21 Shigella sonnei Proteobacteria; Gammaproteobacteria; Enterobacterales; Enterobacteriaceae; Shigella MTGDNTLIHSHGINRRDFMKLCAALAATMGLSSKAAAEMAESVTNPQRPPVIWIGAQECT WP_000145410.1 MULTISPECIES: hydrogenase-2 small chain [Proteobacteria] Length: 372\nNumber of predicted TMHs: 1\nExp number of AAs in TMHs: 23.72226\nExp number, first 60 AAs: 0.22038\nTotal prob of N-in: 0.12994\noutside 1 331\nTMhelix 332 354\ninside 355 372

26864 GCF_001261255.1_8289_7_85 Shigella sonnei Proteobacteria; Gammaproteobacteria; Enterobacterales; Enterobacteriaceae; Shigella MNNEETFYQAMRRQGVTRRSFLKYCSLAATSLGLGAGMAPKIAWALENKPRIPVVWIHGL WP_005139779.1 hydrogenase [Shigella sonnei] Length: 372\nNumber of predicted TMHs: 2\nExp number of AAs in

TMHs: 37.02628\nExp number, first 60 AAs: 19.39684\nTotal prob of N-in: 0.94313\nPOSSIBLE N-term signal sequence\ninside 1 19\nTMhelix 20 39\noutside 40 325\nTMhelix 326 348\ninside 349 372

26865 GCF_001261355.1_9870_6_22 Shigella sonnei Proteobacteria; Gammaproteobacteria; Enterobacterales; Enterobacteriaceae; Shigella MNNEETFYQAMRRQGVTRRSFLKYCSLAATSLGLGAGMAPKIAWALENKPRIPVVWIHGL WP_005139779.1 hydrogenase [Shigella sonnei] Length: 372\nNumber of predicted TMHs: 2\nExp number of AAs in TMHs: 37.02628\nExp number, first 60 AAs: 19.39684\nTotal prob of N-in: 0.94313\nPOSSIBLE N-term signal sequence\ninside 1 19\nTMhelix 20 39\noutside 40 325\nTMhelix 326 348\ninside 349 372

26866 GCF_001261655.1_5417_3_5Shigella sonnei Proteobacteria; Gammaproteobacteria; Enterobacterales; Enterobacteriaceae; Shigella MTGDNTLIHSHGINRRDFMKLCAALAATMGLSSKAAAEMAESVTNPQRPPVIWIGAQECT WP_000145410.1 MULTISPECIES: hydrogenase-2 small chain [Proteobacteria] Length: 372\nNumber of predicted TMHs: 1\nExp number of AAs in TMHs: 23.72226\nExp number, first 60 AAs: 0.22038\nTotal prob of N-in: 0.12994\noutside 1 331\nTMhelix 332 354\ninside 355 372

26867 GCF_001261695.1_12830_1_38 Shigella sonnei Proteobacteria; Gammaproteobacteria; Enterobacterales; Enterobacteriaceae; Shigella MTGDNTLIHSHGINRRDFMKLCAALAATMGLSSKAAAEMAESVTNPQRPPVIWIGAQECT WP_000145410.1 MULTISPECIES: hydrogenase-2 small chain [Proteobacteria] Length: 372\nNumber of predicted TMHs: 1\nExp number of AAs in TMHs: 23.72226\nExp number, first 60 AAs: 0.22038\nTotal prob of N-in: 0.12994\noutside 1 331\nTMhelix 332 354\ninside 355 372

26868 GCF_001261655.1_5417_3_5Shigella sonnei Proteobacteria; Gammaproteobacteria; Enterobacterales; Enterobacteriaceae; Shigella MNNEETFYQAMRRQGVTRRSFLKYCSLAATSLGLGAGMAPKIAWALENKPRIPVVWIHGL WP_005139779.1 hydrogenase [Shigella sonnei] Length: 372\nNumber of predicted TMHs: 2\nExp number of AAs in TMHs: 37.02628\nExp number, first 60 AAs: 19.39684\nTotal prob of N-in: 0.94313\nPOSSIBLE N-term signal sequence\ninside 1 19\nTMhelix 20 39\noutside 40 325\nTMhelix 326 348\ninside 349 372

26869 GCF_001261695.1_12830_1_38 Shigella sonnei Proteobacteria; Gammaproteobacteria; Enterobacterales; Enterobacteriaceae; Shigella MNNEETFYQAMRRQGVTRRSFLKYCSLAATSLGLGAGMAPKIAWALENKPRIPVVWIHGL WP_005139779.1 hydrogenase [Shigella sonnei] Length: 372\nNumber of predicted TMHs: 2\nExp number of AAs in TMHs: 37.02628\nExp number, first 60 AAs: 19.39684\nTotal prob of N-in: 0.94313\nPOSSIBLE N-term signal sequence\ninside 1 19\nTMhelix 20 39\noutside 40 325\nTMhelix 326 348\ninside 349 372

26870 GCF_001261395.1_H14146070601 Shigella sonnei Proteobacteria; Gammaproteobacteria; Enterobacterales; Enterobacteriaceae; Shigella MNNEETFYQAMRRQGVTRRSFLKYCSLAATSLGLGAGMAPKIAWALENKPRIPVVWIHGL WP_052962775.1 hydrogenase [Shigella sonnei] Length: 372\nNumber of predicted TMHs: 2\nExp number of AAs in TMHs: 37.09248\nExp number, first 60 AAs: 19.39296\nTotal prob of N-in: 0.94316\nPOSSIBLE N-term signal sequence\ninside 1 19\nTMhelix 20 39\noutside 40 325\nTMhelix 326 348\ninside 349 372

26871 GCF_001261435.1_Sg_201402995_ATTCCT_L001Shigella sonnei Proteobacteria; Gammaproteobacteria; Enterobacterales; Enterobacteriaceae; Shigella MNNEETFYQAMRRQGVTRRSFLKYCSLAATSLGLGAGMAPKIAWALENKPRIPVVWIHGL WP_005139779.1 hydrogenase [Shigella sonnei] Length: 372\nNumber of predicted TMHs: 2\nExp number of AAs in TMHs: 37.02628\nExp number, first 60 AAs: 19.39684\nTotal prob of N-in: 0.94313\nPOSSIBLE N-term signal sequence\ninside 1 19\nTMhelix 20 39\noutside 40 325\nTMhelix 326 348\ninside 349 372

26872 GCF_001443165.1_ASM144316v1 Shigella sonnei Proteobacteria; Gammaproteobacteria; Enterobacterales; Enterobacteriaceae; Shigella MTGDNTLIHSHGINRRDFMKLCAALAATMGLSSKAAAEMAESVTNPQRPPVIWIGAQECT WP_000145410.1 MULTISPECIES: hydrogenase-2 small chain [Proteobacteria] Length: 372\nNumber of predicted TMHs: 1\nExp number of AAs in TMHs: 23.72226\nExp number, first 60 AAs: 0.22038\nTotal prob of N-in: 0.12994\noutside 1 331\nTMhelix 332 354\ninside 355 372

26873 GCF_001687685.1_ASM168768v1 Shigella sonnei Proteobacteria; Gammaproteobacteria; Enterobacterales; Enterobacteriaceae; Shigella MTGDNTLIHSHGINRRDFMKLCAALAATMGLSSKAAAEMAESVTNPQRPPVIWIGAQECT WP_000145410.1 MULTISPECIES: hydrogenase-2 small chain [Proteobacteria] Length: 372\nNumber of predicted TMHs:

1\nExp number of AAs in TMHs: 23.72226\nExp number, first 60 AAs: 0.22038\nTotal prob of N-in: 0.12994\noutside 1 331\nTMhelix 332 354\ninside 355 372

26874 GCF_001443165.1_ASM144316v1 Shigella sonnei Proteobacteria; Gammaproteobacteria; Enterobacterales; Enterobacteriaceae; Shigella
MNNEETFYQAMRRQGVTRRSFLKYCSLAATSLGLGAGMAPKIAWALENKPRIPVVWIHGL WP_005139779.1
hydrogenase [Shigella sonnei] Length: 372\nNumber of predicted TMHs: 2\nExp number of AAs in TMHs: 37.02628\nExp number, first 60 AAs: 19.39684\nTotal prob of N-in: 0.94313\nPOSSIBLE N-term signal sequence\ninside 1 19\nTMhelix 20 39\noutside 40 325\nTMhelix 326 348\ninside 349 372

26875 GCF_001687685.1_ASM168768v1 Shigella sonnei Proteobacteria; Gammaproteobacteria; Enterobacterales; Enterobacteriaceae; Shigella
MNNEETFYQAMRRQGVTRRSFLKYCSLAATSLGLGAGMAPKIAWALENKPRIPVVWIHGL WP_005139779.1
hydrogenase [Shigella sonnei] Length: 372\nNumber of predicted TMHs: 2\nExp number of AAs in TMHs: 37.02628\nExp number, first 60 AAs: 19.39684\nTotal prob of N-in: 0.94313\nPOSSIBLE N-term signal sequence\ninside 1 19\nTMhelix 20 39\noutside 40 325\nTMhelix 326 348\ninside 349 372

26876 GCF_001261595.1_6350_1_20 Shigella sonnei Proteobacteria; Gammaproteobacteria; Enterobacterales; Enterobacteriaceae; Shigella
MNNEETFYQAMRRQGVTRRSFLKYCSLAATSLGLGAGMAPKIAWALENKPRIPVVWIHGL WP_005139779.1
hydrogenase [Shigella sonnei] Length: 372\nNumber of predicted TMHs: 2\nExp number of AAs in TMHs: 37.02628\nExp number, first 60 AAs: 19.39684\nTotal prob of N-in: 0.94313\nPOSSIBLE N-term signal sequence\ninside 1 19\nTMhelix 20 39\noutside 40 325\nTMhelix 326 348\ninside 349 372

26877 GCF_001261455.1_Sg_201405193_CATTTT_L001Shigella sonnei Proteobacteria; Gammaproteobacteria; Enterobacterales; Enterobacteriaceae; Shigella
MNNEETFYQAMRRQGVTRRSFLKYCSLAATSLGLGAGMAPKIAWALENKPRIPVVWIHGL WP_005139779.1
hydrogenase [Shigella sonnei] Length: 372\nNumber of predicted TMHs: 2\nExp number of AAs in TMHs: 37.02628\nExp number, first 60 AAs: 19.39684\nTotal prob of N-in: 0.94313\nPOSSIBLE N-term signal sequence\ninside 1 19\nTMhelix 20 39\noutside 40 325\nTMhelix 326 348\ninside 349 372

26878 GCF_001687755.1_ASM168775v1 Shigella sonnei Proteobacteria; Gammaproteobacteria; Enterobacterales; Enterobacteriaceae; Shigella
MTGDNTLIHSHGINRRDFMKLCAALAATMGLSSKAAAEMAESVTNPQRPPVIWIGAQECT WP_000145410.1
MULTISPECIES: hydrogenase-2 small chain [Proteobacteria] Length: 372\nNumber of predicted TMHs: 1\nExp number of AAs in TMHs: 23.72226\nExp number, first 60 AAs: 0.22038\nTotal prob of N-in: 0.12994\noutside 1 331\nTMhelix 332 354\ninside 355 372

26879 GCF_001687795.1_ASM168779v1 Shigella sonnei Proteobacteria; Gammaproteobacteria; Enterobacterales; Enterobacteriaceae; Shigella
MTGDNTLIHSHGINRRDFMKLCAALAATMGLSSKAAAEMAESVTNPQRPPVIWIGAQECT WP_000145410.1
MULTISPECIES: hydrogenase-2 small chain [Proteobacteria] Length: 372\nNumber of predicted TMHs: 1\nExp number of AAs in TMHs: 23.72226\nExp number, first 60 AAs: 0.22038\nTotal prob of N-in: 0.12994\noutside 1 331\nTMhelix 332 354\ninside 355 372

26880 GCF_001687825.1_ASM168782v1 Shigella sonnei Proteobacteria; Gammaproteobacteria; Enterobacterales; Enterobacteriaceae; Shigella
MTGDNTLIHSHGINRRDFMKLCAALAATMGLSSKAAAEMAESVTNPQRPPVIWIGAQECT WP_000145410.1
MULTISPECIES: hydrogenase-2 small chain [Proteobacteria] Length: 372\nNumber of predicted TMHs: 1\nExp number of AAs in TMHs: 23.72226\nExp number, first 60 AAs: 0.22038\nTotal prob of N-in: 0.12994\noutside 1 331\nTMhelix 332 354\ninside 355 372

26881 GCF_001687875.1_ASM168787v1 Shigella sonnei Proteobacteria; Gammaproteobacteria; Enterobacterales; Enterobacteriaceae; Shigella
MTGDNTLIHSHGINRRDFMKLCAALAATMGLSSKAAAEMAESVTNPQRPPVIWIGAQECT WP_000145410.1
MULTISPECIES: hydrogenase-2 small chain [Proteobacteria] Length: 372\nNumber of predicted TMHs: 1\nExp number of AAs in TMHs: 23.72226\nExp number, first 60 AAs: 0.22038\nTotal prob of N-in: 0.12994\noutside 1 331\nTMhelix 332 354\ninside 355 372

26882 GCF_001687925.1_ASM168792v1 Shigella sonnei Proteobacteria; Gammaproteobacteria; Enterobacterales; Enterobacteriaceae; Shigella

MTGDNTLIHSHGINRRDFMKLCAALAATMGLSSKAAAEMAESVTNPQRPPVIWIGAQECT WP_000145410.1 MULTISPECIES: hydrogenase-2 small chain [Proteobacteria] Length: 372\nNumber of predicted TMHs: 1\nExp number of AAs in TMHs: 23.72226\nExp number, first 60 AAs: 0.22038\nTotal prob of N-in: 0.12994\noutside 1 331\nTMhelix 332 354\ninside 355 372

26883 GCF_001687965.1_ASM168796v1 Shigella sonnei Proteobacteria; Gammaproteobacteria; Enterobacterales; Enterobacteriaceae; Shigella MTGDNTLIHSHGINRRDFMKLCAALAATMGLSSKAAAEMAESVTNPQRPPVIWIGAQECT WP_000145410.1 MULTISPECIES: hydrogenase-2 small chain [Proteobacteria] Length: 372\nNumber of predicted TMHs: 1\nExp number of AAs in TMHs: 23.72226\nExp number, first 60 AAs: 0.22038\nTotal prob of N-in: 0.12994\noutside 1 331\nTMhelix 332 354\ninside 355 372

26884 GCF_001687995.1_ASM168799v1 Shigella sonnei Proteobacteria; Gammaproteobacteria; Enterobacterales; Enterobacteriaceae; Shigella MTGDNTLIHSHGINRRDFMKLCAALAATMGLSSKAAAEMAESVTNPQRPPVIWIGAQECT WP_000145410.1 MULTISPECIES: hydrogenase-2 small chain [Proteobacteria] Length: 372\nNumber of predicted TMHs: 1\nExp number of AAs in TMHs: 23.72226\nExp number, first 60 AAs: 0.22038\nTotal prob of N-in: 0.12994\noutside 1 331\nTMhelix 332 354\ninside 355 372

26885 GCF_001688025.1_ASM168802v1 Shigella sonnei Proteobacteria; Gammaproteobacteria; Enterobacterales; Enterobacteriaceae; Shigella MTGDNTLIHSHGINRRDFMKLCAALAATMGLSSKAAAEMAESVTNPQRPPVIWIGAQECT WP_000145410.1 MULTISPECIES: hydrogenase-2 small chain [Proteobacteria] Length: 372\nNumber of predicted TMHs: 1\nExp number of AAs in TMHs: 23.72226\nExp number, first 60 AAs: 0.22038\nTotal prob of N-in: 0.12994\noutside 1 331\nTMhelix 332 354\ninside 355 372

26886 GCF_001688025.1_ASM168802v1 Shigella sonnei Proteobacteria; Gammaproteobacteria; Enterobacterales; Enterobacteriaceae; Shigella MNNEETFYQAMRRQGVTRRSFLKYCSLAATSLGLGAGMAPKIAWALENKPRIPVVWIHGL WP_005139779.1 hydrogenase [Shigella sonnei] Length: 372\nNumber of predicted TMHs: 2\nExp number of AAs in TMHs: 37.02628\nExp number, first 60 AAs: 19.39684\nTotal prob of N-in: 0.94313\nPOSSIBLE N-term signal sequence\ninside 1 19\nTMhelix 20 39\noutside 40 325\nTMhelix 326 348\ninside 349 372

26887 GCF_001687755.1_ASM168775v1 Shigella sonnei Proteobacteria; Gammaproteobacteria; Enterobacterales; Enterobacteriaceae; Shigella MNNEETFYQAMRRQGVTRRSFLKYCSLAATSLGLGAGMAPKIAWALENKPRIPVVWIHGL WP_005139779.1 hydrogenase [Shigella sonnei] Length: 372\nNumber of predicted TMHs: 2\nExp number of AAs in TMHs: 37.02628\nExp number, first 60 AAs: 19.39684\nTotal prob of N-in: 0.94313\nPOSSIBLE N-term signal sequence\ninside 1 19\nTMhelix 20 39\noutside 40 325\nTMhelix 326 348\ninside 349 372

26888 GCF_001687795.1_ASM168779v1 Shigella sonnei Proteobacteria; Gammaproteobacteria; Enterobacterales; Enterobacteriaceae; Shigella MNNEETFYQAMRRQGVTRRSFLKYCSLAATSLGLGAGMAPKIAWALENKPRIPVVWIHGL WP_005139779.1 hydrogenase [Shigella sonnei] Length: 372\nNumber of predicted TMHs: 2\nExp number of AAs in TMHs: 37.02628\nExp number, first 60 AAs: 19.39684\nTotal prob of N-in: 0.94313\nPOSSIBLE N-term signal sequence\ninside 1 19\nTMhelix 20 39\noutside 40 325\nTMhelix 326 348\ninside 349 372

26889 GCF_001687825.1_ASM168782v1 Shigella sonnei Proteobacteria; Gammaproteobacteria; Enterobacterales; Enterobacteriaceae; Shigella MNNEETFYQAMRRQGVTRRSFLKYCSLAATSLGLGAGMAPKIAWALENKPRIPVVWIHGL WP_005139779.1 hydrogenase [Shigella sonnei] Length: 372\nNumber of predicted TMHs: 2\nExp number of AAs in TMHs: 37.02628\nExp number, first 60 AAs: 19.39684\nTotal prob of N-in: 0.94313\nPOSSIBLE N-term signal sequence\ninside 1 19\nTMhelix 20 39\noutside 40 325\nTMhelix 326 348\ninside 349 372

26890 GCF_001688055.1_ASM168805v1 Shigella sonnei Proteobacteria; Gammaproteobacteria; Enterobacterales; Enterobacteriaceae; Shigella MTGDNTLIHSHGINRRDFMKLCAALAATMGLSSKAAAEMAESVTNPQRPPVIWIGAQECT WP_000145410.1 MULTISPECIES: hydrogenase-2 small chain [Proteobacteria] Length: 372\nNumber of predicted TMHs: 1\nExp number of AAs in TMHs: 23.72226\nExp number, first 60 AAs: 0.22038\nTotal prob of N-in: 0.12994\noutside 1 331\nTMhelix 332 354\ninside 355 372

26891 GCF_001688105.1_ASM168810v1 Shigella sonnei Proteobacteria; Gammaproteobacteria; Enterobacterales; Enterobacteriaceae; Shigella
MTGDNTLIHSHGINRRDFMKLCAALAATMGLSSKAAAEMAESVTNPQRPPVIWIGAQECT WP_000145410.1
MULTISPECIES: hydrogenase-2 small chain [Proteobacteria] Length: 372\nNumber of predicted TMHs: 1\nExp number of AAs in TMHs: 23.72226\nExp number, first 60 AAs: 0.22038\nTotal prob of N-in: 0.12994\noutside 1 331\nTMhelix 332 354\ninside 355 372

26892 GCF_001688105.1_ASM168810v1 Shigella sonnei Proteobacteria; Gammaproteobacteria; Enterobacterales; Enterobacteriaceae; Shigella
MNNEETFYQAMRRQGVTRRSFLKYCSLAATSLGLGAGMAPKIAWALENKPRIPVVWIHGL WP_005139779.1
hydrogenase [Shigella sonnei] Length: 372\nNumber of predicted TMHs: 2\nExp number of AAs in TMHs: 37.02628\nExp number, first 60 AAs: 19.39684\nTotal prob of N-in: 0.94313\nPOSSIBLE N-term signal sequence\ninside 1 19\nTMhelix 20 39\noutside 40 325\nTMhelix 326 348\ninside 349 372

26893 GCF_001688055.1_ASM168805v1 Shigella sonnei Proteobacteria; Gammaproteobacteria; Enterobacterales; Enterobacteriaceae; Shigella
MNNEETFYQAMRRQGVTRRSFLKYCSLAATSLGLGAGMAPKIAWALENKPRIPVVWIHGL WP_005139779.1
hydrogenase [Shigella sonnei] Length: 372\nNumber of predicted TMHs: 2\nExp number of AAs in TMHs: 37.02628\nExp number, first 60 AAs: 19.39684\nTotal prob of N-in: 0.94313\nPOSSIBLE N-term signal sequence\ninside 1 19\nTMhelix 20 39\noutside 40 325\nTMhelix 326 348\ninside 349 372

26894 GCF_001687925.1_ASM168792v1 Shigella sonnei Proteobacteria; Gammaproteobacteria; Enterobacterales; Enterobacteriaceae; Shigella
MNNEETFYQAMRRQGVTRRSFLKYCSLAATSLGLGAGMAPKIAWALENKPRIPVVWIHGL WP_005139779.1
hydrogenase [Shigella sonnei] Length: 372\nNumber of predicted TMHs: 2\nExp number of AAs in TMHs: 37.02628\nExp number, first 60 AAs: 19.39684\nTotal prob of N-in: 0.94313\nPOSSIBLE N-term signal sequence\ninside 1 19\nTMhelix 20 39\noutside 40 325\nTMhelix 326 348\ninside 349 372

26895 GCF_001687875.1_ASM168787v1 Shigella sonnei Proteobacteria; Gammaproteobacteria; Enterobacterales; Enterobacteriaceae; Shigella
MNNEETFYQAMRRQGVTRRSFLKYCSLAATSLGLGAGMAPKIAWALENKPRIPVVWIHGL WP_005139779.1
hydrogenase [Shigella sonnei] Length: 372\nNumber of predicted TMHs: 2\nExp number of AAs in TMHs: 37.02628\nExp number, first 60 AAs: 19.39684\nTotal prob of N-in: 0.94313\nPOSSIBLE N-term signal sequence\ninside 1 19\nTMhelix 20 39\noutside 40 325\nTMhelix 326 348\ninside 349 372

26896 GCF_001687965.1_ASM168796v1 Shigella sonnei Proteobacteria; Gammaproteobacteria; Enterobacterales; Enterobacteriaceae; Shigella
MNNEETFYQAMRRQGVTRRSFLKYCSLAATSLGLGAGMAPKIAWALENKPRIPVVWIHGL WP_005139779.1
hydrogenase [Shigella sonnei] Length: 372\nNumber of predicted TMHs: 2\nExp number of AAs in TMHs: 37.02628\nExp number, first 60 AAs: 19.39684\nTotal prob of N-in: 0.94313\nPOSSIBLE N-term signal sequence\ninside 1 19\nTMhelix 20 39\noutside 40 325\nTMhelix 326 348\ninside 349 372

26897 GCF_001688115.1_ASM168811v1 Shigella sonnei Proteobacteria; Gammaproteobacteria; Enterobacterales; Enterobacteriaceae; Shigella
MTGDNTLIHSHGINRRDFMKLCAALAATMGLSSKAAAEMAESVTNPQRPPVIWIGAQECT WP_000145410.1
MULTISPECIES: hydrogenase-2 small chain [Proteobacteria] Length: 372\nNumber of predicted TMHs: 1\nExp number of AAs in TMHs: 23.72226\nExp number, first 60 AAs: 0.22038\nTotal prob of N-in: 0.12994\noutside 1 331\nTMhelix 332 354\ninside 355 372

26898 GCF_001688145.1_ASM168814v1 Shigella sonnei Proteobacteria; Gammaproteobacteria; Enterobacterales; Enterobacteriaceae; Shigella
MTGDNTLIHSHGINRRDFMKLCAALAATMGLSSKAAAEMAESVTNPQRPPVIWIGAQECT WP_000145410.1
MULTISPECIES: hydrogenase-2 small chain [Proteobacteria] Length: 372\nNumber of predicted TMHs: 1\nExp number of AAs in TMHs: 23.72226\nExp number, first 60 AAs: 0.22038\nTotal prob of N-in: 0.12994\noutside 1 331\nTMhelix 332 354\ninside 355 372

26899 GCF_001688145.1_ASM168814v1 Shigella sonnei Proteobacteria; Gammaproteobacteria; Enterobacterales; Enterobacteriaceae; Shigella
MNNEETFYQAMRRQGVTRRSFLKYCSLAATSLGLGAGMAPKIAWALENKPRIPVVWIHGL WP_005139779.1
hydrogenase [Shigella sonnei] Length: 372\nNumber of predicted TMHs: 2\nExp number of AAs in

TMHs: 37.02628\nExp number, first 60 AAs: 19.39684\nTotal prob of N-in: 0.94313\nPOSSIBLE N-term signal sequence\ninside 1 19\nTMhelix 20 39\noutside 40 325\nTMhelix 326 348\ninside 349 372

26900 GCF_001688115.1_ASM168811v1 Shigella sonnei Proteobacteria; Gammaproteobacteria; Enterobacterales; Enterobacteriaceae; Shigella
MNNEETFYQAMRRQGVTRRSFLKYCSLAATSLGLGAGMAPKIAWALENKPRIPVVWIHGL WP_005139779.1
hydrogenase [Shigella sonnei] Length: 372\nNumber of predicted TMHs: 2\nExp number of AAs in TMHs: 37.02628\nExp number, first 60 AAs: 19.39684\nTotal prob of N-in: 0.94313\nPOSSIBLE N-term signal sequence\ninside 1 19\nTMhelix 20 39\noutside 40 325\nTMhelix 326 348\ninside 349 372

26901 GCF_001687995.1_ASM168799v1 Shigella sonnei Proteobacteria; Gammaproteobacteria; Enterobacterales; Enterobacteriaceae; Shigella
MNNEETFYQAMRRQGVTRRSFLKYCSLAATSLGLGAGMAPKIAWALENKPRIPVVWIHGL WP_005139779.1
hydrogenase [Shigella sonnei] Length: 372\nNumber of predicted TMHs: 2\nExp number of AAs in TMHs: 37.02628\nExp number, first 60 AAs: 19.39684\nTotal prob of N-in: 0.94313\nPOSSIBLE N-term signal sequence\ninside 1 19\nTMhelix 20 39\noutside 40 325\nTMhelix 326 348\ninside 349 372

26902 GCF_001688235.1_ASM168823v1 Shigella sonnei Proteobacteria; Gammaproteobacteria; Enterobacterales; Enterobacteriaceae; Shigella
MTGDNTLIHSHGINRRDFMKLCAALAATMGLSSKAAAEMAESVTNPQRPPVIWIGAQECT WP_000145410.1
MULTISPECIES: hydrogenase-2 small chain [Proteobacteria] Length: 372\nNumber of predicted TMHs: 1\nExp number of AAs in TMHs: 23.72226\nExp number, first 60 AAs: 0.22038\nTotal prob of N-in: 0.12994\noutside 1 331\nTMhelix 332 354\ninside 355 372

26903 GCF_001688195.1_ASM168819v1 Shigella sonnei Proteobacteria; Gammaproteobacteria; Enterobacterales; Enterobacteriaceae; Shigella
MTGDNTLIHSHGINRRDFMKLCAALAATMGLSSKAAAEMAESVTNPQRPPVIWIGAQECT WP_000145410.1
MULTISPECIES: hydrogenase-2 small chain [Proteobacteria] Length: 372\nNumber of predicted TMHs: 1\nExp number of AAs in TMHs: 23.72226\nExp number, first 60 AAs: 0.22038\nTotal prob of N-in: 0.12994\noutside 1 331\nTMhelix 332 354\ninside 355 372

26904 GCF_001688275.1_ASM168827v1 Shigella sonnei Proteobacteria; Gammaproteobacteria; Enterobacterales; Enterobacteriaceae; Shigella
MTGDNTLIHSHGINRRDFMKLCAALAATMGLSSKAAAEMAESVTNPQRPPVIWIGAQECT WP_000145410.1
MULTISPECIES: hydrogenase-2 small chain [Proteobacteria] Length: 372\nNumber of predicted TMHs: 1\nExp number of AAs in TMHs: 23.72226\nExp number, first 60 AAs: 0.22038\nTotal prob of N-in: 0.12994\noutside 1 331\nTMhelix 332 354\ninside 355 372

26905 GCF_001688305.1_ASM168830v1 Shigella sonnei Proteobacteria; Gammaproteobacteria; Enterobacterales; Enterobacteriaceae; Shigella
MTGDNTLIHSHGINRRDFMKLCAALAATMGLSSKAAAEMAESVTNPQRPPVIWIGAQECT WP_000145410.1
MULTISPECIES: hydrogenase-2 small chain [Proteobacteria] Length: 372\nNumber of predicted TMHs: 1\nExp number of AAs in TMHs: 23.72226\nExp number, first 60 AAs: 0.22038\nTotal prob of N-in: 0.12994\noutside 1 331\nTMhelix 332 354\ninside 355 372

26906 GCF_001688335.1_ASM168833v1 Shigella sonnei Proteobacteria; Gammaproteobacteria; Enterobacterales; Enterobacteriaceae; Shigella
MTGDNTLIHSHGINRRDFMKLCAALAATMGLSSKAAAEMAESVTNPQRPPVIWIGAQECT WP_000145410.1
MULTISPECIES: hydrogenase-2 small chain [Proteobacteria] Length: 372\nNumber of predicted TMHs: 1\nExp number of AAs in TMHs: 23.72226\nExp number, first 60 AAs: 0.22038\nTotal prob of N-in: 0.12994\noutside 1 331\nTMhelix 332 354\ninside 355 372

26907 GCF_001688385.1_ASM168838v1 Shigella sonnei Proteobacteria; Gammaproteobacteria; Enterobacterales; Enterobacteriaceae; Shigella
MTGDNTLIHSHGINRRDFMKLCAALAATMGLSSKAAAEMAESVTNPQRPPVIWIGAQECT WP_000145410.1
MULTISPECIES: hydrogenase-2 small chain [Proteobacteria] Length: 372\nNumber of predicted TMHs: 1\nExp number of AAs in TMHs: 23.72226\nExp number, first 60 AAs: 0.22038\nTotal prob of N-in: 0.12994\noutside 1 331\nTMhelix 332 354\ninside 355 372

26908 GCF_001688395.1_ASM168839v1 Shigella sonnei Proteobacteria; Gammaproteobacteria; Enterobacterales; Enterobacteriaceae; Shigella

MTGDNTLIHSHGINRRDFMKLCAALAATMGLSSKAAAEMAESVTNPQRPPVIWIGAQECT WP_000145410.1 MULTISPECIES: hydrogenase-2 small chain [Proteobacteria] Length: 372\nNumber of predicted TMHs: 1\nExp number of AAs in TMHs: 23.72226\nExp number, first 60 AAs: 0.22038\nTotal prob of N-in: 0.12994\noutside 1 331\nTMhelix 332 354\ninside 355 372

26909 GCF_001688445.1_ASM168844v1 Shigella sonnei Proteobacteria; Gammaproteobacteria; Enterobacterales; Enterobacteriaceae; Shigella MTGDNTLIHSHGINRRDFMKLCAALAATMGLSSKAAAEMAESVTNPQRPPVIWIGAQECT WP_000145410.1 MULTISPECIES: hydrogenase-2 small chain [Proteobacteria] Length: 372\nNumber of predicted TMHs: 1\nExp number of AAs in TMHs: 23.72226\nExp number, first 60 AAs: 0.22038\nTotal prob of N-in: 0.12994\noutside 1 331\nTMhelix 332 354\ninside 355 372

26910 GCF_001688445.1_ASM168844v1 Shigella sonnei Proteobacteria; Gammaproteobacteria; Enterobacterales; Enterobacteriaceae; Shigella MNNEETFYQAMRRQGVTRRSFLKYCSLAATSLGLGAGMAPKIAWALENKPRIPVVWIHGL WP_005139779.1 hydrogenase [Shigella sonnei] Length: 372\nNumber of predicted TMHs: 2\nExp number of AAs in TMHs: 37.02628\nExp number, first 60 AAs: 19.39684\nTotal prob of N-in: 0.94313\nPOSSIBLE N-term signal sequence\ninside 1 19\nTMhelix 20 39\noutside 40 325\nTMhelix 326 348\ninside 349 372

26911 GCF_001688235.1_ASM168823v1 Shigella sonnei Proteobacteria; Gammaproteobacteria; Enterobacterales; Enterobacteriaceae; Shigella MNNEETFYQAMRRQGVTRRSFLKYCSLAATSLGLGAGMAPKIAWALENKPRIPVVWIHGL WP_005139779.1 hydrogenase [Shigella sonnei] Length: 372\nNumber of predicted TMHs: 2\nExp number of AAs in TMHs: 37.02628\nExp number, first 60 AAs: 19.39684\nTotal prob of N-in: 0.94313\nPOSSIBLE N-term signal sequence\ninside 1 19\nTMhelix 20 39\noutside 40 325\nTMhelix 326 348\ninside 349 372

26912 GCF_001688195.1_ASM168819v1 Shigella sonnei Proteobacteria; Gammaproteobacteria; Enterobacterales; Enterobacteriaceae; Shigella MNNEETFYQAMRRQGVTRRSFLKYCSLAATSLGLGAGMAPKIAWALENKPRIPVVWIHGL WP_005139779.1 hydrogenase [Shigella sonnei] Length: 372\nNumber of predicted TMHs: 2\nExp number of AAs in TMHs: 37.02628\nExp number, first 60 AAs: 19.39684\nTotal prob of N-in: 0.94313\nPOSSIBLE N-term signal sequence\ninside 1 19\nTMhelix 20 39\noutside 40 325\nTMhelix 326 348\ninside 349 372

26913 GCF_001688275.1_ASM168827v1 Shigella sonnei Proteobacteria; Gammaproteobacteria; Enterobacterales; Enterobacteriaceae; Shigella MNNEETFYQAMRRQGVTRRSFLKYCSLAATSLGLGAGMAPKIAWALENKPRIPVVWIHGL WP_005139779.1 hydrogenase [Shigella sonnei] Length: 372\nNumber of predicted TMHs: 2\nExp number of AAs in TMHs: 37.02628\nExp number, first 60 AAs: 19.39684\nTotal prob of N-in: 0.94313\nPOSSIBLE N-term signal sequence\ninside 1 19\nTMhelix 20 39\noutside 40 325\nTMhelix 326 348\ninside 349 372

26914 GCF_001688485.1_ASM168848v1 Shigella sonnei Proteobacteria; Gammaproteobacteria; Enterobacterales; Enterobacteriaceae; Shigella MTGDNTLIHSHGINRRDFMKLCAALAATMGLSSKAAAEMAESVTNPQRPPVIWIGAQECT WP_000145410.1 MULTISPECIES: hydrogenase-2 small chain [Proteobacteria] Length: 372\nNumber of predicted TMHs: 1\nExp number of AAs in TMHs: 23.72226\nExp number, first 60 AAs: 0.22038\nTotal prob of N-in: 0.12994\noutside 1 331\nTMhelix 332 354\ninside 355 372

26915 GCF_001688545.1_ASM168854v1 Shigella sonnei Proteobacteria; Gammaproteobacteria; Enterobacterales; Enterobacteriaceae; Shigella MTGDNTLIHSHGINRRDFMKLCAALAATMGLSSKAAAEMAESVTNPQRPPVIWIGAQECT WP_000145410.1 MULTISPECIES: hydrogenase-2 small chain [Proteobacteria] Length: 372\nNumber of predicted TMHs: 1\nExp number of AAs in TMHs: 23.72226\nExp number, first 60 AAs: 0.22038\nTotal prob of N-in: 0.12994\noutside 1 331\nTMhelix 332 354\ninside 355 372

26916 GCF_001688485.1_ASM168848v1 Shigella sonnei Proteobacteria; Gammaproteobacteria; Enterobacterales; Enterobacteriaceae; Shigella MNNEETFYQAMRRQGVTRRSFLKYCSLAATSLGLGAGMAPKIAWALENKPRIPVVWIHGL WP_005139779.1 hydrogenase [Shigella sonnei] Length: 372\nNumber of predicted TMHs: 2\nExp number of AAs in TMHs: 37.02628\nExp number, first 60 AAs: 19.39684\nTotal prob of N-in: 0.94313\nPOSSIBLE N-term signal sequence\ninside 1 19\nTMhelix 20 39\noutside 40 325\nTMhelix 326 348\ninside 349 372

26917 GCF_001688545.1_ASM168854v1 Shigella sonnei Proteobacteria; Gammaproteobacteria; Enterobacterales; Enterobacteriaceae; Shigella
MNNEETFYQAMRRQGVTRRSFLKYCSLAATSLGLGAGMAPKIAWALENKPRIPVVWIHGL WP_005139779.1
hydrogenase [Shigella sonnei] Length: 372\nNumber of predicted TMHs: 2\nExp number of AAs in TMHs: 37.02628\nExp number, first 60 AAs: 19.39684\nTotal prob of N-in: 0.94313\nPOSSIBLE N-term signal sequence\ninside 1 19\nTMhelix 20 39\noutside 40 325\nTMhelix 326 348\ninside 349 372

26918 GCF_001688305.1_ASM168830v1 Shigella sonnei Proteobacteria; Gammaproteobacteria; Enterobacterales; Enterobacteriaceae; Shigella
MNNEETFYQAMRRQGVTRRSFLKYCSLAATSLGLGAGMAPKIAWALENKPRIPVVWIHGL WP_005139779.1
hydrogenase [Shigella sonnei] Length: 372\nNumber of predicted TMHs: 2\nExp number of AAs in TMHs: 37.02628\nExp number, first 60 AAs: 19.39684\nTotal prob of N-in: 0.94313\nPOSSIBLE N-term signal sequence\ninside 1 19\nTMhelix 20 39\noutside 40 325\nTMhelix 326 348\ninside 349 372

26919 GCF_001688335.1_ASM168833v1 Shigella sonnei Proteobacteria; Gammaproteobacteria; Enterobacterales; Enterobacteriaceae; Shigella
MNNEETFYQAMRRQGVTRRSFLKYCSLAATSLGLGAGMAPKIAWALENKPRIPVVWIHGL WP_005139779.1
hydrogenase [Shigella sonnei] Length: 372\nNumber of predicted TMHs: 2\nExp number of AAs in TMHs: 37.02628\nExp number, first 60 AAs: 19.39684\nTotal prob of N-in: 0.94313\nPOSSIBLE N-term signal sequence\ninside 1 19\nTMhelix 20 39\noutside 40 325\nTMhelix 326 348\ninside 349 372

26920 GCF_001688385.1_ASM168838v1 Shigella sonnei Proteobacteria; Gammaproteobacteria; Enterobacterales; Enterobacteriaceae; Shigella
MNNEETFYQAMRRQGVTRRSFLKYCSLAATSLGLGAGMAPKIAWALENKPRIPVVWIHGL WP_005139779.1
hydrogenase [Shigella sonnei] Length: 372\nNumber of predicted TMHs: 2\nExp number of AAs in TMHs: 37.02628\nExp number, first 60 AAs: 19.39684\nTotal prob of N-in: 0.94313\nPOSSIBLE N-term signal sequence\ninside 1 19\nTMhelix 20 39\noutside 40 325\nTMhelix 326 348\ninside 349 372

26921 GCF_001688585.1_ASM168858v1 Shigella sonnei Proteobacteria; Gammaproteobacteria; Enterobacterales; Enterobacteriaceae; Shigella
MTGDNTLIHSHGINRRDFMKLCAALAATMGLSSKAAAEMAESVTNPQRPPVIWIGAQECT WP_000145410.1
MULTISPECIES: hydrogenase-2 small chain [Proteobacteria] Length: 372\nNumber of predicted TMHs: 1\nExp number of AAs in TMHs: 23.72226\nExp number, first 60 AAs: 0.22038\nTotal prob of N-in: 0.12994\noutside 1 331\nTMhelix 332 354\ninside 355 372

26922 GCF_001688605.1_ASM168860v1 Shigella sonnei Proteobacteria; Gammaproteobacteria; Enterobacterales; Enterobacteriaceae; Shigella
MTGDNTLIHSHGINRRDFMKLCAALAATMGLSSKAAAEMAESVTNPQRPPVIWIGAQECT WP_000145410.1
MULTISPECIES: hydrogenase-2 small chain [Proteobacteria] Length: 372\nNumber of predicted TMHs: 1\nExp number of AAs in TMHs: 23.72226\nExp number, first 60 AAs: 0.22038\nTotal prob of N-in: 0.12994\noutside 1 331\nTMhelix 332 354\ninside 355 372

26923 GCF_001688605.1_ASM168860v1 Shigella sonnei Proteobacteria; Gammaproteobacteria; Enterobacterales; Enterobacteriaceae; Shigella
MNNEETFYQAMRRQGVTRRSFLKYCSLAATSLGLGAGMAPKIAWALENKPRIPVVWIHGL WP_005139779.1
hydrogenase [Shigella sonnei] Length: 372\nNumber of predicted TMHs: 2\nExp number of AAs in TMHs: 37.02628\nExp number, first 60 AAs: 19.39684\nTotal prob of N-in: 0.94313\nPOSSIBLE N-term signal sequence\ninside 1 19\nTMhelix 20 39\noutside 40 325\nTMhelix 326 348\ninside 349 372

26924 GCF_001688395.1_ASM168839v1 Shigella sonnei Proteobacteria; Gammaproteobacteria; Enterobacterales; Enterobacteriaceae; Shigella
MNNEETFYQAMRRQGVTRRSFLKYCSLAATSLGLGAGMAPKIAWALENKPRIPVVWIHGL WP_005139779.1
hydrogenase [Shigella sonnei] Length: 372\nNumber of predicted TMHs: 2\nExp number of AAs in TMHs: 37.02628\nExp number, first 60 AAs: 19.39684\nTotal prob of N-in: 0.94313\nPOSSIBLE N-term signal sequence\ninside 1 19\nTMhelix 20 39\noutside 40 325\nTMhelix 326 348\ninside 349 372

26925 GCF_001688785.1_ASM168878v1 Shigella sonnei Proteobacteria; Gammaproteobacteria; Enterobacterales; Enterobacteriaceae; Shigella
MTGDNTLIHSHGINRRDFMKLCAALAATMGLSSKAAAEMAESVTNPQRPPVIWIGAQECT WP_000145410.1
MULTISPECIES: hydrogenase-2 small chain [Proteobacteria] Length: 372\nNumber of predicted TMHs:

1\nExp number of AAs in TMHs: 23.72226\nExp number, first 60 AAs: 0.22038\nTotal prob of N-in: 0.12994\noutside 1 331\nTMhelix 332 354\ninside 355 372

26926 GCF_001688755.1_ASM168875v1 Shigella sonnei Proteobacteria; Gammaproteobacteria; Enterobacterales; Enterobacteriaceae; Shigella
MTGDNTLIHSHGINRRDFMKLCAALAATMGLSSKAAAEMAESVTNPQRPPVIWIGAQECT WP_000145410.1
MULTISPECIES: hydrogenase-2 small chain [Proteobacteria] Length: 372\nNumber of predicted TMHs: 1\nExp number of AAs in TMHs: 23.72226\nExp number, first 60 AAs: 0.22038\nTotal prob of N-in: 0.12994\noutside 1 331\nTMhelix 332 354\ninside 355 372

26927 GCF_001688865.1_ASM168886v1 Shigella sonnei Proteobacteria; Gammaproteobacteria; Enterobacterales; Enterobacteriaceae; Shigella
MTGDNTLIHSHGINRRDFMKLCAALAATMGLSSKAAAEMAESVTNPQRPPVIWIGAQECT WP_000145410.1
MULTISPECIES: hydrogenase-2 small chain [Proteobacteria] Length: 372\nNumber of predicted TMHs: 1\nExp number of AAs in TMHs: 23.72226\nExp number, first 60 AAs: 0.22038\nTotal prob of N-in: 0.12994\noutside 1 331\nTMhelix 332 354\ninside 355 372

26928 GCF_001688885.1_ASM168888v1 Shigella sonnei Proteobacteria; Gammaproteobacteria; Enterobacterales; Enterobacteriaceae; Shigella
MTGDNTLIHSHGINRRDFMKLCAALAATMGLSSKAAAEMAESVTNPQRPPVIWIGAQECT WP_000145410.1
MULTISPECIES: hydrogenase-2 small chain [Proteobacteria] Length: 372\nNumber of predicted TMHs: 1\nExp number of AAs in TMHs: 23.72226\nExp number, first 60 AAs: 0.22038\nTotal prob of N-in: 0.12994\noutside 1 331\nTMhelix 332 354\ninside 355 372

26929 GCF_001688975.1_ASM168897v1 Shigella sonnei Proteobacteria; Gammaproteobacteria; Enterobacterales; Enterobacteriaceae; Shigella
MTGDNTLIHSHGINRRDFMKLCAALAATMGLSSKAAAEMAESVTNPQRPPVIWIGAQECT WP_000145410.1
MULTISPECIES: hydrogenase-2 small chain [Proteobacteria] Length: 372\nNumber of predicted TMHs: 1\nExp number of AAs in TMHs: 23.72226\nExp number, first 60 AAs: 0.22038\nTotal prob of N-in: 0.12994\noutside 1 331\nTMhelix 332 354\ninside 355 372

26930 GCF_001688995.1_ASM168899v1 Shigella sonnei Proteobacteria; Gammaproteobacteria; Enterobacterales; Enterobacteriaceae; Shigella
MTGDNTLIHSHGINRRDFMKLCAALAATMGLSSKAAAEMAESVTNPQRPPVIWIGAQECT WP_000145410.1
MULTISPECIES: hydrogenase-2 small chain [Proteobacteria] Length: 372\nNumber of predicted TMHs: 1\nExp number of AAs in TMHs: 23.72226\nExp number, first 60 AAs: 0.22038\nTotal prob of N-in: 0.12994\noutside 1 331\nTMhelix 332 354\ninside 355 372

26931 GCF_001689015.1_ASM168901v1 Shigella sonnei Proteobacteria; Gammaproteobacteria; Enterobacterales; Enterobacteriaceae; Shigella
MTGDNTLIHSHGINRRDFMKLCAALAATMGLSSKAAAEMAESVTNPQRPPVIWIGAQECT WP_000145410.1
MULTISPECIES: hydrogenase-2 small chain [Proteobacteria] Length: 372\nNumber of predicted TMHs: 1\nExp number of AAs in TMHs: 23.72226\nExp number, first 60 AAs: 0.22038\nTotal prob of N-in: 0.12994\noutside 1 331\nTMhelix 332 354\ninside 355 372

26932 GCF_001689085.1_ASM168908v1 Shigella sonnei Proteobacteria; Gammaproteobacteria; Enterobacterales; Enterobacteriaceae; Shigella
MTGDNTLIHSHGINRRDFMKLCAALAATMGLSSKAAAEMAESVTNPQRPPVIWIGAQECT WP_000145410.1
MULTISPECIES: hydrogenase-2 small chain [Proteobacteria] Length: 372\nNumber of predicted TMHs: 1\nExp number of AAs in TMHs: 23.72226\nExp number, first 60 AAs: 0.22038\nTotal prob of N-in: 0.12994\noutside 1 331\nTMhelix 332 354\ninside 355 372

26933 GCF_001689085.1_ASM168908v1 Shigella sonnei Proteobacteria; Gammaproteobacteria; Enterobacterales; Enterobacteriaceae; Shigella
MNNEETFYQAMRRQGVTRRSFLKYCSLAATSLGLGAGMAPKIAWALENKPRIPVVWIHGL WP_005139779.1
hydrogenase [Shigella sonnei] Length: 372\nNumber of predicted TMHs: 2\nExp number of AAs in TMHs: 37.02628\nExp number, first 60 AAs: 19.39684\nTotal prob of N-in: 0.94313\nPOSSIBLE N-term signal sequence\ninside 1 19\nTMhelix 20 39\noutside 40 325\nTMhelix 326 348\ninside 349 372

26934 GCF_001688785.1_ASM168878v1 Shigella sonnei Proteobacteria; Gammaproteobacteria; Enterobacterales; Enterobacteriaceae; Shigella

MNNEETFYQAMRRQGVTRRSFLKYCSLAATSLGLGAGMAPKIAWALENKPRIPVVWIHGL WP_005139779.1 hydrogenase [Shigella sonnei] Length: 372\nNumber of predicted TMHs: 2\nExp number of AAs in TMHs: 37.02628\nExp number, first 60 AAs: 19.39684\nTotal prob of N-in: 0.94313\nPOSSIBLE N-term signal sequence\ninside 1 19\nTMhelix 20 39\noutside 40 325\nTMhelix 326 348\ninside 349 372

26935 GCF_001688755.1_ASM168875v1 Shigella sonnei Proteobacteria; Gammaproteobacteria; Enterobacterales; Enterobacteriaceae; Shigella MNNEETFYQAMRRQGVTRRSFLKYCSLAATSLGLGAGMAPKIAWALENKPRIPVVWIHGL WP_005139779.1 hydrogenase [Shigella sonnei] Length: 372\nNumber of predicted TMHs: 2\nExp number of AAs in TMHs: 37.02628\nExp number, first 60 AAs: 19.39684\nTotal prob of N-in: 0.94313\nPOSSIBLE N-term signal sequence\ninside 1 19\nTMhelix 20 39\noutside 40 325\nTMhelix 326 348\ninside 349 372

26936 GCF_001688865.1_ASM168886v1 Shigella sonnei Proteobacteria; Gammaproteobacteria; Enterobacterales; Enterobacteriaceae; Shigella MNNEETFYQAMRRQGVTRRSFLKYCSLAATSLGLGAGMAPKIAWALENKPRIPVVWIHGL WP_005139779.1 hydrogenase [Shigella sonnei] Length: 372\nNumber of predicted TMHs: 2\nExp number of AAs in TMHs: 37.02628\nExp number, first 60 AAs: 19.39684\nTotal prob of N-in: 0.94313\nPOSSIBLE N-term signal sequence\ninside 1 19\nTMhelix 20 39\noutside 40 325\nTMhelix 326 348\ninside 349 372

26937 GCF_001689105.1_ASM168910v1 Shigella sonnei Proteobacteria; Gammaproteobacteria; Enterobacterales; Enterobacteriaceae; Shigella MTGDNTLIHSHGINRRDFMKLCAALAATMGLSSKAAAEMAESVTNPQRPPVIWIGAQECT WP_000145410.1 MULTISPECIES: hydrogenase-2 small chain [Proteobacteria] Length: 372\nNumber of predicted TMHs: 1\nExp number of AAs in TMHs: 23.72226\nExp number, first 60 AAs: 0.22038\nTotal prob of N-in: 0.12994\noutside 1 331\nTMhelix 332 354\ninside 355 372

26938 GCF_001689105.1_ASM168910v1 Shigella sonnei Proteobacteria; Gammaproteobacteria; Enterobacterales; Enterobacteriaceae; Shigella MNNEETFYQAMRRQGVTRRSFLKYCSLAATSLGLGAGMAPKIAWALENKPRIPVVWIHGL WP_065555529.1 hydrogenase [Shigella sonnei] Length: 372\nNumber of predicted TMHs: 2\nExp number of AAs in TMHs: 36.85533\nExp number, first 60 AAs: 19.42236\nTotal prob of N-in: 0.94304\nPOSSIBLE N-term signal sequence\ninside 1 19\nTMhelix 20 39\noutside 40 325\nTMhelix 326 348\ninside 349 372

26939 GCF_001688885.1_ASM168888v1 Shigella sonnei Proteobacteria; Gammaproteobacteria; Enterobacterales; Enterobacteriaceae; Shigella MNNEETFYQAMRRQGVTRRSFLKYCSLAATSLGLGAGMAPKIAWALENKPRIPVVWIHGL WP_005139779.1 hydrogenase [Shigella sonnei] Length: 372\nNumber of predicted TMHs: 2\nExp number of AAs in TMHs: 37.02628\nExp number, first 60 AAs: 19.39684\nTotal prob of N-in: 0.94313\nPOSSIBLE N-term signal sequence\ninside 1 19\nTMhelix 20 39\noutside 40 325\nTMhelix 326 348\ninside 349 372

26940 GCF_001689185.1_ASM168918v1 Shigella sonnei Proteobacteria; Gammaproteobacteria; Enterobacterales; Enterobacteriaceae; Shigella MTGDNTLIHSHGINRRDFMKLCAALAATMGLSSKAAAEMAESVTNPQRPPVIWIGAQECT WP_000145410.1 MULTISPECIES: hydrogenase-2 small chain [Proteobacteria] Length: 372\nNumber of predicted TMHs: 1\nExp number of AAs in TMHs: 23.72226\nExp number, first 60 AAs: 0.22038\nTotal prob of N-in: 0.12994\noutside 1 331\nTMhelix 332 354\ninside 355 372

26941 GCF_001689185.1_ASM168918v1 Shigella sonnei Proteobacteria; Gammaproteobacteria; Enterobacterales; Enterobacteriaceae; Shigella MNNEETFYQAMRRQGVTRRSFLKYCSLAATSLGLGAGMAPKIAWALENKPRIPVVWIHGL WP_005139779.1 hydrogenase [Shigella sonnei] Length: 372\nNumber of predicted TMHs: 2\nExp number of AAs in TMHs: 37.02628\nExp number, first 60 AAs: 19.39684\nTotal prob of N-in: 0.94313\nPOSSIBLE N-term signal sequence\ninside 1 19\nTMhelix 20 39\noutside 40 325\nTMhelix 326 348\ninside 349 372

26942 GCF_001688975.1_ASM168897v1 Shigella sonnei Proteobacteria; Gammaproteobacteria; Enterobacterales; Enterobacteriaceae; Shigella MNNEETFYQAMRRQGVTRRSFLKYCSLAATSLGLGAGMAPKIAWALENKPRIPVVWIHGL WP_005139779.1 hydrogenase [Shigella sonnei] Length: 372\nNumber of predicted TMHs: 2\nExp number of AAs in TMHs: 37.02628\nExp number, first 60 AAs: 19.39684\nTotal prob of N-in: 0.94313\nPOSSIBLE N-term signal sequence\ninside 1 19\nTMhelix 20 39\noutside 40 325\nTMhelix 326 348\ninside 349 372

26943 GCF_001688995.1_ASM168899v1 Shigella sonnei Proteobacteria; Gammaproteobacteria; Enterobacterales; Enterobacteriaceae; Shigella
MNNEETFYQAMRRQGVTRRSFLKYCSLAATSLGLGAGMAPKIAWALENKPRIPVVWIHGL WP_005139779.1
hydrogenase [Shigella sonnei] Length: 372\nNumber of predicted TMHs: 2\nExp number of AAs in TMHs: 37.02628\nExp number, first 60 AAs: 19.39684\nTotal prob of N-in: 0.94313\nPOSSIBLE N-term signal sequence\ninside 1 19\nTMhelix 20 39\noutside 40 325\nTMhelix 326 348\ninside 349 372

26944 GCF_001689195.1_ASM168919v1 Shigella sonnei Proteobacteria; Gammaproteobacteria; Enterobacterales; Enterobacteriaceae; Shigella
MTGDNTLIHSHGINRRDFMKLCAALAATMGLSSKAAAEMAESVTNPQRPPVIWIGAQECT WP_000145410.1
MULTISPECIES: hydrogenase-2 small chain [Proteobacteria] Length: 372\nNumber of predicted TMHs: 1\nExp number of AAs in TMHs: 23.72226\nExp number, first 60 AAs: 0.22038\nTotal prob of N-in: 0.12994\noutside 1 331\nTMhelix 332 354\ninside 355 372

26945 GCF_001689225.1_ASM168922v1 Shigella sonnei Proteobacteria; Gammaproteobacteria; Enterobacterales; Enterobacteriaceae; Shigella
MTGDNTLIHSHGINRRDFMKLCAALAATMGLSSKAAAEMAESVTNPQRPPVIWIGAQECT WP_000145410.1
MULTISPECIES: hydrogenase-2 small chain [Proteobacteria] Length: 372\nNumber of predicted TMHs: 1\nExp number of AAs in TMHs: 23.72226\nExp number, first 60 AAs: 0.22038\nTotal prob of N-in: 0.12994\noutside 1 331\nTMhelix 332 354\ninside 355 372

26946 GCF_001689195.1_ASM168919v1 Shigella sonnei Proteobacteria; Gammaproteobacteria; Enterobacterales; Enterobacteriaceae; Shigella
MNNEETFYQAMRRQGVTRRSFLKYCSLAATSLGLGAGMAPKIAWALENKPRIPVVWIHGL WP_005139779.1
hydrogenase [Shigella sonnei] Length: 372\nNumber of predicted TMHs: 2\nExp number of AAs in TMHs: 37.02628\nExp number, first 60 AAs: 19.39684\nTotal prob of N-in: 0.94313\nPOSSIBLE N-term signal sequence\ninside 1 19\nTMhelix 20 39\noutside 40 325\nTMhelix 326 348\ninside 349 372

26947 GCF_001689225.1_ASM168922v1 Shigella sonnei Proteobacteria; Gammaproteobacteria; Enterobacterales; Enterobacteriaceae; Shigella
MNNEETFYQAMRRQGVTRRSFLKYCSLAATSLGLGAGMAPKIAWALENKPRIPVVWIHGL WP_005139779.1
hydrogenase [Shigella sonnei] Length: 372\nNumber of predicted TMHs: 2\nExp number of AAs in TMHs: 37.02628\nExp number, first 60 AAs: 19.39684\nTotal prob of N-in: 0.94313\nPOSSIBLE N-term signal sequence\ninside 1 19\nTMhelix 20 39\noutside 40 325\nTMhelix 326 348\ninside 349 372

26948 GCF_001689015.1_ASM168901v1 Shigella sonnei Proteobacteria; Gammaproteobacteria; Enterobacterales; Enterobacteriaceae; Shigella
MNNEETFYQAMRRQGVTRRSFLKYCSLAATSLGLGAGMAPKIAWALENKPRIPVVWIHGL WP_005139779.1
hydrogenase [Shigella sonnei] Length: 372\nNumber of predicted TMHs: 2\nExp number of AAs in TMHs: 37.02628\nExp number, first 60 AAs: 19.39684\nTotal prob of N-in: 0.94313\nPOSSIBLE N-term signal sequence\ninside 1 19\nTMhelix 20 39\noutside 40 325\nTMhelix 326 348\ninside 349 372

26949 GCF_001689285.1_ASM168928v1 Shigella sonnei Proteobacteria; Gammaproteobacteria; Enterobacterales; Enterobacteriaceae; Shigella
MTGDNTLIHSHGINRRDFMKLCAALAATMGLSSKAAAEMAESVTNPQRPPVIWIGAQECT WP_000145410.1
MULTISPECIES: hydrogenase-2 small chain [Proteobacteria] Length: 372\nNumber of predicted TMHs: 1\nExp number of AAs in TMHs: 23.72226\nExp number, first 60 AAs: 0.22038\nTotal prob of N-in: 0.12994\noutside 1 331\nTMhelix 332 354\ninside 355 372

26950 GCF_001689295.1_ASM168929v1 Shigella sonnei Proteobacteria; Gammaproteobacteria; Enterobacterales; Enterobacteriaceae; Shigella
MTGDNTLIHSHGINRRDFMKLCAALAATMGLSSKAAAEMAESVTNPQRPPVIWIGAQECT WP_000145410.1
MULTISPECIES: hydrogenase-2 small chain [Proteobacteria] Length: 372\nNumber of predicted TMHs: 1\nExp number of AAs in TMHs: 23.72226\nExp number, first 60 AAs: 0.22038\nTotal prob of N-in: 0.12994\noutside 1 331\nTMhelix 332 354\ninside 355 372

26951 GCF_001689345.1_ASM168934v1 Shigella sonnei Proteobacteria; Gammaproteobacteria; Enterobacterales; Enterobacteriaceae; Shigella
MTGDNTLIHSHGINRRDFMKLCAALAATMGLSSKAAAEMAESVTNPQRPPVIWIGAQECT WP_000145410.1
MULTISPECIES: hydrogenase-2 small chain [Proteobacteria] Length: 372\nNumber of predicted TMHs:

1\nExp number of AAs in TMHs: 23.72226\nExp number, first 60 AAs: 0.22038\nTotal prob of N-in: 0.12994\noutside 1 331\nTMhelix 332 354\ninside 355 372

26952 GCF_001689365.1_ASM168936v1 Shigella sonnei Proteobacteria; Gammaproteobacteria; Enterobacterales; Enterobacteriaceae; Shigella MTGDNTLIHSHGINRRDFMKLCAALAATMGLSSKAAAEMAESVTNPQRPPVIWIGAQECT WP_000145410.1 MULTISPECIES: hydrogenase-2 small chain [Proteobacteria] Length: 372\nNumber of predicted TMHs: 1\nExp number of AAs in TMHs: 23.72226\nExp number, first 60 AAs: 0.22038\nTotal prob of N-in: 0.12994\noutside 1 331\nTMhelix 332 354\ninside 355 372

26953 GCF_001933245.2_ASM193324v2 Shigella sonnei Proteobacteria; Gammaproteobacteria; Enterobacterales; Enterobacteriaceae; Shigella MTGDNTLIHSHGINRRDFMKLCAALAATMGLSSKAAAEMAESVTNPQRPPVIWIGAQECT WP_000145410.1 MULTISPECIES: hydrogenase-2 small chain [Proteobacteria] Length: 372\nNumber of predicted TMHs: 1\nExp number of AAs in TMHs: 23.72226\nExp number, first 60 AAs: 0.22038\nTotal prob of N-in: 0.12994\noutside 1 331\nTMhelix 332 354\ninside 355 372

26954 GCF_001255475.1_8290_5_7Shigella sonnei Proteobacteria; Gammaproteobacteria; Enterobacterales; Enterobacteriaceae; Shigella MNNEETFYQAMRRQGVTRRSFLKYCSLAATSLGLGAGMAPKIAWALENKPRIPVVWIHGL WP_005139779.1 hydrogenase [Shigella sonnei] Length: 372\nNumber of predicted TMHs: 2\nExp number of AAs in TMHs: 37.02628\nExp number, first 60 AAs: 19.39684\nTotal prob of N-in: 0.94313\nPOSSIBLE N-term signal sequence\ninside 1 19\nTMhelix 20 39\noutside 40 325\nTMhelix 326 348\ninside 349 372

26955 GCF_001687935.1_ASM168793v1 Shigella sonnei Proteobacteria; Gammaproteobacteria; Enterobacterales; Enterobacteriaceae; Shigella MTGDNTLIHSHGINRRDFMKLCAALAATMGLSSKAAAEMAESVTNPQRPPVIWIGAQECT WP_000145410.1 MULTISPECIES: hydrogenase-2 small chain [Proteobacteria] Length: 372\nNumber of predicted TMHs: 1\nExp number of AAs in TMHs: 23.72226\nExp number, first 60 AAs: 0.22038\nTotal prob of N-in: 0.12994\noutside 1 331\nTMhelix 332 354\ninside 355 372

26956 GCF_001687985.1_ASM168798v1 Shigella sonnei Proteobacteria; Gammaproteobacteria; Enterobacterales; Enterobacteriaceae; Shigella MTGDNTLIHSHGINRRDFMKLCAALAATMGLSSKAAAEMAESVTNPQRPPVIWIGAQECT WP_000145410.1 MULTISPECIES: hydrogenase-2 small chain [Proteobacteria] Length: 372\nNumber of predicted TMHs: 1\nExp number of AAs in TMHs: 23.72226\nExp number, first 60 AAs: 0.22038\nTotal prob of N-in: 0.12994\noutside 1 331\nTMhelix 332 354\ninside 355 372

26957 GCF_001688045.1_ASM168804v1 Shigella sonnei Proteobacteria; Gammaproteobacteria; Enterobacterales; Enterobacteriaceae; Shigella MTGDNTLIHSHGINRRDFMKLCAALAATMGLSSKAAAEMAESVTNPQRPPVIWIGAQECT WP_000145410.1 MULTISPECIES: hydrogenase-2 small chain [Proteobacteria] Length: 372\nNumber of predicted TMHs: 1\nExp number of AAs in TMHs: 23.72226\nExp number, first 60 AAs: 0.22038\nTotal prob of N-in: 0.12994\noutside 1 331\nTMhelix 332 354\ninside 355 372

26958 GCF_001688075.1_ASM168807v1 Shigella sonnei Proteobacteria; Gammaproteobacteria; Enterobacterales; Enterobacteriaceae; Shigella MTGDNTLIHSHGINRRDFMKLCAALAATMGLSSKAAAEMAESVTNPQRPPVIWIGAQECT WP_000145410.1 MULTISPECIES: hydrogenase-2 small chain [Proteobacteria] Length: 372\nNumber of predicted TMHs: 1\nExp number of AAs in TMHs: 23.72226\nExp number, first 60 AAs: 0.22038\nTotal prob of N-in: 0.12994\noutside 1 331\nTMhelix 332 354\ninside 355 372

26959 GCF_001688135.1_ASM168813v1 Shigella sonnei Proteobacteria; Gammaproteobacteria; Enterobacterales; Enterobacteriaceae; Shigella MTGDNTLIHSHGINRRDFMKLCAALAATMGLSSKAAAEMAESVTNPQRPPVIWIGAQECT WP_000145410.1 MULTISPECIES: hydrogenase-2 small chain [Proteobacteria] Length: 372\nNumber of predicted TMHs: 1\nExp number of AAs in TMHs: 23.72226\nExp number, first 60 AAs: 0.22038\nTotal prob of N-in: 0.12994\noutside 1 331\nTMhelix 332 354\ninside 355 372

26960 GCF_001688185.1_ASM168818v1 Shigella sonnei Proteobacteria; Gammaproteobacteria; Enterobacterales; Enterobacteriaceae; Shigella MTGDNTLIHSHGINRRDFMKLCAALAATMGLSSKAAAEMAESVTNPQRPPVIWIGAQECT WP_000145410.1

MULTISPECIES: hydrogenase-2 small chain [Proteobacteria] Length: 372\nNumber of predicted TMHs: 1\nExp number of AAs in TMHs: 23.72226\nExp number, first 60 AAs: 0.22038\nTotal prob of N-in: 0.12994\noutside 1 331\nTMhelix 332 354\ninside 355 372

26961 GCF_001687865.1_ASM168786v1 Shigella sonnei Proteobacteria; Gammaproteobacteria; Enterobacterales; Enterobacteriaceae; Shigella MNNEETFYQAMRRQGVTRRSFLKYCSLAATSLGLGAGMAPKIAWALENKPRIPVVWIHGL WP_005139779.1 hydrogenase [Shigella sonnei] Length: 372\nNumber of predicted TMHs: 2\nExp number of AAs in TMHs: 37.02628\nExp number, first 60 AAs: 19.39684\nTotal prob of N-in: 0.94313\nPOSSIBLE N-term signal sequence\ninside 1 19\nTMhelix 20 39\noutside 40 325\nTMhelix 326 348\ninside 349 372

26962 GCF_001688265.1_ASM168826v1 Shigella sonnei Proteobacteria; Gammaproteobacteria; Enterobacterales; Enterobacteriaceae; Shigella MTGDNTLIHSHGINRRDFMKLCAALAATMGLSSKAAAEMAESVTNPQRPPVIWIGAQECT WP_000145410.1 MULTISPECIES: hydrogenase-2 small chain [Proteobacteria] Length: 372\nNumber of predicted TMHs: 1\nExp number of AAs in TMHs: 23.72226\nExp number, first 60 AAs: 0.22038\nTotal prob of N-in: 0.12994\noutside 1 331\nTMhelix 332 354\ninside 355 372

26963 GCF_001688475.1_ASM168847v1 Shigella sonnei Proteobacteria; Gammaproteobacteria; Enterobacterales; Enterobacteriaceae; Shigella MTGDNTLIHSHGINRRDFMKLCAALAATMGLSSKAAAEMAESVTNPQRPPVIWIGAQECT WP_000145410.1 MULTISPECIES: hydrogenase-2 small chain [Proteobacteria] Length: 372\nNumber of predicted TMHs: 1\nExp number of AAs in TMHs: 23.72226\nExp number, first 60 AAs: 0.22038\nTotal prob of N-in: 0.12994\noutside 1 331\nTMhelix 332 354\ninside 355 372

26964 GCF_001688265.1_ASM168826v1 Shigella sonnei Proteobacteria; Gammaproteobacteria; Enterobacterales; Enterobacteriaceae; Shigella MNNEETFYQAMRRQGVTRRSFLKYCSLAATSLGLGAGMAPKIAWALENKPRIPVVWIHGL WP_005139779.1 hydrogenase [Shigella sonnei] Length: 372\nNumber of predicted TMHs: 2\nExp number of AAs in TMHs: 37.02628\nExp number, first 60 AAs: 19.39684\nTotal prob of N-in: 0.94313\nPOSSIBLE N-term signal sequence\ninside 1 19\nTMhelix 20 39\noutside 40 325\nTMhelix 326 348\ninside 349 372

26965 GCF_001687935.1_ASM168793v1 Shigella sonnei Proteobacteria; Gammaproteobacteria; Enterobacterales; Enterobacteriaceae; Shigella MNNEETFYQAMRRQGVTRRSFLKYCSLAATSLGLGAGMAPKIAWALENKPRIPVVWIHGL WP_005139779.1 hydrogenase [Shigella sonnei] Length: 372\nNumber of predicted TMHs: 2\nExp number of AAs in TMHs: 37.02628\nExp number, first 60 AAs: 19.39684\nTotal prob of N-in: 0.94313\nPOSSIBLE N-term signal sequence\ninside 1 19\nTMhelix 20 39\noutside 40 325\nTMhelix 326 348\ninside 349 372

26966 GCF_001687985.1_ASM168798v1 Shigella sonnei Proteobacteria; Gammaproteobacteria; Enterobacterales; Enterobacteriaceae; Shigella MNNEETFYQAMRRQGVTRRSFLKYCSLAATSLGLGAGMAPKIAWALENKPRIPVVWIHGL WP_005139779.1 hydrogenase [Shigella sonnei] Length: 372\nNumber of predicted TMHs: 2\nExp number of AAs in TMHs: 37.02628\nExp number, first 60 AAs: 19.39684\nTotal prob of N-in: 0.94313\nPOSSIBLE N-term signal sequence\ninside 1 19\nTMhelix 20 39\noutside 40 325\nTMhelix 326 348\ninside 349 372

26967 GCF_001688325.1_ASM168832v1 Shigella sonnei Proteobacteria; Gammaproteobacteria; Enterobacterales; Enterobacteriaceae; Shigella MTGDNTLIHSHGINRRDFMKLCAALAATMGLSSKAAAEMAESVTNPQRPPVIWIGAQECT WP_000145410.1 MULTISPECIES: hydrogenase-2 small chain [Proteobacteria] Length: 372\nNumber of predicted TMHs: 1\nExp number of AAs in TMHs: 23.72226\nExp number, first 60 AAs: 0.22038\nTotal prob of N-in: 0.12994\noutside 1 331\nTMhelix 332 354\ninside 355 372

26968 GCF_001688365.1_ASM168836v1 Shigella sonnei Proteobacteria; Gammaproteobacteria; Enterobacterales; Enterobacteriaceae; Shigella MTGDNTLIHSHGINRRDFMKLCAALAATMGLSSKAAAEMAESVTNPQRPPVIWIGAQECT WP_000145410.1 MULTISPECIES: hydrogenase-2 small chain [Proteobacteria] Length: 372\nNumber of predicted TMHs: 1\nExp number of AAs in TMHs: 23.72226\nExp number, first 60 AAs: 0.22038\nTotal prob of N-in: 0.12994\noutside 1 331\nTMhelix 332 354\ninside 355 372

26969 GCF_001688365.1_ASM168836v1 Shigella sonnei Proteobacteria; Gammaproteobacteria; Enterobacterales; Enterobacteriaceae; Shigella
MNNEETFYQAMRRQGVTRRSFLKYCSLAATSLGLGAGMAPKIAWALENKPRIPVVWIHGL WP_005139779.1
hydrogenase [Shigella sonnei] Length: 372\nNumber of predicted TMHs: 2\nExp number of AAs in TMHs: 37.02628\nExp number, first 60 AAs: 19.39684\nTotal prob of N-in: 0.94313\nPOSSIBLE N-term signal sequence\ninside 1 19\nTMhelix 20 39\noutside 40 325\nTMhelix 326 348\ninside 349 372

26970 GCF_001688325.1_ASM168832v1 Shigella sonnei Proteobacteria; Gammaproteobacteria; Enterobacterales; Enterobacteriaceae; Shigella
MNNEETFYQAMRRQGVTRRSFLKYCSLAATSLGLGAGMAPKIAWALENKPRIPVVWIHGL WP_005139779.1
hydrogenase [Shigella sonnei] Length: 372\nNumber of predicted TMHs: 2\nExp number of AAs in TMHs: 37.02628\nExp number, first 60 AAs: 19.39684\nTotal prob of N-in: 0.94313\nPOSSIBLE N-term signal sequence\ninside 1 19\nTMhelix 20 39\noutside 40 325\nTMhelix 326 348\ninside 349 372

26971 GCF_001688045.1_ASM168804v1 Shigella sonnei Proteobacteria; Gammaproteobacteria; Enterobacterales; Enterobacteriaceae; Shigella
MNNEETFYQAMRRQGVTRRSFLKYCSLAATSLGLGAGMAPKIAWALENKPRIPVVWIHGL WP_005139779.1
hydrogenase [Shigella sonnei] Length: 372\nNumber of predicted TMHs: 2\nExp number of AAs in TMHs: 37.02628\nExp number, first 60 AAs: 19.39684\nTotal prob of N-in: 0.94313\nPOSSIBLE N-term signal sequence\ninside 1 19\nTMhelix 20 39\noutside 40 325\nTMhelix 326 348\ninside 349 372

26972 GCF_001688185.1_ASM168818v1 Shigella sonnei Proteobacteria; Gammaproteobacteria; Enterobacterales; Enterobacteriaceae; Shigella
MNNEETFYQAMRRQGVTRRSFLKYCSLAATSLGLGAGMAPKIAWALENKPRIPVVWIHGL WP_005139779.1
hydrogenase [Shigella sonnei] Length: 372\nNumber of predicted TMHs: 2\nExp number of AAs in TMHs: 37.02628\nExp number, first 60 AAs: 19.39684\nTotal prob of N-in: 0.94313\nPOSSIBLE N-term signal sequence\ninside 1 19\nTMhelix 20 39\noutside 40 325\nTMhelix 326 348\ninside 349 372

26973 GCF_001688135.1_ASM168813v1 Shigella sonnei Proteobacteria; Gammaproteobacteria; Enterobacterales; Enterobacteriaceae; Shigella
MNNEETFYQAMRRQGVTRRSFLKYCSLAATSLGLGAGMAPKIAWALENKPRIPVVWIHGL WP_005139779.1
hydrogenase [Shigella sonnei] Length: 372\nNumber of predicted TMHs: 2\nExp number of AAs in TMHs: 37.02628\nExp number, first 60 AAs: 19.39684\nTotal prob of N-in: 0.94313\nPOSSIBLE N-term signal sequence\ninside 1 19\nTMhelix 20 39\noutside 40 325\nTMhelix 326 348\ninside 349 372

26974 GCF_001688415.1_ASM168841v1 Shigella sonnei Proteobacteria; Gammaproteobacteria; Enterobacterales; Enterobacteriaceae; Shigella
MTGDNTLIHSHGINRRDFMKLCAALAATMGLSSKAAAEMAESVTNPQRPPVIWIGAQECT WP_000145410.1
MULTISPECIES: hydrogenase-2 small chain [Proteobacteria] Length: 372\nNumber of predicted TMHs: 1\nExp number of AAs in TMHs: 23.72226\nExp number, first 60 AAs: 0.22038\nTotal prob of N-in: 0.12994\noutside 1 331\nTMhelix 332 354\ninside 355 372

26975 GCF_001688415.1_ASM168841v1 Shigella sonnei Proteobacteria; Gammaproteobacteria; Enterobacterales; Enterobacteriaceae; Shigella
MNNEETFYQAMRRQGVTRRSFLKYCSLAATSLGLGAGMAPKIAWALENKPRIPVVWIHGL WP_005139779.1
hydrogenase [Shigella sonnei] Length: 372\nNumber of predicted TMHs: 2\nExp number of AAs in TMHs: 37.02628\nExp number, first 60 AAs: 19.39684\nTotal prob of N-in: 0.94313\nPOSSIBLE N-term signal sequence\ninside 1 19\nTMhelix 20 39\noutside 40 325\nTMhelix 326 348\ninside 349 372

26976 GCF_001688075.1_ASM168807v1 Shigella sonnei Proteobacteria; Gammaproteobacteria; Enterobacterales; Enterobacteriaceae; Shigella
MNNEETFYQAMRRQGVTRRSFLKYCSLAATSLGLGAGMAPKIAWALENKPRIPVVWIHGL WP_005139779.1
hydrogenase [Shigella sonnei] Length: 372\nNumber of predicted TMHs: 2\nExp number of AAs in TMHs: 37.02628\nExp number, first 60 AAs: 19.39684\nTotal prob of N-in: 0.94313\nPOSSIBLE N-term signal sequence\ninside 1 19\nTMhelix 20 39\noutside 40 325\nTMhelix 326 348\ninside 349 372

26977 GCF_001688455.1_ASM168845v1 Shigella sonnei Proteobacteria; Gammaproteobacteria; Enterobacterales; Enterobacteriaceae; Shigella
MTGDNTLIHSHGINRRDFMKLCAALAATMGLSSKAAAEMAESVTNPQRPPVIWIGAQECT WP_000145410.1
MULTISPECIES: hydrogenase-2 small chain [Proteobacteria] Length: 372\nNumber of predicted TMHs:

1\nExp number of AAs in TMHs: 23.72226\nExp number, first 60 AAs: 0.22038\nTotal prob of N-in: 0.12994\noutside 1 331\nTMhelix 332 354\ninside 355 372

26978 GCF_001688555.1_ASM168855v1 Shigella sonnei Proteobacteria; Gammaproteobacteria; Enterobacterales; Enterobacteriaceae; Shigella
MTGDNTLIHSHGINRRDFMKLCAALAATMGLSSKAAAEMAESVTNPQRPPVIWIGAQECT WP_000145410.1
MULTISPECIES: hydrogenase-2 small chain [Proteobacteria] Length: 372\nNumber of predicted TMHs: 1\nExp number of AAs in TMHs: 23.72226\nExp number, first 60 AAs: 0.22038\nTotal prob of N-in: 0.12994\noutside 1 331\nTMhelix 332 354\ninside 355 372

26979 GCF_001688745.1_ASM168874v1 Shigella sonnei Proteobacteria; Gammaproteobacteria; Enterobacterales; Enterobacteriaceae; Shigella
MTGDNTLIHSHGINRRDFMKLCAALAATMGLSSKAAAEMAESVTNPQRPPVIWIGAQECT WP_000145410.1
MULTISPECIES: hydrogenase-2 small chain [Proteobacteria] Length: 372\nNumber of predicted TMHs: 1\nExp number of AAs in TMHs: 23.72226\nExp number, first 60 AAs: 0.22038\nTotal prob of N-in: 0.12994\noutside 1 331\nTMhelix 332 354\ninside 355 372

26980 GCF_001688775.1_ASM168877v1 Shigella sonnei Proteobacteria; Gammaproteobacteria; Enterobacterales; Enterobacteriaceae; Shigella
MTGDNTLIHSHGINRRDFMKLCAALAATMGLSSKAAAEMAESVTNPQRPPVIWIGAQECT WP_000145410.1
MULTISPECIES: hydrogenase-2 small chain [Proteobacteria] Length: 372\nNumber of predicted TMHs: 1\nExp number of AAs in TMHs: 23.72226\nExp number, first 60 AAs: 0.22038\nTotal prob of N-in: 0.12994\noutside 1 331\nTMhelix 332 354\ninside 355 372

26981 GCF_001688875.1_ASM168887v1 Shigella sonnei Proteobacteria; Gammaproteobacteria; Enterobacterales; Enterobacteriaceae; Shigella
MTGDNTLIHSHGINRRDFMKLCAALAATMGLSSKAAAEMAESVTNPQRPPVIWIGAQECT WP_000145410.1
MULTISPECIES: hydrogenase-2 small chain [Proteobacteria] Length: 372\nNumber of predicted TMHs: 1\nExp number of AAs in TMHs: 23.72226\nExp number, first 60 AAs: 0.22038\nTotal prob of N-in: 0.12994\noutside 1 331\nTMhelix 332 354\ninside 355 372

26982 GCF_001689005.1_ASM168900v1 Shigella sonnei Proteobacteria; Gammaproteobacteria; Enterobacterales; Enterobacteriaceae; Shigella
MTGDNTLIHSHGINRRDFMKLCAALAATMGLSSKAAAEMAESVTNPQRPPVIWIGAQECT WP_000145410.1
MULTISPECIES: hydrogenase-2 small chain [Proteobacteria] Length: 372\nNumber of predicted TMHs: 1\nExp number of AAs in TMHs: 23.72226\nExp number, first 60 AAs: 0.22038\nTotal prob of N-in: 0.12994\noutside 1 331\nTMhelix 332 354\ninside 355 372

26983 GCF_001688895.1_ASM168889v1 Shigella sonnei Proteobacteria; Gammaproteobacteria; Enterobacterales; Enterobacteriaceae; Shigella
MTGDNTLIHSHGINRRDFMKLCAALAATMGLSSKAAAEMAESVTNPQRPPVIWIGAQECT WP_000145410.1
MULTISPECIES: hydrogenase-2 small chain [Proteobacteria] Length: 372\nNumber of predicted TMHs: 1\nExp number of AAs in TMHs: 23.72226\nExp number, first 60 AAs: 0.22038\nTotal prob of N-in: 0.12994\noutside 1 331\nTMhelix 332 354\ninside 355 372

26984 GCF_001688895.1_ASM168889v1 Shigella sonnei Proteobacteria; Gammaproteobacteria; Enterobacterales; Enterobacteriaceae; Shigella
MNNEETFYQAMRRQGVTRRSFLKYCSLAATSLGLGAGMAPKIAWALENKPRIPVVWIHGL WP_005139779.1
hydrogenase [Shigella sonnei] Length: 372\nNumber of predicted TMHs: 2\nExp number of AAs in TMHs: 37.02628\nExp number, first 60 AAs: 19.39684\nTotal prob of N-in: 0.94313\nPOSSIBLE N-term signal sequence\ninside 1 19\nTMhelix 20 39\noutside 40 325\nTMhelix 326 348\ninside 349 372

26985 GCF_001689215.1_ASM168921v1 Shigella sonnei Proteobacteria; Gammaproteobacteria; Enterobacterales; Enterobacteriaceae; Shigella
MTGDNTLIHSHGINRRDFMKLCAALAATMGLSSKAAAEMAESVTNPQRPPVIWIGAQECT WP_000145410.1
MULTISPECIES: hydrogenase-2 small chain [Proteobacteria] Length: 372\nNumber of predicted TMHs: 1\nExp number of AAs in TMHs: 23.72226\nExp number, first 60 AAs: 0.22038\nTotal prob of N-in: 0.12994\noutside 1 331\nTMhelix 332 354\ninside 355 372

26986 GCF_001688455.1_ASM168845v1 Shigella sonnei Proteobacteria; Gammaproteobacteria; Enterobacterales; Enterobacteriaceae; Shigella

MNNEETFYQAMRRQGVTRRSFLKYCSLAATSLGLGAGMAPKIAWALENKPRIPVVWIHGL	WP_005139779.1	hydrogenase [Shigella sonnei]	Length: 372\nNumber of predicted TMHs:  2\nExp number of AAs in TMHs: 37.02628\nExp number, first 60 AAs:  19.39684\nTotal prob of N-in:   0.94313\nPOSSIBLE N-term signal sequence\ninside   1   19\nTMhelix   20   39\noutside   40   325\nTMhelix   326   348\ninside   349   372

26987	GCF_001688475.1_ASM168847v1	Shigella sonnei	Proteobacteria; Gammaproteobacteria; Enterobacterales; Enterobacteriaceae; Shigella	MNNEETFYQAMRRQGVTRRSFLKYCSLAATSLGLGAGMAPKIAWALENKPRIPVVWIHGL	WP_005139779.1	hydrogenase [Shigella sonnei]	Length: 372\nNumber of predicted TMHs:  2\nExp number of AAs in TMHs: 37.02628\nExp number, first 60 AAs:  19.39684\nTotal prob of N-in:   0.94313\nPOSSIBLE N-term signal sequence\ninside   1   19\nTMhelix   20   39\noutside   40   325\nTMhelix   326   348\ninside   349   372

26988	GCF_001689095.1_ASM168909v1	Shigella sonnei	Proteobacteria; Gammaproteobacteria; Enterobacterales; Enterobacteriaceae; Shigella	MTGDNTLIHSHGINRRDFMKLCAALAATMGLSSKAAAEMAESVTNPQRPPVIWIGAQECT	WP_000145410.1	MULTISPECIES: hydrogenase-2 small chain [Proteobacteria]	Length: 372\nNumber of predicted TMHs:  1\nExp number of AAs in TMHs: 23.72226\nExp number, first 60 AAs:  0.22038\nTotal prob of N-in:   0.12994\noutside   1   331\nTMhelix   332   354\ninside   355   372

26989	GCF_001689095.1_ASM168909v1	Shigella sonnei	Proteobacteria; Gammaproteobacteria; Enterobacterales; Enterobacteriaceae; Shigella	MNNEETFYQAMRRQGVTRRSFLKYCSLAATSLGLGAGMAPKIAWALENKPRIPVVWIHGL	WP_005139779.1	hydrogenase [Shigella sonnei]	Length: 372\nNumber of predicted TMHs:  2\nExp number of AAs in TMHs: 37.02628\nExp number, first 60 AAs:  19.39684\nTotal prob of N-in:   0.94313\nPOSSIBLE N-term signal sequence\ninside   1   19\nTMhelix   20   39\noutside   40   325\nTMhelix   326   348\ninside   349   372

26990	GCF_001688745.1_ASM168874v1	Shigella sonnei	Proteobacteria; Gammaproteobacteria; Enterobacterales; Enterobacteriaceae; Shigella	MNNEETFYQAMRRQGVTRRSFLKYCSLAATSLGLGAGMAPKIAWALENKPRIPVVWIHGL	WP_005139779.1	hydrogenase [Shigella sonnei]	Length: 372\nNumber of predicted TMHs:  2\nExp number of AAs in TMHs: 37.02628\nExp number, first 60 AAs:  19.39684\nTotal prob of N-in:   0.94313\nPOSSIBLE N-term signal sequence\ninside   1   19\nTMhelix   20   39\noutside   40   325\nTMhelix   326   348\ninside   349   372

26991	GCF_001688775.1_ASM168877v1	Shigella sonnei	Proteobacteria; Gammaproteobacteria; Enterobacterales; Enterobacteriaceae; Shigella	MNNEETFYQAMRRQGVTRRSFLKYCSLAATSLGLGAGMAPKIAWALENKPRIPVVWIHGL	WP_005139779.1	hydrogenase [Shigella sonnei]	Length: 372\nNumber of predicted TMHs:  2\nExp number of AAs in TMHs: 37.02628\nExp number, first 60 AAs:  19.39684\nTotal prob of N-in:   0.94313\nPOSSIBLE N-term signal sequence\ninside   1   19\nTMhelix   20   39\noutside   40   325\nTMhelix   326   348\ninside   349   372

26992	GCF_001689145.1_ASM168914v1	Shigella sonnei	Proteobacteria; Gammaproteobacteria; Enterobacterales; Enterobacteriaceae; Shigella	MTGDNTLIHSHGINRRDFMKLCAALAATMGLSSKAAAEMAESVTNPQRPPVIWIGAQECT	WP_000145410.1	MULTISPECIES: hydrogenase-2 small chain [Proteobacteria]	Length: 372\nNumber of predicted TMHs:  1\nExp number of AAs in TMHs: 23.72226\nExp number, first 60 AAs:  0.22038\nTotal prob of N-in:   0.12994\noutside   1   331\nTMhelix   332   354\ninside   355   372

26993	GCF_001689145.1_ASM168914v1	Shigella sonnei	Proteobacteria; Gammaproteobacteria; Enterobacterales; Enterobacteriaceae; Shigella	MNNEETFYQAMRRQGVTRRSFLKYCSLAATSLGLGAGMAPKIAWALENKPRIPVVWIHGL	WP_005139779.1	hydrogenase [Shigella sonnei]	Length: 372\nNumber of predicted TMHs:  2\nExp number of AAs in TMHs: 37.02628\nExp number, first 60 AAs:  19.39684\nTotal prob of N-in:   0.94313\nPOSSIBLE N-term signal sequence\ninside   1   19\nTMhelix   20   39\noutside   40   325\nTMhelix   326   348\ninside   349   372

26994	GCF_001689215.1_ASM168921v1	Shigella sonnei	Proteobacteria; Gammaproteobacteria; Enterobacterales; Enterobacteriaceae; Shigella	MNNEETFYQAMRRQGVTRRSFLKYCSLAATSLGLGAGMAPKIAWALENKPRIPVVWIHGL	WP_005139779.1	hydrogenase [Shigella sonnei]	Length: 372\nNumber of predicted TMHs:  2\nExp number of AAs in TMHs: 37.02628\nExp number, first 60 AAs:  19.39684\nTotal prob of N-in:   0.94313\nPOSSIBLE N-term signal sequence\ninside   1   19\nTMhelix   20   39\noutside   40   325\nTMhelix   326   348\ninside   349   372

26995 GCF_001688875.1_ASM168887v1 Shigella sonnei Proteobacteria; Gammaproteobacteria; Enterobacterales; Enterobacteriaceae; Shigella
MNNEETFYQAMRRQGVTRRSFLKYCSLAATSLGLGAGMAPKIAWALENKPRIPVVWIHGL WP_005139779.1
hydrogenase [Shigella sonnei] Length: 372\nNumber of predicted TMHs: 2\nExp number of AAs in TMHs: 37.02628\nExp number, first 60 AAs: 19.39684\nTotal prob of N-in: 0.94313\nPOSSIBLE N-term signal sequence\ninside 1 19\nTMhelix 20 39\noutside 40 325\nTMhelix 326 348\ninside 349 372

26996 GCF_001689265.1_ASM168926v1 Shigella sonnei Proteobacteria; Gammaproteobacteria; Enterobacterales; Enterobacteriaceae; Shigella
MTGDNTLIHSHGINRRDFMKLCAALAATMGLSSKAAAEMAESVTNPQRPPVIWIGAQECT WP_000145410.1
MULTISPECIES: hydrogenase-2 small chain [Proteobacteria] Length: 372\nNumber of predicted TMHs: 1\nExp number of AAs in TMHs: 23.72226\nExp number, first 60 AAs: 0.22038\nTotal prob of N-in: 0.12994\noutside 1 331\nTMhelix 332 354\ninside 355 372

26997 GCF_001689325.1_ASM168932v1 Shigella sonnei Proteobacteria; Gammaproteobacteria; Enterobacterales; Enterobacteriaceae; Shigella
MTGDNTLIHSHGINRRDFMKLCAALAATMGLSSKAAAEMAESVTNPQRPPVIWIGAQECT WP_000145410.1
MULTISPECIES: hydrogenase-2 small chain [Proteobacteria] Length: 372\nNumber of predicted TMHs: 1\nExp number of AAs in TMHs: 23.72226\nExp number, first 60 AAs: 0.22038\nTotal prob of N-in: 0.12994\noutside 1 331\nTMhelix 332 354\ninside 355 372

26998 GCF_001689325.1_ASM168932v1 Shigella sonnei Proteobacteria; Gammaproteobacteria; Enterobacterales; Enterobacteriaceae; Shigella
MNNEETFYQAMRRQGVTRRSFLKYCSLAATSLGLGAGMAPKIAWALENKPRIPVVWIHGL WP_005139779.1
hydrogenase [Shigella sonnei] Length: 372\nNumber of predicted TMHs: 2\nExp number of AAs in TMHs: 37.02628\nExp number, first 60 AAs: 19.39684\nTotal prob of N-in: 0.94313\nPOSSIBLE N-term signal sequence\ninside 1 19\nTMhelix 20 39\noutside 40 325\nTMhelix 326 348\ninside 349 372

26999 GCF_001689265.1_ASM168926v1 Shigella sonnei Proteobacteria; Gammaproteobacteria; Enterobacterales; Enterobacteriaceae; Shigella
MNNEETFYQAMRRQGVTRRSFLKYCSLAATSLGLGAGMAPKIAWALENKPRIPVVWIHGL WP_005139779.1
hydrogenase [Shigella sonnei] Length: 372\nNumber of predicted TMHs: 2\nExp number of AAs in TMHs: 37.02628\nExp number, first 60 AAs: 19.39684\nTotal prob of N-in: 0.94313\nPOSSIBLE N-term signal sequence\ninside 1 19\nTMhelix 20 39\noutside 40 325\nTMhelix 326 348\ninside 349 372

27000 GCF_001689005.1_ASM168900v1 Shigella sonnei Proteobacteria; Gammaproteobacteria; Enterobacterales; Enterobacteriaceae; Shigella
MNNEETFYQAMRRQGVTRRSFLKYCSLAATSLGLGAGMAPKIAWALENKPRIPVVWIHGL WP_005139779.1
hydrogenase [Shigella sonnei] Length: 372\nNumber of predicted TMHs: 2\nExp number of AAs in TMHs: 37.02628\nExp number, first 60 AAs: 19.39684\nTotal prob of N-in: 0.94313\nPOSSIBLE N-term signal sequence\ninside 1 19\nTMhelix 20 39\noutside 40 325\nTMhelix 326 348\ninside 349 372

27001 GCF_001933295.2_ASM193329v2 Shigella sonnei Proteobacteria; Gammaproteobacteria; Enterobacterales; Enterobacteriaceae; Shigella
MTGDNTLIHSHGINRRDFMKLCAALAATMGLSSKAAAEMAESVTNPQRPPVIWIGAQECT WP_000145410.1
MULTISPECIES: hydrogenase-2 small chain [Proteobacteria] Length: 372\nNumber of predicted TMHs: 1\nExp number of AAs in TMHs: 23.72226\nExp number, first 60 AAs: 0.22038\nTotal prob of N-in: 0.12994\noutside 1 331\nTMhelix 332 354\ninside 355 372

27002 GCF_001689355.1_ASM168935v1 Shigella sonnei Proteobacteria; Gammaproteobacteria; Enterobacterales; Enterobacteriaceae; Shigella
MTGDNTLIHSHGINRRDFMKLCAALAATMGLSSKAAAEMAESVTNPQRPPVIWIGAQECT WP_000145410.1
MULTISPECIES: hydrogenase-2 small chain [Proteobacteria] Length: 372\nNumber of predicted TMHs: 1\nExp number of AAs in TMHs: 23.72226\nExp number, first 60 AAs: 0.22038\nTotal prob of N-in: 0.12994\noutside 1 331\nTMhelix 332 354\ninside 355 372

27003 GCF_900103855.1_IMG-taxon_2654588181_annotated_assembly Shigella sonnei Proteobacteria; Gammaproteobacteria; Enterobacterales; Enterobacteriaceae; Shigella
MTGDNTLIHSHGINRRDFMKLCAALAATMGLSSKAAAEMAESVTNPQRPPVIWIGAQECT WP_000145410.1
MULTISPECIES: hydrogenase-2 small chain [Proteobacteria] Length: 372\nNumber of predicted TMHs:

1\nExp number of AAs in TMHs: 23.72226\nExp number, first 60 AAs: 0.22038\nTotal prob of N-in: 0.12994\noutside 1 331\nTMhelix 332 354\ninside 355 372

27004 GCF_900157495.1_9789_6_60 Shigella sonnei Proteobacteria; Gammaproteobacteria; Enterobacterales; Enterobacteriaceae; Shigella
MTGDNTLIHSHGINRRDFMKLCAALAATMGLSSKAAAEMAESVTNPQRPPVIWIGAQECT WP_000145410.1
MULTISPECIES: hydrogenase-2 small chain [Proteobacteria] Length: 372\nNumber of predicted TMHs: 1\nExp number of AAs in TMHs: 23.72226\nExp number, first 60 AAs: 0.22038\nTotal prob of N-in: 0.12994\noutside 1 331\nTMhelix 332 354\ninside 355 372

27005 GCF_900157505.1_9789_6_74 Shigella sonnei Proteobacteria; Gammaproteobacteria; Enterobacterales; Enterobacteriaceae; Shigella
MTGDNTLIHSHGINRRDFMKLCAALAATMGLSSKAAAEMAESVTNPQRPPVIWIGAQECT WP_000145410.1
MULTISPECIES: hydrogenase-2 small chain [Proteobacteria] Length: 372\nNumber of predicted TMHs: 1\nExp number of AAs in TMHs: 23.72226\nExp number, first 60 AAs: 0.22038\nTotal prob of N-in: 0.12994\noutside 1 331\nTMhelix 332 354\ninside 355 372

27006 GCF_900157535.1_9789_6_20 Shigella sonnei Proteobacteria; Gammaproteobacteria; Enterobacterales; Enterobacteriaceae; Shigella
MTGDNTLIHSHGINRRDFMKLCAALAATMGLSSKAAAEMAESVTNPQRPPVIWIGAQECT WP_000145410.1
MULTISPECIES: hydrogenase-2 small chain [Proteobacteria] Length: 372\nNumber of predicted TMHs: 1\nExp number of AAs in TMHs: 23.72226\nExp number, first 60 AAs: 0.22038\nTotal prob of N-in: 0.12994\noutside 1 331\nTMhelix 332 354\ninside 355 372

27007 GCF_900157555.1_9789_6_58 Shigella sonnei Proteobacteria; Gammaproteobacteria; Enterobacterales; Enterobacteriaceae; Shigella
MTGDNTLIHSHGINRRDFMKLCAALAATMGLSSKAAAEMAESVTNPQRPPVIWIGAQECT WP_000145410.1
MULTISPECIES: hydrogenase-2 small chain [Proteobacteria] Length: 372\nNumber of predicted TMHs: 1\nExp number of AAs in TMHs: 23.72226\nExp number, first 60 AAs: 0.22038\nTotal prob of N-in: 0.12994\noutside 1 331\nTMhelix 332 354\ninside 355 372

27008 GCF_001689355.1_ASM168935v1 Shigella sonnei Proteobacteria; Gammaproteobacteria; Enterobacterales; Enterobacteriaceae; Shigella
MNNEETFYQAMRRQGVTRRSFLKYCSLAATSLGLGAGMAPKIAWALENKPRIPVVWIHGL WP_005139779.1
hydrogenase [Shigella sonnei] Length: 372\nNumber of predicted TMHs: 2\nExp number of AAs in TMHs: 37.02628\nExp number, first 60 AAs: 19.39684\nTotal prob of N-in: 0.94313\nPOSSIBLE N-term signal sequence\ninside 1 19\nTMhelix 20 39\noutside 40 325\nTMhelix 326 348\ninside 349 372

27009 GCF_900103855.1_IMG-taxon_2654588181_annotated_assembly Shigella sonnei Proteobacteria; Gammaproteobacteria; Enterobacterales; Enterobacteriaceae; Shigella
MNNEETFYQAMRRQGVTRRSFLKYCSLAATSLGLGAGMAPKIAWALENKPRIPVVWIHGL WP_001058323.1
MULTISPECIES: hydrogenase-1 small chain [Proteobacteria] Length: 372\nNumber of predicted TMHs: 2\nExp number of AAs in TMHs: 37.2608499999999\nExp number, first 60 AAs: 19.52923\nTotal prob of N-in: 0.94351\nPOSSIBLE N-term signal sequence\ninside 1 19\nTMhelix 20 39\noutside 40 325\nTMhelix 326 348\ninside 349 372

27010 GCF_001933295.2_ASM193329v2 Shigella sonnei Proteobacteria; Gammaproteobacteria; Enterobacterales; Enterobacteriaceae; Shigella
MNNEETFYQAMRRQGVTRRSFLKYCSLAATSLGLGAGMAPKIAWALENKPRIPVVWIHGL WP_005139779.1
hydrogenase [Shigella sonnei] Length: 372\nNumber of predicted TMHs: 2\nExp number of AAs in TMHs: 37.02628\nExp number, first 60 AAs: 19.39684\nTotal prob of N-in: 0.94313\nPOSSIBLE N-term signal sequence\ninside 1 19\nTMhelix 20 39\noutside 40 325\nTMhelix 326 348\ninside 349 372

27011 GCF_900157585.1_9789_6_65 Shigella sonnei Proteobacteria; Gammaproteobacteria; Enterobacterales; Enterobacteriaceae; Shigella
MTGDNTLIHSHGINRRDFMKLCAALAATMGLSSKAAAEMAESVTNPQRPPVIWIGAQECT WP_000145410.1
MULTISPECIES: hydrogenase-2 small chain [Proteobacteria] Length: 372\nNumber of predicted TMHs: 1\nExp number of AAs in TMHs: 23.72226\nExp number, first 60 AAs: 0.22038\nTotal prob of N-in: 0.12994\noutside 1 331\nTMhelix 332 354\ninside 355 372

27012 GCF_900157615.1_9789_6_29 Shigella sonnei Proteobacteria; Gammaproteobacteria; Enterobacterales; Enterobacteriaceae; Shigella
MTGDNTLIHSHGINRRDFMKLCAALAATMGLSSKAAAEMAESVTNPQRPPVIWIGAQECT WP_000145410.1
MULTISPECIES: hydrogenase-2 small chain [Proteobacteria] Length: 372\nNumber of predicted TMHs: 1\nExp number of AAs in TMHs: 23.72226\nExp number, first 60 AAs: 0.22038\nTotal prob of N-in: 0.12994\noutside 1 331\nTMhelix 332 354\ninside 355 372

27013 GCF_900157585.1_9789_6_65 Shigella sonnei Proteobacteria; Gammaproteobacteria; Enterobacterales; Enterobacteriaceae; Shigella
MNNEETFYQAMRRQGVTRRSFLKYCSLAATSLGLGAGMAPKIAWALENKPRIPVVWIHGL WP_005139779.1
hydrogenase [Shigella sonnei] Length: 372\nNumber of predicted TMHs: 2\nExp number of AAs in TMHs: 37.02628\nExp number, first 60 AAs: 19.39684\nTotal prob of N-in: 0.94313\nPOSSIBLE N-term signal sequence\ninside 1 19\nTMhelix 20 39\noutside 40 325\nTMhelix 326 348\ninside 349 372

27014 GCF_900157615.1_9789_6_29 Shigella sonnei Proteobacteria; Gammaproteobacteria; Enterobacterales; Enterobacteriaceae; Shigella
MNNEETFYQAMRRQGVTRRSFLKYCSLAATSLGLGAGMAPKIAWALENKPRIPVVWIHGL WP_005139779.1
hydrogenase [Shigella sonnei] Length: 372\nNumber of predicted TMHs: 2\nExp number of AAs in TMHs: 37.02628\nExp number, first 60 AAs: 19.39684\nTotal prob of N-in: 0.94313\nPOSSIBLE N-term signal sequence\ninside 1 19\nTMhelix 20 39\noutside 40 325\nTMhelix 326 348\ninside 349 372

27015 GCF_900157495.1_9789_6_60 Shigella sonnei Proteobacteria; Gammaproteobacteria; Enterobacterales; Enterobacteriaceae; Shigella
MNNEETFYQAMRRQGVTRRSFLKYCSLAATSLGLGAGMAPKIAWALENKPRIPVVWIHGL WP_005139779.1
hydrogenase [Shigella sonnei] Length: 372\nNumber of predicted TMHs: 2\nExp number of AAs in TMHs: 37.02628\nExp number, first 60 AAs: 19.39684\nTotal prob of N-in: 0.94313\nPOSSIBLE N-term signal sequence\ninside 1 19\nTMhelix 20 39\noutside 40 325\nTMhelix 326 348\ninside 349 372

27016 GCF_900157635.1_9789_6_62 Shigella sonnei Proteobacteria; Gammaproteobacteria; Enterobacterales; Enterobacteriaceae; Shigella
MTGDNTLIHSHGINRRDFMKLCAALAATMGLSSKAAAEMAESVTNPQRPPVIWIGAQECT WP_000145410.1
MULTISPECIES: hydrogenase-2 small chain [Proteobacteria] Length: 372\nNumber of predicted TMHs: 1\nExp number of AAs in TMHs: 23.72226\nExp number, first 60 AAs: 0.22038\nTotal prob of N-in: 0.12994\noutside 1 331\nTMhelix 332 354\ninside 355 372

27017 GCF_900157635.1_9789_6_62 Shigella sonnei Proteobacteria; Gammaproteobacteria; Enterobacterales; Enterobacteriaceae; Shigella
MNNEETFYQAMRRQGVTRRSFLKYCSLAATSLGLGAGMAPKIAWALENKPRIPVVWIHGL WP_005139779.1
hydrogenase [Shigella sonnei] Length: 372\nNumber of predicted TMHs: 2\nExp number of AAs in TMHs: 37.02628\nExp number, first 60 AAs: 19.39684\nTotal prob of N-in: 0.94313\nPOSSIBLE N-term signal sequence\ninside 1 19\nTMhelix 20 39\noutside 40 325\nTMhelix 326 348\ninside 349 372

27018 GCF_900157505.1_9789_6_74 Shigella sonnei Proteobacteria; Gammaproteobacteria; Enterobacterales; Enterobacteriaceae; Shigella
MNNEETFYQAMRRQGVTRRSFLKYCSLAATSLGLGAGMAPKIAWALENKPRIPVVWIHGL WP_005139779.1
hydrogenase [Shigella sonnei] Length: 372\nNumber of predicted TMHs: 2\nExp number of AAs in TMHs: 37.02628\nExp number, first 60 AAs: 19.39684\nTotal prob of N-in: 0.94313\nPOSSIBLE N-term signal sequence\ninside 1 19\nTMhelix 20 39\noutside 40 325\nTMhelix 326 348\ninside 349 372

27019 GCF_900157665.1_9789_6_55 Shigella sonnei Proteobacteria; Gammaproteobacteria; Enterobacterales; Enterobacteriaceae; Shigella
MTGDNTLIHSHGINRRDFMKLCAALAATMGLSSKAAAEMAESVTNPQRPPVIWIGAQECT WP_000145410.1
MULTISPECIES: hydrogenase-2 small chain [Proteobacteria] Length: 372\nNumber of predicted TMHs: 1\nExp number of AAs in TMHs: 23.72226\nExp number, first 60 AAs: 0.22038\nTotal prob of N-in: 0.12994\noutside 1 331\nTMhelix 332 354\ninside 355 372

27020 GCF_900157695.1_9789_6_84 Shigella sonnei Proteobacteria; Gammaproteobacteria; Enterobacterales; Enterobacteriaceae; Shigella
MTGDNTLIHSHGINRRDFMKLCAALAATMGLSSKAAAEMAESVTNPQRPPVIWIGAQECT WP_000145410.1
MULTISPECIES: hydrogenase-2 small chain [Proteobacteria] Length: 372\nNumber of predicted TMHs:

1\nExp number of AAs in TMHs: 23.72226\nExp number, first 60 AAs: 0.22038\nTotal prob of N-in: 0.12994\noutside 1 331\nTMhelix 332 354\ninside 355 372

27021 GCF_900157715.1_9789_6_49 Shigella sonnei Proteobacteria; Gammaproteobacteria; Enterobacterales; Enterobacteriaceae; Shigella
MTGDNTLIHSHGINRRDFMKLCAALAATMGLSSKAAAEMAESVTNPQRPPVIWIGAQECT WP_000145410.1
MULTISPECIES: hydrogenase-2 small chain [Proteobacteria] Length: 372\nNumber of predicted TMHs: 1\nExp number of AAs in TMHs: 23.72226\nExp number, first 60 AAs: 0.22038\nTotal prob of N-in: 0.12994\noutside 1 331\nTMhelix 332 354\ninside 355 372

27022 GCF_900157665.1_9789_6_55 Shigella sonnei Proteobacteria; Gammaproteobacteria; Enterobacterales; Enterobacteriaceae; Shigella
MNNEETFYQAMRRQGVTRRSFLKYCSLAATSLGLGAGMAPKIAWALENKPRIPVVWIHGL WP_005139779.1
hydrogenase [Shigella sonnei] Length: 372\nNumber of predicted TMHs: 2\nExp number of AAs in TMHs: 37.02628\nExp number, first 60 AAs: 19.39684\nTotal prob of N-in: 0.94313\nPOSSIBLE N-term signal sequence\ninside 1 19\nTMhelix 20 39\noutside 40 325\nTMhelix 326 348\ninside 349 372

27023 GCF_900157695.1_9789_6_84 Shigella sonnei Proteobacteria; Gammaproteobacteria; Enterobacterales; Enterobacteriaceae; Shigella
MNNEETFYQAMRRQGVTRRSFLKYCSLAATSLGLGAGMAPKIAWALENKPRIPVVWIHGL WP_005139779.1
hydrogenase [Shigella sonnei] Length: 372\nNumber of predicted TMHs: 2\nExp number of AAs in TMHs: 37.02628\nExp number, first 60 AAs: 19.39684\nTotal prob of N-in: 0.94313\nPOSSIBLE N-term signal sequence\ninside 1 19\nTMhelix 20 39\noutside 40 325\nTMhelix 326 348\ninside 349 372

27024 GCF_900157715.1_9789_6_49 Shigella sonnei Proteobacteria; Gammaproteobacteria; Enterobacterales; Enterobacteriaceae; Shigella
MNNEETFYQAMRRQGVTRRSFLKYCSLAATSLGLGAGMAPKIAWALENKPRIPVVWIHGL WP_005139779.1
hydrogenase [Shigella sonnei] Length: 372\nNumber of predicted TMHs: 2\nExp number of AAs in TMHs: 37.02628\nExp number, first 60 AAs: 19.39684\nTotal prob of N-in: 0.94313\nPOSSIBLE N-term signal sequence\ninside 1 19\nTMhelix 20 39\noutside 40 325\nTMhelix 326 348\ninside 349 372

27025 GCF_900157535.1_9789_6_20 Shigella sonnei Proteobacteria; Gammaproteobacteria; Enterobacterales; Enterobacteriaceae; Shigella
MNNEETFYQAMRRQGVTRRSFLKYCSLAATSLGLGAGMAPKIAWALENKPRIPVVWIHGL WP_005139779.1
hydrogenase [Shigella sonnei] Length: 372\nNumber of predicted TMHs: 2\nExp number of AAs in TMHs: 37.02628\nExp number, first 60 AAs: 19.39684\nTotal prob of N-in: 0.94313\nPOSSIBLE N-term signal sequence\ninside 1 19\nTMhelix 20 39\noutside 40 325\nTMhelix 326 348\ninside 349 372

27026 GCF_900157555.1_9789_6_58 Shigella sonnei Proteobacteria; Gammaproteobacteria; Enterobacterales; Enterobacteriaceae; Shigella
MNNEETFYQAMRRQGVTRRSFLKYCSLAATSLGLGAGMAPKIAWALENKPRIPVVWIHGL WP_005139779.1
hydrogenase [Shigella sonnei] Length: 372\nNumber of predicted TMHs: 2\nExp number of AAs in TMHs: 37.02628\nExp number, first 60 AAs: 19.39684\nTotal prob of N-in: 0.94313\nPOSSIBLE N-term signal sequence\ninside 1 19\nTMhelix 20 39\noutside 40 325\nTMhelix 326 348\ninside 349 372

27027 GCF_900157745.1_9789_6_30 Shigella sonnei Proteobacteria; Gammaproteobacteria; Enterobacterales; Enterobacteriaceae; Shigella
MTGDNTLIHSHGINRRDFMKLCAALAATMGLSSKAAAEMAESVTNPQRPPVIWIGAQECT WP_000145410.1
MULTISPECIES: hydrogenase-2 small chain [Proteobacteria] Length: 372\nNumber of predicted TMHs: 1\nExp number of AAs in TMHs: 23.72226\nExp number, first 60 AAs: 0.22038\nTotal prob of N-in: 0.12994\noutside 1 331\nTMhelix 332 354\ninside 355 372

27028 GCF_900157775.1_9789_6_78 Shigella sonnei Proteobacteria; Gammaproteobacteria; Enterobacterales; Enterobacteriaceae; Shigella
MTGDNTLIHSHGINRRDFMKLCAALAATMGLSSKAAAEMAESVTNPQRPPVIWIGAQECT WP_000145410.1
MULTISPECIES: hydrogenase-2 small chain [Proteobacteria] Length: 372\nNumber of predicted TMHs: 1\nExp number of AAs in TMHs: 23.72226\nExp number, first 60 AAs: 0.22038\nTotal prob of N-in: 0.12994\noutside 1 331\nTMhelix 332 354\ninside 355 372

27029 GCF_900157825.1_9789_6_73 Shigella sonnei Proteobacteria; Gammaproteobacteria; Enterobacterales; Enterobacteriaceae; Shigella

MTGDNTLIHSHGINRRDFMKLCAALAATMGLSSKAAAEMAESVTNPQRPPVIWIGAQECT WP_000145410.1 MULTISPECIES: hydrogenase-2 small chain [Proteobacteria] Length: 372\nNumber of predicted TMHs: 1\nExp number of AAs in TMHs: 23.72226\nExp number, first 60 AAs: 0.22038\nTotal prob of N-in: 0.12994\noutside 1 331\nTMhelix 332 354\ninside 355 372

27030 GCF_900157795.1_9789_6_67 Shigella sonnei Proteobacteria; Gammaproteobacteria; Enterobacterales; Enterobacteriaceae; Shigella MTGDNTLIHSHGINRRDFMKLCAALAATMGLSSKAAAEMAESVTNPQRPPVIWIGAQECT WP_000145410.1 MULTISPECIES: hydrogenase-2 small chain [Proteobacteria] Length: 372\nNumber of predicted TMHs: 1\nExp number of AAs in TMHs: 23.72226\nExp number, first 60 AAs: 0.22038\nTotal prob of N-in: 0.12994\noutside 1 331\nTMhelix 332 354\ninside 355 372

27031 GCF_900157845.1_9789_6_82 Shigella sonnei Proteobacteria; Gammaproteobacteria; Enterobacterales; Enterobacteriaceae; Shigella MTGDNTLIHSHGINRRDFMKLCAALAATMGLSSKAAAEMAESVTNPQRPPVIWIGAQECT WP_000145410.1 MULTISPECIES: hydrogenase-2 small chain [Proteobacteria] Length: 372\nNumber of predicted TMHs: 1\nExp number of AAs in TMHs: 23.72226\nExp number, first 60 AAs: 0.22038\nTotal prob of N-in: 0.12994\noutside 1 331\nTMhelix 332 354\ninside 355 372

27032 GCF_900157885.1_9789_6_13 Shigella sonnei Proteobacteria; Gammaproteobacteria; Enterobacterales; Enterobacteriaceae; Shigella MTGDNTLIHSHGINRRDFMKLCAALAATMGLSSKAAAEMAESVTNPQRPPVIWIGAQECT WP_000145410.1 MULTISPECIES: hydrogenase-2 small chain [Proteobacteria] Length: 372\nNumber of predicted TMHs: 1\nExp number of AAs in TMHs: 23.72226\nExp number, first 60 AAs: 0.22038\nTotal prob of N-in: 0.12994\noutside 1 331\nTMhelix 332 354\ninside 355 372

27033 GCF_900157895.1_9789_6_34 Shigella sonnei Proteobacteria; Gammaproteobacteria; Enterobacterales; Enterobacteriaceae; Shigella MTGDNTLIHSHGINRRDFMKLCAALAATMGLSSKAAAEMAESVTNPQRPPVIWIGAQECT WP_000145410.1 MULTISPECIES: hydrogenase-2 small chain [Proteobacteria] Length: 372\nNumber of predicted TMHs: 1\nExp number of AAs in TMHs: 23.72226\nExp number, first 60 AAs: 0.22038\nTotal prob of N-in: 0.12994\noutside 1 331\nTMhelix 332 354\ninside 355 372

27034 GCF_900157745.1_9789_6_30 Shigella sonnei Proteobacteria; Gammaproteobacteria; Enterobacterales; Enterobacteriaceae; Shigella MNNEETFYQAMRRQGVTRRSFLKYCSLAATSLGLGAGMAPKIAWALENKPRIPVVWIHGL WP_005139779.1 hydrogenase [Shigella sonnei] Length: 372\nNumber of predicted TMHs: 2\nExp number of AAs in TMHs: 37.02628\nExp number, first 60 AAs: 19.39684\nTotal prob of N-in: 0.94313\nPOSSIBLE N-term signal sequence\ninside 1 19\nTMhelix 20 39\noutside 40 325\nTMhelix 326 348\ninside 349 372

27035 GCF_900157775.1_9789_6_78 Shigella sonnei Proteobacteria; Gammaproteobacteria; Enterobacterales; Enterobacteriaceae; Shigella MNNEETFYQAMRRQGVTRRSFLKYCSLAATSLGLGAGMAPKIAWALENKPRIPVVWIHGL WP_005139779.1 hydrogenase [Shigella sonnei] Length: 372\nNumber of predicted TMHs: 2\nExp number of AAs in TMHs: 37.02628\nExp number, first 60 AAs: 19.39684\nTotal prob of N-in: 0.94313\nPOSSIBLE N-term signal sequence\ninside 1 19\nTMhelix 20 39\noutside 40 325\nTMhelix 326 348\ninside 349 372

27036 GCF_900157915.1_9789_6_9Shigella sonnei Proteobacteria; Gammaproteobacteria; Enterobacterales; Enterobacteriaceae; Shigella MTGDNTLIHSHGINRRDFMKLCAALAATMGLSSKAAAEMAESVTNPQRPPVIWIGAQECT WP_000145410.1 MULTISPECIES: hydrogenase-2 small chain [Proteobacteria] Length: 372\nNumber of predicted TMHs: 1\nExp number of AAs in TMHs: 23.72226\nExp number, first 60 AAs: 0.22038\nTotal prob of N-in: 0.12994\noutside 1 331\nTMhelix 332 354\ninside 355 372

27037 GCF_900157945.1_9789_6_85 Shigella sonnei Proteobacteria; Gammaproteobacteria; Enterobacterales; Enterobacteriaceae; Shigella MTGDNTLIHSHGINRRDFMKLCAALAATMGLSSKAAAEMAESVTNPQRPPVIWIGAQECT WP_000145410.1 MULTISPECIES: hydrogenase-2 small chain [Proteobacteria] Length: 372\nNumber of predicted TMHs: 1\nExp number of AAs in TMHs: 23.72226\nExp number, first 60 AAs: 0.22038\nTotal prob of N-in: 0.12994\noutside 1 331\nTMhelix 332 354\ninside 355 372

27038 GCF_900157965.1_9789_6_87 Shigella sonnei Proteobacteria; Gammaproteobacteria; Enterobacterales; Enterobacteriaceae; Shigella
MTGDNTLIHSHGINRRDFMKLCAALAATMGLSSKAAAEMAESVTNPQRPPVIWIGAQECT WP_000145410.1
MULTISPECIES: hydrogenase-2 small chain [Proteobacteria] Length: 372\nNumber of predicted TMHs: 1\nExp number of AAs in TMHs: 23.72226\nExp number, first 60 AAs: 0.22038\nTotal prob of N-in: 0.12994\noutside 1 331\nTMhelix 332 354\ninside 355 372

27039 GCF_900157995.1_9789_6_94 Shigella sonnei Proteobacteria; Gammaproteobacteria; Enterobacterales; Enterobacteriaceae; Shigella
MTGDNTLIHSHGINRRDFMKLCAALAATMGLSSKAAAEMAESVTNPQRPPVIWIGAQECT WP_000145410.1
MULTISPECIES: hydrogenase-2 small chain [Proteobacteria] Length: 372\nNumber of predicted TMHs: 1\nExp number of AAs in TMHs: 23.72226\nExp number, first 60 AAs: 0.22038\nTotal prob of N-in: 0.12994\noutside 1 331\nTMhelix 332 354\ninside 355 372

27040 GCF_900157915.1_9789_6_9Shigella sonnei Proteobacteria; Gammaproteobacteria; Enterobacterales; Enterobacteriaceae; Shigella MNNEETFYQAMRRQGVTRRSFLKYCSLAATSLGLGAGMAPKIAWALENKPRIPVVWIHGL
WP_005139779.1 hydrogenase [Shigella sonnei] Length: 372\nNumber of predicted TMHs: 2\nExp number of AAs in TMHs: 37.02628\nExp number, first 60 AAs: 19.39684\nTotal prob of N-in: 0.94313\nPOSSIBLE N-term signal sequence\ninside 1 19\nTMhelix 20 39\noutside 40 325\nTMhelix 326 348\ninside 349 372

27041 GCF_900157945.1_9789_6_85 Shigella sonnei Proteobacteria; Gammaproteobacteria; Enterobacterales; Enterobacteriaceae; Shigella
MNNEETFYQAMRRQGVTRRSFLKYCSLAATSLGLGAGMAPKIAWALENKPRIPVVWIHGL WP_005139779.1
hydrogenase [Shigella sonnei] Length: 372\nNumber of predicted TMHs: 2\nExp number of AAs in TMHs: 37.02628\nExp number, first 60 AAs: 19.39684\nTotal prob of N-in: 0.94313\nPOSSIBLE N-term signal sequence\ninside 1 19\nTMhelix 20 39\noutside 40 325\nTMhelix 326 348\ninside 349 372

27042 GCF_900157795.1_9789_6_67 Shigella sonnei Proteobacteria; Gammaproteobacteria; Enterobacterales; Enterobacteriaceae; Shigella
MNNEETFYQAMRRQGVTRRSFLKYCSLAATSLGLGAGMAPKIAWALENKPRIPVVWIHGL WP_005139779.1
hydrogenase [Shigella sonnei] Length: 372\nNumber of predicted TMHs: 2\nExp number of AAs in TMHs: 37.02628\nExp number, first 60 AAs: 19.39684\nTotal prob of N-in: 0.94313\nPOSSIBLE N-term signal sequence\ninside 1 19\nTMhelix 20 39\noutside 40 325\nTMhelix 326 348\ninside 349 372

27043 GCF_900158395.1_9803_4_37 Shigella sonnei Proteobacteria; Gammaproteobacteria; Enterobacterales; Enterobacteriaceae; Shigella
MTGDNTLIHSHGINRRDFMKLCAALAATMGLSSKAAAEMAESVTNPQRPPVIWIGAQECT WP_000145410.1
MULTISPECIES: hydrogenase-2 small chain [Proteobacteria] Length: 372\nNumber of predicted TMHs: 1\nExp number of AAs in TMHs: 23.72226\nExp number, first 60 AAs: 0.22038\nTotal prob of N-in: 0.12994\noutside 1 331\nTMhelix 332 354\ninside 355 372

27044 GCF_900157825.1_9789_6_73 Shigella sonnei Proteobacteria; Gammaproteobacteria; Enterobacterales; Enterobacteriaceae; Shigella
MNNEETFYQAMRRQGVTRRSFLKYCSLAATSLGLGAGMAPKIAWALENKPRIPVVWIHGL WP_005139779.1
hydrogenase [Shigella sonnei] Length: 372\nNumber of predicted TMHs: 2\nExp number of AAs in TMHs: 37.02628\nExp number, first 60 AAs: 19.39684\nTotal prob of N-in: 0.94313\nPOSSIBLE N-term signal sequence\ninside 1 19\nTMhelix 20 39\noutside 40 325\nTMhelix 326 348\ninside 349 372

27045 GCF_900158025.1_9789_6_90 Shigella sonnei Proteobacteria; Gammaproteobacteria; Enterobacterales; Enterobacteriaceae; Shigella
MTGDNTLIHSHGINRRDFMKLCAALAATMGLSSKAAAEMAESVTNPQRPPVIWIGAQECT WP_000145410.1
MULTISPECIES: hydrogenase-2 small chain [Proteobacteria] Length: 372\nNumber of predicted TMHs: 1\nExp number of AAs in TMHs: 23.72226\nExp number, first 60 AAs: 0.22038\nTotal prob of N-in: 0.12994\noutside 1 331\nTMhelix 332 354\ninside 355 372

27046 GCF_900158025.1_9789_6_90 Shigella sonnei Proteobacteria; Gammaproteobacteria; Enterobacterales; Enterobacteriaceae; Shigella
MNNEETFYQAMRRQGVTRRSFLKYCSLAATSLGLGAGMAPKIAWALENKPRIPVVWIHGL WP_005139779.1
hydrogenase [Shigella sonnei] Length: 372\nNumber of predicted TMHs: 2\nExp number of AAs in TMHs: 37.02628\nExp number, first 60 AAs: 19.39684\nTotal prob of N-in: 0.94313\nPOSSIBLE N-term signal sequence\ninside 1 19\nTMhelix 20 39\noutside 40 325\nTMhelix 326 348\ninside 349 372

27047 GCF_900157845.1_9789_6_82 Shigella sonnei Proteobacteria; Gammaproteobacteria; Enterobacterales; Enterobacteriaceae; Shigella MNNEETFYQAMRRQGVTRRSFLKYCSLAATSLGLGAGMAPKIAWALENKPRIPVVWIHGL WP_005139779.1 hydrogenase [Shigella sonnei] Length: 372\nNumber of predicted TMHs: 2\nExp number of AAs in TMHs: 37.02628\nExp number, first 60 AAs: 19.39684\nTotal prob of N-in: 0.94313\nPOSSIBLE N-term signal sequence\ninside 1 19\nTMhelix 20 39\noutside 40 325\nTMhelix 326 348\ninside 349 372

27048 GCF_900158045.1_9789_6_91 Shigella sonnei Proteobacteria; Gammaproteobacteria; Enterobacterales; Enterobacteriaceae; Shigella MTGDNTLIHSHGINRRDFMKLCAALAATMGLSSKAAAEMAESVTNPQRPPVIWIGAQECT WP_000145410.1 MULTISPECIES: hydrogenase-2 small chain [Proteobacteria] Length: 372\nNumber of predicted TMHs: 1\nExp number of AAs in TMHs: 23.72226\nExp number, first 60 AAs: 0.22038\nTotal prob of N-in: 0.12994\noutside 1 331\nTMhelix 332 354\ninside 355 372

27049 GCF_900158045.1_9789_6_91 Shigella sonnei Proteobacteria; Gammaproteobacteria; Enterobacterales; Enterobacteriaceae; Shigella MNNEETFYQAMRRQGVTRRSFLKYCSLAATSLGLGAGMAPKIAWALENKPRIPVVWIHGL WP_005139779.1 hydrogenase [Shigella sonnei] Length: 372\nNumber of predicted TMHs: 2\nExp number of AAs in TMHs: 37.02628\nExp number, first 60 AAs: 19.39684\nTotal prob of N-in: 0.94313\nPOSSIBLE N-term signal sequence\ninside 1 19\nTMhelix 20 39\noutside 40 325\nTMhelix 326 348\ninside 349 372

27050 GCF_900157895.1_9789_6_34 Shigella sonnei Proteobacteria; Gammaproteobacteria; Enterobacterales; Enterobacteriaceae; Shigella MNNEETFYQAMRRQGVTRRSFLKYCSLAATSLGLGAGMAPKIAWALENKPRIPVVWIHGL WP_005139779.1 hydrogenase [Shigella sonnei] Length: 372\nNumber of predicted TMHs: 2\nExp number of AAs in TMHs: 37.02628\nExp number, first 60 AAs: 19.39684\nTotal prob of N-in: 0.94313\nPOSSIBLE N-term signal sequence\ninside 1 19\nTMhelix 20 39\noutside 40 325\nTMhelix 326 348\ninside 349 372

27051 GCF_900157885.1_9789_6_13 Shigella sonnei Proteobacteria; Gammaproteobacteria; Enterobacterales; Enterobacteriaceae; Shigella MNNEETFYQAMRRQGVTRRSFLKYCSLAATSLGLGAGMAPKIAWALENKPRIPVVWIHGL WP_005139779.1 hydrogenase [Shigella sonnei] Length: 372\nNumber of predicted TMHs: 2\nExp number of AAs in TMHs: 37.02628\nExp number, first 60 AAs: 19.39684\nTotal prob of N-in: 0.94313\nPOSSIBLE N-term signal sequence\ninside 1 19\nTMhelix 20 39\noutside 40 325\nTMhelix 326 348\ninside 349 372

27052 GCF_900158075.1_9803_4_3Shigella sonnei Proteobacteria; Gammaproteobacteria; Enterobacterales; Enterobacteriaceae; Shigella MTGDNTLIHSHGINRRDFMKLCAALAATMGLSSKAAAEMAESVTNPQRPPVIWIGAQECT WP_000145410.1 MULTISPECIES: hydrogenase-2 small chain [Proteobacteria] Length: 372\nNumber of predicted TMHs: 1\nExp number of AAs in TMHs: 23.72226\nExp number, first 60 AAs: 0.22038\nTotal prob of N-in: 0.12994\noutside 1 331\nTMhelix 332 354\ninside 355 372

27053 GCF_900158125.1_9803_4_8Shigella sonnei Proteobacteria; Gammaproteobacteria; Enterobacterales; Enterobacteriaceae; Shigella MTGDNTLIHSHGINRRDFMKLCAALAATMGLSSKAAAEMAESVTNPQRPPVIWIGAQECT WP_000145410.1 MULTISPECIES: hydrogenase-2 small chain [Proteobacteria] Length: 372\nNumber of predicted TMHs: 1\nExp number of AAs in TMHs: 23.72226\nExp number, first 60 AAs: 0.22038\nTotal prob of N-in: 0.12994\noutside 1 331\nTMhelix 332 354\ninside 355 372

27054 GCF_900158095.1_9803_4_5Shigella sonnei Proteobacteria; Gammaproteobacteria; Enterobacterales; Enterobacteriaceae; Shigella MTGDNTLIHSHGINRRDFMKLCAALAATMGLSSKAAAEMAESVTNPQRPPVIWIGAQECT WP_000145410.1 MULTISPECIES: hydrogenase-2 small chain [Proteobacteria] Length: 372\nNumber of predicted TMHs: 1\nExp number of AAs in TMHs: 23.72226\nExp number, first 60 AAs: 0.22038\nTotal prob of N-in: 0.12994\noutside 1 331\nTMhelix 332 354\ninside 355 372

27055 GCF_900158145.1_9803_4_12 Shigella sonnei Proteobacteria; Gammaproteobacteria; Enterobacterales; Enterobacteriaceae; Shigella MTGDNTLIHSHGINRRDFMKLCAALAATMGLSSKAAAEMAESVTNPQRPPVIWIGAQECT WP_000145410.1 MULTISPECIES: hydrogenase-2 small chain [Proteobacteria] Length: 372\nNumber of predicted TMHs: 1\nExp number of AAs in TMHs: 23.72226\nExp number, first 60 AAs: 0.22038\nTotal prob of N-in: 0.12994\noutside 1 331\nTMhelix 332 354\ninside 355 372

27056	GCF_900158175.1_9803_4_14	Shigella sonnei	Proteobacteria; Gammaproteobacteria; Enterobacterales; Enterobacteriaceae; Shigella
MTGDNTLIHSHGINRRDFMKLCAALAATMGLSSKAAAEMAESVTNPQRPPVIWIGAQECT	WP_000145410.1
MULTISPECIES: hydrogenase-2 small chain [Proteobacteria]	Length: 372\nNumber of predicted TMHs: 1\nExp number of AAs in TMHs: 23.72226\nExp number, first 60 AAs: 0.22038\nTotal prob of N-in: 0.12994\noutside 1 331\nTMhelix 332 354\ninside 355 372

27057	GCF_900158075.1_9803_4_3Shigella sonnei	Proteobacteria; Gammaproteobacteria; Enterobacterales; Enterobacteriaceae; Shigella	MNNEETFYQAMRRQGVTRRSFLKYCSLAATSLGLGAGMAPKIAWALENKPRIPVVWIHGL
WP_005139779.1	hydrogenase [Shigella sonnei]	Length: 372\nNumber of predicted TMHs: 2\nExp number of AAs in TMHs: 37.02628\nExp number, first 60 AAs: 19.39684\nTotal prob of N-in: 0.94313\nPOSSIBLE N-term signal sequence\ninside 1 19\nTMhelix 20 39\noutside 40 325\nTMhelix 326 348\ninside 349 372

27058	GCF_900158605.1_9803_4_60	Shigella sonnei	Proteobacteria; Gammaproteobacteria; Enterobacterales; Enterobacteriaceae; Shigella
MTGDNTLIHSHGINRRDFMKLCAALAATMGLSSKAAAEMAESVTNPQRPPVIWIGAQECT	WP_000145410.1
MULTISPECIES: hydrogenase-2 small chain [Proteobacteria]	Length: 372\nNumber of predicted TMHs: 1\nExp number of AAs in TMHs: 23.72226\nExp number, first 60 AAs: 0.22038\nTotal prob of N-in: 0.12994\noutside 1 331\nTMhelix 332 354\ninside 355 372

27059	GCF_900157965.1_9789_6_87	Shigella sonnei	Proteobacteria; Gammaproteobacteria; Enterobacterales; Enterobacteriaceae; Shigella
MNNEETFYQAMRRQGVTRRSFLKYCSLAATSLGLGAGMAPKIAWALENKPRIPVVWIHGL	WP_005139779.1
hydrogenase [Shigella sonnei]	Length: 372\nNumber of predicted TMHs: 2\nExp number of AAs in TMHs: 37.02628\nExp number, first 60 AAs: 19.39684\nTotal prob of N-in: 0.94313\nPOSSIBLE N-term signal sequence\ninside 1 19\nTMhelix 20 39\noutside 40 325\nTMhelix 326 348\ninside 349 372

27060	GCF_900157995.1_9789_6_94	Shigella sonnei	Proteobacteria; Gammaproteobacteria; Enterobacterales; Enterobacteriaceae; Shigella
MNNEETFYQAMRRQGVTRRSFLKYCSLAATSLGLGAGMAPKIAWALENKPRIPVVWIHGL	WP_005139779.1
hydrogenase [Shigella sonnei]	Length: 372\nNumber of predicted TMHs: 2\nExp number of AAs in TMHs: 37.02628\nExp number, first 60 AAs: 19.39684\nTotal prob of N-in: 0.94313\nPOSSIBLE N-term signal sequence\ninside 1 19\nTMhelix 20 39\noutside 40 325\nTMhelix 326 348\ninside 349 372

27061	GCF_900158205.1_9803_4_17	Shigella sonnei	Proteobacteria; Gammaproteobacteria; Enterobacterales; Enterobacteriaceae; Shigella
MTGDNTLIHSHGINRRDFMKLCAALAATMGLSSKAAAEMAESVTNPQRPPVIWIGAQECT	WP_000145410.1
MULTISPECIES: hydrogenase-2 small chain [Proteobacteria]	Length: 372\nNumber of predicted TMHs: 1\nExp number of AAs in TMHs: 23.72226\nExp number, first 60 AAs: 0.22038\nTotal prob of N-in: 0.12994\noutside 1 331\nTMhelix 332 354\ninside 355 372

27062	GCF_900158225.1_9803_4_18	Shigella sonnei	Proteobacteria; Gammaproteobacteria; Enterobacterales; Enterobacteriaceae; Shigella
MTGDNTLIHSHGINRRDFMKLCAALAATMGLSSKAAAEMAESVTNPQRPPVIWIGAQECT	WP_000145410.1
MULTISPECIES: hydrogenase-2 small chain [Proteobacteria]	Length: 372\nNumber of predicted TMHs: 1\nExp number of AAs in TMHs: 23.72226\nExp number, first 60 AAs: 0.22038\nTotal prob of N-in: 0.12994\noutside 1 331\nTMhelix 332 354\ninside 355 372

27063	GCF_900158275.1_9803_4_24	Shigella sonnei	Proteobacteria; Gammaproteobacteria; Enterobacterales; Enterobacteriaceae; Shigella
MTGDNTLIHSHGINRRDFMKLCAALAATMGLSSKAAAEMAESVTNPQRPPVIWIGAQECT	WP_000145410.1
MULTISPECIES: hydrogenase-2 small chain [Proteobacteria]	Length: 372\nNumber of predicted TMHs: 1\nExp number of AAs in TMHs: 23.72226\nExp number, first 60 AAs: 0.22038\nTotal prob of N-in: 0.12994\noutside 1 331\nTMhelix 332 354\ninside 355 372

27064	GCF_900158255.1_9803_4_22	Shigella sonnei	Proteobacteria; Gammaproteobacteria; Enterobacterales; Enterobacteriaceae; Shigella
MTGDNTLIHSHGINRRDFMKLCAALAATMGLSSKAAAEMAESVTNPQRPPVIWIGAQECT	WP_000145410.1
MULTISPECIES: hydrogenase-2 small chain [Proteobacteria]	Length: 372\nNumber of predicted TMHs: 1\nExp number of AAs in TMHs: 23.72226\nExp number, first 60 AAs: 0.22038\nTotal prob of N-in: 0.12994\noutside 1 331\nTMhelix 332 354\ninside 355 372

27065 GCF_900158205.1_9803_4_17 Shigella sonnei Proteobacteria; Gammaproteobacteria; Enterobacterales; Enterobacteriaceae; Shigella MNNEETFYQAMRRQGVTRRSFLKYCSLAATSLGLGAGMAPKIAWALENKPRIPVVWIHGL WP_005139779.1 hydrogenase [Shigella sonnei] Length: 372\nNumber of predicted TMHs: 2\nExp number of AAs in TMHs: 37.02628\nExp number, first 60 AAs: 19.39684\nTotal prob of N-in: 0.94313\nPOSSIBLE N-term signal sequence\ninside 1 19\nTMhelix 20 39\noutside 40 325\nTMhelix 326 348\ninside 349 372

27066 GCF_900158225.1_9803_4_18 Shigella sonnei Proteobacteria; Gammaproteobacteria; Enterobacterales; Enterobacteriaceae; Shigella MNNEETFYQAMRRQGVTRRSFLKYCSLAATSLGLGAGMAPKIAWALENKPRIPVVWIHGL WP_005139779.1 hydrogenase [Shigella sonnei] Length: 372\nNumber of predicted TMHs: 2\nExp number of AAs in TMHs: 37.02628\nExp number, first 60 AAs: 19.39684\nTotal prob of N-in: 0.94313\nPOSSIBLE N-term signal sequence\ninside 1 19\nTMhelix 20 39\noutside 40 325\nTMhelix 326 348\ninside 349 372

27067 GCF_900158095.1_9803_4_5Shigella sonnei Proteobacteria; Gammaproteobacteria; Enterobacterales; Enterobacteriaceae; Shigella MNNEETFYQAMRRQGVTRRSFLKYCSLAATSLGLGAGMAPKIAWALENKPRIPVVWIHGL WP_005139779.1 hydrogenase [Shigella sonnei] Length: 372\nNumber of predicted TMHs: 2\nExp number of AAs in TMHs: 37.02628\nExp number, first 60 AAs: 19.39684\nTotal prob of N-in: 0.94313\nPOSSIBLE N-term signal sequence\ninside 1 19\nTMhelix 20 39\noutside 40 325\nTMhelix 326 348\ninside 349 372

27068 GCF_900158125.1_9803_4_8Shigella sonnei Proteobacteria; Gammaproteobacteria; Enterobacterales; Enterobacteriaceae; Shigella MNNEETFYQAMRRQGVTRRSFLKYCSLAATSLGLGAGMAPKIAWALENKPRIPVVWIHGL WP_005139779.1 hydrogenase [Shigella sonnei] Length: 372\nNumber of predicted TMHs: 2\nExp number of AAs in TMHs: 37.02628\nExp number, first 60 AAs: 19.39684\nTotal prob of N-in: 0.94313\nPOSSIBLE N-term signal sequence\ninside 1 19\nTMhelix 20 39\noutside 40 325\nTMhelix 326 348\ninside 349 372

27069 GCF_900158285.1_9803_4_25 Shigella sonnei Proteobacteria; Gammaproteobacteria; Enterobacterales; Enterobacteriaceae; Shigella MTGDNTLIHSHGINRRDFMKLCAALAATMGLSSKAAAEMAESVTNPQRPPVIWIGAQECT WP_000145410.1 MULTISPECIES: hydrogenase-2 small chain [Proteobacteria] Length: 372\nNumber of predicted TMHs: 1\nExp number of AAs in TMHs: 23.72226\nExp number, first 60 AAs: 0.22038\nTotal prob of N-in: 0.12994\noutside 1 331\nTMhelix 332 354\ninside 355 372

27070 GCF_900158295.1_9803_4_27 Shigella sonnei Proteobacteria; Gammaproteobacteria; Enterobacterales; Enterobacteriaceae; Shigella MTGDNTLIHSHGINRRDFMKLCAALAATMGLSSKAAAEMAESVTNPQRPPVIWIGAQECT WP_000145410.1 MULTISPECIES: hydrogenase-2 small chain [Proteobacteria] Length: 372\nNumber of predicted TMHs: 1\nExp number of AAs in TMHs: 23.72226\nExp number, first 60 AAs: 0.22038\nTotal prob of N-in: 0.12994\noutside 1 331\nTMhelix 332 354\ninside 355 372

27071 GCF_900158285.1_9803_4_25 Shigella sonnei Proteobacteria; Gammaproteobacteria; Enterobacterales; Enterobacteriaceae; Shigella MNNEETFYQAMRRQGVTRRSFLKYCSLAATSLGLGAGMAPKIAWALENKPRIPVVWIHGL WP_005139779.1 hydrogenase [Shigella sonnei] Length: 372\nNumber of predicted TMHs: 2\nExp number of AAs in TMHs: 37.02628\nExp number, first 60 AAs: 19.39684\nTotal prob of N-in: 0.94313\nPOSSIBLE N-term signal sequence\ninside 1 19\nTMhelix 20 39\noutside 40 325\nTMhelix 326 348\ninside 349 372

27072 GCF_900158295.1_9803_4_27 Shigella sonnei Proteobacteria; Gammaproteobacteria; Enterobacterales; Enterobacteriaceae; Shigella MNNEETFYQAMRRQGVTRRSFLKYCSLAATSLGLGAGMAPKIAWALENKPRIPVVWIHGL WP_005139779.1 hydrogenase [Shigella sonnei] Length: 372\nNumber of predicted TMHs: 2\nExp number of AAs in TMHs: 37.02628\nExp number, first 60 AAs: 19.39684\nTotal prob of N-in: 0.94313\nPOSSIBLE N-term signal sequence\ninside 1 19\nTMhelix 20 39\noutside 40 325\nTMhelix 326 348\ninside 349 372

27073 GCF_900158145.1_9803_4_12 Shigella sonnei Proteobacteria; Gammaproteobacteria; Enterobacterales; Enterobacteriaceae; Shigella MNNEETFYQAMRRQGVTRRSFLKYCSLAATSLGLGAGMAPKIAWALENKPRIPVVWIHGL WP_005139779.1 hydrogenase [Shigella sonnei] Length: 372\nNumber of predicted TMHs: 2\nExp number of AAs in TMHs: 37.02628\nExp number, first 60 AAs: 19.39684\nTotal prob of N-in: 0.94313\nPOSSIBLE N-term signal sequence\ninside 1 19\nTMhelix 20 39\noutside 40 325\nTMhelix 326 348\ninside 349 372

27074 GCF_900158315.1_9803_4_29 Shigella sonnei Proteobacteria; Gammaproteobacteria; Enterobacterales; Enterobacteriaceae; Shigella
MTGDNTLIHSHGINRRDFMKLCAALAATMGLSSKAAAEMAESVTNPQRPPVIWIGAQECT WP_000145410.1
MULTISPECIES: hydrogenase-2 small chain [Proteobacteria] Length: 372\nNumber of predicted TMHs: 1\nExp number of AAs in TMHs: 23.72226\nExp number, first 60 AAs: 0.22038\nTotal prob of N-in: 0.12994\noutside 1 331\nTMhelix 332 354\ninside 355 372

27075 GCF_900158315.1_9803_4_29 Shigella sonnei Proteobacteria; Gammaproteobacteria; Enterobacterales; Enterobacteriaceae; Shigella
MNNEETFYQAMRRQGVTRRSFLKYCSLAATSLGLGAGMAPKIAWALENKPRIPVVWIHGL WP_005139779.1
hydrogenase [Shigella sonnei] Length: 372\nNumber of predicted TMHs: 2\nExp number of AAs in TMHs: 37.02628\nExp number, first 60 AAs: 19.39684\nTotal prob of N-in: 0.94313\nPOSSIBLE N-term signal sequence\ninside 1 19\nTMhelix 20 39\noutside 40 325\nTMhelix 326 348\ninside 349 372

27076 GCF_900158175.1_9803_4_14 Shigella sonnei Proteobacteria; Gammaproteobacteria; Enterobacterales; Enterobacteriaceae; Shigella
MNNEETFYQAMRRQGVTRRSFLKYCSLAATSLGLGAGMAPKIAWALENKPRIPVVWIHGL WP_005139779.1
hydrogenase [Shigella sonnei] Length: 372\nNumber of predicted TMHs: 2\nExp number of AAs in TMHs: 37.02628\nExp number, first 60 AAs: 19.39684\nTotal prob of N-in: 0.94313\nPOSSIBLE N-term signal sequence\ninside 1 19\nTMhelix 20 39\noutside 40 325\nTMhelix 326 348\ninside 349 372

27077 GCF_900158345.1_9803_4_33 Shigella sonnei Proteobacteria; Gammaproteobacteria; Enterobacterales; Enterobacteriaceae; Shigella
MTGDNTLIHSHGINRRDFMKLCAALAATMGLSSKAAAEMAESVTNPQRPPVIWIGAQECT WP_000145410.1
MULTISPECIES: hydrogenase-2 small chain [Proteobacteria] Length: 372\nNumber of predicted TMHs: 1\nExp number of AAs in TMHs: 23.72226\nExp number, first 60 AAs: 0.22038\nTotal prob of N-in: 0.12994\noutside 1 331\nTMhelix 332 354\ninside 355 372

27078 GCF_900158375.1_9803_4_35 Shigella sonnei Proteobacteria; Gammaproteobacteria; Enterobacterales; Enterobacteriaceae; Shigella
MTGDNTLIHSHGINRRDFMKLCAALAATMGLSSKAAAEMAESVTNPQRPPVIWIGAQECT WP_000145410.1
MULTISPECIES: hydrogenase-2 small chain [Proteobacteria] Length: 372\nNumber of predicted TMHs: 1\nExp number of AAs in TMHs: 23.72226\nExp number, first 60 AAs: 0.22038\nTotal prob of N-in: 0.12994\noutside 1 331\nTMhelix 332 354\ninside 355 372

27079 GCF_900158345.1_9803_4_33 Shigella sonnei Proteobacteria; Gammaproteobacteria; Enterobacterales; Enterobacteriaceae; Shigella
MNNEETFYQAMRRQGVTRRSFLKYCSLAATSLGLGAGMAPKIAWALENKPRIPVVWIHGL WP_005139779.1
hydrogenase [Shigella sonnei] Length: 372\nNumber of predicted TMHs: 2\nExp number of AAs in TMHs: 37.02628\nExp number, first 60 AAs: 19.39684\nTotal prob of N-in: 0.94313\nPOSSIBLE N-term signal sequence\ninside 1 19\nTMhelix 20 39\noutside 40 325\nTMhelix 326 348\ninside 349 372

27080 GCF_900158375.1_9803_4_35 Shigella sonnei Proteobacteria; Gammaproteobacteria; Enterobacterales; Enterobacteriaceae; Shigella
MNNEETFYQAMRRQGVTRRSFLKYCSLAATSLGLGAGMAPKIAWALENKPRIPVVWIHGL WP_005139779.1
hydrogenase [Shigella sonnei] Length: 372\nNumber of predicted TMHs: 2\nExp number of AAs in TMHs: 37.02628\nExp number, first 60 AAs: 19.39684\nTotal prob of N-in: 0.94313\nPOSSIBLE N-term signal sequence\ninside 1 19\nTMhelix 20 39\noutside 40 325\nTMhelix 326 348\ninside 349 372

27081 GCF_900158395.1_9803_4_37 Shigella sonnei Proteobacteria; Gammaproteobacteria; Enterobacterales; Enterobacteriaceae; Shigella
MNNEETFYQAMRRQGVTRRSFLKYCSLAATSLGLGAGMAPKIAWALENKPRIPVVWIHGL WP_005139779.1
hydrogenase [Shigella sonnei] Length: 372\nNumber of predicted TMHs: 2\nExp number of AAs in TMHs: 37.02628\nExp number, first 60 AAs: 19.39684\nTotal prob of N-in: 0.94313\nPOSSIBLE N-term signal sequence\ninside 1 19\nTMhelix 20 39\noutside 40 325\nTMhelix 326 348\ninside 349 372

27082 GCF_900158275.1_9803_4_24 Shigella sonnei Proteobacteria; Gammaproteobacteria; Enterobacterales; Enterobacteriaceae; Shigella
MNNEETFYQAMRRQGVTRRSFLKYCSLAATSLGLGAGMAPKIAWALENKPRIPVVWIHGL WP_005139779.1
hydrogenase [Shigella sonnei] Length: 372\nNumber of predicted TMHs: 2\nExp number of AAs in

TMHs: 37.02628\nExp number, first 60 AAs: 19.39684\nTotal prob of N-in: 0.94313\nPOSSIBLE N-term signal sequence\ninside 1 19\nTMhelix 20 39\noutside 40 325\nTMhelix 326 348\ninside 349 372

27083 GCF_900158425.1_9803_4_40 Shigella sonnei Proteobacteria; Gammaproteobacteria; Enterobacterales; Enterobacteriaceae; Shigella
MTGDNTLIHSHGINRRDFMKLCAALAATMGLSSKAAAEMAESVTNPQRPPVIWIGAQECT WP_000145410.1
MULTISPECIES: hydrogenase-2 small chain [Proteobacteria] Length: 372\nNumber of predicted TMHs: 1\nExp number of AAs in TMHs: 23.72226\nExp number, first 60 AAs: 0.22038\nTotal prob of N-in: 0.12994\noutside 1 331\nTMhelix 332 354\ninside 355 372

27084 GCF_900158425.1_9803_4_40 Shigella sonnei Proteobacteria; Gammaproteobacteria; Enterobacterales; Enterobacteriaceae; Shigella
MNNEETFYQAMRRQGVTRRSFLKYCSLAATSLGLGAGMAPKIAWALENKPRIPVVWIHGL WP_005139779.1
hydrogenase [Shigella sonnei] Length: 372\nNumber of predicted TMHs: 2\nExp number of AAs in TMHs: 37.02628\nExp number, first 60 AAs: 19.39684\nTotal prob of N-in: 0.94313\nPOSSIBLE N-term signal sequence\ninside 1 19\nTMhelix 20 39\noutside 40 325\nTMhelix 326 348\ninside 349 372

27085 GCF_900158255.1_9803_4_22 Shigella sonnei Proteobacteria; Gammaproteobacteria; Enterobacterales; Enterobacteriaceae; Shigella
MNNEETFYQAMRRQGVTRRSFLKYCSLAATSLGLGAGMAPKIAWALENKPRIPVVWIHGL WP_005139779.1
hydrogenase [Shigella sonnei] Length: 372\nNumber of predicted TMHs: 2\nExp number of AAs in TMHs: 37.02628\nExp number, first 60 AAs: 19.39684\nTotal prob of N-in: 0.94313\nPOSSIBLE N-term signal sequence\ninside 1 19\nTMhelix 20 39\noutside 40 325\nTMhelix 326 348\ninside 349 372

27086 GCF_900158445.1_9803_4_42 Shigella sonnei Proteobacteria; Gammaproteobacteria; Enterobacterales; Enterobacteriaceae; Shigella
MTGDNTLIHSHGINRRDFMKLCAALAATMGLSSKAAAEMAESVTNPQRPPVIWIGAQECT WP_000145410.1
MULTISPECIES: hydrogenase-2 small chain [Proteobacteria] Length: 372\nNumber of predicted TMHs: 1\nExp number of AAs in TMHs: 23.72226\nExp number, first 60 AAs: 0.22038\nTotal prob of N-in: 0.12994\noutside 1 331\nTMhelix 332 354\ninside 355 372

27087 GCF_900158445.1_9803_4_42 Shigella sonnei Proteobacteria; Gammaproteobacteria; Enterobacterales; Enterobacteriaceae; Shigella
MNNEETFYQAMRRQGVTRRSFLKYCSLAATSLGLGAGMAPKIAWALENKPRIPVVWIHGL WP_005139779.1
hydrogenase [Shigella sonnei] Length: 372\nNumber of predicted TMHs: 2\nExp number of AAs in TMHs: 37.02628\nExp number, first 60 AAs: 19.39684\nTotal prob of N-in: 0.94313\nPOSSIBLE N-term signal sequence\ninside 1 19\nTMhelix 20 39\noutside 40 325\nTMhelix 326 348\ninside 349 372

27088 GCF_900158555.1_9803_4_54 Shigella sonnei Proteobacteria; Gammaproteobacteria; Enterobacterales; Enterobacteriaceae; Shigella
MTGDNTLIHSHGINRRDFMKLCAALAATMGLSSKAAAEMAESVTNPQRPPVIWIGAQECT WP_000145410.1
MULTISPECIES: hydrogenase-2 small chain [Proteobacteria] Length: 372\nNumber of predicted TMHs: 1\nExp number of AAs in TMHs: 23.72226\nExp number, first 60 AAs: 0.22038\nTotal prob of N-in: 0.12994\noutside 1 331\nTMhelix 332 354\ninside 355 372

27089 GCF_900158575.1_9803_4_57 Shigella sonnei Proteobacteria; Gammaproteobacteria; Enterobacterales; Enterobacteriaceae; Shigella
MTGDNTLIHSHGINRRDFMKLCAALAATMGLSSKAAAEMAESVTNPQRPPVIWIGAQECT WP_000145410.1
MULTISPECIES: hydrogenase-2 small chain [Proteobacteria] Length: 372\nNumber of predicted TMHs: 1\nExp number of AAs in TMHs: 23.72226\nExp number, first 60 AAs: 0.22038\nTotal prob of N-in: 0.12994\noutside 1 331\nTMhelix 332 354\ninside 355 372

27090 GCF_900158525.1_9803_4_51 Shigella sonnei Proteobacteria; Gammaproteobacteria; Enterobacterales; Enterobacteriaceae; Shigella
MTGDNTLIHSHGINRRDFMKLCAALAATMGLSSKAAAEMAESVTNPQRPPVIWIGAQECT WP_000145410.1
MULTISPECIES: hydrogenase-2 small chain [Proteobacteria] Length: 372\nNumber of predicted TMHs: 1\nExp number of AAs in TMHs: 23.72226\nExp number, first 60 AAs: 0.22038\nTotal prob of N-in: 0.12994\noutside 1 331\nTMhelix 332 354\ninside 355 372

27091 GCF_900158475.1_9803_4_46 Shigella sonnei Proteobacteria; Gammaproteobacteria; Enterobacterales; Enterobacteriaceae; Shigella

MTGDNTLIHSHGINRRDFMKLCAALAATMGLSSKAAAEMAESVTNPQRPPVIWIGAQECT WP_000145410.1 MULTISPECIES: hydrogenase-2 small chain [Proteobacteria] Length: 372\nNumber of predicted TMHs: 1\nExp number of AAs in TMHs: 23.72226\nExp number, first 60 AAs: 0.22038\nTotal prob of N-in: 0.12994\noutside 1 331\nTMhelix 332 354\ninside 355 372

27092 GCF_900158725.1_9803_4_72 Shigella sonnei Proteobacteria; Gammaproteobacteria; Enterobacterales; Enterobacteriaceae; Shigella MTGDNTLIHSHGINRRDFMKLCAALAATMGLSSKAAAEMAESVTNPQRPPVIWIGAQECT WP_000145410.1 MULTISPECIES: hydrogenase-2 small chain [Proteobacteria] Length: 372\nNumber of predicted TMHs: 1\nExp number of AAs in TMHs: 23.72226\nExp number, first 60 AAs: 0.22038\nTotal prob of N-in: 0.12994\noutside 1 331\nTMhelix 332 354\ninside 355 372

27093 GCF_900158495.1_9803_4_48 Shigella sonnei Proteobacteria; Gammaproteobacteria; Enterobacterales; Enterobacteriaceae; Shigella MTGDNTLIHSHGINRRDFMKLCAALAATMGLSSKAAAEMAESVTNPQRPPVIWIGAQECT WP_000145410.1 MULTISPECIES: hydrogenase-2 small chain [Proteobacteria] Length: 372\nNumber of predicted TMHs: 1\nExp number of AAs in TMHs: 23.72226\nExp number, first 60 AAs: 0.22038\nTotal prob of N-in: 0.12994\noutside 1 331\nTMhelix 332 354\ninside 355 372

27094 GCF_900158705.1_9803_4_70 Shigella sonnei Proteobacteria; Gammaproteobacteria; Enterobacterales; Enterobacteriaceae; Shigella MTGDNTLIHSHGINRRDFMKLCAALAATMGLSSKAAAEMAESVTNPQRPPVIWIGAQECT WP_000145410.1 MULTISPECIES: hydrogenase-2 small chain [Proteobacteria] Length: 372\nNumber of predicted TMHs: 1\nExp number of AAs in TMHs: 23.72226\nExp number, first 60 AAs: 0.22038\nTotal prob of N-in: 0.12994\noutside 1 331\nTMhelix 332 354\ninside 355 372

27095 GCF_900158675.1_9803_4_67 Shigella sonnei Proteobacteria; Gammaproteobacteria; Enterobacterales; Enterobacteriaceae; Shigella MTGDNTLIHSHGINRRDFMKLCAALAATMGLSSKAAAEMAESVTNPQRPPVIWIGAQECT WP_000145410.1 MULTISPECIES: hydrogenase-2 small chain [Proteobacteria] Length: 372\nNumber of predicted TMHs: 1\nExp number of AAs in TMHs: 23.72226\nExp number, first 60 AAs: 0.22038\nTotal prob of N-in: 0.12994\noutside 1 331\nTMhelix 332 354\ninside 355 372

27096 GCF_900158685.1_9803_4_68 Shigella sonnei Proteobacteria; Gammaproteobacteria; Enterobacterales; Enterobacteriaceae; Shigella MTGDNTLIHSHGINRRDFMKLCAALAATMGLSSKAAAEMAESVTNPQRPPVIWIGAQECT WP_000145410.1 MULTISPECIES: hydrogenase-2 small chain [Proteobacteria] Length: 372\nNumber of predicted TMHs: 1\nExp number of AAs in TMHs: 23.72226\nExp number, first 60 AAs: 0.22038\nTotal prob of N-in: 0.12994\noutside 1 331\nTMhelix 332 354\ninside 355 372

27097 GCF_900158655.1_9803_4_65 Shigella sonnei Proteobacteria; Gammaproteobacteria; Enterobacterales; Enterobacteriaceae; Shigella MTGDNTLIHSHGINRRDFMKLCAALAATMGLSSKAAAEMAESVTNPQRPPVIWIGAQECT WP_000145410.1 MULTISPECIES: hydrogenase-2 small chain [Proteobacteria] Length: 372\nNumber of predicted TMHs: 1\nExp number of AAs in TMHs: 23.72226\nExp number, first 60 AAs: 0.22038\nTotal prob of N-in: 0.12994\noutside 1 331\nTMhelix 332 354\ninside 355 372

27098 GCF_900158625.1_9803_4_62 Shigella sonnei Proteobacteria; Gammaproteobacteria; Enterobacterales; Enterobacteriaceae; Shigella MTGDNTLIHSHGINRRDFMKLCAALAATMGLSSKAAAEMAESVTNPQRPPVIWIGAQECT WP_000145410.1 MULTISPECIES: hydrogenase-2 small chain [Proteobacteria] Length: 372\nNumber of predicted TMHs: 1\nExp number of AAs in TMHs: 23.72226\nExp number, first 60 AAs: 0.22038\nTotal prob of N-in: 0.12994\noutside 1 331\nTMhelix 332 354\ninside 355 372

27099 GCF_900158725.1_9803_4_72 Shigella sonnei Proteobacteria; Gammaproteobacteria; Enterobacterales; Enterobacteriaceae; Shigella MNNEETFYQAMRRQGVTRRSFLKYCSLAATSLGLGAGMAPKIAWALENKPRIPVVWIHGL WP_005139779.1 hydrogenase [Shigella sonnei] Length: 372\nNumber of predicted TMHs: 2\nExp number of AAs in TMHs: 37.02628\nExp number, first 60 AAs: 19.39684\nTotal prob of N-in: 0.94313\nPOSSIBLE N-term signal sequence\ninside 1 19\nTMhelix 20 39\noutside 40 325\nTMhelix 326 348\ninside 349 372

27100 GCF_900158835.1_9803_4_83 Shigella sonnei Proteobacteria; Gammaproteobacteria; Enterobacterales; Enterobacteriaceae; Shigella
MTGDNTLIHSHGINRRDFMKLCAALAATMGLSSKAAAEMAESVTNPQRPPVIWIGAQECT WP_000145410.1
MULTISPECIES: hydrogenase-2 small chain [Proteobacteria] Length: 372\nNumber of predicted TMHs: 1\nExp number of AAs in TMHs: 23.72226\nExp number, first 60 AAs: 0.22038\nTotal prob of N-in: 0.12994\noutside 1 331\nTMhelix 332 354\ninside 355 372

27101 GCF_900158525.1_9803_4_51 Shigella sonnei Proteobacteria; Gammaproteobacteria; Enterobacterales; Enterobacteriaceae; Shigella
MNNEETFYQAMRRQGVTRRSFLKYCSLAATSLGLGAGMAPKIAWALENKPRIPVVWIHGL WP_005139779.1
hydrogenase [Shigella sonnei] Length: 372\nNumber of predicted TMHs: 2\nExp number of AAs in TMHs: 37.02628\nExp number, first 60 AAs: 19.39684\nTotal prob of N-in: 0.94313\nPOSSIBLE N-term signal sequence\ninside 1 19\nTMhelix 20 39\noutside 40 325\nTMhelix 326 348\ninside 349 372

27102 GCF_900158625.1_9803_4_62 Shigella sonnei Proteobacteria; Gammaproteobacteria; Enterobacterales; Enterobacteriaceae; Shigella
MNNEETFYQAMRRQGVTRRSFLKYCSLAATSLGLGAGMAPKIAWALENKPRIPVVWIHGL WP_005139779.1
hydrogenase [Shigella sonnei] Length: 372\nNumber of predicted TMHs: 2\nExp number of AAs in TMHs: 37.02628\nExp number, first 60 AAs: 19.39684\nTotal prob of N-in: 0.94313\nPOSSIBLE N-term signal sequence\ninside 1 19\nTMhelix 20 39\noutside 40 325\nTMhelix 326 348\ninside 349 372

27103 GCF_900158575.1_9803_4_57 Shigella sonnei Proteobacteria; Gammaproteobacteria; Enterobacterales; Enterobacteriaceae; Shigella
MNNEETFYQAMRRQGVTRRSFLKYCSLAATSLGLGAGMAPKIAWALENKPRIPVVWIHGL WP_005139779.1
hydrogenase [Shigella sonnei] Length: 372\nNumber of predicted TMHs: 2\nExp number of AAs in TMHs: 37.02628\nExp number, first 60 AAs: 19.39684\nTotal prob of N-in: 0.94313\nPOSSIBLE N-term signal sequence\ninside 1 19\nTMhelix 20 39\noutside 40 325\nTMhelix 326 348\ninside 349 372

27104 GCF_900158475.1_9803_4_46 Shigella sonnei Proteobacteria; Gammaproteobacteria; Enterobacterales; Enterobacteriaceae; Shigella
MNNEETFYQAMRRQGVTRRSFLKYCSLAATSLGLGAGMAPKIAWALENKPRIPVVWIHGL WP_005139779.1
hydrogenase [Shigella sonnei] Length: 372\nNumber of predicted TMHs: 2\nExp number of AAs in TMHs: 37.02628\nExp number, first 60 AAs: 19.39684\nTotal prob of N-in: 0.94313\nPOSSIBLE N-term signal sequence\ninside 1 19\nTMhelix 20 39\noutside 40 325\nTMhelix 326 348\ninside 349 372

27105 GCF_900158555.1_9803_4_54 Shigella sonnei Proteobacteria; Gammaproteobacteria; Enterobacterales; Enterobacteriaceae; Shigella
MNNEETFYQAMRRQGVTRRSFLKYCSLAATSLGLGAGMAPKIAWALENKPRIPVVWIHGL WP_005139779.1
hydrogenase [Shigella sonnei] Length: 372\nNumber of predicted TMHs: 2\nExp number of AAs in TMHs: 37.02628\nExp number, first 60 AAs: 19.39684\nTotal prob of N-in: 0.94313\nPOSSIBLE N-term signal sequence\ninside 1 19\nTMhelix 20 39\noutside 40 325\nTMhelix 326 348\ninside 349 372

27106 GCF_900158675.1_9803_4_67 Shigella sonnei Proteobacteria; Gammaproteobacteria; Enterobacterales; Enterobacteriaceae; Shigella
MNNEETFYQAMRRQGVTRRSFLKYCSLAATSLGLGAGMAPKIAWALENKPRIPVVWIHGL WP_005139779.1
hydrogenase [Shigella sonnei] Length: 372\nNumber of predicted TMHs: 2\nExp number of AAs in TMHs: 37.02628\nExp number, first 60 AAs: 19.39684\nTotal prob of N-in: 0.94313\nPOSSIBLE N-term signal sequence\ninside 1 19\nTMhelix 20 39\noutside 40 325\nTMhelix 326 348\ninside 349 372

27107 GCF_900158705.1_9803_4_70 Shigella sonnei Proteobacteria; Gammaproteobacteria; Enterobacterales; Enterobacteriaceae; Shigella
MNNEETFYQAMRRQGVTRRSFLKYCSLAATSLGLGAGMAPKIAWALENKPRIPVVWIHGL WP_005139779.1
hydrogenase [Shigella sonnei] Length: 372\nNumber of predicted TMHs: 2\nExp number of AAs in TMHs: 37.02628\nExp number, first 60 AAs: 19.39684\nTotal prob of N-in: 0.94313\nPOSSIBLE N-term signal sequence\ninside 1 19\nTMhelix 20 39\noutside 40 325\nTMhelix 326 348\ninside 349 372

27108 GCF_900158605.1_9803_4_60 Shigella sonnei Proteobacteria; Gammaproteobacteria; Enterobacterales; Enterobacteriaceae; Shigella
MNNEETFYQAMRRQGVTRRSFLKYCSLAATSLGLGAGMAPKIAWALENKPRIPVVWIHGL WP_005139779.1
hydrogenase [Shigella sonnei] Length: 372\nNumber of predicted TMHs: 2\nExp number of AAs in

TMHs: 37.02628\nExp number, first 60 AAs: 19.39684\nTotal prob of N-in: 0.94313\nPOSSIBLE N-term signal sequence\ninside 1 19\nTMhelix 20 39\noutside 40 325\nTMhelix 326 348\ninside 349 372

27109 GCF_900158685.1_9803_4_68 Shigella sonnei Proteobacteria; Gammaproteobacteria; Enterobacterales; Enterobacteriaceae; Shigella
MNNEETFYQAMRRQGVTRRSFLKYCSLAATSLGLGAGMAPKIAWALENKPRIPVVWIHGL WP_005139779.1
hydrogenase [Shigella sonnei] Length: 372\nNumber of predicted TMHs: 2\nExp number of AAs in TMHs: 37.02628\nExp number, first 60 AAs: 19.39684\nTotal prob of N-in: 0.94313\nPOSSIBLE N-term signal sequence\ninside 1 19\nTMhelix 20 39\noutside 40 325\nTMhelix 326 348\ninside 349 372

27110 GCF_900158495.1_9803_4_48 Shigella sonnei Proteobacteria; Gammaproteobacteria; Enterobacterales; Enterobacteriaceae; Shigella
MNNEETFYQAMRRQGVTRRSFLKYCSLAATSLGLGAGMAPKIAWALENKPRIPVVWIHGL WP_005139779.1
hydrogenase [Shigella sonnei] Length: 372\nNumber of predicted TMHs: 2\nExp number of AAs in TMHs: 37.02628\nExp number, first 60 AAs: 19.39684\nTotal prob of N-in: 0.94313\nPOSSIBLE N-term signal sequence\ninside 1 19\nTMhelix 20 39\noutside 40 325\nTMhelix 326 348\ninside 349 372

27111 GCF_900158655.1_9803_4_65 Shigella sonnei Proteobacteria; Gammaproteobacteria; Enterobacterales; Enterobacteriaceae; Shigella
MNNEETFYQAMRRQGVTRRSFLKYCSLAATSLGLGAGMAPKIAWALENKPRIPVVWIHGL WP_005139779.1
hydrogenase [Shigella sonnei] Length: 372\nNumber of predicted TMHs: 2\nExp number of AAs in TMHs: 37.02628\nExp number, first 60 AAs: 19.39684\nTotal prob of N-in: 0.94313\nPOSSIBLE N-term signal sequence\ninside 1 19\nTMhelix 20 39\noutside 40 325\nTMhelix 326 348\ninside 349 372

27112 GCF_900158785.1_9803_4_78 Shigella sonnei Proteobacteria; Gammaproteobacteria; Enterobacterales; Enterobacteriaceae; Shigella
MTGDNTLIHSHGINRRDFMKLCAALAATMGLSSKAAAEMAESVTNPQRPPVIWIGAQECT WP_000145410.1
MULTISPECIES: hydrogenase-2 small chain [Proteobacteria] Length: 372\nNumber of predicted TMHs: 1\nExp number of AAs in TMHs: 23.72226\nExp number, first 60 AAs: 0.22038\nTotal prob of N-in: 0.12994\noutside 1 331\nTMhelix 332 354\ninside 355 372

27113 GCF_900158755.1_9803_4_75 Shigella sonnei Proteobacteria; Gammaproteobacteria; Enterobacterales; Enterobacteriaceae; Shigella
MTGDNTLIHSHGINRRDFMKLCAALAATMGLSSKAAAEMAESVTNPQRPPVIWIGAQECT WP_000145410.1
MULTISPECIES: hydrogenase-2 small chain [Proteobacteria] Length: 372\nNumber of predicted TMHs: 1\nExp number of AAs in TMHs: 23.72226\nExp number, first 60 AAs: 0.22038\nTotal prob of N-in: 0.12994\noutside 1 331\nTMhelix 332 354\ninside 355 372

27114 GCF_900158805.1_9803_4_80 Shigella sonnei Proteobacteria; Gammaproteobacteria; Enterobacterales; Enterobacteriaceae; Shigella
MTGDNTLIHSHGINRRDFMKLCAALAATMGLSSKAAAEMAESVTNPQRPPVIWIGAQECT WP_000145410.1
MULTISPECIES: hydrogenase-2 small chain [Proteobacteria] Length: 372\nNumber of predicted TMHs: 1\nExp number of AAs in TMHs: 23.72226\nExp number, first 60 AAs: 0.22038\nTotal prob of N-in: 0.12994\noutside 1 331\nTMhelix 332 354\ninside 355 372

27115 GCF_900158885.1_9803_4_89 Shigella sonnei Proteobacteria; Gammaproteobacteria; Enterobacterales; Enterobacteriaceae; Shigella
MTGDNTLIHSHGINRRDFMKLCAALAATMGLSSKAAAEMAESVTNPQRPPVIWIGAQECT WP_000145410.1
MULTISPECIES: hydrogenase-2 small chain [Proteobacteria] Length: 372\nNumber of predicted TMHs: 1\nExp number of AAs in TMHs: 23.72226\nExp number, first 60 AAs: 0.22038\nTotal prob of N-in: 0.12994\noutside 1 331\nTMhelix 332 354\ninside 355 372

27116 GCF_900158935.1_9803_4_94 Shigella sonnei Proteobacteria; Gammaproteobacteria; Enterobacterales; Enterobacteriaceae; Shigella
MTGDNTLIHSHGINRRDFMKLCAALAATMGLSSKAAAEMAESVTNPQRPPVIWIGAQECT WP_000145410.1
MULTISPECIES: hydrogenase-2 small chain [Proteobacteria] Length: 372\nNumber of predicted TMHs: 1\nExp number of AAs in TMHs: 23.72226\nExp number, first 60 AAs: 0.22038\nTotal prob of N-in: 0.12994\noutside 1 331\nTMhelix 332 354\ninside 355 372

27117 GCF_900158855.1_9803_4_85 Shigella sonnei Proteobacteria; Gammaproteobacteria; Enterobacterales; Enterobacteriaceae; Shigella

MTGDNTLIHSHGINRRDFMKLCAALAATMGLSSKAAAEMAESVTNPQRPPVIWIGAQECT WP_000145410.1 MULTISPECIES: hydrogenase-2 small chain [Proteobacteria] Length: 372\nNumber of predicted TMHs: 1\nExp number of AAs in TMHs: 23.72226\nExp number, first 60 AAs: 0.22038\nTotal prob of N-in: 0.12994\noutside 1 331\nTMhelix 332 354\ninside 355 372

27118 GCF_900158915.1_9803_4_92 Shigella sonnei Proteobacteria; Gammaproteobacteria; Enterobacterales; Enterobacteriaceae; Shigella MTGDNTLIHSHGINRRDFMKLCAALAATMGLSSKAAAEMAESVTNPQRPPVIWIGAQECT WP_000145410.1 MULTISPECIES: hydrogenase-2 small chain [Proteobacteria] Length: 372\nNumber of predicted TMHs: 1\nExp number of AAs in TMHs: 23.72226\nExp number, first 60 AAs: 0.22038\nTotal prob of N-in: 0.12994\noutside 1 331\nTMhelix 332 354\ninside 355 372

27119 GCF_900158965.1_16703_8_1 Shigella sonnei Proteobacteria; Gammaproteobacteria; Enterobacterales; Enterobacteriaceae; Shigella MTGDNTLIHSHGINRRDFMKLCAALAATMGLSSKAAAEMAESVTNPQRPPVIWIGAQECT WP_000145410.1 MULTISPECIES: hydrogenase-2 small chain [Proteobacteria] Length: 372\nNumber of predicted TMHs: 1\nExp number of AAs in TMHs: 23.72226\nExp number, first 60 AAs: 0.22038\nTotal prob of N-in: 0.12994\noutside 1 331\nTMhelix 332 354\ninside 355 372

27120 GCF_900158985.1_16703_8_3 Shigella sonnei Proteobacteria; Gammaproteobacteria; Enterobacterales; Enterobacteriaceae; Shigella MTGDNTLIHSHGINRRDFMKLCAALAATMGLSSKAAAEMAESVTNPQRPPVIWIGAQECT WP_000145410.1 MULTISPECIES: hydrogenase-2 small chain [Proteobacteria] Length: 372\nNumber of predicted TMHs: 1\nExp number of AAs in TMHs: 23.72226\nExp number, first 60 AAs: 0.22038\nTotal prob of N-in: 0.12994\noutside 1 331\nTMhelix 332 354\ninside 355 372

27121 GCF_900159015.1_16703_8_7 Shigella sonnei Proteobacteria; Gammaproteobacteria; Enterobacterales; Enterobacteriaceae; Shigella MTGDNTLIHSHGINRRDFMKLCAALAATMGLSSKAAAEMAESVTNPQRPPVIWIGAQECT WP_000145410.1 MULTISPECIES: hydrogenase-2 small chain [Proteobacteria] Length: 372\nNumber of predicted TMHs: 1\nExp number of AAs in TMHs: 23.72226\nExp number, first 60 AAs: 0.22038\nTotal prob of N-in: 0.12994\noutside 1 331\nTMhelix 332 354\ninside 355 372

27122 GCF_900159045.1_16703_8_10 Shigella sonnei Proteobacteria; Gammaproteobacteria; Enterobacterales; Enterobacteriaceae; Shigella MTGDNTLIHSHGINRRDFMKLCAALAATMGLSSKAAAEMAESVTNPQRPPVIWIGAQECT WP_000145410.1 MULTISPECIES: hydrogenase-2 small chain [Proteobacteria] Length: 372\nNumber of predicted TMHs: 1\nExp number of AAs in TMHs: 23.72226\nExp number, first 60 AAs: 0.22038\nTotal prob of N-in: 0.12994\noutside 1 331\nTMhelix 332 354\ninside 355 372

27123 GCF_900158965.1_16703_8_1 Shigella sonnei Proteobacteria; Gammaproteobacteria; Enterobacterales; Enterobacteriaceae; Shigella MNNEETFYQAMRRQGVTRRSFLKYCSLAATSLGLGAGMAPKIAWALENKPRIPVVWIHGL WP_005139779.1 hydrogenase [Shigella sonnei] Length: 372\nNumber of predicted TMHs: 2\nExp number of AAs in TMHs: 37.02628\nExp number, first 60 AAs: 19.39684\nTotal prob of N-in: 0.94313\nPOSSIBLE N-term signal sequence\ninside 1 19\nTMhelix 20 39\noutside 40 325\nTMhelix 326 348\ninside 349 372

27124 GCF_900158915.1_9803_4_92 Shigella sonnei Proteobacteria; Gammaproteobacteria; Enterobacterales; Enterobacteriaceae; Shigella MNNEETFYQAMRRQGVTRRSFLKYCSLAATSLGLGAGMAPKIAWALENKPRIPVVWIHGL WP_005139779.1 hydrogenase [Shigella sonnei] Length: 372\nNumber of predicted TMHs: 2\nExp number of AAs in TMHs: 37.02628\nExp number, first 60 AAs: 19.39684\nTotal prob of N-in: 0.94313\nPOSSIBLE N-term signal sequence\ninside 1 19\nTMhelix 20 39\noutside 40 325\nTMhelix 326 348\ninside 349 372

27125 GCF_900158835.1_9803_4_83 Shigella sonnei Proteobacteria; Gammaproteobacteria; Enterobacterales; Enterobacteriaceae; Shigella MNNEETFYQAMRRQGVTRRSFLKYCSLAATSLGLGAGMAPKIAWALENKPRIPVVWIHGL WP_005139779.1 hydrogenase [Shigella sonnei] Length: 372\nNumber of predicted TMHs: 2\nExp number of AAs in TMHs: 37.02628\nExp number, first 60 AAs: 19.39684\nTotal prob of N-in: 0.94313\nPOSSIBLE N-term signal sequence\ninside 1 19\nTMhelix 20 39\noutside 40 325\nTMhelix 326 348\ninside 349 372

27126 GCF_900158855.1_9803_4_85 Shigella sonnei Proteobacteria; Gammaproteobacteria; Enterobacterales; Enterobacteriaceae; Shigella
MNNEETFYQAMRRQGVTRRSFLKYCSLAATSLGLGAGMAPKIAWALENKPRIPVVWIHGL WP_005139779.1
hydrogenase [Shigella sonnei] Length: 372\nNumber of predicted TMHs: 2\nExp number of AAs in TMHs: 37.02628\nExp number, first 60 AAs: 19.39684\nTotal prob of N-in: 0.94313\nPOSSIBLE N-term signal sequence\ninside 1 19\nTMhelix 20 39\noutside 40 325\nTMhelix 326 348\ninside 349 372

27127 GCF_900158885.1_9803_4_89 Shigella sonnei Proteobacteria; Gammaproteobacteria; Enterobacterales; Enterobacteriaceae; Shigella
MNNEETFYQAMRRQGVTRRSFLKYCSLAATSLGLGAGMAPKIAWALENKPRIPVVWIHGL WP_005139779.1
hydrogenase [Shigella sonnei] Length: 372\nNumber of predicted TMHs: 2\nExp number of AAs in TMHs: 37.02628\nExp number, first 60 AAs: 19.39684\nTotal prob of N-in: 0.94313\nPOSSIBLE N-term signal sequence\ninside 1 19\nTMhelix 20 39\noutside 40 325\nTMhelix 326 348\ninside 349 372

27128 GCF_900158755.1_9803_4_75 Shigella sonnei Proteobacteria; Gammaproteobacteria; Enterobacterales; Enterobacteriaceae; Shigella
MNNEETFYQAMRRQGVTRRSFLKYCSLAATSLGLGAGMAPKIAWALENKPRIPVVWIHGL WP_005139779.1
hydrogenase [Shigella sonnei] Length: 372\nNumber of predicted TMHs: 2\nExp number of AAs in TMHs: 37.02628\nExp number, first 60 AAs: 19.39684\nTotal prob of N-in: 0.94313\nPOSSIBLE N-term signal sequence\ninside 1 19\nTMhelix 20 39\noutside 40 325\nTMhelix 326 348\ninside 349 372

27129 GCF_900158935.1_9803_4_94 Shigella sonnei Proteobacteria; Gammaproteobacteria; Enterobacterales; Enterobacteriaceae; Shigella
MNNEETFYQAMRRQGVTRRSFLKYCSLAATSLGLGAGMAPKIAWALENKPRIPVVWIHGL WP_005139779.1
hydrogenase [Shigella sonnei] Length: 372\nNumber of predicted TMHs: 2\nExp number of AAs in TMHs: 37.02628\nExp number, first 60 AAs: 19.39684\nTotal prob of N-in: 0.94313\nPOSSIBLE N-term signal sequence\ninside 1 19\nTMhelix 20 39\noutside 40 325\nTMhelix 326 348\ninside 349 372

27130 GCF_900158785.1_9803_4_78 Shigella sonnei Proteobacteria; Gammaproteobacteria; Enterobacterales; Enterobacteriaceae; Shigella
MNNEETFYQAMRRQGVTRRSFLKYCSLAATSLGLGAGMAPKIAWALENKPRIPVVWIHGL WP_005139779.1
hydrogenase [Shigella sonnei] Length: 372\nNumber of predicted TMHs: 2\nExp number of AAs in TMHs: 37.02628\nExp number, first 60 AAs: 19.39684\nTotal prob of N-in: 0.94313\nPOSSIBLE N-term signal sequence\ninside 1 19\nTMhelix 20 39\noutside 40 325\nTMhelix 326 348\ninside 349 372

27131 GCF_900158805.1_9803_4_80 Shigella sonnei Proteobacteria; Gammaproteobacteria; Enterobacterales; Enterobacteriaceae; Shigella
MNNEETFYQAMRRQGVTRRSFLKYCSLAATSLGLGAGMAPKIAWALENKPRIPVVWIHGL WP_005139779.1
hydrogenase [Shigella sonnei] Length: 372\nNumber of predicted TMHs: 2\nExp number of AAs in TMHs: 37.02628\nExp number, first 60 AAs: 19.39684\nTotal prob of N-in: 0.94313\nPOSSIBLE N-term signal sequence\ninside 1 19\nTMhelix 20 39\noutside 40 325\nTMhelix 326 348\ninside 349 372

27132 GCF_900158985.1_16703_8_3 Shigella sonnei Proteobacteria; Gammaproteobacteria; Enterobacterales; Enterobacteriaceae; Shigella
MNNEETFYQAMRRQGVTRRSFLKYCSLAATSLGLGAGMAPKIAWALENKPRIPVVWIHGL WP_005139779.1
hydrogenase [Shigella sonnei] Length: 372\nNumber of predicted TMHs: 2\nExp number of AAs in TMHs: 37.02628\nExp number, first 60 AAs: 19.39684\nTotal prob of N-in: 0.94313\nPOSSIBLE N-term signal sequence\ninside 1 19\nTMhelix 20 39\noutside 40 325\nTMhelix 326 348\ninside 349 372

27133 GCF_900159015.1_16703_8_7 Shigella sonnei Proteobacteria; Gammaproteobacteria; Enterobacterales; Enterobacteriaceae; Shigella
MNNEETFYQAMRRQGVTRRSFLKYCSLAATSLGLGAGMAPKIAWALENKPRIPVVWIHGL WP_005139779.1
hydrogenase [Shigella sonnei] Length: 372\nNumber of predicted TMHs: 2\nExp number of AAs in TMHs: 37.02628\nExp number, first 60 AAs: 19.39684\nTotal prob of N-in: 0.94313\nPOSSIBLE N-term signal sequence\ninside 1 19\nTMhelix 20 39\noutside 40 325\nTMhelix 326 348\ninside 349 372

27134 GCF_900159045.1_16703_8_10 Shigella sonnei Proteobacteria; Gammaproteobacteria; Enterobacterales; Enterobacteriaceae; Shigella
MNNEETFYQAMRRQGVTRRSFLKYCSLAATSLGLGAGMAPKIAWALENKPRIPVVWIHGL WP_005139779.1
hydrogenase [Shigella sonnei] Length: 372\nNumber of predicted TMHs: 2\nExp number of AAs in

TMHs: 37.02628\nExp number, first 60 AAs: 19.39684\nTotal prob of N-in: 0.94313\nPOSSIBLE N-term signal sequence\ninside 1 19\nTMhelix 20 39\noutside 40 325\nTMhelix 326 348\ninside 349 372

27135 GCF_900159065.1_16703_8_12 Shigella sonnei Proteobacteria; Gammaproteobacteria; Enterobacterales; Enterobacteriaceae; Shigella MTGDNTLIHSHGINRRDFMKLCAALAATMGLSSKAAAEMAESVTNPQRPPVIWIGAQECT WP_000145410.1 MULTISPECIES: hydrogenase-2 small chain [Proteobacteria] Length: 372\nNumber of predicted TMHs: 1\nExp number of AAs in TMHs: 23.72226\nExp number, first 60 AAs: 0.22038\nTotal prob of N-in: 0.12994\noutside 1 331\nTMhelix 332 354\ninside 355 372

27136 GCF_900159105.1_16703_8_16 Shigella sonnei Proteobacteria; Gammaproteobacteria; Enterobacterales; Enterobacteriaceae; Shigella MTGDNTLIHSHGINRRDFMKLCAALAATMGLSSKAAAEMAESVTNPQRPPVIWIGAQECT WP_000145410.1 MULTISPECIES: hydrogenase-2 small chain [Proteobacteria] Length: 372\nNumber of predicted TMHs: 1\nExp number of AAs in TMHs: 23.72226\nExp number, first 60 AAs: 0.22038\nTotal prob of N-in: 0.12994\noutside 1 331\nTMhelix 332 354\ninside 355 372

27137 GCF_900159075.1_16703_8_13 Shigella sonnei Proteobacteria; Gammaproteobacteria; Enterobacterales; Enterobacteriaceae; Shigella MTGDNTLIHSHGINRRDFMKLCAALAATMGLSSKAAAEMAESVTNPQRPPVIWIGAQECT WP_000145410.1 MULTISPECIES: hydrogenase-2 small chain [Proteobacteria] Length: 372\nNumber of predicted TMHs: 1\nExp number of AAs in TMHs: 23.72226\nExp number, first 60 AAs: 0.22038\nTotal prob of N-in: 0.12994\noutside 1 331\nTMhelix 332 354\ninside 355 372

27138 GCF_900159085.1_16703_8_14 Shigella sonnei Proteobacteria; Gammaproteobacteria; Enterobacterales; Enterobacteriaceae; Shigella MTGDNTLIHSHGINRRDFMKLCAALAATMGLSSKAAAEMAESVTNPQRPPVIWIGAQECT WP_000145410.1 MULTISPECIES: hydrogenase-2 small chain [Proteobacteria] Length: 372\nNumber of predicted TMHs: 1\nExp number of AAs in TMHs: 23.72226\nExp number, first 60 AAs: 0.22038\nTotal prob of N-in: 0.12994\noutside 1 331\nTMhelix 332 354\ninside 355 372

27139 GCF_900159135.1_16703_8_20 Shigella sonnei Proteobacteria; Gammaproteobacteria; Enterobacterales; Enterobacteriaceae; Shigella MTGDNTLIHSHGINRRDFMKLCAALAATMGLSSKAAAEMAESVTNPQRPPVIWIGAQECT WP_000145410.1 MULTISPECIES: hydrogenase-2 small chain [Proteobacteria] Length: 372\nNumber of predicted TMHs: 1\nExp number of AAs in TMHs: 23.72226\nExp number, first 60 AAs: 0.22038\nTotal prob of N-in: 0.12994\noutside 1 331\nTMhelix 332 354\ninside 355 372

27140 GCF_900159185.1_16703_8_25 Shigella sonnei Proteobacteria; Gammaproteobacteria; Enterobacterales; Enterobacteriaceae; Shigella MTGDNTLIHSHGINRRDFMKLCAALAATMGLSSKAAAEMAESVTNPQRPPVIWIGAQECT WP_000145410.1 MULTISPECIES: hydrogenase-2 small chain [Proteobacteria] Length: 372\nNumber of predicted TMHs: 1\nExp number of AAs in TMHs: 23.72226\nExp number, first 60 AAs: 0.22038\nTotal prob of N-in: 0.12994\noutside 1 331\nTMhelix 332 354\ninside 355 372

27141 GCF_900159165.1_16703_8_23 Shigella sonnei Proteobacteria; Gammaproteobacteria; Enterobacterales; Enterobacteriaceae; Shigella MTGDNTLIHSHGINRRDFMKLCAALAATMGLSSKAAAEMAESVTNPQRPPVIWIGAQECT WP_000145410.1 MULTISPECIES: hydrogenase-2 small chain [Proteobacteria] Length: 372\nNumber of predicted TMHs: 1\nExp number of AAs in TMHs: 23.72226\nExp number, first 60 AAs: 0.22038\nTotal prob of N-in: 0.12994\noutside 1 331\nTMhelix 332 354\ninside 355 372

27142 GCF_900159065.1_16703_8_12 Shigella sonnei Proteobacteria; Gammaproteobacteria; Enterobacterales; Enterobacteriaceae; Shigella MNNEETFYQAMRRQGVTRRSFLKYCSLAATSLGLGAGMAPKIAWALENKPRIPVVWIHGL WP_005139779.1 hydrogenase [Shigella sonnei] Length: 372\nNumber of predicted TMHs: 2\nExp number of AAs in TMHs: 37.02628\nExp number, first 60 AAs: 19.39684\nTotal prob of N-in: 0.94313\nPOSSIBLE N-term signal sequence\ninside 1 19\nTMhelix 20 39\noutside 40 325\nTMhelix 326 348\ninside 349 372

27143 GCF_900159295.1_16703_8_37 Shigella sonnei Proteobacteria; Gammaproteobacteria; Enterobacterales; Enterobacteriaceae; Shigella

MTGDNTLIHSHGINRRDFMKLCAALAATMGLSSKAAAEMAESVTNPQRPPVIWIGAQECT WP_000145410.1 MULTISPECIES: hydrogenase-2 small chain [Proteobacteria] Length: 372\nNumber of predicted TMHs: 1\nExp number of AAs in TMHs: 23.72226\nExp number, first 60 AAs: 0.22038\nTotal prob of N-in: 0.12994\noutside 1 331\nTMhelix 332 354\ninside 355 372

27144 GCF_900159105.1_16703_8_16 Shigella sonnei Proteobacteria; Gammaproteobacteria; Enterobacterales; Enterobacteriaceae; Shigella MNNEETFYQAMRRQGVTRRSFLKYCSLAATSLGLGAGMAPKIAWALENKPRIPVVWIHGL WP_005139779.1 hydrogenase [Shigella sonnei] Length: 372\nNumber of predicted TMHs: 2\nExp number of AAs in TMHs: 37.02628\nExp number, first 60 AAs: 19.39684\nTotal prob of N-in: 0.94313\nPOSSIBLE N-term signal sequence\ninside 1 19\nTMhelix 20 39\noutside 40 325\nTMhelix 326 348\ninside 349 372

27145 GCF_900159185.1_16703_8_25 Shigella sonnei Proteobacteria; Gammaproteobacteria; Enterobacterales; Enterobacteriaceae; Shigella MNNEETFYQAMRRQGVTRRSFLKYCSLAATSLGLGAGMAPKIAWALENKPRIPVVWIHGL WP_005139779.1 hydrogenase [Shigella sonnei] Length: 372\nNumber of predicted TMHs: 2\nExp number of AAs in TMHs: 37.02628\nExp number, first 60 AAs: 19.39684\nTotal prob of N-in: 0.94313\nPOSSIBLE N-term signal sequence\ninside 1 19\nTMhelix 20 39\noutside 40 325\nTMhelix 326 348\ninside 349 372

27146 GCF_900159085.1_16703_8_14 Shigella sonnei Proteobacteria; Gammaproteobacteria; Enterobacterales; Enterobacteriaceae; Shigella MNNEETFYQAMRRQGVTRRSFLKYCSLAATSLGLGAGMAPKIAWALENKPRIPVVWIHGL WP_005139779.1 hydrogenase [Shigella sonnei] Length: 372\nNumber of predicted TMHs: 2\nExp number of AAs in TMHs: 37.02628\nExp number, first 60 AAs: 19.39684\nTotal prob of N-in: 0.94313\nPOSSIBLE N-term signal sequence\ninside 1 19\nTMhelix 20 39\noutside 40 325\nTMhelix 326 348\ninside 349 372

27147 GCF_900159075.1_16703_8_13 Shigella sonnei Proteobacteria; Gammaproteobacteria; Enterobacterales; Enterobacteriaceae; Shigella MNNEETFYQAMRRQGVTRRSFLKYCSLAATSLGLGAGMAPKIAWALENKPRIPVVWIHGL WP_005139779.1 hydrogenase [Shigella sonnei] Length: 372\nNumber of predicted TMHs: 2\nExp number of AAs in TMHs: 37.02628\nExp number, first 60 AAs: 19.39684\nTotal prob of N-in: 0.94313\nPOSSIBLE N-term signal sequence\ninside 1 19\nTMhelix 20 39\noutside 40 325\nTMhelix 326 348\ninside 349 372

27148 GCF_900159135.1_16703_8_20 Shigella sonnei Proteobacteria; Gammaproteobacteria; Enterobacterales; Enterobacteriaceae; Shigella MNNEETFYQAMRRQGVTRRSFLKYCSLAATSLGLGAGMAPKIAWALENKPRIPVVWIHGL WP_005139779.1 hydrogenase [Shigella sonnei] Length: 372\nNumber of predicted TMHs: 2\nExp number of AAs in TMHs: 37.02628\nExp number, first 60 AAs: 19.39684\nTotal prob of N-in: 0.94313\nPOSSIBLE N-term signal sequence\ninside 1 19\nTMhelix 20 39\noutside 40 325\nTMhelix 326 348\ninside 349 372

27149 GCF_900159165.1_16703_8_23 Shigella sonnei Proteobacteria; Gammaproteobacteria; Enterobacterales; Enterobacteriaceae; Shigella MNNEETFYQAMRRQGVTRRSFLKYCSLAATSLGLGAGMAPKIAWALENKPRIPVVWIHGL WP_005139779.1 hydrogenase [Shigella sonnei] Length: 372\nNumber of predicted TMHs: 2\nExp number of AAs in TMHs: 37.02628\nExp number, first 60 AAs: 19.39684\nTotal prob of N-in: 0.94313\nPOSSIBLE N-term signal sequence\ninside 1 19\nTMhelix 20 39\noutside 40 325\nTMhelix 326 348\ninside 349 372

27150 GCF_900159325.1_16703_8_43 Shigella sonnei Proteobacteria; Gammaproteobacteria; Enterobacterales; Enterobacteriaceae; Shigella MTGDNTLIHSHGINRRDFMKLCAALAATMGLSSKAAAEMAESVTNPQRPPVIWIGAQECT WP_000145410.1 MULTISPECIES: hydrogenase-2 small chain [Proteobacteria] Length: 372\nNumber of predicted TMHs: 1\nExp number of AAs in TMHs: 23.72226\nExp number, first 60 AAs: 0.22038\nTotal prob of N-in: 0.12994\noutside 1 331\nTMhelix 332 354\ninside 355 372

27151 GCF_900159485.1_8489_1_70 Shigella sonnei Proteobacteria; Gammaproteobacteria; Enterobacterales; Enterobacteriaceae; Shigella MTGDNTLIHSHGINRRDFMKLCAALAATMGLSSKAAAEMAESVTNPQRPPVIWIGAQECT WP_000145410.1 MULTISPECIES: hydrogenase-2 small chain [Proteobacteria] Length: 372\nNumber of predicted TMHs: 1\nExp number of AAs in TMHs: 23.72226\nExp number, first 60 AAs: 0.22038\nTotal prob of N-in: 0.12994\noutside 1 331\nTMhelix 332 354\ninside 355 372

27152 GCF_900159265.1_16703_8_33 Shigella sonnei Proteobacteria; Gammaproteobacteria; Enterobacterales; Enterobacteriaceae; Shigella
MTGDNTLIHSHGINRRDFMKLCAALAATMGLSSKAAAEMAESVTNPQRPPVIWIGAQECT WP_000145410.1
MULTISPECIES: hydrogenase-2 small chain [Proteobacteria] Length: 372\nNumber of predicted TMHs: 1\nExp number of AAs in TMHs: 23.72226\nExp number, first 60 AAs: 0.22038\nTotal prob of N-in: 0.12994\noutside 1 331\nTMhelix 332 354\ninside 355 372

27153 GCF_900159425.1_8489_1_67 Shigella sonnei Proteobacteria; Gammaproteobacteria; Enterobacterales; Enterobacteriaceae; Shigella
MTGDNTLIHSHGINRRDFMKLCAALAATMGLSSKAAAEMAESVTNPQRPPVIWIGAQECT WP_000145410.1
MULTISPECIES: hydrogenase-2 small chain [Proteobacteria] Length: 372\nNumber of predicted TMHs: 1\nExp number of AAs in TMHs: 23.72226\nExp number, first 60 AAs: 0.22038\nTotal prob of N-in: 0.12994\noutside 1 331\nTMhelix 332 354\ninside 355 372

27154 GCF_900159405.1_8489_1_63 Shigella sonnei Proteobacteria; Gammaproteobacteria; Enterobacterales; Enterobacteriaceae; Shigella
MTGDNTLIHSHGINRRDFMKLCAALAATMGLSSKAAAEMAESVTNPQRPPVIWIGAQECT WP_000145410.1
MULTISPECIES: hydrogenase-2 small chain [Proteobacteria] Length: 372\nNumber of predicted TMHs: 1\nExp number of AAs in TMHs: 23.72226\nExp number, first 60 AAs: 0.22038\nTotal prob of N-in: 0.12994\noutside 1 331\nTMhelix 332 354\ninside 355 372

27155 GCF_900159345.1_16703_8_49 Shigella sonnei Proteobacteria; Gammaproteobacteria; Enterobacterales; Enterobacteriaceae; Shigella
MTGDNTLIHSHGINRRDFMKLCAALAATMGLSSKAAAEMAESVTNPQRPPVIWIGAQECT WP_000145410.1
MULTISPECIES: hydrogenase-2 small chain [Proteobacteria] Length: 372\nNumber of predicted TMHs: 1\nExp number of AAs in TMHs: 23.72226\nExp number, first 60 AAs: 0.22038\nTotal prob of N-in: 0.12994\noutside 1 331\nTMhelix 332 354\ninside 355 372

27156 GCF_900159445.1_8489_1_65 Shigella sonnei Proteobacteria; Gammaproteobacteria; Enterobacterales; Enterobacteriaceae; Shigella
MTGDNTLIHSHGINRRDFMKLCAALAATMGLSSKAAAEMAESVTNPQRPPVIWIGAQECT WP_000145410.1
MULTISPECIES: hydrogenase-2 small chain [Proteobacteria] Length: 372\nNumber of predicted TMHs: 1\nExp number of AAs in TMHs: 23.72226\nExp number, first 60 AAs: 0.22038\nTotal prob of N-in: 0.12994\noutside 1 331\nTMhelix 332 354\ninside 355 372

27157 GCF_900159215.1_16703_8_28 Shigella sonnei Proteobacteria; Gammaproteobacteria; Enterobacterales; Enterobacteriaceae; Shigella
MTGDNTLIHSHGINRRDFMKLCAALAATMGLSSKAAAEMAESVTNPQRPPVIWIGAQECT WP_000145410.1
MULTISPECIES: hydrogenase-2 small chain [Proteobacteria] Length: 372\nNumber of predicted TMHs: 1\nExp number of AAs in TMHs: 23.72226\nExp number, first 60 AAs: 0.22038\nTotal prob of N-in: 0.12994\noutside 1 331\nTMhelix 332 354\ninside 355 372

27158 GCF_900159375.1_8489_1_60 Shigella sonnei Proteobacteria; Gammaproteobacteria; Enterobacterales; Enterobacteriaceae; Shigella
MTGDNTLIHSHGINRRDFMKLCAALAATMGLSSKAAAEMAESVTNPQRPPVIWIGAQECT WP_000145410.1
MULTISPECIES: hydrogenase-2 small chain [Proteobacteria] Length: 372\nNumber of predicted TMHs: 1\nExp number of AAs in TMHs: 23.72226\nExp number, first 60 AAs: 0.22038\nTotal prob of N-in: 0.12994\noutside 1 331\nTMhelix 332 354\ninside 355 372

27159 GCF_900159245.1_16703_8_31 Shigella sonnei Proteobacteria; Gammaproteobacteria; Enterobacterales; Enterobacteriaceae; Shigella
MTGDNTLIHSHGINRRDFMKLCAALAATMGLSSKAAAEMAESVTNPQRPPVIWIGAQECT WP_000145410.1
MULTISPECIES: hydrogenase-2 small chain [Proteobacteria] Length: 372\nNumber of predicted TMHs: 1\nExp number of AAs in TMHs: 23.72226\nExp number, first 60 AAs: 0.22038\nTotal prob of N-in: 0.12994\noutside 1 331\nTMhelix 332 354\ninside 355 372

27160 GCF_900159475.1_8489_1_71 Shigella sonnei Proteobacteria; Gammaproteobacteria; Enterobacterales; Enterobacteriaceae; Shigella
MTGDNTLIHSHGINRRDFMKLCAALAATMGLSSKAAAEMAESVTNPQRPPVIWIGAQECT WP_000145410.1
MULTISPECIES: hydrogenase-2 small chain [Proteobacteria] Length: 372\nNumber of predicted TMHs:

1\nExp number of AAs in TMHs: 23.72226\nExp number, first 60 AAs: 0.22038\nTotal prob of N-in: 0.12994\noutside 1 331\nTMhelix 332 354\ninside 355 372

27161 GCF_900159375.1_8489_1_60 Shigella sonnei Proteobacteria; Gammaproteobacteria; Enterobacterales; Enterobacteriaceae; Shigella
MNNEETFYQAMRRQGVTRRSFLKYCSLAATSLGLGAGMAPKIAWALENKPRIPVVWIHGL WP_005139779.1 hydrogenase [Shigella sonnei] Length: 372\nNumber of predicted TMHs: 2\nExp number of AAs in TMHs: 37.02628\nExp number, first 60 AAs: 19.39684\nTotal prob of N-in: 0.94313\nPOSSIBLE N-term signal sequence\ninside 1 19\nTMhelix 20 39\noutside 40 325\nTMhelix 326 348\ninside 349 372

27162 GCF_900159445.1_8489_1_65 Shigella sonnei Proteobacteria; Gammaproteobacteria; Enterobacterales; Enterobacteriaceae; Shigella
MNNEETFYQAMRRQGVTRRSFLKYCSLAATSLGLGAGMAPKIAWALENKPRIPVVWIHGL WP_005139779.1 hydrogenase [Shigella sonnei] Length: 372\nNumber of predicted TMHs: 2\nExp number of AAs in TMHs: 37.02628\nExp number, first 60 AAs: 19.39684\nTotal prob of N-in: 0.94313\nPOSSIBLE N-term signal sequence\ninside 1 19\nTMhelix 20 39\noutside 40 325\nTMhelix 326 348\ninside 349 372

27163 GCF_900159405.1_8489_1_63 Shigella sonnei Proteobacteria; Gammaproteobacteria; Enterobacterales; Enterobacteriaceae; Shigella
MNNEETFYQAMRRQGVTRRSFLKYCSLAATSLGLGAGMAPKIAWALENKPRIPVVWIHGL WP_005139779.1 hydrogenase [Shigella sonnei] Length: 372\nNumber of predicted TMHs: 2\nExp number of AAs in TMHs: 37.02628\nExp number, first 60 AAs: 19.39684\nTotal prob of N-in: 0.94313\nPOSSIBLE N-term signal sequence\ninside 1 19\nTMhelix 20 39\noutside 40 325\nTMhelix 326 348\ninside 349 372

27164 GCF_900159325.1_16703_8_43 Shigella sonnei Proteobacteria; Gammaproteobacteria; Enterobacterales; Enterobacteriaceae; Shigella
MNNEETFYQAMRRQGVTRRSFLKYCSLAATSLGLGAGMAPKIAWALENKPRIPVVWIHGL WP_005139779.1 hydrogenase [Shigella sonnei] Length: 372\nNumber of predicted TMHs: 2\nExp number of AAs in TMHs: 37.02628\nExp number, first 60 AAs: 19.39684\nTotal prob of N-in: 0.94313\nPOSSIBLE N-term signal sequence\ninside 1 19\nTMhelix 20 39\noutside 40 325\nTMhelix 326 348\ninside 349 372

27165 GCF_900159425.1_8489_1_67 Shigella sonnei Proteobacteria; Gammaproteobacteria; Enterobacterales; Enterobacteriaceae; Shigella
MNNEETFYQAMRRQGVTRRSFLKYCSLAATSLGLGAGMAPKIAWALENKPRIPVVWIHGL WP_005139779.1 hydrogenase [Shigella sonnei] Length: 372\nNumber of predicted TMHs: 2\nExp number of AAs in TMHs: 37.02628\nExp number, first 60 AAs: 19.39684\nTotal prob of N-in: 0.94313\nPOSSIBLE N-term signal sequence\ninside 1 19\nTMhelix 20 39\noutside 40 325\nTMhelix 326 348\ninside 349 372

27166 GCF_900159485.1_8489_1_70 Shigella sonnei Proteobacteria; Gammaproteobacteria; Enterobacterales; Enterobacteriaceae; Shigella
MNNEETFYQAMRRQGVTRRSFLKYCSLAATSLGLGAGMAPKIAWALENKPRIPVVWIHGL WP_005139779.1 hydrogenase [Shigella sonnei] Length: 372\nNumber of predicted TMHs: 2\nExp number of AAs in TMHs: 37.02628\nExp number, first 60 AAs: 19.39684\nTotal prob of N-in: 0.94313\nPOSSIBLE N-term signal sequence\ninside 1 19\nTMhelix 20 39\noutside 40 325\nTMhelix 326 348\ninside 349 372

27167 GCF_900159295.1_16703_8_37 Shigella sonnei Proteobacteria; Gammaproteobacteria; Enterobacterales; Enterobacteriaceae; Shigella
MNNEETFYQAMRRQGVTRRSFLKYCSLAATSLGLGAGMAPKIAWALENKPRIPVVWIHGL WP_005139779.1 hydrogenase [Shigella sonnei] Length: 372\nNumber of predicted TMHs: 2\nExp number of AAs in TMHs: 37.02628\nExp number, first 60 AAs: 19.39684\nTotal prob of N-in: 0.94313\nPOSSIBLE N-term signal sequence\ninside 1 19\nTMhelix 20 39\noutside 40 325\nTMhelix 326 348\ninside 349 372

27168 GCF_900159215.1_16703_8_28 Shigella sonnei Proteobacteria; Gammaproteobacteria; Enterobacterales; Enterobacteriaceae; Shigella
MNNEETFYQAMRRQGVTRRSFLKYCSLAATSLGLGAGMAPKIAWALENKPRIPVVWIHGL WP_005139779.1 hydrogenase [Shigella sonnei] Length: 372\nNumber of predicted TMHs: 2\nExp number of AAs in TMHs: 37.02628\nExp number, first 60 AAs: 19.39684\nTotal prob of N-in: 0.94313\nPOSSIBLE N-term signal sequence\ninside 1 19\nTMhelix 20 39\noutside 40 325\nTMhelix 326 348\ninside 349 372

27169 GCF_900159265.1_16703_8_33 Shigella sonnei Proteobacteria; Gammaproteobacteria; Enterobacterales; Enterobacteriaceae; Shigella

MNNEETFYQAMRRQGVTRRSFLKYCSLAATSLGLGAGMAPKIAWALENKPRIPVVWIHGL	WP_005139779.1	hydrogenase [Shigella sonnei]	Length: 372\nNumber of predicted TMHs: 2\nExp number of AAs in TMHs: 37.02628\nExp number, first 60 AAs: 19.39684\nTotal prob of N-in: 0.94313\nPOSSIBLE N-term signal sequence\ninside 1 19\nTMhelix 20 39\noutside 40 325\nTMhelix 326 348\ninside 349 372

27170	GCF_900159345.1_16703_8_49	Shigella sonnei	Proteobacteria; Gammaproteobacteria; Enterobacterales; Enterobacteriaceae; Shigella	MNNEETFYQAMRRQGVTRRSFLKYCSLAATSLGLGAGMAPKIAWALENKPRIPVVWIHGL	WP_005139779.1	hydrogenase [Shigella sonnei]	Length: 372\nNumber of predicted TMHs: 2\nExp number of AAs in TMHs: 37.02628\nExp number, first 60 AAs: 19.39684\nTotal prob of N-in: 0.94313\nPOSSIBLE N-term signal sequence\ninside 1 19\nTMhelix 20 39\noutside 40 325\nTMhelix 326 348\ninside 349 372

27171	GCF_900159475.1_8489_1_71	Shigella sonnei	Proteobacteria; Gammaproteobacteria; Enterobacterales; Enterobacteriaceae; Shigella	MNNEETFYQAMRRQGVTRRSFLKYCSLAATSLGLGAGMAPKIAWALENKPRIPVVWIHGL	WP_005139779.1	hydrogenase [Shigella sonnei]	Length: 372\nNumber of predicted TMHs: 2\nExp number of AAs in TMHs: 37.02628\nExp number, first 60 AAs: 19.39684\nTotal prob of N-in: 0.94313\nPOSSIBLE N-term signal sequence\ninside 1 19\nTMhelix 20 39\noutside 40 325\nTMhelix 326 348\ninside 349 372

27172	GCF_900159245.1_16703_8_31	Shigella sonnei	Proteobacteria; Gammaproteobacteria; Enterobacterales; Enterobacteriaceae; Shigella	MNNEETFYQAMRRQGVTRRSFLKYCSLAATSLGLGAGMAPKIAWALENKPRIPVVWIHGL	WP_005139779.1	hydrogenase [Shigella sonnei]	Length: 372\nNumber of predicted TMHs: 2\nExp number of AAs in TMHs: 37.02628\nExp number, first 60 AAs: 19.39684\nTotal prob of N-in: 0.94313\nPOSSIBLE N-term signal sequence\ninside 1 19\nTMhelix 20 39\noutside 40 325\nTMhelix 326 348\ninside 349 372

27173	GCF_900159535.1_8489_1_77	Shigella sonnei	Proteobacteria; Gammaproteobacteria; Enterobacterales; Enterobacteriaceae; Shigella	MTGDNTLIHSHGINRRDFMKLCAALAATMGLSSKAAAEMAESVTNPQRPPVIWIGAQECT	WP_000145410.1	MULTISPECIES: hydrogenase-2 small chain [Proteobacteria]	Length: 372\nNumber of predicted TMHs: 1\nExp number of AAs in TMHs: 23.72226\nExp number, first 60 AAs: 0.22038\nTotal prob of N-in: 0.12994\noutside 1 331\nTMhelix 332 354\ninside 355 372

27174	GCF_900159515.1_8489_1_75	Shigella sonnei	Proteobacteria; Gammaproteobacteria; Enterobacterales; Enterobacteriaceae; Shigella	MTGDNTLIHSHGINRRDFMKLCAALAATMGLSSKAAAEMAESVTNPQRPPVIWIGAQECT	WP_000145410.1	MULTISPECIES: hydrogenase-2 small chain [Proteobacteria]	Length: 372\nNumber of predicted TMHs: 1\nExp number of AAs in TMHs: 23.72226\nExp number, first 60 AAs: 0.22038\nTotal prob of N-in: 0.12994\noutside 1 331\nTMhelix 332 354\ninside 355 372

27175	GCF_900159565.1_8489_1_80	Shigella sonnei	Proteobacteria; Gammaproteobacteria; Enterobacterales; Enterobacteriaceae; Shigella	MTGDNTLIHSHGINRRDFMKLCAALAATMGLSSKAAAEMAESVTNPQRPPVIWIGAQECT	WP_000145410.1	MULTISPECIES: hydrogenase-2 small chain [Proteobacteria]	Length: 372\nNumber of predicted TMHs: 1\nExp number of AAs in TMHs: 23.72226\nExp number, first 60 AAs: 0.22038\nTotal prob of N-in: 0.12994\noutside 1 331\nTMhelix 332 354\ninside 355 372

27176	GCF_900159635.1_8489_1_88	Shigella sonnei	Proteobacteria; Gammaproteobacteria; Enterobacterales; Enterobacteriaceae; Shigella	MTGDNTLIHSHGINRRDFMKLCAALAATMGLSSKAAAEMAESVTNPQRPPVIWIGAQECT	WP_000145410.1	MULTISPECIES: hydrogenase-2 small chain [Proteobacteria]	Length: 372\nNumber of predicted TMHs: 1\nExp number of AAs in TMHs: 23.72226\nExp number, first 60 AAs: 0.22038\nTotal prob of N-in: 0.12994\noutside 1 331\nTMhelix 332 354\ninside 355 372

27177	GCF_900159615.1_8489_1_85	Shigella sonnei	Proteobacteria; Gammaproteobacteria; Enterobacterales; Enterobacteriaceae; Shigella	MTGDNTLIHSHGINRRDFMKLCAALAATMGLSSKAAAEMAESVTNPQRPPVIWIGAQECT	WP_000145410.1	MULTISPECIES: hydrogenase-2 small chain [Proteobacteria]	Length: 372\nNumber of predicted TMHs: 1\nExp number of AAs in TMHs: 23.72226\nExp number, first 60 AAs: 0.22038\nTotal prob of N-in: 0.12994\noutside 1 331\nTMhelix 332 354\ninside 355 372

27178 GCF_900159585.1_8489_1_83 Shigella sonnei Proteobacteria; Gammaproteobacteria; Enterobacterales; Enterobacteriaceae; Shigella MTGDNTLIHSHGINRRDFMKLCAALAATMGLSSKAAAEMAESVTNPQRPPVIWIGAQECT WP_000145410.1 MULTISPECIES: hydrogenase-2 small chain [Proteobacteria] Length: 372\nNumber of predicted TMHs: 1\nExp number of AAs in TMHs: 23.72226\nExp number, first 60 AAs: 0.22038\nTotal prob of N-in: 0.12994\noutside 1 331\nTMhelix 332 354\ninside 355 372

27179 GCF_900159655.1_8489_1_90 Shigella sonnei Proteobacteria; Gammaproteobacteria; Enterobacterales; Enterobacteriaceae; Shigella MTGDNTLIHSHGINRRDFMKLCAALAATMGLSSKAAAEMAESVTNPQRPPVIWIGAQECT WP_000145410.1 MULTISPECIES: hydrogenase-2 small chain [Proteobacteria] Length: 372\nNumber of predicted TMHs: 1\nExp number of AAs in TMHs: 23.72226\nExp number, first 60 AAs: 0.22038\nTotal prob of N-in: 0.12994\noutside 1 331\nTMhelix 332 354\ninside 355 372

27180 GCF_900159685.1_8489_1_93 Shigella sonnei Proteobacteria; Gammaproteobacteria; Enterobacterales; Enterobacteriaceae; Shigella MTGDNTLIHSHGINRRDFMKLCAALAATMGLSSKAAAEMAESVTNPQRPPVIWIGAQECT WP_000145410.1 MULTISPECIES: hydrogenase-2 small chain [Proteobacteria] Length: 372\nNumber of predicted TMHs: 1\nExp number of AAs in TMHs: 23.72226\nExp number, first 60 AAs: 0.22038\nTotal prob of N-in: 0.12994\noutside 1 331\nTMhelix 332 354\ninside 355 372

27181 GCF_900159705.1_8489_1_95 Shigella sonnei Proteobacteria; Gammaproteobacteria; Enterobacterales; Enterobacteriaceae; Shigella MTGDNTLIHSHGINRRDFMKLCAALAATMGLSSKAAAEMAESVTNPQRPPVIWIGAQECT WP_000145410.1 MULTISPECIES: hydrogenase-2 small chain [Proteobacteria] Length: 372\nNumber of predicted TMHs: 1\nExp number of AAs in TMHs: 23.72226\nExp number, first 60 AAs: 0.22038\nTotal prob of N-in: 0.12994\noutside 1 331\nTMhelix 332 354\ninside 355 372

27182 GCF_900159735.1_8403_8_3Shigella sonnei Proteobacteria; Gammaproteobacteria; Enterobacterales; Enterobacteriaceae; Shigella MTGDNTLIHSHGINRRDFMKLCAALAATMGLSSKAAAEMAESVTNPQRPPVIWIGAQECT WP_000145410.1 MULTISPECIES: hydrogenase-2 small chain [Proteobacteria] Length: 372\nNumber of predicted TMHs: 1\nExp number of AAs in TMHs: 23.72226\nExp number, first 60 AAs: 0.22038\nTotal prob of N-in: 0.12994\noutside 1 331\nTMhelix 332 354\ninside 355 372

27183 GCF_900159755.1_8403_8_5Shigella sonnei Proteobacteria; Gammaproteobacteria; Enterobacterales; Enterobacteriaceae; Shigella MTGDNTLIHSHGINRRDFMKLCAALAATMGLSSKAAAEMAESVTNPQRPPVIWIGAQECT WP_000145410.1 MULTISPECIES: hydrogenase-2 small chain [Proteobacteria] Length: 372\nNumber of predicted TMHs: 1\nExp number of AAs in TMHs: 23.72226\nExp number, first 60 AAs: 0.22038\nTotal prob of N-in: 0.12994\noutside 1 331\nTMhelix 332 354\ninside 355 372

27184 GCF_900159785.1_8403_8_8Shigella sonnei Proteobacteria; Gammaproteobacteria; Enterobacterales; Enterobacteriaceae; Shigella MTGDNTLIHSHGINRRDFMKLCAALAATMGLSSKAAAEMAESVTNPQRPPVIWIGAQECT WP_000145410.1 MULTISPECIES: hydrogenase-2 small chain [Proteobacteria] Length: 372\nNumber of predicted TMHs: 1\nExp number of AAs in TMHs: 23.72226\nExp number, first 60 AAs: 0.22038\nTotal prob of N-in: 0.12994\noutside 1 331\nTMhelix 332 354\ninside 355 372

27185 GCF_900159685.1_8489_1_93 Shigella sonnei Proteobacteria; Gammaproteobacteria; Enterobacterales; Enterobacteriaceae; Shigella MNNEETFYQAMRRQGVTRRSFLKYCSLAATSLGLGAGMAPKIAWALENKPRIPVVWIHGL WP_005139779.1 hydrogenase [Shigella sonnei] Length: 372\nNumber of predicted TMHs: 2\nExp number of AAs in TMHs: 37.02628\nExp number, first 60 AAs: 19.39684\nTotal prob of N-in: 0.94313\nPOSSIBLE N-term signal sequence\ninside 1 19\nTMhelix 20 39\noutside 40 325\nTMhelix 326 348\ninside 349 372

27186 GCF_900159785.1_8403_8_8Shigella sonnei Proteobacteria; Gammaproteobacteria; Enterobacterales; Enterobacteriaceae; Shigella MNNEETFYQAMRRQGVTRRSFLKYCSLAATSLGLGAGMAPKIAWALENKPRIPVVWIHGL WP_005139779.1 hydrogenase [Shigella sonnei] Length: 372\nNumber of predicted TMHs: 2\nExp number of AAs in TMHs: 37.02628\nExp number, first 60 AAs: 19.39684\nTotal prob of N-in: 0.94313\nPOSSIBLE N-term signal sequence\ninside 1 19\nTMhelix 20 39\noutside 40 325\nTMhelix 326 348\ninside 349 372

27187 GCF_900159935.1_8403_8_23 Shigella sonnei Proteobacteria; Gammaproteobacteria; Enterobacterales; Enterobacteriaceae; Shigella

MTGDNTLIHSHGINRRDFMKLCAALAATMGLSSKAAAEMAESVTNPQRPPVIWIGAQECT WP_000145410.1 MULTISPECIES: hydrogenase-2 small chain [Proteobacteria] Length: 372\nNumber of predicted TMHs: 1\nExp number of AAs in TMHs: 23.72226\nExp number, first 60 AAs: 0.22038\nTotal prob of N-in: 0.12994\noutside 1 331\nTMhelix 332 354\ninside 355 372

27188 GCF_900159515.1_8489_1_75 Shigella sonnei Proteobacteria; Gammaproteobacteria; Enterobacterales; Enterobacteriaceae; Shigella MNNEETFYQAMRRQGVTRRSFLKYCSLAATSLGLGAGMAPKIAWALENKPRIPVVWIHGL WP_005139779.1 hydrogenase [Shigella sonnei] Length: 372\nNumber of predicted TMHs: 2\nExp number of AAs in TMHs: 37.02628\nExp number, first 60 AAs: 19.39684\nTotal prob of N-in: 0.94313\nPOSSIBLE N-term signal sequence\ninside 1 19\nTMhelix 20 39\noutside 40 325\nTMhelix 326 348\ninside 349 372

27189 GCF_900159535.1_8489_1_77 Shigella sonnei Proteobacteria; Gammaproteobacteria; Enterobacterales; Enterobacteriaceae; Shigella MNNEETFYQAMRRQGVTRRSFLKYCSLAATSLGLGAGMAPKIAWALENKPRIPVVWIHGL WP_005139779.1 hydrogenase [Shigella sonnei] Length: 372\nNumber of predicted TMHs: 2\nExp number of AAs in TMHs: 37.02628\nExp number, first 60 AAs: 19.39684\nTotal prob of N-in: 0.94313\nPOSSIBLE N-term signal sequence\ninside 1 19\nTMhelix 20 39\noutside 40 325\nTMhelix 326 348\ninside 349 372

27190 GCF_900159565.1_8489_1_80 Shigella sonnei Proteobacteria; Gammaproteobacteria; Enterobacterales; Enterobacteriaceae; Shigella MNNEETFYQAMRRQGVTRRSFLKYCSLAATSLGLGAGMAPKIAWALENKPRIPVVWIHGL WP_005139779.1 hydrogenase [Shigella sonnei] Length: 372\nNumber of predicted TMHs: 2\nExp number of AAs in TMHs: 37.02628\nExp number, first 60 AAs: 19.39684\nTotal prob of N-in: 0.94313\nPOSSIBLE N-term signal sequence\ninside 1 19\nTMhelix 20 39\noutside 40 325\nTMhelix 326 348\ninside 349 372

27191 GCF_900159635.1_8489_1_88 Shigella sonnei Proteobacteria; Gammaproteobacteria; Enterobacterales; Enterobacteriaceae; Shigella MNNEETFYQAMRRQGVTRRSFLKYCSLAATSLGLGAGMAPKIAWALENKPRIPVVWIHGL WP_005139779.1 hydrogenase [Shigella sonnei] Length: 372\nNumber of predicted TMHs: 2\nExp number of AAs in TMHs: 37.02628\nExp number, first 60 AAs: 19.39684\nTotal prob of N-in: 0.94313\nPOSSIBLE N-term signal sequence\ninside 1 19\nTMhelix 20 39\noutside 40 325\nTMhelix 326 348\ninside 349 372

27192 GCF_900159615.1_8489_1_85 Shigella sonnei Proteobacteria; Gammaproteobacteria; Enterobacterales; Enterobacteriaceae; Shigella MNNEETFYQAMRRQGVTRRSFLKYCSLAATSLGLGAGMAPKIAWALENKPRIPVVWIHGL WP_005139779.1 hydrogenase [Shigella sonnei] Length: 372\nNumber of predicted TMHs: 2\nExp number of AAs in TMHs: 37.02628\nExp number, first 60 AAs: 19.39684\nTotal prob of N-in: 0.94313\nPOSSIBLE N-term signal sequence\ninside 1 19\nTMhelix 20 39\noutside 40 325\nTMhelix 326 348\ninside 349 372

27193 GCF_900159585.1_8489_1_83 Shigella sonnei Proteobacteria; Gammaproteobacteria; Enterobacterales; Enterobacteriaceae; Shigella MNNEETFYQAMRRQGVTRRSFLKYCSLAATSLGLGAGMAPKIAWALENKPRIPVVWIHGL WP_005139779.1 hydrogenase [Shigella sonnei] Length: 372\nNumber of predicted TMHs: 2\nExp number of AAs in TMHs: 37.02628\nExp number, first 60 AAs: 19.39684\nTotal prob of N-in: 0.94313\nPOSSIBLE N-term signal sequence\ninside 1 19\nTMhelix 20 39\noutside 40 325\nTMhelix 326 348\ninside 349 372

27194 GCF_900159655.1_8489_1_90 Shigella sonnei Proteobacteria; Gammaproteobacteria; Enterobacterales; Enterobacteriaceae; Shigella MNNEETFYQAMRRQGVTRRSFLKYCSLAATSLGLGAGMAPKIAWALENKPRIPVVWIHGL WP_005139779.1 hydrogenase [Shigella sonnei] Length: 372\nNumber of predicted TMHs: 2\nExp number of AAs in TMHs: 37.02628\nExp number, first 60 AAs: 19.39684\nTotal prob of N-in: 0.94313\nPOSSIBLE N-term signal sequence\ninside 1 19\nTMhelix 20 39\noutside 40 325\nTMhelix 326 348\ninside 349 372

27195 GCF_900159735.1_8403_8_3Shigella sonnei Proteobacteria; Gammaproteobacteria; Enterobacterales; Enterobacteriaceae; Shigella MNNEETFYQAMRRQGVTRRSFLKYCSLAATSLGLGAGMAPKIAWALENKPRIPVVWIHGL WP_005139779.1 hydrogenase [Shigella sonnei] Length: 372\nNumber of predicted TMHs: 2\nExp number of AAs in TMHs: 37.02628\nExp number, first 60 AAs: 19.39684\nTotal prob of N-in: 0.94313\nPOSSIBLE N-term signal sequence\ninside 1 19\nTMhelix 20 39\noutside 40 325\nTMhelix 326 348\ninside 349 372

27196 GCF_900159705.1_8489_1_95 Shigella sonnei Proteobacteria; Gammaproteobacteria; Enterobacterales; Enterobacteriaceae; Shigella MNNEETFYQAMRRQGVTRRSFLKYCSLAATSLGLGAGMAPKIAWALENKPRIPVVWIHGL WP_005139779.1 hydrogenase [Shigella sonnei] Length: 372\nNumber of predicted TMHs: 2\nExp number of AAs in TMHs: 37.02628\nExp number, first 60 AAs: 19.39684\nTotal prob of N-in: 0.94313\nPOSSIBLE N-term signal sequence\ninside 1 19\nTMhelix 20 39\noutside 40 325\nTMhelix 326 348\ninside 349 372

27197 GCF_900159755.1_8403_8_5Shigella sonnei Proteobacteria; Gammaproteobacteria; Enterobacterales; Enterobacteriaceae; Shigella MNNEETFYQAMRRQGVTRRSFLKYCSLAATSLGLGAGMAPKIAWALENKPRIPVVWIHGL WP_005139779.1 hydrogenase [Shigella sonnei] Length: 372\nNumber of predicted TMHs: 2\nExp number of AAs in TMHs: 37.02628\nExp number, first 60 AAs: 19.39684\nTotal prob of N-in: 0.94313\nPOSSIBLE N-term signal sequence\ninside 1 19\nTMhelix 20 39\noutside 40 325\nTMhelix 326 348\ninside 349 372

27198 GCF_900159805.1_8403_8_10 Shigella sonnei Proteobacteria; Gammaproteobacteria; Enterobacterales; Enterobacteriaceae; Shigella MTGDNTLIHSHGINRRDFMKLCAALAATMGLSSKAAAEMAESVTNPQRPPVIWIGAQECT WP_000145410.1 MULTISPECIES: hydrogenase-2 small chain [Proteobacteria] Length: 372\nNumber of predicted TMHs: 1\nExp number of AAs in TMHs: 23.72226\nExp number, first 60 AAs: 0.22038\nTotal prob of N-in: 0.12994\noutside 1 331\nTMhelix 332 354\ninside 355 372

27199 GCF_900159825.1_8403_8_12 Shigella sonnei Proteobacteria; Gammaproteobacteria; Enterobacterales; Enterobacteriaceae; Shigella MTGDNTLIHSHGINRRDFMKLCAALAATMGLSSKAAAEMAESVTNPQRPPVIWIGAQECT WP_000145410.1 MULTISPECIES: hydrogenase-2 small chain [Proteobacteria] Length: 372\nNumber of predicted TMHs: 1\nExp number of AAs in TMHs: 23.72226\nExp number, first 60 AAs: 0.22038\nTotal prob of N-in: 0.12994\noutside 1 331\nTMhelix 332 354\ninside 355 372

27200 GCF_900159855.1_8403_8_14 Shigella sonnei Proteobacteria; Gammaproteobacteria; Enterobacterales; Enterobacteriaceae; Shigella MTGDNTLIHSHGINRRDFMKLCAALAATMGLSSKAAAEMAESVTNPQRPPVIWIGAQECT WP_000145410.1 MULTISPECIES: hydrogenase-2 small chain [Proteobacteria] Length: 372\nNumber of predicted TMHs: 1\nExp number of AAs in TMHs: 23.72226\nExp number, first 60 AAs: 0.22038\nTotal prob of N-in: 0.12994\noutside 1 331\nTMhelix 332 354\ninside 355 372

27201 GCF_900159895.1_8403_8_19 Shigella sonnei Proteobacteria; Gammaproteobacteria; Enterobacterales; Enterobacteriaceae; Shigella MTGDNTLIHSHGINRRDFMKLCAALAATMGLSSKAAAEMAESVTNPQRPPVIWIGAQECT WP_000145410.1 MULTISPECIES: hydrogenase-2 small chain [Proteobacteria] Length: 372\nNumber of predicted TMHs: 1\nExp number of AAs in TMHs: 23.72226\nExp number, first 60 AAs: 0.22038\nTotal prob of N-in: 0.12994\noutside 1 331\nTMhelix 332 354\ninside 355 372

27202 GCF_900159875.1_8403_8_17 Shigella sonnei Proteobacteria; Gammaproteobacteria; Enterobacterales; Enterobacteriaceae; Shigella MTGDNTLIHSHGINRRDFMKLCAALAATMGLSSKAAAEMAESVTNPQRPPVIWIGAQECT WP_000145410.1 MULTISPECIES: hydrogenase-2 small chain [Proteobacteria] Length: 372\nNumber of predicted TMHs: 1\nExp number of AAs in TMHs: 23.72226\nExp number, first 60 AAs: 0.22038\nTotal prob of N-in: 0.12994\noutside 1 331\nTMhelix 332 354\ninside 355 372

27203 GCF_900159905.1_8403_8_20 Shigella sonnei Proteobacteria; Gammaproteobacteria; Enterobacterales; Enterobacteriaceae; Shigella MTGDNTLIHSHGINRRDFMKLCAALAATMGLSSKAAAEMAESVTNPQRPPVIWIGAQECT WP_000145410.1 MULTISPECIES: hydrogenase-2 small chain [Proteobacteria] Length: 372\nNumber of predicted TMHs: 1\nExp number of AAs in TMHs: 23.72226\nExp number, first 60 AAs: 0.22038\nTotal prob of N-in: 0.12994\noutside 1 331\nTMhelix 332 354\ninside 355 372

27204 GCF_900159955.1_8403_8_25 Shigella sonnei Proteobacteria; Gammaproteobacteria; Enterobacterales; Enterobacteriaceae; Shigella MTGDNTLIHSHGINRRDFMKLCAALAATMGLSSKAAAEMAESVTNPQRPPVIWIGAQECT WP_000145410.1 MULTISPECIES: hydrogenase-2 small chain [Proteobacteria] Length: 372\nNumber of predicted TMHs: 1\nExp number of AAs in TMHs: 23.72226\nExp number, first 60 AAs: 0.22038\nTotal prob of N-in: 0.12994\noutside 1 331\nTMhelix 332 354\ninside 355 372

27205 GCF_900159985.1_8403_8_28 Shigella sonnei Proteobacteria; Gammaproteobacteria; Enterobacterales; Enterobacteriaceae; Shigella
MTGDNTLIHSHGINRRDFMKLCAALAATMGLSSKAAAEMAESVTNPQRPPVIWIGAQECT WP_000145410.1
MULTISPECIES: hydrogenase-2 small chain [Proteobacteria] Length: 372\nNumber of predicted TMHs: 1\nExp number of AAs in TMHs: 23.72226\nExp number, first 60 AAs: 0.22038\nTotal prob of N-in: 0.12994\noutside 1 331\nTMhelix 332 354\ninside 355 372

27206 GCF_900160005.1_8403_8_35 Shigella sonnei Proteobacteria; Gammaproteobacteria; Enterobacterales; Enterobacteriaceae; Shigella
MTGDNTLIHSHGINRRDFMKLCAALAATMGLSSKAAAEMAESVTNPQRPPVIWIGAQECT WP_000145410.1
MULTISPECIES: hydrogenase-2 small chain [Proteobacteria] Length: 372\nNumber of predicted TMHs: 1\nExp number of AAs in TMHs: 23.72226\nExp number, first 60 AAs: 0.22038\nTotal prob of N-in: 0.12994\noutside 1 331\nTMhelix 332 354\ninside 355 372

27207 GCF_900160035.1_8403_8_31 Shigella sonnei Proteobacteria; Gammaproteobacteria; Enterobacterales; Enterobacteriaceae; Shigella
MTGDNTLIHSHGINRRDFMKLCAALAATMGLSSKAAAEMAESVTNPQRPPVIWIGAQECT WP_000145410.1
MULTISPECIES: hydrogenase-2 small chain [Proteobacteria] Length: 372\nNumber of predicted TMHs: 1\nExp number of AAs in TMHs: 23.72226\nExp number, first 60 AAs: 0.22038\nTotal prob of N-in: 0.12994\noutside 1 331\nTMhelix 332 354\ninside 355 372

27208 GCF_900160055.1_8403_8_36 Shigella sonnei Proteobacteria; Gammaproteobacteria; Enterobacterales; Enterobacteriaceae; Shigella
MTGDNTLIHSHGINRRDFMKLCAALAATMGLSSKAAAEMAESVTNPQRPPVIWIGAQECT WP_000145410.1
MULTISPECIES: hydrogenase-2 small chain [Proteobacteria] Length: 372\nNumber of predicted TMHs: 1\nExp number of AAs in TMHs: 23.72226\nExp number, first 60 AAs: 0.22038\nTotal prob of N-in: 0.12994\noutside 1 331\nTMhelix 332 354\ninside 355 372

27209 GCF_900159985.1_8403_8_28 Shigella sonnei Proteobacteria; Gammaproteobacteria; Enterobacterales; Enterobacteriaceae; Shigella
MNNEETFYQAMRRQGVTRRSFLKYCSLAATSLGLGAGMAPKIAWALENKPRIPVVWIHGL WP_005139779.1
hydrogenase [Shigella sonnei] Length: 372\nNumber of predicted TMHs: 2\nExp number of AAs in TMHs: 37.02628\nExp number, first 60 AAs: 19.39684\nTotal prob of N-in: 0.94313\nPOSSIBLE N-term signal sequence\ninside 1 19\nTMhelix 20 39\noutside 40 325\nTMhelix 326 348\ninside 349 372

27210 GCF_900160035.1_8403_8_31 Shigella sonnei Proteobacteria; Gammaproteobacteria; Enterobacterales; Enterobacteriaceae; Shigella
MNNEETFYQAMRRQGVTRRSFLKYCSLAATSLGLGAGMAPKIAWALENKPRIPVVWIHGL WP_005139779.1
hydrogenase [Shigella sonnei] Length: 372\nNumber of predicted TMHs: 2\nExp number of AAs in TMHs: 37.02628\nExp number, first 60 AAs: 19.39684\nTotal prob of N-in: 0.94313\nPOSSIBLE N-term signal sequence\ninside 1 19\nTMhelix 20 39\noutside 40 325\nTMhelix 326 348\ninside 349 372

27211 GCF_900160005.1_8403_8_35 Shigella sonnei Proteobacteria; Gammaproteobacteria; Enterobacterales; Enterobacteriaceae; Shigella
MNNEETFYQAMRRQGVTRRSFLKYCSLAATSLGLGAGMAPKIAWALENKPRIPVVWIHGL WP_005139779.1
hydrogenase [Shigella sonnei] Length: 372\nNumber of predicted TMHs: 2\nExp number of AAs in TMHs: 37.02628\nExp number, first 60 AAs: 19.39684\nTotal prob of N-in: 0.94313\nPOSSIBLE N-term signal sequence\ninside 1 19\nTMhelix 20 39\noutside 40 325\nTMhelix 326 348\ninside 349 372

27212 GCF_900159805.1_8403_8_10 Shigella sonnei Proteobacteria; Gammaproteobacteria; Enterobacterales; Enterobacteriaceae; Shigella
MNNEETFYQAMRRQGVTRRSFLKYCSLAATSLGLGAGMAPKIAWALENKPRIPVVWIHGL WP_005139779.1
hydrogenase [Shigella sonnei] Length: 372\nNumber of predicted TMHs: 2\nExp number of AAs in TMHs: 37.02628\nExp number, first 60 AAs: 19.39684\nTotal prob of N-in: 0.94313\nPOSSIBLE N-term signal sequence\ninside 1 19\nTMhelix 20 39\noutside 40 325\nTMhelix 326 348\ninside 349 372

27213 GCF_900159875.1_8403_8_17 Shigella sonnei Proteobacteria; Gammaproteobacteria; Enterobacterales; Enterobacteriaceae; Shigella
MNNEETFYQAMRRQGVTRRSFLKYCSLAATSLGLGAGMAPKIAWALENKPRIPVVWIHGL WP_005139779.1
hydrogenase [Shigella sonnei] Length: 372\nNumber of predicted TMHs: 2\nExp number of AAs in

TMHs: 37.02628\nExp number, first 60 AAs: 19.39684\nTotal prob of N-in: 0.94313\nPOSSIBLE N-term signal sequence\ninside 1 19\nTMhelix 20 39\noutside 40 325\nTMhelix 326 348\ninside 349 372

27214 GCF_900159955.1_8403_8_25 Shigella sonnei Proteobacteria; Gammaproteobacteria; Enterobacterales; Enterobacteriaceae; Shigella MNNEETFYQAMRRQGVTRRSFLKYCSLAATSLGLGAGMAPKIAWALENKPRIPVVWIHGL WP_005139779.1 hydrogenase [Shigella sonnei] Length: 372\nNumber of predicted TMHs: 2\nExp number of AAs in TMHs: 37.02628\nExp number, first 60 AAs: 19.39684\nTotal prob of N-in: 0.94313\nPOSSIBLE N-term signal sequence\ninside 1 19\nTMhelix 20 39\noutside 40 325\nTMhelix 326 348\ninside 349 372

27215 GCF_900159855.1_8403_8_14 Shigella sonnei Proteobacteria; Gammaproteobacteria; Enterobacterales; Enterobacteriaceae; Shigella MNNEETFYQAMRRQGVTRRSFLKYCSLAATSLGLGAGMAPKIAWALENKPRIPVVWIHGL WP_005139779.1 hydrogenase [Shigella sonnei] Length: 372\nNumber of predicted TMHs: 2\nExp number of AAs in TMHs: 37.02628\nExp number, first 60 AAs: 19.39684\nTotal prob of N-in: 0.94313\nPOSSIBLE N-term signal sequence\ninside 1 19\nTMhelix 20 39\noutside 40 325\nTMhelix 326 348\ninside 349 372

27216 GCF_900159895.1_8403_8_19 Shigella sonnei Proteobacteria; Gammaproteobacteria; Enterobacterales; Enterobacteriaceae; Shigella MNNEETFYQAMRRQGVTRRSFLKYCSLAATSLGLGAGMAPKIAWALENKPRIPVVWIHGL WP_005139779.1 hydrogenase [Shigella sonnei] Length: 372\nNumber of predicted TMHs: 2\nExp number of AAs in TMHs: 37.02628\nExp number, first 60 AAs: 19.39684\nTotal prob of N-in: 0.94313\nPOSSIBLE N-term signal sequence\ninside 1 19\nTMhelix 20 39\noutside 40 325\nTMhelix 326 348\ninside 349 372

27217 GCF_900159825.1_8403_8_12 Shigella sonnei Proteobacteria; Gammaproteobacteria; Enterobacterales; Enterobacteriaceae; Shigella MNNEETFYQAMRRQGVTRRSFLKYCSLAATSLGLGAGMAPKIAWALENKPRIPVVWIHGL WP_005139779.1 hydrogenase [Shigella sonnei] Length: 372\nNumber of predicted TMHs: 2\nExp number of AAs in TMHs: 37.02628\nExp number, first 60 AAs: 19.39684\nTotal prob of N-in: 0.94313\nPOSSIBLE N-term signal sequence\ninside 1 19\nTMhelix 20 39\noutside 40 325\nTMhelix 326 348\ninside 349 372

27218 GCF_900159935.1_8403_8_23 Shigella sonnei Proteobacteria; Gammaproteobacteria; Enterobacterales; Enterobacteriaceae; Shigella MNNEETFYQAMRRQGVTRRSFLKYCSLAATSLGLGAGMAPKIAWALENKPRIPVVWIHGL WP_005139779.1 hydrogenase [Shigella sonnei] Length: 372\nNumber of predicted TMHs: 2\nExp number of AAs in TMHs: 37.02628\nExp number, first 60 AAs: 19.39684\nTotal prob of N-in: 0.94313\nPOSSIBLE N-term signal sequence\ninside 1 19\nTMhelix 20 39\noutside 40 325\nTMhelix 326 348\ninside 349 372

27219 GCF_900159905.1_8403_8_20 Shigella sonnei Proteobacteria; Gammaproteobacteria; Enterobacterales; Enterobacteriaceae; Shigella MNNEETFYQAMRRQGVTRRSFLKYCSLAATSLGLGAGMAPKIAWALENKPRIPVVWIHGL WP_005139779.1 hydrogenase [Shigella sonnei] Length: 372\nNumber of predicted TMHs: 2\nExp number of AAs in TMHs: 37.02628\nExp number, first 60 AAs: 19.39684\nTotal prob of N-in: 0.94313\nPOSSIBLE N-term signal sequence\ninside 1 19\nTMhelix 20 39\noutside 40 325\nTMhelix 326 348\ninside 349 372

27220 GCF_900160055.1_8403_8_36 Shigella sonnei Proteobacteria; Gammaproteobacteria; Enterobacterales; Enterobacteriaceae; Shigella MNNEETFYQAMRRQGVTRRSFLKYCSLAATSLGLGAGMAPKIAWALENKPRIPVVWIHGL WP_005139779.1 hydrogenase [Shigella sonnei] Length: 372\nNumber of predicted TMHs: 2\nExp number of AAs in TMHs: 37.02628\nExp number, first 60 AAs: 19.39684\nTotal prob of N-in: 0.94313\nPOSSIBLE N-term signal sequence\ninside 1 19\nTMhelix 20 39\noutside 40 325\nTMhelix 326 348\ninside 349 372

27221 GCF_900160075.1_8403_8_38 Shigella sonnei Proteobacteria; Gammaproteobacteria; Enterobacterales; Enterobacteriaceae; Shigella MTGDNTLIHSHGINRRDFMKLCAALAATMGLSSKAAAEMAESVTNPQRPPVIWIGAQECT WP_000145410.1 MULTISPECIES: hydrogenase-2 small chain [Proteobacteria] Length: 372\nNumber of predicted TMHs: 1\nExp number of AAs in TMHs: 23.72226\nExp number, first 60 AAs: 0.22038\nTotal prob of N-in: 0.12994\noutside 1 331\nTMhelix 332 354\ninside 355 372

27222 GCF_900160105.1_8403_8_41 Shigella sonnei Proteobacteria; Gammaproteobacteria; Enterobacterales; Enterobacteriaceae; Shigella

MTGDNTLIHSHGINRRDFMKLCAALAATMGLSSKAAAEMAESVTNPQRPPVIWIGAQECT WP_000145410.1 MULTISPECIES: hydrogenase-2 small chain [Proteobacteria] Length: 372\nNumber of predicted TMHs: 1\nExp number of AAs in TMHs: 23.72226\nExp number, first 60 AAs: 0.22038\nTotal prob of N-in: 0.12994\noutside 1 331\nTMhelix 332 354\ninside 355 372

27223 GCF_900160125.1_8403_8_43 Shigella sonnei Proteobacteria; Gammaproteobacteria; Enterobacterales; Enterobacteriaceae; Shigella MTGDNTLIHSHGINRRDFMKLCAALAATMGLSSKAAAEMAESVTNPQRPPVIWIGAQECT WP_000145410.1 MULTISPECIES: hydrogenase-2 small chain [Proteobacteria] Length: 372\nNumber of predicted TMHs: 1\nExp number of AAs in TMHs: 23.72226\nExp number, first 60 AAs: 0.22038\nTotal prob of N-in: 0.12994\noutside 1 331\nTMhelix 332 354\ninside 355 372

27224 GCF_900160155.1_8403_8_46 Shigella sonnei Proteobacteria; Gammaproteobacteria; Enterobacterales; Enterobacteriaceae; Shigella MTGDNTLIHSHGINRRDFMKLCAALAATMGLSSKAAAEMAESVTNPQRPPVIWIGAQECT WP_000145410.1 MULTISPECIES: hydrogenase-2 small chain [Proteobacteria] Length: 372\nNumber of predicted TMHs: 1\nExp number of AAs in TMHs: 23.72226\nExp number, first 60 AAs: 0.22038\nTotal prob of N-in: 0.12994\noutside 1 331\nTMhelix 332 354\ninside 355 372

27225 GCF_900160175.1_8403_8_47 Shigella sonnei Proteobacteria; Gammaproteobacteria; Enterobacterales; Enterobacteriaceae; Shigella MTGDNTLIHSHGINRRDFMKLCAALAATMGLSSKAAAEMAESVTNPQRPPVIWIGAQECT WP_000145410.1 MULTISPECIES: hydrogenase-2 small chain [Proteobacteria] Length: 372\nNumber of predicted TMHs: 1\nExp number of AAs in TMHs: 23.72226\nExp number, first 60 AAs: 0.22038\nTotal prob of N-in: 0.12994\noutside 1 331\nTMhelix 332 354\ninside 355 372

27226 GCF_900160205.1_8403_8_51 Shigella sonnei Proteobacteria; Gammaproteobacteria; Enterobacterales; Enterobacteriaceae; Shigella MTGDNTLIHSHGINRRDFMKLCAALAATMGLSSKAAAEMAESVTNPQRPPVIWIGAQECT WP_000145410.1 MULTISPECIES: hydrogenase-2 small chain [Proteobacteria] Length: 372\nNumber of predicted TMHs: 1\nExp number of AAs in TMHs: 23.72226\nExp number, first 60 AAs: 0.22038\nTotal prob of N-in: 0.12994\noutside 1 331\nTMhelix 332 354\ninside 355 372

27227 GCF_900160225.1_8403_8_53 Shigella sonnei Proteobacteria; Gammaproteobacteria; Enterobacterales; Enterobacteriaceae; Shigella MTGDNTLIHSHGINRRDFMKLCAALAATMGLSSKAAAEMAESVTNPQRPPVIWIGAQECT WP_000145410.1 MULTISPECIES: hydrogenase-2 small chain [Proteobacteria] Length: 372\nNumber of predicted TMHs: 1\nExp number of AAs in TMHs: 23.72226\nExp number, first 60 AAs: 0.22038\nTotal prob of N-in: 0.12994\noutside 1 331\nTMhelix 332 354\ninside 355 372

27228 GCF_900160255.1_8403_8_56 Shigella sonnei Proteobacteria; Gammaproteobacteria; Enterobacterales; Enterobacteriaceae; Shigella MTGDNTLIHSHGINRRDFMKLCAALAATMGLSSKAAAEMAESVTNPQRPPVIWIGAQECT WP_000145410.1 MULTISPECIES: hydrogenase-2 small chain [Proteobacteria] Length: 372\nNumber of predicted TMHs: 1\nExp number of AAs in TMHs: 23.72226\nExp number, first 60 AAs: 0.22038\nTotal prob of N-in: 0.12994\noutside 1 331\nTMhelix 332 354\ninside 355 372

27229 GCF_900160275.1_8403_8_58 Shigella sonnei Proteobacteria; Gammaproteobacteria; Enterobacterales; Enterobacteriaceae; Shigella MTGDNTLIHSHGINRRDFMKLCAALAATMGLSSKAAAEMAESVTNPQRPPVIWIGAQECT WP_000145410.1 MULTISPECIES: hydrogenase-2 small chain [Proteobacteria] Length: 372\nNumber of predicted TMHs: 1\nExp number of AAs in TMHs: 23.72226\nExp number, first 60 AAs: 0.22038\nTotal prob of N-in: 0.12994\noutside 1 331\nTMhelix 332 354\ninside 355 372

27230 GCF_900160295.1_8403_8_59 Shigella sonnei Proteobacteria; Gammaproteobacteria; Enterobacterales; Enterobacteriaceae; Shigella MTGDNTLIHSHGINRRDFMKLCAALAATMGLSSKAAAEMAESVTNPQRPPVIWIGAQECT WP_000145410.1 MULTISPECIES: hydrogenase-2 small chain [Proteobacteria] Length: 372\nNumber of predicted TMHs: 1\nExp number of AAs in TMHs: 23.72226\nExp number, first 60 AAs: 0.22038\nTotal prob of N-in: 0.12994\noutside 1 331\nTMhelix 332 354\ninside 355 372

27231 GCF_900160325.1_8403_8_63 Shigella sonnei Proteobacteria; Gammaproteobacteria; Enterobacterales; Enterobacteriaceae; Shigella
MTGDNTLIHSHGINRRDFMKLCAALAATMGLSSKAAAEMAESVTNPQRPPVIWIGAQECT WP_000145410.1
MULTISPECIES: hydrogenase-2 small chain [Proteobacteria] Length: 372\nNumber of predicted TMHs: 1\nExp number of AAs in TMHs: 23.72226\nExp number, first 60 AAs: 0.22038\nTotal prob of N-in: 0.12994\noutside 1 331\nTMhelix 332 354\ninside 355 372

27232 GCF_900160315.1_8403_8_62 Shigella sonnei Proteobacteria; Gammaproteobacteria; Enterobacterales; Enterobacteriaceae; Shigella
MTGDNTLIHSHGINRRDFMKLCAALAATMGLSSKAAAEMAESVTNPQRPPVIWIGAQECT WP_000145410.1
MULTISPECIES: hydrogenase-2 small chain [Proteobacteria] Length: 372\nNumber of predicted TMHs: 1\nExp number of AAs in TMHs: 23.72226\nExp number, first 60 AAs: 0.22038\nTotal prob of N-in: 0.12994\noutside 1 331\nTMhelix 332 354\ninside 355 372

27233 GCF_900160255.1_8403_8_56 Shigella sonnei Proteobacteria; Gammaproteobacteria; Enterobacterales; Enterobacteriaceae; Shigella
MNNEETFYQAMRRQGVTRRSFLKYCSLAATSLGLGAGMAPKIAWALENKPRIPVVWIHGL WP_005139779.1
hydrogenase [Shigella sonnei] Length: 372\nNumber of predicted TMHs: 2\nExp number of AAs in TMHs: 37.02628\nExp number, first 60 AAs: 19.39684\nTotal prob of N-in: 0.94313\nPOSSIBLE N-term signal sequence\ninside 1 19\nTMhelix 20 39\noutside 40 325\nTMhelix 326 348\ninside 349 372

27234 GCF_900160275.1_8403_8_58 Shigella sonnei Proteobacteria; Gammaproteobacteria; Enterobacterales; Enterobacteriaceae; Shigella
MNNEETFYQAMRRQGVTRRSFLKYCSLAATSLGLGAGMAPKIAWALENKPRIPVVWIHGL WP_005139779.1
hydrogenase [Shigella sonnei] Length: 372\nNumber of predicted TMHs: 2\nExp number of AAs in TMHs: 37.02628\nExp number, first 60 AAs: 19.39684\nTotal prob of N-in: 0.94313\nPOSSIBLE N-term signal sequence\ninside 1 19\nTMhelix 20 39\noutside 40 325\nTMhelix 326 348\ninside 349 372

27235 GCF_900160075.1_8403_8_38 Shigella sonnei Proteobacteria; Gammaproteobacteria; Enterobacterales; Enterobacteriaceae; Shigella
MNNEETFYQAMRRQGVTRRSFLKYCSLAATSLGLGAGMAPKIAWALENKPRIPVVWIHGL WP_005139779.1
hydrogenase [Shigella sonnei] Length: 372\nNumber of predicted TMHs: 2\nExp number of AAs in TMHs: 37.02628\nExp number, first 60 AAs: 19.39684\nTotal prob of N-in: 0.94313\nPOSSIBLE N-term signal sequence\ninside 1 19\nTMhelix 20 39\noutside 40 325\nTMhelix 326 348\ninside 349 372

27236 GCF_900160155.1_8403_8_46 Shigella sonnei Proteobacteria; Gammaproteobacteria; Enterobacterales; Enterobacteriaceae; Shigella
MNNEETFYQAMRRQGVTRRSFLKYCSLAATSLGLGAGMAPKIAWALENKPRIPVVWIHGL WP_005139779.1
hydrogenase [Shigella sonnei] Length: 372\nNumber of predicted TMHs: 2\nExp number of AAs in TMHs: 37.02628\nExp number, first 60 AAs: 19.39684\nTotal prob of N-in: 0.94313\nPOSSIBLE N-term signal sequence\ninside 1 19\nTMhelix 20 39\noutside 40 325\nTMhelix 326 348\ninside 349 372

27237 GCF_900160105.1_8403_8_41 Shigella sonnei Proteobacteria; Gammaproteobacteria; Enterobacterales; Enterobacteriaceae; Shigella
MNNEETFYQAMRRQGVTRRSFLKYCSLAATSLGLGAGMAPKIAWALENKPRIPVVWIHGL WP_005139779.1
hydrogenase [Shigella sonnei] Length: 372\nNumber of predicted TMHs: 2\nExp number of AAs in TMHs: 37.02628\nExp number, first 60 AAs: 19.39684\nTotal prob of N-in: 0.94313\nPOSSIBLE N-term signal sequence\ninside 1 19\nTMhelix 20 39\noutside 40 325\nTMhelix 326 348\ninside 349 372

27238 GCF_900160125.1_8403_8_43 Shigella sonnei Proteobacteria; Gammaproteobacteria; Enterobacterales; Enterobacteriaceae; Shigella
MNNEETFYQAMRRQGVTRRSFLKYCSLAATSLGLGAGMAPKIAWALENKPRIPVVWIHGL WP_005139779.1
hydrogenase [Shigella sonnei] Length: 372\nNumber of predicted TMHs: 2\nExp number of AAs in TMHs: 37.02628\nExp number, first 60 AAs: 19.39684\nTotal prob of N-in: 0.94313\nPOSSIBLE N-term signal sequence\ninside 1 19\nTMhelix 20 39\noutside 40 325\nTMhelix 326 348\ninside 349 372

27239 GCF_900160175.1_8403_8_47 Shigella sonnei Proteobacteria; Gammaproteobacteria; Enterobacterales; Enterobacteriaceae; Shigella
MNNEETFYQAMRRQGVTRRSFLKYCSLAATSLGLGAGMAPKIAWALENKPRIPVVWIHGL WP_005139779.1
hydrogenase [Shigella sonnei] Length: 372\nNumber of predicted TMHs: 2\nExp number of AAs in

TMHs: 37.02628\nExp number, first 60 AAs: 19.39684\nTotal prob of N-in: 0.94313\nPOSSIBLE N-term signal sequence\ninside 1 19\nTMhelix 20 39\noutside 40 325\nTMhelix 326 348\ninside 349 372

27240 GCF_900160205.1_8403_8_51 Shigella sonnei Proteobacteria; Gammaproteobacteria; Enterobacterales; Enterobacteriaceae; Shigella
MNNEETFYQAMRRQGVTRRSFLKYCSLAATSLGLGAGMAPKIAWALENKPRIPVVWIHGL WP_005139779.1
hydrogenase [Shigella sonnei] Length: 372\nNumber of predicted TMHs: 2\nExp number of AAs in TMHs: 37.02628\nExp number, first 60 AAs: 19.39684\nTotal prob of N-in: 0.94313\nPOSSIBLE N-term signal sequence\ninside 1 19\nTMhelix 20 39\noutside 40 325\nTMhelix 326 348\ninside 349 372

27241 GCF_900160225.1_8403_8_53 Shigella sonnei Proteobacteria; Gammaproteobacteria; Enterobacterales; Enterobacteriaceae; Shigella
MNNEETFYQAMRRQGVTRRSFLKYCSLAATSLGLGAGMAPKIAWALENKPRIPVVWIHGL WP_005139779.1
hydrogenase [Shigella sonnei] Length: 372\nNumber of predicted TMHs: 2\nExp number of AAs in TMHs: 37.02628\nExp number, first 60 AAs: 19.39684\nTotal prob of N-in: 0.94313\nPOSSIBLE N-term signal sequence\ninside 1 19\nTMhelix 20 39\noutside 40 325\nTMhelix 326 348\ninside 349 372

27242 GCF_900160295.1_8403_8_59 Shigella sonnei Proteobacteria; Gammaproteobacteria; Enterobacterales; Enterobacteriaceae; Shigella
MNNEETFYQAMRRQGVTRRSFLKYCSLAATSLGLGAGMAPKIAWALENKPRIPVVWIHGL WP_005139779.1
hydrogenase [Shigella sonnei] Length: 372\nNumber of predicted TMHs: 2\nExp number of AAs in TMHs: 37.02628\nExp number, first 60 AAs: 19.39684\nTotal prob of N-in: 0.94313\nPOSSIBLE N-term signal sequence\ninside 1 19\nTMhelix 20 39\noutside 40 325\nTMhelix 326 348\ninside 349 372

27243 GCF_900160325.1_8403_8_63 Shigella sonnei Proteobacteria; Gammaproteobacteria; Enterobacterales; Enterobacteriaceae; Shigella
MNNEETFYQAMRRQGVTRRSFLKYCSLAATSLGLGAGMAPKIAWALENKPRIPVVWIHGL WP_005139779.1
hydrogenase [Shigella sonnei] Length: 372\nNumber of predicted TMHs: 2\nExp number of AAs in TMHs: 37.02628\nExp number, first 60 AAs: 19.39684\nTotal prob of N-in: 0.94313\nPOSSIBLE N-term signal sequence\ninside 1 19\nTMhelix 20 39\noutside 40 325\nTMhelix 326 348\ninside 349 372

27244 GCF_900160315.1_8403_8_62 Shigella sonnei Proteobacteria; Gammaproteobacteria; Enterobacterales; Enterobacteriaceae; Shigella
MNNEETFYQAMRRQGVTRRSFLKYCSLAATSLGLGAGMAPKIAWALENKPRIPVVWIHGL WP_005139779.1
hydrogenase [Shigella sonnei] Length: 372\nNumber of predicted TMHs: 2\nExp number of AAs in TMHs: 37.02628\nExp number, first 60 AAs: 19.39684\nTotal prob of N-in: 0.94313\nPOSSIBLE N-term signal sequence\ninside 1 19\nTMhelix 20 39\noutside 40 325\nTMhelix 326 348\ninside 349 372

27245 GCF_900160345.1_8403_8_65 Shigella sonnei Proteobacteria; Gammaproteobacteria; Enterobacterales; Enterobacteriaceae; Shigella
MTGDNTLIHSHGINRRDFMKLCAALAATMGLSSKAAAEMAESVTNPQRPPVIWIGAQECT WP_000145410.1
MULTISPECIES: hydrogenase-2 small chain [Proteobacteria] Length: 372\nNumber of predicted TMHs: 1\nExp number of AAs in TMHs: 23.72226\nExp number, first 60 AAs: 0.22038\nTotal prob of N-in: 0.12994\noutside 1 331\nTMhelix 332 354\ninside 355 372

27246 GCF_900160365.1_8403_8_67 Shigella sonnei Proteobacteria; Gammaproteobacteria; Enterobacterales; Enterobacteriaceae; Shigella
MTGDNTLIHSHGINRRDFMKLCAALAATMGLSSKAAAEMAESVTNPQRPPVIWIGAQECT WP_000145410.1
MULTISPECIES: hydrogenase-2 small chain [Proteobacteria] Length: 372\nNumber of predicted TMHs: 1\nExp number of AAs in TMHs: 23.72226\nExp number, first 60 AAs: 0.22038\nTotal prob of N-in: 0.12994\noutside 1 331\nTMhelix 332 354\ninside 355 372

27247 GCF_900160395.1_8403_8_70 Shigella sonnei Proteobacteria; Gammaproteobacteria; Enterobacterales; Enterobacteriaceae; Shigella
MTGDNTLIHSHGINRRDFMKLCAALAATMGLSSKAAAEMAESVTNPQRPPVIWIGAQECT WP_000145410.1
MULTISPECIES: hydrogenase-2 small chain [Proteobacteria] Length: 372\nNumber of predicted TMHs: 1\nExp number of AAs in TMHs: 23.72226\nExp number, first 60 AAs: 0.22038\nTotal prob of N-in: 0.12994\noutside 1 331\nTMhelix 332 354\ninside 355 372

27248 GCF_900160415.1_8403_8_72 Shigella sonnei Proteobacteria; Gammaproteobacteria; Enterobacterales; Enterobacteriaceae; Shigella

MTGDNTLIHSHGINRRDFMKLCAALAATMGLSSKAAAEMAESVTNPQRPPVIWIGAQECT WP_000145410.1 MULTISPECIES: hydrogenase-2 small chain [Proteobacteria] Length: 372\nNumber of predicted TMHs: 1\nExp number of AAs in TMHs: 23.72226\nExp number, first 60 AAs: 0.22038\nTotal prob of N-in: 0.12994\noutside 1 331\nTMhelix 332 354\ninside 355 372

27249 GCF_900160445.1_8403_8_75 Shigella sonnei Proteobacteria; Gammaproteobacteria; Enterobacterales; Enterobacteriaceae; Shigella MTGDNTLIHSHGINRRDFMKLCAALAATMGLSSKAAAEMAESVTNPQRPPVIWIGAQECT WP_000145410.1 MULTISPECIES: hydrogenase-2 small chain [Proteobacteria] Length: 372\nNumber of predicted TMHs: 1\nExp number of AAs in TMHs: 23.72226\nExp number, first 60 AAs: 0.22038\nTotal prob of N-in: 0.12994\noutside 1 331\nTMhelix 332 354\ninside 355 372

27250 GCF_900160465.1_8403_8_78 Shigella sonnei Proteobacteria; Gammaproteobacteria; Enterobacterales; Enterobacteriaceae; Shigella MTGDNTLIHSHGINRRDFMKLCAALAATMGLSSKAAAEMAESVTNPQRPPVIWIGAQECT WP_000145410.1 MULTISPECIES: hydrogenase-2 small chain [Proteobacteria] Length: 372\nNumber of predicted TMHs: 1\nExp number of AAs in TMHs: 23.72226\nExp number, first 60 AAs: 0.22038\nTotal prob of N-in: 0.12994\noutside 1 331\nTMhelix 332 354\ninside 355 372

27251 GCF_900160565.1_8403_8_89 Shigella sonnei Proteobacteria; Gammaproteobacteria; Enterobacterales; Enterobacteriaceae; Shigella MTGDNTLIHSHGINRRDFMKLCAALAATMGLSSKAAAEMAESVTNPQRPPVIWIGAQECT WP_000145410.1 MULTISPECIES: hydrogenase-2 small chain [Proteobacteria] Length: 372\nNumber of predicted TMHs: 1\nExp number of AAs in TMHs: 23.72226\nExp number, first 60 AAs: 0.22038\nTotal prob of N-in: 0.12994\noutside 1 331\nTMhelix 332 354\ninside 355 372

27252 GCF_900160495.1_8403_8_82 Shigella sonnei Proteobacteria; Gammaproteobacteria; Enterobacterales; Enterobacteriaceae; Shigella MTGDNTLIHSHGINRRDFMKLCAALAATMGLSSKAAAEMAESVTNPQRPPVIWIGAQECT WP_000145410.1 MULTISPECIES: hydrogenase-2 small chain [Proteobacteria] Length: 372\nNumber of predicted TMHs: 1\nExp number of AAs in TMHs: 23.72226\nExp number, first 60 AAs: 0.22038\nTotal prob of N-in: 0.12994\noutside 1 331\nTMhelix 332 354\ninside 355 372

27253 GCF_900160535.1_8403_8_81 Shigella sonnei Proteobacteria; Gammaproteobacteria; Enterobacterales; Enterobacteriaceae; Shigella MTGDNTLIHSHGINRRDFMKLCAALAATMGLSSKAAAEMAESVTNPQRPPVIWIGAQECT WP_000145410.1 MULTISPECIES: hydrogenase-2 small chain [Proteobacteria] Length: 372\nNumber of predicted TMHs: 1\nExp number of AAs in TMHs: 23.72226\nExp number, first 60 AAs: 0.22038\nTotal prob of N-in: 0.12994\noutside 1 331\nTMhelix 332 354\ninside 355 372

27254 GCF_900160515.1_8403_8_83 Shigella sonnei Proteobacteria; Gammaproteobacteria; Enterobacterales; Enterobacteriaceae; Shigella MTGDNTLIHSHGINRRDFMKLCAALAATMGLSSKAAAEMAESVTNPQRPPVIWIGAQECT WP_000145410.1 MULTISPECIES: hydrogenase-2 small chain [Proteobacteria] Length: 372\nNumber of predicted TMHs: 1\nExp number of AAs in TMHs: 23.72226\nExp number, first 60 AAs: 0.22038\nTotal prob of N-in: 0.12994\noutside 1 331\nTMhelix 332 354\ninside 355 372

27255 GCF_900160585.1_8403_8_91 Shigella sonnei Proteobacteria; Gammaproteobacteria; Enterobacterales; Enterobacteriaceae; Shigella MTGDNTLIHSHGINRRDFMKLCAALAATMGLSSKAAAEMAESVTNPQRPPVIWIGAQECT WP_000145410.1 MULTISPECIES: hydrogenase-2 small chain [Proteobacteria] Length: 372\nNumber of predicted TMHs: 1\nExp number of AAs in TMHs: 23.72226\nExp number, first 60 AAs: 0.22038\nTotal prob of N-in: 0.12994\noutside 1 331\nTMhelix 332 354\ninside 355 372

27256 GCF_900160615.1_8403_8_94 Shigella sonnei Proteobacteria; Gammaproteobacteria; Enterobacterales; Enterobacteriaceae; Shigella MTGDNTLIHSHGINRRDFMKLCAALAATMGLSSKAAAEMAESVTNPQRPPVIWIGAQECT WP_000145410.1 MULTISPECIES: hydrogenase-2 small chain [Proteobacteria] Length: 372\nNumber of predicted TMHs: 1\nExp number of AAs in TMHs: 23.72226\nExp number, first 60 AAs: 0.22038\nTotal prob of N-in: 0.12994\noutside 1 331\nTMhelix 332 354\ninside 355 372

27257	GCF_900160495.1_8403_8_82	Shigella sonnei	Proteobacteria; Gammaproteobacteria; Enterobacterales; Enterobacteriaceae; Shigella
MNNEETFYQAMRRQGVTRRSFLKYCSLAATSLGLGAGMAPKIAWALENKPRIPVVWIHGL	WP_005139779.1
hydrogenase [Shigella sonnei]	Length: 372\nNumber of predicted TMHs: 2\nExp number of AAs in TMHs: 37.02628\nExp number, first 60 AAs: 19.39684\nTotal prob of N-in: 0.94313\nPOSSIBLE N-term signal sequence\ninside 1 19\nTMhelix 20 39\noutside 40 325\nTMhelix 326 348\ninside 349 372

27258	GCF_900160615.1_8403_8_94	Shigella sonnei	Proteobacteria; Gammaproteobacteria; Enterobacterales; Enterobacteriaceae; Shigella
MNNEETFYQAMRRQGVTRRSFLKYCSLAATSLGLGAGMAPKIAWALENKPRIPVVWIHGL	WP_005139779.1
hydrogenase [Shigella sonnei]	Length: 372\nNumber of predicted TMHs: 2\nExp number of AAs in TMHs: 37.02628\nExp number, first 60 AAs: 19.39684\nTotal prob of N-in: 0.94313\nPOSSIBLE N-term signal sequence\ninside 1 19\nTMhelix 20 39\noutside 40 325\nTMhelix 326 348\ninside 349 372

27259	GCF_900160345.1_8403_8_65	Shigella sonnei	Proteobacteria; Gammaproteobacteria; Enterobacterales; Enterobacteriaceae; Shigella
MNNEETFYQAMRRQGVTRRSFLKYCSLAATSLGLGAGMAPKIAWALENKPRIPVVWIHGL	WP_005139779.1
hydrogenase [Shigella sonnei]	Length: 372\nNumber of predicted TMHs: 2\nExp number of AAs in TMHs: 37.02628\nExp number, first 60 AAs: 19.39684\nTotal prob of N-in: 0.94313\nPOSSIBLE N-term signal sequence\ninside 1 19\nTMhelix 20 39\noutside 40 325\nTMhelix 326 348\ninside 349 372

27260	GCF_900160365.1_8403_8_67	Shigella sonnei	Proteobacteria; Gammaproteobacteria; Enterobacterales; Enterobacteriaceae; Shigella
MNNEETFYQAMRRQGVTRRSFLKYCSLAATSLGLGAGMAPKIAWALENKPRIPVVWIHGL	WP_005139779.1
hydrogenase [Shigella sonnei]	Length: 372\nNumber of predicted TMHs: 2\nExp number of AAs in TMHs: 37.02628\nExp number, first 60 AAs: 19.39684\nTotal prob of N-in: 0.94313\nPOSSIBLE N-term signal sequence\ninside 1 19\nTMhelix 20 39\noutside 40 325\nTMhelix 326 348\ninside 349 372

27261	GCF_900160395.1_8403_8_70	Shigella sonnei	Proteobacteria; Gammaproteobacteria; Enterobacterales; Enterobacteriaceae; Shigella
MNNEETFYQAMRRQGVTRRSFLKYCSLAATSLGLGAGMAPKIAWALENKPRIPVVWIHGL	WP_005139779.1
hydrogenase [Shigella sonnei]	Length: 372\nNumber of predicted TMHs: 2\nExp number of AAs in TMHs: 37.02628\nExp number, first 60 AAs: 19.39684\nTotal prob of N-in: 0.94313\nPOSSIBLE N-term signal sequence\ninside 1 19\nTMhelix 20 39\noutside 40 325\nTMhelix 326 348\ninside 349 372

27262	GCF_900160415.1_8403_8_72	Shigella sonnei	Proteobacteria; Gammaproteobacteria; Enterobacterales; Enterobacteriaceae; Shigella
MNNEETFYQAMRRQGVTRRSFLKYCSLAATSLGLGAGMAPKIAWALENKPRIPVVWIHGL	WP_005139779.1
hydrogenase [Shigella sonnei]	Length: 372\nNumber of predicted TMHs: 2\nExp number of AAs in TMHs: 37.02628\nExp number, first 60 AAs: 19.39684\nTotal prob of N-in: 0.94313\nPOSSIBLE N-term signal sequence\ninside 1 19\nTMhelix 20 39\noutside 40 325\nTMhelix 326 348\ninside 349 372

27263	GCF_900160445.1_8403_8_75	Shigella sonnei	Proteobacteria; Gammaproteobacteria; Enterobacterales; Enterobacteriaceae; Shigella
MNNEETFYQAMRRQGVTRRSFLKYCSLAATSLGLGAGMAPKIAWALENKPRIPVVWIHGL	WP_005139779.1
hydrogenase [Shigella sonnei]	Length: 372\nNumber of predicted TMHs: 2\nExp number of AAs in TMHs: 37.02628\nExp number, first 60 AAs: 19.39684\nTotal prob of N-in: 0.94313\nPOSSIBLE N-term signal sequence\ninside 1 19\nTMhelix 20 39\noutside 40 325\nTMhelix 326 348\ninside 349 372

27264	GCF_900160465.1_8403_8_78	Shigella sonnei	Proteobacteria; Gammaproteobacteria; Enterobacterales; Enterobacteriaceae; Shigella
MNNEETFYQAMRRQGVTRRSFLKYCSLAATSLGLGAGMAPKIAWALENKPRIPVVWIHGL	WP_005139779.1
hydrogenase [Shigella sonnei]	Length: 372\nNumber of predicted TMHs: 2\nExp number of AAs in TMHs: 37.02628\nExp number, first 60 AAs: 19.39684\nTotal prob of N-in: 0.94313\nPOSSIBLE N-term signal sequence\ninside 1 19\nTMhelix 20 39\noutside 40 325\nTMhelix 326 348\ninside 349 372

27265	GCF_900160515.1_8403_8_83	Shigella sonnei	Proteobacteria; Gammaproteobacteria; Enterobacterales; Enterobacteriaceae; Shigella
MNNEETFYQAMRRQGVTRRSFLKYCSLAATSLGLGAGMAPKIAWALENKPRIPVVWIHGL	WP_005139779.1
hydrogenase [Shigella sonnei]	Length: 372\nNumber of predicted TMHs: 2\nExp number of AAs in

TMHs: 37.02628\nExp number, first 60 AAs: 19.39684\nTotal prob of N-in: 0.94313\nPOSSIBLE N-term signal sequence\ninside 1 19\nTMhelix 20 39\noutside 40 325\nTMhelix 326 348\ninside 349 372

27266 GCF_900160535.1_8403_8_81 Shigella sonnei Proteobacteria; Gammaproteobacteria; Enterobacterales; Enterobacteriaceae; Shigella
MNNEETFYQAMRRQGVTRRSFLKYCSLAATSLGLGAGMAPKIAWALENKPRIPVVWIHGL WP_005139779.1
hydrogenase [Shigella sonnei] Length: 372\nNumber of predicted TMHs: 2\nExp number of AAs in TMHs: 37.02628\nExp number, first 60 AAs: 19.39684\nTotal prob of N-in: 0.94313\nPOSSIBLE N-term signal sequence\ninside 1 19\nTMhelix 20 39\noutside 40 325\nTMhelix 326 348\ninside 349 372

27267 GCF_900160565.1_8403_8_89 Shigella sonnei Proteobacteria; Gammaproteobacteria; Enterobacterales; Enterobacteriaceae; Shigella
MNNEETFYQAMRRQGVTRRSFLKYCSLAATSLGLGAGMAPKIAWALENKPRIPVVWIHGL WP_005139779.1
hydrogenase [Shigella sonnei] Length: 372\nNumber of predicted TMHs: 2\nExp number of AAs in TMHs: 37.02628\nExp number, first 60 AAs: 19.39684\nTotal prob of N-in: 0.94313\nPOSSIBLE N-term signal sequence\ninside 1 19\nTMhelix 20 39\noutside 40 325\nTMhelix 326 348\ninside 349 372

27268 GCF_900160585.1_8403_8_91 Shigella sonnei Proteobacteria; Gammaproteobacteria; Enterobacterales; Enterobacteriaceae; Shigella
MNNEETFYQAMRRQGVTRRSFLKYCSLAATSLGLGAGMAPKIAWALENKPRIPVVWIHGL WP_005139779.1
hydrogenase [Shigella sonnei] Length: 372\nNumber of predicted TMHs: 2\nExp number of AAs in TMHs: 37.02628\nExp number, first 60 AAs: 19.39684\nTotal prob of N-in: 0.94313\nPOSSIBLE N-term signal sequence\ninside 1 19\nTMhelix 20 39\noutside 40 325\nTMhelix 326 348\ninside 349 372

27269 GCF_900160635.1_8403_8_96 Shigella sonnei Proteobacteria; Gammaproteobacteria; Enterobacterales; Enterobacteriaceae; Shigella
MTGDNTLIHSHGINRRDFMKLCAALAATMGLSSKAAAEMAESVTNPQRPPVIWIGAQECT WP_000145410.1
MULTISPECIES: hydrogenase-2 small chain [Proteobacteria] Length: 372\nNumber of predicted TMHs: 1\nExp number of AAs in TMHs: 23.72226\nExp number, first 60 AAs: 0.22038\nTotal prob of N-in: 0.12994\noutside 1 331\nTMhelix 332 354\ninside 355 372

27270 GCF_900162765.1_16703_8_4 Shigella sonnei Proteobacteria; Gammaproteobacteria; Enterobacterales; Enterobacteriaceae; Shigella
MTGDNTLIHSHGINRRDFMKLCAALAATMGLSSKAAAEMAESVTNPQRPPVIWIGAQECT WP_000145410.1
MULTISPECIES: hydrogenase-2 small chain [Proteobacteria] Length: 372\nNumber of predicted TMHs: 1\nExp number of AAs in TMHs: 23.72226\nExp number, first 60 AAs: 0.22038\nTotal prob of N-in: 0.12994\noutside 1 331\nTMhelix 332 354\ninside 355 372

27271 GCF_900162785.1_9803_4_45 Shigella sonnei Proteobacteria; Gammaproteobacteria; Enterobacterales; Enterobacteriaceae; Shigella
MTGDNTLIHSHGINRRDFMKLCAALAATMGLSSKAAAEMAESVTNPQRPPVIWIGAQECT WP_000145410.1
MULTISPECIES: hydrogenase-2 small chain [Proteobacteria] Length: 372\nNumber of predicted TMHs: 1\nExp number of AAs in TMHs: 23.72226\nExp number, first 60 AAs: 0.22038\nTotal prob of N-in: 0.12994\noutside 1 331\nTMhelix 332 354\ninside 355 372

27272 GCF_900160635.1_8403_8_96 Shigella sonnei Proteobacteria; Gammaproteobacteria; Enterobacterales; Enterobacteriaceae; Shigella
MNNEETFYQAMRRQGVTRRSFLKYCSLAATSLGLGAGMAPKIAWALENKPRIPVVWIHGL WP_005139779.1
hydrogenase [Shigella sonnei] Length: 372\nNumber of predicted TMHs: 2\nExp number of AAs in TMHs: 37.02628\nExp number, first 60 AAs: 19.39684\nTotal prob of N-in: 0.94313\nPOSSIBLE N-term signal sequence\ninside 1 19\nTMhelix 20 39\noutside 40 325\nTMhelix 326 348\ninside 349 372

27273 GCF_900162785.1_9803_4_45 Shigella sonnei Proteobacteria; Gammaproteobacteria; Enterobacterales; Enterobacteriaceae; Shigella
MNNEETFYQAMRRQGVTRRSFLKYCSLAATSLGLGAGMAPKIAWALENKPRIPVVWIHGL WP_005139779.1
hydrogenase [Shigella sonnei] Length: 372\nNumber of predicted TMHs: 2\nExp number of AAs in TMHs: 37.02628\nExp number, first 60 AAs: 19.39684\nTotal prob of N-in: 0.94313\nPOSSIBLE N-term signal sequence\ninside 1 19\nTMhelix 20 39\noutside 40 325\nTMhelix 326 348\ninside 349 372

27274 GCF_900162765.1_16703_8_4 Shigella sonnei Proteobacteria; Gammaproteobacteria; Enterobacterales; Enterobacteriaceae; Shigella

MNNEETFYQAMRRQGVTRRSFLKYCSLAATSLGLGAGMAPKIAWALENKPRIPVVWIHGL WP_005139779.1 hydrogenase [Shigella sonnei] Length: 372\nNumber of predicted TMHs: 2\nExp number of AAs in TMHs: 37.02628\nExp number, first 60 AAs: 19.39684\nTotal prob of N-in: 0.94313\nPOSSIBLE N-term signal sequence\ninside 1 19\nTMhelix 20 39\noutside 40 325\nTMhelix 326 348\ninside 349 372

27275 GCF_001184135.1_ASM118413v1 Shigella sonnei Proteobacteria; Gammaproteobacteria; Enterobacterales; Enterobacteriaceae; Shigella MTGDNTLIHSHGINRRDFMKLCAALAATMGLSSKAAAEMAESVTNPQRPPVIWIGAQECT WP_000145410.1 MULTISPECIES: hydrogenase-2 small chain [Proteobacteria] Length: 372\nNumber of predicted TMHs: 1\nExp number of AAs in TMHs: 23.72226\nExp number, first 60 AAs: 0.22038\nTotal prob of N-in: 0.12994\noutside 1 331\nTMhelix 332 354\ninside 355 372

27276 GCF_001245775.1_8290_4_13 Shigella sonnei Proteobacteria; Gammaproteobacteria; Enterobacterales; Enterobacteriaceae; Shigella MTGDNTLIHSHGINRRDFMKLCAALAATMGLSSKAAAEMAESVTNPQRPPVIWIGAQECT WP_000145410.1 MULTISPECIES: hydrogenase-2 small chain [Proteobacteria] Length: 372\nNumber of predicted TMHs: 1\nExp number of AAs in TMHs: 23.72226\nExp number, first 60 AAs: 0.22038\nTotal prob of N-in: 0.12994\noutside 1 331\nTMhelix 332 354\ninside 355 372

27277 GCF_001246155.1_8290_4_93 Shigella sonnei Proteobacteria; Gammaproteobacteria; Enterobacterales; Enterobacteriaceae; Shigella MTGDNTLIHSHGINRRDFMKLCAALAATMGLSSKAAAEMAESVTNPQRPPVIWIGAQECT WP_000145410.1 MULTISPECIES: hydrogenase-2 small chain [Proteobacteria] Length: 372\nNumber of predicted TMHs: 1\nExp number of AAs in TMHs: 23.72226\nExp number, first 60 AAs: 0.22038\nTotal prob of N-in: 0.12994\noutside 1 331\nTMhelix 332 354\ninside 355 372

27278 GCF_001246455.1_12971_8_24 Shigella sonnei Proteobacteria; Gammaproteobacteria; Enterobacterales; Enterobacteriaceae; Shigella MTGDNTLIHSHGINRRDFMKLCAALAATMGLSSKAAAEMAESVTNPQRPPVIWIGAQECT WP_000145410.1 MULTISPECIES: hydrogenase-2 small chain [Proteobacteria] Length: 372\nNumber of predicted TMHs: 1\nExp number of AAs in TMHs: 23.72226\nExp number, first 60 AAs: 0.22038\nTotal prob of N-in: 0.12994\noutside 1 331\nTMhelix 332 354\ninside 355 372

27279 GCF_001246775.1_8289_7_43 Shigella sonnei Proteobacteria; Gammaproteobacteria; Enterobacterales; Enterobacteriaceae; Shigella MTGDNTLIHSHGINRRDFMKLCAALAATMGLSSKAAAEMAESVTNPQRPPVIWIGAQECT WP_000145410.1 MULTISPECIES: hydrogenase-2 small chain [Proteobacteria] Length: 372\nNumber of predicted TMHs: 1\nExp number of AAs in TMHs: 23.72226\nExp number, first 60 AAs: 0.22038\nTotal prob of N-in: 0.12994\noutside 1 331\nTMhelix 332 354\ninside 355 372

27280 GCF_001247045.1_Sg_201311953_TACAGC_L001 Shigella sonnei Proteobacteria; Gammaproteobacteria; Enterobacterales; Enterobacteriaceae; Shigella MTGDNTLIHSHGINRRDFMKLCAALAATMGLSSKAAAEMAESVTNPQRPPVIWIGAQECT WP_000145410.1 MULTISPECIES: hydrogenase-2 small chain [Proteobacteria] Length: 372\nNumber of predicted TMHs: 1\nExp number of AAs in TMHs: 23.72226\nExp number, first 60 AAs: 0.22038\nTotal prob of N-in: 0.12994\noutside 1 331\nTMhelix 332 354\ninside 355 372

27281 GCF_001247045.1_Sg_201311953_TACAGC_L001 Shigella sonnei Proteobacteria; Gammaproteobacteria; Enterobacterales; Enterobacteriaceae; Shigella MNNEETFYQAMRRQGVTRRSFLKYCSLAATSLGLGAGMAPKIAWALENKPRIPVVWIHGL WP_005139779.1 hydrogenase [Shigella sonnei] Length: 372\nNumber of predicted TMHs: 2\nExp number of AAs in TMHs: 37.02628\nExp number, first 60 AAs: 19.39684\nTotal prob of N-in: 0.94313\nPOSSIBLE N-term signal sequence\ninside 1 19\nTMhelix 20 39\noutside 40 325\nTMhelix 326 348\ninside 349 372

27282 GCF_001247645.1_H14132053701 Shigella sonnei Proteobacteria; Gammaproteobacteria; Enterobacterales; Enterobacteriaceae; Shigella MTGDNTLIHSHGINRRDFMKLCAALAATMGLSSKAAAEMAESVTNPQRPPVIWIGAQECT WP_000145410.1 MULTISPECIES: hydrogenase-2 small chain [Proteobacteria] Length: 372\nNumber of predicted TMHs: 1\nExp number of AAs in TMHs: 23.72226\nExp number, first 60 AAs: 0.22038\nTotal prob of N-in: 0.12994\noutside 1 331\nTMhelix 332 354\ninside 355 372

27283	GCF_001247735.1_5236_8_1Shigella sonnei	Proteobacteria; Gammaproteobacteria; Enterobacterales; Enterobacteriaceae; Shigella	MTGDNTLIHSHGINRRDFMKLCAALAATMGLSSKAAAEMAESVTNPQRPPVIWIGAQECT	WP_000145410.1	MULTISPECIES: hydrogenase-2 small chain [Proteobacteria]	Length: 372\nNumber of predicted TMHs:  1\nExp number of AAs in TMHs: 23.72226\nExp number, first 60 AAs:  0.22038\nTotal prob of N-in:  0.12994\noutside   1   331\nTMhelix   332   354\ninside   355   372

27284	GCF_001247645.1_H14132053701	Shigella sonnei	Proteobacteria; Gammaproteobacteria; Enterobacterales; Enterobacteriaceae; Shigella	MNNEETFYQAMRRQGVTRRSFLKYCSLAATSLGLGAGMAPKIAWALENKPRIPVVWIHGL	WP_052962775.1	hydrogenase [Shigella sonnei]	Length: 372\nNumber of predicted TMHs:  2\nExp number of AAs in TMHs: 37.09248\nExp number, first 60 AAs:  19.39296\nTotal prob of N-in:   0.94316\nPOSSIBLE N-term signal sequence\ninside   1   19\nTMhelix   20   39\noutside   40   325\nTMhelix   326   348\ninside   349   372

27285	GCF_001247735.1_5236_8_1Shigella sonnei	Proteobacteria; Gammaproteobacteria; Enterobacterales; Enterobacteriaceae; Shigella	MNNEETFYQAMRRQGVTRRSFLKYCSLAATSLGLGAGMAPKIAWALENKPRIPVVWIHGL	WP_005139779.1	hydrogenase [Shigella sonnei]	Length: 372\nNumber of predicted TMHs:  2\nExp number of AAs in TMHs: 37.02628\nExp number, first 60 AAs:  19.39684\nTotal prob of N-in:   0.94313\nPOSSIBLE N-term signal sequence\ninside   1   19\nTMhelix   20   39\noutside   40   325\nTMhelix   326   348\ninside   349   372

27286	GCF_001184135.1_ASM118413v1	Shigella sonnei	Proteobacteria; Gammaproteobacteria; Enterobacterales; Enterobacteriaceae; Shigella	MNNEETFYQAMRRQGVTRRSFLKYCSLAATSLGLGAGMAPKIAWALENKPRIPVVWIHGL	WP_005139779.1	hydrogenase [Shigella sonnei]	Length: 372\nNumber of predicted TMHs:  2\nExp number of AAs in TMHs: 37.02628\nExp number, first 60 AAs:  19.39684\nTotal prob of N-in:   0.94313\nPOSSIBLE N-term signal sequence\ninside   1   19\nTMhelix   20   39\noutside   40   325\nTMhelix   326   348\ninside   349   372

27287	GCF_001246155.1_8290_4_93	Shigella sonnei	Proteobacteria; Gammaproteobacteria; Enterobacterales; Enterobacteriaceae; Shigella	MNNEETFYQAMRRQGVTRRSFLKYCSLAATSLGLGAGMAPKIAWALENKPRIPVVWIHGL	WP_005139779.1	hydrogenase [Shigella sonnei]	Length: 372\nNumber of predicted TMHs:  2\nExp number of AAs in TMHs: 37.02628\nExp number, first 60 AAs:  19.39684\nTotal prob of N-in:   0.94313\nPOSSIBLE N-term signal sequence\ninside   1   19\nTMhelix   20   39\noutside   40   325\nTMhelix   326   348\ninside   349   372

27288	GCF_001245775.1_8290_4_13	Shigella sonnei	Proteobacteria; Gammaproteobacteria; Enterobacterales; Enterobacteriaceae; Shigella	MNNEETFYQAMRRQGVTRRSFLKYCSLAATSLGLGAGMAPKIAWALENKPRIPVVWIHGL	WP_005139779.1	hydrogenase [Shigella sonnei]	Length: 372\nNumber of predicted TMHs:  2\nExp number of AAs in TMHs: 37.02628\nExp number, first 60 AAs:  19.39684\nTotal prob of N-in:   0.94313\nPOSSIBLE N-term signal sequence\ninside   1   19\nTMhelix   20   39\noutside   40   325\nTMhelix   326   348\ninside   349   372

27289	GCF_001246455.1_12971_8_24	Shigella sonnei	Proteobacteria; Gammaproteobacteria; Enterobacterales; Enterobacteriaceae; Shigella	MNNEETFYQAMRRQGVTRRSFLKYCSLAATSLGLGAGMAPKIAWALENKPRIPVVWIHGL	WP_005139779.1	hydrogenase [Shigella sonnei]	Length: 372\nNumber of predicted TMHs:  2\nExp number of AAs in TMHs: 37.02628\nExp number, first 60 AAs:  19.39684\nTotal prob of N-in:   0.94313\nPOSSIBLE N-term signal sequence\ninside   1   19\nTMhelix   20   39\noutside   40   325\nTMhelix   326   348\ninside   349   372

27290	GCF_001247785.1_H14144051401	Shigella sonnei	Proteobacteria; Gammaproteobacteria; Enterobacterales; Enterobacteriaceae; Shigella	MTGDNTLIHSHGINRRDFMKLCAALAATMGLSSKAAAEMAESVTNPQRPPVIWIGAQECT	WP_000145410.1	MULTISPECIES: hydrogenase-2 small chain [Proteobacteria]	Length: 372\nNumber of predicted TMHs:  1\nExp number of AAs in TMHs: 23.72226\nExp number, first 60 AAs:  0.22038\nTotal prob of N-in:   0.12994\noutside   1   331\nTMhelix   332   354\ninside   355   372

27291	GCF_001247785.1_H14144051401	Shigella sonnei	Proteobacteria; Gammaproteobacteria; Enterobacterales; Enterobacteriaceae; Shigella	MNNEETFYQAMRRQGVTRRSFLKYCSLAATSLGLGAGMAPKIAWALENKPRIPVVWIHGL	WP_005139779.1	hydrogenase [Shigella sonnei]	Length: 372\nNumber of predicted TMHs:  2\nExp number of AAs in TMHs: 37.02628\nExp number, first 60 AAs:  19.39684\nTotal prob of N-in:   0.94313\nPOSSIBLE N-term signal sequence\ninside   1   19\nTMhelix   20   39\noutside   40   325\nTMhelix   326   348\ninside   349   372

27292 GCF_001246775.1_8289_7_43 Shigella sonnei Proteobacteria; Gammaproteobacteria; Enterobacterales; Enterobacteriaceae; Shigella MNNEETFYQAMRRQGVTRRSFLKYCSLAATSLGLGAGMAPKIAWALENKPRIPVVWIHGL WP_005139779.1 hydrogenase [Shigella sonnei] Length: 372\nNumber of predicted TMHs: 2\nExp number of AAs in TMHs: 37.02628\nExp number, first 60 AAs: 19.39684\nTotal prob of N-in: 0.94313\nPOSSIBLE N-term signal sequence\ninside 1 19\nTMhelix 20 39\noutside 40 325\nTMhelix 326 348\ninside 349 372

27293 GCF_001247865.1_Sg_201403955_CACCGG_L001 Shigella sonnei Proteobacteria; Gammaproteobacteria; Enterobacterales; Enterobacteriaceae; Shigella MTGDNTLIHSHGINRRDFMKLCAALAATMGLSSKAAAEMAESVTNPQRPPVIWIGAQECT WP_000145410.1 MULTISPECIES: hydrogenase-2 small chain [Proteobacteria] Length: 372\nNumber of predicted TMHs: 1\nExp number of AAs in TMHs: 23.72226\nExp number, first 60 AAs: 0.22038\nTotal prob of N-in: 0.12994\noutside 1 331\nTMhelix 332 354\ninside 355 372

27294 GCF_001247945.1_Sg_201404332_CAGGCG_L001 Shigella sonnei Proteobacteria; Gammaproteobacteria; Enterobacterales; Enterobacteriaceae; Shigella MTGDNTLIHSHGINRRDFMKLCAALAATMGLSSKAAAEMAESVTNPQRPPVIWIGAQECT WP_000145410.1 MULTISPECIES: hydrogenase-2 small chain [Proteobacteria] Length: 372\nNumber of predicted TMHs: 1\nExp number of AAs in TMHs: 23.72226\nExp number, first 60 AAs: 0.22038\nTotal prob of N-in: 0.12994\noutside 1 331\nTMhelix 332 354\ninside 355 372

27295 GCF_001248085.1_12850_1_45 Shigella sonnei Proteobacteria; Gammaproteobacteria; Enterobacterales; Enterobacteriaceae; Shigella MTGDNTLIHSHGINRRDFMKLCAALAATMGLSSKAAAEMAESVTNPQRPPVIWIGAQECT WP_000145410.1 MULTISPECIES: hydrogenase-2 small chain [Proteobacteria] Length: 372\nNumber of predicted TMHs: 1\nExp number of AAs in TMHs: 23.72226\nExp number, first 60 AAs: 0.22038\nTotal prob of N-in: 0.12994\noutside 1 331\nTMhelix 332 354\ninside 355 372

27296 GCF_001248175.1_5417_1_7Shigella sonnei Proteobacteria; Gammaproteobacteria; Enterobacterales; Enterobacteriaceae; Shigella MTGDNTLIHSHGINRRDFMKLCAALAATMGLSSKAAAEMAESVTNPQRPPVIWIGAQECT WP_000145410.1 MULTISPECIES: hydrogenase-2 small chain [Proteobacteria] Length: 372\nNumber of predicted TMHs: 1\nExp number of AAs in TMHs: 23.72226\nExp number, first 60 AAs: 0.22038\nTotal prob of N-in: 0.12994\noutside 1 331\nTMhelix 332 354\ninside 355 372

27297 GCF_001248315.1_5236_3_11 Shigella sonnei Proteobacteria; Gammaproteobacteria; Enterobacterales; Enterobacteriaceae; Shigella MTGDNTLIHSHGINRRDFMKLCAALAATMGLSSKAAAEMAESVTNPQRPPVIWIGAQECT WP_000145410.1 MULTISPECIES: hydrogenase-2 small chain [Proteobacteria] Length: 372\nNumber of predicted TMHs: 1\nExp number of AAs in TMHs: 23.72226\nExp number, first 60 AAs: 0.22038\nTotal prob of N-in: 0.12994\noutside 1 331\nTMhelix 332 354\ninside 355 372

27298 GCF_001248275.1_12971_8_29 Shigella sonnei Proteobacteria; Gammaproteobacteria; Enterobacterales; Enterobacteriaceae; Shigella MTGDNTLIHSHGINRRDFMKLCAALAATMGLSSKAAAEMAESVTNPQRPPVIWIGAQECT WP_000145410.1 MULTISPECIES: hydrogenase-2 small chain [Proteobacteria] Length: 372\nNumber of predicted TMHs: 1\nExp number of AAs in TMHs: 23.72226\nExp number, first 60 AAs: 0.22038\nTotal prob of N-in: 0.12994\noutside 1 331\nTMhelix 332 354\ninside 355 372

27299 GCF_001248405.1_6496_8_11 Shigella sonnei Proteobacteria; Gammaproteobacteria; Enterobacterales; Enterobacteriaceae; Shigella MTGDNTLIHSHGINRRDFMKLCAALAATMGLSSKAAAEMAESVTNPQRPPVIWIGAQECT WP_000145410.1 MULTISPECIES: hydrogenase-2 small chain [Proteobacteria] Length: 372\nNumber of predicted TMHs: 1\nExp number of AAs in TMHs: 23.72226\nExp number, first 60 AAs: 0.22038\nTotal prob of N-in: 0.12994\noutside 1 331\nTMhelix 332 354\ninside 355 372

27300 GCF_001248485.1_5417_1_6Shigella sonnei Proteobacteria; Gammaproteobacteria; Enterobacterales; Enterobacteriaceae; Shigella MTGDNTLIHSHGINRRDFMKLCAALAATMGLSSKAAAEMAESVTNPQRPPVIWIGAQECT WP_000145410.1 MULTISPECIES: hydrogenase-2 small chain [Proteobacteria] Length: 372\nNumber of predicted TMHs: 1\nExp number of AAs in TMHs: 23.72226\nExp number, first 60 AAs: 0.22038\nTotal prob of N-in: 0.12994\noutside 1 331\nTMhelix 332 354\ninside 355 372

27301 GCF_001248545.1_6551_1_12 Shigella sonnei Proteobacteria; Gammaproteobacteria; Enterobacterales; Enterobacteriaceae; Shigella MTGDNTLIHSHGINRRDFMKLCAALAATMGLSSKAAAEMAESVTNPQRPPVIWIGAQECT WP_000145410.1 MULTISPECIES: hydrogenase-2 small chain [Proteobacteria] Length: 372\nNumber of predicted TMHs: 1\nExp number of AAs in TMHs: 23.72226\nExp number, first 60 AAs: 0.22038\nTotal prob of N-in: 0.12994\noutside 1 331\nTMhelix 332 354\ninside 355 372

27302 GCF_001248605.1_8489_2_90 Shigella sonnei Proteobacteria; Gammaproteobacteria; Enterobacterales; Enterobacteriaceae; Shigella MTGDNTLIHSHGINRRDFMKLCAALAATMGLSSKAAAEMAESVTNPQRPPVIWIGAQECT WP_000145410.1 MULTISPECIES: hydrogenase-2 small chain [Proteobacteria] Length: 372\nNumber of predicted TMHs: 1\nExp number of AAs in TMHs: 23.72226\nExp number, first 60 AAs: 0.22038\nTotal prob of N-in: 0.12994\noutside 1 331\nTMhelix 332 354\ninside 355 372

27303 GCF_001248485.1_5417_1_6Shigella sonnei Proteobacteria; Gammaproteobacteria; Enterobacterales; Enterobacteriaceae; Shigella MNNEETFYQAMRRQGVTRRSFLKYCSLAATSLGLGAGMAPKIAWALENKPRIPVVWIHGL WP_005139779.1 hydrogenase [Shigella sonnei] Length: 372\nNumber of predicted TMHs: 2\nExp number of AAs in TMHs: 37.02628\nExp number, first 60 AAs: 19.39684\nTotal prob of N-in: 0.94313\nPOSSIBLE N-term signal sequence\ninside 1 19\nTMhelix 20 39\noutside 40 325\nTMhelix 326 348\ninside 349 372

27304 GCF_001248545.1_6551_1_12 Shigella sonnei Proteobacteria; Gammaproteobacteria; Enterobacterales; Enterobacteriaceae; Shigella MNNEETFYQAMRRQGVTRRSFLKYCSLAATSLGLGAGMAPKIAWALENKPRIPVVWIHGL WP_005139779.1 hydrogenase [Shigella sonnei] Length: 372\nNumber of predicted TMHs: 2\nExp number of AAs in TMHs: 37.02628\nExp number, first 60 AAs: 19.39684\nTotal prob of N-in: 0.94313\nPOSSIBLE N-term signal sequence\ninside 1 19\nTMhelix 20 39\noutside 40 325\nTMhelix 326 348\ninside 349 372

27305 GCF_001248605.1_8489_2_90 Shigella sonnei Proteobacteria; Gammaproteobacteria; Enterobacterales; Enterobacteriaceae; Shigella MNNEETFYQAMRRQGVTRRSFLKYCSLAATSLGLGAGMAPKIAWALENKPRIPVVWIHGL WP_005139779.1 hydrogenase [Shigella sonnei] Length: 372\nNumber of predicted TMHs: 2\nExp number of AAs in TMHs: 37.02628\nExp number, first 60 AAs: 19.39684\nTotal prob of N-in: 0.94313\nPOSSIBLE N-term signal sequence\ninside 1 19\nTMhelix 20 39\noutside 40 325\nTMhelix 326 348\ninside 349 372

27306 GCF_001247865.1_Sg_201403955_CACCGG_L001 Shigella sonnei Proteobacteria; Gammaproteobacteria; Enterobacterales; Enterobacteriaceae; Shigella MNNEETFYQAMRRQGVTRRSFLKYCSLAATSLGLGAGMAPKIAWALENKPRIPVVWIHGL WP_005139779.1 hydrogenase [Shigella sonnei] Length: 372\nNumber of predicted TMHs: 2\nExp number of AAs in TMHs: 37.02628\nExp number, first 60 AAs: 19.39684\nTotal prob of N-in: 0.94313\nPOSSIBLE N-term signal sequence\ninside 1 19\nTMhelix 20 39\noutside 40 325\nTMhelix 326 348\ninside 349 372

27307 GCF_001248085.1_12850_1_45 Shigella sonnei Proteobacteria; Gammaproteobacteria; Enterobacterales; Enterobacteriaceae; Shigella MNNEETFYQAMRRQGVTRRSFLKYCSLAATSLGLGAGMAPKIAWALENKPRIPVVWIHGL WP_005139779.1 hydrogenase [Shigella sonnei] Length: 372\nNumber of predicted TMHs: 2\nExp number of AAs in TMHs: 37.02628\nExp number, first 60 AAs: 19.39684\nTotal prob of N-in: 0.94313\nPOSSIBLE N-term signal sequence\ninside 1 19\nTMhelix 20 39\noutside 40 325\nTMhelix 326 348\ninside 349 372

27308 GCF_001247945.1_Sg_201404332_CAGGCG_L001 Shigella sonnei Proteobacteria; Gammaproteobacteria; Enterobacterales; Enterobacteriaceae; Shigella MNNEETFYQAMRRQGVTRRSFLKYCSLAATSLGLGAGMAPKIAWALENKPRIPVVWIHGL WP_005139779.1 hydrogenase [Shigella sonnei] Length: 372\nNumber of predicted TMHs: 2\nExp number of AAs in TMHs: 37.02628\nExp number, first 60 AAs: 19.39684\nTotal prob of N-in: 0.94313\nPOSSIBLE N-term signal sequence\ninside 1 19\nTMhelix 20 39\noutside 40 325\nTMhelix 326 348\ninside 349 372

27309 GCF_001248725.1_9870_6_5Shigella sonnei Proteobacteria; Gammaproteobacteria; Enterobacterales; Enterobacteriaceae; Shigella MTGDNTLIHSHGINRRDFMKLCAALAATMGLSSKAAAEMAESVTNPQRPPVIWIGAQECT WP_000145410.1 MULTISPECIES: hydrogenase-2 small chain [Proteobacteria] Length: 372\nNumber of predicted TMHs: 1\nExp number of AAs in TMHs: 23.72226\nExp number, first 60 AAs: 0.22038\nTotal prob of N-in: 0.12994\noutside 1 331\nTMhelix 332 354\ninside 355 372

27310 GCF_001248725.1_9870_6_5Shigella sonnei Proteobacteria; Gammaproteobacteria; Enterobacterales; Enterobacteriaceae; Shigella MNNEETFYQAMRRQGVTRRSFLKYCSLAATSLGLGAGMAPKIAWALENKPRIPVVWIHGL WP_005139779.1 hydrogenase [Shigella sonnei] Length: 372\nNumber of predicted TMHs: 2\nExp number of AAs in TMHs: 37.02628\nExp number, first 60 AAs: 19.39684\nTotal prob of N-in: 0.94313\nPOSSIBLE N-term signal sequence\ninside 1 19\nTMhelix 20 39\noutside 40 325\nTMhelix 326 348\ninside 349 372

27311 GCF_001248175.1_5417_1_7Shigella sonnei Proteobacteria; Gammaproteobacteria; Enterobacterales; Enterobacteriaceae; Shigella MNNEETFYQAMRRQGVTRRSFLKYCSLAATSLGLGAGMAPKIAWALENKPRIPVVWIHGL WP_005139779.1 hydrogenase [Shigella sonnei] Length: 372\nNumber of predicted TMHs: 2\nExp number of AAs in TMHs: 37.02628\nExp number, first 60 AAs: 19.39684\nTotal prob of N-in: 0.94313\nPOSSIBLE N-term signal sequence\ninside 1 19\nTMhelix 20 39\noutside 40 325\nTMhelix 326 348\ninside 349 372

27312 GCF_001248315.1_5236_3_11 Shigella sonnei Proteobacteria; Gammaproteobacteria; Enterobacterales; Enterobacteriaceae; Shigella MNNEETFYQAMRRQGVTRRSFLKYCSLAATSLGLGAGMAPKIAWALENKPRIPVVWIHGL WP_005139779.1 hydrogenase [Shigella sonnei] Length: 372\nNumber of predicted TMHs: 2\nExp number of AAs in TMHs: 37.02628\nExp number, first 60 AAs: 19.39684\nTotal prob of N-in: 0.94313\nPOSSIBLE N-term signal sequence\ninside 1 19\nTMhelix 20 39\noutside 40 325\nTMhelix 326 348\ninside 349 372

27313 GCF_001248405.1_6496_8_11 Shigella sonnei Proteobacteria; Gammaproteobacteria; Enterobacterales; Enterobacteriaceae; Shigella MNNEETFYQAMRRQGVTRRSFLKYCSLAATSLGLGAGMAPKIAWALENKPRIPVVWIHGL WP_005139779.1 hydrogenase [Shigella sonnei] Length: 372\nNumber of predicted TMHs: 2\nExp number of AAs in TMHs: 37.02628\nExp number, first 60 AAs: 19.39684\nTotal prob of N-in: 0.94313\nPOSSIBLE N-term signal sequence\ninside 1 19\nTMhelix 20 39\noutside 40 325\nTMhelix 326 348\ninside 349 372

27314 GCF_001248275.1_12971_8_29 Shigella sonnei Proteobacteria; Gammaproteobacteria; Enterobacterales; Enterobacteriaceae; Shigella MNNEETFYQAMRRQGVTRRSFLKYCSLAATSLGLGAGMAPKIAWALENKPRIPVVWIHGL WP_005139779.1 hydrogenase [Shigella sonnei] Length: 372\nNumber of predicted TMHs: 2\nExp number of AAs in TMHs: 37.02628\nExp number, first 60 AAs: 19.39684\nTotal prob of N-in: 0.94313\nPOSSIBLE N-term signal sequence\ninside 1 19\nTMhelix 20 39\noutside 40 325\nTMhelix 326 348\ninside 349 372

27315 GCF_001248765.1_12971_8_40 Shigella sonnei Proteobacteria; Gammaproteobacteria; Enterobacterales; Enterobacteriaceae; Shigella MTGDNTLIHSHGINRRDFMKLCAALAATMGLSSKAAAEMAESVTNPQRPPVIWIGAQECT WP_000145410.1 MULTISPECIES: hydrogenase-2 small chain [Proteobacteria] Length: 372\nNumber of predicted TMHs: 1\nExp number of AAs in TMHs: 23.72226\nExp number, first 60 AAs: 0.22038\nTotal prob of N-in: 0.12994\noutside 1 331\nTMhelix 332 354\ninside 355 372

27316 GCF_001248665.1_H14148037701 Shigella sonnei Proteobacteria; Gammaproteobacteria; Enterobacterales; Enterobacteriaceae; Shigella MNNEETFYQAMRRQGVTRRSFLKYCSLAATSLGLGAGMAPKIAWALENKPRIPVVWIHGL WP_005139779.1 hydrogenase [Shigella sonnei] Length: 372\nNumber of predicted TMHs: 2\nExp number of AAs in TMHs: 37.02628\nExp number, first 60 AAs: 19.39684\nTotal prob of N-in: 0.94313\nPOSSIBLE N-term signal sequence\ninside 1 19\nTMhelix 20 39\noutside 40 325\nTMhelix 326 348\ninside 349 372

27317 GCF_001248825.1_12971_8_34 Shigella sonnei Proteobacteria; Gammaproteobacteria; Enterobacterales; Enterobacteriaceae; Shigella MTGDNTLIHSHGINRRDFMKLCAALAATMGLSSKAAAEMAESVTNPQRPPVIWIGAQECT WP_000145410.1 MULTISPECIES: hydrogenase-2 small chain [Proteobacteria] Length: 372\nNumber of predicted TMHs: 1\nExp number of AAs in TMHs: 23.72226\nExp number, first 60 AAs: 0.22038\nTotal prob of N-in: 0.12994\noutside 1 331\nTMhelix 332 354\ninside 355 372

27318 GCF_001248925.1_5236_3_12 Shigella sonnei Proteobacteria; Gammaproteobacteria; Enterobacterales; Enterobacteriaceae; Shigella MTGDNTLIHSHGINRRDFMKLCAALAATMGLSSKAAAEMAESVTNPQRPPVIWIGAQECT WP_000145410.1 MULTISPECIES: hydrogenase-2 small chain [Proteobacteria] Length: 372\nNumber of predicted TMHs: 1\nExp number of AAs in TMHs: 23.72226\nExp number, first 60 AAs: 0.22038\nTotal prob of N-in: 0.12994\noutside 1 331\nTMhelix 332 354\ninside 355 372

27319 GCF_001248965.1_5417_1_11 Shigella sonnei Proteobacteria; Gammaproteobacteria; Enterobacterales; Enterobacteriaceae; Shigella
MTGDNTLIHSHGINRRDFMKLCAALAATMGLSSKAAAEMAESVTNPQRPPVIWIGAQECT WP_000145410.1
MULTISPECIES: hydrogenase-2 small chain [Proteobacteria] Length: 372\nNumber of predicted TMHs: 1\nExp number of AAs in TMHs: 23.72226\nExp number, first 60 AAs: 0.22038\nTotal prob of N-in: 0.12994\noutside 1 331\nTMhelix 332 354\ninside 355 372

27320 GCF_001249005.1_7655_8_14 Shigella sonnei Proteobacteria; Gammaproteobacteria; Enterobacterales; Enterobacteriaceae; Shigella
MTGDNTLIHSHGINRRDFMKLCAALAATMGLSSKAAAEMAESVTNPQRPPVIWIGAQECT WP_000145410.1
MULTISPECIES: hydrogenase-2 small chain [Proteobacteria] Length: 372\nNumber of predicted TMHs: 1\nExp number of AAs in TMHs: 23.72226\nExp number, first 60 AAs: 0.22038\nTotal prob of N-in: 0.12994\noutside 1 331\nTMhelix 332 354\ninside 355 372

27321 GCF_001249045.1_5236_5_8Shigella sonnei Proteobacteria; Gammaproteobacteria; Enterobacterales; Enterobacteriaceae; Shigella MTGDNTLIHSHGINRRDFMKLCAALAATMGLSSKAAAEMAESVTNPQRPPVIWIGAQECT
WP_000145410.1 MULTISPECIES: hydrogenase-2 small chain [Proteobacteria] Length: 372\nNumber of predicted TMHs: 1\nExp number of AAs in TMHs: 23.72226\nExp number, first 60 AAs: 0.22038\nTotal prob of N-in: 0.12994\noutside 1 331\nTMhelix 332 354\ninside 355 372

27322 GCF_001249175.1_5236_3_7Shigella sonnei Proteobacteria; Gammaproteobacteria; Enterobacterales; Enterobacteriaceae; Shigella MTGDNTLIHSHGINRRDFMKLCAALAATMGLSSKAAAEMAESVTNPQRPPVIWIGAQECT
WP_000145410.1 MULTISPECIES: hydrogenase-2 small chain [Proteobacteria] Length: 372\nNumber of predicted TMHs: 1\nExp number of AAs in TMHs: 23.72226\nExp number, first 60 AAs: 0.22038\nTotal prob of N-in: 0.12994\noutside 1 331\nTMhelix 332 354\ninside 355 372

27323 GCF_001249135.1_8289_7_88 Shigella sonnei Proteobacteria; Gammaproteobacteria; Enterobacterales; Enterobacteriaceae; Shigella
MTGDNTLIHSHGINRRDFMKLCAALAATMGLSSKAAAEMAESVTNPQRPPVIWIGAQECT WP_000145410.1
MULTISPECIES: hydrogenase-2 small chain [Proteobacteria] Length: 372\nNumber of predicted TMHs: 1\nExp number of AAs in TMHs: 23.72226\nExp number, first 60 AAs: 0.22038\nTotal prob of N-in: 0.12994\noutside 1 331\nTMhelix 332 354\ninside 355 372

27324 GCF_001249255.1_11822_8_61 Shigella sonnei Proteobacteria; Gammaproteobacteria; Enterobacterales; Enterobacteriaceae; Shigella
MTGDNTLIHSHGINRRDFMKLCAALAATMGLSSKAAAEMAESVTNPQRPPVIWIGAQECT WP_000145410.1
MULTISPECIES: hydrogenase-2 small chain [Proteobacteria] Length: 372\nNumber of predicted TMHs: 1\nExp number of AAs in TMHs: 23.72226\nExp number, first 60 AAs: 0.22038\nTotal prob of N-in: 0.12994\noutside 1 331\nTMhelix 332 354\ninside 355 372

27325 GCF_001249135.1_8289_7_88 Shigella sonnei Proteobacteria; Gammaproteobacteria; Enterobacterales; Enterobacteriaceae; Shigella
MNNEETFYQAMRRQGVTRRSFLKYCSLAATSLGLGAGMAPKIAWALENKPRIPVVWIHGL WP_005139779.1
hydrogenase [Shigella sonnei] Length: 372\nNumber of predicted TMHs: 2\nExp number of AAs in TMHs: 37.02628\nExp number, first 60 AAs: 19.39684\nTotal prob of N-in: 0.94313\nPOSSIBLE N-term signal sequence\ninside 1 19\nTMhelix 20 39\noutside 40 325\nTMhelix 326 348\ninside 349 372

27326 GCF_001249045.1_5236_5_8Shigella sonnei Proteobacteria; Gammaproteobacteria; Enterobacterales; Enterobacteriaceae; Shigella MNNEETFYQAMRRQGVTRRSFLKYCSLAATSLGLGAGMAPKIAWALENKPRIPVVWIHGL
WP_005139779.1 hydrogenase [Shigella sonnei] Length: 372\nNumber of predicted TMHs: 2\nExp number of AAs in TMHs: 37.02628\nExp number, first 60 AAs: 19.39684\nTotal prob of N-in: 0.94313\nPOSSIBLE N-term signal sequence\ninside 1 19\nTMhelix 20 39\noutside 40 325\nTMhelix 326 348\ninside 349 372

27327 GCF_001249175.1_5236_3_7Shigella sonnei Proteobacteria; Gammaproteobacteria; Enterobacterales; Enterobacteriaceae; Shigella MNNEETFYQAMRRQGVTRRSFLKYCSLAATSLGLGAGMAPKIAWALENKPRIPVVWIHGL
WP_005139779.1 hydrogenase [Shigella sonnei] Length: 372\nNumber of predicted TMHs: 2\nExp number of AAs in TMHs: 37.02628\nExp number, first 60 AAs: 19.39684\nTotal prob of N-in: 0.94313\nPOSSIBLE N-term signal sequence\ninside 1 19\nTMhelix 20 39\noutside 40 325\nTMhelix 326 348\ninside 349 372

27328 GCF_001248765.1_12971_8_40 Shigella sonnei Proteobacteria; Gammaproteobacteria; Enterobacterales; Enterobacteriaceae; Shigella

MNNEETFYQAMRRQGVTRRSFLKYCSLAATSLGLGAGMAPKIAWALENKPRIPVVWIHGL WP_005139779.1 hydrogenase [Shigella sonnei] Length: 372\nNumber of predicted TMHs: 2\nExp number of AAs in TMHs: 37.02628\nExp number, first 60 AAs: 19.39684\nTotal prob of N-in: 0.94313\nPOSSIBLE N-term signal sequence\ninside 1 19\nTMhelix 20 39\noutside 40 325\nTMhelix 326 348\ninside 349 372

27329 GCF_001249295.1_Sg_201400049_GCCAAT_L001 Shigella sonnei Proteobacteria; Gammaproteobacteria; Enterobacterales; Enterobacteriaceae; Shigella MTGDNTLIHSHGINRRDFMKLCAALAATMGLSSKAAAEMAESVTNPQRPPVIWIGAQECT WP_000145410.1 MULTISPECIES: hydrogenase-2 small chain [Proteobacteria] Length: 372\nNumber of predicted TMHs: 1\nExp number of AAs in TMHs: 23.72226\nExp number, first 60 AAs: 0.22038\nTotal prob of N-in: 0.12994\noutside 1 331\nTMhelix 332 354\ninside 355 372

27330 GCF_001249375.1_9870_6_31 Shigella sonnei Proteobacteria; Gammaproteobacteria; Enterobacterales; Enterobacteriaceae; Shigella MTGDNTLIHSHGINRRDFMKLCAALAATMGLSSKAAAEMAESVTNPQRPPVIWIGAQECT WP_000145410.1 MULTISPECIES: hydrogenase-2 small chain [Proteobacteria] Length: 372\nNumber of predicted TMHs: 1\nExp number of AAs in TMHs: 23.72226\nExp number, first 60 AAs: 0.22038\nTotal prob of N-in: 0.12994\noutside 1 331\nTMhelix 332 354\ninside 355 372

27331 GCF_001249295.1_Sg_201400049_GCCAAT_L001 Shigella sonnei Proteobacteria; Gammaproteobacteria; Enterobacterales; Enterobacteriaceae; Shigella MNNEETFYQAMRRQGVTRRSFLKYCSLAATSLGLGAGMAPKIAWALENKPRIPVVWIHGL WP_005139779.1 hydrogenase [Shigella sonnei] Length: 372\nNumber of predicted TMHs: 2\nExp number of AAs in TMHs: 37.02628\nExp number, first 60 AAs: 19.39684\nTotal prob of N-in: 0.94313\nPOSSIBLE N-term signal sequence\ninside 1 19\nTMhelix 20 39\noutside 40 325\nTMhelix 326 348\ninside 349 372

27332 GCF_001249375.1_9870_6_31 Shigella sonnei Proteobacteria; Gammaproteobacteria; Enterobacterales; Enterobacteriaceae; Shigella MNNEETFYQAMRRQGVTRRSFLKYCSLAATSLGLGAGMAPKIAWALENKPRIPVVWIHGL WP_005139779.1 hydrogenase [Shigella sonnei] Length: 372\nNumber of predicted TMHs: 2\nExp number of AAs in TMHs: 37.02628\nExp number, first 60 AAs: 19.39684\nTotal prob of N-in: 0.94313\nPOSSIBLE N-term signal sequence\ninside 1 19\nTMhelix 20 39\noutside 40 325\nTMhelix 326 348\ninside 349 372

27333 GCF_001248925.1_5236_3_12 Shigella sonnei Proteobacteria; Gammaproteobacteria; Enterobacterales; Enterobacteriaceae; Shigella MNNEETFYQAMRRQGVTRRSFLKYCSLAATSLGLGAGMAPKIAWALENKPRIPVVWIHGL WP_005139779.1 hydrogenase [Shigella sonnei] Length: 372\nNumber of predicted TMHs: 2\nExp number of AAs in TMHs: 37.02628\nExp number, first 60 AAs: 19.39684\nTotal prob of N-in: 0.94313\nPOSSIBLE N-term signal sequence\ninside 1 19\nTMhelix 20 39\noutside 40 325\nTMhelix 326 348\ninside 349 372

27334 GCF_001248825.1_12971_8_34 Shigella sonnei Proteobacteria; Gammaproteobacteria; Enterobacterales; Enterobacteriaceae; Shigella MNNEETFYQAMRRQGVTRRSFLKYCSLAATSLGLGAGMAPKIAWALENKPRIPVVWIHGL WP_005139779.1 hydrogenase [Shigella sonnei] Length: 372\nNumber of predicted TMHs: 2\nExp number of AAs in TMHs: 37.02628\nExp number, first 60 AAs: 19.39684\nTotal prob of N-in: 0.94313\nPOSSIBLE N-term signal sequence\ninside 1 19\nTMhelix 20 39\noutside 40 325\nTMhelix 326 348\ninside 349 372

27335 GCF_001249435.1_7353_1_19 Shigella sonnei Proteobacteria; Gammaproteobacteria; Enterobacterales; Enterobacteriaceae; Shigella MTGDNTLIHSHGINRRDFMKLCAALAATMGLSSKAAAEMAESVTNPQRPPVIWIGAQECT WP_000145410.1 MULTISPECIES: hydrogenase-2 small chain [Proteobacteria] Length: 372\nNumber of predicted TMHs: 1\nExp number of AAs in TMHs: 23.72226\nExp number, first 60 AAs: 0.22038\nTotal prob of N-in: 0.12994\noutside 1 331\nTMhelix 332 354\ninside 355 372

27336 GCF_001249435.1_7353_1_19 Shigella sonnei Proteobacteria; Gammaproteobacteria; Enterobacterales; Enterobacteriaceae; Shigella MNNEETFYQAMRRQGVTRRSFLKYCSLAATSLGLGAGMAPKIAWALENKPRIPVVWIHGL WP_005139779.1 hydrogenase [Shigella sonnei] Length: 372\nNumber of predicted TMHs: 2\nExp number of AAs in TMHs: 37.02628\nExp number, first 60 AAs: 19.39684\nTotal prob of N-in: 0.94313\nPOSSIBLE N-term signal sequence\ninside 1 19\nTMhelix 20 39\noutside 40 325\nTMhelix 326 348\ninside 349 372

27337 GCF_001248965.1_5417_1_11 Shigella sonnei Proteobacteria; Gammaproteobacteria; Enterobacterales; Enterobacteriaceae; Shigella
MNNEETFYQAMRRQGVTRRSFLKYCSLAATSLGLGAGMAPKIAWALENKPRIPVVWIHGL WP_005139779.1
hydrogenase [Shigella sonnei] Length: 372\nNumber of predicted TMHs: 2\nExp number of AAs in TMHs: 37.02628\nExp number, first 60 AAs: 19.39684\nTotal prob of N-in: 0.94313\nPOSSIBLE N-term signal sequence\ninside 1 19\nTMhelix 20 39\noutside 40 325\nTMhelix 326 348\ninside 349 372

27338 GCF_001249005.1_7655_8_14 Shigella sonnei Proteobacteria; Gammaproteobacteria; Enterobacterales; Enterobacteriaceae; Shigella
MNNEETFYQAMRRQGVTRRSFLKYCSLAATSLGLGAGMAPKIAWALENKPRIPVVWIHGL WP_005139779.1
hydrogenase [Shigella sonnei] Length: 372\nNumber of predicted TMHs: 2\nExp number of AAs in TMHs: 37.02628\nExp number, first 60 AAs: 19.39684\nTotal prob of N-in: 0.94313\nPOSSIBLE N-term signal sequence\ninside 1 19\nTMhelix 20 39\noutside 40 325\nTMhelix 326 348\ninside 349 372

27339 GCF_001249255.1_11822_8_61 Shigella sonnei Proteobacteria; Gammaproteobacteria; Enterobacterales; Enterobacteriaceae; Shigella
MNNEETFYQAMRRQGVTRRSFLKYCSLAATSLGLGAGMAPKIAWALENKPRIPVVWIHGL WP_005139779.1
hydrogenase [Shigella sonnei] Length: 372\nNumber of predicted TMHs: 2\nExp number of AAs in TMHs: 37.02628\nExp number, first 60 AAs: 19.39684\nTotal prob of N-in: 0.94313\nPOSSIBLE N-term signal sequence\ninside 1 19\nTMhelix 20 39\noutside 40 325\nTMhelix 326 348\ninside 349 372

27340 GCF_001249475.1_6246_5_5Shigella sonnei Proteobacteria; Gammaproteobacteria; Enterobacterales; Enterobacteriaceae; Shigella MTGDNTLIHSHGINRRDFMKLCAALAATMGLSSKAAAEMAESVTNPQRPPVIWIGAQECT
WP_000145410.1 MULTISPECIES: hydrogenase-2 small chain [Proteobacteria] Length: 372\nNumber of predicted TMHs: 1\nExp number of AAs in TMHs: 23.72226\nExp number, first 60 AAs: 0.22038\nTotal prob of N-in: 0.12994\noutside 1 331\nTMhelix 332 354\ninside 355 372

27341 GCF_001249555.1_Sg_201401054_AGTCAA_L001 Shigella sonnei Proteobacteria; Gammaproteobacteria; Enterobacterales; Enterobacteriaceae; Shigella
MTGDNTLIHSHGINRRDFMKLCAALAATMGLSSKAAAEMAESVTNPQRPPVIWIGAQECT WP_000145410.1
MULTISPECIES: hydrogenase-2 small chain [Proteobacteria] Length: 372\nNumber of predicted TMHs: 1\nExp number of AAs in TMHs: 23.72226\nExp number, first 60 AAs: 0.22038\nTotal prob of N-in: 0.12994\noutside 1 331\nTMhelix 332 354\ninside 355 372

27342 GCF_001249595.1_12971_8_38 Shigella sonnei Proteobacteria; Gammaproteobacteria; Enterobacterales; Enterobacteriaceae; Shigella
MTGDNTLIHSHGINRRDFMKLCAALAATMGLSSKAAAEMAESVTNPQRPPVIWIGAQECT WP_000145410.1
MULTISPECIES: hydrogenase-2 small chain [Proteobacteria] Length: 372\nNumber of predicted TMHs: 1\nExp number of AAs in TMHs: 23.72226\nExp number, first 60 AAs: 0.22038\nTotal prob of N-in: 0.12994\noutside 1 331\nTMhelix 332 354\ninside 355 372

27343 GCF_001249655.1_8289_7_60 Shigella sonnei Proteobacteria; Gammaproteobacteria; Enterobacterales; Enterobacteriaceae; Shigella
MTGDNTLIHSHGINRRDFMKLCAALAATMGLSSKAAAEMAESVTNPQRPPVIWIGAQECT WP_000145410.1
MULTISPECIES: hydrogenase-2 small chain [Proteobacteria] Length: 372\nNumber of predicted TMHs: 1\nExp number of AAs in TMHs: 23.72226\nExp number, first 60 AAs: 0.22038\nTotal prob of N-in: 0.12994\noutside 1 331\nTMhelix 332 354\ninside 355 372

27344 GCF_001249695.1_8289_7_82 Shigella sonnei Proteobacteria; Gammaproteobacteria; Enterobacterales; Enterobacteriaceae; Shigella
MTGDNTLIHSHGINRRDFMKLCAALAATMGLSSKAAAEMAESVTNPQRPPVIWIGAQECT WP_000145410.1
MULTISPECIES: hydrogenase-2 small chain [Proteobacteria] Length: 372\nNumber of predicted TMHs: 1\nExp number of AAs in TMHs: 23.72226\nExp number, first 60 AAs: 0.22038\nTotal prob of N-in: 0.12994\noutside 1 331\nTMhelix 332 354\ninside 355 372

27345 GCF_001249775.1_H14132053901 Shigella sonnei Proteobacteria; Gammaproteobacteria; Enterobacterales; Enterobacteriaceae; Shigella
MTGDNTLIHSHGINRRDFMKLCAALAATMGLSSKAAAEMAESVTNPQRPPVIWIGAQECT WP_000145410.1
MULTISPECIES: hydrogenase-2 small chain [Proteobacteria] Length: 372\nNumber of predicted TMHs: 1\nExp number of AAs in TMHs: 23.72226\nExp number, first 60 AAs: 0.22038\nTotal prob of N-in: 0.12994\noutside 1 331\nTMhelix 332 354\ninside 355 372

27346 GCF_001249855.1_11822_8_58 Shigella sonnei Proteobacteria; Gammaproteobacteria; Enterobacterales; Enterobacteriaceae; Shigella MTGDNTLIHSHGINRRDFMKLCAALAATMGLSSKAAAEMAESVTNPQRPPVIWIGAQECT WP_000145410.1 MULTISPECIES: hydrogenase-2 small chain [Proteobacteria] Length: 372\nNumber of predicted TMHs: 1\nExp number of AAs in TMHs: 23.72226\nExp number, first 60 AAs: 0.22038\nTotal prob of N-in: 0.12994\noutside 1 331\nTMhelix 332 354\ninside 355 372

27347 GCF_001249915.1_5236_6_8Shigella sonnei Proteobacteria; Gammaproteobacteria; Enterobacterales; Enterobacteriaceae; Shigella MTGDNTLIHSHGINRRDFMKLCAALAATMGLSSKAAAEMAESVTNPQRPPVIWIGAQECT WP_000145410.1 MULTISPECIES: hydrogenase-2 small chain [Proteobacteria] Length: 372\nNumber of predicted TMHs: 1\nExp number of AAs in TMHs: 23.72226\nExp number, first 60 AAs: 0.22038\nTotal prob of N-in: 0.12994\noutside 1 331\nTMhelix 332 354\ninside 355 372

27348 GCF_001249955.1_8290_5_9Shigella sonnei Proteobacteria; Gammaproteobacteria; Enterobacterales; Enterobacteriaceae; Shigella MTGDNTLIHSHGINRRDFMKLCAALAATMGLSSKAAAEMAESVTNPQRPPVIWIGAQECT WP_000145410.1 MULTISPECIES: hydrogenase-2 small chain [Proteobacteria] Length: 372\nNumber of predicted TMHs: 1\nExp number of AAs in TMHs: 23.72226\nExp number, first 60 AAs: 0.22038\nTotal prob of N-in: 0.12994\noutside 1 331\nTMhelix 332 354\ninside 355 372

27349 GCF_001249915.1_5236_6_8Shigella sonnei Proteobacteria; Gammaproteobacteria; Enterobacterales; Enterobacteriaceae; Shigella MNNEETFYQAMRRQGVTRRSFLKYCSLAATSLGLGAGMAPKIAWALENKPRIPVVWIHGL WP_005139779.1 hydrogenase [Shigella sonnei] Length: 372\nNumber of predicted TMHs: 2\nExp number of AAs in TMHs: 37.02628\nExp number, first 60 AAs: 19.39684\nTotal prob of N-in: 0.94313\nPOSSIBLE N-term signal sequence\ninside 1 19\nTMhelix 20 39\noutside 40 325\nTMhelix 326 348\ninside 349 372

27350 GCF_001249955.1_8290_5_9Shigella sonnei Proteobacteria; Gammaproteobacteria; Enterobacterales; Enterobacteriaceae; Shigella MNNEETFYQAMRRQGVTRRSFLKYCSLAATSLGLGAGMAPKIAWALENKPRIPVVWIHGL WP_005139779.1 hydrogenase [Shigella sonnei] Length: 372\nNumber of predicted TMHs: 2\nExp number of AAs in TMHs: 37.02628\nExp number, first 60 AAs: 19.39684\nTotal prob of N-in: 0.94313\nPOSSIBLE N-term signal sequence\ninside 1 19\nTMhelix 20 39\noutside 40 325\nTMhelix 326 348\ninside 349 372

27351 GCF_001250135.1_9870_6_24 Shigella sonnei Proteobacteria; Gammaproteobacteria; Enterobacterales; Enterobacteriaceae; Shigella MTGDNTLIHSHGINRRDFMKLCAALAATMGLSSKAAAEMAESVTNPQRPPVIWIGAQECT WP_000145410.1 MULTISPECIES: hydrogenase-2 small chain [Proteobacteria] Length: 372\nNumber of predicted TMHs: 1\nExp number of AAs in TMHs: 23.72226\nExp number, first 60 AAs: 0.22038\nTotal prob of N-in: 0.12994\noutside 1 331\nTMhelix 332 354\ninside 355 372

27352 GCF_001249475.1_6246_5_5Shigella sonnei Proteobacteria; Gammaproteobacteria; Enterobacterales; Enterobacteriaceae; Shigella MNNEETFYQAMRRQGVTRRSFLKYCSLAATSLGLGAGMAPKIAWALENKPRIPVVWIHGL WP_005139779.1 hydrogenase [Shigella sonnei] Length: 372\nNumber of predicted TMHs: 2\nExp number of AAs in TMHs: 37.02628\nExp number, first 60 AAs: 19.39684\nTotal prob of N-in: 0.94313\nPOSSIBLE N-term signal sequence\ninside 1 19\nTMhelix 20 39\noutside 40 325\nTMhelix 326 348\ninside 349 372

27353 GCF_001250015.1_12971_8_6 Shigella sonnei Proteobacteria; Gammaproteobacteria; Enterobacterales; Enterobacteriaceae; Shigella MTGDNTLIHSHGINRRDFMKLCAALAATMGLSSKAAAEMAESVTNPQRPPVIWIGAQECT WP_000145410.1 MULTISPECIES: hydrogenase-2 small chain [Proteobacteria] Length: 372\nNumber of predicted TMHs: 1\nExp number of AAs in TMHs: 23.72226\nExp number, first 60 AAs: 0.22038\nTotal prob of N-in: 0.12994\noutside 1 331\nTMhelix 332 354\ninside 355 372

27354 GCF_001250095.1_12971_8_71 Shigella sonnei Proteobacteria; Gammaproteobacteria; Enterobacterales; Enterobacteriaceae; Shigella MTGDNTLIHSHGINRRDFMKLCAALAATMGLSSKAAAEMAESVTNPQRPPVIWIGAQECT WP_000145410.1 MULTISPECIES: hydrogenase-2 small chain [Proteobacteria] Length: 372\nNumber of predicted TMHs: 1\nExp number of AAs in TMHs: 23.72226\nExp number, first 60 AAs: 0.22038\nTotal prob of N-in: 0.12994\noutside 1 331\nTMhelix 332 354\ninside 355 372

27355 GCF_001250015.1_12971_8_6 Shigella sonnei Proteobacteria; Gammaproteobacteria; Enterobacterales; Enterobacteriaceae; Shigella MNNEETFYQAMRRQGVTRRSFLKYCSLAATSLGLGAGMAPKIAWALENKPRIPVVWIHGL WP_005139779.1

hydrogenase [Shigella sonnei] Length: 372\nNumber of predicted TMHs: 2\nExp number of AAs in TMHs: 37.02628\nExp number, first 60 AAs: 19.39684\nTotal prob of N-in: 0.94313\nPOSSIBLE N-term signal sequence\ninside 1 19\nTMhelix 20 39\noutside 40 325\nTMhelix 326 348\ninside 349 372

27356 GCF_001250095.1_12971_8_71 Shigella sonnei Proteobacteria; Gammaproteobacteria; Enterobacterales; Enterobacteriaceae; Shigella
MNNEETFYQAMRRQGVTRRSFLKYCSLAATSLGLGAGMAPKIAWALENKPRIPVVWIHGL WP_005139779.1
hydrogenase [Shigella sonnei] Length: 372\nNumber of predicted TMHs: 2\nExp number of AAs in TMHs: 37.02628\nExp number, first 60 AAs: 19.39684\nTotal prob of N-in: 0.94313\nPOSSIBLE N-term signal sequence\ninside 1 19\nTMhelix 20 39\noutside 40 325\nTMhelix 326 348\ninside 349 372

27357 GCF_001249555.1_Sg_201401054_AGTCAA_L001 Shigella sonnei Proteobacteria; Gammaproteobacteria; Enterobacterales; Enterobacteriaceae; Shigella
MNNEETFYQAMRRQGVTRRSFLKYCSLAATSLGLGAGMAPKIAWALENKPRIPVVWIHGL WP_005139779.1
hydrogenase [Shigella sonnei] Length: 372\nNumber of predicted TMHs: 2\nExp number of AAs in TMHs: 37.02628\nExp number, first 60 AAs: 19.39684\nTotal prob of N-in: 0.94313\nPOSSIBLE N-term signal sequence\ninside 1 19\nTMhelix 20 39\noutside 40 325\nTMhelix 326 348\ninside 349 372

27358 GCF_001249595.1_12971_8_38 Shigella sonnei Proteobacteria; Gammaproteobacteria; Enterobacterales; Enterobacteriaceae; Shigella
MNNEETFYQAMRRQGVTRRSFLKYCSLAATSLGLGAGMAPKIAWALENKPRIPVVWIHGL WP_005139779.1
hydrogenase [Shigella sonnei] Length: 372\nNumber of predicted TMHs: 2\nExp number of AAs in TMHs: 37.02628\nExp number, first 60 AAs: 19.39684\nTotal prob of N-in: 0.94313\nPOSSIBLE N-term signal sequence\ninside 1 19\nTMhelix 20 39\noutside 40 325\nTMhelix 326 348\ninside 349 372

27359 GCF_001250135.1_9870_6_24 Shigella sonnei Proteobacteria; Gammaproteobacteria; Enterobacterales; Enterobacteriaceae; Shigella
MNNEETFYQAMRRQGVTRRSFLKYCSLAATSLGLGAGMAPKIAWALENKPRIPVVWIHGL WP_005139779.1
hydrogenase [Shigella sonnei] Length: 372\nNumber of predicted TMHs: 2\nExp number of AAs in TMHs: 37.02628\nExp number, first 60 AAs: 19.39684\nTotal prob of N-in: 0.94313\nPOSSIBLE N-term signal sequence\ninside 1 19\nTMhelix 20 39\noutside 40 325\nTMhelix 326 348\ninside 349 372

27360 GCF_001249655.1_8289_7_60 Shigella sonnei Proteobacteria; Gammaproteobacteria; Enterobacterales; Enterobacteriaceae; Shigella
MNNEETFYQAMRRQGVTRRSFLKYCSLAATSLGLGAGMAPKIAWALENKPRIPVVWIHGL WP_005139779.1
hydrogenase [Shigella sonnei] Length: 372\nNumber of predicted TMHs: 2\nExp number of AAs in TMHs: 37.02628\nExp number, first 60 AAs: 19.39684\nTotal prob of N-in: 0.94313\nPOSSIBLE N-term signal sequence\ninside 1 19\nTMhelix 20 39\noutside 40 325\nTMhelix 326 348\ninside 349 372

27361 GCF_001249695.1_8289_7_82 Shigella sonnei Proteobacteria; Gammaproteobacteria; Enterobacterales; Enterobacteriaceae; Shigella
MNNEETFYQAMRRQGVTRRSFLKYCSLAATSLGLGAGMAPKIAWALENKPRIPVVWIHGL WP_005139779.1
hydrogenase [Shigella sonnei] Length: 372\nNumber of predicted TMHs: 2\nExp number of AAs in TMHs: 37.02628\nExp number, first 60 AAs: 19.39684\nTotal prob of N-in: 0.94313\nPOSSIBLE N-term signal sequence\ninside 1 19\nTMhelix 20 39\noutside 40 325\nTMhelix 326 348\ninside 349 372

27362 GCF_001249855.1_11822_8_58 Shigella sonnei Proteobacteria; Gammaproteobacteria; Enterobacterales; Enterobacteriaceae; Shigella
MNNEETFYQAMRRQGVTRRSFLKYCSLAATSLGLGAGMAPKIAWALENKPRIPVVWIHGL WP_005139779.1
hydrogenase [Shigella sonnei] Length: 372\nNumber of predicted TMHs: 2\nExp number of AAs in TMHs: 37.02628\nExp number, first 60 AAs: 19.39684\nTotal prob of N-in: 0.94313\nPOSSIBLE N-term signal sequence\ninside 1 19\nTMhelix 20 39\noutside 40 325\nTMhelix 326 348\ninside 349 372

27363 GCF_001250215.1_12971_8_31 Shigella sonnei Proteobacteria; Gammaproteobacteria; Enterobacterales; Enterobacteriaceae; Shigella
MTGDNTLIHSHGINRRDFMKLCAALAATMGLSSKAAAEMAESVTNPQRPPVIWIGAQECT WP_000145410.1
MULTISPECIES: hydrogenase-2 small chain [Proteobacteria] Length: 372\nNumber of predicted TMHs: 1\nExp number of AAs in TMHs: 23.72226\nExp number, first 60 AAs: 0.22038\nTotal prob of N-in: 0.12994\noutside 1 331\nTMhelix 332 354\ninside 355 372

27364 GCF_001250215.1_12971_8_31 Shigella sonnei Proteobacteria; Gammaproteobacteria; Enterobacterales; Enterobacteriaceae; Shigella MNNEETFYQAMRRQGVTRRSFLKYCSLAATSLGLGAGMAPKIAWALENKPRIPVVWIHGL WP_005139779.1 hydrogenase [Shigella sonnei] Length: 372\nNumber of predicted TMHs: 2\nExp number of AAs in TMHs: 37.02628\nExp number, first 60 AAs: 19.39684\nTotal prob of N-in: 0.94313\nPOSSIBLE N-term signal sequence\ninside 1 19\nTMhelix 20 39\noutside 40 325\nTMhelix 326 348\ninside 349 372

27365 GCF_001249775.1_H14132053901 Shigella sonnei Proteobacteria; Gammaproteobacteria; Enterobacterales; Enterobacteriaceae; Shigella MNNEETFYQAMRRQGVTRRSFLKYCSLAATSLGLGAGMAPKIAWALENKPRIPVVWIHGL WP_052962775.1 hydrogenase [Shigella sonnei] Length: 372\nNumber of predicted TMHs: 2\nExp number of AAs in TMHs: 37.09248\nExp number, first 60 AAs: 19.39296\nTotal prob of N-in: 0.94316\nPOSSIBLE N-term signal sequence\ninside 1 19\nTMhelix 20 39\noutside 40 325\nTMhelix 326 348\ninside 349 372

27366 GCF_001250295.1_8289_7_74 Shigella sonnei Proteobacteria; Gammaproteobacteria; Enterobacterales; Enterobacteriaceae; Shigella MTGDNTLIHSHGINRRDFMKLCAALAATMGLSSKAAAEMAESVTNPQRPPVIWIGAQECT WP_000145410.1 MULTISPECIES: hydrogenase-2 small chain [Proteobacteria] Length: 372\nNumber of predicted TMHs: 1\nExp number of AAs in TMHs: 23.72226\nExp number, first 60 AAs: 0.22038\nTotal prob of N-in: 0.12994\noutside 1 331\nTMhelix 332 354\ninside 355 372

27367 GCF_001250355.1_8290_5_2Shigella sonnei Proteobacteria; Gammaproteobacteria; Enterobacterales; Enterobacteriaceae; Shigella MTGDNTLIHSHGINRRDFMKLCAALAATMGLSSKAAAEMAESVTNPQRPPVIWIGAQECT WP_000145410.1 MULTISPECIES: hydrogenase-2 small chain [Proteobacteria] Length: 372\nNumber of predicted TMHs: 1\nExp number of AAs in TMHs: 23.72226\nExp number, first 60 AAs: 0.22038\nTotal prob of N-in: 0.12994\noutside 1 331\nTMhelix 332 354\ninside 355 372

27368 GCF_001250395.1_8290_4_16 Shigella sonnei Proteobacteria; Gammaproteobacteria; Enterobacterales; Enterobacteriaceae; Shigella MTGDNTLIHSHGINRRDFMKLCAALAATMGLSSKAAAEMAESVTNPQRPPVIWIGAQECT WP_000145410.1 MULTISPECIES: hydrogenase-2 small chain [Proteobacteria] Length: 372\nNumber of predicted TMHs: 1\nExp number of AAs in TMHs: 23.72226\nExp number, first 60 AAs: 0.22038\nTotal prob of N-in: 0.12994\noutside 1 331\nTMhelix 332 354\ninside 355 372

27369 GCF_001250495.1_5236_7_4Shigella sonnei Proteobacteria; Gammaproteobacteria; Enterobacterales; Enterobacteriaceae; Shigella MTGDNTLIHSHGINRRDFMKLCAALAATMGLSSKAAAEMAESVTNPQRPPVIWIGAQECT WP_000145410.1 MULTISPECIES: hydrogenase-2 small chain [Proteobacteria] Length: 372\nNumber of predicted TMHs: 1\nExp number of AAs in TMHs: 23.72226\nExp number, first 60 AAs: 0.22038\nTotal prob of N-in: 0.12994\noutside 1 331\nTMhelix 332 354\ninside 355 372

27370 GCF_001250535.1_H14122052001 Shigella sonnei Proteobacteria; Gammaproteobacteria; Enterobacterales; Enterobacteriaceae; Shigella MTGDNTLIHSHGINRRDFMKLCAALAATMGLSSKAAAEMAESVTNPQRPPVIWIGAQECT WP_000145410.1 MULTISPECIES: hydrogenase-2 small chain [Proteobacteria] Length: 372\nNumber of predicted TMHs: 1\nExp number of AAs in TMHs: 23.72226\nExp number, first 60 AAs: 0.22038\nTotal prob of N-in: 0.12994\noutside 1 331\nTMhelix 332 354\ninside 355 372

27371 GCF_001250595.1_Sg_201311626_GCGCTA_L001 Shigella sonnei Proteobacteria; Gammaproteobacteria; Enterobacterales; Enterobacteriaceae; Shigella MTGDNTLIHSHGINRRDFMKLCAALAATMGLSSKAAAEMAESVTNPQRPPVIWIGAQECT WP_000145410.1 MULTISPECIES: hydrogenase-2 small chain [Proteobacteria] Length: 372\nNumber of predicted TMHs: 1\nExp number of AAs in TMHs: 23.72226\nExp number, first 60 AAs: 0.22038\nTotal prob of N-in: 0.12994\noutside 1 331\nTMhelix 332 354\ninside 355 372

27372 GCF_001250675.1_9870_7_7Shigella sonnei Proteobacteria; Gammaproteobacteria; Enterobacterales; Enterobacteriaceae; Shigella MTGDNTLIHSHGINRRDFMKLCAALAATMGLSSKAAAEMAESVTNPQRPPVIWIGAQECT WP_000145410.1 MULTISPECIES: hydrogenase-2 small chain [Proteobacteria] Length: 372\nNumber of predicted TMHs: 1\nExp number of AAs in TMHs: 23.72226\nExp number, first 60 AAs: 0.22038\nTotal prob of N-in: 0.12994\noutside 1 331\nTMhelix 332 354\ninside 355 372

27373 GCF_001250715.1_5236_1_10 Shigella sonnei Proteobacteria; Gammaproteobacteria; Enterobacterales; Enterobacteriaceae; Shigella MTGDNTLIHSHGINRRDFMKLCAALAATMGLSSKAAAEMAESVTNPQRPPVIWIGAQECT WP_000145410.1 MULTISPECIES: hydrogenase-2 small chain [Proteobacteria] Length: 372\nNumber of predicted TMHs: 1\nExp number of AAs in TMHs: 23.72226\nExp number, first 60 AAs: 0.22038\nTotal prob of N-in: 0.12994\noutside 1 331\nTMhelix 332 354\ninside 355 372

27374 GCF_001250775.1_8289_7_69 Shigella sonnei Proteobacteria; Gammaproteobacteria; Enterobacterales; Enterobacteriaceae; Shigella MTGDNTLIHSHGINRRDFMKLCAALAATMGLSSKAAAEMAESVTNPQRPPVIWIGAQECT WP_000145410.1 MULTISPECIES: hydrogenase-2 small chain [Proteobacteria] Length: 372\nNumber of predicted TMHs: 1\nExp number of AAs in TMHs: 23.72226\nExp number, first 60 AAs: 0.22038\nTotal prob of N-in: 0.12994\noutside 1 331\nTMhelix 332 354\ninside 355 372

27375 GCF_001250715.1_5236_1_10 Shigella sonnei Proteobacteria; Gammaproteobacteria; Enterobacterales; Enterobacteriaceae; Shigella MNNEETFYQAMRRQGVTRRSFLKYCSLAATSLGLGAGMAPKIAWALENKPRIPVVWIHGL WP_005139779.1 hydrogenase [Shigella sonnei] Length: 372\nNumber of predicted TMHs: 2\nExp number of AAs in TMHs: 37.02628\nExp number, first 60 AAs: 19.39684\nTotal prob of N-in: 0.94313\nPOSSIBLE N-term signal sequence\ninside 1 19\nTMhelix 20 39\noutside 40 325\nTMhelix 326 348\ninside 349 372

27376 GCF_001250775.1_8289_7_69 Shigella sonnei Proteobacteria; Gammaproteobacteria; Enterobacterales; Enterobacteriaceae; Shigella MNNEETFYQAMRRQGVTRRSFLKYCSLAATSLGLGAGMAPKIAWALENKPRIPVVWIHGL WP_005139779.1 hydrogenase [Shigella sonnei] Length: 372\nNumber of predicted TMHs: 2\nExp number of AAs in TMHs: 37.02628\nExp number, first 60 AAs: 19.39684\nTotal prob of N-in: 0.94313\nPOSSIBLE N-term signal sequence\ninside 1 19\nTMhelix 20 39\noutside 40 325\nTMhelix 326 348\ninside 349 372

27377 GCF_001250295.1_8289_7_74 Shigella sonnei Proteobacteria; Gammaproteobacteria; Enterobacterales; Enterobacteriaceae; Shigella MNNEETFYQAMRRQGVTRRSFLKYCSLAATSLGLGAGMAPKIAWALENKPRIPVVWIHGL WP_005139779.1 hydrogenase [Shigella sonnei] Length: 372\nNumber of predicted TMHs: 2\nExp number of AAs in TMHs: 37.02628\nExp number, first 60 AAs: 19.39684\nTotal prob of N-in: 0.94313\nPOSSIBLE N-term signal sequence\ninside 1 19\nTMhelix 20 39\noutside 40 325\nTMhelix 326 348\ninside 349 372

27378 GCF_001250835.1_5236_1_9Shigella sonnei Proteobacteria; Gammaproteobacteria; Enterobacterales; Enterobacteriaceae; Shigella MTGDNTLIHSHGINRRDFMKLCAALAATMGLSSKAAAEMAESVTNPQRPPVIWIGAQECT WP_000145410.1 MULTISPECIES: hydrogenase-2 small chain [Proteobacteria] Length: 372\nNumber of predicted TMHs: 1\nExp number of AAs in TMHs: 23.72226\nExp number, first 60 AAs: 0.22038\nTotal prob of N-in: 0.12994\noutside 1 331\nTMhelix 332 354\ninside 355 372

27379 GCF_001250835.1_5236_1_9Shigella sonnei Proteobacteria; Gammaproteobacteria; Enterobacterales; Enterobacteriaceae; Shigella MNNEETFYQAMRRQGVTRRSFLKYCSLAATSLGLGAGMAPKIAWALENKPRIPVVWIHGL WP_005139779.1 hydrogenase [Shigella sonnei] Length: 372\nNumber of predicted TMHs: 2\nExp number of AAs in TMHs: 37.02628\nExp number, first 60 AAs: 19.39684\nTotal prob of N-in: 0.94313\nPOSSIBLE N-term signal sequence\ninside 1 19\nTMhelix 20 39\noutside 40 325\nTMhelix 326 348\ninside 349 372

27380 GCF_001250355.1_8290_5_2Shigella sonnei Proteobacteria; Gammaproteobacteria; Enterobacterales; Enterobacteriaceae; Shigella MNNEETFYQAMRRQGVTRRSFLKYCSLAATSLGLGAGMAPKIAWALENKPRIPVVWIHGL WP_005139779.1 hydrogenase [Shigella sonnei] Length: 372\nNumber of predicted TMHs: 2\nExp number of AAs in TMHs: 37.02628\nExp number, first 60 AAs: 19.39684\nTotal prob of N-in: 0.94313\nPOSSIBLE N-term signal sequence\ninside 1 19\nTMhelix 20 39\noutside 40 325\nTMhelix 326 348\ninside 349 372

27381 GCF_001250395.1_8290_4_16 Shigella sonnei Proteobacteria; Gammaproteobacteria; Enterobacterales; Enterobacteriaceae; Shigella MNNEETFYQAMRRQGVTRRSFLKYCSLAATSLGLGAGMAPKIAWALENKPRIPVVWIHGL WP_005139779.1 hydrogenase [Shigella sonnei] Length: 372\nNumber of predicted TMHs: 2\nExp number of AAs in TMHs: 37.02628\nExp number, first 60 AAs: 19.39684\nTotal prob of N-in: 0.94313\nPOSSIBLE N-term signal sequence\ninside 1 19\nTMhelix 20 39\noutside 40 325\nTMhelix 326 348\ninside 349 372

27382	GCF_001250895.1_8489_2_93	Shigella sonnei	Proteobacteria; Gammaproteobacteria; Enterobacterales; Enterobacteriaceae; Shigella	MTGDNTLIHSHGINRRDFMKLCAALAATMGLSSKAAAEMAESVTNPQRPPVIWIGAQECT	WP_000145410.1	MULTISPECIES: hydrogenase-2 small chain [Proteobacteria]	Length: 372\nNumber of predicted TMHs: 1\nExp number of AAs in TMHs: 23.72226\nExp number, first 60 AAs: 0.22038\nTotal prob of N-in: 0.12994\noutside 1 331\nTMhelix 332 354\ninside 355 372

27383	GCF_001250895.1_8489_2_93	Shigella sonnei	Proteobacteria; Gammaproteobacteria; Enterobacterales; Enterobacteriaceae; Shigella	MNNEETFYQAMRRQGVTRRSFLKYCSLAATSLGLGAGMAPKIAWALENKPRIPVVWIHGL	WP_005139779.1	hydrogenase [Shigella sonnei]	Length: 372\nNumber of predicted TMHs: 2\nExp number of AAs in TMHs: 37.02628\nExp number, first 60 AAs: 19.39684\nTotal prob of N-in: 0.94313\nPOSSIBLE N-term signal sequence\ninside 1 19\nTMhelix 20 39\noutside 40 325\nTMhelix 326 348\ninside 349 372

27384	GCF_001250495.1_5236_7_4Shigella sonnei	Proteobacteria; Gammaproteobacteria; Enterobacterales; Enterobacteriaceae; Shigella	MNNEETFYQAMRRQGVTRRSFLKYCSLAATSLGLGAGMAPKIAWALENKPRIPVVWIHGL	WP_005139779.1	hydrogenase [Shigella sonnei]	Length: 372\nNumber of predicted TMHs: 2\nExp number of AAs in TMHs: 37.02628\nExp number, first 60 AAs: 19.39684\nTotal prob of N-in: 0.94313\nPOSSIBLE N-term signal sequence\ninside 1 19\nTMhelix 20 39\noutside 40 325\nTMhelix 326 348\ninside 349 372

27385	GCF_001250595.1_Sg_201311626_GCGCTA_L001	Shigella sonnei	Proteobacteria; Gammaproteobacteria; Enterobacterales; Enterobacteriaceae; Shigella	MNNEETFYQAMRRQGVTRRSFLKYCSLAATSLGLGAGMAPKIAWALENKPRIPVVWIHGL	WP_005139779.1	hydrogenase [Shigella sonnei]	Length: 372\nNumber of predicted TMHs: 2\nExp number of AAs in TMHs: 37.02628\nExp number, first 60 AAs: 19.39684\nTotal prob of N-in: 0.94313\nPOSSIBLE N-term signal sequence\ninside 1 19\nTMhelix 20 39\noutside 40 325\nTMhelix 326 348\ninside 349 372

27386	GCF_001250915.1_8289_7_65	Shigella sonnei	Proteobacteria; Gammaproteobacteria; Enterobacterales; Enterobacteriaceae; Shigella	MTGDNTLIHSHGINRRDFMKLCAALAATMGLSSKAAAEMAESVTNPQRPPVIWIGAQECT	WP_000145410.1	MULTISPECIES: hydrogenase-2 small chain [Proteobacteria]	Length: 372\nNumber of predicted TMHs: 1\nExp number of AAs in TMHs: 23.72226\nExp number, first 60 AAs: 0.22038\nTotal prob of N-in: 0.12994\noutside 1 331\nTMhelix 332 354\ninside 355 372

27387	GCF_001250915.1_8289_7_65	Shigella sonnei	Proteobacteria; Gammaproteobacteria; Enterobacterales; Enterobacteriaceae; Shigella	MNNEETFYQAMRRQGVTRRSFLKYCSLAATSLGLGAGMAPKIAWALENKPRIPVVWIHGL	WP_005139779.1	hydrogenase [Shigella sonnei]	Length: 372\nNumber of predicted TMHs: 2\nExp number of AAs in TMHs: 37.02628\nExp number, first 60 AAs: 19.39684\nTotal prob of N-in: 0.94313\nPOSSIBLE N-term signal sequence\ninside 1 19\nTMhelix 20 39\noutside 40 325\nTMhelix 326 348\ninside 349 372

27388	GCF_001250535.1_H14122052001	Shigella sonnei	Proteobacteria; Gammaproteobacteria; Enterobacterales; Enterobacteriaceae; Shigella	MNNEETFYQAMRRQGVTRRSFLKYCSLAATSLGLGAGMAPKIAWALENKPRIPVVWIHGL	WP_052962775.1	hydrogenase [Shigella sonnei]	Length: 372\nNumber of predicted TMHs: 2\nExp number of AAs in TMHs: 37.09248\nExp number, first 60 AAs: 19.39296\nTotal prob of N-in: 0.94316\nPOSSIBLE N-term signal sequence\ninside 1 19\nTMhelix 20 39\noutside 40 325\nTMhelix 326 348\ninside 349 372

27389	GCF_001250675.1_9870_7_7Shigella sonnei	Proteobacteria; Gammaproteobacteria; Enterobacterales; Enterobacteriaceae; Shigella	MNNEETFYQAMRRQGVTRRSFLKYCSLAATSLGLGAGMAPKIAWALENKPRIPVVWIHGL	WP_005139779.1	hydrogenase [Shigella sonnei]	Length: 372\nNumber of predicted TMHs: 2\nExp number of AAs in TMHs: 37.02628\nExp number, first 60 AAs: 19.39684\nTotal prob of N-in: 0.94313\nPOSSIBLE N-term signal sequence\ninside 1 19\nTMhelix 20 39\noutside 40 325\nTMhelix 326 348\ninside 349 372

27390	GCF_001250975.1_8290_4_56	Shigella sonnei	Proteobacteria; Gammaproteobacteria; Enterobacterales; Enterobacteriaceae; Shigella	MTGDNTLIHSHGINRRDFMKLCAALAATMGLSSKAAAEMAESVTNPQRPPVIWIGAQECT	WP_000145410.1	MULTISPECIES: hydrogenase-2 small chain [Proteobacteria]	Length: 372\nNumber of predicted TMHs: 1\nExp number of AAs in TMHs: 23.72226\nExp number, first 60 AAs: 0.22038\nTotal prob of N-in: 0.12994\noutside 1 331\nTMhelix 332 354\ninside 355 372

27391 GCF_001251115.1_8289_7_39 Shigella sonnei Proteobacteria; Gammaproteobacteria; Enterobacterales; Enterobacteriaceae; Shigella
MTGDNTLIHSHGINRRDFMKLCAALAATMGLSSKAAAEMAESVTNPQRPPVIWIGAQECT WP_000145410.1
MULTISPECIES: hydrogenase-2 small chain [Proteobacteria] Length: 372\nNumber of predicted TMHs: 1\nExp number of AAs in TMHs: 23.72226\nExp number, first 60 AAs: 0.22038\nTotal prob of N-in: 0.12994\noutside 1 331\nTMhelix 332 354\ninside 355 372

27392 GCF_001251035.1_Sg_201400752_GATCAG_L001 Shigella sonnei Proteobacteria; Gammaproteobacteria; Enterobacterales; Enterobacteriaceae; Shigella
MTGDNTLIHSHGINRRDFMKLCAALAATMGLSSKAAAEMAESVTNPQRPPVIWIGAQECT WP_000145410.1
MULTISPECIES: hydrogenase-2 small chain [Proteobacteria] Length: 372\nNumber of predicted TMHs: 1\nExp number of AAs in TMHs: 23.72226\nExp number, first 60 AAs: 0.22038\nTotal prob of N-in: 0.12994\noutside 1 331\nTMhelix 332 354\ninside 355 372

27393 GCF_001251175.1_H14046047401 Shigella sonnei Proteobacteria; Gammaproteobacteria; Enterobacterales; Enterobacteriaceae; Shigella
MTGDNTLIHSHGINRRDFMKLCAALAATMGLSSKAAAEMAESVTNPQRPPVIWIGAQECT WP_000145410.1
MULTISPECIES: hydrogenase-2 small chain [Proteobacteria] Length: 372\nNumber of predicted TMHs: 1\nExp number of AAs in TMHs: 23.72226\nExp number, first 60 AAs: 0.22038\nTotal prob of N-in: 0.12994\noutside 1 331\nTMhelix 332 354\ninside 355 372

27394 GCF_001251215.1_6350_1_6Shigella sonnei Proteobacteria; Gammaproteobacteria; Enterobacterales; Enterobacteriaceae; Shigella MTGDNTLIHSHGINRRDFMKLCAALAATMGLSSKAAAEMAESVTNPQRPPVIWIGAQECT
WP_000145410.1 MULTISPECIES: hydrogenase-2 small chain [Proteobacteria] Length: 372\nNumber of predicted TMHs: 1\nExp number of AAs in TMHs: 23.72226\nExp number, first 60 AAs: 0.22038\nTotal prob of N-in: 0.12994\noutside 1 331\nTMhelix 332 354\ninside 355 372

27395 GCF_001251275.1_9870_7_12 Shigella sonnei Proteobacteria; Gammaproteobacteria; Enterobacterales; Enterobacteriaceae; Shigella
MTGDNTLIHSHGINRRDFMKLCAALAATMGLSSKAAAEMAESVTNPQRPPVIWIGAQECT WP_000145410.1
MULTISPECIES: hydrogenase-2 small chain [Proteobacteria] Length: 372\nNumber of predicted TMHs: 1\nExp number of AAs in TMHs: 23.72226\nExp number, first 60 AAs: 0.22038\nTotal prob of N-in: 0.12994\noutside 1 331\nTMhelix 332 354\ninside 355 372

27396 GCF_001251315.1_Sg_201403906_CAACTA_L001 Shigella sonnei Proteobacteria; Gammaproteobacteria; Enterobacterales; Enterobacteriaceae; Shigella
MTGDNTLIHSHGINRRDFMKLCAALAATMGLSSKAAAEMAESVTNPQRPPVIWIGAQECT WP_000145410.1
MULTISPECIES: hydrogenase-2 small chain [Proteobacteria] Length: 372\nNumber of predicted TMHs: 1\nExp number of AAs in TMHs: 23.72226\nExp number, first 60 AAs: 0.22038\nTotal prob of N-in: 0.12994\noutside 1 331\nTMhelix 332 354\ninside 355 372

27397 GCF_001251375.1_8290_4_10 Shigella sonnei Proteobacteria; Gammaproteobacteria; Enterobacterales; Enterobacteriaceae; Shigella
MTGDNTLIHSHGINRRDFMKLCAALAATMGLSSKAAAEMAESVTNPQRPPVIWIGAQECT WP_000145410.1
MULTISPECIES: hydrogenase-2 small chain [Proteobacteria] Length: 372\nNumber of predicted TMHs: 1\nExp number of AAs in TMHs: 23.72226\nExp number, first 60 AAs: 0.22038\nTotal prob of N-in: 0.12994\noutside 1 331\nTMhelix 332 354\ninside 355 372

27398 GCF_001251515.1_8489_2_83 Shigella sonnei Proteobacteria; Gammaproteobacteria; Enterobacterales; Enterobacteriaceae; Shigella
MTGDNTLIHSHGINRRDFMKLCAALAATMGLSSKAAAEMAESVTNPQRPPVIWIGAQECT WP_000145410.1
MULTISPECIES: hydrogenase-2 small chain [Proteobacteria] Length: 372\nNumber of predicted TMHs: 1\nExp number of AAs in TMHs: 23.72226\nExp number, first 60 AAs: 0.22038\nTotal prob of N-in: 0.12994\noutside 1 331\nTMhelix 332 354\ninside 355 372

27399 GCF_001251375.1_8290_4_10 Shigella sonnei Proteobacteria; Gammaproteobacteria; Enterobacterales; Enterobacteriaceae; Shigella
MNNEETFYQAMRRQGVTRRSFLKYCSLAATSLGLGAGMAPKIAWALENKPRIPVVWIHGL WP_005139779.1
hydrogenase [Shigella sonnei] Length: 372\nNumber of predicted TMHs: 2\nExp number of AAs in TMHs: 37.02628\nExp number, first 60 AAs: 19.39684\nTotal prob of N-in: 0.94313\nPOSSIBLE N-term signal sequence\ninside 1 19\nTMhelix 20 39\noutside 40 325\nTMhelix 326 348\ninside 349 372

27400 GCF_001251515.1_8489_2_83 Shigella sonnei Proteobacteria; Gammaproteobacteria; Enterobacterales; Enterobacteriaceae; Shigella MNNEETFYQAMRRQGVTRRSFLKYCSLAATSLGLGAGMAPKIAWALENKPRIPVVWIHGL WP_005139779.1 hydrogenase [Shigella sonnei] Length: 372\nNumber of predicted TMHs: 2\nExp number of AAs in TMHs: 37.02628\nExp number, first 60 AAs: 19.39684\nTotal prob of N-in: 0.94313\nPOSSIBLE N-term signal sequence\ninside 1 19\nTMhelix 20 39\noutside 40 325\nTMhelix 326 348\ninside 349 372

27401 GCF_001250975.1_8290_4_56 Shigella sonnei Proteobacteria; Gammaproteobacteria; Enterobacterales; Enterobacteriaceae; Shigella MNNEETFYQAMRRQGVTRRSFLKYCSLAATSLGLGAGMAPKIAWALENKPRIPVVWIHGL WP_005139779.1 hydrogenase [Shigella sonnei] Length: 372\nNumber of predicted TMHs: 2\nExp number of AAs in TMHs: 37.02628\nExp number, first 60 AAs: 19.39684\nTotal prob of N-in: 0.94313\nPOSSIBLE N-term signal sequence\ninside 1 19\nTMhelix 20 39\noutside 40 325\nTMhelix 326 348\ninside 349 372

27402 GCF_001251555.1_8289_7_45 Shigella sonnei Proteobacteria; Gammaproteobacteria; Enterobacterales; Enterobacteriaceae; Shigella MTGDNTLIHSHGINRRDFMKLCAALAATMGLSSKAAAEMAESVTNPQRPPVIWIGAQECT WP_000145410.1 MULTISPECIES: hydrogenase-2 small chain [Proteobacteria] Length: 372\nNumber of predicted TMHs: 1\nExp number of AAs in TMHs: 23.72226\nExp number, first 60 AAs: 0.22038\nTotal prob of N-in: 0.12994\noutside 1 331\nTMhelix 332 354\ninside 355 372

27403 GCF_001251555.1_8289_7_45 Shigella sonnei Proteobacteria; Gammaproteobacteria; Enterobacterales; Enterobacteriaceae; Shigella MNNEETFYQAMRRQGVTRRSFLKYCSLAATSLGLGAGMAPKIAWALENKPRIPVVWIHGL WP_005139779.1 hydrogenase [Shigella sonnei] Length: 372\nNumber of predicted TMHs: 2\nExp number of AAs in TMHs: 37.02628\nExp number, first 60 AAs: 19.39684\nTotal prob of N-in: 0.94313\nPOSSIBLE N-term signal sequence\ninside 1 19\nTMhelix 20 39\noutside 40 325\nTMhelix 326 348\ninside 349 372

27404 GCF_001251215.1_6350_1_6Shigella sonnei Proteobacteria; Gammaproteobacteria; Enterobacterales; Enterobacteriaceae; Shigella MNNEETFYQAMRRQGVTRRSFLKYCSLAATSLGLGAGMAPKIAWALENKPRIPVVWIHGL WP_005139779.1 hydrogenase [Shigella sonnei] Length: 372\nNumber of predicted TMHs: 2\nExp number of AAs in TMHs: 37.02628\nExp number, first 60 AAs: 19.39684\nTotal prob of N-in: 0.94313\nPOSSIBLE N-term signal sequence\ninside 1 19\nTMhelix 20 39\noutside 40 325\nTMhelix 326 348\ninside 349 372

27405 GCF_001251175.1_H14046047401 Shigella sonnei Proteobacteria; Gammaproteobacteria; Enterobacterales; Enterobacteriaceae; Shigella MNNEETFYQAMRRQGVTRRSFLKYCSLAATSLGLGAGMAPKIAWALENKPRIPVVWIHGL WP_005139779.1 hydrogenase [Shigella sonnei] Length: 372\nNumber of predicted TMHs: 2\nExp number of AAs in TMHs: 37.02628\nExp number, first 60 AAs: 19.39684\nTotal prob of N-in: 0.94313\nPOSSIBLE N-term signal sequence\ninside 1 19\nTMhelix 20 39\noutside 40 325\nTMhelix 326 348\ninside 349 372

27406 GCF_001251035.1_Sg_201400752_GATCAG_L001 Shigella sonnei Proteobacteria; Gammaproteobacteria; Enterobacterales; Enterobacteriaceae; Shigella MNNEETFYQAMRRQGVTRRSFLKYCSLAATSLGLGAGMAPKIAWALENKPRIPVVWIHGL WP_005139779.1 hydrogenase [Shigella sonnei] Length: 372\nNumber of predicted TMHs: 2\nExp number of AAs in TMHs: 37.02628\nExp number, first 60 AAs: 19.39684\nTotal prob of N-in: 0.94313\nPOSSIBLE N-term signal sequence\ninside 1 19\nTMhelix 20 39\noutside 40 325\nTMhelix 326 348\ninside 349 372

27407 GCF_001251115.1_8289_7_39 Shigella sonnei Proteobacteria; Gammaproteobacteria; Enterobacterales; Enterobacteriaceae; Shigella MNNEETFYQAMRRQGVTRRSFLKYCSLAATSLGLGAGMAPKIAWALENKPRIPVVWIHGL WP_005139779.1 hydrogenase [Shigella sonnei] Length: 372\nNumber of predicted TMHs: 2\nExp number of AAs in TMHs: 37.02628\nExp number, first 60 AAs: 19.39684\nTotal prob of N-in: 0.94313\nPOSSIBLE N-term signal sequence\ninside 1 19\nTMhelix 20 39\noutside 40 325\nTMhelix 326 348\ninside 349 372

27408 GCF_001251635.1_8290_5_8Shigella sonnei Proteobacteria; Gammaproteobacteria; Enterobacterales; Enterobacteriaceae; Shigella MTGDNTLIHSHGINRRDFMKLCAALAATMGLSSKAAAEMAESVTNPQRPPVIWIGAQECT WP_000145410.1 MULTISPECIES: hydrogenase-2 small chain [Proteobacteria] Length: 372\nNumber of predicted TMHs: 1\nExp number of AAs in TMHs: 23.72226\nExp number, first 60 AAs: 0.22038\nTotal prob of N-in: 0.12994\noutside 1 331\nTMhelix 332 354\ninside 355 372

27409 GCF_001251635.1_8290_5_8Shigella sonnei Proteobacteria; Gammaproteobacteria; Enterobacterales; Enterobacteriaceae; Shigella MNNEETFYQAMRRQGVTRRSFLKYCSLAATSLGLGAGMAPKIAWALENKPRIPVVWIHGL WP_005139779.1 hydrogenase [Shigella sonnei] Length: 372\nNumber of predicted TMHs: 2\nExp number of AAs in TMHs: 37.02628\nExp number, first 60 AAs: 19.39684\nTotal prob of N-in: 0.94313\nPOSSIBLE N-term signal sequence\ninside 1 19\nTMhelix 20 39\noutside 40 325\nTMhelix 326 348\ninside 349 372

27410 GCF_001251275.1_9870_7_12 Shigella sonnei Proteobacteria; Gammaproteobacteria; Enterobacterales; Enterobacteriaceae; Shigella MNNEETFYQAMRRQGVTRRSFLKYCSLAATSLGLGAGMAPKIAWALENKPRIPVVWIHGL WP_005139779.1 hydrogenase [Shigella sonnei] Length: 372\nNumber of predicted TMHs: 2\nExp number of AAs in TMHs: 37.02628\nExp number, first 60 AAs: 19.39684\nTotal prob of N-in: 0.94313\nPOSSIBLE N-term signal sequence\ninside 1 19\nTMhelix 20 39\noutside 40 325\nTMhelix 326 348\ninside 349 372

27411 GCF_001251315.1_Sg_201403906_CAACTA_L001 Shigella sonnei Proteobacteria; Gammaproteobacteria; Enterobacterales; Enterobacteriaceae; Shigella MNNEETFYQAMRRQGVTRRSFLKYCSLAATSLGLGAGMAPKIAWALENKPRIPVVWIHGL WP_005139779.1 hydrogenase [Shigella sonnei] Length: 372\nNumber of predicted TMHs: 2\nExp number of AAs in TMHs: 37.02628\nExp number, first 60 AAs: 19.39684\nTotal prob of N-in: 0.94313\nPOSSIBLE N-term signal sequence\ninside 1 19\nTMhelix 20 39\noutside 40 325\nTMhelix 326 348\ninside 349 372

27412 GCF_001251675.1_5236_2_4Shigella sonnei Proteobacteria; Gammaproteobacteria; Enterobacterales; Enterobacteriaceae; Shigella MTGDNTLIHSHGINRRDFMKLCAALAATMGLSSKAAAEMAESVTNPQRPPVIWIGAQECT WP_000145410.1 MULTISPECIES: hydrogenase-2 small chain [Proteobacteria] Length: 372\nNumber of predicted TMHs: 1\nExp number of AAs in TMHs: 23.72226\nExp number, first 60 AAs: 0.22038\nTotal prob of N-in: 0.12994\noutside 1 331\nTMhelix 332 354\ninside 355 372

27413 GCF_001251735.1_11822_8_62 Shigella sonnei Proteobacteria; Gammaproteobacteria; Enterobacterales; Enterobacteriaceae; Shigella MTGDNTLIHSHGINRRDFMKLCAALAATMGLSSKAAAEMAESVTNPQRPPVIWIGAQECT WP_000145410.1 MULTISPECIES: hydrogenase-2 small chain [Proteobacteria] Length: 372\nNumber of predicted TMHs: 1\nExp number of AAs in TMHs: 23.72226\nExp number, first 60 AAs: 0.22038\nTotal prob of N-in: 0.12994\noutside 1 331\nTMhelix 332 354\ninside 355 372

27414 GCF_001251795.1_12971_8_60 Shigella sonnei Proteobacteria; Gammaproteobacteria; Enterobacterales; Enterobacteriaceae; Shigella MTGDNTLIHSHGINRRDFMKLCAALAATMGLSSKAAAEMAESVTNPQRPPVIWIGAQECT WP_000145410.1 MULTISPECIES: hydrogenase-2 small chain [Proteobacteria] Length: 372\nNumber of predicted TMHs: 1\nExp number of AAs in TMHs: 23.72226\nExp number, first 60 AAs: 0.22038\nTotal prob of N-in: 0.12994\noutside 1 331\nTMhelix 332 354\ninside 355 372

27415 GCF_001251855.1_5417_2_1Shigella sonnei Proteobacteria; Gammaproteobacteria; Enterobacterales; Enterobacteriaceae; Shigella MTGDNTLIHSHGINRRDFMKLCAALAATMGLSSKAAAEMAESVTNPQRPPVIWIGAQECT WP_000145410.1 MULTISPECIES: hydrogenase-2 small chain [Proteobacteria] Length: 372\nNumber of predicted TMHs: 1\nExp number of AAs in TMHs: 23.72226\nExp number, first 60 AAs: 0.22038\nTotal prob of N-in: 0.12994\noutside 1 331\nTMhelix 332 354\ninside 355 372

27416 GCF_001251895.1_8290_5_3Shigella sonnei Proteobacteria; Gammaproteobacteria; Enterobacterales; Enterobacteriaceae; Shigella MTGDNTLIHSHGINRRDFMKLCAALAATMGLSSKAAAEMAESVTNPQRPPVIWIGAQECT WP_000145410.1 MULTISPECIES: hydrogenase-2 small chain [Proteobacteria] Length: 372\nNumber of predicted TMHs: 1\nExp number of AAs in TMHs: 23.72226\nExp number, first 60 AAs: 0.22038\nTotal prob of N-in: 0.12994\noutside 1 331\nTMhelix 332 354\ninside 355 372

27417 GCF_001251915.1_Sg_201401742_GTTTCG_L001 Shigella sonnei Proteobacteria; Gammaproteobacteria; Enterobacterales; Enterobacteriaceae; Shigella MTGDNTLIHSHGINRRDFMKLCAALAATMGLSSKAAAEMAESVTNPQRPPVIWIGAQECT WP_000145410.1 MULTISPECIES: hydrogenase-2 small chain [Proteobacteria] Length: 372\nNumber of predicted TMHs: 1\nExp number of AAs in TMHs: 23.72226\nExp number, first 60 AAs: 0.22038\nTotal prob of N-in: 0.12994\noutside 1 331\nTMhelix 332 354\ninside 355 372

27418 GCF_001252015.1_H14142025201 Shigella sonnei Proteobacteria; Gammaproteobacteria; Enterobacterales; Enterobacteriaceae; Shigella

MTGDNTLIHSHGINRRDFMKLCAALAATMGLSSKAAAEMAESVTNPQRPPVIWIGAQECT WP_000145410.1 MULTISPECIES: hydrogenase-2 small chain [Proteobacteria] Length: 372\nNumber of predicted TMHs: 1\nExp number of AAs in TMHs: 23.72226\nExp number, first 60 AAs: 0.22038\nTotal prob of N-in: 0.12994\noutside 1 331\nTMhelix 332 354\ninside 355 372

27419 GCF_001252115.1_9870_7_1Shigella sonnei Proteobacteria; Gammaproteobacteria; Enterobacterales; Enterobacteriaceae; Shigella MTGDNTLIHSHGINRRDFMKLCAALAATMGLSSKAAAEMAESVTNPQRPPVIWIGAQECT WP_000145410.1 MULTISPECIES: hydrogenase-2 small chain [Proteobacteria] Length: 372\nNumber of predicted TMHs: 1\nExp number of AAs in TMHs: 23.72226\nExp number, first 60 AAs: 0.22038\nTotal prob of N-in: 0.12994\noutside 1 331\nTMhelix 332 354\ninside 355 372

27420 GCF_001252155.1_12830_1_35 Shigella sonnei Proteobacteria; Gammaproteobacteria; Enterobacterales; Enterobacteriaceae; Shigella MTGDNTLIHSHGINRRDFMKLCAALAATMGLSSKAAAEMAESVTNPQRPPVIWIGAQECT WP_000145410.1 MULTISPECIES: hydrogenase-2 small chain [Proteobacteria] Length: 372\nNumber of predicted TMHs: 1\nExp number of AAs in TMHs: 23.72226\nExp number, first 60 AAs: 0.22038\nTotal prob of N-in: 0.12994\noutside 1 331\nTMhelix 332 354\ninside 355 372

27421 GCF_001252155.1_12830_1_35 Shigella sonnei Proteobacteria; Gammaproteobacteria; Enterobacterales; Enterobacteriaceae; Shigella MNNEETFYQAMRRQGVTRRSFLKYCSLAATSLGLGAGMAPKIAWALENKPRIPVVWIHGL WP_005139779.1 hydrogenase [Shigella sonnei] Length: 372\nNumber of predicted TMHs: 2\nExp number of AAs in TMHs: 37.02628\nExp number, first 60 AAs: 19.39684\nTotal prob of N-in: 0.94313\nPOSSIBLE N-term signal sequence\ninside 1 19\nTMhelix 20 39\noutside 40 325\nTMhelix 326 348\ninside 349 372

27422 GCF_001252115.1_9870_7_1Shigella sonnei Proteobacteria; Gammaproteobacteria; Enterobacterales; Enterobacteriaceae; Shigella MNNEETFYQAMRRQGVTRRSFLKYCSLAATSLGLGAGMAPKIAWALENKPRIPVVWIHGL WP_005139779.1 hydrogenase [Shigella sonnei] Length: 372\nNumber of predicted TMHs: 2\nExp number of AAs in TMHs: 37.02628\nExp number, first 60 AAs: 19.39684\nTotal prob of N-in: 0.94313\nPOSSIBLE N-term signal sequence\ninside 1 19\nTMhelix 20 39\noutside 40 325\nTMhelix 326 348\ninside 349 372

27423 GCF_001252615.1_8489_2_87 Shigella sonnei Proteobacteria; Gammaproteobacteria; Enterobacterales; Enterobacteriaceae; Shigella MTGDNTLIHSHGINRRDFMKLCAALAATMGLSSKAAAEMAESVTNPQRPPVIWIGAQECT WP_000145410.1 MULTISPECIES: hydrogenase-2 small chain [Proteobacteria] Length: 372\nNumber of predicted TMHs: 1\nExp number of AAs in TMHs: 23.72226\nExp number, first 60 AAs: 0.22038\nTotal prob of N-in: 0.12994\noutside 1 331\nTMhelix 332 354\ninside 355 372

27424 GCF_001251675.1_5236_2_4Shigella sonnei Proteobacteria; Gammaproteobacteria; Enterobacterales; Enterobacteriaceae; Shigella MNNEETFYQAMRRQGVTRRSFLKYCSLAATSLGLGAGMAPKIAWALENKPRIPVVWIHGL WP_005139779.1 hydrogenase [Shigella sonnei] Length: 372\nNumber of predicted TMHs: 2\nExp number of AAs in TMHs: 37.02628\nExp number, first 60 AAs: 19.39684\nTotal prob of N-in: 0.94313\nPOSSIBLE N-term signal sequence\ninside 1 19\nTMhelix 20 39\noutside 40 325\nTMhelix 326 348\ninside 349 372

27425 GCF_001252235.1_12971_8_58 Shigella sonnei Proteobacteria; Gammaproteobacteria; Enterobacterales; Enterobacteriaceae; Shigella MTGDNTLIHSHGINRRDFMKLCAALAATMGLSSKAAAEMAESVTNPQRPPVIWIGAQECT WP_000145410.1 MULTISPECIES: hydrogenase-2 small chain [Proteobacteria] Length: 372\nNumber of predicted TMHs: 1\nExp number of AAs in TMHs: 23.72226\nExp number, first 60 AAs: 0.22038\nTotal prob of N-in: 0.12994\noutside 1 331\nTMhelix 332 354\ninside 355 372

27426 GCF_001252235.1_12971_8_58 Shigella sonnei Proteobacteria; Gammaproteobacteria; Enterobacterales; Enterobacteriaceae; Shigella MNNEETFYQAMRRQGVTRRSFLKYCSLAATSLGLGAGMAPKIAWALENKPRIPVVWIHGL WP_005139779.1 hydrogenase [Shigella sonnei] Length: 372\nNumber of predicted TMHs: 2\nExp number of AAs in TMHs: 37.02628\nExp number, first 60 AAs: 19.39684\nTotal prob of N-in: 0.94313\nPOSSIBLE N-term signal sequence\ninside 1 19\nTMhelix 20 39\noutside 40 325\nTMhelix 326 348\ninside 349 372

27427 GCF_001251735.1_11822_8_62 Shigella sonnei Proteobacteria; Gammaproteobacteria; Enterobacterales; Enterobacteriaceae; Shigella MNNEETFYQAMRRQGVTRRSFLKYCSLAATSLGLGAGMAPKIAWALENKPRIPVVWIHGL WP_005139779.1

hydrogenase [Shigella sonnei] Length: 372\nNumber of predicted TMHs: 2\nExp number of AAs in TMHs: 37.02628\nExp number, first 60 AAs: 19.39684\nTotal prob of N-in: 0.94313\nPOSSIBLE N-term signal sequence\ninside 1 19\nTMhelix 20 39\noutside 40 325\nTMhelix 326 348\ninside 349 372

27428 GCF_001252275.1_8290_4_54 Shigella sonnei Proteobacteria; Gammaproteobacteria; Enterobacterales; Enterobacteriaceae; Shigella MTGDNTLIHSHGINRRDFMKLCAALAATMGLSSKAAAEMAESVTNPQRPPVIWIGAQECT WP_000145410.1 MULTISPECIES: hydrogenase-2 small chain [Proteobacteria] Length: 372\nNumber of predicted TMHs: 1\nExp number of AAs in TMHs: 23.72226\nExp number, first 60 AAs: 0.22038\nTotal prob of N-in: 0.12994\noutside 1 331\nTMhelix 332 354\ninside 355 372

27429 GCF_001252275.1_8290_4_54 Shigella sonnei Proteobacteria; Gammaproteobacteria; Enterobacterales; Enterobacteriaceae; Shigella MNNEETFYQAMRRQGVTRRSFLKYCSLAATSLGLGAGMAPKIAWALENKPRIPVVWIHGL WP_005139779.1 hydrogenase [Shigella sonnei] Length: 372\nNumber of predicted TMHs: 2\nExp number of AAs in TMHs: 37.02628\nExp number, first 60 AAs: 19.39684\nTotal prob of N-in: 0.94313\nPOSSIBLE N-term signal sequence\ninside 1 19\nTMhelix 20 39\noutside 40 325\nTMhelix 326 348\ninside 349 372

27430 GCF_001251795.1_12971_8_60 Shigella sonnei Proteobacteria; Gammaproteobacteria; Enterobacterales; Enterobacteriaceae; Shigella MNNEETFYQAMRRQGVTRRSFLKYCSLAATSLGLGAGMAPKIAWALENKPRIPVVWIHGL WP_005139779.1 hydrogenase [Shigella sonnei] Length: 372\nNumber of predicted TMHs: 2\nExp number of AAs in TMHs: 37.02628\nExp number, first 60 AAs: 19.39684\nTotal prob of N-in: 0.94313\nPOSSIBLE N-term signal sequence\ninside 1 19\nTMhelix 20 39\noutside 40 325\nTMhelix 326 348\ninside 349 372

27431 GCF_001251895.1_8290_5_3Shigella sonnei Proteobacteria; Gammaproteobacteria; Enterobacterales; Enterobacteriaceae; Shigella MNNEETFYQAMRRQGVTRRSFLKYCSLAATSLGLGAGMAPKIAWALENKPRIPVVWIHGL WP_005139779.1 hydrogenase [Shigella sonnei] Length: 372\nNumber of predicted TMHs: 2\nExp number of AAs in TMHs: 37.02628\nExp number, first 60 AAs: 19.39684\nTotal prob of N-in: 0.94313\nPOSSIBLE N-term signal sequence\ninside 1 19\nTMhelix 20 39\noutside 40 325\nTMhelix 326 348\ninside 349 372

27432 GCF_001251855.1_5417_2_1Shigella sonnei Proteobacteria; Gammaproteobacteria; Enterobacterales; Enterobacteriaceae; Shigella MNNEETFYQAMRRQGVTRRSFLKYCSLAATSLGLGAGMAPKIAWALENKPRIPVVWIHGL WP_005139779.1 hydrogenase [Shigella sonnei] Length: 372\nNumber of predicted TMHs: 2\nExp number of AAs in TMHs: 37.02628\nExp number, first 60 AAs: 19.39684\nTotal prob of N-in: 0.94313\nPOSSIBLE N-term signal sequence\ninside 1 19\nTMhelix 20 39\noutside 40 325\nTMhelix 326 348\ninside 349 372

27433 GCF_001251915.1_Sg_201401742_GTTTCG_L001 Shigella sonnei Proteobacteria; Gammaproteobacteria; Enterobacterales; Enterobacteriaceae; Shigella MNNEETFYQAMRRQGVTRRSFLKYCSLAATSLGLGAGMAPKIAWALENKPRIPVVWIHGL WP_005139779.1 hydrogenase [Shigella sonnei] Length: 372\nNumber of predicted TMHs: 2\nExp number of AAs in TMHs: 37.02628\nExp number, first 60 AAs: 19.39684\nTotal prob of N-in: 0.94313\nPOSSIBLE N-term signal sequence\ninside 1 19\nTMhelix 20 39\noutside 40 325\nTMhelix 326 348\ninside 349 372

27434 GCF_001252335.1_12844_1_55 Shigella sonnei Proteobacteria; Gammaproteobacteria; Enterobacterales; Enterobacteriaceae; Shigella MTGDNTLIHSHGINRRDFMKLCAALAATMGLSSKAAAEMAESVTNPQRPPVIWIGAQECT WP_000145410.1 MULTISPECIES: hydrogenase-2 small chain [Proteobacteria] Length: 372\nNumber of predicted TMHs: 1\nExp number of AAs in TMHs: 23.72226\nExp number, first 60 AAs: 0.22038\nTotal prob of N-in: 0.12994\noutside 1 331\nTMhelix 332 354\ninside 355 372

27435 GCF_001252375.1_8290_4_51 Shigella sonnei Proteobacteria; Gammaproteobacteria; Enterobacterales; Enterobacteriaceae; Shigella MTGDNTLIHSHGINRRDFMKLCAALAATMGLSSKAAAEMAESVTNPQRPPVIWIGAQECT WP_000145410.1 MULTISPECIES: hydrogenase-2 small chain [Proteobacteria] Length: 372\nNumber of predicted TMHs: 1\nExp number of AAs in TMHs: 23.72226\nExp number, first 60 AAs: 0.22038\nTotal prob of N-in: 0.12994\noutside 1 331\nTMhelix 332 354\ninside 355 372

27436 GCF_001252435.1_H14128036201 Shigella sonnei Proteobacteria; Gammaproteobacteria; Enterobacterales; Enterobacteriaceae; Shigella MTGDNTLIHSHGINRRDFMKLCAALAATMGLSSKAAAEMAESVTNPQRPPVIWIGAQECT WP_000145410.1

MULTISPECIES: hydrogenase-2 small chain [Proteobacteria] Length: 372\nNumber of predicted TMHs: 1\nExp number of AAs in TMHs: 23.72226\nExp number, first 60 AAs: 0.22038\nTotal prob of N-in: 0.12994\noutside 1 331\nTMhelix 332 354\ninside 355 372

27437 GCF_001252475.1_8290_4_7Shigella sonnei Proteobacteria; Gammaproteobacteria; Enterobacterales; Enterobacteriaceae; Shigella MTGDNTLIHSHGINRRDFMKLCAALAATMGLSSKAAAEMAESVTNPQRPPVIWIGAQECT WP_000145410.1 MULTISPECIES: hydrogenase-2 small chain [Proteobacteria] Length: 372\nNumber of predicted TMHs: 1\nExp number of AAs in TMHs: 23.72226\nExp number, first 60 AAs: 0.22038\nTotal prob of N-in: 0.12994\noutside 1 331\nTMhelix 332 354\ninside 355 372

27438 GCF_001252535.1_5417_2_5Shigella sonnei Proteobacteria; Gammaproteobacteria; Enterobacterales; Enterobacteriaceae; Shigella MTGDNTLIHSHGINRRDFMKLCAALAATMGLSSKAAAEMAESVTNPQRPPVIWIGAQECT WP_000145410.1 MULTISPECIES: hydrogenase-2 small chain [Proteobacteria] Length: 372\nNumber of predicted TMHs: 1\nExp number of AAs in TMHs: 23.72226\nExp number, first 60 AAs: 0.22038\nTotal prob of N-in: 0.12994\noutside 1 331\nTMhelix 332 354\ninside 355 372

27439 GCF_001252695.1_H14030023601 Shigella sonnei Proteobacteria; Gammaproteobacteria; Enterobacterales; Enterobacteriaceae; Shigella MTGDNTLIHSHGINRRDFMKLCAALAATMGLSSKAAAEMAESVTNPQRPPVIWIGAQECT WP_000145410.1 MULTISPECIES: hydrogenase-2 small chain [Proteobacteria] Length: 372\nNumber of predicted TMHs: 1\nExp number of AAs in TMHs: 23.72226\nExp number, first 60 AAs: 0.22038\nTotal prob of N-in: 0.12994\noutside 1 331\nTMhelix 332 354\ninside 355 372

27440 GCF_001252735.1_H14128036301 Shigella sonnei Proteobacteria; Gammaproteobacteria; Enterobacterales; Enterobacteriaceae; Shigella MTGDNTLIHSHGINRRDFMKLCAALAATMGLSSKAAAEMAESVTNPQRPPVIWIGAQECT WP_000145410.1 MULTISPECIES: hydrogenase-2 small chain [Proteobacteria] Length: 372\nNumber of predicted TMHs: 1\nExp number of AAs in TMHs: 23.72226\nExp number, first 60 AAs: 0.22038\nTotal prob of N-in: 0.12994\noutside 1 331\nTMhelix 332 354\ninside 355 372

27441 GCF_001252335.1_12844_1_55 Shigella sonnei Proteobacteria; Gammaproteobacteria; Enterobacterales; Enterobacteriaceae; Shigella MNNEETFYQAMRRQGVTRRSFLKYCSLAATSLGLGAGMAPKIAWALENKPRIPVVWIHGL WP_005139779.1 hydrogenase [Shigella sonnei] Length: 372\nNumber of predicted TMHs: 2\nExp number of AAs in TMHs: 37.02628\nExp number, first 60 AAs: 19.39684\nTotal prob of N-in: 0.94313\nPOSSIBLE N-term signal sequence\ninside 1 19\nTMhelix 20 39\noutside 40 325\nTMhelix 326 348\ninside 349 372

27442 GCF_001252815.1_12844_1_52 Shigella sonnei Proteobacteria; Gammaproteobacteria; Enterobacterales; Enterobacteriaceae; Shigella MTGDNTLIHSHGINRRDFMKLCAALAATMGLSSKAAAEMAESVTNPQRPPVIWIGAQECT WP_000145410.1 MULTISPECIES: hydrogenase-2 small chain [Proteobacteria] Length: 372\nNumber of predicted TMHs: 1\nExp number of AAs in TMHs: 23.72226\nExp number, first 60 AAs: 0.22038\nTotal prob of N-in: 0.12994\noutside 1 331\nTMhelix 332 354\ninside 355 372

27443 GCF_001252915.1_6496_8_6Shigella sonnei Proteobacteria; Gammaproteobacteria; Enterobacterales; Enterobacteriaceae; Shigella MTGDNTLIHSHGINRRDFMKLCAALAATMGLSSKAAAEMAESVTNPQRPPVIWIGAQECT WP_000145410.1 MULTISPECIES: hydrogenase-2 small chain [Proteobacteria] Length: 372\nNumber of predicted TMHs: 1\nExp number of AAs in TMHs: 23.72226\nExp number, first 60 AAs: 0.22038\nTotal prob of N-in: 0.12994\noutside 1 331\nTMhelix 332 354\ninside 355 372

27444 GCF_001252815.1_12844_1_52 Shigella sonnei Proteobacteria; Gammaproteobacteria; Enterobacterales; Enterobacteriaceae; Shigella MNNEETFYQAMRRQGVTRRSFLKYCSLAATSLGLGAGMAPKIAWALENKPRIPVVWIHGL WP_005139779.1 hydrogenase [Shigella sonnei] Length: 372\nNumber of predicted TMHs: 2\nExp number of AAs in TMHs: 37.02628\nExp number, first 60 AAs: 19.39684\nTotal prob of N-in: 0.94313\nPOSSIBLE N-term signal sequence\ninside 1 19\nTMhelix 20 39\noutside 40 325\nTMhelix 326 348\ninside 349 372

27445 GCF_001254015.1_H14088028501 Shigella sonnei Proteobacteria; Gammaproteobacteria; Enterobacterales; Enterobacteriaceae; Shigella MTGDNTLIHSHGINRRDFMKLCAALAATMGLSSKAAAEMAESVTNPQRPPVIWIGAQECT WP_000145410.1 MULTISPECIES: hydrogenase-2 small chain [Proteobacteria] Length: 372\nNumber of predicted TMHs:

1\nExp number of AAs in TMHs: 23.72226\nExp number, first 60 AAs: 0.22038\nTotal prob of N-in: 0.12994\noutside 1 331\nTMhelix 332 354\ninside 355 372

27446 GCF_001252915.1_6496_8_6Shigella sonnei Proteobacteria; Gammaproteobacteria; Enterobacterales; Enterobacteriaceae; Shigella MNNEETFYQAMRRQGVTRRSFLKYCSLAATSLGLGAGMAPKIAWALENKPRIPVVWIHGL WP_005139779.1 hydrogenase [Shigella sonnei] Length: 372\nNumber of predicted TMHs: 2\nExp number of AAs in TMHs: 37.02628\nExp number, first 60 AAs: 19.39684\nTotal prob of N-in: 0.94313\nPOSSIBLE N-term signal sequence\ninside 1 19\nTMhelix 20 39\noutside 40 325\nTMhelix 326 348\ninside 349 372

27447 GCF_001252375.1_8290_4_51 Shigella sonnei Proteobacteria; Gammaproteobacteria; Enterobacterales; Enterobacteriaceae; Shigella MNNEETFYQAMRRQGVTRRSFLKYCSLAATSLGLGAGMAPKIAWALENKPRIPVVWIHGL WP_005139779.1 hydrogenase [Shigella sonnei] Length: 372\nNumber of predicted TMHs: 2\nExp number of AAs in TMHs: 37.02628\nExp number, first 60 AAs: 19.39684\nTotal prob of N-in: 0.94313\nPOSSIBLE N-term signal sequence\ninside 1 19\nTMhelix 20 39\noutside 40 325\nTMhelix 326 348\ninside 349 372

27448 GCF_001252935.1_6350_1_1Shigella sonnei Proteobacteria; Gammaproteobacteria; Enterobacterales; Enterobacteriaceae; Shigella MTGDNTLIHSHGINRRDFMKLCAALAATMGLSSKAAAEMAESVTNPQRPPVIWIGAQECT WP_000145410.1 MULTISPECIES: hydrogenase-2 small chain [Proteobacteria] Length: 372\nNumber of predicted TMHs: 1\nExp number of AAs in TMHs: 23.72226\nExp number, first 60 AAs: 0.22038\nTotal prob of N-in: 0.12994\noutside 1 331\nTMhelix 332 354\ninside 355 372

27449 GCF_001252975.1_8290_5_15 Shigella sonnei Proteobacteria; Gammaproteobacteria; Enterobacterales; Enterobacteriaceae; Shigella MTGDNTLIHSHGINRRDFMKLCAALAATMGLSSKAAAEMAESVTNPQRPPVIWIGAQECT WP_000145410.1 MULTISPECIES: hydrogenase-2 small chain [Proteobacteria] Length: 372\nNumber of predicted TMHs: 1\nExp number of AAs in TMHs: 23.72226\nExp number, first 60 AAs: 0.22038\nTotal prob of N-in: 0.12994\noutside 1 331\nTMhelix 332 354\ninside 355 372

27450 GCF_001252935.1_6350_1_1Shigella sonnei Proteobacteria; Gammaproteobacteria; Enterobacterales; Enterobacteriaceae; Shigella MNNEETFYQAMRRQGVTRRSFLKYCSLAATSLGLGAGMAPKIAWALENKPRIPVVWIHGL WP_005139779.1 hydrogenase [Shigella sonnei] Length: 372\nNumber of predicted TMHs: 2\nExp number of AAs in TMHs: 37.02628\nExp number, first 60 AAs: 19.39684\nTotal prob of N-in: 0.94313\nPOSSIBLE N-term signal sequence\ninside 1 19\nTMhelix 20 39\noutside 40 325\nTMhelix 326 348\ninside 349 372

27451 GCF_001252975.1_8290_5_15 Shigella sonnei Proteobacteria; Gammaproteobacteria; Enterobacterales; Enterobacteriaceae; Shigella MNNEETFYQAMRRQGVTRRSFLKYCSLAATSLGLGAGMAPKIAWALENKPRIPVVWIHGL WP_005139779.1 hydrogenase [Shigella sonnei] Length: 372\nNumber of predicted TMHs: 2\nExp number of AAs in TMHs: 37.02628\nExp number, first 60 AAs: 19.39684\nTotal prob of N-in: 0.94313\nPOSSIBLE N-term signal sequence\ninside 1 19\nTMhelix 20 39\noutside 40 325\nTMhelix 326 348\ninside 349 372

27452 GCF_001252695.1_H14030023601 Shigella sonnei Proteobacteria; Gammaproteobacteria; Enterobacterales; Enterobacteriaceae; Shigella MNNEETFYQAMRRQGVTRRSFLKYCSLAATSLGLGAGMAPKIAWALENKPRIPVVWIHGL WP_005139779.1 hydrogenase [Shigella sonnei] Length: 372\nNumber of predicted TMHs: 2\nExp number of AAs in TMHs: 37.02628\nExp number, first 60 AAs: 19.39684\nTotal prob of N-in: 0.94313\nPOSSIBLE N-term signal sequence\ninside 1 19\nTMhelix 20 39\noutside 40 325\nTMhelix 326 348\ninside 349 372

27453 GCF_001252475.1_8290_4_7Shigella sonnei Proteobacteria; Gammaproteobacteria; Enterobacterales; Enterobacteriaceae; Shigella MNNEETFYQAMRRQGVTRRSFLKYCSLAATSLGLGAGMAPKIAWALENKPRIPVVWIHGL WP_005139779.1 hydrogenase [Shigella sonnei] Length: 372\nNumber of predicted TMHs: 2\nExp number of AAs in TMHs: 37.02628\nExp number, first 60 AAs: 19.39684\nTotal prob of N-in: 0.94313\nPOSSIBLE N-term signal sequence\ninside 1 19\nTMhelix 20 39\noutside 40 325\nTMhelix 326 348\ninside 349 372

27454 GCF_001252615.1_8489_2_87 Shigella sonnei Proteobacteria; Gammaproteobacteria; Enterobacterales; Enterobacteriaceae; Shigella MNNEETFYQAMRRQGVTRRSFLKYCSLAATSLGLGAGMAPKIAWALENKPRIPVVWIHGL WP_005139779.1 hydrogenase [Shigella sonnei] Length: 372\nNumber of predicted TMHs: 2\nExp number of AAs in TMHs: 37.02628\nExp number, first 60 AAs: 19.39684\nTotal prob of N-in: 0.94313\nPOSSIBLE N-term signal sequence\ninside 1 19\nTMhelix 20 39\noutside 40 325\nTMhelix 326 348\ninside 349 372

27455 GCF_001252535.1_5417_2_5Shigella sonnei Proteobacteria; Gammaproteobacteria; Enterobacterales; Enterobacteriaceae; Shigella MNNEETFYQAMRRQGVTRRSFLKYCSLAATSLGLGAGMAPKIAWALENKPRIPVVWIHGL WP_005139779.1 hydrogenase [Shigella sonnei] Length: 372\nNumber of predicted TMHs: 2\nExp number of AAs in TMHs: 37.02628\nExp number, first 60 AAs: 19.39684\nTotal prob of N-in: 0.94313\nPOSSIBLE N-term signal sequence\ninside 1 19\nTMhelix 20 39\noutside 40 325\nTMhelix 326 348\ninside 349 372

27456 GCF_001252435.1_H14128036201 Shigella sonnei Proteobacteria; Gammaproteobacteria; Enterobacterales; Enterobacteriaceae; Shigella MNNEETFYQAMRRQGVTRRSFLKYCSLAATSLGLGAGMAPKIAWALENKPRIPVVWIHGL WP_052962775.1 hydrogenase [Shigella sonnei] Length: 372\nNumber of predicted TMHs: 2\nExp number of AAs in TMHs: 37.09248\nExp number, first 60 AAs: 19.39296\nTotal prob of N-in: 0.94316\nPOSSIBLE N-term signal sequence\ninside 1 19\nTMhelix 20 39\noutside 40 325\nTMhelix 326 348\ninside 349 372

27457 GCF_001253015.1_7353_1_18 Shigella sonnei Proteobacteria; Gammaproteobacteria; Enterobacterales; Enterobacteriaceae; Shigella MTGDNTLIHSHGINRRDFMKLCAALAATMGLSSKAAAEMAESVTNPQRPPVIWIGAQECT WP_000145410.1 MULTISPECIES: hydrogenase-2 small chain [Proteobacteria] Length: 372\nNumber of predicted TMHs: 1\nExp number of AAs in TMHs: 23.72226\nExp number, first 60 AAs: 0.22038\nTotal prob of N-in: 0.12994\noutside 1 331\nTMhelix 332 354\ninside 355 372

27458 GCF_001253115.1_5236_7_6Shigella sonnei Proteobacteria; Gammaproteobacteria; Enterobacterales; Enterobacteriaceae; Shigella MTGDNTLIHSHGINRRDFMKLCAALAATMGLSSKAAAEMAESVTNPQRPPVIWIGAQECT WP_000145410.1 MULTISPECIES: hydrogenase-2 small chain [Proteobacteria] Length: 372\nNumber of predicted TMHs: 1\nExp number of AAs in TMHs: 23.72226\nExp number, first 60 AAs: 0.22038\nTotal prob of N-in: 0.12994\noutside 1 331\nTMhelix 332 354\ninside 355 372

27459 GCF_001253195.1_12844_1_61 Shigella sonnei Proteobacteria; Gammaproteobacteria; Enterobacterales; Enterobacteriaceae; Shigella MTGDNTLIHSHGINRRDFMKLCAALAATMGLSSKAAAEMAESVTNPQRPPVIWIGAQECT WP_000145410.1 MULTISPECIES: hydrogenase-2 small chain [Proteobacteria] Length: 372\nNumber of predicted TMHs: 1\nExp number of AAs in TMHs: 23.72226\nExp number, first 60 AAs: 0.22038\nTotal prob of N-in: 0.12994\noutside 1 331\nTMhelix 332 354\ninside 355 372

27460 GCF_001253335.1_Sg_201312520_TCCCGA_L001 Shigella sonnei Proteobacteria; Gammaproteobacteria; Enterobacterales; Enterobacteriaceae; Shigella MTGDNTLIHSHGINRRDFMKLCAALAATMGLSSKAAAEMAESVTNPQRPPVIWIGAQECT WP_000145410.1 MULTISPECIES: hydrogenase-2 small chain [Proteobacteria] Length: 372\nNumber of predicted TMHs: 1\nExp number of AAs in TMHs: 23.72226\nExp number, first 60 AAs: 0.22038\nTotal prob of N-in: 0.12994\noutside 1 331\nTMhelix 332 354\ninside 355 372

27461 GCF_001253375.1_H14132054201 Shigella sonnei Proteobacteria; Gammaproteobacteria; Enterobacterales; Enterobacteriaceae; Shigella MTGDNTLIHSHGINRRDFMKLCAALAATMGLSSKAAAEMAESVTNPQRPPVIWIGAQECT WP_000145410.1 MULTISPECIES: hydrogenase-2 small chain [Proteobacteria] Length: 372\nNumber of predicted TMHs: 1\nExp number of AAs in TMHs: 23.72226\nExp number, first 60 AAs: 0.22038\nTotal prob of N-in: 0.12994\noutside 1 331\nTMhelix 332 354\ninside 355 372

27462 GCF_001253475.1_H14112033401 Shigella sonnei Proteobacteria; Gammaproteobacteria; Enterobacterales; Enterobacteriaceae; Shigella MTGDNTLIHSHGINRRDFMKLCAALAATMGLSSKAAAEMAESVTNPQRPPVIWIGAQECT WP_000145410.1 MULTISPECIES: hydrogenase-2 small chain [Proteobacteria] Length: 372\nNumber of predicted TMHs: 1\nExp number of AAs in TMHs: 23.72226\nExp number, first 60 AAs: 0.22038\nTotal prob of N-in: 0.12994\noutside 1 331\nTMhelix 332 354\ninside 355 372

27463 GCF_001253535.1_Sg_201400063_CAGATC_L001 Shigella sonnei Proteobacteria; Gammaproteobacteria; Enterobacterales; Enterobacteriaceae; Shigella MTGDNTLIHSHGINRRDFMKLCAALAATMGLSSKAAAEMAESVTNPQRPPVIWIGAQECT WP_000145410.1 MULTISPECIES: hydrogenase-2 small chain [Proteobacteria] Length: 372\nNumber of predicted TMHs: 1\nExp number of AAs in TMHs: 23.72226\nExp number, first 60 AAs: 0.22038\nTotal prob of N-in: 0.12994\noutside 1 331\nTMhelix 332 354\ninside 355 372

27464 GCF_001253655.1_12830_1_36 Shigella sonnei Proteobacteria; Gammaproteobacteria; Enterobacterales; Enterobacteriaceae; Shigella
MTGDNTLIHSHGINRRDFMKLCAALAATMGLSSKAAAEMAESVTNPQRPPVIWIGAQECT WP_000145410.1
MULTISPECIES: hydrogenase-2 small chain [Proteobacteria] Length: 372\nNumber of predicted TMHs: 1\nExp number of AAs in TMHs: 23.72226\nExp number, first 60 AAs: 0.22038\nTotal prob of N-in: 0.12994\noutside 1 331\nTMhelix 332 354\ninside 355 372

27465 GCF_001253735.1_12971_8_73 Shigella sonnei Proteobacteria; Gammaproteobacteria; Enterobacterales; Enterobacteriaceae; Shigella
MTGDNTLIHSHGINRRDFMKLCAALAATMGLSSKAAAEMAESVTNPQRPPVIWIGAQECT WP_000145410.1
MULTISPECIES: hydrogenase-2 small chain [Proteobacteria] Length: 372\nNumber of predicted TMHs: 1\nExp number of AAs in TMHs: 23.72226\nExp number, first 60 AAs: 0.22038\nTotal prob of N-in: 0.12994\noutside 1 331\nTMhelix 332 354\ninside 355 372

27466 GCF_001253795.1_Sg_201401124_ATGTCA_L001 Shigella sonnei Proteobacteria; Gammaproteobacteria; Enterobacterales; Enterobacteriaceae; Shigella
MTGDNTLIHSHGINRRDFMKLCAALAATMGLSSKAAAEMAESVTNPQRPPVIWIGAQECT WP_000145410.1
MULTISPECIES: hydrogenase-2 small chain [Proteobacteria] Length: 372\nNumber of predicted TMHs: 1\nExp number of AAs in TMHs: 23.72226\nExp number, first 60 AAs: 0.22038\nTotal prob of N-in: 0.12994\noutside 1 331\nTMhelix 332 354\ninside 355 372

27467 GCF_001253655.1_12830_1_36 Shigella sonnei Proteobacteria; Gammaproteobacteria; Enterobacterales; Enterobacteriaceae; Shigella
MNNEETFYQAMRRQGVTRRSFLKYCSLAATSLGLGAGMAPKIAWALENKPRIPVVWIHGL WP_053008205.1
hydrogenase [Shigella sonnei] Length: 372\nNumber of predicted TMHs: 2\nExp number of AAs in TMHs: 37.06004\nExp number, first 60 AAs: 19.39728\nTotal prob of N-in: 0.94317\nPOSSIBLE N-term signal sequence\ninside 1 19\nTMhelix 20 39\noutside 40 325\nTMhelix 326 348\ninside 349 372

27468 GCF_001253795.1_Sg_201401124_ATGTCA_L001 Shigella sonnei Proteobacteria; Gammaproteobacteria; Enterobacterales; Enterobacteriaceae; Shigella
MNNEETFYQAMRRQGVTRRSFLKYCSLAATSLGLGAGMAPKIAWALENKPRIPVVWIHGL WP_005139779.1
hydrogenase [Shigella sonnei] Length: 372\nNumber of predicted TMHs: 2\nExp number of AAs in TMHs: 37.02628\nExp number, first 60 AAs: 19.39684\nTotal prob of N-in: 0.94313\nPOSSIBLE N-term signal sequence\ninside 1 19\nTMhelix 20 39\noutside 40 325\nTMhelix 326 348\ninside 349 372

27469 GCF_001253735.1_12971_8_73 Shigella sonnei Proteobacteria; Gammaproteobacteria; Enterobacterales; Enterobacteriaceae; Shigella
MNNEETFYQAMRRQGVTRRSFLKYCSLAATSLGLGAGMAPKIAWALENKPRIPVVWIHGL WP_052962775.1
hydrogenase [Shigella sonnei] Length: 372\nNumber of predicted TMHs: 2\nExp number of AAs in TMHs: 37.09248\nExp number, first 60 AAs: 19.39296\nTotal prob of N-in: 0.94316\nPOSSIBLE N-term signal sequence\ninside 1 19\nTMhelix 20 39\noutside 40 325\nTMhelix 326 348\ninside 349 372

27470 GCF_001253015.1_7353_1_18 Shigella sonnei Proteobacteria; Gammaproteobacteria; Enterobacterales; Enterobacteriaceae; Shigella
MNNEETFYQAMRRQGVTRRSFLKYCSLAATSLGLGAGMAPKIAWALENKPRIPVVWIHGL WP_005139779.1
hydrogenase [Shigella sonnei] Length: 372\nNumber of predicted TMHs: 2\nExp number of AAs in TMHs: 37.02628\nExp number, first 60 AAs: 19.39684\nTotal prob of N-in: 0.94313\nPOSSIBLE N-term signal sequence\ninside 1 19\nTMhelix 20 39\noutside 40 325\nTMhelix 326 348\ninside 349 372

27471 GCF_001253875.1_8290_4_3Shigella sonnei Proteobacteria; Gammaproteobacteria; Enterobacterales; Enterobacteriaceae; Shigella MTGDNTLIHSHGINRRDFMKLCAALAATMGLSSKAAAEMAESVTNPQRPPVIWIGAQECT
WP_000145410.1 MULTISPECIES: hydrogenase-2 small chain [Proteobacteria] Length: 372\nNumber of predicted TMHs: 1\nExp number of AAs in TMHs: 23.72226\nExp number, first 60 AAs: 0.22038\nTotal prob of N-in: 0.12994\noutside 1 331\nTMhelix 332 354\ninside 355 372

27472 GCF_001253875.1_8290_4_3Shigella sonnei Proteobacteria; Gammaproteobacteria; Enterobacterales; Enterobacteriaceae; Shigella MNNEETFYQAMRRQGVTRRSFLKYCSLAATSLGLGAGMAPKIAWALENKPRIPVVWIHGL
WP_005139779.1 hydrogenase [Shigella sonnei] Length: 372\nNumber of predicted TMHs: 2\nExp number of AAs in TMHs: 37.02628\nExp number, first 60 AAs: 19.39684\nTotal prob of N-in: 0.94313\nPOSSIBLE N-term signal sequence\ninside 1 19\nTMhelix 20 39\noutside 40 325\nTMhelix 326 348\ninside 349 372

27473 GCF_001253115.1_5236_7_6Shigella sonnei Proteobacteria; Gammaproteobacteria; Enterobacterales; Enterobacteriaceae; Shigella MNNEETFYQAMRRQGVTRRSFLKYCSLAATSLGLGAGMAPKIAWALENKPRIPVVWIHGL WP_005139779.1 hydrogenase [Shigella sonnei] Length: 372\nNumber of predicted TMHs: 2\nExp number of AAs in TMHs: 37.02628\nExp number, first 60 AAs: 19.39684\nTotal prob of N-in: 0.94313\nPOSSIBLE N-term signal sequence\ninside 1 19\nTMhelix 20 39\noutside 40 325\nTMhelix 326 348\ninside 349 372

27474 GCF_001253195.1_12844_1_61 Shigella sonnei Proteobacteria; Gammaproteobacteria; Enterobacterales; Enterobacteriaceae; Shigella MNNEETFYQAMRRQGVTRRSFLKYCSLAATSLGLGAGMAPKIAWALENKPRIPVVWIHGL WP_005139779.1 hydrogenase [Shigella sonnei] Length: 372\nNumber of predicted TMHs: 2\nExp number of AAs in TMHs: 37.02628\nExp number, first 60 AAs: 19.39684\nTotal prob of N-in: 0.94313\nPOSSIBLE N-term signal sequence\ninside 1 19\nTMhelix 20 39\noutside 40 325\nTMhelix 326 348\ninside 349 372

27475 GCF_001253475.1_H14112033401 Shigella sonnei Proteobacteria; Gammaproteobacteria; Enterobacterales; Enterobacteriaceae; Shigella MNNEETFYQAMRRQGVTRRSFLKYCSLAATSLGLGAGMAPKIAWALENKPRIPVVWIHGL WP_005139779.1 hydrogenase [Shigella sonnei] Length: 372\nNumber of predicted TMHs: 2\nExp number of AAs in TMHs: 37.02628\nExp number, first 60 AAs: 19.39684\nTotal prob of N-in: 0.94313\nPOSSIBLE N-term signal sequence\ninside 1 19\nTMhelix 20 39\noutside 40 325\nTMhelix 326 348\ninside 349 372

27476 GCF_001253375.1_H14132054201 Shigella sonnei Proteobacteria; Gammaproteobacteria; Enterobacterales; Enterobacteriaceae; Shigella MNNEETFYQAMRRQGVTRRSFLKYCSLAATSLGLGAGMAPKIAWALENKPRIPVVWIHGL WP_005139779.1 hydrogenase [Shigella sonnei] Length: 372\nNumber of predicted TMHs: 2\nExp number of AAs in TMHs: 37.02628\nExp number, first 60 AAs: 19.39684\nTotal prob of N-in: 0.94313\nPOSSIBLE N-term signal sequence\ninside 1 19\nTMhelix 20 39\noutside 40 325\nTMhelix 326 348\ninside 349 372

27477 GCF_001253335.1_Sg_201312520_TCCCGA_L001 Shigella sonnei Proteobacteria; Gammaproteobacteria; Enterobacterales; Enterobacteriaceae; Shigella MNNEETFYQAMRRQGVTRRSFLKYCSLAATSLGLGAGMAPKIAWALENKPRIPVVWIHGL WP_005139779.1 hydrogenase [Shigella sonnei] Length: 372\nNumber of predicted TMHs: 2\nExp number of AAs in TMHs: 37.02628\nExp number, first 60 AAs: 19.39684\nTotal prob of N-in: 0.94313\nPOSSIBLE N-term signal sequence\ninside 1 19\nTMhelix 20 39\noutside 40 325\nTMhelix 326 348\ninside 349 372

27478 GCF_001253915.1_5236_1_2Shigella sonnei Proteobacteria; Gammaproteobacteria; Enterobacterales; Enterobacteriaceae; Shigella MTGDNTLIHSHGINRRDFMKLCAALAATMGLSSKAAAEMAESVTNPQRPPVIWIGAQECT WP_000145410.1 MULTISPECIES: hydrogenase-2 small chain [Proteobacteria] Length: 372\nNumber of predicted TMHs: 1\nExp number of AAs in TMHs: 23.72226\nExp number, first 60 AAs: 0.22038\nTotal prob of N-in: 0.12994\noutside 1 331\nTMhelix 332 354\ninside 355 372

27479 GCF_001253915.1_5236_1_2Shigella sonnei Proteobacteria; Gammaproteobacteria; Enterobacterales; Enterobacteriaceae; Shigella MNNEETFYQAMRRQGVTRRSFLKYCSLAATSLGLGAGMAPKIAWALENKPRIPVVWIHGL WP_005139779.1 hydrogenase [Shigella sonnei] Length: 372\nNumber of predicted TMHs: 2\nExp number of AAs in TMHs: 37.02628\nExp number, first 60 AAs: 19.39684\nTotal prob of N-in: 0.94313\nPOSSIBLE N-term signal sequence\ninside 1 19\nTMhelix 20 39\noutside 40 325\nTMhelix 326 348\ninside 349 372

27480 GCF_001253535.1_Sg_201400063_CAGATC_L001 Shigella sonnei Proteobacteria; Gammaproteobacteria; Enterobacterales; Enterobacteriaceae; Shigella MNNEETFYQAMRRQGVTRRSFLKYCSLAATSLGLGAGMAPKIAWALENKPRIPVVWIHGL WP_005139779.1 hydrogenase [Shigella sonnei] Length: 372\nNumber of predicted TMHs: 2\nExp number of AAs in TMHs: 37.02628\nExp number, first 60 AAs: 19.39684\nTotal prob of N-in: 0.94313\nPOSSIBLE N-term signal sequence\ninside 1 19\nTMhelix 20 39\noutside 40 325\nTMhelix 326 348\ninside 349 372

27481 GCF_001253975.1_5236_1_12 Shigella sonnei Proteobacteria; Gammaproteobacteria; Enterobacterales; Enterobacteriaceae; Shigella MTGDNTLIHSHGINRRDFMKLCAALAATMGLSSKAAAEMAESVTNPQRPPVIWIGAQECT WP_000145410.1 MULTISPECIES: hydrogenase-2 small chain [Proteobacteria] Length: 372\nNumber of predicted TMHs: 1\nExp number of AAs in TMHs: 23.72226\nExp number, first 60 AAs: 0.22038\nTotal prob of N-in: 0.12994\noutside 1 331\nTMhelix 332 354\ninside 355 372

27482 GCF_001254115.1_5236_3_10 Shigella sonnei Proteobacteria; Gammaproteobacteria; Enterobacterales; Enterobacteriaceae; Shigella
MTGDNTLIHSHGINRRDFMKLCAALAATMGLSSKAAAEMAESVTNPQRPPVIWIGAQECT WP_000145410.1
MULTISPECIES: hydrogenase-2 small chain [Proteobacteria] Length: 372\nNumber of predicted TMHs: 1\nExp number of AAs in TMHs: 23.72226\nExp number, first 60 AAs: 0.22038\nTotal prob of N-in: 0.12994\noutside 1 331\nTMhelix 332 354\ninside 355 372

27483 GCF_001254135.1_6350_1_21 Shigella sonnei Proteobacteria; Gammaproteobacteria; Enterobacterales; Enterobacteriaceae; Shigella
MTGDNTLIHSHGINRRDFMKLCAALAATMGLSSKAAAEMAESVTNPQRPPVIWIGAQECT WP_000145410.1
MULTISPECIES: hydrogenase-2 small chain [Proteobacteria] Length: 372\nNumber of predicted TMHs: 1\nExp number of AAs in TMHs: 23.72226\nExp number, first 60 AAs: 0.22038\nTotal prob of N-in: 0.12994\noutside 1 331\nTMhelix 332 354\ninside 355 372

27484 GCF_001254175.1_8290_4_52 Shigella sonnei Proteobacteria; Gammaproteobacteria; Enterobacterales; Enterobacteriaceae; Shigella
MTGDNTLIHSHGINRRDFMKLCAALAATMGLSSKAAAEMAESVTNPQRPPVIWIGAQECT WP_000145410.1
MULTISPECIES: hydrogenase-2 small chain [Proteobacteria] Length: 372\nNumber of predicted TMHs: 1\nExp number of AAs in TMHs: 23.72226\nExp number, first 60 AAs: 0.22038\nTotal prob of N-in: 0.12994\noutside 1 331\nTMhelix 332 354\ninside 355 372

27485 GCF_001254235.1_8290_5_11 Shigella sonnei Proteobacteria; Gammaproteobacteria; Enterobacterales; Enterobacteriaceae; Shigella
MTGDNTLIHSHGINRRDFMKLCAALAATMGLSSKAAAEMAESVTNPQRPPVIWIGAQECT WP_000145410.1
MULTISPECIES: hydrogenase-2 small chain [Proteobacteria] Length: 372\nNumber of predicted TMHs: 1\nExp number of AAs in TMHs: 23.72226\nExp number, first 60 AAs: 0.22038\nTotal prob of N-in: 0.12994\noutside 1 331\nTMhelix 332 354\ninside 355 372

27486 GCF_001254295.1_5008_8_12 Shigella sonnei Proteobacteria; Gammaproteobacteria; Enterobacterales; Enterobacteriaceae; Shigella
MTGDNTLIHSHGINRRDFMKLCAALAATMGLSSKAAAEMAESVTNPQRPPVIWIGAQECT WP_000145410.1
MULTISPECIES: hydrogenase-2 small chain [Proteobacteria] Length: 372\nNumber of predicted TMHs: 1\nExp number of AAs in TMHs: 23.72226\nExp number, first 60 AAs: 0.22038\nTotal prob of N-in: 0.12994\noutside 1 331\nTMhelix 332 354\ninside 355 372

27487 GCF_001254415.1_8290_4_15 Shigella sonnei Proteobacteria; Gammaproteobacteria; Enterobacterales; Enterobacteriaceae; Shigella
MTGDNTLIHSHGINRRDFMKLCAALAATMGLSSKAAAEMAESVTNPQRPPVIWIGAQECT WP_000145410.1
MULTISPECIES: hydrogenase-2 small chain [Proteobacteria] Length: 372\nNumber of predicted TMHs: 1\nExp number of AAs in TMHs: 23.72226\nExp number, first 60 AAs: 0.22038\nTotal prob of N-in: 0.12994\noutside 1 331\nTMhelix 332 354\ninside 355 372

27488 GCF_001254415.1_8290_4_15 Shigella sonnei Proteobacteria; Gammaproteobacteria; Enterobacterales; Enterobacteriaceae; Shigella
MNNEETFYQAMRRQGVTRRSFLKYCSLAATSLGLGAGMAPKIAWALENKPRIPVVWIHGL WP_005139779.1
hydrogenase [Shigella sonnei] Length: 372\nNumber of predicted TMHs: 2\nExp number of AAs in TMHs: 37.02628\nExp number, first 60 AAs: 19.39684\nTotal prob of N-in: 0.94313\nPOSSIBLE N-term signal sequence\ninside 1 19\nTMhelix 20 39\noutside 40 325\nTMhelix 326 348\ninside 349 372

27489 GCF_001254795.1_12971_8_11 Shigella sonnei Proteobacteria; Gammaproteobacteria; Enterobacterales; Enterobacteriaceae; Shigella
MTGDNTLIHSHGINRRDFMKLCAALAATMGLSSKAAAEMAESVTNPQRPPVIWIGAQECT WP_000145410.1
MULTISPECIES: hydrogenase-2 small chain [Proteobacteria] Length: 372\nNumber of predicted TMHs: 1\nExp number of AAs in TMHs: 23.72226\nExp number, first 60 AAs: 0.22038\nTotal prob of N-in: 0.12994\noutside 1 331\nTMhelix 332 354\ninside 355 372

27490 GCF_001253975.1_5236_1_12 Shigella sonnei Proteobacteria; Gammaproteobacteria; Enterobacterales; Enterobacteriaceae; Shigella
MNNEETFYQAMRRQGVTRRSFLKYCSLAATSLGLGAGMAPKIAWALENKPRIPVVWIHGL WP_005139779.1
hydrogenase [Shigella sonnei] Length: 372\nNumber of predicted TMHs: 2\nExp number of AAs in

TMHs: 37.02628\nExp number, first 60 AAs: 19.39684\nTotal prob of N-in: 0.94313\nPOSSIBLE N-term signal sequence\ninside 1 19\nTMhelix 20 39\noutside 40 325\nTMhelix 326 348\ninside 349 372

27491 GCF_001254475.1_Sg_201400046_ACAGTG_L001 Shigella sonnei Proteobacteria; Gammaproteobacteria; Enterobacterales; Enterobacteriaceae; Shigella MTGDNTLIHSHGINRRDFMKLCAALAATMGLSSKAAAEMAESVTNPQRPPVIWIGAQECT WP_000145410.1 MULTISPECIES: hydrogenase-2 small chain [Proteobacteria] Length: 372\nNumber of predicted TMHs: 1\nExp number of AAs in TMHs: 23.72226\nExp number, first 60 AAs: 0.22038\nTotal prob of N-in: 0.12994\noutside 1 331\nTMhelix 332 354\ninside 355 372

27492 GCF_001254555.1_8289_7_79 Shigella sonnei Proteobacteria; Gammaproteobacteria; Enterobacterales; Enterobacteriaceae; Shigella MTGDNTLIHSHGINRRDFMKLCAALAATMGLSSKAAAEMAESVTNPQRPPVIWIGAQECT WP_000145410.1 MULTISPECIES: hydrogenase-2 small chain [Proteobacteria] Length: 372\nNumber of predicted TMHs: 1\nExp number of AAs in TMHs: 23.72226\nExp number, first 60 AAs: 0.22038\nTotal prob of N-in: 0.12994\noutside 1 331\nTMhelix 332 354\ninside 355 372

27493 GCF_001254615.1_5008_7_3Shigella sonnei Proteobacteria; Gammaproteobacteria; Enterobacterales; Enterobacteriaceae; Shigella MTGDNTLIHSHGINRRDFMKLCAALAATMGLSSKAAAEMAESVTNPQRPPVIWIGAQECT WP_000145410.1 MULTISPECIES: hydrogenase-2 small chain [Proteobacteria] Length: 372\nNumber of predicted TMHs: 1\nExp number of AAs in TMHs: 23.72226\nExp number, first 60 AAs: 0.22038\nTotal prob of N-in: 0.12994\noutside 1 331\nTMhelix 332 354\ninside 355 372

27494 GCF_001254475.1_Sg_201400046_ACAGTG_L001 Shigella sonnei Proteobacteria; Gammaproteobacteria; Enterobacterales; Enterobacteriaceae; Shigella MNNEETFYQAMRRQGVTRRSFLKYCSLAATSLGLGAGMAPKIAWALENKPRIPVVWIHGL WP_005139779.1 hydrogenase [Shigella sonnei] Length: 372\nNumber of predicted TMHs: 2\nExp number of AAs in TMHs: 37.02628\nExp number, first 60 AAs: 19.39684\nTotal prob of N-in: 0.94313\nPOSSIBLE N-term signal sequence\ninside 1 19\nTMhelix 20 39\noutside 40 325\nTMhelix 326 348\ninside 349 372

27495 GCF_001254615.1_5008_7_3Shigella sonnei Proteobacteria; Gammaproteobacteria; Enterobacterales; Enterobacteriaceae; Shigella MNNEETFYQAMRRQGVTRRSFLKYCSLAATSLGLGAGMAPKIAWALENKPRIPVVWIHGL WP_005139779.1 hydrogenase [Shigella sonnei] Length: 372\nNumber of predicted TMHs: 2\nExp number of AAs in TMHs: 37.02628\nExp number, first 60 AAs: 19.39684\nTotal prob of N-in: 0.94313\nPOSSIBLE N-term signal sequence\ninside 1 19\nTMhelix 20 39\noutside 40 325\nTMhelix 326 348\ninside 349 372

27496 GCF_001254555.1_8289_7_79 Shigella sonnei Proteobacteria; Gammaproteobacteria; Enterobacterales; Enterobacteriaceae; Shigella MNNEETFYQAMRRQGVTRRSFLKYCSLAATSLGLGAGMAPKIAWALENKPRIPVVWIHGL WP_005139779.1 hydrogenase [Shigella sonnei] Length: 372\nNumber of predicted TMHs: 2\nExp number of AAs in TMHs: 37.02628\nExp number, first 60 AAs: 19.39684\nTotal prob of N-in: 0.94313\nPOSSIBLE N-term signal sequence\ninside 1 19\nTMhelix 20 39\noutside 40 325\nTMhelix 326 348\ninside 349 372

27497 GCF_001254135.1_6350_1_21 Shigella sonnei Proteobacteria; Gammaproteobacteria; Enterobacterales; Enterobacteriaceae; Shigella MNNEETFYQAMRRQGVTRRSFLKYCSLAATSLGLGAGMAPKIAWALENKPRIPVVWIHGL WP_005139779.1 hydrogenase [Shigella sonnei] Length: 372\nNumber of predicted TMHs: 2\nExp number of AAs in TMHs: 37.02628\nExp number, first 60 AAs: 19.39684\nTotal prob of N-in: 0.94313\nPOSSIBLE N-term signal sequence\ninside 1 19\nTMhelix 20 39\noutside 40 325\nTMhelix 326 348\ninside 349 372

27498 GCF_001254115.1_5236_3_10 Shigella sonnei Proteobacteria; Gammaproteobacteria; Enterobacterales; Enterobacteriaceae; Shigella MNNEETFYQAMRRQGVTRRSFLKYCSLAATSLGLGAGMAPKIAWALENKPRIPVVWIHGL WP_005139779.1 hydrogenase [Shigella sonnei] Length: 372\nNumber of predicted TMHs: 2\nExp number of AAs in TMHs: 37.02628\nExp number, first 60 AAs: 19.39684\nTotal prob of N-in: 0.94313\nPOSSIBLE N-term signal sequence\ninside 1 19\nTMhelix 20 39\noutside 40 325\nTMhelix 326 348\ninside 349 372

27499 GCF_001254235.1_8290_5_11 Shigella sonnei Proteobacteria; Gammaproteobacteria; Enterobacterales; Enterobacteriaceae; Shigella MNNEETFYQAMRRQGVTRRSFLKYCSLAATSLGLGAGMAPKIAWALENKPRIPVVWIHGL WP_005139779.1 hydrogenase [Shigella sonnei] Length: 372\nNumber of predicted TMHs: 2\nExp number of AAs in TMHs:

TMHs: 37.02628\nExp number, first 60 AAs: 19.39684\nTotal prob of N-in: 0.94313\nPOSSIBLE N-term signal sequence\ninside 1 19\nTMhelix 20 39\noutside 40 325\nTMhelix 326 348\ninside 349 372

27500 GCF_001254175.1_8290_4_52 Shigella sonnei Proteobacteria; Gammaproteobacteria; Enterobacterales; Enterobacteriaceae; Shigella
MNNEETFYQAMRRQGVTRRSFLKYCSLAATSLGLGAGMAPKIAWALENKPRIPVVWIHGL WP_005139779.1
hydrogenase [Shigella sonnei] Length: 372\nNumber of predicted TMHs: 2\nExp number of AAs in TMHs: 37.02628\nExp number, first 60 AAs: 19.39684\nTotal prob of N-in: 0.94313\nPOSSIBLE N-term signal sequence\ninside 1 19\nTMhelix 20 39\noutside 40 325\nTMhelix 326 348\ninside 349 372

27501 GCF_001254755.1_9870_6_39 Shigella sonnei Proteobacteria; Gammaproteobacteria; Enterobacterales; Enterobacteriaceae; Shigella
MTGDNTLIHSHGINRRDFMKLCAALAATMGLSSKAAAEMAESVTNPQRPPVIWIGAQECT WP_000145410.1
MULTISPECIES: hydrogenase-2 small chain [Proteobacteria] Length: 372\nNumber of predicted TMHs: 1\nExp number of AAs in TMHs: 23.72226\nExp number, first 60 AAs: 0.22038\nTotal prob of N-in: 0.12994\noutside 1 331\nTMhelix 332 354\ninside 355 372

27502 GCF_001254755.1_9870_6_39 Shigella sonnei Proteobacteria; Gammaproteobacteria; Enterobacterales; Enterobacteriaceae; Shigella
MNNEETFYQAMRRQGVTRRSFLKYCSLAATSLGLGAGMAPKIAWALENKPRIPVVWIHGL WP_005139779.1
hydrogenase [Shigella sonnei] Length: 372\nNumber of predicted TMHs: 2\nExp number of AAs in TMHs: 37.02628\nExp number, first 60 AAs: 19.39684\nTotal prob of N-in: 0.94313\nPOSSIBLE N-term signal sequence\ninside 1 19\nTMhelix 20 39\noutside 40 325\nTMhelix 326 348\ninside 349 372

27503 GCF_001254295.1_5008_8_12 Shigella sonnei Proteobacteria; Gammaproteobacteria; Enterobacterales; Enterobacteriaceae; Shigella
MNNEETFYQAMRRQGVTRRSFLKYCSLAATSLGLGAGMAPKIAWALENKPRIPVVWIHGL WP_005139779.1
hydrogenase [Shigella sonnei] Length: 372\nNumber of predicted TMHs: 2\nExp number of AAs in TMHs: 37.02628\nExp number, first 60 AAs: 19.39684\nTotal prob of N-in: 0.94313\nPOSSIBLE N-term signal sequence\ninside 1 19\nTMhelix 20 39\noutside 40 325\nTMhelix 326 348\ninside 349 372

27504 GCF_001254935.1_12971_8_76 Shigella sonnei Proteobacteria; Gammaproteobacteria; Enterobacterales; Enterobacteriaceae; Shigella
MTGDNTLIHSHGINRRDFMKLCAALAATMGLSSKAAAEMAESVTNPQRPPVIWIGAQECT WP_000145410.1
MULTISPECIES: hydrogenase-2 small chain [Proteobacteria] Length: 372\nNumber of predicted TMHs: 1\nExp number of AAs in TMHs: 23.72226\nExp number, first 60 AAs: 0.22038\nTotal prob of N-in: 0.12994\noutside 1 331\nTMhelix 332 354\ninside 355 372

27505 GCF_001254995.1_8289_7_68 Shigella sonnei Proteobacteria; Gammaproteobacteria; Enterobacterales; Enterobacteriaceae; Shigella
MTGDNTLIHSHGINRRDFMKLCAALAATMGLSSKAAAEMAESVTNPQRPPVIWIGAQECT WP_000145410.1
MULTISPECIES: hydrogenase-2 small chain [Proteobacteria] Length: 372\nNumber of predicted TMHs: 1\nExp number of AAs in TMHs: 23.72226\nExp number, first 60 AAs: 0.22038\nTotal prob of N-in: 0.12994\noutside 1 331\nTMhelix 332 354\ninside 355 372

27506 GCF_001255055.1_8289_7_49 Shigella sonnei Proteobacteria; Gammaproteobacteria; Enterobacterales; Enterobacteriaceae; Shigella
MTGDNTLIHSHGINRRDFMKLCAALAATMGLSSKAAAEMAESVTNPQRPPVIWIGAQECT WP_000145410.1
MULTISPECIES: hydrogenase-2 small chain [Proteobacteria] Length: 372\nNumber of predicted TMHs: 1\nExp number of AAs in TMHs: 23.72226\nExp number, first 60 AAs: 0.22038\nTotal prob of N-in: 0.12994\noutside 1 331\nTMhelix 332 354\ninside 355 372

27507 GCF_001255095.1_6350_1_13 Shigella sonnei Proteobacteria; Gammaproteobacteria; Enterobacterales; Enterobacteriaceae; Shigella
MTGDNTLIHSHGINRRDFMKLCAALAATMGLSSKAAAEMAESVTNPQRPPVIWIGAQECT WP_000145410.1
MULTISPECIES: hydrogenase-2 small chain [Proteobacteria] Length: 372\nNumber of predicted TMHs: 1\nExp number of AAs in TMHs: 23.72226\nExp number, first 60 AAs: 0.22038\nTotal prob of N-in: 0.12994\noutside 1 331\nTMhelix 332 354\ninside 355 372

27508 GCF_001255175.1_5008_7_5Shigella sonnei Proteobacteria; Gammaproteobacteria; Enterobacterales; Enterobacteriaceae; Shigella MTGDNTLIHSHGINRRDFMKLCAALAATMGLSSKAAAEMAESVTNPQRPPVIWIGAQECT

WP_000145410.1 MULTISPECIES: hydrogenase-2 small chain [Proteobacteria] Length: 372\nNumber of predicted TMHs: 1\nExp number of AAs in TMHs: 23.72226\nExp number, first 60 AAs: 0.22038\nTotal prob of N-in: 0.12994\noutside 1 331\nTMhelix 332 354\ninside 355 372

27509 GCF_001255235.1_8290_4_12 Shigella sonnei Proteobacteria; Gammaproteobacteria; Enterobacterales; Enterobacteriaceae; Shigella MTGDNTLIHSHGINRRDFMKLCAALAATMGLSSKAAAEMAESVTNPQRPPVIWIGAQECT WP_000145410.1 MULTISPECIES: hydrogenase-2 small chain [Proteobacteria] Length: 372\nNumber of predicted TMHs: 1\nExp number of AAs in TMHs: 23.72226\nExp number, first 60 AAs: 0.22038\nTotal prob of N-in: 0.12994\noutside 1 331\nTMhelix 332 354\ninside 355 372

27510 GCF_001255315.1_8290_4_36 Shigella sonnei Proteobacteria; Gammaproteobacteria; Enterobacterales; Enterobacteriaceae; Shigella MTGDNTLIHSHGINRRDFMKLCAALAATMGLSSKAAAEMAESVTNPQRPPVIWIGAQECT WP_000145410.1 MULTISPECIES: hydrogenase-2 small chain [Proteobacteria] Length: 372\nNumber of predicted TMHs: 1\nExp number of AAs in TMHs: 23.72226\nExp number, first 60 AAs: 0.22038\nTotal prob of N-in: 0.12994\noutside 1 331\nTMhelix 332 354\ninside 355 372

27511 GCF_001255315.1_8290_4_36 Shigella sonnei Proteobacteria; Gammaproteobacteria; Enterobacterales; Enterobacteriaceae; Shigella MNNEETFYQAMRRQGVTRRSFLKYCSLAATSLGLGAGMAPKIAWALENKPRIPVVWIHGL WP_005139779.1 hydrogenase [Shigella sonnei] Length: 372\nNumber of predicted TMHs: 2\nExp number of AAs in TMHs: 37.02628\nExp number, first 60 AAs: 19.39684\nTotal prob of N-in: 0.94313\nPOSSIBLE N-term signal sequence\ninside 1 19\nTMhelix 20 39\noutside 40 325\nTMhelix 326 348\ninside 349 372

27512 GCF_001254795.1_12971_8_11 Shigella sonnei Proteobacteria; Gammaproteobacteria; Enterobacterales; Enterobacteriaceae; Shigella MNNEETFYQAMRRQGVTRRSFLKYCSLAATSLGLGAGMAPKIAWALENKPRIPVVWIHGL WP_005139779.1 hydrogenase [Shigella sonnei] Length: 372\nNumber of predicted TMHs: 2\nExp number of AAs in TMHs: 37.02628\nExp number, first 60 AAs: 19.39684\nTotal prob of N-in: 0.94313\nPOSSIBLE N-term signal sequence\ninside 1 19\nTMhelix 20 39\noutside 40 325\nTMhelix 326 348\ninside 349 372

27513 GCF_001255395.1_8290_5_6Shigella sonnei Proteobacteria; Gammaproteobacteria; Enterobacterales; Enterobacteriaceae; Shigella MTGDNTLIHSHGINRRDFMKLCAALAATMGLSSKAAAEMAESVTNPQRPPVIWIGAQECT WP_000145410.1 MULTISPECIES: hydrogenase-2 small chain [Proteobacteria] Length: 372\nNumber of predicted TMHs: 1\nExp number of AAs in TMHs: 23.72226\nExp number, first 60 AAs: 0.22038\nTotal prob of N-in: 0.12994\noutside 1 331\nTMhelix 332 354\ninside 355 372

27514 GCF_001255355.1_Sg_201312549_TCGGCA_L001 Shigella sonnei Proteobacteria; Gammaproteobacteria; Enterobacterales; Enterobacteriaceae; Shigella MTGDNTLIHSHGINRRDFMKLCAALAATMGLSSKAAAEMAESVTNPQRPPVIWIGAQECT WP_000145410.1 MULTISPECIES: hydrogenase-2 small chain [Proteobacteria] Length: 372\nNumber of predicted TMHs: 1\nExp number of AAs in TMHs: 23.72226\nExp number, first 60 AAs: 0.22038\nTotal prob of N-in: 0.12994\noutside 1 331\nTMhelix 332 354\ninside 355 372

27515 GCF_001255475.1_8290_5_7Shigella sonnei Proteobacteria; Gammaproteobacteria; Enterobacterales; Enterobacteriaceae; Shigella MTGDNTLIHSHGINRRDFMKLCAALAATMGLSSKAAAEMAESVTNPQRPPVIWIGAQECT WP_000145410.1 MULTISPECIES: hydrogenase-2 small chain [Proteobacteria] Length: 372\nNumber of predicted TMHs: 1\nExp number of AAs in TMHs: 23.72226\nExp number, first 60 AAs: 0.22038\nTotal prob of N-in: 0.12994\noutside 1 331\nTMhelix 332 354\ninside 355 372

27516 GCF_001255355.1_Sg_201312549_TCGGCA_L001 Shigella sonnei Proteobacteria; Gammaproteobacteria; Enterobacterales; Enterobacteriaceae; Shigella MNNEETFYQAMRRQGVTRRSFLKYCSLAATSLGLGAGMAPKIAWALENKPRIPVVWIHGL WP_005139779.1 hydrogenase [Shigella sonnei] Length: 372\nNumber of predicted TMHs: 2\nExp number of AAs in TMHs: 37.02628\nExp number, first 60 AAs: 19.39684\nTotal prob of N-in: 0.94313\nPOSSIBLE N-term signal sequence\ninside 1 19\nTMhelix 20 39\noutside 40 325\nTMhelix 326 348\ninside 349 372

27517 GCF_001255395.1_8290_5_6Shigella sonnei Proteobacteria; Gammaproteobacteria; Enterobacterales; Enterobacteriaceae; Shigella MNNEETFYQAMRRQGVTRRSFLKYCSLAATSLGLGAGMAPKIAWALENKPRIPVVWIHGL WP_005139779.1 hydrogenase [Shigella sonnei] Length: 372\nNumber of predicted TMHs: 2\nExp

number of AAs in TMHs: 37.02628\nExp number, first 60 AAs: 19.39684\nTotal prob of N-in: 0.94313\nPOSSIBLE N-term signal sequence\ninside 1 19\nTMhelix 20 39\noutside 40 325\nTMhelix 326 348\ninside 349 372

27518 GCF_001254995.1_8289_7_68 Shigella sonnei Proteobacteria; Gammaproteobacteria; Enterobacterales; Enterobacteriaceae; Shigella MNNEETFYQAMRRQGVTRRSFLKYCSLAATSLGLGAGMAPKIAWALENKPRIPVVWIHGL WP_005139779.1 hydrogenase [Shigella sonnei] Length: 372\nNumber of predicted TMHs: 2\nExp number of AAs in TMHs: 37.02628\nExp number, first 60 AAs: 19.39684\nTotal prob of N-in: 0.94313\nPOSSIBLE N-term signal sequence\ninside 1 19\nTMhelix 20 39\noutside 40 325\nTMhelix 326 348\ninside 349 372

27519 GCF_001255055.1_8289_7_49 Shigella sonnei Proteobacteria; Gammaproteobacteria; Enterobacterales; Enterobacteriaceae; Shigella MNNEETFYQAMRRQGVTRRSFLKYCSLAATSLGLGAGMAPKIAWALENKPRIPVVWIHGL WP_005139779.1 hydrogenase [Shigella sonnei] Length: 372\nNumber of predicted TMHs: 2\nExp number of AAs in TMHs: 37.02628\nExp number, first 60 AAs: 19.39684\nTotal prob of N-in: 0.94313\nPOSSIBLE N-term signal sequence\ninside 1 19\nTMhelix 20 39\noutside 40 325\nTMhelix 326 348\ninside 349 372

27520 GCF_001255095.1_6350_1_13 Shigella sonnei Proteobacteria; Gammaproteobacteria; Enterobacterales; Enterobacteriaceae; Shigella MNNEETFYQAMRRQGVTRRSFLKYCSLAATSLGLGAGMAPKIAWALENKPRIPVVWIHGL WP_005139779.1 hydrogenase [Shigella sonnei] Length: 372\nNumber of predicted TMHs: 2\nExp number of AAs in TMHs: 37.02628\nExp number, first 60 AAs: 19.39684\nTotal prob of N-in: 0.94313\nPOSSIBLE N-term signal sequence\ninside 1 19\nTMhelix 20 39\noutside 40 325\nTMhelix 326 348\ninside 349 372

27521 GCF_001254935.1_12971_8_76 Shigella sonnei Proteobacteria; Gammaproteobacteria; Enterobacterales; Enterobacteriaceae; Shigella MNNEETFYQAMRRQGVTRRSFLKYCSLAATSLGLGAGMAPKIAWALENKPRIPVVWIHGL WP_052962775.1 hydrogenase [Shigella sonnei] Length: 372\nNumber of predicted TMHs: 2\nExp number of AAs in TMHs: 37.09248\nExp number, first 60 AAs: 19.39296\nTotal prob of N-in: 0.94316\nPOSSIBLE N-term signal sequence\ninside 1 19\nTMhelix 20 39\noutside 40 325\nTMhelix 326 348\ninside 349 372

27522 GCF_001255175.1_5008_7_5Shigella sonnei Proteobacteria; Gammaproteobacteria; Enterobacterales; Enterobacteriaceae; Shigella MNNEETFYQAMRRQGVTRRSFLKYCSLAATSLGLGAGMAPKIAWALENKPRIPVVWIHGL WP_005139779.1 hydrogenase [Shigella sonnei] Length: 372\nNumber of predicted TMHs: 2\nExp number of AAs in TMHs: 37.02628\nExp number, first 60 AAs: 19.39684\nTotal prob of N-in: 0.94313\nPOSSIBLE N-term signal sequence\ninside 1 19\nTMhelix 20 39\noutside 40 325\nTMhelix 326 348\ninside 349 372

27523 GCF_001255535.1_5417_2_8Shigella sonnei Proteobacteria; Gammaproteobacteria; Enterobacterales; Enterobacteriaceae; Shigella MTGDNTLIHSHGINRRDFMKLCAALAATMGLSSKAAAEMAESVTNPQRPPVIWIGAQECT WP_000145410.1 MULTISPECIES: hydrogenase-2 small chain [Proteobacteria] Length: 372\nNumber of predicted TMHs: 1\nExp number of AAs in TMHs: 23.72226\nExp number, first 60 AAs: 0.22038\nTotal prob of N-in: 0.12994\noutside 1 331\nTMhelix 332 354\ninside 355 372

27524 GCF_001255535.1_5417_2_8Shigella sonnei Proteobacteria; Gammaproteobacteria; Enterobacterales; Enterobacteriaceae; Shigella MNNEETFYQAMRRQGVTRRSFLKYCSLAATSLGLGAGMAPKIAWALENKPRIPVVWIHGL WP_005139779.1 hydrogenase [Shigella sonnei] Length: 372\nNumber of predicted TMHs: 2\nExp number of AAs in TMHs: 37.02628\nExp number, first 60 AAs: 19.39684\nTotal prob of N-in: 0.94313\nPOSSIBLE N-term signal sequence\ninside 1 19\nTMhelix 20 39\noutside 40 325\nTMhelix 326 348\ninside 349 372

27525 GCF_001255235.1_8290_4_12 Shigella sonnei Proteobacteria; Gammaproteobacteria; Enterobacterales; Enterobacteriaceae; Shigella MNNEETFYQAMRRQGVTRRSFLKYCSLAATSLGLGAGMAPKIAWALENKPRIPVVWIHGL WP_005139779.1 hydrogenase [Shigella sonnei] Length: 372\nNumber of predicted TMHs: 2\nExp number of AAs in TMHs: 37.02628\nExp number, first 60 AAs: 19.39684\nTotal prob of N-in: 0.94313\nPOSSIBLE N-term signal sequence\ninside 1 19\nTMhelix 20 39\noutside 40 325\nTMhelix 326 348\ninside 349 372

27526 GCF_001255615.1_5008_8_4Shigella sonnei Proteobacteria; Gammaproteobacteria; Enterobacterales; Enterobacteriaceae; Shigella MTGDNTLIHSHGINRRDFMKLCAALAATMGLSSKAAAEMAESVTNPQRPPVIWIGAQECT WP_000145410.1 MULTISPECIES: hydrogenase-2 small chain [Proteobacteria] Length: 372\nNumber of predicted TMHs: 1\nExp number of AAs in TMHs: 23.72226\nExp number, first 60 AAs: 0.22038\nTotal prob of N-in: 0.12994\noutside 1 331\nTMhelix 332 354\ninside 355 372

27527 GCF_001255655.1_5236_3_3Shigella sonnei Proteobacteria; Gammaproteobacteria; Enterobacterales; Enterobacteriaceae; Shigella MTGDNTLIHSHGINRRDFMKLCAALAATMGLSSKAAAEMAESVTNPQRPPVIWIGAQECT WP_000145410.1 MULTISPECIES: hydrogenase-2 small chain [Proteobacteria] Length: 372\nNumber of predicted TMHs: 1\nExp number of AAs in TMHs: 23.72226\nExp number, first 60 AAs: 0.22038\nTotal prob of N-in: 0.12994\noutside 1 331\nTMhelix 332 354\ninside 355 372

27528 GCF_001255715.1_5236_3_9Shigella sonnei Proteobacteria; Gammaproteobacteria; Enterobacterales; Enterobacteriaceae; Shigella MTGDNTLIHSHGINRRDFMKLCAALAATMGLSSKAAAEMAESVTNPQRPPVIWIGAQECT WP_000145410.1 MULTISPECIES: hydrogenase-2 small chain [Proteobacteria] Length: 372\nNumber of predicted TMHs: 1\nExp number of AAs in TMHs: 23.72226\nExp number, first 60 AAs: 0.22038\nTotal prob of N-in: 0.12994\noutside 1 331\nTMhelix 332 354\ninside 355 372

27529 GCF_001255775.1_12971_8_45 Shigella sonnei Proteobacteria; Gammaproteobacteria; Enterobacterales; Enterobacteriaceae; Shigella MTGDNTLIHSHGINRRDFMKLCAALAATMGLSSKAAAEMAESVTNPQRPPVIWIGAQECT WP_000145410.1 MULTISPECIES: hydrogenase-2 small chain [Proteobacteria] Length: 372\nNumber of predicted TMHs: 1\nExp number of AAs in TMHs: 23.72226\nExp number, first 60 AAs: 0.22038\nTotal prob of N-in: 0.12994\noutside 1 331\nTMhelix 332 354\ninside 355 372

27530 GCF_001255855.1_H14094030901 Shigella sonnei Proteobacteria; Gammaproteobacteria; Enterobacterales; Enterobacteriaceae; Shigella MTGDNTLIHSHGINRRDFMKLCAALAATMGLSSKAAAEMAESVTNPQRPPVIWIGAQECT WP_000145410.1 MULTISPECIES: hydrogenase-2 small chain [Proteobacteria] Length: 372\nNumber of predicted TMHs: 1\nExp number of AAs in TMHs: 23.72226\nExp number, first 60 AAs: 0.22038\nTotal prob of N-in: 0.12994\noutside 1 331\nTMhelix 332 354\ninside 355 372

27531 GCF_001255935.1_12844_1_53 Shigella sonnei Proteobacteria; Gammaproteobacteria; Enterobacterales; Enterobacteriaceae; Shigella MTGDNTLIHSHGINRRDFMKLCAALAATMGLSSKAAAEMAESVTNPQRPPVIWIGAQECT WP_000145410.1 MULTISPECIES: hydrogenase-2 small chain [Proteobacteria] Length: 372\nNumber of predicted TMHs: 1\nExp number of AAs in TMHs: 23.72226\nExp number, first 60 AAs: 0.22038\nTotal prob of N-in: 0.12994\noutside 1 331\nTMhelix 332 354\ninside 355 372

27532 GCF_001255975.1_7655_8_4Shigella sonnei Proteobacteria; Gammaproteobacteria; Enterobacterales; Enterobacteriaceae; Shigella MTGDNTLIHSHGINRRDFMKLCAALAATMGLSSKAAAEMAESVTNPQRPPVIWIGAQECT WP_000145410.1 MULTISPECIES: hydrogenase-2 small chain [Proteobacteria] Length: 372\nNumber of predicted TMHs: 1\nExp number of AAs in TMHs: 23.72226\nExp number, first 60 AAs: 0.22038\nTotal prob of N-in: 0.12994\noutside 1 331\nTMhelix 332 354\ninside 355 372

27533 GCF_001256055.1_9870_6_63 Shigella sonnei Proteobacteria; Gammaproteobacteria; Enterobacterales; Enterobacteriaceae; Shigella MTGDNTLIHSHGINRRDFMKLCAALAATMGLSSKAAAEMAESVTNPQRPPVIWIGAQECT WP_000145410.1 MULTISPECIES: hydrogenase-2 small chain [Proteobacteria] Length: 372\nNumber of predicted TMHs: 1\nExp number of AAs in TMHs: 23.72226\nExp number, first 60 AAs: 0.22038\nTotal prob of N-in: 0.12994\noutside 1 331\nTMhelix 332 354\ninside 355 372

27534 GCF_001256095.1_8290_4_2Shigella sonnei Proteobacteria; Gammaproteobacteria; Enterobacterales; Enterobacteriaceae; Shigella MTGDNTLIHSHGINRRDFMKLCAALAATMGLSSKAAAEMAESVTNPQRPPVIWIGAQECT WP_000145410.1 MULTISPECIES: hydrogenase-2 small chain [Proteobacteria] Length: 372\nNumber of predicted TMHs: 1\nExp number of AAs in TMHs: 23.72226\nExp number, first 60 AAs: 0.22038\nTotal prob of N-in: 0.12994\noutside 1 331\nTMhelix 332 354\ninside 355 372

27535 GCF_001256155.1_H14118090301 Shigella sonnei Proteobacteria; Gammaproteobacteria; Enterobacterales; Enterobacteriaceae; Shigella MTGDNTLIHSHGINRRDFMKLCAALAATMGLSSKAAAEMAESVTNPQRPPVIWIGAQECT WP_000145410.1 MULTISPECIES: hydrogenase-2 small chain [Proteobacteria] Length: 372\nNumber of predicted TMHs: 1\nExp number of AAs in TMHs: 23.72226\nExp number, first 60 AAs: 0.22038\nTotal prob of N-in: 0.12994\noutside 1 331\nTMhelix 332 354\ninside 355 372

27536 GCF_001256055.1_9870_6_63 Shigella sonnei Proteobacteria; Gammaproteobacteria; Enterobacterales; Enterobacteriaceae; Shigella

MNNEETFYQAMRRQGVTRRSFLKYCSLAATSLGLGAGMAPKIAWALENKPRIPVVWIHGL WP_005139779.1 hydrogenase [Shigella sonnei] Length: 372\nNumber of predicted TMHs: 2\nExp number of AAs in TMHs: 37.02628\nExp number, first 60 AAs: 19.39684\nTotal prob of N-in: 0.94313\nPOSSIBLE N-term signal sequence\ninside 1 19\nTMhelix 20 39\noutside 40 325\nTMhelix 326 348\ninside 349 372

27537 GCF_001256095.1_8290_4_2Shigella sonnei Proteobacteria; Gammaproteobacteria; Enterobacterales; Enterobacteriaceae; Shigella MNNEETFYQAMRRQGVTRRSFLKYCSLAATSLGLGAGMAPKIAWALENKPRIPVVWIHGL WP_005139779.1 hydrogenase [Shigella sonnei] Length: 372\nNumber of predicted TMHs: 2\nExp number of AAs in TMHs: 37.02628\nExp number, first 60 AAs: 19.39684\nTotal prob of N-in: 0.94313\nPOSSIBLE N-term signal sequence\ninside 1 19\nTMhelix 20 39\noutside 40 325\nTMhelix 326 348\ninside 349 372

27538 GCF_001256155.1_H14118090301 Shigella sonnei Proteobacteria; Gammaproteobacteria; Enterobacterales; Enterobacteriaceae; Shigella MNNEETFYQAMRRQGVTRRSFLKYCSLAATSLGLGAGMAPKIAWALENKPRIPVVWIHGL WP_052962775.1 hydrogenase [Shigella sonnei] Length: 372\nNumber of predicted TMHs: 2\nExp number of AAs in TMHs: 37.09248\nExp number, first 60 AAs: 19.39296\nTotal prob of N-in: 0.94316\nPOSSIBLE N-term signal sequence\ninside 1 19\nTMhelix 20 39\noutside 40 325\nTMhelix 326 348\ninside 349 372

27539 GCF_001255715.1_5236_3_9Shigella sonnei Proteobacteria; Gammaproteobacteria; Enterobacterales; Enterobacteriaceae; Shigella MNNEETFYQAMRRQGVTRRSFLKYCSLAATSLGLGAGMAPKIAWALENKPRIPVVWIHGL WP_005139779.1 hydrogenase [Shigella sonnei] Length: 372\nNumber of predicted TMHs: 2\nExp number of AAs in TMHs: 37.02628\nExp number, first 60 AAs: 19.39684\nTotal prob of N-in: 0.94313\nPOSSIBLE N-term signal sequence\ninside 1 19\nTMhelix 20 39\noutside 40 325\nTMhelix 326 348\ninside 349 372

27540 GCF_001255655.1_5236_3_3Shigella sonnei Proteobacteria; Gammaproteobacteria; Enterobacterales; Enterobacteriaceae; Shigella MNNEETFYQAMRRQGVTRRSFLKYCSLAATSLGLGAGMAPKIAWALENKPRIPVVWIHGL WP_005139779.1 hydrogenase [Shigella sonnei] Length: 372\nNumber of predicted TMHs: 2\nExp number of AAs in TMHs: 37.02628\nExp number, first 60 AAs: 19.39684\nTotal prob of N-in: 0.94313\nPOSSIBLE N-term signal sequence\ninside 1 19\nTMhelix 20 39\noutside 40 325\nTMhelix 326 348\ninside 349 372

27541 GCF_001255615.1_5008_8_4Shigella sonnei Proteobacteria; Gammaproteobacteria; Enterobacterales; Enterobacteriaceae; Shigella MNNEETFYQAMRRQGVTRRSFLKYCSLAATSLGLGAGMAPKIAWALENKPRIPVVWIHGL WP_005139779.1 hydrogenase [Shigella sonnei] Length: 372\nNumber of predicted TMHs: 2\nExp number of AAs in TMHs: 37.02628\nExp number, first 60 AAs: 19.39684\nTotal prob of N-in: 0.94313\nPOSSIBLE N-term signal sequence\ninside 1 19\nTMhelix 20 39\noutside 40 325\nTMhelix 326 348\ninside 349 372

27542 GCF_001255775.1_12971_8_45 Shigella sonnei Proteobacteria; Gammaproteobacteria; Enterobacterales; Enterobacteriaceae; Shigella MNNEETFYQAMRRQGVTRRSFLKYCSLAATSLGLGAGMAPKIAWALENKPRIPVVWIHGL WP_005139779.1 hydrogenase [Shigella sonnei] Length: 372\nNumber of predicted TMHs: 2\nExp number of AAs in TMHs: 37.02628\nExp number, first 60 AAs: 19.39684\nTotal prob of N-in: 0.94313\nPOSSIBLE N-term signal sequence\ninside 1 19\nTMhelix 20 39\noutside 40 325\nTMhelix 326 348\ninside 349 372

27543 GCF_001256195.1_7655_8_5Shigella sonnei Proteobacteria; Gammaproteobacteria; Enterobacterales; Enterobacteriaceae; Shigella MTGDNTLIHSHGINRRDFMKLCAALAATMGLSSKAAAEMAESVTNPQRPPVIWIGAQECT WP_000145410.1 MULTISPECIES: hydrogenase-2 small chain [Proteobacteria] Length: 372\nNumber of predicted TMHs: 1\nExp number of AAs in TMHs: 23.72226\nExp number, first 60 AAs: 0.22038\nTotal prob of N-in: 0.12994\noutside 1 331\nTMhelix 332 354\ninside 355 372

27544 GCF_001256195.1_7655_8_5Shigella sonnei Proteobacteria; Gammaproteobacteria; Enterobacterales; Enterobacteriaceae; Shigella MNNEETFYQAMRRQGVTRRSFLKYCSLAATSLGLGAGMAPKIAWALENKPRIPVVWIHGL WP_005139779.1 hydrogenase [Shigella sonnei] Length: 372\nNumber of predicted TMHs: 2\nExp number of AAs in TMHs: 37.02628\nExp number, first 60 AAs: 19.39684\nTotal prob of N-in: 0.94313\nPOSSIBLE N-term signal sequence\ninside 1 19\nTMhelix 20 39\noutside 40 325\nTMhelix 326 348\ninside 349 372

27545 GCF_001255855.1_H14094030901 Shigella sonnei Proteobacteria; Gammaproteobacteria; Enterobacterales; Enterobacteriaceae; Shigella MNNEETFYQAMRRQGVTRRSFLKYCSLAATSLGLGAGMAPKIAWALENKPRIPVVWIHGL WP_005139779.1 hydrogenase [Shigella sonnei] Length: 372\nNumber of predicted TMHs: 2\nExp number of AAs in TMHs: 37.02628\nExp number, first 60 AAs: 19.39684\nTotal prob of N-in: 0.94313\nPOSSIBLE N-term signal sequence\ninside 1 19\nTMhelix 20 39\noutside 40 325\nTMhelix 326 348\ninside 349 372

27546 GCF_001255935.1_12844_1_53 Shigella sonnei Proteobacteria; Gammaproteobacteria; Enterobacterales; Enterobacteriaceae; Shigella MNNEETFYQAMRRQGVTRRSFLKYCSLAATSLGLGAGMAPKIAWALENKPRIPVVWIHGL WP_005139779.1 hydrogenase [Shigella sonnei] Length: 372\nNumber of predicted TMHs: 2\nExp number of AAs in TMHs: 37.02628\nExp number, first 60 AAs: 19.39684\nTotal prob of N-in: 0.94313\nPOSSIBLE N-term signal sequence\ninside 1 19\nTMhelix 20 39\noutside 40 325\nTMhelix 326 348\ninside 349 372

27547 GCF_001255975.1_7655_8_4Shigella sonnei Proteobacteria; Gammaproteobacteria; Enterobacterales; Enterobacteriaceae; Shigella MNNEETFYQAMRRQGVTRRSFLKYCSLAATSLGLGAGMAPKIAWALENKPRIPVVWIHGL WP_005139779.1 hydrogenase [Shigella sonnei] Length: 372\nNumber of predicted TMHs: 2\nExp number of AAs in TMHs: 37.02628\nExp number, first 60 AAs: 19.39684\nTotal prob of N-in: 0.94313\nPOSSIBLE N-term signal sequence\ninside 1 19\nTMhelix 20 39\noutside 40 325\nTMhelix 326 348\ninside 349 372

27548 GCF_001256255.1_7655_8_10 Shigella sonnei Proteobacteria; Gammaproteobacteria; Enterobacterales; Enterobacteriaceae; Shigella MTGDNTLIHSHGINRRDFMKLCAALAATMGLSSKAAAEMAESVTNPQRPPVIWIGAQECT WP_000145410.1 MULTISPECIES: hydrogenase-2 small chain [Proteobacteria] Length: 372\nNumber of predicted TMHs: 1\nExp number of AAs in TMHs: 23.72226\nExp number, first 60 AAs: 0.22038\nTotal prob of N-in: 0.12994\noutside 1 331\nTMhelix 332 354\ninside 355 372

27549 GCF_001256275.1_6350_1_17 Shigella sonnei Proteobacteria; Gammaproteobacteria; Enterobacterales; Enterobacteriaceae; Shigella MTGDNTLIHSHGINRRDFMKLCAALAATMGLSSKAAAEMAESVTNPQRPPVIWIGAQECT WP_000145410.1 MULTISPECIES: hydrogenase-2 small chain [Proteobacteria] Length: 372\nNumber of predicted TMHs: 1\nExp number of AAs in TMHs: 23.72226\nExp number, first 60 AAs: 0.22038\nTotal prob of N-in: 0.12994\noutside 1 331\nTMhelix 332 354\ninside 355 372

27550 GCF_001256335.1_H14072036301 Shigella sonnei Proteobacteria; Gammaproteobacteria; Enterobacterales; Enterobacteriaceae; Shigella MTGDNTLIHSHGINRRDFMKLCAALAATMGLSSKAAAEMAESVTNPQRPPVIWIGAQECT WP_000145410.1 MULTISPECIES: hydrogenase-2 small chain [Proteobacteria] Length: 372\nNumber of predicted TMHs: 1\nExp number of AAs in TMHs: 23.72226\nExp number, first 60 AAs: 0.22038\nTotal prob of N-in: 0.12994\noutside 1 331\nTMhelix 332 354\ninside 355 372

27551 GCF_001256395.1_12971_8_25 Shigella sonnei Proteobacteria; Gammaproteobacteria; Enterobacterales; Enterobacteriaceae; Shigella MTGDNTLIHSHGINRRDFMKLCAALAATMGLSSKAAAEMAESVTNPQRPPVIWIGAQECT WP_000145410.1 MULTISPECIES: hydrogenase-2 small chain [Proteobacteria] Length: 372\nNumber of predicted TMHs: 1\nExp number of AAs in TMHs: 23.72226\nExp number, first 60 AAs: 0.22038\nTotal prob of N-in: 0.12994\noutside 1 331\nTMhelix 332 354\ninside 355 372

27552 GCF_001256455.1_Sg_201400526_ATCACG_L001 Shigella sonnei Proteobacteria; Gammaproteobacteria; Enterobacterales; Enterobacteriaceae; Shigella MTGDNTLIHSHGINRRDFMKLCAALAATMGLSSKAAAEMAESVTNPQRPPVIWIGAQECT WP_000145410.1 MULTISPECIES: hydrogenase-2 small chain [Proteobacteria] Length: 372\nNumber of predicted TMHs: 1\nExp number of AAs in TMHs: 23.72226\nExp number, first 60 AAs: 0.22038\nTotal prob of N-in: 0.12994\noutside 1 331\nTMhelix 332 354\ninside 355 372

27553 GCF_001256515.1_5236_7_3Shigella sonnei Proteobacteria; Gammaproteobacteria; Enterobacterales; Enterobacteriaceae; Shigella MTGDNTLIHSHGINRRDFMKLCAALAATMGLSSKAAAEMAESVTNPQRPPVIWIGAQECT WP_000145410.1 MULTISPECIES: hydrogenase-2 small chain [Proteobacteria] Length: 372\nNumber of predicted TMHs: 1\nExp number of AAs in TMHs: 23.72226\nExp number, first 60 AAs: 0.22038\nTotal prob of N-in: 0.12994\noutside 1 331\nTMhelix 332 354\ninside 355 372

27554 GCF_001256575.1_8489_2_89 Shigella sonnei Proteobacteria; Gammaproteobacteria; Enterobacterales; Enterobacteriaceae; Shigella MTGDNTLIHSHGINRRDFMKLCAALAATMGLSSKAAAEMAESVTNPQRPPVIWIGAQECT WP_000145410.1 MULTISPECIES: hydrogenase-2 small chain [Proteobacteria] Length: 372\nNumber of predicted TMHs: 1\nExp number of AAs in TMHs: 23.72226\nExp number, first 60 AAs: 0.22038\nTotal prob of N-in: 0.12994\noutside 1 331\nTMhelix 332 354\ninside 355 372

27555 GCF_001256715.1_5236_6_3Shigella sonnei Proteobacteria; Gammaproteobacteria; Enterobacterales; Enterobacteriaceae; Shigella MTGDNTLIHSHGINRRDFMKLCAALAATMGLSSKAAAEMAESVTNPQRPPVIWIGAQECT WP_000145410.1 MULTISPECIES: hydrogenase-2 small chain [Proteobacteria] Length: 372\nNumber of predicted TMHs: 1\nExp number of AAs in TMHs: 23.72226\nExp number, first 60 AAs: 0.22038\nTotal prob of N-in: 0.12994\noutside 1 331\nTMhelix 332 354\ninside 355 372

27556 GCF_001256715.1_5236_6_3Shigella sonnei Proteobacteria; Gammaproteobacteria; Enterobacterales; Enterobacteriaceae; Shigella MNNEETFYQAMRRQGVTRRSFLKYCSLAATSLGLGAGMAPKIAWALENKPRIPVVWIHGL WP_005139779.1 hydrogenase [Shigella sonnei] Length: 372\nNumber of predicted TMHs: 2\nExp number of AAs in TMHs: 37.02628\nExp number, first 60 AAs: 19.39684\nTotal prob of N-in: 0.94313\nPOSSIBLE N-term signal sequence\ninside 1 19\nTMhelix 20 39\noutside 40 325\nTMhelix 326 348\ninside 349 372

27557 GCF_001256275.1_6350_1_17 Shigella sonnei Proteobacteria; Gammaproteobacteria; Enterobacterales; Enterobacteriaceae; Shigella MNNEETFYQAMRRQGVTRRSFLKYCSLAATSLGLGAGMAPKIAWALENKPRIPVVWIHGL WP_005139779.1 hydrogenase [Shigella sonnei] Length: 372\nNumber of predicted TMHs: 2\nExp number of AAs in TMHs: 37.02628\nExp number, first 60 AAs: 19.39684\nTotal prob of N-in: 0.94313\nPOSSIBLE N-term signal sequence\ninside 1 19\nTMhelix 20 39\noutside 40 325\nTMhelix 326 348\ninside 349 372

27558 GCF_001256855.1_8289_7_91 Shigella sonnei Proteobacteria; Gammaproteobacteria; Enterobacterales; Enterobacteriaceae; Shigella MTGDNTLIHSHGINRRDFMKLCAALAATMGLSSKAAAEMAESVTNPQRPPVIWIGAQECT WP_000145410.1 MULTISPECIES: hydrogenase-2 small chain [Proteobacteria] Length: 372\nNumber of predicted TMHs: 1\nExp number of AAs in TMHs: 23.72226\nExp number, first 60 AAs: 0.22038\nTotal prob of N-in: 0.12994\noutside 1 331\nTMhelix 332 354\ninside 355 372

27559 GCF_001256755.1_6551_1_4Shigella sonnei Proteobacteria; Gammaproteobacteria; Enterobacterales; Enterobacteriaceae; Shigella MTGDNTLIHSHGINRRDFMKLCAALAATMGLSSKAAAEMAESVTNPQRPPVIWIGAQECT WP_000145410.1 MULTISPECIES: hydrogenase-2 small chain [Proteobacteria] Length: 372\nNumber of predicted TMHs: 1\nExp number of AAs in TMHs: 23.72226\nExp number, first 60 AAs: 0.22038\nTotal prob of N-in: 0.12994\noutside 1 331\nTMhelix 332 354\ninside 355 372

27560 GCF_001256895.1_9870_6_38 Shigella sonnei Proteobacteria; Gammaproteobacteria; Enterobacterales; Enterobacteriaceae; Shigella MTGDNTLIHSHGINRRDFMKLCAALAATMGLSSKAAAEMAESVTNPQRPPVIWIGAQECT WP_000145410.1 MULTISPECIES: hydrogenase-2 small chain [Proteobacteria] Length: 372\nNumber of predicted TMHs: 1\nExp number of AAs in TMHs: 23.72226\nExp number, first 60 AAs: 0.22038\nTotal prob of N-in: 0.12994\noutside 1 331\nTMhelix 332 354\ninside 355 372

27561 GCF_001256755.1_6551_1_4Shigella sonnei Proteobacteria; Gammaproteobacteria; Enterobacterales; Enterobacteriaceae; Shigella MNNEETFYQAMRRQGVTRRSFLKYCSLAATSLGLGAGMAPKIAWALENKPRIPVVWIHGL WP_005139779.1 hydrogenase [Shigella sonnei] Length: 372\nNumber of predicted TMHs: 2\nExp number of AAs in TMHs: 37.02628\nExp number, first 60 AAs: 19.39684\nTotal prob of N-in: 0.94313\nPOSSIBLE N-term signal sequence\ninside 1 19\nTMhelix 20 39\noutside 40 325\nTMhelix 326 348\ninside 349 372

27562 GCF_001256855.1_8289_7_91 Shigella sonnei Proteobacteria; Gammaproteobacteria; Enterobacterales; Enterobacteriaceae; Shigella MNNEETFYQAMRRQGVTRRSFLKYCSLAATSLGLGAGMAPKIAWALENKPRIPVVWIHGL WP_005139779.1 hydrogenase [Shigella sonnei] Length: 372\nNumber of predicted TMHs: 2\nExp number of AAs in TMHs: 37.02628\nExp number, first 60 AAs: 19.39684\nTotal prob of N-in: 0.94313\nPOSSIBLE N-term signal sequence\ninside 1 19\nTMhelix 20 39\noutside 40 325\nTMhelix 326 348\ninside 349 372

27563 GCF_001256895.1_9870_6_38 Shigella sonnei Proteobacteria; Gammaproteobacteria; Enterobacterales; Enterobacteriaceae; Shigella MNNEETFYQAMRRQGVTRRSFLKYCSLAATSLGLGAGMAPKIAWALENKPRIPVVWIHGL WP_005139779.1 hydrogenase [Shigella sonnei] Length: 372\nNumber of predicted TMHs: 2\nExp number of AAs in TMHs: 37.02628\nExp number, first 60 AAs: 19.39684\nTotal prob of N-in: 0.94313\nPOSSIBLE N-term signal sequence\ninside 1 19\nTMhelix 20 39\noutside 40 325\nTMhelix 326 348\ninside 349 372

27564 GCF_001256255.1_7655_8_10 Shigella sonnei Proteobacteria; Gammaproteobacteria; Enterobacterales; Enterobacteriaceae; Shigella

MNNEETFYQAMRRQGVTRRSFLKYCSLAATSLGLGAGMAPKIAWALENKPRIPVVWIHGL WP_005139779.1 hydrogenase [Shigella sonnei] Length: 372\nNumber of predicted TMHs: 2\nExp number of AAs in TMHs: 37.02628\nExp number, first 60 AAs: 19.39684\nTotal prob of N-in: 0.94313\nPOSSIBLE N-term signal sequence\ninside 1 19\nTMhelix 20 39\noutside 40 325\nTMhelix 326 348\ninside 349 372

27565 GCF_001256955.1_12971_8_10 Shigella sonnei Proteobacteria; Gammaproteobacteria; Enterobacterales; Enterobacteriaceae; Shigella MTGDNTLIHSHGINRRDFMKLCAALAATMGLSSKAAAEMAESVTNPQRPPVIWIGAQECT WP_000145410.1 MULTISPECIES: hydrogenase-2 small chain [Proteobacteria] Length: 372\nNumber of predicted TMHs: 1\nExp number of AAs in TMHs: 23.72226\nExp number, first 60 AAs: 0.22038\nTotal prob of N-in: 0.12994\noutside 1 331\nTMhelix 332 354\ninside 355 372

27566 GCF_001256955.1_12971_8_10 Shigella sonnei Proteobacteria; Gammaproteobacteria; Enterobacterales; Enterobacteriaceae; Shigella MNNEETFYQAMRRQGVTRRSFLKYCSLAATSLGLGAGMAPKIAWALENKPRIPVVWIHGL WP_005139779.1 hydrogenase [Shigella sonnei] Length: 372\nNumber of predicted TMHs: 2\nExp number of AAs in TMHs: 37.02628\nExp number, first 60 AAs: 19.39684\nTotal prob of N-in: 0.94313\nPOSSIBLE N-term signal sequence\ninside 1 19\nTMhelix 20 39\noutside 40 325\nTMhelix 326 348\ninside 349 372

27567 GCF_001256335.1_H14072036301 Shigella sonnei Proteobacteria; Gammaproteobacteria; Enterobacterales; Enterobacteriaceae; Shigella MNNEETFYQAMRRQGVTRRSFLKYCSLAATSLGLGAGMAPKIAWALENKPRIPVVWIHGL WP_005139779.1 hydrogenase [Shigella sonnei] Length: 372\nNumber of predicted TMHs: 2\nExp number of AAs in TMHs: 37.02628\nExp number, first 60 AAs: 19.39684\nTotal prob of N-in: 0.94313\nPOSSIBLE N-term signal sequence\ninside 1 19\nTMhelix 20 39\noutside 40 325\nTMhelix 326 348\ninside 349 372

27568 GCF_001256455.1_Sg_201400526_ATCACG_L001 Shigella sonnei Proteobacteria; Gammaproteobacteria; Enterobacterales; Enterobacteriaceae; Shigella MNNEETFYQAMRRQGVTRRSFLKYCSLAATSLGLGAGMAPKIAWALENKPRIPVVWIHGL WP_005139779.1 hydrogenase [Shigella sonnei] Length: 372\nNumber of predicted TMHs: 2\nExp number of AAs in TMHs: 37.02628\nExp number, first 60 AAs: 19.39684\nTotal prob of N-in: 0.94313\nPOSSIBLE N-term signal sequence\ninside 1 19\nTMhelix 20 39\noutside 40 325\nTMhelix 326 348\ninside 349 372

27569 GCF_001256395.1_12971_8_25 Shigella sonnei Proteobacteria; Gammaproteobacteria; Enterobacterales; Enterobacteriaceae; Shigella MNNEETFYQAMRRQGVTRRSFLKYCSLAATSLGLGAGMAPKIAWALENKPRIPVVWIHGL WP_005139779.1 hydrogenase [Shigella sonnei] Length: 372\nNumber of predicted TMHs: 2\nExp number of AAs in TMHs: 37.02628\nExp number, first 60 AAs: 19.39684\nTotal prob of N-in: 0.94313\nPOSSIBLE N-term signal sequence\ninside 1 19\nTMhelix 20 39\noutside 40 325\nTMhelix 326 348\ninside 349 372

27570 GCF_001256515.1_5236_7_3Shigella sonnei Proteobacteria; Gammaproteobacteria; Enterobacterales; Enterobacteriaceae; Shigella MNNEETFYQAMRRQGVTRRSFLKYCSLAATSLGLGAGMAPKIAWALENKPRIPVVWIHGL WP_005139779.1 hydrogenase [Shigella sonnei] Length: 372\nNumber of predicted TMHs: 2\nExp number of AAs in TMHs: 37.02628\nExp number, first 60 AAs: 19.39684\nTotal prob of N-in: 0.94313\nPOSSIBLE N-term signal sequence\ninside 1 19\nTMhelix 20 39\noutside 40 325\nTMhelix 326 348\ninside 349 372

27571 GCF_001256575.1_8489_2_89 Shigella sonnei Proteobacteria; Gammaproteobacteria; Enterobacterales; Enterobacteriaceae; Shigella MNNEETFYQAMRRQGVTRRSFLKYCSLAATSLGLGAGMAPKIAWALENKPRIPVVWIHGL WP_005139779.1 hydrogenase [Shigella sonnei] Length: 372\nNumber of predicted TMHs: 2\nExp number of AAs in TMHs: 37.02628\nExp number, first 60 AAs: 19.39684\nTotal prob of N-in: 0.94313\nPOSSIBLE N-term signal sequence\ninside 1 19\nTMhelix 20 39\noutside 40 325\nTMhelix 326 348\ninside 349 372

27572 GCF_001257015.1_12971_8_74 Shigella sonnei Proteobacteria; Gammaproteobacteria; Enterobacterales; Enterobacteriaceae; Shigella MTGDNTLIHSHGINRRDFMKLCAALAATMGLSSKAAAEMAESVTNPQRPPVIWIGAQECT WP_000145410.1 MULTISPECIES: hydrogenase-2 small chain [Proteobacteria] Length: 372\nNumber of predicted TMHs: 1\nExp number of AAs in TMHs: 23.72226\nExp number, first 60 AAs: 0.22038\nTotal prob of N-in: 0.12994\noutside 1 331\nTMhelix 332 354\ninside 355 372

27573	GCF_001257115.1_7655_8_7Shigella sonnei	Proteobacteria; Gammaproteobacteria; Enterobacterales; Enterobacteriaceae; Shigella	MTGDNTLIHSHGINRRDFMKLCAALAATMGLSSKAAAEMAESVTNPQRPPVIWIGAQECT	WP_000145410.1	MULTISPECIES: hydrogenase-2 small chain [Proteobacteria]	Length: 372\nNumber of predicted TMHs: 1\nExp number of AAs in TMHs: 23.72226\nExp number, first 60 AAs: 0.22038\nTotal prob of N-in: 0.12994\noutside 1 331\nTMhelix 332 354\ninside 355 372

27574	GCF_001257235.1_6551_1_7Shigella sonnei	Proteobacteria; Gammaproteobacteria; Enterobacterales; Enterobacteriaceae; Shigella	MTGDNTLIHSHGINRRDFMKLCAALAATMGLSSKAAAEMAESVTNPQRPPVIWIGAQECT	WP_000145410.1	MULTISPECIES: hydrogenase-2 small chain [Proteobacteria]	Length: 372\nNumber of predicted TMHs: 1\nExp number of AAs in TMHs: 23.72226\nExp number, first 60 AAs: 0.22038\nTotal prob of N-in: 0.12994\noutside 1 331\nTMhelix 332 354\ninside 355 372

27575	GCF_001257155.1_12971_8_67	Shigella sonnei	Proteobacteria; Gammaproteobacteria; Enterobacterales; Enterobacteriaceae; Shigella	MTGDNTLIHSHGINRRDFMKLCAALAATMGLSSKAAAEMAESVTNPQRPPVIWIGAQECT	WP_000145410.1	MULTISPECIES: hydrogenase-2 small chain [Proteobacteria]	Length: 372\nNumber of predicted TMHs: 1\nExp number of AAs in TMHs: 23.72226\nExp number, first 60 AAs: 0.22038\nTotal prob of N-in: 0.12994\noutside 1 331\nTMhelix 332 354\ninside 355 372

27576	GCF_001257295.1_Sg_201405079_CATGGC_L001	Shigella sonnei	Proteobacteria; Gammaproteobacteria; Enterobacterales; Enterobacteriaceae; Shigella	MTGDNTLIHSHGINRRDFMKLCAALAATMGLSSKAAAEMAESVTNPQRPPVIWIGAQECT	WP_000145410.1	MULTISPECIES: hydrogenase-2 small chain [Proteobacteria]	Length: 372\nNumber of predicted TMHs: 1\nExp number of AAs in TMHs: 23.72226\nExp number, first 60 AAs: 0.22038\nTotal prob of N-in: 0.12994\noutside 1 331\nTMhelix 332 354\ninside 355 372

27577	GCF_001257315.1_12971_8_70	Shigella sonnei	Proteobacteria; Gammaproteobacteria; Enterobacterales; Enterobacteriaceae; Shigella	MTGDNTLIHSHGINRRDFMKLCAALAATMGLSSKAAAEMAESVTNPQRPPVIWIGAQECT	WP_000145410.1	MULTISPECIES: hydrogenase-2 small chain [Proteobacteria]	Length: 372\nNumber of predicted TMHs: 1\nExp number of AAs in TMHs: 23.72226\nExp number, first 60 AAs: 0.22038\nTotal prob of N-in: 0.12994\noutside 1 331\nTMhelix 332 354\ninside 355 372

27578	GCF_001257355.1_9870_6_2Shigella sonnei	Proteobacteria; Gammaproteobacteria; Enterobacterales; Enterobacteriaceae; Shigella	MTGDNTLIHSHGINRRDFMKLCAALAATMGLSSKAAAEMAESVTNPQRPPVIWIGAQECT	WP_000145410.1	MULTISPECIES: hydrogenase-2 small chain [Proteobacteria]	Length: 372\nNumber of predicted TMHs: 1\nExp number of AAs in TMHs: 23.72226\nExp number, first 60 AAs: 0.22038\nTotal prob of N-in: 0.12994\noutside 1 331\nTMhelix 332 354\ninside 355 372

27579	GCF_001257395.1_H14138024801	Shigella sonnei	Proteobacteria; Gammaproteobacteria; Enterobacterales; Enterobacteriaceae; Shigella	MTGDNTLIHSHGINRRDFMKLCAALAATMGLSSKAAAEMAESVTNPQRPPVIWIGAQECT	WP_000145410.1	MULTISPECIES: hydrogenase-2 small chain [Proteobacteria]	Length: 372\nNumber of predicted TMHs: 1\nExp number of AAs in TMHs: 23.72226\nExp number, first 60 AAs: 0.22038\nTotal prob of N-in: 0.12994\noutside 1 331\nTMhelix 332 354\ninside 355 372

27580	GCF_001257395.1_H14138024801	Shigella sonnei	Proteobacteria; Gammaproteobacteria; Enterobacterales; Enterobacteriaceae; Shigella	MNNEETFYQAMRRQGVTRRSFLKYCSLAATSLGLGAGMAPKIAWALENKPRIPVVWIHGL	WP_052962775.1	hydrogenase [Shigella sonnei]	Length: 372\nNumber of predicted TMHs: 2\nExp number of AAs in TMHs: 37.09248\nExp number, first 60 AAs: 19.39296\nTotal prob of N-in: 0.94316\nPOSSIBLE N-term signal sequence\ninside 1 19\nTMhelix 20 39\noutside 40 325\nTMhelix 326 348\ninside 349 372

27581	GCF_001257015.1_12971_8_74	Shigella sonnei	Proteobacteria; Gammaproteobacteria; Enterobacterales; Enterobacteriaceae; Shigella	MNNEETFYQAMRRQGVTRRSFLKYCSLAATSLGLGAGMAPKIAWALENKPRIPVVWIHGL	WP_005139779.1	hydrogenase [Shigella sonnei]	Length: 372\nNumber of predicted TMHs: 2\nExp number of AAs in TMHs: 37.02628\nExp number, first 60 AAs: 19.39684\nTotal prob of N-in: 0.94313\nPOSSIBLE N-term signal sequence\ninside 1 19\nTMhelix 20 39\noutside 40 325\nTMhelix 326 348\ninside 349 372

27582 GCF_001257435.1_5236_1_8Shigella sonnei Proteobacteria; Gammaproteobacteria; Enterobacterales; Enterobacteriaceae; Shigella MTGDNTLIHSHGINRRDFMKLCAALAATMGLSSKAAAEMAESVTNPQRPPVIWIGAQECT WP_000145410.1 MULTISPECIES: hydrogenase-2 small chain [Proteobacteria] Length: 372\nNumber of predicted TMHs: 1\nExp number of AAs in TMHs: 23.72226\nExp number, first 60 AAs: 0.22038\nTotal prob of N-in: 0.12994\noutside 1 331\nTMhelix 332 354\ninside 355 372

27583 GCF_001257435.1_5236_1_8Shigella sonnei Proteobacteria; Gammaproteobacteria; Enterobacterales; Enterobacteriaceae; Shigella MNNEETFYQAMRRQGVTRRSFLKYCSLAATSLGLGAGMAPKIAWALENKPRIPVVWIHGL WP_005139779.1 hydrogenase [Shigella sonnei] Length: 372\nNumber of predicted TMHs: 2\nExp number of AAs in TMHs: 37.02628\nExp number, first 60 AAs: 19.39684\nTotal prob of N-in: 0.94313\nPOSSIBLE N-term signal sequence\ninside 1 19\nTMhelix 20 39\noutside 40 325\nTMhelix 326 348\ninside 349 372

27584 GCF_001257115.1_7655_8_7Shigella sonnei Proteobacteria; Gammaproteobacteria; Enterobacterales; Enterobacteriaceae; Shigella MNNEETFYQAMRRQGVTRRSFLKYCSLAATSLGLGAGMAPKIAWALENKPRIPVVWIHGL WP_005139779.1 hydrogenase [Shigella sonnei] Length: 372\nNumber of predicted TMHs: 2\nExp number of AAs in TMHs: 37.02628\nExp number, first 60 AAs: 19.39684\nTotal prob of N-in: 0.94313\nPOSSIBLE N-term signal sequence\ninside 1 19\nTMhelix 20 39\noutside 40 325\nTMhelix 326 348\ninside 349 372

27585 GCF_001257645.1_8289_7_44 Shigella sonnei Proteobacteria; Gammaproteobacteria; Enterobacterales; Enterobacteriaceae; Shigella MTGDNTLIHSHGINRRDFMKLCAALAATMGLSSKAAAEMAESVTNPQRPPVIWIGAQECT WP_000145410.1 MULTISPECIES: hydrogenase-2 small chain [Proteobacteria] Length: 372\nNumber of predicted TMHs: 1\nExp number of AAs in TMHs: 23.72226\nExp number, first 60 AAs: 0.22038\nTotal prob of N-in: 0.12994\noutside 1 331\nTMhelix 332 354\ninside 355 372

27586 GCF_001257735.1_12971_8_19 Shigella sonnei Proteobacteria; Gammaproteobacteria; Enterobacterales; Enterobacteriaceae; Shigella MTGDNTLIHSHGINRRDFMKLCAALAATMGLSSKAAAEMAESVTNPQRPPVIWIGAQECT WP_000145410.1 MULTISPECIES: hydrogenase-2 small chain [Proteobacteria] Length: 372\nNumber of predicted TMHs: 1\nExp number of AAs in TMHs: 23.72226\nExp number, first 60 AAs: 0.22038\nTotal prob of N-in: 0.12994\noutside 1 331\nTMhelix 332 354\ninside 355 372

27587 GCF_001257735.1_12971_8_19 Shigella sonnei Proteobacteria; Gammaproteobacteria; Enterobacterales; Enterobacteriaceae; Shigella MNNEETFYQAMRRQGVTRRSFLKYCSLAATSLGLGAGMAPKIAWALENKPRIPVVWIHGL WP_005139779.1 hydrogenase [Shigella sonnei] Length: 372\nNumber of predicted TMHs: 2\nExp number of AAs in TMHs: 37.02628\nExp number, first 60 AAs: 19.39684\nTotal prob of N-in: 0.94313\nPOSSIBLE N-term signal sequence\ninside 1 19\nTMhelix 20 39\noutside 40 325\nTMhelix 326 348\ninside 349 372

27588 GCF_001257645.1_8289_7_44 Shigella sonnei Proteobacteria; Gammaproteobacteria; Enterobacterales; Enterobacteriaceae; Shigella MNNEETFYQAMRRQGVTRRSFLKYCSLAATSLGLGAGMAPKIAWALENKPRIPVVWIHGL WP_005139779.1 hydrogenase [Shigella sonnei] Length: 372\nNumber of predicted TMHs: 2\nExp number of AAs in TMHs: 37.02628\nExp number, first 60 AAs: 19.39684\nTotal prob of N-in: 0.94313\nPOSSIBLE N-term signal sequence\ninside 1 19\nTMhelix 20 39\noutside 40 325\nTMhelix 326 348\ninside 349 372

27589 GCF_001257235.1_6551_1_7Shigella sonnei Proteobacteria; Gammaproteobacteria; Enterobacterales; Enterobacteriaceae; Shigella MNNEETFYQAMRRQGVTRRSFLKYCSLAATSLGLGAGMAPKIAWALENKPRIPVVWIHGL WP_005139779.1 hydrogenase [Shigella sonnei] Length: 372\nNumber of predicted TMHs: 2\nExp number of AAs in TMHs: 37.02628\nExp number, first 60 AAs: 19.39684\nTotal prob of N-in: 0.94313\nPOSSIBLE N-term signal sequence\ninside 1 19\nTMhelix 20 39\noutside 40 325\nTMhelix 326 348\ninside 349 372

27590 GCF_001257155.1_12971_8_67 Shigella sonnei Proteobacteria; Gammaproteobacteria; Enterobacterales; Enterobacteriaceae; Shigella MNNEETFYQAMRRQGVTRRSFLKYCSLAATSLGLGAGMAPKIAWALENKPRIPVVWIHGL WP_005139779.1 hydrogenase [Shigella sonnei] Length: 372\nNumber of predicted TMHs: 2\nExp number of AAs in TMHs: 37.02628\nExp number, first 60 AAs: 19.39684\nTotal prob of N-in: 0.94313\nPOSSIBLE N-term signal sequence\ninside 1 19\nTMhelix 20 39\noutside 40 325\nTMhelix 326 348\ninside 349 372

27591 GCF_001257775.1_8289_7_48 Shigella sonnei Proteobacteria; Gammaproteobacteria; Enterobacterales; Enterobacteriaceae; Shigella

MTGDNTLIHSHGINRRDFMKLCAALAATMGLSSKAAAEMAESVTNPQRPPVIWIGAQECT WP_000145410.1 MULTISPECIES: hydrogenase-2 small chain [Proteobacteria] Length: 372\nNumber of predicted TMHs: 1\nExp number of AAs in TMHs: 23.72226\nExp number, first 60 AAs: 0.22038\nTotal prob of N-in: 0.12994\noutside 1 331\nTMhelix 332 354\ninside 355 372

27592 GCF_001257775.1_8289_7_48 Shigella sonnei Proteobacteria; Gammaproteobacteria; Enterobacterales; Enterobacteriaceae; Shigella MNNEETFYQAMRRQGVTRRSFLKYCSLAATSLGLGAGMAPKIAWALENKPRIPVVWIHGL WP_005139779.1 hydrogenase [Shigella sonnei] Length: 372\nNumber of predicted TMHs: 2\nExp number of AAs in TMHs: 37.02628\nExp number, first 60 AAs: 19.39684\nTotal prob of N-in: 0.94313\nPOSSIBLE N-term signal sequence\ninside 1 19\nTMhelix 20 39\noutside 40 325\nTMhelix 326 348\ninside 349 372

27593 GCF_001257295.1_Sg_201405079_CATGGC_L001 Shigella sonnei Proteobacteria; Gammaproteobacteria; Enterobacterales; Enterobacteriaceae; Shigella MNNEETFYQAMRRQGVTRRSFLKYCSLAATSLGLGAGMAPKIAWALENKPRIPVVWIHGL WP_005139779.1 hydrogenase [Shigella sonnei] Length: 372\nNumber of predicted TMHs: 2\nExp number of AAs in TMHs: 37.02628\nExp number, first 60 AAs: 19.39684\nTotal prob of N-in: 0.94313\nPOSSIBLE N-term signal sequence\ninside 1 19\nTMhelix 20 39\noutside 40 325\nTMhelix 326 348\ninside 349 372

27594 GCF_001257315.1_12971_8_70 Shigella sonnei Proteobacteria; Gammaproteobacteria; Enterobacterales; Enterobacteriaceae; Shigella MNNEETFYQAMRRQGVTRRSFLKYCSLAATSLGLGAGMAPKIAWALENKPRIPVVWIHGL WP_005139779.1 hydrogenase [Shigella sonnei] Length: 372\nNumber of predicted TMHs: 2\nExp number of AAs in TMHs: 37.02628\nExp number, first 60 AAs: 19.39684\nTotal prob of N-in: 0.94313\nPOSSIBLE N-term signal sequence\ninside 1 19\nTMhelix 20 39\noutside 40 325\nTMhelix 326 348\ninside 349 372

27595 GCF_001257355.1_9870_6_2Shigella sonnei Proteobacteria; Gammaproteobacteria; Enterobacterales; Enterobacteriaceae; Shigella MNNEETFYQAMRRQGVTRRSFLKYCSLAATSLGLGAGMAPKIAWALENKPRIPVVWIHGL WP_005139779.1 hydrogenase [Shigella sonnei] Length: 372\nNumber of predicted TMHs: 2\nExp number of AAs in TMHs: 37.02628\nExp number, first 60 AAs: 19.39684\nTotal prob of N-in: 0.94313\nPOSSIBLE N-term signal sequence\ninside 1 19\nTMhelix 20 39\noutside 40 325\nTMhelix 326 348\ninside 349 372

27596 GCF_001257855.1_H14126044201 Shigella sonnei Proteobacteria; Gammaproteobacteria; Enterobacterales; Enterobacteriaceae; Shigella MTGDNTLIHSHGINRRDFMKLCAALAATMGLSSKAAAEMAESVTNPQRPPVIWIGAQECT WP_000145410.1 MULTISPECIES: hydrogenase-2 small chain [Proteobacteria] Length: 372\nNumber of predicted TMHs: 1\nExp number of AAs in TMHs: 23.72226\nExp number, first 60 AAs: 0.22038\nTotal prob of N-in: 0.12994\noutside 1 331\nTMhelix 332 354\ninside 355 372

27597 GCF_001257915.1_8290_4_50 Shigella sonnei Proteobacteria; Gammaproteobacteria; Enterobacterales; Enterobacteriaceae; Shigella MTGDNTLIHSHGINRRDFMKLCAALAATMGLSSKAAAEMAESVTNPQRPPVIWIGAQECT WP_000145410.1 MULTISPECIES: hydrogenase-2 small chain [Proteobacteria] Length: 372\nNumber of predicted TMHs: 1\nExp number of AAs in TMHs: 23.72226\nExp number, first 60 AAs: 0.22038\nTotal prob of N-in: 0.12994\noutside 1 331\nTMhelix 332 354\ninside 355 372

27598 GCF_001257995.1_Sg_201403834_CAAAAG_L001 Shigella sonnei Proteobacteria; Gammaproteobacteria; Enterobacterales; Enterobacteriaceae; Shigella MTGDNTLIHSHGINRRDFMKLCAALAATMGLSSKAAAEMAESVTNPQRPPVIWIGAQECT WP_000145410.1 MULTISPECIES: hydrogenase-2 small chain [Proteobacteria] Length: 372\nNumber of predicted TMHs: 1\nExp number of AAs in TMHs: 23.72226\nExp number, first 60 AAs: 0.22038\nTotal prob of N-in: 0.12994\noutside 1 331\nTMhelix 332 354\ninside 355 372

27599 GCF_001258035.1_5236_7_1Shigella sonnei Proteobacteria; Gammaproteobacteria; Enterobacterales; Enterobacteriaceae; Shigella MTGDNTLIHSHGINRRDFMKLCAALAATMGLSSKAAAEMAESVTNPQRPPVIWIGAQECT WP_000145410.1 MULTISPECIES: hydrogenase-2 small chain [Proteobacteria] Length: 372\nNumber of predicted TMHs: 1\nExp number of AAs in TMHs: 23.72226\nExp number, first 60 AAs: 0.22038\nTotal prob of N-in: 0.12994\noutside 1 331\nTMhelix 332 354\ninside 355 372

27600 GCF_001258115.1_8289_7_77 Shigella sonnei Proteobacteria; Gammaproteobacteria; Enterobacterales; Enterobacteriaceae; Shigella

MTGDNTLIHSHGINRRDFMKLCAALAATMGLSSKAAAEMAESVTNPQRPPVIWIGAQECT WP_000145410.1 MULTISPECIES: hydrogenase-2 small chain [Proteobacteria] Length: 372\nNumber of predicted TMHs: 1\nExp number of AAs in TMHs: 23.72226\nExp number, first 60 AAs: 0.22038\nTotal prob of N-in: 0.12994\noutside 1 331\nTMhelix 332 354\ninside 355 372

27601 GCF_001258155.1_12971_8_72 Shigella sonnei Proteobacteria; Gammaproteobacteria; Enterobacterales; Enterobacteriaceae; Shigella MTGDNTLIHSHGINRRDFMKLCAALAATMGLSSKAAAEMAESVTNPQRPPVIWIGAQECT WP_000145410.1 MULTISPECIES: hydrogenase-2 small chain [Proteobacteria] Length: 372\nNumber of predicted TMHs: 1\nExp number of AAs in TMHs: 23.72226\nExp number, first 60 AAs: 0.22038\nTotal prob of N-in: 0.12994\noutside 1 331\nTMhelix 332 354\ninside 355 372

27602 GCF_001258215.1_Sg_201400633_TTAGGC_L001 Shigella sonnei Proteobacteria; Gammaproteobacteria; Enterobacterales; Enterobacteriaceae; Shigella MTGDNTLIHSHGINRRDFMKLCAALAATMGLSSKAAAEMAESVTNPQRPPVIWIGAQECT WP_000145410.1 MULTISPECIES: hydrogenase-2 small chain [Proteobacteria] Length: 372\nNumber of predicted TMHs: 1\nExp number of AAs in TMHs: 23.72226\nExp number, first 60 AAs: 0.22038\nTotal prob of N-in: 0.12994\noutside 1 331\nTMhelix 332 354\ninside 355 372

27603 GCF_001258275.1_8289_7_46 Shigella sonnei Proteobacteria; Gammaproteobacteria; Enterobacterales; Enterobacteriaceae; Shigella MTGDNTLIHSHGINRRDFMKLCAALAATMGLSSKAAAEMAESVTNPQRPPVIWIGAQECT WP_000145410.1 MULTISPECIES: hydrogenase-2 small chain [Proteobacteria] Length: 372\nNumber of predicted TMHs: 1\nExp number of AAs in TMHs: 23.72226\nExp number, first 60 AAs: 0.22038\nTotal prob of N-in: 0.12994\noutside 1 331\nTMhelix 332 354\ninside 355 372

27604 GCF_001258275.1_8289_7_46 Shigella sonnei Proteobacteria; Gammaproteobacteria; Enterobacterales; Enterobacteriaceae; Shigella MNNEETFYQAMRRQGVTRRSFLKYCSLAATSLGLGAGMAPKIAWALENKPRIPVVWIHGL WP_005139779.1 hydrogenase [Shigella sonnei] Length: 372\nNumber of predicted TMHs: 2\nExp number of AAs in TMHs: 37.02628\nExp number, first 60 AAs: 19.39684\nTotal prob of N-in: 0.94313\nPOSSIBLE N-term signal sequence\ninside 1 19\nTMhelix 20 39\noutside 40 325\nTMhelix 326 348\ninside 349 372

27605 GCF_001257915.1_8290_4_50 Shigella sonnei Proteobacteria; Gammaproteobacteria; Enterobacterales; Enterobacteriaceae; Shigella MNNEETFYQAMRRQGVTRRSFLKYCSLAATSLGLGAGMAPKIAWALENKPRIPVVWIHGL WP_005139779.1 hydrogenase [Shigella sonnei] Length: 372\nNumber of predicted TMHs: 2\nExp number of AAs in TMHs: 37.02628\nExp number, first 60 AAs: 19.39684\nTotal prob of N-in: 0.94313\nPOSSIBLE N-term signal sequence\ninside 1 19\nTMhelix 20 39\noutside 40 325\nTMhelix 326 348\ninside 349 372

27606 GCF_001258315.1_8290_4_43 Shigella sonnei Proteobacteria; Gammaproteobacteria; Enterobacterales; Enterobacteriaceae; Shigella MTGDNTLIHSHGINRRDFMKLCAALAATMGLSSKAAAEMAESVTNPQRPPVIWIGAQECT WP_000145410.1 MULTISPECIES: hydrogenase-2 small chain [Proteobacteria] Length: 372\nNumber of predicted TMHs: 1\nExp number of AAs in TMHs: 23.72226\nExp number, first 60 AAs: 0.22038\nTotal prob of N-in: 0.12994\noutside 1 331\nTMhelix 332 354\ninside 355 372

27607 GCF_001258375.1_8289_7_36 Shigella sonnei Proteobacteria; Gammaproteobacteria; Enterobacterales; Enterobacteriaceae; Shigella MTGDNTLIHSHGINRRDFMKLCAALAATMGLSSKAAAEMAESVTNPQRPPVIWIGAQECT WP_000145410.1 MULTISPECIES: hydrogenase-2 small chain [Proteobacteria] Length: 372\nNumber of predicted TMHs: 1\nExp number of AAs in TMHs: 23.72226\nExp number, first 60 AAs: 0.22038\nTotal prob of N-in: 0.12994\noutside 1 331\nTMhelix 332 354\ninside 355 372

27608 GCF_001258315.1_8290_4_43 Shigella sonnei Proteobacteria; Gammaproteobacteria; Enterobacterales; Enterobacteriaceae; Shigella MNNEETFYQAMRRQGVTRRSFLKYCSLAATSLGLGAGMAPKIAWALENKPRIPVVWIHGL WP_005139779.1 hydrogenase [Shigella sonnei] Length: 372\nNumber of predicted TMHs: 2\nExp number of AAs in TMHs: 37.02628\nExp number, first 60 AAs: 19.39684\nTotal prob of N-in: 0.94313\nPOSSIBLE N-term signal sequence\ninside 1 19\nTMhelix 20 39\noutside 40 325\nTMhelix 326 348\ninside 349 372

27609 GCF_001258375.1_8289_7_36 Shigella sonnei Proteobacteria; Gammaproteobacteria; Enterobacterales; Enterobacteriaceae; Shigella
MNNEETFYQAMRRQGVTRRSFLKYCSLAATSLGLGAGMAPKIAWALENKPRIPVVWIHGL WP_005139779.1
hydrogenase [Shigella sonnei] Length: 372\nNumber of predicted TMHs: 2\nExp number of AAs in TMHs: 37.02628\nExp number, first 60 AAs: 19.39684\nTotal prob of N-in: 0.94313\nPOSSIBLE N-term signal sequence\ninside 1 19\nTMhelix 20 39\noutside 40 325\nTMhelix 326 348\ninside 349 372

27610 GCF_001257995.1_Sg_201403834_CAAAAG_L001 Shigella sonnei Proteobacteria; Gammaproteobacteria; Enterobacterales; Enterobacteriaceae; Shigella
MNNEETFYQAMRRQGVTRRSFLKYCSLAATSLGLGAGMAPKIAWALENKPRIPVVWIHGL WP_005139779.1
hydrogenase [Shigella sonnei] Length: 372\nNumber of predicted TMHs: 2\nExp number of AAs in TMHs: 37.02628\nExp number, first 60 AAs: 19.39684\nTotal prob of N-in: 0.94313\nPOSSIBLE N-term signal sequence\ninside 1 19\nTMhelix 20 39\noutside 40 325\nTMhelix 326 348\ninside 349 372

27611 GCF_001258415.1_5417_1_8Shigella sonnei Proteobacteria; Gammaproteobacteria; Enterobacterales; Enterobacteriaceae; Shigella MTGDNTLIHSHGINRRDFMKLCAALAATMGLSSKAAAEMAESVTNPQRPPVIWIGAQECT
WP_000145410.1 MULTISPECIES: hydrogenase-2 small chain [Proteobacteria] Length: 372\nNumber of predicted TMHs: 1\nExp number of AAs in TMHs: 23.72226\nExp number, first 60 AAs: 0.22038\nTotal prob of N-in: 0.12994\noutside 1 331\nTMhelix 332 354\ninside 355 372

27612 GCF_001258415.1_5417_1_8Shigella sonnei Proteobacteria; Gammaproteobacteria; Enterobacterales; Enterobacteriaceae; Shigella MNNEETFYQAMRRQGVTRRSFLKYCSLAATSLGLGAGMAPKIAWALENKPRIPVVWIHGL
WP_005139779.1 hydrogenase [Shigella sonnei] Length: 372\nNumber of predicted TMHs: 2\nExp number of AAs in TMHs: 37.02628\nExp number, first 60 AAs: 19.39684\nTotal prob of N-in: 0.94313\nPOSSIBLE N-term signal sequence\ninside 1 19\nTMhelix 20 39\noutside 40 325\nTMhelix 326 348\ninside 349 372

27613 GCF_001258035.1_5236_7_1Shigella sonnei Proteobacteria; Gammaproteobacteria; Enterobacterales; Enterobacteriaceae; Shigella MNNEETFYQAMRRQGVTRRSFLKYCSLAATSLGLGAGMAPKIAWALENKPRIPVVWIHGL
WP_005139779.1 hydrogenase [Shigella sonnei] Length: 372\nNumber of predicted TMHs: 2\nExp number of AAs in TMHs: 37.02628\nExp number, first 60 AAs: 19.39684\nTotal prob of N-in: 0.94313\nPOSSIBLE N-term signal sequence\ninside 1 19\nTMhelix 20 39\noutside 40 325\nTMhelix 326 348\ninside 349 372

27614 GCF_001258155.1_12971_8_72 Shigella sonnei Proteobacteria; Gammaproteobacteria; Enterobacterales; Enterobacteriaceae; Shigella
MNNEETFYQAMRRQGVTRRSFLKYCSLAATSLGLGAGMAPKIAWALENKPRIPVVWIHGL WP_005139779.1
hydrogenase [Shigella sonnei] Length: 372\nNumber of predicted TMHs: 2\nExp number of AAs in TMHs: 37.02628\nExp number, first 60 AAs: 19.39684\nTotal prob of N-in: 0.94313\nPOSSIBLE N-term signal sequence\ninside 1 19\nTMhelix 20 39\noutside 40 325\nTMhelix 326 348\ninside 349 372

27615 GCF_001258115.1_8289_7_77 Shigella sonnei Proteobacteria; Gammaproteobacteria; Enterobacterales; Enterobacteriaceae; Shigella
MNNEETFYQAMRRQGVTRRSFLKYCSLAATSLGLGAGMAPKIAWALENKPRIPVVWIHGL WP_005139779.1
hydrogenase [Shigella sonnei] Length: 372\nNumber of predicted TMHs: 2\nExp number of AAs in TMHs: 37.02628\nExp number, first 60 AAs: 19.39684\nTotal prob of N-in: 0.94313\nPOSSIBLE N-term signal sequence\ninside 1 19\nTMhelix 20 39\noutside 40 325\nTMhelix 326 348\ninside 349 372

27616 GCF_001258455.1_7353_1_21 Shigella sonnei Proteobacteria; Gammaproteobacteria; Enterobacterales; Enterobacteriaceae; Shigella
MTGDNTLIHSHGINRRDFMKLCAALAATMGLSSKAAAEMAESVTNPQRPPVIWIGAQECT WP_000145410.1
MULTISPECIES: hydrogenase-2 small chain [Proteobacteria] Length: 372\nNumber of predicted TMHs: 1\nExp number of AAs in TMHs: 23.72226\nExp number, first 60 AAs: 0.22038\nTotal prob of N-in: 0.12994\noutside 1 331\nTMhelix 332 354\ninside 355 372

27617 GCF_001258575.1_8289_7_73 Shigella sonnei Proteobacteria; Gammaproteobacteria; Enterobacterales; Enterobacteriaceae; Shigella
MTGDNTLIHSHGINRRDFMKLCAALAATMGLSSKAAAEMAESVTNPQRPPVIWIGAQECT WP_000145410.1
MULTISPECIES: hydrogenase-2 small chain [Proteobacteria] Length: 372\nNumber of predicted TMHs: 1\nExp number of AAs in TMHs: 23.72226\nExp number, first 60 AAs: 0.22038\nTotal prob of N-in: 0.12994\noutside 1 331\nTMhelix 332 354\ninside 355 372

27618 GCF_001258455.1_7353_1_21 Shigella sonnei Proteobacteria; Gammaproteobacteria; Enterobacterales; Enterobacteriaceae; Shigella MNNEETFYQAMRRQGVTRRSFLKYCSLAATSLGLGAGMAPKIAWALENKPRIPVVWIHGL WP_005139779.1 hydrogenase [Shigella sonnei] Length: 372\nNumber of predicted TMHs: 2\nExp number of AAs in TMHs: 37.02628\nExp number, first 60 AAs: 19.39684\nTotal prob of N-in: 0.94313\nPOSSIBLE N-term signal sequence\ninside 1 19\nTMhelix 20 39\noutside 40 325\nTMhelix 326 348\ninside 349 372

27619 GCF_001258575.1_8289_7_73 Shigella sonnei Proteobacteria; Gammaproteobacteria; Enterobacterales; Enterobacteriaceae; Shigella MNNEETFYQAMRRQGVTRRSFLKYCSLAATSLGLGAGMAPKIAWALENKPRIPVVWIHGL WP_005139779.1 hydrogenase [Shigella sonnei] Length: 372\nNumber of predicted TMHs: 2\nExp number of AAs in TMHs: 37.02628\nExp number, first 60 AAs: 19.39684\nTotal prob of N-in: 0.94313\nPOSSIBLE N-term signal sequence\ninside 1 19\nTMhelix 20 39\noutside 40 325\nTMhelix 326 348\ninside 349 372

27620 GCF_001258215.1_Sg_201400633_TTAGGC_L001 Shigella sonnei Proteobacteria; Gammaproteobacteria; Enterobacterales; Enterobacteriaceae; Shigella MNNEETFYQAMRRQGVTRRSFLKYCSLAATSLGLGAGMAPKIAWALENKPRIPVVWIHGL WP_005139779.1 hydrogenase [Shigella sonnei] Length: 372\nNumber of predicted TMHs: 2\nExp number of AAs in TMHs: 37.02628\nExp number, first 60 AAs: 19.39684\nTotal prob of N-in: 0.94313\nPOSSIBLE N-term signal sequence\ninside 1 19\nTMhelix 20 39\noutside 40 325\nTMhelix 326 348\ninside 349 372

27621 GCF_001258615.1_12971_8_77 Shigella sonnei Proteobacteria; Gammaproteobacteria; Enterobacterales; Enterobacteriaceae; Shigella MTGDNTLIHSHGINRRDFMKLCAALAATMGLSSKAAAEMAESVTNPQRPPVIWIGAQECT WP_000145410.1 MULTISPECIES: hydrogenase-2 small chain [Proteobacteria] Length: 372\nNumber of predicted TMHs: 1\nExp number of AAs in TMHs: 23.72226\nExp number, first 60 AAs: 0.22038\nTotal prob of N-in: 0.12994\noutside 1 331\nTMhelix 332 354\ninside 355 372

27622 GCF_001258675.1_12850_1_43 Shigella sonnei Proteobacteria; Gammaproteobacteria; Enterobacterales; Enterobacteriaceae; Shigella MTGDNTLIHSHGINRRDFMKLCAALAATMGLSSKAAAEMAESVTNPQRPPVIWIGAQECT WP_000145410.1 MULTISPECIES: hydrogenase-2 small chain [Proteobacteria] Length: 372\nNumber of predicted TMHs: 1\nExp number of AAs in TMHs: 23.72226\nExp number, first 60 AAs: 0.22038\nTotal prob of N-in: 0.12994\noutside 1 331\nTMhelix 332 354\ninside 355 372

27623 GCF_001258715.1_7353_1_23 Shigella sonnei Proteobacteria; Gammaproteobacteria; Enterobacterales; Enterobacteriaceae; Shigella MTGDNTLIHSHGINRRDFMKLCAALAATMGLSSKAAAEMAESVTNPQRPPVIWIGAQECT WP_000145410.1 MULTISPECIES: hydrogenase-2 small chain [Proteobacteria] Length: 372\nNumber of predicted TMHs: 1\nExp number of AAs in TMHs: 23.72226\nExp number, first 60 AAs: 0.22038\nTotal prob of N-in: 0.12994\noutside 1 331\nTMhelix 332 354\ninside 355 372

27624 GCF_001258795.1_5236_6_2Shigella sonnei Proteobacteria; Gammaproteobacteria; Enterobacterales; Enterobacteriaceae; Shigella MTGDNTLIHSHGINRRDFMKLCAALAATMGLSSKAAAEMAESVTNPQRPPVIWIGAQECT WP_000145410.1 MULTISPECIES: hydrogenase-2 small chain [Proteobacteria] Length: 372\nNumber of predicted TMHs: 1\nExp number of AAs in TMHs: 23.72226\nExp number, first 60 AAs: 0.22038\nTotal prob of N-in: 0.12994\noutside 1 331\nTMhelix 332 354\ninside 355 372

27625 GCF_001258915.1_5417_3_7Shigella sonnei Proteobacteria; Gammaproteobacteria; Enterobacterales; Enterobacteriaceae; Shigella MTGDNTLIHSHGINRRDFMKLCAALAATMGLSSKAAAEMAESVTNPQRPPVIWIGAQECT WP_000145410.1 MULTISPECIES: hydrogenase-2 small chain [Proteobacteria] Length: 372\nNumber of predicted TMHs: 1\nExp number of AAs in TMHs: 23.72226\nExp number, first 60 AAs: 0.22038\nTotal prob of N-in: 0.12994\noutside 1 331\nTMhelix 332 354\ninside 355 372

27626 GCF_001258855.1_5236_5_6Shigella sonnei Proteobacteria; Gammaproteobacteria; Enterobacterales; Enterobacteriaceae; Shigella MTGDNTLIHSHGINRRDFMKLCAALAATMGLSSKAAAEMAESVTNPQRPPVIWIGAQECT WP_000145410.1 MULTISPECIES: hydrogenase-2 small chain [Proteobacteria] Length: 372\nNumber of predicted TMHs: 1\nExp number of AAs in TMHs: 23.72226\nExp number, first 60 AAs: 0.22038\nTotal prob of N-in: 0.12994\noutside 1 331\nTMhelix 332 354\ninside 355 372

27627	GCF_001258955.1_5236_6_10	Shigella sonnei	Proteobacteria; Gammaproteobacteria; Enterobacterales; Enterobacteriaceae; Shigella
MTGDNTLIHSHGINRRDFMKLCAALAATMGLSSKAAAEMAESVTNPQRPPVIWIGAQECT	WP_000145410.1
MULTISPECIES: hydrogenase-2 small chain [Proteobacteria]	Length: 372\nNumber of predicted TMHs: 1\nExp number of AAs in TMHs: 23.72226\nExp number, first 60 AAs: 0.22038\nTotal prob of N-in: 0.12994\noutside 1 331\nTMhelix 332 354\ninside 355 372

27628	GCF_001259075.1_Sg_201405302_CGGAAT_L001	Shigella sonnei	Proteobacteria; Gammaproteobacteria; Enterobacterales; Enterobacteriaceae; Shigella
MTGDNTLIHSHGINRRDFMKLCAALAATMGLSSKAAAEMAESVTNPQRPPVIWIGAQECT	WP_000145410.1
MULTISPECIES: hydrogenase-2 small chain [Proteobacteria]	Length: 372\nNumber of predicted TMHs: 1\nExp number of AAs in TMHs: 23.72226\nExp number, first 60 AAs: 0.22038\nTotal prob of N-in: 0.12994\noutside 1 331\nTMhelix 332 354\ninside 355 372

27629	GCF_001259155.1_5236_7_2Shigella sonnei	Proteobacteria; Gammaproteobacteria; Enterobacterales; Enterobacteriaceae; Shigella	MTGDNTLIHSHGINRRDFMKLCAALAATMGLSSKAAAEMAESVTNPQRPPVIWIGAQECT
WP_000145410.1	MULTISPECIES: hydrogenase-2 small chain [Proteobacteria]	Length: 372\nNumber of predicted TMHs: 1\nExp number of AAs in TMHs: 23.72226\nExp number, first 60 AAs: 0.22038\nTotal prob of N-in: 0.12994\noutside 1 331\nTMhelix 332 354\ninside 355 372

27630	GCF_001259075.1_Sg_201405302_CGGAAT_L001	Shigella sonnei	Proteobacteria; Gammaproteobacteria; Enterobacterales; Enterobacteriaceae; Shigella
MNNEETFYQAMRRQGVTRRSFLKYCSLAATSLGLGAGMAPKIAWALENKPRIPVVWIHGL	WP_005139779.1
hydrogenase [Shigella sonnei]	Length: 372\nNumber of predicted TMHs: 2\nExp number of AAs in TMHs: 37.02628\nExp number, first 60 AAs: 19.39684\nTotal prob of N-in: 0.94313\nPOSSIBLE N-term signal sequence\ninside 1 19\nTMhelix 20 39\noutside 40 325\nTMhelix 326 348\ninside 349 372

27631	GCF_001259155.1_5236_7_2Shigella sonnei	Proteobacteria; Gammaproteobacteria; Enterobacterales; Enterobacteriaceae; Shigella	MNNEETFYQAMRRQGVTRRSFLKYCSLAATSLGLGAGMAPKIAWALENKPRIPVVWIHGL
WP_005139779.1	hydrogenase [Shigella sonnei]	Length: 372\nNumber of predicted TMHs: 2\nExp number of AAs in TMHs: 37.02628\nExp number, first 60 AAs: 19.39684\nTotal prob of N-in: 0.94313\nPOSSIBLE N-term signal sequence\ninside 1 19\nTMhelix 20 39\noutside 40 325\nTMhelix 326 348\ninside 349 372

27632	GCF_001258615.1_12971_8_77	Shigella sonnei	Proteobacteria; Gammaproteobacteria; Enterobacterales; Enterobacteriaceae; Shigella
MNNEETFYQAMRRQGVTRRSFLKYCSLAATSLGLGAGMAPKIAWALENKPRIPVVWIHGL	WP_005139779.1
hydrogenase [Shigella sonnei]	Length: 372\nNumber of predicted TMHs: 2\nExp number of AAs in TMHs: 37.02628\nExp number, first 60 AAs: 19.39684\nTotal prob of N-in: 0.94313\nPOSSIBLE N-term signal sequence\ninside 1 19\nTMhelix 20 39\noutside 40 325\nTMhelix 326 348\ninside 349 372

27633	GCF_001258675.1_12850_1_43	Shigella sonnei	Proteobacteria; Gammaproteobacteria; Enterobacterales; Enterobacteriaceae; Shigella
MNNEETFYQAMRRQGVTRRSFLKYCSLAATSLGLGAGMAPKIAWALENKPRIPVVWIHGL	WP_005139779.1
hydrogenase [Shigella sonnei]	Length: 372\nNumber of predicted TMHs: 2\nExp number of AAs in TMHs: 37.02628\nExp number, first 60 AAs: 19.39684\nTotal prob of N-in: 0.94313\nPOSSIBLE N-term signal sequence\ninside 1 19\nTMhelix 20 39\noutside 40 325\nTMhelix 326 348\ninside 349 372

27634	GCF_001259195.1_5236_3_8Shigella sonnei	Proteobacteria; Gammaproteobacteria; Enterobacterales; Enterobacteriaceae; Shigella	MTGDNTLIHSHGINRRDFMKLCAALAATMGLSSKAAAEMAESVTNPQRPPVIWIGAQECT
WP_000145410.1	MULTISPECIES: hydrogenase-2 small chain [Proteobacteria]	Length: 372\nNumber of predicted TMHs: 1\nExp number of AAs in TMHs: 23.72226\nExp number, first 60 AAs: 0.22038\nTotal prob of N-in: 0.12994\noutside 1 331\nTMhelix 332 354\ninside 355 372

27635	GCF_001259275.1_12971_8_55	Shigella sonnei	Proteobacteria; Gammaproteobacteria; Enterobacterales; Enterobacteriaceae; Shigella
MTGDNTLIHSHGINRRDFMKLCAALAATMGLSSKAAAEMAESVTNPQRPPVIWIGAQECT	WP_000145410.1
MULTISPECIES: hydrogenase-2 small chain [Proteobacteria]	Length: 372\nNumber of predicted TMHs: 1\nExp number of AAs in TMHs: 23.72226\nExp number, first 60 AAs: 0.22038\nTotal prob of N-in: 0.12994\noutside 1 331\nTMhelix 332 354\ninside 355 372

27636 GCF_001259195.1_5236_3_8Shigella sonnei Proteobacteria; Gammaproteobacteria; Enterobacterales; Enterobacteriaceae; Shigella MNNEETFYQAMRRQGVTRRSFLKYCSLAATSLGLGAGMAPKIAWALENKPRIPVVWIHGL WP_005139779.1 hydrogenase [Shigella sonnei] Length: 372\nNumber of predicted TMHs: 2\nExp number of AAs in TMHs: 37.02628\nExp number, first 60 AAs: 19.39684\nTotal prob of N-in: 0.94313\nPOSSIBLE N-term signal sequence\ninside 1 19\nTMhelix 20 39\noutside 40 325\nTMhelix 326 348\ninside 349 372

27637 GCF_001259275.1_12971_8_55 Shigella sonnei Proteobacteria; Gammaproteobacteria; Enterobacterales; Enterobacteriaceae; Shigella MNNEETFYQAMRRQGVTRRSFLKYCSLAATSLGLGAGMAPKIAWALENKPRIPVVWIHGL WP_005139779.1 hydrogenase [Shigella sonnei] Length: 372\nNumber of predicted TMHs: 2\nExp number of AAs in TMHs: 37.02628\nExp number, first 60 AAs: 19.39684\nTotal prob of N-in: 0.94313\nPOSSIBLE N-term signal sequence\ninside 1 19\nTMhelix 20 39\noutside 40 325\nTMhelix 326 348\ninside 349 372

27638 GCF_001258715.1_7353_1_23 Shigella sonnei Proteobacteria; Gammaproteobacteria; Enterobacterales; Enterobacteriaceae; Shigella MNNEETFYQAMRRQGVTRRSFLKYCSLAATSLGLGAGMAPKIAWALENKPRIPVVWIHGL WP_005139779.1 hydrogenase [Shigella sonnei] Length: 372\nNumber of predicted TMHs: 2\nExp number of AAs in TMHs: 37.02628\nExp number, first 60 AAs: 19.39684\nTotal prob of N-in: 0.94313\nPOSSIBLE N-term signal sequence\ninside 1 19\nTMhelix 20 39\noutside 40 325\nTMhelix 326 348\ninside 349 372

27639 GCF_001258795.1_5236_6_2Shigella sonnei Proteobacteria; Gammaproteobacteria; Enterobacterales; Enterobacteriaceae; Shigella MNNEETFYQAMRRQGVTRRSFLKYCSLAATSLGLGAGMAPKIAWALENKPRIPVVWIHGL WP_005139779.1 hydrogenase [Shigella sonnei] Length: 372\nNumber of predicted TMHs: 2\nExp number of AAs in TMHs: 37.02628\nExp number, first 60 AAs: 19.39684\nTotal prob of N-in: 0.94313\nPOSSIBLE N-term signal sequence\ninside 1 19\nTMhelix 20 39\noutside 40 325\nTMhelix 326 348\ninside 349 372

27640 GCF_001259335.1_5008_8_10 Shigella sonnei Proteobacteria; Gammaproteobacteria; Enterobacterales; Enterobacteriaceae; Shigella MTGDNTLIHSHGINRRDFMKLCAALAATMGLSSKAAAEMAESVTNPQRPPVIWIGAQECT WP_000145410.1 MULTISPECIES: hydrogenase-2 small chain [Proteobacteria] Length: 372\nNumber of predicted TMHs: 1\nExp number of AAs in TMHs: 23.72226\nExp number, first 60 AAs: 0.22038\nTotal prob of N-in: 0.12994\noutside 1 331\nTMhelix 332 354\ninside 355 372

27641 GCF_001259335.1_5008_8_10 Shigella sonnei Proteobacteria; Gammaproteobacteria; Enterobacterales; Enterobacteriaceae; Shigella MNNEETFYQAMRRQGVTRRSFLKYCSLAATSLGLGAGMAPKIAWALENKPRIPVVWIHGL WP_005139779.1 hydrogenase [Shigella sonnei] Length: 372\nNumber of predicted TMHs: 2\nExp number of AAs in TMHs: 37.02628\nExp number, first 60 AAs: 19.39684\nTotal prob of N-in: 0.94313\nPOSSIBLE N-term signal sequence\ninside 1 19\nTMhelix 20 39\noutside 40 325\nTMhelix 326 348\ninside 349 372

27642 GCF_001258855.1_5236_5_6Shigella sonnei Proteobacteria; Gammaproteobacteria; Enterobacterales; Enterobacteriaceae; Shigella MNNEETFYQAMRRQGVTRRSFLKYCSLAATSLGLGAGMAPKIAWALENKPRIPVVWIHGL WP_005139779.1 hydrogenase [Shigella sonnei] Length: 372\nNumber of predicted TMHs: 2\nExp number of AAs in TMHs: 37.02628\nExp number, first 60 AAs: 19.39684\nTotal prob of N-in: 0.94313\nPOSSIBLE N-term signal sequence\ninside 1 19\nTMhelix 20 39\noutside 40 325\nTMhelix 326 348\ninside 349 372

27643 GCF_001258915.1_5417_3_7Shigella sonnei Proteobacteria; Gammaproteobacteria; Enterobacterales; Enterobacteriaceae; Shigella MNNEETFYQAMRRQGVTRRSFLKYCSLAATSLGLGAGMAPKIAWALENKPRIPVVWIHGL WP_005139779.1 hydrogenase [Shigella sonnei] Length: 372\nNumber of predicted TMHs: 2\nExp number of AAs in TMHs: 37.02628\nExp number, first 60 AAs: 19.39684\nTotal prob of N-in: 0.94313\nPOSSIBLE N-term signal sequence\ninside 1 19\nTMhelix 20 39\noutside 40 325\nTMhelix 326 348\ninside 349 372

27644 GCF_001258955.1_5236_6_10 Shigella sonnei Proteobacteria; Gammaproteobacteria; Enterobacterales; Enterobacteriaceae; Shigella MNNEETFYQAMRRQGVTRRSFLKYCSLAATSLGLGAGMAPKIAWALENKPRIPVVWIHGL WP_005139779.1 hydrogenase [Shigella sonnei] Length: 372\nNumber of predicted TMHs: 2\nExp number of AAs in TMHs: 37.02628\nExp number, first 60 AAs: 19.39684\nTotal prob of N-in: 0.94313\nPOSSIBLE N-term signal sequence\ninside 1 19\nTMhelix 20 39\noutside 40 325\nTMhelix 326 348\ninside 349 372

27645 GCF_001259415.1_8290_5_13 Shigella sonnei Proteobacteria; Gammaproteobacteria; Enterobacterales; Enterobacteriaceae; Shigella

MTGDNTLIHSHGINRRDFMKLCAALAATMGLSSKAAAEMAESVTNPQRPPVIWIGAQECT WP_000145410.1 MULTISPECIES: hydrogenase-2 small chain [Proteobacteria] Length: 372\nNumber of predicted TMHs: 1\nExp number of AAs in TMHs: 23.72226\nExp number, first 60 AAs: 0.22038\nTotal prob of N-in: 0.12994\noutside 1 331\nTMhelix 332 354\ninside 355 372

27646 GCF_001259455.1_12971_8_8 Shigella sonnei Proteobacteria; Gammaproteobacteria; Enterobacterales; Enterobacteriaceae; Shigella MTGDNTLIHSHGINRRDFMKLCAALAATMGLSSKAAAEMAESVTNPQRPPVIWIGAQECT WP_000145410.1 MULTISPECIES: hydrogenase-2 small chain [Proteobacteria] Length: 372\nNumber of predicted TMHs: 1\nExp number of AAs in TMHs: 23.72226\nExp number, first 60 AAs: 0.22038\nTotal prob of N-in: 0.12994\noutside 1 331\nTMhelix 332 354\ninside 355 372

27647 GCF_001259535.1_Sg_201405267_CCAACA_L001 Shigella sonnei Proteobacteria; Gammaproteobacteria; Enterobacterales; Enterobacteriaceae; Shigella MTGDNTLIHSHGINRRDFMKLCAALAATMGLSSKAAAEMAESVTNPQRPPVIWIGAQECT WP_000145410.1 MULTISPECIES: hydrogenase-2 small chain [Proteobacteria] Length: 372\nNumber of predicted TMHs: 1\nExp number of AAs in TMHs: 23.72226\nExp number, first 60 AAs: 0.22038\nTotal prob of N-in: 0.12994\noutside 1 331\nTMhelix 332 354\ninside 355 372

27648 GCF_001259675.1_8290_4_46 Shigella sonnei Proteobacteria; Gammaproteobacteria; Enterobacterales; Enterobacteriaceae; Shigella MTGDNTLIHSHGINRRDFMKLCAALAATMGLSSKAAAEMAESVTNPQRPPVIWIGAQECT WP_000145410.1 MULTISPECIES: hydrogenase-2 small chain [Proteobacteria] Length: 372\nNumber of predicted TMHs: 1\nExp number of AAs in TMHs: 23.72226\nExp number, first 60 AAs: 0.22038\nTotal prob of N-in: 0.12994\noutside 1 331\nTMhelix 332 354\ninside 355 372

27649 GCF_001259735.1_12971_8_48 Shigella sonnei Proteobacteria; Gammaproteobacteria; Enterobacterales; Enterobacteriaceae; Shigella MTGDNTLIHSHGINRRDFMKLCAALAATMGLSSKAAAEMAESVTNPQRPPVIWIGAQECT WP_000145410.1 MULTISPECIES: hydrogenase-2 small chain [Proteobacteria] Length: 372\nNumber of predicted TMHs: 1\nExp number of AAs in TMHs: 23.72226\nExp number, first 60 AAs: 0.22038\nTotal prob of N-in: 0.12994\noutside 1 331\nTMhelix 332 354\ninside 355 372

27650 GCF_001259815.1_5236_5_3Shigella sonnei Proteobacteria; Gammaproteobacteria; Enterobacterales; Enterobacteriaceae; Shigella MTGDNTLIHSHGINRRDFMKLCAALAATMGLSSKAAAEMAESVTNPQRPPVIWIGAQECT WP_000145410.1 MULTISPECIES: hydrogenase-2 small chain [Proteobacteria] Length: 372\nNumber of predicted TMHs: 1\nExp number of AAs in TMHs: 23.72226\nExp number, first 60 AAs: 0.22038\nTotal prob of N-in: 0.12994\noutside 1 331\nTMhelix 332 354\ninside 355 372

27651 GCF_001259855.1_8289_7_93 Shigella sonnei Proteobacteria; Gammaproteobacteria; Enterobacterales; Enterobacteriaceae; Shigella MTGDNTLIHSHGINRRDFMKLCAALAATMGLSSKAAAEMAESVTNPQRPPVIWIGAQECT WP_000145410.1 MULTISPECIES: hydrogenase-2 small chain [Proteobacteria] Length: 372\nNumber of predicted TMHs: 1\nExp number of AAs in TMHs: 23.72226\nExp number, first 60 AAs: 0.22038\nTotal prob of N-in: 0.12994\noutside 1 331\nTMhelix 332 354\ninside 355 372

27652 GCF_001260035.1_12971_8_20 Shigella sonnei Proteobacteria; Gammaproteobacteria; Enterobacterales; Enterobacteriaceae; Shigella MTGDNTLIHSHGINRRDFMKLCAALAATMGLSSKAAAEMAESVTNPQRPPVIWIGAQECT WP_000145410.1 MULTISPECIES: hydrogenase-2 small chain [Proteobacteria] Length: 372\nNumber of predicted TMHs: 1\nExp number of AAs in TMHs: 23.72226\nExp number, first 60 AAs: 0.22038\nTotal prob of N-in: 0.12994\noutside 1 331\nTMhelix 332 354\ninside 355 372

27653 GCF_001259975.1_8290_4_30 Shigella sonnei Proteobacteria; Gammaproteobacteria; Enterobacterales; Enterobacteriaceae; Shigella MTGDNTLIHSHGINRRDFMKLCAALAATMGLSSKAAAEMAESVTNPQRPPVIWIGAQECT WP_000145410.1 MULTISPECIES: hydrogenase-2 small chain [Proteobacteria] Length: 372\nNumber of predicted TMHs: 1\nExp number of AAs in TMHs: 23.72226\nExp number, first 60 AAs: 0.22038\nTotal prob of N-in: 0.12994\noutside 1 331\nTMhelix 332 354\ninside 355 372

27654 GCF_001260035.1_12971_8_20 Shigella sonnei Proteobacteria; Gammaproteobacteria; Enterobacterales; Enterobacteriaceae; Shigella MNNEETFYQAMRRQGVTRRSFLKYCSLAATSLGLGAGMAPKIAWALENKPRIPVVWIHGL WP_005139779.1 hydrogenase [Shigella sonnei] Length: 372\nNumber of predicted TMHs: 2\nExp number of AAs in TMHs: 37.02628\nExp number, first 60 AAs: 19.39684\nTotal prob of N-in: 0.94313\nPOSSIBLE N-term signal sequence\ninside 1 19\nTMhelix 20 39\noutside 40 325\nTMhelix 326 348\ninside 349 372

27655 GCF_001259975.1_8290_4_30 Shigella sonnei Proteobacteria; Gammaproteobacteria; Enterobacterales; Enterobacteriaceae; Shigella MNNEETFYQAMRRQGVTRRSFLKYCSLAATSLGLGAGMAPKIAWALENKPRIPVVWIHGL WP_005139779.1 hydrogenase [Shigella sonnei] Length: 372\nNumber of predicted TMHs: 2\nExp number of AAs in TMHs: 37.02628\nExp number, first 60 AAs: 19.39684\nTotal prob of N-in: 0.94313\nPOSSIBLE N-term signal sequence\ninside 1 19\nTMhelix 20 39\noutside 40 325\nTMhelix 326 348\ninside 349 372

27656 GCF_001259415.1_8290_5_13 Shigella sonnei Proteobacteria; Gammaproteobacteria; Enterobacterales; Enterobacteriaceae; Shigella MNNEETFYQAMRRQGVTRRSFLKYCSLAATSLGLGAGMAPKIAWALENKPRIPVVWIHGL WP_005139779.1 hydrogenase [Shigella sonnei] Length: 372\nNumber of predicted TMHs: 2\nExp number of AAs in TMHs: 37.02628\nExp number, first 60 AAs: 19.39684\nTotal prob of N-in: 0.94313\nPOSSIBLE N-term signal sequence\ninside 1 19\nTMhelix 20 39\noutside 40 325\nTMhelix 326 348\ninside 349 372

27657 GCF_001260135.1_5236_8_3Shigella sonnei Proteobacteria; Gammaproteobacteria; Enterobacterales; Enterobacteriaceae; Shigella MTGDNTLIHSHGINRRDFMKLCAALAATMGLSSKAAAEMAESVTNPQRPPVIWIGAQECT WP_000145410.1 MULTISPECIES: hydrogenase-2 small chain [Proteobacteria] Length: 372\nNumber of predicted TMHs: 1\nExp number of AAs in TMHs: 23.72226\nExp number, first 60 AAs: 0.22038\nTotal prob of N-in: 0.12994\noutside 1 331\nTMhelix 332 354\ninside 355 372

27658 GCF_001260135.1_5236_8_3Shigella sonnei Proteobacteria; Gammaproteobacteria; Enterobacterales; Enterobacteriaceae; Shigella MNNEETFYQAMRRQGVTRRSFLKYCSLAATSLGLGAGMAPKIAWALENKPRIPVVWIHGL WP_005139779.1 hydrogenase [Shigella sonnei] Length: 372\nNumber of predicted TMHs: 2\nExp number of AAs in TMHs: 37.02628\nExp number, first 60 AAs: 19.39684\nTotal prob of N-in: 0.94313\nPOSSIBLE N-term signal sequence\ninside 1 19\nTMhelix 20 39\noutside 40 325\nTMhelix 326 348\ninside 349 372

27659 GCF_001259535.1_Sg_201405267_CCAACA_L001 Shigella sonnei Proteobacteria; Gammaproteobacteria; Enterobacterales; Enterobacteriaceae; Shigella MNNEETFYQAMRRQGVTRRSFLKYCSLAATSLGLGAGMAPKIAWALENKPRIPVVWIHGL WP_005139779.1 hydrogenase [Shigella sonnei] Length: 372\nNumber of predicted TMHs: 2\nExp number of AAs in TMHs: 37.02628\nExp number, first 60 AAs: 19.39684\nTotal prob of N-in: 0.94313\nPOSSIBLE N-term signal sequence\ninside 1 19\nTMhelix 20 39\noutside 40 325\nTMhelix 326 348\ninside 349 372

27660 GCF_001259455.1_12971_8_8 Shigella sonnei Proteobacteria; Gammaproteobacteria; Enterobacterales; Enterobacteriaceae; Shigella MNNEETFYQAMRRQGVTRRSFLKYCSLAATSLGLGAGMAPKIAWALENKPRIPVVWIHGL WP_005139779.1 hydrogenase [Shigella sonnei] Length: 372\nNumber of predicted TMHs: 2\nExp number of AAs in TMHs: 37.02628\nExp number, first 60 AAs: 19.39684\nTotal prob of N-in: 0.94313\nPOSSIBLE N-term signal sequence\ninside 1 19\nTMhelix 20 39\noutside 40 325\nTMhelix 326 348\ninside 349 372

27661 GCF_001260215.1_12844_1_57 Shigella sonnei Proteobacteria; Gammaproteobacteria; Enterobacterales; Enterobacteriaceae; Shigella MTGDNTLIHSHGINRRDFMKLCAALAATMGLSSKAAAEMAESVTNPQRPPVIWIGAQECT WP_000145410.1 MULTISPECIES: hydrogenase-2 small chain [Proteobacteria] Length: 372\nNumber of predicted TMHs: 1\nExp number of AAs in TMHs: 23.72226\nExp number, first 60 AAs: 0.22038\nTotal prob of N-in: 0.12994\noutside 1 331\nTMhelix 332 354\ninside 355 372

27662 GCF_001260215.1_12844_1_57 Shigella sonnei Proteobacteria; Gammaproteobacteria; Enterobacterales; Enterobacteriaceae; Shigella MNNEETFYQAMRRQGVTRRSFLKYCSLAATSLGLGAGMAPKIAWALENKPRIPVVWIHGL WP_005139779.1 hydrogenase [Shigella sonnei] Length: 372\nNumber of predicted TMHs: 2\nExp number of AAs in TMHs: 37.02628\nExp number, first 60 AAs: 19.39684\nTotal prob of N-in: 0.94313\nPOSSIBLE N-term signal sequence\ninside 1 19\nTMhelix 20 39\noutside 40 325\nTMhelix 326 348\ninside 349 372

27663 GCF_001259675.1_8290_4_46 Shigella sonnei Proteobacteria; Gammaproteobacteria; Enterobacterales; Enterobacteriaceae; Shigella MNNEETFYQAMRRQGVTRRSFLKYCSLAATSLGLGAGMAPKIAWALENKPRIPVVWIHGL WP_005139779.1 hydrogenase [Shigella sonnei] Length: 372\nNumber of predicted TMHs: 2\nExp number of AAs in TMHs: 37.02628\nExp number, first 60 AAs: 19.39684\nTotal prob of N-in: 0.94313\nPOSSIBLE N-term signal sequence\ninside 1 19\nTMhelix 20 39\noutside 40 325\nTMhelix 326 348\ninside 349 372

27664 GCF_001259735.1_12971_8_48 Shigella sonnei Proteobacteria; Gammaproteobacteria; Enterobacterales; Enterobacteriaceae; Shigella MNNEETFYQAMRRQGVTRRSFLKYCSLAATSLGLGAGMAPKIAWALENKPRIPVVWIHGL WP_005139779.1 hydrogenase [Shigella sonnei] Length: 372\nNumber of predicted TMHs: 2\nExp number of AAs in TMHs: 37.02628\nExp number, first 60 AAs: 19.39684\nTotal prob of N-in: 0.94313\nPOSSIBLE N-term signal sequence\ninside 1 19\nTMhelix 20 39\noutside 40 325\nTMhelix 326 348\ninside 349 372

27665 GCF_001260255.1_8290_4_32 Shigella sonnei Proteobacteria; Gammaproteobacteria; Enterobacterales; Enterobacteriaceae; Shigella MTGDNTLIHSHGINRRDFMKLCAALAATMGLSSKAAAEMAESVTNPQRPPVIWIGAQECT WP_000145410.1 MULTISPECIES: hydrogenase-2 small chain [Proteobacteria] Length: 372\nNumber of predicted TMHs: 1\nExp number of AAs in TMHs: 23.72226\nExp number, first 60 AAs: 0.22038\nTotal prob of N-in: 0.12994\noutside 1 331\nTMhelix 332 354\ninside 355 372

27666 GCF_001260255.1_8290_4_32 Shigella sonnei Proteobacteria; Gammaproteobacteria; Enterobacterales; Enterobacteriaceae; Shigella MNNEETFYQAMRRQGVTRRSFLKYCSLAATSLGLGAGMAPKIAWALENKPRIPVVWIHGL WP_005139779.1 hydrogenase [Shigella sonnei] Length: 372\nNumber of predicted TMHs: 2\nExp number of AAs in TMHs: 37.02628\nExp number, first 60 AAs: 19.39684\nTotal prob of N-in: 0.94313\nPOSSIBLE N-term signal sequence\ninside 1 19\nTMhelix 20 39\noutside 40 325\nTMhelix 326 348\ninside 349 372

27667 GCF_001259815.1_5236_5_3Shigella sonnei Proteobacteria; Gammaproteobacteria; Enterobacterales; Enterobacteriaceae; Shigella MNNEETFYQAMRRQGVTRRSFLKYCSLAATSLGLGAGMAPKIAWALENKPRIPVVWIHGL WP_005139779.1 hydrogenase [Shigella sonnei] Length: 372\nNumber of predicted TMHs: 2\nExp number of AAs in TMHs: 37.02628\nExp number, first 60 AAs: 19.39684\nTotal prob of N-in: 0.94313\nPOSSIBLE N-term signal sequence\ninside 1 19\nTMhelix 20 39\noutside 40 325\nTMhelix 326 348\ninside 349 372

27668 GCF_001259855.1_8289_7_93 Shigella sonnei Proteobacteria; Gammaproteobacteria; Enterobacterales; Enterobacteriaceae; Shigella MNNEETFYQAMRRQGVTRRSFLKYCSLAATSLGLGAGMAPKIAWALENKPRIPVVWIHGL WP_005139779.1 hydrogenase [Shigella sonnei] Length: 372\nNumber of predicted TMHs: 2\nExp number of AAs in TMHs: 37.02628\nExp number, first 60 AAs: 19.39684\nTotal prob of N-in: 0.94313\nPOSSIBLE N-term signal sequence\ninside 1 19\nTMhelix 20 39\noutside 40 325\nTMhelix 326 348\ninside 349 372

27669 GCF_001260355.1_8290_5_4Shigella sonnei Proteobacteria; Gammaproteobacteria; Enterobacterales; Enterobacteriaceae; Shigella MTGDNTLIHSHGINRRDFMKLCAALAATMGLSSKAAAEMAESVTNPQRPPVIWIGAQECT WP_000145410.1 MULTISPECIES: hydrogenase-2 small chain [Proteobacteria] Length: 372\nNumber of predicted TMHs: 1\nExp number of AAs in TMHs: 23.72226\nExp number, first 60 AAs: 0.22038\nTotal prob of N-in: 0.12994\noutside 1 331\nTMhelix 332 354\ninside 355 372

27670 GCF_001260395.1_8289_7_35 Shigella sonnei Proteobacteria; Gammaproteobacteria; Enterobacterales; Enterobacteriaceae; Shigella MTGDNTLIHSHGINRRDFMKLCAALAATMGLSSKAAAEMAESVTNPQRPPVIWIGAQECT WP_000145410.1 MULTISPECIES: hydrogenase-2 small chain [Proteobacteria] Length: 372\nNumber of predicted TMHs: 1\nExp number of AAs in TMHs: 23.72226\nExp number, first 60 AAs: 0.22038\nTotal prob of N-in: 0.12994\noutside 1 331\nTMhelix 332 354\ninside 355 372

27671 GCF_001260475.1_5236_2_8Shigella sonnei Proteobacteria; Gammaproteobacteria; Enterobacterales; Enterobacteriaceae; Shigella MTGDNTLIHSHGINRRDFMKLCAALAATMGLSSKAAAEMAESVTNPQRPPVIWIGAQECT WP_000145410.1 MULTISPECIES: hydrogenase-2 small chain [Proteobacteria] Length: 372\nNumber of predicted TMHs: 1\nExp number of AAs in TMHs: 23.72226\nExp number, first 60 AAs: 0.22038\nTotal prob of N-in: 0.12994\noutside 1 331\nTMhelix 332 354\ninside 355 372

27672 GCF_001260515.1_Sg_201403956_CACGAT_L001 Shigella sonnei Proteobacteria; Gammaproteobacteria; Enterobacterales; Enterobacteriaceae; Shigella MTGDNTLIHSHGINRRDFMKLCAALAATMGLSSKAAAEMAESVTNPQRPPVIWIGAQECT WP_000145410.1 MULTISPECIES: hydrogenase-2 small chain [Proteobacteria] Length: 372\nNumber of predicted TMHs: 1\nExp number of AAs in TMHs: 23.72226\nExp number, first 60 AAs: 0.22038\nTotal prob of N-in: 0.12994\noutside 1 331\nTMhelix 332 354\ninside 355 372

27673 GCF_001260575.1_5236_2_3Shigella sonnei Proteobacteria; Gammaproteobacteria; Enterobacterales; Enterobacteriaceae; Shigella MTGDNTLIHSHGINRRDFMKLCAALAATMGLSSKAAAEMAESVTNPQRPPVIWIGAQECT WP_000145410.1 MULTISPECIES: hydrogenase-2 small chain [Proteobacteria] Length: 372\nNumber of predicted TMHs: 1\nExp number of AAs in TMHs: 23.72226\nExp number, first 60 AAs: 0.22038\nTotal prob of N-in: 0.12994\noutside 1 331\nTMhelix 332 354\ninside 355 372

27674 GCF_001260595.1_8289_7_63 Shigella sonnei Proteobacteria; Gammaproteobacteria; Enterobacterales; Enterobacteriaceae; Shigella MTGDNTLIHSHGINRRDFMKLCAALAATMGLSSKAAAEMAESVTNPQRPPVIWIGAQECT WP_000145410.1 MULTISPECIES: hydrogenase-2 small chain [Proteobacteria] Length: 372\nNumber of predicted TMHs: 1\nExp number of AAs in TMHs: 23.72226\nExp number, first 60 AAs: 0.22038\nTotal prob of N-in: 0.12994\noutside 1 331\nTMhelix 332 354\ninside 355 372

27675 GCF_001260735.1_8289_7_67 Shigella sonnei Proteobacteria; Gammaproteobacteria; Enterobacterales; Enterobacteriaceae; Shigella MTGDNTLIHSHGINRRDFMKLCAALAATMGLSSKAAAEMAESVTNPQRPPVIWIGAQECT WP_000145410.1 MULTISPECIES: hydrogenase-2 small chain [Proteobacteria] Length: 372\nNumber of predicted TMHs: 1\nExp number of AAs in TMHs: 23.72226\nExp number, first 60 AAs: 0.22038\nTotal prob of N-in: 0.12994\noutside 1 331\nTMhelix 332 354\ninside 355 372

27676 GCF_001260655.1_8290_4_1Shigella sonnei Proteobacteria; Gammaproteobacteria; Enterobacterales; Enterobacteriaceae; Shigella MTGDNTLIHSHGINRRDFMKLCAALAATMGLSSKAAAEMAESVTNPQRPPVIWIGAQECT WP_000145410.1 MULTISPECIES: hydrogenase-2 small chain [Proteobacteria] Length: 372\nNumber of predicted TMHs: 1\nExp number of AAs in TMHs: 23.72226\nExp number, first 60 AAs: 0.22038\nTotal prob of N-in: 0.12994\noutside 1 331\nTMhelix 332 354\ninside 355 372

27677 GCF_001260775.1_5417_1_3Shigella sonnei Proteobacteria; Gammaproteobacteria; Enterobacterales; Enterobacteriaceae; Shigella MTGDNTLIHSHGINRRDFMKLCAALAATMGLSSKAAAEMAESVTNPQRPPVIWIGAQECT WP_000145410.1 MULTISPECIES: hydrogenase-2 small chain [Proteobacteria] Length: 372\nNumber of predicted TMHs: 1\nExp number of AAs in TMHs: 23.72226\nExp number, first 60 AAs: 0.22038\nTotal prob of N-in: 0.12994\noutside 1 331\nTMhelix 332 354\ninside 355 372

27678 GCF_001260655.1_8290_4_1Shigella sonnei Proteobacteria; Gammaproteobacteria; Enterobacterales; Enterobacteriaceae; Shigella MNNEETFYQAMRRQGVTRRSFLKYCSLAATSLGLGAGMAPKIAWALENKPRIPVVWIHGL WP_005139779.1 hydrogenase [Shigella sonnei] Length: 372\nNumber of predicted TMHs: 2\nExp number of AAs in TMHs: 37.02628\nExp number, first 60 AAs: 19.39684\nTotal prob of N-in: 0.94313\nPOSSIBLE N-term signal sequence\ninside 1 19\nTMhelix 20 39\noutside 40 325\nTMhelix 326 348\ninside 349 372

27679 GCF_001260775.1_5417_1_3Shigella sonnei Proteobacteria; Gammaproteobacteria; Enterobacterales; Enterobacteriaceae; Shigella MNNEETFYQAMRRQGVTRRSFLKYCSLAATSLGLGAGMAPKIAWALENKPRIPVVWIHGL WP_005139779.1 hydrogenase [Shigella sonnei] Length: 372\nNumber of predicted TMHs: 2\nExp number of AAs in TMHs: 37.02628\nExp number, first 60 AAs: 19.39684\nTotal prob of N-in: 0.94313\nPOSSIBLE N-term signal sequence\ninside 1 19\nTMhelix 20 39\noutside 40 325\nTMhelix 326 348\ninside 349 372

27680 GCF_001260355.1_8290_5_4Shigella sonnei Proteobacteria; Gammaproteobacteria; Enterobacterales; Enterobacteriaceae; Shigella MNNEETFYQAMRRQGVTRRSFLKYCSLAATSLGLGAGMAPKIAWALENKPRIPVVWIHGL WP_005139779.1 hydrogenase [Shigella sonnei] Length: 372\nNumber of predicted TMHs: 2\nExp number of AAs in TMHs: 37.02628\nExp number, first 60 AAs: 19.39684\nTotal prob of N-in: 0.94313\nPOSSIBLE N-term signal sequence\ninside 1 19\nTMhelix 20 39\noutside 40 325\nTMhelix 326 348\ninside 349 372

27681 GCF_001260575.1_5236_2_3Shigella sonnei Proteobacteria; Gammaproteobacteria; Enterobacterales; Enterobacteriaceae; Shigella MNNEETFYQAMRRQGVTRRSFLKYCSLAATSLGLGAGMAPKIAWALENKPRIPVVWIHGL WP_005139779.1 hydrogenase [Shigella sonnei] Length: 372\nNumber of predicted TMHs: 2\nExp

number of AAs in TMHs: 37.02628\nExp number, first 60 AAs: 19.39684\nTotal prob of N-in: 0.94313\nPOSSIBLE N-term signal sequence\ninside 1 19\nTMhelix 20 39\noutside 40 325\nTMhelix 326 348\ninside 349 372

27682 GCF_001260395.1_8289_7_35 Shigella sonnei Proteobacteria; Gammaproteobacteria; Enterobacterales; Enterobacteriaceae; Shigella MNNEETFYQAMRRQGVTRRSFLKYCSLAATSLGLGAGMAPKIAWALENKPRIPVVWIHGL WP_005139779.1 hydrogenase [Shigella sonnei] Length: 372\nNumber of predicted TMHs: 2\nExp number of AAs in TMHs: 37.02628\nExp number, first 60 AAs: 19.39684\nTotal prob of N-in: 0.94313\nPOSSIBLE N-term signal sequence\ninside 1 19\nTMhelix 20 39\noutside 40 325\nTMhelix 326 348\ninside 349 372

27683 GCF_001260475.1_5236_2_8Shigella sonnei Proteobacteria; Gammaproteobacteria; Enterobacterales; Enterobacteriaceae; Shigella MNNEETFYQAMRRQGVTRRSFLKYCSLAATSLGLGAGMAPKIAWALENKPRIPVVWIHGL WP_005139779.1 hydrogenase [Shigella sonnei] Length: 372\nNumber of predicted TMHs: 2\nExp number of AAs in TMHs: 37.02628\nExp number, first 60 AAs: 19.39684\nTotal prob of N-in: 0.94313\nPOSSIBLE N-term signal sequence\ninside 1 19\nTMhelix 20 39\noutside 40 325\nTMhelix 326 348\ninside 349 372

27684 GCF_001260835.1_12971_8_35 Shigella sonnei Proteobacteria; Gammaproteobacteria; Enterobacterales; Enterobacteriaceae; Shigella MTGDNTLIHSHGINRRDFMKLCAALAATMGLSSKAAAEMAESVTNPQRPPVIWIGAQECT WP_000145410.1 MULTISPECIES: hydrogenase-2 small chain [Proteobacteria] Length: 372\nNumber of predicted TMHs: 1\nExp number of AAs in TMHs: 23.72226\nExp number, first 60 AAs: 0.22038\nTotal prob of N-in: 0.12994\noutside 1 331\nTMhelix 332 354\ninside 355 372

27685 GCF_001260835.1_12971_8_35 Shigella sonnei Proteobacteria; Gammaproteobacteria; Enterobacterales; Enterobacteriaceae; Shigella MNNEETFYQAMRRQGVTRRSFLKYCSLAATSLGLGAGMAPKIAWALENKPRIPVVWIHGL WP_005139779.1 hydrogenase [Shigella sonnei] Length: 372\nNumber of predicted TMHs: 2\nExp number of AAs in TMHs: 37.02628\nExp number, first 60 AAs: 19.39684\nTotal prob of N-in: 0.94313\nPOSSIBLE N-term signal sequence\ninside 1 19\nTMhelix 20 39\noutside 40 325\nTMhelix 326 348\ninside 349 372

27686 GCF_001260515.1_Sg_201403956_CACGAT_L001 Shigella sonnei Proteobacteria; Gammaproteobacteria; Enterobacterales; Enterobacteriaceae; Shigella MNNEETFYQAMRRQGVTRRSFLKYCSLAATSLGLGAGMAPKIAWALENKPRIPVVWIHGL WP_005139779.1 hydrogenase [Shigella sonnei] Length: 372\nNumber of predicted TMHs: 2\nExp number of AAs in TMHs: 37.02628\nExp number, first 60 AAs: 19.39684\nTotal prob of N-in: 0.94313\nPOSSIBLE N-term signal sequence\ninside 1 19\nTMhelix 20 39\noutside 40 325\nTMhelix 326 348\ninside 349 372

27687 GCF_001260875.1_9870_6_8Shigella sonnei Proteobacteria; Gammaproteobacteria; Enterobacterales; Enterobacteriaceae; Shigella MTGDNTLIHSHGINRRDFMKLCAALAATMGLSSKAAAEMAESVTNPQRPPVIWIGAQECT WP_000145410.1 MULTISPECIES: hydrogenase-2 small chain [Proteobacteria] Length: 372\nNumber of predicted TMHs: 1\nExp number of AAs in TMHs: 23.72226\nExp number, first 60 AAs: 0.22038\nTotal prob of N-in: 0.12994\noutside 1 331\nTMhelix 332 354\ninside 355 372

27688 GCF_001260955.1_8290_4_19 Shigella sonnei Proteobacteria; Gammaproteobacteria; Enterobacterales; Enterobacteriaceae; Shigella MTGDNTLIHSHGINRRDFMKLCAALAATMGLSSKAAAEMAESVTNPQRPPVIWIGAQECT WP_000145410.1 MULTISPECIES: hydrogenase-2 small chain [Proteobacteria] Length: 372\nNumber of predicted TMHs: 1\nExp number of AAs in TMHs: 23.72226\nExp number, first 60 AAs: 0.22038\nTotal prob of N-in: 0.12994\noutside 1 331\nTMhelix 332 354\ninside 355 372

27689 GCF_001260875.1_9870_6_8Shigella sonnei Proteobacteria; Gammaproteobacteria; Enterobacterales; Enterobacteriaceae; Shigella MNNEETFYQAMRRQGVTRRSFLKYCSLAATSLGLGAGMAPKIAWALENKPRIPVVWIHGL WP_005139779.1 hydrogenase [Shigella sonnei] Length: 372\nNumber of predicted TMHs: 2\nExp number of AAs in TMHs: 37.02628\nExp number, first 60 AAs: 19.39684\nTotal prob of N-in: 0.94313\nPOSSIBLE N-term signal sequence\ninside 1 19\nTMhelix 20 39\noutside 40 325\nTMhelix 326 348\ninside 349 372

27690 GCF_001260955.1_8290_4_19 Shigella sonnei Proteobacteria; Gammaproteobacteria; Enterobacterales; Enterobacteriaceae; Shigella MNNEETFYQAMRRQGVTRRSFLKYCSLAATSLGLGAGMAPKIAWALENKPRIPVVWIHGL WP_005139779.1 hydrogenase [Shigella sonnei] Length: 372\nNumber of predicted TMHs: 2\nExp number of AAs in

TMHs: 37.02628\nExp number, first 60 AAs: 19.39684\nTotal prob of N-in: 0.94313\nPOSSIBLE N-term signal sequence\ninside 1 19\nTMhelix 20 39\noutside 40 325\nTMhelix 326 348\ninside 349 372

27691 GCF_001260595.1_8289_7_63 Shigella sonnei Proteobacteria; Gammaproteobacteria; Enterobacterales; Enterobacteriaceae; Shigella MNNEETFYQAMRRQGVTRRSFLKYCSLAATSLGLGAGMAPKIAWALENKPRIPVVWIHGL WP_005139779.1 hydrogenase [Shigella sonnei] Length: 372\nNumber of predicted TMHs: 2\nExp number of AAs in TMHs: 37.02628\nExp number, first 60 AAs: 19.39684\nTotal prob of N-in: 0.94313\nPOSSIBLE N-term signal sequence\ninside 1 19\nTMhelix 20 39\noutside 40 325\nTMhelix 326 348\ninside 349 372

27692 GCF_001260735.1_8289_7_67 Shigella sonnei Proteobacteria; Gammaproteobacteria; Enterobacterales; Enterobacteriaceae; Shigella MNNEETFYQAMRRQGVTRRSFLKYCSLAATSLGLGAGMAPKIAWALENKPRIPVVWIHGL WP_005139779.1 hydrogenase [Shigella sonnei] Length: 372\nNumber of predicted TMHs: 2\nExp number of AAs in TMHs: 37.02628\nExp number, first 60 AAs: 19.39684\nTotal prob of N-in: 0.94313\nPOSSIBLE N-term signal sequence\ninside 1 19\nTMhelix 20 39\noutside 40 325\nTMhelix 326 348\ninside 349 372

27693 GCF_001261015.1_5236_3_5Shigella sonnei Proteobacteria; Gammaproteobacteria; Enterobacterales; Enterobacteriaceae; Shigella MTGDNTLIHSHGINRRDFMKLCAALAATMGLSSKAAAEMAESVTNPQRPPVIWIGAQECT WP_000145410.1 MULTISPECIES: hydrogenase-2 small chain [Proteobacteria] Length: 372\nNumber of predicted TMHs: 1\nExp number of AAs in TMHs: 23.72226\nExp number, first 60 AAs: 0.22038\nTotal prob of N-in: 0.12994\noutside 1 331\nTMhelix 332 354\ninside 355 372

27694 GCF_001261115.1_9870_6_32 Shigella sonnei Proteobacteria; Gammaproteobacteria; Enterobacterales; Enterobacteriaceae; Shigella MTGDNTLIHSHGINRRDFMKLCAALAATMGLSSKAAAEMAESVTNPQRPPVIWIGAQECT WP_000145410.1 MULTISPECIES: hydrogenase-2 small chain [Proteobacteria] Length: 372\nNumber of predicted TMHs: 1\nExp number of AAs in TMHs: 23.72226\nExp number, first 60 AAs: 0.22038\nTotal prob of N-in: 0.12994\noutside 1 331\nTMhelix 332 354\ninside 355 372

27695 GCF_001261235.1_12971_8_49 Shigella sonnei Proteobacteria; Gammaproteobacteria; Enterobacterales; Enterobacteriaceae; Shigella MTGDNTLIHSHGINRRDFMKLCAALAATMGLSSKAAAEMAESVTNPQRPPVIWIGAQECT WP_000145410.1 MULTISPECIES: hydrogenase-2 small chain [Proteobacteria] Length: 372\nNumber of predicted TMHs: 1\nExp number of AAs in TMHs: 23.72226\nExp number, first 60 AAs: 0.22038\nTotal prob of N-in: 0.12994\noutside 1 331\nTMhelix 332 354\ninside 355 372

27696 GCF_001261175.1_12971_8_68 Shigella sonnei Proteobacteria; Gammaproteobacteria; Enterobacterales; Enterobacteriaceae; Shigella MTGDNTLIHSHGINRRDFMKLCAALAATMGLSSKAAAEMAESVTNPQRPPVIWIGAQECT WP_000145410.1 MULTISPECIES: hydrogenase-2 small chain [Proteobacteria] Length: 372\nNumber of predicted TMHs: 1\nExp number of AAs in TMHs: 23.72226\nExp number, first 60 AAs: 0.22038\nTotal prob of N-in: 0.12994\noutside 1 331\nTMhelix 332 354\ninside 355 372

27697 GCF_001261295.1_12971_8_59 Shigella sonnei Proteobacteria; Gammaproteobacteria; Enterobacterales; Enterobacteriaceae; Shigella MTGDNTLIHSHGINRRDFMKLCAALAATMGLSSKAAAEMAESVTNPQRPPVIWIGAQECT WP_000145410.1 MULTISPECIES: hydrogenase-2 small chain [Proteobacteria] Length: 372\nNumber of predicted TMHs: 1\nExp number of AAs in TMHs: 23.72226\nExp number, first 60 AAs: 0.22038\nTotal prob of N-in: 0.12994\noutside 1 331\nTMhelix 332 354\ninside 355 372

27698 GCF_001261375.1_5008_7_1Shigella sonnei Proteobacteria; Gammaproteobacteria; Enterobacterales; Enterobacteriaceae; Shigella MTGDNTLIHSHGINRRDFMKLCAALAATMGLSSKAAAEMAESVTNPQRPPVIWIGAQECT WP_000145410.1 MULTISPECIES: hydrogenase-2 small chain [Proteobacteria] Length: 372\nNumber of predicted TMHs: 1\nExp number of AAs in TMHs: 23.72226\nExp number, first 60 AAs: 0.22038\nTotal prob of N-in: 0.12994\noutside 1 331\nTMhelix 332 354\ninside 355 372

27699 GCF_001261415.1_8290_4_95 Shigella sonnei Proteobacteria; Gammaproteobacteria; Enterobacterales; Enterobacteriaceae; Shigella MTGDNTLIHSHGINRRDFMKLCAALAATMGLSSKAAAEMAESVTNPQRPPVIWIGAQECT WP_000145410.1 MULTISPECIES: hydrogenase-2 small chain [Proteobacteria] Length: 372\nNumber of predicted TMHs:

1\nExp number of AAs in TMHs: 23.72226\nExp number, first 60 AAs: 0.22038\nTotal prob of N-in: 0.12994\noutside 1 331\nTMhelix 332 354\ninside 355 372

27700 GCF_001261575.1_12844_1_60 Shigella sonnei Proteobacteria; Gammaproteobacteria; Enterobacterales; Enterobacteriaceae; Shigella
MTGDNTLIHSHGINRRDFMKLCAALAATMGLSSKAAAEMAESVTNPQRPPVIWIGAQECT WP_000145410.1
MULTISPECIES: hydrogenase-2 small chain [Proteobacteria] Length: 372\nNumber of predicted TMHs: 1\nExp number of AAs in TMHs: 23.72226\nExp number, first 60 AAs: 0.22038\nTotal prob of N-in: 0.12994\noutside 1 331\nTMhelix 332 354\ninside 355 372

27701 GCF_001261575.1_12844_1_60 Shigella sonnei Proteobacteria; Gammaproteobacteria; Enterobacterales; Enterobacteriaceae; Shigella
MNNEETFYQAMRRQGVTRRSFLKYCSLAATSLGLGAGMAPKIAWALENKPRIPVVWIHGL WP_005139779.1
hydrogenase [Shigella sonnei] Length: 372\nNumber of predicted TMHs: 2\nExp number of AAs in TMHs: 37.02628\nExp number, first 60 AAs: 19.39684\nTotal prob of N-in: 0.94313\nPOSSIBLE N-term signal sequence\ninside 1 19\nTMhelix 20 39\noutside 40 325\nTMhelix 326 348\ninside 349 372

27702 GCF_001261015.1_5236_3_5Shigella sonnei Proteobacteria; Gammaproteobacteria; Enterobacterales; Enterobacteriaceae; Shigella MNNEETFYQAMRRQGVTRRSFLKYCSLAATSLGLGAGMAPKIAWALENKPRIPVVWIHGL
WP_005139779.1 hydrogenase [Shigella sonnei] Length: 372\nNumber of predicted TMHs: 2\nExp number of AAs in TMHs: 37.02628\nExp number, first 60 AAs: 19.39684\nTotal prob of N-in: 0.94313\nPOSSIBLE N-term signal sequence\ninside 1 19\nTMhelix 20 39\noutside 40 325\nTMhelix 326 348\ninside 349 372

27703 GCF_001261115.1_9870_6_32 Shigella sonnei Proteobacteria; Gammaproteobacteria; Enterobacterales; Enterobacteriaceae; Shigella
MNNEETFYQAMRRQGVTRRSFLKYCSLAATSLGLGAGMAPKIAWALENKPRIPVVWIHGL WP_005139779.1
hydrogenase [Shigella sonnei] Length: 372\nNumber of predicted TMHs: 2\nExp number of AAs in TMHs: 37.02628\nExp number, first 60 AAs: 19.39684\nTotal prob of N-in: 0.94313\nPOSSIBLE N-term signal sequence\ninside 1 19\nTMhelix 20 39\noutside 40 325\nTMhelix 326 348\ninside 349 372

27704 GCF_001261235.1_12971_8_49 Shigella sonnei Proteobacteria; Gammaproteobacteria; Enterobacterales; Enterobacteriaceae; Shigella
MNNEETFYQAMRRQGVTRRSFLKYCSLAATSLGLGAGMAPKIAWALENKPRIPVVWIHGL WP_005139779.1
hydrogenase [Shigella sonnei] Length: 372\nNumber of predicted TMHs: 2\nExp number of AAs in TMHs: 37.02628\nExp number, first 60 AAs: 19.39684\nTotal prob of N-in: 0.94313\nPOSSIBLE N-term signal sequence\ninside 1 19\nTMhelix 20 39\noutside 40 325\nTMhelix 326 348\ninside 349 372

27705 GCF_001261175.1_12971_8_68 Shigella sonnei Proteobacteria; Gammaproteobacteria; Enterobacterales; Enterobacteriaceae; Shigella
MNNEETFYQAMRRQGVTRRSFLKYCSLAATSLGLGAGMAPKIAWALENKPRIPVVWIHGL WP_005139779.1
hydrogenase [Shigella sonnei] Length: 372\nNumber of predicted TMHs: 2\nExp number of AAs in TMHs: 37.02628\nExp number, first 60 AAs: 19.39684\nTotal prob of N-in: 0.94313\nPOSSIBLE N-term signal sequence\ninside 1 19\nTMhelix 20 39\noutside 40 325\nTMhelix 326 348\ninside 349 372

27706 GCF_001261635.1_12850_1_49 Shigella sonnei Proteobacteria; Gammaproteobacteria; Enterobacterales; Enterobacteriaceae; Shigella
MTGDNTLIHSHGINRRDFMKLCAALAATMGLSSKAAAEMAESVTNPQRPPVIWIGAQECT WP_000145410.1
MULTISPECIES: hydrogenase-2 small chain [Proteobacteria] Length: 372\nNumber of predicted TMHs: 1\nExp number of AAs in TMHs: 23.72226\nExp number, first 60 AAs: 0.22038\nTotal prob of N-in: 0.12994\noutside 1 331\nTMhelix 332 354\ninside 355 372

27707 GCF_001261675.1_8290_5_14 Shigella sonnei Proteobacteria; Gammaproteobacteria; Enterobacterales; Enterobacteriaceae; Shigella
MTGDNTLIHSHGINRRDFMKLCAALAATMGLSSKAAAEMAESVTNPQRPPVIWIGAQECT WP_000145410.1
MULTISPECIES: hydrogenase-2 small chain [Proteobacteria] Length: 372\nNumber of predicted TMHs: 1\nExp number of AAs in TMHs: 23.72226\nExp number, first 60 AAs: 0.22038\nTotal prob of N-in: 0.12994\noutside 1 331\nTMhelix 332 354\ninside 355 372

27708 GCF_001261635.1_12850_1_49 Shigella sonnei Proteobacteria; Gammaproteobacteria; Enterobacterales; Enterobacteriaceae; Shigella
MNNEETFYQAMRRQGVTRRSFLKYCSLAATSLGLGAGMAPKIAWALENKPRIPVVWIHGL WP_005139779.1

hydrogenase [Shigella sonnei] Length: 372\nNumber of predicted TMHs: 2\nExp number of AAs in TMHs: 37.02628\nExp number, first 60 AAs: 19.39684\nTotal prob of N-in: 0.94313\nPOSSIBLE N-term signal sequence\ninside 1 19\nTMhelix 20 39\noutside 40 325\nTMhelix 326 348\ninside 349 372

27709 GCF_001261675.1_8290_5_14 Shigella sonnei Proteobacteria; Gammaproteobacteria; Enterobacterales; Enterobacteriaceae; Shigella MNNEETFYQAMRRQGVTRRSFLKYCSLAATSLGLGAGMAPKIAWALENKPRIPVVWIHGL WP_005139779.1 hydrogenase [Shigella sonnei] Length: 372\nNumber of predicted TMHs: 2\nExp number of AAs in TMHs: 37.02628\nExp number, first 60 AAs: 19.39684\nTotal prob of N-in: 0.94313\nPOSSIBLE N-term signal sequence\ninside 1 19\nTMhelix 20 39\noutside 40 325\nTMhelix 326 348\ninside 349 372

27710 GCF_001261295.1_12971_8_59 Shigella sonnei Proteobacteria; Gammaproteobacteria; Enterobacterales; Enterobacteriaceae; Shigella MNNEETFYQAMRRQGVTRRSFLKYCSLAATSLGLGAGMAPKIAWALENKPRIPVVWIHGL WP_005139779.1 hydrogenase [Shigella sonnei] Length: 372\nNumber of predicted TMHs: 2\nExp number of AAs in TMHs: 37.02628\nExp number, first 60 AAs: 19.39684\nTotal prob of N-in: 0.94313\nPOSSIBLE N-term signal sequence\ninside 1 19\nTMhelix 20 39\noutside 40 325\nTMhelix 326 348\ninside 349 372

27711 GCF_001262135.1_8290_5_1Shigella sonnei Proteobacteria; Gammaproteobacteria; Enterobacterales; Enterobacteriaceae; Shigella MTGDNTLIHSHGINRRDFMKLCAALAATMGLSSKAAAEMAESVTNPQRPPVIWIGAQECT WP_000145410.1 MULTISPECIES: hydrogenase-2 small chain [Proteobacteria] Length: 372\nNumber of predicted TMHs: 1\nExp number of AAs in TMHs: 23.72226\nExp number, first 60 AAs: 0.22038\nTotal prob of N-in: 0.12994\noutside 1 331\nTMhelix 332 354\ninside 355 372

27712 GCF_001262135.1_8290_5_1Shigella sonnei Proteobacteria; Gammaproteobacteria; Enterobacterales; Enterobacteriaceae; Shigella MNNEETFYQAMRRQGVTRRSFLKYCSLAATSLGLGAGMAPKIAWALENKPRIPVVWIHGL WP_005139779.1 hydrogenase [Shigella sonnei] Length: 372\nNumber of predicted TMHs: 2\nExp number of AAs in TMHs: 37.02628\nExp number, first 60 AAs: 19.39684\nTotal prob of N-in: 0.94313\nPOSSIBLE N-term signal sequence\ninside 1 19\nTMhelix 20 39\noutside 40 325\nTMhelix 326 348\ninside 349 372

27713 GCF_001261415.1_8290_4_95 Shigella sonnei Proteobacteria; Gammaproteobacteria; Enterobacterales; Enterobacteriaceae; Shigella MNNEETFYQAMRRQGVTRRSFLKYCSLAATSLGLGAGMAPKIAWALENKPRIPVVWIHGL WP_005139779.1 hydrogenase [Shigella sonnei] Length: 372\nNumber of predicted TMHs: 2\nExp number of AAs in TMHs: 37.02628\nExp number, first 60 AAs: 19.39684\nTotal prob of N-in: 0.94313\nPOSSIBLE N-term signal sequence\ninside 1 19\nTMhelix 20 39\noutside 40 325\nTMhelix 326 348\ninside 349 372

27714 GCF_001687705.1_ASM168770v1 Shigella sonnei Proteobacteria; Gammaproteobacteria; Enterobacterales; Enterobacteriaceae; Shigella MTGDNTLIHSHGINRRDFMKLCAALAATMGLSSKAAAEMAESVTNPQRPPVIWIGAQECT WP_000145410.1 MULTISPECIES: hydrogenase-2 small chain [Proteobacteria] Length: 372\nNumber of predicted TMHs: 1\nExp number of AAs in TMHs: 23.72226\nExp number, first 60 AAs: 0.22038\nTotal prob of N-in: 0.12994\noutside 1 331\nTMhelix 332 354\ninside 355 372

27715 GCF_001687785.1_ASM168778v1 Shigella sonnei Proteobacteria; Gammaproteobacteria; Enterobacterales; Enterobacteriaceae; Shigella MTGDNTLIHSHGINRRDFMKLCAALAATMGLSSKAAAEMAESVTNPQRPPVIWIGAQECT WP_000145410.1 MULTISPECIES: hydrogenase-2 small chain [Proteobacteria] Length: 372\nNumber of predicted TMHs: 1\nExp number of AAs in TMHs: 23.72226\nExp number, first 60 AAs: 0.22038\nTotal prob of N-in: 0.12994\noutside 1 331\nTMhelix 332 354\ninside 355 372

27716 GCF_001687705.1_ASM168770v1 Shigella sonnei Proteobacteria; Gammaproteobacteria; Enterobacterales; Enterobacteriaceae; Shigella MNNEETFYQAMRRQGVTRRSFLKYCSLAATSLGLGAGMAPKIAWALENKPRIPVVWIHGL WP_005139779.1 hydrogenase [Shigella sonnei] Length: 372\nNumber of predicted TMHs: 2\nExp number of AAs in TMHs: 37.02628\nExp number, first 60 AAs: 19.39684\nTotal prob of N-in: 0.94313\nPOSSIBLE N-term signal sequence\ninside 1 19\nTMhelix 20 39\noutside 40 325\nTMhelix 326 348\ninside 349 372

27717 GCF_001687785.1_ASM168778v1 Shigella sonnei Proteobacteria; Gammaproteobacteria; Enterobacterales; Enterobacteriaceae; Shigella MNNEETFYQAMRRQGVTRRSFLKYCSLAATSLGLGAGMAPKIAWALENKPRIPVVWIHGL WP_005139779.1

hydrogenase [Shigella sonnei] Length: 372\nNumber of predicted TMHs: 2\nExp number of AAs in TMHs: 37.02628\nExp number, first 60 AAs: 19.39684\nTotal prob of N-in: 0.94313\nPOSSIBLE N-term signal sequence\ninside 1 19\nTMhelix 20 39\noutside 40 325\nTMhelix 326 348\ninside 349 372

27718 GCF_001261375.1_5008_7_1Shigella sonnei Proteobacteria; Gammaproteobacteria; Enterobacterales; Enterobacteriaceae; Shigella MNNEETFYQAMRRQGVTRRSFLKYCSLAATSLGLGAGMAPKIAWALENKPRIPVVWIHGL WP_005139779.1 hydrogenase [Shigella sonnei] Length: 372\nNumber of predicted TMHs: 2\nExp number of AAs in TMHs: 37.02628\nExp number, first 60 AAs: 19.39684\nTotal prob of N-in: 0.94313\nPOSSIBLE N-term signal sequence\ninside 1 19\nTMhelix 20 39\noutside 40 325\nTMhelix 326 348\ninside 349 372

27719 GCF_001687865.1_ASM168786v1 Shigella sonnei Proteobacteria; Gammaproteobacteria; Enterobacterales; Enterobacteriaceae; Shigella MTGDNTLIHSHGINRRDFMKLCAALAATMGLSSKAAAEMAESVTNPQRPPVIWIGAQECT WP_000145410.1 MULTISPECIES: hydrogenase-2 small chain [Proteobacteria] Length: 372\nNumber of predicted TMHs: 1\nExp number of AAs in TMHs: 23.72226\nExp number, first 60 AAs: 0.22038\nTotal prob of N-in: 0.12994\noutside 1 331\nTMhelix 332 354\ninside 355 372

27720 GCF_001722135.1_ASM172213v1 Shigella sp. FC130 Proteobacteria; Gammaproteobacteria; Enterobacterales; Enterobacteriaceae; Shigella MALGNNVFSHHGINRRDFMKLCAALSATMGLSGKAAAEMSEAMASPERPPVIWIGAQECT WP_069368630.1 MULTISPECIES: hydrogenase 2 small subunit [Shigella] Length: 374\nNumber of predicted TMHs: 1\nExp number of AAs in TMHs: 21.24001\nExp number, first 60 AAs: 0.08735\nTotal prob of N-in: 0.01872\noutside 1 333\nTMhelix 334 356\ninside 357 374

27721 GCF_001742985.1_ASM174298v1 Shigella sp. FC1655 Proteobacteria; Gammaproteobacteria; Enterobacterales; Enterobacteriaceae; Shigella MALGNNVFSHHGINRRDFMKLCAALSATMGLSGKAAAEMSEAMASPERPPVIWIGAQECT WP_069368630.1 MULTISPECIES: hydrogenase 2 small subunit [Shigella] Length: 374\nNumber of predicted TMHs: 1\nExp number of AAs in TMHs: 21.24001\nExp number, first 60 AAs: 0.08735\nTotal prob of N-in: 0.01872\noutside 1 333\nTMhelix 334 356\ninside 357 374

27722 GCF_001730415.1_ASM173041v1 Shigella sp. FC2175 Proteobacteria; Gammaproteobacteria; Enterobacterales; Enterobacteriaceae; Shigella MNNEETFYQAMRRQGVTRRSFLKYCSLAATSLGLGAGMAPKIAWALENKPRIPVVWIHGL WP_001058323.1 MULTISPECIES: hydrogenase-1 small chain [Proteobacteria] Length: 372\nNumber of predicted TMHs: 2\nExp number of AAs in TMHs: 37.2608499999999\nExp number, first 60 AAs: 19.52923\nTotal prob of N-in: 0.94351\nPOSSIBLE N-term signal sequence\ninside 1 19\nTMhelix 20 39\noutside 40 325\nTMhelix 326 348\ninside 349 372

27723 GCF_001722155.1_ASM172215v1 Shigella sp. FC569 Proteobacteria; Gammaproteobacteria; Enterobacterales; Enterobacteriaceae; Shigella MNNEETFYQAMRRQGVTRRSFLKYCSLAATSLGLGAGMAPKIAWALENKPRIPVVWIHGL WP_001058323.1 MULTISPECIES: hydrogenase-1 small chain [Proteobacteria] Length: 372\nNumber of predicted TMHs: 2\nExp number of AAs in TMHs: 37.2608499999999\nExp number, first 60 AAs: 19.52923\nTotal prob of N-in: 0.94351\nPOSSIBLE N-term signal sequence\ninside 1 19\nTMhelix 20 39\noutside 40 325\nTMhelix 326 348\ninside 349 372

27724 GCF_001722155.1_ASM172215v1 Shigella sp. FC569 Proteobacteria; Gammaproteobacteria; Enterobacterales; Enterobacteriaceae; Shigella MTGDNTLIHSHGINRRDFMKLCAALAATMGLSSKAAAEMAESVTNPQRPPVIWIGAQECT WP_000145410.1 MULTISPECIES: hydrogenase-2 small chain [Proteobacteria] Length: 372\nNumber of predicted TMHs: 1\nExp number of AAs in TMHs: 23.72226\nExp number, first 60 AAs: 0.22038\nTotal prob of N-in: 0.12994\noutside 1 331\nTMhelix 332 354\ninside 355 372

27725 GCF_001596115.1_ASM159611v1 Shigella sp. PAMC 28760 Proteobacteria; Gammaproteobacteria; Enterobacterales; Enterobacteriaceae; Shigella MTGDNTLIHSHGINRRDFMKLCAALAATMGLSSKAAAEMAESVTNPQRPPVIWIGAQECT WP_000145410.1 MULTISPECIES: hydrogenase-2 small chain [Proteobacteria] Length: 372\nNumber of predicted TMHs: 1\nExp number of AAs in TMHs: 23.72226\nExp number, first 60 AAs: 0.22038\nTotal prob of N-in: 0.12994\noutside 1 331\nTMhelix 332 354\ninside 355 372

27726	GCF_001596115.1_ASM159611v1	Shigella sp. PAMC 28760	Proteobacteria; Gammaproteobacteria; Enterobacterales; Enterobacteriaceae; Shigella
MNNEETFYQAMRRQGVTRRSFLKYCSLAATSLGLGAGMAPKIAWALENKPRIPVVWIHGL	WP_001058323.1
MULTISPECIES: hydrogenase-1 small chain [Proteobacteria]	Length: 372\nNumber of predicted TMHs: 2\nExp number of AAs in TMHs: 37.2608499999999\nExp number, first 60 AAs: 19.52923\nTotal prob of N-in: 0.94351\nPOSSIBLE N-term signal sequence\ninside 1 19\nTMhelix 20 39\noutside 40 325\nTMhelix 326 348\ninside 349 372

27727	GCF_000815685.1_ASM81568v1	Shigella boydii 08-0009	Proteobacteria; Gammaproteobacteria; Enterobacterales; Enterobacteriaceae; Shigella; Shigella boydii
MTGDNTLIHSHGINRRDFMKLCAALAATMGLSSKAAAEMAESVTNPQRPPVIWIGAQECT	WP_000145410.1
MULTISPECIES: hydrogenase-2 small chain [Proteobacteria]	Length: 372\nNumber of predicted TMHs: 1\nExp number of AAs in TMHs: 23.72226\nExp number, first 60 AAs: 0.22038\nTotal prob of N-in: 0.12994\noutside 1 331\nTMhelix 332 354\ninside 355 372

27728	GCF_000815685.1_ASM81568v1	Shigella boydii 08-0009	Proteobacteria; Gammaproteobacteria; Enterobacterales; Enterobacteriaceae; Shigella; Shigella boydii
MNNEETFYQAMRRQGVTRRSFLKYCSLAATSLGLGAGMAPKIAWALENKPRIPVVWIHGL	WP_001058323.1
MULTISPECIES: hydrogenase-1 small chain [Proteobacteria]	Length: 372\nNumber of predicted TMHs: 2\nExp number of AAs in TMHs: 37.2608499999999\nExp number, first 60 AAs: 19.52923\nTotal prob of N-in: 0.94351\nPOSSIBLE N-term signal sequence\ninside 1 19\nTMhelix 20 39\noutside 40 325\nTMhelix 326 348\ninside 349 372

27729	GCF_000815705.1_ASM81570v1	Shigella boydii 08-0280	Proteobacteria; Gammaproteobacteria; Enterobacterales; Enterobacteriaceae; Shigella; Shigella boydii
MTGDNTLIHSHGINRRDFMKLCAALAATMGLSSKAAAEMAESVTNPQRPPVIWIGAQECT	WP_000145410.1
MULTISPECIES: hydrogenase-2 small chain [Proteobacteria]	Length: 372\nNumber of predicted TMHs: 1\nExp number of AAs in TMHs: 23.72226\nExp number, first 60 AAs: 0.22038\nTotal prob of N-in: 0.12994\noutside 1 331\nTMhelix 332 354\ninside 355 372

27730	GCF_000815705.1_ASM81570v1	Shigella boydii 08-0280	Proteobacteria; Gammaproteobacteria; Enterobacterales; Enterobacteriaceae; Shigella; Shigella boydii
MNNEETFYQAMRRQGVTRRSFLKYCSLAATSLGLGAGMAPKIAWALENKPRIPVVWIHGL	WP_001058323.1
MULTISPECIES: hydrogenase-1 small chain [Proteobacteria]	Length: 372\nNumber of predicted TMHs: 2\nExp number of AAs in TMHs: 37.2608499999999\nExp number, first 60 AAs: 19.52923\nTotal prob of N-in: 0.94351\nPOSSIBLE N-term signal sequence\ninside 1 19\nTMhelix 20 39\noutside 40 325\nTMhelix 326 348\ninside 349 372

27731	GCF_000815725.1_ASM81572v1	Shigella boydii 08-2671	Proteobacteria; Gammaproteobacteria; Enterobacterales; Enterobacteriaceae; Shigella; Shigella boydii
MNNEETFYQAMRRQGVTRRSFLKYCSLAATSLGLGAGMAPKIAWALENKPRIPVVWIHGL	WP_001058323.1
MULTISPECIES: hydrogenase-1 small chain [Proteobacteria]	Length: 372\nNumber of predicted TMHs: 2\nExp number of AAs in TMHs: 37.2608499999999\nExp number, first 60 AAs: 19.52923\nTotal prob of N-in: 0.94351\nPOSSIBLE N-term signal sequence\ninside 1 19\nTMhelix 20 39\noutside 40 325\nTMhelix 326 348\ninside 349 372

27732	GCF_000815725.1_ASM81572v1	Shigella boydii 08-2671	Proteobacteria; Gammaproteobacteria; Enterobacterales; Enterobacteriaceae; Shigella; Shigella boydii
MTGDNTLIHSHGINRRDFMKLCAALAATMGLSSKAAAEMAESVTNPQRPPVIWIGAQECT	WP_000145410.1
MULTISPECIES: hydrogenase-2 small chain [Proteobacteria]	Length: 372\nNumber of predicted TMHs: 1\nExp number of AAs in TMHs: 23.72226\nExp number, first 60 AAs: 0.22038\nTotal prob of N-in: 0.12994\noutside 1 331\nTMhelix 332 354\ninside 355 372

27733	GCF_000815735.1_ASM81573v1	Shigella boydii 08-2675	Proteobacteria; Gammaproteobacteria; Enterobacterales; Enterobacteriaceae; Shigella; Shigella boydii
MTGDNTLIHSHGINRRDFMKLCAALAATMGLSSKAAAEMAESVTNPQRPPVIWIGAQECT	WP_000145410.1
MULTISPECIES: hydrogenase-2 small chain [Proteobacteria]	Length: 372\nNumber of predicted TMHs: 1\nExp number of AAs in TMHs: 23.72226\nExp number, first 60 AAs: 0.22038\nTotal prob of N-in: 0.12994\noutside 1 331\nTMhelix 332 354\ninside 355 372

27734 GCF_000815735.1_ASM81573v1 Shigella boydii 08-2675 Proteobacteria; Gammaproteobacteria; Enterobacterales; Enterobacteriaceae; Shigella; Shigella boydii
MNNEETFYQAMRRQGVTRRSFLKYCSLAATSLGLGAGMAPKIAWALENKPRIPVVWIHGL WP_001058323.1
MULTISPECIES: hydrogenase-1 small chain [Proteobacteria] Length: 372\nNumber of predicted TMHs: 2\nExp number of AAs in TMHs: 37.2608499999999\nExp number, first 60 AAs: 19.52923\nTotal prob of N-in: 0.94351\nPOSSIBLE N-term signal sequence\ninside 1 19\nTMhelix 20 39\noutside 40 325\nTMhelix 326 348\ninside 349 372

27735 GCF_000815765.1_ASM81576v1 Shigella boydii 08-6341 Proteobacteria; Gammaproteobacteria; Enterobacterales; Enterobacteriaceae; Shigella; Shigella boydii
MTGDNTLIHSHGINRRDFMKLCAALAATMGLSSKAAAEMAESVTNPQRPPVIWIGAQECT WP_000145410.1
MULTISPECIES: hydrogenase-2 small chain [Proteobacteria] Length: 372\nNumber of predicted TMHs: 1\nExp number of AAs in TMHs: 23.72226\nExp number, first 60 AAs: 0.22038\nTotal prob of N-in: 0.12994\noutside 1 331\nTMhelix 332 354\ninside 355 372

27736 GCF_000815765.1_ASM81576v1 Shigella boydii 08-6341 Proteobacteria; Gammaproteobacteria; Enterobacterales; Enterobacteriaceae; Shigella; Shigella boydii
MNNEETFYQAMRRQGVTRRSFLKYCSLAATSLGLGAGMAPKIAWALENKPRIPVVWIHGL WP_001058323.1
MULTISPECIES: hydrogenase-1 small chain [Proteobacteria] Length: 372\nNumber of predicted TMHs: 2\nExp number of AAs in TMHs: 37.2608499999999\nExp number, first 60 AAs: 19.52923\nTotal prob of N-in: 0.94351\nPOSSIBLE N-term signal sequence\ninside 1 19\nTMhelix 20 39\noutside 40 325\nTMhelix 326 348\ninside 349 372

27737 GCF_000815785.1_ASM81578v1 Shigella boydii 09-0344 Proteobacteria; Gammaproteobacteria; Enterobacterales; Enterobacteriaceae; Shigella; Shigella boydii
MNNEETFYQAMRRQGVTRRSFLKYCSLAATSLGLGAGMAPKIAWALENKPRIPVVWIHGL WP_001058323.1
MULTISPECIES: hydrogenase-1 small chain [Proteobacteria] Length: 372\nNumber of predicted TMHs: 2\nExp number of AAs in TMHs: 37.2608499999999\nExp number, first 60 AAs: 19.52923\nTotal prob of N-in: 0.94351\nPOSSIBLE N-term signal sequence\ninside 1 19\nTMhelix 20 39\noutside 40 325\nTMhelix 326 348\ninside 349 372

27738 GCF_000815785.1_ASM81578v1 Shigella boydii 09-0344 Proteobacteria; Gammaproteobacteria; Enterobacterales; Enterobacteriaceae; Shigella; Shigella boydii
MTGDNTLIHSHGINRRDFMKLCAALAATMGLSSKAAAEMAESVTNPQRPPVIWIGAQECT WP_000145410.1
MULTISPECIES: hydrogenase-2 small chain [Proteobacteria] Length: 372\nNumber of predicted TMHs: 1\nExp number of AAs in TMHs: 23.72226\nExp number, first 60 AAs: 0.22038\nTotal prob of N-in: 0.12994\noutside 1 331\nTMhelix 332 354\ninside 355 372

27739 GCF_000268185.1_ASM26818v1 Shigella boydii 4444-74 Proteobacteria; Gammaproteobacteria; Enterobacterales; Enterobacteriaceae; Shigella; Shigella boydii
MTGDNTLIHSHGINRRDFMKLCAALAATMGLSSKAAAEMAESVTNPQRPPVIWIGAQECT WP_000145410.1
MULTISPECIES: hydrogenase-2 small chain [Proteobacteria] Length: 372\nNumber of predicted TMHs: 1\nExp number of AAs in TMHs: 23.72226\nExp number, first 60 AAs: 0.22038\nTotal prob of N-in: 0.12994\noutside 1 331\nTMhelix 332 354\ninside 355 372

27740 GCF_000268185.1_ASM26818v1 Shigella boydii 4444-74 Proteobacteria; Gammaproteobacteria; Enterobacterales; Enterobacteriaceae; Shigella; Shigella boydii
MNNEETFYQAMRRQGVTRRSFLKYCSLAATSLGLGAGMAPKIAWALENKPRIPVVWIHGL WP_001058323.1
MULTISPECIES: hydrogenase-1 small chain [Proteobacteria] Length: 372\nNumber of predicted TMHs: 2\nExp number of AAs in TMHs: 37.2608499999999\nExp number, first 60 AAs: 19.52923\nTotal prob of N-in: 0.94351\nPOSSIBLE N-term signal sequence\ninside 1 19\nTMhelix 20 39\noutside 40 325\nTMhelix 326 348\ninside 349 372

27741 GCF_000020185.1_ASM2018v1 Shigella boydii CDC 3083-94 Proteobacteria; Gammaproteobacteria; Enterobacterales; Enterobacteriaceae; Shigella; Shigella boydii
MNNEETFYQAMRRQGVTRRSFLKYCSLAATSLGLGAGMAPKIAWALENKPRIPVVWIHGL WP_001058323.1
MULTISPECIES: hydrogenase-1 small chain [Proteobacteria] Length: 372\nNumber of predicted TMHs: 2\nExp number of AAs in TMHs: 37.2608499999999\nExp number, first 60 AAs: 19.52923\nTotal prob of N-in: 0.94351\nPOSSIBLE N-term signal sequence\ninside 1 19\nTMhelix 20 39\noutside 40 325\nTMhelix 326 348\ninside 349 372

27742	GCF_000020185.1_ASM2018v1	Shigella boydii CDC 3083-94	Proteobacteria; Gammaproteobacteria; Enterobacterales; Enterobacteriaceae; Shigella; Shigella boydii
MTGDNTLIHSHGINRRDFMKLCAALAATMGLSSKAAAEMAESVTNPQRPPVIWIGAQECT	WP_000145410.1
MULTISPECIES: hydrogenase-2 small chain [Proteobacteria]	Length: 372\nNumber of predicted TMHs: 1\nExp number of AAs in TMHs: 23.72226\nExp number, first 60 AAs: 0.22038\nTotal prob of N-in: 0.12994\noutside 1 331\nTMhelix 332 354\ninside 355 372

27743	GCF_000815585.1_ASM81558v1	Shigella boydii S6614	Proteobacteria; Gammaproteobacteria; Enterobacterales; Enterobacteriaceae; Shigella; Shigella boydii
MNNEETFYQAMRRQGVTRRSFLKYCSLAATSLGLGAGMAPKIAWALENKPRIPVVWIHGL	WP_001058323.1
MULTISPECIES: hydrogenase-1 small chain [Proteobacteria]	Length: 372\nNumber of predicted TMHs: 2\nExp number of AAs in TMHs: 37.2608499999999\nExp number, first 60 AAs: 19.52923\nTotal prob of N-in: 0.94351\nPOSSIBLE N-term signal sequence\ninside 1 19\nTMhelix 20 39\noutside 40 325\nTMhelix 326 348\ninside 349 372

27744	GCF_000815585.1_ASM81558v1	Shigella boydii S6614	Proteobacteria; Gammaproteobacteria; Enterobacterales; Enterobacteriaceae; Shigella; Shigella boydii
MTGDNTLIHSHGINRRDFMKLCAALAATMGLSSKAAAEMAESVTNPQRPPVIWIGAQECT	WP_000145410.1
MULTISPECIES: hydrogenase-2 small chain [Proteobacteria]	Length: 372\nNumber of predicted TMHs: 1\nExp number of AAs in TMHs: 23.72226\nExp number, first 60 AAs: 0.22038\nTotal prob of N-in: 0.12994\noutside 1 331\nTMhelix 332 354\ninside 355 372

27745	GCF_000815645.1_ASM81564v1	Shigella boydii S7334	Proteobacteria; Gammaproteobacteria; Enterobacterales; Enterobacteriaceae; Shigella; Shigella boydii
MTGDNTLIHSHGINRRDFMKLCAALAATMGLSSKAAAEMAESVTNPQRPPVIWIGAQECT	WP_000145410.1
MULTISPECIES: hydrogenase-2 small chain [Proteobacteria]	Length: 372\nNumber of predicted TMHs: 1\nExp number of AAs in TMHs: 23.72226\nExp number, first 60 AAs: 0.22038\nTotal prob of N-in: 0.12994\noutside 1 331\nTMhelix 332 354\ninside 355 372

27746	GCF_000815645.1_ASM81564v1	Shigella boydii S7334	Proteobacteria; Gammaproteobacteria; Enterobacterales; Enterobacteriaceae; Shigella; Shigella boydii
MNNEETFYQAMRRQGVTRRSFLKYCSLAATSLGLGAGMAPKIAWALENKPRIPVVWIHGL	WP_001058323.1
MULTISPECIES: hydrogenase-1 small chain [Proteobacteria]	Length: 372\nNumber of predicted TMHs: 2\nExp number of AAs in TMHs: 37.2608499999999\nExp number, first 60 AAs: 19.52923\nTotal prob of N-in: 0.94351\nPOSSIBLE N-term signal sequence\ninside 1 19\nTMhelix 20 39\noutside 40 325\nTMhelix 326 348\ninside 349 372

27747	GCF_000012025.1_ASM1202v1	Shigella boydii Sb227	Proteobacteria; Gammaproteobacteria; Enterobacterales; Enterobacteriaceae; Shigella; Shigella boydii
MTGDNTLIHSHGINRRDFMKLCAALAATMGLSSKAAAEMAESVTNPQRPPVIWIGAQECT	WP_000145410.1
MULTISPECIES: hydrogenase-2 small chain [Proteobacteria]	Length: 372\nNumber of predicted TMHs: 1\nExp number of AAs in TMHs: 23.72226\nExp number, first 60 AAs: 0.22038\nTotal prob of N-in: 0.12994\noutside 1 331\nTMhelix 332 354\ninside 355 372

27748	GCF_000012025.1_ASM1202v1	Shigella boydii Sb227	Proteobacteria; Gammaproteobacteria; Enterobacterales; Enterobacteriaceae; Shigella; Shigella boydii
MNNEETFYQAMRRQGVTRRSFLKYCSLAATSLGLGAGMAPKIAWALENKPRIPVVWIHGL	WP_001058323.1
MULTISPECIES: hydrogenase-1 small chain [Proteobacteria]	Length: 372\nNumber of predicted TMHs: 2\nExp number of AAs in TMHs: 37.2608499999999\nExp number, first 60 AAs: 19.52923\nTotal prob of N-in: 0.94351\nPOSSIBLE N-term signal sequence\ninside 1 19\nTMhelix 20 39\noutside 40 325\nTMhelix 326 348\ninside 349 372

27749	GCF_000268105.1_ASM26810v1	Shigella dysenteriae 225-75	Proteobacteria; Gammaproteobacteria; Enterobacterales; Enterobacteriaceae; Shigella; Shigella dysenteriae
MTGDNTLIHSHGINRRDFMKLCAALAATMGLSSKAAAEMAESVTNPQRPPVIWIGAQECT	WP_000145410.1
MULTISPECIES: hydrogenase-2 small chain [Proteobacteria]	Length: 372\nNumber of predicted TMHs: 1\nExp number of AAs in TMHs: 23.72226\nExp number, first 60 AAs: 0.22038\nTotal prob of N-in: 0.12994\noutside 1 331\nTMhelix 332 354\ninside 355 372

27750	GCF_000268105.1_ASM26810v1	Shigella dysenteriae 225-75	Proteobacteria; Gammaproteobacteria; Enterobacterales; Enterobacteriaceae; Shigella; Shigella dysenteriae

MNNEETFYQAMRRQGVTRRSFLKYCSLAATSLGLGAGMAPKIAWALENKPRIPVVWIHGL WP_001058323.1 MULTISPECIES: hydrogenase-1 small chain [Proteobacteria] Length: 372\nNumber of predicted TMHs: 2\nExp number of AAs in TMHs: 37.2608499999999\nExp number, first 60 AAs: 19.52923\nTotal prob of N-in: 0.94351\nPOSSIBLE N-term signal sequence\ninside 1 19\nTMhelix 20 39\noutside 40 325\nTMhelix 326 348\ninside 349 372

27751 GCF_001017215.1_ASM101721v1 Shigella dysenteriae 4 Proteobacteria; Gammaproteobacteria; Enterobacterales; Enterobacteriaceae; Shigella; Shigella dysenteriae MTGDNTLIHSHGINRRDFMKLCAALAATMGLSSKAAAEMAESVTNPQRPPVIWIGAQECT WP_000145410.1 MULTISPECIES: hydrogenase-2 small chain [Proteobacteria] Length: 372\nNumber of predicted TMHs: 1\nExp number of AAs in TMHs: 23.72226\nExp number, first 60 AAs: 0.22038\nTotal prob of N-in: 0.12994\noutside 1 331\nTMhelix 332 354\ninside 355 372

27752 GCF_000193895.1_ASM19389v2 Shigella dysenteriae CDC 74-1112 Proteobacteria; Gammaproteobacteria; Enterobacterales; Enterobacteriaceae; Shigella; Shigella dysenteriae MTGDNTLIHSHGINRRDFMKLCAALAATMGLSSKAAAEMAESVTNPQRPPVIWIGAQECT WP_000145410.1 MULTISPECIES: hydrogenase-2 small chain [Proteobacteria] Length: 372\nNumber of predicted TMHs: 1\nExp number of AAs in TMHs: 23.72226\nExp number, first 60 AAs: 0.22038\nTotal prob of N-in: 0.12994\noutside 1 331\nTMhelix 332 354\ninside 355 372

27753 GCF_000815515.1_ASM81551v1 Shigella dysenteriae S6554 Proteobacteria; Gammaproteobacteria; Enterobacterales; Enterobacteriaceae; Shigella; Shigella dysenteriae MNNEETFYQAMRRQGVTRRSFLKYCSLAATSLGLGAGMAPKIAWALENKPRIPVVWIHGL WP_001058323.1 MULTISPECIES: hydrogenase-1 small chain [Proteobacteria] Length: 372\nNumber of predicted TMHs: 2\nExp number of AAs in TMHs: 37.2608499999999\nExp number, first 60 AAs: 19.52923\nTotal prob of N-in: 0.94351\nPOSSIBLE N-term signal sequence\ninside 1 19\nTMhelix 20 39\noutside 40 325\nTMhelix 326 348\ninside 349 372

27754 GCF_000815515.1_ASM81551v1 Shigella dysenteriae S6554 Proteobacteria; Gammaproteobacteria; Enterobacterales; Enterobacteriaceae; Shigella; Shigella dysenteriae MTGDNTLIHSHGINRRDFMKLCAALAATMGLSSKAAAEMAESVTNPQRPPVIWIGAQECT WP_000145410.1 MULTISPECIES: hydrogenase-2 small chain [Proteobacteria] Length: 372\nNumber of predicted TMHs: 1\nExp number of AAs in TMHs: 23.72226\nExp number, first 60 AAs: 0.22038\nTotal prob of N-in: 0.12994\noutside 1 331\nTMhelix 332 354\ninside 355 372

27755 GCF_000012005.1_ASM1200v1 Shigella dysenteriae Sd197 Proteobacteria; Gammaproteobacteria; Enterobacterales; Enterobacteriaceae; Shigella; Shigella dysenteriae MNNEETFYQAMRRQGVTRRSFLKYCSLAATSLGLGAGMAPKIAWALENKPRIPVVWIHGL YP_402611.1 hydrogenase-1 small subunit [Shigella dysenteriae Sd197] Length: 372\nNumber of predicted TMHs: 2\nExp number of AAs in TMHs: 37.2608499999999\nExp number, first 60 AAs: 19.52923\nTotal prob of N-in: 0.94351\nPOSSIBLE N-term signal sequence\ninside 1 19\nTMhelix 20 39\noutside 40 325\nTMhelix 326 348\ninside 349 372

27756 GCF_000012005.1_ASM1200v1 Shigella dysenteriae Sd197 Proteobacteria; Gammaproteobacteria; Enterobacterales; Enterobacteriaceae; Shigella; Shigella dysenteriae MTGDNTLIHSHGINRRDFMKLCAALAATMGLSSKAAAETAESVTNPQRPPVIWIGAQECT YP_404583.1 hydrogenase 2 small subunit [Shigella dysenteriae Sd197] Length: 372\nNumber of predicted TMHs: 1\nExp number of AAs in TMHs: 23.58948\nExp number, first 60 AAs: 0.08017\nTotal prob of N-in: 0.12395\noutside 1 331\nTMhelix 332 354\ninside 355 372

27757 GCF_000467245.1_Shigella_dysenteriae_strain_SD1D Shigella dysenteriae SD1D Proteobacteria; Gammaproteobacteria; Enterobacterales; Enterobacteriaceae; Shigella; Shigella dysenteriae MTGDNTLIHSHGINRRDFMKLCAALAATMGLSSKAAAEMAESVTNPQRPPVIWIGAQECT WP_000145410.1 MULTISPECIES: hydrogenase-2 small chain [Proteobacteria] Length: 372\nNumber of predicted TMHs: 1\nExp number of AAs in TMHs: 23.72226\nExp number, first 60 AAs: 0.22038\nTotal prob of N-in: 0.12994\noutside 1 331\nTMhelix 332 354\ninside 355 372

27758 GCF_000467245.1_Shigella_dysenteriae_strain_SD1D Shigella dysenteriae SD1D Proteobacteria; Gammaproteobacteria; Enterobacterales; Enterobacteriaceae; Shigella; Shigella dysenteriae MNNEETFYQAMRRQGVTRRSFLKYCSLAATSLGLGAGMAPKIAWALENKPRIPVVWIHGL WP_001058323.1 MULTISPECIES: hydrogenase-1 small chain [Proteobacteria] Length: 372\nNumber of predicted TMHs:

2\nExp number of AAs in TMHs: 37.2608499999999\nExp number, first 60 AAs: 19.52923\nTotal prob of N-in: 0.94351\nPOSSIBLE N-term signal sequence\ninside 1 19\nTMhelix 20 39\noutside 40 325\nTMhelix 326 348\ninside 349 372

27759 GCF_000499065.1_XNQ_WAC Shigella dysenteriae WRSd3 Proteobacteria; Gammaproteobacteria; Enterobacterales; Enterobacteriaceae; Shigella; Shigella dysenteriae MTGDNTLIHSHGINRRDFMKLCAALAATMGLSSKAAAETAESVTNPQRPPVIWIGAQECT WP_000145424.1 hydrogenase 2 small subunit [Shigella dysenteriae] Length: 372\nNumber of predicted TMHs: 1\nExp number of AAs in TMHs: 23.58948\nExp number, first 60 AAs: 0.08017\nTotal prob of N-in: 0.12395\noutside 1 331\nTMhelix 332 354\ninside 355 372

27760 GCF_000499065.1_XNQ_WAC Shigella dysenteriae WRSd3 Proteobacteria; Gammaproteobacteria; Enterobacterales; Enterobacteriaceae; Shigella; Shigella dysenteriae MNNEETFYQAMRRQGVTRRSFLKYCSLAATSLGLGAGMAPKIAWALENKPRIPVVWIHGL WP_001058323.1 MULTISPECIES: hydrogenase-1 small chain [Proteobacteria] Length: 372\nNumber of predicted TMHs: 2\nExp number of AAs in TMHs: 37.2608499999999\nExp number, first 60 AAs: 19.52923\nTotal prob of N-in: 0.94351\nPOSSIBLE N-term signal sequence\ninside 1 19\nTMhelix 20 39\noutside 40 325\nTMhelix 326 348\ninside 349 372

27761 GCF_000499085.1_RRH_AFF Shigella dysenteriae WRSd5 Proteobacteria; Gammaproteobacteria; Enterobacterales; Enterobacteriaceae; Shigella; Shigella dysenteriae MNNEETFYQAMRRQGVTRRSFLKYCSLAATSLGLGAGMAPKIAWALENKPRIPVVWIHGL WP_001058323.1 MULTISPECIES: hydrogenase-1 small chain [Proteobacteria] Length: 372\nNumber of predicted TMHs: 2\nExp number of AAs in TMHs: 37.2608499999999\nExp number, first 60 AAs: 19.52923\nTotal prob of N-in: 0.94351\nPOSSIBLE N-term signal sequence\ninside 1 19\nTMhelix 20 39\noutside 40 325\nTMhelix 326 348\ninside 349 372

27762 GCF_000499085.1_RRH_AFF Shigella dysenteriae WRSd5 Proteobacteria; Gammaproteobacteria; Enterobacterales; Enterobacteriaceae; Shigella; Shigella dysenteriae MTGDNTLIHSHGINRRDFMKLCAALAATMGLSSKAAAETAESVTNPQRPPVIWIGAQECT WP_000145424.1 hydrogenase 2 small subunit [Shigella dysenteriae] Length: 372\nNumber of predicted TMHs: 1\nExp number of AAs in TMHs: 23.58948\nExp number, first 60 AAs: 0.08017\nTotal prob of N-in: 0.12395\noutside 1 331\nTMhelix 332 354\ninside 355 372

27763 GCF_000566045.1_ASM56604v1 Shigella flexneri 1997005 Proteobacteria; Gammaproteobacteria; Enterobacterales; Enterobacteriaceae; Shigella; Shigella flexneri MNNEETFYQAMRRQGVTRRSFLKYCSLAATSLGLGAGMAPKIAWALENKPRIPVVWIHGL WP_001058323.1 MULTISPECIES: hydrogenase-1 small chain [Proteobacteria] Length: 372\nNumber of predicted TMHs: 2\nExp number of AAs in TMHs: 37.2608499999999\nExp number, first 60 AAs: 19.52923\nTotal prob of N-in: 0.94351\nPOSSIBLE N-term signal sequence\ninside 1 19\nTMhelix 20 39\noutside 40 325\nTMhelix 326 348\ninside 349 372

27764 GCF_000566065.1_ASM56606v1 Shigella flexneri 2000019 Proteobacteria; Gammaproteobacteria; Enterobacterales; Enterobacteriaceae; Shigella; Shigella flexneri MNNEETFYQAMRRQGVTRRSFLKYCSLAATSLGLGAGMAPKIAWALENKPRIPVVWIHGL WP_001058323.1 MULTISPECIES: hydrogenase-1 small chain [Proteobacteria] Length: 372\nNumber of predicted TMHs: 2\nExp number of AAs in TMHs: 37.2608499999999\nExp number, first 60 AAs: 19.52923\nTotal prob of N-in: 0.94351\nPOSSIBLE N-term signal sequence\ninside 1 19\nTMhelix 20 39\noutside 40 325\nTMhelix 326 348\ninside 349 372

27765 GCF_000566065.1_ASM56606v1 Shigella flexneri 2000019 Proteobacteria; Gammaproteobacteria; Enterobacterales; Enterobacteriaceae; Shigella; Shigella flexneri MTGDNTLIHSHGINRRDFMKLCAALAATMGLSSKAAAEMAESVTNPQRPPVIWIGAQECT WP_000145421.1 MULTISPECIES: hydrogenase 2 small subunit [Shigella] Length: 372\nNumber of predicted TMHs: 1\nExp number of AAs in TMHs: 22.90256\nExp number, first 60 AAs: 0.2236\nTotal prob of N-in: 0.09847\noutside 1 331\nTMhelix 332 354\ninside 355 372

27766 GCF_000566105.1_ASM56610v1 Shigella flexneri 2001004 Proteobacteria; Gammaproteobacteria; Enterobacterales; Enterobacteriaceae; Shigella; Shigella flexneri MTGDNTLIHSHGINRRDFMKLCAALAATMGLSSKAAAEMAESVTNPQRPPVIWIGAQECT WP_000145421.1 MULTISPECIES: hydrogenase 2 small subunit [Shigella] Length: 372\nNumber of predicted TMHs: 1\nExp

number of AAs in TMHs: 22.90256\nExp number, first 60 AAs:  0.2236\nTotal prob of N-in:   0.09847\noutside   1   331\nTMhelix   332   354\ninside   355   372

27767 GCF_000566105.1_ASM56610v1 Shigella flexneri 2001004 Proteobacteria; Gammaproteobacteria; Enterobacterales; Enterobacteriaceae; Shigella; Shigella flexneri MNNEETFYQAMRRQGVTRRSFLKYCSLAATSLGLGAGMAPKIAWALENKPRIPVVWIHGL WP_001058323.1 MULTISPECIES: hydrogenase-1 small chain [Proteobacteria] Length: 372\nNumber of predicted TMHs: 2\nExp number of AAs in TMHs: 37.2608499999999\nExp number, first 60 AAs:  19.52923\nTotal prob of N-in: 0.94351\nPOSSIBLE N-term signal sequence\ninside   1   19\nTMhelix   20   39\noutside   40   325\nTMhelix   326   348\ninside   349   372

27768 GCF_000566085.1_ASM56608v1 Shigella flexneri 2001020 Proteobacteria; Gammaproteobacteria; Enterobacterales; Enterobacteriaceae; Shigella; Shigella flexneri MNNEETFYQAMRRQGVTRRSFLKYCSLAATSLGLGAGMAPKIAWALENKPRIPVVWIHGL WP_001058323.1 MULTISPECIES: hydrogenase-1 small chain [Proteobacteria] Length: 372\nNumber of predicted TMHs: 2\nExp number of AAs in TMHs: 37.2608499999999\nExp number, first 60 AAs:  19.52923\nTotal prob of N-in: 0.94351\nPOSSIBLE N-term signal sequence\ninside   1   19\nTMhelix   20   39\noutside   40   325\nTMhelix   326   348\ninside   349   372

27769 GCF_000566545.1_ASM56654v1 Shigella flexneri 2001025 Proteobacteria; Gammaproteobacteria; Enterobacterales; Enterobacteriaceae; Shigella; Shigella flexneri MTGDNTLIHSHGINRRDFMKLCAALAATMGLSSKAAAEMAESVTNPQRPPVIWIGAQECT WP_000145421.1 MULTISPECIES: hydrogenase 2 small subunit [Shigella] Length: 372\nNumber of predicted TMHs:  1\nExp number of AAs in TMHs: 22.90256\nExp number, first 60 AAs:  0.2236\nTotal prob of N-in:   0.09847\noutside   1   331\nTMhelix   332   354\ninside   355   372

27770 GCF_000566545.1_ASM56654v1 Shigella flexneri 2001025 Proteobacteria; Gammaproteobacteria; Enterobacterales; Enterobacteriaceae; Shigella; Shigella flexneri MNNEETFYQAMRRQGVTRRSFLKYCSLAATSLGLGAGMAPKIAWALENKPRIPVVWIHGL WP_001058323.1 MULTISPECIES: hydrogenase-1 small chain [Proteobacteria] Length: 372\nNumber of predicted TMHs: 2\nExp number of AAs in TMHs: 37.2608499999999\nExp number, first 60 AAs:  19.52923\nTotal prob of N-in: 0.94351\nPOSSIBLE N-term signal sequence\ninside   1   19\nTMhelix   20   39\noutside   40   325\nTMhelix   326   348\ninside   349   372

27771 GCF_000566565.1_ASM56656v1 Shigella flexneri 2001027 Proteobacteria; Gammaproteobacteria; Enterobacterales; Enterobacteriaceae; Shigella; Shigella flexneri MNNEETFYQAMRRQGVTRRSFLKYCSLAATSLGLGAGMAPKIAWALENKPRIPVVWIHGL WP_001058323.1 MULTISPECIES: hydrogenase-1 small chain [Proteobacteria] Length: 372\nNumber of predicted TMHs: 2\nExp number of AAs in TMHs: 37.2608499999999\nExp number, first 60 AAs:  19.52923\nTotal prob of N-in: 0.94351\nPOSSIBLE N-term signal sequence\ninside   1   19\nTMhelix   20   39\noutside   40   325\nTMhelix   326   348\ninside   349   372

27772 GCF_000566565.1_ASM56656v1 Shigella flexneri 2001027 Proteobacteria; Gammaproteobacteria; Enterobacterales; Enterobacteriaceae; Shigella; Shigella flexneri MTGDNTLIHSHGINRRDFMKLCAALAATMGLSSKAAAEMAESVTNPQRPPVIWIGAQECT WP_000145421.1 MULTISPECIES: hydrogenase 2 small subunit [Shigella] Length: 372\nNumber of predicted TMHs:  1\nExp number of AAs in TMHs: 22.90256\nExp number, first 60 AAs:  0.2236\nTotal prob of N-in:   0.09847\noutside   1   331\nTMhelix   332   354\ninside   355   372

27773 GCF_000566125.1_ASM56612v1 Shigella flexneri 2001042 Proteobacteria; Gammaproteobacteria; Enterobacterales; Enterobacteriaceae; Shigella; Shigella flexneri MNNEETFYQAMRRQGVTRRSFLKYCSLAATSLGLGAGMAPKIAWALENKPRIPVVWIHGL WP_001058323.1 MULTISPECIES: hydrogenase-1 small chain [Proteobacteria] Length: 372\nNumber of predicted TMHs: 2\nExp number of AAs in TMHs: 37.2608499999999\nExp number, first 60 AAs:  19.52923\nTotal prob of N-in: 0.94351\nPOSSIBLE N-term signal sequence\ninside   1   19\nTMhelix   20   39\noutside   40   325\nTMhelix   326   348\ninside   349   372

27774 GCF_000566125.1_ASM56612v1 Shigella flexneri 2001042 Proteobacteria; Gammaproteobacteria; Enterobacterales; Enterobacteriaceae; Shigella; Shigella flexneri MTGDNTLIHSHGINRRDFMKLCAALAATMGLSSKAAAEMAESVTNPQRPPVIWIGAQECT WP_000145421.1 MULTISPECIES: hydrogenase 2 small subunit [Shigella] Length: 372\nNumber of predicted TMHs:  1\nExp

number of AAs in TMHs: 22.90256\nExp number, first 60 AAs: 0.2236\nTotal prob of N-in: 0.09847\noutside 1 331\nTMhelix 332 354\ninside 355 372

27775 GCF_000566145.1_ASM56614v1 Shigella flexneri 2001044 Proteobacteria; Gammaproteobacteria; Enterobacterales; Enterobacteriaceae; Shigella; Shigella flexneri
MTGDNTLIHSHGINRRDFMKLCAALAATMGLSSKAAAEMAESVTNPQRPPVIWIGAQECT WP_000145421.1
MULTISPECIES: hydrogenase 2 small subunit [Shigella] Length: 372\nNumber of predicted TMHs: 1\nExp number of AAs in TMHs: 22.90256\nExp number, first 60 AAs: 0.2236\nTotal prob of N-in: 0.09847\noutside 1 331\nTMhelix 332 354\ninside 355 372

27776 GCF_000566585.1_ASM56658v1 Shigella flexneri 2001048 Proteobacteria; Gammaproteobacteria; Enterobacterales; Enterobacteriaceae; Shigella; Shigella flexneri
MTGDNTLIHSHGINRRDFMKLCAALAATMGLSSKAAAEMAESVTNPQRPPVIWIGAQECT WP_000145421.1
MULTISPECIES: hydrogenase 2 small subunit [Shigella] Length: 372\nNumber of predicted TMHs: 1\nExp number of AAs in TMHs: 22.90256\nExp number, first 60 AAs: 0.2236\nTotal prob of N-in: 0.09847\noutside 1 331\nTMhelix 332 354\ninside 355 372

27777 GCF_000565945.1_ASM56594v1 Shigella flexneri 2002007 Proteobacteria; Gammaproteobacteria; Enterobacterales; Enterobacteriaceae; Shigella; Shigella flexneri
MNNEETFYQAMRRQGVTRRSFLKYCSLAATSLGLGAGMAPKIAWALENKPRIPVVWIHGL WP_001058323.1
MULTISPECIES: hydrogenase-1 small chain [Proteobacteria] Length: 372\nNumber of predicted TMHs: 2\nExp number of AAs in TMHs: 37.2608499999999\nExp number, first 60 AAs: 19.52923\nTotal prob of N-in: 0.94351\nPOSSIBLE N-term signal sequence\ninside 1 19\nTMhelix 20 39\noutside 40 325\nTMhelix 326 348\ninside 349 372

27778 GCF_000565945.1_ASM56594v1 Shigella flexneri 2002007 Proteobacteria; Gammaproteobacteria; Enterobacterales; Enterobacteriaceae; Shigella; Shigella flexneri
MTGDNTLIHSHGINRRDFMKLCAALAATMGLSSKAAAEMAESVTNPQRPPVIWIGAQECT WP_000145421.1
MULTISPECIES: hydrogenase 2 small subunit [Shigella] Length: 372\nNumber of predicted TMHs: 1\nExp number of AAs in TMHs: 22.90256\nExp number, first 60 AAs: 0.2236\nTotal prob of N-in: 0.09847\noutside 1 331\nTMhelix 332 354\ninside 355 372

27779 GCF_000022245.1_ASM2224v1 Shigella flexneri 2002017 Proteobacteria; Gammaproteobacteria; Enterobacterales; Enterobacteriaceae; Shigella; Shigella flexneri
MNNEETFYQAMRRQGVTRRSFLKYCSLAATSLGLGAGMAPKIAWALENKPRIPVVWIHGL WP_001058323.1
MULTISPECIES: hydrogenase-1 small chain [Proteobacteria] Length: 372\nNumber of predicted TMHs: 2\nExp number of AAs in TMHs: 37.2608499999999\nExp number, first 60 AAs: 19.52923\nTotal prob of N-in: 0.94351\nPOSSIBLE N-term signal sequence\ninside 1 19\nTMhelix 20 39\noutside 40 325\nTMhelix 326 348\ninside 349 372

27780 GCF_000022245.1_ASM2224v1 Shigella flexneri 2002017 Proteobacteria; Gammaproteobacteria; Enterobacterales; Enterobacteriaceae; Shigella; Shigella flexneri
MTGDNTLIHSHGINRRDFMKLCAALAATMGLSSKAAAEMAESVTNPQRPPVIWIGAQECT WP_000145421.1
MULTISPECIES: hydrogenase 2 small subunit [Shigella] Length: 372\nNumber of predicted TMHs: 1\nExp number of AAs in TMHs: 22.90256\nExp number, first 60 AAs: 0.2236\nTotal prob of N-in: 0.09847\noutside 1 331\nTMhelix 332 354\ninside 355 372

27781 GCF_000566185.1_ASM56618v1 Shigella flexneri 2002028 Proteobacteria; Gammaproteobacteria; Enterobacterales; Enterobacteriaceae; Shigella; Shigella flexneri
MTGDNTLIHSHGINRRDFMKLCAALAATMGLSSKAAAEMAESVTNPQRPPVIWIGAQECT WP_000145421.1
MULTISPECIES: hydrogenase 2 small subunit [Shigella] Length: 372\nNumber of predicted TMHs: 1\nExp number of AAs in TMHs: 22.90256\nExp number, first 60 AAs: 0.2236\nTotal prob of N-in: 0.09847\noutside 1 331\nTMhelix 332 354\ninside 355 372

27782 GCF_000566605.1_ASM56660v1 Shigella flexneri 2002035 Proteobacteria; Gammaproteobacteria; Enterobacterales; Enterobacteriaceae; Shigella; Shigella flexneri
MTGDNTLIHSHGINRRDFMKLCAALAATMGLSSKAAAEMAESVTNPQRPPVIWIGAQECT WP_000145421.1
MULTISPECIES: hydrogenase 2 small subunit [Shigella] Length: 372\nNumber of predicted TMHs: 1\nExp number of AAs in TMHs: 22.90256\nExp number, first 60 AAs: 0.2236\nTotal prob of N-in: 0.09847\noutside 1 331\nTMhelix 332 354\ninside 355 372

27783 GCF_000566605.1_ASM56660v1 Shigella flexneri 2002035 Proteobacteria; Gammaproteobacteria; Enterobacterales; Enterobacteriaceae; Shigella; Shigella flexneri
MNNEETFYQAMRRQGVTRRSFLKYCSLAATSLGLGAGMAPKIAWALENKPRIPVVWIHGL WP_001058323.1
MULTISPECIES: hydrogenase-1 small chain [Proteobacteria] Length: 372\nNumber of predicted TMHs: 2\nExp number of AAs in TMHs: 37.2608499999999\nExp number, first 60 AAs: 19.52923\nTotal prob of N-in: 0.94351\nPOSSIBLE N-term signal sequence\ninside 1 19\nTMhelix 20 39\noutside 40 325\nTMhelix 326 348\ninside 349 372

27784 GCF_000566205.1_ASM56620v1 Shigella flexneri 2002069 Proteobacteria; Gammaproteobacteria; Enterobacterales; Enterobacteriaceae; Shigella; Shigella flexneri
MNNEETFYQAMRRQGVTRRSFLKYCSLAATSLGLGAGMAPKIAWALENKPRIPVVWIHGL WP_001058323.1
MULTISPECIES: hydrogenase-1 small chain [Proteobacteria] Length: 372\nNumber of predicted TMHs: 2\nExp number of AAs in TMHs: 37.2608499999999\nExp number, first 60 AAs: 19.52923\nTotal prob of N-in: 0.94351\nPOSSIBLE N-term signal sequence\ninside 1 19\nTMhelix 20 39\noutside 40 325\nTMhelix 326 348\ninside 349 372

27785 GCF_000566205.1_ASM56620v1 Shigella flexneri 2002069 Proteobacteria; Gammaproteobacteria; Enterobacterales; Enterobacteriaceae; Shigella; Shigella flexneri
MTGDNTLIHSHGINRRDFMKLCAALAATMGLSSKAAAEMAESVTNPQRPPVIWIGAQECT WP_000145421.1
MULTISPECIES: hydrogenase 2 small subunit [Shigella] Length: 372\nNumber of predicted TMHs: 1\nExp number of AAs in TMHs: 22.90256\nExp number, first 60 AAs: 0.2236\nTotal prob of N-in: 0.09847\noutside 1 331\nTMhelix 332 354\ninside 355 372

27786 GCF_000566225.1_ASM56622v1 Shigella flexneri 2002091 Proteobacteria; Gammaproteobacteria; Enterobacterales; Enterobacteriaceae; Shigella; Shigella flexneri
MNNEETFYQAMRRQGVTRRSFLKYCSLAATSLGLGAGMAPKIAWALENKPRIPVVWIHGL WP_025746233.1
hydrogenase [Shigella flexneri] Length: 372\nNumber of predicted TMHs: 2\nExp number of AAs in TMHs: 38.0705399999999\nExp number, first 60 AAs: 19.53132\nTotal prob of N-in: 0.94350\nPOSSIBLE N-term signal sequence\ninside 1 19\nTMhelix 20 39\noutside 40 325\nTMhelix 326 348\ninside 349 372

27787 GCF_000566225.1_ASM56622v1 Shigella flexneri 2002091 Proteobacteria; Gammaproteobacteria; Enterobacterales; Enterobacteriaceae; Shigella; Shigella flexneri
MTGDNTLIHSHGINRRDFMKLCAALAATMGLSSKAAAEMAESVTNPQRPPVIWIGAQECT WP_000145421.1
MULTISPECIES: hydrogenase 2 small subunit [Shigella] Length: 372\nNumber of predicted TMHs: 1\nExp number of AAs in TMHs: 22.90256\nExp number, first 60 AAs: 0.2236\nTotal prob of N-in: 0.09847\noutside 1 331\nTMhelix 332 354\ninside 355 372

27788 GCF_000565885.1_ASM56588v1 Shigella flexneri 2002103 Proteobacteria; Gammaproteobacteria; Enterobacterales; Enterobacteriaceae; Shigella; Shigella flexneri
MNNEETFYQAMRRQGVTRRSFLKYCSLAATSLGLGAGMAPKIAWALENKPRIPVVWIHGL WP_001058323.1
MULTISPECIES: hydrogenase-1 small chain [Proteobacteria] Length: 372\nNumber of predicted TMHs: 2\nExp number of AAs in TMHs: 37.2608499999999\nExp number, first 60 AAs: 19.52923\nTotal prob of N-in: 0.94351\nPOSSIBLE N-term signal sequence\ninside 1 19\nTMhelix 20 39\noutside 40 325\nTMhelix 326 348\ninside 349 372

27789 GCF_000565885.1_ASM56588v1 Shigella flexneri 2002103 Proteobacteria; Gammaproteobacteria; Enterobacterales; Enterobacteriaceae; Shigella; Shigella flexneri
MTGDNTLIHSHGINRRDFMKLCAALAATMGLSSKAAAEMAESVTNPQRPPVIWIGAQECT WP_000145421.1
MULTISPECIES: hydrogenase 2 small subunit [Shigella] Length: 372\nNumber of predicted TMHs: 1\nExp number of AAs in TMHs: 22.90256\nExp number, first 60 AAs: 0.2236\nTotal prob of N-in: 0.09847\noutside 1 331\nTMhelix 332 354\ninside 355 372

27790 GCF_000566245.1_ASM56624v1 Shigella flexneri 2002106 Proteobacteria; Gammaproteobacteria; Enterobacterales; Enterobacteriaceae; Shigella; Shigella flexneri
MTGDNTLIHSHGINRRDFMKLCAALAATMGLSSKAAAEMAESVTNPQRPPVIWIGAQECT WP_000145421.1
MULTISPECIES: hydrogenase 2 small subunit [Shigella] Length: 372\nNumber of predicted TMHs: 1\nExp number of AAs in TMHs: 22.90256\nExp number, first 60 AAs: 0.2236\nTotal prob of N-in: 0.09847\noutside 1 331\nTMhelix 332 354\ninside 355 372

27791 GCF_000566245.1_ASM56624v1 Shigella flexneri 2002106 Proteobacteria; Gammaproteobacteria; Enterobacterales; Enterobacteriaceae; Shigella; Shigella flexneri

MNNEETFYQAMRRQGVTRRSFLKYCSLAATSLGLGAGMAPKIAWALENKPRIPVVWIHGL	WP_001058323.1	MULTISPECIES: hydrogenase-1 small chain [Proteobacteria]	Length: 372\nNumber of predicted TMHs: 2\nExp number of AAs in TMHs: 37.2608499999999\nExp number, first 60 AAs: 19.52923\nTotal prob of N-in: 0.94351\nPOSSIBLE N-term signal sequence\ninside 1 19\nTMhelix 20 39\noutside 40 325\nTMhelix 326 348\ninside 349 372

27792	GCF_000566265.1_ASM56626v1	Shigella flexneri 2002110	Proteobacteria; Gammaproteobacteria; Enterobacterales; Enterobacteriaceae; Shigella; Shigella flexneri	MTGDNTLIHSHGINRRDFMKLCAALAATMGLSSKAAAEMAESVTNPQRPPVIWIGAQECT	WP_000145421.1	MULTISPECIES: hydrogenase 2 small subunit [Shigella]	Length: 372\nNumber of predicted TMHs: 1\nExp number of AAs in TMHs: 22.90256\nExp number, first 60 AAs: 0.2236\nTotal prob of N-in: 0.09847\noutside 1 331\nTMhelix 332 354\ninside 355 372

27793	GCF_000566625.1_ASM56662v1	Shigella flexneri 2002127	Proteobacteria; Gammaproteobacteria; Enterobacterales; Enterobacteriaceae; Shigella; Shigella flexneri	MNNEETFYQAMRRQGVTRRSFLKYCSLAATSLGLGAGMAPKIAWALENKPRIPVVWIHGL	WP_001058323.1	MULTISPECIES: hydrogenase-1 small chain [Proteobacteria]	Length: 372\nNumber of predicted TMHs: 2\nExp number of AAs in TMHs: 37.2608499999999\nExp number, first 60 AAs: 19.52923\nTotal prob of N-in: 0.94351\nPOSSIBLE N-term signal sequence\ninside 1 19\nTMhelix 20 39\noutside 40 325\nTMhelix 326 348\ninside 349 372

27794	GCF_000566625.1_ASM56662v1	Shigella flexneri 2002127	Proteobacteria; Gammaproteobacteria; Enterobacterales; Enterobacteriaceae; Shigella; Shigella flexneri	MTGDNTLIHSHGINRRDFMKLCAALAATMGLSSKAAAEMAESVTNPQRPPVIWIGAQECT	WP_000145421.1	MULTISPECIES: hydrogenase 2 small subunit [Shigella]	Length: 372\nNumber of predicted TMHs: 1\nExp number of AAs in TMHs: 22.90256\nExp number, first 60 AAs: 0.2236\nTotal prob of N-in: 0.09847\noutside 1 331\nTMhelix 332 354\ninside 355 372

27795	GCF_000566285.1_ASM56628v1	Shigella flexneri 2002140	Proteobacteria; Gammaproteobacteria; Enterobacterales; Enterobacteriaceae; Shigella; Shigella flexneri	MNNEETFYQAMRRQGVTRRSFLKYCSLAATSLGLGAGMAPKIAWALENKPRIPVVWIHGL	WP_001058323.1	MULTISPECIES: hydrogenase-1 small chain [Proteobacteria]	Length: 372\nNumber of predicted TMHs: 2\nExp number of AAs in TMHs: 37.2608499999999\nExp number, first 60 AAs: 19.52923\nTotal prob of N-in: 0.94351\nPOSSIBLE N-term signal sequence\ninside 1 19\nTMhelix 20 39\noutside 40 325\nTMhelix 326 348\ninside 349 372

27796	GCF_000566285.1_ASM56628v1	Shigella flexneri 2002140	Proteobacteria; Gammaproteobacteria; Enterobacterales; Enterobacteriaceae; Shigella; Shigella flexneri	MTGDNTLIHSHGINRRDFMKLCAALAATMGLSSKAAAEMAESVTNPQRPPVIWIGAQECT	WP_000145421.1	MULTISPECIES: hydrogenase 2 small subunit [Shigella]	Length: 372\nNumber of predicted TMHs: 1\nExp number of AAs in TMHs: 22.90256\nExp number, first 60 AAs: 0.2236\nTotal prob of N-in: 0.09847\noutside 1 331\nTMhelix 332 354\ninside 355 372

27797	GCF_000566645.1_ASM56664v1	Shigella flexneri 2002141	Proteobacteria; Gammaproteobacteria; Enterobacterales; Enterobacteriaceae; Shigella; Shigella flexneri	MNNEETFYQAMRRQGVTRRSFLKYCSLAATSLGLGAGMAPKIAWALENKPRIPVVWIHGL	WP_001058323.1	MULTISPECIES: hydrogenase-1 small chain [Proteobacteria]	Length: 372\nNumber of predicted TMHs: 2\nExp number of AAs in TMHs: 37.2608499999999\nExp number, first 60 AAs: 19.52923\nTotal prob of N-in: 0.94351\nPOSSIBLE N-term signal sequence\ninside 1 19\nTMhelix 20 39\noutside 40 325\nTMhelix 326 348\ninside 349 372

27798	GCF_000566645.1_ASM56664v1	Shigella flexneri 2002141	Proteobacteria; Gammaproteobacteria; Enterobacterales; Enterobacteriaceae; Shigella; Shigella flexneri	MTGDNTLIHSHGINRRDFMKLCAALAATMGLSSKAAAEMAESVTNPQRPPVIWIGAQECT	WP_000145421.1	MULTISPECIES: hydrogenase 2 small subunit [Shigella]	Length: 372\nNumber of predicted TMHs: 1\nExp number of AAs in TMHs: 22.90256\nExp number, first 60 AAs: 0.2236\nTotal prob of N-in: 0.09847\noutside 1 331\nTMhelix 332 354\ninside 355 372

27799	GCF_000566665.1_ASM56666v1	Shigella flexneri 2002142	Proteobacteria; Gammaproteobacteria; Enterobacterales; Enterobacteriaceae; Shigella; Shigella flexneri	MTGDNTLIHSHGINRRDFMKLCAALAATMGLSSKAAAEMAESVTNPQRPPVIWIGAQECT	WP_000145421.1

MULTISPECIES: hydrogenase 2 small subunit [Shigella] Length: 372\nNumber of predicted TMHs: 1\nExp number of AAs in TMHs: 22.90256\nExp number, first 60 AAs: 0.2236\nTotal prob of N-in: 0.09847\noutside 1 331\nTMhelix 332 354\ninside 355 372

27800 GCF_000566665.1_ASM56666v1 Shigella flexneri 2002142 Proteobacteria; Gammaproteobacteria; Enterobacterales; Enterobacteriaceae; Shigella; Shigella flexneri MNNEETFYQAMRRQGVTRRSFLKYCSLAATSLGLGAGMAPKIAWALENKPRIPVVWIHGL WP_001058323.1 MULTISPECIES: hydrogenase-1 small chain [Proteobacteria] Length: 372\nNumber of predicted TMHs: 2\nExp number of AAs in TMHs: 37.2608499999999\nExp number, first 60 AAs: 19.52923\nTotal prob of N-in: 0.94351\nPOSSIBLE N-term signal sequence\ninside 1 19\nTMhelix 20 39\noutside 40 325\nTMhelix 326 348\ninside 349 372

27801 GCF_000566305.1_ASM56630v1 Shigella flexneri 2003035 Proteobacteria; Gammaproteobacteria; Enterobacterales; Enterobacteriaceae; Shigella; Shigella flexneri MTGDNTLIHSHGINRRDFMKLCAALAATMGLSSKAAAEMAESVTNPQRPPVIWIGAQECT WP_000145421.1 MULTISPECIES: hydrogenase 2 small subunit [Shigella] Length: 372\nNumber of predicted TMHs: 1\nExp number of AAs in TMHs: 22.90256\nExp number, first 60 AAs: 0.2236\nTotal prob of N-in: 0.09847\noutside 1 331\nTMhelix 332 354\ninside 355 372

27802 GCF_000743955.1_ASM74395v1 Shigella flexneri 2003036 Proteobacteria; Gammaproteobacteria; Enterobacterales; Enterobacteriaceae; Shigella; Shigella flexneri MNNEETFYQAMRRQGVTRRSFLKYCSLAATSLGLGAGMAPKIAWALENKPRIPVVWIHGL WP_001058323.1 MULTISPECIES: hydrogenase-1 small chain [Proteobacteria] Length: 372\nNumber of predicted TMHs: 2\nExp number of AAs in TMHs: 37.2608499999999\nExp number, first 60 AAs: 19.52923\nTotal prob of N-in: 0.94351\nPOSSIBLE N-term signal sequence\ninside 1 19\nTMhelix 20 39\noutside 40 325\nTMhelix 326 348\ninside 349 372

27803 GCF_000743955.1_ASM74395v1 Shigella flexneri 2003036 Proteobacteria; Gammaproteobacteria; Enterobacterales; Enterobacteriaceae; Shigella; Shigella flexneri MTGDNTLIHSHGINRRDFMKLCAALAATMGLSSKAAAEMAESVTNPQRPPVIWIGAQECT WP_000145421.1 MULTISPECIES: hydrogenase 2 small subunit [Shigella] Length: 372\nNumber of predicted TMHs: 1\nExp number of AAs in TMHs: 22.90256\nExp number, first 60 AAs: 0.2236\nTotal prob of N-in: 0.09847\noutside 1 331\nTMhelix 332 354\ninside 355 372

27804 GCF_000565805.1_ASM56580v1 Shigella flexneri 2003055 Proteobacteria; Gammaproteobacteria; Enterobacterales; Enterobacteriaceae; Shigella; Shigella flexneri MTGDNTLIHSHGINRRDFMKLCAALAATMGLSSKAAAEMAESVTNPQRPPVIWIGAQECT WP_000145421.1 MULTISPECIES: hydrogenase 2 small subunit [Shigella] Length: 372\nNumber of predicted TMHs: 1\nExp number of AAs in TMHs: 22.90256\nExp number, first 60 AAs: 0.2236\nTotal prob of N-in: 0.09847\noutside 1 331\nTMhelix 332 354\ninside 355 372

27805 GCF_000565805.1_ASM56580v1 Shigella flexneri 2003055 Proteobacteria; Gammaproteobacteria; Enterobacterales; Enterobacteriaceae; Shigella; Shigella flexneri MNNEETFYQAMRRQGVTRRSFLKYCSLAATSLGLGAGMAPKIAWALENKPRIPVVWIHGL WP_001058323.1 MULTISPECIES: hydrogenase-1 small chain [Proteobacteria] Length: 372\nNumber of predicted TMHs: 2\nExp number of AAs in TMHs: 37.2608499999999\nExp number, first 60 AAs: 19.52923\nTotal prob of N-in: 0.94351\nPOSSIBLE N-term signal sequence\ninside 1 19\nTMhelix 20 39\noutside 40 325\nTMhelix 326 348\ninside 349 372

27806 GCF_000566345.1_ASM56634v1 Shigella flexneri 2005025 Proteobacteria; Gammaproteobacteria; Enterobacterales; Enterobacteriaceae; Shigella; Shigella flexneri MTGDNTLIHSHGINRRDFMKLCAALAATMGLSSKAAAEMAESVTNPQRPPVIWIGAQECT WP_000145421.1 MULTISPECIES: hydrogenase 2 small subunit [Shigella] Length: 372\nNumber of predicted TMHs: 1\nExp number of AAs in TMHs: 22.90256\nExp number, first 60 AAs: 0.2236\nTotal prob of N-in: 0.09847\noutside 1 331\nTMhelix 332 354\ninside 355 372

27807 GCF_000566365.1_ASM56636v1 Shigella flexneri 2005051 Proteobacteria; Gammaproteobacteria; Enterobacterales; Enterobacteriaceae; Shigella; Shigella flexneri MNNEETFYQAMRRQGVTRRSFLKYCSLAATSLGLGAGMAPKIAWALENKPRIPVVWIHGL WP_001058323.1 MULTISPECIES: hydrogenase-1 small chain [Proteobacteria] Length: 372\nNumber of predicted TMHs: 2\nExp number of AAs in TMHs: 37.2608499999999\nExp number, first 60 AAs: 19.52923\nTotal prob of N-in:

0.94351\nPOSSIBLE N-term signal sequence\ninside 1 19\nTMhelix 20 39\noutside 40 325\nTMhelix 326 348\ninside 349 372

27808 GCF_000566365.1_ASM56636v1 Shigella flexneri 2005051 Proteobacteria; Gammaproteobacteria; Enterobacterales; Enterobacteriaceae; Shigella; Shigella flexneri
MTGDNTLIHSHGINRRDFMKLCAALAATMGLSSKAAAEMAESVTNPQRPPVIWIGAQECT WP_000145421.1
MULTISPECIES: hydrogenase 2 small subunit [Shigella] Length: 372\nNumber of predicted TMHs: 1\nExp number of AAs in TMHs: 22.90256\nExp number, first 60 AAs: 0.2236\nTotal prob of N-in: 0.09847\noutside 1 331\nTMhelix 332 354\ninside 355 372

27809 GCF_000565645.1_ASM56564v1 Shigella flexneri 2005AH264 Proteobacteria; Gammaproteobacteria; Enterobacterales; Enterobacteriaceae; Shigella; Shigella flexneri
MNNEETFYQAMRRQGVTRRSFLKYCSLAATSLGLGAGMAPKIAWALENKPRIPVVWIHGL WP_001058323.1
MULTISPECIES: hydrogenase-1 small chain [Proteobacteria] Length: 372\nNumber of predicted TMHs: 2\nExp number of AAs in TMHs: 37.2608499999999\nExp number, first 60 AAs: 19.52923\nTotal prob of N-in: 0.94351\nPOSSIBLE N-term signal sequence\ninside 1 19\nTMhelix 20 39\noutside 40 325\nTMhelix 326 348\ninside 349 372

27810 GCF_000565645.1_ASM56564v1 Shigella flexneri 2005AH264 Proteobacteria; Gammaproteobacteria; Enterobacterales; Enterobacteriaceae; Shigella; Shigella flexneri
MTGDNTLIHSHGINRRDFMKLCAALAATMGLSSKAAAEMAESVTNPQRPPVIWIGAQECT WP_000145421.1
MULTISPECIES: hydrogenase 2 small subunit [Shigella] Length: 372\nNumber of predicted TMHs: 1\nExp number of AAs in TMHs: 22.90256\nExp number, first 60 AAs: 0.2236\nTotal prob of N-in: 0.09847\noutside 1 331\nTMhelix 332 354\ninside 355 372

27811 GCF_000566405.1_ASM56640v1 Shigella flexneri 2005GS061 Proteobacteria; Gammaproteobacteria; Enterobacterales; Enterobacteriaceae; Shigella; Shigella flexneri
MNNEETFYQAMRRQGVTRRSFLKYCSLAATSLGLGAGMAPKIAWALENKPRIPVVWIHGL WP_001058323.1
MULTISPECIES: hydrogenase-1 small chain [Proteobacteria] Length: 372\nNumber of predicted TMHs: 2\nExp number of AAs in TMHs: 37.2608499999999\nExp number, first 60 AAs: 19.52923\nTotal prob of N-in: 0.94351\nPOSSIBLE N-term signal sequence\ninside 1 19\nTMhelix 20 39\noutside 40 325\nTMhelix 326 348\ninside 349 372

27812 GCF_000566405.1_ASM56640v1 Shigella flexneri 2005GS061 Proteobacteria; Gammaproteobacteria; Enterobacterales; Enterobacteriaceae; Shigella; Shigella flexneri
MTGDNTLIHSHGINRRDFMKLCAALAATMGLSSKAAAEMAESVTNPQRPPVIWIGAQECT WP_000145421.1
MULTISPECIES: hydrogenase 2 small subunit [Shigella] Length: 372\nNumber of predicted TMHs: 1\nExp number of AAs in TMHs: 22.90256\nExp number, first 60 AAs: 0.2236\nTotal prob of N-in: 0.09847\noutside 1 331\nTMhelix 332 354\ninside 355 372

27813 GCF_000213495.1_ASM21349v2 Shigella flexneri 2930-71 Proteobacteria; Gammaproteobacteria; Enterobacterales; Enterobacteriaceae; Shigella; Shigella flexneri
MTGDNTLIHSHGINRRDFMKLCAALAATMGLSSKAAAEMAESVTNPQRPPVIWIGAQECT WP_000145421.1
MULTISPECIES: hydrogenase 2 small subunit [Shigella] Length: 372\nNumber of predicted TMHs: 1\nExp number of AAs in TMHs: 22.90256\nExp number, first 60 AAs: 0.2236\nTotal prob of N-in: 0.09847\noutside 1 331\nTMhelix 332 354\ninside 355 372

27814 GCF_000213495.1_ASM21349v2 Shigella flexneri 2930-71 Proteobacteria; Gammaproteobacteria; Enterobacterales; Enterobacteriaceae; Shigella; Shigella flexneri
MNNEETFYQAMRRQGVTRRSFLKYCSLAATSLGLGAGMAPKIAWALENKPRIPVVWIHGL WP_001058323.1
MULTISPECIES: hydrogenase-1 small chain [Proteobacteria] Length: 372\nNumber of predicted TMHs: 2\nExp number of AAs in TMHs: 37.2608499999999\nExp number, first 60 AAs: 19.52923\nTotal prob of N-in: 0.94351\nPOSSIBLE N-term signal sequence\ninside 1 19\nTMhelix 20 39\noutside 40 325\nTMhelix 326 348\ninside 349 372

27815 GCF_001017355.1_ASM101735v1 Shigella flexneri 2a Proteobacteria; Gammaproteobacteria; Enterobacterales; Enterobacteriaceae; Shigella; Shigella flexneri
MNNEETFYQAMRRQGVTRRSFLKYCSLAATSLGLGAGMAPKIAWALENKPRIPVVWIHGL WP_001058323.1
MULTISPECIES: hydrogenase-1 small chain [Proteobacteria] Length: 372\nNumber of predicted TMHs: 2\nExp number of AAs in TMHs: 37.2608499999999\nExp number, first 60 AAs: 19.52923\nTotal prob of N-in:

0.94351\nPOSSIBLE N-term signal sequence\ninside 1 19\nTMhelix 20 39\noutside 40 325\nTMhelix 326 348\ninside 349 372

27816 GCF_001017355.1_ASM101735v1 Shigella flexneri 2a Proteobacteria; Gammaproteobacteria; Enterobacterales; Enterobacteriaceae; Shigella; Shigella flexneri
MTGDNTLIHSHGINRRDFMKLCAALAATMGLSSKAAAEMAESVTNPQRPPVIWIGAQECT WP_000145421.1
MULTISPECIES: hydrogenase 2 small subunit [Shigella] Length: 372\nNumber of predicted TMHs: 1\nExp number of AAs in TMHs: 22.90256\nExp number, first 60 AAs: 0.2236\nTotal prob of N-in: 0.09847\noutside 1 331\nTMhelix 332 354\ninside 355 372

27817 GCF_001184125.1_ASM118412v1 Shigella flexneri 2a Proteobacteria; Gammaproteobacteria; Enterobacterales; Enterobacteriaceae; Shigella; Shigella flexneri
MTGDNTLIHSHGINRRDFMKLCAALAATMGLSSKAAAEMAESVTNPQRPPVIWIGAQECT WP_000145421.1
MULTISPECIES: hydrogenase 2 small subunit [Shigella] Length: 372\nNumber of predicted TMHs: 1\nExp number of AAs in TMHs: 22.90256\nExp number, first 60 AAs: 0.2236\nTotal prob of N-in: 0.09847\noutside 1 331\nTMhelix 332 354\ninside 355 372

27818 GCF_001078805.1_14928_1_79 Shigella flexneri 2a Proteobacteria; Gammaproteobacteria; Enterobacterales; Enterobacteriaceae; Shigella; Shigella flexneri
MTGDNTLIHSHGINRRDFMKLCAALAATMGLSSKAAAEMAESVTNPQRPPVIWIGAQECT WP_000145421.1
MULTISPECIES: hydrogenase 2 small subunit [Shigella] Length: 372\nNumber of predicted TMHs: 1\nExp number of AAs in TMHs: 22.90256\nExp number, first 60 AAs: 0.2236\nTotal prob of N-in: 0.09847\noutside 1 331\nTMhelix 332 354\ninside 355 372

27819 GCF_001017225.1_ASM101722v1 Shigella flexneri 2a Proteobacteria; Gammaproteobacteria; Enterobacterales; Enterobacteriaceae; Shigella; Shigella flexneri
MNNEETFYQAMRRQGVTRRSFLKYCSLAATSLGLGAGMAPKIAWALENKPRIPVVWIHGL WP_001058323.1
MULTISPECIES: hydrogenase-1 small chain [Proteobacteria] Length: 372\nNumber of predicted TMHs: 2\nExp number of AAs in TMHs: 37.2608499999999\nExp number, first 60 AAs: 19.52923\nTotal prob of N-in: 0.94351\nPOSSIBLE N-term signal sequence\ninside 1 19\nTMhelix 20 39\noutside 40 325\nTMhelix 326 348\ninside 349 372

27820 GCF_001017285.1_ASM101728v1 Shigella flexneri 2a Proteobacteria; Gammaproteobacteria; Enterobacterales; Enterobacteriaceae; Shigella; Shigella flexneri
MNNEETFYQAMRRQGVTRRSFLKYCSLAATSLGLGAGMAPKIAWALENKPRIPVVWIHGL WP_001058323.1
MULTISPECIES: hydrogenase-1 small chain [Proteobacteria] Length: 372\nNumber of predicted TMHs: 2\nExp number of AAs in TMHs: 37.2608499999999\nExp number, first 60 AAs: 19.52923\nTotal prob of N-in: 0.94351\nPOSSIBLE N-term signal sequence\ninside 1 19\nTMhelix 20 39\noutside 40 325\nTMhelix 326 348\ninside 349 372

27821 GCF_001017305.1_ASM101730v1 Shigella flexneri 2a Proteobacteria; Gammaproteobacteria; Enterobacterales; Enterobacteriaceae; Shigella; Shigella flexneri
MNNEETFYQAMRRQGVTRRSFLKYCSLAATSLGLGAGMAPKIAWALENKPRIPVVWIHGL WP_001058323.1
MULTISPECIES: hydrogenase-1 small chain [Proteobacteria] Length: 372\nNumber of predicted TMHs: 2\nExp number of AAs in TMHs: 37.2608499999999\nExp number, first 60 AAs: 19.52923\nTotal prob of N-in: 0.94351\nPOSSIBLE N-term signal sequence\ninside 1 19\nTMhelix 20 39\noutside 40 325\nTMhelix 326 348\ninside 349 372

27822 GCF_001078805.1_14928_1_79 Shigella flexneri 2a Proteobacteria; Gammaproteobacteria; Enterobacterales; Enterobacteriaceae; Shigella; Shigella flexneri
MNNEETFYQAMRRQGVTRRSFLKYCSLAATSLGLGAGMAPKIAWALENKPRIPVVWIHGL WP_001058323.1
MULTISPECIES: hydrogenase-1 small chain [Proteobacteria] Length: 372\nNumber of predicted TMHs: 2\nExp number of AAs in TMHs: 37.2608499999999\nExp number, first 60 AAs: 19.52923\nTotal prob of N-in: 0.94351\nPOSSIBLE N-term signal sequence\ninside 1 19\nTMhelix 20 39\noutside 40 325\nTMhelix 326 348\ninside 349 372

27823 GCF_001184125.1_ASM118412v1 Shigella flexneri 2a Proteobacteria; Gammaproteobacteria; Enterobacterales; Enterobacteriaceae; Shigella; Shigella flexneri
MNNEETFYQAMRRQGVTRRSFLKYCSLAATSLGLGAGMAPKIAWALENKPRIPVVWIHGL WP_001058323.1
MULTISPECIES: hydrogenase-1 small chain [Proteobacteria] Length: 372\nNumber of predicted TMHs: 2\nExp number of AAs in TMHs: 37.2608499999999\nExp number, first 60 AAs: 19.52923\nTotal prob of N-in:

0.94351\nPOSSIBLE N-term signal sequence\ninside 1 19\nTMhelix 20 39\noutside 40 325\nTMhelix 326 348\ninside 349 372

27824 GCF_001017165.1_ASM101716v1 Shigella flexneri 2a Proteobacteria; Gammaproteobacteria; Enterobacterales; Enterobacteriaceae; Shigella; Shigella flexneri
MTGDNTLIHSHGINRRDFMKLCAALAATMGLSSKAAAEMAESVTNPQRPPVIWIGAQECT WP_000145421.1
MULTISPECIES: hydrogenase 2 small subunit [Shigella] Length: 372\nNumber of predicted TMHs: 1\nExp number of AAs in TMHs: 22.90256\nExp number, first 60 AAs: 0.2236\nTotal prob of N-in: 0.09847\noutside 1 331\nTMhelix 332 354\ninside 355 372

27825 GCF_001017165.1_ASM101716v1 Shigella flexneri 2a Proteobacteria; Gammaproteobacteria; Enterobacterales; Enterobacteriaceae; Shigella; Shigella flexneri
MNNEETFYQAMRRQGVTRRSFLKYCSLAATSLGLGAGMAPKIAWALENKPRIPVVWIHGL WP_001058323.1
MULTISPECIES: hydrogenase-1 small chain [Proteobacteria] Length: 372\nNumber of predicted TMHs: 2\nExp number of AAs in TMHs: 37.2608499999999\nExp number, first 60 AAs: 19.52923\nTotal prob of N-in: 0.94351\nPOSSIBLE N-term signal sequence\ninside 1 19\nTMhelix 20 39\noutside 40 325\nTMhelix 326 348\ninside 349 372

27826 GCF_001017285.1_ASM101728v1 Shigella flexneri 2a Proteobacteria; Gammaproteobacteria; Enterobacterales; Enterobacteriaceae; Shigella; Shigella flexneri
MTGDNTLIHSHGINRRDFMKLCAALAATMGLSSKAAAEMAESVTNPQRPPVIWIGAQECT WP_000145421.1
MULTISPECIES: hydrogenase 2 small subunit [Shigella] Length: 372\nNumber of predicted TMHs: 1\nExp number of AAs in TMHs: 22.90256\nExp number, first 60 AAs: 0.2236\nTotal prob of N-in: 0.09847\noutside 1 331\nTMhelix 332 354\ninside 355 372

27827 GCF_001017155.1_ASM101715v1 Shigella flexneri 2a Proteobacteria; Gammaproteobacteria; Enterobacterales; Enterobacteriaceae; Shigella; Shigella flexneri
MTGDNTLIHSHGINRRDFMKLCAALAATMGLSSKAAAEMAESVTNPQRPPVIWIGAQECT WP_000145421.1
MULTISPECIES: hydrogenase 2 small subunit [Shigella] Length: 372\nNumber of predicted TMHs: 1\nExp number of AAs in TMHs: 22.90256\nExp number, first 60 AAs: 0.2236\nTotal prob of N-in: 0.09847\noutside 1 331\nTMhelix 332 354\ninside 355 372

27828 GCF_001017225.1_ASM101722v1 Shigella flexneri 2a Proteobacteria; Gammaproteobacteria; Enterobacterales; Enterobacteriaceae; Shigella; Shigella flexneri
MTGDNTLIHSHGINRRDFMKLCAALAATMGLSSKAAAEMAESVTNPQRPPVIWIGAQECT WP_000145421.1
MULTISPECIES: hydrogenase 2 small subunit [Shigella] Length: 372\nNumber of predicted TMHs: 1\nExp number of AAs in TMHs: 22.90256\nExp number, first 60 AAs: 0.2236\nTotal prob of N-in: 0.09847\noutside 1 331\nTMhelix 332 354\ninside 355 372

27829 GCF_001017155.1_ASM101715v1 Shigella flexneri 2a Proteobacteria; Gammaproteobacteria; Enterobacterales; Enterobacteriaceae; Shigella; Shigella flexneri
MNNEETFYQAMRRQGVTRRSFLKYCSLAATSLGLGAGMAPKIAWALENKPRIPVVWIHGL WP_001058323.1
MULTISPECIES: hydrogenase-1 small chain [Proteobacteria] Length: 372\nNumber of predicted TMHs: 2\nExp number of AAs in TMHs: 37.2608499999999\nExp number, first 60 AAs: 19.52923\nTotal prob of N-in: 0.94351\nPOSSIBLE N-term signal sequence\ninside 1 19\nTMhelix 20 39\noutside 40 325\nTMhelix 326 348\ninside 349 372

27830 GCF_001017305.1_ASM101730v1 Shigella flexneri 2a Proteobacteria; Gammaproteobacteria; Enterobacterales; Enterobacteriaceae; Shigella; Shigella flexneri
MTGDNTLIHSHGINRRDFMKLCAALAATMGLSSKAAAEMAESVTNPQRPPVIWIGAQECT WP_000145421.1
MULTISPECIES: hydrogenase 2 small subunit [Shigella] Length: 372\nNumber of predicted TMHs: 1\nExp number of AAs in TMHs: 22.90256\nExp number, first 60 AAs: 0.2236\nTotal prob of N-in: 0.09847\noutside 1 331\nTMhelix 332 354\ninside 355 372

27831 GCF_001017295.1_ASM101729v1 Shigella flexneri 2a Proteobacteria; Gammaproteobacteria; Enterobacterales; Enterobacteriaceae; Shigella; Shigella flexneri
MTGDNTLIHSHGINRRDFMKLCAALAATMGLSSKAAAEMAESVTNPQRPPVIWIGAQECT WP_000145421.1
MULTISPECIES: hydrogenase 2 small subunit [Shigella] Length: 372\nNumber of predicted TMHs: 1\nExp number of AAs in TMHs: 22.90256\nExp number, first 60 AAs: 0.2236\nTotal prob of N-in: 0.09847\noutside 1 331\nTMhelix 332 354\ninside 355 372

27832 GCF_001017295.1_ASM101729v1 Shigella flexneri 2a Proteobacteria; Gammaproteobacteria; Enterobacterales; Enterobacteriaceae; Shigella; Shigella flexneri
MNNEETFYQAMRRQGVTRRSFLKYCSLAATSLGLGAGMAPKIAWALENKPRIPVVWIHGL WP_001058323.1
MULTISPECIES: hydrogenase-1 small chain [Proteobacteria] Length: 372\nNumber of predicted TMHs: 2\nExp number of AAs in TMHs: 37.2608499999999\nExp number, first 60 AAs: 19.52923\nTotal prob of N-in: 0.94351\nPOSSIBLE N-term signal sequence\ninside 1 19\nTMhelix 20 39\noutside 40 325\nTMhelix 326 348\ninside 349 372

27833 GCF_001184075.1_ASM118407v1 Shigella flexneri 2a Proteobacteria; Gammaproteobacteria; Enterobacterales; Enterobacteriaceae; Shigella; Shigella flexneri
MTGDNTLIHSHGINRRDFMKLCAALAATMGLSSKAAAEMAESVTNPQRPPVIWIGAQECT WP_000145421.1
MULTISPECIES: hydrogenase 2 small subunit [Shigella] Length: 372\nNumber of predicted TMHs: 1\nExp number of AAs in TMHs: 22.90256\nExp number, first 60 AAs: 0.2236\nTotal prob of N-in: 0.09847\noutside 1 331\nTMhelix 332 354\ninside 355 372

27834 GCF_001017345.1_ASM101734v1 Shigella flexneri 2a Proteobacteria; Gammaproteobacteria; Enterobacterales; Enterobacteriaceae; Shigella; Shigella flexneri
MTGDNTLIHSHGINRRDFMKLCAALAATMGLSSKAAAEMAESVTNPQRPPVIWIGAQECT WP_000145421.1
MULTISPECIES: hydrogenase 2 small subunit [Shigella] Length: 372\nNumber of predicted TMHs: 1\nExp number of AAs in TMHs: 22.90256\nExp number, first 60 AAs: 0.2236\nTotal prob of N-in: 0.09847\noutside 1 331\nTMhelix 332 354\ninside 355 372

27835 GCF_001184075.1_ASM118407v1 Shigella flexneri 2a Proteobacteria; Gammaproteobacteria; Enterobacterales; Enterobacteriaceae; Shigella; Shigella flexneri
MNNEETFYQAMRRQGVTRRSFLKYCSLAATSLGLGAGMAPKIAWALENKPRIPVVWIHGL WP_001058323.1
MULTISPECIES: hydrogenase-1 small chain [Proteobacteria] Length: 372\nNumber of predicted TMHs: 2\nExp number of AAs in TMHs: 37.2608499999999\nExp number, first 60 AAs: 19.52923\nTotal prob of N-in: 0.94351\nPOSSIBLE N-term signal sequence\ninside 1 19\nTMhelix 20 39\noutside 40 325\nTMhelix 326 348\ninside 349 372

27836 GCF_001017405.1_ASM101740v1 Shigella flexneri 2a Proteobacteria; Gammaproteobacteria; Enterobacterales; Enterobacteriaceae; Shigella; Shigella flexneri
MNNEETFYQAMRRQGVTRRSFLKYCSLAATSLGLGAGMAPKIAWALENKPRIPVVWIHGL WP_001058323.1
MULTISPECIES: hydrogenase-1 small chain [Proteobacteria] Length: 372\nNumber of predicted TMHs: 2\nExp number of AAs in TMHs: 37.2608499999999\nExp number, first 60 AAs: 19.52923\nTotal prob of N-in: 0.94351\nPOSSIBLE N-term signal sequence\ninside 1 19\nTMhelix 20 39\noutside 40 325\nTMhelix 326 348\ninside 349 372

27837 GCF_001017405.1_ASM101740v1 Shigella flexneri 2a Proteobacteria; Gammaproteobacteria; Enterobacterales; Enterobacteriaceae; Shigella; Shigella flexneri
MTGDNTLIHSHGINRRDFMKLCAALAATMGLSSKAAAEMAESVTNPQRPPVIWIGAQECT WP_000145421.1
MULTISPECIES: hydrogenase 2 small subunit [Shigella] Length: 372\nNumber of predicted TMHs: 1\nExp number of AAs in TMHs: 22.90256\nExp number, first 60 AAs: 0.2236\nTotal prob of N-in: 0.09847\noutside 1 331\nTMhelix 332 354\ninside 355 372

27838 GCF_001017235.1_ASM101723v1 Shigella flexneri 2a Proteobacteria; Gammaproteobacteria; Enterobacterales; Enterobacteriaceae; Shigella; Shigella flexneri
MNNEETFYQAMRRQGVTRRSFLKYCSLAATSLGLGAGMAPKIAWALENKPRIPVVWIHGL WP_001058323.1
MULTISPECIES: hydrogenase-1 small chain [Proteobacteria] Length: 372\nNumber of predicted TMHs: 2\nExp number of AAs in TMHs: 37.2608499999999\nExp number, first 60 AAs: 19.52923\nTotal prob of N-in: 0.94351\nPOSSIBLE N-term signal sequence\ninside 1 19\nTMhelix 20 39\noutside 40 325\nTMhelix 326 348\ninside 349 372

27839 GCF_001017345.1_ASM101734v1 Shigella flexneri 2a Proteobacteria; Gammaproteobacteria; Enterobacterales; Enterobacteriaceae; Shigella; Shigella flexneri
MNNEETFYQAMRRQGVTRRSFLKYCSLAATSLGLGAGMAPKIAWALENKPRIPVVWIHGL WP_001058323.1
MULTISPECIES: hydrogenase-1 small chain [Proteobacteria] Length: 372\nNumber of predicted TMHs: 2\nExp number of AAs in TMHs: 37.2608499999999\nExp number, first 60 AAs: 19.52923\nTotal prob of N-in: 0.94351\nPOSSIBLE N-term signal sequence\ninside 1 19\nTMhelix 20 39\noutside 40 325\nTMhelix 326 348\ninside 349 372

27840 GCF_001017235.1_ASM101723v1 Shigella flexneri 2a Proteobacteria; Gammaproteobacteria; Enterobacterales; Enterobacteriaceae; Shigella; Shigella flexneri
MTGDNTLIHSHGINRRDFMKLCAALAATMGLSSKAAAEMAESVTNPQRPPVIWIGAQECT WP_000145421.1
MULTISPECIES: hydrogenase 2 small subunit [Shigella] Length: 372\nNumber of predicted TMHs: 1\nExp number of AAs in TMHs: 22.90256\nExp number, first 60 AAs: 0.2236\nTotal prob of N-in: 0.09847\noutside 1 331\nTMhelix 332 354\ninside 355 372

27841 GCF_000213475.1_ASM21347v2 Shigella flexneri 4343-70 Proteobacteria; Gammaproteobacteria; Enterobacterales; Enterobacteriaceae; Shigella; Shigella flexneri
MTGDNTLIHSHGINRRDFMKLCAALAATMGLSSKAAAEMAESVTNPQRPPVIWIGAQECT WP_000145421.1
MULTISPECIES: hydrogenase 2 small subunit [Shigella] Length: 372\nNumber of predicted TMHs: 1\nExp number of AAs in TMHs: 22.90256\nExp number, first 60 AAs: 0.2236\nTotal prob of N-in: 0.09847\noutside 1 331\nTMhelix 332 354\ninside 355 372

27842 GCF_000213475.1_ASM21347v2 Shigella flexneri 4343-70 Proteobacteria; Gammaproteobacteria; Enterobacterales; Enterobacteriaceae; Shigella; Shigella flexneri
MNNEETFYQAMRRQGVTRRSFLKYCSLAATSLGLGAGMAPKIAWALENKPRIPVVWIHGL WP_001058323.1
MULTISPECIES: hydrogenase-1 small chain [Proteobacteria] Length: 372\nNumber of predicted TMHs: 2\nExp number of AAs in TMHs: 37.2608499999999\nExp number, first 60 AAs: 19.52923\nTotal prob of N-in: 0.94351\nPOSSIBLE N-term signal sequence\ninside 1 19\nTMhelix 20 39\noutside 40 325\nTMhelix 326 348\ninside 349 372

27843 GCF_000565965.1_ASM56596v1 Shigella flexneri 51575 Proteobacteria; Gammaproteobacteria; Enterobacterales; Enterobacteriaceae; Shigella; Shigella flexneri
MNNEETFYQAMRRQGVTRRSFLKYCSLAATSLGLGAGMAPKIAWALENKPRIPVVWIHGL WP_025758242.1
hydrogenase [Shigella flexneri] Length: 372\nNumber of predicted TMHs: 2\nExp number of AAs in TMHs: 37.0188199999999\nExp number, first 60 AAs: 19.52908\nTotal prob of N-in: 0.94338\nPOSSIBLE N-term signal sequence\ninside 1 19\nTMhelix 20 39\noutside 40 325\nTMhelix 326 348\ninside 349 372

27844 GCF_000565965.1_ASM56596v1 Shigella flexneri 51575 Proteobacteria; Gammaproteobacteria; Enterobacterales; Enterobacteriaceae; Shigella; Shigella flexneri
MTGDNTLIHSHGINRRDFMKLCAALAATMGLSSKAAAEMAESVTNPQRPPVIWIGAQECT WP_000145413.1
MULTISPECIES: hydrogenase 2 small subunit [Shigella] Length: 372\nNumber of predicted TMHs: 1\nExp number of AAs in TMHs: 23.72614\nExp number, first 60 AAs: 0.22038\nTotal prob of N-in: 0.13004\noutside 1 331\nTMhelix 332 354\ninside 355 372

27845 GCF_000565985.1_ASM56598v1 Shigella flexneri 51576 Proteobacteria; Gammaproteobacteria; Enterobacterales; Enterobacteriaceae; Shigella; Shigella flexneri
MTGDNTLIHSHGINRRDFMKLCAALAATMGLSSKAAAEMAESVTNPQRPPVIWIGAQECT WP_000145421.1
MULTISPECIES: hydrogenase 2 small subunit [Shigella] Length: 372\nNumber of predicted TMHs: 1\nExp number of AAs in TMHs: 22.90256\nExp number, first 60 AAs: 0.2236\nTotal prob of N-in: 0.09847\noutside 1 331\nTMhelix 332 354\ninside 355 372

27846 GCF_000566025.1_ASM56602v1 Shigella flexneri 51581 Proteobacteria; Gammaproteobacteria; Enterobacterales; Enterobacteriaceae; Shigella; Shigella flexneri
MTGDNTLIHSHGINRRDFMKLCAALAATMGLSSKAAAEMAESVTNPQRPPVIWIGAQECT WP_000145421.1
MULTISPECIES: hydrogenase 2 small subunit [Shigella] Length: 372\nNumber of predicted TMHs: 1\nExp number of AAs in TMHs: 22.90256\nExp number, first 60 AAs: 0.2236\nTotal prob of N-in: 0.09847\noutside 1 331\nTMhelix 332 354\ninside 355 372

27847 GCF_000566025.1_ASM56602v1 Shigella flexneri 51581 Proteobacteria; Gammaproteobacteria; Enterobacterales; Enterobacteriaceae; Shigella; Shigella flexneri
MNNEETFYQAMRRQGVTRRSFLKYCSLAATSLGLGAGMAPKIAWALENKPRIPVVWIHGL WP_001058323.1
MULTISPECIES: hydrogenase-1 small chain [Proteobacteria] Length: 372\nNumber of predicted TMHs: 2\nExp number of AAs in TMHs: 37.2608499999999\nExp number, first 60 AAs: 19.52923\nTotal prob of N-in: 0.94351\nPOSSIBLE N-term signal sequence\ninside 1 19\nTMhelix 20 39\noutside 40 325\nTMhelix 326 348\ninside 349 372

27848 GCF_000267985.1_ASM26798v1 Shigella flexneri CCH060 Proteobacteria; Gammaproteobacteria; Enterobacterales; Enterobacteriaceae; Shigella; Shigella flexneri
MNNEETFYQAMRRQGVTRRSFLKYCSLAATSLGLGAGMAPKIAWALENKPRIPVVWIHGL WP_001058323.1

MULTISPECIES: hydrogenase-1 small chain [Proteobacteria] Length: 372\nNumber of predicted TMHs: 2\nExp number of AAs in TMHs: 37.2608499999999\nExp number, first 60 AAs: 19.52923\nTotal prob of N-in: 0.94351\nPOSSIBLE N-term signal sequence\ninside 1 19\nTMhelix 20 39\noutside 40 325\nTMhelix 326 348\ninside 349 372

27849 GCF_000267985.1_ASM26798v1 Shigella flexneri CCH060 Proteobacteria; Gammaproteobacteria; Enterobacterales; Enterobacteriaceae; Shigella; Shigella flexneri
MTGDNTLIHSHGINRRDFMKLCAALAATMGLSSKAAAEMAESVTNPQRPPVIWIGAQECT WP_000145410.1
MULTISPECIES: hydrogenase-2 small chain [Proteobacteria] Length: 372\nNumber of predicted TMHs: 1\nExp number of AAs in TMHs: 23.72226\nExp number, first 60 AAs: 0.22038\nTotal prob of N-in: 0.12994\noutside 1 331\nTMhelix 332 354\ninside 355 372

27850 GCF_000193935.1_ASM19393v2 Shigella flexneri CDC 796-83 Proteobacteria; Gammaproteobacteria; Enterobacterales; Enterobacteriaceae; Shigella; Shigella flexneri
MNNEETFYQAMRRQGVTRRSFLKYCSLAATSLGLGAGMAPKIAWALENKPRIPVVWIHGL WP_001058323.1
MULTISPECIES: hydrogenase-1 small chain [Proteobacteria] Length: 372\nNumber of predicted TMHs: 2\nExp number of AAs in TMHs: 37.2608499999999\nExp number, first 60 AAs: 19.52923\nTotal prob of N-in: 0.94351\nPOSSIBLE N-term signal sequence\ninside 1 19\nTMhelix 20 39\noutside 40 325\nTMhelix 326 348\ninside 349 372

27851 GCF_000193935.1_ASM19393v2 Shigella flexneri CDC 796-83 Proteobacteria; Gammaproteobacteria; Enterobacterales; Enterobacteriaceae; Shigella; Shigella flexneri
MTGDNTLIHSHGINRRDFMKLCAALAATMGLSSKAAAEMAESVTNPQRPPVIWIGAQECT WP_000145409.1
hydrogenase-2 small chain [Shigella flexneri] Length: 372\nNumber of predicted TMHs: 1\nExp number of AAs in TMHs: 23.57417\nExp number, first 60 AAs: 0.22323\nTotal prob of N-in: 0.11155\noutside 1 331\nTMhelix 332 354\ninside 355 372

27852 GCF_000815805.1_ASM81580v1 Shigella flexneri MT1457 Proteobacteria; Gammaproteobacteria; Enterobacterales; Enterobacteriaceae; Shigella; Shigella flexneri
MTGDNTLIHSHGINRRDFMKLCAALAATMGLSSKAAAEMAESVTNPQRPPVIWIGAQECT WP_000145413.1
MULTISPECIES: hydrogenase 2 small subunit [Shigella] Length: 372\nNumber of predicted TMHs: 1\nExp number of AAs in TMHs: 23.72614\nExp number, first 60 AAs: 0.22038\nTotal prob of N-in: 0.13004\noutside 1 331\nTMhelix 332 354\ninside 355 372

27853 GCF_000815805.1_ASM81580v1 Shigella flexneri MT1457 Proteobacteria; Gammaproteobacteria; Enterobacterales; Enterobacteriaceae; Shigella; Shigella flexneri
MNNEETFYQAMRRQGVTRRSFLKYCSLAATSLGLGAGMAPKIAWALENKPRIPVVWIHGL WP_001058323.1
MULTISPECIES: hydrogenase-1 small chain [Proteobacteria] Length: 372\nNumber of predicted TMHs: 2\nExp number of AAs in TMHs: 37.2608499999999\nExp number, first 60 AAs: 19.52923\nTotal prob of N-in: 0.94351\nPOSSIBLE N-term signal sequence\ninside 1 19\nTMhelix 20 39\noutside 40 325\nTMhelix 326 348\ninside 349 372

27854 GCF_000815485.1_ASM81548v1 Shigella flexneri S5717 Proteobacteria; Gammaproteobacteria; Enterobacterales; Enterobacteriaceae; Shigella; Shigella flexneri
MNNEETFYQAMRRQGVTRRSFLKYCSLAATSLGLGAGMAPKIAWALENKPRIPVVWIHGL WP_001058323.1
MULTISPECIES: hydrogenase-1 small chain [Proteobacteria] Length: 372\nNumber of predicted TMHs: 2\nExp number of AAs in TMHs: 37.2608499999999\nExp number, first 60 AAs: 19.52923\nTotal prob of N-in: 0.94351\nPOSSIBLE N-term signal sequence\ninside 1 19\nTMhelix 20 39\noutside 40 325\nTMhelix 326 348\ninside 349 372

27855 GCF_000815485.1_ASM81548v1 Shigella flexneri S5717 Proteobacteria; Gammaproteobacteria; Enterobacterales; Enterobacteriaceae; Shigella; Shigella flexneri
MTGDNTLIHSHGINRRDFMKLCAALAATMGLSSKAAAEMAESVTNPQRPPVIWIGAQECT WP_000145421.1
MULTISPECIES: hydrogenase 2 small subunit [Shigella] Length: 372\nNumber of predicted TMHs: 1\nExp number of AAs in TMHs: 22.90256\nExp number, first 60 AAs: 0.2236\nTotal prob of N-in: 0.09847\noutside 1 331\nTMhelix 332 354\ninside 355 372

27856 GCF_000815565.1_ASM81556v1 Shigella flexneri S6585 Proteobacteria; Gammaproteobacteria; Enterobacterales; Enterobacteriaceae; Shigella; Shigella flexneri
MNNEETFYQAMRRQGVTRRSFLKYCSLAATSLGLGAGMAPKIAWALENKPRIPVVWIHGL WP_001058323.1
MULTISPECIES: hydrogenase-1 small chain [Proteobacteria] Length: 372\nNumber of predicted TMHs:

2\nExp number of AAs in TMHs: 37.2608499999999\nExp number, first 60 AAs:  19.52923\nTotal prob of N-in: 0.94351\nPOSSIBLE N-term signal sequence\ninside   1   19\nTMhelix   20   39\noutside   40   325\nTMhelix   326   348\ninside   349   372

27857 GCF_000815565.1_ASM81556v1 Shigella flexneri S6585 Proteobacteria; Gammaproteobacteria; Enterobacterales; Enterobacteriaceae; Shigella; Shigella flexneri MTGDNTLIHSHGINRRDFMKLCAALAATMGLSSKAAAEMAESVTNPQRPPVIWIGAQECT WP_000145421.1 MULTISPECIES: hydrogenase 2 small subunit [Shigella] Length: 372\nNumber of predicted TMHs:  1\nExp number of AAs in TMHs: 22.90256\nExp number, first 60 AAs:  0.2236\nTotal prob of N-in:   0.09847\noutside   1   331\nTMhelix   332   354\ninside   355   372

27858 GCF_000815595.1_ASM81559v1 Shigella flexneri S6678 Proteobacteria; Gammaproteobacteria; Enterobacterales; Enterobacteriaceae; Shigella; Shigella flexneri MNNEETFYQAMRRQGVTRRSFLKYCSLAATSLGLGAGMAPKIAWALENKPRIPVVWIHGL WP_001058323.1 MULTISPECIES: hydrogenase-1 small chain [Proteobacteria] Length: 372\nNumber of predicted TMHs: 2\nExp number of AAs in TMHs: 37.2608499999999\nExp number, first 60 AAs:  19.52923\nTotal prob of N-in: 0.94351\nPOSSIBLE N-term signal sequence\ninside   1   19\nTMhelix   20   39\noutside   40   325\nTMhelix   326   348\ninside   349   372

27859 GCF_000815595.1_ASM81559v1 Shigella flexneri S6678 Proteobacteria; Gammaproteobacteria; Enterobacterales; Enterobacteriaceae; Shigella; Shigella flexneri MTGDNTLIHSHGINRRDFMKLCAALAATMGLSSKAAAEMAESVTNPQRPPVIWIGAQECT WP_000145421.1 MULTISPECIES: hydrogenase 2 small subunit [Shigella] Length: 372\nNumber of predicted TMHs:  1\nExp number of AAs in TMHs: 22.90256\nExp number, first 60 AAs:  0.2236\nTotal prob of N-in:   0.09847\noutside   1   331\nTMhelix   332   354\ninside   355   372

27860 GCF_000815625.1_ASM81562v1 Shigella flexneri S6764 Proteobacteria; Gammaproteobacteria; Enterobacterales; Enterobacteriaceae; Shigella; Shigella flexneri MNNEETFYQAMRRQGVTRRSFLKYCSLAATSLGLGAGMAPKIAWALENKPRIPVVWIHGL WP_001058323.1 MULTISPECIES: hydrogenase-1 small chain [Proteobacteria] Length: 372\nNumber of predicted TMHs: 2\nExp number of AAs in TMHs: 37.2608499999999\nExp number, first 60 AAs:  19.52923\nTotal prob of N-in: 0.94351\nPOSSIBLE N-term signal sequence\ninside   1   19\nTMhelix   20   39\noutside   40   325\nTMhelix   326   348\ninside   349   372

27861 GCF_000815625.1_ASM81562v1 Shigella flexneri S6764 Proteobacteria; Gammaproteobacteria; Enterobacterales; Enterobacteriaceae; Shigella; Shigella flexneri MTGDNTLIHSHGINRRDFMKLCAALAATMGLSSKAAAEMAESVTNPQRPPVIWIGAQECT WP_000145421.1 MULTISPECIES: hydrogenase 2 small subunit [Shigella] Length: 372\nNumber of predicted TMHs:  1\nExp number of AAs in TMHs: 22.90256\nExp number, first 60 AAs:  0.2236\nTotal prob of N-in:   0.09847\noutside   1   331\nTMhelix   332   354\ninside   355   372

27862 GCF_000815665.1_ASM81566v1 Shigella flexneri S7737 Proteobacteria; Gammaproteobacteria; Enterobacterales; Enterobacteriaceae; Shigella; Shigella flexneri MNNEETFYQAMRRQGVTRRSFLKYCSLAATSLGLGAGMAPKIAWALENKPRIPVVWIHGL WP_001058323.1 MULTISPECIES: hydrogenase-1 small chain [Proteobacteria] Length: 372\nNumber of predicted TMHs: 2\nExp number of AAs in TMHs: 37.2608499999999\nExp number, first 60 AAs:  19.52923\nTotal prob of N-in: 0.94351\nPOSSIBLE N-term signal sequence\ninside   1   19\nTMhelix   20   39\noutside   40   325\nTMhelix   326   348\ninside   349   372

27863 GCF_000815665.1_ASM81566v1 Shigella flexneri S7737 Proteobacteria; Gammaproteobacteria; Enterobacterales; Enterobacteriaceae; Shigella; Shigella flexneri MTGDNTLIHSHGINRRDFMKLCAALAATMGLSSKAAAEMAESVTNPQRPPVIWIGAQECT WP_000145421.1 MULTISPECIES: hydrogenase 2 small subunit [Shigella] Length: 372\nNumber of predicted TMHs:  1\nExp number of AAs in TMHs: 22.90256\nExp number, first 60 AAs:  0.2236\nTotal prob of N-in:   0.09847\noutside   1   331\nTMhelix   332   354\ninside   355   372

27864 GCF_000565905.1_ASM56590v1 Shigella flexneri Shi05SX04 Proteobacteria; Gammaproteobacteria; Enterobacterales; Enterobacteriaceae; Shigella; Shigella flexneri MNNEETFYQAMRRQGVTRRSFLKYCSLAATSLGLGAGMAPKIAWALENKPRIPVVWIHGL WP_001058323.1 MULTISPECIES: hydrogenase-1 small chain [Proteobacteria] Length: 372\nNumber of predicted TMHs: 2\nExp number of AAs in TMHs: 37.2608499999999\nExp number, first 60 AAs:  19.52923\nTotal prob of N-in:

0.94351\nPOSSIBLE N-term signal sequence\ninside 1 19\nTMhelix 20 39\noutside 40 325\nTMhelix 326 348\ninside 349 372

27865 GCF_000565905.1_ASM56590v1 Shigella flexneri Shi05SX04 Proteobacteria; Gammaproteobacteria; Enterobacterales; Enterobacteriaceae; Shigella; Shigella flexneri MTGDNTLIHSHGINRRDFMKLCAALAATMGLSSKAAAEMAESVTNPQRPPVIWIGAQECT WP_000145421.1 MULTISPECIES: hydrogenase 2 small subunit [Shigella] Length: 372\nNumber of predicted TMHs: 1\nExp number of AAs in TMHs: 22.90256\nExp number, first 60 AAs: 0.2236\nTotal prob of N-in: 0.09847\noutside 1 331\nTMhelix 332 354\ninside 355 372

27866 GCF_000565745.1_ASM56574v1 Shigella flexneri Shi06AH028 Proteobacteria; Gammaproteobacteria; Enterobacterales; Enterobacteriaceae; Shigella; Shigella flexneri MTGDNTLIHSHGINRRDFMKLCAALAATMGLSSKAAAEMAESVTNPQRPPVIWIGAQECT WP_000145421.1 MULTISPECIES: hydrogenase 2 small subunit [Shigella] Length: 372\nNumber of predicted TMHs: 1\nExp number of AAs in TMHs: 22.90256\nExp number, first 60 AAs: 0.2236\nTotal prob of N-in: 0.09847\noutside 1 331\nTMhelix 332 354\ninside 355 372

27867 GCF_000565745.1_ASM56574v1 Shigella flexneri Shi06AH028 Proteobacteria; Gammaproteobacteria; Enterobacterales; Enterobacteriaceae; Shigella; Shigella flexneri MNNEETFYQAMRRQGVTRRSFLKYCSLAATSLGLGAGMAPKIAWALENKPRIPVVWIHGL WP_001058323.1 MULTISPECIES: hydrogenase-1 small chain [Proteobacteria] Length: 372\nNumber of predicted TMHs: 2\nExp number of AAs in TMHs: 37.2608499999999\nExp number, first 60 AAs: 19.52923\nTotal prob of N-in: 0.94351\nPOSSIBLE N-term signal sequence\ninside 1 19\nTMhelix 20 39\noutside 40 325\nTMhelix 326 348\ninside 349 372

27868 GCF_000566525.1_ASM56652v1 Shigella flexneri Shi06AH091 Proteobacteria; Gammaproteobacteria; Enterobacterales; Enterobacteriaceae; Shigella; Shigella flexneri MTGDNTLIHSHGINRRDFMKLCAALAATMGLSSKAAAEMAESVTNPQRPPVIWIGAQECT WP_000145421.1 MULTISPECIES: hydrogenase 2 small subunit [Shigella] Length: 372\nNumber of predicted TMHs: 1\nExp number of AAs in TMHs: 22.90256\nExp number, first 60 AAs: 0.2236\nTotal prob of N-in: 0.09847\noutside 1 331\nTMhelix 332 354\ninside 355 372

27869 GCF_000566525.1_ASM56652v1 Shigella flexneri Shi06AH091 Proteobacteria; Gammaproteobacteria; Enterobacterales; Enterobacteriaceae; Shigella; Shigella flexneri MNNEETFYQAMRRQGVTRRSFLKYCSLAATSLGLGAGMAPKIAWALENKPRIPVVWIHGL WP_001058323.1 MULTISPECIES: hydrogenase-1 small chain [Proteobacteria] Length: 372\nNumber of predicted TMHs: 2\nExp number of AAs in TMHs: 37.2608499999999\nExp number, first 60 AAs: 19.52923\nTotal prob of N-in: 0.94351\nPOSSIBLE N-term signal sequence\ninside 1 19\nTMhelix 20 39\noutside 40 325\nTMhelix 326 348\ninside 349 372

27870 GCF_000565545.1_ASM56554v1 Shigella flexneri Shi06AH116 Proteobacteria; Gammaproteobacteria; Enterobacterales; Enterobacteriaceae; Shigella; Shigella flexneri MNNEETFYQAMRRQGVTRRSFLKYCSLAATSLGLGAGMAPKIAWALENKPRIPVVWIHGL WP_001058323.1 MULTISPECIES: hydrogenase-1 small chain [Proteobacteria] Length: 372\nNumber of predicted TMHs: 2\nExp number of AAs in TMHs: 37.2608499999999\nExp number, first 60 AAs: 19.52923\nTotal prob of N-in: 0.94351\nPOSSIBLE N-term signal sequence\ninside 1 19\nTMhelix 20 39\noutside 40 325\nTMhelix 326 348\ninside 349 372

27871 GCF_000565545.1_ASM56554v1 Shigella flexneri Shi06AH116 Proteobacteria; Gammaproteobacteria; Enterobacterales; Enterobacteriaceae; Shigella; Shigella flexneri MTGDNTLIHSHGINRRDFMKLCAALAATMGLSSKAAAEMAESVTNPQRPPVIWIGAQECT WP_000145421.1 MULTISPECIES: hydrogenase 2 small subunit [Shigella] Length: 372\nNumber of predicted TMHs: 1\nExp number of AAs in TMHs: 22.90256\nExp number, first 60 AAs: 0.2236\nTotal prob of N-in: 0.09847\noutside 1 331\nTMhelix 332 354\ninside 355 372

27872 GCF_000565825.1_ASM56582v1 Shigella flexneri Shi06AH130 Proteobacteria; Gammaproteobacteria; Enterobacterales; Enterobacteriaceae; Shigella; Shigella flexneri MNNEETFYQAMRRQGVTRRSFLKYCSLAATSLGLGAGMAPKIAWALENKPRIPVVWIHGL WP_001058323.1 MULTISPECIES: hydrogenase-1 small chain [Proteobacteria] Length: 372\nNumber of predicted TMHs: 2\nExp number of AAs in TMHs: 37.2608499999999\nExp number, first 60 AAs: 19.52923\nTotal prob of N-in:

0.94351\nPOSSIBLE N-term signal sequence\ninside 1 19\nTMhelix 20 39\noutside 40 325\nTMhelix 326 348\ninside 349 372

27873 GCF_000565825.1_ASM56582v1 Shigella flexneri Shi06AH130 Proteobacteria; Gammaproteobacteria; Enterobacterales; Enterobacteriaceae; Shigella; Shigella flexneri MTGDNTLIHSHGINRRDFMKLCAALAATMGLSSKAAAEMAESVTNPQRPPVIWIGAQECT WP_000145421.1 MULTISPECIES: hydrogenase 2 small subunit [Shigella] Length: 372\nNumber of predicted TMHs: 1\nExp number of AAs in TMHs: 22.90256\nExp number, first 60 AAs: 0.2236\nTotal prob of N-in: 0.09847\noutside 1 331\nTMhelix 332 354\ninside 355 372

27874 GCF_000565665.1_ASM56566v1 Shigella flexneri Shi06AH135 Proteobacteria; Gammaproteobacteria; Enterobacterales; Enterobacteriaceae; Shigella; Shigella flexneri MTGDNTLIHSHGINRRDFMKLCAALAATMGLSSKAAAEMAESVTNPQRPPVIWIGAQECT WP_000145421.1 MULTISPECIES: hydrogenase 2 small subunit [Shigella] Length: 372\nNumber of predicted TMHs: 1\nExp number of AAs in TMHs: 22.90256\nExp number, first 60 AAs: 0.2236\nTotal prob of N-in: 0.09847\noutside 1 331\nTMhelix 332 354\ninside 355 372

27875 GCF_000565665.1_ASM56566v1 Shigella flexneri Shi06AH135 Proteobacteria; Gammaproteobacteria; Enterobacterales; Enterobacteriaceae; Shigella; Shigella flexneri MNNEETFYQAMRRQGVTRRSFLKYCSLAATSLGLGAGMAPKIAWALENKPRIPVVWIHGL WP_001058323.1 MULTISPECIES: hydrogenase-1 small chain [Proteobacteria] Length: 372\nNumber of predicted TMHs: 2\nExp number of AAs in TMHs: 37.2608499999999\nExp number, first 60 AAs: 19.52923\nTotal prob of N-in: 0.94351\nPOSSIBLE N-term signal sequence\ninside 1 19\nTMhelix 20 39\noutside 40 325\nTMhelix 326 348\ninside 349 372

27876 GCF_000565505.1_ASM56550v1 Shigella flexneri Shi06AH66 Proteobacteria; Gammaproteobacteria; Enterobacterales; Enterobacteriaceae; Shigella; Shigella flexneri MNNEETFYQAMRRQGVTRRSFLKYCSLAATSLGLGAGMAPKIAWALENKPRIPVVWIHGL WP_001058323.1 MULTISPECIES: hydrogenase-1 small chain [Proteobacteria] Length: 372\nNumber of predicted TMHs: 2\nExp number of AAs in TMHs: 37.2608499999999\nExp number, first 60 AAs: 19.52923\nTotal prob of N-in: 0.94351\nPOSSIBLE N-term signal sequence\ninside 1 19\nTMhelix 20 39\noutside 40 325\nTMhelix 326 348\ninside 349 372

27877 GCF_000565505.1_ASM56550v1 Shigella flexneri Shi06AH66 Proteobacteria; Gammaproteobacteria; Enterobacterales; Enterobacteriaceae; Shigella; Shigella flexneri MTGDNTLIHSHGINRRDFMKLCAALAATMGLSSKAAAEMAESVTNPQRPPVIWIGAQECT WP_000145421.1 MULTISPECIES: hydrogenase 2 small subunit [Shigella] Length: 372\nNumber of predicted TMHs: 1\nExp number of AAs in TMHs: 22.90256\nExp number, first 60 AAs: 0.2236\nTotal prob of N-in: 0.09847\noutside 1 331\nTMhelix 332 354\ninside 355 372

27878 GCF_000566425.1_ASM56642v1 Shigella flexneri Shi06GS02 Proteobacteria; Gammaproteobacteria; Enterobacterales; Enterobacteriaceae; Shigella; Shigella flexneri MNNEETFYQAMRRQGVTRRSFLKYCSLAATSLGLGAGMAPKIAWALENKPRIPVVWIHGL WP_001058323.1 MULTISPECIES: hydrogenase-1 small chain [Proteobacteria] Length: 372\nNumber of predicted TMHs: 2\nExp number of AAs in TMHs: 37.2608499999999\nExp number, first 60 AAs: 19.52923\nTotal prob of N-in: 0.94351\nPOSSIBLE N-term signal sequence\ninside 1 19\nTMhelix 20 39\noutside 40 325\nTMhelix 326 348\ninside 349 372

27879 GCF_000565565.1_ASM56556v1 Shigella flexneri Shi06GS07 Proteobacteria; Gammaproteobacteria; Enterobacterales; Enterobacteriaceae; Shigella; Shigella flexneri MNNEETFYQAMRRQGVTRRSFLKYCSLAATSLGLGAGMAPKIAWALENKPRIPVVWIHGL WP_001058323.1 MULTISPECIES: hydrogenase-1 small chain [Proteobacteria] Length: 372\nNumber of predicted TMHs: 2\nExp number of AAs in TMHs: 37.2608499999999\nExp number, first 60 AAs: 19.52923\nTotal prob of N-in: 0.94351\nPOSSIBLE N-term signal sequence\ninside 1 19\nTMhelix 20 39\noutside 40 325\nTMhelix 326 348\ninside 349 372

27880 GCF_000565565.1_ASM56556v1 Shigella flexneri Shi06GS07 Proteobacteria; Gammaproteobacteria; Enterobacterales; Enterobacteriaceae; Shigella; Shigella flexneri MTGDNTLIHSHGINRRDFMKLCAALAATMGLSSKAAAEMAESVTNPQRPPVIWIGAQECT WP_000145421.1 MULTISPECIES: hydrogenase 2 small subunit [Shigella] Length: 372\nNumber of predicted TMHs: 1\nExp

number of AAs in TMHs: 22.90256\nExp number, first 60 AAs:  0.2236\nTotal prob of N-in:   0.09847\noutside   1   331\nTMhelix   332   354\ninside   355   372

27881 GCF_000565785.1_ASM56578v1 Shigella flexneri Shi06GS37 Proteobacteria; Gammaproteobacteria; Enterobacterales; Enterobacteriaceae; Shigella; Shigella flexneri MNNEETFYQAMRRQGVTRRSFLKYCSLAATSLGLGAGMAPKIAWALENKPRIPVVWIHGL WP_001058323.1 MULTISPECIES: hydrogenase-1 small chain [Proteobacteria] Length: 372\nNumber of predicted TMHs: 2\nExp number of AAs in TMHs: 37.2608499999999\nExp number, first 60 AAs:  19.52923\nTotal prob of N-in: 0.94351\nPOSSIBLE N-term signal sequence\ninside   1   19\nTMhelix   20   39\noutside   40   325\nTMhelix   326   348\ninside   349   372

27882 GCF_000565785.1_ASM56578v1 Shigella flexneri Shi06GS37 Proteobacteria; Gammaproteobacteria; Enterobacterales; Enterobacteriaceae; Shigella; Shigella flexneri MTGDNTLIHSHGINRRDFMKLCAALAATMGLSSKAAAEMAESVTNPQRPPVIWIGAQECT WP_000145421.1 MULTISPECIES: hydrogenase 2 small subunit [Shigella] Length: 372\nNumber of predicted TMHs:  1\nExp number of AAs in TMHs: 22.90256\nExp number, first 60 AAs:  0.2236\nTotal prob of N-in:   0.09847\noutside   1   331\nTMhelix   332   354\ninside   355   372

27883 GCF_000565525.1_ASM56552v1 Shigella flexneri Shi06GS43 Proteobacteria; Gammaproteobacteria; Enterobacterales; Enterobacteriaceae; Shigella; Shigella flexneri MNNEETFYQAMRRQGVTRRSFLKYCSLAATSLGLGAGMAPKIAWALENKPRIPVVWIHGL WP_001058323.1 MULTISPECIES: hydrogenase-1 small chain [Proteobacteria] Length: 372\nNumber of predicted TMHs: 2\nExp number of AAs in TMHs: 37.2608499999999\nExp number, first 60 AAs:  19.52923\nTotal prob of N-in: 0.94351\nPOSSIBLE N-term signal sequence\ninside   1   19\nTMhelix   20   39\noutside   40   325\nTMhelix   326   348\ninside   349   372

27884 GCF_000565525.1_ASM56552v1 Shigella flexneri Shi06GS43 Proteobacteria; Gammaproteobacteria; Enterobacterales; Enterobacteriaceae; Shigella; Shigella flexneri MTGDNTLIHSHGINRRDFMKLCAALAATMGLSSKAAAEMAESVTNPQRPPVIWIGAQECT WP_000145421.1 MULTISPECIES: hydrogenase 2 small subunit [Shigella] Length: 372\nNumber of predicted TMHs:  1\nExp number of AAs in TMHs: 22.90256\nExp number, first 60 AAs:  0.2236\nTotal prob of N-in:   0.09847\noutside   1   331\nTMhelix   332   354\ninside   355   372

27885 GCF_000565585.1_ASM56558v1 Shigella flexneri Shi06GS48 Proteobacteria; Gammaproteobacteria; Enterobacterales; Enterobacteriaceae; Shigella; Shigella flexneri MTGDNTLIHSHGINRRDFMKLCAALAATMGLSSKAAAEMAESVTNPQRPPVIWIGAQECT WP_000145421.1 MULTISPECIES: hydrogenase 2 small subunit [Shigella] Length: 372\nNumber of predicted TMHs:  1\nExp number of AAs in TMHs: 22.90256\nExp number, first 60 AAs:  0.2236\nTotal prob of N-in:   0.09847\noutside   1   331\nTMhelix   332   354\ninside   355   372

27886 GCF_000565585.1_ASM56558v1 Shigella flexneri Shi06GS48 Proteobacteria; Gammaproteobacteria; Enterobacterales; Enterobacteriaceae; Shigella; Shigella flexneri MNNEETFYQAMRRQGVTRRSFLKYCSLAATSLGLGAGMAPKIAWALENKPRIPVVWIHGL WP_001058323.1 MULTISPECIES: hydrogenase-1 small chain [Proteobacteria] Length: 372\nNumber of predicted TMHs: 2\nExp number of AAs in TMHs: 37.2608499999999\nExp number, first 60 AAs:  19.52923\nTotal prob of N-in: 0.94351\nPOSSIBLE N-term signal sequence\ninside   1   19\nTMhelix   20   39\noutside   40   325\nTMhelix   326   348\ninside   349   372

27887 GCF_000565845.1_ASM56584v1 Shigella flexneri Shi06GS55 Proteobacteria; Gammaproteobacteria; Enterobacterales; Enterobacteriaceae; Shigella; Shigella flexneri MTGDNTLIHSHGINRRDFMKLCAALAATMGLSSKAAAEMAESVTNPQRPPVIWIGAQECT WP_000145421.1 MULTISPECIES: hydrogenase 2 small subunit [Shigella] Length: 372\nNumber of predicted TMHs:  1\nExp number of AAs in TMHs: 22.90256\nExp number, first 60 AAs:  0.2236\nTotal prob of N-in:   0.09847\noutside   1   331\nTMhelix   332   354\ninside   355   372

27888 GCF_000565845.1_ASM56584v1 Shigella flexneri Shi06GS55 Proteobacteria; Gammaproteobacteria; Enterobacterales; Enterobacteriaceae; Shigella; Shigella flexneri MNNEETFYQAMRRQGVTRRSFLKYCSLAATSLGLGAGMAPKIAWALENKPRIPVVWIHGL WP_001058323.1 MULTISPECIES: hydrogenase-1 small chain [Proteobacteria] Length: 372\nNumber of predicted TMHs: 2\nExp number of AAs in TMHs: 37.2608499999999\nExp number, first 60 AAs:  19.52923\nTotal prob of N-in:

0.94351\nPOSSIBLE N-term signal sequence\ninside 1 19\nTMhelix 20 39\noutside 40 325\nTMhelix 326 348\ninside 349 372

27889 GCF_000743995.1_ASM74399v1 Shigella flexneri Shi06HN006 Proteobacteria; Gammaproteobacteria; Enterobacterales; Enterobacteriaceae; Shigella; Shigella flexneri
MTGDNTLIHSHGINRRDFMKLCAALAATMGLSSKAAAEMAESVTNPQRPPVIWIGAQECT WP_000145421.1
MULTISPECIES: hydrogenase 2 small subunit [Shigella] Length: 372\nNumber of predicted TMHs: 1\nExp number of AAs in TMHs: 22.90256\nExp number, first 60 AAs: 0.2236\nTotal prob of N-in: 0.09847\noutside 1 331\nTMhelix 332 354\ninside 355 372

27890 GCF_000743995.1_ASM74399v1 Shigella flexneri Shi06HN006 Proteobacteria; Gammaproteobacteria; Enterobacterales; Enterobacteriaceae; Shigella; Shigella flexneri
MNNEETFYQAMRRQGVTRRSFLKYCSLAATSLGLGAGMAPKIAWALENKPRIPVVWIHGL WP_001058323.1
MULTISPECIES: hydrogenase-1 small chain [Proteobacteria] Length: 372\nNumber of predicted TMHs: 2\nExp number of AAs in TMHs: 37.2608499999999\nExp number, first 60 AAs: 19.52923\nTotal prob of N-in: 0.94351\nPOSSIBLE N-term signal sequence\ninside 1 19\nTMhelix 20 39\noutside 40 325\nTMhelix 326 348\ninside 349 372

27891 GCF_000565705.1_ASM56570v1 Shigella flexneri Shi06HN016 Proteobacteria; Gammaproteobacteria; Enterobacterales; Enterobacteriaceae; Shigella; Shigella flexneri
MNNEETFYQAMRRQGVTRRSFLKYCSLAATSLGLGAGMAPKIAWALENKPRIPVVWIHGL WP_001058323.1
MULTISPECIES: hydrogenase-1 small chain [Proteobacteria] Length: 372\nNumber of predicted TMHs: 2\nExp number of AAs in TMHs: 37.2608499999999\nExp number, first 60 AAs: 19.52923\nTotal prob of N-in: 0.94351\nPOSSIBLE N-term signal sequence\ninside 1 19\nTMhelix 20 39\noutside 40 325\nTMhelix 326 348\ninside 349 372

27892 GCF_000565705.1_ASM56570v1 Shigella flexneri Shi06HN016 Proteobacteria; Gammaproteobacteria; Enterobacterales; Enterobacteriaceae; Shigella; Shigella flexneri
MTGDNTLIHSHGINRRDFMKLCAALAATMGLSSKAAAEMAESVTNPQRPPVIWIGAQECT WP_000145421.1
MULTISPECIES: hydrogenase 2 small subunit [Shigella] Length: 372\nNumber of predicted TMHs: 1\nExp number of AAs in TMHs: 22.90256\nExp number, first 60 AAs: 0.2236\nTotal prob of N-in: 0.09847\noutside 1 331\nTMhelix 332 354\ninside 355 372

27893 GCF_000565625.1_ASM56562v1 Shigella flexneri Shi06HN023 Proteobacteria; Gammaproteobacteria; Enterobacterales; Enterobacteriaceae; Shigella; Shigella flexneri
MTGDNTLIHSHGINRRDFMKLCAALAATMGLSSKAAAEMAESVTNPQRPPVIWIGAQECT WP_000145421.1
MULTISPECIES: hydrogenase 2 small subunit [Shigella] Length: 372\nNumber of predicted TMHs: 1\nExp number of AAs in TMHs: 22.90256\nExp number, first 60 AAs: 0.2236\nTotal prob of N-in: 0.09847\noutside 1 331\nTMhelix 332 354\ninside 355 372

27894 GCF_000565625.1_ASM56562v1 Shigella flexneri Shi06HN023 Proteobacteria; Gammaproteobacteria; Enterobacterales; Enterobacteriaceae; Shigella; Shigella flexneri
MNNEETFYQAMRRQGVTRRSFLKYCSLAATSLGLGAGMAPKIAWALENKPRIPVVWIHGL WP_001058323.1
MULTISPECIES: hydrogenase-1 small chain [Proteobacteria] Length: 372\nNumber of predicted TMHs: 2\nExp number of AAs in TMHs: 37.2608499999999\nExp number, first 60 AAs: 19.52923\nTotal prob of N-in: 0.94351\nPOSSIBLE N-term signal sequence\ninside 1 19\nTMhelix 20 39\noutside 40 325\nTMhelix 326 348\ninside 349 372

27895 GCF_000566505.1_ASM56650v1 Shigella flexneri Shi06HN091 Proteobacteria; Gammaproteobacteria; Enterobacterales; Enterobacteriaceae; Shigella; Shigella flexneri
MTGDNTLIHSHGINRRDFMKLCAALAATMGLSSKAAAEMAESVTNPQRPPVIWIGAQECT WP_000145421.1
MULTISPECIES: hydrogenase 2 small subunit [Shigella] Length: 372\nNumber of predicted TMHs: 1\nExp number of AAs in TMHs: 22.90256\nExp number, first 60 AAs: 0.2236\nTotal prob of N-in: 0.09847\noutside 1 331\nTMhelix 332 354\ninside 355 372

27896 GCF_000566505.1_ASM56650v1 Shigella flexneri Shi06HN091 Proteobacteria; Gammaproteobacteria; Enterobacterales; Enterobacteriaceae; Shigella; Shigella flexneri
MNNEETFYQAMRRQGVTRRSFLKYCSLAATSLGLGAGMAPKIAWALENKPRIPVVWIHGL WP_001058323.1
MULTISPECIES: hydrogenase-1 small chain [Proteobacteria] Length: 372\nNumber of predicted TMHs: 2\nExp number of AAs in TMHs: 37.2608499999999\nExp number, first 60 AAs: 19.52923\nTotal prob of N-in:

0.94351\nPOSSIBLE N-term signal sequence\ninside 1 19\nTMhelix 20 39\noutside 40 325\nTMhelix 326 348\ninside 349 372

27897 GCF_000565605.1_ASM56560v1 Shigella flexneri Shi06HN118 Proteobacteria; Gammaproteobacteria; Enterobacterales; Enterobacteriaceae; Shigella; Shigella flexneri MNNEETFYQAMRRQGVTRRSFLKYCSLAATSLGLGAGMAPKIAWALENKPRIPVVWIHGL WP_001058323.1 MULTISPECIES: hydrogenase-1 small chain [Proteobacteria] Length: 372\nNumber of predicted TMHs: 2\nExp number of AAs in TMHs: 37.2608499999999\nExp number, first 60 AAs: 19.52923\nTotal prob of N-in: 0.94351\nPOSSIBLE N-term signal sequence\ninside 1 19\nTMhelix 20 39\noutside 40 325\nTMhelix 326 348\ninside 349 372

27898 GCF_000565605.1_ASM56560v1 Shigella flexneri Shi06HN118 Proteobacteria; Gammaproteobacteria; Enterobacterales; Enterobacteriaceae; Shigella; Shigella flexneri MTGDNTLIHSHGINRRDFMKLCAALAATMGLSSKAAAEMAESVTNPQRPPVIWIGAQECT WP_000145421.1 MULTISPECIES: hydrogenase 2 small subunit [Shigella] Length: 372\nNumber of predicted TMHs: 1\nExp number of AAs in TMHs: 22.90256\nExp number, first 60 AAs: 0.2236\nTotal prob of N-in: 0.09847\noutside 1 331\nTMhelix 332 354\ninside 355 372

27899 GCF_000566445.1_ASM56644v1 Shigella flexneri Shi06HN159 Proteobacteria; Gammaproteobacteria; Enterobacterales; Enterobacteriaceae; Shigella; Shigella flexneri MTGDNTLIHSHGINRRDFMKLCAALAATMGLSSKAAAEMAESVTNPQRPPVIWIGAQECT WP_000145421.1 MULTISPECIES: hydrogenase 2 small subunit [Shigella] Length: 372\nNumber of predicted TMHs: 1\nExp number of AAs in TMHs: 22.90256\nExp number, first 60 AAs: 0.2236\nTotal prob of N-in: 0.09847\noutside 1 331\nTMhelix 332 354\ninside 355 372

27900 GCF_000566445.1_ASM56644v1 Shigella flexneri Shi06HN159 Proteobacteria; Gammaproteobacteria; Enterobacterales; Enterobacteriaceae; Shigella; Shigella flexneri MNNEETFYQAMRRQGVTRRSFLKYCSLAATSLGLGAGMAPKIAWALENKPRIPVVWIHGL WP_001058323.1 MULTISPECIES: hydrogenase-1 small chain [Proteobacteria] Length: 372\nNumber of predicted TMHs: 2\nExp number of AAs in TMHs: 37.2608499999999\nExp number, first 60 AAs: 19.52923\nTotal prob of N-in: 0.94351\nPOSSIBLE N-term signal sequence\ninside 1 19\nTMhelix 20 39\noutside 40 325\nTMhelix 326 348\ninside 349 372

27901 GCF_000565765.1_ASM56576v1 Shigella flexneri Shi06HN244 Proteobacteria; Gammaproteobacteria; Enterobacterales; Enterobacteriaceae; Shigella; Shigella flexneri MNNEETFYQAMRRQGVTRRSFLKYCSLAATSLGLGAGMAPKIAWALENKPRIPVVWIHGL WP_001058323.1 MULTISPECIES: hydrogenase-1 small chain [Proteobacteria] Length: 372\nNumber of predicted TMHs: 2\nExp number of AAs in TMHs: 37.2608499999999\nExp number, first 60 AAs: 19.52923\nTotal prob of N-in: 0.94351\nPOSSIBLE N-term signal sequence\ninside 1 19\nTMhelix 20 39\noutside 40 325\nTMhelix 326 348\ninside 349 372

27902 GCF_000565765.1_ASM56576v1 Shigella flexneri Shi06HN244 Proteobacteria; Gammaproteobacteria; Enterobacterales; Enterobacteriaceae; Shigella; Shigella flexneri MTGDNTLIHSHGINRRDFMKLCAALAATMGLSSKAAAEMAESVTNPQRPPVIWIGAQECT WP_000145421.1 MULTISPECIES: hydrogenase 2 small subunit [Shigella] Length: 372\nNumber of predicted TMHs: 1\nExp number of AAs in TMHs: 22.90256\nExp number, first 60 AAs: 0.2236\nTotal prob of N-in: 0.09847\noutside 1 331\nTMhelix 332 354\ninside 355 372

27903 GCF_000565725.1_ASM56572v1 Shigella flexneri Shi06HN250 Proteobacteria; Gammaproteobacteria; Enterobacterales; Enterobacteriaceae; Shigella; Shigella flexneri MNNEETFYQAMRRQGVTRRSFLKYCSLAATSLGLGAGMAPKIAWALENKPRIPVVWIHGL WP_001058323.1 MULTISPECIES: hydrogenase-1 small chain [Proteobacteria] Length: 372\nNumber of predicted TMHs: 2\nExp number of AAs in TMHs: 37.2608499999999\nExp number, first 60 AAs: 19.52923\nTotal prob of N-in: 0.94351\nPOSSIBLE N-term signal sequence\ninside 1 19\nTMhelix 20 39\noutside 40 325\nTMhelix 326 348\ninside 349 372

27904 GCF_000565725.1_ASM56572v1 Shigella flexneri Shi06HN250 Proteobacteria; Gammaproteobacteria; Enterobacterales; Enterobacteriaceae; Shigella; Shigella flexneri MTGDNTLIHSHGINRRDFMKLCAALAATMGLSSKAAAEMAESVTNPQRPPVIWIGAQECT WP_000145421.1 MULTISPECIES: hydrogenase 2 small subunit [Shigella] Length: 372\nNumber of predicted TMHs: 1\nExp

number of AAs in TMHs: 22.90256\nExp number, first 60 AAs: 0.2236\nTotal prob of N-in: 0.09847\noutside 1 331\nTMhelix 332 354\ninside 355 372

27905 GCF_000566465.1_ASM56646v1 Shigella flexneri Shi06HN344 Proteobacteria; Gammaproteobacteria; Enterobacterales; Enterobacteriaceae; Shigella; Shigella flexneri MTGDNTLIHSHGINRRDFMKLCAALAATMGLSSKAAAEMAESVTNPQRPPVIWIGAQECT WP_000145421.1 MULTISPECIES: hydrogenase 2 small subunit [Shigella] Length: 372\nNumber of predicted TMHs: 1\nExp number of AAs in TMHs: 22.90256\nExp number, first 60 AAs: 0.2236\nTotal prob of N-in: 0.09847\noutside 1 331\nTMhelix 332 354\ninside 355 372

27906 GCF_000565685.1_ASM56568v1 Shigella flexneri Shi06HN378 Proteobacteria; Gammaproteobacteria; Enterobacterales; Enterobacteriaceae; Shigella; Shigella flexneri MNNEETFYQAMRRQGVTRRSFLKYCSLAATSLGLGAGMAPKIAWALENKPRIPVVWIHGL WP_001058323.1 MULTISPECIES: hydrogenase-1 small chain [Proteobacteria] Length: 372\nNumber of predicted TMHs: 2\nExp number of AAs in TMHs: 37.2608499999999\nExp number, first 60 AAs: 19.52923\nTotal prob of N-in: 0.94351\nPOSSIBLE N-term signal sequence\ninside 1 19\nTMhelix 20 39\noutside 40 325\nTMhelix 326 348\ninside 349 372

27907 GCF_000565685.1_ASM56568v1 Shigella flexneri Shi06HN378 Proteobacteria; Gammaproteobacteria; Enterobacterales; Enterobacteriaceae; Shigella; Shigella flexneri MTGDNTLIHSHGINRRDFMKLCAALAATMGLSSKAAAEMAESVTNPQRPPVIWIGAQECT WP_000145421.1 MULTISPECIES: hydrogenase 2 small subunit [Shigella] Length: 372\nNumber of predicted TMHs: 1\nExp number of AAs in TMHs: 22.90256\nExp number, first 60 AAs: 0.2236\nTotal prob of N-in: 0.09847\noutside 1 331\nTMhelix 332 354\ninside 355 372

27908 GCF_000565865.1_ASM56586v1 Shigella flexneri Shi06SX36 Proteobacteria; Gammaproteobacteria; Enterobacterales; Enterobacteriaceae; Shigella; Shigella flexneri MNNEETFYQAMRRQGVTRRSFLKYCSLAATSLGLGAGMAPKIAWALENKPRIPVVWIHGL WP_001058323.1 MULTISPECIES: hydrogenase-1 small chain [Proteobacteria] Length: 372\nNumber of predicted TMHs: 2\nExp number of AAs in TMHs: 37.2608499999999\nExp number, first 60 AAs: 19.52923\nTotal prob of N-in: 0.94351\nPOSSIBLE N-term signal sequence\ninside 1 19\nTMhelix 20 39\noutside 40 325\nTMhelix 326 348\ninside 349 372

27909 GCF_000565865.1_ASM56586v1 Shigella flexneri Shi06SX36 Proteobacteria; Gammaproteobacteria; Enterobacterales; Enterobacteriaceae; Shigella; Shigella flexneri MTGDNTLIHSHGINRRDFMKLCAALAATMGLSSKAAAEMAESVTNPQRPPVIWIGAQECT WP_000145421.1 MULTISPECIES: hydrogenase 2 small subunit [Shigella] Length: 372\nNumber of predicted TMHs: 1\nExp number of AAs in TMHs: 22.90256\nExp number, first 60 AAs: 0.2236\nTotal prob of N-in: 0.09847\noutside 1 331\nTMhelix 332 354\ninside 355 372

27910 GCF_000565925.1_ASM56592v1 Shigella flexneri Shi06SX53 Proteobacteria; Gammaproteobacteria; Enterobacterales; Enterobacteriaceae; Shigella; Shigella flexneri MTGDNTLIHSHGINRRDFMKLCAALAATMGLSSKAAAEMAESVTNPQRPPVIWIGAQECT WP_000145421.1 MULTISPECIES: hydrogenase 2 small subunit [Shigella] Length: 372\nNumber of predicted TMHs: 1\nExp number of AAs in TMHs: 22.90256\nExp number, first 60 AAs: 0.2236\nTotal prob of N-in: 0.09847\noutside 1 331\nTMhelix 332 354\ninside 355 372

27911 GCF_000565925.1_ASM56592v1 Shigella flexneri Shi06SX53 Proteobacteria; Gammaproteobacteria; Enterobacterales; Enterobacteriaceae; Shigella; Shigella flexneri MNNEETFYQAMRRQGVTRRSFLKYCSLAATSLGLGAGMAPKIAWALENKPRIPVVWIHGL WP_001058323.1 MULTISPECIES: hydrogenase-1 small chain [Proteobacteria] Length: 372\nNumber of predicted TMHs: 2\nExp number of AAs in TMHs: 37.2608499999999\nExp number, first 60 AAs: 19.52923\nTotal prob of N-in: 0.94351\nPOSSIBLE N-term signal sequence\ninside 1 19\nTMhelix 20 39\noutside 40 325\nTMhelix 326 348\ninside 349 372

27912 GCF_000213695.1_ASM21369v2 Shigella flexneri VA-6 Proteobacteria; Gammaproteobacteria; Enterobacterales; Enterobacteriaceae; Shigella; Shigella flexneri MNNEETFYQAMRRQGVTRRSFLKYCSLAATSLGLGAGMAPKIAWALENKPRIPVVWIHGL WP_001058326.1 hydrogenase [Shigella flexneri] Length: 372\nNumber of predicted TMHs: 2\nExp number of AAs in TMHs: 37.0215599999999\nExp number, first 60 AAs: 19.52998\nTotal prob of N-in: 0.94338\nPOSSIBLE N-term signal sequence\ninside 1 19\nTMhelix 20 39\noutside 40 325\nTMhelix 326 348\ninside 349 372

27913	GCF_000213695.1_ASM21369v2	Shigella flexneri VA-6	Proteobacteria; Gammaproteobacteria; Enterobacterales; Enterobacteriaceae; Shigella; Shigella flexneri
	MTGDNTLIHSHGINRRDFMKLCAALAATMGLSSKAAAEMAESVTNPQRPPVIWIGAQECT	WP_000145413.1
	MULTISPECIES: hydrogenase 2 small subunit [Shigella]	Length: 372\nNumber of predicted TMHs: 1\nExp number of AAs in TMHs: 23.72614\nExp number, first 60 AAs: 0.22038\nTotal prob of N-in: 0.13004\noutside 1 331\nTMhelix 332 354\ninside 355 372

27914	GCF_001017365.1_ASM101736v1	Shigella flexneri Y	Proteobacteria; Gammaproteobacteria; Enterobacterales; Enterobacteriaceae; Shigella; Shigella flexneri
	MNNEETFYQAMRRQGVTRRSFLKYCSLAATSLGLGAGMAPKIAWALENKPRIPVVWIHGL	WP_001058323.1
	MULTISPECIES: hydrogenase-1 small chain [Proteobacteria]	Length: 372\nNumber of predicted TMHs: 2\nExp number of AAs in TMHs: 37.2608499999999\nExp number, first 60 AAs: 19.52923\nTotal prob of N-in: 0.94351\nPOSSIBLE N-term signal sequence\ninside 1 19\nTMhelix 20 39\noutside 40 325\nTMhelix 326 348\ninside 349 372

27915	GCF_001184145.1_ASM118414v1	Shigella flexneri Y	Proteobacteria; Gammaproteobacteria; Enterobacterales; Enterobacteriaceae; Shigella; Shigella flexneri
	MTGDNTLIHSHGINRRDFMKLCAALAATMGLSSKAAAEMAESVTNPQRPPVIWIGAQECT	WP_000145421.1
	MULTISPECIES: hydrogenase 2 small subunit [Shigella]	Length: 372\nNumber of predicted TMHs: 1\nExp number of AAs in TMHs: 22.90256\nExp number, first 60 AAs: 0.2236\nTotal prob of N-in: 0.09847\noutside 1 331\nTMhelix 332 354\ninside 355 372

27916	GCF_001184145.1_ASM118414v1	Shigella flexneri Y	Proteobacteria; Gammaproteobacteria; Enterobacterales; Enterobacteriaceae; Shigella; Shigella flexneri
	MNNEETFYQAMRRQGVTRRSFLKYCSLAATSLGLGAGMAPKIAWALENKPRIPVVWIHGL	WP_001058323.1
	MULTISPECIES: hydrogenase-1 small chain [Proteobacteria]	Length: 372\nNumber of predicted TMHs: 2\nExp number of AAs in TMHs: 37.2608499999999\nExp number, first 60 AAs: 19.52923\nTotal prob of N-in: 0.94351\nPOSSIBLE N-term signal sequence\ninside 1 19\nTMhelix 20 39\noutside 40 325\nTMhelix 326 348\ninside 349 372

27917	GCF_001017365.1_ASM101736v1	Shigella flexneri Y	Proteobacteria; Gammaproteobacteria; Enterobacterales; Enterobacteriaceae; Shigella; Shigella flexneri
	MTGDNTLIHSHGINRRDFMKLCAALAATMGLSSKAAAEMAESVTNPQRPPVIWIGAQECT	WP_000145421.1
	MULTISPECIES: hydrogenase 2 small subunit [Shigella]	Length: 372\nNumber of predicted TMHs: 1\nExp number of AAs in TMHs: 22.90256\nExp number, first 60 AAs: 0.2236\nTotal prob of N-in: 0.09847\noutside 1 331\nTMhelix 332 354\ninside 355 372

27918	GCF_000183785.1_ASM18378v2	Shigella flexneri 2a str. 2457T	Proteobacteria; Gammaproteobacteria; Enterobacterales; Enterobacteriaceae; Shigella; Shigella flexneri; Shigella flexneri 2a
	MTGDNTLIHSHGINRRDFMKLCAALAATMGLSSKAAAEMAESVTNPQRPPVIWIGAQECT	WP_000145421.1
	MULTISPECIES: hydrogenase 2 small subunit [Shigella]	Length: 372\nNumber of predicted TMHs: 1\nExp number of AAs in TMHs: 22.90256\nExp number, first 60 AAs: 0.2236\nTotal prob of N-in: 0.09847\noutside 1 331\nTMhelix 332 354\ninside 355 372

27919	GCF_000183785.1_ASM18378v2	Shigella flexneri 2a str. 2457T	Proteobacteria; Gammaproteobacteria; Enterobacterales; Enterobacteriaceae; Shigella; Shigella flexneri; Shigella flexneri 2a
	MNNEETFYQAMRRQGVTRRSFLKYCSLAATSLGLGAGMAPKIAWALENKPRIPVVWIHGL	WP_001058323.1
	MULTISPECIES: hydrogenase-1 small chain [Proteobacteria]	Length: 372\nNumber of predicted TMHs: 2\nExp number of AAs in TMHs: 37.2608499999999\nExp number, first 60 AAs: 19.52923\nTotal prob of N-in: 0.94351\nPOSSIBLE N-term signal sequence\ninside 1 19\nTMhelix 20 39\noutside 40 325\nTMhelix 326 348\ninside 349 372

27920	GCF_000007405.1_ASM740v1	Shigella flexneri 2a str. 2457T	Proteobacteria; Gammaproteobacteria; Enterobacterales; Enterobacteriaceae; Shigella; Shigella flexneri; Shigella flexneri 2a
	MNNEETFYQAMRRQGVTRRSFLKYCSLAATSLGLGAGMAPKIAWALENKPRIPVVWIHGL	WP_001058323.1
	MULTISPECIES: hydrogenase-1 small chain [Proteobacteria]	Length: 372\nNumber of predicted TMHs: 2\nExp number of AAs in TMHs: 37.2608499999999\nExp number, first 60 AAs: 19.52923\nTotal prob of N-in: 0.94351\nPOSSIBLE N-term signal sequence\ninside 1 19\nTMhelix 20 39\noutside 40 325\nTMhelix 326 348\ninside 349 372

27921 GCF_000007405.1_ASM740v1 Shigella flexneri 2a str. 2457T Proteobacteria; Gammaproteobacteria; Enterobacterales; Enterobacteriaceae; Shigella; Shigella flexneri; Shigella flexneri 2a
MTGDNTLIHSHGINRRDFMKLCAALAATMGLSSKAAAEMAESVTNPQRPPVIWIGAQECT WP_000145421.1
MULTISPECIES: hydrogenase 2 small subunit [Shigella] Length: 372\nNumber of predicted TMHs: 1\nExp number of AAs in TMHs: 22.90256\nExp number, first 60 AAs: 0.2236\nTotal prob of N-in: 0.09847\noutside 1 331\nTMhelix 332 354\ninside 355 372

27922 GCF_000006925.2_ASM692v2 Shigella flexneri 2a str. 301 Proteobacteria; Gammaproteobacteria; Enterobacterales; Enterobacteriaceae; Shigella; Shigella flexneri; Shigella flexneri 2a
MTGDNTLIHSHGINRRDFMKLCAALAATMGLSSKAAAEMAESVTNPQRPPVIWIGAQECT NP_708815.1
hydrogenase 2 small subunit [Shigella flexneri 2a str. 301] Length: 372\nNumber of predicted TMHs: 1\nExp number of AAs in TMHs: 22.90256\nExp number, first 60 AAs: 0.2236\nTotal prob of N-in: 0.09847\noutside 1 331\nTMhelix 332 354\ninside 355 372

27923 GCF_000013585.1_ASM1358v1 Shigella flexneri 5 str. 8401 Proteobacteria; Gammaproteobacteria; Enterobacterales; Enterobacteriaceae; Shigella; Shigella flexneri; Shigella flexneri 5
MNNEETFYQAMRRQGVTRRSFLKYCSLAATSLGLGAGMAPKIAWALENKPRIPVVWSHGL WP_001058327.1
hydrogenase [Shigella flexneri] Length: 372\nNumber of predicted TMHs: 2\nExp number of AAs in TMHs: 37.2451399999999\nExp number, first 60 AAs: 19.52414\nTotal prob of N-in: 0.94348\nPOSSIBLE N-term signal sequence\ninside 1 19\nTMhelix 20 39\noutside 40 325\nTMhelix 326 348\ninside 349 372

27924 GCF_000013585.1_ASM1358v1 Shigella flexneri 5 str. 8401 Proteobacteria; Gammaproteobacteria; Enterobacterales; Enterobacteriaceae; Shigella; Shigella flexneri; Shigella flexneri 5
MTGDNTLIHSHGINRRDFMKLCAALAATMGLSSKAAAEMAESVTNPQRPPVIWIGAQECT WP_000145413.1
MULTISPECIES: hydrogenase 2 small subunit [Shigella] Length: 372\nNumber of predicted TMHs: 1\nExp number of AAs in TMHs: 23.72614\nExp number, first 60 AAs: 0.22038\nTotal prob of N-in: 0.13004\noutside 1 331\nTMhelix 332 354\ninside 355 372

27925 GCF_000252895.1_ASM25289v1 Shigella flexneri 5a str. M90T Proteobacteria; Gammaproteobacteria; Enterobacterales; Enterobacteriaceae; Shigella; Shigella flexneri; Shigella flexneri 5a
MTGDNTLIHSHGINRRDFMKLCAALAATMGLSSKAAAEMAESVTNPQRPPVIWIGAQECT WP_000145413.1
MULTISPECIES: hydrogenase 2 small subunit [Shigella] Length: 372\nNumber of predicted TMHs: 1\nExp number of AAs in TMHs: 23.72614\nExp number, first 60 AAs: 0.22038\nTotal prob of N-in: 0.13004\noutside 1 331\nTMhelix 332 354\ninside 355 372

27926 GCF_000252895.1_ASM25289v1 Shigella flexneri 5a str. M90T Proteobacteria; Gammaproteobacteria; Enterobacterales; Enterobacteriaceae; Shigella; Shigella flexneri; Shigella flexneri 5a
MNNEETFYQAMRRQGVTRRSFLKYCSLAATSLGLGAGMAPKIAWALENKPRIPVVWSHGL WP_001058327.1
hydrogenase [Shigella flexneri] Length: 372\nNumber of predicted TMHs: 2\nExp number of AAs in TMHs: 37.2451399999999\nExp number, first 60 AAs: 19.52414\nTotal prob of N-in: 0.94348\nPOSSIBLE N-term signal sequence\ninside 1 19\nTMhelix 20 39\noutside 40 325\nTMhelix 326 348\ninside 349 372

27927 GCF_000467205.1_Shigella_sonnei_strain_1DT-1Shigella sonnei 1DT-1 Proteobacteria; Gammaproteobacteria; Enterobacterales; Enterobacteriaceae; Shigella; Shigella sonnei
MNNEETFYQAMRRQGVTRRSFLKYCSLAATSLGLGAGMAPKIAWALENKPRIPVVWIHGL WP_001058323.1
MULTISPECIES: hydrogenase-1 small chain [Proteobacteria] Length: 372\nNumber of predicted TMHs: 2\nExp number of AAs in TMHs: 37.2608499999999\nExp number, first 60 AAs: 19.52923\nTotal prob of N-in: 0.94351\nPOSSIBLE N-term signal sequence\ninside 1 19\nTMhelix 20 39\noutside 40 325\nTMhelix 326 348\ninside 349 372

27928 GCF_000467205.1_Shigella_sonnei_strain_1DT-1Shigella sonnei 1DT-1 Proteobacteria; Gammaproteobacteria; Enterobacterales; Enterobacteriaceae; Shigella; Shigella sonnei
MTGDNTLIHSHGINRRDFMKLCAALAATMGLSSKAAAEMAESVTNPQRPPVIWIGAQECT WP_000145410.1
MULTISPECIES: hydrogenase-2 small chain [Proteobacteria] Length: 372\nNumber of predicted TMHs: 1\nExp number of AAs in TMHs: 23.72226\nExp number, first 60 AAs: 0.22038\nTotal prob of N-in: 0.12994\noutside 1 331\nTMhelix 332 354\ninside 355 372

27929 GCF_000268005.1_ASM26800v1 Shigella sonnei 3233-85 Proteobacteria; Gammaproteobacteria; Enterobacterales; Enterobacteriaceae; Shigella; Shigella sonnei
MNNEETFYQAMRRQGVTRRSFLKYCSLAATSLGLGAGMAPKIAWALENKPRIPVVWIHGL WP_005139779.1
hydrogenase [Shigella sonnei] Length: 372\nNumber of predicted TMHs: 2\nExp number of AAs in

TMHs: 37.02628\nExp number, first 60 AAs: 19.39684\nTotal prob of N-in: 0.94313\nPOSSIBLE N-term signal sequence\ninside 1 19\nTMhelix 20 39\noutside 40 325\nTMhelix 326 348\ninside 349 372

27930 GCF_000268005.1_ASM26800v1 Shigella sonnei 3233-85 Proteobacteria; Gammaproteobacteria; Enterobacterales; Enterobacteriaceae; Shigella; Shigella sonnei
MTGDNTLIHSHGINRRDFMKLCAALAATMGLSSKAAAEMAESVTNPQRPPVIWIGAQECT WP_000145410.1
MULTISPECIES: hydrogenase-2 small chain [Proteobacteria] Length: 372\nNumber of predicted TMHs: 1\nExp number of AAs in TMHs: 23.72226\nExp number, first 60 AAs: 0.22038\nTotal prob of N-in: 0.12994\noutside 1 331\nTMhelix 332 354\ninside 355 372

27931 GCF_000268225.1_ASM26822v1 Shigella sonnei 4822-66 Proteobacteria; Gammaproteobacteria; Enterobacterales; Enterobacteriaceae; Shigella; Shigella sonnei
MTGDNTLIHSHGINRRDFMKLCAALAATMGLSSKAAAEMAESVTNPQRPPVIWIGAQECT WP_000145410.1
MULTISPECIES: hydrogenase-2 small chain [Proteobacteria] Length: 372\nNumber of predicted TMHs: 1\nExp number of AAs in TMHs: 23.72226\nExp number, first 60 AAs: 0.22038\nTotal prob of N-in: 0.12994\noutside 1 331\nTMhelix 332 354\ninside 355 372

27932 GCF_000268225.1_ASM26822v1 Shigella sonnei 4822-66 Proteobacteria; Gammaproteobacteria; Enterobacterales; Enterobacteriaceae; Shigella; Shigella sonnei
MNNEETFYQAMRRQGVTRRSFLKYCSLAATSLGLGAGMAPKIAWALENKPRIPVVWIHGL WP_005139779.1
hydrogenase [Shigella sonnei] Length: 372\nNumber of predicted TMHs: 2\nExp number of AAs in TMHs: 37.02628\nExp number, first 60 AAs: 19.39684\nTotal prob of N-in: 0.94313\nPOSSIBLE N-term signal sequence\ninside 1 19\nTMhelix 20 39\noutside 40 325\nTMhelix 326 348\ninside 349 372

27933 GCF_000283715.1_ASM28371v1 Shigella sonnei 53G Proteobacteria; Gammaproteobacteria; Enterobacterales; Enterobacteriaceae; Shigella; Shigella sonnei
MNNEETFYQAMRRQGVTRRSFLKYCSLAATSLGLGAGMAPKIAWALENKPRIPVVWIHGL WP_005139779.1
hydrogenase [Shigella sonnei] Length: 372\nNumber of predicted TMHs: 2\nExp number of AAs in TMHs: 37.02628\nExp number, first 60 AAs: 19.39684\nTotal prob of N-in: 0.94313\nPOSSIBLE N-term signal sequence\ninside 1 19\nTMhelix 20 39\noutside 40 325\nTMhelix 326 348\ninside 349 372

27934 GCF_000283715.1_ASM28371v1 Shigella sonnei 53G Proteobacteria; Gammaproteobacteria; Enterobacterales; Enterobacteriaceae; Shigella; Shigella sonnei
MTGDNTLIHSHGINRRDFMKLCAALAATMGLSSKAAAEMAESVTNPQRPPVIWIGAQECT WP_000145410.1
MULTISPECIES: hydrogenase-2 small chain [Proteobacteria] Length: 372\nNumber of predicted TMHs: 1\nExp number of AAs in TMHs: 23.72226\nExp number, first 60 AAs: 0.22038\nTotal prob of N-in: 0.12994\noutside 1 331\nTMhelix 332 354\ninside 355 372

27935 GCF_000815505.1_ASM81550v1 Shigella sonnei S6513 Proteobacteria; Gammaproteobacteria; Enterobacterales; Enterobacteriaceae; Shigella; Shigella sonnei
MNNEETFYQAMRRQGVTRRSFLKYCSLAATSLGLGAGMAPKIAWALENKPRIPVVWIHGL WP_005139779.1
hydrogenase [Shigella sonnei] Length: 372\nNumber of predicted TMHs: 2\nExp number of AAs in TMHs: 37.02628\nExp number, first 60 AAs: 19.39684\nTotal prob of N-in: 0.94313\nPOSSIBLE N-term signal sequence\ninside 1 19\nTMhelix 20 39\noutside 40 325\nTMhelix 326 348\ninside 349 372

27936 GCF_000815505.1_ASM81550v1 Shigella sonnei S6513 Proteobacteria; Gammaproteobacteria; Enterobacterales; Enterobacteriaceae; Shigella; Shigella sonnei
MTGDNTLIHSHGINRRDFMKLCAALAATMGLSSKAAAEMAESVTNPQRPPVIWIGAQECT WP_000145410.1
MULTISPECIES: hydrogenase-2 small chain [Proteobacteria] Length: 372\nNumber of predicted TMHs: 1\nExp number of AAs in TMHs: 23.72226\nExp number, first 60 AAs: 0.22038\nTotal prob of N-in: 0.12994\noutside 1 331\nTMhelix 332 354\ninside 355 372

27937 GCF_000092525.1_ASM9252v1 Shigella sonnei Ss046 Proteobacteria; Gammaproteobacteria; Enterobacterales; Enterobacteriaceae; Shigella; Shigella sonnei
MTGDNTLIHSHGINRRDFMKLCAALAATMGLSSKAAAEMAESVTNPQRPPVIWIGAQECT WP_000145410.1
MULTISPECIES: hydrogenase-2 small chain [Proteobacteria] Length: 372\nNumber of predicted TMHs: 1\nExp number of AAs in TMHs: 23.72226\nExp number, first 60 AAs: 0.22038\nTotal prob of N-in: 0.12994\noutside 1 331\nTMhelix 332 354\ninside 355 372

27938 GCF_000092525.1_ASM9252v1 Shigella sonnei Ss046 Proteobacteria; Gammaproteobacteria; Enterobacterales; Enterobacteriaceae; Shigella; Shigella sonnei

MNNEETFYQAMRRQGVTRRSFLKYCSLAATSLGLGAGMAPKIAWALENKPRIPVVWIHGL WP_005139779.1 hydrogenase [Shigella sonnei] Length: 372\nNumber of predicted TMHs: 2\nExp number of AAs in TMHs: 37.02628\nExp number, first 60 AAs: 19.39684\nTotal prob of N-in: 0.94313\nPOSSIBLE N-term signal sequence\ninside 1 19\nTMhelix 20 39\noutside 40 325\nTMhelix 326 348\ninside 349 372

27939 GCF_000281815.1_ASM28181v1 Shigella sonnei str. Moseley Proteobacteria; Gammaproteobacteria; Enterobacterales; Enterobacteriaceae; Shigella; Shigella sonnei MNNEETFYQAMRRQGVTRRSFLKYCSLAATSLGLGAGMAPKIAWALENKPRIPVVWIHGL WP_005139779.1 hydrogenase [Shigella sonnei] Length: 372\nNumber of predicted TMHs: 2\nExp number of AAs in TMHs: 37.02628\nExp number, first 60 AAs: 19.39684\nTotal prob of N-in: 0.94313\nPOSSIBLE N-term signal sequence\ninside 1 19\nTMhelix 20 39\noutside 40 325\nTMhelix 326 348\ninside 349 372

27940 GCF_000281815.1_ASM28181v1 Shigella sonnei str. Moseley Proteobacteria; Gammaproteobacteria; Enterobacterales; Enterobacteriaceae; Shigella; Shigella sonnei MTGDNTLIHSHGINRRDFMKLCAALAATMGLSSKAAAEMAESVTNPQRPPVIWIGAQECT WP_000145410.1 MULTISPECIES: hydrogenase-2 small chain [Proteobacteria] Length: 372\nNumber of predicted TMHs: 1\nExp number of AAs in TMHs: 23.72226\nExp number, first 60 AAs: 0.22038\nTotal prob of N-in: 0.12994\noutside 1 331\nTMhelix 332 354\ninside 355 372

27941 GCF_000211975.1_ASM21197v2 Shigella boydii 5216-82 Proteobacteria; Gammaproteobacteria; Enterobacterales; Enterobacteriaceae; Shigella; Shigella sp. SF-2015 MTGDNTLIHSHGINRRDFMKLCAALAATMGLSSKAAAEMAESVTNPQRPPVIWIGAQECT WP_000145410.1 MULTISPECIES: hydrogenase-2 small chain [Proteobacteria] Length: 372\nNumber of predicted TMHs: 1\nExp number of AAs in TMHs: 23.72226\nExp number, first 60 AAs: 0.22038\nTotal prob of N-in: 0.12994\noutside 1 331\nTMhelix 332 354\ninside 355 372

27942 GCF_000193915.1_ASM19391v2 Shigella boydii ATCC 9905 Proteobacteria; Gammaproteobacteria; Enterobacterales; Enterobacteriaceae; Shigella; Shigella sp. SF-2015 MNNEETFYQAMRRQGVTRRSFLKYCSLAATSLGLGAGMAPKIAWALENKPRIPVVWIHGL WP_024258461.1 hydrogenase [Shigella sp. SF-2015] Length: 372\nNumber of predicted TMHs: 2\nExp number of AAs in TMHs: 37.42235\nExp number, first 60 AAs: 19.53031\nTotal prob of N-in: 0.94351\nPOSSIBLE N-term signal sequence\ninside 1 19\nTMhelix 20 39\noutside 40 325\nTMhelix 326 348\ninside 349 372

27943 GCF_000193915.1_ASM19391v2 Shigella boydii ATCC 9905 Proteobacteria; Gammaproteobacteria; Enterobacterales; Enterobacteriaceae; Shigella; Shigella sp. SF-2015 MAGDNTLIHSHGINRRDFMKLCAALAATMGLSSKAAAEMAESVTNPQRPPVIWIGAQECT WP_000896631.1 hydrogenase-2 small chain [Shigella sp. SF-2015] Length: 372\nNumber of predicted TMHs: 1\nExp number of AAs in TMHs: 23.29146\nExp number, first 60 AAs: 0.17789\nTotal prob of N-in: 0.10465\noutside 1 331\nTMhelix 332 354\ninside 355 372

27944 GCF_000168075.1_ASM16807v1 Shigella dysenteriae 1012 Proteobacteria; Gammaproteobacteria; Enterobacterales; Enterobacteriaceae; Shigella; Shigella sp. SF-2015 MNNEETFYQAMRRQGVTRRSFLKYCSLAATSLGLGAGMAPKIAWALENKPRIPVVWIHGL WP_004999783.1 MULTISPECIES: hydrogenase [Enterobacteriaceae] Length: 372\nNumber of predicted TMHs: 2\nExp number of AAs in TMHs: 37.2630799999999\nExp number, first 60 AAs: 19.52922\nTotal prob of N-in: 0.94351\nPOSSIBLE N-term signal sequence\ninside 1 19\nTMhelix 20 39\noutside 40 325\nTMhelix 326 348\ninside 349 372

27945 GCF_000168075.1_ASM16807v1 Shigella dysenteriae 1012 Proteobacteria; Gammaproteobacteria; Enterobacterales; Enterobacteriaceae; Shigella; Shigella sp. SF-2015 MTGDNTLIHSHGINRRDFMKLCAALAATMGLSSKAAAEMAESVTNPQRPPVIWIGAQECT WP_000145410.1 MULTISPECIES: hydrogenase-2 small chain [Proteobacteria] Length: 372\nNumber of predicted TMHs: 1\nExp number of AAs in TMHs: 23.72226\nExp number, first 60 AAs: 0.22038\nTotal prob of N-in: 0.12994\noutside 1 331\nTMhelix 332 354\ninside 355 372

27946 GCF_000815495.1_ASM81549v1 Shigella dysenteriae S6205 Proteobacteria; Gammaproteobacteria; Enterobacterales; Enterobacteriaceae; Shigella; Shigella sp. SF-2015 MNNEETFYQAMRRQGVTRRSFLKYCSLAATSLGLGAGMAPKIAWALENKPRIPVVWIHGL WP_004999783.1 MULTISPECIES: hydrogenase [Enterobacteriaceae] Length: 372\nNumber of predicted TMHs: 2\nExp number of AAs in TMHs: 37.2630799999999\nExp number, first 60 AAs: 19.52922\nTotal prob of N-in:

0.94351\nPOSSIBLE N-term signal sequence\ninside 1 19\nTMhelix 20 39\noutside 40 325\nTMhelix 326 348\ninside 349 372

27947 GCF_000815495.1_ASM81549v1 Shigella dysenteriae S6205 Proteobacteria; Gammaproteobacteria; Enterobacterales; Enterobacteriaceae; Shigella; Shigella sp. SF-2015 MTGDNTLIHSHGINRRDFMKLCAALAATMGLSSKAAAEMAESVTNPQRPPVIWIGAQECT WP_000145410.1 MULTISPECIES: hydrogenase-2 small chain [Proteobacteria] Length: 372\nNumber of predicted TMHs: 1\nExp number of AAs in TMHs: 23.72226\nExp number, first 60 AAs: 0.22038\nTotal prob of N-in: 0.12994\noutside 1 331\nTMhelix 332 354\ninside 355 372

27948 GCF_001188495.1_ASM118849v1 Trabulsiella odontotermitis Proteobacteria; Gammaproteobacteria; Enterobacterales; Enterobacteriaceae; Trabulsiella MEGVNALISSHGVSRRDFMKLCTALAATMGLSSEAAAAIADSVSSPQRPPVIWIGAQECT WP_049850194.1 hydrogenase 2 small subunit [Trabulsiella odontotermitis] Length: 372\nNumber of predicted TMHs: 1\nExp number of AAs in TMHs: 20.85045\nExp number, first 60 AAs: 0.54192\nTotal prob of N-in: 0.04835\noutside 1 331\nTMhelix 332 354\ninside 355 372

27949 GCF_001297785.1_ASM129778v1 Trabulsiella odontotermitis Proteobacteria; Gammaproteobacteria; Enterobacterales; Enterobacteriaceae; Trabulsiella MEGVNALISSHGVSRRDFMKLCTALAATMGLSSEAAAAIAESVSSPQRPPVVWIGAQECT WP_054178753.1 hydrogenase 2 small subunit [Trabulsiella odontotermitis] Length: 372\nNumber of predicted TMHs: 1\nExp number of AAs in TMHs: 20.13817\nExp number, first 60 AAs: 0.4614\nTotal prob of N-in: 0.05744\noutside 1 331\nTMhelix 332 354\ninside 355 372

27950 GCF_001297765.1_ASM129776v1 Trabulsiella odontotermitis Proteobacteria; Gammaproteobacteria; Enterobacterales; Enterobacteriaceae; Trabulsiella MEGVNALISSHGVSRRDFMKLCTALAATMGLSSEAAAAIAESVSSPQRPPVVWIGAQECT WP_054178753.1 hydrogenase 2 small subunit [Trabulsiella odontotermitis] Length: 372\nNumber of predicted TMHs: 1\nExp number of AAs in TMHs: 20.13817\nExp number, first 60 AAs: 0.4614\nTotal prob of N-in: 0.05744\noutside 1 331\nTMhelix 332 354\ninside 355 372

27951 GCF_001188485.1_ASM118848v1 Trabulsiella odontotermitis Proteobacteria; Gammaproteobacteria; Enterobacterales; Enterobacteriaceae; Trabulsiella MEGVNALISSHGVSRRDFMKLCTALAATMGLSSEAAAAIADSVSSPQRPPVIWIGAQECT WP_049850194.1 hydrogenase 2 small subunit [Trabulsiella odontotermitis] Length: 372\nNumber of predicted TMHs: 1\nExp number of AAs in TMHs: 20.85045\nExp number, first 60 AAs: 0.54192\nTotal prob of N-in: 0.04835\noutside 1 331\nTMhelix 332 354\ninside 355 372

27952 GCF_001297845.1_ASM129784v1 Trabulsiella odontotermitis Proteobacteria; Gammaproteobacteria; Enterobacterales; Enterobacteriaceae; Trabulsiella MEGVNALISSHGVSRRDFMKLCTALAATMGLSSEAAAAIAESVSSPQRPPVVWIGAQECT WP_054178753.1 hydrogenase 2 small subunit [Trabulsiella odontotermitis] Length: 372\nNumber of predicted TMHs: 1\nExp number of AAs in TMHs: 20.13817\nExp number, first 60 AAs: 0.4614\nTotal prob of N-in: 0.05744\noutside 1 331\nTMhelix 332 354\ninside 355 372

27953 GCF_001297775.1_ASM129777v1 Trabulsiella odontotermitis Proteobacteria; Gammaproteobacteria; Enterobacterales; Enterobacteriaceae; Trabulsiella MEGVNALISSHGVSRRDFMKLCTALAATMGLSSEAAAAIAESVSSPQRPPVVWIGAQECT WP_054178753.1 hydrogenase 2 small subunit [Trabulsiella odontotermitis] Length: 372\nNumber of predicted TMHs: 1\nExp number of AAs in TMHs: 20.13817\nExp number, first 60 AAs: 0.4614\nTotal prob of N-in: 0.05744\noutside 1 331\nTMhelix 332 354\ninside 355 372

27954 GCF_001297855.1_ASM129785v1 Trabulsiella odontotermitis Proteobacteria; Gammaproteobacteria; Enterobacterales; Enterobacteriaceae; Trabulsiella MEGVNALISSHGVSRRDFMKLCTALAATMGLSSEAAAAIAESVSSPQRPPVVWIGAQECT WP_054178753.1 hydrogenase 2 small subunit [Trabulsiella odontotermitis] Length: 372\nNumber of predicted TMHs: 1\nExp number of AAs in TMHs: 20.13817\nExp number, first 60 AAs: 0.4614\nTotal prob of N-in: 0.05744\noutside 1 331\nTMhelix 332 354\ninside 355 372

27955 GCF_000734965.1_ASM73496v1 Trabulsiella guamensis ATCC 49490 Proteobacteria; Gammaproteobacteria; Enterobacterales; Enterobacteriaceae; Trabulsiella; Trabulsiella guamensis

MEGVNALITSHGVSRRDFMKLCTALAATMGLSSEAAAAIAESVSSPQRPPVIWIGAQECT WP_038154568.1 hydrogenase 2 small subunit [Trabulsiella guamensis] Length: 372\nNumber of predicted TMHs: 1\nExp number of AAs in TMHs: 21.25787\nExp number, first 60 AAs: 0.7061\nTotal prob of N-in: 0.12035\noutside 1 331\nTMhelix 332 354\ninside 355 372

27956 GCF_000330845.1_ASM33084v1 Enterobacteriaceae bacterium strain FGI 57 Proteobacteria; Gammaproteobacteria; Enterobacterales; Enterobacteriaceae; unclassified Enterobacteriaceae; unclassified Enterobacteriaceae (miscellaneous) MTGDNTLSTSHGVNRRDFMKLCAALAATMGLSSKAAAEMAESITRPQRPPVIWIGAQECT WP_015963621.1 hydrogenase (NiFe) small subunit HydA [Enterobacteriaceae bacterium strain FGI 57] Length: 372\nNumber of predicted TMHs: 1\nExp number of AAs in TMHs: 27.69452\nExp number, first 60 AAs: 0.41067\nTotal prob of N-in: 0.28488\noutside 1 331\nTMhelix 332 354\ninside 355 372

27957 GCF_000239335.1_ASM23933v1 Yokenella regensburgei ATCC 43003 Proteobacteria; Gammaproteobacteria; Enterobacterales; Enterobacteriaceae; Yokenella; Yokenella regensburgei MDNQSIFPQGVNRRDFLKLCTALAATMGLTPGAAAEMANAVTSPQRPPVIWIGAQECTGC WP_006819760.1 hydrogenase 2 small subunit [Yokenella regensburgei] Length: 375\nNumber of predicted TMHs: 1\nExp number of AAs in TMHs: 19.82817\nExp number, first 60 AAs: 0.89054\nTotal prob of N-in: 0.05616\noutside 1 329\nTMhelix 330 352\ninside 353 375

27958 GCF_001484765.1_ASM148476v1 Erwinia teleogrylli Proteobacteria; Gammaproteobacteria; Enterobacterales; Erwiniaceae; Erwinia MSEGPNTFLNHGISRRDFMTLCAALSASMGLSSTASAQLTEAMASPARPPVIWIGAQECT WP_058912695.1 hydrogenase 2 small subunit [Erwinia teleogrylli] Length: 376\nNumber of predicted TMHs: 1\nExp number of AAs in TMHs: 19.66396\nExp number, first 60 AAs: 0.33887\nTotal prob of N-in: 0.04862\noutside 1 332\nTMhelix 333 355\ninside 356 376

27959 GCF_002163605.1_ASM216360v1 Tatumella citrea Proteobacteria; Gammaproteobacteria; Enterobacterales; Erwiniaceae; Tatumella MIDMLNMISRRRILQGMGALAATTLLPSGILPAFADTPANSDFNDISRLLTGRNTLSAEF WP_087488285.1 hypothetical protein [Tatumella citrea] Length: 167\nNumber of predicted TMHs: 2\nExp number of AAs in TMHs: 40.36885\nExp number, first 60 AAs: 22.17399\nTotal prob of N-in: 0.97263\nPOSSIBLE N-term signal sequence\ninside 1 12\nTMhelix 13 35\noutside 36 105\nTMhelix 106 128\ninside 129 167

27960 GCF_002163585.1_ASM216358v1 Tatumella citrea Proteobacteria; Gammaproteobacteria; Enterobacterales; Erwiniaceae; Tatumella MIDMLNMISRRRILQGMGALAATTLLPSGILPAFADTPANSDFNDISRLLTGRNTLSAEF WP_087488285.1 hypothetical protein [Tatumella citrea] Length: 167\nNumber of predicted TMHs: 2\nExp number of AAs in TMHs: 40.36885\nExp number, first 60 AAs: 22.17399\nTotal prob of N-in: 0.97263\nPOSSIBLE N-term signal sequence\ninside 1 12\nTMhelix 13 35\noutside 36 105\nTMhelix 106 128\ninside 129 167

27961 GCF_000757425.2_Tmorbiroseiassem5 Tatumella morbirosei Proteobacteria; Gammaproteobacteria; Enterobacterales; Erwiniaceae; Tatumella MIDMLNMISRRRILQGMGALAATTLLPSGILPAFADTPANSDFNDISRLLTGRTTLSPEF WP_038020607.1 hypothetical protein [Tatumella morbirosei] Length: 167\nNumber of predicted TMHs: 2\nExp number of AAs in TMHs: 38.857\nExp number, first 60 AAs: 21.9113\nTotal prob of N-in: 0.96012\nPOSSIBLE N-term signal sequence\ninside 1 12\nTMhelix 13 35\noutside 36 105\nTMhelix 106 128\ninside 129 167

27962 GCF_001761405.1_ASM176140v1 Edwardsiella hoshinae Proteobacteria; Gammaproteobacteria; Enterobacterales; Hafniaceae; Edwardsiella MNNEDTFYQAMRRQGVTRRSFLKYCSLAATSLGLGAGMAPKIAWALENKPRIPVVWIHGL WP_024524425.1 hydrogenase [Edwardsiella hoshinae] Length: 372\nNumber of predicted TMHs: 2\nExp number of AAs in TMHs: 39.01093\nExp number, first 60 AAs: 19.72184\nTotal prob of N-in: 0.95251\nPOSSIBLE N-term signal sequence\ninside 1 19\nTMhelix 20 39\noutside 40 325\nTMhelix 326 348\ninside 349 372

27963 GCF_001761405.1_ASM176140v1 Edwardsiella hoshinae Proteobacteria; Gammaproteobacteria; Enterobacterales; Hafniaceae; Edwardsiella MVEDSFLSAHGINRRDFMKLCAGMAATLGLSSNAAAEMAAAITSPQRPPVIWIGAQECTG WP_024522509.1 hydrogenase 2 small subunit [Edwardsiella hoshinae] Length: 374\nNumber of predicted TMHs: 1\nExp number of AAs in TMHs: 21.62924\nExp number, first 60 AAs: 2.31877\nTotal prob of N-in: 0.10501\noutside 1 331\nTMhelix 332 354\ninside 355 374

27964	GCF_002076875.1_ASM207687v1	Edwardsiella ictaluri	Proteobacteria; Gammaproteobacteria; Enterobacterales; Hafniaceae; Edwardsiella
MVEDSFLSAHGINRRDFMKLCAGMAATLGLSSNAAAEMAAAITSPQRPPVIWIGAQECTG	WP_049640271.1
hydrogenase 2 small subunit [Edwardsiella ictaluri]	Length: 374\nNumber of predicted TMHs: 1\nExp number of AAs in TMHs: 20.14577\nExp number, first 60 AAs: 2.31859\nTotal prob of N-in: 0.10532\noutside 1 331\nTMhelix 332 354\ninside 355 374

27965	GCF_002076875.1_ASM207687v1	Edwardsiella ictaluri	Proteobacteria; Gammaproteobacteria; Enterobacterales; Hafniaceae; Edwardsiella
MNNEDTFYQAMRRQGVTRRSFLKYCSLAATSLGLGAGMAPKIAWALENKPRIPVVWIHGL	WP_015872619.1
hydrogenase [Edwardsiella ictaluri]	Length: 373\nNumber of predicted TMHs: 2\nExp number of AAs in TMHs: 38.57269\nExp number, first 60 AAs: 19.72695\nTotal prob of N-in: 0.95270\nPOSSIBLE N-term signal sequence\ninside 1 19\nTMhelix 20 39\noutside 40 325\nTMhelix 326 348\ninside 349 373

27966	GCF_002075835.1_ASM207583v1	Edwardsiella piscicida	Proteobacteria; Gammaproteobacteria; Enterobacterales; Hafniaceae; Edwardsiella
MNNEDTFYQAMRRQGVTRRSFLKYCSLAATSLGLGAGMAPKIAWALENKPRIPVVWIHGL	WP_015462312.1
MULTISPECIES: hydrogenase [Edwardsiella]	Length: 373\nNumber of predicted TMHs: 2\nExp number of AAs in TMHs: 38.3616\nExp number, first 60 AAs: 19.72644\nTotal prob of N-in: 0.95270\nPOSSIBLE N-term signal sequence\ninside 1 19\nTMhelix 20 39\noutside 40 325\nTMhelix 326 348\ninside 349 373

27967	GCF_001896205.1_ASM189620v1	Edwardsiella piscicida	Proteobacteria; Gammaproteobacteria; Enterobacterales; Hafniaceae; Edwardsiella
MNNEDTFYQAMRRQGVTRRSFLKYCSLAATSLGLGAGMAPKIAWALENKPRIPVVWIHGL	WP_015462312.1
MULTISPECIES: hydrogenase [Edwardsiella]	Length: 373\nNumber of predicted TMHs: 2\nExp number of AAs in TMHs: 38.3616\nExp number, first 60 AAs: 19.72644\nTotal prob of N-in: 0.95270\nPOSSIBLE N-term signal sequence\ninside 1 19\nTMhelix 20 39\noutside 40 325\nTMhelix 326 348\ninside 349 373

27968	GCF_000711155.1_ASM71115v1	Edwardsiella piscicida	Proteobacteria; Gammaproteobacteria; Enterobacterales; Hafniaceae; Edwardsiella
MVEDSFLSAHGINRRDFMKLCAGMAATLGLSSNAAAEMAAAITSPQRPPVIWIGAQECTG	WP_012849197.1
MULTISPECIES: hydrogenase 2 small subunit [Edwardsiella]Length: 374\nNumber of predicted TMHs: 1\nExp number of AAs in TMHs: 21.63218\nExp number, first 60 AAs: 2.31899\nTotal prob of N-in: 0.10509\noutside 1 331\nTMhelix 332 354\ninside 355 374

27969	GCF_000711155.1_ASM71115v1	Edwardsiella piscicida	Proteobacteria; Gammaproteobacteria; Enterobacterales; Hafniaceae; Edwardsiella
MNNEDTFYQAMRRQGVTRRSFLKYCSLAATSLGLGAGMAPKIAWALENKPRIPVVWIHGL	WP_015462312.1
MULTISPECIES: hydrogenase [Edwardsiella]	Length: 373\nNumber of predicted TMHs: 2\nExp number of AAs in TMHs: 38.3616\nExp number, first 60 AAs: 19.72644\nTotal prob of N-in: 0.95270\nPOSSIBLE N-term signal sequence\ninside 1 19\nTMhelix 20 39\noutside 40 325\nTMhelix 326 348\ninside 349 373

27970	GCF_001896205.1_ASM189620v1	Edwardsiella piscicida	Proteobacteria; Gammaproteobacteria; Enterobacterales; Hafniaceae; Edwardsiella
MVEDSFLSAHGINRRDFMKLCAGMAATLGLSSNAAAEMAAAITSPQRPPVIWIGAQECTG	WP_012849197.1
MULTISPECIES: hydrogenase 2 small subunit [Edwardsiella]Length: 374\nNumber of predicted TMHs: 1\nExp number of AAs in TMHs: 21.63218\nExp number, first 60 AAs: 2.31899\nTotal prob of N-in: 0.10509\noutside 1 331\nTMhelix 332 354\ninside 355 374

27971	GCF_001729265.1_ASM172926v1	Edwardsiella piscicida	Proteobacteria; Gammaproteobacteria; Enterobacterales; Hafniaceae; Edwardsiella
MVEDSFLSAHGINRRDFMKLCAGMAATLGLSSNAAAEMAAAITSPQRPPVIWIGAQECTG	WP_012849197.1
MULTISPECIES: hydrogenase 2 small subunit [Edwardsiella]Length: 374\nNumber of predicted TMHs: 1\nExp number of AAs in TMHs: 21.63218\nExp number, first 60 AAs: 2.31899\nTotal prob of N-in: 0.10509\noutside 1 331\nTMhelix 332 354\ninside 355 374

27972	GCF_000711175.1_ASM71117v1	Edwardsiella piscicida	Proteobacteria; Gammaproteobacteria; Enterobacterales; Hafniaceae; Edwardsiella
MVEDSFLSAHGINRRDFMKLCAGMAATLGLSSNAAAEMAAAITSPQRPPVIWIGAQECTG	WP_012849197.1
MULTISPECIES: hydrogenase 2 small subunit [Edwardsiella]Length: 374\nNumber of predicted TMHs: 1\nExp

number of AAs in TMHs: 21.63218\nExp number, first 60 AAs: 2.31899\nTotal prob of N-in: 0.10509\noutside 1 331\nTMhelix 332 354\ninside 355 374

27973 GCF_000711175.1_ASM71117v1 Edwardsiella piscicida Proteobacteria; Gammaproteobacteria; Enterobacterales; Hafniaceae; Edwardsiella
MNNEDTFYQAMRRQGVTRRSFLKYCSLAATSLGLGAGMAPKIAWALENKPRIPVVWIHGL WP_034165849.1
MULTISPECIES: hydrogenase [Edwardsiella] Length: 373\nNumber of predicted TMHs: 2\nExp number of AAs in TMHs: 38.5762099999999\nExp number, first 60 AAs: 19.72682\nTotal prob of N-in: 0.95270\nPOSSIBLE N-term signal sequence\ninside 1 19\nTMhelix 20 39\noutside 40 325\nTMhelix 326 348\ninside 349 373

27974 GCF_002075835.1_ASM207583v1 Edwardsiella piscicida Proteobacteria; Gammaproteobacteria; Enterobacterales; Hafniaceae; Edwardsiella
MVEDSFLSAHGINRRDFMKLCAGMAATLGLSSNAAAEMAAAITSPQRPPVIWIGAQECTG WP_012849197.1
MULTISPECIES: hydrogenase 2 small subunit [Edwardsiella] Length: 374\nNumber of predicted TMHs: 1\nExp number of AAs in TMHs: 21.63218\nExp number, first 60 AAs: 2.31899\nTotal prob of N-in: 0.10509\noutside 1 331\nTMhelix 332 354\ninside 355 374

27975 GCF_001729265.1_ASM172926v1 Edwardsiella piscicida Proteobacteria; Gammaproteobacteria; Enterobacterales; Hafniaceae; Edwardsiella
MNNEDTFYQAMRRQGVTRRSFLKYCSLAATSLGLGAGMAPKIAWALENKPRIPVVWIHGL WP_015462312.1
MULTISPECIES: hydrogenase [Edwardsiella] Length: 373\nNumber of predicted TMHs: 2\nExp number of AAs in TMHs: 38.3616\nExp number, first 60 AAs: 19.72644\nTotal prob of N-in: 0.95270\nPOSSIBLE N-term signal sequence\ninside 1 19\nTMhelix 20 39\noutside 40 325\nTMhelix 326 348\ninside 349 373

27976 GCF_000800725.1_ASM80072v2 Edwardsiella sp. EA181011 Proteobacteria; Gammaproteobacteria; Enterobacterales; Hafniaceae; Edwardsiella
MNNEDTFYQAMRRQGVTRRSFLKYCSLAATSLGLGAGMAPKIAWALENKPRIPVVWIHGL WP_034165849.1
MULTISPECIES: hydrogenase [Edwardsiella] Length: 373\nNumber of predicted TMHs: 2\nExp number of AAs in TMHs: 38.5762099999999\nExp number, first 60 AAs: 19.72682\nTotal prob of N-in: 0.95270\nPOSSIBLE N-term signal sequence\ninside 1 19\nTMhelix 20 39\noutside 40 325\nTMhelix 326 348\ninside 349 373

27977 GCF_000800725.1_ASM80072v2 Edwardsiella sp. EA181011 Proteobacteria; Gammaproteobacteria; Enterobacterales; Hafniaceae; Edwardsiella
MVEDSFLSAHGINRRDFMKLCAGMAATLGLSSNAAAEMAAAITSPQRPPVIWIGAQECTG WP_012849197.1
MULTISPECIES: hydrogenase 2 small subunit [Edwardsiella] Length: 374\nNumber of predicted TMHs: 1\nExp number of AAs in TMHs: 21.63218\nExp number, first 60 AAs: 2.31899\nTotal prob of N-in: 0.10509\noutside 1 331\nTMhelix 332 354\ninside 355 374

27978 GCF_001186215.1_ASM118621v1 Edwardsiella sp. LADL05-105 Proteobacteria; Gammaproteobacteria; Enterobacterales; Hafniaceae; Edwardsiella
MNNEDTFYQAMRRQGVTRRSFLKYCSLAATSLGLGAGMAPKIAWALENKPRIPVVWIHGL WP_034165849.1
MULTISPECIES: hydrogenase [Edwardsiella] Length: 373\nNumber of predicted TMHs: 2\nExp number of AAs in TMHs: 38.5762099999999\nExp number, first 60 AAs: 19.72682\nTotal prob of N-in: 0.95270\nPOSSIBLE N-term signal sequence\ninside 1 19\nTMhelix 20 39\noutside 40 325\nTMhelix 326 348\ninside 349 373

27979 GCF_001186215.1_ASM118621v1 Edwardsiella sp. LADL05-105 Proteobacteria; Gammaproteobacteria; Enterobacterales; Hafniaceae; Edwardsiella
MVEDSFLSAHGINRRDFMKLCAGMAATLGLSSNAAAEMAAAITSPQRPPVIWIGAQECTG WP_012849197.1
MULTISPECIES: hydrogenase 2 small subunit [Edwardsiella] Length: 374\nNumber of predicted TMHs: 1\nExp number of AAs in TMHs: 21.63218\nExp number, first 60 AAs: 2.31899\nTotal prob of N-in: 0.10509\noutside 1 331\nTMhelix 332 354\ninside 355 374

27980 GCF_001011055.1_ASM101105v1 Edwardsiella tarda Proteobacteria; Gammaproteobacteria; Enterobacterales; Hafniaceae; Edwardsiella
MNNEDTFYQAMRRQGVTRRSFLKYCSLAATSLGLGAGMAPKIAWALENKPRIPVVWIHGL WP_005282286.1
hydrogenase [Edwardsiella tarda] Length: 372\nNumber of predicted TMHs: 2\nExp number of AAs in TMHs: 39.00806\nExp number, first 60 AAs: 19.72168\nTotal prob of N-in: 0.95251\nPOSSIBLE N-term signal sequence\ninside 1 19\nTMhelix 20 39\noutside 40 325\nTMhelix 326 348\ninside 349 372

27981 GCF_001011055.1_ASM101105v1 Edwardsiella tarda Proteobacteria; Gammaproteobacteria; Enterobacterales; Hafniaceae; Edwardsiella

MVEDSFLSAHGINRRDFMKLCAGMAATLGLSSNAAAEMAAAITSPQRPPVIWIGAQECTG WP_005288186.1
hydrogenase 2 small subunit [Edwardsiella tarda]Length: 374\nNumber of predicted TMHs: 1\nExp number of AAs in TMHs: 21.63044\nExp number, first 60 AAs: 2.3188\nTotal prob of N-in: 0.10504\noutside 1 331\nTMhelix 332 354\ninside 355 374

27982 GCF_002006945.1_ASM200694v1 Edwardsiella tarda Proteobacteria; Gammaproteobacteria; Enterobacterales; Hafniaceae; Edwardsiella
MNNEDTFYQAMRRQGVTRRSFLKYCSLAATSLGLGAGMAPKIAWALENKPRIPVVWIHGL WP_005282286.1
hydrogenase [Edwardsiella tarda] Length: 372\nNumber of predicted TMHs: 2\nExp number of AAs in TMHs: 39.00806\nExp number, first 60 AAs: 19.72168\nTotal prob of N-in: 0.95251\nPOSSIBLE N-term signal sequence\ninside 1 19\nTMhelix 20 39\noutside 40 325\nTMhelix 326 348\ninside 349 372

27983 GCF_002006945.1_ASM200694v1 Edwardsiella tarda Proteobacteria; Gammaproteobacteria; Enterobacterales; Hafniaceae; Edwardsiella
MVEDSFLSAHGINRRDFMKLCAGMAATLGLSSNAAAEMAAAITSPQRPPVIWIGAQECTG WP_005288186.1
hydrogenase 2 small subunit [Edwardsiella tarda]Length: 374\nNumber of predicted TMHs: 1\nExp number of AAs in TMHs: 21.63044\nExp number, first 60 AAs: 2.3188\nTotal prob of N-in: 0.10504\noutside 1 331\nTMhelix 332 354\ninside 355 374

27984 GCF_000804515.1_ASM80451v1 Edwardsiella tarda Proteobacteria; Gammaproteobacteria; Enterobacterales; Hafniaceae; Edwardsiella
MNNEDTFYQAMRRQGVTRRSFLKYCSLAATSLGLGAGMAPKIAWALENKPRIPVVWIHGL WP_015462312.1
MULTISPECIES: hydrogenase [Edwardsiella] Length: 373\nNumber of predicted TMHs: 2\nExp number of AAs in TMHs: 38.3616\nExp number, first 60 AAs: 19.72644\nTotal prob of N-in: 0.95270\nPOSSIBLE N-term signal sequence\ninside 1 19\nTMhelix 20 39\noutside 40 325\nTMhelix 326 348\ninside 349 373

27985 GCF_001700615.1_ASM170061v1 Edwardsiella tarda Proteobacteria; Gammaproteobacteria; Enterobacterales; Hafniaceae; Edwardsiella
MVEDSFLSAHGINRRDFMKLCAGMAATLGLSSNAAAEMAAAITSPQRPPVIWIGAQECTG WP_005288186.1
hydrogenase 2 small subunit [Edwardsiella tarda]Length: 374\nNumber of predicted TMHs: 1\nExp number of AAs in TMHs: 21.63044\nExp number, first 60 AAs: 2.3188\nTotal prob of N-in: 0.10504\noutside 1 331\nTMhelix 332 354\ninside 355 374

27986 GCF_000804515.1_ASM80451v1 Edwardsiella tarda Proteobacteria; Gammaproteobacteria; Enterobacterales; Hafniaceae; Edwardsiella
MVEDSFLSAHGINRRDFMKLCAGMAATLGLSSNAAAEMAAAITSPQRPPVIWIGAQECTG WP_012849197.1
MULTISPECIES: hydrogenase 2 small subunit [Edwardsiella]Length: 374\nNumber of predicted TMHs: 1\nExp number of AAs in TMHs: 21.63218\nExp number, first 60 AAs: 2.31899\nTotal prob of N-in: 0.10509\noutside 1 331\nTMhelix 332 354\ninside 355 374

27987 GCF_001700615.1_ASM170061v1 Edwardsiella tarda Proteobacteria; Gammaproteobacteria; Enterobacterales; Hafniaceae; Edwardsiella
MNNEDTFYQAMRRQGVTRRSFLKYCSLAATSLGLGAGMAPKIAWALENKPRIPVVWIHGL WP_005282286.1
hydrogenase [Edwardsiella tarda] Length: 372\nNumber of predicted TMHs: 2\nExp number of AAs in TMHs: 39.00806\nExp number, first 60 AAs: 19.72168\nTotal prob of N-in: 0.95251\nPOSSIBLE N-term signal sequence\ninside 1 19\nTMhelix 20 39\noutside 40 325\nTMhelix 326 348\ninside 349 372

27988 GCF_000804595.1_ASM80459v1 Edwardsiella anguillarum ET070829 Proteobacteria; Gammaproteobacteria; Enterobacterales; Hafniaceae; Edwardsiella; Edwardsiella anguillarum
MNNEDTFYQAMRRQGVTRRSFLKYCSLAATSLGLGAGMAPKIAWALENKPRIPVVWIHGL WP_034165849.1
MULTISPECIES: hydrogenase [Edwardsiella] Length: 373\nNumber of predicted TMHs: 2\nExp number of AAs in TMHs: 38.5762099999999\nExp number, first 60 AAs: 19.72682\nTotal prob of N-in: 0.95270\nPOSSIBLE N-term signal sequence\ninside 1 19\nTMhelix 20 39\noutside 40 325\nTMhelix 326 348\ninside 349 373

27989 GCF_000804595.1_ASM80459v1 Edwardsiella anguillarum ET070829 Proteobacteria; Gammaproteobacteria; Enterobacterales; Hafniaceae; Edwardsiella; Edwardsiella anguillarum
MVEDSFLSAHGINRRDFMKLCAGMAATLGLSSNAAAEMAAAITSPQRPPVIWIGAQECTG WP_012849197.1
MULTISPECIES: hydrogenase 2 small subunit [Edwardsiella]Length: 374\nNumber of predicted TMHs: 1\nExp number of AAs in TMHs: 21.63218\nExp number, first 60 AAs: 2.31899\nTotal prob of N-in: 0.10509\noutside 1 331\nTMhelix 332 354\ninside 355 374

27990 GCF_000804575.1_ASM80457v1 Edwardsiella anguillarum ET080729 Proteobacteria; Gammaproteobacteria; Enterobacterales; Hafniaceae; Edwardsiella; Edwardsiella anguillarum
MVEDSFLSAHGINRRDFMKLCAGMAATLGLSSNAAAEMAAAITSPQRPPVIWIGAQECTG WP_012849197.1
MULTISPECIES: hydrogenase 2 small subunit [Edwardsiella] Length: 374\nNumber of predicted TMHs: 1\nExp number of AAs in TMHs: 21.63218\nExp number, first 60 AAs: 2.31899\nTotal prob of N-in: 0.10509\noutside 1 331\nTMhelix 332 354\ninside 355 374

27991 GCF_000804575.1_ASM80457v1 Edwardsiella anguillarum ET080729 Proteobacteria; Gammaproteobacteria; Enterobacterales; Hafniaceae; Edwardsiella; Edwardsiella anguillarum
MNNEDTFYQAMRRQGVTRRSFLKYCSLAATSLGLGAGMAPKIAWALENKPRIPVVWIHGL WP_034165849.1
MULTISPECIES: hydrogenase [Edwardsiella] Length: 373\nNumber of predicted TMHs: 2\nExp number of AAs in TMHs: 38.5762099999999\nExp number, first 60 AAs: 19.72682\nTotal prob of N-in: 0.95270\nPOSSIBLE N-term signal sequence\ninside 1 19\nTMhelix 20 39\noutside 40 325\nTMhelix 326 348\ninside 349 373

27992 GCF_000264765.2_ASM26476v2 Edwardsiella anguillarum ET080813 Proteobacteria; Gammaproteobacteria; Enterobacterales; Hafniaceae; Edwardsiella; Edwardsiella anguillarum
MVEDSFLSAHGINRRDFMKLCAGMAATLGLSSNAAAEMAAAITSPQRPPVIWIGAQECTG WP_012849197.1
MULTISPECIES: hydrogenase 2 small subunit [Edwardsiella] Length: 374\nNumber of predicted TMHs: 1\nExp number of AAs in TMHs: 21.63218\nExp number, first 60 AAs: 2.31899\nTotal prob of N-in: 0.10509\noutside 1 331\nTMhelix 332 354\ninside 355 374

27993 GCF_000264765.2_ASM26476v2 Edwardsiella anguillarum ET080813 Proteobacteria; Gammaproteobacteria; Enterobacterales; Hafniaceae; Edwardsiella; Edwardsiella anguillarum
MNNEDTFYQAMRRQGVTRRSFLKYCSLAATSLGLGAGMAPKIAWALENKPRIPVVWIHGL WP_034165849.1
MULTISPECIES: hydrogenase [Edwardsiella] Length: 373\nNumber of predicted TMHs: 2\nExp number of AAs in TMHs: 38.5762099999999\nExp number, first 60 AAs: 19.72682\nTotal prob of N-in: 0.95270\nPOSSIBLE N-term signal sequence\ninside 1 19\nTMhelix 20 39\noutside 40 325\nTMhelix 326 348\ninside 349 373

27994 GCF_000804615.1_ASM80461v1 Edwardsiella anguillarum ET081126R Proteobacteria; Gammaproteobacteria; Enterobacterales; Hafniaceae; Edwardsiella; Edwardsiella anguillarum
MVEDSFLSAHGINRRDFMKLCAGMAATLGLSSNAAAEMAAAITSPQRPPVIWIGAQECTG WP_012849197.1
MULTISPECIES: hydrogenase 2 small subunit [Edwardsiella] Length: 374\nNumber of predicted TMHs: 1\nExp number of AAs in TMHs: 21.63218\nExp number, first 60 AAs: 2.31899\nTotal prob of N-in: 0.10509\noutside 1 331\nTMhelix 332 354\ninside 355 374

27995 GCF_000804615.1_ASM80461v1 Edwardsiella anguillarum ET081126R Proteobacteria; Gammaproteobacteria; Enterobacterales; Hafniaceae; Edwardsiella; Edwardsiella anguillarum
MNNEDTFYQAMRRQGVTRRSFLKYCSLAATSLGLGAGMAPKIAWALENKPRIPVVWIHGL WP_034165849.1
MULTISPECIES: hydrogenase [Edwardsiella] Length: 373\nNumber of predicted TMHs: 2\nExp number of AAs in TMHs: 38.5762099999999\nExp number, first 60 AAs: 19.72682\nTotal prob of N-in: 0.95270\nPOSSIBLE N-term signal sequence\ninside 1 19\nTMhelix 20 39\noutside 40 325\nTMhelix 326 348\ninside 349 373

27996 GCF_000474215.1_ASM47421v1 Edwardsiella hoshinae NBRC 105699 = ATCC 33379
Proteobacteria; Gammaproteobacteria; Enterobacterales; Hafniaceae; Edwardsiella; Edwardsiella hoshinae
MVEDSFLSAHGINRRDFMKLCAGMAATLGLSSNAAAEMAAAITSPQRPPVIWIGAQECTG WP_024522509.1
hydrogenase 2 small subunit [Edwardsiella hoshinae] Length: 374\nNumber of predicted TMHs: 1\nExp number of AAs in TMHs: 21.62924\nExp number, first 60 AAs: 2.31877\nTotal prob of N-in: 0.10501\noutside 1 331\nTMhelix 332 354\ninside 355 374

27997 GCF_000474215.1_ASM47421v1 Edwardsiella hoshinae NBRC 105699 = ATCC 33379
Proteobacteria; Gammaproteobacteria; Enterobacterales; Hafniaceae; Edwardsiella; Edwardsiella hoshinae
MNNEDTFYQAMRRQGVTRRSFLKYCSLAATSLGLGAGMAPKIAWALENKPRIPVVWIHGL WP_024524425.1
hydrogenase [Edwardsiella hoshinae] Length: 372\nNumber of predicted TMHs: 2\nExp number of AAs in TMHs: 39.01093\nExp number, first 60 AAs: 19.72184\nTotal prob of N-in: 0.95251\nPOSSIBLE N-term signal sequence\ninside 1 19\nTMhelix 20 39\noutside 40 325\nTMhelix 326 348\ninside 349 372

27998 GCF_000022885.2_ASM2288v2 Edwardsiella ictaluri 93-146 Proteobacteria; Gammaproteobacteria; Enterobacterales; Hafniaceae; Edwardsiella; Edwardsiella ictaluri
MVEDSFLSAHGINRRDFMKLCAGMAATLGLSSNAAAEMAAAITSPQRPPVIWIGAQECTG WP_015871935.1
hydrogenase 2 small subunit [Edwardsiella ictaluri] Length: 374\nNumber of predicted TMHs: 1\nExp

number of AAs in TMHs: 20.14542\nExp number, first 60 AAs: 2.31859\nTotal prob of N-in: 0.10530\noutside 1 331\nTMhelix 332 354\ninside 355 374

27999 GCF_000022885.2_ASM2288v2 Edwardsiella ictaluri 93-146 Proteobacteria; Gammaproteobacteria; Enterobacterales; Hafniaceae; Edwardsiella; Edwardsiella ictaluri
MNNEDTFYQAMRRQGVTRRSFLKYCSLAATSLGLGAGMAPKIAWALENKPRIPVVWIHGL WP_015872619.1
hydrogenase [Edwardsiella ictaluri] Length: 373\nNumber of predicted TMHs: 2\nExp number of AAs in TMHs: 38.57269\nExp number, first 60 AAs: 19.72695\nTotal prob of N-in: 0.95270\nPOSSIBLE N-term signal sequence\ninside 1 19\nTMhelix 20 39\noutside 40 325\nTMhelix 326 348\ninside 349 373

28000 GCF_000264785.1_ASM26478v1 Edwardsiella ictaluri ATCC 33202 Proteobacteria; Gammaproteobacteria; Enterobacterales; Hafniaceae; Edwardsiella; Edwardsiella ictaluri
MNNEDTFYQAMRRQGVTRRSFLKYCSLAATSLGLGAGMAPKIAWALENKPRIPVVWIHGL WP_015872619.1
hydrogenase [Edwardsiella ictaluri] Length: 373\nNumber of predicted TMHs: 2\nExp number of AAs in TMHs: 38.57269\nExp number, first 60 AAs: 19.72695\nTotal prob of N-in: 0.95270\nPOSSIBLE N-term signal sequence\ninside 1 19\nTMhelix 20 39\noutside 40 325\nTMhelix 326 348\ninside 349 373

28001 GCF_000264785.1_ASM26478v1 Edwardsiella ictaluri ATCC 33202 Proteobacteria; Gammaproteobacteria; Enterobacterales; Hafniaceae; Edwardsiella; Edwardsiella ictaluri
MVEDSFLSAHGINRRDFMKLCAGMAATLGLSSNAAAEMAAAITSPQRPPVIWIGAQECTG WP_015871935.1
hydrogenase 2 small subunit [Edwardsiella ictaluri] Length: 374\nNumber of predicted TMHs: 1\nExp number of AAs in TMHs: 20.14542\nExp number, first 60 AAs: 2.31859\nTotal prob of N-in: 0.10530\noutside 1 331\nTMhelix 332 354\ninside 355 374

28002 GCF_000348565.1_ASM34856v1 Edwardsiella piscicida C07-087 Proteobacteria; Gammaproteobacteria; Enterobacterales; Hafniaceae; Edwardsiella; Edwardsiella piscicida
MNNEDTFYQAMRRQGVTRRSFLKYCSLAATSLGLGAGMAPKIAWALENKPRIPVVWIHGL WP_015462312.1
MULTISPECIES: hydrogenase [Edwardsiella] Length: 373\nNumber of predicted TMHs: 2\nExp number of AAs in TMHs: 38.3616\nExp number, first 60 AAs: 19.72644\nTotal prob of N-in: 0.95270\nPOSSIBLE N-term signal sequence\ninside 1 19\nTMhelix 20 39\noutside 40 325\nTMhelix 326 348\ninside 349 373

28003 GCF_000348565.1_ASM34856v1 Edwardsiella piscicida C07-087 Proteobacteria; Gammaproteobacteria; Enterobacterales; Hafniaceae; Edwardsiella; Edwardsiella piscicida
MVEDSFLSAHGINRRDFMKLCAGMAATLGLSSNAAAEMAAAITSPQRPPVIWIGAQECTG WP_012849197.1
MULTISPECIES: hydrogenase 2 small subunit [Edwardsiella]Length: 374\nNumber of predicted TMHs: 1\nExp number of AAs in TMHs: 21.63218\nExp number, first 60 AAs: 2.31899\nTotal prob of N-in: 0.10509\noutside 1 331\nTMhelix 332 354\ninside 355 374

28004 GCF_000264805.1_ASM26480v1 Edwardsiella tarda ATCC 15947 = NBRC 105688 Proteobacteria; Gammaproteobacteria; Enterobacterales; Hafniaceae; Edwardsiella; Edwardsiella tarda
MVEDSFLSAHGINRRDFMKLCAGMAATLGLSSNAAAEMAAAITSPQRPPVIWIGAQECTG WP_005288186.1
hydrogenase 2 small subunit [Edwardsiella tarda]Length: 374\nNumber of predicted TMHs: 1\nExp number of AAs in TMHs: 21.63044\nExp number, first 60 AAs: 2.3188\nTotal prob of N-in: 0.10504\noutside 1 331\nTMhelix 332 354\ninside 355 374

28005 GCF_000264805.1_ASM26480v1 Edwardsiella tarda ATCC 15947 = NBRC 105688 Proteobacteria; Gammaproteobacteria; Enterobacterales; Hafniaceae; Edwardsiella; Edwardsiella tarda
MNNEDTFYQAMRRQGVTRRSFLKYCSLAATSLGLGAGMAPKIAWALENKPRIPVVWIHGL WP_005282286.1
hydrogenase [Edwardsiella tarda] Length: 372\nNumber of predicted TMHs: 2\nExp number of AAs in TMHs: 39.00806\nExp number, first 60 AAs: 19.72168\nTotal prob of N-in: 0.95251\nPOSSIBLE N-term signal sequence\ninside 1 19\nTMhelix 20 39\noutside 40 325\nTMhelix 326 348\ninside 349 372

28006 GCF_000341505.1_ASM34150v1 Edwardsiella tarda ATCC 15947 = NBRC 105688 Proteobacteria; Gammaproteobacteria; Enterobacterales; Hafniaceae; Edwardsiella; Edwardsiella tarda
MVEDSFLSAHGINRRDFMKLCAGMAATLGLSSNAAAEMAAAITSPQRPPVIWIGAQECTG WP_005288186.1
hydrogenase 2 small subunit [Edwardsiella tarda]Length: 374\nNumber of predicted TMHs: 1\nExp number of AAs in TMHs: 21.63044\nExp number, first 60 AAs: 2.3188\nTotal prob of N-in: 0.10504\noutside 1 331\nTMhelix 332 354\ninside 355 374

28007 GCF_000341505.1_ASM34150v1 Edwardsiella tarda ATCC 15947 = NBRC 105688 Proteobacteria; Gammaproteobacteria; Enterobacterales; Hafniaceae; Edwardsiella; Edwardsiella tarda

MNNEDTFYQAMRRQGVTRRSFLKYCSLAATSLGLGAGMAPKIAWALENKPRIPVVWIHGL WP_005282286.1 hydrogenase [Edwardsiella tarda] Length: 372\nNumber of predicted TMHs: 2\nExp number of AAs in TMHs: 39.00806\nExp number, first 60 AAs: 19.72168\nTotal prob of N-in: 0.95251\nPOSSIBLE N-term signal sequence\ninside 1 19\nTMhelix 20 39\noutside 40 325\nTMhelix 326 348\ninside 349 372

28008 GCF_000163955.1_ASM16395v1 Edwardsiella tarda ATCC 23685 Proteobacteria; Gammaproteobacteria; Enterobacterales; Hafniaceae; Edwardsiella; Edwardsiella tarda MVEDSFLSAHGINRRDFMKLCAGMAATLGLSSNAAAEMAAAITSPQRPPVIWIGAQECTG WP_005288186.1 hydrogenase 2 small subunit [Edwardsiella tarda]Length: 374\nNumber of predicted TMHs: 1\nExp number of AAs in TMHs: 21.63044\nExp number, first 60 AAs: 2.3188\nTotal prob of N-in: 0.10504\noutside 1 331\nTMhelix 332 354\ninside 355 374

28009 GCF_000163955.1_ASM16395v1 Edwardsiella tarda ATCC 23685 Proteobacteria; Gammaproteobacteria; Enterobacterales; Hafniaceae; Edwardsiella; Edwardsiella tarda MNNEDTFYQAMRRQGVTRRSFLKYCSLAATSLGLGAGMAPKIAWALENKPRIPVVWIHGL WP_005282286.1 hydrogenase [Edwardsiella tarda] Length: 372\nNumber of predicted TMHs: 2\nExp number of AAs in TMHs: 39.00806\nExp number, first 60 AAs: 19.72168\nTotal prob of N-in: 0.95251\nPOSSIBLE N-term signal sequence\ninside 1 19\nTMhelix 20 39\noutside 40 325\nTMhelix 326 348\ninside 349 372

28010 GCF_000264825.1_ASM26482v1 Edwardsiella tarda DT Proteobacteria; Gammaproteobacteria; Enterobacterales; Hafniaceae; Edwardsiella; Edwardsiella tarda MNNEDTFYQAMRRQGVTRRSFLKYCSLAATSLGLGAGMAPKIAWALENKPRIPVVWIHGL WP_005282286.1 hydrogenase [Edwardsiella tarda] Length: 372\nNumber of predicted TMHs: 2\nExp number of AAs in TMHs: 39.00806\nExp number, first 60 AAs: 19.72168\nTotal prob of N-in: 0.95251\nPOSSIBLE N-term signal sequence\ninside 1 19\nTMhelix 20 39\noutside 40 325\nTMhelix 326 348\ninside 349 372

28011 GCF_000264825.1_ASM26482v1 Edwardsiella tarda DT Proteobacteria; Gammaproteobacteria; Enterobacterales; Hafniaceae; Edwardsiella; Edwardsiella tarda MVEDSFLSAHGINRRDFMKLCAGMAATLGLSSNAAAEMAAAITSPQRPPVIWIGAQECTG WP_005288186.1 hydrogenase 2 small subunit [Edwardsiella tarda]Length: 374\nNumber of predicted TMHs: 1\nExp number of AAs in TMHs: 21.63044\nExp number, first 60 AAs: 2.3188\nTotal prob of N-in: 0.10504\noutside 1 331\nTMhelix 332 354\ninside 355 374

28012 GCF_000020865.1_ASM2086v1 Edwardsiella tarda EIB202 Proteobacteria; Gammaproteobacteria; Enterobacterales; Hafniaceae; Edwardsiella; Edwardsiella tarda MVEDSFLSAHGINRRDFMKLCAGMAATLGLSSNAAAEMAAAITSPQRPPVIWIGAQECTG WP_012849197.1 MULTISPECIES: hydrogenase 2 small subunit [Edwardsiella] Length: 374\nNumber of predicted TMHs: 1\nExp number of AAs in TMHs: 21.63218\nExp number, first 60 AAs: 2.31899\nTotal prob of N-in: 0.10509\noutside 1 331\nTMhelix 332 354\ninside 355 374

28013 GCF_000020865.1_ASM2086v1 Edwardsiella tarda EIB202 Proteobacteria; Gammaproteobacteria; Enterobacterales; Hafniaceae; Edwardsiella; Edwardsiella tarda MNNEDTFYQAMRRQGVTRRSFLKYCSLAATSLGLGAGMAPKIAWALENKPRIPVVWIHGL WP_015462312.1 MULTISPECIES: hydrogenase [Edwardsiella] Length: 373\nNumber of predicted TMHs: 2\nExp number of AAs in TMHs: 38.3616\nExp number, first 60 AAs: 19.72644\nTotal prob of N-in: 0.95270\nPOSSIBLE N-term signal sequence\ninside 1 19\nTMhelix 20 39\noutside 40 325\nTMhelix 326 348\ninside 349 373

28014 GCF_000146305.1_ASM14630v1 Edwardsiella tarda FL6-60 Proteobacteria; Gammaproteobacteria; Enterobacterales; Hafniaceae; Edwardsiella; Edwardsiella tarda MVEDSFLSAHGINRRDFMKLCAGMAATLGLSSNAAAEMAAAITSPQRPPVIWIGAQECTG WP_012849197.1 MULTISPECIES: hydrogenase 2 small subunit [Edwardsiella] Length: 374\nNumber of predicted TMHs: 1\nExp number of AAs in TMHs: 21.63218\nExp number, first 60 AAs: 2.31899\nTotal prob of N-in: 0.10509\noutside 1 331\nTMhelix 332 354\ninside 355 374

28015 GCF_000146305.1_ASM14630v1 Edwardsiella tarda FL6-60 Proteobacteria; Gammaproteobacteria; Enterobacterales; Hafniaceae; Edwardsiella; Edwardsiella tarda MNNEDTFYQAMRRQGVTRRSFLKYCSLAATSLGLGAGMAPKIAWALENKPRIPVVWIHGL WP_015462312.1 MULTISPECIES: hydrogenase [Edwardsiella] Length: 373\nNumber of predicted TMHs: 2\nExp number of AAs in TMHs: 38.3616\nExp number, first 60 AAs: 19.72644\nTotal prob of N-in: 0.95270\nPOSSIBLE N-term signal sequence\ninside 1 19\nTMhelix 20 39\noutside 40 325\nTMhelix 326 348\ninside 349 373

28016 GCF_000818795.1_ASM81879v1 Hafnia alvei Proteobacteria; Gammaproteobacteria; Enterobacterales; Hafniaceae; Hafnia MAEDNFLSAHGVNRRDFMKLCAGMAATLGLSSNAAAEMAAAITNPQRPPVIWIGAQECTG WP_040044698.1 hydrogenase 2 small subunit [Hafnia alvei] Length: 384\nNumber of predicted TMHs: 1\nExp number of AAs in TMHs: 22.00032\nExp number, first 60 AAs: 2.80789\nTotal prob of N-in: 0.13211\noutside 1 331\nTMhelix 332 354\ninside 355 384

28017 GCF_000818805.2_ASM81880v2 Hafnia alvei Proteobacteria; Gammaproteobacteria; Enterobacterales; Hafniaceae; Hafnia MAEDNFLSAHGVNRRDFMKLCAGMAATLGLSSNAAAEMAAAITNPQRPPVIWIGAQECTG WP_046360240.1 hydrogenase 2 small subunit [Hafnia alvei] Length: 384\nNumber of predicted TMHs: 1\nExp number of AAs in TMHs: 21.98363\nExp number, first 60 AAs: 2.80775\nTotal prob of N-in: 0.13205\noutside 1 331\nTMhelix 332 354\ninside 355 384

28018 GCF_001636255.1_ASM163625v1 Hafnia alvei Proteobacteria; Gammaproteobacteria; Enterobacterales; Hafniaceae; Hafnia MAEDNFLSAHGVNRRDFMKLCAGMAATLGLSSNAAAEMAAAITNPQRPPVIWIGAQECTG WP_040044698.1 hydrogenase 2 small subunit [Hafnia alvei] Length: 384\nNumber of predicted TMHs: 1\nExp number of AAs in TMHs: 22.00032\nExp number, first 60 AAs: 2.80789\nTotal prob of N-in: 0.13211\noutside 1 331\nTMhelix 332 354\ninside 355 384

28019 GCF_900095695.1_GB001_A1 Hafnia alvei Proteobacteria; Gammaproteobacteria; Enterobacterales; Hafniaceae; Hafnia MAEDNFLSAHGVNRRDFMKLCAGMAATLGLSSNAAAEMAAAITNPQRPPVIWIGAQECTG WP_040044698.1 hydrogenase 2 small subunit [Hafnia alvei] Length: 384\nNumber of predicted TMHs: 1\nExp number of AAs in TMHs: 22.00032\nExp number, first 60 AAs: 2.80789\nTotal prob of N-in: 0.13211\noutside 1 331\nTMhelix 332 354\ninside 355 384

28020 GCF_000980975.1_ASM98097v1 Hafnia alvei Proteobacteria; Gammaproteobacteria; Enterobacterales; Hafniaceae; Hafnia MAEDNFLSAHGVNRRDFMKLCAGMAATLGLSSNAAAEMAAAITNPQRPPVIWIGAQECTG WP_043494122.1 hydrogenase 2 small subunit [Hafnia alvei] Length: 384\nNumber of predicted TMHs: 1\nExp number of AAs in TMHs: 22.00049\nExp number, first 60 AAs: 2.80791\nTotal prob of N-in: 0.13212\noutside 1 331\nTMhelix 332 354\ninside 355 384

28021 GCF_000980985.1_ASM98098v1 Hafnia alvei Proteobacteria; Gammaproteobacteria; Enterobacterales; Hafniaceae; Hafnia MAEDNFLSAHGVNRRDFMKLCAGMAATLGLSSNAAAEMAAAITNPQRPPVIWIGAQECTG WP_046459282.1 MULTISPECIES: hydrogenase 2 small subunit [Hafniaceae] Length: 384\nNumber of predicted TMHs: 1\nExp number of AAs in TMHs: 22.00049\nExp number, first 60 AAs: 2.80791\nTotal prob of N-in: 0.13212\noutside 1 331\nTMhelix 332 354\ninside 355 384

28022 GCF_000735375.1_GHAL_DRAFTv1 Hafnia alvei ATCC 13337 Proteobacteria; Gammaproteobacteria; Enterobacterales; Hafniaceae; Hafnia; Hafnia alvei MAEDNFLSAHGVNRRDFMKLCAGMAATLGLSSNAAAEMAAAITNPQRPPVIWIGAQECTG WP_043494122.1 hydrogenase 2 small subunit [Hafnia alvei] Length: 384\nNumber of predicted TMHs: 1\nExp number of AAs in TMHs: 22.00049\nExp number, first 60 AAs: 2.80791\nTotal prob of N-in: 0.13212\noutside 1 331\nTMhelix 332 354\ninside 355 384

28023 GCF_000597785.2_ASM59778v2 Hafnia alvei FB1 Proteobacteria; Gammaproteobacteria; Enterobacterales; Hafniaceae; Hafnia; Hafnia alvei MAEDNFLSAHGVNRRDFMKLCAGMAATLGLSSNAAAEMAAAITNPQRPPVIWIGAQECTG WP_025797933.1 hydrogenase 2 small subunit [Hafnia alvei] Length: 384\nNumber of predicted TMHs: 1\nExp number of AAs in TMHs: 21.98365\nExp number, first 60 AAs: 2.80775\nTotal prob of N-in: 0.13206\noutside 1 331\nTMhelix 332 354\ninside 355 384

28024 GCF_001586165.1_ASM158616v1 Obesumbacterium proteus Proteobacteria; Gammaproteobacteria; Enterobacterales; Hafniaceae; Obesumbacterium MIDIPSRFTRRRLLQGMGVLAATTLVPTGVFPAYASSAISNDFITISIFLTGREKLSAGY WP_061552847.1 hypothetical protein [Obesumbacterium proteus] Length: 167\nNumber of predicted TMHs: 2\nExp number of AAs in TMHs: 40.75495\nExp number, first 60 AAs: 22.47043\nTotal prob of N-in: 0.98581\nPOSSIBLE N-term signal sequence\ninside 1 12\nTMhelix 13 35\noutside 36 103\nTMhelix 104 126\ninside 127 167

28025 GCF_001586165.1_ASM158616v1 Obesumbacterium proteus Proteobacteria; Gammaproteobacteria; Enterobacterales; Hafniaceae; Obesumbacterium MAEDNFLSAHGVNRRDFMKLCAGMAATLGLSSNAAAEMAAAITNPQRPPVIWIGAQECTG WP_046459282.1 MULTISPECIES: hydrogenase 2 small subunit [Hafniaceae] Length: 384\nNumber of predicted TMHs: 1\nExp

number of AAs in TMHs: 22.00049\nExp number, first 60 AAs: 2.80791\nTotal prob of N-in: 0.13212\noutside 1 331\nTMhelix 332 354\ninside 355 384

28026 GCF_001655035.1_Opr12841_DRAFTv1 Obesumbacterium proteus ATCC 12841 Proteobacteria; Gammaproteobacteria; Enterobacterales; Hafniaceae; Obesumbacterium; Obesumbacterium proteus MAEDNFLSAHGVNRRDFMKLCAGMAATLGLSSNAAAEMAAAITNPQRPPVIWIGAQECTG WP_046459282.1 MULTISPECIES: hydrogenase 2 small subunit [Hafniaceae] Length: 384\nNumber of predicted TMHs: 1\nExp number of AAs in TMHs: 22.00049\nExp number, first 60 AAs: 2.80791\nTotal prob of N-in: 0.13212\noutside 1 331\nTMhelix 332 354\ninside 355 384

28027 GCF_001655035.1_Opr12841_DRAFTv1 Obesumbacterium proteus ATCC 12841 Proteobacteria; Gammaproteobacteria; Enterobacterales; Hafniaceae; Obesumbacterium; Obesumbacterium proteus MIDIPSRFTRRRLLQGMGVLAATTLVPTGVFPAYASSAISNDFITISIFLTGREKLSAGY WP_061552847.1 hypothetical protein [Obesumbacterium proteus] Length: 167\nNumber of predicted TMHs: 2\nExp number of AAs in TMHs: 40.75495\nExp number, first 60 AAs: 22.47043\nTotal prob of N-in: 0.98581\nPOSSIBLE N-term signal sequence\ninside 1 12\nTMhelix 13 35\noutside 36 103\nTMhelix 104 126\ninside 127 167

28028 GCF_001654855.1_Cmy19692_DRAFTv1 Cosenzaea myxofaciens ATCC 19692 Proteobacteria; Gammaproteobacteria; Enterobacterales; Morganellaceae; Cosenzaea; Cosenzaea myxofaciens MALGNNVFSQHGINRRDFMKLCAALSATMGLSGKAAAQMSQAMASPERPPVIWIGAQECT WP_066750637.1 hydrogenase 2 small subunit [Cosenzaea myxofaciens] Length: 375\nNumber of predicted TMHs: 1\nExp number of AAs in TMHs: 21.6093\nExp number, first 60 AAs: 0.53064\nTotal prob of N-in: 0.05337\noutside 1 333\nTMhelix 334 356\ninside 357 375

28029 GCF_001263435.1_ASM126343v1 Morganella morganii Proteobacteria; Gammaproteobacteria; Enterobacterales; Morganellaceae; Morganella MIDDNTLITSDMTSEGIDRRSFLKLCAALAATMGLSANASAEIAASVSRKDCPPVIWISA WP_046892967.1 hydrogenase 2 small subunit [Morganella morganii] Length: 377\nNumber of predicted TMHs: 1\nExp number of AAs in TMHs: 26.47066\nExp number, first 60 AAs: 2.01431\nTotal prob of N-in: 0.22925\noutside 1 336\nTMhelix 337 359\ninside 360 377

28030 GCF_001066005.1_ASM106600v1 Morganella morganii Proteobacteria; Gammaproteobacteria; Enterobacterales; Morganellaceae; Morganella MIEDNTFITSDITSEGIDRRSFLKLCAALAATMGLSANASAEIAASVSRKDCPPVIWISA WP_024474542.1 MULTISPECIES: hydrogenase 2 small subunit [Morganella] Length: 377\nNumber of predicted TMHs: 1\nExp number of AAs in TMHs: 24.35345\nExp number, first 60 AAs: 0.88171\nTotal prob of N-in: 0.14103\noutside 1 336\nTMhelix 337 359\ninside 360 377

28031 GCF_000770295.1_ASM77029v1 Morganella morganii Proteobacteria; Gammaproteobacteria; Enterobacterales; Morganellaceae; Morganella MIEDNTFITSDITSEGIDRRSFLKLCAALAATMGLSANASAEIAASVSRKDCPPVIWISA WP_004240464.1 Uptake hydrogenase small subunit precursor [Morganella morganii] Length: 377\nNumber of predicted TMHs: 1\nExp number of AAs in TMHs: 24.59041\nExp number, first 60 AAs: 0.8813\nTotal prob of N-in: 0.14248\noutside 1 336\nTMhelix 337 359\ninside 360 377

28032 GCF_000747035.1_ASM74703v1 Morganella morganii Proteobacteria; Gammaproteobacteria; Enterobacterales; Morganellaceae; Morganella MIEDNTFITSDITSEGIDRRSFLKLCAALAATMGLSANASAEIAASVSRKDCPPVIWISA WP_032097959.1 hydrogenase 2 small subunit [Morganella morganii] Length: 377\nNumber of predicted TMHs: 1\nExp number of AAs in TMHs: 24.35208\nExp number, first 60 AAs: 0.88157\nTotal prob of N-in: 0.14101\noutside 1 336\nTMhelix 337 359\ninside 360 377

28033 GCF_002077705.1_ASM207770v1 Morganella morganii Proteobacteria; Gammaproteobacteria; Enterobacterales; Morganellaceae; Morganella MIEDNTFITSDITSEGIDRRSFLKLCAALAATMGLSANASAEIAASVSRKDCPPVIWISA WP_081120125.1 hydrogenase 2 small subunit [Morganella morganii] Length: 377\nNumber of predicted TMHs: 1\nExp number of AAs in TMHs: 24.35419\nExp number, first 60 AAs: 0.8818\nTotal prob of N-in: 0.14104\noutside 1 336\nTMhelix 337 359\ninside 360 377

28034 GCF_001653675.1_ASM165367v1 Morganella morganii Proteobacteria; Gammaproteobacteria; Enterobacterales; Morganellaceae; Morganella

MIEDNTFITSDITSEGIDRRSFLKLCAALAATMGLSANASAEIAASVSRKDCPPVIWISA WP_064483459.1 hydrogenase 2 small subunit [Morganella morganii] Length: 377\nNumber of predicted TMHs: 1\nExp number of AAs in TMHs: 24.35436\nExp number, first 60 AAs: 0.88204\nTotal prob of N-in: 0.14105\noutside 1 336\nTMhelix 337 359\ninside 360 377

28035 GCF_000966695.1_ASM96669v1 Morganella morganii Proteobacteria; Gammaproteobacteria; Enterobacterales; Morganellaceae; Morganella
MIEDNTFITSDITSEGIDRRSFLKLCAALAATMGLSANASAEIAASVSRKDCPPVIWISA WP_004240464.1 Uptake hydrogenase small subunit precursor [Morganella morganii] Length: 377\nNumber of predicted TMHs: 1\nExp number of AAs in TMHs: 24.59041\nExp number, first 60 AAs: 0.8813\nTotal prob of N-in: 0.14248\noutside 1 336\nTMhelix 337 359\ninside 360 377

28036 GCF_002029935.1_ASM202993v1 Morganella morganii Proteobacteria; Gammaproteobacteria; Enterobacterales; Morganellaceae; Morganella
MIEDNTFITSDITSEGIDRRSFLKLCAALAATMGLSANASAEIAASVSRKDCPPVIWISA WP_032097959.1 hydrogenase 2 small subunit [Morganella morganii] Length: 377\nNumber of predicted TMHs: 1\nExp number of AAs in TMHs: 24.35208\nExp number, first 60 AAs: 0.88157\nTotal prob of N-in: 0.14101\noutside 1 336\nTMhelix 337 359\ninside 360 377

28037 GCF_001558895.1_ASM155889v1 Morganella morganii Proteobacteria; Gammaproteobacteria; Enterobacterales; Morganellaceae; Morganella
MIEDNTFITSDITSEGIDRRSFLKLCAALAATMGLSANASAEIAASVSRKDCPPVIWISA WP_004236377.1 hydrogenase 2 small subunit [Morganella morganii] Length: 377\nNumber of predicted TMHs: 1\nExp number of AAs in TMHs: 24.3561\nExp number, first 60 AAs: 0.88157\nTotal prob of N-in: 0.14113\noutside 1 336\nTMhelix 337 359\ninside 360 377

28038 GCF_001066745.1_ASM106674v1 Morganella morganii Proteobacteria; Gammaproteobacteria; Enterobacterales; Morganellaceae; Morganella
MIEDNTFITSDITSEGIDRRSFLKLCAALAATMGLSANASAEIAASVSRKDCPPVIWISA WP_004240464.1 Uptake hydrogenase small subunit precursor [Morganella morganii] Length: 377\nNumber of predicted TMHs: 1\nExp number of AAs in TMHs: 24.59041\nExp number, first 60 AAs: 0.8813\nTotal prob of N-in: 0.14248\noutside 1 336\nTMhelix 337 359\ninside 360 377

28039 GCF_001274995.1_ASM127499v1 Morganella morganii Proteobacteria; Gammaproteobacteria; Enterobacterales; Morganellaceae; Morganella
MIDDNTLITSDMTSEGIDRRSFLKLCAALAATMGLSANASAEIAASVSRKDCPPVIWISA WP_046892967.1 hydrogenase 2 small subunit [Morganella morganii] Length: 377\nNumber of predicted TMHs: 1\nExp number of AAs in TMHs: 26.47066\nExp number, first 60 AAs: 2.01431\nTotal prob of N-in: 0.22925\noutside 1 336\nTMhelix 337 359\ninside 360 377

28040 GCF_900142745.1_IMG-taxon_2654588210_annotated_assembly Morganella morganii
Proteobacteria; Gammaproteobacteria; Enterobacterales; Morganellaceae; Morganella
MIEDNTFITSDITSEGIDRRSFLKLCAALAATMGLSANASAEIAASVSRKDCPPVIWISA WP_004236377.1 hydrogenase 2 small subunit [Morganella morganii] Length: 377\nNumber of predicted TMHs: 1\nExp number of AAs in TMHs: 24.3561\nExp number, first 60 AAs: 0.88157\nTotal prob of N-in: 0.14113\noutside 1 336\nTMhelix 337 359\ninside 360 377

28041 GCF_001006565.1_ASM100656v1 Morganella morganii Proteobacteria; Gammaproteobacteria; Enterobacterales; Morganellaceae; Morganella
MIDDNTLITSDMTSEGIDRRSFLKLCAALAATMGLSANASAEIAASVSRKDCPPVIWISA WP_046892967.1 hydrogenase 2 small subunit [Morganella morganii] Length: 377\nNumber of predicted TMHs: 1\nExp number of AAs in TMHs: 26.47066\nExp number, first 60 AAs: 2.01431\nTotal prob of N-in: 0.22925\noutside 1 336\nTMhelix 337 359\ninside 360 377

28042 GCF_000783955.1_ASM78395v1 Morganella morganii Proteobacteria; Gammaproteobacteria; Enterobacterales; Morganellaceae; Morganella
MIEDNTFITSDITSEGIDRRSFLKLCAALAATMGLSANASAEIAASVSRKDCPPVIWISA WP_036412934.1 hydrogenase 2 small subunit [Morganella morganii] Length: 377\nNumber of predicted TMHs: 1\nExp number of AAs in TMHs: 24.59719\nExp number, first 60 AAs: 0.88201\nTotal prob of N-in: 0.14259\noutside 1 336\nTMhelix 337 359\ninside 360 377

28043 GCF_002180575.1_ASM218057v1 Morganella morganii Proteobacteria; Gammaproteobacteria; Enterobacterales; Morganellaceae; Morganella
MIEDNTFITSDITSEGIDRRSFLKLCAALAATMGLSANASAEIAASVSRKDCPPVIWISA WP_081120125.1 hydrogenase 2 small subunit [Morganella morganii] Length: 377\nNumber of predicted TMHs: 1\nExp number of AAs in TMHs: 24.35419\nExp number, first 60 AAs: 0.8818\nTotal prob of N-in: 0.14104\noutside 1 336\nTMhelix 337 359\ninside 360 377

28044 GCF_002077675.1_ASM207767v1 Morganella morganii Proteobacteria; Gammaproteobacteria; Enterobacterales; Morganellaceae; Morganella
MIEDNTFITSDITSEGIDRRSFLKLCAALAATMGLSANASAEIAASVSRKDCPPVIWISA WP_081120125.1 hydrogenase 2 small subunit [Morganella morganii] Length: 377\nNumber of predicted TMHs: 1\nExp number of AAs in TMHs: 24.35419\nExp number, first 60 AAs: 0.8818\nTotal prob of N-in: 0.14104\noutside 1 336\nTMhelix 337 359\ninside 360 377

28045 GCF_001676225.1_ASM167622v1 Morganella psychrotolerans Proteobacteria; Gammaproteobacteria; Enterobacterales; Morganellaceae; Morganella
MIEDNTFITSEGINRRGFLKLCAALAATMGLSANASAEIAASVSRKDCPPVIWISAQECT WP_067401458.1 hydrogenase 2 small subunit [Morganella psychrotolerans] Length: 373\nNumber of predicted TMHs: 1\nExp number of AAs in TMHs: 25.39115\nExp number, first 60 AAs: 1.59255\nTotal prob of N-in: 0.18277\noutside 1 332\nTMhelix 333 355\ninside 356 373

28046 GCF_001676155.1_ASM167615v1 Morganella psychrotolerans Proteobacteria; Gammaproteobacteria; Enterobacterales; Morganellaceae; Morganella
MIEDNTFITSEGINRRGFLKLCAALAATMGLSSNASAEIAASVSRKDCPPVIWISAQECT WP_067367128.1 hydrogenase 2 small subunit [Morganella psychrotolerans] Length: 373\nNumber of predicted TMHs: 1\nExp number of AAs in TMHs: 24.47032\nExp number, first 60 AAs: 0.61117\nTotal prob of N-in: 0.14537\noutside 1 332\nTMhelix 333 355\ninside 356 373

28047 GCF_001676055.1_ASM167605v1 Morganella psychrotolerans Proteobacteria; Gammaproteobacteria; Enterobacterales; Morganellaceae; Morganella
MIEDNTFITSEGINRRGFLKLCAALAATMGLSANASAEIAASVSRKDCPPVIWISAQECT WP_067425327.1 hydrogenase 2 small subunit [Morganella psychrotolerans] Length: 373\nNumber of predicted TMHs: 1\nExp number of AAs in TMHs: 25.29862\nExp number, first 60 AAs: 1.60767\nTotal prob of N-in: 0.16730\noutside 1 332\nTMhelix 333 355\ninside 356 373

28048 GCF_000512315.2_GS_De_Novo_Assembly Morganella sp. EGD-HP17 Proteobacteria; Gammaproteobacteria; Enterobacterales; Morganellaceae; Morganella
MIEDNTFITSDITSEGIDRRSFLKLCAALAATMGLSANASAEIAASVSRKDCPPVIWISA WP_036416587.1 hydrogenase 2 small subunit [Morganella sp. EGD-HP17] Length: 377\nNumber of predicted TMHs: 1\nExp number of AAs in TMHs: 24.59037\nExp number, first 60 AAs: 0.88129\nTotal prob of N-in: 0.14248\noutside 1 336\nTMhelix 337 359\ninside 360 377

28049 GCF_001807955.1_ASM180795v1 Morganella sp. HMSC11D09 Proteobacteria; Gammaproteobacteria; Enterobacterales; Morganellaceae; Morganella
MIEDNTFITSDITSEGIDRRSFLKLCAALAATMGLSANASAEIAASVSRKDCPPVIWISA WP_024474542.1 MULTISPECIES: hydrogenase 2 small subunit [Morganella] Length: 377\nNumber of predicted TMHs: 1\nExp number of AAs in TMHs: 24.35345\nExp number, first 60 AAs: 0.88171\nTotal prob of N-in: 0.14103\noutside 1 336\nTMhelix 337 359\ninside 360 377

28050 GCF_000633515.1_de_novo Morganella morganii H1r Proteobacteria; Gammaproteobacteria; Enterobacterales; Morganellaceae; Morganella; Morganella morganii
MIEDNTFITSDITSEGINRRSFLKLCAALAATMGLSANTSAEIAASVSRKDCPPVIWISA WP_025151798.1 hydrogenase 2 small subunit [Morganella morganii] Length: 377\nNumber of predicted TMHs: 1\nExp number of AAs in TMHs: 24.87262\nExp number, first 60 AAs: 0.74549\nTotal prob of N-in: 0.16168\noutside 1 336\nTMhelix 337 359\ninside 360 377

28051 GCF_000307755.2_ASM30775v2 Morganella morganii SC01 Proteobacteria; Gammaproteobacteria; Enterobacterales; Morganellaceae; Morganella; Morganella morganii
MIEDNTFITSDITSEGIDRRSFLKLCAALAATMGLSANASAEIAASVSRKDCPPVIWISA WP_004240464.1 Uptake hydrogenase small subunit precursor [Morganella morganii] Length: 377\nNumber of predicted TMHs: 1\nExp

number of AAs in TMHs: 24.59041\nExp number, first 60 AAs:  0.8813\nTotal prob of N-in:   0.14248\noutside   1   336\nTMhelix   337   359\ninside   360   377

28052	GCF_002184465.1_ASM218446v1	Morganella morganii subsp. morganii	Proteobacteria; Gammaproteobacteria; Enterobacterales; Morganellaceae; Morganella; Morganella morganii	MIEDNTFITSDITSEGIDRRSFLKLCAALAATMGLSANASAEIAASVSRKDCPPVIWISA	WP_004236377.1	hydrogenase 2 small subunit [Morganella morganii]	Length: 377\nNumber of predicted TMHs:  1\nExp number of AAs in TMHs: 24.3561\nExp number, first 60 AAs:  0.88157\nTotal prob of N-in:   0.14113\noutside   1   336\nTMhelix   337   359\ninside   360   377

28053	GCF_002185325.1_ASM218532v1	Morganella morganii subsp. morganii	Proteobacteria; Gammaproteobacteria; Enterobacterales; Morganellaceae; Morganella; Morganella morganii	MIEDNTFITSDITSEGIDRRSFLKLCAALAATMGLSANASAEIAASVSRKDCPPVIWISA	WP_087826282.1	hydrogenase 2 small subunit [Morganella morganii]	Length: 377\nNumber of predicted TMHs:  1\nExp number of AAs in TMHs: 24.72224\nExp number, first 60 AAs:  0.88822\nTotal prob of N-in:   0.13143\noutside   1   336\nTMhelix   337   359\ninside   360   377

28054	GCF_000286435.2_ASM28643v2	Morganella morganii subsp. morganii KT	Proteobacteria; Gammaproteobacteria; Enterobacterales; Morganellaceae; Morganella; Morganella morganii; Morganella morganii subsp. morganii	MIEDNTFITSDITSEGIDRRSFLKLCAALAATMGLSANASAEIAASVSRKDCPPVIWISA	WP_004236377.1	hydrogenase 2 small subunit [Morganella morganii]	Length: 377\nNumber of predicted TMHs:  1\nExp number of AAs in TMHs: 24.3561\nExp number, first 60 AAs:  0.88157\nTotal prob of N-in:   0.14113\noutside   1   336\nTMhelix   337   359\ninside   360   377

28055	GCF_001598895.1_ASM159889v1	Morganella morganii subsp. morganii NBRC 3848	Proteobacteria; Gammaproteobacteria; Enterobacterales; Morganellaceae; Morganella; Morganella morganii; Morganella morganii subsp. morganii	MIEDNTFITSDITSEGIDRRSFLKLCAALAATMGLSANASAEIAASVSRKDCPPVIWISA	WP_004236377.1	hydrogenase 2 small subunit [Morganella morganii]	Length: 377\nNumber of predicted TMHs:  1\nExp number of AAs in TMHs: 24.3561\nExp number, first 60 AAs:  0.88157\nTotal prob of N-in:   0.14113\noutside   1   336\nTMhelix   337   359\ninside   360   377

28056	GCF_001060835.1_ASM106083v1	Proteus mirabilis	Proteobacteria; Gammaproteobacteria; Enterobacterales; Morganellaceae; Proteus	MALGNNVFSQHGINRRDFMKLCAALSATMGLSGKAAAQMSQAMTSPERPPVIWIGAQECT	WP_004245185.1	MULTISPECIES: hydrogenase 2 small subunit [Proteus]	Length: 374\nNumber of predicted TMHs:  1\nExp number of AAs in TMHs: 21.84144\nExp number, first 60 AAs:  0.50836\nTotal prob of N-in:   0.03961\noutside   1   333\nTMhelix   334   356\ninside   357   374

28057	GCF_001060325.1_ASM106032v1	Proteus mirabilis	Proteobacteria; Gammaproteobacteria; Enterobacterales; Morganellaceae; Proteus	MALGNNVFSQHGINRRDFMKLCAALSATMGLSGKAAAQMSQAMTSPERPPVIWIGAQECT	WP_004245185.1	MULTISPECIES: hydrogenase 2 small subunit [Proteus]	Length: 374\nNumber of predicted TMHs:  1\nExp number of AAs in TMHs: 21.84144\nExp number, first 60 AAs:  0.50836\nTotal prob of N-in:   0.03961\noutside   1   333\nTMhelix   334   356\ninside   357   374

28058	GCF_001064375.1_ASM106437v1	Proteus mirabilis	Proteobacteria; Gammaproteobacteria; Enterobacterales; Morganellaceae; Proteus	MALGNNVFSQHGINRRDFMKLCAALSATMGLSGKAAAQMSQAMTSPERPPVIWIGAQECT	WP_004245185.1	MULTISPECIES: hydrogenase 2 small subunit [Proteus]	Length: 374\nNumber of predicted TMHs:  1\nExp number of AAs in TMHs: 21.84144\nExp number, first 60 AAs:  0.50836\nTotal prob of N-in:   0.03961\noutside   1   333\nTMhelix   334   356\ninside   357   374

28059	GCF_001076705.1_ASM107670v1	Proteus mirabilis	Proteobacteria; Gammaproteobacteria; Enterobacterales; Morganellaceae; Proteus	MALGNNVFSQHGINRRDFMKLCAALSATMGLSGKAAAQMSQAMTSPERPPVIWIGAQECT	WP_004245185.1	MULTISPECIES: hydrogenase 2 small subunit [Proteus]	Length: 374\nNumber of predicted TMHs:  1\nExp number of AAs in TMHs: 21.84144\nExp number, first 60 AAs:  0.50836\nTotal prob of N-in:   0.03961\noutside   1   333\nTMhelix   334   356\ninside   357   374

28060	GCF_001062135.1_ASM106213v1	Proteus mirabilis	Proteobacteria; Gammaproteobacteria; Enterobacterales; Morganellaceae; Proteus

MALGNNVFSQHGINRRDFMKLCAALSATMGLSGKAAAQMSQAMTSPERPPVIWIGAQECT WP_004245185.1 MULTISPECIES: hydrogenase 2 small subunit [Proteus] Length: 374\nNumber of predicted TMHs: 1\nExp number of AAs in TMHs: 21.84144\nExp number, first 60 AAs: 0.50836\nTotal prob of N-in: 0.03961\noutside 1 333\nTMhelix 334 356\ninside 357 374

28061 GCF_000972665.1_ASM97266v1 Proteus mirabilis Proteobacteria; Gammaproteobacteria; Enterobacterales; Morganellaceae; Proteus MALGNNVFSQHGINRRDFMKLCAALSATMGLSGKAAAQMSQAMTSPERPPVIWIGAQECT WP_004245185.1 MULTISPECIES: hydrogenase 2 small subunit [Proteus] Length: 374\nNumber of predicted TMHs: 1\nExp number of AAs in TMHs: 21.84144\nExp number, first 60 AAs: 0.50836\nTotal prob of N-in: 0.03961\noutside 1 333\nTMhelix 334 356\ninside 357 374

28062 GCF_001077105.1_ASM107710v1 Proteus mirabilis Proteobacteria; Gammaproteobacteria; Enterobacterales; Morganellaceae; Proteus MALGNNVFSQHGINRRDFMKLCAALSATMGLSGKAAAQMSQAMTSPERPPVIWIGAQECT WP_004245185.1 MULTISPECIES: hydrogenase 2 small subunit [Proteus] Length: 374\nNumber of predicted TMHs: 1\nExp number of AAs in TMHs: 21.84144\nExp number, first 60 AAs: 0.50836\nTotal prob of N-in: 0.03961\noutside 1 333\nTMhelix 334 356\ninside 357 374

28063 GCF_000770765.1_ASM77076v1 Proteus mirabilis Proteobacteria; Gammaproteobacteria; Enterobacterales; Morganellaceae; Proteus MALGNNVFSQHGINRRDFMKLCAALSATMGLSGKAAAQMSQAMTSPERPPVIWIGAQECT WP_004245185.1 MULTISPECIES: hydrogenase 2 small subunit [Proteus] Length: 374\nNumber of predicted TMHs: 1\nExp number of AAs in TMHs: 21.84144\nExp number, first 60 AAs: 0.50836\nTotal prob of N-in: 0.03961\noutside 1 333\nTMhelix 334 356\ninside 357 374

28064 GCF_001619795.1_ASM161979v1 Proteus mirabilis Proteobacteria; Gammaproteobacteria; Enterobacterales; Morganellaceae; Proteus MALGNNVFSQHGINRRDFMKLCAALSATMGLSGKAAAQMSQAMTSPERPPVIWIGAQECT WP_004245185.1 MULTISPECIES: hydrogenase 2 small subunit [Proteus] Length: 374\nNumber of predicted TMHs: 1\nExp number of AAs in TMHs: 21.84144\nExp number, first 60 AAs: 0.50836\nTotal prob of N-in: 0.03961\noutside 1 333\nTMhelix 334 356\ninside 357 374

28065 GCF_002184635.1_ASM218463v1 Proteus mirabilis Proteobacteria; Gammaproteobacteria; Enterobacterales; Morganellaceae; Proteus MALGNNVFSHHGINRRDFMKLCAALSATMGLSGKAAAEMSEAMVSTERPPVIWIGAQECT WP_036932958.1 hydrogenase 2 small subunit [Proteus vulgaris] Length: 374\nNumber of predicted TMHs: 1\nExp number of AAs in TMHs: 21.23761\nExp number, first 60 AAs: 0.07839\nTotal prob of N-in: 0.01866\noutside 1 333\nTMhelix 334 356\ninside 357 374

28066 GCF_001617295.1_ASM161729v1 Proteus mirabilis Proteobacteria; Gammaproteobacteria; Enterobacterales; Morganellaceae; Proteus MALGNNVFSQHGINRRDFMKLCAALSATMGLSGKAAAQMSQAMTSPERPPVIWIGAQECT WP_004245185.1 MULTISPECIES: hydrogenase 2 small subunit [Proteus] Length: 374\nNumber of predicted TMHs: 1\nExp number of AAs in TMHs: 21.84144\nExp number, first 60 AAs: 0.50836\nTotal prob of N-in: 0.03961\noutside 1 333\nTMhelix 334 356\ninside 357 374

28067 GCF_001867325.1_ASM186732v1 Proteus mirabilis Proteobacteria; Gammaproteobacteria; Enterobacterales; Morganellaceae; Proteus MALGNNVFSQHGINRRDFMKLCAALSATMGLSGKAAAQMSQAMTSPERPPVIWIGAQECT WP_004245185.1 MULTISPECIES: hydrogenase 2 small subunit [Proteus] Length: 374\nNumber of predicted TMHs: 1\nExp number of AAs in TMHs: 21.84144\nExp number, first 60 AAs: 0.50836\nTotal prob of N-in: 0.03961\noutside 1 333\nTMhelix 334 356\ninside 357 374

28068 GCF_002206145.1_ASM220614v1 Proteus mirabilis Proteobacteria; Gammaproteobacteria; Enterobacterales; Morganellaceae; Proteus MALGNNVFSHHGINRRDFMKLCAALSATMGLSGKAAAEMSEAMASPERPPVIWIGAQECT WP_075674248.1 hydrogenase 2 small subunit [Proteus sp. H24] Length: 374\nNumber of predicted TMHs: 1\nExp number of AAs in TMHs: 21.24004\nExp number, first 60 AAs: 0.08735\nTotal prob of N-in: 0.01872\noutside 1 333\nTMhelix 334 356\ninside 357 374

28069 GCF_001463025.1_ASM146302v1 Proteus mirabilis Proteobacteria; Gammaproteobacteria; Enterobacterales; Morganellaceae; Proteus
MALGNNVFSQHGINRRDFMKLCAALSATMGLSGKAAAQMSEAMTSPERPPVIWIGAQECT WP_060557833.1
hydrogenase 2 small subunit [Proteus mirabilis] Length: 374\nNumber of predicted TMHs: 1\nExp number of AAs in TMHs: 21.96902\nExp number, first 60 AAs: 0.29882\nTotal prob of N-in: 0.03012\noutside 1 333\nTMhelix 334 356\ninside 357 374

28070 GCF_001060355.1_ASM106035v1 Proteus mirabilis Proteobacteria; Gammaproteobacteria; Enterobacterales; Morganellaceae; Proteus
MALGNNVFSQHGINRRDFMKLCAALSATMGLSGKAAAQMSQAMTSPERPPVIWIGAQECT WP_004245185.1
MULTISPECIES: hydrogenase 2 small subunit [Proteus] Length: 374\nNumber of predicted TMHs: 1\nExp number of AAs in TMHs: 21.84144\nExp number, first 60 AAs: 0.50836\nTotal prob of N-in: 0.03961\noutside 1 333\nTMhelix 334 356\ninside 357 374

28071 GCF_001653835.1_ASM165383v1 Proteus mirabilis Proteobacteria; Gammaproteobacteria; Enterobacterales; Morganellaceae; Proteus
MALGNNVFSQHGINRRDFMKLCAALSATMGLSGKAAAQMSQAMTSPERPPVIWIGAQECT WP_004245185.1
MULTISPECIES: hydrogenase 2 small subunit [Proteus] Length: 374\nNumber of predicted TMHs: 1\nExp number of AAs in TMHs: 21.84144\nExp number, first 60 AAs: 0.50836\nTotal prob of N-in: 0.03961\noutside 1 333\nTMhelix 334 356\ninside 357 374

28072 GCF_001640165.1_ASM164016v1 Proteus mirabilis Proteobacteria; Gammaproteobacteria; Enterobacterales; Morganellaceae; Proteus
MALGNNVFSQHGINRRDFMKLCAALSATMGLSGKAAAQMSQAMTSPERPPVIWIGAQECT WP_004245185.1
MULTISPECIES: hydrogenase 2 small subunit [Proteus] Length: 374\nNumber of predicted TMHs: 1\nExp number of AAs in TMHs: 21.84144\nExp number, first 60 AAs: 0.50836\nTotal prob of N-in: 0.03961\noutside 1 333\nTMhelix 334 356\ninside 357 374

28073 GCF_000783465.1_ASM78346v1 Proteus mirabilis Proteobacteria; Gammaproteobacteria; Enterobacterales; Morganellaceae; Proteus
MALGNNVFSQHGINRRDFMKLCAALSATMGLSGKAAAQMSQAMTSPERPPVIWIGAQECT WP_004245185.1
MULTISPECIES: hydrogenase 2 small subunit [Proteus] Length: 374\nNumber of predicted TMHs: 1\nExp number of AAs in TMHs: 21.84144\nExp number, first 60 AAs: 0.50836\nTotal prob of N-in: 0.03961\noutside 1 333\nTMhelix 334 356\ninside 357 374

28074 GCF_001858185.1_ASM185818v1 Proteus mirabilis Proteobacteria; Gammaproteobacteria; Enterobacterales; Morganellaceae; Proteus
MALGNNVFSQHGINRRDFMKLCAALSATMGLSGKAAAQMSQAMTSPERPPVIWIGAQECT WP_004245185.1
MULTISPECIES: hydrogenase 2 small subunit [Proteus] Length: 374\nNumber of predicted TMHs: 1\nExp number of AAs in TMHs: 21.84144\nExp number, first 60 AAs: 0.50836\nTotal prob of N-in: 0.03961\noutside 1 333\nTMhelix 334 356\ninside 357 374

28075 GCF_001553525.1_ASM155352v1 Proteus mirabilis Proteobacteria; Gammaproteobacteria; Enterobacterales; Morganellaceae; Proteus
MALGNNVFSQHGINRRDFMKLCAALSATMGLSGKAAAQMSQAMTSPERSPVIWIGAQECT WP_036908378.1
hydrogenase 2 small subunit [Proteus mirabilis] Length: 374\nNumber of predicted TMHs: 1\nExp number of AAs in TMHs: 21.81625\nExp number, first 60 AAs: 0.48316\nTotal prob of N-in: 0.03842\noutside 1 333\nTMhelix 334 356\ninside 357 374

28076 GCF_001061595.1_ASM106159v1 Proteus mirabilis Proteobacteria; Gammaproteobacteria; Enterobacterales; Morganellaceae; Proteus
MALGNNVFSQHGINRRDFMKLCAALSATMGLSGKAAAQMSQAMTSPERPPVIWIGAQECT WP_004245185.1
MULTISPECIES: hydrogenase 2 small subunit [Proteus] Length: 374\nNumber of predicted TMHs: 1\nExp number of AAs in TMHs: 21.84144\nExp number, first 60 AAs: 0.50836\nTotal prob of N-in: 0.03961\noutside 1 333\nTMhelix 334 356\ninside 357 374

28077 GCF_002180115.1_ASM218011v1 Proteus mirabilis Proteobacteria; Gammaproteobacteria; Enterobacterales; Morganellaceae; Proteus
MALGNNVFSQHGINRRDFMKLCAALSATMGLSGKAAAQMSQAMTSPERPPVIWIGAQECT WP_004245185.1
MULTISPECIES: hydrogenase 2 small subunit [Proteus] Length: 374\nNumber of predicted TMHs: 1\nExp

number of AAs in TMHs: 21.84144\nExp number, first 60 AAs: 0.50836\nTotal prob of N-in: 0.03961\noutside 1 333\nTMhelix 334 356\ninside 357 374

28078 GCF_002197405.1_ASM219740v1 Proteus mirabilis Proteobacteria; Gammaproteobacteria; Enterobacterales; Morganellaceae; Proteus
MALGNNVFSQHGINRRDFMKLCAALSATMGLSGKAAAQMSQAMTSPERPPVIWIGAQECT WP_004245185.1
MULTISPECIES: hydrogenase 2 small subunit [Proteus] Length: 374\nNumber of predicted TMHs: 1\nExp number of AAs in TMHs: 21.84144\nExp number, first 60 AAs: 0.50836\nTotal prob of N-in: 0.03961\noutside 1 333\nTMhelix 334 356\ninside 357 374

28079 GCF_001062655.1_ASM106265v1 Proteus mirabilis Proteobacteria; Gammaproteobacteria; Enterobacterales; Morganellaceae; Proteus
MALGNNVFSQHGINRRDFMKLCAALSATMGLSGKAAAQMSQAMTSPERPPVIWIGAQECT WP_004245185.1
MULTISPECIES: hydrogenase 2 small subunit [Proteus] Length: 374\nNumber of predicted TMHs: 1\nExp number of AAs in TMHs: 21.84144\nExp number, first 60 AAs: 0.50836\nTotal prob of N-in: 0.03961\noutside 1 333\nTMhelix 334 356\ninside 357 374

28080 GCF_001062385.1_ASM106238v1 Proteus mirabilis Proteobacteria; Gammaproteobacteria; Enterobacterales; Morganellaceae; Proteus
MALGNNVFSQHGINRRDFMKLCAALSATMGLSGKAAAQMSQAMTSPERPPVIWIGAQECT WP_004245185.1
MULTISPECIES: hydrogenase 2 small subunit [Proteus] Length: 374\nNumber of predicted TMHs: 1\nExp number of AAs in TMHs: 21.84144\nExp number, first 60 AAs: 0.50836\nTotal prob of N-in: 0.03961\noutside 1 333\nTMhelix 334 356\ninside 357 374

28081 GCF_001060765.1_ASM106076v1 Proteus mirabilis Proteobacteria; Gammaproteobacteria; Enterobacterales; Morganellaceae; Proteus
MALGNNVFSQHGINRRDFMKLCAALSATMGLSGKAAAQMSQAMTSPERPPVIWIGAQECT WP_004245185.1
MULTISPECIES: hydrogenase 2 small subunit [Proteus] Length: 374\nNumber of predicted TMHs: 1\nExp number of AAs in TMHs: 21.84144\nExp number, first 60 AAs: 0.50836\nTotal prob of N-in: 0.03961\noutside 1 333\nTMhelix 334 356\ninside 357 374

28082 GCF_000783575.1_ASM78357v1 Proteus mirabilis Proteobacteria; Gammaproteobacteria; Enterobacterales; Morganellaceae; Proteus
MALGNNVFSQHGINRRDFMKLCAALSATMGLSGKAAAQMSQAMTSPERPPVIWIGAQECT WP_004245185.1
MULTISPECIES: hydrogenase 2 small subunit [Proteus] Length: 374\nNumber of predicted TMHs: 1\nExp number of AAs in TMHs: 21.84144\nExp number, first 60 AAs: 0.50836\nTotal prob of N-in: 0.03961\noutside 1 333\nTMhelix 334 356\ninside 357 374

28083 GCF_000783875.1_ASM78387v1 Proteus mirabilis Proteobacteria; Gammaproteobacteria; Enterobacterales; Morganellaceae; Proteus
MALGNNVFSQHGINRRDFMKLCAALSATMGLSGKAAAQMSQAMTSPERPPVIWIGAQECT WP_004245185.1
MULTISPECIES: hydrogenase 2 small subunit [Proteus] Length: 374\nNumber of predicted TMHs: 1\nExp number of AAs in TMHs: 21.84144\nExp number, first 60 AAs: 0.50836\nTotal prob of N-in: 0.03961\noutside 1 333\nTMhelix 334 356\ninside 357 374

28084 GCF_001640985.1_ASM164098v1 Proteus mirabilis Proteobacteria; Gammaproteobacteria; Enterobacterales; Morganellaceae; Proteus
MALGNNVFSQHGINRRDFMKLCAALSATMGLSGKAAAQMSQAMTSPERPPVIWIGAQECT WP_004245185.1
MULTISPECIES: hydrogenase 2 small subunit [Proteus] Length: 374\nNumber of predicted TMHs: 1\nExp number of AAs in TMHs: 21.84144\nExp number, first 60 AAs: 0.50836\nTotal prob of N-in: 0.03961\noutside 1 333\nTMhelix 334 356\ninside 357 374

28085 GCF_001063205.1_ASM106320v1 Proteus mirabilis Proteobacteria; Gammaproteobacteria; Enterobacterales; Morganellaceae; Proteus
MALGNNVFSQHGINRRDFMKLCAALSATMGLSGKAAAQMSQAMTSPERPPVIWIGAQECT WP_004245185.1
MULTISPECIES: hydrogenase 2 small subunit [Proteus] Length: 374\nNumber of predicted TMHs: 1\nExp number of AAs in TMHs: 21.84144\nExp number, first 60 AAs: 0.50836\nTotal prob of N-in: 0.03961\noutside 1 333\nTMhelix 334 356\ninside 357 374

28086 GCF_001062745.1_ASM106274v1 Proteus mirabilis Proteobacteria; Gammaproteobacteria; Enterobacterales; Morganellaceae; Proteus

MALGNNVFSQHGINRRDFMKLCAALSATMGLSGKAAAQMSQAMTSPERPPVIWIGAQECT WP_004245185.1 MULTISPECIES: hydrogenase 2 small subunit [Proteus] Length: 374\nNumber of predicted TMHs: 1\nExp number of AAs in TMHs: 21.84144\nExp number, first 60 AAs: 0.50836\nTotal prob of N-in: 0.03961\noutside 1 333\nTMhelix 334 356\ninside 357 374

28087 GCF_001653845.1_ASM165384v1 Proteus mirabilis Proteobacteria; Gammaproteobacteria; Enterobacterales; Morganellaceae; Proteus MALGNNVFSQHGINRRDFMKLCAALSATMGLSGKAAAQMSQAMTSPERPPVIWIGAQECT WP_004245185.1 MULTISPECIES: hydrogenase 2 small subunit [Proteus] Length: 374\nNumber of predicted TMHs: 1\nExp number of AAs in TMHs: 21.84144\nExp number, first 60 AAs: 0.50836\nTotal prob of N-in: 0.03961\noutside 1 333\nTMhelix 334 356\ninside 357 374

28088 GCF_001643755.1_ASM164375v1 Proteus mirabilis Proteobacteria; Gammaproteobacteria; Enterobacterales; Morganellaceae; Proteus MALGNNVFSQHGINRRDFMKLCAALSATMGLSGKAAAQMSQAMTSPERPPVIWIGAQECT WP_004245185.1 MULTISPECIES: hydrogenase 2 small subunit [Proteus] Length: 374\nNumber of predicted TMHs: 1\nExp number of AAs in TMHs: 21.84144\nExp number, first 60 AAs: 0.50836\nTotal prob of N-in: 0.03961\noutside 1 333\nTMhelix 334 356\ninside 357 374

28089 GCF_002055685.1_ASM205568v1 Proteus mirabilis Proteobacteria; Gammaproteobacteria; Enterobacterales; Morganellaceae; Proteus MALGNNVFSQHGINRRDFMKLCAALSATMGLSGKAAAQMSQAMTSPERPPVIWIGAQECT WP_004245185.1 MULTISPECIES: hydrogenase 2 small subunit [Proteus] Length: 374\nNumber of predicted TMHs: 1\nExp number of AAs in TMHs: 21.84144\nExp number, first 60 AAs: 0.50836\nTotal prob of N-in: 0.03961\noutside 1 333\nTMhelix 334 356\ninside 357 374

28090 GCF_001448445.1_ASM144844v1 Proteus mirabilis Proteobacteria; Gammaproteobacteria; Enterobacterales; Morganellaceae; Proteus MALGNNVFSQHGINRRDFMKLCAALSATMGLSGKAAAQMSQAMTSPERPPVIWIGAQECT WP_004245185.1 MULTISPECIES: hydrogenase 2 small subunit [Proteus] Length: 374\nNumber of predicted TMHs: 1\nExp number of AAs in TMHs: 21.84144\nExp number, first 60 AAs: 0.50836\nTotal prob of N-in: 0.03961\noutside 1 333\nTMhelix 334 356\ninside 357 374

28091 GCF_000783595.1_ASM78359v1 Proteus mirabilis Proteobacteria; Gammaproteobacteria; Enterobacterales; Morganellaceae; Proteus MALGNNVFSQHGINRRDFMKLCAALSATMGLSGKAAAQMSQAMTSPERPPVIWIGAQECT WP_004245185.1 MULTISPECIES: hydrogenase 2 small subunit [Proteus] Length: 374\nNumber of predicted TMHs: 1\nExp number of AAs in TMHs: 21.84144\nExp number, first 60 AAs: 0.50836\nTotal prob of N-in: 0.03961\noutside 1 333\nTMhelix 334 356\ninside 357 374

28092 GCF_001066855.1_ASM106685v1 Proteus mirabilis Proteobacteria; Gammaproteobacteria; Enterobacterales; Morganellaceae; Proteus MALGNNVFSQHGINRRDFMKLCAALSATMGLSGKAAAQMSQAMTSPERPPVIWIGAQECT WP_004245185.1 MULTISPECIES: hydrogenase 2 small subunit [Proteus] Length: 374\nNumber of predicted TMHs: 1\nExp number of AAs in TMHs: 21.84144\nExp number, first 60 AAs: 0.50836\nTotal prob of N-in: 0.03961\noutside 1 333\nTMhelix 334 356\ninside 357 374

28093 GCF_001063035.1_ASM106303v1 Proteus mirabilis Proteobacteria; Gammaproteobacteria; Enterobacterales; Morganellaceae; Proteus MALGNNVFSQHGINRRDFMKLCAALSATMGLSGKAAAQMSQAMTSPERPPVIWIGAQECT WP_004245185.1 MULTISPECIES: hydrogenase 2 small subunit [Proteus] Length: 374\nNumber of predicted TMHs: 1\nExp number of AAs in TMHs: 21.84144\nExp number, first 60 AAs: 0.50836\nTotal prob of N-in: 0.03961\noutside 1 333\nTMhelix 334 356\ninside 357 374

28094 GCF_900101725.1_IMG-taxon_2654588119_annotated_assembly Proteus mirabilis Proteobacteria; Gammaproteobacteria; Enterobacterales; Morganellaceae; Proteus MALGNNVFSQHGINRRDFMKLCAALSATMGLSGKAAAQMSQAMTSPERPPVIWIGAQECT WP_004245185.1 MULTISPECIES: hydrogenase 2 small subunit [Proteus] Length: 374\nNumber of predicted TMHs: 1\nExp number of AAs in TMHs: 21.84144\nExp number, first 60 AAs: 0.50836\nTotal prob of N-in: 0.03961\noutside 1 333\nTMhelix 334 356\ninside 357 374

28095 GCF_001604705.1_ASM160470v1 Proteus mirabilis Proteobacteria; Gammaproteobacteria; Enterobacterales; Morganellaceae; Proteus
MALGNNVFSQHGINRRDFMKLCAALSATMGLSGKAAAQMSQAMTSPERPPVIWIGAQECT WP_004245185.1
MULTISPECIES: hydrogenase 2 small subunit [Proteus] Length: 374\nNumber of predicted TMHs: 1\nExp number of AAs in TMHs: 21.84144\nExp number, first 60 AAs: 0.50836\nTotal prob of N-in: 0.03961\noutside 1 333\nTMhelix 334 356\ninside 357 374

28096 GCF_001617305.1_ASM161730v1 Proteus mirabilis Proteobacteria; Gammaproteobacteria; Enterobacterales; Morganellaceae; Proteus
MALGNNVFSQHGINRRDFMKLCAALSATMGLSGKAAAQMSQAMTSPERPPVIWIGAQECT WP_004245185.1
MULTISPECIES: hydrogenase 2 small subunit [Proteus] Length: 374\nNumber of predicted TMHs: 1\nExp number of AAs in TMHs: 21.84144\nExp number, first 60 AAs: 0.50836\nTotal prob of N-in: 0.03961\noutside 1 333\nTMhelix 334 356\ninside 357 374

28097 GCF_002180235.1_ASM218023v1 Proteus mirabilis Proteobacteria; Gammaproteobacteria; Enterobacterales; Morganellaceae; Proteus
MALGNNVFSQHGINRRDFMKLCAALSATMGLSGKAAAQMSQAMTSPERPPVIWIGAQECT WP_004245185.1
MULTISPECIES: hydrogenase 2 small subunit [Proteus] Length: 374\nNumber of predicted TMHs: 1\nExp number of AAs in TMHs: 21.84144\nExp number, first 60 AAs: 0.50836\nTotal prob of N-in: 0.03961\noutside 1 333\nTMhelix 334 356\ninside 357 374

28098 GCF_900113495.1_IMG-taxon_2623620510_annotated_assembly Proteus mirabilis Proteobacteria; Gammaproteobacteria; Enterobacterales; Morganellaceae; Proteus
MALGNNVFSQHGINRRDFMKLCAALSATMGLSGKAAAQMSQAMTSPERPPVIWIGAQECT WP_004245185.1
MULTISPECIES: hydrogenase 2 small subunit [Proteus] Length: 374\nNumber of predicted TMHs: 1\nExp number of AAs in TMHs: 21.84144\nExp number, first 60 AAs: 0.50836\nTotal prob of N-in: 0.03961\noutside 1 333\nTMhelix 334 356\ninside 357 374

28099 GCF_001281545.1_ASM128154v1 Proteus mirabilis Proteobacteria; Gammaproteobacteria; Enterobacterales; Morganellaceae; Proteus
MALGNNVFSQHGINRRDFMKLCAALSATMGLSGKAAAQMSQAMTSPERPPVIWIGAQECT WP_004245185.1
MULTISPECIES: hydrogenase 2 small subunit [Proteus] Length: 374\nNumber of predicted TMHs: 1\nExp number of AAs in TMHs: 21.84144\nExp number, first 60 AAs: 0.50836\nTotal prob of N-in: 0.03961\noutside 1 333\nTMhelix 334 356\ninside 357 374

28100 GCF_001076755.1_ASM107675v1 Proteus mirabilis Proteobacteria; Gammaproteobacteria; Enterobacterales; Morganellaceae; Proteus
MALGNNVFSQHGINRRDFMKLCAALSATMGLSGKAAAQMSQAMTSPERPPVIWIGAQECT WP_004245185.1
MULTISPECIES: hydrogenase 2 small subunit [Proteus] Length: 374\nNumber of predicted TMHs: 1\nExp number of AAs in TMHs: 21.84144\nExp number, first 60 AAs: 0.50836\nTotal prob of N-in: 0.03961\noutside 1 333\nTMhelix 334 356\ninside 357 374

28101 GCF_001076715.1_ASM107671v1 Proteus mirabilis Proteobacteria; Gammaproteobacteria; Enterobacterales; Morganellaceae; Proteus
MALGNNVFSQHGINRRDFMKLCAALSATMGLSGKAAAQMSQAMTSPERPPVIWIGAQECT WP_004245185.1
MULTISPECIES: hydrogenase 2 small subunit [Proteus] Length: 374\nNumber of predicted TMHs: 1\nExp number of AAs in TMHs: 21.84144\nExp number, first 60 AAs: 0.50836\nTotal prob of N-in: 0.03961\noutside 1 333\nTMhelix 334 356\ninside 357 374

28102 GCF_000784015.1_ASM78401v1 Proteus mirabilis Proteobacteria; Gammaproteobacteria; Enterobacterales; Morganellaceae; Proteus
MALGNNVFSQHGINRRDFMKLCAALSATMGLSGKAAAQMSQAMTSPERSPVIWIGAQECT WP_036908378.1
hydrogenase 2 small subunit [Proteus mirabilis] Length: 374\nNumber of predicted TMHs: 1\nExp number of AAs in TMHs: 21.81625\nExp number, first 60 AAs: 0.48316\nTotal prob of N-in: 0.03842\noutside 1 333\nTMhelix 334 356\ninside 357 374

28103 GCF_001065915.1_ASM106591v1 Proteus mirabilis Proteobacteria; Gammaproteobacteria; Enterobacterales; Morganellaceae; Proteus
MALGNNVFSQHGINRRDFMKLCAALSATMGLSGKAAAQMSQAMTSPERPPVIWIGAQECT WP_004245185.1
MULTISPECIES: hydrogenase 2 small subunit [Proteus] Length: 374\nNumber of predicted TMHs: 1\nExp

number of AAs in TMHs: 21.84144\nExp number, first 60 AAs: 0.50836\nTotal prob of N-in: 0.03961\noutside 1 333\nTMhelix 334 356\ninside 357 374

28104 GCF_001065085.1_ASM106508v1 Proteus mirabilis Proteobacteria; Gammaproteobacteria; Enterobacterales; Morganellaceae; Proteus
MALGNNVFSQHGINRRDFMKLCAALSATMGLSGKAAAQMSQAMTSPERPPVIWIGAQECT WP_004245185.1
MULTISPECIES: hydrogenase 2 small subunit [Proteus] Length: 374\nNumber of predicted TMHs: 1\nExp number of AAs in TMHs: 21.84144\nExp number, first 60 AAs: 0.50836\nTotal prob of N-in: 0.03961\noutside 1 333\nTMhelix 334 356\ninside 357 374

28105 GCF_001063575.1_ASM106357v1 Proteus mirabilis Proteobacteria; Gammaproteobacteria; Enterobacterales; Morganellaceae; Proteus
MALGNNVFSQHGINRRDFMKLCAALSATMGLSGKAAAQMSQAMTSPERPPVIWIGAQECT WP_004245185.1
MULTISPECIES: hydrogenase 2 small subunit [Proteus] Length: 374\nNumber of predicted TMHs: 1\nExp number of AAs in TMHs: 21.84144\nExp number, first 60 AAs: 0.50836\nTotal prob of N-in: 0.03961\noutside 1 333\nTMhelix 334 356\ninside 357 374

28106 GCF_001064105.1_ASM106410v1 Proteus mirabilis Proteobacteria; Gammaproteobacteria; Enterobacterales; Morganellaceae; Proteus
MALGNNVFSQHGINRRDFMKLCAALSATMGLSGKAAAQMSQAMTSPERPPVIWIGAQECT WP_004245185.1
MULTISPECIES: hydrogenase 2 small subunit [Proteus] Length: 374\nNumber of predicted TMHs: 1\nExp number of AAs in TMHs: 21.84144\nExp number, first 60 AAs: 0.50836\nTotal prob of N-in: 0.03961\noutside 1 333\nTMhelix 334 356\ninside 357 374

28107 GCF_000755485.1_ASM75548v1 Proteus mirabilis Proteobacteria; Gammaproteobacteria; Enterobacterales; Morganellaceae; Proteus
MALGNNVFSQHGINRRDFMKLCAALSATMGLSGKAAAQMSQAMTSPERSPVIWIGAQECT WP_036908378.1
hydrogenase 2 small subunit [Proteus mirabilis] Length: 374\nNumber of predicted TMHs: 1\nExp number of AAs in TMHs: 21.81625\nExp number, first 60 AAs: 0.48316\nTotal prob of N-in: 0.03842\noutside 1 333\nTMhelix 334 356\ninside 357 374

28108 GCF_001462385.1_ASM146238v1 Proteus mirabilis Proteobacteria; Gammaproteobacteria; Enterobacterales; Morganellaceae; Proteus
MALGNNVFSQHGINRRDFMKLCAALSATMGLSGKAAAQMSQAMTSPERPPVIWIGAQECT WP_004245185.1
MULTISPECIES: hydrogenase 2 small subunit [Proteus] Length: 374\nNumber of predicted TMHs: 1\nExp number of AAs in TMHs: 21.84144\nExp number, first 60 AAs: 0.50836\nTotal prob of N-in: 0.03961\noutside 1 333\nTMhelix 334 356\ninside 357 374

28109 GCF_001060285.1_ASM106028v1 Proteus mirabilis Proteobacteria; Gammaproteobacteria; Enterobacterales; Morganellaceae; Proteus
MALGNNVFSQHGINRRDFMKLCAALSATMGLSGKAAAQMSQAMTSPERSPVIWIGAQECT WP_036908378.1
hydrogenase 2 small subunit [Proteus mirabilis] Length: 374\nNumber of predicted TMHs: 1\nExp number of AAs in TMHs: 21.81625\nExp number, first 60 AAs: 0.48316\nTotal prob of N-in: 0.03842\noutside 1 333\nTMhelix 334 356\ninside 357 374

28110 GCF_001462475.1_ASM146247v1 Proteus mirabilis Proteobacteria; Gammaproteobacteria; Enterobacterales; Morganellaceae; Proteus
MALGNNVFSQHGINRRDFMKLCAALSATMGLSGKAAAQMSQAMTSPERPPVIWIGAQECT WP_004245185.1
MULTISPECIES: hydrogenase 2 small subunit [Proteus] Length: 374\nNumber of predicted TMHs: 1\nExp number of AAs in TMHs: 21.84144\nExp number, first 60 AAs: 0.50836\nTotal prob of N-in: 0.03961\noutside 1 333\nTMhelix 334 356\ninside 357 374

28111 GCF_001076695.1_ASM107669v1 Proteus mirabilis Proteobacteria; Gammaproteobacteria; Enterobacterales; Morganellaceae; Proteus
MALGNNVFSQHGINRRDFMKLCAALSATMGLSGKAAAQMLQAMTSPERPPVIWIGAQECT WP_049221622.1
hydrogenase 2 small subunit [Proteus mirabilis] Length: 374\nNumber of predicted TMHs: 1\nExp number of AAs in TMHs: 23.04552\nExp number, first 60 AAs: 1.71241\nTotal prob of N-in: 0.09538\noutside 1 333\nTMhelix 334 356\ninside 357 374

28112 GCF_001939795.1_ASM193979v1 Proteus sp. H24 Proteobacteria; Gammaproteobacteria; Enterobacterales; Morganellaceae; Proteus

MALGNNVFSHHGINRRDFMKLCAALSATMGLSGKAAAEMSEAMASPERPPVIWIGAQECT WP_075674248.1 hydrogenase 2 small subunit [Proteus sp. H24] Length: 374\nNumber of predicted TMHs: 1\nExp number of AAs in TMHs: 21.24004\nExp number, first 60 AAs: 0.08735\nTotal prob of N-in: 0.01872\noutside 1 333\nTMhelix 334 356\ninside 357 374

28113 GCF_001808035.1_ASM180803v1 Proteus sp. HMSC14B05 Proteobacteria; Gammaproteobacteria; Enterobacterales; Morganellaceae; Proteus MALGNNVFSQHGINRRDFMKLCAALSATMGLSGKAAAQMSQAMTSPERPPVIWIGAQECT WP_004245185.1 MULTISPECIES: hydrogenase 2 small subunit [Proteus] Length: 374\nNumber of predicted TMHs: 1\nExp number of AAs in TMHs: 21.84144\nExp number, first 60 AAs: 0.50836\nTotal prob of N-in: 0.03961\noutside 1 333\nTMhelix 334 356\ninside 357 374

28114 GCF_001281565.1_ASM128156v1 Proteus vulgaris Proteobacteria; Gammaproteobacteria; Enterobacterales; Morganellaceae; Proteus MALGNNVFSQHGINRRDFMKLCAALSATMGLSGKAAAQMSQAMTSPERPPVIWIGAQECT WP_004245185.1 MULTISPECIES: hydrogenase 2 small subunit [Proteus] Length: 374\nNumber of predicted TMHs: 1\nExp number of AAs in TMHs: 21.84144\nExp number, first 60 AAs: 0.50836\nTotal prob of N-in: 0.03961\noutside 1 333\nTMhelix 334 356\ninside 357 374

28115 GCF_000754995.1_PVA Proteus vulgaris Proteobacteria; Gammaproteobacteria; Enterobacterales; Morganellaceae; Proteus MALGNNVFSHHGINRRDFMKLCAALSATMGLSGKAAAEMSEAMVSTERPPVIWIGAQECT WP_036932958.1 hydrogenase 2 small subunit [Proteus vulgaris] Length: 374\nNumber of predicted TMHs: 1\nExp number of AAs in TMHs: 21.23761\nExp number, first 60 AAs: 0.07839\nTotal prob of N-in: 0.01866\noutside 1 333\nTMhelix 334 356\ninside 357 374

28116 GCF_001049975.1_PRJEB9439_assembly_1 Proteus vulgaris Proteobacteria; Gammaproteobacteria; Enterobacterales; Morganellaceae; Proteus MALGNNVFSHHGINRRDFMKLCAALSATMGLSGKAAAEMSEAMASSERPPVIWIGAQECT WP_072069443.1 hydrogenase 2 small subunit [Proteus vulgaris] Length: 374\nNumber of predicted TMHs: 1\nExp number of AAs in TMHs: 21.23488\nExp number, first 60 AAs: 0.08218\nTotal prob of N-in: 0.01845\noutside 1 333\nTMhelix 334 356\ninside 357 374

28117 GCF_001049955.1_PRJEB9438_assembly_1 Proteus vulgaris Proteobacteria; Gammaproteobacteria; Enterobacterales; Morganellaceae; Proteus MALGNNVFSHHGINRRDFMKLCAALSATMGLSGKAAAEMSEAMVSPERPPVIWIGAQECT WP_072063712.1 hydrogenase 2 small subunit [Proteus vulgaris] Length: 374\nNumber of predicted TMHs: 1\nExp number of AAs in TMHs: 21.23569\nExp number, first 60 AAs: 0.083\nTotal prob of N-in: 0.01850\noutside 1 333\nTMhelix 334 356\ninside 357 374

28118 GCF_001654965.1_Pha700826_DRAFTv1 Proteus hauseri ATCC 700826Proteobacteria; Gammaproteobacteria; Enterobacterales; Morganellaceae; Proteus; Proteus hauseri MALGNNVFSHHGINRRDFMKLCAALSATMGLSGKAAAEMSEAMASSERPPVIWIGAQECT WP_064718971.1 hydrogenase 2 small subunit [Proteus hauseri] Length: 375\nNumber of predicted TMHs: 1\nExp number of AAs in TMHs: 21.15373\nExp number, first 60 AAs: 0.08208\nTotal prob of N-in: 0.01907\noutside 1 333\nTMhelix 334 356\ninside 357 375

28119 GCF_000497855.1_version_3_of_Proteus_hauseri_genome Proteus hauseri ZMd44 Proteobacteria; Gammaproteobacteria; Enterobacterales; Morganellaceae; Proteus; Proteus hauseri MALGNNVFSHHGINRRDFMKLCAALSATMGLSGKAAAEMSEAMASPERPPVIWIGAQECT WP_023580919.1 hydrogenase 2 small subunit [Proteus hauseri] Length: 374\nNumber of predicted TMHs: 1\nExp number of AAs in TMHs: 21.24006\nExp number, first 60 AAs: 0.08735\nTotal prob of N-in: 0.01872\noutside 1 333\nTMhelix 334 356\ninside 357 374

28120 GCF_000160755.1_ASM16075v1 Proteus mirabilis ATCC 29906Proteobacteria; Gammaproteobacteria; Enterobacterales; Morganellaceae; Proteus; Proteus mirabilis MALGNNVFSQHGINRRDFMKLCAALSATMGLSGKAAAQMSQAMTSPERPPVIWIGAQECT WP_004245185.1 MULTISPECIES: hydrogenase 2 small subunit [Proteus] Length: 374\nNumber of predicted TMHs: 1\nExp number of AAs in TMHs: 21.84144\nExp number, first 60 AAs: 0.50836\nTotal prob of N-in: 0.03961\noutside 1 333\nTMhelix 334 356\ninside 357 374

28121 GCF_000444425.1_ASM44442v1 Proteus mirabilis BB2000 Proteobacteria; Gammaproteobacteria; Enterobacterales; Morganellaceae; Proteus; Proteus mirabilis
MALGNNVFSQHGINRRDFMKLCAALSATMGLSGKAAAQMSQAMTSPERPPVIWIGAQECT WP_004245185.1
MULTISPECIES: hydrogenase 2 small subunit [Proteus] Length: 374\nNumber of predicted TMHs: 1\nExp number of AAs in TMHs: 21.84144\nExp number, first 60 AAs: 0.50836\nTotal prob of N-in: 0.03961\noutside 1 333\nTMhelix 334 356\ninside 357 374

28122 GCF_000313255.1_Version_1_for_Proteus_mirabilis_C05028_genome Proteus mirabilis C05028
Proteobacteria; Gammaproteobacteria; Enterobacterales; Morganellaceae; Proteus; Proteus mirabilis
MALGNNVFSQHGINRRDFMKLCAALSATMGLSGKAAAQMSQAMTSPERPPVIWIGAQECT WP_004245185.1
MULTISPECIES: hydrogenase 2 small subunit [Proteus] Length: 374\nNumber of predicted TMHs: 1\nExp number of AAs in TMHs: 21.84144\nExp number, first 60 AAs: 0.50836\nTotal prob of N-in: 0.03961\noutside 1 333\nTMhelix 334 356\ninside 357 374

28123 GCF_000069965.1_ASM6996v1 Proteus mirabilis HI4320 Proteobacteria; Gammaproteobacteria; Enterobacterales; Morganellaceae; Proteus; Proteus mirabilis
MALGNNVFSQHGINRRDFMKLCAALSATMGLSGKAAAQMSQAMTSPERPPVIWIGAQECT WP_004245185.1
MULTISPECIES: hydrogenase 2 small subunit [Proteus] Length: 374\nNumber of predicted TMHs: 1\nExp number of AAs in TMHs: 21.84144\nExp number, first 60 AAs: 0.50836\nTotal prob of N-in: 0.03961\noutside 1 333\nTMhelix 334 356\ninside 357 374

28124 GCF_000372565.1_PM_version_1 Proteus mirabilis PR03 Proteobacteria; Gammaproteobacteria; Enterobacterales; Morganellaceae; Proteus; Proteus mirabilis
MALGNNVFSQHGINRRDFMKLCAALSATMGLSGKAAAQMSQAMTSPERPPVIWIGAQECT WP_004245185.1
MULTISPECIES: hydrogenase 2 small subunit [Proteus] Length: 374\nNumber of predicted TMHs: 1\nExp number of AAs in TMHs: 21.84144\nExp number, first 60 AAs: 0.50836\nTotal prob of N-in: 0.03961\noutside 1 333\nTMhelix 334 356\ninside 357 374

28125 GCF_000297835.1_Prot_mira_WGLW4_V1 Proteus mirabilis WGLW4 Proteobacteria; Gammaproteobacteria; Enterobacterales; Morganellaceae; Proteus; Proteus mirabilis
MALGNNVFSQHGINRRDFMKLCAALSATMGLSGKAAAQMSQAMTSPERPPVIWIGAQECT WP_004245185.1
MULTISPECIES: hydrogenase 2 small subunit [Proteus] Length: 374\nNumber of predicted TMHs: 1\nExp number of AAs in TMHs: 21.84144\nExp number, first 60 AAs: 0.50836\nTotal prob of N-in: 0.03961\noutside 1 333\nTMhelix 334 356\ninside 357 374

28126 GCF_000297815.1_Prot_mira_WGLW6_V1 Proteus mirabilis WGLW6 Proteobacteria; Gammaproteobacteria; Enterobacterales; Morganellaceae; Proteus; Proteus mirabilis
MALGNNVFSQHGINRRDFMKLCAALSATMGLSGKAAAQMSQAMTSPERPPVIWIGAQECT WP_004245185.1
MULTISPECIES: hydrogenase 2 small subunit [Proteus] Length: 374\nNumber of predicted TMHs: 1\nExp number of AAs in TMHs: 21.84144\nExp number, first 60 AAs: 0.50836\nTotal prob of N-in: 0.03961\noutside 1 333\nTMhelix 334 356\ninside 357 374

28127 GCF_900061445.1_Providencia_heimbachae Providencia heimbachae Proteobacteria; Gammaproteobacteria; Enterobacterales; Morganellaceae; Providencia
MAGDNSIISPHGVNRRDFMKLCTALAATMGLSSNAAAQMADAVSDPGRPPVIWIGAQECT WP_068440982.1
hydrogenase 2 small subunit [Providencia heimbachae] Length: 372\nNumber of predicted TMHs: 1\nExp number of AAs in TMHs: 22.20978\nExp number, first 60 AAs: 0.6395\nTotal prob of N-in: 0.08809\noutside 1 332\nTMhelix 333 355\ninside 356 372

28128 GCF_000805715.1_Providencia_rettgeri_CCBH11880_NDM-1 Providencia rettgeri
Proteobacteria; Gammaproteobacteria; Enterobacterales; Morganellaceae; Providencia
MIGDNSIISSHGINRRDFMKLCTALAATMGLSANAAAQIAESISDPSRPPVIWIGAQECT WP_042846126.1 hydrogenase 2 small subunit [Providencia rettgeri] Length: 372\nNumber of predicted TMHs: 1\nExp number of AAs in TMHs: 23.57993\nExp number, first 60 AAs: 2.65908\nTotal prob of N-in: 0.14478\noutside 1 332\nTMhelix 333 355\ninside 356 372

28129 GCF_001874625.1_ASM187462v1 Providencia rettgeri Proteobacteria; Gammaproteobacteria; Enterobacterales; Morganellaceae; Providencia
MIGDNSIISSHGINRRDFMKLCTALAATMGLSANAAAQIAESISDPSRPPVIWIGAQECT WP_042846126.1 hydrogenase 2 small subunit [Providencia rettgeri] Length: 372\nNumber of predicted TMHs: 1\nExp number of AAs in TMHs:

23.57993\nExp number, first 60 AAs: 2.65908\nTotal prob of N-in: 0.14478\noutside 1 332\nTMhelix 333 355\ninside 356 372

28130 GCF_001049815.1_Providencia_rettgeri_H1736 Providencia rettgeri Proteobacteria; Gammaproteobacteria; Enterobacterales; Morganellaceae; Providencia MIGDNSIISSHGINRRDFMKLCTALAATMGLSANAAAQIAESISDPSRPPVIWIGAQECT WP_042846126.1 hydrogenase 2 small subunit [Providencia rettgeri] Length: 372\nNumber of predicted TMHs: 1\nExp number of AAs in TMHs: 23.57993\nExp number, first 60 AAs: 2.65908\nTotal prob of N-in: 0.14478\noutside 1 332\nTMhelix 333 355\ninside 356 372

28131 GCF_000527335.1_Pal20592v1.0 Providencia alcalifaciens 205/92 Proteobacteria; Gammaproteobacteria; Enterobacterales; Morganellaceae; Providencia; Providencia alcalifaciens MAGDNTIHSPHGINRRDFMKLCTALAATMGLSGKAAAQMADAVSDPERPPVIWIGAQECT WP_006662936.1 hydrogenase 2 small subunit [Providencia alcalifaciens] Length: 372\nNumber of predicted TMHs: 1\nExp number of AAs in TMHs: 23.09916\nExp number, first 60 AAs: 0.49604\nTotal prob of N-in: 0.06612\noutside 1 332\nTMhelix 333 355\ninside 356 372

28132 GCF_000314875.2_ASM31487v2 Providencia alcalifaciens Dmel2 Proteobacteria; Gammaproteobacteria; Enterobacterales; Morganellaceae; Providencia; Providencia alcalifaciens MAGDNTIHSPHGINRRDFMKLCTALAATMGLSGKAAAQMADAVSDPERPPVIWIGAQECT WP_006662936.1 hydrogenase 2 small subunit [Providencia alcalifaciens] Length: 372\nNumber of predicted TMHs: 1\nExp number of AAs in TMHs: 23.09916\nExp number, first 60 AAs: 0.49604\nTotal prob of N-in: 0.06612\noutside 1 332\nTMhelix 333 355\ninside 356 372

28133 GCF_000173415.1_ASM17341v1 Providencia alcalifaciens DSM 30120 Proteobacteria; Gammaproteobacteria; Enterobacterales; Morganellaceae; Providencia; Providencia alcalifaciens MAGDNTIHSPHGINRRDFMKLCTALAATMGLSGKAAAQMADAVSDPERPPVIWIGAQECT WP_006659159.1 hydrogenase 2 small subunit [Providencia alcalifaciens] Length: 372\nNumber of predicted TMHs: 1\nExp number of AAs in TMHs: 23.09904\nExp number, first 60 AAs: 0.49604\nTotal prob of N-in: 0.06612\noutside 1 332\nTMhelix 333 355\ninside 356 372

28134 GCF_000517805.1_PalF902004v1.0 Providencia alcalifaciens F90-2004 Proteobacteria; Gammaproteobacteria; Enterobacterales; Morganellaceae; Providencia; Providencia alcalifaciens MAGDNTIHSPHGINRRDFMKLCTALAATMGLSGKAAAQMADAVSDPERPPVIWIGAQECT WP_006662936.1 hydrogenase 2 small subunit [Providencia alcalifaciens] Length: 372\nNumber of predicted TMHs: 1\nExp number of AAs in TMHs: 23.09916\nExp number, first 60 AAs: 0.49604\nTotal prob of N-in: 0.06612\noutside 1 332\nTMhelix 333 355\ninside 356 372

28135 GCF_000527275.1_PalPAL1v1.0 Providencia alcalifaciens PAL-1 Proteobacteria; Gammaproteobacteria; Enterobacterales; Morganellaceae; Providencia; Providencia alcalifaciens MAGDNSILSPHGINRRDFMKLCTALAATMGLSGKAAAQMADAVSDPERPPVIWIGAQECT WP_036956217.1 hydrogenase 2 small subunit [Providencia alcalifaciens] Length: 372\nNumber of predicted TMHs: 1\nExp number of AAs in TMHs: 22.62342\nExp number, first 60 AAs: 0.18757\nTotal prob of N-in: 0.02923\noutside 1 332\nTMhelix 333 355\ninside 356 372

28136 GCF_000527255.1_PalPAL2v1.0 Providencia alcalifaciens PAL-2 Proteobacteria; Gammaproteobacteria; Enterobacterales; Morganellaceae; Providencia; Providencia alcalifaciens MAGDNTIHSPHGINRRDFMKLCTALAATMGLSGKAAAQMADAVSDPERPPVIWIGAQECT WP_006662936.1 hydrogenase 2 small subunit [Providencia alcalifaciens] Length: 372\nNumber of predicted TMHs: 1\nExp number of AAs in TMHs: 23.09916\nExp number, first 60 AAs: 0.49604\nTotal prob of N-in: 0.06612\noutside 1 332\nTMhelix 333 355\ninside 356 372

28137 GCF_000517745.1_PalPAL3v1.0 Providencia alcalifaciens PAL-3 Proteobacteria; Gammaproteobacteria; Enterobacterales; Morganellaceae; Providencia; Providencia alcalifaciens MAGDNSILSPHGINRRDFMKLCTALAATMGLSGKAAAQMADAVSDPERPPVIWIGAQECT WP_036956217.1 hydrogenase 2 small subunit [Providencia alcalifaciens] Length: 372\nNumber of predicted TMHs: 1\nExp number of AAs in TMHs: 22.62342\nExp number, first 60 AAs: 0.18757\nTotal prob of N-in: 0.02923\noutside 1 332\nTMhelix 333 355\ninside 356 372

28138 GCF_000527315.1_PalR901475v1.0 Providencia alcalifaciens R90-1475 Proteobacteria; Gammaproteobacteria; Enterobacterales; Morganellaceae; Providencia; Providencia alcalifaciens

MAGDNTIHSPHGINRRDFMKLCTALAATMGLSGKAAAQMADAVSDPERPPVIWIGAQECT WP_036968787.1 hydrogenase 2 small subunit [Providencia alcalifaciens] Length: 372\nNumber of predicted TMHs: 1\nExp number of AAs in TMHs: 23.11788\nExp number, first 60 AAs: 0.49611\nTotal prob of N-in: 0.06654\noutside 1 332\nTMhelix 333 355\ninside 356 372

28139 GCF_000527295.1_PalRIMD1656011v1.0 Providencia alcalifaciens RIMD 1656011 Proteobacteria; Gammaproteobacteria; Enterobacterales; Morganellaceae; Providencia; Providencia alcalifaciens MAGDNTIHSPHGINRRDFMKLCTALAATMGLSGKAAAQMADAVSDPERPPVIWIGAQECT WP_006659159.1 hydrogenase 2 small subunit [Providencia alcalifaciens] Length: 372\nNumber of predicted TMHs: 1\nExp number of AAs in TMHs: 23.09904\nExp number, first 60 AAs: 0.49604\nTotal prob of N-in: 0.06612\noutside 1 332\nTMhelix 333 355\ninside 356 372

28140 GCF_000314855.2_ASM31485v2 Providencia burhodogranariea DSM 19968 Proteobacteria; Gammaproteobacteria; Enterobacterales; Morganellaceae; Providencia; Providencia burhodogranariea MTGDSAFVSPHGINRRDFMKLCTALAATMGLSSKAAAQMADAISDTDRPPVIWIGAQECT WP_008912427.1 hydrogenase 2 small subunit [Providencia burhodogranariea] Length: 372\nNumber of predicted TMHs: 1\nExp number of AAs in TMHs: 20.38902\nExp number, first 60 AAs: 0.14001\nTotal prob of N-in: 0.01623\noutside 1 332\nTMhelix 333 355\ninside 356 372

28141 GCF_001655055.1_Phe35613_DRAFTv1 Providencia heimbachae ATCC 35613 Proteobacteria; Gammaproteobacteria; Enterobacterales; Morganellaceae; Providencia; Providencia heimbachae MAGDNSIISPHGVNRRDFMKLCTALAATMGLSSNAAAQMADAVSDPGRPPVIWIGAQECT WP_068908148.1 hydrogenase 2 small subunit [Providencia heimbachae] Length: 372\nNumber of predicted TMHs: 1\nExp number of AAs in TMHs: 20.46673\nExp number, first 60 AAs: 0.64382\nTotal prob of N-in: 0.09022\noutside 1 332\nTMhelix 333 355\ninside 356 372

28142 GCF_000314835.2_ASM31483v2 Providencia rettgeri Dmel1 Proteobacteria; Gammaproteobacteria; Enterobacterales; Morganellaceae; Providencia; Providencia rettgeri MIGDNSIISSHGINRRDFMKLCTALAATMGLSGKAAAQIAESISDPARPPVIWIGAQECT WP_004908437.1 hydrogenase 2 small subunit [Providencia rettgeri] Length: 372\nNumber of predicted TMHs: 1\nExp number of AAs in TMHs: 21.45764\nExp number, first 60 AAs: 0.53706\nTotal prob of N-in: 0.04172\noutside 1 332\nTMhelix 333 355\ninside 356 372

28143 GCF_000158055.1_ASM15805v1 Providencia rettgeri DSM 1131 Proteobacteria; Gammaproteobacteria; Enterobacterales; Morganellaceae; Providencia; Providencia rettgeri MIGDNSIISSHGINRRDFMKLCTALAATMGLSGKAAAQIAESISDPARPPVIWIGAQECT WP_004257951.1 hydrogenase 2 small subunit [Providencia rettgeri] Length: 372\nNumber of predicted TMHs: 1\nExp number of AAs in TMHs: 21.45583\nExp number, first 60 AAs: 0.53694\nTotal prob of N-in: 0.04170\noutside 1 332\nTMhelix 333 355\ninside 356 372

28144 GCF_000156395.1_ASM15639v1 Providencia rustigianii DSM 4541 Proteobacteria; Gammaproteobacteria; Enterobacterales; Morganellaceae; Providencia; Providencia rustigianii MAGDNSIISPHGINRRDFMKLCTALAATMGLSGKAAAQMADAVSDPERPPVIWIGAQECT WP_006814155.1 hydrogenase 2 small subunit [Providencia rustigianii] Length: 372\nNumber of predicted TMHs: 1\nExp number of AAs in TMHs: 22.74518\nExp number, first 60 AAs: 0.28996\nTotal prob of N-in: 0.04072\noutside 1 332\nTMhelix 333 355\ninside 356 372

28145 GCF_000314895.2_ASM31489v2 Providencia sneebia DSM 19967 Proteobacteria; Gammaproteobacteria; Enterobacterales; Morganellaceae; Providencia; Providencia sneebia MAGENTIISPHGISRRDFMKLCTALAATMGLSSKAAAQMADAVSDPGRPPVIWIGAQECT WP_008916678.1 hydrogenase 2 small subunit [Providencia sneebia] Length: 372\nNumber of predicted TMHs: 1\nExp number of AAs in TMHs: 22.07262\nExp number, first 60 AAs: 0.17465\nTotal prob of N-in: 0.02087\noutside 1 332\nTMhelix 333 355\ninside 356 372

28146 GCF_000696465.1_ASM69646v1 Pectobacterium atrosepticum Proteobacteria; Gammaproteobacteria; Enterobacterales; Pectobacteriaceae; Pectobacterium MTEENKMFFREGVNRRDFMKLCAALAATMGLSSRAAADIASSVSASARPPVIWIGAQECT WP_011092815.1 hydrogenase 2 small subunit [Pectobacterium atrosepticum] Length: 377\nNumber of predicted TMHs: 1\nExp number of AAs in TMHs: 20.77346\nExp number, first 60 AAs: 0.28964\nTotal prob of N-in: 0.09443\noutside 1 333\nTMhelix 334 356\ninside 357 377

28147	GCF_000749905.1_ASM74990v1	Pectobacterium atrosepticum	Proteobacteria; Gammaproteobacteria; Enterobacterales; Pectobacteriaceae; Pectobacterium
	MTEENKMFFREGVNRRDFMKLCAALAATMGLSSRAAADIASSVSASARPPVIWIGAQECT	WP_011092815.1
	hydrogenase 2 small subunit [Pectobacterium atrosepticum]	Length: 377\nNumber of predicted TMHs: 1\nExp number of AAs in TMHs: 20.77346\nExp number, first 60 AAs: 0.28964\nTotal prob of N-in: 0.09443\noutside 1 333\nTMhelix 334 356\ninside 357 377

28148	GCF_000749965.1_ASM74996v1	Pectobacterium atrosepticum	Proteobacteria; Gammaproteobacteria; Enterobacterales; Pectobacteriaceae; Pectobacterium
	MTEENKMFFREGVNRRDFMKLCAALAATMGLGSRAAADIASSVSASARPPVIWIGAQECT	WP_039296234.1
	hydrogenase 2 small subunit [Pectobacterium atrosepticum]	Length: 377\nNumber of predicted TMHs: 1\nExp number of AAs in TMHs: 21.05543\nExp number, first 60 AAs: 0.57228\nTotal prob of N-in: 0.10695\noutside 1 333\nTMhelix 334 356\ninside 357 377

28149	GCF_000740965.1_ASM74096v1	Pectobacterium atrosepticum	Proteobacteria; Gammaproteobacteria; Enterobacterales; Pectobacteriaceae; Pectobacterium
	MTEENKMFFREGVNRRDFMKLCAALAATMGLSSRAAADIASSVSASARPPVIWIGAQECT	WP_011092815.1
	hydrogenase 2 small subunit [Pectobacterium atrosepticum]	Length: 377\nNumber of predicted TMHs: 1\nExp number of AAs in TMHs: 20.77346\nExp number, first 60 AAs: 0.28964\nTotal prob of N-in: 0.09443\noutside 1 333\nTMhelix 334 356\ninside 357 377

28150	GCF_000389755.1_Pa6276v1.0	Pectobacterium atrosepticum CFBP 6276	Proteobacteria; Gammaproteobacteria; Enterobacterales; Pectobacteriaceae; Pectobacterium; Pectobacterium atrosepticum
	MTEENKMFFREGVNRRDFMKLCAALAATMGLGSRAAADIASSVSASARPPVIWIGAQECT	WP_039296234.1
	hydrogenase 2 small subunit [Pectobacterium atrosepticum]	Length: 377\nNumber of predicted TMHs: 1\nExp number of AAs in TMHs: 21.05543\nExp number, first 60 AAs: 0.57228\nTotal prob of N-in: 0.10695\noutside 1 333\nTMhelix 334 356\ninside 357 377

28151	GCF_001038685.1_ASM103868v1	Pectobacterium atrosepticum ICMP 1526	Proteobacteria; Gammaproteobacteria; Enterobacterales; Pectobacteriaceae; Pectobacterium; Pectobacterium atrosepticum
	MTEENKMFFREGVNRRDFMKLCAALAATMGLSSRAAADIASSVSASARPPVIWIGAQECT	WP_011092815.1
	hydrogenase 2 small subunit [Pectobacterium atrosepticum]	Length: 377\nNumber of predicted TMHs: 1\nExp number of AAs in TMHs: 20.77346\nExp number, first 60 AAs: 0.28964\nTotal prob of N-in: 0.09443\noutside 1 333\nTMhelix 334 356\ninside 357 377

28152	GCF_000011605.1_ASM1160v1	Pectobacterium atrosepticum SCRI1043	Proteobacteria; Gammaproteobacteria; Enterobacterales; Pectobacteriaceae; Pectobacterium; Pectobacterium atrosepticum
	MTEENKMFFREGVNRRDFMKLCAALAATMGLSSRAAADIASSVSASARPPVIWIGAQECT	WP_011092815.1
	hydrogenase 2 small subunit [Pectobacterium atrosepticum]	Length: 377\nNumber of predicted TMHs: 1\nExp number of AAs in TMHs: 20.77346\nExp number, first 60 AAs: 0.28964\nTotal prob of N-in: 0.09443\noutside 1 333\nTMhelix 334 356\ninside 357 377

28153	GCF_002093895.1_ASM209389v1	Plesiomonas shigelloides	Proteobacteria; Gammaproteobacteria; Enterobacterales; unclassified Enterobacterales; Plesiomonas
	MAMNNSFLDRLGVSRRDFMKLCTGLAASMGLSANAATKIAAAVSDPARPPVIWIGAQECT	WP_010863703.1
	hydrogenase 2 small subunit [Plesiomonas shigelloides]	Length: 381\nNumber of predicted TMHs: 1\nExp number of AAs in TMHs: 20.33985\nExp number, first 60 AAs: 2.29085\nTotal prob of N-in: 0.11522\noutside 1 333\nTMhelix 334 356\ninside 357 381

28154	GCF_000813415.1_ASM81341v1	Plesiomonas shigelloides	Proteobacteria; Gammaproteobacteria; Enterobacterales; unclassified Enterobacterales; Plesiomonas
	MAMNNSFLDRLGVSRRDFMKLCTGLAASMGLSANAATKIAAAVSDPARPPVIWIGAQECT	WP_010863703.1
	hydrogenase 2 small subunit [Plesiomonas shigelloides]	Length: 381\nNumber of predicted TMHs: 1\nExp number of AAs in TMHs: 20.33985\nExp number, first 60 AAs: 2.29085\nTotal prob of N-in: 0.11522\noutside 1 333\nTMhelix 334 356\ninside 357 381

28155	GCF_900087055.1_31289_E01	Plesiomonas shigelloides	Proteobacteria; Gammaproteobacteria; Enterobacterales; unclassified Enterobacterales; Plesiomonas
	MAMNNSFLDRLGVSRRDFMKLCTGLAASMGLSANAATKIAAAVSDPARPPVIWIGAQECT	WP_010863703.1
	hydrogenase 2 small subunit [Plesiomonas shigelloides]	Length: 381\nNumber of predicted TMHs: 1\nExp

number of AAs in TMHs: 20.33985\nExp number, first 60 AAs: 2.29085\nTotal prob of N-in: 0.11522\noutside 1 333\nTMhelix 334 356\ninside 357 381

28156 GCF_000800945.1_ZOR0011.1 Plesiomonas sp. ZOR0011 Proteobacteria; Gammaproteobacteria; Enterobacterales; unclassified Enterobacterales; Plesiomonas
MAMNNSFLDRLGVSRRDFMKLCTGLAASMGLSANAATKIAAAVSDPARPPVIWIGAQECT WP_047706871.1
hydrogenase 2 small subunit [Plesiomonas sp. ZOR0011] Length: 381\nNumber of predicted TMHs: 1\nExp number of AAs in TMHs: 20.34013\nExp number, first 60 AAs: 2.29084\nTotal prob of N-in: 0.11523\noutside 1 333\nTMhelix 334 356\ninside 357 381

28157 GCF_000392595.1_PleShi1.0 Plesiomonas shigelloides 302-73 Proteobacteria; Gammaproteobacteria; Enterobacterales; unclassified Enterobacterales; Plesiomonas; Plesiomonas shigelloides
MAMNNSFLDRLGVSRRDFMKLCTGLAASMGLSANAATKIAAAVSDPARPPVIWIGAQECT WP_010863703.1
hydrogenase 2 small subunit [Plesiomonas shigelloides] Length: 381\nNumber of predicted TMHs: 1\nExp number of AAs in TMHs: 20.33985\nExp number, first 60 AAs: 2.29085\nTotal prob of N-in: 0.11522\noutside 1 333\nTMhelix 334 356\ninside 357 381

28158 GCF_001880985.1_ASM188098v1 Serratia fonticola Proteobacteria; Gammaproteobacteria; Enterobacterales; Yersiniaceae; Serratia
MIRENQTFFRDGVNRRDFMKLCAALAATMGLSTKAAAEIADSVGSPQRPPVIWIGAQECT WP_071680561.1
hydrogenase 2 small subunit [Serratia fonticola] Length: 377\nNumber of predicted TMHs: 1\nExp number of AAs in TMHs: 19.73066\nExp number, first 60 AAs: 0.69249\nTotal prob of N-in: 0.07499\noutside 1 333\nTMhelix 334 356\ninside 357 377

28159 GCF_000734885.1_SerGri_SPAdes Serratia grimesii Proteobacteria; Gammaproteobacteria; Enterobacterales; Yersiniaceae; Serratia
MSKRRTTLTALALIVSLGLSSAPVLADKGGNGNGGGNGHGNSGNHGNNGNHGNNGNHGNK WP_037422402.1
membrane protein [Serratia grimesii] Length: 160\nNumber of predicted TMHs: 2\nExp number of AAs in TMHs: 33.04158\nExp number, first 60 AAs: 16.06517\nTotal prob of N-in: 0.94244\nPOSSIBLE N-term signal sequence\ninside 1 6\nTMhelix 7 26\noutside 27 135\nTMhelix 136 158\ninside 159 160

28160 GCF_000975245.1_ASM97524v1 Serratia liquefaciens Proteobacteria; Gammaproteobacteria; Enterobacterales; Yersiniaceae; Serratia
MSKRRTTFTVLALIVSLGLSSAPALADKGGNGNGGGNGHGNSGNHGNNGNNGKHGNKVKD WP_046373790.1
membrane protein [Serratia liquefaciens] Length: 157\nNumber of predicted TMHs: 2\nExp number of AAs in TMHs: 36.25847\nExp number, first 60 AAs: 17.0716\nTotal prob of N-in: 0.96941\nPOSSIBLE N-term signal sequence\ninside 1 6\nTMhelix 7 26\noutside 27 132\nTMhelix 133 155\ninside 156 157

28161 GCF_001540785.1_12082_3_83 Serratia marcescens Proteobacteria; Gammaproteobacteria; Enterobacterales; Yersiniaceae; Serratia
MNKRRTTLTVLALIASLGLSSAPALADKGGNGNGNGNGHGNSGNHGNSGNHGNNGNKGNK WP_033646351.1
MULTISPECIES: membrane protein [Enterobacterales] Length: 161\nNumber of predicted TMHs: 2\nExp number of AAs in TMHs: 31.11104\nExp number, first 60 AAs: 14.67675\nTotal prob of N-in: 0.95023\nPOSSIBLE N-term signal sequence\ninside 1 6\nTMhelix 7 26\noutside 27 141\nTMhelix 142 160\ninside 161 161

28162 GCF_001063325.1_ASM106332v1 Serratia marcescens Proteobacteria; Gammaproteobacteria; Enterobacterales; Yersiniaceae; Serratia
MNKRRTTLTVLALIASLGLSSAPALADKGGNGNGNGNGHGNSGNHGNNGNNGNHGNNGNH WP_049209162.1
membrane protein [Serratia marcescens] Length: 167\nNumber of predicted TMHs: 2\nExp number of AAs in TMHs: 30.02384\nExp number, first 60 AAs: 14.11185\nTotal prob of N-in: 0.94170\nPOSSIBLE N-term signal sequence\ninside 1 6\nTMhelix 7 26\noutside 27 147\nTMhelix 148 166\ninside 167 167

28163 GCF_001065845.1_ASM106584v1 Serratia marcescens Proteobacteria; Gammaproteobacteria; Enterobacterales; Yersiniaceae; Serratia
MNKRRTTLTVLALIASLGLSSAPALADKGGNGNGNGNGHGNSGNHGNNGNHGNNGNNGNH WP_004941297.1
MULTISPECIES: hypothetical protein [Serratia] Length: 167\nNumber of predicted TMHs: 2\nExp number of AAs in TMHs: 30.02384\nExp number, first 60 AAs: 14.11185\nTotal prob of N-in: 0.94170\nPOSSIBLE N-term signal sequence\ninside 1 6\nTMhelix 7 26\noutside 27 147\nTMhelix 148 166\ninside 167 167

28164 GCF_001065935.1_ASM106593v1 Serratia marcescens Proteobacteria; Gammaproteobacteria; Enterobacterales; Yersiniaceae; Serratia

MNKRRTTLTVLALIASLGLSSAPALADKGGNGNGNGNGHGNSGNHGNNGNHGNNGNNGNH	WP_004941297.1	MULTISPECIES: hypothetical protein [Serratia]	Length: 167\nNumber of predicted TMHs:  2\nExp number of AAs in TMHs: 30.02384\nExp number, first 60 AAs:  14.11185\nTotal prob of N-in:   0.94170\nPOSSIBLE N-term signal sequence\ninside   1   6\nTMhelix   7   26\noutside   27   147\nTMhelix   148   166\ninside   167   167

28165	GCF_001064715.1_ASM106471v1	Serratia marcescens	Proteobacteria; Gammaproteobacteria; Enterobacterales; Yersiniaceae; Serratia	MNKRRTTLTVLALIASLGLSSAPALADKGGNGNGNGNGHGNSGNHGNNGNNGNHGNNGNH	WP_049209162.1	membrane protein [Serratia marcescens]	Length: 167\nNumber of predicted TMHs:  2\nExp number of AAs in TMHs: 30.02384\nExp number, first 60 AAs:  14.11185\nTotal prob of N-in:   0.94170\nPOSSIBLE N-term signal sequence\ninside   1   6\nTMhelix   7   26\noutside   27   147\nTMhelix   148   166\ninside   167   167

28166	GCF_001536365.1_12082_3_76	Serratia marcescens	Proteobacteria; Gammaproteobacteria; Enterobacterales; Yersiniaceae; Serratia	MNKRRTTLTVLALIASLGLSSAPALADKGGNGNGNGNGHGNSGNHGNSGNHGNNGNKGNK	WP_033646351.1	MULTISPECIES: membrane protein [Enterobacterales]	Length: 161\nNumber of predicted TMHs:  2\nExp number of AAs in TMHs: 31.11104\nExp number, first 60 AAs:  14.67675\nTotal prob of N-in:   0.95023\nPOSSIBLE N-term signal sequence\ninside   1   6\nTMhelix   7   26\noutside   27   141\nTMhelix   142   160\ninside   161   161

28167	GCF_001536465.1_12082_2_70	Serratia marcescens	Proteobacteria; Gammaproteobacteria; Enterobacterales; Yersiniaceae; Serratia	MNKRRTTLTVLALIASLGLSSAPALADKGGNGNGNGNGHGNSGNHGNSGNHGNNGNKGNK	WP_033646351.1	MULTISPECIES: membrane protein [Enterobacterales]	Length: 161\nNumber of predicted TMHs:  2\nExp number of AAs in TMHs: 31.11104\nExp number, first 60 AAs:  14.67675\nTotal prob of N-in:   0.95023\nPOSSIBLE N-term signal sequence\ninside   1   6\nTMhelix   7   26\noutside   27   141\nTMhelix   142   160\ninside   161   161

28168	GCF_001536385.1_12082_3_28	Serratia marcescens	Proteobacteria; Gammaproteobacteria; Enterobacterales; Yersiniaceae; Serratia	MNKRRTTLTVLALIASLGLSSAPALADKGGNGNGNGNGHGNSGNHGNSGNHGNNGNKGNK	WP_033646351.1	MULTISPECIES: membrane protein [Enterobacterales]	Length: 161\nNumber of predicted TMHs:  2\nExp number of AAs in TMHs: 31.11104\nExp number, first 60 AAs:  14.67675\nTotal prob of N-in:   0.95023\nPOSSIBLE N-term signal sequence\ninside   1   6\nTMhelix   7   26\noutside   27   141\nTMhelix   142   160\ninside   161   161

28169	GCF_001536525.1_12082_3_49	Serratia marcescens	Proteobacteria; Gammaproteobacteria; Enterobacterales; Yersiniaceae; Serratia	MNKRRTTLTVLALIASLGLSSAPALADKGGNGNGNGNGHGNSGNHGNSGNHGNNGNKGNK	WP_033646351.1	MULTISPECIES: membrane protein [Enterobacterales]	Length: 161\nNumber of predicted TMHs:  2\nExp number of AAs in TMHs: 31.11104\nExp number, first 60 AAs:  14.67675\nTotal prob of N-in:   0.95023\nPOSSIBLE N-term signal sequence\ninside   1   6\nTMhelix   7   26\noutside   27   141\nTMhelix   142   160\ninside   161   161

28170	GCF_001536505.1_12082_2_64	Serratia marcescens	Proteobacteria; Gammaproteobacteria; Enterobacterales; Yersiniaceae; Serratia	MNKRRTTLTVLALIASLGLSSAPALADKGGNGNGNGNGHGNSGNHGNSGNHGNNGNKGNK	WP_033646351.1	MULTISPECIES: membrane protein [Enterobacterales]	Length: 161\nNumber of predicted TMHs:  2\nExp number of AAs in TMHs: 31.11104\nExp number, first 60 AAs:  14.67675\nTotal prob of N-in:   0.95023\nPOSSIBLE N-term signal sequence\ninside   1   6\nTMhelix   7   26\noutside   27   141\nTMhelix   142   160\ninside   161   161

28171	GCF_001536565.1_12082_5_49	Serratia marcescens	Proteobacteria; Gammaproteobacteria; Enterobacterales; Yersiniaceae; Serratia	MNKRRTTLTVLALIASLGLSSAPALADKGGNGNGNGNGHGNSGNHGNSGNHGNNGNKGNK	WP_033646351.1	MULTISPECIES: membrane protein [Enterobacterales]	Length: 161\nNumber of predicted TMHs:  2\nExp number of AAs in TMHs: 31.11104\nExp number, first 60 AAs:  14.67675\nTotal prob of N-in:   0.95023\nPOSSIBLE N-term signal sequence\ninside   1   6\nTMhelix   7   26\noutside   27   141\nTMhelix   142   160\ninside   161   161

28172	GCF_001536625.1_12082_3_56	Serratia marcescens	Proteobacteria; Gammaproteobacteria; Enterobacterales; Yersiniaceae; Serratia	MNKRRTTLTVLALIASLGLSSAPALADKGGNGNGNGNGHGNSGNHGNSGNHGNNGNKGNK	WP_033646351.1	MULTISPECIES: membrane protein [Enterobacterales]	Length: 161\nNumber of predicted TMHs:  2\nExp number of AAs in TMHs: 31.11104\nExp number, first 60 AAs:  14.67675\nTotal prob of N-in:   0.95023\nPOSSIBLE N-term signal sequence\ninside   1   6\nTMhelix   7   26\noutside   27   141\nTMhelix   142   160\ninside   161   161

28173 GCF_001536725.1_12082_3_14 Serratia marcescens Proteobacteria; Gammaproteobacteria; Enterobacterales; Yersiniaceae; Serratia
MNKRRTTLTVLALIASLGLSSAPALADKGGNGNGNGNGHGNSGNHGNSGNHGNNGNKGNK WP_033646351.1
MULTISPECIES: membrane protein [Enterobacterales] Length: 161\nNumber of predicted TMHs: 2\nExp number of AAs in TMHs: 31.11104\nExp number, first 60 AAs: 14.67675\nTotal prob of N-in: 0.95023\nPOSSIBLE N-term signal sequence\ninside 1 6\nTMhelix 7 26\noutside 27 141\nTMhelix 142 160\ninside 161 161

28174 GCF_001536605.1_12045_8_19 Serratia marcescens Proteobacteria; Gammaproteobacteria; Enterobacterales; Yersiniaceae; Serratia
MNKRRTTLTVLALIASLGLSSAPALADKGGNGNGNGNGHGNSGNNGNHGNNGNHGNKGNK WP_060419303.1
MULTISPECIES: hypothetical protein [Serratia] Length: 164\nNumber of predicted TMHs: 2\nExp number of AAs in TMHs: 30.51677\nExp number, first 60 AAs: 14.42464\nTotal prob of N-in: 0.94579\nPOSSIBLE N-term signal sequence\ninside 1 6\nTMhelix 7 26\noutside 27 144\nTMhelix 145 163\ninside 164 164

28175 GCF_001536765.1_12082_2_66 Serratia marcescens Proteobacteria; Gammaproteobacteria; Enterobacterales; Yersiniaceae; Serratia
MNKRRTTLTVLALIASLGLSSAPALADKGGNGNGNGNGHGNSGNHGNSGNHGNNGNKGNK WP_033646351.1
MULTISPECIES: membrane protein [Enterobacterales] Length: 161\nNumber of predicted TMHs: 2\nExp number of AAs in TMHs: 31.11104\nExp number, first 60 AAs: 14.67675\nTotal prob of N-in: 0.95023\nPOSSIBLE N-term signal sequence\ninside 1 6\nTMhelix 7 26\noutside 27 141\nTMhelix 142 160\ninside 161 161

28176 GCF_001536665.1_12082_3_57 Serratia marcescens Proteobacteria; Gammaproteobacteria; Enterobacterales; Yersiniaceae; Serratia
MNKRRTTLTVLALIASLGLSSAPALADKGGNGNGNGNGHGNSGNHGNSGNHGNNGNKGNK WP_060427691.1
hypothetical protein [Serratia marcescens] Length: 161\nNumber of predicted TMHs: 2\nExp number of AAs in TMHs: 31.15272\nExp number, first 60 AAs: 14.60251\nTotal prob of N-in: 0.95045\nPOSSIBLE N-term signal sequence\ninside 1 6\nTMhelix 7 26\noutside 27 141\nTMhelix 142 160\ninside 161 161

28177 GCF_001536685.1_12082_2_86 Serratia marcescens Proteobacteria; Gammaproteobacteria; Enterobacterales; Yersiniaceae; Serratia
MNKRRTTLTVLALIASLGLSSAPALADKGGNGNGNGNGHGNSGNHGNSGNHGNNGNKGNK WP_060427691.1
hypothetical protein [Serratia marcescens] Length: 161\nNumber of predicted TMHs: 2\nExp number of AAs in TMHs: 31.15272\nExp number, first 60 AAs: 14.60251\nTotal prob of N-in: 0.95045\nPOSSIBLE N-term signal sequence\ninside 1 6\nTMhelix 7 26\noutside 27 141\nTMhelix 142 160\ninside 161 161

28178 GCF_001536845.1_12082_3_30 Serratia marcescens Proteobacteria; Gammaproteobacteria; Enterobacterales; Yersiniaceae; Serratia
MNKRRTTLTVLALIASLGLSSAPALADKGGNGNGNGNGHGNSGNHGNSGNHGNNGNKGNK WP_033646351.1
MULTISPECIES: membrane protein [Enterobacterales] Length: 161\nNumber of predicted TMHs: 2\nExp number of AAs in TMHs: 31.11104\nExp number, first 60 AAs: 14.67675\nTotal prob of N-in: 0.95023\nPOSSIBLE N-term signal sequence\ninside 1 6\nTMhelix 7 26\noutside 27 141\nTMhelix 142 160\ninside 161 161

28179 GCF_001536945.1_12082_3_63 Serratia marcescens Proteobacteria; Gammaproteobacteria; Enterobacterales; Yersiniaceae; Serratia
MNKRRTTLTVLALIASLGLSSAPALADKGGNGNGNGNGHGNSGNHGNSGNHGNNGNKGNK WP_033646351.1
MULTISPECIES: membrane protein [Enterobacterales] Length: 161\nNumber of predicted TMHs: 2\nExp number of AAs in TMHs: 31.11104\nExp number, first 60 AAs: 14.67675\nTotal prob of N-in: 0.95023\nPOSSIBLE N-term signal sequence\ninside 1 6\nTMhelix 7 26\noutside 27 141\nTMhelix 142 160\ninside 161 161

28180 GCF_001536985.1_12082_2_79 Serratia marcescens Proteobacteria; Gammaproteobacteria; Enterobacterales; Yersiniaceae; Serratia
MNKRRTTLTVLALIASLGLSSAPALADKGGNGNGNGNGHGNSGNHGNSGNHGNNGNKGNK WP_033646351.1
MULTISPECIES: membrane protein [Enterobacterales] Length: 161\nNumber of predicted TMHs: 2\nExp number of AAs in TMHs: 31.11104\nExp number, first 60 AAs: 14.67675\nTotal prob of N-in: 0.95023\nPOSSIBLE N-term signal sequence\ninside 1 6\nTMhelix 7 26\noutside 27 141\nTMhelix 142 160\ninside 161 161

28181 GCF_001537005.1_12082_3_48 Serratia marcescens Proteobacteria; Gammaproteobacteria; Enterobacterales; Yersiniaceae; Serratia
MNKRRTTLTVLALIASLGLSSAPALADKGGNGNGNGNGHGNSGNHGNSGNHGNNGNKGNK WP_033646351.1
MULTISPECIES: membrane protein [Enterobacterales] Length: 161\nNumber of predicted TMHs: 2\nExp

number of AAs in TMHs: 31.11104\nExp number, first 60 AAs: 14.67675\nTotal prob of N-in: 0.95023\nPOSSIBLE N-term signal sequence\ninside 1 6\nTMhelix 7 26\noutside 27 141\nTMhelix 142 160\ninside 161 161

28182 GCF_001537145.1_12082_2_28 Serratia marcescens Proteobacteria; Gammaproteobacteria; Enterobacterales; Yersiniaceae; Serratia
MNKRRTTLTVLALIASLGLSSAPALADKGGNGNGNGNGHGNSGNHGNSGNHGNNGNKGNK WP_033646351.1
MULTISPECIES: membrane protein [Enterobacterales] Length: 161\nNumber of predicted TMHs: 2\nExp number of AAs in TMHs: 31.11104\nExp number, first 60 AAs: 14.67675\nTotal prob of N-in: 0.95023\nPOSSIBLE N-term signal sequence\ninside 1 6\nTMhelix 7 26\noutside 27 141\nTMhelix 142 160\ninside 161 161

28183 GCF_001537185.1_12082_3_26 Serratia marcescens Proteobacteria; Gammaproteobacteria; Enterobacterales; Yersiniaceae; Serratia
MNKRRTTLTVLALIASLGLSSAPALADKGGNGNGNGNGHGNSGNHGNSGNHGNNGNKGNK WP_033646351.1
MULTISPECIES: membrane protein [Enterobacterales] Length: 161\nNumber of predicted TMHs: 2\nExp number of AAs in TMHs: 31.11104\nExp number, first 60 AAs: 14.67675\nTotal prob of N-in: 0.95023\nPOSSIBLE N-term signal sequence\ninside 1 6\nTMhelix 7 26\noutside 27 141\nTMhelix 142 160\ninside 161 161

28184 GCF_001537105.1_12045_8_17 Serratia marcescens Proteobacteria; Gammaproteobacteria; Enterobacterales; Yersiniaceae; Serratia
MNKRRTTLTVLALIASLGLSSAPALADKGGNGNGNGNGHGNSGNHGNSGNHGNNGNKGNK WP_033646351.1
MULTISPECIES: membrane protein [Enterobacterales] Length: 161\nNumber of predicted TMHs: 2\nExp number of AAs in TMHs: 31.11104\nExp number, first 60 AAs: 14.67675\nTotal prob of N-in: 0.95023\nPOSSIBLE N-term signal sequence\ninside 1 6\nTMhelix 7 26\noutside 27 141\nTMhelix 142 160\ninside 161 161

28185 GCF_001537085.1_12082_3_93 Serratia marcescens Proteobacteria; Gammaproteobacteria; Enterobacterales; Yersiniaceae; Serratia
MNKRRTTLTVLALIASLGLSSAPALADKGGNGNGNGNGNGHGNSGNNGNHGNNGNHGNKG WP_060432922.1
hypothetical protein [Serratia marcescens] Length: 163\nNumber of predicted TMHs: 2\nExp number of AAs in TMHs: 30.71713\nExp number, first 60 AAs: 14.46723\nTotal prob of N-in: 0.94731\nPOSSIBLE N-term signal sequence\ninside 1 6\nTMhelix 7 26\noutside 27 143\nTMhelix 144 162\ninside 163 163

28186 GCF_001537045.1_12082_5_37 Serratia marcescens Proteobacteria; Gammaproteobacteria; Enterobacterales; Yersiniaceae; Serratia
MNKRRTTLTVLALIASLGLSSAPALADKGGNGNGNGNGHGNSGNHGNSGNHGNNGNKGNK WP_060427691.1
hypothetical protein [Serratia marcescens] Length: 161\nNumber of predicted TMHs: 2\nExp number of AAs in TMHs: 31.15272\nExp number, first 60 AAs: 14.60251\nTotal prob of N-in: 0.95045\nPOSSIBLE N-term signal sequence\ninside 1 6\nTMhelix 7 26\noutside 27 141\nTMhelix 142 160\ninside 161 161

28187 GCF_001537205.1_12082_2_37 Serratia marcescens Proteobacteria; Gammaproteobacteria; Enterobacterales; Yersiniaceae; Serratia
MNKRRTTLTVLALIASLGLSSAPALADKGGNGNGNGNGHGNSGNNGNHGNNGNHGNKGNK WP_060419303.1
MULTISPECIES: hypothetical protein [Serratia] Length: 164\nNumber of predicted TMHs: 2\nExp number of AAs in TMHs: 30.51677\nExp number, first 60 AAs: 14.42464\nTotal prob of N-in: 0.94579\nPOSSIBLE N-term signal sequence\ninside 1 6\nTMhelix 7 26\noutside 27 144\nTMhelix 145 163\ninside 164 164

28188 GCF_001537245.1_12082_2_77 Serratia marcescens Proteobacteria; Gammaproteobacteria; Enterobacterales; Yersiniaceae; Serratia
MNKRRTTLTVLALIASLGLSSAPALADKGGNGNGNGNGHGNSGNNGNHGNNGNHGNKGNK WP_060419303.1
MULTISPECIES: hypothetical protein [Serratia] Length: 164\nNumber of predicted TMHs: 2\nExp number of AAs in TMHs: 30.51677\nExp number, first 60 AAs: 14.42464\nTotal prob of N-in: 0.94579\nPOSSIBLE N-term signal sequence\ninside 1 6\nTMhelix 7 26\noutside 27 144\nTMhelix 145 163\ninside 164 164

28189 GCF_001537725.1_12082_3_7 Serratia marcescens Proteobacteria; Gammaproteobacteria; Enterobacterales; Yersiniaceae; Serratia
MNKRRTTLTVLALIASLGLSSAPALADKGGNGNGNGNGHGNSGNHGNNGNHGNNGNNGNH WP_004941297.1
MULTISPECIES: hypothetical protein [Serratia] Length: 167\nNumber of predicted TMHs: 2\nExp number of AAs in TMHs: 30.02384\nExp number, first 60 AAs: 14.11185\nTotal prob of N-in: 0.94170\nPOSSIBLE N-term signal sequence\ninside 1 6\nTMhelix 7 26\noutside 27 147\nTMhelix 148 166\ninside 167 167

28190 GCF_001537665.1_12082_2_85 Serratia marcescens Proteobacteria; Gammaproteobacteria; Enterobacterales; Yersiniaceae; Serratia

MNKRRTTLTVLALIASLGLSSAPALADKGGNGNGNGNGHGNSGNHGNSGNHGNNGNKGNK WP_033646351.1 MULTISPECIES: membrane protein [Enterobacterales] Length: 161\nNumber of predicted TMHs: 2\nExp number of AAs in TMHs: 31.11104\nExp number, first 60 AAs: 14.67675\nTotal prob of N-in: 0.95023\nPOSSIBLE N-term signal sequence\ninside 1 6\nTMhelix 7 26\noutside 27 141\nTMhelix 142 160\ninside 161 161

28191 GCF_001537705.1_12082_3_82 Serratia marcescens Proteobacteria; Gammaproteobacteria; Enterobacterales; Yersiniaceae; Serratia MNKRRTTLTVLALIASLGLSSAPALADKGGNGNGNGNGHGNSGNHGNSGNHGNNGNKGNK WP_033646351.1 MULTISPECIES: membrane protein [Enterobacterales] Length: 161\nNumber of predicted TMHs: 2\nExp number of AAs in TMHs: 31.11104\nExp number, first 60 AAs: 14.67675\nTotal prob of N-in: 0.95023\nPOSSIBLE N-term signal sequence\ninside 1 6\nTMhelix 7 26\noutside 27 141\nTMhelix 142 160\ninside 161 161

28192 GCF_001537765.1_12082_2_29 Serratia marcescens Proteobacteria; Gammaproteobacteria; Enterobacterales; Yersiniaceae; Serratia MNKRRTTLTVLALIASLGLSSAPALADKGGNGNGNGNGHGNSGNHGNSGNHGNNGNKGNK WP_033646351.1 MULTISPECIES: membrane protein [Enterobacterales] Length: 161\nNumber of predicted TMHs: 2\nExp number of AAs in TMHs: 31.11104\nExp number, first 60 AAs: 14.67675\nTotal prob of N-in: 0.95023\nPOSSIBLE N-term signal sequence\ninside 1 6\nTMhelix 7 26\noutside 27 141\nTMhelix 142 160\ninside 161 161

28193 GCF_001537565.1_12045_8_25 Serratia marcescens Proteobacteria; Gammaproteobacteria; Enterobacterales; Yersiniaceae; Serratia MNKRRTTLTVLALIASLGLSSAPALADKGGNGNGNGNGHGNSGNHGNSGNHGNNGNNGNK WP_060424250.1 hypothetical protein [Serratia marcescens] Length: 164\nNumber of predicted TMHs: 2\nExp number of AAs in TMHs: 30.64504\nExp number, first 60 AAs: 14.33456\nTotal prob of N-in: 0.94646\nPOSSIBLE N-term signal sequence\ninside 1 6\nTMhelix 7 26\noutside 27 144\nTMhelix 145 163\ninside 164 164

28194 GCF_001537805.1_12082_5_51 Serratia marcescens Proteobacteria; Gammaproteobacteria; Enterobacterales; Yersiniaceae; Serratia MNKRRTTLTVLALIASLGLSSAPALADKGGNGNGNGNGHGNSGNHGNSGNHGNNGNKGNK WP_033646351.1 MULTISPECIES: membrane protein [Enterobacterales] Length: 161\nNumber of predicted TMHs: 2\nExp number of AAs in TMHs: 31.11104\nExp number, first 60 AAs: 14.67675\nTotal prob of N-in: 0.95023\nPOSSIBLE N-term signal sequence\ninside 1 6\nTMhelix 7 26\noutside 27 141\nTMhelix 142 160\ninside 161 161

28195 GCF_001537825.1_12045_8_21 Serratia marcescens Proteobacteria; Gammaproteobacteria; Enterobacterales; Yersiniaceae; Serratia MNKRRTTLTVLALIASLGLSSAPALADKGGNGNGNGNGHGNSGNHGNSGNHGNNGNKGNK WP_033646351.1 MULTISPECIES: membrane protein [Enterobacterales] Length: 161\nNumber of predicted TMHs: 2\nExp number of AAs in TMHs: 31.11104\nExp number, first 60 AAs: 14.67675\nTotal prob of N-in: 0.95023\nPOSSIBLE N-term signal sequence\ninside 1 6\nTMhelix 7 26\noutside 27 141\nTMhelix 142 160\ninside 161 161

28196 GCF_001537865.1_12082_5_46 Serratia marcescens Proteobacteria; Gammaproteobacteria; Enterobacterales; Yersiniaceae; Serratia MNKRRTTLTVLALIASLGLSSAPALADKGGNGNGNGNGHGNSGNHGNSGNHGNNGNKGNK WP_033646351.1 MULTISPECIES: membrane protein [Enterobacterales] Length: 161\nNumber of predicted TMHs: 2\nExp number of AAs in TMHs: 31.11104\nExp number, first 60 AAs: 14.67675\nTotal prob of N-in: 0.95023\nPOSSIBLE N-term signal sequence\ninside 1 6\nTMhelix 7 26\noutside 27 141\nTMhelix 142 160\ninside 161 161

28197 GCF_001537905.1_12082_5_34 Serratia marcescens Proteobacteria; Gammaproteobacteria; Enterobacterales; Yersiniaceae; Serratia MNKRRTTLTVLALIASLGLSSAPALADKGGNGNGNGNGHGNSGNHGNSGNHGNNGNKGNK WP_033646351.1 MULTISPECIES: membrane protein [Enterobacterales] Length: 161\nNumber of predicted TMHs: 2\nExp number of AAs in TMHs: 31.11104\nExp number, first 60 AAs: 14.67675\nTotal prob of N-in: 0.95023\nPOSSIBLE N-term signal sequence\ninside 1 6\nTMhelix 7 26\noutside 27 141\nTMhelix 142 160\ninside 161 161

28198 GCF_001537925.1_12082_3_89 Serratia marcescens Proteobacteria; Gammaproteobacteria; Enterobacterales; Yersiniaceae; Serratia MNKRRTTLTVLALIASLGLSSAPALADKGGNGNGNGNGHGNSGNHGNSGNHGNNGNKGNK WP_033646351.1 MULTISPECIES: membrane protein [Enterobacterales] Length: 161\nNumber of predicted TMHs: 2\nExp number of AAs in TMHs: 31.11104\nExp number, first 60 AAs: 14.67675\nTotal prob of N-in: 0.95023\nPOSSIBLE N-term signal sequence\ninside 1 6\nTMhelix 7 26\noutside 27 141\nTMhelix 142 160\ninside 161 161

28199	GCF_001538325.1_12082_5_42	Serratia marcescens	Proteobacteria; Gammaproteobacteria; Enterobacterales; Yersiniaceae; Serratia
MNKRRTTLTVLALIASLGLSSAPALADKGGNGNGNGNGHGNSGNHGNSGNHGNNGNKGNK	WP_033646351.1
MULTISPECIES: membrane protein [Enterobacterales]	Length: 161\nNumber of predicted TMHs:  2\nExp number of AAs in TMHs: 31.11104\nExp number, first 60 AAs:  14.67675\nTotal prob of N-in:   0.95023\nPOSSIBLE N-term signal sequence\ninside   1   6\nTMhelix   7   26\noutside   27   141\nTMhelix   142   160\ninside   161   161

28200	GCF_001538365.1_12082_2_91	Serratia marcescens	Proteobacteria; Gammaproteobacteria; Enterobacterales; Yersiniaceae; Serratia
MNKRRTTLTVLALIASLGLSSAPALADKGGNGNGNGNGHGNSGNHGNSGNHGNNGNKGNK	WP_033646351.1
MULTISPECIES: membrane protein [Enterobacterales]	Length: 161\nNumber of predicted TMHs:  2\nExp number of AAs in TMHs: 31.11104\nExp number, first 60 AAs:  14.67675\nTotal prob of N-in:   0.95023\nPOSSIBLE N-term signal sequence\ninside   1   6\nTMhelix   7   26\noutside   27   141\nTMhelix   142   160\ninside   161   161

28201	GCF_001538275.1_12082_5_36	Serratia marcescens	Proteobacteria; Gammaproteobacteria; Enterobacterales; Yersiniaceae; Serratia
MNKRRTTLTVLALIASLGLSSAPALADKGGNGNGNGNGHGNSGNHGNSGNHGNNGNKGNK	WP_033646351.1
MULTISPECIES: membrane protein [Enterobacterales]	Length: 161\nNumber of predicted TMHs:  2\nExp number of AAs in TMHs: 31.11104\nExp number, first 60 AAs:  14.67675\nTotal prob of N-in:   0.95023\nPOSSIBLE N-term signal sequence\ninside   1   6\nTMhelix   7   26\noutside   27   141\nTMhelix   142   160\ninside   161   161

28202	GCF_001538385.1_12082_5_30	Serratia marcescens	Proteobacteria; Gammaproteobacteria; Enterobacterales; Yersiniaceae; Serratia
MNKRRTTLTVLALIASLGLSSAPALADKGGNGNGNGNGHGNSGNHGNSGNHGNNGNKGNK	WP_033646351.1
MULTISPECIES: membrane protein [Enterobacterales]	Length: 161\nNumber of predicted TMHs:  2\nExp number of AAs in TMHs: 31.11104\nExp number, first 60 AAs:  14.67675\nTotal prob of N-in:   0.95023\nPOSSIBLE N-term signal sequence\ninside   1   6\nTMhelix   7   26\noutside   27   141\nTMhelix   142   160\ninside   161   161

28203	GCF_001538425.1_12082_2_74	Serratia marcescens	Proteobacteria; Gammaproteobacteria; Enterobacterales; Yersiniaceae; Serratia
MNKRRTTLTVLALIASLGLSSAPALADKGGNGNGNGNGHGNSGNHGNSGNHGNNGNKGNK	WP_033646351.1
MULTISPECIES: membrane protein [Enterobacterales]	Length: 161\nNumber of predicted TMHs:  2\nExp number of AAs in TMHs: 31.11104\nExp number, first 60 AAs:  14.67675\nTotal prob of N-in:   0.95023\nPOSSIBLE N-term signal sequence\ninside   1   6\nTMhelix   7   26\noutside   27   141\nTMhelix   142   160\ninside   161   161

28204	GCF_001538115.1_12082_3_68	Serratia marcescens	Proteobacteria; Gammaproteobacteria; Enterobacterales; Yersiniaceae; Serratia
MNKRRTTLTVLALIASLGLSSAPALADKGGNGNGNGNGHGNSGNHGNSGNHGNNGNKGNK	WP_060427691.1
hypothetical protein [Serratia marcescens]	Length: 161\nNumber of predicted TMHs:  2\nExp number of AAs in TMHs: 31.15272\nExp number, first 60 AAs:  14.60251\nTotal prob of N-in:   0.95045\nPOSSIBLE N-term signal sequence\ninside   1   6\nTMhelix   7   26\noutside   27   141\nTMhelix   142   160\ninside   161   161

28205	GCF_001538705.1_12045_8_23	Serratia marcescens	Proteobacteria; Gammaproteobacteria; Enterobacterales; Yersiniaceae; Serratia
MNKRRTTLTVLALIASLGLSSAPALADKGGNGNGNGNGHGNSGNHGNSGNHGNNGNKGNK	WP_033646351.1
MULTISPECIES: membrane protein [Enterobacterales]	Length: 161\nNumber of predicted TMHs:  2\nExp number of AAs in TMHs: 31.11104\nExp number, first 60 AAs:  14.67675\nTotal prob of N-in:   0.95023\nPOSSIBLE N-term signal sequence\ninside   1   6\nTMhelix   7   26\noutside   27   141\nTMhelix   142   160\ninside   161   161

28206	GCF_001538805.1_12082_2_55	Serratia marcescens	Proteobacteria; Gammaproteobacteria; Enterobacterales; Yersiniaceae; Serratia
MNKRRTTLTVLALIASLGLSSAPALADKGGNGNGNGNGHGNSGNHGNSGNHGNNGNKGNK	WP_033646351.1
MULTISPECIES: membrane protein [Enterobacterales]	Length: 161\nNumber of predicted TMHs:  2\nExp number of AAs in TMHs: 31.11104\nExp number, first 60 AAs:  14.67675\nTotal prob of N-in:   0.95023\nPOSSIBLE N-term signal sequence\ninside   1   6\nTMhelix   7   26\noutside   27   141\nTMhelix   142   160\ninside   161   161

28207	GCF_001538845.1_12082_3_90	Serratia marcescens	Proteobacteria; Gammaproteobacteria; Enterobacterales; Yersiniaceae; Serratia
MNKRRTTLTVLALIASLGLSSAPALADKGGNGNGNGNGHGNSGNHGNSGNHGNNGNKGNK	WP_033646351.1
MULTISPECIES: membrane protein [Enterobacterales]	Length: 161\nNumber of predicted TMHs:  2\nExp

number of AAs in TMHs: 31.11104\nExp number, first 60 AAs: 14.67675\nTotal prob of N-in: 0.95023\nPOSSIBLE N-term signal sequence\ninside 1 6\nTMhelix 7 26\noutside 27 141\nTMhelix 142 160\ninside 161 161

28208 GCF_001538525.1_12082_2_33 Serratia marcescens Proteobacteria; Gammaproteobacteria; Enterobacterales; Yersiniaceae; Serratia MNKRRTTLTVLALIASLGLSSAPALADKGGNGNGNGNGHGNSGNHGNSGNHGNNGNKGNK WP_060427691.1 hypothetical protein [Serratia marcescens] Length: 161\nNumber of predicted TMHs: 2\nExp number of AAs in TMHs: 31.15272\nExp number, first 60 AAs: 14.60251\nTotal prob of N-in: 0.95045\nPOSSIBLE N-term signal sequence\ninside 1 6\nTMhelix 7 26\noutside 27 141\nTMhelix 142 160\ninside 161 161

28209 GCF_001538885.1_12082_3_32 Serratia marcescens Proteobacteria; Gammaproteobacteria; Enterobacterales; Yersiniaceae; Serratia MNKRRTTLTVLALIASLGLSSAPALADKGGNGNGNGNGHGNSGNHGNSGNHGNNGNKGNK WP_033646351.1 MULTISPECIES: membrane protein [Enterobacterales] Length: 161\nNumber of predicted TMHs: 2\nExp number of AAs in TMHs: 31.11104\nExp number, first 60 AAs: 14.67675\nTotal prob of N-in: 0.95023\nPOSSIBLE N-term signal sequence\ninside 1 6\nTMhelix 7 26\noutside 27 141\nTMhelix 142 160\ninside 161 161

28210 GCF_001538905.1_12082_2_23 Serratia marcescens Proteobacteria; Gammaproteobacteria; Enterobacterales; Yersiniaceae; Serratia MNKRRTTLTVLALIASLGLSSAPALADKGGNGNGNGNGHGNSGNHGNSGNHGNNGNKGNK WP_033646351.1 MULTISPECIES: membrane protein [Enterobacterales] Length: 161\nNumber of predicted TMHs: 2\nExp number of AAs in TMHs: 31.11104\nExp number, first 60 AAs: 14.67675\nTotal prob of N-in: 0.95023\nPOSSIBLE N-term signal sequence\ninside 1 6\nTMhelix 7 26\noutside 27 141\nTMhelix 142 160\ninside 161 161

28211 GCF_001538945.1_12082_3_34 Serratia marcescens Proteobacteria; Gammaproteobacteria; Enterobacterales; Yersiniaceae; Serratia MNKRRTTLTVLALIASLGLSSAPALADKGGNGNGNGNGHGNSGNHGNSGNHGNNGNKGNK WP_033646351.1 MULTISPECIES: membrane protein [Enterobacterales] Length: 161\nNumber of predicted TMHs: 2\nExp number of AAs in TMHs: 31.11104\nExp number, first 60 AAs: 14.67675\nTotal prob of N-in: 0.95023\nPOSSIBLE N-term signal sequence\ninside 1 6\nTMhelix 7 26\noutside 27 141\nTMhelix 142 160\ninside 161 161

28212 GCF_001538985.1_12045_8_27 Serratia marcescens Proteobacteria; Gammaproteobacteria; Enterobacterales; Yersiniaceae; Serratia MNKRRTTLTVLALIASLGLSSAPALADKGGNGNGNGNGHGNSGNHGNSGNHGNNGNKGNK WP_033646351.1 MULTISPECIES: membrane protein [Enterobacterales] Length: 161\nNumber of predicted TMHs: 2\nExp number of AAs in TMHs: 31.11104\nExp number, first 60 AAs: 14.67675\nTotal prob of N-in: 0.95023\nPOSSIBLE N-term signal sequence\ninside 1 6\nTMhelix 7 26\noutside 27 141\nTMhelix 142 160\ninside 161 161

28213 GCF_001539045.1_12082_3_36 Serratia marcescens Proteobacteria; Gammaproteobacteria; Enterobacterales; Yersiniaceae; Serratia MNKRRTTLTVLALIASLGLSSAPALADKGGNGNGNGNGHGNSGNHGNNGNHGNNGNNGNH WP_004941297.1 MULTISPECIES: hypothetical protein [Serratia] Length: 167\nNumber of predicted TMHs: 2\nExp number of AAs in TMHs: 30.02384\nExp number, first 60 AAs: 14.11185\nTotal prob of N-in: 0.94170\nPOSSIBLE N-term signal sequence\ninside 1 6\nTMhelix 7 26\noutside 27 147\nTMhelix 148 166\ninside 167 167

28214 GCF_001539145.1_12082_3_16 Serratia marcescens Proteobacteria; Gammaproteobacteria; Enterobacterales; Yersiniaceae; Serratia MNKRRTTLTVLALIASLGLSSAPALADKGGNGNGNGNGHGNSGNHGNSGNHGNNGNKGNK WP_033646351.1 MULTISPECIES: membrane protein [Enterobacterales] Length: 161\nNumber of predicted TMHs: 2\nExp number of AAs in TMHs: 31.11104\nExp number, first 60 AAs: 14.67675\nTotal prob of N-in: 0.95023\nPOSSIBLE N-term signal sequence\ninside 1 6\nTMhelix 7 26\noutside 27 141\nTMhelix 142 160\ninside 161 161

28215 GCF_001539225.1_12045_8_7 Serratia marcescens Proteobacteria; Gammaproteobacteria; Enterobacterales; Yersiniaceae; Serratia MNKRRTTLTVLALIASLGLSSAPALADKGGNGNGNGNGHGNSGNHGNSGNHGNNGNKGNK WP_033646351.1 MULTISPECIES: membrane protein [Enterobacterales] Length: 161\nNumber of predicted TMHs: 2\nExp number of AAs in TMHs: 31.11104\nExp number, first 60 AAs: 14.67675\nTotal prob of N-in: 0.95023\nPOSSIBLE N-term signal sequence\ninside 1 6\nTMhelix 7 26\noutside 27 141\nTMhelix 142 160\ninside 161 161

28216 GCF_001539185.1_12045_8_4 Serratia marcescens Proteobacteria; Gammaproteobacteria; Enterobacterales; Yersiniaceae; Serratia

MNKRRTTLTVLALIASLGLSSAPALADKGGNGNGNGNGHGNSGNHGNSGNHGNNGNKGNK WP_033646351.1 MULTISPECIES: membrane protein [Enterobacterales] Length: 161\nNumber of predicted TMHs: 2\nExp number of AAs in TMHs: 31.11104\nExp number, first 60 AAs: 14.67675\nTotal prob of N-in: 0.95023\nPOSSIBLE N-term signal sequence\ninside 1 6\nTMhelix 7 26\noutside 27 141\nTMhelix 142 160\ninside 161 161

28217 GCF_001539065.1_12082_5_48 Serratia marcescens Proteobacteria; Gammaproteobacteria; Enterobacterales; Yersiniaceae; Serratia MNKRRTTLTVLALIASLGLSSAPALADKGGNGNGNGNGHGNSGNNGNHGNNGNHGNKGNK WP_060419303.1 MULTISPECIES: hypothetical protein [Serratia] Length: 164\nNumber of predicted TMHs: 2\nExp number of AAs in TMHs: 30.51677\nExp number, first 60 AAs: 14.42464\nTotal prob of N-in: 0.94579\nPOSSIBLE N-term signal sequence\ninside 1 6\nTMhelix 7 26\noutside 27 144\nTMhelix 145 163\ninside 164 164

28218 GCF_001539005.1_12082_5_33 Serratia marcescens Proteobacteria; Gammaproteobacteria; Enterobacterales; Yersiniaceae; Serratia MNKRRTTLTVLALIASLGLSSAPALADKGGNGNGNGNGHGNSGNNGNHGNNGNHGNKGNK WP_060419303.1 MULTISPECIES: hypothetical protein [Serratia] Length: 164\nNumber of predicted TMHs: 2\nExp number of AAs in TMHs: 30.51677\nExp number, first 60 AAs: 14.42464\nTotal prob of N-in: 0.94579\nPOSSIBLE N-term signal sequence\ninside 1 6\nTMhelix 7 26\noutside 27 144\nTMhelix 145 163\ninside 164 164

28219 GCF_001539325.1_12082_3_53 Serratia marcescens Proteobacteria; Gammaproteobacteria; Enterobacterales; Yersiniaceae; Serratia MNKRRTTLTVLALIASLGLSSAPALADKGGNGNGNGNGHGNSGNHGNSGNHGNNGNKGNK WP_033646351.1 MULTISPECIES: membrane protein [Enterobacterales] Length: 161\nNumber of predicted TMHs: 2\nExp number of AAs in TMHs: 31.11104\nExp number, first 60 AAs: 14.67675\nTotal prob of N-in: 0.95023\nPOSSIBLE N-term signal sequence\ninside 1 6\nTMhelix 7 26\noutside 27 141\nTMhelix 142 160\ninside 161 161

28220 GCF_001539105.1_12045_8_10 Serratia marcescens Proteobacteria; Gammaproteobacteria; Enterobacterales; Yersiniaceae; Serratia MNKRRTTLTVLALIASLGLSSAPALADKGGNGNGNGNGHGNSGNHGNSGNHGNNGNKGNK WP_060427691.1 hypothetical protein [Serratia marcescens] Length: 161\nNumber of predicted TMHs: 2\nExp number of AAs in TMHs: 31.15272\nExp number, first 60 AAs: 14.60251\nTotal prob of N-in: 0.95045\nPOSSIBLE N-term signal sequence\ninside 1 6\nTMhelix 7 26\noutside 27 141\nTMhelix 142 160\ninside 161 161

28221 GCF_001539585.1_12082_2_69 Serratia marcescens Proteobacteria; Gammaproteobacteria; Enterobacterales; Yersiniaceae; Serratia MNKRRTTLTVLALIASLGLSSAPALADKGGNGNGNGNGHGNSGNHGNSGNHGNNGNKGNK WP_033646351.1 MULTISPECIES: membrane protein [Enterobacterales] Length: 161\nNumber of predicted TMHs: 2\nExp number of AAs in TMHs: 31.11104\nExp number, first 60 AAs: 14.67675\nTotal prob of N-in: 0.95023\nPOSSIBLE N-term signal sequence\ninside 1 6\nTMhelix 7 26\noutside 27 141\nTMhelix 142 160\ninside 161 161

28222 GCF_001539605.1_12082_3_80 Serratia marcescens Proteobacteria; Gammaproteobacteria; Enterobacterales; Yersiniaceae; Serratia MNKRRTTLTVLALIASLGLSSAPALADKGGNGNGNGNGHGNSGNHGNSGNHGNNGNKGNK WP_033646351.1 MULTISPECIES: membrane protein [Enterobacterales] Length: 161\nNumber of predicted TMHs: 2\nExp number of AAs in TMHs: 31.11104\nExp number, first 60 AAs: 14.67675\nTotal prob of N-in: 0.95023\nPOSSIBLE N-term signal sequence\ninside 1 6\nTMhelix 7 26\noutside 27 141\nTMhelix 142 160\ninside 161 161

28223 GCF_001539345.1_12082_2_88 Serratia marcescens Proteobacteria; Gammaproteobacteria; Enterobacterales; Yersiniaceae; Serratia MNKRRTTLTVLALIASLGLSSAPALADKGGNGNGNGNGHGNSGNHGNSGNHGNNGNKGNK WP_060427691.1 hypothetical protein [Serratia marcescens] Length: 161\nNumber of predicted TMHs: 2\nExp number of AAs in TMHs: 31.15272\nExp number, first 60 AAs: 14.60251\nTotal prob of N-in: 0.95045\nPOSSIBLE N-term signal sequence\ninside 1 6\nTMhelix 7 26\noutside 27 141\nTMhelix 142 160\ninside 161 161

28224 GCF_001539685.1_12082_3_79 Serratia marcescens Proteobacteria; Gammaproteobacteria; Enterobacterales; Yersiniaceae; Serratia MNKRRTTLTVLALIASLGLSSAPALADKGGNGNGNGNGHGNSGNHGNSGNHGNNGNKGNK WP_033646351.1 MULTISPECIES: membrane protein [Enterobacterales] Length: 161\nNumber of predicted TMHs: 2\nExp number of AAs in TMHs: 31.11104\nExp number, first 60 AAs: 14.67675\nTotal prob of N-in: 0.95023\nPOSSIBLE N-term signal sequence\ninside 1 6\nTMhelix 7 26\noutside 27 141\nTMhelix 142 160\ninside 161 161

28225 GCF_001539705.1_12082_2_51 Serratia marcescens Proteobacteria; Gammaproteobacteria; Enterobacterales; Yersiniaceae; Serratia
MNKRRTTLTVLALIASLGLSSAPALADKGGNGNGNGNGHGNSGNHGNSGNHGNNGNKGNK WP_033646351.1
MULTISPECIES: membrane protein [Enterobacterales] Length: 161\nNumber of predicted TMHs: 2\nExp number of AAs in TMHs: 31.11104\nExp number, first 60 AAs: 14.67675\nTotal prob of N-in: 0.95023\nPOSSIBLE N-term signal sequence\ninside 1 6\nTMhelix 7 26\noutside 27 141\nTMhelix 142 160\ninside 161 161

28226 GCF_001539745.1_12082_2_93 Serratia marcescens Proteobacteria; Gammaproteobacteria; Enterobacterales; Yersiniaceae; Serratia
MNKRRTTLTVLALIASLGLSSAPALADKGGNGNGNGNGHGNSGNHGNSGNHGNNGNKGNK WP_033646351.1
MULTISPECIES: membrane protein [Enterobacterales] Length: 161\nNumber of predicted TMHs: 2\nExp number of AAs in TMHs: 31.11104\nExp number, first 60 AAs: 14.67675\nTotal prob of N-in: 0.95023\nPOSSIBLE N-term signal sequence\ninside 1 6\nTMhelix 7 26\noutside 27 141\nTMhelix 142 160\ninside 161 161

28227 GCF_001539785.1_12082_2_36 Serratia marcescens Proteobacteria; Gammaproteobacteria; Enterobacterales; Yersiniaceae; Serratia
MNKRRTTLTVLALIASLGLSSAPALADKGGNGNGNGNGHGNSGNHGNSGNHGNNGNKGNK WP_033646351.1
MULTISPECIES: membrane protein [Enterobacterales] Length: 161\nNumber of predicted TMHs: 2\nExp number of AAs in TMHs: 31.11104\nExp number, first 60 AAs: 14.67675\nTotal prob of N-in: 0.95023\nPOSSIBLE N-term signal sequence\ninside 1 6\nTMhelix 7 26\noutside 27 141\nTMhelix 142 160\ninside 161 161

28228 GCF_001539805.1_12082_2_42 Serratia marcescens Proteobacteria; Gammaproteobacteria; Enterobacterales; Yersiniaceae; Serratia
MNKRRTTLTVLALIASLGLSSAPALADKGGNGNGNGNGHGNSGNHGNSGNHGNNGNKGNK WP_033646351.1
MULTISPECIES: membrane protein [Enterobacterales] Length: 161\nNumber of predicted TMHs: 2\nExp number of AAs in TMHs: 31.11104\nExp number, first 60 AAs: 14.67675\nTotal prob of N-in: 0.95023\nPOSSIBLE N-term signal sequence\ninside 1 6\nTMhelix 7 26\noutside 27 141\nTMhelix 142 160\ninside 161 161

28229 GCF_001539845.1_12082_2_73 Serratia marcescens Proteobacteria; Gammaproteobacteria; Enterobacterales; Yersiniaceae; Serratia
MNKRRTTLTVLALIASLGLSSAPALADKGGNGNGNGNGHGNSGNHGNSGNHGNNGNKGNK WP_033646351.1
MULTISPECIES: membrane protein [Enterobacterales] Length: 161\nNumber of predicted TMHs: 2\nExp number of AAs in TMHs: 31.11104\nExp number, first 60 AAs: 14.67675\nTotal prob of N-in: 0.95023\nPOSSIBLE N-term signal sequence\ninside 1 6\nTMhelix 7 26\noutside 27 141\nTMhelix 142 160\ninside 161 161

28230 GCF_001539865.1_12082_3_11 Serratia marcescens Proteobacteria; Gammaproteobacteria; Enterobacterales; Yersiniaceae; Serratia
MNKRRTTLTVLALIASLGLSSAPALADKGGNGNGNGNGHGNSGNHGNSGNHGNNGNKGNK WP_033646351.1
MULTISPECIES: membrane protein [Enterobacterales] Length: 161\nNumber of predicted TMHs: 2\nExp number of AAs in TMHs: 31.11104\nExp number, first 60 AAs: 14.67675\nTotal prob of N-in: 0.95023\nPOSSIBLE N-term signal sequence\ninside 1 6\nTMhelix 7 26\noutside 27 141\nTMhelix 142 160\ninside 161 161

28231 GCF_001539905.1_12082_3_58 Serratia marcescens Proteobacteria; Gammaproteobacteria; Enterobacterales; Yersiniaceae; Serratia
MNKRRTTLTVLALIASLGLSSAPALADKGGNGNGNGNGHGNSGNHGNSGNHGNNGNKGNK WP_033646351.1
MULTISPECIES: membrane protein [Enterobacterales] Length: 161\nNumber of predicted TMHs: 2\nExp number of AAs in TMHs: 31.11104\nExp number, first 60 AAs: 14.67675\nTotal prob of N-in: 0.95023\nPOSSIBLE N-term signal sequence\ninside 1 6\nTMhelix 7 26\noutside 27 141\nTMhelix 142 160\ninside 161 161

28232 GCF_001539945.1_12082_3_60 Serratia marcescens Proteobacteria; Gammaproteobacteria; Enterobacterales; Yersiniaceae; Serratia
MNKRRTTLTVLALIASLGLSSAPALADKGGNGNGNGNGHGNSGNHGNSGNHGNNGNKGNK WP_033646351.1
MULTISPECIES: membrane protein [Enterobacterales] Length: 161\nNumber of predicted TMHs: 2\nExp number of AAs in TMHs: 31.11104\nExp number, first 60 AAs: 14.67675\nTotal prob of N-in: 0.95023\nPOSSIBLE N-term signal sequence\ninside 1 6\nTMhelix 7 26\noutside 27 141\nTMhelix 142 160\ninside 161 161

28233 GCF_001061145.1_ASM106114v1 Serratia marcescens Proteobacteria; Gammaproteobacteria; Enterobacterales; Yersiniaceae; Serratia
MNKRRTTLTVLALIASLGLSSAPALADKGGNGNGNGNGHGNSGNHGNNGNHGNNGNNGNN WP_048323073.1
membrane protein [Serratia marcescens] Length: 170\nNumber of predicted TMHs: 2\nExp number of

AAs in TMHs: 29.49026\nExp number, first 60 AAs: 13.84277\nTotal prob of N-in: 0.93726\nPOSSIBLE N-term signal sequence\ninside 1 6\nTMhelix 7 26\noutside 27 150\nTMhelix 151 169\ninside 170 170

28234 GCF_001540005.1_12082_3_55 Serratia marcescens Proteobacteria; Gammaproteobacteria; Enterobacterales; Yersiniaceae; Serratia MNKRRTTLTVLALIASLGLSSAPALADKGGNGNGNGNGHGNSGNHGNSGNHGNNGNKGNK WP_033646351.1 MULTISPECIES: membrane protein [Enterobacterales] Length: 161\nNumber of predicted TMHs: 2\nExp number of AAs in TMHs: 31.11104\nExp number, first 60 AAs: 14.67675\nTotal prob of N-in: 0.95023\nPOSSIBLE N-term signal sequence\ninside 1 6\nTMhelix 7 26\noutside 27 141\nTMhelix 142 160\ninside 161 161

28235 GCF_001061225.1_ASM106122v1 Serratia marcescens Proteobacteria; Gammaproteobacteria; Enterobacterales; Yersiniaceae; Serratia MNKRRTTLTVLALIASLGLSSAPALADKGGNGNGNGNGHGNSGNHGNSGNHGNNGNKGNK WP_033646351.1 MULTISPECIES: membrane protein [Enterobacterales] Length: 161\nNumber of predicted TMHs: 2\nExp number of AAs in TMHs: 31.11104\nExp number, first 60 AAs: 14.67675\nTotal prob of N-in: 0.95023\nPOSSIBLE N-term signal sequence\ninside 1 6\nTMhelix 7 26\noutside 27 141\nTMhelix 142 160\ninside 161 161

28236 GCF_001539965.1_12082_2_38 Serratia marcescens Proteobacteria; Gammaproteobacteria; Enterobacterales; Yersiniaceae; Serratia MNKRRTTLTVLALIASLGLSSAPALADKGGNGNGNGNGHGNSGNHGNSGNHGNNGNKGNK WP_033646351.1 MULTISPECIES: membrane protein [Enterobacterales] Length: 161\nNumber of predicted TMHs: 2\nExp number of AAs in TMHs: 31.11104\nExp number, first 60 AAs: 14.67675\nTotal prob of N-in: 0.95023\nPOSSIBLE N-term signal sequence\ninside 1 6\nTMhelix 7 26\noutside 27 141\nTMhelix 142 160\ninside 161 161

28237 GCF_001540065.1_12082_3_84 Serratia marcescens Proteobacteria; Gammaproteobacteria; Enterobacterales; Yersiniaceae; Serratia MNKRRTTLTVLALIASLGLSSAPALADKGGNGNGNGNGHGNSGNHGNSGNHGNNGNKGNK WP_033646351.1 MULTISPECIES: membrane protein [Enterobacterales] Length: 161\nNumber of predicted TMHs: 2\nExp number of AAs in TMHs: 31.11104\nExp number, first 60 AAs: 14.67675\nTotal prob of N-in: 0.95023\nPOSSIBLE N-term signal sequence\ninside 1 6\nTMhelix 7 26\noutside 27 141\nTMhelix 142 160\ninside 161 161

28238 GCF_001540105.1_12082_3_10 Serratia marcescens Proteobacteria; Gammaproteobacteria; Enterobacterales; Yersiniaceae; Serratia MNKRRTTLTVLALIASLGLSSAPALADKGGNGNGNGNGHGNSGNHGNSGNHGNNGNKGNK WP_033646351.1 MULTISPECIES: membrane protein [Enterobacterales] Length: 161\nNumber of predicted TMHs: 2\nExp number of AAs in TMHs: 31.11104\nExp number, first 60 AAs: 14.67675\nTotal prob of N-in: 0.95023\nPOSSIBLE N-term signal sequence\ninside 1 6\nTMhelix 7 26\noutside 27 141\nTMhelix 142 160\ninside 161 161

28239 GCF_001540425.1_12082_2_62 Serratia marcescens Proteobacteria; Gammaproteobacteria; Enterobacterales; Yersiniaceae; Serratia MNKRRTTLTVLALIASLGLSSAPALADKGGNGNGNGNGHGNSGNHGNSGNHGNNGNKGNK WP_033646351.1 MULTISPECIES: membrane protein [Enterobacterales] Length: 161\nNumber of predicted TMHs: 2\nExp number of AAs in TMHs: 31.11104\nExp number, first 60 AAs: 14.67675\nTotal prob of N-in: 0.95023\nPOSSIBLE N-term signal sequence\ninside 1 6\nTMhelix 7 26\noutside 27 141\nTMhelix 142 160\ninside 161 161

28240 GCF_001540385.1_12082_2_84 Serratia marcescens Proteobacteria; Gammaproteobacteria; Enterobacterales; Yersiniaceae; Serratia MNKRRTTLTVLALIASLGLSSAPALADKGGNGNGNGNGHGNSGNHGNSGNHGNNGNKGNK WP_033646351.1 MULTISPECIES: membrane protein [Enterobacterales] Length: 161\nNumber of predicted TMHs: 2\nExp number of AAs in TMHs: 31.11104\nExp number, first 60 AAs: 14.67675\nTotal prob of N-in: 0.95023\nPOSSIBLE N-term signal sequence\ninside 1 6\nTMhelix 7 26\noutside 27 141\nTMhelix 142 160\ninside 161 161

28241 GCF_001540485.1_12082_3_23 Serratia marcescens Proteobacteria; Gammaproteobacteria; Enterobacterales; Yersiniaceae; Serratia MNKRRTTLTVLALIASLGLSSAPALADKGGNGNGNGNGHGNSGNHGNSGNHGNNGNKGNK WP_033646351.1 MULTISPECIES: membrane protein [Enterobacterales] Length: 161\nNumber of predicted TMHs: 2\nExp number of AAs in TMHs: 31.11104\nExp number, first 60 AAs: 14.67675\nTotal prob of N-in: 0.95023\nPOSSIBLE N-term signal sequence\ninside 1 6\nTMhelix 7 26\noutside 27 141\nTMhelix 142 160\ninside 161 161

28242 GCF_001540465.1_12082_5_38 Serratia marcescens Proteobacteria; Gammaproteobacteria; Enterobacterales; Yersiniaceae; Serratia

MNKRRTTLTVLALIASLGLSSAPALADKGGNGNGNGNGHGNSGNNGNHGNNGNHGNKGNK WP_060419303.1 MULTISPECIES: hypothetical protein [Serratia] Length: 164\nNumber of predicted TMHs: 2\nExp number of AAs in TMHs: 30.51677\nExp number, first 60 AAs: 14.42464\nTotal prob of N-in: 0.94579\nPOSSIBLE N-term signal sequence\ninside 1 6\nTMhelix 7 26\noutside 27 144\nTMhelix 145 163\ninside 164 164

28243 GCF_001540525.1_12082_2_76 Serratia marcescens Proteobacteria; Gammaproteobacteria; Enterobacterales; Yersiniaceae; Serratia MNKRRTTLTVLALIASLGLSSAPALADKGGNGNGNGNGHGNSGNHGNSGNHGNNGNKGNK WP_033646351.1 MULTISPECIES: membrane protein [Enterobacterales] Length: 161\nNumber of predicted TMHs: 2\nExp number of AAs in TMHs: 31.11104\nExp number, first 60 AAs: 14.67675\nTotal prob of N-in: 0.95023\nPOSSIBLE N-term signal sequence\ninside 1 6\nTMhelix 7 26\noutside 27 141\nTMhelix 142 160\ninside 161 161

28244 GCF_001540565.1_12082_3_94 Serratia marcescens Proteobacteria; Gammaproteobacteria; Enterobacterales; Yersiniaceae; Serratia MNKRRTTLTVLALIASLGLSSAPALADKGGNGNGNGNGHGNSGNHGNSGNHGNNGNKGNK WP_033646351.1 MULTISPECIES: membrane protein [Enterobacterales] Length: 161\nNumber of predicted TMHs: 2\nExp number of AAs in TMHs: 31.11104\nExp number, first 60 AAs: 14.67675\nTotal prob of N-in: 0.95023\nPOSSIBLE N-term signal sequence\ninside 1 6\nTMhelix 7 26\noutside 27 141\nTMhelix 142 160\ninside 161 161

28245 GCF_001540585.1_12082_3_85 Serratia marcescens Proteobacteria; Gammaproteobacteria; Enterobacterales; Yersiniaceae; Serratia MNKRRTTLTVLALIASLGLSSAPALADKGGNGNGNGNGHGNSGNHGNSGNHGNNGNKGNK WP_033646351.1 MULTISPECIES: membrane protein [Enterobacterales] Length: 161\nNumber of predicted TMHs: 2\nExp number of AAs in TMHs: 31.11104\nExp number, first 60 AAs: 14.67675\nTotal prob of N-in: 0.95023\nPOSSIBLE N-term signal sequence\ninside 1 6\nTMhelix 7 26\noutside 27 141\nTMhelix 142 160\ninside 161 161

28246 GCF_001540625.1_12082_2_89 Serratia marcescens Proteobacteria; Gammaproteobacteria; Enterobacterales; Yersiniaceae; Serratia MNKRRTTLTVLALIASLGLSSAPALADKGGNGNGNGNGHGNSGNHGNSGNHGNNGNKGNK WP_033646351.1 MULTISPECIES: membrane protein [Enterobacterales] Length: 161\nNumber of predicted TMHs: 2\nExp number of AAs in TMHs: 31.11104\nExp number, first 60 AAs: 14.67675\nTotal prob of N-in: 0.95023\nPOSSIBLE N-term signal sequence\ninside 1 6\nTMhelix 7 26\noutside 27 141\nTMhelix 142 160\ninside 161 161

28247 GCF_001540665.1_12045_8_12 Serratia marcescens Proteobacteria; Gammaproteobacteria; Enterobacterales; Yersiniaceae; Serratia MNKRRTTLTVLALIASLGLSSAPALADKGGNGNGNGNGHGNSGNHGNSGNHGNNGNKGNK WP_033646351.1 MULTISPECIES: membrane protein [Enterobacterales] Length: 161\nNumber of predicted TMHs: 2\nExp number of AAs in TMHs: 31.11104\nExp number, first 60 AAs: 14.67675\nTotal prob of N-in: 0.95023\nPOSSIBLE N-term signal sequence\ninside 1 6\nTMhelix 7 26\noutside 27 141\nTMhelix 142 160\ninside 161 161

28248 GCF_001540685.1_12082_3_6 Serratia marcescens Proteobacteria; Gammaproteobacteria; Enterobacterales; Yersiniaceae; Serratia MNKRRTTLTVLALIASLGLSSAPALADKGGNGNGNGNGHGNSGNHGNSGNHGNNGNKGNK WP_033646351.1 MULTISPECIES: membrane protein [Enterobacterales] Length: 161\nNumber of predicted TMHs: 2\nExp number of AAs in TMHs: 31.11104\nExp number, first 60 AAs: 14.67675\nTotal prob of N-in: 0.95023\nPOSSIBLE N-term signal sequence\ninside 1 6\nTMhelix 7 26\noutside 27 141\nTMhelix 142 160\ninside 161 161

28249 GCF_001540725.1_12082_2_52 Serratia marcescens Proteobacteria; Gammaproteobacteria; Enterobacterales; Yersiniaceae; Serratia MNKRRTTLTVLALIASLGLSSAPALADKGGNGNGNGNGHGNSGNHGNSGNHGNNGNKGNK WP_033646351.1 MULTISPECIES: membrane protein [Enterobacterales] Length: 161\nNumber of predicted TMHs: 2\nExp number of AAs in TMHs: 31.11104\nExp number, first 60 AAs: 14.67675\nTotal prob of N-in: 0.95023\nPOSSIBLE N-term signal sequence\ninside 1 6\nTMhelix 7 26\noutside 27 141\nTMhelix 142 160\ninside 161 161

28250 GCF_001564475.1_12082_3_81 Serratia marcescens Proteobacteria; Gammaproteobacteria; Enterobacterales; Yersiniaceae; Serratia MNKRRTTLTVLALIASLGLSSAPALADKGGNGNGNGNGHGNSGNHGNSGNHGNNGNKGNK WP_033646351.1 MULTISPECIES: membrane protein [Enterobacterales] Length: 161\nNumber of predicted TMHs: 2\nExp number of AAs in TMHs: 31.11104\nExp number, first 60 AAs: 14.67675\nTotal prob of N-in: 0.95023\nPOSSIBLE N-term signal sequence\ninside 1 6\nTMhelix 7 26\noutside 27 141\nTMhelix 142 160\ninside 161 161

28251 GCF_001540825.1_12082_3_54 Serratia marcescens Proteobacteria; Gammaproteobacteria; Enterobacterales; Yersiniaceae; Serratia
MNKRRTTLTVLALIASLGLSSAPALADKGGNGNGNGNGHGNSGNHGNSGNHGNNGNKGNK WP_033646351.1
MULTISPECIES: membrane protein [Enterobacterales] Length: 161\nNumber of predicted TMHs: 2\nExp number of AAs in TMHs: 31.11104\nExp number, first 60 AAs: 14.67675\nTotal prob of N-in: 0.95023\nPOSSIBLE N-term signal sequence\ninside 1 6\nTMhelix 7 26\noutside 27 141\nTMhelix 142 160\ninside 161 161

28252 GCF_001064335.1_ASM106433v1 Serratia marcescens Proteobacteria; Gammaproteobacteria; Enterobacterales; Yersiniaceae; Serratia
MNKRRTTLTVLALIASLGLSSAPALADKGGNGNGNGNGHGNSGNHGNNGNHGNNGNNGNH WP_004941297.1
MULTISPECIES: hypothetical protein [Serratia] Length: 167\nNumber of predicted TMHs: 2\nExp number of AAs in TMHs: 30.02384\nExp number, first 60 AAs: 14.11185\nTotal prob of N-in: 0.94170\nPOSSIBLE N-term signal sequence\ninside 1 6\nTMhelix 7 26\noutside 27 147\nTMhelix 148 166\ninside 167 167

28253 GCF_001643155.1_ASM164315v1 Serratia marcescens Proteobacteria; Gammaproteobacteria; Enterobacterales; Yersiniaceae; Serratia
MNKRRTTLTVLALIASLGLSSAPALADKGGNGNGHGNSGNHGNSGNHGNNGNKGNKDKGG WP_063989634.1
hypothetical protein [Serratia marcescens] Length: 157\nNumber of predicted TMHs: 2\nExp number of AAs in TMHs: 31.94676\nExp number, first 60 AAs: 15.13538\nTotal prob of N-in: 0.95672\nPOSSIBLE N-term signal sequence\ninside 1 6\nTMhelix 7 26\noutside 27 137\nTMhelix 138 156\ninside 157 157

28254 GCF_001540745.1_12082_3_4 Serratia marcescens Proteobacteria; Gammaproteobacteria; Enterobacterales; Yersiniaceae; Serratia
MNKRRTTLTVLALIASLGLSSAPALADKGGNGNGNGNGHGNSGNHGNSGNHGNNGNKGNK WP_060427691.1
hypothetical protein [Serratia marcescens] Length: 161\nNumber of predicted TMHs: 2\nExp number of AAs in TMHs: 31.15272\nExp number, first 60 AAs: 14.60251\nTotal prob of N-in: 0.95045\nPOSSIBLE N-term signal sequence\ninside 1 6\nTMhelix 7 26\noutside 27 141\nTMhelix 142 160\ninside 161 161

28255 GCF_002108655.1_ASM210865v1 Serratia marcescens Proteobacteria; Gammaproteobacteria; Enterobacterales; Yersiniaceae; Serratia
MNKRRTTLTVLALIASLGLSSAPALADKGGNGNGNGNGHGNSGNHGNNGNHGNNGNNGNH WP_004941297.1
MULTISPECIES: hypothetical protein [Serratia] Length: 167\nNumber of predicted TMHs: 2\nExp number of AAs in TMHs: 30.02384\nExp number, first 60 AAs: 14.11185\nTotal prob of N-in: 0.94170\nPOSSIBLE N-term signal sequence\ninside 1 6\nTMhelix 7 26\noutside 27 147\nTMhelix 148 166\ninside 167 167

28256 GCF_002029225.1_ASM202922v1 Serratia marcescens Proteobacteria; Gammaproteobacteria; Enterobacterales; Yersiniaceae; Serratia
MNKRRTTLTVLALIASLGLSSAPALADKGGNGNGNGNGHGNSGNHGNNGNHGNNGNHGNK WP_033640576.1
MULTISPECIES: membrane protein [Serratia] Length: 164\nNumber of predicted TMHs: 2\nExp number of AAs in TMHs: 30.54778\nExp number, first 60 AAs: 14.38033\nTotal prob of N-in: 0.94595\nPOSSIBLE N-term signal sequence\ninside 1 6\nTMhelix 7 26\noutside 27 144\nTMhelix 145 163\ninside 164 164

28257 GCF_002152845.1_ASM215284v1 Serratia marcescens Proteobacteria; Gammaproteobacteria; Enterobacterales; Yersiniaceae; Serratia
MNKRRTTLTVLALIASLGLSSAPALADKGGNGNGNGNGHGNSGNHGNSGNHGNNGNKGNK WP_033646351.1
MULTISPECIES: membrane protein [Enterobacterales] Length: 161\nNumber of predicted TMHs: 2\nExp number of AAs in TMHs: 31.11104\nExp number, first 60 AAs: 14.67675\nTotal prob of N-in: 0.95023\nPOSSIBLE N-term signal sequence\ninside 1 6\nTMhelix 7 26\noutside 27 141\nTMhelix 142 160\ninside 161 161

28258 GCF_002029205.1_ASM202920v1 Serratia marcescens Proteobacteria; Gammaproteobacteria; Enterobacterales; Yersiniaceae; Serratia
MNKRRTTLTVLALIASLGLSSAPALADKGGNGNGNGNGNGHGNSGNHGNHGNNGNHGNKG WP_079451327.1
hypothetical protein [Serratia marcescens] Length: 163\nNumber of predicted TMHs: 2\nExp number of AAs in TMHs: 30.71713\nExp number, first 60 AAs: 14.46723\nTotal prob of N-in: 0.94731\nPOSSIBLE N-term signal sequence\ninside 1 6\nTMhelix 7 26\noutside 27 143\nTMhelix 144 162\ninside 163 163

28259 GCF_002205475.1_ASM220547v1 Serratia marcescens Proteobacteria; Gammaproteobacteria; Enterobacterales; Yersiniaceae; Serratia
MNKRRTTLTVLALIASLGLSSAPALADKGGNGNGNGNGHGNSGNHGNNGNHGNNGNNGNH WP_004941297.1
MULTISPECIES: hypothetical protein [Serratia] Length: 167\nNumber of predicted TMHs: 2\nExp number of

AAs in TMHs: 30.02384\nExp number, first 60 AAs:  14.11185\nTotal prob of N-in:   0.94170\nPOSSIBLE N-term signal sequence\ninside   1   6\nTMhelix   7   26\noutside   27   147\nTMhelix   148   166\ninside   167   167

28260 GCF_001060335.1_ASM106033v1 Serratia marcescens Proteobacteria; Gammaproteobacteria; Enterobacterales; Yersiniaceae; Serratia
MNKRRTTLTVLALIASLGLSSAPALADKGGNGNGNGNGHGNSGNNGNHGNNGNHGNKGNK WP_049200367.1
MULTISPECIES: membrane protein [Enterobacterales] Length: 164\nNumber of predicted TMHs:  2\nExp number of AAs in TMHs: 30.51526\nExp number, first 60 AAs:  14.42673\nTotal prob of N-in:   0.94579\nPOSSIBLE N-term signal sequence\ninside   1   6\nTMhelix   7   26\noutside   27   144\nTMhelix   145   163\ninside   164   164

28261 GCF_001034405.1_Serr_marc_UCI88_V1 Serratia marcescens Proteobacteria; Gammaproteobacteria; Enterobacterales; Yersiniaceae; Serratia
MNKRRTTLTVLALIASLGLSSAPALADKGGNGNGNGNGHGNSGNHGNSGNHGNNGNKGNK WP_033646351.1
MULTISPECIES: membrane protein [Enterobacterales] Length: 161\nNumber of predicted TMHs:  2\nExp number of AAs in TMHs: 31.11104\nExp number, first 60 AAs:  14.67675\nTotal prob of N-in:   0.95023\nPOSSIBLE N-term signal sequence\ninside   1   6\nTMhelix   7   26\noutside   27   141\nTMhelix   142   160\ninside   161   161

28262 GCF_001060585.1_ASM106058v1 Serratia marcescens Proteobacteria; Gammaproteobacteria; Enterobacterales; Yersiniaceae; Serratia
MNKRRTTLTVLALIASLGLSSAPALADKGGNGNGNGNGHGNSGNHGNSGNHGNNGNKGNK WP_033646351.1
MULTISPECIES: membrane protein [Enterobacterales] Length: 161\nNumber of predicted TMHs:  2\nExp number of AAs in TMHs: 31.11104\nExp number, first 60 AAs:  14.67675\nTotal prob of N-in:   0.95023\nPOSSIBLE N-term signal sequence\ninside   1   6\nTMhelix   7   26\noutside   27   141\nTMhelix   142   160\ninside   161   161

28263 GCF_000783975.1_ASM78397v1 Serratia marcescens Proteobacteria; Gammaproteobacteria; Enterobacterales; Yersiniaceae; Serratia
MNKRRTTLTVLALIASLGLSSAPAMADKGGNGNGNGHGNSGNHGNNGNHGNNGNHGNKDK WP_047576109.1
membrane protein [Serratia marcescens] Length: 162\nNumber of predicted TMHs:  2\nExp number of AAs in TMHs: 31.23828\nExp number, first 60 AAs:  14.93725\nTotal prob of N-in:   0.95239\nPOSSIBLE N-term signal sequence\ninside   1   6\nTMhelix   7   26\noutside   27   142\nTMhelix   143   161\ninside   162   162

28264 GCF_001007555.1_2015-04-28 Serratia marcescens Proteobacteria; Gammaproteobacteria; Enterobacterales; Yersiniaceae; Serratia
MNKRRTTLSVLALIASLGLSSAPAMADKGGNGNGNGHGNSGNHGNNGNHGNNGNHGNKDK WP_046898556.1
membrane protein [Serratia marcescens] Length: 162\nNumber of predicted TMHs:  2\nExp number of AAs in TMHs: 32.71622\nExp number, first 60 AAs:  16.20553\nTotal prob of N-in:   0.95416\nPOSSIBLE N-term signal sequence\ninside   1   6\nTMhelix   7   26\noutside   27   142\nTMhelix   143   161\ninside   162   162

28265 GCF_000751195.1_Serratia_assembly Serratia marcescens Proteobacteria; Gammaproteobacteria; Enterobacterales; Yersiniaceae; Serratia
MNKRRTTLTVLALIASLGLSSAPALADKGGNGNGNGNGHGNSGNHGNNGNHGNNGNHGNK WP_033640576.1
MULTISPECIES: membrane protein [Serratia] Length: 164\nNumber of predicted TMHs:  2\nExp number of AAs in TMHs: 30.54778\nExp number, first 60 AAs:  14.38033\nTotal prob of N-in:   0.94595\nPOSSIBLE N-term signal sequence\ninside   1   6\nTMhelix   7   26\noutside   27   144\nTMhelix   145   163\ninside   164   164

28266 GCF_001063125.1_ASM106312v1 Serratia marcescens Proteobacteria; Gammaproteobacteria; Enterobacterales; Yersiniaceae; Serratia
MNKRRTTLTVLALIASLGLSSAPALADKGGNGNGNGNGHGNSGNHGNNGNHGNNGNNGNH WP_004941297.1
MULTISPECIES: hypothetical protein [Serratia] Length: 167\nNumber of predicted TMHs:  2\nExp number of AAs in TMHs: 30.02384\nExp number, first 60 AAs:  14.11185\nTotal prob of N-in:   0.94170\nPOSSIBLE N-term signal sequence\ninside   1   6\nTMhelix   7   26\noutside   27   147\nTMhelix   148   166\ninside   167   167

28267 GCF_001062235.1_ASM106223v1 Serratia marcescens Proteobacteria; Gammaproteobacteria; Enterobacterales; Yersiniaceae; Serratia
MNKRRTTLTVLALIASLGLSSAPALADKGGNGNGNGNGHGNSGNHGNNGNHGNNGNNGNN WP_048323073.1
membrane protein [Serratia marcescens] Length: 170\nNumber of predicted TMHs:  2\nExp number of AAs in TMHs: 29.49026\nExp number, first 60 AAs:  13.84277\nTotal prob of N-in:   0.93726\nPOSSIBLE N-term signal sequence\ninside   1   6\nTMhelix   7   26\noutside   27   150\nTMhelix   151   169\ninside   170   170

28268 GCF_001063175.1_ASM106317v1 Serratia marcescens Proteobacteria; Gammaproteobacteria; Enterobacterales; Yersiniaceae; Serratia

MNKRRTTLTVLALIASLGLSSAPALADKGGNGNGNGNGHGNSGNHGNNGNHGNNGNNGNH WP_004941297.1 MULTISPECIES: hypothetical protein [Serratia] Length: 167\nNumber of predicted TMHs: 2\nExp number of AAs in TMHs: 30.02384\nExp number, first 60 AAs: 14.11185\nTotal prob of N-in: 0.94170\nPOSSIBLE N-term signal sequence\ninside 1 6\nTMhelix 7 26\noutside 27 147\nTMhelix 148 166\ninside 167 167

28269 GCF_001064455.1_ASM106445v1 Serratia marcescens Proteobacteria; Gammaproteobacteria; Enterobacterales; Yersiniaceae; Serratia MNKRRTTLTVLALIASLGLSSAPALADKGGNGNGNGNGHGNSGNHGNNGNHGNNGNNGNH WP_004941297.1 MULTISPECIES: hypothetical protein [Serratia] Length: 167\nNumber of predicted TMHs: 2\nExp number of AAs in TMHs: 30.02384\nExp number, first 60 AAs: 14.11185\nTotal prob of N-in: 0.94170\nPOSSIBLE N-term signal sequence\ninside 1 6\nTMhelix 7 26\noutside 27 147\nTMhelix 148 166\ninside 167 167

28270 GCF_001538505.1_12082_3_45 Serratia marcescens Proteobacteria; Gammaproteobacteria; Enterobacterales; Yersiniaceae; Serratia MNKRRTTLTVLALIASLGLSSAPALADKGGNGNGNGNGHGNSGNHGNSGNHGNNGNKGNK WP_033646351.1 MULTISPECIES: membrane protein [Enterobacterales] Length: 161\nNumber of predicted TMHs: 2\nExp number of AAs in TMHs: 31.11104\nExp number, first 60 AAs: 14.67675\nTotal prob of N-in: 0.95023\nPOSSIBLE N-term signal sequence\ninside 1 6\nTMhelix 7 26\noutside 27 141\nTMhelix 142 160\ninside 161 161

28271 GCF_001536585.1_12082_5_39 Serratia marcescens Proteobacteria; Gammaproteobacteria; Enterobacterales; Yersiniaceae; Serratia MNKRRTTLTVLALIASLGLSSAPALADKGGNGNGNGNGHGNSGNHGNSGNHGNNGNKGNK WP_033646351.1 MULTISPECIES: membrane protein [Enterobacterales] Length: 161\nNumber of predicted TMHs: 2\nExp number of AAs in TMHs: 31.11104\nExp number, first 60 AAs: 14.67675\nTotal prob of N-in: 0.95023\nPOSSIBLE N-term signal sequence\ninside 1 6\nTMhelix 7 26\noutside 27 141\nTMhelix 142 160\ninside 161 161

28272 GCF_000783915.1_ASM78391v1 Serratia marcescens Proteobacteria; Gammaproteobacteria; Enterobacterales; Yersiniaceae; Serratia MNKRRTTLTVLALIASLGLSSAPALADKGGNGNGNGNGHGNSGNHGNSGNHGNNGNNGNN WP_047567898.1 membrane protein [Serratia marcescens] Length: 170\nNumber of predicted TMHs: 2\nExp number of AAs in TMHs: 29.59643\nExp number, first 60 AAs: 13.79423\nTotal prob of N-in: 0.93785\nPOSSIBLE N-term signal sequence\ninside 1 6\nTMhelix 7 26\noutside 27 150\nTMhelix 151 169\ninside 170 170

28273 GCF_001034395.1_Serr_marc_UCI87_V1 Serratia marcescens Proteobacteria; Gammaproteobacteria; Enterobacterales; Yersiniaceae; Serratia MNKRRTTLTVLALIASLGLSSAPALADKGGNGNGNGNGHGNSGNHGNNGNHGNNGNNGNN WP_048323073.1 membrane protein [Serratia marcescens] Length: 170\nNumber of predicted TMHs: 2\nExp number of AAs in TMHs: 29.49026\nExp number, first 60 AAs: 13.84277\nTotal prob of N-in: 0.93726\nPOSSIBLE N-term signal sequence\ninside 1 6\nTMhelix 7 26\noutside 27 150\nTMhelix 151 169\ninside 170 170

28274 GCF_000743395.1_SAS_1 Serratia marcescens Proteobacteria; Gammaproteobacteria; Enterobacterales; Yersiniaceae; Serratia MNKRRTTLTVLALIASLGLSSAPALADKGGNGNGNGNGHGNSGNHGNNGNHGNNGNHGNK WP_033640576.1 MULTISPECIES: membrane protein [Serratia] Length: 164\nNumber of predicted TMHs: 2\nExp number of AAs in TMHs: 30.54778\nExp number, first 60 AAs: 14.38033\nTotal prob of N-in: 0.94595\nPOSSIBLE N-term signal sequence\ninside 1 6\nTMhelix 7 26\noutside 27 144\nTMhelix 145 163\ninside 164 164

28275 GCF_001060655.1_ASM106065v1 Serratia marcescens Proteobacteria; Gammaproteobacteria; Enterobacterales; Yersiniaceae; Serratia MNKRRTTLTVLALIASLGLSSAPALADKGGNGNGNGNGHGNSGNHGNNGNHGNNGNHGNK WP_033640576.1 MULTISPECIES: membrane protein [Serratia] Length: 164\nNumber of predicted TMHs: 2\nExp number of AAs in TMHs: 30.54778\nExp number, first 60 AAs: 14.38033\nTotal prob of N-in: 0.94595\nPOSSIBLE N-term signal sequence\ninside 1 6\nTMhelix 7 26\noutside 27 144\nTMhelix 145 163\ninside 164 164

28276 GCF_001061195.1_ASM106119v1 Serratia marcescens Proteobacteria; Gammaproteobacteria; Enterobacterales; Yersiniaceae; Serratia MNKRRTTLTVLALIASLGLSSAPALADKGGNGNGNGNGHGNSGNHGNSGNHGNNGNKGNK WP_033646351.1 MULTISPECIES: membrane protein [Enterobacterales] Length: 161\nNumber of predicted TMHs: 2\nExp number of AAs in TMHs: 31.11104\nExp number, first 60 AAs: 14.67675\nTotal prob of N-in: 0.95023\nPOSSIBLE N-term signal sequence\ninside 1 6\nTMhelix 7 26\noutside 27 141\nTMhelix 142 160\ninside 161 161

28277	GCF_001062285.1_ASM106228v1	Serratia marcescens	Proteobacteria; Gammaproteobacteria; Enterobacterales; Yersiniaceae; Serratia
MNKRRTTLTVLALIASLGLSSAPALADKGGNGNGNGNGHGNSGNHGNSGNHGNNGNKGNK	WP_033646351.1
MULTISPECIES: membrane protein [Enterobacterales]	Length: 161\nNumber of predicted TMHs:  2\nExp number of AAs in TMHs: 31.11104\nExp number, first 60 AAs:  14.67675\nTotal prob of N-in:   0.95023\nPOSSIBLE N-term signal sequence\ninside   1   6\nTMhelix   7   26\noutside   27   141\nTMhelix   142   160\ninside   161   161

28278	GCF_001063145.1_ASM106314v1	Serratia marcescens	Proteobacteria; Gammaproteobacteria; Enterobacterales; Yersiniaceae; Serratia
MNKRRTTLTVLALIASLGLSSAPALADKGGNGNGNGNGHGNSGNHGNNGNHGNNGNNGNH	WP_004941297.1
MULTISPECIES: hypothetical protein [Serratia]	Length: 167\nNumber of predicted TMHs:  2\nExp number of AAs in TMHs: 30.02384\nExp number, first 60 AAs:  14.11185\nTotal prob of N-in:   0.94170\nPOSSIBLE N-term signal sequence\ninside   1   6\nTMhelix   7   26\noutside   27   147\nTMhelix   148   166\ninside   167   167

28279	GCF_001536645.1_12082_3_73	Serratia marcescens	Proteobacteria; Gammaproteobacteria; Enterobacterales; Yersiniaceae; Serratia
MNKRRTTLTVLALIASLGLSSAPALADKGGNGNGNGNGHGNSGNHGNSGNHGNNGNKGNK	WP_033646351.1
MULTISPECIES: membrane protein [Enterobacterales]	Length: 161\nNumber of predicted TMHs:  2\nExp number of AAs in TMHs: 31.11104\nExp number, first 60 AAs:  14.67675\nTotal prob of N-in:   0.95023\nPOSSIBLE N-term signal sequence\ninside   1   6\nTMhelix   7   26\noutside   27   141\nTMhelix   142   160\ninside   161   161

28280	GCF_001536705.1_12082_3_61	Serratia marcescens	Proteobacteria; Gammaproteobacteria; Enterobacterales; Yersiniaceae; Serratia
MNKRRTTLTVLALIASLGLSSAPALADKGGNGNGNGNGHGNSGNHGNSGNHGNNGNKGNK	WP_033646351.1
MULTISPECIES: membrane protein [Enterobacterales]	Length: 161\nNumber of predicted TMHs:  2\nExp number of AAs in TMHs: 31.11104\nExp number, first 60 AAs:  14.67675\nTotal prob of N-in:   0.95023\nPOSSIBLE N-term signal sequence\ninside   1   6\nTMhelix   7   26\noutside   27   141\nTMhelix   142   160\ninside   161   161

28281	GCF_001536745.1_12082_3_86	Serratia marcescens	Proteobacteria; Gammaproteobacteria; Enterobacterales; Yersiniaceae; Serratia
MNKRRTTLTVLALIASLGLSSAPALADKGGNGNGNGNGHGNSGNHGNSGNHGNNGNKGNK	WP_033646351.1
MULTISPECIES: membrane protein [Enterobacterales]	Length: 161\nNumber of predicted TMHs:  2\nExp number of AAs in TMHs: 31.11104\nExp number, first 60 AAs:  14.67675\nTotal prob of N-in:   0.95023\nPOSSIBLE N-term signal sequence\ninside   1   6\nTMhelix   7   26\noutside   27   141\nTMhelix   142   160\ninside   161   161

28282	GCF_001536895.1_12082_3_47	Serratia marcescens	Proteobacteria; Gammaproteobacteria; Enterobacterales; Yersiniaceae; Serratia
MNKRRTTLTVLALIASLGLSSAPALADKGGNGNGNGNGHGNSGNHGNSGNHGNNGNKGNK	WP_033646351.1
MULTISPECIES: membrane protein [Enterobacterales]	Length: 161\nNumber of predicted TMHs:  2\nExp number of AAs in TMHs: 31.11104\nExp number, first 60 AAs:  14.67675\nTotal prob of N-in:   0.95023\nPOSSIBLE N-term signal sequence\ninside   1   6\nTMhelix   7   26\noutside   27   141\nTMhelix   142   160\ninside   161   161

28283	GCF_001536485.1_12082_2_24	Serratia marcescens	Proteobacteria; Gammaproteobacteria; Enterobacterales; Yersiniaceae; Serratia
MNKRRTTLTVLALIASLGLSSAPALADKGGNGNGNGNGHGNSGNHGNSGNHGNNGNKGNK	WP_033646351.1
MULTISPECIES: membrane protein [Enterobacterales]	Length: 161\nNumber of predicted TMHs:  2\nExp number of AAs in TMHs: 31.11104\nExp number, first 60 AAs:  14.67675\nTotal prob of N-in:   0.95023\nPOSSIBLE N-term signal sequence\ninside   1   6\nTMhelix   7   26\noutside   27   141\nTMhelix   142   160\ninside   161   161

28284	GCF_001536965.1_12082_3_42	Serratia marcescens	Proteobacteria; Gammaproteobacteria; Enterobacterales; Yersiniaceae; Serratia
MNKRRTTLTVLALIASLGLSSAPALADKGGNGNGNGNGHGNSGNHGNSGNHGNNGNKGNK	WP_033646351.1
MULTISPECIES: membrane protein [Enterobacterales]	Length: 161\nNumber of predicted TMHs:  2\nExp number of AAs in TMHs: 31.11104\nExp number, first 60 AAs:  14.67675\nTotal prob of N-in:   0.95023\nPOSSIBLE N-term signal sequence\ninside   1   6\nTMhelix   7   26\noutside   27   141\nTMhelix   142   160\ninside   161   161

28285	GCF_001537025.1_12082_2_54	Serratia marcescens	Proteobacteria; Gammaproteobacteria; Enterobacterales; Yersiniaceae; Serratia
MNKRRTTLTVLALIASLGLSSAPALADKGGNGNGNGNGHGNSGNHGNSGNHGNNGNKGNK	WP_033646351.1
MULTISPECIES: membrane protein [Enterobacterales]	Length: 161\nNumber of predicted TMHs:  2\nExp

number of AAs in TMHs: 31.11104\nExp number, first 60 AAs: 14.67675\nTotal prob of N-in: 0.95023\nPOSSIBLE N-term signal sequence\ninside 1 6\nTMhelix 7 26\noutside 27 141\nTMhelix 142 160\ninside 161 161

28286 GCF_001537165.1_12082_5_44 Serratia marcescens Proteobacteria; Gammaproteobacteria; Enterobacterales; Yersiniaceae; Serratia
MNKRRTTLTVLALIASLGLSSAPALADKGGNGNGNGNGHGNSGNHGNSGNHGNNGNKGNK WP_033646351.1
MULTISPECIES: membrane protein [Enterobacterales] Length: 161\nNumber of predicted TMHs: 2\nExp number of AAs in TMHs: 31.11104\nExp number, first 60 AAs: 14.67675\nTotal prob of N-in: 0.95023\nPOSSIBLE N-term signal sequence\ninside 1 6\nTMhelix 7 26\noutside 27 141\nTMhelix 142 160\ninside 161 161

28287 GCF_001537225.1_12082_2_72 Serratia marcescens Proteobacteria; Gammaproteobacteria; Enterobacterales; Yersiniaceae; Serratia
MNKRRTTLTVLALIASLGLSSAPALADKGGNGNGNGNGHGNSGNHGNSGNHGNNGNKGNK WP_033646351.1
MULTISPECIES: membrane protein [Enterobacterales] Length: 161\nNumber of predicted TMHs: 2\nExp number of AAs in TMHs: 31.11104\nExp number, first 60 AAs: 14.67675\nTotal prob of N-in: 0.95023\nPOSSIBLE N-term signal sequence\ninside 1 6\nTMhelix 7 26\noutside 27 141\nTMhelix 142 160\ninside 161 161

28288 GCF_001537125.1_12082_3_20 Serratia marcescens Proteobacteria; Gammaproteobacteria; Enterobacterales; Yersiniaceae; Serratia
MNKRRTTLTVLALIASLGLSSAPALADKGGNGNGNGNGHGNSGNHGNSGNHGNNGNKGNK WP_033646351.1
MULTISPECIES: membrane protein [Enterobacterales] Length: 161\nNumber of predicted TMHs: 2\nExp number of AAs in TMHs: 31.11104\nExp number, first 60 AAs: 14.67675\nTotal prob of N-in: 0.95023\nPOSSIBLE N-term signal sequence\ninside 1 6\nTMhelix 7 26\noutside 27 141\nTMhelix 142 160\ninside 161 161

28289 GCF_001537305.1_12082_5_41 Serratia marcescens Proteobacteria; Gammaproteobacteria; Enterobacterales; Yersiniaceae; Serratia
MNKRRTTLTVLALIASLGLSSAPALADKGGNGNGNGNGHGNSGNHGNSGNHGNNGNKGNK WP_033646351.1
MULTISPECIES: membrane protein [Enterobacterales] Length: 161\nNumber of predicted TMHs: 2\nExp number of AAs in TMHs: 31.11104\nExp number, first 60 AAs: 14.67675\nTotal prob of N-in: 0.95023\nPOSSIBLE N-term signal sequence\ninside 1 6\nTMhelix 7 26\noutside 27 141\nTMhelix 142 160\ninside 161 161

28290 GCF_001537365.1_12082_3_5 Serratia marcescens Proteobacteria; Gammaproteobacteria; Enterobacterales; Yersiniaceae; Serratia
MNKRRTTLTVLALIASLGLSSAPALADKGGNGNGNGNGHGNSGNHGNSGNHGNNGNKGNK WP_033646351.1
MULTISPECIES: membrane protein [Enterobacterales] Length: 161\nNumber of predicted TMHs: 2\nExp number of AAs in TMHs: 31.11104\nExp number, first 60 AAs: 14.67675\nTotal prob of N-in: 0.95023\nPOSSIBLE N-term signal sequence\ninside 1 6\nTMhelix 7 26\noutside 27 141\nTMhelix 142 160\ninside 161 161

28291 GCF_001537485.1_12082_2_75 Serratia marcescens Proteobacteria; Gammaproteobacteria; Enterobacterales; Yersiniaceae; Serratia
MNKRRTTLTVLALIASLGLSSAPALADKGGNGNGNGNGHGNSGNHGNSGNHGNNGNKGNK WP_033646351.1
MULTISPECIES: membrane protein [Enterobacterales] Length: 161\nNumber of predicted TMHs: 2\nExp number of AAs in TMHs: 31.11104\nExp number, first 60 AAs: 14.67675\nTotal prob of N-in: 0.95023\nPOSSIBLE N-term signal sequence\ninside 1 6\nTMhelix 7 26\noutside 27 141\nTMhelix 142 160\ninside 161 161

28292 GCF_001537525.1_12082_2_25 Serratia marcescens Proteobacteria; Gammaproteobacteria; Enterobacterales; Yersiniaceae; Serratia
MNKRRTTLTVLALIASLGLSSAPALADKGGNGNGNGNGHGNSGNHGNSGNHGNNGNKGNK WP_033646351.1
MULTISPECIES: membrane protein [Enterobacterales] Length: 161\nNumber of predicted TMHs: 2\nExp number of AAs in TMHs: 31.11104\nExp number, first 60 AAs: 14.67675\nTotal prob of N-in: 0.95023\nPOSSIBLE N-term signal sequence\ninside 1 6\nTMhelix 7 26\noutside 27 141\nTMhelix 142 160\ninside 161 161

28293 GCF_001537345.1_12082_2_78 Serratia marcescens Proteobacteria; Gammaproteobacteria; Enterobacterales; Yersiniaceae; Serratia
MNKRRTTLTVLALIASLGLSSAPALADKGGNGNGNGNGHGNSGNHGNSGNHGNNGNKGNK WP_060427691.1
hypothetical protein [Serratia marcescens] Length: 161\nNumber of predicted TMHs: 2\nExp number of AAs in TMHs: 31.15272\nExp number, first 60 AAs: 14.60251\nTotal prob of N-in: 0.95045\nPOSSIBLE N-term signal sequence\ninside 1 6\nTMhelix 7 26\noutside 27 141\nTMhelix 142 160\ninside 161 161

28294 GCF_001537585.1_12082_3_71 Serratia marcescens Proteobacteria; Gammaproteobacteria; Enterobacterales; Yersiniaceae; Serratia

MNKRRTTLTVLALIASLGLSSAPALADKGGNGNGNGNGHGNSGNHGNSGNHGNNGNKGNK WP_033646351.1 MULTISPECIES: membrane protein [Enterobacterales] Length: 161\nNumber of predicted TMHs: 2\nExp number of AAs in TMHs: 31.11104\nExp number, first 60 AAs: 14.67675\nTotal prob of N-in: 0.95023\nPOSSIBLE N-term signal sequence\ninside 1 6\nTMhelix 7 26\noutside 27 141\nTMhelix 142 160\ninside 161 161

28295 GCF_001538195.1_12045_8_14 Serratia marcescens Proteobacteria; Gammaproteobacteria; Enterobacterales; Yersiniaceae; Serratia MNKRRTTLTVLALIASLGLSSAPALADKGGNGNGNGNGHGNSGNHGNSGNHGNNGNKGNK WP_033646351.1 MULTISPECIES: membrane protein [Enterobacterales] Length: 161\nNumber of predicted TMHs: 2\nExp number of AAs in TMHs: 31.11104\nExp number, first 60 AAs: 14.67675\nTotal prob of N-in: 0.95023\nPOSSIBLE N-term signal sequence\ninside 1 6\nTMhelix 7 26\noutside 27 141\nTMhelix 142 160\ninside 161 161

28296 GCF_001538345.1_12082_2_92 Serratia marcescens Proteobacteria; Gammaproteobacteria; Enterobacterales; Yersiniaceae; Serratia MNKRRTTLTVLALIASLGLSSAPALADKGGNGNGNGNGHGNSGNHGNSGNHGNNGNKGNK WP_033646351.1 MULTISPECIES: membrane protein [Enterobacterales] Length: 161\nNumber of predicted TMHs: 2\nExp number of AAs in TMHs: 31.11104\nExp number, first 60 AAs: 14.67675\nTotal prob of N-in: 0.95023\nPOSSIBLE N-term signal sequence\ninside 1 6\nTMhelix 7 26\noutside 27 141\nTMhelix 142 160\ninside 161 161

28297 GCF_001538405.1_12082_3_65 Serratia marcescens Proteobacteria; Gammaproteobacteria; Enterobacterales; Yersiniaceae; Serratia MNKRRTTLTVLALIASLGLSSAPALADKGGNGNGNGNGHGNSGNHGNSGNHGNNGNKGNK WP_033646351.1 MULTISPECIES: membrane protein [Enterobacterales] Length: 161\nNumber of predicted TMHs: 2\nExp number of AAs in TMHs: 31.11104\nExp number, first 60 AAs: 14.67675\nTotal prob of N-in: 0.95023\nPOSSIBLE N-term signal sequence\ninside 1 6\nTMhelix 7 26\noutside 27 141\nTMhelix 142 160\ninside 161 161

28298 GCF_001538445.1_12082_2_56 Serratia marcescens Proteobacteria; Gammaproteobacteria; Enterobacterales; Yersiniaceae; Serratia MNKRRTTLTVLALIASLGLSSAPALADKGGNGNGNGNGHGNSGNHGNSGNHGNNGNKGNK WP_033646351.1 MULTISPECIES: membrane protein [Enterobacterales] Length: 161\nNumber of predicted TMHs: 2\nExp number of AAs in TMHs: 31.11104\nExp number, first 60 AAs: 14.67675\nTotal prob of N-in: 0.95023\nPOSSIBLE N-term signal sequence\ninside 1 6\nTMhelix 7 26\noutside 27 141\nTMhelix 142 160\ninside 161 161

28299 GCF_001538465.1_12045_8_5 Serratia marcescens Proteobacteria; Gammaproteobacteria; Enterobacterales; Yersiniaceae; Serratia MNKRRTTLTVLALIASLGLSSAPALADKGGNGNGNGNGHGNSGNHGNSGNHGNNGNKGNK WP_033646351.1 MULTISPECIES: membrane protein [Enterobacterales] Length: 161\nNumber of predicted TMHs: 2\nExp number of AAs in TMHs: 31.11104\nExp number, first 60 AAs: 14.67675\nTotal prob of N-in: 0.95023\nPOSSIBLE N-term signal sequence\ninside 1 6\nTMhelix 7 26\noutside 27 141\nTMhelix 142 160\ninside 161 161

28300 GCF_001538565.1_12082_3_96 Serratia marcescens Proteobacteria; Gammaproteobacteria; Enterobacterales; Yersiniaceae; Serratia MNKRRTTLTVLALIASLGLSSAPALADKGGNGNGNGNGHGNSGNHGNSGNHGNNGNKGNK WP_033646351.1 MULTISPECIES: membrane protein [Enterobacterales] Length: 161\nNumber of predicted TMHs: 2\nExp number of AAs in TMHs: 31.11104\nExp number, first 60 AAs: 14.67675\nTotal prob of N-in: 0.95023\nPOSSIBLE N-term signal sequence\ninside 1 6\nTMhelix 7 26\noutside 27 141\nTMhelix 142 160\ninside 161 161

28301 GCF_001538625.1_12082_2_44 Serratia marcescens Proteobacteria; Gammaproteobacteria; Enterobacterales; Yersiniaceae; Serratia MNKRRTTLTVLALIASLGLSSAPALADKGGNGNGNGNGHGNSGNHGNSGNHGNNGNKGNK WP_033646351.1 MULTISPECIES: membrane protein [Enterobacterales] Length: 161\nNumber of predicted TMHs: 2\nExp number of AAs in TMHs: 31.11104\nExp number, first 60 AAs: 14.67675\nTotal prob of N-in: 0.95023\nPOSSIBLE N-term signal sequence\ninside 1 6\nTMhelix 7 26\noutside 27 141\nTMhelix 142 160\ninside 161 161

28302 GCF_001539125.1_12045_8_6 Serratia marcescens Proteobacteria; Gammaproteobacteria; Enterobacterales; Yersiniaceae; Serratia MNKRRTTLTVLALIASLGLSSAPALADKGGNGNGNGNGHGNSGNHGNSGNHGNNGNKGNK WP_033646351.1 MULTISPECIES: membrane protein [Enterobacterales] Length: 161\nNumber of predicted TMHs: 2\nExp number of AAs in TMHs: 31.11104\nExp number, first 60 AAs: 14.67675\nTotal prob of N-in: 0.95023\nPOSSIBLE N-term signal sequence\ninside 1 6\nTMhelix 7 26\noutside 27 141\nTMhelix 142 160\ninside 161 161

28303 GCF_001539205.1_12082_3_59 Serratia marcescens Proteobacteria; Gammaproteobacteria; Enterobacterales; Yersiniaceae; Serratia
MNKRRTTLTVLALIASLGLSSAPALADKGGNGNGNGNGHGNSGNHGNSGNHGNNGNKGNK WP_033646351.1
MULTISPECIES: membrane protein [Enterobacterales] Length: 161\nNumber of predicted TMHs: 2\nExp number of AAs in TMHs: 31.11104\nExp number, first 60 AAs: 14.67675\nTotal prob of N-in: 0.95023\nPOSSIBLE N-term signal sequence\ninside 1 6\nTMhelix 7 26\noutside 27 141\nTMhelix 142 160\ninside 161 161

28304 GCF_001538925.1_12082_3_9 Serratia marcescens Proteobacteria; Gammaproteobacteria; Enterobacterales; Yersiniaceae; Serratia
MNKRRTTLTVLALIASLGLSSAPALADKGGNGNGNGNGHGNSGNHGNSGNHGNNGNKGNK WP_060427691.1
hypothetical protein [Serratia marcescens] Length: 161\nNumber of predicted TMHs: 2\nExp number of AAs in TMHs: 31.15272\nExp number, first 60 AAs: 14.60251\nTotal prob of N-in: 0.95045\nPOSSIBLE N-term signal sequence\ninside 1 6\nTMhelix 7 26\noutside 27 141\nTMhelix 142 160\ninside 161 161

28305 GCF_001539265.1_12082_2_61 Serratia marcescens Proteobacteria; Gammaproteobacteria; Enterobacterales; Yersiniaceae; Serratia
MNKRRTTLTVLALIASLGLSSAPALADKGGNGNGNGNGHGNSGNHGNSGNHGNNGNKGNK WP_033646351.1
MULTISPECIES: membrane protein [Enterobacterales] Length: 161\nNumber of predicted TMHs: 2\nExp number of AAs in TMHs: 31.11104\nExp number, first 60 AAs: 14.67675\nTotal prob of N-in: 0.95023\nPOSSIBLE N-term signal sequence\ninside 1 6\nTMhelix 7 26\noutside 27 141\nTMhelix 142 160\ninside 161 161

28306 GCF_001539245.1_12082_3_24 Serratia marcescens Proteobacteria; Gammaproteobacteria; Enterobacterales; Yersiniaceae; Serratia
MNKRRTTLTVLALIASLGLSSAPALADKGGNGNGNGNGHGNSGNHGNSGNHGNNGNKGNK WP_033646351.1
MULTISPECIES: membrane protein [Enterobacterales] Length: 161\nNumber of predicted TMHs: 2\nExp number of AAs in TMHs: 31.11104\nExp number, first 60 AAs: 14.67675\nTotal prob of N-in: 0.95023\nPOSSIBLE N-term signal sequence\ninside 1 6\nTMhelix 7 26\noutside 27 141\nTMhelix 142 160\ninside 161 161

28307 GCF_001539365.1_12082_2_58 Serratia marcescens Proteobacteria; Gammaproteobacteria; Enterobacterales; Yersiniaceae; Serratia
MNKRRTTLTVLALIASLGLSSAPALADKGGNGNGNGNGHGNSGNHGNSGNHGNNGNKGNK WP_033646351.1
MULTISPECIES: membrane protein [Enterobacterales] Length: 161\nNumber of predicted TMHs: 2\nExp number of AAs in TMHs: 31.11104\nExp number, first 60 AAs: 14.67675\nTotal prob of N-in: 0.95023\nPOSSIBLE N-term signal sequence\ninside 1 6\nTMhelix 7 26\noutside 27 141\nTMhelix 142 160\ninside 161 161

28308 GCF_001539425.1_12082_2_81 Serratia marcescens Proteobacteria; Gammaproteobacteria; Enterobacterales; Yersiniaceae; Serratia
MNKRRTTLTVLALIASLGLSSAPALADKGGNGNGNGNGHGNSGNHGNSGNHGNNGNKGNK WP_033646351.1
MULTISPECIES: membrane protein [Enterobacterales] Length: 161\nNumber of predicted TMHs: 2\nExp number of AAs in TMHs: 31.11104\nExp number, first 60 AAs: 14.67675\nTotal prob of N-in: 0.95023\nPOSSIBLE N-term signal sequence\ninside 1 6\nTMhelix 7 26\noutside 27 141\nTMhelix 142 160\ninside 161 161

28309 GCF_001536425.1_12082_3_92 Serratia marcescens Proteobacteria; Gammaproteobacteria; Enterobacterales; Yersiniaceae; Serratia
MNKRRTTLTVLALIASLGLSSAPALADKGGNGNGNGNGHGNSGNHGNSGNHGNNGNKGNK WP_033646351.1
MULTISPECIES: membrane protein [Enterobacterales] Length: 161\nNumber of predicted TMHs: 2\nExp number of AAs in TMHs: 31.11104\nExp number, first 60 AAs: 14.67675\nTotal prob of N-in: 0.95023\nPOSSIBLE N-term signal sequence\ninside 1 6\nTMhelix 7 26\noutside 27 141\nTMhelix 142 160\ninside 161 161

28310 GCF_001539305.1_12045_8_2 Serratia marcescens Proteobacteria; Gammaproteobacteria; Enterobacterales; Yersiniaceae; Serratia
MNKRRTTLTVLALIASLGLSSAPALADKGGNGNGNGNGHGNSGNNGNHGNNGNHGNKGNK WP_060419303.1
MULTISPECIES: hypothetical protein [Serratia] Length: 164\nNumber of predicted TMHs: 2\nExp number of AAs in TMHs: 30.51677\nExp number, first 60 AAs: 14.42464\nTotal prob of N-in: 0.94579\nPOSSIBLE N-term signal sequence\ninside 1 6\nTMhelix 7 26\noutside 27 144\nTMhelix 145 163\ninside 164 164

28311 GCF_001539925.1_12082_2_80 Serratia marcescens Proteobacteria; Gammaproteobacteria; Enterobacterales; Yersiniaceae; Serratia
MNKRRTTLTVLALIASLGLSSAPALADKGGNGNGNGNGHGNSGNHGNSGNHGNNGNKGNK WP_033646351.1
MULTISPECIES: membrane protein [Enterobacterales] Length: 161\nNumber of predicted TMHs: 2\nExp

number of AAs in TMHs: 31.11104\nExp number, first 60 AAs: 14.67675\nTotal prob of N-in: 0.95023\nPOSSIBLE N-term signal sequence\ninside 1 6\nTMhelix 7 26\noutside 27 141\nTMhelix 142 160\ninside 161 161

28312 GCF_001539825.1_12082_3_74 Serratia marcescens Proteobacteria; Gammaproteobacteria; Enterobacterales; Yersiniaceae; Serratia
MNKRRTTLTVLALIASLGLSSAPALADKGGNGNGNGNGHGNSGNNGNHGNNGNHGNKGNK WP_060419303.1
MULTISPECIES: hypothetical protein [Serratia] Length: 164\nNumber of predicted TMHs: 2\nExp number of AAs in TMHs: 30.51677\nExp number, first 60 AAs: 14.42464\nTotal prob of N-in: 0.94579\nPOSSIBLE N-term signal sequence\ninside 1 6\nTMhelix 7 26\noutside 27 144\nTMhelix 145 163\ninside 164 164

28313 GCF_001539985.1_12045_8_24 Serratia marcescens Proteobacteria; Gammaproteobacteria; Enterobacterales; Yersiniaceae; Serratia
MNKRRTTLTVLALIASLGLSSAPALADKGGNGNGNGNGHGNSGNHGNNGNHGNNGNHGNK WP_033640576.1
MULTISPECIES: membrane protein [Serratia] Length: 164\nNumber of predicted TMHs: 2\nExp number of AAs in TMHs: 30.54778\nExp number, first 60 AAs: 14.38033\nTotal prob of N-in: 0.94595\nPOSSIBLE N-term signal sequence\ninside 1 6\nTMhelix 7 26\noutside 27 144\nTMhelix 145 163\ninside 164 164

28314 GCF_001540045.1_12082_3_77 Serratia marcescens Proteobacteria; Gammaproteobacteria; Enterobacterales; Yersiniaceae; Serratia
MNKRRTTLTVLALIASLGLSSAPALADKGGNGNGNGNGHGNSGNHGNSGNHGNNGNKGNK WP_033646351.1
MULTISPECIES: membrane protein [Enterobacterales] Length: 161\nNumber of predicted TMHs: 2\nExp number of AAs in TMHs: 31.11104\nExp number, first 60 AAs: 14.67675\nTotal prob of N-in: 0.95023\nPOSSIBLE N-term signal sequence\ninside 1 6\nTMhelix 7 26\noutside 27 141\nTMhelix 142 160\ninside 161 161

28315 GCF_001540085.1_12082_5_40 Serratia marcescens Proteobacteria; Gammaproteobacteria; Enterobacterales; Yersiniaceae; Serratia
MNKRRTTLTVLALIASLGLSSAPALADKGGNGNGNGNGHGNSGNHGNSGNHGNNGNKGNK WP_033646351.1
MULTISPECIES: membrane protein [Enterobacterales] Length: 161\nNumber of predicted TMHs: 2\nExp number of AAs in TMHs: 31.11104\nExp number, first 60 AAs: 14.67675\nTotal prob of N-in: 0.95023\nPOSSIBLE N-term signal sequence\ninside 1 6\nTMhelix 7 26\noutside 27 141\nTMhelix 142 160\ninside 161 161

28316 GCF_001540025.1_12082_3_40 Serratia marcescens Proteobacteria; Gammaproteobacteria; Enterobacterales; Yersiniaceae; Serratia
MNKRRTTLTVLALIASLGLSSAPALADKGGNGNGNGNGHGNSGNNGNHGNNGNHGNKGNK WP_060419303.1
MULTISPECIES: hypothetical protein [Serratia] Length: 164\nNumber of predicted TMHs: 2\nExp number of AAs in TMHs: 30.51677\nExp number, first 60 AAs: 14.42464\nTotal prob of N-in: 0.94579\nPOSSIBLE N-term signal sequence\ninside 1 6\nTMhelix 7 26\noutside 27 144\nTMhelix 145 163\ninside 164 164

28317 GCF_001540145.1_12082_3_33 Serratia marcescens Proteobacteria; Gammaproteobacteria; Enterobacterales; Yersiniaceae; Serratia
MNKRRTTLTVLALIASLGLSSAPALADKGGNGNGNGNGHGNSGNHGNSGNHGNNGNKGNK WP_033646351.1
MULTISPECIES: membrane protein [Enterobacterales] Length: 161\nNumber of predicted TMHs: 2\nExp number of AAs in TMHs: 31.11104\nExp number, first 60 AAs: 14.67675\nTotal prob of N-in: 0.95023\nPOSSIBLE N-term signal sequence\ninside 1 6\nTMhelix 7 26\noutside 27 141\nTMhelix 142 160\ninside 161 161

28318 GCF_001540185.1_12082_2_41 Serratia marcescens Proteobacteria; Gammaproteobacteria; Enterobacterales; Yersiniaceae; Serratia
MNKRRTTLTVLALIASLGLSSAPALADKGGNGNGNGNGHGNSGNHGNSGNHGNNGNKGNK WP_033646351.1
MULTISPECIES: membrane protein [Enterobacterales] Length: 161\nNumber of predicted TMHs: 2\nExp number of AAs in TMHs: 31.11104\nExp number, first 60 AAs: 14.67675\nTotal prob of N-in: 0.95023\nPOSSIBLE N-term signal sequence\ninside 1 6\nTMhelix 7 26\noutside 27 141\nTMhelix 142 160\ninside 161 161

28319 GCF_001540645.1_12045_8_13 Serratia marcescens Proteobacteria; Gammaproteobacteria; Enterobacterales; Yersiniaceae; Serratia
MNKRRTTLTVLALIASLGLSSAPALADKGGNGNGNGNGHGNSGNHGNSGNHGNNGNKGNK WP_033646351.1
MULTISPECIES: membrane protein [Enterobacterales] Length: 161\nNumber of predicted TMHs: 2\nExp number of AAs in TMHs: 31.11104\nExp number, first 60 AAs: 14.67675\nTotal prob of N-in: 0.95023\nPOSSIBLE N-term signal sequence\ninside 1 6\nTMhelix 7 26\noutside 27 141\nTMhelix 142 160\ninside 161 161

28320 GCF_001540705.1_12082_3_70 Serratia marcescens Proteobacteria; Gammaproteobacteria; Enterobacterales; Yersiniaceae; Serratia

MNKRRTTLTVLALIASLGLSSAPALADKGGNGNGNGNGHGNSGNHGNSGNHGNNGNKGNK WP_033646351.1 MULTISPECIES: membrane protein [Enterobacterales] Length: 161\nNumber of predicted TMHs: 2\nExp number of AAs in TMHs: 31.11104\nExp number, first 60 AAs: 14.67675\nTotal prob of N-in: 0.95023\nPOSSIBLE N-term signal sequence\ninside 1 6\nTMhelix 7 26\noutside 27 141\nTMhelix 142 160\ninside 161 161

28321 GCF_001540765.1_12082_2_96 Serratia marcescens Proteobacteria; Gammaproteobacteria; Enterobacterales; Yersiniaceae; Serratia MNKRRTTLTVLALIASLGLSSAPALADKGGNGNGNGNGHGNSGNHGNSGNHGNNGNKGNK WP_033646351.1 MULTISPECIES: membrane protein [Enterobacterales] Length: 161\nNumber of predicted TMHs: 2\nExp number of AAs in TMHs: 31.11104\nExp number, first 60 AAs: 14.67675\nTotal prob of N-in: 0.95023\nPOSSIBLE N-term signal sequence\ninside 1 6\nTMhelix 7 26\noutside 27 141\nTMhelix 142 160\ninside 161 161

28322 GCF_001536405.1_12082_3_29 Serratia marcescens Proteobacteria; Gammaproteobacteria; Enterobacterales; Yersiniaceae; Serratia MNKRRTTLTVLALIASLGLSSAPALADKGGNGNGNGNGHGNSGNHGNSGNHGNNGNKGNK WP_033646351.1 MULTISPECIES: membrane protein [Enterobacterales] Length: 161\nNumber of predicted TMHs: 2\nExp number of AAs in TMHs: 31.11104\nExp number, first 60 AAs: 14.67675\nTotal prob of N-in: 0.95023\nPOSSIBLE N-term signal sequence\ninside 1 6\nTMhelix 7 26\noutside 27 141\nTMhelix 142 160\ninside 161 161

28323 GCF_001540805.1_12082_2_45 Serratia marcescens Proteobacteria; Gammaproteobacteria; Enterobacterales; Yersiniaceae; Serratia MNKRRTTLTVLALIASLGLSSAPALADKGGNGNGNGNGHGNSGNHGNSGNHGNNGNKGNK WP_033646351.1 MULTISPECIES: membrane protein [Enterobacterales] Length: 161\nNumber of predicted TMHs: 2\nExp number of AAs in TMHs: 31.11104\nExp number, first 60 AAs: 14.67675\nTotal prob of N-in: 0.95023\nPOSSIBLE N-term signal sequence\ninside 1 6\nTMhelix 7 26\noutside 27 141\nTMhelix 142 160\ninside 161 161

28324 GCF_001853455.1_ASM185345v1 Serratia marcescens Proteobacteria; Gammaproteobacteria; Enterobacterales; Yersiniaceae; Serratia MNKRRTTLTVLALIASLGLSSAPALADKGGNGNGNGNGHGNSGNNGNHGNNGNHGNKGNK WP_060419303.1 MULTISPECIES: hypothetical protein [Serratia] Length: 164\nNumber of predicted TMHs: 2\nExp number of AAs in TMHs: 30.51677\nExp number, first 60 AAs: 14.42464\nTotal prob of N-in: 0.94579\nPOSSIBLE N-term signal sequence\ninside 1 6\nTMhelix 7 26\noutside 27 144\nTMhelix 145 163\ninside 164 164

28325 GCF_001940505.1_ASM194050v1 Serratia marcescens Proteobacteria; Gammaproteobacteria; Enterobacterales; Yersiniaceae; Serratia MNKRRTTLTVLALIASLGLSSAPALADKGGNGNGNGNGHGNSGNHGNNGNHGNNGNNGNH WP_004941297.1 MULTISPECIES: hypothetical protein [Serratia] Length: 167\nNumber of predicted TMHs: 2\nExp number of AAs in TMHs: 30.02384\nExp number, first 60 AAs: 14.11185\nTotal prob of N-in: 0.94170\nPOSSIBLE N-term signal sequence\ninside 1 6\nTMhelix 7 26\noutside 27 147\nTMhelix 148 166\ninside 167 167

28326 GCF_001594385.1_ASM159438v1 Serratia marcescens Proteobacteria; Gammaproteobacteria; Enterobacterales; Yersiniaceae; Serratia MNKRRTTLTVLALIASLGLSSAPALADKGGNGNGNGNGHGNSGNHGNHGNHGNNGNNGNH WP_061872057.1 hypothetical protein [Serratia marcescens] Length: 167\nNumber of predicted TMHs: 2\nExp number of AAs in TMHs: 30.02384\nExp number, first 60 AAs: 14.11185\nTotal prob of N-in: 0.94170\nPOSSIBLE N-term signal sequence\ninside 1 6\nTMhelix 7 26\noutside 27 147\nTMhelix 148 166\ninside 167 167

28327 GCF_001064345.1_ASM106434v1 Serratia marcescens Proteobacteria; Gammaproteobacteria; Enterobacterales; Yersiniaceae; Serratia MNKRRTTLTVLALIASLGLSSAPALADKGGNGNGNGNGHGNSGNHGNNGNHGNNGNNGNH WP_004941297.1 MULTISPECIES: hypothetical protein [Serratia] Length: 167\nNumber of predicted TMHs: 2\nExp number of AAs in TMHs: 30.02384\nExp number, first 60 AAs: 14.11185\nTotal prob of N-in: 0.94170\nPOSSIBLE N-term signal sequence\ninside 1 6\nTMhelix 7 26\noutside 27 147\nTMhelix 148 166\ninside 167 167

28328 GCF_001022215.1_ASM102221v1 Serratia marcescens Proteobacteria; Gammaproteobacteria; Enterobacterales; Yersiniaceae; Serratia MNKRRTTLTVLALIASLGLSSAPALADKGGNGNGNGNGHGNSGNHGNNGNHGNNGNNGNH WP_004941297.1 MULTISPECIES: hypothetical protein [Serratia] Length: 167\nNumber of predicted TMHs: 2\nExp number of AAs in TMHs: 30.02384\nExp number, first 60 AAs: 14.11185\nTotal prob of N-in: 0.94170\nPOSSIBLE N-term signal sequence\ninside 1 6\nTMhelix 7 26\noutside 27 147\nTMhelix 148 166\ninside 167 167

28329	GCF_001060625.1_ASM106062v1	Serratia marcescens	Proteobacteria; Gammaproteobacteria; Enterobacterales; Yersiniaceae; Serratia
	MNKRRTTLTVLALIASLGLSSAPALADKGGNGNGNGNGHGNSGNHGNNGNHGNNGNHGNK	WP_033640576.1
	MULTISPECIES: membrane protein [Serratia]	Length: 164\nNumber of predicted TMHs:  2\nExp number of AAs in TMHs: 30.54778\nExp number, first 60 AAs:  14.38033\nTotal prob of N-in:   0.94595\nPOSSIBLE N-term signal sequence\ninside   1   6\nTMhelix   7   26\noutside   27   144\nTMhelix   145   163\ninside   164   164

28330	GCF_001536345.1_12082_2_67	Serratia marcescens	Proteobacteria; Gammaproteobacteria; Enterobacterales; Yersiniaceae; Serratia
	MNKRRTTLTVLALIASLGLSSAPALADKGGNGNGNGNGHGNSGNHGNSGNHGNNGNKGNK	WP_033646351.1
	MULTISPECIES: membrane protein [Enterobacterales]	Length: 161\nNumber of predicted TMHs:  2\nExp number of AAs in TMHs: 31.11104\nExp number, first 60 AAs:  14.67675\nTotal prob of N-in:   0.95023\nPOSSIBLE N-term signal sequence\ninside   1   6\nTMhelix   7   26\noutside   27   141\nTMhelix   142   160\ninside   161   161

28331	GCF_001537645.1_12082_2_39	Serratia marcescens	Proteobacteria; Gammaproteobacteria; Enterobacterales; Yersiniaceae; Serratia
	MNKRRTTLTVLALIASLGLSSAPALADKGGNGNGNGNGHGNSGNHGNSGNHGNNGNKGNK	WP_033646351.1
	MULTISPECIES: membrane protein [Enterobacterales]	Length: 161\nNumber of predicted TMHs:  2\nExp number of AAs in TMHs: 31.11104\nExp number, first 60 AAs:  14.67675\nTotal prob of N-in:   0.95023\nPOSSIBLE N-term signal sequence\ninside   1   6\nTMhelix   7   26\noutside   27   141\nTMhelix   142   160\ninside   161   161

28332	GCF_001537425.1_12082_2_48	Serratia marcescens	Proteobacteria; Gammaproteobacteria; Enterobacterales; Yersiniaceae; Serratia
	MNKRRTTLTVLALIASLGLSSAPALADKGGNGNGNGNGHGNSGNHGNSGNHGNNGNKGNK	WP_060427691.1
	hypothetical protein [Serratia marcescens]	Length: 161\nNumber of predicted TMHs:  2\nExp number of AAs in TMHs: 31.15272\nExp number, first 60 AAs:  14.60251\nTotal prob of N-in:   0.95045\nPOSSIBLE N-term signal sequence\ninside   1   6\nTMhelix   7   26\noutside   27   141\nTMhelix   142   160\ninside   161   161

28333	GCF_001537685.1_12082_5_32	Serratia marcescens	Proteobacteria; Gammaproteobacteria; Enterobacterales; Yersiniaceae; Serratia
	MNKRRTTLTVLALIASLGLSSAPALADKGGNGNGNGNGHGNSGNHGNSGNHGNNGNKGNK	WP_033646351.1
	MULTISPECIES: membrane protein [Enterobacterales]	Length: 161\nNumber of predicted TMHs:  2\nExp number of AAs in TMHs: 31.11104\nExp number, first 60 AAs:  14.67675\nTotal prob of N-in:   0.95023\nPOSSIBLE N-term signal sequence\ninside   1   6\nTMhelix   7   26\noutside   27   141\nTMhelix   142   160\ninside   161   161

28334	GCF_001537845.1_12082_2_32	Serratia marcescens	Proteobacteria; Gammaproteobacteria; Enterobacterales; Yersiniaceae; Serratia
	MNKRRTTLTVLALIASLGLSSAPALADKGGNGNGNGNGHGNSGNHGNSGNHGNNGNKGNK	WP_033646351.1
	MULTISPECIES: membrane protein [Enterobacterales]	Length: 161\nNumber of predicted TMHs:  2\nExp number of AAs in TMHs: 31.11104\nExp number, first 60 AAs:  14.67675\nTotal prob of N-in:   0.95023\nPOSSIBLE N-term signal sequence\ninside   1   6\nTMhelix   7   26\noutside   27   141\nTMhelix   142   160\ninside   161   161

28335	GCF_001537785.1_12045_8_1	Serratia marcescens	Proteobacteria; Gammaproteobacteria; Enterobacterales; Yersiniaceae; Serratia
	MNKRRTTLTVLALIASLGLSSAPALADKGGNGNGNGNGHGNSGNHGNSGNHGNNGNKGNK	WP_033646351.1
	MULTISPECIES: membrane protein [Enterobacterales]	Length: 161\nNumber of predicted TMHs:  2\nExp number of AAs in TMHs: 31.11104\nExp number, first 60 AAs:  14.67675\nTotal prob of N-in:   0.95023\nPOSSIBLE N-term signal sequence\ninside   1   6\nTMhelix   7   26\noutside   27   141\nTMhelix   142   160\ninside   161   161

28336	GCF_001537885.1_12082_2_34	Serratia marcescens	Proteobacteria; Gammaproteobacteria; Enterobacterales; Yersiniaceae; Serratia
	MNKRRTTLTVLALIASLGLSSAPALADKGGNGNGNGNGHGNSGNHGNSGNHGNNGNKGNK	WP_033646351.1
	MULTISPECIES: membrane protein [Enterobacterales]	Length: 161\nNumber of predicted TMHs:  2\nExp number of AAs in TMHs: 31.11104\nExp number, first 60 AAs:  14.67675\nTotal prob of N-in:   0.95023\nPOSSIBLE N-term signal sequence\ninside   1   6\nTMhelix   7   26\noutside   27   141\nTMhelix   142   160\ninside   161   161

28337	GCF_001538005.1_12082_3_46	Serratia marcescens	Proteobacteria; Gammaproteobacteria; Enterobacterales; Yersiniaceae; Serratia
	MNKRRTTLTVLALIASLGLSSAPALADKGGNGNGNGNGHGNSGNHGNSGNHGNNGNKGNK	WP_033646351.1
	MULTISPECIES: membrane protein [Enterobacterales]	Length: 161\nNumber of predicted TMHs:  2\nExp

number of AAs in TMHs: 31.11104\nExp number, first 60 AAs:  14.67675\nTotal prob of N-in:   0.95023\nPOSSIBLE N-term signal sequence\ninside   1   6\nTMhelix   7   26\noutside   27   141\nTMhelix   142   160\ninside   161   161

28338 GCF_001537745.1_12082_3_95 Serratia marcescens Proteobacteria; Gammaproteobacteria; Enterobacterales; Yersiniaceae; Serratia
MNKRRTTLTVLALIASLGLSSAPALADKGGNGNGNGNGHGNSGNNGNHGNNGNHGNKGNK WP_060419303.1
MULTISPECIES: hypothetical protein [Serratia] Length: 164\nNumber of predicted TMHs:  2\nExp number of AAs in TMHs: 30.51677\nExp number, first 60 AAs:  14.42464\nTotal prob of N-in:   0.94579\nPOSSIBLE N-term signal sequence\ninside   1   6\nTMhelix   7   26\noutside   27   144\nTMhelix   145   163\ninside   164   164

28339 GCF_001538725.1_12082_2_60 Serratia marcescens Proteobacteria; Gammaproteobacteria; Enterobacterales; Yersiniaceae; Serratia
MNKRRTTLTVLALIASLGLSSAPALADKGGNGNGNGNGHGNSGNHGNSGNHGNNGNKGNK WP_033646351.1
MULTISPECIES: membrane protein [Enterobacterales] Length: 161\nNumber of predicted TMHs:  2\nExp number of AAs in TMHs: 31.11104\nExp number, first 60 AAs:  14.67675\nTotal prob of N-in:   0.95023\nPOSSIBLE N-term signal sequence\ninside   1   6\nTMhelix   7   26\noutside   27   141\nTMhelix   142   160\ninside   161   161

28340 GCF_001538765.1_12082_2_35 Serratia marcescens Proteobacteria; Gammaproteobacteria; Enterobacterales; Yersiniaceae; Serratia
MNKRRTTLTVLALIASLGLSSAPALADKGGNGNGNGNGHGNSGNHGNSGNHGNNGNKGNK WP_033646351.1
MULTISPECIES: membrane protein [Enterobacterales] Length: 161\nNumber of predicted TMHs:  2\nExp number of AAs in TMHs: 31.11104\nExp number, first 60 AAs:  14.67675\nTotal prob of N-in:   0.95023\nPOSSIBLE N-term signal sequence\ninside   1   6\nTMhelix   7   26\noutside   27   141\nTMhelix   142   160\ninside   161   161

28341 GCF_001538825.1_12082_2_87 Serratia marcescens Proteobacteria; Gammaproteobacteria; Enterobacterales; Yersiniaceae; Serratia
MNKRRTTLTVLALIASLGLSSAPALADKGGNGNGNGNGHGNSGNHGNSGNHGNNGNKGNK WP_033646351.1
MULTISPECIES: membrane protein [Enterobacterales] Length: 161\nNumber of predicted TMHs:  2\nExp number of AAs in TMHs: 31.11104\nExp number, first 60 AAs:  14.67675\nTotal prob of N-in:   0.95023\nPOSSIBLE N-term signal sequence\ninside   1   6\nTMhelix   7   26\noutside   27   141\nTMhelix   142   160\ninside   161   161

28342 GCF_001538865.1_12082_3_87 Serratia marcescens Proteobacteria; Gammaproteobacteria; Enterobacterales; Yersiniaceae; Serratia
MNKRRTTLTVLALIASLGLSSAPALADKGGNGNGNGNGHGNSGNHGNSGNHGNNGNKGNK WP_033646351.1
MULTISPECIES: membrane protein [Enterobacterales] Length: 161\nNumber of predicted TMHs:  2\nExp number of AAs in TMHs: 31.11104\nExp number, first 60 AAs:  14.67675\nTotal prob of N-in:   0.95023\nPOSSIBLE N-term signal sequence\ninside   1   6\nTMhelix   7   26\noutside   27   141\nTMhelix   142   160\ninside   161   161

28343 GCF_001538965.1_12082_3_39 Serratia marcescens Proteobacteria; Gammaproteobacteria; Enterobacterales; Yersiniaceae; Serratia
MNKRRTTLTVLALIASLGLSSAPALADKGGNGNGNGNGHGNSGNHGNSGNHGNNGNKGNK WP_033646351.1
MULTISPECIES: membrane protein [Enterobacterales] Length: 161\nNumber of predicted TMHs:  2\nExp number of AAs in TMHs: 31.11104\nExp number, first 60 AAs:  14.67675\nTotal prob of N-in:   0.95023\nPOSSIBLE N-term signal sequence\ninside   1   6\nTMhelix   7   26\noutside   27   141\nTMhelix   142   160\ninside   161   161

28344 GCF_001538665.1_12082_5_47 Serratia marcescens Proteobacteria; Gammaproteobacteria; Enterobacterales; Yersiniaceae; Serratia
MNKRRTTLTVLALIASLGLSSAPALADKGGNGNGNGNGHGNSGNHGNSGNHGNNGNKGNK WP_060427691.1
hypothetical protein [Serratia marcescens] Length: 161\nNumber of predicted TMHs:  2\nExp number of AAs in TMHs: 31.15272\nExp number, first 60 AAs:  14.60251\nTotal prob of N-in:   0.95045\nPOSSIBLE N-term signal sequence\ninside   1   6\nTMhelix   7   26\noutside   27   141\nTMhelix   142   160\ninside   161   161

28345 GCF_001539025.1_12082_2_27 Serratia marcescens Proteobacteria; Gammaproteobacteria; Enterobacterales; Yersiniaceae; Serratia
MNKRRTTLTVLALIASLGLSSAPALADKGGNGNGNGNGHGNSGNHGNSGNHGNNGNKGNK WP_033646351.1
MULTISPECIES: membrane protein [Enterobacterales] Length: 161\nNumber of predicted TMHs:  2\nExp number of AAs in TMHs: 31.11104\nExp number, first 60 AAs:  14.67675\nTotal prob of N-in:   0.95023\nPOSSIBLE N-term signal sequence\ninside   1   6\nTMhelix   7   26\noutside   27   141\nTMhelix   142   160\ninside   161   161

28346 GCF_001539085.1_12082_3_12 Serratia marcescens Proteobacteria; Gammaproteobacteria; Enterobacterales; Yersiniaceae; Serratia

MNKRRTTLTVLALIASLGLSSAPALADKGGNGNGNGNGHGNSGNHGNSGNHGNNGNKGNK WP_033646351.1 MULTISPECIES: membrane protein [Enterobacterales] Length: 161\nNumber of predicted TMHs: 2\nExp number of AAs in TMHs: 31.11104\nExp number, first 60 AAs: 14.67675\nTotal prob of N-in: 0.95023\nPOSSIBLE N-term signal sequence\ninside 1 6\nTMhelix 7 26\noutside 27 141\nTMhelix 142 160\ninside 161 161

28347 GCF_001539525.1_12082_5_52 Serratia marcescens Proteobacteria; Gammaproteobacteria; Enterobacterales; Yersiniaceae; Serratia MNKRRTTLTVLALIASLGLSSAPALADKGGNGNGNGNGHGNSGNHGNSGNHGNNGNKGNK WP_033646351.1 MULTISPECIES: membrane protein [Enterobacterales] Length: 161\nNumber of predicted TMHs: 2\nExp number of AAs in TMHs: 31.11104\nExp number, first 60 AAs: 14.67675\nTotal prob of N-in: 0.95023\nPOSSIBLE N-term signal sequence\ninside 1 6\nTMhelix 7 26\noutside 27 141\nTMhelix 142 160\ninside 161 161

28348 GCF_001539565.1_12082_3_1 Serratia marcescens Proteobacteria; Gammaproteobacteria; Enterobacterales; Yersiniaceae; Serratia MNKRRTTLTVLALIASLGLSSAPALADKGGNGNGNGNGHGNSGNHGNSGNHGNNGNKGNK WP_033646351.1 MULTISPECIES: membrane protein [Enterobacterales] Length: 161\nNumber of predicted TMHs: 2\nExp number of AAs in TMHs: 31.11104\nExp number, first 60 AAs: 14.67675\nTotal prob of N-in: 0.95023\nPOSSIBLE N-term signal sequence\ninside 1 6\nTMhelix 7 26\noutside 27 141\nTMhelix 142 160\ninside 161 161

28349 GCF_001539625.1_12082_3_44 Serratia marcescens Proteobacteria; Gammaproteobacteria; Enterobacterales; Yersiniaceae; Serratia MNKRRTTLTVLALIASLGLSSAPALADKGGNGNGNGNGHGNSGNHGNSGNHGNNGNKGNK WP_033646351.1 MULTISPECIES: membrane protein [Enterobacterales] Length: 161\nNumber of predicted TMHs: 2\nExp number of AAs in TMHs: 31.11104\nExp number, first 60 AAs: 14.67675\nTotal prob of N-in: 0.95023\nPOSSIBLE N-term signal sequence\ninside 1 6\nTMhelix 7 26\noutside 27 141\nTMhelix 142 160\ninside 161 161

28350 GCF_001539665.1_12082_3_51 Serratia marcescens Proteobacteria; Gammaproteobacteria; Enterobacterales; Yersiniaceae; Serratia MNKRRTTLTVLALIASLGLSSAPALADKGGNGNGNGNGHGNSGNHGNSGNHGNNGNKGNK WP_033646351.1 MULTISPECIES: membrane protein [Enterobacterales] Length: 161\nNumber of predicted TMHs: 2\nExp number of AAs in TMHs: 31.11104\nExp number, first 60 AAs: 14.67675\nTotal prob of N-in: 0.95023\nPOSSIBLE N-term signal sequence\ninside 1 6\nTMhelix 7 26\noutside 27 141\nTMhelix 142 160\ninside 161 161

28351 GCF_001539725.1_12082_2_47 Serratia marcescens Proteobacteria; Gammaproteobacteria; Enterobacterales; Yersiniaceae; Serratia MNKRRTTLTVLALIASLGLSSAPALADKGGNGNGNGNGHGNSGNHGNSGNHGNNGNKGNK WP_033646351.1 MULTISPECIES: membrane protein [Enterobacterales] Length: 161\nNumber of predicted TMHs: 2\nExp number of AAs in TMHs: 31.11104\nExp number, first 60 AAs: 14.67675\nTotal prob of N-in: 0.95023\nPOSSIBLE N-term signal sequence\ninside 1 6\nTMhelix 7 26\noutside 27 141\nTMhelix 142 160\ninside 161 161

28352 GCF_001539465.1_12082_5_50 Serratia marcescens Proteobacteria; Gammaproteobacteria; Enterobacterales; Yersiniaceae; Serratia MNKRRTTLTVLALIASLGLSSAPALADKGGNGNGNGNGHGNSGNHGNSGNHGNNGNKGNK WP_060427691.1 hypothetical protein [Serratia marcescens] Length: 161\nNumber of predicted TMHs: 2\nExp number of AAs in TMHs: 31.15272\nExp number, first 60 AAs: 14.60251\nTotal prob of N-in: 0.95045\nPOSSIBLE N-term signal sequence\ninside 1 6\nTMhelix 7 26\noutside 27 141\nTMhelix 142 160\ninside 161 161

28353 GCF_001539765.1_12082_2_65 Serratia marcescens Proteobacteria; Gammaproteobacteria; Enterobacterales; Yersiniaceae; Serratia MNKRRTTLTVLALIASLGLSSAPALADKGGNGNGNGNGHGNSGNHGNSGNHGNNGNKGNK WP_033646351.1 MULTISPECIES: membrane protein [Enterobacterales] Length: 161\nNumber of predicted TMHs: 2\nExp number of AAs in TMHs: 31.11104\nExp number, first 60 AAs: 14.67675\nTotal prob of N-in: 0.95023\nPOSSIBLE N-term signal sequence\ninside 1 6\nTMhelix 7 26\noutside 27 141\nTMhelix 142 160\ninside 161 161

28354 GCF_001539885.1_12082_2_71 Serratia marcescens Proteobacteria; Gammaproteobacteria; Enterobacterales; Yersiniaceae; Serratia MNKRRTTLTVLALIASLGLSSAPALADKGGNGNGNGNGHGNSGNHGNSGNHGNNGNKGNK WP_033646351.1 MULTISPECIES: membrane protein [Enterobacterales] Length: 161\nNumber of predicted TMHs: 2\nExp number of AAs in TMHs: 31.11104\nExp number, first 60 AAs: 14.67675\nTotal prob of N-in: 0.95023\nPOSSIBLE N-term signal sequence\ninside 1 6\nTMhelix 7 26\noutside 27 141\nTMhelix 142 160\ninside 161 161

28355 GCF_001076625.1_ASM107662v1 Serratia marcescens Proteobacteria; Gammaproteobacteria; Enterobacterales; Yersiniaceae; Serratia
MNKRRTTLTVLALIASLGLSSAPALADKGGNGNGNGNGHGNSGNHGNNGNHGNNGNNGNH WP_004941297.1
MULTISPECIES: hypothetical protein [Serratia] Length: 167\nNumber of predicted TMHs: 2\nExp number of AAs in TMHs: 30.02384\nExp number, first 60 AAs: 14.11185\nTotal prob of N-in: 0.94170\nPOSSIBLE N-term signal sequence\ninside 1 6\nTMhelix 7 26\noutside 27 147\nTMhelix 148 166\ninside 167 167

28356 GCF_001067015.1_ASM106701v1 Serratia marcescens Proteobacteria; Gammaproteobacteria; Enterobacterales; Yersiniaceae; Serratia
MNKRRTTLTVLALIASLGLSSAPALADKGGNGNGNGNGHGNSGNHGNSGNHGNNGNKGNK WP_033646351.1
MULTISPECIES: membrane protein [Enterobacterales] Length: 161\nNumber of predicted TMHs: 2\nExp number of AAs in TMHs: 31.11104\nExp number, first 60 AAs: 14.67675\nTotal prob of N-in: 0.95023\nPOSSIBLE N-term signal sequence\ninside 1 6\nTMhelix 7 26\noutside 27 141\nTMhelix 142 160\ninside 161 161

28357 GCF_001066015.1_ASM106601v1 Serratia marcescens Proteobacteria; Gammaproteobacteria; Enterobacterales; Yersiniaceae; Serratia
MNKRRTTLTVLALIASLGLSSAPALADKGGNGNGNGNGHGNSGNHGNSGNHGNNGNKGNK WP_033646351.1
MULTISPECIES: membrane protein [Enterobacterales] Length: 161\nNumber of predicted TMHs: 2\nExp number of AAs in TMHs: 31.11104\nExp number, first 60 AAs: 14.67675\nTotal prob of N-in: 0.95023\nPOSSIBLE N-term signal sequence\ninside 1 6\nTMhelix 7 26\noutside 27 141\nTMhelix 142 160\ninside 161 161

28358 GCF_001540245.1_12082_3_22 Serratia marcescens Proteobacteria; Gammaproteobacteria; Enterobacterales; Yersiniaceae; Serratia
MNKRRTTLTVLALIASLGLSSAPALADKGGNGNGNGNGHGNSGNHGNSGNHGNNGNKGNK WP_033646351.1
MULTISPECIES: membrane protein [Enterobacterales] Length: 161\nNumber of predicted TMHs: 2\nExp number of AAs in TMHs: 31.11104\nExp number, first 60 AAs: 14.67675\nTotal prob of N-in: 0.95023\nPOSSIBLE N-term signal sequence\ninside 1 6\nTMhelix 7 26\noutside 27 141\nTMhelix 142 160\ninside 161 161

28359 GCF_001540305.1_12082_2_94 Serratia marcescens Proteobacteria; Gammaproteobacteria; Enterobacterales; Yersiniaceae; Serratia
MNKRRTTLTVLALIASLGLSSAPALADKGGNGNGNGNGHGNSGNHGNSGNHGNNGNKGNK WP_033646351.1
MULTISPECIES: membrane protein [Enterobacterales] Length: 161\nNumber of predicted TMHs: 2\nExp number of AAs in TMHs: 31.11104\nExp number, first 60 AAs: 14.67675\nTotal prob of N-in: 0.95023\nPOSSIBLE N-term signal sequence\ninside 1 6\nTMhelix 7 26\noutside 27 141\nTMhelix 142 160\ninside 161 161

28360 GCF_001063375.1_ASM106337v1 Serratia marcescens Proteobacteria; Gammaproteobacteria; Enterobacterales; Yersiniaceae; Serratia
MNKRRTTLTVLALIASLGLSSAPALADKGGNGNGNGNGHGNSGNHGNNGNNGNHGNNGNH WP_049209162.1
membrane protein [Serratia marcescens] Length: 167\nNumber of predicted TMHs: 2\nExp number of AAs in TMHs: 30.02384\nExp number, first 60 AAs: 14.11185\nTotal prob of N-in: 0.94170\nPOSSIBLE N-term signal sequence\ninside 1 6\nTMhelix 7 26\noutside 27 147\nTMhelix 148 166\ninside 167 167

28361 GCF_001064855.1_ASM106485v1 Serratia marcescens Proteobacteria; Gammaproteobacteria; Enterobacterales; Yersiniaceae; Serratia
MNKRRTTLTVLALIASLGLSSAPALADKGGNGNGNGNGHGNSGNHGNNGNNGNHGNNGNH WP_049209162.1
membrane protein [Serratia marcescens] Length: 167\nNumber of predicted TMHs: 2\nExp number of AAs in TMHs: 30.02384\nExp number, first 60 AAs: 14.11185\nTotal prob of N-in: 0.94170\nPOSSIBLE N-term signal sequence\ninside 1 6\nTMhelix 7 26\noutside 27 147\nTMhelix 148 166\ninside 167 167

28362 GCF_001064725.1_ASM106472v1 Serratia marcescens Proteobacteria; Gammaproteobacteria; Enterobacterales; Yersiniaceae; Serratia
MNKRRTTLTVLALIASLGLSSAPALADKGGNGNGNGNGHGNSGNHGNNGNNGNHGNNGNH WP_049209162.1
membrane protein [Serratia marcescens] Length: 167\nNumber of predicted TMHs: 2\nExp number of AAs in TMHs: 30.02384\nExp number, first 60 AAs: 14.11185\nTotal prob of N-in: 0.94170\nPOSSIBLE N-term signal sequence\ninside 1 6\nTMhelix 7 26\noutside 27 147\nTMhelix 148 166\ninside 167 167

28363 GCF_001540445.1_12082_3_72 Serratia marcescens Proteobacteria; Gammaproteobacteria; Enterobacterales; Yersiniaceae; Serratia
MNKRRTTLTVLALIASLGLSSAPALADKGGNGNGNGNGHGNSGNHGNSGNHGNNGNKGNK WP_033646351.1
MULTISPECIES: membrane protein [Enterobacterales] Length: 161\nNumber of predicted TMHs: 2\nExp

number of AAs in TMHs: 31.11104\nExp number, first 60 AAs: 14.67675\nTotal prob of N-in: 0.95023\nPOSSIBLE N-term signal sequence\ninside 1 6\nTMhelix 7 26\noutside 27 141\nTMhelix 142 160\ninside 161 161

28364 GCF_001540345.1_12082_3_67 Serratia marcescens Proteobacteria; Gammaproteobacteria; Enterobacterales; Yersiniaceae; Serratia
MNKRRTTLTVLALIASLGLSSAPALADKGGNGNGNGNGHGNSGNHGNSGNHGNNGNKGNK WP_033646351.1
MULTISPECIES: membrane protein [Enterobacterales] Length: 161\nNumber of predicted TMHs: 2\nExp number of AAs in TMHs: 31.11104\nExp number, first 60 AAs: 14.67675\nTotal prob of N-in: 0.95023\nPOSSIBLE N-term signal sequence\ninside 1 6\nTMhelix 7 26\noutside 27 141\nTMhelix 142 160\ninside 161 161

28365 GCF_001294565.1_ASM129456v1 Serratia marcescens Proteobacteria; Gammaproteobacteria; Enterobacterales; Yersiniaceae; Serratia
MNKRRTTLTVLALIASLGLSSAPALADKGGNGNGNGNGHGNSGNHGNSGNHGNNGNKGNK WP_033646351.1
MULTISPECIES: membrane protein [Enterobacterales] Length: 161\nNumber of predicted TMHs: 2\nExp number of AAs in TMHs: 31.11104\nExp number, first 60 AAs: 14.67675\nTotal prob of N-in: 0.95023\nPOSSIBLE N-term signal sequence\ninside 1 6\nTMhelix 7 26\noutside 27 141\nTMhelix 142 160\ninside 161 161

28366 GCF_001536545.1_12082_3_88 Serratia marcescens Proteobacteria; Gammaproteobacteria; Enterobacterales; Yersiniaceae; Serratia
MNKRRTTLTVLALIASLGLSSAPALADKGGNGNGNGNGHGNSGNHGNSGNHGNNGNKGNK WP_033646351.1
MULTISPECIES: membrane protein [Enterobacterales] Length: 161\nNumber of predicted TMHs: 2\nExp number of AAs in TMHs: 31.11104\nExp number, first 60 AAs: 14.67675\nTotal prob of N-in: 0.95023\nPOSSIBLE N-term signal sequence\ninside 1 6\nTMhelix 7 26\noutside 27 141\nTMhelix 142 160\ninside 161 161

28367 GCF_001536445.1_12082_2_63 Serratia marcescens Proteobacteria; Gammaproteobacteria; Enterobacterales; Yersiniaceae; Serratia
MNKRRTTLTVLALIASLGLSSAPALADKGGNGNGNGNGHGNSGNHGNSGNHGNNGNKGNK WP_033646351.1
MULTISPECIES: membrane protein [Enterobacterales] Length: 161\nNumber of predicted TMHs: 2\nExp number of AAs in TMHs: 31.11104\nExp number, first 60 AAs: 14.67675\nTotal prob of N-in: 0.95023\nPOSSIBLE N-term signal sequence\ninside 1 6\nTMhelix 7 26\noutside 27 141\nTMhelix 142 160\ninside 161 161

28368 GCF_001537065.1_12082_3_13 Serratia marcescens Proteobacteria; Gammaproteobacteria; Enterobacterales; Yersiniaceae; Serratia
MNKRRTTLTVLALIASLGLSSAPALADKGGNGNGNGNGHGNSGNHGNSGNHGNNGNKGNK WP_033646351.1
MULTISPECIES: membrane protein [Enterobacterales] Length: 161\nNumber of predicted TMHs: 2\nExp number of AAs in TMHs: 31.11104\nExp number, first 60 AAs: 14.67675\nTotal prob of N-in: 0.95023\nPOSSIBLE N-term signal sequence\ninside 1 6\nTMhelix 7 26\noutside 27 141\nTMhelix 142 160\ninside 161 161

28369 GCF_002153355.1_ASM215335v1 Serratia marcescens Proteobacteria; Gammaproteobacteria; Enterobacterales; Yersiniaceae; Serratia
MNKRRTTLTVLALIASLGLSSAPALADKGGNGNGNGNGHGNSGNHGNSGNHGNNDNKGNK WP_086579703.1
hypothetical protein [Serratia marcescens] Length: 170\nNumber of predicted TMHs: 2\nExp number of AAs in TMHs: 29.54105\nExp number, first 60 AAs: 13.87393\nTotal prob of N-in: 0.93752\nPOSSIBLE N-term signal sequence\ninside 1 6\nTMhelix 7 26\noutside 27 150\nTMhelix 151 169\ninside 170 170

28370 GCF_002118055.1_ASM211805v1 Serratia marcescens Proteobacteria; Gammaproteobacteria; Enterobacterales; Yersiniaceae; Serratia
MNKRRTTLTVLALIASLGLSSAPALADKGGNGNGNGNGNGNGHGNSGNHGNNGNHGNNGN WP_055312290.1
hypothetical protein [Serratia marcescens] Length: 171\nNumber of predicted TMHs: 2\nExp number of AAs in TMHs: 29.30784\nExp number, first 60 AAs: 13.75093\nTotal prob of N-in: 0.93571\nPOSSIBLE N-term signal sequence\ninside 1 6\nTMhelix 7 26\noutside 27 151\nTMhelix 152 170\ninside 171 171

28371 GCF_002094145.1_ASM209414v1 Serratia marcescens Proteobacteria; Gammaproteobacteria; Enterobacterales; Yersiniaceae; Serratia
MNKRRTTLTVLALIASLGLSSAPALADKGGNGNGNGNGHGNSGNHGNNGNHGNNGNNGNH WP_033633261.1
MULTISPECIES: membrane protein [Serratia] Length: 176\nNumber of predicted TMHs: 2\nExp number of AAs in TMHs: 28.40352\nExp number, first 60 AAs: 13.30908\nTotal prob of N-in: 0.92780\nPOSSIBLE N-term signal sequence\ninside 1 6\nTMhelix 7 26\noutside 27 156\nTMhelix 157 175\ninside 176 176

28372 GCF_900108835.1_SM1978 Serratia marcescens Proteobacteria; Gammaproteobacteria; Enterobacterales; Yersiniaceae; Serratia

MNKRRTTLTVLALIASLGLSSAPALADKGGNGNGNGNGHGNSGNHGNNGNHGNNGNNGNH WP_004941297.1 MULTISPECIES: hypothetical protein [Serratia] Length: 167\nNumber of predicted TMHs: 2\nExp number of AAs in TMHs: 30.02384\nExp number, first 60 AAs: 14.11185\nTotal prob of N-in: 0.94170\nPOSSIBLE N-term signal sequence\ninside 1 6\nTMhelix 7 26\noutside 27 147\nTMhelix 148 166\ninside 167 167

28373 GCF_002104105.1_ASM210410v1 Serratia marcescens Proteobacteria; Gammaproteobacteria; Enterobacterales; Yersiniaceae; Serratia MNKRRTTLTVLALIASLGLSSAPALADKGGNGNGNGNGHGNSGNHGNSGNHGNNGNKGNK WP_033646351.1 MULTISPECIES: membrane protein [Enterobacterales] Length: 161\nNumber of predicted TMHs: 2\nExp number of AAs in TMHs: 31.11104\nExp number, first 60 AAs: 14.67675\nTotal prob of N-in: 0.95023\nPOSSIBLE N-term signal sequence\ninside 1 6\nTMhelix 7 26\noutside 27 141\nTMhelix 142 160\ninside 161 161

28374 GCF_001540605.1_12082_2_83 Serratia marcescens Proteobacteria; Gammaproteobacteria; Enterobacterales; Yersiniaceae; Serratia MNKRRTTLTVLALIASLGLSSAPALADKGGNGNGNGNGHGNSGNHGNSGNHGNNGNKGNK WP_033646351.1 MULTISPECIES: membrane protein [Enterobacterales] Length: 161\nNumber of predicted TMHs: 2\nExp number of AAs in TMHs: 31.11104\nExp number, first 60 AAs: 14.67675\nTotal prob of N-in: 0.95023\nPOSSIBLE N-term signal sequence\ninside 1 6\nTMhelix 7 26\noutside 27 141\nTMhelix 142 160\ninside 161 161

28375 GCF_001540545.1_12082_3_66 Serratia marcescens Proteobacteria; Gammaproteobacteria; Enterobacterales; Yersiniaceae; Serratia MNKRRTTLTVLALIASLGLSSAPALADKGGNGNGNGNGHGNSGNHGNSGNHGNNGNKGNK WP_033646351.1 MULTISPECIES: membrane protein [Enterobacterales] Length: 161\nNumber of predicted TMHs: 2\nExp number of AAs in TMHs: 31.11104\nExp number, first 60 AAs: 14.67675\nTotal prob of N-in: 0.95023\nPOSSIBLE N-term signal sequence\ninside 1 6\nTMhelix 7 26\noutside 27 141\nTMhelix 142 160\ninside 161 161

28376 GCF_001540505.1_12082_3_78 Serratia marcescens Proteobacteria; Gammaproteobacteria; Enterobacterales; Yersiniaceae; Serratia MNKRRTTLTVLALIASLGLSSAPALADKGGNGNGNGNGHGNSGNHGNSGNHGNNGNKGNK WP_033646351.1 MULTISPECIES: membrane protein [Enterobacterales] Length: 161\nNumber of predicted TMHs: 2\nExp number of AAs in TMHs: 31.11104\nExp number, first 60 AAs: 14.67675\nTotal prob of N-in: 0.95023\nPOSSIBLE N-term signal sequence\ninside 1 6\nTMhelix 7 26\noutside 27 141\nTMhelix 142 160\ninside 161 161

28377 GCF_001540405.1_12082_5_45 Serratia marcescens Proteobacteria; Gammaproteobacteria; Enterobacterales; Yersiniaceae; Serratia MNKRRTTLTVLALIASLGLSSAPALADKGGNGNGNGNGHGNSGNHGNSGNHGNNGNKGNK WP_033646351.1 MULTISPECIES: membrane protein [Enterobacterales] Length: 161\nNumber of predicted TMHs: 2\nExp number of AAs in TMHs: 31.11104\nExp number, first 60 AAs: 14.67675\nTotal prob of N-in: 0.95023\nPOSSIBLE N-term signal sequence\ninside 1 6\nTMhelix 7 26\noutside 27 141\nTMhelix 142 160\ninside 161 161

28378 GCF_002104115.1_ASM210411v1 Serratia marcescens Proteobacteria; Gammaproteobacteria; Enterobacterales; Yersiniaceae; Serratia MNKRRTTLTVLALIASLGLSSAPALADKGGNGNGNGNGHGNSGNHGNSGNHGNNGNKGNK WP_033646351.1 MULTISPECIES: membrane protein [Enterobacterales] Length: 161\nNumber of predicted TMHs: 2\nExp number of AAs in TMHs: 31.11104\nExp number, first 60 AAs: 14.67675\nTotal prob of N-in: 0.95023\nPOSSIBLE N-term signal sequence\ninside 1 6\nTMhelix 7 26\noutside 27 141\nTMhelix 142 160\ninside 161 161

28379 GCF_000734475.1_ASM73447v1 Serratia marcescens Proteobacteria; Gammaproteobacteria; Enterobacterales; Yersiniaceae; Serratia MNKRRTTLTVLALIASLGLSSAPALADKGGNGNGNGNGHGNSGNHGNSGNHGNNGNKGNK WP_033646351.1 MULTISPECIES: membrane protein [Enterobacterales] Length: 161\nNumber of predicted TMHs: 2\nExp number of AAs in TMHs: 31.11104\nExp number, first 60 AAs: 14.67675\nTotal prob of N-in: 0.95023\nPOSSIBLE N-term signal sequence\ninside 1 6\nTMhelix 7 26\noutside 27 141\nTMhelix 142 160\ninside 161 161

28380 GCF_000805875.1_SmRM66262_v1.0 Serratia marcescens Proteobacteria; Gammaproteobacteria; Enterobacterales; Yersiniaceae; Serratia MNKRRTTLTVLALIASLGLSSAPALADKGGNGNGNGNGHGNSGNHGNSGNHGNNGNKGNK WP_033646351.1 MULTISPECIES: membrane protein [Enterobacterales] Length: 161\nNumber of predicted TMHs: 2\nExp number of AAs in TMHs: 31.11104\nExp number, first 60 AAs: 14.67675\nTotal prob of N-in: 0.95023\nPOSSIBLE N-term signal sequence\ninside 1 6\nTMhelix 7 26\noutside 27 141\nTMhelix 142 160\ninside 161 161

28381 GCF_000739215.1_ASM73921v1 Serratia marcescens Proteobacteria; Gammaproteobacteria; Enterobacterales; Yersiniaceae; Serratia
MNKRRTTLTVLALIASLGLSSAPALADKGGNGNGNGNGHGNSGNHGNSGNHGNNGNKGNK WP_033646351.1
MULTISPECIES: membrane protein [Enterobacterales] Length: 161\nNumber of predicted TMHs: 2\nExp number of AAs in TMHs: 31.11104\nExp number, first 60 AAs: 14.67675\nTotal prob of N-in: 0.95023\nPOSSIBLE N-term signal sequence\ninside 1 6\nTMhelix 7 26\noutside 27 141\nTMhelix 142 160\ninside 161 161

28382 GCF_000738535.1_ASM73853v1 Serratia marcescens Proteobacteria; Gammaproteobacteria; Enterobacterales; Yersiniaceae; Serratia
MNKRRTTLTVLALIASLGLSSAPALADKGGNGNGNGNGHGNSGNHGNNGNHGNNGNHGNK WP_033640576.1
MULTISPECIES: membrane protein [Serratia] Length: 164\nNumber of predicted TMHs: 2\nExp number of AAs in TMHs: 30.54778\nExp number, first 60 AAs: 14.38033\nTotal prob of N-in: 0.94595\nPOSSIBLE N-term signal sequence\ninside 1 6\nTMhelix 7 26\noutside 27 144\nTMhelix 145 163\ninside 164 164

28383 GCF_000695485.1_smgenome Serratia marcescens Proteobacteria; Gammaproteobacteria; Enterobacterales; Yersiniaceae; Serratia
MNKRRTTLTVLALIASLGLSSAPALADKGGNGNGNGNGHGNSGNHGNNGNHGNNGNHGNK WP_016928034.1
membrane protein [Serratia marcescens] Length: 164\nNumber of predicted TMHs: 2\nExp number of AAs in TMHs: 30.54438\nExp number, first 60 AAs: 14.38759\nTotal prob of N-in: 0.94593\nPOSSIBLE N-term signal sequence\ninside 1 6\nTMhelix 7 26\noutside 27 144\nTMhelix 145 163\ninside 164 164

28384 GCF_001030265.1_Serr_marc_BWH57_V1 Serratia marcescens Proteobacteria; Gammaproteobacteria; Enterobacterales; Yersiniaceae; Serratia
MNKRRTTLTVLALIASLGLSSAPALADKGGNGNGNGNGHGNSGNHGNNGNHGNNGNNGNH WP_004941297.1
MULTISPECIES: hypothetical protein [Serratia] Length: 167\nNumber of predicted TMHs: 2\nExp number of AAs in TMHs: 30.02384\nExp number, first 60 AAs: 14.11185\nTotal prob of N-in: 0.94170\nPOSSIBLE N-term signal sequence\ninside 1 6\nTMhelix 7 26\noutside 27 147\nTMhelix 148 166\ninside 167 167

28385 GCF_001065325.1_ASM106532v1 Serratia marcescens Proteobacteria; Gammaproteobacteria; Enterobacterales; Yersiniaceae; Serratia
MNKRRTTLTVLALIASLGLSSAPALADKGGNGNGNGNGHGNSGNHGNHGNNGNHGNKGNK WP_049300632.1
membrane protein [Serratia marcescens] Length: 161\nNumber of predicted TMHs: 2\nExp number of AAs in TMHs: 31.06334\nExp number, first 60 AAs: 14.63729\nTotal prob of N-in: 0.95002\nPOSSIBLE N-term signal sequence\ninside 1 6\nTMhelix 7 26\noutside 27 141\nTMhelix 142 160\ninside 161 161

28386 GCF_001034375.1_Serr_marc_BWH56_V1 Serratia marcescens Proteobacteria; Gammaproteobacteria; Enterobacterales; Yersiniaceae; Serratia
MNKRRTTLTVLALIASLGLSSAPALADKGGNGNGNGNGHGNSGNHGNNGNHGNNGNNGNH WP_004941297.1
MULTISPECIES: hypothetical protein [Serratia] Length: 167\nNumber of predicted TMHs: 2\nExp number of AAs in TMHs: 30.02384\nExp number, first 60 AAs: 14.11185\nTotal prob of N-in: 0.94170\nPOSSIBLE N-term signal sequence\ninside 1 6\nTMhelix 7 26\noutside 27 147\nTMhelix 148 166\ninside 167 167

28387 GCF_001066945.1_ASM106694v1 Serratia marcescens Proteobacteria; Gammaproteobacteria; Enterobacterales; Yersiniaceae; Serratia
MNKRRTTLTVLALIASLGLSSAPALADKGGNGNGNGNGHGNSGNHGNSGNHGNNGNKGNK WP_033646351.1
MULTISPECIES: membrane protein [Enterobacterales] Length: 161\nNumber of predicted TMHs: 2\nExp number of AAs in TMHs: 31.11104\nExp number, first 60 AAs: 14.67675\nTotal prob of N-in: 0.95023\nPOSSIBLE N-term signal sequence\ninside 1 6\nTMhelix 7 26\noutside 27 141\nTMhelix 142 160\ninside 161 161

28388 GCF_001280365.1_ASM128036v1 Serratia marcescens Proteobacteria; Gammaproteobacteria; Enterobacterales; Yersiniaceae; Serratia
MNKRRTTLTVLALIASLGLSSAPAMADKGGNGNGNGNGHGNSGNHGNNGNHGNNGNHGNK WP_033642812.1
MULTISPECIES: membrane protein [Serratia] Length: 164\nNumber of predicted TMHs: 2\nExp number of AAs in TMHs: 30.89657\nExp number, first 60 AAs: 14.76607\nTotal prob of N-in: 0.94978\nPOSSIBLE N-term signal sequence\ninside 1 6\nTMhelix 7 26\noutside 27 144\nTMhelix 145 163\ninside 164 164

28389 GCF_001067375.1_ASM106737v1 Serratia marcescens Proteobacteria; Gammaproteobacteria; Enterobacterales; Yersiniaceae; Serratia
MNKRRTTLTVLALIASLGLSSAPALADKGGNGNGNGNGHGNSGNHGNSGNHGNNGNKGNK WP_033646351.1
MULTISPECIES: membrane protein [Enterobacterales] Length: 161\nNumber of predicted TMHs: 2\nExp

number of AAs in TMHs: 31.11104\nExp number, first 60 AAs: 14.67675\nTotal prob of N-in: 0.95023\nPOSSIBLE N-term signal sequence\ninside 1 6\nTMhelix 7 26\noutside 27 141\nTMhelix 142 160\ninside 161 161

28390 GCF_001536325.1_12082_3_52 Serratia marcescens Proteobacteria; Gammaproteobacteria; Enterobacterales; Yersiniaceae; Serratia
MNKRRTTLTVLALIASLGLSSAPALADKGGNGNGNGNGHGNSGNHGNSGNHGNNGNKGNK WP_033646351.1
MULTISPECIES: membrane protein [Enterobacterales] Length: 161\nNumber of predicted TMHs: 2\nExp number of AAs in TMHs: 31.11104\nExp number, first 60 AAs: 14.67675\nTotal prob of N-in: 0.95023\nPOSSIBLE N-term signal sequence\ninside 1 6\nTMhelix 7 26\noutside 27 141\nTMhelix 142 160\ninside 161 161

28391 GCF_001068085.1_ASM106808v1 Serratia marcescens Proteobacteria; Gammaproteobacteria; Enterobacterales; Yersiniaceae; Serratia
MNKRRTTLTVLALIASLGLSSAPALADKGGNGNGNGNGHGNSGNNGNHGNNGNHGNKGNK WP_049200367.1
MULTISPECIES: membrane protein [Enterobacterales] Length: 164\nNumber of predicted TMHs: 2\nExp number of AAs in TMHs: 30.51526\nExp number, first 60 AAs: 14.42673\nTotal prob of N-in: 0.94579\nPOSSIBLE N-term signal sequence\ninside 1 6\nTMhelix 7 26\noutside 27 144\nTMhelix 145 163\ninside 164 164

28392 GCF_001417865.2_ASM141786v2 Serratia marcescens Proteobacteria; Gammaproteobacteria; Enterobacterales; Yersiniaceae; Serratia
MNKRRTTLTVLALIASLGLSSAPALADKGGNGNGNGNGHGNSGNHGNNGNHGNNGNNGNH WP_033633261.1
MULTISPECIES: membrane protein [Serratia] Length: 176\nNumber of predicted TMHs: 2\nExp number of AAs in TMHs: 28.40352\nExp number, first 60 AAs: 13.30908\nTotal prob of N-in: 0.92780\nPOSSIBLE N-term signal sequence\ninside 1 6\nTMhelix 7 26\noutside 27 156\nTMhelix 157 175\ninside 176 176

28393 GCF_001064835.1_ASM106483v1 Serratia marcescens Proteobacteria; Gammaproteobacteria; Enterobacterales; Yersiniaceae; Serratia
MNKRRTTLTVLALIASLGLSSAPALADKGGNGNGNGNGHGNSGNHGNNGNNGNHGNNGNH WP_049209162.1
membrane protein [Serratia marcescens] Length: 167\nNumber of predicted TMHs: 2\nExp number of AAs in TMHs: 30.02384\nExp number, first 60 AAs: 14.11185\nTotal prob of N-in: 0.94170\nPOSSIBLE N-term signal sequence\ninside 1 6\nTMhelix 7 26\noutside 27 147\nTMhelix 148 166\ninside 167 167

28394 GCF_001064975.1_ASM106497v1 Serratia marcescens Proteobacteria; Gammaproteobacteria; Enterobacterales; Yersiniaceae; Serratia
MNKRRTTLTVLALIASLGLSSAPALADKGGNGNGNGNGHGNSGNHGNNGNNGNHGNNGNH WP_049209162.1
membrane protein [Serratia marcescens] Length: 167\nNumber of predicted TMHs: 2\nExp number of AAs in TMHs: 30.02384\nExp number, first 60 AAs: 14.11185\nTotal prob of N-in: 0.94170\nPOSSIBLE N-term signal sequence\ninside 1 6\nTMhelix 7 26\noutside 27 147\nTMhelix 148 166\ninside 167 167

28395 GCF_001537325.1_12082_2_53 Serratia marcescens Proteobacteria; Gammaproteobacteria; Enterobacterales; Yersiniaceae; Serratia
MNKRRTTLTVLALIASLGLSSAPALADKGGNGNGNGNGHGNSGNHGNSGNHGNNGNKGNK WP_033646351.1
MULTISPECIES: membrane protein [Enterobacterales] Length: 161\nNumber of predicted TMHs: 2\nExp number of AAs in TMHs: 31.11104\nExp number, first 60 AAs: 14.67675\nTotal prob of N-in: 0.95023\nPOSSIBLE N-term signal sequence\ninside 1 6\nTMhelix 7 26\noutside 27 141\nTMhelix 142 160\ninside 161 161

28396 GCF_001537405.1_12045_8_26 Serratia marcescens Proteobacteria; Gammaproteobacteria; Enterobacterales; Yersiniaceae; Serratia
MNKRRTTLTVLALIASLGLSSAPALADKGGNGNGNGNGHGNSGNHGNSGNHGNNGNKGNK WP_033646351.1
MULTISPECIES: membrane protein [Enterobacterales] Length: 161\nNumber of predicted TMHs: 2\nExp number of AAs in TMHs: 31.11104\nExp number, first 60 AAs: 14.67675\nTotal prob of N-in: 0.95023\nPOSSIBLE N-term signal sequence\ninside 1 6\nTMhelix 7 26\noutside 27 141\nTMhelix 142 160\ninside 161 161

28397 GCF_001537445.1_12045_8_9 Serratia marcescens Proteobacteria; Gammaproteobacteria; Enterobacterales; Yersiniaceae; Serratia
MNKRRTTLTVLALIASLGLSSAPALADKGGNGNGNGNGHGNSGNHGNSGNHGNNGNKGNK WP_033646351.1
MULTISPECIES: membrane protein [Enterobacterales] Length: 161\nNumber of predicted TMHs: 2\nExp number of AAs in TMHs: 31.11104\nExp number, first 60 AAs: 14.67675\nTotal prob of N-in: 0.95023\nPOSSIBLE N-term signal sequence\ninside 1 6\nTMhelix 7 26\noutside 27 141\nTMhelix 142 160\ninside 161 161

28398 GCF_001537465.1_12082_2_82 Serratia marcescens Proteobacteria; Gammaproteobacteria; Enterobacterales; Yersiniaceae; Serratia

MNKRRTTLTVLALIASLGLSSAPALADKGGNGNGNGNGHGNSGNHGNSGNHGNNGNKGNK WP_033646351.1 MULTISPECIES: membrane protein [Enterobacterales] Length: 161\nNumber of predicted TMHs: 2\nExp number of AAs in TMHs: 31.11104\nExp number, first 60 AAs: 14.67675\nTotal prob of N-in: 0.95023\nPOSSIBLE N-term signal sequence\ninside 1 6\nTMhelix 7 26\noutside 27 141\nTMhelix 142 160\ninside 161 161

28399 GCF_001537505.1_12082_5_35 Serratia marcescens Proteobacteria; Gammaproteobacteria; Enterobacterales; Yersiniaceae; Serratia MNKRRTTLTVLALIASLGLSSAPALADKGGNGNGNGNGHGNSGNHGNSGNHGNNGNKGNK WP_033646351.1 MULTISPECIES: membrane protein [Enterobacterales] Length: 161\nNumber of predicted TMHs: 2\nExp number of AAs in TMHs: 31.11104\nExp number, first 60 AAs: 14.67675\nTotal prob of N-in: 0.95023\nPOSSIBLE N-term signal sequence\ninside 1 6\nTMhelix 7 26\noutside 27 141\nTMhelix 142 160\ninside 161 161

28400 GCF_001537285.1_12045_8_16 Serratia marcescens Proteobacteria; Gammaproteobacteria; Enterobacterales; Yersiniaceae; Serratia MHNTNKRRTTLTVLALIASLGLSSAPALADKGGNGNGNGNGHGNSGNHGNSGNHGNNGNK WP_060418187.1 hypothetical protein [Serratia marcescens] Length: 164\nNumber of predicted TMHs: 1\nExp number of AAs in TMHs: 24.4976\nExp number, first 60 AAs: 8.86012\nTotal prob of N-in: 0.94445\ninside 1 144\nTMhelix 145 163\noutside 164 164

28401 GCF_001537545.1_12082_3_18 Serratia marcescens Proteobacteria; Gammaproteobacteria; Enterobacterales; Yersiniaceae; Serratia MNKRRTTLTVLALIASLGLSSAPALADKGGNGNGNGNGHGNSGNHGNSGNHGNNGNKGNK WP_033646351.1 MULTISPECIES: membrane protein [Enterobacterales] Length: 161\nNumber of predicted TMHs: 2\nExp number of AAs in TMHs: 31.11104\nExp number, first 60 AAs: 14.67675\nTotal prob of N-in: 0.95023\nPOSSIBLE N-term signal sequence\ninside 1 6\nTMhelix 7 26\noutside 27 141\nTMhelix 142 160\ninside 161 161

28402 GCF_001537605.1_12082_2_59 Serratia marcescens Proteobacteria; Gammaproteobacteria; Enterobacterales; Yersiniaceae; Serratia MNKRRTTLTVLALIASLGLSSAPALADKGGNGNGNGNGHGNSGNHGNSGNHGNNGNKGNK WP_033646351.1 MULTISPECIES: membrane protein [Enterobacterales] Length: 161\nNumber of predicted TMHs: 2\nExp number of AAs in TMHs: 31.11104\nExp number, first 60 AAs: 14.67675\nTotal prob of N-in: 0.95023\nPOSSIBLE N-term signal sequence\ninside 1 6\nTMhelix 7 26\noutside 27 141\nTMhelix 142 160\ninside 161 161

28403 GCF_001537625.1_12082_3_27 Serratia marcescens Proteobacteria; Gammaproteobacteria; Enterobacterales; Yersiniaceae; Serratia MNKRRTTLTVLALIASLGLSSAPALADKGGNGNGNGNGHGNSGNHGNSGNHGNNGNKGNK WP_033646351.1 MULTISPECIES: membrane protein [Enterobacterales] Length: 161\nNumber of predicted TMHs: 2\nExp number of AAs in TMHs: 31.11104\nExp number, first 60 AAs: 14.67675\nTotal prob of N-in: 0.95023\nPOSSIBLE N-term signal sequence\ninside 1 6\nTMhelix 7 26\noutside 27 141\nTMhelix 142 160\ninside 161 161

28404 GCF_001538545.1_12045_8_11 Serratia marcescens Proteobacteria; Gammaproteobacteria; Enterobacterales; Yersiniaceae; Serratia MNKRRTTLTVLALIASLGLSSAPALADKGGNGNGNGNGHGNSGNHGNSGNHGNNGNKGNK WP_033646351.1 MULTISPECIES: membrane protein [Enterobacterales] Length: 161\nNumber of predicted TMHs: 2\nExp number of AAs in TMHs: 31.11104\nExp number, first 60 AAs: 14.67675\nTotal prob of N-in: 0.95023\nPOSSIBLE N-term signal sequence\ninside 1 6\nTMhelix 7 26\noutside 27 141\nTMhelix 142 160\ninside 161 161

28405 GCF_001538585.1_12082_3_75 Serratia marcescens Proteobacteria; Gammaproteobacteria; Enterobacterales; Yersiniaceae; Serratia MNKRRTTLTVLALIASLGLSSAPALADKGGNGNGNGNGHGNSGNHGNSGNHGNNGNKGNK WP_033646351.1 MULTISPECIES: membrane protein [Enterobacterales] Length: 161\nNumber of predicted TMHs: 2\nExp number of AAs in TMHs: 31.11104\nExp number, first 60 AAs: 14.67675\nTotal prob of N-in: 0.95023\nPOSSIBLE N-term signal sequence\ninside 1 6\nTMhelix 7 26\noutside 27 141\nTMhelix 142 160\ninside 161 161

28406 GCF_001538485.1_12045_8_15 Serratia marcescens Proteobacteria; Gammaproteobacteria; Enterobacterales; Yersiniaceae; Serratia MNKRRTTLTVLALIASLGLSSAPALADKGGNGNGNGNGHGNSGNNGNHGNNGNHGNKGNK WP_060419303.1 MULTISPECIES: hypothetical protein [Serratia] Length: 164\nNumber of predicted TMHs: 2\nExp number of AAs in TMHs: 30.51677\nExp number, first 60 AAs: 14.42464\nTotal prob of N-in: 0.94579\nPOSSIBLE N-term signal sequence\ninside 1 6\nTMhelix 7 26\noutside 27 144\nTMhelix 145 163\ninside 164 164

28407 GCF_001932655.1_ASM193265v1 Serratia marcescens Proteobacteria; Gammaproteobacteria; Enterobacterales; Yersiniaceae; Serratia
MNKRRTTLTVLALIASLGLSSAPALADKGGNGNGNGNGHGNSGNNGNHGNNGNHGNKGNK WP_060419303.1
MULTISPECIES: hypothetical protein [Serratia] Length: 164\nNumber of predicted TMHs: 2\nExp number of AAs in TMHs: 30.51677\nExp number, first 60 AAs: 14.42464\nTotal prob of N-in: 0.94579\nPOSSIBLE N-term signal sequence\ninside 1 6\nTMhelix 7 26\noutside 27 144\nTMhelix 145 163\ninside 164 164

28408 GCF_001975745.1_ASM197574v1 Serratia marcescens Proteobacteria; Gammaproteobacteria; Enterobacterales; Yersiniaceae; Serratia
MNKRRTTLTVLALIASLGLSSAPALADKGGNGNGNGNGHGNSGNHGNNGNHGNNGNNGNH WP_004941297.1
MULTISPECIES: hypothetical protein [Serratia] Length: 167\nNumber of predicted TMHs: 2\nExp number of AAs in TMHs: 30.02384\nExp number, first 60 AAs: 14.11185\nTotal prob of N-in: 0.94170\nPOSSIBLE N-term signal sequence\ninside 1 6\nTMhelix 7 26\noutside 27 147\nTMhelix 148 166\ninside 167 167

28409 GCF_002104095.1_ASM210409v1 Serratia marcescens Proteobacteria; Gammaproteobacteria; Enterobacterales; Yersiniaceae; Serratia
MNKRRTTLTVLALIASLGLSSAPALADKGGNGNGNGNGHGNSGNHGNSGNHGNNGNKGNK WP_033646351.1
MULTISPECIES: membrane protein [Enterobacterales] Length: 161\nNumber of predicted TMHs: 2\nExp number of AAs in TMHs: 31.11104\nExp number, first 60 AAs: 14.67675\nTotal prob of N-in: 0.95023\nPOSSIBLE N-term signal sequence\ninside 1 6\nTMhelix 7 26\noutside 27 141\nTMhelix 142 160\ninside 161 161

28410 GCF_001538605.1_12082_2_30 Serratia marcescens Proteobacteria; Gammaproteobacteria; Enterobacterales; Yersiniaceae; Serratia
MNKRRTTLTVLALIASLGLSSAPALADKGGNGNGNGNGHGNSGNHGNSGNHGNNGNKGNK WP_033646351.1
MULTISPECIES: membrane protein [Enterobacterales] Length: 161\nNumber of predicted TMHs: 2\nExp number of AAs in TMHs: 31.11104\nExp number, first 60 AAs: 14.67675\nTotal prob of N-in: 0.95023\nPOSSIBLE N-term signal sequence\ninside 1 6\nTMhelix 7 26\noutside 27 141\nTMhelix 142 160\ninside 161 161

28411 GCF_001538645.1_12082_3_15 Serratia marcescens Proteobacteria; Gammaproteobacteria; Enterobacterales; Yersiniaceae; Serratia
MNKRRTTLTVLALIASLGLSSAPALADKGGNGNGNGNGHGNSGNHGNSGNHGNNGNKGNK WP_033646351.1
MULTISPECIES: membrane protein [Enterobacterales] Length: 161\nNumber of predicted TMHs: 2\nExp number of AAs in TMHs: 31.11104\nExp number, first 60 AAs: 14.67675\nTotal prob of N-in: 0.95023\nPOSSIBLE N-term signal sequence\ninside 1 6\nTMhelix 7 26\noutside 27 141\nTMhelix 142 160\ninside 161 161

28412 GCF_001538785.1_12045_8_18 Serratia marcescens Proteobacteria; Gammaproteobacteria; Enterobacterales; Yersiniaceae; Serratia
MNKRRTTLTVLALIASLGLSSAPALADKGGNGNGNGNGHGNSGNHGNNGNHGNNGNNGNH WP_004941297.1
MULTISPECIES: hypothetical protein [Serratia] Length: 167\nNumber of predicted TMHs: 2\nExp number of AAs in TMHs: 30.02384\nExp number, first 60 AAs: 14.11185\nTotal prob of N-in: 0.94170\nPOSSIBLE N-term signal sequence\ninside 1 6\nTMhelix 7 26\noutside 27 147\nTMhelix 148 166\ninside 167 167

28413 GCF_001538685.1_12082_3_64 Serratia marcescens Proteobacteria; Gammaproteobacteria; Enterobacterales; Yersiniaceae; Serratia
MNKRRTTLTVLALIASLGLSSAPALADKGGNGNGNGNGHGNSGNHGNSGNHGNNGNKGNK WP_033646351.1
MULTISPECIES: membrane protein [Enterobacterales] Length: 161\nNumber of predicted TMHs: 2\nExp number of AAs in TMHs: 31.11104\nExp number, first 60 AAs: 14.67675\nTotal prob of N-in: 0.95023\nPOSSIBLE N-term signal sequence\ninside 1 6\nTMhelix 7 26\noutside 27 141\nTMhelix 142 160\ninside 161 161

28414 GCF_001538745.1_12082_3_35 Serratia marcescens Proteobacteria; Gammaproteobacteria; Enterobacterales; Yersiniaceae; Serratia
MNKRRTTLTVLALIASLGLSSAPALADKGGNGNGNGNGHGNSGNHGNSGNHGNNGNKGNK WP_033646351.1
MULTISPECIES: membrane protein [Enterobacterales] Length: 161\nNumber of predicted TMHs: 2\nExp number of AAs in TMHs: 31.11104\nExp number, first 60 AAs: 14.67675\nTotal prob of N-in: 0.95023\nPOSSIBLE N-term signal sequence\ninside 1 6\nTMhelix 7 26\noutside 27 141\nTMhelix 142 160\ninside 161 161

28415 GCF_001539385.1_12082_3_19 Serratia marcescens Proteobacteria; Gammaproteobacteria; Enterobacterales; Yersiniaceae; Serratia
MNKRRTTLTVLALIASLGLSSAPALADKGGNGNGNGNGHGNSGNHGNSGNHGNNGNKGNK WP_033646351.1
MULTISPECIES: membrane protein [Enterobacterales] Length: 161\nNumber of predicted TMHs: 2\nExp

number of AAs in TMHs: 31.11104\nExp number, first 60 AAs: 14.67675\nTotal prob of N-in: 0.95023\nPOSSIBLE N-term signal sequence\ninside 1 6\nTMhelix 7 26\noutside 27 141\nTMhelix 142 160\ninside 161 161

28416 GCF_001539285.1_12082_3_37 Serratia marcescens Proteobacteria; Gammaproteobacteria; Enterobacterales; Yersiniaceae; Serratia MNKRRTTLTVLALIASLGLSSAPALADKGGNGNGNGNGHGNSGNHGNSGNHGNNGNKGNK WP_060427691.1 hypothetical protein [Serratia marcescens] Length: 161\nNumber of predicted TMHs: 2\nExp number of AAs in TMHs: 31.15272\nExp number, first 60 AAs: 14.60251\nTotal prob of N-in: 0.95045\nPOSSIBLE N-term signal sequence\ninside 1 6\nTMhelix 7 26\noutside 27 141\nTMhelix 142 160\ninside 161 161

28417 GCF_001539405.1_12082_3_38 Serratia marcescens Proteobacteria; Gammaproteobacteria; Enterobacterales; Yersiniaceae; Serratia MNKRRTTLTVLALIASLGLSSAPALADKGGNGNGNGNGHGNSGNHGNSGNHGNNGNKGNK WP_033646351.1 MULTISPECIES: membrane protein [Enterobacterales] Length: 161\nNumber of predicted TMHs: 2\nExp number of AAs in TMHs: 31.11104\nExp number, first 60 AAs: 14.67675\nTotal prob of N-in: 0.95023\nPOSSIBLE N-term signal sequence\ninside 1 6\nTMhelix 7 26\noutside 27 141\nTMhelix 142 160\ninside 161 161

28418 GCF_001539485.1_12045_8_3 Serratia marcescens Proteobacteria; Gammaproteobacteria; Enterobacterales; Yersiniaceae; Serratia MNKRRTTLTVLALIASLGLSSAPALADKGGNGNGNGNGHGNSGNHGNSGNHGNNGNKGNK WP_033646351.1 MULTISPECIES: membrane protein [Enterobacterales] Length: 161\nNumber of predicted TMHs: 2\nExp number of AAs in TMHs: 31.11104\nExp number, first 60 AAs: 14.67675\nTotal prob of N-in: 0.95023\nPOSSIBLE N-term signal sequence\ninside 1 6\nTMhelix 7 26\noutside 27 141\nTMhelix 142 160\ninside 161 161

28419 GCF_001539505.1_12082_2_40 Serratia marcescens Proteobacteria; Gammaproteobacteria; Enterobacterales; Yersiniaceae; Serratia MNKRRTTLTVLALIASLGLSSAPALADKGGNGNGNGNGHGNSGNHGNSGNHGNNGNKGNK WP_033646351.1 MULTISPECIES: membrane protein [Enterobacterales] Length: 161\nNumber of predicted TMHs: 2\nExp number of AAs in TMHs: 31.11104\nExp number, first 60 AAs: 14.67675\nTotal prob of N-in: 0.95023\nPOSSIBLE N-term signal sequence\ninside 1 6\nTMhelix 7 26\noutside 27 141\nTMhelix 142 160\ninside 161 161

28420 GCF_001539445.1_12082_3_62 Serratia marcescens Proteobacteria; Gammaproteobacteria; Enterobacterales; Yersiniaceae; Serratia MNKRRTTLTVLALIASLGLSSAPALADKGGNGNGNGNGHGNSGNHGNNGNHGNNGNHGNK WP_033640576.1 MULTISPECIES: membrane protein [Serratia] Length: 164\nNumber of predicted TMHs: 2\nExp number of AAs in TMHs: 30.54778\nExp number, first 60 AAs: 14.38033\nTotal prob of N-in: 0.94595\nPOSSIBLE N-term signal sequence\ninside 1 6\nTMhelix 7 26\noutside 27 144\nTMhelix 145 163\ninside 164 164

28421 GCF_001539545.1_12082_3_50 Serratia marcescens Proteobacteria; Gammaproteobacteria; Enterobacterales; Yersiniaceae; Serratia MNKRRTTLTVLALIASLGLSSAPALADKGGNGNGNGNGHGNSGNHGNSGNHGNNGNKGNK WP_033646351.1 MULTISPECIES: membrane protein [Enterobacterales] Length: 161\nNumber of predicted TMHs: 2\nExp number of AAs in TMHs: 31.11104\nExp number, first 60 AAs: 14.67675\nTotal prob of N-in: 0.95023\nPOSSIBLE N-term signal sequence\ninside 1 6\nTMhelix 7 26\noutside 27 141\nTMhelix 142 160\ninside 161 161

28422 GCF_001539645.1_12082_2_46 Serratia marcescens Proteobacteria; Gammaproteobacteria; Enterobacterales; Yersiniaceae; Serratia MNKRRTTLTVLALIASLGLSSAPALADKGGNGNGNGNGHGNSGNHGNSGNHGNNGNKGNK WP_033646351.1 MULTISPECIES: membrane protein [Enterobacterales] Length: 161\nNumber of predicted TMHs: 2\nExp number of AAs in TMHs: 31.11104\nExp number, first 60 AAs: 14.67675\nTotal prob of N-in: 0.95023\nPOSSIBLE N-term signal sequence\ninside 1 6\nTMhelix 7 26\noutside 27 141\nTMhelix 142 160\ninside 161 161

28423 GCF_001540125.1_12045_8_28 Serratia marcescens Proteobacteria; Gammaproteobacteria; Enterobacterales; Yersiniaceae; Serratia MNKRRTTLTVLALIASLGLSSAPALADKGGNGNGNGNGHGNSGNHGNSGNHGNNGNKGNK WP_033646351.1 MULTISPECIES: membrane protein [Enterobacterales] Length: 161\nNumber of predicted TMHs: 2\nExp number of AAs in TMHs: 31.11104\nExp number, first 60 AAs: 14.67675\nTotal prob of N-in: 0.95023\nPOSSIBLE N-term signal sequence\ninside 1 6\nTMhelix 7 26\noutside 27 141\nTMhelix 142 160\ninside 161 161

28424 GCF_001540165.1_12082_3_91 Serratia marcescens Proteobacteria; Gammaproteobacteria; Enterobacterales; Yersiniaceae; Serratia

MNKRRTTLTVLALIASLGLSSAPALADKGGNGNGNGNGHGNSGNHGNSGNHGNNGNKGNK WP_033646351.1 MULTISPECIES: membrane protein [Enterobacterales] Length: 161\nNumber of predicted TMHs: 2\nExp number of AAs in TMHs: 31.11104\nExp number, first 60 AAs: 14.67675\nTotal prob of N-in: 0.95023\nPOSSIBLE N-term signal sequence\ninside 1 6\nTMhelix 7 26\noutside 27 141\nTMhelix 142 160\ninside 161 161

28425 GCF_001540225.1_12045_8_22 Serratia marcescens Proteobacteria; Gammaproteobacteria; Enterobacterales; Yersiniaceae; Serratia MNKRRTTLTVLALIASLGLSSAPALADKGGNGNGNGNGHGNSGNHGNSGNHGNNGNKGNK WP_033646351.1 MULTISPECIES: membrane protein [Enterobacterales] Length: 161\nNumber of predicted TMHs: 2\nExp number of AAs in TMHs: 31.11104\nExp number, first 60 AAs: 14.67675\nTotal prob of N-in: 0.95023\nPOSSIBLE N-term signal sequence\ninside 1 6\nTMhelix 7 26\noutside 27 141\nTMhelix 142 160\ninside 161 161

28426 GCF_001540265.1_12082_2_57 Serratia marcescens Proteobacteria; Gammaproteobacteria; Enterobacterales; Yersiniaceae; Serratia MNKRRTTLTVLALIASLGLSSAPALADKGGNGNGNGNGHGNSGNHGNSGNHGNNGNKGNK WP_033646351.1 MULTISPECIES: membrane protein [Enterobacterales] Length: 161\nNumber of predicted TMHs: 2\nExp number of AAs in TMHs: 31.11104\nExp number, first 60 AAs: 14.67675\nTotal prob of N-in: 0.95023\nPOSSIBLE N-term signal sequence\ninside 1 6\nTMhelix 7 26\noutside 27 141\nTMhelix 142 160\ninside 161 161

28427 GCF_001540285.1_12082_3_21 Serratia marcescens Proteobacteria; Gammaproteobacteria; Enterobacterales; Yersiniaceae; Serratia MNKRRTTLTVLALIASLGLSSAPALADKGGNGNGNGNGHGNSGNHGNSGNHGNNGNKGNK WP_033646351.1 MULTISPECIES: membrane protein [Enterobacterales] Length: 161\nNumber of predicted TMHs: 2\nExp number of AAs in TMHs: 31.11104\nExp number, first 60 AAs: 14.67675\nTotal prob of N-in: 0.95023\nPOSSIBLE N-term signal sequence\ninside 1 6\nTMhelix 7 26\noutside 27 141\nTMhelix 142 160\ninside 161 161

28428 GCF_001540205.1_12082_2_50 Serratia marcescens Proteobacteria; Gammaproteobacteria; Enterobacterales; Yersiniaceae; Serratia MNKRRTTLTVLALIASLGLSSAPALADKGGNGNGNGNGHGNSGNNGNHGNNGNHGNKGNK WP_060419303.1 MULTISPECIES: hypothetical protein [Serratia] Length: 164\nNumber of predicted TMHs: 2\nExp number of AAs in TMHs: 30.51677\nExp number, first 60 AAs: 14.42464\nTotal prob of N-in: 0.94579\nPOSSIBLE N-term signal sequence\ninside 1 6\nTMhelix 7 26\noutside 27 144\nTMhelix 145 163\ninside 164 164

28429 GCF_001540325.1_12082_2_95 Serratia marcescens Proteobacteria; Gammaproteobacteria; Enterobacterales; Yersiniaceae; Serratia MNKRRTTLTVLALIASLGLSSAPALADKGGNGNGNGNGHGNSGNHGNSGNHGNNGNKGNK WP_033646351.1 MULTISPECIES: membrane protein [Enterobacterales] Length: 161\nNumber of predicted TMHs: 2\nExp number of AAs in TMHs: 31.11104\nExp number, first 60 AAs: 14.67675\nTotal prob of N-in: 0.95023\nPOSSIBLE N-term signal sequence\ninside 1 6\nTMhelix 7 26\noutside 27 141\nTMhelix 142 160\ninside 161 161

28430 GCF_001540365.1_12082_2_68 Serratia marcescens Proteobacteria; Gammaproteobacteria; Enterobacterales; Yersiniaceae; Serratia MNKRRTTLTVLALIASLGLSSAPALADKGGNGNGNGNGHGNSGNHGNSGNHGNNGNKGNK WP_033646351.1 MULTISPECIES: membrane protein [Enterobacterales] Length: 161\nNumber of predicted TMHs: 2\nExp number of AAs in TMHs: 31.11104\nExp number, first 60 AAs: 14.67675\nTotal prob of N-in: 0.95023\nPOSSIBLE N-term signal sequence\ninside 1 6\nTMhelix 7 26\noutside 27 141\nTMhelix 142 160\ninside 161 161

28431 GCF_001889685.1_ASM188968v1 Serratia marcescens Proteobacteria; Gammaproteobacteria; Enterobacterales; Yersiniaceae; Serratia MNKRRTTLTVLALIASLGLSSAPAMADKGGNGNGNGNGHGNSGNHGNNGNHGNNGNHGNK WP_033642812.1 MULTISPECIES: membrane protein [Serratia] Length: 164\nNumber of predicted TMHs: 2\nExp number of AAs in TMHs: 30.89657\nExp number, first 60 AAs: 14.76607\nTotal prob of N-in: 0.94978\nPOSSIBLE N-term signal sequence\ninside 1 6\nTMhelix 7 26\noutside 27 144\nTMhelix 145 163\ninside 164 164

28432 GCF_001853495.1_ASM185349v1 Serratia marcescens Proteobacteria; Gammaproteobacteria; Enterobacterales; Yersiniaceae; Serratia MNKRRTTLTVLALIASLGLSSAPALADKGGNGNGNGNGHGNSGNNGNHGNNGNHGNKGNK WP_060419303.1 MULTISPECIES: hypothetical protein [Serratia] Length: 164\nNumber of predicted TMHs: 2\nExp number of AAs in TMHs: 30.51677\nExp number, first 60 AAs: 14.42464\nTotal prob of N-in: 0.94579\nPOSSIBLE N-term signal sequence\ninside 1 6\nTMhelix 7 26\noutside 27 144\nTMhelix 145 163\ninside 164 164

28433	GCF_001672055.1_ASM167205v1	Serratia marcescens	Proteobacteria; Gammaproteobacteria; Enterobacterales; Yersiniaceae; Serratia
MNKRRTTLTVLALIASLGLSSAPALADKGGNGNGNGNGHGNSGNHGNNGNHGNNGNNGNH	WP_033633261.1
MULTISPECIES: membrane protein [Serratia]	Length: 176\nNumber of predicted TMHs:  2\nExp number of AAs in TMHs: 28.40352\nExp number, first 60 AAs:  13.30908\nTotal prob of N-in:   0.92780\nPOSSIBLE N-term signal sequence\ninside   1   6\nTMhelix   7   26\noutside   27   156\nTMhelix   157   175\ninside   176   176

28434	GCF_002185265.1_ASM218526v1	Serratia nematodiphila	Proteobacteria; Gammaproteobacteria; Enterobacterales; Yersiniaceae; Serratia
MNKRRTTLTVLALIASLGLSSAPALADKGGNGNGNGNGHGNSGNNGNHGNNGNHGNKGNK	WP_087763533.1
hypothetical protein [Serratia nematodiphila]	Length: 164\nNumber of predicted TMHs:  2\nExp number of AAs in TMHs: 30.5153\nExp number, first 60 AAs:  14.42673\nTotal prob of N-in:   0.94579\nPOSSIBLE N-term signal sequence\ninside   1   6\nTMhelix   7   26\noutside   27   144\nTMhelix   145   163\ninside   164   164

28435	GCF_900101535.1_IMG-taxon_2596583578_annotated_assembly	Serratia nematodiphila
Proteobacteria; Gammaproteobacteria; Enterobacterales; Yersiniaceae; Serratia
MNKRRTTLTVLALIASLGLSSAPALADKGGNGNGNGNGHGNSGNHGNNGNHGNNGNNGNH	WP_033633261.1
MULTISPECIES: membrane protein [Serratia]	Length: 176\nNumber of predicted TMHs:  2\nExp number of AAs in TMHs: 28.40352\nExp number, first 60 AAs:  13.30908\nTotal prob of N-in:   0.92780\nPOSSIBLE N-term signal sequence\ninside   1   6\nTMhelix   7   26\noutside   27   156\nTMhelix   157   175\ninside   176   176

28436	GCF_002082115.1_ASM208211v1	Serratia nematodiphila	Proteobacteria; Gammaproteobacteria; Enterobacterales; Yersiniaceae; Serratia
MNKRRTTLTVLALIASLGLSSAPALADKGGNGNGNGNGNGHGNSGNHGNNGNHGNNGNNG	WP_082997693.1
hypothetical protein [Serratia nematodiphila]	Length: 169\nNumber of predicted TMHs:  2\nExp number of AAs in TMHs: 29.66631\nExp number, first 60 AAs:  13.93018\nTotal prob of N-in:   0.93875\nPOSSIBLE N-term signal sequence\ninside   1   6\nTMhelix   7   26\noutside   27   149\nTMhelix   150   168\ninside   169   169

28437	GCF_001590765.1_ASM159076v1	Serratia plymuthica	Proteobacteria; Gammaproteobacteria; Enterobacterales; Yersiniaceae; Serratia
MKGVLEKCTVEDFNYISATLDSYVSFTNDKRRKTLLSAYQNNPALHAELISLIDTQIKYF	WP_020438466.1	hypothetical protein [Serratia plymuthica]	Length: 249\nNumber of predicted TMHs:  2\nExp number of AAs in TMHs: 39.37829\nExp number, first 60 AAs:  0.01949\nTotal prob of N-in:   0.09376\noutside   1   157\nTMhelix   158   180\ninside   181   200\nTMhelix   201   220\noutside   221   249

28438	GCF_001537385.1_12082_3_2	Serratia sp. 2880STDY5682894	Proteobacteria; Gammaproteobacteria; Enterobacterales; Yersiniaceae; Serratia
MNKRRTTLTVLALIASLGLSSAPALADKGGNGNGNGNGHGNSGNNGNHGNNGNHGNKGNK	WP_060419303.1
MULTISPECIES: hypothetical protein [Serratia]	Length: 164\nNumber of predicted TMHs:  2\nExp number of AAs in TMHs: 30.51677\nExp number, first 60 AAs:  14.42464\nTotal prob of N-in:   0.94579\nPOSSIBLE N-term signal sequence\ninside   1   6\nTMhelix   7   26\noutside   27   144\nTMhelix   145   163\ninside   164   164

28439	GCF_001537265.1_12082_3_3	Serratia sp. 2880STDY5682895	Proteobacteria; Gammaproteobacteria; Enterobacterales; Yersiniaceae; Serratia
MNKRRTTLTVLALIASLGLSSAPALADKGGNGNGNGNGHGNSGNHGNSGNHGNNGNKGNK	WP_033646351.1
MULTISPECIES: membrane protein [Enterobacterales]	Length: 161\nNumber of predicted TMHs:  2\nExp number of AAs in TMHs: 31.11104\nExp number, first 60 AAs:  14.67675\nTotal prob of N-in:   0.95023\nPOSSIBLE N-term signal sequence\ninside   1   6\nTMhelix   7   26\noutside   27   141\nTMhelix   142   160\ninside   161   161

28440	GCF_001076875.1_ASM107687v1	Serratia sp. 506_PEND	Proteobacteria; Gammaproteobacteria; Enterobacterales; Yersiniaceae; Serratia
MNKRRTTLTVLALIASLGLSSAPALADKGGNGNGNGNGHGNSGNHGNNGNHGNNGNHGNK	WP_033640576.1
MULTISPECIES: membrane protein [Serratia]	Length: 164\nNumber of predicted TMHs:  2\nExp number of AAs in TMHs: 30.54778\nExp number, first 60 AAs:  14.38033\nTotal prob of N-in:   0.94595\nPOSSIBLE N-term signal sequence\ninside   1   6\nTMhelix   7   26\noutside   27   144\nTMhelix   145   163\ninside   164   164

28441	GCF_000214195.1_ASM21419v1	Serratia sp. AS12	Proteobacteria; Gammaproteobacteria; Enterobacterales; Yersiniaceae; Serratia
MNKRRTTLTALALIVSLGLSSAPAFADKGGNGHGNGNGGGHGNSQGNGNGHGNGNSGDHG	WP_013812568.1
MULTISPECIES: membrane protein [Serratia]	Length: 167\nNumber of predicted TMHs:  2\nExp number of

AAs in TMHs: 33.54956\nExp number, first 60 AAs: 15.01343\nTotal prob of N-in: 0.96357\nPOSSIBLE N-term signal sequence\ninside 1 6\nTMhelix 7 26\noutside 27 142\nTMhelix 143 165\ninside 166 167

28442 GCF_000214805.1_ASM21480v1 Serratia sp. AS13 Proteobacteria; Gammaproteobacteria; Enterobacterales; Yersiniaceae; Serratia MNKRRTTLTALALIVSLGLSSAPAFADKGGNGHGNGNGGGHGNSQGNGNGHGNGNSGDHG WP_013812568.1 MULTISPECIES: membrane protein [Serratia] Length: 167\nNumber of predicted TMHs: 2\nExp number of AAs in TMHs: 33.54956\nExp number, first 60 AAs: 15.01343\nTotal prob of N-in: 0.96357\nPOSSIBLE N-term signal sequence\ninside 1 6\nTMhelix 7 26\noutside 27 142\nTMhelix 143 165\ninside 166 167

28443 GCF_000695995.1_ASM69599v1 Serratia sp. FS14 Proteobacteria; Gammaproteobacteria; Enterobacterales; Yersiniaceae; Serratia MNKRRTTLTVLALIASLGLSSAPALADKGGNGNGNGNGHGNSGNHGNNGNHGNNGNNGNH WP_004941297.1 MULTISPECIES: hypothetical protein [Serratia] Length: 167\nNumber of predicted TMHs: 2\nExp number of AAs in TMHs: 30.02384\nExp number, first 60 AAs: 14.11185\nTotal prob of N-in: 0.94170\nPOSSIBLE N-term signal sequence\ninside 1 6\nTMhelix 7 26\noutside 27 147\nTMhelix 148 166\ninside 167 167

28444 GCF_001808215.1_ASM180821v1 Serratia sp. HMSC15F11 Proteobacteria; Gammaproteobacteria; Enterobacterales; Yersiniaceae; Serratia MNKRRTTLTVLALIASLGLSSAPALADKGGNGNGNGNGHGNSGNHGNNGNHGNNGNNGNH WP_004941297.1 MULTISPECIES: hypothetical protein [Serratia] Length: 167\nNumber of predicted TMHs: 2\nExp number of AAs in TMHs: 30.02384\nExp number, first 60 AAs: 14.11185\nTotal prob of N-in: 0.94170\nPOSSIBLE N-term signal sequence\ninside 1 6\nTMhelix 7 26\noutside 27 147\nTMhelix 148 166\ninside 167 167

28445 GCF_001714765.1_SerISTD041.0 Serratia sp. ISTD04 Proteobacteria; Gammaproteobacteria; Enterobacterales; Yersiniaceae; Serratia MNKRRTTLTVLALIASLGLSSAPAMADKGGNGNGNGNGHGNSGNHGNNGNHGNNGNHGNK WP_033642812.1 MULTISPECIES: membrane protein [Serratia] Length: 164\nNumber of predicted TMHs: 2\nExp number of AAs in TMHs: 30.89657\nExp number, first 60 AAs: 14.76607\nTotal prob of N-in: 0.94978\nPOSSIBLE N-term signal sequence\ninside 1 6\nTMhelix 7 26\noutside 27 144\nTMhelix 145 163\ninside 164 164

28446 GCF_001422565.1_Leaf50 Serratia sp. Leaf50 Proteobacteria; Gammaproteobacteria; Enterobacterales; Yersiniaceae; Serratia MIDANSNLTRRRLLQGMGVLAAATLIPTGVFPAYASSAISNDFITISIFLTGREKLSANY WP_055770949.1 hypothetical protein [Serratia sp. Leaf50] Length: 167\nNumber of predicted TMHs: 2\nExp number of AAs in TMHs: 41.47245\nExp number, first 60 AAs: 22.67645\nTotal prob of N-in: 0.99317\nPOSSIBLE N-term signal sequence\ninside 1 12\nTMhelix 13 35\noutside 36 104\nTMhelix 105 127\ninside 128 167

28447 GCF_001984565.1_ASM198456v1 Serratia sp. S119 Proteobacteria; Gammaproteobacteria; Enterobacterales; Yersiniaceae; Serratia MNKRRTTLTVLALIASLGLSSAPALADKGGNGNGNGNGHGNSGNHGNSGNHGNNVNKGNK WP_077039006.1 hypothetical protein [Serratia sp. S119] Length: 161\nNumber of predicted TMHs: 2\nExp number of AAs in TMHs: 31.11104\nExp number, first 60 AAs: 14.67675\nTotal prob of N-in: 0.95023\nPOSSIBLE N-term signal sequence\ninside 1 6\nTMhelix 7 26\noutside 27 141\nTMhelix 142 160\ninside 161 161

28448 GCF_000347995.1_ASM34799v1 Serratia sp. S4 Proteobacteria; Gammaproteobacteria; Enterobacterales; Yersiniaceae; Serratia MSKRRTTLTVLALIVSMGLSSAPAFADKGGNGNGNGHGNSGNHGNNGNHGNKAKGNESDQ WP_017892578.1 membrane protein [Serratia sp. S4] Length: 152\nNumber of predicted TMHs: 2\nExp number of AAs in TMHs: 36.38921\nExp number, first 60 AAs: 18.00995\nTotal prob of N-in: 0.97278\nPOSSIBLE N-term signal sequence\ninside 1 6\nTMhelix 7 26\noutside 27 127\nTMhelix 128 150\ninside 151 152

28449 GCF_000747565.1_ASM74756v1 Serratia sp. SCBI Proteobacteria; Gammaproteobacteria; Enterobacterales; Yersiniaceae; Serratia MNKRRTTLTVLALIASLGLSSAPAMADKGGNGNGNGHGNSGNHGNNGNHGNKDKGDYRDY WP_042784306.1 membrane protein [Serratia sp. SCBI] Length: 156\nNumber of predicted TMHs: 2\nExp number of AAs in TMHs: 32.20711\nExp number, first 60 AAs: 15.4275\nTotal prob of N-in: 0.95951\nPOSSIBLE N-term signal sequence\ninside 1 6\nTMhelix 7 26\noutside 27 136\nTMhelix 137 155\ninside 156 156

28450 GCF_001011075.1_ASM101107v1 Serratia sp. TEL Proteobacteria; Gammaproteobacteria; Enterobacterales; Yersiniaceae; Serratia MNKRRTTLTVLALIASLGLSSAPAMADKGGNGNGNGNGHGNSGNHGNNGNHGNKDKGDYR WP_047026237.1

membrane protein [Serratia sp. TEL] Length: 158\nNumber of predicted TMHs: 2\nExp number of AAs in TMHs: 31.88626\nExp number, first 60 AAs: 15.26199\nTotal prob of N-in: 0.95719\nPOSSIBLE N-term signal sequence\ninside 1 6\nTMhelix 7 26\noutside 27 138\nTMhelix 139 157\ninside 158 158

28451 GCF_000988045.1_GFC_24 Serratia ureilytica Proteobacteria; Gammaproteobacteria; Enterobacterales; Yersiniaceae; Serratia MNKRRTTLTVLALIASLGLSSAPAMADKGGNGNGNRHGNSGNHGNNGNHGNNGNHGNKDK WP_046686926.1 membrane protein [Serratia ureilytica] Length: 162\nNumber of predicted TMHs: 2\nExp number of AAs in TMHs: 30.41441\nExp number, first 60 AAs: 14.76836\nTotal prob of N-in: 0.94981\nPOSSIBLE N-term signal sequence\ninside 1 6\nTMhelix 7 26\noutside 27 142\nTMhelix 143 161\ninside 162 162

28452 GCF_001590885.1_ASM159088v1 Serratia ficaria NBRC 102596 Proteobacteria; Gammaproteobacteria; Enterobacterales; Yersiniaceae; Serratia; Serratia ficaria MSKRRTTLTVLALIASLGLSSAPALADKGGNGNGNGGGNGHGNSGNHGNSGDHDKGNKGN WP_061795424.1 hypothetical protein [Serratia ficaria] Length: 161\nNumber of predicted TMHs: 2\nExp number of AAs in TMHs: 31.67648\nExp number, first 60 AAs: 14.05019\nTotal prob of N-in: 0.93632\nPOSSIBLE N-term signal sequence\ninside 1 6\nTMhelix 7 26\noutside 27 141\nTMhelix 142 160\ninside 161 161

28453 GCF_001590905.1_ASM159090v1 Serratia grimesii NBRC 13537 Proteobacteria; Gammaproteobacteria; Enterobacterales; Yersiniaceae; Serratia; Serratia grimesii MSKRRTTLTALALIVSLGLSSAPVLADKGGNGNGGGNGHGNSGNHGNNGNHGNNGNHGNK WP_061807603.1 hypothetical protein [Serratia grimesii] Length: 160\nNumber of predicted TMHs: 2\nExp number of AAs in TMHs: 33.04249\nExp number, first 60 AAs: 16.06267\nTotal prob of N-in: 0.94245\nPOSSIBLE N-term signal sequence\ninside 1 6\nTMhelix 7 26\noutside 27 135\nTMhelix 136 158\ninside 159 160

28454 GCF_000422085.1_ASM42208v1 Serratia liquefaciens ATCC 27592 Proteobacteria; Gammaproteobacteria; Enterobacterales; Yersiniaceae; Serratia; Serratia liquefaciens MSKRRTTFTVLALIVSLGLSSAPALADKGGNGNGGGNGHGNSGNHGNNGNNGKHGNKAKD WP_020826508.1 membrane protein [Serratia liquefaciens] Length: 157\nNumber of predicted TMHs: 2\nExp number of AAs in TMHs: 36.25849\nExp number, first 60 AAs: 17.0716\nTotal prob of N-in: 0.96941\nPOSSIBLE N-term signal sequence\ninside 1 6\nTMhelix 7 26\noutside 27 132\nTMhelix 133 155\ninside 156 157

28455 GCF_000699165.1_ASM69916v1 Serratia liquefaciens FK01 Proteobacteria; Gammaproteobacteria; Enterobacterales; Yersiniaceae; Serratia; Serratia liquefaciens MSKRRTAFTVLALIVSLGLSSAPALADKGGNGNGHGNSGNHGNNGNNGKHGNKAKDNDGD WP_044550180.1 membrane protein [Serratia liquefaciens] Length: 153\nNumber of predicted TMHs: 2\nExp number of AAs in TMHs: 38.82143\nExp number, first 60 AAs: 18.81863\nTotal prob of N-in: 0.98666\nPOSSIBLE N-term signal sequence\ninside 1 6\nTMhelix 7 26\noutside 27 128\nTMhelix 129 151\ninside 152 153

28456 GCF_000418935.1_S_marcescensAB42556419-isolate1-1.0Serratia marcescens AB42556419-isolate1 Proteobacteria; Gammaproteobacteria; Enterobacterales; Yersiniaceae; Serratia; Serratia marcescens MNKRRTTLTVLALIASLGLSSAPALADKGGNGNGNGNGHGNSGNHGNNGNHGNNGNNGNH WP_004941297.1 MULTISPECIES: hypothetical protein [Serratia] Length: 167\nNumber of predicted TMHs: 2\nExp number of AAs in TMHs: 30.02384\nExp number, first 60 AAs: 14.11185\nTotal prob of N-in: 0.94170\nPOSSIBLE N-term signal sequence\ninside 1 6\nTMhelix 7 26\noutside 27 147\nTMhelix 148 166\ninside 167 167

28457 GCF_000521925.1_Serr_marc_BIDMC_44_V1 Serratia marcescens BIDMC 44 Proteobacteria; Gammaproteobacteria; Enterobacterales; Yersiniaceae; Serratia; Serratia marcescens MNKRRTTLTVLALIASLGLSSAPALADKGGNGNGNGNGHGNSGNHGNSGNHGNNDNKDNK WP_033638251.1 membrane protein [Serratia marcescens] Length: 173\nNumber of predicted TMHs: 2\nExp number of AAs in TMHs: 28.99941\nExp number, first 60 AAs: 13.60587\nTotal prob of N-in: 0.93289\nPOSSIBLE N-term signal sequence\ninside 1 6\nTMhelix 7 26\noutside 27 153\nTMhelix 154 172\ninside 173 173

28458 GCF_000521905.1_Serr_marc_BIDMC_50_V1 Serratia marcescens BIDMC 50 Proteobacteria; Gammaproteobacteria; Enterobacterales; Yersiniaceae; Serratia; Serratia marcescens MNKRRTTLTVLALIASLGLSSAPAMADKGGNGNGNGNGHGNSGNHGNNGNHGNNGNHGNK WP_033642812.1 MULTISPECIES: membrane protein [Serratia] Length: 164\nNumber of predicted TMHs: 2\nExp number of AAs in TMHs: 30.89657\nExp number, first 60 AAs: 14.76607\nTotal prob of N-in: 0.94978\nPOSSIBLE N-term signal sequence\ninside 1 6\nTMhelix 7 26\noutside 27 144\nTMhelix 145 163\ninside 164 164

28459 GCF_000633715.1_Serr_marc_BIDMC_80_V1 Serratia marcescens BIDMC 80 Proteobacteria; Gammaproteobacteria; Enterobacterales; Yersiniaceae; Serratia; Serratia marcescens

MNKRRTTLTVLALIASLGLSSAPALADKGGNGNGNGNGHGNSGNHGNSGNHGNNGNKGNK WP_033646351.1 MULTISPECIES: membrane protein [Enterobacterales] Length: 161\nNumber of predicted TMHs: 2\nExp number of AAs in TMHs: 31.11104\nExp number, first 60 AAs: 14.67675\nTotal prob of N-in: 0.95023\nPOSSIBLE N-term signal sequence\ninside 1 6\nTMhelix 7 26\noutside 27 141\nTMhelix 142 160\ninside 161 161

28460 GCF_000633695.1_Serr_marc_BIDMC_81_V1 Serratia marcescens BIDMC 81 Proteobacteria; Gammaproteobacteria; Enterobacterales; Yersiniaceae; Serratia; Serratia marcescens MNKRRTTLTVLALIASLGLSSAPAMADKGGNGNGNGNGHGNSGNHGNNGNHGNHGNKDKG WP_033652092.1 membrane protein [Serratia marcescens] Length: 161\nNumber of predicted TMHs: 2\nExp number of AAs in TMHs: 31.39885\nExp number, first 60 AAs: 15.01586\nTotal prob of N-in: 0.95359\nPOSSIBLE N-term signal sequence\ninside 1 6\nTMhelix 7 26\noutside 27 141\nTMhelix 142 160\ninside 161 161

28461 GCF_000465615.2_GS_De_Novo_Assembly Serratia marcescens EGD-HP20 Proteobacteria; Gammaproteobacteria; Enterobacterales; Yersiniaceae; Serratia; Serratia marcescens MNKRRTTLTVLALIASLGLSSAPALADKGGNGNGNGNGHGNSGNHGNNGNHGNNGNNGNH WP_004941297.1 MULTISPECIES: hypothetical protein [Serratia] Length: 167\nNumber of predicted TMHs: 2\nExp number of AAs in TMHs: 30.02384\nExp number, first 60 AAs: 14.11185\nTotal prob of N-in: 0.94170\nPOSSIBLE N-term signal sequence\ninside 1 6\nTMhelix 7 26\noutside 27 147\nTMhelix 148 166\ninside 167 167

28462 GCF_000442455.1_S.marcescens_LCT-SM166 Serratia marcescens LCT-SM166 Proteobacteria; Gammaproteobacteria; Enterobacterales; Yersiniaceae; Serratia; Serratia marcescens MNKRRTTLTVLALIASLGLSSAPALADKGGNGNGNGNGHGNSGNHGNNGNHGNNGNHGNK WP_016928034.1 membrane protein [Serratia marcescens] Length: 164\nNumber of predicted TMHs: 2\nExp number of AAs in TMHs: 30.54438\nExp number, first 60 AAs: 14.38759\nTotal prob of N-in: 0.94593\nPOSSIBLE N-term signal sequence\ninside 1 6\nTMhelix 7 26\noutside 27 144\nTMhelix 145 163\ninside 164 164

28463 GCF_000264275.1_ASM26427v1 Serratia marcescens LCT-SM213 Proteobacteria; Gammaproteobacteria; Enterobacterales; Yersiniaceae; Serratia; Serratia marcescens MNKRRTTLTVLALIASLGLSSAPALADKGGNGNGNGNGHGNSGNHGNNGNHGNNGNHGNK WP_016928034.1 membrane protein [Serratia marcescens] Length: 164\nNumber of predicted TMHs: 2\nExp number of AAs in TMHs: 30.54438\nExp number, first 60 AAs: 14.38759\nTotal prob of N-in: 0.94593\nPOSSIBLE N-term signal sequence\ninside 1 6\nTMhelix 7 26\noutside 27 144\nTMhelix 145 163\ninside 164 164

28464 GCF_000442375.1_S.marcescens_LCT-SM262 Serratia marcescens LCT-SM262 Proteobacteria; Gammaproteobacteria; Enterobacterales; Yersiniaceae; Serratia; Serratia marcescens MNKRRTTLTVLALIASLGLSSAPALADKGGNGNGNGNGHGNSGNHGNNGNHGNNGNHGNK WP_016928034.1 membrane protein [Serratia marcescens] Length: 164\nNumber of predicted TMHs: 2\nExp number of AAs in TMHs: 30.54438\nExp number, first 60 AAs: 14.38759\nTotal prob of N-in: 0.94593\nPOSSIBLE N-term signal sequence\ninside 1 6\nTMhelix 7 26\noutside 27 144\nTMhelix 145 163\ninside 164 164

28465 GCF_000418915.1_S_marcescensMC458-1.0 Serratia marcescens MC458 Proteobacteria; Gammaproteobacteria; Enterobacterales; Yersiniaceae; Serratia; Serratia marcescens MNKRRTTLTVLALIASLGLSSAPALADKGGNGNGNGNGHGNSGNHGNNGNHGNNGNNGNH WP_004941297.1 MULTISPECIES: hypothetical protein [Serratia] Length: 167\nNumber of predicted TMHs: 2\nExp number of AAs in TMHs: 30.02384\nExp number, first 60 AAs: 14.11185\nTotal prob of N-in: 0.94170\nPOSSIBLE N-term signal sequence\ninside 1 6\nTMhelix 7 26\noutside 27 147\nTMhelix 148 166\ninside 167 167

28466 GCF_000418895.1_S_marcescensMC459-1.0 Serratia marcescens MC459 Proteobacteria; Gammaproteobacteria; Enterobacterales; Yersiniaceae; Serratia; Serratia marcescens MNKRRTTLTVLALIASLGLSSAPALADKGGNGNGNGNGHGNSGNHGNNGNHGNNGNNGNH WP_004941297.1 MULTISPECIES: hypothetical protein [Serratia] Length: 167\nNumber of predicted TMHs: 2\nExp number of AAs in TMHs: 30.02384\nExp number, first 60 AAs: 14.11185\nTotal prob of N-in: 0.94170\nPOSSIBLE N-term signal sequence\ninside 1 6\nTMhelix 7 26\noutside 27 147\nTMhelix 148 166\ninside 167 167

28467 GCF_000418875.1_S_marcescensMC460-1.0 Serratia marcescens MC460 Proteobacteria; Gammaproteobacteria; Enterobacterales; Yersiniaceae; Serratia; Serratia marcescens MNKRRTTLTVLALIASLGLSSAPALADKGGNGNGNGNGHGNSGNHGNNGNHGNNGNNGNH WP_004941297.1 MULTISPECIES: hypothetical protein [Serratia] Length: 167\nNumber of predicted TMHs: 2\nExp number of AAs in TMHs: 30.02384\nExp number, first 60 AAs: 14.11185\nTotal prob of N-in: 0.94170\nPOSSIBLE N-term signal sequence\ninside 1 6\nTMhelix 7 26\noutside 27 147\nTMhelix 148 166\ninside 167 167

28468 GCF_000418855.2_S_marcescensMC6000-1.0 Serratia marcescens MC6000 Proteobacteria; Gammaproteobacteria; Enterobacterales; Yersiniaceae; Serratia; Serratia marcescens MNKRRTTLTVLALIASLGLSSAPALADKGGNGNGNGNGHGNSGNHGNNGNHGNNGNNGNH WP_004941297.1 MULTISPECIES: hypothetical protein [Serratia] Length: 167\nNumber of predicted TMHs: 2\nExp number of AAs in TMHs: 30.02384\nExp number, first 60 AAs: 14.11185\nTotal prob of N-in: 0.94170\nPOSSIBLE N-term signal sequence\ninside 1 6\nTMhelix 7 26\noutside 27 147\nTMhelix 148 166\ninside 167 167

28469 GCF_000418835.1_S_marcescensMC6001-1.0 Serratia marcescens MC6001 Proteobacteria; Gammaproteobacteria; Enterobacterales; Yersiniaceae; Serratia; Serratia marcescens MNKRRTTLTVLALIASLGLSSAPALADKGGNGNGNGNGHGNSGNHGNNGNHGNNGNNGNH WP_004941297.1 MULTISPECIES: hypothetical protein [Serratia] Length: 167\nNumber of predicted TMHs: 2\nExp number of AAs in TMHs: 30.02384\nExp number, first 60 AAs: 14.11185\nTotal prob of N-in: 0.94170\nPOSSIBLE N-term signal sequence\ninside 1 6\nTMhelix 7 26\noutside 27 147\nTMhelix 148 166\ninside 167 167

28470 GCF_000418815.1_S_marcescensMC620-1.0 Serratia marcescens MC620 Proteobacteria; Gammaproteobacteria; Enterobacterales; Yersiniaceae; Serratia; Serratia marcescens MNKRRTTLTVLALIASLGLSSAPALADKGGNGNGNGNGHGNSGNHGNNGNHGNNGNNGNH WP_004941297.1 MULTISPECIES: hypothetical protein [Serratia] Length: 167\nNumber of predicted TMHs: 2\nExp number of AAs in TMHs: 30.02384\nExp number, first 60 AAs: 14.11185\nTotal prob of N-in: 0.94170\nPOSSIBLE N-term signal sequence\ninside 1 6\nTMhelix 7 26\noutside 27 147\nTMhelix 148 166\ninside 167 167

28471 GCF_900029885.1_Sm_SMB2099 Serratia marcescens SMB2099 Proteobacteria; Gammaproteobacteria; Enterobacterales; Yersiniaceae; Serratia; Serratia marcescens MNKRRTTLTVLALIASLGLSSAPALADKGGNGNGNGNGHGNSGNHGNSGNHGNNGNKGNK WP_033646351.1 MULTISPECIES: membrane protein [Enterobacterales] Length: 161\nNumber of predicted TMHs: 2\nExp number of AAs in TMHs: 31.11104\nExp number, first 60 AAs: 14.67675\nTotal prob of N-in: 0.95023\nPOSSIBLE N-term signal sequence\ninside 1 6\nTMhelix 7 26\noutside 27 141\nTMhelix 142 160\ninside 161 161

28472 GCF_001317205.2_56_S29v.02 Serratia marcescens subsp. marcescens Proteobacteria; Gammaproteobacteria; Enterobacterales; Yersiniaceae; Serratia; Serratia marcescens MNKRRTTLTVLALIASLGLSSAPALADKGGNGNGNGNGHGNSGNHGNHGNNGNHGNNGNN WP_055316240.1 hypothetical protein [Serratia marcescens] Length: 170\nNumber of predicted TMHs: 2\nExp number of AAs in TMHs: 29.49026\nExp number, first 60 AAs: 13.84277\nTotal prob of N-in: 0.93726\nPOSSIBLE N-term signal sequence\ninside 1 6\nTMhelix 7 26\noutside 27 150\nTMhelix 151 169\ninside 170 170

28473 GCF_001908035.1_ASM190803v1 Serratia marcescens subsp. marcescens Proteobacteria; Gammaproteobacteria; Enterobacterales; Yersiniaceae; Serratia; Serratia marcescens MNKRRTTLTVLALIASLGLSSAPALADKGGNGNGNGNGHGNSGNHGNSGNHGNNGNKGNK WP_033646351.1 MULTISPECIES: membrane protein [Enterobacterales] Length: 161\nNumber of predicted TMHs: 2\nExp number of AAs in TMHs: 31.11104\nExp number, first 60 AAs: 14.67675\nTotal prob of N-in: 0.95023\nPOSSIBLE N-term signal sequence\ninside 1 6\nTMhelix 7 26\noutside 27 141\nTMhelix 142 160\ninside 161 161

28474 GCF_001051865.1_ASM105186v1 Serratia marcescens subsp. marcescens Proteobacteria; Gammaproteobacteria; Enterobacterales; Yersiniaceae; Serratia; Serratia marcescens MNKRRTTLTVLALIASLGLSSAPALADKGGNGNGNGNGHGNSGNHGNNGNNGNHGNNGNH WP_049209162.1 membrane protein [Serratia marcescens] Length: 167\nNumber of predicted TMHs: 2\nExp number of AAs in TMHs: 30.02384\nExp number, first 60 AAs: 14.11185\nTotal prob of N-in: 0.94170\nPOSSIBLE N-term signal sequence\ninside 1 6\nTMhelix 7 26\noutside 27 147\nTMhelix 148 166\ninside 167 167

28475 GCF_001317425.2_59_S30v.02 Serratia marcescens subsp. marcescens Proteobacteria; Gammaproteobacteria; Enterobacterales; Yersiniaceae; Serratia; Serratia marcescens MNKRRTTLTVLALIASLGLSSAPALADKGGNGNGNGNGHGNSGNHGNHGNNGNHGNNGNN WP_055316240.1 hypothetical protein [Serratia marcescens] Length: 170\nNumber of predicted TMHs: 2\nExp number of AAs in TMHs: 29.49026\nExp number, first 60 AAs: 13.84277\nTotal prob of N-in: 0.93726\nPOSSIBLE N-term signal sequence\ninside 1 6\nTMhelix 7 26\noutside 27 150\nTMhelix 151 169\ninside 170 170

28476 GCF_001065405.1_ASM106540v1 Serratia marcescens subsp. marcescens Proteobacteria; Gammaproteobacteria; Enterobacterales; Yersiniaceae; Serratia; Serratia marcescens MNKRRTTLTVLALIASLGLSSAPALADKGGNGNGNGNGHGNSGNHGNNGNHGNNGNNGNN WP_048323073.1 membrane protein [Serratia marcescens] Length: 170\nNumber of predicted TMHs: 2\nExp number of

AAs in TMHs: 29.49026\nExp number, first 60 AAs: 13.84277\nTotal prob of N-in: 0.93726\nPOSSIBLE N-term signal sequence\ninside 1 6\nTMhelix 7 26\noutside 27 150\nTMhelix 151 169\ninside 170 170

28477 GCF_001908015.1_ASM190801v1 Serratia marcescens subsp. marcescens Proteobacteria; Gammaproteobacteria; Enterobacterales; Yersiniaceae; Serratia; Serratia marcescens MNKRRTTLTVLALIASLGLSSAPALADKGGNGNGNGNGHGNSGNHGNSGNHGNNGNKGNK WP_033646351.1 MULTISPECIES: membrane protein [Enterobacterales] Length: 161\nNumber of predicted TMHs: 2\nExp number of AAs in TMHs: 31.11104\nExp number, first 60 AAs: 14.67675\nTotal prob of N-in: 0.95023\nPOSSIBLE N-term signal sequence\ninside 1 6\nTMhelix 7 26\noutside 27 141\nTMhelix 142 160\ninside 161 161

28478 GCF_000342205.1_SerMar_1.0 Serratia marcescens VGH107 Proteobacteria; Gammaproteobacteria; Enterobacterales; Yersiniaceae; Serratia; Serratia marcescens MNKRRTTLTVLALIASLGLSSAPALADKGGNGNGNGNGHGNSGNHGNNGNHGNNGNNGNH WP_004941297.1 MULTISPECIES: hypothetical protein [Serratia] Length: 167\nNumber of predicted TMHs: 2\nExp number of AAs in TMHs: 30.02384\nExp number, first 60 AAs: 14.11185\nTotal prob of N-in: 0.94170\nPOSSIBLE N-term signal sequence\ninside 1 6\nTMhelix 7 26\noutside 27 147\nTMhelix 148 166\ninside 167 167

28479 GCF_000336425.1_ASM33642v1 Serratia marcescens WW4 Proteobacteria; Gammaproteobacteria; Enterobacterales; Yersiniaceae; Serratia; Serratia marcescens MNKRRTTLTVLALIASLGLSSAPALADKGGNGNGNGNGHGNSGNHGNNGNHGNNGNNGNH WP_004941297.1 MULTISPECIES: hypothetical protein [Serratia] Length: 167\nNumber of predicted TMHs: 2\nExp number of AAs in TMHs: 30.02384\nExp number, first 60 AAs: 14.11185\nTotal prob of N-in: 0.94170\nPOSSIBLE N-term signal sequence\ninside 1 6\nTMhelix 7 26\noutside 27 147\nTMhelix 148 166\ninside 167 167

28480 GCF_000735445.1_GSMA_DRAFTv1 Serratia marcescens subsp. marcescens ATCC 13880 Proteobacteria; Gammaproteobacteria; Enterobacterales; Yersiniaceae; Serratia; Serratia marcescens; Serratia marcescens subsp. marcescens MNKRRTTLTVLALIASLGLSSAPALADKGGNGNGNGNGHGNSGNHGNNGNHGNNGNHGNK WP_033640576.1 MULTISPECIES: membrane protein [Serratia] Length: 164\nNumber of predicted TMHs: 2\nExp number of AAs in TMHs: 30.54778\nExp number, first 60 AAs: 14.38033\nTotal prob of N-in: 0.94595\nPOSSIBLE N-term signal sequence\ninside 1 6\nTMhelix 7 26\noutside 27 144\nTMhelix 145 163\ninside 164 164

28481 GCF_000513215.1_DB11 Serratia marcescens subsp. marcescens Db11 Proteobacteria; Gammaproteobacteria; Enterobacterales; Yersiniaceae; Serratia; Serratia marcescens; Serratia marcescens subsp. marcescens MNKRRTTLTVLALIASLGLSSAPALADKGGNGNGNGNGHGNSGNNGNHGNNGNHGNKGNK WP_025302560.1 membrane protein [Serratia marcescens] Length: 161\nNumber of predicted TMHs: 2\nExp number of AAs in TMHs: 31.05977\nExp number, first 60 AAs: 14.64678\nTotal prob of N-in: 0.94999\nPOSSIBLE N-term signal sequence\ninside 1 6\nTMhelix 7 26\noutside 27 141\nTMhelix 142 160\ninside 161 161

28482 GCF_000738675.1_ASM73867v1 Serratia nematodiphila DZ0503SBS1 Proteobacteria; Gammaproteobacteria; Enterobacterales; Yersiniaceae; Serratia; Serratia nematodiphila MNKRRTTLTVLALIASLGLSSAPALADKGGNGNGNGNGHGNSGNHGNNGNHGNNGNNGNH WP_033633261.1 MULTISPECIES: membrane protein [Serratia] Length: 176\nNumber of predicted TMHs: 2\nExp number of AAs in TMHs: 28.40352\nExp number, first 60 AAs: 13.30908\nTotal prob of N-in: 0.92780\nPOSSIBLE N-term signal sequence\ninside 1 6\nTMhelix 7 26\noutside 27 156\nTMhelix 157 175\ninside 176 176

28483 GCF_000214235.1_ASM21423v1 Serratia plymuthica AS9 Proteobacteria; Gammaproteobacteria; Enterobacterales; Yersiniaceae; Serratia; Serratia plymuthica MNKRRTTLTALALIVSLGLSSAPAFADKGGNGHGNGNGGGHGNSQGNGNGHGNGNSGDHG WP_013812568.1 MULTISPECIES: membrane protein [Serratia] Length: 167\nNumber of predicted TMHs: 2\nExp number of AAs in TMHs: 33.54956\nExp number, first 60 AAs: 15.01343\nTotal prob of N-in: 0.96357\nPOSSIBLE N-term signal sequence\ninside 1 6\nTMhelix 7 26\noutside 27 142\nTMhelix 143 165\ninside 166 167

28484 GCF_000438825.1_ASM43882v1 Serratia plymuthica S13 Proteobacteria; Gammaproteobacteria; Enterobacterales; Yersiniaceae; Serratia; Serratia plymuthica MKGVLEKCTVEDFNYISATLDSYVSFTNDKRRKTLLSAYQNNPALHAELISLIDTQIKYF WP_020438466.1 hypothetical protein [Serratia plymuthica] Length: 249\nNumber of predicted TMHs: 2\nExp number of AAs in TMHs: 39.37829\nExp number, first 60 AAs: 0.01949\nTotal prob of N-in: 0.09376\noutside 1 157\nTMhelix 158 180\ninside 181 200\nTMhelix 201 220\noutside 221 249

28485 GCF_000018085.1_ASM1808v1 Serratia proteamaculans 568 Proteobacteria; Gammaproteobacteria; Enterobacterales; Yersiniaceae; Serratia; Serratia proteamaculans MSKRRTTLTVLALIVSMGLSSAPAFADKGGNGNGNGHGNSGNHGNSGNHGNNGNHGNKAK WP_012006495.1 membrane protein [Serratia proteamaculans] Length: 158\nNumber of predicted TMHs: 2\nExp number of AAs in TMHs: 35.71623\nExp number, first 60 AAs: 17.72829\nTotal prob of N-in: 0.96723\nPOSSIBLE N-term signal sequence\ninside 1 6\nTMhelix 7 26\noutside 27 133\nTMhelix 134 156\ninside 157 158

28486 GCF_001139945.1_8016_2_90 Yersinia aldovae Proteobacteria; Gammaproteobacteria; Enterobacterales; Yersiniaceae; Yersinia MLSQHGVNRRDFMKLCAALAATMGLSSKAVAEIANTVSSPQRPPVIWIGAQECTGCTESL WP_049604143.1 hydrogenase 2 small subunit [Yersinia aldovae] Length: 371\nNumber of predicted TMHs: 1\nExp number of AAs in TMHs: 17.71025\nExp number, first 60 AAs: 0.20937\nTotal prob of N-in: 0.02429\noutside 1 327\nTMhelix 328 350\ninside 351 371

28487 GCF_001091225.1_8016_2_89 Yersinia aldovae Proteobacteria; Gammaproteobacteria; Enterobacterales; Yersiniaceae; Yersinia MLSQHGVNRRDFMKLCAALAATMGLSSKAAAEIANTVSSPQRPPVIWIGAQECTGCTESL WP_004703028.1 hydrogenase 2 small subunit [Yersinia aldovae] Length: 371\nNumber of predicted TMHs: 1\nExp number of AAs in TMHs: 17.64702\nExp number, first 60 AAs: 0.1462\nTotal prob of N-in: 0.02153\noutside 1 327\nTMhelix 328 350\ninside 351 371

28488 GCF_001122605.1_8016_2_88 Yersinia aldovae Proteobacteria; Gammaproteobacteria; Enterobacterales; Yersiniaceae; Yersinia MIGEKNMLSQHGVNRRDFMKLCAALAATMGLSSKAVAEIANTVSSPQRPPVIWIGAQECT WP_072082296.1 hydrogenase 2 small subunit [Yersinia aldovae] Length: 377\nNumber of predicted TMHs: 1\nExp number of AAs in TMHs: 18.20867\nExp number, first 60 AAs: 0.73101\nTotal prob of N-in: 0.03938\noutside 1 333\nTMhelix 334 356\ninside 357 377

28489 GCF_001164505.1_8016_2_87 Yersinia aldovae Proteobacteria; Gammaproteobacteria; Enterobacterales; Yersiniaceae; Yersinia MLSQHGVNRRDFMKLCAALAATMGLSSKAVAEIANTVSSPQRPPVIWIGAQECTGCTESL WP_049688611.1 hydrogenase 2 small subunit [Yersinia aldovae] Length: 371\nNumber of predicted TMHs: 1\nExp number of AAs in TMHs: 17.71634\nExp number, first 60 AAs: 0.20936\nTotal prob of N-in: 0.02452\noutside 1 327\nTMhelix 328 350\ninside 351 371

28490 GCF_001047675.1_ASM104767v1 Yersinia aleksiciae Proteobacteria; Gammaproteobacteria; Enterobacterales; Yersiniaceae; Yersinia MTGEKNMLSQHGVNRRDFMKLCAALAATMGLSSKAAAEIASTVSSPQRPPVIWIGAQECT WP_071840476.1 MULTISPECIES: hydrogenase 2 small subunit [Yersinia] Length: 377\nNumber of predicted TMHs: 1\nExp number of AAs in TMHs: 18.66557\nExp number, first 60 AAs: 1.14034\nTotal prob of N-in: 0.06014\noutside 1 333\nTMhelix 334 356\ninside 357 377

28491 GCF_001319845.1_4821_6_8Yersinia aleksiciae Proteobacteria; Gammaproteobacteria; Enterobacterales; Yersiniaceae; Yersinia MTGEKNMLSQHGVNRRDFMKLCAALAATMGLSSKAAAEIASTVSSPQRPPVIWIGAQECT WP_071840476.1 MULTISPECIES: hydrogenase 2 small subunit [Yersinia] Length: 377\nNumber of predicted TMHs: 1\nExp number of AAs in TMHs: 18.66557\nExp number, first 60 AAs: 1.14034\nTotal prob of N-in: 0.06014\noutside 1 333\nTMhelix 334 356\ninside 357 377

28492 GCF_001131685.1_5139_5_11 Yersinia bercovieri Proteobacteria; Gammaproteobacteria; Enterobacterales; Yersiniaceae; Yersinia MVGEKNMLSQHGVNRRDFMKLCAALAATMGLSSKAAAEIANTVSSPQRPPVIWIGAQECT WP_072083829.1 MULTISPECIES: hydrogenase 2 small subunit [Yersinia] Length: 377\nNumber of predicted TMHs: 1\nExp number of AAs in TMHs: 17.97676\nExp number, first 60 AAs: 0.48377\nTotal prob of N-in: 0.02854\noutside 1 333\nTMhelix 334 356\ninside 357 377

28493 GCF_001319545.1_4821_6_4Yersinia bercovieri Proteobacteria; Gammaproteobacteria; Enterobacterales; Yersiniaceae; Yersinia MVGEKNMLSQHGVNRRDFMKLCAALAATMGLSSKAAAEIANTVSSPQRPPVIWIGAQECT WP_072083829.1 MULTISPECIES: hydrogenase 2 small subunit [Yersinia] Length: 377\nNumber of predicted TMHs: 1\nExp number of AAs in TMHs: 17.97676\nExp number, first 60 AAs: 0.48377\nTotal prob of N-in: 0.02854\noutside 1 333\nTMhelix 334 356\ninside 357 377

28494 GCF_001102905.1_4976_7_9Yersinia enterocolitica Proteobacteria; Gammaproteobacteria; Enterobacterales; Yersiniaceae; Yersinia
MLSQHGVNRRDFMKLCAALAATMGLSSKAAAEIANTVSSPQRPPVIWIGAQECTGCTESL WP_019079071.1
hydrogenase 2 small subunit [Yersinia enterocolitica] Length: 371\nNumber of predicted TMHs: 1\nExp number of AAs in TMHs: 18.80245\nExp number, first 60 AAs: 0.14643\nTotal prob of N-in: 0.01944\noutside 1 327\nTMhelix 328 350\ninside 351 371

28495 GCF_001218005.1_5103_7_11 Yersinia enterocolitica Proteobacteria; Gammaproteobacteria; Enterobacterales; Yersiniaceae; Yersinia
MLSQHGVNRRDFMKLCAALAATMGLSSKAAAEIANTVSSPQRPPVIWIGAQECTGCTESL WP_019079071.1
hydrogenase 2 small subunit [Yersinia enterocolitica] Length: 371\nNumber of predicted TMHs: 1\nExp number of AAs in TMHs: 18.80245\nExp number, first 60 AAs: 0.14643\nTotal prob of N-in: 0.01944\noutside 1 327\nTMhelix 328 350\ninside 351 371

28496 GCF_001218525.1_5116_2_6Yersinia enterocolitica Proteobacteria; Gammaproteobacteria; Enterobacterales; Yersiniaceae; Yersinia
MLSQHGVNRRDFMKLCAALAATMGLSSKAAAEIANTVSSPQRPPVIWIGAQECTGCTESL WP_005157252.1
hydrogenase 2 small subunit [Yersinia enterocolitica] Length: 371\nNumber of predicted TMHs: 1\nExp number of AAs in TMHs: 18.80217\nExp number, first 60 AAs: 0.14643\nTotal prob of N-in: 0.01943\noutside 1 327\nTMhelix 328 350\ninside 351 371

28497 GCF_001182245.1_5116_3_6Yersinia enterocolitica Proteobacteria; Gammaproteobacteria; Enterobacterales; Yersiniaceae; Yersinia
MLSQHGVNRRDFMKLCAALAATMGLSSKAAAEIANTVSSPQRPPVIWIGAQECTGCTESL WP_005157252.1
hydrogenase 2 small subunit [Yersinia enterocolitica] Length: 371\nNumber of predicted TMHs: 1\nExp number of AAs in TMHs: 18.80217\nExp number, first 60 AAs: 0.14643\nTotal prob of N-in: 0.01943\noutside 1 327\nTMhelix 328 350\ninside 351 371

28498 GCF_001182185.1_4976_7_1Yersinia enterocolitica Proteobacteria; Gammaproteobacteria; Enterobacterales; Yersiniaceae; Yersinia
MLSQHGVNRRDFMKLCAALAATMGLSSKAAAEIANTVSSPQRPPVIWIGAQECTGCTESL WP_005157252.1
hydrogenase 2 small subunit [Yersinia enterocolitica] Length: 371\nNumber of predicted TMHs: 1\nExp number of AAs in TMHs: 18.80217\nExp number, first 60 AAs: 0.14643\nTotal prob of N-in: 0.01943\noutside 1 327\nTMhelix 328 350\ninside 351 371

28499 GCF_001182145.1_4976_7_5Yersinia enterocolitica Proteobacteria; Gammaproteobacteria; Enterobacterales; Yersiniaceae; Yersinia
MLSQHGVNRRDFMKLCAALAATMGLSSKAAAEIANTVSSPQRPPVIWIGAQECTGCTESL WP_005157252.1
hydrogenase 2 small subunit [Yersinia enterocolitica] Length: 371\nNumber of predicted TMHs: 1\nExp number of AAs in TMHs: 18.80217\nExp number, first 60 AAs: 0.14643\nTotal prob of N-in: 0.01943\noutside 1 327\nTMhelix 328 350\ninside 351 371

28500 GCF_001174965.1_5749_7_1Yersinia enterocolitica Proteobacteria; Gammaproteobacteria; Enterobacterales; Yersiniaceae; Yersinia
MLSQHGVNRRDFMKLCAALAATMGLSSKAAAEIANTVSSPQRPPVIWIGAQECTGCTESL WP_019079071.1
hydrogenase 2 small subunit [Yersinia enterocolitica] Length: 371\nNumber of predicted TMHs: 1\nExp number of AAs in TMHs: 18.80245\nExp number, first 60 AAs: 0.14643\nTotal prob of N-in: 0.01944\noutside 1 327\nTMhelix 328 350\ninside 351 371

28501 GCF_001182125.1_5116_3_9Yersinia enterocolitica Proteobacteria; Gammaproteobacteria; Enterobacterales; Yersiniaceae; Yersinia
MLSQHGVNRRDFMKLCAALAATMGLSSKAAAEIANTVSSPQRPPVIWIGAQECTGCTESL WP_005157252.1
hydrogenase 2 small subunit [Yersinia enterocolitica] Length: 371\nNumber of predicted TMHs: 1\nExp number of AAs in TMHs: 18.80217\nExp number, first 60 AAs: 0.14643\nTotal prob of N-in: 0.01943\noutside 1 327\nTMhelix 328 350\ninside 351 371

28502 GCF_001182085.1_5749_7_4Yersinia enterocolitica Proteobacteria; Gammaproteobacteria; Enterobacterales; Yersiniaceae; Yersinia
MLSQHGVNRRDFMKLCAALAATMGLSSKAAAEIANTVSSPQRPPVIWIGAQECTGCTESL WP_019079071.1
hydrogenase 2 small subunit [Yersinia enterocolitica] Length: 371\nNumber of predicted TMHs: 1\nExp

number of AAs in TMHs: 18.80245\nExp number, first 60 AAs: 0.14643\nTotal prob of N-in: 0.01944\noutside 1 327\nTMhelix 328 350\ninside 351 371

28503 GCF_001150845.1_4976_3_6Yersinia enterocolitica Proteobacteria; Gammaproteobacteria; Enterobacterales; Yersiniaceae; Yersinia
MLSQHGVNRRDFMKLCAALAATMGLSSKAAAEIANTVSSPQRPPVIWIGAQECTGCTESL WP_005157252.1
hydrogenase 2 small subunit [Yersinia enterocolitica] Length: 371\nNumber of predicted TMHs: 1\nExp number of AAs in TMHs: 18.80217\nExp number, first 60 AAs: 0.14643\nTotal prob of N-in: 0.01943\noutside 1 327\nTMhelix 328 350\ninside 351 371

28504 GCF_001170365.1_5103_7_9Yersinia enterocolitica Proteobacteria; Gammaproteobacteria; Enterobacterales; Yersiniaceae; Yersinia
MLSQHGVNRRDFMKLCAALAATMGLSSKAAAEIANTVSSPQRPPVIWIGAQECTGCTESL WP_005157252.1
hydrogenase 2 small subunit [Yersinia enterocolitica] Length: 371\nNumber of predicted TMHs: 1\nExp number of AAs in TMHs: 18.80217\nExp number, first 60 AAs: 0.14643\nTotal prob of N-in: 0.01943\noutside 1 327\nTMhelix 328 350\ninside 351 371

28505 GCF_001149865.1_4976_6_10 Yersinia enterocolitica Proteobacteria; Gammaproteobacteria; Enterobacterales; Yersiniaceae; Yersinia
MLSQHGVNRRDFMKLCAALAATMGLSSKAAAEIANTVSSPQRPPVIWIGAQECTGCTESL WP_005157252.1
hydrogenase 2 small subunit [Yersinia enterocolitica] Length: 371\nNumber of predicted TMHs: 1\nExp number of AAs in TMHs: 18.80217\nExp number, first 60 AAs: 0.14643\nTotal prob of N-in: 0.01943\noutside 1 327\nTMhelix 328 350\ninside 351 371

28506 GCF_001135745.1_8016_2_62 Yersinia enterocolitica Proteobacteria; Gammaproteobacteria; Enterobacterales; Yersiniaceae; Yersinia
MLSQHGVNRRDFMKLCAALAATMGLSSKAAAEIANTVSSPQRPPVIWIGAQECTGCTESL WP_019079071.1
hydrogenase 2 small subunit [Yersinia enterocolitica] Length: 371\nNumber of predicted TMHs: 1\nExp number of AAs in TMHs: 18.80245\nExp number, first 60 AAs: 0.14643\nTotal prob of N-in: 0.01944\noutside 1 327\nTMhelix 328 350\ninside 351 371

28507 GCF_001136265.1_4840_6_6Yersinia enterocolitica Proteobacteria; Gammaproteobacteria; Enterobacterales; Yersiniaceae; Yersinia
MLSQHGVNRRDFMKLCAALAATMGLSSKAAAEIANTVSSPQRPPVIWIGAQECTGCTESL WP_005157252.1
hydrogenase 2 small subunit [Yersinia enterocolitica] Length: 371\nNumber of predicted TMHs: 1\nExp number of AAs in TMHs: 18.80217\nExp number, first 60 AAs: 0.14643\nTotal prob of N-in: 0.01943\noutside 1 327\nTMhelix 328 350\ninside 351 371

28508 GCF_001160345.1_8016_2_61 Yersinia enterocolitica Proteobacteria; Gammaproteobacteria; Enterobacterales; Yersiniaceae; Yersinia
MLSQHGVNRRDFMKLCAALAATMGLSSKAAAEIANTVSSPQRPPVIWIGAQECTGCTESL WP_019079071.1
hydrogenase 2 small subunit [Yersinia enterocolitica] Length: 371\nNumber of predicted TMHs: 1\nExp number of AAs in TMHs: 18.80245\nExp number, first 60 AAs: 0.14643\nTotal prob of N-in: 0.01944\noutside 1 327\nTMhelix 328 350\ninside 351 371

28509 GCF_000754975.1_YEA Yersinia enterocolitica Proteobacteria; Gammaproteobacteria; Enterobacterales; Yersiniaceae; Yersinia
MLSQHGVNRRDFMKLCAALAATMGLSSKAAAEIANTVSSPQRPPVIWIGAQECTGCTESL WP_005157252.1
hydrogenase 2 small subunit [Yersinia enterocolitica] Length: 371\nNumber of predicted TMHs: 1\nExp number of AAs in TMHs: 18.80217\nExp number, first 60 AAs: 0.14643\nTotal prob of N-in: 0.01943\noutside 1 327\nTMhelix 328 350\ninside 351 371

28510 GCF_001097785.1_4840_7_8Yersinia enterocolitica Proteobacteria; Gammaproteobacteria; Enterobacterales; Yersiniaceae; Yersinia
MLSQHGVNRRDFMKLCAALAATMGLSSKAAAEIANTVSSPQRPPVIWIGAQECTGCTESL WP_005157252.1
hydrogenase 2 small subunit [Yersinia enterocolitica] Length: 371\nNumber of predicted TMHs: 1\nExp number of AAs in TMHs: 18.80217\nExp number, first 60 AAs: 0.14643\nTotal prob of N-in: 0.01943\noutside 1 327\nTMhelix 328 350\ninside 351 371

28511 GCF_001105265.1_8016_2_79 Yersinia enterocolitica Proteobacteria; Gammaproteobacteria; Enterobacterales; Yersiniaceae; Yersinia

MLSQHGVNRRDFMKLCAALAATMGLSSKAAAEIANTVSSPQRPPVIWIGAQECTGCTESL WP_019079071.1 hydrogenase 2 small subunit [Yersinia enterocolitica] Length: 371\nNumber of predicted TMHs: 1\nExp number of AAs in TMHs: 18.80245\nExp number, first 60 AAs: 0.14643\nTotal prob of N-in: 0.01944\noutside 1 327\nTMhelix 328 350\ninside 351 371

28512 GCF_001124705.1_8016_2_74 Yersinia enterocolitica Proteobacteria; Gammaproteobacteria; Enterobacterales; Yersiniaceae; Yersinia MLSQHGVNRRDFMKLCAALAATMGLSSKAAAEIANTVSSPQRPPVIWIGAQECTGCTESL WP_005157252.1 hydrogenase 2 small subunit [Yersinia enterocolitica] Length: 371\nNumber of predicted TMHs: 1\nExp number of AAs in TMHs: 18.80217\nExp number, first 60 AAs: 0.14643\nTotal prob of N-in: 0.01943\noutside 1 327\nTMhelix 328 350\ninside 351 371

28513 GCF_001118285.1_4976_3_10 Yersinia enterocolitica Proteobacteria; Gammaproteobacteria; Enterobacterales; Yersiniaceae; Yersinia MLSQHGVNRRDFMKLCAALAATMGLSSKAAAEIANTVSSPQRPPVIWIGAQECTGCTESL WP_005157252.1 hydrogenase 2 small subunit [Yersinia enterocolitica] Length: 371\nNumber of predicted TMHs: 1\nExp number of AAs in TMHs: 18.80217\nExp number, first 60 AAs: 0.14643\nTotal prob of N-in: 0.01943\noutside 1 327\nTMhelix 328 350\ninside 351 371

28514 GCF_000834735.1_ASM83473v1 Yersinia enterocolitica Proteobacteria; Gammaproteobacteria; Enterobacterales; Yersiniaceae; Yersinia MLSQHGVNRRDFMKLCAALAATMGLSSKAAAEIANTVSSPQRPPVIWIGAQECTGCTESL WP_005157252.1 hydrogenase 2 small subunit [Yersinia enterocolitica] Length: 371\nNumber of predicted TMHs: 1\nExp number of AAs in TMHs: 18.80217\nExp number, first 60 AAs: 0.14643\nTotal prob of N-in: 0.01943\noutside 1 327\nTMhelix 328 350\ninside 351 371

28515 GCF_001087805.1_8016_2_69 Yersinia enterocolitica Proteobacteria; Gammaproteobacteria; Enterobacterales; Yersiniaceae; Yersinia MLSQHGVNRRDFMKLCAALAATMGLSSKAAAEIANTVSSPQRPPVIWIGAQECTGCTESL WP_005157252.1 hydrogenase 2 small subunit [Yersinia enterocolitica] Length: 371\nNumber of predicted TMHs: 1\nExp number of AAs in TMHs: 18.80217\nExp number, first 60 AAs: 0.14643\nTotal prob of N-in: 0.01943\noutside 1 327\nTMhelix 328 350\ninside 351 371

28516 GCF_001086365.1_4976_3_8Yersinia enterocolitica Proteobacteria; Gammaproteobacteria; Enterobacterales; Yersiniaceae; Yersinia MLSQHGVNRRDFMKLCAALAATMGLSSKAAAEIANTVSSPQRPPVIWIGAQECTGCTESL WP_005157252.1 hydrogenase 2 small subunit [Yersinia enterocolitica] Length: 371\nNumber of predicted TMHs: 1\nExp number of AAs in TMHs: 18.80217\nExp number, first 60 AAs: 0.14643\nTotal prob of N-in: 0.01943\noutside 1 327\nTMhelix 328 350\ninside 351 371

28517 GCF_000755055.1_YEG_1 Yersinia enterocolitica Proteobacteria; Gammaproteobacteria; Enterobacterales; Yersiniaceae; Yersinia MLSQHGVNRRDFMKLCAALAATMGLSSKAAAEIANTVSSPQRPPVIWIGAQECTGCTESL WP_005173957.1 hydrogenase 2 small subunit [Yersinia enterocolitica] Length: 371\nNumber of predicted TMHs: 1\nExp number of AAs in TMHs: 18.61508\nExp number, first 60 AAs: 0.14605\nTotal prob of N-in: 0.02327\noutside 1 327\nTMhelix 328 350\ninside 351 371

28518 GCF_001087205.1_5116_2_3Yersinia enterocolitica Proteobacteria; Gammaproteobacteria; Enterobacterales; Yersiniaceae; Yersinia MLSQHGVNRRDFMKLCAALAATMGLSSKAAAEIANTVSSPQRPPVIWIGAQECTGCTESL WP_019079071.1 hydrogenase 2 small subunit [Yersinia enterocolitica] Length: 371\nNumber of predicted TMHs: 1\nExp number of AAs in TMHs: 18.80245\nExp number, first 60 AAs: 0.14643\nTotal prob of N-in: 0.01944\noutside 1 327\nTMhelix 328 350\ninside 351 371

28519 GCF_001090445.1_4976_6_8Yersinia enterocolitica Proteobacteria; Gammaproteobacteria; Enterobacterales; Yersiniaceae; Yersinia MLSQHGVNRRDFMKLCAALAATMGLSSKAAAEIANTVSSPQRPPVIWIGAQECTGCTESL WP_005157252.1 hydrogenase 2 small subunit [Yersinia enterocolitica] Length: 371\nNumber of predicted TMHs: 1\nExp number of AAs in TMHs: 18.80217\nExp number, first 60 AAs: 0.14643\nTotal prob of N-in: 0.01943\noutside 1 327\nTMhelix 328 350\ninside 351 371

28520 GCF_001084885.1_8016_2_75 Yersinia enterocolitica Proteobacteria; Gammaproteobacteria; Enterobacterales; Yersiniaceae; Yersinia
MLSQHGVNRRDFMKLCAALAATMGLSSKAAAEIANTVSSPQRPPVIWIGAQECTGCTESL WP_005173957.1
hydrogenase 2 small subunit [Yersinia enterocolitica] Length: 371\nNumber of predicted TMHs: 1\nExp number of AAs in TMHs: 18.61508\nExp number, first 60 AAs: 0.14605\nTotal prob of N-in: 0.02327\noutside 1 327\nTMhelix 328 350\ninside 351 371

28521 GCF_001107025.1_8016_2_72 Yersinia enterocolitica Proteobacteria; Gammaproteobacteria; Enterobacterales; Yersiniaceae; Yersinia
MLSQHGVNRRDFMKLCAALAATMGLSSKAAAEIANTVSSPQRPPVIWIGAQECTGCTESL WP_005157252.1
hydrogenase 2 small subunit [Yersinia enterocolitica] Length: 371\nNumber of predicted TMHs: 1\nExp number of AAs in TMHs: 18.80217\nExp number, first 60 AAs: 0.14643\nTotal prob of N-in: 0.01943\noutside 1 327\nTMhelix 328 350\ninside 351 371

28522 GCF_900079955.1_IP35471 Yersinia enterocolitica Proteobacteria; Gammaproteobacteria; Enterobacterales; Yersiniaceae; Yersinia
MLSQHGVNRRDFMKLCAALAATMGLSSKAAAEIANTVSSPQRPPVIWIGAQECTGCTESL WP_005157252.1
hydrogenase 2 small subunit [Yersinia enterocolitica] Length: 371\nNumber of predicted TMHs: 1\nExp number of AAs in TMHs: 18.80217\nExp number, first 60 AAs: 0.14643\nTotal prob of N-in: 0.01943\noutside 1 327\nTMhelix 328 350\ninside 351 371

28523 GCF_002082245.1_ASM208224v1 Yersinia enterocolitica Proteobacteria; Gammaproteobacteria; Enterobacterales; Yersiniaceae; Yersinia
MLSQHGVNRRDFMKLCAALAATMGLSSKAAAEIANMVGSPQRPPVIWIGAQECTGCTESL WP_083162457.1
hydrogenase 2 small subunit [Yersinia enterocolitica] Length: 371\nNumber of predicted TMHs: 1\nExp number of AAs in TMHs: 18.89409\nExp number, first 60 AAs: 0.43102\nTotal prob of N-in: 0.04471\noutside 1 327\nTMhelix 328 350\ninside 351 371

28524 GCF_002082275.1_ASM208227v1 Yersinia enterocolitica Proteobacteria; Gammaproteobacteria; Enterobacterales; Yersiniaceae; Yersinia
MLSQHGVNRRDFMKLCAALAATMGLSSKAAAEIANTVSSPQRPPVIWIGAQECTGCTESL WP_019079071.1
hydrogenase 2 small subunit [Yersinia enterocolitica] Length: 371\nNumber of predicted TMHs: 1\nExp number of AAs in TMHs: 18.80245\nExp number, first 60 AAs: 0.14643\nTotal prob of N-in: 0.01944\noutside 1 327\nTMhelix 328 350\ninside 351 371

28525 GCF_900079935.1_IP35472 Yersinia enterocolitica Proteobacteria; Gammaproteobacteria; Enterobacterales; Yersiniaceae; Yersinia
MLSQHGVNRRDFMKLCAALAATMGLSSKAAAEIANTVSSPQRPPVIWIGAQECTGCTESL WP_005157252.1
hydrogenase 2 small subunit [Yersinia enterocolitica] Length: 371\nNumber of predicted TMHs: 1\nExp number of AAs in TMHs: 18.80217\nExp number, first 60 AAs: 0.14643\nTotal prob of N-in: 0.01943\noutside 1 327\nTMhelix 328 350\ninside 351 371

28526 GCF_900079905.1_IP35478 Yersinia enterocolitica Proteobacteria; Gammaproteobacteria; Enterobacterales; Yersiniaceae; Yersinia
MLSQHGVNRRDFMKLCAALAATMGLSSKAAAEIANTVSSPQRPPVIWIGAQECTGCTESL WP_005157252.1
hydrogenase 2 small subunit [Yersinia enterocolitica] Length: 371\nNumber of predicted TMHs: 1\nExp number of AAs in TMHs: 18.80217\nExp number, first 60 AAs: 0.14643\nTotal prob of N-in: 0.01943\noutside 1 327\nTMhelix 328 350\ninside 351 371

28527 GCF_900079885.1_IP134 Yersinia enterocolitica Proteobacteria; Gammaproteobacteria; Enterobacterales; Yersiniaceae; Yersinia
MLSQHGVNRRDFMKLCAALAATMGLSSKAAAEIANTVSSPQRPPVIWIGAQECTGCTESL WP_005157252.1
hydrogenase 2 small subunit [Yersinia enterocolitica] Length: 371\nNumber of predicted TMHs: 1\nExp number of AAs in TMHs: 18.80217\nExp number, first 60 AAs: 0.14643\nTotal prob of N-in: 0.01943\noutside 1 327\nTMhelix 328 350\ninside 351 371

28528 GCF_900079855.1_IP35466 Yersinia enterocolitica Proteobacteria; Gammaproteobacteria; Enterobacterales; Yersiniaceae; Yersinia
MLSQHGVNRRDFMKLCAALAATMGLSSKAAAEIANTVSSPQRPPVIWIGAQECTGCTESL WP_005157252.1
hydrogenase 2 small subunit [Yersinia enterocolitica] Length: 371\nNumber of predicted TMHs: 1\nExp

number of AAs in TMHs: 18.80217\nExp number, first 60 AAs: 0.14643\nTotal prob of N-in: 0.01943\noutside 1 327\nTMhelix 328 350\ninside 351 371

28529 GCF_900079825.1_IP35464 Yersinia enterocolitica Proteobacteria; Gammaproteobacteria; Enterobacterales; Yersiniaceae; Yersinia
MLSQHGVNRRDFMKLCAALAATMGLSSKAAAEIANTVSSPQRPPVIWIGAQECTGCTESL WP_005157252.1
hydrogenase 2 small subunit [Yersinia enterocolitica] Length: 371\nNumber of predicted TMHs: 1\nExp number of AAs in TMHs: 18.80217\nExp number, first 60 AAs: 0.14643\nTotal prob of N-in: 0.01943\noutside 1 327\nTMhelix 328 350\ninside 351 371

28530 GCF_001319955.1_5139_5_3Yersinia enterocolitica Proteobacteria; Gammaproteobacteria; Enterobacterales; Yersiniaceae; Yersinia
MVGEKNMLSQHGVNRRDFMKLCAALAATMGLSSKAAAEIANTVSSPQRPPVIWIGAQECT WP_072083829.1
MULTISPECIES: hydrogenase 2 small subunit [Yersinia] Length: 377\nNumber of predicted TMHs: 1\nExp number of AAs in TMHs: 17.97676\nExp number, first 60 AAs: 0.48377\nTotal prob of N-in: 0.02854\noutside 1 333\nTMhelix 334 356\ninside 357 377

28531 GCF_001983255.1_ASM198325v1 Yersinia enterocolitica Proteobacteria; Gammaproteobacteria; Enterobacterales; Yersiniaceae; Yersinia
MLSQHGVNRRDFMKLCAALAATMGLSSKAAAEIANTVSSPQRPPVIWIGAQECTGCTESL WP_019079071.1
hydrogenase 2 small subunit [Yersinia enterocolitica] Length: 371\nNumber of predicted TMHs: 1\nExp number of AAs in TMHs: 18.80245\nExp number, first 60 AAs: 0.14643\nTotal prob of N-in: 0.01944\noutside 1 327\nTMhelix 328 350\ninside 351 371

28532 GCF_001319465.1_5116_3_2Yersinia enterocolitica Proteobacteria; Gammaproteobacteria; Enterobacterales; Yersiniaceae; Yersinia
MLSQHGVNRRDFMKLCAALAATMGLSSKAAAEIANTVSSPQRPPVIWIGAQECTGCTESL WP_019079071.1
hydrogenase 2 small subunit [Yersinia enterocolitica] Length: 371\nNumber of predicted TMHs: 1\nExp number of AAs in TMHs: 18.80245\nExp number, first 60 AAs: 0.14643\nTotal prob of N-in: 0.01944\noutside 1 327\nTMhelix 328 350\ninside 351 371

28533 GCF_001354595.1_8016_2_71 Yersinia enterocolitica Proteobacteria; Gammaproteobacteria; Enterobacterales; Yersiniaceae; Yersinia
MLSQHGVNRRDFMKLCAALAATMGLSSKAAAEIANTVSSPQRPPVIWIGAQECTGCTESL WP_005157252.1
hydrogenase 2 small subunit [Yersinia enterocolitica] Length: 371\nNumber of predicted TMHs: 1\nExp number of AAs in TMHs: 18.80217\nExp number, first 60 AAs: 0.14643\nTotal prob of N-in: 0.01943\noutside 1 327\nTMhelix 328 350\ninside 351 371

28534 GCF_001319405.1_4976_7_6Yersinia enterocolitica Proteobacteria; Gammaproteobacteria; Enterobacterales; Yersiniaceae; Yersinia
MLSQHGVNRRDFMKLCAALAATMGLSSKAAAEIANTVSSPQRPPVIWIGAQECTGCTESL WP_019079071.1
hydrogenase 2 small subunit [Yersinia enterocolitica] Length: 371\nNumber of predicted TMHs: 1\nExp number of AAs in TMHs: 18.80245\nExp number, first 60 AAs: 0.14643\nTotal prob of N-in: 0.01944\noutside 1 327\nTMhelix 328 350\ninside 351 371

28535 GCF_001319365.1_5103_7_6Yersinia enterocolitica Proteobacteria; Gammaproteobacteria; Enterobacterales; Yersiniaceae; Yersinia
MLSQHGVNRRDFMKLCAALAATMGLSSKAAAEIANTVSSPQRPPVIWIGAQECTGCTESL WP_005157252.1
hydrogenase 2 small subunit [Yersinia enterocolitica] Length: 371\nNumber of predicted TMHs: 1\nExp number of AAs in TMHs: 18.80217\nExp number, first 60 AAs: 0.14643\nTotal prob of N-in: 0.01943\noutside 1 327\nTMhelix 328 350\ninside 351 371

28536 GCF_001319305.1_4976_6_11 Yersinia enterocolitica Proteobacteria; Gammaproteobacteria; Enterobacterales; Yersiniaceae; Yersinia
MLSQHGVNRRDFMKLCAALAATMGLSSKAAAEIANTVSSPQRPPVIWIGAQECTGCTESL WP_005157252.1
hydrogenase 2 small subunit [Yersinia enterocolitica] Length: 371\nNumber of predicted TMHs: 1\nExp number of AAs in TMHs: 18.80217\nExp number, first 60 AAs: 0.14643\nTotal prob of N-in: 0.01943\noutside 1 327\nTMhelix 328 350\ninside 351 371

28537 GCF_001223065.1_4976_2_7Yersinia enterocolitica Proteobacteria; Gammaproteobacteria; Enterobacterales; Yersiniaceae; Yersinia

MLSQHGVNRRDFMKLCAALAATMGLSSKAAAEIANTVSSPQRPPVIWIGAQECTGCTESL WP_019079071.1 hydrogenase 2 small subunit [Yersinia enterocolitica] Length: 371\nNumber of predicted TMHs: 1\nExp number of AAs in TMHs: 18.80245\nExp number, first 60 AAs: 0.14643\nTotal prob of N-in: 0.01944\noutside 1 327\nTMhelix 328 350\ninside 351 371

28538 GCF_001223425.1_5749_7_2Yersinia enterocolitica Proteobacteria; Gammaproteobacteria; Enterobacterales; Yersiniaceae; Yersinia MLSQHGVNRRDFMKLCAALAATMGLSSKAAAEIANTVSSPQRPPVIWIGAQECTGCTESL WP_005157252.1 hydrogenase 2 small subunit [Yersinia enterocolitica] Length: 371\nNumber of predicted TMHs: 1\nExp number of AAs in TMHs: 18.80217\nExp number, first 60 AAs: 0.14643\nTotal prob of N-in: 0.01943\noutside 1 327\nTMhelix 328 350\ninside 351 371

28539 GCF_001222745.1_4976_1_10 Yersinia enterocolitica Proteobacteria; Gammaproteobacteria; Enterobacterales; Yersiniaceae; Yersinia MLSQHGVNRRDFMKLCAALAATMGLSSKAAAEIANTVSSPQRPPVIWIGAQECTGCTESL WP_005157252.1 hydrogenase 2 small subunit [Yersinia enterocolitica] Length: 371\nNumber of predicted TMHs: 1\nExp number of AAs in TMHs: 18.80217\nExp number, first 60 AAs: 0.14643\nTotal prob of N-in: 0.01943\noutside 1 327\nTMhelix 328 350\ninside 351 371

28540 GCF_001222305.1_5103_7_2Yersinia enterocolitica Proteobacteria; Gammaproteobacteria; Enterobacterales; Yersiniaceae; Yersinia MLSQHGVNRRDFMKLCAALAATMGLSSKAAAEIANTVSSPQRPPVIWIGAQECTGCTESL WP_005157252.1 hydrogenase 2 small subunit [Yersinia enterocolitica] Length: 371\nNumber of predicted TMHs: 1\nExp number of AAs in TMHs: 18.80217\nExp number, first 60 AAs: 0.14643\nTotal prob of N-in: 0.01943\noutside 1 327\nTMhelix 328 350\ninside 351 371

28541 GCF_001222285.1_4976_3_2Yersinia enterocolitica Proteobacteria; Gammaproteobacteria; Enterobacterales; Yersiniaceae; Yersinia MLSQHGVNRRDFMKLCAALAATMGLSSKAAAEIANTVSSPQRPPVIWIGAQECTGCTESL WP_005157252.1 hydrogenase 2 small subunit [Yersinia enterocolitica] Length: 371\nNumber of predicted TMHs: 1\nExp number of AAs in TMHs: 18.80217\nExp number, first 60 AAs: 0.14643\nTotal prob of N-in: 0.01943\noutside 1 327\nTMhelix 328 350\ninside 351 371

28542 GCF_001221145.1_4976_2_10 Yersinia enterocolitica Proteobacteria; Gammaproteobacteria; Enterobacterales; Yersiniaceae; Yersinia MLSQHGVNRRDFMKLCAALAATMGLSSKAAAEIANTVSSPQRPPVIWIGAQECTGCTESL WP_019079071.1 hydrogenase 2 small subunit [Yersinia enterocolitica] Length: 371\nNumber of predicted TMHs: 1\nExp number of AAs in TMHs: 18.80245\nExp number, first 60 AAs: 0.14643\nTotal prob of N-in: 0.01944\noutside 1 327\nTMhelix 328 350\ninside 351 371

28543 GCF_001222105.1_5116_2_1Yersinia enterocolitica Proteobacteria; Gammaproteobacteria; Enterobacterales; Yersiniaceae; Yersinia MLSQHGVNRRDFMKLCAALAATMGLSSKAAAEIANTVSSPQRPPVIWIGAQECTGCTESL WP_005157252.1 hydrogenase 2 small subunit [Yersinia enterocolitica] Length: 371\nNumber of predicted TMHs: 1\nExp number of AAs in TMHs: 18.80217\nExp number, first 60 AAs: 0.14643\nTotal prob of N-in: 0.01943\noutside 1 327\nTMhelix 328 350\ninside 351 371

28544 GCF_001220745.1_4976_2_3Yersinia enterocolitica Proteobacteria; Gammaproteobacteria; Enterobacterales; Yersiniaceae; Yersinia MLSQHGVNRRDFMKLCAALAATMGLSSKAAAEIANTVSSPQRPPVIWIGAQECTGCTESL WP_019079071.1 hydrogenase 2 small subunit [Yersinia enterocolitica] Length: 371\nNumber of predicted TMHs: 1\nExp number of AAs in TMHs: 18.80245\nExp number, first 60 AAs: 0.14643\nTotal prob of N-in: 0.01944\noutside 1 327\nTMhelix 328 350\ninside 351 371

28545 GCF_001220485.1_5116_2_9Yersinia enterocolitica Proteobacteria; Gammaproteobacteria; Enterobacterales; Yersiniaceae; Yersinia MLSQHGVNRRDFMKLCAALAATMGLSSKAAAEIANTVSSPQRPPVIWIGAQECTGCTESL WP_019079071.1 hydrogenase 2 small subunit [Yersinia enterocolitica] Length: 371\nNumber of predicted TMHs: 1\nExp number of AAs in TMHs: 18.80245\nExp number, first 60 AAs: 0.14643\nTotal prob of N-in: 0.01944\noutside 1 327\nTMhelix 328 350\ninside 351 371

28546 GCF_001220085.1_8016_2_67 Yersinia enterocolitica Proteobacteria; Gammaproteobacteria; Enterobacterales; Yersiniaceae; Yersinia
MLSQHGVNRRDFMKLCAALAATMGLSSKAAAEIANTVSSPQRPPVIWIGAQECTGCTESL WP_005157252.1
hydrogenase 2 small subunit [Yersinia enterocolitica] Length: 371\nNumber of predicted TMHs: 1\nExp number of AAs in TMHs: 18.80217\nExp number, first 60 AAs: 0.14643\nTotal prob of N-in: 0.01943\noutside 1 327\nTMhelix 328 350\ninside 351 371

28547 GCF_001219605.1_5116_2_4Yersinia enterocolitica Proteobacteria; Gammaproteobacteria; Enterobacterales; Yersiniaceae; Yersinia
MLSQHGVNRRDFMKLCAALAATMGLSSKAAAEIANTVSSPQRPPVIWIGAQECTGCTESL WP_019079071.1
hydrogenase 2 small subunit [Yersinia enterocolitica] Length: 371\nNumber of predicted TMHs: 1\nExp number of AAs in TMHs: 18.80245\nExp number, first 60 AAs: 0.14643\nTotal prob of N-in: 0.01944\noutside 1 327\nTMhelix 328 350\ninside 351 371

28548 GCF_001219785.1_4976_1_8Yersinia enterocolitica Proteobacteria; Gammaproteobacteria; Enterobacterales; Yersiniaceae; Yersinia
MLSQHGVNRRDFMKLCAALAATMGLSSKAAAEIANTVSSPQRPPVIWIGAQECTGCTESL WP_005157252.1
hydrogenase 2 small subunit [Yersinia enterocolitica] Length: 371\nNumber of predicted TMHs: 1\nExp number of AAs in TMHs: 18.80217\nExp number, first 60 AAs: 0.14643\nTotal prob of N-in: 0.01943\noutside 1 327\nTMhelix 328 350\ninside 351 371

28549 GCF_001218905.1_5116_2_2Yersinia enterocolitica Proteobacteria; Gammaproteobacteria; Enterobacterales; Yersiniaceae; Yersinia
MLSQHGVNRRDFMKLCAALAATMGLSSKAAAEIANTVSSPQRPPVIWIGAQECTGCTESL WP_019079071.1
hydrogenase 2 small subunit [Yersinia enterocolitica] Length: 371\nNumber of predicted TMHs: 1\nExp number of AAs in TMHs: 18.80245\nExp number, first 60 AAs: 0.14643\nTotal prob of N-in: 0.01944\noutside 1 327\nTMhelix 328 350\ninside 351 371

28550 GCF_001182365.1_8016_2_77 Yersinia enterocolitica Proteobacteria; Gammaproteobacteria; Enterobacterales; Yersiniaceae; Yersinia
MLSQHGVNRRDFMKLCAALAATMGLSSKAAAEIANTVSSPQRPPVIWIGAQECTGCTESL WP_019079071.1
hydrogenase 2 small subunit [Yersinia enterocolitica] Length: 371\nNumber of predicted TMHs: 1\nExp number of AAs in TMHs: 18.80245\nExp number, first 60 AAs: 0.14643\nTotal prob of N-in: 0.01944\noutside 1 327\nTMhelix 328 350\ninside 351 371

28551 GCF_001219345.1_4976_1_3Yersinia enterocolitica Proteobacteria; Gammaproteobacteria; Enterobacterales; Yersiniaceae; Yersinia
MLSQHGVNRRDFMKLCAALAATMGLSSKAAAEIANTVSSPQRPPVIWIGAQECTGCTESL WP_005157252.1
hydrogenase 2 small subunit [Yersinia enterocolitica] Length: 371\nNumber of predicted TMHs: 1\nExp number of AAs in TMHs: 18.80217\nExp number, first 60 AAs: 0.14643\nTotal prob of N-in: 0.01943\noutside 1 327\nTMhelix 328 350\ninside 351 371

28552 GCF_001217685.1_5103_7_4Yersinia enterocolitica Proteobacteria; Gammaproteobacteria; Enterobacterales; Yersiniaceae; Yersinia
MLSQHGVNRRDFMKLCAALAATMGLSSKAAAEIANTVSSPQRPPVIWIGAQECTGCTESL WP_019079071.1
hydrogenase 2 small subunit [Yersinia enterocolitica] Length: 371\nNumber of predicted TMHs: 1\nExp number of AAs in TMHs: 18.80245\nExp number, first 60 AAs: 0.14643\nTotal prob of N-in: 0.01944\noutside 1 327\nTMhelix 328 350\ninside 351 371

28553 GCF_900079835.1_IP35459 Yersinia enterocolitica Proteobacteria; Gammaproteobacteria; Enterobacterales; Yersiniaceae; Yersinia
MLSQHGVNRRDFMKLCAALAATMGLSSKAAAEIANTVSSPQRPPVIWIGAQECTGCTESL WP_005157252.1
hydrogenase 2 small subunit [Yersinia enterocolitica] Length: 371\nNumber of predicted TMHs: 1\nExp number of AAs in TMHs: 18.80217\nExp number, first 60 AAs: 0.14643\nTotal prob of N-in: 0.01943\noutside 1 327\nTMhelix 328 350\ninside 351 371

28554 GCF_001995305.1_ASM199530v1 Yersinia enterocolitica Proteobacteria; Gammaproteobacteria; Enterobacterales; Yersiniaceae; Yersinia
MLSQHGVNRRDFMKLCAALAATMGLSSKAAAEIANTVSSPQRPPVIWIGAQECTGCTESL WP_050077148.1
MULTISPECIES: hydrogenase 2 small subunit [Yersinia] Length: 371\nNumber of predicted TMHs: 1\nExp

number of AAs in TMHs: 18.80844\nExp number, first 60 AAs:  0.14641\nTotal prob of N-in:   0.01969\noutside   1   327\nTMhelix   328   350\ninside   351   371

28555 GCF_001086025.1_8016_2_70 Yersinia enterocolitica Proteobacteria; Gammaproteobacteria; Enterobacterales; Yersiniaceae; Yersinia
MLSQHGVNRRDFMKLCAALAATMGLSSKAAAEIANTVSSPQRPPVIWIGAQECTGCTESL WP_005157252.1
hydrogenase 2 small subunit [Yersinia enterocolitica] Length: 371\nNumber of predicted TMHs:  1\nExp number of AAs in TMHs: 18.80217\nExp number, first 60 AAs:  0.14643\nTotal prob of N-in:   0.01943\noutside   1   327\nTMhelix   328   350\ninside   351   371

28556 GCF_001220045.1_4976_3_1Yersinia enterocolitica Proteobacteria; Gammaproteobacteria; Enterobacterales; Yersiniaceae; Yersinia
MLSQHGVNRRDFMKLCAALAATMGLSSKAAAEIANTVSSPQRPPVIWIGAQECTGCTESL WP_005157252.1
hydrogenase 2 small subunit [Yersinia enterocolitica] Length: 371\nNumber of predicted TMHs:  1\nExp number of AAs in TMHs: 18.80217\nExp number, first 60 AAs:  0.14643\nTotal prob of N-in:   0.01943\noutside   1   327\nTMhelix   328   350\ninside   351   371

28557 GCF_001220145.1_5116_2_8Yersinia enterocolitica Proteobacteria; Gammaproteobacteria; Enterobacterales; Yersiniaceae; Yersinia
MLSQHGVNRRDFMKLCAALAATMGLSSKAAAEIANTVSSPQRPPVIWIGAQECTGCTESL WP_019079071.1
hydrogenase 2 small subunit [Yersinia enterocolitica] Length: 371\nNumber of predicted TMHs:  1\nExp number of AAs in TMHs: 18.80245\nExp number, first 60 AAs:  0.14643\nTotal prob of N-in:   0.01944\noutside   1   327\nTMhelix   328   350\ninside   351   371

28558 GCF_001220185.1_4976_1_11 Yersinia enterocolitica Proteobacteria; Gammaproteobacteria; Enterobacterales; Yersiniaceae; Yersinia
MLSQHGVNRRDFMKLCAALAATMGLSSKAAAEIANTVSSPQRPPVIWIGAQECTGCTESL WP_005157252.1
hydrogenase 2 small subunit [Yersinia enterocolitica] Length: 371\nNumber of predicted TMHs:  1\nExp number of AAs in TMHs: 18.80217\nExp number, first 60 AAs:  0.14643\nTotal prob of N-in:   0.01943\noutside   1   327\nTMhelix   328   350\ninside   351   371

28559 GCF_001220565.1_5103_7_10 Yersinia enterocolitica Proteobacteria; Gammaproteobacteria; Enterobacterales; Yersiniaceae; Yersinia
MLSQHGVNRRDFMKLCAALAATMGLSSKAAAEIANTVSSPQRPPVIWIGAQECTGCTESL WP_005157252.1
hydrogenase 2 small subunit [Yersinia enterocolitica] Length: 371\nNumber of predicted TMHs:  1\nExp number of AAs in TMHs: 18.80217\nExp number, first 60 AAs:  0.14643\nTotal prob of N-in:   0.01943\noutside   1   327\nTMhelix   328   350\ninside   351   371

28560 GCF_001220825.1_4976_2_9Yersinia enterocolitica Proteobacteria; Gammaproteobacteria; Enterobacterales; Yersiniaceae; Yersinia
MLSQHGVNRRDFMKLCAALAATMGLSSKAAAEIANTVSSPQRPPVIWIGAQECTGCTESL WP_019079071.1
hydrogenase 2 small subunit [Yersinia enterocolitica] Length: 371\nNumber of predicted TMHs:  1\nExp number of AAs in TMHs: 18.80245\nExp number, first 60 AAs:  0.14643\nTotal prob of N-in:   0.01944\noutside   1   327\nTMhelix   328   350\ninside   351   371

28561 GCF_001221085.1_5116_2_5Yersinia enterocolitica Proteobacteria; Gammaproteobacteria; Enterobacterales; Yersiniaceae; Yersinia
MLSQHGVNRRDFMKLCAALAATMGLSSKAAAEIANTVSSPQRPPVIWIGAQECTGCTESL WP_005157252.1
hydrogenase 2 small subunit [Yersinia enterocolitica] Length: 371\nNumber of predicted TMHs:  1\nExp number of AAs in TMHs: 18.80217\nExp number, first 60 AAs:  0.14643\nTotal prob of N-in:   0.01943\noutside   1   327\nTMhelix   328   350\ninside   351   371

28562 GCF_001221865.1_5116_2_11 Yersinia enterocolitica Proteobacteria; Gammaproteobacteria; Enterobacterales; Yersiniaceae; Yersinia
MLSQHGVNRRDFMKLCAALAATMGLSSKAAAEIANTVSSPQRPPVIWIGAQECTGCTESL WP_005157252.1
hydrogenase 2 small subunit [Yersinia enterocolitica] Length: 371\nNumber of predicted TMHs:  1\nExp number of AAs in TMHs: 18.80217\nExp number, first 60 AAs:  0.14643\nTotal prob of N-in:   0.01943\noutside   1   327\nTMhelix   328   350\ninside   351   371

28563 GCF_001222445.1_4976_6_3Yersinia enterocolitica Proteobacteria; Gammaproteobacteria; Enterobacterales; Yersiniaceae; Yersinia

MLSQHGVNRRDFMKLCAALAATMGLSSKAAAEIANTVSSPQRPPVIWIGAQECTGCTESL WP_005157252.1 hydrogenase 2 small subunit [Yersinia enterocolitica] Length: 371\nNumber of predicted TMHs: 1\nExp number of AAs in TMHs: 18.80217\nExp number, first 60 AAs: 0.14643\nTotal prob of N-in: 0.01943\noutside 1 327\nTMhelix 328 350\ninside 351 371

28564 GCF_001222265.1_4976_3_3Yersinia enterocolitica Proteobacteria; Gammaproteobacteria; Enterobacterales; Yersiniaceae; Yersinia MLSQHGVNRRDFMKLCAALAATMGLSSKAAAEIANTVSSPQRPPVIWIGAQECTGCTESL WP_019079071.1 hydrogenase 2 small subunit [Yersinia enterocolitica] Length: 371\nNumber of predicted TMHs: 1\nExp number of AAs in TMHs: 18.80245\nExp number, first 60 AAs: 0.14643\nTotal prob of N-in: 0.01944\noutside 1 327\nTMhelix 328 350\ninside 351 371

28565 GCF_001222805.1_4976_1_1Yersinia enterocolitica Proteobacteria; Gammaproteobacteria; Enterobacterales; Yersiniaceae; Yersinia MLSQHGVNRRDFMKLCAALAATMGLSSKAAAEIANTVSSPQRPPVIWIGAQECTGCTESL WP_019079071.1 hydrogenase 2 small subunit [Yersinia enterocolitica] Length: 371\nNumber of predicted TMHs: 1\nExp number of AAs in TMHs: 18.80245\nExp number, first 60 AAs: 0.14643\nTotal prob of N-in: 0.01944\noutside 1 327\nTMhelix 328 350\ninside 351 371

28566 GCF_001222985.1_4976_2_5Yersinia enterocolitica Proteobacteria; Gammaproteobacteria; Enterobacterales; Yersiniaceae; Yersinia MLSQHGVNRRDFMKLCAALAATMGLSSKAAAEIANTVSSPQRPPVIWIGAQECTGCTESL WP_005157252.1 hydrogenase 2 small subunit [Yersinia enterocolitica] Length: 371\nNumber of predicted TMHs: 1\nExp number of AAs in TMHs: 18.80217\nExp number, first 60 AAs: 0.14643\nTotal prob of N-in: 0.01943\noutside 1 327\nTMhelix 328 350\ninside 351 371

28567 GCF_001223145.1_4976_1_5Yersinia enterocolitica Proteobacteria; Gammaproteobacteria; Enterobacterales; Yersiniaceae; Yersinia MLSQHGVNRRDFMKLCAALAATMGLSSKAAAEIANTVSSPQRPPVIWIGAQECTGCTESL WP_019079071.1 hydrogenase 2 small subunit [Yersinia enterocolitica] Length: 371\nNumber of predicted TMHs: 1\nExp number of AAs in TMHs: 18.80245\nExp number, first 60 AAs: 0.14643\nTotal prob of N-in: 0.01944\noutside 1 327\nTMhelix 328 350\ninside 351 371

28568 GCF_001305635.1_ASM130563v1 Yersinia enterocolitica Proteobacteria; Gammaproteobacteria; Enterobacterales; Yersiniaceae; Yersinia MLSQHGVNRRDFMKLCAALAATMGLSSKAAAEIANTVSSPQRPPVIWIGAQECTGCTESL WP_005157252.1 hydrogenase 2 small subunit [Yersinia enterocolitica] Length: 371\nNumber of predicted TMHs: 1\nExp number of AAs in TMHs: 18.80217\nExp number, first 60 AAs: 0.14643\nTotal prob of N-in: 0.01943\noutside 1 327\nTMhelix 328 350\ninside 351 371

28569 GCF_001304755.1_ASM130475v1 Yersinia enterocolitica Proteobacteria; Gammaproteobacteria; Enterobacterales; Yersiniaceae; Yersinia MLSQHGVNRRDFMKLCAALAATMGLSSKAAAEIANTVSSPQRPPVIWIGAQECTGCTESL WP_019079071.1 hydrogenase 2 small subunit [Yersinia enterocolitica] Length: 371\nNumber of predicted TMHs: 1\nExp number of AAs in TMHs: 18.80245\nExp number, first 60 AAs: 0.14643\nTotal prob of N-in: 0.01944\noutside 1 327\nTMhelix 328 350\ninside 351 371

28570 GCF_001319325.1_4976_6_9Yersinia enterocolitica Proteobacteria; Gammaproteobacteria; Enterobacterales; Yersiniaceae; Yersinia MLSQHGVNRRDFMKLCAALAATMGLSSKAAAEIANTVSSPQRPPVIWIGAQECTGCTESL WP_019079071.1 hydrogenase 2 small subunit [Yersinia enterocolitica] Length: 371\nNumber of predicted TMHs: 1\nExp number of AAs in TMHs: 18.80245\nExp number, first 60 AAs: 0.14643\nTotal prob of N-in: 0.01944\noutside 1 327\nTMhelix 328 350\ninside 351 371

28571 GCF_001319345.1_4976_6_5Yersinia enterocolitica Proteobacteria; Gammaproteobacteria; Enterobacterales; Yersiniaceae; Yersinia MLSQHGVNRRDFMKLCAALAATMGLSSKAAAEIANTVSSPQRPPVIWIGAQECTGCTESL WP_019079071.1 hydrogenase 2 small subunit [Yersinia enterocolitica] Length: 371\nNumber of predicted TMHs: 1\nExp number of AAs in TMHs: 18.80245\nExp number, first 60 AAs: 0.14643\nTotal prob of N-in: 0.01944\noutside 1 327\nTMhelix 328 350\ninside 351 371

28572	GCF_001319385.1_5103_7_8Yersinia enterocolitica	Proteobacteria; Gammaproteobacteria; Enterobacterales; Yersiniaceae; Yersinia
MLSQHGVNRRDFMKLCAALAATMGLSSKAAAEIANTVSSPQRPPVIWIGAQECTGCTESL	WP_005157252.1
hydrogenase 2 small subunit [Yersinia enterocolitica]	Length: 371\nNumber of predicted TMHs: 1\nExp number of AAs in TMHs: 18.80217\nExp number, first 60 AAs: 0.14643\nTotal prob of N-in: 0.01943\noutside 1 327\nTMhelix 328 350\ninside 351 371

28573	GCF_001319425.1_4976_7_4Yersinia enterocolitica	Proteobacteria; Gammaproteobacteria; Enterobacterales; Yersiniaceae; Yersinia
MLSQHGVNRRDFMKLCAALAATMGLSSKAAAEIANTVSSPQRPPVIWIGAQECTGCTESL	WP_005157252.1
hydrogenase 2 small subunit [Yersinia enterocolitica]	Length: 371\nNumber of predicted TMHs: 1\nExp number of AAs in TMHs: 18.80217\nExp number, first 60 AAs: 0.14643\nTotal prob of N-in: 0.01943\noutside 1 327\nTMhelix 328 350\ninside 351 371

28574	GCF_001319445.1_4976_7_2Yersinia enterocolitica	Proteobacteria; Gammaproteobacteria; Enterobacterales; Yersiniaceae; Yersinia
MLSQHGVNRRDFMKLCAALAATMGLSSKAAAEIANTVSSPQRPPVIWIGAQECTGCTESL	WP_005157252.1
hydrogenase 2 small subunit [Yersinia enterocolitica]	Length: 371\nNumber of predicted TMHs: 1\nExp number of AAs in TMHs: 18.80217\nExp number, first 60 AAs: 0.14643\nTotal prob of N-in: 0.01943\noutside 1 327\nTMhelix 328 350\ninside 351 371

28575	GCF_001354575.1_8016_2_65	Yersinia enterocolitica	Proteobacteria; Gammaproteobacteria; Enterobacterales; Yersiniaceae; Yersinia
MLSQHGVNRRDFMKLCAALAATMGLSSKAAAEIANTVSSPQRPPVIWIGAQECTGCTESL	WP_005157252.1
hydrogenase 2 small subunit [Yersinia enterocolitica]	Length: 371\nNumber of predicted TMHs: 1\nExp number of AAs in TMHs: 18.80217\nExp number, first 60 AAs: 0.14643\nTotal prob of N-in: 0.01943\noutside 1 327\nTMhelix 328 350\ninside 351 371

28576	GCF_001319505.1_4976_3_5Yersinia enterocolitica	Proteobacteria; Gammaproteobacteria; Enterobacterales; Yersiniaceae; Yersinia
MLSQHGVNRRDFMKLCAALAATMGLSSKAAAEIANTVSSPQRPPVIWIGAQECTGCTESL	WP_019079071.1
hydrogenase 2 small subunit [Yersinia enterocolitica]	Length: 371\nNumber of predicted TMHs: 1\nExp number of AAs in TMHs: 18.80245\nExp number, first 60 AAs: 0.14643\nTotal prob of N-in: 0.01944\noutside 1 327\nTMhelix 328 350\ninside 351 371

28577	GCF_001354615.1_8016_2_73	Yersinia enterocolitica	Proteobacteria; Gammaproteobacteria; Enterobacterales; Yersiniaceae; Yersinia
MLSQHGVNRRDFMKLCAALAATMGLSSKAAAEIANTVSSPQRPPVIWIGAQECTGCTESL	WP_005157252.1
hydrogenase 2 small subunit [Yersinia enterocolitica]	Length: 371\nNumber of predicted TMHs: 1\nExp number of AAs in TMHs: 18.80217\nExp number, first 60 AAs: 0.14643\nTotal prob of N-in: 0.01943\noutside 1 327\nTMhelix 328 350\ninside 351 371

28578	GCF_001219385.1_5103_7_3Yersinia enterocolitica	Proteobacteria; Gammaproteobacteria; Enterobacterales; Yersiniaceae; Yersinia
MLSQHGVNRRDFMKLCAALAATMGLSSKAAAEIANTVSSPQRPPVIWIGAQECTGCTESL	WP_019079071.1
hydrogenase 2 small subunit [Yersinia enterocolitica]	Length: 371\nNumber of predicted TMHs: 1\nExp number of AAs in TMHs: 18.80245\nExp number, first 60 AAs: 0.14643\nTotal prob of N-in: 0.01944\noutside 1 327\nTMhelix 328 350\ninside 351 371

28579	GCF_001219625.1_4976_2_8Yersinia enterocolitica	Proteobacteria; Gammaproteobacteria; Enterobacterales; Yersiniaceae; Yersinia
MLSQHGVNRRDFMKLCAALAATMGLSSKAAAEIANTVSSPQRPPVIWIGAQECTGCTESL	WP_019079071.1
hydrogenase 2 small subunit [Yersinia enterocolitica]	Length: 371\nNumber of predicted TMHs: 1\nExp number of AAs in TMHs: 18.80245\nExp number, first 60 AAs: 0.14643\nTotal prob of N-in: 0.01944\noutside 1 327\nTMhelix 328 350\ninside 351 371

28580	GCF_001995075.1_ASM199507v1	Yersinia enterocolitica	Proteobacteria; Gammaproteobacteria; Enterobacterales; Yersiniaceae; Yersinia
MLSQHGVNRRDFMKLCAALAATMGLSSKAAAEIANTVSSPQRPPVIWIGAQECTGCTESL	WP_050077148.1
MULTISPECIES: hydrogenase 2 small subunit [Yersinia]	Length: 371\nNumber of predicted TMHs: 1\nExp

number of AAs in TMHs: 18.80844\nExp number, first 60 AAs: 0.14641\nTotal prob of N-in: 0.01969\noutside 1 327\nTMhelix 328 350\ninside 351 371

28581 GCF_900079815.1_IP35475 Yersinia enterocolitica Proteobacteria; Gammaproteobacteria; Enterobacterales; Yersiniaceae; Yersinia
MLSQHGVNRRDFMKLCAALAATMGLSSKAAAEIANTVSSPQRPPVIWIGAQECTGCTESL WP_005157252.1
hydrogenase 2 small subunit [Yersinia enterocolitica] Length: 371\nNumber of predicted TMHs: 1\nExp number of AAs in TMHs: 18.80217\nExp number, first 60 AAs: 0.14643\nTotal prob of N-in: 0.01943\noutside 1 327\nTMhelix 328 350\ninside 351 371

28582 GCF_001219325.1_4976_1_7Yersinia enterocolitica Proteobacteria; Gammaproteobacteria; Enterobacterales; Yersiniaceae; Yersinia
MLSQHGVNRRDFMKLCAALAATMGLSSKAAAEIANTVSSPQRPPVIWIGAQECTGCTESL WP_005157252.1
hydrogenase 2 small subunit [Yersinia enterocolitica] Length: 371\nNumber of predicted TMHs: 1\nExp number of AAs in TMHs: 18.80217\nExp number, first 60 AAs: 0.14643\nTotal prob of N-in: 0.01943\noutside 1 327\nTMhelix 328 350\ninside 351 371

28583 GCF_001219245.1_4976_6_2Yersinia enterocolitica Proteobacteria; Gammaproteobacteria; Enterobacterales; Yersiniaceae; Yersinia
MLSQHGVNRRDFMKLCAALAATMGLSSKAAAEIANTVSSPQRPPVIWIGAQECTGCTESL WP_005157252.1
hydrogenase 2 small subunit [Yersinia enterocolitica] Length: 371\nNumber of predicted TMHs: 1\nExp number of AAs in TMHs: 18.80217\nExp number, first 60 AAs: 0.14643\nTotal prob of N-in: 0.01943\noutside 1 327\nTMhelix 328 350\ninside 351 371

28584 GCF_001218705.1_4976_1_9Yersinia enterocolitica Proteobacteria; Gammaproteobacteria; Enterobacterales; Yersiniaceae; Yersinia
MLSQHGVNRRDFMKLCAALAATMGLSSKAAAEIANTVSSPQRPPVIWIGAQECTGCTESL WP_019079071.1
hydrogenase 2 small subunit [Yersinia enterocolitica] Length: 371\nNumber of predicted TMHs: 1\nExp number of AAs in TMHs: 18.80245\nExp number, first 60 AAs: 0.14643\nTotal prob of N-in: 0.01944\noutside 1 327\nTMhelix 328 350\ninside 351 371

28585 GCF_001218165.1_4976_2_2Yersinia enterocolitica Proteobacteria; Gammaproteobacteria; Enterobacterales; Yersiniaceae; Yersinia
MLSQHGVNRRDFMKLCAALAATMGLSSKAAAEIANTVSSPQRPPVIWIGAQECTGCTESL WP_005157252.1
hydrogenase 2 small subunit [Yersinia enterocolitica] Length: 371\nNumber of predicted TMHs: 1\nExp number of AAs in TMHs: 18.80217\nExp number, first 60 AAs: 0.14643\nTotal prob of N-in: 0.01943\noutside 1 327\nTMhelix 328 350\ninside 351 371

28586 GCF_001182345.1_8016_2_78 Yersinia enterocolitica Proteobacteria; Gammaproteobacteria; Enterobacterales; Yersiniaceae; Yersinia
MLSQHGVNRRDFMKLCAALAATMGLSSKAAAEIANTVSSPQRPPVIWIGAQECTGCTESL WP_019079071.1
hydrogenase 2 small subunit [Yersinia enterocolitica] Length: 371\nNumber of predicted TMHs: 1\nExp number of AAs in TMHs: 18.80245\nExp number, first 60 AAs: 0.14643\nTotal prob of N-in: 0.01944\noutside 1 327\nTMhelix 328 350\ninside 351 371

28587 GCF_001217745.1_4976_6_4Yersinia enterocolitica Proteobacteria; Gammaproteobacteria; Enterobacterales; Yersiniaceae; Yersinia
MLSQHGVNRRDFMKLCAALAATMGLSSKAAAEIANTVSSPQRPPVIWIGAQECTGCTESL WP_019079071.1
hydrogenase 2 small subunit [Yersinia enterocolitica] Length: 371\nNumber of predicted TMHs: 1\nExp number of AAs in TMHs: 18.80245\nExp number, first 60 AAs: 0.14643\nTotal prob of N-in: 0.01944\noutside 1 327\nTMhelix 328 350\ninside 351 371

28588 GCF_001182965.1_8016_2_76 Yersinia enterocolitica Proteobacteria; Gammaproteobacteria; Enterobacterales; Yersiniaceae; Yersinia
MLSQHGVNRRDFMKLCAALAATMGLSSKAAAEIANTVSSPQRPPVIWIGAQECTGCTESL WP_005173957.1
hydrogenase 2 small subunit [Yersinia enterocolitica] Length: 371\nNumber of predicted TMHs: 1\nExp number of AAs in TMHs: 18.61508\nExp number, first 60 AAs: 0.14605\nTotal prob of N-in: 0.02327\noutside 1 327\nTMhelix 328 350\ninside 351 371

28589 GCF_001217565.1_4976_1_4Yersinia enterocolitica Proteobacteria; Gammaproteobacteria; Enterobacterales; Yersiniaceae; Yersinia

MLSQHGVNRRDFMKLCAALAATMGLSSKAAAEIANTVSSPQRPPVIWIGAQECTGCTESL WP_019079071.1 hydrogenase 2 small subunit [Yersinia enterocolitica] Length: 371\nNumber of predicted TMHs: 1\nExp number of AAs in TMHs: 18.80245\nExp number, first 60 AAs: 0.14643\nTotal prob of N-in: 0.01944\noutside 1 327\nTMhelix 328 350\ninside 351 371

28590 GCF_001171185.1_5116_3_5Yersinia enterocolitica Proteobacteria; Gammaproteobacteria; Enterobacterales; Yersiniaceae; Yersinia MLSQHGVNRRDFMKLCAALAATMGLSSKAAAEIANTVSSPQRPPVIWIGAQECTGCTESL WP_019079071.1 hydrogenase 2 small subunit [Yersinia enterocolitica] Length: 371\nNumber of predicted TMHs: 1\nExp number of AAs in TMHs: 18.80245\nExp number, first 60 AAs: 0.14643\nTotal prob of N-in: 0.01944\noutside 1 327\nTMhelix 328 350\ninside 351 371

28591 GCF_001182225.1_5116_3_3Yersinia enterocolitica Proteobacteria; Gammaproteobacteria; Enterobacterales; Yersiniaceae; Yersinia MLSQHGVNRRDFMKLCAALAATMGLSSKAAAEIANTVSSPQRPPVIWIGAQECTGCTESL WP_005157252.1 hydrogenase 2 small subunit [Yersinia enterocolitica] Length: 371\nNumber of predicted TMHs: 1\nExp number of AAs in TMHs: 18.80217\nExp number, first 60 AAs: 0.14643\nTotal prob of N-in: 0.01943\noutside 1 327\nTMhelix 328 350\ninside 351 371

28592 GCF_001182165.1_4976_7_3Yersinia enterocolitica Proteobacteria; Gammaproteobacteria; Enterobacterales; Yersiniaceae; Yersinia MLSQHGVNRRDFMKLCAALAATMGLSSKAAAEIANTVSSPQRPPVIWIGAQECTGCTESL WP_005157252.1 hydrogenase 2 small subunit [Yersinia enterocolitica] Length: 371\nNumber of predicted TMHs: 1\nExp number of AAs in TMHs: 18.80217\nExp number, first 60 AAs: 0.14643\nTotal prob of N-in: 0.01943\noutside 1 327\nTMhelix 328 350\ninside 351 371

28593 GCF_001182205.1_5116_3_1Yersinia enterocolitica Proteobacteria; Gammaproteobacteria; Enterobacterales; Yersiniaceae; Yersinia MLSQHGVNRRDFMKLCAALAATMGLSSKAAAEIANTVSSPQRPPVIWIGAQECTGCTESL WP_005157252.1 hydrogenase 2 small subunit [Yersinia enterocolitica] Length: 371\nNumber of predicted TMHs: 1\nExp number of AAs in TMHs: 18.80217\nExp number, first 60 AAs: 0.14643\nTotal prob of N-in: 0.01943\noutside 1 327\nTMhelix 328 350\ninside 351 371

28594 GCF_001174985.1_5749_7_3Yersinia enterocolitica Proteobacteria; Gammaproteobacteria; Enterobacterales; Yersiniaceae; Yersinia MLSQHGVNRRDFMKLCAALAATMGLSSKAAAEIANTVSSPQRPPVIWIGAQECTGCTESL WP_019079071.1 hydrogenase 2 small subunit [Yersinia enterocolitica] Length: 371\nNumber of predicted TMHs: 1\nExp number of AAs in TMHs: 18.80245\nExp number, first 60 AAs: 0.14643\nTotal prob of N-in: 0.01944\noutside 1 327\nTMhelix 328 350\ninside 351 371

28595 GCF_001168925.1_5116_3_7Yersinia enterocolitica Proteobacteria; Gammaproteobacteria; Enterobacterales; Yersiniaceae; Yersinia MLSQHGVNRRDFMKLCAALAATMGLSSKAAAEIANTVSSPQRPPVIWIGAQECTGCTESL WP_019079071.1 hydrogenase 2 small subunit [Yersinia enterocolitica] Length: 371\nNumber of predicted TMHs: 1\nExp number of AAs in TMHs: 18.80245\nExp number, first 60 AAs: 0.14643\nTotal prob of N-in: 0.01944\noutside 1 327\nTMhelix 328 350\ninside 351 371

28596 GCF_001174945.1_5749_7_5Yersinia enterocolitica Proteobacteria; Gammaproteobacteria; Enterobacterales; Yersiniaceae; Yersinia MLSQHGVNRRDFMKLCAALAATMGLSSKAAAEIANTVSSPQRPPVIWIGAQECTGCTESL WP_005157252.1 hydrogenase 2 small subunit [Yersinia enterocolitica] Length: 371\nNumber of predicted TMHs: 1\nExp number of AAs in TMHs: 18.80217\nExp number, first 60 AAs: 0.14643\nTotal prob of N-in: 0.01943\noutside 1 327\nTMhelix 328 350\ninside 351 371

28597 GCF_001125285.1_5116_3_4Yersinia enterocolitica Proteobacteria; Gammaproteobacteria; Enterobacterales; Yersiniaceae; Yersinia MLSQHGVNRRDFMKLCAALAATMGLSSKAAAEIDNTVSSPQRPPVIWIGAQECTGCTESL WP_050148956.1 hydrogenase 2 small subunit [Yersinia enterocolitica] Length: 371\nNumber of predicted TMHs: 1\nExp number of AAs in TMHs: 18.69135\nExp number, first 60 AAs: 0.03535\nTotal prob of N-in: 0.01462\noutside 1 327\nTMhelix 328 350\ninside 351 371

28598 GCF_001151605.1_4976_7_8Yersinia enterocolitica Proteobacteria; Gammaproteobacteria; Enterobacterales; Yersiniaceae; Yersinia
MLSQHGVNRRDFMKLCAALAATMGLSSKAAAEIANTVSSPQRPPVIWIGAQECTGCTESL WP_019079071.1
hydrogenase 2 small subunit [Yersinia enterocolitica] Length: 371\nNumber of predicted TMHs: 1\nExp number of AAs in TMHs: 18.80245\nExp number, first 60 AAs: 0.14643\nTotal prob of N-in: 0.01944\noutside 1 327\nTMhelix 328 350\ninside 351 371

28599 GCF_001106265.1_5116_3_10 Yersinia enterocolitica Proteobacteria; Gammaproteobacteria; Enterobacterales; Yersiniaceae; Yersinia
MLSQHGVNRRDFMKLCAALAATMGLSSKAAAEIASKVSAPQRPPVIWIGAQECTGCTESL WP_050143758.1
MULTISPECIES: hydrogenase 2 small subunit [Yersinia] Length: 371\nNumber of predicted TMHs: 1\nExp number of AAs in TMHs: 16.08592\nExp number, first 60 AAs: 0.22113\nTotal prob of N-in: 0.02765\noutside 1 327\nTMhelix 328 350\ninside 351 371

28600 GCF_001146765.1_8016_2_63 Yersinia enterocolitica Proteobacteria; Gammaproteobacteria; Enterobacterales; Yersiniaceae; Yersinia
MLSQHGVNRRDFMKLCAALAATMGLSSKAAAEIANTVSSPQRPPVIWIGAQECTGCTESL WP_019079071.1
hydrogenase 2 small subunit [Yersinia enterocolitica] Length: 371\nNumber of predicted TMHs: 1\nExp number of AAs in TMHs: 18.80245\nExp number, first 60 AAs: 0.14643\nTotal prob of N-in: 0.01944\noutside 1 327\nTMhelix 328 350\ninside 351 371

28601 GCF_001139785.1_8016_2_64 Yersinia enterocolitica Proteobacteria; Gammaproteobacteria; Enterobacterales; Yersiniaceae; Yersinia
MLSQHGVNRRDFMKLCAALAATMGLSSKAAAEIANTVSSPQRPPVIWIGAQECTGCTESL WP_005157252.1
hydrogenase 2 small subunit [Yersinia enterocolitica] Length: 371\nNumber of predicted TMHs: 1\nExp number of AAs in TMHs: 18.80217\nExp number, first 60 AAs: 0.14643\nTotal prob of N-in: 0.01943\noutside 1 327\nTMhelix 328 350\ninside 351 371

28602 GCF_001135865.1_4976_3_4Yersinia enterocolitica Proteobacteria; Gammaproteobacteria; Enterobacterales; Yersiniaceae; Yersinia
MLSQHGVNRRDFMKLCAALAATMGLSSKAAAEIANTVSSPQRPPVIWIGAQECTGCTESL WP_005157252.1
hydrogenase 2 small subunit [Yersinia enterocolitica] Length: 371\nNumber of predicted TMHs: 1\nExp number of AAs in TMHs: 18.80217\nExp number, first 60 AAs: 0.14643\nTotal prob of N-in: 0.01943\noutside 1 327\nTMhelix 328 350\ninside 351 371

28603 GCF_001125325.1_5103_7_5Yersinia enterocolitica Proteobacteria; Gammaproteobacteria; Enterobacterales; Yersiniaceae; Yersinia
MLSQHGVNRRDFMKLCAALAATMGLSSKAAAEIANTVSSPQRPPVIWIGAQECTGCTESL WP_005157252.1
hydrogenase 2 small subunit [Yersinia enterocolitica] Length: 371\nNumber of predicted TMHs: 1\nExp number of AAs in TMHs: 18.80217\nExp number, first 60 AAs: 0.14643\nTotal prob of N-in: 0.01943\noutside 1 327\nTMhelix 328 350\ninside 351 371

28604 GCF_001121425.1_8016_2_68 Yersinia enterocolitica Proteobacteria; Gammaproteobacteria; Enterobacterales; Yersiniaceae; Yersinia
MLSQHGVNRRDFMKLCAALAATMGLSSKAAAEIANTVSSPQRPPVIWIGAQECTGCTESL WP_005157252.1
hydrogenase 2 small subunit [Yersinia enterocolitica] Length: 371\nNumber of predicted TMHs: 1\nExp number of AAs in TMHs: 18.80217\nExp number, first 60 AAs: 0.14643\nTotal prob of N-in: 0.01943\noutside 1 327\nTMhelix 328 350\ninside 351 371

28605 GCF_001120245.1_4976_3_9Yersinia enterocolitica Proteobacteria; Gammaproteobacteria; Enterobacterales; Yersiniaceae; Yersinia
MLSQHGVNRRDFMKLCAALAATMGLSSKAAAEIANTVSSPQRPPVIWIGAQECTGCTESL WP_005157252.1
hydrogenase 2 small subunit [Yersinia enterocolitica] Length: 371\nNumber of predicted TMHs: 1\nExp number of AAs in TMHs: 18.80217\nExp number, first 60 AAs: 0.14643\nTotal prob of N-in: 0.01943\noutside 1 327\nTMhelix 328 350\ninside 351 371

28606 GCF_001116445.1_4976_3_7Yersinia enterocolitica Proteobacteria; Gammaproteobacteria; Enterobacterales; Yersiniaceae; Yersinia
MLSQHGVNRRDFMKLCAALAATMGLSSKAAAEIANTVSSPQRPPVIWIGAQECTGCTESL WP_005157252.1
hydrogenase 2 small subunit [Yersinia enterocolitica] Length: 371\nNumber of predicted TMHs: 1\nExp

number of AAs in TMHs: 18.80217\nExp number, first 60 AAs: 0.14643\nTotal prob of N-in: 0.01943\noutside 1 327\nTMhelix 328 350\ninside 351 371

28607 GCF_001103745.1_4976_7_7Yersinia enterocolitica Proteobacteria; Gammaproteobacteria; Enterobacterales; Yersiniaceae; Yersinia
MLSQHGVNRRDFMKLCAALAATMGLSSKAAAEIANTVSSPQRPPVIWIGAQECTGCTESL WP_019079071.1
hydrogenase 2 small subunit [Yersinia enterocolitica] Length: 371\nNumber of predicted TMHs: 1\nExp number of AAs in TMHs: 18.80245\nExp number, first 60 AAs: 0.14643\nTotal prob of N-in: 0.01944\noutside 1 327\nTMhelix 328 350\ninside 351 371

28608 GCF_001102185.1_4976_6_1Yersinia enterocolitica Proteobacteria; Gammaproteobacteria; Enterobacterales; Yersiniaceae; Yersinia
MLSQHGVNRRDFMKLCAALAATMGLSSKAAAEIANTVSSPQRPPVIWIGAQECTGCTESL WP_005157252.1
hydrogenase 2 small subunit [Yersinia enterocolitica] Length: 371\nNumber of predicted TMHs: 1\nExp number of AAs in TMHs: 18.80217\nExp number, first 60 AAs: 0.14643\nTotal prob of N-in: 0.01943\noutside 1 327\nTMhelix 328 350\ninside 351 371

28609 GCF_001090885.1_5116_3_8Yersinia enterocolitica Proteobacteria; Gammaproteobacteria; Enterobacterales; Yersiniaceae; Yersinia
MLSQHGVNRRDFMKLCAALAATMGLSSKAAAEIANTVSSPQRPPVIWIGAQECTGCTESL WP_005157252.1
hydrogenase 2 small subunit [Yersinia enterocolitica] Length: 371\nNumber of predicted TMHs: 1\nExp number of AAs in TMHs: 18.80217\nExp number, first 60 AAs: 0.14643\nTotal prob of N-in: 0.01943\noutside 1 327\nTMhelix 328 350\ninside 351 371

28610 GCF_001098085.1_4976_6_6Yersinia enterocolitica Proteobacteria; Gammaproteobacteria; Enterobacterales; Yersiniaceae; Yersinia
MLSQHGVNRRDFMKLCAALAATMGLSSKAAAEIANTVSSPQRPPVIWIGAQECTGCTESL WP_005157252.1
hydrogenase 2 small subunit [Yersinia enterocolitica] Length: 371\nNumber of predicted TMHs: 1\nExp number of AAs in TMHs: 18.80217\nExp number, first 60 AAs: 0.14643\nTotal prob of N-in: 0.01943\noutside 1 327\nTMhelix 328 350\ninside 351 371

28611 GCF_001086525.1_4976_6_7Yersinia enterocolitica Proteobacteria; Gammaproteobacteria; Enterobacterales; Yersiniaceae; Yersinia
MLSQHGVNRRDFMKLCAALAATMGLSSKAAAEIANTVSSPQRPPVIWIGAQECTGCTESL WP_019079071.1
hydrogenase 2 small subunit [Yersinia enterocolitica] Length: 371\nNumber of predicted TMHs: 1\nExp number of AAs in TMHs: 18.80245\nExp number, first 60 AAs: 0.14643\nTotal prob of N-in: 0.01944\noutside 1 327\nTMhelix 328 350\ninside 351 371

28612 GCF_001090225.1_8016_2_66 Yersinia enterocolitica Proteobacteria; Gammaproteobacteria; Enterobacterales; Yersiniaceae; Yersinia
MLSQHGVNRRDFMKLCAALAATMGLSSKAAAEIANTVSSPQRPPVIWIGAQECTGCTESL WP_005157252.1
hydrogenase 2 small subunit [Yersinia enterocolitica] Length: 371\nNumber of predicted TMHs: 1\nExp number of AAs in TMHs: 18.80217\nExp number, first 60 AAs: 0.14643\nTotal prob of N-in: 0.01943\noutside 1 327\nTMhelix 328 350\ninside 351 371

28613 GCF_000754985.1_YEE Yersinia enterocolitica Proteobacteria; Gammaproteobacteria; Enterobacterales; Yersiniaceae; Yersinia
MLSQHGVNRRDFMKLCAALAATMGLSSKAAAEIANTVSSPQRPPVIWIGAQECTGCTESL WP_005157252.1
hydrogenase 2 small subunit [Yersinia enterocolitica] Length: 371\nNumber of predicted TMHs: 1\nExp number of AAs in TMHs: 18.80217\nExp number, first 60 AAs: 0.14643\nTotal prob of N-in: 0.01943\noutside 1 327\nTMhelix 328 350\ninside 351 371

28614 GCF_000755045.1_YEF Yersinia enterocolitica Proteobacteria; Gammaproteobacteria; Enterobacterales; Yersiniaceae; Yersinia
MLSQHGVNRRDFMKLCAALAATMGLSSKAAAEIANTVSSPQRPPVIWIGAQECTGCTESL WP_019079071.1
hydrogenase 2 small subunit [Yersinia enterocolitica] Length: 371\nNumber of predicted TMHs: 1\nExp number of AAs in TMHs: 18.80245\nExp number, first 60 AAs: 0.14643\nTotal prob of N-in: 0.01944\noutside 1 327\nTMhelix 328 350\ninside 351 371

28615 GCF_000834795.1_ASM83479v1 Yersinia enterocolitica Proteobacteria; Gammaproteobacteria; Enterobacterales; Yersiniaceae; Yersinia

MLSQHGVNRRDFMKLCAALAATMGLSSKAAAEIANTVSSPQRPPVIWIGAQECTGCTESL WP_005173957.1 hydrogenase 2 small subunit [Yersinia enterocolitica] Length: 371\nNumber of predicted TMHs: 1\nExp number of AAs in TMHs: 18.61508\nExp number, first 60 AAs: 0.14605\nTotal prob of N-in: 0.02327\noutside 1 327\nTMhelix 328 350\ninside 351 371

28616 GCF_000834195.1_ASM83419v1 Yersinia enterocolitica Proteobacteria; Gammaproteobacteria; Enterobacterales; Yersiniaceae; Yersinia MLSQHGVNRRDFMKLCAALAATMGLSSKAAAEIANTVSSPQRPPVIWIGAQECTGCTESL WP_005173957.1 hydrogenase 2 small subunit [Yersinia enterocolitica] Length: 371\nNumber of predicted TMHs: 1\nExp number of AAs in TMHs: 18.61508\nExp number, first 60 AAs: 0.14605\nTotal prob of N-in: 0.02327\noutside 1 327\nTMhelix 328 350\ninside 351 371

28617 GCF_001085265.1_5103_7_7Yersinia enterocolitica Proteobacteria; Gammaproteobacteria; Enterobacterales; Yersiniaceae; Yersinia MLSQHGVNRRDFMKLCAALAATMGLSSKAAAEIANTVSSPQRPPVIWIGAQECTGCTESL WP_019079071.1 hydrogenase 2 small subunit [Yersinia enterocolitica] Length: 371\nNumber of predicted TMHs: 1\nExp number of AAs in TMHs: 18.80245\nExp number, first 60 AAs: 0.14643\nTotal prob of N-in: 0.01944\noutside 1 327\nTMhelix 328 350\ninside 351 371

28618 GCF_000987925.1_ASM98792v1 Yersinia enterocolitica Proteobacteria; Gammaproteobacteria; Enterobacterales; Yersiniaceae; Yersinia MLSQHGVNRRDFMKLCAALAATMGLSSKAAAEIANTVSSPQRPPVIWIGAQECTGCTESL WP_019079071.1 hydrogenase 2 small subunit [Yersinia enterocolitica] Length: 371\nNumber of predicted TMHs: 1\nExp number of AAs in TMHs: 18.80245\nExp number, first 60 AAs: 0.14643\nTotal prob of N-in: 0.01944\noutside 1 327\nTMhelix 328 350\ninside 351 371

28619 GCF_001219745.1_4976_1_6Yersinia enterocolitica Proteobacteria; Gammaproteobacteria; Enterobacterales; Yersiniaceae; Yersinia MLSQHGVNRRDFMKLCAALAATMGLSSKAAAEIANTVSSPQRPPVIWIGAQECTGCTESL WP_005157252.1 hydrogenase 2 small subunit [Yersinia enterocolitica] Length: 371\nNumber of predicted TMHs: 1\nExp number of AAs in TMHs: 18.80217\nExp number, first 60 AAs: 0.14643\nTotal prob of N-in: 0.01943\noutside 1 327\nTMhelix 328 350\ninside 351 371

28620 GCF_900079865.1_IP35467 Yersinia enterocolitica Proteobacteria; Gammaproteobacteria; Enterobacterales; Yersiniaceae; Yersinia MLSQHGVNRRDFMKLCAALAATMGLSSKAAAEIANTVSSPQRPPVIWIGAQECTGCTESL WP_005157252.1 hydrogenase 2 small subunit [Yersinia enterocolitica] Length: 371\nNumber of predicted TMHs: 1\nExp number of AAs in TMHs: 18.80217\nExp number, first 60 AAs: 0.14643\nTotal prob of N-in: 0.01943\noutside 1 327\nTMhelix 328 350\ninside 351 371

28621 GCF_900079895.1_IP35474 Yersinia enterocolitica Proteobacteria; Gammaproteobacteria; Enterobacterales; Yersiniaceae; Yersinia MLSQHGVNRRDFMKLCAALAATMGLSSKAAAEIANTVSSPQRPPVIWIGAQECTGCTESL WP_005157252.1 hydrogenase 2 small subunit [Yersinia enterocolitica] Length: 371\nNumber of predicted TMHs: 1\nExp number of AAs in TMHs: 18.80217\nExp number, first 60 AAs: 0.14643\nTotal prob of N-in: 0.01943\noutside 1 327\nTMhelix 328 350\ninside 351 371

28622 GCF_900079925.1_IP35465 Yersinia enterocolitica Proteobacteria; Gammaproteobacteria; Enterobacterales; Yersiniaceae; Yersinia MLSQHGVNRRDFMKLCAALAATMGLSSKAAAEIANTVSSPQRPPVIWIGAQECTGCTESL WP_005157252.1 hydrogenase 2 small subunit [Yersinia enterocolitica] Length: 371\nNumber of predicted TMHs: 1\nExp number of AAs in TMHs: 18.80217\nExp number, first 60 AAs: 0.14643\nTotal prob of N-in: 0.01943\noutside 1 327\nTMhelix 328 350\ninside 351 371

28623 GCF_900079915.1_IP35470 Yersinia enterocolitica Proteobacteria; Gammaproteobacteria; Enterobacterales; Yersiniaceae; Yersinia MLSQHGVNRRDFMKLCAALAATMGLSSKAAAEIANTVSSPQRPPVIWIGAQECTGCTESL WP_005157252.1 hydrogenase 2 small subunit [Yersinia enterocolitica] Length: 371\nNumber of predicted TMHs: 1\nExp number of AAs in TMHs: 18.80217\nExp number, first 60 AAs: 0.14643\nTotal prob of N-in: 0.01943\noutside 1 327\nTMhelix 328 350\ninside 351 371

28624 GCF_900079945.1_IP35462 Yersinia enterocolitica Proteobacteria; Gammaproteobacteria; Enterobacterales; Yersiniaceae; Yersinia
MLSQHGVNRRDFMKLCAALAATMGLSSKAAAEIANTVSSPQRPPVIWIGAQECTGCTESL WP_005157252.1
hydrogenase 2 small subunit [Yersinia enterocolitica] Length: 371\nNumber of predicted TMHs: 1\nExp number of AAs in TMHs: 18.80217\nExp number, first 60 AAs: 0.14643\nTotal prob of N-in: 0.01943\noutside 1 327\nTMhelix 328 350\ninside 351 371

28625 GCF_900080125.1_IP35477 Yersinia enterocolitica Proteobacteria; Gammaproteobacteria; Enterobacterales; Yersiniaceae; Yersinia
MLSQHGVNRRDFMKLCAALAATMGLSSKAAAEIANTVSSPQRPPVIWIGAQECTGCTESL WP_005157252.1
hydrogenase 2 small subunit [Yersinia enterocolitica] Length: 371\nNumber of predicted TMHs: 1\nExp number of AAs in TMHs: 18.80217\nExp number, first 60 AAs: 0.14643\nTotal prob of N-in: 0.01943\noutside 1 327\nTMhelix 328 350\ninside 351 371

28626 GCF_002083285.1_ASM208328v1 Yersinia enterocolitica Proteobacteria; Gammaproteobacteria; Enterobacterales; Yersiniaceae; Yersinia
MLSQHGVNRRDFMKLCAALAATMGLSSKAAAEIANTVSSPQRPPVIWIGAQECTGCTESL WP_005173957.1
hydrogenase 2 small subunit [Yersinia enterocolitica] Length: 371\nNumber of predicted TMHs: 1\nExp number of AAs in TMHs: 18.61508\nExp number, first 60 AAs: 0.14605\nTotal prob of N-in: 0.02327\noutside 1 327\nTMhelix 328 350\ninside 351 371

28627 GCF_900079875.1_IP35463 Yersinia enterocolitica Proteobacteria; Gammaproteobacteria; Enterobacterales; Yersiniaceae; Yersinia
MLSQHGVNRRDFMKLCAALAATMGLSSKAAAEIANTVSSPQRPPVIWIGAQECTGCTESL WP_005157252.1
hydrogenase 2 small subunit [Yersinia enterocolitica] Length: 371\nNumber of predicted TMHs: 1\nExp number of AAs in TMHs: 18.80217\nExp number, first 60 AAs: 0.14643\nTotal prob of N-in: 0.01943\noutside 1 327\nTMhelix 328 350\ninside 351 371

28628 GCF_001319585.1_4821_6_7Yersinia frederiksenii Proteobacteria; Gammaproteobacteria; Enterobacterales; Yersiniaceae; Yersinia
MLSQHGVNRRDFMKLCAALAATMGLSSKAAAEIASTVSSPQRPPVIWIGAQECTGCTESL WP_004711888.1
hydrogenase 2 small subunit [Yersinia frederiksenii] Length: 371\nNumber of predicted TMHs: 1\nExp number of AAs in TMHs: 19.47103\nExp number, first 60 AAs: 0.16503\nTotal prob of N-in: 0.02370\noutside 1 327\nTMhelix 328 350\ninside 351 371

28629 GCF_001089645.1_4821_6_9Yersinia frederiksenii Proteobacteria; Gammaproteobacteria; Enterobacterales; Yersiniaceae; Yersinia
MLSQHGVNRRDFMKLCAALAATMGLSSKAAAEIASTVSSPQRPPVIWIGAQECTGCTESL WP_050123536.1
hydrogenase 2 small subunit [Yersinia frederiksenii] Length: 371\nNumber of predicted TMHs: 1\nExp number of AAs in TMHs: 20.13947\nExp number, first 60 AAs: 0.16438\nTotal prob of N-in: 0.02557\noutside 1 327\nTMhelix 328 350\ninside 351 371

28630 GCF_001319525.1_4840_7_4Yersinia frederiksenii Proteobacteria; Gammaproteobacteria; Enterobacterales; Yersiniaceae; Yersinia
MLSQHGVNRRDFMKLCAALAATMGLSSKAAAEIASTVSSPQRPPVIWIGAQECTGCTESL WP_050123536.1
hydrogenase 2 small subunit [Yersinia frederiksenii] Length: 371\nNumber of predicted TMHs: 1\nExp number of AAs in TMHs: 20.13947\nExp number, first 60 AAs: 0.16438\nTotal prob of N-in: 0.02557\noutside 1 327\nTMhelix 328 350\ninside 351 371

28631 GCF_002188925.1_ASM218892v1 Yersinia frederiksenii Proteobacteria; Gammaproteobacteria; Enterobacterales; Yersiniaceae; Yersinia
MLSQHGVNRRDFMKLCAALAATMGLSSKAAAEIAATVSSPQRPPVIWIGAQECTGCTEAL WP_050097866.1
hydrogenase 2 small subunit [Yersinia frederiksenii] Length: 371\nNumber of predicted TMHs: 1\nExp number of AAs in TMHs: 19.35295\nExp number, first 60 AAs: 0.23099\nTotal prob of N-in: 0.04591\noutside 1 327\nTMhelix 328 350\ninside 351 371

28632 GCF_002188975.1_ASM218897v1 Yersinia frederiksenii Proteobacteria; Gammaproteobacteria; Enterobacterales; Yersiniaceae; Yersinia
MLSQHGVNRRDFMKLCAALAATMGLSSKAAAEIASTVSSPQRPPVIWIGAQECTGCTESL WP_050123536.1
hydrogenase 2 small subunit [Yersinia frederiksenii] Length: 371\nNumber of predicted TMHs: 1\nExp

number of AAs in TMHs: 20.13947\nExp number, first 60 AAs: 0.16438\nTotal prob of N-in: 0.02557\noutside 1 327\nTMhelix 328 350\ninside 351 371

28633 GCF_001157485.1_4821_6_3Yersinia frederiksenii Proteobacteria; Gammaproteobacteria; Enterobacterales; Yersiniaceae; Yersinia
MLSQHGVNRRDFMKLCAALAATMGLSSKAAAEIASTVSSPQRPPVIWIGAQECTGCTESL WP_050296612.1
hydrogenase 2 small subunit [Yersinia frederiksenii] Length: 371\nNumber of predicted TMHs: 1\nExp number of AAs in TMHs: 20.13821\nExp number, first 60 AAs: 0.16439\nTotal prob of N-in: 0.02551\noutside 1 327\nTMhelix 328 350\ninside 351 371

28634 GCF_001147425.1_4821_5_9Yersinia frederiksenii Proteobacteria; Gammaproteobacteria; Enterobacterales; Yersiniaceae; Yersinia
MLSQHGVNRRDFMKLCAALAATMGLSSKAAAEIASTVSSPQRPPVIWIGAQECTGCTESL WP_050123536.1
hydrogenase 2 small subunit [Yersinia frederiksenii] Length: 371\nNumber of predicted TMHs: 1\nExp number of AAs in TMHs: 20.13947\nExp number, first 60 AAs: 0.16438\nTotal prob of N-in: 0.02557\noutside 1 327\nTMhelix 328 350\ninside 351 371

28635 GCF_002192925.1_ASM219292v1 Yersinia frederiksenii Proteobacteria; Gammaproteobacteria; Enterobacterales; Yersiniaceae; Yersinia
MLSQHGVNRRDFMKLCAALAATMGLSSKAAAEIADTVGSPQRPPVIWIGAQECTGCTESL WP_049610771.1
MULTISPECIES: hydrogenase 2 small subunit [Yersinia] Length: 371\nNumber of predicted TMHs: 1\nExp number of AAs in TMHs: 17.72326\nExp number, first 60 AAs: 0.1354\nTotal prob of N-in: 0.02052\noutside 1 327\nTMhelix 328 350\ninside 351 371

28636 GCF_001135485.1_5139_5_6Yersinia frederiksenii Proteobacteria; Gammaproteobacteria; Enterobacterales; Yersiniaceae; Yersinia
MLSQHGVNRRDFMKLCAALAATMGLSSKAAAEIADTVGSPQRPPVIWIGAQECTGCTESL WP_049610771.1
MULTISPECIES: hydrogenase 2 small subunit [Yersinia] Length: 371\nNumber of predicted TMHs: 1\nExp number of AAs in TMHs: 17.72326\nExp number, first 60 AAs: 0.1354\nTotal prob of N-in: 0.02052\noutside 1 327\nTMhelix 328 350\ninside 351 371

28637 GCF_001105765.1_4821_5_3Yersinia frederiksenii Proteobacteria; Gammaproteobacteria; Enterobacterales; Yersiniaceae; Yersinia
MLSQHGVNRRDFMKLCAALAATMGLSSKAAAEIAATVSSPQRPPVIWIGAQECTGCTEAL WP_050097866.1
hydrogenase 2 small subunit [Yersinia frederiksenii] Length: 371\nNumber of predicted TMHs: 1\nExp number of AAs in TMHs: 19.35295\nExp number, first 60 AAs: 0.23099\nTotal prob of N-in: 0.04591\noutside 1 327\nTMhelix 328 350\ninside 351 371

28638 GCF_001090165.1_4821_6_2Yersinia frederiksenii Proteobacteria; Gammaproteobacteria; Enterobacterales; Yersiniaceae; Yersinia
MLSQHGVNRRDFMKLCAALAATMGLSSKAAAEIANTVSSPQRPPVIWIGAQECTGCTESL WP_050108507.1
hydrogenase 2 small subunit [Yersinia frederiksenii] Length: 371\nNumber of predicted TMHs: 1\nExp number of AAs in TMHs: 20.12094\nExp number, first 60 AAs: 0.14584\nTotal prob of N-in: 0.02476\noutside 1 327\nTMhelix 328 350\ninside 351 371

28639 GCF_001151185.1_4821_8_8Yersinia frederiksenii Proteobacteria; Gammaproteobacteria; Enterobacterales; Yersiniaceae; Yersinia
MLSQHGVNRRDFMKLCAALAATMGLSSKAAAEIADTVGSPQRPPVIWIGAQECTGCTESL WP_049610771.1
MULTISPECIES: hydrogenase 2 small subunit [Yersinia] Length: 371\nNumber of predicted TMHs: 1\nExp number of AAs in TMHs: 17.72326\nExp number, first 60 AAs: 0.1354\nTotal prob of N-in: 0.02052\noutside 1 327\nTMhelix 328 350\ninside 351 371

28640 GCF_001319565.1_4821_6_6Yersinia frederiksenii Proteobacteria; Gammaproteobacteria; Enterobacterales; Yersiniaceae; Yersinia
MLSQHGVNRRDFMKLCAALAATMGLSSKAAAEIADTVGSPQRPPVIWIGAQECTGCTESL WP_049610771.1
MULTISPECIES: hydrogenase 2 small subunit [Yersinia] Length: 371\nNumber of predicted TMHs: 1\nExp number of AAs in TMHs: 17.72326\nExp number, first 60 AAs: 0.1354\nTotal prob of N-in: 0.02052\noutside 1 327\nTMhelix 328 350\ninside 351 371

28641 GCF_001218185.1_4821_8_1Yersinia frederiksenii Proteobacteria; Gammaproteobacteria; Enterobacterales; Yersiniaceae; Yersinia

MLSQHGVNRRDFMKLCAALAATMGLSSKAAAEIADTVGSPQRPPVIWIGAQECTGCTESL WP_049610771.1 MULTISPECIES: hydrogenase 2 small subunit [Yersinia] Length: 371\nNumber of predicted TMHs: 1\nExp number of AAs in TMHs: 17.72326\nExp number, first 60 AAs: 0.1354\nTotal prob of N-in: 0.02052\noutside 1 327\nTMhelix 328 350\ninside 351 371

28642 GCF_001319985.1_4821_5_10 Yersinia frederiksenii Proteobacteria; Gammaproteobacteria; Enterobacterales; Yersiniaceae; Yersinia MLSQHGVNRRDFMKLCAALAATMGLSSKAAAEIADTVGSPQRPPVIWIGAQECTGCTESL WP_049610771.1 MULTISPECIES: hydrogenase 2 small subunit [Yersinia] Length: 371\nNumber of predicted TMHs: 1\nExp number of AAs in TMHs: 17.72326\nExp number, first 60 AAs: 0.1354\nTotal prob of N-in: 0.02052\noutside 1 327\nTMhelix 328 350\ninside 351 371

28643 GCF_001158365.1_5139_5_1Yersinia frederiksenii Proteobacteria; Gammaproteobacteria; Enterobacterales; Yersiniaceae; Yersinia MLSQHGVNRRDFMKLCAALAATMGLSSKAAAEIASTVSSPQRPPVIWIGAQECTGCTESL WP_004711888.1 hydrogenase 2 small subunit [Yersinia frederiksenii] Length: 371\nNumber of predicted TMHs: 1\nExp number of AAs in TMHs: 19.47103\nExp number, first 60 AAs: 0.16503\nTotal prob of N-in: 0.02370\noutside 1 327\nTMhelix 328 350\ninside 351 371

28644 GCF_001222845.1_4840_8_10 Yersinia frederiksenii Proteobacteria; Gammaproteobacteria; Enterobacterales; Yersiniaceae; Yersinia MLSQHGVNRRDFMKLCAALAATMGLSSKAAAEIADTVGSPQRPPVIWIGAQECTGCTESL WP_049610771.1 MULTISPECIES: hydrogenase 2 small subunit [Yersinia] Length: 371\nNumber of predicted TMHs: 1\nExp number of AAs in TMHs: 17.72326\nExp number, first 60 AAs: 0.1354\nTotal prob of N-in: 0.02052\noutside 1 327\nTMhelix 328 350\ninside 351 371

28645 GCF_001121225.1_4821_8_11 Yersinia frederiksenii Proteobacteria; Gammaproteobacteria; Enterobacterales; Yersiniaceae; Yersinia MLSQHGVNRRDFMKLCAALAATMGLSSKAAAEIASTVSSPQRPPVIWIGAQECTGCTESL WP_050123536.1 hydrogenase 2 small subunit [Yersinia frederiksenii] Length: 371\nNumber of predicted TMHs: 1\nExp number of AAs in TMHs: 20.13947\nExp number, first 60 AAs: 0.16438\nTotal prob of N-in: 0.02557\noutside 1 327\nTMhelix 328 350\ninside 351 371

28646 GCF_001142745.1_4821_6_5Yersinia frederiksenii Proteobacteria; Gammaproteobacteria; Enterobacterales; Yersiniaceae; Yersinia MLSQHGVNRRDFMKLCAALAATMGLSSKAAAEIASTVSSPQRPPVIWIGAQECTGCTESL WP_004711888.1 hydrogenase 2 small subunit [Yersinia frederiksenii] Length: 371\nNumber of predicted TMHs: 1\nExp number of AAs in TMHs: 19.47103\nExp number, first 60 AAs: 0.16503\nTotal prob of N-in: 0.02370\noutside 1 327\nTMhelix 328 350\ninside 351 371

28647 GCF_001147045.1_4821_5_5Yersinia frederiksenii Proteobacteria; Gammaproteobacteria; Enterobacterales; Yersiniaceae; Yersinia MLSQHGVNRRDFMKLCAALAATMGLSSKAAAEIAATVSSPQRPPVIWIGAQECTGCTEAL WP_050097866.1 hydrogenase 2 small subunit [Yersinia frederiksenii] Length: 371\nNumber of predicted TMHs: 1\nExp number of AAs in TMHs: 19.35295\nExp number, first 60 AAs: 0.23099\nTotal prob of N-in: 0.04591\noutside 1 327\nTMhelix 328 350\ninside 351 371

28648 GCF_001155565.1_4840_6_2Yersinia frederiksenii Proteobacteria; Gammaproteobacteria; Enterobacterales; Yersiniaceae; Yersinia MLSQHGVNRRDFMKLCAALAATMGLSSKAAAEIAATVSSPQRPPVIWIGAQECTGCTEAL WP_050097866.1 hydrogenase 2 small subunit [Yersinia frederiksenii] Length: 371\nNumber of predicted TMHs: 1\nExp number of AAs in TMHs: 19.35295\nExp number, first 60 AAs: 0.23099\nTotal prob of N-in: 0.04591\noutside 1 327\nTMhelix 328 350\ninside 351 371

28649 GCF_001098625.1_5139_5_4Yersinia frederiksenii Proteobacteria; Gammaproteobacteria; Enterobacterales; Yersiniaceae; Yersinia MLSQHGVNRRDFMKLCAALAATMGLSSKAAAEIASTVSSPQRPPVIWIGAQECTGCTESL WP_050123536.1 hydrogenase 2 small subunit [Yersinia frederiksenii] Length: 371\nNumber of predicted TMHs: 1\nExp number of AAs in TMHs: 20.13947\nExp number, first 60 AAs: 0.16438\nTotal prob of N-in: 0.02557\noutside 1 327\nTMhelix 328 350\ninside 351 371

28650 GCF_001150525.1_4840_7_2Yersinia frederiksenii Proteobacteria; Gammaproteobacteria; Enterobacterales; Yersiniaceae; Yersinia
MLSQHGVNRRDFMKLCAALAATMGLSSKAAAEIANTVGSPQRPPVIWIGAQECTGCTESL WP_050086543.1
MULTISPECIES: hydrogenase 2 small subunit [Yersinia] Length: 371\nNumber of predicted TMHs: 1\nExp number of AAs in TMHs: 19.43469\nExp number, first 60 AAs: 0.16653\nTotal prob of N-in: 0.02906\noutside 1 327\nTMhelix 328 350\ninside 351 371

28651 GCF_001171065.1_4840_6_7Yersinia frederiksenii Proteobacteria; Gammaproteobacteria; Enterobacterales; Yersiniaceae; Yersinia
MLSQHGVNRRDFMKLCAALAATMGLSSKAAAEIASTVSSPQRPPVIWIGAQECTGCTESL WP_050123536.1
hydrogenase 2 small subunit [Yersinia frederiksenii] Length: 371\nNumber of predicted TMHs: 1\nExp number of AAs in TMHs: 20.13947\nExp number, first 60 AAs: 0.16438\nTotal prob of N-in: 0.02557\noutside 1 327\nTMhelix 328 350\ninside 351 371

28652 GCF_001088505.1_4840_7_1Yersinia intermedia Proteobacteria; Gammaproteobacteria; Enterobacterales; Yersiniaceae; Yersinia MLSQHGVNRRDFMKLCAALAATMGLSSKAAAEIANTVGSPQRPPVIWIGAQECTGCTESL
WP_005188113.1 MULTISPECIES: hydrogenase 2 small subunit [Yersinia] Length: 371\nNumber of predicted TMHs: 1\nExp number of AAs in TMHs: 19.43348\nExp number, first 60 AAs: 0.16654\nTotal prob of N-in: 0.02901\noutside 1 327\nTMhelix 328 350\ninside 351 371

28653 GCF_001116865.1_4840_6_5Yersinia intermedia Proteobacteria; Gammaproteobacteria; Enterobacterales; Yersiniaceae; Yersinia MLSQHGVNRRDFMKLCAALAATMGLSSKAAAEIANTVSSPQRPPVIWIGAQECTGCTESL
WP_050077148.1 MULTISPECIES: hydrogenase 2 small subunit [Yersinia] Length: 371\nNumber of predicted TMHs: 1\nExp number of AAs in TMHs: 18.80844\nExp number, first 60 AAs: 0.14641\nTotal prob of N-in: 0.01969\noutside 1 327\nTMhelix 328 350\ninside 351 371

28654 GCF_002188935.1_ASM218893v1 Yersinia intermedia Proteobacteria; Gammaproteobacteria; Enterobacterales; Yersiniaceae; Yersinia
MLSQHGVNRRDFMKLCAALAATMGLSSKAAAEIANTVGSPQRPPVIWIGAQECTGCTESL WP_050086543.1
MULTISPECIES: hydrogenase 2 small subunit [Yersinia] Length: 371\nNumber of predicted TMHs: 1\nExp number of AAs in TMHs: 19.43469\nExp number, first 60 AAs: 0.16653\nTotal prob of N-in: 0.02906\noutside 1 327\nTMhelix 328 350\ninside 351 371

28655 GCF_001140765.1_4821_5_11 Yersinia intermedia Proteobacteria; Gammaproteobacteria; Enterobacterales; Yersiniaceae; Yersinia
MLSQHGVNRRDFMKLCAALAATMGLSSKAAAEIANTVGSPQRPPVIWIGAQECTGCTESL WP_050086543.1
MULTISPECIES: hydrogenase 2 small subunit [Yersinia] Length: 371\nNumber of predicted TMHs: 1\nExp number of AAs in TMHs: 19.43469\nExp number, first 60 AAs: 0.16653\nTotal prob of N-in: 0.02906\noutside 1 327\nTMhelix 328 350\ninside 351 371

28656 GCF_001319645.1_4821_8_6Yersinia intermedia Proteobacteria; Gammaproteobacteria; Enterobacterales; Yersiniaceae; Yersinia MLSQHGVNRRDFMKLCAALAATMGLSSKAAAEIANTVGSPQRPPVIWIGAQECTGCTESL
WP_050086543.1 MULTISPECIES: hydrogenase 2 small subunit [Yersinia] Length: 371\nNumber of predicted TMHs: 1\nExp number of AAs in TMHs: 19.43469\nExp number, first 60 AAs: 0.16653\nTotal prob of N-in: 0.02906\noutside 1 327\nTMhelix 328 350\ninside 351 371

28657 GCF_000834515.1_ASM83451v1 Yersinia intermedia Proteobacteria; Gammaproteobacteria; Enterobacterales; Yersiniaceae; Yersinia
MLSQHGVNRRDFMKLCAALAATMGLSSKAAAEIANTVGSPQRPPVIWIGAQECTGCTESL WP_005188113.1
MULTISPECIES: hydrogenase 2 small subunit [Yersinia] Length: 371\nNumber of predicted TMHs: 1\nExp number of AAs in TMHs: 19.43348\nExp number, first 60 AAs: 0.16654\nTotal prob of N-in: 0.02901\noutside 1 327\nTMhelix 328 350\ninside 351 371

28658 GCF_001167085.1_4821_8_7Yersinia intermedia Proteobacteria; Gammaproteobacteria; Enterobacterales; Yersiniaceae; Yersinia MLSQHGVNRRDFMKLCAALAATMGLSSKAAAEIANTVGSPQRPPVIWIGAQECTGCTESL
WP_050086543.1 MULTISPECIES: hydrogenase 2 small subunit [Yersinia] Length: 371\nNumber of predicted TMHs: 1\nExp number of AAs in TMHs: 19.43469\nExp number, first 60 AAs: 0.16653\nTotal prob of N-in: 0.02906\noutside 1 327\nTMhelix 328 350\ninside 351 371

28659 GCF_001218045.1_4821_8_5Yersinia intermedia Proteobacteria; Gammaproteobacteria; Enterobacterales; Yersiniaceae; Yersinia MLSQHGVNRRDFMKLCAALAATMGLSSKAAAEIANTVGSPQRPPVIWIGAQECTGCTESL

WP_005188113.1 MULTISPECIES: hydrogenase 2 small subunit [Yersinia] Length: 371\nNumber of predicted TMHs: 1\nExp number of AAs in TMHs: 19.43348\nExp number, first 60 AAs: 0.16654\nTotal prob of N-in: 0.02901\noutside 1 327\nTMhelix 328 350\ninside 351 371

28660 GCF_001157385.1_5139_5_8Yersinia intermedia Proteobacteria; Gammaproteobacteria; Enterobacterales; Yersiniaceae; Yersinia MLSQHGVNRRDFMKLCAALAATMGLSSKAAAEIANTVGSPQRPPVIWIGAQECTGCTESL WP_005188113.1 MULTISPECIES: hydrogenase 2 small subunit [Yersinia] Length: 371\nNumber of predicted TMHs: 1\nExp number of AAs in TMHs: 19.43348\nExp number, first 60 AAs: 0.16654\nTotal prob of N-in: 0.02901\noutside 1 327\nTMhelix 328 350\ninside 351 371

28661 GCF_001244675.1_5139_5_2Yersinia intermedia Proteobacteria; Gammaproteobacteria; Enterobacterales; Yersiniaceae; Yersinia MLSQHGVNRRDFMKLCAALAATMGLSSKAAAEIANTVGSPQRPPVIWIGAQECTGCTESL WP_053011723.1 hydrogenase 2 small subunit [Yersinia intermedia] Length: 371\nNumber of predicted TMHs: 1\nExp number of AAs in TMHs: 19.4359\nExp number, first 60 AAs: 0.16653\nTotal prob of N-in: 0.02910\noutside 1 327\nTMhelix 328 350\ninside 351 371

28662 GCF_001221165.1_4821_7_8Yersinia intermedia Proteobacteria; Gammaproteobacteria; Enterobacterales; Yersiniaceae; Yersinia MLSQHGVNRRDFMKLCAALAATMGLSSKAAAEIANTVGSPQRPPVIWIGAQECTGCTESL WP_050086543.1 MULTISPECIES: hydrogenase 2 small subunit [Yersinia] Length: 371\nNumber of predicted TMHs: 1\nExp number of AAs in TMHs: 19.43469\nExp number, first 60 AAs: 0.16653\nTotal prob of N-in: 0.02906\noutside 1 327\nTMhelix 328 350\ninside 351 371

28663 GCF_001319925.1_4821_5_1Yersinia intermedia Proteobacteria; Gammaproteobacteria; Enterobacterales; Yersiniaceae; Yersinia MLSQHGVNRRDFMKLCAALAATMGLSSKAAAEIANTVGSPQRPPVIWIGAQECTGCTESL WP_050086543.1 MULTISPECIES: hydrogenase 2 small subunit [Yersinia] Length: 371\nNumber of predicted TMHs: 1\nExp number of AAs in TMHs: 19.43469\nExp number, first 60 AAs: 0.16653\nTotal prob of N-in: 0.02906\noutside 1 327\nTMhelix 328 350\ninside 351 371

28664 GCF_001104645.1_4821_5_8Yersinia intermedia Proteobacteria; Gammaproteobacteria; Enterobacterales; Yersiniaceae; Yersinia MLSQHGVNRRDFMKLCAALAATMGLSSKAAAEIANTVGSPQRPPVIWIGAQECTGCTESL WP_005188113.1 MULTISPECIES: hydrogenase 2 small subunit [Yersinia] Length: 371\nNumber of predicted TMHs: 1\nExp number of AAs in TMHs: 19.43348\nExp number, first 60 AAs: 0.16654\nTotal prob of N-in: 0.02901\noutside 1 327\nTMhelix 328 350\ninside 351 371

28665 GCF_001153385.1_4821_5_2Yersinia intermedia Proteobacteria; Gammaproteobacteria; Enterobacterales; Yersiniaceae; Yersinia MLSQHGVNRRDFMKLCAALAATMGLSSKAAAEIANTVGSPQRPPVIWIGAQECTGCTESL WP_005188113.1 MULTISPECIES: hydrogenase 2 small subunit [Yersinia] Length: 371\nNumber of predicted TMHs: 1\nExp number of AAs in TMHs: 19.43348\nExp number, first 60 AAs: 0.16654\nTotal prob of N-in: 0.02901\noutside 1 327\nTMhelix 328 350\ninside 351 371

28666 GCF_001166625.1_4821_6_10 Yersinia intermedia Proteobacteria; Gammaproteobacteria; Enterobacterales; Yersiniaceae; Yersinia MLSQHGVNRRDFMKLCAALAATMGLSSKAAAEIANTVGSPQRPPVIWIGAQECTGCTESL WP_005188113.1 MULTISPECIES: hydrogenase 2 small subunit [Yersinia] Length: 371\nNumber of predicted TMHs: 1\nExp number of AAs in TMHs: 19.43348\nExp number, first 60 AAs: 0.16654\nTotal prob of N-in: 0.02901\noutside 1 327\nTMhelix 328 350\ninside 351 371

28667 GCF_001244565.1_5139_2_5Yersinia intermedia Proteobacteria; Gammaproteobacteria; Enterobacterales; Yersiniaceae; Yersinia MLSQHGVNRRDFMKLCAALAATMGLSSKAAAEIADTVGSPQRPPVIWIGAQECTGCTESL WP_049610771.1 MULTISPECIES: hydrogenase 2 small subunit [Yersinia] Length: 371\nNumber of predicted TMHs: 1\nExp number of AAs in TMHs: 17.72326\nExp number, first 60 AAs: 0.1354\nTotal prob of N-in: 0.02052\noutside 1 327\nTMhelix 328 350\ninside 351 371

28668 GCF_002188995.1_ASM218899v1 Yersinia intermedia Proteobacteria; Gammaproteobacteria; Enterobacterales; Yersiniaceae; Yersinia MLSQHGVNRRDFMKLCAALAATMGLSSKAAAEIANTVGSPQRPPVIWIGAQECTGCTESL WP_005188113.1 MULTISPECIES: hydrogenase 2 small subunit [Yersinia] Length: 371\nNumber of predicted TMHs: 1\nExp number of AAs in TMHs: 19.43348\nExp number, first 60 AAs: 0.16654\nTotal prob of N-in: 0.02901\noutside 1 327\nTMhelix 328 350\ninside 351 371

28669 GCF_001085865.1_4821_5_6Yersinia kristensenii Proteobacteria; Gammaproteobacteria; Enterobacterales; Yersiniaceae; Yersinia

MTGEKNMLSQHGVNRRDFMKLCAALAATMGLSSKAAAEIASIVSSPQRPPVIWIGAQECT WP_072076348.1 hydrogenase 2 small subunit [Yersinia kristensenii] Length: 377\nNumber of predicted TMHs: 1\nExp number of AAs in TMHs: 21.8128\nExp number, first 60 AAs: 3.25632\nTotal prob of N-in: 0.15090\noutside 1 333\nTMhelix 334 356\ninside 357 377

28670 GCF_000834865.1_ASM83486v1 Yersinia kristensenii Proteobacteria; Gammaproteobacteria; Enterobacterales; Yersiniaceae; Yersinia MTGEKNMLSQHGVNRRDFMKLCAALAATMGLSSKAAAEIASTVSSPQRPPVIWIGAQECT WP_071841774.1 MULTISPECIES: hydrogenase 2 small subunit [Yersinia] Length: 377\nNumber of predicted TMHs: 1\nExp number of AAs in TMHs: 19.63724\nExp number, first 60 AAs: 1.13949\nTotal prob of N-in: 0.05911\noutside 1 333\nTMhelix 334 356\ninside 357 377

28671 GCF_002188895.1_ASM218889v1 Yersinia kristensenii Proteobacteria; Gammaproteobacteria; Enterobacterales; Yersiniaceae; Yersinia MLSQHGVNRRDFMKLCAALAATMGLSSKAAAEIANTVTSPQRPPVIWIGAQECTGCTESL WP_087768347.1 hydrogenase 2 small subunit [Yersinia kristensenii] Length: 371\nNumber of predicted TMHs: 1\nExp number of AAs in TMHs: 19.03378\nExp number, first 60 AAs: 0.15058\nTotal prob of N-in: 0.02082\noutside 1 327\nTMhelix 328 350\ninside 351 371

28672 GCF_001134585.1_4821_5_7Yersinia kristensenii Proteobacteria; Gammaproteobacteria; Enterobacterales; Yersiniaceae; Yersinia MTGEKNMLSQHGVNRRDFMKLCAALAATMGLSSKAAAEIASIVSSPQRPPVIWIGAQECT WP_072076348.1 hydrogenase 2 small subunit [Yersinia kristensenii] Length: 377\nNumber of predicted TMHs: 1\nExp number of AAs in TMHs: 21.8128\nExp number, first 60 AAs: 3.25632\nTotal prob of N-in: 0.15090\noutside 1 333\nTMhelix 334 356\ninside 357 377

28673 GCF_001115185.1_5139_3_1Yersinia kristensenii Proteobacteria; Gammaproteobacteria; Enterobacterales; Yersiniaceae; Yersinia MTGEKNMLSQHGVNRRDFMKLCAALAATMGLSSKAAAEIASTVSSPQRPPVIWIGAQECT WP_071840476.1 MULTISPECIES: hydrogenase 2 small subunit [Yersinia] Length: 377\nNumber of predicted TMHs: 1\nExp number of AAs in TMHs: 18.66557\nExp number, first 60 AAs: 1.14034\nTotal prob of N-in: 0.06014\noutside 1 333\nTMhelix 334 356\ninside 357 377

28674 GCF_001319625.1_4821_8_9Yersinia kristensenii Proteobacteria; Gammaproteobacteria; Enterobacterales; Yersiniaceae; Yersinia MTGEKNMLSQHGVNRRDFMKLCAALAATMGLSSKAAAEIASIVSSPQRPPVIWIGAQECT WP_072076348.1 hydrogenase 2 small subunit [Yersinia kristensenii] Length: 377\nNumber of predicted TMHs: 1\nExp number of AAs in TMHs: 21.8128\nExp number, first 60 AAs: 3.25632\nTotal prob of N-in: 0.15090\noutside 1 333\nTMhelix 334 356\ninside 357 377

28675 GCF_001144805.1_4821_5_12 Yersinia kristensenii Proteobacteria; Gammaproteobacteria; Enterobacterales; Yersiniaceae; Yersinia MTGEKNMLSQHGVNRRDFMKLCAALAATMGLSSKAAAEIASIVSSPQRPPVIWIGAQECT WP_072076348.1 hydrogenase 2 small subunit [Yersinia kristensenii] Length: 377\nNumber of predicted TMHs: 1\nExp number of AAs in TMHs: 21.8128\nExp number, first 60 AAs: 3.25632\nTotal prob of N-in: 0.15090\noutside 1 333\nTMhelix 334 356\ninside 357 377

28676 GCF_001158505.1_5139_5_5Yersinia kristensenii Proteobacteria; Gammaproteobacteria; Enterobacterales; Yersiniaceae; Yersinia MTGEKNMLSQHGVNRRDFMKLCAALAATMGLSSKAAAEIASTVSSPQRPPVIWIGAQECT WP_072089171.1 hydrogenase 2 small subunit [Yersinia kristensenii] Length: 377\nNumber of predicted TMHs: 1\nExp number of AAs in TMHs: 19.69673\nExp number, first 60 AAs: 1.13954\nTotal prob of N-in: 0.05909\noutside 1 333\nTMhelix 334 356\ninside 357 377

28677 GCF_002188915.1_ASM218891v1 Yersinia kristensenii Proteobacteria; Gammaproteobacteria; Enterobacterales; Yersiniaceae; Yersinia MTGEKNMLSQHGVNRRDFMKLCAALAATMGLSSKAAAEIASIVSSPQRPPVIWIGAQECT WP_087795668.1 hydrogenase 2 small subunit [Yersinia kristensenii] Length: 377\nNumber of predicted TMHs: 1\nExp number of AAs in TMHs: 21.82261\nExp number, first 60 AAs: 3.25499\nTotal prob of N-in: 0.15131\noutside 1 333\nTMhelix 334 356\ninside 357 377

28678	GCF_001153365.1_5139_5_10	Yersinia kristensenii	Proteobacteria; Gammaproteobacteria; Enterobacterales; Yersiniaceae; Yersinia
MTGEKNMLSQHGVNRRDFMKLCAALAATMGLSSKAAAEIASIVSSPQRPPVIWIGAQECT	WP_072076348.1
hydrogenase 2 small subunit [Yersinia kristensenii]	Length: 377\nNumber of predicted TMHs: 1\nExp number of AAs in TMHs: 21.8128\nExp number, first 60 AAs: 3.25632\nTotal prob of N-in: 0.15090\noutside 1 333\nTMhelix 334 356\ninside 357 377

28679	GCF_001091065.1_4821_5_4Yersinia kristensenii	Proteobacteria; Gammaproteobacteria; Enterobacterales; Yersiniaceae; Yersinia
MTGEKNMLSQHGVNRRDFMKLCAALAATMGLSSKAAAEIASIVSSPQRPPVIWIGAQECT	WP_072076348.1
hydrogenase 2 small subunit [Yersinia kristensenii]	Length: 377\nNumber of predicted TMHs: 1\nExp number of AAs in TMHs: 21.8128\nExp number, first 60 AAs: 3.25632\nTotal prob of N-in: 0.15090\noutside 1 333\nTMhelix 334 356\ninside 357 377

28680	GCF_001244605.1_5139_2_4Yersinia kristensenii	Proteobacteria; Gammaproteobacteria; Enterobacterales; Yersiniaceae; Yersinia
MTGEKNMLSQHGVNRRDFMKLCAALAATMGLSSKAAAEIASTVSSPQRPPVIWIGAQECT	WP_071841774.1
MULTISPECIES: hydrogenase 2 small subunit [Yersinia]	Length: 377\nNumber of predicted TMHs: 1\nExp number of AAs in TMHs: 19.63724\nExp number, first 60 AAs: 1.13949\nTotal prob of N-in: 0.05911\noutside 1 333\nTMhelix 334 356\ninside 357 377

28681	GCF_000750355.1_ASM75035v1	Yersinia kristensenii	Proteobacteria; Gammaproteobacteria; Enterobacterales; Yersiniaceae; Yersinia
MTGEKNMLSQHGVNRRDFMKLCAALAATMGLSSKAAAEIASTVSSPQRPPVIWIGAQECT	WP_071841774.1
MULTISPECIES: hydrogenase 2 small subunit [Yersinia]	Length: 377\nNumber of predicted TMHs: 1\nExp number of AAs in TMHs: 19.63724\nExp number, first 60 AAs: 1.13949\nTotal prob of N-in: 0.05911\noutside 1 333\nTMhelix 334 356\ninside 357 377

28682	GCF_001142305.1_4840_6_3Yersinia massiliensis	Proteobacteria; Gammaproteobacteria; Enterobacterales; Yersiniaceae; Yersinia
MLSQHGVNRRDFMKLCAALAATMGLSSKAAAEIADTVGSPQRPPVIWIGAQECTGCTESL	WP_049610771.1
MULTISPECIES: hydrogenase 2 small subunit [Yersinia]	Length: 371\nNumber of predicted TMHs: 1\nExp number of AAs in TMHs: 17.72326\nExp number, first 60 AAs: 0.1354\nTotal prob of N-in: 0.02052\noutside 1 327\nTMhelix 328 350\ninside 351 371

28683	GCF_001222925.1_4821_7_3Yersinia mollaretii	Proteobacteria; Gammaproteobacteria; Enterobacterales; Yersiniaceae; Yersinia	MLSQHGVNRRDFMKLCAALAATMGLSSKAAAEIANTVSSPQRPPVIWIGAQECTGCTESL
WP_004877360.1	hydrogenase 2 small subunit [Yersinia mollaretii] Length: 371\nNumber of predicted TMHs: 1\nExp number of AAs in TMHs: 17.39821\nExp number, first 60 AAs: 0.14649\nTotal prob of N-in: 0.02220\noutside 1 327\nTMhelix 328 350\ninside 351 371

28684	GCF_001218645.1_4821_7_4Yersinia mollaretii	Proteobacteria; Gammaproteobacteria; Enterobacterales; Yersiniaceae; Yersinia	MLSQHGVNRRDFMKLCAALAATMGLSSKAAAEIANTVSSPQRPPVIWIGAQECTGCTESL
WP_049679370.1	hydrogenase 2 small subunit [Yersinia mollaretii] Length: 371\nNumber of predicted TMHs: 1\nExp number of AAs in TMHs: 16.2166\nExp number, first 60 AAs: 0.14616\nTotal prob of N-in: 0.02405\noutside 1 327\nTMhelix 328 350\ninside 351 371

28685	GCF_001108385.1_4840_7_3Yersinia mollaretii	Proteobacteria; Gammaproteobacteria; Enterobacterales; Yersiniaceae; Yersinia	MLSQHGVNRRDFMKLCAALAATMGLSSKAAAEIANTVSSPQRPSVIWIGAQECTGCTESL
WP_049648612.1	hydrogenase 2 small subunit [Yersinia mollaretii] Length: 371\nNumber of predicted TMHs: 1\nExp number of AAs in TMHs: 16.21007\nExp number, first 60 AAs: 0.1396\nTotal prob of N-in: 0.02375\noutside 1 327\nTMhelix 328 350\ninside 351 371

28686	GCF_001220765.1_4821_7_10	Yersinia mollaretii	Proteobacteria; Gammaproteobacteria; Enterobacterales; Yersiniaceae; Yersinia
MLSQHGVNRRDFMKLCAALAATMGLSSKAAAEIANTVSSPQRPPVIWIGAQECTGCTESL	WP_004877360.1
hydrogenase 2 small subunit [Yersinia mollaretii] Length: 371\nNumber of predicted TMHs: 1\nExp number of AAs in TMHs: 17.39821\nExp number, first 60 AAs: 0.14649\nTotal prob of N-in: 0.02220\noutside 1 327\nTMhelix 328 350\ninside 351 371

28687 GCF_001149645.1_4821_8_10 Yersinia mollaretii Proteobacteria; Gammaproteobacteria; Enterobacterales; Yersiniaceae; Yersinia MLSQHGVNRRDFMKLCAALAATMGLSSKAAAEIANTVSSPQRPPVIWIGAQECTGCTESL WP_049612764.1 hydrogenase 2 small subunit [Yersinia mollaretii] Length: 371\nNumber of predicted TMHs: 1\nExp number of AAs in TMHs: 16.01105\nExp number, first 60 AAs: 0.14611\nTotal prob of N-in: 0.02430\noutside 1 327\nTMhelix 328 350\ninside 351 371

28688 GCF_001127065.1_5139_5_7Yersinia mollaretii Proteobacteria; Gammaproteobacteria; Enterobacterales; Yersiniaceae; Yersinia MLSQHGVNRRDFMKLCAALAATMGLSSKAAAEIANTVSSPQRPPVIWIGAQECTGCTESL WP_049679370.1 hydrogenase 2 small subunit [Yersinia mollaretii] Length: 371\nNumber of predicted TMHs: 1\nExp number of AAs in TMHs: 16.2166\nExp number, first 60 AAs: 0.14616\nTotal prob of N-in: 0.02405\noutside 1 327\nTMhelix 328 350\ninside 351 371

28689 GCF_001319605.1_4821_6_11 Yersinia mollaretii Proteobacteria; Gammaproteobacteria; Enterobacterales; Yersiniaceae; Yersinia MLSQHGVNRRDFMKLCAALAATMGLSSKAAAEIASKVSAPQRPPVIWIGAQECTGCTESL WP_050143758.1 MULTISPECIES: hydrogenase 2 small subunit [Yersinia] Length: 371\nNumber of predicted TMHs: 1\nExp number of AAs in TMHs: 16.08592\nExp number, first 60 AAs: 0.22113\nTotal prob of N-in: 0.02765\noutside 1 327\nTMhelix 328 350\ninside 351 371

28690 GCF_001220845.1_4840_8_8Yersinia mollaretii Proteobacteria; Gammaproteobacteria; Enterobacterales; Yersiniaceae; Yersinia MLSQHGVNRRDFMKLCAALAATMGLSSKAAAEIASKVSAPQRPPVIWIGAQECTGCTESL WP_050143758.1 MULTISPECIES: hydrogenase 2 small subunit [Yersinia] Length: 371\nNumber of predicted TMHs: 1\nExp number of AAs in TMHs: 16.08592\nExp number, first 60 AAs: 0.22113\nTotal prob of N-in: 0.02765\noutside 1 327\nTMhelix 328 350\ninside 351 371

28691 GCF_001244635.1_8016_2_92 Yersinia pekkanenii Proteobacteria; Gammaproteobacteria; Enterobacterales; Yersiniaceae; Yersinia MLSQHGVNRRDFMKLCAALAATMGLSSRAAAEIANTVSSPQRPPVIWIGAQECTGCTESL WP_049611356.1 hydrogenase 2 small subunit [Yersinia pekkanenii] Length: 371\nNumber of predicted TMHs: 1\nExp number of AAs in TMHs: 18.84792\nExp number, first 60 AAs: 0.17222\nTotal prob of N-in: 0.02206\noutside 1 327\nTMhelix 328 350\ninside 351 371

28692 GCF_001152565.1_4821_6_12 Yersinia pekkanenii Proteobacteria; Gammaproteobacteria; Enterobacterales; Yersiniaceae; Yersinia MLSQHGVNRRDFMKLCAALAATMGLSSRAAAEIANTVSSPQRPPVIWIGAQECTGCTESL WP_049611356.1 hydrogenase 2 small subunit [Yersinia pekkanenii] Length: 371\nNumber of predicted TMHs: 1\nExp number of AAs in TMHs: 18.84792\nExp number, first 60 AAs: 0.17222\nTotal prob of N-in: 0.02206\noutside 1 327\nTMhelix 328 350\ninside 351 371

28693 GCF_001217805.1_4840_8_6Yersinia rohdei Proteobacteria; Gammaproteobacteria; Enterobacterales; Yersiniaceae; Yersinia MLSQHGVNRRDFMKLCAALAATMGLSSKAAAEIATTVSSPARPPVIWIGAQECTGCTESL WP_004716851.1 hydrogenase 2 small subunit [Yersinia rohdei] Length: 371\nNumber of predicted TMHs: 1\nExp number of AAs in TMHs: 19.43087\nExp number, first 60 AAs: 0.19417\nTotal prob of N-in: 0.02626\noutside 1 327\nTMhelix 328 350\ninside 351 371

28694 GCF_001152185.1_5139_5_9Yersinia rohdei Proteobacteria; Gammaproteobacteria; Enterobacterales; Yersiniaceae; Yersinia MLSQHGVNRRDFMKLCAALAATMGLSSKAAAEIATTVSSPARPPVIWIGAQECTGCTESL WP_004716851.1 hydrogenase 2 small subunit [Yersinia rohdei] Length: 371\nNumber of predicted TMHs: 1\nExp number of AAs in TMHs: 19.43087\nExp number, first 60 AAs: 0.19417\nTotal prob of N-in: 0.02626\noutside 1 327\nTMhelix 328 350\ninside 351 371

28695 GCF_001320025.1_4840_6_1Yersinia rohdei Proteobacteria; Gammaproteobacteria; Enterobacterales; Yersiniaceae; Yersinia MLSQHGVNRRDFMKLCAALAATMGLSSKAAAEIATTVSSPARPPVIWIGAQECTGCTESL WP_004716851.1 hydrogenase 2 small subunit [Yersinia rohdei] Length: 371\nNumber of predicted TMHs: 1\nExp number of AAs in TMHs: 19.43087\nExp number, first 60 AAs: 0.19417\nTotal prob of N-in: 0.02626\noutside 1 327\nTMhelix 328 350\ninside 351 371

28696 GCF_001223385.1_4821_7_2Yersinia rohdei Proteobacteria; Gammaproteobacteria; Enterobacterales; Yersiniaceae; Yersinia MLSQHGVNRRDFMKLCAALAATMGLSSKAAAEIATTVSSPARPPVIWIGAQECTGCTESL WP_004716851.1 hydrogenase 2 small subunit [Yersinia rohdei] Length: 371\nNumber of predicted TMHs:

1\nExp number of AAs in TMHs: 19.43087\nExp number, first 60 AAs: 0.19417\nTotal prob of N-in: 0.02626\noutside 1 327\nTMhelix 328 350\ninside 351 371

28697 GCF_000834455.1_ASM83445v1 Yersinia rohdei Proteobacteria; Gammaproteobacteria; Enterobacterales; Yersiniaceae; Yersinia MLSQHGVNRRDFMKLCAALAATMGLSSKAAAEIATTVSSPARPPVIWIGAQECTGCTESL WP_004716851.1 hydrogenase 2 small subunit [Yersinia rohdei] Length: 371\nNumber of predicted TMHs: 1\nExp number of AAs in TMHs: 19.43087\nExp number, first 60 AAs: 0.19417\nTotal prob of N-in: 0.02626\noutside 1 327\nTMhelix 328 350\ninside 351 371

28698 GCF_002073315.1_ASM207331v1 Yersinia sp. FDAARGOS_228 Proteobacteria; Gammaproteobacteria; Enterobacterales; Yersiniaceae; Yersinia MLSQHGVNRRDFMKLCAALAATMGLSSKAAAEIANTVGSPQRPPVIWIGAQECTGCTESL WP_005188113.1 MULTISPECIES: hydrogenase 2 small subunit [Yersinia] Length: 371\nNumber of predicted TMHs: 1\nExp number of AAs in TMHs: 19.43348\nExp number, first 60 AAs: 0.16654\nTotal prob of N-in: 0.02901\noutside 1 327\nTMhelix 328 350\ninside 351 371

28699 GCF_000834395.1_ASM83439v1 Yersinia aldovae 670-83 Proteobacteria; Gammaproteobacteria; Enterobacterales; Yersiniaceae; Yersinia; Yersinia aldovae MTGEKNMLSQHGVNRRDFMKLCAALAATMGLSSKAVAEIANTVSSPQRPPVIWIGAQECT WP_071841507.1 hydrogenase 2 small subunit [Yersinia aldovae] Length: 377\nNumber of predicted TMHs: 1\nExp number of AAs in TMHs: 18.55284\nExp number, first 60 AAs: 1.0683\nTotal prob of N-in: 0.05755\noutside 1 333\nTMhelix 334 356\ninside 357 377

28700 GCF_000173735.1_ASM17373v1 Yersinia aldovae ATCC 35236 Proteobacteria; Gammaproteobacteria; Enterobacterales; Yersiniaceae; Yersinia; Yersinia aldovae MLSQHGVNRRDFMKLCAALAATMGLSSKAAAEIANTVSSPQRPPVIWIGAQECTGCTESL WP_004703028.1 hydrogenase 2 small subunit [Yersinia aldovae] Length: 371\nNumber of predicted TMHs: 1\nExp number of AAs in TMHs: 17.64702\nExp number, first 60 AAs: 0.1462\nTotal prob of N-in: 0.02153\noutside 1 327\nTMhelix 328 350\ninside 351 371

28701 GCF_000167975.1_ASM16797v1 Yersinia bercovieri ATCC 43970 Proteobacteria; Gammaproteobacteria; Enterobacterales; Yersiniaceae; Yersinia; Yersinia bercovieri MLSQHGVNRRDFMKLCAALAATMGLSSKAAAEIANTVSSPQRPPVIWIGAQECTGCTESL WP_005275305.1 hydrogenase 2 small subunit [Yersinia bercovieri]Length: 371\nNumber of predicted TMHs: 1\nExp number of AAs in TMHs: 17.6704\nExp number, first 60 AAs: 0.14651\nTotal prob of N-in: 0.02137\noutside 1 327\nTMhelix 328 350\ninside 351 371

28702 GCF_000009345.1_ASM934v1 Yersinia enterocolitica subsp. enterocolitica 8081Proteobacteria; Gammaproteobacteria; Enterobacterales; Yersiniaceae; Yersinia; Yersinia enterocolitica; Yersinia enterocolitica subsp. enterocolitica MLSQHGVNRRDFMKLCAALAATMGLSSKAAAEIANTVSSPQRPPVIWIGAQECTGCTESL YP_001007770.1 hydrogenase 2 small subunit [Yersinia enterocolitica subsp. enterocolitica 8081] Length: 371\nNumber of predicted TMHs: 1\nExp number of AAs in TMHs: 18.61508\nExp number, first 60 AAs: 0.14605\nTotal prob of N-in: 0.02327\noutside 1 327\nTMhelix 328 350\ninside 351 371

28703 GCF_001598735.1_ASM159873v1 Yersinia enterocolitica subsp. enterocolitica NBRC 105693 Proteobacteria; Gammaproteobacteria; Enterobacterales; Yersiniaceae; Yersinia; Yersinia enterocolitica; Yersinia enterocolitica subsp. enterocolitica MLSQHGVNRRDFMKLCAALAATMGLSSKAAAEIANTVSSPQRPPVIWIGAQECTGCTESL WP_005173957.1 hydrogenase 2 small subunit [Yersinia enterocolitica] Length: 371\nNumber of predicted TMHs: 1\nExp number of AAs in TMHs: 18.61508\nExp number, first 60 AAs: 0.14605\nTotal prob of N-in: 0.02327\noutside 1 327\nTMhelix 328 350\ninside 351 371

28704 GCF_000297175.1_ASM29717v1 Yersinia enterocolitica subsp. enterocolitica WA-314 Proteobacteria; Gammaproteobacteria; Enterobacterales; Yersiniaceae; Yersinia; Yersinia enterocolitica; Yersinia enterocolitica subsp. enterocolitica MLSQHGVNRRDFMKLCAALAATMGLSSKAAAEIANTVSSPQRPPVIWIGAQECTGCTESL WP_005173957.1 hydrogenase 2 small subunit [Yersinia enterocolitica] Length: 371\nNumber of predicted TMHs: 1\nExp number of AAs in TMHs: 18.61508\nExp number, first 60 AAs: 0.14605\nTotal prob of N-in: 0.02327\noutside 1 327\nTMhelix 328 350\ninside 351 371

28705 GCF_000192105.1_ASM19210v1 Yersinia enterocolitica subsp. palearctica 105.5R(r) Proteobacteria; Gammaproteobacteria; Enterobacterales; Yersiniaceae; Yersinia; Yersinia enterocolitica; Yersinia enterocolitica subsp. palearctica MLSQHGVNRRDFMKLCAALAATMGLSSKAAAEIANTVSSPQRPPVIWIGAQECTGCTESL WP_005157252.1 hydrogenase 2 small subunit [Yersinia enterocolitica] Length: 371\nNumber of predicted TMHs: 1\nExp number of AAs in TMHs: 18.80217\nExp number, first 60 AAs: 0.14643\nTotal prob of N-in: 0.01943\noutside 1 327\nTMhelix 328 350\ninside 351 371

28706 GCF_000284935.1_ASM28493v1 Yersinia enterocolitica subsp. palearctica 556/8265 Proteobacteria; Gammaproteobacteria; Enterobacterales; Yersiniaceae; Yersinia; Yersinia enterocolitica; Yersinia enterocolitica subsp. palearctica MLSQHGVNRRDFMKLCAALAATMGLSSKAAAEIANTVSSPQRPPVIWIGAQECTGCTESL WP_005157252.1 hydrogenase 2 small subunit [Yersinia enterocolitica] Length: 371\nNumber of predicted TMHs: 1\nExp number of AAs in TMHs: 18.80217\nExp number, first 60 AAs: 0.14643\nTotal prob of N-in: 0.01943\noutside 1 327\nTMhelix 328 350\ninside 351 371

28707 GCF_000284955.1_ASM28495v1 Yersinia enterocolitica subsp. palearctica 647/5307 Proteobacteria; Gammaproteobacteria; Enterobacterales; Yersiniaceae; Yersinia; Yersinia enterocolitica; Yersinia enterocolitica subsp. palearctica MLSQHGVNRRDFMKLCAALAATMGLSSKAAAEIANTVSSPQRPPVIWIGAQECTGCTESL WP_005157252.1 hydrogenase 2 small subunit [Yersinia enterocolitica] Length: 371\nNumber of predicted TMHs: 1\nExp number of AAs in TMHs: 18.80217\nExp number, first 60 AAs: 0.14643\nTotal prob of N-in: 0.01943\noutside 1 327\nTMhelix 328 350\ninside 351 371

28708 GCF_000230775.1_ASM23077v2 Yersinia enterocolitica subsp. palearctica PhRBD_Ye1 Proteobacteria; Gammaproteobacteria; Enterobacterales; Yersiniaceae; Yersinia; Yersinia enterocolitica; Yersinia enterocolitica subsp. palearctica MLSQHGVNRRDFMKLCAALAATMGLSSKAAAEIANTVSSPQRPPVIWIGAQECTGCTESL WP_005157252.1 hydrogenase 2 small subunit [Yersinia enterocolitica] Length: 371\nNumber of predicted TMHs: 1\nExp number of AAs in TMHs: 18.80217\nExp number, first 60 AAs: 0.14643\nTotal prob of N-in: 0.01943\noutside 1 327\nTMhelix 328 350\ninside 351 371

28709 GCF_000253175.1_ASM25317v1 Yersinia enterocolitica subsp. palearctica Y11 Proteobacteria; Gammaproteobacteria; Enterobacterales; Yersiniaceae; Yersinia; Yersinia enterocolitica; Yersinia enterocolitica subsp. palearctica MLSQHGVNRRDFMKLCAALAATMGLSSKAAAEIANTVSSPQRPPVIWIGAQECTGCTESL WP_005157252.1 hydrogenase 2 small subunit [Yersinia enterocolitica] Length: 371\nNumber of predicted TMHs: 1\nExp number of AAs in TMHs: 18.80217\nExp number, first 60 AAs: 0.14643\nTotal prob of N-in: 0.01943\noutside 1 327\nTMhelix 328 350\ninside 351 371

28710 GCF_000401935.1_YE-149_1.0 Yersinia enterocolitica subsp. palearctica YE-149 Proteobacteria; Gammaproteobacteria; Enterobacterales; Yersiniaceae; Yersinia; Yersinia enterocolitica; Yersinia enterocolitica subsp. palearctica MLSQHGVNRRDFMKLCAALAATMGLSSKAAAEIANTVSSPQRPPVIWIGAQECTGCTESL WP_005157252.1 hydrogenase 2 small subunit [Yersinia enterocolitica] Length: 371\nNumber of predicted TMHs: 1\nExp number of AAs in TMHs: 18.80217\nExp number, first 60 AAs: 0.14643\nTotal prob of N-in: 0.01943\noutside 1 327\nTMhelix 328 350\ninside 351 371

28711 GCF_000401995.1_YE-150_1.0 Yersinia enterocolitica subsp. palearctica YE-150 Proteobacteria; Gammaproteobacteria; Enterobacterales; Yersiniaceae; Yersinia; Yersinia enterocolitica; Yersinia enterocolitica subsp. palearctica MLSQHGVNRRDFMKLCAALAATMGLSSKAAAEIANTVSSPQRPPVIWIGAQECTGCTESL WP_005157252.1 hydrogenase 2 small subunit [Yersinia enterocolitica] Length: 371\nNumber of predicted TMHs: 1\nExp number of AAs in TMHs: 18.80217\nExp number, first 60 AAs: 0.14643\nTotal prob of N-in: 0.01943\noutside 1 327\nTMhelix 328 350\ninside 351 371

28712 GCF_000401975.1_YE-P1_1.0 Yersinia enterocolitica subsp. palearctica YE-P1 Proteobacteria; Gammaproteobacteria; Enterobacterales; Yersiniaceae; Yersinia; Yersinia enterocolitica; Yersinia enterocolitica subsp. palearctica MLSQHGVNRRDFMKLCAALAATMGLSSKAAAEIANTVSSPQRPPVIWIGAQECTGCTESL WP_005157252.1 hydrogenase 2 small subunit [Yersinia enterocolitica] Length: 371\nNumber of predicted TMHs: 1\nExp number of AAs in TMHs: 18.80217\nExp number, first 60 AAs: 0.14643\nTotal prob of N-in: 0.01943\noutside 1 327\nTMhelix 328 350\ninside 351 371

28713 GCF_000401955.1_YE-P4_1.0 Yersinia enterocolitica subsp. palearctica YE-P4 Proteobacteria; Gammaproteobacteria; Enterobacterales; Yersiniaceae; Yersinia; Yersinia enterocolitica; Yersinia enterocolitica subsp. palearctica MLSQHGVNRRDFMKLCAALAATMGLSSKAAAEIANTVSSPQRPPVIWIGAQECTGCTESL WP_005157252.1 hydrogenase 2 small subunit [Yersinia enterocolitica] Length: 371\nNumber of predicted TMHs: 1\nExp number of AAs in TMHs: 18.80217\nExp number, first 60 AAs: 0.14643\nTotal prob of N-in: 0.01943\noutside 1 327\nTMhelix 328 350\ninside 351 371

28714 GCF_000284975.1_ASM28497v1 Yersinia enterocolitica subsp. palearctica YO527 Proteobacteria; Gammaproteobacteria; Enterobacterales; Yersiniaceae; Yersinia; Yersinia enterocolitica; Yersinia enterocolitica subsp. palearctica MLSQHGVNRRDFMKLCAALAATMGLSSKAAAEIANTVSSPQRPPVIWIGAQECTGCTESL WP_005157252.1 hydrogenase 2 small subunit [Yersinia enterocolitica] Length: 371\nNumber of predicted TMHs: 1\nExp number of AAs in TMHs: 18.80217\nExp number, first 60 AAs: 0.14643\nTotal prob of N-in: 0.01943\noutside 1 327\nTMhelix 328 350\ninside 351 371

28715 GCF_000330605.1_ASM33060v1 Yersinia enterocolitica IP 10393 Proteobacteria; Gammaproteobacteria; Enterobacterales; Yersiniaceae; Yersinia; Yersinia enterocolitica; Yersinia enterocolitica (type O MLSQHGVNRRDFMKLCAALAATMGLSSKAAAEIANTVSSPQRPPVIWIGAQECTGCTESL WP_005157252.1 hydrogenase 2 small subunit [Yersinia enterocolitica] Length: 371\nNumber of predicted TMHs: 1\nExp number of AAs in TMHs: 18.80217\nExp number, first 60 AAs: 0.14643\nTotal prob of N-in: 0.01943\noutside 1 327\nTMhelix 328 350\ninside 351 371

28716 GCF_000285015.1_ASM28501v1 Yersinia enterocolitica IP2222 Proteobacteria; Gammaproteobacteria; Enterobacterales; Yersiniaceae; Yersinia; Yersinia enterocolitica; Yersinia enterocolitica (type O MLSQHGVNRRDFMKLCAALAATMGLSSKAAAEIANTVSSPQRPPVIWIGAQECTGCTESL WP_019079071.1 hydrogenase 2 small subunit [Yersinia enterocolitica] Length: 371\nNumber of predicted TMHs: 1\nExp number of AAs in TMHs: 18.80245\nExp number, first 60 AAs: 0.14643\nTotal prob of N-in: 0.01944\noutside 1 327\nTMhelix 328 350\ninside 351 371

28717 GCF_000284995.1_ASM28499v1 Yersinia enterocolitica NFO Proteobacteria; Gammaproteobacteria; Enterobacterales; Yersiniaceae; Yersinia; Yersinia enterocolitica; Yersinia enterocolitica (type O MLSQHGVNRRDFMKLCAALAATMGLSSKAAAEIANTVSSPQRPPVIWIGAQECTGCTESL WP_019079071.1 hydrogenase 2 small subunit [Yersinia enterocolitica] Length: 371\nNumber of predicted TMHs: 1\nExp number of AAs in TMHs: 18.80245\nExp number, first 60 AAs: 0.14643\nTotal prob of N-in: 0.01944\noutside 1 327\nTMhelix 328 350\ninside 351 371

28718 GCF_001050795.1_ASM105079v1 Yersinia enterocolitica (type O:3) str. YE12/03 Proteobacteria; Gammaproteobacteria; Enterobacterales; Yersiniaceae; Yersinia; Yersinia enterocolitica; Yersinia enterocolitica (type O MLSQHGVNRRDFMKLCAALAATMGLSSKAAAEIANTVSSPQRPPVIWIGAQECTGCTESL WP_005157252.1 hydrogenase 2 small subunit [Yersinia enterocolitica] Length: 371\nNumber of predicted TMHs: 1\nExp number of AAs in TMHs: 18.80217\nExp number, first 60 AAs: 0.14643\nTotal prob of N-in: 0.01943\noutside 1 327\nTMhelix 328 350\ninside 351 371

28719 GCF_001050775.1_ASM105077v1 Yersinia enterocolitica (type O:5,27) str. YE149/02 Proteobacteria; Gammaproteobacteria; Enterobacterales; Yersiniaceae; Yersinia; Yersinia enterocolitica; Yersinia enterocolitica (type O MLSQHGVNRRDFMKLCAALAATMGLSSKAAAEIANTVSSPQRPPVIWIGAQECTGCTESL WP_005157252.1 hydrogenase 2 small subunit [Yersinia enterocolitica] Length: 371\nNumber of predicted TMHs: 1\nExp number of AAs in TMHs: 18.80217\nExp number, first 60 AAs: 0.14643\nTotal prob of N-in: 0.01943\noutside 1 327\nTMhelix 328 350\ninside 351 371

28720 GCF_000968115.1_ASM96811v1 Yersinia enterocolitica (type O:5) str. YE53/03 Proteobacteria; Gammaproteobacteria; Enterobacterales; Yersiniaceae; Yersinia; Yersinia enterocolitica; Yersinia enterocolitica (type O MLSQHGVNRRDFMKLCAALAATMGLSSKAAAEIANTVSSPQRPPVIWIGAQECTGCTESL WP_019079071.1 hydrogenase 2 small subunit [Yersinia enterocolitica] Length: 371\nNumber of predicted TMHs: 1\nExp number of AAs in TMHs: 18.80245\nExp number, first 60 AAs: 0.14643\nTotal prob of N-in: 0.01944\noutside 1 327\nTMhelix 328 350\ninside 351 371

28721 GCF_001050735.1_ASM105073v1 Yersinia enterocolitica (type O:9) str. YE212/02 Proteobacteria; Gammaproteobacteria; Enterobacterales; Yersiniaceae; Yersinia; Yersinia enterocolitica; Yersinia enterocolitica (type O MLSQHGVNRRDFMKLCAALAATMGLSSKAAAEIANTVSSPQRPPVIWIGAQECTGCTESL WP_005157252.1 hydrogenase 2 small subunit [Yersinia enterocolitica] Length: 371\nNumber of predicted TMHs: 1\nExp number of AAs in TMHs: 18.80217\nExp number, first 60 AAs: 0.14643\nTotal prob of N-in: 0.01943\noutside 1 327\nTMhelix 328 350\ninside 351 371

28722 GCF_001050755.1_ASM105075v1 Yersinia enterocolitica (type O:9) str. YE56/03 Proteobacteria; Gammaproteobacteria; Enterobacterales; Yersiniaceae; Yersinia; Yersinia enterocolitica; Yersinia enterocolitica (type O MLSQHGVNRRDFMKLCAALAATMGLSSKAAAEIANTVSSPQRPPVIWIGAQECTGCTESL WP_005157252.1 hydrogenase 2 small subunit [Yersinia enterocolitica] Length: 371\nNumber of predicted TMHs: 1\nExp

number of AAs in TMHs: 18.80217\nExp number, first 60 AAs: 0.14643\nTotal prob of N-in: 0.01943\noutside 1 327\nTMhelix 328 350\ninside 351 371

28723 GCF_000168015.1_ASM16801v1 Yersinia frederiksenii ATCC 33641 Proteobacteria; Gammaproteobacteria; Enterobacterales; Yersiniaceae; Yersinia; Yersinia frederiksenii MLSQHGVNRRDFMKLCAALAATMGLSSKAAAEIASTVSSPQRPPVIWIGAQECTGCTESL WP_004711888.1 hydrogenase 2 small subunit [Yersinia frederiksenii] Length: 371\nNumber of predicted TMHs: 1\nExp number of AAs in TMHs: 19.47103\nExp number, first 60 AAs: 0.16503\nTotal prob of N-in: 0.02370\noutside 1 327\nTMhelix 328 350\ninside 351 371

28724 GCF_000754805.1_YFB Yersinia frederiksenii ATCC 33641 Proteobacteria; Gammaproteobacteria; Enterobacterales; Yersiniaceae; Yersinia; Yersinia frederiksenii MLSQHGVNRRDFMKLCAALAATMGLSSKAAAEIASTVSSPQRPPVIWIGAQECTGCTESL WP_004711888.1 hydrogenase 2 small subunit [Yersinia frederiksenii] Length: 371\nNumber of predicted TMHs: 1\nExp number of AAs in TMHs: 19.47103\nExp number, first 60 AAs: 0.16503\nTotal prob of N-in: 0.02370\noutside 1 327\nTMhelix 328 350\ninside 351 371

28725 GCF_000834215.1_ASM83421v1 Yersinia frederiksenii Y225 Proteobacteria; Gammaproteobacteria; Enterobacterales; Yersiniaceae; Yersinia; Yersinia frederiksenii MTGEKNMLSQHGVNRRDFMKLCAALAATMGLSSKAAAEIASTVSSPQRPPVIWIGAQECT WP_071841774.1 MULTISPECIES: hydrogenase 2 small subunit [Yersinia] Length: 377\nNumber of predicted TMHs: 1\nExp number of AAs in TMHs: 19.63724\nExp number, first 60 AAs: 1.13949\nTotal prob of N-in: 0.05911\noutside 1 333\nTMhelix 334 356\ninside 357 377

28726 GCF_000168035.1_ASM16803v1 Yersinia intermedia ATCC 29909 Proteobacteria; Gammaproteobacteria; Enterobacterales; Yersiniaceae; Yersinia; Yersinia intermedia MLSQHGVNRRDFMKLCAALAATMGLSSKAAAEIANTVGSPQRPPVIWIGAQECTGCTESL WP_005188113.1 MULTISPECIES: hydrogenase 2 small subunit [Yersinia] Length: 371\nNumber of predicted TMHs: 1\nExp number of AAs in TMHs: 19.43348\nExp number, first 60 AAs: 0.16654\nTotal prob of N-in: 0.02901\noutside 1 327\nTMhelix 328 350\ninside 351 371

28727 GCF_000167995.1_ASM16799v1 Yersinia mollaretii ATCC 43969 Proteobacteria; Gammaproteobacteria; Enterobacterales; Yersiniaceae; Yersinia; Yersinia mollaretii MLSQHGVNRRDFMKLCAALAATMGLSSKAAAEIANTVSSPQRPPVIWIGAQECTGCTESL WP_004877360.1 hydrogenase 2 small subunit [Yersinia mollaretii] Length: 371\nNumber of predicted TMHs: 1\nExp number of AAs in TMHs: 17.39821\nExp number, first 60 AAs: 0.14649\nTotal prob of N-in: 0.02220\noutside 1 327\nTMhelix 328 350\ninside 351 371

28728 GCF_000173775.1_ASM17377v1 Yersinia rohdei ATCC 43380 Proteobacteria; Gammaproteobacteria; Enterobacterales; Yersiniaceae; Yersinia; Yersinia rohdei MLSQHGVNRRDFMKLCAALAATMGLSSKAAAEIATTVSSPARPPVIWIGAQECTGCTESL WP_004716851.1 hydrogenase 2 small subunit [Yersinia rohdei] Length: 371\nNumber of predicted TMHs: 1\nExp number of AAs in TMHs: 19.43087\nExp number, first 60 AAs: 0.19417\nTotal prob of N-in: 0.02626\noutside 1 327\nTMhelix 328 350\ninside 351 371

28729 GCF_001467615.1_ASM146761v1 Legionella erythra Proteobacteria; Gammaproteobacteria; Legionellales; Legionellaceae; Legionella MDKNIRIYAETAHALGLPAFFQPELNVLTIKLGRRSYFFHAAITPFNQGASIYLAKHKYL WP_058527013.1 hypothetical protein [Legionella erythra] Length: 360\nNumber of predicted TMHs: 1\nExp number of AAs in TMHs: 21.90514\nExp number, first 60 AAs: 1.76555\nTotal prob of N-in: 0.22936\noutside 1 334\nTMhelix 335 357\ninside 358 360

28730 GCF_000746145.1_ASM74614v1 Methylobacter sp. BBA5.1 Proteobacteria; Gammaproteobacteria; Methylococcales; Methylococcaceae; Methylobacter MSDMREINSEIIDAHGRRSVILQTIPSHQGLMKINKLLLIVVFILMTVVFILGFLLFPAH WP_020157341.1 MULTISPECIES: hypothetical protein [Methylobacter] Length: 204\nNumber of predicted TMHs: 2\nExp number of AAs in TMHs: 44.18268\nExp number, first 60 AAs: 22.42089\nTotal prob of N-in: 0.80733\nPOSSIBLE N-term signal sequence\ninside 1 36\nTMhelix 37 59\noutside 60 163\nTMhelix 164 186\ninside 187 204

28731 GCF_000745375.1_ASM74537v1 Methylobacter whittenburyi Proteobacteria; Gammaproteobacteria; Methylococcales; Methylococcaceae; Methylobacter MSDMREINSEIIDAHGRRSVILQTIPSHQGLMKINKLLLIVVFILMTVVFILGFLLFPAH WP_036295385.1 hypothetical protein [Methylobacter whittenburyi] Length: 204\nNumber of predicted TMHs: 2\nExp number of AAs in TMHs: 44.15254\nExp number, first 60 AAs: 22.41751\nTotal prob of N-in: 0.88818\nPOSSIBLE N-term signal sequence\ninside 1 36\nTMhelix 37 59\noutside 60 163\nTMhelix 164 186\ninside 187 204

28732 GCF_000383855.1_ASM38385v1 Methylobacter marinus A45 Proteobacteria; Gammaproteobacteria; Methylococcales; Methylococcaceae; Methylobacter; Methylobacter marinus MSDMREINSEIIDAHGRRSVILQTIPSHQGLMKINKLLLIVVFILMTVVFILGFLLFPAH WP_020157341.1 MULTISPECIES: hypothetical protein [Methylobacter] Length: 204\nNumber of predicted TMHs: 2\nExp number of AAs in TMHs: 44.18268\nExp number, first 60 AAs: 22.42089\nTotal prob of N-in: 0.80733\nPOSSIBLE N-term signal sequence\ninside 1 36\nTMhelix 37 59\noutside 60 163\nTMhelix 164 186\ninside 187 204

28733 GCF_002005105.1_14B Methylocaldum sp. 14B Proteobacteria; Gammaproteobacteria; Methylococcales; Methylococcaceae; Methylocaldum MVIRQESVHRFSGLASSGRELFFARRSFIHLFRPAAVLLLTFLLAACGSPLNNPYPAGDK WP_077728929.1 peptide ABC transporter substrate-binding protein [Methylocaldum sp. 14B] Length: 746\nNumber of predicted TMHs: 1\nExp number of AAs in TMHs: 38.99376\nExp number, first 60 AAs: 16.97743\nTotal prob of N-in: 0.80265\nPOSSIBLE N-term signal sequence\noutside 1 714\nTMhelix 715 737\ninside 738 746

28734 GCF_002127725.1_ASM212772v1 Methylocaldum sp. SAD2 Proteobacteria; Gammaproteobacteria; Methylococcales; Methylococcaceae; Methylocaldum MAIRQESVHRFSGPASSGRELFFARRSFIHLFRPAAVLLLTFLLAACGSPLNNPYPAGDK WP_086135304.1 peptide ABC transporter substrate-binding protein [Methylocaldum sp. SAD2] Length: 746\nNumber of predicted TMHs: 2\nExp number of AAs in TMHs: 39.28717\nExp number, first 60 AAs: 17.27293\nTotal prob of N-in: 0.81699\nPOSSIBLE N-term signal sequence\ninside 1 28\nTMhelix 29 51\noutside 52 714\nTMhelix 715 737\ninside 738 746

28735 GCF_000427385.1_ASM42738v1 Methylocaldum szegediense O-12 Proteobacteria; Gammaproteobacteria; Methylococcales; Methylococcaceae; Methylocaldum; Methylocaldum szegediense MLANHQIQPPQPITLLLRHRRKALTAFLIALSAGAITALFEKPLYEAHAALLFNLGREYQ WP_026610324.1 hypothetical protein [Methylocaldum szegediense] Length: 470\nNumber of predicted TMHs: 1\nExp number of AAs in TMHs: 32.27667\nExp number, first 60 AAs: 12.62692\nTotal prob of N-in: 0.63136\nPOSSIBLE N-term signal sequence\noutside 1 423\nTMhelix 424 441\ninside 442 470

28736 GCF_001644015.1_ASM164401v1 Methylomonas lenta Proteobacteria; Gammaproteobacteria; Methylococcales; Methylococcaceae; Methylomonas MQRGRRTTVLGLLFSTLVWAASEYSDYLAALSAKESSNNPNSMNQYGFLGRYQMGESALI WP_083960596.1 hypothetical protein [Methylomonas lenta] Length: 297\nNumber of predicted TMHs: 2\nExp number of AAs in TMHs: 63.07311\nExp number, first 60 AAs: 7.95913\nTotal prob of N-in: 0.92874\ninside 1 241\nTMhelix 242 264\noutside 265 273\nTMhelix 274 296\ninside 297 297

28737 GCF_000963695.1_Methcu1.0 Methylococcaceae bacterium Sn10-6 Proteobacteria; Gammaproteobacteria; Methylococcales; Methylococcaceae; unclassified Methylococcaceae MDRRSFIKSIGLGAVATSAPLTATTAKAASSVYKIAIVGGGFAGASIAKYLKLWGQNQVE WP_052700213.1 hypothetical protein [Methylococcaceae bacterium Sn10-6] Length: 262\nNumber of predicted TMHs: 1\nExp number of AAs in TMHs: 27.54693\nExp number, first 60 AAs: 5.3737\nTotal prob of N-in: 0.56671\noutside 1 223\nTMhelix 224 246\ninside 247 262

28738 GCF_000421465.1_ASM42146v1 Methylohalobius crimeensis 10Ki Proteobacteria; Gammaproteobacteria; Methylococcales; Methylothermaceae; Methylohalobius; Methylohalobius crimeensis MVKSRRGYFKPKWTLTALTVVAVGLFLLLARWQVHRAEEKQSLLDMNERRRQASAMVLSG WP_022948069.1 hypothetical protein [Methylohalobius crimeensis] Length: 250\nNumber of predicted TMHs: 2\nExp number of AAs in TMHs: 42.08076\nExp number, first 60 AAs: 21.51863\nTotal prob of N-in: 0.91137\nPOSSIBLE N-term signal sequence\ninside 1 12\nTMhelix 13 35\noutside 36 211\nTMhelix 212 234\ninside 235 250

28739 GCF_000711955.1_ASM71195v1 Nevskia soli DSM 19509 Proteobacteria; Gammaproteobacteria; Nevskiales; Sinobacteraceae; Nevskia; Nevskia soli MSGKSLLQKLFDGRRRSLLAGGMLLALLAAPLPSLAAWEATLSKAVTPDASAGMAPVPAA WP_084183403.1 hypothetical protein [Nevskia soli] Length: 1106\nNumber of predicted TMHs: 1\nExp number of AAs in TMHs: 32.9258299999999\nExp number, first 60 AAs: 12.27761\nTotal prob of N-in: 0.56879\nPOSSIBLE N-term signal sequence\noutside 1 1076\nTMhelix 1077 1099\ninside 1100 1106

28740 GCF_001683815.1_ASM168381v1 Alcanivorax sp. CP2C Proteobacteria; Gammaproteobacteria; Oceanospirillales; Alcanivoracaceae; Alcanivorax
MPNRNARRSLLGLALTLLVIALCLNLAWWQWQRAGEKREWLADQAAKAQAAPVSVTAALD WP_067611404.1 hypothetical protein [Alcanivorax sp. CP2C] Length: 248\nNumber of predicted TMHs: 2\nExp number of AAs in TMHs: 42.33485\nExp number, first 60 AAs: 22.37799\nTotal prob of N-in: 0.94593\nPOSSIBLE N-term signal sequence\ninside 1 8\nTMhelix 9 31\noutside 32 216\nTMhelix 217 236\ninside 237 248

28741 GCF_001635515.1_ASM163551v1 Alcanivorax sp. HI0003 Proteobacteria; Gammaproteobacteria; Oceanospirillales; Alcanivoracaceae; Alcanivorax
MSNTENRRPVLSWIAFLATLAVVAICLRLSWWQLERAEEKRLWLATQEEKARQPAAGLPA WP_063508475.1 MULTISPECIES: hypothetical protein [Alcanivorax] Length: 250\nNumber of predicted TMHs: 2\nExp number of AAs in TMHs: 42.45417\nExp number, first 60 AAs: 21.55813\nTotal prob of N-in: 0.86094\nPOSSIBLE N-term signal sequence\ninside 1 12\nTMhelix 13 32\noutside 33 219\nTMhelix 220 239\ninside 240 250

28742 GCF_001635525.1_ASM163552v1 Alcanivorax sp. HI0007 Proteobacteria; Gammaproteobacteria; Oceanospirillales; Alcanivoracaceae; Alcanivorax
MSNTENRRPVLSWIAFLATLAVVAICLRLSWWQLERAEEKRLWLATQEEKARQPAAGLPA WP_063508475.1 MULTISPECIES: hypothetical protein [Alcanivorax] Length: 250\nNumber of predicted TMHs: 2\nExp number of AAs in TMHs: 42.45417\nExp number, first 60 AAs: 21.55813\nTotal prob of N-in: 0.86094\nPOSSIBLE N-term signal sequence\ninside 1 12\nTMhelix 13 32\noutside 33 219\nTMhelix 220 239\ninside 240 250

28743 GCF_001634835.1_ASM163483v1 Alcanivorax sp. HI0033 Proteobacteria; Gammaproteobacteria; Oceanospirillales; Alcanivoracaceae; Alcanivorax
MSNTENRRPVLSWIAFLATLAVVAICLRLSWWQLERAEEKRLWLATQEEKARQPAAGLPA WP_063508475.1 MULTISPECIES: hypothetical protein [Alcanivorax] Length: 250\nNumber of predicted TMHs: 2\nExp number of AAs in TMHs: 42.45417\nExp number, first 60 AAs: 21.55813\nTotal prob of N-in: 0.86094\nPOSSIBLE N-term signal sequence\ninside 1 12\nTMhelix 13 32\noutside 33 219\nTMhelix 220 239\ninside 240 250

28744 GCF_001602235.1_ASM160223v1 Alcanivorax sp. NBRC 102024 Proteobacteria; Gammaproteobacteria; Oceanospirillales; Alcanivoracaceae; Alcanivorax
MSNTENRRPVLSWIAFLATLAVVAICLRLSWWQLERAEEKRLWLATQEEKARQPAAGLPA WP_062817430.1 hypothetical protein [Alcanivorax sp. NBRC 102024] Length: 250\nNumber of predicted TMHs: 2\nExp number of AAs in TMHs: 42.45477\nExp number, first 60 AAs: 21.55817\nTotal prob of N-in: 0.86092\nPOSSIBLE N-term signal sequence\ninside 1 12\nTMhelix 13 32\noutside 33 219\nTMhelix 220 239\ninside 240 250

28745 GCF_000012985.1_ASM1298v1 Hahella chejuensis KCTC 2396 Proteobacteria; Gammaproteobacteria; Oceanospirillales; Hahellaceae; Hahella; Hahella chejuensis
MNIEQALNAFYKESGFELETGKRPAFVEVFVGCLLIPLPNVETRRKYIKYHDLHHVITGY WP_011396790.1 hypothetical protein [Hahella chejuensis] Length: 199\nNumber of predicted TMHs: 1\nExp number of AAs in TMHs: 43.23952\nExp number, first 60 AAs: 2.42433\nTotal prob of N-in: 0.24615\noutside 1 156\nTMhelix 157 179\ninside 180 199

28746 GCF_001639695.1_ASM163969v1 Marinomonas aquimarina Proteobacteria; Gammaproteobacteria; Oceanospirillales; Oceanospirillaceae; Marinomonas
MNATAILNKILPIVSPNMHKTRRKALSACVLSLAQGSLCTVTSIGRGIQSNAYEKHRIKR WP_067213518.1 hypothetical protein [Marinomonas aquimarina] Length: 393\nNumber of predicted TMHs: 1\nExp number of AAs in TMHs: 24.56661\nExp number, first 60 AAs: 3.34039\nTotal prob of N-in: 0.23077\noutside 1 320\nTMhelix 321 343\ninside 344 393

28747 GCF_900089755.1_MaquimarinaCECT5080_SPAdes_Prokka Marinomonas aquimarina Proteobacteria; Gammaproteobacteria; Oceanospirillales; Oceanospirillaceae; Marinomonas
MNATAILNKILPIVSPNMHKTRRKALSACVLSLAQGSLCTVTSIGRGIQSNAYEKHRIKR WP_067213518.1 hypothetical protein [Marinomonas aquimarina] Length: 393\nNumber of predicted TMHs: 1\nExp number of AAs in TMHs: 24.56661\nExp number, first 60 AAs: 3.34039\nTotal prob of N-in: 0.23077\noutside 1 320\nTMhelix 321 343\ninside 344 393

28748 GCF_900129155.1_IMG-taxon_2582581270_annotated_assembly Marinomonas polaris DSM 16579 Proteobacteria; Gammaproteobacteria; Oceanospirillales; Oceanospirillaceae; Marinomonas; Marinomonas polaris MNATAILNKILPIVSPNMHKTRRNALSVCVLSLAQGNLCTVTSIGRGIQSKAYEKHRIKR WP_084122697.1 hypothetical protein [Marinomonas polaris] Length: 287\nNumber of predicted TMHs: 1\nExp number of AAs in TMHs: 24.67225\nExp number, first 60 AAs: 2.62615\nTotal prob of N-in: 0.28343\noutside 1 214\nTMhelix 215 237\ninside 238 287

28749 GCF_001709345.1_ASM170934v1 Terasakiispira papahanaumokuakeensis Proteobacteria; Gammaproteobacteria; Oceanospirillales; unclassified Oceanospirillales; Terasakiispira
MAKTNRLPADARIWITGASSGIGAAVAQRLLRQGYRLVLSGRRREPLDAIAQTAPASQTL WP_068999571.1
short-chain dehydrogenase [Terasakiispira papahanaumokuakeensis] Length: 257\nNumber of predicted TMHs: 1\nExp number of AAs in TMHs: 27.42007\nExp number, first 60 AAs: 5.70122\nTotal prob of N-in: 0.42959\noutside 1 227\nTMhelix 228 250\ninside 251 257

28750 GCF_000017245.1_ASM1724v1 Actinobacillus succinogenes 130Z Proteobacteria; Gammaproteobacteria; Pasteurellales; Pasteurellaceae; Actinobacillus; Actinobacillus succinogenes
MQRHDGFVSALNSVSRRDFMKLCTALAATMGLSSKAGAEMTNALTNPQRPPVIWIGAQEC WP_012073014.1
hydrogenase 2 small subunit [Actinobacillus succinogenes] Length: 381\nNumber of predicted TMHs: 1\nExp number of AAs in TMHs: 20.61645\nExp number, first 60 AAs: 0.03339\nTotal prob of N-in: 0.02135\noutside 1 334\nTMhelix 335 357\ninside 358 381

28751 GCF_001262035.1_ASM126203v1 Aggregatibacter aphrophilus Proteobacteria; Gammaproteobacteria; Pasteurellales; Pasteurellaceae; Aggregatibacter
MQRSDGLFSALAEVSRRDFMKLCTALAATMGLSSKAGAEMTAALIEPKRPPVLWIGAQEC WP_050693696.1
hydrogenase 2 small subunit [Aggregatibacter aphrophilus]Length: 388\nNumber of predicted TMHs: 1\nExp number of AAs in TMHs: 18.82995\nExp number, first 60 AAs: 0.06305\nTotal prob of N-in: 0.01889\noutside 1 335\nTMhelix 336 358\ninside 359 388

28752 GCF_001680765.1_ASM168076v1 Aggregatibacter aphrophilus Proteobacteria; Gammaproteobacteria; Pasteurellales; Pasteurellaceae; Aggregatibacter
MQKSDGLFSALAEVSRRDFMKLCTALAATMGLSSKAGAEMTAALTEPKRPPVLWIGAQEC WP_005703197.1
hydrogenase 2 small subunit [Aggregatibacter aphrophilus]Length: 388\nNumber of predicted TMHs: 1\nExp number of AAs in TMHs: 18.77495\nExp number, first 60 AAs: 0.03719\nTotal prob of N-in: 0.01612\noutside 1 335\nTMhelix 336 358\ninside 359 388

28753 GCF_002083125.1_ASM208312v1 Aggregatibacter aphrophilus Proteobacteria; Gammaproteobacteria; Pasteurellales; Pasteurellaceae; Aggregatibacter
MQRSDGLFSALADVSRRDFMKLCTALAATMGLSSKAGAEMTEALTAPARPPVLWIGAQEC WP_083015466.1
hydrogenase 2 small subunit [Aggregatibacter aphrophilus]Length: 387\nNumber of predicted TMHs: 1\nExp number of AAs in TMHs: 18.71446\nExp number, first 60 AAs: 0.01028\nTotal prob of N-in: 0.01294\noutside 1 335\nTMhelix 336 358\ninside 359 387

28754 GCF_001680805.1_ASM168080v1 Aggregatibacter aphrophilus Proteobacteria; Gammaproteobacteria; Pasteurellales; Pasteurellaceae; Aggregatibacter
MQRCDGLFSALAEVSRRDFMKLCTALAATMGLSSKAGAEMTAALTEPKRPPVLWIGAQEC WP_005700513.1
hydrogenase 2 small subunit [Aggregatibacter aphrophilus]Length: 388\nNumber of predicted TMHs: 1\nExp number of AAs in TMHs: 18.89369\nExp number, first 60 AAs: 0.05449\nTotal prob of N-in: 0.02474\noutside 1 335\nTMhelix 336 358\ninside 359 388

28755 GCF_001059425.1_ASM105942v1 Aggregatibacter segnis Proteobacteria; Gammaproteobacteria; Pasteurellales; Pasteurellaceae; Aggregatibacter
MQRSDGLFSALADVSRRDFMKLCTALAATMGLSSKAGAQMTEALTAPARPPVLWIGAQEC WP_048750393.1
hydrogenase 2 small subunit [Aggregatibacter segnis] Length: 382\nNumber of predicted TMHs: 1\nExp number of AAs in TMHs: 19.75945\nExp number, first 60 AAs: 0.02593\nTotal prob of N-in: 0.01528\noutside 1 335\nTMhelix 336 358\ninside 359 382

28756 GCF_000259915.1_ASM25991v1 Aggregatibacter actinomycetemcomitans RhAA1 Proteobacteria; Gammaproteobacteria; Pasteurellales; Pasteurellaceae; Aggregatibacter; Aggregatibacter actinomycetemcomitans
MQRSDGLFSALADVSRRDFMKLCTALAATMGLSSKAGAEMTAALTAPARPPVVWIGAQEC WP_005576248.1
hydrogenase 2 small subunit [Aggregatibacter actinomycetemcomitans] Length: 382\nNumber of predicted TMHs: 1\nExp number of AAs in TMHs: 20.26456\nExp number, first 60 AAs: 0.04434\nTotal prob of N-in: 0.02109\noutside 1 335\nTMhelix 336 358\ninside 359 382

28757 GCF_001190115.1_ASM119011v1 Aggregatibacter actinomycetemcomitans RhAA1 Proteobacteria; Gammaproteobacteria; Pasteurellales; Pasteurellaceae; Aggregatibacter; Aggregatibacter actinomycetemcomitans
MQRSDGLFSALADVSRRDFMKLCTALAATMGLSSKAGAEMTAALTAPARPPVVWIGAQEC WP_005576248.1
hydrogenase 2 small subunit [Aggregatibacter actinomycetemcomitans] Length: 382\nNumber of

predicted TMHs: 1\nExp number of AAs in TMHs: 20.26456\nExp number, first 60 AAs: 0.04434\nTotal prob of N-in: 0.02109\noutside 1 335\nTMhelix 336 358\ninside 359 382

28758 GCF_001596315.1_ANH9776v1.0 Aggregatibacter actinomycetemcomitans serotype e str. ANH9776 Proteobacteria; Gammaproteobacteria; Pasteurellales; Pasteurellaceae; Aggregatibacter; Aggregatibacter actinomycetemcomitans MQRSDGLFSALADVSRRDFMKLCTALAATMGLSSKAGAEMTAALTAPARPPVVWIGAQEC WP_005556985.1 hydrogenase 2 small subunit [Aggregatibacter actinomycetemcomitans] Length: 382\nNumber of predicted TMHs: 1\nExp number of AAs in TMHs: 20.38217\nExp number, first 60 AAs: 0.04445\nTotal prob of N-in: 0.02587\noutside 1 335\nTMhelix 336 358\ninside 359 382

28759 GCF_001596425.1_SA3096v1.0 Aggregatibacter actinomycetemcomitans serotype e str. SA3096 Proteobacteria; Gammaproteobacteria; Pasteurellales; Pasteurellaceae; Aggregatibacter; Aggregatibacter actinomycetemcomitans MQRSDGLFSALADVSRRDFMKLCTALAATMGLSSKAGAEMTAALTAPARPPVVWIGAQEC WP_005556985.1 hydrogenase 2 small subunit [Aggregatibacter actinomycetemcomitans] Length: 382\nNumber of predicted TMHs: 1\nExp number of AAs in TMHs: 20.38217\nExp number, first 60 AAs: 0.04445\nTotal prob of N-in: 0.02587\noutside 1 335\nTMhelix 336 358\ninside 359 382

28760 GCF_000226755.1_ASM22675v2 Aggregatibacter actinomycetemcomitans serotype e str. SC1083 Proteobacteria; Gammaproteobacteria; Pasteurellales; Pasteurellaceae; Aggregatibacter; Aggregatibacter actinomycetemcomitans MQRSDGLFSALADVSRRDFMKLCTALAATMGLSSKAGAEMTAALTAPARPPVVWIGAQEC WP_005556985.1 hydrogenase 2 small subunit [Aggregatibacter actinomycetemcomitans] Length: 382\nNumber of predicted TMHs: 1\nExp number of AAs in TMHs: 20.38217\nExp number, first 60 AAs: 0.04445\nTotal prob of N-in: 0.02587\noutside 1 335\nTMhelix 336 358\ninside 359 382

28761 GCF_001596235.1_SC936v1.0 Aggregatibacter actinomycetemcomitans serotype e str. SC936 Proteobacteria; Gammaproteobacteria; Pasteurellales; Pasteurellaceae; Aggregatibacter; Aggregatibacter actinomycetemcomitans MQRSDGLFSALADVSRRDFMKLCTALAATMGLSSKAGAEMTAALTAPARPPVVWIGAQEC WP_061886967.1 hydrogenase 2 small subunit [Aggregatibacter actinomycetemcomitans] Length: 382\nNumber of predicted TMHs: 1\nExp number of AAs in TMHs: 20.38253\nExp number, first 60 AAs: 0.04445\nTotal prob of N-in: 0.02589\noutside 1 335\nTMhelix 336 358\ninside 359 382

28762 GCF_000226495.2_ASM22649v3 Aggregatibacter aphrophilus ATCC 33389 Proteobacteria; Gammaproteobacteria; Pasteurellales; Pasteurellaceae; Aggregatibacter; Aggregatibacter aphrophilus MQKSDGLFSALAEVSRRDFMKLCTALAATMGLSSKAGAEMTAALTEPKRPPVLWIGAQEC WP_005703197.1 hydrogenase 2 small subunit [Aggregatibacter aphrophilus]Length: 388\nNumber of predicted TMHs: 1\nExp number of AAs in TMHs: 18.77495\nExp number, first 60 AAs: 0.03719\nTotal prob of N-in: 0.01612\noutside 1 335\nTMhelix 336 358\ninside 359 388

28763 GCF_000231255.1_Aggr_aphr_F0387_V1 Aggregatibacter aphrophilus F0387 Proteobacteria; Gammaproteobacteria; Pasteurellales; Pasteurellaceae; Aggregatibacter; Aggregatibacter aphrophilus MQRCDGLFSALAEVSRRDFMKLCTALAATMGLSSKAGAEMTAALTEPKRPPVLWIGAQEC WP_005700513.1 hydrogenase 2 small subunit [Aggregatibacter aphrophilus]Length: 388\nNumber of predicted TMHs: 1\nExp number of AAs in TMHs: 18.89369\nExp number, first 60 AAs: 0.05449\nTotal prob of N-in: 0.02474\noutside 1 335\nTMhelix 336 358\ninside 359 388

28764 GCF_000022985.1_ASM2298v1 Aggregatibacter aphrophilus NJ8700 Proteobacteria; Gammaproteobacteria; Pasteurellales; Pasteurellaceae; Aggregatibacter; Aggregatibacter aphrophilus MQRCDGLFSALAEVSRRDFMKLCTALAATMGLSSKAGAEMTAALTEPKRPPVLWIGAQEC WP_005700513.1 hydrogenase 2 small subunit [Aggregatibacter aphrophilus]Length: 388\nNumber of predicted TMHs: 1\nExp number of AAs in TMHs: 18.89369\nExp number, first 60 AAs: 0.05449\nTotal prob of N-in: 0.02474\noutside 1 335\nTMhelix 336 358\ninside 359 388

28765 GCF_001188835.1_ASM118883v1 Aggregatibacter aphrophilus NJ8700 Proteobacteria; Gammaproteobacteria; Pasteurellales; Pasteurellaceae; Aggregatibacter; Aggregatibacter aphrophilus MQRCDGLFSALAEVSRRDFMKLCTALAATMGLSSKAGAEMTAALTEPKRPPVLWIGAQEC WP_005700513.1 hydrogenase 2 small subunit [Aggregatibacter aphrophilus]Length: 388\nNumber of predicted TMHs: 1\nExp number of AAs in TMHs: 18.89369\nExp number, first 60 AAs: 0.05449\nTotal prob of N-in: 0.02474\noutside 1 335\nTMhelix 336 358\ninside 359 388

28766 GCF_000185305.1_ASM18530v1 Aggregatibacter segnis ATCC 33393 Proteobacteria; Gammaproteobacteria; Pasteurellales; Pasteurellaceae; Aggregatibacter; Aggregatibacter segnis

MQRSDGLFSALADVSRRDFMKLCTALAATMGLSSKASAQMTEALTAPARPPVLWIGAQEC WP_006718272.1 hydrogenase 2 small subunit [Aggregatibacter segnis] Length: 382\nNumber of predicted TMHs: 1\nExp number of AAs in TMHs: 19.72503\nExp number, first 60 AAs: 0.01396\nTotal prob of N-in: 0.01545\noutside 1 335\nTMhelix 336 358\ninside 359 382

28767 GCF_000466335.1_ASM46633v1 Aggregatibacter sp. oral taxon 458 str. W10330 Proteobacteria; Gammaproteobacteria; Pasteurellales; Pasteurellaceae; Aggregatibacter; Aggregatibacter sp. oral taxon 458 MQRSDGLFSALADVSRRDFMKLCTALAATMGLSSKAGAEMTNALTAPARPPVLWIGAQEC WP_033002376.1 hydrogenase 2 small subunit [Aggregatibacter sp. oral taxon 458] Length: 382\nNumber of predicted TMHs: 1\nExp number of AAs in TMHs: 20.24146\nExp number, first 60 AAs: 0.01746\nTotal prob of N-in: 0.01288\noutside 1 335\nTMhelix 336 358\ninside 359 382

28768 GCF_001679045.1_ASM167904v1 Haemophilus haemolyticus Proteobacteria; Gammaproteobacteria; Pasteurellales; Pasteurellaceae; Haemophilus MNNPDGVFSALADVSRRDFMKLCTALAATMGLNSKAGAEMTAAMTSPARPPVLWIGAQEC WP_005645504.1 hydrogenase 2 small subunit [Haemophilus haemolyticus] Length: 381\nNumber of predicted TMHs: 1\nExp number of AAs in TMHs: 18.19451\nExp number, first 60 AAs: 0.02615\nTotal prob of N-in: 0.00939\noutside 1 333\nTMhelix 334 356\ninside 357 381

28769 GCF_001008215.1_ASM100821v1 Haemophilus haemolyticus Proteobacteria; Gammaproteobacteria; Pasteurellales; Pasteurellaceae; Haemophilus MNNPDGVFSALADVSRRDFMKLCTALAATMGLNSKAGAEMTAAMTSPARPPVLWIGAQEC WP_046949274.1 hydrogenase 2 small subunit [Haemophilus haemolyticus] Length: 381\nNumber of predicted TMHs: 1\nExp number of AAs in TMHs: 18.12462\nExp number, first 60 AAs: 0.02612\nTotal prob of N-in: 0.00902\noutside 1 333\nTMhelix 334 356\ninside 357 381

28770 GCF_001059125.1_ASM105912v1 Haemophilus influenzae Proteobacteria; Gammaproteobacteria; Pasteurellales; Pasteurellaceae; Haemophilus MQRSDGLFSALAEVSRRDFMKLCTALAATMGLSSKAGAEMTAALTEPKRPPVLWIGAQEC WP_005697985.1 MULTISPECIES: hydrogenase 2 small subunit [Haemophilus] Length: 382\nNumber of predicted TMHs: 1\nExp number of AAs in TMHs: 20.08151\nExp number, first 60 AAs: 0.03967\nTotal prob of N-in: 0.01472\noutside 1 335\nTMhelix 336 358\ninside 359 382

28771 GCF_001058575.1_ASM105857v1 Haemophilus influenzae Proteobacteria; Gammaproteobacteria; Pasteurellales; Pasteurellaceae; Haemophilus MQRSDGLFSALAEVSRRDFMKLCTALAATMGLSSKAGAEMTAALTEPKRPPVLWIGAQEC WP_005697985.1 MULTISPECIES: hydrogenase 2 small subunit [Haemophilus] Length: 382\nNumber of predicted TMHs: 1\nExp number of AAs in TMHs: 20.08151\nExp number, first 60 AAs: 0.03967\nTotal prob of N-in: 0.01472\noutside 1 335\nTMhelix 336 358\ninside 359 382

28772 GCF_001054455.1_ASM105445v1 Haemophilus influenzae Proteobacteria; Gammaproteobacteria; Pasteurellales; Pasteurellaceae; Haemophilus MQRSDGLFSALAEVSRRDFMKLCTALAATMGLSSKAGAEMTAALTEPKRPPVLWIGAQEC WP_005697985.1 MULTISPECIES: hydrogenase 2 small subunit [Haemophilus] Length: 382\nNumber of predicted TMHs: 1\nExp number of AAs in TMHs: 20.08151\nExp number, first 60 AAs: 0.03967\nTotal prob of N-in: 0.01472\noutside 1 335\nTMhelix 336 358\ninside 359 382

28773 GCF_001679455.1_ASM167945v1 Haemophilus parainfluenzae Proteobacteria; Gammaproteobacteria; Pasteurellales; Pasteurellaceae; Haemophilus MQRSDGLFSALAEVSRRDFMKLCTALAATMGLSSKAGAEMTAALTEPKRPPVLWIGAQEC WP_005697985.1 MULTISPECIES: hydrogenase 2 small subunit [Haemophilus] Length: 382\nNumber of predicted TMHs: 1\nExp number of AAs in TMHs: 20.08151\nExp number, first 60 AAs: 0.03967\nTotal prob of N-in: 0.01472\noutside 1 335\nTMhelix 336 358\ninside 359 382

28774 GCF_001679405.1_ASM167940v1 Haemophilus parainfluenzae Proteobacteria; Gammaproteobacteria; Pasteurellales; Pasteurellaceae; Haemophilus MQRSDGLFSALAEVSRRDFMKLCTALAATMGLSSKAGAEMTAALTEPKRPPVLWIGAQEC WP_065242640.1 hydrogenase 2 small subunit [Haemophilus parainfluenzae] Length: 382\nNumber of predicted TMHs: 1\nExp number of AAs in TMHs: 19.94247\nExp number, first 60 AAs: 0.03969\nTotal prob of N-in: 0.01286\noutside 1 335\nTMhelix 336 358\ninside 359 382

28775 GCF_001055595.1_ASM105559v1 Haemophilus parainfluenzae Proteobacteria; Gammaproteobacteria; Pasteurellales; Pasteurellaceae; Haemophilus
MQRSDGLFSALAEVSRRDFMKLCTALAATMGLSSKAGAEMTAALTEPKRPPVLWIGAQEC WP_049371969.1
hydrogenase 2 small subunit [Haemophilus parainfluenzae] Length: 382\nNumber of predicted TMHs: 1\nExp number of AAs in TMHs: 20.26374\nExp number, first 60 AAs: 0.03967\nTotal prob of N-in: 0.01386\noutside 1 335\nTMhelix 336 358\ninside 359 382

28776 GCF_001949885.1_ASM194988v1 Haemophilus parainfluenzae Proteobacteria; Gammaproteobacteria; Pasteurellales; Pasteurellaceae; Haemophilus
MQRSDGLFSALAEVSRRDFMKLCTALAATMGLSSKAGAEMTAALTEPKRPPVLWIGAQEC WP_075875963.1
hydrogenase 2 small subunit [Haemophilus parainfluenzae] Length: 382\nNumber of predicted TMHs: 1\nExp number of AAs in TMHs: 18.64643\nExp number, first 60 AAs: 0.03966\nTotal prob of N-in: 0.01388\noutside 1 335\nTMhelix 336 358\ninside 359 382

28777 GCF_001053915.1_ASM105391v1 Haemophilus parainfluenzae Proteobacteria; Gammaproteobacteria; Pasteurellales; Pasteurellaceae; Haemophilus
MQRSDGLFSALAEVSRRDFMKLCTALAATMGLSSKAGAEMTAALTEPKRPPVLWIGAQEC WP_005697985.1
MULTISPECIES: hydrogenase 2 small subunit [Haemophilus] Length: 382\nNumber of predicted TMHs: 1\nExp number of AAs in TMHs: 20.08151\nExp number, first 60 AAs: 0.03967\nTotal prob of N-in: 0.01472\noutside 1 335\nTMhelix 336 358\ninside 359 382

28778 GCF_001279145.1_ASM127914v1 Haemophilus parainfluenzae Proteobacteria; Gammaproteobacteria; Pasteurellales; Pasteurellaceae; Haemophilus
MQRSDGLFSALAEVSRRDFMKLCTALAATMGLSSKAGAEMTAALTEPKRPPVLWIGAQEC WP_005697985.1
MULTISPECIES: hydrogenase 2 small subunit [Haemophilus] Length: 382\nNumber of predicted TMHs: 1\nExp number of AAs in TMHs: 20.08151\nExp number, first 60 AAs: 0.03967\nTotal prob of N-in: 0.01472\noutside 1 335\nTMhelix 336 358\ninside 359 382

28779 GCF_001058435.1_ASM105843v1 Haemophilus parainfluenzae Proteobacteria; Gammaproteobacteria; Pasteurellales; Pasteurellaceae; Haemophilus
MQRSDGLFSALAEVSRRDFMKLCTALAATMGLSSKAGAEMTAALTEPKRPPVLWIGAQEC WP_005697985.1
MULTISPECIES: hydrogenase 2 small subunit [Haemophilus] Length: 382\nNumber of predicted TMHs: 1\nExp number of AAs in TMHs: 20.08151\nExp number, first 60 AAs: 0.03967\nTotal prob of N-in: 0.01472\noutside 1 335\nTMhelix 336 358\ninside 359 382

28780 GCF_001054475.1_ASM105447v1 Haemophilus parainfluenzae Proteobacteria; Gammaproteobacteria; Pasteurellales; Pasteurellaceae; Haemophilus
MQRSDGLFSALAEVSRRDFMKLCTALAATMGLSSKAGAEMTAALTEPKRPPVLWIGAQEC WP_005697985.1
MULTISPECIES: hydrogenase 2 small subunit [Haemophilus] Length: 382\nNumber of predicted TMHs: 1\nExp number of AAs in TMHs: 20.08151\nExp number, first 60 AAs: 0.03967\nTotal prob of N-in: 0.01472\noutside 1 335\nTMhelix 336 358\ninside 359 382

28781 GCF_001057005.1_ASM105700v1 Haemophilus parainfluenzae Proteobacteria; Gammaproteobacteria; Pasteurellales; Pasteurellaceae; Haemophilus
MQRSDGLFSALAEVSRRDFMKLCTALAATMGLSSKAGAEMTAALTEPKRPPVLWIGAQEC WP_049371969.1
hydrogenase 2 small subunit [Haemophilus parainfluenzae] Length: 382\nNumber of predicted TMHs: 1\nExp number of AAs in TMHs: 20.26374\nExp number, first 60 AAs: 0.03967\nTotal prob of N-in: 0.01386\noutside 1 335\nTMhelix 336 358\ninside 359 382

28782 GCF_001053535.1_ASM105353v1 Haemophilus parainfluenzae Proteobacteria; Gammaproteobacteria; Pasteurellales; Pasteurellaceae; Haemophilus
MQRSDGLFSALAEVSRRDFMKLCTALAATMGLSSKAGAEMTAALTEPKRPPVLWIGAQEC WP_049371969.1
hydrogenase 2 small subunit [Haemophilus parainfluenzae] Length: 382\nNumber of predicted TMHs: 1\nExp number of AAs in TMHs: 20.26374\nExp number, first 60 AAs: 0.03967\nTotal prob of N-in: 0.01386\noutside 1 335\nTMhelix 336 358\ninside 359 382

28783 GCF_001679325.1_ASM167932v1 Haemophilus parainfluenzae Proteobacteria; Gammaproteobacteria; Pasteurellales; Pasteurellaceae; Haemophilus
MQRSDGLFSALAEVSRRDFMKLCTALAATMGLSSKAGAEMTAALTEPKRPPVLWIGAQEC WP_005697985.1
MULTISPECIES: hydrogenase 2 small subunit [Haemophilus] Length: 382\nNumber of predicted TMHs:

1\nExp number of AAs in TMHs: 20.08151\nExp number, first 60 AAs: 0.03967\nTotal prob of N-in: 0.01472\noutside 1 335\nTMhelix 336 358\ninside 359 382

28784 GCF_001680775.1_ASM168077v1 Haemophilus parainfluenzae Proteobacteria; Gammaproteobacteria; Pasteurellales; Pasteurellaceae; Haemophilus
MQRSDGLFSALAEVSRRDFMKLCTALAATMGLSSKAGAEMTAALTEPKRPPVLWIGAQEY WP_065286329.1
hydrogenase 2 small subunit [Haemophilus parainfluenzae] Length: 382\nNumber of predicted TMHs: 1\nExp number of AAs in TMHs: 19.98911\nExp number, first 60 AAs: 0.03967\nTotal prob of N-in: 0.01491\noutside 1 335\nTMhelix 336 358\ninside 359 382

28785 GCF_001949895.1_ASM194989v1 Haemophilus parainfluenzae Proteobacteria; Gammaproteobacteria; Pasteurellales; Pasteurellaceae; Haemophilus
MQRSDGLFSALAEVSRRDFMKLCTALAATMGLSSKAGAEMTAALTEPKRPPVLWIGAQEC WP_075875963.1
hydrogenase 2 small subunit [Haemophilus parainfluenzae] Length: 382\nNumber of predicted TMHs: 1\nExp number of AAs in TMHs: 18.64643\nExp number, first 60 AAs: 0.03966\nTotal prob of N-in: 0.01388\noutside 1 335\nTMhelix 336 358\ninside 359 382

28786 GCF_001055095.1_ASM105509v1 Haemophilus parainfluenzae Proteobacteria; Gammaproteobacteria; Pasteurellales; Pasteurellaceae; Haemophilus
MQRSDGLFSALAEVSRRDFMKLCTALAATMGLSSKAGAEMTTALTEPKRPPVLWIGAQEC WP_049369733.1
hydrogenase 2 small subunit [Haemophilus parainfluenzae] Length: 382\nNumber of predicted TMHs: 1\nExp number of AAs in TMHs: 19.93856\nExp number, first 60 AAs: 0.02577\nTotal prob of N-in: 0.01432\noutside 1 335\nTMhelix 336 358\ninside 359 382

28787 GCF_001055885.1_ASM105588v1 Haemophilus parainfluenzae Proteobacteria; Gammaproteobacteria; Pasteurellales; Pasteurellaceae; Haemophilus
MQRSDGLFSALAEVSRRDFMKLCTALAATMGLSSKAGAEMTAALTEPKRPPVLWIGAQEC WP_049365049.1
MULTISPECIES: hydrogenase 2 small subunit [Haemophilus] Length: 382\nNumber of predicted TMHs: 1\nExp number of AAs in TMHs: 19.98526\nExp number, first 60 AAs: 0.03967\nTotal prob of N-in: 0.01483\noutside 1 335\nTMhelix 336 358\ninside 359 382

28788 GCF_001059815.1_ASM105981v1 Haemophilus parainfluenzae Proteobacteria; Gammaproteobacteria; Pasteurellales; Pasteurellaceae; Haemophilus
MQRSDGLFSALAEVSRRDFIKLCTALAATMGLSSKAGAEMTAALTEPKRPPVLWIGAQEC WP_049383969.1
hydrogenase 2 small subunit [Haemophilus parainfluenzae] Length: 383\nNumber of predicted TMHs: 1\nExp number of AAs in TMHs: 19.92284\nExp number, first 60 AAs: 0.03046\nTotal prob of N-in: 0.01444\noutside 1 335\nTMhelix 336 358\ninside 359 383

28789 GCF_001053575.1_ASM105357v1 Haemophilus parainfluenzae Proteobacteria; Gammaproteobacteria; Pasteurellales; Pasteurellaceae; Haemophilus
MQRSDGLFSALAEVSRRDFMKLCTALAATMGLSSKAGAEMTAALTEPKRPPVLWIGAQEC WP_005697985.1
MULTISPECIES: hydrogenase 2 small subunit [Haemophilus] Length: 382\nNumber of predicted TMHs: 1\nExp number of AAs in TMHs: 20.08151\nExp number, first 60 AAs: 0.03967\nTotal prob of N-in: 0.01472\noutside 1 335\nTMhelix 336 358\ninside 359 382

28790 GCF_001053035.1_ASM105303v1 Haemophilus parainfluenzae Proteobacteria; Gammaproteobacteria; Pasteurellales; Pasteurellaceae; Haemophilus
MQRSDGLFSALVEVSRRDFMKLCTALAATMGLSSKASAEMTTALTEPKRPPVLWIGAQEC WP_049355647.1
hydrogenase 2 small subunit [Haemophilus parainfluenzae] Length: 382\nNumber of predicted TMHs: 1\nExp number of AAs in TMHs: 20.05377\nExp number, first 60 AAs: 0.01189\nTotal prob of N-in: 0.01348\noutside 1 335\nTMhelix 336 358\ninside 359 382

28791 GCF_001752465.1_ASM175246v1 Haemophilus quentini Proteobacteria; Gammaproteobacteria; Pasteurellales; Pasteurellaceae; Haemophilus
MNNLDGVFSALADVSRRDFMKLCTALAATMGLNSKAGAEMTAAMTSPARPPVLWIGAQEC WP_005641824.1
MULTISPECIES: hydrogenase 2 small subunit [Haemophilus] Length: 381\nNumber of predicted TMHs: 1\nExp number of AAs in TMHs: 18.24867\nExp number, first 60 AAs: 0.02404\nTotal prob of N-in: 0.00866\noutside 1 333\nTMhelix 334 356\ninside 357 381

28792 GCF_001752475.1_ASM175247v1 Haemophilus quentini Proteobacteria; Gammaproteobacteria; Pasteurellales; Pasteurellaceae; Haemophilus

MNNLDGVFSALADVSRRDFMKLCTALAATMGLNSKAGAEMTAAMTSPARPPVLWIGAQEC WP_005641824.1 MULTISPECIES: hydrogenase 2 small subunit [Haemophilus] Length: 381\nNumber of predicted TMHs: 1\nExp number of AAs in TMHs: 18.24867\nExp number, first 60 AAs: 0.02404\nTotal prob of N-in: 0.00866\noutside 1 333\nTMhelix 334 356\ninside 357 381

28793 GCF_001702075.1_ASM170207v2 Haemophilus quentini Proteobacteria; Gammaproteobacteria; Pasteurellales; Pasteurellaceae; Haemophilus MNNLDGVFSALADVSRRDFMKLCTALAATMGLNSKAGAEMTAAMTSPARPPVLWIGAQEC WP_005641824.1 MULTISPECIES: hydrogenase 2 small subunit [Haemophilus] Length: 381\nNumber of predicted TMHs: 1\nExp number of AAs in TMHs: 18.24867\nExp number, first 60 AAs: 0.02404\nTotal prob of N-in: 0.00866\noutside 1 333\nTMhelix 334 356\ninside 357 381

28794 GCF_001276515.1_ASM127651v1 Haemophilus sp. C1 Proteobacteria; Gammaproteobacteria; Pasteurellales; Pasteurellaceae; Haemophilus MNNPDGVFSALADVSRRDFMKLCTALAATMGLNSKAGAEMTAAMTSPARPPVLWIGAQEC WP_053465944.1 hydrogenase 2 small subunit [Haemophilus sp. C1] Length: 381\nNumber of predicted TMHs: 1\nExp number of AAs in TMHs: 18.17194\nExp number, first 60 AAs: 0.02615\nTotal prob of N-in: 0.00935\noutside 1 333\nTMhelix 334 356\ninside 357 381

28795 GCF_001679485.1_ASM167948v1 Haemophilus sp. CCUG 60358 Proteobacteria; Gammaproteobacteria; Pasteurellales; Pasteurellaceae; Haemophilus MQRSDGLFSALAEVSRRDFMKLCTALAATMGLSSKAGAEMTAALTEPKRPPVLWIGAQEC WP_065243663.1 hydrogenase 2 small subunit [Haemophilus sp. CCUG 60358] Length: 382\nNumber of predicted TMHs: 1\nExp number of AAs in TMHs: 19.98847\nExp number, first 60 AAs: 0.03967\nTotal prob of N-in: 0.01489\noutside 1 335\nTMhelix 336 358\ninside 359 382

28796 GCF_001679495.1_ASM167949v1 Haemophilus sp. CCUG 66565 Proteobacteria; Gammaproteobacteria; Pasteurellales; Pasteurellaceae; Haemophilus MNNPDGVFSALADVSRRDFMKLCTALAATMGLNSKAGAEMTAAMTSPARPPVLWIGAQEC WP_065264789.1 hydrogenase 2 small subunit [Haemophilus sp. CCUG 66565] Length: 381\nNumber of predicted TMHs: 1\nExp number of AAs in TMHs: 19.0493\nExp number, first 60 AAs: 0.02601\nTotal prob of N-in: 0.00894\noutside 1 333\nTMhelix 334 356\ninside 357 381

28797 GCF_001810345.1_ASM181034v1 Haemophilus sp. HMSC061E01 Proteobacteria; Gammaproteobacteria; Pasteurellales; Pasteurellaceae; Haemophilus MQRSDGLFSALAEVSRRDFMKLCTALAATMGLSSKAGAEMTAALTEPKRPPVLWIGAQEC WP_005697985.1 MULTISPECIES: hydrogenase 2 small subunit [Haemophilus] Length: 382\nNumber of predicted TMHs: 1\nExp number of AAs in TMHs: 20.08151\nExp number, first 60 AAs: 0.03967\nTotal prob of N-in: 0.01472\noutside 1 335\nTMhelix 336 358\ninside 359 382

28798 GCF_001810545.1_ASM181054v1 Haemophilus sp. HMSC066D02 Proteobacteria; Gammaproteobacteria; Pasteurellales; Pasteurellaceae; Haemophilus MQRSDGLFSALAEVSRRDFMKLCTALAATMGLSSKAGAEMTAALTEPKRPPVLWIGAQEC WP_049380266.1 MULTISPECIES: hydrogenase 2 small subunit [Haemophilus] Length: 382\nNumber of predicted TMHs: 1\nExp number of AAs in TMHs: 19.98847\nExp number, first 60 AAs: 0.03967\nTotal prob of N-in: 0.01489\noutside 1 335\nTMhelix 336 358\ninside 359 382

28799 GCF_001811025.1_ASM181102v1 Haemophilus sp. HMSC066D03 Proteobacteria; Gammaproteobacteria; Pasteurellales; Pasteurellaceae; Haemophilus MQRSDGLFSALAEVSRRDFMKLCTALAATMGLSSKAGAEMTAALTEPKRPPVLWIGAQEC WP_049380266.1 MULTISPECIES: hydrogenase 2 small subunit [Haemophilus] Length: 382\nNumber of predicted TMHs: 1\nExp number of AAs in TMHs: 19.98847\nExp number, first 60 AAs: 0.03967\nTotal prob of N-in: 0.01489\noutside 1 335\nTMhelix 336 358\ninside 359 382

28800 GCF_001815355.1_ASM181535v1 Haemophilus sp. HMSC068C11 Proteobacteria; Gammaproteobacteria; Pasteurellales; Pasteurellaceae; Haemophilus MQRSDGLFSALAEVSRRDFMKLCTALAATMGLSSKAGAEMTAALTEPKRPPVLWIGAQEC WP_070775114.1 hydrogenase 2 small subunit [Haemophilus sp. HMSC068C11] Length: 383\nNumber of predicted TMHs: 1\nExp number of AAs in TMHs: 19.93204\nExp number, first 60 AAs: 0.03967\nTotal prob of N-in: 0.01483\noutside 1 335\nTMhelix 336 358\ninside 359 383

28801 GCF_001814055.1_ASM181405v1 Haemophilus sp. HMSC073C03 Proteobacteria; Gammaproteobacteria; Pasteurellales; Pasteurellaceae; Haemophilus
MQRSDGLFSALAEVSRRDFMKLCTALAATMGLSSKAGAEMTAALTEPKRPPVLWIGAQEC WP_005697985.1
MULTISPECIES: hydrogenase 2 small subunit [Haemophilus] Length: 382\nNumber of predicted TMHs: 1\nExp number of AAs in TMHs: 20.08151\nExp number, first 60 AAs: 0.03967\nTotal prob of N-in: 0.01472\noutside 1 335\nTMhelix 336 358\ninside 359 382

28802 GCF_001838615.1_ASM183861v1 Haemophilus sp. HMSC61B11 Proteobacteria; Gammaproteobacteria; Pasteurellales; Pasteurellaceae; Haemophilus
MQRSDGLFSALAEVSRRDFMKLCTALAATMGLSSKAGAEMTAALTEPKRPPVLWIGAQEC WP_005697985.1
MULTISPECIES: hydrogenase 2 small subunit [Haemophilus] Length: 382\nNumber of predicted TMHs: 1\nExp number of AAs in TMHs: 20.08151\nExp number, first 60 AAs: 0.03967\nTotal prob of N-in: 0.01472\noutside 1 335\nTMhelix 336 358\ninside 359 382

28803 GCF_001838635.1_ASM183863v1 Haemophilus sp. HMSC71H05 Proteobacteria; Gammaproteobacteria; Pasteurellales; Pasteurellaceae; Haemophilus
MQRSDGLFSALAEVSRRDFMKLCTALAATMGLSSKAGAEMTAALTEPKRPPVLWIGAQEC WP_049365049.1
MULTISPECIES: hydrogenase 2 small subunit [Haemophilus] Length: 382\nNumber of predicted TMHs: 1\nExp number of AAs in TMHs: 19.98526\nExp number, first 60 AAs: 0.03967\nTotal prob of N-in: 0.01483\noutside 1 335\nTMhelix 336 358\ninside 359 382

28804 GCF_000262285.1_ASM26228v1 Haemophilus haemolyticus HK386 Proteobacteria; Gammaproteobacteria; Pasteurellales; Pasteurellaceae; Haemophilus; Haemophilus haemolyticus
MNNPDGVFSALADVSRRDFMKLCTALAATMGLNSKAGAEMTAAMTSPARPPVLWIGAQEC WP_005645504.1
hydrogenase 2 small subunit [Haemophilus haemolyticus] Length: 381\nNumber of predicted TMHs: 1\nExp number of AAs in TMHs: 18.19451\nExp number, first 60 AAs: 0.02615\nTotal prob of N-in: 0.00939\noutside 1 333\nTMhelix 334 356\ninside 357 381

28805 GCF_000222025.1_ASM22202v2 Haemophilus haemolyticus M19501 Proteobacteria; Gammaproteobacteria; Pasteurellales; Pasteurellaceae; Haemophilus; Haemophilus haemolyticus
MNNLDGVFSALADVSRRDFMKLCTALAATMGLNSKAGAEMTAAMTSPARPPVLWIGAQEC WP_005631853.1
hydrogenase 2 small subunit [Haemophilus haemolyticus] Length: 381\nNumber of predicted TMHs: 1\nExp number of AAs in TMHs: 18.16781\nExp number, first 60 AAs: 0.02404\nTotal prob of N-in: 0.00869\noutside 1 333\nTMhelix 334 356\ninside 357 381

28806 GCF_000222085.1_ASM22208v2 Haemophilus haemolyticus M21639 Proteobacteria; Gammaproteobacteria; Pasteurellales; Pasteurellaceae; Haemophilus; Haemophilus haemolyticus
MNNLDGVFSALADVSRRDFMKLCTALAATMGLNSKAGAEMTAAMTSPARPPVLWIGAQEC WP_005641824.1
MULTISPECIES: hydrogenase 2 small subunit [Haemophilus] Length: 381\nNumber of predicted TMHs: 1\nExp number of AAs in TMHs: 18.24867\nExp number, first 60 AAs: 0.02404\nTotal prob of N-in: 0.00866\noutside 1 333\nTMhelix 334 356\ninside 357 381

28807 GCF_000746485.1_ASM74648v1 Haemophilus parainfluenzae ATCC 33392 Proteobacteria; Gammaproteobacteria; Pasteurellales; Pasteurellaceae; Haemophilus; Haemophilus parainfluenzae
MQRSDGLFSALAEVSRRDFMKLCTALAATMGLSSKAGAEMTAALTEPKRPPVLWIGAQEC WP_005695040.1
hydrogenase 2 small subunit [Haemophilus parainfluenzae] Length: 382\nNumber of predicted TMHs: 1\nExp number of AAs in TMHs: 20.09879\nExp number, first 60 AAs: 0.03965\nTotal prob of N-in: 0.01748\noutside 1 335\nTMhelix 336 358\ninside 359 382

28808 GCF_000191405.1_ASM19140v1 Haemophilus parainfluenzae ATCC 33392 Proteobacteria; Gammaproteobacteria; Pasteurellales; Pasteurellaceae; Haemophilus; Haemophilus parainfluenzae
MQRSDGLFSALAEVSRRDFMKLCTALAATMGLSSKAGAEMTAALTEPKRPPVLWIGAQEC WP_005695040.1
hydrogenase 2 small subunit [Haemophilus parainfluenzae] Length: 382\nNumber of predicted TMHs: 1\nExp number of AAs in TMHs: 20.09879\nExp number, first 60 AAs: 0.03965\nTotal prob of N-in: 0.01748\noutside 1 335\nTMhelix 336 358\ninside 359 382

28809 GCF_000261285.1_HparainfluenzaeHK2019v1.0 Haemophilus parainfluenzae HK2019 Proteobacteria; Gammaproteobacteria; Pasteurellales; Pasteurellaceae; Haemophilus; Haemophilus parainfluenzae
MQRSDGLFSALAEVSRRDFMKLCTALAATMGLSSKAGAEMTAALTEPKRPPVLWIGAQEC WP_005697985.1
MULTISPECIES: hydrogenase 2 small subunit [Haemophilus] Length: 382\nNumber of predicted TMHs:

1\nExp number of AAs in TMHs: 20.08151\nExp number, first 60 AAs: 0.03967\nTotal prob of N-in: 0.01472\noutside 1 335\nTMhelix 336 358\ninside 359 382

28810 GCF_000259485.1_HparainfluenzaeHK262v1.0 Haemophilus parainfluenzae HK262 Proteobacteria; Gammaproteobacteria; Pasteurellales; Pasteurellaceae; Haemophilus; Haemophilus parainfluenzae MQRSDGLFSALAEVSRRDFMKLCTALAATMGLSSKAGAEMTAALTEPKRPPVLWIGAQEC WP_005697985.1 MULTISPECIES: hydrogenase 2 small subunit [Haemophilus] Length: 382\nNumber of predicted TMHs: 1\nExp number of AAs in TMHs: 20.08151\nExp number, first 60 AAs: 0.03967\nTotal prob of N-in: 0.01472\noutside 1 335\nTMhelix 336 358\ninside 359 382

28811 GCF_000210895.1_ASM21089v1 Haemophilus parainfluenzae T3T1 Proteobacteria; Gammaproteobacteria; Pasteurellales; Pasteurellaceae; Haemophilus; Haemophilus parainfluenzae MQRSDGLFSALAEVSRRDFMKLCTALAATMGLSSKAGAEMTAALTEPKRPPVLWIGAQEC WP_014064535.1 hydrogenase 2 small subunit [Haemophilus parainfluenzae] Length: 382\nNumber of predicted TMHs: 1\nExp number of AAs in TMHs: 20.08105\nExp number, first 60 AAs: 0.03967\nTotal prob of N-in: 0.01471\noutside 1 335\nTMhelix 336 358\ninside 359 382

28812 GCF_000940515.1_Mannheimia_massilioguelmaensis_MG13T Mannheimia massilioguelmaensis Proteobacteria; Gammaproteobacteria; Pasteurellales; Pasteurellaceae; Mannheimia MQRTDGLLSALNDSMADVSRRDFMKLCTALAATMGLSSKASAEEMTHALTNPQRPPVIWI WP_044469494.1 hydrogenase 2 small subunit [Mannheimia massilioguelmaensis] Length: 390\nNumber of predicted TMHs: 1\nExp number of AAs in TMHs: 20.56313\nExp number, first 60 AAs: 0.00314\nTotal prob of N-in: 0.01634\noutside 1 337\nTMhelix 338 360\ninside 361 390

28813 GCF_000262245.1_ASM26224v1 Pasteurella bettyae CCUG 2042 Proteobacteria; Gammaproteobacteria; Pasteurellales; Pasteurellaceae; Pasteurella; Pasteurella bettyae MQRTDGLLSALNDSMADVSRRDFMKLCTALAATMGLNSKASAEEMTHALTNPQRPPVIWI WP_005760791.1 hydrogenase 2 small subunit [Pasteurella bettyae] Length: 390\nNumber of predicted TMHs: 1\nExp number of AAs in TMHs: 20.56151\nExp number, first 60 AAs: 0.00151\nTotal prob of N-in: 0.01626\noutside 1 337\nTMhelix 338 360\ninside 361 390

28814 GCF_002000425.1_ASM200042v1 Rodentibacter ratti Proteobacteria; Gammaproteobacteria; Pasteurellales; Pasteurellaceae; Rodentibacter MGNQDNVFSTLADISRRDFMKLCSALAATMGLSSKAGAEMTAAMTNPARPPVIWIGAQEC WP_077474212.1 MULTISPECIES: hydrogenase 2 small subunit [Rodentibacter] Length: 376\nNumber of predicted TMHs: 1\nExp number of AAs in TMHs: 18.75431\nExp number, first 60 AAs: 0.16849\nTotal prob of N-in: 0.02918\noutside 1 335\nTMhelix 336 358\ninside 359 376

28815 GCF_002000545.1_ASM200054v1 Rodentibacter trehalosifermentans Proteobacteria; Gammaproteobacteria; Pasteurellales; Pasteurellaceae; Rodentibacter MGNQDNVFSTLADISRRDFMKLCSALAATMGLSSKAGAEMTAAMTNPARPPVIWIGAQEC WP_077474212.1 MULTISPECIES: hydrogenase 2 small subunit [Rodentibacter] Length: 376\nNumber of predicted TMHs: 1\nExp number of AAs in TMHs: 18.75431\nExp number, first 60 AAs: 0.16849\nTotal prob of N-in: 0.02918\noutside 1 335\nTMhelix 336 358\ninside 359 376

28816 GCF_001999265.1_ASM199926v1 Rodentibacter trehalosifermentans Proteobacteria; Gammaproteobacteria; Pasteurellales; Pasteurellaceae; Rodentibacter MGNQDNVFSTLADISRRDFMKLCSALAATMGLSSKAGAEMTAAMTNPARPPVIWIGAQEC WP_077421846.1 hydrogenase 2 small subunit [Rodentibacter trehalosifermentans] Length: 376\nNumber of predicted TMHs: 1\nExp number of AAs in TMHs: 20.26832\nExp number, first 60 AAs: 0.16911\nTotal prob of N-in: 0.02391\noutside 1 335\nTMhelix 336 358\ninside 359 376

28817 GCF_002002485.1_ASM200248v1 Pasteurellaceae bacterium 15-036681 Proteobacteria; Gammaproteobacteria; Pasteurellales; Pasteurellaceae; unclassified Pasteurellaceae; unclassified Pasteurellaceae (miscellaneous) MQRYDSLFSTFTDVSRRDFMKLCTALAATMGLSSKAGAEMTQALTSPQRPPVIWIGAQEC WP_077558482.1 hydrogenase 2 small subunit [Pasteurellaceae bacterium 15-036681] Length: 376\nNumber of predicted TMHs: 1\nExp number of AAs in TMHs: 16.93332\nExp number, first 60 AAs: 0.04754\nTotal prob of N-in: 0.06185\noutside 1 337\nTMhelix 338 360\ninside 361 376

28818 GCF_000974525.1_ASM97452v1 Pseudomonas abietaniphila Proteobacteria; Gammaproteobacteria; Pseudomonadales; Pseudomonadaceae; Pseudomonas

MSLSRPVSASLPSSQPPGARRRLLCAGLLGSLLTVGFATQVQAEGKDRWVTDTLSTYVRS WP_062381708.1 peptide-binding protein [Pseudomonas abietaniphila] Length: 225\nNumber of predicted TMHs: 1\nExp number of AAs in TMHs: 24.90346\nExp number, first 60 AAs: 2.6889\nTotal prob of N-in: 0.18906\noutside 1 192\nTMhelix 193 215\ninside 216 225

28819 GCF_000876035.1_ASM87603v1 Pseudomonas abietaniphila Proteobacteria; Gammaproteobacteria; Pseudomonadales; Pseudomonadaceae; Pseudomonas MSLSRPVSASLPSSQPPGARRRLLCAGLLGSLLTVGFATQVQAEGKDRWVTDTLSTYVRS WP_062381708.1 peptide-binding protein [Pseudomonas abietaniphila] Length: 225\nNumber of predicted TMHs: 1\nExp number of AAs in TMHs: 24.90346\nExp number, first 60 AAs: 2.6889\nTotal prob of N-in: 0.18906\noutside 1 192\nTMhelix 193 215\ninside 216 225

28820 GCF_900100795.1_IMG-taxon_2667527211_annotated_assembly Pseudomonas abietaniphila Proteobacteria; Gammaproteobacteria; Pseudomonadales; Pseudomonadaceae; Pseudomonas MSLSRPVSASLPSSLLPGARRRLLSAGLLGTLLTVGVVAQAQAEGKDRWVNDTLSTYVRS WP_074751114.1 peptide-binding protein [Pseudomonas abietaniphila] Length: 225\nNumber of predicted TMHs: 1\nExp number of AAs in TMHs: 27.62505\nExp number, first 60 AAs: 5.40251\nTotal prob of N-in: 0.21335\noutside 1 192\nTMhelix 193 215\ninside 216 225

28821 GCF_000820515.1_ASM82051v1 Pseudomonas batumici Proteobacteria; Gammaproteobacteria; Pseudomonadales; Pseudomonadaceae; Pseudomonas MTHTPLTRGALRHTRRPLMFLASSALCLGGWVHNAWAETAAAVPVAPANVAPQAPLGSYS WP_040070684.1 cellulose synthase BcsB subunit [Pseudomonas batumici] Length: 768\nNumber of predicted TMHs: 1\nExp number of AAs in TMHs: 25.70837\nExp number, first 60 AAs: 3.07318\nTotal prob of N-in: 0.15462\noutside 1 736\nTMhelix 737 759\ninside 760 768

28822 GCF_000759445.1_ASM75944v1 Pseudomonas luteaProteobacteria; Gammaproteobacteria; Pseudomonadales; Pseudomonadaceae; Pseudomonas MPIPASYASLRLNTLPGARRRLLSAGLLTTFLTAGFAADDALAESPGKDRWVSDTLSTYV WP_052075064.1 peptide-binding protein [Pseudomonas lutea] Length: 227\nNumber of predicted TMHs: 1\nExp number of AAs in TMHs: 28.29192\nExp number, first 60 AAs: 6.05987\nTotal prob of N-in: 0.34736\noutside 1 194\nTMhelix 195 217\ninside 218 227

28823 GCF_002157495.1_ASM215749v1 Pseudomonas sp. 1239 Proteobacteria; Gammaproteobacteria; Pseudomonadales; Pseudomonadaceae; Pseudomonas MPPSFQLRARRSLIPGLLGSLVILCGPAHAEEGSGNERWVSDSLSTYVRSGPTDGHRIVG WP_054916521.1 MULTISPECIES: peptide-binding protein [Pseudomonas] Length: 215\nNumber of predicted TMHs: 1\nExp number of AAs in TMHs: 23.55908\nExp number, first 60 AAs: 1.33747\nTotal prob of N-in: 0.24943\noutside 1 182\nTMhelix 183 205\ninside 206 215

28824 GCF_002080045.1_ASM208004v1 Pseudomonas sp. Bc-h Proteobacteria; Gammaproteobacteria; Pseudomonadales; Pseudomonadaceae; Pseudomonas MSLTRPLSVSASQSSLLPGARRRLLSAGLVGTLLTVGIASQAQAEGKDRWVTDTLSTYVR WP_081564362.1 peptide-binding protein [Pseudomonas sp. Bc-h] Length: 226\nNumber of predicted TMHs: 1\nExp number of AAs in TMHs: 23.86131\nExp number, first 60 AAs: 1.6624\nTotal prob of N-in: 0.11059\noutside 1 193\nTMhelix 194 216\ninside 217 226

28825 GCF_000282495.1_GM79.fsa Pseudomonas sp. GM79 Proteobacteria; Gammaproteobacteria; Pseudomonadales; Pseudomonadaceae; Pseudomonas MKPSTRRTFLPRRSLILITCSLVGWSGWALSAAAQINATEPAMVPVTALPGTDAPGNSYT WP_081499302.1 cellulose synthase regulator BcsB [Pseudomonas sp. GM79] Length: 767\nNumber of predicted TMHs: 1\nExp number of AAs in TMHs: 35.21641\nExp number, first 60 AAs: 12.37491\nTotal prob of N-in: 0.56814\nPOSSIBLE N-term signal sequence\noutside 1 734\nTMhelix 735 757\ninside 758 767

28826 GCF_001466905.1_ASM146690v1 Pseudomonas sp. ICMP 3272 Proteobacteria; Gammaproteobacteria; Pseudomonadales; Pseudomonadaceae; Pseudomonas MSKRRRVLRLLIVATLASLLQACDIDLYTNLGEREANAMLAVLLRDGIPASRKVQDNGQL WP_057412399.1 MULTISPECIES: EscJ/YscJ/HrcJ family type III secretion inner membrane ring protein [Pseudomonas] Length: 259\nNumber of predicted TMHs: 1\nExp number of AAs in TMHs: 33.56925\nExp number, first 60 AAs: 8.63141\nTotal prob of N-in: 0.58822\noutside 1 217\nTMhelix 218 240\ninside 241 259

28827 GCF_002018915.1_ASM201891v1 Pseudomonas sp. MF6396 Proteobacteria; Gammaproteobacteria; Pseudomonadales; Pseudomonadaceae; Pseudomonas
MPPSFQLRARRSLIPGLLGSLVILCGPAHAEEGSGNERWVSDSLSTYVRSGPTDGHRIVG WP_054916521.1 MULTISPECIES: peptide-binding protein [Pseudomonas] Length: 215\nNumber of predicted TMHs: 1\nExp number of AAs in TMHs: 23.55908\nExp number, first 60 AAs: 1.33747\nTotal prob of N-in: 0.24943\noutside 1 182\nTMhelix 183 205\ninside 206 215

28828 GCF_001320045.1_ASM132004v1 Pseudomonas sp. NBRC 111117 Proteobacteria; Gammaproteobacteria; Pseudomonadales; Pseudomonadaceae; Pseudomonas
MPPSFQLRARRSLIPGLLGSLVILCGPAHAEEGSGNERWVSDSLSTYVRSGPTDGHRIVG WP_054916521.1 MULTISPECIES: peptide-binding protein [Pseudomonas] Length: 215\nNumber of predicted TMHs: 1\nExp number of AAs in TMHs: 23.55908\nExp number, first 60 AAs: 1.33747\nTotal prob of N-in: 0.24943\noutside 1 182\nTMhelix 183 205\ninside 206 215

28829 GCF_001547895.1_ASM154789v1 Pseudomonas sp. Os17 Proteobacteria; Gammaproteobacteria; Pseudomonadales; Pseudomonadaceae; Pseudomonas
MQKGMPRRALLKGSVMALGALVAGSLGGCLTSQLYESQASNEVYTETVSQFYVTADAKSF WP_060840229.1 hypothetical protein [Pseudomonas sp. Os17] Length: 219\nNumber of predicted TMHs: 2\nExp number of AAs in TMHs: 35.45664\nExp number, first 60 AAs: 13.81897\nTotal prob of N-in: 0.74895\nPOSSIBLE N-term signal sequence\ninside 1 12\nTMhelix 13 35\noutside 36 190\nTMhelix 191 213\ninside 214 219

28830 GCF_000633395.1_de_novo Pseudomonas sp. PH1b Proteobacteria; Gammaproteobacteria; Pseudomonadales; Pseudomonadaceae; Pseudomonas
MQKGIPRRGLLKGTVVALGAVVAGSLGGCLTSQLYEERGSDEVYTETVGQFYITADEQSF WP_029978019.1 hypothetical protein [Pseudomonas sp. PH1b] Length: 219\nNumber of predicted TMHs: 1\nExp number of AAs in TMHs: 33.93338\nExp number, first 60 AAs: 12.94791\nTotal prob of N-in: 0.71507\nPOSSIBLE N-term signal sequence\ninside 1 184\nTMhelix 185 207\noutside 208 219

28831 GCF_000801235.1_V1 Pseudomonas sp. SHC52 Proteobacteria; Gammaproteobacteria; Pseudomonadales; Pseudomonadaceae; Pseudomonas
MNAKHFSGAGQRDNRRSLVILASTLLALSGGAAFADTAPADTGNGYNLTLKQLGRAYPMN WP_041021757.1 cellulose synthase BcsB subunit [Pseudomonas sp. SHC52] Length: 751\nNumber of predicted TMHs: 1\nExp number of AAs in TMHs: 35.25302\nExp number, first 60 AAs: 12.53898\nTotal prob of N-in: 0.60020\nPOSSIBLE N-term signal sequence\noutside 1 718\nTMhelix 719 741\ninside 742 751

28832 GCF_001547915.1_ASM154791v1 Pseudomonas sp. St29 Proteobacteria; Gammaproteobacteria; Pseudomonadales; Pseudomonadaceae; Pseudomonas
MQKGMPRRALLKGSVMALGALVAGSLGGCLTSQLYESQASNEVYTETVSQFYVTADAKSF WP_060844157.1 hypothetical protein [Pseudomonas sp. St29] Length: 219\nNumber of predicted TMHs: 2\nExp number of AAs in TMHs: 35.45852\nExp number, first 60 AAs: 13.8195\nTotal prob of N-in: 0.74897\nPOSSIBLE N-term signal sequence\ninside 1 12\nTMhelix 13 35\noutside 36 190\nTMhelix 191 213\ninside 214 219

28833 GCF_000795625.1_AZPAE13872 Pseudomonas aeruginosa Proteobacteria; Gammaproteobacteria; Pseudomonadales; Pseudomonadaceae; Pseudomonas; Pseudomonas aeruginosa group
MFRPLSVFIGTRYTRAKRRSHFVSFISLTSMIGLALGVLVMIVVLSVMNGFDREMRTRIL WP_043106916.1 lipoprotein-releasing system transmembrane subunit LolC, partial [Pseudomonas aeruginosa] Length: 313\nNumber of predicted TMHs: 2\nExp number of AAs in TMHs: 47.69245\nExp number, first 60 AAs: 23.32288\nTotal prob of N-in: 0.41319\nPOSSIBLE N-term signal sequence\ninside 1 20\nTMhelix 21 43\noutside 44 274\nTMhelix 275 297\ninside 298 313

28834 GCF_000412555.1_C_1426 Pseudomonas aeruginosa str. C 1426 Proteobacteria; Gammaproteobacteria; Pseudomonadales; Pseudomonadaceae; Pseudomonas; Pseudomonas aeruginosa group; Pseudomonas aeruginosa
MSASPSPRQLLHEALASLRGLGRRSLLALLGIVMGCSSVIAMLNIGSNATDEAMSIFKDMWP_049878540.1 ABC transporter permease, partial [Pseudomonas aeruginosa] Length: 316\nNumber of predicted TMHs: 2\nExp number of AAs in TMHs: 48.75428\nExp number, first 60 AAs: 21.53429\nTotal prob of N-in: 0.27756\nPOSSIBLE N-term signal sequence\noutside 1 25\nTMhelix 26 48\ninside 49 272\nTMhelix 273 295\noutside 296 316

28835 GCF_000412495.1_PA_17_SmallColonyVariant Pseudomonas aeruginosa str. PA 17SCV Proteobacteria; Gammaproteobacteria; Pseudomonadales; Pseudomonadaceae; Pseudomonas; Pseudomonas aeruginosa group; Pseudomonas aeruginosa; Pseudomonas aeruginosa str. PA 17
MFRPLSVFIGTRYTRAKRRSHFVSFISLTSMIGLALGVLVMIVVLSVMNGFDREMRTRIL WP_043174684.1 lipoprotein-

releasing system transmembrane subunit LolC, partial [Pseudomonas aeruginosa]	Length: 312\nNumber of predicted TMHs:  2\nExp number of AAs in TMHs: 47.16645\nExp number, first 60 AAs:  23.28881\nTotal prob of N-in:  0.37259\nPOSSIBLE N-term signal sequence\noutside   1   24\nTMhelix   25   47\ninside   48   273\nTMhelix   274   296\noutside   297   312

28836	GCF_001411965.1_ASM141196v1	Pseudomonas corrugata	Proteobacteria; Gammaproteobacteria; Pseudomonadales; Pseudomonadaceae; Pseudomonas; Pseudomonas fluorescens group	MNVKHSSGVGQRENRRSLVILASTLLALSGGAAFADTAPADTSDGYTLTLKQLGRAYPMN	WP_024776312.1	cellulose synthase BcsB subunit [Pseudomonas corrugata]  Length: 751\nNumber of predicted TMHs:  1\nExp number of AAs in TMHs: 34.13377\nExp number, first 60 AAs:  11.47394\nTotal prob of N-in:   0.54857\nPOSSIBLE N-term signal sequence\noutside   1   718\nTMhelix   719   741\ninside   742   751

28837	GCF_001708425.1_ASM170842v1	Pseudomonas corrugata	Proteobacteria; Gammaproteobacteria; Pseudomonadales; Pseudomonadaceae; Pseudomonas; Pseudomonas fluorescens group	MNVKHSSGVGQRENRRSLVILASTLLALSGGAAFADTAPADTSDGYTLTLKQLGRAYPMN	WP_055135416.1	cellulose synthase BcsB subunit [Pseudomonas corrugata]  Length: 751\nNumber of predicted TMHs:  1\nExp number of AAs in TMHs: 34.13387\nExp number, first 60 AAs:  11.47386\nTotal prob of N-in:   0.54858\nPOSSIBLE N-term signal sequence\noutside   1   718\nTMhelix   719   741\ninside   742   751

28838	GCF_001269905.1_ASM126990v1	Pseudomonas corrugata	Proteobacteria; Gammaproteobacteria; Pseudomonadales; Pseudomonadaceae; Pseudomonas; Pseudomonas fluorescens group	MNVKHSSGVGQRENRRSLVILASTLLALSGGAAFADTAPADTSDGYTLTLKQLGRAYPMN	WP_024776312.1	cellulose synthase BcsB subunit [Pseudomonas corrugata]  Length: 751\nNumber of predicted TMHs:  1\nExp number of AAs in TMHs: 34.13377\nExp number, first 60 AAs:  11.47394\nTotal prob of N-in:   0.54857\nPOSSIBLE N-term signal sequence\noutside   1   718\nTMhelix   719   741\ninside   742   751

28839	GCF_001412065.1_ASM141206v1	Pseudomonas corrugata	Proteobacteria; Gammaproteobacteria; Pseudomonadales; Pseudomonadaceae; Pseudomonas; Pseudomonas fluorescens group	MNVKHSSGVGQRENRRSLVILASTLLALSGGAAFADTAPADTSDGYTLTLKQLGRAYPMN	WP_055135416.1	cellulose synthase BcsB subunit [Pseudomonas corrugata]  Length: 751\nNumber of predicted TMHs:  1\nExp number of AAs in TMHs: 34.13387\nExp number, first 60 AAs:  11.47386\nTotal prob of N-in:   0.54858\nPOSSIBLE N-term signal sequence\noutside   1   718\nTMhelix   719   741\ninside   742   751

28840	GCF_001412195.1_ASM141219v1	Pseudomonas corrugata	Proteobacteria; Gammaproteobacteria; Pseudomonadales; Pseudomonadaceae; Pseudomonas; Pseudomonas fluorescens group	MNVKHSSGVGQRENRRSLVILASTLLALSGGAAFADTAPADTSDGYTLTLKQLGRAYPMN	WP_055135416.1	cellulose synthase BcsB subunit [Pseudomonas corrugata]  Length: 751\nNumber of predicted TMHs:  1\nExp number of AAs in TMHs: 34.13387\nExp number, first 60 AAs:  11.47386\nTotal prob of N-in:   0.54858\nPOSSIBLE N-term signal sequence\noutside   1   718\nTMhelix   719   741\ninside   742   751

28841	GCF_900106085.1_IMG-taxon_2639763195_annotated_assembly	Pseudomonas corrugata	Proteobacteria; Gammaproteobacteria; Pseudomonadales; Pseudomonadaceae; Pseudomonas; Pseudomonas fluorescens group	MNVKHSSGVGQRENRRSLVILASTLLALSGGAAFADTAPADTSDGYTLTLKQLGRAYPMN	WP_024776312.1	cellulose synthase BcsB subunit [Pseudomonas corrugata]  Length: 751\nNumber of predicted TMHs:  1\nExp number of AAs in TMHs: 34.13377\nExp number, first 60 AAs:  11.47394\nTotal prob of N-in:   0.54857\nPOSSIBLE N-term signal sequence\noutside   1   718\nTMhelix   719   741\ninside   742   751

28842	GCF_001625455.1_ASM162545v1	Pseudomonas fluorescens	Proteobacteria; Gammaproteobacteria; Pseudomonadales; Pseudomonadaceae; Pseudomonas; Pseudomonas fluorescens group	MKPSTRRTFLPRRSLILITCSLVGWSGWALSAAAQISATEPAMMPVTALPGTDAPSNSYT	WP_081235347.1	cellulose synthase regulator BcsB [Pseudomonas fluorescens]	Length: 767\nNumber of predicted TMHs:  2\nExp number of AAs in TMHs: 39.57567\nExp number, first 60 AAs:  16.73599\nTotal prob of N-in:   0.75033\nPOSSIBLE N-term signal sequence\ninside   1   12\nTMhelix   13   35\noutside   36   734\nTMhelix   735   757\ninside   758   767

28843	GCF_001269485.1_ASM126948v1	Pseudomonas protegens	Proteobacteria; Gammaproteobacteria; Pseudomonadales; Pseudomonadaceae; Pseudomonas; Pseudomonas fluorescens group	MQKRTPRRALLKGTVMALGAAVAGSLGGCLTSRLYESRASNEVYTETVSQFYITADAKSFWP_053152798.1	hypothetical protein [Pseudomonas protegens]	Length: 219\nNumber of predicted TMHs:  1\nExp number of AAs in TMHs: 31.59185\nExp number, first 60 AAs:  11.36731\nTotal prob of N-in:   0.76688\nPOSSIBLE N-term signal sequence\ninside   1   184\nTMhelix   185   207\noutside   208   219

28844	GCF_002006545.1_ASM200654v1	Pseudomonas protegens	Proteobacteria; Gammaproteobacteria; Pseudomonadales; Pseudomonadaceae; Pseudomonas; Pseudomonas fluorescens group	MQKRTPRRALLKGTVMALGAAVAGSLGGCLTSRLYESRASNEVYTETVSQFYITADAKSFWP_077934522.1	hypothetical protein [Pseudomonas protegens]	Length: 219\nNumber of predicted TMHs: 1\nExp number of AAs in TMHs: 31.79975\nExp number, first 60 AAs: 11.50763\nTotal prob of N-in: 0.76814\nPOSSIBLE N-term signal sequence\ninside 1 184\nTMhelix 185 207\noutside 208 219

28845	GCF_001269495.1_ASM126949v1	Pseudomonas protegens	Proteobacteria; Gammaproteobacteria; Pseudomonadales; Pseudomonadaceae; Pseudomonas; Pseudomonas fluorescens group	MQKGRPRRALLKGTVMALGAAVAGSLGGCLTSKLYESRASNEVYTETVSQFYLTADAKSF	WP_053159530.1	hypothetical protein [Pseudomonas protegens]	Length: 219\nNumber of predicted TMHs: 1\nExp number of AAs in TMHs: 28.16201\nExp number, first 60 AAs: 8.08636\nTotal prob of N-in: 0.67073\ninside 1 184\nTMhelix 185 207\noutside 208 219

28846	GCF_000522485.1_Pco1	Pseudomonas corrugata CFBP 5454	Proteobacteria; Gammaproteobacteria; Pseudomonadales; Pseudomonadaceae; Pseudomonas; Pseudomonas fluorescens group; Pseudomonas corrugata	MNVKHSSGVGQRENRRSLVILASTLLALSGGAAFADTAPADTSDGYTLTLKQLGRAYPMN	WP_024776312.1	cellulose synthase BcsB subunit [Pseudomonas corrugata]	Length: 751\nNumber of predicted TMHs: 1\nExp number of AAs in TMHs: 34.13377\nExp number, first 60 AAs: 11.47394\nTotal prob of N-in: 0.54857\nPOSSIBLE N-term signal sequence\noutside 1 718\nTMhelix 719 741\ninside 742 751

28847	GCF_000828695.1_ASM82869v1	Pseudomonas protegens Cab57	Proteobacteria; Gammaproteobacteria; Pseudomonadales; Pseudomonadaceae; Pseudomonas; Pseudomonas fluorescens group; Pseudomonas protegens	MQKGRPRRALLKGTVMALGAAVAGSLGGCLTSKLYESRASNEVYTETVSQFYLTADAKSF	WP_041119189.1	hypothetical protein [Pseudomonas protegens]	Length: 219\nNumber of predicted TMHs: 1\nExp number of AAs in TMHs: 30.4417\nExp number, first 60 AAs: 8.56013\nTotal prob of N-in: 0.67913\ninside 1 184\nTMhelix 185 207\noutside 208 219

28848	GCF_000276565.1_ASM27656v1	Pseudomonas tolaasii PMS117	Proteobacteria; Gammaproteobacteria; Pseudomonadales; Pseudomonadaceae; Pseudomonas; Pseudomonas fluorescens group; Pseudomonas tolaasii	MSRRTTLLLLLALSAGHAQAANLTASVDRSRLNSGETVELTVESSDVTQFGKPDLSPLDA	WP_016971764.1	protein BatD, partial [Pseudomonas tolaasii]	Length: 472\nNumber of predicted TMHs: 1\nExp number of AAs in TMHs: 23.20248\nExp number, first 60 AAs: 0.69115\nTotal prob of N-in: 0.03683\noutside 1 408\nTMhelix 409 431\ninside 432 472

28849	GCF_001467305.1_ASM146730v1	Pseudomonas putida	Proteobacteria; Gammaproteobacteria; Pseudomonadales; Pseudomonadaceae; Pseudomonas; Pseudomonas putida group	MPPSFQLRARRSLIPGLLGSLVILCGPAHAEEGSGNERWVSDSLSTYVRSGPTDGHRIVG WP_058540537.1	peptide-binding protein [Pseudomonas putida]	Length: 215\nNumber of predicted TMHs: 1\nExp number of AAs in TMHs: 23.55908\nExp number, first 60 AAs: 1.33747\nTotal prob of N-in: 0.24943\noutside 1 182\nTMhelix 183 205\ninside 206 215

28850	GCF_000878325.1_UASWS0946_1.0	Pseudomonas putida	Proteobacteria; Gammaproteobacteria; Pseudomonadales; Pseudomonadaceae; Pseudomonas; Pseudomonas putida group	MPPSFQLRARRSLIPGLLGSLVILCGPAHAEEGSGNERWVSDSLSTYVRSGPTDGHRIVG WP_043861144.1	peptide-binding protein [Pseudomonas putida]	Length: 215\nNumber of predicted TMHs: 1\nExp number of AAs in TMHs: 23.55908\nExp number, first 60 AAs: 1.33747\nTotal prob of N-in: 0.24943\noutside 1 182\nTMhelix 183 205\ninside 206 215

28851	GCF_001976065.1_ASM197606v1	Pseudomonas putida	Proteobacteria; Gammaproteobacteria; Pseudomonadales; Pseudomonadaceae; Pseudomonas; Pseudomonas putida group	MNSKSFAFGSLRAPRRALVRMTCALLAMGGGATAFAETVPVDAAPGANSYSVTLKQLGRN	WP_077186208.1	cellulose synthase regulator BcsB [Pseudomonas putida]	Length: 757\nNumber of predicted TMHs: 1\nExp number of AAs in TMHs: 26.03203\nExp number, first 60 AAs: 3.22969\nTotal prob of N-in: 0.15300\noutside 1 724\nTMhelix 725 747\ninside 748 757

28852	GCF_001064225.1_ASM106422v1	Pseudomonas stutzeri	Proteobacteria; Gammaproteobacteria; Pseudomonadales; Pseudomonadaceae; Pseudomonas; Pseudomonas stutzeri group; Pseudomonas stutzeri subgroup	MFRPISVFIGSRYTRAKRRNHFISFISLTSLIGLALGVLAMIMVLSVMNGFQREMSARIL	WP_049325590.1	multidrug ABC transporter substrate-binding protein, partial [Pseudomonas stutzeri]	Length: 323\nNumber of predicted TMHs:

2\nExp number of AAs in TMHs: 47.57917\nExp number, first 60 AAs: 23.36115\nTotal prob of N-in: 0.96077\nPOSSIBLE N-term signal sequence\ninside 1 20\nTMhelix 21 43\noutside 44 273\nTMhelix 274 296\ninside 297 323

28853 GCF_000341615.1_PstNF13_1.0 Pseudomonas stutzeri NF13 Proteobacteria; Gammaproteobacteria; Pseudomonadales; Pseudomonadaceae; Pseudomonas; Pseudomonas stutzeri group; Pseudomonas stutzeri subgroup; Pseudomonas stutzeri MNATHAYGRRQVLLGSASLGVAAMLPAVGIGQTPDTVGLLNVPATDIGTYHREILFETTR WP_003298078.1 hypothetical protein [Pseudomonas stutzeri] Length: 209\nNumber of predicted TMHs: 1\nExp number of AAs in TMHs: 44.35717\nExp number, first 60 AAs: 16.92885\nTotal prob of N-in: 0.97522\nPOSSIBLE N-term signal sequence\ninside 1 180\nTMhelix 181 203\noutside 204 209

28854 GCF_001698815.1_ASM169881v1 Pseudomonas syringae Proteobacteria; Gammaproteobacteria; Pseudomonadales; Pseudomonadaceae; Pseudomonas; Pseudomonas syringae group; Pseudomonas syringae group genomosp. 1 MSMSRHLSAFLSRLPQLLNQPRRAIVGAGLLGGLLALCGPTHAAETKPNERWVSDSLNTF WP_065832530.1 peptide-binding protein [Pseudomonas syringae] Length: 228\nNumber of predicted TMHs: 1\nExp number of AAs in TMHs: 26.44853\nExp number, first 60 AAs: 4.34135\nTotal prob of N-in: 0.35490\noutside 1 195\nTMhelix 196 218\ninside 219 228

28855 GCF_001467105.1_ASM146710v1 Pseudomonas syringae ICMP 19498 Proteobacteria; Gammaproteobacteria; Pseudomonadales; Pseudomonadaceae; Pseudomonas; Pseudomonas syringae group; Pseudomonas syringae group genomosp. 1; Pseudomonas syringae MSKRRRVLRLLIVATLASLLQACDIDLYTNLGEREANAMLAVLLRDGIPASRKVQDNGQL WP_057412399.1 MULTISPECIES: EscJ/YscJ/HrcJ family type III secretion inner membrane ring protein [Pseudomonas] Length: 259\nNumber of predicted TMHs: 1\nExp number of AAs in TMHs: 33.56925\nExp number, first 60 AAs: 8.63141\nTotal prob of N-in: 0.58822\noutside 1 217\nTMhelix 218 240\ninside 241 259

28856 GCF_001467115.1_ASM146711v1 Pseudomonas syringae ICMP 19499 Proteobacteria; Gammaproteobacteria; Pseudomonadales; Pseudomonadaceae; Pseudomonas; Pseudomonas syringae group; Pseudomonas syringae group genomosp. 1; Pseudomonas syringae MSKRRRVLRLLIVATLASLLQACDIDLYTNLGEREANAMLAVLLRDGIPASRKVQDNGQL WP_057412399.1 MULTISPECIES: EscJ/YscJ/HrcJ family type III secretion inner membrane ring protein [Pseudomonas] Length: 259\nNumber of predicted TMHs: 1\nExp number of AAs in TMHs: 33.56925\nExp number, first 60 AAs: 8.63141\nTotal prob of N-in: 0.58822\noutside 1 217\nTMhelix 218 240\ninside 241 259

28857 GCF_001538245.1_ASM153824v1 Pseudomonas syringae pv. castaneae Proteobacteria; Gammaproteobacteria; Pseudomonadales; Pseudomonadaceae; Pseudomonas; Pseudomonas syringae group; Pseudomonas syringae group genomosp. 1; Pseudomonas syringae; Pseudomonas syringae group pathovars incertae sedis MSKRRRVLRLLIVATLASLLQACDIDLYTNLGEREANAMLAVLLRDGIPASRKVQDNGQL WP_057412399.1 MULTISPECIES: EscJ/YscJ/HrcJ family type III secretion inner membrane ring protein [Pseudomonas] Length: 259\nNumber of predicted TMHs: 1\nExp number of AAs in TMHs: 33.56925\nExp number, first 60 AAs: 8.63141\nTotal prob of N-in: 0.58822\noutside 1 217\nTMhelix 218 240\ninside 241 259

28858 GCF_001400285.1_PcsICMP9419 Pseudomonas syringae pv. castaneae Proteobacteria; Gammaproteobacteria; Pseudomonadales; Pseudomonadaceae; Pseudomonas; Pseudomonas syringae group; Pseudomonas syringae group genomosp. 1; Pseudomonas syringae; Pseudomonas syringae group pathovars incertae sedis MSKRRRVLRLLIVATLASLLQACDIDLYTNLGEREANAMLAVLLRDGIPASRKVQDNGQL WP_057412399.1 MULTISPECIES: EscJ/YscJ/HrcJ family type III secretion inner membrane ring protein [Pseudomonas] Length: 259\nNumber of predicted TMHs: 1\nExp number of AAs in TMHs: 33.56925\nExp number, first 60 AAs: 8.63141\nTotal prob of N-in: 0.58822\noutside 1 217\nTMhelix 218 240\ninside 241 259

28859 GCF_001537945.1_ASM153794v1 Pseudomonas syringae pv. cerasicola Proteobacteria; Gammaproteobacteria; Pseudomonadales; Pseudomonadaceae; Pseudomonas; Pseudomonas syringae group; Pseudomonas syringae group genomosp. 1; Pseudomonas syringae; Pseudomonas syringae group pathovars incertae sedis MPKRRRVLRLLIVATLASLLQACDIDLYTNLGEREANAMLAVLLRDGIPASRKVQDNGQL WP_005733361.1 MULTISPECIES: EscJ/YscJ/HrcJ family type III secretion inner membrane ring protein [Pseudomonas syringae group] Length: 259\nNumber of predicted TMHs: 1\nExp number of AAs in TMHs: 31.46284\nExp number, first 60 AAs: 7.15654999999999\nTotal prob of N-in: 0.48729\noutside 1 217\nTMhelix 218 240\ninside 241 259

28860 GCF_001400215.1_PceICMP17524 Pseudomonas syringae pv. cerasicola Proteobacteria; Gammaproteobacteria; Pseudomonadales; Pseudomonadaceae; Pseudomonas; Pseudomonas syringae group; Pseudomonas syringae group genomosp. 1; Pseudomonas syringae; Pseudomonas syringae group pathovars incertae sedis MPKRRRVLRLLIVATLASLLQACDIDLYTNLGEREANAMLAVLLRDGIPASRKVQDNGQL WP_005733361.1 MULTISPECIES:

EscJ/YscJ/HrcJ family type III secretion inner membrane ring protein [Pseudomonas syringae group] Length: 259\nNumber of predicted TMHs: 1\nExp number of AAs in TMHs: 31.46284\nExp number, first 60 AAs: 7.15654999999999\nTotal prob of N-in: 0.48729\noutside 1 217\nTMhelix 218 240\ninside 241 259

28861 GCF_001400795.1_PcuICMP11894 Pseudomonas syringae pv. cunninghamiae Proteobacteria; Gammaproteobacteria; Pseudomonadales; Pseudomonadaceae; Pseudomonas; Pseudomonas syringae group; Pseudomonas syringae group genomosp. 1; Pseudomonas syringae; Pseudomonas syringae group pathovars incertae sedis MPKRRRVLRLLIVATLASLLQACDIDLYTNLGEREANAMLAVLLRDGIPASRKVQDNGQL WP_002553574.1 MULTISPECIES: EscJ/YscJ/HrcJ family type III secretion inner membrane ring protein [Pseudomonas syringae group] Length: 259\nNumber of predicted TMHs: 1\nExp number of AAs in TMHs: 31.58284\nExp number, first 60 AAs: 7.2383\nTotal prob of N-in: 0.48165\noutside 1 217\nTMhelix 218 240\ninside 241 259

28862 GCF_001400275.1_PdaICMP9757 Pseudomonas syringae pv. daphniphylli Proteobacteria; Gammaproteobacteria; Pseudomonadales; Pseudomonadaceae; Pseudomonas; Pseudomonas syringae group; Pseudomonas syringae group genomosp. 1; Pseudomonas syringae; Pseudomonas syringae group pathovars incertae sedis MPKRRRVLRLLIVATLASLLQACDIDLYTNLGEREANAMLAVLLRDGIPASRKVQDNGQL WP_002553574.1 MULTISPECIES: EscJ/YscJ/HrcJ family type III secretion inner membrane ring protein [Pseudomonas syringae group] Length: 259\nNumber of predicted TMHs: 1\nExp number of AAs in TMHs: 31.58284\nExp number, first 60 AAs: 7.2383\nTotal prob of N-in: 0.48165\noutside 1 217\nTMhelix 218 240\ninside 241 259

28863 GCF_001538255.1_ASM153825v1 Pseudomonas syringae pv. daphniphylli Proteobacteria; Gammaproteobacteria; Pseudomonadales; Pseudomonadaceae; Pseudomonas; Pseudomonas syringae group; Pseudomonas syringae group genomosp. 1; Pseudomonas syringae; Pseudomonas syringae group pathovars incertae sedis MPKRRRVLRLLIVATLASLLQACDIDLYTNLGEREANAMLAVLLRDGIPASRKVQDNGQL WP_002553574.1 MULTISPECIES: EscJ/YscJ/HrcJ family type III secretion inner membrane ring protein [Pseudomonas syringae group] Length: 259\nNumber of predicted TMHs: 1\nExp number of AAs in TMHs: 31.58284\nExp number, first 60 AAs: 7.2383\nTotal prob of N-in: 0.48165\noutside 1 217\nTMhelix 218 240\ninside 241 259

28864 GCF_000302795.1_Pav013_1.0 Pseudomonas syringae pv. avellanae str. ISPaVe013 Proteobacteria; Gammaproteobacteria; Pseudomonadales; Pseudomonadaceae; Pseudomonas; Pseudomonas syringae group; Pseudomonas syringae group genomosp. 1; Pseudomonas syringae; Pseudomonas syringae pv. syringae MSKRRRVLRLLIVATLASLLQACDIDLYTNLGEREANAMLAVLLRDGIPASRKVQDNGQL WP_003410081.1 EscJ/YscJ/HrcJ family type III secretion inner membrane ring protein [Pseudomonas syringae] Length: 259\nNumber of predicted TMHs: 1\nExp number of AAs in TMHs: 33.3785799999999\nExp number, first 60 AAs: 9.04202999999999\nTotal prob of N-in: 0.57849\noutside 1 217\nTMhelix 218 240\ninside 241 259

28865 GCF_000972195.1_ASM97219v1 Pseudomonas syringae pv. syringae CRAFRU12 Proteobacteria; Gammaproteobacteria; Pseudomonadales; Pseudomonadaceae; Pseudomonas; Pseudomonas syringae group; Pseudomonas syringae group genomosp. 1; Pseudomonas syringae; Pseudomonas syringae pv. syringae MSKRRRVLRLLIVATLASLLQACDIDLYTNLGEREANAMLAVLLRDGIPASRKVQDNGQL WP_003410081.1 EscJ/YscJ/HrcJ family type III secretion inner membrane ring protein [Pseudomonas syringae] Length: 259\nNumber of predicted TMHs: 1\nExp number of AAs in TMHs: 33.3785799999999\nExp number, first 60 AAs: 9.04202999999999\nTotal prob of N-in: 0.57849\noutside 1 217\nTMhelix 218 240\ninside 241 259

28866 GCF_001400115.1_PamICMP3918 Pseudomonas amygdali Proteobacteria; Gammaproteobacteria; Pseudomonadales; Pseudomonadaceae; Pseudomonas; Pseudomonas syringae group; Pseudomonas syringae group genomosp. 2 MPKRRRVLRLLIVATLASLLQACDIDLYTNLGEREANAMLAVLLRDGIPASRKVQDNGQL WP_004660817.1 MULTISPECIES: EscJ/YscJ/HrcJ family type III secretion inner membrane ring protein [Pseudomonas syringae group genomosp. 2] Length: 259\nNumber of predicted TMHs: 1\nExp number of AAs in TMHs: 31.46243\nExp number, first 60 AAs: 7.15651999999999\nTotal prob of N-in: 0.48728\noutside 1 217\nTMhelix 218 240\ninside 241 259

28867 GCF_000935645.1_PSAVPseNe107-G1 Pseudomonas amygdali Proteobacteria; Gammaproteobacteria; Pseudomonadales; Pseudomonadaceae; Pseudomonas; Pseudomonas syringae group; Pseudomonas syringae group genomosp. 2 MPKRRRVLRLLIVATLASLLQACDIDLYTNLGEREANAMLAVLLRDGIPASRKVQDNGQL WP_002553574.1 MULTISPECIES: EscJ/YscJ/HrcJ family type III secretion inner membrane ring protein [Pseudomonas syringae group] Length: 259\nNumber of predicted TMHs: 1\nExp number of AAs in TMHs: 31.58284\nExp number, first 60 AAs: 7.2383\nTotal prob of N-in: 0.48165\noutside 1 217\nTMhelix 218 240\ninside 241 259

28868 GCF_000935675.1_PMCFBP3225-G1 Pseudomonas meliae Proteobacteria; Gammaproteobacteria; Pseudomonadales; Pseudomonadaceae; Pseudomonas; Pseudomonas syringae group; Pseudomonas syringae group genomosp. 2	MPKRRRVLRLLIVATLASLLQACDIDLYTNLGEREANAMLAVLLRDGIPASRKVQDNGQLWP_005733361.1	MULTISPECIES: EscJ/YscJ/HrcJ family type III secretion inner membrane ring protein [Pseudomonas syringae group]	Length: 259\nNumber of predicted TMHs: 1\nExp number of AAs in TMHs: 31.46284\nExp number, first 60 AAs: 7.15654999999999\nTotal prob of N-in: 0.48729\noutside 1 217\nTMhelix 218 240\ninside 241 259

28869	GCF_001293725.1_Psv4352	Pseudomonas savastanoi	Proteobacteria; Gammaproteobacteria; Pseudomonadales; Pseudomonadaceae; Pseudomonas; Pseudomonas syringae group; Pseudomonas syringae group genomosp. 2	MPKRRRVLRLLIVATLASLLQACDIDLYTNLGEREANAMLAVLLRDGIPASRKVQDNGQLWP_002553574.1	MULTISPECIES: EscJ/YscJ/HrcJ family type III secretion inner membrane ring protein [Pseudomonas syringae group]	Length: 259\nNumber of predicted TMHs: 1\nExp number of AAs in TMHs: 31.58284\nExp number, first 60 AAs: 7.2383\nTotal prob of N-in: 0.48165\noutside 1 217\nTMhelix 218 240\ninside 241 259

28870	GCF_001537985.1_ASM153798v1	Pseudomonas amygdali pv. aesculi	Proteobacteria; Gammaproteobacteria; Pseudomonadales; Pseudomonadaceae; Pseudomonas; Pseudomonas syringae group; Pseudomonas syringae group genomosp. 2; Pseudomonas amygdali	MPKRRRVLRLLIVATLASLLQACDIDLYTNLGEREANAMLAVLLRDGIPASRKVQDNGQLWP_005733361.1	MULTISPECIES: EscJ/YscJ/HrcJ family type III secretion inner membrane ring protein [Pseudomonas syringae group]	Length: 259\nNumber of predicted TMHs: 1\nExp number of AAs in TMHs: 31.46284\nExp number, first 60 AAs: 7.15654999999999\nTotal prob of N-in: 0.48729\noutside 1 217\nTMhelix 218 240\ninside 241 259

28871	GCF_001538185.1_ASM153818v1	Pseudomonas amygdali pv. aesculi	Proteobacteria; Gammaproteobacteria; Pseudomonadales; Pseudomonadaceae; Pseudomonas; Pseudomonas syringae group; Pseudomonas syringae group genomosp. 2; Pseudomonas amygdali	MPKRRRVLRLLIVATLASLLQACDIDLYTNLGEREANAMLAVLLRDGIPASRKVQDNGQLWP_005733361.1	MULTISPECIES: EscJ/YscJ/HrcJ family type III secretion inner membrane ring protein [Pseudomonas syringae group]	Length: 259\nNumber of predicted TMHs: 1\nExp number of AAs in TMHs: 31.46284\nExp number, first 60 AAs: 7.15654999999999\nTotal prob of N-in: 0.48729\noutside 1 217\nTMhelix 218 240\ninside 241 259

28872	GCF_001538085.1_ASM153808v1	Pseudomonas amygdali pv. aesculi	Proteobacteria; Gammaproteobacteria; Pseudomonadales; Pseudomonadaceae; Pseudomonas; Pseudomonas syringae group; Pseudomonas syringae group genomosp. 2; Pseudomonas amygdali	MPKRRRVLRLLIVATLASLLQACDIDLYTNLGEREANAMLAVLLRDGIPASRKVQDNGQLWP_005733361.1	MULTISPECIES: EscJ/YscJ/HrcJ family type III secretion inner membrane ring protein [Pseudomonas syringae group]	Length: 259\nNumber of predicted TMHs: 1\nExp number of AAs in TMHs: 31.46284\nExp number, first 60 AAs: 7.15654999999999\nTotal prob of N-in: 0.48729\noutside 1 217\nTMhelix 218 240\ninside 241 259

28873	GCF_001537965.1_ASM153796v1	Pseudomonas amygdali pv. aesculi	Proteobacteria; Gammaproteobacteria; Pseudomonadales; Pseudomonadaceae; Pseudomonas; Pseudomonas syringae group; Pseudomonas syringae group genomosp. 2; Pseudomonas amygdali	MPKRRRVLRLLIVATLASLLQACDIDLYTNLGEREANAMLAVLLRDGIPASRKVQDNGQLWP_005733361.1	MULTISPECIES: EscJ/YscJ/HrcJ family type III secretion inner membrane ring protein [Pseudomonas syringae group]	Length: 259\nNumber of predicted TMHs: 1\nExp number of AAs in TMHs: 31.46284\nExp number, first 60 AAs: 7.15654999999999\nTotal prob of N-in: 0.48729\noutside 1 217\nTMhelix 218 240\ninside 241 259

28874	GCF_001538045.1_ASM153804v1	Pseudomonas amygdali pv. aesculi	Proteobacteria; Gammaproteobacteria; Pseudomonadales; Pseudomonadaceae; Pseudomonas; Pseudomonas syringae group; Pseudomonas syringae group genomosp. 2; Pseudomonas amygdali	MPKRRRVLRLLIVATLASLLQACDIDLYTNLGEREANAMLAVLLRDGIPASRKVQDNGQLWP_005733361.1	MULTISPECIES: EscJ/YscJ/HrcJ family type III secretion inner membrane ring protein [Pseudomonas syringae group]	Length: 259\nNumber of predicted TMHs: 1\nExp number of AAs in TMHs: 31.46284\nExp number, first 60 AAs: 7.15654999999999\nTotal prob of N-in: 0.48729\noutside 1 217\nTMhelix 218 240\ninside 241 259

28875	GCF_001537955.1_ASM153795v1	Pseudomonas amygdali pv. aesculi	Proteobacteria; Gammaproteobacteria; Pseudomonadales; Pseudomonadaceae; Pseudomonas; Pseudomonas syringae group; Pseudomonas syringae group genomosp. 2; Pseudomonas amygdali	MPKRRRVLRLLIVATLASLLQACDIDLYTNLGEREANAMLAVLLRDGIPASRKVQDNGQLWP_005733361.1	MULTISPECIES: EscJ/YscJ/HrcJ family type III secretion inner membrane ring protein [Pseudomonas syringae group]	Length: 259\nNumber of predicted TMHs: 1\nExp number of AAs in TMHs: 31.46284\nExp number, first 60 AAs: 7.15654999999999\nTotal prob of N-in: 0.48729\noutside 1 217\nTMhelix 218 240\ninside 241 259

28876 GCF_001400755.1_PccICMP5710 Pseudomonas amygdali pv. ciccaronei Proteobacteria; Gammaproteobacteria; Pseudomonadales; Pseudomonadaceae; Pseudomonas; Pseudomonas syringae group; Pseudomonas syringae group genomosp. 2; Pseudomonas amygdali MPKRRRVLRLLIVATLASLLQACDIDLYTNLGEREANAMLAVLLRDGIPASRKVQDNGQL WP_002553574.1 MULTISPECIES: EscJ/YscJ/HrcJ family type III secretion inner membrane ring protein [Pseudomonas syringae group] Length: 259\nNumber of predicted TMHs: 1\nExp number of AAs in TMHs: 31.58284\nExp number, first 60 AAs: 7.2383\nTotal prob of N-in: 0.48165\noutside 1 217\nTMhelix 218 240\ninside 241 259

28877 GCF_000935665.1_PDCFBP4219-G1 Pseudomonas amygdali pv. dendropanacis Proteobacteria; Gammaproteobacteria; Pseudomonadales; Pseudomonadaceae; Pseudomonas; Pseudomonas syringae group; Pseudomonas syringae group genomosp. 2; Pseudomonas amygdali MPKRRRVLRLLIVATLASLLQACDIDLYTNLGEREANAMLAVLLRDGIPASRKVQDNGQL WP_002553574.1 MULTISPECIES: EscJ/YscJ/HrcJ family type III secretion inner membrane ring protein [Pseudomonas syringae group] Length: 259\nNumber of predicted TMHs: 1\nExp number of AAs in TMHs: 31.58284\nExp number, first 60 AAs: 7.2383\nTotal prob of N-in: 0.48165\noutside 1 217\nTMhelix 218 240\ninside 241 259

28878 GCF_001538055.1_ASM153805v1 Pseudomonas amygdali pv. eriobotryae Proteobacteria; Gammaproteobacteria; Pseudomonadales; Pseudomonadaceae; Pseudomonas; Pseudomonas syringae group; Pseudomonas syringae group genomosp. 2; Pseudomonas amygdali MPKRRRVLRLLIVATLASLLQACDIDLYTNLGEREANAMLAVLLRDGIPASRKVQDNGQL WP_002553574.1 MULTISPECIES: EscJ/YscJ/HrcJ family type III secretion inner membrane ring protein [Pseudomonas syringae group] Length: 259\nNumber of predicted TMHs: 1\nExp number of AAs in TMHs: 31.58284\nExp number, first 60 AAs: 7.2383\nTotal prob of N-in: 0.48165\noutside 1 217\nTMhelix 218 240\ninside 241 259

28879 GCF_001400395.1_PhiICMP9623 Pseudomonas amygdali pv. hibisci Proteobacteria; Gammaproteobacteria; Pseudomonadales; Pseudomonadaceae; Pseudomonas; Pseudomonas syringae group; Pseudomonas syringae group genomosp. 2; Pseudomonas amygdali MPKRRRVLRLLIVATLASLLQACDIDLYTNLGEREANAMLAVLLRDGIPASRKVQDNGQL WP_005733361.1 MULTISPECIES: EscJ/YscJ/HrcJ family type III secretion inner membrane ring protein [Pseudomonas syringae group] Length: 259\nNumber of predicted TMHs: 1\nExp number of AAs in TMHs: 31.46284\nExp number, first 60 AAs: 7.15654999999999\nTotal prob of N-in: 0.48729\noutside 1 217\nTMhelix 218 240\ninside 241 259

28880 GCF_001006445.1_PSl_v01 Pseudomonas amygdali pv. lachrymans Proteobacteria; Gammaproteobacteria; Pseudomonadales; Pseudomonadaceae; Pseudomonas; Pseudomonas syringae group; Pseudomonas syringae group genomosp. 2; Pseudomonas amygdali MPKRRRVLRLLIVATLASLLQTCDIDLYTNLGEREANAMLAVLLRDGIPASRKVQDNGQL WP_005743513.1 EscJ/YscJ/HrcJ family type III secretion inner membrane ring protein [Pseudomonas amygdali] Length: 259\nNumber of predicted TMHs: 1\nExp number of AAs in TMHs: 30.06245\nExp number, first 60 AAs: 4.73138\nTotal prob of N-in: 0.42541\noutside 1 217\nTMhelix 218 240\ninside 241 259

28881 GCF_001293845.1_PlaYM7902 Pseudomonas amygdali pv. lachrymans Proteobacteria; Gammaproteobacteria; Pseudomonadales; Pseudomonadaceae; Pseudomonas; Pseudomonas syringae group; Pseudomonas syringae group genomosp. 2; Pseudomonas amygdali MPKRRRVLRLLIVATLASLLQTCDIDLYTNLGEREANAMLAVLLRDGIPASRKVQDNGQL WP_005743513.1 EscJ/YscJ/HrcJ family type III secretion inner membrane ring protein [Pseudomonas amygdali] Length: 259\nNumber of predicted TMHs: 1\nExp number of AAs in TMHs: 30.06245\nExp number, first 60 AAs: 4.73138\nTotal prob of N-in: 0.42541\noutside 1 217\nTMhelix 218 240\ninside 241 259

28882 GCF_002068135.1_ASM206813v1 Pseudomonas amygdali pv. lachrymans Proteobacteria; Gammaproteobacteria; Pseudomonadales; Pseudomonadaceae; Pseudomonas; Pseudomonas syringae group; Pseudomonas syringae group genomosp. 2; Pseudomonas amygdali MPKRRRVLRLLIVATLASLLQTCDIDLYTNLGEREANAMLAVLLRDGIPASRKVQDNGQL WP_005743513.1 EscJ/YscJ/HrcJ family type III secretion inner membrane ring protein [Pseudomonas amygdali] Length: 259\nNumber of predicted TMHs: 1\nExp number of AAs in TMHs: 30.06245\nExp number, first 60 AAs: 4.73138\nTotal prob of N-in: 0.42541\noutside 1 217\nTMhelix 218 240\ninside 241 259

28883 GCF_001400875.1_PmeICMP5711 Pseudomonas amygdali pv. mellea Proteobacteria; Gammaproteobacteria; Pseudomonadales; Pseudomonadaceae; Pseudomonas; Pseudomonas syringae group; Pseudomonas syringae group genomosp. 2; Pseudomonas amygdali MPKRRRVLRLLIVATLASLLQACDIDLYTNLGEREANAMLAVLLRDGIPASRKVQDNGQL WP_004660817.1 MULTISPECIES: EscJ/YscJ/HrcJ family type III secretion inner membrane ring protein [Pseudomonas syringae group genomosp. 2]

Length: 259\nNumber of predicted TMHs: 1\nExp number of AAs in TMHs: 31.46243\nExp number, first 60 AAs: 7.15651999999999\nTotal prob of N-in: 0.48728\noutside 1 217\nTMhelix 218 240\ninside 241 259

28884 GCF_001293975.1_PmeN6801 Pseudomonas amygdali pv. mellea Proteobacteria; Gammaproteobacteria; Pseudomonadales; Pseudomonadaceae; Pseudomonas; Pseudomonas syringae group; Pseudomonas syringae group genomosp. 2; Pseudomonas amygdali MPKRRRVLRLLIVATLASLLQACDIDLYTNLGEREANAMLAVLLRDGIPASRKVQDNGQL WP_004660817.1 MULTISPECIES: EscJ/YscJ/HrcJ family type III secretion inner membrane ring protein [Pseudomonas syringae group genomosp. 2] Length: 259\nNumber of predicted TMHs: 1\nExp number of AAs in TMHs: 31.46243\nExp number, first 60 AAs: 7.15651999999999\nTotal prob of N-in: 0.48728\noutside 1 217\nTMhelix 218 240\ninside 241 259

28885 GCF_001294215.1_PmpFTRS_U7805 Pseudomonas amygdali pv. morsprunorum Proteobacteria; Gammaproteobacteria; Pseudomonadales; Pseudomonadaceae; Pseudomonas; Pseudomonas syringae group; Pseudomonas syringae group genomosp. 2; Pseudomonas amygdali MPKRRRVLRLLIVATLASLLQACDIDLYTNLGEREANAMLAVLLRDGIPASRKVQDNGQL WP_005733361.1 MULTISPECIES: EscJ/YscJ/HrcJ family type III secretion inner membrane ring protein [Pseudomonas syringae group] Length: 259\nNumber of predicted TMHs: 1\nExp number of AAs in TMHs: 31.46284\nExp number, first 60 AAs: 7.15654999999999\nTotal prob of N-in: 0.48729\noutside 1 217\nTMhelix 218 240\ninside 241 259

28886 GCF_001535785.1_ASM153578v1 Pseudomonas amygdali pv. morsprunorum Proteobacteria; Gammaproteobacteria; Pseudomonadales; Pseudomonadaceae; Pseudomonas; Pseudomonas syringae group; Pseudomonas syringae group genomosp. 2; Pseudomonas amygdali MSKRRRVLRLLIVATLASLLQACDIDLYTNLGEREANAMLAVLLRDGIPASRKVQDNGQL WP_057412399.1 MULTISPECIES: EscJ/YscJ/HrcJ family type III secretion inner membrane ring protein [Pseudomonas] Length: 259\nNumber of predicted TMHs: 1\nExp number of AAs in TMHs: 33.56925\nExp number, first 60 AAs: 8.63141\nTotal prob of N-in: 0.58822\noutside 1 217\nTMhelix 218 240\ninside 241 259

28887 GCF_001535735.1_ASM153573v1 Pseudomonas amygdali pv. morsprunorum Proteobacteria; Gammaproteobacteria; Pseudomonadales; Pseudomonadaceae; Pseudomonas; Pseudomonas syringae group; Pseudomonas syringae group genomosp. 2; Pseudomonas amygdali MSKRRRVLRLLIVATLASLLQACDIDLYTNLGEREANAMLAVLLRDGIPASRKVQDNGQL WP_057412399.1 MULTISPECIES: EscJ/YscJ/HrcJ family type III secretion inner membrane ring protein [Pseudomonas] Length: 259\nNumber of predicted TMHs: 1\nExp number of AAs in TMHs: 33.56925\nExp number, first 60 AAs: 8.63141\nTotal prob of N-in: 0.58822\noutside 1 217\nTMhelix 218 240\ninside 241 259

28888 GCF_001293985.1_PmyAZ84488 Pseudomonas amygdali pv. myricae Proteobacteria; Gammaproteobacteria; Pseudomonadales; Pseudomonadaceae; Pseudomonas; Pseudomonas syringae group; Pseudomonas syringae group genomosp. 2; Pseudomonas amygdali MSKRRRVLRLLIVATLASLLQACDIDLYTNLGEREANAMLAVLLRDGIPASRKVQDNGQL WP_054072647.1 EscJ/YscJ/HrcJ family type III secretion inner membrane ring protein [Pseudomonas amygdali] Length: 259\nNumber of predicted TMHs: 1\nExp number of AAs in TMHs: 33.56895\nExp number, first 60 AAs: 8.6315\nTotal prob of N-in: 0.58821\noutside 1 217\nTMhelix 218 240\ninside 241 259

28889 GCF_001535805.1_ASM153580v1 Pseudomonas amygdali pv. myricae Proteobacteria; Gammaproteobacteria; Pseudomonadales; Pseudomonadaceae; Pseudomonas; Pseudomonas syringae group; Pseudomonas syringae group genomosp. 2; Pseudomonas amygdali MSKRRRVLRLLIVATLASLLQACDIDLYTNLGEREANAMLAVLLRDGIPASRKVQDNGQL WP_054072647.1 EscJ/YscJ/HrcJ family type III secretion inner membrane ring protein [Pseudomonas amygdali] Length: 259\nNumber of predicted TMHs: 1\nExp number of AAs in TMHs: 33.56895\nExp number, first 60 AAs: 8.6315\nTotal prob of N-in: 0.58821\noutside 1 217\nTMhelix 218 240\ninside 241 259

28890 GCF_001400555.1_PmyICMP7118 Pseudomonas amygdali pv. myricae Proteobacteria; Gammaproteobacteria; Pseudomonadales; Pseudomonadaceae; Pseudomonas; Pseudomonas syringae group; Pseudomonas syringae group genomosp. 2; Pseudomonas amygdali MSKRRRVLRLLIVATLASLLQACDIDLYTNLGEREANAMLAVLLRDGIPASRKVQDNGQL WP_054072647.1 EscJ/YscJ/HrcJ family type III secretion inner membrane ring protein [Pseudomonas amygdali] Length: 259\nNumber of predicted TMHs: 1\nExp number of AAs in TMHs: 33.56895\nExp number, first 60 AAs: 8.6315\nTotal prob of N-in: 0.58821\noutside 1 217\nTMhelix 218 240\ninside 241 259

28891 GCF_001400415.1_PhoICMP7840 Pseudomonas amygdali pv. photiniae Proteobacteria; Gammaproteobacteria; Pseudomonadales; Pseudomonadaceae; Pseudomonas; Pseudomonas syringae group;

Pseudomonas syringae group genomosp. 2; Pseudomonas amygdali
MSKRRRVLRLLIVATLASLLQACDIDLYTNLGEREANAMLAVLLRDGIPASRKVQDNGQL WP_057412399.1 MULTISPECIES: EscJ/YscJ/HrcJ family type III secretion inner membrane ring protein [Pseudomonas] Length: 259\nNumber of predicted TMHs: 1\nExp number of AAs in TMHs: 33.56925\nExp number, first 60 AAs: 8.63141\nTotal prob of N-in: 0.58822\noutside 1 217\nTMhelix 218 240\ninside 241 259

28892 GCF_001294115.1_PseHC_1 Pseudomonas amygdali pv. sesami Proteobacteria; Gammaproteobacteria; Pseudomonadales; Pseudomonadaceae; Pseudomonas; Pseudomonas syringae group; Pseudomonas syringae group genomosp. 2; Pseudomonas amygdali MPKRRRVLRLLIVATLASLLQACDIDLYTNLGEREANAMLAVLLRDGIPASRKVQDNGQL WP_054077311.1 EscJ/YscJ/HrcJ family type III secretion inner membrane ring protein [Pseudomonas amygdali] Length: 259\nNumber of predicted TMHs: 1\nExp number of AAs in TMHs: 30.61995\nExp number, first 60 AAs: 7.21721\nTotal prob of N-in: 0.44727\noutside 1 217\nTMhelix 218 240\ninside 241 259

28893 GCF_001401265.1_PseICMP763 Pseudomonas amygdali pv. sesami Proteobacteria; Gammaproteobacteria; Pseudomonadales; Pseudomonadaceae; Pseudomonas; Pseudomonas syringae group; Pseudomonas syringae group genomosp. 2; Pseudomonas amygdali
MPKRRRVLRLLIVATLASLLQACDIDLYTNLGEREANAMLAVLLRDGIPASRKVQDNGQL WP_054077311.1 EscJ/YscJ/HrcJ family type III secretion inner membrane ring protein [Pseudomonas amygdali] Length: 259\nNumber of predicted TMHs: 1\nExp number of AAs in TMHs: 30.61995\nExp number, first 60 AAs: 7.21721\nTotal prob of N-in: 0.44727\noutside 1 217\nTMhelix 218 240\ninside 241 259

28894 GCF_000934645.1_ASM93464v1 Pseudomonas amygdali pv. tabaci Proteobacteria; Gammaproteobacteria; Pseudomonadales; Pseudomonadaceae; Pseudomonas; Pseudomonas syringae group; Pseudomonas syringae group genomosp. 2; Pseudomonas amygdali
MPKRRRVLRLLIVATLASLLQACDIDLYTNLGEREANAMLAVLLRDGIPASRKVQDNGQL WP_005777666.1 EscJ/YscJ/HrcJ family type III secretion inner membrane ring protein [Pseudomonas amygdali] Length: 259\nNumber of predicted TMHs: 1\nExp number of AAs in TMHs: 31.78523\nExp number, first 60 AAs: 6.9796\nTotal prob of N-in: 0.50005\noutside 1 217\nTMhelix 218 240\ninside 241 259

28895 GCF_001401095.1_PtaICMP2835 Pseudomonas amygdali pv. tabaci Proteobacteria; Gammaproteobacteria; Pseudomonadales; Pseudomonadaceae; Pseudomonas; Pseudomonas syringae group; Pseudomonas syringae group genomosp. 2; Pseudomonas amygdali
MPKRRRVLRLLIVATLASLLQACDIDLYTNLGEREANAMLAVLLRDGIPASRKVQDNGQL WP_005777666.1 EscJ/YscJ/HrcJ family type III secretion inner membrane ring protein [Pseudomonas amygdali] Length: 259\nNumber of predicted TMHs: 1\nExp number of AAs in TMHs: 31.78523\nExp number, first 60 AAs: 6.9796\nTotal prob of N-in: 0.50005\noutside 1 217\nTMhelix 218 240\ninside 241 259

28896 GCF_001401165.1_PumICMP3962 Pseudomonas amygdali pv. ulmi Proteobacteria; Gammaproteobacteria; Pseudomonadales; Pseudomonadaceae; Pseudomonas; Pseudomonas syringae group; Pseudomonas syringae group genomosp. 2; Pseudomonas amygdali
MPKRRRVLRLLIVATLASLLQACDIDLYTNLGEREANAMLAVLLRDGIPASRKVQDNGQL WP_057430882.1 EscJ/YscJ/HrcJ family type III secretion inner membrane ring protein [Pseudomonas amygdali] Length: 259\nNumber of predicted TMHs: 1\nExp number of AAs in TMHs: 32.00344\nExp number, first 60 AAs: 6.85934999999999\nTotal prob of N-in: 0.50874\noutside 1 217\nTMhelix 218 240\ninside 241 259

28897 GCF_001535915.1_ASM153591v1 Pseudomonas amygdali pv. ulmi Proteobacteria; Gammaproteobacteria; Pseudomonadales; Pseudomonadaceae; Pseudomonas; Pseudomonas syringae group; Pseudomonas syringae group genomosp. 2; Pseudomonas amygdali
MPKRRRVLRLLIVATLASLLQACDIDLYTNLGEREANAMLAVLLRDGIPASRKVQDNGQL WP_057430882.1 EscJ/YscJ/HrcJ family type III secretion inner membrane ring protein [Pseudomonas amygdali] Length: 259\nNumber of predicted TMHs: 1\nExp number of AAs in TMHs: 32.00344\nExp number, first 60 AAs: 6.85934999999999\nTotal prob of N-in: 0.50874\noutside 1 217\nTMhelix 218 240\ninside 241 259

28898 GCF_000145685.1_ASM14568v1 Pseudomonas amygdali pv. aesculi str. 0893_23 Proteobacteria; Gammaproteobacteria; Pseudomonadales; Pseudomonadaceae; Pseudomonas; Pseudomonas syringae group; Pseudomonas syringae group genomosp. 2; Pseudomonas amygdali; Pseudomonas amygdali pv. aesculi
MPKRRRVLRLLIVATLASLLQACDIDLYTNLGEREANAMLAVLLRDGIPASRKVQDNGQL WP_005733361.1 MULTISPECIES: EscJ/YscJ/HrcJ family type III secretion inner membrane ring protein [Pseudomonas syringae group] Length: 259\nNumber of predicted TMHs: 1\nExp number of AAs in TMHs: 31.46284\nExp number, first 60 AAs: 7.15654999999999\nTotal prob of N-in: 0.48729\noutside 1 217\nTMhelix 218 240\ninside 241 259

28899 GCF_000163275.1_ASM16327v1 Pseudomonas amygdali pv. aesculi str. 2250 Proteobacteria; Gammaproteobacteria; Pseudomonadales; Pseudomonadaceae; Pseudomonas; Pseudomonas syringae group; Pseudomonas syringae group genomosp. 2; Pseudomonas amygdali; Pseudomonas amygdali pv. aesculi MPKRRRVLRLLIVATLASLLQACDIDLYTNLGEREANAMLAVLLRDGIPASRKVQDNGQL WP_005733361.1 MULTISPECIES: EscJ/YscJ/HrcJ family type III secretion inner membrane ring protein [Pseudomonas syringae group] Length: 259\nNumber of predicted TMHs: 1\nExp number of AAs in TMHs: 31.46284\nExp number, first 60 AAs: 7.15654999999999\nTotal prob of N-in: 0.48729\noutside 1 217\nTMhelix 218 240\ninside 241 259

28900 GCF_000163255.1_ASM16325v1 Pseudomonas amygdali pv. aesculi str. NCPPB 3681 Proteobacteria; Gammaproteobacteria; Pseudomonadales; Pseudomonadaceae; Pseudomonas; Pseudomonas syringae group; Pseudomonas syringae group genomosp. 2; Pseudomonas amygdali; Pseudomonas amygdali pv. aesculi MPKRRRVLRLLIVATLASLLQACDIDLYTNLGEREANAMLAVLLRDGIPASRKVQDNGQL WP_005733361.1 MULTISPECIES: EscJ/YscJ/HrcJ family type III secretion inner membrane ring protein [Pseudomonas syringae group] Length: 259\nNumber of predicted TMHs: 1\nExp number of AAs in TMHs: 31.46284\nExp number, first 60 AAs: 7.15654999999999\nTotal prob of N-in: 0.48729\noutside 1 217\nTMhelix 218 240\ninside 241 259

28901 GCF_000146005.1_ASM14600v1 Pseudomonas amygdali pv. lachrymans str. M301315 Proteobacteria; Gammaproteobacteria; Pseudomonadales; Pseudomonadaceae; Pseudomonas; Pseudomonas syringae group; Pseudomonas syringae group genomosp. 2; Pseudomonas amygdali; Pseudomonas amygdali pv. lachrymans MPKRRRVLRLLIVATLASLLQTCDIDLYTNLGEREANAMLAVLLRDGIPASRKVQDNGQL WP_005743513.1 EscJ/YscJ/HrcJ family type III secretion inner membrane ring protein [Pseudomonas amygdali] Length: 259\nNumber of predicted TMHs: 1\nExp number of AAs in TMHs: 30.06245\nExp number, first 60 AAs: 4.73138\nTotal prob of N-in: 0.42541\noutside 1 217\nTMhelix 218 240\ninside 241 259

28902 GCF_000275945.1_ASM27594v1 Pseudomonas amygdali pv. tabaci str. 6605 Proteobacteria; Gammaproteobacteria; Pseudomonadales; Pseudomonadaceae; Pseudomonas; Pseudomonas syringae group; Pseudomonas syringae group genomosp. 2; Pseudomonas amygdali; Pseudomonas amygdali pv. tabaci MPKRRRVLRLLIVATLASLLQACDIDLYTNLGEREANAMLAVLLRDGIPASRKVQDNGQL WP_005777666.1 EscJ/YscJ/HrcJ family type III secretion inner membrane ring protein [Pseudomonas amygdali] Length: 259\nNumber of predicted TMHs: 1\nExp number of AAs in TMHs: 31.78523\nExp number, first 60 AAs: 6.9796\nTotal prob of N-in: 0.50005\noutside 1 217\nTMhelix 218 240\ninside 241 259

28903 GCF_001006455.1_PSt_v01 Pseudomonas amygdali pv. tabaci str. ATCC 11528 Proteobacteria; Gammaproteobacteria; Pseudomonadales; Pseudomonadaceae; Pseudomonas; Pseudomonas syringae group; Pseudomonas syringae group genomosp. 2; Pseudomonas amygdali; Pseudomonas amygdali pv. tabaci MPKRRRVLRLLIVATLASLLQACDIDLYTNLGEREANAMLAVLLRDGIPASRKVQDNGQL WP_005777666.1 EscJ/YscJ/HrcJ family type III secretion inner membrane ring protein [Pseudomonas amygdali] Length: 259\nNumber of predicted TMHs: 1\nExp number of AAs in TMHs: 31.78523\nExp number, first 60 AAs: 6.9796\nTotal prob of N-in: 0.50005\noutside 1 217\nTMhelix 218 240\ninside 241 259

28904 GCF_000145945.1_ASM14594v1 Pseudomonas amygdali pv. tabaci str. ATCC 11528 Proteobacteria; Gammaproteobacteria; Pseudomonadales; Pseudomonadaceae; Pseudomonas; Pseudomonas syringae group; Pseudomonas syringae group genomosp. 2; Pseudomonas amygdali; Pseudomonas amygdali pv. tabaci MPKRRRVLRLLIVATLASLLQACDIDLYTNLGEREANAMLAVLLRDGIPASRKVQDNGQL WP_005777666.1 EscJ/YscJ/HrcJ family type III secretion inner membrane ring protein [Pseudomonas amygdali] Length: 259\nNumber of predicted TMHs: 1\nExp number of AAs in TMHs: 31.78523\nExp number, first 60 AAs: 6.9796\nTotal prob of N-in: 0.50005\noutside 1 217\nTMhelix 218 240\ninside 241 259

28905 GCF_001483545.1_PfrICMP7711 Pseudomonas savastanoi pv. fraxini Proteobacteria; Gammaproteobacteria; Pseudomonadales; Pseudomonadaceae; Pseudomonas; Pseudomonas syringae group; Pseudomonas syringae group genomosp. 2; Pseudomonas savastanoi MPKRRRVLRLLIVATLASLLQACDIDLYTNLGEREANAMLAVLLRDGIPASRKVQDNGQL WP_002553574.1 MULTISPECIES: EscJ/YscJ/HrcJ family type III secretion inner membrane ring protein [Pseudomonas syringae group] Length: 259\nNumber of predicted TMHs: 1\nExp number of AAs in TMHs: 31.58284\nExp number, first 60 AAs: 7.2383\nTotal prob of N-in: 0.48165\noutside 1 217\nTMhelix 218 240\ninside 241 259

28906 GCF_001538155.1_ASM153815v1 Pseudomonas savastanoi pv. fraxini Proteobacteria; Gammaproteobacteria; Pseudomonadales; Pseudomonadaceae; Pseudomonas; Pseudomonas syringae group; Pseudomonas syringae group genomosp. 2; Pseudomonas savastanoi MPKRRRVLRLLIVATLASLLQACDIDLYTNLGEREANAMLAVLLRDGIPASRKVQDNGQL WP_002553574.1 MULTISPECIES: EscJ/YscJ/HrcJ family type III secretion inner membrane ring protein [Pseudomonas syringae group] Length:

259\nNumber of predicted TMHs: 1\nExp number of AAs in TMHs: 31.58284\nExp number, first 60 AAs: 7.2383\nTotal prob of N-in: 0.48165\noutside 1 217\nTMhelix 218 240\ninside 241 259

28907 GCF_001294275.1_PgyUnB647 Pseudomonas savastanoi pv. glycinea Proteobacteria; Gammaproteobacteria; Pseudomonadales; Pseudomonadaceae; Pseudomonas; Pseudomonas syringae group; Pseudomonas syringae group genomosp. 2; Pseudomonas savastanoi MPKRRRVLRLLIVATLASLLQACDIDLYTNLGEREANAMLAVLLRDGIPASRKVQDNGQL WP_004660817.1 MULTISPECIES: EscJ/YscJ/HrcJ family type III secretion inner membrane ring protein [Pseudomonas syringae group genomosp. 2] Length: 259\nNumber of predicted TMHs: 1\nExp number of AAs in TMHs: 31.46243\nExp number, first 60 AAs: 7.15651999999999\nTotal prob of N-in: 0.48728\noutside 1 217\nTMhelix 218 240\ninside 241 259

28908 GCF_001293765.1_PgyLN10 Pseudomonas savastanoi pv. glycinea Proteobacteria; Gammaproteobacteria; Pseudomonadales; Pseudomonadaceae; Pseudomonas; Pseudomonas syringae group; Pseudomonas syringae group genomosp. 2; Pseudomonas savastanoiMPKRRRVLRLLIVATLASLLQACDIDLYTNLGEREANAMLAVLLRDGIPASRKVQDNGQL WP_004660817.1 MULTISPECIES: EscJ/YscJ/HrcJ family type III secretion inner membrane ring protein [Pseudomonas syringae group genomosp. 2] Length: 259\nNumber of predicted TMHs: 1\nExp number of AAs in TMHs: 31.46243\nExp number, first 60 AAs: 7.15651999999999\nTotal prob of N-in: 0.48728\noutside 1 217\nTMhelix 218 240\ninside 241 259

28909 GCF_001294195.1_PgyKN44 Pseudomonas savastanoi pv. glycinea Proteobacteria; Gammaproteobacteria; Pseudomonadales; Pseudomonadaceae; Pseudomonas; Pseudomonas syringae group; Pseudomonas syringae group genomosp. 2; Pseudomonas savastanoiMPKRRRVLRLLIVATLASLLQACDIDLYTNLGEREANAMLAVLLRDGIPASRKVQDNGQL WP_004660817.1 MULTISPECIES: EscJ/YscJ/HrcJ family type III secretion inner membrane ring protein [Pseudomonas syringae group genomosp. 2] Length: 259\nNumber of predicted TMHs: 1\nExp number of AAs in TMHs: 31.46243\nExp number, first 60 AAs: 7.15651999999999\nTotal prob of N-in: 0.48728\noutside 1 217\nTMhelix 218 240\ninside 241 259

28910 GCF_001294265.1_Pseudomonas_syringae_pv._glycinea_BR1 Pseudomonas savastanoi pv. glycinea Proteobacteria; Gammaproteobacteria; Pseudomonadales; Pseudomonadaceae; Pseudomonas; Pseudomonas syringae group; Pseudomonas syringae group genomosp. 2; Pseudomonas savastanoi MPKRRRVLRLLIVATLASLLQACDIDLYTNLGEREANAMLAVLLRDGIPASRKVQDNGQL WP_004660817.1 MULTISPECIES: EscJ/YscJ/HrcJ family type III secretion inner membrane ring protein [Pseudomonas syringae group genomosp. 2] Length: 259\nNumber of predicted TMHs: 1\nExp number of AAs in TMHs: 31.46243\nExp number, first 60 AAs: 7.15651999999999\nTotal prob of N-in: 0.48728\noutside 1 217\nTMhelix 218 240\ninside 241 259

28911 GCF_001400375.1_PgyICMP2189 Pseudomonas savastanoi pv. glycinea Proteobacteria; Gammaproteobacteria; Pseudomonadales; Pseudomonadaceae; Pseudomonas; Pseudomonas syringae group; Pseudomonas syringae group genomosp. 2; Pseudomonas savastanoi MPKRRRVLRLLIVATLASLLQACDIDLYTNLGEREANAMLAVLLRDGIPASRKVQDNGQL WP_004660817.1 MULTISPECIES: EscJ/YscJ/HrcJ family type III secretion inner membrane ring protein [Pseudomonas syringae group genomosp. 2] Length: 259\nNumber of predicted TMHs: 1\nExp number of AAs in TMHs: 31.46243\nExp number, first 60 AAs: 7.15651999999999\nTotal prob of N-in: 0.48728\noutside 1 217\nTMhelix 218 240\ninside 241 259

28912 GCF_001535815.1_ASM153581v1 Pseudomonas savastanoi pv. nerii Proteobacteria; Gammaproteobacteria; Pseudomonadales; Pseudomonadaceae; Pseudomonas; Pseudomonas syringae group; Pseudomonas syringae group genomosp. 2; Pseudomonas savastanoi MPKRRRVLRLLIVATLASLLQACDIDLYTNLGEREANAMLAVLLRDGIPASRKVQDNGQL WP_002553574.1 MULTISPECIES: EscJ/YscJ/HrcJ family type III secretion inner membrane ring protein [Pseudomonas syringae group] Length: 259\nNumber of predicted TMHs: 1\nExp number of AAs in TMHs: 31.58284\nExp number, first 60 AAs: 7.2383\nTotal prob of N-in: 0.48165\noutside 1 217\nTMhelix 218 240\ninside 241 259

28913 GCF_001400575.1_PneICMP16943 Pseudomonas savastanoi pv. nerii Proteobacteria; Gammaproteobacteria; Pseudomonadales; Pseudomonadaceae; Pseudomonas; Pseudomonas syringae group; Pseudomonas syringae group genomosp. 2; Pseudomonas savastanoi MPKRRRVLRLLIVATLASLLQACDIDLYTNLGEREANAMLAVLLRDGIPASRKVQDNGQL WP_002553574.1 MULTISPECIES: EscJ/YscJ/HrcJ family type III secretion inner membrane ring protein [Pseudomonas syringae group] Length: 259\nNumber of predicted TMHs: 1\nExp number of AAs in TMHs: 31.58284\nExp number, first 60 AAs: 7.2383\nTotal prob of N-in: 0.48165\noutside 1 217\nTMhelix 218 240\ninside 241 259

28914 GCF_001400605.1_PphICMP2740 Pseudomonas savastanoi pv. phaseolicola Proteobacteria; Gammaproteobacteria; Pseudomonadales; Pseudomonadaceae; Pseudomonas; Pseudomonas syringae group;

Pseudomonas syringae group genomosp. 2; Pseudomonas savastanoi
MPKRRRVLRLLIVATLASLLQACDIDLYTNLGEREANAMLAVLLRDGIPASRKVQDNGQL WP_004660817.1 MULTISPECIES: EscJ/YscJ/HrcJ family type III secretion inner membrane ring protein [Pseudomonas syringae group genomosp. 2]
Length: 259\nNumber of predicted TMHs: 1\nExp number of AAs in TMHs: 31.46243\nExp number, first 60 AAs: 7.15651999999999\nTotal prob of N-in: 0.48728\noutside 1 217\nTMhelix 218 240\ninside 241 259

28915 GCF_001294025.1_Pph1302A Pseudomonas savastanoi pv. phaseolicola Proteobacteria; Gammaproteobacteria; Pseudomonadales; Pseudomonadaceae; Pseudomonas; Pseudomonas syringae group; Pseudomonas syringae group genomosp. 2; Pseudomonas savastanoi
MPKRRRVLRLLIVATLASLLQACDIDLYTNLGEREANAMLAVLLRDGIPASRKVQDNGQL WP_004660817.1 MULTISPECIES: EscJ/YscJ/HrcJ family type III secretion inner membrane ring protein [Pseudomonas syringae group genomosp. 2]
Length: 259\nNumber of predicted TMHs: 1\nExp number of AAs in TMHs: 31.46243\nExp number, first 60 AAs: 7.15651999999999\nTotal prob of N-in: 0.48728\noutside 1 217\nTMhelix 218 240\ninside 241 259

28916 GCF_001294065.1_PphY5_2 Pseudomonas savastanoi pv. phaseolicola Proteobacteria; Gammaproteobacteria; Pseudomonadales; Pseudomonadaceae; Pseudomonas; Pseudomonas syringae group; Pseudomonas syringae group genomosp. 2; Pseudomonas savastanoi
MPKRRRVLRLLIVATLASLLQACDIDLYTNLGEREANAMLAVLLRDGIPASRKVQDNGQL WP_004660817.1 MULTISPECIES: EscJ/YscJ/HrcJ family type III secretion inner membrane ring protein [Pseudomonas syringae group genomosp. 2]
Length: 259\nNumber of predicted TMHs: 1\nExp number of AAs in TMHs: 31.46243\nExp number, first 60 AAs: 7.15651999999999\nTotal prob of N-in: 0.48728\noutside 1 217\nTMhelix 218 240\ninside 241 259

28917 GCF_001294105.1_PphNPS3121 Pseudomonas savastanoi pv. phaseolicola Proteobacteria; Gammaproteobacteria; Pseudomonadales; Pseudomonadaceae; Pseudomonas; Pseudomonas syringae group; Pseudomonas syringae group genomosp. 2; Pseudomonas savastanoi
MPKRRRVLRLLIVATLASLLQACDIDLYTNLGEREANAMLAVLLRDGIPASRKVQDNGQL WP_004660817.1 MULTISPECIES: EscJ/YscJ/HrcJ family type III secretion inner membrane ring protein [Pseudomonas syringae group genomosp. 2]
Length: 259\nNumber of predicted TMHs: 1\nExp number of AAs in TMHs: 31.46243\nExp number, first 60 AAs: 7.15651999999999\nTotal prob of N-in: 0.48728\noutside 1 217\nTMhelix 218 240\ninside 241 259

28918 GCF_001294035.1_PphHB10Y Pseudomonas savastanoi pv. phaseolicola Proteobacteria; Gammaproteobacteria; Pseudomonadales; Pseudomonadaceae; Pseudomonas; Pseudomonas syringae group; Pseudomonas syringae group genomosp. 2; Pseudomonas savastanoi
MPKRRRVLRLLIVATLASLLQACDIDLYTNLGEREANAMLAVLLRDGIPASRKVQDNGQL WP_004660817.1 MULTISPECIES: EscJ/YscJ/HrcJ family type III secretion inner membrane ring protein [Pseudomonas syringae group genomosp. 2]
Length: 259\nNumber of predicted TMHs: 1\nExp number of AAs in TMHs: 31.46243\nExp number, first 60 AAs: 7.15651999999999\nTotal prob of N-in: 0.48728\noutside 1 217\nTMhelix 218 240\ninside 241 259

28919 GCF_001400935.1_PreICMP16945 Pseudomonas savastanoi pv. retacarpa Proteobacteria; Gammaproteobacteria; Pseudomonadales; Pseudomonadaceae; Pseudomonas; Pseudomonas syringae group; Pseudomonas syringae group genomosp. 2; Pseudomonas savastanoi
MPKRRRVLRLLIVATLASLLQACDIDLYTNLGEREANAMLAVLLRDGIPASRKVQDNGQL WP_002553574.1 MULTISPECIES: EscJ/YscJ/HrcJ family type III secretion inner membrane ring protein [Pseudomonas syringae group] Length: 259\nNumber of predicted TMHs: 1\nExp number of AAs in TMHs: 31.58284\nExp number, first 60 AAs: 7.2383\nTotal prob of N-in: 0.48165\noutside 1 217\nTMhelix 218 240\ninside 241 259

28920 GCF_002115545.1_ASM211554v1 Pseudomonas savastanoi pv. retacarpa Proteobacteria; Gammaproteobacteria; Pseudomonadales; Pseudomonadaceae; Pseudomonas; Pseudomonas syringae group; Pseudomonas syringae group genomosp. 2; Pseudomonas savastanoi
MPKRRRVLRLLIVATLASLLQACDIDLYTNLGEREANAMLAVLLRDGIPASRKVQDNGQL WP_002553574.1 MULTISPECIES: EscJ/YscJ/HrcJ family type III secretion inner membrane ring protein [Pseudomonas syringae group] Length: 259\nNumber of predicted TMHs: 1\nExp number of AAs in TMHs: 31.58284\nExp number, first 60 AAs: 7.2383\nTotal prob of N-in: 0.48165\noutside 1 217\nTMhelix 218 240\ninside 241 259

28921 GCF_001401285.1_PsvICMP4352 Pseudomonas savastanoi pv. savastanoi Proteobacteria; Gammaproteobacteria; Pseudomonadales; Pseudomonadaceae; Pseudomonas; Pseudomonas syringae group; Pseudomonas syringae group genomosp. 2; Pseudomonas savastanoi
MPKRRRVLRLLIVATLASLLQACDIDLYTNLGEREANAMLAVLLRDGIPASRKVQDNGQL WP_002553574.1 MULTISPECIES: EscJ/YscJ/HrcJ family type III secretion inner membrane ring protein [Pseudomonas syringae group] Length: 259\nNumber of predicted TMHs: 1\nExp number of AAs in TMHs: 31.58284\nExp number, first 60 AAs: 7.2383\nTotal prob of N-in: 0.48165\noutside 1 217\nTMhelix 218 240\ninside 241 259

28922 GCF_000751155.1_PSV1.0 Pseudomonas savastanoi pv. savastanoi Proteobacteria; Gammaproteobacteria; Pseudomonadales; Pseudomonadaceae; Pseudomonas; Pseudomonas syringae group; Pseudomonas syringae group genomosp. 2; Pseudomonas savastanoi
MPKRRRVLRLLIVATLASLLQACDIDLYTNLGEREANAMLAVLLRDGIPASRKVQDNGQLWP_002553574.1 MULTISPECIES: EscJ/YscJ/HrcJ family type III secretion inner membrane ring protein [Pseudomonas syringae group] Length: 259\nNumber of predicted TMHs: 1\nExp number of AAs in TMHs: 31.58284\nExp number, first 60 AAs: 7.2383\nTotal prob of N-in: 0.48165\noutside 1 217\nTMhelix 218 240\ninside 241 259

28923 GCF_000935695.1_PSAVPseNe107-G1 Pseudomonas savastanoi pv. savastanoi Proteobacteria; Gammaproteobacteria; Pseudomonadales; Pseudomonadaceae; Pseudomonas; Pseudomonas syringae group; Pseudomonas syringae group genomosp. 2; Pseudomonas savastanoi
MPKRRRVLRLLIVATLASLLQACDIDLYTNLGEREANAMLAVLLRDGIPASRKVQDNGQLWP_002553574.1 MULTISPECIES: EscJ/YscJ/HrcJ family type III secretion inner membrane ring protein [Pseudomonas syringae group] Length: 259\nNumber of predicted TMHs: 1\nExp number of AAs in TMHs: 31.58284\nExp number, first 60 AAs: 7.2383\nTotal prob of N-in: 0.48165\noutside 1 217\nTMhelix 218 240\ninside 241 259

28924 GCF_000732035.1_PSV1.0 Pseudomonas savastanoi pv. savastanoi Proteobacteria; Gammaproteobacteria; Pseudomonadales; Pseudomonadaceae; Pseudomonas; Pseudomonas syringae group; Pseudomonas syringae group genomosp. 2; Pseudomonas savastanoi
MPKRRRVLRLLIVATLASLLQACDIDLYTNLGEREANAMLAVLLRDGIPASRKVQDNGQLWP_002553574.1 MULTISPECIES: EscJ/YscJ/HrcJ family type III secretion inner membrane ring protein [Pseudomonas syringae group] Length: 259\nNumber of predicted TMHs: 1\nExp number of AAs in TMHs: 31.58284\nExp number, first 60 AAs: 7.2383\nTotal prob of N-in: 0.48165\noutside 1 217\nTMhelix 218 240\ninside 241 259

28925 GCF_000187045.1_ASM18704v2 Pseudomonas savastanoi pv. glycinea str. B076 Proteobacteria; Gammaproteobacteria; Pseudomonadales; Pseudomonadaceae; Pseudomonas; Pseudomonas syringae group; Pseudomonas syringae group genomosp. 2; Pseudomonas savastanoi; Pseudomonas savastanoi pv. glycinea
MPKRRRVLRLLIVATLASLLQACDIDLYTNLGEREANAMLAVLLRDGIPASRKVQDNGQLWP_004660817.1 MULTISPECIES: EscJ/YscJ/HrcJ family type III secretion inner membrane ring protein [Pseudomonas syringae group genomosp. 2] Length: 259\nNumber of predicted TMHs: 1\nExp number of AAs in TMHs: 31.46243\nExp number, first 60 AAs: 7.15651999999999\nTotal prob of N-in: 0.48728\noutside 1 217\nTMhelix 218 240\ninside 241 259

28926 GCF_000187065.1_ASM18706v2 Pseudomonas savastanoi pv. glycinea str. race 4 Proteobacteria; Gammaproteobacteria; Pseudomonadales; Pseudomonadaceae; Pseudomonas; Pseudomonas syringae group; Pseudomonas syringae group genomosp. 2; Pseudomonas savastanoi; Pseudomonas savastanoi pv. glycinea
MPKRRRVLRLLIVATLASLLQACDIDLYTNLGEREANAMLAVLLRDGIPASRKVQDNGQLWP_004660817.1 MULTISPECIES: EscJ/YscJ/HrcJ family type III secretion inner membrane ring protein [Pseudomonas syringae group genomosp. 2] Length: 259\nNumber of predicted TMHs: 1\nExp number of AAs in TMHs: 31.46243\nExp number, first 60 AAs: 7.15651999999999\nTotal prob of N-in: 0.48728\noutside 1 217\nTMhelix 218 240\ninside 241 259

28927 GCF_000012205.1_ASM1220v1 Pseudomonas savastanoi pv. phaseolicola 1448A Proteobacteria; Gammaproteobacteria; Pseudomonadales; Pseudomonadaceae; Pseudomonas; Pseudomonas syringae group; Pseudomonas syringae group genomosp. 2; Pseudomonas savastanoi; Pseudomonas savastanoi pv. phaseolicola
MPKRRRVLRLLIVATLASLLQACDIDLYTNLGEREANAMLAVLLRDGIPASRKVQDNGQLWP_004660817.1 MULTISPECIES: EscJ/YscJ/HrcJ family type III secretion inner membrane ring protein [Pseudomonas syringae group genomosp. 2] Length: 259\nNumber of predicted TMHs: 1\nExp number of AAs in TMHs: 31.46243\nExp number, first 60 AAs: 7.15651999999999\nTotal prob of N-in: 0.48728\noutside 1 217\nTMhelix 218 240\ninside 241 259

28928 GCF_000225805.1_ASM22580v1 Pseudomonas savastanoi pv. phaseolicola 1644R Proteobacteria; Gammaproteobacteria; Pseudomonadales; Pseudomonadaceae; Pseudomonas; Pseudomonas syringae group; Pseudomonas syringae group genomosp. 2; Pseudomonas savastanoi; Pseudomonas savastanoi pv. phaseolicola
MPKRRRVLRLLIVATLASLLQACDIDLYTNLGEREANAMLAVLLRDGIPASRKVQDNGQLWP_004660817.1 MULTISPECIES: EscJ/YscJ/HrcJ family type III secretion inner membrane ring protein [Pseudomonas syringae group genomosp. 2] Length: 259\nNumber of predicted TMHs: 1\nExp number of AAs in TMHs: 31.46243\nExp number, first 60 AAs: 7.15651999999999\nTotal prob of N-in: 0.48728\noutside 1 217\nTMhelix 218 240\ninside 241 259

28929 GCF_000164015.3_ASM16401v3 Pseudomonas savastanoi pv. savastanoi NCPPB 3335 Proteobacteria; Gammaproteobacteria; Pseudomonadales; Pseudomonadaceae; Pseudomonas; Pseudomonas syringae group; Pseudomonas syringae group genomosp. 2; Pseudomonas savastanoi; Pseudomonas savastanoi pv. savastanoi MPKRRRVLRLLIVATLASLLQACDIDLYTNLGEREANAMLAVLLRDGIPASRKVQDNGQLWP_002553574.1 MULTISPECIES: EscJ/YscJ/HrcJ family type III secretion inner membrane ring protein [Pseudomonas syringae group] Length: 259\nNumber of predicted TMHs: 1\nExp number of AAs in TMHs: 31.58284\nExp number, first 60 AAs: 7.2383\nTotal prob of N-in: 0.48165\noutside 1 217\nTMhelix 218 240\ninside 241 259

28930 GCF_000245015.1_ASM24501v1 Beggiatoa alba B18LD Proteobacteria; Gammaproteobacteria; Thiotrichales; Thiotrichaceae; Beggiatoa; Beggiatoa alba
MDSKLTIGDSLRAQGMSRRGFLQFCTVAAGMMGLSSSLIPQIARALEKAQRPSVIWLSFQ WP_002684417.1
Ni/Fe hydrogenase [Beggiatoa alba] Length: 371\nNumber of predicted TMHs: 2\nExp number of AAs in TMHs: 51.4432999999999\nExp number, first 60 AAs: 20.56714\nTotal prob of N-in: 0.55518\nPOSSIBLE N-term signal sequence\ninside 1 20\nTMhelix 21 43\noutside 44 319\nTMhelix 320 342\ninside 343 371

28931 GCF_000621325.1_ASM62132v1 Thiothrix lacustris DSM 21227 Proteobacteria; Gammaproteobacteria; Thiotrichales; Thiotrichaceae; Thiothrix; Thiothrix lacustris
MRTTQHEPHEETLGERLRSQGISRRGFLKFCGLLASSMALAPAMVPKIAAALEQAKRPSV WP_038141922.1
Ni/Fe hydrogenase [Thiothrix lacustris] Length: 370\nNumber of predicted TMHs: 1\nExp number of AAs in TMHs: 72.5891499999999\nExp number, first 60 AAs: 10.62907\nTotal prob of N-in: 0.82258\nPOSSIBLE N-term signal sequence\noutside 1 320\nTMhelix 321 343\ninside 344 370

28932 GCF_001051235.1_ASM105123v2 Candidatus Competibacter denitrificans Run_A_D11
Proteobacteria; Gammaproteobacteria; unclassified Gammaproteobacteria; Competibacteraceae; Candidatus Competibacter; Candidatus Competibacter denitrificans
MKSFFSRWLPDPHSLRTHKSLRFLGDHLHDPRLWHFSRRSTVRGLTAGAFFAFVPVPWQM WP_048669916.1
hypothetical protein [Candidatus Competibacter denitrificans] Length: 172\nNumber of predicted TMHs: 2\nExp number of AAs in TMHs: 63.50229\nExp number, first 60 AAs: 14.17881\nTotal prob of N-in: 0.19584\nPOSSIBLE N-term signal sequence\ninside 1 47\nTMhelix 48 70\noutside 71 135\nTMhelix 136 158\ninside 159 172

28933 GCF_001247655.1_ASM124765v1 Gallaecimonas pentaromativorans Proteobacteria; Gammaproteobacteria; unclassified Gammaproteobacteria; Gallaecimonas
MMKKATPTFPGRRRRWLRLTALTLALALLGALLWRWQTLAQAGLEVERQQIGVMARALTS WP_083445970.1
hypothetical protein [Gallaecimonas pentaromativorans] Length: 186\nNumber of predicted TMHs: 2\nExp number of AAs in TMHs: 39.57771\nExp number, first 60 AAs: 19.92365\nTotal prob of N-in: 0.67766\nPOSSIBLE N-term signal sequence\noutside 1 14\nTMhelix 15 34\ninside 35 164\nTMhelix 165 184\noutside 185 186

28934 GCF_000299915.1_ASM29991v1 Gallaecimonas xiamenensis 3-C-1 Proteobacteria; Gammaproteobacteria; unclassified Gammaproteobacteria; Gallaecimonas; Gallaecimonas xiamenensis
MMKKATPTFPGRRRRWLRLVALALALSLLGVLLWRWQNLAQAGLQVERQQISVMARALTS WP_008484916.1
putative SerB-cotransposed membrane protein [Gallaecimonas xiamenensis] Length: 186\nNumber of predicted TMHs: 2\nExp number of AAs in TMHs: 39.28387\nExp number, first 60 AAs: 19.77852\nTotal prob of N-in: 0.80252\nPOSSIBLE N-term signal sequence\noutside 1 14\nTMhelix 15 34\ninside 35 164\nTMhelix 165 184\noutside 185 186

28935 GCF_000828615.1_ASM82861v1 Thiolapillus brandeum Proteobacteria; Gammaproteobacteria; unclassified Gammaproteobacteria; Thiolapillus
MTDIHQQHKSSVGEIKLSSILMTISKHRRSFLLVLGASVLLGIIFAMSAPVRHNYVTTIQ WP_041066896.1 hypothetical protein [Thiolapillus brandeum] Length: 342\nNumber of predicted TMHs: 2\nExp number of AAs in TMHs: 43.40701\nExp number, first 60 AAs: 20.69148\nTotal prob of N-in: 0.99672\nPOSSIBLE N-term signal sequence\ninside 1 29\nTMhelix 30 49\noutside 50 308\nTMhelix 309 331\ninside 332 342

28936 GCF_000198515.1_ASM19851v1 gamma proteobacterium HdN1 Proteobacteria; Gammaproteobacteria; unclassified Gammaproteobacteria; unclassified Gammaproteobacteria (miscellaneous)
MTDILTNTRRPTLQQLQQQAYSVGQKALPFARSTSTLALGVFILSTLASTDVLAELPKTV WP_013261885.1 hypothetical protein [gamma proteobacterium HdN1] Length: 155\nNumber of predicted TMHs: 2\nExp number of AAs in TMHs: 56.36342\nExp number, first 60 AAs: 11.48675\nTotal prob of N-in: 0.95730\nPOSSIBLE N-term signal sequence\ninside 1 82\nTMhelix 83 105\noutside 106 119\nTMhelix 120 142\ninside 143 155

28937 GCF_000497835.1_L18_assembly_1.0 gamma proteobacterium L18 Proteobacteria; Gammaproteobacteria; unclassified Gammaproteobacteria; unclassified Gammaproteobacteria (miscellaneous)
MTLKLSSSRRPFMRLASSVLCLATLVHNAYAAAPAPEATAVVTPVADSAPAGSYSRSITL WP_027980621.1 cellulose synthase BcsB subunit [gamma proteobacterium L18] Length: 764\nNumber of predicted TMHs: 1\nExp number of AAs in TMHs: 24.81588\nExp number, first 60 AAs: 2.08156\nTotal prob of N-in: 0.09774\noutside 1 732\nTMhelix 733 755\ninside 756 764

28938 GCF_001676085.1_ASM167608v1 Photobacterium aquimaris Proteobacteria; Gammaproteobacteria; Vibrionales; Vibrionaceae; Photobacterium
MDSHAALFELGQKRLAELRKTKPLREGSFQKKLEKNGITRRDFMIWSATITSLLALPLPF WP_060997517.1 hydrogenase [Photobacterium aquimaris] Length: 378\nNumber of predicted TMHs: 1\nExp number of AAs in TMHs: 24.7955\nExp number, first 60 AAs: 4.86911\nTotal prob of N-in: 0.22840\noutside 1 350\nTMhelix 351 373\ninside 374 378

28939 GCF_001676065.1_ASM167606v1 Photobacterium aquimaris Proteobacteria; Gammaproteobacteria; Vibrionales; Vibrionaceae; Photobacterium
MDSHAALFELGQKRLAELRKTKPLREGSFQKKLEKNGITRRDFMIWSATITSLLALPLPF WP_065168551.1 hydrogenase [Photobacterium aquimaris] Length: 378\nNumber of predicted TMHs: 1\nExp number of AAs in TMHs: 25.03809\nExp number, first 60 AAs: 4.90855\nTotal prob of N-in: 0.22850\noutside 1 350\nTMhelix 351 373\ninside 374 378

28940 GCF_900185565.1_P.aquimaris_CECT9191_Spades_Prokka_EMBL Photobacterium aquimaris Proteobacteria; Gammaproteobacteria; Vibrionales; Vibrionaceae; Photobacterium
MDSHAALFELGQKRLAELRKTKPLREGSFQKKLEKNGITRRDFMIWSATITSLLALPLPF WP_060997517.1 hydrogenase [Photobacterium aquimaris] Length: 378\nNumber of predicted TMHs: 1\nExp number of AAs in TMHs: 24.7955\nExp number, first 60 AAs: 4.86911\nTotal prob of N-in: 0.22840\noutside 1 350\nTMhelix 351 373\ninside 374 378

28941 GCF_001558115.1_ASM155811v1 Photobacterium aquimaris Proteobacteria; Gammaproteobacteria; Vibrionales; Vibrionaceae; Photobacterium
MDSHAALFELGQKRLAELRKTKPLREGSFQKKLEKNGITRRDFMIWSATITSLLALPLPF WP_060997517.1 hydrogenase [Photobacterium aquimaris] Length: 378\nNumber of predicted TMHs: 1\nExp number of AAs in TMHs: 24.7955\nExp number, first 60 AAs: 4.86911\nTotal prob of N-in: 0.22840\noutside 1 350\nTMhelix 351 373\ninside 374 378

28942 GCF_000950265.1_CFSAN029431_01.0 Photobacterium iliopiscarium Proteobacteria; Gammaproteobacteria; Vibrionales; Vibrionaceae; Photobacterium
MDSHAALFELGQKRLAELRKMKPLREGSFQKKLEQNGITRRDFMIWSATITSLLALPLPF WP_045036997.1 hydrogenase [Photobacterium iliopiscarium] Length: 378\nNumber of predicted TMHs: 1\nExp number of AAs in TMHs: 23.40825\nExp number, first 60 AAs: 3.32225\nTotal prob of N-in: 0.15738\noutside 1 350\nTMhelix 351 373\ninside 374 378

28943 GCF_000949935.1_CFSAN029430_01.0 Photobacterium iliopiscarium Proteobacteria; Gammaproteobacteria; Vibrionales; Vibrionaceae; Photobacterium
MDSHAALFELGQKRLAELRKMKPLREGSFQKKLEQNGITRRDFMIWSATITSLLALPLPF WP_045036997.1 hydrogenase [Photobacterium iliopiscarium] Length: 378\nNumber of predicted TMHs: 1\nExp number of AAs in TMHs: 23.40825\nExp number, first 60 AAs: 3.32225\nTotal prob of N-in: 0.15738\noutside 1 350\nTMhelix 351 373\ninside 374 378

28944 GCF_000950135.1_CFSAN029417_01.0 Photobacterium kishitanii Proteobacteria; Gammaproteobacteria; Vibrionales; Vibrionaceae; Photobacterium
MDSHAALFELGQKRLAELRKTKPLREGSFQKKLEKNGITRRDFMIWSATITSLLALPLPF WP_036794300.1 MULTISPECIES: hydrogenase [Photobacterium] Length: 378\nNumber of predicted TMHs: 1\nExp number of AAs in TMHs: 25.24052\nExp number, first 60 AAs: 4.73518\nTotal prob of N-in: 0.22727\noutside 1 350\nTMhelix 351 373\ninside 374 378

28945 GCF_000949965.1_CFSAN029419_01.0 Photobacterium kishitanii Proteobacteria; Gammaproteobacteria; Vibrionales; Vibrionaceae; Photobacterium
MDSHAALFELGQKRLAELRKTKPLREGSFQKKLEKNGITRRDFMIWSATITSLLALPLPF WP_036794300.1 MULTISPECIES: hydrogenase [Photobacterium] Length: 378\nNumber of predicted TMHs: 1\nExp number of AAs in TMHs: 25.24052\nExp number, first 60 AAs: 4.73518\nTotal prob of N-in: 0.22727\noutside 1 350\nTMhelix 351 373\ninside 374 378

28946 GCF_001455895.1_ASM145589v1 Photobacterium kishitanii Proteobacteria; Gammaproteobacteria; Vibrionales; Vibrionaceae; Photobacterium
MDSHAALFELGQKRLAELRKTKPLREGSFQKKLEKNGITRRDFMIWSATITSLLALPLPF WP_036794300.1 MULTISPECIES: hydrogenase [Photobacterium] Length: 378\nNumber of predicted TMHs: 1\nExp number of AAs in TMHs: 25.24052\nExp number, first 60 AAs: 4.73518\nTotal prob of N-in: 0.22727\noutside 1 350\nTMhelix 351 373\ninside 374 378

28947 GCF_000950335.1_CFSAN029420_01.0 Photobacterium kishitanii Proteobacteria; Gammaproteobacteria; Vibrionales; Vibrionaceae; Photobacterium

MDSHAALFELGQKRLAELRKTKPLREGSFQKKLEKNGITRRDFMIWSATITSLLALPLPF WP_036794300.1 MULTISPECIES: hydrogenase [Photobacterium] Length: 378\nNumber of predicted TMHs: 1\nExp number of AAs in TMHs: 25.24052\nExp number, first 60 AAs: 4.73518\nTotal prob of N-in: 0.22727\noutside 1 350\nTMhelix 351 373\ninside 374 378

28948 GCF_000950095.1_CFSAN029432_01.0 Photobacterium kishitanii Proteobacteria; Gammaproteobacteria; Vibrionales; Vibrionaceae; Photobacterium MDSHAALFELGQKRLAELRKTKPLREGSFQKKLEKNGITRRDFMIWSATITSLLALPLPF WP_036794300.1 MULTISPECIES: hydrogenase [Photobacterium] Length: 378\nNumber of predicted TMHs: 1\nExp number of AAs in TMHs: 25.24052\nExp number, first 60 AAs: 4.73518\nTotal prob of N-in: 0.22727\noutside 1 350\nTMhelix 351 373\ninside 374 378

28949 GCF_001676235.1_ASM167623v1 Photobacterium kishitanii Proteobacteria; Gammaproteobacteria; Vibrionales; Vibrionaceae; Photobacterium MDSHAALFELGQKRLAELRKTKPLREGSFQKKLEKNGITRRDFMIWSATITSLLALPLPF WP_036794300.1 MULTISPECIES: hydrogenase [Photobacterium] Length: 378\nNumber of predicted TMHs: 1\nExp number of AAs in TMHs: 25.24052\nExp number, first 60 AAs: 4.73518\nTotal prob of N-in: 0.22727\noutside 1 350\nTMhelix 351 373\ninside 374 378

28950 GCF_001676215.1_ASM167621v1 Photobacterium kishitanii Proteobacteria; Gammaproteobacteria; Vibrionales; Vibrionaceae; Photobacterium MDSHAALFELGQKRLAELRKTKPLREGSFQKKLEKNGITRRDFMIWSATITSLLALPLPF WP_036794300.1 MULTISPECIES: hydrogenase [Photobacterium] Length: 378\nNumber of predicted TMHs: 1\nExp number of AAs in TMHs: 25.24052\nExp number, first 60 AAs: 4.73518\nTotal prob of N-in: 0.22727\noutside 1 350\nTMhelix 351 373\ninside 374 378

28951 GCF_000950075.1_CFSAN029418_01.0 Photobacterium kishitanii Proteobacteria; Gammaproteobacteria; Vibrionales; Vibrionaceae; Photobacterium MDSHAALFELGQKRLAELRKTKPLREGSFQKKLEKNGITRRDFMIWSATITSLLALPLPF WP_036794300.1 MULTISPECIES: hydrogenase [Photobacterium] Length: 378\nNumber of predicted TMHs: 1\nExp number of AAs in TMHs: 25.24052\nExp number, first 60 AAs: 4.73518\nTotal prob of N-in: 0.22727\noutside 1 350\nTMhelix 351 373\ninside 374 378

28952 GCF_000949955.1_CFSAN029438_01.0 Photobacterium phosphoreum Proteobacteria; Gammaproteobacteria; Vibrionales; Vibrionaceae; Photobacterium MDSHAALFELGQKRLAELRKTKPLREGSFQKKLEQNGFTRRDFMIWSATITSLLALPLPF WP_045031529.1 hydrogenase [Photobacterium phosphoreum] Length: 378\nNumber of predicted TMHs: 1\nExp number of AAs in TMHs: 22.44152\nExp number, first 60 AAs: 3.66781\nTotal prob of N-in: 0.15231\noutside 1 350\nTMhelix 351 373\ninside 374 378

28953 GCF_001676275.1_ASM167627v1 Photobacterium phosphoreum Proteobacteria; Gammaproteobacteria; Vibrionales; Vibrionaceae; Photobacterium MDSHAALFELGQKRLAELRKTKPLREGSFQKKLEQNGFTRRDFMIWSATITSLLALPLPF WP_045031529.1 hydrogenase [Photobacterium phosphoreum] Length: 378\nNumber of predicted TMHs: 1\nExp number of AAs in TMHs: 22.44152\nExp number, first 60 AAs: 3.66781\nTotal prob of N-in: 0.15231\noutside 1 350\nTMhelix 351 373\ninside 374 378

28954 GCF_001676135.1_ASM167613v1 Photobacterium phosphoreum Proteobacteria; Gammaproteobacteria; Vibrionales; Vibrionaceae; Photobacterium MDSHAALFELGQKRLAELRKTKPLREGSFQKKLEQNGFTRRDFMIWSATITSLLALPLPF WP_045031529.1 hydrogenase [Photobacterium phosphoreum] Length: 378\nNumber of predicted TMHs: 1\nExp number of AAs in TMHs: 22.44152\nExp number, first 60 AAs: 3.66781\nTotal prob of N-in: 0.15231\noutside 1 350\nTMhelix 351 373\ninside 374 378

28955 GCF_001676195.1_ASM167619v1 Photobacterium phosphoreum Proteobacteria; Gammaproteobacteria; Vibrionales; Vibrionaceae; Photobacterium MDSHAALFELGQKRLAELRKTKPLREGSFQKKLEQNGFTRRDFMIWSATITSLLALPLPF WP_045031529.1 hydrogenase [Photobacterium phosphoreum] Length: 378\nNumber of predicted TMHs: 1\nExp number of AAs in TMHs: 22.44152\nExp number, first 60 AAs: 3.66781\nTotal prob of N-in: 0.15231\noutside 1 350\nTMhelix 351 373\ninside 374 378

28956 GCF_900166965.1_P.piscicola_NCCB100098T_SPAdes_Prokka_EMBL Photobacterium piscicola Proteobacteria; Gammaproteobacteria; Vibrionales; Vibrionaceae; Photobacterium MDSHAALFELGQKRLAELRKMKPLREGSFQKKLEKNGITRRDFMIWSATITSLLALPLPF WP_080157639.1 hydrogenase [Photobacterium piscicola] Length: 378\nNumber of predicted TMHs: 1\nExp number of AAs in TMHs: 24.189\nExp number, first 60 AAs: 4.76643\nTotal prob of N-in: 0.23035\noutside 1 350\nTMhelix 351 373\ninside 374 378

28957 GCF_900185615.1_P.malacitana_CECT9190T_SPAdes_Prokka_EMBL Photobacterium sp. CECT 9190 Proteobacteria; Gammaproteobacteria; Vibrionales; Vibrionaceae; Photobacterium MDSHAALFELGQKRLAELRKTKPLREGSFQKKLEKNGITRRDFMIWSATITSLLALPLPF WP_065168551.1 hydrogenase [Photobacterium aquimaris] Length: 378\nNumber of predicted TMHs: 1\nExp number of AAs in TMHs: 25.03809\nExp number, first 60 AAs: 4.90855\nTotal prob of N-in: 0.22850\noutside 1 350\nTMhelix 351 373\ninside 374 378

28958 GCF_900185625.1_P.andalusiensis_CECT9192T_SPAdes_Prokka_EMBL Photobacterium sp. CECT 9192 Proteobacteria; Gammaproteobacteria; Vibrionales; Vibrionaceae; Photobacterium MDSHAALFELGQKRLAELRKTKPLREGSFQKKLEKNGITRRDFMIWSATITSLLALPLPF WP_060997517.1 hydrogenase [Photobacterium aquimaris] Length: 378\nNumber of predicted TMHs: 1\nExp number of AAs in TMHs: 24.7955\nExp number, first 60 AAs: 4.86911\nTotal prob of N-in: 0.22840\noutside 1 350\nTMhelix 351 373\ninside 374 378

28959 GCF_900166975.1_P.torugnensis_CECT9189T_SPAdes_Prokka_EMBL.gz Photobacterium sp. H01100410B Proteobacteria; Gammaproteobacteria; Vibrionales; Vibrionaceae; Photobacterium MDSHAALFELGQKRLAELRNTKPLREGSFQKKLEQNGFTRRDFMIWSATITSLLALPLPF WP_080174681.1 hydrogenase [Photobacterium sp. H01100410B] Length: 378\nNumber of predicted TMHs: 1\nExp number of AAs in TMHs: 23.14172\nExp number, first 60 AAs: 3.83728\nTotal prob of N-in: 0.15380\noutside 1 350\nTMhelix 351 373\ninside 374 378

28960 GCF_000613045.2_BDW_PRJEB5262 Photobacterium phosphoreum ANT-2200 Proteobacteria; Gammaproteobacteria; Vibrionales; Vibrionaceae; Photobacterium; Photobacterium phosphoreum MDSHAALFELGQKRLAELRKTKPLREGSFQKKLEKNGITRRDFMIWSATITSLLALPLPF WP_036794300.1 MULTISPECIES: hydrogenase [Photobacterium] Length: 378\nNumber of predicted TMHs: 1\nExp number of AAs in TMHs: 25.24052\nExp number, first 60 AAs: 4.73518\nTotal prob of N-in: 0.22727\noutside 1 350\nTMhelix 351 373\ninside 374 378

28961 GCF_000967545.1_S2757 Vibrio galatheae Proteobacteria; Gammaproteobacteria; Vibrionales; Vibrionaceae; Vibrio MNNTRRRLLAGLSLLPIASLARISTAEEIDDIPECESGSLENSEWQVVWDETSVLLFPIE WP_052706166.1 hypothetical protein [Vibrio galatheae] Length: 344\nNumber of predicted TMHs: 1\nExp number of AAs in TMHs: 19.2654\nExp number, first 60 AAs: 0.25076\nTotal prob of N-in: 0.21480\noutside 1 311\nTMhelix 312 334\ninside 335 344

28962 GCF_002157735.1_ASM215773v1 Vibrio sp. CA-1004 Proteobacteria; Gammaproteobacteria; Vibrionales; Vibrionaceae; Vibrio MDTHNALFQLGKSRLKKLKQDPSLKSSSNLIDKLAEHGISRRDFMKWSAALTSILALPSP WP_086981138.1 hydrogenase [Vibrio sp. CA-1004] Length: 379\nNumber of predicted TMHs: 1\nExp number of AAs in TMHs: 19.99979\nExp number, first 60 AAs: 0.46496\nTotal prob of N-in: 0.15315\noutside 1 350\nTMhelix 351 373\ninside 374 379

28963 GCF_001006405.1_ASM100640v1 Vibrio parahaemolyticus Proteobacteria; Gammaproteobacteria; Vibrionales; Vibrionaceae; Vibrio; Vibrio harveyi group MTGDNTLINSHGVNRRDFMKLCAALAATMGLSSKAAAEMAESVSRPQRPPVVWIGAQECT WP_042998143.1 MULTISPECIES: hydrogenase 2 small subunit [Gammaproteobacteria] Length: 372\nNumber of predicted TMHs: 1\nExp number of AAs in TMHs: 27.41754\nExp number, first 60 AAs: 0.28297\nTotal prob of N-in: 0.25533\noutside 1 331\nTMhelix 332 354\ninside 355 372

28964 GCF_000978665.1_ASM97866v1 Vibrio parahaemolyticus Proteobacteria; Gammaproteobacteria; Vibrionales; Vibrionaceae; Vibrio; Vibrio harveyi group MTGDNTLINSHGVNRRDFMKLCAALAATMGLSSKAAAEMAESVSRPQRPPVVWIGAQECT WP_042998143.1 MULTISPECIES: hydrogenase 2 small subunit [Gammaproteobacteria] Length: 372\nNumber of predicted TMHs: 1\nExp number of AAs in TMHs: 27.41754\nExp number, first 60 AAs: 0.28297\nTotal prob of N-in: 0.25533\noutside 1 331\nTMhelix 332 354\ninside 355 372

28965 GCF_001632775.1_ASM163277v1 Dokdonella koreensis DS-123 Proteobacteria; Gammaproteobacteria; Xanthomonadales; Rhodanobacteraceae; Dokdonella; Dokdonella koreensis MQQTRRSIVGFVLGLAAVLAPAAMAAGGGALQASNTSLSDQGSLQNGAKLFFNYCSGCHS WP_067651820.1

ubiquinol cytochrome C oxidoreductase [Dokdonella koreensis] Length: 250\nNumber of predicted TMHs: 2\nExp number of AAs in TMHs: 42.7331799999999\nExp number, first 60 AAs: 22.38995\nTotal prob of N-in: 0.99531\nPOSSIBLE N-term signal sequence\ninside 1 6\nTMhelix 7 29\noutside 30 221\nTMhelix 222 241\ninside 242 250

28966 GCF_900100425.1_IMG-taxon_2623620556_annotated_assembly Dyella jiangningensis Proteobacteria; Gammaproteobacteria; Xanthomonadales; Rhodanobacteraceae; Dyella MSMKRIHFSRRAVLLAALAALAGLFVVFVLTSSRWRMGRVDLTSDKLYTLTPGTVHIVEG WP_074551242.1 ABC transporter [Dyella jiangningensis] Length: 632\nNumber of predicted TMHs: 2\nExp number of AAs in TMHs: 40.53489\nExp number, first 60 AAs: 20.59524\nTotal prob of N-in: 0.99989\nPOSSIBLE N-term signal sequence\ninside 1 12\nTMhelix 13 35\noutside 36 601\nTMhelix 602 621\ninside 622 632

28967 GCF_001641285.1_ASM164128v1 Dyella thiooxydans Proteobacteria; Gammaproteobacteria; Xanthomonadales; Rhodanobacteraceae; Dyella MDIVNADRRRFVCVLAGAIAAGVAGVPSFAEQTQPIGRRWPLWSIRRDARTIYLTAETPP WP_063670164.1 TraB/GumN family protein [Dyella thiooxydans] Length: 299\nNumber of predicted TMHs: 2\nExp number of AAs in TMHs: 34.09217\nExp number, first 60 AAs: 18.01114\nTotal prob of N-in: 0.93419\nPOSSIBLE N-term signal sequence\ninside 1 10\nTMhelix 11 30\noutside 31 266\nTMhelix 267 289\ninside 290 299

28968 GCF_000264355.1_Rh24678 Rhodanobacter denitrificans Proteobacteria; Gammaproteobacteria; Xanthomonadales; Rhodanobacteraceae; Rhodanobacter MLRHPDSSGTRWPRRSLLALGMLWTAPNTLLGLLLGVAAVSFGARMRWQPRELALVVRRW WP_043292041.1 hypothetical protein [Rhodanobacter denitrificans] Length: 152\nNumber of predicted TMHs: 2\nExp number of AAs in TMHs: 44.42985\nExp number, first 60 AAs: 22.93277\nTotal prob of N-in: 0.54254\nPOSSIBLE N-term signal sequence\ninside 1 19\nTMhelix 20 42\noutside 43 109\nTMhelix 110 132\ninside 133 152

28969 GCF_000230695.2_ASM23069v3 Rhodanobacter denitrificans Proteobacteria; Gammaproteobacteria; Xanthomonadales; Rhodanobacteraceae; Rhodanobacter MLRHPDSSGTRWPRRSLLALGMLWTAPNTLLGLLLGVAAVSFGARMRWQPRELALVVRRW WP_015448274.1 hypothetical protein [Rhodanobacter denitrificans] Length: 158\nNumber of predicted TMHs: 2\nExp number of AAs in TMHs: 44.19229\nExp number, first 60 AAs: 22.938\nTotal prob of N-in: 0.56471\nPOSSIBLE N-term signal sequence\ninside 1 19\nTMhelix 20 42\noutside 43 109\nTMhelix 110 132\ninside 133 158

28970 GCF_002001125.1_ASM200112v1 Rhodanobacter sp. B04 Proteobacteria; Gammaproteobacteria; Xanthomonadales; Rhodanobacteraceae; Rhodanobacter MTRRTLLPALALLAAVAFATPASPAHAETGDSSPSAGTHSAKLEDFLGSLHLRTGNIPIA WP_077554286.1 hypothetical protein [Rhodanobacter sp. B04] Length: 312\nNumber of predicted TMHs: 1\nExp number of AAs in TMHs: 40.90922\nExp number, first 60 AAs: 6.40113999999999\nTotal prob of N-in: 0.54218\noutside 1 279\nTMhelix 280 302\ninside 303 312

28971 GCF_002001045.1_ASM200104v1 Rhodanobacter sp. C01 Proteobacteria; Gammaproteobacteria; Xanthomonadales; Rhodanobacteraceae; Rhodanobacter MRHLRLPRVLSTPLRLTRRTALYGGLLLLVLVFVPLIVASSRWLHASRIDLTTDKLYTLT WP_077441289.1 ABC transporter [Rhodanobacter sp. C01] Length: 638\nNumber of predicted TMHs: 2\nExp number of AAs in TMHs: 40.88274\nExp number, first 60 AAs: 21.02807\nTotal prob of N-in: 0.99966\nPOSSIBLE N-term signal sequence\ninside 1 20\nTMhelix 21 43\noutside 44 608\nTMhelix 609 628\ninside 629 638

28972 GCF_002001105.1_ASM200110v1 Rhodanobacter sp. C06 Proteobacteria; Gammaproteobacteria; Xanthomonadales; Rhodanobacteraceae; Rhodanobacter MARLRLPAVFTTPLRWNRRATLYAALVLLTLVFVSLIVSSGHWLRTRRIDLTADRLYTLS WP_077482847.1 ABC transporter [Rhodanobacter sp. C06] Length: 637\nNumber of predicted TMHs: 2\nExp number of AAs in TMHs: 42.15202\nExp number, first 60 AAs: 22.27236\nTotal prob of N-in: 0.99954\nPOSSIBLE N-term signal sequence\ninside 1 20\nTMhelix 21 43\noutside 44 607\nTMhelix 608 627\ninside 628 637

28973 GCF_900142825.1_IMG-taxon_2690315657_annotated_assembly Rhodanobacter sp. OK091 Proteobacteria; Gammaproteobacteria; Xanthomonadales; Rhodanobacteraceae; Rhodanobacter MTSLIEKTIDRRGFLRMSSGVALSAGALALLGGHSALAAALPAVAAHDIGILNVEVALEY WP_072761292.1 ferritin [Rhodanobacter sp. OK091] Length: 193\nNumber of predicted TMHs: 2\nExp number of AAs in TMHs: 39.62091\nExp number, first 60 AAs: 22.91473\nTotal prob of N-in: 0.97660\nPOSSIBLE N-term signal sequence\ninside 1 20\nTMhelix 21 43\noutside 44 131\nTMhelix 132 154\ninside 155 193

28974 GCF_000427505.1_ASM42750v1 Rhodanobacter sp. OR444 Proteobacteria; Gammaproteobacteria; Xanthomonadales; Rhodanobacteraceae; Rhodanobacter
MPRSRLHRLLATPLHLTRRGALYGALALLLLLFVPLIVASSRWLHASRVDLTTDKLYTLT WP_027494017.1 ABC transporter [Rhodanobacter sp. OR444] Length: 638\nNumber of predicted TMHs: 2\nExp number of AAs in TMHs: 40.59727\nExp number, first 60 AAs: 20.76642\nTotal prob of N-in: 0.99881\nPOSSIBLE N-term signal sequence\ninside 1 19\nTMhelix 20 39\noutside 40 608\nTMhelix 609 628\ninside 629 638

28975 GCF_000473345.1_ASM47334v1 Rhodanobacter sp. OR87 Proteobacteria; Gammaproteobacteria; Xanthomonadales; Rhodanobacteraceae; Rhodanobacter
MLRHPDSSGTRWPRRSPLALGMLWTVPNTLLGLLLGVVGVAFGARLRWQRRELALVVRRW WP_027484742.1 hypothetical protein [Rhodanobacter sp. OR87] Length: 158\nNumber of predicted TMHs: 2\nExp number of AAs in TMHs: 46.22787\nExp number, first 60 AAs: 23.30831\nTotal prob of N-in: 0.18051\nPOSSIBLE N-term signal sequence\noutside 1 19\nTMhelix 20 42\ninside 43 109\nTMhelix 110 132\noutside 133 158

28976 GCF_000427545.1_ASM42754v1 Rhodanobacter sp. OR92 Proteobacteria; Gammaproteobacteria; Xanthomonadales; Rhodanobacteraceae; Rhodanobacter
MLRHPDSSGTRWPRRSLLALGMLWTAPNTLLGLLLGVAAVSFGARMRWQPRELALVVRRW WP_027490105.1 MULTISPECIES: hypothetical protein [Rhodanobacter] Length: 158\nNumber of predicted TMHs: 2\nExp number of AAs in TMHs: 45.48161\nExp number, first 60 AAs: 23.03149\nTotal prob of N-in: 0.52956\nPOSSIBLE N-term signal sequence\ninside 1 19\nTMhelix 20 42\noutside 43 109\nTMhelix 110 132\ninside 133 158

28977 GCF_001428385.1_Soil772 Rhodanobacter sp. Soil772 Proteobacteria; Gammaproteobacteria; Xanthomonadales; Rhodanobacteraceae; Rhodanobacter
MRLDAWLPKRRAWLLCLLLPLTAGAGTPADYAYRYALDTAGSSAAWRVELTPAVYAASTP WP_056388158.1 hypothetical protein [Rhodanobacter sp. Soil772]Length: 464\nNumber of predicted TMHs: 1\nExp number of AAs in TMHs: 34.96844\nExp number, first 60 AAs: 12.4281\nTotal prob of N-in: 0.61362\nPOSSIBLE N-term signal sequence\noutside 1 431\nTMhelix 432 454\ninside 455 464

28978 GCF_001182895.1_ASM118289v1 Dyella-like sp. DHo Proteobacteria; Gammaproteobacteria; Xanthomonadales; Rhodanobacteraceae; unclassified Rhodanobacteraceae
MESPSLSQDQPTPTPRRRRHLLRLAACLAVAAAGVASAFAQEKTPVATPAAGQVQSDPVK WP_082190363.1 cytochrome C [Dyella-like sp. DHo] Length: 497\nNumber of predicted TMHs: 2\nExp number of AAs in TMHs: 38.9458499999999\nExp number, first 60 AAs: 17.91395\nTotal prob of N-in: 0.89552\nPOSSIBLE N-term signal sequence\ninside 1 20\nTMhelix 21 40\noutside 41 467\nTMhelix 468 490\ninside 491 497

28979 GCF_001482195.1_ASM148219v1 Luteimonas abyssi Proteobacteria; Gammaproteobacteria; Xanthomonadales; Xanthomonadaceae; Luteimonas
MTGPAAPAVDARPRRPVLRIVFAAALAAAFVGFVALGIWQVQRLAWKQDLIERVDARIHA WP_058835255.1 hypothetical protein [Luteimonas abyssi] Length: 267\nNumber of predicted TMHs: 2\nExp number of AAs in TMHs: 43.89214\nExp number, first 60 AAs: 22.07863\nTotal prob of N-in: 0.96604\nPOSSIBLE N-term signal sequence\ninside 1 19\nTMhelix 20 42\noutside 43 217\nTMhelix 218 240\ninside 241 267

28980 GCF_000731095.1_ASM73109v1 Lysobacter antibioticus Proteobacteria; Gammaproteobacteria; Xanthomonadales; Xanthomonadaceae; Lysobacter
MRLGPGRARRQGQVRHRARSRRTAGADVGRMGCVSGRRRRAALAAYAIALLAMCVFAGTA WP_081931066.1 hypothetical protein [Lysobacter antibioticus] Length: 286\nNumber of predicted TMHs: 2\nExp number of AAs in TMHs: 40.90567\nExp number, first 60 AAs: 19.47043\nTotal prob of N-in: 0.99750\nPOSSIBLE N-term signal sequence\ninside 1 39\nTMhelix 40 62\noutside 63 257\nTMhelix 258 277\ninside 278 286

28981 GCF_000731095.1_ASM73109v1 Lysobacter antibioticus Proteobacteria; Gammaproteobacteria; Xanthomonadales; Xanthomonadaceae; Lysobacter
MSRRGTLLFGWTLALAAIALFANLGLWQGRRAVEKQAMLDAAAAVLNERTPRPLAAADDP WP_031371299.1 hypothetical protein [Lysobacter antibioticus] Length: 239\nNumber of predicted TMHs: 2\nExp number of AAs in TMHs: 45.77492\nExp number, first 60 AAs: 22.61365\nTotal prob of N-in: 0.82499\nPOSSIBLE N-term signal sequence\ninside 1 6\nTMhelix 7 29\noutside 30 214\nTMhelix 215 234\ninside 235 239

28982 GCF_001442785.1_ASM144278v1 Lysobacter capsici Proteobacteria; Gammaproteobacteria; Xanthomonadales; Xanthomonadaceae; Lysobacter
MSRRGTLLFGWILALATIAVFASLGRWQSHRAVEKQAMLDAAAAVLHDRRPKPLALADDP WP_057920519.1 hypothetical protein [Lysobacter capsici] Length: 246\nNumber of predicted TMHs: 2\nExp number of

AAs in TMHs: 42.60352\nExp number, first 60 AAs: 21.51965\nTotal prob of N-in: 0.96581\nPOSSIBLE N-term signal sequence\ninside 1 6\nTMhelix 7 29\noutside 30 214\nTMhelix 215 234\ninside 235 246

28983 GCF_000987985.1_ASM98798v1 Lysobacter capsici Proteobacteria; Gammaproteobacteria; Xanthomonadales; Xanthomonadaceae; Lysobacter
MSRRGTLLFGWILALATIAVFASLGRWQSHRAVEKQAMLDAAAAVLHDRRPKPLALADDP WP_046659779.1
hypothetical protein [Lysobacter capsici] Length: 246\nNumber of predicted TMHs: 2\nExp number of AAs in TMHs: 41.89857\nExp number, first 60 AAs: 21.51933\nTotal prob of N-in: 0.97821\nPOSSIBLE N-term signal sequence\ninside 1 6\nTMhelix 7 29\noutside 30 214\nTMhelix 215 234\ninside 235 246

28984 GCF_002024615.1_ASM202461v1 Lysobacter enzymogenes Proteobacteria; Gammaproteobacteria; Xanthomonadales; Xanthomonadaceae; Lysobacter
MSRRGNLLLGWTLALAAIALFANLGLWQARRAVEKQAMLDAAARVLGERVPQPLSVADDP WP_057945932.1
hypothetical protein [Lysobacter enzymogenes] Length: 243\nNumber of predicted TMHs: 2\nExp number of AAs in TMHs: 43.47799\nExp number, first 60 AAs: 22.72454\nTotal prob of N-in: 0.96028\nPOSSIBLE N-term signal sequence\ninside 1 6\nTMhelix 7 29\noutside 30 214\nTMhelix 215 234\ninside 235 243

28985 GCF_001442515.1_ASM144251v1 Lysobacter enzymogenes Proteobacteria; Gammaproteobacteria; Xanthomonadales; Xanthomonadaceae; Lysobacter
MSRRGNLLLGWTLALAAIALFANLGLWQARRAVEKQAMLDAAARVLGERVPQPLSVADDP WP_057945932.1
hypothetical protein [Lysobacter enzymogenes] Length: 243\nNumber of predicted TMHs: 2\nExp number of AAs in TMHs: 43.47799\nExp number, first 60 AAs: 22.72454\nTotal prob of N-in: 0.96028\nPOSSIBLE N-term signal sequence\ninside 1 6\nTMhelix 7 29\noutside 30 214\nTMhelix 215 234\ninside 235 243

28986 GCF_900106525.1_IMG-taxon_2675903697_annotated_assembly Lysobacter enzymogenes
Proteobacteria; Gammaproteobacteria; Xanthomonadales; Xanthomonadaceae; Lysobacter
MSRRGTLLLGWTLALATIALFANLGLWQARRAVEKQAMLDAAARVLGERVPQPLSRAADP WP_074872556.1
hypothetical protein [Lysobacter enzymogenes] Length: 243\nNumber of predicted TMHs: 2\nExp number of AAs in TMHs: 43.39872\nExp number, first 60 AAs: 22.75195\nTotal prob of N-in: 0.95340\nPOSSIBLE N-term signal sequence\ninside 1 6\nTMhelix 7 29\noutside 30 214\nTMhelix 215 234\ninside 235 243

28987 GCF_001427065.1_Root559 Lysobacter sp. Root559 Proteobacteria; Gammaproteobacteria; Xanthomonadales; Xanthomonadaceae; Lysobacter
MSDRRHAAVEQARGRRPLLLAVLASSLLVACAGFLALGGWQVQRMAWKHDLIARVEARLT WP_055909325.1
hypothetical protein [Lysobacter sp. Root559] Length: 267\nNumber of predicted TMHs: 2\nExp number of AAs in TMHs: 44.37713\nExp number, first 60 AAs: 22.56303\nTotal prob of N-in: 0.98509\nPOSSIBLE N-term signal sequence\ninside 1 19\nTMhelix 20 42\noutside 43 219\nTMhelix 220 242\ninside 243 267

28988 GCF_001427065.1_Root559 Lysobacter sp. Root559 Proteobacteria; Gammaproteobacteria; Xanthomonadales; Xanthomonadaceae; Lysobacter
MSRRRNLVVGWTLALLTIATFANLGLWQSRRAIEKQAMLDAAARVLGERRPGALALAADP WP_055901957.1
hypothetical protein [Lysobacter sp. Root559] Length: 239\nNumber of predicted TMHs: 2\nExp number of AAs in TMHs: 42.28955\nExp number, first 60 AAs: 22.45163\nTotal prob of N-in: 0.99766\nPOSSIBLE N-term signal sequence\ninside 1 6\nTMhelix 7 29\noutside 30 214\nTMhelix 215 234\ninside 235 239

28989 GCF_001427505.1_Root604 Lysobacter sp. Root604 Proteobacteria; Gammaproteobacteria; Xanthomonadales; Xanthomonadaceae; Lysobacter
MSRRRNLIVGWTLALLTIAAFVNLGLWQSRRAVEKQAMLDASARVVGERRPGSLALAADP WP_056181154.1
hypothetical protein [Lysobacter sp. Root604] Length: 239\nNumber of predicted TMHs: 2\nExp number of AAs in TMHs: 42.86084\nExp number, first 60 AAs: 22.5007\nTotal prob of N-in: 0.99726\nPOSSIBLE N-term signal sequence\ninside 1 6\nTMhelix 7 29\noutside 30 214\nTMhelix 215 234\ninside 235 239

28990 GCF_001427505.1_Root604 Lysobacter sp. Root604 Proteobacteria; Gammaproteobacteria; Xanthomonadales; Xanthomonadaceae; Lysobacter
MSDERHAAAVAPAHGRRPLLLAVFAASLAVALIGFLALGVWQVQRMGWKHALIARVEARL WP_056178414.1
hypothetical protein [Lysobacter sp. Root604] Length: 268\nNumber of predicted TMHs: 2\nExp number of AAs in TMHs: 45.21831\nExp number, first 60 AAs: 22.77502\nTotal prob of N-in: 0.89840\nPOSSIBLE N-term signal sequence\ninside 1 18\nTMhelix 19 41\noutside 42 220\nTMhelix 221 243\ninside 244 268

28991 GCF_001427715.1_Root667 Lysobacter sp. Root667 Proteobacteria; Gammaproteobacteria; Xanthomonadales; Xanthomonadaceae; Lysobacter

MSRRRNLVVGWTLALLTIATFANLGLWQSRRAVEKQAMLDAAARVLGERRPGALALAADP WP_056100861.1 hypothetical protein [Lysobacter sp. Root667] Length: 239\nNumber of predicted TMHs: 2\nExp number of AAs in TMHs: 42.28209\nExp number, first 60 AAs: 22.43552\nTotal prob of N-in: 0.99748\nPOSSIBLE N-term signal sequence\ninside 1 6\nTMhelix 7 29\noutside 30 214\nTMhelix 215 234\ninside 235 239

28992 GCF_001428685.1_Root76 Lysobacter sp. Root76 Proteobacteria; Gammaproteobacteria; Xanthomonadales; Xanthomonadaceae; Lysobacter MSRRRNLVVGWTLALLTIAAFANLGLWQSRRAVEKQAMLDAAARVLGERRPGALALAADP WP_056306103.1 MULTISPECIES: hypothetical protein [Lysobacter] Length: 239\nNumber of predicted TMHs: 2\nExp number of AAs in TMHs: 42.28512\nExp number, first 60 AAs: 22.43945\nTotal prob of N-in: 0.99775\nPOSSIBLE N-term signal sequence\ninside 1 6\nTMhelix 7 29\noutside 30 214\nTMhelix 215 234\ninside 235 239

28993 GCF_001429785.1_Root916 Lysobacter sp. Root916 Proteobacteria; Gammaproteobacteria; Xanthomonadales; Xanthomonadaceae; Lysobacter MSRRRNLIVGWTLALLTIAAFVNLGLWQSRRAVEKQAMLDASARVLGERRPGSLALAADP WP_057164304.1 hypothetical protein [Lysobacter sp. Root916] Length: 239\nNumber of predicted TMHs: 2\nExp number of AAs in TMHs: 42.89752\nExp number, first 60 AAs: 22.50051\nTotal prob of N-in: 0.99725\nPOSSIBLE N-term signal sequence\ninside 1 6\nTMhelix 7 29\noutside 30 214\nTMhelix 215 234\ninside 235 239

28994 GCF_001429785.1_Root916 Lysobacter sp. Root916 Proteobacteria; Gammaproteobacteria; Xanthomonadales; Xanthomonadaceae; Lysobacter MSDERHTAAVAPAHGRRPLLLAVFAASLAVALIGFLALGVWQVQRMGWKHALIARVEARL WP_057163649.1 hypothetical protein [Lysobacter sp. Root916] Length: 268\nNumber of predicted TMHs: 2\nExp number of AAs in TMHs: 45.19269\nExp number, first 60 AAs: 22.78069\nTotal prob of N-in: 0.90954\nPOSSIBLE N-term signal sequence\ninside 1 18\nTMhelix 19 41\noutside 42 220\nTMhelix 221 243\ninside 244 268

28995 GCF_001429315.1_Root96 Lysobacter sp. Root96 Proteobacteria; Gammaproteobacteria; Xanthomonadales; Xanthomonadaceae; Lysobacter MSRRRNLVVGWTLALLTIAAFANLGLWQSRRAVEKQAMLDAAARVLGERRPGALALAADP WP_056306103.1 MULTISPECIES: hypothetical protein [Lysobacter] Length: 239\nNumber of predicted TMHs: 2\nExp number of AAs in TMHs: 42.28512\nExp number, first 60 AAs: 22.43945\nTotal prob of N-in: 0.99775\nPOSSIBLE N-term signal sequence\ninside 1 6\nTMhelix 7 29\noutside 30 214\nTMhelix 215 234\ninside 235 239

28996 GCF_001429345.1_Root983 Lysobacter sp. Root983 Proteobacteria; Gammaproteobacteria; Xanthomonadales; Xanthomonadaceae; Lysobacter MSRRRNLIVGWTLALLTIAAFVNLGLWQSRRAVEKQAMLDASARVLGERRPGSLALAADP WP_057161099.1 hypothetical protein [Lysobacter sp. Root983] Length: 239\nNumber of predicted TMHs: 2\nExp number of AAs in TMHs: 42.86128\nExp number, first 60 AAs: 22.50072\nTotal prob of N-in: 0.99726\nPOSSIBLE N-term signal sequence\ninside 1 6\nTMhelix 7 29\noutside 30 214\nTMhelix 215 234\ninside 235 239

28997 GCF_001429345.1_Root983 Lysobacter sp. Root983 Proteobacteria; Gammaproteobacteria; Xanthomonadales; Xanthomonadaceae; Lysobacter MSDERHAAAAAPARGRRPLLLAVFAASLAVALIGFLALGVWQVQRMGWKHALIARVEARL WP_057160327.1 hypothetical protein [Lysobacter sp. Root983] Length: 268\nNumber of predicted TMHs: 2\nExp number of AAs in TMHs: 44.99883\nExp number, first 60 AAs: 22.81043\nTotal prob of N-in: 0.97015\nPOSSIBLE N-term signal sequence\ninside 1 18\nTMhelix 19 41\noutside 42 220\nTMhelix 221 243\ninside 244 268

28998 GCF_000426005.1_ASM42600v1 Lysobacter sp. URHA0019 Proteobacteria; Gammaproteobacteria; Xanthomonadales; Xanthomonadaceae; Lysobacter MSRRGTLLLGWTFALLTAAGFARLGVWQSDRAVEKERLLANVASVLKDRRAVPLSAANDA WP_027084088.1 hypothetical protein [Lysobacter sp. URHA0019] Length: 238\nNumber of predicted TMHs: 2\nExp number of AAs in TMHs: 42.47302\nExp number, first 60 AAs: 20.90589\nTotal prob of N-in: 0.89578\nPOSSIBLE N-term signal sequence\ninside 1 4\nTMhelix 5 27\noutside 28 213\nTMhelix 214 233\ninside 234 238

28999 GCF_000768335.1_ASM76833v1 Lysobacter arseniciresistens ZS79 Proteobacteria; Gammaproteobacteria; Xanthomonadales; Xanthomonadaceae; Lysobacter; Lysobacter arseniciresistens MAPVQRRTAMQAQPHVVIVGGGFAGLWATRGLARAPVRITLVDRRNHHLFQPLLYQVATA WP_052101578.1 pyridine nucleotide-disulfide oxidoreductase [Lysobacter arseniciresistens] Length: 443\nNumber of predicted TMHs: 1\nExp number of AAs in TMHs: 28.98714\nExp number, first 60 AAs: 5.32222\nTotal prob of N-in: 0.34021\noutside 1 383\nTMhelix 384 401\ninside 402 443

29000 GCF_000604005.2_ASM60400v2 Lysobacter capsici AZ78 Proteobacteria; Gammaproteobacteria; Xanthomonadales; Xanthomonadaceae; Lysobacter; Lysobacter capsici MSRRGTLLFGWILALATIAVFASLGRWQSHRAVEKQAMLDAAAAVLHDRRPKPLALADDP WP_036112499.1 hypothetical protein [Lysobacter capsici] Length: 246\nNumber of predicted TMHs: 2\nExp number of AAs in TMHs: 42.18005\nExp number, first 60 AAs: 21.5197\nTotal prob of N-in: 0.97232\nPOSSIBLE N-term signal sequence\ninside 1 6\nTMhelix 7 29\noutside 30 214\nTMhelix 215 234\ninside 235 246

29001 GCF_000768345.1_ASM76834v1 Lysobacter concretionis Ko07 = DSM 16239 Proteobacteria; Gammaproteobacteria; Xanthomonadales; Xanthomonadaceae; Lysobacter; Lysobacter concretionis MSRRATLLIGWLLALATIAVFTSLGFWQLRRAVEKQAMLDAAVQVVTERRAAPLAAAART WP_036194832.1 hypothetical protein [Lysobacter concretionis] Length: 243\nNumber of predicted TMHs: 2\nExp number of AAs in TMHs: 43.21483\nExp number, first 60 AAs: 22.82041\nTotal prob of N-in: 0.99038\nPOSSIBLE N-term signal sequence\ninside 1 6\nTMhelix 7 29\noutside 30 212\nTMhelix 213 232\ninside 233 243

29002 GCF_900167055.1_IMG-taxon_2568526003_annotated_assembly Lysobacter spongiicola DSM 21749 Proteobacteria; Gammaproteobacteria; Xanthomonadales; Xanthomonadaceae; Lysobacter; Lysobacter spongiicola MSRRTTLVLGWTLAIVGVALFVALGLWQSRRALEKEAVLAETASTVAGRAAQPLSAASDP WP_078758809.1 hypothetical protein [Lysobacter spongiicola] Length: 246\nNumber of predicted TMHs: 2\nExp number of AAs in TMHs: 43.43748\nExp number, first 60 AAs: 22.15113\nTotal prob of N-in: 0.93809\nPOSSIBLE N-term signal sequence\ninside 1 6\nTMhelix 7 29\noutside 30 214\nTMhelix 215 234\ninside 235 246

29003 GCF_001431405.1_ASM143140v1 Pseudoxanthomonas dokdonensis Proteobacteria; Gammaproteobacteria; Xanthomonadales; Xanthomonadaceae; Pseudoxanthomonas MTAGAPRRGRRRGLLALFASALLLAFAGFTALGVWQLQRLAWKHDLIARVDARIHAPAVA WP_057657592.1 hypothetical protein [Pseudoxanthomonas dokdonensis] Length: 257\nNumber of predicted TMHs: 2\nExp number of AAs in TMHs: 45.31914\nExp number, first 60 AAs: 22.88377\nTotal prob of N-in: 0.96974\nPOSSIBLE N-term signal sequence\ninside 1 12\nTMhelix 13 35\noutside 36 216\nTMhelix 217 239\ninside 240 257

29004 GCF_001700965.1_ASM170096v1 Stenotrophomonas nitritireducens Proteobacteria; Gammaproteobacteria; Xanthomonadales; Xanthomonadaceae; Stenotrophomonas MKIINLQDRRELLPAIALAHVQAFGTLLPNWTLAQAEAELRAQQRDGIPCTWLAEDDSGW WP_062169329.1 MULTISPECIES: N-acetyltransferase [Stenotrophomonas] Length: 155\nNumber of predicted TMHs: 1\nExp number of AAs in TMHs: 29.51751\nExp number, first 60 AAs: 8.02971\nTotal prob of N-in: 0.84697\ninside 1 94\nTMhelix 95 117\noutside 118 155

29005 GCF_002205885.1_ASM220588v1 Stenotrophomonas pavanii Proteobacteria; Gammaproteobacteria; Xanthomonadales; Xanthomonadaceae; Stenotrophomonas MDRRELLKMIVAATGTAMVGLPALVHGKVPAAGAMPAFSDADIGMLDEIAETILPRTRTP WP_049467359.1 twin-arginine translocation pathway signal [Stenotrophomonas maltophilia] Length: 191\nNumber of predicted TMHs: 1\nExp number of AAs in TMHs: 20.52951\nExp number, first 60 AAs: 3.49352\nTotal prob of N-in: 0.72346\ninside 1 149\nTMhelix 150 172\noutside 173 191

29006 GCF_001431565.1_ASM143156v1 Stenotrophomonas pavanii Proteobacteria; Gammaproteobacteria; Xanthomonadales; Xanthomonadaceae; Stenotrophomonas MDRRELLKMIVAATGTAMVGLPALVHGKVPAAGAMPAFSDADIGMLDEIAETILPRTRTP WP_057494184.1 twin-arginine translocation pathway signal [Stenotrophomonas pavanii] Length: 191\nNumber of predicted TMHs: 1\nExp number of AAs in TMHs: 20.52218\nExp number, first 60 AAs: 3.49142\nTotal prob of N-in: 0.72338\ninside 1 149\nTMhelix 150 172\noutside 173 191

29007 GCF_900101175.1_IMG-taxon_2623620583_annotated_assembly Stenotrophomonas pavanii Proteobacteria; Gammaproteobacteria; Xanthomonadales; Xanthomonadaceae; Stenotrophomonas MDRRELLKMIVAATGTAMVGLPALVHGKVPATGAMPAFSDADIGMLDEIAETILPRTRTP WP_074726554.1 Twin-arginine translocation pathway signal [Stenotrophomonas pavanii] Length: 191\nNumber of predicted TMHs: 1\nExp number of AAs in TMHs: 20.58681\nExp number, first 60 AAs: 3.47678\nTotal prob of N-in: 0.71877\ninside 1 149\nTMhelix 150 172\noutside 173 191

29008 GCF_001580555.1_ASM158055v1 Stenotrophomonas sp. DDT-1 Proteobacteria; Gammaproteobacteria; Xanthomonadales; Xanthomonadaceae; Stenotrophomonas MDRRELLKMIVAATGAAMVGLPALVQGKAPAAGAMVPFSDADIGTLDEIAETILPRTRTP WP_061479189.1 Twin-arginine translocation pathway signal [Stenotrophomonas sp. DDT-1] Length: 191\nNumber of

predicted TMHs:  1\nExp number of AAs in TMHs: 19.68696\nExp number, first 60 AAs:  2.97036\nTotal prob of N-in: 0.74776\ninside 1 149\nTMhelix 150 172\noutside 173 191

29009 GCF_001580555.1_ASM158055v1 Stenotrophomonas sp. DDT-1 Proteobacteria; Gammaproteobacteria; Xanthomonadales; Xanthomonadaceae; Stenotrophomonas MRPVRAWTRRAALPVLLLALGACAKQGDMVGDAGATAAEAAIASPEGAFLAYEHDVQVQL WP_061480680.1 hypothetical protein [Stenotrophomonas sp. DDT-1] Length: 270\nNumber of predicted TMHs:  1\nExp number of AAs in TMHs: 22.94304\nExp number, first 60 AAs:  0.2824\nTotal prob of N-in:  0.08079\noutside 1 233\nTMhelix 234 256\ninside 257 270

29010 GCF_001807895.1_ASM180789v1 Stenotrophomonas sp. HMSC10F07 Proteobacteria; Gammaproteobacteria; Xanthomonadales; Xanthomonadaceae; Stenotrophomonas MRPVRAWTRRAALPVLLLALGACAKQGDMVGDAGATAAEAAIASPEGAFLAYEHDVQVQL WP_053497069.1 MULTISPECIES: hypothetical protein [Stenotrophomonas] Length: 271\nNumber of predicted TMHs:  1\nExp number of AAs in TMHs: 22.8986\nExp number, first 60 AAs:  0.29597\nTotal prob of N-in:  0.05695\noutside 1 233\nTMhelix 234 256\ninside 257 271

29011 GCF_001807895.1_ASM180789v1 Stenotrophomonas sp. HMSC10F07 Proteobacteria; Gammaproteobacteria; Xanthomonadales; Xanthomonadaceae; Stenotrophomonas MDRRELLKMIVAATGAAMVGLPALVHGKAPAAGMMAAFSDAEIGTLDEIAETILPRTRTP WP_005415638.1 MULTISPECIES: hypothetical protein [Proteobacteria] Length: 191\nNumber of predicted TMHs:  1\nExp number of AAs in TMHs: 26.26404\nExp number, first 60 AAs:  9.53785\nTotal prob of N-in:  0.59430\ninside 1 149\nTMhelix 150 172\noutside 173 191

29012 GCF_001975525.1_ASM197552v1 Stenotrophomonas sp. KAs 5-3 Proteobacteria; Gammaproteobacteria; Xanthomonadales; Xanthomonadaceae; Stenotrophomonas MDRRELLKMIVAATGAAMVGLPALVHGKVPAAGAMPAFSDADIGMLDEIAETILPRTRTP WP_005412559.1 MULTISPECIES: hypothetical protein [Stenotrophomonas] Length: 191\nNumber of predicted TMHs:  1\nExp number of AAs in TMHs: 22.08369\nExp number, first 60 AAs:  5.28303\nTotal prob of N-in:  0.70356\ninside 1 149\nTMhelix 150 172\noutside 173 191

29013 GCF_001562215.1_ASM156221v1 Stenotrophomonas sp. KCTC 12332 Proteobacteria; Gammaproteobacteria; Xanthomonadales; Xanthomonadaceae; Stenotrophomonas MKIINLQDRRELLPAIALAHVQAFGTLLPNWTLAQAEAELRAQQRDGIPCTWLAEDDSGW WP_062169329.1 MULTISPECIES: N-acetyltransferase [Stenotrophomonas] Length: 155\nNumber of predicted TMHs:  1\nExp number of AAs in TMHs: 29.51751\nExp number, first 60 AAs:  8.02971\nTotal prob of N-in:  0.84697\ninside 1 94\nTMhelix 95 117\noutside 118 155

29014 GCF_000632045.1_Willow_Endophyte_RIT309 Stenotrophomonas sp. RIT309 Proteobacteria; Gammaproteobacteria; Xanthomonadales; Xanthomonadaceae; Stenotrophomonas MTQTGRRSFLVVSSVLALASSAAAGAADSAAGQFTEWTGDSTVSRQMTLRQLGFRQPLVL WP_032978925.1 membrane protein [Stenotrophomonas sp. RIT309] Length: 733\nNumber of predicted TMHs:  1\nExp number of AAs in TMHs: 26.10562\nExp number, first 60 AAs:  5.83459\nTotal prob of N-in:  0.28306\noutside 1 699\nTMhelix 700 717\ninside 718 733

29015 GCF_002192255.1_ASM219225v1 Stenotrophomonas sp. WZN-1 Proteobacteria; Gammaproteobacteria; Xanthomonadales; Xanthomonadaceae; Stenotrophomonas MDRRELLKMIVAATGAAMVGLPALAQGKAAAVPVKTLFSDADVGTLDEIAETILPRTRTP WP_087921816.1 Twin-arginine translocation pathway signal [Stenotrophomonas sp. WZN-1] Length: 191\nNumber of predicted TMHs:  1\nExp number of AAs in TMHs: 20.42429\nExp number, first 60 AAs:  2.82062\nTotal prob of N-in: 0.70745\ninside 1 149\nTMhelix 150 172\noutside 173 191

29016 GCF_001075985.1_ASM107598v1 Stenotrophomonas maltophilia Proteobacteria; Gammaproteobacteria; Xanthomonadales; Xanthomonadaceae; Stenotrophomonas; Stenotrophomonas maltophilia group MDRRELLKMIVAATGAAMVGLPALAQGNAAAAPVKTLFSDADIGTLDEIAETILPRTRTP WP_049404954.1 twin-arginine translocation pathway signal [Stenotrophomonas maltophilia] Length: 191\nNumber of predicted TMHs: 1\nExp number of AAs in TMHs: 22.01925\nExp number, first 60 AAs:  4.63872\nTotal prob of N-in:  0.67790\ninside 1 149\nTMhelix 150 172\noutside 173 191

29017 GCF_001072195.1_ASM107219v1 Stenotrophomonas maltophilia Proteobacteria; Gammaproteobacteria; Xanthomonadales; Xanthomonadaceae; Stenotrophomonas; Stenotrophomonas maltophilia group

MDRRELLKMIVAATGAAMVGLPALVHGKVPAAGAMPAFSDADIGMLDEIAETILPQTRTP WP_049403241.1 twin-arginine translocation pathway signal [Stenotrophomonas maltophilia] Length: 191\nNumber of predicted TMHs: 1\nExp number of AAs in TMHs: 22.06874\nExp number, first 60 AAs: 5.27759\nTotal prob of N-in: 0.70349\ninside 1 149\nTMhelix 150 172\noutside 173 191

29018 GCF_001073045.1_ASM107304v1 Stenotrophomonas maltophilia Proteobacteria; Gammaproteobacteria; Xanthomonadales; Xanthomonadaceae; Stenotrophomonas; Stenotrophomonas maltophilia group MDRRELLKMIVAATGAAMVGLPALVHGQAPAGATKIHFSDADVGTLDEIAETILPRTRTP WP_049434537.1 twin-arginine translocation pathway signal [Stenotrophomonas maltophilia] Length: 191\nNumber of predicted TMHs: 1\nExp number of AAs in TMHs: 22.50442\nExp number, first 60 AAs: 5.58558\nTotal prob of N-in: 0.69433\ninside 1 149\nTMhelix 150 172\noutside 173 191

29019 GCF_001069435.1_ASM106943v1 Stenotrophomonas maltophilia Proteobacteria; Gammaproteobacteria; Xanthomonadales; Xanthomonadaceae; Stenotrophomonas; Stenotrophomonas maltophilia group MDRRELLKMIVAATGAAMVGLPALVHGKAPAAGAMAAFSDADIGTLDEIAETILPRTRTP WP_049438573.1 twin-arginine translocation pathway signal [Stenotrophomonas maltophilia] Length: 191\nNumber of predicted TMHs: 1\nExp number of AAs in TMHs: 22.84658\nExp number, first 60 AAs: 5.7273\nTotal prob of N-in: 0.66404\ninside 1 149\nTMhelix 150 172\noutside 173 191

29020 GCF_001069235.1_ASM106923v1 Stenotrophomonas maltophilia Proteobacteria; Gammaproteobacteria; Xanthomonadales; Xanthomonadaceae; Stenotrophomonas; Stenotrophomonas maltophilia group MDRRELLKMIVAATGAAMVGLPALAQGNAAAAPVKTLFSDADIGTLDEIAETILPRTRTP WP_049404954.1 twin-arginine translocation pathway signal [Stenotrophomonas maltophilia] Length: 191\nNumber of predicted TMHs: 1\nExp number of AAs in TMHs: 22.01925\nExp number, first 60 AAs: 4.63872\nTotal prob of N-in: 0.67790\ninside 1 149\nTMhelix 150 172\noutside 173 191

29021 GCF_001075145.1_ASM107514v1 Stenotrophomonas maltophilia Proteobacteria; Gammaproteobacteria; Xanthomonadales; Xanthomonadaceae; Stenotrophomonas; Stenotrophomonas maltophilia group MDRRELLKMIVAATGAAMVGLPALVHGKVPAAGAMPAFSDADIGMLDEIAETILPRTRTP WP_005412559.1 MULTISPECIES: hypothetical protein [Stenotrophomonas] Length: 191\nNumber of predicted TMHs: 1\nExp number of AAs in TMHs: 22.08369\nExp number, first 60 AAs: 5.28303\nTotal prob of N-in: 0.70356\ninside 1 149\nTMhelix 150 172\noutside 173 191

29022 GCF_001069685.1_ASM106968v1 Stenotrophomonas maltophilia Proteobacteria; Gammaproteobacteria; Xanthomonadales; Xanthomonadaceae; Stenotrophomonas; Stenotrophomonas maltophilia group MDRRELLKMIVAATGAAMVGLPALAQGNAAAAPVKALFSDADIGTLDEIAETILPRTRTP WP_046429410.1 twin-arginine translocation pathway signal [Stenotrophomonas maltophilia] Length: 191\nNumber of predicted TMHs: 1\nExp number of AAs in TMHs: 21.48701\nExp number, first 60 AAs: 4.22916\nTotal prob of N-in: 0.69390\ninside 1 149\nTMhelix 150 172\noutside 173 191

29023 GCF_001676375.1_ASM167637v1 Stenotrophomonas maltophilia Proteobacteria; Gammaproteobacteria; Xanthomonadales; Xanthomonadaceae; Stenotrophomonas; Stenotrophomonas maltophilia group MDRRELLKMIVAATGAAMVGLPALVHGKVPAAGAMPAFSDADIGMLDEIAETILPRTRTP WP_005412559.1 MULTISPECIES: hypothetical protein [Stenotrophomonas] Length: 191\nNumber of predicted TMHs: 1\nExp number of AAs in TMHs: 22.08369\nExp number, first 60 AAs: 5.28303\nTotal prob of N-in: 0.70356\ninside 1 149\nTMhelix 150 172\noutside 173 191

29024 GCF_002189545.1_ASM218954v1 Stenotrophomonas maltophilia Proteobacteria; Gammaproteobacteria; Xanthomonadales; Xanthomonadaceae; Stenotrophomonas; Stenotrophomonas maltophilia group MDRRELLKMIVAATGAAMVGLPALAQGNAAAAPVKTLFSDADIGTLDEIAETILPRTRTP WP_049404954.1 twin-arginine translocation pathway signal [Stenotrophomonas maltophilia] Length: 191\nNumber of predicted TMHs: 1\nExp number of AAs in TMHs: 22.01925\nExp number, first 60 AAs: 4.63872\nTotal prob of N-in: 0.67790\ninside 1 149\nTMhelix 150 172\noutside 173 191

29025 GCF_001676295.1_ASM167629v1 Stenotrophomonas maltophilia Proteobacteria; Gammaproteobacteria; Xanthomonadales; Xanthomonadaceae; Stenotrophomonas; Stenotrophomonas maltophilia group MDRRELLKMIVSATGAAMVGLPALVHGKVPAAGMMAAFSDAEIGTLDEIAETILPRTRTP WP_065196568.1 Twin-arginine translocation pathway signal [Stenotrophomonas maltophilia] Length: 191\nNumber of predicted TMHs: 1\nExp number of AAs in TMHs: 22.19258\nExp number, first 60 AAs: 5.23944\nTotal prob of N-in: 0.67472\ninside 1 149\nTMhelix 150 172\noutside 173 191

29026	GCF_001069565.1_ASM106956v1	Stenotrophomonas maltophilia	Proteobacteria; Gammaproteobacteria; Xanthomonadales; Xanthomonadaceae; Stenotrophomonas; Stenotrophomonas maltophilia group	MDRRELLKMIVAATGAAMVGLPALAQGNAAAAPVKTLFSDADIGTLDEIAETILPRTRTP	WP_049404954.1	twin-arginine translocation pathway signal [Stenotrophomonas maltophilia]	Length: 191\nNumber of predicted TMHs: 1\nExp number of AAs in TMHs: 22.01925\nExp number, first 60 AAs: 4.63872\nTotal prob of N-in: 0.67790\ninside 1 149\nTMhelix 150 172\noutside 173 191

29027	GCF_001676395.1_ASM167639v1	Stenotrophomonas maltophilia	Proteobacteria; Gammaproteobacteria; Xanthomonadales; Xanthomonadaceae; Stenotrophomonas; Stenotrophomonas maltophilia group	MDRRELLKMIVSATGAAMVGLPALVHGKAPAAGMMAAFSDAEIGTLDEIAETILPRTRTP	WP_065186014.1	Twin-arginine translocation pathway signal [Stenotrophomonas maltophilia]	Length: 191\nNumber of predicted TMHs: 1\nExp number of AAs in TMHs: 21.77718\nExp number, first 60 AAs: 4.87292\nTotal prob of N-in: 0.68173\ninside 1 149\nTMhelix 150 172\noutside 173 191

29028	GCF_001542395.1_ASM154239v1	Stenotrophomonas maltophilia	Proteobacteria; Gammaproteobacteria; Xanthomonadales; Xanthomonadaceae; Stenotrophomonas; Stenotrophomonas maltophilia group	MDRRELLKMIVAATGAAMVGLPALVHGKVPAAGAMPAFSDADIGMLDEIAETILPRTRTP	WP_005412559.1	MULTISPECIES: hypothetical protein [Stenotrophomonas]	Length: 191\nNumber of predicted TMHs: 1\nExp number of AAs in TMHs: 22.08369\nExp number, first 60 AAs: 5.28303\nTotal prob of N-in: 0.70356\ninside 1 149\nTMhelix 150 172\noutside 173 191

29029	GCF_001499715.1_BurA1	Stenotrophomonas maltophilia	Proteobacteria; Gammaproteobacteria; Xanthomonadales; Xanthomonadaceae; Stenotrophomonas; Stenotrophomonas maltophilia group	MDRRELLKMIVAATGAAMVGLPALVQGRAPAAGAMPAFTDADIGTLDEIAETILPRTRTP	WP_058980795.1	Twin-arginine translocation pathway signal [Stenotrophomonas maltophilia]	Length: 191\nNumber of predicted TMHs: 1\nExp number of AAs in TMHs: 21.06326\nExp number, first 60 AAs: 4.59546\nTotal prob of N-in: 0.72998\ninside 1 149\nTMhelix 150 172\noutside 173 191

29030	GCF_001676295.1_ASM167629v1	Stenotrophomonas maltophilia	Proteobacteria; Gammaproteobacteria; Xanthomonadales; Xanthomonadaceae; Stenotrophomonas; Stenotrophomonas maltophilia group	MRPVRAWTRRAVLPVLLLALGACAKQGDMVGDAGATAAEAAIASPEGAFLAYEHDVQVQL	WP_065177445.1	hypothetical protein [Stenotrophomonas maltophilia]	Length: 271\nNumber of predicted TMHs: 1\nExp number of AAs in TMHs: 23.07034\nExp number, first 60 AAs: 0.46424\nTotal prob of N-in: 0.06963\noutside 1 233\nTMhelix 234 256\ninside 257 271

29031	GCF_001068915.1_ASM106891v1	Stenotrophomonas maltophilia	Proteobacteria; Gammaproteobacteria; Xanthomonadales; Xanthomonadaceae; Stenotrophomonas; Stenotrophomonas maltophilia group	MDRRELLKMIVAATGAAMVGLPALVHGKVPAAGAMPAFSDADIGMLDEIAETILPRTRTP	WP_005412559.1	MULTISPECIES: hypothetical protein [Stenotrophomonas]	Length: 191\nNumber of predicted TMHs: 1\nExp number of AAs in TMHs: 22.08369\nExp number, first 60 AAs: 5.28303\nTotal prob of N-in: 0.70356\ninside 1 149\nTMhelix 150 172\noutside 173 191

29032	GCF_001286925.1_Stenotrophomonas_maltophilia_mecca_isolate	Stenotrophomonas maltophilia	Proteobacteria; Gammaproteobacteria; Xanthomonadales; Xanthomonadaceae; Stenotrophomonas; Stenotrophomonas maltophilia group	MDRRELLKMIVAATGAAMVGLPVLVQGKAPAAGAMAAFSDAEISTLDEIAETILPRTRTP	WP_072168650.1	Twin-arginine translocation pathway signal [Stenotrophomonas maltophilia]	Length: 191\nNumber of predicted TMHs: 1\nExp number of AAs in TMHs: 21.90848\nExp number, first 60 AAs: 5.11694\nTotal prob of N-in: 0.69279\ninside 1 149\nTMhelix 150 172\noutside 173 191

29033	GCF_001286925.1_Stenotrophomonas_maltophilia_mecca_isolate	Stenotrophomonas maltophilia	Proteobacteria; Gammaproteobacteria; Xanthomonadales; Xanthomonadaceae; Stenotrophomonas; Stenotrophomonas maltophilia group	MRPVRAWTRRAALPVLLLALGACAKQGDMVGDAGATAAEAAIASPEGAFLAYEHDVQVQL	WP_072166824.1	hypothetical protein [Stenotrophomonas maltophilia]	Length: 270\nNumber of predicted TMHs: 1\nExp number of AAs in TMHs: 22.86251\nExp number, first 60 AAs: 0.28612\nTotal prob of N-in: 0.07493\noutside 1 233\nTMhelix 234 256\ninside 257 270

29034	GCF_001499695.1_BurE1	Stenotrophomonas maltophilia	Proteobacteria; Gammaproteobacteria; Xanthomonadales; Xanthomonadaceae; Stenotrophomonas; Stenotrophomonas maltophilia group	MDRRELLKMIVAATGAAMVGLPALVHGKVPAAGAMPAFSDADIGMLDEIAETILPRTRTP	WP_005412559.1	MULTISPECIES: hypothetical protein [Stenotrophomonas]	Length: 191\nNumber of predicted TMHs: 1\nExp

number of AAs in TMHs: 22.08369\nExp number, first 60 AAs: 5.28303\nTotal prob of N-in: 0.70356\ninside 1 149\nTMhelix 150 172\noutside 173 191

29035 GCF_001073355.1_ASM107335v1 Stenotrophomonas maltophilia Proteobacteria; Gammaproteobacteria; Xanthomonadales; Xanthomonadaceae; Stenotrophomonas; Stenotrophomonas maltophilia group MDRRELLKMIVAATGAAMVGLPALVHGKVPAAGAMPAFSDADIGMLDEIAETILPRTRTP WP_012479395.1 membrane protein [Stenotrophomonas maltophilia] Length: 191\nNumber of predicted TMHs: 1\nExp number of AAs in TMHs: 22.08345\nExp number, first 60 AAs: 5.28293\nTotal prob of N-in: 0.70356\ninside 1 149\nTMhelix 150 172\noutside 173 191

29036 GCF_001431665.1_ASM143166v1 Stenotrophomonas maltophilia Proteobacteria; Gammaproteobacteria; Xanthomonadales; Xanthomonadaceae; Stenotrophomonas; Stenotrophomonas maltophilia group MDRRELLKMIVAATGAAMVGLPALAQGQGRAASAAVHFSDADIGMLDEIAETILPRTRTP WP_057496275.1 twin-arginine translocation pathway signal [Stenotrophomonas maltophilia] Length: 191\nNumber of predicted TMHs: 1\nExp number of AAs in TMHs: 18.71657\nExp number, first 60 AAs: 1.74745\nTotal prob of N-in: 0.74861\ninside 1 149\nTMhelix 150 172\noutside 173 191

29037 GCF_001676385.1_ASM167638v1 Stenotrophomonas maltophilia Proteobacteria; Gammaproteobacteria; Xanthomonadales; Xanthomonadaceae; Stenotrophomonas; Stenotrophomonas maltophilia group MDRRELLKMIVAATGAAMVGLPALVQGQVPAAGAMPAFSDADIGMLDEIAETILPRTRTP WP_065198061.1 Twin-arginine translocation pathway signal [Stenotrophomonas maltophilia] Length: 191\nNumber of predicted TMHs: 1\nExp number of AAs in TMHs: 21.3475\nExp number, first 60 AAs: 5.22098\nTotal prob of N-in: 0.77710\ninside 1 149\nTMhelix 150 172\noutside 173 191

29038 GCF_001651505.1_ISMMS7v1 Stenotrophomonas maltophilia Proteobacteria; Gammaproteobacteria; Xanthomonadales; Xanthomonadaceae; Stenotrophomonas; Stenotrophomonas maltophilia group MDRRELLKMIVAATGTAMVGLPALVHGKVPAAGAMPAFSDADIGMLDEIAETILPRTRTP WP_064237859.1 Twin-arginine translocation pathway signal [Stenotrophomonas maltophilia] Length: 191\nNumber of predicted TMHs: 1\nExp number of AAs in TMHs: 20.52934\nExp number, first 60 AAs: 3.49346\nTotal prob of N-in: 0.72346\ninside 1 149\nTMhelix 150 172\noutside 173 191

29039 GCF_001275085.1_ISMMS4v1 Stenotrophomonas maltophilia Proteobacteria; Gammaproteobacteria; Xanthomonadales; Xanthomonadaceae; Stenotrophomonas; Stenotrophomonas maltophilia group MDRRELLKMIVAATGAAMVGLPALVHGKVPAAGAMPAFSDADIGMLDEIAETILPRTRTP WP_005412559.1 MULTISPECIES: hypothetical protein [Stenotrophomonas] Length: 191\nNumber of predicted TMHs: 1\nExp number of AAs in TMHs: 22.08369\nExp number, first 60 AAs: 5.28303\nTotal prob of N-in: 0.70356\ninside 1 149\nTMhelix 150 172\noutside 173 191

29040 GCF_001069295.1_ASM106929v1 Stenotrophomonas maltophilia Proteobacteria; Gammaproteobacteria; Xanthomonadales; Xanthomonadaceae; Stenotrophomonas; Stenotrophomonas maltophilia group MDRRELLKMIVAATGAAMVGLPALVHGKVPAAGAMPAFSDADIGMLDEIAETILPRTRTP WP_005412559.1 MULTISPECIES: hypothetical protein [Stenotrophomonas] Length: 191\nNumber of predicted TMHs: 1\nExp number of AAs in TMHs: 22.08369\nExp number, first 60 AAs: 5.28303\nTotal prob of N-in: 0.70356\ninside 1 149\nTMhelix 150 172\noutside 173 191

29041 GCF_001069595.1_ASM106959v1 Stenotrophomonas maltophilia Proteobacteria; Gammaproteobacteria; Xanthomonadales; Xanthomonadaceae; Stenotrophomonas; Stenotrophomonas maltophilia group MDRRELLKMIVAATGAAMVGLPALAQGNAAAAPVKTLFSDADIGTLDEIAETILPRTRTP WP_049404954.1 twin-arginine translocation pathway signal [Stenotrophomonas maltophilia] Length: 191\nNumber of predicted TMHs: 1\nExp number of AAs in TMHs: 22.01925\nExp number, first 60 AAs: 4.63872\nTotal prob of N-in: 0.67790\ninside 1 149\nTMhelix 150 172\noutside 173 191

29042 GCF_001071065.1_ASM107106v1 Stenotrophomonas maltophilia Proteobacteria; Gammaproteobacteria; Xanthomonadales; Xanthomonadaceae; Stenotrophomonas; Stenotrophomonas maltophilia group MDRRELLKMIVAATGAAMVGLPALVHGQAPAGATKIHFSDADVGTLDEIAETILPRTRTP WP_049434537.1 twin-arginine translocation pathway signal [Stenotrophomonas maltophilia] Length: 191\nNumber of predicted TMHs: 1\nExp number of AAs in TMHs: 22.50442\nExp number, first 60 AAs: 5.58558\nTotal prob of N-in: 0.69433\ninside 1 149\nTMhelix 150 172\noutside 173 191

29043 GCF_001071045.1_ASM107104v1 Stenotrophomonas maltophilia Proteobacteria; Gammaproteobacteria; Xanthomonadales; Xanthomonadaceae; Stenotrophomonas; Stenotrophomonas maltophilia group

MDRRELLKMIVAATGAAMVGLPALVHGQAPAGATKIHFSDADVGTLDEIAETILPRTRTP WP_049434537.1 twin-arginine translocation pathway signal [Stenotrophomonas maltophilia] Length: 191\nNumber of predicted TMHs: 1\nExp number of AAs in TMHs: 22.50442\nExp number, first 60 AAs: 5.58558\nTotal prob of N-in: 0.69433\ninside 1 149\nTMhelix 150 172\noutside 173 191

29044 GCF_002189565.1_ASM218956v1 Stenotrophomonas maltophilia Proteobacteria; Gammaproteobacteria; Xanthomonadales; Xanthomonadaceae; Stenotrophomonas; Stenotrophomonas maltophilia group MDRRELLKMIVAATGAAMVGLPALVHGKVPAAGAMPAFSDADIGMLDEIAEIILPRTRTP WP_087802949.1 Twin-arginine translocation pathway signal [Stenotrophomonas maltophilia] Length: 191\nNumber of predicted TMHs: 1\nExp number of AAs in TMHs: 22.21156\nExp number, first 60 AAs: 5.0856\nTotal prob of N-in: 0.71312\ninside 1 149\nTMhelix 150 172\noutside 173 191

29045 GCF_002205215.1_ASM220521v1 Stenotrophomonas maltophilia Proteobacteria; Gammaproteobacteria; Xanthomonadales; Xanthomonadaceae; Stenotrophomonas; Stenotrophomonas maltophilia group MDRRELLKMIVAATGAAMVGLPALVQGKAPAAGAMAAFSDAEISTLDEIAETILPRTRTP WP_088496559.1 Twin-arginine translocation pathway signal [Stenotrophomonas maltophilia] Length: 191\nNumber of predicted TMHs: 1\nExp number of AAs in TMHs: 20.16074\nExp number, first 60 AAs: 3.37567\nTotal prob of N-in: 0.72933\ninside 1 149\nTMhelix 150 172\noutside 173 191

29046 GCF_002205215.1_ASM220521v1 Stenotrophomonas maltophilia Proteobacteria; Gammaproteobacteria; Xanthomonadales; Xanthomonadaceae; Stenotrophomonas; Stenotrophomonas maltophilia group MRPVRAWTRRAALPVLLLALGACAKQGDMVGDAGATAAEAAIASPEGAFLAYEHDVQVQL WP_088496929.1 hypothetical protein [Stenotrophomonas maltophilia] Length: 270\nNumber of predicted TMHs: 1\nExp number of AAs in TMHs: 22.94304\nExp number, first 60 AAs: 0.2824\nTotal prob of N-in: 0.08079\noutside 1 233\nTMhelix 234 256\ninside 257 270

29047 GCF_001074705.1_ASM107470v1 Stenotrophomonas maltophilia Proteobacteria; Gammaproteobacteria; Xanthomonadales; Xanthomonadaceae; Stenotrophomonas; Stenotrophomonas maltophilia group MDRRELLKMIVAATGAAMVGLPALVHGKVPAAGAMPAFSDADIGMLDEIAETILPRTRTP WP_005412559.1 MULTISPECIES: hypothetical protein [Stenotrophomonas] Length: 191\nNumber of predicted TMHs: 1\nExp number of AAs in TMHs: 22.08369\nExp number, first 60 AAs: 5.28303\nTotal prob of N-in: 0.70356\ninside 1 149\nTMhelix 150 172\noutside 173 191

29048 GCF_000978875.1_ASM97887v1 Stenotrophomonas maltophilia Proteobacteria; Gammaproteobacteria; Xanthomonadales; Xanthomonadaceae; Stenotrophomonas; Stenotrophomonas maltophilia group MDRRELLKMIVAATGAAMVGLPALAQGNAAAAPVKALFSDADIGTLDEIAETILPRTRTP WP_046429410.1 twin-arginine translocation pathway signal [Stenotrophomonas maltophilia] Length: 191\nNumber of predicted TMHs: 1\nExp number of AAs in TMHs: 21.48701\nExp number, first 60 AAs: 4.22916\nTotal prob of N-in: 0.69390\ninside 1 149\nTMhelix 150 172\noutside 173 191

29049 GCF_002205225.1_ASM220522v1 Stenotrophomonas maltophilia Proteobacteria; Gammaproteobacteria; Xanthomonadales; Xanthomonadaceae; Stenotrophomonas; Stenotrophomonas maltophilia group MDRRELLKMIVAATGAAMVGLPALAQGNAAAAPVKTLFSDADIGTLDEIAETILPRTRTP WP_049404954.1 twin-arginine translocation pathway signal [Stenotrophomonas maltophilia] Length: 191\nNumber of predicted TMHs: 1\nExp number of AAs in TMHs: 22.01925\nExp number, first 60 AAs: 4.63872\nTotal prob of N-in: 0.67790\ninside 1 149\nTMhelix 150 172\noutside 173 191

29050 GCF_002205275.1_ASM220527v1 Stenotrophomonas maltophilia Proteobacteria; Gammaproteobacteria; Xanthomonadales; Xanthomonadaceae; Stenotrophomonas; Stenotrophomonas maltophilia group MDRRELLKMIVAATGAAMVGLPALAQGNAAAAPVKALFSDADIGTLDEIAETILPRTRTP WP_046429410.1 twin-arginine translocation pathway signal [Stenotrophomonas maltophilia] Length: 191\nNumber of predicted TMHs: 1\nExp number of AAs in TMHs: 21.48701\nExp number, first 60 AAs: 4.22916\nTotal prob of N-in: 0.69390\ninside 1 149\nTMhelix 150 172\noutside 173 191

29051 GCF_002208885.1_ASM220888v1 Stenotrophomonas maltophilia Proteobacteria; Gammaproteobacteria; Xanthomonadales; Xanthomonadaceae; Stenotrophomonas; Stenotrophomonas maltophilia group MDRRELLKMIVAATGAAMVGLPALVHGKVPAAGAMPAFSDADIGMLDEIAETILPRTRTP WP_005412559.1 MULTISPECIES: hypothetical protein [Stenotrophomonas] Length: 191\nNumber of predicted TMHs: 1\nExp number of AAs in TMHs: 22.08369\nExp number, first 60 AAs: 5.28303\nTotal prob of N-in: 0.70356\ninside 1 149\nTMhelix 150 172\noutside 173 191

29052 GCF_001274655.1_ASM127465v1 Stenotrophomonas maltophilia Proteobacteria; Gammaproteobacteria; Xanthomonadales; Xanthomonadaceae; Stenotrophomonas; Stenotrophomonas maltophilia group MDRRELLKMIVAATGAAMVGLPALVQGQAPAGATKIHFSDADVGTLDEIAETILPRTRTP WP_053442649.1 twin-arginine translocation pathway signal [Stenotrophomonas maltophilia] Length: 191\nNumber of predicted TMHs: 1\nExp number of AAs in TMHs: 20.2318\nExp number, first 60 AAs: 3.42144\nTotal prob of N-in: 0.74184\ninside 1 149\nTMhelix 150 172\noutside 173 191

29053 GCF_001275095.1_ISMMS6v1 Stenotrophomonas maltophilia Proteobacteria; Gammaproteobacteria; Xanthomonadales; Xanthomonadaceae; Stenotrophomonas; Stenotrophomonas maltophilia group MDRRELLKMIVAATGTAMVGLPALVHGKVPAAGAMPAFSDADIGMLDEIAETILPRTRTP WP_053462412.1 twin-arginine translocation pathway signal [Stenotrophomonas maltophilia] Length: 191\nNumber of predicted TMHs: 1\nExp number of AAs in TMHs: 20.52951\nExp number, first 60 AAs: 3.49352\nTotal prob of N-in: 0.72346\ninside 1 149\nTMhelix 150 172\noutside 173 191

29054 GCF_002205295.1_ASM220529v1 Stenotrophomonas maltophilia Proteobacteria; Gammaproteobacteria; Xanthomonadales; Xanthomonadaceae; Stenotrophomonas; Stenotrophomonas maltophilia group MDRRELLKMIVAATGAAMVGLPALVHGKVPAAGAMPAFSDADIGMLDEIAETILPRTRTP WP_005412559.1 MULTISPECIES: hypothetical protein [Stenotrophomonas] Length: 191\nNumber of predicted TMHs: 1\nExp number of AAs in TMHs: 22.08369\nExp number, first 60 AAs: 5.28303\nTotal prob of N-in: 0.70356\ninside 1 149\nTMhelix 150 172\noutside 173 191

29055 GCF_001276305.1_ASM127630v1 Stenotrophomonas maltophilia Proteobacteria; Gammaproteobacteria; Xanthomonadales; Xanthomonadaceae; Stenotrophomonas; Stenotrophomonas maltophilia group MDRRELLKMIVAATGAAMVGLPALAQGNAAAAPVKALFSDADIGTLDEIAETILPRTRTP WP_046429410.1 twin-arginine translocation pathway signal [Stenotrophomonas maltophilia] Length: 191\nNumber of predicted TMHs: 1\nExp number of AAs in TMHs: 21.48701\nExp number, first 60 AAs: 4.22916\nTotal prob of N-in: 0.69390\ninside 1 149\nTMhelix 150 172\noutside 173 191

29056 GCF_001275075.1_ISMMS5v1 Stenotrophomonas maltophilia Proteobacteria; Gammaproteobacteria; Xanthomonadales; Xanthomonadaceae; Stenotrophomonas; Stenotrophomonas maltophilia group MDRRELLKMIVAATGAAMVGLPALAQGNAAAAPVKTLFSDADIGTLDEIAETILPRTRTP WP_049404954.1 twin-arginine translocation pathway signal [Stenotrophomonas maltophilia] Length: 191\nNumber of predicted TMHs: 1\nExp number of AAs in TMHs: 22.01925\nExp number, first 60 AAs: 4.63872\nTotal prob of N-in: 0.67790\ninside 1 149\nTMhelix 150 172\noutside 173 191

29057 GCF_001276185.1_ASM127618v1 Stenotrophomonas maltophilia Proteobacteria; Gammaproteobacteria; Xanthomonadales; Xanthomonadaceae; Stenotrophomonas; Stenotrophomonas maltophilia group MDRRELLKMIVAATGTAMVGLPALVHGKVPAAGAMPAFSDADIGMLDEIAETILPRTRTP WP_053505877.1 twin-arginine translocation pathway signal [Stenotrophomonas maltophilia] Length: 191\nNumber of predicted TMHs: 1\nExp number of AAs in TMHs: 20.20849\nExp number, first 60 AAs: 3.60413\nTotal prob of N-in: 0.71158\ninside 1 149\nTMhelix 150 172\noutside 173 191

29058 GCF_001274675.1_ASM127467v1 Stenotrophomonas maltophilia Proteobacteria; Gammaproteobacteria; Xanthomonadales; Xanthomonadaceae; Stenotrophomonas; Stenotrophomonas maltophilia group MDRRELLKMIVAATGAAMVGLPALVQGQAPAGATKIHFSDADVGTLDEIAETILPRTRTP WP_053442649.1 twin-arginine translocation pathway signal [Stenotrophomonas maltophilia] Length: 191\nNumber of predicted TMHs: 1\nExp number of AAs in TMHs: 20.2318\nExp number, first 60 AAs: 3.42144\nTotal prob of N-in: 0.74184\ninside 1 149\nTMhelix 150 172\noutside 173 191

29059 GCF_001297005.1_SmOC94scaf_imr500 Stenotrophomonas maltophilia Proteobacteria; Gammaproteobacteria; Xanthomonadales; Xanthomonadaceae; Stenotrophomonas; Stenotrophomonas maltophilia group MDRRELLKMIVAATGAAMVGLPALVQGQAPAPAGATKIHFSDADVGTLDEIAETILPRTR WP_054171204.1 twin-arginine translocation pathway signal [Stenotrophomonas maltophilia] Length: 193\nNumber of predicted TMHs: 1\nExp number of AAs in TMHs: 19.02731\nExp number, first 60 AAs: 2.21787\nTotal prob of N-in: 0.75526\ninside 1 151\nTMhelix 152 174\noutside 175 193

29060 GCF_001075445.1_ASM107544v1 Stenotrophomonas maltophilia Proteobacteria; Gammaproteobacteria; Xanthomonadales; Xanthomonadaceae; Stenotrophomonas; Stenotrophomonas maltophilia group MDRRELLKMIVAATGAAMVGLPALAQGNAAAAPVKTLFSDADIGTLDEIAETILPRTRTP WP_049404954.1 twin-arginine translocation pathway signal [Stenotrophomonas maltophilia] Length: 191\nNumber of predicted TMHs:

1\nExp number of AAs in TMHs: 22.01925\nExp number, first 60 AAs: 4.63872\nTotal prob of N-in: 0.67790\ninside 1 149\nTMhelix 150 172\noutside 173 191

29061 GCF_001595975.1_ASM159597v1 Stenotrophomonas maltophilia Proteobacteria; Gammaproteobacteria; Xanthomonadales; Xanthomonadaceae; Stenotrophomonas; Stenotrophomonas maltophilia group MDRRELLKMIVAATGAAMVGLPALVHGKVPAAGAMPAFSDADIGMLDEIAETILPRTRTP WP_062606679.1 Twin-arginine translocation pathway signal [Stenotrophomonas maltophilia] Length: 191\nNumber of predicted TMHs: 1\nExp number of AAs in TMHs: 22.25413\nExp number, first 60 AAs: 5.3498\nTotal prob of N-in: 0.70477\ninside 1 149\nTMhelix 150 172\noutside 173 191

29062 GCF_001997185.1_ASM199718v1 Stenotrophomonas maltophilia Proteobacteria; Gammaproteobacteria; Xanthomonadales; Xanthomonadaceae; Stenotrophomonas; Stenotrophomonas maltophilia group MDRRELLKMIVAATGAAMVGLPALVHGKVPAAGAMPAFSDADIGMLDEIAETILPRTRTP WP_005412559.1 MULTISPECIES: hypothetical protein [Stenotrophomonas] Length: 191\nNumber of predicted TMHs: 1\nExp number of AAs in TMHs: 22.08369\nExp number, first 60 AAs: 5.28303\nTotal prob of N-in: 0.70356\ninside 1 149\nTMhelix 150 172\noutside 173 191

29063 GCF_001676395.1_ASM167639v1 Stenotrophomonas maltophilia Proteobacteria; Gammaproteobacteria; Xanthomonadales; Xanthomonadaceae; Stenotrophomonas; Stenotrophomonas maltophilia group MRPVRAWTRRAALPVLLLALGACAKQGDMVGDAGATAAEAAIASPEGAFLAYEHDVQVQL WP_065186376.1 hypothetical protein [Stenotrophomonas maltophilia] Length: 271\nNumber of predicted TMHs: 1\nExp number of AAs in TMHs: 22.9003\nExp number, first 60 AAs: 0.29444\nTotal prob of N-in: 0.06107\noutside 1 233\nTMhelix 234 256\ninside 257 271

29064 GCF_001071195.1_ASM107119v1 Stenotrophomonas maltophilia Proteobacteria; Gammaproteobacteria; Xanthomonadales; Xanthomonadaceae; Stenotrophomonas; Stenotrophomonas maltophilia group MDRRELLKMIVAATGAAMVGLPALVHGKVPAAGAMPAFSDADIGMLDEIAETILPRTRTP WP_005408476.1 membrane protein [Stenotrophomonas maltophilia] Length: 191\nNumber of predicted TMHs: 1\nExp number of AAs in TMHs: 22.20941\nExp number, first 60 AAs: 5.35942\nTotal prob of N-in: 0.70371\ninside 1 149\nTMhelix 150 172\noutside 173 191

29065 GCF_001071285.1_ASM107128v1 Stenotrophomonas maltophilia Proteobacteria; Gammaproteobacteria; Xanthomonadales; Xanthomonadaceae; Stenotrophomonas; Stenotrophomonas maltophilia group MDRRELLKMIVAATGAAMVGLPALVHGKVPAAGAMPAFSDADIGMLDEIAETILPRTRTP WP_012479395.1 membrane protein [Stenotrophomonas maltophilia] Length: 191\nNumber of predicted TMHs: 1\nExp number of AAs in TMHs: 22.08345\nExp number, first 60 AAs: 5.28293\nTotal prob of N-in: 0.70356\ninside 1 149\nTMhelix 150 172\noutside 173 191

29066 GCF_001075175.1_ASM107517v1 Stenotrophomonas maltophilia Proteobacteria; Gammaproteobacteria; Xanthomonadales; Xanthomonadaceae; Stenotrophomonas; Stenotrophomonas maltophilia group MDRRELLKMIVAATGAAMVGLPALVHGKVPAAGAMPAFSDADIGMLDEIAETILPRTRTP WP_005412559.1 MULTISPECIES: hypothetical protein [Stenotrophomonas] Length: 191\nNumber of predicted TMHs: 1\nExp number of AAs in TMHs: 22.08369\nExp number, first 60 AAs: 5.28303\nTotal prob of N-in: 0.70356\ninside 1 149\nTMhelix 150 172\noutside 173 191

29067 GCF_001072535.1_ASM107253v1 Stenotrophomonas maltophilia Proteobacteria; Gammaproteobacteria; Xanthomonadales; Xanthomonadaceae; Stenotrophomonas; Stenotrophomonas maltophilia group MDRRELLKMIVAATGAAMVGLPALVHGKVPAAGAMPAFSDADIGMLDEIAETILPQTRTP WP_049403241.1 twin-arginine translocation pathway signal [Stenotrophomonas maltophilia] Length: 191\nNumber of predicted TMHs: 1\nExp number of AAs in TMHs: 22.06874\nExp number, first 60 AAs: 5.27759\nTotal prob of N-in: 0.70349\ninside 1 149\nTMhelix 150 172\noutside 173 191

29068 GCF_001074595.1_ASM107459v1 Stenotrophomonas maltophilia Proteobacteria; Gammaproteobacteria; Xanthomonadales; Xanthomonadaceae; Stenotrophomonas; Stenotrophomonas maltophilia group MDRRELLKMIVAATGAAMVGLPALVHGKVPAAGAMPAFSDADIGMLDEIAETILPRTRTP WP_005412559.1 MULTISPECIES: hypothetical protein [Stenotrophomonas] Length: 191\nNumber of predicted TMHs: 1\nExp number of AAs in TMHs: 22.08369\nExp number, first 60 AAs: 5.28303\nTotal prob of N-in: 0.70356\ninside 1 149\nTMhelix 150 172\noutside 173 191

29069 GCF_001070325.1_ASM107032v1 Stenotrophomonas maltophilia Proteobacteria; Gammaproteobacteria; Xanthomonadales; Xanthomonadaceae; Stenotrophomonas; Stenotrophomonas maltophilia group

MDRRELLKMIVAATGTAMVGLPALVHGKVPAAGAMPAFSDADIGMLDEIAETILPRTRTP WP_049467359.1 twin-arginine translocation pathway signal [Stenotrophomonas maltophilia] Length: 191\nNumber of predicted TMHs: 1\nExp number of AAs in TMHs: 20.52951\nExp number, first 60 AAs: 3.49352\nTotal prob of N-in: 0.72346\ninside 1 149\nTMhelix 150 172\noutside 173 191

29070 GCF_001071115.1_ASM107111v1 Stenotrophomonas maltophilia Proteobacteria; Gammaproteobacteria; Xanthomonadales; Xanthomonadaceae; Stenotrophomonas; Stenotrophomonas maltophilia group MDRRELLKMIVAATGAAMVGLPALAQGNAAAAPVKTLFSDADIGTLDEIAETILPRTRTP WP_014646300.1 membrane protein [Stenotrophomonas maltophilia] Length: 191\nNumber of predicted TMHs: 1\nExp number of AAs in TMHs: 22.03151\nExp number, first 60 AAs: 4.64253\nTotal prob of N-in: 0.67798\ninside 1 149\nTMhelix 150 172\noutside 173 191

29071 GCF_000742995.1_ASM74299v1 Stenotrophomonas maltophilia Proteobacteria; Gammaproteobacteria; Xanthomonadales; Xanthomonadaceae; Stenotrophomonas; Stenotrophomonas maltophilia group MDRRELLKMIVAATGAAMVGLPALVHGKVPAAGAMPAFSDADIGMLDEIAETILPRTRTP WP_005412559.1 MULTISPECIES: hypothetical protein [Stenotrophomonas] Length: 191\nNumber of predicted TMHs: 1\nExp number of AAs in TMHs: 22.08369\nExp number, first 60 AAs: 5.28303\nTotal prob of N-in: 0.70356\ninside 1 149\nTMhelix 150 172\noutside 173 191

29072 GCF_001069005.1_ASM106900v1 Stenotrophomonas maltophilia Proteobacteria; Gammaproteobacteria; Xanthomonadales; Xanthomonadaceae; Stenotrophomonas; Stenotrophomonas maltophilia group MDRRELLKMIVAATGAAMVGLPALVHGKVPAAGAMPAFSDADIGMLDEIAETILPRTRTP WP_005412559.1 MULTISPECIES: hypothetical protein [Stenotrophomonas] Length: 191\nNumber of predicted TMHs: 1\nExp number of AAs in TMHs: 22.08369\nExp number, first 60 AAs: 5.28303\nTotal prob of N-in: 0.70356\ninside 1 149\nTMhelix 150 172\noutside 173 191

29073 GCF_001276345.1_ASM127634v1 Stenotrophomonas maltophilia Proteobacteria; Gammaproteobacteria; Xanthomonadales; Xanthomonadaceae; Stenotrophomonas; Stenotrophomonas maltophilia group MDRRELLKMIVAATGAAMVGLPALVHGKAPAAGMMAAFSDAEIGTLDEIAETILPRTRTP WP_005415638.1 MULTISPECIES: hypothetical protein [Proteobacteria] Length: 191\nNumber of predicted TMHs: 1\nExp number of AAs in TMHs: 26.26404\nExp number, first 60 AAs: 9.53785\nTotal prob of N-in: 0.59430\ninside 1 149\nTMhelix 150 172\noutside 173 191

29074 GCF_001676445.1_ASM167644v1 Stenotrophomonas maltophilia Proteobacteria; Gammaproteobacteria; Xanthomonadales; Xanthomonadaceae; Stenotrophomonas; Stenotrophomonas maltophilia group MRPVRAWTRRAVLPVLLLALGACAKQGDMVGDAGATAAEAAIASPEGAFLAYEHDVQVQL WP_065177445.1 hypothetical protein [Stenotrophomonas maltophilia] Length: 271\nNumber of predicted TMHs: 1\nExp number of AAs in TMHs: 23.07034\nExp number, first 60 AAs: 0.46424\nTotal prob of N-in: 0.06963\noutside 1 233\nTMhelix 234 256\ninside 257 271

29075 GCF_001276345.1_ASM127634v1 Stenotrophomonas maltophilia Proteobacteria; Gammaproteobacteria; Xanthomonadales; Xanthomonadaceae; Stenotrophomonas; Stenotrophomonas maltophilia group MRPVRAWTRRAALPVLLLALGACAKQGDMVGDAGATAAEAAIASPEGAFLAYEHDVQVQL WP_053497069.1 MULTISPECIES: hypothetical protein [Stenotrophomonas] Length: 271\nNumber of predicted TMHs: 1\nExp number of AAs in TMHs: 22.8986\nExp number, first 60 AAs: 0.29597\nTotal prob of N-in: 0.05695\noutside 1 233\nTMhelix 234 256\ninside 257 271

29076 GCF_002205155.1_ASM220515v1 Stenotrophomonas maltophilia Proteobacteria; Gammaproteobacteria; Xanthomonadales; Xanthomonadaceae; Stenotrophomonas; Stenotrophomonas maltophilia group MDRRELLKMIVAATGAAMVGLPALVHGKVPAAGAMPAFSDADIGMLDEIAETILPRTRTP WP_005412559.1 MULTISPECIES: hypothetical protein [Stenotrophomonas] Length: 191\nNumber of predicted TMHs: 1\nExp number of AAs in TMHs: 22.08369\nExp number, first 60 AAs: 5.28303\nTotal prob of N-in: 0.70356\ninside 1 149\nTMhelix 150 172\noutside 173 191

29077 GCF_002205175.1_ASM220517v1 Stenotrophomonas maltophilia Proteobacteria; Gammaproteobacteria; Xanthomonadales; Xanthomonadaceae; Stenotrophomonas; Stenotrophomonas maltophilia group MDRRELLKMIVAATGAAMVGLPALVHGKVPAAGAMPAFSDADIGMLDEIAETILPRTRTP WP_005412559.1 MULTISPECIES: hypothetical protein [Stenotrophomonas] Length: 191\nNumber of predicted TMHs: 1\nExp number of AAs in TMHs: 22.08369\nExp number, first 60 AAs: 5.28303\nTotal prob of N-in: 0.70356\ninside 1 149\nTMhelix 150 172\noutside 173 191

29078 GCF_001619675.1_ASM161967v1 Stenotrophomonas maltophilia Proteobacteria; Gammaproteobacteria; Xanthomonadales; Xanthomonadaceae; Stenotrophomonas; Stenotrophomonas maltophilia group MDRRELLKMIVAATGAAMVGLPALAQGNAAAAPVKTLFSDADIGTLDEIAETILPRTRTP WP_014646300.1 membrane protein [Stenotrophomonas maltophilia] Length: 191\nNumber of predicted TMHs: 1\nExp number of AAs in TMHs: 22.03151\nExp number, first 60 AAs: 4.64253\nTotal prob of N-in: 0.67798\ninside 1 149\nTMhelix 150 172\noutside 173 191

29079 GCF_000287935.1_Sten_malt_Ab55555_V1 Stenotrophomonas maltophilia Ab55555 Proteobacteria; Gammaproteobacteria; Xanthomonadales; Xanthomonadaceae; Stenotrophomonas; Stenotrophomonas maltophilia group; Stenotrophomonas maltophilia MDRRELLKMIVAATGAAMVGLPALVHGKVPAAGAMPAFSDADIGMLDEIAETILPRTRTP WP_005408476.1 membrane protein [Stenotrophomonas maltophilia] Length: 191\nNumber of predicted TMHs: 1\nExp number of AAs in TMHs: 22.20941\nExp number, first 60 AAs: 5.35942\nTotal prob of N-in: 0.70371\ninside 1 149\nTMhelix 150 172\noutside 173 191

29080 GCF_000382065.1_ASM38206v1 Stenotrophomonas maltophilia ATCC 19867 Proteobacteria; Gammaproteobacteria; Xanthomonadales; Xanthomonadaceae; Stenotrophomonas; Stenotrophomonas maltophilia group; Stenotrophomonas maltophilia MDRRELLKMIVAATGAAMVGLPALVQGQAPAAGAMPAFTDADIGTLDEIAETILPRTRTP WP_019659369.1 membrane protein [Stenotrophomonas maltophilia] Length: 191\nNumber of predicted TMHs: 1\nExp number of AAs in TMHs: 20.56127\nExp number, first 60 AAs: 4.29189\nTotal prob of N-in: 0.76983\ninside 1 149\nTMhelix 150 172\noutside 173 191

29081 GCF_000346445.1_ASM34644v1 Stenotrophomonas maltophilia AU12-09 Proteobacteria; Gammaproteobacteria; Xanthomonadales; Xanthomonadaceae; Stenotrophomonas; Stenotrophomonas maltophilia group; Stenotrophomonas maltophilia MRPVRAWTRRAALPVLLLALGACAKQGDMVGDAGATAAEAAIASPEGAFLAYEHDVQVQL WP_005415830.1 MULTISPECIES: hypothetical protein [Proteobacteria] Length: 271\nNumber of predicted TMHs: 1\nExp number of AAs in TMHs: 22.9003\nExp number, first 60 AAs: 0.29444\nTotal prob of N-in: 0.06107\noutside 1 233\nTMhelix 234 256\ninside 257 271

29082 GCF_000346445.1_ASM34644v1 Stenotrophomonas maltophilia AU12-09 Proteobacteria; Gammaproteobacteria; Xanthomonadales; Xanthomonadaceae; Stenotrophomonas; Stenotrophomonas maltophilia group; Stenotrophomonas maltophilia MDRRELLKMIVAATGAAMVGLPALVHGKAPAAGMMAAFSDAEIGTLDEIAETILPRTRTP WP_005415638.1 MULTISPECIES: hypothetical protein [Proteobacteria] Length: 191\nNumber of predicted TMHs: 1\nExp number of AAs in TMHs: 26.26404\nExp number, first 60 AAs: 9.53785\nTotal prob of N-in: 0.59430\ninside 1 149\nTMhelix 150 172\noutside 173 191

29083 GCF_000284595.1_ASM28459v1 Stenotrophomonas maltophilia D457 Proteobacteria; Gammaproteobacteria; Xanthomonadales; Xanthomonadaceae; Stenotrophomonas; Stenotrophomonas maltophilia group; Stenotrophomonas maltophilia MDRRELLKMIVAATGAAMVGLPALAQGNAAAAPVKTLFSDADIGTLDEIAETILPRTRTP WP_014646300.1 membrane protein [Stenotrophomonas maltophilia] Length: 191\nNumber of predicted TMHs: 1\nExp number of AAs in TMHs: 22.03151\nExp number, first 60 AAs: 4.64253\nTotal prob of N-in: 0.67798\ninside 1 149\nTMhelix 150 172\noutside 173 191

29084 GCF_000344215.1_EPM1_1.0 Stenotrophomonas maltophilia EPM1 Proteobacteria; Gammaproteobacteria; Xanthomonadales; Xanthomonadaceae; Stenotrophomonas; Stenotrophomonas maltophilia group; Stenotrophomonas maltophilia MDRRELLKMIVAATGAAMVGLPALVHGKVPAAGAMPAFSDADIGMLDEIAETILPRTRTP WP_005412559.1 MULTISPECIES: hypothetical protein [Stenotrophomonas] Length: 191\nNumber of predicted TMHs: 1\nExp number of AAs in TMHs: 22.08369\nExp number, first 60 AAs: 5.28303\nTotal prob of N-in: 0.70356\ninside 1 149\nTMhelix 150 172\noutside 173 191

29085 GCF_000072485.1_ASM7248v1 Stenotrophomonas maltophilia K279a Proteobacteria; Gammaproteobacteria; Xanthomonadales; Xanthomonadaceae; Stenotrophomonas; Stenotrophomonas maltophilia group; Stenotrophomonas maltophilia MDRRELLKMIVAATGAAMVGLPALVHGKVPAAGAMPAFSDADIGMLDEIAETILPRTRTP WP_012479395.1 membrane protein [Stenotrophomonas maltophilia] Length: 191\nNumber of predicted TMHs: 1\nExp number of AAs in TMHs: 22.08345\nExp number, first 60 AAs: 5.28293\nTotal prob of N-in: 0.70356\ninside 1 149\nTMhelix 150 172\noutside 173 191

29086 GCF_000455685.1_ASM45568v1 Stenotrophomonas maltophilia MF89 Proteobacteria; Gammaproteobacteria; Xanthomonadales; Xanthomonadaceae; Stenotrophomonas; Stenotrophomonas maltophilia group; Stenotrophomonas maltophilia MDRRELLKMIVAATGAAMVGLPALVQGQAPAAGAMPAFTDADIGTLDEIAETILPRTRTP WP_021203063.1 membrane protein [Stenotrophomonas maltophilia] Length: 191\nNumber of

predicted TMHs:  1\nExp number of AAs in TMHs: 20.56092\nExp number, first 60 AAs:  4.29176\nTotal prob of N-in: 0.76983\ninside   1   149\nTMhelix   150   172\noutside   173   191

29087 GCF_000597745.1_XMT-16JAN14 Stenotrophomonas maltophilia MTCC 434 Proteobacteria; Gammaproteobacteria; Xanthomonadales; Xanthomonadaceae; Stenotrophomonas; Stenotrophomonas maltophilia group; Stenotrophomonas maltophilia MDRRELLKMIVAATGAAMVGLPALVHGKVPAAGAMPAFSDADIGMLDEIAETILPRTRTP WP_005412559.1 MULTISPECIES: hypothetical protein [Stenotrophomonas] Length: 191\nNumber of predicted TMHs:  1\nExp number of AAs in TMHs: 22.08369\nExp number, first 60 AAs:  5.28303\nTotal prob of N-in: 0.70356\ninside   1   149\nTMhelix   150   172\noutside   173   191

29088 GCF_001591205.1_ASM159120v1 Stenotrophomonas maltophilia NBRC 14161 Proteobacteria; Gammaproteobacteria; Xanthomonadales; Xanthomonadaceae; Stenotrophomonas; Stenotrophomonas maltophilia group; Stenotrophomonas maltophilia MDRRELLKMIVAATGAAMVGLPALVHGKVPAAGAMPAFSDADIGMLDEIAETILPRTRTP WP_005412559.1 MULTISPECIES: hypothetical protein [Stenotrophomonas] Length: 191\nNumber of predicted TMHs:  1\nExp number of AAs in TMHs: 22.08369\nExp number, first 60 AAs:  5.28303\nTotal prob of N-in: 0.70356\ninside   1   149\nTMhelix   150   172\noutside   173   191

29089 GCF_000237025.1_ASM23702v2 Stenotrophomonas maltophilia RR-10 Proteobacteria; Gammaproteobacteria; Xanthomonadales; Xanthomonadaceae; Stenotrophomonas; Stenotrophomonas maltophilia group; Stenotrophomonas maltophilia MDRRELLKMIVAATGAAMVGLPALVQGQAPAGATKIHFSDADVGTLDEIAETILPRTRTP WP_019337066.1 membrane protein [Stenotrophomonas maltophilia] Length: 191\nNumber of predicted TMHs:  1\nExp number of AAs in TMHs: 20.23153\nExp number, first 60 AAs:  3.42135\nTotal prob of N-in: 0.74184\ninside   1   149\nTMhelix   150   172\noutside   173   191

29090 GCF_000499565.1_Stmalt0377 Stenotrophomonas maltophilia stmalt0377 Proteobacteria; Gammaproteobacteria; Xanthomonadales; Xanthomonadaceae; Stenotrophomonas; Stenotrophomonas maltophilia group; Stenotrophomonas maltophilia MDRRELLKMIVAATGAAMVGLPALVHGKVPAAGAMPAFSDADIGMLDEIAETILPRTRTP WP_005412559.1 MULTISPECIES: hypothetical protein [Stenotrophomonas] Length: 191\nNumber of predicted TMHs:  1\nExp number of AAs in TMHs: 22.08369\nExp number, first 60 AAs:  5.28303\nTotal prob of N-in: 0.70356\ninside   1   149\nTMhelix   150   172\noutside   173   191

29091 GCF_000455625.1_Stmalt0435 Stenotrophomonas maltophilia stmalt0435 Proteobacteria; Gammaproteobacteria; Xanthomonadales; Xanthomonadaceae; Stenotrophomonas; Stenotrophomonas maltophilia group; Stenotrophomonas maltophilia MDRRELLKMIVAATGAAMVGLPALVHGKVPAAGAMPAFSDADIGMLDEIAETILPRTRTP WP_005412559.1 MULTISPECIES: hypothetical protein [Stenotrophomonas] Length: 191\nNumber of predicted TMHs:  1\nExp number of AAs in TMHs: 22.08369\nExp number, first 60 AAs:  5.28303\nTotal prob of N-in: 0.70356\ninside   1   149\nTMhelix   150   172\noutside   173   191

29092 GCF_000877275.1_ASM87727v1 Stenotrophomonas maltophilia WJ66 Proteobacteria; Gammaproteobacteria; Xanthomonadales; Xanthomonadaceae; Stenotrophomonas; Stenotrophomonas maltophilia group; Stenotrophomonas maltophilia MTQTGRRTFLLVSTALALASGAAIASAADTAAGQFAEWTGDSTVTRQMTLRQLGFRQPLV WP_043033836.1 membrane protein [Stenotrophomonas maltophilia] Length: 733\nNumber of predicted TMHs:  1\nExp number of AAs in TMHs: 35.41193\nExp number, first 60 AAs:  15.6288\nTotal prob of N-in: 0.74935\nPOSSIBLE N-term signal sequence\noutside   1   700\nTMhelix   701   718\ninside   719   733

29093 GCF_000877275.1_ASM87727v1 Stenotrophomonas maltophilia WJ66 Proteobacteria; Gammaproteobacteria; Xanthomonadales; Xanthomonadaceae; Stenotrophomonas; Stenotrophomonas maltophilia group; Stenotrophomonas maltophilia MDRRELLKMIVAATGAAMVGLPALVHGKVPAAGAMPAFSDADIGMLDEIAETILPRTRTP WP_005412559.1 MULTISPECIES: hypothetical protein [Stenotrophomonas] Length: 191\nNumber of predicted TMHs:  1\nExp number of AAs in TMHs: 22.08369\nExp number, first 60 AAs:  5.28303\nTotal prob of N-in: 0.70356\ninside   1   149\nTMhelix   150   172\noutside   173   191

29094 GCF_000962995.1_Xal-FIJ080-G1 Xanthomonas albilineans Proteobacteria; Gammaproteobacteria; Xanthomonadales; Xanthomonadaceae; Xanthomonas MNRRRRALRCGLLVLLAWSTLAGAAPVSDPTPLAYRSAAEEARFHALTAELRCVQCQNQS WP_045765622.1 cytochrome c biogenesis protein [Xanthomonas albilineans] Length: 151\nNumber of predicted TMHs: 2\nExp number of AAs in TMHs: 42.07513\nExp number, first 60 AAs:  19.85387\nTotal prob of N-in:   0.98732\nPOSSIBLE N-term signal sequence\ninside   1   6\nTMhelix   7   26\noutside   27   110\nTMhelix   111   133\ninside   134   151

29095 GCF_000962925.1_Xal-PNG_130-G1 Xanthomonas albilineans Proteobacteria; Gammaproteobacteria; Xanthomonadales; Xanthomonadaceae; Xanthomonas

MNRRRRALRCGLLVLLAWSTLAGAAPVSDPTPLAYRSAAEEARFHALTAELRCVQCQNQS WP_045765622.1 cytochrome c biogenesis protein [Xanthomonas albilineans] Length: 151\nNumber of predicted TMHs: 2\nExp number of AAs in TMHs: 42.07513\nExp number, first 60 AAs: 19.85387\nTotal prob of N-in: 0.98732\nPOSSIBLE N-term signal sequence\ninside 1 6\nTMhelix 7 26\noutside 27 110\nTMhelix 111 133\ninside 134 151

29096 GCF_000963115.1_Xal-REU174-G1 Xanthomonas albilineans Proteobacteria; Gammaproteobacteria; Xanthomonadales; Xanthomonadaceae; Xanthomonas MNRRRRALRCGLLVLLAWSTVAGAAPVSDPTPLAYRSAAEEARFHALTAELRCVQCQNQS WP_045757957.1 cytochrome c biogenesis protein [Xanthomonas albilineans] Length: 151\nNumber of predicted TMHs: 2\nExp number of AAs in TMHs: 42.08172\nExp number, first 60 AAs: 19.86081\nTotal prob of N-in: 0.98692\nPOSSIBLE N-term signal sequence\ninside 1 6\nTMhelix 7 26\noutside 27 110\nTMhelix 111 133\ninside 134 151

29097 GCF_000963055.1_Xal-HVO005-G1 Xanthomonas albilineans Proteobacteria; Gammaproteobacteria; Xanthomonadales; Xanthomonadaceae; Xanthomonas MNRRRRALRCGLLVLLAWSTVAGAAPVSDPTPLAYRSAAEEARFHALTAELRCVQCQNQS WP_045760628.1 cytochrome c biogenesis protein [Xanthomonas albilineans] Length: 151\nNumber of predicted TMHs: 2\nExp number of AAs in TMHs: 41.97025\nExp number, first 60 AAs: 19.87538\nTotal prob of N-in: 0.98697\nPOSSIBLE N-term signal sequence\ninside 1 6\nTMhelix 7 26\noutside 27 110\nTMhelix 111 133\ninside 134 151

29098 GCF_000963135.1_Xal-REU209-G1 Xanthomonas albilineans Proteobacteria; Gammaproteobacteria; Xanthomonadales; Xanthomonadaceae; Xanthomonas MNRRRRALRCGLLVLLAWSTVAGAAPVSDPTPLAYRSAAEEARFHALTAELRCVQCQNQS WP_045760628.1 cytochrome c biogenesis protein [Xanthomonas albilineans] Length: 151\nNumber of predicted TMHs: 2\nExp number of AAs in TMHs: 41.97025\nExp number, first 60 AAs: 19.87538\nTotal prob of N-in: 0.98697\nPOSSIBLE N-term signal sequence\ninside 1 6\nTMhelix 7 26\noutside 27 110\nTMhelix 111 133\ninside 134 151

29099 GCF_000963065.1_Xal-GAB266-G1 Xanthomonas albilineans Proteobacteria; Gammaproteobacteria; Xanthomonadales; Xanthomonadaceae; Xanthomonas MNRRRRALRCGLLVLLAWSTVAGAAPVSDPTPLAYRSAAEEARFHALTAELRCVQCQNQS WP_045760628.1 cytochrome c biogenesis protein [Xanthomonas albilineans] Length: 151\nNumber of predicted TMHs: 2\nExp number of AAs in TMHs: 41.97025\nExp number, first 60 AAs: 19.87538\nTotal prob of N-in: 0.98697\nPOSSIBLE N-term signal sequence\ninside 1 6\nTMhelix 7 26\noutside 27 110\nTMhelix 111 133\ninside 134 151

29100 GCF_000963155.1_Xal-Xa23R1-G1 Xanthomonas albilineans Proteobacteria; Gammaproteobacteria; Xanthomonadales; Xanthomonadaceae; Xanthomonas MNRRRRALRCGLLVLLAWSTLAGAAPVSDPTPLAYRSAAEEARFHALTAELRCVQCQNQS WP_045765622.1 cytochrome c biogenesis protein [Xanthomonas albilineans] Length: 151\nNumber of predicted TMHs: 2\nExp number of AAs in TMHs: 42.07513\nExp number, first 60 AAs: 19.85387\nTotal prob of N-in: 0.98732\nPOSSIBLE N-term signal sequence\ninside 1 6\nTMhelix 7 26\noutside 27 110\nTMhelix 111 133\ninside 134 151

29101 GCF_000963145.1_Xal-USA048-G1 Xanthomonas albilineans Proteobacteria; Gammaproteobacteria; Xanthomonadales; Xanthomonadaceae; Xanthomonas MNRRRRALRCGLLVLLAWSTLAGAAPVSDPTPLAYRSAAEEARFHALTAELRCVQCQNQS WP_045765622.1 cytochrome c biogenesis protein [Xanthomonas albilineans] Length: 151\nNumber of predicted TMHs: 2\nExp number of AAs in TMHs: 42.07513\nExp number, first 60 AAs: 19.85387\nTotal prob of N-in: 0.98732\nPOSSIBLE N-term signal sequence\ninside 1 6\nTMhelix 7 26\noutside 27 110\nTMhelix 111 133\ninside 134 151

29102 GCF_000962945.1_Xal-LKA070-G1 Xanthomonas albilineans Proteobacteria; Gammaproteobacteria; Xanthomonadales; Xanthomonadaceae; Xanthomonas MNRRRRALRCGLLVLLAWSTVAGAAPVSDPTPLAYRSAAEEARFHALTAELRCVQCQNQS WP_045760628.1 cytochrome c biogenesis protein [Xanthomonas albilineans] Length: 151\nNumber of predicted TMHs: 2\nExp number of AAs in TMHs: 41.97025\nExp number, first 60 AAs: 19.87538\nTotal prob of N-in: 0.98697\nPOSSIBLE N-term signal sequence\ninside 1 6\nTMhelix 7 26\noutside 27 110\nTMhelix 111 133\ninside 134 151

29103 GCF_000962915.1_Xal-HVO_082-G1 Xanthomonas albilineans Proteobacteria; Gammaproteobacteria; Xanthomonadales; Xanthomonadaceae; Xanthomonas MNRRRRALRCGLLVLLAWSTVAGAAPVSDPTPLAYRSAAEEARFHALTAELRCVQCQNQS WP_045760628.1 cytochrome c biogenesis protein [Xanthomonas albilineans] Length: 151\nNumber of predicted TMHs: 2\nExp number of AAs in TMHs: 41.97025\nExp number, first 60 AAs: 19.87538\nTotal prob of N-in: 0.98697\nPOSSIBLE N-term signal sequence\ninside 1 6\nTMhelix 7 26\noutside 27 110\nTMhelix 111 133\ninside 134 151

29104 GCF_001277045.1_XANT002-SEQ-2-ASM-1 Xanthomonas oryzae Proteobacteria; Gammaproteobacteria; Xanthomonadales; Xanthomonadaceae; Xanthomonas
MVIRATRLLHLLDALRGRRQPVAGAQLATALGVSLRTLYRDIATLRAQGADILGDPGVGYWP_029217638.1 transcriptional regulator [Xanthomonas oryzae] Length: 231\nNumber of predicted TMHs: 1\nExp number of AAs in TMHs: 16.88277\nExp number, first 60 AAs: 0.38088\nTotal prob of N-in: 0.59248\ninside 1 167\nTMhelix 168 185\noutside 186 231

29105 GCF_000815185.1_ASM81518v1 Xanthomonas sacchari Proteobacteria; Gammaproteobacteria; Xanthomonadales; Xanthomonadaceae; Xanthomonas
MSQHRRPALRLGLLALLAWSTLAGAEPVSDPTPLAYRSAAEEARFHALTAELRCVQCQNQ WP_052250939.1 cytochrome c biogenesis protein [Xanthomonas sacchari] Length: 152\nNumber of predicted TMHs: 1\nExp number of AAs in TMHs: 33.26602\nExp number, first 60 AAs: 12.58762\nTotal prob of N-in: 0.75411\nPOSSIBLE N-term signal sequence\noutside 1 111\nTMhelix 112 131\ninside 132 152

29106 GCF_000831625.1_Xsac-LMG_476-G1 Xanthomonas sacchari Proteobacteria; Gammaproteobacteria; Xanthomonadales; Xanthomonadaceae; Xanthomonas
MSQHRRPALRLGLLALLAWSTLAGAEPVSDPTPLAYRSAAEEARFHALTAELRCVQCQNQ WP_017907310.1 MULTISPECIES: cytochrome c biogenesis protein [Xanthomonas] Length: 152\nNumber of predicted TMHs: 1\nExp number of AAs in TMHs: 33.5356\nExp number, first 60 AAs: 12.3579\nTotal prob of N-in: 0.75555\nPOSSIBLE N-term signal sequence\noutside 1 111\nTMhelix 112 134\ninside 135 152

29107 GCF_000364685.1_SHU166 Xanthomonas sp. SHU166 Proteobacteria; Gammaproteobacteria; Xanthomonadales; Xanthomonadaceae; Xanthomonas
MSQHRRPALRLGLLALLAWSTLAGAEPVSDPTPLAYRSAAEEARFHALTAELRCVQCQNQ WP_017907310.1 MULTISPECIES: cytochrome c-type biogenesis protein CcmH [Xanthomonas] Length: 152\nNumber of predicted TMHs: 1\nExp number of AAs in TMHs: 33.5356\nExp number, first 60 AAs: 12.3579\nTotal prob of N-in: 0.75555\nPOSSIBLE N-term signal sequence\noutside 1 111\nTMhelix 112 134\ninside 135 152

29108 GCF_000364665.1_SHU199 Xanthomonas sp. SHU199 Proteobacteria; Gammaproteobacteria; Xanthomonadales; Xanthomonadaceae; Xanthomonas
MSQHRRPALRLGLLALLAWSTLAGAEPVSDPTPLAYRSAAEEARFHALTAELRCVQCQNQ WP_017907310.1 MULTISPECIES: cytochrome c-type biogenesis protein CcmH [Xanthomonas] Length: 152\nNumber of predicted TMHs: 1\nExp number of AAs in TMHs: 33.5356\nExp number, first 60 AAs: 12.3579\nTotal prob of N-in: 0.75555\nPOSSIBLE N-term signal sequence\noutside 1 111\nTMhelix 112 134\ninside 135 152

29109 GCF_000364645.1_SHU308 Xanthomonas sp. SHU308 Proteobacteria; Gammaproteobacteria; Xanthomonadales; Xanthomonadaceae; Xanthomonas
MSQHRRPALRLGLLALLAWSTLAGAEPVSDPTPLAYRSAAEEARFHALTAELRCVQCQNQ WP_017907310.1 MULTISPECIES: cytochrome c-type biogenesis protein CcmH [Xanthomonas] Length: 152\nNumber of predicted TMHs: 1\nExp number of AAs in TMHs: 33.5356\nExp number, first 60 AAs: 12.3579\nTotal prob of N-in: 0.75555\nPOSSIBLE N-term signal sequence\noutside 1 111\nTMhelix 112 134\ninside 135 152

29110 GCF_001455815.1_ASM145581v1 Xanthomonas translucens Proteobacteria; Gammaproteobacteria; Xanthomonadales; Xanthomonadaceae; Xanthomonas
MAIRATRLLHLLDALRGRRRPVAGAQLATALGVSLRTLYRDIATLRVQGADILGDPGVGY WP_039007220.1 transcriptional regulator [Xanthomonas translucens] Length: 231\nNumber of predicted TMHs: 1\nExp number of AAs in TMHs: 20.84011\nExp number, first 60 AAs: 0.88569\nTotal prob of N-in: 0.39505\noutside 1 166\nTMhelix 167 186\ninside 187 231

29111 GCF_000212755.1_ASM21275v2 Xanthomonas oryzae X11-5A Proteobacteria; Gammaproteobacteria; Xanthomonadales; Xanthomonadaceae; Xanthomonas; Xanthomonas oryzae
MVIRATRLLHLLDALRGRRQPVAGAQLATALGVSLRTLYRDIATLRAQGADILGDPGVGYWP_029217638.1 transcriptional regulator [Xanthomonas oryzae] Length: 231\nNumber of predicted TMHs: 1\nExp number of AAs in TMHs: 16.88277\nExp number, first 60 AAs: 0.38088\nTotal prob of N-in: 0.59248\ninside 1 167\nTMhelix 168 185\noutside 186 231

29112 GCF_000212775.1_ASM21277v2 Xanthomonas oryzae X8-1A Proteobacteria; Gammaproteobacteria; Xanthomonadales; Xanthomonadaceae; Xanthomonas; Xanthomonas oryzae
MVIRATRLLHLLDALRGRRRPVAGAQLATALGVSLRTLYRDIATLRAQGADILGDPGVGY WP_029561485.1 DNA-binding protein [Xanthomonas oryzae] Length: 231\nNumber of predicted TMHs: 1\nExp number of AAs in TMHs:

16.98802\nExp number, first 60 AAs: 0.53234\nTotal prob of N-in: 0.65795\ninside 1 167\nTMhelix 168 185\noutside 186 231

29113 GCF_000225975.1_ASM22597v2 Xanthomonas sacchari NCPPB 4393 Proteobacteria; Gammaproteobacteria; Xanthomonadales; Xanthomonadaceae; Xanthomonas; Xanthomonas sacchari MSQPRRPALRLGLLALLAWSTLAGAEPVSDPTPLAYRSAAEEARFHALTAELRCVQCQNQ WP_050946593.1 cytochrome c-type biogenesis protein CcmH [Xanthomonas sacchari] Length: 152\nNumber of predicted TMHs: 1\nExp number of AAs in TMHs: 31.60173\nExp number, first 60 AAs: 10.43971\nTotal prob of N-in: 0.63814\nPOSSIBLE N-term signal sequence\noutside 1 111\nTMhelix 112 134\ninside 135 152

29114 GCF_000807145.1_Xtc-CFBP2541-G1 Xanthomonas translucens pv. cerealis Proteobacteria; Gammaproteobacteria; Xanthomonadales; Xanthomonadaceae; Xanthomonas; Xanthomonas translucens MAIRATRLLHLLDALRGRRRPVAGAQLATALGVSLRTLYRDIATLRVQGADILGDPGVGY WP_039007220.1 transcriptional regulator [Xanthomonas translucens] Length: 231\nNumber of predicted TMHs: 1\nExp number of AAs in TMHs: 20.84011\nExp number, first 60 AAs: 0.88569\nTotal prob of N-in: 0.39505\noutside 1 166\nTMhelix 167 186\ninside 187 231

29115 GCF_001707115.1_XTC_AA Xanthomonas translucens pv. cerealis Proteobacteria; Gammaproteobacteria; Xanthomonadales; Xanthomonadaceae; Xanthomonas; Xanthomonas translucens MAIRATRLLHLLDALRGRRRPVAGAQLATALGVSLRTLYRDIATLRVQGADILGDPGVGY WP_039007220.1 transcriptional regulator [Xanthomonas translucens] Length: 231\nNumber of predicted TMHs: 1\nExp number of AAs in TMHs: 20.84011\nExp number, first 60 AAs: 0.88569\nTotal prob of N-in: 0.39505\noutside 1 166\nTMhelix 167 186\ninside 187 231

29116 GCF_001707285.1_XTG_AA Xanthomonas translucens pv. poae Proteobacteria; Gammaproteobacteria; Xanthomonadales; Xanthomonadaceae; Xanthomonas; Xanthomonas translucens MAIRATRLLHLLDALRGRRRPVAGAQLATALGVSLRTLYRDIATLRAQGADILGDPGVGY WP_064538702.1 transcriptional regulator [Xanthomonas translucens] Length: 231\nNumber of predicted TMHs: 1\nExp number of AAs in TMHs: 21.13389\nExp number, first 60 AAs: 0.91977\nTotal prob of N-in: 0.41860\noutside 1 166\nTMhelix 167 186\ninside 187 231

29117 GCF_001707435.1_XTP_AA Xanthomonas translucens pv. poae Proteobacteria; Gammaproteobacteria; Xanthomonadales; Xanthomonadaceae; Xanthomonas; Xanthomonas translucens MAIRATRLLHLLDALRGRRRPVAGAQLATALGVSLRTLYRDIATLRAQGADILGDPGVGY WP_064538702.1 transcriptional regulator [Xanthomonas translucens] Length: 231\nNumber of predicted TMHs: 1\nExp number of AAs in TMHs: 21.13389\nExp number, first 60 AAs: 0.91977\nTotal prob of N-in: 0.41860\noutside 1 166\nTMhelix 167 186\ninside 187 231

29118 GCF_001659965.1_ASM165996v1 Xanthomonas translucens pv. poae Proteobacteria; Gammaproteobacteria; Xanthomonadales; Xanthomonadaceae; Xanthomonas; Xanthomonas translucens MAIRATRLLHLLDALRGRRRPVAGAQLATALGVSLRTLYRDIATLRAQGADILGDPGVGY WP_064538702.1 transcriptional regulator [Xanthomonas translucens] Length: 231\nNumber of predicted TMHs: 1\nExp number of AAs in TMHs: 21.13389\nExp number, first 60 AAs: 0.91977\nTotal prob of N-in: 0.41860\noutside 1 166\nTMhelix 167 186\ninside 187 231

29119 GCF_000284115.1_ASM28411v1 Phycisphaera mikurensis NBRC 102666 PVC group; Planctomycetes; Phycisphaerae; Phycisphaerales; Phycisphaeraceae; Phycisphaera; Phycisphaera mikurensis MPVTPSSPRSAPSAARRTAVLLAASCASFGGAGSAFADEVTVTGATLFEEFFKVAANASD WP_014438486.1 PEP-CTERM sorting domain-containing protein [Phycisphaera mikurensis] Length: 1060\nNumber of predicted TMHs: 1\nExp number of AAs in TMHs: 18.5353\nExp number, first 60 AAs: 2.04571\nTotal prob of N-in: 0.09377\noutside 1 1031\nTMhelix 1032 1054\ninside 1055 1060

29120 GCF_000255705.1_ASM25570v1 Zavarzinella formosa DSM 19928 PVC group; Planctomycetes; Planctomycetia; Planctomycetales; Gemmataceae; Zavarzinella; Zavarzinella formosa MSPVNLPKRTLARAIARRAVLLSLANVGMGILATAGVAAAFQAPVLLLPVGAYFLLPGKR WP_020469605.1 hypothetical protein [Zavarzinella formosa] Length: 576\nNumber of predicted TMHs: 2\nExp number of AAs in TMHs: 42.24883\nExp number, first 60 AAs: 24.0221\nTotal prob of N-in: 0.98577\nPOSSIBLE N-term signal sequence\ninside 1 19\nTMhelix 20 42\noutside 43 557\nTMhelix 558 575\ninside 576 576

29121 GCF_900129635.1_IMG-taxon_2695420965_annotated_assembly Singulisphaera sp. GP187 PVC group; Planctomycetes; Planctomycetia; Planctomycetales; Isosphaeraceae; Singulisphaera

MPIVLLAAWASFAGYGLWIASIKRRSPLEGMLLGLLLGPVGCLVEASRRERSAEEVEEEQ WP_074307161.1 hypothetical protein [Singulisphaera sp. GP187] Length: 154\nNumber of predicted TMHs: 2\nExp number of AAs in TMHs: 40.37635\nExp number, first 60 AAs: 18.75332\nTotal prob of N-in: 0.73059\nPOSSIBLE N-term signal sequence\ninside 1 4\nTMhelix 5 22\noutside 23 129\nTMhelix 130 152\ninside 153 154

29122 GCF_900129635.1_IMG-taxon_2695420965_annotated_assembly Singulisphaera sp. GP187 PVC group; Planctomycetes; Planctomycetia; Planctomycetales; Isosphaeraceae; Singulisphaera MQVRFPVILAASEPRRLRGRRAALGPRAGVILLLLGVLPSPPHGPAGLAFASIPSPREEG WP_074311720.1 bacteriocin/lantibiotic ABC transporter [Singulisphaera sp. GP187] Length: 276\nNumber of predicted TMHs: 1\nExp number of AAs in TMHs: 25.75046\nExp number, first 60 AAs: 3.06048\nTotal prob of N-in: 0.15294\noutside 1 239\nTMhelix 240 262\ninside 263 276

29123 GCF_900129635.1_IMG-taxon_2695420965_annotated_assembly Singulisphaera sp. GP187 PVC group; Planctomycetes; Planctomycetia; Planctomycetales; Isosphaeraceae; Singulisphaera MDDVPAGPAYPEPRRPPLSVVIPVHNGGLDFERCLRRLRESDQADFELIVVDDGSADNSA WP_074307323.1 glycosyl transferase [Singulisphaera sp. GP187] Length: 391\nNumber of predicted TMHs: 2\nExp number of AAs in TMHs: 47.23213\nExp number, first 60 AAs: 0.00216\nTotal prob of N-in: 0.10343\noutside 1 276\nTMhelix 277 299\ninside 300 319\nTMhelix 320 342\noutside 343 391

29124 GCF_001707835.1_ASM170783v1 Planctopirus sp. JC280 PVC group; Planctomycetes; Planctomycetia; Planctomycetales; Planctomycetaceae; Planctopirus MSSSPPSLAAASIPPNSRKGEIRRRWFRRRTAVGLPPGTITPHEQRPARMELISYSPGDY WP_068853454.1 magnesium and cobalt transport protein CorA [Planctopirus sp. JC280] Length: 381\nNumber of predicted TMHs: 2\nExp number of AAs in TMHs: 43.9639\nExp number, first 60 AAs: 0.00212\nTotal prob of N-in: 0.66696\ninside 1 310\nTMhelix 311 333\noutside 334 342\nTMhelix 343 365\ninside 366 381

29125 GCF_000092105.1_ASM9210v1 Planctopirus limnophila DSM 3776 PVC group; Planctomycetes; Planctomycetia; Planctomycetales; Planctomycetaceae; Planctopirus; Planctopirus limnophila MSSSSIDVAAASIPANSRKGEIRRRWFRRRTAVGLQPGTITPHEQRPARMELISYSLSDY WP_013112430.1 magnesium and cobalt transport protein CorA [Planctopirus limnophila] Length: 382\nNumber of predicted TMHs: 2\nExp number of AAs in TMHs: 43.86001\nExp number, first 60 AAs: 0.00617\nTotal prob of N-in: 0.90991\ninside 1 310\nTMhelix 311 333\noutside 334 342\nTMhelix 343 365\ninside 366 382

29126 GCF_000165715.2_ASM16571v3 Rubinisphaera brasiliensis DSM 5305 PVC group; Planctomycetes; Planctomycetia; Planctomycetales; Planctomycetaceae; Rubinisphaera; Rubinisphaera brasiliensis MKNVSIFRAARPRRGLISSAMVCLLFSGCMGQKHSLPSFSWKQSSETSEKVEAAAEKTTP WP_013628474.1 hypothetical protein [Rubinisphaera brasiliensis] Length: 548\nNumber of predicted TMHs: 1\nExp number of AAs in TMHs: 19.87908\nExp number, first 60 AAs: 1.2282\nTotal prob of N-in: 0.07181\noutside 1 528\nTMhelix 529 546\ninside 547 548

29127 GCF_000255655.1_ASM25565v1 Schlesneria paludicola DSM 18645 PVC group; Planctomycetes; Planctomycetia; Planctomycetales; Planctomycetaceae; Schlesneria; Schlesneria paludicola MPSVSHGRRNVLGLVTLVLACLCVAGWVSSIRQEVCLYLTQKDRTPFMVVSRDHYLCVVV WP_010585528.1 hypothetical protein [Schlesneria paludicola] Length: 215\nNumber of predicted TMHs: 2\nExp number of AAs in TMHs: 43.01919\nExp number, first 60 AAs: 20.87203\nTotal prob of N-in: 0.10348\nPOSSIBLE N-term signal sequence\noutside 1 9\nTMhelix 10 32\ninside 33 176\nTMhelix 177 199\noutside 200 215

29128 GCF_000019665.1_ASM1966v1 Methylacidiphilum infernorum V4 PVC group; Verrucomicrobia; Methylacidiphilae; Methylacidiphilales; Methylacidiphilaceae; Methylacidiphilum; Methylacidiphilum infernorum MEQDRQTFYSLLQQQGISRRSFIKFCTLSAFSLGLGVEVIPEIVHAFETKPRIPVLWLHG WP_048810203.1 uptake hydrogenase small subunit [Methylacidiphilum infernorum] Length: 382\nNumber of predicted TMHs: 1\nExp number of AAs in TMHs: 26.8925399999999\nExp number, first 60 AAs: 9.28213999999999\nTotal prob of N-in: 0.47787\noutside 1 327\nTMhelix 328 350\ninside 351 382

29129 GCF_000171235.2_ASM17123v2 Didymococcus colitermitum TAV2 PVC group; Verrucomicrobia; Opitutae; Opitutales; Opitutaceae; Didymococcus; Didymococcus colitermitum MSVRSPSRRQVIATNSTCCTSTGMWKRSNRPPPPPRLHTYTNLRSLYFTLAGSFPCRPPS WP_081722086.1 hypothetical protein [Didymococcus colitermitum] Length: 382\nNumber of predicted TMHs: 2\nExp number of AAs in TMHs: 55.64986\nExp number, first 60 AAs: 0.00199\nTotal prob of N-in: 0.95798\ninside 1 89\nTMhelix 90 112\noutside 113 354\nTMhelix 355 377\ninside 378 382

29130 GCF_000019965.1_ASM1996v1 Opitutus terrae PB90-1 PVC group; Verrucomicrobia; Opitutae; Opitutales; Opitutaceae; Opitutus; Opitutus terrae
MTLPRSRACLSAVNVSQPHTTLLRRLTNPFSARQRRGVLACGVRLSVLGFAFAALSLPAA WP_083767733.1 hypothetical protein [Opitutus terrae] Length: 1919\nNumber of predicted TMHs: 2\nExp number of AAs in TMHs: 42.6056999999998\nExp number, first 60 AAs: 15.26192\nTotal prob of N-in: 0.78714\nPOSSIBLE N-term signal sequence\ninside 1 41\nTMhelix 42 64\noutside 65 1891\nTMhelix 1892 1909\ninside 1910 1919

29131 GCF_000243495.2_ASM24349v3 Opitutaceae bacterium TAV1 PVC group; Verrucomicrobia; Opitutae; Opitutales; Opitutaceae; unclassified Opitutaceae
MHTNKSILAGSGSRRRLLLALAATALAASSASAQTWIMGDGATLTLLSSGSLTSISPVAI WP_007362680.1 HAF repeat/PEP-CTERM domain-containing protein [Opitutaceae bacterium TAV1] Length: 409\nNumber of predicted TMHs: 2\nExp number of AAs in TMHs: 37.4584599999999\nExp number, first 60 AAs: 17.87411\nTotal prob of N-in: 0.80809\nPOSSIBLE N-term signal sequence\ninside 1 16\nTMhelix 17 39\noutside 40 376\nTMhelix 377 399\ninside 400 409

29132 GCF_000242935.2_ASM24293v3 Opitutaceae bacterium TAV5 PVC group; Verrucomicrobia; Opitutae; Opitutales; Opitutaceae; unclassified Opitutaceae
MTLLPAAARANRPDHRLFPRRRPCLLARGAGLATLLAGFPFAMARAADAPRYTYTVLATP WP_009513422.1 HAF repeat-containing protein [Opitutaceae bacterium TAV5] Length: 596\nNumber of predicted TMHs: 1\nExp number of AAs in TMHs: 35.10324\nExp number, first 60 AAs: 15.306\nTotal prob of N-in: 0.72515\nPOSSIBLE N-term signal sequence\noutside 1 562\nTMhelix 563 585\ninside 586 596

29133 GCF_000173075.1_ASM17307v1 Chthoniobacter flavus Ellin428 PVC group; Verrucomicrobia; Spartobacteria; Chthoniobacterales; Chthoniobacteraceae; Chthoniobacter; Chthoniobacter flavus
MPSRRPPLLRACALLLLSLTATFFAGCGNDPNLQPEHEKRADGSPWQVRYSYMPDDPRSL WP_006980818.1 ABC transporter substrate-binding protein [Chthoniobacter flavus] Length: 662\nNumber of predicted TMHs: 1\nExp number of AAs in TMHs: 23.81619\nExp number, first 60 AAs: 5.09376\nTotal prob of N-in: 0.25577\noutside 1 636\nTMhelix 637 656\ninside 657 662

29134 GCF_001683795.1_ASM168379v1 Akkermansia glycaniphila PVC group; Verrucomicrobia; Verrucomicrobiae; Verrucomicrobiales; Akkermansiaceae; Akkermansia
MKPSFSRLARRTAIAALLIFAVQASAEITTRENSVFSINFYNNGDTFQNQISSTMDWSET WP_067774360.1 PEP-CTERM sorting domain-containing protein [Akkermansia glycaniphila] Length: 322\nNumber of predicted TMHs: 1\nExp number of AAs in TMHs: 19.22085\nExp number, first 60 AAs: 2.34144\nTotal prob of N-in: 0.14111\noutside 1 297\nTMhelix 298 317\ninside 318 322

29135 GCF_900097105.1_WK001 Akkermansia glycaniphila PVC group; Verrucomicrobia; Verrucomicrobiae; Verrucomicrobiales; Akkermansiaceae; Akkermansia
MKPSFSRLARRTAIAALLIFAVQASAEITTRENSVFSINFYNNGDTFQNQISSTMDWSET WP_067774360.1 PEP-CTERM sorting domain-containing protein [Akkermansia glycaniphila] Length: 322\nNumber of predicted TMHs: 1\nExp number of AAs in TMHs: 19.22085\nExp number, first 60 AAs: 2.34144\nTotal prob of N-in: 0.14111\noutside 1 297\nTMhelix 298 317\ninside 318 322

29136 GCF_000304275.1_gls454017v1.0 Leptospira meyeri serovar Hardjo str. Went 5 Spirochaetes; Spirochaetia; Leptospirales; Leptospiraceae; Leptospira; Leptospira meyeri; Leptospira meyeri serovar Hardjo
MNIHSLFYEDEYGSVMAISRKNLISRRTVVKTSAFGAVLLFLSTWNCKPSPGSVITGRGI WP_004786587.1 hypothetical protein [Leptospira meyeri] Length: 203\nNumber of predicted TMHs: 1\nExp number of AAs in TMHs: 20.01486\nExp number, first 60 AAs: 1.95522\nTotal prob of N-in: 0.73279\ninside 1 160\nTMhelix 161 180\noutside 181 203

29137 GCF_000347075.1_gls454196v01 Leptospira meyeri serovar Semaranga str. Veldrot Semarang 173 Spirochaetes; Spirochaetia; Leptospirales; Leptospiraceae; Leptospira; Leptospira meyeri; Leptospira meyeri serovar Semaranga MNIHSLFYEDEYGSVMAISRKNLISRRTVVKTSAFGAVLLFLSTWNCKPSSGSVITGRGI WP_035983057.1 hypothetical protein [Leptospira meyeri] Length: 203\nNumber of predicted TMHs: 1\nExp number of AAs in TMHs: 20.81871\nExp number, first 60 AAs: 2.65065\nTotal prob of N-in: 0.70650\ninside 1 160\nTMhelix 161 180\noutside 181 203

29138 GCF_000332455.1_gls454199v02 Leptospira vanthielii serovar Holland str. Waz Holland = ATCC 700522 Spirochaetes; Spirochaetia; Leptospirales; Leptospiraceae; Leptospira; Leptospira vanthielii; Leptospira vanthielii serovar Holland MAISRKNLISRRTVVKTSAFGAVFLFLSTWNCKPSTTSVIAGRGIDCKFLDPEAQEFFSH WP_002985403.1 hypothetical protein [Leptospira vanthielii] Length: 188\nNumber of predicted TMHs: 1\nExp number of

AAs in TMHs: 21.78985\nExp number, first 60 AAs: 1.50152\nTotal prob of N-in: 0.93929\ninside 1 145\nTMhelix 146 165\noutside 166 188

29139 GCF_000332515.2_gls454195v02 Leptospira wolbachii serovar Codice str. CDC Spirochaetes; Spirochaetia; Leptospirales; Leptospiraceae; Leptospira; Leptospira wolbachii; Leptospira wolbachii serovar Codice MAISRKNLISRRTVVKTSAFGAVFLFLSTWKCKPTTSVIVGRGLSCKFLDPEAQEFFSHL WP_015681490.1 hypothetical protein [Leptospira wolbachii] Length: 187\nNumber of predicted TMHs: 1\nExp number of AAs in TMHs: 21.6434\nExp number, first 60 AAs: 1.3902\nTotal prob of N-in: 0.93485\ninside 1 144\nTMhelix 145 164\noutside 165 187

29140 GCF_000507245.1_ASM50724v1 Salinispira pacifica Spirochaetes; Spirochaetia; Spirochaetales; Spirochaetaceae; Salinispira MDTQPVNNPQPDEISLVDIIVVFIKRRRLLVASASLAIFLSILVLYIFPLFNVYLYDPGT WP_024269372.1 hypothetical protein [Salinispira pacifica] Length: 293\nNumber of predicted TMHs: 2\nExp number of AAs in TMHs: 44.53888\nExp number, first 60 AAs: 22.64385\nTotal prob of N-in: 0.35903\nPOSSIBLE N-term signal sequence\noutside 1 33\nTMhelix 34 56\ninside 57 247\nTMhelix 248 270\noutside 271 293

29141 GCF_000165795.1_ASM16579v1 Aminomonas paucivorans DSM 12260 Synergistetes; Synergistia; Synergistales; Synergistaceae; Aminomonas; Aminomonas paucivorans MTMRRSTRRAWVAFAGGLALWGTAFLTGTLPVSATPRSTTYPYPINSPADLFAAGSGLAQ WP_040345071.1 hypothetical protein [Aminomonas paucivorans] Length: 567\nNumber of predicted TMHs: 2\nExp number of AAs in TMHs: 40.2564799999999\nExp number, first 60 AAs: 22.27557\nTotal prob of N-in: 0.99334\nPOSSIBLE N-term signal sequence\ninside 1 11\nTMhelix 12 34\noutside 35 540\nTMhelix 541 563\ninside 564 567

29142 GCF_002159945.1_ASM215994v1 Cloacibacillus sp. An23 Synergistetes; Synergistia; Synergistales; Synergistaceae; Cloacibacillus MGNDNVPRDGMYAWNDDYSEIDLVDILSSLWKRRKLIAAVTAACLVCAAGYLALSPRVYE WP_087364264.1 hypothetical protein [Cloacibacillus sp. An23] Length: 437\nNumber of predicted TMHs: 2\nExp number of AAs in TMHs: 41.86002\nExp number, first 60 AAs: 20.19452\nTotal prob of N-in: 0.60690\nPOSSIBLE N-term signal sequence\ninside 1 35\nTMhelix 36 58\noutside 59 403\nTMhelix 404 426\ninside 427 437

29143 GCF_000299415.1_ASM29941v1 Candidatus Microthrix parvicella Bio17-1 Terrabacteria group; Actinobacteria; Acidimicrobiia; Acidimicrobiales; Microthrixaceae; Candidatus Microthrix; Candidatus Microthrix parvicella MSYWYVQFEFRLGGVMGSATSRPVGRGVALSRRGCVGVVAAVAVLALGLVAVVGVASPAS WP_081582096.1 hypothetical protein [Candidatus Microthrix parvicella] Length: 956\nNumber of predicted TMHs: 2\nExp number of AAs in TMHs: 44.7976199999999\nExp number, first 60 AAs: 22.33231\nTotal prob of N-in: 0.97994\nPOSSIBLE N-term signal sequence\ninside 1 33\nTMhelix 34 56\noutside 57 923\nTMhelix 924 946\ninside 947 956

29144 GCF_000455525.1_M.parvicella.RN1 Candidatus Microthrix parvicella RN1 Terrabacteria group; Actinobacteria; Acidimicrobiia; Acidimicrobiales; Microthrixaceae; Candidatus Microthrix; Candidatus Microthrix parvicella MGSATSRPVGRGVALSRRGCVGVVAAVAVLALGLVAVVGVASPASAAAGSTGWSQVSAGD WP_012224248.1 exported hypothetical protein [Candidatus Microthrix parvicella] Length: 941\nNumber of predicted TMHs: 2\nExp number of AAs in TMHs: 44.9075299999999\nExp number, first 60 AAs: 22.38448\nTotal prob of N-in: 0.99843\nPOSSIBLE N-term signal sequence\ninside 1 19\nTMhelix 20 42\noutside 43 908\nTMhelix 909 931\ninside 932 941

29145 GCF_000466165.1_ASM46616v1 Actinobaculum sp. oral taxon 183 str. F0552 Terrabacteria group; Actinobacteria; Actinobacteria; Actinomycetales; Actinomycetaceae; Actinobaculum; Actinobaculum sp. oral taxon 183 MMMWLWPRWAVAALLVGGLGACFALWRTTRGDNSKAWVDWTRRALMIVVVASMGMTPSVEWP_021602770.1 VWA domain-containing protein [Actinobaculum sp. oral taxon 183] Length: 326\nNumber of predicted TMHs: 2\nExp number of AAs in TMHs: 45.3627\nExp number, first 60 AAs: 24.51126\nTotal prob of N-in: 0.08129\nPOSSIBLE N-term signal sequence\noutside 1 3\nTMhelix 4 26\ninside 27 293\nTMhelix 294 316\noutside 317 326

29146 GCF_900141725.1_IMG-taxon_2651870314_annotated_assembly Actinomyces denticolens Terrabacteria group; Actinobacteria; Actinobacteria; Actinomycetales; Actinomycetaceae; Actinomyces MRQPPQRRRTLLTALAIALLALTGLVAPLRPAPALALLDPGYNPIHLTLLGDSYSAGNGA WP_073453776.1 hypothetical protein [Actinomyces denticolens] Length: 877\nNumber of predicted TMHs: 2\nExp number of AAs in TMHs: 40.74275\nExp number, first 60 AAs: 20.00504\nTotal prob of N-in: 0.98549\nPOSSIBLE N-term signal sequence\ninside 1 11\nTMhelix 12 29\noutside 30 846\nTMhelix 847 869\ninside 870 877

29147 GCF_001746855.1_ASM174685v1 Actinomyces hongkongensis Terrabacteria group; Actinobacteria; Actinobacteria; Actinomycetales; Actinomycetaceae; Actinomyces

MPATRSRRRALARLVPALLAALLACSLAPLRADAADIRRVGDILSAIGGTVYIHDNPDWS WP_009743428.1 MULTISPECIES: pilus assembly protein [Actinomyces] Length: 769\nNumber of predicted TMHs: 1\nExp number of AAs in TMHs: 37.02313\nExp number, first 60 AAs: 14.86808\nTotal prob of N-in: 0.72647\nPOSSIBLE N-term signal sequence\noutside 1 736\nTMhelix 737 759\ninside 760 769

29148 GCF_001746855.1_ASM174685v1 Actinomyces hongkongensis Terrabacteria group; Actinobacteria; Actinobacteria; Actinomycetales; Actinomycetaceae; Actinomyces MHTTTRRAVVAGCAVFALFAAPMAALAEPAGPGAPQSGSAQSGAADAPQSEPAAAQGAQG WP_009743508.1 MULTISPECIES: bifunctional metallophosphatase/5-nucleotidase [Actinomyces] Length: 765\nNumber of predicted TMHs: 2\nExp number of AAs in TMHs: 41.53601\nExp number, first 60 AAs: 19.57571\nTotal prob of N-in: 0.96746\nPOSSIBLE N-term signal sequence\ninside 1 6\nTMhelix 7 26\noutside 27 736\nTMhelix 737 759\ninside 760 765

29149 GCF_001907235.1_ASM190723v1 Actinomyces hordeovulneris Terrabacteria group; Actinobacteria; Actinobacteria; Actinomycetales; Actinomycetaceae; Actinomyces MTRVWQRRAVLGLLVGALSLNGLAALAAPTPTPSPEIPSAVAGTARSSGEPLVIVFDISG WP_084542572.1 hypothetical protein [Actinomyces hordeovulneris] Length: 817\nNumber of predicted TMHs: 2\nExp number of AAs in TMHs: 40.8464399999999\nExp number, first 60 AAs: 19.18728\nTotal prob of N-in: 0.90958\nPOSSIBLE N-term signal sequence\ninside 1 8\nTMhelix 9 28\noutside 29 783\nTMhelix 784 806\ninside 807 817

29150 GCF_001907235.1_ASM190723v1 Actinomyces hordeovulneris Terrabacteria group; Actinobacteria; Actinobacteria; Actinomycetales; Actinomycetaceae; Actinomyces MKPSARRTILASTLSVLLVAGGLTTGTAALAVENPDAPPTVSSEASESETAQPGSGDESE WP_084542603.1 hypothetical protein [Actinomyces hordeovulneris] Length: 1337\nNumber of predicted TMHs: 2\nExp number of AAs in TMHs: 37.53764\nExp number, first 60 AAs: 16.68719\nTotal prob of N-in: 0.76241\nPOSSIBLE N-term signal sequence\ninside 1 8\nTMhelix 9 31\noutside 32 1307\nTMhelix 1308 1330\ninside 1331 1337

29151 GCF_001457875.1_Actinomyces_ihumii Actinomyces ihumiiTerrabacteria group; Actinobacteria; Actinobacteria; Actinomycetales; Actinomycetaceae; Actinomyces MRESSSPPGRCLFTRRGAIIAACACAGALCLAGTACAADPAAPEAEKPYAHPSAARVVAQ WP_058236945.1 hypothetical protein [Actinomyces ihumii] Length: 637\nNumber of predicted TMHs: 1\nExp number of AAs in TMHs: 30.53116\nExp number, first 60 AAs: 8.46308\nTotal prob of N-in: 0.37869\noutside 1 608\nTMhelix 609 631\ninside 632 637

29152 GCF_001457875.1_Actinomyces_ihumii Actinomyces ihumiiTerrabacteria group; Actinobacteria; Actinobacteria; Actinomycetales; Actinomycetaceae; Actinomyces MKRSQPRSRSAAFFDLDKTILATSTSVALRAPLREAGLLTRRGALLGLLIQLPYLIKGAD WP_058237475.1 hypothetical protein [Actinomyces ihumii] Length: 267\nNumber of predicted TMHs: 1\nExp number of AAs in TMHs: 19.7489\nExp number, first 60 AAs: 0.10076\nTotal prob of N-in: 0.67289\ninside 1 238\nTMhelix 239 261\noutside 262 267

29153 GCF_001262055.1_ASM126205v1 Actinomyces meyeri Terrabacteria group; Actinobacteria; Actinobacteria; Actinomycetales; Actinomycetaceae; Actinomyces MVRAAFFDLDKTILDTSSNVALSGPFIEAGLMNRRTAVASVLVQLPYLLAGADESRMEQM WP_082175956.1 hydrolase [Actinomyces meyeri] Length: 279\nNumber of predicted TMHs: 1\nExp number of AAs in TMHs: 16.02729\nExp number, first 60 AAs: 0.02801\nTotal prob of N-in: 0.01235\noutside 1 235\nTMhelix 236 254\ninside 255 279

29154 GCF_900105015.1_IMG-taxon_2634166153_annotated_assembly Actinomyces meyeri Terrabacteria group; Actinobacteria; Actinobacteria; Actinomycetales; Actinomycetaceae; Actinomyces MVRAAFFDLDKTILDTSSNVALSGPFIEAGLMNRRTAVASVLVQLPYLLAGADESRMEQM WP_070778599.1 MULTISPECIES: hydrolase [Actinomyces] Length: 279\nNumber of predicted TMHs: 1\nExp number of AAs in TMHs: 16.02741\nExp number, first 60 AAs: 0.02813\nTotal prob of N-in: 0.01236\noutside 1 235\nTMhelix 236 254\ninside 255 279

29155 GCF_001956415.1_ASM195641v1 Actinomyces naeslundii Terrabacteria group; Actinobacteria; Actinobacteria; Actinomycetales; Actinomycetaceae; Actinomyces MLRRRTVLSRLRKLVAGAAVLTCAAAGLGGIERPSAVAAPGGLPADALPAVFAQGGSSRY WP_076252647.1 fimbrial protein [Actinomyces naeslundii] Length: 953\nNumber of predicted TMHs: 1\nExp number of AAs in TMHs: 29.48508\nExp number, first 60 AAs: 7.96407\nTotal prob of N-in: 0.40402\noutside 1 922\nTMhelix 923 945\ninside 946 953

29156 GCF_001937475.1_ASM193747v1 Actinomyces naeslundii Terrabacteria group; Actinobacteria; Actinobacteria; Actinomycetales; Actinomycetaceae; Actinomyces
MLRRRTVLSRLRKLVAGAAVLTCAAAALGGIERPSAVAAPGGLPADALPAVFAQGGTSRY WP_075413273.1
fimbrial protein [Actinomyces naeslundii] Length: 953\nNumber of predicted TMHs: 1\nExp number of AAs in TMHs: 31.60165\nExp number, first 60 AAs: 10.09274\nTotal prob of N-in: 0.51245\nPOSSIBLE N-term signal sequence\noutside 1 922\nTMhelix 923 945\ninside 946 953

29157 GCF_001956355.1_ASM195635v1 Actinomyces naeslundii Terrabacteria group; Actinobacteria; Actinobacteria; Actinomycetales; Actinomycetaceae; Actinomyces
MLRRRTVLSRLRKLVAGAAVLTCAAAALGGIERPSAVAAPGGLPADALPAVFAQGGTSRY WP_075413273.1
fimbrial protein [Actinomyces naeslundii] Length: 953\nNumber of predicted TMHs: 1\nExp number of AAs in TMHs: 31.60165\nExp number, first 60 AAs: 10.09274\nTotal prob of N-in: 0.51245\nPOSSIBLE N-term signal sequence\noutside 1 922\nTMhelix 923 945\ninside 946 953

29158 GCF_001937615.1_ASM193761v1 Actinomyces naeslundii Terrabacteria group; Actinobacteria; Actinobacteria; Actinomycetales; Actinomycetaceae; Actinomyces
MLRRRTVLSRLRKLVAGAAVLTCAAAALGGIERPSAVAAPGGLPADALPAVFAQGGTSRY WP_075416240.1
fimbrial protein [Actinomyces naeslundii] Length: 953\nNumber of predicted TMHs: 1\nExp number of AAs in TMHs: 31.6123\nExp number, first 60 AAs: 10.09274\nTotal prob of N-in: 0.51245\nPOSSIBLE N-term signal sequence\noutside 1 922\nTMhelix 923 945\ninside 946 953

29159 GCF_001937605.1_ASM193760v1 Actinomyces naeslundii Terrabacteria group; Actinobacteria; Actinobacteria; Actinomycetales; Actinomycetaceae; Actinomyces
MLRRRTVLSRLRKLVAGAAVLTCAAAALGGIERPSAVAAPGGLPADALPAVFAQGGTSRY WP_075408720.1
fimbrial protein [Actinomyces naeslundii] Length: 953\nNumber of predicted TMHs: 1\nExp number of AAs in TMHs: 31.60165\nExp number, first 60 AAs: 10.09274\nTotal prob of N-in: 0.51245\nPOSSIBLE N-term signal sequence\noutside 1 922\nTMhelix 923 945\ninside 946 953

29160 GCF_001937735.1_ASM193773v1 Actinomyces naeslundii Terrabacteria group; Actinobacteria; Actinobacteria; Actinomycetales; Actinomycetaceae; Actinomyces
MLRRRTVLSRLRKLVAGAAVLTCAAAALGGIERPSAVAAPGGLPADALPAVFAQGGSSRY WP_075407058.1
fimbrial protein [Actinomyces naeslundii] Length: 953\nNumber of predicted TMHs: 1\nExp number of AAs in TMHs: 31.60073\nExp number, first 60 AAs: 10.09199\nTotal prob of N-in: 0.51244\nPOSSIBLE N-term signal sequence\noutside 1 922\nTMhelix 923 945\ninside 946 953

29161 GCF_001956445.1_ASM195644v1 Actinomyces naeslundii Terrabacteria group; Actinobacteria; Actinobacteria; Actinomycetales; Actinomycetaceae; Actinomyces
MLRRRTVLSRLRKLVAGAAVLTCAAAGLGGIERPSAVAAPGGLPADALPAVFAQGGSSRY WP_076134956.1
fimbrial protein [Actinomyces naeslundii] Length: 953\nNumber of predicted TMHs: 1\nExp number of AAs in TMHs: 29.48509\nExp number, first 60 AAs: 7.96407\nTotal prob of N-in: 0.40402\noutside 1 922\nTMhelix 923 945\ninside 946 953

29162 GCF_001956365.1_ASM195636v1 Actinomyces naeslundii Terrabacteria group; Actinobacteria; Actinobacteria; Actinomycetales; Actinomycetaceae; Actinomyces
MLRRRTVLSRLRKLVAGAAVLTCAAAGLGGIERPSAVAAPGGLPADALPAVFAQGGSSRY WP_076132099.1
fimbrial protein [Actinomyces naeslundii] Length: 953\nNumber of predicted TMHs: 1\nExp number of AAs in TMHs: 29.48509\nExp number, first 60 AAs: 7.96408\nTotal prob of N-in: 0.40402\noutside 1 922\nTMhelix 923 945\ninside 946 953

29163 GCF_001937595.1_ASM193759v1 Actinomyces naeslundii Terrabacteria group; Actinobacteria; Actinobacteria; Actinomycetales; Actinomycetaceae; Actinomyces
MLRRRTVLSRLRKLVAGAAVLTCAAAALGGIERPSAVAAPGGLPADALPAVFAQGGSSRY WP_075407058.1
fimbrial protein [Actinomyces naeslundii] Length: 953\nNumber of predicted TMHs: 1\nExp number of AAs in TMHs: 31.60073\nExp number, first 60 AAs: 10.09199\nTotal prob of N-in: 0.51244\nPOSSIBLE N-term signal sequence\noutside 1 922\nTMhelix 923 945\ninside 946 953

29164 GCF_001907295.1_ASM190729v1 Actinomyces nasicola Terrabacteria group; Actinobacteria; Actinobacteria; Actinomycetales; Actinomycetaceae; Actinomyces
MRPMTERNLSPPYPRRAFGRRHRRRSLVRRLRTRIAALAALTLMLTTAMFTTAAMADEPA WP_083603136.1
hypothetical protein [Actinomyces nasicola] Length: 1309\nNumber of predicted TMHs: 2\nExp number of

AAs in TMHs: 42.06741\nExp number, first 60 AAs:  22.00269\nTotal prob of N-in:   0.99498\nPOSSIBLE N-term signal sequence\ninside   1   33\nTMhelix   34   56\noutside   57  1283\nTMhelix   1284  1303\ninside   1304  1309

29165 GCF_001937725.1_ASM193772v1 Actinomyces oris Terrabacteria group; Actinobacteria; Actinobacteria; Actinomycetales; Actinomycetaceae; Actinomyces
MFPLPRRRAILPRLRKLVAGAAALTCVAAALVGTEQPSAGAAPRGLPADALPAVFAQGGT WP_075392615.1
fimbrial protein [Actinomyces oris] Length: 956\nNumber of predicted TMHs:  1\nExp number of AAs in TMHs: 24.29367\nExp number, first 60 AAs:  2.70988\nTotal prob of N-in:   0.13565\noutside   1   925\nTMhelix   926   948\ninside   949   956

29166 GCF_001937425.1_ASM193742v1 Actinomyces oris Terrabacteria group; Actinobacteria; Actinobacteria; Actinomycetales; Actinomycetaceae; Actinomyces
MLRRRAILPRLRKLVAGAAALTCVAAALVGTEQPSAGAAPRGLPADALPAVFAQGGTSWY WP_075401950.1
fimbrial protein [Actinomyces oris] Length: 953\nNumber of predicted TMHs:  1\nExp number of AAs in TMHs: 27.25057\nExp number, first 60 AAs:  5.95578\nTotal prob of N-in:   0.30227\noutside   1   921\nTMhelix   922   944\ninside   945   953

29167 GCF_001937385.1_ASM193738v1 Actinomyces oris Terrabacteria group; Actinobacteria; Actinobacteria; Actinomycetales; Actinomycetaceae; Actinomyces
MLRRRAILPRLRKLVAGAAALTCVAAALVGTEQPSAGAAPRGLPADALPAVFAQGGTSWY WP_070661474.1
MULTISPECIES: fimbrial protein [Actinomyces] Length: 953\nNumber of predicted TMHs:  1\nExp number of AAs in TMHs: 27.25057\nExp number, first 60 AAs:  5.95578\nTotal prob of N-in:   0.30227\noutside   1   921\nTMhelix   922   944\ninside   945   953

29168 GCF_001937535.1_ASM193753v1 Actinomyces oris Terrabacteria group; Actinobacteria; Actinobacteria; Actinomycetales; Actinomycetaceae; Actinomyces
MLRRRAILPRLRKLVAGAAALTCVAAALVGTEQPSAGAAPRGLPADALPAVFAQGGTSWY WP_075374766.1
fimbrial protein [Actinomyces oris] Length: 953\nNumber of predicted TMHs:  1\nExp number of AAs in TMHs: 27.2658\nExp number, first 60 AAs:  5.95576\nTotal prob of N-in:   0.30227\noutside   1   921\nTMhelix   922   944\ninside   945   953

29169 GCF_001937715.1_ASM193771v1 Actinomyces oris Terrabacteria group; Actinobacteria; Actinobacteria; Actinomycetales; Actinomycetaceae; Actinomyces
MLRRRAILPRLRKLVAGAAALTCVAAALVGTEQPSAGAAPRGLPADALPAVFAQGGTSWY WP_075378877.1
fimbrial protein [Actinomyces oris] Length: 953\nNumber of predicted TMHs:  1\nExp number of AAs in TMHs: 27.2656\nExp number, first 60 AAs:  5.95581999999999\nTotal prob of N-in:   0.30226\noutside   1   921\nTMhelix   922   944\ninside   945   953

29170 GCF_001937485.1_ASM193748v1 Actinomyces oris Terrabacteria group; Actinobacteria; Actinobacteria; Actinomycetales; Actinomycetaceae; Actinomyces
MFPLPRRRAILPRLRKLVAGAAALTCVAAALVGTEQPSAGAAPRGLPADALPAVFAQGGT WP_075371541.1
fimbrial protein [Actinomyces oris] Length: 956\nNumber of predicted TMHs:  1\nExp number of AAs in TMHs: 24.29323\nExp number, first 60 AAs:  2.7099\nTotal prob of N-in:   0.13564\noutside   1   925\nTMhelix   926   948\ninside   949   956

29171 GCF_001937445.1_ASM193744v1 Actinomyces oris Terrabacteria group; Actinobacteria; Actinobacteria; Actinomycetales; Actinomycetaceae; Actinomyces
MLRRRAILPRLRKLVAGAAALTCVAAALVGTEQPSAGAAPRGLPADALPAVFAQGGTSWY WP_075390447.1
fimbrial protein [Actinomyces oris] Length: 953\nNumber of predicted TMHs:  1\nExp number of AAs in TMHs: 27.26579\nExp number, first 60 AAs:  5.95576\nTotal prob of N-in:   0.30227\noutside   1   921\nTMhelix   922   944\ninside   945   953

29172 GCF_001937415.1_ASM193741v1 Actinomyces oris Terrabacteria group; Actinobacteria; Actinobacteria; Actinomycetales; Actinomycetaceae; Actinomyces
MLRRRAILPRLRKLVAGAAALTCVAAALVGTEQPSAGAAPRGLPADALPAVFAQGGTSWY WP_070661474.1
MULTISPECIES: fimbrial protein [Actinomyces] Length: 953\nNumber of predicted TMHs:  1\nExp number of AAs in TMHs: 27.25057\nExp number, first 60 AAs:  5.95578\nTotal prob of N-in:   0.30227\noutside   1   921\nTMhelix   922   944\ninside   945   953

29173 GCF_000820725.1_Actinomyces_polynesiensis Actinomyces polynesiensis Terrabacteria group; Actinobacteria; Actinobacteria; Actinomycetales; Actinomycetaceae; Actinomyces

MDEPRGRFARLLPPEQGASRRTMLLSFGLWPVAMLLTLRGHGIGLGGLGLVAIPLCVTAF WP_043534360.1 hypothetical protein [Actinomyces polynesiensis]Length: 155\nNumber of predicted TMHs: 2\nExp number of AAs in TMHs: 47.79644\nExp number, first 60 AAs: 22.74902\nTotal prob of N-in: 0.63516\nPOSSIBLE N-term signal sequence\ninside 1 37\nTMhelix 38 60\noutside 61 84\nTMhelix 85 107\ninside 108 155

29174 GCF_001553565.1_ASM155356v1 Actinomyces radicidentis Terrabacteria group; Actinobacteria; Actinobacteria; Actinomycetales; Actinomycetaceae; Actinomyces MSTGTPYAAPWSPGAPAPAPKGRRGPVVLMVVGGVLVIIAVIAFVVSIVSVTRAATSLER WP_067940768.1 hypothetical protein [Actinomyces radicidentis] Length: 199\nNumber of predicted TMHs: 2\nExp number of AAs in TMHs: 59.66404\nExp number, first 60 AAs: 22.87323\nTotal prob of N-in: 0.17905\nPOSSIBLE N-term signal sequence\ninside 1 26\nTMhelix 27 49\noutside 50 155\nTMhelix 156 178\ninside 179 199

29175 GCF_001813985.1_ASM181398v1 Actinomyces sp. HMSC062G12 Terrabacteria group; Actinobacteria; Actinobacteria; Actinomycetales; Actinomycetaceae; Actinomyces MVRAAFFDLDKTILDTSSNVALSGPFIEAGLMNRRTAVASVLVQLPYLLAGADESRMEQM WP_070778599.1 MULTISPECIES: hydrolase [Actinomyces] Length: 279\nNumber of predicted TMHs: 1\nExp number of AAs in TMHs: 16.02741\nExp number, first 60 AAs: 0.02813\nTotal prob of N-in: 0.01236\noutside 1 235\nTMhelix 236 254\ninside 255 279

29176 GCF_001814535.1_ASM181453v1 Actinomyces sp. HMSC065F12 Terrabacteria group; Actinobacteria; Actinobacteria; Actinomycetales; Actinomycetaceae; Actinomyces MRESSSPPGRCLFTRRGAIIAACTCAGALFLAGTACAADPAAPEAEKPYAHPSAARVVAQ WP_070725188.1 hypothetical protein [Actinomyces sp. HMSC065F12] Length: 625\nNumber of predicted TMHs: 1\nExp number of AAs in TMHs: 35.79845\nExp number, first 60 AAs: 13.73061\nTotal prob of N-in: 0.61852\nPOSSIBLE N-term signal sequence\noutside 1 596\nTMhelix 597 619\ninside 620 625

29177 GCF_001814535.1_ASM181453v1 Actinomyces sp. HMSC065F12 Terrabacteria group; Actinobacteria; Actinobacteria; Actinomycetales; Actinomycetaceae; Actinomyces MKRSQPRSRSAAFFDLDKTILATSTSVALRAPLREAGLLTRRGALLGLLIQLPYLIKGAD WP_070725291.1 hypothetical protein [Actinomyces sp. HMSC065F12] Length: 267\nNumber of predicted TMHs: 1\nExp number of AAs in TMHs: 19.24274\nExp number, first 60 AAs: 0.10453\nTotal prob of N-in: 0.64051\ninside 1 238\nTMhelix 239 261\noutside 262 267

29178 GCF_001812965.1_ASM181296v1 Actinomyces sp. HMSC075C01 Terrabacteria group; Actinobacteria; Actinobacteria; Actinomycetales; Actinomycetaceae; Actinomyces MLRRRAILPRLRKLVAGAAALTCVAAALVGTEQPSAGAAPRGLPADALPAVFAQGGTSWY WP_070661474.1 MULTISPECIES: fimbrial protein [Actinomyces] Length: 953\nNumber of predicted TMHs: 1\nExp number of AAs in TMHs: 27.25057\nExp number, first 60 AAs: 5.95578\nTotal prob of N-in: 0.30227\noutside 1 921\nTMhelix 922 944\ninside 945 953

29179 GCF_001807465.1_ASM180746v1 Actinomyces sp. HMSC08A09 Terrabacteria group; Actinobacteria; Actinobacteria; Actinomycetales; Actinomycetaceae; Actinomyces MLRRRAILPRLRKLVAGAAALTCVAAALVGTEQPSAGAAPRGLPADALPAVFAQGGTSWY WP_070511598.1 fimbrial protein [Actinomyces sp. HMSC08A09] Length: 953\nNumber of predicted TMHs: 1\nExp number of AAs in TMHs: 27.16086\nExp number, first 60 AAs: 5.95555999999999\nTotal prob of N-in: 0.30230\noutside 1 921\nTMhelix 922 944\ninside 945 953

29180 GCF_000278725.1_AspICM47v1.0 Actinomyces sp. ICM47 Terrabacteria group; Actinobacteria; Actinobacteria; Actinomycetales; Actinomycetaceae; Actinomyces MVRAAFFDLDKTILDTSSNVALSGPFIEAGLMSRRTAITSFLVQLPYLLSGADESRMGQM WP_009649116.1 hydrolase [Actinomyces sp. ICM47] Length: 256\nNumber of predicted TMHs: 1\nExp number of AAs in TMHs: 18.30293\nExp number, first 60 AAs: 0.0403\nTotal prob of N-in: 0.02201\noutside 1 235\nTMhelix 236 254\ninside 255 256

29181 GCF_900155605.1_PRJEB18271 Actinomyces sp. Marseille-P2818 Terrabacteria group; Actinobacteria; Actinobacteria; Actinomycetales; Actinomycetaceae; Actinomyces MLTRTRRAVIATGAALALFATPLAAFADPAGEATSTTEASPEATAPTPAAEATASASQAT WP_077209678.1 bifunctional metallophosphatase/5-nucleotidase [Actinomyces sp. Marseille-P2818] Length: 767\nNumber of predicted TMHs: 2\nExp number of AAs in TMHs: 32.65226\nExp number, first 60 AAs: 15.0263\nTotal prob of N-in: 0.74396\nPOSSIBLE N-term signal sequence\ninside 1 6\nTMhelix 7 26\noutside 27 744\nTMhelix 745 762\ninside 763 767

29182 GCF_000308055.1_ASM30805v1 Actinomyces sp. ph3 Terrabacteria group; Actinobacteria; Actinobacteria; Actinomycetales; Actinomycetaceae; Actinomyces
MRASISRRSHVLLSIAACALMISASLFGATPTHAVRNQAEIREFFMILSDGNVSHSITVI WP_019126397.1 hypothetical protein [Actinomyces sp. ph3] Length: 329\nNumber of predicted TMHs: 2\nExp number of AAs in TMHs: 42.7606\nExp number, first 60 AAs: 22.03899\nTotal prob of N-in: 0.97975\nPOSSIBLE N-term signal sequence\ninside 1 12\nTMhelix 13 35\noutside 36 257\nTMhelix 258 280\ninside 281 329

29183 GCF_000308055.1_ASM30805v1 Actinomyces sp. ph3 Terrabacteria group; Actinobacteria; Actinobacteria; Actinomycetales; Actinomycetaceae; Actinomyces
MAFLSFTRQHRRRRVIFTALLSAAIACGFLGVSSTRLRAQADSYGSDLILLKDAAHVQMS WP_019126250.1 hypothetical protein [Actinomyces sp. ph3] Length: 341\nNumber of predicted TMHs: 2\nExp number of AAs in TMHs: 42.77012\nExp number, first 60 AAs: 20.81858\nTotal prob of N-in: 0.89717\nPOSSIBLE N-term signal sequence\ninside 1 12\nTMhelix 13 35\noutside 36 312\nTMhelix 313 335\ninside 336 341

29184 GCF_000308055.1_ASM30805v1 Actinomyces sp. ph3 Terrabacteria group; Actinobacteria; Actinobacteria; Actinomycetales; Actinomycetaceae; Actinomyces
MSHLTARRAKRRRTLLATFVSASVALGIVGASASHHQADAAPNDGADLIVMKDAAHVKME WP_019126251.1 hypothetical protein [Actinomyces sp. ph3] Length: 344\nNumber of predicted TMHs: 2\nExp number of AAs in TMHs: 38.28528\nExp number, first 60 AAs: 19.05414\nTotal prob of N-in: 0.89483\nPOSSIBLE N-term signal sequence\ninside 1 12\nTMhelix 13 35\noutside 36 315\nTMhelix 316 338\ninside 339 344

29185 GCF_000758755.1_04_NF40_HMP1627v01 Actinomyces sp. S6-Spd3 Terrabacteria group; Actinobacteria; Actinobacteria; Actinomycetales; Actinomycetaceae; Actinomyces
MLTRTRRAVIATGAALALFATPLAAFADPAGEATSTTEASPEATAPTSAAEATASANQAT WP_034501857.1 bifunctional metallophosphatase/5-nucleotidase [Actinomyces sp. S6-Spd3] Length: 767\nNumber of predicted TMHs: 2\nExp number of AAs in TMHs: 33.33585\nExp number, first 60 AAs: 15.02517\nTotal prob of N-in: 0.74397\nPOSSIBLE N-term signal sequence\ninside 1 6\nTMhelix 7 26\noutside 27 744\nTMhelix 745 762\ninside 763 767

29186 GCF_001856685.1_ASM185668v1 Actinomyces sp. VUL4_3 Terrabacteria group; Actinobacteria; Actinobacteria; Actinomycetales; Actinomycetaceae; Actinomyces
MKDTKKASRRSLVNFAAIGSTVAVALAMLVPTYFGLGANARTATAVEGVELTVPDISMAV WP_071163387.1 hypothetical protein [Actinomyces sp. VUL4_3] Length: 2413\nNumber of predicted TMHs: 2\nExp number of AAs in TMHs: 42.87408\nExp number, first 60 AAs: 22.62855\nTotal prob of N-in: 0.99985\nPOSSIBLE N-term signal sequence\ninside 1 11\nTMhelix 12 34\noutside 35 2380\nTMhelix 2381 2403\ninside 2404 2413

29187 GCF_001856685.1_ASM185668v1 Actinomyces sp. VUL4_3 Terrabacteria group; Actinobacteria; Actinobacteria; Actinomycetales; Actinomycetaceae; Actinomyces
MSSFYTAEAAGRRRNTLRRGMAGLLAAGMAVTLGAFGVNFPSDSAQAYTVETVPGPAIDK WP_071164834.1 hypothetical protein [Actinomyces sp. VUL4_3] Length: 1570\nNumber of predicted TMHs: 2\nExp number of AAs in TMHs: 40.51249\nExp number, first 60 AAs: 19.82357\nTotal prob of N-in: 0.99211\nPOSSIBLE N-term signal sequence\ninside 1 20\nTMhelix 21 38\noutside 39 1532\nTMhelix 1533 1555\ninside 1556 1570

29188 GCF_001687305.1_ASM168730v1 Actinomyces sp. VUL7 Terrabacteria group; Actinobacteria; Actinobacteria; Actinomycetales; Actinomycetaceae; Actinomyces
MLTRRNALSALALGVVGVSIAPTALADGVSVDVSTQDGPYTHEVVIPVVLRSPGGDPDKP WP_067783620.1 hypothetical protein [Actinomyces sp. VUL7] Length: 233\nNumber of predicted TMHs: 1\nExp number of AAs in TMHs: 35.49347\nExp number, first 60 AAs: 16.66611\nTotal prob of N-in: 0.89815\nPOSSIBLE N-term signal sequence\ninside 1 205\nTMhelix 206 225\noutside 226 233

29189 GCF_000364865.1_ASM36486v1 Actinomyces cardiffensis F0333 Terrabacteria group; Actinobacteria; Actinobacteria; Actinomycetales; Actinomycetaceae; Actinomyces; Actinomyces cardiffensis
MKRHRRGLVAVMFSAILLSSPCVASYPRVTHAALADTPAEATANAPAGAEETPAKALMDA WP_005963289.1 thermitase [Actinomyces cardiffensis] Length: 477\nNumber of predicted TMHs: 2\nExp number of AAs in TMHs: 41.87437\nExp number, first 60 AAs: 18.41127\nTotal prob of N-in: 0.87066\nPOSSIBLE N-term signal sequence\ninside 1 6\nTMhelix 7 29\noutside 30 450\nTMhelix 451 473\ninside 474 477

29190 GCF_000429225.1_ASM42922v1 Actinomyces dentalis DSM 19115 Terrabacteria group; Actinobacteria; Actinobacteria; Actinomycetales; Actinomycetaceae; Actinomyces; Actinomyces dentalis
MDARRSHMVVLRGVGVLLVLLLGSVLALVQPASAAPALIATESPRPPDNVQADHTGMIDD WP_026410447.1 hypothetical protein [Actinomyces dentalis] Length: 238\nNumber of predicted TMHs: 2\nExp number of

AAs in TMHs: 46.35323\nExp number, first 60 AAs:  21.89874\nTotal prob of N-in:   0.98396\nPOSSIBLE N-term signal sequence\ninside   1   12\nTMhelix   13   35\noutside   36   203\nTMhelix   204   226\ninside   227   238

29191 GCF_000411155.1_Acti_euro_ACS-120-V-COL10B_V1 Actinomyces europaeus ACS-120-V-Col10b Terrabacteria group; Actinobacteria; Actinobacteria; Actinomycetales; Actinomycetaceae; Actinomyces; Actinomyces europaeus MGCVRNVHRRSMFSKMRTTTFGASVCTLAIALTGVGQAFATPPVEALAGEQVIATNGVGG WP_083938950.1 hypothetical protein [Actinomyces europaeus] Length: 967\nNumber of predicted TMHs: 2\nExp number of AAs in TMHs: 41.99042\nExp number, first 60 AAs:  20.63804\nTotal prob of N-in:   0.99464\nPOSSIBLE N-term signal sequence\ninside   1   20\nTMhelix   21   40\noutside   41   936\nTMhelix   937   959\ninside   960   967

29192 GCF_000411155.1_Acti_euro_ACS-120-V-COL10B_V1 Actinomyces europaeus ACS-120-V-Col10b Terrabacteria group; Actinobacteria; Actinobacteria; Actinomycetales; Actinomycetaceae; Actinomyces; Actinomyces europaeus MKTERRRVVAASTALFAALTLSAPAALAAEEPWQILGQGQLFVAKSGEYYGVCAGDDQLR WP_016443822.1 hypothetical protein [Actinomyces europaeus] Length: 376\nNumber of predicted TMHs: 2\nExp number of AAs in TMHs: 42.34834\nExp number, first 60 AAs:  20.23947\nTotal prob of N-in:   0.90624\nPOSSIBLE N-term signal sequence\ninside   1   6\nTMhelix   7   29\noutside   30   346\nTMhelix   347   369\ninside   370   376

29193 GCF_000411155.1_Acti_euro_ACS-120-V-COL10B_V1 Actinomyces europaeus ACS-120-V-Col10b Terrabacteria group; Actinobacteria; Actinobacteria; Actinomycetales; Actinomycetaceae; Actinomyces; Actinomyces europaeus MQRIFAFHTQRRKLMRRSVKTLAAGAIATAALVVSPLAAFAEDPVSESEANPVNAASSEA WP_083938905.1 hypothetical protein [Actinomyces europaeus] Length: 806\nNumber of predicted TMHs: 2\nExp number of AAs in TMHs: 42.12195\nExp number, first 60 AAs:  20.01997\nTotal prob of N-in:   0.98343\nPOSSIBLE N-term signal sequence\ninside   1   20\nTMhelix   21   40\noutside   41   778\nTMhelix   779   801\ninside   802   806

29194 GCF_000429245.1_ASM42924v1 Actinomyces georgiae DSM 6843 Terrabacteria group; Actinobacteria; Actinobacteria; Actinomycetales; Actinomycetaceae; Actinomyces; Actinomyces georgiae MHTTTRRAVVAGCAAFALFAAPMTALAEPADPGSPQSGGAQSGAADAPQSDPAATQGAQG WP_026430636.1 bifunctional metallophosphatase/5-nucleotidase [Actinomyces georgiae] Length: 765\nNumber of predicted TMHs:  2\nExp number of AAs in TMHs: 39.20111\nExp number, first 60 AAs:  17.397\nTotal prob of N-in: 0.86895\nPOSSIBLE N-term signal sequence\ninside   1   6\nTMhelix   7   26\noutside   27   736\nTMhelix   737 759\ninside   760   765

29195 GCF_000277685.1_AgeorgiaeF0490v1.0 Actinomyces georgiae F0490 Terrabacteria group; Actinobacteria; Actinobacteria; Actinomycetales; Actinomycetaceae; Actinomyces; Actinomyces georgiae MHTTTRRAVVAGCAAFALFAAPMTALAEPADPGSPQSGGAQSGAADAPQSDPAATQGAQG WP_005868763.1 bifunctional metallophosphatase/5-nucleotidase [Actinomyces georgiae] Length: 765\nNumber of predicted TMHs:  2\nExp number of AAs in TMHs: 39.20111\nExp number, first 60 AAs:  17.397\nTotal prob of N-in: 0.86895\nPOSSIBLE N-term signal sequence\ninside   1   6\nTMhelix   7   26\noutside   27   736\nTMhelix   737 759\ninside   760   765

29196 GCF_000466205.2_ASM46620v2 Actinomyces johnsonii F0510 Terrabacteria group; Actinobacteria; Actinobacteria; Actinomycetales; Actinomycetaceae; Actinomyces; Actinomyces johnsonii MLRRRTVVSRLRKLVASAAVLTCAAAAFAGIERPPAAAAPPGLPSDALPAVFAQGGTSRY WP_021607699.1 hypothetical protein [Actinomyces johnsonii] Length: 953\nNumber of predicted TMHs:  1\nExp number of AAs in TMHs: 35.53717\nExp number, first 60 AAs:  14.16833\nTotal prob of N-in:   0.72667\nPOSSIBLE N-term signal sequence\noutside   1   921\nTMhelix   922   944\ninside   945   953

29197 GCF_000466245.1_ASM46624v1 Actinomyces johnsonii F0542 Terrabacteria group; Actinobacteria; Actinobacteria; Actinomycetales; Actinomycetaceae; Actinomyces; Actinomyces johnsonii MLRRRTVVSRLRKLVASAAVLTCAAAAFAGIERPSAAAAPPGLPSDALPAVFAQGGTSRY WP_021608244.1 hypothetical protein [Actinomyces johnsonii] Length: 953\nNumber of predicted TMHs:  1\nExp number of AAs in TMHs: 35.31399\nExp number, first 60 AAs:  13.95463\nTotal prob of N-in:   0.71490\nPOSSIBLE N-term signal sequence\noutside   1   921\nTMhelix   922   944\ninside   945   953

29198 GCF_000269805.1_ASM26980v1 Actinomyces massiliensis 4401292 Terrabacteria group; Actinobacteria; Actinobacteria; Actinomycetales; Actinomycetaceae; Actinomyces; Actinomyces massiliensis MRHRRPLIRHLTAAGAVALVAGSALLAPTAAADQPILTCAGSGFMCMGTGHFGTVSASYT WP_008733546.1 hypothetical protein [Actinomyces massiliensis] Length: 381\nNumber of predicted TMHs:  2\nExp number of AAs in TMHs: 40.55178\nExp number, first 60 AAs:  18.88437\nTotal prob of N-in:   0.86434\nPOSSIBLE N-term signal sequence\ninside   1   6\nTMhelix   7   29\noutside   30   352\nTMhelix   353   375\ninside   376   381

29199 GCF_000296275.1_AmassiliensisF0489v1.0 Actinomyces massiliensis F0489 Terrabacteria group; Actinobacteria; Actinobacteria; Actinomycetales; Actinomycetaceae; Actinomyces; Actinomyces massiliensis MRHRRPLIRHLTAAGAVALVAGSALLAPTAAADQPILTCAGSGFMCMGTGHFGTVSASYT WP_008733546.1 hypothetical protein [Actinomyces massiliensis] Length: 381\nNumber of predicted TMHs: 2\nExp number of AAs in TMHs: 40.55178\nExp number, first 60 AAs: 18.88437\nTotal prob of N-in: 0.86434\nPOSSIBLE N-term signal sequence\ninside 1 6\nTMhelix 7 29\noutside 30 352\nTMhelix 353 375\ninside 376 381

29200 GCF_000163415.1_ASM16341v1 Actinomyces odontolyticus F0309 Terrabacteria group; Actinobacteria; Actinobacteria; Actinomycetales; Actinomycetaceae; Actinomyces; Actinomyces odontolyticus MVRAAFFDLDKTILDTSSNVALSGPFIEAGLMNRRTALASVLVQLPYLLAGADESRMEQM WP_003795913.1 hydrolase [Actinomyces odontolyticus] Length: 256\nNumber of predicted TMHs: 1\nExp number of AAs in TMHs: 15.21451\nExp number, first 60 AAs: 0.03492\nTotal prob of N-in: 0.01232\noutside 1 235\nTMhelix 236 254\ninside 255 256

29201 GCF_000428685.1_ASM42868v1 Actinomyces slackii ATCC 49928 Terrabacteria group; Actinobacteria; Actinobacteria; Actinomycetales; Actinomycetaceae; Actinomyces; Actinomyces slackii MIVPLQPTRSHRTPRRRALVSAACLLAASLSVSAPALASEGQSATSMRDSSVTAQALAEK WP_034515529.1 LPXTG cell wall anchor domain-containing protein [Actinomyces slackii] Length: 399\nNumber of predicted TMHs: 1\nExp number of AAs in TMHs: 28.99899\nExp number, first 60 AAs: 9.53129\nTotal prob of N-in: 0.46790\noutside 1 374\nTMhelix 375 393\ninside 394 399

29202 GCF_000195595.1_ASM19559v1 Actinomyces sp. oral taxon 170 str. F0386 Terrabacteria group; Actinobacteria; Actinobacteria; Actinomycetales; Actinomycetaceae; Actinomyces; Actinomyces sp. oral taxon 170 MIPRSTQVLSPESTNPSTIRSRRGLLARAGALTSVLALSTAGMAASAAHAVEETPAPAPA WP_050793000.1 hypothetical protein [Actinomyces sp. oral taxon 170] Length: 536\nNumber of predicted TMHs: 1\nExp number of AAs in TMHs: 26.44752\nExp number, first 60 AAs: 4.76956\nTotal prob of N-in: 0.22186\noutside 1 509\nTMhelix 510 532\ninside 533 536

29203 GCF_000186685.1_ASM18668v1 Actinomyces sp. oral taxon 178 str. F0338 Terrabacteria group; Actinobacteria; Actinobacteria; Actinomycetales; Actinomycetaceae; Actinomyces; Actinomyces sp. oral taxon 178 MPATRSRRRALARLVPALLAALLACSLAPLRADAADIRRVGDILSAIGGTVYIHDNPDWS WP_009743428.1 MULTISPECIES: pilus assembly protein [Actinomyces] Length: 769\nNumber of predicted TMHs: 1\nExp number of AAs in TMHs: 37.02313\nExp number, first 60 AAs: 14.86808\nTotal prob of N-in: 0.72647\nPOSSIBLE N-term signal sequence\noutside 1 736\nTMhelix 737 759\ninside 760 769

29204 GCF_000186685.1_ASM18668v1 Actinomyces sp. oral taxon 178 str. F0338 Terrabacteria group; Actinobacteria; Actinobacteria; Actinomycetales; Actinomycetaceae; Actinomyces; Actinomyces sp. oral taxon 178 MHTTTRRAVVAGCAVFALFAAPMAALAEPAGPGAPQSGSAQSGAADAPQSEPAAAQGAQG WP_009743508.1 MULTISPECIES: bifunctional metallophosphatase/5-nucleotidase [Actinomyces] Length: 765\nNumber of predicted TMHs: 2\nExp number of AAs in TMHs: 41.53601\nExp number, first 60 AAs: 19.57571\nTotal prob of N-in: 0.96746\nPOSSIBLE N-term signal sequence\ninside 1 6\nTMhelix 7 26\noutside 27 736\nTMhelix 737 759\ninside 760 765

29205 GCF_000185285.1_ASM18528v1 Actinomyces sp. oral taxon 180 str. F0310 Terrabacteria group; Actinobacteria; Actinobacteria; Actinomycetales; Actinomycetaceae; Actinomyces; Actinomyces sp. oral taxon 180 MVFRPAVNLVLLVLIVLLGAALVFLTARRATRRSFIDILRRSMILAVVVVMGAGPSIPGE WP_009213069.1 MULTISPECIES: hypothetical protein [Actinobacteria] Length: 344\nNumber of predicted TMHs: 2\nExp number of AAs in TMHs: 46.33274\nExp number, first 60 AAs: 26.28713\nTotal prob of N-in: 0.39658\nPOSSIBLE N-term signal sequence\noutside 1 4\nTMhelix 5 27\ninside 28 310\nTMhelix 311 330\noutside 331 344

29206 GCF_000185285.1_ASM18528v1 Actinomyces sp. oral taxon 180 str. F0310 Terrabacteria group; Actinobacteria; Actinobacteria; Actinomycetales; Actinomycetaceae; Actinomyces; Actinomyces sp. oral taxon 180 MARAAFFDLDKTILDTSSNVALSGPFIEAGLMSRRAAVASVLVHLPYLLSGADESRMQQM WP_009212972.1 hydrolase [Actinomyces sp. oral taxon 180] Length: 260\nNumber of predicted TMHs: 1\nExp number of AAs in TMHs: 18.32419\nExp number, first 60 AAs: 0.06986\nTotal prob of N-in: 0.01564\noutside 1 231\nTMhelix 232 254\ninside 255 260

29207 GCF_000318335.2_ASM31833v2 Actinomyces sp. oral taxon 181 str. F0379 Terrabacteria group; Actinobacteria; Actinobacteria; Actinomycetales; Actinomycetaceae; Actinomyces; Actinomyces sp. oral taxon 181 MSHLTARRAKRRRTLLATFVSASVALGIVGASASHHQADAAPNDGADLIVMKDAAHVKME WP_009409131.1 hypothetical protein [Actinomyces sp. oral taxon 181] Length: 353\nNumber of predicted TMHs: 2\nExp

number of AAs in TMHs: 38.2668\nExp number, first 60 AAs:  19.05335\nTotal prob of N-in:   0.89440\nPOSSIBLE N-term signal sequence\ninside   1   12\nTMhelix   13   35\noutside   36   324\nTMhelix   325   347\ninside   348   353

29208 GCF_000318335.2_ASM31833v2 Actinomyces sp. oral taxon 181 str. F0379 Terrabacteria group; Actinobacteria; Actinobacteria; Actinomycetales; Actinomycetaceae; Actinomyces; Actinomyces sp. oral taxon 181 MAFLSFTRQRRRRRVIFTALLSAAIACGFLGVSSTQLRAQADSYDGDLVLLKDAAHVQMS WP_009409132.1 repeat-containing protein [Actinomyces sp. oral taxon 181]Length: 341\nNumber of predicted TMHs:  2\nExp number of AAs in TMHs: 42.7219499999999\nExp number, first 60 AAs:  20.68403\nTotal prob of N-in: 0.94932\nPOSSIBLE N-term signal sequence\ninside   1   12\nTMhelix   13   35\noutside   36   312\nTMhelix   313 335\ninside   336   341

29209 GCF_000239715.1_Actinomyces_sp_F0330_V1 Actinomyces sp. oral taxon 849 str. F0330 Terrabacteria group; Actinobacteria; Actinobacteria; Actinomycetales; Actinomycetaceae; Actinomyces; Actinomyces sp. oral taxon 849 MLRRRTVVSRLRKLVASAAVLTCAAAAFAGIERPSAAAAPPGLPSDALPAVFAQGGTSRY WP_009233255.1 hypothetical protein [Actinomyces sp. oral taxon 849] Length: 953\nNumber of predicted TMHs:  1\nExp number of AAs in TMHs: 35.31399\nExp number, first 60 AAs:  13.95463\nTotal prob of N-in:   0.71490\nPOSSIBLE N-term signal sequence\noutside   1   921\nTMhelix   922   944\ninside   945   953

29210 GCF_000466305.1_ASM46630v1 Actinomyces sp. oral taxon 877 str. F0543 Terrabacteria group; Actinobacteria; Actinobacteria; Actinomycetales; Actinomycetaceae; Actinomyces; Actinomyces sp. oral taxon 877 MHTTTRRAVVAGCAVFALFAAPMAALAEPAGPGAPQSGGAQSGAADAPQSEPAAAQGAQG WP_021613375.1 bifunctional metallophosphatase/5-nucleotidase [Actinomyces sp. oral taxon 877] Length: 765\nNumber of predicted TMHs:  2\nExp number of AAs in TMHs: 41.60768\nExp number, first 60 AAs:  19.64738\nTotal prob of N-in: 0.96820\nPOSSIBLE N-term signal sequence\ninside   1   6\nTMhelix   7   26\noutside   27   736\nTMhelix   737 759\ninside   760   765

29211 GCF_000429105.1_ASM42910v1 Actinomyces suimastitidis DSM 15538 Terrabacteria group; Actinobacteria; Actinobacteria; Actinomycetales; Actinomycetaceae; Actinomyces; Actinomyces suimastitidis MLPAPRRRTCFAPMRLVAFLTAALLTATACIFAGPTAYAQDAWPTVLKSGKINIFDLDAT WP_051259476.1 hypothetical protein [Actinomyces suimastitidis] Length: 1443\nNumber of predicted TMHs:  2\nExp number of AAs in TMHs: 43.84744\nExp number, first 60 AAs:  22.51818\nTotal prob of N-in:   0.98892\nPOSSIBLE N-term signal sequence\ninside 1   15\nTMhelix   16   38\noutside   39  1414\nTMhelix   1415  1437\ninside   1438  1443

29212 GCF_000295095.1_ASM29509v1 Actinomyces timonensis DSM 23838 Terrabacteria group; Actinobacteria; Actinobacteria; Actinomycetales; Actinomycetaceae; Actinomyces; Actinomyces timonensis MTEPIAAPPPPEERWQNLARQEEQARRRRNLILTGAGAAGVGVIGLGALSMCGGPTTAPN WP_017177339.1 hypothetical protein [Actinomyces timonensis] Length: 231\nNumber of predicted TMHs:  2\nExp number of AAs in TMHs: 43.54767\nExp number, first 60 AAs:  21.23586\nTotal prob of N-in:   0.94816\nPOSSIBLE N-term signal sequence\ninside   1   29\nTMhelix   30   52\noutside   53   189\nTMhelix   190   212\ninside   213   231

29213 GCF_000420425.1_ASM42042v1 Actinomyces vaccimaxillae DSM 15804 Terrabacteria group; Actinobacteria; Actinobacteria; Actinomycetales; Actinomycetaceae; Actinomyces; Actinomyces vaccimaxillae MLLCWVRVFWCWFDQYGLDFCMSRSLWNGRRRLVAVAGAVAVMGGFVVAGPAVAAPSVLR WP_083940714.1 hypothetical protein [Actinomyces vaccimaxillae]Length: 1330\nNumber of predicted TMHs:  2\nExp number of AAs in TMHs: 43.4889\nExp number, first 60 AAs:  22.45803\nTotal prob of N-in:   0.98290\nPOSSIBLE N-term signal sequence\ninside   1   32\nTMhelix   33   55\noutside   56  1300\nTMhelix   1301  1323\ninside   1324  1330

29214 GCF_000411135.1_Acti_scha_FB123-CNA2_V1 Actinotignum schaalii FB123-CNA-2 Terrabacteria group; Actinobacteria; Actinobacteria; Actinomycetales; Actinomycetaceae; Actinotignum; Actinotignum schaalii MNRHLMMPLASRRSLIALLCTVLLLSLGWGVLPQASAAPAAGTLTNVSVSITGDDTYGDY WP_016442126.1 hypothetical protein [Actinotignum schaalii] Length: 1678\nNumber of predicted TMHs:  2\nExp number of AAs in TMHs: 37.8282699999999\nExp number, first 60 AAs:  19.25508\nTotal prob of N-in:   0.91211\nPOSSIBLE N-term signal sequence\ninside   1   12\nTMhelix   13   32\noutside   33  1653\nTMhelix   1654  1671\ninside   1672  1678

29215 GCF_000612055.1_ASM61205v1 Trueperella pyogenes Terrabacteria group; Actinobacteria; Actinobacteria; Actinomycetales; Actinomycetaceae; Trueperella MNNTTRPGRRRATLALLSVAAVAVVNTITPFAVADSNSDLVDQLLGERFGDDFNRGPSTP WP_080712419.1 hypothetical protein [Trueperella pyogenes] Length: 951\nNumber of predicted TMHs:  2\nExp number of AAs in TMHs: 45.42229\nExp number, first 60 AAs:  22.4851\nTotal prob of N-in:   0.99425\nPOSSIBLE N-term signal sequence\ninside   1   11\nTMhelix   12   34\noutside   35   921\nTMhelix   922   944\ninside   945   951

29216	GCF_001281085.1_ASM128108v1	Trueperella pyogenes	Terrabacteria group; Actinobacteria; Actinobacteria; Actinomycetales; Actinomycetaceae; Trueperella
MNNTTRPGRRRATLALLSVAAVAVVNTITPFAVADSNLDSAYDEILRERFGGDDDFNRGP	WP_080999149.1
hypothetical protein [Trueperella pyogenes]	Length: 926\nNumber of predicted TMHs:  2\nExp number of AAs in TMHs: 45.02005\nExp number, first 60 AAs:  22.18977\nTotal prob of N-in:   0.99222\nPOSSIBLE N-term signal sequence\ninside   1   11\nTMhelix   12   34\noutside   35   896\nTMhelix   897   919\ninside   920   926

29217	GCF_000599565.1_TruePyoMS2391.0	Trueperella pyogenes MS249	Terrabacteria group; Actinobacteria; Actinobacteria; Actinomycetales; Actinomycetaceae; Trueperella; Trueperella pyogenes
MNNTTRPGRRRATLALLSVAAVAVVNTITPFAVADSNHRSSYEDILKDRFDDDFAPRPSTWP_080689132.1	hypothetical protein [Trueperella pyogenes]	Length: 938\nNumber of predicted TMHs:  2\nExp number of AAs in TMHs: 45.0878999999999\nExp number, first 60 AAs:  22.21234\nTotal prob of N-in:   0.99315\nPOSSIBLE N-term signal sequence\ninside   1   11\nTMhelix   12   34\noutside   35   908\nTMhelix   909   931\ninside   932   938

29218	GCF_000816065.1_ASM81606v1	Trueperella pyogenes TP8	Terrabacteria group; Actinobacteria; Actinobacteria; Actinomycetales; Actinomycetaceae; Trueperella; Trueperella pyogenes
MNNTTRPGRRRATLALLSVAAVAVVNTITPFAVADSNLDSAYDEILRERFGGDDDFNRGP	WP_080754043.1
hypothetical protein [Trueperella pyogenes]	Length: 940\nNumber of predicted TMHs:  2\nExp number of AAs in TMHs: 45.1085599999999\nExp number, first 60 AAs:  22.18984\nTotal prob of N-in:   0.99222\nPOSSIBLE N-term signal sequence\ninside   1   11\nTMhelix   12   34\noutside   35   910\nTMhelix   911   933\ninside   934   940

29219	GCF_000763095.1_ASM76309v1	Actinopolyspora erythraea	Terrabacteria group; Actinobacteria; Actinobacteria; Actinopolysporales; Actinopolysporaceae; Actinopolyspora
MTQSEHPPENHERRGAAFFDLDRTIIARSSTLAFSKPFFREGLINRRAVLKSTYAQFMFM WP_043569318.1	inhibition of morphological differentiation protein [Actinopolyspora erythraea]	Length: 274\nNumber of predicted TMHs:  1\nExp number of AAs in TMHs: 22.76849\nExp number, first 60 AAs:  0.04212\nTotal prob of N-in:   0.00762\noutside   1   245\nTMhelix   246   268\ninside   269   274

29220	GCF_000371785.1_ASM37178v1	Actinopolyspora halophila DSM 43834	Terrabacteria group; Actinobacteria; Actinobacteria; Actinopolysporales; Actinopolysporaceae; Actinopolyspora; Actinopolyspora halophila
MTLHRAMRHRRTPMRRRWVQAGLIGVILAALTVVIIGVASSRSPDSENRAYELEQQLRCP	WP_085945762.1
cytochrome C biosynthesis protein [Actinopolyspora halophila]	Length: 174\nNumber of predicted TMHs:  2\nExp number of AAs in TMHs: 41.26669\nExp number, first 60 AAs:  20.59755\nTotal prob of N-in:   0.99999\nPOSSIBLE N-term signal sequence\ninside   1   20\nTMhelix   21   40\noutside   41   117\nTMhelix   118   137\ninside   138   174

29221	GCF_002108035.1_Bado703Bv1	Bifidobacterium adolescentis	Terrabacteria group; Actinobacteria; Actinobacteria; Bifidobacteriales; Bifidobacteriaceae; Bifidobacterium
MNGIFTTMDGDKGARRRGIISAVAAAAMLLPLAFAPTAMAADPDYPGGIKGEYNPLGINA	WP_085346658.1
hypothetical protein [Bifidobacterium adolescentis]	Length: 818\nNumber of predicted TMHs:  2\nExp number of AAs in TMHs: 44.11243\nExp number, first 60 AAs:  21.75671\nTotal prob of N-in:   0.98446\nPOSSIBLE N-term signal sequence\ninside   1   18\nTMhelix   19   41\noutside   42   792\nTMhelix   793   815\ninside   816   818

29222	GCF_001417815.1_ASM141781v1	Bifidobacterium aesculapii	Terrabacteria group; Actinobacteria; Actinobacteria; Bifidobacteriales; Bifidobacteriaceae; Bifidobacterium
MKRRLRTPRRAVVAAAALTLAAAPLSACGEPPTTVMQPEGPTIAIGVAADEPGLSRWHDG	WP_055427006.1
hypothetical protein [Bifidobacterium aesculapii]	Length: 279\nNumber of predicted TMHs:  1\nExp number of AAs in TMHs: 15.7199\nExp number, first 60 AAs:  1.39298\nTotal prob of N-in:   0.82848\ninside   1   229\nTMhelix   230   249\noutside   250   279

29223	GCF_002075855.1_Bbif1899B	Bifidobacterium catenulatum	Terrabacteria group; Actinobacteria; Actinobacteria; Bifidobacteriales; Bifidobacteriaceae; Bifidobacterium
MNGIFTTMDGDKGARRRGIISAVAAAAMLLPLAFAPTAMAADPDYPGGIKGEYNPLGINA	WP_080788544.1
hypothetical protein [Bifidobacterium catenulatum]	Length: 823\nNumber of predicted TMHs:  2\nExp number of AAs in TMHs: 44.11243\nExp number, first 60 AAs:  21.75671\nTotal prob of N-in:   0.98446\nPOSSIBLE N-term signal sequence\ninside   1   18\nTMhelix   19   41\noutside   42   797\nTMhelix   798   820\ninside   821   823

29224	GCF_000741135.1_Bifcho	Bifidobacterium choerinum	Terrabacteria group; Actinobacteria; Actinobacteria; Bifidobacteriales; Bifidobacteriaceae; Bifidobacterium
MTTGREQMMGRRERYCSARHGALFAPRIRRDLDRRDPVTAAFAIAGWACYIIAALLLAWV	WP_024540489.1	class C sortase [Bifidobacterium choerinum]	Length: 334\nNumber of predicted TMHs:  2\nExp number of AAs in TMHs:

45.6432699999999\nExp number, first 60 AAs: 21.36985\nTotal prob of N-in: 0.96879\nPOSSIBLE N-term signal sequence\ninside 1 37\nTMhelix 38 60\noutside 61 294\nTMhelix 295 317\ninside 318 334

29225 GCF_000738005.1_ASM73800v1 Bifidobacterium crudilactis Terrabacteria group; Actinobacteria; Actinobacteria; Bifidobacteriales; Bifidobacteriaceae; Bifidobacterium
MGRLYAFVAFFSVSRGSESRRGVLAGIAFALSILMLLTGVMVALPAEQAEADSLELGPDG WP_084674512.1 hypothetical protein [Bifidobacterium crudilactis] Length: 1013\nNumber of predicted TMHs: 2\nExp number of AAs in TMHs: 44.9151299999999\nExp number, first 60 AAs: 22.50662\nTotal prob of N-in: 0.99062\nPOSSIBLE N-term signal sequence\ninside 1 21\nTMhelix 22 44\noutside 45 973\nTMhelix 974 996\ninside 997 1013

29226 GCF_000741705.1_Bifpsy Bifidobacterium psychraerophilum Terrabacteria group; Actinobacteria; Actinobacteria; Bifidobacteriales; Bifidobacteriaceae; Bifidobacterium
MDVKRRRFISRTAAVLVAAAMTMTAGLVTTSAQAATGSDASITGGMPVVITELAVKTSNG WP_033496003.1 hypothetical protein [Bifidobacterium psychraerophilum] Length: 1740\nNumber of predicted TMHs: 2\nExp number of AAs in TMHs: 43.60284\nExp number, first 60 AAs: 21.86564\nTotal prob of N-in: 0.98168\nPOSSIBLE N-term signal sequence\ninside 1 12\nTMhelix 13 35\noutside 36 1712\nTMhelix 1713 1735\ninside 1736 1740

29227 GCF_000741335.1_Bifpul Bifidobacterium pullorum Terrabacteria group; Actinobacteria; Actinobacteria; Bifidobacteriales; Bifidobacteriaceae; Bifidobacterium
MRKVMDTLFNTTDERGASKRRGIVAALAAAAMLLPLAFSPTAMAADPDYPGGIKGEYNPL WP_081883253.1 hypothetical protein [Bifidobacterium pullorum] Length: 847\nNumber of predicted TMHs: 2\nExp number of AAs in TMHs: 44.21133\nExp number, first 60 AAs: 21.85569\nTotal prob of N-in: 0.98297\nPOSSIBLE N-term signal sequence\ninside 1 20\nTMhelix 21 43\noutside 44 821\nTMhelix 822 844\ninside 845 847

29228 GCF_000424225.1_ASM42422v1 Bifidobacterium sp. AGR2158 Terrabacteria group; Actinobacteria; Actinobacteria; Bifidobacteriales; Bifidobacteriaceae; Bifidobacterium
MLAQGRSPIPMHSNDDARRAPIWRRVTAGVGAMVMCASLGVAGVHTAFADTTSSTAVATA WP_081664073.1 carbon-nitrogen hydrolase family protein [Bifidobacterium sp. AGR2158] Length: 951\nNumber of predicted TMHs: 1\nExp number of AAs in TMHs: 38.4570299999999\nExp number, first 60 AAs: 14.43514\nTotal prob of N-in: 0.60774\nPOSSIBLE N-term signal sequence\noutside 1 921\nTMhelix 922 944\ninside 945 951

29229 GCF_000424225.1_ASM42422v1 Bifidobacterium sp. AGR2158 Terrabacteria group; Actinobacteria; Actinobacteria; Bifidobacteriales; Bifidobacteriaceae; Bifidobacterium
MTTGREQMMGRRERHCSARHGAPFAPRIRRDLDRRDPVTAAFAIAGWVCYIIAALLLAWV WP_051198764.1 class C sortase [Bifidobacterium sp. AGR2158] Length: 334\nNumber of predicted TMHs: 2\nExp number of AAs in TMHs: 45.7468799999999\nExp number, first 60 AAs: 21.33837\nTotal prob of N-in: 0.97165\nPOSSIBLE N-term signal sequence\ninside 1 37\nTMhelix 38 60\noutside 61 294\nTMhelix 295 317\ninside 318 334

29230 GCF_000741785.1_Bifste Bifidobacterium stellenboschense Terrabacteria group; Actinobacteria; Actinobacteria; Bifidobacteriales; Bifidobacteriaceae; Bifidobacterium
MLSRPPTSDSRESGAESAKNLRKGTKGTRMTASSRRAAIATFAAVAALGAGLVAPTAYAA WP_084686092.1 hypothetical protein [Bifidobacterium stellenboschense] Length: 623\nNumber of predicted TMHs: 2\nExp number of AAs in TMHs: 44.08332\nExp number, first 60 AAs: 20.98507\nTotal prob of N-in: 0.91631\nPOSSIBLE N-term signal sequence\ninside 1 37\nTMhelix 38 60\noutside 61 595\nTMhelix 596 618\ninside 619 623

29231 GCF_000741495.1_Bifthelum Bifidobacterium thermophilum Terrabacteria group; Actinobacteria; Actinobacteria; Bifidobacteriales; Bifidobacteriaceae; Bifidobacterium
MPGFSSRKGNEVKQPSRRGIIAGLSAALAGALCLTAVPGAFAADSPDVSALSVHGGAQRI WP_081885647.1 alkaline phosphatase [Bifidobacterium thermophilum] Length: 688\nNumber of predicted TMHs: 2\nExp number of AAs in TMHs: 43.64897\nExp number, first 60 AAs: 21.97014\nTotal prob of N-in: 0.96418\nPOSSIBLE N-term signal sequence\ninside 1 19\nTMhelix 20 42\noutside 43 657\nTMhelix 658 680\ninside 681 688

29232 GCF_001263945.1_ASM126394v1 Bifidobacterium animalis subsp. animalis MCC 0483 Terrabacteria group; Actinobacteria; Actinobacteria; Bifidobacteriales; Bifidobacteriaceae; Bifidobacterium; Bifidobacterium animalis; Bifidobacterium animalis subsp. animalis
MVRRRTIVLRALWGAPVVALLAMIIMIMTPRANAVENAFAINAPRGSVAVWIDVESNEAQ WP_052826618.1 hypothetical protein [Bifidobacterium animalis] Length: 235\nNumber of predicted TMHs: 2\nExp number of AAs in TMHs: 45.46195\nExp number, first 60 AAs: 22.60568\nTotal prob of N-in: 0.99672\nPOSSIBLE N-term signal sequence\ninside 1 6\nTMhelix 7 29\noutside 30 202\nTMhelix 203 225\ninside 226 235

29233 GCF_000741165.1_Bifbia Bifidobacterium biavatii DSM 23969 Terrabacteria group; Actinobacteria; Actinobacteria; Bifidobacteriales; Bifidobacteriaceae; Bifidobacterium; Bifidobacterium biavatii
MSKEERRGSAVRGWPRRAAVIVAGLTAVGMLAGPVTAASADQPSTADTTQVLVPTAGYRL WP_051924085.1
adhesin isopeptide-forming adherence domain-containing protein [Bifidobacterium biavatii] Length: 1278\nNumber of predicted TMHs: 2\nExp number of AAs in TMHs: 39.8290299999999\nExp number, first 60 AAs: 18.22633\nTotal prob of N-in: 0.88418\nPOSSIBLE N-term signal sequence\ninside 1 12\nTMhelix 13 35\noutside 36 1207\nTMhelix 1208 1230\ninside 1231 1278

29234 GCF_000741165.1_Bifbia Bifidobacterium biavatii DSM 23969 Terrabacteria group; Actinobacteria; Actinobacteria; Bifidobacteriales; Bifidobacteriaceae; Bifidobacterium; Bifidobacterium biavatii
MRFGLNRRKDRSKNGAKPQRFHTFDRRKTVASLVVAALSLTCAIPAQAQNIDISSIVNST WP_051923592.1 cell surface protein [Bifidobacterium biavatii] Length: 2888\nNumber of predicted TMHs: 1\nExp number of AAs in TMHs: 32.9665099999999\nExp number, first 60 AAs: 10.97572\nTotal prob of N-in: 0.54754\nPOSSIBLE N-term signal sequence\noutside 1 2854\nTMhelix 2855 2874\ninside 2875 2888

29235 GCF_000771645.1_DSM-23969 Bifidobacterium biavatii DSM 23969 Terrabacteria group; Actinobacteria; Actinobacteria; Bifidobacteriales; Bifidobacteriaceae; Bifidobacterium; Bifidobacterium biavatii
MRFGLNRRKDRSKNGAKPQRFHTFDRRKTVASLVVAALSLTCAIPAQAQNIDISSIVNST WP_051923592.1 cell surface protein [Bifidobacterium biavatii] Length: 2888\nNumber of predicted TMHs: 1\nExp number of AAs in TMHs: 32.9665099999999\nExp number, first 60 AAs: 10.97572\nTotal prob of N-in: 0.54754\nPOSSIBLE N-term signal sequence\noutside 1 2854\nTMhelix 2855 2874\ninside 2875 2888

29236 GCF_000771645.1_DSM-23969 Bifidobacterium biavatii DSM 23969 Terrabacteria group; Actinobacteria; Actinobacteria; Bifidobacteriales; Bifidobacteriaceae; Bifidobacterium; Bifidobacterium biavatii
MSKEERRGSAVRGWPRRAAVIVAGLTAVGMLAGPVTAASADQPSTADTTQVLVPTAGYRL WP_051924085.1
adhesin isopeptide-forming adherence domain-containing protein [Bifidobacterium biavatii] Length: 1278\nNumber of predicted TMHs: 2\nExp number of AAs in TMHs: 39.8290299999999\nExp number, first 60 AAs: 18.22633\nTotal prob of N-in: 0.88418\nPOSSIBLE N-term signal sequence\ninside 1 12\nTMhelix 13 35\noutside 36 1207\nTMhelix 1208 1230\ninside 1231 1278

29237 GCF_000771085.1_DSM-19703 Bifidobacterium bombi DSM 19703 Terrabacteria group; Actinobacteria; Actinobacteria; Bifidobacteriales; Bifidobacteriaceae; Bifidobacterium; Bifidobacterium bombi
MPAMANKRGNRRTPIRSWLKWLVALVAALSTLLIGGVPAATAIQPGKYQFLGLSQSVGGV WP_044087456.1
hypothetical protein [Bifidobacterium bombi] Length: 2571\nNumber of predicted TMHs: 2\nExp number of AAs in TMHs: 45.45986\nExp number, first 60 AAs: 22.89119\nTotal prob of N-in: 0.99996\nPOSSIBLE N-term signal sequence\ninside 1 20\nTMhelix 21 43\noutside 44 2529\nTMhelix 2530 2552\ninside 2553 2571

29238 GCF_000771665.1_DSM-23973 Bifidobacterium callitrichos DSM 23973 Terrabacteria group; Actinobacteria; Actinobacteria; Bifidobacteriales; Bifidobacteriaceae; Bifidobacterium; Bifidobacterium callitrichos
MARRAVVSAVAALGACGLALAVPAGAMAADSSSSSLGVAGVSGESSVTAPASGSTGSGDA WP_052119059.1
hypothetical protein [Bifidobacterium callitrichos] Length: 320\nNumber of predicted TMHs: 2\nExp number of AAs in TMHs: 44.45868\nExp number, first 60 AAs: 22.35804\nTotal prob of N-in: 0.99011\nPOSSIBLE N-term signal sequence\ninside 1 6\nTMhelix 7 29\noutside 30 285\nTMhelix 286 308\ninside 309 320

29239 GCF_000741175.1_Bifcal Bifidobacterium callitrichos DSM 23973 Terrabacteria group; Actinobacteria; Actinobacteria; Bifidobacteriales; Bifidobacteriaceae; Bifidobacterium; Bifidobacterium callitrichos
MARRAVVSAVAALGACGLALAVPAGAMAADSSSSSLGVAGVSGESSVTAPASGSTGSGDA WP_052119059.1
hypothetical protein [Bifidobacterium callitrichos] Length: 320\nNumber of predicted TMHs: 2\nExp number of AAs in TMHs: 44.45868\nExp number, first 60 AAs: 22.35804\nTotal prob of N-in: 0.99011\nPOSSIBLE N-term signal sequence\ninside 1 6\nTMhelix 7 29\noutside 30 285\nTMhelix 286 308\ninside 309 320

29240 GCF_000771425.1_DSM-20434 Bifidobacterium choerinum DSM 20434 Terrabacteria group; Actinobacteria; Actinobacteria; Bifidobacteriales; Bifidobacteriaceae; Bifidobacterium; Bifidobacterium choerinum
MTTGREQMMGRRERYCSARHGALFAPRIRRDLDRRDPVTAAFAIAGWACYIIAALLLAWV WP_024540489.1 class C sortase [Bifidobacterium choerinum] Length: 334\nNumber of predicted TMHs: 2\nExp number of AAs in TMHs: 45.6432699999999\nExp number, first 60 AAs: 21.36985\nTotal prob of N-in: 0.96879\nPOSSIBLE N-term signal sequence\ninside 1 37\nTMhelix 38 60\noutside 61 294\nTMhelix 295 317\ninside 318 334

29241 GCF_000484675.1_ASM48467v1 Bifidobacterium choerinum DSM 20434 Terrabacteria group; Actinobacteria; Actinobacteria; Bifidobacteriales; Bifidobacteriaceae; Bifidobacterium; Bifidobacterium choerinum
MTTGREQMMGRRERYCSARHGALFAPRIRRDLDRRDPVTAAFAIAGWACYIIAALLLAWV WP_024540489.1 class

C sortase [Bifidobacterium choerinum] Length: 334\nNumber of predicted TMHs: 2\nExp number of AAs in TMHs: 45.6432699999999\nExp number, first 60 AAs: 21.36985\nTotal prob of N-in: 0.96879\nPOSSIBLE N-term signal sequence\ninside 1 37\nTMhelix 38 60\noutside 61 294\nTMhelix 295 317\ninside 318 334

29242 GCF_000741205.1_Bifgalcum Bifidobacterium gallicum DSM 20093 = LMG 11596 Terrabacteria group; Actinobacteria; Actinobacteria; Bifidobacteriales; Bifidobacteriaceae; Bifidobacterium; Bifidobacterium gallicum MTKTTMARRAWLVTAAMMSTAMAFCGLTAPTTFADESTGSIKSGQNSVTVTYEHEDKPVA WP_006294973.1 hypothetical protein [Bifidobacterium gallicum] Length: 390\nNumber of predicted TMHs: 2\nExp number of AAs in TMHs: 43.68915\nExp number, first 60 AAs: 21.79015\nTotal prob of N-in: 0.96462\nPOSSIBLE N-term signal sequence\ninside 1 11\nTMhelix 12 34\noutside 35 357\nTMhelix 358 380\ninside 381 390

29243 GCF_000173375.1_ASM17337v1 Bifidobacterium gallicum DSM 20093 = LMG 11596 Terrabacteria group; Actinobacteria; Actinobacteria; Bifidobacteriales; Bifidobacteriaceae; Bifidobacterium; Bifidobacterium gallicum MTKTTMARRAWLVTAAMMSTAMAFCGLTAPTTFADESTGSIKSGQNSVTVTYEHEDKPVA WP_006294973.1 hypothetical protein [Bifidobacterium gallicum] Length: 390\nNumber of predicted TMHs: 2\nExp number of AAs in TMHs: 43.68915\nExp number, first 60 AAs: 21.79015\nTotal prob of N-in: 0.96462\nPOSSIBLE N-term signal sequence\ninside 1 11\nTMhelix 12 34\noutside 35 357\nTMhelix 358 380\ninside 381 390

29244 GCF_000771165.1_DSM-20093 Bifidobacterium gallicum DSM 20093 = LMG 11596 Terrabacteria group; Actinobacteria; Actinobacteria; Bifidobacteriales; Bifidobacteriaceae; Bifidobacterium; Bifidobacterium gallicum MTKTTMARRAWLVTAAMMSTAMAFCGLTAPTTFADESTGSIKSGQNSVTVTYEHEDKPVA WP_006294973.1 hypothetical protein [Bifidobacterium gallicum] Length: 390\nNumber of predicted TMHs: 2\nExp number of AAs in TMHs: 43.68915\nExp number, first 60 AAs: 21.79015\nTotal prob of N-in: 0.96462\nPOSSIBLE N-term signal sequence\ninside 1 11\nTMhelix 12 34\noutside 35 357\nTMhelix 358 380\ninside 381 390

29245 GCF_000771545.1_DSM-21854 Bifidobacterium kashiwanohense JCM 15439 = DSM 21854 Terrabacteria group; Actinobacteria; Actinobacteria; Bifidobacteriales; Bifidobacteriaceae; Bifidobacterium; Bifidobacterium kashiwanohense MNGIFTTMDGDKGARRRGIVSAVAAAAMLLPLAFAPTAMAADPDYPGGIKGEYNPLGINA WP_033501893.1 hypothetical protein [Bifidobacterium kashiwanohense] Length: 818\nNumber of predicted TMHs: 2\nExp number of AAs in TMHs: 44.04916\nExp number, first 60 AAs: 21.69345\nTotal prob of N-in: 0.98135\nPOSSIBLE N-term signal sequence\ninside 1 18\nTMhelix 19 41\noutside 42 792\nTMhelix 793 815\ninside 816 818

29246 GCF_001042615.1_ASM104261v1 Bifidobacterium kashiwanohense JCM 15439 = DSM 21854 Terrabacteria group; Actinobacteria; Actinobacteria; Bifidobacteriales; Bifidobacteriaceae; Bifidobacterium; Bifidobacterium kashiwanohense MNGIFTTMDGDKGARRRGIVSAVAAAAMLLPLAFAPTAMAADPDYPGGIKGEYNPLGINA WP_033501893.1 hypothetical protein [Bifidobacterium kashiwanohense] Length: 818\nNumber of predicted TMHs: 2\nExp number of AAs in TMHs: 44.04916\nExp number, first 60 AAs: 21.69345\nTotal prob of N-in: 0.98135\nPOSSIBLE N-term signal sequence\ninside 1 18\nTMhelix 19 41\noutside 42 792\nTMhelix 793 815\ninside 816 818

29247 GCF_000800475.2_ASM80047v2 Bifidobacterium pseudolongum PV8-2 Terrabacteria group; Actinobacteria; Actinobacteria; Bifidobacteriales; Bifidobacteriaceae; Bifidobacterium; Bifidobacterium pseudolongum MDASADARDDRRTAMIALTGPVWMVVAACGVAAAVWWLPAPHAATVRLQDVRAGAPHGDD WP_081277489.1 hypothetical protein [Bifidobacterium pseudolongum] Length: 211\nNumber of predicted TMHs: 2\nExp number of AAs in TMHs: 50.92317\nExp number, first 60 AAs: 22.6514\nTotal prob of N-in: 0.30904\nPOSSIBLE N-term signal sequence\noutside 1 14\nTMhelix 15 37\ninside 38 183\nTMhelix 184 206\noutside 207 211

29248 GCF_000741295.1_Bifpse_sub.glo Bifidobacterium pseudolongum subsp. globosum Terrabacteria group; Actinobacteria; Actinobacteria; Bifidobacteriales; Bifidobacteriaceae; Bifidobacterium; Bifidobacterium pseudolongum MDASADDRDDRRTAMIALTGPVWMVVAACGAAAAVWWLPAPHAATVRLQDARAGAPHGDD WP_080699286.1 hypothetical protein [Bifidobacterium pseudolongum] Length: 211\nNumber of predicted TMHs: 2\nExp number of AAs in TMHs: 49.81793\nExp number, first 60 AAs: 22.70895\nTotal prob of N-in: 0.23760\nPOSSIBLE N-term signal sequence\noutside 1 14\nTMhelix 15 37\ninside 38 183\nTMhelix 184 206\noutside 207 211

29249 GCF_000771145.1_DSM-20092 Bifidobacterium pseudolongum subsp. globosum DSM 20092 Terrabacteria group; Actinobacteria; Actinobacteria; Bifidobacteriales; Bifidobacteriaceae; Bifidobacterium; Bifidobacterium pseudolongum; Bifidobacterium pseudolongum subsp. globosum MDASADDRDDRRTAMIALTGPVWMVVAACGAAAAVWWLPAPHAATVRLQDARAGAPHGDD WP_080699286.1 hypothetical protein [Bifidobacterium pseudolongum] Length: 211\nNumber of predicted TMHs: 2\nExp number of AAs in TMHs: 49.81793\nExp number, first 60 AAs:  22.70895\nTotal prob of N-in:   0.23760\nPOSSIBLE N-term signal sequence\noutside   1   14\nTMhelix   15   37\ninside   38   183\nTMhelix   184   206\noutside   207   211

29250 GCF_000687595.1_ASM68759v1 Bifidobacterium pseudolongum subsp. globosum DSM 20092 Terrabacteria group; Actinobacteria; Actinobacteria; Bifidobacteriales; Bifidobacteriaceae; Bifidobacterium; Bifidobacterium pseudolongum; Bifidobacterium pseudolongum subsp. globosum MDASADDRDDRRTAMIALTGPVWMVVAACGAAAAVWWLPAPHAATVRLQDARAGAPHGDD WP_080699286.1 hypothetical protein [Bifidobacterium pseudolongum] Length: 211\nNumber of predicted TMHs:  2\nExp number of AAs in TMHs: 49.81793\nExp number, first 60 AAs:  22.70895\nTotal prob of N-in:   0.23760\nPOSSIBLE N-term signal sequence\noutside   1   14\nTMhelix   15   37\ninside   38   183\nTMhelix   184   206\noutside   207   211

29251 GCF_000771225.1_DSM-20099 Bifidobacterium pseudolongum subsp. pseudolongum DSM 20099 Terrabacteria group; Actinobacteria; Actinobacteria; Bifidobacteriales; Bifidobacteriaceae; Bifidobacterium; Bifidobacterium pseudolongum; Bifidobacterium pseudolongum subsp. pseudolongum MPRELRGALGRPDRENRAKGEEPMRMKHGSDRRPTISVKPRLLAAFTAVAMMFATAVPSL WP_080745829.1 hypothetical protein [Bifidobacterium pseudolongum] Length: 1475\nNumber of predicted TMHs:  1\nExp number of AAs in TMHs: 27.34236\nExp number, first 60 AAs:  5.58909\nTotal prob of N-in:   0.29206\noutside   1   1447\nTMhelix   1448  1470\ninside   1471  1475

29252 GCF_000771565.1_DSM-22366 Bifidobacterium psychraerophilum DSM 22366 Terrabacteria group; Actinobacteria; Actinobacteria; Bifidobacteriales; Bifidobacteriaceae; Bifidobacterium; Bifidobacterium psychraerophilum MDVKRRRFISRTAAVLVAAAMTMTAGLVTTSAQAATGSDASITGGMPVVITELAVKTSNG WP_033496003.1 hypothetical protein [Bifidobacterium psychraerophilum] Length: 1740\nNumber of predicted TMHs:  2\nExp number of AAs in TMHs: 43.60284\nExp number, first 60 AAs:  21.86564\nTotal prob of N-in:   0.98168\nPOSSIBLE N-term signal sequence\ninside   1   12\nTMhelix   13   35\noutside   36  1712\nTMhelix   1713  1735\ninside   1736  1740

29253 GCF_000263595.1_ASM26359v1 Gardnerella vaginalis 1500E Terrabacteria group; Actinobacteria; Actinobacteria; Bifidobacteriales; Bifidobacteriaceae; Gardnerella; Gardnerella vaginalis MSMQQRKRDFRKCTRRALFAFFVASATVLATFAIAPIAPSNASSEISNKISTKTENKAET WP_004128748.1 hypothetical protein [Gardnerella vaginalis] Length: 958\nNumber of predicted TMHs:  2\nExp number of AAs in TMHs: 44.81562\nExp number, first 60 AAs:  21.9107\nTotal prob of N-in:   0.99979\nPOSSIBLE N-term signal sequence\ninside   1   16\nTMhelix   17   39\noutside   40   910\nTMhelix   911   933\ninside   934   958

29254 GCF_000263655.1_ASM26365v1 Gardnerella vaginalis 6119V5 Terrabacteria group; Actinobacteria; Actinobacteria; Bifidobacteriales; Bifidobacteriaceae; Gardnerella; Gardnerella vaginalis MSMQQRKRDFRKCTRRALFAFFVAAATVLATFAIAPIAPSNASSETSTKAATETKADTKA WP_004136222.1 hypothetical protein [Gardnerella vaginalis] Length: 976\nNumber of predicted TMHs:  2\nExp number of AAs in TMHs: 44.45261\nExp number, first 60 AAs:  21.88622\nTotal prob of N-in:   0.99990\nPOSSIBLE N-term signal sequence\ninside   1   16\nTMhelix   17   39\noutside   40   904\nTMhelix   905   927\ninside   928   976

29255 GCF_000717315.1_ASM71731v1 Actinospica acidiphila Terrabacteria group; Actinobacteria; Actinobacteria; Catenulisporales; Actinospicaceae; Actinospica MLRGVENHLSPRTAAFFDLDKTVIAKSSTLTFSKSFYQGGLINRRAVLRTAYSQFVFLAG WP_033274879.1 MULTISPECIES: inhibition of morphological differentiation protein [Actinobacteria] Length: 277\nNumber of predicted TMHs:  1\nExp number of AAs in TMHs: 21.81221\nExp number, first 60 AAs:  0.02078\nTotal prob of N-in:   0.00383\noutside   1   247\nTMhelix   248   270\ninside   271   277

29256 GCF_000717315.1_ASM71731v1 Actinospica acidiphila Terrabacteria group; Actinobacteria; Actinobacteria; Catenulisporales; Actinospicaceae; Actinospica MRHARRRVVRRVTRLAAVGGLLLGGTMVTHAVASEPGTPDAVPFSTASPADASGPGAGLV WP_033275253.1 protease [Actinospica acidiphila] Length: 457\nNumber of predicted TMHs:  2\nExp number of AAs in TMHs: 38.82844\nExp number, first 60 AAs:  21.8309\nTotal prob of N-in:   0.97535\nPOSSIBLE N-term signal sequence\ninside   1   11\nTMhelix   12   34\noutside   35   418\nTMhelix   419   441\ninside   442   457

29257 GCF_000504285.1_ASM50428v1 Actinospica robiniae DSM 44927 Terrabacteria group; Actinobacteria; Actinobacteria; Catenulisporales; Actinospicaceae; Actinospica; Actinospica robiniae MARLWIRPPRARRPATGSRRSTVSTLKKVGAAAASTALALGTGFAAPTAAAGGTVLADAS WP_084316449.1 hypothetical protein [Actinospica robiniae] Length: 485\nNumber of predicted TMHs:  1\nExp number of AAs in TMHs: 30.83547\nExp number, first 60 AAs:  8.01693\nTotal prob of N-in:   0.36382\noutside   1   450\nTMhelix   451   473\ninside   474   485

29258 GCF_000504285.1_ASM50428v1 Actinospica robiniae DSM 44927 Terrabacteria group; Actinobacteria; Actinobacteria; Catenulisporales; Actinospicaceae; Actinospica; Actinospica robiniae MATRRTLVTSVCAAVLAGCALSFAAPAASASSVELKITESWSGTLPQRVEPGQTVTLLTS WP_034263843.1 hypothetical protein [Actinospica robiniae] Length: 296\nNumber of predicted TMHs: 2\nExp number of AAs in TMHs: 42.79354\nExp number, first 60 AAs: 21.45553\nTotal prob of N-in: 0.97260\nPOSSIBLE N-term signal sequence\ninside 1 6\nTMhelix 7 29\noutside 30 265\nTMhelix 266 288\ninside 289 296

29259 GCF_000504285.1_ASM50428v1 Actinospica robiniae DSM 44927 Terrabacteria group; Actinobacteria; Actinobacteria; Catenulisporales; Actinospicaceae; Actinospica; Actinospica robiniae MRTAAFFDLDKTIIAKSSTLAFGRSFYQGGLINRRAVLKTAYAQFVYLAGGADHDQMERM WP_034266494.1 inhibition of morphological differentiation protein [Actinospica robiniae] Length: 266\nNumber of predicted TMHs: 1\nExp number of AAs in TMHs: 20.93182\nExp number, first 60 AAs: 0.11197\nTotal prob of N-in: 0.05509\noutside 1 237\nTMhelix 238 257\ninside 258 266

29260 GCF_000024025.1_ASM2402v1 Catenulispora acidiphila DSM 44928 Terrabacteria group; Actinobacteria; Actinobacteria; Catenulisporales; Catenulisporaceae; Catenulispora; Catenulispora acidiphila MSTKLTSGARRTAVLLATAGVALIAAASAAEAHVTVNPNTEPQGGYAKVSFRVPNEEANA WP_012787270.1 nuclear export factor GLE1 [Catenulispora acidiphila] Length: 253\nNumber of predicted TMHs: 2\nExp number of AAs in TMHs: 41.51837\nExp number, first 60 AAs: 20.32248\nTotal prob of N-in: 0.96276\nPOSSIBLE N-term signal sequence\ninside 1 12\nTMhelix 13 35\noutside 36 222\nTMhelix 223 245\ninside 246 253

29261 GCF_000024025.1_ASM2402v1 Catenulispora acidiphila DSM 44928 Terrabacteria group; Actinobacteria; Actinobacteria; Catenulisporales; Catenulisporaceae; Catenulispora; Catenulispora acidiphila MDEHADARADGAGRPAAFFDLDKTIIAKSSTLAFGRSFYNGGLINRRTVLRTAYAQFVYL WP_012784304.1 haloacid dehalogenase [Catenulispora acidiphila] Length: 281\nNumber of predicted TMHs: 1\nExp number of AAs in TMHs: 20.07884\nExp number, first 60 AAs: 0.13227\nTotal prob of N-in: 0.14637\noutside 1 247\nTMhelix 248 270\ninside 271 281

29262 GCF_000024025.1_ASM2402v1 Catenulispora acidiphila DSM 44928 Terrabacteria group; Actinobacteria; Actinobacteria; Catenulisporales; Catenulisporaceae; Catenulispora; Catenulispora acidiphila MPNPNPRRRAFALAAALAGAVALSAPAAAASGPAQARARSSFSQTSDAQAAPASAAASGK WP_015793958.1 ABC transporter substrate-binding protein [Catenulispora acidiphila] Length: 629\nNumber of predicted TMHs: 2\nExp number of AAs in TMHs: 44.9360299999999\nExp number, first 60 AAs: 18.86601\nTotal prob of N-in: 0.71812\nPOSSIBLE N-term signal sequence\ninside 1 11\nTMhelix 12 34\noutside 35 594\nTMhelix 595 617\ninside 618 629

29263 GCF_900156035.1_IMG-taxon_2681813557_annotated_assembly Corynebacterium afermentans Terrabacteria group; Actinobacteria; Actinobacteria; Corynebacteriales; Corynebacteriaceae; Corynebacterium MRPAPFSPRTSRRAALLAVPAALLLAAPIAPVAHAQLPDGMSPEEVQSMIPGEISVPAGE WP_082855643.1 hypothetical protein [Corynebacterium afermentans] Length: 376\nNumber of predicted TMHs: 2\nExp number of AAs in TMHs: 39.93319\nExp number, first 60 AAs: 17.68082\nTotal prob of N-in: 0.80899\nPOSSIBLE N-term signal sequence\ninside 1 12\nTMhelix 13 35\noutside 36 346\nTMhelix 347 369\ninside 370 376

29264 GCF_000732945.1_ASM73294v1 Corynebacterium atypicum Terrabacteria group; Actinobacteria; Actinobacteria; Corynebacteriales; Corynebacteriaceae; Corynebacterium MADQHPSERCYGGRGRRGLFGFAIAVLAGVLSTVLAIPQAAAHDSVIDADPGVGAAVAEF WP_051866792.1 hypothetical protein [Corynebacterium atypicum] Length: 218\nNumber of predicted TMHs: 2\nExp number of AAs in TMHs: 45.38686\nExp number, first 60 AAs: 22.51125\nTotal prob of N-in: 0.99927\nPOSSIBLE N-term signal sequence\ninside 1 19\nTMhelix 20 42\noutside 43 177\nTMhelix 178 200\ninside 201 218

29265 GCF_900078305.1_PRJEB13138 Corynebacterium bouchesdurhonense Terrabacteria group; Actinobacteria; Actinobacteria; Corynebacteriales; Corynebacteriaceae; Corynebacterium MFESKAARRSTLIAAALSTALIAPSLAPAATAQSSNADLPSSSTGSSNDSATSKPSNAAV WP_066523952.1 hypothetical protein [Corynebacterium bouchesdurhonense] Length: 293\nNumber of predicted TMHs: 2\nExp number of AAs in TMHs: 44.09596\nExp number, first 60 AAs: 3.08509\nTotal prob of N-in: 0.83503\ninside 1 193\nTMhelix 194 216\noutside 217 269\nTMhelix 270 292\ninside 293 293

29266 GCF_000732145.1_ASM73214v1 Corynebacterium crenatum Terrabacteria group; Actinobacteria; Actinobacteria; Corynebacteriales; Corynebacteriaceae; Corynebacterium MGLPFKKAGPLSRRKAVFSALGAAALIGAALPTIPTAQAQTPTGYGFDATASIGEEPEFS WP_003856086.1 MULTISPECIES: hypothetical protein [Corynebacterium] Length: 919\nNumber of predicted TMHs: 1\nExp number of AAs in

TMHs: 33.37495\nExp number, first 60 AAs: 11.01947\nTotal prob of N-in: 0.50897\nPOSSIBLE N-term signal sequence\noutside 1 894\nTMhelix 895 917\ninside 918 919

29267 GCF_001021025.1_ASM102102v1 Corynebacterium epidermidicanis Terrabacteria group; Actinobacteria; Actinobacteria; Corynebacteriales; Corynebacteriaceae; Corynebacterium MFLPMSRRTLIATVTATSLIFGSLTAPAQAASFPGDVGRIILGAGADQTQSTVSWRAKGF WP_083984450.1 hypothetical protein [Corynebacterium epidermidicanis] Length: 618\nNumber of predicted TMHs: 1\nExp number of AAs in TMHs: 23.91348\nExp number, first 60 AAs: 2.34284\nTotal prob of N-in: 0.10858\noutside 1 580\nTMhelix 581 603\ninside 604 618

29268 GCF_001912755.1_ASM191275v1 Corynebacterium glutamicum Terrabacteria group; Actinobacteria; Actinobacteria; Corynebacteriales; Corynebacteriaceae; Corynebacterium MGLPFKKAGPLSRRKAVFSALGAAALIGAALPTIPTAQAQTPTGYGFDATASISEEPEFS WP_074508171.1 hypothetical protein [Corynebacterium glutamicum] Length: 919\nNumber of predicted TMHs: 1\nExp number of AAs in TMHs: 33.36375\nExp number, first 60 AAs: 11.01921\nTotal prob of N-in: 0.50896\nPOSSIBLE N-term signal sequence\noutside 1 894\nTMhelix 895 917\ninside 918 919

29269 GCF_001912735.1_ASM191273v1 Corynebacterium glutamicum Terrabacteria group; Actinobacteria; Actinobacteria; Corynebacteriales; Corynebacteriaceae; Corynebacterium MGLPFKKAGPLSRRKAVFSALGAAALIGAALPTIPTAQAQTPTGYGFDATASIGEEPEFS WP_003856086.1 MULTISPECIES: hypothetical protein [Corynebacterium] Length: 919\nNumber of predicted TMHs: 1\nExp number of AAs in TMHs: 33.37495\nExp number, first 60 AAs: 11.01947\nTotal prob of N-in: 0.50897\nPOSSIBLE N-term signal sequence\noutside 1 894\nTMhelix 895 917\ninside 918 919

29270 GCF_002003445.1_ASM200344v1 Corynebacterium glutamicum Terrabacteria group; Actinobacteria; Actinobacteria; Corynebacteriales; Corynebacteriaceae; Corynebacterium MGFPFKKAGPLSRRKAVFSALGAAALIGAALPAIPTAQAQTGTGYGFDATASISEEPKFT WP_077312225.1 hypothetical protein [Corynebacterium glutamicum] Length: 903\nNumber of predicted TMHs: 1\nExp number of AAs in TMHs: 36.67884\nExp number, first 60 AAs: 14.23033\nTotal prob of N-in: 0.65672\nPOSSIBLE N-term signal sequence\noutside 1 878\nTMhelix 879 901\ninside 902 903

29271 GCF_001912155.1_ASM191215v1 Corynebacterium glutamicum Terrabacteria group; Actinobacteria; Actinobacteria; Corynebacteriales; Corynebacteriaceae; Corynebacterium MGLPFKKAGPLSRRKAVFSALGAAALIGAALPTIPTAQAQTPTGYGFDATASIGEEPEFS WP_003856086.1 MULTISPECIES: hypothetical protein [Corynebacterium] Length: 919\nNumber of predicted TMHs: 1\nExp number of AAs in TMHs: 33.37495\nExp number, first 60 AAs: 11.01947\nTotal prob of N-in: 0.50897\nPOSSIBLE N-term signal sequence\noutside 1 894\nTMhelix 895 917\ninside 918 919

29272 GCF_000403725.1_GD7 Corynebacterium ihumii Terrabacteria group; Actinobacteria; Actinobacteria; Corynebacteriales; Corynebacteriaceae; Corynebacterium MTRSRRTAVVAAVLSAALVAPFTSPLQGGAVEASDFDTRYTTTNFWDVDHAIVQGLTLEP WP_034996857.1 hypothetical protein [Corynebacterium ihumii] Length: 748\nNumber of predicted TMHs: 2\nExp number of AAs in TMHs: 40.30832\nExp number, first 60 AAs: 18.45986\nTotal prob of N-in: 0.86830\nPOSSIBLE N-term signal sequence\ninside 1 6\nTMhelix 7 29\noutside 30 689\nTMhelix 690 712\ninside 713 748

29273 GCF_000403725.1_GD7 Corynebacterium ihumii Terrabacteria group; Actinobacteria; Actinobacteria; Corynebacteriales; Corynebacteriaceae; Corynebacterium MRPAPFSPRTSRRAALLAVPAALLLAAPIAPVAHAQLPDGMSPEEVQSMIPGEISVPAGE WP_051106395.1 hypothetical protein [Corynebacterium ihumii] Length: 382\nNumber of predicted TMHs: 2\nExp number of AAs in TMHs: 39.61497\nExp number, first 60 AAs: 17.67833\nTotal prob of N-in: 0.80886\nPOSSIBLE N-term signal sequence\ninside 1 12\nTMhelix 13 35\noutside 36 352\nTMhelix 353 375\ninside 376 382

29274 GCF_000739455.1_ASM73945v1 Corynebacterium imitans Terrabacteria group; Actinobacteria; Actinobacteria; Corynebacteriales; Corynebacteriaceae; Corynebacterium MHTVARRAAVFAAAAALAAPPVAGAAEVVQQGSQIVVADASSCAIAYNDRAENISYTAAH WP_038588689.1 hypothetical protein [Corynebacterium imitans] Length: 297\nNumber of predicted TMHs: 1\nExp number of AAs in TMHs: 24.4953\nExp number, first 60 AAs: 3.32174\nTotal prob of N-in: 0.68966\ninside 1 260\nTMhelix 261 283\noutside 284 297

29275 GCF_000739455.1_ASM73945v1 Corynebacterium imitans Terrabacteria group; Actinobacteria; Actinobacteria; Corynebacteriales; Corynebacteriaceae; Corynebacterium

MHTSSRRRALRSTTVASALALSLTVAAPGAGADEDTNGEVFGSGPDASAEFSEQYAAEHG WP_051904793.1 hypothetical protein [Corynebacterium imitans] Length: 480\nNumber of predicted TMHs: 1\nExp number of AAs in TMHs: 23.80377\nExp number, first 60 AAs: 2.23932\nTotal prob of N-in: 0.45982\noutside 1 444\nTMhelix 445 467\ninside 468 480

29276 GCF_000738185.1_Cj14566 Corynebacterium jeikeium Terrabacteria group; Actinobacteria; Actinobacteria; Corynebacteriales; Corynebacteriaceae; Corynebacterium MKLGNCSPLRLNVVSLEYEVPLIMFLRSRRAMIAVAAASSLVLTATAVPQSPFSPVAAQA WP_080719734.1 hypothetical protein [Corynebacterium jeikeium] Length: 960\nNumber of predicted TMHs: 1\nExp number of AAs in TMHs: 24.72671\nExp number, first 60 AAs: 3.72801\nTotal prob of N-in: 0.18657\noutside 1 931\nTMhelix 932 954\ninside 955 960

29277 GCF_000738325.1_Cj47446 Corynebacterium jeikeium Terrabacteria group; Actinobacteria; Actinobacteria; Corynebacteriales; Corynebacteriaceae; Corynebacterium MMHLRSRRAMIAVAASASMILTASAVPNSPVSPSVAQAATANKTQDTNLPLKCTLFAAGG WP_051894215.1 hypothetical protein [Corynebacterium jeikeium] Length: 929\nNumber of predicted TMHs: 1\nExp number of AAs in TMHs: 22.03769\nExp number, first 60 AAs: 1.0221\nTotal prob of N-in: 0.05103\noutside 1 900\nTMhelix 901 923\ninside 924 929

29278 GCF_000738305.1_Cj38002 Corynebacterium jeikeium Terrabacteria group; Actinobacteria; Actinobacteria; Corynebacteriales; Corynebacteriaceae; Corynebacterium MMFLRSRRAMIAVAAASSLVLTATAVPQSPFSPVAAQAAMANKTQDTTLPLKCNISAGGG WP_051894060.1 hypothetical protein [Corynebacterium jeikeium] Length: 927\nNumber of predicted TMHs: 1\nExp number of AAs in TMHs: 24.16114\nExp number, first 60 AAs: 2.97994\nTotal prob of N-in: 0.14205\noutside 1 898\nTMhelix 899 921\ninside 922 927

29279 GCF_000738195.1_Cj16348 Corynebacterium jeikeium Terrabacteria group; Actinobacteria; Actinobacteria; Corynebacteriales; Corynebacteriaceae; Corynebacterium MMFLRSRRAMIAVAAASSLVLTATAVPQSPFSPVAAQAATANKTQDTTLPLKCNISAGGG WP_035002726.1 hypothetical protein [Corynebacterium jeikeium] Length: 927\nNumber of predicted TMHs: 1\nExp number of AAs in TMHs: 22.82794\nExp number, first 60 AAs: 1.83154\nTotal prob of N-in: 0.09062\noutside 1 898\nTMhelix 899 921\ninside 922 927

29280 GCF_900100745.1_CJ Corynebacterium jeikeium Terrabacteria group; Actinobacteria; Actinobacteria; Corynebacteriales; Corynebacteriaceae; Corynebacterium MTSQGFATQGRRSLVAAFAISAGLLGSAHAAVGVADAARVAGLNDQPGVSRDYGADTLSI WP_071057585.1 adhesin [Corynebacterium jeikeium] Length: 279\nNumber of predicted TMHs: 2\nExp number of AAs in TMHs: 44.37943\nExp number, first 60 AAs: 21.64447\nTotal prob of N-in: 0.96175\nPOSSIBLE N-term signal sequence\ninside 1 12\nTMhelix 13 35\noutside 36 248\nTMhelix 249 271\ninside 272 279

29281 GCF_001059385.1_ASM105938v1 Corynebacterium jeikeium Terrabacteria group; Actinobacteria; Actinobacteria; Corynebacteriales; Corynebacteriaceae; Corynebacterium MLPAHSVPKAHATQPPRTRRAAMALLAFLTVLALVPTLAVDAQAKDKDSVGQHGGSGQRR WP_049049856.1 hypothetical protein [Corynebacterium jeikeium] Length: 652\nNumber of predicted TMHs: 2\nExp number of AAs in TMHs: 42.10599\nExp number, first 60 AAs: 19.64436\nTotal prob of N-in: 0.96249\nPOSSIBLE N-term signal sequence\ninside 1 20\nTMhelix 21 40\noutside 41 624\nTMhelix 625 647\ninside 648 652

29282 GCF_000738255.1_Cj21382 Corynebacterium jeikeium Terrabacteria group; Actinobacteria; Actinobacteria; Corynebacteriales; Corynebacteriaceae; Corynebacterium MMFLRSRRAMIAVAAASSLVLTATAVPQSPFSPVAAQAATANKTQDTNLDYRCNVFAGGG WP_051879522.1 hypothetical protein [Corynebacterium jeikeium] Length: 929\nNumber of predicted TMHs: 1\nExp number of AAs in TMHs: 22.86218\nExp number, first 60 AAs: 1.83778\nTotal prob of N-in: 0.09163\noutside 1 900\nTMhelix 901 923\ninside 924 929

29283 GCF_000738265.1_Cj37130 Corynebacterium jeikeium Terrabacteria group; Actinobacteria; Actinobacteria; Corynebacteriales; Corynebacteriaceae; Corynebacterium MMFLRSRRAMIAVAAASSLVLTATAVPQSPLSPVAARAATANKTQDTKLPIKCTVFAAGG WP_034981692.1 hypothetical protein [Corynebacterium jeikeium] Length: 938\nNumber of predicted TMHs: 1\nExp number of AAs in TMHs: 22.83202\nExp number, first 60 AAs: 1.81359\nTotal prob of N-in: 0.08948\noutside 1 909\nTMhelix 910 932\ninside 933 938

29284 GCF_000738355.1_Cj30952 Corynebacterium jeikeium Terrabacteria group; Actinobacteria; Actinobacteria; Corynebacteriales; Corynebacteriaceae; Corynebacterium
MSSHVSAVQGRRRLVTTIAISAGLLGGAHAAVGVADAAKVAGLNDQPGVSRDYGTETLSI WP_035003110.1 hypothetical protein [Corynebacterium jeikeium] Length: 279\nNumber of predicted TMHs: 2\nExp number of AAs in TMHs: 41.04341\nExp number, first 60 AAs: 18.33485\nTotal prob of N-in: 0.79827\nPOSSIBLE N-term signal sequence\ninside 1 12\nTMhelix 13 35\noutside 36 248\nTMhelix 249 271\ninside 272 279

29285 GCF_000738245.1_Cj30184 Corynebacterium jeikeium Terrabacteria group; Actinobacteria; Actinobacteria; Corynebacteriales; Corynebacteriaceae; Corynebacterium
MMFLRSRRAMIAVAAASSLVLTATAVPQSPFSPVAAQAATANKTQDTTLPLHCSLNAGGN WP_043014821.1 hypothetical protein [Corynebacterium jeikeium] Length: 572\nNumber of predicted TMHs: 1\nExp number of AAs in TMHs: 22.83715\nExp number, first 60 AAs: 1.83015\nTotal prob of N-in: 0.09141\noutside 1 543\nTMhelix 544 566\ninside 567 572

29286 GCF_001059385.1_ASM105938v1 Corynebacterium jeikeium Terrabacteria group; Actinobacteria; Actinobacteria; Corynebacteriales; Corynebacteriaceae; Corynebacterium
MMHLRSRRAMIAVAVSASMILAVSAVPNSPVSPSVAQAATANKTHNTSLPLKCNLSARGG WP_080971943.1 hypothetical protein [Corynebacterium jeikeium] Length: 1305\nNumber of predicted TMHs: 1\nExp number of AAs in TMHs: 26.89344\nExp number, first 60 AAs: 4.32729\nTotal prob of N-in: 0.21249\noutside 1 1275\nTMhelix 1276 1298\ninside 1299 1305

29287 GCF_001412085.1_ASM141208v1 Corynebacterium lowii Terrabacteria group; Actinobacteria; Actinobacteria; Corynebacteriales; Corynebacteriaceae; Corynebacterium
MTFPGPFSRRALLRPLSLAAVVSTLALGTTLGVGLPAQADVLPLSPTAEDSKDAWVNPAV WP_055174723.1 hypothetical protein [Corynebacterium lowii] Length: 917\nNumber of predicted TMHs: 1\nExp number of AAs in TMHs: 34.34201\nExp number, first 60 AAs: 11.8421\nTotal prob of N-in: 0.52141\nPOSSIBLE N-term signal sequence\noutside 1 876\nTMhelix 877 899\ninside 900 917

29288 GCF_001020985.1_ASM102098v1 Corynebacterium mustelae Terrabacteria group; Actinobacteria; Actinobacteria; Corynebacteriales; Corynebacteriaceae; Corynebacterium
MRKLTRRTPLRSVAAVVLAAACLATSAVPVAAGDLNQVVSADEAIAPAGELIEFRSGHAD WP_047261629.1 hypothetical protein [Corynebacterium mustelae] Length: 313\nNumber of predicted TMHs: 2\nExp number of AAs in TMHs: 42.85202\nExp number, first 60 AAs: 19.47672\nTotal prob of N-in: 0.95187\nPOSSIBLE N-term signal sequence\ninside 1 12\nTMhelix 13 32\noutside 33 280\nTMhelix 281 303\ninside 304 313

29289 GCF_001020985.1_ASM102098v1 Corynebacterium mustelae Terrabacteria group; Actinobacteria; Actinobacteria; Corynebacteriales; Corynebacteriaceae; Corynebacterium
MGIFNRRRGAILAAISIASLLPIHSVAWAQDAPTGSLSWGIRASFDNYTSGATFIEDGAT WP_047261182.1 hypothetical protein [Corynebacterium mustelae] Length: 730\nNumber of predicted TMHs: 1\nExp number of AAs in TMHs: 31.3099\nExp number, first 60 AAs: 8.82205\nTotal prob of N-in: 0.43447\noutside 1 695\nTMhelix 696 718\ninside 719 730

29290 GCF_001412105.1_ASM141210v1 Corynebacterium oculi Terrabacteria group; Actinobacteria; Actinobacteria; Corynebacteriales; Corynebacteriaceae; Corynebacterium
MRLTGFPSRRALLRPLSLAAITGTVALCTVLPPLTPPSARADVLPLSPTSETSREAWMNP WP_055122566.1 hypothetical protein [Corynebacterium oculi] Length: 867\nNumber of predicted TMHs: 1\nExp number of AAs in TMHs: 34.72794\nExp number, first 60 AAs: 12.11675\nTotal prob of N-in: 0.53621\nPOSSIBLE N-term signal sequence\noutside 1 834\nTMhelix 835 857\ninside 858 867

29291 GCF_900092335.1_PRJEB14666 Corynebacterium phoceense Terrabacteria group; Actinobacteria; Actinobacteria; Corynebacteriales; Corynebacteriaceae; Corynebacterium
MNFSPRRALVAAAVSVSLAVTATPVATAQSSLPSLPSGLVSVNGTGIENGAFTGAIQKDG WP_068801324.1 hypothetical protein [Corynebacterium phoceense] Length: 284\nNumber of predicted TMHs: 1\nExp number of AAs in TMHs: 26.17242\nExp number, first 60 AAs: 5.01523\nTotal prob of N-in: 0.59282\noutside 1 245\nTMhelix 246 268\ninside 269 284

29292 GCF_001053555.1_ASM105355v1 Corynebacterium propinquum Terrabacteria group; Actinobacteria; Actinobacteria; Corynebacteriales; Corynebacteriaceae; Corynebacterium
MNESKPGRRALVRYGSTCCAAVLATGMVTGTTAFGIGALNSAAPHAAATPLIHAQQEEPT WP_049167940.1 D-alanyl-D-alanine carboxypeptidase [Corynebacterium propinquum] Length: 459\nNumber of predicted TMHs: 2\nExp

number of AAs in TMHs: 43.39179\nExp number, first 60 AAs: 20.47679\nTotal prob of N-in: 0.91643\nPOSSIBLE N-term signal sequence\ninside 1 11\nTMhelix 12 34\noutside 35 433\nTMhelix 434 456\ninside 457 459

29293 GCF_001052555.1_ASM105255v1 Corynebacterium propinquum Terrabacteria group; Actinobacteria; Actinobacteria; Corynebacteriales; Corynebacteriaceae; Corynebacterium MNESKPGRRALVRYGSTCCAAVLATGLVTGTTAFGIGTLNSAAPHAAAAPLIHAQQEEPT WP_049150213.1 D-alanyl-D-alanine carboxypeptidase [Corynebacterium propinquum] Length: 459\nNumber of predicted TMHs: 2\nExp number of AAs in TMHs: 43.27689\nExp number, first 60 AAs: 20.36391\nTotal prob of N-in: 0.91075\nPOSSIBLE N-term signal sequence\ninside 1 11\nTMhelix 12 34\noutside 35 433\nTMhelix 434 456\ninside 457 459

29294 GCF_001586235.1_ASM158623v1 Corynebacterium simulans Terrabacteria group; Actinobacteria; Actinobacteria; Corynebacteriales; Corynebacteriaceae; Corynebacterium MSIAVPGPNPADVAERRRTLRKYQWGVTGLLILAAVIFFACSWWQHQDANAPAWVGFVRA WP_062042481.1 DUF445 domain-containing protein [Corynebacterium simulans] Length: 428\nNumber of predicted TMHs: 2\nExp number of AAs in TMHs: 53.7136500000001\nExp number, first 60 AAs: 24.15165\nTotal prob of N-in: 0.50482\nPOSSIBLE N-term signal sequence\ninside 1 24\nTMhelix 25 47\noutside 48 404\nTMhelix 405 427\ninside 428 428

29295 GCF_001579275.1_ASM157927v1 Corynebacterium simulans Terrabacteria group; Actinobacteria; Actinobacteria; Corynebacteriales; Corynebacteriaceae; Corynebacterium MSIAVPGPNPADVAERRRTLRKYQWGVTGLLILAAVIFFACSWWQHQDATAPAWVGFVRA WP_061920275.1 MULTISPECIES: DUF445 domain-containing protein [Corynebacterium] Length: 428\nNumber of predicted TMHs: 2\nExp number of AAs in TMHs: 53.5800100000001\nExp number, first 60 AAs: 24.17271\nTotal prob of N-in: 0.49808\nPOSSIBLE N-term signal sequence\ninside 1 24\nTMhelix 25 47\noutside 48 404\nTMhelix 405 427\ninside 428 428

29296 GCF_001586215.1_ASM158621v1 Corynebacterium simulans Terrabacteria group; Actinobacteria; Actinobacteria; Corynebacteriales; Corynebacteriaceae; Corynebacterium MSIAVPGPNPADVAERRRTLRKYQWGVTGLLILAAVIFFACSWWQHQDANAPAWVGFVRA WP_062036085.1 DUF445 domain-containing protein [Corynebacterium simulans] Length: 428\nNumber of predicted TMHs: 2\nExp number of AAs in TMHs: 53.7142000000001\nExp number, first 60 AAs: 24.15164\nTotal prob of N-in: 0.50482\nPOSSIBLE N-term signal sequence\ninside 1 24\nTMhelix 25 47\noutside 48 404\nTMhelix 405 427\ninside 428 428

29297 GCF_001563605.1_ASM156360v1 Corynebacterium sp. CMW7794 Terrabacteria group; Actinobacteria; Actinobacteria; Corynebacteriales; Corynebacteriaceae; Corynebacterium MNFSPRRALVAAAVSVSLAVTATPAATAQSSLPSLPSGLVSVNGTGIENGAFTGAIQKDG WP_066490205.1 MULTISPECIES: hypothetical protein [Corynebacterium] Length: 284\nNumber of predicted TMHs: 1\nExp number of AAs in TMHs: 24.87519\nExp number, first 60 AAs: 3.69521\nTotal prob of N-in: 0.56645\noutside 1 245\nTMhelix 246 268\ninside 269 284

29298 GCF_001942345.1_ASM194234v1 Corynebacterium sp. CNJ-954Terrabacteria group; Actinobacteria; Actinobacteria; Corynebacteriales; Corynebacteriaceae; Corynebacterium MTAHTLPRRRTLRAPALATALATTLTIGLTGPLTAPAVAETGLNNVEKDDWLPLTADPGA WP_075811183.1 D-alanyl-D-alanine carboxypeptidase [Corynebacterium sp. CNJ-954] Length: 443\nNumber of predicted TMHs: 1\nExp number of AAs in TMHs: 33.15399\nExp number, first 60 AAs: 12.39296\nTotal prob of N-in: 0.56932\nPOSSIBLE N-term signal sequence\noutside 1 418\nTMhelix 419 438\ninside 439 443

29299 GCF_001552965.1_ASM155296v1 Corynebacterium sp. DNF00584 Terrabacteria group; Actinobacteria; Actinobacteria; Corynebacteriales; Corynebacteriaceae; Corynebacterium MNFSPRRALVAAAVSVSLAVTATPAATAQSSLPSLPSGLVSVNGTGIENGAFTGAIQKDG WP_066490205.1 MULTISPECIES: hypothetical protein [Corynebacterium] Length: 284\nNumber of predicted TMHs: 1\nExp number of AAs in TMHs: 24.87519\nExp number, first 60 AAs: 3.69521\nTotal prob of N-in: 0.56645\noutside 1 245\nTMhelix 246 268\ninside 269 284

29300 GCF_001665955.1_ASM166595v1 Corynebacterium sp. EPI-003-04-2554_SCH2473622 Terrabacteria group; Actinobacteria; Actinobacteria; Corynebacteriales; Corynebacteriaceae; Corynebacterium MNESKPGRRAIVRYSSTCCANLLVVGMATGAATLGIGAFGSASPHAAATPHTEAAPLLHA WP_064834573.1 D-alanyl-D-alanine carboxypeptidase [Corynebacterium sp. EPI-003-04-2554_SCH2473622] Length: 467\nNumber of predicted TMHs: 2\nExp number of AAs in TMHs: 44.48775\nExp number, first 60 AAs: 21.59742\nTotal prob of N-in:

0.98117\nPOSSIBLE N-term signal sequence\ninside 1 19\nTMhelix 20 42\noutside 43 441\nTMhelix 442 464\ninside 465 467

29301 GCF_001838175.1_ASM183817v1 Corynebacterium sp. HMSC034A01 Terrabacteria group; Actinobacteria; Actinobacteria; Corynebacteriales; Corynebacteriaceae; Corynebacterium MAKSVRSRRTAIAAGALSIALVAPFVSPVTDAPTAQAVEASAFDTRYTTDNFWNANEAVV WP_070844356.1 hypothetical protein [Corynebacterium sp. HMSC034A01] Length: 782\nNumber of predicted TMHs: 1\nExp number of AAs in TMHs: 41.96303\nExp number, first 60 AAs: 10.79534\nTotal prob of N-in: 0.58881\nPOSSIBLE N-term signal sequence\noutside 1 725\nTMhelix 726 748\ninside 749 782

29302 GCF_001835805.1_ASM183580v1 Corynebacterium sp. HMSC034B08 Terrabacteria group; Actinobacteria; Actinobacteria; Corynebacteriales; Corynebacteriaceae; Corynebacterium MRFPSRRNALLCATTAGSLLLAGSVCAPAHAQSSLSSSSSSGSSGSSRPSSPSNESYTFP WP_070816867.1 hypothetical protein [Corynebacterium sp. HMSC034B08] Length: 500\nNumber of predicted TMHs: 1\nExp number of AAs in TMHs: 20.74084\nExp number, first 60 AAs: 2.89969\nTotal prob of N-in: 0.15869\noutside 1 455\nTMhelix 456 478\ninside 479 500

29303 GCF_001835905.1_ASM183590v1 Corynebacterium sp. HMSC034E11 Terrabacteria group; Actinobacteria; Actinobacteria; Corynebacteriales; Corynebacteriaceae; Corynebacterium MRFPSRRNALLCATTAGSLLLAGSVCAPAHAQSSLSSSSSSGSSGSSRPSSPSNESYTFP WP_070819865.1 hypothetical protein [Corynebacterium sp. HMSC034E11] Length: 500\nNumber of predicted TMHs: 1\nExp number of AAs in TMHs: 22.62238\nExp number, first 60 AAs: 2.87033\nTotal prob of N-in: 0.16754\noutside 1 455\nTMhelix 456 478\ninside 479 500

29304 GCF_001836185.1_ASM183618v1 Corynebacterium sp. HMSC036D03 Terrabacteria group; Actinobacteria; Actinobacteria; Corynebacteriales; Corynebacteriaceae; Corynebacterium MSIAVPGPNPADVAERRRTLRKYQWGVTGLLILAAVIFFACSWWQHQDANAPAWVGFVRA WP_070496988.1 MULTISPECIES: DUF445 domain-containing protein [Corynebacterium] Length: 428\nNumber of predicted TMHs: 2\nExp number of AAs in TMHs: 53.7142000000001\nExp number, first 60 AAs: 24.15164\nTotal prob of N-in: 0.50482\nPOSSIBLE N-term signal sequence\ninside 1 24\nTMhelix 25 47\noutside 48 404\nTMhelix 405 427\ninside 428 428

29305 GCF_001836265.1_ASM183626v1 Corynebacterium sp. HMSC036E10 Terrabacteria group; Actinobacteria; Actinobacteria; Corynebacteriales; Corynebacteriaceae; Corynebacterium MRFSSRRNALLCATTAGSLLLAGSVCAPAHAQSSLSSSSSSGSSGSSRPSSPSNESYTFP WP_070568623.1 MULTISPECIES: hypothetical protein [Corynebacterium] Length: 500\nNumber of predicted TMHs: 1\nExp number of AAs in TMHs: 23.44927\nExp number, first 60 AAs: 4.02226\nTotal prob of N-in: 0.22351\noutside 1 455\nTMhelix 456 478\ninside 479 500

29306 GCF_001815935.1_ASM181593v1 Corynebacterium sp. HMSC04H06 Terrabacteria group; Actinobacteria; Actinobacteria; Corynebacteriales; Corynebacteriaceae; Corynebacterium MSAVSRRRWFPATVAATMTFSALSVPTALAQETPKSPVVINEVESNGDSVGDWVELANTD WP_070769551.1 hypothetical protein [Corynebacterium sp. HMSC04H06] Length: 530\nNumber of predicted TMHs: 1\nExp number of AAs in TMHs: 21.62788\nExp number, first 60 AAs: 2.09467\nTotal prob of N-in: 0.17109\noutside 1 487\nTMhelix 488 510\ninside 511 530

29307 GCF_001815935.1_ASM181593v1 Corynebacterium sp. HMSC04H06 Terrabacteria group; Actinobacteria; Actinobacteria; Corynebacteriales; Corynebacteriaceae; Corynebacterium MSSPQLSSPQSSRRVARRRGTLVLAAVTSTVSAGLAVPAASAAPAGDNVVISEVYGGGGN WP_070768625.1 nuclease [Corynebacterium sp. HMSC04H06] Length: 911\nNumber of predicted TMHs: 2\nExp number of AAs in TMHs: 37.69841\nExp number, first 60 AAs: 18.75258\nTotal prob of N-in: 0.84184\nPOSSIBLE N-term signal sequence\ninside 1 19\nTMhelix 20 42\noutside 43 868\nTMhelix 869 891\ninside 892 911

29308 GCF_001810365.1_ASM181036v1 Corynebacterium sp. HMSC055D05 Terrabacteria group; Actinobacteria; Actinobacteria; Corynebacteriales; Corynebacteriaceae; Corynebacterium MRFSSRRNALLCATTAGSLLLAGSVCAPAHAQSSLSSSSSSGSSGSSRPSSPSNESYTFP WP_070568623.1 MULTISPECIES: hypothetical protein [Corynebacterium] Length: 500\nNumber of predicted TMHs: 1\nExp number of AAs in TMHs: 23.44927\nExp number, first 60 AAs: 4.02226\nTotal prob of N-in: 0.22351\noutside 1 455\nTMhelix 456 478\ninside 479 500

29309 GCF_001807155.1_ASM180715v1 Corynebacterium sp. HMSC05C01 Terrabacteria group; Actinobacteria; Actinobacteria; Corynebacteriales; Corynebacteriaceae; Corynebacterium
MRFPSRRNALLCATTAGSLLLAGSVCAPAHAQSSLSSSSSSGSSGSSRPSSPSNESYTFP WP_070614106.1 MULTISPECIES: hypothetical protein [Corynebacterium] Length: 500\nNumber of predicted TMHs: 1\nExp number of AAs in TMHs: 22.30507\nExp number, first 60 AAs: 2.893\nTotal prob of N-in: 0.16070\noutside 1 455\nTMhelix 456 478\ninside 479 500

29310 GCF_001807215.1_ASM180721v1 Corynebacterium sp. HMSC05D08 Terrabacteria group; Actinobacteria; Actinobacteria; Corynebacteriales; Corynebacteriaceae; Corynebacterium
MTSRRGTFLAALVTASLIPLAPPALAIPATDGHLTWGIRASFNNYTGGATLVKDGATRKG WP_070508929.1 hypothetical protein [Corynebacterium sp. HMSC05D08] Length: 655\nNumber of predicted TMHs: 2\nExp number of AAs in TMHs: 37.09915\nExp number, first 60 AAs: 17.62274\nTotal prob of N-in: 0.78739\nPOSSIBLE N-term signal sequence\ninside 1 6\nTMhelix 7 29\noutside 30 616\nTMhelix 617 639\ninside 640 655

29311 GCF_001807215.1_ASM180721v1 Corynebacterium sp. HMSC05D08 Terrabacteria group; Actinobacteria; Actinobacteria; Corynebacteriales; Corynebacteriaceae; Corynebacterium
MTIAVPGPNPADVAERRRTLRKYQWGVTGLLILAAVIFFACSWWQHHNPDAPTWVGYVRA WP_049151243.1
MULTISPECIES: DUF445 domain-containing protein [Corynebacterium] Length: 428\nNumber of predicted TMHs: 2\nExp number of AAs in TMHs: 52.76767\nExp number, first 60 AAs: 23.54962\nTotal prob of N-in: 0.78331\nPOSSIBLE N-term signal sequence\ninside 1 24\nTMhelix 25 47\noutside 48 404\nTMhelix 405 427\ninside 428 428

29312 GCF_001807225.1_ASM180722v1 Corynebacterium sp. HMSC05E07 Terrabacteria group; Actinobacteria; Actinobacteria; Corynebacteriales; Corynebacteriaceae; Corynebacterium
MISWHRRSCLGMATVIAAGLVAVPTATAQELNVESSRCLREDCFSAFSSKADETEEYKPS WP_070537220.1 hypothetical protein [Corynebacterium sp. HMSC05E07] Length: 175\nNumber of predicted TMHs: 1\nExp number of AAs in TMHs: 24.27834\nExp number, first 60 AAs: 3.61639\nTotal prob of N-in: 0.83106\ninside 1 138\nTMhelix 139 161\noutside 162 175

29313 GCF_001814975.1_ASM181497v1 Corynebacterium sp. HMSC065A05 Terrabacteria group; Actinobacteria; Actinobacteria; Corynebacteriales; Corynebacteriaceae; Corynebacterium
MNFSPRRALVAAAVSVSLAVTATPAATAQSSLPSLPSGLVSVNGTGIENGAFTGAIQKDG WP_066490205.1
MULTISPECIES: hypothetical protein [Corynebacterium] Length: 284\nNumber of predicted TMHs: 1\nExp number of AAs in TMHs: 24.87519\nExp number, first 60 AAs: 3.69521\nTotal prob of N-in: 0.56645\noutside 1 245\nTMhelix 246 268\ninside 269 284

29314 GCF_001811185.1_ASM181118v1 Corynebacterium sp. HMSC067D03 Terrabacteria group; Actinobacteria; Actinobacteria; Corynebacteriales; Corynebacteriaceae; Corynebacterium
MRFPSRRNALLCATTAGSLLLAGSVCAPAHAQSSLSSSSSSGSSGSSRPSSPSNESYTFP WP_070614106.1 MULTISPECIES: hypothetical protein [Corynebacterium] Length: 500\nNumber of predicted TMHs: 1\nExp number of AAs in TMHs: 22.30507\nExp number, first 60 AAs: 2.893\nTotal prob of N-in: 0.16070\noutside 1 455\nTMhelix 456 478\ninside 479 500

29315 GCF_001807295.1_ASM180729v1 Corynebacterium sp. HMSC06C06 Terrabacteria group; Actinobacteria; Actinobacteria; Corynebacteriales; Corynebacteriaceae; Corynebacterium
MTSRRGTLLAALVTASLIPLAPPALAIPATDGHLTWGIRASFNNYTGGATLVKDGATRKG WP_070420471.1 hypothetical protein [Corynebacterium sp. HMSC06C06] Length: 653\nNumber of predicted TMHs: 2\nExp number of AAs in TMHs: 37.05362\nExp number, first 60 AAs: 17.51134\nTotal prob of N-in: 0.78184\nPOSSIBLE N-term signal sequence\ninside 1 6\nTMhelix 7 29\noutside 30 614\nTMhelix 615 637\ninside 638 653

29316 GCF_001807295.1_ASM180729v1 Corynebacterium sp. HMSC06C06 Terrabacteria group; Actinobacteria; Actinobacteria; Corynebacteriales; Corynebacteriaceae; Corynebacterium
MTIAVPGPNPADVAERRRTLRKYQWGVTGLLILAAVIFFACSWWQHHNPDAPTWVGYVRA WP_070421008.1
DUF445 domain-containing protein [Corynebacterium sp. HMSC06C06] Length: 428\nNumber of predicted TMHs: 2\nExp number of AAs in TMHs: 52.76767\nExp number, first 60 AAs: 23.54962\nTotal prob of N-in: 0.78331\nPOSSIBLE N-term signal sequence\ninside 1 24\nTMhelix 25 47\noutside 48 404\nTMhelix 405 427\ninside 428 428

29317 GCF_001807315.1_ASM180731v1 Corynebacterium sp. HMSC06D04 Terrabacteria group; Actinobacteria; Actinobacteria; Corynebacteriales; Corynebacteriaceae; Corynebacterium
MSIAVPGPNPADVAERRRTLRKYQWGVTGLLILAAVIFFACSWWQHQDANAPAWVGFVRA WP_070432509.1

DUF445 domain-containing protein [Corynebacterium sp. HMSC06D04] Length: 428\nNumber of predicted TMHs: 2\nExp number of AAs in TMHs: 53.7137200000001\nExp number, first 60 AAs: 24.15164\nTotal prob of N-in: 0.50482\nPOSSIBLE N-term signal sequence\ninside 1 24\nTMhelix 25 47\noutside 48 404\nTMhelix 405 427\ninside 428 428

29318 GCF_001807365.1_ASM180736v1 Corynebacterium sp. HMSC06G04 Terrabacteria group; Actinobacteria; Actinobacteria; Corynebacteriales; Corynebacteriaceae; Corynebacterium MSIAVPGPNPADVAERRRTLRKYQWGVTGLLILAAVIFFACSWWQHQDANAPAWVGFVRA WP_070496988.1 MULTISPECIES: DUF445 domain-containing protein [Corynebacterium] Length: 428\nNumber of predicted TMHs: 2\nExp number of AAs in TMHs: 53.7142000000001\nExp number, first 60 AAs: 24.15164\nTotal prob of N-in: 0.50482\nPOSSIBLE N-term signal sequence\ninside 1 24\nTMhelix 25 47\noutside 48 404\nTMhelix 405 427\ninside 428 428

29319 GCF_001837615.1_ASM183761v1 Corynebacterium sp. HMSC070H05 Terrabacteria group; Actinobacteria; Actinobacteria; Corynebacteriales; Corynebacteriaceae; Corynebacterium MRFSSRRNALLCATTAGSLLLAGSVCAPAHAQSSLSSSSSSGSSGSSRPSSPSNESYTFP WP_070568623.1 MULTISPECIES: hypothetical protein [Corynebacterium] Length: 500\nNumber of predicted TMHs: 1\nExp number of AAs in TMHs: 23.44927\nExp number, first 60 AAs: 4.02226\nTotal prob of N-in: 0.22351\noutside 1 455\nTMhelix 456 478\ninside 479 500

29320 GCF_001814825.1_ASM181482v1 Corynebacterium sp. HMSC071B10 Terrabacteria group; Actinobacteria; Actinobacteria; Corynebacteriales; Corynebacteriaceae; Corynebacterium MHTVARRAAVLAAAAALAAPPVAGAAEVVQQGSQIVVADASSCAIAYNDRAENISYTAAH WP_070764635.1 hypothetical protein [Corynebacterium sp. HMSC071B10] Length: 297\nNumber of predicted TMHs: 1\nExp number of AAs in TMHs: 23.65279\nExp number, first 60 AAs: 2.80289\nTotal prob of N-in: 0.64048\ninside 1 260\nTMhelix 261 283\noutside 284 297

29321 GCF_001814825.1_ASM181482v1 Corynebacterium sp. HMSC071B10 Terrabacteria group; Actinobacteria; Actinobacteria; Corynebacteriales; Corynebacteriaceae; Corynebacterium MHTSPRRRALRSTTIASAVALSLTVASPGAGADEDSNEEVFGSGPDASAEFSEQYAAEHG WP_070763091.1 hypothetical protein [Corynebacterium sp. HMSC071B10] Length: 480\nNumber of predicted TMHs: 1\nExp number of AAs in TMHs: 21.75226\nExp number, first 60 AAs: 0.54788\nTotal prob of N-in: 0.39527\noutside 1 444\nTMhelix 445 467\ninside 468 480

29322 GCF_001809185.1_ASM180918v1 Corynebacterium sp. HMSC072G08 Terrabacteria group; Actinobacteria; Actinobacteria; Corynebacteriales; Corynebacteriaceae; Corynebacterium MNFSPRRALVAAAVSVSLAVTATPAATAQSSLPSLPSGLVSVNGTGIENGAFTGAIQKDG WP_066490205.1 MULTISPECIES: hypothetical protein [Corynebacterium] Length: 284\nNumber of predicted TMHs: 1\nExp number of AAs in TMHs: 24.87519\nExp number, first 60 AAs: 3.69521\nTotal prob of N-in: 0.56645\noutside 1 245\nTMhelix 246 268\ninside 269 284

29323 GCF_001831515.1_ASM183151v1 Corynebacterium sp. HMSC074A01 Terrabacteria group; Actinobacteria; Actinobacteria; Corynebacteriales; Corynebacteriaceae; Corynebacterium MHTSPRRRALRSTTIASAVALSLTVASPGAGADEDSNEEVFGSGPDASAEFSEQYAAEHG WP_070791085.1 hypothetical protein [Corynebacterium sp. HMSC074A01] Length: 480\nNumber of predicted TMHs: 1\nExp number of AAs in TMHs: 21.75227\nExp number, first 60 AAs: 0.54788\nTotal prob of N-in: 0.39527\noutside 1 444\nTMhelix 445 467\ninside 468 480

29324 GCF_001831515.1_ASM183151v1 Corynebacterium sp. HMSC074A01 Terrabacteria group; Actinobacteria; Actinobacteria; Corynebacteriales; Corynebacteriaceae; Corynebacterium MGSTGRRTTVRRGVTLAACALGASLVIVGVEPIAAPQALPTAEAAPSRFSRSDAERTNIT WP_070790698.1 hypothetical protein [Corynebacterium sp. HMSC074A01] Length: 981\nNumber of predicted TMHs: 2\nExp number of AAs in TMHs: 43.48519\nExp number, first 60 AAs: 21.83677\nTotal prob of N-in: 0.98500\nPOSSIBLE N-term signal sequence\ninside 1 12\nTMhelix 13 35\noutside 36 942\nTMhelix 943 965\ninside 966 981

29325 GCF_001831515.1_ASM183151v1 Corynebacterium sp. HMSC074A01 Terrabacteria group; Actinobacteria; Actinobacteria; Corynebacteriales; Corynebacteriaceae; Corynebacterium MHTVARRAAVLAAAAALAAPPVAGAAEVVQQGSQIVVADASSCAIAYNDRAENISYTAAH WP_070789886.1 hypothetical protein [Corynebacterium sp. HMSC074A01] Length: 297\nNumber of predicted TMHs: 1\nExp number of AAs in TMHs: 23.66432\nExp number, first 60 AAs: 2.80383\nTotal prob of N-in: 0.64056\ninside 1 260\nTMhelix 261 283\noutside 284 297

29326 GCF_001815635.1_ASM181563v1 Corynebacterium sp. HMSC075D04 Terrabacteria group; Actinobacteria; Actinobacteria; Corynebacteriales; Corynebacteriaceae; Corynebacterium
MRFPSRRNALLCATTAGSLLLAGSVCAPAHAQSSLSSSSSSGSSGSSRPSSPSNESYTFP WP_083316703.1 hypothetical protein [Corynebacterium sp. HMSC075D04] Length: 500\nNumber of predicted TMHs: 1\nExp number of AAs in TMHs: 22.3051\nExp number, first 60 AAs: 2.893\nTotal prob of N-in: 0.16070\noutside 1 455\nTMhelix 456 478\ninside 479 500

29327 GCF_001813815.1_ASM181381v1 Corynebacterium sp. HMSC076D02 Terrabacteria group; Actinobacteria; Actinobacteria; Corynebacteriales; Corynebacteriaceae; Corynebacterium
MSIAVPGPNPADVAERRRTLRKYQWGVTGLLILAAVIFFACSWWQHQDANAPAWVGFVRA WP_070518604.1 MULTISPECIES: DUF445 domain-containing protein [Corynebacterium] Length: 428\nNumber of predicted TMHs: 2\nExp number of AAs in TMHs: 53.7136500000001\nExp number, first 60 AAs: 24.15165\nTotal prob of N-in: 0.50482\nPOSSIBLE N-term signal sequence\ninside 1 24\nTMhelix 25 47\noutside 48 404\nTMhelix 405 427\ninside 428 428

29328 GCF_001810505.1_ASM181050v1 Corynebacterium sp. HMSC077B05 Terrabacteria group; Actinobacteria; Actinobacteria; Corynebacteriales; Corynebacteriaceae; Corynebacterium
MNFSPRRALVAAAVSVSLAVTATPAATAQSSLPSLPSGLVSVNGTGIENGAFTGAIQKDG WP_066490205.1 MULTISPECIES: hypothetical protein [Corynebacterium] Length: 284\nNumber of predicted TMHs: 1\nExp number of AAs in TMHs: 24.87519\nExp number, first 60 AAs: 3.69521\nTotal prob of N-in: 0.56645\noutside 1 245\nTMhelix 246 268\ninside 269 284

29329 GCF_001808545.1_ASM180854v1 Corynebacterium sp. HMSC077D03 Terrabacteria group; Actinobacteria; Actinobacteria; Corynebacteriales; Corynebacteriaceae; Corynebacterium
MSIAVPGPNPADVAERRRTLRKYQWGVTGLLILAAVIFFACSWWQHQDANAPAWVGFVRA WP_070518604.1 MULTISPECIES: DUF445 domain-containing protein [Corynebacterium] Length: 428\nNumber of predicted TMHs: 2\nExp number of AAs in TMHs: 53.7136500000001\nExp number, first 60 AAs: 24.15165\nTotal prob of N-in: 0.50482\nPOSSIBLE N-term signal sequence\ninside 1 24\nTMhelix 25 47\noutside 48 404\nTMhelix 405 427\ninside 428 428

29330 GCF_001814575.1_ASM181457v1 Corynebacterium sp. HMSC077D10 Terrabacteria group; Actinobacteria; Actinobacteria; Corynebacteriales; Corynebacteriaceae; Corynebacterium
MNFSPRRALVAAAVSVSLAVTATPAATAQSSLPSLPSGLVSVNGTGIENGAFTGAIQKDG WP_066490205.1 MULTISPECIES: hypothetical protein [Corynebacterium] Length: 284\nNumber of predicted TMHs: 1\nExp number of AAs in TMHs: 24.87519\nExp number, first 60 AAs: 3.69521\nTotal prob of N-in: 0.56645\noutside 1 245\nTMhelix 246 268\ninside 269 284

29331 GCF_001807495.1_ASM180749v1 Corynebacterium sp. HMSC08C04 Terrabacteria group; Actinobacteria; Actinobacteria; Corynebacteriales; Corynebacteriaceae; Corynebacterium
MSIAVPGPNPADVAERRRTLRKYQWGVTGLLILAAVIFFACSWWQHQDATAPAWVGFVRA WP_061920275.1 MULTISPECIES: DUF445 domain-containing protein [Corynebacterium] Length: 428\nNumber of predicted TMHs: 2\nExp number of AAs in TMHs: 53.5800100000001\nExp number, first 60 AAs: 24.17271\nTotal prob of N-in: 0.49808\nPOSSIBLE N-term signal sequence\ninside 1 24\nTMhelix 25 47\noutside 48 404\nTMhelix 405 427\ninside 428 428

29332 GCF_001807545.1_ASM180754v1 Corynebacterium sp. HMSC08F01 Terrabacteria group; Actinobacteria; Actinobacteria; Corynebacteriales; Corynebacteriaceae; Corynebacterium
MRFPSRRNALLCATTAGSLLLAGSVCAPAHAQSSLSSSSSSGSSGSSRPSSPSNESYTFP WP_083292036.1 hypothetical protein [Corynebacterium sp. HMSC08F01] Length: 500\nNumber of predicted TMHs: 1\nExp number of AAs in TMHs: 22.30359\nExp number, first 60 AAs: 2.89311\nTotal prob of N-in: 0.16068\noutside 1 455\nTMhelix 456 478\ninside 479 500

29333 GCF_001806995.1_ASM180699v1 Corynebacterium sp. HMSC11D10 Terrabacteria group; Actinobacteria; Actinobacteria; Corynebacteriales; Corynebacteriaceae; Corynebacterium
MRFSSRRNALLCATTAGSLLLAGSVCAPAHAQSSLSSSSSSGSSGSSRPSSPSNESYTFP WP_070452682.1 hypothetical protein [Corynebacterium sp. HMSC11D10] Length: 500\nNumber of predicted TMHs: 1\nExp number of AAs in TMHs: 23.44912\nExp number, first 60 AAs: 4.02228\nTotal prob of N-in: 0.22351\noutside 1 455\nTMhelix 456 478\ninside 479 500

29334 GCF_001806865.1_ASM180686v1 Corynebacterium sp. HMSC28B08 Terrabacteria group; Actinobacteria; Actinobacteria; Corynebacteriales; Corynebacteriaceae; Corynebacterium

MAQKRKLSARRATLATTSVLCLTALGMAGSFGLEGVGPEGFGPATANAQESPTVLKNGHI WP_070433763.1 hypothetical protein [Corynebacterium sp. HMSC28B08] Length: 607\nNumber of predicted TMHs: 2\nExp number of AAs in TMHs: 41.7209\nExp number, first 60 AAs: 19.22362\nTotal prob of N-in: 0.90321\nPOSSIBLE N-term signal sequence\ninside 1 11\nTMhelix 12 31\noutside 32 571\nTMhelix 572 594\ninside 595 607

29335 GCF_000477955.1_Cory_sp_KPL1989_V1 Corynebacterium sp. KPL1989 Terrabacteria group; Actinobacteria; Actinobacteria; Corynebacteriales; Corynebacteriaceae; Corynebacterium MGQTARQLTSRKRRRPIATTLALVTAFVGLPQLFVPQQSYAYDVLETHREIRNVQTTFPDWP_023019151.1 MULTISPECIES: LPXTG cell wall anchor domain-containing protein [Corynebacterium] Length: 707\nNumber of predicted TMHs: 2\nExp number of AAs in TMHs: 39.44537\nExp number, first 60 AAs: 18.43098\nTotal prob of N-in: 0.91695\nPOSSIBLE N-term signal sequence\ninside 1 16\nTMhelix 17 35\noutside 36 677\nTMhelix 678 700\ninside 701 707

29336 GCF_000477955.1_Cory_sp_KPL1989_V1 Corynebacterium sp. KPL1989 Terrabacteria group; Actinobacteria; Actinobacteria; Corynebacteriales; Corynebacteriaceae; Corynebacterium MNESKPGRRAIVRYGSTCCANLLVVGMATGATTLGIGAFGSATPHAAATPHAEAAPLLHA WP_023019056.1 MULTISPECIES: hypothetical protein [Corynebacterium] Length: 467\nNumber of predicted TMHs: 2\nExp number of AAs in TMHs: 44.4092\nExp number, first 60 AAs: 21.51669\nTotal prob of N-in: 0.97067\nPOSSIBLE N-term signal sequence\ninside 1 19\nTMhelix 20 42\noutside 43 441\nTMhelix 442 464\ninside 465 467

29337 GCF_000477935.1_Cory_sp_KPL1995_V1 Corynebacterium sp. KPL1995 Terrabacteria group; Actinobacteria; Actinobacteria; Corynebacteriales; Corynebacteriaceae; Corynebacterium MNESKPGRRAIVRYGSTCCANLLVVGMATGATTLGIGAFGSATPHAAATPHAEAAPLLHA WP_023019056.1 MULTISPECIES: hypothetical protein [Corynebacterium] Length: 467\nNumber of predicted TMHs: 2\nExp number of AAs in TMHs: 44.4092\nExp number, first 60 AAs: 21.51669\nTotal prob of N-in: 0.97067\nPOSSIBLE N-term signal sequence\ninside 1 19\nTMhelix 20 42\noutside 43 441\nTMhelix 442 464\ninside 465 467

29338 GCF_000477935.1_Cory_sp_KPL1995_V1 Corynebacterium sp. KPL1995 Terrabacteria group; Actinobacteria; Actinobacteria; Corynebacteriales; Corynebacteriaceae; Corynebacterium MGQTARQLTSRKRRRPIATTLALVTAFVGLPQLFVPQQSYAYDVLETHREIRNVQTTFPDWP_023019151.1 MULTISPECIES: LPXTG cell wall anchor domain-containing protein [Corynebacterium] Length: 707\nNumber of predicted TMHs: 2\nExp number of AAs in TMHs: 39.44537\nExp number, first 60 AAs: 18.43098\nTotal prob of N-in: 0.91695\nPOSSIBLE N-term signal sequence\ninside 1 16\nTMhelix 17 35\noutside 36 677\nTMhelix 678 700\ninside 701 707

29339 GCF_900099485.1_PRJEB15651 Corynebacterium sp. MC3 Terrabacteria group; Actinobacteria; Actinobacteria; Corynebacteriales; Corynebacteriaceae; Corynebacterium MRNFGASRAARRRALIAGTIICSLALNDVPVLAADPHPAVEQAQSANDAIAAGRVTSLEQ WP_070975208.1 hypothetical protein [Corynebacterium sp. MC3] Length: 1007\nNumber of predicted TMHs: 1\nExp number of AAs in TMHs: 25.89845\nExp number, first 60 AAs: 3.35723\nTotal prob of N-in: 0.15923\noutside 1 971\nTMhelix 972 994\ninside 995 1007

29340 GCF_001875735.1_ASM187573v1 Corynebacterium sp. NML120713 Terrabacteria group; Actinobacteria; Actinobacteria; Corynebacteriales; Corynebacteriaceae; Corynebacterium MRNFGASRAARRRALIAGTIICSLALNDAPALAADPHPAVEQAQTENDAIAAGRVTSLEQ WP_071573543.1 hypothetical protein [Corynebacterium sp. NML120713] Length: 991\nNumber of predicted TMHs: 1\nExp number of AAs in TMHs: 24.34471\nExp number, first 60 AAs: 1.70925\nTotal prob of N-in: 0.08254\noutside 1 955\nTMhelix 956 978\ninside 979 991

29341 GCF_001875665.1_ASM187566v1 Corynebacterium sp. NML130628 Terrabacteria group; Actinobacteria; Actinobacteria; Corynebacteriales; Corynebacteriaceae; Corynebacterium MRKIGASRAARRRAMIAGTVICSLALNTVSANAVTPHYAVNPESAAQQLDGNQNWRDANG WP_083584111.1 hypothetical protein [Corynebacterium sp. NML130628] Length: 876\nNumber of predicted TMHs: 1\nExp number of AAs in TMHs: 34.92409\nExp number, first 60 AAs: 12.93009\nTotal prob of N-in: 0.59410\nPOSSIBLE N-term signal sequence\noutside 1 841\nTMhelix 842 864\ninside 865 876

29342 GCF_001875725.1_ASM187572v1 Corynebacterium sp. NML140438 Terrabacteria group; Actinobacteria; Actinobacteria; Corynebacteriales; Corynebacteriaceae; Corynebacterium MRNVGPVPAAHRRALIAGTIICSLALNTVHANATTPHYATNPERAAEQLGENQNWRDSKG WP_071567683.1 hypothetical protein [Corynebacterium sp. NML140438] Length: 903\nNumber of predicted TMHs: 1\nExp number of AAs in TMHs: 24.63452\nExp number, first 60 AAs: 2.099\nTotal prob of N-in: 0.09935\noutside 1 867\nTMhelix 868 890\ninside 891 903

29343 GCF_001767255.1_ASM176725v1 Corynebacterium sp. NML98-0116 Terrabacteria group; Actinobacteria; Actinobacteria; Corynebacteriales; Corynebacteriaceae; Corynebacterium MRNFGASRAARRRALIAGTIICSLALNDVPVLAADPHPAVEQAQSANDAIAAGRVTSLEQ WP_070362059.1 hypothetical protein [Corynebacterium sp. NML98-0116] Length: 927\nNumber of predicted TMHs: 1\nExp number of AAs in TMHs: 25.89954\nExp number, first 60 AAs: 3.35576\nTotal prob of N-in: 0.15963\noutside 1 891\nTMhelix 892 914\ninside 915 927

29344 GCF_001941345.1_ASM194134v1 Corynebacterium stationis Terrabacteria group; Actinobacteria; Actinobacteria; Corynebacteriales; Corynebacteriaceae; Corynebacterium MAKVPRRTIFRPLALMMAMALAIGWTAWPALAQDSSESAEAATEGSKIALIMDASDSMSE WP_075723495.1 VWA domain-containing protein [Corynebacterium stationis] Length: 664\nNumber of predicted TMHs: 2\nExp number of AAs in TMHs: 42.41753\nExp number, first 60 AAs: 19.70338\nTotal prob of N-in: 0.95191\nPOSSIBLE N-term signal sequence\ninside 1 12\nTMhelix 13 32\noutside 33 635\nTMhelix 636 658\ninside 659 664

29345 GCF_001643075.1_ASM164307v1 Corynebacterium stationis Terrabacteria group; Actinobacteria; Actinobacteria; Corynebacteriales; Corynebacteriaceae; Corynebacterium MVKVPRRTTFRQLVLMMAMALAIGWTAWPALAQDSSESAEAATEGSKIALIMDASDSMNE WP_066839588.1 VWA domain-containing protein [Corynebacterium stationis] Length: 664\nNumber of predicted TMHs: 2\nExp number of AAs in TMHs: 41.86115\nExp number, first 60 AAs: 19.14711\nTotal prob of N-in: 0.97736\nPOSSIBLE N-term signal sequence\ninside 1 12\nTMhelix 13 32\noutside 33 635\nTMhelix 636 658\ninside 659 664

29346 GCF_001053405.1_ASM105340v1 Corynebacterium striatum Terrabacteria group; Actinobacteria; Actinobacteria; Corynebacteriales; Corynebacteriaceae; Corynebacterium MTIAVPGPNPADVAERRRTLRKYQWGVTGLLILAAVIFFACSWWQHHNPDAPTWVGYVRA WP_049151243.1 MULTISPECIES: DUF445 domain-containing protein [Corynebacterium] Length: 428\nNumber of predicted TMHs: 2\nExp number of AAs in TMHs: 52.76767\nExp number, first 60 AAs: 23.54962\nTotal prob of N-in: 0.78331\nPOSSIBLE N-term signal sequence\ninside 1 24\nTMhelix 25 47\noutside 48 404\nTMhelix 405 427\ninside 428 428

29347 GCF_001055405.1_ASM105540v1 Corynebacterium striatum Terrabacteria group; Actinobacteria; Actinobacteria; Corynebacteriales; Corynebacteriaceae; Corynebacterium MTIAVPGPNPADVAERRRTLRKYQWGVTGLLILAAVIFFACSWWQHHNPDAPTWVGYVRA WP_049062446.1 DUF445 domain-containing protein [Corynebacterium striatum] Length: 428\nNumber of predicted TMHs: 2\nExp number of AAs in TMHs: 52.76731\nExp number, first 60 AAs: 23.54963\nTotal prob of N-in: 0.78331\nPOSSIBLE N-term signal sequence\ninside 1 24\nTMhelix 25 47\noutside 48 404\nTMhelix 405 427\ninside 428 428

29348 GCF_001053405.1_ASM105340v1 Corynebacterium striatum Terrabacteria group; Actinobacteria; Actinobacteria; Corynebacteriales; Corynebacteriaceae; Corynebacterium MTSRRGTFLAALVTASLIPLAPPALAIPATDGHLTWGIRASFNNYTGGATLVKDGATRKG WP_049151632.1 hypothetical protein [Corynebacterium striatum] Length: 665\nNumber of predicted TMHs: 2\nExp number of AAs in TMHs: 37.10191\nExp number, first 60 AAs: 17.62673\nTotal prob of N-in: 0.78737\nPOSSIBLE N-term signal sequence\ninside 1 6\nTMhelix 7 29\noutside 30 626\nTMhelix 627 649\ninside 650 665

29349 GCF_001053435.1_ASM105343v1 Corynebacterium striatum Terrabacteria group; Actinobacteria; Actinobacteria; Corynebacteriales; Corynebacteriaceae; Corynebacterium MTIAVPGPNPADVAERRRTLRKYQWGVTGLLILAAVIFFACSWWQHHNPDAPTWVGYVRA WP_049151243.1 MULTISPECIES: DUF445 domain-containing protein [Corynebacterium] Length: 428\nNumber of predicted TMHs: 2\nExp number of AAs in TMHs: 52.76767\nExp number, first 60 AAs: 23.54962\nTotal prob of N-in: 0.78331\nPOSSIBLE N-term signal sequence\ninside 1 24\nTMhelix 25 47\noutside 48 404\nTMhelix 405 427\ninside 428 428

29350 GCF_000988225.1_ASM98822v1 Corynebacterium striatum Terrabacteria group; Actinobacteria; Actinobacteria; Corynebacteriales; Corynebacteriaceae; Corynebacterium MTSRRGTLLAALVTASLIPLAPPALAIPATDGHLTWGIRASFNNYTGGATLVKDGATRKG WP_046646201.1 hypothetical protein [Corynebacterium striatum] Length: 673\nNumber of predicted TMHs: 2\nExp number of AAs in TMHs: 37.05714\nExp number, first 60 AAs: 17.51593\nTotal prob of N-in: 0.78181\nPOSSIBLE N-term signal sequence\ninside 1 6\nTMhelix 7 29\noutside 30 634\nTMhelix 635 657\ninside 658 673

29351 GCF_001058455.1_ASM105845v1 Corynebacterium striatum Terrabacteria group; Actinobacteria; Actinobacteria; Corynebacteriales; Corynebacteriaceae; Corynebacterium MTSRRGTFLAALVTASLIPLAPPALAIPATDGHLTWGIRASFNNYTGGATLVKDGATRKG WP_049064233.1 hypothetical protein [Corynebacterium striatum]	Length: 661\nNumber of predicted TMHs:  2\nExp number of AAs in TMHs: 37.19204\nExp number, first 60 AAs:  17.6393\nTotal prob of N-in:   0.78729\nPOSSIBLE N-term signal sequence\ninside   1   6\nTMhelix   7   29\noutside   30   622\nTMhelix   623   645\ninside   646   661

29352	GCF_001076975.1_ASM107697v1	Corynebacterium striatum	Terrabacteria group; Actinobacteria; Actinobacteria; Corynebacteriales; Corynebacteriaceae; Corynebacterium	MTSRRGTFLAALVTASLIPLAPPALAIPATDGHLTWGIRASFNNYTGGATLVKDGATRKG	WP_049146658.1	hypothetical protein [Corynebacterium striatum]	Length: 661\nNumber of predicted TMHs:  2\nExp number of AAs in TMHs: 37.19209\nExp number, first 60 AAs:  17.6393\nTotal prob of N-in:   0.78729\nPOSSIBLE N-term signal sequence\ninside   1   6\nTMhelix   7   29\noutside   30   622\nTMhelix   623   645\ninside   646   661

29353	GCF_000988225.1_ASM98822v1	Corynebacterium striatum	Terrabacteria group; Actinobacteria; Actinobacteria; Corynebacteriales; Corynebacteriaceae; Corynebacterium	MTIAVPGPNPADVAERRRTLRKYQWGVTGLLILAAVIFFACSWWQHHNPDAPTWVGYVRA	WP_046645688.1	DUF445 domain-containing protein [Corynebacterium striatum]	Length: 428\nNumber of predicted TMHs: 2\nExp number of AAs in TMHs: 52.76767\nExp number, first 60 AAs:  23.54962\nTotal prob of N-in:   0.78331\nPOSSIBLE N-term signal sequence\ninside   1   24\nTMhelix   25   47\noutside   48   404\nTMhelix   405   427\ninside   428   428

29354	GCF_001053435.1_ASM105343v1	Corynebacterium striatum	Terrabacteria group; Actinobacteria; Actinobacteria; Corynebacteriales; Corynebacteriaceae; Corynebacterium	MTSRRGTFLAALVTASLIPLAPPALAIPATDGHLTWGIRASFNNYTGGATLVKDGATRKG	WP_049151632.1	hypothetical protein [Corynebacterium striatum]	Length: 665\nNumber of predicted TMHs:  2\nExp number of AAs in TMHs: 37.10191\nExp number, first 60 AAs:  17.62673\nTotal prob of N-in:   0.78737\nPOSSIBLE N-term signal sequence\ninside   1   6\nTMhelix   7   29\noutside   30   626\nTMhelix   627   649\ninside   650   665

29355	GCF_001076975.1_ASM107697v1	Corynebacterium striatum	Terrabacteria group; Actinobacteria; Actinobacteria; Corynebacteriales; Corynebacteriaceae; Corynebacterium	MTIAVPGPNPADVAERRRTLRKYQWGVTGLLILAAVIFFACSWWQYHNPDAPTWVGYVRA	WP_049146110.1	DUF445 domain-containing protein [Corynebacterium striatum]	Length: 428\nNumber of predicted TMHs: 2\nExp number of AAs in TMHs: 51.9667500000001\nExp number, first 60 AAs:  23.77035\nTotal prob of N-in: 0.85104\nPOSSIBLE N-term signal sequence\ninside   1   24\nTMhelix   25   47\noutside   48   404\nTMhelix   405   427\ninside   428   428

29356	GCF_001058455.1_ASM105845v1	Corynebacterium striatum	Terrabacteria group; Actinobacteria; Actinobacteria; Corynebacteriales; Corynebacteriaceae; Corynebacterium	MTIAVPGPNPADVAERRRTLRKYQWGVTGLLILAAVIFFACSWWQHHNPDAPTWVGYVRA	WP_049192460.1	DUF445 domain-containing protein [Corynebacterium striatum]	Length: 428\nNumber of predicted TMHs: 2\nExp number of AAs in TMHs: 52.76767\nExp number, first 60 AAs:  23.54962\nTotal prob of N-in:   0.78331\nPOSSIBLE N-term signal sequence\ninside   1   24\nTMhelix   25   47\noutside   48   404\nTMhelix   405   427\ninside   428   428

29357	GCF_001059665.1_ASM105966v1	Corynebacterium striatum	Terrabacteria group; Actinobacteria; Actinobacteria; Corynebacteriales; Corynebacteriaceae; Corynebacterium	MTSRRGTFLAALVTASLIPLAPPALAIPATDGHLTWGIRASFNNYTGGATLVKDGATRKG	WP_049160753.1	hypothetical protein [Corynebacterium striatum]	Length: 653\nNumber of predicted TMHs:  2\nExp number of AAs in TMHs: 37.18925\nExp number, first 60 AAs:  17.6374\nTotal prob of N-in:   0.78731\nPOSSIBLE N-term signal sequence\ninside   1   6\nTMhelix   7   29\noutside   30   614\nTMhelix   615   637\ninside   638   653

29358	GCF_001059955.1_ASM105995v1	Corynebacterium striatum	Terrabacteria group; Actinobacteria; Actinobacteria; Corynebacteriales; Corynebacteriaceae; Corynebacterium	MTIAVPGPNPADVAERRRTLRKYQWGVTGLLILAAVIFFACSWWQHHNPDAPTWVGYVRA	WP_049160355.1	DUF445 domain-containing protein [Corynebacterium striatum]	Length: 428\nNumber of predicted TMHs: 2\nExp number of AAs in TMHs: 52.76767\nExp number, first 60 AAs:  23.54962\nTotal prob of N-in:   0.78331\nPOSSIBLE N-term signal sequence\ninside   1   24\nTMhelix   25   47\noutside   48   404\nTMhelix   405   427\ninside   428   428

29359	GCF_001055405.1_ASM105540v1	Corynebacterium striatum	Terrabacteria group; Actinobacteria; Actinobacteria; Corynebacteriales; Corynebacteriaceae; Corynebacterium	MTSRRGTFLAALVTASLIPLAPPALAIPATDGHLTWGIRASFNNYTGGATLVKDGATRKG	WP_049064233.1	hypothetical protein [Corynebacterium striatum]	Length: 661\nNumber of predicted TMHs:  2\nExp number of AAs in TMHs: 37.19204\nExp number, first 60 AAs:  17.6393\nTotal prob of N-in:   0.78729\nPOSSIBLE N-term signal sequence\ninside   1   6\nTMhelix   7   29\noutside   30   622\nTMhelix   623   645\ninside   646   661

29360 GCF_002156805.1_ASM215680v1 Corynebacterium striatum Terrabacteria group; Actinobacteria; Actinobacteria; Corynebacteriales; Corynebacteriaceae; Corynebacterium MTIAVPGPNPADVAERRRTLRKYQWGVTSLLILAAVIFFACSWWQHHNPDAPTWVGYVRA WP_086892183.1 DUF445 domain-containing protein [Corynebacterium striatum] Length: 428\nNumber of predicted TMHs: 2\nExp number of AAs in TMHs: 52.26671\nExp number, first 60 AAs: 23.38147\nTotal prob of N-in: 0.80419\nPOSSIBLE N-term signal sequence\ninside 1 24\nTMhelix 25 47\noutside 48 404\nTMhelix 405 427\ninside 428 428

29361 GCF_001059665.1_ASM105966v1 Corynebacterium striatum Terrabacteria group; Actinobacteria; Actinobacteria; Corynebacteriales; Corynebacteriaceae; Corynebacterium MTIAVPGPNPADVAERRRTLRKYQWGVTGLLILAAVIFFACSWWQHHNPDAPTWVGYVRA WP_049160355.1 DUF445 domain-containing protein [Corynebacterium striatum] Length: 428\nNumber of predicted TMHs: 2\nExp number of AAs in TMHs: 52.76767\nExp number, first 60 AAs: 23.54962\nTotal prob of N-in: 0.78331\nPOSSIBLE N-term signal sequence\ninside 1 24\nTMhelix 25 47\noutside 48 404\nTMhelix 405 427\ninside 428 428

29362 GCF_900105305.1_IMG-taxon_2636416018_annotated_assembly Corynebacterium timonense Terrabacteria group; Actinobacteria; Actinobacteria; Corynebacteriales; Corynebacteriaceae; Corynebacterium MKIAYKRRGAIALAVCLTLSPLNPVLSPVAAAVERPLSLSWGVRASFNNYSKGPTHMLGG WP_019193688.1 hypothetical protein [Corynebacterium timonense] Length: 700\nNumber of predicted TMHs: 1\nExp number of AAs in TMHs: 33.0897699999999\nExp number, first 60 AAs: 10.66706\nTotal prob of N-in: 0.57095\nPOSSIBLE N-term signal sequence\noutside 1 661\nTMhelix 662 684\ninside 685 700

29363 GCF_900105305.1_IMG-taxon_2636416018_annotated_assembly Corynebacterium timonense Terrabacteria group; Actinobacteria; Actinobacteria; Corynebacteriales; Corynebacteriaceae; Corynebacterium MKAINRRFLARRGTVLTATAATAALTLSMVSPAGAEEGSSLKGSSVGSSKVTTGTVEAGSWP_081582885.1 lipase [Corynebacterium timonense] Length: 541\nNumber of predicted TMHs: 2\nExp number of AAs in TMHs: 38.3442499999999\nExp number, first 60 AAs: 16.25429\nTotal prob of N-in: 0.73553\nPOSSIBLE N-term signal sequence\ninside 1 12\nTMhelix 13 35\noutside 36 509\nTMhelix 510 532\ninside 533 541

29364 GCF_001302345.1_ASM130234v1 Corynebacterium ulcerans Terrabacteria group; Actinobacteria; Actinobacteria; Corynebacteriales; Corynebacteriaceae; Corynebacterium MDSKDTGVDAAGMGARNTLFVRSEKHARRAPIIAFVTAAALFIIAAAGPALMISRIKPIP WP_054314932.1 hypothetical protein [Corynebacterium ulcerans] Length: 395\nNumber of predicted TMHs: 2\nExp number of AAs in TMHs: 43.50942\nExp number, first 60 AAs: 22.4666\nTotal prob of N-in: 0.60803\nPOSSIBLE N-term signal sequence\noutside 1 31\nTMhelix 32 54\ninside 55 363\nTMhelix 364 386\noutside 387 395

29365 GCF_001021065.1_ASM102106v1 Corynebacterium uterequi Terrabacteria group; Actinobacteria; Actinobacteria; Corynebacteriales; Corynebacteriaceae; Corynebacterium MISSLRRRTALTGVAALTALTALTALAGATGDVAAQPVDPALEQNVTSAEPVIVGEPVIIPAG WP_052844124.1 hypothetical protein [Corynebacterium uterequi] Length: 286\nNumber of predicted TMHs: 1\nExp number of AAs in TMHs: 29.06309\nExp number, first 60 AAs: 6.03143\nTotal prob of N-in: 0.35103\noutside 1 258\nTMhelix 259 281\ninside 282 286

29366 GCF_001639025.1_ASM163902v1 Corynebacterium afermentans subsp. afermentans Terrabacteria group; Actinobacteria; Actinobacteria; Corynebacteriales; Corynebacteriaceae; Corynebacterium; Corynebacterium afermentans MRPAPFSPRTSRRAALLAVPAALLLAAPIAPVAHAQLPDGMSPEEVQSMIPGEISVPAGE WP_082855643.1 hypothetical protein [Corynebacterium afermentans] Length: 376\nNumber of predicted TMHs: 2\nExp number of AAs in TMHs: 39.93319\nExp number, first 60 AAs: 17.68082\nTotal prob of N-in: 0.80899\nPOSSIBLE N-term signal sequence\ninside 1 12\nTMhelix 13 35\noutside 36 346\nTMhelix 347 369\ninside 370 376

29367 GCF_001941445.1_ASM194144v1 Corynebacterium aquilae DSM 44791 Terrabacteria group; Actinobacteria; Actinobacteria; Corynebacteriales; Corynebacteriaceae; Corynebacterium; Corynebacterium aquilae MLKTRRRLIAGLCSCALLAGAVTPAHADIPNEFDKSLDILFFEALESGSSASGNDPDSPE WP_075727961.1 hypothetical protein [Corynebacterium aquilae] Length: 171\nNumber of predicted TMHs: 1\nExp number of AAs in TMHs: 25.05072\nExp number, first 60 AAs: 2.34231\nTotal prob of N-in: 0.95987\ninside 1 134\nTMhelix 135 157\noutside 158 171

29368 GCF_000550785.1_ASM55078v1 Corynebacterium casei LMG S-19264 Terrabacteria group; Actinobacteria; Actinobacteria; Corynebacteriales; Corynebacteriaceae; Corynebacterium; Corynebacterium casei MNAISNYTSALNQRAQLIQARRRRFLTAVVASSLAAATLSMSVPVVSAQDVEPTELDGMA WP_006821438.1 D-alanyl-D-alanine carboxypeptidase [Corynebacterium casei] Length: 496\nNumber of predicted TMHs: 2\nExp

number of AAs in TMHs: 40.36576\nExp number, first 60 AAs: 17.47733\nTotal prob of N-in: 0.78513\nPOSSIBLE N-term signal sequence\ninside 1 24\nTMhelix 25 47\noutside 48 460\nTMhelix 461 483\ninside 484 496

29369 GCF_000234765.1_ASM23476v1 Corynebacterium casei UCMA 3821 Terrabacteria group; Actinobacteria; Actinobacteria; Corynebacteriales; Corynebacteriaceae; Corynebacterium; Corynebacterium casei MNAISNYTSALNQRAQLIQARRRRFLTAVVASSLAAATLSMSVPVVSAQDVEPTELDGMA WP_006821438.1 D-alanyl-D-alanine carboxypeptidase [Corynebacterium casei] Length: 496\nNumber of predicted TMHs: 2\nExp number of AAs in TMHs: 40.36576\nExp number, first 60 AAs: 17.47733\nTotal prob of N-in: 0.78513\nPOSSIBLE N-term signal sequence\ninside 1 24\nTMhelix 25 47\noutside 48 460\nTMhelix 461 483\ninside 484 496

29370 GCF_000380545.1_contig Corynebacterium crenatum MT Terrabacteria group; Actinobacteria; Actinobacteria; Corynebacteriales; Corynebacteriaceae; Corynebacterium; Corynebacterium crenatum MGLPFKKAGPLSRRKAVFSALGAAALIGAALPTIPTAQAQTPTGYGFDATASIGEEPEFS WP_003856086.1 MULTISPECIES: hypothetical protein [Corynebacterium] Length: 919\nNumber of predicted TMHs: 1\nExp number of AAs in TMHs: 33.37495\nExp number, first 60 AAs: 11.01947\nTotal prob of N-in: 0.50897\nPOSSIBLE N-term signal sequence\noutside 1 894\nTMhelix 895 917\ninside 918 919

29371 GCF_000372245.1_ASM37224v1 Corynebacterium doosanense CAU 212 = DSM 45436 Terrabacteria group; Actinobacteria; Actinobacteria; Corynebacteriales; Corynebacteriaceae; Corynebacterium; Corynebacterium doosanense MNFSTRPSRRSPLRTGAAVAAAAGLLTLGAPIAIAHDAVIGGDPADGATVSEFPESVTLE WP_018023019.1 hypothetical protein [Corynebacterium doosanense] Length: 206\nNumber of predicted TMHs: 2\nExp number of AAs in TMHs: 43.97473\nExp number, first 60 AAs: 22.3808\nTotal prob of N-in: 0.98699\nPOSSIBLE N-term signal sequence\ninside 1 12\nTMhelix 13 35\noutside 36 164\nTMhelix 165 187\ninside 188 206

29372 GCF_000767055.1_ASM76705v1 Corynebacterium doosanense CAU 212 = DSM 45436 Terrabacteria group; Actinobacteria; Actinobacteria; Corynebacteriales; Corynebacteriaceae; Corynebacterium; Corynebacterium doosanense MNFSTRPSRRSPLRTGAAVAAAAGLLTLGAPIAIAHDAVIGGDPADGATVSEFPESVTLE WP_018023019.1 hypothetical protein [Corynebacterium doosanense] Length: 206\nNumber of predicted TMHs: 2\nExp number of AAs in TMHs: 43.97473\nExp number, first 60 AAs: 22.3808\nTotal prob of N-in: 0.98699\nPOSSIBLE N-term signal sequence\ninside 1 12\nTMhelix 13 35\noutside 36 164\nTMhelix 165 187\ninside 188 206

29373 GCF_000318135.1_ASM31813v1 Corynebacterium durum F0235 Terrabacteria group; Actinobacteria; Actinobacteria; Corynebacteriales; Corynebacteriaceae; Corynebacterium; Corynebacterium durum MGFLSLFRHNHRVATLKPMDDDHFRIGDNDRRRALESLTSLAASGHLTVDEFDERSAAAA WP_040359116.1 hypothetical protein [Corynebacterium durum] Length: 201\nNumber of predicted TMHs: 1\nExp number of AAs in TMHs: 37.33378\nExp number, first 60 AAs: 0.00035\nTotal prob of N-in: 0.67113\noutside 1 139\nTMhelix 140 162\ninside 163 201

29374 GCF_000318135.1_ASM31813v1 Corynebacterium durum F0235 Terrabacteria group; Actinobacteria; Actinobacteria; Corynebacteriales; Corynebacteriaceae; Corynebacterium; Corynebacterium durum MRLSFHQSPLLPTPATRRPIVTTVGTAAVVTLAALLTPPASSAFTVTADTEKQLCRVDAN WP_006062599.1 hypothetical protein [Corynebacterium durum] Length: 258\nNumber of predicted TMHs: 1\nExp number of AAs in TMHs: 24.52819\nExp number, first 60 AAs: 5.71495\nTotal prob of N-in: 0.20196\noutside 1 227\nTMhelix 228 247\ninside 248 258

29375 GCF_000160795.1_ASM16079v1 Corynebacterium efficiens YS-314 Terrabacteria group; Actinobacteria; Actinobacteria; Corynebacteriales; Corynebacteriaceae; Corynebacterium; Corynebacterium efficiens MKTLKRRGLLAKAGALLAVAALAFSASPSYAQTNDVPETANVVITKLEQPTVAGSVASGQ WP_006769188.1 type 1 fimbrial protein [Corynebacterium efficiens] Length: 498\nNumber of predicted TMHs: 1\nExp number of AAs in TMHs: 28.90576\nExp number, first 60 AAs: 6.94614\nTotal prob of N-in: 0.34164\noutside 1 467\nTMhelix 468 490\ninside 491 498

29376 GCF_000011305.1_ASM1130v1 Corynebacterium efficiens YS-314 Terrabacteria group; Actinobacteria; Actinobacteria; Corynebacteriales; Corynebacteriaceae; Corynebacterium; Corynebacterium efficiens MKTLKRRGLLAKAGALLAVAALAFSASPSYAQTNDVPETANVVITKLEQPTVAGSVASGQ WP_006769188.1 type 1 fimbrial protein [Corynebacterium efficiens] Length: 498\nNumber of predicted TMHs: 1\nExp number of AAs in TMHs: 28.90576\nExp number, first 60 AAs: 6.94614\nTotal prob of N-in: 0.34164\noutside 1 467\nTMhelix 468 490\ninside 491 498

29377 GCF_000525655.1_ASM52565v1 Corynebacterium falsenii DSM 44353 Terrabacteria group; Actinobacteria; Actinobacteria; Corynebacteriales; Corynebacteriaceae; Corynebacterium; Corynebacterium falsenii

MTNNTPSRRTFLTGLGLGAGAGALAAGTGVWAATDRAGATPSDTVPFYGMHQAGIDTSQQ WP_025402979.1 deferrochelatase/peroxidase EfeB [Corynebacterium falsenii] Length: 418\nNumber of predicted TMHs: 2\nExp number of AAs in TMHs: 39.46359\nExp number, first 60 AAs: 21.78232\nTotal prob of N-in: 0.89324\nPOSSIBLE N-term signal sequence\ninside 1 11\nTMhelix 12 34\noutside 35 355\nTMhelix 356 378\ninside 379 418

29378 GCF_000525655.1_ASM52565v1 Corynebacterium falsenii DSM 44353 Terrabacteria group; Actinobacteria; Actinobacteria; Corynebacteriales; Corynebacteriaceae; Corynebacterium; Corynebacterium falsenii MTHETPEPTPTKRRGVLPKSTLALGIGLLIVAFVAGLVATMNVKNFSEGYDVQANKQYYL WP_025403387.1 hypothetical protein [Corynebacterium falsenii] Length: 179\nNumber of predicted TMHs: 2\nExp number of AAs in TMHs: 44.41195\nExp number, first 60 AAs: 22.01817\nTotal prob of N-in: 0.89322\nPOSSIBLE N-term signal sequence\ninside 1 20\nTMhelix 21 43\noutside 44 145\nTMhelix 146 168\ninside 169 179

29379 GCF_001941485.1_ASM194148v1 Corynebacterium frankenforstense DSM 45800 Terrabacteria group; Actinobacteria; Actinobacteria; Corynebacteriales; Corynebacteriaceae; Corynebacterium; Corynebacterium frankenforstense MTPRRGILKAAALAAAMLLMAPSAAVGQETADPAATAGESTASLVPDADTAGTAGTADGA WP_075663971.1 hypothetical protein [Corynebacterium frankenforstense] Length: 1159\nNumber of predicted TMHs: 1\nExp number of AAs in TMHs: 25.44342\nExp number, first 60 AAs: 4.82024\nTotal prob of N-in: 0.23741\noutside 1 1135\nTMhelix 1136 1155\ninside 1156 1159

29380 GCF_001941485.1_ASM194148v1 Corynebacterium frankenforstense DSM 45800 Terrabacteria group; Actinobacteria; Actinobacteria; Corynebacteriales; Corynebacteriaceae; Corynebacterium; Corynebacterium frankenforstense MGLVGQPGTRRRAALATLAAVLVLLLVGDVAPRVVLYLMYDLEVGVSRSVRSVDPDARVV WP_075664640.1 hypothetical protein [Corynebacterium frankenforstense] Length: 411\nNumber of predicted TMHs: 2\nExp number of AAs in TMHs: 43.75991\nExp number, first 60 AAs: 20.97276\nTotal prob of N-in: 0.94191\nPOSSIBLE N-term signal sequence\ninside 1 12\nTMhelix 13 35\noutside 36 377\nTMhelix 378 400\ninside 401 411

29381 GCF_000010225.1_ASM1022v1 Corynebacterium glutamicum R Terrabacteria group; Actinobacteria; Actinobacteria; Corynebacteriales; Corynebacteriaceae; Corynebacterium; Corynebacterium glutamicum MGLPFKKAGPLSRRKAVFSALGAAALIGAALPTIPTAQAQTPTGYGFDATASIGEEPEFS WP_044027174.1 hypothetical protein [Corynebacterium glutamicum] Length: 919\nNumber of predicted TMHs: 1\nExp number of AAs in TMHs: 33.36401\nExp number, first 60 AAs: 11.01947\nTotal prob of N-in: 0.50897\nPOSSIBLE N-term signal sequence\noutside 1 894\nTMhelix 895 917\ninside 918 919

29382 GCF_000224315.1_ASM22431v2 Corynebacterium glutamicum S9114 Terrabacteria group; Actinobacteria; Actinobacteria; Corynebacteriales; Corynebacteriaceae; Corynebacterium; Corynebacterium glutamicum MGLPFKKAGPLSRRKAVFSALGAAALIGAALPTIPTAQAQTPTGYGFDATASIGEEPEFS WP_003856086.1 MULTISPECIES: hypothetical protein [Corynebacterium] Length: 919\nNumber of predicted TMHs: 1\nExp number of AAs in TMHs: 33.37495\nExp number, first 60 AAs: 11.01947\nTotal prob of N-in: 0.50897\nPOSSIBLE N-term signal sequence\noutside 1 894\nTMhelix 895 917\ninside 918 919

29383 GCF_000404145.1_ASM40414v1 Corynebacterium glutamicum SCgG1 Terrabacteria group; Actinobacteria; Actinobacteria; Corynebacteriales; Corynebacteriaceae; Corynebacterium; Corynebacterium glutamicum MGLPFKKAGPLSRRKAVFSALGAAALIGAALPTIPTAQAQTPTGYGFDATASIGEEPEFS WP_003856086.1 MULTISPECIES: hypothetical protein [Corynebacterium] Length: 919\nNumber of predicted TMHs: 1\nExp number of AAs in TMHs: 33.37495\nExp number, first 60 AAs: 11.01947\nTotal prob of N-in: 0.50897\nPOSSIBLE N-term signal sequence\noutside 1 894\nTMhelix 895 917\ninside 918 919

29384 GCF_000404185.1_ASM40418v1 Corynebacterium glutamicum SCgG2 Terrabacteria group; Actinobacteria; Actinobacteria; Corynebacteriales; Corynebacteriaceae; Corynebacterium; Corynebacterium glutamicum MGLPFKKAGPLSRRKAVFSALGAAALIGAALPTIPTAQAQTPTGYGFDATASIGEEPEFS WP_003856086.1 MULTISPECIES: hypothetical protein [Corynebacterium] Length: 919\nNumber of predicted TMHs: 1\nExp number of AAs in TMHs: 33.37495\nExp number, first 60 AAs: 11.01947\nTotal prob of N-in: 0.50897\nPOSSIBLE N-term signal sequence\noutside 1 894\nTMhelix 895 917\ninside 918 919

29385 GCF_000417765.1_ASM41776v1 Corynebacterium glutamicum Z188 Terrabacteria group; Actinobacteria; Actinobacteria; Corynebacteriales; Corynebacteriaceae; Corynebacterium; Corynebacterium glutamicum MGLPFKKAGPLSRRKAVFSALGAAALIGAALPTIPTAQAQTPTGYGFDATASIGEEPEFS WP_003856086.1 MULTISPECIES: hypothetical protein [Corynebacterium] Length: 919\nNumber of predicted TMHs: 1\nExp number of AAs in TMHs: 33.37495\nExp number, first 60 AAs: 11.01947\nTotal prob of N-in: 0.50897\nPOSSIBLE N-term signal sequence\noutside 1 894\nTMhelix 895 917\ninside 918 919

29386 GCF_000626675.1_ASM62667v1 Corynebacterium glyciniphilum AJ 3170 Terrabacteria group; Actinobacteria; Actinobacteria; Corynebacteriales; Corynebacteriaceae; Corynebacterium; Corynebacterium glyciniphilum MTRFQRLQRFHRSPRRPAVGRVLLSALLLSTMPIPSSDSQPRCDHPAFGDAIPVDVQLPH WP_081803761.1 hypothetical protein [Corynebacterium glyciniphilum] Length: 379\nNumber of predicted TMHs: 1\nExp number of AAs in TMHs: 24.31638\nExp number, first 60 AAs: 0.42114\nTotal prob of N-in: 0.17441\noutside 1 355\nTMhelix 356 378\ninside 379 379

29387 GCF_000626675.1_ASM62667v1 Corynebacterium glyciniphilum AJ 3170 Terrabacteria group; Actinobacteria; Actinobacteria; Corynebacteriales; Corynebacteriaceae; Corynebacterium; Corynebacterium glyciniphilum MTAHTLPRRRTLRATTLATALAATLTISLTGPLTAPAVAETGLNNIEKDDWLPLTVDPGA WP_038546560.1 D-alanyl-D-alanine carboxypeptidase [Corynebacterium glyciniphilum] Length: 440\nNumber of predicted TMHs: 1\nExp number of AAs in TMHs: 34.43\nExp number, first 60 AAs: 13.72465\nTotal prob of N-in: 0.63297\nPOSSIBLE N-term signal sequence\noutside 1 415\nTMhelix 416 435\ninside 436 440

29388 GCF_000375365.1_ASM37536v1 Corynebacterium mastitidis DSM 44356 Terrabacteria group; Actinobacteria; Actinobacteria; Corynebacteriales; Corynebacteriaceae; Corynebacterium; Corynebacterium mastitidis MRRQGKRRCRVAARIARRGAIALSLGIAGGLISPGVALAHDSVIAATPEDGGTVEEFPRD WP_026166134.1 copper resistance protein [Corynebacterium mastitidis] Length: 193\nNumber of predicted TMHs: 2\nExp number of AAs in TMHs: 42.53094\nExp number, first 60 AAs: 20.18822\nTotal prob of N-in: 0.99965\nPOSSIBLE N-term signal sequence\ninside 1 19\nTMhelix 20 39\noutside 40 165\nTMhelix 166 188\ninside 189 193

29389 GCF_000375525.1_ASM37552v1 Corynebacterium propinquum DSM 44285 Terrabacteria group; Actinobacteria; Actinobacteria; Corynebacteriales; Corynebacteriaceae; Corynebacterium; Corynebacterium propinquum MNESKPGRRALVRYGSTCCAAVLATGLVTGTTAFGIGTLNSAAPHAAAAPLIHAQQEEPT WP_018121428.1 hypothetical protein [Corynebacterium propinquum] Length: 459\nNumber of predicted TMHs: 2\nExp number of AAs in TMHs: 43.30554\nExp number, first 60 AAs: 20.39469\nTotal prob of N-in: 0.91193\nPOSSIBLE N-term signal sequence\ninside 1 11\nTMhelix 12 34\noutside 35 433\nTMhelix 434 456\ninside 457 459

29390 GCF_000466825.2_ASM46682v2 Corynebacterium pseudodiphtheriticum 090104 Terrabacteria group; Actinobacteria; Actinobacteria; Corynebacteriales; Corynebacteriaceae; Corynebacterium; Corynebacterium pseudodiphtheriticum MNESKPGRRAIVRYGSTCCANLLVVGMATGAATLGIGAFGSATPHAAATPHAEAAPLLHA WP_021351856.1 hypothetical protein [Corynebacterium pseudodiphtheriticum] Length: 467\nNumber of predicted TMHs: 2\nExp number of AAs in TMHs: 44.73665\nExp number, first 60 AAs: 21.79333\nTotal prob of N-in: 0.98221\nPOSSIBLE N-term signal sequence\ninside 1 19\nTMhelix 20 42\noutside 43 441\nTMhelix 442 464\ninside 465 467

29391 GCF_000688415.1_ASM68841v1 Corynebacterium pseudodiphtheriticum DSM 44287 Terrabacteria group; Actinobacteria; Actinobacteria; Corynebacteriales; Corynebacteriaceae; Corynebacterium; Corynebacterium pseudodiphtheriticum MNESKPGRRAIVRYSSTCCANLLVVGMATGAATLGIGAFGSASPHAAATPHTEAAPLLHA WP_027017577.1 D-alanyl-D-alanine carboxypeptidase [Corynebacterium pseudodiphtheriticum] Length: 467\nNumber of predicted TMHs: 2\nExp number of AAs in TMHs: 44.48778\nExp number, first 60 AAs: 21.59742\nTotal prob of N-in: 0.98117\nPOSSIBLE N-term signal sequence\ninside 1 19\nTMhelix 20 42\noutside 43 441\nTMhelix 442 464\ninside 465 467

29392 GCF_000159135.1_ASM15913v1 Corynebacterium striatum ATCC 6940 Terrabacteria group; Actinobacteria; Actinobacteria; Corynebacteriales; Corynebacteriaceae; Corynebacterium; Corynebacterium striatum MTIAVPGPNPADVAERRRTLRKYQWGVTGLLILAAVIFFACSWWQHHNPDAPTWVGYVRA WP_005531173.1 DUF445 domain-containing protein [Corynebacterium striatum] Length: 428\nNumber of predicted TMHs: 2\nExp number of AAs in TMHs: 52.76767\nExp number, first 60 AAs: 23.54962\nTotal prob of N-in: 0.78331\nPOSSIBLE N-term signal sequence\ninside 1 24\nTMhelix 25 47\noutside 48 404\nTMhelix 405 427\ninside 428 428

29393 GCF_000159135.1_ASM15913v1 Corynebacterium striatum ATCC 6940 Terrabacteria group; Actinobacteria; Actinobacteria; Corynebacteriales; Corynebacteriaceae; Corynebacterium; Corynebacterium striatum MTSRRGTFLAALVTASLIPLAPPALAIPATDGHLTWGIRASFNNYTGGATLVKDGATRKG WP_005531760.1 hypothetical protein [Corynebacterium striatum] Length: 643\nNumber of predicted TMHs: 2\nExp number of AAs in TMHs: 37.17762\nExp number, first 60 AAs: 17.61764\nTotal prob of N-in: 0.78742\nPOSSIBLE N-term signal sequence\ninside 1 6\nTMhelix 7 29\noutside 30 604\nTMhelix 605 627\ninside 628 643

29394 GCF_000312345.1_ASM31234v1 Corynebacterium timonense 5401744 Terrabacteria group; Actinobacteria; Actinobacteria; Corynebacteriales; Corynebacteriaceae; Corynebacterium; Corynebacterium timonense MKAINRRFLARRGTVLTATAATAALTLSMVSPAGAEEGSSLKGSSVGSSKVTTGTVEAGS WP_081582885.1 lipase

[Corynebacterium timonense] Length: 541\nNumber of predicted TMHs: 2\nExp number of AAs in TMHs: 38.3442499999999\nExp number, first 60 AAs: 16.25429\nTotal prob of N-in: 0.73553\nPOSSIBLE N-term signal sequence\ninside 1 12\nTMhelix 13 35\noutside 36 509\nTMhelix 510 532\ninside 533 541

29395 GCF_000312345.1_ASM31234v1 Corynebacterium timonense 5401744 Terrabacteria group; Actinobacteria; Actinobacteria; Corynebacteriales; Corynebacteriaceae; Corynebacterium; Corynebacterium timonense MKIAYKRRGAIALAVCLTLSPLNPVLSPVAAAVERPLSLSWGVRASFNNYSKGPTHMLGG WP_019193688.1 hypothetical protein [Corynebacterium timonense] Length: 700\nNumber of predicted TMHs: 1\nExp number of AAs in TMHs: 33.0897699999999\nExp number, first 60 AAs: 10.66706\nTotal prob of N-in: 0.57095\nPOSSIBLE N-term signal sequence\noutside 1 661\nTMhelix 662 684\ninside 685 700

29396 GCF_000333435.1_ASM33343v1 Corynebacterium-like bacterium B27 Terrabacteria group; Actinobacteria; Actinobacteria; Corynebacteriales; Corynebacteriaceae; unclassified Corynebacteriaceae MALSAHQGGGRRRTLILASAALLTAASIGVPAAAYADITDNGGATRNDGDKAAALRASIV WP_031284473.1 alkaline phosphatase [Corynebacterium-like bacterium B27] Length: 631\nNumber of predicted TMHs: 2\nExp number of AAs in TMHs: 41.3861999999999\nExp number, first 60 AAs: 19.12163\nTotal prob of N-in: 0.84576\nPOSSIBLE N-term signal sequence\ninside 1 12\nTMhelix 13 35\noutside 36 601\nTMhelix 602 624\ninside 625 631

29397 GCF_000333435.1_ASM33343v1 Corynebacterium-like bacterium B27 Terrabacteria group; Actinobacteria; Actinobacteria; Corynebacteriales; Corynebacteriaceae; unclassified Corynebacteriaceae MMHTIDIARRAAVAVAIGLLVALAAVLGAAAPAALAHGGPFQLTVSPDGAGGLIVNAAYT WP_051048134.1 hypothetical protein [Corynebacterium-like bacterium B27] Length: 201\nNumber of predicted TMHs: 2\nExp number of AAs in TMHs: 45.23913\nExp number, first 60 AAs: 22.78555\nTotal prob of N-in: 0.99996\nPOSSIBLE N-term signal sequence\ninside 1 12\nTMhelix 13 35\noutside 36 169\nTMhelix 170 192\ninside 193 201

29398 GCF_001722515.1_ASM172251v1 Dietzia alimentaria Terrabacteria group; Actinobacteria; Actinobacteria; Corynebacteriales; Dietziaceae; Dietzia MNSPLPPAADTPSPDPVVAPARPVSTKRRRKAVGALVIAGGLVGAGAVATAVIPEPQVAT WP_082923770.1 MULTISPECIES: cytochrome C [Dietzia] Length: 294\nNumber of predicted TMHs: 2\nExp number of AAs in TMHs: 41.55666\nExp number, first 60 AAs: 21.33987\nTotal prob of N-in: 0.94497\nPOSSIBLE N-term signal sequence\ninside 1 31\nTMhelix 32 54\noutside 55 272\nTMhelix 273 292\ninside 293 294

29399 GCF_001643175.1_ASM164317v1 Dietzia cinnamea Terrabacteria group; Actinobacteria; Actinobacteria; Corynebacteriales; Dietziaceae; Dietzia MTAGTVSARRARRAVFARQGVRVVAAAAAVFSVLTAVLILLACFVNDQRIDRDMGSATAT WP_007630985.1 hypothetical protein [Dietzia cinnamea] Length: 160\nNumber of predicted TMHs: 2\nExp number of AAs in TMHs: 45.12231\nExp number, first 60 AAs: 22.95898\nTotal prob of N-in: 0.85240\nPOSSIBLE N-term signal sequence\ninside 1 19\nTMhelix 20 42\noutside 43 117\nTMhelix 118 140\ninside 141 160

29400 GCF_001658045.1_ASM165804v1 Dietzia sp. 111N12-1 Terrabacteria group; Actinobacteria; Actinobacteria; Corynebacteriales; Dietziaceae; Dietzia MNSPLPPAADTPSPDPVVAPARPVSTKRRRKAVGALVIAGGLVGAGAVATAVIPEPQVAT WP_082923770.1 MULTISPECIES: cytochrome C [Dietzia] Length: 294\nNumber of predicted TMHs: 2\nExp number of AAs in TMHs: 41.55666\nExp number, first 60 AAs: 21.33987\nTotal prob of N-in: 0.94497\nPOSSIBLE N-term signal sequence\ninside 1 31\nTMhelix 32 54\noutside 55 272\nTMhelix 273 292\ninside 293 294

29401 GCF_001658045.1_ASM165804v1 Dietzia sp. 111N12-1 Terrabacteria group; Actinobacteria; Actinobacteria; Corynebacteriales; Dietziaceae; Dietzia MTSTRRRPVQRAAGLVTAALVALAPLAAAAPVTAVEPAPAVPLTEPPPAAVLPSTTSPSY WP_067712163.1 penicillin-binding protein [Dietzia sp. 111N12-1] Length: 465\nNumber of predicted TMHs: 2\nExp number of AAs in TMHs: 43.2672799999999\nExp number, first 60 AAs: 22.51758\nTotal prob of N-in: 0.99484\nPOSSIBLE N-term signal sequence\ninside 1 12\nTMhelix 13 35\noutside 36 439\nTMhelix 440 462\ninside 463 465

29402 GCF_001815995.1_ASM181599v1 Dietzia sp. HMSC21D01 Terrabacteria group; Actinobacteria; Actinobacteria; Corynebacteriales; Dietziaceae; Dietzia MTAGTVSARRARRAVLTRQGVRVVAAAAAVFSVLTAVLILLACFVNDQRIDRDMGSATAT WP_061228443.1 MULTISPECIES: hypothetical protein [Dietzia] Length: 160\nNumber of predicted TMHs: 2\nExp number of AAs in TMHs: 43.93491\nExp number, first 60 AAs: 22.95451\nTotal prob of N-in: 0.87106\nPOSSIBLE N-term signal sequence\ninside 1 19\nTMhelix 20 42\noutside 43 117\nTMhelix 118 140\ninside 141 160

29403 GCF_001651825.1_ASM165182v1 Dietzia timorensis Terrabacteria group; Actinobacteria; Actinobacteria; Corynebacteriales; Dietziaceae; Dietzia MKSPQLPQEDQSGKDSPSKARRRTPFRRRKATGALVIAAGLLGAGAFAAAVTPEPQVATA

WP_067473214.1 cytochrome C [Dietzia timorensis] Length: 293\nNumber of predicted TMHs: 2\nExp number of AAs in TMHs: 41.9526\nExp number, first 60 AAs: 21.87616\nTotal prob of N-in: 0.99122\nPOSSIBLE N-term signal sequence\ninside 1 30\nTMhelix 31 53\noutside 54 270\nTMhelix 271 290\ninside 291 293

29404 GCF_001659785.1_ASM165978v1 Dietzia timorensis Terrabacteria group; Actinobacteria; Actinobacteria; Corynebacteriales; Dietziaceae; Dietzia MKSPQLPQEDQSGKDSPSKARRRTPFRRRKATGALVIAAGLLGAGAFAAAVTPEPQVATA WP_067473214.1 cytochrome C [Dietzia timorensis] Length: 293\nNumber of predicted TMHs: 2\nExp number of AAs in TMHs: 41.9526\nExp number, first 60 AAs: 21.87616\nTotal prob of N-in: 0.99122\nPOSSIBLE N-term signal sequence\ninside 1 30\nTMhelix 31 53\noutside 54 270\nTMhelix 271 290\ninside 291 293

29405 GCF_001571065.1_ASM157106v1 Dietzia cinnamea NBRC 102147 Terrabacteria group; Actinobacteria; Actinobacteria; Corynebacteriales; Dietziaceae; Dietzia; Dietzia cinnamea MTAGTVSARRARRAVLTRQGVRVVAAAAAVFSVLTAVLILLACFVNDQRIDRDMGSATAT WP_061228443.1 MULTISPECIES: hypothetical protein [Dietzia] Length: 160\nNumber of predicted TMHs: 2\nExp number of AAs in TMHs: 43.93491\nExp number, first 60 AAs: 22.95451\nTotal prob of N-in: 0.87106\nPOSSIBLE N-term signal sequence\ninside 1 19\nTMhelix 20 42\noutside 43 117\nTMhelix 118 140\ninside 141 160

29406 GCF_000186325.1_ASM18632v1 Dietzia cinnamea P4 Terrabacteria group; Actinobacteria; Actinobacteria; Corynebacteriales; Dietziaceae; Dietzia; Dietzia cinnamea MTAGTVSARRARRAVFARQGVRVVAAAAAVFSVLTAVLILLACFVNDQRIDRDMGSATAT WP_007630985.1 hypothetical protein [Dietzia cinnamea] Length: 160\nNumber of predicted TMHs: 2\nExp number of AAs in TMHs: 45.12231\nExp number, first 60 AAs: 22.95898\nTotal prob of N-in: 0.85240\nPOSSIBLE N-term signal sequence\ninside 1 19\nTMhelix 20 42\noutside 43 117\nTMhelix 118 140\ninside 141 160

29407 GCF_002043085.1_ASM204308v1 Gordonia sp. i37 Terrabacteria group; Actinobacteria; Actinobacteria; Corynebacteriales; Gordoniaceae; Gordonia MVRRRAALVLAVAGTVVALMIGLSATASAHVKVAGIDATEGGYGVLTFRVPTESDTASTT WP_079930690.1 hypothetical protein [Gordonia sp. i37] Length: 232\nNumber of predicted TMHs: 2\nExp number of AAs in TMHs: 45.06376\nExp number, first 60 AAs: 22.4045\nTotal prob of N-in: 0.99946\nPOSSIBLE N-term signal sequence\ninside 1 6\nTMhelix 7 29\noutside 30 201\nTMhelix 202 224\ninside 225 232

29408 GCF_000347295.2_ASM34729v2 Gordonia sp. NB4-1Y Terrabacteria group; Actinobacteria; Actinobacteria; Corynebacteriales; Gordoniaceae; Gordonia MRRRDPATRRGVILTAVAVIALAPSVLPSVPAQAITPPQVSPGALVRGATVGPPEPTEQRWP_053777677.1 type VII secretion-associated serine protease mycosin [Gordonia sp. NB4-1Y] Length: 467\nNumber of predicted TMHs: 2\nExp number of AAs in TMHs: 45.2644299999999\nExp number, first 60 AAs: 20.68148\nTotal prob of N-in: 0.96518\nPOSSIBLE N-term signal sequence\ninside 1 12\nTMhelix 13 35\noutside 36 433\nTMhelix 434 456\ninside 457 467

29409 GCF_001305675.1_ASM130567v1 Gordonia sp. QH-11 Terrabacteria group; Actinobacteria; Actinobacteria; Corynebacteriales; Gordoniaceae; Gordonia MTSTSSPRIAAFFDLDKTVIARSSALAFTRPFFEGGLLTRRAMLKSAIAQLQFLLTSAEA WP_062391465.1 inhibition of morphological differentiation protein [Gordonia sp. QH-11] Length: 272\nNumber of predicted TMHs: 1\nExp number of AAs in TMHs: 21.48059\nExp number, first 60 AAs: 0.05216\nTotal prob of N-in: 0.07420\noutside 1 237\nTMhelix 238 260\ninside 261 272

29410 GCF_000472025.1_Gordonia_amicalis_CCMA-559 Gordonia amicalis CCMA-559 Terrabacteria group; Actinobacteria; Actinobacteria; Corynebacteriales; Gordoniaceae; Gordonia; Gordonia amicalis MNEPTAPAVDESPTVKFGRDESLEATDSSAEQPAERRSVLHHLWQVLSWLLLIGAVAIMC WP_024500549.1 S26 family signal peptidase [Gordonia amicalis] Length: 222\nNumber of predicted TMHs: 2\nExp number of AAs in TMHs: 43.29195\nExp number, first 60 AAs: 15.99377\nTotal prob of N-in: 0.16830\nPOSSIBLE N-term signal sequence\noutside 1 42\nTMhelix 43 65\ninside 66 180\nTMhelix 181 200\noutside 201 222

29411 GCF_001485495.1_ASM148549v1 Gordonia desulfuricans NBRC 100010 Terrabacteria group; Actinobacteria; Actinobacteria; Corynebacteriales; Gordoniaceae; Gordonia; Gordonia desulfuricans MRRRDPATRRGVILTAVAVIALAPSVLPSVPAQAITPPQVSPGALVRGATVGPPEPTEQRWP_059036185.1 type VII secretion-associated serine protease mycosin [Gordonia desulfuricans] Length: 467\nNumber of predicted TMHs: 2\nExp number of AAs in TMHs: 44.22433\nExp number, first 60 AAs: 20.92026\nTotal prob of N-in: 0.96492\nPOSSIBLE N-term signal sequence\ninside 1 12\nTMhelix 13 35\noutside 36 433\nTMhelix 434 456\ninside 457 467

29412 GCF_000241305.1_ASM24130v2 Gordonia effusa NBRC 100432 Terrabacteria group; Actinobacteria; Actinobacteria; Corynebacteriales; Gordoniaceae; Gordonia; Gordonia effusa
MTNESAPWGAPRIAAFFDLDKTVIARSSALAFTRPFYAEGLINRRSVLKSTYAQLMFLLT WP_007318184.1 hypothetical protein [Gordonia effusa] Length: 274\nNumber of predicted TMHs: 1\nExp number of AAs in TMHs: 22.62305\nExp number, first 60 AAs: 0.02705\nTotal prob of N-in: 0.06474\noutside 1 243\nTMhelix 244 266\ninside 267 274

29413 GCF_001592365.1_ASM159236v1 Gordonia hydrophobica NBRC 16057 Terrabacteria group; Actinobacteria; Actinobacteria; Corynebacteriales; Gordoniaceae; Gordonia; Gordonia hydrophobica
MTSTSSTPIAAFFDLDKTVIARSSALIFTRPFFDGGLLTRRAMFKSAIAQLQFLLTSAEA WP_066169164.1 inhibition of morphological differentiation protein [Gordonia hydrophobica] Length: 272\nNumber of predicted TMHs: 1\nExp number of AAs in TMHs: 19.3192\nExp number, first 60 AAs: 0.40484\nTotal prob of N-in: 0.25959\noutside 1 237\nTMhelix 238 260\ninside 261 272

29414 GCF_000192435.1_ASM19243v1 Gordonia neofelifaecis NRRL B-59395 Terrabacteria group; Actinobacteria; Actinobacteria; Corynebacteriales; Gordoniaceae; Gordonia; Gordonia neofelifaecis
MTSTSSRPIAAFFDLDKTVIARSSALAFTRPFFEGGLLTRRAMLRSAVAQLQFLLTSAEA WP_009679440.1 haloacid dehalogenase [Gordonia neofelifaecis] Length: 272\nNumber of predicted TMHs: 1\nExp number of AAs in TMHs: 17.2786\nExp number, first 60 AAs: 0.05157\nTotal prob of N-in: 0.07902\noutside 1 236\nTMhelix 237 259\ninside 260 272

29415 GCF_000385355.1_ASM38535v1 Gordonia polyisoprenivorans HW436 Terrabacteria group; Actinobacteria; Actinobacteria; Corynebacteriales; Gordoniaceae; Gordonia; Gordonia polyisoprenivorans
MVRRRAALVLAVAGTAVALMIGLSGTASAHVKVSGIDATEGGYGVLTFRVPTESDTASTT WP_026919933.1 hypothetical protein [Gordonia polyisoprenivorans] Length: 232\nNumber of predicted TMHs: 2\nExp number of AAs in TMHs: 44.94222\nExp number, first 60 AAs: 22.28068\nTotal prob of N-in: 0.99855\nPOSSIBLE N-term signal sequence\ninside 1 6\nTMhelix 7 29\noutside 30 201\nTMhelix 202 224\ninside 225 232

29416 GCF_000241325.1_ASM24132v2 Gordonia polyisoprenivorans NBRC 16320 = JCM 10675 Terrabacteria group; Actinobacteria; Actinobacteria; Corynebacteriales; Gordoniaceae; Gordonia; Gordonia polyisoprenivorans
MVRRRAALVLAVAGTAVALMIGLSGTASAHVKVSGIDATEGGYGVLTFRVPTESDTASTT WP_035726910.1 hypothetical protein [Gordonia polyisoprenivorans] Length: 232\nNumber of predicted TMHs: 2\nExp number of AAs in TMHs: 44.94222\nExp number, first 60 AAs: 22.28068\nTotal prob of N-in: 0.99855\nPOSSIBLE N-term signal sequence\ninside 1 6\nTMhelix 7 29\noutside 30 201\nTMhelix 202 224\ninside 225 232

29417 GCF_000247715.1_ASM24771v1 Gordonia polyisoprenivorans VH2 Terrabacteria group; Actinobacteria; Actinobacteria; Corynebacteriales; Gordoniaceae; Gordonia; Gordonia polyisoprenivorans
MVRRRAALVLAVAGTVVALMIGLSATASAHVKVSGIDATEGGYGVLTFRVPTESDTASTT WP_014361013.1 hypothetical protein [Gordonia polyisoprenivorans] Length: 232\nNumber of predicted TMHs: 2\nExp number of AAs in TMHs: 45.06925\nExp number, first 60 AAs: 22.40535\nTotal prob of N-in: 0.99953\nPOSSIBLE N-term signal sequence\ninside 1 6\nTMhelix 7 29\noutside 30 201\nTMhelix 202 224\ninside 225 232

29418 GCF_000298195.1_ASM29819v1 Gordonia rhizosphera NBRC 16068 Terrabacteria group; Actinobacteria; Actinobacteria; Corynebacteriales; Gordoniaceae; Gordonia; Gordonia rhizosphera
MSTTTVPTRAARRTAIRLGLIVAVTLGLAVLSPGSACACTCAPRQAAAVVKDASAVILGT WP_006339621.1 hypothetical protein [Gordonia rhizosphera] Length: 200\nNumber of predicted TMHs: 2\nExp number of AAs in TMHs: 44.12997\nExp number, first 60 AAs: 21.13537\nTotal prob of N-in: 0.80807\nPOSSIBLE N-term signal sequence\ninside 1 19\nTMhelix 20 42\noutside 43 164\nTMhelix 165 187\ninside 188 200

29419 GCF_000334455.1_ASM33445v1 Gordonia soli NBRC 108243 Terrabacteria group; Actinobacteria; Actinobacteria; Corynebacteriales; Gordoniaceae; Gordonia; Gordonia soli
MTHRIEPRQRRRRGPLAALLLAVLTVGALLVSAGIGAATPSGGSDSGGSSSEGSQSGGSD WP_051989624.1 hypothetical protein [Gordonia soli] Length: 404\nNumber of predicted TMHs: 2\nExp number of AAs in TMHs: 44.88044\nExp number, first 60 AAs: 22.45981\nTotal prob of N-in: 0.99911\nPOSSIBLE N-term signal sequence\ninside 1 15\nTMhelix 16 38\noutside 39 373\nTMhelix 374 396\ninside 397 404

29420 GCF_000334455.1_ASM33445v1 Gordonia soli NBRC 108243 Terrabacteria group; Actinobacteria; Actinobacteria; Corynebacteriales; Gordoniaceae; Gordonia; Gordonia soli
MKSTDRKIAAFFDLDKTVIAKSSTLAFSRPFFDEGLINRRSVLKSSYAQFLFLVTAADHD WP_007618975.1 hypothetical protein [Gordonia soli] Length: 266\nNumber of predicted TMHs: 1\nExp number of AAs in TMHs: 18.32106\nExp number, first 60 AAs: 0.00618\nTotal prob of N-in: 0.60377\ninside 1 239\nTMhelix 240 262\noutside 263 266

29421 GCF_002086115.1_ASM208611v1 Mycobacterium bacteremicum Terrabacteria group; Actinobacteria; Actinobacteria; Corynebacteriales; Mycobacteriaceae; Mycobacterium MTRGSTVINDRVRERLIERAALSAPGVVARRTLVPGGNLPAVHLAGQSGVDIQIAASWPV WP_083061113.1 hypothetical protein [Mycobacterium bacteremicum] Length: 236\nNumber of predicted TMHs: 2\nExp number of AAs in TMHs: 44.45423\nExp number, first 60 AAs: 0.00172\nTotal prob of N-in: 0.99985\ninside 1 128\nTMhelix 129 151\noutside 152 174\nTMhelix 175 197\ninside 198 236

29422 GCF_002101555.1_ASM210155v1 Mycobacterium canariasense Terrabacteria group; Actinobacteria; Actinobacteria; Corynebacteriales; Mycobacteriaceae; Mycobacterium MWGTDSRRAEAVAELLHNGRDSRWRLRRCGSNSRVPRHRRSVLTHLHILHAAAEPTEQEQ WP_036439627.1 MULTISPECIES: hypothetical protein [Mycobacterium] Length: 179\nNumber of predicted TMHs: 2\nExp number of AAs in TMHs: 42.06263\nExp number, first 60 AAs: 0.00181\nTotal prob of N-in: 0.87116\ninside 1 66\nTMhelix 67 89\noutside 90 142\nTMhelix 143 162\ninside 163 179

29423 GCF_001570445.1_ASM157044v1 Mycobacterium canariasense Terrabacteria group; Actinobacteria; Actinobacteria; Corynebacteriales; Mycobacteriaceae; Mycobacterium MWGTDSRRAEAVAELLHNGRDSRWRLRRCGSNSRVPRHRRSVLTHLHILHAAAEPTEQEQ WP_036439627.1 MULTISPECIES: hypothetical protein [Mycobacterium] Length: 179\nNumber of predicted TMHs: 2\nExp number of AAs in TMHs: 42.06263\nExp number, first 60 AAs: 0.00181\nTotal prob of N-in: 0.87116\ninside 1 66\nTMhelix 67 89\noutside 90 142\nTMhelix 143 162\ninside 163 179

29424 GCF_001667965.1_ASM166796v1 Mycobacterium conceptionense Terrabacteria group; Actinobacteria; Actinobacteria; Corynebacteriales; Mycobacteriaceae; Mycobacterium MPQHTSAQRRRTTVALFAVATLVFAPACAAEHSPSSSAPAAPQSDAGFAPSAPAPEAGPS WP_064899274.1 hypothetical protein [Mycobacterium conceptionense] Length: 330\nNumber of predicted TMHs: 2\nExp number of AAs in TMHs: 37.64732\nExp number, first 60 AAs: 14.14843\nTotal prob of N-in: 0.77356\nPOSSIBLE N-term signal sequence\ninside 1 12\nTMhelix 13 30\noutside 31 261\nTMhelix 262 284\ninside 285 330

29425 GCF_001667235.1_ASM166723v1 Mycobacterium conceptionense Terrabacteria group; Actinobacteria; Actinobacteria; Corynebacteriales; Mycobacteriaceae; Mycobacterium MPQHTSAQRRRTTVALFAVATLVFAPACAAEHSPSSSAPAAPQSDAGFAPSAPAPEAGPS WP_064899274.1 hypothetical protein [Mycobacterium conceptionense] Length: 330\nNumber of predicted TMHs: 2\nExp number of AAs in TMHs: 37.64732\nExp number, first 60 AAs: 14.14843\nTotal prob of N-in: 0.77356\nPOSSIBLE N-term signal sequence\ninside 1 12\nTMhelix 13 30\noutside 31 261\nTMhelix 262 284\ninside 285 330

29426 GCF_001954125.1_ASM195412v1 Mycobacterium conceptionense Terrabacteria group; Actinobacteria; Actinobacteria; Corynebacteriales; Mycobacteriaceae; Mycobacterium MPQHTSAQRRRTTVALFAVATLVFAPACAAEHSPSSSAPAAPQSDAGFAPAAPAPEAGPS WP_047037357.1 MULTISPECIES: hypothetical protein [Mycobacterium] Length: 330\nNumber of predicted TMHs: 2\nExp number of AAs in TMHs: 37.49221\nExp number, first 60 AAs: 13.96769\nTotal prob of N-in: 0.77513\nPOSSIBLE N-term signal sequence\ninside 1 12\nTMhelix 13 30\noutside 31 261\nTMhelix 262 284\ninside 285 330

29427 GCF_002102065.1_ASM210206v1 Mycobacterium conceptionense Terrabacteria group; Actinobacteria; Actinobacteria; Corynebacteriales; Mycobacteriaceae; Mycobacterium MPQHTSAQRRRTTVALFAVATLVFAPACAAEHSPSSSAPAAPQSDAGFAPSAPAPEAGPS WP_085140301.1 hypothetical protein [Mycobacterium conceptionense] Length: 330\nNumber of predicted TMHs: 2\nExp number of AAs in TMHs: 37.79226\nExp number, first 60 AAs: 14.31476\nTotal prob of N-in: 0.77064\nPOSSIBLE N-term signal sequence\ninside 1 12\nTMhelix 13 30\noutside 31 261\nTMhelix 262 284\ninside 285 330

29428 GCF_001667225.1_ASM166722v1 Mycobacterium conceptionense Terrabacteria group; Actinobacteria; Actinobacteria; Corynebacteriales; Mycobacteriaceae; Mycobacterium MPQHTSAQRRRTTVALFAVATLVFAPACAAEHSPSSSAPAAPQSDAGFAPSAPAPEAGPS WP_064899274.1 hypothetical protein [Mycobacterium conceptionense] Length: 330\nNumber of predicted TMHs: 2\nExp number of AAs in TMHs: 37.64732\nExp number, first 60 AAs: 14.14843\nTotal prob of N-in: 0.77356\nPOSSIBLE N-term signal sequence\ninside 1 12\nTMhelix 13 30\noutside 31 261\nTMhelix 262 284\ninside 285 330

29429 GCF_001673265.1_ASM167326v1 Mycobacterium conceptionense Terrabacteria group; Actinobacteria; Actinobacteria; Corynebacteriales; Mycobacteriaceae; Mycobacterium MPQHTSAQRRRTTVALFAVATLVFAPACAAEHSPSSSAPAAPQSDAGFAPAAPAPEAGPS WP_065064343.1 hypothetical protein [Mycobacterium conceptionense] Length: 330\nNumber of predicted TMHs: 2\nExp

number of AAs in TMHs: 37.49221\nExp number, first 60 AAs: 13.96769\nTotal prob of N-in: 0.77513\nPOSSIBLE N-term signal sequence\ninside 1 12\nTMhelix 13 30\noutside 31 261\nTMhelix 262 284\ninside 285 330

29430 GCF_001954235.1_ASM195423v1 Mycobacterium conceptionense Terrabacteria group; Actinobacteria; Actinobacteria; Corynebacteriales; Mycobacteriaceae; Mycobacterium MPQHTSAQRRRTTVALFAVATLVFAPACAAEHSPSSSAPAAPQSDAGFAPAAPAPEAGPS WP_047037357.1 MULTISPECIES: hypothetical protein [Mycobacterium] Length: 330\nNumber of predicted TMHs: 2\nExp number of AAs in TMHs: 37.49221\nExp number, first 60 AAs: 13.96769\nTotal prob of N-in: 0.77513\nPOSSIBLE N-term signal sequence\ninside 1 12\nTMhelix 13 30\noutside 31 261\nTMhelix 262 284\ninside 285 330

29431 GCF_001077745.1_ASM107774v1 Mycobacterium conceptionense Terrabacteria group; Actinobacteria; Actinobacteria; Corynebacteriales; Mycobacteriaceae; Mycobacterium MPQHTSAQRRRTTVALFAVATLVFAPACAAEHSPSSSAPAAPQSDAGFAPAAPAPEAGPS WP_047037357.1 MULTISPECIES: hypothetical protein [Mycobacterium] Length: 330\nNumber of predicted TMHs: 2\nExp number of AAs in TMHs: 37.49221\nExp number, first 60 AAs: 13.96769\nTotal prob of N-in: 0.77513\nPOSSIBLE N-term signal sequence\ninside 1 12\nTMhelix 13 30\noutside 31 261\nTMhelix 262 284\ninside 285 330

29432 GCF_001665335.1_ASM166533v1 Mycobacterium conceptionense Terrabacteria group; Actinobacteria; Actinobacteria; Corynebacteriales; Mycobacteriaceae; Mycobacterium MPQHTSAQRRRTTVALFAVATLVFAPACAAEHSPSSSAPAAPQSDAGFAPSAPAPEAGPS WP_064961509.1 hypothetical protein [Mycobacterium conceptionense] Length: 330\nNumber of predicted TMHs: 2\nExp number of AAs in TMHs: 37.65141\nExp number, first 60 AAs: 14.14837\nTotal prob of N-in: 0.77360\nPOSSIBLE N-term signal sequence\ninside 1 12\nTMhelix 13 30\noutside 31 261\nTMhelix 262 284\ninside 285 330

29433 GCF_001667305.1_ASM166730v1 Mycobacterium conceptionense Terrabacteria group; Actinobacteria; Actinobacteria; Corynebacteriales; Mycobacteriaceae; Mycobacterium MPQHTSAQRRRTTVALFAVATLVFAPACAAEHSPSSSAPAAPQSDAGFAPSAPAPEAGPS WP_064899274.1 hypothetical protein [Mycobacterium conceptionense] Length: 330\nNumber of predicted TMHs: 2\nExp number of AAs in TMHs: 37.64732\nExp number, first 60 AAs: 14.14843\nTotal prob of N-in: 0.77356\nPOSSIBLE N-term signal sequence\ninside 1 12\nTMhelix 13 30\noutside 31 261\nTMhelix 262 284\ninside 285 330

29434 GCF_001954205.1_ASM195420v1 Mycobacterium conceptionense Terrabacteria group; Actinobacteria; Actinobacteria; Corynebacteriales; Mycobacteriaceae; Mycobacterium MPQHTSAQRRRTTVALFAVATLVFAPACAAEHSPSSSAPAAPQSDAGFAPAAPAPEAGPS WP_076216568.1 hypothetical protein [Mycobacterium conceptionense] Length: 330\nNumber of predicted TMHs: 2\nExp number of AAs in TMHs: 37.64723\nExp number, first 60 AAs: 14.14842\nTotal prob of N-in: 0.77356\nPOSSIBLE N-term signal sequence\ninside 1 12\nTMhelix 13 30\noutside 31 261\nTMhelix 262 284\ninside 285 330

29435 GCF_001077745.1_ASM107774v1 Mycobacterium conceptionense Terrabacteria group; Actinobacteria; Actinobacteria; Corynebacteriales; Mycobacteriaceae; Mycobacterium MSGPLGLSIGTTNLVAARVGNQPVSRRAVLTLSTDRTPQVGVPASGSGVTLGGFVERVGD WP_048895863.1 molecular chaperone, partial [Mycobacterium conceptionense] Length: 488\nNumber of predicted TMHs: 1\nExp number of AAs in TMHs: 23.59491\nExp number, first 60 AAs: 0.00464\nTotal prob of N-in: 0.00383\noutside 1 427\nTMhelix 428 450\ninside 451 488

29436 GCF_002102105.1_ASM210210v1 Mycobacterium confluentis Terrabacteria group; Actinobacteria; Actinobacteria; Corynebacteriales; Mycobacteriaceae; Mycobacterium MTSGQRSSRWAGVPLRDRPARRRDALIAAGVQLLGGVAGPAITVRAVCRHAGLTERYFYE WP_085156310.1 TetR family transcriptional regulator [Mycobacterium confluentis] Length: 194\nNumber of predicted TMHs: 2\nExp number of AAs in TMHs: 38.87978\nExp number, first 60 AAs: 20.16028\nTotal prob of N-in: 0.90460\nPOSSIBLE N-term signal sequence\ninside 1 24\nTMhelix 25 47\noutside 48 145\nTMhelix 146 168\ninside 169 194

29437 GCF_002102095.1_ASM210209v1 Mycobacterium conspicuum Terrabacteria group; Actinobacteria; Actinobacteria; Corynebacteriales; Mycobacteriaceae; Mycobacterium MTRRRYVTRLGPLLAVLALAATWTAPLASAHAARVSCQPADNAVLAAGPAQVSATFNEQL WP_085235053.1 copper resistance protein CopC [Mycobacterium conspicuum] Length: 178\nNumber of predicted TMHs: 2\nExp number of AAs in TMHs: 43.71219\nExp number, first 60 AAs: 20.88373\nTotal prob of N-in: 0.98387\nPOSSIBLE N-term signal sequence\ninside 1 12\nTMhelix 13 35\noutside 36 146\nTMhelix 147 169\ninside 170 178

29438 GCF_002086605.1_ASM208660v1 Mycobacterium elephantis Terrabacteria group; Actinobacteria; Actinobacteria; Corynebacteriales; Mycobacteriaceae; Mycobacterium

MPSPPPTLSRRRVLFGAAALTLLGVTAAGCGSPPPPEVDELLAQLNRARADSQLASDAAT WP_083042553.1 hypothetical protein [Mycobacterium elephantis] Length: 165\nNumber of predicted TMHs: 2\nExp number of AAs in TMHs: 35.68521\nExp number, first 60 AAs: 17.35725\nTotal prob of N-in: 0.85200\nPOSSIBLE N-term signal sequence\ninside 1 12\nTMhelix 13 35\noutside 36 137\nTMhelix 138 160\ninside 161 165

29439 GCF_001005175.1_ASM100517v1 Mycobacterium elephantis Terrabacteria group; Actinobacteria; Actinobacteria; Corynebacteriales; Mycobacteriaceae; Mycobacterium MPSPPPTLSRRRVLFGAAALTLLGVTAAGCGSPPPPEVDELLAQLNRARADSQLASDAAA WP_046753310.1 MULTISPECIES: membrane protein [Mycobacterium] Length: 165\nNumber of predicted TMHs: 2\nExp number of AAs in TMHs: 35.68644\nExp number, first 60 AAs: 17.35768\nTotal prob of N-in: 0.85201\nPOSSIBLE N-term signal sequence\ninside 1 12\nTMhelix 13 35\noutside 36 137\nTMhelix 138 160\ninside 161 165

29440 GCF_002101585.1_ASM210158v1 Mycobacterium engbaekii Terrabacteria group; Actinobacteria; Actinobacteria; Corynebacteriales; Mycobacteriaceae; Mycobacterium MPSPPSAIDRRRVLTGAGMLALLAVVTPACGSAPTPPAISELEAQRQRALRDSALAAAAA WP_085130694.1 hypothetical protein [Mycobacterium engbaekii] Length: 167\nNumber of predicted TMHs: 2\nExp number of AAs in TMHs: 34.67801\nExp number, first 60 AAs: 18.31742\nTotal prob of N-in: 0.83641\nPOSSIBLE N-term signal sequence\ninside 1 12\nTMhelix 13 35\noutside 36 141\nTMhelix 142 164\ninside 165 167

29441 GCF_000723385.1_PRJEB5746_assembly_1 Mycobacterium farcinogenes Terrabacteria group; Actinobacteria; Actinobacteria; Corynebacteriales; Mycobacteriaceae; Mycobacterium MPQHTSAQRRRTTVALFAVATLVFAPACAAEHSPSSSAPAAPQSDAGFAPSAPAPEAGPS WP_036393700.1 hypothetical protein [Mycobacterium farcinogenes] Length: 330\nNumber of predicted TMHs: 2\nExp number of AAs in TMHs: 37.64712\nExp number, first 60 AAs: 14.14842\nTotal prob of N-in: 0.77356\nPOSSIBLE N-term signal sequence\ninside 1 12\nTMhelix 13 30\noutside 31 261\nTMhelix 262 284\ninside 285 330

29442 GCF_001722335.1_ASM172233v1 Mycobacterium flavescens Terrabacteria group; Actinobacteria; Actinobacteria; Corynebacteriales; Mycobacteriaceae; Mycobacterium MPAADNPTRTAAFFDLDKTVIAKSSTLAFSKPFFDQGLMNRRSVLKSTYAQFLFLMSGAD WP_069415385.1 inhibition of morphological differentiation protein [Mycobacterium flavescens] Length: 269\nNumber of predicted TMHs: 1\nExp number of AAs in TMHs: 21.62925\nExp number, first 60 AAs: 0.03719\nTotal prob of N-in: 0.14821\noutside 1 241\nTMhelix 242 264\ninside 265 269

29443 GCF_001673255.1_ASM167325v1 Mycobacterium gordonae Terrabacteria group; Actinobacteria; Actinobacteria; Corynebacteriales; Mycobacteriaceae; Mycobacterium MFPGFDALPEALRPVARTRREHVHPVASPPTQTLVDCGVYVDGQRLPGKFTPAEALAKAR WP_065044213.1 magnesium and cobalt transport protein CorA [Mycobacterium gordonae] Length: 366\nNumber of predicted TMHs: 2\nExp number of AAs in TMHs: 42.5335\nExp number, first 60 AAs: 0.00172\nTotal prob of N-in: 0.67098\ninside 1 308\nTMhelix 309 328\noutside 329 337\nTMhelix 338 360\ninside 361 366

29444 GCF_001673475.1_ASM167347v1 Mycobacterium gordonae Terrabacteria group; Actinobacteria; Actinobacteria; Corynebacteriales; Mycobacteriaceae; Mycobacterium MFPGFDALPEALRPVARTRRQHVHPVASPPTQTLVDCGVYVDGQRLPGKFTPAEALAKAR WP_065161838.1 magnesium and cobalt transport protein CorA [Mycobacterium gordonae] Length: 366\nNumber of predicted TMHs: 2\nExp number of AAs in TMHs: 42.53405\nExp number, first 60 AAs: 0.00232\nTotal prob of N-in: 0.67097\ninside 1 308\nTMhelix 309 328\noutside 329 337\nTMhelix 338 360\ninside 361 366

29445 GCF_001417955.2_ASM141795v2 Mycobacterium gordonae Terrabacteria group; Actinobacteria; Actinobacteria; Corynebacteriales; Mycobacteriaceae; Mycobacterium MFPGFDALPEALRPVARTRREHVHPVANPPAQTLVDCGVYVDGHRLPGKFSPAEALAKVR WP_055579670.1 magnesium transporter CorA [Mycobacterium gordonae] Length: 366\nNumber of predicted TMHs: 2\nExp number of AAs in TMHs: 42.43736\nExp number, first 60 AAs: 0.00096\nTotal prob of N-in: 0.68267\ninside 1 308\nTMhelix 309 328\noutside 329 337\nTMhelix 338 360\ninside 361 366

29446 GCF_001021485.1_ASM102148v1 Mycobacterium haemophilum Terrabacteria group; Actinobacteria; Actinobacteria; Corynebacteriales; Mycobacteriaceae; Mycobacterium MPRSGTGSKRARRGCVPRALVAVALLTSGAFAGLPPAYAISPPTIDPGAVPPDGPPGPAA WP_047314210.1 type VII secretion-associated serine protease mycosin [Mycobacterium haemophilum] Length: 591\nNumber of predicted TMHs: 2\nExp number of AAs in TMHs: 43.906\nExp number, first 60 AAs: 20.08392\nTotal prob of N-in: 0.93656\nPOSSIBLE N-term signal sequence\ninside 1 19\nTMhelix 20 42\noutside 43 556\nTMhelix 557 579\ninside 580 591

29447 GCF_001021415.1_ASM102141v1 Mycobacterium haemophilum Terrabacteria group; Actinobacteria; Actinobacteria; Corynebacteriales; Mycobacteriaceae; Mycobacterium MPRSGTGSKRARRGCVPRALVAVALLTSGAFAGLPPAYAISPPTIDPGAVPPDGPPGPAA WP_047314210.1 type VII secretion-associated serine protease mycosin [Mycobacterium haemophilum] Length: 591\nNumber of predicted TMHs: 2\nExp number of AAs in TMHs: 43.906\nExp number, first 60 AAs: 20.08392\nTotal prob of N-in: 0.93656\nPOSSIBLE N-term signal sequence\ninside 1 19\nTMhelix 20 42\noutside 43 556\nTMhelix 557 579\ninside 580 591

29448 GCF_001021405.1_ASM102140v1 Mycobacterium haemophilum Terrabacteria group; Actinobacteria; Actinobacteria; Corynebacteriales; Mycobacteriaceae; Mycobacterium MPRSGTGSKRARRGCVPRALVAVALLTSGAFAGLPPAYAISPPTIDPGAVPPDGPPGPAA WP_047314210.1 type VII secretion-associated serine protease mycosin [Mycobacterium haemophilum] Length: 591\nNumber of predicted TMHs: 2\nExp number of AAs in TMHs: 43.906\nExp number, first 60 AAs: 20.08392\nTotal prob of N-in: 0.93656\nPOSSIBLE N-term signal sequence\ninside 1 19\nTMhelix 20 42\noutside 43 556\nTMhelix 557 579\ninside 580 591

29449 GCF_001021435.1_ASM102143v1 Mycobacterium haemophilum Terrabacteria group; Actinobacteria; Actinobacteria; Corynebacteriales; Mycobacteriaceae; Mycobacterium MPRSGTGSKRARRGCVPRALVAVALLTSGAFAGLPPAYAISPPTIDPGAVPPDGPPGPAA WP_047314210.1 type VII secretion-associated serine protease mycosin [Mycobacterium haemophilum] Length: 591\nNumber of predicted TMHs: 2\nExp number of AAs in TMHs: 43.906\nExp number, first 60 AAs: 20.08392\nTotal prob of N-in: 0.93656\nPOSSIBLE N-term signal sequence\ninside 1 19\nTMhelix 20 42\noutside 43 556\nTMhelix 557 579\ninside 580 591

29450 GCF_001953995.1_ASM195399v1 Mycobacterium heraklionense Terrabacteria group; Actinobacteria; Actinobacteria; Corynebacteriales; Mycobacteriaceae; Mycobacterium MPGPLSAIDRRRVLTGAGMLALLAVAAPACGSVPTAPAIDGLEAQREAAQHDSALAAAAA WP_076048688.1 hypothetical protein [Mycobacterium heraklionense] Length: 169\nNumber of predicted TMHs: 2\nExp number of AAs in TMHs: 37.71154\nExp number, first 60 AAs: 19.67711\nTotal prob of N-in: 0.85700\nPOSSIBLE N-term signal sequence\ninside 1 12\nTMhelix 13 35\noutside 36 143\nTMhelix 144 166\ninside 167 169

29451 GCF_001021505.1_ASM102150v1 Mycobacterium heraklionense Terrabacteria group; Actinobacteria; Actinobacteria; Corynebacteriales; Mycobacteriaceae; Mycobacterium MPGPLSVIDRRRVLTGAGMFALLALAAPACGSAPTTPVVDDLEAQRQLALHDSELATAAA WP_047318566.1 hypothetical protein [Mycobacterium heraklionense] Length: 169\nNumber of predicted TMHs: 2\nExp number of AAs in TMHs: 41.71377\nExp number, first 60 AAs: 20.37173\nTotal prob of N-in: 0.92016\nPOSSIBLE N-term signal sequence\ninside 1 12\nTMhelix 13 35\noutside 36 144\nTMhelix 145 167\ninside 168 169

29452 GCF_001673575.1_ASM167357v1 Mycobacterium heraklionense Terrabacteria group; Actinobacteria; Actinobacteria; Corynebacteriales; Mycobacteriaceae; Mycobacterium MPGPLSAIDRRRMLTGAGMLALLAVAAPACGSAPTAPLVDDLEAQRQLALRDSELATAAA WP_082980945.1 hypothetical protein [Mycobacterium heraklionense] Length: 169\nNumber of predicted TMHs: 2\nExp number of AAs in TMHs: 38.96919\nExp number, first 60 AAs: 18.96961\nTotal prob of N-in: 0.82803\nPOSSIBLE N-term signal sequence\ninside 1 12\nTMhelix 13 35\noutside 36 143\nTMhelix 144 166\ninside 167 169

29453 GCF_001672655.1_ASM167265v1 Mycobacterium heraklionense Terrabacteria group; Actinobacteria; Actinobacteria; Corynebacteriales; Mycobacteriaceae; Mycobacterium MPGPLSAIDRRRVLTGAGMLALLAVAAPACGSGPTAPAVDDLEAQREAAQHDSALAAAAA WP_082961219.1 hypothetical protein [Mycobacterium heraklionense] Length: 169\nNumber of predicted TMHs: 2\nExp number of AAs in TMHs: 36.65862\nExp number, first 60 AAs: 17.86862\nTotal prob of N-in: 0.83739\nPOSSIBLE N-term signal sequence\ninside 1 12\nTMhelix 13 35\noutside 36 143\nTMhelix 144 166\ninside 167 169

29454 GCF_002086255.1_ASM208625v1 Mycobacterium insubricum Terrabacteria group; Actinobacteria; Actinobacteria; Corynebacteriales; Mycobacteriaceae; Mycobacterium MNSGQRRGRWSGVPLSDRPSRRRDALLAAGIAALGAPDGPALTVRAICREARLTERYFYE WP_083033625.1 TetR family transcriptional regulator [Mycobacterium insubricum] Length: 193\nNumber of predicted TMHs: 1\nExp number of AAs in TMHs: 22.50786\nExp number, first 60 AAs: 4.80379\nTotal prob of N-in: 0.48342\noutside 1 145\nTMhelix 146 168\ninside 169 193

29455 GCF_001672775.1_ASM167277v1 Mycobacterium kyorinense Terrabacteria group; Actinobacteria; Actinobacteria; Corynebacteriales; Mycobacteriaceae; Mycobacterium

MPPTVPVVTRRRVLAGGAALAVLGITASACSPQPDQPPPVDELEAELTLARHDSELAAAA WP_065015780.1 hypothetical protein [Mycobacterium kyorinense] Length: 172\nNumber of predicted TMHs: 1\nExp number of AAs in TMHs: 31.56968\nExp number, first 60 AAs: 9.06798\nTotal prob of N-in: 0.88929\ninside 1 146\nTMhelix 147 169\noutside 170 172

29456 GCF_002007745.1_ASM200774v1 Mycobacterium litorale Terrabacteria group; Actinobacteria; Actinobacteria; Corynebacteriales; Mycobacteriaceae; Mycobacterium MVAGGVADRRCARSRRDRRPRALGRPDRRRAAVTHRNRRLVAVTAAVLLTGVHCAPPAYA WP_078020650.1 type VII secretion-associated serine protease mycosin [Mycobacterium litorale] Length: 469\nNumber of predicted TMHs: 2\nExp number of AAs in TMHs: 36.67971\nExp number, first 60 AAs: 16.58259\nTotal prob of N-in: 0.77497\nPOSSIBLE N-term signal sequence\ninside 1 38\nTMhelix 39 61\noutside 62 434\nTMhelix 435 457\ninside 458 469

29457 GCF_002102265.1_ASM210226v1 Mycobacterium longobardum Terrabacteria group; Actinobacteria; Actinobacteria; Corynebacteriales; Mycobacteriaceae; Mycobacterium MPGPLSAIDRRRVLTGAGMLALLAVVAPACGSAPAPPAVDDLEAQRQLAQRDSELAAAAR WP_085266655.1 hypothetical protein [Mycobacterium longobardum] Length: 170\nNumber of predicted TMHs: 2\nExp number of AAs in TMHs: 40.38309\nExp number, first 60 AAs: 20.71172\nTotal prob of N-in: 0.91772\nPOSSIBLE N-term signal sequence\ninside 1 12\nTMhelix 13 35\noutside 36 145\nTMhelix 146 168\ninside 169 170

29458 GCF_002086395.1_ASM208639v1 Mycobacterium moriokaenseTerrabacteria group; Actinobacteria; Actinobacteria; Corynebacteriales; Mycobacteriaceae; Mycobacterium MASPPGPVRTAAFFDLDKTVIAKSSTLAFSKPFFDQGLITRRTVLKSAYAQFLFLMSGAD WP_083155272.1 inhibition of morphological differentiation protein [Mycobacterium moriokaense]Length: 269\nNumber of predicted TMHs: 1\nExp number of AAs in TMHs: 21.64539\nExp number, first 60 AAs: 0.04664\nTotal prob of N-in: 0.01762\noutside 1 241\nTMhelix 242 264\ninside 265 269

29459 GCF_002086415.1_ASM208641v1 Mycobacterium noviomagense Terrabacteria group; Actinobacteria; Actinobacteria; Corynebacteriales; Mycobacteriaceae; Mycobacterium MPSAAPDVTRRRVLVGGAALAAVALVGSACSSPPPPPNVDDLESQLKLARHDSELAAAAA WP_083087896.1 hypothetical protein [Mycobacterium noviomagense] Length: 173\nNumber of predicted TMHs: 2\nExp number of AAs in TMHs: 34.70344\nExp number, first 60 AAs: 16.96688\nTotal prob of N-in: 0.91955\nPOSSIBLE N-term signal sequence\ninside 1 12\nTMhelix 13 35\noutside 36 144\nTMhelix 145 167\ninside 168 173

29460 GCF_002086815.1_ASM208681v1 Mycobacterium parafortuitum Terrabacteria group; Actinobacteria; Actinobacteria; Corynebacteriales; Mycobacteriaceae; Mycobacterium MWHVPSRPPTLSRRRVLAGAAVLAALGVAGAGCGTPPPPADLDDLTTALDRSRSDSDLAA WP_083145723.1 hypothetical protein [Mycobacterium parafortuitum] Length: 177\nNumber of predicted TMHs: 2\nExp number of AAs in TMHs: 36.26924\nExp number, first 60 AAs: 15.29187\nTotal prob of N-in: 0.73671\nPOSSIBLE N-term signal sequence\ninside 1 15\nTMhelix 16 38\noutside 39 151\nTMhelix 152 174\ninside 175 177

29461 GCF_002086695.1_ASM208669v1 Mycobacterium rhodesiae Terrabacteria group; Actinobacteria; Actinobacteria; Corynebacteriales; Mycobacteriaceae; Mycobacterium MHVRPEPATGLNSRTAAFFDLDKTVIAKSSTLAFSKPFFDQGLLNRRAVLKSSYAQFLFL WP_083120947.1 inhibition of morphological differentiation protein [Mycobacterium rhodesiae] Length: 274\nNumber of predicted TMHs: 1\nExp number of AAs in TMHs: 21.44177\nExp number, first 60 AAs: 0.01193\nTotal prob of N-in: 0.04980\noutside 1 246\nTMhelix 247 269\ninside 270 274

29462 GCF_002013645.1_ASM201364v1 Mycobacterium salmoniphilum Terrabacteria group; Actinobacteria; Actinobacteria; Corynebacteriales; Mycobacteriaceae; Mycobacterium MTIDGTQSRRSWLPWATTIAVLPIATTACAGGTSSHLSQGADDAPATTGAPTYTQAPTAP WP_078323872.1 hypothetical protein [Mycobacterium salmoniphilum] Length: 323\nNumber of predicted TMHs: 1\nExp number of AAs in TMHs: 37.20016\nExp number, first 60 AAs: 13.30723\nTotal prob of N-in: 0.64982\nPOSSIBLE N-term signal sequence\noutside 1 276\nTMhelix 277 299\ninside 300 323

29463 GCF_002013645.1_ASM201364v1 Mycobacterium salmoniphilum Terrabacteria group; Actinobacteria; Actinobacteria; Corynebacteriales; Mycobacteriaceae; Mycobacterium MIKHLTRRATVALVVAAAAMSPAAYPAVADTATPLNGLFALAPGSCAEGRATGSYFRMIL WP_078324820.1 hypothetical protein [Mycobacterium salmoniphilum] Length: 328\nNumber of predicted TMHs: 1\nExp number of AAs in TMHs: 37.14882\nExp number, first 60 AAs: 14.77973\nTotal prob of N-in: 0.73657\nPOSSIBLE N-term signal sequence\noutside 1 296\nTMhelix 297 319\ninside 320 328

29464 GCF_002013685.1_ASM201368v1 Mycobacterium salmoniphilum Terrabacteria group; Actinobacteria; Actinobacteria; Corynebacteriales; Mycobacteriaceae; Mycobacterium
MIKHLTRRATVALVVAAAAMSPAAYPAVADTATPLNGLFALAPGSCAEGRATGSYFRMIL WP_078324820.1
hypothetical protein [Mycobacterium salmoniphilum] Length: 328\nNumber of predicted TMHs: 1\nExp number of AAs in TMHs: 37.14882\nExp number, first 60 AAs: 14.77973\nTotal prob of N-in: 0.73657\nPOSSIBLE N-term signal sequence\noutside 1 296\nTMhelix 297 319\ninside 320 328

29465 GCF_001021425.1_ASM102142v1 Mycobacterium senegalense Terrabacteria group; Actinobacteria; Actinobacteria; Corynebacteriales; Mycobacteriaceae; Mycobacterium
MPQHTSAQRRRTTVALFAVATLVFAPACAAEHSPSSSAPAAPQSDAGFAPAAPAPEAGPS WP_047037357.1
MULTISPECIES: hypothetical protein [Mycobacterium] Length: 330\nNumber of predicted TMHs: 2\nExp number of AAs in TMHs: 37.49221\nExp number, first 60 AAs: 13.96769\nTotal prob of N-in: 0.77513\nPOSSIBLE N-term signal sequence\ninside 1 12\nTMhelix 13 30\noutside 31 261\nTMhelix 262 284\ninside 285 330

29466 GCF_001012795.1_ASM101279v1 Mycobacterium senegalense Terrabacteria group; Actinobacteria; Actinobacteria; Corynebacteriales; Mycobacteriaceae; Mycobacterium
MPQHTSAQRRRTTVALFAVATLVFAPACAAEHSPSSSAPAAPQSDAGFAPAAPAPEAGPS WP_047037357.1
MULTISPECIES: hypothetical protein [Mycobacterium] Length: 330\nNumber of predicted TMHs: 2\nExp number of AAs in TMHs: 37.49221\nExp number, first 60 AAs: 13.96769\nTotal prob of N-in: 0.77513\nPOSSIBLE N-term signal sequence\ninside 1 12\nTMhelix 13 30\noutside 31 261\nTMhelix 262 284\ninside 285 330

29467 GCF_002086755.1_ASM208675v1 Mycobacterium shinjukuenseTerrabacteria group; Actinobacteria; Actinobacteria; Corynebacteriales; Mycobacteriaceae; Mycobacterium
MARRAAVVGTAAGCLAVFAVPAAAPRATAGEPSPTPFVQVRIDQVTPDVVTTTSEPVVTV WP_083049427.1
hypothetical protein [Mycobacterium shinjukuense] Length: 793\nNumber of predicted TMHs: 2\nExp number of AAs in TMHs: 38.7079199999999\nExp number, first 60 AAs: 18.70723\nTotal prob of N-in: 0.89499\nPOSSIBLE N-term signal sequence\ninside 1 6\nTMhelix 7 29\noutside 30 731\nTMhelix 732 754\ninside 755 793

29468 GCF_001667945.1_ASM166794v1 Mycobacterium sinense Terrabacteria group; Actinobacteria; Actinobacteria; Corynebacteriales; Mycobacteriaceae; Mycobacterium
MRSPRPVGSRTRTAAFLDLDHTVIAKSSALAFSKPFMDQGLINRRTVLKSSYAQFLMLLS WP_064922776.1 inhibition of morphological differentiation protein [Mycobacterium sinense] Length: 272\nNumber of predicted TMHs: 1\nExp number of AAs in TMHs: 21.72556\nExp number, first 60 AAs: 0.01879\nTotal prob of N-in: 0.05333\noutside 1 244\nTMhelix 245 267\ninside 268 272

29469 GCF_001673565.1_ASM167356v1 Mycobacterium sinense Terrabacteria group; Actinobacteria; Actinobacteria; Corynebacteriales; Mycobacteriaceae; Mycobacterium
MRSPGQAGSRTRTAAFLDLDHTVIAKSSALAFSKPFMDQGLINRRTVLKSSYAQFLMLLS WP_065024600.1 inhibition of morphological differentiation protein [Mycobacterium sinense] Length: 272\nNumber of predicted TMHs: 1\nExp number of AAs in TMHs: 21.70116\nExp number, first 60 AAs: 0.02232\nTotal prob of N-in: 0.13772\noutside 1 244\nTMhelix 245 267\ninside 268 272

29470 GCF_001667395.1_ASM166739v1 Mycobacterium sinense Terrabacteria group; Actinobacteria; Actinobacteria; Corynebacteriales; Mycobacteriaceae; Mycobacterium
MRSPGQAGSRTRTAAFLDLDHTVIAKSSALAFSKPFMDQGLINRRTVLKSSYAQFLMLLS WP_064856216.1 inhibition of morphological differentiation protein [Mycobacterium sinense] Length: 272\nNumber of predicted TMHs: 1\nExp number of AAs in TMHs: 21.69067\nExp number, first 60 AAs: 0.01947\nTotal prob of N-in: 0.09107\noutside 1 244\nTMhelix 245 267\ninside 268 272

29471 GCF_001672735.1_ASM167273v1 Mycobacterium sinense Terrabacteria group; Actinobacteria; Actinobacteria; Corynebacteriales; Mycobacteriaceae; Mycobacterium
MRSPRPVGSRTRTAAFLDLDHTVIAKSSALAFSKPFMDQGLINRRTVLKSSYAQFLMLLS WP_064922776.1 inhibition of morphological differentiation protein [Mycobacterium sinense] Length: 272\nNumber of predicted TMHs: 1\nExp number of AAs in TMHs: 21.72556\nExp number, first 60 AAs: 0.01879\nTotal prob of N-in: 0.05333\noutside 1 244\nTMhelix 245 267\ninside 268 272

29472 GCF_001667375.1_ASM166737v1 Mycobacterium sinense Terrabacteria group; Actinobacteria; Actinobacteria; Corynebacteriales; Mycobacteriaceae; Mycobacterium
MRSPGQAGSRTRTAAFLDLDHTVIAKSSALAFSKPFMDQGLINRRTVLKSSYAQFLMLLS WP_064856216.1 inhibition of morphological differentiation protein [Mycobacterium sinense] Length: 272\nNumber of predicted TMHs: 1\nExp

number of AAs in TMHs: 21.69067\nExp number, first 60 AAs: 0.01947\nTotal prob of N-in: 0.09107\noutside 1 244\nTMhelix 245 267\ninside 268 272

29473 GCF_001665235.1_ASM166523v1 Mycobacterium sp. 1100029.7 Terrabacteria group; Actinobacteria; Actinobacteria; Corynebacteriales; Mycobacteriaceae; Mycobacterium MRRRAVVGWAGMLLVTMLLTATLTAQVASAHATRVSADPAADAVLSVGPARATATFNERL WP_066814184.1 copper resistance protein CopC [Mycobacterium sp. 1100029.7] Length: 170\nNumber of predicted TMHs: 2\nExp number of AAs in TMHs: 42.82088\nExp number, first 60 AAs: 22.40832\nTotal prob of N-in: 0.99844\nPOSSIBLE N-term signal sequence\ninside 1 4\nTMhelix 5 27\noutside 28 146\nTMhelix 147 166\ninside 167 170

29474 GCF_001668575.1_ASM166857v1 Mycobacterium sp. 1164985.4 Terrabacteria group; Actinobacteria; Actinobacteria; Corynebacteriales; Mycobacteriaceae; Mycobacterium MAHRPSAGRPAPTAPTAPRRSFAESLAGADSAADAERRRALRRMKVVALSFLIGATVVFL WP_067289482.1 DUF445 domain-containing protein [Mycobacterium sp. 1164985.4] Length: 446\nNumber of predicted TMHs: 2\nExp number of AAs in TMHs: 57.9346000000001\nExp number, first 60 AAs: 14.69917\nTotal prob of N-in: 0.99800\nPOSSIBLE N-term signal sequence\ninside 1 45\nTMhelix 46 68\noutside 69 419\nTMhelix 420 442\ninside 443 446

29475 GCF_001673415.1_ASM167341v1 Mycobacterium sp. 1165196.3 Terrabacteria group; Actinobacteria; Actinobacteria; Corynebacteriales; Mycobacteriaceae; Mycobacterium MVPGLRLVAGGFLVSGDASDDPVAAVDALVAAAGPGLDPPAIQRRDAVLVTGPWLAGVTA WP_067179189.1 hypothetical protein, partial [Mycobacterium sp. 1165196.3] Length: 393\nNumber of predicted TMHs: 2\nExp number of AAs in TMHs: 43.87335\nExp number, first 60 AAs: 0.0275\nTotal prob of N-in: 0.01374\noutside 1 340\nTMhelix 341 363\ninside 364 369\nTMhelix 370 392\noutside 393 393

29476 GCF_001673405.1_ASM167340v1 Mycobacterium sp. 1245111.1 Terrabacteria group; Actinobacteria; Actinobacteria; Corynebacteriales; Mycobacteriaceae; Mycobacterium MGRRRRAAVAALAVLPWFTAPPANAVSPPPVDESRLPPPAPAHPPQPTVQREICAVSSVI WP_067342238.1 type VII secretion-associated serine protease mycosin [Mycobacterium sp. 1245111.1] Length: 447\nNumber of predicted TMHs: 1\nExp number of AAs in TMHs: 26.03048\nExp number, first 60 AAs: 3.94878\nTotal prob of N-in: 0.21968\noutside 1 405\nTMhelix 406 428\ninside 429 447

29477 GCF_001673615.1_ASM167361v1 Mycobacterium sp. 1245499.0 Terrabacteria group; Actinobacteria; Actinobacteria; Corynebacteriales; Mycobacteriaceae; Mycobacterium MVPVLRLVAGGFLVSGDASDDPVAAVDALVAAAGPGLDPPAIQRRDAVLVTGPWMAGVTA WP_067022955.1 hypothetical protein, partial [Mycobacterium sp. 1245499.0] Length: 393\nNumber of predicted TMHs: 2\nExp number of AAs in TMHs: 43.85088\nExp number, first 60 AAs: 0.02478\nTotal prob of N-in: 0.01761\noutside 1 340\nTMhelix 341 363\ninside 364 369\nTMhelix 370 392\noutside 393 393

29478 GCF_001669335.1_ASM166933v1 Mycobacterium sp. 1245852.3 Terrabacteria group; Actinobacteria; Actinobacteria; Corynebacteriales; Mycobacteriaceae; Mycobacterium MVPGLRLVAGGFLVSGDASDDPVAAVDALVAAAGPGLDPPAIQRRDAVLVTGPWLAGVTA WP_067138430.1 hypothetical protein, partial [Mycobacterium sp. 1245852.3] Length: 393\nNumber of predicted TMHs: 2\nExp number of AAs in TMHs: 43.86338\nExp number, first 60 AAs: 0.02651\nTotal prob of N-in: 0.01366\noutside 1 340\nTMhelix 341 363\ninside 364 369\nTMhelix 370 392\noutside 393 393

29479 GCF_001668615.1_ASM166861v1 Mycobacterium sp. 1274761.0 Terrabacteria group; Actinobacteria; Actinobacteria; Corynebacteriales; Mycobacteriaceae; Mycobacterium MPSALPVISRRRMLFGTAALAVVGVTGAACGGKPAPPQDLDRLTAQLDRARSDSQLASDA WP_066975116.1 hypothetical protein [Mycobacterium sp. 1274761.0] Length: 161\nNumber of predicted TMHs: 2\nExp number of AAs in TMHs: 34.22633\nExp number, first 60 AAs: 17.58948\nTotal prob of N-in: 0.79772\nPOSSIBLE N-term signal sequence\ninside 1 12\nTMhelix 13 32\noutside 33 137\nTMhelix 138 160\ninside 161 161

29480 GCF_001665365.1_ASM166536v1 Mycobacterium sp. 1477680.9 Terrabacteria group; Actinobacteria; Actinobacteria; Corynebacteriales; Mycobacteriaceae; Mycobacterium MPSPPPTLSRRRVLFGAAALTLLGVTAAGCGSPPPPEVDELLAQLNRARADSQLASDAAA WP_046753310.1 MULTISPECIES: membrane protein [Mycobacterium] Length: 165\nNumber of predicted TMHs: 2\nExp number of AAs in TMHs: 35.68644\nExp number, first 60 AAs: 17.35768\nTotal prob of N-in: 0.85201\nPOSSIBLE N-term signal sequence\ninside 1 12\nTMhelix 13 35\noutside 36 137\nTMhelix 138 160\ninside 161 165

29481 GCF_001665255.1_ASM166525v1 Mycobacterium sp. 1482268.1 Terrabacteria group; Actinobacteria; Actinobacteria; Corynebacteriales; Mycobacteriaceae; Mycobacterium
MRFPPRAPRRALIAVAGALTIGLACAGPAGAHVRAEADNPAPGSFSVVTFRVPGESETGA WP_067074559.1
nuclear export factor GLE1 [Mycobacterium sp. 1482268.1] Length: 216\nNumber of predicted TMHs: 2\nExp number of AAs in TMHs: 40.6698999999999\nExp number, first 60 AAs: 19.63461\nTotal prob of N-in: 0.90994\nPOSSIBLE N-term signal sequence\ninside 1 12\nTMhelix 13 35\noutside 36 189\nTMhelix 190 212\ninside 213 216

29482 GCF_001667535.1_ASM166753v1 Mycobacterium sp. 852002-10318_SCH4845932 Terrabacteria group; Actinobacteria; Actinobacteria; Corynebacteriales; Mycobacteriaceae; Mycobacterium
MPSPPPTLSRRRVLFGAAALTLLGVTAAGCGSPPPPEVDELLAQLNRARADSQLASDAAA WP_046753310.1
MULTISPECIES: membrane protein [Mycobacterium] Length: 165\nNumber of predicted TMHs: 2\nExp number of AAs in TMHs: 35.68644\nExp number, first 60 AAs: 17.35768\nTotal prob of N-in: 0.85201\nPOSSIBLE N-term signal sequence\ninside 1 12\nTMhelix 13 35\noutside 36 137\nTMhelix 138 160\ninside 161 165

29483 GCF_001665535.1_ASM166553v1 Mycobacterium sp. 852002-51961_SCH5331710 Terrabacteria group; Actinobacteria; Actinobacteria; Corynebacteriales; Mycobacteriaceae; Mycobacterium
MAHRPSAGRPAPTAPTAPRRSFAESLAGADSAADAERRRALRRMKVVALSFLIGATVVFL WP_067088552.1
MULTISPECIES: DUF445 domain-containing protein [Mycobacterium]Length: 446\nNumber of predicted TMHs: 2\nExp number of AAs in TMHs: 59.8128700000001\nExp number, first 60 AAs: 14.72976\nTotal prob of N-in: 0.99783\nPOSSIBLE N-term signal sequence\ninside 1 45\nTMhelix 46 68\noutside 69 419\nTMhelix 420 442\ninside 443 446

29484 GCF_001665685.1_ASM166568v1 Mycobacterium sp. 852013-50091_SCH5140682 Terrabacteria group; Actinobacteria; Actinobacteria; Corynebacteriales; Mycobacteriaceae; Mycobacterium
MPSALPKISRRRALVSAAALAVLGVAAAGCADPPPSPDLADLAAQLDRARADSKLASAAA WP_064943920.1
hypothetical protein [Mycobacterium sp. 852013-50091_SCH5140682] Length: 174\nNumber of predicted TMHs: 2\nExp number of AAs in TMHs: 35.72083\nExp number, first 60 AAs: 15.50477\nTotal prob of N-in: 0.91109\nPOSSIBLE N-term signal sequence\ninside 1 12\nTMhelix 13 31\noutside 32 149\nTMhelix 150 172\ninside 173 174

29485 GCF_001665755.1_ASM166575v1 Mycobacterium sp. 852014-51730_SCH5271717 Terrabacteria group; Actinobacteria; Actinobacteria; Corynebacteriales; Mycobacteriaceae; Mycobacterium
MPSPPPTLSRRRVLFGAAALTLLGVTAAGCGSPPPPEVDELLAQLNRARADSQLASDAAA WP_064892568.1
hypothetical protein [Mycobacterium sp. 852014-51730_SCH5271717] Length: 165\nNumber of predicted TMHs: 2\nExp number of AAs in TMHs: 35.68956\nExp number, first 60 AAs: 17.35854\nTotal prob of N-in: 0.85202\nPOSSIBLE N-term signal sequence\ninside 1 12\nTMhelix 13 35\noutside 36 137\nTMhelix 138 160\ninside 161 165

29486 GCF_002013545.1_ASM201354v1 Mycobacterium sp. 96-892 Terrabacteria group; Actinobacteria; Actinobacteria; Corynebacteriales; Mycobacteriaceae; Mycobacterium
MTIDSTQSRRGWLPWATTIAVVLIAATACAGGTSSRLSRGADDAPAAPAASTYTQAPAAP WP_078308829.1
MULTISPECIES: hypothetical protein [Mycobacterium] Length: 314\nNumber of predicted TMHs: 2\nExp number of AAs in TMHs: 44.98378\nExp number, first 60 AAs: 21.82869\nTotal prob of N-in: 0.95713\nPOSSIBLE N-term signal sequence\ninside 1 11\nTMhelix 12 34\noutside 35 267\nTMhelix 268 290\ninside 291 314

29487 GCF_002013545.1_ASM201354v1 Mycobacterium sp. 96-892 Terrabacteria group; Actinobacteria; Actinobacteria; Corynebacteriales; Mycobacteriaceae; Mycobacterium
MIKHLTRRATVALVVAAAAMSPAAYPAVADTATPLNGLFALAPGSCAEGRATGSYFRMIL WP_078312905.1
MULTISPECIES: hypothetical protein [Mycobacterium] Length: 328\nNumber of predicted TMHs: 1\nExp number of AAs in TMHs: 37.14407\nExp number, first 60 AAs: 14.80443\nTotal prob of N-in: 0.73615\nPOSSIBLE N-term signal sequence\noutside 1 296\nTMhelix 297 319\ninside 320 328

29488 GCF_900157365.1_PRJEB19151 Mycobacterium sp. AB215 Terrabacteria group; Actinobacteria; Actinobacteria; Corynebacteriales; Mycobacteriaceae; Mycobacterium
MRRRAVLAWAGVMLAAVALTAAYTAPVASAHATRLSADPADNAALTTGPDRVSATFNERL WP_077078633.1
copper resistance protein CopC [Mycobacterium sp. AB215] Length: 169\nNumber of predicted TMHs: 2\nExp number of AAs in TMHs: 44.8848\nExp number, first 60 AAs: 22.37707\nTotal prob of N-in: 0.99808\nPOSSIBLE N-term signal sequence\ninside 1 4\nTMhelix 5 27\noutside 28 142\nTMhelix 143 165\ninside 166 169

29489 GCF_001667505.1_ASM166750v1 Mycobacterium sp. ACS1612 Terrabacteria group; Actinobacteria; Actinobacteria; Corynebacteriales; Mycobacteriaceae; Mycobacterium
MASQTTPTPGKPIRTAAFFDLDKTVIAKSSTLAFSKPFFDQGLINRRAVLKSTYAQFLFL WP_067796236.1 inhibition of morphological differentiation protein [Mycobacterium sp. ACS1612] Length: 274\nNumber of predicted TMHs: 1\nExp number of AAs in TMHs: 21.59376\nExp number, first 60 AAs: 0.03489\nTotal prob of N-in: 0.06796\noutside 1 246\nTMhelix 247 269\ninside 270 274

29490 GCF_002013375.1_ASM201337v1 Mycobacterium sp. D16Q13 Terrabacteria group; Actinobacteria; Actinobacteria; Corynebacteriales; Mycobacteriaceae; Mycobacterium
MTIDSTQSRRGWLPWATTIAVVLIAATACAGGTSSRLSRGADDAPAAPAASTYTQAPAAP WP_078308829.1 MULTISPECIES: hypothetical protein [Mycobacterium] Length: 314\nNumber of predicted TMHs: 2\nExp number of AAs in TMHs: 44.98378\nExp number, first 60 AAs: 21.82869\nTotal prob of N-in: 0.95713\nPOSSIBLE N-term signal sequence\ninside 1 11\nTMhelix 12 34\noutside 35 267\nTMhelix 268 290\ninside 291 314

29491 GCF_002013375.1_ASM201337v1 Mycobacterium sp. D16Q13 Terrabacteria group; Actinobacteria; Actinobacteria; Corynebacteriales; Mycobacteriaceae; Mycobacterium
MIKHLTRRATVALVVAAAAMSPAAYPAVADTATPLNGLFALAPGSCAEGRATGSYFRMIL WP_078312905.1 MULTISPECIES: hypothetical protein [Mycobacterium] Length: 328\nNumber of predicted TMHs: 1\nExp number of AAs in TMHs: 37.14407\nExp number, first 60 AAs: 14.80443\nTotal prob of N-in: 0.73615\nPOSSIBLE N-term signal sequence\noutside 1 296\nTMhelix 297 319\ninside 320 328

29492 GCF_002013415.1_ASM201341v1 Mycobacterium sp. D16Q14 Terrabacteria group; Actinobacteria; Actinobacteria; Corynebacteriales; Mycobacteriaceae; Mycobacterium
MIKHLTRRATIALVVAAAAMSPAAYPAVADTATPLNGLFALAPGSCAEGRATGSYFRMIL WP_078341856.1 hypothetical protein [Mycobacterium sp. D16Q14] Length: 328\nNumber of predicted TMHs: 1\nExp number of AAs in TMHs: 38.04666\nExp number, first 60 AAs: 15.66895\nTotal prob of N-in: 0.77230\nPOSSIBLE N-term signal sequence\noutside 1 296\nTMhelix 297 319\ninside 320 328

29493 GCF_002013745.1_ASM201374v1 Mycobacterium sp. D16Q16 Terrabacteria group; Actinobacteria; Actinobacteria; Corynebacteriales; Mycobacteriaceae; Mycobacterium
MIKHLTRRATVALVVAAAAMSPAAYPAVADTATPLNGLFALAPGSCAEGRATGSYFRMIL WP_078312905.1 MULTISPECIES: hypothetical protein [Mycobacterium] Length: 328\nNumber of predicted TMHs: 1\nExp number of AAs in TMHs: 37.14407\nExp number, first 60 AAs: 14.80443\nTotal prob of N-in: 0.73615\nPOSSIBLE N-term signal sequence\noutside 1 296\nTMhelix 297 319\ninside 320 328

29494 GCF_002013745.1_ASM201374v1 Mycobacterium sp. D16Q16 Terrabacteria group; Actinobacteria; Actinobacteria; Corynebacteriales; Mycobacteriaceae; Mycobacterium
MTIDSTQSRRGWLPWATTIAVVLIAATACAGGTSSRLSRGADDAPAAPAASTYTQAPAAP WP_078308829.1 MULTISPECIES: hypothetical protein [Mycobacterium] Length: 314\nNumber of predicted TMHs: 2\nExp number of AAs in TMHs: 44.98378\nExp number, first 60 AAs: 21.82869\nTotal prob of N-in: 0.95713\nPOSSIBLE N-term signal sequence\ninside 1 11\nTMhelix 12 34\noutside 35 267\nTMhelix 268 290\ninside 291 314

29495 GCF_002013785.1_ASM201378v1 Mycobacterium sp. D16Q20 Terrabacteria group; Actinobacteria; Actinobacteria; Corynebacteriales; Mycobacteriaceae; Mycobacterium
MIKHLTRRATIALVVAAAAMSPAAYPAVADTATPLNGLFALAPGSCAEGRATGSYFRMIL WP_078292814.1 MULTISPECIES: hypothetical protein [Mycobacterium] Length: 328\nNumber of predicted TMHs: 1\nExp number of AAs in TMHs: 38.04253\nExp number, first 60 AAs: 15.67632\nTotal prob of N-in: 0.77221\nPOSSIBLE N-term signal sequence\noutside 1 296\nTMhelix 297 319\ninside 320 328

29496 GCF_002013465.1_ASM201346v1 Mycobacterium sp. D16R12 Terrabacteria group; Actinobacteria; Actinobacteria; Corynebacteriales; Mycobacteriaceae; Mycobacterium
MIKHLTRRATVALVVAAAAMSPAAYPAMADTATPLNGLFALAPGSCAEGRATGSYFRMIL WP_078303742.1 hypothetical protein [Mycobacterium sp. D16R12] Length: 328\nNumber of predicted TMHs: 1\nExp number of AAs in TMHs: 37.48192\nExp number, first 60 AAs: 15.01848\nTotal prob of N-in: 0.75053\nPOSSIBLE N-term signal sequence\noutside 1 296\nTMhelix 297 319\ninside 320 328

29497 GCF_002013775.1_ASM201377v1 Mycobacterium sp. D16R18 Terrabacteria group; Actinobacteria; Actinobacteria; Corynebacteriales; Mycobacteriaceae; Mycobacterium
MTIDDTQSRRSWLPWATTIAVLLIATTACAGGTSSHLSQGADDAPAATGAPTYTQAPAAP WP_078359203.1 hypothetical protein [Mycobacterium sp. D16R18] Length: 315\nNumber of predicted TMHs: 2\nExp

number of AAs in TMHs: 44.0938\nExp number, first 60 AAs: 21.19393\nTotal prob of N-in: 0.94569\nPOSSIBLE N-term signal sequence\ninside 1 11\nTMhelix 12 34\noutside 35 268\nTMhelix 269 291\ninside 292 315

29498 GCF_002013775.1_ASM201377v1 Mycobacterium sp. D16R18 Terrabacteria group; Actinobacteria; Actinobacteria; Corynebacteriales; Mycobacteriaceae; Mycobacterium MIKHLTRRATVALVVAAAAMSPAAYPAVADTATPLNGLFALAPGSCAEGRATGSYFRMIL WP_078361061.1 hypothetical protein [Mycobacterium sp. D16R18] Length: 328\nNumber of predicted TMHs: 1\nExp number of AAs in TMHs: 37.27563\nExp number, first 60 AAs: 14.80821\nTotal prob of N-in: 0.73397\nPOSSIBLE N-term signal sequence\noutside 1 296\nTMhelix 297 319\ninside 320 328

29499 GCF_002013385.1_ASM201338v1 Mycobacterium sp. D16R24 Terrabacteria group; Actinobacteria; Actinobacteria; Corynebacteriales; Mycobacteriaceae; Mycobacterium MIKHLTRRATIALVVAAAAMSPAAYPAVADTATPLNGLFALAPGSCAEGRATGSYFRMIL WP_078292814.1 MULTISPECIES: hypothetical protein [Mycobacterium] Length: 328\nNumber of predicted TMHs: 1\nExp number of AAs in TMHs: 38.04253\nExp number, first 60 AAs: 15.67632\nTotal prob of N-in: 0.77221\nPOSSIBLE N-term signal sequence\noutside 1 296\nTMhelix 297 319\ninside 320 328

29500 GCF_002013425.1_ASM201342v1 Mycobacterium sp. D17A2 Terrabacteria group; Actinobacteria; Actinobacteria; Corynebacteriales; Mycobacteriaceae; Mycobacterium MIKHLTRRATIALVVAAAAMSPAAYPAVADTATPLNGLFALAPGSCAEGRATGSYFRMIL WP_078292814.1 MULTISPECIES: hypothetical protein [Mycobacterium] Length: 328\nNumber of predicted TMHs: 1\nExp number of AAs in TMHs: 38.04253\nExp number, first 60 AAs: 15.67632\nTotal prob of N-in: 0.77221\nPOSSIBLE N-term signal sequence\noutside 1 296\nTMhelix 297 319\ninside 320 328

29501 GCF_001666865.1_ASM166686v1 Mycobacterium sp. E136 Terrabacteria group; Actinobacteria; Actinobacteria; Corynebacteriales; Mycobacteriaceae; Mycobacterium MAHRPSAGRPAPTAPTAPRRSFAESLAGADSAADAERRRALRRMKVVALSFLIGATVVFL WP_067088552.1 MULTISPECIES: DUF445 domain-containing protein [Mycobacterium]Length: 446\nNumber of predicted TMHs: 2\nExp number of AAs in TMHs: 59.8128700000001\nExp number, first 60 AAs: 14.72976\nTotal prob of N-in: 0.99783\nPOSSIBLE N-term signal sequence\ninside 1 45\nTMhelix 46 68\noutside 69 419\nTMhelix 420 442\ninside 443 446

29502 GCF_001021385.1_ASM102138v1 Mycobacterium sp. EPa45 Terrabacteria group; Actinobacteria; Actinobacteria; Corynebacteriales; Mycobacteriaceae; Mycobacterium MHVRPGPTTGAPSRTAAFFDLDKTVIAKSSTLAFSKPFFDQGLLNRRAVLKSSYAQFLFL WP_047333843.1 inhibition of morphological differentiation protein [Mycobacterium sp. EPa45] Length: 274\nNumber of predicted TMHs: 1\nExp number of AAs in TMHs: 21.4451\nExp number, first 60 AAs: 0.01307\nTotal prob of N-in: 0.05165\noutside 1 246\nTMhelix 247 269\ninside 270 274

29503 GCF_001545925.1_ASM154592v1 Mycobacterium sp. NAZ190054 Terrabacteria group; Actinobacteria; Actinobacteria; Corynebacteriales; Mycobacteriaceae; Mycobacterium MGCSRGCARRSPVASPACAPSFMSPLSPPVRTGMAARALPAAAGRWLLLRIPVGTVWSEE WP_082753821.1 CPBP family intramembrane metalloprotease [Mycobacterium sp. NAZ190054] Length: 171\nNumber of predicted TMHs: 2\nExp number of AAs in TMHs: 37.93473\nExp number, first 60 AAs: 0.04385\nTotal prob of N-in: 0.96351\ninside 1 102\nTMhelix 103 122\noutside 123 125\nTMhelix 126 148\ninside 149 171

29504 GCF_001545925.1_ASM154592v1 Mycobacterium sp. NAZ190054 Terrabacteria group; Actinobacteria; Actinobacteria; Corynebacteriales; Mycobacteriaceae; Mycobacterium MPGWQDVRVPCQPPTVSRRRVLIGAAALAALGVGATACGAAPPPPDLDDLTTALDRARSD WP_067958698.1 hypothetical protein [Mycobacterium sp. NAZ190054] Length: 177\nNumber of predicted TMHs: 2\nExp number of AAs in TMHs: 43.9776\nExp number, first 60 AAs: 22.1328\nTotal prob of N-in: 0.98984\nPOSSIBLE N-term signal sequence\ninside 1 20\nTMhelix 21 43\noutside 44 151\nTMhelix 152 174\ninside 175 177

29505 GCF_001611855.1_ASM161185v1 Mycobacterium sp. QIA-37 Terrabacteria group; Actinobacteria; Actinobacteria; Corynebacteriales; Mycobacteriaceae; Mycobacterium MIKHLTRRATVALVVAAAAMSPAAYPAVADTATPLNGLFALTPGSCAEGRATGSYFRMIL WP_064407838.1 hypothetical protein [Mycobacterium sp. QIA-37] Length: 329\nNumber of predicted TMHs: 1\nExp number of AAs in TMHs: 37.0585\nExp number, first 60 AAs: 14.7336\nTotal prob of N-in: 0.73308\nPOSSIBLE N-term signal sequence\noutside 1 297\nTMhelix 298 320\ninside 321 329

29506 GCF_000972925.1_ASM97292v1 Mycobacterium sp. UM_Kg1 Terrabacteria group; Actinobacteria; Actinobacteria; Corynebacteriales; Mycobacteriaceae; Mycobacterium MPSPPSAIDRRRVLTGAGMLALLAVVAPACGSAPTPPAVGELEAQRRLALRDSALAAAAA WP_046318871.1 hypothetical protein [Mycobacterium sp. UM_Kg1] Length: 166\nNumber of predicted TMHs: 2\nExp number of AAs in TMHs: 37.63294\nExp number, first 60 AAs: 19.92755\nTotal prob of N-in: 0.90478\nPOSSIBLE N-term signal sequence\ninside 1 12\nTMhelix 13 35\noutside 36 140\nTMhelix 141 163\ninside 164 166

29507 GCF_000455145.1_Mycobacterium_sp_UM_WWY Mycobacterium sp. UM_WWY Terrabacteria group; Actinobacteria; Actinobacteria; Corynebacteriales; Mycobacteriaceae; Mycobacterium MPSALPKISRRRALVSAAAVAVLGVTAAGCADPPPSPDLADLAAQLDRARADSKLASDAA WP_036426178.1 hypothetical protein [Mycobacterium sp. UM_WWY] Length: 171\nNumber of predicted TMHs: 2\nExp number of AAs in TMHs: 33.84544\nExp number, first 60 AAs: 13.72738\nTotal prob of N-in: 0.87874\nPOSSIBLE N-term signal sequence\ninside 1 12\nTMhelix 13 31\noutside 32 146\nTMhelix 147 169\ninside 170 171

29508 GCF_001886515.1_ASM188651v1 Mycobacterium sp. WY10 Terrabacteria group; Actinobacteria; Actinobacteria; Corynebacteriales; Mycobacteriaceae; Mycobacterium MPVRPEPTPGPPSRTAAFFDLDKTVIAKSSTLAFSKPFFDQGLLNRRAVLKSSYAQFLFL WP_083542983.1 inhibition of morphological differentiation protein [Mycobacterium sp. WY10] Length: 274\nNumber of predicted TMHs: 1\nExp number of AAs in TMHs: 21.41819\nExp number, first 60 AAs: 0.00939\nTotal prob of N-in: 0.01714\noutside 1 246\nTMhelix 247 269\ninside 270 274

29509 GCF_002086795.1_ASM208679v1 Mycobacterium tusciae Terrabacteria group; Actinobacteria; Actinobacteria; Corynebacteriales; Mycobacteriaceae; Mycobacterium MRPASEPVRTAAFFDLDKTVIAKSSTLAFSKPFFDQGLINRRTVLKSAYAQFLFLMSGAD WP_083126251.1 inhibition of morphological differentiation protein [Mycobacterium tusciae] Length: 269\nNumber of predicted TMHs: 1\nExp number of AAs in TMHs: 21.62138\nExp number, first 60 AAs: 0.03467\nTotal prob of N-in: 0.05696\noutside 1 241\nTMhelix 242 264\ninside 265 269

29510 GCF_000559085.1_ASM55908v2 Mycobacterium aromaticivorans JS19b1 = JCM 16368 Terrabacteria group; Actinobacteria; Actinobacteria; Corynebacteriales; Mycobacteriaceae; Mycobacterium; Mycobacterium aromaticivorans MHVRPEPATGLNSRTAAFFDLDKTVIAKSSTLAFSKPFFDQGLLNRRAVLKSSYAQFLFL WP_051660360.1 inhibition of morphological differentiation protein [Mycobacterium aromaticivorans] Length: 274\nNumber of predicted TMHs: 1\nExp number of AAs in TMHs: 21.44144\nExp number, first 60 AAs: 0.01193\nTotal prob of N-in: 0.04980\noutside 1 246\nTMhelix 247 269\ninside 270 274

29511 GCF_002089735.1_ASM208973v1 Mycobacterium avium subsp. hominissuis Terrabacteria group; Actinobacteria; Actinobacteria; Corynebacteriales; Mycobacteriaceae; Mycobacterium; Mycobacterium avium complex (MAC); Mycobacterium avium MDKTIIAKSSTLAFSKPFFAQGLLNRRAVLKSSYAQFIYLLSGADHDQMDRMRAHMTNMC WP_084023994.1 inhibition of morphological differentiation protein [Mycobacterium avium] Length: 254\nNumber of predicted TMHs: 1\nExp number of AAs in TMHs: 22.37707\nExp number, first 60 AAs: 0.07236\nTotal prob of N-in: 0.17718\noutside 1 226\nTMhelix 227 249\ninside 250 254

29512 GCF_002089815.1_ASM208981v1 Mycobacterium avium subsp. hominissuis Terrabacteria group; Actinobacteria; Actinobacteria; Corynebacteriales; Mycobacteriaceae; Mycobacterium; Mycobacterium avium complex (MAC); Mycobacterium avium MDKTIIAKSSTLAFSKPFFAQGLLNRRAVLKSSYAQFIYLLSGADHDQMDRMRAHMTNMC WP_084023994.1 inhibition of morphological differentiation protein [Mycobacterium avium] Length: 254\nNumber of predicted TMHs: 1\nExp number of AAs in TMHs: 22.37707\nExp number, first 60 AAs: 0.07236\nTotal prob of N-in: 0.17718\noutside 1 226\nTMhelix 227 249\ninside 250 254

29513 GCF_002089275.1_ASM208927v1 Mycobacterium avium subsp. hominissuis Terrabacteria group; Actinobacteria; Actinobacteria; Corynebacteriales; Mycobacteriaceae; Mycobacterium; Mycobacterium avium complex (MAC); Mycobacterium avium MDKTIIAKSSTLAFSKPFFAQGLLNRRAVLKSSYAQFIYLLSGADHDQMDRMRAHMTNMC WP_084023994.1 inhibition of morphological differentiation protein [Mycobacterium avium] Length: 254\nNumber of predicted TMHs: 1\nExp number of AAs in TMHs: 22.37707\nExp number, first 60 AAs: 0.07236\nTotal prob of N-in: 0.17718\noutside 1 226\nTMhelix 227 249\ninside 250 254

29514 GCF_002089215.1_ASM208921v1 Mycobacterium avium subsp. hominissuis Terrabacteria group; Actinobacteria; Actinobacteria; Corynebacteriales; Mycobacteriaceae; Mycobacterium; Mycobacterium avium complex (MAC); Mycobacterium avium MDKTIIAKSSTLAFSKPFFAQGLLNRRAVLKSSYAQFIYLLSGADHDQMDRMRAHMTNMC WP_084023994.1 inhibition of morphological differentiation protein [Mycobacterium avium] Length:

254\nNumber of predicted TMHs:  1\nExp number of AAs in TMHs: 22.37707\nExp number, first 60 AAs:  0.07236\nTotal prob of N-in:   0.17718\noutside   1   226\nTMhelix   227   249\ninside   250   254

29515 GCF_002089555.1_ASM208955v1 Mycobacterium avium subsp. hominissuis Terrabacteria group; Actinobacteria; Actinobacteria; Corynebacteriales; Mycobacteriaceae; Mycobacterium; Mycobacterium avium complex (MAC); Mycobacterium avium MDKTIIAKSSTLAFSKPFFAQGLLNRRAVLKSSYAQFIYLLSGADHDQMDRMRAHMTNMC WP_084023994.1 inhibition of morphological differentiation protein [Mycobacterium avium] Length: 254\nNumber of predicted TMHs:  1\nExp number of AAs in TMHs: 22.37707\nExp number, first 60 AAs:  0.07236\nTotal prob of N-in:   0.17718\noutside   1   226\nTMhelix   227   249\ninside   250   254

29516 GCF_002089975.1_ASM208997v1 Mycobacterium avium subsp. hominissuis Terrabacteria group; Actinobacteria; Actinobacteria; Corynebacteriales; Mycobacteriaceae; Mycobacterium; Mycobacterium avium complex (MAC); Mycobacterium avium MDKTIIAKSSTLAFSKPFFAQGLLNRRAVLKSSYAQFIYLLSGADHDQMDRMRAHMTNMC WP_084023994.1 inhibition of morphological differentiation protein [Mycobacterium avium] Length: 254\nNumber of predicted TMHs:  1\nExp number of AAs in TMHs: 22.37707\nExp number, first 60 AAs:  0.07236\nTotal prob of N-in:   0.17718\noutside   1   226\nTMhelix   227   249\ninside   250   254

29517 GCF_002089875.1_ASM208987v1 Mycobacterium avium subsp. hominissuis Terrabacteria group; Actinobacteria; Actinobacteria; Corynebacteriales; Mycobacteriaceae; Mycobacterium; Mycobacterium avium complex (MAC); Mycobacterium avium MDKTIIAKSSTLAFSKPFFAQGLLNRRAVLKSSYAQFIYLLSGADHDQMDRMRAHMTNMC WP_084023994.1 inhibition of morphological differentiation protein [Mycobacterium avium] Length: 254\nNumber of predicted TMHs:  1\nExp number of AAs in TMHs: 22.37707\nExp number, first 60 AAs:  0.07236\nTotal prob of N-in:   0.17718\noutside   1   226\nTMhelix   227   249\ninside   250   254

29518 GCF_002089695.1_ASM208969v1 Mycobacterium avium subsp. hominissuis Terrabacteria group; Actinobacteria; Actinobacteria; Corynebacteriales; Mycobacteriaceae; Mycobacterium; Mycobacterium avium complex (MAC); Mycobacterium avium MDKTIIAKSSTLAFSKPFFAQGLLNRRAVLKSSYAQFIYLLSGADHDQMDRMRAHMTNMC WP_084023994.1 inhibition of morphological differentiation protein [Mycobacterium avium] Length: 254\nNumber of predicted TMHs:  1\nExp number of AAs in TMHs: 22.37707\nExp number, first 60 AAs:  0.07236\nTotal prob of N-in:   0.17718\noutside   1   226\nTMhelix   227   249\ninside   250   254

29519 GCF_000240505.1_ASM24050v2 Mycobacterium avium subsp. avium Env 77 Terrabacteria group; Actinobacteria; Actinobacteria; Corynebacteriales; Mycobacteriaceae; Mycobacterium; Mycobacterium avium complex (MAC); Mycobacterium avium; Mycobacterium avium subsp. avium MRSPKQPGSRTRTAAFLDLDHTVIAKSSALAFSKPFMDQGLINRRTVLKSSYAQFLMLLS WP_052312168.1 MULTISPECIES: inhibition of morphological differentiation protein [Mycobacterium] Length: 272\nNumber of predicted TMHs:  1\nExp number of AAs in TMHs: 21.73312\nExp number, first 60 AAs:  0.0228\nTotal prob of N-in:   0.06677\noutside   1   244\nTMhelix   245   267\ninside   268   272

29520 GCF_001853945.1_ASM185394v1 Mycobacterium chelonae Terrabacteria group; Actinobacteria; Actinobacteria; Corynebacteriales; Mycobacteriaceae; Mycobacterium; Mycobacterium chelonae group MIKHLTRRATVALVVAAAAMSPAAYPAVADTATPLNGLFALTPGSCAEGRATGSYFRMIL WP_030094292.1 MULTISPECIES: hypothetical protein [Mycobacterium chelonae group] Length: 329\nNumber of predicted TMHs:  1\nExp number of AAs in TMHs: 37.05801\nExp number, first 60 AAs:  14.73407\nTotal prob of N-in: 0.73307\nPOSSIBLE N-term signal sequence\noutside   1   297\nTMhelix   298   320\ninside   321   329

29521 GCF_002013485.1_ASM201348v1 Mycobacterium chelonae Terrabacteria group; Actinobacteria; Actinobacteria; Corynebacteriales; Mycobacteriaceae; Mycobacterium; Mycobacterium chelonae group MIKHLTRRATVALVVAAAAMSPAAYPAVADTATPLNGLFALTPGSCAEGRATGSYFRMIL WP_030094292.1 MULTISPECIES: hypothetical protein [Mycobacterium chelonae group] Length: 329\nNumber of predicted TMHs:  1\nExp number of AAs in TMHs: 37.05801\nExp number, first 60 AAs:  14.73407\nTotal prob of N-in: 0.73307\nPOSSIBLE N-term signal sequence\noutside   1   297\nTMhelix   298   320\ninside   321   329

29522 GCF_002013535.1_ASM201353v1 Mycobacterium chelonae Terrabacteria group; Actinobacteria; Actinobacteria; Corynebacteriales; Mycobacteriaceae; Mycobacterium; Mycobacterium chelonae group MIKHLTRRATVALVVAAAAMSPAAYPAVADTATPLNGLFALTPGSCAEGRATGSYFRMIL WP_030094292.1 MULTISPECIES: hypothetical protein [Mycobacterium chelonae group] Length: 329\nNumber of predicted TMHs:  1\nExp number of AAs in TMHs: 37.05801\nExp number, first 60 AAs:  14.73407\nTotal prob of N-in: 0.73307\nPOSSIBLE N-term signal sequence\noutside   1   297\nTMhelix   298   320\ninside   321   329

29523 GCF_002013635.1_ASM201363v1 Mycobacterium chelonae Terrabacteria group; Actinobacteria; Actinobacteria; Corynebacteriales; Mycobacteriaceae; Mycobacterium; Mycobacterium chelonae group MIKHLTRRATVALVVAAAAMSPAAYPAVADTATPLNGLFALTPGSCAEGRATGSYFRMIL WP_030094292.1 MULTISPECIES: hypothetical protein [Mycobacterium chelonae group] Length: 329\nNumber of predicted TMHs: 1\nExp number of AAs in TMHs: 37.05801\nExp number, first 60 AAs: 14.73407\nTotal prob of N-in: 0.73307\nPOSSIBLE N-term signal sequence\noutside 1 297\nTMhelix 298 320\ninside 321 329

29524 GCF_001853745.1_ASM185374v1 Mycobacterium chelonae Terrabacteria group; Actinobacteria; Actinobacteria; Corynebacteriales; Mycobacteriaceae; Mycobacterium; Mycobacterium chelonae group MIKHLTRRATVALVVAAAAMSPAAYPAVADTATPLNGLFALTPGSCAEGRATGSYFRMIL WP_030094292.1 MULTISPECIES: hypothetical protein [Mycobacterium chelonae group] Length: 329\nNumber of predicted TMHs: 1\nExp number of AAs in TMHs: 37.05801\nExp number, first 60 AAs: 14.73407\nTotal prob of N-in: 0.73307\nPOSSIBLE N-term signal sequence\noutside 1 297\nTMhelix 298 320\ninside 321 329

29525 GCF_002013595.1_ASM201359v1 Mycobacterium chelonae Terrabacteria group; Actinobacteria; Actinobacteria; Corynebacteriales; Mycobacteriaceae; Mycobacterium; Mycobacterium chelonae group MIKHLTRRATVALVVAAAAMSPAAYPAVADTATPLNGLFALTPGSCAEGRATGSYFRMIL WP_078321647.1 hypothetical protein [Mycobacterium chelonae] Length: 328\nNumber of predicted TMHs: 1\nExp number of AAs in TMHs: 37.05882\nExp number, first 60 AAs: 14.7315\nTotal prob of N-in: 0.73338\nPOSSIBLE N-term signal sequence\noutside 1 296\nTMhelix 297 319\ninside 320 328

29526 GCF_002102055.1_ASM210205v1 Mycobacterium chelonae Terrabacteria group; Actinobacteria; Actinobacteria; Corynebacteriales; Mycobacteriaceae; Mycobacterium; Mycobacterium chelonae group MIKHLTRRATVALVVAAAAMSPAAYPAVADTATPLNGLFALTPGSCAEGRATGSYFRMIL WP_046252588.1 hypothetical protein [Mycobacterium chelonae] Length: 329\nNumber of predicted TMHs: 1\nExp number of AAs in TMHs: 37.06635\nExp number, first 60 AAs: 14.74582\nTotal prob of N-in: 0.73289\nPOSSIBLE N-term signal sequence\noutside 1 297\nTMhelix 298 320\ninside 321 329

29527 GCF_002013855.1_ASM201385v1 Mycobacterium chelonae Terrabacteria group; Actinobacteria; Actinobacteria; Corynebacteriales; Mycobacteriaceae; Mycobacterium; Mycobacterium chelonae group MIKHLTRRATVALVVAAAAMSPAAYPAVADTATPLNGLFALTPGSCAEGRATGSYFRMIL WP_030094292.1 MULTISPECIES: hypothetical protein [Mycobacterium chelonae group] Length: 329\nNumber of predicted TMHs: 1\nExp number of AAs in TMHs: 37.05801\nExp number, first 60 AAs: 14.73407\nTotal prob of N-in: 0.73307\nPOSSIBLE N-term signal sequence\noutside 1 297\nTMhelix 298 320\ninside 321 329

29528 GCF_001853895.1_ASM185389v1 Mycobacterium chelonae Terrabacteria group; Actinobacteria; Actinobacteria; Corynebacteriales; Mycobacteriaceae; Mycobacterium; Mycobacterium chelonae group MIKHLTRRATVALVVAAAAMSPAAYPAVADTATPLNGLFALTPGSCAEGRATGSYFRMIL WP_030094292.1 MULTISPECIES: hypothetical protein [Mycobacterium chelonae group] Length: 329\nNumber of predicted TMHs: 1\nExp number of AAs in TMHs: 37.05801\nExp number, first 60 AAs: 14.73407\nTotal prob of N-in: 0.73307\nPOSSIBLE N-term signal sequence\noutside 1 297\nTMhelix 298 320\ninside 321 329

29529 GCF_001853985.1_ASM185398v1 Mycobacterium chelonae Terrabacteria group; Actinobacteria; Actinobacteria; Corynebacteriales; Mycobacteriaceae; Mycobacterium; Mycobacterium chelonae group MIKHLTRRATVALVVAAAAMSPAAYPAVADTATPLNGLFALTPGSCAEGRATGSYFRMIL WP_030094292.1 MULTISPECIES: hypothetical protein [Mycobacterium chelonae group] Length: 329\nNumber of predicted TMHs: 1\nExp number of AAs in TMHs: 37.05801\nExp number, first 60 AAs: 14.73407\nTotal prob of N-in: 0.73307\nPOSSIBLE N-term signal sequence\noutside 1 297\nTMhelix 298 320\ninside 321 329

29530 GCF_001854015.1_ASM185401v1 Mycobacterium chelonae Terrabacteria group; Actinobacteria; Actinobacteria; Corynebacteriales; Mycobacteriaceae; Mycobacterium; Mycobacterium chelonae group MIKHLTRRATVALVVAAAAMSPAAYPAVADTATPLNGLFALTPGSCAEGRATGSYFRMIL WP_030094292.1 MULTISPECIES: hypothetical protein [Mycobacterium chelonae group] Length: 329\nNumber of predicted TMHs: 1\nExp number of AAs in TMHs: 37.05801\nExp number, first 60 AAs: 14.73407\nTotal prob of N-in: 0.73307\nPOSSIBLE N-term signal sequence\noutside 1 297\nTMhelix 298 320\ninside 321 329

29531 GCF_000971805.1_ASM97180v1 Mycobacterium chelonae Terrabacteria group; Actinobacteria; Actinobacteria; Corynebacteriales; Mycobacteriaceae; Mycobacterium; Mycobacterium chelonae group MIKHLTRRATVALVVAAAAMSPAAYPAVADTATPLNGLFALTPGSCAEGRATGSYFRMIL WP_046252588.1 hypothetical protein [Mycobacterium chelonae] Length: 329\nNumber of predicted TMHs: 1\nExp number of

AAs in TMHs: 37.06635\nExp number, first 60 AAs: 14.74582\nTotal prob of N-in: 0.73289\nPOSSIBLE N-term signal sequence\noutside 1 297\nTMhelix 298 320\ninside 321 329

29532 GCF_001853585.1_ASM185358v1 Mycobacterium chelonae Terrabacteria group; Actinobacteria; Actinobacteria; Corynebacteriales; Mycobacteriaceae; Mycobacterium; Mycobacterium chelonae group MIKHLTRRATVALVVAAAAMSPAAYPAVADTATPLNGLFALTPGSCAEGRATGSYFRMIL WP_030094292.1 MULTISPECIES: hypothetical protein [Mycobacterium chelonae group] Length: 329\nNumber of predicted TMHs: 1\nExp number of AAs in TMHs: 37.05801\nExp number, first 60 AAs: 14.73407\nTotal prob of N-in: 0.73307\nPOSSIBLE N-term signal sequence\noutside 1 297\nTMhelix 298 320\ninside 321 329

29533 GCF_001853615.1_ASM185361v1 Mycobacterium chelonae Terrabacteria group; Actinobacteria; Actinobacteria; Corynebacteriales; Mycobacteriaceae; Mycobacterium; Mycobacterium chelonae group MIKHLTRRATVALVVAAAAMSPAAYPAVADTATPLNGLFALTPGSCAEGRATGSYFRMIL WP_030094292.1 MULTISPECIES: hypothetical protein [Mycobacterium chelonae group] Length: 329\nNumber of predicted TMHs: 1\nExp number of AAs in TMHs: 37.05801\nExp number, first 60 AAs: 14.73407\nTotal prob of N-in: 0.73307\nPOSSIBLE N-term signal sequence\noutside 1 297\nTMhelix 298 320\ninside 321 329

29534 GCF_001853775.1_ASM185377v1 Mycobacterium chelonae Terrabacteria group; Actinobacteria; Actinobacteria; Corynebacteriales; Mycobacteriaceae; Mycobacterium; Mycobacterium chelonae group MIKHLTRRATVALVVAAAAMSPAAYPAVADTATPLNGLFALTPGSCAEGRATGSYFRMIL WP_030094292.1 MULTISPECIES: hypothetical protein [Mycobacterium chelonae group] Length: 329\nNumber of predicted TMHs: 1\nExp number of AAs in TMHs: 37.05801\nExp number, first 60 AAs: 14.73407\nTotal prob of N-in: 0.73307\nPOSSIBLE N-term signal sequence\noutside 1 297\nTMhelix 298 320\ninside 321 329

29535 GCF_002013455.1_ASM201345v1 Mycobacterium chelonae Terrabacteria group; Actinobacteria; Actinobacteria; Corynebacteriales; Mycobacteriaceae; Mycobacterium; Mycobacterium chelonae group MIKHLTRRATVALVVAAAAMSPAAYPAVADTATPLNGLFALTPGSCAEGRATGSYFRMIL WP_030094292.1 MULTISPECIES: hypothetical protein [Mycobacterium chelonae group] Length: 329\nNumber of predicted TMHs: 1\nExp number of AAs in TMHs: 37.05801\nExp number, first 60 AAs: 14.73407\nTotal prob of N-in: 0.73307\nPOSSIBLE N-term signal sequence\noutside 1 297\nTMhelix 298 320\ninside 321 329

29536 GCF_002013515.1_ASM201351v1 Mycobacterium chelonae Terrabacteria group; Actinobacteria; Actinobacteria; Corynebacteriales; Mycobacteriaceae; Mycobacterium; Mycobacterium chelonae group MIKHLTRRATVALVVAAAAMSPAAYPAVADTATPLNGLFALTPGSCAEGRATGSYFRMIL WP_030094292.1 MULTISPECIES: hypothetical protein [Mycobacterium chelonae group] Length: 329\nNumber of predicted TMHs: 1\nExp number of AAs in TMHs: 37.05801\nExp number, first 60 AAs: 14.73407\nTotal prob of N-in: 0.73307\nPOSSIBLE N-term signal sequence\noutside 1 297\nTMhelix 298 320\ninside 321 329

29537 GCF_002013675.1_ASM201367v1 Mycobacterium chelonae Terrabacteria group; Actinobacteria; Actinobacteria; Corynebacteriales; Mycobacteriaceae; Mycobacterium; Mycobacterium chelonae group MIKHLTRRATVALVVAAAAMSPAAYPAVADTATPLNGLFALTPGSCAEGRATGSYFRMIL WP_030094292.1 MULTISPECIES: hypothetical protein [Mycobacterium chelonae group] Length: 329\nNumber of predicted TMHs: 1\nExp number of AAs in TMHs: 37.05801\nExp number, first 60 AAs: 14.73407\nTotal prob of N-in: 0.73307\nPOSSIBLE N-term signal sequence\noutside 1 297\nTMhelix 298 320\ninside 321 329

29538 GCF_001853665.1_ASM185366v1 Mycobacterium chelonae Terrabacteria group; Actinobacteria; Actinobacteria; Corynebacteriales; Mycobacteriaceae; Mycobacterium; Mycobacterium chelonae group MIKHLTRRATVALVVAAAAMSPAAYPAVADTATPLNGLFALTPGSCAEGRATGSYFRMIL WP_030094292.1 MULTISPECIES: hypothetical protein [Mycobacterium chelonae group] Length: 329\nNumber of predicted TMHs: 1\nExp number of AAs in TMHs: 37.05801\nExp number, first 60 AAs: 14.73407\nTotal prob of N-in: 0.73307\nPOSSIBLE N-term signal sequence\noutside 1 297\nTMhelix 298 320\ninside 321 329

29539 GCF_002013815.1_ASM201381v1 Mycobacterium chelonae Terrabacteria group; Actinobacteria; Actinobacteria; Corynebacteriales; Mycobacteriaceae; Mycobacterium; Mycobacterium chelonae group MIKHLTRRATVALVVAAAAMSPAAYPAVADTATPLNGLFALTPGSCAEGRATGSYFRMIL WP_030094292.1 MULTISPECIES: hypothetical protein [Mycobacterium chelonae group] Length: 329\nNumber of predicted TMHs: 1\nExp number of AAs in TMHs: 37.05801\nExp number, first 60 AAs: 14.73407\nTotal prob of N-in: 0.73307\nPOSSIBLE N-term signal sequence\noutside 1 297\nTMhelix 298 320\ninside 321 329

29540 GCF_002013825.1_ASM201382v1 Mycobacterium chelonae Terrabacteria group; Actinobacteria; Actinobacteria; Corynebacteriales; Mycobacteriaceae; Mycobacterium; Mycobacterium chelonae group

MIKHLTRRATVALVVAAAAMSPAAYPAVADTATPLNGLFALTPGSCAEGRATGSYFRMIL WP_030094292.1 MULTISPECIES: hypothetical protein [Mycobacterium chelonae group] Length: 329\nNumber of predicted TMHs: 1\nExp number of AAs in TMHs: 37.05801\nExp number, first 60 AAs: 14.73407\nTotal prob of N-in: 0.73307\nPOSSIBLE N-term signal sequence\noutside 1 297\nTMhelix 298 320\ninside 321 329

29541 GCF_001853995.1_ASM185399v1 Mycobacterium chelonae Terrabacteria group; Actinobacteria; Actinobacteria; Corynebacteriales; Mycobacteriaceae; Mycobacterium; Mycobacterium chelonae group MIKHLTRRATVALVVAAAAMSPAAYPAVADTATPLNGLFALTPGSCAEGRATGSYFRMIL WP_030094292.1 MULTISPECIES: hypothetical protein [Mycobacterium chelonae group] Length: 329\nNumber of predicted TMHs: 1\nExp number of AAs in TMHs: 37.05801\nExp number, first 60 AAs: 14.73407\nTotal prob of N-in: 0.73307\nPOSSIBLE N-term signal sequence\noutside 1 297\nTMhelix 298 320\ninside 321 329

29542 GCF_001854035.1_ASM185403v1 Mycobacterium chelonae Terrabacteria group; Actinobacteria; Actinobacteria; Corynebacteriales; Mycobacteriaceae; Mycobacterium; Mycobacterium chelonae group MIKHLTRRATVALVVAAAAMSPAAYPAVADTATPLNGLFALTPGSCAEGRATGSYFRMIL WP_030094292.1 MULTISPECIES: hypothetical protein [Mycobacterium chelonae group] Length: 329\nNumber of predicted TMHs: 1\nExp number of AAs in TMHs: 37.05801\nExp number, first 60 AAs: 14.73407\nTotal prob of N-in: 0.73307\nPOSSIBLE N-term signal sequence\noutside 1 297\nTMhelix 298 320\ninside 321 329

29543 GCF_001942555.1_ASM194255v1 Mycobacterium chelonae Terrabacteria group; Actinobacteria; Actinobacteria; Corynebacteriales; Mycobacteriaceae; Mycobacterium; Mycobacterium chelonae group MIKHLTRRATVALVVAAAAMSPAAYPAVADTATPLNGLFALTPGSCAEGRATGSYFRMIL WP_030094292.1 MULTISPECIES: hypothetical protein [Mycobacterium chelonae group] Length: 329\nNumber of predicted TMHs: 1\nExp number of AAs in TMHs: 37.05801\nExp number, first 60 AAs: 14.73407\nTotal prob of N-in: 0.73307\nPOSSIBLE N-term signal sequence\noutside 1 297\nTMhelix 298 320\ninside 321 329

29544 GCF_002013305.1_ASM201330v1 Mycobacterium chelonae Terrabacteria group; Actinobacteria; Actinobacteria; Corynebacteriales; Mycobacteriaceae; Mycobacterium; Mycobacterium chelonae group MIKHLTRRATVALVVAAAAMSPAAYPAVADTATPLNGLFALTPGSCAEGRATGSYFRMIL WP_030094292.1 MULTISPECIES: hypothetical protein [Mycobacterium chelonae group] Length: 329\nNumber of predicted TMHs: 1\nExp number of AAs in TMHs: 37.05801\nExp number, first 60 AAs: 14.73407\nTotal prob of N-in: 0.73307\nPOSSIBLE N-term signal sequence\noutside 1 297\nTMhelix 298 320\ninside 321 329

29545 GCF_001853545.1_ASM185354v1 Mycobacterium chelonae Terrabacteria group; Actinobacteria; Actinobacteria; Corynebacteriales; Mycobacteriaceae; Mycobacterium; Mycobacterium chelonae group MIKHLTRRATVALVVAAAAMSPAAYPAVADTATPLNGLFALTPGSCAEGRATGSYFRMIL WP_030094292.1 MULTISPECIES: hypothetical protein [Mycobacterium chelonae group] Length: 329\nNumber of predicted TMHs: 1\nExp number of AAs in TMHs: 37.05801\nExp number, first 60 AAs: 14.73407\nTotal prob of N-in: 0.73307\nPOSSIBLE N-term signal sequence\noutside 1 297\nTMhelix 298 320\ninside 321 329

29546 GCF_001853925.1_ASM185392v1 Mycobacterium chelonae Terrabacteria group; Actinobacteria; Actinobacteria; Corynebacteriales; Mycobacteriaceae; Mycobacterium; Mycobacterium chelonae group MIKHLTRRATVALVVAAAAMSPAAYPAVADTATPLNGLFALTPGSCAEGRATGSYFRMIL WP_030094292.1 MULTISPECIES: hypothetical protein [Mycobacterium chelonae group] Length: 329\nNumber of predicted TMHs: 1\nExp number of AAs in TMHs: 37.05801\nExp number, first 60 AAs: 14.73407\nTotal prob of N-in: 0.73307\nPOSSIBLE N-term signal sequence\noutside 1 297\nTMhelix 298 320\ninside 321 329

29547 GCF_001853965.1_ASM185396v1 Mycobacterium chelonae Terrabacteria group; Actinobacteria; Actinobacteria; Corynebacteriales; Mycobacteriaceae; Mycobacterium; Mycobacterium chelonae group MIKHLTRRATVALVVAAAAMSPAAYPAVADTATPLNGLFALTPGSCAEGRATGSYFRMIL WP_030094292.1 MULTISPECIES: hypothetical protein [Mycobacterium chelonae group] Length: 329\nNumber of predicted TMHs: 1\nExp number of AAs in TMHs: 37.05801\nExp number, first 60 AAs: 14.73407\nTotal prob of N-in: 0.73307\nPOSSIBLE N-term signal sequence\noutside 1 297\nTMhelix 298 320\ninside 321 329

29548 GCF_001853885.1_ASM185388v1 Mycobacterium chelonae Terrabacteria group; Actinobacteria; Actinobacteria; Corynebacteriales; Mycobacteriaceae; Mycobacterium; Mycobacterium chelonae group MIKHLTRRATVALVVAAAAMSPAAYPAVADTATPLNGLFTLTPGSCAEGRATGSYFRMIL WP_070949002.1 hypothetical protein [Mycobacterium chelonae] Length: 329\nNumber of predicted TMHs: 1\nExp number of AAs in TMHs: 36.955\nExp number, first 60 AAs: 14.62982\nTotal prob of N-in: 0.72947\nPOSSIBLE N-term signal sequence\noutside 1 297\nTMhelix 298 320\ninside 321 329

29549 GCF_001853805.1_ASM185380v1 Mycobacterium chelonae Terrabacteria group; Actinobacteria; Actinobacteria; Corynebacteriales; Mycobacteriaceae; Mycobacterium; Mycobacterium chelonae group
MIKHLTRRATVALVVAAAAMSPAAYPAVADTATPLNGLFALTPGSCAEGRATGSYFRMIL WP_070941957.1
hypothetical protein [Mycobacterium chelonae] Length: 329\nNumber of predicted TMHs: 1\nExp number of AAs in TMHs: 37.0581\nExp number, first 60 AAs: 14.73385\nTotal prob of N-in: 0.73307\nPOSSIBLE N-term signal sequence\noutside 1 297\nTMhelix 298 320\ninside 321 329

29550 GCF_001853605.1_ASM185360v1 Mycobacterium chelonae Terrabacteria group; Actinobacteria; Actinobacteria; Corynebacteriales; Mycobacteriaceae; Mycobacterium; Mycobacterium chelonae group
MIKHLTRRATVALVVAAAAMSPAAYPAVADTATPLNGLFALTPGSCAEGRATGSYFRMIL WP_030094292.1
MULTISPECIES: hypothetical protein [Mycobacterium chelonae group] Length: 329\nNumber of predicted TMHs: 1\nExp number of AAs in TMHs: 37.05801\nExp number, first 60 AAs: 14.73407\nTotal prob of N-in: 0.73307\nPOSSIBLE N-term signal sequence\noutside 1 297\nTMhelix 298 320\ninside 321 329

29551 GCF_001853865.1_ASM185386v1 Mycobacterium chelonae Terrabacteria group; Actinobacteria; Actinobacteria; Corynebacteriales; Mycobacteriaceae; Mycobacterium; Mycobacterium chelonae group
MIKHLTRRATVALVVAAAAMSPAAYPAVADTATPLNGLFALTPGSCAEGRATGSYFRMIL WP_030094292.1
MULTISPECIES: hypothetical protein [Mycobacterium chelonae group] Length: 329\nNumber of predicted TMHs: 1\nExp number of AAs in TMHs: 37.05801\nExp number, first 60 AAs: 14.73407\nTotal prob of N-in: 0.73307\nPOSSIBLE N-term signal sequence\noutside 1 297\nTMhelix 298 320\ninside 321 329

29552 GCF_001942575.1_ASM194257v1 Mycobacterium chelonae Terrabacteria group; Actinobacteria; Actinobacteria; Corynebacteriales; Mycobacteriaceae; Mycobacterium; Mycobacterium chelonae group
MIKHLTRRATVALVVAAAAMSPAAYPAVADTATPLNGLFALTPGSCAEGRATGSYFRMIL WP_030094292.1
MULTISPECIES: hypothetical protein [Mycobacterium chelonae group] Length: 329\nNumber of predicted TMHs: 1\nExp number of AAs in TMHs: 37.05801\nExp number, first 60 AAs: 14.73407\nTotal prob of N-in: 0.73307\nPOSSIBLE N-term signal sequence\noutside 1 297\nTMhelix 298 320\ninside 321 329

29553 GCF_002013725.1_ASM201372v1 Mycobacterium chelonae Terrabacteria group; Actinobacteria; Actinobacteria; Corynebacteriales; Mycobacteriaceae; Mycobacterium; Mycobacterium chelonae group
MIKHLTRRATVALVVAAAAMSPAAYPAVADTATPLNGLFALTPGSCAEGRATGSYFRMIL WP_030094292.1
MULTISPECIES: hypothetical protein [Mycobacterium chelonae group] Length: 329\nNumber of predicted TMHs: 1\nExp number of AAs in TMHs: 37.05801\nExp number, first 60 AAs: 14.73407\nTotal prob of N-in: 0.73307\nPOSSIBLE N-term signal sequence\noutside 1 297\nTMhelix 298 320\ninside 321 329

29554 GCF_001942545.1_ASM194254v1 Mycobacterium chelonae Terrabacteria group; Actinobacteria; Actinobacteria; Corynebacteriales; Mycobacteriaceae; Mycobacterium; Mycobacterium chelonae group
MIKHLTRRATVALVVAAAAMSPAAYPAVADTATPLNGLFALTPGSCAEGRATGSYFRMIL WP_046252588.1
hypothetical protein [Mycobacterium chelonae] Length: 329\nNumber of predicted TMHs: 1\nExp number of AAs in TMHs: 37.06635\nExp number, first 60 AAs: 14.74582\nTotal prob of N-in: 0.73289\nPOSSIBLE N-term signal sequence\noutside 1 297\nTMhelix 298 320\ninside 321 329

29555 GCF_002013315.1_ASM201331v1 Mycobacterium chelonae Terrabacteria group; Actinobacteria; Actinobacteria; Corynebacteriales; Mycobacteriaceae; Mycobacterium; Mycobacterium chelonae group
MIKHLTRRATVALVVAAAAMSPAAYPAVADTATPLNGLFALTPGSCAEGRATGSYFRMIL WP_030094292.1
MULTISPECIES: hypothetical protein [Mycobacterium chelonae group] Length: 329\nNumber of predicted TMHs: 1\nExp number of AAs in TMHs: 37.05801\nExp number, first 60 AAs: 14.73407\nTotal prob of N-in: 0.73307\nPOSSIBLE N-term signal sequence\noutside 1 297\nTMhelix 298 320\ninside 321 329

29556 GCF_001853535.1_ASM185353v1 Mycobacterium chelonae Terrabacteria group; Actinobacteria; Actinobacteria; Corynebacteriales; Mycobacteriaceae; Mycobacterium; Mycobacterium chelonae group
MIKHLTRRATVALVVAAAAMSPAAYPAVADTATPLNGLFALTPGSCAEGRATGSYFRMIL WP_030094292.1
MULTISPECIES: hypothetical protein [Mycobacterium chelonae group] Length: 329\nNumber of predicted TMHs: 1\nExp number of AAs in TMHs: 37.05801\nExp number, first 60 AAs: 14.73407\nTotal prob of N-in: 0.73307\nPOSSIBLE N-term signal sequence\noutside 1 297\nTMhelix 298 320\ninside 321 329

29557 GCF_001853845.1_ASM185384v1 Mycobacterium chelonae Terrabacteria group; Actinobacteria; Actinobacteria; Corynebacteriales; Mycobacteriaceae; Mycobacterium; Mycobacterium chelonae group
MIKHLTRRATVALVVAAAAMSPAAYPAVADTATPLNGLFALTPGSCAEGRATGSYFRMIL WP_030094292.1
MULTISPECIES: hypothetical protein [Mycobacterium chelonae group] Length: 329\nNumber of

predicted TMHs: 1\nExp number of AAs in TMHs: 37.05801\nExp number, first 60 AAs: 14.73407\nTotal prob of N-in: 0.73307\nPOSSIBLE N-term signal sequence\noutside 1 297\nTMhelix 298 320\ninside 321 329

29558 GCF_001853685.1_ASM185368v1 Mycobacterium chelonae Terrabacteria group; Actinobacteria; Actinobacteria; Corynebacteriales; Mycobacteriaceae; Mycobacterium; Mycobacterium chelonae group MIKHLTRRATVALVVAAAAMSPAAYPAVADTATPLNGLFALTPGSCAEGRATGSYFRMIL WP_030094292.1 MULTISPECIES: hypothetical protein [Mycobacterium chelonae group] Length: 329\nNumber of predicted TMHs: 1\nExp number of AAs in TMHs: 37.05801\nExp number, first 60 AAs: 14.73407\nTotal prob of N-in: 0.73307\nPOSSIBLE N-term signal sequence\noutside 1 297\nTMhelix 298 320\ninside 321 329

29559 GCF_001853715.1_ASM185371v1 Mycobacterium chelonae Terrabacteria group; Actinobacteria; Actinobacteria; Corynebacteriales; Mycobacteriaceae; Mycobacterium; Mycobacterium chelonae group MIKHLTRRATVALVVAAAAMSPAAYPAVADTATPLNGLFALTPGSCAEGRATGSYFRMIL WP_030094292.1 MULTISPECIES: hypothetical protein [Mycobacterium chelonae group] Length: 329\nNumber of predicted TMHs: 1\nExp number of AAs in TMHs: 37.05801\nExp number, first 60 AAs: 14.73407\nTotal prob of N-in: 0.73307\nPOSSIBLE N-term signal sequence\noutside 1 297\nTMhelix 298 320\ninside 321 329

29560 GCF_001853695.1_ASM185369v1 Mycobacterium chelonae Terrabacteria group; Actinobacteria; Actinobacteria; Corynebacteriales; Mycobacteriaceae; Mycobacterium; Mycobacterium chelonae group MIKHLTRRATVALVVAAAAMSPAAYPAVADTATPLNGLFALTPGSCAEGRATGSYFRMIL WP_030094292.1 MULTISPECIES: hypothetical protein [Mycobacterium chelonae group] Length: 329\nNumber of predicted TMHs: 1\nExp number of AAs in TMHs: 37.05801\nExp number, first 60 AAs: 14.73407\nTotal prob of N-in: 0.73307\nPOSSIBLE N-term signal sequence\noutside 1 297\nTMhelix 298 320\ninside 321 329

29561 GCF_002013895.1_ASM201389v1 Mycobacterium franklinii Terrabacteria group; Actinobacteria; Actinobacteria; Corynebacteriales; Mycobacteriaceae; Mycobacterium; Mycobacterium chelonae group MIKHLTRRATVALVVAAAAMSPATYPAVADTATPLNGLFALTPGSCAEGRATGSYFRMIL WP_078334177.1 hypothetical protein [Mycobacterium franklinii] Length: 329\nNumber of predicted TMHs: 1\nExp number of AAs in TMHs: 34.88048\nExp number, first 60 AAs: 11.69834\nTotal prob of N-in: 0.64184\nPOSSIBLE N-term signal sequence\noutside 1 297\nTMhelix 298 320\ninside 321 329

29562 GCF_002013295.1_ASM201329v1 Mycobacterium franklinii Terrabacteria group; Actinobacteria; Actinobacteria; Corynebacteriales; Mycobacteriaceae; Mycobacterium; Mycobacterium chelonae group MIESTDTVSAPRTAAFFDLDKTVIAKSSTLAFSKPFFDQGLINRRAVLKSSYAQFFFLLS WP_070939071.1 inhibition of morphological differentiation protein [Mycobacterium franklinii] Length: 272\nNumber of predicted TMHs: 1\nExp number of AAs in TMHs: 20.94357\nExp number, first 60 AAs: 0.03048\nTotal prob of N-in: 0.02931\noutside 1 244\nTMhelix 245 267\ninside 268 272

29563 GCF_002086225.1_ASM208622v1 Mycobacterium franklinii Terrabacteria group; Actinobacteria; Actinobacteria; Corynebacteriales; Mycobacteriaceae; Mycobacterium; Mycobacterium chelonae group MTVDSTQSRRSWLPWATTIAVVVIATTACAGGTSTHLSRGTDDAPAATGASTYPQAPAAP WP_078335677.1 hypothetical protein [Mycobacterium franklinii] Length: 314\nNumber of predicted TMHs: 2\nExp number of AAs in TMHs: 44.40492\nExp number, first 60 AAs: 21.27241\nTotal prob of N-in: 0.86630\nPOSSIBLE N-term signal sequence\ninside 1 11\nTMhelix 12 34\noutside 35 269\nTMhelix 270 292\ninside 293 314

29564 GCF_002013715.1_ASM201371v1 Mycobacterium franklinii Terrabacteria group; Actinobacteria; Actinobacteria; Corynebacteriales; Mycobacteriaceae; Mycobacterium; Mycobacterium chelonae group MIESTDTVSAPRTAAFFDLDKTVIAKSSTLAFSKPFFDQGLINRRAVLKSSYAQFFFLLS WP_070939071.1 inhibition of morphological differentiation protein [Mycobacterium franklinii] Length: 272\nNumber of predicted TMHs: 1\nExp number of AAs in TMHs: 20.94357\nExp number, first 60 AAs: 0.03048\nTotal prob of N-in: 0.02931\noutside 1 244\nTMhelix 245 267\ninside 268 272

29565 GCF_002013715.1_ASM201371v1 Mycobacterium franklinii Terrabacteria group; Actinobacteria; Actinobacteria; Corynebacteriales; Mycobacteriaceae; Mycobacterium; Mycobacterium chelonae group MIKHLTRRATVALVVAAAAMSPAAYPAVADTATPLNGLFALTPGSCAEGRATGSYFRMIL WP_078282577.1 hypothetical protein [Mycobacterium franklinii] Length: 329\nNumber of predicted TMHs: 1\nExp number of AAs in TMHs: 37.07163\nExp number, first 60 AAs: 14.55215\nTotal prob of N-in: 0.73812\nPOSSIBLE N-term signal sequence\noutside 1 297\nTMhelix 298 320\ninside 321 329

29566 GCF_001853795.1_ASM185379v1 Mycobacterium franklinii Terrabacteria group; Actinobacteria; Actinobacteria; Corynebacteriales; Mycobacteriaceae; Mycobacterium; Mycobacterium chelonae group

MIESTDTVSAPRTAAFFDLDKTVIAKSSTLAFSKPFFDQGLINRRAVLKSSYAQFFFLLS WP_070939071.1 inhibition of morphological differentiation protein [Mycobacterium franklinii] Length: 272\nNumber of predicted TMHs: 1\nExp number of AAs in TMHs: 20.94357\nExp number, first 60 AAs: 0.03048\nTotal prob of N-in: 0.02931\noutside 1 244\nTMhelix 245 267\ninside 268 272

29567 GCF_001853795.1_ASM185379v1 Mycobacterium franklinii Terrabacteria group; Actinobacteria; Actinobacteria; Corynebacteriales; Mycobacteriaceae; Mycobacterium; Mycobacterium chelonae group MIKHLTRRATVALVVAAAAMSPAAYPAVADTATPLNGLFALTPGSCAEGRATGSYFRMIL WP_070936631.1 hypothetical protein [Mycobacterium franklinii] Length: 329\nNumber of predicted TMHs: 1\nExp number of AAs in TMHs: 37.2019\nExp number, first 60 AAs: 14.67614\nTotal prob of N-in: 0.73605\nPOSSIBLE N-term signal sequence\noutside 1 297\nTMhelix 298 320\ninside 321 329

29568 GCF_002086225.1_ASM208622v1 Mycobacterium franklinii Terrabacteria group; Actinobacteria; Actinobacteria; Corynebacteriales; Mycobacteriaceae; Mycobacterium; Mycobacterium chelonae group MIKHLTRRATVALVVAAAAMSPATYPAVADTATPLNGLFALTPGSCAEGRATGSYFRMIL WP_078334177.1 hypothetical protein [Mycobacterium franklinii] Length: 329\nNumber of predicted TMHs: 1\nExp number of AAs in TMHs: 34.88048\nExp number, first 60 AAs: 11.69834\nTotal prob of N-in: 0.64184\nPOSSIBLE N-term signal sequence\noutside 1 297\nTMhelix 298 320\ninside 321 329

29569 GCF_002013895.1_ASM201389v1 Mycobacterium franklinii Terrabacteria group; Actinobacteria; Actinobacteria; Corynebacteriales; Mycobacteriaceae; Mycobacterium; Mycobacterium chelonae group MTVDSTQSRRSWLPWATTIAVVVIATTACAGGTSTHLSRGTDDAPAATGASTYPQAPAAP WP_078335677.1 hypothetical protein [Mycobacterium franklinii] Length: 314\nNumber of predicted TMHs: 2\nExp number of AAs in TMHs: 44.40492\nExp number, first 60 AAs: 21.27241\nTotal prob of N-in: 0.86630\nPOSSIBLE N-term signal sequence\ninside 1 11\nTMhelix 12 34\noutside 35 269\nTMhelix 270 292\ninside 293 314

29570 GCF_002013295.1_ASM201329v1 Mycobacterium franklinii Terrabacteria group; Actinobacteria; Actinobacteria; Corynebacteriales; Mycobacteriaceae; Mycobacterium; Mycobacterium chelonae group MIKHLTRRATVALVVAAAAMSPAAYPAVADTATPLNGLFALTPGSCAEGRATGSYFRMIL WP_078282577.1 hypothetical protein [Mycobacterium franklinii] Length: 329\nNumber of predicted TMHs: 1\nExp number of AAs in TMHs: 37.07163\nExp number, first 60 AAs: 14.55215\nTotal prob of N-in: 0.73812\nPOSSIBLE N-term signal sequence\noutside 1 297\nTMhelix 298 320\ninside 321 329

29571 GCF_001605725.1_ASM160572v1 Mycobacterium immunogenum Terrabacteria group; Actinobacteria; Actinobacteria; Corynebacteriales; Mycobacteriaceae; Mycobacterium; Mycobacterium chelonae group MIKHLTRRATVALVVAAAAMSPAAYPAMADTATPLNGLFALAPGSCAEGRATGSYFRMIL WP_043079207.1 hypothetical protein [Mycobacterium immunogenum] Length: 331\nNumber of predicted TMHs: 1\nExp number of AAs in TMHs: 37.23212\nExp number, first 60 AAs: 14.87724\nTotal prob of N-in: 0.75569\nPOSSIBLE N-term signal sequence\noutside 1 299\nTMhelix 300 322\ninside 323 331

29572 GCF_001296335.1_ASM129633v1 Mycobacterium immunogenum Terrabacteria group; Actinobacteria; Actinobacteria; Corynebacteriales; Mycobacteriaceae; Mycobacterium; Mycobacterium chelonae group MIKHLTRRATVALVVAAAAMSPAAYPAMADTATPLNGLFALAPGSCAEGRATGSYFRMIL WP_043079207.1 hypothetical protein [Mycobacterium immunogenum] Length: 331\nNumber of predicted TMHs: 1\nExp number of AAs in TMHs: 37.23212\nExp number, first 60 AAs: 14.87724\nTotal prob of N-in: 0.75569\nPOSSIBLE N-term signal sequence\noutside 1 299\nTMhelix 300 322\ninside 323 331

29573 GCF_001677135.1_ASM167713v1 Mycobacterium immunogenum Terrabacteria group; Actinobacteria; Actinobacteria; Corynebacteriales; Mycobacteriaceae; Mycobacterium; Mycobacterium chelonae group MIKHLTRRATVALVVAAAAMSPAAYPAMADTATPLNGLFALAPGSCAEGRATGSYFRMIL WP_043079207.1 hypothetical protein [Mycobacterium immunogenum] Length: 331\nNumber of predicted TMHs: 1\nExp number of AAs in TMHs: 37.23212\nExp number, first 60 AAs: 14.87724\nTotal prob of N-in: 0.75569\nPOSSIBLE N-term signal sequence\noutside 1 299\nTMhelix 300 322\ninside 323 331

29574 GCF_002101665.1_ASM210166v1 Mycobacterium immunogenum Terrabacteria group; Actinobacteria; Actinobacteria; Corynebacteriales; Mycobacteriaceae; Mycobacterium; Mycobacterium chelonae group MIKHLTRRATVALVVAAAAMSPAAYPAMADTATPLNGLFALAPGSCAEGRATGSYFRMIL WP_043079207.1 hypothetical protein [Mycobacterium immunogenum] Length: 331\nNumber of predicted TMHs: 1\nExp number of AAs in TMHs: 37.23212\nExp number, first 60 AAs: 14.87724\nTotal prob of N-in: 0.75569\nPOSSIBLE N-term signal sequence\noutside 1 299\nTMhelix 300 322\ninside 323 331

29575 GCF_001296415.1_ASM129641v1 Mycobacterium immunogenum Terrabacteria group; Actinobacteria; Actinobacteria; Corynebacteriales; Mycobacteriaceae; Mycobacterium; Mycobacterium chelonae group MIKHLTRRATVALVVAAAAMSPAAYPAMADTATPLNGLFALAPGSCAEGRATGSYFRMIL WP_043079207.1 hypothetical protein [Mycobacterium immunogenum] Length: 331\nNumber of predicted TMHs: 1\nExp number of AAs in TMHs: 37.23212\nExp number, first 60 AAs: 14.87724\nTotal prob of N-in: 0.75569\nPOSSIBLE N-term signal sequence\noutside 1 299\nTMhelix 300 322\ninside 323 331

29576 GCF_001296245.1_ASM129624v1 Mycobacterium immunogenum Terrabacteria group; Actinobacteria; Actinobacteria; Corynebacteriales; Mycobacteriaceae; Mycobacterium; Mycobacterium chelonae group MTTRTSVPQRRQAMRSAATRLAAATAAGLIVAALPLSTPLASAEPGFTPPDTSGCPYKVV WP_043076567.1 penicillin-binding protein [Mycobacterium immunogenum] Length: 428\nNumber of predicted TMHs: 2\nExp number of AAs in TMHs: 43.211\nExp number, first 60 AAs: 21.69159\nTotal prob of N-in: 0.99257\nPOSSIBLE N-term signal sequence\ninside 1 20\nTMhelix 21 43\noutside 44 397\nTMhelix 398 420\ninside 421 428

29577 GCF_001296345.1_ASM129634v1 Mycobacterium immunogenum Terrabacteria group; Actinobacteria; Actinobacteria; Corynebacteriales; Mycobacteriaceae; Mycobacterium; Mycobacterium chelonae group MIKHLTRRATVALVVAAAAMSPAAYPAMADTATPLNGLFALAPGSCAEGRATGSYFRMIL WP_043079207.1 hypothetical protein [Mycobacterium immunogenum] Length: 331\nNumber of predicted TMHs: 1\nExp number of AAs in TMHs: 37.23212\nExp number, first 60 AAs: 14.87724\nTotal prob of N-in: 0.75569\nPOSSIBLE N-term signal sequence\noutside 1 299\nTMhelix 300 322\ninside 323 331

29578 GCF_001296245.1_ASM129624v1 Mycobacterium immunogenum Terrabacteria group; Actinobacteria; Actinobacteria; Corynebacteriales; Mycobacteriaceae; Mycobacterium; Mycobacterium chelonae group MIKHLTRRATVALVVAAAAMSPAAYPAMADTATPLNGLFALAPGSCAEGRATGSYFRMIL WP_043079207.1 hypothetical protein [Mycobacterium immunogenum] Length: 331\nNumber of predicted TMHs: 1\nExp number of AAs in TMHs: 37.23212\nExp number, first 60 AAs: 14.87724\nTotal prob of N-in: 0.75569\nPOSSIBLE N-term signal sequence\noutside 1 299\nTMhelix 300 322\ninside 323 331

29579 GCF_001296425.1_ASM129642v1 Mycobacterium immunogenum Terrabacteria group; Actinobacteria; Actinobacteria; Corynebacteriales; Mycobacteriaceae; Mycobacterium; Mycobacterium chelonae group MTTRTSVPQRRQAMRSAATRLAAATAAGLIVAALPLSTPLASAEPGFTPPDTSGCPYKVV WP_043076567.1 penicillin-binding protein [Mycobacterium immunogenum] Length: 428\nNumber of predicted TMHs: 2\nExp number of AAs in TMHs: 43.211\nExp number, first 60 AAs: 21.69159\nTotal prob of N-in: 0.99257\nPOSSIBLE N-term signal sequence\ninside 1 20\nTMhelix 21 43\noutside 44 397\nTMhelix 398 420\ninside 421 428

29580 GCF_001296425.1_ASM129642v1 Mycobacterium immunogenum Terrabacteria group; Actinobacteria; Actinobacteria; Corynebacteriales; Mycobacteriaceae; Mycobacterium; Mycobacterium chelonae group MIKHLTRRATVALVVAAAAMSPAAYPAMADTATPLNGLFALAPGSCAEGRATGSYFRMIL WP_043079207.1 hypothetical protein [Mycobacterium immunogenum] Length: 331\nNumber of predicted TMHs: 1\nExp number of AAs in TMHs: 37.23212\nExp number, first 60 AAs: 14.87724\nTotal prob of N-in: 0.75569\nPOSSIBLE N-term signal sequence\noutside 1 299\nTMhelix 300 322\ninside 323 331

29581 GCF_001296405.1_ASM129640v1 Mycobacterium immunogenum Terrabacteria group; Actinobacteria; Actinobacteria; Corynebacteriales; Mycobacteriaceae; Mycobacterium; Mycobacterium chelonae group MIKHLTRRATVALVVAAAAMSPAAYPAMADTATPLNGLFALAPGSCAEGRATGSYFRMIL WP_043079207.1 hypothetical protein [Mycobacterium immunogenum] Length: 331\nNumber of predicted TMHs: 1\nExp number of AAs in TMHs: 37.23212\nExp number, first 60 AAs: 14.87724\nTotal prob of N-in: 0.75569\nPOSSIBLE N-term signal sequence\noutside 1 299\nTMhelix 300 322\ninside 323 331

29582 GCF_001655155.1_ASM165515v1 Mycobacterium immunogenum Terrabacteria group; Actinobacteria; Actinobacteria; Corynebacteriales; Mycobacteriaceae; Mycobacterium; Mycobacterium chelonae group MTTRTSVPQRRQAMRSAATRLAAATAAGLIVAALPLSTPLASAEPGFTPPDTSGCPYKVV WP_064627889.1 penicillin-binding protein [Mycobacterium immunogenum] Length: 428\nNumber of predicted TMHs: 2\nExp number of AAs in TMHs: 43.21104\nExp number, first 60 AAs: 21.69159\nTotal prob of N-in: 0.99257\nPOSSIBLE N-term signal sequence\ninside 1 20\nTMhelix 21 43\noutside 44 397\nTMhelix 398 420\ninside 421 428

29583 GCF_002013555.1_ASM201355v1 Mycobacterium immunogenum Terrabacteria group; Actinobacteria; Actinobacteria; Corynebacteriales; Mycobacteriaceae; Mycobacterium; Mycobacterium chelonae group MIKHLTRRATVALVVAAAAMSPAAYPAMADTATPLNGLFALAPGSCAEGRATGSYFRMIL WP_043079207.1 hypothetical protein [Mycobacterium immunogenum] Length: 331\nNumber of predicted TMHs: 1\nExp

number of AAs in TMHs: 37.23212\nExp number, first 60 AAs: 14.87724\nTotal prob of N-in: 0.75569\nPOSSIBLE N-term signal sequence\noutside 1 299\nTMhelix 300 322\ninside 323 331

29584 GCF_001655155.1_ASM165515v1 Mycobacterium immunogenum Terrabacteria group; Actinobacteria; Actinobacteria; Corynebacteriales; Mycobacteriaceae; Mycobacterium; Mycobacterium chelonae group MIKHLTRRATVALVVAAAAMSPAAYPAVADTATPLNGLFALAPGSCAEGRATGSYFRMIL WP_064632136.1 hypothetical protein [Mycobacterium immunogenum] Length: 331\nNumber of predicted TMHs: 1\nExp number of AAs in TMHs: 37.22268\nExp number, first 60 AAs: 14.85817\nTotal prob of N-in: 0.73601\nPOSSIBLE N-term signal sequence\noutside 1 299\nTMhelix 300 322\ninside 323 331

29585 GCF_001296345.1_ASM129634v1 Mycobacterium immunogenum Terrabacteria group; Actinobacteria; Actinobacteria; Corynebacteriales; Mycobacteriaceae; Mycobacterium; Mycobacterium chelonae group MTTRTSVPQRRQAMRSAATRLAAATAAGLIVAALPLSTPLASAEPGFTPPDTSGCPYKVV WP_043076567.1 penicillin-binding protein [Mycobacterium immunogenum] Length: 428\nNumber of predicted TMHs: 2\nExp number of AAs in TMHs: 43.211\nExp number, first 60 AAs: 21.69159\nTotal prob of N-in: 0.99257\nPOSSIBLE N-term signal sequence\ninside 1 20\nTMhelix 21 43\noutside 44 397\nTMhelix 398 420\ninside 421 428

29586 GCF_001296325.1_ASM129632v1 Mycobacterium immunogenum Terrabacteria group; Actinobacteria; Actinobacteria; Corynebacteriales; Mycobacteriaceae; Mycobacterium; Mycobacterium chelonae group MTTRTSVPQRRQAMRSAATRLAAATAAGLIVAALPLSTPLASAEPGFTPPDTSGCPYKVV WP_043076567.1 penicillin-binding protein [Mycobacterium immunogenum] Length: 428\nNumber of predicted TMHs: 2\nExp number of AAs in TMHs: 43.211\nExp number, first 60 AAs: 21.69159\nTotal prob of N-in: 0.99257\nPOSSIBLE N-term signal sequence\ninside 1 20\nTMhelix 21 43\noutside 44 397\nTMhelix 398 420\ninside 421 428

29587 GCF_001296265.1_ASM129626v1 Mycobacterium immunogenum Terrabacteria group; Actinobacteria; Actinobacteria; Corynebacteriales; Mycobacteriaceae; Mycobacterium; Mycobacterium chelonae group MTTRTSVPQRRQAMRSAATRLAAATAAGLIVAALPLSTPLASAEPGFTPPDTSGCPYKVV WP_043076567.1 penicillin-binding protein [Mycobacterium immunogenum] Length: 428\nNumber of predicted TMHs: 2\nExp number of AAs in TMHs: 43.211\nExp number, first 60 AAs: 21.69159\nTotal prob of N-in: 0.99257\nPOSSIBLE N-term signal sequence\ninside 1 20\nTMhelix 21 43\noutside 44 397\nTMhelix 398 420\ninside 421 428

29588 GCF_001296405.1_ASM129640v1 Mycobacterium immunogenum Terrabacteria group; Actinobacteria; Actinobacteria; Corynebacteriales; Mycobacteriaceae; Mycobacterium; Mycobacterium chelonae group MTTRTSVPQRRQAMRSAATRLAAATAAGLIVAALPLSTPLASAEPGFTPPDTSGCPYKVV WP_043076567.1 penicillin-binding protein [Mycobacterium immunogenum] Length: 428\nNumber of predicted TMHs: 2\nExp number of AAs in TMHs: 43.211\nExp number, first 60 AAs: 21.69159\nTotal prob of N-in: 0.99257\nPOSSIBLE N-term signal sequence\ninside 1 20\nTMhelix 21 43\noutside 44 397\nTMhelix 398 420\ninside 421 428

29589 GCF_001296325.1_ASM129632v1 Mycobacterium immunogenum Terrabacteria group; Actinobacteria; Actinobacteria; Corynebacteriales; Mycobacteriaceae; Mycobacterium; Mycobacterium chelonae group MIKHLTRRATVALVVAAAAMSPAAYPAMADTATPLNGLFALAPGSCAEGRATGSYFRMIL WP_043079207.1 hypothetical protein [Mycobacterium immunogenum] Length: 331\nNumber of predicted TMHs: 1\nExp number of AAs in TMHs: 37.23212\nExp number, first 60 AAs: 14.87724\nTotal prob of N-in: 0.75569\nPOSSIBLE N-term signal sequence\noutside 1 299\nTMhelix 300 322\ninside 323 331

29590 GCF_001296265.1_ASM129626v1 Mycobacterium immunogenum Terrabacteria group; Actinobacteria; Actinobacteria; Corynebacteriales; Mycobacteriaceae; Mycobacterium; Mycobacterium chelonae group MIKHLTRRATVALVVAAAAMSPAAYPAMADTATPLNGLFALAPGSCAEGRATGSYFRMIL WP_043079207.1 hypothetical protein [Mycobacterium immunogenum] Length: 331\nNumber of predicted TMHs: 1\nExp number of AAs in TMHs: 37.23212\nExp number, first 60 AAs: 14.87724\nTotal prob of N-in: 0.75569\nPOSSIBLE N-term signal sequence\noutside 1 299\nTMhelix 300 322\ninside 323 331

29591 GCF_001296335.1_ASM129633v1 Mycobacterium immunogenum Terrabacteria group; Actinobacteria; Actinobacteria; Corynebacteriales; Mycobacteriaceae; Mycobacterium; Mycobacterium chelonae group MTTRTSVPQRRQAMRSAATRLAAATAAGLIVAALPLSTPLASAEPGFTPPDTSGCPYKVV WP_043076567.1 penicillin-binding protein [Mycobacterium immunogenum] Length: 428\nNumber of predicted TMHs: 2\nExp number of AAs in TMHs: 43.211\nExp number, first 60 AAs: 21.69159\nTotal prob of N-in: 0.99257\nPOSSIBLE N-term signal sequence\ninside 1 20\nTMhelix 21 43\noutside 44 397\nTMhelix 398 420\ninside 421 428

29592 GCF_001296385.1_ASM129638v1 Mycobacterium immunogenum Terrabacteria group; Actinobacteria; Actinobacteria; Corynebacteriales; Mycobacteriaceae; Mycobacterium; Mycobacterium chelonae group

MTTRTSVPQRRQAMRSAATRLAAATAAGLIVAALPLSTPLASAEPGFTPPDTSGCPYKVV WP_043076567.1 penicillin-binding protein [Mycobacterium immunogenum] Length: 428\nNumber of predicted TMHs: 2\nExp number of AAs in TMHs: 43.211\nExp number, first 60 AAs: 21.69159\nTotal prob of N-in: 0.99257\nPOSSIBLE N-term signal sequence\ninside 1 20\nTMhelix 21 43\noutside 44 397\nTMhelix 398 420\ninside 421 428

29593 GCF_001296415.1_ASM129641v1 Mycobacterium immunogenum Terrabacteria group; Actinobacteria; Actinobacteria; Corynebacteriales; Mycobacteriaceae; Mycobacterium; Mycobacterium chelonae group MTTRTSVPQRRQAMRSAATRLAAATAAGLIVAALPLSTPLASAEPGFTPPDTSGCPYKVV WP_043076567.1 penicillin-binding protein [Mycobacterium immunogenum] Length: 428\nNumber of predicted TMHs: 2\nExp number of AAs in TMHs: 43.211\nExp number, first 60 AAs: 21.69159\nTotal prob of N-in: 0.99257\nPOSSIBLE N-term signal sequence\ninside 1 20\nTMhelix 21 43\noutside 44 397\nTMhelix 398 420\ninside 421 428

29594 GCF_001296255.1_ASM129625v1 Mycobacterium immunogenum Terrabacteria group; Actinobacteria; Actinobacteria; Corynebacteriales; Mycobacteriaceae; Mycobacterium; Mycobacterium chelonae group MTTRTSVPQRRQAMRSAATRLAAATAAGLIVAALPLSTPLASAEPGFTPPDTSGCPYKVV WP_043076567.1 penicillin-binding protein [Mycobacterium immunogenum] Length: 428\nNumber of predicted TMHs: 2\nExp number of AAs in TMHs: 43.211\nExp number, first 60 AAs: 21.69159\nTotal prob of N-in: 0.99257\nPOSSIBLE N-term signal sequence\ninside 1 20\nTMhelix 21 43\noutside 44 397\nTMhelix 398 420\ninside 421 428

29595 GCF_001296255.1_ASM129625v1 Mycobacterium immunogenum Terrabacteria group; Actinobacteria; Actinobacteria; Corynebacteriales; Mycobacteriaceae; Mycobacterium; Mycobacterium chelonae group MIKHLTRRATVALVVAAAAMSPAAYPAMADTATPLNGLFALAPGSCAEGRATGSYFRMIL WP_043079207.1 hypothetical protein [Mycobacterium immunogenum] Length: 331\nNumber of predicted TMHs: 1\nExp number of AAs in TMHs: 37.23212\nExp number, first 60 AAs: 14.87724\nTotal prob of N-in: 0.75569\nPOSSIBLE N-term signal sequence\noutside 1 299\nTMhelix 300 322\ninside 323 331

29596 GCF_001296275.1_ASM129627v1 Mycobacterium immunogenum Terrabacteria group; Actinobacteria; Actinobacteria; Corynebacteriales; Mycobacteriaceae; Mycobacterium; Mycobacterium chelonae group MTTRTSVPQRRQAMRSAATRLAAATAAGLIVAALPLSTPLASAEPGFTPPDTSGCPYKVV WP_043076567.1 penicillin-binding protein [Mycobacterium immunogenum] Length: 428\nNumber of predicted TMHs: 2\nExp number of AAs in TMHs: 43.211\nExp number, first 60 AAs: 21.69159\nTotal prob of N-in: 0.99257\nPOSSIBLE N-term signal sequence\ninside 1 20\nTMhelix 21 43\noutside 44 397\nTMhelix 398 420\ninside 421 428

29597 GCF_001296385.1_ASM129638v1 Mycobacterium immunogenum Terrabacteria group; Actinobacteria; Actinobacteria; Corynebacteriales; Mycobacteriaceae; Mycobacterium; Mycobacterium chelonae group MIKHLTRRATVALVVAAAAMSPAAYPAMADTATPLNGLFALAPGSCAEGRATGSYFRMIL WP_043079207.1 hypothetical protein [Mycobacterium immunogenum] Length: 331\nNumber of predicted TMHs: 1\nExp number of AAs in TMHs: 37.23212\nExp number, first 60 AAs: 14.87724\nTotal prob of N-in: 0.75569\nPOSSIBLE N-term signal sequence\noutside 1 299\nTMhelix 300 322\ninside 323 331

29598 GCF_001296275.1_ASM129627v1 Mycobacterium immunogenum Terrabacteria group; Actinobacteria; Actinobacteria; Corynebacteriales; Mycobacteriaceae; Mycobacterium; Mycobacterium chelonae group MIKHLTRRATVALVVAAAAMSPAAYPAMADTATPLNGLFALAPGSCAEGRATGSYFRMIL WP_043079207.1 hypothetical protein [Mycobacterium immunogenum] Length: 331\nNumber of predicted TMHs: 1\nExp number of AAs in TMHs: 37.23212\nExp number, first 60 AAs: 14.87724\nTotal prob of N-in: 0.75569\nPOSSIBLE N-term signal sequence\noutside 1 299\nTMhelix 300 322\ninside 323 331

29599 GCF_000878425.1_ASM87842v1 Mycobacterium immunogenum Terrabacteria group; Actinobacteria; Actinobacteria; Corynebacteriales; Mycobacteriaceae; Mycobacterium; Mycobacterium chelonae group MIKHLTRRATVALVVAAAAMSPAAYPAMADTATPLNGLFALAPGSCAEGRATGSYFRMIL WP_043079207.1 hypothetical protein [Mycobacterium immunogenum] Length: 331\nNumber of predicted TMHs: 1\nExp number of AAs in TMHs: 37.23212\nExp number, first 60 AAs: 14.87724\nTotal prob of N-in: 0.75569\nPOSSIBLE N-term signal sequence\noutside 1 299\nTMhelix 300 322\ninside 323 331

29600 GCF_002101665.1_ASM210166v1 Mycobacterium immunogenum Terrabacteria group; Actinobacteria; Actinobacteria; Corynebacteriales; Mycobacteriaceae; Mycobacterium; Mycobacterium chelonae group MTTRTSVPQRRQAMRSAATRLAAATAAGLIVAALPLSTPLASAEPGFTPPDTSGCPYKVV WP_043076567.1 penicillin-binding protein [Mycobacterium immunogenum] Length: 428\nNumber of predicted TMHs: 2\nExp number of AAs in TMHs: 43.211\nExp number, first 60 AAs: 21.69159\nTotal prob of N-in: 0.99257\nPOSSIBLE N-term signal sequence\ninside 1 20\nTMhelix 21 43\noutside 44 397\nTMhelix 398 420\ninside 421 428

29601 GCF_001677135.1_ASM167713v1 Mycobacterium immunogenum Terrabacteria group; Actinobacteria; Actinobacteria; Corynebacteriales; Mycobacteriaceae; Mycobacterium; Mycobacterium chelonae group MTTRTSVPQRRQAMRSAATRLAAATAAGLIVAALPLSTPLASAEPGFTPPDTSGCPYKVV WP_043076567.1 penicillin-binding protein [Mycobacterium immunogenum] Length: 428\nNumber of predicted TMHs: 2\nExp number of AAs in TMHs: 43.211\nExp number, first 60 AAs: 21.69159\nTotal prob of N-in: 0.99257\nPOSSIBLE N-term signal sequence\ninside 1 20\nTMhelix 21 43\noutside 44 397\nTMhelix 398 420\ninside 421 428

29602 GCF_000878425.1_ASM87842v1 Mycobacterium immunogenum Terrabacteria group; Actinobacteria; Actinobacteria; Corynebacteriales; Mycobacteriaceae; Mycobacterium; Mycobacterium chelonae group MTTRTSVPQRRQAMRSAATRLAAATAAGLIVAALPLSTPLASAEPGFTPPDTSGCPYKVV WP_043076567.1 penicillin-binding protein [Mycobacterium immunogenum] Length: 428\nNumber of predicted TMHs: 2\nExp number of AAs in TMHs: 43.211\nExp number, first 60 AAs: 21.69159\nTotal prob of N-in: 0.99257\nPOSSIBLE N-term signal sequence\ninside 1 20\nTMhelix 21 43\noutside 44 397\nTMhelix 398 420\ninside 421 428

29603 GCF_001605725.1_ASM160572v1 Mycobacterium immunogenum Terrabacteria group; Actinobacteria; Actinobacteria; Corynebacteriales; Mycobacteriaceae; Mycobacterium; Mycobacterium chelonae group MTTRTSVPQRRQAMRSAATRLAAATAAGLIVAALPLSTPLASAEPGFTPPDTSGCPYKVV WP_043076567.1 penicillin-binding protein [Mycobacterium immunogenum] Length: 428\nNumber of predicted TMHs: 2\nExp number of AAs in TMHs: 43.211\nExp number, first 60 AAs: 21.69159\nTotal prob of N-in: 0.99257\nPOSSIBLE N-term signal sequence\ninside 1 20\nTMhelix 21 43\noutside 44 397\nTMhelix 398 420\ninside 421 428

29604 GCF_001853765.1_ASM185376v1 Mycobacterium saopaulense Terrabacteria group; Actinobacteria; Actinobacteria; Corynebacteriales; Mycobacteriaceae; Mycobacterium; Mycobacterium chelonae group MIKHLTRRATVALVVAAAAMSPAAYPAVADTATPLNGLFALTPGSCAEGRATGSYFRMIL WP_070913086.1 hypothetical protein [Mycobacterium saopaulense] Length: 330\nNumber of predicted TMHs: 1\nExp number of AAs in TMHs: 37.0896\nExp number, first 60 AAs: 14.50762\nTotal prob of N-in: 0.73854\nPOSSIBLE N-term signal sequence\noutside 1 298\nTMhelix 299 321\ninside 322 330

29605 GCF_001456355.1_ASM145635v1 Mycobacterium saopaulense Terrabacteria group; Actinobacteria; Actinobacteria; Corynebacteriales; Mycobacteriaceae; Mycobacterium; Mycobacterium chelonae group MIKHLTRRATVALVVAAAAMSPAAYPAVADTATPLNGLFALTPGSCAEGRATGSYFRMIL WP_070913086.1 hypothetical protein [Mycobacterium saopaulense] Length: 330\nNumber of predicted TMHs: 1\nExp number of AAs in TMHs: 37.0896\nExp number, first 60 AAs: 14.50762\nTotal prob of N-in: 0.73854\nPOSSIBLE N-term signal sequence\noutside 1 298\nTMhelix 299 321\ninside 322 330

29606 GCF_002086715.1_ASM208671v1 Mycobacterium saopaulense Terrabacteria group; Actinobacteria; Actinobacteria; Corynebacteriales; Mycobacteriaceae; Mycobacterium; Mycobacterium chelonae group MIKHLTRRATVALVVAAAAMSPAAYPAVADTATPLNGLFALTPGSCAEGRATGSYFRMIL WP_070913086.1 hypothetical protein [Mycobacterium saopaulense] Length: 330\nNumber of predicted TMHs: 1\nExp number of AAs in TMHs: 37.0896\nExp number, first 60 AAs: 14.50762\nTotal prob of N-in: 0.73854\nPOSSIBLE N-term signal sequence\noutside 1 298\nTMhelix 299 321\ninside 322 330

29607 GCF_001853625.1_ASM185362v1 Mycobacterium saopaulense Terrabacteria group; Actinobacteria; Actinobacteria; Corynebacteriales; Mycobacteriaceae; Mycobacterium; Mycobacterium chelonae group MIKHLTRRATVALVVAAAAMSPAAYPAVADTATPLNGLFALTPGSCAEGRATGSYFRMIL WP_070913086.1 hypothetical protein [Mycobacterium saopaulense] Length: 330\nNumber of predicted TMHs: 1\nExp number of AAs in TMHs: 37.0896\nExp number, first 60 AAs: 14.50762\nTotal prob of N-in: 0.73854\nPOSSIBLE N-term signal sequence\noutside 1 298\nTMhelix 299 321\ninside 322 330

29608 GCF_001440005.1_ASM144000v1 Mycobacterium sp. H001 Terrabacteria group; Actinobacteria; Actinobacteria; Corynebacteriales; Mycobacteriaceae; Mycobacterium; Mycobacterium chelonae group MIKHLTRRATVALVVAAAAMSPAAYPAVADTATPLNGLFALTPGSCAEGRATGSYFRMIL WP_030094292.1 MULTISPECIES: hypothetical protein [Mycobacterium chelonae group] Length: 329\nNumber of predicted TMHs: 1\nExp number of AAs in TMHs: 37.05801\nExp number, first 60 AAs: 14.73407\nTotal prob of N-in: 0.73307\nPOSSIBLE N-term signal sequence\noutside 1 297\nTMhelix 298 320\ninside 321 329

29609 GCF_001440085.1_ASM144008v1 Mycobacterium sp. H002 Terrabacteria group; Actinobacteria; Actinobacteria; Corynebacteriales; Mycobacteriaceae; Mycobacterium; Mycobacterium chelonae group MIKHLTRRATVALVVAAAAMSPAAYPAVADTATPLNGLFALTPGSCAEGRATGSYFRMIL WP_030094292.1 MULTISPECIES: hypothetical protein [Mycobacterium chelonae group] Length: 329\nNumber of

predicted TMHs: 1\nExp number of AAs in TMHs: 37.05801\nExp number, first 60 AAs: 14.73407\nTotal prob of N-in: 0.73307\nPOSSIBLE N-term signal sequence\noutside 1 297\nTMhelix 298 320\ninside 321 329

29610 GCF_001440135.1_ASM144013v1 Mycobacterium sp. H003 Terrabacteria group; Actinobacteria; Actinobacteria; Corynebacteriales; Mycobacteriaceae; Mycobacterium; Mycobacterium chelonae group MIKHLTRRATVALVVAAAAMSPAAYPAVADTATPLNGLFALTPGSCAEGRATGSYFRMIL WP_030094292.1 MULTISPECIES: hypothetical protein [Mycobacterium chelonae group] Length: 329\nNumber of predicted TMHs: 1\nExp number of AAs in TMHs: 37.05801\nExp number, first 60 AAs: 14.73407\nTotal prob of N-in: 0.73307\nPOSSIBLE N-term signal sequence\noutside 1 297\nTMhelix 298 320\ninside 321 329

29611 GCF_001440125.1_ASM144012v1 Mycobacterium sp. H054 Terrabacteria group; Actinobacteria; Actinobacteria; Corynebacteriales; Mycobacteriaceae; Mycobacterium; Mycobacterium chelonae group MIKHLTRRATVALVVAAAAMSPAAYPAVADTATPLNGLFALTPGSCAEGRATGSYFRMIL WP_030094292.1 MULTISPECIES: hypothetical protein [Mycobacterium chelonae group] Length: 329\nNumber of predicted TMHs: 1\nExp number of AAs in TMHs: 37.05801\nExp number, first 60 AAs: 14.73407\nTotal prob of N-in: 0.73307\nPOSSIBLE N-term signal sequence\noutside 1 297\nTMhelix 298 320\ninside 321 329

29612 GCF_001440155.1_ASM144015v1 Mycobacterium sp. H063 Terrabacteria group; Actinobacteria; Actinobacteria; Corynebacteriales; Mycobacteriaceae; Mycobacterium; Mycobacterium chelonae group MIKHLTRRATVALVVAAAAMSPAAYPAVADTATPLNGLFALTPGSCAEGRATGSYFRMIL WP_030094292.1 MULTISPECIES: hypothetical protein [Mycobacterium chelonae group] Length: 329\nNumber of predicted TMHs: 1\nExp number of AAs in TMHs: 37.05801\nExp number, first 60 AAs: 14.73407\nTotal prob of N-in: 0.73307\nPOSSIBLE N-term signal sequence\noutside 1 297\nTMhelix 298 320\ninside 321 329

29613 GCF_001440185.1_ASM144018v1 Mycobacterium sp. H070 Terrabacteria group; Actinobacteria; Actinobacteria; Corynebacteriales; Mycobacteriaceae; Mycobacterium; Mycobacterium chelonae group MIKHLTRRATVALVVAAAAMSPAAYPAVADTATPLNGLFALTPGSCAEGRATGSYFRMIL WP_030094292.1 MULTISPECIES: hypothetical protein [Mycobacterium chelonae group] Length: 329\nNumber of predicted TMHs: 1\nExp number of AAs in TMHs: 37.05801\nExp number, first 60 AAs: 14.73407\nTotal prob of N-in: 0.73307\nPOSSIBLE N-term signal sequence\noutside 1 297\nTMhelix 298 320\ninside 321 329

29614 GCF_001440105.1_ASM144010v1 Mycobacterium sp. H072 Terrabacteria group; Actinobacteria; Actinobacteria; Corynebacteriales; Mycobacteriaceae; Mycobacterium; Mycobacterium chelonae group MIKHLTRRATVALVVAAAAMSPAAYPAVADTATPLNGLFALTPGSCAEGRATGSYFRMIL WP_030094292.1 MULTISPECIES: hypothetical protein [Mycobacterium chelonae group] Length: 329\nNumber of predicted TMHs: 1\nExp number of AAs in TMHs: 37.05801\nExp number, first 60 AAs: 14.73407\nTotal prob of N-in: 0.73307\nPOSSIBLE N-term signal sequence\noutside 1 297\nTMhelix 298 320\ninside 321 329

29615 GCF_001440205.1_ASM144020v1 Mycobacterium sp. H079 Terrabacteria group; Actinobacteria; Actinobacteria; Corynebacteriales; Mycobacteriaceae; Mycobacterium; Mycobacterium chelonae group MIKHLTRRATVALVVAAAAMSPAAYPAVADTATPLNGLFALTPGSCAEGRATGSYFRMIL WP_030094292.1 MULTISPECIES: hypothetical protein [Mycobacterium chelonae group] Length: 329\nNumber of predicted TMHs: 1\nExp number of AAs in TMHs: 37.05801\nExp number, first 60 AAs: 14.73407\nTotal prob of N-in: 0.73307\nPOSSIBLE N-term signal sequence\noutside 1 297\nTMhelix 298 320\ninside 321 329

29616 GCF_001440225.1_ASM144022v1 Mycobacterium sp. H092 Terrabacteria group; Actinobacteria; Actinobacteria; Corynebacteriales; Mycobacteriaceae; Mycobacterium; Mycobacterium chelonae group MIKHLTRRATVALVVAAAAMSPAAYPAVADTATPLNGLFALTPGSCAEGRATGSYFRMIL WP_030094292.1 MULTISPECIES: hypothetical protein [Mycobacterium chelonae group] Length: 329\nNumber of predicted TMHs: 1\nExp number of AAs in TMHs: 37.05801\nExp number, first 60 AAs: 14.73407\nTotal prob of N-in: 0.73307\nPOSSIBLE N-term signal sequence\noutside 1 297\nTMhelix 298 320\ninside 321 329

29617 GCF_001440245.1_ASM144024v1 Mycobacterium sp. H101 Terrabacteria group; Actinobacteria; Actinobacteria; Corynebacteriales; Mycobacteriaceae; Mycobacterium; Mycobacterium chelonae group MIKHLTRRATVALVVAAAAMSPAAYPAVADTATPLNGLFALTPGSCAEGRATGSYFRMIL WP_030094292.1 MULTISPECIES: hypothetical protein [Mycobacterium chelonae group] Length: 329\nNumber of predicted TMHs: 1\nExp number of AAs in TMHs: 37.05801\nExp number, first 60 AAs: 14.73407\nTotal prob of N-in: 0.73307\nPOSSIBLE N-term signal sequence\noutside 1 297\nTMhelix 298 320\ninside 321 329

29618 GCF_001440265.1_ASM144026v1 Mycobacterium sp. H110 Terrabacteria group; Actinobacteria; Actinobacteria; Corynebacteriales; Mycobacteriaceae; Mycobacterium; Mycobacterium chelonae group

MIKHLTRRATVALVVAAAAMSPAAYPAVADTATPLNGLFALTPGSCAEGRATGSYFRMIL WP_030094292.1 MULTISPECIES: hypothetical protein [Mycobacterium chelonae group] Length: 329\nNumber of predicted TMHs: 1\nExp number of AAs in TMHs: 37.05801\nExp number, first 60 AAs: 14.73407\nTotal prob of N-in: 0.73307\nPOSSIBLE N-term signal sequence\noutside 1 297\nTMhelix 298 320\ninside 321 329

29619 GCF_001440275.1_ASM144027v1 Mycobacterium sp. HXVII Terrabacteria group; Actinobacteria; Actinobacteria; Corynebacteriales; Mycobacteriaceae; Mycobacterium; Mycobacterium chelonae group MIKHLTRRATVALVVAAAAMSPAAYPAVADTATPLNGLFALTPGSCAEGRATGSYFRMIL WP_030094292.1 MULTISPECIES: hypothetical protein [Mycobacterium chelonae group] Length: 329\nNumber of predicted TMHs: 1\nExp number of AAs in TMHs: 37.05801\nExp number, first 60 AAs: 14.73407\nTotal prob of N-in: 0.73307\nPOSSIBLE N-term signal sequence\noutside 1 297\nTMhelix 298 320\ninside 321 329

29620 GCF_001440305.1_ASM144030v1 Mycobacterium sp. HXXIII Terrabacteria group; Actinobacteria; Actinobacteria; Corynebacteriales; Mycobacteriaceae; Mycobacterium; Mycobacterium chelonae group MIKHLTRRATVALVVAAAAMSPAAYPAVADTATPLNGLFALTPGSCAEGRATGSYFRMIL WP_030094292.1 MULTISPECIES: hypothetical protein [Mycobacterium chelonae group] Length: 329\nNumber of predicted TMHs: 1\nExp number of AAs in TMHs: 37.05801\nExp number, first 60 AAs: 14.73407\nTotal prob of N-in: 0.73307\nPOSSIBLE N-term signal sequence\noutside 1 297\nTMhelix 298 320\ninside 321 329

29621 GCF_001216305.1_7396_7_35 Mycobacterium abscessus Terrabacteria group; Actinobacteria; Actinobacteria; Corynebacteriales; Mycobacteriaceae; Mycobacterium; Mycobacterium chelonae group; Mycobacterium abscessus subgroup MIKHLTRRATVALVVAAAAISPAAYPAMADTATPLNGLFALTPGSCAEGRATGSYFRMIL WP_005073213.1 hypothetical protein [Mycobacterium abscessus] Length: 330\nNumber of predicted TMHs: 2\nExp number of AAs in TMHs: 39.3416199999999\nExp number, first 60 AAs: 17.07688\nTotal prob of N-in: 0.84431\nPOSSIBLE N-term signal sequence\ninside 1 11\nTMhelix 12 29\noutside 30 298\nTMhelix 299 321\ninside 322 330

29622 GCF_002142845.1_ASM214284v1 Mycobacterium abscessus Terrabacteria group; Actinobacteria; Actinobacteria; Corynebacteriales; Mycobacteriaceae; Mycobacterium; Mycobacterium chelonae group; Mycobacterium abscessus subgroup MIKHLTRRATVALVVAAAAISPAAYPAMADTATPLNGLFALTPGSCAEGRATGSYFRMIL WP_005073213.1 hypothetical protein [Mycobacterium abscessus] Length: 330\nNumber of predicted TMHs: 2\nExp number of AAs in TMHs: 39.3416199999999\nExp number, first 60 AAs: 17.07688\nTotal prob of N-in: 0.84431\nPOSSIBLE N-term signal sequence\ninside 1 11\nTMhelix 12 29\noutside 30 298\nTMhelix 299 321\ninside 322 330

29623 GCF_001606215.1_ASM160621v1 Mycobacterium abscessus Terrabacteria group; Actinobacteria; Actinobacteria; Corynebacteriales; Mycobacteriaceae; Mycobacterium; Mycobacterium chelonae group; Mycobacterium abscessus subgroup MIKHLTRRATVALVVAAAAISPAAYPAVADTATPLNGLFALTPGSCAEGRATGSYFRMIL WP_005063674.1 hypothetical protein [Mycobacterium abscessus] Length: 330\nNumber of predicted TMHs: 1\nExp number of AAs in TMHs: 39.25259\nExp number, first 60 AAs: 16.98187\nTotal prob of N-in: 0.83210\nPOSSIBLE N-term signal sequence\noutside 1 298\nTMhelix 299 321\ninside 322 330

29624 GCF_001606275.1_ASM160627v1 Mycobacterium abscessus Terrabacteria group; Actinobacteria; Actinobacteria; Corynebacteriales; Mycobacteriaceae; Mycobacterium; Mycobacterium chelonae group; Mycobacterium abscessus subgroup MIKHLTRRATVALVVAAAAISPAAYPAVADTATPLNGLFALTPGSCAEGRATGSYFRMIL WP_005063674.1 hypothetical protein [Mycobacterium abscessus] Length: 330\nNumber of predicted TMHs: 1\nExp number of AAs in TMHs: 39.25259\nExp number, first 60 AAs: 16.98187\nTotal prob of N-in: 0.83210\nPOSSIBLE N-term signal sequence\noutside 1 298\nTMhelix 299 321\ninside 322 330

29625 GCF_001610615.1_ASM161061v1 Mycobacterium abscessus Terrabacteria group; Actinobacteria; Actinobacteria; Corynebacteriales; Mycobacteriaceae; Mycobacterium; Mycobacterium chelonae group; Mycobacterium abscessus subgroup MIKHLTRRATVALVVAAAAISPAAYPAVADTATPLNGLFALTPGSCAEGRATGSYFRMIL WP_005063674.1 hypothetical protein [Mycobacterium abscessus] Length: 330\nNumber of predicted TMHs: 1\nExp number of AAs in TMHs: 39.25259\nExp number, first 60 AAs: 16.98187\nTotal prob of N-in: 0.83210\nPOSSIBLE N-term signal sequence\noutside 1 298\nTMhelix 299 321\ninside 322 330

29626 GCF_001606235.1_ASM160623v1 Mycobacterium abscessus Terrabacteria group; Actinobacteria; Actinobacteria; Corynebacteriales; Mycobacteriaceae; Mycobacterium; Mycobacterium chelonae group; Mycobacterium abscessus subgroup MIKHLTRRATVALVVAAAAISPAAYPAMADTATPLNGLFALTPGSCAEGRATGSYFRMIL WP_005073213.1 hypothetical protein [Mycobacterium abscessus] Length: 330\nNumber of predicted TMHs: 2\nExp number of AAs in TMHs: 39.3416199999999\nExp number, first 60 AAs: 17.07688\nTotal prob of N-in:

0.84431\nPOSSIBLE N-term signal sequence\ninside 1 11\nTMhelix 12 29\noutside 30 298\nTMhelix 299 321\ninside 322 330

29627 GCF_001217405.1_7396_8_57 Mycobacterium abscessus Terrabacteria group; Actinobacteria; Actinobacteria; Corynebacteriales; Mycobacteriaceae; Mycobacterium; Mycobacterium chelonae group; Mycobacterium abscessus subgroup MIKHLTRRATVALVVAAAAISPAAYPAVADTATPLNGLFALTPGSCAEGRATGSYFRMIL WP_005063674.1 hypothetical protein [Mycobacterium abscessus] Length: 330\nNumber of predicted TMHs: 1\nExp number of AAs in TMHs: 39.25259\nExp number, first 60 AAs: 16.98187\nTotal prob of N-in: 0.83210\nPOSSIBLE N-term signal sequence\noutside 1 298\nTMhelix 299 321\ninside 322 330

29628 GCF_001606195.1_ASM160619v1 Mycobacterium abscessus Terrabacteria group; Actinobacteria; Actinobacteria; Corynebacteriales; Mycobacteriaceae; Mycobacterium; Mycobacterium chelonae group; Mycobacterium abscessus subgroup MIKHLTRRATVALVVAAAAISPAAYPAMADTATPLNGLFALTPGSCAEGRATGSYFRMIL WP_005073213.1 hypothetical protein [Mycobacterium abscessus] Length: 330\nNumber of predicted TMHs: 2\nExp number of AAs in TMHs: 39.3416199999999\nExp number, first 60 AAs: 17.07688\nTotal prob of N-in: 0.84431\nPOSSIBLE N-term signal sequence\ninside 1 11\nTMhelix 12 29\noutside 30 298\nTMhelix 299 321\ninside 322 330

29629 GCF_001350895.1_7396_8_79 Mycobacterium abscessus Terrabacteria group; Actinobacteria; Actinobacteria; Corynebacteriales; Mycobacteriaceae; Mycobacterium; Mycobacterium chelonae group; Mycobacterium abscessus subgroup MIKHLTRRATVALVVAAAAISPAAYPAMADTATPLNGLFALTPGSCAEGRATGSYFRMIL WP_005073213.1 hypothetical protein [Mycobacterium abscessus] Length: 330\nNumber of predicted TMHs: 2\nExp number of AAs in TMHs: 39.3416199999999\nExp number, first 60 AAs: 17.07688\nTotal prob of N-in: 0.84431\nPOSSIBLE N-term signal sequence\ninside 1 11\nTMhelix 12 29\noutside 30 298\nTMhelix 299 321\ninside 322 330

29630 GCF_001217225.1_7396_7_33 Mycobacterium abscessus Terrabacteria group; Actinobacteria; Actinobacteria; Corynebacteriales; Mycobacteriaceae; Mycobacterium; Mycobacterium chelonae group; Mycobacterium abscessus subgroup MIKHLTRRATVALVVAAAAISPAAYPAVADTATPLNGLFALTPGSCAEGRATGSYFRMIL WP_005063674.1 hypothetical protein [Mycobacterium abscessus] Length: 330\nNumber of predicted TMHs: 1\nExp number of AAs in TMHs: 39.25259\nExp number, first 60 AAs: 16.98187\nTotal prob of N-in: 0.83210\nPOSSIBLE N-term signal sequence\noutside 1 298\nTMhelix 299 321\ninside 322 330

29631 GCF_001217105.1_7396_8_96 Mycobacterium abscessus Terrabacteria group; Actinobacteria; Actinobacteria; Corynebacteriales; Mycobacteriaceae; Mycobacterium; Mycobacterium chelonae group; Mycobacterium abscessus subgroup MIKHLTRRATVALVVAAAAISPAAYPAVADTATPLNGLFALTPGSCAEGRATGSYFRMIL WP_005063674.1 hypothetical protein [Mycobacterium abscessus] Length: 330\nNumber of predicted TMHs: 1\nExp number of AAs in TMHs: 39.25259\nExp number, first 60 AAs: 16.98187\nTotal prob of N-in: 0.83210\nPOSSIBLE N-term signal sequence\noutside 1 298\nTMhelix 299 321\ninside 322 330

29632 GCF_001216805.1_7396_8_88 Mycobacterium abscessus Terrabacteria group; Actinobacteria; Actinobacteria; Corynebacteriales; Mycobacteriaceae; Mycobacterium; Mycobacterium chelonae group; Mycobacterium abscessus subgroup MIKHLTRRATVALVVAAAAISPAAYPAVADTATPLNGLFALTPGSCAEGRATGSYFRMIL WP_005063674.1 hypothetical protein [Mycobacterium abscessus] Length: 330\nNumber of predicted TMHs: 1\nExp number of AAs in TMHs: 39.25259\nExp number, first 60 AAs: 16.98187\nTotal prob of N-in: 0.83210\nPOSSIBLE N-term signal sequence\noutside 1 298\nTMhelix 299 321\ninside 322 330

29633 GCF_001216905.1_7396_8_72 Mycobacterium abscessus Terrabacteria group; Actinobacteria; Actinobacteria; Corynebacteriales; Mycobacteriaceae; Mycobacterium; Mycobacterium chelonae group; Mycobacterium abscessus subgroup MIKHLTRRATVALVVAAAAISPAAYPAVADTATPLNGLFALTPGSCAEGRATGSYFRMIL WP_005063674.1 hypothetical protein [Mycobacterium abscessus] Length: 330\nNumber of predicted TMHs: 1\nExp number of AAs in TMHs: 39.25259\nExp number, first 60 AAs: 16.98187\nTotal prob of N-in: 0.83210\nPOSSIBLE N-term signal sequence\noutside 1 298\nTMhelix 299 321\ninside 322 330

29634 GCF_001216245.1_7520_7_28 Mycobacterium abscessus Terrabacteria group; Actinobacteria; Actinobacteria; Corynebacteriales; Mycobacteriaceae; Mycobacterium; Mycobacterium chelonae group; Mycobacterium abscessus subgroup MIKHLTRRATVALVVAAAAISPAAYPAVADTATPLNGLFALTPGSCAEGRATGSYFRMIL WP_005063674.1 hypothetical protein [Mycobacterium abscessus] Length: 330\nNumber of predicted TMHs: 1\nExp number of AAs in TMHs: 39.25259\nExp number, first 60 AAs: 16.98187\nTotal prob of N-in: 0.83210\nPOSSIBLE N-term signal sequence\noutside 1 298\nTMhelix 299 321\ninside 322 330

29635 GCF_001216185.1_7396_8_53 Mycobacterium abscessus Terrabacteria group; Actinobacteria; Actinobacteria; Corynebacteriales; Mycobacteriaceae; Mycobacterium; Mycobacterium chelonae group; Mycobacterium abscessus subgroup MIKHLTRRATVALVVAAAAISPAAYPAVADTATPLNGLFALTPGSCAEGRATGSYFRMIL WP_005063674.1 hypothetical protein [Mycobacterium abscessus] Length: 330\nNumber of predicted TMHs: 1\nExp number of AAs in TMHs: 39.25259\nExp number, first 60 AAs: 16.98187\nTotal prob of N-in: 0.83210\nPOSSIBLE N-term signal sequence\noutside 1 298\nTMhelix 299 321\ninside 322 330

29636 GCF_001216045.1_7396_8_89 Mycobacterium abscessus Terrabacteria group; Actinobacteria; Actinobacteria; Corynebacteriales; Mycobacteriaceae; Mycobacterium; Mycobacterium chelonae group; Mycobacterium abscessus subgroup MIKHLTRRATVALVVAAAAISPAAYPAVADTATPLNGLFALTPGSCAEGRATGSYFRMIL WP_005063674.1 hypothetical protein [Mycobacterium abscessus] Length: 330\nNumber of predicted TMHs: 1\nExp number of AAs in TMHs: 39.25259\nExp number, first 60 AAs: 16.98187\nTotal prob of N-in: 0.83210\nPOSSIBLE N-term signal sequence\noutside 1 298\nTMhelix 299 321\ninside 322 330

29637 GCF_001216005.1_7396_7_39 Mycobacterium abscessus Terrabacteria group; Actinobacteria; Actinobacteria; Corynebacteriales; Mycobacteriaceae; Mycobacterium; Mycobacterium chelonae group; Mycobacterium abscessus subgroup MIKHLTRRATVALVVAAAAISPAAYPAVADTATPLNGLFALTPGSCAEGRATGSYFRMIL WP_005063674.1 hypothetical protein [Mycobacterium abscessus] Length: 330\nNumber of predicted TMHs: 1\nExp number of AAs in TMHs: 39.25259\nExp number, first 60 AAs: 16.98187\nTotal prob of N-in: 0.83210\nPOSSIBLE N-term signal sequence\noutside 1 298\nTMhelix 299 321\ninside 322 330

29638 GCF_001216145.1_7396_8_65 Mycobacterium abscessus Terrabacteria group; Actinobacteria; Actinobacteria; Corynebacteriales; Mycobacteriaceae; Mycobacterium; Mycobacterium chelonae group; Mycobacterium abscessus subgroup MIKHLTRRATVALVVAAAAISPAAYPAMADTATPLNGLFALTPGSCAEGRATGSYFRMIL WP_005073213.1 hypothetical protein [Mycobacterium abscessus] Length: 330\nNumber of predicted TMHs: 2\nExp number of AAs in TMHs: 39.3416199999999\nExp number, first 60 AAs: 17.07688\nTotal prob of N-in: 0.84431\nPOSSIBLE N-term signal sequence\ninside 1 11\nTMhelix 12 29\noutside 30 298\nTMhelix 299 321\ninside 322 330

29639 GCF_001216065.1_7396_8_90 Mycobacterium abscessus Terrabacteria group; Actinobacteria; Actinobacteria; Corynebacteriales; Mycobacteriaceae; Mycobacterium; Mycobacterium chelonae group; Mycobacterium abscessus subgroup MIKHLTRRATVALVVAAAAISPAAYPAMADTATPLNGLFALTPGSCAEGRATGSYFRMIL WP_005073213.1 hypothetical protein [Mycobacterium abscessus] Length: 330\nNumber of predicted TMHs: 2\nExp number of AAs in TMHs: 39.3416199999999\nExp number, first 60 AAs: 17.07688\nTotal prob of N-in: 0.84431\nPOSSIBLE N-term signal sequence\ninside 1 11\nTMhelix 12 29\noutside 30 298\nTMhelix 299 321\ninside 322 330

29640 GCF_001215345.1_7396_8_52 Mycobacterium abscessus Terrabacteria group; Actinobacteria; Actinobacteria; Corynebacteriales; Mycobacteriaceae; Mycobacterium; Mycobacterium chelonae group; Mycobacterium abscessus subgroup MIKHLTRRATVALVVAAAAISPAAYPAVADTATPLNGLFALTPGSCAEGRATGSYFRMIL WP_005063674.1 hypothetical protein [Mycobacterium abscessus] Length: 330\nNumber of predicted TMHs: 1\nExp number of AAs in TMHs: 39.25259\nExp number, first 60 AAs: 16.98187\nTotal prob of N-in: 0.83210\nPOSSIBLE N-term signal sequence\noutside 1 298\nTMhelix 299 321\ninside 322 330

29641 GCF_001215305.1_7396_8_78 Mycobacterium abscessus Terrabacteria group; Actinobacteria; Actinobacteria; Corynebacteriales; Mycobacteriaceae; Mycobacterium; Mycobacterium chelonae group; Mycobacterium abscessus subgroup MIKHLTRRATVALVVAAAAISPAAYPAVADTATPLNGLFALTPGSCAEGRATGSYFRMIL WP_005063674.1 hypothetical protein [Mycobacterium abscessus] Length: 330\nNumber of predicted TMHs: 1\nExp number of AAs in TMHs: 39.25259\nExp number, first 60 AAs: 16.98187\nTotal prob of N-in: 0.83210\nPOSSIBLE N-term signal sequence\noutside 1 298\nTMhelix 299 321\ninside 322 330

29642 GCF_001215445.1_7396_7_27 Mycobacterium abscessus Terrabacteria group; Actinobacteria; Actinobacteria; Corynebacteriales; Mycobacteriaceae; Mycobacterium; Mycobacterium chelonae group; Mycobacterium abscessus subgroup MIKHLTRRATVALVVAAAAISPAAYPAMADTATPLNGLFALTPGSCAEGRATGSYFRMIL WP_005073213.1 hypothetical protein [Mycobacterium abscessus] Length: 330\nNumber of predicted TMHs: 2\nExp number of AAs in TMHs: 39.3416199999999\nExp number, first 60 AAs: 17.07688\nTotal prob of N-in: 0.84431\nPOSSIBLE N-term signal sequence\ninside 1 11\nTMhelix 12 29\noutside 30 298\nTMhelix 299 321\ninside 322 330

29643 GCF_001215225.1_7520_7_30 Mycobacterium abscessus Terrabacteria group; Actinobacteria; Actinobacteria; Corynebacteriales; Mycobacteriaceae; Mycobacterium; Mycobacterium chelonae group; Mycobacterium abscessus subgroup MIKHLTRRATVALVVAAAAISPAAYPAVADTATPLNGLFALTPGSCAEGRATGSYFRMIL WP_005063674.1 hypothetical protein [Mycobacterium abscessus] Length: 330\nNumber of predicted TMHs: 1\nExp number of AAs in TMHs: 39.25259\nExp number, first 60 AAs: 16.98187\nTotal prob of N-in: 0.83210\nPOSSIBLE N-term signal sequence\noutside 1 298\nTMhelix 299 321\ninside 322 330

29644 GCF_001215205.1_7396_8_87 Mycobacterium abscessus Terrabacteria group; Actinobacteria; Actinobacteria; Corynebacteriales; Mycobacteriaceae; Mycobacterium; Mycobacterium chelonae group; Mycobacterium abscessus subgroup MIKHLTRRATVALVVAAAAISPAAYPAVADTATPLNGLFALTPGSCAEGRATGSYFRMIL WP_005063674.1 hypothetical protein [Mycobacterium abscessus] Length: 330\nNumber of predicted TMHs: 1\nExp number of AAs in TMHs: 39.25259\nExp number, first 60 AAs: 16.98187\nTotal prob of N-in: 0.83210\nPOSSIBLE N-term signal sequence\noutside 1 298\nTMhelix 299 321\ninside 322 330

29645 GCF_001215065.1_7396_7_41 Mycobacterium abscessus Terrabacteria group; Actinobacteria; Actinobacteria; Corynebacteriales; Mycobacteriaceae; Mycobacterium; Mycobacterium chelonae group; Mycobacterium abscessus subgroup MIKHLTRRATVALVVAAAAISPAAYPAVADTATPLNGLFALTPGSCAEGRATGSYFRMIL WP_005063674.1 hypothetical protein [Mycobacterium abscessus] Length: 330\nNumber of predicted TMHs: 1\nExp number of AAs in TMHs: 39.25259\nExp number, first 60 AAs: 16.98187\nTotal prob of N-in: 0.83210\nPOSSIBLE N-term signal sequence\noutside 1 298\nTMhelix 299 321\ninside 322 330

29646 GCF_001215025.1_7396_8_75 Mycobacterium abscessus Terrabacteria group; Actinobacteria; Actinobacteria; Corynebacteriales; Mycobacteriaceae; Mycobacterium; Mycobacterium chelonae group; Mycobacterium abscessus subgroup MIKHLTRRATVALVVAAAAISPAAYPAVADTATPLNGLFALTPGSCAEGRATGSYFRMIL WP_005063674.1 hypothetical protein [Mycobacterium abscessus] Length: 330\nNumber of predicted TMHs: 1\nExp number of AAs in TMHs: 39.25259\nExp number, first 60 AAs: 16.98187\nTotal prob of N-in: 0.83210\nPOSSIBLE N-term signal sequence\noutside 1 298\nTMhelix 299 321\ninside 322 330

29647 GCF_001214645.1_7396_7_24 Mycobacterium abscessus Terrabacteria group; Actinobacteria; Actinobacteria; Corynebacteriales; Mycobacteriaceae; Mycobacterium; Mycobacterium chelonae group; Mycobacterium abscessus subgroup MIKHLTRRATVALVVAAAAISPAAYPAVADTATPLNGLFALTPGSCAEGRATGSYFRMIL WP_005063674.1 hypothetical protein [Mycobacterium abscessus] Length: 330\nNumber of predicted TMHs: 1\nExp number of AAs in TMHs: 39.25259\nExp number, first 60 AAs: 16.98187\nTotal prob of N-in: 0.83210\nPOSSIBLE N-term signal sequence\noutside 1 298\nTMhelix 299 321\ninside 322 330

29648 GCF_001214505.1_7396_8_76 Mycobacterium abscessus Terrabacteria group; Actinobacteria; Actinobacteria; Corynebacteriales; Mycobacteriaceae; Mycobacterium; Mycobacterium chelonae group; Mycobacterium abscessus subgroup MIKHLTRRATVALVVAAAAISPAAYPAVADTATPLNGLFALTPGSCAEGRATGSYFRMIL WP_005063674.1 hypothetical protein [Mycobacterium abscessus] Length: 330\nNumber of predicted TMHs: 1\nExp number of AAs in TMHs: 39.25259\nExp number, first 60 AAs: 16.98187\nTotal prob of N-in: 0.83210\nPOSSIBLE N-term signal sequence\noutside 1 298\nTMhelix 299 321\ninside 322 330

29649 GCF_001214465.1_7396_7_36 Mycobacterium abscessus Terrabacteria group; Actinobacteria; Actinobacteria; Corynebacteriales; Mycobacteriaceae; Mycobacterium; Mycobacterium chelonae group; Mycobacterium abscessus subgroup MIKHLTRRATVALVVAAAAISPAAYPAVADTATPLNGLFALTPGSCAEGRATGSYFRMIL WP_005063674.1 hypothetical protein [Mycobacterium abscessus] Length: 330\nNumber of predicted TMHs: 1\nExp number of AAs in TMHs: 39.25259\nExp number, first 60 AAs: 16.98187\nTotal prob of N-in: 0.83210\nPOSSIBLE N-term signal sequence\noutside 1 298\nTMhelix 299 321\ninside 322 330

29650 GCF_001214605.1_7520_7_51 Mycobacterium abscessus Terrabacteria group; Actinobacteria; Actinobacteria; Corynebacteriales; Mycobacteriaceae; Mycobacterium; Mycobacterium chelonae group; Mycobacterium abscessus subgroup MIKHLTRRATVALVVAAAAISPAAYPAVADTATPLNGLFALTPGSCAEGRATGSYFRMIL WP_005063674.1 hypothetical protein [Mycobacterium abscessus] Length: 330\nNumber of predicted TMHs: 1\nExp number of AAs in TMHs: 39.25259\nExp number, first 60 AAs: 16.98187\nTotal prob of N-in: 0.83210\nPOSSIBLE N-term signal sequence\noutside 1 298\nTMhelix 299 321\ninside 322 330

29651 GCF_001214565.1_7396_7_3Mycobacterium abscessus Terrabacteria group; Actinobacteria; Actinobacteria; Corynebacteriales; Mycobacteriaceae; Mycobacterium; Mycobacterium chelonae group; Mycobacterium abscessus subgroup MIKHLTRRATVALVVAAAAISPAAYPAVADTATPLNGLFALTPGSCAEGRATGSYFRMIL WP_005063674.1 hypothetical protein [Mycobacterium abscessus] Length: 330\nNumber of predicted TMHs: 1\nExp number of AAs in TMHs: 39.25259\nExp number, first 60 AAs: 16.98187\nTotal prob of N-in: 0.83210\nPOSSIBLE N-term signal sequence\noutside 1 298\nTMhelix 299 321\ninside 322 330

29652 GCF_001214405.1_7396_7_45 Mycobacterium abscessus Terrabacteria group; Actinobacteria; Actinobacteria; Corynebacteriales; Mycobacteriaceae; Mycobacterium; Mycobacterium chelonae group; Mycobacterium abscessus subgroup MIKHLTRRATVALVVAAAAISPAAYPAVADTATPLNGLFALTPGSCAEGRATGSYFRMIL WP_005063674.1 hypothetical protein [Mycobacterium abscessus] Length: 330\nNumber of predicted TMHs: 1\nExp number of AAs in TMHs: 39.25259\nExp number, first 60 AAs: 16.98187\nTotal prob of N-in: 0.83210\nPOSSIBLE N-term signal sequence\noutside 1 298\nTMhelix 299 321\ninside 322 330

29653 GCF_001214085.1_7396_6_94 Mycobacterium abscessus Terrabacteria group; Actinobacteria; Actinobacteria; Corynebacteriales; Mycobacteriaceae; Mycobacterium; Mycobacterium chelonae group; Mycobacterium abscessus subgroup MIKHLTRRATVALVVAAAAISPAAYPAMADTATPLNGLFALTPGSCAEGRATGSYFRMIL WP_005073213.1 hypothetical protein [Mycobacterium abscessus] Length: 330\nNumber of predicted TMHs: 2\nExp number of AAs in TMHs: 39.3416199999999\nExp number, first 60 AAs: 17.07688\nTotal prob of N-in: 0.84431\nPOSSIBLE N-term signal sequence\ninside 1 11\nTMhelix 12 29\noutside 30 298\nTMhelix 299 321\ninside 322 330

29654 GCF_001214045.1_7396_7_16 Mycobacterium abscessus Terrabacteria group; Actinobacteria; Actinobacteria; Corynebacteriales; Mycobacteriaceae; Mycobacterium; Mycobacterium chelonae group; Mycobacterium abscessus subgroup MIKHLTRRATVALVVAAAAISPAAYPAMADTATPLNGLFALTPGSCAEGRATGSYFRMIL WP_005073213.1 hypothetical protein [Mycobacterium abscessus] Length: 330\nNumber of predicted TMHs: 2\nExp number of AAs in TMHs: 39.3416199999999\nExp number, first 60 AAs: 17.07688\nTotal prob of N-in: 0.84431\nPOSSIBLE N-term signal sequence\ninside 1 11\nTMhelix 12 29\noutside 30 298\nTMhelix 299 321\ninside 322 330

29655 GCF_001213805.1_7396_6_73 Mycobacterium abscessus Terrabacteria group; Actinobacteria; Actinobacteria; Corynebacteriales; Mycobacteriaceae; Mycobacterium; Mycobacterium chelonae group; Mycobacterium abscessus subgroup MIKHLTRRATVALVVAAAAISPAAYPAVADTATPLNGLFALTPGSCAEGRATGSYFRMIL WP_005063674.1 hypothetical protein [Mycobacterium abscessus] Length: 330\nNumber of predicted TMHs: 1\nExp number of AAs in TMHs: 39.25259\nExp number, first 60 AAs: 16.98187\nTotal prob of N-in: 0.83210\nPOSSIBLE N-term signal sequence\noutside 1 298\nTMhelix 299 321\ninside 322 330

29656 GCF_001213985.1_7396_6_62 Mycobacterium abscessus Terrabacteria group; Actinobacteria; Actinobacteria; Corynebacteriales; Mycobacteriaceae; Mycobacterium; Mycobacterium chelonae group; Mycobacterium abscessus subgroup MIKHLTRRATVALVVAAAAISPAAYPAMADTATPLNGLFALTPGSCAEGRATGSYFRMIL WP_005073213.1 hypothetical protein [Mycobacterium abscessus] Length: 330\nNumber of predicted TMHs: 2\nExp number of AAs in TMHs: 39.3416199999999\nExp number, first 60 AAs: 17.07688\nTotal prob of N-in: 0.84431\nPOSSIBLE N-term signal sequence\ninside 1 11\nTMhelix 12 29\noutside 30 298\nTMhelix 299 321\ninside 322 330

29657 GCF_001213945.1_7396_6_56 Mycobacterium abscessus Terrabacteria group; Actinobacteria; Actinobacteria; Corynebacteriales; Mycobacteriaceae; Mycobacterium; Mycobacterium chelonae group; Mycobacterium abscessus subgroup MIKHLTRRATVALVVAAAAISPAAYPAMADTATPLNGLFALTPGSCAEGRATGSYFRMIL WP_005073213.1 hypothetical protein [Mycobacterium abscessus] Length: 330\nNumber of predicted TMHs: 2\nExp number of AAs in TMHs: 39.3416199999999\nExp number, first 60 AAs: 17.07688\nTotal prob of N-in: 0.84431\nPOSSIBLE N-term signal sequence\ninside 1 11\nTMhelix 12 29\noutside 30 298\nTMhelix 299 321\ninside 322 330

29658 GCF_001213885.1_7396_7_14 Mycobacterium abscessus Terrabacteria group; Actinobacteria; Actinobacteria; Corynebacteriales; Mycobacteriaceae; Mycobacterium; Mycobacterium chelonae group; Mycobacterium abscessus subgroup MIKHLTRRATVALVVAAAAISPAAYPAMADTATPLNGLFALTPGSCAEGRATGSYFRMIL WP_005073213.1 hypothetical protein [Mycobacterium abscessus] Length: 330\nNumber of predicted TMHs: 2\nExp number of AAs in TMHs: 39.3416199999999\nExp number, first 60 AAs: 17.07688\nTotal prob of N-in: 0.84431\nPOSSIBLE N-term signal sequence\ninside 1 11\nTMhelix 12 29\noutside 30 298\nTMhelix 299 321\ninside 322 330

29659 GCF_001213425.1_7396_7_10 Mycobacterium abscessus Terrabacteria group; Actinobacteria; Actinobacteria; Corynebacteriales; Mycobacteriaceae; Mycobacterium; Mycobacterium chelonae group; Mycobacterium abscessus subgroup MIKHLTRRATVALVVAAAAISPAAYPAVADTATPLNGLFALTPGSCAEGRATGSYFRMIL WP_005063674.1 hypothetical protein [Mycobacterium abscessus] Length: 330\nNumber of predicted TMHs: 1\nExp number of AAs in TMHs: 39.25259\nExp number, first 60 AAs: 16.98187\nTotal prob of N-in: 0.83210\nPOSSIBLE N-term signal sequence\noutside 1 298\nTMhelix 299 321\ninside 322 330

29660 GCF_001213405.1_7396_6_87 Mycobacterium abscessus Terrabacteria group; Actinobacteria; Actinobacteria; Corynebacteriales; Mycobacteriaceae; Mycobacterium; Mycobacterium chelonae group; Mycobacterium abscessus subgroup MIKHLTRRATVALVVAAAAISPAAYPAVADTATPLNGLFALTPGSCAEGRATGSYFRMIL WP_005063674.1 hypothetical protein [Mycobacterium abscessus] Length: 330\nNumber of predicted TMHs: 1\nExp number of AAs in TMHs: 39.25259\nExp number, first 60 AAs: 16.98187\nTotal prob of N-in: 0.83210\nPOSSIBLE N-term signal sequence\noutside 1 298\nTMhelix 299 321\ninside 322 330

29661 GCF_001213365.1_7396_7_47 Mycobacterium abscessus Terrabacteria group; Actinobacteria; Actinobacteria; Corynebacteriales; Mycobacteriaceae; Mycobacterium; Mycobacterium chelonae group; Mycobacterium abscessus subgroup MIKHLTRRATVALVVAAAAISPAAYPAVADTATPLNGLFALTPGSCAEGRATGSYFRMIL WP_005063674.1 hypothetical protein [Mycobacterium abscessus] Length: 330\nNumber of predicted TMHs: 1\nExp number of AAs in TMHs: 39.25259\nExp number, first 60 AAs: 16.98187\nTotal prob of N-in: 0.83210\nPOSSIBLE N-term signal sequence\noutside 1 298\nTMhelix 299 321\ninside 322 330

29662 GCF_001213685.1_7396_6_80 Mycobacterium abscessus Terrabacteria group; Actinobacteria; Actinobacteria; Corynebacteriales; Mycobacteriaceae; Mycobacterium; Mycobacterium chelonae group; Mycobacterium abscessus subgroup MIKHLTRRATVALVVAAAAISPAAYPAMADTATPLNGLFALTPGSCAEGRATGSYFRMIL WP_005073213.1 hypothetical protein [Mycobacterium abscessus] Length: 330\nNumber of predicted TMHs: 2\nExp number of AAs in TMHs: 39.3416199999999\nExp number, first 60 AAs: 17.07688\nTotal prob of N-in: 0.84431\nPOSSIBLE N-term signal sequence\ninside 1 11\nTMhelix 12 29\noutside 30 298\nTMhelix 299 321\ninside 322 330

29663 GCF_001213245.1_7396_6_75 Mycobacterium abscessus Terrabacteria group; Actinobacteria; Actinobacteria; Corynebacteriales; Mycobacteriaceae; Mycobacterium; Mycobacterium chelonae group; Mycobacterium abscessus subgroup MIKHLTRRATVALVVAAAAISPAAYPAVADTATPLNGLFALTPGSCAEGRATGSYFRMIL WP_005063674.1 hypothetical protein [Mycobacterium abscessus] Length: 330\nNumber of predicted TMHs: 1\nExp number of AAs in TMHs: 39.25259\nExp number, first 60 AAs: 16.98187\nTotal prob of N-in: 0.83210\nPOSSIBLE N-term signal sequence\noutside 1 298\nTMhelix 299 321\ninside 322 330

29664 GCF_001213305.1_7396_6_63 Mycobacterium abscessus Terrabacteria group; Actinobacteria; Actinobacteria; Corynebacteriales; Mycobacteriaceae; Mycobacterium; Mycobacterium chelonae group; Mycobacterium abscessus subgroup MIKHLTRRATVALVVAAAAISPAAYPAMADTATPLNGLFALTPGSCAEGRATGSYFRMIL WP_005073213.1 hypothetical protein [Mycobacterium abscessus] Length: 330\nNumber of predicted TMHs: 2\nExp number of AAs in TMHs: 39.3416199999999\nExp number, first 60 AAs: 17.07688\nTotal prob of N-in: 0.84431\nPOSSIBLE N-term signal sequence\ninside 1 11\nTMhelix 12 29\noutside 30 298\nTMhelix 299 321\ninside 322 330

29665 GCF_001213165.1_7396_6_70 Mycobacterium abscessus Terrabacteria group; Actinobacteria; Actinobacteria; Corynebacteriales; Mycobacteriaceae; Mycobacterium; Mycobacterium chelonae group; Mycobacterium abscessus subgroup MIKHLTRRATVALVVAAAAISPAAYPAMADTATPLNGLFALTPGSCAEGRATGSYFRMIL WP_005073213.1 hypothetical protein [Mycobacterium abscessus] Length: 330\nNumber of predicted TMHs: 2\nExp number of AAs in TMHs: 39.3416199999999\nExp number, first 60 AAs: 17.07688\nTotal prob of N-in: 0.84431\nPOSSIBLE N-term signal sequence\ninside 1 11\nTMhelix 12 29\noutside 30 298\nTMhelix 299 321\ninside 322 330

29666 GCF_001212865.1_7396_6_77 Mycobacterium abscessus Terrabacteria group; Actinobacteria; Actinobacteria; Corynebacteriales; Mycobacteriaceae; Mycobacterium; Mycobacterium chelonae group; Mycobacterium abscessus subgroup MIKHLTRRATVALVVAAAAISPAAYPAMADTATPLNGLFALTPGSCAEGRATGSYFRMIL WP_005073213.1 hypothetical protein [Mycobacterium abscessus] Length: 330\nNumber of predicted TMHs: 2\nExp number of AAs in TMHs: 39.3416199999999\nExp number, first 60 AAs: 17.07688\nTotal prob of N-in: 0.84431\nPOSSIBLE N-term signal sequence\ninside 1 11\nTMhelix 12 29\noutside 30 298\nTMhelix 299 321\ninside 322 330

29667 GCF_001213745.1_7396_6_89 Mycobacterium abscessus Terrabacteria group; Actinobacteria; Actinobacteria; Corynebacteriales; Mycobacteriaceae; Mycobacterium; Mycobacterium chelonae group; Mycobacterium abscessus subgroup MIKHLTRRATVALVVAAAAISPAAYPAMADTATPLNGLFALTPGSCAEGRATGSYFRMIL WP_005073213.1 hypothetical protein [Mycobacterium abscessus] Length: 330\nNumber of predicted TMHs: 2\nExp number of AAs in TMHs: 39.3416199999999\nExp number, first 60 AAs: 17.07688\nTotal prob of N-in: 0.84431\nPOSSIBLE N-term signal sequence\ninside 1 11\nTMhelix 12 29\noutside 30 298\nTMhelix 299 321\ninside 322 330

29668 GCF_001213965.1_7396_6_78 Mycobacterium abscessus Terrabacteria group; Actinobacteria; Actinobacteria; Corynebacteriales; Mycobacteriaceae; Mycobacterium; Mycobacterium chelonae group; Mycobacterium abscessus subgroup MIKHLTRRATVALVVAAAAISPAAYPAMADTATPLNGLFALTPGSCAEGRATGSYFRMIL WP_005073213.1 hypothetical protein [Mycobacterium abscessus] Length: 330\nNumber of predicted TMHs: 2\nExp number of AAs in TMHs: 39.3416199999999\nExp number, first 60 AAs: 17.07688\nTotal prob of N-in: 0.84431\nPOSSIBLE N-term signal sequence\ninside 1 11\nTMhelix 12 29\noutside 30 298\nTMhelix 299 321\ninside 322 330

29669 GCF_001606295.1_ASM160629v1 Mycobacterium abscessus Terrabacteria group; Actinobacteria; Actinobacteria; Corynebacteriales; Mycobacteriaceae; Mycobacterium; Mycobacterium chelonae group; Mycobacterium abscessus subgroup MIKHLTRRATVALVVAAAAISPAAYPAMADTATPLNGLFALTPGSCAEGRATGSYFRMIL WP_005073213.1 hypothetical protein [Mycobacterium abscessus] Length: 330\nNumber of predicted TMHs: 2\nExp number of AAs in TMHs: 39.3416199999999\nExp number, first 60 AAs: 17.07688\nTotal prob of N-in: 0.84431\nPOSSIBLE N-term signal sequence\ninside 1 11\nTMhelix 12 29\noutside 30 298\nTMhelix 299 321\ninside 322 330

29670 GCF_001606335.1_ASM160633v1 Mycobacterium abscessus Terrabacteria group; Actinobacteria; Actinobacteria; Corynebacteriales; Mycobacteriaceae; Mycobacterium; Mycobacterium chelonae group; Mycobacterium abscessus subgroup MIKHLTRRATVALVVAAAAISPAAYPAMADTATPLNGLFALTPGSCAEGRATGSYFRMIL WP_005073213.1 hypothetical protein [Mycobacterium abscessus] Length: 330\nNumber of predicted TMHs: 2\nExp number of AAs in TMHs: 39.3416199999999\nExp number, first 60 AAs: 17.07688\nTotal prob of N-in: 0.84431\nPOSSIBLE N-term signal sequence\ninside 1 11\nTMhelix 12 29\noutside 30 298\nTMhelix 299 321\ninside 322 330

29671 GCF_002142915.1_ASM214291v1 Mycobacterium abscessus Terrabacteria group; Actinobacteria; Actinobacteria; Corynebacteriales; Mycobacteriaceae; Mycobacterium; Mycobacterium chelonae group; Mycobacterium abscessus subgroup MIKHLTRRATVALVVAAAAISPAAYPAVADTATPLNGLFALTPGSCAEGRATGSYFRMIL WP_005063674.1 hypothetical protein [Mycobacterium abscessus] Length: 330\nNumber of predicted TMHs: 1\nExp number of AAs in TMHs: 39.25259\nExp number, first 60 AAs: 16.98187\nTotal prob of N-in: 0.83210\nPOSSIBLE N-term signal sequence\noutside 1 298\nTMhelix 299 321\ninside 322 330

29672 GCF_001213785.1_7396_7_20 Mycobacterium abscessus Terrabacteria group; Actinobacteria; Actinobacteria; Corynebacteriales; Mycobacteriaceae; Mycobacterium; Mycobacterium chelonae group; Mycobacterium abscessus subgroup MIKHLTRRATVALVVAAAAISPAAYPAMADTATPLNGLFALTPGSCAEGRATGSYFRMIL WP_005073213.1 hypothetical protein [Mycobacterium abscessus] Length: 330\nNumber of predicted TMHs: 2\nExp number of AAs in TMHs: 39.3416199999999\nExp number, first 60 AAs: 17.07688\nTotal prob of N-in: 0.84431\nPOSSIBLE N-term signal sequence\ninside 1 11\nTMhelix 12 29\noutside 30 298\nTMhelix 299 321\ninside 322 330

29673 GCF_001213825.1_7396_6_68 Mycobacterium abscessus Terrabacteria group; Actinobacteria; Actinobacteria; Corynebacteriales; Mycobacteriaceae; Mycobacterium; Mycobacterium chelonae group; Mycobacterium abscessus subgroup MIKHLTRRATVALVVAAAAISPAAYPAMADTATPLNGLFALTPGSCAEGRATGSYFRMIL WP_005073213.1 hypothetical protein [Mycobacterium abscessus] Length: 330\nNumber of predicted TMHs: 2\nExp number of AAs in TMHs: 39.3416199999999\nExp number, first 60 AAs: 17.07688\nTotal prob of N-in: 0.84431\nPOSSIBLE N-term signal sequence\ninside 1 11\nTMhelix 12 29\noutside 30 298\nTMhelix 299 321\ninside 322 330

29674 GCF_001215005.1_7396_8_54 Mycobacterium abscessus Terrabacteria group; Actinobacteria; Actinobacteria; Corynebacteriales; Mycobacteriaceae; Mycobacterium; Mycobacterium chelonae group; Mycobacterium abscessus subgroup MIKHLTRRATVALVVAAAAISPAAYPAMADTATPLNGLFALTPGSCAEGRATGSYFRMIL WP_005073213.1 hypothetical protein [Mycobacterium abscessus] Length: 330\nNumber of predicted TMHs: 2\nExp number of AAs in TMHs: 39.3416199999999\nExp number, first 60 AAs: 17.07688\nTotal prob of N-in: 0.84431\nPOSSIBLE N-term signal sequence\ninside 1 11\nTMhelix 12 29\noutside 30 298\nTMhelix 299 321\ninside 322 330

29675 GCF_001215045.1_7396_7_29 Mycobacterium abscessus Terrabacteria group; Actinobacteria; Actinobacteria; Corynebacteriales; Mycobacteriaceae; Mycobacterium; Mycobacterium chelonae group; Mycobacterium abscessus subgroup MIKHLTRRATVALVVAAAAISPAAYPAMADTATPLNGLFALTPGSCAEGRATGSYFRMIL WP_005073213.1 hypothetical protein [Mycobacterium abscessus] Length: 330\nNumber of predicted TMHs: 2\nExp number of AAs in TMHs: 39.3416199999999\nExp number, first 60 AAs: 17.07688\nTotal prob of N-in:

0.84431\nPOSSIBLE N-term signal sequence\ninside 1 11\nTMhelix 12 29\noutside 30 298\nTMhelix 299 321\ninside 322 330

29676 GCF_001212805.1_7396_6_90 Mycobacterium abscessus Terrabacteria group; Actinobacteria; Actinobacteria; Corynebacteriales; Mycobacteriaceae; Mycobacterium; Mycobacterium chelonae group; Mycobacterium abscessus subgroup MIKHLTRRATVALVVAAAAISPAAYPAMADTATPLNGLFALTPGSCAEGRATGSYFRMIL WP_005073213.1 hypothetical protein [Mycobacterium abscessus] Length: 330\nNumber of predicted TMHs: 2\nExp number of AAs in TMHs: 39.3416199999999\nExp number, first 60 AAs: 17.07688\nTotal prob of N-in: 0.84431\nPOSSIBLE N-term signal sequence\ninside 1 11\nTMhelix 12 29\noutside 30 298\nTMhelix 299 321\ninside 322 330

29677 GCF_001214725.1_7396_7_30 Mycobacterium abscessus Terrabacteria group; Actinobacteria; Actinobacteria; Corynebacteriales; Mycobacteriaceae; Mycobacterium; Mycobacterium chelonae group; Mycobacterium abscessus subgroup MIKHLTRRATVALVVAAAAISPAAYPAVADTATPLNGLFALTPGSCAEGRATGSYFRMIL WP_005063674.1 hypothetical protein [Mycobacterium abscessus] Length: 330\nNumber of predicted TMHs: 1\nExp number of AAs in TMHs: 39.25259\nExp number, first 60 AAs: 16.98187\nTotal prob of N-in: 0.83210\nPOSSIBLE N-term signal sequence\noutside 1 298\nTMhelix 299 321\ninside 322 330

29678 GCF_001214825.1_7396_8_71 Mycobacterium abscessus Terrabacteria group; Actinobacteria; Actinobacteria; Corynebacteriales; Mycobacteriaceae; Mycobacterium; Mycobacterium chelonae group; Mycobacterium abscessus subgroup MIKHLTRRATVALVVAAAAISPAAYPAVADTATPLNGLFALTPGSCAEGRATGSYFRMIL WP_005063674.1 hypothetical protein [Mycobacterium abscessus] Length: 330\nNumber of predicted TMHs: 1\nExp number of AAs in TMHs: 39.25259\nExp number, first 60 AAs: 16.98187\nTotal prob of N-in: 0.83210\nPOSSIBLE N-term signal sequence\noutside 1 298\nTMhelix 299 321\ninside 322 330

29679 GCF_001212845.1_7396_6_50 Mycobacterium abscessus Terrabacteria group; Actinobacteria; Actinobacteria; Corynebacteriales; Mycobacteriaceae; Mycobacterium; Mycobacterium chelonae group; Mycobacterium abscessus subgroup MIKHLTRRATVALVVAAAAISPAAYPAMADTATPLNGLFALTPGSCAEGRATGSYFRMIL WP_005073213.1 hypothetical protein [Mycobacterium abscessus] Length: 330\nNumber of predicted TMHs: 2\nExp number of AAs in TMHs: 39.3416199999999\nExp number, first 60 AAs: 17.07688\nTotal prob of N-in: 0.84431\nPOSSIBLE N-term signal sequence\ninside 1 11\nTMhelix 12 29\noutside 30 298\nTMhelix 299 321\ninside 322 330

29680 GCF_001215165.1_7396_8_92 Mycobacterium abscessus Terrabacteria group; Actinobacteria; Actinobacteria; Corynebacteriales; Mycobacteriaceae; Mycobacterium; Mycobacterium chelonae group; Mycobacterium abscessus subgroup MIKHLTRRATVALVVAAAAISPAAYPAMADTATPLNGLFALTPGSCAEGRATGSYFRMIL WP_005073213.1 hypothetical protein [Mycobacterium abscessus] Length: 330\nNumber of predicted TMHs: 2\nExp number of AAs in TMHs: 39.3416199999999\nExp number, first 60 AAs: 17.07688\nTotal prob of N-in: 0.84431\nPOSSIBLE N-term signal sequence\ninside 1 11\nTMhelix 12 29\noutside 30 298\nTMhelix 299 321\ninside 322 330

29681 GCF_001212885.1_7396_7_12 Mycobacterium abscessus Terrabacteria group; Actinobacteria; Actinobacteria; Corynebacteriales; Mycobacteriaceae; Mycobacterium; Mycobacterium chelonae group; Mycobacterium abscessus subgroup MIKHLTRRATVALVVAAAAISPAAYPAMADTATPLNGLFALTPGSCAEGRATGSYFRMIL WP_005073213.1 hypothetical protein [Mycobacterium abscessus] Length: 330\nNumber of predicted TMHs: 2\nExp number of AAs in TMHs: 39.3416199999999\nExp number, first 60 AAs: 17.07688\nTotal prob of N-in: 0.84431\nPOSSIBLE N-term signal sequence\ninside 1 11\nTMhelix 12 29\noutside 30 298\nTMhelix 299 321\ninside 322 330

29682 GCF_001215325.1_7396_7_32 Mycobacterium abscessus Terrabacteria group; Actinobacteria; Actinobacteria; Corynebacteriales; Mycobacteriaceae; Mycobacterium; Mycobacterium chelonae group; Mycobacterium abscessus subgroup MIKHLTRRATVALVVAAAAISPAAYPAVADTATPLNGLFALTPGSCAEGRATGSYFRMIL WP_005063674.1 hypothetical protein [Mycobacterium abscessus] Length: 330\nNumber of predicted TMHs: 1\nExp number of AAs in TMHs: 39.25259\nExp number, first 60 AAs: 16.98187\nTotal prob of N-in: 0.83210\nPOSSIBLE N-term signal sequence\noutside 1 298\nTMhelix 299 321\ninside 322 330

29683 GCF_001215905.1_7396_7_48 Mycobacterium abscessus Terrabacteria group; Actinobacteria; Actinobacteria; Corynebacteriales; Mycobacteriaceae; Mycobacterium; Mycobacterium chelonae group; Mycobacterium abscessus subgroup MIKHLTRRATVALVVAAAAISPAAYPAVADTATPLNGLFALTPGSCAEGRATGSYFRMIL WP_005063674.1 hypothetical protein [Mycobacterium abscessus] Length: 330\nNumber of predicted TMHs:

1\nExp number of AAs in TMHs: 39.25259\nExp number, first 60 AAs: 16.98187\nTotal prob of N-in: 0.83210\nPOSSIBLE N-term signal sequence\noutside 1 298\nTMhelix 299 321\ninside 322 330

29684 GCF_001217165.1_7396_7_26 Mycobacterium abscessus Terrabacteria group; Actinobacteria; Actinobacteria; Corynebacteriales; Mycobacteriaceae; Mycobacterium; Mycobacterium chelonae group; Mycobacterium abscessus subgroup MIKHLTRRATVALVVAAAAISPAAYPAVADTATPLNGLFALTPGSCAEGRATGSYFRMIL WP_005063674.1 hypothetical protein [Mycobacterium abscessus] Length: 330\nNumber of predicted TMHs: 1\nExp number of AAs in TMHs: 39.25259\nExp number, first 60 AAs: 16.98187\nTotal prob of N-in: 0.83210\nPOSSIBLE N-term signal sequence\noutside 1 298\nTMhelix 299 321\ninside 322 330

29685 GCF_001217285.1_7396_8_66 Mycobacterium abscessus Terrabacteria group; Actinobacteria; Actinobacteria; Corynebacteriales; Mycobacteriaceae; Mycobacterium; Mycobacterium chelonae group; Mycobacterium abscessus subgroup MIKHLTRRATVALVVAAAAISPAAYPAVADTATPLNGLFALTPGSCAEGRATGSYFRMIL WP_005063674.1 hypothetical protein [Mycobacterium abscessus] Length: 330\nNumber of predicted TMHs: 1\nExp number of AAs in TMHs: 39.25259\nExp number, first 60 AAs: 16.98187\nTotal prob of N-in: 0.83210\nPOSSIBLE N-term signal sequence\noutside 1 298\nTMhelix 299 321\ninside 322 330

29686 GCF_001217365.1_7396_8_50 Mycobacterium abscessus Terrabacteria group; Actinobacteria; Actinobacteria; Corynebacteriales; Mycobacteriaceae; Mycobacterium; Mycobacterium chelonae group; Mycobacterium abscessus subgroup MIKHLTRRATVALVVAAAAISPAAYPAVADTATPLNGLFALTPGSCAEGRATGSYFRMIL WP_005063674.1 hypothetical protein [Mycobacterium abscessus] Length: 330\nNumber of predicted TMHs: 1\nExp number of AAs in TMHs: 39.25259\nExp number, first 60 AAs: 16.98187\nTotal prob of N-in: 0.83210\nPOSSIBLE N-term signal sequence\noutside 1 298\nTMhelix 299 321\ninside 322 330

29687 GCF_001217465.1_7396_8_86 Mycobacterium abscessus Terrabacteria group; Actinobacteria; Actinobacteria; Corynebacteriales; Mycobacteriaceae; Mycobacterium; Mycobacterium chelonae group; Mycobacterium abscessus subgroup MIKHLTRRATVALVVAAAAISPAAYPAVADTATPLNGLFALTPGSCAEGRATGSYFRMIL WP_005063674.1 hypothetical protein [Mycobacterium abscessus] Length: 330\nNumber of predicted TMHs: 1\nExp number of AAs in TMHs: 39.25259\nExp number, first 60 AAs: 16.98187\nTotal prob of N-in: 0.83210\nPOSSIBLE N-term signal sequence\noutside 1 298\nTMhelix 299 321\ninside 322 330

29688 GCF_001213665.1_7396_6_81 Mycobacterium abscessus Terrabacteria group; Actinobacteria; Actinobacteria; Corynebacteriales; Mycobacteriaceae; Mycobacterium; Mycobacterium chelonae group; Mycobacterium abscessus subgroup MIKHLTRRATVALVVAAAAISPAAYPAMADTATPLNGLFALTPGSCAEGRATGSYFRMIL WP_005073213.1 hypothetical protein [Mycobacterium abscessus] Length: 330\nNumber of predicted TMHs: 2\nExp number of AAs in TMHs: 39.3416199999999\nExp number, first 60 AAs: 17.07688\nTotal prob of N-in: 0.84431\nPOSSIBLE N-term signal sequence\ninside 1 11\nTMhelix 12 29\noutside 30 298\nTMhelix 299 321\ninside 322 330

29689 GCF_001215625.1_7396_7_6Mycobacterium abscessus Terrabacteria group; Actinobacteria; Actinobacteria; Corynebacteriales; Mycobacteriaceae; Mycobacterium; Mycobacterium chelonae group; Mycobacterium abscessus subgroup MIKHLTRRATVALVVAAAAISPAAYPAMADTATPLNGLFALTPGSCAEGRATGSYFRMIL WP_005073213.1 hypothetical protein [Mycobacterium abscessus] Length: 330\nNumber of predicted TMHs: 2\nExp number of AAs in TMHs: 39.3416199999999\nExp number, first 60 AAs: 17.07688\nTotal prob of N-in: 0.84431\nPOSSIBLE N-term signal sequence\ninside 1 11\nTMhelix 12 29\noutside 30 298\nTMhelix 299 321\ninside 322 330

29690 GCF_001215965.1_7396_8_67 Mycobacterium abscessus Terrabacteria group; Actinobacteria; Actinobacteria; Corynebacteriales; Mycobacteriaceae; Mycobacterium; Mycobacterium chelonae group; Mycobacterium abscessus subgroup MIKHLTRRATVALVVAAAAISPAAYPAMADTATPLNGLFALTPGSCAEGRATGSYFRMIL WP_005073213.1 hypothetical protein [Mycobacterium abscessus] Length: 330\nNumber of predicted TMHs: 2\nExp number of AAs in TMHs: 39.3416199999999\nExp number, first 60 AAs: 17.07688\nTotal prob of N-in: 0.84431\nPOSSIBLE N-term signal sequence\ninside 1 11\nTMhelix 12 29\noutside 30 298\nTMhelix 299 321\ninside 322 330

29691 GCF_001216025.1_7396_8_51 Mycobacterium abscessus Terrabacteria group; Actinobacteria; Actinobacteria; Corynebacteriales; Mycobacteriaceae; Mycobacterium; Mycobacterium chelonae group; Mycobacterium abscessus subgroup MIKHLTRRATVALVVAAAAISPAAYPAMADTATPLNGLFALTPGSCAEGRATGSYFRMIL WP_005073213.1 hypothetical protein [Mycobacterium abscessus] Length: 330\nNumber of predicted TMHs: 2\nExp number of AAs in TMHs: 39.3416199999999\nExp number, first 60 AAs: 17.07688\nTotal prob of N-in: 0.84431\nPOSSIBLE N-term signal sequence\ninside 1 11\nTMhelix 12 29\noutside 30 298\nTMhelix 299 321\ninside 322 330

29692 GCF_001215705.1_7520_7_34 Mycobacterium abscessus Terrabacteria group; Actinobacteria; Actinobacteria; Corynebacteriales; Mycobacteriaceae; Mycobacterium; Mycobacterium chelonae group; Mycobacterium abscessus subgroup MIKHLTRRATVALVVAAAAISPAAYPAVADTATPLNGLFALTPGSCAEGRATGSYFRMIL WP_005063674.1 hypothetical protein [Mycobacterium abscessus] Length: 330\nNumber of predicted TMHs: 1\nExp number of AAs in TMHs: 39.25259\nExp number, first 60 AAs: 16.98187\nTotal prob of N-in: 0.83210\nPOSSIBLE N-term signal sequence\noutside 1 298\nTMhelix 299 321\ninside 322 330

29693 GCF_001215465.1_7396_8_83 Mycobacterium abscessus Terrabacteria group; Actinobacteria; Actinobacteria; Corynebacteriales; Mycobacteriaceae; Mycobacterium; Mycobacterium chelonae group; Mycobacterium abscessus subgroup MIKHLTRRATVALVVAAAAISPAAYPAMADTATPLNGLFALTPGSCAEGRATGSYFRMIL WP_005073213.1 hypothetical protein [Mycobacterium abscessus] Length: 330\nNumber of predicted TMHs: 2\nExp number of AAs in TMHs: 39.3416199999999\nExp number, first 60 AAs: 17.07688\nTotal prob of N-in: 0.84431\nPOSSIBLE N-term signal sequence\ninside 1 11\nTMhelix 12 29\noutside 30 298\nTMhelix 299 321\ninside 322 330

29694 GCF_001214105.1_7396_7_17 Mycobacterium abscessus Terrabacteria group; Actinobacteria; Actinobacteria; Corynebacteriales; Mycobacteriaceae; Mycobacterium; Mycobacterium chelonae group; Mycobacterium abscessus subgroup MIKHLTRRATVALVVAAAAISPAAYPAMADTATPLNGLFALTPGSCAEGRATGSYFRMIL WP_005073213.1 hypothetical protein [Mycobacterium abscessus] Length: 330\nNumber of predicted TMHs: 2\nExp number of AAs in TMHs: 39.3416199999999\nExp number, first 60 AAs: 17.07688\nTotal prob of N-in: 0.84431\nPOSSIBLE N-term signal sequence\ninside 1 11\nTMhelix 12 29\noutside 30 298\nTMhelix 299 321\ninside 322 330

29695 GCF_001214145.1_7396_7_22 Mycobacterium abscessus Terrabacteria group; Actinobacteria; Actinobacteria; Corynebacteriales; Mycobacteriaceae; Mycobacterium; Mycobacterium chelonae group; Mycobacterium abscessus subgroup MIKHLTRRATVALVVAAAAISPAAYPAVADTATPLNGLFALTPGSCAEGRATGSYFRMIL WP_005063674.1 hypothetical protein [Mycobacterium abscessus] Length: 330\nNumber of predicted TMHs: 1\nExp number of AAs in TMHs: 39.25259\nExp number, first 60 AAs: 16.98187\nTotal prob of N-in: 0.83210\nPOSSIBLE N-term signal sequence\noutside 1 298\nTMhelix 299 321\ninside 322 330

29696 GCF_001214885.1_7396_7_46 Mycobacterium abscessus Terrabacteria group; Actinobacteria; Actinobacteria; Corynebacteriales; Mycobacteriaceae; Mycobacterium; Mycobacterium chelonae group; Mycobacterium abscessus subgroup MIKHLTRRATVALVVAAAAISPAAYPAVADTATPLNGLFALTPGSCAEGRATGSYFRMIL WP_005063674.1 hypothetical protein [Mycobacterium abscessus] Length: 330\nNumber of predicted TMHs: 1\nExp number of AAs in TMHs: 39.25259\nExp number, first 60 AAs: 16.98187\nTotal prob of N-in: 0.83210\nPOSSIBLE N-term signal sequence\noutside 1 298\nTMhelix 299 321\ninside 322 330

29697 GCF_001214945.1_7520_7_29 Mycobacterium abscessus Terrabacteria group; Actinobacteria; Actinobacteria; Corynebacteriales; Mycobacteriaceae; Mycobacterium; Mycobacterium chelonae group; Mycobacterium abscessus subgroup MIKHLTRRATVALVVAAAAISPAAYPAVADTATPLNGLFALTPGSCAEGRATGSYFRMIL WP_005063674.1 hypothetical protein [Mycobacterium abscessus] Length: 330\nNumber of predicted TMHs: 1\nExp number of AAs in TMHs: 39.25259\nExp number, first 60 AAs: 16.98187\nTotal prob of N-in: 0.83210\nPOSSIBLE N-term signal sequence\noutside 1 298\nTMhelix 299 321\ninside 322 330

29698 GCF_001213145.1_7396_6_82 Mycobacterium abscessus Terrabacteria group; Actinobacteria; Actinobacteria; Corynebacteriales; Mycobacteriaceae; Mycobacterium; Mycobacterium chelonae group; Mycobacterium abscessus subgroup MIKHLTRRATVALVVAAAAISPAAYPAVADTATPLNGLFALTPGSCAEGRATGSYFRMIL WP_005063674.1 hypothetical protein [Mycobacterium abscessus] Length: 330\nNumber of predicted TMHs: 1\nExp number of AAs in TMHs: 39.25259\nExp number, first 60 AAs: 16.98187\nTotal prob of N-in: 0.83210\nPOSSIBLE N-term signal sequence\noutside 1 298\nTMhelix 299 321\ninside 322 330

29699 GCF_001217525.1_7396_8_82 Mycobacterium abscessus Terrabacteria group; Actinobacteria; Actinobacteria; Corynebacteriales; Mycobacteriaceae; Mycobacterium; Mycobacterium chelonae group; Mycobacterium abscessus subgroup MIKHLTRRATVALVVAAAAISPAAYPAMADTATPLNGLFALTPGSCAEGRATGSYFRMIL WP_005073213.1 hypothetical protein [Mycobacterium abscessus] Length: 330\nNumber of predicted TMHs: 2\nExp number of AAs in TMHs: 39.3416199999999\nExp number, first 60 AAs: 17.07688\nTotal prob of N-in: 0.84431\nPOSSIBLE N-term signal sequence\ninside 1 11\nTMhelix 12 29\noutside 30 298\nTMhelix 299 321\ninside 322 330

29700 GCF_001213065.1_7396_6_57 Mycobacterium abscessus Terrabacteria group; Actinobacteria; Actinobacteria; Corynebacteriales; Mycobacteriaceae; Mycobacterium; Mycobacterium chelonae group; Mycobacterium

abscessus subgroup	MIKHLTRRATVALVVAAAAISPAAYPAVADTATPLNGLFALTPGSCAEGRATGSYFRMIL
WP_005063674.1	hypothetical protein [Mycobacterium abscessus] Length: 330\nNumber of predicted TMHs: 1\nExp number of AAs in TMHs: 39.25259\nExp number, first 60 AAs: 16.98187\nTotal prob of N-in: 0.83210\nPOSSIBLE N-term signal sequence\noutside 1 298\nTMhelix 299 321\ninside 322 330

29701	GCF_001610635.1_ASM161063v1	Mycobacterium abscessus	Terrabacteria group; Actinobacteria; Actinobacteria; Corynebacteriales; Mycobacteriaceae; Mycobacterium; Mycobacterium chelonae group; Mycobacterium abscessus subgroup	MIKHLTRRATVALVVAAAAISPAAYPAVADTATPLNGLFALTPGSCAEGRATGSYFRMIL
WP_005063674.1	hypothetical protein [Mycobacterium abscessus] Length: 330\nNumber of predicted TMHs: 1\nExp number of AAs in TMHs: 39.25259\nExp number, first 60 AAs: 16.98187\nTotal prob of N-in: 0.83210\nPOSSIBLE N-term signal sequence\noutside 1 298\nTMhelix 299 321\ninside 322 330

29702	GCF_001606255.1_ASM160625v1	Mycobacterium abscessus	Terrabacteria group; Actinobacteria; Actinobacteria; Corynebacteriales; Mycobacteriaceae; Mycobacterium; Mycobacterium chelonae group; Mycobacterium abscessus subgroup	MIKHLTRRATVALVVAAAAISPAAYPAVADTATPLNGLFALTPGSCAEGRATGSYFRMIL
WP_005063674.1	hypothetical protein [Mycobacterium abscessus] Length: 330\nNumber of predicted TMHs: 1\nExp number of AAs in TMHs: 39.25259\nExp number, first 60 AAs: 16.98187\nTotal prob of N-in: 0.83210\nPOSSIBLE N-term signal sequence\noutside 1 298\nTMhelix 299 321\ninside 322 330

29703	GCF_001677155.1_ASM167715v1	Mycobacterium abscessus	Terrabacteria group; Actinobacteria; Actinobacteria; Corynebacteriales; Mycobacteriaceae; Mycobacterium; Mycobacterium chelonae group; Mycobacterium abscessus subgroup	MIKHLTRRATVALVVAAAAISPAAYPAMADTATPLNGLFALTPGSCAEGRATGSYFRMIL
WP_005073213.1	hypothetical protein [Mycobacterium abscessus] Length: 330\nNumber of predicted TMHs: 2\nExp number of AAs in TMHs: 39.3416199999999\nExp number, first 60 AAs: 17.07688\nTotal prob of N-in: 0.84431\nPOSSIBLE N-term signal sequence\ninside 1 11\nTMhelix 12 29\noutside 30 298\nTMhelix 299 321\ninside 322 330

29704	GCF_001212945.1_7396_8_49	Mycobacterium abscessus	Terrabacteria group; Actinobacteria; Actinobacteria; Corynebacteriales; Mycobacteriaceae; Mycobacterium; Mycobacterium chelonae group; Mycobacterium abscessus subgroup	MIKHLTRRATVALVVAAAAISPAAYPAVADTATPLNGLFALTPGSCAEGRATGSYFRMIL
WP_005063674.1	hypothetical protein [Mycobacterium abscessus] Length: 330\nNumber of predicted TMHs: 1\nExp number of AAs in TMHs: 39.25259\nExp number, first 60 AAs: 16.98187\nTotal prob of N-in: 0.83210\nPOSSIBLE N-term signal sequence\noutside 1 298\nTMhelix 299 321\ninside 322 330

29705	GCF_001213225.1_7396_6_88	Mycobacterium abscessus	Terrabacteria group; Actinobacteria; Actinobacteria; Corynebacteriales; Mycobacteriaceae; Mycobacterium; Mycobacterium chelonae group; Mycobacterium abscessus subgroup	MIKHLTRRATVALVVAAAAISPAAYPAMADTATPLNGLFALTPGSCAEGRATGSYFRMIL
WP_005073213.1	hypothetical protein [Mycobacterium abscessus] Length: 330\nNumber of predicted TMHs: 2\nExp number of AAs in TMHs: 39.3416199999999\nExp number, first 60 AAs: 17.07688\nTotal prob of N-in: 0.84431\nPOSSIBLE N-term signal sequence\ninside 1 11\nTMhelix 12 29\noutside 30 298\nTMhelix 299 321\ninside 322 330

29706	GCF_001610675.1_ASM161067v1	Mycobacterium abscessus	Terrabacteria group; Actinobacteria; Actinobacteria; Corynebacteriales; Mycobacteriaceae; Mycobacterium; Mycobacterium chelonae group; Mycobacterium abscessus subgroup	MIKHLTRRATVALVVAAAAISPAAYPAVADTATPLNGLFALTPGSCAEGRATGSYFRMIL
WP_005063674.1	hypothetical protein [Mycobacterium abscessus] Length: 330\nNumber of predicted TMHs: 1\nExp number of AAs in TMHs: 39.25259\nExp number, first 60 AAs: 16.98187\nTotal prob of N-in: 0.83210\nPOSSIBLE N-term signal sequence\noutside 1 298\nTMhelix 299 321\ninside 322 330

29707	GCF_001213005.1_7396_6_51	Mycobacterium abscessus	Terrabacteria group; Actinobacteria; Actinobacteria; Corynebacteriales; Mycobacteriaceae; Mycobacterium; Mycobacterium chelonae group; Mycobacterium abscessus subgroup	MIKHLTRRATVALVVAAAAISPAAYPAMADTATPLNGLFALTPGSCAEGRATGSYFRMIL
WP_005073213.1	hypothetical protein [Mycobacterium abscessus] Length: 330\nNumber of predicted TMHs: 2\nExp number of AAs in TMHs: 39.3416199999999\nExp number, first 60 AAs: 17.07688\nTotal prob of N-in: 0.84431\nPOSSIBLE N-term signal sequence\ninside 1 11\nTMhelix 12 29\noutside 30 298\nTMhelix 299 321\ninside 322 330

29708	GCF_001677195.1_ASM167719v1	Mycobacterium abscessus	Terrabacteria group; Actinobacteria; Actinobacteria; Corynebacteriales; Mycobacteriaceae; Mycobacterium; Mycobacterium chelonae group; Mycobacterium abscessus subgroup	MIKHLTRRATVALVVAAAAISPAAYPAMADTATPLNGLFALTPGSCAEGRATGSYFRMIL
WP_005073213.1	hypothetical protein [Mycobacterium abscessus] Length: 330\nNumber of predicted TMHs:

2\nExp number of AAs in TMHs: 39.3416199999999\nExp number, first 60 AAs: 17.07688\nTotal prob of N-in: 0.84431\nPOSSIBLE N-term signal sequence\ninside 1 11\nTMhelix 12 29\noutside 30 298\nTMhelix 299 321\ninside 322 330

29709 GCF_001213285.1_7396_7_21 Mycobacterium abscessus Terrabacteria group; Actinobacteria; Actinobacteria; Corynebacteriales; Mycobacteriaceae; Mycobacterium; Mycobacterium chelonae group; Mycobacterium abscessus subgroup MIKHLTRRATVALVVAAAAISPAAYPAMADTATPLNGLFALTPGSCAEGRATGSYFRMIL WP_005073213.1 hypothetical protein [Mycobacterium abscessus] Length: 330\nNumber of predicted TMHs: 2\nExp number of AAs in TMHs: 39.3416199999999\nExp number, first 60 AAs: 17.07688\nTotal prob of N-in: 0.84431\nPOSSIBLE N-term signal sequence\ninside 1 11\nTMhelix 12 29\noutside 30 298\nTMhelix 299 321\ninside 322 330

29710 GCF_001216885.1_7396_8_73 Mycobacterium abscessus Terrabacteria group; Actinobacteria; Actinobacteria; Corynebacteriales; Mycobacteriaceae; Mycobacterium; Mycobacterium chelonae group; Mycobacterium abscessus subgroup MIKHLTRRATVALVVAAAAISPAAYPAVADTATPLNGLFALTPGSCAEGRATGSYFRMIL WP_005063674.1 hypothetical protein [Mycobacterium abscessus] Length: 330\nNumber of predicted TMHs: 1\nExp number of AAs in TMHs: 39.25259\nExp number, first 60 AAs: 16.98187\nTotal prob of N-in: 0.83210\nPOSSIBLE N-term signal sequence\noutside 1 298\nTMhelix 299 321\ninside 322 330

29711 GCF_001216825.1_7396_7_25 Mycobacterium abscessus Terrabacteria group; Actinobacteria; Actinobacteria; Corynebacteriales; Mycobacteriaceae; Mycobacterium; Mycobacterium chelonae group; Mycobacterium abscessus subgroup MIKHLTRRATVALVVAAAAISPAAYPAVADTATPLNGLFALTPGSCAEGRATGSYFRMIL WP_005063674.1 hypothetical protein [Mycobacterium abscessus] Length: 330\nNumber of predicted TMHs: 1\nExp number of AAs in TMHs: 39.25259\nExp number, first 60 AAs: 16.98187\nTotal prob of N-in: 0.83210\nPOSSIBLE N-term signal sequence\noutside 1 298\nTMhelix 299 321\ninside 322 330

29712 GCF_001217125.1_7396_7_42 Mycobacterium abscessus Terrabacteria group; Actinobacteria; Actinobacteria; Corynebacteriales; Mycobacteriaceae; Mycobacterium; Mycobacterium chelonae group; Mycobacterium abscessus subgroup MIKHLTRRATVALVVAAAAISPAAYPAMADTATPLNGLFALTPGSCAEGRATGSYFRMIL WP_005073213.1 hypothetical protein [Mycobacterium abscessus] Length: 330\nNumber of predicted TMHs: 2\nExp number of AAs in TMHs: 39.3416199999999\nExp number, first 60 AAs: 17.07688\nTotal prob of N-in: 0.84431\nPOSSIBLE N-term signal sequence\ninside 1 11\nTMhelix 12 29\noutside 30 298\nTMhelix 299 321\ninside 322 330

29713 GCF_001217065.1_7520_7_38 Mycobacterium abscessus Terrabacteria group; Actinobacteria; Actinobacteria; Corynebacteriales; Mycobacteriaceae; Mycobacterium; Mycobacterium chelonae group; Mycobacterium abscessus subgroup MIKHLTRRATVALVVAAAAISPAAYPAVADTATPLNGLFALTPGSCAEGRATGSYFRMIL WP_005063674.1 hypothetical protein [Mycobacterium abscessus] Length: 330\nNumber of predicted TMHs: 1\nExp number of AAs in TMHs: 39.25259\nExp number, first 60 AAs: 16.98187\nTotal prob of N-in: 0.83210\nPOSSIBLE N-term signal sequence\noutside 1 298\nTMhelix 299 321\ninside 322 330

29714 GCF_001216925.1_7396_7_40 Mycobacterium abscessus Terrabacteria group; Actinobacteria; Actinobacteria; Corynebacteriales; Mycobacteriaceae; Mycobacterium; Mycobacterium chelonae group; Mycobacterium abscessus subgroup MIKHLTRRATVALVVAAAAISPAAYPAVADTATPLNGLFALTPGSCAEGRATGSYFRMIL WP_005063674.1 hypothetical protein [Mycobacterium abscessus] Length: 330\nNumber of predicted TMHs: 1\nExp number of AAs in TMHs: 39.25259\nExp number, first 60 AAs: 16.98187\nTotal prob of N-in: 0.83210\nPOSSIBLE N-term signal sequence\noutside 1 298\nTMhelix 299 321\ninside 322 330

29715 GCF_001216685.1_7520_7_39 Mycobacterium abscessus Terrabacteria group; Actinobacteria; Actinobacteria; Corynebacteriales; Mycobacteriaceae; Mycobacterium; Mycobacterium chelonae group; Mycobacterium abscessus subgroup MIKHLTRRATVALVVAAAAISPAAYPAVADTATPLNGLFALTPGSCAEGRATGSYFRMIL WP_005063674.1 hypothetical protein [Mycobacterium abscessus] Length: 330\nNumber of predicted TMHs: 1\nExp number of AAs in TMHs: 39.25259\nExp number, first 60 AAs: 16.98187\nTotal prob of N-in: 0.83210\nPOSSIBLE N-term signal sequence\noutside 1 298\nTMhelix 299 321\ninside 322 330

29716 GCF_001215565.1_7396_8_58 Mycobacterium abscessus Terrabacteria group; Actinobacteria; Actinobacteria; Corynebacteriales; Mycobacteriaceae; Mycobacterium; Mycobacterium chelonae group; Mycobacterium abscessus subgroup MIKHLTRRATVALVVAAAAISPAAYPAMADTATPLNGLFALTPGSCAEGRATGSYFRMIL WP_005073213.1 hypothetical protein [Mycobacterium abscessus] Length: 330\nNumber of predicted TMHs: 2\nExp number of AAs in TMHs: 39.3416199999999\nExp number, first 60 AAs: 17.07688\nTotal prob of N-in:

0.84431\nPOSSIBLE N-term signal sequence\ninside 1 11\nTMhelix 12 29\noutside 30 298\nTMhelix 299 321\ninside 322 330

29717 GCF_001213865.1_7396_6_93 Mycobacterium abscessus Terrabacteria group; Actinobacteria; Actinobacteria; Corynebacteriales; Mycobacteriaceae; Mycobacterium; Mycobacterium chelonae group; Mycobacterium abscessus subgroup MIKHLTRRATVALVVAAAAISPAAYPAVADTATPLNGLFALTPGSCAEGRATGSYFRMIL WP_005063674.1 hypothetical protein [Mycobacterium abscessus] Length: 330\nNumber of predicted TMHs: 1\nExp number of AAs in TMHs: 39.25259\nExp number, first 60 AAs: 16.98187\nTotal prob of N-in: 0.83210\nPOSSIBLE N-term signal sequence\noutside 1 298\nTMhelix 299 321\ninside 322 330

29718 GCF_001214065.1_7396_7_15 Mycobacterium abscessus Terrabacteria group; Actinobacteria; Actinobacteria; Corynebacteriales; Mycobacteriaceae; Mycobacterium; Mycobacterium chelonae group; Mycobacterium abscessus subgroup MIKHLTRRATVALVVAAAAISPAAYPAMADTATPLNGLFALTPGSCAEGRATGSYFRMIL WP_005073213.1 hypothetical protein [Mycobacterium abscessus] Length: 330\nNumber of predicted TMHs: 2\nExp number of AAs in TMHs: 39.3416199999999\nExp number, first 60 AAs: 17.07688\nTotal prob of N-in: 0.84431\nPOSSIBLE N-term signal sequence\ninside 1 11\nTMhelix 12 29\noutside 30 298\nTMhelix 299 321\ninside 322 330

29719 GCF_001213905.1_7396_6_76 Mycobacterium abscessus Terrabacteria group; Actinobacteria; Actinobacteria; Corynebacteriales; Mycobacteriaceae; Mycobacterium; Mycobacterium chelonae group; Mycobacterium abscessus subgroup MIKHLTRRATVALVVAAAAISPAAYPAVADTATPLNGLFALTPGSCAEGRATGSYFRMIL WP_005063674.1 hypothetical protein [Mycobacterium abscessus] Length: 330\nNumber of predicted TMHs: 1\nExp number of AAs in TMHs: 39.25259\nExp number, first 60 AAs: 16.98187\nTotal prob of N-in: 0.83210\nPOSSIBLE N-term signal sequence\noutside 1 298\nTMhelix 299 321\ninside 322 330

29720 GCF_001215285.1_7396_8_61 Mycobacterium abscessus Terrabacteria group; Actinobacteria; Actinobacteria; Corynebacteriales; Mycobacteriaceae; Mycobacterium; Mycobacterium chelonae group; Mycobacterium abscessus subgroup MIKHLTRRATVALVVAAAAISPAAYPAVADTATPLNGLFALTPGSCAEGRATGSYFRMIL WP_005063674.1 hypothetical protein [Mycobacterium abscessus] Length: 330\nNumber of predicted TMHs: 1\nExp number of AAs in TMHs: 39.25259\nExp number, first 60 AAs: 16.98187\nTotal prob of N-in: 0.83210\nPOSSIBLE N-term signal sequence\noutside 1 298\nTMhelix 299 321\ninside 322 330

29721 GCF_001215385.1_7396_7_34 Mycobacterium abscessus Terrabacteria group; Actinobacteria; Actinobacteria; Corynebacteriales; Mycobacteriaceae; Mycobacterium; Mycobacterium chelonae group; Mycobacterium abscessus subgroup MIKHLTRRATVALVVAAAAISPAAYPAMADTATPLNGLFALTPGSCAEGRATGSYFRMIL WP_005073213.1 hypothetical protein [Mycobacterium abscessus] Length: 330\nNumber of predicted TMHs: 2\nExp number of AAs in TMHs: 39.3416199999999\nExp number, first 60 AAs: 17.07688\nTotal prob of N-in: 0.84431\nPOSSIBLE N-term signal sequence\ninside 1 11\nTMhelix 12 29\noutside 30 298\nTMhelix 299 321\ninside 322 330

29722 GCF_001214185.1_7396_6_67 Mycobacterium abscessus Terrabacteria group; Actinobacteria; Actinobacteria; Corynebacteriales; Mycobacteriaceae; Mycobacterium; Mycobacterium chelonae group; Mycobacterium abscessus subgroup MIKHLTRRATVALVVAAAAISPAAYPAMADTATPLNGLFALTPGSCAEGRATGSYFRMIL WP_005073213.1 hypothetical protein [Mycobacterium abscessus] Length: 330\nNumber of predicted TMHs: 2\nExp number of AAs in TMHs: 39.3416199999999\nExp number, first 60 AAs: 17.07688\nTotal prob of N-in: 0.84431\nPOSSIBLE N-term signal sequence\ninside 1 11\nTMhelix 12 29\noutside 30 298\nTMhelix 299 321\ninside 322 330

29723 GCF_001214225.1_7520_7_40 Mycobacterium abscessus Terrabacteria group; Actinobacteria; Actinobacteria; Corynebacteriales; Mycobacteriaceae; Mycobacterium; Mycobacterium chelonae group; Mycobacterium abscessus subgroup MIKHLTRRATVALVVAAAAISPAAYPAMADTATPLNGLFALTPGSCAEGRATGSYFRMIL WP_005073213.1 hypothetical protein [Mycobacterium abscessus] Length: 330\nNumber of predicted TMHs: 2\nExp number of AAs in TMHs: 39.3416199999999\nExp number, first 60 AAs: 17.07688\nTotal prob of N-in: 0.84431\nPOSSIBLE N-term signal sequence\ninside 1 11\nTMhelix 12 29\noutside 30 298\nTMhelix 299 321\ninside 322 330

29724 GCF_001215245.1_7396_7_7Mycobacterium abscessus Terrabacteria group; Actinobacteria; Actinobacteria; Corynebacteriales; Mycobacteriaceae; Mycobacterium; Mycobacterium chelonae group; Mycobacterium abscessus subgroup MIKHLTRRATVALVVAAAAISPAAYPAVADTATPLNGLFALTPGSCAEGRATGSYFRMIL WP_005063674.1 hypothetical protein [Mycobacterium abscessus] Length: 330\nNumber of predicted TMHs: 1\nExp number of AAs in TMHs:

39.25259\nExp number, first 60 AAs: 16.98187\nTotal prob of N-in: 0.83210\nPOSSIBLE N-term signal sequence\noutside 1 298\nTMhelix 299 321\ninside 322 330

29725 GCF_000069185.1_ASM6918v1 Mycobacterium abscessus Terrabacteria group; Actinobacteria; Actinobacteria; Corynebacteriales; Mycobacteriaceae; Mycobacterium; Mycobacterium chelonae group; Mycobacterium abscessus subgroup MIKHLTRRATVALVVAAAAISPAAYPAMADTATPLNGLFALTPGSCAEGRATGSYFRMIL YP_001701589.1 Hypothetical protein MAB_0841 [Mycobacterium abscessus] Length: 330\nNumber of predicted TMHs: 2\nExp number of AAs in TMHs: 39.3416199999999\nExp number, first 60 AAs: 17.07688\nTotal prob of N-in: 0.84431\nPOSSIBLE N-term signal sequence\ninside 1 11\nTMhelix 12 29\noutside 30 298\nTMhelix 299 321\ninside 322 330

29726 GCF_001213445.1_7396_6_86 Mycobacterium abscessus Terrabacteria group; Actinobacteria; Actinobacteria; Corynebacteriales; Mycobacteriaceae; Mycobacterium; Mycobacterium chelonae group; Mycobacterium abscessus subgroup MIKHLTRRATVALVVAAAAISPAAYPAMADTATPLNGLFALTPGSCAEGRATGSYFRMIL WP_005073213.1 hypothetical protein [Mycobacterium abscessus] Length: 330\nNumber of predicted TMHs: 2\nExp number of AAs in TMHs: 39.3416199999999\nExp number, first 60 AAs: 17.07688\nTotal prob of N-in: 0.84431\nPOSSIBLE N-term signal sequence\ninside 1 11\nTMhelix 12 29\noutside 30 298\nTMhelix 299 321\ninside 322 330

29727 GCF_001213385.1_7396_6_60 Mycobacterium abscessus Terrabacteria group; Actinobacteria; Actinobacteria; Corynebacteriales; Mycobacteriaceae; Mycobacterium; Mycobacterium chelonae group; Mycobacterium abscessus subgroup MIKHLTRRATVALVVAAAAISPAAYPAMADTATPLNGLFALTPGSCAEGRATGSYFRMIL WP_005073213.1 hypothetical protein [Mycobacterium abscessus] Length: 330\nNumber of predicted TMHs: 2\nExp number of AAs in TMHs: 39.3416199999999\nExp number, first 60 AAs: 17.07688\nTotal prob of N-in: 0.84431\nPOSSIBLE N-term signal sequence\ninside 1 11\nTMhelix 12 29\noutside 30 298\nTMhelix 299 321\ninside 322 330

29728 GCF_001213185.1_7396_6_54 Mycobacterium abscessus Terrabacteria group; Actinobacteria; Actinobacteria; Corynebacteriales; Mycobacteriaceae; Mycobacterium; Mycobacterium chelonae group; Mycobacterium abscessus subgroup MIKHLTRRATVALVVAAAAISPAAYPAVADTATPLNGLFALTPGSCAEGRATGSYFRMIL WP_005063674.1 hypothetical protein [Mycobacterium abscessus] Length: 330\nNumber of predicted TMHs: 1\nExp number of AAs in TMHs: 39.25259\nExp number, first 60 AAs: 16.98187\nTotal prob of N-in: 0.83210\nPOSSIBLE N-term signal sequence\noutside 1 298\nTMhelix 299 321\ninside 322 330

29729 GCF_002142755.1_ASM214275v1 Mycobacterium abscessus Terrabacteria group; Actinobacteria; Actinobacteria; Corynebacteriales; Mycobacteriaceae; Mycobacterium; Mycobacterium chelonae group; Mycobacterium abscessus subgroup MIKHLTRRATVALVVAAAAISPAAYPAVADTATPLNGLFALTPGSCAEGRATGSYFRMIL WP_005063674.1 hypothetical protein [Mycobacterium abscessus] Length: 330\nNumber of predicted TMHs: 1\nExp number of AAs in TMHs: 39.25259\nExp number, first 60 AAs: 16.98187\nTotal prob of N-in: 0.83210\nPOSSIBLE N-term signal sequence\noutside 1 298\nTMhelix 299 321\ninside 322 330

29730 GCF_002142775.1_ASM214277v1 Mycobacterium abscessus Terrabacteria group; Actinobacteria; Actinobacteria; Corynebacteriales; Mycobacteriaceae; Mycobacterium; Mycobacterium chelonae group; Mycobacterium abscessus subgroup MIKHLTRRATVALVVAAAAISPAAYPAMADTATPLNGLFALTPGSCAEGRATGSYFRMIL WP_005073213.1 hypothetical protein [Mycobacterium abscessus] Length: 330\nNumber of predicted TMHs: 2\nExp number of AAs in TMHs: 39.3416199999999\nExp number, first 60 AAs: 17.07688\nTotal prob of N-in: 0.84431\nPOSSIBLE N-term signal sequence\ninside 1 11\nTMhelix 12 29\noutside 30 298\nTMhelix 299 321\ninside 322 330

29731 GCF_001213505.1_7396_6_95 Mycobacterium abscessus Terrabacteria group; Actinobacteria; Actinobacteria; Corynebacteriales; Mycobacteriaceae; Mycobacterium; Mycobacterium chelonae group; Mycobacterium abscessus subgroup MIKHLTRRATVALVVAAAAISPAAYPAMADTATPLNGLFALTPGSCAEGRATGSYFRMIL WP_005073213.1 hypothetical protein [Mycobacterium abscessus] Length: 330\nNumber of predicted TMHs: 2\nExp number of AAs in TMHs: 39.3416199999999\nExp number, first 60 AAs: 17.07688\nTotal prob of N-in: 0.84431\nPOSSIBLE N-term signal sequence\ninside 1 11\nTMhelix 12 29\noutside 30 298\nTMhelix 299 321\ninside 322 330

29732 GCF_001216165.1_7396_8_69 Mycobacterium abscessus Terrabacteria group; Actinobacteria; Actinobacteria; Corynebacteriales; Mycobacteriaceae; Mycobacterium; Mycobacterium chelonae group; Mycobacterium abscessus subgroup MIKHLTRRATVALVVAAAAISPAAYPAVADTATPLNGLFALTPGSCAEGRATGSYFRMIL WP_005063674.1 hypothetical protein [Mycobacterium abscessus] Length: 330\nNumber of predicted TMHs:

1\nExp number of AAs in TMHs: 39.25259\nExp number, first 60 AAs: 16.98187\nTotal prob of N-in: 0.83210\nPOSSIBLE N-term signal sequence\noutside 1 298\nTMhelix 299 321\ninside 322 330

29733 GCF_001214525.1_7396_8_68 Mycobacterium abscessus Terrabacteria group; Actinobacteria; Actinobacteria; Corynebacteriales; Mycobacteriaceae; Mycobacterium; Mycobacterium chelonae group; Mycobacterium abscessus subgroup MIKHLTRRATVALVVAAAAISPAAYPAVADTATPLNGLFALTPGSCAEGRATGSYFRMIL WP_005063674.1 hypothetical protein [Mycobacterium abscessus] Length: 330\nNumber of predicted TMHs: 1\nExp number of AAs in TMHs: 39.25259\nExp number, first 60 AAs: 16.98187\nTotal prob of N-in: 0.83210\nPOSSIBLE N-term signal sequence\noutside 1 298\nTMhelix 299 321\ninside 322 330

29734 GCF_001216265.1_7396_8_94 Mycobacterium abscessus Terrabacteria group; Actinobacteria; Actinobacteria; Corynebacteriales; Mycobacteriaceae; Mycobacterium; Mycobacterium chelonae group; Mycobacterium abscessus subgroup MIKHLTRRATVALVVAAAAISPAAYPAMADTATPLNGLFALTPGSCAEGRATGSYFRMIL WP_005073213.1 hypothetical protein [Mycobacterium abscessus] Length: 330\nNumber of predicted TMHs: 2\nExp number of AAs in TMHs: 39.3416199999999\nExp number, first 60 AAs: 17.07688\nTotal prob of N-in: 0.84431\nPOSSIBLE N-term signal sequence\ninside 1 11\nTMhelix 12 29\noutside 30 298\nTMhelix 299 321\ninside 322 330

29735 GCF_001216225.1_7396_8_55 Mycobacterium abscessus Terrabacteria group; Actinobacteria; Actinobacteria; Corynebacteriales; Mycobacteriaceae; Mycobacterium; Mycobacterium chelonae group; Mycobacterium abscessus subgroup MIKHLTRRATVALVVAAAAISPAAYPAVADTATPLNGLFALTPGSCAEGRATGSYFRMIL WP_005063674.1 hypothetical protein [Mycobacterium abscessus] Length: 330\nNumber of predicted TMHs: 1\nExp number of AAs in TMHs: 39.25259\nExp number, first 60 AAs: 16.98187\nTotal prob of N-in: 0.83210\nPOSSIBLE N-term signal sequence\noutside 1 298\nTMhelix 299 321\ninside 322 330

29736 GCF_001216745.1_7520_7_46 Mycobacterium abscessus Terrabacteria group; Actinobacteria; Actinobacteria; Corynebacteriales; Mycobacteriaceae; Mycobacterium; Mycobacterium chelonae group; Mycobacterium abscessus subgroup MIKHLTRRATVALVVAAAAISPAAYPAMADTATPLNGLFALTPGSCAEGRATGSYFRMIL WP_005073213.1 hypothetical protein [Mycobacterium abscessus] Length: 330\nNumber of predicted TMHs: 2\nExp number of AAs in TMHs: 39.3416199999999\nExp number, first 60 AAs: 17.07688\nTotal prob of N-in: 0.84431\nPOSSIBLE N-term signal sequence\ninside 1 11\nTMhelix 12 29\noutside 30 298\nTMhelix 299 321\ninside 322 330

29737 GCF_001214445.1_7396_8_62 Mycobacterium abscessus Terrabacteria group; Actinobacteria; Actinobacteria; Corynebacteriales; Mycobacteriaceae; Mycobacterium; Mycobacterium chelonae group; Mycobacterium abscessus subgroup MIKHLTRRATVALVVAAAAISPAAYPAVADTATPLNGLFALTPGSCAEGRATGSYFRMIL WP_005063674.1 hypothetical protein [Mycobacterium abscessus] Length: 330\nNumber of predicted TMHs: 1\nExp number of AAs in TMHs: 39.25259\nExp number, first 60 AAs: 16.98187\nTotal prob of N-in: 0.83210\nPOSSIBLE N-term signal sequence\noutside 1 298\nTMhelix 299 321\ninside 322 330

29738 GCF_001214625.1_7396_7_37 Mycobacterium abscessus Terrabacteria group; Actinobacteria; Actinobacteria; Corynebacteriales; Mycobacteriaceae; Mycobacterium; Mycobacterium chelonae group; Mycobacterium abscessus subgroup MIKHLTRRATVALVVAAAAISPAAYPAMADTATPLNGLFALTPGSCAEGRATGSYFRMIL WP_005073213.1 hypothetical protein [Mycobacterium abscessus] Length: 330\nNumber of predicted TMHs: 2\nExp number of AAs in TMHs: 39.3416199999999\nExp number, first 60 AAs: 17.07688\nTotal prob of N-in: 0.84431\nPOSSIBLE N-term signal sequence\ninside 1 11\nTMhelix 12 29\noutside 30 298\nTMhelix 299 321\ninside 322 330

29739 GCF_001214345.1_7396_8_95 Mycobacterium abscessus Terrabacteria group; Actinobacteria; Actinobacteria; Corynebacteriales; Mycobacteriaceae; Mycobacterium; Mycobacterium chelonae group; Mycobacterium abscessus subgroup MIKHLTRRATVALVVAAAAISPAAYPAVADTATPLNGLFALTPGSCAEGRATGSYFRMIL WP_005063674.1 hypothetical protein [Mycobacterium abscessus] Length: 330\nNumber of predicted TMHs: 1\nExp number of AAs in TMHs: 39.25259\nExp number, first 60 AAs: 16.98187\nTotal prob of N-in: 0.83210\nPOSSIBLE N-term signal sequence\noutside 1 298\nTMhelix 299 321\ninside 322 330

29740 GCF_001214585.1_7396_8_85 Mycobacterium abscessus Terrabacteria group; Actinobacteria; Actinobacteria; Corynebacteriales; Mycobacteriaceae; Mycobacterium; Mycobacterium chelonae group; Mycobacterium abscessus subgroup MIKHLTRRATVALVVAAAAISPAAYPAMADTATPLNGLFALTPGSCAEGRATGSYFRMIL WP_005073213.1 hypothetical protein [Mycobacterium abscessus] Length: 330\nNumber of predicted TMHs: 2\nExp number of AAs in TMHs: 39.3416199999999\nExp number, first 60 AAs: 17.07688\nTotal prob of N-in:

0.84431\nPOSSIBLE N-term signal sequence\ninside 1 11\nTMhelix 12 29\noutside 30 298\nTMhelix 299 321\ninside 322 330

29741 GCF_001214285.1_7396_7_43 Mycobacterium abscessus Terrabacteria group; Actinobacteria; Actinobacteria; Corynebacteriales; Mycobacteriaceae; Mycobacterium; Mycobacterium chelonae group; Mycobacterium abscessus subgroup MIKHLTRRATVALVVAAAAISPAAYPAVADTATPLNGLFALTPGSCAEGRATGSYFRMIL WP_005063674.1 hypothetical protein [Mycobacterium abscessus] Length: 330\nNumber of predicted TMHs: 1\nExp number of AAs in TMHs: 39.25259\nExp number, first 60 AAs: 16.98187\nTotal prob of N-in: 0.83210\nPOSSIBLE N-term signal sequence\noutside 1 298\nTMhelix 299 321\ninside 322 330

29742 GCF_001214005.1_7396_7_11 Mycobacterium abscessus Terrabacteria group; Actinobacteria; Actinobacteria; Corynebacteriales; Mycobacteriaceae; Mycobacterium; Mycobacterium chelonae group; Mycobacterium abscessus subgroup MIKHLTRRATVALVVAAAAISPAAYPAVADTATPLNGLFALTPGSCAEGRATGSYFRMIL WP_005063674.1 hypothetical protein [Mycobacterium abscessus] Length: 330\nNumber of predicted TMHs: 1\nExp number of AAs in TMHs: 39.25259\nExp number, first 60 AAs: 16.98187\nTotal prob of N-in: 0.83210\nPOSSIBLE N-term signal sequence\noutside 1 298\nTMhelix 299 321\ninside 322 330

29743 GCF_001213605.1_7396_6_84 Mycobacterium abscessus Terrabacteria group; Actinobacteria; Actinobacteria; Corynebacteriales; Mycobacteriaceae; Mycobacterium; Mycobacterium chelonae group; Mycobacterium abscessus subgroup MIKHLTRRATVALVVAAAAISPAAYPAMADTATPLNGLFALTPGSCAEGRATGSYFRMIL WP_005073213.1 hypothetical protein [Mycobacterium abscessus] Length: 330\nNumber of predicted TMHs: 2\nExp number of AAs in TMHs: 39.3416199999999\nExp number, first 60 AAs: 17.07688\nTotal prob of N-in: 0.84431\nPOSSIBLE N-term signal sequence\ninside 1 11\nTMhelix 12 29\noutside 30 298\nTMhelix 299 321\ninside 322 330

29744 GCF_001213565.1_7396_6_52 Mycobacterium abscessus Terrabacteria group; Actinobacteria; Actinobacteria; Corynebacteriales; Mycobacteriaceae; Mycobacterium; Mycobacterium chelonae group; Mycobacterium abscessus subgroup MIKHLTRRATVALVVAAAAISPAAYPAMADTATPLNGLFALTPGSCAEGRATGSYFRMIL WP_005073213.1 hypothetical protein [Mycobacterium abscessus] Length: 330\nNumber of predicted TMHs: 2\nExp number of AAs in TMHs: 39.3416199999999\nExp number, first 60 AAs: 17.07688\nTotal prob of N-in: 0.84431\nPOSSIBLE N-term signal sequence\ninside 1 11\nTMhelix 12 29\noutside 30 298\nTMhelix 299 321\ninside 322 330

29745 GCF_001216525.1_7520_7_44 Mycobacterium abscessus Terrabacteria group; Actinobacteria; Actinobacteria; Corynebacteriales; Mycobacteriaceae; Mycobacterium; Mycobacterium chelonae group; Mycobacterium abscessus subgroup MIKHLTRRATVALVVAAAAISPAAYPAVADTATPLNGLFALTPGSCAEGRATGSYFRMIL WP_005063674.1 hypothetical protein [Mycobacterium abscessus] Length: 330\nNumber of predicted TMHs: 1\nExp number of AAs in TMHs: 39.25259\nExp number, first 60 AAs: 16.98187\nTotal prob of N-in: 0.83210\nPOSSIBLE N-term signal sequence\noutside 1 298\nTMhelix 299 321\ninside 322 330

29746 GCF_001213325.1_7396_6_72 Mycobacterium abscessus Terrabacteria group; Actinobacteria; Actinobacteria; Corynebacteriales; Mycobacteriaceae; Mycobacterium; Mycobacterium chelonae group; Mycobacterium abscessus subgroup MIKHLTRRATVALVVAAAAISPAAYPAVADTATPLNGLFALTPGSCAEGRATGSYFRMIL WP_005063674.1 hypothetical protein [Mycobacterium abscessus] Length: 330\nNumber of predicted TMHs: 1\nExp number of AAs in TMHs: 39.25259\nExp number, first 60 AAs: 16.98187\nTotal prob of N-in: 0.83210\nPOSSIBLE N-term signal sequence\noutside 1 298\nTMhelix 299 321\ninside 322 330

29747 GCF_001216365.1_7396_8_60 Mycobacterium abscessus Terrabacteria group; Actinobacteria; Actinobacteria; Corynebacteriales; Mycobacteriaceae; Mycobacterium; Mycobacterium chelonae group; Mycobacterium abscessus subgroup MIKHLTRRATVALVVAAAAISPAAYPAVADTATPLNGLFALTPGSCAEGRATGSYFRMIL WP_005063674.1 hypothetical protein [Mycobacterium abscessus] Length: 330\nNumber of predicted TMHs: 1\nExp number of AAs in TMHs: 39.25259\nExp number, first 60 AAs: 16.98187\nTotal prob of N-in: 0.83210\nPOSSIBLE N-term signal sequence\noutside 1 298\nTMhelix 299 321\ninside 322 330

29748 GCF_002142855.1_ASM214285v1 Mycobacterium abscessus Terrabacteria group; Actinobacteria; Actinobacteria; Corynebacteriales; Mycobacteriaceae; Mycobacterium; Mycobacterium chelonae group; Mycobacterium abscessus subgroup MIKHLTRRATVALVVAAAAISPAAYPAVADTATPLNGLFALTPGSCAEGRATGSYFRMIL WP_005063674.1 hypothetical protein [Mycobacterium abscessus] Length: 330\nNumber of predicted TMHs: 1\nExp number of AAs in TMHs: 39.25259\nExp number, first 60 AAs: 16.98187\nTotal prob of N-in: 0.83210\nPOSSIBLE N-term signal sequence\noutside 1 298\nTMhelix 299 321\ninside 322 330

29749 GCF_002142835.1_ASM214283v1 Mycobacterium abscessus Terrabacteria group; Actinobacteria; Actinobacteria; Corynebacteriales; Mycobacteriaceae; Mycobacterium; Mycobacterium chelonae group; Mycobacterium abscessus subgroup MIKHLTRRATVALVVAAAAISPAAYPAVADTATPLNGLFALTPGSCAEGRATGSYFRMIL WP_005063674.1 hypothetical protein [Mycobacterium abscessus] Length: 330\nNumber of predicted TMHs: 1\nExp number of AAs in TMHs: 39.25259\nExp number, first 60 AAs: 16.98187\nTotal prob of N-in: 0.83210\nPOSSIBLE N-term signal sequence\noutside 1 298\nTMhelix 299 321\ninside 322 330

29750 GCF_001216865.1_7396_8_64 Mycobacterium abscessus Terrabacteria group; Actinobacteria; Actinobacteria; Corynebacteriales; Mycobacteriaceae; Mycobacterium; Mycobacterium chelonae group; Mycobacterium abscessus subgroup MIKHLTRRATVALVVAAAAISPAAYPAVADTATPLNGLFALTPGSCAEGRATGSYFRMIL WP_005063674.1 hypothetical protein [Mycobacterium abscessus] Length: 330\nNumber of predicted TMHs: 1\nExp number of AAs in TMHs: 39.25259\nExp number, first 60 AAs: 16.98187\nTotal prob of N-in: 0.83210\nPOSSIBLE N-term signal sequence\noutside 1 298\nTMhelix 299 321\ninside 322 330

29751 GCF_001217325.1_7396_8_81 Mycobacterium abscessus Terrabacteria group; Actinobacteria; Actinobacteria; Corynebacteriales; Mycobacteriaceae; Mycobacterium; Mycobacterium chelonae group; Mycobacterium abscessus subgroup MIKHLTRRATVALVVAAAAISPAAYPAVADTATPLNGLFALTPGSCAEGRATGSYFRMIL WP_052618647.1 hypothetical protein [Mycobacterium abscessus] Length: 330\nNumber of predicted TMHs: 1\nExp number of AAs in TMHs: 38.94211\nExp number, first 60 AAs: 17.07574\nTotal prob of N-in: 0.83166\nPOSSIBLE N-term signal sequence\noutside 1 298\nTMhelix 299 321\ninside 322 330

29752 GCF_002142785.1_ASM214278v1 Mycobacterium abscessus Terrabacteria group; Actinobacteria; Actinobacteria; Corynebacteriales; Mycobacteriaceae; Mycobacterium; Mycobacterium chelonae group; Mycobacterium abscessus subgroup MIKHLTRRATVALVVAAAAISPAAYPAVADTATPLNGLFALTPGSCAEGRATGSYFRMIL WP_005063674.1 hypothetical protein [Mycobacterium abscessus] Length: 330\nNumber of predicted TMHs: 1\nExp number of AAs in TMHs: 39.25259\nExp number, first 60 AAs: 16.98187\nTotal prob of N-in: 0.83210\nPOSSIBLE N-term signal sequence\noutside 1 298\nTMhelix 299 321\ninside 322 330

29753 GCF_002142765.1_ASM214276v1 Mycobacterium abscessus Terrabacteria group; Actinobacteria; Actinobacteria; Corynebacteriales; Mycobacteriaceae; Mycobacterium; Mycobacterium chelonae group; Mycobacterium abscessus subgroup MIKHLTRRATVALVVAAAAISPAAYPAVADTATPLNGLFALTPGSCAEGRATGSYFRMIL WP_005063674.1 hypothetical protein [Mycobacterium abscessus] Length: 330\nNumber of predicted TMHs: 1\nExp number of AAs in TMHs: 39.25259\nExp number, first 60 AAs: 16.98187\nTotal prob of N-in: 0.83210\nPOSSIBLE N-term signal sequence\noutside 1 298\nTMhelix 299 321\ninside 322 330

29754 GCF_001215265.1_7396_8_84 Mycobacterium abscessus Terrabacteria group; Actinobacteria; Actinobacteria; Corynebacteriales; Mycobacteriaceae; Mycobacterium; Mycobacterium chelonae group; Mycobacterium abscessus subgroup MIKHLTRRATVALVVAAAAISPAAYPAMADTATPLNGLFALTPGSCAEGRATGSYFRMIL WP_005073213.1 hypothetical protein [Mycobacterium abscessus] Length: 330\nNumber of predicted TMHs: 2\nExp number of AAs in TMHs: 39.3416199999999\nExp number, first 60 AAs: 17.07688\nTotal prob of N-in: 0.84431\nPOSSIBLE N-term signal sequence\ninside 1 11\nTMhelix 12 29\noutside 30 298\nTMhelix 299 321\ninside 322 330

29755 GCF_001215605.1_7520_7_26 Mycobacterium abscessus Terrabacteria group; Actinobacteria; Actinobacteria; Corynebacteriales; Mycobacteriaceae; Mycobacterium; Mycobacterium chelonae group; Mycobacterium abscessus subgroup MIKHLTRRATVALVVAAAAISPAAYPAVADTATPLNGLFALTPGSCAEGRATGSYFRMIL WP_005063674.1 hypothetical protein [Mycobacterium abscessus] Length: 330\nNumber of predicted TMHs: 1\nExp number of AAs in TMHs: 39.25259\nExp number, first 60 AAs: 16.98187\nTotal prob of N-in: 0.83210\nPOSSIBLE N-term signal sequence\noutside 1 298\nTMhelix 299 321\ninside 322 330

29756 GCF_001215485.1_7396_7_9Mycobacterium abscessus Terrabacteria group; Actinobacteria; Actinobacteria; Corynebacteriales; Mycobacteriaceae; Mycobacterium; Mycobacterium chelonae group; Mycobacterium abscessus subgroup MIKHLTRRATVALVVAAAAISPAAYPAVADTATPLNGLFALTPGSCAEGRATGSYFRMIL WP_005063674.1 hypothetical protein [Mycobacterium abscessus] Length: 330\nNumber of predicted TMHs: 1\nExp number of AAs in TMHs: 39.25259\nExp number, first 60 AAs: 16.98187\nTotal prob of N-in: 0.83210\nPOSSIBLE N-term signal sequence\noutside 1 298\nTMhelix 299 321\ninside 322 330

29757 GCF_001215845.1_7396_7_28 Mycobacterium abscessus Terrabacteria group; Actinobacteria; Actinobacteria; Corynebacteriales; Mycobacteriaceae; Mycobacterium; Mycobacterium chelonae group; Mycobacterium abscessus subgroup MIKHLTRRATVALVVAAAAISPAAYPAMADTATPLNGLFALTPGSCAEGRATGSYFRMIL WP_005073213.1 hypothetical protein [Mycobacterium abscessus] Length: 330\nNumber of predicted TMHs:

2\nExp number of AAs in TMHs: 39.3416199999999\nExp number, first 60 AAs: 17.07688\nTotal prob of N-in: 0.84431\nPOSSIBLE N-term signal sequence\ninside 1 11\nTMhelix 12 29\noutside 30 298\nTMhelix 299 321\ninside 322 330

29758 GCF_001215945.1_7520_7_50 Mycobacterium abscessus Terrabacteria group; Actinobacteria; Actinobacteria; Corynebacteriales; Mycobacteriaceae; Mycobacterium; Mycobacterium chelonae group; Mycobacterium abscessus subgroup MIKHLTRRATVALVVAAAAISPAAYPAMADTATPLNGLFALTPGSCAEGRATGSYFRMIL WP_005073213.1 hypothetical protein [Mycobacterium abscessus] Length: 330\nNumber of predicted TMHs: 2\nExp number of AAs in TMHs: 39.3416199999999\nExp number, first 60 AAs: 17.07688\nTotal prob of N-in: 0.84431\nPOSSIBLE N-term signal sequence\ninside 1 11\nTMhelix 12 29\noutside 30 298\nTMhelix 299 321\ninside 322 330

29759 GCF_001215645.1_7396_7_38 Mycobacterium abscessus Terrabacteria group; Actinobacteria; Actinobacteria; Corynebacteriales; Mycobacteriaceae; Mycobacterium; Mycobacterium chelonae group; Mycobacterium abscessus subgroup MIKHLTRRATVALVVAAAAISPAAYPAVADTATPLNGLFALTPGSCAEGRATGSYFRMIL WP_005063674.1 hypothetical protein [Mycobacterium abscessus] Length: 330\nNumber of predicted TMHs: 1\nExp number of AAs in TMHs: 39.25259\nExp number, first 60 AAs: 16.98187\nTotal prob of N-in: 0.83210\nPOSSIBLE N-term signal sequence\noutside 1 298\nTMhelix 299 321\ninside 322 330

29760 GCF_001215745.1_7520_7_37 Mycobacterium abscessus Terrabacteria group; Actinobacteria; Actinobacteria; Corynebacteriales; Mycobacteriaceae; Mycobacterium; Mycobacterium chelonae group; Mycobacterium abscessus subgroup MIKHLTRRATVALVVAAAAISPAAYPAVADTATPLNGLFALTPGSCAEGRATGSYFRMIL WP_005063674.1 hypothetical protein [Mycobacterium abscessus] Length: 330\nNumber of predicted TMHs: 1\nExp number of AAs in TMHs: 39.25259\nExp number, first 60 AAs: 16.98187\nTotal prob of N-in: 0.83210\nPOSSIBLE N-term signal sequence\noutside 1 298\nTMhelix 299 321\ninside 322 330

29761 GCF_001216405.1_7396_8_63 Mycobacterium abscessus Terrabacteria group; Actinobacteria; Actinobacteria; Corynebacteriales; Mycobacteriaceae; Mycobacterium; Mycobacterium chelonae group; Mycobacterium abscessus subgroup MIKHLTRRATVALVVAAAAISPAAYPAVADTATPLNGLFALTPGSCAEGRATGSYFRMIL WP_005063674.1 hypothetical protein [Mycobacterium abscessus] Length: 330\nNumber of predicted TMHs: 1\nExp number of AAs in TMHs: 39.25259\nExp number, first 60 AAs: 16.98187\nTotal prob of N-in: 0.83210\nPOSSIBLE N-term signal sequence\noutside 1 298\nTMhelix 299 321\ninside 322 330

29762 GCF_001216665.1_7396_8_59 Mycobacterium abscessus Terrabacteria group; Actinobacteria; Actinobacteria; Corynebacteriales; Mycobacteriaceae; Mycobacterium; Mycobacterium chelonae group; Mycobacterium abscessus subgroup MIKHLTRRATVALVVAAAAISPAAYPAVADTATPLNGLFALTPGSCAEGRATGSYFRMIL WP_005063674.1 hypothetical protein [Mycobacterium abscessus] Length: 330\nNumber of predicted TMHs: 1\nExp number of AAs in TMHs: 39.25259\nExp number, first 60 AAs: 16.98187\nTotal prob of N-in: 0.83210\nPOSSIBLE N-term signal sequence\noutside 1 298\nTMhelix 299 321\ninside 322 330

29763 GCF_001216625.1_7396_8_56 Mycobacterium abscessus Terrabacteria group; Actinobacteria; Actinobacteria; Corynebacteriales; Mycobacteriaceae; Mycobacterium; Mycobacterium chelonae group; Mycobacterium abscessus subgroup MIKHLTRRATVALVVAAAAISPAAYPAVADTATPLNGLFALTPGSCAEGRATGSYFRMIL WP_005063674.1 hypothetical protein [Mycobacterium abscessus] Length: 330\nNumber of predicted TMHs: 1\nExp number of AAs in TMHs: 39.25259\nExp number, first 60 AAs: 16.98187\nTotal prob of N-in: 0.83210\nPOSSIBLE N-term signal sequence\noutside 1 298\nTMhelix 299 321\ninside 322 330

29764 GCF_001216965.1_7396_8_91 Mycobacterium abscessus Terrabacteria group; Actinobacteria; Actinobacteria; Corynebacteriales; Mycobacteriaceae; Mycobacterium; Mycobacterium chelonae group; Mycobacterium abscessus subgroup MIKHLTRRATVALVVAAAAISPAAYPAMADTATPLNGLFALTPGSCAEGRATGSYFRMIL WP_005073213.1 hypothetical protein [Mycobacterium abscessus] Length: 330\nNumber of predicted TMHs: 2\nExp number of AAs in TMHs: 39.3416199999999\nExp number, first 60 AAs: 17.07688\nTotal prob of N-in: 0.84431\nPOSSIBLE N-term signal sequence\ninside 1 11\nTMhelix 12 29\noutside 30 298\nTMhelix 299 321\ninside 322 330

29765 GCF_001217145.1_7396_7_31 Mycobacterium abscessus Terrabacteria group; Actinobacteria; Actinobacteria; Corynebacteriales; Mycobacteriaceae; Mycobacterium; Mycobacterium chelonae group; Mycobacterium abscessus subgroup MIKHLTRRATVALVVAAAAISPAAYPAMADTATPLNGLFALTPGSCAEGRATGSYFRMIL WP_005073213.1 hypothetical protein [Mycobacterium abscessus] Length: 330\nNumber of predicted TMHs: 2\nExp number of AAs in TMHs: 39.3416199999999\nExp number, first 60 AAs: 17.07688\nTotal prob of N-in:

0.84431\nPOSSIBLE N-term signal sequence\ninside 1 11\nTMhelix 12 29\noutside 30 298\nTMhelix 299 321\ninside 322 330

29766 GCF_001216725.1_7396_8_70 Mycobacterium abscessus Terrabacteria group; Actinobacteria; Actinobacteria; Corynebacteriales; Mycobacteriaceae; Mycobacterium; Mycobacterium chelonae group; Mycobacterium abscessus subgroup MIKHLTRRATVALVVAAAAISPAAYPAVADTATPLNGLFALTPGSCAEGRATGSYFRMIL WP_005063674.1 hypothetical protein [Mycobacterium abscessus] Length: 330\nNumber of predicted TMHs: 1\nExp number of AAs in TMHs: 39.25259\nExp number, first 60 AAs: 16.98187\nTotal prob of N-in: 0.83210\nPOSSIBLE N-term signal sequence\noutside 1 298\nTMhelix 299 321\ninside 322 330

29767 GCF_001213125.1_7396_6_65 Mycobacterium abscessus Terrabacteria group; Actinobacteria; Actinobacteria; Corynebacteriales; Mycobacteriaceae; Mycobacterium; Mycobacterium chelonae group; Mycobacterium abscessus subgroup MIKHLTRRATVALVVAAAAISPAAYPAVADTATPLNGLFALTPGSCAEGRATGSYFRMIL WP_005063674.1 hypothetical protein [Mycobacterium abscessus] Length: 330\nNumber of predicted TMHs: 1\nExp number of AAs in TMHs: 39.25259\nExp number, first 60 AAs: 16.98187\nTotal prob of N-in: 0.83210\nPOSSIBLE N-term signal sequence\noutside 1 298\nTMhelix 299 321\ninside 322 330

29768 GCF_001212965.1_7396_6_71 Mycobacterium abscessus Terrabacteria group; Actinobacteria; Actinobacteria; Corynebacteriales; Mycobacteriaceae; Mycobacterium; Mycobacterium chelonae group; Mycobacterium abscessus subgroup MIKHLTRRATVALVVAAAAISPAAYPAVADTATPLNGLFALTPGSCAEGRATGSYFRMIL WP_005063674.1 hypothetical protein [Mycobacterium abscessus] Length: 330\nNumber of predicted TMHs: 1\nExp number of AAs in TMHs: 39.25259\nExp number, first 60 AAs: 16.98187\nTotal prob of N-in: 0.83210\nPOSSIBLE N-term signal sequence\noutside 1 298\nTMhelix 299 321\ninside 322 330

29769 GCF_002142865.1_ASM214286v1 Mycobacterium abscessus Terrabacteria group; Actinobacteria; Actinobacteria; Corynebacteriales; Mycobacteriaceae; Mycobacterium; Mycobacterium chelonae group; Mycobacterium abscessus subgroup MIKHLTRRATVALVVAAAAISPAAYPAMADTATPLNGLFALTPGSCAEGRATGSYFRMIL WP_005073213.1 hypothetical protein [Mycobacterium abscessus] Length: 330\nNumber of predicted TMHs: 2\nExp number of AAs in TMHs: 39.3416199999999\nExp number, first 60 AAs: 17.07688\nTotal prob of N-in: 0.84431\nPOSSIBLE N-term signal sequence\ninside 1 11\nTMhelix 12 29\noutside 30 298\nTMhelix 299 321\ninside 322 330

29770 GCF_001677215.1_ASM167721v1 Mycobacterium abscessus Terrabacteria group; Actinobacteria; Actinobacteria; Corynebacteriales; Mycobacteriaceae; Mycobacterium; Mycobacterium chelonae group; Mycobacterium abscessus subgroup MIKHLTRRATVALVVAAAAISPAAYPAVADTATPLNGLFALTPGSCAEGRATGSYFRMIL WP_005063674.1 hypothetical protein [Mycobacterium abscessus] Length: 330\nNumber of predicted TMHs: 1\nExp number of AAs in TMHs: 39.25259\nExp number, first 60 AAs: 16.98187\nTotal prob of N-in: 0.83210\nPOSSIBLE N-term signal sequence\noutside 1 298\nTMhelix 299 321\ninside 322 330

29771 GCF_001677095.1_ASM167709v1 Mycobacterium abscessus Terrabacteria group; Actinobacteria; Actinobacteria; Corynebacteriales; Mycobacteriaceae; Mycobacterium; Mycobacterium chelonae group; Mycobacterium abscessus subgroup MIKHLTRRATVALVVAAAAISPAAYPAVADTATPLNGLFALTPGSCAEGRATGSYFRMIL WP_005063674.1 hypothetical protein [Mycobacterium abscessus] Length: 330\nNumber of predicted TMHs: 1\nExp number of AAs in TMHs: 39.25259\nExp number, first 60 AAs: 16.98187\nTotal prob of N-in: 0.83210\nPOSSIBLE N-term signal sequence\noutside 1 298\nTMhelix 299 321\ninside 322 330

29772 GCF_001677175.1_ASM167717v1 Mycobacterium abscessus Terrabacteria group; Actinobacteria; Actinobacteria; Corynebacteriales; Mycobacteriaceae; Mycobacterium; Mycobacterium chelonae group; Mycobacterium abscessus subgroup MIKHLTRRATVALVVAAAAISPAAYPAVADTATPLNGLFALTPGSCAEGRATGSYFRMIL WP_005063674.1 hypothetical protein [Mycobacterium abscessus] Length: 330\nNumber of predicted TMHs: 1\nExp number of AAs in TMHs: 39.25259\nExp number, first 60 AAs: 16.98187\nTotal prob of N-in: 0.83210\nPOSSIBLE N-term signal sequence\noutside 1 298\nTMhelix 299 321\ninside 322 330

29773 GCF_001606315.1_ASM160631v1 Mycobacterium abscessus Terrabacteria group; Actinobacteria; Actinobacteria; Corynebacteriales; Mycobacteriaceae; Mycobacterium; Mycobacterium chelonae group; Mycobacterium abscessus subgroup MIKHLTRRATVALVVAAAAISPAAYPAMADTATPLNGLFALTPGSCAEGRATGSYFRMIL WP_005073213.1 hypothetical protein [Mycobacterium abscessus] Length: 330\nNumber of predicted TMHs: 2\nExp number of AAs in TMHs: 39.3416199999999\nExp number, first 60 AAs: 17.07688\nTotal prob of N-in: 0.84431\nPOSSIBLE N-term signal sequence\ninside 1 11\nTMhelix 12 29\noutside 30 298\nTMhelix 299 321\ninside 322 330

29774 GCF_001610655.1_ASM161065v1 Mycobacterium abscessus Terrabacteria group; Actinobacteria; Actinobacteria; Corynebacteriales; Mycobacteriaceae; Mycobacterium; Mycobacterium chelonae group; Mycobacterium abscessus subgroup MIKHLTRRATVALVVAAAAISPAAYPAMADTATPLNGLFALTPGSCAEGRATGSYFRMIL WP_005073213.1 hypothetical protein [Mycobacterium abscessus] Length: 330\nNumber of predicted TMHs: 2\nExp number of AAs in TMHs: 39.3416199999999\nExp number, first 60 AAs: 17.07688\nTotal prob of N-in: 0.84431\nPOSSIBLE N-term signal sequence\ninside 1 11\nTMhelix 12 29\noutside 30 298\nTMhelix 299 321\ninside 322 330

29775 GCF_001212905.1_7396_6_61 Mycobacterium abscessus Terrabacteria group; Actinobacteria; Actinobacteria; Corynebacteriales; Mycobacteriaceae; Mycobacterium; Mycobacterium chelonae group; Mycobacterium abscessus subgroup MIKHLTRRATVALVVAAAAISPAAYPAMADTATPLNGLFALTPGSCAEGRATGSYFRMIL WP_005073213.1 hypothetical protein [Mycobacterium abscessus] Length: 330\nNumber of predicted TMHs: 2\nExp number of AAs in TMHs: 39.3416199999999\nExp number, first 60 AAs: 17.07688\nTotal prob of N-in: 0.84431\nPOSSIBLE N-term signal sequence\ninside 1 11\nTMhelix 12 29\noutside 30 298\nTMhelix 299 321\ninside 322 330

29776 GCF_001212985.1_7396_6_58 Mycobacterium abscessus Terrabacteria group; Actinobacteria; Actinobacteria; Corynebacteriales; Mycobacteriaceae; Mycobacterium; Mycobacterium chelonae group; Mycobacterium abscessus subgroup MIKHLTRRATVALVVAAAAISPAAYPAMADTATPLNGLFALTPGSCAEGRATGSYFRMIL WP_005073213.1 hypothetical protein [Mycobacterium abscessus] Length: 330\nNumber of predicted TMHs: 2\nExp number of AAs in TMHs: 39.3416199999999\nExp number, first 60 AAs: 17.07688\nTotal prob of N-in: 0.84431\nPOSSIBLE N-term signal sequence\ninside 1 11\nTMhelix 12 29\noutside 30 298\nTMhelix 299 321\ninside 322 330

29777 GCF_001212825.1_7396_6_66 Mycobacterium abscessus Terrabacteria group; Actinobacteria; Actinobacteria; Corynebacteriales; Mycobacteriaceae; Mycobacterium; Mycobacterium chelonae group; Mycobacterium abscessus subgroup MIKHLTRRATVALVVAAAAISPAAYPAVADTATPLNGLFALTPGSCAEGRATGSYFRMIL WP_005063674.1 hypothetical protein [Mycobacterium abscessus] Length: 330\nNumber of predicted TMHs: 1\nExp number of AAs in TMHs: 39.25259\nExp number, first 60 AAs: 16.98187\nTotal prob of N-in: 0.83210\nPOSSIBLE N-term signal sequence\noutside 1 298\nTMhelix 299 321\ninside 322 330

29778 GCF_001213025.1_7396_6_91 Mycobacterium abscessus Terrabacteria group; Actinobacteria; Actinobacteria; Corynebacteriales; Mycobacteriaceae; Mycobacterium; Mycobacterium chelonae group; Mycobacterium abscessus subgroup MIKHLTRRATVALVVAAAAISPAAYPAVADTATPLNGLFALTPGSCAEGRATGSYFRMIL WP_005063674.1 hypothetical protein [Mycobacterium abscessus] Length: 330\nNumber of predicted TMHs: 1\nExp number of AAs in TMHs: 39.25259\nExp number, first 60 AAs: 16.98187\nTotal prob of N-in: 0.83210\nPOSSIBLE N-term signal sequence\noutside 1 298\nTMhelix 299 321\ninside 322 330

29779 GCF_001213205.1_7396_6_49 Mycobacterium abscessus Terrabacteria group; Actinobacteria; Actinobacteria; Corynebacteriales; Mycobacteriaceae; Mycobacterium; Mycobacterium chelonae group; Mycobacterium abscessus subgroup MIKHLTRRATVALVVAAAAISPAAYPAMADTATPLNGLFALTPGSCAEGRATGSYFRMIL WP_005073213.1 hypothetical protein [Mycobacterium abscessus] Length: 330\nNumber of predicted TMHs: 2\nExp number of AAs in TMHs: 39.3416199999999\nExp number, first 60 AAs: 17.07688\nTotal prob of N-in: 0.84431\nPOSSIBLE N-term signal sequence\ninside 1 11\nTMhelix 12 29\noutside 30 298\nTMhelix 299 321\ninside 322 330

29780 GCF_001213705.1_7396_7_19 Mycobacterium abscessus Terrabacteria group; Actinobacteria; Actinobacteria; Corynebacteriales; Mycobacteriaceae; Mycobacterium; Mycobacterium chelonae group; Mycobacterium abscessus subgroup MIKHLTRRATVALVVAAAAISPAAYPAMADTATPLNGLFALTPGSCAEGRATGSYFRMIL WP_005073213.1 hypothetical protein [Mycobacterium abscessus] Length: 330\nNumber of predicted TMHs: 2\nExp number of AAs in TMHs: 39.3416199999999\nExp number, first 60 AAs: 17.07688\nTotal prob of N-in: 0.84431\nPOSSIBLE N-term signal sequence\ninside 1 11\nTMhelix 12 29\noutside 30 298\nTMhelix 299 321\ninside 322 330

29781 GCF_001213465.1_7396_7_2Mycobacterium abscessus Terrabacteria group; Actinobacteria; Actinobacteria; Corynebacteriales; Mycobacteriaceae; Mycobacterium; Mycobacterium chelonae group; Mycobacterium abscessus subgroup MIKHLTRRATVALVVAAAAISPAAYPAVADTATPLNGLFALTPGSCAEGRATGSYFRMIL WP_005063674.1 hypothetical protein [Mycobacterium abscessus] Length: 330\nNumber of predicted TMHs: 1\nExp number of AAs in TMHs: 39.25259\nExp number, first 60 AAs: 16.98187\nTotal prob of N-in: 0.83210\nPOSSIBLE N-term signal sequence\noutside 1 298\nTMhelix 299 321\ninside 322 330

29782 GCF_001213725.1_7396_6_59 Mycobacterium abscessus Terrabacteria group; Actinobacteria; Actinobacteria; Corynebacteriales; Mycobacteriaceae; Mycobacterium; Mycobacterium chelonae group; Mycobacterium abscessus subgroup MIKHLTRRATVALVVAAAAISPAAYPAMADTATPLNGLFALTPGSCAEGRATGSYFRMIL WP_005073213.1 hypothetical protein [Mycobacterium abscessus] Length: 330\nNumber of predicted TMHs: 2\nExp number of AAs in TMHs: 39.3416199999999\nExp number, first 60 AAs: 17.07688\nTotal prob of N-in: 0.84431\nPOSSIBLE N-term signal sequence\ninside 1 11\nTMhelix 12 29\noutside 30 298\nTMhelix 299 321\ninside 322 330

29783 GCF_001213545.1_7396_6_64 Mycobacterium abscessus Terrabacteria group; Actinobacteria; Actinobacteria; Corynebacteriales; Mycobacteriaceae; Mycobacterium; Mycobacterium chelonae group; Mycobacterium abscessus subgroup MIKHLTRRATVALVVAAAAISPAAYPAVADTATPLNGLFALTPGSCAEGRATGSYFRMIL WP_005063674.1 hypothetical protein [Mycobacterium abscessus] Length: 330\nNumber of predicted TMHs: 1\nExp number of AAs in TMHs: 39.25259\nExp number, first 60 AAs: 16.98187\nTotal prob of N-in: 0.83210\nPOSSIBLE N-term signal sequence\noutside 1 298\nTMhelix 299 321\ninside 322 330

29784 GCF_001213625.1_7396_6_96 Mycobacterium abscessus Terrabacteria group; Actinobacteria; Actinobacteria; Corynebacteriales; Mycobacteriaceae; Mycobacterium; Mycobacterium chelonae group; Mycobacterium abscessus subgroup MIKHLTRRATVALVVAAAAISPAAYPAVADTATPLNGLFALTPGSCAEGRATGSYFRMIL WP_005063674.1 hypothetical protein [Mycobacterium abscessus] Length: 330\nNumber of predicted TMHs: 1\nExp number of AAs in TMHs: 39.25259\nExp number, first 60 AAs: 16.98187\nTotal prob of N-in: 0.83210\nPOSSIBLE N-term signal sequence\noutside 1 298\nTMhelix 299 321\ninside 322 330

29785 GCF_001213585.1_7396_6_85 Mycobacterium abscessus Terrabacteria group; Actinobacteria; Actinobacteria; Corynebacteriales; Mycobacteriaceae; Mycobacterium; Mycobacterium chelonae group; Mycobacterium abscessus subgroup MIKHLTRRATVALVVAAAAISPAAYPAVADTATPLNGLFALTPGSCAEGRATGSYFRMIL WP_005063674.1 hypothetical protein [Mycobacterium abscessus] Length: 330\nNumber of predicted TMHs: 1\nExp number of AAs in TMHs: 39.25259\nExp number, first 60 AAs: 16.98187\nTotal prob of N-in: 0.83210\nPOSSIBLE N-term signal sequence\noutside 1 298\nTMhelix 299 321\ninside 322 330

29786 GCF_001213845.1_7396_6_92 Mycobacterium abscessus Terrabacteria group; Actinobacteria; Actinobacteria; Corynebacteriales; Mycobacteriaceae; Mycobacterium; Mycobacterium chelonae group; Mycobacterium abscessus subgroup MIKHLTRRATVALVVAAAAISPAAYPAMADTATPLNGLFALTPGSCAEGRATGSYFRMIL WP_005073213.1 hypothetical protein [Mycobacterium abscessus] Length: 330\nNumber of predicted TMHs: 2\nExp number of AAs in TMHs: 39.3416199999999\nExp number, first 60 AAs: 17.07688\nTotal prob of N-in: 0.84431\nPOSSIBLE N-term signal sequence\ninside 1 11\nTMhelix 12 29\noutside 30 298\nTMhelix 299 321\ninside 322 330

29787 GCF_001214025.1_7396_6_55 Mycobacterium abscessus Terrabacteria group; Actinobacteria; Actinobacteria; Corynebacteriales; Mycobacteriaceae; Mycobacterium; Mycobacterium chelonae group; Mycobacterium abscessus subgroup MIKHLTRRATVALVVAAAAISPAAYPAVADTATPLNGLFALTPGSCAEGRATGSYFRMIL WP_005063674.1 hypothetical protein [Mycobacterium abscessus] Length: 330\nNumber of predicted TMHs: 1\nExp number of AAs in TMHs: 39.25259\nExp number, first 60 AAs: 16.98187\nTotal prob of N-in: 0.83210\nPOSSIBLE N-term signal sequence\noutside 1 298\nTMhelix 299 321\ninside 322 330

29788 GCF_001214125.1_7396_6_74 Mycobacterium abscessus Terrabacteria group; Actinobacteria; Actinobacteria; Corynebacteriales; Mycobacteriaceae; Mycobacterium; Mycobacterium chelonae group; Mycobacterium abscessus subgroup MIKHLTRRATVALVVAAAAISPAAYPAVADTATPLNGLFALTPGSCAEGRATGSYFRMIL WP_005063674.1 hypothetical protein [Mycobacterium abscessus] Length: 330\nNumber of predicted TMHs: 1\nExp number of AAs in TMHs: 39.25259\nExp number, first 60 AAs: 16.98187\nTotal prob of N-in: 0.83210\nPOSSIBLE N-term signal sequence\noutside 1 298\nTMhelix 299 321\ninside 322 330

29789 GCF_001214165.1_7396_7_1Mycobacterium abscessus Terrabacteria group; Actinobacteria; Actinobacteria; Corynebacteriales; Mycobacteriaceae; Mycobacterium; Mycobacterium chelonae group; Mycobacterium abscessus subgroup MIKHLTRRATVALVVAAAAISPAAYPAVADTATPLNGLFALTPGSCAEGRATGSYFRMIL WP_005063674.1 hypothetical protein [Mycobacterium abscessus] Length: 330\nNumber of predicted TMHs: 1\nExp number of AAs in TMHs: 39.25259\nExp number, first 60 AAs: 16.98187\nTotal prob of N-in: 0.83210\nPOSSIBLE N-term signal sequence\noutside 1 298\nTMhelix 299 321\ninside 322 330

29790 GCF_001214205.1_7396_7_4Mycobacterium abscessus Terrabacteria group; Actinobacteria; Actinobacteria; Corynebacteriales; Mycobacteriaceae; Mycobacterium; Mycobacterium chelonae group; Mycobacterium abscessus subgroup MIKHLTRRATVALVVAAAAISPAAYPAVADTATPLNGLFALTPGSCAEGRATGSYFRMIL WP_005063674.1 hypothetical protein [Mycobacterium abscessus] Length: 330\nNumber of predicted TMHs: 1\nExp number of AAs in TMHs: 39.25259\nExp number, first 60 AAs: 16.98187\nTotal prob of N-in: 0.83210\nPOSSIBLE N-term signal sequence\noutside 1 298\nTMhelix 299 321\ninside 322 330

29791 GCF_001214265.1_7396_8_77 Mycobacterium abscessus Terrabacteria group; Actinobacteria; Actinobacteria; Corynebacteriales; Mycobacteriaceae; Mycobacterium; Mycobacterium chelonae group; Mycobacterium abscessus subgroup MIKHLTRRATVALVVAAAAISPAAYPAVADTATPLNGLFALTPGSCAEGRATGSYFRMIL WP_005063674.1 hypothetical protein [Mycobacterium abscessus] Length: 330\nNumber of predicted TMHs: 1\nExp number of AAs in TMHs: 39.25259\nExp number, first 60 AAs: 16.98187\nTotal prob of N-in: 0.83210\nPOSSIBLE N-term signal sequence\noutside 1 298\nTMhelix 299 321\ninside 322 330

29792 GCF_001214305.1_7396_7_44 Mycobacterium abscessus Terrabacteria group; Actinobacteria; Actinobacteria; Corynebacteriales; Mycobacteriaceae; Mycobacterium; Mycobacterium chelonae group; Mycobacterium abscessus subgroup MIKHLTRRATVALVVAAAAISPAAYPAVADTATPLNGLFALTPGSCAEGRATGSYFRMIL WP_005063674.1 hypothetical protein [Mycobacterium abscessus] Length: 330\nNumber of predicted TMHs: 1\nExp number of AAs in TMHs: 39.25259\nExp number, first 60 AAs: 16.98187\nTotal prob of N-in: 0.83210\nPOSSIBLE N-term signal sequence\noutside 1 298\nTMhelix 299 321\ninside 322 330

29793 GCF_001214925.1_7520_7_41 Mycobacterium abscessus Terrabacteria group; Actinobacteria; Actinobacteria; Corynebacteriales; Mycobacteriaceae; Mycobacterium; Mycobacterium chelonae group; Mycobacterium abscessus subgroup MIKHLTRRATVALVVAAAAISPAAYPAMADTATPLNGLFALTPGSCAEGRATGSYFRMIL WP_005073213.1 hypothetical protein [Mycobacterium abscessus] Length: 330\nNumber of predicted TMHs: 2\nExp number of AAs in TMHs: 39.3416199999999\nExp number, first 60 AAs: 17.07688\nTotal prob of N-in: 0.84431\nPOSSIBLE N-term signal sequence\ninside 1 11\nTMhelix 12 29\noutside 30 298\nTMhelix 299 321\ninside 322 330

29794 GCF_001214665.1_7396_8_74 Mycobacterium abscessus Terrabacteria group; Actinobacteria; Actinobacteria; Corynebacteriales; Mycobacteriaceae; Mycobacterium; Mycobacterium chelonae group; Mycobacterium abscessus subgroup MIKHLTRRATVALVVAAAAISPAAYPAVADTATPLNGLFALTPGSCAEGRATGSYFRMIL WP_005063674.1 hypothetical protein [Mycobacterium abscessus] Length: 330\nNumber of predicted TMHs: 1\nExp number of AAs in TMHs: 39.25259\nExp number, first 60 AAs: 16.98187\nTotal prob of N-in: 0.83210\nPOSSIBLE N-term signal sequence\noutside 1 298\nTMhelix 299 321\ninside 322 330

29795 GCF_001214985.1_7396_7_23 Mycobacterium abscessus Terrabacteria group; Actinobacteria; Actinobacteria; Corynebacteriales; Mycobacteriaceae; Mycobacterium; Mycobacterium chelonae group; Mycobacterium abscessus subgroup MIKHLTRRATVALVVAAAAISPAAYPAMADTATPLNGLFALTPGSCAEGRATGSYFRMIL WP_005073213.1 hypothetical protein [Mycobacterium abscessus] Length: 330\nNumber of predicted TMHs: 2\nExp number of AAs in TMHs: 39.3416199999999\nExp number, first 60 AAs: 17.07688\nTotal prob of N-in: 0.84431\nPOSSIBLE N-term signal sequence\ninside 1 11\nTMhelix 12 29\noutside 30 298\nTMhelix 299 321\ninside 322 330

29796 GCF_001214685.1_7396_8_93 Mycobacterium abscessus Terrabacteria group; Actinobacteria; Actinobacteria; Corynebacteriales; Mycobacteriaceae; Mycobacterium; Mycobacterium chelonae group; Mycobacterium abscessus subgroup MIKHLTRRATVALVVAAAAISPAAYPAVADTATPLNGLFALTPGSCAEGRATGSYFRMIL WP_005063674.1 hypothetical protein [Mycobacterium abscessus] Length: 330\nNumber of predicted TMHs: 1\nExp number of AAs in TMHs: 39.25259\nExp number, first 60 AAs: 16.98187\nTotal prob of N-in: 0.83210\nPOSSIBLE N-term signal sequence\noutside 1 298\nTMhelix 299 321\ninside 322 330

29797 GCF_001214805.1_7520_7_25 Mycobacterium abscessus Terrabacteria group; Actinobacteria; Actinobacteria; Corynebacteriales; Mycobacteriaceae; Mycobacterium; Mycobacterium chelonae group; Mycobacterium abscessus subgroup MIKHLTRRATVALVVAAAAISPAAYPAVADTATPLNGLFALTPGSCAEGRATGSYFRMIL WP_005063674.1 hypothetical protein [Mycobacterium abscessus] Length: 330\nNumber of predicted TMHs: 1\nExp number of AAs in TMHs: 39.25259\nExp number, first 60 AAs: 16.98187\nTotal prob of N-in: 0.83210\nPOSSIBLE N-term signal sequence\noutside 1 298\nTMhelix 299 321\ninside 322 330

29798 GCF_001214845.1_7396_8_80 Mycobacterium abscessus Terrabacteria group; Actinobacteria; Actinobacteria; Corynebacteriales; Mycobacteriaceae; Mycobacterium; Mycobacterium chelonae group; Mycobacterium abscessus subgroup MIKHLTRRATVALVVAAAAISPAAYPAVADTATPLNGLFALTPGSCAEGRATGSYFRMIL WP_005063674.1 hypothetical protein [Mycobacterium abscessus] Length: 330\nNumber of predicted TMHs: 1\nExp number of AAs in TMHs: 39.25259\nExp number, first 60 AAs: 16.98187\nTotal prob of N-in: 0.83210\nPOSSIBLE N-term signal sequence\noutside 1 298\nTMhelix 299 321\ninside 322 330

29799 GCF_000270845.1_ASM27084v1 Mycobacterium abscessus 3A-0122-R Terrabacteria group; Actinobacteria; Actinobacteria; Corynebacteriales; Mycobacteriaceae; Mycobacterium; Mycobacterium chelonae group; Mycobacterium abscessus subgroup; Mycobacterium abscessus
MIKHLTRRATVALVVAAAAISPAAYPAMADTATPLNGLFALTPGSCAEGRATGSYFRMIL WP_005073213.1
hypothetical protein [Mycobacterium abscessus] Length: 330\nNumber of predicted TMHs: 2\nExp number of AAs in TMHs: 39.3416199999999\nExp number, first 60 AAs: 17.07688\nTotal prob of N-in: 0.84431\nPOSSIBLE N-term signal sequence\ninside 1 11\nTMhelix 12 29\noutside 30 298\nTMhelix 299 321\ninside 322 330

29800 GCF_000270705.1_ASM27070v1 Mycobacterium abscessus 3A-0731 Terrabacteria group; Actinobacteria; Actinobacteria; Corynebacteriales; Mycobacteriaceae; Mycobacterium; Mycobacterium chelonae group; Mycobacterium abscessus subgroup; Mycobacterium abscessus
MIKHLTRRATVALVVAAAAISPAAYPAMADTATPLNGLFALTPGSCAEGRATGSYFRMIL WP_005073213.1
hypothetical protein [Mycobacterium abscessus] Length: 330\nNumber of predicted TMHs: 2\nExp number of AAs in TMHs: 39.3416199999999\nExp number, first 60 AAs: 17.07688\nTotal prob of N-in: 0.84431\nPOSSIBLE N-term signal sequence\ninside 1 11\nTMhelix 12 29\noutside 30 298\nTMhelix 299 321\ninside 322 330

29801 GCF_000271105.1_ASM27110v1 Mycobacterium abscessus 3A-0810-R Terrabacteria group; Actinobacteria; Actinobacteria; Corynebacteriales; Mycobacteriaceae; Mycobacterium; Mycobacterium chelonae group; Mycobacterium abscessus subgroup; Mycobacterium abscessus
MIKHLTRRATVALVVAAAAISPAAYPAMADTATPLNGLFALTPGSCAEGRATGSYFRMIL WP_005073213.1
hypothetical protein [Mycobacterium abscessus] Length: 330\nNumber of predicted TMHs: 2\nExp number of AAs in TMHs: 39.3416199999999\nExp number, first 60 AAs: 17.07688\nTotal prob of N-in: 0.84431\nPOSSIBLE N-term signal sequence\ninside 1 11\nTMhelix 12 29\noutside 30 298\nTMhelix 299 321\ninside 322 330

29802 GCF_000270565.1_ASM27056v1 Mycobacterium abscessus 3A-0930-R Terrabacteria group; Actinobacteria; Actinobacteria; Corynebacteriales; Mycobacteriaceae; Mycobacterium; Mycobacterium chelonae group; Mycobacterium abscessus subgroup; Mycobacterium abscessus
MIKHLTRRATVALVVAAAAISPAAYPAMADTATPLNGLFALTPGSCAEGRATGSYFRMIL WP_005073213.1
hypothetical protein [Mycobacterium abscessus] Length: 330\nNumber of predicted TMHs: 2\nExp number of AAs in TMHs: 39.3416199999999\nExp number, first 60 AAs: 17.07688\nTotal prob of N-in: 0.84431\nPOSSIBLE N-term signal sequence\ninside 1 11\nTMhelix 12 29\noutside 30 298\nTMhelix 299 321\ninside 322 330

29803 GCF_000231155.1_ASM23115v2 Mycobacterium abscessus 47J26 Terrabacteria group; Actinobacteria; Actinobacteria; Corynebacteriales; Mycobacteriaceae; Mycobacterium; Mycobacterium chelonae group; Mycobacterium abscessus subgroup; Mycobacterium abscessus
MIKHLTRRATVALVVAAAAISPAAYPAVADTATPLNGLFALTPGSCAEGRATGSYFRMIL WP_005063674.1 hypothetical protein [Mycobacterium abscessus] Length: 330\nNumber of predicted TMHs: 1\nExp number of AAs in TMHs: 39.25259\nExp number, first 60 AAs: 16.98187\nTotal prob of N-in: 0.83210\nPOSSIBLE N-term signal sequence\noutside 1 298\nTMhelix 299 321\ninside 322 330

29804 GCF_000270685.2_ASM27068v2 Mycobacterium abscessus 4S-0116-R Terrabacteria group; Actinobacteria; Actinobacteria; Corynebacteriales; Mycobacteriaceae; Mycobacterium; Mycobacterium chelonae group; Mycobacterium abscessus subgroup; Mycobacterium abscessus
MIKHLTRRATVALVVAAAAISPAAYPAMADTATPLNGLFALTPGSCAEGRATGSYFRMIL WP_005073213.1
hypothetical protein [Mycobacterium abscessus] Length: 330\nNumber of predicted TMHs: 2\nExp number of AAs in TMHs: 39.3416199999999\nExp number, first 60 AAs: 17.07688\nTotal prob of N-in: 0.84431\nPOSSIBLE N-term signal sequence\ninside 1 11\nTMhelix 12 29\noutside 30 298\nTMhelix 299 321\ninside 322 330

29805 GCF_000270625.1_ASM27062v1 Mycobacterium abscessus 4S-0116-S Terrabacteria group; Actinobacteria; Actinobacteria; Corynebacteriales; Mycobacteriaceae; Mycobacterium; Mycobacterium chelonae group; Mycobacterium abscessus subgroup; Mycobacterium abscessus
MIKHLTRRATVALVVAAAAISPAAYPAMADTATPLNGLFALTPGSCAEGRATGSYFRMIL WP_005073213.1
hypothetical protein [Mycobacterium abscessus] Length: 330\nNumber of predicted TMHs: 2\nExp number of AAs in TMHs: 39.3416199999999\nExp number, first 60 AAs: 17.07688\nTotal prob of N-in: 0.84431\nPOSSIBLE N-term signal sequence\ninside 1 11\nTMhelix 12 29\noutside 30 298\nTMhelix 299 321\ninside 322 330

29806 GCF_000271265.1_ASM27126v1 Mycobacterium abscessus 4S-0726-RB Terrabacteria group; Actinobacteria; Actinobacteria; Corynebacteriales; Mycobacteriaceae; Mycobacterium; Mycobacterium chelonae group; Mycobacterium abscessus subgroup; Mycobacterium abscessus
MIKHLTRRATVALVVAAAAISPAAYPAMADTATPLNGLFALTPGSCAEGRATGSYFRMIL WP_005073213.1
hypothetical protein [Mycobacterium abscessus] Length: 330\nNumber of predicted TMHs: 2\nExp number of

AAs in TMHs: 39.3416199999999\nExp number, first 60 AAs: 17.07688\nTotal prob of N-in: 0.84431\nPOSSIBLE N-term signal sequence\ninside 1 11\nTMhelix 12 29\noutside 30 298\nTMhelix 299 321\ninside 322 330

29807 GCF_000271205.1_ASM27120v1 Mycobacterium abscessus 5S-0304 Terrabacteria group; Actinobacteria; Actinobacteria; Corynebacteriales; Mycobacteriaceae; Mycobacterium; Mycobacterium chelonae group; Mycobacterium abscessus subgroup; Mycobacterium abscessus
MIKHLTRRATVALVVAAAAISPAAYPAVADTATPLNGLFALTPGSCAEGRATGSYFRMIL WP_005063674.1 hypothetical protein [Mycobacterium abscessus] Length: 330\nNumber of predicted TMHs: 1\nExp number of AAs in TMHs: 39.25259\nExp number, first 60 AAs: 16.98187\nTotal prob of N-in: 0.83210\nPOSSIBLE N-term signal sequence\noutside 1 298\nTMhelix 299 321\ninside 322 330

29808 GCF_000271005.1_ASM27100v1 Mycobacterium abscessus 5S-0817 Terrabacteria group; Actinobacteria; Actinobacteria; Corynebacteriales; Mycobacteriaceae; Mycobacterium; Mycobacterium chelonae group; Mycobacterium abscessus subgroup; Mycobacterium abscessus
MIKHLTRRATVALVVAAAAISPAAYPAVADTATPLNGLFALTPGSCAEGRATGSYFRMIL WP_005063674.1 hypothetical protein [Mycobacterium abscessus] Length: 330\nNumber of predicted TMHs: 1\nExp number of AAs in TMHs: 39.25259\nExp number, first 60 AAs: 16.98187\nTotal prob of N-in: 0.83210\nPOSSIBLE N-term signal sequence\noutside 1 298\nTMhelix 299 321\ninside 322 330

29809 GCF_000271025.1_ASM27102v1 Mycobacterium abscessus 5S-1212 Terrabacteria group; Actinobacteria; Actinobacteria; Corynebacteriales; Mycobacteriaceae; Mycobacterium; Mycobacterium chelonae group; Mycobacterium abscessus subgroup; Mycobacterium abscessus
MIKHLTRRATVALVVAAAAISPAAYPAVADTATPLNGLFALTPGSCAEGRATGSYFRMIL WP_005063674.1 hypothetical protein [Mycobacterium abscessus] Length: 330\nNumber of predicted TMHs: 1\nExp number of AAs in TMHs: 39.25259\nExp number, first 60 AAs: 16.98187\nTotal prob of N-in: 0.83210\nPOSSIBLE N-term signal sequence\noutside 1 298\nTMhelix 299 321\ninside 322 330

29810 GCF_000271045.1_ASM27104v1 Mycobacterium abscessus 5S-1215 Terrabacteria group; Actinobacteria; Actinobacteria; Corynebacteriales; Mycobacteriaceae; Mycobacterium; Mycobacterium chelonae group; Mycobacterium abscessus subgroup; Mycobacterium abscessus
MIKHLTRRATVALVVAAAAISPAAYPAVADTATPLNGLFALTPGSCAEGRATGSYFRMIL WP_005063674.1 hypothetical protein [Mycobacterium abscessus] Length: 330\nNumber of predicted TMHs: 1\nExp number of AAs in TMHs: 39.25259\nExp number, first 60 AAs: 16.98187\nTotal prob of N-in: 0.83210\nPOSSIBLE N-term signal sequence\noutside 1 298\nTMhelix 299 321\ninside 322 330

29811 GCF_000270985.1_ASM27098v1 Mycobacterium abscessus 6G-0125-R Terrabacteria group; Actinobacteria; Actinobacteria; Corynebacteriales; Mycobacteriaceae; Mycobacterium; Mycobacterium chelonae group; Mycobacterium abscessus subgroup; Mycobacterium abscessus
MIKHLTRRATVALVVAAAAISPAAYPAMADTATPLNGLFALTPGSCAEGRATGSYFRMIL WP_005073213.1 hypothetical protein [Mycobacterium abscessus] Length: 330\nNumber of predicted TMHs: 2\nExp number of AAs in TMHs: 39.3416199999999\nExp number, first 60 AAs: 17.07688\nTotal prob of N-in: 0.84431\nPOSSIBLE N-term signal sequence\ninside 1 11\nTMhelix 12 29\noutside 30 298\nTMhelix 299 321\ninside 322 330

29812 GCF_000270865.1_ASM27086v1 Mycobacterium abscessus 6G-0125-S Terrabacteria group; Actinobacteria; Actinobacteria; Corynebacteriales; Mycobacteriaceae; Mycobacterium; Mycobacterium chelonae group; Mycobacterium abscessus subgroup; Mycobacterium abscessus
MIKHLTRRATVALVVAAAAISPAAYPAMADTATPLNGLFALTPGSCAEGRATGSYFRMIL WP_005073213.1 hypothetical protein [Mycobacterium abscessus] Length: 330\nNumber of predicted TMHs: 2\nExp number of AAs in TMHs: 39.3416199999999\nExp number, first 60 AAs: 17.07688\nTotal prob of N-in: 0.84431\nPOSSIBLE N-term signal sequence\ninside 1 11\nTMhelix 12 29\noutside 30 298\nTMhelix 299 321\ninside 322 330

29813 GCF_000270605.1_ASM27060v1 Mycobacterium abscessus 6G-0728-R Terrabacteria group; Actinobacteria; Actinobacteria; Corynebacteriales; Mycobacteriaceae; Mycobacterium; Mycobacterium chelonae group; Mycobacterium abscessus subgroup; Mycobacterium abscessus
MIKHLTRRATVALVVAAAAISPAAYPAMADTATPLNGLFALTPGSCAEGRATGSYFRMIL WP_005073213.1 hypothetical protein [Mycobacterium abscessus] Length: 330\nNumber of predicted TMHs: 2\nExp number of AAs in TMHs: 39.3416199999999\nExp number, first 60 AAs: 17.07688\nTotal prob of N-in: 0.84431\nPOSSIBLE N-term signal sequence\ninside 1 11\nTMhelix 12 29\noutside 30 298\nTMhelix 299 321\ninside 322 330

29814 GCF_000271185.1_ASM27118v1 Mycobacterium abscessus 6G-0728-S Terrabacteria group; Actinobacteria; Actinobacteria; Corynebacteriales; Mycobacteriaceae; Mycobacterium; Mycobacterium chelonae group;

Mycobacterium abscessus subgroup; Mycobacterium abscessus
MIKHLTRRATVALVVAAAAISPAAYPAMADTATPLNGLFALTPGSCAEGRATGSYFRMIL WP_005073213.1 hypothetical protein [Mycobacterium abscessus] Length: 330\nNumber of predicted TMHs: 2\nExp number of AAs in TMHs: 39.3416199999999\nExp number, first 60 AAs: 17.07688\nTotal prob of N-in: 0.84431\nPOSSIBLE N-term signal sequence\ninside 1 11\nTMhelix 12 29\noutside 30 298\nTMhelix 299 321\ninside 322 330

29815 GCF_000332605.1_SOAPdenovo_v1.06 Mycobacterium abscessus 9808 Terrabacteria group; Actinobacteria; Actinobacteria; Corynebacteriales; Mycobacteriaceae; Mycobacterium; Mycobacterium chelonae group; Mycobacterium abscessus subgroup; Mycobacterium abscessus
MIKHLTRRATVALVVAAAAISPAAYPAMADTATPLNGLFALTPGSCAEGRATGSYFRMIL WP_005073213.1 hypothetical protein [Mycobacterium abscessus] Length: 330\nNumber of predicted TMHs: 2\nExp number of AAs in TMHs: 39.3416199999999\nExp number, first 60 AAs: 17.07688\nTotal prob of N-in: 0.84431\nPOSSIBLE N-term signal sequence\ninside 1 11\nTMhelix 12 29\noutside 30 298\nTMhelix 299 321\ninside 322 330

29816 GCF_000333695.1_ASM33369v1 Mycobacterium abscessus CF Terrabacteria group; Actinobacteria; Actinobacteria; Corynebacteriales; Mycobacteriaceae; Mycobacterium; Mycobacterium chelonae group; Mycobacterium abscessus subgroup; Mycobacterium abscessus
MIKHLTRRATVALVVAAAAISPAAYPAMADTATPLNGLFALTPGSCAEGRATGSYFRMIL WP_005073213.1 hypothetical protein [Mycobacterium abscessus] Length: 330\nNumber of predicted TMHs: 2\nExp number of AAs in TMHs: 39.3416199999999\nExp number, first 60 AAs: 17.07688\nTotal prob of N-in: 0.84431\nPOSSIBLE N-term signal sequence\ninside 1 11\nTMhelix 12 29\noutside 30 298\nTMhelix 299 321\ninside 322 330

29817 GCF_000260595.1_ASM26059v1 Mycobacterium abscessus M115 Terrabacteria group; Actinobacteria; Actinobacteria; Corynebacteriales; Mycobacteriaceae; Mycobacterium; Mycobacterium chelonae group; Mycobacterium abscessus subgroup; Mycobacterium abscessus
MIKHLTRRATVALVVAAAAISPAAYPAVADTATPLNGLFALTPGSCAEGRATGSYFRMIL WP_005063674.1 hypothetical protein [Mycobacterium abscessus] Length: 330\nNumber of predicted TMHs: 1\nExp number of AAs in TMHs: 39.25259\nExp number, first 60 AAs: 16.98187\nTotal prob of N-in: 0.83210\nPOSSIBLE N-term signal sequence\noutside 1 298\nTMhelix 299 321\ninside 322 330

29818 GCF_000280595.1_ASM28059v1 Mycobacterium abscessus M139 Terrabacteria group; Actinobacteria; Actinobacteria; Corynebacteriales; Mycobacteriaceae; Mycobacterium; Mycobacterium chelonae group; Mycobacterium abscessus subgroup; Mycobacterium abscessus
MIKHLTRRATVALVVAAAAISPAAYPAMADTATPLNGLFALTPGSCAEGRATGSYFRMIL WP_005073213.1 hypothetical protein [Mycobacterium abscessus] Length: 330\nNumber of predicted TMHs: 2\nExp number of AAs in TMHs: 39.3416199999999\nExp number, first 60 AAs: 17.07688\nTotal prob of N-in: 0.84431\nPOSSIBLE N-term signal sequence\ninside 1 11\nTMhelix 12 29\noutside 30 298\nTMhelix 299 321\ninside 322 330

29819 GCF_000280615.1_ASM28061v1 Mycobacterium abscessus M152 Terrabacteria group; Actinobacteria; Actinobacteria; Corynebacteriales; Mycobacteriaceae; Mycobacterium; Mycobacterium chelonae group; Mycobacterium abscessus subgroup; Mycobacterium abscessus
MIKHLTRRATVALVVAAAAISPAAYPAMADTATPLNGLFALTPGSCAEGRATGSYFRMIL WP_005073213.1 hypothetical protein [Mycobacterium abscessus] Length: 330\nNumber of predicted TMHs: 2\nExp number of AAs in TMHs: 39.3416199999999\nExp number, first 60 AAs: 17.07688\nTotal prob of N-in: 0.84431\nPOSSIBLE N-term signal sequence\ninside 1 11\nTMhelix 12 29\noutside 30 298\nTMhelix 299 321\ninside 322 330

29820 GCF_000260575.2_ASM26057v2 Mycobacterium abscessus M154 Terrabacteria group; Actinobacteria; Actinobacteria; Corynebacteriales; Mycobacteriaceae; Mycobacterium; Mycobacterium chelonae group; Mycobacterium abscessus subgroup; Mycobacterium abscessus
MIKHLTRRATVALVVAAAAISPAAYPAVADTATPLNGLFALTPGSCAEGRATGSYFRMIL WP_005063674.1 hypothetical protein [Mycobacterium abscessus] Length: 330\nNumber of predicted TMHs: 1\nExp number of AAs in TMHs: 39.25259\nExp number, first 60 AAs: 16.98187\nTotal prob of N-in: 0.83210\nPOSSIBLE N-term signal sequence\noutside 1 298\nTMhelix 299 321\ninside 322 330

29821 GCF_000261105.1_ASM26110v1 Mycobacterium abscessus M159 Terrabacteria group; Actinobacteria; Actinobacteria; Corynebacteriales; Mycobacteriaceae; Mycobacterium; Mycobacterium chelonae group; Mycobacterium abscessus subgroup; Mycobacterium abscessus
MIKHLTRRATVALVVAAAAISPAAYPAVADTATPLNGLFALTPGSCAEGRATGSYFRMIL WP_005063674.1 hypothetical protein [Mycobacterium abscessus] Length: 330\nNumber of predicted TMHs: 1\nExp number of AAs in TMHs: 39.25259\nExp number, first 60 AAs: 16.98187\nTotal prob of N-in: 0.83210\nPOSSIBLE N-term signal sequence\noutside 1 298\nTMhelix 299 321\ninside 322 330

29822 GCF_000261125.1_ASM26112v1 Mycobacterium abscessus M172 Terrabacteria group; Actinobacteria; Actinobacteria; Corynebacteriales; Mycobacteriaceae; Mycobacterium; Mycobacterium chelonae group; Mycobacterium abscessus subgroup; Mycobacterium abscessus
MIKHLTRRATVALVVAAAAISPAAYPAVADTATPLNGLFALTPGSCAEGRATGSYFRMIL WP_005063674.1 hypothetical protein [Mycobacterium abscessus] Length: 330\nNumber of predicted TMHs: 1\nExp number of AAs in TMHs: 39.25259\nExp number, first 60 AAs: 16.98187\nTotal prob of N-in: 0.83210\nPOSSIBLE N-term signal sequence\noutside 1 298\nTMhelix 299 321\ninside 322 330

29823 GCF_000260615.2_M24_version02 Mycobacterium abscessus M24 Terrabacteria group; Actinobacteria; Actinobacteria; Corynebacteriales; Mycobacteriaceae; Mycobacterium; Mycobacterium chelonae group; Mycobacterium abscessus subgroup; Mycobacterium abscessus
MIKHLTRRATVALVVAAAAISPAAYPAMADTATPLNGLFALTPGSCAEGRATGSYFRMIL WP_005073213.1
hypothetical protein [Mycobacterium abscessus] Length: 330\nNumber of predicted TMHs: 2\nExp number of AAs in TMHs: 39.3416199999999\nExp number, first 60 AAs: 17.07688\nTotal prob of N-in: 0.84431\nPOSSIBLE N-term signal sequence\ninside 1 11\nTMhelix 12 29\noutside 30 298\nTMhelix 299 321\ninside 322 330

29824 GCF_000257225.2_ASM25722v2 Mycobacterium abscessus M93 Terrabacteria group; Actinobacteria; Actinobacteria; Corynebacteriales; Mycobacteriaceae; Mycobacterium; Mycobacterium chelonae group; Mycobacterium abscessus subgroup; Mycobacterium abscessus
MIKHLTRRATVALVVAAAAISPAAYPAMADTATPLNGLFALTPGSCAEGRATGSYFRMIL WP_005073213.1
hypothetical protein [Mycobacterium abscessus] Length: 330\nNumber of predicted TMHs: 2\nExp number of AAs in TMHs: 39.3416199999999\nExp number, first 60 AAs: 17.07688\nTotal prob of N-in: 0.84431\nPOSSIBLE N-term signal sequence\ninside 1 11\nTMhelix 12 29\noutside 30 298\nTMhelix 299 321\ninside 322 330

29825 GCF_000715335.1_denovo_assembly Mycobacterium abscessus MAB_082312_2273 Terrabacteria group; Actinobacteria; Actinobacteria; Corynebacteriales; Mycobacteriaceae; Mycobacterium; Mycobacterium chelonae group; Mycobacterium abscessus subgroup; Mycobacterium abscessus
MIKHLTRRATVALVVAAAAMSPAAYPAVADTATPLNGLFALTPGSCAEGRATGSYFRMIL WP_030094292.1
MULTISPECIES: hypothetical protein [Mycobacterium chelonae group] Length: 329\nNumber of predicted TMHs: 1\nExp number of AAs in TMHs: 37.05801\nExp number, first 60 AAs: 14.73407\nTotal prob of N-in: 0.73307\nPOSSIBLE N-term signal sequence\noutside 1 297\nTMhelix 298 320\ninside 321 329

29826 GCF_000523915.1_ASM52391v1 Mycobacterium abscessus MAB_091912_2455 Terrabacteria group; Actinobacteria; Actinobacteria; Corynebacteriales; Mycobacteriaceae; Mycobacterium; Mycobacterium chelonae group; Mycobacterium abscessus subgroup; Mycobacterium abscessus
MIKHLTRRATVALVVAAAAISPAAYPAVADTATPLNGLFALTPGSCAEGRATGSYFRMIL WP_005063674.1 hypothetical protein [Mycobacterium abscessus] Length: 330\nNumber of predicted TMHs: 1\nExp number of AAs in TMHs: 39.25259\nExp number, first 60 AAs: 16.98187\nTotal prob of N-in: 0.83210\nPOSSIBLE N-term signal sequence\noutside 1 298\nTMhelix 299 321\ninside 322 330

29827 GCF_000523955.1_ASM52395v1 Mycobacterium abscessus MAB_110811_1470 Terrabacteria group; Actinobacteria; Actinobacteria; Corynebacteriales; Mycobacteriaceae; Mycobacterium; Mycobacterium chelonae group; Mycobacterium abscessus subgroup; Mycobacterium abscessus
MIKHLTRRATVALVVAAAAISPAAYPAMADTATPLNGLFALTPGSCAEGRATGSYFRMIL WP_005073213.1
hypothetical protein [Mycobacterium abscessus] Length: 330\nNumber of predicted TMHs: 2\nExp number of AAs in TMHs: 39.3416199999999\nExp number, first 60 AAs: 17.07688\nTotal prob of N-in: 0.84431\nPOSSIBLE N-term signal sequence\ninside 1 11\nTMhelix 12 29\noutside 30 298\nTMhelix 299 321\ninside 322 330

29828 GCF_000523935.1_ASM52393v1 Mycobacterium abscessus MAB_110811_2726 Terrabacteria group; Actinobacteria; Actinobacteria; Corynebacteriales; Mycobacteriaceae; Mycobacterium; Mycobacterium chelonae group; Mycobacterium abscessus subgroup; Mycobacterium abscessus
MIKHLTRRATVALVVAAAAISPAAYPAMADTATPLNGLFALTPGSCAEGRATGSYFRMIL WP_005073213.1
hypothetical protein [Mycobacterium abscessus] Length: 330\nNumber of predicted TMHs: 2\nExp number of AAs in TMHs: 39.3416199999999\nExp number, first 60 AAs: 17.07688\nTotal prob of N-in: 0.84431\nPOSSIBLE N-term signal sequence\ninside 1 11\nTMhelix 12 29\noutside 30 298\nTMhelix 299 321\ninside 322 330

29829 GCF_900139935.1_10660_1_37 Mycobacterium abscessus subsp. abscessus Terrabacteria group; Actinobacteria; Actinobacteria; Corynebacteriales; Mycobacteriaceae; Mycobacterium; Mycobacterium chelonae group; Mycobacterium abscessus subgroup; Mycobacterium abscessus
MIKHLTRRATVALVVAAAAISPAAYPAMADTATPLNGLFALTPGSCAEGRATGSYFRMIL WP_005073213.1
hypothetical protein [Mycobacterium abscessus] Length: 330\nNumber of predicted TMHs: 2\nExp number of

AAs in TMHs: 39.3416199999999\nExp number, first 60 AAs: 17.07688\nTotal prob of N-in: 0.84431\nPOSSIBLE N-term signal sequence\ninside 1 11\nTMhelix 12 29\noutside 30 298\nTMhelix 299 321\ninside 322 330

29830 GCF_000770215.1_ASM77021v1 Mycobacterium abscessus subsp. abscessus Terrabacteria group; Actinobacteria; Actinobacteria; Corynebacteriales; Mycobacteriaceae; Mycobacterium; Mycobacterium chelonae group; Mycobacterium abscessus subgroup; Mycobacterium abscessus MIKHLTRRATVALVVAAAAISPAAYPAVADTATPLNGLFALTPGSCAEGRATGSYFRMIL WP_005063674.1 hypothetical protein [Mycobacterium abscessus] Length: 330\nNumber of predicted TMHs: 1\nExp number of AAs in TMHs: 39.25259\nExp number, first 60 AAs: 16.98187\nTotal prob of N-in: 0.83210\nPOSSIBLE N-term signal sequence\noutside 1 298\nTMhelix 299 321\ninside 322 330

29831 GCF_001942565.1_ASM194256v1 Mycobacterium abscessus subsp. abscessus Terrabacteria group; Actinobacteria; Actinobacteria; Corynebacteriales; Mycobacteriaceae; Mycobacterium; Mycobacterium chelonae group; Mycobacterium abscessus subgroup; Mycobacterium abscessus MIKHLTRRATVALVVAAAAISPAAYPAMADTATPLNGLFALTPGSCAEGRATGSYFRMIL WP_005073213.1 hypothetical protein [Mycobacterium abscessus] Length: 330\nNumber of predicted TMHs: 2\nExp number of AAs in TMHs: 39.3416199999999\nExp number, first 60 AAs: 17.07688\nTotal prob of N-in: 0.84431\nPOSSIBLE N-term signal sequence\ninside 1 11\nTMhelix 12 29\noutside 30 298\nTMhelix 299 321\ninside 322 330

29832 GCF_900130245.1_10071_6_72 Mycobacterium abscessus subsp. abscessus Terrabacteria group; Actinobacteria; Actinobacteria; Corynebacteriales; Mycobacteriaceae; Mycobacterium; Mycobacterium chelonae group; Mycobacterium abscessus subgroup; Mycobacterium abscessus MIKHLTRRATVALVVAAAAISPAAYPAMADTATPLNGLFALTPGSCAEGRATGSYFRMIL WP_005073213.1 hypothetical protein [Mycobacterium abscessus] Length: 330\nNumber of predicted TMHs: 2\nExp number of AAs in TMHs: 39.3416199999999\nExp number, first 60 AAs: 17.07688\nTotal prob of N-in: 0.84431\nPOSSIBLE N-term signal sequence\ninside 1 11\nTMhelix 12 29\noutside 30 298\nTMhelix 299 321\ninside 322 330

29833 GCF_900130255.1_10071_6_76 Mycobacterium abscessus subsp. abscessus Terrabacteria group; Actinobacteria; Actinobacteria; Corynebacteriales; Mycobacteriaceae; Mycobacterium; Mycobacterium chelonae group; Mycobacterium abscessus subgroup; Mycobacterium abscessus MIKHLTRRATVALVVAAAAISPAAYPAMADTATPLNGLFALTPGSCAEGRATGSYFRMIL WP_005073213.1 hypothetical protein [Mycobacterium abscessus] Length: 330\nNumber of predicted TMHs: 2\nExp number of AAs in TMHs: 39.3416199999999\nExp number, first 60 AAs: 17.07688\nTotal prob of N-in: 0.84431\nPOSSIBLE N-term signal sequence\ninside 1 11\nTMhelix 12 29\noutside 30 298\nTMhelix 299 321\ninside 322 330

29834 GCF_900130425.1_10625_5_78 Mycobacterium abscessus subsp. abscessus Terrabacteria group; Actinobacteria; Actinobacteria; Corynebacteriales; Mycobacteriaceae; Mycobacterium; Mycobacterium chelonae group; Mycobacterium abscessus subgroup; Mycobacterium abscessus MIKHLTRRATVALVVAAAAISPAAYPAMADTATPLNGLFALTPGSCAEGRATGSYFRMIL WP_005073213.1 hypothetical protein [Mycobacterium abscessus] Length: 330\nNumber of predicted TMHs: 2\nExp number of AAs in TMHs: 39.3416199999999\nExp number, first 60 AAs: 17.07688\nTotal prob of N-in: 0.84431\nPOSSIBLE N-term signal sequence\ninside 1 11\nTMhelix 12 29\noutside 30 298\nTMhelix 299 321\ninside 322 330

29835 GCF_900130445.1_10625_5_81 Mycobacterium abscessus subsp. abscessus Terrabacteria group; Actinobacteria; Actinobacteria; Corynebacteriales; Mycobacteriaceae; Mycobacterium; Mycobacterium chelonae group; Mycobacterium abscessus subgroup; Mycobacterium abscessus MIKHLTRRATVALVVAAAAISPAAYPAVADTATPLNGLFALTPGSCAEGRATGSYFRMIL WP_005063674.1 hypothetical protein [Mycobacterium abscessus] Length: 330\nNumber of predicted TMHs: 1\nExp number of AAs in TMHs: 39.25259\nExp number, first 60 AAs: 16.98187\nTotal prob of N-in: 0.83210\nPOSSIBLE N-term signal sequence\noutside 1 298\nTMhelix 299 321\ninside 322 330

29836 GCF_900130475.1_10625_5_85 Mycobacterium abscessus subsp. abscessus Terrabacteria group; Actinobacteria; Actinobacteria; Corynebacteriales; Mycobacteriaceae; Mycobacterium; Mycobacterium chelonae group; Mycobacterium abscessus subgroup; Mycobacterium abscessus MIKHLTRRATVALVVAAAAISPAAYPAVADTATPLNGLFALTPGSCAEGRATGSYFRMIL WP_005063674.1 hypothetical protein [Mycobacterium abscessus] Length: 330\nNumber of predicted TMHs: 1\nExp number of AAs in TMHs: 39.25259\nExp number, first 60 AAs: 16.98187\nTotal prob of N-in: 0.83210\nPOSSIBLE N-term signal sequence\noutside 1 298\nTMhelix 299 321\ninside 322 330

29837 GCF_900130505.1_10665_3_3 Mycobacterium abscessus subsp. abscessus Terrabacteria group; Actinobacteria; Actinobacteria; Corynebacteriales; Mycobacteriaceae; Mycobacterium; Mycobacterium chelonae group;

Mycobacterium abscessus subgroup; Mycobacterium abscessus
MIKHLTRRATVALVVAAAAISPAAYPAMADTATPLNGLFALTPGSCAEGRATGSYFRMIL WP_005073213.1
hypothetical protein [Mycobacterium abscessus] Length: 330\nNumber of predicted TMHs: 2\nExp number of AAs in TMHs: 39.3416199999999\nExp number, first 60 AAs: 17.07688\nTotal prob of N-in: 0.84431\nPOSSIBLE N-term signal sequence\ninside 1 11\nTMhelix 12 29\noutside 30 298\nTMhelix 299 321\ninside 322 330

29838 GCF_900130495.1_10665_3_2 Mycobacterium abscessus subsp. abscessus Terrabacteria group; Actinobacteria; Actinobacteria; Corynebacteriales; Mycobacteriaceae; Mycobacterium; Mycobacterium chelonae group; Mycobacterium abscessus subgroup; Mycobacterium abscessus
MIKHLTRRATVALVVAAAAISPAAYPAMADTATPLNGLFALTPGSCAEGRATGSYFRMIL WP_005073213.1
hypothetical protein [Mycobacterium abscessus] Length: 330\nNumber of predicted TMHs: 2\nExp number of AAs in TMHs: 39.3416199999999\nExp number, first 60 AAs: 17.07688\nTotal prob of N-in: 0.84431\nPOSSIBLE N-term signal sequence\ninside 1 11\nTMhelix 12 29\noutside 30 298\nTMhelix 299 321\ninside 322 330

29839 GCF_900130565.1_11861_7_25 Mycobacterium abscessus subsp. abscessus Terrabacteria group; Actinobacteria; Actinobacteria; Corynebacteriales; Mycobacteriaceae; Mycobacterium; Mycobacterium chelonae group; Mycobacterium abscessus subgroup; Mycobacterium abscessus
MIKHLTRRATVALVVAAAAISPAAYPAMADTATPLNGLFALTPGSCAEGRATGSYFRMIL WP_074356318.1
hypothetical protein [Mycobacterium abscessus] Length: 330\nNumber of predicted TMHs: 2\nExp number of AAs in TMHs: 39.34118\nExp number, first 60 AAs: 17.07668\nTotal prob of N-in: 0.84431\nPOSSIBLE N-term signal sequence\ninside 1 11\nTMhelix 12 29\noutside 30 298\nTMhelix 299 321\ninside 322 330

29840 GCF_900130845.1_10665_3_35 Mycobacterium abscessus subsp. abscessus Terrabacteria group; Actinobacteria; Actinobacteria; Corynebacteriales; Mycobacteriaceae; Mycobacterium; Mycobacterium chelonae group; Mycobacterium abscessus subgroup; Mycobacterium abscessus
MIKHLTRRATVALVVAAAAISPAAYPAMADTATPLNGLFALTPGSCAEGRATGSYFRMIL WP_005073213.1
hypothetical protein [Mycobacterium abscessus] Length: 330\nNumber of predicted TMHs: 2\nExp number of AAs in TMHs: 39.3416199999999\nExp number, first 60 AAs: 17.07688\nTotal prob of N-in: 0.84431\nPOSSIBLE N-term signal sequence\ninside 1 11\nTMhelix 12 29\noutside 30 298\nTMhelix 299 321\ninside 322 330

29841 GCF_900130895.1_10665_3_42 Mycobacterium abscessus subsp. abscessus Terrabacteria group; Actinobacteria; Actinobacteria; Corynebacteriales; Mycobacteriaceae; Mycobacterium; Mycobacterium chelonae group; Mycobacterium abscessus subgroup; Mycobacterium abscessus
MIKHLTRRATVALVVAAAAISPAAYPAMADTATPLNGLFALTPGSCAEGRATGSYFRMIL WP_005073213.1
hypothetical protein [Mycobacterium abscessus] Length: 330\nNumber of predicted TMHs: 2\nExp number of AAs in TMHs: 39.3416199999999\nExp number, first 60 AAs: 17.07688\nTotal prob of N-in: 0.84431\nPOSSIBLE N-term signal sequence\ninside 1 11\nTMhelix 12 29\noutside 30 298\nTMhelix 299 321\ninside 322 330

29842 GCF_900130935.1_10665_3_47 Mycobacterium abscessus subsp. abscessus Terrabacteria group; Actinobacteria; Actinobacteria; Corynebacteriales; Mycobacteriaceae; Mycobacterium; Mycobacterium chelonae group; Mycobacterium abscessus subgroup; Mycobacterium abscessus
MIKHLTRRATVALVVAAAAISPAAYPAMADTATPLNGLFALTPGSCAEGRATGSYFRMIL WP_005073213.1
hypothetical protein [Mycobacterium abscessus] Length: 330\nNumber of predicted TMHs: 2\nExp number of AAs in TMHs: 39.3416199999999\nExp number, first 60 AAs: 17.07688\nTotal prob of N-in: 0.84431\nPOSSIBLE N-term signal sequence\ninside 1 11\nTMhelix 12 29\noutside 30 298\nTMhelix 299 321\ninside 322 330

29843 GCF_900130915.1_10665_3_44 Mycobacterium abscessus subsp. abscessus Terrabacteria group; Actinobacteria; Actinobacteria; Corynebacteriales; Mycobacteriaceae; Mycobacterium; Mycobacterium chelonae group; Mycobacterium abscessus subgroup; Mycobacterium abscessus
MIKHLTRRATVALVVAAAAISPAAYPAMADTATPLNGLFALTPGSCAEGRATGSYFRMIL WP_005073213.1
hypothetical protein [Mycobacterium abscessus] Length: 330\nNumber of predicted TMHs: 2\nExp number of AAs in TMHs: 39.3416199999999\nExp number, first 60 AAs: 17.07688\nTotal prob of N-in: 0.84431\nPOSSIBLE N-term signal sequence\ninside 1 11\nTMhelix 12 29\noutside 30 298\nTMhelix 299 321\ninside 322 330

29844 GCF_900130955.1_10665_3_46 Mycobacterium abscessus subsp. abscessus Terrabacteria group; Actinobacteria; Actinobacteria; Corynebacteriales; Mycobacteriaceae; Mycobacterium; Mycobacterium chelonae group; Mycobacterium abscessus subgroup; Mycobacterium abscessus
MIKHLTRRATVALVVAAAAISPAAYPAMADTATPLNGLFALTPGSCAEGRATGSYFRMIL WP_005073213.1
hypothetical protein [Mycobacterium abscessus] Length: 330\nNumber of predicted TMHs: 2\nExp number of AAs in TMHs: 39.3416199999999\nExp number, first 60 AAs: 17.07688\nTotal prob of N-in: 0.84431\nPOSSIBLE N-term signal sequence\ninside 1 11\nTMhelix 12 29\noutside 30 298\nTMhelix 299 321\ninside 322 330

29845 GCF_900131085.1_10665_3_52 Mycobacterium abscessus subsp. abscessus Terrabacteria group; Actinobacteria; Actinobacteria; Corynebacteriales; Mycobacteriaceae; Mycobacterium; Mycobacterium chelonae group; Mycobacterium abscessus subgroup; Mycobacterium abscessus
MIKHLTRRATVALVVAAAAISPAAYPAMADTATPLNGLFALTPGSCAEGRATGSYFRMIL WP_005073213.1 hypothetical protein [Mycobacterium abscessus] Length: 330\nNumber of predicted TMHs: 2\nExp number of AAs in TMHs: 39.3416199999999\nExp number, first 60 AAs: 17.07688\nTotal prob of N-in: 0.84431\nPOSSIBLE N-term signal sequence\ninside 1 11\nTMhelix 12 29\noutside 30 298\nTMhelix 299 321\ninside 322 330

29846 GCF_900131155.1_10665_3_60 Mycobacterium abscessus subsp. abscessus Terrabacteria group; Actinobacteria; Actinobacteria; Corynebacteriales; Mycobacteriaceae; Mycobacterium; Mycobacterium chelonae group; Mycobacterium abscessus subgroup; Mycobacterium abscessus
MIKHLTRRATVALVVAAAAISPAAYPAMADTATPLNGLFALTPGSCAEGRATGSYFRMIL WP_005073213.1 hypothetical protein [Mycobacterium abscessus] Length: 330\nNumber of predicted TMHs: 2\nExp number of AAs in TMHs: 39.3416199999999\nExp number, first 60 AAs: 17.07688\nTotal prob of N-in: 0.84431\nPOSSIBLE N-term signal sequence\ninside 1 11\nTMhelix 12 29\noutside 30 298\nTMhelix 299 321\ninside 322 330

29847 GCF_900131165.1_10665_3_61 Mycobacterium abscessus subsp. abscessus Terrabacteria group; Actinobacteria; Actinobacteria; Corynebacteriales; Mycobacteriaceae; Mycobacterium; Mycobacterium chelonae group; Mycobacterium abscessus subgroup; Mycobacterium abscessus
MIKHLTRRATVALVVAAAAISPAAYPAVADTATPLNGLFALTPGSCAEGRATGSYFRMIL WP_005063674.1 hypothetical protein [Mycobacterium abscessus] Length: 330\nNumber of predicted TMHs: 1\nExp number of AAs in TMHs: 39.25259\nExp number, first 60 AAs: 16.98187\nTotal prob of N-in: 0.83210\nPOSSIBLE N-term signal sequence\noutside 1 298\nTMhelix 299 321\ninside 322 330

29848 GCF_900131275.1_10665_3_72 Mycobacterium abscessus subsp. abscessus Terrabacteria group; Actinobacteria; Actinobacteria; Corynebacteriales; Mycobacteriaceae; Mycobacterium; Mycobacterium chelonae group; Mycobacterium abscessus subgroup; Mycobacterium abscessus
MIKHLTRRATVALVVAAAAISPAAYPAMADTATPLNGLFALTPGSCAEGRATGSYFRMIL WP_005073213.1 hypothetical protein [Mycobacterium abscessus] Length: 330\nNumber of predicted TMHs: 2\nExp number of AAs in TMHs: 39.3416199999999\nExp number, first 60 AAs: 17.07688\nTotal prob of N-in: 0.84431\nPOSSIBLE N-term signal sequence\ninside 1 11\nTMhelix 12 29\noutside 30 298\nTMhelix 299 321\ninside 322 330

29849 GCF_900131335.1_10665_3_80 Mycobacterium abscessus subsp. abscessus Terrabacteria group; Actinobacteria; Actinobacteria; Corynebacteriales; Mycobacteriaceae; Mycobacterium; Mycobacterium chelonae group; Mycobacterium abscessus subgroup; Mycobacterium abscessus
MIKHLTRRATVALVVAAAAISPAAYPAMADTATPLNGLFALTPGSCAEGRATGSYFRMIL WP_005073213.1 hypothetical protein [Mycobacterium abscessus] Length: 330\nNumber of predicted TMHs: 2\nExp number of AAs in TMHs: 39.3416199999999\nExp number, first 60 AAs: 17.07688\nTotal prob of N-in: 0.84431\nPOSSIBLE N-term signal sequence\ninside 1 11\nTMhelix 12 29\noutside 30 298\nTMhelix 299 321\ninside 322 330

29850 GCF_900131345.1_10665_3_81 Mycobacterium abscessus subsp. abscessus Terrabacteria group; Actinobacteria; Actinobacteria; Corynebacteriales; Mycobacteriaceae; Mycobacterium; Mycobacterium chelonae group; Mycobacterium abscessus subgroup; Mycobacterium abscessus
MIKHLTRRATVALVVAAAAISPAAYPAMADTATPLNGLFALTPGSCAEGRATGSYFRMIL WP_005073213.1 hypothetical protein [Mycobacterium abscessus] Length: 330\nNumber of predicted TMHs: 2\nExp number of AAs in TMHs: 39.3416199999999\nExp number, first 60 AAs: 17.07688\nTotal prob of N-in: 0.84431\nPOSSIBLE N-term signal sequence\ninside 1 11\nTMhelix 12 29\noutside 30 298\nTMhelix 299 321\ninside 322 330

29851 GCF_900131365.1_10665_3_83 Mycobacterium abscessus subsp. abscessus Terrabacteria group; Actinobacteria; Actinobacteria; Corynebacteriales; Mycobacteriaceae; Mycobacterium; Mycobacterium chelonae group; Mycobacterium abscessus subgroup; Mycobacterium abscessus
MIKHLTRRATVALVVAAAAISPAAYPAMADTATPLNGLFALTPGSCAEGRATGSYFRMIL WP_005073213.1 hypothetical protein [Mycobacterium abscessus] Length: 330\nNumber of predicted TMHs: 2\nExp number of AAs in TMHs: 39.3416199999999\nExp number, first 60 AAs: 17.07688\nTotal prob of N-in: 0.84431\nPOSSIBLE N-term signal sequence\ninside 1 11\nTMhelix 12 29\noutside 30 298\nTMhelix 299 321\ninside 322 330

29852 GCF_900131575.1_10665_4_14 Mycobacterium abscessus subsp. abscessus Terrabacteria group; Actinobacteria; Actinobacteria; Corynebacteriales; Mycobacteriaceae; Mycobacterium; Mycobacterium chelonae group; Mycobacterium abscessus subgroup; Mycobacterium abscessus
MIKHLTRRATVALVVAAAAISPAAYPAMADTATPLNGLFALTPGSCAEGRATGSYFRMIL WP_005073213.1 hypothetical protein [Mycobacterium abscessus] Length: 330\nNumber of predicted TMHs: 2\nExp number of

AAs in TMHs: 39.3416199999999\nExp number, first 60 AAs: 17.07688\nTotal prob of N-in: 0.84431\nPOSSIBLE N-term signal sequence\ninside 1 11\nTMhelix 12 29\noutside 30 298\nTMhelix 299 321\ninside 322 330

29853 GCF_900131645.1_10665_4_22 Mycobacterium abscessus subsp. abscessus Terrabacteria group; Actinobacteria; Actinobacteria; Corynebacteriales; Mycobacteriaceae; Mycobacterium; Mycobacterium chelonae group; Mycobacterium abscessus subgroup; Mycobacterium abscessus
MIKHLTRRATVALVVAAAAISPAAYPAMADTATPLNGLFALTPGSCAEGRATGSYFRMIL WP_005073213.1
hypothetical protein [Mycobacterium abscessus] Length: 330\nNumber of predicted TMHs: 2\nExp number of AAs in TMHs: 39.3416199999999\nExp number, first 60 AAs: 17.07688\nTotal prob of N-in: 0.84431\nPOSSIBLE N-term signal sequence\ninside 1 11\nTMhelix 12 29\noutside 30 298\nTMhelix 299 321\ninside 322 330

29854 GCF_900131665.1_10665_4_25 Mycobacterium abscessus subsp. abscessus Terrabacteria group; Actinobacteria; Actinobacteria; Corynebacteriales; Mycobacteriaceae; Mycobacterium; Mycobacterium chelonae group; Mycobacterium abscessus subgroup; Mycobacterium abscessus
MIKHLTRRATVALVVAAAAISPAAYPAMADTATPLNGLFALTPGSCAEGRATGSYFRMIL WP_005073213.1
hypothetical protein [Mycobacterium abscessus] Length: 330\nNumber of predicted TMHs: 2\nExp number of AAs in TMHs: 39.3416199999999\nExp number, first 60 AAs: 17.07688\nTotal prob of N-in: 0.84431\nPOSSIBLE N-term signal sequence\ninside 1 11\nTMhelix 12 29\noutside 30 298\nTMhelix 299 321\ninside 322 330

29855 GCF_900131715.1_10665_4_30 Mycobacterium abscessus subsp. abscessus Terrabacteria group; Actinobacteria; Actinobacteria; Corynebacteriales; Mycobacteriaceae; Mycobacterium; Mycobacterium chelonae group; Mycobacterium abscessus subgroup; Mycobacterium abscessus
MIKHLTRRATVALVVAAAAISPAAYPAMADTATPLNGLFALTPGSCAEGRATGSYFRMIL WP_074250269.1
hypothetical protein [Mycobacterium abscessus] Length: 330\nNumber of predicted TMHs: 2\nExp number of AAs in TMHs: 39.3415199999999\nExp number, first 60 AAs: 17.07715\nTotal prob of N-in: 0.84431\nPOSSIBLE N-term signal sequence\ninside 1 11\nTMhelix 12 29\noutside 30 298\nTMhelix 299 321\ninside 322 330

29856 GCF_900131925.1_10665_4_38 Mycobacterium abscessus subsp. abscessus Terrabacteria group; Actinobacteria; Actinobacteria; Corynebacteriales; Mycobacteriaceae; Mycobacterium; Mycobacterium chelonae group; Mycobacterium abscessus subgroup; Mycobacterium abscessus
MIKHLTRRATVALVVAAAAISPAAYPAMADTATPLNGLFALTPGSCAEGRATGSYFRMIL WP_005073213.1
hypothetical protein [Mycobacterium abscessus] Length: 330\nNumber of predicted TMHs: 2\nExp number of AAs in TMHs: 39.3416199999999\nExp number, first 60 AAs: 17.07688\nTotal prob of N-in: 0.84431\nPOSSIBLE N-term signal sequence\ninside 1 11\nTMhelix 12 29\noutside 30 298\nTMhelix 299 321\ninside 322 330

29857 GCF_900131805.1_11861_7_30 Mycobacterium abscessus subsp. abscessus Terrabacteria group; Actinobacteria; Actinobacteria; Corynebacteriales; Mycobacteriaceae; Mycobacterium; Mycobacterium chelonae group; Mycobacterium abscessus subgroup; Mycobacterium abscessus
MIKHLTRRATVALVVAAAAISPAAYPAMADTATPLNGLFALTPGSCAEGRATGSYFRMIL WP_074340136.1
hypothetical protein [Mycobacterium abscessus] Length: 330\nNumber of predicted TMHs: 2\nExp number of AAs in TMHs: 39.3416799999999\nExp number, first 60 AAs: 17.07671\nTotal prob of N-in: 0.84432\nPOSSIBLE N-term signal sequence\ninside 1 11\nTMhelix 12 29\noutside 30 298\nTMhelix 299 321\ninside 322 330

29858 GCF_900131965.1_10665_4_41 Mycobacterium abscessus subsp. abscessus Terrabacteria group; Actinobacteria; Actinobacteria; Corynebacteriales; Mycobacteriaceae; Mycobacterium; Mycobacterium chelonae group; Mycobacterium abscessus subgroup; Mycobacterium abscessus
MIKHLTRRATVALVVAAAAISPAAYPAMADTATPLNGLFALTPGSCAEGRATGSYFRMIL WP_005073213.1
hypothetical protein [Mycobacterium abscessus] Length: 330\nNumber of predicted TMHs: 2\nExp number of AAs in TMHs: 39.3416199999999\nExp number, first 60 AAs: 17.07688\nTotal prob of N-in: 0.84431\nPOSSIBLE N-term signal sequence\ninside 1 11\nTMhelix 12 29\noutside 30 298\nTMhelix 299 321\ninside 322 330

29859 GCF_900132135.1_10665_4_59 Mycobacterium abscessus subsp. abscessus Terrabacteria group; Actinobacteria; Actinobacteria; Corynebacteriales; Mycobacteriaceae; Mycobacterium; Mycobacterium chelonae group; Mycobacterium abscessus subgroup; Mycobacterium abscessus
MIKHLTRRATVALVVAAAAISPAAYPAMADTATPLNGLFALTPGSCAEGRATGSYFRMIL WP_005073213.1
hypothetical protein [Mycobacterium abscessus] Length: 330\nNumber of predicted TMHs: 2\nExp number of AAs in TMHs: 39.3416199999999\nExp number, first 60 AAs: 17.07688\nTotal prob of N-in: 0.84431\nPOSSIBLE N-term signal sequence\ninside 1 11\nTMhelix 12 29\noutside 30 298\nTMhelix 299 321\ninside 322 330

29860 GCF_900132015.1_10665_4_44 Mycobacterium abscessus subsp. abscessus Terrabacteria group; Actinobacteria; Actinobacteria; Corynebacteriales; Mycobacteriaceae; Mycobacterium; Mycobacterium chelonae group;

Mycobacterium abscessus subgroup; Mycobacterium abscessus
MIKHLTRRATVALVVAAAAISPAAYPAMADTATPLNGLFALTPGSCAEGRATGSYFRMIL WP_074250269.1
hypothetical protein [Mycobacterium abscessus] Length: 330\nNumber of predicted TMHs: 2\nExp number of AAs in TMHs: 39.3415199999999\nExp number, first 60 AAs: 17.07715\nTotal prob of N-in: 0.84431\nPOSSIBLE N-term signal sequence\ninside 1 11\nTMhelix 12 29\noutside 30 298\nTMhelix 299 321\ninside 322 330

29861 GCF_900132365.1_10665_4_68 Mycobacterium abscessus subsp. abscessus Terrabacteria group; Actinobacteria; Actinobacteria; Corynebacteriales; Mycobacteriaceae; Mycobacterium; Mycobacterium chelonae group; Mycobacterium abscessus subgroup; Mycobacterium abscessus
MIKHLTRRATVALVVAAAAISPAAYPAMADTATPLNGLFALTPGSCAEGRATGSYFRMIL WP_005073213.1
hypothetical protein [Mycobacterium abscessus] Length: 330\nNumber of predicted TMHs: 2\nExp number of AAs in TMHs: 39.3416199999999\nExp number, first 60 AAs: 17.07688\nTotal prob of N-in: 0.84431\nPOSSIBLE N-term signal sequence\ninside 1 11\nTMhelix 12 29\noutside 30 298\nTMhelix 299 321\ninside 322 330

29862 GCF_900132415.1_10665_4_73 Mycobacterium abscessus subsp. abscessus Terrabacteria group; Actinobacteria; Actinobacteria; Corynebacteriales; Mycobacteriaceae; Mycobacterium; Mycobacterium chelonae group; Mycobacterium abscessus subgroup; Mycobacterium abscessus
MIKHLTRRATVALVVAAAAISPAAYPAMADTATPLNGLFALTPGSCAEGRATGSYFRMIL WP_005073213.1
hypothetical protein [Mycobacterium abscessus] Length: 330\nNumber of predicted TMHs: 2\nExp number of AAs in TMHs: 39.3416199999999\nExp number, first 60 AAs: 17.07688\nTotal prob of N-in: 0.84431\nPOSSIBLE N-term signal sequence\ninside 1 11\nTMhelix 12 29\noutside 30 298\nTMhelix 299 321\ninside 322 330

29863 GCF_900132435.1_10665_4_75 Mycobacterium abscessus subsp. abscessus Terrabacteria group; Actinobacteria; Actinobacteria; Corynebacteriales; Mycobacteriaceae; Mycobacterium; Mycobacterium chelonae group; Mycobacterium abscessus subgroup; Mycobacterium abscessus
MIKHLTRRATVALVVAAAAISPAAYPAMADTATPLNGLFALTPGSCAEGRATGSYFRMIL WP_005073213.1
hypothetical protein [Mycobacterium abscessus] Length: 330\nNumber of predicted TMHs: 2\nExp number of AAs in TMHs: 39.3416199999999\nExp number, first 60 AAs: 17.07688\nTotal prob of N-in: 0.84431\nPOSSIBLE N-term signal sequence\ninside 1 11\nTMhelix 12 29\noutside 30 298\nTMhelix 299 321\ninside 322 330

29864 GCF_900132665.1_10250_1_15 Mycobacterium abscessus subsp. abscessus Terrabacteria group; Actinobacteria; Actinobacteria; Corynebacteriales; Mycobacteriaceae; Mycobacterium; Mycobacterium chelonae group; Mycobacterium abscessus subgroup; Mycobacterium abscessus
MIKHLTRRATVALVVAAAAISPAAYPAMADTATPLNGLFALTPGSCAEGRATGSYFRMIL WP_005073213.1
hypothetical protein [Mycobacterium abscessus] Length: 330\nNumber of predicted TMHs: 2\nExp number of AAs in TMHs: 39.3416199999999\nExp number, first 60 AAs: 17.07688\nTotal prob of N-in: 0.84431\nPOSSIBLE N-term signal sequence\ninside 1 11\nTMhelix 12 29\noutside 30 298\nTMhelix 299 321\ninside 322 330

29865 GCF_900132675.1_10250_1_17 Mycobacterium abscessus subsp. abscessus Terrabacteria group; Actinobacteria; Actinobacteria; Corynebacteriales; Mycobacteriaceae; Mycobacterium; Mycobacterium chelonae group; Mycobacterium abscessus subgroup; Mycobacterium abscessus
MIKHLTRRATVALVVAAAAISPAAYPAMADTATPLNGLFALTPGSCAEGRATGSYFRMIL WP_005073213.1
hypothetical protein [Mycobacterium abscessus] Length: 330\nNumber of predicted TMHs: 2\nExp number of AAs in TMHs: 39.3416199999999\nExp number, first 60 AAs: 17.07688\nTotal prob of N-in: 0.84431\nPOSSIBLE N-term signal sequence\ninside 1 11\nTMhelix 12 29\noutside 30 298\nTMhelix 299 321\ninside 322 330

29866 GCF_900132515.1_11861_7_33 Mycobacterium abscessus subsp. abscessus Terrabacteria group; Actinobacteria; Actinobacteria; Corynebacteriales; Mycobacteriaceae; Mycobacterium; Mycobacterium chelonae group; Mycobacterium abscessus subgroup; Mycobacterium abscessus
MIKHLTRRATVALVVAAAAISPAAYPAMADTATPLNGLFALTPGSCAEGRATGSYFRMIL WP_074340136.1
hypothetical protein [Mycobacterium abscessus] Length: 330\nNumber of predicted TMHs: 2\nExp number of AAs in TMHs: 39.3416799999999\nExp number, first 60 AAs: 17.07671\nTotal prob of N-in: 0.84432\nPOSSIBLE N-term signal sequence\ninside 1 11\nTMhelix 12 29\noutside 30 298\nTMhelix 299 321\ninside 322 330

29867 GCF_900132845.1_10250_1_8 Mycobacterium abscessus subsp. abscessus Terrabacteria group; Actinobacteria; Actinobacteria; Corynebacteriales; Mycobacteriaceae; Mycobacterium; Mycobacterium chelonae group; Mycobacterium abscessus subgroup; Mycobacterium abscessus
MIKHLTRRATVALVVAAAAISPAAYPAMADTATPLNGLFALTPGSCAEGRATGSYFRMIL WP_005073213.1
hypothetical protein [Mycobacterium abscessus] Length: 330\nNumber of predicted TMHs: 2\nExp number of AAs in TMHs: 39.3416199999999\nExp number, first 60 AAs: 17.07688\nTotal prob of N-in: 0.84431\nPOSSIBLE N-term signal sequence\ninside 1 11\nTMhelix 12 29\noutside 30 298\nTMhelix 299 321\ninside 322 330

29868 GCF_900132855.1_10665_4_85 Mycobacterium abscessus subsp. abscessus Terrabacteria group; Actinobacteria; Actinobacteria; Corynebacteriales; Mycobacteriaceae; Mycobacterium; Mycobacterium chelonae group; Mycobacterium abscessus subgroup; Mycobacterium abscessus
MIKHLTRRATVALVVAAAAISPAAYPAMADTATPLNGLFALTPGSCAEGRATGSYFRMIL WP_005073213.1
hypothetical protein [Mycobacterium abscessus] Length: 330\nNumber of predicted TMHs: 2\nExp number of AAs in TMHs: 39.3416199999999\nExp number, first 60 AAs: 17.07688\nTotal prob of N-in: 0.84431\nPOSSIBLE N-term signal sequence\ninside 1 11\nTMhelix 12 29\noutside 30 298\nTMhelix 299 321\ninside 322 330

29869 GCF_900133005.1_11893_6_62 Mycobacterium abscessus subsp. abscessus Terrabacteria group; Actinobacteria; Actinobacteria; Corynebacteriales; Mycobacteriaceae; Mycobacterium; Mycobacterium chelonae group; Mycobacterium abscessus subgroup; Mycobacterium abscessus
MIKHLTRRATVALVVAAAAISPAAYPAMADTATPLNGLFALTPGSCAEGRATGSYFRMIL WP_005073213.1
hypothetical protein [Mycobacterium abscessus] Length: 330\nNumber of predicted TMHs: 2\nExp number of AAs in TMHs: 39.3416199999999\nExp number, first 60 AAs: 17.07688\nTotal prob of N-in: 0.84431\nPOSSIBLE N-term signal sequence\ninside 1 11\nTMhelix 12 29\noutside 30 298\nTMhelix 299 321\ninside 322 330

29870 GCF_900133175.1_12082_5_71 Mycobacterium abscessus subsp. abscessus Terrabacteria group; Actinobacteria; Actinobacteria; Corynebacteriales; Mycobacteriaceae; Mycobacterium; Mycobacterium chelonae group; Mycobacterium abscessus subgroup; Mycobacterium abscessus
MIKHLTRRATVALVVAAAAISPAAYPAMADTATPLNGLFALTPGSCAEGRATGSYFRMIL WP_005073213.1
hypothetical protein [Mycobacterium abscessus] Length: 330\nNumber of predicted TMHs: 2\nExp number of AAs in TMHs: 39.3416199999999\nExp number, first 60 AAs: 17.07688\nTotal prob of N-in: 0.84431\nPOSSIBLE N-term signal sequence\ninside 1 11\nTMhelix 12 29\noutside 30 298\nTMhelix 299 321\ninside 322 330

29871 GCF_900133215.1_12163_2_83 Mycobacterium abscessus subsp. abscessus Terrabacteria group; Actinobacteria; Actinobacteria; Corynebacteriales; Mycobacteriaceae; Mycobacterium; Mycobacterium chelonae group; Mycobacterium abscessus subgroup; Mycobacterium abscessus
MIKHLTRRATVALVVAAAAISPAAYPAMADTATPLNGLFALTPGSCAEGRATGSYFRMIL WP_005073213.1
hypothetical protein [Mycobacterium abscessus] Length: 330\nNumber of predicted TMHs: 2\nExp number of AAs in TMHs: 39.3416199999999\nExp number, first 60 AAs: 17.07688\nTotal prob of N-in: 0.84431\nPOSSIBLE N-term signal sequence\ninside 1 11\nTMhelix 12 29\noutside 30 298\nTMhelix 299 321\ninside 322 330

29872 GCF_900133295.1_10250_1_34 Mycobacterium abscessus subsp. abscessus Terrabacteria group; Actinobacteria; Actinobacteria; Corynebacteriales; Mycobacteriaceae; Mycobacterium; Mycobacterium chelonae group; Mycobacterium abscessus subgroup; Mycobacterium abscessus
MIKHLTRRATVALVVAAAAISPAAYPAMADTATPLNGLFALTPGSCAEGRATGSYFRMIL WP_005073213.1
hypothetical protein [Mycobacterium abscessus] Length: 330\nNumber of predicted TMHs: 2\nExp number of AAs in TMHs: 39.3416199999999\nExp number, first 60 AAs: 17.07688\nTotal prob of N-in: 0.84431\nPOSSIBLE N-term signal sequence\ninside 1 11\nTMhelix 12 29\noutside 30 298\nTMhelix 299 321\ninside 322 330

29873 GCF_900133435.1_10250_1_51 Mycobacterium abscessus subsp. abscessus Terrabacteria group; Actinobacteria; Actinobacteria; Corynebacteriales; Mycobacteriaceae; Mycobacterium; Mycobacterium chelonae group; Mycobacterium abscessus subgroup; Mycobacterium abscessus
MIKHLTRRATVALVVAAAAISPAAYPAMADTATPLNGLFALTPGSCAEGRATGSYFRMIL WP_005073213.1
hypothetical protein [Mycobacterium abscessus] Length: 330\nNumber of predicted TMHs: 2\nExp number of AAs in TMHs: 39.3416199999999\nExp number, first 60 AAs: 17.07688\nTotal prob of N-in: 0.84431\nPOSSIBLE N-term signal sequence\ninside 1 11\nTMhelix 12 29\noutside 30 298\nTMhelix 299 321\ninside 322 330

29874 GCF_900133465.1_10396_8_2 Mycobacterium abscessus subsp. abscessus Terrabacteria group; Actinobacteria; Actinobacteria; Corynebacteriales; Mycobacteriaceae; Mycobacterium; Mycobacterium chelonae group; Mycobacterium abscessus subgroup; Mycobacterium abscessus
MIKHLTRRATVALVVAAAAISPAAYPAMADTATPLNGLFALTPGSCAEGRATGSYFRMIL WP_005073213.1
hypothetical protein [Mycobacterium abscessus] Length: 330\nNumber of predicted TMHs: 2\nExp number of AAs in TMHs: 39.3416199999999\nExp number, first 60 AAs: 17.07688\nTotal prob of N-in: 0.84431\nPOSSIBLE N-term signal sequence\ninside 1 11\nTMhelix 12 29\noutside 30 298\nTMhelix 299 321\ninside 322 330

29875 GCF_900133325.1_10250_1_39 Mycobacterium abscessus subsp. abscessus Terrabacteria group; Actinobacteria; Actinobacteria; Corynebacteriales; Mycobacteriaceae; Mycobacterium; Mycobacterium chelonae group; Mycobacterium abscessus subgroup; Mycobacterium abscessus
MIKHLTRRATVALVVAAAAISPAAYPAMADTATPLNGLFALTPGSCAEGRATGSYFRMIL WP_074250269.1
hypothetical protein [Mycobacterium abscessus] Length: 330\nNumber of predicted TMHs: 2\nExp number of

AAs in TMHs: 39.3415199999999\nExp number, first 60 AAs: 17.07715\nTotal prob of N-in: 0.84431\nPOSSIBLE N-term signal sequence\ninside 1 11\nTMhelix 12 29\noutside 30 298\nTMhelix 299 321\ninside 322 330

29876 GCF_900133505.1_10625_4_74 Mycobacterium abscessus subsp. abscessus Terrabacteria group; Actinobacteria; Actinobacteria; Corynebacteriales; Mycobacteriaceae; Mycobacterium; Mycobacterium chelonae group; Mycobacterium abscessus subgroup; Mycobacterium abscessus
MIKHLTRRATVALVVAAAAISPAAYPAMADTATPLNGLFALTPGSCAEGRATGSYFRMIL WP_005073213.1
hypothetical protein [Mycobacterium abscessus] Length: 330\nNumber of predicted TMHs: 2\nExp number of AAs in TMHs: 39.3416199999999\nExp number, first 60 AAs: 17.07688\nTotal prob of N-in: 0.84431\nPOSSIBLE N-term signal sequence\ninside 1 11\nTMhelix 12 29\noutside 30 298\nTMhelix 299 321\ninside 322 330

29877 GCF_900133515.1_11861_7_47 Mycobacterium abscessus subsp. abscessus Terrabacteria group; Actinobacteria; Actinobacteria; Corynebacteriales; Mycobacteriaceae; Mycobacterium; Mycobacterium chelonae group; Mycobacterium abscessus subgroup; Mycobacterium abscessus
MIKHLTRRATVALVVAAAAISPAAYPAMADTATPLNGLFALTPGSCAEGRATGSYFRMIL WP_005073213.1
hypothetical protein [Mycobacterium abscessus] Length: 330\nNumber of predicted TMHs: 2\nExp number of AAs in TMHs: 39.3416199999999\nExp number, first 60 AAs: 17.07688\nTotal prob of N-in: 0.84431\nPOSSIBLE N-term signal sequence\ninside 1 11\nTMhelix 12 29\noutside 30 298\nTMhelix 299 321\ninside 322 330

29878 GCF_900133645.1_12163_1_14 Mycobacterium abscessus subsp. abscessus Terrabacteria group; Actinobacteria; Actinobacteria; Corynebacteriales; Mycobacteriaceae; Mycobacterium; Mycobacterium chelonae group; Mycobacterium abscessus subgroup; Mycobacterium abscessus
MIKHLTRRATVALVVAAAAISPAAYPAMADTATPLNGLFALTPGSCAEGRATGSYFRMIL WP_005073213.1
hypothetical protein [Mycobacterium abscessus] Length: 330\nNumber of predicted TMHs: 2\nExp number of AAs in TMHs: 39.3416199999999\nExp number, first 60 AAs: 17.07688\nTotal prob of N-in: 0.84431\nPOSSIBLE N-term signal sequence\ninside 1 11\nTMhelix 12 29\noutside 30 298\nTMhelix 299 321\ninside 322 330

29879 GCF_900133665.1_12163_1_13 Mycobacterium abscessus subsp. abscessus Terrabacteria group; Actinobacteria; Actinobacteria; Corynebacteriales; Mycobacteriaceae; Mycobacterium; Mycobacterium chelonae group; Mycobacterium abscessus subgroup; Mycobacterium abscessus
MIKHLTRRATVALVVAAAAISPAAYPAMADTATPLNGLFALTPGSCAEGRATGSYFRMIL WP_005073213.1
hypothetical protein [Mycobacterium abscessus] Length: 330\nNumber of predicted TMHs: 2\nExp number of AAs in TMHs: 39.3416199999999\nExp number, first 60 AAs: 17.07688\nTotal prob of N-in: 0.84431\nPOSSIBLE N-term signal sequence\ninside 1 11\nTMhelix 12 29\noutside 30 298\nTMhelix 299 321\ninside 322 330

29880 GCF_900133705.1_12163_1_20 Mycobacterium abscessus subsp. abscessus Terrabacteria group; Actinobacteria; Actinobacteria; Corynebacteriales; Mycobacteriaceae; Mycobacterium; Mycobacterium chelonae group; Mycobacterium abscessus subgroup; Mycobacterium abscessus
MIKHLTRRATVALVVAAAAISPAAYPAMADTATPLNGLFALTPGSCAEGRATGSYFRMIL WP_005073213.1
hypothetical protein [Mycobacterium abscessus] Length: 330\nNumber of predicted TMHs: 2\nExp number of AAs in TMHs: 39.3416199999999\nExp number, first 60 AAs: 17.07688\nTotal prob of N-in: 0.84431\nPOSSIBLE N-term signal sequence\ninside 1 11\nTMhelix 12 29\noutside 30 298\nTMhelix 299 321\ninside 322 330

29881 GCF_900133815.1_12163_1_7 Mycobacterium abscessus subsp. abscessus Terrabacteria group; Actinobacteria; Actinobacteria; Corynebacteriales; Mycobacteriaceae; Mycobacterium; Mycobacterium chelonae group; Mycobacterium abscessus subgroup; Mycobacterium abscessus
MIKHLTRRATVALVVAAAAISPAAYPAMADTATPLNGLFALTPGSCAEGRATGSYFRMIL WP_005073213.1
hypothetical protein [Mycobacterium abscessus] Length: 330\nNumber of predicted TMHs: 2\nExp number of AAs in TMHs: 39.3416199999999\nExp number, first 60 AAs: 17.07688\nTotal prob of N-in: 0.84431\nPOSSIBLE N-term signal sequence\ninside 1 11\nTMhelix 12 29\noutside 30 298\nTMhelix 299 321\ninside 322 330

29882 GCF_900133835.1_12163_1_9 Mycobacterium abscessus subsp. abscessus Terrabacteria group; Actinobacteria; Actinobacteria; Corynebacteriales; Mycobacteriaceae; Mycobacterium; Mycobacterium chelonae group; Mycobacterium abscessus subgroup; Mycobacterium abscessus
MIKHLTRRATVALVVAAAAISPAAYPAMADTATPLNGLFALTPGSCAEGRATGSYFRMIL WP_005073213.1
hypothetical protein [Mycobacterium abscessus] Length: 330\nNumber of predicted TMHs: 2\nExp number of AAs in TMHs: 39.3416199999999\nExp number, first 60 AAs: 17.07688\nTotal prob of N-in: 0.84431\nPOSSIBLE N-term signal sequence\ninside 1 11\nTMhelix 12 29\noutside 30 298\nTMhelix 299 321\ninside 322 330

29883 GCF_900133855.1_10250_1_54 Mycobacterium abscessus subsp. abscessus Terrabacteria group; Actinobacteria; Actinobacteria; Corynebacteriales; Mycobacteriaceae; Mycobacterium; Mycobacterium chelonae group;

Mycobacterium abscessus subgroup; Mycobacterium abscessus
MIKHLTRRATVALVVAAAAISPAAYPAMADTATPLNGLFALTPGSCAEGRATGSYFRMIL WP_005073213.1
hypothetical protein [Mycobacterium abscessus] Length: 330\nNumber of predicted TMHs: 2\nExp number of AAs in TMHs: 39.3416199999999\nExp number, first 60 AAs: 17.07688\nTotal prob of N-in: 0.84431\nPOSSIBLE N-term signal sequence\ninside 1 11\nTMhelix 12 29\noutside 30 298\nTMhelix 299 321\ninside 322 330

29884 GCF_900133905.1_10250_1_60 Mycobacterium abscessus subsp. abscessus Terrabacteria group; Actinobacteria; Actinobacteria; Corynebacteriales; Mycobacteriaceae; Mycobacterium; Mycobacterium chelonae group; Mycobacterium abscessus subgroup; Mycobacterium abscessus
MIKHLTRRATVALVVAAAAISPAAYPAMADTATPLNGLFALTPGSCAEGRATGSYFRMIL WP_005073213.1
hypothetical protein [Mycobacterium abscessus] Length: 330\nNumber of predicted TMHs: 2\nExp number of AAs in TMHs: 39.3416199999999\nExp number, first 60 AAs: 17.07688\nTotal prob of N-in: 0.84431\nPOSSIBLE N-term signal sequence\ninside 1 11\nTMhelix 12 29\noutside 30 298\nTMhelix 299 321\ninside 322 330

29885 GCF_900133895.1_10250_1_58 Mycobacterium abscessus subsp. abscessus Terrabacteria group; Actinobacteria; Actinobacteria; Corynebacteriales; Mycobacteriaceae; Mycobacterium; Mycobacterium chelonae group; Mycobacterium abscessus subgroup; Mycobacterium abscessus
MIKHLTRRATVALVVAAAAISPAAYPAMADTATPLNGLFALTPGSCAEGRATGSYFRMIL WP_005073213.1
hypothetical protein [Mycobacterium abscessus] Length: 330\nNumber of predicted TMHs: 2\nExp number of AAs in TMHs: 39.3416199999999\nExp number, first 60 AAs: 17.07688\nTotal prob of N-in: 0.84431\nPOSSIBLE N-term signal sequence\ninside 1 11\nTMhelix 12 29\noutside 30 298\nTMhelix 299 321\ninside 322 330

29886 GCF_900133945.1_10250_1_69 Mycobacterium abscessus subsp. abscessus Terrabacteria group; Actinobacteria; Actinobacteria; Corynebacteriales; Mycobacteriaceae; Mycobacterium; Mycobacterium chelonae group; Mycobacterium abscessus subgroup; Mycobacterium abscessus
MIKHLTRRATVALVVAAAAISPAAYPAMADTATPLNGLFALTPGSCAEGRATGSYFRMIL WP_005073213.1
hypothetical protein [Mycobacterium abscessus] Length: 330\nNumber of predicted TMHs: 2\nExp number of AAs in TMHs: 39.3416199999999\nExp number, first 60 AAs: 17.07688\nTotal prob of N-in: 0.84431\nPOSSIBLE N-term signal sequence\ninside 1 11\nTMhelix 12 29\noutside 30 298\nTMhelix 299 321\ninside 322 330

29887 GCF_900134215.1_11861_7_90 Mycobacterium abscessus subsp. abscessus Terrabacteria group; Actinobacteria; Actinobacteria; Corynebacteriales; Mycobacteriaceae; Mycobacterium; Mycobacterium chelonae group; Mycobacterium abscessus subgroup; Mycobacterium abscessus
MIKHLTRRATVALVVAAAAISPAAYPAMADTATPLNGLFALTPGSCAEGRATGSYFRMIL WP_005073213.1
hypothetical protein [Mycobacterium abscessus] Length: 330\nNumber of predicted TMHs: 2\nExp number of AAs in TMHs: 39.3416199999999\nExp number, first 60 AAs: 17.07688\nTotal prob of N-in: 0.84431\nPOSSIBLE N-term signal sequence\ninside 1 11\nTMhelix 12 29\noutside 30 298\nTMhelix 299 321\ninside 322 330

29888 GCF_900134185.1_11861_7_34 Mycobacterium abscessus subsp. abscessus Terrabacteria group; Actinobacteria; Actinobacteria; Corynebacteriales; Mycobacteriaceae; Mycobacterium; Mycobacterium chelonae group; Mycobacterium abscessus subgroup; Mycobacterium abscessus
MIKHLTRRATVALVVAAAAISPAAYPAMADTATPLNGLFALTPGSCAEGRATGSYFRMIL WP_074340136.1
hypothetical protein [Mycobacterium abscessus] Length: 330\nNumber of predicted TMHs: 2\nExp number of AAs in TMHs: 39.3416799999999\nExp number, first 60 AAs: 17.07671\nTotal prob of N-in: 0.84432\nPOSSIBLE N-term signal sequence\ninside 1 11\nTMhelix 12 29\noutside 30 298\nTMhelix 299 321\ninside 322 330

29889 GCF_900134385.1_12082_5_78 Mycobacterium abscessus subsp. abscessus Terrabacteria group; Actinobacteria; Actinobacteria; Corynebacteriales; Mycobacteriaceae; Mycobacterium; Mycobacterium chelonae group; Mycobacterium abscessus subgroup; Mycobacterium abscessus
MIKHLTRRATVALVVAAAAISPAAYPAMADTATPLNGLFALTPGSCAEGRATGSYFRMIL WP_005073213.1
hypothetical protein [Mycobacterium abscessus] Length: 330\nNumber of predicted TMHs: 2\nExp number of AAs in TMHs: 39.3416199999999\nExp number, first 60 AAs: 17.07688\nTotal prob of N-in: 0.84431\nPOSSIBLE N-term signal sequence\ninside 1 11\nTMhelix 12 29\noutside 30 298\nTMhelix 299 321\ninside 322 330

29890 GCF_900134485.1_12163_1_38 Mycobacterium abscessus subsp. abscessus Terrabacteria group; Actinobacteria; Actinobacteria; Corynebacteriales; Mycobacteriaceae; Mycobacterium; Mycobacterium chelonae group; Mycobacterium abscessus subgroup; Mycobacterium abscessus
MIKHLTRRATVALVVAAAAISPAAYPAMADTATPLNGLFALTPGSCAEGRATGSYFRMIL WP_005073213.1
hypothetical protein [Mycobacterium abscessus] Length: 330\nNumber of predicted TMHs: 2\nExp number of AAs in TMHs: 39.3416199999999\nExp number, first 60 AAs: 17.07688\nTotal prob of N-in: 0.84431\nPOSSIBLE N-term signal sequence\ninside 1 11\nTMhelix 12 29\noutside 30 298\nTMhelix 299 321\ninside 322 330

29891 GCF_900134505.1_12163_1_39 Mycobacterium abscessus subsp. abscessus Terrabacteria group; Actinobacteria; Actinobacteria; Corynebacteriales; Mycobacteriaceae; Mycobacterium; Mycobacterium chelonae group; Mycobacterium abscessus subgroup; Mycobacterium abscessus
MIKHLTRRATVALVVAAAAISPAAYPAMADTATPLNGLFALTPGSCAEGRATGSYFRMIL WP_005073213.1
hypothetical protein [Mycobacterium abscessus] Length: 330\nNumber of predicted TMHs: 2\nExp number of AAs in TMHs: 39.3416199999999\nExp number, first 60 AAs: 17.07688\nTotal prob of N-in: 0.84431\nPOSSIBLE N-term signal sequence\ninside 1 11\nTMhelix 12 29\noutside 30 298\nTMhelix 299 321\ninside 322 330

29892 GCF_900134525.1_12163_1_41 Mycobacterium abscessus subsp. abscessus Terrabacteria group; Actinobacteria; Actinobacteria; Corynebacteriales; Mycobacteriaceae; Mycobacterium; Mycobacterium chelonae group; Mycobacterium abscessus subgroup; Mycobacterium abscessus
MIKHLTRRATVALVVAAAAISPAAYPAMADTATPLNGLFALTPGSCAEGRATGSYFRMIL WP_005073213.1
hypothetical protein [Mycobacterium abscessus] Length: 330\nNumber of predicted TMHs: 2\nExp number of AAs in TMHs: 39.3416199999999\nExp number, first 60 AAs: 17.07688\nTotal prob of N-in: 0.84431\nPOSSIBLE N-term signal sequence\ninside 1 11\nTMhelix 12 29\noutside 30 298\nTMhelix 299 321\ninside 322 330

29893 GCF_900134595.1_12163_1_51 Mycobacterium abscessus subsp. abscessus Terrabacteria group; Actinobacteria; Actinobacteria; Corynebacteriales; Mycobacteriaceae; Mycobacterium; Mycobacterium chelonae group; Mycobacterium abscessus subgroup; Mycobacterium abscessus
MIKHLTRRATVALVVAAAAISPAAYPAMADTATPLNGLFALTPGSCAEGRATGSYFRMIL WP_005073213.1
hypothetical protein [Mycobacterium abscessus] Length: 330\nNumber of predicted TMHs: 2\nExp number of AAs in TMHs: 39.3416199999999\nExp number, first 60 AAs: 17.07688\nTotal prob of N-in: 0.84431\nPOSSIBLE N-term signal sequence\ninside 1 11\nTMhelix 12 29\noutside 30 298\nTMhelix 299 321\ninside 322 330

29894 GCF_900134615.1_12163_1_50 Mycobacterium abscessus subsp. abscessus Terrabacteria group; Actinobacteria; Actinobacteria; Corynebacteriales; Mycobacteriaceae; Mycobacterium; Mycobacterium chelonae group; Mycobacterium abscessus subgroup; Mycobacterium abscessus
MIKHLTRRATVALVVAAAAISPAAYPAMADTATPLNGLFALTPGSCAEGRATGSYFRMIL WP_005073213.1
hypothetical protein [Mycobacterium abscessus] Length: 330\nNumber of predicted TMHs: 2\nExp number of AAs in TMHs: 39.3416199999999\nExp number, first 60 AAs: 17.07688\nTotal prob of N-in: 0.84431\nPOSSIBLE N-term signal sequence\ninside 1 11\nTMhelix 12 29\noutside 30 298\nTMhelix 299 321\ninside 322 330

29895 GCF_900134715.1_10208_3_11 Mycobacterium abscessus subsp. abscessus Terrabacteria group; Actinobacteria; Actinobacteria; Corynebacteriales; Mycobacteriaceae; Mycobacterium; Mycobacterium chelonae group; Mycobacterium abscessus subgroup; Mycobacterium abscessus
MIKHLTRRATVALVVAAAAISPAAYPAMADTATPLNGLFALTPGSCAEGRATGSYFRMIL WP_005073213.1
hypothetical protein [Mycobacterium abscessus] Length: 330\nNumber of predicted TMHs: 2\nExp number of AAs in TMHs: 39.3416199999999\nExp number, first 60 AAs: 17.07688\nTotal prob of N-in: 0.84431\nPOSSIBLE N-term signal sequence\ninside 1 11\nTMhelix 12 29\noutside 30 298\nTMhelix 299 321\ninside 322 330

29896 GCF_900134695.1_10208_3_1 Mycobacterium abscessus subsp. abscessus Terrabacteria group; Actinobacteria; Actinobacteria; Corynebacteriales; Mycobacteriaceae; Mycobacterium; Mycobacterium chelonae group; Mycobacterium abscessus subgroup; Mycobacterium abscessus
MIKHLTRRATVALVVAAAAISPAAYPAMADTATPLNGLFALTPGSCAEGRATGSYFRMIL WP_005073213.1
hypothetical protein [Mycobacterium abscessus] Length: 330\nNumber of predicted TMHs: 2\nExp number of AAs in TMHs: 39.3416199999999\nExp number, first 60 AAs: 17.07688\nTotal prob of N-in: 0.84431\nPOSSIBLE N-term signal sequence\ninside 1 11\nTMhelix 12 29\noutside 30 298\nTMhelix 299 321\ninside 322 330

29897 GCF_900134745.1_10208_3_12 Mycobacterium abscessus subsp. abscessus Terrabacteria group; Actinobacteria; Actinobacteria; Corynebacteriales; Mycobacteriaceae; Mycobacterium; Mycobacterium chelonae group; Mycobacterium abscessus subgroup; Mycobacterium abscessus
MIKHLTRRATVALVVAAAAISPAAYPAMADTATPLNGLFALTPGSCAEGRATGSYFRMIL WP_005073213.1
hypothetical protein [Mycobacterium abscessus] Length: 330\nNumber of predicted TMHs: 2\nExp number of AAs in TMHs: 39.3416199999999\nExp number, first 60 AAs: 17.07688\nTotal prob of N-in: 0.84431\nPOSSIBLE N-term signal sequence\ninside 1 11\nTMhelix 12 29\noutside 30 298\nTMhelix 299 321\ninside 322 330

29898 GCF_900134725.1_10208_3_14 Mycobacterium abscessus subsp. abscessus Terrabacteria group; Actinobacteria; Actinobacteria; Corynebacteriales; Mycobacteriaceae; Mycobacterium; Mycobacterium chelonae group; Mycobacterium abscessus subgroup; Mycobacterium abscessus
MIKHLTRRATVALVVAAAAISPAAYPAMADTATPLNGLFALTPGSCAEGRATGSYFRMIL WP_005073213.1
hypothetical protein [Mycobacterium abscessus] Length: 330\nNumber of predicted TMHs: 2\nExp number of

AAs in TMHs: 39.3416199999999\nExp number, first 60 AAs: 17.07688\nTotal prob of N-in: 0.84431\nPOSSIBLE N-term signal sequence\ninside 1 11\nTMhelix 12 29\noutside 30 298\nTMhelix 299 321\ninside 322 330

29899 GCF_900134805.1_10208_3_19 Mycobacterium abscessus subsp. abscessus Terrabacteria group; Actinobacteria; Actinobacteria; Corynebacteriales; Mycobacteriaceae; Mycobacterium; Mycobacterium chelonae group; Mycobacterium abscessus subgroup; Mycobacterium abscessus
MIKHLTRRATVALVVAAAAISPAAYPAMADTATPLNGLFALTPGSCAEGRATGSYFRMIL WP_005073213.1
hypothetical protein [Mycobacterium abscessus] Length: 330\nNumber of predicted TMHs: 2\nExp number of AAs in TMHs: 39.3416199999999\nExp number, first 60 AAs: 17.07688\nTotal prob of N-in: 0.84431\nPOSSIBLE N-term signal sequence\ninside 1 11\nTMhelix 12 29\noutside 30 298\nTMhelix 299 321\ninside 322 330

29900 GCF_900134975.1_10208_3_42 Mycobacterium abscessus subsp. abscessus Terrabacteria group; Actinobacteria; Actinobacteria; Corynebacteriales; Mycobacteriaceae; Mycobacterium; Mycobacterium chelonae group; Mycobacterium abscessus subgroup; Mycobacterium abscessus
MIKHLTRRATVALVVAAAAISPAAYPAMADTATPLNGLFALTPGSCAEGRATGSYFRMIL WP_005073213.1
hypothetical protein [Mycobacterium abscessus] Length: 330\nNumber of predicted TMHs: 2\nExp number of AAs in TMHs: 39.3416199999999\nExp number, first 60 AAs: 17.07688\nTotal prob of N-in: 0.84431\nPOSSIBLE N-term signal sequence\ninside 1 11\nTMhelix 12 29\noutside 30 298\nTMhelix 299 321\ninside 322 330

29901 GCF_900134945.1_10208_3_36 Mycobacterium abscessus subsp. abscessus Terrabacteria group; Actinobacteria; Actinobacteria; Corynebacteriales; Mycobacteriaceae; Mycobacterium; Mycobacterium chelonae group; Mycobacterium abscessus subgroup; Mycobacterium abscessus
MIKHLTRRATVALVVAAAAISPAAYPAMADTATPLNGLFALTPGSCAEGRATGSYFRMIL WP_005073213.1
hypothetical protein [Mycobacterium abscessus] Length: 330\nNumber of predicted TMHs: 2\nExp number of AAs in TMHs: 39.3416199999999\nExp number, first 60 AAs: 17.07688\nTotal prob of N-in: 0.84431\nPOSSIBLE N-term signal sequence\ninside 1 11\nTMhelix 12 29\noutside 30 298\nTMhelix 299 321\ninside 322 330

29902 GCF_900135005.1_10208_3_5 Mycobacterium abscessus subsp. abscessus Terrabacteria group; Actinobacteria; Actinobacteria; Corynebacteriales; Mycobacteriaceae; Mycobacterium; Mycobacterium chelonae group; Mycobacterium abscessus subgroup; Mycobacterium abscessus
MIKHLTRRATVALVVAAAAISPAAYPAMADTATPLNGLFALTPGSCAEGRATGSYFRMIL WP_005073213.1
hypothetical protein [Mycobacterium abscessus] Length: 330\nNumber of predicted TMHs: 2\nExp number of AAs in TMHs: 39.3416199999999\nExp number, first 60 AAs: 17.07688\nTotal prob of N-in: 0.84431\nPOSSIBLE N-term signal sequence\ninside 1 11\nTMhelix 12 29\noutside 30 298\nTMhelix 299 321\ninside 322 330

29903 GCF_900135025.1_10208_3_8 Mycobacterium abscessus subsp. abscessus Terrabacteria group; Actinobacteria; Actinobacteria; Corynebacteriales; Mycobacteriaceae; Mycobacterium; Mycobacterium chelonae group; Mycobacterium abscessus subgroup; Mycobacterium abscessus
MIKHLTRRATVALVVAAAAISPAAYPAMADTATPLNGLFALTPGSCAEGRATGSYFRMIL WP_005073213.1
hypothetical protein [Mycobacterium abscessus] Length: 330\nNumber of predicted TMHs: 2\nExp number of AAs in TMHs: 39.3416199999999\nExp number, first 60 AAs: 17.07688\nTotal prob of N-in: 0.84431\nPOSSIBLE N-term signal sequence\ninside 1 11\nTMhelix 12 29\noutside 30 298\nTMhelix 299 321\ninside 322 330

29904 GCF_900135045.1_10250_1_81 Mycobacterium abscessus subsp. abscessus Terrabacteria group; Actinobacteria; Actinobacteria; Corynebacteriales; Mycobacteriaceae; Mycobacterium; Mycobacterium chelonae group; Mycobacterium abscessus subgroup; Mycobacterium abscessus
MIKHLTRRATVALVVAAAAISPAAYPAMADTATPLNGLFALTPGSCAEGRATGSYFRMIL WP_005073213.1
hypothetical protein [Mycobacterium abscessus] Length: 330\nNumber of predicted TMHs: 2\nExp number of AAs in TMHs: 39.3416199999999\nExp number, first 60 AAs: 17.07688\nTotal prob of N-in: 0.84431\nPOSSIBLE N-term signal sequence\ninside 1 11\nTMhelix 12 29\noutside 30 298\nTMhelix 299 321\ninside 322 330

29905 GCF_900135125.1_10625_4_23 Mycobacterium abscessus subsp. abscessus Terrabacteria group; Actinobacteria; Actinobacteria; Corynebacteriales; Mycobacteriaceae; Mycobacterium; Mycobacterium chelonae group; Mycobacterium abscessus subgroup; Mycobacterium abscessus
MIKHLTRRATVALVVAAAAISPAAYPAMADTATPLNGLFALTPGSCAEGRATGSYFRMIL WP_005073213.1
hypothetical protein [Mycobacterium abscessus] Length: 330\nNumber of predicted TMHs: 2\nExp number of AAs in TMHs: 39.3416199999999\nExp number, first 60 AAs: 17.07688\nTotal prob of N-in: 0.84431\nPOSSIBLE N-term signal sequence\ninside 1 11\nTMhelix 12 29\noutside 30 298\nTMhelix 299 321\ninside 322 330

29906 GCF_900135145.1_10625_4_24 Mycobacterium abscessus subsp. abscessus Terrabacteria group; Actinobacteria; Actinobacteria; Corynebacteriales; Mycobacteriaceae; Mycobacterium; Mycobacterium chelonae group;

Mycobacterium abscessus subgroup; Mycobacterium abscessus
MIKHLTRRATVALVVAAAAISPAAYPAMADTATPLNGLFALTPGSCAEGRATGSYFRMIL WP_005073213.1
hypothetical protein [Mycobacterium abscessus] Length: 330\nNumber of predicted TMHs: 2\nExp number of AAs in TMHs: 39.3416199999999\nExp number, first 60 AAs: 17.07688\nTotal prob of N-in: 0.84431\nPOSSIBLE N-term signal sequence\ninside 1 11\nTMhelix 12 29\noutside 30 298\nTMhelix 299 321\ninside 322 330

29907 GCF_900135165.1_10625_4_34 Mycobacterium abscessus subsp. abscessus Terrabacteria group; Actinobacteria; Actinobacteria; Corynebacteriales; Mycobacteriaceae; Mycobacterium; Mycobacterium chelonae group; Mycobacterium abscessus subgroup; Mycobacterium abscessus
MIKHLTRRATVALVVAAAAISPAAYPAMADTATPLNGLFALTPGSCAEGRATGSYFRMIL WP_005073213.1
hypothetical protein [Mycobacterium abscessus] Length: 330\nNumber of predicted TMHs: 2\nExp number of AAs in TMHs: 39.3416199999999\nExp number, first 60 AAs: 17.07688\nTotal prob of N-in: 0.84431\nPOSSIBLE N-term signal sequence\ninside 1 11\nTMhelix 12 29\noutside 30 298\nTMhelix 299 321\ninside 322 330

29908 GCF_900135185.1_10625_4_21 Mycobacterium abscessus subsp. abscessus Terrabacteria group; Actinobacteria; Actinobacteria; Corynebacteriales; Mycobacteriaceae; Mycobacterium; Mycobacterium chelonae group; Mycobacterium abscessus subgroup; Mycobacterium abscessus
MIKHLTRRATVALVVAAAAISPAAYPAMADTATPLNGLFALTPGSCAEGRATGSYFRMIL WP_005073213.1
hypothetical protein [Mycobacterium abscessus] Length: 330\nNumber of predicted TMHs: 2\nExp number of AAs in TMHs: 39.3416199999999\nExp number, first 60 AAs: 17.07688\nTotal prob of N-in: 0.84431\nPOSSIBLE N-term signal sequence\ninside 1 11\nTMhelix 12 29\noutside 30 298\nTMhelix 299 321\ninside 322 330

29909 GCF_900135205.1_10625_4_33 Mycobacterium abscessus subsp. abscessus Terrabacteria group; Actinobacteria; Actinobacteria; Corynebacteriales; Mycobacteriaceae; Mycobacterium; Mycobacterium chelonae group; Mycobacterium abscessus subgroup; Mycobacterium abscessus
MIKHLTRRATVALVVAAAAISPAAYPAMADTATPLNGLFALTPGSCAEGRATGSYFRMIL WP_005073213.1
hypothetical protein [Mycobacterium abscessus] Length: 330\nNumber of predicted TMHs: 2\nExp number of AAs in TMHs: 39.3416199999999\nExp number, first 60 AAs: 17.07688\nTotal prob of N-in: 0.84431\nPOSSIBLE N-term signal sequence\ninside 1 11\nTMhelix 12 29\noutside 30 298\nTMhelix 299 321\ninside 322 330

29910 GCF_900135335.1_10625_4_47 Mycobacterium abscessus subsp. abscessus Terrabacteria group; Actinobacteria; Actinobacteria; Corynebacteriales; Mycobacteriaceae; Mycobacterium; Mycobacterium chelonae group; Mycobacterium abscessus subgroup; Mycobacterium abscessus
MIKHLTRRATVALVVAAAAISPAAYPAMADTATPLNGLFALTPGSCAEGRATGSYFRMIL WP_005073213.1
hypothetical protein [Mycobacterium abscessus] Length: 330\nNumber of predicted TMHs: 2\nExp number of AAs in TMHs: 39.3416199999999\nExp number, first 60 AAs: 17.07688\nTotal prob of N-in: 0.84431\nPOSSIBLE N-term signal sequence\ninside 1 11\nTMhelix 12 29\noutside 30 298\nTMhelix 299 321\ninside 322 330

29911 GCF_900135355.1_10625_4_49 Mycobacterium abscessus subsp. abscessus Terrabacteria group; Actinobacteria; Actinobacteria; Corynebacteriales; Mycobacteriaceae; Mycobacterium; Mycobacterium chelonae group; Mycobacterium abscessus subgroup; Mycobacterium abscessus
MIKHLTRRATVALVVAAAAISPAAYPAMADTATPLNGLFALTPGSCAEGRATGSYFRMIL WP_005073213.1
hypothetical protein [Mycobacterium abscessus] Length: 330\nNumber of predicted TMHs: 2\nExp number of AAs in TMHs: 39.3416199999999\nExp number, first 60 AAs: 17.07688\nTotal prob of N-in: 0.84431\nPOSSIBLE N-term signal sequence\ninside 1 11\nTMhelix 12 29\noutside 30 298\nTMhelix 299 321\ninside 322 330

29912 GCF_900135375.1_10625_4_54 Mycobacterium abscessus subsp. abscessus Terrabacteria group; Actinobacteria; Actinobacteria; Corynebacteriales; Mycobacteriaceae; Mycobacterium; Mycobacterium chelonae group; Mycobacterium abscessus subgroup; Mycobacterium abscessus
MIKHLTRRATVALVVAAAAISPAAYPAMADTATPLNGLFALTPGSCAEGRATGSYFRMIL WP_005073213.1
hypothetical protein [Mycobacterium abscessus] Length: 330\nNumber of predicted TMHs: 2\nExp number of AAs in TMHs: 39.3416199999999\nExp number, first 60 AAs: 17.07688\nTotal prob of N-in: 0.84431\nPOSSIBLE N-term signal sequence\ninside 1 11\nTMhelix 12 29\noutside 30 298\nTMhelix 299 321\ninside 322 330

29913 GCF_900135395.1_10625_4_56 Mycobacterium abscessus subsp. abscessus Terrabacteria group; Actinobacteria; Actinobacteria; Corynebacteriales; Mycobacteriaceae; Mycobacterium; Mycobacterium chelonae group; Mycobacterium abscessus subgroup; Mycobacterium abscessus
MIKHLTRRATVALVVAAAAISPAAYPAMADTATPLNGLFALTPGSCAEGRATGSYFRMIL WP_005073213.1
hypothetical protein [Mycobacterium abscessus] Length: 330\nNumber of predicted TMHs: 2\nExp number of AAs in TMHs: 39.3416199999999\nExp number, first 60 AAs: 17.07688\nTotal prob of N-in: 0.84431\nPOSSIBLE N-term signal sequence\ninside 1 11\nTMhelix 12 29\noutside 30 298\nTMhelix 299 321\ninside 322 330

29914 GCF_900135435.1_10625_4_61 Mycobacterium abscessus subsp. abscessus Terrabacteria group; Actinobacteria; Actinobacteria; Corynebacteriales; Mycobacteriaceae; Mycobacterium; Mycobacterium chelonae group; Mycobacterium abscessus subgroup; Mycobacterium abscessus
MIKHLTRRATVALVVAAAAISPAAYPAMADTATPLNGLFALTPGSCAEGRATGSYFRMIL WP_005073213.1
hypothetical protein [Mycobacterium abscessus] Length: 330\nNumber of predicted TMHs: 2\nExp number of AAs in TMHs: 39.3416199999999\nExp number, first 60 AAs: 17.07688\nTotal prob of N-in: 0.84431\nPOSSIBLE N-term signal sequence\ninside 1 11\nTMhelix 12 29\noutside 30 298\nTMhelix 299 321\ninside 322 330

29915 GCF_900135415.1_10625_4_62 Mycobacterium abscessus subsp. abscessus Terrabacteria group; Actinobacteria; Actinobacteria; Corynebacteriales; Mycobacteriaceae; Mycobacterium; Mycobacterium chelonae group; Mycobacterium abscessus subgroup; Mycobacterium abscessus
MIKHLTRRATVALVVAAAAISPAAYPAMADTATPLNGLFALTPGSCAEGRATGSYFRMIL WP_005073213.1
hypothetical protein [Mycobacterium abscessus] Length: 330\nNumber of predicted TMHs: 2\nExp number of AAs in TMHs: 39.3416199999999\nExp number, first 60 AAs: 17.07688\nTotal prob of N-in: 0.84431\nPOSSIBLE N-term signal sequence\ninside 1 11\nTMhelix 12 29\noutside 30 298\nTMhelix 299 321\ninside 322 330

29916 GCF_900135555.1_10625_4_72 Mycobacterium abscessus subsp. abscessus Terrabacteria group; Actinobacteria; Actinobacteria; Corynebacteriales; Mycobacteriaceae; Mycobacterium; Mycobacterium chelonae group; Mycobacterium abscessus subgroup; Mycobacterium abscessus
MIKHLTRRATVALVVAAAAISPAAYPAMADTATPLNGLFALTPGSCAEGRATGSYFRMIL WP_005073213.1
hypothetical protein [Mycobacterium abscessus] Length: 330\nNumber of predicted TMHs: 2\nExp number of AAs in TMHs: 39.3416199999999\nExp number, first 60 AAs: 17.07688\nTotal prob of N-in: 0.84431\nPOSSIBLE N-term signal sequence\ninside 1 11\nTMhelix 12 29\noutside 30 298\nTMhelix 299 321\ninside 322 330

29917 GCF_900135585.1_10625_5_15 Mycobacterium abscessus subsp. abscessus Terrabacteria group; Actinobacteria; Actinobacteria; Corynebacteriales; Mycobacteriaceae; Mycobacterium; Mycobacterium chelonae group; Mycobacterium abscessus subgroup; Mycobacterium abscessus
MIKHLTRRATVALVVAAAAISPAAYPAMADTATPLNGLFALTPGSCAEGRATGSYFRMIL WP_005073213.1
hypothetical protein [Mycobacterium abscessus] Length: 330\nNumber of predicted TMHs: 2\nExp number of AAs in TMHs: 39.3416199999999\nExp number, first 60 AAs: 17.07688\nTotal prob of N-in: 0.84431\nPOSSIBLE N-term signal sequence\ninside 1 11\nTMhelix 12 29\noutside 30 298\nTMhelix 299 321\ninside 322 330

29918 GCF_900135575.1_10625_5_16 Mycobacterium abscessus subsp. abscessus Terrabacteria group; Actinobacteria; Actinobacteria; Corynebacteriales; Mycobacteriaceae; Mycobacterium; Mycobacterium chelonae group; Mycobacterium abscessus subgroup; Mycobacterium abscessus
MIKHLTRRATVALVVAAAAISPAAYPAMADTATPLNGLFALTPGSCAEGRATGSYFRMIL WP_005073213.1
hypothetical protein [Mycobacterium abscessus] Length: 330\nNumber of predicted TMHs: 2\nExp number of AAs in TMHs: 39.3416199999999\nExp number, first 60 AAs: 17.07688\nTotal prob of N-in: 0.84431\nPOSSIBLE N-term signal sequence\ninside 1 11\nTMhelix 12 29\noutside 30 298\nTMhelix 299 321\ninside 322 330

29919 GCF_900135605.1_10625_5_20 Mycobacterium abscessus subsp. abscessus Terrabacteria group; Actinobacteria; Actinobacteria; Corynebacteriales; Mycobacteriaceae; Mycobacterium; Mycobacterium chelonae group; Mycobacterium abscessus subgroup; Mycobacterium abscessus
MIKHLTRRATVALVVAAAAISPAAYPAMADTATPLNGLFALTPGSCAEGRATGSYFRMIL WP_005073213.1
hypothetical protein [Mycobacterium abscessus] Length: 330\nNumber of predicted TMHs: 2\nExp number of AAs in TMHs: 39.3416199999999\nExp number, first 60 AAs: 17.07688\nTotal prob of N-in: 0.84431\nPOSSIBLE N-term signal sequence\ninside 1 11\nTMhelix 12 29\noutside 30 298\nTMhelix 299 321\ninside 322 330

29920 GCF_900135625.1_10625_5_19 Mycobacterium abscessus subsp. abscessus Terrabacteria group; Actinobacteria; Actinobacteria; Corynebacteriales; Mycobacteriaceae; Mycobacterium; Mycobacterium chelonae group; Mycobacterium abscessus subgroup; Mycobacterium abscessus
MIKHLTRRATVALVVAAAAISPAAYPAMADTATPLNGLFALTPGSCAEGRATGSYFRMIL WP_005073213.1
hypothetical protein [Mycobacterium abscessus] Length: 330\nNumber of predicted TMHs: 2\nExp number of AAs in TMHs: 39.3416199999999\nExp number, first 60 AAs: 17.07688\nTotal prob of N-in: 0.84431\nPOSSIBLE N-term signal sequence\ninside 1 11\nTMhelix 12 29\noutside 30 298\nTMhelix 299 321\ninside 322 330

29921 GCF_900135645.1_10665_2_59 Mycobacterium abscessus subsp. abscessus Terrabacteria group; Actinobacteria; Actinobacteria; Corynebacteriales; Mycobacteriaceae; Mycobacterium; Mycobacterium chelonae group; Mycobacterium abscessus subgroup; Mycobacterium abscessus
MIKHLTRRATVALVVAAAAISPAAYPAMADTATPLNGLFALTPGSCAEGRATGSYFRMIL WP_005073213.1
hypothetical protein [Mycobacterium abscessus] Length: 330\nNumber of predicted TMHs: 2\nExp number of

AAs in TMHs: 39.3416199999999\nExp number, first 60 AAs: 17.07688\nTotal prob of N-in: 0.84431\nPOSSIBLE N-term signal sequence\ninside 1 11\nTMhelix 12 29\noutside 30 298\nTMhelix 299 321\ninside 322 330

29922 GCF_900135855.1_11893_6_23 Mycobacterium abscessus subsp. abscessus Terrabacteria group; Actinobacteria; Actinobacteria; Corynebacteriales; Mycobacteriaceae; Mycobacterium; Mycobacterium chelonae group; Mycobacterium abscessus subgroup; Mycobacterium abscessus
MIKHLTRRATVALVVAAAAISPAAYPAMADTATPLNGLFALTPGSCAEGRATGSYFRMIL WP_005073213.1
hypothetical protein [Mycobacterium abscessus] Length: 330\nNumber of predicted TMHs: 2\nExp number of AAs in TMHs: 39.3416199999999\nExp number, first 60 AAs: 17.07688\nTotal prob of N-in: 0.84431\nPOSSIBLE N-term signal sequence\ninside 1 11\nTMhelix 12 29\noutside 30 298\nTMhelix 299 321\ninside 322 330

29923 GCF_900135935.1_11893_6_39 Mycobacterium abscessus subsp. abscessus Terrabacteria group; Actinobacteria; Actinobacteria; Corynebacteriales; Mycobacteriaceae; Mycobacterium; Mycobacterium chelonae group; Mycobacterium abscessus subgroup; Mycobacterium abscessus
MIKHLTRRATVALVVAAAAISPAAYPAMADTATPLNGLFALTPGSCAEGRATGSYFRMIL WP_005073213.1
hypothetical protein [Mycobacterium abscessus] Length: 330\nNumber of predicted TMHs: 2\nExp number of AAs in TMHs: 39.3416199999999\nExp number, first 60 AAs: 17.07688\nTotal prob of N-in: 0.84431\nPOSSIBLE N-term signal sequence\ninside 1 11\nTMhelix 12 29\noutside 30 298\nTMhelix 299 321\ninside 322 330

29924 GCF_900135765.1_11861_7_73 Mycobacterium abscessus subsp. abscessus Terrabacteria group; Actinobacteria; Actinobacteria; Corynebacteriales; Mycobacteriaceae; Mycobacterium; Mycobacterium chelonae group; Mycobacterium abscessus subgroup; Mycobacterium abscessus
MPRRRTILSALASFSMVIGLLVVSTPTAWAAPFPVVDPAKLPPSQPPAPPGDMRLNGAPL WP_074362414.1 type VII secretion-associated serine protease mycosin [Mycobacterium abscessus] Length: 470\nNumber of predicted TMHs: 2\nExp number of AAs in TMHs: 44.7482799999999\nExp number, first 60 AAs: 22.65577\nTotal prob of N-in: 0.99532\nPOSSIBLE N-term signal sequence\ninside 1 6\nTMhelix 7 29\noutside 30 441\nTMhelix 442 464\ninside 465 470

29925 GCF_900136095.1_12163_1_66 Mycobacterium abscessus subsp. abscessus Terrabacteria group; Actinobacteria; Actinobacteria; Corynebacteriales; Mycobacteriaceae; Mycobacterium; Mycobacterium chelonae group; Mycobacterium abscessus subgroup; Mycobacterium abscessus
MIKHLTRRATVALVVAAAAISPAAYPAMADTATPLNGLFALTPGSCAEGRATGSYFRMIL WP_005073213.1
hypothetical protein [Mycobacterium abscessus] Length: 330\nNumber of predicted TMHs: 2\nExp number of AAs in TMHs: 39.3416199999999\nExp number, first 60 AAs: 17.07688\nTotal prob of N-in: 0.84431\nPOSSIBLE N-term signal sequence\ninside 1 11\nTMhelix 12 29\noutside 30 298\nTMhelix 299 321\ninside 322 330

29926 GCF_900136135.1_12163_1_71 Mycobacterium abscessus subsp. abscessus Terrabacteria group; Actinobacteria; Actinobacteria; Corynebacteriales; Mycobacteriaceae; Mycobacterium; Mycobacterium chelonae group; Mycobacterium abscessus subgroup; Mycobacterium abscessus
MIKHLTRRATVALVVAAAAISPAAYPAMADTATPLNGLFALTPGSCAEGRATGSYFRMIL WP_005073213.1
hypothetical protein [Mycobacterium abscessus] Length: 330\nNumber of predicted TMHs: 2\nExp number of AAs in TMHs: 39.3416199999999\nExp number, first 60 AAs: 17.07688\nTotal prob of N-in: 0.84431\nPOSSIBLE N-term signal sequence\ninside 1 11\nTMhelix 12 29\noutside 30 298\nTMhelix 299 321\ninside 322 330

29927 GCF_900136115.1_12163_1_70 Mycobacterium abscessus subsp. abscessus Terrabacteria group; Actinobacteria; Actinobacteria; Corynebacteriales; Mycobacteriaceae; Mycobacterium; Mycobacterium chelonae group; Mycobacterium abscessus subgroup; Mycobacterium abscessus
MIKHLTRRATVALVVAAAAISPAAYPAMADTATPLNGLFALTPGSCAEGRATGSYFRMIL WP_005073213.1
hypothetical protein [Mycobacterium abscessus] Length: 330\nNumber of predicted TMHs: 2\nExp number of AAs in TMHs: 39.3416199999999\nExp number, first 60 AAs: 17.07688\nTotal prob of N-in: 0.84431\nPOSSIBLE N-term signal sequence\ninside 1 11\nTMhelix 12 29\noutside 30 298\nTMhelix 299 321\ninside 322 330

29928 GCF_900136145.1_12163_1_72 Mycobacterium abscessus subsp. abscessus Terrabacteria group; Actinobacteria; Actinobacteria; Corynebacteriales; Mycobacteriaceae; Mycobacterium; Mycobacterium chelonae group; Mycobacterium abscessus subgroup; Mycobacterium abscessus
MIKHLTRRATVALVVAAAAISPAAYPAMADTATPLNGLFALTPGSCAEGRATGSYFRMIL WP_005073213.1
hypothetical protein [Mycobacterium abscessus] Length: 330\nNumber of predicted TMHs: 2\nExp number of AAs in TMHs: 39.3416199999999\nExp number, first 60 AAs: 17.07688\nTotal prob of N-in: 0.84431\nPOSSIBLE N-term signal sequence\ninside 1 11\nTMhelix 12 29\noutside 30 298\nTMhelix 299 321\ninside 322 330

29929 GCF_900136215.1_12163_1_76 Mycobacterium abscessus subsp. abscessus Terrabacteria group; Actinobacteria; Actinobacteria; Corynebacteriales; Mycobacteriaceae; Mycobacterium; Mycobacterium chelonae group; Mycobacterium abscessus subgroup; Mycobacterium abscessus
MIKHLTRRATVALVVAAAAISPAAYPAMADTATPLNGLFALTPGSCAEGRATGSYFRMIL WP_005073213.1
hypothetical protein [Mycobacterium abscessus] Length: 330\nNumber of predicted TMHs: 2\nExp number of AAs in TMHs: 39.3416199999999\nExp number, first 60 AAs: 17.07688\nTotal prob of N-in: 0.84431\nPOSSIBLE N-term signal sequence\ninside 1 11\nTMhelix 12 29\noutside 30 298\nTMhelix 299 321\ninside 322 330

29930 GCF_900136355.1_12163_1_90 Mycobacterium abscessus subsp. abscessus Terrabacteria group; Actinobacteria; Actinobacteria; Corynebacteriales; Mycobacteriaceae; Mycobacterium; Mycobacterium chelonae group; Mycobacterium abscessus subgroup; Mycobacterium abscessus
MIKHLTRRATVALVVAAAAISPAAYPAMADTATPLNGLFALTPGSCAEGRATGSYFRMIL WP_005073213.1
hypothetical protein [Mycobacterium abscessus] Length: 330\nNumber of predicted TMHs: 2\nExp number of AAs in TMHs: 39.3416199999999\nExp number, first 60 AAs: 17.07688\nTotal prob of N-in: 0.84431\nPOSSIBLE N-term signal sequence\ninside 1 11\nTMhelix 12 29\noutside 30 298\nTMhelix 299 321\ninside 322 330

29931 GCF_900136415.1_10208_3_44 Mycobacterium abscessus subsp. abscessus Terrabacteria group; Actinobacteria; Actinobacteria; Corynebacteriales; Mycobacteriaceae; Mycobacterium; Mycobacterium chelonae group; Mycobacterium abscessus subgroup; Mycobacterium abscessus
MIKHLTRRATVALVVAAAAISPAAYPAMADTATPLNGLFALTPGSCAEGRATGSYFRMIL WP_005073213.1
hypothetical protein [Mycobacterium abscessus] Length: 330\nNumber of predicted TMHs: 2\nExp number of AAs in TMHs: 39.3416199999999\nExp number, first 60 AAs: 17.07688\nTotal prob of N-in: 0.84431\nPOSSIBLE N-term signal sequence\ninside 1 11\nTMhelix 12 29\noutside 30 298\nTMhelix 299 321\ninside 322 330

29932 GCF_900136435.1_10208_3_66 Mycobacterium abscessus subsp. abscessus Terrabacteria group; Actinobacteria; Actinobacteria; Corynebacteriales; Mycobacteriaceae; Mycobacterium; Mycobacterium chelonae group; Mycobacterium abscessus subgroup; Mycobacterium abscessus
MIKHLTRRATVALVVAAAAISPAAYPAMADTATPLNGLFALTPGSCAEGRATGSYFRMIL WP_005073213.1
hypothetical protein [Mycobacterium abscessus] Length: 330\nNumber of predicted TMHs: 2\nExp number of AAs in TMHs: 39.3416199999999\nExp number, first 60 AAs: 17.07688\nTotal prob of N-in: 0.84431\nPOSSIBLE N-term signal sequence\ninside 1 11\nTMhelix 12 29\noutside 30 298\nTMhelix 299 321\ninside 322 330

29933 GCF_900136535.1_10208_3_79 Mycobacterium abscessus subsp. abscessus Terrabacteria group; Actinobacteria; Actinobacteria; Corynebacteriales; Mycobacteriaceae; Mycobacterium; Mycobacterium chelonae group; Mycobacterium abscessus subgroup; Mycobacterium abscessus
MIKHLTRRATVALVVAAAAISPAAYPAMADTATPLNGLFALTPGSCAEGRATGSYFRMIL WP_005073213.1
hypothetical protein [Mycobacterium abscessus] Length: 330\nNumber of predicted TMHs: 2\nExp number of AAs in TMHs: 39.3416199999999\nExp number, first 60 AAs: 17.07688\nTotal prob of N-in: 0.84431\nPOSSIBLE N-term signal sequence\ninside 1 11\nTMhelix 12 29\noutside 30 298\nTMhelix 299 321\ninside 322 330

29934 GCF_900136575.1_10208_3_80 Mycobacterium abscessus subsp. abscessus Terrabacteria group; Actinobacteria; Actinobacteria; Corynebacteriales; Mycobacteriaceae; Mycobacterium; Mycobacterium chelonae group; Mycobacterium abscessus subgroup; Mycobacterium abscessus
MIKHLTRRATVALVVAAAAISPAAYPAMADTATPLNGLFALTPGSCAEGRATGSYFRMIL WP_005073213.1
hypothetical protein [Mycobacterium abscessus] Length: 330\nNumber of predicted TMHs: 2\nExp number of AAs in TMHs: 39.3416199999999\nExp number, first 60 AAs: 17.07688\nTotal prob of N-in: 0.84431\nPOSSIBLE N-term signal sequence\ninside 1 11\nTMhelix 12 29\noutside 30 298\nTMhelix 299 321\ninside 322 330

29935 GCF_900136595.1_10625_4_30 Mycobacterium abscessus subsp. abscessus Terrabacteria group; Actinobacteria; Actinobacteria; Corynebacteriales; Mycobacteriaceae; Mycobacterium; Mycobacterium chelonae group; Mycobacterium abscessus subgroup; Mycobacterium abscessus
MIKHLTRRATVALVVAAAAISPAAYPAMADTATPLNGLFALTPGSCAEGRATGSYFRMIL WP_005073213.1
hypothetical protein [Mycobacterium abscessus] Length: 330\nNumber of predicted TMHs: 2\nExp number of AAs in TMHs: 39.3416199999999\nExp number, first 60 AAs: 17.07688\nTotal prob of N-in: 0.84431\nPOSSIBLE N-term signal sequence\ninside 1 11\nTMhelix 12 29\noutside 30 298\nTMhelix 299 321\ninside 322 330

29936 GCF_900136605.1_10208_3_84 Mycobacterium abscessus subsp. abscessus Terrabacteria group; Actinobacteria; Actinobacteria; Corynebacteriales; Mycobacteriaceae; Mycobacterium; Mycobacterium chelonae group; Mycobacterium abscessus subgroup; Mycobacterium abscessus
MIKHLTRRATVALVVAAAAISPAAYPAMADTATPLNGLFALTPGSCAEGRATGSYFRMIL WP_005073213.1
hypothetical protein [Mycobacterium abscessus] Length: 330\nNumber of predicted TMHs: 2\nExp number of

AAs in TMHs: 39.3416199999999\nExp number, first 60 AAs: 17.07688\nTotal prob of N-in: 0.84431\nPOSSIBLE N-term signal sequence\ninside 1 11\nTMhelix 12 29\noutside 30 298\nTMhelix 299 321\ninside 322 330

29937 GCF_900136775.1_10625_5_35 Mycobacterium abscessus subsp. abscessus Terrabacteria group; Actinobacteria; Actinobacteria; Corynebacteriales; Mycobacteriaceae; Mycobacterium; Mycobacterium chelonae group; Mycobacterium abscessus subgroup; Mycobacterium abscessus
MIKHLTRRATVALVVAAAAISPAAYPAMADTATPLNGLFALTPGSCAEGRATGSYFRMIL WP_005073213.1
hypothetical protein [Mycobacterium abscessus] Length: 330\nNumber of predicted TMHs: 2\nExp number of AAs in TMHs: 39.3416199999999\nExp number, first 60 AAs: 17.07688\nTotal prob of N-in: 0.84431\nPOSSIBLE N-term signal sequence\ninside 1 11\nTMhelix 12 29\noutside 30 298\nTMhelix 299 321\ninside 322 330

29938 GCF_900136795.1_10625_5_36 Mycobacterium abscessus subsp. abscessus Terrabacteria group; Actinobacteria; Actinobacteria; Corynebacteriales; Mycobacteriaceae; Mycobacterium; Mycobacterium chelonae group; Mycobacterium abscessus subgroup; Mycobacterium abscessus
MIKHLTRRATVALVVAAAAISPAAYPAMADTATPLNGLFALTPGSCAEGRATGSYFRMIL WP_005073213.1
hypothetical protein [Mycobacterium abscessus] Length: 330\nNumber of predicted TMHs: 2\nExp number of AAs in TMHs: 39.3416199999999\nExp number, first 60 AAs: 17.07688\nTotal prob of N-in: 0.84431\nPOSSIBLE N-term signal sequence\ninside 1 11\nTMhelix 12 29\noutside 30 298\nTMhelix 299 321\ninside 322 330

29939 GCF_900136815.1_10625_5_37 Mycobacterium abscessus subsp. abscessus Terrabacteria group; Actinobacteria; Actinobacteria; Corynebacteriales; Mycobacteriaceae; Mycobacterium; Mycobacterium chelonae group; Mycobacterium abscessus subgroup; Mycobacterium abscessus
MIKHLTRRATVALVVAAAAISPAAYPAMADTATPLNGLFALTPGSCAEGRATGSYFRMIL WP_005073213.1
hypothetical protein [Mycobacterium abscessus] Length: 330\nNumber of predicted TMHs: 2\nExp number of AAs in TMHs: 39.3416199999999\nExp number, first 60 AAs: 17.07688\nTotal prob of N-in: 0.84431\nPOSSIBLE N-term signal sequence\ninside 1 11\nTMhelix 12 29\noutside 30 298\nTMhelix 299 321\ninside 322 330

29940 GCF_900136845.1_10625_5_41 Mycobacterium abscessus subsp. abscessus Terrabacteria group; Actinobacteria; Actinobacteria; Corynebacteriales; Mycobacteriaceae; Mycobacterium; Mycobacterium chelonae group; Mycobacterium abscessus subgroup; Mycobacterium abscessus
MTKHLTRRATVALVVAAAAISPAAYPAMADTATPLNGLFALTPGSCAEGRATGSYFRMIL WP_074319467.1
hypothetical protein [Mycobacterium abscessus] Length: 330\nNumber of predicted TMHs: 2\nExp number of AAs in TMHs: 40.2609399999999\nExp number, first 60 AAs: 17.99061\nTotal prob of N-in: 0.88931\nPOSSIBLE N-term signal sequence\ninside 1 11\nTMhelix 12 29\noutside 30 298\nTMhelix 299 321\ninside 322 330

29941 GCF_900136985.1_11861_7_74 Mycobacterium abscessus subsp. abscessus Terrabacteria group; Actinobacteria; Actinobacteria; Corynebacteriales; Mycobacteriaceae; Mycobacterium; Mycobacterium chelonae group; Mycobacterium abscessus subgroup; Mycobacterium abscessus
MIKHLTRRATVALVVAAAAISPAAYPAMADTATPLNGLFALTPGSCAEGRATGSYFRMIL WP_005073213.1
hypothetical protein [Mycobacterium abscessus] Length: 330\nNumber of predicted TMHs: 2\nExp number of AAs in TMHs: 39.3416199999999\nExp number, first 60 AAs: 17.07688\nTotal prob of N-in: 0.84431\nPOSSIBLE N-term signal sequence\ninside 1 11\nTMhelix 12 29\noutside 30 298\nTMhelix 299 321\ninside 322 330

29942 GCF_900136995.1_11861_7_75 Mycobacterium abscessus subsp. abscessus Terrabacteria group; Actinobacteria; Actinobacteria; Corynebacteriales; Mycobacteriaceae; Mycobacterium; Mycobacterium chelonae group; Mycobacterium abscessus subgroup; Mycobacterium abscessus
MIKHLTRRATVALVVAAAAISPAAYPAMADTATPLNGLFALTPGSCAEGRATGSYFRMIL WP_005073213.1
hypothetical protein [Mycobacterium abscessus] Length: 330\nNumber of predicted TMHs: 2\nExp number of AAs in TMHs: 39.3416199999999\nExp number, first 60 AAs: 17.07688\nTotal prob of N-in: 0.84431\nPOSSIBLE N-term signal sequence\ninside 1 11\nTMhelix 12 29\noutside 30 298\nTMhelix 299 321\ninside 322 330

29943 GCF_900137065.1_11861_8_14 Mycobacterium abscessus subsp. abscessus Terrabacteria group; Actinobacteria; Actinobacteria; Corynebacteriales; Mycobacteriaceae; Mycobacterium; Mycobacterium chelonae group; Mycobacterium abscessus subgroup; Mycobacterium abscessus
MIKHLTRRATVALVVAAAAISPAAYPAMADTATPLNGLFALTPGSCAEGRATGSYFRMIL WP_005073213.1
hypothetical protein [Mycobacterium abscessus] Length: 330\nNumber of predicted TMHs: 2\nExp number of AAs in TMHs: 39.3416199999999\nExp number, first 60 AAs: 17.07688\nTotal prob of N-in: 0.84431\nPOSSIBLE N-term signal sequence\ninside 1 11\nTMhelix 12 29\noutside 30 298\nTMhelix 299 321\ninside 322 330

29944 GCF_900137225.1_12082_5_69 Mycobacterium abscessus subsp. abscessus Terrabacteria group; Actinobacteria; Actinobacteria; Corynebacteriales; Mycobacteriaceae; Mycobacterium; Mycobacterium chelonae group;

Mycobacterium abscessus subgroup; Mycobacterium abscessus
MIKHLTRRATVALVVAAAAISPAAYPAMADTATPLNGLFALTPGSCAEGRATGSYFRMIL WP_005073213.1
hypothetical protein [Mycobacterium abscessus] Length: 330\nNumber of predicted TMHs: 2\nExp number of AAs in TMHs: 39.3416199999999\nExp number, first 60 AAs: 17.07688\nTotal prob of N-in: 0.84431\nPOSSIBLE N-term signal sequence\ninside 1 11\nTMhelix 12 29\noutside 30 298\nTMhelix 299 321\ninside 322 330

29945 GCF_900137205.1_12082_5_66 Mycobacterium abscessus subsp. abscessus Terrabacteria group; Actinobacteria; Actinobacteria; Corynebacteriales; Mycobacteriaceae; Mycobacterium; Mycobacterium chelonae group; Mycobacterium abscessus subgroup; Mycobacterium abscessus
MIKHLTRRATVALVVAAAAISPAAYPAMADTATPLNGLFALTPGSCAEGRATGSYFRMIL WP_005073213.1
hypothetical protein [Mycobacterium abscessus] Length: 330\nNumber of predicted TMHs: 2\nExp number of AAs in TMHs: 39.3416199999999\nExp number, first 60 AAs: 17.07688\nTotal prob of N-in: 0.84431\nPOSSIBLE N-term signal sequence\ninside 1 11\nTMhelix 12 29\noutside 30 298\nTMhelix 299 321\ninside 322 330

29946 GCF_900137265.1_12082_5_82 Mycobacterium abscessus subsp. abscessus Terrabacteria group; Actinobacteria; Actinobacteria; Corynebacteriales; Mycobacteriaceae; Mycobacterium; Mycobacterium chelonae group; Mycobacterium abscessus subgroup; Mycobacterium abscessus
MIKHLTRRATVALVVAAAAISPAAYPAMADTATPLNGLFALTPGSCAEGRATGSYFRMIL WP_005073213.1
hypothetical protein [Mycobacterium abscessus] Length: 330\nNumber of predicted TMHs: 2\nExp number of AAs in TMHs: 39.3416199999999\nExp number, first 60 AAs: 17.07688\nTotal prob of N-in: 0.84431\nPOSSIBLE N-term signal sequence\ninside 1 11\nTMhelix 12 29\noutside 30 298\nTMhelix 299 321\ninside 322 330

29947 GCF_900137285.1_12163_1_93 Mycobacterium abscessus subsp. abscessus Terrabacteria group; Actinobacteria; Actinobacteria; Corynebacteriales; Mycobacteriaceae; Mycobacterium; Mycobacterium chelonae group; Mycobacterium abscessus subgroup; Mycobacterium abscessus
MIKHLTRRATVALVVAAAAISPAAYPAMADTATPLNGLFALTPGSCAEGRATGSYFRMIL WP_005073213.1
hypothetical protein [Mycobacterium abscessus] Length: 330\nNumber of predicted TMHs: 2\nExp number of AAs in TMHs: 39.3416199999999\nExp number, first 60 AAs: 17.07688\nTotal prob of N-in: 0.84431\nPOSSIBLE N-term signal sequence\ninside 1 11\nTMhelix 12 29\noutside 30 298\nTMhelix 299 321\ninside 322 330

29948 GCF_900137315.1_12163_1_95 Mycobacterium abscessus subsp. abscessus Terrabacteria group; Actinobacteria; Actinobacteria; Corynebacteriales; Mycobacteriaceae; Mycobacterium; Mycobacterium chelonae group; Mycobacterium abscessus subgroup; Mycobacterium abscessus
MIKHLTRRATVALVVAAAAISPAAYPAMADTATPLNGLFALTPGSCAEGRATGSYFRMIL WP_005073213.1
hypothetical protein [Mycobacterium abscessus] Length: 330\nNumber of predicted TMHs: 2\nExp number of AAs in TMHs: 39.3416199999999\nExp number, first 60 AAs: 17.07688\nTotal prob of N-in: 0.84431\nPOSSIBLE N-term signal sequence\ninside 1 11\nTMhelix 12 29\noutside 30 298\nTMhelix 299 321\ninside 322 330

29949 GCF_900137405.1_12163_2_17 Mycobacterium abscessus subsp. abscessus Terrabacteria group; Actinobacteria; Actinobacteria; Corynebacteriales; Mycobacteriaceae; Mycobacterium; Mycobacterium chelonae group; Mycobacterium abscessus subgroup; Mycobacterium abscessus
MIKHLTRRATVALVVAAAAISPAAYPAMADTATPLNGLFALTPGSCAEGRATGSYFRMIL WP_005073213.1
hypothetical protein [Mycobacterium abscessus] Length: 330\nNumber of predicted TMHs: 2\nExp number of AAs in TMHs: 39.3416199999999\nExp number, first 60 AAs: 17.07688\nTotal prob of N-in: 0.84431\nPOSSIBLE N-term signal sequence\ninside 1 11\nTMhelix 12 29\noutside 30 298\nTMhelix 299 321\ninside 322 330

29950 GCF_900137435.1_12163_2_19 Mycobacterium abscessus subsp. abscessus Terrabacteria group; Actinobacteria; Actinobacteria; Corynebacteriales; Mycobacteriaceae; Mycobacterium; Mycobacterium chelonae group; Mycobacterium abscessus subgroup; Mycobacterium abscessus
MIKHLTRRATVALVVAAAAISPAAYPAMADTATPLNGLFALTPGSCAEGRATGSYFRMIL WP_005073213.1
hypothetical protein [Mycobacterium abscessus] Length: 330\nNumber of predicted TMHs: 2\nExp number of AAs in TMHs: 39.3416199999999\nExp number, first 60 AAs: 17.07688\nTotal prob of N-in: 0.84431\nPOSSIBLE N-term signal sequence\ninside 1 11\nTMhelix 12 29\noutside 30 298\nTMhelix 299 321\ninside 322 330

29951 GCF_900137455.1_12163_2_2 Mycobacterium abscessus subsp. abscessus Terrabacteria group; Actinobacteria; Actinobacteria; Corynebacteriales; Mycobacteriaceae; Mycobacterium; Mycobacterium chelonae group; Mycobacterium abscessus subgroup; Mycobacterium abscessus
MIKHLTRRATVALVVAAAAISPAAYPAMADTATPLNGLFALTPGSCAEGRATGSYFRMIL WP_005073213.1
hypothetical protein [Mycobacterium abscessus] Length: 330\nNumber of predicted TMHs: 2\nExp number of AAs in TMHs: 39.3416199999999\nExp number, first 60 AAs: 17.07688\nTotal prob of N-in: 0.84431\nPOSSIBLE N-term signal sequence\ninside 1 11\nTMhelix 12 29\noutside 30 298\nTMhelix 299 321\ninside 322 330

29952 GCF_900137495.1_12163_2_24 Mycobacterium abscessus subsp. abscessus Terrabacteria group; Actinobacteria; Actinobacteria; Corynebacteriales; Mycobacteriaceae; Mycobacterium; Mycobacterium chelonae group; Mycobacterium abscessus subgroup; Mycobacterium abscessus
MIKHLTRRATVALVVAAAAISPAAYPAMADTATPLNGLFALTPGSCAEGRATGSYFRMIL WP_005073213.1
hypothetical protein [Mycobacterium abscessus] Length: 330\nNumber of predicted TMHs: 2\nExp number of AAs in TMHs: 39.3416199999999\nExp number, first 60 AAs: 17.07688\nTotal prob of N-in: 0.84431\nPOSSIBLE N-term signal sequence\ninside 1 11\nTMhelix 12 29\noutside 30 298\nTMhelix 299 321\ninside 322 330

29953 GCF_900137505.1_12163_2_26 Mycobacterium abscessus subsp. abscessus Terrabacteria group; Actinobacteria; Actinobacteria; Corynebacteriales; Mycobacteriaceae; Mycobacterium; Mycobacterium chelonae group; Mycobacterium abscessus subgroup; Mycobacterium abscessus
MIKHLTRRATVALVVAAAAISPAAYPAMADTATPLNGLFALTPGSCAEGRATGSYFRMIL WP_005073213.1
hypothetical protein [Mycobacterium abscessus] Length: 330\nNumber of predicted TMHs: 2\nExp number of AAs in TMHs: 39.3416199999999\nExp number, first 60 AAs: 17.07688\nTotal prob of N-in: 0.84431\nPOSSIBLE N-term signal sequence\ninside 1 11\nTMhelix 12 29\noutside 30 298\nTMhelix 299 321\ninside 322 330

29954 GCF_900137525.1_12163_2_27 Mycobacterium abscessus subsp. abscessus Terrabacteria group; Actinobacteria; Actinobacteria; Corynebacteriales; Mycobacteriaceae; Mycobacterium; Mycobacterium chelonae group; Mycobacterium abscessus subgroup; Mycobacterium abscessus
MIKHLTRRATVALVVAAAAISPAAYPAMADTATPLNGLFALTPGSCAEGRATGSYFRMIL WP_005073213.1
hypothetical protein [Mycobacterium abscessus] Length: 330\nNumber of predicted TMHs: 2\nExp number of AAs in TMHs: 39.3416199999999\nExp number, first 60 AAs: 17.07688\nTotal prob of N-in: 0.84431\nPOSSIBLE N-term signal sequence\ninside 1 11\nTMhelix 12 29\noutside 30 298\nTMhelix 299 321\ninside 322 330

29955 GCF_900137645.1_12163_2_4 Mycobacterium abscessus subsp. abscessus Terrabacteria group; Actinobacteria; Actinobacteria; Corynebacteriales; Mycobacteriaceae; Mycobacterium; Mycobacterium chelonae group; Mycobacterium abscessus subgroup; Mycobacterium abscessus
MIKHLTRRATVALVVAAAAISPAAYPAMADTATPLNGLFALTPGSCAEGRATGSYFRMIL WP_005073213.1
hypothetical protein [Mycobacterium abscessus] Length: 330\nNumber of predicted TMHs: 2\nExp number of AAs in TMHs: 39.3416199999999\nExp number, first 60 AAs: 17.07688\nTotal prob of N-in: 0.84431\nPOSSIBLE N-term signal sequence\ninside 1 11\nTMhelix 12 29\noutside 30 298\nTMhelix 299 321\ninside 322 330

29956 GCF_900137685.1_12163_2_44 Mycobacterium abscessus subsp. abscessus Terrabacteria group; Actinobacteria; Actinobacteria; Corynebacteriales; Mycobacteriaceae; Mycobacterium; Mycobacterium chelonae group; Mycobacterium abscessus subgroup; Mycobacterium abscessus
MIKHLTRRATVALVVAAAAISPAAYPAMADTATPLNGLFALTPGSCAEGRATGSYFRMIL WP_005073213.1
hypothetical protein [Mycobacterium abscessus] Length: 330\nNumber of predicted TMHs: 2\nExp number of AAs in TMHs: 39.3416199999999\nExp number, first 60 AAs: 17.07688\nTotal prob of N-in: 0.84431\nPOSSIBLE N-term signal sequence\ninside 1 11\nTMhelix 12 29\noutside 30 298\nTMhelix 299 321\ninside 322 330

29957 GCF_900137665.1_12163_2_38 Mycobacterium abscessus subsp. abscessus Terrabacteria group; Actinobacteria; Actinobacteria; Corynebacteriales; Mycobacteriaceae; Mycobacterium; Mycobacterium chelonae group; Mycobacterium abscessus subgroup; Mycobacterium abscessus
MIKHLTRRATVALVVAAAAISPAAYPAMADTATPLNGLFALTPGSCAEGRATGSYFRMIL WP_005073213.1
hypothetical protein [Mycobacterium abscessus] Length: 330\nNumber of predicted TMHs: 2\nExp number of AAs in TMHs: 39.3416199999999\nExp number, first 60 AAs: 17.07688\nTotal prob of N-in: 0.84431\nPOSSIBLE N-term signal sequence\ninside 1 11\nTMhelix 12 29\noutside 30 298\nTMhelix 299 321\ninside 322 330

29958 GCF_900137745.1_12163_2_51 Mycobacterium abscessus subsp. abscessus Terrabacteria group; Actinobacteria; Actinobacteria; Corynebacteriales; Mycobacteriaceae; Mycobacterium; Mycobacterium chelonae group; Mycobacterium abscessus subgroup; Mycobacterium abscessus
MIKHLTRRATVALVVAAAAISPAAYPAMADTATPLNGLFALTPGSCAEGRATGSYFRMIL WP_005073213.1
hypothetical protein [Mycobacterium abscessus] Length: 330\nNumber of predicted TMHs: 2\nExp number of AAs in TMHs: 39.3416199999999\nExp number, first 60 AAs: 17.07688\nTotal prob of N-in: 0.84431\nPOSSIBLE N-term signal sequence\ninside 1 11\nTMhelix 12 29\noutside 30 298\nTMhelix 299 321\ninside 322 330

29959 GCF_900137755.1_12163_2_49 Mycobacterium abscessus subsp. abscessus Terrabacteria group; Actinobacteria; Actinobacteria; Corynebacteriales; Mycobacteriaceae; Mycobacterium; Mycobacterium chelonae group; Mycobacterium abscessus subgroup; Mycobacterium abscessus
MIKHLTRRATVALVVAAAAISPAAYPAMADTATPLNGLFALTPGSCAEGRATGSYFRMIL WP_005073213.1
hypothetical protein [Mycobacterium abscessus] Length: 330\nNumber of predicted TMHs: 2\nExp number of

AAs in TMHs: 39.3416199999999\nExp number, first 60 AAs: 17.07688\nTotal prob of N-in: 0.84431\nPOSSIBLE N-term signal sequence\ninside 1 11\nTMhelix 12 29\noutside 30 298\nTMhelix 299 321\ninside 322 330

29960 GCF_900137955.1_10665_2_79 Mycobacterium abscessus subsp. abscessus Terrabacteria group; Actinobacteria; Actinobacteria; Corynebacteriales; Mycobacteriaceae; Mycobacterium; Mycobacterium chelonae group; Mycobacterium abscessus subgroup; Mycobacterium abscessus
MIKHLTRRATVALVVAAAAISPAAYPAMADTATPLNGLFALTPGSCAEGRATGSYFRMIL WP_005073213.1
hypothetical protein [Mycobacterium abscessus] Length: 330\nNumber of predicted TMHs: 2\nExp number of AAs in TMHs: 39.3416199999999\nExp number, first 60 AAs: 17.07688\nTotal prob of N-in: 0.84431\nPOSSIBLE N-term signal sequence\ninside 1 11\nTMhelix 12 29\noutside 30 298\nTMhelix 299 321\ninside 322 330

29961 GCF_900137975.1_12163_2_89 Mycobacterium abscessus subsp. abscessus Terrabacteria group; Actinobacteria; Actinobacteria; Corynebacteriales; Mycobacteriaceae; Mycobacterium; Mycobacterium chelonae group; Mycobacterium abscessus subgroup; Mycobacterium abscessus
MIKHLTRRATVALVVAAAAISPAAYPAMADTATPLNGLFALTPGSCAEGRATGSYFRMIL WP_005073213.1
hypothetical protein [Mycobacterium abscessus] Length: 330\nNumber of predicted TMHs: 2\nExp number of AAs in TMHs: 39.3416199999999\nExp number, first 60 AAs: 17.07688\nTotal prob of N-in: 0.84431\nPOSSIBLE N-term signal sequence\ninside 1 11\nTMhelix 12 29\noutside 30 298\nTMhelix 299 321\ninside 322 330

29962 GCF_900137995.1_11893_6_11 Mycobacterium abscessus subsp. abscessus Terrabacteria group; Actinobacteria; Actinobacteria; Corynebacteriales; Mycobacteriaceae; Mycobacterium; Mycobacterium chelonae group; Mycobacterium abscessus subgroup; Mycobacterium abscessus
MIKHLTRRATVALVVAAAAISPAAYPAMADTATPLNGLFALTPGSCAEGRATGSYFRMIL WP_005073213.1
hypothetical protein [Mycobacterium abscessus] Length: 330\nNumber of predicted TMHs: 2\nExp number of AAs in TMHs: 39.3416199999999\nExp number, first 60 AAs: 17.07688\nTotal prob of N-in: 0.84431\nPOSSIBLE N-term signal sequence\ninside 1 11\nTMhelix 12 29\noutside 30 298\nTMhelix 299 321\ninside 322 330

29963 GCF_900138115.1_12045_8_47 Mycobacterium abscessus subsp. abscessus Terrabacteria group; Actinobacteria; Actinobacteria; Corynebacteriales; Mycobacteriaceae; Mycobacterium; Mycobacterium chelonae group; Mycobacterium abscessus subgroup; Mycobacterium abscessus
MIKHLTRRATVALVVAAAAISPAAYPAMADTATPLNGLFALTPGSCAEGRATGSYFRMIL WP_005073213.1
hypothetical protein [Mycobacterium abscessus] Length: 330\nNumber of predicted TMHs: 2\nExp number of AAs in TMHs: 39.3416199999999\nExp number, first 60 AAs: 17.07688\nTotal prob of N-in: 0.84431\nPOSSIBLE N-term signal sequence\ninside 1 11\nTMhelix 12 29\noutside 30 298\nTMhelix 299 321\ninside 322 330

29964 GCF_900138145.1_12082_5_80 Mycobacterium abscessus subsp. abscessus Terrabacteria group; Actinobacteria; Actinobacteria; Corynebacteriales; Mycobacteriaceae; Mycobacterium; Mycobacterium chelonae group; Mycobacterium abscessus subgroup; Mycobacterium abscessus
MIKHLTRRATVALVVAAAAISPAAYPAMADTATPLNGLFALTPGSCAEGRATGSYFRMIL WP_005073213.1
hypothetical protein [Mycobacterium abscessus] Length: 330\nNumber of predicted TMHs: 2\nExp number of AAs in TMHs: 39.3416199999999\nExp number, first 60 AAs: 17.07688\nTotal prob of N-in: 0.84431\nPOSSIBLE N-term signal sequence\ninside 1 11\nTMhelix 12 29\noutside 30 298\nTMhelix 299 321\ninside 322 330

29965 GCF_900138185.1_12082_5_89 Mycobacterium abscessus subsp. abscessus Terrabacteria group; Actinobacteria; Actinobacteria; Corynebacteriales; Mycobacteriaceae; Mycobacterium; Mycobacterium chelonae group; Mycobacterium abscessus subgroup; Mycobacterium abscessus
MIKHLTRRATVALVVAAAAISPAAYPAMADTATPLNGLFALTPGSCAEGRATGSYFRMIL WP_005073213.1
hypothetical protein [Mycobacterium abscessus] Length: 330\nNumber of predicted TMHs: 2\nExp number of AAs in TMHs: 39.3416199999999\nExp number, first 60 AAs: 17.07688\nTotal prob of N-in: 0.84431\nPOSSIBLE N-term signal sequence\ninside 1 11\nTMhelix 12 29\noutside 30 298\nTMhelix 299 321\ninside 322 330

29966 GCF_900138205.1_12082_5_88 Mycobacterium abscessus subsp. abscessus Terrabacteria group; Actinobacteria; Actinobacteria; Corynebacteriales; Mycobacteriaceae; Mycobacterium; Mycobacterium chelonae group; Mycobacterium abscessus subgroup; Mycobacterium abscessus
MIKHLTRRATVALVVAAAAISPAAYPAMADTATPLNGLFALTPGSCAEGRATGSYFRMIL WP_005073213.1
hypothetical protein [Mycobacterium abscessus] Length: 330\nNumber of predicted TMHs: 2\nExp number of AAs in TMHs: 39.3416199999999\nExp number, first 60 AAs: 17.07688\nTotal prob of N-in: 0.84431\nPOSSIBLE N-term signal sequence\ninside 1 11\nTMhelix 12 29\noutside 30 298\nTMhelix 299 321\ninside 322 330

29967 GCF_900138215.1_12082_5_91 Mycobacterium abscessus subsp. abscessus Terrabacteria group; Actinobacteria; Actinobacteria; Corynebacteriales; Mycobacteriaceae; Mycobacterium; Mycobacterium chelonae group;

Mycobacterium abscessus subgroup; Mycobacterium abscessus
MIKHLTRRATVALVVAAAAISPAAYPAMADTATPLNGLFALTPGSCAEGRATGSYFRMIL WP_005073213.1
hypothetical protein [Mycobacterium abscessus] Length: 330\nNumber of predicted TMHs: 2\nExp number of AAs in TMHs: 39.3416199999999\nExp number, first 60 AAs: 17.07688\nTotal prob of N-in: 0.84431\nPOSSIBLE N-term signal sequence\ninside 1 11\nTMhelix 12 29\noutside 30 298\nTMhelix 299 321\ninside 322 330

29968 GCF_900138235.1_12082_5_95 Mycobacterium abscessus subsp. abscessus Terrabacteria group; Actinobacteria; Actinobacteria; Corynebacteriales; Mycobacteriaceae; Mycobacterium; Mycobacterium chelonae group; Mycobacterium abscessus subgroup; Mycobacterium abscessus
MIKHLTRRATVALVVAAAAISPAAYPAMADTATPLNGLFALTPGSCAEGRATGSYFRMIL WP_005073213.1
hypothetical protein [Mycobacterium abscessus] Length: 330\nNumber of predicted TMHs: 2\nExp number of AAs in TMHs: 39.3416199999999\nExp number, first 60 AAs: 17.07688\nTotal prob of N-in: 0.84431\nPOSSIBLE N-term signal sequence\ninside 1 11\nTMhelix 12 29\noutside 30 298\nTMhelix 299 321\ninside 322 330

29969 GCF_900138255.1_12082_5_92 Mycobacterium abscessus subsp. abscessus Terrabacteria group; Actinobacteria; Actinobacteria; Corynebacteriales; Mycobacteriaceae; Mycobacterium; Mycobacterium chelonae group; Mycobacterium abscessus subgroup; Mycobacterium abscessus
MIKHLTRRATVALVVAAAAISPAAYPAMADTATPLNGLFALTPGSCAEGRATGSYFRMIL WP_005073213.1
hypothetical protein [Mycobacterium abscessus] Length: 330\nNumber of predicted TMHs: 2\nExp number of AAs in TMHs: 39.3416199999999\nExp number, first 60 AAs: 17.07688\nTotal prob of N-in: 0.84431\nPOSSIBLE N-term signal sequence\ninside 1 11\nTMhelix 12 29\noutside 30 298\nTMhelix 299 321\ninside 322 330

29970 GCF_900138275.1_12163_2_66 Mycobacterium abscessus subsp. abscessus Terrabacteria group; Actinobacteria; Actinobacteria; Corynebacteriales; Mycobacteriaceae; Mycobacterium; Mycobacterium chelonae group; Mycobacterium abscessus subgroup; Mycobacterium abscessus
MIKHLTRRATVALVVAAAAISPAAYPAMADTATPLNGLFALTPGSCAEGRATGSYFRMIL WP_005073213.1
hypothetical protein [Mycobacterium abscessus] Length: 330\nNumber of predicted TMHs: 2\nExp number of AAs in TMHs: 39.3416199999999\nExp number, first 60 AAs: 17.07688\nTotal prob of N-in: 0.84431\nPOSSIBLE N-term signal sequence\ninside 1 11\nTMhelix 12 29\noutside 30 298\nTMhelix 299 321\ninside 322 330

29971 GCF_900138935.1_10208_3_61 Mycobacterium abscessus subsp. abscessus Terrabacteria group; Actinobacteria; Actinobacteria; Corynebacteriales; Mycobacteriaceae; Mycobacterium; Mycobacterium chelonae group; Mycobacterium abscessus subgroup; Mycobacterium abscessus
MIKHLTRRATVALVVAAAAISPAAYPAMADTATPLNGLFALTPGSCAEGRATGSYFRMIL WP_005073213.1
hypothetical protein [Mycobacterium abscessus] Length: 330\nNumber of predicted TMHs: 2\nExp number of AAs in TMHs: 39.3416199999999\nExp number, first 60 AAs: 17.07688\nTotal prob of N-in: 0.84431\nPOSSIBLE N-term signal sequence\ninside 1 11\nTMhelix 12 29\noutside 30 298\nTMhelix 299 321\ninside 322 330

29972 GCF_900138975.1_10208_3_65 Mycobacterium abscessus subsp. abscessus Terrabacteria group; Actinobacteria; Actinobacteria; Corynebacteriales; Mycobacteriaceae; Mycobacterium; Mycobacterium chelonae group; Mycobacterium abscessus subgroup; Mycobacterium abscessus
MIKHLTRRATVALVVAAAAISPAAYPAMADTATPLNGLFALTPGSCAEGRATGSYFRMIL WP_005073213.1
hypothetical protein [Mycobacterium abscessus] Length: 330\nNumber of predicted TMHs: 2\nExp number of AAs in TMHs: 39.3416199999999\nExp number, first 60 AAs: 17.07688\nTotal prob of N-in: 0.84431\nPOSSIBLE N-term signal sequence\ninside 1 11\nTMhelix 12 29\noutside 30 298\nTMhelix 299 321\ninside 322 330

29973 GCF_900138925.1_10208_3_62 Mycobacterium abscessus subsp. abscessus Terrabacteria group; Actinobacteria; Actinobacteria; Corynebacteriales; Mycobacteriaceae; Mycobacterium; Mycobacterium chelonae group; Mycobacterium abscessus subgroup; Mycobacterium abscessus
MIKHLTRRATVALVVAAAAISPAAYPAMADTATPLNGLFALTPGSCAEGRATGSYFRMIL WP_074258304.1
hypothetical protein [Mycobacterium abscessus] Length: 330\nNumber of predicted TMHs: 2\nExp number of AAs in TMHs: 39.34165\nExp number, first 60 AAs: 17.07689\nTotal prob of N-in: 0.84431\nPOSSIBLE N-term signal sequence\ninside 1 11\nTMhelix 12 29\noutside 30 298\nTMhelix 299 321\ninside 322 330

29974 GCF_900139025.1_10465_1_49 Mycobacterium abscessus subsp. abscessus Terrabacteria group; Actinobacteria; Actinobacteria; Corynebacteriales; Mycobacteriaceae; Mycobacterium; Mycobacterium chelonae group; Mycobacterium abscessus subgroup; Mycobacterium abscessus
MIKHLTRRATVALVVAAAAISPAAYPAMADTATPLNGLFALTPGSCAEGRATGSYFRMIL WP_005073213.1
hypothetical protein [Mycobacterium abscessus] Length: 330\nNumber of predicted TMHs: 2\nExp number of AAs in TMHs: 39.3416199999999\nExp number, first 60 AAs: 17.07688\nTotal prob of N-in: 0.84431\nPOSSIBLE N-term signal sequence\ninside 1 11\nTMhelix 12 29\noutside 30 298\nTMhelix 299 321\ninside 322 330

29975 GCF_900139115.1_10465_1_57 Mycobacterium abscessus subsp. abscessus Terrabacteria group; Actinobacteria; Actinobacteria; Corynebacteriales; Mycobacteriaceae; Mycobacterium; Mycobacterium chelonae group; Mycobacterium abscessus subgroup; Mycobacterium abscessus
MIKHLTRRATVALVVAAAAISPAAYPAMADTATPLNGLFALTPGSCAEGRATGSYFRMIL WP_005073213.1
hypothetical protein [Mycobacterium abscessus] Length: 330\nNumber of predicted TMHs: 2\nExp number of AAs in TMHs: 39.3416199999999\nExp number, first 60 AAs: 17.07688\nTotal prob of N-in: 0.84431\nPOSSIBLE N-term signal sequence\ninside 1 11\nTMhelix 12 29\noutside 30 298\nTMhelix 299 321\ninside 322 330

29976 GCF_900139135.1_10465_1_58 Mycobacterium abscessus subsp. abscessus Terrabacteria group; Actinobacteria; Actinobacteria; Corynebacteriales; Mycobacteriaceae; Mycobacterium; Mycobacterium chelonae group; Mycobacterium abscessus subgroup; Mycobacterium abscessus
MIKHLTRRATVALVVAAAAISPAAYPAMADTATPLNGLFALTPGSCAEGRATGSYFRMIL WP_005073213.1
hypothetical protein [Mycobacterium abscessus] Length: 330\nNumber of predicted TMHs: 2\nExp number of AAs in TMHs: 39.3416199999999\nExp number, first 60 AAs: 17.07688\nTotal prob of N-in: 0.84431\nPOSSIBLE N-term signal sequence\ninside 1 11\nTMhelix 12 29\noutside 30 298\nTMhelix 299 321\ninside 322 330

29977 GCF_900139185.1_10465_1_61 Mycobacterium abscessus subsp. abscessus Terrabacteria group; Actinobacteria; Actinobacteria; Corynebacteriales; Mycobacteriaceae; Mycobacterium; Mycobacterium chelonae group; Mycobacterium abscessus subgroup; Mycobacterium abscessus
MIKHLTRRATVALVVAAAAISPAAYPAMADTATPLNGLFALTPGSCAEGRATGSYFRMIL WP_005073213.1
hypothetical protein [Mycobacterium abscessus] Length: 330\nNumber of predicted TMHs: 2\nExp number of AAs in TMHs: 39.3416199999999\nExp number, first 60 AAs: 17.07688\nTotal prob of N-in: 0.84431\nPOSSIBLE N-term signal sequence\ninside 1 11\nTMhelix 12 29\noutside 30 298\nTMhelix 299 321\ninside 322 330

29978 GCF_900139155.1_10465_1_64 Mycobacterium abscessus subsp. abscessus Terrabacteria group; Actinobacteria; Actinobacteria; Corynebacteriales; Mycobacteriaceae; Mycobacterium; Mycobacterium chelonae group; Mycobacterium abscessus subgroup; Mycobacterium abscessus
MIKHLTRRATVALVVAAAAISPAAYPAMADTATPLNGLFALTPGSCAEGRATGSYFRMIL WP_074343781.1
hypothetical protein [Mycobacterium abscessus] Length: 330\nNumber of predicted TMHs: 2\nExp number of AAs in TMHs: 39.3417499999999\nExp number, first 60 AAs: 17.07664\nTotal prob of N-in: 0.84432\nPOSSIBLE N-term signal sequence\ninside 1 11\nTMhelix 12 29\noutside 30 298\nTMhelix 299 321\ninside 322 330

29979 GCF_900139245.1_10465_1_70 Mycobacterium abscessus subsp. abscessus Terrabacteria group; Actinobacteria; Actinobacteria; Corynebacteriales; Mycobacteriaceae; Mycobacterium; Mycobacterium chelonae group; Mycobacterium abscessus subgroup; Mycobacterium abscessus
MIKHLTRRATVALVVAAAAISPAAYPAMADTATPLNGLFALTPGSCAEGRATGSYFRMIL WP_005073213.1
hypothetical protein [Mycobacterium abscessus] Length: 330\nNumber of predicted TMHs: 2\nExp number of AAs in TMHs: 39.3416199999999\nExp number, first 60 AAs: 17.07688\nTotal prob of N-in: 0.84431\nPOSSIBLE N-term signal sequence\ninside 1 11\nTMhelix 12 29\noutside 30 298\nTMhelix 299 321\ninside 322 330

29980 GCF_900139345.1_10625_4_10 Mycobacterium abscessus subsp. abscessus Terrabacteria group; Actinobacteria; Actinobacteria; Corynebacteriales; Mycobacteriaceae; Mycobacterium; Mycobacterium chelonae group; Mycobacterium abscessus subgroup; Mycobacterium abscessus
MIKHLTRRATVALVVAAAAISPAAYPAMADTATPLNGLFALTPGSCAEGRATGSYFRMIL WP_005073213.1
hypothetical protein [Mycobacterium abscessus] Length: 330\nNumber of predicted TMHs: 2\nExp number of AAs in TMHs: 39.3416199999999\nExp number, first 60 AAs: 17.07688\nTotal prob of N-in: 0.84431\nPOSSIBLE N-term signal sequence\ninside 1 11\nTMhelix 12 29\noutside 30 298\nTMhelix 299 321\ninside 322 330

29981 GCF_900139395.1_10625_4_16 Mycobacterium abscessus subsp. abscessus Terrabacteria group; Actinobacteria; Actinobacteria; Corynebacteriales; Mycobacteriaceae; Mycobacterium; Mycobacterium chelonae group; Mycobacterium abscessus subgroup; Mycobacterium abscessus
MIKHLTRRATVALVVAAAAISPAAYPAMADTATPLNGLFALTPGSCAEGRATGSYFRMIL WP_005073213.1
hypothetical protein [Mycobacterium abscessus] Length: 330\nNumber of predicted TMHs: 2\nExp number of AAs in TMHs: 39.3416199999999\nExp number, first 60 AAs: 17.07688\nTotal prob of N-in: 0.84431\nPOSSIBLE N-term signal sequence\ninside 1 11\nTMhelix 12 29\noutside 30 298\nTMhelix 299 321\ninside 322 330

29982 GCF_900139415.1_10625_4_18 Mycobacterium abscessus subsp. abscessus Terrabacteria group; Actinobacteria; Actinobacteria; Corynebacteriales; Mycobacteriaceae; Mycobacterium; Mycobacterium chelonae group; Mycobacterium abscessus subgroup; Mycobacterium abscessus
MIKHLTRRATVALVVAAAAISPAAYPAMADTATPLNGLFALTPGSCAEGRATGSYFRMIL WP_005073213.1
hypothetical protein [Mycobacterium abscessus] Length: 330\nNumber of predicted TMHs: 2\nExp number of

AAs in TMHs: 39.3416199999999\nExp number, first 60 AAs:  17.07688\nTotal prob of N-in:   0.84431\nPOSSIBLE N-term signal sequence\ninside   1   11\nTMhelix   12   29\noutside   30   298\nTMhelix   299   321\ninside   322   330

29983 GCF_900139435.1_10625_4_85 Mycobacterium abscessus subsp. abscessus Terrabacteria group; Actinobacteria; Actinobacteria; Corynebacteriales; Mycobacteriaceae; Mycobacterium; Mycobacterium chelonae group; Mycobacterium abscessus subgroup; Mycobacterium abscessus
MIKHLTRRATVALVVAAAAISPAAYPAMADTATPLNGLFALTPGSCAEGRATGSYFRMIL WP_074250269.1
hypothetical protein [Mycobacterium abscessus] Length: 330\nNumber of predicted TMHs:  2\nExp number of AAs in TMHs: 39.3415199999999\nExp number, first 60 AAs:  17.07715\nTotal prob of N-in:   0.84431\nPOSSIBLE N-term signal sequence\ninside   1   11\nTMhelix   12   29\noutside   30   298\nTMhelix   299   321\ninside   322   330

29984 GCF_900139455.1_10625_4_7 Mycobacterium abscessus subsp. abscessus Terrabacteria group; Actinobacteria; Actinobacteria; Corynebacteriales; Mycobacteriaceae; Mycobacterium; Mycobacterium chelonae group; Mycobacterium abscessus subgroup; Mycobacterium abscessus
MIKHLTRRATVALVVAAAAISPAAYPAMADTATPLNGLFALTPGSCAEGRATGSYFRMIL WP_005073213.1
hypothetical protein [Mycobacterium abscessus] Length: 330\nNumber of predicted TMHs:  2\nExp number of AAs in TMHs: 39.3416199999999\nExp number, first 60 AAs:  17.07688\nTotal prob of N-in:   0.84431\nPOSSIBLE N-term signal sequence\ninside   1   11\nTMhelix   12   29\noutside   30   298\nTMhelix   299   321\ninside   322   330

29985 GCF_900139705.1_10660_1_12 Mycobacterium abscessus subsp. abscessus Terrabacteria group; Actinobacteria; Actinobacteria; Corynebacteriales; Mycobacteriaceae; Mycobacterium; Mycobacterium chelonae group; Mycobacterium abscessus subgroup; Mycobacterium abscessus
MIKHLTRRATVALVVAAAAISPAAYPAMADTATPLNGLFALTPGSCAEGRATGSYFRMIL WP_005073213.1
hypothetical protein [Mycobacterium abscessus] Length: 330\nNumber of predicted TMHs:  2\nExp number of AAs in TMHs: 39.3416199999999\nExp number, first 60 AAs:  17.07688\nTotal prob of N-in:   0.84431\nPOSSIBLE N-term signal sequence\ninside   1   11\nTMhelix   12   29\noutside   30   298\nTMhelix   299   321\ninside   322   330

29986 GCF_900139735.1_10660_1_17 Mycobacterium abscessus subsp. abscessus Terrabacteria group; Actinobacteria; Actinobacteria; Corynebacteriales; Mycobacteriaceae; Mycobacterium; Mycobacterium chelonae group; Mycobacterium abscessus subgroup; Mycobacterium abscessus
MIKHLTRRATVALVVAAAAISPAAYPAMADTATPLNGLFALTPGSCAEGRATGSYFRMIL WP_005073213.1
hypothetical protein [Mycobacterium abscessus] Length: 330\nNumber of predicted TMHs:  2\nExp number of AAs in TMHs: 39.3416199999999\nExp number, first 60 AAs:  17.07688\nTotal prob of N-in:   0.84431\nPOSSIBLE N-term signal sequence\ninside   1   11\nTMhelix   12   29\noutside   30   298\nTMhelix   299   321\ninside   322   330

29987 GCF_900139895.1_10660_1_35 Mycobacterium abscessus subsp. abscessus Terrabacteria group; Actinobacteria; Actinobacteria; Corynebacteriales; Mycobacteriaceae; Mycobacterium; Mycobacterium chelonae group; Mycobacterium abscessus subgroup; Mycobacterium abscessus
MIKHLTRRATVALVVAAAAISPAAYPAMADTATPLNGLFALTPGSCAEGRATGSYFRMIL WP_005073213.1
hypothetical protein [Mycobacterium abscessus] Length: 330\nNumber of predicted TMHs:  2\nExp number of AAs in TMHs: 39.3416199999999\nExp number, first 60 AAs:  17.07688\nTotal prob of N-in:   0.84431\nPOSSIBLE N-term signal sequence\ninside   1   11\nTMhelix   12   29\noutside   30   298\nTMhelix   299   321\ninside   322   330

29988 GCF_900139955.1_10660_1_38 Mycobacterium abscessus subsp. abscessus Terrabacteria group; Actinobacteria; Actinobacteria; Corynebacteriales; Mycobacteriaceae; Mycobacterium; Mycobacterium chelonae group; Mycobacterium abscessus subgroup; Mycobacterium abscessus
MIKHLTRRATVALVVAAAAISPAAYPAMADTATPLNGLFALTPGSCAEGRATGSYFRMIL WP_005073213.1
hypothetical protein [Mycobacterium abscessus] Length: 330\nNumber of predicted TMHs:  2\nExp number of AAs in TMHs: 39.3416199999999\nExp number, first 60 AAs:  17.07688\nTotal prob of N-in:   0.84431\nPOSSIBLE N-term signal sequence\ninside   1   11\nTMhelix   12   29\noutside   30   298\nTMhelix   299   321\ninside   322   330

29989 GCF_900140005.1_10660_1_53 Mycobacterium abscessus subsp. abscessus Terrabacteria group; Actinobacteria; Actinobacteria; Corynebacteriales; Mycobacteriaceae; Mycobacterium; Mycobacterium chelonae group; Mycobacterium abscessus subgroup; Mycobacterium abscessus
MIKHLTRRATVALVVAAAAISPAAYPAMADTATPLNGLFALTPGSCAEGRATGSYFRMIL WP_005073213.1
hypothetical protein [Mycobacterium abscessus] Length: 330\nNumber of predicted TMHs:  2\nExp number of AAs in TMHs: 39.3416199999999\nExp number, first 60 AAs:  17.07688\nTotal prob of N-in:   0.84431\nPOSSIBLE N-term signal sequence\ninside   1   11\nTMhelix   12   29\noutside   30   298\nTMhelix   299   321\ninside   322   330

29990 GCF_900140105.1_10665_2_67 Mycobacterium abscessus subsp. abscessus Terrabacteria group; Actinobacteria; Actinobacteria; Corynebacteriales; Mycobacteriaceae; Mycobacterium; Mycobacterium chelonae group;

Mycobacterium abscessus subgroup; Mycobacterium abscessus
MIKHLTRRATVALVVAAAAISPAAYPAMADTATPLNGLFALTPGSCAEGRATGSYFRMIL WP_005073213.1
hypothetical protein [Mycobacterium abscessus] Length: 330\nNumber of predicted TMHs: 2\nExp number of AAs in TMHs: 39.3416199999999\nExp number, first 60 AAs: 17.07688\nTotal prob of N-in: 0.84431\nPOSSIBLE N-term signal sequence\ninside 1 11\nTMhelix 12 29\noutside 30 298\nTMhelix 299 321\ninside 322 330

29991 GCF_900140125.1_10665_2_64 Mycobacterium abscessus subsp. abscessus Terrabacteria group; Actinobacteria; Actinobacteria; Corynebacteriales; Mycobacteriaceae; Mycobacterium; Mycobacterium chelonae group; Mycobacterium abscessus subgroup; Mycobacterium abscessus
MIKHLTRRATVALVVAAAAISPAAYPAMADTATPLNGLFALTPGSCAEGRATGSYFRMIL WP_005073213.1
hypothetical protein [Mycobacterium abscessus] Length: 330\nNumber of predicted TMHs: 2\nExp number of AAs in TMHs: 39.3416199999999\nExp number, first 60 AAs: 17.07688\nTotal prob of N-in: 0.84431\nPOSSIBLE N-term signal sequence\ninside 1 11\nTMhelix 12 29\noutside 30 298\nTMhelix 299 321\ninside 322 330

29992 GCF_900140145.1_10665_2_61 Mycobacterium abscessus subsp. abscessus Terrabacteria group; Actinobacteria; Actinobacteria; Corynebacteriales; Mycobacteriaceae; Mycobacterium; Mycobacterium chelonae group; Mycobacterium abscessus subgroup; Mycobacterium abscessus
MIKHLTRRATVALVVAAAAISPAAYPAMADTATPLNGLFALTPGSCAEGRATGSYFRMIL WP_005073213.1
hypothetical protein [Mycobacterium abscessus] Length: 330\nNumber of predicted TMHs: 2\nExp number of AAs in TMHs: 39.3416199999999\nExp number, first 60 AAs: 17.07688\nTotal prob of N-in: 0.84431\nPOSSIBLE N-term signal sequence\ninside 1 11\nTMhelix 12 29\noutside 30 298\nTMhelix 299 321\ninside 322 330

29993 GCF_900140165.1_10665_2_68 Mycobacterium abscessus subsp. abscessus Terrabacteria group; Actinobacteria; Actinobacteria; Corynebacteriales; Mycobacteriaceae; Mycobacterium; Mycobacterium chelonae group; Mycobacterium abscessus subgroup; Mycobacterium abscessus
MIKHLTRRATVALVVAAAAISPAAYPAMADTATPLNGLFALTPGSCAEGRATGSYFRMIL WP_005073213.1
hypothetical protein [Mycobacterium abscessus] Length: 330\nNumber of predicted TMHs: 2\nExp number of AAs in TMHs: 39.3416199999999\nExp number, first 60 AAs: 17.07688\nTotal prob of N-in: 0.84431\nPOSSIBLE N-term signal sequence\ninside 1 11\nTMhelix 12 29\noutside 30 298\nTMhelix 299 321\ninside 322 330

29994 GCF_900140375.1_10665_3_78 Mycobacterium abscessus subsp. abscessus Terrabacteria group; Actinobacteria; Actinobacteria; Corynebacteriales; Mycobacteriaceae; Mycobacterium; Mycobacterium chelonae group; Mycobacterium abscessus subgroup; Mycobacterium abscessus
MIKHLTRRATVALVVAAAAISPAAYPAMADTATPLNGLFALTPGSCAEGRATGSYFRMIL WP_005073213.1
hypothetical protein [Mycobacterium abscessus] Length: 330\nNumber of predicted TMHs: 2\nExp number of AAs in TMHs: 39.3416199999999\nExp number, first 60 AAs: 17.07688\nTotal prob of N-in: 0.84431\nPOSSIBLE N-term signal sequence\ninside 1 11\nTMhelix 12 29\noutside 30 298\nTMhelix 299 321\ninside 322 330

29995 GCF_900140395.1_10665_4_24 Mycobacterium abscessus subsp. abscessus Terrabacteria group; Actinobacteria; Actinobacteria; Corynebacteriales; Mycobacteriaceae; Mycobacterium; Mycobacterium chelonae group; Mycobacterium abscessus subgroup; Mycobacterium abscessus
MIKHLTRRATVALVVAAAAISPAAYPAMADTATPLNGLFALTPGSCAEGRATGSYFRMIL WP_005073213.1
hypothetical protein [Mycobacterium abscessus] Length: 330\nNumber of predicted TMHs: 2\nExp number of AAs in TMHs: 39.3416199999999\nExp number, first 60 AAs: 17.07688\nTotal prob of N-in: 0.84431\nPOSSIBLE N-term signal sequence\ninside 1 11\nTMhelix 12 29\noutside 30 298\nTMhelix 299 321\ninside 322 330

29996 GCF_900140325.1_10665_3_26 Mycobacterium abscessus subsp. abscessus Terrabacteria group; Actinobacteria; Actinobacteria; Corynebacteriales; Mycobacteriaceae; Mycobacterium; Mycobacterium chelonae group; Mycobacterium abscessus subgroup; Mycobacterium abscessus
MIKHLTRRATVALVVAAAAISPAAYPAMADTATPLNGLFALTPGSCAEGRATGSYFRMIL WP_074250269.1
hypothetical protein [Mycobacterium abscessus] Length: 330\nNumber of predicted TMHs: 2\nExp number of AAs in TMHs: 39.3415199999999\nExp number, first 60 AAs: 17.07715\nTotal prob of N-in: 0.84431\nPOSSIBLE N-term signal sequence\ninside 1 11\nTMhelix 12 29\noutside 30 298\nTMhelix 299 321\ninside 322 330

29997 GCF_900140345.1_10665_3_25 Mycobacterium abscessus subsp. abscessus Terrabacteria group; Actinobacteria; Actinobacteria; Corynebacteriales; Mycobacteriaceae; Mycobacterium; Mycobacterium chelonae group; Mycobacterium abscessus subgroup; Mycobacterium abscessus
MIKHLTRRATVALVVAAAAISPAAYPAMADTATPLNGLFALTPGSCAEGRATGSYFRMIL WP_074250269.1
hypothetical protein [Mycobacterium abscessus] Length: 330\nNumber of predicted TMHs: 2\nExp number of AAs in TMHs: 39.3415199999999\nExp number, first 60 AAs: 17.07715\nTotal prob of N-in: 0.84431\nPOSSIBLE N-term signal sequence\ninside 1 11\nTMhelix 12 29\noutside 30 298\nTMhelix 299 321\ninside 322 330

29998 GCF_900140415.1_10665_4_74 Mycobacterium abscessus subsp. abscessus Terrabacteria group; Actinobacteria; Actinobacteria; Corynebacteriales; Mycobacteriaceae; Mycobacterium; Mycobacterium chelonae group; Mycobacterium abscessus subgroup; Mycobacterium abscessus
MIKHLTRRATVALVVAAAAISPAAYPAMADTATPLNGLFALTPGSCAEGRATGSYFRMIL WP_079642101.1
hypothetical protein [Mycobacterium abscessus] Length: 330\nNumber of predicted TMHs: 2\nExp number of AAs in TMHs: 39.34159\nExp number, first 60 AAs: 17.07688\nTotal prob of N-in: 0.84431\nPOSSIBLE N-term signal sequence\ninside 1 11\nTMhelix 12 29\noutside 30 298\nTMhelix 299 321\ninside 322 330

29999 GCF_900140785.1_10702_1_41 Mycobacterium abscessus subsp. abscessus Terrabacteria group; Actinobacteria; Actinobacteria; Corynebacteriales; Mycobacteriaceae; Mycobacterium; Mycobacterium chelonae group; Mycobacterium abscessus subgroup; Mycobacterium abscessus
MIKHLTRRATVALVVAAAAISPAAYPAMADTATPLNGLFALTPGSCAEGRATGSYFRMIL WP_005073213.1
hypothetical protein [Mycobacterium abscessus] Length: 330\nNumber of predicted TMHs: 2\nExp number of AAs in TMHs: 39.3416199999999\nExp number, first 60 AAs: 17.07688\nTotal prob of N-in: 0.84431\nPOSSIBLE N-term signal sequence\ninside 1 11\nTMhelix 12 29\noutside 30 298\nTMhelix 299 321\ninside 322 330

30000 GCF_900140905.1_10702_1_53 Mycobacterium abscessus subsp. abscessus Terrabacteria group; Actinobacteria; Actinobacteria; Corynebacteriales; Mycobacteriaceae; Mycobacterium; Mycobacterium chelonae group; Mycobacterium abscessus subgroup; Mycobacterium abscessus
MIKHLTRRATVALVVAAAAISPAAYPAMADTATPLNGLFALTPGSCAEGRATGSYFRMIL WP_005073213.1
hypothetical protein [Mycobacterium abscessus] Length: 330\nNumber of predicted TMHs: 2\nExp number of AAs in TMHs: 39.3416199999999\nExp number, first 60 AAs: 17.07688\nTotal prob of N-in: 0.84431\nPOSSIBLE N-term signal sequence\ninside 1 11\nTMhelix 12 29\noutside 30 298\nTMhelix 299 321\ninside 322 330

30001 GCF_900140925.1_10702_1_55 Mycobacterium abscessus subsp. abscessus Terrabacteria group; Actinobacteria; Actinobacteria; Corynebacteriales; Mycobacteriaceae; Mycobacterium; Mycobacterium chelonae group; Mycobacterium abscessus subgroup; Mycobacterium abscessus
MIKHLTRRATVALVVAAAAISPAAYPAMADTATPLNGLFALTPGSCAEGRATGSYFRMIL WP_005073213.1
hypothetical protein [Mycobacterium abscessus] Length: 330\nNumber of predicted TMHs: 2\nExp number of AAs in TMHs: 39.3416199999999\nExp number, first 60 AAs: 17.07688\nTotal prob of N-in: 0.84431\nPOSSIBLE N-term signal sequence\ninside 1 11\nTMhelix 12 29\noutside 30 298\nTMhelix 299 321\ninside 322 330

30002 GCF_900140935.1_10702_1_60 Mycobacterium abscessus subsp. abscessus Terrabacteria group; Actinobacteria; Actinobacteria; Corynebacteriales; Mycobacteriaceae; Mycobacterium; Mycobacterium chelonae group; Mycobacterium abscessus subgroup; Mycobacterium abscessus
MIKHLTRRATVALVVAAAAISPAAYPAMADTATPLNGLFALTPGSCAEGRATGSYFRMIL WP_005073213.1
hypothetical protein [Mycobacterium abscessus] Length: 330\nNumber of predicted TMHs: 2\nExp number of AAs in TMHs: 39.3416199999999\nExp number, first 60 AAs: 17.07688\nTotal prob of N-in: 0.84431\nPOSSIBLE N-term signal sequence\ninside 1 11\nTMhelix 12 29\noutside 30 298\nTMhelix 299 321\ninside 322 330

30003 GCF_900140885.1_10702_1_51 Mycobacterium abscessus subsp. abscessus Terrabacteria group; Actinobacteria; Actinobacteria; Corynebacteriales; Mycobacteriaceae; Mycobacterium; Mycobacterium chelonae group; Mycobacterium abscessus subgroup; Mycobacterium abscessus
MIKHLTRRATVALVVAAAAISPAAYPAVADTATPLNGLFALTPGSCAEGRATGSYFRMIL WP_079583063.1 hypothetical protein [Mycobacterium abscessus] Length: 330\nNumber of predicted TMHs: 1\nExp number of AAs in TMHs: 39.25299\nExp number, first 60 AAs: 16.98088\nTotal prob of N-in: 0.83210\nPOSSIBLE N-term signal sequence\noutside 1 298\nTMhelix 299 321\ninside 322 330

30004 GCF_900141025.1_10702_1_69 Mycobacterium abscessus subsp. abscessus Terrabacteria group; Actinobacteria; Actinobacteria; Corynebacteriales; Mycobacteriaceae; Mycobacterium; Mycobacterium chelonae group; Mycobacterium abscessus subgroup; Mycobacterium abscessus
MIKHLTRRATVALVVAAAAISPAAYPAMADTATPLNGLFALTPGSCAEGRATGSYFRMIL WP_005073213.1
hypothetical protein [Mycobacterium abscessus] Length: 330\nNumber of predicted TMHs: 2\nExp number of AAs in TMHs: 39.3416199999999\nExp number, first 60 AAs: 17.07688\nTotal prob of N-in: 0.84431\nPOSSIBLE N-term signal sequence\ninside 1 11\nTMhelix 12 29\noutside 30 298\nTMhelix 299 321\ninside 322 330

30005 GCF_900141525.1_12082_5_83 Mycobacterium abscessus subsp. abscessus Terrabacteria group; Actinobacteria; Actinobacteria; Corynebacteriales; Mycobacteriaceae; Mycobacterium; Mycobacterium chelonae group; Mycobacterium abscessus subgroup; Mycobacterium abscessus
MIKHLTRRATVALVVAAAAISPAAYPAMADTATPLNGLFALTPGSCAEGRATGSYFRMIL WP_005073213.1
hypothetical protein [Mycobacterium abscessus] Length: 330\nNumber of predicted TMHs: 2\nExp number of

AAs in TMHs: 39.3416199999999\nExp number, first 60 AAs: 17.07688\nTotal prob of N-in: 0.84431\nPOSSIBLE N-term signal sequence\ninside 1 11\nTMhelix 12 29\noutside 30 298\nTMhelix 299 321\ninside 322 330

30006 GCF_900141535.1_12163_2_68 Mycobacterium abscessus subsp. abscessus Terrabacteria group; Actinobacteria; Actinobacteria; Corynebacteriales; Mycobacteriaceae; Mycobacterium; Mycobacterium chelonae group; Mycobacterium abscessus subgroup; Mycobacterium abscessus
MIKHLTRRATVALVVAAAAISPAAYPAMADTATPLNGLFALTPGSCAEGRATGSYFRMIL WP_005073213.1
hypothetical protein [Mycobacterium abscessus] Length: 330\nNumber of predicted TMHs: 2\nExp number of AAs in TMHs: 39.3416199999999\nExp number, first 60 AAs: 17.07688\nTotal prob of N-in: 0.84431\nPOSSIBLE N-term signal sequence\ninside 1 11\nTMhelix 12 29\noutside 30 298\nTMhelix 299 321\ninside 322 330

30007 GCF_900141555.1_12082_5_74 Mycobacterium abscessus subsp. abscessus Terrabacteria group; Actinobacteria; Actinobacteria; Corynebacteriales; Mycobacteriaceae; Mycobacterium; Mycobacterium chelonae group; Mycobacterium abscessus subgroup; Mycobacterium abscessus
MIKHLTRRATVALVVAAAAISPAAYPAMADTATPLNGLFALTPGSCAEGRATGSYFRMIL WP_005073213.1
hypothetical protein [Mycobacterium abscessus] Length: 330\nNumber of predicted TMHs: 2\nExp number of AAs in TMHs: 39.3416199999999\nExp number, first 60 AAs: 17.07688\nTotal prob of N-in: 0.84431\nPOSSIBLE N-term signal sequence\ninside 1 11\nTMhelix 12 29\noutside 30 298\nTMhelix 299 321\ninside 322 330

30008 GCF_900141575.1_12163_2_69 Mycobacterium abscessus subsp. abscessus Terrabacteria group; Actinobacteria; Actinobacteria; Corynebacteriales; Mycobacteriaceae; Mycobacterium; Mycobacterium chelonae group; Mycobacterium abscessus subgroup; Mycobacterium abscessus
MIKHLTRRATVALVVAAAAISPAAYPAMADTATPLNGLFALTPGSCAEGRATGSYFRMIL WP_005073213.1
hypothetical protein [Mycobacterium abscessus] Length: 330\nNumber of predicted TMHs: 2\nExp number of AAs in TMHs: 39.3416199999999\nExp number, first 60 AAs: 17.07688\nTotal prob of N-in: 0.84431\nPOSSIBLE N-term signal sequence\ninside 1 11\nTMhelix 12 29\noutside 30 298\nTMhelix 299 321\ninside 322 330

30009 GCF_900141585.1_12163_2_70 Mycobacterium abscessus subsp. abscessus Terrabacteria group; Actinobacteria; Actinobacteria; Corynebacteriales; Mycobacteriaceae; Mycobacterium; Mycobacterium chelonae group; Mycobacterium abscessus subgroup; Mycobacterium abscessus
MIKHLTRRATVALVVAAAAISPAAYPAMADTATPLNGLFALTPGSCAEGRATGSYFRMIL WP_005073213.1
hypothetical protein [Mycobacterium abscessus] Length: 330\nNumber of predicted TMHs: 2\nExp number of AAs in TMHs: 39.3416199999999\nExp number, first 60 AAs: 17.07688\nTotal prob of N-in: 0.84431\nPOSSIBLE N-term signal sequence\ninside 1 11\nTMhelix 12 29\noutside 30 298\nTMhelix 299 321\ninside 322 330

30010 GCF_900138175.1_12082_5_87 Mycobacterium abscessus subsp. abscessus Terrabacteria group; Actinobacteria; Actinobacteria; Corynebacteriales; Mycobacteriaceae; Mycobacterium; Mycobacterium chelonae group; Mycobacterium abscessus subgroup; Mycobacterium abscessus
MIKHLTRRATVALVVAAAAISPAAYPAMADTATPLNGLFALTPGSCAEGRATGSYFRMIL WP_005073213.1
hypothetical protein [Mycobacterium abscessus] Length: 330\nNumber of predicted TMHs: 2\nExp number of AAs in TMHs: 39.3416199999999\nExp number, first 60 AAs: 17.07688\nTotal prob of N-in: 0.84431\nPOSSIBLE N-term signal sequence\ninside 1 11\nTMhelix 12 29\noutside 30 298\nTMhelix 299 321\ninside 322 330

30011 GCF_002072635.1_ASM207263v1 Mycobacterium abscessus subsp. abscessus Terrabacteria group; Actinobacteria; Actinobacteria; Corynebacteriales; Mycobacteriaceae; Mycobacterium; Mycobacterium chelonae group; Mycobacterium abscessus subgroup; Mycobacterium abscessus
MIKHLTRRATVALVVAAAAISPAAYPAMADTATPLNGLFALTPGSCAEGRATGSYFRMIL WP_005073213.1
hypothetical protein [Mycobacterium abscessus] Length: 330\nNumber of predicted TMHs: 2\nExp number of AAs in TMHs: 39.3416199999999\nExp number, first 60 AAs: 17.07688\nTotal prob of N-in: 0.84431\nPOSSIBLE N-term signal sequence\ninside 1 11\nTMhelix 12 29\noutside 30 298\nTMhelix 299 321\ninside 322 330

30012 GCF_900130415.1_10625_5_77 Mycobacterium abscessus subsp. abscessus Terrabacteria group; Actinobacteria; Actinobacteria; Corynebacteriales; Mycobacteriaceae; Mycobacterium; Mycobacterium chelonae group; Mycobacterium abscessus subgroup; Mycobacterium abscessus
MIKHLTRRATVALVVAAAAISPAAYPAMADTATPLNGLFALTPGSCAEGRATGSYFRMIL WP_005073213.1
hypothetical protein [Mycobacterium abscessus] Length: 330\nNumber of predicted TMHs: 2\nExp number of AAs in TMHs: 39.3416199999999\nExp number, first 60 AAs: 17.07688\nTotal prob of N-in: 0.84431\nPOSSIBLE N-term signal sequence\ninside 1 11\nTMhelix 12 29\noutside 30 298\nTMhelix 299 321\ninside 322 330

30013 GCF_900130555.1_10665_3_6 Mycobacterium abscessus subsp. abscessus Terrabacteria group; Actinobacteria; Actinobacteria; Corynebacteriales; Mycobacteriaceae; Mycobacterium; Mycobacterium chelonae group;

Mycobacterium abscessus subgroup; Mycobacterium abscessus
MIKHLTRRATVALVVAAAAISPAAYPAMADTATPLNGLFALTPGSCAEGRATGSYFRMIL WP_005073213.1
hypothetical protein [Mycobacterium abscessus] Length: 330\nNumber of predicted TMHs: 2\nExp number of AAs in TMHs: 39.3416199999999\nExp number, first 60 AAs: 17.07688\nTotal prob of N-in: 0.84431\nPOSSIBLE N-term signal sequence\ninside 1 11\nTMhelix 12 29\noutside 30 298\nTMhelix 299 321\ninside 322 330

30014 GCF_900130925.1_10665_3_45 Mycobacterium abscessus subsp. abscessus Terrabacteria group; Actinobacteria; Actinobacteria; Corynebacteriales; Mycobacteriaceae; Mycobacterium; Mycobacterium chelonae group; Mycobacterium abscessus subgroup; Mycobacterium abscessus
MIKHLTRRATVALVVAAAAISPAAYPAMADTATPLNGLFALTPGSCAEGRATGSYFRMIL WP_005073213.1
hypothetical protein [Mycobacterium abscessus] Length: 330\nNumber of predicted TMHs: 2\nExp number of AAs in TMHs: 39.3416199999999\nExp number, first 60 AAs: 17.07688\nTotal prob of N-in: 0.84431\nPOSSIBLE N-term signal sequence\ninside 1 11\nTMhelix 12 29\noutside 30 298\nTMhelix 299 321\ninside 322 330

30015 GCF_900131355.1_10665_3_82 Mycobacterium abscessus subsp. abscessus Terrabacteria group; Actinobacteria; Actinobacteria; Corynebacteriales; Mycobacteriaceae; Mycobacterium; Mycobacterium chelonae group; Mycobacterium abscessus subgroup; Mycobacterium abscessus
MIKHLTRRATVALVVAAAAISPAAYPAMADTATPLNGLFALTPGSCAEGRATGSYFRMIL WP_005073213.1
hypothetical protein [Mycobacterium abscessus] Length: 330\nNumber of predicted TMHs: 2\nExp number of AAs in TMHs: 39.3416199999999\nExp number, first 60 AAs: 17.07688\nTotal prob of N-in: 0.84431\nPOSSIBLE N-term signal sequence\ninside 1 11\nTMhelix 12 29\noutside 30 298\nTMhelix 299 321\ninside 322 330

30016 GCF_900131375.1_10665_3_84 Mycobacterium abscessus subsp. abscessus Terrabacteria group; Actinobacteria; Actinobacteria; Corynebacteriales; Mycobacteriaceae; Mycobacterium; Mycobacterium chelonae group; Mycobacterium abscessus subgroup; Mycobacterium abscessus
MIKHLTRRATVALVVAAAAISPAAYPAMADTATPLNGLFALTPGSCAEGRATGSYFRMIL WP_005073213.1
hypothetical protein [Mycobacterium abscessus] Length: 330\nNumber of predicted TMHs: 2\nExp number of AAs in TMHs: 39.3416199999999\nExp number, first 60 AAs: 17.07688\nTotal prob of N-in: 0.84431\nPOSSIBLE N-term signal sequence\ninside 1 11\nTMhelix 12 29\noutside 30 298\nTMhelix 299 321\ninside 322 330

30017 GCF_900131675.1_10665_4_26 Mycobacterium abscessus subsp. abscessus Terrabacteria group; Actinobacteria; Actinobacteria; Corynebacteriales; Mycobacteriaceae; Mycobacterium; Mycobacterium chelonae group; Mycobacterium abscessus subgroup; Mycobacterium abscessus
MIKHLTRRATVALVVAAAAISPAAYPAMADTATPLNGLFALTPGSCAEGRATGSYFRMIL WP_005073213.1
hypothetical protein [Mycobacterium abscessus] Length: 330\nNumber of predicted TMHs: 2\nExp number of AAs in TMHs: 39.3416199999999\nExp number, first 60 AAs: 17.07688\nTotal prob of N-in: 0.84431\nPOSSIBLE N-term signal sequence\ninside 1 11\nTMhelix 12 29\noutside 30 298\nTMhelix 299 321\ninside 322 330

30018 GCF_900131475.1_11861_7_29 Mycobacterium abscessus subsp. abscessus Terrabacteria group; Actinobacteria; Actinobacteria; Corynebacteriales; Mycobacteriaceae; Mycobacterium; Mycobacterium chelonae group; Mycobacterium abscessus subgroup; Mycobacterium abscessus
MIKHLTRRATVALVVAAAAISPAAYPAMADTATPLNGLFALTPGSCAEGRATGSYFRMIL WP_074340136.1
hypothetical protein [Mycobacterium abscessus] Length: 330\nNumber of predicted TMHs: 2\nExp number of AAs in TMHs: 39.3416799999999\nExp number, first 60 AAs: 17.07671\nTotal prob of N-in: 0.84432\nPOSSIBLE N-term signal sequence\ninside 1 11\nTMhelix 12 29\noutside 30 298\nTMhelix 299 321\ninside 322 330

30019 GCF_900132155.1_10665_4_60 Mycobacterium abscessus subsp. abscessus Terrabacteria group; Actinobacteria; Actinobacteria; Corynebacteriales; Mycobacteriaceae; Mycobacterium; Mycobacterium chelonae group; Mycobacterium abscessus subgroup; Mycobacterium abscessus
MIKHLTRRATVALVVAAAAISPAAYPAMADTATPLNGLFALTPGSCAEGRATGSYFRMIL WP_005073213.1
hypothetical protein [Mycobacterium abscessus] Length: 330\nNumber of predicted TMHs: 2\nExp number of AAs in TMHs: 39.3416199999999\nExp number, first 60 AAs: 17.07688\nTotal prob of N-in: 0.84431\nPOSSIBLE N-term signal sequence\ninside 1 11\nTMhelix 12 29\noutside 30 298\nTMhelix 299 321\ninside 322 330

30020 GCF_900131975.1_10665_4_43 Mycobacterium abscessus subsp. abscessus Terrabacteria group; Actinobacteria; Actinobacteria; Corynebacteriales; Mycobacteriaceae; Mycobacterium; Mycobacterium chelonae group; Mycobacterium abscessus subgroup; Mycobacterium abscessus
MIKHLTRRATVALVVAAAAISPAAYPAMADTATPLNGLFALTPGSCAEGRATGSYFRMIL WP_074250269.1
hypothetical protein [Mycobacterium abscessus] Length: 330\nNumber of predicted TMHs: 2\nExp number of AAs in TMHs: 39.3415199999999\nExp number, first 60 AAs: 17.07715\nTotal prob of N-in: 0.84431\nPOSSIBLE N-term signal sequence\ninside 1 11\nTMhelix 12 29\noutside 30 298\nTMhelix 299 321\ninside 322 330

30021 GCF_900132585.1_10250_1_11 Mycobacterium abscessus subsp. abscessus Terrabacteria group; Actinobacteria; Actinobacteria; Corynebacteriales; Mycobacteriaceae; Mycobacterium; Mycobacterium chelonae group; Mycobacterium abscessus subgroup; Mycobacterium abscessus
MIKHLTRRATVALVVAAAAISPAAYPAMADTATPLNGLFALTPGSCAEGRATGSYFRMIL WP_005073213.1
hypothetical protein [Mycobacterium abscessus] Length: 330\nNumber of predicted TMHs: 2\nExp number of AAs in TMHs: 39.3416199999999\nExp number, first 60 AAs: 17.07688\nTotal prob of N-in: 0.84431\nPOSSIBLE N-term signal sequence\ninside 1 11\nTMhelix 12 29\noutside 30 298\nTMhelix 299 321\ninside 322 330

30022 GCF_900132605.1_11893_6_50 Mycobacterium abscessus subsp. abscessus Terrabacteria group; Actinobacteria; Actinobacteria; Corynebacteriales; Mycobacteriaceae; Mycobacterium; Mycobacterium chelonae group; Mycobacterium abscessus subgroup; Mycobacterium abscessus
MIKHLTRRATVALVVAAAAISPAAYPAMADTATPLNGLFALTPGSCAEGRATGSYFRMIL WP_005073213.1
hypothetical protein [Mycobacterium abscessus] Length: 330\nNumber of predicted TMHs: 2\nExp number of AAs in TMHs: 39.3416199999999\nExp number, first 60 AAs: 17.07688\nTotal prob of N-in: 0.84431\nPOSSIBLE N-term signal sequence\ninside 1 11\nTMhelix 12 29\noutside 30 298\nTMhelix 299 321\ninside 322 330

30023 GCF_900132655.1_10250_1_13 Mycobacterium abscessus subsp. abscessus Terrabacteria group; Actinobacteria; Actinobacteria; Corynebacteriales; Mycobacteriaceae; Mycobacterium; Mycobacterium chelonae group; Mycobacterium abscessus subgroup; Mycobacterium abscessus
MIKHLTRRATVALVVAAAAISPAAYPAMADTATPLNGLFALTPGSCAEGRATGSYFRMIL WP_005073213.1
hypothetical protein [Mycobacterium abscessus] Length: 330\nNumber of predicted TMHs: 2\nExp number of AAs in TMHs: 39.3416199999999\nExp number, first 60 AAs: 17.07688\nTotal prob of N-in: 0.84431\nPOSSIBLE N-term signal sequence\ninside 1 11\nTMhelix 12 29\noutside 30 298\nTMhelix 299 321\ninside 322 330

30024 GCF_900132685.1_10250_1_18 Mycobacterium abscessus subsp. abscessus Terrabacteria group; Actinobacteria; Actinobacteria; Corynebacteriales; Mycobacteriaceae; Mycobacterium; Mycobacterium chelonae group; Mycobacterium abscessus subgroup; Mycobacterium abscessus
MIKHLTRRATVALVVAAAAISPAAYPAMADTATPLNGLFALTPGSCAEGRATGSYFRMIL WP_005073213.1
hypothetical protein [Mycobacterium abscessus] Length: 330\nNumber of predicted TMHs: 2\nExp number of AAs in TMHs: 39.3416199999999\nExp number, first 60 AAs: 17.07688\nTotal prob of N-in: 0.84431\nPOSSIBLE N-term signal sequence\ninside 1 11\nTMhelix 12 29\noutside 30 298\nTMhelix 299 321\ninside 322 330

30025 GCF_900133035.1_11893_6_63 Mycobacterium abscessus subsp. abscessus Terrabacteria group; Actinobacteria; Actinobacteria; Corynebacteriales; Mycobacteriaceae; Mycobacterium; Mycobacterium chelonae group; Mycobacterium abscessus subgroup; Mycobacterium abscessus
MIKHLTRRATVALVVAAAAISPAAYPAMADTATPLNGLFALTPGSCAEGRATGSYFRMIL WP_005073213.1
hypothetical protein [Mycobacterium abscessus] Length: 330\nNumber of predicted TMHs: 2\nExp number of AAs in TMHs: 39.3416199999999\nExp number, first 60 AAs: 17.07688\nTotal prob of N-in: 0.84431\nPOSSIBLE N-term signal sequence\ninside 1 11\nTMhelix 12 29\noutside 30 298\nTMhelix 299 321\ninside 322 330

30026 GCF_900133085.1_11893_6_69 Mycobacterium abscessus subsp. abscessus Terrabacteria group; Actinobacteria; Actinobacteria; Corynebacteriales; Mycobacteriaceae; Mycobacterium; Mycobacterium chelonae group; Mycobacterium abscessus subgroup; Mycobacterium abscessus
MIKHLTRRATVALVVAAAAISPAAYPAMADTATPLNGLFALTPGSCAEGRATGSYFRMIL WP_005073213.1
hypothetical protein [Mycobacterium abscessus] Length: 330\nNumber of predicted TMHs: 2\nExp number of AAs in TMHs: 39.3416199999999\nExp number, first 60 AAs: 17.07688\nTotal prob of N-in: 0.84431\nPOSSIBLE N-term signal sequence\ninside 1 11\nTMhelix 12 29\noutside 30 298\nTMhelix 299 321\ninside 322 330

30027 GCF_900133125.1_12082_5_53 Mycobacterium abscessus subsp. abscessus Terrabacteria group; Actinobacteria; Actinobacteria; Corynebacteriales; Mycobacteriaceae; Mycobacterium; Mycobacterium chelonae group; Mycobacterium abscessus subgroup; Mycobacterium abscessus
MIKHLTRRATVALVVAAAAISPAAYPAMADTATPLNGLFALTPGSCAEGRATGSYFRMIL WP_005073213.1
hypothetical protein [Mycobacterium abscessus] Length: 330\nNumber of predicted TMHs: 2\nExp number of AAs in TMHs: 39.3416199999999\nExp number, first 60 AAs: 17.07688\nTotal prob of N-in: 0.84431\nPOSSIBLE N-term signal sequence\ninside 1 11\nTMhelix 12 29\noutside 30 298\nTMhelix 299 321\ninside 322 330

30028 GCF_900133655.1_12163_1_10 Mycobacterium abscessus subsp. abscessus Terrabacteria group; Actinobacteria; Actinobacteria; Corynebacteriales; Mycobacteriaceae; Mycobacterium; Mycobacterium chelonae group; Mycobacterium abscessus subgroup; Mycobacterium abscessus
MIKHLTRRATVALVVAAAAISPAAYPAMADTATPLNGLFALTPGSCAEGRATGSYFRMIL WP_005073213.1
hypothetical protein [Mycobacterium abscessus] Length: 330\nNumber of predicted TMHs: 2\nExp number of

AAs in TMHs: 39.3416199999999\nExp number, first 60 AAs: 17.07688\nTotal prob of N-in: 0.84431\nPOSSIBLE N-term signal sequence\ninside 1 11\nTMhelix 12 29\noutside 30 298\nTMhelix 299 321\ninside 322 330

30029 GCF_900133685.1_12163_1_15 Mycobacterium abscessus subsp. abscessus Terrabacteria group; Actinobacteria; Actinobacteria; Corynebacteriales; Mycobacteriaceae; Mycobacterium; Mycobacterium chelonae group; Mycobacterium abscessus subgroup; Mycobacterium abscessus
MIKHLTRRATVALVVAAAAISPAAYPAMADTATPLNGLFALTPGSCAEGRATGSYFRMIL WP_005073213.1
hypothetical protein [Mycobacterium abscessus] Length: 330\nNumber of predicted TMHs: 2\nExp number of AAs in TMHs: 39.3416199999999\nExp number, first 60 AAs: 17.07688\nTotal prob of N-in: 0.84431\nPOSSIBLE N-term signal sequence\ninside 1 11\nTMhelix 12 29\noutside 30 298\nTMhelix 299 321\ninside 322 330

30030 GCF_900133715.1_12163_1_19 Mycobacterium abscessus subsp. abscessus Terrabacteria group; Actinobacteria; Actinobacteria; Corynebacteriales; Mycobacteriaceae; Mycobacterium; Mycobacterium chelonae group; Mycobacterium abscessus subgroup; Mycobacterium abscessus
MIKHLTRRATVALVVAAAAISPAAYPAMADTATPLNGLFALTPGSCAEGRATGSYFRMIL WP_005073213.1
hypothetical protein [Mycobacterium abscessus] Length: 330\nNumber of predicted TMHs: 2\nExp number of AAs in TMHs: 39.3416199999999\nExp number, first 60 AAs: 17.07688\nTotal prob of N-in: 0.84431\nPOSSIBLE N-term signal sequence\ninside 1 11\nTMhelix 12 29\noutside 30 298\nTMhelix 299 321\ninside 322 330

30031 GCF_900134015.1_10396_8_8 Mycobacterium abscessus subsp. abscessus Terrabacteria group; Actinobacteria; Actinobacteria; Corynebacteriales; Mycobacteriaceae; Mycobacterium; Mycobacterium chelonae group; Mycobacterium abscessus subgroup; Mycobacterium abscessus
MIKHLTRRATVALVVAAAAISPAAYPAMADTATPLNGLFALTPGSCAEGRATGSYFRMIL WP_005073213.1
hypothetical protein [Mycobacterium abscessus] Length: 330\nNumber of predicted TMHs: 2\nExp number of AAs in TMHs: 39.3416199999999\nExp number, first 60 AAs: 17.07688\nTotal prob of N-in: 0.84431\nPOSSIBLE N-term signal sequence\ninside 1 11\nTMhelix 12 29\noutside 30 298\nTMhelix 299 321\ninside 322 330

30032 GCF_900133975.1_10250_1_70 Mycobacterium abscessus subsp. abscessus Terrabacteria group; Actinobacteria; Actinobacteria; Corynebacteriales; Mycobacteriaceae; Mycobacterium; Mycobacterium chelonae group; Mycobacterium abscessus subgroup; Mycobacterium abscessus
MIKHLTRRATVALVVAAAAISPAAYPAMADTATPLNGLFALTPGSCAEGRATGSYFRMIL WP_005073213.1
hypothetical protein [Mycobacterium abscessus] Length: 330\nNumber of predicted TMHs: 2\nExp number of AAs in TMHs: 39.3416199999999\nExp number, first 60 AAs: 17.07688\nTotal prob of N-in: 0.84431\nPOSSIBLE N-term signal sequence\ninside 1 11\nTMhelix 12 29\noutside 30 298\nTMhelix 299 321\ninside 322 330

30033 GCF_900134045.1_10250_1_78 Mycobacterium abscessus subsp. abscessus Terrabacteria group; Actinobacteria; Actinobacteria; Corynebacteriales; Mycobacteriaceae; Mycobacterium; Mycobacterium chelonae group; Mycobacterium abscessus subgroup; Mycobacterium abscessus
MIKHLTRRATVALVVAAAAISPAAYPAMADTATPLNGLFALTPGSCAEGRATGSYFRMIL WP_005073213.1
hypothetical protein [Mycobacterium abscessus] Length: 330\nNumber of predicted TMHs: 2\nExp number of AAs in TMHs: 39.3416199999999\nExp number, first 60 AAs: 17.07688\nTotal prob of N-in: 0.84431\nPOSSIBLE N-term signal sequence\ninside 1 11\nTMhelix 12 29\noutside 30 298\nTMhelix 299 321\ninside 322 330

30034 GCF_900134095.1_10625_4_35 Mycobacterium abscessus subsp. abscessus Terrabacteria group; Actinobacteria; Actinobacteria; Corynebacteriales; Mycobacteriaceae; Mycobacterium; Mycobacterium chelonae group; Mycobacterium abscessus subgroup; Mycobacterium abscessus
MIKHLTRRATVALVVAAAAISPAAYPAMADTATPLNGLFALTPGSCAEGRATGSYFRMIL WP_005073213.1
hypothetical protein [Mycobacterium abscessus] Length: 330\nNumber of predicted TMHs: 2\nExp number of AAs in TMHs: 39.3416199999999\nExp number, first 60 AAs: 17.07688\nTotal prob of N-in: 0.84431\nPOSSIBLE N-term signal sequence\ninside 1 11\nTMhelix 12 29\noutside 30 298\nTMhelix 299 321\ninside 322 330

30035 GCF_900134125.1_10625_4_73 Mycobacterium abscessus subsp. abscessus Terrabacteria group; Actinobacteria; Actinobacteria; Corynebacteriales; Mycobacteriaceae; Mycobacterium; Mycobacterium chelonae group; Mycobacterium abscessus subgroup; Mycobacterium abscessus
MIKHLTRRATVALVVAAAAISPAAYPAMADTATPLNGLFALTPGSCAEGRATGSYFRMIL WP_005073213.1
hypothetical protein [Mycobacterium abscessus] Length: 330\nNumber of predicted TMHs: 2\nExp number of AAs in TMHs: 39.3416199999999\nExp number, first 60 AAs: 17.07688\nTotal prob of N-in: 0.84431\nPOSSIBLE N-term signal sequence\ninside 1 11\nTMhelix 12 29\noutside 30 298\nTMhelix 299 321\ninside 322 330

30036 GCF_900134465.1_12163_1_37 Mycobacterium abscessus subsp. abscessus Terrabacteria group; Actinobacteria; Actinobacteria; Corynebacteriales; Mycobacteriaceae; Mycobacterium; Mycobacterium chelonae group;

Mycobacterium abscessus subgroup; Mycobacterium abscessus
MIKHLTRRATVALVVAAAAISPAAYPAMADTATPLNGLFALTPGSCAEGRATGSYFRMIL WP_005073213.1
hypothetical protein [Mycobacterium abscessus] Length: 330\nNumber of predicted TMHs: 2\nExp number of AAs in TMHs: 39.3416199999999\nExp number, first 60 AAs: 17.07688\nTotal prob of N-in: 0.84431\nPOSSIBLE N-term signal sequence\ninside 1 11\nTMhelix 12 29\noutside 30 298\nTMhelix 299 321\ninside 322 330

30037 GCF_900134515.1_12163_1_40 Mycobacterium abscessus subsp. abscessus Terrabacteria group; Actinobacteria; Actinobacteria; Corynebacteriales; Mycobacteriaceae; Mycobacterium; Mycobacterium chelonae group; Mycobacterium abscessus subgroup; Mycobacterium abscessus
MIKHLTRRATVALVVAAAAISPAAYPAMADTATPLNGLFALTPGSCAEGRATGSYFRMIL WP_005073213.1
hypothetical protein [Mycobacterium abscessus] Length: 330\nNumber of predicted TMHs: 2\nExp number of AAs in TMHs: 39.3416199999999\nExp number, first 60 AAs: 17.07688\nTotal prob of N-in: 0.84431\nPOSSIBLE N-term signal sequence\ninside 1 11\nTMhelix 12 29\noutside 30 298\nTMhelix 299 321\ninside 322 330

30038 GCF_900134585.1_12163_1_46 Mycobacterium abscessus subsp. abscessus Terrabacteria group; Actinobacteria; Actinobacteria; Corynebacteriales; Mycobacteriaceae; Mycobacterium; Mycobacterium chelonae group; Mycobacterium abscessus subgroup; Mycobacterium abscessus
MIKHLTRRATVALVVAAAAISPAAYPAMADTATPLNGLFALTPGSCAEGRATGSYFRMIL WP_005073213.1
hypothetical protein [Mycobacterium abscessus] Length: 330\nNumber of predicted TMHs: 2\nExp number of AAs in TMHs: 39.3416199999999\nExp number, first 60 AAs: 17.07688\nTotal prob of N-in: 0.84431\nPOSSIBLE N-term signal sequence\ninside 1 11\nTMhelix 12 29\noutside 30 298\nTMhelix 299 321\ninside 322 330

30039 GCF_900134555.1_12163_1_48 Mycobacterium abscessus subsp. abscessus Terrabacteria group; Actinobacteria; Actinobacteria; Corynebacteriales; Mycobacteriaceae; Mycobacterium; Mycobacterium chelonae group; Mycobacterium abscessus subgroup; Mycobacterium abscessus
MIKHLTRRATVALVVAAAAISPAAYPAMADTATPLNGLFALTPGSCAEGRATGSYFRMIL WP_005073213.1
hypothetical protein [Mycobacterium abscessus] Length: 330\nNumber of predicted TMHs: 2\nExp number of AAs in TMHs: 39.3416199999999\nExp number, first 60 AAs: 17.07688\nTotal prob of N-in: 0.84431\nPOSSIBLE N-term signal sequence\ninside 1 11\nTMhelix 12 29\noutside 30 298\nTMhelix 299 321\ninside 322 330

30040 GCF_900134955.1_10208_3_39 Mycobacterium abscessus subsp. abscessus Terrabacteria group; Actinobacteria; Actinobacteria; Corynebacteriales; Mycobacteriaceae; Mycobacterium; Mycobacterium chelonae group; Mycobacterium abscessus subgroup; Mycobacterium abscessus
MIKHLTRRATVALVVAAAAISPAAYPAMADTATPLNGLFALTPGSCAEGRATGSYFRMIL WP_005073213.1
hypothetical protein [Mycobacterium abscessus] Length: 330\nNumber of predicted TMHs: 2\nExp number of AAs in TMHs: 39.3416199999999\nExp number, first 60 AAs: 17.07688\nTotal prob of N-in: 0.84431\nPOSSIBLE N-term signal sequence\ninside 1 11\nTMhelix 12 29\noutside 30 298\nTMhelix 299 321\ninside 322 330

30041 GCF_900134985.1_10208_3_4 Mycobacterium abscessus subsp. abscessus Terrabacteria group; Actinobacteria; Actinobacteria; Corynebacteriales; Mycobacteriaceae; Mycobacterium; Mycobacterium chelonae group; Mycobacterium abscessus subgroup; Mycobacterium abscessus
MIKHLTRRATVALVVAAAAISPAAYPAMADTATPLNGLFALTPGSCAEGRATGSYFRMIL WP_005073213.1
hypothetical protein [Mycobacterium abscessus] Length: 330\nNumber of predicted TMHs: 2\nExp number of AAs in TMHs: 39.3416199999999\nExp number, first 60 AAs: 17.07688\nTotal prob of N-in: 0.84431\nPOSSIBLE N-term signal sequence\ninside 1 11\nTMhelix 12 29\noutside 30 298\nTMhelix 299 321\ninside 322 330

30042 GCF_900135015.1_10208_3_6 Mycobacterium abscessus subsp. abscessus Terrabacteria group; Actinobacteria; Actinobacteria; Corynebacteriales; Mycobacteriaceae; Mycobacterium; Mycobacterium chelonae group; Mycobacterium abscessus subgroup; Mycobacterium abscessus
MIKHLTRRATVALVVAAAAISPAAYPAMADTATPLNGLFALTPGSCAEGRATGSYFRMIL WP_005073213.1
hypothetical protein [Mycobacterium abscessus] Length: 330\nNumber of predicted TMHs: 2\nExp number of AAs in TMHs: 39.3416199999999\nExp number, first 60 AAs: 17.07688\nTotal prob of N-in: 0.84431\nPOSSIBLE N-term signal sequence\ninside 1 11\nTMhelix 12 29\noutside 30 298\nTMhelix 299 321\ninside 322 330

30043 GCF_900135155.1_10625_4_20 Mycobacterium abscessus subsp. abscessus Terrabacteria group; Actinobacteria; Actinobacteria; Corynebacteriales; Mycobacteriaceae; Mycobacterium; Mycobacterium chelonae group; Mycobacterium abscessus subgroup; Mycobacterium abscessus
MIKHLTRRATVALVVAAAAISPAAYPAMADTATPLNGLFALTPGSCAEGRATGSYFRMIL WP_005073213.1
hypothetical protein [Mycobacterium abscessus] Length: 330\nNumber of predicted TMHs: 2\nExp number of AAs in TMHs: 39.3416199999999\nExp number, first 60 AAs: 17.07688\nTotal prob of N-in: 0.84431\nPOSSIBLE N-term signal sequence\ninside 1 11\nTMhelix 12 29\noutside 30 298\nTMhelix 299 321\ninside 322 330

30044 GCF_900135195.1_10625_4_25 Mycobacterium abscessus subsp. abscessus Terrabacteria group; Actinobacteria; Actinobacteria; Corynebacteriales; Mycobacteriaceae; Mycobacterium; Mycobacterium chelonae group; Mycobacterium abscessus subgroup; Mycobacterium abscessus
MIKHLTRRATVALVVAAAAISPAAYPAMADTATPLNGLFALTPGSCAEGRATGSYFRMIL WP_005073213.1
hypothetical protein [Mycobacterium abscessus] Length: 330\nNumber of predicted TMHs: 2\nExp number of AAs in TMHs: 39.3416199999999\nExp number, first 60 AAs: 17.07688\nTotal prob of N-in: 0.84431\nPOSSIBLE N-term signal sequence\ninside 1 11\nTMhelix 12 29\noutside 30 298\nTMhelix 299 321\ninside 322 330

30045 GCF_900135245.1_10625_4_38 Mycobacterium abscessus subsp. abscessus Terrabacteria group; Actinobacteria; Actinobacteria; Corynebacteriales; Mycobacteriaceae; Mycobacterium; Mycobacterium chelonae group; Mycobacterium abscessus subgroup; Mycobacterium abscessus
MIKHLTRRATVALVVAAAAISPAAYPAMADTATPLNGLFALTPGSCAEGRATGSYFRMIL WP_005073213.1
hypothetical protein [Mycobacterium abscessus] Length: 330\nNumber of predicted TMHs: 2\nExp number of AAs in TMHs: 39.3416199999999\nExp number, first 60 AAs: 17.07688\nTotal prob of N-in: 0.84431\nPOSSIBLE N-term signal sequence\ninside 1 11\nTMhelix 12 29\noutside 30 298\nTMhelix 299 321\ninside 322 330

30046 GCF_900135505.1_10625_4_69 Mycobacterium abscessus subsp. abscessus Terrabacteria group; Actinobacteria; Actinobacteria; Corynebacteriales; Mycobacteriaceae; Mycobacterium; Mycobacterium chelonae group; Mycobacterium abscessus subgroup; Mycobacterium abscessus
MIKHLTRRATVALVVAAAAISPAAYPAMADTATPLNGLFALTPGSCAEGRATGSYFRMIL WP_005073213.1
hypothetical protein [Mycobacterium abscessus] Length: 330\nNumber of predicted TMHs: 2\nExp number of AAs in TMHs: 39.3416199999999\nExp number, first 60 AAs: 17.07688\nTotal prob of N-in: 0.84431\nPOSSIBLE N-term signal sequence\ninside 1 11\nTMhelix 12 29\noutside 30 298\nTMhelix 299 321\ninside 322 330

30047 GCF_900135525.1_10625_4_71 Mycobacterium abscessus subsp. abscessus Terrabacteria group; Actinobacteria; Actinobacteria; Corynebacteriales; Mycobacteriaceae; Mycobacterium; Mycobacterium chelonae group; Mycobacterium abscessus subgroup; Mycobacterium abscessus
MIKHLTRRATVALVVAAAAISPAAYPAMADTATPLNGLFALTPGSCAEGRATGSYFRMIL WP_005073213.1
hypothetical protein [Mycobacterium abscessus] Length: 330\nNumber of predicted TMHs: 2\nExp number of AAs in TMHs: 39.3416199999999\nExp number, first 60 AAs: 17.07688\nTotal prob of N-in: 0.84431\nPOSSIBLE N-term signal sequence\ninside 1 11\nTMhelix 12 29\noutside 30 298\nTMhelix 299 321\ninside 322 330

30048 GCF_900135545.1_10625_5_10 Mycobacterium abscessus subsp. abscessus Terrabacteria group; Actinobacteria; Actinobacteria; Corynebacteriales; Mycobacteriaceae; Mycobacterium; Mycobacterium chelonae group; Mycobacterium abscessus subgroup; Mycobacterium abscessus
MIKHLTRRATVALVVAAAAISPAAYPAMADTATPLNGLFALTPGSCAEGRATGSYFRMIL WP_005073213.1
hypothetical protein [Mycobacterium abscessus] Length: 330\nNumber of predicted TMHs: 2\nExp number of AAs in TMHs: 39.3416199999999\nExp number, first 60 AAs: 17.07688\nTotal prob of N-in: 0.84431\nPOSSIBLE N-term signal sequence\ninside 1 11\nTMhelix 12 29\noutside 30 298\nTMhelix 299 321\ninside 322 330

30049 GCF_900135565.1_10625_5_11 Mycobacterium abscessus subsp. abscessus Terrabacteria group; Actinobacteria; Actinobacteria; Corynebacteriales; Mycobacteriaceae; Mycobacterium; Mycobacterium chelonae group; Mycobacterium abscessus subgroup; Mycobacterium abscessus
MIKHLTRRATVALVVAAAAISPAAYPAMADTATPLNGLFALTPGSCAEGRATGSYFRMIL WP_005073213.1
hypothetical protein [Mycobacterium abscessus] Length: 330\nNumber of predicted TMHs: 2\nExp number of AAs in TMHs: 39.3416199999999\nExp number, first 60 AAs: 17.07688\nTotal prob of N-in: 0.84431\nPOSSIBLE N-term signal sequence\ninside 1 11\nTMhelix 12 29\noutside 30 298\nTMhelix 299 321\ninside 322 330

30050 GCF_900135595.1_10625_5_17 Mycobacterium abscessus subsp. abscessus Terrabacteria group; Actinobacteria; Actinobacteria; Corynebacteriales; Mycobacteriaceae; Mycobacterium; Mycobacterium chelonae group; Mycobacterium abscessus subgroup; Mycobacterium abscessus
MIKHLTRRATVALVVAAAAISPAAYPAMADTATPLNGLFALTPGSCAEGRATGSYFRMIL WP_005073213.1
hypothetical protein [Mycobacterium abscessus] Length: 330\nNumber of predicted TMHs: 2\nExp number of AAs in TMHs: 39.3416199999999\nExp number, first 60 AAs: 17.07688\nTotal prob of N-in: 0.84431\nPOSSIBLE N-term signal sequence\ninside 1 11\nTMhelix 12 29\noutside 30 298\nTMhelix 299 321\ninside 322 330

30051 GCF_900135945.1_12082_5_61 Mycobacterium abscessus subsp. abscessus Terrabacteria group; Actinobacteria; Actinobacteria; Corynebacteriales; Mycobacteriaceae; Mycobacterium; Mycobacterium chelonae group; Mycobacterium abscessus subgroup; Mycobacterium abscessus
MIKHLTRRATVALVVAAAAISPAAYPAMADTATPLNGLFALTPGSCAEGRATGSYFRMIL WP_005073213.1
hypothetical protein [Mycobacterium abscessus] Length: 330\nNumber of predicted TMHs: 2\nExp number of

AAs in TMHs: 39.3416199999999\nExp number, first 60 AAs: 17.07688\nTotal prob of N-in: 0.84431\nPOSSIBLE N-term signal sequence\ninside 1 11\nTMhelix 12 29\noutside 30 298\nTMhelix 299 321\ninside 322 330

30052 GCF_900136035.1_12163_1_59 Mycobacterium abscessus subsp. abscessus Terrabacteria group; Actinobacteria; Actinobacteria; Corynebacteriales; Mycobacteriaceae; Mycobacterium; Mycobacterium chelonae group; Mycobacterium abscessus subgroup; Mycobacterium abscessus
MIKHLTRRATVALVVAAAAISPAAYPAMADTATPLNGLFALTPGSCAEGRATGSYFRMIL WP_005073213.1
hypothetical protein [Mycobacterium abscessus] Length: 330\nNumber of predicted TMHs: 2\nExp number of AAs in TMHs: 39.3416199999999\nExp number, first 60 AAs: 17.07688\nTotal prob of N-in: 0.84431\nPOSSIBLE N-term signal sequence\ninside 1 11\nTMhelix 12 29\noutside 30 298\nTMhelix 299 321\ninside 322 330

30053 GCF_900136015.1_12163_1_57 Mycobacterium abscessus subsp. abscessus Terrabacteria group; Actinobacteria; Actinobacteria; Corynebacteriales; Mycobacteriaceae; Mycobacterium; Mycobacterium chelonae group; Mycobacterium abscessus subgroup; Mycobacterium abscessus
MIKHLTRRATVALVVAAAAISPAAYPAMADTATPLNGLFALTPGSCAEGRATGSYFRMIL WP_005073213.1
hypothetical protein [Mycobacterium abscessus] Length: 330\nNumber of predicted TMHs: 2\nExp number of AAs in TMHs: 39.3416199999999\nExp number, first 60 AAs: 17.07688\nTotal prob of N-in: 0.84431\nPOSSIBLE N-term signal sequence\ninside 1 11\nTMhelix 12 29\noutside 30 298\nTMhelix 299 321\ninside 322 330

30054 GCF_900136055.1_12163_1_60 Mycobacterium abscessus subsp. abscessus Terrabacteria group; Actinobacteria; Actinobacteria; Corynebacteriales; Mycobacteriaceae; Mycobacterium; Mycobacterium chelonae group; Mycobacterium abscessus subgroup; Mycobacterium abscessus
MIKHLTRRATVALVVAAAAISPAAYPAMADTATPLNGLFALTPGSCAEGRATGSYFRMIL WP_005073213.1
hypothetical protein [Mycobacterium abscessus] Length: 330\nNumber of predicted TMHs: 2\nExp number of AAs in TMHs: 39.3416199999999\nExp number, first 60 AAs: 17.07688\nTotal prob of N-in: 0.84431\nPOSSIBLE N-term signal sequence\ninside 1 11\nTMhelix 12 29\noutside 30 298\nTMhelix 299 321\ninside 322 330

30055 GCF_900136475.1_10208_3_69 Mycobacterium abscessus subsp. abscessus Terrabacteria group; Actinobacteria; Actinobacteria; Corynebacteriales; Mycobacteriaceae; Mycobacterium; Mycobacterium chelonae group; Mycobacterium abscessus subgroup; Mycobacterium abscessus
MIKHLTRRATVALVVAAAAISPAAYPAMADTATPLNGLFALTPGSCAEGRATGSYFRMIL WP_005073213.1
hypothetical protein [Mycobacterium abscessus] Length: 330\nNumber of predicted TMHs: 2\nExp number of AAs in TMHs: 39.3416199999999\nExp number, first 60 AAs: 17.07688\nTotal prob of N-in: 0.84431\nPOSSIBLE N-term signal sequence\ninside 1 11\nTMhelix 12 29\noutside 30 298\nTMhelix 299 321\ninside 322 330

30056 GCF_900136515.1_10208_3_75 Mycobacterium abscessus subsp. abscessus Terrabacteria group; Actinobacteria; Actinobacteria; Corynebacteriales; Mycobacteriaceae; Mycobacterium; Mycobacterium chelonae group; Mycobacterium abscessus subgroup; Mycobacterium abscessus
MIKHLTRRATVALVVAAAAISPAAYPAMADTATPLNGLFALTPGSCAEGRATGSYFRMIL WP_005073213.1
hypothetical protein [Mycobacterium abscessus] Length: 330\nNumber of predicted TMHs: 2\nExp number of AAs in TMHs: 39.3416199999999\nExp number, first 60 AAs: 17.07688\nTotal prob of N-in: 0.84431\nPOSSIBLE N-term signal sequence\ninside 1 11\nTMhelix 12 29\noutside 30 298\nTMhelix 299 321\ninside 322 330

30057 GCF_900136545.1_10208_3_81 Mycobacterium abscessus subsp. abscessus Terrabacteria group; Actinobacteria; Actinobacteria; Corynebacteriales; Mycobacteriaceae; Mycobacterium; Mycobacterium chelonae group; Mycobacterium abscessus subgroup; Mycobacterium abscessus
MIKHLTRRATVALVVAAAAISPAAYPAMADTATPLNGLFALTPGSCAEGRATGSYFRMIL WP_005073213.1
hypothetical protein [Mycobacterium abscessus] Length: 330\nNumber of predicted TMHs: 2\nExp number of AAs in TMHs: 39.3416199999999\nExp number, first 60 AAs: 17.07688\nTotal prob of N-in: 0.84431\nPOSSIBLE N-term signal sequence\ninside 1 11\nTMhelix 12 29\noutside 30 298\nTMhelix 299 321\ninside 322 330

30058 GCF_900136885.1_10625_5_56 Mycobacterium abscessus subsp. abscessus Terrabacteria group; Actinobacteria; Actinobacteria; Corynebacteriales; Mycobacteriaceae; Mycobacterium; Mycobacterium chelonae group; Mycobacterium abscessus subgroup; Mycobacterium abscessus
MIKHLTRRATVALVVAAAAISPAAYPAMADTATPLNGLFALTPGSCAEGRATGSYFRMIL WP_005073213.1
hypothetical protein [Mycobacterium abscessus] Length: 330\nNumber of predicted TMHs: 2\nExp number of AAs in TMHs: 39.3416199999999\nExp number, first 60 AAs: 17.07688\nTotal prob of N-in: 0.84431\nPOSSIBLE N-term signal sequence\ninside 1 11\nTMhelix 12 29\noutside 30 298\nTMhelix 299 321\ninside 322 330

30059 GCF_900136905.1_10625_5_58 Mycobacterium abscessus subsp. abscessus Terrabacteria group; Actinobacteria; Actinobacteria; Corynebacteriales; Mycobacteriaceae; Mycobacterium; Mycobacterium chelonae group;

Mycobacterium abscessus subgroup; Mycobacterium abscessus
MIKHLTRRATVALVVAAAAISPAAYPAMADTATPLNGLFALTPGSCAEGRATGSYFRMIL WP_005073213.1
hypothetical protein [Mycobacterium abscessus] Length: 330\nNumber of predicted TMHs: 2\nExp number of AAs in TMHs: 39.3416199999999\nExp number, first 60 AAs: 17.07688\nTotal prob of N-in: 0.84431\nPOSSIBLE N-term signal sequence\ninside 1 11\nTMhelix 12 29\noutside 30 298\nTMhelix 299 321\ninside 322 330

30060 GCF_900136935.1_11861_7_14 Mycobacterium abscessus subsp. abscessus Terrabacteria group; Actinobacteria; Actinobacteria; Corynebacteriales; Mycobacteriaceae; Mycobacterium; Mycobacterium chelonae group; Mycobacterium abscessus subgroup; Mycobacterium abscessus
MIKHLTRRATVALVVAAAAISPAAYPAMADTATPLNGLFALTPGSCAEGRATGSYFRMIL WP_005073213.1
hypothetical protein [Mycobacterium abscessus] Length: 330\nNumber of predicted TMHs: 2\nExp number of AAs in TMHs: 39.3416199999999\nExp number, first 60 AAs: 17.07688\nTotal prob of N-in: 0.84431\nPOSSIBLE N-term signal sequence\ninside 1 11\nTMhelix 12 29\noutside 30 298\nTMhelix 299 321\ninside 322 330

30061 GCF_900137005.1_11861_7_76 Mycobacterium abscessus subsp. abscessus Terrabacteria group; Actinobacteria; Actinobacteria; Corynebacteriales; Mycobacteriaceae; Mycobacterium; Mycobacterium chelonae group; Mycobacterium abscessus subgroup; Mycobacterium abscessus
MIKHLTRRATVALVVAAAAISPAAYPAMADTATPLNGLFALTPGSCAEGRATGSYFRMIL WP_005073213.1
hypothetical protein [Mycobacterium abscessus] Length: 330\nNumber of predicted TMHs: 2\nExp number of AAs in TMHs: 39.3416199999999\nExp number, first 60 AAs: 17.07688\nTotal prob of N-in: 0.84431\nPOSSIBLE N-term signal sequence\ninside 1 11\nTMhelix 12 29\noutside 30 298\nTMhelix 299 321\ninside 322 330

30062 GCF_900137035.1_11861_7_93 Mycobacterium abscessus subsp. abscessus Terrabacteria group; Actinobacteria; Actinobacteria; Corynebacteriales; Mycobacteriaceae; Mycobacterium; Mycobacterium chelonae group; Mycobacterium abscessus subgroup; Mycobacterium abscessus
MIKHLTRRATVALVVAAAAISPAAYPAMADTATPLNGLFALTPGSCAEGRATGSYFRMIL WP_005073213.1
hypothetical protein [Mycobacterium abscessus] Length: 330\nNumber of predicted TMHs: 2\nExp number of AAs in TMHs: 39.3416199999999\nExp number, first 60 AAs: 17.07688\nTotal prob of N-in: 0.84431\nPOSSIBLE N-term signal sequence\ninside 1 11\nTMhelix 12 29\noutside 30 298\nTMhelix 299 321\ninside 322 330

30063 GCF_900137325.1_12163_2_1 Mycobacterium abscessus subsp. abscessus Terrabacteria group; Actinobacteria; Actinobacteria; Corynebacteriales; Mycobacteriaceae; Mycobacterium; Mycobacterium chelonae group; Mycobacterium abscessus subgroup; Mycobacterium abscessus
MIKHLTRRATVALVVAAAAISPAAYPAMADTATPLNGLFALTPGSCAEGRATGSYFRMIL WP_005073213.1
hypothetical protein [Mycobacterium abscessus] Length: 330\nNumber of predicted TMHs: 2\nExp number of AAs in TMHs: 39.3416199999999\nExp number, first 60 AAs: 17.07688\nTotal prob of N-in: 0.84431\nPOSSIBLE N-term signal sequence\ninside 1 11\nTMhelix 12 29\noutside 30 298\nTMhelix 299 321\ninside 322 330

30064 GCF_900137345.1_12163_2_10 Mycobacterium abscessus subsp. abscessus Terrabacteria group; Actinobacteria; Actinobacteria; Corynebacteriales; Mycobacteriaceae; Mycobacterium; Mycobacterium chelonae group; Mycobacterium abscessus subgroup; Mycobacterium abscessus
MIKHLTRRATVALVVAAAAISPAAYPAMADTATPLNGLFALTPGSCAEGRATGSYFRMIL WP_005073213.1
hypothetical protein [Mycobacterium abscessus] Length: 330\nNumber of predicted TMHs: 2\nExp number of AAs in TMHs: 39.3416199999999\nExp number, first 60 AAs: 17.07688\nTotal prob of N-in: 0.84431\nPOSSIBLE N-term signal sequence\ninside 1 11\nTMhelix 12 29\noutside 30 298\nTMhelix 299 321\ninside 322 330

30065 GCF_900137375.1_12163_2_13 Mycobacterium abscessus subsp. abscessus Terrabacteria group; Actinobacteria; Actinobacteria; Corynebacteriales; Mycobacteriaceae; Mycobacterium; Mycobacterium chelonae group; Mycobacterium abscessus subgroup; Mycobacterium abscessus
MIKHLTRRATVALVVAAAAISPAAYPAMADTATPLNGLFALTPGSCAEGRATGSYFRMIL WP_005073213.1
hypothetical protein [Mycobacterium abscessus] Length: 330\nNumber of predicted TMHs: 2\nExp number of AAs in TMHs: 39.3416199999999\nExp number, first 60 AAs: 17.07688\nTotal prob of N-in: 0.84431\nPOSSIBLE N-term signal sequence\ninside 1 11\nTMhelix 12 29\noutside 30 298\nTMhelix 299 321\ninside 322 330

30066 GCF_900137655.1_12163_2_42 Mycobacterium abscessus subsp. abscessus Terrabacteria group; Actinobacteria; Actinobacteria; Corynebacteriales; Mycobacteriaceae; Mycobacterium; Mycobacterium chelonae group; Mycobacterium abscessus subgroup; Mycobacterium abscessus
MIKHLTRRATVALVVAAAAISPAAYPAMADTATPLNGLFALTPGSCAEGRATGSYFRMIL WP_005073213.1
hypothetical protein [Mycobacterium abscessus] Length: 330\nNumber of predicted TMHs: 2\nExp number of AAs in TMHs: 39.3416199999999\nExp number, first 60 AAs: 17.07688\nTotal prob of N-in: 0.84431\nPOSSIBLE N-term signal sequence\ninside 1 11\nTMhelix 12 29\noutside 30 298\nTMhelix 299 321\ninside 322 330

30067 GCF_900137675.1_12163_2_40 Mycobacterium abscessus subsp. abscessus Terrabacteria group; Actinobacteria; Actinobacteria; Corynebacteriales; Mycobacteriaceae; Mycobacterium; Mycobacterium chelonae group; Mycobacterium abscessus subgroup; Mycobacterium abscessus
MIKHLTRRATVALVVAAAAISPAAYPAMADTATPLNGLFALTPGSCAEGRATGSYFRMIL WP_005073213.1
hypothetical protein [Mycobacterium abscessus] Length: 330\nNumber of predicted TMHs: 2\nExp number of AAs in TMHs: 39.3416199999999\nExp number, first 60 AAs: 17.07688\nTotal prob of N-in: 0.84431\nPOSSIBLE N-term signal sequence\ninside 1 11\nTMhelix 12 29\noutside 30 298\nTMhelix 299 321\ninside 322 330

30068 GCF_900137735.1_12163_2_50 Mycobacterium abscessus subsp. abscessus Terrabacteria group; Actinobacteria; Actinobacteria; Corynebacteriales; Mycobacteriaceae; Mycobacterium; Mycobacterium chelonae group; Mycobacterium abscessus subgroup; Mycobacterium abscessus
MIKHLTRRATVALVVAAAAISPAAYPAMADTATPLNGLFALTPGSCAEGRATGSYFRMIL WP_005073213.1
hypothetical protein [Mycobacterium abscessus] Length: 330\nNumber of predicted TMHs: 2\nExp number of AAs in TMHs: 39.3416199999999\nExp number, first 60 AAs: 17.07688\nTotal prob of N-in: 0.84431\nPOSSIBLE N-term signal sequence\ninside 1 11\nTMhelix 12 29\noutside 30 298\nTMhelix 299 321\ninside 322 330

30069 GCF_900137715.1_12163_2_47 Mycobacterium abscessus subsp. abscessus Terrabacteria group; Actinobacteria; Actinobacteria; Corynebacteriales; Mycobacteriaceae; Mycobacterium; Mycobacterium chelonae group; Mycobacterium abscessus subgroup; Mycobacterium abscessus
MIKHLTRRATVALVVAAAAISPAAYPAMADTATPLNGLFALTPGSCAEGRATGSYFRMIL WP_005073213.1
hypothetical protein [Mycobacterium abscessus] Length: 330\nNumber of predicted TMHs: 2\nExp number of AAs in TMHs: 39.3416199999999\nExp number, first 60 AAs: 17.07688\nTotal prob of N-in: 0.84431\nPOSSIBLE N-term signal sequence\ninside 1 11\nTMhelix 12 29\noutside 30 298\nTMhelix 299 321\ninside 322 330

30070 GCF_900138055.1_12045_8_37 Mycobacterium abscessus subsp. abscessus Terrabacteria group; Actinobacteria; Actinobacteria; Corynebacteriales; Mycobacteriaceae; Mycobacterium; Mycobacterium chelonae group; Mycobacterium abscessus subgroup; Mycobacterium abscessus
MIKHLTRRATVALVVAAAAISPAAYPAMADTATPLNGLFALTPGSCAEGRATGSYFRMIL WP_005073213.1
hypothetical protein [Mycobacterium abscessus] Length: 330\nNumber of predicted TMHs: 2\nExp number of AAs in TMHs: 39.3416199999999\nExp number, first 60 AAs: 17.07688\nTotal prob of N-in: 0.84431\nPOSSIBLE N-term signal sequence\ninside 1 11\nTMhelix 12 29\noutside 30 298\nTMhelix 299 321\ninside 322 330

30071 GCF_900138075.1_12045_8_38 Mycobacterium abscessus subsp. abscessus Terrabacteria group; Actinobacteria; Actinobacteria; Corynebacteriales; Mycobacteriaceae; Mycobacterium; Mycobacterium chelonae group; Mycobacterium abscessus subgroup; Mycobacterium abscessus
MIKHLTRRATVALVVAAAAISPAAYPAMADTATPLNGLFALTPGSCAEGRATGSYFRMIL WP_005073213.1
hypothetical protein [Mycobacterium abscessus] Length: 330\nNumber of predicted TMHs: 2\nExp number of AAs in TMHs: 39.3416199999999\nExp number, first 60 AAs: 17.07688\nTotal prob of N-in: 0.84431\nPOSSIBLE N-term signal sequence\ninside 1 11\nTMhelix 12 29\noutside 30 298\nTMhelix 299 321\ninside 322 330

30072 GCF_900138105.1_12045_8_39 Mycobacterium abscessus subsp. abscessus Terrabacteria group; Actinobacteria; Actinobacteria; Corynebacteriales; Mycobacteriaceae; Mycobacterium; Mycobacterium chelonae group; Mycobacterium abscessus subgroup; Mycobacterium abscessus
MIKHLTRRATVALVVAAAAISPAAYPAMADTATPLNGLFALTPGSCAEGRATGSYFRMIL WP_005073213.1
hypothetical protein [Mycobacterium abscessus] Length: 330\nNumber of predicted TMHs: 2\nExp number of AAs in TMHs: 39.3416199999999\nExp number, first 60 AAs: 17.07688\nTotal prob of N-in: 0.84431\nPOSSIBLE N-term signal sequence\ninside 1 11\nTMhelix 12 29\noutside 30 298\nTMhelix 299 321\ninside 322 330

30073 GCF_900138125.1_12045_8_40 Mycobacterium abscessus subsp. abscessus Terrabacteria group; Actinobacteria; Actinobacteria; Corynebacteriales; Mycobacteriaceae; Mycobacterium; Mycobacterium chelonae group; Mycobacterium abscessus subgroup; Mycobacterium abscessus
MIKHLTRRATVALVVAAAAISPAAYPAMADTATPLNGLFALTPGSCAEGRATGSYFRMIL WP_005073213.1
hypothetical protein [Mycobacterium abscessus] Length: 330\nNumber of predicted TMHs: 2\nExp number of AAs in TMHs: 39.3416199999999\nExp number, first 60 AAs: 17.07688\nTotal prob of N-in: 0.84431\nPOSSIBLE N-term signal sequence\ninside 1 11\nTMhelix 12 29\noutside 30 298\nTMhelix 299 321\ninside 322 330

30074 GCF_900138755.1_10060_7_2 Mycobacterium abscessus subsp. abscessus Terrabacteria group; Actinobacteria; Actinobacteria; Corynebacteriales; Mycobacteriaceae; Mycobacterium; Mycobacterium chelonae group; Mycobacterium abscessus subgroup; Mycobacterium abscessus
MIKHLTRRATVALVVAAAAISPAAYPAMADTATPLNGLFALTPGSCAEGRATGSYFRMIL WP_005073213.1
hypothetical protein [Mycobacterium abscessus] Length: 330\nNumber of predicted TMHs: 2\nExp number of

AAs in TMHs: 39.3416199999999\nExp number, first 60 AAs: 17.07688\nTotal prob of N-in: 0.84431\nPOSSIBLE N-term signal sequence\ninside 1 11\nTMhelix 12 29\noutside 30 298\nTMhelix 299 321\ninside 322 330

30075 GCF_900138775.1_10208_3_20 Mycobacterium abscessus subsp. abscessus Terrabacteria group; Actinobacteria; Actinobacteria; Corynebacteriales; Mycobacteriaceae; Mycobacterium; Mycobacterium chelonae group; Mycobacterium abscessus subgroup; Mycobacterium abscessus
MIKHLTRRATVALVVAAAAISPAAYPAMADTATPLNGLFALTPGSCAEGRATGSYFRMIL WP_005073213.1
hypothetical protein [Mycobacterium abscessus] Length: 330\nNumber of predicted TMHs: 2\nExp number of AAs in TMHs: 39.3416199999999\nExp number, first 60 AAs: 17.07688\nTotal prob of N-in: 0.84431\nPOSSIBLE N-term signal sequence\ninside 1 11\nTMhelix 12 29\noutside 30 298\nTMhelix 299 321\ninside 322 330

30076 GCF_900138895.1_10208_3_59 Mycobacterium abscessus subsp. abscessus Terrabacteria group; Actinobacteria; Actinobacteria; Corynebacteriales; Mycobacteriaceae; Mycobacterium; Mycobacterium chelonae group; Mycobacterium abscessus subgroup; Mycobacterium abscessus
MIKHLTRRATVALVVAAAAISPAAYPAMADTATPLNGLFALTPGSCAEGRATGSYFRMIL WP_005073213.1
hypothetical protein [Mycobacterium abscessus] Length: 330\nNumber of predicted TMHs: 2\nExp number of AAs in TMHs: 39.3416199999999\nExp number, first 60 AAs: 17.07688\nTotal prob of N-in: 0.84431\nPOSSIBLE N-term signal sequence\ninside 1 11\nTMhelix 12 29\noutside 30 298\nTMhelix 299 321\ninside 322 330

30077 GCF_900138945.1_10208_3_63 Mycobacterium abscessus subsp. abscessus Terrabacteria group; Actinobacteria; Actinobacteria; Corynebacteriales; Mycobacteriaceae; Mycobacterium; Mycobacterium chelonae group; Mycobacterium abscessus subgroup; Mycobacterium abscessus
MIKHLTRRATVALVVAAAAISPAAYPAMADTATPLNGLFALTPGSCAEGRATGSYFRMIL WP_005073213.1
hypothetical protein [Mycobacterium abscessus] Length: 330\nNumber of predicted TMHs: 2\nExp number of AAs in TMHs: 39.3416199999999\nExp number, first 60 AAs: 17.07688\nTotal prob of N-in: 0.84431\nPOSSIBLE N-term signal sequence\ninside 1 11\nTMhelix 12 29\noutside 30 298\nTMhelix 299 321\ninside 322 330

30078 GCF_900138965.1_10208_3_64 Mycobacterium abscessus subsp. abscessus Terrabacteria group; Actinobacteria; Actinobacteria; Corynebacteriales; Mycobacteriaceae; Mycobacterium; Mycobacterium chelonae group; Mycobacterium abscessus subgroup; Mycobacterium abscessus
MIKHLTRRATVALVVAAAAISPAAYPAMADTATPLNGLFALTPGSCAEGRATGSYFRMIL WP_005073213.1
hypothetical protein [Mycobacterium abscessus] Length: 330\nNumber of predicted TMHs: 2\nExp number of AAs in TMHs: 39.3416199999999\nExp number, first 60 AAs: 17.07688\nTotal prob of N-in: 0.84431\nPOSSIBLE N-term signal sequence\ninside 1 11\nTMhelix 12 29\noutside 30 298\nTMhelix 299 321\ninside 322 330

30079 GCF_900139265.1_10465_1_73 Mycobacterium abscessus subsp. abscessus Terrabacteria group; Actinobacteria; Actinobacteria; Corynebacteriales; Mycobacteriaceae; Mycobacterium; Mycobacterium chelonae group; Mycobacterium abscessus subgroup; Mycobacterium abscessus
MIKHLTRRATVALVVAAAAISPAAYPAMADTATPLNGLFALTPGSCAEGRATGSYFRMIL WP_005073213.1
hypothetical protein [Mycobacterium abscessus] Length: 330\nNumber of predicted TMHs: 2\nExp number of AAs in TMHs: 39.3416199999999\nExp number, first 60 AAs: 17.07688\nTotal prob of N-in: 0.84431\nPOSSIBLE N-term signal sequence\ninside 1 11\nTMhelix 12 29\noutside 30 298\nTMhelix 299 321\ninside 322 330

30080 GCF_900139285.1_10465_1_72 Mycobacterium abscessus subsp. abscessus Terrabacteria group; Actinobacteria; Actinobacteria; Corynebacteriales; Mycobacteriaceae; Mycobacterium; Mycobacterium chelonae group; Mycobacterium abscessus subgroup; Mycobacterium abscessus
MIKHLTRRATVALVVAAAAISPAAYPAMADTATPLNGLFALTPGSCAEGRATGSYFRMIL WP_005073213.1
hypothetical protein [Mycobacterium abscessus] Length: 330\nNumber of predicted TMHs: 2\nExp number of AAs in TMHs: 39.3416199999999\nExp number, first 60 AAs: 17.07688\nTotal prob of N-in: 0.84431\nPOSSIBLE N-term signal sequence\ninside 1 11\nTMhelix 12 29\noutside 30 298\nTMhelix 299 321\ninside 322 330

30081 GCF_900139315.1_10625_4_1 Mycobacterium abscessus subsp. abscessus Terrabacteria group; Actinobacteria; Actinobacteria; Corynebacteriales; Mycobacteriaceae; Mycobacterium; Mycobacterium chelonae group; Mycobacterium abscessus subgroup; Mycobacterium abscessus
MIKHLTRRATVALVVAAAAISPAAYPAMADTATPLNGLFALTPGSCAEGRATGSYFRMIL WP_005073213.1
hypothetical protein [Mycobacterium abscessus] Length: 330\nNumber of predicted TMHs: 2\nExp number of AAs in TMHs: 39.3416199999999\nExp number, first 60 AAs: 17.07688\nTotal prob of N-in: 0.84431\nPOSSIBLE N-term signal sequence\ninside 1 11\nTMhelix 12 29\noutside 30 298\nTMhelix 299 321\ninside 322 330

30082 GCF_900139785.1_10660_1_24 Mycobacterium abscessus subsp. abscessus Terrabacteria group; Actinobacteria; Actinobacteria; Corynebacteriales; Mycobacteriaceae; Mycobacterium; Mycobacterium chelonae group;

Mycobacterium abscessus subgroup; Mycobacterium abscessus
MIKHLTRRATVALVVAAAAISPAAYPAMADTATPLNGLFALTPGSCAEGRATGSYFRMIL WP_005073213.1
hypothetical protein [Mycobacterium abscessus] Length: 330\nNumber of predicted TMHs: 2\nExp number of AAs in TMHs: 39.3416199999999\nExp number, first 60 AAs: 17.07688\nTotal prob of N-in: 0.84431\nPOSSIBLE N-term signal sequence\ninside 1 11\nTMhelix 12 29\noutside 30 298\nTMhelix 299 321\ninside 322 330

30083 GCF_900139855.1_10660_1_3 Mycobacterium abscessus subsp. abscessus Terrabacteria group; Actinobacteria; Actinobacteria; Corynebacteriales; Mycobacteriaceae; Mycobacterium; Mycobacterium chelonae group; Mycobacterium abscessus subgroup; Mycobacterium abscessus
MIKHLTRRATVALVVAAAAISPAAYPAMADTATPLNGLFALTPGSCAEGRATGSYFRMIL WP_005073213.1
hypothetical protein [Mycobacterium abscessus] Length: 330\nNumber of predicted TMHs: 2\nExp number of AAs in TMHs: 39.3416199999999\nExp number, first 60 AAs: 17.07688\nTotal prob of N-in: 0.84431\nPOSSIBLE N-term signal sequence\ninside 1 11\nTMhelix 12 29\noutside 30 298\nTMhelix 299 321\ninside 322 330

30084 GCF_900139615.1_10625_5_55 Mycobacterium abscessus subsp. abscessus Terrabacteria group; Actinobacteria; Actinobacteria; Corynebacteriales; Mycobacteriaceae; Mycobacterium; Mycobacterium chelonae group; Mycobacterium abscessus subgroup; Mycobacterium abscessus
MTKHLTRRATVALVVAAAAISPAAYPAMADTATPLNGLFALTPGSCAEGRATGSYFRMIL WP_074319467.1
hypothetical protein [Mycobacterium abscessus] Length: 330\nNumber of predicted TMHs: 2\nExp number of AAs in TMHs: 40.2609399999999\nExp number, first 60 AAs: 17.99061\nTotal prob of N-in: 0.88931\nPOSSIBLE N-term signal sequence\ninside 1 11\nTMhelix 12 29\noutside 30 298\nTMhelix 299 321\ninside 322 330

30085 GCF_900139875.1_10660_1_32 Mycobacterium abscessus subsp. abscessus Terrabacteria group; Actinobacteria; Actinobacteria; Corynebacteriales; Mycobacteriaceae; Mycobacterium; Mycobacterium chelonae group; Mycobacterium abscessus subgroup; Mycobacterium abscessus
MIKHLTRRATVALVVAAAAISPAAYPAMADTATPLNGLFALTPGSCAEGRATGSYFRMIL WP_005073213.1
hypothetical protein [Mycobacterium abscessus] Length: 330\nNumber of predicted TMHs: 2\nExp number of AAs in TMHs: 39.3416199999999\nExp number, first 60 AAs: 17.07688\nTotal prob of N-in: 0.84431\nPOSSIBLE N-term signal sequence\ninside 1 11\nTMhelix 12 29\noutside 30 298\nTMhelix 299 321\ninside 322 330

30086 GCF_900140185.1_10665_2_71 Mycobacterium abscessus subsp. abscessus Terrabacteria group; Actinobacteria; Actinobacteria; Corynebacteriales; Mycobacteriaceae; Mycobacterium; Mycobacterium chelonae group; Mycobacterium abscessus subgroup; Mycobacterium abscessus
MIKHLTRRATVALVVAAAAISPAAYPAMADTATPLNGLFALTPGSCAEGRATGSYFRMIL WP_005073213.1
hypothetical protein [Mycobacterium abscessus] Length: 330\nNumber of predicted TMHs: 2\nExp number of AAs in TMHs: 39.3416199999999\nExp number, first 60 AAs: 17.07688\nTotal prob of N-in: 0.84431\nPOSSIBLE N-term signal sequence\ninside 1 11\nTMhelix 12 29\noutside 30 298\nTMhelix 299 321\ninside 322 330

30087 GCF_900140255.1_10665_2_91 Mycobacterium abscessus subsp. abscessus Terrabacteria group; Actinobacteria; Actinobacteria; Corynebacteriales; Mycobacteriaceae; Mycobacterium; Mycobacterium chelonae group; Mycobacterium abscessus subgroup; Mycobacterium abscessus
MIKHLTRRATVALVVAAAAISPAAYPAMADTATPLNGLFALTPGSCAEGRATGSYFRMIL WP_005073213.1
hypothetical protein [Mycobacterium abscessus] Length: 330\nNumber of predicted TMHs: 2\nExp number of AAs in TMHs: 39.3416199999999\nExp number, first 60 AAs: 17.07688\nTotal prob of N-in: 0.84431\nPOSSIBLE N-term signal sequence\ninside 1 11\nTMhelix 12 29\noutside 30 298\nTMhelix 299 321\ninside 322 330

30088 GCF_900140665.1_10702_1_30 Mycobacterium abscessus subsp. abscessus Terrabacteria group; Actinobacteria; Actinobacteria; Corynebacteriales; Mycobacteriaceae; Mycobacterium; Mycobacterium chelonae group; Mycobacterium abscessus subgroup; Mycobacterium abscessus
MIKHLTRRATVALVVAAAAISPAAYPAMADTATPLNGLFALTPGSCAEGRATGSYFRMIL WP_005073213.1
hypothetical protein [Mycobacterium abscessus] Length: 330\nNumber of predicted TMHs: 2\nExp number of AAs in TMHs: 39.3416199999999\nExp number, first 60 AAs: 17.07688\nTotal prob of N-in: 0.84431\nPOSSIBLE N-term signal sequence\ninside 1 11\nTMhelix 12 29\noutside 30 298\nTMhelix 299 321\ninside 322 330

30089 GCF_900140705.1_10702_1_34 Mycobacterium abscessus subsp. abscessus Terrabacteria group; Actinobacteria; Actinobacteria; Corynebacteriales; Mycobacteriaceae; Mycobacterium; Mycobacterium chelonae group; Mycobacterium abscessus subgroup; Mycobacterium abscessus
MIKHLTRRATVALVVAAAAISPAAYPAMADTATPLNGLFALTPGSCAEGRATGSYFRMIL WP_005073213.1
hypothetical protein [Mycobacterium abscessus] Length: 330\nNumber of predicted TMHs: 2\nExp number of AAs in TMHs: 39.3416199999999\nExp number, first 60 AAs: 17.07688\nTotal prob of N-in: 0.84431\nPOSSIBLE N-term signal sequence\ninside 1 11\nTMhelix 12 29\noutside 30 298\nTMhelix 299 321\ninside 322 330

30090 GCF_900140875.1_10702_1_54 Mycobacterium abscessus subsp. abscessus Terrabacteria group; Actinobacteria; Actinobacteria; Corynebacteriales; Mycobacteriaceae; Mycobacterium; Mycobacterium chelonae group; Mycobacterium abscessus subgroup; Mycobacterium abscessus
MIKHLTRRATVALVVAAAAISPAAYPAMADTATPLNGLFALTPGSCAEGRATGSYFRMIL WP_005073213.1
hypothetical protein [Mycobacterium abscessus] Length: 330\nNumber of predicted TMHs: 2\nExp number of AAs in TMHs: 39.3416199999999\nExp number, first 60 AAs: 17.07688\nTotal prob of N-in: 0.84431\nPOSSIBLE N-term signal sequence\ninside 1 11\nTMhelix 12 29\noutside 30 298\nTMhelix 299 321\ninside 322 330

30091 GCF_900140895.1_10702_1_56 Mycobacterium abscessus subsp. abscessus Terrabacteria group; Actinobacteria; Actinobacteria; Corynebacteriales; Mycobacteriaceae; Mycobacterium; Mycobacterium chelonae group; Mycobacterium abscessus subgroup; Mycobacterium abscessus
MIKHLTRRATVALVVAAAAISPAAYPAMADTATPLNGLFALTPGSCAEGRATGSYFRMIL WP_005073213.1
hypothetical protein [Mycobacterium abscessus] Length: 330\nNumber of predicted TMHs: 2\nExp number of AAs in TMHs: 39.3416199999999\nExp number, first 60 AAs: 17.07688\nTotal prob of N-in: 0.84431\nPOSSIBLE N-term signal sequence\ninside 1 11\nTMhelix 12 29\noutside 30 298\nTMhelix 299 321\ninside 322 330

30092 GCF_900140945.1_10702_1_59 Mycobacterium abscessus subsp. abscessus Terrabacteria group; Actinobacteria; Actinobacteria; Corynebacteriales; Mycobacteriaceae; Mycobacterium; Mycobacterium chelonae group; Mycobacterium abscessus subgroup; Mycobacterium abscessus
MIKHLTRRATVALVVAAAAISPAAYPAMADTATPLNGLFALTPGSCAEGRATGSYFRMIL WP_005073213.1
hypothetical protein [Mycobacterium abscessus] Length: 330\nNumber of predicted TMHs: 2\nExp number of AAs in TMHs: 39.3416199999999\nExp number, first 60 AAs: 17.07688\nTotal prob of N-in: 0.84431\nPOSSIBLE N-term signal sequence\ninside 1 11\nTMhelix 12 29\noutside 30 298\nTMhelix 299 321\ninside 322 330

30093 GCF_900141545.1_12082_5_73 Mycobacterium abscessus subsp. abscessus Terrabacteria group; Actinobacteria; Actinobacteria; Corynebacteriales; Mycobacteriaceae; Mycobacterium; Mycobacterium chelonae group; Mycobacterium abscessus subgroup; Mycobacterium abscessus
MIKHLTRRATVALVVAAAAISPAAYPAMADTATPLNGLFALTPGSCAEGRATGSYFRMIL WP_005073213.1
hypothetical protein [Mycobacterium abscessus] Length: 330\nNumber of predicted TMHs: 2\nExp number of AAs in TMHs: 39.3416199999999\nExp number, first 60 AAs: 17.07688\nTotal prob of N-in: 0.84431\nPOSSIBLE N-term signal sequence\ninside 1 11\nTMhelix 12 29\noutside 30 298\nTMhelix 299 321\ninside 322 330

30094 GCF_900141565.1_12163_2_67 Mycobacterium abscessus subsp. abscessus Terrabacteria group; Actinobacteria; Actinobacteria; Corynebacteriales; Mycobacteriaceae; Mycobacterium; Mycobacterium chelonae group; Mycobacterium abscessus subgroup; Mycobacterium abscessus
MIKHLTRRATVALVVAAAAISPAAYPAMADTATPLNGLFALTPGSCAEGRATGSYFRMIL WP_005073213.1
hypothetical protein [Mycobacterium abscessus] Length: 330\nNumber of predicted TMHs: 2\nExp number of AAs in TMHs: 39.3416199999999\nExp number, first 60 AAs: 17.07688\nTotal prob of N-in: 0.84431\nPOSSIBLE N-term signal sequence\ninside 1 11\nTMhelix 12 29\noutside 30 298\nTMhelix 299 321\ninside 322 330

30095 GCF_900141595.1_12082_5_96 Mycobacterium abscessus subsp. abscessus Terrabacteria group; Actinobacteria; Actinobacteria; Corynebacteriales; Mycobacteriaceae; Mycobacterium; Mycobacterium chelonae group; Mycobacterium abscessus subgroup; Mycobacterium abscessus
MIKHLTRRATVALVVAAAAISPAAYPAMADTATPLNGLFALTPGSCAEGRATGSYFRMIL WP_005073213.1
hypothetical protein [Mycobacterium abscessus] Length: 330\nNumber of predicted TMHs: 2\nExp number of AAs in TMHs: 39.3416199999999\nExp number, first 60 AAs: 17.07688\nTotal prob of N-in: 0.84431\nPOSSIBLE N-term signal sequence\ninside 1 11\nTMhelix 12 29\noutside 30 298\nTMhelix 299 321\ninside 322 330

30096 GCF_900141615.1_12163_2_71 Mycobacterium abscessus subsp. abscessus Terrabacteria group; Actinobacteria; Actinobacteria; Corynebacteriales; Mycobacteriaceae; Mycobacterium; Mycobacterium chelonae group; Mycobacterium abscessus subgroup; Mycobacterium abscessus
MIKHLTRRATVALVVAAAAISPAAYPAMADTATPLNGLFALTPGSCAEGRATGSYFRMIL WP_005073213.1
hypothetical protein [Mycobacterium abscessus] Length: 330\nNumber of predicted TMHs: 2\nExp number of AAs in TMHs: 39.3416199999999\nExp number, first 60 AAs: 17.07688\nTotal prob of N-in: 0.84431\nPOSSIBLE N-term signal sequence\ninside 1 11\nTMhelix 12 29\noutside 30 298\nTMhelix 299 321\ninside 322 330

30097 GCF_900141675.1_12163_2_95 Mycobacterium abscessus subsp. abscessus Terrabacteria group; Actinobacteria; Actinobacteria; Corynebacteriales; Mycobacteriaceae; Mycobacterium; Mycobacterium chelonae group; Mycobacterium abscessus subgroup; Mycobacterium abscessus
MIKHLTRRATVALVVAAAAISPAAYPAMADTATPLNGLFALTPGSCAEGRATGSYFRMIL WP_005073213.1
hypothetical protein [Mycobacterium abscessus] Length: 330\nNumber of predicted TMHs: 2\nExp number of

AAs in TMHs: 39.3416199999999\nExp number, first 60 AAs: 17.07688\nTotal prob of N-in: 0.84431\nPOSSIBLE N-term signal sequence\ninside 1 11\nTMhelix 12 29\noutside 30 298\nTMhelix 299 321\ninside 322 330

30098 GCF_900134145.1_10625_4_77 Mycobacterium abscessus subsp. abscessus Terrabacteria group; Actinobacteria; Actinobacteria; Corynebacteriales; Mycobacteriaceae; Mycobacterium; Mycobacterium chelonae group; Mycobacterium abscessus subgroup; Mycobacterium abscessus
MIKHLTRRATVALVVAAAAISPAAYPAMADTATPLNGLFALTPGSCAEGRATGSYFRMIL WP_005073213.1
hypothetical protein [Mycobacterium abscessus] Length: 330\nNumber of predicted TMHs: 2\nExp number of AAs in TMHs: 39.3416199999999\nExp number, first 60 AAs: 17.07688\nTotal prob of N-in: 0.84431\nPOSSIBLE N-term signal sequence\ninside 1 11\nTMhelix 12 29\noutside 30 298\nTMhelix 299 321\ninside 322 330

30099 GCF_900141455.1_11893_6_58 Mycobacterium abscessus subsp. abscessus Terrabacteria group; Actinobacteria; Actinobacteria; Corynebacteriales; Mycobacteriaceae; Mycobacterium; Mycobacterium chelonae group; Mycobacterium abscessus subgroup; Mycobacterium abscessus
MIKHLTRRATVALVVAAAAISPAAYPAMADTATPLNGLFALTPGSCAEGRATGSYFRMIL WP_005073213.1
hypothetical protein [Mycobacterium abscessus] Length: 330\nNumber of predicted TMHs: 2\nExp number of AAs in TMHs: 39.3416199999999\nExp number, first 60 AAs: 17.07688\nTotal prob of N-in: 0.84431\nPOSSIBLE N-term signal sequence\ninside 1 11\nTMhelix 12 29\noutside 30 298\nTMhelix 299 321\ninside 322 330

30100 GCF_900134055.1_10396_8_7 Mycobacterium abscessus subsp. abscessus Terrabacteria group; Actinobacteria; Actinobacteria; Corynebacteriales; Mycobacteriaceae; Mycobacterium; Mycobacterium chelonae group; Mycobacterium abscessus subgroup; Mycobacterium abscessus
MIKHLTRRATVALVVAAAAISPAAYPAMADTATPLNGLFALTPGSCAEGRATGSYFRMIL WP_005073213.1
hypothetical protein [Mycobacterium abscessus] Length: 330\nNumber of predicted TMHs: 2\nExp number of AAs in TMHs: 39.3416199999999\nExp number, first 60 AAs: 17.07688\nTotal prob of N-in: 0.84431\nPOSSIBLE N-term signal sequence\ninside 1 11\nTMhelix 12 29\noutside 30 298\nTMhelix 299 321\ninside 322 330

30101 GCF_900134065.1_10396_8_6 Mycobacterium abscessus subsp. abscessus Terrabacteria group; Actinobacteria; Actinobacteria; Corynebacteriales; Mycobacteriaceae; Mycobacterium; Mycobacterium chelonae group; Mycobacterium abscessus subgroup; Mycobacterium abscessus
MIKHLTRRATVALVVAAAAISPAAYPAMADTATPLNGLFALTPGSCAEGRATGSYFRMIL WP_005073213.1
hypothetical protein [Mycobacterium abscessus] Length: 330\nNumber of predicted TMHs: 2\nExp number of AAs in TMHs: 39.3416199999999\nExp number, first 60 AAs: 17.07688\nTotal prob of N-in: 0.84431\nPOSSIBLE N-term signal sequence\ninside 1 11\nTMhelix 12 29\noutside 30 298\nTMhelix 299 321\ninside 322 330

30102 GCF_900134085.1_10625_4_52 Mycobacterium abscessus subsp. abscessus Terrabacteria group; Actinobacteria; Actinobacteria; Corynebacteriales; Mycobacteriaceae; Mycobacterium; Mycobacterium chelonae group; Mycobacterium abscessus subgroup; Mycobacterium abscessus
MIKHLTRRATVALVVAAAAISPAAYPAMADTATPLNGLFALTPGSCAEGRATGSYFRMIL WP_005073213.1
hypothetical protein [Mycobacterium abscessus] Length: 330\nNumber of predicted TMHs: 2\nExp number of AAs in TMHs: 39.3416199999999\nExp number, first 60 AAs: 17.07688\nTotal prob of N-in: 0.84431\nPOSSIBLE N-term signal sequence\ninside 1 11\nTMhelix 12 29\noutside 30 298\nTMhelix 299 321\ninside 322 330

30103 GCF_900134115.1_10625_4_53 Mycobacterium abscessus subsp. abscessus Terrabacteria group; Actinobacteria; Actinobacteria; Corynebacteriales; Mycobacteriaceae; Mycobacterium; Mycobacterium chelonae group; Mycobacterium abscessus subgroup; Mycobacterium abscessus
MIKHLTRRATVALVVAAAAISPAAYPAMADTATPLNGLFALTPGSCAEGRATGSYFRMIL WP_005073213.1
hypothetical protein [Mycobacterium abscessus] Length: 330\nNumber of predicted TMHs: 2\nExp number of AAs in TMHs: 39.3416199999999\nExp number, first 60 AAs: 17.07688\nTotal prob of N-in: 0.84431\nPOSSIBLE N-term signal sequence\ninside 1 11\nTMhelix 12 29\noutside 30 298\nTMhelix 299 321\ninside 322 330

30104 GCF_900132325.1_10665_4_65 Mycobacterium abscessus subsp. abscessus Terrabacteria group; Actinobacteria; Actinobacteria; Corynebacteriales; Mycobacteriaceae; Mycobacterium; Mycobacterium chelonae group; Mycobacterium abscessus subgroup; Mycobacterium abscessus
MIKHLTRRATVALVVAAAAISPAAYPAMADTATPLNGLFALTPGSCAEGRATGSYFRMIL WP_005073213.1
hypothetical protein [Mycobacterium abscessus] Length: 330\nNumber of predicted TMHs: 2\nExp number of AAs in TMHs: 39.3416199999999\nExp number, first 60 AAs: 17.07688\nTotal prob of N-in: 0.84431\nPOSSIBLE N-term signal sequence\ninside 1 11\nTMhelix 12 29\noutside 30 298\nTMhelix 299 321\ninside 322 330

30105 GCF_900132255.1_11893_6_49 Mycobacterium abscessus subsp. abscessus Terrabacteria group; Actinobacteria; Actinobacteria; Corynebacteriales; Mycobacteriaceae; Mycobacterium; Mycobacterium chelonae group;

Mycobacterium abscessus subgroup; Mycobacterium abscessus
MIKHLTRRATVALVVAAAAISPAAYPAMADTATPLNGLFALTPGSCAEGRATGSYFRMIL WP_005073213.1
hypothetical protein [Mycobacterium abscessus] Length: 330\nNumber of predicted TMHs: 2\nExp number of AAs in TMHs: 39.3416199999999\nExp number, first 60 AAs: 17.07688\nTotal prob of N-in: 0.84431\nPOSSIBLE N-term signal sequence\ninside 1 11\nTMhelix 12 29\noutside 30 298\nTMhelix 299 321\ninside 322 330

30106 GCF_900134165.1_10625_4_79 Mycobacterium abscessus subsp. abscessus Terrabacteria group; Actinobacteria; Actinobacteria; Corynebacteriales; Mycobacteriaceae; Mycobacterium; Mycobacterium chelonae group; Mycobacterium abscessus subgroup; Mycobacterium abscessus
MIKHLTRRATVALVVAAAAISPAAYPAMADTATPLNGLFALTPGSCAEGRATGSYFRMIL WP_005073213.1
hypothetical protein [Mycobacterium abscessus] Length: 330\nNumber of predicted TMHs: 2\nExp number of AAs in TMHs: 39.3416199999999\nExp number, first 60 AAs: 17.07688\nTotal prob of N-in: 0.84431\nPOSSIBLE N-term signal sequence\ninside 1 11\nTMhelix 12 29\noutside 30 298\nTMhelix 299 321\ninside 322 330

30107 GCF_900134345.1_12082_5_59 Mycobacterium abscessus subsp. abscessus Terrabacteria group; Actinobacteria; Actinobacteria; Corynebacteriales; Mycobacteriaceae; Mycobacterium; Mycobacterium chelonae group; Mycobacterium abscessus subgroup; Mycobacterium abscessus
MIKHLTRRATVALVVAAAAISPAAYPAMADTATPLNGLFALTPGSCAEGRATGSYFRMIL WP_005073213.1
hypothetical protein [Mycobacterium abscessus] Length: 330\nNumber of predicted TMHs: 2\nExp number of AAs in TMHs: 39.3416199999999\nExp number, first 60 AAs: 17.07688\nTotal prob of N-in: 0.84431\nPOSSIBLE N-term signal sequence\ninside 1 11\nTMhelix 12 29\noutside 30 298\nTMhelix 299 321\ninside 322 330

30108 GCF_900134365.1_12082_5_72 Mycobacterium abscessus subsp. abscessus Terrabacteria group; Actinobacteria; Actinobacteria; Corynebacteriales; Mycobacteriaceae; Mycobacterium; Mycobacterium chelonae group; Mycobacterium abscessus subgroup; Mycobacterium abscessus
MIKHLTRRATVALVVAAAAISPAAYPAMADTATPLNGLFALTPGSCAEGRATGSYFRMIL WP_005073213.1
hypothetical protein [Mycobacterium abscessus] Length: 330\nNumber of predicted TMHs: 2\nExp number of AAs in TMHs: 39.3416199999999\nExp number, first 60 AAs: 17.07688\nTotal prob of N-in: 0.84431\nPOSSIBLE N-term signal sequence\ninside 1 11\nTMhelix 12 29\noutside 30 298\nTMhelix 299 321\ninside 322 330

30109 GCF_900134395.1_12163_1_28 Mycobacterium abscessus subsp. abscessus Terrabacteria group; Actinobacteria; Actinobacteria; Corynebacteriales; Mycobacteriaceae; Mycobacterium; Mycobacterium chelonae group; Mycobacterium abscessus subgroup; Mycobacterium abscessus
MIKHLTRRATVALVVAAAAISPAAYPAMADTATPLNGLFALTPGSCAEGRATGSYFRMIL WP_005073213.1
hypothetical protein [Mycobacterium abscessus] Length: 330\nNumber of predicted TMHs: 2\nExp number of AAs in TMHs: 39.3416199999999\nExp number, first 60 AAs: 17.07688\nTotal prob of N-in: 0.84431\nPOSSIBLE N-term signal sequence\ninside 1 11\nTMhelix 12 29\noutside 30 298\nTMhelix 299 321\ninside 322 330

30110 GCF_900134455.1_12163_1_30 Mycobacterium abscessus subsp. abscessus Terrabacteria group; Actinobacteria; Actinobacteria; Corynebacteriales; Mycobacteriaceae; Mycobacterium; Mycobacterium chelonae group; Mycobacterium abscessus subgroup; Mycobacterium abscessus
MIKHLTRRATVALVVAAAAISPAAYPAMADTATPLNGLFALTPGSCAEGRATGSYFRMIL WP_005073213.1
hypothetical protein [Mycobacterium abscessus] Length: 330\nNumber of predicted TMHs: 2\nExp number of AAs in TMHs: 39.3416199999999\nExp number, first 60 AAs: 17.07688\nTotal prob of N-in: 0.84431\nPOSSIBLE N-term signal sequence\ninside 1 11\nTMhelix 12 29\noutside 30 298\nTMhelix 299 321\ninside 322 330

30111 GCF_900132115.1_10665_4_56 Mycobacterium abscessus subsp. abscessus Terrabacteria group; Actinobacteria; Actinobacteria; Corynebacteriales; Mycobacteriaceae; Mycobacterium; Mycobacterium chelonae group; Mycobacterium abscessus subgroup; Mycobacterium abscessus
MIKHLTRRATVALVVAAAAISPAAYPAMADTATPLNGLFALTPGSCAEGRATGSYFRMIL WP_005073213.1
hypothetical protein [Mycobacterium abscessus] Length: 330\nNumber of predicted TMHs: 2\nExp number of AAs in TMHs: 39.3416199999999\nExp number, first 60 AAs: 17.07688\nTotal prob of N-in: 0.84431\nPOSSIBLE N-term signal sequence\ninside 1 11\nTMhelix 12 29\noutside 30 298\nTMhelix 299 321\ninside 322 330

30112 GCF_900134665.1_12163_2_64 Mycobacterium abscessus subsp. abscessus Terrabacteria group; Actinobacteria; Actinobacteria; Corynebacteriales; Mycobacteriaceae; Mycobacterium; Mycobacterium chelonae group; Mycobacterium abscessus subgroup; Mycobacterium abscessus
MIKHLTRRATVALVVAAAAISPAAYPAMADTATPLNGLFALTPGSCAEGRATGSYFRMIL WP_005073213.1
hypothetical protein [Mycobacterium abscessus] Length: 330\nNumber of predicted TMHs: 2\nExp number of AAs in TMHs: 39.3416199999999\nExp number, first 60 AAs: 17.07688\nTotal prob of N-in: 0.84431\nPOSSIBLE N-term signal sequence\ninside 1 11\nTMhelix 12 29\noutside 30 298\nTMhelix 299 321\ninside 322 330

30113 GCF_900134675.1_12163_2_82 Mycobacterium abscessus subsp. abscessus Terrabacteria group; Actinobacteria; Actinobacteria; Corynebacteriales; Mycobacteriaceae; Mycobacterium; Mycobacterium chelonae group; Mycobacterium abscessus subgroup; Mycobacterium abscessus
MIKHLTRRATVALVVAAAAISPAAYPAMADTATPLNGLFALTPGSCAEGRATGSYFRMIL WP_005073213.1
hypothetical protein [Mycobacterium abscessus] Length: 330\nNumber of predicted TMHs: 2\nExp number of AAs in TMHs: 39.3416199999999\nExp number, first 60 AAs: 17.07688\nTotal prob of N-in: 0.84431\nPOSSIBLE N-term signal sequence\ninside 1 11\nTMhelix 12 29\noutside 30 298\nTMhelix 299 321\ninside 322 330

30114 GCF_900134815.1_10208_3_25 Mycobacterium abscessus subsp. abscessus Terrabacteria group; Actinobacteria; Actinobacteria; Corynebacteriales; Mycobacteriaceae; Mycobacterium; Mycobacterium chelonae group; Mycobacterium abscessus subgroup; Mycobacterium abscessus
MIKHLTRRATVALVVAAAAISPAAYPAMADTATPLNGLFALTPGSCAEGRATGSYFRMIL WP_005073213.1
hypothetical protein [Mycobacterium abscessus] Length: 330\nNumber of predicted TMHs: 2\nExp number of AAs in TMHs: 39.3416199999999\nExp number, first 60 AAs: 17.07688\nTotal prob of N-in: 0.84431\nPOSSIBLE N-term signal sequence\ninside 1 11\nTMhelix 12 29\noutside 30 298\nTMhelix 299 321\ninside 322 330

30115 GCF_900134875.1_10208_3_29 Mycobacterium abscessus subsp. abscessus Terrabacteria group; Actinobacteria; Actinobacteria; Corynebacteriales; Mycobacteriaceae; Mycobacterium; Mycobacterium chelonae group; Mycobacterium abscessus subgroup; Mycobacterium abscessus
MIKHLTRRATVALVVAAAAISPAAYPAMADTATPLNGLFALTPGSCAEGRATGSYFRMIL WP_005073213.1
hypothetical protein [Mycobacterium abscessus] Length: 330\nNumber of predicted TMHs: 2\nExp number of AAs in TMHs: 39.3416199999999\nExp number, first 60 AAs: 17.07688\nTotal prob of N-in: 0.84431\nPOSSIBLE N-term signal sequence\ninside 1 11\nTMhelix 12 29\noutside 30 298\nTMhelix 299 321\ninside 322 330

30116 GCF_900134855.1_10208_3_31 Mycobacterium abscessus subsp. abscessus Terrabacteria group; Actinobacteria; Actinobacteria; Corynebacteriales; Mycobacteriaceae; Mycobacterium; Mycobacterium chelonae group; Mycobacterium abscessus subgroup; Mycobacterium abscessus
MIKHLTRRATVALVVAAAAISPAAYPAMADTATPLNGLFALTPGSCAEGRATGSYFRMIL WP_005073213.1
hypothetical protein [Mycobacterium abscessus] Length: 330\nNumber of predicted TMHs: 2\nExp number of AAs in TMHs: 39.3416199999999\nExp number, first 60 AAs: 17.07688\nTotal prob of N-in: 0.84431\nPOSSIBLE N-term signal sequence\ninside 1 11\nTMhelix 12 29\noutside 30 298\nTMhelix 299 321\ninside 322 330

30117 GCF_900134895.1_10208_3_30 Mycobacterium abscessus subsp. abscessus Terrabacteria group; Actinobacteria; Actinobacteria; Corynebacteriales; Mycobacteriaceae; Mycobacterium; Mycobacterium chelonae group; Mycobacterium abscessus subgroup; Mycobacterium abscessus
MIKHLTRRATVALVVAAAAISPAAYPAMADTATPLNGLFALTPGSCAEGRATGSYFRMIL WP_005073213.1
hypothetical protein [Mycobacterium abscessus] Length: 330\nNumber of predicted TMHs: 2\nExp number of AAs in TMHs: 39.3416199999999\nExp number, first 60 AAs: 17.07688\nTotal prob of N-in: 0.84431\nPOSSIBLE N-term signal sequence\ninside 1 11\nTMhelix 12 29\noutside 30 298\nTMhelix 299 321\ninside 322 330

30118 GCF_900131985.1_10665_4_47 Mycobacterium abscessus subsp. abscessus Terrabacteria group; Actinobacteria; Actinobacteria; Corynebacteriales; Mycobacteriaceae; Mycobacterium; Mycobacterium chelonae group; Mycobacterium abscessus subgroup; Mycobacterium abscessus
MIKHLTRRATVALVVAAAAISPAAYPAMADTATPLNGLFALTPGSCAEGRATGSYFRMIL WP_005073213.1
hypothetical protein [Mycobacterium abscessus] Length: 330\nNumber of predicted TMHs: 2\nExp number of AAs in TMHs: 39.3416199999999\nExp number, first 60 AAs: 17.07688\nTotal prob of N-in: 0.84431\nPOSSIBLE N-term signal sequence\ninside 1 11\nTMhelix 12 29\noutside 30 298\nTMhelix 299 321\ninside 322 330

30119 GCF_900135055.1_10208_3_9 Mycobacterium abscessus subsp. abscessus Terrabacteria group; Actinobacteria; Actinobacteria; Corynebacteriales; Mycobacteriaceae; Mycobacterium; Mycobacterium chelonae group; Mycobacterium abscessus subgroup; Mycobacterium abscessus
MIKHLTRRATVALVVAAAAISPAAYPAMADTATPLNGLFALTPGSCAEGRATGSYFRMIL WP_005073213.1
hypothetical protein [Mycobacterium abscessus] Length: 330\nNumber of predicted TMHs: 2\nExp number of AAs in TMHs: 39.3416199999999\nExp number, first 60 AAs: 17.07688\nTotal prob of N-in: 0.84431\nPOSSIBLE N-term signal sequence\ninside 1 11\nTMhelix 12 29\noutside 30 298\nTMhelix 299 321\ninside 322 330

30120 GCF_900135075.1_10250_1_82 Mycobacterium abscessus subsp. abscessus Terrabacteria group; Actinobacteria; Actinobacteria; Corynebacteriales; Mycobacteriaceae; Mycobacterium; Mycobacterium chelonae group; Mycobacterium abscessus subgroup; Mycobacterium abscessus
MIKHLTRRATVALVVAAAAISPAAYPAMADTATPLNGLFALTPGSCAEGRATGSYFRMIL WP_005073213.1
hypothetical protein [Mycobacterium abscessus] Length: 330\nNumber of predicted TMHs: 2\nExp number of

AAs in TMHs: 39.3416199999999\nExp number, first 60 AAs: 17.07688\nTotal prob of N-in: 0.84431\nPOSSIBLE N-term signal sequence\ninside 1 11\nTMhelix 12 29\noutside 30 298\nTMhelix 299 321\ninside 322 330

30121 GCF_900135095.1_10250_1_85 Mycobacterium abscessus subsp. abscessus Terrabacteria group; Actinobacteria; Actinobacteria; Corynebacteriales; Mycobacteriaceae; Mycobacterium; Mycobacterium chelonae group; Mycobacterium abscessus subgroup; Mycobacterium abscessus
MIKHLTRRATVALVVAAAAISPAAYPAMADTATPLNGLFALTPGSCAEGRATGSYFRMIL WP_005073213.1
hypothetical protein [Mycobacterium abscessus] Length: 330\nNumber of predicted TMHs: 2\nExp number of AAs in TMHs: 39.3416199999999\nExp number, first 60 AAs: 17.07688\nTotal prob of N-in: 0.84431\nPOSSIBLE N-term signal sequence\ninside 1 11\nTMhelix 12 29\noutside 30 298\nTMhelix 299 321\ninside 322 330

30122 GCF_900135115.1_10625_4_19 Mycobacterium abscessus subsp. abscessus Terrabacteria group; Actinobacteria; Actinobacteria; Corynebacteriales; Mycobacteriaceae; Mycobacterium; Mycobacterium chelonae group; Mycobacterium abscessus subgroup; Mycobacterium abscessus
MIKHLTRRATVALVVAAAAISPAAYPAMADTATPLNGLFALTPGSCAEGRATGSYFRMIL WP_005073213.1
hypothetical protein [Mycobacterium abscessus] Length: 330\nNumber of predicted TMHs: 2\nExp number of AAs in TMHs: 39.3416199999999\nExp number, first 60 AAs: 17.07688\nTotal prob of N-in: 0.84431\nPOSSIBLE N-term signal sequence\ninside 1 11\nTMhelix 12 29\noutside 30 298\nTMhelix 299 321\ninside 322 330

30123 GCF_900135215.1_10625_4_28 Mycobacterium abscessus subsp. abscessus Terrabacteria group; Actinobacteria; Actinobacteria; Corynebacteriales; Mycobacteriaceae; Mycobacterium; Mycobacterium chelonae group; Mycobacterium abscessus subgroup; Mycobacterium abscessus
MIKHLTRRATVALVVAAAAISPAAYPAMADTATPLNGLFALTPGSCAEGRATGSYFRMIL WP_005073213.1
hypothetical protein [Mycobacterium abscessus] Length: 330\nNumber of predicted TMHs: 2\nExp number of AAs in TMHs: 39.3416199999999\nExp number, first 60 AAs: 17.07688\nTotal prob of N-in: 0.84431\nPOSSIBLE N-term signal sequence\ninside 1 11\nTMhelix 12 29\noutside 30 298\nTMhelix 299 321\ninside 322 330

30124 GCF_900135255.1_10625_4_40 Mycobacterium abscessus subsp. abscessus Terrabacteria group; Actinobacteria; Actinobacteria; Corynebacteriales; Mycobacteriaceae; Mycobacterium; Mycobacterium chelonae group; Mycobacterium abscessus subgroup; Mycobacterium abscessus
MIKHLTRRATVALVVAAAAISPAAYPAMADTATPLNGLFALTPGSCAEGRATGSYFRMIL WP_005073213.1
hypothetical protein [Mycobacterium abscessus] Length: 330\nNumber of predicted TMHs: 2\nExp number of AAs in TMHs: 39.3416199999999\nExp number, first 60 AAs: 17.07688\nTotal prob of N-in: 0.84431\nPOSSIBLE N-term signal sequence\ninside 1 11\nTMhelix 12 29\noutside 30 298\nTMhelix 299 321\ninside 322 330

30125 GCF_900135275.1_10625_4_41 Mycobacterium abscessus subsp. abscessus Terrabacteria group; Actinobacteria; Actinobacteria; Corynebacteriales; Mycobacteriaceae; Mycobacterium; Mycobacterium chelonae group; Mycobacterium abscessus subgroup; Mycobacterium abscessus
MIKHLTRRATVALVVAAAAISPAAYPAMADTATPLNGLFALTPGSCAEGRATGSYFRMIL WP_005073213.1
hypothetical protein [Mycobacterium abscessus] Length: 330\nNumber of predicted TMHs: 2\nExp number of AAs in TMHs: 39.3416199999999\nExp number, first 60 AAs: 17.07688\nTotal prob of N-in: 0.84431\nPOSSIBLE N-term signal sequence\ninside 1 11\nTMhelix 12 29\noutside 30 298\nTMhelix 299 321\ninside 322 330

30126 GCF_900135315.1_10625_4_45 Mycobacterium abscessus subsp. abscessus Terrabacteria group; Actinobacteria; Actinobacteria; Corynebacteriales; Mycobacteriaceae; Mycobacterium; Mycobacterium chelonae group; Mycobacterium abscessus subgroup; Mycobacterium abscessus
MIKHLTRRATVALVVAAAAISPAAYPAMADTATPLNGLFALTPGSCAEGRATGSYFRMIL WP_005073213.1
hypothetical protein [Mycobacterium abscessus] Length: 330\nNumber of predicted TMHs: 2\nExp number of AAs in TMHs: 39.3416199999999\nExp number, first 60 AAs: 17.07688\nTotal prob of N-in: 0.84431\nPOSSIBLE N-term signal sequence\ninside 1 11\nTMhelix 12 29\noutside 30 298\nTMhelix 299 321\ninside 322 330

30127 GCF_900135305.1_10625_4_44 Mycobacterium abscessus subsp. abscessus Terrabacteria group; Actinobacteria; Actinobacteria; Corynebacteriales; Mycobacteriaceae; Mycobacterium; Mycobacterium chelonae group; Mycobacterium abscessus subgroup; Mycobacterium abscessus
MIKHLTRRATVALVVAAAAISPAAYPAMADTATPLNGLFALTPGSCAEGRATGSYFRMIL WP_005073213.1
hypothetical protein [Mycobacterium abscessus] Length: 330\nNumber of predicted TMHs: 2\nExp number of AAs in TMHs: 39.3416199999999\nExp number, first 60 AAs: 17.07688\nTotal prob of N-in: 0.84431\nPOSSIBLE N-term signal sequence\ninside 1 11\nTMhelix 12 29\noutside 30 298\nTMhelix 299 321\ninside 322 330

30128 GCF_900131765.1_10665_4_8 Mycobacterium abscessus subsp. abscessus Terrabacteria group; Actinobacteria; Actinobacteria; Corynebacteriales; Mycobacteriaceae; Mycobacterium; Mycobacterium chelonae group;

Mycobacterium abscessus subgroup; Mycobacterium abscessus
MIKHLTRRATVALVVAAAAISPAAYPAMADTATPLNGLFALTPGSCAEGRATGSYFRMIL WP_005073213.1
hypothetical protein [Mycobacterium abscessus] Length: 330\nNumber of predicted TMHs: 2\nExp number of AAs in TMHs: 39.3416199999999\nExp number, first 60 AAs: 17.07688\nTotal prob of N-in: 0.84431\nPOSSIBLE N-term signal sequence\ninside 1 11\nTMhelix 12 29\noutside 30 298\nTMhelix 299 321\ninside 322 330

30129 GCF_900135465.1_10625_4_64 Mycobacterium abscessus subsp. abscessus Terrabacteria group; Actinobacteria; Actinobacteria; Corynebacteriales; Mycobacteriaceae; Mycobacterium; Mycobacterium chelonae group; Mycobacterium abscessus subgroup; Mycobacterium abscessus
MIKHLTRRATVALVVAAAAISPAAYPAMADTATPLNGLFALTPGSCAEGRATGSYFRMIL WP_005073213.1
hypothetical protein [Mycobacterium abscessus] Length: 330\nNumber of predicted TMHs: 2\nExp number of AAs in TMHs: 39.3416199999999\nExp number, first 60 AAs: 17.07688\nTotal prob of N-in: 0.84431\nPOSSIBLE N-term signal sequence\ninside 1 11\nTMhelix 12 29\noutside 30 298\nTMhelix 299 321\ninside 322 330

30130 GCF_900131705.1_10665_4_28 Mycobacterium abscessus subsp. abscessus Terrabacteria group; Actinobacteria; Actinobacteria; Corynebacteriales; Mycobacteriaceae; Mycobacterium; Mycobacterium chelonae group; Mycobacterium abscessus subgroup; Mycobacterium abscessus
MIKHLTRRATVALVVAAAAISPAAYPAMADTATPLNGLFALTPGSCAEGRATGSYFRMIL WP_005073213.1
hypothetical protein [Mycobacterium abscessus] Length: 330\nNumber of predicted TMHs: 2\nExp number of AAs in TMHs: 39.3416199999999\nExp number, first 60 AAs: 17.07688\nTotal prob of N-in: 0.84431\nPOSSIBLE N-term signal sequence\ninside 1 11\nTMhelix 12 29\noutside 30 298\nTMhelix 299 321\ninside 322 330

30131 GCF_900135485.1_10625_4_66 Mycobacterium abscessus subsp. abscessus Terrabacteria group; Actinobacteria; Actinobacteria; Corynebacteriales; Mycobacteriaceae; Mycobacterium; Mycobacterium chelonae group; Mycobacterium abscessus subgroup; Mycobacterium abscessus
MIKHLTRRATVALVVAAAAISPAAYPAMADTATPLNGLFALTPGSCAEGRATGSYFRMIL WP_005073213.1
hypothetical protein [Mycobacterium abscessus] Length: 330\nNumber of predicted TMHs: 2\nExp number of AAs in TMHs: 39.3416199999999\nExp number, first 60 AAs: 17.07688\nTotal prob of N-in: 0.84431\nPOSSIBLE N-term signal sequence\ninside 1 11\nTMhelix 12 29\noutside 30 298\nTMhelix 299 321\ninside 322 330

30132 GCF_900135495.1_10625_4_65 Mycobacterium abscessus subsp. abscessus Terrabacteria group; Actinobacteria; Actinobacteria; Corynebacteriales; Mycobacteriaceae; Mycobacterium; Mycobacterium chelonae group; Mycobacterium abscessus subgroup; Mycobacterium abscessus
MIKHLTRRATVALVVAAAAISPAAYPAMADTATPLNGLFALTPGSCAEGRATGSYFRMIL WP_005073213.1
hypothetical protein [Mycobacterium abscessus] Length: 330\nNumber of predicted TMHs: 2\nExp number of AAs in TMHs: 39.3416199999999\nExp number, first 60 AAs: 17.07688\nTotal prob of N-in: 0.84431\nPOSSIBLE N-term signal sequence\ninside 1 11\nTMhelix 12 29\noutside 30 298\nTMhelix 299 321\ninside 322 330

30133 GCF_900135515.1_10625_4_67 Mycobacterium abscessus subsp. abscessus Terrabacteria group; Actinobacteria; Actinobacteria; Corynebacteriales; Mycobacteriaceae; Mycobacterium; Mycobacterium chelonae group; Mycobacterium abscessus subgroup; Mycobacterium abscessus
MIKHLTRRATVALVVAAAAISPAAYPAMADTATPLNGLFALTPGSCAEGRATGSYFRMIL WP_005073213.1
hypothetical protein [Mycobacterium abscessus] Length: 330\nNumber of predicted TMHs: 2\nExp number of AAs in TMHs: 39.3416199999999\nExp number, first 60 AAs: 17.07688\nTotal prob of N-in: 0.84431\nPOSSIBLE N-term signal sequence\ninside 1 11\nTMhelix 12 29\noutside 30 298\nTMhelix 299 321\ninside 322 330

30134 GCF_900135535.1_10625_4_70 Mycobacterium abscessus subsp. abscessus Terrabacteria group; Actinobacteria; Actinobacteria; Corynebacteriales; Mycobacteriaceae; Mycobacterium; Mycobacterium chelonae group; Mycobacterium abscessus subgroup; Mycobacterium abscessus
MIKHLTRRATVALVVAAAAISPAAYPAMADTATPLNGLFALTPGSCAEGRATGSYFRMIL WP_005073213.1
hypothetical protein [Mycobacterium abscessus] Length: 330\nNumber of predicted TMHs: 2\nExp number of AAs in TMHs: 39.3416199999999\nExp number, first 60 AAs: 17.07688\nTotal prob of N-in: 0.84431\nPOSSIBLE N-term signal sequence\ninside 1 11\nTMhelix 12 29\noutside 30 298\nTMhelix 299 321\ninside 322 330

30135 GCF_900135665.1_11861_7_26 Mycobacterium abscessus subsp. abscessus Terrabacteria group; Actinobacteria; Actinobacteria; Corynebacteriales; Mycobacteriaceae; Mycobacterium; Mycobacterium chelonae group; Mycobacterium abscessus subgroup; Mycobacterium abscessus
MIKHLTRRATVALVVAAAAISPAAYPAMADTATPLNGLFALTPGSCAEGRATGSYFRMIL WP_005073213.1
hypothetical protein [Mycobacterium abscessus] Length: 330\nNumber of predicted TMHs: 2\nExp number of AAs in TMHs: 39.3416199999999\nExp number, first 60 AAs: 17.07688\nTotal prob of N-in: 0.84431\nPOSSIBLE N-term signal sequence\ninside 1 11\nTMhelix 12 29\noutside 30 298\nTMhelix 299 321\ninside 322 330

30136 GCF_900135765.1_11861_7_73 Mycobacterium abscessus subsp. abscessus Terrabacteria group; Actinobacteria; Actinobacteria; Corynebacteriales; Mycobacteriaceae; Mycobacterium; Mycobacterium chelonae group; Mycobacterium abscessus subgroup; Mycobacterium abscessus
MIKHLTRRATVALVVAAAAISPAAYPAMADTATPLNGLFALTPGSCAEGRATGSYFRMIL WP_005073213.1 hypothetical protein [Mycobacterium abscessus] Length: 330\nNumber of predicted TMHs: 2\nExp number of AAs in TMHs: 39.3416199999999\nExp number, first 60 AAs: 17.07688\nTotal prob of N-in: 0.84431\nPOSSIBLE N-term signal sequence\ninside 1 11\nTMhelix 12 29\noutside 30 298\nTMhelix 299 321\ninside 322 330

30137 GCF_900135775.1_11861_7_80 Mycobacterium abscessus subsp. abscessus Terrabacteria group; Actinobacteria; Actinobacteria; Corynebacteriales; Mycobacteriaceae; Mycobacterium; Mycobacterium chelonae group; Mycobacterium abscessus subgroup; Mycobacterium abscessus
MIKHLTRRATVALVVAAAAISPAAYPAMADTATPLNGLFALTPGSCAEGRATGSYFRMIL WP_005073213.1 hypothetical protein [Mycobacterium abscessus] Length: 330\nNumber of predicted TMHs: 2\nExp number of AAs in TMHs: 39.3416199999999\nExp number, first 60 AAs: 17.07688\nTotal prob of N-in: 0.84431\nPOSSIBLE N-term signal sequence\ninside 1 11\nTMhelix 12 29\noutside 30 298\nTMhelix 299 321\ninside 322 330

30138 GCF_900131595.1_10665_4_15 Mycobacterium abscessus subsp. abscessus Terrabacteria group; Actinobacteria; Actinobacteria; Corynebacteriales; Mycobacteriaceae; Mycobacterium; Mycobacterium chelonae group; Mycobacterium abscessus subgroup; Mycobacterium abscessus
MIKHLTRRATVALVVAAAAISPAAYPAMADTATPLNGLFALTPGSCAEGRATGSYFRMIL WP_005073213.1 hypothetical protein [Mycobacterium abscessus] Length: 330\nNumber of predicted TMHs: 2\nExp number of AAs in TMHs: 39.3416199999999\nExp number, first 60 AAs: 17.07688\nTotal prob of N-in: 0.84431\nPOSSIBLE N-term signal sequence\ninside 1 11\nTMhelix 12 29\noutside 30 298\nTMhelix 299 321\ninside 322 330

30139 GCF_900135675.1_11861_7_27 Mycobacterium abscessus subsp. abscessus Terrabacteria group; Actinobacteria; Actinobacteria; Corynebacteriales; Mycobacteriaceae; Mycobacterium; Mycobacterium chelonae group; Mycobacterium abscessus subgroup; Mycobacterium abscessus
MIKHLTRRATVALVVAAAAISPAAYPAMADTATPLNGLFALTPGSCTEGRATGSYFRMILWP_079655361.1 hypothetical protein [Mycobacterium abscessus] Length: 330\nNumber of predicted TMHs: 2\nExp number of AAs in TMHs: 39.3376499999999\nExp number, first 60 AAs: 17.0732\nTotal prob of N-in: 0.84430\nPOSSIBLE N-term signal sequence\ninside 1 11\nTMhelix 12 29\noutside 30 298\nTMhelix 299 321\ninside 322 330

30140 GCF_900135825.1_11861_8_17 Mycobacterium abscessus subsp. abscessus Terrabacteria group; Actinobacteria; Actinobacteria; Corynebacteriales; Mycobacteriaceae; Mycobacterium; Mycobacterium chelonae group; Mycobacterium abscessus subgroup; Mycobacterium abscessus
MIKHLTRRATVALVVAAAAISPAAYPAMADTATPLNGLFALTPGSCAEGRATGSYFRMIL WP_005073213.1 hypothetical protein [Mycobacterium abscessus] Length: 330\nNumber of predicted TMHs: 2\nExp number of AAs in TMHs: 39.3416199999999\nExp number, first 60 AAs: 17.07688\nTotal prob of N-in: 0.84431\nPOSSIBLE N-term signal sequence\ninside 1 11\nTMhelix 12 29\noutside 30 298\nTMhelix 299 321\ninside 322 330

30141 GCF_900131415.1_10665_4_3 Mycobacterium abscessus subsp. abscessus Terrabacteria group; Actinobacteria; Actinobacteria; Corynebacteriales; Mycobacteriaceae; Mycobacterium; Mycobacterium chelonae group; Mycobacterium abscessus subgroup; Mycobacterium abscessus
MIKHLTRRATVALVVAAAAISPAAYPAMADTATPLNGLFALTPGSCAEGRATGSYFRMIL WP_005073213.1 hypothetical protein [Mycobacterium abscessus] Length: 330\nNumber of predicted TMHs: 2\nExp number of AAs in TMHs: 39.3416199999999\nExp number, first 60 AAs: 17.07688\nTotal prob of N-in: 0.84431\nPOSSIBLE N-term signal sequence\ninside 1 11\nTMhelix 12 29\noutside 30 298\nTMhelix 299 321\ninside 322 330

30142 GCF_900135725.1_11861_7_72 Mycobacterium abscessus subsp. abscessus Terrabacteria group; Actinobacteria; Actinobacteria; Corynebacteriales; Mycobacteriaceae; Mycobacterium; Mycobacterium chelonae group; Mycobacterium abscessus subgroup; Mycobacterium abscessus
MPRRRTILSALASFSMVIGLLVVSTPTAWAAPFPVVDPAKLPPSQPPAPPGDMRLNGAPL WP_074362414.1 type VII secretion-associated serine protease mycosin [Mycobacterium abscessus] Length: 470\nNumber of predicted TMHs: 2\nExp number of AAs in TMHs: 44.7482799999999\nExp number, first 60 AAs: 22.65577\nTotal prob of N-in: 0.99532\nPOSSIBLE N-term signal sequence\ninside 1 6\nTMhelix 7 29\noutside 30 441\nTMhelix 442 464\ninside 465 470

30143 GCF_900131395.1_10665_3_85 Mycobacterium abscessus subsp. abscessus Terrabacteria group; Actinobacteria; Actinobacteria; Corynebacteriales; Mycobacteriaceae; Mycobacterium; Mycobacterium chelonae group; Mycobacterium abscessus subgroup; Mycobacterium abscessus
MIKHLTRRATVALVVAAAAISPAAYPAMADTATPLNGLFALTPGSCAEGRATGSYFRMIL WP_005073213.1

hypothetical protein [Mycobacterium abscessus] Length: 330\nNumber of predicted TMHs: 2\nExp number of AAs in TMHs: 39.3416199999999\nExp number, first 60 AAs: 17.07688\nTotal prob of N-in: 0.84431\nPOSSIBLE N-term signal sequence\ninside 1 11\nTMhelix 12 29\noutside 30 298\nTMhelix 299 321\ninside 322 330

30144 GCF_900136045.1_12163_1_61 Mycobacterium abscessus subsp. abscessus Terrabacteria group; Actinobacteria; Actinobacteria; Corynebacteriales; Mycobacteriaceae; Mycobacterium; Mycobacterium chelonae group; Mycobacterium abscessus subgroup; Mycobacterium abscessus
MIKHLTRRATVALVVAAAAISPAAYPAMADTATPLNGLFALTPGSCAEGRATGSYFRMIL WP_005073213.1
hypothetical protein [Mycobacterium abscessus] Length: 330\nNumber of predicted TMHs: 2\nExp number of AAs in TMHs: 39.3416199999999\nExp number, first 60 AAs: 17.07688\nTotal prob of N-in: 0.84431\nPOSSIBLE N-term signal sequence\ninside 1 11\nTMhelix 12 29\noutside 30 298\nTMhelix 299 321\ninside 322 330

30145 GCF_900136065.1_12163_1_63 Mycobacterium abscessus subsp. abscessus Terrabacteria group; Actinobacteria; Actinobacteria; Corynebacteriales; Mycobacteriaceae; Mycobacterium; Mycobacterium chelonae group; Mycobacterium abscessus subgroup; Mycobacterium abscessus
MIKHLTRRATVALVVAAAAISPAAYPAMADTATPLNGLFALTPGSCAEGRATGSYFRMIL WP_005073213.1
hypothetical protein [Mycobacterium abscessus] Length: 330\nNumber of predicted TMHs: 2\nExp number of AAs in TMHs: 39.3416199999999\nExp number, first 60 AAs: 17.07688\nTotal prob of N-in: 0.84431\nPOSSIBLE N-term signal sequence\ninside 1 11\nTMhelix 12 29\noutside 30 298\nTMhelix 299 321\ninside 322 330

30146 GCF_900136075.1_12163_1_64 Mycobacterium abscessus subsp. abscessus Terrabacteria group; Actinobacteria; Actinobacteria; Corynebacteriales; Mycobacteriaceae; Mycobacterium; Mycobacterium chelonae group; Mycobacterium abscessus subgroup; Mycobacterium abscessus
MIKHLTRRATVALVVAAAAISPAAYPAMADTATPLNGLFALTPGSCAEGRATGSYFRMIL WP_005073213.1
hypothetical protein [Mycobacterium abscessus] Length: 330\nNumber of predicted TMHs: 2\nExp number of AAs in TMHs: 39.3416199999999\nExp number, first 60 AAs: 17.07688\nTotal prob of N-in: 0.84431\nPOSSIBLE N-term signal sequence\ninside 1 11\nTMhelix 12 29\noutside 30 298\nTMhelix 299 321\ninside 322 330

30147 GCF_900136245.1_12163_1_80 Mycobacterium abscessus subsp. abscessus Terrabacteria group; Actinobacteria; Actinobacteria; Corynebacteriales; Mycobacteriaceae; Mycobacterium; Mycobacterium chelonae group; Mycobacterium abscessus subgroup; Mycobacterium abscessus
MIKHLTRRATVALVVAAAAISPAAYPAMADTATPLNGLFALTPGSCAEGRATGSYFRMIL WP_005073213.1
hypothetical protein [Mycobacterium abscessus] Length: 330\nNumber of predicted TMHs: 2\nExp number of AAs in TMHs: 39.3416199999999\nExp number, first 60 AAs: 17.07688\nTotal prob of N-in: 0.84431\nPOSSIBLE N-term signal sequence\ninside 1 11\nTMhelix 12 29\noutside 30 298\nTMhelix 299 321\ninside 322 330

30148 GCF_900136255.1_12163_1_81 Mycobacterium abscessus subsp. abscessus Terrabacteria group; Actinobacteria; Actinobacteria; Corynebacteriales; Mycobacteriaceae; Mycobacterium; Mycobacterium chelonae group; Mycobacterium abscessus subgroup; Mycobacterium abscessus
MIKHLTRRATVALVVAAAAISPAAYPAMADTATPLNGLFALTPGSCAEGRATGSYFRMIL WP_005073213.1
hypothetical protein [Mycobacterium abscessus] Length: 330\nNumber of predicted TMHs: 2\nExp number of AAs in TMHs: 39.3416199999999\nExp number, first 60 AAs: 17.07688\nTotal prob of N-in: 0.84431\nPOSSIBLE N-term signal sequence\ninside 1 11\nTMhelix 12 29\noutside 30 298\nTMhelix 299 321\ninside 322 330

30149 GCF_900136445.1_10208_3_68 Mycobacterium abscessus subsp. abscessus Terrabacteria group; Actinobacteria; Actinobacteria; Corynebacteriales; Mycobacteriaceae; Mycobacterium; Mycobacterium chelonae group; Mycobacterium abscessus subgroup; Mycobacterium abscessus
MIKHLTRRATVALVVAAAAISPAAYPAMADTATPLNGLFALTPGSCAEGRATGSYFRMIL WP_005073213.1
hypothetical protein [Mycobacterium abscessus] Length: 330\nNumber of predicted TMHs: 2\nExp number of AAs in TMHs: 39.3416199999999\nExp number, first 60 AAs: 17.07688\nTotal prob of N-in: 0.84431\nPOSSIBLE N-term signal sequence\ninside 1 11\nTMhelix 12 29\noutside 30 298\nTMhelix 299 321\ninside 322 330

30150 GCF_900136485.1_10208_3_76 Mycobacterium abscessus subsp. abscessus Terrabacteria group; Actinobacteria; Actinobacteria; Corynebacteriales; Mycobacteriaceae; Mycobacterium; Mycobacterium chelonae group; Mycobacterium abscessus subgroup; Mycobacterium abscessus
MIKHLTRRATVALVVAAAAISPAAYPAMADTATPLNGLFALTPGSCAEGRATGSYFRMIL WP_005073213.1
hypothetical protein [Mycobacterium abscessus] Length: 330\nNumber of predicted TMHs: 2\nExp number of AAs in TMHs: 39.3416199999999\nExp number, first 60 AAs: 17.07688\nTotal prob of N-in: 0.84431\nPOSSIBLE N-term signal sequence\ninside 1 11\nTMhelix 12 29\noutside 30 298\nTMhelix 299 321\ninside 322 330

30151 GCF_900136505.1_10208_3_77 Mycobacterium abscessus subsp. abscessus Terrabacteria group; Actinobacteria; Actinobacteria; Corynebacteriales; Mycobacteriaceae; Mycobacterium; Mycobacterium chelonae group; Mycobacterium abscessus subgroup; Mycobacterium abscessus
MIKHLTRRATVALVVAAAAISPAAYPAMADTATPLNGLFALTPGSCAEGRATGSYFRMIL WP_005073213.1
hypothetical protein [Mycobacterium abscessus] Length: 330\nNumber of predicted TMHs: 2\nExp number of AAs in TMHs: 39.3416199999999\nExp number, first 60 AAs: 17.07688\nTotal prob of N-in: 0.84431\nPOSSIBLE N-term signal sequence\ninside 1 11\nTMhelix 12 29\noutside 30 298\nTMhelix 299 321\ninside 322 330

30152 GCF_900136555.1_10208_3_78 Mycobacterium abscessus subsp. abscessus Terrabacteria group; Actinobacteria; Actinobacteria; Corynebacteriales; Mycobacteriaceae; Mycobacterium; Mycobacterium chelonae group; Mycobacterium abscessus subgroup; Mycobacterium abscessus
MIKHLTRRATVALVVAAAAISPAAYPAMADTATPLNGLFALTPGSCAEGRATGSYFRMIL WP_005073213.1
hypothetical protein [Mycobacterium abscessus] Length: 330\nNumber of predicted TMHs: 2\nExp number of AAs in TMHs: 39.3416199999999\nExp number, first 60 AAs: 17.07688\nTotal prob of N-in: 0.84431\nPOSSIBLE N-term signal sequence\ninside 1 11\nTMhelix 12 29\noutside 30 298\nTMhelix 299 321\ninside 322 330

30153 GCF_900131245.1_10665_3_69 Mycobacterium abscessus subsp. abscessus Terrabacteria group; Actinobacteria; Actinobacteria; Corynebacteriales; Mycobacteriaceae; Mycobacterium; Mycobacterium chelonae group; Mycobacterium abscessus subgroup; Mycobacterium abscessus
MIKHLTRRATVALVVAAAAISPAAYPAMADTATPLNGLFALTPGSCAEGRATGSYFRMIL WP_005073213.1
hypothetical protein [Mycobacterium abscessus] Length: 330\nNumber of predicted TMHs: 2\nExp number of AAs in TMHs: 39.3416199999999\nExp number, first 60 AAs: 17.07688\nTotal prob of N-in: 0.84431\nPOSSIBLE N-term signal sequence\ninside 1 11\nTMhelix 12 29\noutside 30 298\nTMhelix 299 321\ninside 322 330

30154 GCF_900136625.1_10625_4_31 Mycobacterium abscessus subsp. abscessus Terrabacteria group; Actinobacteria; Actinobacteria; Corynebacteriales; Mycobacteriaceae; Mycobacterium; Mycobacterium chelonae group; Mycobacterium abscessus subgroup; Mycobacterium abscessus
MIKHLTRRATVALVVAAAAISPAAYPAMADTATPLNGLFALTPGSCAEGRATGSYFRMIL WP_005073213.1
hypothetical protein [Mycobacterium abscessus] Length: 330\nNumber of predicted TMHs: 2\nExp number of AAs in TMHs: 39.3416199999999\nExp number, first 60 AAs: 17.07688\nTotal prob of N-in: 0.84431\nPOSSIBLE N-term signal sequence\ninside 1 11\nTMhelix 12 29\noutside 30 298\nTMhelix 299 321\ninside 322 330

30155 GCF_900136645.1_10625_5_12 Mycobacterium abscessus subsp. abscessus Terrabacteria group; Actinobacteria; Actinobacteria; Corynebacteriales; Mycobacteriaceae; Mycobacterium; Mycobacterium chelonae group; Mycobacterium abscessus subgroup; Mycobacterium abscessus
MIKHLTRRATVALVVAAAAISPAAYPAMADTATPLNGLFALTPGSCAEGRATGSYFRMIL WP_005073213.1
hypothetical protein [Mycobacterium abscessus] Length: 330\nNumber of predicted TMHs: 2\nExp number of AAs in TMHs: 39.3416199999999\nExp number, first 60 AAs: 17.07688\nTotal prob of N-in: 0.84431\nPOSSIBLE N-term signal sequence\ninside 1 11\nTMhelix 12 29\noutside 30 298\nTMhelix 299 321\ninside 322 330

30156 GCF_900136675.1_10625_5_13 Mycobacterium abscessus subsp. abscessus Terrabacteria group; Actinobacteria; Actinobacteria; Corynebacteriales; Mycobacteriaceae; Mycobacterium; Mycobacterium chelonae group; Mycobacterium abscessus subgroup; Mycobacterium abscessus
MIKHLTRRATVALVVAAAAISPAAYPAMADTATPLNGLFALTPGSCAEGRATGSYFRMIL WP_005073213.1
hypothetical protein [Mycobacterium abscessus] Length: 330\nNumber of predicted TMHs: 2\nExp number of AAs in TMHs: 39.3416199999999\nExp number, first 60 AAs: 17.07688\nTotal prob of N-in: 0.84431\nPOSSIBLE N-term signal sequence\ninside 1 11\nTMhelix 12 29\noutside 30 298\nTMhelix 299 321\ninside 322 330

30157 GCF_900136695.1_10625_5_25 Mycobacterium abscessus subsp. abscessus Terrabacteria group; Actinobacteria; Actinobacteria; Corynebacteriales; Mycobacteriaceae; Mycobacterium; Mycobacterium chelonae group; Mycobacterium abscessus subgroup; Mycobacterium abscessus
MIKHLTRRATVALVVAAAAISPAAYPAMADTATPLNGLFALTPGSCAEGRATGSYFRMIL WP_005073213.1
hypothetical protein [Mycobacterium abscessus] Length: 330\nNumber of predicted TMHs: 2\nExp number of AAs in TMHs: 39.3416199999999\nExp number, first 60 AAs: 17.07688\nTotal prob of N-in: 0.84431\nPOSSIBLE N-term signal sequence\ninside 1 11\nTMhelix 12 29\noutside 30 298\nTMhelix 299 321\ninside 322 330

30158 GCF_900136745.1_10625_5_32 Mycobacterium abscessus subsp. abscessus Terrabacteria group; Actinobacteria; Actinobacteria; Corynebacteriales; Mycobacteriaceae; Mycobacterium; Mycobacterium chelonae group; Mycobacterium abscessus subgroup; Mycobacterium abscessus
MIKHLTRRATVALVVAAAAISPAAYPAMADTATPLNGLFALTPGSCAEGRATGSYFRMIL WP_005073213.1
hypothetical protein [Mycobacterium abscessus] Length: 330\nNumber of predicted TMHs: 2\nExp number of

AAs in TMHs: 39.3416199999999\nExp number, first 60 AAs: 17.07688\nTotal prob of N-in: 0.84431\nPOSSIBLE N-term signal sequence\ninside 1 11\nTMhelix 12 29\noutside 30 298\nTMhelix 299 321\ninside 322 330

30159 GCF_900136865.1_10625_5_57 Mycobacterium abscessus subsp. abscessus Terrabacteria group; Actinobacteria; Actinobacteria; Corynebacteriales; Mycobacteriaceae; Mycobacterium; Mycobacterium chelonae group; Mycobacterium abscessus subgroup; Mycobacterium abscessus MIKHLTRRATVALVVAAAAISPAAYPAMADTATPLNGLFALTPGSCAEGRATGSYFRMIL WP_005073213.1 hypothetical protein [Mycobacterium abscessus] Length: 330\nNumber of predicted TMHs: 2\nExp number of AAs in TMHs: 39.3416199999999\nExp number, first 60 AAs: 17.07688\nTotal prob of N-in: 0.84431\nPOSSIBLE N-term signal sequence\ninside 1 11\nTMhelix 12 29\noutside 30 298\nTMhelix 299 321\ninside 322 330

30160 GCF_900136895.1_10625_5_59 Mycobacterium abscessus subsp. abscessus Terrabacteria group; Actinobacteria; Actinobacteria; Corynebacteriales; Mycobacteriaceae; Mycobacterium; Mycobacterium chelonae group; Mycobacterium abscessus subgroup; Mycobacterium abscessus MIKHLTRRATVALVVAAAAISPAAYPAMADTATPLNGLFALTPGSCAEGRATGSYFRMIL WP_005073213.1 hypothetical protein [Mycobacterium abscessus] Length: 330\nNumber of predicted TMHs: 2\nExp number of AAs in TMHs: 39.3416199999999\nExp number, first 60 AAs: 17.07688\nTotal prob of N-in: 0.84431\nPOSSIBLE N-term signal sequence\ninside 1 11\nTMhelix 12 29\noutside 30 298\nTMhelix 299 321\ninside 322 330

30161 GCF_900136915.1_11861_7_15 Mycobacterium abscessus subsp. abscessus Terrabacteria group; Actinobacteria; Actinobacteria; Corynebacteriales; Mycobacteriaceae; Mycobacterium; Mycobacterium chelonae group; Mycobacterium abscessus subgroup; Mycobacterium abscessus MIKHLTRRATVALVVAAAAISPAAYPAMADTATPLNGLFALTPGSCAEGRATGSYFRMIL WP_005073213.1 hypothetical protein [Mycobacterium abscessus] Length: 330\nNumber of predicted TMHs: 2\nExp number of AAs in TMHs: 39.3416199999999\nExp number, first 60 AAs: 17.07688\nTotal prob of N-in: 0.84431\nPOSSIBLE N-term signal sequence\ninside 1 11\nTMhelix 12 29\noutside 30 298\nTMhelix 299 321\ninside 322 330

30162 GCF_900136945.1_10665_2_78 Mycobacterium abscessus subsp. abscessus Terrabacteria group; Actinobacteria; Actinobacteria; Corynebacteriales; Mycobacteriaceae; Mycobacterium; Mycobacterium chelonae group; Mycobacterium abscessus subgroup; Mycobacterium abscessus MIKHLTRRATVALVVAAAAISPAAYPAMADTATPLNGLFALTPGSCAEGRATGSYFRMIL WP_005073213.1 hypothetical protein [Mycobacterium abscessus] Length: 330\nNumber of predicted TMHs: 2\nExp number of AAs in TMHs: 39.3416199999999\nExp number, first 60 AAs: 17.07688\nTotal prob of N-in: 0.84431\nPOSSIBLE N-term signal sequence\ninside 1 11\nTMhelix 12 29\noutside 30 298\nTMhelix 299 321\ninside 322 330

30163 GCF_900136925.1_10660_1_22 Mycobacterium abscessus subsp. abscessus Terrabacteria group; Actinobacteria; Actinobacteria; Corynebacteriales; Mycobacteriaceae; Mycobacterium; Mycobacterium chelonae group; Mycobacterium abscessus subgroup; Mycobacterium abscessus MIKHLTRRATVALVVAAAAISPAAYPAMADTATPLNGLFALTPGSCAEGRATGSYFRMIL WP_005073213.1 hypothetical protein [Mycobacterium abscessus] Length: 330\nNumber of predicted TMHs: 2\nExp number of AAs in TMHs: 39.3416199999999\nExp number, first 60 AAs: 17.07688\nTotal prob of N-in: 0.84431\nPOSSIBLE N-term signal sequence\ninside 1 11\nTMhelix 12 29\noutside 30 298\nTMhelix 299 321\ninside 322 330

30164 GCF_900136985.1_11861_7_74 Mycobacterium abscessus subsp. abscessus Terrabacteria group; Actinobacteria; Actinobacteria; Corynebacteriales; Mycobacteriaceae; Mycobacterium; Mycobacterium chelonae group; Mycobacterium abscessus subgroup; Mycobacterium abscessus MPRRRTILSALASFSMVIGLLVVSTPTAWAAPFPVVDPAKLPPSQPPAPPGDMRLNGAPL WP_074362414.1 type VII secretion-associated serine protease mycosin [Mycobacterium abscessus] Length: 470\nNumber of predicted TMHs: 2\nExp number of AAs in TMHs: 44.7482799999999\nExp number, first 60 AAs: 22.65577\nTotal prob of N-in: 0.99532\nPOSSIBLE N-term signal sequence\ninside 1 6\nTMhelix 7 29\noutside 30 441\nTMhelix 442 464\ninside 465 470

30165 GCF_900136995.1_11861_7_75 Mycobacterium abscessus subsp. abscessus Terrabacteria group; Actinobacteria; Actinobacteria; Corynebacteriales; Mycobacteriaceae; Mycobacterium; Mycobacterium chelonae group; Mycobacterium abscessus subgroup; Mycobacterium abscessus MPRRRTILSALASFSMVIGLLVVSTPTAWAAPFPVVDPAKLPPSQPPAPPGDMRLNGAPL WP_074362414.1 type VII secretion-associated serine protease mycosin [Mycobacterium abscessus] Length: 470\nNumber of predicted TMHs: 2\nExp number of AAs in TMHs: 44.7482799999999\nExp number, first 60 AAs: 22.65577\nTotal prob of N-in: 0.99532\nPOSSIBLE N-term signal sequence\ninside 1 6\nTMhelix 7 29\noutside 30 441\nTMhelix 442 464\ninside 465 470

30166 GCF_900137115.1_11893_6_3 Mycobacterium abscessus subsp. abscessus Terrabacteria group; Actinobacteria; Actinobacteria; Corynebacteriales; Mycobacteriaceae; Mycobacterium; Mycobacterium chelonae group; Mycobacterium abscessus subgroup; Mycobacterium abscessus
MIKHLTRRATVALVVAAAAISPAAYPAMADTATPLNGLFALTPGSCAEGRATGSYFRMIL WP_005073213.1
hypothetical protein [Mycobacterium abscessus] Length: 330\nNumber of predicted TMHs: 2\nExp number of AAs in TMHs: 39.3416199999999\nExp number, first 60 AAs: 17.07688\nTotal prob of N-in: 0.84431\nPOSSIBLE N-term signal sequence\ninside 1 11\nTMhelix 12 29\noutside 30 298\nTMhelix 299 321\ninside 322 330

30167 GCF_900137175.1_12078_1_62 Mycobacterium abscessus subsp. abscessus Terrabacteria group; Actinobacteria; Actinobacteria; Corynebacteriales; Mycobacteriaceae; Mycobacterium; Mycobacterium chelonae group; Mycobacterium abscessus subgroup; Mycobacterium abscessus
MIKHLTRRATVALVVAAAAISPAAYPAMADTATPLNGLFALTPGSCAEGRATGSYFRMIL WP_005073213.1
hypothetical protein [Mycobacterium abscessus] Length: 330\nNumber of predicted TMHs: 2\nExp number of AAs in TMHs: 39.3416199999999\nExp number, first 60 AAs: 17.07688\nTotal prob of N-in: 0.84431\nPOSSIBLE N-term signal sequence\ninside 1 11\nTMhelix 12 29\noutside 30 298\nTMhelix 299 321\ninside 322 330

30168 GCF_900137155.1_12078_1_68 Mycobacterium abscessus subsp. abscessus Terrabacteria group; Actinobacteria; Actinobacteria; Corynebacteriales; Mycobacteriaceae; Mycobacterium; Mycobacterium chelonae group; Mycobacterium abscessus subgroup; Mycobacterium abscessus
MIKHLTRRATVALVVAAAAISPAAYPAMADTATPLNGLFALTPGSCAEGRATGSYFRMIL WP_005073213.1
hypothetical protein [Mycobacterium abscessus] Length: 330\nNumber of predicted TMHs: 2\nExp number of AAs in TMHs: 39.3416199999999\nExp number, first 60 AAs: 17.07688\nTotal prob of N-in: 0.84431\nPOSSIBLE N-term signal sequence\ninside 1 11\nTMhelix 12 29\noutside 30 298\nTMhelix 299 321\ninside 322 330

30169 GCF_900137365.1_12163_2_11 Mycobacterium abscessus subsp. abscessus Terrabacteria group; Actinobacteria; Actinobacteria; Corynebacteriales; Mycobacteriaceae; Mycobacterium; Mycobacterium chelonae group; Mycobacterium abscessus subgroup; Mycobacterium abscessus
MIKHLTRRATVALVVAAAAISPAAYPAMADTATPLNGLFALTPGSCAEGRATGSYFRMIL WP_005073213.1
hypothetical protein [Mycobacterium abscessus] Length: 330\nNumber of predicted TMHs: 2\nExp number of AAs in TMHs: 39.3416199999999\nExp number, first 60 AAs: 17.07688\nTotal prob of N-in: 0.84431\nPOSSIBLE N-term signal sequence\ninside 1 11\nTMhelix 12 29\noutside 30 298\nTMhelix 299 321\ninside 322 330

30170 GCF_900137355.1_12163_2_14 Mycobacterium abscessus subsp. abscessus Terrabacteria group; Actinobacteria; Actinobacteria; Corynebacteriales; Mycobacteriaceae; Mycobacterium; Mycobacterium chelonae group; Mycobacterium abscessus subgroup; Mycobacterium abscessus
MIKHLTRRATVALVVAAAAISPAAYPAMADTATPLNGLFALTPGSCAEGRATGSYFRMIL WP_005073213.1
hypothetical protein [Mycobacterium abscessus] Length: 330\nNumber of predicted TMHs: 2\nExp number of AAs in TMHs: 39.3416199999999\nExp number, first 60 AAs: 17.07688\nTotal prob of N-in: 0.84431\nPOSSIBLE N-term signal sequence\ninside 1 11\nTMhelix 12 29\noutside 30 298\nTMhelix 299 321\ninside 322 330

30171 GCF_900137385.1_12163_2_15 Mycobacterium abscessus subsp. abscessus Terrabacteria group; Actinobacteria; Actinobacteria; Corynebacteriales; Mycobacteriaceae; Mycobacterium; Mycobacterium chelonae group; Mycobacterium abscessus subgroup; Mycobacterium abscessus
MIKHLTRRATVALVVAAAAISPAAYPAMADTATPLNGLFALTPGSCAEGRATGSYFRMIL WP_005073213.1
hypothetical protein [Mycobacterium abscessus] Length: 330\nNumber of predicted TMHs: 2\nExp number of AAs in TMHs: 39.3416199999999\nExp number, first 60 AAs: 17.07688\nTotal prob of N-in: 0.84431\nPOSSIBLE N-term signal sequence\ninside 1 11\nTMhelix 12 29\noutside 30 298\nTMhelix 299 321\ninside 322 330

30172 GCF_900137415.1_12163_2_18 Mycobacterium abscessus subsp. abscessus Terrabacteria group; Actinobacteria; Actinobacteria; Corynebacteriales; Mycobacteriaceae; Mycobacterium; Mycobacterium chelonae group; Mycobacterium abscessus subgroup; Mycobacterium abscessus
MIKHLTRRATVALVVAAAAISPAAYPAMADTATPLNGLFALTPGSCAEGRATGSYFRMIL WP_005073213.1
hypothetical protein [Mycobacterium abscessus] Length: 330\nNumber of predicted TMHs: 2\nExp number of AAs in TMHs: 39.3416199999999\nExp number, first 60 AAs: 17.07688\nTotal prob of N-in: 0.84431\nPOSSIBLE N-term signal sequence\ninside 1 11\nTMhelix 12 29\noutside 30 298\nTMhelix 299 321\ninside 322 330

30173 GCF_900137545.1_12163_2_3 Mycobacterium abscessus subsp. abscessus Terrabacteria group; Actinobacteria; Actinobacteria; Corynebacteriales; Mycobacteriaceae; Mycobacterium; Mycobacterium chelonae group; Mycobacterium abscessus subgroup; Mycobacterium abscessus
MIKHLTRRATVALVVAAAAISPAAYPAMADTATPLNGLFALTPGSCAEGRATGSYFRMIL WP_005073213.1
hypothetical protein [Mycobacterium abscessus] Length: 330\nNumber of predicted TMHs: 2\nExp number of

AAs in TMHs: 39.3416199999999\nExp number, first 60 AAs: 17.07688\nTotal prob of N-in: 0.84431\nPOSSIBLE N-term signal sequence\ninside 1 11\nTMhelix 12 29\noutside 30 298\nTMhelix 299 321\ninside 322 330

30174 GCF_900131185.1_10665_3_65 Mycobacterium abscessus subsp. abscessus Terrabacteria group; Actinobacteria; Actinobacteria; Corynebacteriales; Mycobacteriaceae; Mycobacterium; Mycobacterium chelonae group; Mycobacterium abscessus subgroup; Mycobacterium abscessus
MIKHLTRRATVALVVAAAAISPAAYPAMADTATPLNGLFALTPGSCAEGRATGSYFRMIL WP_005073213.1
hypothetical protein [Mycobacterium abscessus] Length: 330\nNumber of predicted TMHs: 2\nExp number of AAs in TMHs: 39.3416199999999\nExp number, first 60 AAs: 17.07688\nTotal prob of N-in: 0.84431\nPOSSIBLE N-term signal sequence\ninside 1 11\nTMhelix 12 29\noutside 30 298\nTMhelix 299 321\ninside 322 330

30175 GCF_900137565.1_12163_2_33 Mycobacterium abscessus subsp. abscessus Terrabacteria group; Actinobacteria; Actinobacteria; Corynebacteriales; Mycobacteriaceae; Mycobacterium; Mycobacterium chelonae group; Mycobacterium abscessus subgroup; Mycobacterium abscessus
MIKHLTRRATVALVVAAAAISPAAYPAMADTATPLNGLFALTPGSCAEGRATGSYFRMIL WP_005073213.1
hypothetical protein [Mycobacterium abscessus] Length: 330\nNumber of predicted TMHs: 2\nExp number of AAs in TMHs: 39.3416199999999\nExp number, first 60 AAs: 17.07688\nTotal prob of N-in: 0.84431\nPOSSIBLE N-term signal sequence\ninside 1 11\nTMhelix 12 29\noutside 30 298\nTMhelix 299 321\ninside 322 330

30176 GCF_900137595.1_12163_2_32 Mycobacterium abscessus subsp. abscessus Terrabacteria group; Actinobacteria; Actinobacteria; Corynebacteriales; Mycobacteriaceae; Mycobacterium; Mycobacterium chelonae group; Mycobacterium abscessus subgroup; Mycobacterium abscessus
MIKHLTRRATVALVVAAAAISPAAYPAMADTATPLNGLFALTPGSCAEGRATGSYFRMIL WP_005073213.1
hypothetical protein [Mycobacterium abscessus] Length: 330\nNumber of predicted TMHs: 2\nExp number of AAs in TMHs: 39.3416199999999\nExp number, first 60 AAs: 17.07688\nTotal prob of N-in: 0.84431\nPOSSIBLE N-term signal sequence\ninside 1 11\nTMhelix 12 29\noutside 30 298\nTMhelix 299 321\ninside 322 330

30177 GCF_900137585.1_12163_2_34 Mycobacterium abscessus subsp. abscessus Terrabacteria group; Actinobacteria; Actinobacteria; Corynebacteriales; Mycobacteriaceae; Mycobacterium; Mycobacterium chelonae group; Mycobacterium abscessus subgroup; Mycobacterium abscessus
MIKHLTRRATVALVVAAAAISPAAYPAMADTATPLNGLFALTPGSCAEGRATGSYFRMIL WP_005073213.1
hypothetical protein [Mycobacterium abscessus] Length: 330\nNumber of predicted TMHs: 2\nExp number of AAs in TMHs: 39.3416199999999\nExp number, first 60 AAs: 17.07688\nTotal prob of N-in: 0.84431\nPOSSIBLE N-term signal sequence\ninside 1 11\nTMhelix 12 29\noutside 30 298\nTMhelix 299 321\ninside 322 330

30178 GCF_900137615.1_12163_2_41 Mycobacterium abscessus subsp. abscessus Terrabacteria group; Actinobacteria; Actinobacteria; Corynebacteriales; Mycobacteriaceae; Mycobacterium; Mycobacterium chelonae group; Mycobacterium abscessus subgroup; Mycobacterium abscessus
MIKHLTRRATVALVVAAAAISPAAYPAMADTATPLNGLFALTPGSCAEGRATGSYFRMIL WP_005073213.1
hypothetical protein [Mycobacterium abscessus] Length: 330\nNumber of predicted TMHs: 2\nExp number of AAs in TMHs: 39.3416199999999\nExp number, first 60 AAs: 17.07688\nTotal prob of N-in: 0.84431\nPOSSIBLE N-term signal sequence\ninside 1 11\nTMhelix 12 29\noutside 30 298\nTMhelix 299 321\ninside 322 330

30179 GCF_900137785.1_12163_2_55 Mycobacterium abscessus subsp. abscessus Terrabacteria group; Actinobacteria; Actinobacteria; Corynebacteriales; Mycobacteriaceae; Mycobacterium; Mycobacterium chelonae group; Mycobacterium abscessus subgroup; Mycobacterium abscessus
MIKHLTRRATVALVVAAAAISPAAYPAMADTATPLNGLFALTPGSCAEGRATGSYFRMIL WP_005073213.1
hypothetical protein [Mycobacterium abscessus] Length: 330\nNumber of predicted TMHs: 2\nExp number of AAs in TMHs: 39.3416199999999\nExp number, first 60 AAs: 17.07688\nTotal prob of N-in: 0.84431\nPOSSIBLE N-term signal sequence\ninside 1 11\nTMhelix 12 29\noutside 30 298\nTMhelix 299 321\ninside 322 330

30180 GCF_900137815.1_12163_2_56 Mycobacterium abscessus subsp. abscessus Terrabacteria group; Actinobacteria; Actinobacteria; Corynebacteriales; Mycobacteriaceae; Mycobacterium; Mycobacterium chelonae group; Mycobacterium abscessus subgroup; Mycobacterium abscessus
MIKHLTRRATVALVVAAAAISPAAYPAMADTATPLNGLFALTPGSCAEGRATGSYFRMIL WP_005073213.1
hypothetical protein [Mycobacterium abscessus] Length: 330\nNumber of predicted TMHs: 2\nExp number of AAs in TMHs: 39.3416199999999\nExp number, first 60 AAs: 17.07688\nTotal prob of N-in: 0.84431\nPOSSIBLE N-term signal sequence\ninside 1 11\nTMhelix 12 29\noutside 30 298\nTMhelix 299 321\ninside 322 330

30181 GCF_900137805.1_12163_2_54 Mycobacterium abscessus subsp. abscessus Terrabacteria group; Actinobacteria; Actinobacteria; Corynebacteriales; Mycobacteriaceae; Mycobacterium; Mycobacterium chelonae group;

Mycobacterium abscessus subgroup; Mycobacterium abscessus
MIKHLTRRATVALVVAAAAISPAAYPAMADTATPLNGLFALTPGSCAEGRATGSYFRMIL WP_005073213.1
hypothetical protein [Mycobacterium abscessus] Length: 330\nNumber of predicted TMHs: 2\nExp number of AAs in TMHs: 39.3416199999999\nExp number, first 60 AAs: 17.07688\nTotal prob of N-in: 0.84431\nPOSSIBLE N-term signal sequence\ninside 1 11\nTMhelix 12 29\noutside 30 298\nTMhelix 299 321\ninside 322 330

30182 GCF_900137895.1_12163_2_87 Mycobacterium abscessus subsp. abscessus Terrabacteria group; Actinobacteria; Actinobacteria; Corynebacteriales; Mycobacteriaceae; Mycobacterium; Mycobacterium chelonae group; Mycobacterium abscessus subgroup; Mycobacterium abscessus
MIKHLTRRATVALVVAAAAISPAAYPAMADTATPLNGLFALTPGSCAEGRATGSYFRMIL WP_005073213.1
hypothetical protein [Mycobacterium abscessus] Length: 330\nNumber of predicted TMHs: 2\nExp number of AAs in TMHs: 39.3416199999999\nExp number, first 60 AAs: 17.07688\nTotal prob of N-in: 0.84431\nPOSSIBLE N-term signal sequence\ninside 1 11\nTMhelix 12 29\noutside 30 298\nTMhelix 299 321\ninside 322 330

30183 GCF_900137915.1_12163_2_90 Mycobacterium abscessus subsp. abscessus Terrabacteria group; Actinobacteria; Actinobacteria; Corynebacteriales; Mycobacteriaceae; Mycobacterium; Mycobacterium chelonae group; Mycobacterium abscessus subgroup; Mycobacterium abscessus
MIKHLTRRATVALVVAAAAISPAAYPAMADTATPLNGLFALTPGSCAEGRATGSYFRMIL WP_005073213.1
hypothetical protein [Mycobacterium abscessus] Length: 330\nNumber of predicted TMHs: 2\nExp number of AAs in TMHs: 39.3416199999999\nExp number, first 60 AAs: 17.07688\nTotal prob of N-in: 0.84431\nPOSSIBLE N-term signal sequence\ninside 1 11\nTMhelix 12 29\noutside 30 298\nTMhelix 299 321\ninside 322 330

30184 GCF_900138005.1_10665_2_80 Mycobacterium abscessus subsp. abscessus Terrabacteria group; Actinobacteria; Actinobacteria; Corynebacteriales; Mycobacteriaceae; Mycobacterium; Mycobacterium chelonae group; Mycobacterium abscessus subgroup; Mycobacterium abscessus
MIKHLTRRATVALVVAAAAISPAAYPAMADTATPLNGLFALTPGSCAEGRATGSYFRMIL WP_005073213.1
hypothetical protein [Mycobacterium abscessus] Length: 330\nNumber of predicted TMHs: 2\nExp number of AAs in TMHs: 39.3416199999999\nExp number, first 60 AAs: 17.07688\nTotal prob of N-in: 0.84431\nPOSSIBLE N-term signal sequence\ninside 1 11\nTMhelix 12 29\noutside 30 298\nTMhelix 299 321\ninside 322 330

30185 GCF_900138025.1_11893_6_2 Mycobacterium abscessus subsp. abscessus Terrabacteria group; Actinobacteria; Actinobacteria; Corynebacteriales; Mycobacteriaceae; Mycobacterium; Mycobacterium chelonae group; Mycobacterium abscessus subgroup; Mycobacterium abscessus
MIKHLTRRATVALVVAAAAISPAAYPAMADTATPLNGLFALTPGSCAEGRATGSYFRMIL WP_005073213.1
hypothetical protein [Mycobacterium abscessus] Length: 330\nNumber of predicted TMHs: 2\nExp number of AAs in TMHs: 39.3416199999999\nExp number, first 60 AAs: 17.07688\nTotal prob of N-in: 0.84431\nPOSSIBLE N-term signal sequence\ninside 1 11\nTMhelix 12 29\noutside 30 298\nTMhelix 299 321\ninside 322 330

30186 GCF_900138045.1_12045_8_33 Mycobacterium abscessus subsp. abscessus Terrabacteria group; Actinobacteria; Actinobacteria; Corynebacteriales; Mycobacteriaceae; Mycobacterium; Mycobacterium chelonae group; Mycobacterium abscessus subgroup; Mycobacterium abscessus
MIKHLTRRATVALVVAAAAISPAAYPAMADTATPLNGLFALTPGSCAEGRATGSYFRMIL WP_005073213.1
hypothetical protein [Mycobacterium abscessus] Length: 330\nNumber of predicted TMHs: 2\nExp number of AAs in TMHs: 39.3416199999999\nExp number, first 60 AAs: 17.07688\nTotal prob of N-in: 0.84431\nPOSSIBLE N-term signal sequence\ninside 1 11\nTMhelix 12 29\noutside 30 298\nTMhelix 299 321\ninside 322 330

30187 GCF_900138065.1_11893_6_60 Mycobacterium abscessus subsp. abscessus Terrabacteria group; Actinobacteria; Actinobacteria; Corynebacteriales; Mycobacteriaceae; Mycobacterium; Mycobacterium chelonae group; Mycobacterium abscessus subgroup; Mycobacterium abscessus
MIKHLTRRATVALVVAAAAISPAAYPAMADTATPLNGLFALTPGSCAEGRATGSYFRMIL WP_005073213.1
hypothetical protein [Mycobacterium abscessus] Length: 330\nNumber of predicted TMHs: 2\nExp number of AAs in TMHs: 39.3416199999999\nExp number, first 60 AAs: 17.07688\nTotal prob of N-in: 0.84431\nPOSSIBLE N-term signal sequence\ninside 1 11\nTMhelix 12 29\noutside 30 298\nTMhelix 299 321\ninside 322 330

30188 GCF_900138095.1_12045_8_36 Mycobacterium abscessus subsp. abscessus Terrabacteria group; Actinobacteria; Actinobacteria; Corynebacteriales; Mycobacteriaceae; Mycobacterium; Mycobacterium chelonae group; Mycobacterium abscessus subgroup; Mycobacterium abscessus
MIKHLTRRATVALVVAAAAISPAAYPAMADTATPLNGLFALTPGSCAEGRATGSYFRMIL WP_005073213.1
hypothetical protein [Mycobacterium abscessus] Length: 330\nNumber of predicted TMHs: 2\nExp number of AAs in TMHs: 39.3416199999999\nExp number, first 60 AAs: 17.07688\nTotal prob of N-in: 0.84431\nPOSSIBLE N-term signal sequence\ninside 1 11\nTMhelix 12 29\noutside 30 298\nTMhelix 299 321\ninside 322 330

30189 GCF_900131055.1_10665_3_51 Mycobacterium abscessus subsp. abscessus Terrabacteria group; Actinobacteria; Actinobacteria; Corynebacteriales; Mycobacteriaceae; Mycobacterium; Mycobacterium chelonae group; Mycobacterium abscessus subgroup; Mycobacterium abscessus
MIKHLTRRATVALVVAAAAISPAAYPAMADTATPLNGLFALTPGSCAEGRATGSYFRMIL WP_005073213.1
hypothetical protein [Mycobacterium abscessus] Length: 330\nNumber of predicted TMHs: 2\nExp number of AAs in TMHs: 39.3416199999999\nExp number, first 60 AAs: 17.07688\nTotal prob of N-in: 0.84431\nPOSSIBLE N-term signal sequence\ninside 1 11\nTMhelix 12 29\noutside 30 298\nTMhelix 299 321\ninside 322 330

30190 GCF_900130875.1_10665_3_40 Mycobacterium abscessus subsp. abscessus Terrabacteria group; Actinobacteria; Actinobacteria; Corynebacteriales; Mycobacteriaceae; Mycobacterium; Mycobacterium chelonae group; Mycobacterium abscessus subgroup; Mycobacterium abscessus
MIKHLTRRATVALVVAAAAISPAAYPAMADTATPLNGLFALTPGSCAEGRATGSYFRMIL WP_074250269.1
hypothetical protein [Mycobacterium abscessus] Length: 330\nNumber of predicted TMHs: 2\nExp number of AAs in TMHs: 39.3415199999999\nExp number, first 60 AAs: 17.07715\nTotal prob of N-in: 0.84431\nPOSSIBLE N-term signal sequence\ninside 1 11\nTMhelix 12 29\noutside 30 298\nTMhelix 299 321\ninside 322 330

30191 GCF_900131005.1_11861_7_58 Mycobacterium abscessus subsp. abscessus Terrabacteria group; Actinobacteria; Actinobacteria; Corynebacteriales; Mycobacteriaceae; Mycobacterium; Mycobacterium chelonae group; Mycobacterium abscessus subgroup; Mycobacterium abscessus
MIKHLTRRATVALVVAAAAISPAAYPAMADTATPLNGLFALTPGSCAEGRATGSYFRMIL WP_005073213.1
hypothetical protein [Mycobacterium abscessus] Length: 330\nNumber of predicted TMHs: 2\nExp number of AAs in TMHs: 39.3416199999999\nExp number, first 60 AAs: 17.07688\nTotal prob of N-in: 0.84431\nPOSSIBLE N-term signal sequence\ninside 1 11\nTMhelix 12 29\noutside 30 298\nTMhelix 299 321\ninside 322 330

30192 GCF_900138745.1_10208_3_37 Mycobacterium abscessus subsp. abscessus Terrabacteria group; Actinobacteria; Actinobacteria; Corynebacteriales; Mycobacteriaceae; Mycobacterium; Mycobacterium chelonae group; Mycobacterium abscessus subgroup; Mycobacterium abscessus
MIKHLTRRATVALVVAAAAISPAAYPAMADTATPLNGLFALTPGSCAEGRATGSYFRMIL WP_005073213.1
hypothetical protein [Mycobacterium abscessus] Length: 330\nNumber of predicted TMHs: 2\nExp number of AAs in TMHs: 39.3416199999999\nExp number, first 60 AAs: 17.07688\nTotal prob of N-in: 0.84431\nPOSSIBLE N-term signal sequence\ninside 1 11\nTMhelix 12 29\noutside 30 298\nTMhelix 299 321\ninside 322 330

30193 GCF_900138765.1_10208_3_45 Mycobacterium abscessus subsp. abscessus Terrabacteria group; Actinobacteria; Actinobacteria; Corynebacteriales; Mycobacteriaceae; Mycobacterium; Mycobacterium chelonae group; Mycobacterium abscessus subgroup; Mycobacterium abscessus
MIKHLTRRATVALVVAAAAISPAAYPAMADTATPLNGLFALTPGSCAEGRATGSYFRMIL WP_005073213.1
hypothetical protein [Mycobacterium abscessus] Length: 330\nNumber of predicted TMHs: 2\nExp number of AAs in TMHs: 39.3416199999999\nExp number, first 60 AAs: 17.07688\nTotal prob of N-in: 0.84431\nPOSSIBLE N-term signal sequence\ninside 1 11\nTMhelix 12 29\noutside 30 298\nTMhelix 299 321\ninside 322 330

30194 GCF_900138785.1_10208_3_47 Mycobacterium abscessus subsp. abscessus Terrabacteria group; Actinobacteria; Actinobacteria; Corynebacteriales; Mycobacteriaceae; Mycobacterium; Mycobacterium chelonae group; Mycobacterium abscessus subgroup; Mycobacterium abscessus
MIKHLTRRATVALVVAAAAISPAAYPAMADTATPLNGLFALTPGSCAEGRATGSYFRMIL WP_005073213.1
hypothetical protein [Mycobacterium abscessus] Length: 330\nNumber of predicted TMHs: 2\nExp number of AAs in TMHs: 39.3416199999999\nExp number, first 60 AAs: 17.07688\nTotal prob of N-in: 0.84431\nPOSSIBLE N-term signal sequence\ninside 1 11\nTMhelix 12 29\noutside 30 298\nTMhelix 299 321\ninside 322 330

30195 GCF_900130985.1_11861_7_12 Mycobacterium abscessus subsp. abscessus Terrabacteria group; Actinobacteria; Actinobacteria; Corynebacteriales; Mycobacteriaceae; Mycobacterium; Mycobacterium chelonae group; Mycobacterium abscessus subgroup; Mycobacterium abscessus
MIKHLTRRATVALVVAAAAISPAAYPAMADTATPLNGLFALTPGSCAEGRATGSYFRMIL WP_005073213.1
hypothetical protein [Mycobacterium abscessus] Length: 330\nNumber of predicted TMHs: 2\nExp number of AAs in TMHs: 39.3416199999999\nExp number, first 60 AAs: 17.07688\nTotal prob of N-in: 0.84431\nPOSSIBLE N-term signal sequence\ninside 1 11\nTMhelix 12 29\noutside 30 298\nTMhelix 299 321\ninside 322 330

30196 GCF_900138885.1_10208_3_55 Mycobacterium abscessus subsp. abscessus Terrabacteria group; Actinobacteria; Actinobacteria; Corynebacteriales; Mycobacteriaceae; Mycobacterium; Mycobacterium chelonae group; Mycobacterium abscessus subgroup; Mycobacterium abscessus
MIKHLTRRATVALVVAAAAISPAAYPAMADTATPLNGLFALTPGSCAEGRATGSYFRMIL WP_005073213.1
hypothetical protein [Mycobacterium abscessus] Length: 330\nNumber of predicted TMHs: 2\nExp number of

AAs in TMHs: 39.3416199999999\nExp number, first 60 AAs: 17.07688\nTotal prob of N-in: 0.84431\nPOSSIBLE N-term signal sequence\ninside 1 11\nTMhelix 12 29\noutside 30 298\nTMhelix 299 321\ninside 322 330

30197 GCF_900138875.1_10208_3_56 Mycobacterium abscessus subsp. abscessus Terrabacteria group; Actinobacteria; Actinobacteria; Corynebacteriales; Mycobacteriaceae; Mycobacterium; Mycobacterium chelonae group; Mycobacterium abscessus subgroup; Mycobacterium abscessus
MIKHLTRRATVALVVAAAAISPAAYPAMADTATPLNGLFALTPGSCAEGRATGSYFRMIL WP_005073213.1
hypothetical protein [Mycobacterium abscessus] Length: 330\nNumber of predicted TMHs: 2\nExp number of AAs in TMHs: 39.3416199999999\nExp number, first 60 AAs: 17.07688\nTotal prob of N-in: 0.84431\nPOSSIBLE N-term signal sequence\ninside 1 11\nTMhelix 12 29\noutside 30 298\nTMhelix 299 321\ninside 322 330

30198 GCF_900139015.1_10396_8_10 Mycobacterium abscessus subsp. abscessus Terrabacteria group; Actinobacteria; Actinobacteria; Corynebacteriales; Mycobacteriaceae; Mycobacterium; Mycobacterium chelonae group; Mycobacterium abscessus subgroup; Mycobacterium abscessus
MIKHLTRRATVALVVAAAAISPAAYPAMADTATPLNGLFALTPGSCAEGRATGSYFRMIL WP_005073213.1
hypothetical protein [Mycobacterium abscessus] Length: 330\nNumber of predicted TMHs: 2\nExp number of AAs in TMHs: 39.3416199999999\nExp number, first 60 AAs: 17.07688\nTotal prob of N-in: 0.84431\nPOSSIBLE N-term signal sequence\ninside 1 11\nTMhelix 12 29\noutside 30 298\nTMhelix 299 321\ninside 322 330

30199 GCF_900138995.1_10250_1_68 Mycobacterium abscessus subsp. abscessus Terrabacteria group; Actinobacteria; Actinobacteria; Corynebacteriales; Mycobacteriaceae; Mycobacterium; Mycobacterium chelonae group; Mycobacterium abscessus subgroup; Mycobacterium abscessus
MIKHLTRRATVALVVAAAAISPAAYPAMADTATPLNGLFALTPGSCAEGRATGSYFRMIL WP_005073213.1
hypothetical protein [Mycobacterium abscessus] Length: 330\nNumber of predicted TMHs: 2\nExp number of AAs in TMHs: 39.3416199999999\nExp number, first 60 AAs: 17.07688\nTotal prob of N-in: 0.84431\nPOSSIBLE N-term signal sequence\ninside 1 11\nTMhelix 12 29\noutside 30 298\nTMhelix 299 321\ninside 322 330

30200 GCF_900138835.1_10208_3_50 Mycobacterium abscessus subsp. abscessus Terrabacteria group; Actinobacteria; Actinobacteria; Corynebacteriales; Mycobacteriaceae; Mycobacterium; Mycobacterium chelonae group; Mycobacterium abscessus subgroup; Mycobacterium abscessus
MIKHLTRRATVALVVAAAAISPAAYPAMADTATPLNGLFALTPGSCAEGRATGSYFRMIL WP_005073213.1
hypothetical protein [Mycobacterium abscessus] Length: 330\nNumber of predicted TMHs: 2\nExp number of AAs in TMHs: 39.3416199999999\nExp number, first 60 AAs: 17.07688\nTotal prob of N-in: 0.84431\nPOSSIBLE N-term signal sequence\ninside 1 11\nTMhelix 12 29\noutside 30 298\nTMhelix 299 321\ninside 322 330

30201 GCF_900130785.1_10665_3_29 Mycobacterium abscessus subsp. abscessus Terrabacteria group; Actinobacteria; Actinobacteria; Corynebacteriales; Mycobacteriaceae; Mycobacterium; Mycobacterium chelonae group; Mycobacterium abscessus subgroup; Mycobacterium abscessus
MIKHLTRRATVALVVAAAAISPAAYPAMADTATPLNGLFALTPGSCAEGRATGSYFRMIL WP_074250269.1
hypothetical protein [Mycobacterium abscessus] Length: 330\nNumber of predicted TMHs: 2\nExp number of AAs in TMHs: 39.3415199999999\nExp number, first 60 AAs: 17.07715\nTotal prob of N-in: 0.84431\nPOSSIBLE N-term signal sequence\ninside 1 11\nTMhelix 12 29\noutside 30 298\nTMhelix 299 321\ninside 322 330

30202 GCF_900138815.1_10208_3_48 Mycobacterium abscessus subsp. abscessus Terrabacteria group; Actinobacteria; Actinobacteria; Corynebacteriales; Mycobacteriaceae; Mycobacterium; Mycobacterium chelonae group; Mycobacterium abscessus subgroup; Mycobacterium abscessus
MIKHLTRRATVALVVAAAAISPAAYPAMADTATPLNGLFALTPGSCAEGRATGSYFRMIL WP_005073213.1
hypothetical protein [Mycobacterium abscessus] Length: 330\nNumber of predicted TMHs: 2\nExp number of AAs in TMHs: 39.3416199999999\nExp number, first 60 AAs: 17.07688\nTotal prob of N-in: 0.84431\nPOSSIBLE N-term signal sequence\ninside 1 11\nTMhelix 12 29\noutside 30 298\nTMhelix 299 321\ninside 322 330

30203 GCF_900139065.1_10465_1_51 Mycobacterium abscessus subsp. abscessus Terrabacteria group; Actinobacteria; Actinobacteria; Corynebacteriales; Mycobacteriaceae; Mycobacterium; Mycobacterium chelonae group; Mycobacterium abscessus subgroup; Mycobacterium abscessus
MIKHLTRRATVALVVAAAAISPAAYPAMADTATPLNGLFALTPGSCAEGRATGSYFRMIL WP_005073213.1
hypothetical protein [Mycobacterium abscessus] Length: 330\nNumber of predicted TMHs: 2\nExp number of AAs in TMHs: 39.3416199999999\nExp number, first 60 AAs: 17.07688\nTotal prob of N-in: 0.84431\nPOSSIBLE N-term signal sequence\ninside 1 11\nTMhelix 12 29\noutside 30 298\nTMhelix 299 321\ninside 322 330

30204 GCF_900130685.1_11861_7_38 Mycobacterium abscessus subsp. abscessus Terrabacteria group; Actinobacteria; Actinobacteria; Corynebacteriales; Mycobacteriaceae; Mycobacterium; Mycobacterium chelonae group;

Mycobacterium abscessus subgroup; Mycobacterium abscessus
MIKHLTRRATVALVVAAAAISPAAYPAMADTATPLNGLFALTPGSCAEGRATGSYFRMIL WP_005073213.1
hypothetical protein [Mycobacterium abscessus] Length: 330\nNumber of predicted TMHs: 2\nExp number of AAs in TMHs: 39.3416199999999\nExp number, first 60 AAs: 17.07688\nTotal prob of N-in: 0.84431\nPOSSIBLE N-term signal sequence\ninside 1 11\nTMhelix 12 29\noutside 30 298\nTMhelix 299 321\ninside 322 330

30205 GCF_900130645.1_10665_3_19 Mycobacterium abscessus subsp. abscessus Terrabacteria group; Actinobacteria; Actinobacteria; Corynebacteriales; Mycobacteriaceae; Mycobacterium; Mycobacterium chelonae group; Mycobacterium abscessus subgroup; Mycobacterium abscessus
MIKHLTRRATVALVVAAAAISPAAYPAMADTATPLNGLFALTPGSCAEGRATGSYFRMIL WP_005073213.1
hypothetical protein [Mycobacterium abscessus] Length: 330\nNumber of predicted TMHs: 2\nExp number of AAs in TMHs: 39.3416199999999\nExp number, first 60 AAs: 17.07688\nTotal prob of N-in: 0.84431\nPOSSIBLE N-term signal sequence\ninside 1 11\nTMhelix 12 29\noutside 30 298\nTMhelix 299 321\ninside 322 330

30206 GCF_900139045.1_10465_1_48 Mycobacterium abscessus subsp. abscessus Terrabacteria group; Actinobacteria; Actinobacteria; Corynebacteriales; Mycobacteriaceae; Mycobacterium; Mycobacterium chelonae group; Mycobacterium abscessus subgroup; Mycobacterium abscessus
MIKHLTRRATVALVVAAAAISPAAYPAMADTATPLNGLFALTPGSCAEGRATGSYFRMIL WP_005073213.1
hypothetical protein [Mycobacterium abscessus] Length: 330\nNumber of predicted TMHs: 2\nExp number of AAs in TMHs: 39.3416199999999\nExp number, first 60 AAs: 17.07688\nTotal prob of N-in: 0.84431\nPOSSIBLE N-term signal sequence\ninside 1 11\nTMhelix 12 29\noutside 30 298\nTMhelix 299 321\ninside 322 330

30207 GCF_900139105.1_10465_1_56 Mycobacterium abscessus subsp. abscessus Terrabacteria group; Actinobacteria; Actinobacteria; Corynebacteriales; Mycobacteriaceae; Mycobacterium; Mycobacterium chelonae group; Mycobacterium abscessus subgroup; Mycobacterium abscessus
MIKHLTRRATVALVVAAAAISPAAYPAMADTATPLNGLFALTPGSCAEGRATGSYFRMIL WP_005073213.1
hypothetical protein [Mycobacterium abscessus] Length: 330\nNumber of predicted TMHs: 2\nExp number of AAs in TMHs: 39.3416199999999\nExp number, first 60 AAs: 17.07688\nTotal prob of N-in: 0.84431\nPOSSIBLE N-term signal sequence\ninside 1 11\nTMhelix 12 29\noutside 30 298\nTMhelix 299 321\ninside 322 330

30208 GCF_900130525.1_10665_3_5 Mycobacterium abscessus subsp. abscessus Terrabacteria group; Actinobacteria; Actinobacteria; Corynebacteriales; Mycobacteriaceae; Mycobacterium; Mycobacterium chelonae group; Mycobacterium abscessus subgroup; Mycobacterium abscessus
MIKHLTRRATVALVVAAAAISPAAYPAMADTATPLNGLFALTPGSCAEGRATGSYFRMIL WP_074250269.1
hypothetical protein [Mycobacterium abscessus] Length: 330\nNumber of predicted TMHs: 2\nExp number of AAs in TMHs: 39.3415199999999\nExp number, first 60 AAs: 17.07715\nTotal prob of N-in: 0.84431\nPOSSIBLE N-term signal sequence\ninside 1 11\nTMhelix 12 29\noutside 30 298\nTMhelix 299 321\ninside 322 330

30209 GCF_900130595.1_10665_3_10 Mycobacterium abscessus subsp. abscessus Terrabacteria group; Actinobacteria; Actinobacteria; Corynebacteriales; Mycobacteriaceae; Mycobacterium; Mycobacterium chelonae group; Mycobacterium abscessus subgroup; Mycobacterium abscessus
MIKHLTRRATVALVVAAAAISPAAYPAMADTATPLNGLFALTPGSCAEGRATGSYFRMIL WP_005073213.1
hypothetical protein [Mycobacterium abscessus] Length: 330\nNumber of predicted TMHs: 2\nExp number of AAs in TMHs: 39.3416199999999\nExp number, first 60 AAs: 17.07688\nTotal prob of N-in: 0.84431\nPOSSIBLE N-term signal sequence\ninside 1 11\nTMhelix 12 29\noutside 30 298\nTMhelix 299 321\ninside 322 330

30210 GCF_900139085.1_10465_1_55 Mycobacterium abscessus subsp. abscessus Terrabacteria group; Actinobacteria; Actinobacteria; Corynebacteriales; Mycobacteriaceae; Mycobacterium; Mycobacterium chelonae group; Mycobacterium abscessus subgroup; Mycobacterium abscessus
MIKHLTRRATVALVVAAAAISPAAYPAMADTATPLNGLFALTPGSCAEGRATGSYFRMIL WP_005073213.1
hypothetical protein [Mycobacterium abscessus] Length: 330\nNumber of predicted TMHs: 2\nExp number of AAs in TMHs: 39.3416199999999\nExp number, first 60 AAs: 17.07688\nTotal prob of N-in: 0.84431\nPOSSIBLE N-term signal sequence\ninside 1 11\nTMhelix 12 29\noutside 30 298\nTMhelix 299 321\ninside 322 330

30211 GCF_900139175.1_10465_1_66 Mycobacterium abscessus subsp. abscessus Terrabacteria group; Actinobacteria; Actinobacteria; Corynebacteriales; Mycobacteriaceae; Mycobacterium; Mycobacterium chelonae group; Mycobacterium abscessus subgroup; Mycobacterium abscessus
MIKHLTRRATVALVVAAAAISPAAYPAMADTATPLNGLFALTPGSCAEGRATGSYFRMIL WP_005073213.1
hypothetical protein [Mycobacterium abscessus] Length: 330\nNumber of predicted TMHs: 2\nExp number of AAs in TMHs: 39.3416199999999\nExp number, first 60 AAs: 17.07688\nTotal prob of N-in: 0.84431\nPOSSIBLE N-term signal sequence\ninside 1 11\nTMhelix 12 29\noutside 30 298\nTMhelix 299 321\ninside 322 330

30212 GCF_900139205.1_10465_1_65 Mycobacterium abscessus subsp. abscessus Terrabacteria group; Actinobacteria; Actinobacteria; Corynebacteriales; Mycobacteriaceae; Mycobacterium; Mycobacterium chelonae group; Mycobacterium abscessus subgroup; Mycobacterium abscessus
MIKHLTRRATVALVVAAAAISPAAYPAMADTATPLNGLFALTPGSCAEGRATGSYFRMIL WP_005073213.1
hypothetical protein [Mycobacterium abscessus] Length: 330\nNumber of predicted TMHs: 2\nExp number of AAs in TMHs: 39.3416199999999\nExp number, first 60 AAs: 17.07688\nTotal prob of N-in: 0.84431\nPOSSIBLE N-term signal sequence\ninside 1 11\nTMhelix 12 29\noutside 30 298\nTMhelix 299 321\ninside 322 330

30213 GCF_900130395.1_10625_5_75 Mycobacterium abscessus subsp. abscessus Terrabacteria group; Actinobacteria; Actinobacteria; Corynebacteriales; Mycobacteriaceae; Mycobacterium; Mycobacterium chelonae group; Mycobacterium abscessus subgroup; Mycobacterium abscessus
MIKHLTRRATVALVVAAAAISPAAYPAMADTATPLNGLFALTPGSCAEGRATGSYFRMIL WP_005073213.1
hypothetical protein [Mycobacterium abscessus] Length: 330\nNumber of predicted TMHs: 2\nExp number of AAs in TMHs: 39.3416199999999\nExp number, first 60 AAs: 17.07688\nTotal prob of N-in: 0.84431\nPOSSIBLE N-term signal sequence\ninside 1 11\nTMhelix 12 29\noutside 30 298\nTMhelix 299 321\ninside 322 330

30214 GCF_900139275.1_10465_1_74 Mycobacterium abscessus subsp. abscessus Terrabacteria group; Actinobacteria; Actinobacteria; Corynebacteriales; Mycobacteriaceae; Mycobacterium; Mycobacterium chelonae group; Mycobacterium abscessus subgroup; Mycobacterium abscessus
MIKHLTRRATVALVVAAAAISPAAYPAMADTATPLNGLFALTPGSCAEGRATGSYFRMIL WP_005073213.1
hypothetical protein [Mycobacterium abscessus] Length: 330\nNumber of predicted TMHs: 2\nExp number of AAs in TMHs: 39.3416199999999\nExp number, first 60 AAs: 17.07688\nTotal prob of N-in: 0.84431\nPOSSIBLE N-term signal sequence\ninside 1 11\nTMhelix 12 29\noutside 30 298\nTMhelix 299 321\ninside 322 330

30215 GCF_900139325.1_10625_4_11 Mycobacterium abscessus subsp. abscessus Terrabacteria group; Actinobacteria; Actinobacteria; Corynebacteriales; Mycobacteriaceae; Mycobacterium; Mycobacterium chelonae group; Mycobacterium abscessus subgroup; Mycobacterium abscessus
MIKHLTRRATVALVVAAAAISPAAYPAMADTATPLNGLFALTPGSCAEGRATGSYFRMIL WP_005073213.1
hypothetical protein [Mycobacterium abscessus] Length: 330\nNumber of predicted TMHs: 2\nExp number of AAs in TMHs: 39.3416199999999\nExp number, first 60 AAs: 17.07688\nTotal prob of N-in: 0.84431\nPOSSIBLE N-term signal sequence\ninside 1 11\nTMhelix 12 29\noutside 30 298\nTMhelix 299 321\ninside 322 330

30216 GCF_900139255.1_10465_1_71 Mycobacterium abscessus subsp. abscessus Terrabacteria group; Actinobacteria; Actinobacteria; Corynebacteriales; Mycobacteriaceae; Mycobacterium; Mycobacterium chelonae group; Mycobacterium abscessus subgroup; Mycobacterium abscessus
MIKHLTRRATVALVVAAAAISPAAYPAMADTATPLNGLFALTPGSCAEGRATGSYFRMIL WP_005073213.1
hypothetical protein [Mycobacterium abscessus] Length: 330\nNumber of predicted TMHs: 2\nExp number of AAs in TMHs: 39.3416199999999\nExp number, first 60 AAs: 17.07688\nTotal prob of N-in: 0.84431\nPOSSIBLE N-term signal sequence\ninside 1 11\nTMhelix 12 29\noutside 30 298\nTMhelix 299 321\ninside 322 330

30217 GCF_900139295.1_10465_1_75 Mycobacterium abscessus subsp. abscessus Terrabacteria group; Actinobacteria; Actinobacteria; Corynebacteriales; Mycobacteriaceae; Mycobacterium; Mycobacterium chelonae group; Mycobacterium abscessus subgroup; Mycobacterium abscessus
MIKHLTRRATVALVVAAAAISPAAYPAMADTATPLNGLFALTPGSCAEGRATGSYFRMIL WP_005073213.1
hypothetical protein [Mycobacterium abscessus] Length: 330\nNumber of predicted TMHs: 2\nExp number of AAs in TMHs: 39.3416199999999\nExp number, first 60 AAs: 17.07688\nTotal prob of N-in: 0.84431\nPOSSIBLE N-term signal sequence\ninside 1 11\nTMhelix 12 29\noutside 30 298\nTMhelix 299 321\ninside 322 330

30218 GCF_900139305.1_10465_1_76 Mycobacterium abscessus subsp. abscessus Terrabacteria group; Actinobacteria; Actinobacteria; Corynebacteriales; Mycobacteriaceae; Mycobacterium; Mycobacterium chelonae group; Mycobacterium abscessus subgroup; Mycobacterium abscessus
MIKHLTRRATVALVVAAAAISPAAYPAMADTATPLNGLFALTPGSCAEGRATGSYFRMIL WP_005073213.1
hypothetical protein [Mycobacterium abscessus] Length: 330\nNumber of predicted TMHs: 2\nExp number of AAs in TMHs: 39.3416199999999\nExp number, first 60 AAs: 17.07688\nTotal prob of N-in: 0.84431\nPOSSIBLE N-term signal sequence\ninside 1 11\nTMhelix 12 29\noutside 30 298\nTMhelix 299 321\ninside 322 330

30219 GCF_900139365.1_10625_4_13 Mycobacterium abscessus subsp. abscessus Terrabacteria group; Actinobacteria; Actinobacteria; Corynebacteriales; Mycobacteriaceae; Mycobacterium; Mycobacterium chelonae group; Mycobacterium abscessus subgroup; Mycobacterium abscessus
MIKHLTRRATVALVVAAAAISPAAYPAMADTATPLNGLFALTPGSCAEGRATGSYFRMIL WP_005073213.1
hypothetical protein [Mycobacterium abscessus] Length: 330\nNumber of predicted TMHs: 2\nExp number of

AAs in TMHs: 39.3416199999999\nExp number, first 60 AAs: 17.07688\nTotal prob of N-in: 0.84431\nPOSSIBLE N-term signal sequence\ninside 1 11\nTMhelix 12 29\noutside 30 298\nTMhelix 299 321\ninside 322 330

30220 GCF_900139495.1_10625_4_9 Mycobacterium abscessus subsp. abscessus Terrabacteria group; Actinobacteria; Actinobacteria; Corynebacteriales; Mycobacteriaceae; Mycobacterium; Mycobacterium chelonae group; Mycobacterium abscessus subgroup; Mycobacterium abscessus MIKHLTRRATVALVVAAAAISPAAYPAMADTATPLNGLFALTPGSCAEGRATGSYFRMIL WP_005073213.1 hypothetical protein [Mycobacterium abscessus] Length: 330\nNumber of predicted TMHs: 2\nExp number of AAs in TMHs: 39.3416199999999\nExp number, first 60 AAs: 17.07688\nTotal prob of N-in: 0.84431\nPOSSIBLE N-term signal sequence\ninside 1 11\nTMhelix 12 29\noutside 30 298\nTMhelix 299 321\ninside 322 330

30221 GCF_900139475.1_10625_5_3 Mycobacterium abscessus subsp. abscessus Terrabacteria group; Actinobacteria; Actinobacteria; Corynebacteriales; Mycobacteriaceae; Mycobacterium; Mycobacterium chelonae group; Mycobacterium abscessus subgroup; Mycobacterium abscessus MIKHLTRRATVALVVAAAAISPAAYPAMADTATPLNGLFALTPGSCAEGRATGSYFRMIL WP_005073213.1 hypothetical protein [Mycobacterium abscessus] Length: 330\nNumber of predicted TMHs: 2\nExp number of AAs in TMHs: 39.3416199999999\nExp number, first 60 AAs: 17.07688\nTotal prob of N-in: 0.84431\nPOSSIBLE N-term signal sequence\ninside 1 11\nTMhelix 12 29\noutside 30 298\nTMhelix 299 321\ninside 322 330

30222 GCF_900139505.1_10625_5_44 Mycobacterium abscessus subsp. abscessus Terrabacteria group; Actinobacteria; Actinobacteria; Corynebacteriales; Mycobacteriaceae; Mycobacterium; Mycobacterium chelonae group; Mycobacterium abscessus subgroup; Mycobacterium abscessus MIKHLTRRATVALVVAAAAISPAAYPAMADTATPLNGLFALTPGSCAEGRATGSYFRMIL WP_005073213.1 hypothetical protein [Mycobacterium abscessus] Length: 330\nNumber of predicted TMHs: 2\nExp number of AAs in TMHs: 39.3416199999999\nExp number, first 60 AAs: 17.07688\nTotal prob of N-in: 0.84431\nPOSSIBLE N-term signal sequence\ninside 1 11\nTMhelix 12 29\noutside 30 298\nTMhelix 299 321\ninside 322 330

30223 GCF_900139685.1_10660_1_14 Mycobacterium abscessus subsp. abscessus Terrabacteria group; Actinobacteria; Actinobacteria; Corynebacteriales; Mycobacteriaceae; Mycobacterium; Mycobacterium chelonae group; Mycobacterium abscessus subgroup; Mycobacterium abscessus MIKHLTRRATVALVVAAAAISPAAYPAMADTATPLNGLFALTPGSCAEGRATGSYFRMIL WP_005073213.1 hypothetical protein [Mycobacterium abscessus] Length: 330\nNumber of predicted TMHs: 2\nExp number of AAs in TMHs: 39.3416199999999\nExp number, first 60 AAs: 17.07688\nTotal prob of N-in: 0.84431\nPOSSIBLE N-term signal sequence\ninside 1 11\nTMhelix 12 29\noutside 30 298\nTMhelix 299 321\ninside 322 330

30224 GCF_900139805.1_10660_1_25 Mycobacterium abscessus subsp. abscessus Terrabacteria group; Actinobacteria; Actinobacteria; Corynebacteriales; Mycobacteriaceae; Mycobacterium; Mycobacterium chelonae group; Mycobacterium abscessus subgroup; Mycobacterium abscessus MIKHLTRRATVALVVAAAAISPAAYPAMADTATPLNGLFALTPGSCAEGRATGSYFRMIL WP_005073213.1 hypothetical protein [Mycobacterium abscessus] Length: 330\nNumber of predicted TMHs: 2\nExp number of AAs in TMHs: 39.3416199999999\nExp number, first 60 AAs: 17.07688\nTotal prob of N-in: 0.84431\nPOSSIBLE N-term signal sequence\ninside 1 11\nTMhelix 12 29\noutside 30 298\nTMhelix 299 321\ninside 322 330

30225 GCF_900139865.1_10660_1_29 Mycobacterium abscessus subsp. abscessus Terrabacteria group; Actinobacteria; Actinobacteria; Corynebacteriales; Mycobacteriaceae; Mycobacterium; Mycobacterium chelonae group; Mycobacterium abscessus subgroup; Mycobacterium abscessus MIKHLTRRATVALVVAAAAISPAAYPAMADTATPLNGLFALTPGSCAEGRATGSYFRMIL WP_005073213.1 hypothetical protein [Mycobacterium abscessus] Length: 330\nNumber of predicted TMHs: 2\nExp number of AAs in TMHs: 39.3416199999999\nExp number, first 60 AAs: 17.07688\nTotal prob of N-in: 0.84431\nPOSSIBLE N-term signal sequence\ninside 1 11\nTMhelix 12 29\noutside 30 298\nTMhelix 299 321\ninside 322 330

30226 GCF_900139825.1_10660_1_27 Mycobacterium abscessus subsp. abscessus Terrabacteria group; Actinobacteria; Actinobacteria; Corynebacteriales; Mycobacteriaceae; Mycobacterium; Mycobacterium chelonae group; Mycobacterium abscessus subgroup; Mycobacterium abscessus MIKHLTRRATVALVVAAAAISPAAYPAMADTATPLNGLFALTPGSCAEGRATGSYFRMIL WP_005073213.1 hypothetical protein [Mycobacterium abscessus] Length: 330\nNumber of predicted TMHs: 2\nExp number of AAs in TMHs: 39.3416199999999\nExp number, first 60 AAs: 17.07688\nTotal prob of N-in: 0.84431\nPOSSIBLE N-term signal sequence\ninside 1 11\nTMhelix 12 29\noutside 30 298\nTMhelix 299 321\ninside 322 330

30227 GCF_900139885.1_10660_1_33 Mycobacterium abscessus subsp. abscessus Terrabacteria group; Actinobacteria; Actinobacteria; Corynebacteriales; Mycobacteriaceae; Mycobacterium; Mycobacterium chelonae group;

Mycobacterium abscessus subgroup; Mycobacterium abscessus
MIKHLTRRATVALVVAAAAISPAAYPAMADTATPLNGLFALTPGSCAEGRATGSYFRMIL WP_005073213.1
hypothetical protein [Mycobacterium abscessus] Length: 330\nNumber of predicted TMHs: 2\nExp number of AAs in TMHs: 39.3416199999999\nExp number, first 60 AAs: 17.07688\nTotal prob of N-in: 0.84431\nPOSSIBLE N-term signal sequence\ninside 1 11\nTMhelix 12 29\noutside 30 298\nTMhelix 299 321\ninside 322 330

30228 GCF_900140035.1_10660_1_7 Mycobacterium abscessus subsp. abscessus Terrabacteria group; Actinobacteria; Actinobacteria; Corynebacteriales; Mycobacteriaceae; Mycobacterium; Mycobacterium chelonae group; Mycobacterium abscessus subgroup; Mycobacterium abscessus
MIKHLTRRATVALVVAAAAISPAAYPAMADTATPLNGLFALTPGSCAEGRATGSYFRMIL WP_005073213.1
hypothetical protein [Mycobacterium abscessus] Length: 330\nNumber of predicted TMHs: 2\nExp number of AAs in TMHs: 39.3416199999999\nExp number, first 60 AAs: 17.07688\nTotal prob of N-in: 0.84431\nPOSSIBLE N-term signal sequence\ninside 1 11\nTMhelix 12 29\noutside 30 298\nTMhelix 299 321\ninside 322 330

30229 GCF_900140025.1_10660_1_52 Mycobacterium abscessus subsp. abscessus Terrabacteria group; Actinobacteria; Actinobacteria; Corynebacteriales; Mycobacteriaceae; Mycobacterium; Mycobacterium chelonae group; Mycobacterium abscessus subgroup; Mycobacterium abscessus
MIKHLTRRATVALVVAAAAISPAAYPAMADTATPLNGLFALTPGSCAEGRATGSYFRMIL WP_005073213.1
hypothetical protein [Mycobacterium abscessus] Length: 330\nNumber of predicted TMHs: 2\nExp number of AAs in TMHs: 39.3416199999999\nExp number, first 60 AAs: 17.07688\nTotal prob of N-in: 0.84431\nPOSSIBLE N-term signal sequence\ninside 1 11\nTMhelix 12 29\noutside 30 298\nTMhelix 299 321\ninside 322 330

30230 GCF_900140085.1_10665_2_66 Mycobacterium abscessus subsp. abscessus Terrabacteria group; Actinobacteria; Actinobacteria; Corynebacteriales; Mycobacteriaceae; Mycobacterium; Mycobacterium chelonae group; Mycobacterium abscessus subgroup; Mycobacterium abscessus
MIKHLTRRATVALVVAAAAISPAAYPAMADTATPLNGLFALTPGSCAEGRATGSYFRMIL WP_005073213.1
hypothetical protein [Mycobacterium abscessus] Length: 330\nNumber of predicted TMHs: 2\nExp number of AAs in TMHs: 39.3416199999999\nExp number, first 60 AAs: 17.07688\nTotal prob of N-in: 0.84431\nPOSSIBLE N-term signal sequence\ninside 1 11\nTMhelix 12 29\noutside 30 298\nTMhelix 299 321\ninside 322 330

30231 GCF_900140195.1_10665_2_72 Mycobacterium abscessus subsp. abscessus Terrabacteria group; Actinobacteria; Actinobacteria; Corynebacteriales; Mycobacteriaceae; Mycobacterium; Mycobacterium chelonae group; Mycobacterium abscessus subgroup; Mycobacterium abscessus
MIKHLTRRATVALVVAAAAISPAAYPAMADTATPLNGLFALTPGSCAEGRATGSYFRMIL WP_005073213.1
hypothetical protein [Mycobacterium abscessus] Length: 330\nNumber of predicted TMHs: 2\nExp number of AAs in TMHs: 39.3416199999999\nExp number, first 60 AAs: 17.07688\nTotal prob of N-in: 0.84431\nPOSSIBLE N-term signal sequence\ninside 1 11\nTMhelix 12 29\noutside 30 298\nTMhelix 299 321\ninside 322 330

30232 GCF_900140245.1_10665_2_74 Mycobacterium abscessus subsp. abscessus Terrabacteria group; Actinobacteria; Actinobacteria; Corynebacteriales; Mycobacteriaceae; Mycobacterium; Mycobacterium chelonae group; Mycobacterium abscessus subgroup; Mycobacterium abscessus
MIKHLTRRATVALVVAAAAISPAAYPAMADTATPLNGLFALTPGSCAEGRATGSYFRMIL WP_005073213.1
hypothetical protein [Mycobacterium abscessus] Length: 330\nNumber of predicted TMHs: 2\nExp number of AAs in TMHs: 39.3416199999999\nExp number, first 60 AAs: 17.07688\nTotal prob of N-in: 0.84431\nPOSSIBLE N-term signal sequence\ninside 1 11\nTMhelix 12 29\noutside 30 298\nTMhelix 299 321\ninside 322 330

30233 GCF_900140265.1_10665_2_93 Mycobacterium abscessus subsp. abscessus Terrabacteria group; Actinobacteria; Actinobacteria; Corynebacteriales; Mycobacteriaceae; Mycobacterium; Mycobacterium chelonae group; Mycobacterium abscessus subgroup; Mycobacterium abscessus
MIKHLTRRATVALVVAAAAISPAAYPAMADTATPLNGLFALTPGSCAEGRATGSYFRMIL WP_005073213.1
hypothetical protein [Mycobacterium abscessus] Length: 330\nNumber of predicted TMHs: 2\nExp number of AAs in TMHs: 39.3416199999999\nExp number, first 60 AAs: 17.07688\nTotal prob of N-in: 0.84431\nPOSSIBLE N-term signal sequence\ninside 1 11\nTMhelix 12 29\noutside 30 298\nTMhelix 299 321\ninside 322 330

30234 GCF_900140285.1_10665_2_92 Mycobacterium abscessus subsp. abscessus Terrabacteria group; Actinobacteria; Actinobacteria; Corynebacteriales; Mycobacteriaceae; Mycobacterium; Mycobacterium chelonae group; Mycobacterium abscessus subgroup; Mycobacterium abscessus
MIKHLTRRATVALVVAAAAISPAAYPAMADTATPLNGLFALTPGSCAEGRATGSYFRMIL WP_005073213.1
hypothetical protein [Mycobacterium abscessus] Length: 330\nNumber of predicted TMHs: 2\nExp number of AAs in TMHs: 39.3416199999999\nExp number, first 60 AAs: 17.07688\nTotal prob of N-in: 0.84431\nPOSSIBLE N-term signal sequence\ninside 1 11\nTMhelix 12 29\noutside 30 298\nTMhelix 299 321\ninside 322 330

30235 GCF_900140625.1_10702_1_27 Mycobacterium abscessus subsp. abscessus Terrabacteria group; Actinobacteria; Actinobacteria; Corynebacteriales; Mycobacteriaceae; Mycobacterium; Mycobacterium chelonae group; Mycobacterium abscessus subgroup; Mycobacterium abscessus
MIKHLTRRATVALVVAAAAISPAAYPAMADTATPLNGLFALTPGSCAEGRATGSYFRMIL WP_005073213.1
hypothetical protein [Mycobacterium abscessus] Length: 330\nNumber of predicted TMHs: 2\nExp number of AAs in TMHs: 39.3416199999999\nExp number, first 60 AAs: 17.07688\nTotal prob of N-in: 0.84431\nPOSSIBLE N-term signal sequence\ninside 1 11\nTMhelix 12 29\noutside 30 298\nTMhelix 299 321\ninside 322 330

30236 GCF_900140475.1_10702_1_13 Mycobacterium abscessus subsp. abscessus Terrabacteria group; Actinobacteria; Actinobacteria; Corynebacteriales; Mycobacteriaceae; Mycobacterium; Mycobacterium chelonae group; Mycobacterium abscessus subgroup; Mycobacterium abscessus
MIKHLTRRATVALVVAAAAISPAAYPAMADTATPLNGLFALTPGSCAEGRATGSYFRMIL WP_074340136.1
hypothetical protein [Mycobacterium abscessus] Length: 330\nNumber of predicted TMHs: 2\nExp number of AAs in TMHs: 39.3416799999999\nExp number, first 60 AAs: 17.07671\nTotal prob of N-in: 0.84432\nPOSSIBLE N-term signal sequence\ninside 1 11\nTMhelix 12 29\noutside 30 298\nTMhelix 299 321\ninside 322 330

30237 GCF_900140655.1_10702_1_32 Mycobacterium abscessus subsp. abscessus Terrabacteria group; Actinobacteria; Actinobacteria; Corynebacteriales; Mycobacteriaceae; Mycobacterium; Mycobacterium chelonae group; Mycobacterium abscessus subgroup; Mycobacterium abscessus
MIKHLTRRATVALVVAAAAISPAAYPAMADTATPLNGLFALTPGSCAEGRATGSYFRMIL WP_005073213.1
hypothetical protein [Mycobacterium abscessus] Length: 330\nNumber of predicted TMHs: 2\nExp number of AAs in TMHs: 39.3416199999999\nExp number, first 60 AAs: 17.07688\nTotal prob of N-in: 0.84431\nPOSSIBLE N-term signal sequence\ninside 1 11\nTMhelix 12 29\noutside 30 298\nTMhelix 299 321\ninside 322 330

30238 GCF_900140695.1_10702_1_35 Mycobacterium abscessus subsp. abscessus Terrabacteria group; Actinobacteria; Actinobacteria; Corynebacteriales; Mycobacteriaceae; Mycobacterium; Mycobacterium chelonae group; Mycobacterium abscessus subgroup; Mycobacterium abscessus
MIKHLTRRATVALVVAAAAISPAAYPAMADTATPLNGLFALTPGSCAEGRATGSYFRMIL WP_005073213.1
hypothetical protein [Mycobacterium abscessus] Length: 330\nNumber of predicted TMHs: 2\nExp number of AAs in TMHs: 39.3416199999999\nExp number, first 60 AAs: 17.07688\nTotal prob of N-in: 0.84431\nPOSSIBLE N-term signal sequence\ninside 1 11\nTMhelix 12 29\noutside 30 298\nTMhelix 299 321\ninside 322 330

30239 GCF_900140715.1_10702_1_36 Mycobacterium abscessus subsp. abscessus Terrabacteria group; Actinobacteria; Actinobacteria; Corynebacteriales; Mycobacteriaceae; Mycobacterium; Mycobacterium chelonae group; Mycobacterium abscessus subgroup; Mycobacterium abscessus
MIKHLTRRATVALVVAAAAISPAAYPAMADTATPLNGLFALTPGSCAEGRATGSYFRMIL WP_005073213.1
hypothetical protein [Mycobacterium abscessus] Length: 330\nNumber of predicted TMHs: 2\nExp number of AAs in TMHs: 39.3416199999999\nExp number, first 60 AAs: 17.07688\nTotal prob of N-in: 0.84431\nPOSSIBLE N-term signal sequence\ninside 1 11\nTMhelix 12 29\noutside 30 298\nTMhelix 299 321\ninside 322 330

30240 GCF_900140745.1_10702_1_39 Mycobacterium abscessus subsp. abscessus Terrabacteria group; Actinobacteria; Actinobacteria; Corynebacteriales; Mycobacteriaceae; Mycobacterium; Mycobacterium chelonae group; Mycobacterium abscessus subgroup; Mycobacterium abscessus
MIKHLTRRATVALVVAAAAISPAAYPAMADTATPLNGLFALTPGSCAEGRATGSYFRMIL WP_005073213.1
hypothetical protein [Mycobacterium abscessus] Length: 330\nNumber of predicted TMHs: 2\nExp number of AAs in TMHs: 39.3416199999999\nExp number, first 60 AAs: 17.07688\nTotal prob of N-in: 0.84431\nPOSSIBLE N-term signal sequence\ninside 1 11\nTMhelix 12 29\noutside 30 298\nTMhelix 299 321\ninside 322 330

30241 GCF_900141385.1_11893_6_51 Mycobacterium abscessus subsp. abscessus Terrabacteria group; Actinobacteria; Actinobacteria; Corynebacteriales; Mycobacteriaceae; Mycobacterium; Mycobacterium chelonae group; Mycobacterium abscessus subgroup; Mycobacterium abscessus
MIKHLTRRATVALVVAAAAISPAAYPAMADTATPLNGLFALTPGSCAEGRATGSYFRMIL WP_005073213.1
hypothetical protein [Mycobacterium abscessus] Length: 330\nNumber of predicted TMHs: 2\nExp number of AAs in TMHs: 39.3416199999999\nExp number, first 60 AAs: 17.07688\nTotal prob of N-in: 0.84431\nPOSSIBLE N-term signal sequence\ninside 1 11\nTMhelix 12 29\noutside 30 298\nTMhelix 299 321\ninside 322 330

30242 GCF_000770175.1_ASM77017v1 Mycobacterium abscessus subsp. abscessus Terrabacteria group; Actinobacteria; Actinobacteria; Corynebacteriales; Mycobacteriaceae; Mycobacterium; Mycobacterium chelonae group; Mycobacterium abscessus subgroup; Mycobacterium abscessus
MIKHLTRRATVALVVAAAAISPAAYPAVADTATPLNGLFALTPGSCAEGRATGSYFRMIL WP_005063674.1 hypothetical protein [Mycobacterium abscessus] Length: 330\nNumber of predicted TMHs: 1\nExp number of AAs in TMHs:

39.25259\nExp number, first 60 AAs: 16.98187\nTotal prob of N-in: 0.83210\nPOSSIBLE N-term signal sequence\noutside 1 298\nTMhelix 299 321\ninside 322 330

30243 GCF_900141395.1_11893_6_55 Mycobacterium abscessus subsp. abscessus Terrabacteria group; Actinobacteria; Actinobacteria; Corynebacteriales; Mycobacteriaceae; Mycobacterium; Mycobacterium chelonae group; Mycobacterium abscessus subgroup; Mycobacterium abscessus
MIKHLTRRATVALVVAAAAISPAAYPAMADTATPLNGLFALTPGSCAEGRATGSYFRMIL WP_005073213.1
hypothetical protein [Mycobacterium abscessus] Length: 330\nNumber of predicted TMHs: 2\nExp number of AAs in TMHs: 39.3416199999999\nExp number, first 60 AAs: 17.07688\nTotal prob of N-in: 0.84431\nPOSSIBLE N-term signal sequence\ninside 1 11\nTMhelix 12 29\noutside 30 298\nTMhelix 299 321\ninside 322 330

30244 GCF_900141435.1_11893_6_54 Mycobacterium abscessus subsp. abscessus Terrabacteria group; Actinobacteria; Actinobacteria; Corynebacteriales; Mycobacteriaceae; Mycobacterium; Mycobacterium chelonae group; Mycobacterium abscessus subgroup; Mycobacterium abscessus
MIKHLTRRATVALVVAAAAISPAAYPAMADTATPLNGLFALTPGSCAEGRATGSYFRMIL WP_005073213.1
hypothetical protein [Mycobacterium abscessus] Length: 330\nNumber of predicted TMHs: 2\nExp number of AAs in TMHs: 39.3416199999999\nExp number, first 60 AAs: 17.07688\nTotal prob of N-in: 0.84431\nPOSSIBLE N-term signal sequence\ninside 1 11\nTMhelix 12 29\noutside 30 298\nTMhelix 299 321\ninside 322 330

30245 GCF_900132485.1_11861_7_81 Mycobacterium abscessus subsp. abscessus Terrabacteria group; Actinobacteria; Actinobacteria; Corynebacteriales; Mycobacteriaceae; Mycobacterium; Mycobacterium chelonae group; Mycobacterium abscessus subgroup; Mycobacterium abscessus
MIKHLTRRATVALVVAAAAISPAAYPAMADTATPLNGLFALTPGSCAEGRATGSYFRMIL WP_005073213.1
hypothetical protein [Mycobacterium abscessus] Length: 330\nNumber of predicted TMHs: 2\nExp number of AAs in TMHs: 39.3416199999999\nExp number, first 60 AAs: 17.07688\nTotal prob of N-in: 0.84431\nPOSSIBLE N-term signal sequence\ninside 1 11\nTMhelix 12 29\noutside 30 298\nTMhelix 299 321\ninside 322 330

30246 GCF_900141485.1_12045_8_45 Mycobacterium abscessus subsp. abscessus Terrabacteria group; Actinobacteria; Actinobacteria; Corynebacteriales; Mycobacteriaceae; Mycobacterium; Mycobacterium chelonae group; Mycobacterium abscessus subgroup; Mycobacterium abscessus
MIKHLTRRATVALVVAAAAISPAAYPAMADTATPLNGLFALTPGSCAEGRATGSYFRMIL WP_005073213.1
hypothetical protein [Mycobacterium abscessus] Length: 330\nNumber of predicted TMHs: 2\nExp number of AAs in TMHs: 39.3416199999999\nExp number, first 60 AAs: 17.07688\nTotal prob of N-in: 0.84431\nPOSSIBLE N-term signal sequence\ninside 1 11\nTMhelix 12 29\noutside 30 298\nTMhelix 299 321\ninside 322 330

30247 GCF_900141645.1_12163_2_94 Mycobacterium abscessus subsp. abscessus Terrabacteria group; Actinobacteria; Actinobacteria; Corynebacteriales; Mycobacteriaceae; Mycobacterium; Mycobacterium chelonae group; Mycobacterium abscessus subgroup; Mycobacterium abscessus
MIKHLTRRATVALVVAAAAISPAAYPAMADTATPLNGLFALTPGSCAEGRATGSYFRMIL WP_005073213.1
hypothetical protein [Mycobacterium abscessus] Length: 330\nNumber of predicted TMHs: 2\nExp number of AAs in TMHs: 39.3416199999999\nExp number, first 60 AAs: 17.07688\nTotal prob of N-in: 0.84431\nPOSSIBLE N-term signal sequence\ninside 1 11\nTMhelix 12 29\noutside 30 298\nTMhelix 299 321\ninside 322 330

30248 GCF_900141685.1_12163_2_91 Mycobacterium abscessus subsp. abscessus Terrabacteria group; Actinobacteria; Actinobacteria; Corynebacteriales; Mycobacteriaceae; Mycobacterium; Mycobacterium chelonae group; Mycobacterium abscessus subgroup; Mycobacterium abscessus
MIKHLTRRATVALVVAAAAISPAAYPAMADTATPLNGLFALTPGSCAEGRATGSYFRMIL WP_005073213.1
hypothetical protein [Mycobacterium abscessus] Length: 330\nNumber of predicted TMHs: 2\nExp number of AAs in TMHs: 39.3416199999999\nExp number, first 60 AAs: 17.07688\nTotal prob of N-in: 0.84431\nPOSSIBLE N-term signal sequence\ninside 1 11\nTMhelix 12 29\noutside 30 298\nTMhelix 299 321\ninside 322 330

30249 GCF_900141695.1_12163_2_93 Mycobacterium abscessus subsp. abscessus Terrabacteria group; Actinobacteria; Actinobacteria; Corynebacteriales; Mycobacteriaceae; Mycobacterium; Mycobacterium chelonae group; Mycobacterium abscessus subgroup; Mycobacterium abscessus
MIKHLTRRATVALVVAAAAISPAAYPAMADTATPLNGLFALTPGSCAEGRATGSYFRMIL WP_005073213.1
hypothetical protein [Mycobacterium abscessus] Length: 330\nNumber of predicted TMHs: 2\nExp number of AAs in TMHs: 39.3416199999999\nExp number, first 60 AAs: 17.07688\nTotal prob of N-in: 0.84431\nPOSSIBLE N-term signal sequence\ninside 1 11\nTMhelix 12 29\noutside 30 298\nTMhelix 299 321\ninside 322 330

30250 GCF_900133255.1_10250_1_30 Mycobacterium abscessus subsp. abscessus Terrabacteria group; Actinobacteria; Actinobacteria; Corynebacteriales; Mycobacteriaceae; Mycobacterium; Mycobacterium chelonae group;

Mycobacterium abscessus subgroup; Mycobacterium abscessus
MTKHLTRRATVALVVAAAAISPAAYPAMADTATPLNGLFALTPGSCAEGRATGSYFRMIL WP_074319467.1
hypothetical protein [Mycobacterium abscessus] Length: 330\nNumber of predicted TMHs: 2\nExp number of AAs in TMHs: 40.2609399999999\nExp number, first 60 AAs: 17.99061\nTotal prob of N-in: 0.88931\nPOSSIBLE N-term signal sequence\ninside 1 11\nTMhelix 12 29\noutside 30 298\nTMhelix 299 321\ninside 322 330

30251 GCF_900133275.1_10250_1_32 Mycobacterium abscessus subsp. abscessus Terrabacteria group; Actinobacteria; Actinobacteria; Corynebacteriales; Mycobacteriaceae; Mycobacterium; Mycobacterium chelonae group; Mycobacterium abscessus subgroup; Mycobacterium abscessus
MIKHLTRRATVALVVAAAAISPAAYPAMADTATPLNGLFALTPGSCAEGRATGSYFRMIL WP_074250269.1
hypothetical protein [Mycobacterium abscessus] Length: 330\nNumber of predicted TMHs: 2\nExp number of AAs in TMHs: 39.3415199999999\nExp number, first 60 AAs: 17.07715\nTotal prob of N-in: 0.84431\nPOSSIBLE N-term signal sequence\ninside 1 11\nTMhelix 12 29\noutside 30 298\nTMhelix 299 321\ninside 322 330

30252 GCF_900133375.1_10250_1_40 Mycobacterium abscessus subsp. abscessus Terrabacteria group; Actinobacteria; Actinobacteria; Corynebacteriales; Mycobacteriaceae; Mycobacterium; Mycobacterium chelonae group; Mycobacterium abscessus subgroup; Mycobacterium abscessus
MIKHLTRRATVALVVAAAAISPAAYPAMADTATPLNGLFALTPGSCAEGRATGSYFRMIL WP_005073213.1
hypothetical protein [Mycobacterium abscessus] Length: 330\nNumber of predicted TMHs: 2\nExp number of AAs in TMHs: 39.3416199999999\nExp number, first 60 AAs: 17.07688\nTotal prob of N-in: 0.84431\nPOSSIBLE N-term signal sequence\ninside 1 11\nTMhelix 12 29\noutside 30 298\nTMhelix 299 321\ninside 322 330

30253 GCF_900133535.1_11861_7_89 Mycobacterium abscessus subsp. abscessus Terrabacteria group; Actinobacteria; Actinobacteria; Corynebacteriales; Mycobacteriaceae; Mycobacterium; Mycobacterium chelonae group; Mycobacterium abscessus subgroup; Mycobacterium abscessus
MIKHLTRRATVALVVAAAAISPAAYPAMADTATPLNGLFALTPGSCAEGRATGSYFRMIL WP_005073213.1
hypothetical protein [Mycobacterium abscessus] Length: 330\nNumber of predicted TMHs: 2\nExp number of AAs in TMHs: 39.3416199999999\nExp number, first 60 AAs: 17.07688\nTotal prob of N-in: 0.84431\nPOSSIBLE N-term signal sequence\ninside 1 11\nTMhelix 12 29\noutside 30 298\nTMhelix 299 321\ninside 322 330

30254 GCF_900133345.1_10250_1_44 Mycobacterium abscessus subsp. abscessus Terrabacteria group; Actinobacteria; Actinobacteria; Corynebacteriales; Mycobacteriaceae; Mycobacterium; Mycobacterium chelonae group; Mycobacterium abscessus subgroup; Mycobacterium abscessus
MIKHLTRRATVALVVAAAAISPAAYPAMADTATPLNGLFALTPGSCAEGRATGSYFRMIL WP_005073213.1
hypothetical protein [Mycobacterium abscessus] Length: 330\nNumber of predicted TMHs: 2\nExp number of AAs in TMHs: 39.3416199999999\nExp number, first 60 AAs: 17.07688\nTotal prob of N-in: 0.84431\nPOSSIBLE N-term signal sequence\ninside 1 11\nTMhelix 12 29\noutside 30 298\nTMhelix 299 321\ninside 322 330

30255 GCF_900133305.1_10250_1_35 Mycobacterium abscessus subsp. abscessus Terrabacteria group; Actinobacteria; Actinobacteria; Corynebacteriales; Mycobacteriaceae; Mycobacterium; Mycobacterium chelonae group; Mycobacterium abscessus subgroup; Mycobacterium abscessus
MIKHLTRRATVALVVAAAAISPAAYPAMADTATPLNGLFALTPGSCAEGRATGSYFRMIL WP_005073213.1
hypothetical protein [Mycobacterium abscessus] Length: 330\nNumber of predicted TMHs: 2\nExp number of AAs in TMHs: 39.3416199999999\nExp number, first 60 AAs: 17.07688\nTotal prob of N-in: 0.84431\nPOSSIBLE N-term signal sequence\ninside 1 11\nTMhelix 12 29\noutside 30 298\nTMhelix 299 321\ninside 322 330

30256 GCF_900133115.1_11893_6_70 Mycobacterium abscessus subsp. abscessus Terrabacteria group; Actinobacteria; Actinobacteria; Corynebacteriales; Mycobacteriaceae; Mycobacterium; Mycobacterium chelonae group; Mycobacterium abscessus subgroup; Mycobacterium abscessus
MIKHLTRRATVALVVAAAAISPAAYPAMADTATPLNGLFALTPGSCAEGRATGSYFRMIL WP_005073213.1
hypothetical protein [Mycobacterium abscessus] Length: 330\nNumber of predicted TMHs: 2\nExp number of AAs in TMHs: 39.3416199999999\nExp number, first 60 AAs: 17.07688\nTotal prob of N-in: 0.84431\nPOSSIBLE N-term signal sequence\ninside 1 11\nTMhelix 12 29\noutside 30 298\nTMhelix 299 321\ninside 322 330

30257 GCF_900133095.1_11893_6_68 Mycobacterium abscessus subsp. abscessus Terrabacteria group; Actinobacteria; Actinobacteria; Corynebacteriales; Mycobacteriaceae; Mycobacterium; Mycobacterium chelonae group; Mycobacterium abscessus subgroup; Mycobacterium abscessus
MIKHLTRRATVALVVAAAAISPAAYPAMADTATPLNGLFALTPGSCAEGRATGSYFRMIL WP_005073213.1
hypothetical protein [Mycobacterium abscessus] Length: 330\nNumber of predicted TMHs: 2\nExp number of AAs in TMHs: 39.3416199999999\nExp number, first 60 AAs: 17.07688\nTotal prob of N-in: 0.84431\nPOSSIBLE N-term signal sequence\ninside 1 11\nTMhelix 12 29\noutside 30 298\nTMhelix 299 321\ninside 322 330

30258 GCF_900133075.1_11893_6_65 Mycobacterium abscessus subsp. abscessus Terrabacteria group; Actinobacteria; Actinobacteria; Corynebacteriales; Mycobacteriaceae; Mycobacterium; Mycobacterium chelonae group; Mycobacterium abscessus subgroup; Mycobacterium abscessus
MIKHLTRRATVALVVAAAAISPAAYPAMADTATPLNGLFALTPGSCAEGRATGSYFRMIL WP_005073213.1
hypothetical protein [Mycobacterium abscessus] Length: 330\nNumber of predicted TMHs: 2\nExp number of AAs in TMHs: 39.3416199999999\nExp number, first 60 AAs: 17.07688\nTotal prob of N-in: 0.84431\nPOSSIBLE N-term signal sequence\ninside 1 11\nTMhelix 12 29\noutside 30 298\nTMhelix 299 321\ninside 322 330

30259 GCF_900133555.1_11893_6_19 Mycobacterium abscessus subsp. abscessus Terrabacteria group; Actinobacteria; Actinobacteria; Corynebacteriales; Mycobacteriaceae; Mycobacterium; Mycobacterium chelonae group; Mycobacterium abscessus subgroup; Mycobacterium abscessus
MIKHLTRRATVALVVAAAAISPAAYPAMADTATPLNGLFALTPGSCAEGRATGSYFRMIL WP_005073213.1
hypothetical protein [Mycobacterium abscessus] Length: 330\nNumber of predicted TMHs: 2\nExp number of AAs in TMHs: 39.3416199999999\nExp number, first 60 AAs: 17.07688\nTotal prob of N-in: 0.84431\nPOSSIBLE N-term signal sequence\ninside 1 11\nTMhelix 12 29\noutside 30 298\nTMhelix 299 321\ninside 322 330

30260 GCF_900133355.1_10250_1_41 Mycobacterium abscessus subsp. abscessus Terrabacteria group; Actinobacteria; Actinobacteria; Corynebacteriales; Mycobacteriaceae; Mycobacterium; Mycobacterium chelonae group; Mycobacterium abscessus subgroup; Mycobacterium abscessus
MIKHLTRRATVALVVAAAAISPAAYPAMADTATPLNGLFALTPGSCAEGRATGSYFRMIL WP_074250269.1
hypothetical protein [Mycobacterium abscessus] Length: 330\nNumber of predicted TMHs: 2\nExp number of AAs in TMHs: 39.3415199999999\nExp number, first 60 AAs: 17.07715\nTotal prob of N-in: 0.84431\nPOSSIBLE N-term signal sequence\ninside 1 11\nTMhelix 12 29\noutside 30 298\nTMhelix 299 321\ninside 322 330

30261 GCF_900132805.1_10250_1_5 Mycobacterium abscessus subsp. abscessus Terrabacteria group; Actinobacteria; Actinobacteria; Corynebacteriales; Mycobacteriaceae; Mycobacterium; Mycobacterium chelonae group; Mycobacterium abscessus subgroup; Mycobacterium abscessus
MIKHLTRRATVALVVAAAAISPAAYPAMADTATPLNGLFALTPGSCAEGRATGSYFRMIL WP_005073213.1
hypothetical protein [Mycobacterium abscessus] Length: 330\nNumber of predicted TMHs: 2\nExp number of AAs in TMHs: 39.3416199999999\nExp number, first 60 AAs: 17.07688\nTotal prob of N-in: 0.84431\nPOSSIBLE N-term signal sequence\ninside 1 11\nTMhelix 12 29\noutside 30 298\nTMhelix 299 321\ninside 322 330

30262 GCF_900133575.1_12082_5_56 Mycobacterium abscessus subsp. abscessus Terrabacteria group; Actinobacteria; Actinobacteria; Corynebacteriales; Mycobacteriaceae; Mycobacterium; Mycobacterium chelonae group; Mycobacterium abscessus subgroup; Mycobacterium abscessus
MIKHLTRRATVALVVAAAAISPAAYPAMADTATPLNGLFALTPGSCAEGRATGSYFRMIL WP_005073213.1
hypothetical protein [Mycobacterium abscessus] Length: 330\nNumber of predicted TMHs: 2\nExp number of AAs in TMHs: 39.3416199999999\nExp number, first 60 AAs: 17.07688\nTotal prob of N-in: 0.84431\nPOSSIBLE N-term signal sequence\ninside 1 11\nTMhelix 12 29\noutside 30 298\nTMhelix 299 321\ninside 322 330

30263 GCF_900132615.1_10250_1_1 Mycobacterium abscessus subsp. abscessus Terrabacteria group; Actinobacteria; Actinobacteria; Corynebacteriales; Mycobacteriaceae; Mycobacterium; Mycobacterium chelonae group; Mycobacterium abscessus subgroup; Mycobacterium abscessus
MTKHLTRRATVALVVAAAAISPAAYPAMADTATPLNGLFALTPGSCAEGRATGSYFRMIL WP_074319467.1
hypothetical protein [Mycobacterium abscessus] Length: 330\nNumber of predicted TMHs: 2\nExp number of AAs in TMHs: 40.2609399999999\nExp number, first 60 AAs: 17.99061\nTotal prob of N-in: 0.88931\nPOSSIBLE N-term signal sequence\ninside 1 11\nTMhelix 12 29\noutside 30 298\nTMhelix 299 321\ninside 322 330

30264 GCF_900133395.1_10250_1_49 Mycobacterium abscessus subsp. abscessus Terrabacteria group; Actinobacteria; Actinobacteria; Corynebacteriales; Mycobacteriaceae; Mycobacterium; Mycobacterium chelonae group; Mycobacterium abscessus subgroup; Mycobacterium abscessus
MIKHLTRRATVALVVAAAAISPAAYPAMADTATPLNGLFALTPGSCAEGRATGSYFRMIL WP_074250269.1
hypothetical protein [Mycobacterium abscessus] Length: 330\nNumber of predicted TMHs: 2\nExp number of AAs in TMHs: 39.3415199999999\nExp number, first 60 AAs: 17.07715\nTotal prob of N-in: 0.84431\nPOSSIBLE N-term signal sequence\ninside 1 11\nTMhelix 12 29\noutside 30 298\nTMhelix 299 321\ninside 322 330

30265 GCF_900132755.1_10250_1_24 Mycobacterium abscessus subsp. abscessus Terrabacteria group; Actinobacteria; Actinobacteria; Corynebacteriales; Mycobacteriaceae; Mycobacterium; Mycobacterium chelonae group; Mycobacterium abscessus subgroup; Mycobacterium abscessus
MIKHLTRRATVALVVAAAAISPAAYPAMADTATPLNGLFALTPGSCAEGRATGSYFRMIL WP_005073213.1
hypothetical protein [Mycobacterium abscessus] Length: 330\nNumber of predicted TMHs: 2\nExp number of

AAs in TMHs: 39.3416199999999\nExp number, first 60 AAs: 17.07688\nTotal prob of N-in: 0.84431\nPOSSIBLE N-term signal sequence\ninside 1 11\nTMhelix 12 29\noutside 30 298\nTMhelix 299 321\ninside 322 330

30266 GCF_900133805.1_12163_1_27 Mycobacterium abscessus subsp. abscessus Terrabacteria group; Actinobacteria; Actinobacteria; Corynebacteriales; Mycobacteriaceae; Mycobacterium; Mycobacterium chelonae group; Mycobacterium abscessus subgroup; Mycobacterium abscessus
MIKHLTRRATVALVVAAAAISPAAYPAMADTATPLNGLFALTPGSCAEGRATGSYFRMIL WP_005073213.1
hypothetical protein [Mycobacterium abscessus] Length: 330\nNumber of predicted TMHs: 2\nExp number of AAs in TMHs: 39.3416199999999\nExp number, first 60 AAs: 17.07688\nTotal prob of N-in: 0.84431\nPOSSIBLE N-term signal sequence\ninside 1 11\nTMhelix 12 29\noutside 30 298\nTMhelix 299 321\ninside 322 330

30267 GCF_900132735.1_10250_1_21 Mycobacterium abscessus subsp. abscessus Terrabacteria group; Actinobacteria; Actinobacteria; Corynebacteriales; Mycobacteriaceae; Mycobacterium; Mycobacterium chelonae group; Mycobacterium abscessus subgroup; Mycobacterium abscessus
MIKHLTRRATVALVVAAAAISPAAYPAMADTATPLNGLFALTPGSCAEGRATGSYFRMIL WP_005073213.1
hypothetical protein [Mycobacterium abscessus] Length: 330\nNumber of predicted TMHs: 2\nExp number of AAs in TMHs: 39.3416199999999\nExp number, first 60 AAs: 17.07688\nTotal prob of N-in: 0.84431\nPOSSIBLE N-term signal sequence\ninside 1 11\nTMhelix 12 29\noutside 30 298\nTMhelix 299 321\ninside 322 330

30268 GCF_900132595.1_10250_1_10 Mycobacterium abscessus subsp. abscessus Terrabacteria group; Actinobacteria; Actinobacteria; Corynebacteriales; Mycobacteriaceae; Mycobacterium; Mycobacterium chelonae group; Mycobacterium abscessus subgroup; Mycobacterium abscessus
MIKHLTRRATVALVVAAAAISPAAYPAMADTATPLNGLFALTPGSCAEGRATGSYFRMIL WP_005073213.1
hypothetical protein [Mycobacterium abscessus] Length: 330\nNumber of predicted TMHs: 2\nExp number of AAs in TMHs: 39.3416199999999\nExp number, first 60 AAs: 17.07688\nTotal prob of N-in: 0.84431\nPOSSIBLE N-term signal sequence\ninside 1 11\nTMhelix 12 29\noutside 30 298\nTMhelix 299 321\ninside 322 330

30269 GCF_900132575.1_12163_2_76 Mycobacterium abscessus subsp. abscessus Terrabacteria group; Actinobacteria; Actinobacteria; Corynebacteriales; Mycobacteriaceae; Mycobacterium; Mycobacterium chelonae group; Mycobacterium abscessus subgroup; Mycobacterium abscessus
MIKHLTRRATVALVVAAAAISPAAYPAMADTATPLNGLFALTPGSCAEGRATGSYFRMIL WP_005073213.1
hypothetical protein [Mycobacterium abscessus] Length: 330\nNumber of predicted TMHs: 2\nExp number of AAs in TMHs: 39.3416199999999\nExp number, first 60 AAs: 17.07688\nTotal prob of N-in: 0.84431\nPOSSIBLE N-term signal sequence\ninside 1 11\nTMhelix 12 29\noutside 30 298\nTMhelix 299 321\ninside 322 330

30270 GCF_900133965.1_10250_1_73 Mycobacterium abscessus subsp. abscessus Terrabacteria group; Actinobacteria; Actinobacteria; Corynebacteriales; Mycobacteriaceae; Mycobacterium; Mycobacterium chelonae group; Mycobacterium abscessus subgroup; Mycobacterium abscessus
MIKHLTRRATVALVVAAAAISPAAYPAMADTATPLNGLFALTPGSCAEGRATGSYFRMIL WP_005073213.1
hypothetical protein [Mycobacterium abscessus] Length: 330\nNumber of predicted TMHs: 2\nExp number of AAs in TMHs: 39.3416199999999\nExp number, first 60 AAs: 17.07688\nTotal prob of N-in: 0.84431\nPOSSIBLE N-term signal sequence\ninside 1 11\nTMhelix 12 29\noutside 30 298\nTMhelix 299 321\ninside 322 330

30271 GCF_900133985.1_10250_1_74 Mycobacterium abscessus subsp. abscessus Terrabacteria group; Actinobacteria; Actinobacteria; Corynebacteriales; Mycobacteriaceae; Mycobacterium; Mycobacterium chelonae group; Mycobacterium abscessus subgroup; Mycobacterium abscessus
MIKHLTRRATVALVVAAAAISPAAYPAMADTATPLNGLFALTPGSCAEGRATGSYFRMIL WP_005073213.1
hypothetical protein [Mycobacterium abscessus] Length: 330\nNumber of predicted TMHs: 2\nExp number of AAs in TMHs: 39.3416199999999\nExp number, first 60 AAs: 17.07688\nTotal prob of N-in: 0.84431\nPOSSIBLE N-term signal sequence\ninside 1 11\nTMhelix 12 29\noutside 30 298\nTMhelix 299 321\ninside 322 330

30272 GCF_900134005.1_10396_8_11 Mycobacterium abscessus subsp. abscessus Terrabacteria group; Actinobacteria; Actinobacteria; Corynebacteriales; Mycobacteriaceae; Mycobacterium; Mycobacterium chelonae group; Mycobacterium abscessus subgroup; Mycobacterium abscessus
MIKHLTRRATVALVVAAAAISPAAYPAMADTATPLNGLFALTPGSCAEGRATGSYFRMIL WP_005073213.1
hypothetical protein [Mycobacterium abscessus] Length: 330\nNumber of predicted TMHs: 2\nExp number of AAs in TMHs: 39.3416199999999\nExp number, first 60 AAs: 17.07688\nTotal prob of N-in: 0.84431\nPOSSIBLE N-term signal sequence\ninside 1 11\nTMhelix 12 29\noutside 30 298\nTMhelix 299 321\ninside 322 330

30273 GCF_900132555.1_11893_6_41 Mycobacterium abscessus subsp. abscessus Terrabacteria group; Actinobacteria; Actinobacteria; Corynebacteriales; Mycobacteriaceae; Mycobacterium; Mycobacterium chelonae group;

Mycobacterium abscessus subgroup; Mycobacterium abscessus
MIKHLTRRATVALVVAAAAISPAAYPAMADTATPLNGLFALTPGSCAEGRATGSYFRMIL WP_005073213.1
hypothetical protein [Mycobacterium abscessus] Length: 330\nNumber of predicted TMHs: 2\nExp number of AAs in TMHs: 39.3416199999999\nExp number, first 60 AAs: 17.07688\nTotal prob of N-in: 0.84431\nPOSSIBLE N-term signal sequence\ninside 1 11\nTMhelix 12 29\noutside 30 298\nTMhelix 299 321\ninside 322 330

30274 GCF_900132505.1_10665_4_81 Mycobacterium abscessus subsp. abscessus Terrabacteria group; Actinobacteria; Actinobacteria; Corynebacteriales; Mycobacteriaceae; Mycobacterium; Mycobacterium chelonae group; Mycobacterium abscessus subgroup; Mycobacterium abscessus
MIKHLTRRATVALVVAAAAISPAAYPAMADTATPLNGLFALTPGSCAEGRATGSYFRMIL WP_005073213.1
hypothetical protein [Mycobacterium abscessus] Length: 330\nNumber of predicted TMHs: 2\nExp number of AAs in TMHs: 39.3416199999999\nExp number, first 60 AAs: 17.07688\nTotal prob of N-in: 0.84431\nPOSSIBLE N-term signal sequence\ninside 1 11\nTMhelix 12 29\noutside 30 298\nTMhelix 299 321\ninside 322 330

30275 GCF_900136105.1_12163_1_62 Mycobacterium abscessus subsp. abscessus Terrabacteria group; Actinobacteria; Actinobacteria; Corynebacteriales; Mycobacteriaceae; Mycobacterium; Mycobacterium chelonae group; Mycobacterium abscessus subgroup; Mycobacterium abscessus
MIKHLTRRATVALVVAAAAISPAAYPAMADTATPLNGLFALTPGSCAEGRATGSYFRMIL WP_005073213.1
hypothetical protein [Mycobacterium abscessus] Length: 330\nNumber of predicted TMHs: 2\nExp number of AAs in TMHs: 39.3416199999999\nExp number, first 60 AAs: 17.07688\nTotal prob of N-in: 0.84431\nPOSSIBLE N-term signal sequence\ninside 1 11\nTMhelix 12 29\noutside 30 298\nTMhelix 299 321\ninside 322 330

30276 GCF_900132335.1_10665_4_64 Mycobacterium abscessus subsp. abscessus Terrabacteria group; Actinobacteria; Actinobacteria; Corynebacteriales; Mycobacteriaceae; Mycobacterium; Mycobacterium chelonae group; Mycobacterium abscessus subgroup; Mycobacterium abscessus
MIKHLTRRATVALVVAAAAISPAAYPAMADTATPLNGLFALTPGSCAEGRATGSYFRMIL WP_005073213.1
hypothetical protein [Mycobacterium abscessus] Length: 330\nNumber of predicted TMHs: 2\nExp number of AAs in TMHs: 39.3416199999999\nExp number, first 60 AAs: 17.07688\nTotal prob of N-in: 0.84431\nPOSSIBLE N-term signal sequence\ninside 1 11\nTMhelix 12 29\noutside 30 298\nTMhelix 299 321\ninside 322 330

30277 GCF_900141125.1_10702_1_75 Mycobacterium abscessus subsp. abscessus Terrabacteria group; Actinobacteria; Actinobacteria; Corynebacteriales; Mycobacteriaceae; Mycobacterium; Mycobacterium chelonae group; Mycobacterium abscessus subgroup; Mycobacterium abscessus
MIKHLTRRATVALVVAAAAISPAAYPAMADTATPLNGLFALTPGSCAEGRATGSYFRMIL WP_005073213.1
hypothetical protein [Mycobacterium abscessus] Length: 330\nNumber of predicted TMHs: 2\nExp number of AAs in TMHs: 39.3416199999999\nExp number, first 60 AAs: 17.07688\nTotal prob of N-in: 0.84431\nPOSSIBLE N-term signal sequence\ninside 1 11\nTMhelix 12 29\noutside 30 298\nTMhelix 299 321\ninside 322 330

30278 GCF_900140355.1_10665_4_12 Mycobacterium abscessus subsp. abscessus Terrabacteria group; Actinobacteria; Actinobacteria; Corynebacteriales; Mycobacteriaceae; Mycobacterium; Mycobacterium chelonae group; Mycobacterium abscessus subgroup; Mycobacterium abscessus
MIKHLTRRATVALVVAAAAISPAAYPAMADTATPLNGLFALTPGSCAEGRATGSYFRMIL WP_005073213.1
hypothetical protein [Mycobacterium abscessus] Length: 330\nNumber of predicted TMHs: 2\nExp number of AAs in TMHs: 39.3416199999999\nExp number, first 60 AAs: 17.07688\nTotal prob of N-in: 0.84431\nPOSSIBLE N-term signal sequence\ninside 1 11\nTMhelix 12 29\noutside 30 298\nTMhelix 299 321\ninside 322 330

30279 GCF_900136085.1_12163_1_68 Mycobacterium abscessus subsp. abscessus Terrabacteria group; Actinobacteria; Actinobacteria; Corynebacteriales; Mycobacteriaceae; Mycobacterium; Mycobacterium chelonae group; Mycobacterium abscessus subgroup; Mycobacterium abscessus
MIKHLTRRATVALVVAAAAISPAAYPAMADTATPLNGLFALTPGSCAEGRATGSYFRMIL WP_005073213.1
hypothetical protein [Mycobacterium abscessus] Length: 330\nNumber of predicted TMHs: 2\nExp number of AAs in TMHs: 39.3416199999999\nExp number, first 60 AAs: 17.07688\nTotal prob of N-in: 0.84431\nPOSSIBLE N-term signal sequence\ninside 1 11\nTMhelix 12 29\noutside 30 298\nTMhelix 299 321\ninside 322 330

30280 GCF_900140335.1_10665_3_39 Mycobacterium abscessus subsp. abscessus Terrabacteria group; Actinobacteria; Actinobacteria; Corynebacteriales; Mycobacteriaceae; Mycobacterium; Mycobacterium chelonae group; Mycobacterium abscessus subgroup; Mycobacterium abscessus
MIKHLTRRATVALVVAAAAISPAAYPAMADTATPLNGLFALTPGSCAEGRATGSYFRMIL WP_005073213.1
hypothetical protein [Mycobacterium abscessus] Length: 330\nNumber of predicted TMHs: 2\nExp number of AAs in TMHs: 39.3416199999999\nExp number, first 60 AAs: 17.07688\nTotal prob of N-in: 0.84431\nPOSSIBLE N-term signal sequence\ninside 1 11\nTMhelix 12 29\noutside 30 298\nTMhelix 299 321\ninside 322 330

30281 GCF_900141095.1_10702_1_76 Mycobacterium abscessus subsp. abscessus Terrabacteria group; Actinobacteria; Actinobacteria; Corynebacteriales; Mycobacteriaceae; Mycobacterium; Mycobacterium chelonae group; Mycobacterium abscessus subgroup; Mycobacterium abscessus
MIKHLTRRATVALVVAAAAISPAAYPAMADTATPLNGLFALTPGSCAEGRATGSYFRMIL WP_005073213.1
hypothetical protein [Mycobacterium abscessus] Length: 330\nNumber of predicted TMHs: 2\nExp number of AAs in TMHs: 39.3416199999999\nExp number, first 60 AAs: 17.07688\nTotal prob of N-in: 0.84431\nPOSSIBLE N-term signal sequence\ninside 1 11\nTMhelix 12 29\noutside 30 298\nTMhelix 299 321\ninside 322 330

30282 GCF_900139485.1_10625_5_2 Mycobacterium abscessus subsp. abscessus Terrabacteria group; Actinobacteria; Actinobacteria; Corynebacteriales; Mycobacteriaceae; Mycobacterium; Mycobacterium chelonae group; Mycobacterium abscessus subgroup; Mycobacterium abscessus
MIKHLTRRATVALVVAAAAISPAAYPAMADTATPLNGLFALTPGSCAEGRATGSYFRMIL WP_005073213.1
hypothetical protein [Mycobacterium abscessus] Length: 330\nNumber of predicted TMHs: 2\nExp number of AAs in TMHs: 39.3416199999999\nExp number, first 60 AAs: 17.07688\nTotal prob of N-in: 0.84431\nPOSSIBLE N-term signal sequence\ninside 1 11\nTMhelix 12 29\noutside 30 298\nTMhelix 299 321\ninside 322 330

30283 GCF_900139235.1_10465_1_69 Mycobacterium abscessus subsp. abscessus Terrabacteria group; Actinobacteria; Actinobacteria; Corynebacteriales; Mycobacteriaceae; Mycobacterium; Mycobacterium chelonae group; Mycobacterium abscessus subgroup; Mycobacterium abscessus
MIKHLTRRATVALVVAAAAISPAAYPAMADTATPLNGLFALTPGSCAEGRATGSYFRMIL WP_005073213.1
hypothetical protein [Mycobacterium abscessus] Length: 330\nNumber of predicted TMHs: 2\nExp number of AAs in TMHs: 39.3416199999999\nExp number, first 60 AAs: 17.07688\nTotal prob of N-in: 0.84431\nPOSSIBLE N-term signal sequence\ninside 1 11\nTMhelix 12 29\noutside 30 298\nTMhelix 299 321\ninside 322 330

30284 GCF_900139195.1_10465_1_62 Mycobacterium abscessus subsp. abscessus Terrabacteria group; Actinobacteria; Actinobacteria; Corynebacteriales; Mycobacteriaceae; Mycobacterium; Mycobacterium chelonae group; Mycobacterium abscessus subgroup; Mycobacterium abscessus
MIKHLTRRATVALVVAAAAISPAAYPAMADTATPLNGLFALTPGSCAEGRATGSYFRMIL WP_005073213.1
hypothetical protein [Mycobacterium abscessus] Length: 330\nNumber of predicted TMHs: 2\nExp number of AAs in TMHs: 39.3416199999999\nExp number, first 60 AAs: 17.07688\nTotal prob of N-in: 0.84431\nPOSSIBLE N-term signal sequence\ninside 1 11\nTMhelix 12 29\noutside 30 298\nTMhelix 299 321\ninside 322 330

30285 GCF_900139515.1_10625_5_4 Mycobacterium abscessus subsp. abscessus Terrabacteria group; Actinobacteria; Actinobacteria; Corynebacteriales; Mycobacteriaceae; Mycobacterium; Mycobacterium chelonae group; Mycobacterium abscessus subgroup; Mycobacterium abscessus
MIKHLTRRATVALVVAAAAISPAAYPAMADTATPLNGLFALTPGSCAEGRATGSYFRMIL WP_005073213.1
hypothetical protein [Mycobacterium abscessus] Length: 330\nNumber of predicted TMHs: 2\nExp number of AAs in TMHs: 39.3416199999999\nExp number, first 60 AAs: 17.07688\nTotal prob of N-in: 0.84431\nPOSSIBLE N-term signal sequence\ninside 1 11\nTMhelix 12 29\noutside 30 298\nTMhelix 299 321\ninside 322 330

30286 GCF_900139535.1_10625_5_46 Mycobacterium abscessus subsp. abscessus Terrabacteria group; Actinobacteria; Actinobacteria; Corynebacteriales; Mycobacteriaceae; Mycobacterium; Mycobacterium chelonae group; Mycobacterium abscessus subgroup; Mycobacterium abscessus
MIKHLTRRATVALVVAAAAISPAAYPAMADTATPLNGLFALTPGSCAEGRATGSYFRMIL WP_005073213.1
hypothetical protein [Mycobacterium abscessus] Length: 330\nNumber of predicted TMHs: 2\nExp number of AAs in TMHs: 39.3416199999999\nExp number, first 60 AAs: 17.07688\nTotal prob of N-in: 0.84431\nPOSSIBLE N-term signal sequence\ninside 1 11\nTMhelix 12 29\noutside 30 298\nTMhelix 299 321\ninside 322 330

30287 GCF_900139215.1_10465_1_68 Mycobacterium abscessus subsp. abscessus Terrabacteria group; Actinobacteria; Actinobacteria; Corynebacteriales; Mycobacteriaceae; Mycobacterium; Mycobacterium chelonae group; Mycobacterium abscessus subgroup; Mycobacterium abscessus
MIKHLTRRATVALVVAAAAISPAAYPAMADTATPLNGLFALTPGSCAEGRATGSYFRMIL WP_005073213.1
hypothetical protein [Mycobacterium abscessus] Length: 330\nNumber of predicted TMHs: 2\nExp number of AAs in TMHs: 39.3416199999999\nExp number, first 60 AAs: 17.07688\nTotal prob of N-in: 0.84431\nPOSSIBLE N-term signal sequence\ninside 1 11\nTMhelix 12 29\noutside 30 298\nTMhelix 299 321\ninside 322 330

30288 GCF_900138985.1_10250_1_36 Mycobacterium abscessus subsp. abscessus Terrabacteria group; Actinobacteria; Actinobacteria; Corynebacteriales; Mycobacteriaceae; Mycobacterium; Mycobacterium chelonae group; Mycobacterium abscessus subgroup; Mycobacterium abscessus
MIKHLTRRATVALVVAAAAISPAAYPAMADTATPLNGLFALTPGSCAEGRATGSYFRMIL WP_074250269.1
hypothetical protein [Mycobacterium abscessus] Length: 330\nNumber of predicted TMHs: 2\nExp number of

AAs in TMHs: 39.3415199999999\nExp number, first 60 AAs: 17.07715\nTotal prob of N-in: 0.84431\nPOSSIBLE N-term signal sequence\ninside 1 11\nTMhelix 12 29\noutside 30 298\nTMhelix 299 321\ninside 322 330

30289 GCF_900139165.1_10465_1_63 Mycobacterium abscessus subsp. abscessus Terrabacteria group; Actinobacteria; Actinobacteria; Corynebacteriales; Mycobacteriaceae; Mycobacterium; Mycobacterium chelonae group; Mycobacterium abscessus subgroup; Mycobacterium abscessus
MIKHLTRRATVALVVAAAAISPAAYPAMADTATPLNGLFALTPGSCAEGRATGSYFRMIL WP_005073213.1
hypothetical protein [Mycobacterium abscessus] Length: 330\nNumber of predicted TMHs: 2\nExp number of AAs in TMHs: 39.3416199999999\nExp number, first 60 AAs: 17.07688\nTotal prob of N-in: 0.84431\nPOSSIBLE N-term signal sequence\ninside 1 11\nTMhelix 12 29\noutside 30 298\nTMhelix 299 321\ninside 322 330

30290 GCF_900138825.1_10208_3_52 Mycobacterium abscessus subsp. abscessus Terrabacteria group; Actinobacteria; Actinobacteria; Corynebacteriales; Mycobacteriaceae; Mycobacterium; Mycobacterium chelonae group; Mycobacterium abscessus subgroup; Mycobacterium abscessus
MIKHLTRRATVALVVAAAAISPAAYPAMADTATPLNGLFALTPGSCAEGRATGSYFRMIL WP_005073213.1
hypothetical protein [Mycobacterium abscessus] Length: 330\nNumber of predicted TMHs: 2\nExp number of AAs in TMHs: 39.3416199999999\nExp number, first 60 AAs: 17.07688\nTotal prob of N-in: 0.84431\nPOSSIBLE N-term signal sequence\ninside 1 11\nTMhelix 12 29\noutside 30 298\nTMhelix 299 321\ninside 322 330

30291 GCF_900139565.1_10625_5_51 Mycobacterium abscessus subsp. abscessus Terrabacteria group; Actinobacteria; Actinobacteria; Corynebacteriales; Mycobacteriaceae; Mycobacterium; Mycobacterium chelonae group; Mycobacterium abscessus subgroup; Mycobacterium abscessus
MIKHLTRRATVALVVAAAAISPAAYPAMADTATPLNGLFALTPGSCAEGRATGSYFRMIL WP_005073213.1
hypothetical protein [Mycobacterium abscessus] Length: 330\nNumber of predicted TMHs: 2\nExp number of AAs in TMHs: 39.3416199999999\nExp number, first 60 AAs: 17.07688\nTotal prob of N-in: 0.84431\nPOSSIBLE N-term signal sequence\ninside 1 11\nTMhelix 12 29\noutside 30 298\nTMhelix 299 321\ninside 322 330

30292 GCF_900139925.1_10660_1_42 Mycobacterium abscessus subsp. abscessus Terrabacteria group; Actinobacteria; Actinobacteria; Corynebacteriales; Mycobacteriaceae; Mycobacterium; Mycobacterium chelonae group; Mycobacterium abscessus subgroup; Mycobacterium abscessus
MIKHLTRRATVALVVAAAAISPAAYPAMADTATPLNGLFALTPGSCAEGRATGSYFRMIL WP_005073213.1
hypothetical protein [Mycobacterium abscessus] Length: 330\nNumber of predicted TMHs: 2\nExp number of AAs in TMHs: 39.3416199999999\nExp number, first 60 AAs: 17.07688\nTotal prob of N-in: 0.84431\nPOSSIBLE N-term signal sequence\ninside 1 11\nTMhelix 12 29\noutside 30 298\nTMhelix 299 321\ninside 322 330

30293 GCF_900139945.1_10660_1_44 Mycobacterium abscessus subsp. abscessus Terrabacteria group; Actinobacteria; Actinobacteria; Corynebacteriales; Mycobacteriaceae; Mycobacterium; Mycobacterium chelonae group; Mycobacterium abscessus subgroup; Mycobacterium abscessus
MIKHLTRRATVALVVAAAAISPAAYPAMADTATPLNGLFALTPGSCAEGRATGSYFRMIL WP_005073213.1
hypothetical protein [Mycobacterium abscessus] Length: 330\nNumber of predicted TMHs: 2\nExp number of AAs in TMHs: 39.3416199999999\nExp number, first 60 AAs: 17.07688\nTotal prob of N-in: 0.84431\nPOSSIBLE N-term signal sequence\ninside 1 11\nTMhelix 12 29\noutside 30 298\nTMhelix 299 321\ninside 322 330

30294 GCF_900139965.1_10660_1_46 Mycobacterium abscessus subsp. abscessus Terrabacteria group; Actinobacteria; Actinobacteria; Corynebacteriales; Mycobacteriaceae; Mycobacterium; Mycobacterium chelonae group; Mycobacterium abscessus subgroup; Mycobacterium abscessus
MIKHLTRRATVALVVAAAAISPAAYPAMADTATPLNGLFALTPGSCAEGRATGSYFRMIL WP_005073213.1
hypothetical protein [Mycobacterium abscessus] Length: 330\nNumber of predicted TMHs: 2\nExp number of AAs in TMHs: 39.3416199999999\nExp number, first 60 AAs: 17.07688\nTotal prob of N-in: 0.84431\nPOSSIBLE N-term signal sequence\ninside 1 11\nTMhelix 12 29\noutside 30 298\nTMhelix 299 321\ninside 322 330

30295 GCF_900138285.1_12163_2_58 Mycobacterium abscessus subsp. abscessus Terrabacteria group; Actinobacteria; Actinobacteria; Corynebacteriales; Mycobacteriaceae; Mycobacterium; Mycobacterium chelonae group; Mycobacterium abscessus subgroup; Mycobacterium abscessus
MIKHLTRRATVALVVAAAAISPAAYPAMADTATPLNGLFALTPGSCAEGRATGSYFRMIL WP_005073213.1
hypothetical protein [Mycobacterium abscessus] Length: 330\nNumber of predicted TMHs: 2\nExp number of AAs in TMHs: 39.3416199999999\nExp number, first 60 AAs: 17.07688\nTotal prob of N-in: 0.84431\nPOSSIBLE N-term signal sequence\ninside 1 11\nTMhelix 12 29\noutside 30 298\nTMhelix 299 321\ninside 322 330

30296 GCF_900138265.1_12163_2_96 Mycobacterium abscessus subsp. abscessus Terrabacteria group; Actinobacteria; Actinobacteria; Corynebacteriales; Mycobacteriaceae; Mycobacterium; Mycobacterium chelonae group;

Mycobacterium abscessus subgroup; Mycobacterium abscessus
MIKHLTRRATVALVVAAAAISPAAYPAMADTATPLNGLFALTPGSCAEGRATGSYFRMIL WP_005073213.1
hypothetical protein [Mycobacterium abscessus] Length: 330\nNumber of predicted TMHs: 2\nExp number of AAs in TMHs: 39.3416199999999\nExp number, first 60 AAs: 17.07688\nTotal prob of N-in: 0.84431\nPOSSIBLE N-term signal sequence\ninside 1 11\nTMhelix 12 29\noutside 30 298\nTMhelix 299 321\ninside 322 330

30297 GCF_900138245.1_12082_5_94 Mycobacterium abscessus subsp. abscessus Terrabacteria group; Actinobacteria; Actinobacteria; Corynebacteriales; Mycobacteriaceae; Mycobacterium; Mycobacterium chelonae group; Mycobacterium abscessus subgroup; Mycobacterium abscessus
MIKHLTRRATVALVVAAAAISPAAYPAMADTATPLNGLFALTPGSCAEGRATGSYFRMIL WP_005073213.1
hypothetical protein [Mycobacterium abscessus] Length: 330\nNumber of predicted TMHs: 2\nExp number of AAs in TMHs: 39.3416199999999\nExp number, first 60 AAs: 17.07688\nTotal prob of N-in: 0.84431\nPOSSIBLE N-term signal sequence\ninside 1 11\nTMhelix 12 29\noutside 30 298\nTMhelix 299 321\ninside 322 330

30298 GCF_900141005.1_10702_1_65 Mycobacterium abscessus subsp. abscessus Terrabacteria group; Actinobacteria; Actinobacteria; Corynebacteriales; Mycobacteriaceae; Mycobacterium; Mycobacterium chelonae group; Mycobacterium abscessus subgroup; Mycobacterium abscessus
MIKHLTRRATVALVVAAAAISPAAYPAMADTATPLNGLFALTPGSCAEGRATGSYFRMIL WP_074250269.1
hypothetical protein [Mycobacterium abscessus] Length: 330\nNumber of predicted TMHs: 2\nExp number of AAs in TMHs: 39.3415199999999\nExp number, first 60 AAs: 17.07715\nTotal prob of N-in: 0.84431\nPOSSIBLE N-term signal sequence\ninside 1 11\nTMhelix 12 29\noutside 30 298\nTMhelix 299 321\ninside 322 330

30299 GCF_900138225.1_12082_5_93 Mycobacterium abscessus subsp. abscessus Terrabacteria group; Actinobacteria; Actinobacteria; Corynebacteriales; Mycobacteriaceae; Mycobacterium; Mycobacterium chelonae group; Mycobacterium abscessus subgroup; Mycobacterium abscessus
MIKHLTRRATVALVVAAAAISPAAYPAMADTATPLNGLFALTPGSCAEGRATGSYFRMIL WP_005073213.1
hypothetical protein [Mycobacterium abscessus] Length: 330\nNumber of predicted TMHs: 2\nExp number of AAs in TMHs: 39.3416199999999\nExp number, first 60 AAs: 17.07688\nTotal prob of N-in: 0.84431\nPOSSIBLE N-term signal sequence\ninside 1 11\nTMhelix 12 29\noutside 30 298\nTMhelix 299 321\ninside 322 330

30300 GCF_900138195.1_12082_5_90 Mycobacterium abscessus subsp. abscessus Terrabacteria group; Actinobacteria; Actinobacteria; Corynebacteriales; Mycobacteriaceae; Mycobacterium; Mycobacterium chelonae group; Mycobacterium abscessus subgroup; Mycobacterium abscessus
MIKHLTRRATVALVVAAAAISPAAYPAMADTATPLNGLFALTPGSCAEGRATGSYFRMIL WP_005073213.1
hypothetical protein [Mycobacterium abscessus] Length: 330\nNumber of predicted TMHs: 2\nExp number of AAs in TMHs: 39.3416199999999\nExp number, first 60 AAs: 17.07688\nTotal prob of N-in: 0.84431\nPOSSIBLE N-term signal sequence\ninside 1 11\nTMhelix 12 29\noutside 30 298\nTMhelix 299 321\ninside 322 330

30301 GCF_900137865.1_12163_2_7 Mycobacterium abscessus subsp. abscessus Terrabacteria group; Actinobacteria; Actinobacteria; Corynebacteriales; Mycobacteriaceae; Mycobacterium; Mycobacterium chelonae group; Mycobacterium abscessus subgroup; Mycobacterium abscessus
MIKHLTRRATVALVVAAAAISPAAYPAMADTATPLNGLFALTPGSCAEGRATGSYFRMIL WP_005073213.1
hypothetical protein [Mycobacterium abscessus] Length: 330\nNumber of predicted TMHs: 2\nExp number of AAs in TMHs: 39.3416199999999\nExp number, first 60 AAs: 17.07688\nTotal prob of N-in: 0.84431\nPOSSIBLE N-term signal sequence\ninside 1 11\nTMhelix 12 29\noutside 30 298\nTMhelix 299 321\ninside 322 330

30302 GCF_900137845.1_12163_2_65 Mycobacterium abscessus subsp. abscessus Terrabacteria group; Actinobacteria; Actinobacteria; Corynebacteriales; Mycobacteriaceae; Mycobacterium; Mycobacterium chelonae group; Mycobacterium abscessus subgroup; Mycobacterium abscessus
MIKHLTRRATVALVVAAAAISPAAYPAMADTATPLNGLFALTPGSCAEGRATGSYFRMIL WP_005073213.1
hypothetical protein [Mycobacterium abscessus] Length: 330\nNumber of predicted TMHs: 2\nExp number of AAs in TMHs: 39.3416199999999\nExp number, first 60 AAs: 17.07688\nTotal prob of N-in: 0.84431\nPOSSIBLE N-term signal sequence\ninside 1 11\nTMhelix 12 29\noutside 30 298\nTMhelix 299 321\ninside 322 330

30303 GCF_900130265.1_10071_6_80 Mycobacterium abscessus subsp. abscessus Terrabacteria group; Actinobacteria; Actinobacteria; Corynebacteriales; Mycobacteriaceae; Mycobacterium; Mycobacterium chelonae group; Mycobacterium abscessus subgroup; Mycobacterium abscessus
MIKHLTRRATVALVVAAAAISPAAYPAMADTATPLNGLFALTPGSCAEGRATGSYFRMIL WP_005073213.1
hypothetical protein [Mycobacterium abscessus] Length: 330\nNumber of predicted TMHs: 2\nExp number of AAs in TMHs: 39.3416199999999\nExp number, first 60 AAs: 17.07688\nTotal prob of N-in: 0.84431\nPOSSIBLE N-term signal sequence\ninside 1 11\nTMhelix 12 29\noutside 30 298\nTMhelix 299 321\ninside 322 330

30304 GCF_900130315.1_10625_5_63 Mycobacterium abscessus subsp. abscessus Terrabacteria group; Actinobacteria; Actinobacteria; Corynebacteriales; Mycobacteriaceae; Mycobacterium; Mycobacterium chelonae group; Mycobacterium abscessus subgroup; Mycobacterium abscessus
MIKHLTRRATVALVVAAAAISPAAYPAMADTATPLNGLFALTPGSCAEGRATGSYFRMIL WP_005073213.1
hypothetical protein [Mycobacterium abscessus] Length: 330\nNumber of predicted TMHs: 2\nExp number of AAs in TMHs: 39.3416199999999\nExp number, first 60 AAs: 17.07688\nTotal prob of N-in: 0.84431\nPOSSIBLE N-term signal sequence\ninside 1 11\nTMhelix 12 29\noutside 30 298\nTMhelix 299 321\ninside 322 330

30305 GCF_900139815.1_10660_1_26 Mycobacterium abscessus subsp. abscessus Terrabacteria group; Actinobacteria; Actinobacteria; Corynebacteriales; Mycobacteriaceae; Mycobacterium; Mycobacterium chelonae group; Mycobacterium abscessus subgroup; Mycobacterium abscessus
MIKHLTRRATVALVVAAAAISPAAYPAMADTATPLNGLFALTPGSCAEGRATGSYFRMIL WP_074340136.1
hypothetical protein [Mycobacterium abscessus] Length: 330\nNumber of predicted TMHs: 2\nExp number of AAs in TMHs: 39.3416799999999\nExp number, first 60 AAs: 17.07671\nTotal prob of N-in: 0.84432\nPOSSIBLE N-term signal sequence\ninside 1 11\nTMhelix 12 29\noutside 30 298\nTMhelix 299 321\ninside 322 330

30306 GCF_900132835.1_10665_4_82 Mycobacterium abscessus subsp. abscessus Terrabacteria group; Actinobacteria; Actinobacteria; Corynebacteriales; Mycobacteriaceae; Mycobacterium; Mycobacterium chelonae group; Mycobacterium abscessus subgroup; Mycobacterium abscessus
MIKHLTRRATVALVVAAAAISPAAYPAMADTATPLNGLFALTPGSCAEGRATGSYFRMIL WP_005073213.1
hypothetical protein [Mycobacterium abscessus] Length: 330\nNumber of predicted TMHs: 2\nExp number of AAs in TMHs: 39.3416199999999\nExp number, first 60 AAs: 17.07688\nTotal prob of N-in: 0.84431\nPOSSIBLE N-term signal sequence\ninside 1 11\nTMhelix 12 29\noutside 30 298\nTMhelix 299 321\ninside 322 330

30307 GCF_900137825.1_12163_2_57 Mycobacterium abscessus subsp. abscessus Terrabacteria group; Actinobacteria; Actinobacteria; Corynebacteriales; Mycobacteriaceae; Mycobacterium; Mycobacterium chelonae group; Mycobacterium abscessus subgroup; Mycobacterium abscessus
MIKHLTRRATVALVVAAAAISPAAYPAMADTATPLNGLFALTPGSCAEGRATGSYFRMIL WP_005073213.1
hypothetical protein [Mycobacterium abscessus] Length: 330\nNumber of predicted TMHs: 2\nExp number of AAs in TMHs: 39.3416199999999\nExp number, first 60 AAs: 17.07688\nTotal prob of N-in: 0.84431\nPOSSIBLE N-term signal sequence\ninside 1 11\nTMhelix 12 29\noutside 30 298\nTMhelix 299 321\ninside 322 330

30308 GCF_900132775.1_10250_1_3 Mycobacterium abscessus subsp. abscessus Terrabacteria group; Actinobacteria; Actinobacteria; Corynebacteriales; Mycobacteriaceae; Mycobacterium; Mycobacterium chelonae group; Mycobacterium abscessus subgroup; Mycobacterium abscessus
MIKHLTRRATVALVVAAAAISPAAYPAMADTATPLNGLFALTPGSCAEGRATGSYFRMIL WP_005073213.1
hypothetical protein [Mycobacterium abscessus] Length: 330\nNumber of predicted TMHs: 2\nExp number of AAs in TMHs: 39.3416199999999\nExp number, first 60 AAs: 17.07688\nTotal prob of N-in: 0.84431\nPOSSIBLE N-term signal sequence\ninside 1 11\nTMhelix 12 29\noutside 30 298\nTMhelix 299 321\ninside 322 330

30309 GCF_900137795.1_12163_2_53 Mycobacterium abscessus subsp. abscessus Terrabacteria group; Actinobacteria; Actinobacteria; Corynebacteriales; Mycobacteriaceae; Mycobacterium; Mycobacterium chelonae group; Mycobacterium abscessus subgroup; Mycobacterium abscessus
MIKHLTRRATVALVVAAAAISPAAYPAMADTATPLNGLFALTPGSCAEGRATGSYFRMIL WP_005073213.1
hypothetical protein [Mycobacterium abscessus] Length: 330\nNumber of predicted TMHs: 2\nExp number of AAs in TMHs: 39.3416199999999\nExp number, first 60 AAs: 17.07688\nTotal prob of N-in: 0.84431\nPOSSIBLE N-term signal sequence\ninside 1 11\nTMhelix 12 29\noutside 30 298\nTMhelix 299 321\ninside 322 330

30310 GCF_900137765.1_12163_2_48 Mycobacterium abscessus subsp. abscessus Terrabacteria group; Actinobacteria; Actinobacteria; Corynebacteriales; Mycobacteriaceae; Mycobacterium; Mycobacterium chelonae group; Mycobacterium abscessus subgroup; Mycobacterium abscessus
MIKHLTRRATVALVVAAAAISPAAYPAMADTATPLNGLFALTPGSCAEGRATGSYFRMIL WP_005073213.1
hypothetical protein [Mycobacterium abscessus] Length: 330\nNumber of predicted TMHs: 2\nExp number of AAs in TMHs: 39.3416199999999\nExp number, first 60 AAs: 17.07688\nTotal prob of N-in: 0.84431\nPOSSIBLE N-term signal sequence\ninside 1 11\nTMhelix 12 29\noutside 30 298\nTMhelix 299 321\ninside 322 330

30311 GCF_900132445.1_10665_4_77 Mycobacterium abscessus subsp. abscessus Terrabacteria group; Actinobacteria; Actinobacteria; Corynebacteriales; Mycobacteriaceae; Mycobacterium; Mycobacterium chelonae group; Mycobacterium abscessus subgroup; Mycobacterium abscessus
MIKHLTRRATVALVVAAAAISPAAYPAMADTATPLNGLFALTPGSCAEGRATGSYFRMIL WP_074250269.1
hypothetical protein [Mycobacterium abscessus] Length: 330\nNumber of predicted TMHs: 2\nExp number of

AAs in TMHs: 39.3415199999999\nExp number, first 60 AAs:  17.07715\nTotal prob of N-in:   0.84431\nPOSSIBLE N-term signal sequence\ninside   1   11\nTMhelix   12   29\noutside   30   298\nTMhelix   299   321\ninside   322   330

30312 GCF_900140045.1_10660_1_6 Mycobacterium abscessus subsp. abscessus Terrabacteria group; Actinobacteria; Actinobacteria; Corynebacteriales; Mycobacteriaceae; Mycobacterium; Mycobacterium chelonae group; Mycobacterium abscessus subgroup; Mycobacterium abscessus
MIKHLTRRATVALVVAAAAISPAAYPAMADTATPLNGLFALTPGSCAEGRATGSYFRMIL WP_005073213.1
hypothetical protein [Mycobacterium abscessus] Length: 330\nNumber of predicted TMHs:  2\nExp number of AAs in TMHs: 39.3416199999999\nExp number, first 60 AAs:  17.07688\nTotal prob of N-in:   0.84431\nPOSSIBLE N-term signal sequence\ninside   1   11\nTMhelix   12   29\noutside   30   298\nTMhelix   299   321\ninside   322   330

30313 GCF_900137465.1_12163_2_23 Mycobacterium abscessus subsp. abscessus Terrabacteria group; Actinobacteria; Actinobacteria; Corynebacteriales; Mycobacteriaceae; Mycobacterium; Mycobacterium chelonae group; Mycobacterium abscessus subgroup; Mycobacterium abscessus
MIKHLTRRATVALVVAAAAISPAAYPAMADTATPLNGLFALTPGSCAEGRATGSYFRMIL WP_005073213.1
hypothetical protein [Mycobacterium abscessus] Length: 330\nNumber of predicted TMHs:  2\nExp number of AAs in TMHs: 39.3416199999999\nExp number, first 60 AAs:  17.07688\nTotal prob of N-in:   0.84431\nPOSSIBLE N-term signal sequence\ninside   1   11\nTMhelix   12   29\noutside   30   298\nTMhelix   299   321\ninside   322   330

30314 GCF_900137425.1_12163_2_20 Mycobacterium abscessus subsp. abscessus Terrabacteria group; Actinobacteria; Actinobacteria; Corynebacteriales; Mycobacteriaceae; Mycobacterium; Mycobacterium chelonae group; Mycobacterium abscessus subgroup; Mycobacterium abscessus
MIKHLTRRATVALVVAAAAISPAAYPAMADTATPLNGLFALTPGSCAEGRATGSYFRMIL WP_005073213.1
hypothetical protein [Mycobacterium abscessus] Length: 330\nNumber of predicted TMHs:  2\nExp number of AAs in TMHs: 39.3416199999999\nExp number, first 60 AAs:  17.07688\nTotal prob of N-in:   0.84431\nPOSSIBLE N-term signal sequence\ninside   1   11\nTMhelix   12   29\noutside   30   298\nTMhelix   299   321\ninside   322   330

30315 GCF_900137445.1_12163_2_21 Mycobacterium abscessus subsp. abscessus Terrabacteria group; Actinobacteria; Actinobacteria; Corynebacteriales; Mycobacteriaceae; Mycobacterium; Mycobacterium chelonae group; Mycobacterium abscessus subgroup; Mycobacterium abscessus
MIKHLTRRATVALVVAAAAISPAAYPAMADTATPLNGLFALTPGSCAEGRATGSYFRMIL WP_005073213.1
hypothetical protein [Mycobacterium abscessus] Length: 330\nNumber of predicted TMHs:  2\nExp number of AAs in TMHs: 39.3416199999999\nExp number, first 60 AAs:  17.07688\nTotal prob of N-in:   0.84431\nPOSSIBLE N-term signal sequence\ninside   1   11\nTMhelix   12   29\noutside   30   298\nTMhelix   299   321\ninside   322   330

30316 GCF_900137395.1_12163_2_16 Mycobacterium abscessus subsp. abscessus Terrabacteria group; Actinobacteria; Actinobacteria; Corynebacteriales; Mycobacteriaceae; Mycobacterium; Mycobacterium chelonae group; Mycobacterium abscessus subgroup; Mycobacterium abscessus
MIKHLTRRATVALVVAAAAISPAAYPAMADTATPLNGLFALTPGSCAEGRATGSYFRMIL WP_005073213.1
hypothetical protein [Mycobacterium abscessus] Length: 330\nNumber of predicted TMHs:  2\nExp number of AAs in TMHs: 39.3416199999999\nExp number, first 60 AAs:  17.07688\nTotal prob of N-in:   0.84431\nPOSSIBLE N-term signal sequence\ninside   1   11\nTMhelix   12   29\noutside   30   298\nTMhelix   299   321\ninside   322   330

30317 GCF_900137145.1_12078_1_66 Mycobacterium abscessus subsp. abscessus Terrabacteria group; Actinobacteria; Actinobacteria; Corynebacteriales; Mycobacteriaceae; Mycobacterium; Mycobacterium chelonae group; Mycobacterium abscessus subgroup; Mycobacterium abscessus
MIKHLTRRATVALVVAAAAISPAAYPAMADTATPLNGLFALTPGSCAEGRATGSYFRMIL WP_005073213.1
hypothetical protein [Mycobacterium abscessus] Length: 330\nNumber of predicted TMHs:  2\nExp number of AAs in TMHs: 39.3416199999999\nExp number, first 60 AAs:  17.07688\nTotal prob of N-in:   0.84431\nPOSSIBLE N-term signal sequence\ninside   1   11\nTMhelix   12   29\noutside   30   298\nTMhelix   299   321\ninside   322   330

30318 GCF_900137125.1_12045_8_44 Mycobacterium abscessus subsp. abscessus Terrabacteria group; Actinobacteria; Actinobacteria; Corynebacteriales; Mycobacteriaceae; Mycobacterium; Mycobacterium chelonae group; Mycobacterium abscessus subgroup; Mycobacterium abscessus
MIKHLTRRATVALVVAAAAISPAAYPAMADTATPLNGLFALTPGSCAEGRATGSYFRMIL WP_005073213.1
hypothetical protein [Mycobacterium abscessus] Length: 330\nNumber of predicted TMHs:  2\nExp number of AAs in TMHs: 39.3416199999999\nExp number, first 60 AAs:  17.07688\nTotal prob of N-in:   0.84431\nPOSSIBLE N-term signal sequence\ninside   1   11\nTMhelix   12   29\noutside   30   298\nTMhelix   299   321\ninside   322   330

30319 GCF_900136855.1_10625_5_43 Mycobacterium abscessus subsp. abscessus Terrabacteria group; Actinobacteria; Actinobacteria; Corynebacteriales; Mycobacteriaceae; Mycobacterium; Mycobacterium chelonae group;

Mycobacterium abscessus subgroup; Mycobacterium abscessus
MTKHLTRRATVALVVAAAAISPAAYPAMADTATPLNGLFALTPGSCAEGRATGSYFRMIL WP_074319467.1
hypothetical protein [Mycobacterium abscessus] Length: 330\nNumber of predicted TMHs: 2\nExp number of AAs in TMHs: 40.2609399999999\nExp number, first 60 AAs: 17.99061\nTotal prob of N-in: 0.88931\nPOSSIBLE N-term signal sequence\ninside 1 11\nTMhelix 12 29\noutside 30 298\nTMhelix 299 321\ninside 322 330

30320 GCF_900137075.1_11893_6_1 Mycobacterium abscessus subsp. abscessus Terrabacteria group; Actinobacteria; Actinobacteria; Corynebacteriales; Mycobacteriaceae; Mycobacterium; Mycobacterium chelonae group; Mycobacterium abscessus subgroup; Mycobacterium abscessus
MIKHLTRRATVALVVAAAAISPAAYPAMADTATPLNGLFALTPGSCAEGRATGSYFRMIL WP_005073213.1
hypothetical protein [Mycobacterium abscessus] Length: 330\nNumber of predicted TMHs: 2\nExp number of AAs in TMHs: 39.3416199999999\nExp number, first 60 AAs: 17.07688\nTotal prob of N-in: 0.84431\nPOSSIBLE N-term signal sequence\ninside 1 11\nTMhelix 12 29\noutside 30 298\nTMhelix 299 321\ninside 322 330

30321 GCF_900136835.1_10625_5_42 Mycobacterium abscessus subsp. abscessus Terrabacteria group; Actinobacteria; Actinobacteria; Corynebacteriales; Mycobacteriaceae; Mycobacterium; Mycobacterium chelonae group; Mycobacterium abscessus subgroup; Mycobacterium abscessus
MTKHLTRRATVALVVAAAAISPAAYPAMADTATPLNGLFALTPGSCAEGRATGSYFRMIL WP_074319467.1
hypothetical protein [Mycobacterium abscessus] Length: 330\nNumber of predicted TMHs: 2\nExp number of AAs in TMHs: 40.2609399999999\nExp number, first 60 AAs: 17.99061\nTotal prob of N-in: 0.88931\nPOSSIBLE N-term signal sequence\ninside 1 11\nTMhelix 12 29\noutside 30 298\nTMhelix 299 321\ninside 322 330

30322 GCF_900136805.1_10625_5_40 Mycobacterium abscessus subsp. abscessus Terrabacteria group; Actinobacteria; Actinobacteria; Corynebacteriales; Mycobacteriaceae; Mycobacterium; Mycobacterium chelonae group; Mycobacterium abscessus subgroup; Mycobacterium abscessus
MTKHLTRRATVALVVAAAAISPAAYPAMADTATPLNGLFALTPGSCAEGRATGSYFRMIL WP_074319467.1
hypothetical protein [Mycobacterium abscessus] Length: 330\nNumber of predicted TMHs: 2\nExp number of AAs in TMHs: 40.2609399999999\nExp number, first 60 AAs: 17.99061\nTotal prob of N-in: 0.88931\nPOSSIBLE N-term signal sequence\ninside 1 11\nTMhelix 12 29\noutside 30 298\nTMhelix 299 321\ninside 322 330

30323 GCF_900136615.1_10625_4_32 Mycobacterium abscessus subsp. abscessus Terrabacteria group; Actinobacteria; Actinobacteria; Corynebacteriales; Mycobacteriaceae; Mycobacterium; Mycobacterium chelonae group; Mycobacterium abscessus subgroup; Mycobacterium abscessus
MIKHLTRRATVALVVAAAAISPAAYPAMADTATPLNGLFALTPGSCAEGRATGSYFRMIL WP_005073213.1
hypothetical protein [Mycobacterium abscessus] Length: 330\nNumber of predicted TMHs: 2\nExp number of AAs in TMHs: 39.3416199999999\nExp number, first 60 AAs: 17.07688\nTotal prob of N-in: 0.84431\nPOSSIBLE N-term signal sequence\ninside 1 11\nTMhelix 12 29\noutside 30 298\nTMhelix 299 321\ninside 322 330

30324 GCF_900136585.1_10208_3_82 Mycobacterium abscessus subsp. abscessus Terrabacteria group; Actinobacteria; Actinobacteria; Corynebacteriales; Mycobacteriaceae; Mycobacterium; Mycobacterium chelonae group; Mycobacterium abscessus subgroup; Mycobacterium abscessus
MIKHLTRRATVALVVAAAAISPAAYPAMADTATPLNGLFALTPGSCAEGRATGSYFRMIL WP_005073213.1
hypothetical protein [Mycobacterium abscessus] Length: 330\nNumber of predicted TMHs: 2\nExp number of AAs in TMHs: 39.3416199999999\nExp number, first 60 AAs: 17.07688\nTotal prob of N-in: 0.84431\nPOSSIBLE N-term signal sequence\ninside 1 11\nTMhelix 12 29\noutside 30 298\nTMhelix 299 321\ninside 322 330

30325 GCF_900136565.1_10208_3_73 Mycobacterium abscessus subsp. abscessus Terrabacteria group; Actinobacteria; Actinobacteria; Corynebacteriales; Mycobacteriaceae; Mycobacterium; Mycobacterium chelonae group; Mycobacterium abscessus subgroup; Mycobacterium abscessus
MIKHLTRRATVALVVAAAAISPAAYPAMADTATPLNGLFALTPGSCAEGRATGSYFRMIL WP_005073213.1
hypothetical protein [Mycobacterium abscessus] Length: 330\nNumber of predicted TMHs: 2\nExp number of AAs in TMHs: 39.3416199999999\nExp number, first 60 AAs: 17.07688\nTotal prob of N-in: 0.84431\nPOSSIBLE N-term signal sequence\ninside 1 11\nTMhelix 12 29\noutside 30 298\nTMhelix 299 321\ninside 322 330

30326 GCF_900136185.1_12163_1_65 Mycobacterium abscessus subsp. abscessus Terrabacteria group; Actinobacteria; Actinobacteria; Corynebacteriales; Mycobacteriaceae; Mycobacterium; Mycobacterium chelonae group; Mycobacterium abscessus subgroup; Mycobacterium abscessus
MIKHLTRRATVALVVAAAAISPAAYPAMADTATPLNGLFALTPGSCAEGRATGSYFRMIL WP_005073213.1
hypothetical protein [Mycobacterium abscessus] Length: 330\nNumber of predicted TMHs: 2\nExp number of AAs in TMHs: 39.3416199999999\nExp number, first 60 AAs: 17.07688\nTotal prob of N-in: 0.84431\nPOSSIBLE N-term signal sequence\ninside 1 11\nTMhelix 12 29\noutside 30 298\nTMhelix 299 321\ninside 322 330

30327 GCF_900136155.1_12163_1_67 Mycobacterium abscessus subsp. abscessus Terrabacteria group; Actinobacteria; Actinobacteria; Corynebacteriales; Mycobacteriaceae; Mycobacterium; Mycobacterium chelonae group; Mycobacterium abscessus subgroup; Mycobacterium abscessus
MIKHLTRRATVALVVAAAAISPAAYPAMADTATPLNGLFALTPGSCAEGRATGSYFRMIL WP_005073213.1
hypothetical protein [Mycobacterium abscessus] Length: 330\nNumber of predicted TMHs: 2\nExp number of AAs in TMHs: 39.3416199999999\nExp number, first 60 AAs: 17.07688\nTotal prob of N-in: 0.84431\nPOSSIBLE N-term signal sequence\ninside 1 11\nTMhelix 12 29\noutside 30 298\nTMhelix 299 321\ninside 322 330

30328 GCF_900133425.1_10396_8_3 Mycobacterium abscessus subsp. abscessus Terrabacteria group; Actinobacteria; Actinobacteria; Corynebacteriales; Mycobacteriaceae; Mycobacterium; Mycobacterium chelonae group; Mycobacterium abscessus subgroup; Mycobacterium abscessus
MIKHLTRRATVALVVAAAAISPAAYPAMADTATPLNGLFALTPGSCAEGRATGSYFRMIL WP_005073213.1
hypothetical protein [Mycobacterium abscessus] Length: 330\nNumber of predicted TMHs: 2\nExp number of AAs in TMHs: 39.3416199999999\nExp number, first 60 AAs: 17.07688\nTotal prob of N-in: 0.84431\nPOSSIBLE N-term signal sequence\ninside 1 11\nTMhelix 12 29\noutside 30 298\nTMhelix 299 321\ninside 322 330

30329 GCF_900133205.1_10250_1_25 Mycobacterium abscessus subsp. abscessus Terrabacteria group; Actinobacteria; Actinobacteria; Corynebacteriales; Mycobacteriaceae; Mycobacterium; Mycobacterium chelonae group; Mycobacterium abscessus subgroup; Mycobacterium abscessus
MIKHLTRRATVALVVAAAAISPAAYPAMADTATPLNGLFALTPGSCAEGRATGSYFRMIL WP_074250269.1
hypothetical protein [Mycobacterium abscessus] Length: 330\nNumber of predicted TMHs: 2\nExp number of AAs in TMHs: 39.3415199999999\nExp number, first 60 AAs: 17.07715\nTotal prob of N-in: 0.84431\nPOSSIBLE N-term signal sequence\ninside 1 11\nTMhelix 12 29\noutside 30 298\nTMhelix 299 321\ninside 322 330

30330 GCF_900133285.1_10250_1_33 Mycobacterium abscessus subsp. abscessus Terrabacteria group; Actinobacteria; Actinobacteria; Corynebacteriales; Mycobacteriaceae; Mycobacterium; Mycobacterium chelonae group; Mycobacterium abscessus subgroup; Mycobacterium abscessus
MIKHLTRRATVALVVAAAAISPAAYPAMADTATPLNGLFALTPGSCAEGRATGSYFRMIL WP_005073213.1
hypothetical protein [Mycobacterium abscessus] Length: 330\nNumber of predicted TMHs: 2\nExp number of AAs in TMHs: 39.3416199999999\nExp number, first 60 AAs: 17.07688\nTotal prob of N-in: 0.84431\nPOSSIBLE N-term signal sequence\ninside 1 11\nTMhelix 12 29\noutside 30 298\nTMhelix 299 321\ninside 322 330

30331 GCF_900133265.1_10250_1_31 Mycobacterium abscessus subsp. abscessus Terrabacteria group; Actinobacteria; Actinobacteria; Corynebacteriales; Mycobacteriaceae; Mycobacterium; Mycobacterium chelonae group; Mycobacterium abscessus subgroup; Mycobacterium abscessus
MIKHLTRRATVALVVAAAAISPAAYPAMADTATPLNGLFALTPGSCAEGRATGSYFRMIL WP_005073213.1
hypothetical protein [Mycobacterium abscessus] Length: 330\nNumber of predicted TMHs: 2\nExp number of AAs in TMHs: 39.3416199999999\nExp number, first 60 AAs: 17.07688\nTotal prob of N-in: 0.84431\nPOSSIBLE N-term signal sequence\ninside 1 11\nTMhelix 12 29\noutside 30 298\nTMhelix 299 321\ninside 322 330

30332 GCF_900131195.1_10665_3_63 Mycobacterium abscessus subsp. abscessus Terrabacteria group; Actinobacteria; Actinobacteria; Corynebacteriales; Mycobacteriaceae; Mycobacterium; Mycobacterium chelonae group; Mycobacterium abscessus subgroup; Mycobacterium abscessus
MIKHLTRRATVALVVAAAAISPAAYPAMADTATPLNGLFALTPGSCAEGRATGSYFRMIL WP_005073213.1
hypothetical protein [Mycobacterium abscessus] Length: 330\nNumber of predicted TMHs: 2\nExp number of AAs in TMHs: 39.3416199999999\nExp number, first 60 AAs: 17.07688\nTotal prob of N-in: 0.84431\nPOSSIBLE N-term signal sequence\ninside 1 11\nTMhelix 12 29\noutside 30 298\nTMhelix 299 321\ninside 322 330

30333 GCF_900131285.1_10665_3_75 Mycobacterium abscessus subsp. abscessus Terrabacteria group; Actinobacteria; Actinobacteria; Corynebacteriales; Mycobacteriaceae; Mycobacterium; Mycobacterium chelonae group; Mycobacterium abscessus subgroup; Mycobacterium abscessus
MIKHLTRRATVALVVAAAAISPAAYPAMADTATPLNGLFALTPGSCAEGRATGSYFRMIL WP_005073213.1
hypothetical protein [Mycobacterium abscessus] Length: 330\nNumber of predicted TMHs: 2\nExp number of AAs in TMHs: 39.3416199999999\nExp number, first 60 AAs: 17.07688\nTotal prob of N-in: 0.84431\nPOSSIBLE N-term signal sequence\ninside 1 11\nTMhelix 12 29\noutside 30 298\nTMhelix 299 321\ninside 322 330

30334 GCF_900133225.1_10250_1_27 Mycobacterium abscessus subsp. abscessus Terrabacteria group; Actinobacteria; Actinobacteria; Corynebacteriales; Mycobacteriaceae; Mycobacterium; Mycobacterium chelonae group; Mycobacterium abscessus subgroup; Mycobacterium abscessus
MIKHLTRRATVALVVAAAAISPAAYPAMADTATPLNGLFALTPGSCAEGRATGSYFRMIL WP_005073213.1
hypothetical protein [Mycobacterium abscessus] Length: 330\nNumber of predicted TMHs: 2\nExp number of

AAs in TMHs: 39.3416199999999\nExp number, first 60 AAs:  17.07688\nTotal prob of N-in:   0.84431\nPOSSIBLE N-term signal sequence\ninside   1   11\nTMhelix   12   29\noutside   30   298\nTMhelix   299   321\ninside   322   330

30335 GCF_900132005.1_10665_4_42 Mycobacterium abscessus subsp. abscessus Terrabacteria group; Actinobacteria; Actinobacteria; Corynebacteriales; Mycobacteriaceae; Mycobacterium; Mycobacterium chelonae group; Mycobacterium abscessus subgroup; Mycobacterium abscessus
MIKHLTRRATVALVVAAAAISPAAYPAMADTATPLNGLFALTPGSCAEGRATGSYFRMIL WP_074250269.1
hypothetical protein [Mycobacterium abscessus] Length: 330\nNumber of predicted TMHs:  2\nExp number of AAs in TMHs: 39.3415199999999\nExp number, first 60 AAs:  17.07715\nTotal prob of N-in:   0.84431\nPOSSIBLE N-term signal sequence\ninside   1   11\nTMhelix   12   29\noutside   30   298\nTMhelix   299   321\ninside   322   330

30336 GCF_900131915.1_10665_4_36 Mycobacterium abscessus subsp. abscessus Terrabacteria group; Actinobacteria; Actinobacteria; Corynebacteriales; Mycobacteriaceae; Mycobacterium; Mycobacterium chelonae group; Mycobacterium abscessus subgroup; Mycobacterium abscessus
MIKHLTRRATVALVVAAAAISPAAYPAMADTATPLNGLFALTPGSCAEGRATGSYFRMIL WP_005073213.1
hypothetical protein [Mycobacterium abscessus] Length: 330\nNumber of predicted TMHs:  2\nExp number of AAs in TMHs: 39.3416199999999\nExp number, first 60 AAs:  17.07688\nTotal prob of N-in:   0.84431\nPOSSIBLE N-term signal sequence\ninside   1   11\nTMhelix   12   29\noutside   30   298\nTMhelix   299   321\ninside   322   330

30337 GCF_900140405.1_10665_4_83 Mycobacterium abscessus subsp. abscessus Terrabacteria group; Actinobacteria; Actinobacteria; Corynebacteriales; Mycobacteriaceae; Mycobacterium; Mycobacterium chelonae group; Mycobacterium abscessus subgroup; Mycobacterium abscessus
MIKHLTRRATVALVVAAAAISPAAYPAMADTATPLNGLFALTPGSCAEGRATGSYFRMIL WP_005073213.1
hypothetical protein [Mycobacterium abscessus] Length: 330\nNumber of predicted TMHs:  2\nExp number of AAs in TMHs: 39.3416199999999\nExp number, first 60 AAs:  17.07688\nTotal prob of N-in:   0.84431\nPOSSIBLE N-term signal sequence\ninside   1   11\nTMhelix   12   29\noutside   30   298\nTMhelix   299   321\ninside   322   330

30338 GCF_900132625.1_12082_5_70 Mycobacterium abscessus subsp. abscessus Terrabacteria group; Actinobacteria; Actinobacteria; Corynebacteriales; Mycobacteriaceae; Mycobacterium; Mycobacterium chelonae group; Mycobacterium abscessus subgroup; Mycobacterium abscessus
MIKHLTRRATVALVVAAAAISPAAYPAMADTATPLNGLFALTPGSCAEGRATGSYFRMIL WP_074356318.1
hypothetical protein [Mycobacterium abscessus] Length: 330\nNumber of predicted TMHs:  2\nExp number of AAs in TMHs: 39.34118\nExp number, first 60 AAs:  17.07668\nTotal prob of N-in:   0.84431\nPOSSIBLE N-term signal sequence\ninside   1   11\nTMhelix   12   29\noutside   30   298\nTMhelix   299   321\ninside   322   330

30339 GCF_900132295.1_12163_2_73 Mycobacterium abscessus subsp. abscessus Terrabacteria group; Actinobacteria; Actinobacteria; Corynebacteriales; Mycobacteriaceae; Mycobacterium; Mycobacterium chelonae group; Mycobacterium abscessus subgroup; Mycobacterium abscessus
MIKHLTRRATVALVVAAAAISPAAYPAMADTATPLNGLFALTPGSCAEGRATGSYFRMIL WP_005073213.1
hypothetical protein [Mycobacterium abscessus] Length: 330\nNumber of predicted TMHs:  2\nExp number of AAs in TMHs: 39.3416199999999\nExp number, first 60 AAs:  17.07688\nTotal prob of N-in:   0.84431\nPOSSIBLE N-term signal sequence\ninside   1   11\nTMhelix   12   29\noutside   30   298\nTMhelix   299   321\ninside   322   330

30340 GCF_900135805.1_11893_6_21 Mycobacterium abscessus subsp. abscessus Terrabacteria group; Actinobacteria; Actinobacteria; Corynebacteriales; Mycobacteriaceae; Mycobacterium; Mycobacterium chelonae group; Mycobacterium abscessus subgroup; Mycobacterium abscessus
MIKHLTRRATVALVVAAAAISPAAYPAMADTATPLNGLFALTPGSCAEGRATGSYFRMIL WP_005073213.1
hypothetical protein [Mycobacterium abscessus] Length: 330\nNumber of predicted TMHs:  2\nExp number of AAs in TMHs: 39.3416199999999\nExp number, first 60 AAs:  17.07688\nTotal prob of N-in:   0.84431\nPOSSIBLE N-term signal sequence\ninside   1   11\nTMhelix   12   29\noutside   30   298\nTMhelix   299   321\ninside   322   330

30341 GCF_900140435.1_10665_4_9 Mycobacterium abscessus subsp. abscessus Terrabacteria group; Actinobacteria; Actinobacteria; Corynebacteriales; Mycobacteriaceae; Mycobacterium; Mycobacterium chelonae group; Mycobacterium abscessus subgroup; Mycobacterium abscessus
MIKHLTRRATVALVVAAAAISPAAYPAMADTATPLNGLFALTPGSCAEGRATGSYFRMIL WP_005073213.1
hypothetical protein [Mycobacterium abscessus] Length: 330\nNumber of predicted TMHs:  2\nExp number of AAs in TMHs: 39.3416199999999\nExp number, first 60 AAs:  17.07688\nTotal prob of N-in:   0.84431\nPOSSIBLE N-term signal sequence\ninside   1   11\nTMhelix   12   29\noutside   30   298\nTMhelix   299   321\ninside   322   330

30342 GCF_900135785.1_11861_8_12 Mycobacterium abscessus subsp. abscessus Terrabacteria group; Actinobacteria; Actinobacteria; Corynebacteriales; Mycobacteriaceae; Mycobacterium; Mycobacterium chelonae group;

Mycobacterium abscessus subgroup; Mycobacterium abscessus
MIKHLTRRATVALVVAAAAISPAAYPAMADTATPLNGLFALTPGSCAEGRATGSYFRMIL WP_005073213.1
hypothetical protein [Mycobacterium abscessus] Length: 330\nNumber of predicted TMHs: 2\nExp number of AAs in TMHs: 39.3416199999999\nExp number, first 60 AAs: 17.07688\nTotal prob of N-in: 0.84431\nPOSSIBLE N-term signal sequence\ninside 1 11\nTMhelix 12 29\noutside 30 298\nTMhelix 299 321\ninside 322 330

30343 GCF_900135655.1_11861_7_24 Mycobacterium abscessus subsp. abscessus Terrabacteria group; Actinobacteria; Actinobacteria; Corynebacteriales; Mycobacteriaceae; Mycobacterium; Mycobacterium chelonae group; Mycobacterium abscessus subgroup; Mycobacterium abscessus
MIKHLTRRATVALVVAAAAISPAAYPAMADTATPLNGLFALTPGSCAEGRATGSYFRMIL WP_005073213.1
hypothetical protein [Mycobacterium abscessus] Length: 330\nNumber of predicted TMHs: 2\nExp number of AAs in TMHs: 39.3416199999999\nExp number, first 60 AAs: 17.07688\nTotal prob of N-in: 0.84431\nPOSSIBLE N-term signal sequence\ninside 1 11\nTMhelix 12 29\noutside 30 298\nTMhelix 299 321\ninside 322 330

30344 GCF_900135635.1_10625_5_9 Mycobacterium abscessus subsp. abscessus Terrabacteria group; Actinobacteria; Actinobacteria; Corynebacteriales; Mycobacteriaceae; Mycobacterium; Mycobacterium chelonae group; Mycobacterium abscessus subgroup; Mycobacterium abscessus
MIKHLTRRATVALVVAAAAISPAAYPAMADTATPLNGLFALTPGSCAEGRATGSYFRMIL WP_005073213.1
hypothetical protein [Mycobacterium abscessus] Length: 330\nNumber of predicted TMHs: 2\nExp number of AAs in TMHs: 39.3416199999999\nExp number, first 60 AAs: 17.07688\nTotal prob of N-in: 0.84431\nPOSSIBLE N-term signal sequence\ninside 1 11\nTMhelix 12 29\noutside 30 298\nTMhelix 299 321\ninside 322 330

30345 GCF_900135615.1_10625_5_18 Mycobacterium abscessus subsp. abscessus Terrabacteria group; Actinobacteria; Actinobacteria; Corynebacteriales; Mycobacteriaceae; Mycobacterium; Mycobacterium chelonae group; Mycobacterium abscessus subgroup; Mycobacterium abscessus
MIKHLTRRATVALVVAAAAISPAAYPAMADTATPLNGLFALTPGSCAEGRATGSYFRMIL WP_005073213.1
hypothetical protein [Mycobacterium abscessus] Length: 330\nNumber of predicted TMHs: 2\nExp number of AAs in TMHs: 39.3416199999999\nExp number, first 60 AAs: 17.07688\nTotal prob of N-in: 0.84431\nPOSSIBLE N-term signal sequence\ninside 1 11\nTMhelix 12 29\noutside 30 298\nTMhelix 299 321\ninside 322 330

30346 GCF_900135345.1_10625_4_51 Mycobacterium abscessus subsp. abscessus Terrabacteria group; Actinobacteria; Actinobacteria; Corynebacteriales; Mycobacteriaceae; Mycobacterium; Mycobacterium chelonae group; Mycobacterium abscessus subgroup; Mycobacterium abscessus
MIKHLTRRATVALVVAAAAISPAAYPAMADTATPLNGLFALTPGSCAEGRATGSYFRMIL WP_005073213.1
hypothetical protein [Mycobacterium abscessus] Length: 330\nNumber of predicted TMHs: 2\nExp number of AAs in TMHs: 39.3416199999999\nExp number, first 60 AAs: 17.07688\nTotal prob of N-in: 0.84431\nPOSSIBLE N-term signal sequence\ninside 1 11\nTMhelix 12 29\noutside 30 298\nTMhelix 299 321\ninside 322 330

30347 GCF_900135325.1_10625_4_46 Mycobacterium abscessus subsp. abscessus Terrabacteria group; Actinobacteria; Actinobacteria; Corynebacteriales; Mycobacteriaceae; Mycobacterium; Mycobacterium chelonae group; Mycobacterium abscessus subgroup; Mycobacterium abscessus
MIKHLTRRATVALVVAAAAISPAAYPAMADTATPLNGLFALTPGSCAEGRATGSYFRMIL WP_005073213.1
hypothetical protein [Mycobacterium abscessus] Length: 330\nNumber of predicted TMHs: 2\nExp number of AAs in TMHs: 39.3416199999999\nExp number, first 60 AAs: 17.07688\nTotal prob of N-in: 0.84431\nPOSSIBLE N-term signal sequence\ninside 1 11\nTMhelix 12 29\noutside 30 298\nTMhelix 299 321\ninside 322 330

30348 GCF_900135295.1_10625_4_43 Mycobacterium abscessus subsp. abscessus Terrabacteria group; Actinobacteria; Actinobacteria; Corynebacteriales; Mycobacteriaceae; Mycobacterium; Mycobacterium chelonae group; Mycobacterium abscessus subgroup; Mycobacterium abscessus
MIKHLTRRATVALVVAAAAISPAAYPAMADTATPLNGLFALTPGSCAEGRATGSYFRMIL WP_005073213.1
hypothetical protein [Mycobacterium abscessus] Length: 330\nNumber of predicted TMHs: 2\nExp number of AAs in TMHs: 39.3416199999999\nExp number, first 60 AAs: 17.07688\nTotal prob of N-in: 0.84431\nPOSSIBLE N-term signal sequence\ninside 1 11\nTMhelix 12 29\noutside 30 298\nTMhelix 299 321\ninside 322 330

30349 GCF_900135265.1_10625_4_39 Mycobacterium abscessus subsp. abscessus Terrabacteria group; Actinobacteria; Actinobacteria; Corynebacteriales; Mycobacteriaceae; Mycobacterium; Mycobacterium chelonae group; Mycobacterium abscessus subgroup; Mycobacterium abscessus
MIKHLTRRATVALVVAAAAISPAAYPAMADTATPLNGLFALTPGSCAEGRATGSYFRMIL WP_005073213.1
hypothetical protein [Mycobacterium abscessus] Length: 330\nNumber of predicted TMHs: 2\nExp number of AAs in TMHs: 39.3416199999999\nExp number, first 60 AAs: 17.07688\nTotal prob of N-in: 0.84431\nPOSSIBLE N-term signal sequence\ninside 1 11\nTMhelix 12 29\noutside 30 298\nTMhelix 299 321\ninside 322 330

30350 GCF_900134935.1_10208_3_35 Mycobacterium abscessus subsp. abscessus Terrabacteria group; Actinobacteria; Actinobacteria; Corynebacteriales; Mycobacteriaceae; Mycobacterium; Mycobacterium chelonae group; Mycobacterium abscessus subgroup; Mycobacterium abscessus
MIKHLTRRATVALVVAAAAISPAAYPAMADTATPLNGLFALTPGSCAEGRATGSYFRMIL WP_005073213.1
hypothetical protein [Mycobacterium abscessus] Length: 330\nNumber of predicted TMHs: 2\nExp number of AAs in TMHs: 39.3416199999999\nExp number, first 60 AAs: 17.07688\nTotal prob of N-in: 0.84431\nPOSSIBLE N-term signal sequence\ninside 1 11\nTMhelix 12 29\noutside 30 298\nTMhelix 299 321\ninside 322 330

30351 GCF_900134825.1_10208_3_22 Mycobacterium abscessus subsp. abscessus Terrabacteria group; Actinobacteria; Actinobacteria; Corynebacteriales; Mycobacteriaceae; Mycobacterium; Mycobacterium chelonae group; Mycobacterium abscessus subgroup; Mycobacterium abscessus
MIKHLTRRATVALVVAAAAISPAAYPAMADTATPLNGLFALTPGSCAEGRATGSYFRMIL WP_005073213.1
hypothetical protein [Mycobacterium abscessus] Length: 330\nNumber of predicted TMHs: 2\nExp number of AAs in TMHs: 39.3416199999999\nExp number, first 60 AAs: 17.07688\nTotal prob of N-in: 0.84431\nPOSSIBLE N-term signal sequence\ninside 1 11\nTMhelix 12 29\noutside 30 298\nTMhelix 299 321\ninside 322 330

30352 GCF_900134795.1_10208_3_21 Mycobacterium abscessus subsp. abscessus Terrabacteria group; Actinobacteria; Actinobacteria; Corynebacteriales; Mycobacteriaceae; Mycobacterium; Mycobacterium chelonae group; Mycobacterium abscessus subgroup; Mycobacterium abscessus
MIKHLTRRATVALVVAAAAISPAAYPAMADTATPLNGLFALTPGSCAEGRATGSYFRMIL WP_005073213.1
hypothetical protein [Mycobacterium abscessus] Length: 330\nNumber of predicted TMHs: 2\nExp number of AAs in TMHs: 39.3416199999999\nExp number, first 60 AAs: 17.07688\nTotal prob of N-in: 0.84431\nPOSSIBLE N-term signal sequence\ninside 1 11\nTMhelix 12 29\noutside 30 298\nTMhelix 299 321\ninside 322 330

30353 GCF_900134775.1_10208_3_2 Mycobacterium abscessus subsp. abscessus Terrabacteria group; Actinobacteria; Actinobacteria; Corynebacteriales; Mycobacteriaceae; Mycobacterium; Mycobacterium chelonae group; Mycobacterium abscessus subgroup; Mycobacterium abscessus
MIKHLTRRATVALVVAAAAISPAAYPAMADTATPLNGLFALTPGSCAEGRATGSYFRMIL WP_005073213.1
hypothetical protein [Mycobacterium abscessus] Length: 330\nNumber of predicted TMHs: 2\nExp number of AAs in TMHs: 39.3416199999999\nExp number, first 60 AAs: 17.07688\nTotal prob of N-in: 0.84431\nPOSSIBLE N-term signal sequence\ninside 1 11\nTMhelix 12 29\noutside 30 298\nTMhelix 299 321\ninside 322 330

30354 GCF_900141035.1_10702_1_68 Mycobacterium abscessus subsp. abscessus Terrabacteria group; Actinobacteria; Actinobacteria; Corynebacteriales; Mycobacteriaceae; Mycobacterium; Mycobacterium chelonae group; Mycobacterium abscessus subgroup; Mycobacterium abscessus
MIKHLTRRATVALVVAAAAISPAAYPAMADTATPLNGLFALTPGSCAEGRATGSYFRMIL WP_005073213.1
hypothetical protein [Mycobacterium abscessus] Length: 330\nNumber of predicted TMHs: 2\nExp number of AAs in TMHs: 39.3416199999999\nExp number, first 60 AAs: 17.07688\nTotal prob of N-in: 0.84431\nPOSSIBLE N-term signal sequence\ninside 1 11\nTMhelix 12 29\noutside 30 298\nTMhelix 299 321\ninside 322 330

30355 GCF_900140825.1_10702_1_50 Mycobacterium abscessus subsp. abscessus Terrabacteria group; Actinobacteria; Actinobacteria; Corynebacteriales; Mycobacteriaceae; Mycobacterium; Mycobacterium chelonae group; Mycobacterium abscessus subgroup; Mycobacterium abscessus
MIKHLTRRATVALVVAAAAISPAAYPAMADTATPLNGLFALTPGSCAEGRATGSYFRMIL WP_074343781.1
hypothetical protein [Mycobacterium abscessus] Length: 330\nNumber of predicted TMHs: 2\nExp number of AAs in TMHs: 39.3417499999999\nExp number, first 60 AAs: 17.07664\nTotal prob of N-in: 0.84432\nPOSSIBLE N-term signal sequence\ninside 1 11\nTMhelix 12 29\noutside 30 298\nTMhelix 299 321\ninside 322 330

30356 GCF_900134435.1_12163_1_32 Mycobacterium abscessus subsp. abscessus Terrabacteria group; Actinobacteria; Actinobacteria; Corynebacteriales; Mycobacteriaceae; Mycobacterium; Mycobacterium chelonae group; Mycobacterium abscessus subgroup; Mycobacterium abscessus
MIKHLTRRATVALVVAAAAISPAAYPAMADTATPLNGLFALTPGSCAEGRATGSYFRMIL WP_005073213.1
hypothetical protein [Mycobacterium abscessus] Length: 330\nNumber of predicted TMHs: 2\nExp number of AAs in TMHs: 39.3416199999999\nExp number, first 60 AAs: 17.07688\nTotal prob of N-in: 0.84431\nPOSSIBLE N-term signal sequence\ninside 1 11\nTMhelix 12 29\noutside 30 298\nTMhelix 299 321\ninside 322 330

30357 GCF_900134375.1_12082_5_75 Mycobacterium abscessus subsp. abscessus Terrabacteria group; Actinobacteria; Actinobacteria; Corynebacteriales; Mycobacteriaceae; Mycobacterium; Mycobacterium chelonae group; Mycobacterium abscessus subgroup; Mycobacterium abscessus
MIKHLTRRATVALVVAAAAISPAAYPAMADTATPLNGLFALTPGSCAEGRATGSYFRMIL WP_005073213.1
hypothetical protein [Mycobacterium abscessus] Length: 330\nNumber of predicted TMHs: 2\nExp number of

AAs in TMHs: 39.3416199999999\nExp number, first 60 AAs: 17.07688\nTotal prob of N-in: 0.84431\nPOSSIBLE N-term signal sequence\ninside 1 11\nTMhelix 12 29\noutside 30 298\nTMhelix 299 321\ninside 322 330

30358 GCF_900133955.1_10250_1_66 Mycobacterium abscessus subsp. abscessus Terrabacteria group; Actinobacteria; Actinobacteria; Corynebacteriales; Mycobacteriaceae; Mycobacterium; Mycobacterium chelonae group; Mycobacterium abscessus subgroup; Mycobacterium abscessus MIKHLTRRATVALVVAAAAISPAAYPAMADTATPLNGLFALTPGSCAEGRATGSYFRMIL WP_005073213.1 hypothetical protein [Mycobacterium abscessus] Length: 330\nNumber of predicted TMHs: 2\nExp number of AAs in TMHs: 39.3416199999999\nExp number, first 60 AAs: 17.07688\nTotal prob of N-in: 0.84431\nPOSSIBLE N-term signal sequence\ninside 1 11\nTMhelix 12 29\noutside 30 298\nTMhelix 299 321\ninside 322 330

30359 GCF_900133935.1_10250_1_67 Mycobacterium abscessus subsp. abscessus Terrabacteria group; Actinobacteria; Actinobacteria; Corynebacteriales; Mycobacteriaceae; Mycobacterium; Mycobacterium chelonae group; Mycobacterium abscessus subgroup; Mycobacterium abscessus MIKHLTRRATVALVVAAAAISPAAYPAMADTATPLNGLFALTPGSCAEGRATGSYFRMIL WP_005073213.1 hypothetical protein [Mycobacterium abscessus] Length: 330\nNumber of predicted TMHs: 2\nExp number of AAs in TMHs: 39.3416199999999\nExp number, first 60 AAs: 17.07688\nTotal prob of N-in: 0.84431\nPOSSIBLE N-term signal sequence\ninside 1 11\nTMhelix 12 29\noutside 30 298\nTMhelix 299 321\ninside 322 330

30360 GCF_900133915.1_10250_1_62 Mycobacterium abscessus subsp. abscessus Terrabacteria group; Actinobacteria; Actinobacteria; Corynebacteriales; Mycobacteriaceae; Mycobacterium; Mycobacterium chelonae group; Mycobacterium abscessus subgroup; Mycobacterium abscessus MIKHLTRRATVALVVAAAAISPAAYPAMADTATPLNGLFALTPGSCAEGRATGSYFRMIL WP_005073213.1 hypothetical protein [Mycobacterium abscessus] Length: 330\nNumber of predicted TMHs: 2\nExp number of AAs in TMHs: 39.3416199999999\nExp number, first 60 AAs: 17.07688\nTotal prob of N-in: 0.84431\nPOSSIBLE N-term signal sequence\ninside 1 11\nTMhelix 12 29\noutside 30 298\nTMhelix 299 321\ninside 322 330

30361 GCF_900133885.1_10250_1_59 Mycobacterium abscessus subsp. abscessus Terrabacteria group; Actinobacteria; Actinobacteria; Corynebacteriales; Mycobacteriaceae; Mycobacterium; Mycobacterium chelonae group; Mycobacterium abscessus subgroup; Mycobacterium abscessus MIKHLTRRATVALVVAAAAISPAAYPAMADTATPLNGLFALTPGSCAEGRATGSYFRMIL WP_005073213.1 hypothetical protein [Mycobacterium abscessus] Length: 330\nNumber of predicted TMHs: 2\nExp number of AAs in TMHs: 39.3416199999999\nExp number, first 60 AAs: 17.07688\nTotal prob of N-in: 0.84431\nPOSSIBLE N-term signal sequence\ninside 1 11\nTMhelix 12 29\noutside 30 298\nTMhelix 299 321\ninside 322 330

30362 GCF_900133565.1_11893_6_20 Mycobacterium abscessus subsp. abscessus Terrabacteria group; Actinobacteria; Actinobacteria; Corynebacteriales; Mycobacteriaceae; Mycobacterium; Mycobacterium chelonae group; Mycobacterium abscessus subgroup; Mycobacterium abscessus MIKHLTRRATVALVVAAAAISPAAYPAMADTATPLNGLFALTPGSCAEGRATGSYFRMIL WP_005073213.1 hypothetical protein [Mycobacterium abscessus] Length: 330\nNumber of predicted TMHs: 2\nExp number of AAs in TMHs: 39.3416199999999\nExp number, first 60 AAs: 17.07688\nTotal prob of N-in: 0.84431\nPOSSIBLE N-term signal sequence\ninside 1 11\nTMhelix 12 29\noutside 30 298\nTMhelix 299 321\ninside 322 330

30363 GCF_900133525.1_11861_7_84 Mycobacterium abscessus subsp. abscessus Terrabacteria group; Actinobacteria; Actinobacteria; Corynebacteriales; Mycobacteriaceae; Mycobacterium; Mycobacterium chelonae group; Mycobacterium abscessus subgroup; Mycobacterium abscessus MIKHLTRRATVALVVAAAAISPAAYPAMADTATPLNGLFALTPGSCAEGRATGSYFRMIL WP_005073213.1 hypothetical protein [Mycobacterium abscessus] Length: 330\nNumber of predicted TMHs: 2\nExp number of AAs in TMHs: 39.3416199999999\nExp number, first 60 AAs: 17.07688\nTotal prob of N-in: 0.84431\nPOSSIBLE N-term signal sequence\ninside 1 11\nTMhelix 12 29\noutside 30 298\nTMhelix 299 321\ninside 322 330

30364 GCF_900133455.1_11861_7_44 Mycobacterium abscessus subsp. abscessus Terrabacteria group; Actinobacteria; Actinobacteria; Corynebacteriales; Mycobacteriaceae; Mycobacterium; Mycobacterium chelonae group; Mycobacterium abscessus subgroup; Mycobacterium abscessus MIKHLTRRATVALVVAAAAISPAAYPAMADTATPLNGLFALTPGSCAEGRATGSYFRMIL WP_005073213.1 hypothetical protein [Mycobacterium abscessus] Length: 330\nNumber of predicted TMHs: 2\nExp number of AAs in TMHs: 39.3416199999999\nExp number, first 60 AAs: 17.07688\nTotal prob of N-in: 0.84431\nPOSSIBLE N-term signal sequence\ninside 1 11\nTMhelix 12 29\noutside 30 298\nTMhelix 299 321\ninside 322 330

30365 GCF_900139675.1_10660_1_1 Mycobacterium abscessus subsp. abscessus Terrabacteria group; Actinobacteria; Actinobacteria; Corynebacteriales; Mycobacteriaceae; Mycobacterium; Mycobacterium chelonae group;

Mycobacterium abscessus subgroup; Mycobacterium abscessus
MIKHLTRRATVALVVAAAAISPAAYPAMADTATPLNGLFALTPGSCAEGRATGSYFRMIL WP_005073213.1
hypothetical protein [Mycobacterium abscessus] Length: 330\nNumber of predicted TMHs: 2\nExp number of AAs in TMHs: 39.3416199999999\nExp number, first 60 AAs: 17.07688\nTotal prob of N-in: 0.84431\nPOSSIBLE N-term signal sequence\ninside 1 11\nTMhelix 12 29\noutside 30 298\nTMhelix 299 321\ninside 322 330

30366 GCF_900141495.1_12045_8_46 Mycobacterium abscessus subsp. abscessus Terrabacteria group; Actinobacteria; Actinobacteria; Corynebacteriales; Mycobacteriaceae; Mycobacterium; Mycobacterium chelonae group; Mycobacterium abscessus subgroup; Mycobacterium abscessus
MIKHLTRRATVALVVAAAAISPAAYPAMADTATPLNGLFALTPGSCAEGRATGSYFRMIL WP_005073213.1
hypothetical protein [Mycobacterium abscessus] Length: 330\nNumber of predicted TMHs: 2\nExp number of AAs in TMHs: 39.3416199999999\nExp number, first 60 AAs: 17.07688\nTotal prob of N-in: 0.84431\nPOSSIBLE N-term signal sequence\ninside 1 11\nTMhelix 12 29\noutside 30 298\nTMhelix 299 321\ninside 322 330

30367 GCF_900141465.1_12045_8_41 Mycobacterium abscessus subsp. abscessus Terrabacteria group; Actinobacteria; Actinobacteria; Corynebacteriales; Mycobacteriaceae; Mycobacterium; Mycobacterium chelonae group; Mycobacterium abscessus subgroup; Mycobacterium abscessus
MIKHLTRRATVALVVAAAAISPAAYPAMADTATPLNGLFALTPGSCAEGRATGSYFRMIL WP_005073213.1
hypothetical protein [Mycobacterium abscessus] Length: 330\nNumber of predicted TMHs: 2\nExp number of AAs in TMHs: 39.3416199999999\nExp number, first 60 AAs: 17.07688\nTotal prob of N-in: 0.84431\nPOSSIBLE N-term signal sequence\ninside 1 11\nTMhelix 12 29\noutside 30 298\nTMhelix 299 321\ninside 322 330

30368 GCF_900141445.1_11893_6_53 Mycobacterium abscessus subsp. abscessus Terrabacteria group; Actinobacteria; Actinobacteria; Corynebacteriales; Mycobacteriaceae; Mycobacterium; Mycobacterium chelonae group; Mycobacterium abscessus subgroup; Mycobacterium abscessus
MIKHLTRRATVALVVAAAAISPAAYPAMADTATPLNGLFALTPGSCAEGRATGSYFRMIL WP_005073213.1
hypothetical protein [Mycobacterium abscessus] Length: 330\nNumber of predicted TMHs: 2\nExp number of AAs in TMHs: 39.3416199999999\nExp number, first 60 AAs: 17.07688\nTotal prob of N-in: 0.84431\nPOSSIBLE N-term signal sequence\ninside 1 11\nTMhelix 12 29\noutside 30 298\nTMhelix 299 321\ninside 322 330

30369 GCF_900141405.1_11893_6_57 Mycobacterium abscessus subsp. abscessus Terrabacteria group; Actinobacteria; Actinobacteria; Corynebacteriales; Mycobacteriaceae; Mycobacterium; Mycobacterium chelonae group; Mycobacterium abscessus subgroup; Mycobacterium abscessus
MIKHLTRRATVALVVAAAAISPAAYPAMADTATPLNGLFALTPGSCAEGRATGSYFRMIL WP_005073213.1
hypothetical protein [Mycobacterium abscessus] Length: 330\nNumber of predicted TMHs: 2\nExp number of AAs in TMHs: 39.3416199999999\nExp number, first 60 AAs: 17.07688\nTotal prob of N-in: 0.84431\nPOSSIBLE N-term signal sequence\ninside 1 11\nTMhelix 12 29\noutside 30 298\nTMhelix 299 321\ninside 322 330

30370 GCF_900140645.1_10702_1_28 Mycobacterium abscessus subsp. abscessus Terrabacteria group; Actinobacteria; Actinobacteria; Corynebacteriales; Mycobacteriaceae; Mycobacterium; Mycobacterium chelonae group; Mycobacterium abscessus subgroup; Mycobacterium abscessus
MIKHLTRRATVALVVAAAAISPAAYPAMADTATPLNGLFALTPGSCAEGRATGSYFRMIL WP_005073213.1
hypothetical protein [Mycobacterium abscessus] Length: 330\nNumber of predicted TMHs: 2\nExp number of AAs in TMHs: 39.3416199999999\nExp number, first 60 AAs: 17.07688\nTotal prob of N-in: 0.84431\nPOSSIBLE N-term signal sequence\ninside 1 11\nTMhelix 12 29\noutside 30 298\nTMhelix 299 321\ninside 322 330

30371 GCF_900140615.1_10702_1_26 Mycobacterium abscessus subsp. abscessus Terrabacteria group; Actinobacteria; Actinobacteria; Corynebacteriales; Mycobacteriaceae; Mycobacterium; Mycobacterium chelonae group; Mycobacterium abscessus subgroup; Mycobacterium abscessus
MIKHLTRRATVALVVAAAAISPAAYPAMADTATPLNGLFALTPGSCAEGRATGSYFRMIL WP_005073213.1
hypothetical protein [Mycobacterium abscessus] Length: 330\nNumber of predicted TMHs: 2\nExp number of AAs in TMHs: 39.3416199999999\nExp number, first 60 AAs: 17.07688\nTotal prob of N-in: 0.84431\nPOSSIBLE N-term signal sequence\ninside 1 11\nTMhelix 12 29\noutside 30 298\nTMhelix 299 321\ninside 322 330

30372 GCF_900140595.1_10702_1_23 Mycobacterium abscessus subsp. abscessus Terrabacteria group; Actinobacteria; Actinobacteria; Corynebacteriales; Mycobacteriaceae; Mycobacterium; Mycobacterium chelonae group; Mycobacterium abscessus subgroup; Mycobacterium abscessus
MIKHLTRRATVALVVAAAAISPAAYPAMADTATPLNGLFALTPGSCAEGRATGSYFRMIL WP_005073213.1
hypothetical protein [Mycobacterium abscessus] Length: 330\nNumber of predicted TMHs: 2\nExp number of AAs in TMHs: 39.3416199999999\nExp number, first 60 AAs: 17.07688\nTotal prob of N-in: 0.84431\nPOSSIBLE N-term signal sequence\ninside 1 11\nTMhelix 12 29\noutside 30 298\nTMhelix 299 321\ninside 322 330

30373 GCF_900140575.1_10702_1_22 Mycobacterium abscessus subsp. abscessus Terrabacteria group; Actinobacteria; Actinobacteria; Corynebacteriales; Mycobacteriaceae; Mycobacterium; Mycobacterium chelonae group; Mycobacterium abscessus subgroup; Mycobacterium abscessus
MIKHLTRRATVALVVAAAAISPAAYPAMADTATPLNGLFALTPGSCAEGRATGSYFRMIL WP_005073213.1
hypothetical protein [Mycobacterium abscessus] Length: 330\nNumber of predicted TMHs: 2\nExp number of AAs in TMHs: 39.3416199999999\nExp number, first 60 AAs: 17.07688\nTotal prob of N-in: 0.84431\nPOSSIBLE N-term signal sequence\ninside 1 11\nTMhelix 12 29\noutside 30 298\nTMhelix 299 321\ninside 322 330

30374 GCF_900140315.1_10665_3_24 Mycobacterium abscessus subsp. abscessus Terrabacteria group; Actinobacteria; Actinobacteria; Corynebacteriales; Mycobacteriaceae; Mycobacterium; Mycobacterium chelonae group; Mycobacterium abscessus subgroup; Mycobacterium abscessus
MIKHLTRRATVALVVAAAAISPAAYPAMADTATPLNGLFALTPGSCAEGRATGSYFRMIL WP_074250269.1
hypothetical protein [Mycobacterium abscessus] Length: 330\nNumber of predicted TMHs: 2\nExp number of AAs in TMHs: 39.3415199999999\nExp number, first 60 AAs: 17.07715\nTotal prob of N-in: 0.84431\nPOSSIBLE N-term signal sequence\ninside 1 11\nTMhelix 12 29\noutside 30 298\nTMhelix 299 321\ninside 322 330

30375 GCF_900140505.1_10702_1_14 Mycobacterium abscessus subsp. abscessus Terrabacteria group; Actinobacteria; Actinobacteria; Corynebacteriales; Mycobacteriaceae; Mycobacterium; Mycobacterium chelonae group; Mycobacterium abscessus subgroup; Mycobacterium abscessus
MIKHLTRRATVALVVAAAAISPAAYPAMADTATPLNGLFALTPGSCAEGRATGSYFRMIL WP_005073213.1
hypothetical protein [Mycobacterium abscessus] Length: 330\nNumber of predicted TMHs: 2\nExp number of AAs in TMHs: 39.3416199999999\nExp number, first 60 AAs: 17.07688\nTotal prob of N-in: 0.84431\nPOSSIBLE N-term signal sequence\ninside 1 11\nTMhelix 12 29\noutside 30 298\nTMhelix 299 321\ninside 322 330

30376 GCF_900140115.1_10665_2_65 Mycobacterium abscessus subsp. abscessus Terrabacteria group; Actinobacteria; Actinobacteria; Corynebacteriales; Mycobacteriaceae; Mycobacterium; Mycobacterium chelonae group; Mycobacterium abscessus subgroup; Mycobacterium abscessus
MIKHLTRRATVALVVAAAAISPAAYPAMADTATPLNGLFALTPGSCAEGRATGSYFRMIL WP_005073213.1
hypothetical protein [Mycobacterium abscessus] Length: 330\nNumber of predicted TMHs: 2\nExp number of AAs in TMHs: 39.3416199999999\nExp number, first 60 AAs: 17.07688\nTotal prob of N-in: 0.84431\nPOSSIBLE N-term signal sequence\ninside 1 11\nTMhelix 12 29\noutside 30 298\nTMhelix 299 321\ninside 322 330

30377 GCF_900140095.1_10660_1_9 Mycobacterium abscessus subsp. abscessus Terrabacteria group; Actinobacteria; Actinobacteria; Corynebacteriales; Mycobacteriaceae; Mycobacterium; Mycobacterium chelonae group; Mycobacterium abscessus subgroup; Mycobacterium abscessus
MIKHLTRRATVALVVAAAAISPAAYPAMADTATPLNGLFALTPGSCAEGRATGSYFRMIL WP_005073213.1
hypothetical protein [Mycobacterium abscessus] Length: 330\nNumber of predicted TMHs: 2\nExp number of AAs in TMHs: 39.3416199999999\nExp number, first 60 AAs: 17.07688\nTotal prob of N-in: 0.84431\nPOSSIBLE N-term signal sequence\ninside 1 11\nTMhelix 12 29\noutside 30 298\nTMhelix 299 321\ninside 322 330

30378 GCF_900140065.1_10660_1_8 Mycobacterium abscessus subsp. abscessus Terrabacteria group; Actinobacteria; Actinobacteria; Corynebacteriales; Mycobacteriaceae; Mycobacterium; Mycobacterium chelonae group; Mycobacterium abscessus subgroup; Mycobacterium abscessus
MIKHLTRRATVALVVAAAAISPAAYPAMADTATPLNGLFALTPGSCAEGRATGSYFRMIL WP_005073213.1
hypothetical protein [Mycobacterium abscessus] Length: 330\nNumber of predicted TMHs: 2\nExp number of AAs in TMHs: 39.3416199999999\nExp number, first 60 AAs: 17.07688\nTotal prob of N-in: 0.84431\nPOSSIBLE N-term signal sequence\ninside 1 11\nTMhelix 12 29\noutside 30 298\nTMhelix 299 321\ninside 322 330

30379 GCF_900139765.1_10660_1_2 Mycobacterium abscessus subsp. abscessus Terrabacteria group; Actinobacteria; Actinobacteria; Corynebacteriales; Mycobacteriaceae; Mycobacterium; Mycobacterium chelonae group; Mycobacterium abscessus subgroup; Mycobacterium abscessus
MIKHLTRRATVALVVAAAAISPAAYPAMADTATPLNGLFALTPGSCAEGRATGSYFRMIL WP_005073213.1
hypothetical protein [Mycobacterium abscessus] Length: 330\nNumber of predicted TMHs: 2\nExp number of AAs in TMHs: 39.3416199999999\nExp number, first 60 AAs: 17.07688\nTotal prob of N-in: 0.84431\nPOSSIBLE N-term signal sequence\ninside 1 11\nTMhelix 12 29\noutside 30 298\nTMhelix 299 321\ninside 322 330

30380 GCF_900139745.1_10660_1_19 Mycobacterium abscessus subsp. abscessus Terrabacteria group; Actinobacteria; Actinobacteria; Corynebacteriales; Mycobacteriaceae; Mycobacterium; Mycobacterium chelonae group; Mycobacterium abscessus subgroup; Mycobacterium abscessus
MIKHLTRRATVALVVAAAAISPAAYPAMADTATPLNGLFALTPGSCAEGRATGSYFRMIL WP_005073213.1
hypothetical protein [Mycobacterium abscessus] Length: 330\nNumber of predicted TMHs: 2\nExp number of

AAs in TMHs: 39.3416199999999\nExp number, first 60 AAs: 17.07688\nTotal prob of N-in: 0.84431\nPOSSIBLE N-term signal sequence\ninside 1 11\nTMhelix 12 29\noutside 30 298\nTMhelix 299 321\ninside 322 330

30381 GCF_900139725.1_10660_1_13 Mycobacterium abscessus subsp. abscessus Terrabacteria group; Actinobacteria; Actinobacteria; Corynebacteriales; Mycobacteriaceae; Mycobacterium; Mycobacterium chelonae group; Mycobacterium abscessus subgroup; Mycobacterium abscessus MIKHLTRRATVALVVAAAAISPAAYPAMADTATPLNGLFALTPGSCAEGRATGSYFRMIL WP_005073213.1 hypothetical protein [Mycobacterium abscessus] Length: 330\nNumber of predicted TMHs: 2\nExp number of AAs in TMHs: 39.3416199999999\nExp number, first 60 AAs: 17.07688\nTotal prob of N-in: 0.84431\nPOSSIBLE N-term signal sequence\ninside 1 11\nTMhelix 12 29\noutside 30 298\nTMhelix 299 321\ninside 322 330

30382 GCF_900139695.1_10660_1_11 Mycobacterium abscessus subsp. abscessus Terrabacteria group; Actinobacteria; Actinobacteria; Corynebacteriales; Mycobacteriaceae; Mycobacterium; Mycobacterium chelonae group; Mycobacterium abscessus subgroup; Mycobacterium abscessus MIKHLTRRATVALVVAAAAISPAAYPAMADTATPLNGLFALTPGSCAEGRATGSYFRMIL WP_005073213.1 hypothetical protein [Mycobacterium abscessus] Length: 330\nNumber of predicted TMHs: 2\nExp number of AAs in TMHs: 39.3416199999999\nExp number, first 60 AAs: 17.07688\nTotal prob of N-in: 0.84431\nPOSSIBLE N-term signal sequence\ninside 1 11\nTMhelix 12 29\noutside 30 298\nTMhelix 299 321\ninside 322 330

30383 GCF_900139465.1_10625_4_5 Mycobacterium abscessus subsp. abscessus Terrabacteria group; Actinobacteria; Actinobacteria; Corynebacteriales; Mycobacteriaceae; Mycobacterium; Mycobacterium chelonae group; Mycobacterium abscessus subgroup; Mycobacterium abscessus MIKHLTRRATVALVVAAAAISPAAYPAMADTATPLNGLFALTPGSCAEGRATGSYFRMIL WP_005073213.1 hypothetical protein [Mycobacterium abscessus] Length: 330\nNumber of predicted TMHs: 2\nExp number of AAs in TMHs: 39.3416199999999\nExp number, first 60 AAs: 17.07688\nTotal prob of N-in: 0.84431\nPOSSIBLE N-term signal sequence\ninside 1 11\nTMhelix 12 29\noutside 30 298\nTMhelix 299 321\ninside 322 330

30384 GCF_900139425.1_10625_4_8 Mycobacterium abscessus subsp. abscessus Terrabacteria group; Actinobacteria; Actinobacteria; Corynebacteriales; Mycobacteriaceae; Mycobacterium; Mycobacterium chelonae group; Mycobacterium abscessus subgroup; Mycobacterium abscessus MIKHLTRRATVALVVAAAAISPAAYPAMADTATPLNGLFALTPGSCAEGRATGSYFRMIL WP_005073213.1 hypothetical protein [Mycobacterium abscessus] Length: 330\nNumber of predicted TMHs: 2\nExp number of AAs in TMHs: 39.3416199999999\nExp number, first 60 AAs: 17.07688\nTotal prob of N-in: 0.84431\nPOSSIBLE N-term signal sequence\ninside 1 11\nTMhelix 12 29\noutside 30 298\nTMhelix 299 321\ninside 322 330

30385 GCF_900139405.1_10625_4_4 Mycobacterium abscessus subsp. abscessus Terrabacteria group; Actinobacteria; Actinobacteria; Corynebacteriales; Mycobacteriaceae; Mycobacterium; Mycobacterium chelonae group; Mycobacterium abscessus subgroup; Mycobacterium abscessus MIKHLTRRATVALVVAAAAISPAAYPAMADTATPLNGLFALTPGSCAEGRATGSYFRMIL WP_005073213.1 hypothetical protein [Mycobacterium abscessus] Length: 330\nNumber of predicted TMHs: 2\nExp number of AAs in TMHs: 39.3416199999999\nExp number, first 60 AAs: 17.07688\nTotal prob of N-in: 0.84431\nPOSSIBLE N-term signal sequence\ninside 1 11\nTMhelix 12 29\noutside 30 298\nTMhelix 299 321\ninside 322 330

30386 GCF_900139375.1_10625_4_15 Mycobacterium abscessus subsp. abscessus Terrabacteria group; Actinobacteria; Actinobacteria; Corynebacteriales; Mycobacteriaceae; Mycobacterium; Mycobacterium chelonae group; Mycobacterium abscessus subgroup; Mycobacterium abscessus MIKHLTRRATVALVVAAAAISPAAYPAMADTATPLNGLFALTPGSCAEGRATGSYFRMIL WP_005073213.1 hypothetical protein [Mycobacterium abscessus] Length: 330\nNumber of predicted TMHs: 2\nExp number of AAs in TMHs: 39.3416199999999\nExp number, first 60 AAs: 17.07688\nTotal prob of N-in: 0.84431\nPOSSIBLE N-term signal sequence\ninside 1 11\nTMhelix 12 29\noutside 30 298\nTMhelix 299 321\ninside 322 330

30387 GCF_900139335.1_10625_4_14 Mycobacterium abscessus subsp. abscessus Terrabacteria group; Actinobacteria; Actinobacteria; Corynebacteriales; Mycobacteriaceae; Mycobacterium; Mycobacterium chelonae group; Mycobacterium abscessus subgroup; Mycobacterium abscessus MIKHLTRRATVALVVAAAAISPAAYPAMADTATPLNGLFALTPGSCAEGRATGSYFRMIL WP_005073213.1 hypothetical protein [Mycobacterium abscessus] Length: 330\nNumber of predicted TMHs: 2\nExp number of AAs in TMHs: 39.3416199999999\nExp number, first 60 AAs: 17.07688\nTotal prob of N-in: 0.84431\nPOSSIBLE N-term signal sequence\ninside 1 11\nTMhelix 12 29\noutside 30 298\nTMhelix 299 321\ninside 322 330

30388 GCF_900139125.1_10465_1_59 Mycobacterium abscessus subsp. abscessus Terrabacteria group; Actinobacteria; Actinobacteria; Corynebacteriales; Mycobacteriaceae; Mycobacterium; Mycobacterium chelonae group;

Mycobacterium abscessus subgroup; Mycobacterium abscessus
MIKHLTRRATVALVVAAAAISPAAYPAMADTATPLNGLFALTPGSCAEGRATGSYFRMIL WP_005073213.1
hypothetical protein [Mycobacterium abscessus] Length: 330\nNumber of predicted TMHs: 2\nExp number of AAs in TMHs: 39.3416199999999\nExp number, first 60 AAs: 17.07688\nTotal prob of N-in: 0.84431\nPOSSIBLE N-term signal sequence\ninside 1 11\nTMhelix 12 29\noutside 30 298\nTMhelix 299 321\ninside 322 330

30389 GCF_900139075.1_10465_1_54 Mycobacterium abscessus subsp. abscessus Terrabacteria group; Actinobacteria; Actinobacteria; Corynebacteriales; Mycobacteriaceae; Mycobacterium; Mycobacterium chelonae group; Mycobacterium abscessus subgroup; Mycobacterium abscessus
MIKHLTRRATVALVVAAAAISPAAYPAMADTATPLNGLFALTPGSCAEGRATGSYFRMIL WP_005073213.1
hypothetical protein [Mycobacterium abscessus] Length: 330\nNumber of predicted TMHs: 2\nExp number of AAs in TMHs: 39.3416199999999\nExp number, first 60 AAs: 17.07688\nTotal prob of N-in: 0.84431\nPOSSIBLE N-term signal sequence\ninside 1 11\nTMhelix 12 29\noutside 30 298\nTMhelix 299 321\ninside 322 330

30390 GCF_900139055.1_10465_1_52 Mycobacterium abscessus subsp. abscessus Terrabacteria group; Actinobacteria; Actinobacteria; Corynebacteriales; Mycobacteriaceae; Mycobacterium; Mycobacterium chelonae group; Mycobacterium abscessus subgroup; Mycobacterium abscessus
MIKHLTRRATVALVVAAAAISPAAYPAMADTATPLNGLFALTPGSCAEGRATGSYFRMIL WP_005073213.1
hypothetical protein [Mycobacterium abscessus] Length: 330\nNumber of predicted TMHs: 2\nExp number of AAs in TMHs: 39.3416199999999\nExp number, first 60 AAs: 17.07688\nTotal prob of N-in: 0.84431\nPOSSIBLE N-term signal sequence\ninside 1 11\nTMhelix 12 29\noutside 30 298\nTMhelix 299 321\ninside 322 330

30391 GCF_900139035.1_10465_1_50 Mycobacterium abscessus subsp. abscessus Terrabacteria group; Actinobacteria; Actinobacteria; Corynebacteriales; Mycobacteriaceae; Mycobacterium; Mycobacterium chelonae group; Mycobacterium abscessus subgroup; Mycobacterium abscessus
MIKHLTRRATVALVVAAAAISPAAYPAMADTATPLNGLFALTPGSCAEGRATGSYFRMIL WP_005073213.1
hypothetical protein [Mycobacterium abscessus] Length: 330\nNumber of predicted TMHs: 2\nExp number of AAs in TMHs: 39.3416199999999\nExp number, first 60 AAs: 17.07688\nTotal prob of N-in: 0.84431\nPOSSIBLE N-term signal sequence\ninside 1 11\nTMhelix 12 29\noutside 30 298\nTMhelix 299 321\ninside 322 330

30392 GCF_900138035.1_12045_8_35 Mycobacterium abscessus subsp. abscessus Terrabacteria group; Actinobacteria; Actinobacteria; Corynebacteriales; Mycobacteriaceae; Mycobacterium; Mycobacterium chelonae group; Mycobacterium abscessus subgroup; Mycobacterium abscessus
MIKHLTRRATVALVVAAAAISPAAYPAMADTATPLNGLFALTPGSCAEGRATGSYFRMIL WP_005073213.1
hypothetical protein [Mycobacterium abscessus] Length: 330\nNumber of predicted TMHs: 2\nExp number of AAs in TMHs: 39.3416199999999\nExp number, first 60 AAs: 17.07688\nTotal prob of N-in: 0.84431\nPOSSIBLE N-term signal sequence\ninside 1 11\nTMhelix 12 29\noutside 30 298\nTMhelix 299 321\ninside 322 330

30393 GCF_900138015.1_12045_8_34 Mycobacterium abscessus subsp. abscessus Terrabacteria group; Actinobacteria; Actinobacteria; Corynebacteriales; Mycobacteriaceae; Mycobacterium; Mycobacterium chelonae group; Mycobacterium abscessus subgroup; Mycobacterium abscessus
MIKHLTRRATVALVVAAAAISPAAYPAMADTATPLNGLFALTPGSCAEGRATGSYFRMIL WP_005073213.1
hypothetical protein [Mycobacterium abscessus] Length: 330\nNumber of predicted TMHs: 2\nExp number of AAs in TMHs: 39.3416199999999\nExp number, first 60 AAs: 17.07688\nTotal prob of N-in: 0.84431\nPOSSIBLE N-term signal sequence\ninside 1 11\nTMhelix 12 29\noutside 30 298\nTMhelix 299 321\ninside 322 330

30394 GCF_900137965.1_10660_1_4 Mycobacterium abscessus subsp. abscessus Terrabacteria group; Actinobacteria; Actinobacteria; Corynebacteriales; Mycobacteriaceae; Mycobacterium; Mycobacterium chelonae group; Mycobacterium abscessus subgroup; Mycobacterium abscessus
MIKHLTRRATVALVVAAAAISPAAYPAMADTATPLNGLFALTPGSCAEGRATGSYFRMIL WP_005073213.1
hypothetical protein [Mycobacterium abscessus] Length: 330\nNumber of predicted TMHs: 2\nExp number of AAs in TMHs: 39.3416199999999\nExp number, first 60 AAs: 17.07688\nTotal prob of N-in: 0.84431\nPOSSIBLE N-term signal sequence\ninside 1 11\nTMhelix 12 29\noutside 30 298\nTMhelix 299 321\ninside 322 330

30395 GCF_900137925.1_10208_3_83 Mycobacterium abscessus subsp. abscessus Terrabacteria group; Actinobacteria; Actinobacteria; Corynebacteriales; Mycobacteriaceae; Mycobacterium; Mycobacterium chelonae group; Mycobacterium abscessus subgroup; Mycobacterium abscessus
MIKHLTRRATVALVVAAAAISPAAYPAMADTATPLNGLFALTPGSCAEGRATGSYFRMIL WP_005073213.1
hypothetical protein [Mycobacterium abscessus] Length: 330\nNumber of predicted TMHs: 2\nExp number of AAs in TMHs: 39.3416199999999\nExp number, first 60 AAs: 17.07688\nTotal prob of N-in: 0.84431\nPOSSIBLE N-term signal sequence\ninside 1 11\nTMhelix 12 29\noutside 30 298\nTMhelix 299 321\ninside 322 330

30396 GCF_900137575.1_12163_2_35 Mycobacterium abscessus subsp. abscessus Terrabacteria group; Actinobacteria; Actinobacteria; Corynebacteriales; Mycobacteriaceae; Mycobacterium; Mycobacterium chelonae group; Mycobacterium abscessus subgroup; Mycobacterium abscessus
MIKHLTRRATVALVVAAAAISPAAYPAMADTATPLNGLFALTPGSCAEGRATGSYFRMIL WP_005073213.1
hypothetical protein [Mycobacterium abscessus] Length: 330\nNumber of predicted TMHs: 2\nExp number of AAs in TMHs: 39.3416199999999\nExp number, first 60 AAs: 17.07688\nTotal prob of N-in: 0.84431\nPOSSIBLE N-term signal sequence\ninside 1 11\nTMhelix 12 29\noutside 30 298\nTMhelix 299 321\ninside 322 330

30397 GCF_900137605.1_12163_2_36 Mycobacterium abscessus subsp. abscessus Terrabacteria group; Actinobacteria; Actinobacteria; Corynebacteriales; Mycobacteriaceae; Mycobacterium; Mycobacterium chelonae group; Mycobacterium abscessus subgroup; Mycobacterium abscessus
MIKHLTRRATVALVVAAAAISPAAYPAMADTATPLNGLFALTPGSCAEGRATGSYFRMIL WP_005073213.1
hypothetical protein [Mycobacterium abscessus] Length: 330\nNumber of predicted TMHs: 2\nExp number of AAs in TMHs: 39.3416199999999\nExp number, first 60 AAs: 17.07688\nTotal prob of N-in: 0.84431\nPOSSIBLE N-term signal sequence\ninside 1 11\nTMhelix 12 29\noutside 30 298\nTMhelix 299 321\ninside 322 330

30398 GCF_900137515.1_12163_2_28 Mycobacterium abscessus subsp. abscessus Terrabacteria group; Actinobacteria; Actinobacteria; Corynebacteriales; Mycobacteriaceae; Mycobacterium; Mycobacterium chelonae group; Mycobacterium abscessus subgroup; Mycobacterium abscessus
MIKHLTRRATVALVVAAAAISPAAYPAMADTATPLNGLFALTPGSCAEGRATGSYFRMIL WP_005073213.1
hypothetical protein [Mycobacterium abscessus] Length: 330\nNumber of predicted TMHs: 2\nExp number of AAs in TMHs: 39.3416199999999\nExp number, first 60 AAs: 17.07688\nTotal prob of N-in: 0.84431\nPOSSIBLE N-term signal sequence\ninside 1 11\nTMhelix 12 29\noutside 30 298\nTMhelix 299 321\ninside 322 330

30399 GCF_900137305.1_12163_1_96 Mycobacterium abscessus subsp. abscessus Terrabacteria group; Actinobacteria; Actinobacteria; Corynebacteriales; Mycobacteriaceae; Mycobacterium; Mycobacterium chelonae group; Mycobacterium abscessus subgroup; Mycobacterium abscessus
MIKHLTRRATVALVVAAAAISPAAYPAMADTATPLNGLFALTPGSCAEGRATGSYFRMIL WP_005073213.1
hypothetical protein [Mycobacterium abscessus] Length: 330\nNumber of predicted TMHs: 2\nExp number of AAs in TMHs: 39.3416199999999\nExp number, first 60 AAs: 17.07688\nTotal prob of N-in: 0.84431\nPOSSIBLE N-term signal sequence\ninside 1 11\nTMhelix 12 29\noutside 30 298\nTMhelix 299 321\ninside 322 330

30400 GCF_900137275.1_12163_1_94 Mycobacterium abscessus subsp. abscessus Terrabacteria group; Actinobacteria; Actinobacteria; Corynebacteriales; Mycobacteriaceae; Mycobacterium; Mycobacterium chelonae group; Mycobacterium abscessus subgroup; Mycobacterium abscessus
MIKHLTRRATVALVVAAAAISPAAYPAMADTATPLNGLFALTPGSCAEGRATGSYFRMIL WP_005073213.1
hypothetical protein [Mycobacterium abscessus] Length: 330\nNumber of predicted TMHs: 2\nExp number of AAs in TMHs: 39.3416199999999\nExp number, first 60 AAs: 17.07688\nTotal prob of N-in: 0.84431\nPOSSIBLE N-term signal sequence\ninside 1 11\nTMhelix 12 29\noutside 30 298\nTMhelix 299 321\ninside 322 330

30401 GCF_900137255.1_12163_1_91 Mycobacterium abscessus subsp. abscessus Terrabacteria group; Actinobacteria; Actinobacteria; Corynebacteriales; Mycobacteriaceae; Mycobacterium; Mycobacterium chelonae group; Mycobacterium abscessus subgroup; Mycobacterium abscessus
MIKHLTRRATVALVVAAAAISPAAYPAMADTATPLNGLFALTPGSCAEGRATGSYFRMIL WP_005073213.1
hypothetical protein [Mycobacterium abscessus] Length: 330\nNumber of predicted TMHs: 2\nExp number of AAs in TMHs: 39.3416199999999\nExp number, first 60 AAs: 17.07688\nTotal prob of N-in: 0.84431\nPOSSIBLE N-term signal sequence\ninside 1 11\nTMhelix 12 29\noutside 30 298\nTMhelix 299 321\ninside 322 330

30402 GCF_900137235.1_12082_5_68 Mycobacterium abscessus subsp. abscessus Terrabacteria group; Actinobacteria; Actinobacteria; Corynebacteriales; Mycobacteriaceae; Mycobacterium; Mycobacterium chelonae group; Mycobacterium abscessus subgroup; Mycobacterium abscessus
MIKHLTRRATVALVVAAAAISPAAYPAMADTATPLNGLFALTPGSCAEGRATGSYFRMIL WP_005073213.1
hypothetical protein [Mycobacterium abscessus] Length: 330\nNumber of predicted TMHs: 2\nExp number of AAs in TMHs: 39.3416199999999\nExp number, first 60 AAs: 17.07688\nTotal prob of N-in: 0.84431\nPOSSIBLE N-term signal sequence\ninside 1 11\nTMhelix 12 29\noutside 30 298\nTMhelix 299 321\ninside 322 330

30403 GCF_900137215.1_12082_5_86 Mycobacterium abscessus subsp. abscessus Terrabacteria group; Actinobacteria; Actinobacteria; Corynebacteriales; Mycobacteriaceae; Mycobacterium; Mycobacterium chelonae group; Mycobacterium abscessus subgroup; Mycobacterium abscessus
MIKHLTRRATVALVVAAAAISPAAYPAMADTATPLNGLFALTPGSCAEGRATGSYFRMIL WP_005073213.1
hypothetical protein [Mycobacterium abscessus] Length: 330\nNumber of predicted TMHs: 2\nExp number of

AAs in TMHs: 39.3416199999999\nExp number, first 60 AAs: 17.07688\nTotal prob of N-in: 0.84431\nPOSSIBLE N-term signal sequence\ninside 1 11\nTMhelix 12 29\noutside 30 298\nTMhelix 299 321\ninside 322 330

30404 GCF_900137165.1_12078_1_67 Mycobacterium abscessus subsp. abscessus Terrabacteria group; Actinobacteria; Actinobacteria; Corynebacteriales; Mycobacteriaceae; Mycobacterium; Mycobacterium chelonae group; Mycobacterium abscessus subgroup; Mycobacterium abscessus
MIKHLTRRATVALVVAAAAISPAAYPAMADTATPLNGLFALTPGSCAEGRATGSYFRMIL WP_005073213.1
hypothetical protein [Mycobacterium abscessus] Length: 330\nNumber of predicted TMHs: 2\nExp number of AAs in TMHs: 39.3416199999999\nExp number, first 60 AAs: 17.07688\nTotal prob of N-in: 0.84431\nPOSSIBLE N-term signal sequence\ninside 1 11\nTMhelix 12 29\noutside 30 298\nTMhelix 299 321\ninside 322 330

30405 GCF_900136765.1_10625_5_34 Mycobacterium abscessus subsp. abscessus Terrabacteria group; Actinobacteria; Actinobacteria; Corynebacteriales; Mycobacteriaceae; Mycobacterium; Mycobacterium chelonae group; Mycobacterium abscessus subgroup; Mycobacterium abscessus
MIKHLTRRATVALVVAAAAISPAAYPAMADTATPLNGLFALTPGSCAEGRATGSYFRMIL WP_005073213.1
hypothetical protein [Mycobacterium abscessus] Length: 330\nNumber of predicted TMHs: 2\nExp number of AAs in TMHs: 39.3416199999999\nExp number, first 60 AAs: 17.07688\nTotal prob of N-in: 0.84431\nPOSSIBLE N-term signal sequence\ninside 1 11\nTMhelix 12 29\noutside 30 298\nTMhelix 299 321\ninside 322 330

30406 GCF_900136705.1_10625_5_24 Mycobacterium abscessus subsp. abscessus Terrabacteria group; Actinobacteria; Actinobacteria; Corynebacteriales; Mycobacteriaceae; Mycobacterium; Mycobacterium chelonae group; Mycobacterium abscessus subgroup; Mycobacterium abscessus
MIKHLTRRATVALVVAAAAISPAAYPAMADTATPLNGLFALTPGSCAEGRATGSYFRMIL WP_005073213.1
hypothetical protein [Mycobacterium abscessus] Length: 330\nNumber of predicted TMHs: 2\nExp number of AAs in TMHs: 39.3416199999999\nExp number, first 60 AAs: 17.07688\nTotal prob of N-in: 0.84431\nPOSSIBLE N-term signal sequence\ninside 1 11\nTMhelix 12 29\noutside 30 298\nTMhelix 299 321\ninside 322 330

30407 GCF_900136685.1_10625_5_23 Mycobacterium abscessus subsp. abscessus Terrabacteria group; Actinobacteria; Actinobacteria; Corynebacteriales; Mycobacteriaceae; Mycobacterium; Mycobacterium chelonae group; Mycobacterium abscessus subgroup; Mycobacterium abscessus
MIKHLTRRATVALVVAAAAISPAAYPAMADTATPLNGLFALTPGSCAEGRATGSYFRMIL WP_005073213.1
hypothetical protein [Mycobacterium abscessus] Length: 330\nNumber of predicted TMHs: 2\nExp number of AAs in TMHs: 39.3416199999999\nExp number, first 60 AAs: 17.07688\nTotal prob of N-in: 0.84431\nPOSSIBLE N-term signal sequence\ninside 1 11\nTMhelix 12 29\noutside 30 298\nTMhelix 299 321\ninside 322 330

30408 GCF_900136665.1_10625_5_21 Mycobacterium abscessus subsp. abscessus Terrabacteria group; Actinobacteria; Actinobacteria; Corynebacteriales; Mycobacteriaceae; Mycobacterium; Mycobacterium chelonae group; Mycobacterium abscessus subgroup; Mycobacterium abscessus
MIKHLTRRATVALVVAAAAISPAAYPAMADTATPLNGLFALTPGSCAEGRATGSYFRMIL WP_005073213.1
hypothetical protein [Mycobacterium abscessus] Length: 330\nNumber of predicted TMHs: 2\nExp number of AAs in TMHs: 39.3416199999999\nExp number, first 60 AAs: 17.07688\nTotal prob of N-in: 0.84431\nPOSSIBLE N-term signal sequence\ninside 1 11\nTMhelix 12 29\noutside 30 298\nTMhelix 299 321\ninside 322 330

30409 GCF_900136365.1_12163_2_74 Mycobacterium abscessus subsp. abscessus Terrabacteria group; Actinobacteria; Actinobacteria; Corynebacteriales; Mycobacteriaceae; Mycobacterium; Mycobacterium chelonae group; Mycobacterium abscessus subgroup; Mycobacterium abscessus
MIKHLTRRATVALVVAAAAISPAAYPAMADTATPLNGLFALTPGSCAEGRATGSYFRMIL WP_005073213.1
hypothetical protein [Mycobacterium abscessus] Length: 330\nNumber of predicted TMHs: 2\nExp number of AAs in TMHs: 39.3416199999999\nExp number, first 60 AAs: 17.07688\nTotal prob of N-in: 0.84431\nPOSSIBLE N-term signal sequence\ninside 1 11\nTMhelix 12 29\noutside 30 298\nTMhelix 299 321\ninside 322 330

30410 GCF_900136385.1_12163_2_84 Mycobacterium abscessus subsp. abscessus Terrabacteria group; Actinobacteria; Actinobacteria; Corynebacteriales; Mycobacteriaceae; Mycobacterium; Mycobacterium chelonae group; Mycobacterium abscessus subgroup; Mycobacterium abscessus
MIKHLTRRATVALVVAAAAISPAAYPAMADTATPLNGLFALTPGSCAEGRATGSYFRMIL WP_005073213.1
hypothetical protein [Mycobacterium abscessus] Length: 330\nNumber of predicted TMHs: 2\nExp number of AAs in TMHs: 39.3416199999999\nExp number, first 60 AAs: 17.07688\nTotal prob of N-in: 0.84431\nPOSSIBLE N-term signal sequence\ninside 1 11\nTMhelix 12 29\noutside 30 298\nTMhelix 299 321\ninside 322 330

30411 GCF_900136305.1_12163_1_87 Mycobacterium abscessus subsp. abscessus Terrabacteria group; Actinobacteria; Actinobacteria; Corynebacteriales; Mycobacteriaceae; Mycobacterium; Mycobacterium chelonae group;

Mycobacterium abscessus subgroup; Mycobacterium abscessus
MIKHLTRRATVALVVAAAAISPAAYPAMADTATPLNGLFALTPGSCAEGRATGSYFRMIL WP_005073213.1 hypothetical protein [Mycobacterium abscessus] Length: 330\nNumber of predicted TMHs: 2\nExp number of AAs in TMHs: 39.3416199999999\nExp number, first 60 AAs: 17.07688\nTotal prob of N-in: 0.84431\nPOSSIBLE N-term signal sequence\ninside 1 11\nTMhelix 12 29\noutside 30 298\nTMhelix 299 321\ninside 322 330

30412 GCF_900136285.1_12163_1_84 Mycobacterium abscessus subsp. abscessus Terrabacteria group; Actinobacteria; Actinobacteria; Corynebacteriales; Mycobacteriaceae; Mycobacterium; Mycobacterium chelonae group; Mycobacterium abscessus subgroup; Mycobacterium abscessus
MIKHLTRRATVALVVAAAAISPAAYPAMADTATPLNGLFALTPGSCAEGRATGSYFRMIL WP_005073213.1 hypothetical protein [Mycobacterium abscessus] Length: 330\nNumber of predicted TMHs: 2\nExp number of AAs in TMHs: 39.3416199999999\nExp number, first 60 AAs: 17.07688\nTotal prob of N-in: 0.84431\nPOSSIBLE N-term signal sequence\ninside 1 11\nTMhelix 12 29\noutside 30 298\nTMhelix 299 321\ninside 322 330

30413 GCF_900136265.1_12163_1_82 Mycobacterium abscessus subsp. abscessus Terrabacteria group; Actinobacteria; Actinobacteria; Corynebacteriales; Mycobacteriaceae; Mycobacterium; Mycobacterium chelonae group; Mycobacterium abscessus subgroup; Mycobacterium abscessus
MIKHLTRRATVALVVAAAAISPAAYPAMADTATPLNGLFALTPGSCAEGRATGSYFRMIL WP_005073213.1 hypothetical protein [Mycobacterium abscessus] Length: 330\nNumber of predicted TMHs: 2\nExp number of AAs in TMHs: 39.3416199999999\nExp number, first 60 AAs: 17.07688\nTotal prob of N-in: 0.84431\nPOSSIBLE N-term signal sequence\ninside 1 11\nTMhelix 12 29\noutside 30 298\nTMhelix 299 321\ninside 322 330

30414 GCF_900135975.1_12163_1_55 Mycobacterium abscessus subsp. abscessus Terrabacteria group; Actinobacteria; Actinobacteria; Corynebacteriales; Mycobacteriaceae; Mycobacterium; Mycobacterium chelonae group; Mycobacterium abscessus subgroup; Mycobacterium abscessus
MIKHLTRRATVALVVAAAAISPAAYPAMADTATPLNGLFALTPGSCAEGRATGSYFRMIL WP_005073213.1 hypothetical protein [Mycobacterium abscessus] Length: 330\nNumber of predicted TMHs: 2\nExp number of AAs in TMHs: 39.3416199999999\nExp number, first 60 AAs: 17.07688\nTotal prob of N-in: 0.84431\nPOSSIBLE N-term signal sequence\ninside 1 11\nTMhelix 12 29\noutside 30 298\nTMhelix 299 321\ninside 322 330

30415 GCF_900135925.1_12082_5_62 Mycobacterium abscessus subsp. abscessus Terrabacteria group; Actinobacteria; Actinobacteria; Corynebacteriales; Mycobacteriaceae; Mycobacterium; Mycobacterium chelonae group; Mycobacterium abscessus subgroup; Mycobacterium abscessus
MIKHLTRRATVALVVAAAAISPAAYPAMADTATPLNGLFALTPGSCAEGRATGSYFRMIL WP_005073213.1 hypothetical protein [Mycobacterium abscessus] Length: 330\nNumber of predicted TMHs: 2\nExp number of AAs in TMHs: 39.3416199999999\nExp number, first 60 AAs: 17.07688\nTotal prob of N-in: 0.84431\nPOSSIBLE N-term signal sequence\ninside 1 11\nTMhelix 12 29\noutside 30 298\nTMhelix 299 321\ninside 322 330

30416 GCF_900135705.1_11861_7_69 Mycobacterium abscessus subsp. abscessus Terrabacteria group; Actinobacteria; Actinobacteria; Corynebacteriales; Mycobacteriaceae; Mycobacterium; Mycobacterium chelonae group; Mycobacterium abscessus subgroup; Mycobacterium abscessus
MPRRRTILSALASFSMVIGLLVVSTPTAWAAPFPVVDPAKLPPSQPPAPPGDMRLNGAPL WP_074362414.1 type VII secretion-associated serine protease mycosin [Mycobacterium abscessus] Length: 470\nNumber of predicted TMHs: 2\nExp number of AAs in TMHs: 44.7482799999999\nExp number, first 60 AAs: 22.65577\nTotal prob of N-in: 0.99532\nPOSSIBLE N-term signal sequence\ninside 1 6\nTMhelix 7 29\noutside 30 441\nTMhelix 442 464\ninside 465 470

30417 GCF_900135475.1_10625_4_68 Mycobacterium abscessus subsp. abscessus Terrabacteria group; Actinobacteria; Actinobacteria; Corynebacteriales; Mycobacteriaceae; Mycobacterium; Mycobacterium chelonae group; Mycobacterium abscessus subgroup; Mycobacterium abscessus
MIKHLTRRATVALVVAAAAISPAAYPAMADTATPLNGLFALTPGSCAEGRATGSYFRMIL WP_005073213.1 hypothetical protein [Mycobacterium abscessus] Length: 330\nNumber of predicted TMHs: 2\nExp number of AAs in TMHs: 39.3416199999999\nExp number, first 60 AAs: 17.07688\nTotal prob of N-in: 0.84431\nPOSSIBLE N-term signal sequence\ninside 1 11\nTMhelix 12 29\noutside 30 298\nTMhelix 299 321\ninside 322 330

30418 GCF_900135455.1_10625_4_63 Mycobacterium abscessus subsp. abscessus Terrabacteria group; Actinobacteria; Actinobacteria; Corynebacteriales; Mycobacteriaceae; Mycobacterium; Mycobacterium chelonae group; Mycobacterium abscessus subgroup; Mycobacterium abscessus
MIKHLTRRATVALVVAAAAISPAAYPAMADTATPLNGLFALTPGSCAEGRATGSYFRMIL WP_005073213.1 hypothetical protein [Mycobacterium abscessus] Length: 330\nNumber of predicted TMHs: 2\nExp number of

AAs in TMHs: 39.3416199999999\nExp number, first 60 AAs: 17.07688\nTotal prob of N-in: 0.84431\nPOSSIBLE N-term signal sequence\ninside 1 11\nTMhelix 12 29\noutside 30 298\nTMhelix 299 321\ninside 322 330

30419 GCF_900135385.1_10625_4_55 Mycobacterium abscessus subsp. abscessus Terrabacteria group; Actinobacteria; Actinobacteria; Corynebacteriales; Mycobacteriaceae; Mycobacterium; Mycobacterium chelonae group; Mycobacterium abscessus subgroup; Mycobacterium abscessus
MIKHLTRRATVALVVAAAAISPAAYPAMADTATPLNGLFALTPGSCAEGRATGSYFRMIL WP_005073213.1
hypothetical protein [Mycobacterium abscessus] Length: 330\nNumber of predicted TMHs: 2\nExp number of AAs in TMHs: 39.3416199999999\nExp number, first 60 AAs: 17.07688\nTotal prob of N-in: 0.84431\nPOSSIBLE N-term signal sequence\ninside 1 11\nTMhelix 12 29\noutside 30 298\nTMhelix 299 321\ninside 322 330

30420 GCF_900135365.1_10625_4_50 Mycobacterium abscessus subsp. abscessus Terrabacteria group; Actinobacteria; Actinobacteria; Corynebacteriales; Mycobacteriaceae; Mycobacterium; Mycobacterium chelonae group; Mycobacterium abscessus subgroup; Mycobacterium abscessus
MIKHLTRRATVALVVAAAAISPAAYPAMADTATPLNGLFALTPGSCAEGRATGSYFRMIL WP_005073213.1
hypothetical protein [Mycobacterium abscessus] Length: 330\nNumber of predicted TMHs: 2\nExp number of AAs in TMHs: 39.3416199999999\nExp number, first 60 AAs: 17.07688\nTotal prob of N-in: 0.84431\nPOSSIBLE N-term signal sequence\ninside 1 11\nTMhelix 12 29\noutside 30 298\nTMhelix 299 321\ninside 322 330

30421 GCF_900135135.1_10625_4_22 Mycobacterium abscessus subsp. abscessus Terrabacteria group; Actinobacteria; Actinobacteria; Corynebacteriales; Mycobacteriaceae; Mycobacterium; Mycobacterium chelonae group; Mycobacterium abscessus subgroup; Mycobacterium abscessus
MIKHLTRRATVALVVAAAAISPAAYPAMADTATPLNGLFALTPGSCAEGRATGSYFRMIL WP_005073213.1
hypothetical protein [Mycobacterium abscessus] Length: 330\nNumber of predicted TMHs: 2\nExp number of AAs in TMHs: 39.3416199999999\nExp number, first 60 AAs: 17.07688\nTotal prob of N-in: 0.84431\nPOSSIBLE N-term signal sequence\ninside 1 11\nTMhelix 12 29\noutside 30 298\nTMhelix 299 321\ninside 322 330

30422 GCF_900135085.1_10250_1_84 Mycobacterium abscessus subsp. abscessus Terrabacteria group; Actinobacteria; Actinobacteria; Corynebacteriales; Mycobacteriaceae; Mycobacterium; Mycobacterium chelonae group; Mycobacterium abscessus subgroup; Mycobacterium abscessus
MIKHLTRRATVALVVAAAAISPAAYPAMADTATPLNGLFALTPGSCAEGRATGSYFRMIL WP_005073213.1
hypothetical protein [Mycobacterium abscessus] Length: 330\nNumber of predicted TMHs: 2\nExp number of AAs in TMHs: 39.3416199999999\nExp number, first 60 AAs: 17.07688\nTotal prob of N-in: 0.84431\nPOSSIBLE N-term signal sequence\ninside 1 11\nTMhelix 12 29\noutside 30 298\nTMhelix 299 321\ninside 322 330

30423 GCF_900135065.1_10250_1_83 Mycobacterium abscessus subsp. abscessus Terrabacteria group; Actinobacteria; Actinobacteria; Corynebacteriales; Mycobacteriaceae; Mycobacterium; Mycobacterium chelonae group; Mycobacterium abscessus subgroup; Mycobacterium abscessus
MIKHLTRRATVALVVAAAAISPAAYPAMADTATPLNGLFALTPGSCAEGRATGSYFRMIL WP_005073213.1
hypothetical protein [Mycobacterium abscessus] Length: 330\nNumber of predicted TMHs: 2\nExp number of AAs in TMHs: 39.3416199999999\nExp number, first 60 AAs: 17.07688\nTotal prob of N-in: 0.84431\nPOSSIBLE N-term signal sequence\ninside 1 11\nTMhelix 12 29\noutside 30 298\nTMhelix 299 321\ninside 322 330

30424 GCF_900135105.1_10250_1_9 Mycobacterium abscessus subsp. abscessus Terrabacteria group; Actinobacteria; Actinobacteria; Corynebacteriales; Mycobacteriaceae; Mycobacterium; Mycobacterium chelonae group; Mycobacterium abscessus subgroup; Mycobacterium abscessus
MIKHLTRRATVALVVAAAAISPAAYPAMADTATPLNGLFALTPGSCAEGRATGSYFRMIL WP_005073213.1
hypothetical protein [Mycobacterium abscessus] Length: 330\nNumber of predicted TMHs: 2\nExp number of AAs in TMHs: 39.3416199999999\nExp number, first 60 AAs: 17.07688\nTotal prob of N-in: 0.84431\nPOSSIBLE N-term signal sequence\ninside 1 11\nTMhelix 12 29\noutside 30 298\nTMhelix 299 321\ninside 322 330

30425 GCF_900134735.1_10208_3_15 Mycobacterium abscessus subsp. abscessus Terrabacteria group; Actinobacteria; Actinobacteria; Corynebacteriales; Mycobacteriaceae; Mycobacterium; Mycobacterium chelonae group; Mycobacterium abscessus subgroup; Mycobacterium abscessus
MIKHLTRRATVALVVAAAAISPAAYPAMADTATPLNGLFALTPGSCAEGRATGSYFRMIL WP_005073213.1
hypothetical protein [Mycobacterium abscessus] Length: 330\nNumber of predicted TMHs: 2\nExp number of AAs in TMHs: 39.3416199999999\nExp number, first 60 AAs: 17.07688\nTotal prob of N-in: 0.84431\nPOSSIBLE N-term signal sequence\ninside 1 11\nTMhelix 12 29\noutside 30 298\nTMhelix 299 321\ninside 322 330

30426 GCF_900134705.1_10208_3_10 Mycobacterium abscessus subsp. abscessus Terrabacteria group; Actinobacteria; Actinobacteria; Corynebacteriales; Mycobacteriaceae; Mycobacterium; Mycobacterium chelonae group;

Mycobacterium abscessus subgroup; Mycobacterium abscessus
MIKHLTRRATVALVVAAAAISPAAYPAMADTATPLNGLFALTPGSCAEGRATGSYFRMIL WP_005073213.1
hypothetical protein [Mycobacterium abscessus] Length: 330\nNumber of predicted TMHs: 2\nExp number of AAs in TMHs: 39.3416199999999\nExp number, first 60 AAs: 17.07688\nTotal prob of N-in: 0.84431\nPOSSIBLE N-term signal sequence\ninside 1 11\nTMhelix 12 29\noutside 30 298\nTMhelix 299 321\ninside 322 330

30427 GCF_900134625.1_12163_1_52 Mycobacterium abscessus subsp. abscessus Terrabacteria group; Actinobacteria; Actinobacteria; Corynebacteriales; Mycobacteriaceae; Mycobacterium; Mycobacterium chelonae group; Mycobacterium abscessus subgroup; Mycobacterium abscessus
MIKHLTRRATVALVVAAAAISPAAYPAMADTATPLNGLFALTPGSCAEGRATGSYFRMIL WP_005073213.1
hypothetical protein [Mycobacterium abscessus] Length: 330\nNumber of predicted TMHs: 2\nExp number of AAs in TMHs: 39.3416199999999\nExp number, first 60 AAs: 17.07688\nTotal prob of N-in: 0.84431\nPOSSIBLE N-term signal sequence\ninside 1 11\nTMhelix 12 29\noutside 30 298\nTMhelix 299 321\ninside 322 330

30428 GCF_900134645.1_12163_2_59 Mycobacterium abscessus subsp. abscessus Terrabacteria group; Actinobacteria; Actinobacteria; Corynebacteriales; Mycobacteriaceae; Mycobacterium; Mycobacterium chelonae group; Mycobacterium abscessus subgroup; Mycobacterium abscessus
MIKHLTRRATVALVVAAAAISPAAYPAMADTATPLNGLFALTPGSCAEGRATGSYFRMIL WP_005073213.1
hypothetical protein [Mycobacterium abscessus] Length: 330\nNumber of predicted TMHs: 2\nExp number of AAs in TMHs: 39.3416199999999\nExp number, first 60 AAs: 17.07688\nTotal prob of N-in: 0.84431\nPOSSIBLE N-term signal sequence\ninside 1 11\nTMhelix 12 29\noutside 30 298\nTMhelix 299 321\ninside 322 330

30429 GCF_900134605.1_12163_1_53 Mycobacterium abscessus subsp. abscessus Terrabacteria group; Actinobacteria; Actinobacteria; Corynebacteriales; Mycobacteriaceae; Mycobacterium; Mycobacterium chelonae group; Mycobacterium abscessus subgroup; Mycobacterium abscessus
MIKHLTRRATVALVVAAAAISPAAYPAMADTATPLNGLFALTPGSCAEGRATGSYFRMIL WP_005073213.1
hypothetical protein [Mycobacterium abscessus] Length: 330\nNumber of predicted TMHs: 2\nExp number of AAs in TMHs: 39.3416199999999\nExp number, first 60 AAs: 17.07688\nTotal prob of N-in: 0.84431\nPOSSIBLE N-term signal sequence\ninside 1 11\nTMhelix 12 29\noutside 30 298\nTMhelix 299 321\ninside 322 330

30430 GCF_900134355.1_12082_5_60 Mycobacterium abscessus subsp. abscessus Terrabacteria group; Actinobacteria; Actinobacteria; Corynebacteriales; Mycobacteriaceae; Mycobacterium; Mycobacterium chelonae group; Mycobacterium abscessus subgroup; Mycobacterium abscessus
MIKHLTRRATVALVVAAAAISPAAYPAMADTATPLNGLFALTPGSCAEGRATGSYFRMIL WP_005073213.1
hypothetical protein [Mycobacterium abscessus] Length: 330\nNumber of predicted TMHs: 2\nExp number of AAs in TMHs: 39.3416199999999\nExp number, first 60 AAs: 17.07688\nTotal prob of N-in: 0.84431\nPOSSIBLE N-term signal sequence\ninside 1 11\nTMhelix 12 29\noutside 30 298\nTMhelix 299 321\ninside 322 330

30431 GCF_900134155.1_11861_7_48 Mycobacterium abscessus subsp. abscessus Terrabacteria group; Actinobacteria; Actinobacteria; Corynebacteriales; Mycobacteriaceae; Mycobacterium; Mycobacterium chelonae group; Mycobacterium abscessus subgroup; Mycobacterium abscessus
MIKHLTRRATVALVVAAAAISPAAYPAMADTATPLNGLFALTPGSCAEGRATGSYFRMIL WP_005073213.1
hypothetical protein [Mycobacterium abscessus] Length: 330\nNumber of predicted TMHs: 2\nExp number of AAs in TMHs: 39.3416199999999\nExp number, first 60 AAs: 17.07688\nTotal prob of N-in: 0.84431\nPOSSIBLE N-term signal sequence\ninside 1 11\nTMhelix 12 29\noutside 30 298\nTMhelix 299 321\ninside 322 330

30432 GCF_900133845.1_10250_1_57 Mycobacterium abscessus subsp. abscessus Terrabacteria group; Actinobacteria; Actinobacteria; Corynebacteriales; Mycobacteriaceae; Mycobacterium; Mycobacterium chelonae group; Mycobacterium abscessus subgroup; Mycobacterium abscessus
MIKHLTRRATVALVVAAAAISPAAYPAMADTATPLNGLFALTPGSCAEGRATGSYFRMIL WP_005073213.1
hypothetical protein [Mycobacterium abscessus] Length: 330\nNumber of predicted TMHs: 2\nExp number of AAs in TMHs: 39.3416199999999\nExp number, first 60 AAs: 17.07688\nTotal prob of N-in: 0.84431\nPOSSIBLE N-term signal sequence\ninside 1 11\nTMhelix 12 29\noutside 30 298\nTMhelix 299 321\ninside 322 330

30433 GCF_900133775.1_12163_1_22 Mycobacterium abscessus subsp. abscessus Terrabacteria group; Actinobacteria; Actinobacteria; Corynebacteriales; Mycobacteriaceae; Mycobacterium; Mycobacterium chelonae group; Mycobacterium abscessus subgroup; Mycobacterium abscessus
MIKHLTRRATVALVVAAAAISPAAYPAMADTATPLNGLFALTPGSCAEGRATGSYFRMIL WP_005073213.1
hypothetical protein [Mycobacterium abscessus] Length: 330\nNumber of predicted TMHs: 2\nExp number of AAs in TMHs: 39.3416199999999\nExp number, first 60 AAs: 17.07688\nTotal prob of N-in: 0.84431\nPOSSIBLE N-term signal sequence\ninside 1 11\nTMhelix 12 29\noutside 30 298\nTMhelix 299 321\ninside 322 330

30434 GCF_900133405.1_10250_1_48 Mycobacterium abscessus subsp. abscessus Terrabacteria group; Actinobacteria; Actinobacteria; Corynebacteriales; Mycobacteriaceae; Mycobacterium; Mycobacterium chelonae group; Mycobacterium abscessus subgroup; Mycobacterium abscessus
MIKHLTRRATVALVVAAAAISPAAYPAMADTATPLNGLFALTPGSCAEGRATGSYFRMIL WP_005073213.1
hypothetical protein [Mycobacterium abscessus] Length: 330\nNumber of predicted TMHs: 2\nExp number of AAs in TMHs: 39.3416199999999\nExp number, first 60 AAs: 17.07688\nTotal prob of N-in: 0.84431\nPOSSIBLE N-term signal sequence\ninside 1 11\nTMhelix 12 29\noutside 30 298\nTMhelix 299 321\ninside 322 330

30435 GCF_900133385.1_10250_1_45 Mycobacterium abscessus subsp. abscessus Terrabacteria group; Actinobacteria; Actinobacteria; Corynebacteriales; Mycobacteriaceae; Mycobacterium; Mycobacterium chelonae group; Mycobacterium abscessus subgroup; Mycobacterium abscessus
MIKHLTRRATVALVVAAAAISPAAYPAMADTATPLNGLFALTPGSCAEGRATGSYFRMIL WP_005073213.1
hypothetical protein [Mycobacterium abscessus] Length: 330\nNumber of predicted TMHs: 2\nExp number of AAs in TMHs: 39.3416199999999\nExp number, first 60 AAs: 17.07688\nTotal prob of N-in: 0.84431\nPOSSIBLE N-term signal sequence\ninside 1 11\nTMhelix 12 29\noutside 30 298\nTMhelix 299 321\ninside 322 330

30436 GCF_900133365.1_10250_1_42 Mycobacterium abscessus subsp. abscessus Terrabacteria group; Actinobacteria; Actinobacteria; Corynebacteriales; Mycobacteriaceae; Mycobacterium; Mycobacterium chelonae group; Mycobacterium abscessus subgroup; Mycobacterium abscessus
MIKHLTRRATVALVVAAAAISPAAYPAMADTATPLNGLFALTPGSCAEGRATGSYFRMIL WP_005073213.1
hypothetical protein [Mycobacterium abscessus] Length: 330\nNumber of predicted TMHs: 2\nExp number of AAs in TMHs: 39.3416199999999\nExp number, first 60 AAs: 17.07688\nTotal prob of N-in: 0.84431\nPOSSIBLE N-term signal sequence\ninside 1 11\nTMhelix 12 29\noutside 30 298\nTMhelix 299 321\ninside 322 330

30437 GCF_900133315.1_10250_1_38 Mycobacterium abscessus subsp. abscessus Terrabacteria group; Actinobacteria; Actinobacteria; Corynebacteriales; Mycobacteriaceae; Mycobacterium; Mycobacterium chelonae group; Mycobacterium abscessus subgroup; Mycobacterium abscessus
MIKHLTRRATVALVVAAAAISPAAYPAMADTATPLNGLFALTPGSCAEGRATGSYFRMIL WP_005073213.1
hypothetical protein [Mycobacterium abscessus] Length: 330\nNumber of predicted TMHs: 2\nExp number of AAs in TMHs: 39.3416199999999\nExp number, first 60 AAs: 17.07688\nTotal prob of N-in: 0.84431\nPOSSIBLE N-term signal sequence\ninside 1 11\nTMhelix 12 29\noutside 30 298\nTMhelix 299 321\ninside 322 330

30438 GCF_900132995.1_11893_6_18 Mycobacterium abscessus subsp. abscessus Terrabacteria group; Actinobacteria; Actinobacteria; Corynebacteriales; Mycobacteriaceae; Mycobacterium; Mycobacterium chelonae group; Mycobacterium abscessus subgroup; Mycobacterium abscessus
MIKHLTRRATVALVVAAAAISPAAYPAMADTATPLNGLFALTPGSCAEGRATGSYFRMIL WP_005073213.1
hypothetical protein [Mycobacterium abscessus] Length: 330\nNumber of predicted TMHs: 2\nExp number of AAs in TMHs: 39.3416199999999\nExp number, first 60 AAs: 17.07688\nTotal prob of N-in: 0.84431\nPOSSIBLE N-term signal sequence\ninside 1 11\nTMhelix 12 29\noutside 30 298\nTMhelix 299 321\ninside 322 330

30439 GCF_900132865.1_10665_4_84 Mycobacterium abscessus subsp. abscessus Terrabacteria group; Actinobacteria; Actinobacteria; Corynebacteriales; Mycobacteriaceae; Mycobacterium; Mycobacterium chelonae group; Mycobacterium abscessus subgroup; Mycobacterium abscessus
MIKHLTRRATVALVVAAAAISPAAYPAMADTATPLNGLFALTPGSCAEGRATGSYFRMIL WP_005073213.1
hypothetical protein [Mycobacterium abscessus] Length: 330\nNumber of predicted TMHs: 2\nExp number of AAs in TMHs: 39.3416199999999\nExp number, first 60 AAs: 17.07688\nTotal prob of N-in: 0.84431\nPOSSIBLE N-term signal sequence\ninside 1 11\nTMhelix 12 29\noutside 30 298\nTMhelix 299 321\ninside 322 330

30440 GCF_900132545.1_11893_6_42 Mycobacterium abscessus subsp. abscessus Terrabacteria group; Actinobacteria; Actinobacteria; Corynebacteriales; Mycobacteriaceae; Mycobacterium; Mycobacterium chelonae group; Mycobacterium abscessus subgroup; Mycobacterium abscessus
MIKHLTRRATVALVVAAAAISPAAYPAMADTATPLNGLFALTPGSCAEGRATGSYFRMIL WP_005073213.1
hypothetical protein [Mycobacterium abscessus] Length: 330\nNumber of predicted TMHs: 2\nExp number of AAs in TMHs: 39.3416199999999\nExp number, first 60 AAs: 17.07688\nTotal prob of N-in: 0.84431\nPOSSIBLE N-term signal sequence\ninside 1 11\nTMhelix 12 29\noutside 30 298\nTMhelix 299 321\ninside 322 330

30441 GCF_900132315.1_10665_4_63 Mycobacterium abscessus subsp. abscessus Terrabacteria group; Actinobacteria; Actinobacteria; Corynebacteriales; Mycobacteriaceae; Mycobacterium; Mycobacterium chelonae group; Mycobacterium abscessus subgroup; Mycobacterium abscessus
MIKHLTRRATVALVVAAAAISPAAYPAMADTATPLNGLFALTPGSCAEGRATGSYFRMIL WP_074250269.1
hypothetical protein [Mycobacterium abscessus] Length: 330\nNumber of predicted TMHs: 2\nExp number of

AAs in TMHs: 39.3415199999999\nExp number, first 60 AAs: 17.07715\nTotal prob of N-in: 0.84431\nPOSSIBLE N-term signal sequence\ninside 1 11\nTMhelix 12 29\noutside 30 298\nTMhelix 299 321\ninside 322 330

30442 GCF_900132425.1_10665_4_78 Mycobacterium abscessus subsp. abscessus Terrabacteria group; Actinobacteria; Actinobacteria; Corynebacteriales; Mycobacteriaceae; Mycobacterium; Mycobacterium chelonae group; Mycobacterium abscessus subgroup; Mycobacterium abscessus
MIKHLTRRATVALVVAAAAISPAAYPAMADTATPLNGLFALTPGSCAEGRATGSYFRMIL WP_005073213.1
hypothetical protein [Mycobacterium abscessus] Length: 330\nNumber of predicted TMHs: 2\nExp number of AAs in TMHs: 39.3416199999999\nExp number, first 60 AAs: 17.07688\nTotal prob of N-in: 0.84431\nPOSSIBLE N-term signal sequence\ninside 1 11\nTMhelix 12 29\noutside 30 298\nTMhelix 299 321\ninside 322 330

30443 GCF_900131845.1_11861_8_25 Mycobacterium abscessus subsp. abscessus Terrabacteria group; Actinobacteria; Actinobacteria; Corynebacteriales; Mycobacteriaceae; Mycobacterium; Mycobacterium chelonae group; Mycobacterium abscessus subgroup; Mycobacterium abscessus
MIKHLTRRATVALVVAAAAISPAAYPAMADTATPLNGLFALTPGSCAEGRATGSYFRMIL WP_074356318.1
hypothetical protein [Mycobacterium abscessus] Length: 330\nNumber of predicted TMHs: 2\nExp number of AAs in TMHs: 39.34118\nExp number, first 60 AAs: 17.07668\nTotal prob of N-in: 0.84431\nPOSSIBLE N-term signal sequence\ninside 1 11\nTMhelix 12 29\noutside 30 298\nTMhelix 299 321\ninside 322 330

30444 GCF_900131825.1_11861_7_32 Mycobacterium abscessus subsp. abscessus Terrabacteria group; Actinobacteria; Actinobacteria; Corynebacteriales; Mycobacteriaceae; Mycobacterium; Mycobacterium chelonae group; Mycobacterium abscessus subgroup; Mycobacterium abscessus
MIKHLTRRATVALVVAAAAISPAAYPAMADTATPLNGLFALTPGSCAEGRATGSYFRMIL WP_074340136.1
hypothetical protein [Mycobacterium abscessus] Length: 330\nNumber of predicted TMHs: 2\nExp number of AAs in TMHs: 39.3416799999999\nExp number, first 60 AAs: 17.07671\nTotal prob of N-in: 0.84432\nPOSSIBLE N-term signal sequence\ninside 1 11\nTMhelix 12 29\noutside 30 298\nTMhelix 299 321\ninside 322 330

30445 GCF_900132045.1_10665_4_49 Mycobacterium abscessus subsp. abscessus Terrabacteria group; Actinobacteria; Actinobacteria; Corynebacteriales; Mycobacteriaceae; Mycobacterium; Mycobacterium chelonae group; Mycobacterium abscessus subgroup; Mycobacterium abscessus
MIKHLTRRATVALVVAAAAISPAAYPAMADTATPLNGLFALTPGSCAEGRATGSYFRMIL WP_005073213.1
hypothetical protein [Mycobacterium abscessus] Length: 330\nNumber of predicted TMHs: 2\nExp number of AAs in TMHs: 39.3416199999999\nExp number, first 60 AAs: 17.07688\nTotal prob of N-in: 0.84431\nPOSSIBLE N-term signal sequence\ninside 1 11\nTMhelix 12 29\noutside 30 298\nTMhelix 299 321\ninside 322 330

30446 GCF_900131605.1_10665_4_17 Mycobacterium abscessus subsp. abscessus Terrabacteria group; Actinobacteria; Actinobacteria; Corynebacteriales; Mycobacteriaceae; Mycobacterium; Mycobacterium chelonae group; Mycobacterium abscessus subgroup; Mycobacterium abscessus
MIKHLTRRATVALVVAAAAISPAAYPAMADTATPLNGLFALTPGSCAEGRATGSYFRMIL WP_005073213.1
hypothetical protein [Mycobacterium abscessus] Length: 330\nNumber of predicted TMHs: 2\nExp number of AAs in TMHs: 39.3416199999999\nExp number, first 60 AAs: 17.07688\nTotal prob of N-in: 0.84431\nPOSSIBLE N-term signal sequence\ninside 1 11\nTMhelix 12 29\noutside 30 298\nTMhelix 299 321\ninside 322 330

30447 GCF_900131175.1_10665_3_62 Mycobacterium abscessus subsp. abscessus Terrabacteria group; Actinobacteria; Actinobacteria; Corynebacteriales; Mycobacteriaceae; Mycobacterium; Mycobacterium chelonae group; Mycobacterium abscessus subgroup; Mycobacterium abscessus
MIKHLTRRATVALVVAAAAISPAAYPAMADTATPLNGLFALTPGSCAEGRATGSYFRMIL WP_005073213.1
hypothetical protein [Mycobacterium abscessus] Length: 330\nNumber of predicted TMHs: 2\nExp number of AAs in TMHs: 39.3416199999999\nExp number, first 60 AAs: 17.07688\nTotal prob of N-in: 0.84431\nPOSSIBLE N-term signal sequence\ninside 1 11\nTMhelix 12 29\noutside 30 298\nTMhelix 299 321\ninside 322 330

30448 GCF_900131145.1_10665_3_59 Mycobacterium abscessus subsp. abscessus Terrabacteria group; Actinobacteria; Actinobacteria; Corynebacteriales; Mycobacteriaceae; Mycobacterium; Mycobacterium chelonae group; Mycobacterium abscessus subgroup; Mycobacterium abscessus
MIKHLTRRATVALVVAAAAISPAAYPAMADTATPLNGLFALTPGSCAEGRATGSYFRMIL WP_005073213.1
hypothetical protein [Mycobacterium abscessus] Length: 330\nNumber of predicted TMHs: 2\nExp number of AAs in TMHs: 39.3416199999999\nExp number, first 60 AAs: 17.07688\nTotal prob of N-in: 0.84431\nPOSSIBLE N-term signal sequence\ninside 1 11\nTMhelix 12 29\noutside 30 298\nTMhelix 299 321\ninside 322 330

30449 GCF_900131075.1_10665_3_53 Mycobacterium abscessus subsp. abscessus Terrabacteria group; Actinobacteria; Actinobacteria; Corynebacteriales; Mycobacteriaceae; Mycobacterium; Mycobacterium chelonae group;

Mycobacterium abscessus subgroup; Mycobacterium abscessus
MIKHLTRRATVALVVAAAAISPAAYPAVADTATPLNGLFALTPGSCAEGRATGSYFRMIL WP_005063674.1 hypothetical protein [Mycobacterium abscessus] Length: 330\nNumber of predicted TMHs: 1\nExp number of AAs in TMHs: 39.25259\nExp number, first 60 AAs: 16.98187\nTotal prob of N-in: 0.83210\nPOSSIBLE N-term signal sequence\noutside 1 298\nTMhelix 299 321\ninside 322 330

30450 GCF_900130655.1_10665_3_18 Mycobacterium abscessus subsp. abscessus Terrabacteria group; Actinobacteria; Actinobacteria; Corynebacteriales; Mycobacteriaceae; Mycobacterium; Mycobacterium chelonae group; Mycobacterium abscessus subgroup; Mycobacterium abscessus
MIKHLTRRATVALVVAAAAISPAAYPAMADTATPLNGLFALTPGSCAEGRATGSYFRMIL WP_005073213.1 hypothetical protein [Mycobacterium abscessus] Length: 330\nNumber of predicted TMHs: 2\nExp number of AAs in TMHs: 39.3416199999999\nExp number, first 60 AAs: 17.07688\nTotal prob of N-in: 0.84431\nPOSSIBLE N-term signal sequence\ninside 1 11\nTMhelix 12 29\noutside 30 298\nTMhelix 299 321\ninside 322 330

30451 GCF_900130605.1_10665_3_11 Mycobacterium abscessus subsp. abscessus Terrabacteria group; Actinobacteria; Actinobacteria; Corynebacteriales; Mycobacteriaceae; Mycobacterium; Mycobacterium chelonae group; Mycobacterium abscessus subgroup; Mycobacterium abscessus
MIKHLTRRATVALVVAAAAISPAAYPAMADTATPLNGLFALTPGSCAEGRATGSYFRMIL WP_005073213.1 hypothetical protein [Mycobacterium abscessus] Length: 330\nNumber of predicted TMHs: 2\nExp number of AAs in TMHs: 39.3416199999999\nExp number, first 60 AAs: 17.07688\nTotal prob of N-in: 0.84431\nPOSSIBLE N-term signal sequence\ninside 1 11\nTMhelix 12 29\noutside 30 298\nTMhelix 299 321\ninside 322 330

30452 GCF_001942505.1_ASM194250v1 Mycobacterium abscessus subsp. abscessus Terrabacteria group; Actinobacteria; Actinobacteria; Corynebacteriales; Mycobacteriaceae; Mycobacterium; Mycobacterium chelonae group; Mycobacterium abscessus subgroup; Mycobacterium abscessus
MIKHLTRRATVALVVAAAAISPAAYPAMADTATPLNGLFALTPGSCAEGRATGSYFRMIL WP_005073213.1 hypothetical protein [Mycobacterium abscessus] Length: 330\nNumber of predicted TMHs: 2\nExp number of AAs in TMHs: 39.3416199999999\nExp number, first 60 AAs: 17.07688\nTotal prob of N-in: 0.84431\nPOSSIBLE N-term signal sequence\ninside 1 11\nTMhelix 12 29\noutside 30 298\nTMhelix 299 321\ninside 322 330

30453 GCF_900134295.1_11893_6_15 Mycobacterium abscessus subsp. bolletii Terrabacteria group; Actinobacteria; Actinobacteria; Corynebacteriales; Mycobacteriaceae; Mycobacterium; Mycobacterium chelonae group; Mycobacterium abscessus subgroup; Mycobacterium abscessus
MIKHLTRRATVALVVAAAAISPAAYPAMADTATPLNGLFALTPGSCAEGRATGSYFRMIL WP_005073213.1 hypothetical protein [Mycobacterium abscessus] Length: 330\nNumber of predicted TMHs: 2\nExp number of AAs in TMHs: 39.3416199999999\nExp number, first 60 AAs: 17.07688\nTotal prob of N-in: 0.84431\nPOSSIBLE N-term signal sequence\ninside 1 11\nTMhelix 12 29\noutside 30 298\nTMhelix 299 321\ninside 322 330

30454 GCF_900134335.1_12082_5_58 Mycobacterium abscessus subsp. bolletii Terrabacteria group; Actinobacteria; Actinobacteria; Corynebacteriales; Mycobacteriaceae; Mycobacterium; Mycobacterium chelonae group; Mycobacterium abscessus subgroup; Mycobacterium abscessus
MIKHLTRRATVALVVAAAAISPAAYPAMADTATPLNGLFALTPGSCAEGRATGSYFRMIL WP_005073213.1 hypothetical protein [Mycobacterium abscessus] Length: 330\nNumber of predicted TMHs: 2\nExp number of AAs in TMHs: 39.3416199999999\nExp number, first 60 AAs: 17.07688\nTotal prob of N-in: 0.84431\nPOSSIBLE N-term signal sequence\ninside 1 11\nTMhelix 12 29\noutside 30 298\nTMhelix 299 321\ninside 322 330

30455 GCF_900131565.1_10665_4_13 Mycobacterium abscessus subsp. bolletii Terrabacteria group; Actinobacteria; Actinobacteria; Corynebacteriales; Mycobacteriaceae; Mycobacterium; Mycobacterium chelonae group; Mycobacterium abscessus subgroup; Mycobacterium abscessus
MIKHLTRRATVALVVAAAAISPAAYPAMADTATPLNGLFALTPGSCAEGRATGSYFRMIL WP_005073213.1 hypothetical protein [Mycobacterium abscessus] Length: 330\nNumber of predicted TMHs: 2\nExp number of AAs in TMHs: 39.3416199999999\nExp number, first 60 AAs: 17.07688\nTotal prob of N-in: 0.84431\nPOSSIBLE N-term signal sequence\ninside 1 11\nTMhelix 12 29\noutside 30 298\nTMhelix 299 321\ninside 322 330

30456 GCF_900133335.1_10250_1_37 Mycobacterium abscessus subsp. bolletii Terrabacteria group; Actinobacteria; Actinobacteria; Corynebacteriales; Mycobacteriaceae; Mycobacterium; Mycobacterium chelonae group; Mycobacterium abscessus subgroup; Mycobacterium abscessus
MIKHLTRRATVALVVAAAAISPAAYPAMADTATPLNGLFALTPGSCAEGRATGSYFRMIL WP_005073213.1 hypothetical protein [Mycobacterium abscessus] Length: 330\nNumber of predicted TMHs: 2\nExp number of AAs in TMHs: 39.3416199999999\nExp number, first 60 AAs: 17.07688\nTotal prob of N-in: 0.84431\nPOSSIBLE N-term signal sequence\ninside 1 11\nTMhelix 12 29\noutside 30 298\nTMhelix 299 321\ninside 322 330

30457 GCF_900131125.1_10665_3_57 Mycobacterium abscessus subsp. bolletii Terrabacteria group; Actinobacteria; Actinobacteria; Corynebacteriales; Mycobacteriaceae; Mycobacterium; Mycobacterium chelonae group; Mycobacterium abscessus subgroup; Mycobacterium abscessus
MIKHLTRRATVALVVAAAAISPAAYPAMADTATPLNGLFALTPGSCAEGRATGSYFRMIL WP_005073213.1
hypothetical protein [Mycobacterium abscessus] Length: 330\nNumber of predicted TMHs: 2\nExp number of AAs in TMHs: 39.3416199999999\nExp number, first 60 AAs: 17.07688\nTotal prob of N-in: 0.84431\nPOSSIBLE N-term signal sequence\ninside 1 11\nTMhelix 12 29\noutside 30 298\nTMhelix 299 321\ninside 322 330

30458 GCF_900134915.1_10208_3_32 Mycobacterium abscessus subsp. bolletii Terrabacteria group; Actinobacteria; Actinobacteria; Corynebacteriales; Mycobacteriaceae; Mycobacterium; Mycobacterium chelonae group; Mycobacterium abscessus subgroup; Mycobacterium abscessus
MIKHLTRRATVALVVAAAAISPAAYPAVADTATPLNGLFALTPGSCAEGRATGSYFRMIL WP_074330522.1 hypothetical protein [Mycobacterium abscessus] Length: 330\nNumber of predicted TMHs: 1\nExp number of AAs in TMHs: 39.25295\nExp number, first 60 AAs: 16.98073\nTotal prob of N-in: 0.83211\nPOSSIBLE N-term signal sequence\noutside 1 298\nTMhelix 299 321\ninside 322 330

30459 GCF_900135895.1_11893_6_38 Mycobacterium abscessus subsp. bolletii Terrabacteria group; Actinobacteria; Actinobacteria; Corynebacteriales; Mycobacteriaceae; Mycobacterium; Mycobacterium chelonae group; Mycobacterium abscessus subgroup; Mycobacterium abscessus
MIKHLTRRATVALVVAAAAISPAAYPAMADTATPLNGLFALTPGSCAEGRATGSYFRMIL WP_005073213.1
hypothetical protein [Mycobacterium abscessus] Length: 330\nNumber of predicted TMHs: 2\nExp number of AAs in TMHs: 39.3416199999999\nExp number, first 60 AAs: 17.07688\nTotal prob of N-in: 0.84431\nPOSSIBLE N-term signal sequence\ninside 1 11\nTMhelix 12 29\noutside 30 298\nTMhelix 299 321\ninside 322 330

30460 GCF_900137885.1_12163_2_86 Mycobacterium abscessus subsp. bolletii Terrabacteria group; Actinobacteria; Actinobacteria; Corynebacteriales; Mycobacteriaceae; Mycobacterium; Mycobacterium chelonae group; Mycobacterium abscessus subgroup; Mycobacterium abscessus
MIKHLTRRATVALVVAAAAISPAAYPAMADTATPLNGLFALTPGSCAEGRATGSYFRMIL WP_074254038.1
hypothetical protein [Mycobacterium abscessus] Length: 330\nNumber of predicted TMHs: 2\nExp number of AAs in TMHs: 39.3496799999999\nExp number, first 60 AAs: 17.05555\nTotal prob of N-in: 0.84446\nPOSSIBLE N-term signal sequence\ninside 1 11\nTMhelix 12 29\noutside 30 298\nTMhelix 299 321\ninside 322 330

30461 GCF_900132745.1_10250_1_23 Mycobacterium abscessus subsp. bolletii Terrabacteria group; Actinobacteria; Actinobacteria; Corynebacteriales; Mycobacteriaceae; Mycobacterium; Mycobacterium chelonae group; Mycobacterium abscessus subgroup; Mycobacterium abscessus
MIKHLTRRATVALVVAAAAISPAAYPAMADTATPLNGLFALTPGSCAEGRATGSYFRMIL WP_074254038.1
hypothetical protein [Mycobacterium abscessus] Length: 330\nNumber of predicted TMHs: 2\nExp number of AAs in TMHs: 39.3496799999999\nExp number, first 60 AAs: 17.05555\nTotal prob of N-in: 0.84446\nPOSSIBLE N-term signal sequence\ninside 1 11\nTMhelix 12 29\noutside 30 298\nTMhelix 299 321\ninside 322 330

30462 GCF_900140135.1_10665_2_63 Mycobacterium abscessus subsp. bolletii Terrabacteria group; Actinobacteria; Actinobacteria; Corynebacteriales; Mycobacteriaceae; Mycobacterium; Mycobacterium chelonae group; Mycobacterium abscessus subgroup; Mycobacterium abscessus
MIKHLTRRATVALVVAAAAISPAAYPAMADTATPLNGLFALTPGSCAEGRATGSYFRMIL WP_005073213.1
hypothetical protein [Mycobacterium abscessus] Length: 330\nNumber of predicted TMHs: 2\nExp number of AAs in TMHs: 39.3416199999999\nExp number, first 60 AAs: 17.07688\nTotal prob of N-in: 0.84431\nPOSSIBLE N-term signal sequence\ninside 1 11\nTMhelix 12 29\noutside 30 298\nTMhelix 299 321\ninside 322 330

30463 GCF_900133795.1_12163_1_5 Mycobacterium abscessus subsp. bolletii Terrabacteria group; Actinobacteria; Actinobacteria; Corynebacteriales; Mycobacteriaceae; Mycobacterium; Mycobacterium chelonae group; Mycobacterium abscessus subgroup; Mycobacterium abscessus
MIKHLTRRATVALVVAAAAISPAAYPAMADTATPLNGLFALTPGSCAEGRATGSYFRMIL WP_005073213.1
hypothetical protein [Mycobacterium abscessus] Length: 330\nNumber of predicted TMHs: 2\nExp number of AAs in TMHs: 39.3416199999999\nExp number, first 60 AAs: 17.07688\nTotal prob of N-in: 0.84431\nPOSSIBLE N-term signal sequence\ninside 1 11\nTMhelix 12 29\noutside 30 298\nTMhelix 299 321\ninside 322 330

30464 GCF_900137535.1_12163_2_29 Mycobacterium abscessus subsp. bolletii Terrabacteria group; Actinobacteria; Actinobacteria; Corynebacteriales; Mycobacteriaceae; Mycobacterium; Mycobacterium chelonae group; Mycobacterium abscessus subgroup; Mycobacterium abscessus
MIKHLTRRATVALVVAAAAISPAAYPAMADTATPLNGLFALTPGSCAEGRATGSYFRMIL WP_005073213.1
hypothetical protein [Mycobacterium abscessus] Length: 330\nNumber of predicted TMHs: 2\nExp number of

AAs in TMHs: 39.3416199999999\nExp number, first 60 AAs:  17.07688\nTotal prob of N-in:   0.84431\nPOSSIBLE N-term signal sequence\ninside   1   11\nTMhelix   12   29\noutside   30   298\nTMhelix   299   321\ninside   322   330

30465 GCF_900130485.1_10665_3_1 Mycobacterium abscessus subsp. bolletii Terrabacteria group; Actinobacteria; Actinobacteria; Corynebacteriales; Mycobacteriaceae; Mycobacterium; Mycobacterium chelonae group; Mycobacterium abscessus subgroup; Mycobacterium abscessus
MIKHLTRRATVALVVAAAAISPAAYPAMADTATPLNGLFALTPGSCAEGRATGSYFRMIL WP_005073213.1
hypothetical protein [Mycobacterium abscessus] Length: 330\nNumber of predicted TMHs:  2\nExp number of AAs in TMHs: 39.3416199999999\nExp number, first 60 AAs:  17.07688\nTotal prob of N-in:   0.84431\nPOSSIBLE N-term signal sequence\ninside   1   11\nTMhelix   12   29\noutside   30   298\nTMhelix   299   321\ninside   322   330

30466 GCF_900131095.1_10665_3_55 Mycobacterium abscessus subsp. bolletii Terrabacteria group; Actinobacteria; Actinobacteria; Corynebacteriales; Mycobacteriaceae; Mycobacterium; Mycobacterium chelonae group; Mycobacterium abscessus subgroup; Mycobacterium abscessus
MIKHLTRRATVALVVAAAAISPAAYPAMADTATPLNGLFALTPGSCAEGRATGSYFRMIL WP_074254038.1
hypothetical protein [Mycobacterium abscessus] Length: 330\nNumber of predicted TMHs:  2\nExp number of AAs in TMHs: 39.3496799999999\nExp number, first 60 AAs:  17.05555\nTotal prob of N-in:   0.84446\nPOSSIBLE N-term signal sequence\ninside   1   11\nTMhelix   12   29\noutside   30   298\nTMhelix   299   321\ninside   322   330

30467 GCF_900134755.1_10208_3_17 Mycobacterium abscessus subsp. bolletii Terrabacteria group; Actinobacteria; Actinobacteria; Corynebacteriales; Mycobacteriaceae; Mycobacterium; Mycobacterium chelonae group; Mycobacterium abscessus subgroup; Mycobacterium abscessus
MIKHLTRRATVALVVAAAAISPAAYPAMADTATPLNGLFALTPGSCAEGRATGSYFRMIL WP_074254038.1
hypothetical protein [Mycobacterium abscessus] Length: 330\nNumber of predicted TMHs:  2\nExp number of AAs in TMHs: 39.3496799999999\nExp number, first 60 AAs:  17.05555\nTotal prob of N-in:   0.84446\nPOSSIBLE N-term signal sequence\ninside   1   11\nTMhelix   12   29\noutside   30   298\nTMhelix   299   321\ninside   322   330

30468 GCF_900134535.1_12163_1_44 Mycobacterium abscessus subsp. bolletii Terrabacteria group; Actinobacteria; Actinobacteria; Corynebacteriales; Mycobacteriaceae; Mycobacterium; Mycobacterium chelonae group; Mycobacterium abscessus subgroup; Mycobacterium abscessus
MIKHLTRRATVALVVAAAAISPAAYPAMADTATPLNGLFALTPGSCAEGRATGSYFRMIL WP_005073213.1
hypothetical protein [Mycobacterium abscessus] Length: 330\nNumber of predicted TMHs:  2\nExp number of AAs in TMHs: 39.3416199999999\nExp number, first 60 AAs:  17.07688\nTotal prob of N-in:   0.84431\nPOSSIBLE N-term signal sequence\ninside   1   11\nTMhelix   12   29\noutside   30   298\nTMhelix   299   321\ninside   322   330

30469 GCF_900133795.1_12163_1_5 Mycobacterium abscessus subsp. bolletii Terrabacteria group; Actinobacteria; Actinobacteria; Corynebacteriales; Mycobacteriaceae; Mycobacterium; Mycobacterium chelonae group; Mycobacterium abscessus subgroup; Mycobacterium abscessus
MRGGRRRGLLGFIQVLAAIMASLALVWTAPAAQAISPPVIDPGAVAANEPPMPPEELEST WP_079610166.1 type VII secretion-associated serine protease mycosin [Mycobacterium abscessus] Length: 489\nNumber of predicted TMHs:  2\nExp number of AAs in TMHs: 55.05414\nExp number, first 60 AAs:  22.16163\nTotal prob of N-in:   0.98239\nPOSSIBLE N-term signal sequence\ninside   1   6\nTMhelix   7   29\noutside   30   453\nTMhelix   454   473\ninside   474   489

30470 GCF_900140205.1_10665_2_73 Mycobacterium abscessus subsp. bolletii Terrabacteria group; Actinobacteria; Actinobacteria; Corynebacteriales; Mycobacteriaceae; Mycobacterium; Mycobacterium chelonae group; Mycobacterium abscessus subgroup; Mycobacterium abscessus
MIKHLTRRATVALVVAAAAISPAAYPAMADTATPLNGLFALTPGSCAEGRATGSYFRMIL WP_005073213.1
hypothetical protein [Mycobacterium abscessus] Length: 330\nNumber of predicted TMHs:  2\nExp number of AAs in TMHs: 39.3416199999999\nExp number, first 60 AAs:  17.07688\nTotal prob of N-in:   0.84431\nPOSSIBLE N-term signal sequence\ninside   1   11\nTMhelix   12   29\noutside   30   298\nTMhelix   299   321\ninside   322   330

30471 GCF_900139995.1_10660_1_51 Mycobacterium abscessus subsp. bolletii Terrabacteria group; Actinobacteria; Actinobacteria; Corynebacteriales; Mycobacteriaceae; Mycobacterium; Mycobacterium chelonae group; Mycobacterium abscessus subgroup; Mycobacterium abscessus
MIKHLTRRATVALVVAAAAISPAAYPAMADTATPLNGLFALTPGSCAEGRATGSYFRMIL WP_074254038.1
hypothetical protein [Mycobacterium abscessus] Length: 330\nNumber of predicted TMHs:  2\nExp number of AAs in TMHs: 39.3496799999999\nExp number, first 60 AAs:  17.05555\nTotal prob of N-in:   0.84446\nPOSSIBLE N-term signal sequence\ninside   1   11\nTMhelix   12   29\noutside   30   298\nTMhelix   299   321\ninside   322   330

30472 GCF_900132405.1_10665_4_72 Mycobacterium abscessus subsp. bolletii Terrabacteria group; Actinobacteria; Actinobacteria; Corynebacteriales; Mycobacteriaceae; Mycobacterium; Mycobacterium chelonae group;

Mycobacterium abscessus subgroup; Mycobacterium abscessus
MIKHLTRRATVALVVAAAAISPAAYPAMADTATPLNGLFALTPGSCAEGRATGSYFRMIL WP_074254038.1
hypothetical protein [Mycobacterium abscessus] Length: 330\nNumber of predicted TMHs: 2\nExp number of AAs in TMHs: 39.3496799999999\nExp number, first 60 AAs: 17.05555\nTotal prob of N-in: 0.84446\nPOSSIBLE N-term signal sequence\ninside 1 11\nTMhelix 12 29\noutside 30 298\nTMhelix 299 321\ninside 322 330

30473 GCF_002086525.1_ASM208652v1 Mycobacterium abscessus subsp. bolletii Terrabacteria group; Actinobacteria; Actinobacteria; Corynebacteriales; Mycobacteriaceae; Mycobacterium; Mycobacterium chelonae group; Mycobacterium abscessus subgroup; Mycobacterium abscessus
MIKHLTRRATVALVVAAAAISPAAYPAMADTATPLNGLFALTPGSCAEGRATGSYFRMIL WP_005073213.1
hypothetical protein [Mycobacterium abscessus] Length: 330\nNumber of predicted TMHs: 2\nExp number of AAs in TMHs: 39.3416199999999\nExp number, first 60 AAs: 17.07688\nTotal prob of N-in: 0.84431\nPOSSIBLE N-term signal sequence\ninside 1 11\nTMhelix 12 29\noutside 30 298\nTMhelix 299 321\ninside 322 330

30474 GCF_900138865.1_10208_3_54 Mycobacterium abscessus subsp. bolletii Terrabacteria group; Actinobacteria; Actinobacteria; Corynebacteriales; Mycobacteriaceae; Mycobacterium; Mycobacterium chelonae group; Mycobacterium abscessus subgroup; Mycobacterium abscessus
MIKHLTRRATVALVVAAAAISPAAYPAMADTATPLNGLFALTPGSCAEGRATGSYFRMIL WP_005073213.1
hypothetical protein [Mycobacterium abscessus] Length: 330\nNumber of predicted TMHs: 2\nExp number of AAs in TMHs: 39.3416199999999\nExp number, first 60 AAs: 17.07688\nTotal prob of N-in: 0.84431\nPOSSIBLE N-term signal sequence\ninside 1 11\nTMhelix 12 29\noutside 30 298\nTMhelix 299 321\ninside 322 330

30475 GCF_900136205.1_12163_1_77 Mycobacterium abscessus subsp. bolletii Terrabacteria group; Actinobacteria; Actinobacteria; Corynebacteriales; Mycobacteriaceae; Mycobacterium; Mycobacterium chelonae group; Mycobacterium abscessus subgroup; Mycobacterium abscessus
MIKHLTRRATVALVVAAAAISPAAYPAMADTATPLNGLFALTPGSCAEGRATGSYFRMIL WP_074254038.1
hypothetical protein [Mycobacterium abscessus] Length: 330\nNumber of predicted TMHs: 2\nExp number of AAs in TMHs: 39.3496799999999\nExp number, first 60 AAs: 17.05555\nTotal prob of N-in: 0.84446\nPOSSIBLE N-term signal sequence\ninside 1 11\nTMhelix 12 29\noutside 30 298\nTMhelix 299 321\ninside 322 330

30476 GCF_900135035.1_10208_3_7 Mycobacterium abscessus subsp. bolletii Terrabacteria group; Actinobacteria; Actinobacteria; Corynebacteriales; Mycobacteriaceae; Mycobacterium; Mycobacterium chelonae group; Mycobacterium abscessus subgroup; Mycobacterium abscessus
MIKHLTRRATVALVVAAAAISPAAYPAMADTATPLNGLFALTPGSCAEGRATGSYFRMIL WP_074254038.1
hypothetical protein [Mycobacterium abscessus] Length: 330\nNumber of predicted TMHs: 2\nExp number of AAs in TMHs: 39.3496799999999\nExp number, first 60 AAs: 17.05555\nTotal prob of N-in: 0.84446\nPOSSIBLE N-term signal sequence\ninside 1 11\nTMhelix 12 29\noutside 30 298\nTMhelix 299 321\ninside 322 330

30477 GCF_900133635.1_12163_1_12 Mycobacterium abscessus subsp. bolletii Terrabacteria group; Actinobacteria; Actinobacteria; Corynebacteriales; Mycobacteriaceae; Mycobacterium; Mycobacterium chelonae group; Mycobacterium abscessus subgroup; Mycobacterium abscessus
MIKHLTRRATVALVVAAAAISPAAYPAMADTATPLNGLFALTPGSCAEGRATGSYFRMIL WP_005073213.1
hypothetical protein [Mycobacterium abscessus] Length: 330\nNumber of predicted TMHs: 2\nExp number of AAs in TMHs: 39.3416199999999\nExp number, first 60 AAs: 17.07688\nTotal prob of N-in: 0.84431\nPOSSIBLE N-term signal sequence\ninside 1 11\nTMhelix 12 29\noutside 30 298\nTMhelix 299 321\ninside 322 330

30478 GCF_900135225.1_10625_4_37 Mycobacterium abscessus subsp. bolletii Terrabacteria group; Actinobacteria; Actinobacteria; Corynebacteriales; Mycobacteriaceae; Mycobacterium; Mycobacterium chelonae group; Mycobacterium abscessus subgroup; Mycobacterium abscessus
MIKHLTRRATVALVVAAAAISPAAYPAMADTATPLNGLFALTPGSCAEGRATGSYFRMIL WP_005073213.1
hypothetical protein [Mycobacterium abscessus] Length: 330\nNumber of predicted TMHs: 2\nExp number of AAs in TMHs: 39.3416199999999\nExp number, first 60 AAs: 17.07688\nTotal prob of N-in: 0.84431\nPOSSIBLE N-term signal sequence\ninside 1 11\nTMhelix 12 29\noutside 30 298\nTMhelix 299 321\ninside 322 330

30479 GCF_900131955.1_10665_4_40 Mycobacterium abscessus subsp. bolletii Terrabacteria group; Actinobacteria; Actinobacteria; Corynebacteriales; Mycobacteriaceae; Mycobacterium; Mycobacterium chelonae group; Mycobacterium abscessus subgroup; Mycobacterium abscessus
MIKHLTRRATVALVVAAAAISPAAYPAMADTATPLNGLFALTPGSCAEGRATGSYFRMIL WP_074254038.1
hypothetical protein [Mycobacterium abscessus] Length: 330\nNumber of predicted TMHs: 2\nExp number of AAs in TMHs: 39.3496799999999\nExp number, first 60 AAs: 17.05555\nTotal prob of N-in: 0.84446\nPOSSIBLE N-term signal sequence\ninside 1 11\nTMhelix 12 29\noutside 30 298\nTMhelix 299 321\ninside 322 330

30480 GCF_900136235.1_12163_1_79 Mycobacterium abscessus subsp. bolletii Terrabacteria group; Actinobacteria; Actinobacteria; Corynebacteriales; Mycobacteriaceae; Mycobacterium; Mycobacterium chelonae group; Mycobacterium abscessus subgroup; Mycobacterium abscessus
MIKHLTRRATVALVVAAAAISPAAYPAMADTATPLNGLFALTPGSCAEGRATGSYFRMIL WP_074254038.1
hypothetical protein [Mycobacterium abscessus] Length: 330\nNumber of predicted TMHs: 2\nExp number of AAs in TMHs: 39.3496799999999\nExp number, first 60 AAs: 17.05555\nTotal prob of N-in: 0.84446\nPOSSIBLE N-term signal sequence\ninside 1 11\nTMhelix 12 29\noutside 30 298\nTMhelix 299 321\ninside 322 330

30481 GCF_900130815.1_10665_3_32 Mycobacterium abscessus subsp. bolletii Terrabacteria group; Actinobacteria; Actinobacteria; Corynebacteriales; Mycobacteriaceae; Mycobacterium; Mycobacterium chelonae group; Mycobacterium abscessus subgroup; Mycobacterium abscessus
MIKHLTRRATVALVVAAAAISPAAYPAMADTATPLNGLFALTPGSCAEGRATGSYFRMIL WP_074254038.1
hypothetical protein [Mycobacterium abscessus] Length: 330\nNumber of predicted TMHs: 2\nExp number of AAs in TMHs: 39.3496799999999\nExp number, first 60 AAs: 17.05555\nTotal prob of N-in: 0.84446\nPOSSIBLE N-term signal sequence\ninside 1 11\nTMhelix 12 29\noutside 30 298\nTMhelix 299 321\ninside 322 330

30482 GCF_900134845.1_10208_3_28 Mycobacterium abscessus subsp. bolletii Terrabacteria group; Actinobacteria; Actinobacteria; Corynebacteriales; Mycobacteriaceae; Mycobacterium; Mycobacterium chelonae group; Mycobacterium abscessus subgroup; Mycobacterium abscessus
MIKHLTRRATVALVVAAAAISPAAYPAMADTATPLNGLFALTPGSCAEGRATGSYFRMIL WP_074254038.1
hypothetical protein [Mycobacterium abscessus] Length: 330\nNumber of predicted TMHs: 2\nExp number of AAs in TMHs: 39.3496799999999\nExp number, first 60 AAs: 17.05555\nTotal prob of N-in: 0.84446\nPOSSIBLE N-term signal sequence\ninside 1 11\nTMhelix 12 29\noutside 30 298\nTMhelix 299 321\ninside 322 330

30483 GCF_900134285.1_11893_6_14 Mycobacterium abscessus subsp. bolletii Terrabacteria group; Actinobacteria; Actinobacteria; Corynebacteriales; Mycobacteriaceae; Mycobacterium; Mycobacterium chelonae group; Mycobacterium abscessus subgroup; Mycobacterium abscessus
MIKHLTRRATVALVVAAAAISPAAYPAMADTATPLNGLFALTPGSCAEGRATGSYFRMIL WP_005073213.1
hypothetical protein [Mycobacterium abscessus] Length: 330\nNumber of predicted TMHs: 2\nExp number of AAs in TMHs: 39.3416199999999\nExp number, first 60 AAs: 17.07688\nTotal prob of N-in: 0.84431\nPOSSIBLE N-term signal sequence\ninside 1 11\nTMhelix 12 29\noutside 30 298\nTMhelix 299 321\ninside 322 330

30484 GCF_900132985.1_11893_6_12 Mycobacterium abscessus subsp. bolletii Terrabacteria group; Actinobacteria; Actinobacteria; Corynebacteriales; Mycobacteriaceae; Mycobacterium; Mycobacterium chelonae group; Mycobacterium abscessus subgroup; Mycobacterium abscessus
MIKHLTRRATVALVVAAAAISPAAYPAMADTATPLNGLFALTPGSCAEGRATGSYFRMIL WP_005073213.1
hypothetical protein [Mycobacterium abscessus] Length: 330\nNumber of predicted TMHs: 2\nExp number of AAs in TMHs: 39.3416199999999\nExp number, first 60 AAs: 17.07688\nTotal prob of N-in: 0.84431\nPOSSIBLE N-term signal sequence\ninside 1 11\nTMhelix 12 29\noutside 30 298\nTMhelix 299 321\ninside 322 330

30485 GCF_900134995.1_10208_3_40 Mycobacterium abscessus subsp. bolletii Terrabacteria group; Actinobacteria; Actinobacteria; Corynebacteriales; Mycobacteriaceae; Mycobacterium; Mycobacterium chelonae group; Mycobacterium abscessus subgroup; Mycobacterium abscessus
MIKHLTRRATVALVVAAAAISPAAYPAMADTATPLNGLFALTPGSCAEGRATGSYFRMIL WP_074254038.1
hypothetical protein [Mycobacterium abscessus] Length: 330\nNumber of predicted TMHs: 2\nExp number of AAs in TMHs: 39.3496799999999\nExp number, first 60 AAs: 17.05555\nTotal prob of N-in: 0.84446\nPOSSIBLE N-term signal sequence\ninside 1 11\nTMhelix 12 29\noutside 30 298\nTMhelix 299 321\ninside 322 330

30486 GCF_900140075.1_10665_2_62 Mycobacterium abscessus subsp. bolletii Terrabacteria group; Actinobacteria; Actinobacteria; Corynebacteriales; Mycobacteriaceae; Mycobacterium; Mycobacterium chelonae group; Mycobacterium abscessus subgroup; Mycobacterium abscessus
MIKHLTRRATVALVVAAAAISPAAYPAMADTATPLNGLFALTPGSCAEGRATGSYFRMIL WP_074254038.1
hypothetical protein [Mycobacterium abscessus] Length: 330\nNumber of predicted TMHs: 2\nExp number of AAs in TMHs: 39.3496799999999\nExp number, first 60 AAs: 17.05555\nTotal prob of N-in: 0.84446\nPOSSIBLE N-term signal sequence\ninside 1 11\nTMhelix 12 29\noutside 30 298\nTMhelix 299 321\ninside 322 330

30487 GCF_900141605.1_12163_2_72 Mycobacterium abscessus subsp. bolletii Terrabacteria group; Actinobacteria; Actinobacteria; Corynebacteriales; Mycobacteriaceae; Mycobacterium; Mycobacterium chelonae group; Mycobacterium abscessus subgroup; Mycobacterium abscessus
MIKHLTRRATVALVVAAAAISPAAYPAMADTATPLNGLFALTPGSCAEGRATGSYFRMIL WP_005073213.1
hypothetical protein [Mycobacterium abscessus] Length: 330\nNumber of predicted TMHs: 2\nExp number of

AAs in TMHs: 39.3416199999999\nExp number, first 60 AAs: 17.07688\nTotal prob of N-in: 0.84431\nPOSSIBLE N-term signal sequence\ninside 1 11\nTMhelix 12 29\noutside 30 298\nTMhelix 299 321\ninside 322 330

30488 GCF_900134785.1_10208_3_24 Mycobacterium abscessus subsp. bolletii Terrabacteria group; Actinobacteria; Actinobacteria; Corynebacteriales; Mycobacteriaceae; Mycobacterium; Mycobacterium chelonae group; Mycobacterium abscessus subgroup; Mycobacterium abscessus MIKHLTRRATVALVVAAAAISPAAYPAVADTATPLNGLFALTPGSCAEGRATGSYFRMIL WP_074330522.1 hypothetical protein [Mycobacterium abscessus] Length: 330\nNumber of predicted TMHs: 1\nExp number of AAs in TMHs: 39.25295\nExp number, first 60 AAs: 16.98073\nTotal prob of N-in: 0.83211\nPOSSIBLE N-term signal sequence\noutside 1 298\nTMhelix 299 321\ninside 322 330

30489 GCF_900134105.1_10396_8_9 Mycobacterium abscessus subsp. bolletii Terrabacteria group; Actinobacteria; Actinobacteria; Corynebacteriales; Mycobacteriaceae; Mycobacterium; Mycobacterium chelonae group; Mycobacterium abscessus subgroup; Mycobacterium abscessus MIKHLTRRATVALVVAAAAISPAAYPAMADTATPLNGLFALTPGSCAEGRATGSYFRMIL WP_074254038.1 hypothetical protein [Mycobacterium abscessus] Length: 330\nNumber of predicted TMHs: 2\nExp number of AAs in TMHs: 39.3496799999999\nExp number, first 60 AAs: 17.05555\nTotal prob of N-in: 0.84446\nPOSSIBLE N-term signal sequence\ninside 1 11\nTMhelix 12 29\noutside 30 298\nTMhelix 299 321\ninside 322 330

30490 GCF_900133625.1_12163_1_11 Mycobacterium abscessus subsp. bolletii Terrabacteria group; Actinobacteria; Actinobacteria; Corynebacteriales; Mycobacteriaceae; Mycobacterium; Mycobacterium chelonae group; Mycobacterium abscessus subgroup; Mycobacterium abscessus MIKHLTRRATVALVVAAAAISPAAYPAMADTATPLNGLFALTPGSCAEGRATGSYFRMIL WP_005073213.1 hypothetical protein [Mycobacterium abscessus] Length: 330\nNumber of predicted TMHs: 2\nExp number of AAs in TMHs: 39.3416199999999\nExp number, first 60 AAs: 17.07688\nTotal prob of N-in: 0.84431\nPOSSIBLE N-term signal sequence\ninside 1 11\nTMhelix 12 29\noutside 30 298\nTMhelix 299 321\ninside 322 330

30491 GCF_900130615.1_10665_3_15 Mycobacterium abscessus subsp. bolletii Terrabacteria group; Actinobacteria; Actinobacteria; Corynebacteriales; Mycobacteriaceae; Mycobacterium; Mycobacterium chelonae group; Mycobacterium abscessus subgroup; Mycobacterium abscessus MIKHLTRRATVALVVAAAAISPAAYPAMADTATPLNGLFALTPGSCAEGRATGSYFRMIL WP_074254038.1 hypothetical protein [Mycobacterium abscessus] Length: 330\nNumber of predicted TMHs: 2\nExp number of AAs in TMHs: 39.3496799999999\nExp number, first 60 AAs: 17.05555\nTotal prob of N-in: 0.84446\nPOSSIBLE N-term signal sequence\ninside 1 11\nTMhelix 12 29\noutside 30 298\nTMhelix 299 321\ninside 322 330

30492 GCF_900130735.1_10665_3_21 Mycobacterium abscessus subsp. bolletii Terrabacteria group; Actinobacteria; Actinobacteria; Corynebacteriales; Mycobacteriaceae; Mycobacterium; Mycobacterium chelonae group; Mycobacterium abscessus subgroup; Mycobacterium abscessus MIKHLTRRATVALVVAAAAISPAAYPAMADTATPLNGLFALTPGSCAEGRATGSYFRMIL WP_074254038.1 hypothetical protein [Mycobacterium abscessus] Length: 330\nNumber of predicted TMHs: 2\nExp number of AAs in TMHs: 39.3496799999999\nExp number, first 60 AAs: 17.05555\nTotal prob of N-in: 0.84446\nPOSSIBLE N-term signal sequence\ninside 1 11\nTMhelix 12 29\noutside 30 298\nTMhelix 299 321\ninside 322 330

30493 GCF_900137935.1_10660_1_45 Mycobacterium abscessus subsp. bolletii Terrabacteria group; Actinobacteria; Actinobacteria; Corynebacteriales; Mycobacteriaceae; Mycobacterium; Mycobacterium chelonae group; Mycobacterium abscessus subgroup; Mycobacterium abscessus MIKHLTRRATVALVVAAAAISPAAYPAMADTATPLNGLFALTPGSCAEGRATGSYFRMIL WP_005073213.1 hypothetical protein [Mycobacterium abscessus] Length: 330\nNumber of predicted TMHs: 2\nExp number of AAs in TMHs: 39.3416199999999\nExp number, first 60 AAs: 17.07688\nTotal prob of N-in: 0.84431\nPOSSIBLE N-term signal sequence\ninside 1 11\nTMhelix 12 29\noutside 30 298\nTMhelix 299 321\ninside 322 330

30494 GCF_900132785.1_10250_1_4 Mycobacterium abscessus subsp. bolletii Terrabacteria group; Actinobacteria; Actinobacteria; Corynebacteriales; Mycobacteriaceae; Mycobacterium; Mycobacterium chelonae group; Mycobacterium abscessus subgroup; Mycobacterium abscessus MIKHLTRRATVALVVAAAAISPAAYPAMADTATPLNGLFALTPGSCAEGRATGSYFRMIL WP_074254038.1 hypothetical protein [Mycobacterium abscessus] Length: 330\nNumber of predicted TMHs: 2\nExp number of AAs in TMHs: 39.3496799999999\nExp number, first 60 AAs: 17.05555\nTotal prob of N-in: 0.84446\nPOSSIBLE N-term signal sequence\ninside 1 11\nTMhelix 12 29\noutside 30 298\nTMhelix 299 321\ninside 322 330

30495 GCF_900135885.1_11893_6_37 Mycobacterium abscessus subsp. bolletii Terrabacteria group; Actinobacteria; Actinobacteria; Corynebacteriales; Mycobacteriaceae; Mycobacterium; Mycobacterium chelonae group;

Mycobacterium abscessus subgroup; Mycobacterium abscessus
MIKHLTRRATVALVVAAAAISPAAYPAMADTATPLNGLFALTPGSCAEGRATGSYFRMIL WP_005073213.1
hypothetical protein [Mycobacterium abscessus] Length: 330\nNumber of predicted TMHs: 2\nExp number of AAs in TMHs: 39.3416199999999\nExp number, first 60 AAs: 17.07688\nTotal prob of N-in: 0.84431\nPOSSIBLE N-term signal sequence\ninside 1 11\nTMhelix 12 29\noutside 30 298\nTMhelix 299 321\ninside 322 330

30496 GCF_900132715.1_10250_1_2 Mycobacterium abscessus subsp. bolletii Terrabacteria group; Actinobacteria; Actinobacteria; Corynebacteriales; Mycobacteriaceae; Mycobacterium; Mycobacterium chelonae group; Mycobacterium abscessus subgroup; Mycobacterium abscessus
MIKHLTRRATVALVVAAAAISPAAYPAMADTATPLNGLFALTPGSCAEGRATGSYFRMIL WP_005073213.1
hypothetical protein [Mycobacterium abscessus] Length: 330\nNumber of predicted TMHs: 2\nExp number of AAs in TMHs: 39.3416199999999\nExp number, first 60 AAs: 17.07688\nTotal prob of N-in: 0.84431\nPOSSIBLE N-term signal sequence\ninside 1 11\nTMhelix 12 29\noutside 30 298\nTMhelix 299 321\ninside 322 330

30497 GCF_000758405.1_ASM75840v1 Mycobacterium abscessus subsp. bolletii Terrabacteria group; Actinobacteria; Actinobacteria; Corynebacteriales; Mycobacteriaceae; Mycobacterium; Mycobacterium chelonae group; Mycobacterium abscessus subgroup; Mycobacterium abscessus
MIKHLTRRATVALVVAAAAISPAAYPAMADTATPLNGLFALTPGSCAEGRATGSYFRMIL WP_005073213.1
hypothetical protein [Mycobacterium abscessus] Length: 330\nNumber of predicted TMHs: 2\nExp number of AAs in TMHs: 39.3416199999999\nExp number, first 60 AAs: 17.07688\nTotal prob of N-in: 0.84431\nPOSSIBLE N-term signal sequence\ninside 1 11\nTMhelix 12 29\noutside 30 298\nTMhelix 299 321\ninside 322 330

30498 GCF_900132455.1_10665_4_79 Mycobacterium abscessus subsp. bolletii Terrabacteria group; Actinobacteria; Actinobacteria; Corynebacteriales; Mycobacteriaceae; Mycobacterium; Mycobacterium chelonae group; Mycobacterium abscessus subgroup; Mycobacterium abscessus
MIKHLTRRATVALVVAAAAISPAAYPAMADTATPLNGLFALTPGSCAEGRATGSYFRMIL WP_074254038.1
hypothetical protein [Mycobacterium abscessus] Length: 330\nNumber of predicted TMHs: 2\nExp number of AAs in TMHs: 39.3496799999999\nExp number, first 60 AAs: 17.05555\nTotal prob of N-in: 0.84446\nPOSSIBLE N-term signal sequence\ninside 1 11\nTMhelix 12 29\noutside 30 298\nTMhelix 299 321\ninside 322 330

30499 GCF_900132345.1_10665_4_66 Mycobacterium abscessus subsp. bolletii Terrabacteria group; Actinobacteria; Actinobacteria; Corynebacteriales; Mycobacteriaceae; Mycobacterium; Mycobacterium chelonae group; Mycobacterium abscessus subgroup; Mycobacterium abscessus
MIKHLTRRATVALVVAAAAISPAAYPAMADTATPLNGLFALTPGSCAEGRATGSYFRMIL WP_005073213.1
hypothetical protein [Mycobacterium abscessus] Length: 330\nNumber of predicted TMHs: 2\nExp number of AAs in TMHs: 39.3416199999999\nExp number, first 60 AAs: 17.07688\nTotal prob of N-in: 0.84431\nPOSSIBLE N-term signal sequence\ninside 1 11\nTMhelix 12 29\noutside 30 298\nTMhelix 299 321\ninside 322 330

30500 GCF_900131555.1_10665_4_10 Mycobacterium abscessus subsp. bolletii Terrabacteria group; Actinobacteria; Actinobacteria; Corynebacteriales; Mycobacteriaceae; Mycobacterium; Mycobacterium chelonae group; Mycobacterium abscessus subgroup; Mycobacterium abscessus
MIKHLTRRATVALVVAAAAISPAAYPAMADTATPLNGLFALTPGSCAEGRATGSYFRMIL WP_005073213.1
hypothetical protein [Mycobacterium abscessus] Length: 330\nNumber of predicted TMHs: 2\nExp number of AAs in TMHs: 39.3416199999999\nExp number, first 60 AAs: 17.07688\nTotal prob of N-in: 0.84431\nPOSSIBLE N-term signal sequence\ninside 1 11\nTMhelix 12 29\noutside 30 298\nTMhelix 299 321\ninside 322 330

30501 GCF_900139385.1_10625_4_17 Mycobacterium abscessus subsp. bolletii Terrabacteria group; Actinobacteria; Actinobacteria; Corynebacteriales; Mycobacteriaceae; Mycobacterium; Mycobacterium chelonae group; Mycobacterium abscessus subgroup; Mycobacterium abscessus
MIKHLTRRATVALVVAAAAISPAAYPAMADTATPLNGLFALTPGSCAEGRATGSYFRMIL WP_074254038.1
hypothetical protein [Mycobacterium abscessus] Length: 330\nNumber of predicted TMHs: 2\nExp number of AAs in TMHs: 39.3496799999999\nExp number, first 60 AAs: 17.05555\nTotal prob of N-in: 0.84446\nPOSSIBLE N-term signal sequence\ninside 1 11\nTMhelix 12 29\noutside 30 298\nTMhelix 299 321\ninside 322 330

30502 GCF_900134425.1_12163_1_35 Mycobacterium abscessus subsp. bolletii Terrabacteria group; Actinobacteria; Actinobacteria; Corynebacteriales; Mycobacteriaceae; Mycobacterium; Mycobacterium chelonae group; Mycobacterium abscessus subgroup; Mycobacterium abscessus
MIKHLTRRATVALVVAAAAISPAAYPAMADTATPLNGLFALTPGSCAEGRATGSYFRMIL WP_074254038.1
hypothetical protein [Mycobacterium abscessus] Length: 330\nNumber of predicted TMHs: 2\nExp number of AAs in TMHs: 39.3496799999999\nExp number, first 60 AAs: 17.05555\nTotal prob of N-in: 0.84446\nPOSSIBLE N-term signal sequence\ninside 1 11\nTMhelix 12 29\noutside 30 298\nTMhelix 299 321\ninside 322 330

30503 GCF_900131135.1_10665_3_58 Mycobacterium abscessus subsp. bolletii Terrabacteria group; Actinobacteria; Actinobacteria; Corynebacteriales; Mycobacteriaceae; Mycobacterium; Mycobacterium chelonae group; Mycobacterium abscessus subgroup; Mycobacterium abscessus
MIKHLTRRATVALVVAAAAISPAAYPAMADTATPLNGLFALTPGSCAEGRATGSYFRMIL WP_005073213.1
hypothetical protein [Mycobacterium abscessus] Length: 330\nNumber of predicted TMHs: 2\nExp number of AAs in TMHs: 39.3416199999999\nExp number, first 60 AAs: 17.07688\nTotal prob of N-in: 0.84431\nPOSSIBLE N-term signal sequence\ninside 1 11\nTMhelix 12 29\noutside 30 298\nTMhelix 299 321\ninside 322 330

30504 GCF_900131105.1_10665_3_54 Mycobacterium abscessus subsp. bolletii Terrabacteria group; Actinobacteria; Actinobacteria; Corynebacteriales; Mycobacteriaceae; Mycobacterium; Mycobacterium chelonae group; Mycobacterium abscessus subgroup; Mycobacterium abscessus
MIKHLTRRATVALVVAAAAISPAAYPAMADTATPLNGLFALTPGSCAEGRATGSYFRMIL WP_005073213.1
hypothetical protein [Mycobacterium abscessus] Length: 330\nNumber of predicted TMHs: 2\nExp number of AAs in TMHs: 39.3416199999999\nExp number, first 60 AAs: 17.07688\nTotal prob of N-in: 0.84431\nPOSSIBLE N-term signal sequence\ninside 1 11\nTMhelix 12 29\noutside 30 298\nTMhelix 299 321\ninside 322 330

30505 GCF_900134275.1_11893_6_13 Mycobacterium abscessus subsp. bolletii Terrabacteria group; Actinobacteria; Actinobacteria; Corynebacteriales; Mycobacteriaceae; Mycobacterium; Mycobacterium chelonae group; Mycobacterium abscessus subgroup; Mycobacterium abscessus
MIKHLTRRATVALVVAAAAISPAAYPAMADTATPLNGLFALTPGSCAEGRATGSYFRMIL WP_005073213.1
hypothetical protein [Mycobacterium abscessus] Length: 330\nNumber of predicted TMHs: 2\nExp number of AAs in TMHs: 39.3416199999999\nExp number, first 60 AAs: 17.07688\nTotal prob of N-in: 0.84431\nPOSSIBLE N-term signal sequence\ninside 1 11\nTMhelix 12 29\noutside 30 298\nTMhelix 299 321\ninside 322 330

30506 GCF_900140175.1_10665_2_69 Mycobacterium abscessus subsp. bolletii Terrabacteria group; Actinobacteria; Actinobacteria; Corynebacteriales; Mycobacteriaceae; Mycobacterium; Mycobacterium chelonae group; Mycobacterium abscessus subgroup; Mycobacterium abscessus
MIKHLTRRATVALVVAAAAISPAAYPAMADTATPLNGLFALTPGSCAEGRATGSYFRMIL WP_005073213.1
hypothetical protein [Mycobacterium abscessus] Length: 330\nNumber of predicted TMHs: 2\nExp number of AAs in TMHs: 39.3416199999999\nExp number, first 60 AAs: 17.07688\nTotal prob of N-in: 0.84431\nPOSSIBLE N-term signal sequence\ninside 1 11\nTMhelix 12 29\noutside 30 298\nTMhelix 299 321\ninside 322 330

30507 GCF_900135995.1_12163_1_54 Mycobacterium abscessus subsp. bolletii Terrabacteria group; Actinobacteria; Actinobacteria; Corynebacteriales; Mycobacteriaceae; Mycobacterium; Mycobacterium chelonae group; Mycobacterium abscessus subgroup; Mycobacterium abscessus
MIKHLTRRATVALVVAAAAISPAAYPAMADTATPLNGLFALTPGSCAEGRATGSYFRMIL WP_074254038.1
hypothetical protein [Mycobacterium abscessus] Length: 330\nNumber of predicted TMHs: 2\nExp number of AAs in TMHs: 39.3496799999999\nExp number, first 60 AAs: 17.05555\nTotal prob of N-in: 0.84446\nPOSSIBLE N-term signal sequence\ninside 1 11\nTMhelix 12 29\noutside 30 298\nTMhelix 299 321\ninside 322 330

30508 GCF_900132355.1_10665_4_67 Mycobacterium abscessus subsp. bolletii Terrabacteria group; Actinobacteria; Actinobacteria; Corynebacteriales; Mycobacteriaceae; Mycobacterium; Mycobacterium chelonae group; Mycobacterium abscessus subgroup; Mycobacterium abscessus
MIKHLTRRATVALVVAAAAISPAAYPAMADTATPLNGLFALTPGSCAEGRATGSYFRMIL WP_005073213.1
hypothetical protein [Mycobacterium abscessus] Length: 330\nNumber of predicted TMHs: 2\nExp number of AAs in TMHs: 39.3416199999999\nExp number, first 60 AAs: 17.07688\nTotal prob of N-in: 0.84431\nPOSSIBLE N-term signal sequence\ninside 1 11\nTMhelix 12 29\noutside 30 298\nTMhelix 299 321\ninside 322 330

30509 GCF_900130515.1_10665_3_4 Mycobacterium abscessus subsp. bolletii Terrabacteria group; Actinobacteria; Actinobacteria; Corynebacteriales; Mycobacteriaceae; Mycobacterium; Mycobacterium chelonae group; Mycobacterium abscessus subgroup; Mycobacterium abscessus
MIKHLTRRATVALVVAAAAISPAAYPAMADTATPLNGLFALTPGSCAEGRATGSYFRMIL WP_074254038.1
hypothetical protein [Mycobacterium abscessus] Length: 330\nNumber of predicted TMHs: 2\nExp number of AAs in TMHs: 39.3496799999999\nExp number, first 60 AAs: 17.05555\nTotal prob of N-in: 0.84446\nPOSSIBLE N-term signal sequence\ninside 1 11\nTMhelix 12 29\noutside 30 298\nTMhelix 299 321\ninside 322 330

30510 GCF_000770125.1_ASM77012v1 Mycobacterium abscessus subsp. bolletii Terrabacteria group; Actinobacteria; Actinobacteria; Corynebacteriales; Mycobacteriaceae; Mycobacterium; Mycobacterium chelonae group; Mycobacterium abscessus subgroup; Mycobacterium abscessus
MIKHLTRRATVALVVAAAAISPAAYPAMADTATPLNGLFALTPGSCAEGRATGSYFRMIL WP_005073213.1
hypothetical protein [Mycobacterium abscessus] Length: 330\nNumber of predicted TMHs: 2\nExp number of

AAs in TMHs: 39.3416199999999\nExp number, first 60 AAs: 17.07688\nTotal prob of N-in: 0.84431\nPOSSIBLE N-term signal sequence\ninside 1 11\nTMhelix 12 29\noutside 30 298\nTMhelix 299 321\ninside 322 330

30511 GCF_900137535.1_12163_2_29 Mycobacterium abscessus subsp. bolletii Terrabacteria group; Actinobacteria; Actinobacteria; Corynebacteriales; Mycobacteriaceae; Mycobacterium; Mycobacterium chelonae group; Mycobacterium abscessus subgroup; Mycobacterium abscessus MRGGRRRGLLGFIQVLAAIMASLALVWTAPAAQAISPPVIDPGAVAANEPPMPPEELEST WP_079610166.1 type VII secretion-associated serine protease mycosin [Mycobacterium abscessus] Length: 489\nNumber of predicted TMHs: 2\nExp number of AAs in TMHs: 55.05414\nExp number, first 60 AAs: 22.16163\nTotal prob of N-in: 0.98239\nPOSSIBLE N-term signal sequence\ninside 1 6\nTMhelix 7 29\noutside 30 453\nTMhelix 454 473\ninside 474 489

30512 GCF_900133105.1_11893_6_71 Mycobacterium abscessus subsp. bolletii Terrabacteria group; Actinobacteria; Actinobacteria; Corynebacteriales; Mycobacteriaceae; Mycobacterium; Mycobacterium chelonae group; Mycobacterium abscessus subgroup; Mycobacterium abscessus MIKHLTRRATVALVVAAAAISPAAYPAMADTATPLNGLFALTPGSCAEGRATGSYFRMIL WP_005073213.1 hypothetical protein [Mycobacterium abscessus] Length: 330\nNumber of predicted TMHs: 2\nExp number of AAs in TMHs: 39.3416199999999\nExp number, first 60 AAs: 17.07688\nTotal prob of N-in: 0.84431\nPOSSIBLE N-term signal sequence\ninside 1 11\nTMhelix 12 29\noutside 30 298\nTMhelix 299 321\ninside 322 330

30513 GCF_900131215.1_10665_3_64 Mycobacterium abscessus subsp. bolletii Terrabacteria group; Actinobacteria; Actinobacteria; Corynebacteriales; Mycobacteriaceae; Mycobacterium; Mycobacterium chelonae group; Mycobacterium abscessus subgroup; Mycobacterium abscessus MIKHLTRRATVALVVAAAAISPAAYPAMADTATPLNGLFALTPGSCAEGRATGSYFRMIL WP_005073213.1 hypothetical protein [Mycobacterium abscessus] Length: 330\nNumber of predicted TMHs: 2\nExp number of AAs in TMHs: 39.3416199999999\nExp number, first 60 AAs: 17.07688\nTotal prob of N-in: 0.84431\nPOSSIBLE N-term signal sequence\ninside 1 11\nTMhelix 12 29\noutside 30 298\nTMhelix 299 321\ninside 322 330

30514 GCF_900131745.1_10665_4_31 Mycobacterium abscessus subsp. bolletii Terrabacteria group; Actinobacteria; Actinobacteria; Corynebacteriales; Mycobacteriaceae; Mycobacterium; Mycobacterium chelonae group; Mycobacterium abscessus subgroup; Mycobacterium abscessus MIKHLTRRATVALVVAAAAISPAAYPAMADTATPLNGLFALTPGSCAEGRATGSYFRMIL WP_074254038.1 hypothetical protein [Mycobacterium abscessus] Length: 330\nNumber of predicted TMHs: 2\nExp number of AAs in TMHs: 39.3496799999999\nExp number, first 60 AAs: 17.05555\nTotal prob of N-in: 0.84446\nPOSSIBLE N-term signal sequence\ninside 1 11\nTMhelix 12 29\noutside 30 298\nTMhelix 299 321\ninside 322 330

30515 GCF_900131885.1_10665_4_33 Mycobacterium abscessus subsp. bolletii Terrabacteria group; Actinobacteria; Actinobacteria; Corynebacteriales; Mycobacteriaceae; Mycobacterium; Mycobacterium chelonae group; Mycobacterium abscessus subgroup; Mycobacterium abscessus MIKHLTRRATVALVVAAAAISPAAYPAMADTATPLNGLFALTPGSCAEGRATGSYFRMIL WP_005073213.1 hypothetical protein [Mycobacterium abscessus] Length: 330\nNumber of predicted TMHs: 2\nExp number of AAs in TMHs: 39.3416199999999\nExp number, first 60 AAs: 17.07688\nTotal prob of N-in: 0.84431\nPOSSIBLE N-term signal sequence\ninside 1 11\nTMhelix 12 29\noutside 30 298\nTMhelix 299 321\ninside 322 330

30516 GCF_900132305.1_10665_4_62 Mycobacterium abscessus subsp. bolletii Terrabacteria group; Actinobacteria; Actinobacteria; Corynebacteriales; Mycobacteriaceae; Mycobacterium; Mycobacterium chelonae group; Mycobacterium abscessus subgroup; Mycobacterium abscessus MIKHLTRRATVALVVAAAAISPAAYPAMADTATPLNGLFALTPGSCAEGRATGSYFRMIL WP_074254038.1 hypothetical protein [Mycobacterium abscessus] Length: 330\nNumber of predicted TMHs: 2\nExp number of AAs in TMHs: 39.3496799999999\nExp number, first 60 AAs: 17.05555\nTotal prob of N-in: 0.84446\nPOSSIBLE N-term signal sequence\ninside 1 11\nTMhelix 12 29\noutside 30 298\nTMhelix 299 321\ninside 322 330

30517 GCF_900135235.1_10625_4_36 Mycobacterium abscessus subsp. bolletii Terrabacteria group; Actinobacteria; Actinobacteria; Corynebacteriales; Mycobacteriaceae; Mycobacterium; Mycobacterium chelonae group; Mycobacterium abscessus subgroup; Mycobacterium abscessus MIKHLTRRATVALVVAAAAISPAAYPAMADTATPLNGLFALTPGSCAEGRATGSYFRMIL WP_005073213.1 hypothetical protein [Mycobacterium abscessus] Length: 330\nNumber of predicted TMHs: 2\nExp number of AAs in TMHs: 39.3416199999999\nExp number, first 60 AAs: 17.07688\nTotal prob of N-in: 0.84431\nPOSSIBLE N-term signal sequence\ninside 1 11\nTMhelix 12 29\noutside 30 298\nTMhelix 299 321\ninside 322 330

30518 GCF_900134925.1_10208_3_33 Mycobacterium abscessus subsp. bolletii Terrabacteria group; Actinobacteria; Actinobacteria; Corynebacteriales; Mycobacteriaceae; Mycobacterium; Mycobacterium chelonae group;

Mycobacterium abscessus subgroup; Mycobacterium abscessus
MIKHLTRRATVALVVAAAAISPAAYPAVADTATPLNGLFALTPGSCAEGRATGSYFRMIL WP_074330522.1 hypothetical protein [Mycobacterium abscessus] Length: 330\nNumber of predicted TMHs: 1\nExp number of AAs in TMHs: 39.25295\nExp number, first 60 AAs: 16.98073\nTotal prob of N-in: 0.83211\nPOSSIBLE N-term signal sequence\noutside 1 298\nTMhelix 299 321\ninside 322 330

30519 GCF_900134905.1_10208_3_34 Mycobacterium abscessus subsp. bolletii Terrabacteria group; Actinobacteria; Actinobacteria; Corynebacteriales; Mycobacteriaceae; Mycobacterium; Mycobacterium chelonae group; Mycobacterium abscessus subgroup; Mycobacterium abscessus
MIKHLTRRATVALVVAAAAISPAAYPAVADTATPLNGLFALTPGSCAEGRATGSYFRMIL WP_074330522.1 hypothetical protein [Mycobacterium abscessus] Length: 330\nNumber of predicted TMHs: 1\nExp number of AAs in TMHs: 39.25295\nExp number, first 60 AAs: 16.98073\nTotal prob of N-in: 0.83211\nPOSSIBLE N-term signal sequence\noutside 1 298\nTMhelix 299 321\ninside 322 330

30520 GCF_002013835.1_ASM201383v1 Mycobacterium abscessus subsp. bolletii Terrabacteria group; Actinobacteria; Actinobacteria; Corynebacteriales; Mycobacteriaceae; Mycobacterium; Mycobacterium chelonae group; Mycobacterium abscessus subgroup; Mycobacterium abscessus
MIKHLTRRATVALVVAAAAISPAAYPAMADTATPLNGLFALTPGSCAEGRATGSYFRMIL WP_005073213.1 hypothetical protein [Mycobacterium abscessus] Length: 330\nNumber of predicted TMHs: 2\nExp number of AAs in TMHs: 39.3416199999999\nExp number, first 60 AAs: 17.07688\nTotal prob of N-in: 0.84431\nPOSSIBLE N-term signal sequence\ninside 1 11\nTMhelix 12 29\noutside 30 298\nTMhelix 299 321\ninside 322 330

30521 GCF_000758225.1_ASM75822v1 Mycobacterium abscessus subsp. bolletii Terrabacteria group; Actinobacteria; Actinobacteria; Corynebacteriales; Mycobacteriaceae; Mycobacterium; Mycobacterium chelonae group; Mycobacterium abscessus subgroup; Mycobacterium abscessus
MIKHLTRRATVALVVAAAAISPAAYPAVADTATPLNGLFALTPGSCAEGRATGSYFRMIL WP_005063674.1 hypothetical protein [Mycobacterium abscessus] Length: 330\nNumber of predicted TMHs: 1\nExp number of AAs in TMHs: 39.25259\nExp number, first 60 AAs: 16.98187\nTotal prob of N-in: 0.83210\nPOSSIBLE N-term signal sequence\noutside 1 298\nTMhelix 299 321\ninside 322 330

30522 GCF_900139985.1_10660_1_49 Mycobacterium abscessus subsp. bolletii Terrabacteria group; Actinobacteria; Actinobacteria; Corynebacteriales; Mycobacteriaceae; Mycobacterium; Mycobacterium chelonae group; Mycobacterium abscessus subgroup; Mycobacterium abscessus
MIKHLTRRATVALVVAAAAISPAAYPAMADTATPLNGLFALTPGSCAEGRATGSYFRMIL WP_005073213.1 hypothetical protein [Mycobacterium abscessus] Length: 330\nNumber of predicted TMHs: 2\nExp number of AAs in TMHs: 39.3416199999999\nExp number, first 60 AAs: 17.07688\nTotal prob of N-in: 0.84431\nPOSSIBLE N-term signal sequence\ninside 1 11\nTMhelix 12 29\noutside 30 298\nTMhelix 299 321\ninside 322 330

30523 GCF_900132055.1_10665_4_50 Mycobacterium abscessus subsp. bolletii Terrabacteria group; Actinobacteria; Actinobacteria; Corynebacteriales; Mycobacteriaceae; Mycobacterium; Mycobacterium chelonae group; Mycobacterium abscessus subgroup; Mycobacterium abscessus
MIKHLTRRATVALVVAAAAISPAAYPAMADTATPLNGLFALTPGSCAEGRATGSYFRMIL WP_074254038.1 hypothetical protein [Mycobacterium abscessus] Length: 330\nNumber of predicted TMHs: 2\nExp number of AAs in TMHs: 39.3496799999999\nExp number, first 60 AAs: 17.05555\nTotal prob of N-in: 0.84446\nPOSSIBLE N-term signal sequence\ninside 1 11\nTMhelix 12 29\noutside 30 298\nTMhelix 299 321\ninside 322 330

30524 GCF_900131035.1_10665_3_48 Mycobacterium abscessus subsp. bolletii Terrabacteria group; Actinobacteria; Actinobacteria; Corynebacteriales; Mycobacteriaceae; Mycobacterium; Mycobacterium chelonae group; Mycobacterium abscessus subgroup; Mycobacterium abscessus
MIKHLTRRATVALVVAAAAISPAAYPAMADTATPLNGLFALTPGSCAEGRATGSYFRMIL WP_074254038.1 hypothetical protein [Mycobacterium abscessus] Length: 330\nNumber of predicted TMHs: 2\nExp number of AAs in TMHs: 39.3496799999999\nExp number, first 60 AAs: 17.05555\nTotal prob of N-in: 0.84446\nPOSSIBLE N-term signal sequence\ninside 1 11\nTMhelix 12 29\noutside 30 298\nTMhelix 299 321\ninside 322 330

30525 GCF_900132145.1_10665_4_58 Mycobacterium abscessus subsp. bolletii Terrabacteria group; Actinobacteria; Actinobacteria; Corynebacteriales; Mycobacteriaceae; Mycobacterium; Mycobacterium chelonae group; Mycobacterium abscessus subgroup; Mycobacterium abscessus
MIKHLTRRATVALVVAAAAISPAAYPAMADTATPLNGLFALTPGSCAEGRATGSYFRMIL WP_074254038.1 hypothetical protein [Mycobacterium abscessus] Length: 330\nNumber of predicted TMHs: 2\nExp number of AAs in TMHs: 39.3496799999999\nExp number, first 60 AAs: 17.05555\nTotal prob of N-in: 0.84446\nPOSSIBLE N-term signal sequence\ninside 1 11\nTMhelix 12 29\noutside 30 298\nTMhelix 299 321\ninside 322 330

30526 GCF_900137855.1_12163_2_6 Mycobacterium abscessus subsp. bolletii Terrabacteria group; Actinobacteria; Actinobacteria; Corynebacteriales; Mycobacteriaceae; Mycobacterium; Mycobacterium chelonae group; Mycobacterium abscessus subgroup; Mycobacterium abscessus
MIKHLTRRATVALVVAAAAISPAAYPAMADTATPLNGLFALTPGSCAEGRATGSYFRMIL WP_074254038.1
hypothetical protein [Mycobacterium abscessus] Length: 330\nNumber of predicted TMHs: 2\nExp number of AAs in TMHs: 39.3496799999999\nExp number, first 60 AAs: 17.05555\nTotal prob of N-in: 0.84446\nPOSSIBLE N-term signal sequence\ninside 1 11\nTMhelix 12 29\noutside 30 298\nTMhelix 299 321\ninside 322 330

30527 GCF_900132285.1_10665_4_61 Mycobacterium abscessus subsp. bolletii Terrabacteria group; Actinobacteria; Actinobacteria; Corynebacteriales; Mycobacteriaceae; Mycobacterium; Mycobacterium chelonae group; Mycobacterium abscessus subgroup; Mycobacterium abscessus
MIKHLTRRATVALVVAAAAISPAAYPAMADTATPLNGLFALTPGSCAEGRATGSYFRMIL WP_005073213.1
hypothetical protein [Mycobacterium abscessus] Length: 330\nNumber of predicted TMHs: 2\nExp number of AAs in TMHs: 39.3416199999999\nExp number, first 60 AAs: 17.07688\nTotal prob of N-in: 0.84431\nPOSSIBLE N-term signal sequence\ninside 1 11\nTMhelix 12 29\noutside 30 298\nTMhelix 299 321\ninside 322 330

30528 GCF_900130295.1_10625_5_61 Mycobacterium abscessus subsp. bolletii Terrabacteria group; Actinobacteria; Actinobacteria; Corynebacteriales; Mycobacteriaceae; Mycobacterium; Mycobacterium chelonae group; Mycobacterium abscessus subgroup; Mycobacterium abscessus
MIKHLTRRATVALVVAAAAISPAAYPAMADTATPLNGLFALTPGSCAEGRATGSYFRMIL WP_074254038.1
hypothetical protein [Mycobacterium abscessus] Length: 330\nNumber of predicted TMHs: 2\nExp number of AAs in TMHs: 39.3496799999999\nExp number, first 60 AAs: 17.05555\nTotal prob of N-in: 0.84446\nPOSSIBLE N-term signal sequence\ninside 1 11\nTMhelix 12 29\noutside 30 298\nTMhelix 299 321\ninside 322 330

30529 GCF_900133925.1_10250_1_65 Mycobacterium abscessus subsp. bolletii Terrabacteria group; Actinobacteria; Actinobacteria; Corynebacteriales; Mycobacteriaceae; Mycobacterium; Mycobacterium chelonae group; Mycobacterium abscessus subgroup; Mycobacterium abscessus
MIKHLTRRATVALVVAAAAISPAAYPAMADTATPLNGLFALTPGSCAEGRATGSYFRMIL WP_074254038.1
hypothetical protein [Mycobacterium abscessus] Length: 330\nNumber of predicted TMHs: 2\nExp number of AAs in TMHs: 39.3496799999999\nExp number, first 60 AAs: 17.05555\nTotal prob of N-in: 0.84446\nPOSSIBLE N-term signal sequence\ninside 1 11\nTMhelix 12 29\noutside 30 298\nTMhelix 299 321\ninside 322 330

30530 GCF_900133055.1_11893_6_67 Mycobacterium abscessus subsp. bolletii Terrabacteria group; Actinobacteria; Actinobacteria; Corynebacteriales; Mycobacteriaceae; Mycobacterium; Mycobacterium chelonae group; Mycobacterium abscessus subgroup; Mycobacterium abscessus
MIKHLTRRATVALVVAAAAISPAAYPAMADTATPLNGLFALTPGSCAEGRATGSYFRMIL WP_005073213.1
hypothetical protein [Mycobacterium abscessus] Length: 330\nNumber of predicted TMHs: 2\nExp number of AAs in TMHs: 39.3416199999999\nExp number, first 60 AAs: 17.07688\nTotal prob of N-in: 0.84431\nPOSSIBLE N-term signal sequence\ninside 1 11\nTMhelix 12 29\noutside 30 298\nTMhelix 299 321\ninside 322 330

30531 GCF_900137195.1_12082_5_65 Mycobacterium abscessus subsp. bolletii Terrabacteria group; Actinobacteria; Actinobacteria; Corynebacteriales; Mycobacteriaceae; Mycobacterium; Mycobacterium chelonae group; Mycobacterium abscessus subgroup; Mycobacterium abscessus
MIKHLTRRATVALVVAAAAISPAAYPAMADTATPLNGLFALTPGSCAEGRATGSYFRMIL WP_074254038.1
hypothetical protein [Mycobacterium abscessus] Length: 330\nNumber of predicted TMHs: 2\nExp number of AAs in TMHs: 39.3496799999999\nExp number, first 60 AAs: 17.05555\nTotal prob of N-in: 0.84446\nPOSSIBLE N-term signal sequence\ninside 1 11\nTMhelix 12 29\noutside 30 298\nTMhelix 299 321\ninside 322 330

30532 GCF_900137135.1_11893_6_17 Mycobacterium abscessus subsp. bolletii Terrabacteria group; Actinobacteria; Actinobacteria; Corynebacteriales; Mycobacteriaceae; Mycobacterium; Mycobacterium chelonae group; Mycobacterium abscessus subgroup; Mycobacterium abscessus
MIKHLTRRATVALVVAAAAISPAAYPAMADTATPLNGLFALTPGSCAEGRATGSYFRMIL WP_005073213.1
hypothetical protein [Mycobacterium abscessus] Length: 330\nNumber of predicted TMHs: 2\nExp number of AAs in TMHs: 39.3416199999999\nExp number, first 60 AAs: 17.07688\nTotal prob of N-in: 0.84431\nPOSSIBLE N-term signal sequence\ninside 1 11\nTMhelix 12 29\noutside 30 298\nTMhelix 299 321\ninside 322 330

30533 GCF_900133785.1_12163_1_24 Mycobacterium abscessus subsp. bolletii Terrabacteria group; Actinobacteria; Actinobacteria; Corynebacteriales; Mycobacteriaceae; Mycobacterium; Mycobacterium chelonae group; Mycobacterium abscessus subgroup; Mycobacterium abscessus
MIKHLTRRATVALVVAAAAISPAAYPAMADTATPLNGLFALTPGSCAEGRATGSYFRMIL WP_005073213.1
hypothetical protein [Mycobacterium abscessus] Length: 330\nNumber of predicted TMHs: 2\nExp number of

AAs in TMHs: 39.3416199999999\nExp number, first 60 AAs:  17.07688\nTotal prob of N-in:   0.84431\nPOSSIBLE N-term signal sequence\ninside   1   11\nTMhelix   12   29\noutside   30   298\nTMhelix   299   321\ninside   322   330

30534 GCF_900133415.1_10250_1_50 Mycobacterium abscessus subsp. bolletii Terrabacteria group; Actinobacteria; Actinobacteria; Corynebacteriales; Mycobacteriaceae; Mycobacterium; Mycobacterium chelonae group; Mycobacterium abscessus subgroup; Mycobacterium abscessus
MIKHLTRRATVALVVAAAAISPAAYPAMADTATPLNGLFALTPGSCAEGRATGSYFRMIL WP_074254038.1
hypothetical protein [Mycobacterium abscessus] Length: 330\nNumber of predicted TMHs:  2\nExp number of AAs in TMHs: 39.3496799999999\nExp number, first 60 AAs:  17.05555\nTotal prob of N-in:   0.84446\nPOSSIBLE N-term signal sequence\ninside   1   11\nTMhelix   12   29\noutside   30   298\nTMhelix   299   321\ninside   322   330

30535 GCF_900136655.1_10625_5_22 Mycobacterium abscessus subsp. bolletii Terrabacteria group; Actinobacteria; Actinobacteria; Corynebacteriales; Mycobacteriaceae; Mycobacterium; Mycobacterium chelonae group; Mycobacterium abscessus subgroup; Mycobacterium abscessus
MIKHLTRRATVALVVAAAAISPAAYPAMADTATPLNGLFALTPGSCAEGRATGSYFRMIL WP_005073213.1
hypothetical protein [Mycobacterium abscessus] Length: 330\nNumber of predicted TMHs:  2\nExp number of AAs in TMHs: 39.3416199999999\nExp number, first 60 AAs:  17.07688\nTotal prob of N-in:   0.84431\nPOSSIBLE N-term signal sequence\ninside   1   11\nTMhelix   12   29\noutside   30   298\nTMhelix   299   321\ninside   322   330

30536 GCF_900131255.1_10665_3_70 Mycobacterium abscessus subsp. bolletii Terrabacteria group; Actinobacteria; Actinobacteria; Corynebacteriales; Mycobacteriaceae; Mycobacterium; Mycobacterium chelonae group; Mycobacterium abscessus subgroup; Mycobacterium abscessus
MIKHLTRRATVALVVAAAAISPAAYPAMADTATPLNGLFALTPGSCAEGRATGSYFRMIL WP_005073213.1
hypothetical protein [Mycobacterium abscessus] Length: 330\nNumber of predicted TMHs:  2\nExp number of AAs in TMHs: 39.3416199999999\nExp number, first 60 AAs:  17.07688\nTotal prob of N-in:   0.84431\nPOSSIBLE N-term signal sequence\ninside   1   11\nTMhelix   12   29\noutside   30   298\nTMhelix   299   321\ninside   322   330

30537 GCF_900133045.1_11893_6_64 Mycobacterium abscessus subsp. bolletii Terrabacteria group; Actinobacteria; Actinobacteria; Corynebacteriales; Mycobacteriaceae; Mycobacterium; Mycobacterium chelonae group; Mycobacterium abscessus subgroup; Mycobacterium abscessus
MIKHLTRRATVALVVAAAAISPAAYPAMADTATPLNGLFALTPGSCAEGRATGSYFRMIL WP_005073213.1
hypothetical protein [Mycobacterium abscessus] Length: 330\nNumber of predicted TMHs:  2\nExp number of AAs in TMHs: 39.3416199999999\nExp number, first 60 AAs:  17.07688\nTotal prob of N-in:   0.84431\nPOSSIBLE N-term signal sequence\ninside   1   11\nTMhelix   12   29\noutside   30   298\nTMhelix   299   321\ninside   322   330

30538 GCF_900131295.1_10665_3_74 Mycobacterium abscessus subsp. bolletii Terrabacteria group; Actinobacteria; Actinobacteria; Corynebacteriales; Mycobacteriaceae; Mycobacterium; Mycobacterium chelonae group; Mycobacterium abscessus subgroup; Mycobacterium abscessus
MIKHLTRRATVALVVAAAAISPAAYPAMADTATPLNGLFALTPGSCAEGRATGSYFRMIL WP_074254038.1
hypothetical protein [Mycobacterium abscessus] Length: 330\nNumber of predicted TMHs:  2\nExp number of AAs in TMHs: 39.3496799999999\nExp number, first 60 AAs:  17.05555\nTotal prob of N-in:   0.84446\nPOSSIBLE N-term signal sequence\ninside   1   11\nTMhelix   12   29\noutside   30   298\nTMhelix   299   321\ninside   322   330

30539 GCF_900140215.1_10665_2_75 Mycobacterium abscessus subsp. bolletii Terrabacteria group; Actinobacteria; Actinobacteria; Corynebacteriales; Mycobacteriaceae; Mycobacterium; Mycobacterium chelonae group; Mycobacterium abscessus subgroup; Mycobacterium abscessus
MIKHLTRRATVALVVAAAAISPAAYPAMADTATPLNGLFALTPGSCAEGRATGSYFRMIL WP_074254038.1
hypothetical protein [Mycobacterium abscessus] Length: 330\nNumber of predicted TMHs:  2\nExp number of AAs in TMHs: 39.3496799999999\nExp number, first 60 AAs:  17.05555\nTotal prob of N-in:   0.84446\nPOSSIBLE N-term signal sequence\ninside   1   11\nTMhelix   12   29\noutside   30   298\nTMhelix   299   321\ninside   322   330

30540 GCF_900132765.1_10250_1_26 Mycobacterium abscessus subsp. bolletii Terrabacteria group; Actinobacteria; Actinobacteria; Corynebacteriales; Mycobacteriaceae; Mycobacterium; Mycobacterium chelonae group; Mycobacterium abscessus subgroup; Mycobacterium abscessus
MIKHLTRRATVALVVAAAAISPAAYPAMADTATPLNGLFALTPGSCAEGRATGSYFRMIL WP_005073213.1
hypothetical protein [Mycobacterium abscessus] Length: 330\nNumber of predicted TMHs:  2\nExp number of AAs in TMHs: 39.3416199999999\nExp number, first 60 AAs:  17.07688\nTotal prob of N-in:   0.84431\nPOSSIBLE N-term signal sequence\ninside   1   11\nTMhelix   12   29\noutside   30   298\nTMhelix   299   321\ninside   322   330

30541 GCF_900140055.1_10665_2_60 Mycobacterium abscessus subsp. bolletii Terrabacteria group; Actinobacteria; Actinobacteria; Corynebacteriales; Mycobacteriaceae; Mycobacterium; Mycobacterium chelonae group;

Mycobacterium abscessus subgroup; Mycobacterium abscessus
MIKHLTRRATVALVVAAAAISPAAYPAMADTATPLNGLFALTPGSCAEGRATGSYFRMIL WP_005073213.1 hypothetical protein [Mycobacterium abscessus] Length: 330\nNumber of predicted TMHs: 2\nExp number of AAs in TMHs: 39.3416199999999\nExp number, first 60 AAs: 17.07688\nTotal prob of N-in: 0.84431\nPOSSIBLE N-term signal sequence\ninside 1 11\nTMhelix 12 29\noutside 30 298\nTMhelix 299 321\ninside 322 330

30542 GCF_900133875.1_12163_2_61 Mycobacterium abscessus subsp. massiliense Terrabacteria group; Actinobacteria; Actinobacteria; Corynebacteriales; Mycobacteriaceae; Mycobacterium; Mycobacterium chelonae group; Mycobacterium abscessus subgroup; Mycobacterium abscessus
MIKHLTRRATVALVVAAAAISPAAYPAMADTATPLNGLFALTPGSCAEGRATGSYFRMIL WP_074356318.1 hypothetical protein [Mycobacterium abscessus] Length: 330\nNumber of predicted TMHs: 2\nExp number of AAs in TMHs: 39.34118\nExp number, first 60 AAs: 17.07668\nTotal prob of N-in: 0.84431\nPOSSIBLE N-term signal sequence\ninside 1 11\nTMhelix 12 29\noutside 30 298\nTMhelix 299 321\ninside 322 330

30543 GCF_900138305.1_11893_7_42 Mycobacterium abscessus subsp. massiliense Terrabacteria group; Actinobacteria; Actinobacteria; Corynebacteriales; Mycobacteriaceae; Mycobacterium; Mycobacterium chelonae group; Mycobacterium abscessus subgroup; Mycobacterium abscessus
MIKHLTRRATVALVVAAAAISPAAYPAVADTATPLNGLFALTPGSCAEGRATGSYFRMIL WP_005063674.1 hypothetical protein [Mycobacterium abscessus] Length: 330\nNumber of predicted TMHs: 1\nExp number of AAs in TMHs: 39.25259\nExp number, first 60 AAs: 16.98187\nTotal prob of N-in: 0.83210\nPOSSIBLE N-term signal sequence\noutside 1 298\nTMhelix 299 321\ninside 322 330

30544 GCF_900138325.1_11893_7_44 Mycobacterium abscessus subsp. massiliense Terrabacteria group; Actinobacteria; Actinobacteria; Corynebacteriales; Mycobacteriaceae; Mycobacterium; Mycobacterium chelonae group; Mycobacterium abscessus subgroup; Mycobacterium abscessus
MIKHLTRRATVALVVAAAAISPAAYPAVADTATPLNGLFALTPGSCAEGRATGSYFRMIL WP_005063674.1 hypothetical protein [Mycobacterium abscessus] Length: 330\nNumber of predicted TMHs: 1\nExp number of AAs in TMHs: 39.25259\nExp number, first 60 AAs: 16.98187\nTotal prob of N-in: 0.83210\nPOSSIBLE N-term signal sequence\noutside 1 298\nTMhelix 299 321\ninside 322 330

30545 GCF_900138345.1_11893_7_46 Mycobacterium abscessus subsp. massiliense Terrabacteria group; Actinobacteria; Actinobacteria; Corynebacteriales; Mycobacteriaceae; Mycobacterium; Mycobacterium chelonae group; Mycobacterium abscessus subgroup; Mycobacterium abscessus
MIKHLTRRATVALVVAAAAISPAAYPAVADTATPLNGLFALTPGSCAEGRATGSYFRMIL WP_005063674.1 hypothetical protein [Mycobacterium abscessus] Length: 330\nNumber of predicted TMHs: 1\nExp number of AAs in TMHs: 39.25259\nExp number, first 60 AAs: 16.98187\nTotal prob of N-in: 0.83210\nPOSSIBLE N-term signal sequence\noutside 1 298\nTMhelix 299 321\ninside 322 330

30546 GCF_900138365.1_11893_7_50 Mycobacterium abscessus subsp. massiliense Terrabacteria group; Actinobacteria; Actinobacteria; Corynebacteriales; Mycobacteriaceae; Mycobacterium; Mycobacterium chelonae group; Mycobacterium abscessus subgroup; Mycobacterium abscessus
MIKHLTRRATVALVVAAAAISPAAYPAVADTATPLNGLFALTPGSCAEGRATGSYFRMIL WP_005063674.1 hypothetical protein [Mycobacterium abscessus] Length: 330\nNumber of predicted TMHs: 1\nExp number of AAs in TMHs: 39.25259\nExp number, first 60 AAs: 16.98187\nTotal prob of N-in: 0.83210\nPOSSIBLE N-term signal sequence\noutside 1 298\nTMhelix 299 321\ninside 322 330

30547 GCF_900138385.1_11893_7_49 Mycobacterium abscessus subsp. massiliense Terrabacteria group; Actinobacteria; Actinobacteria; Corynebacteriales; Mycobacteriaceae; Mycobacterium; Mycobacterium chelonae group; Mycobacterium abscessus subgroup; Mycobacterium abscessus
MIKHLTRRATVALVVAAAAISPAAYPAVADTATPLNGLFALTPGSCAEGRATGSYFRMIL WP_005063674.1 hypothetical protein [Mycobacterium abscessus] Length: 330\nNumber of predicted TMHs: 1\nExp number of AAs in TMHs: 39.25259\nExp number, first 60 AAs: 16.98187\nTotal prob of N-in: 0.83210\nPOSSIBLE N-term signal sequence\noutside 1 298\nTMhelix 299 321\ninside 322 330

30548 GCF_900138415.1_11893_7_55 Mycobacterium abscessus subsp. massiliense Terrabacteria group; Actinobacteria; Actinobacteria; Corynebacteriales; Mycobacteriaceae; Mycobacterium; Mycobacterium chelonae group; Mycobacterium abscessus subgroup; Mycobacterium abscessus
MIKHLTRRATVALVVAAAAISPAAYPAVADTATPLNGLFALTPGSCAEGRATGSYFRMIL WP_005063674.1 hypothetical protein [Mycobacterium abscessus] Length: 330\nNumber of predicted TMHs: 1\nExp number of AAs in TMHs: 39.25259\nExp number, first 60 AAs: 16.98187\nTotal prob of N-in: 0.83210\nPOSSIBLE N-term signal sequence\noutside 1 298\nTMhelix 299 321\ninside 322 330

30549 GCF_900138435.1_11893_7_56 Mycobacterium abscessus subsp. massiliense Terrabacteria group; Actinobacteria; Actinobacteria; Corynebacteriales; Mycobacteriaceae; Mycobacterium; Mycobacterium chelonae group; Mycobacterium abscessus subgroup; Mycobacterium abscessus
MIKHLTRRATVALVVAAAAISPAAYPAVADTATPLNGLFALTPGSCAEGRATGSYFRMIL WP_005063674.1 hypothetical protein [Mycobacterium abscessus] Length: 330\nNumber of predicted TMHs: 1\nExp number of AAs in TMHs: 39.25259\nExp number, first 60 AAs: 16.98187\nTotal prob of N-in: 0.83210\nPOSSIBLE N-term signal sequence\noutside 1 298\nTMhelix 299 321\ninside 322 330

30550 GCF_900138545.1_20034_1_10 Mycobacterium abscessus subsp. massiliense Terrabacteria group; Actinobacteria; Actinobacteria; Corynebacteriales; Mycobacteriaceae; Mycobacterium; Mycobacterium chelonae group; Mycobacterium abscessus subgroup; Mycobacterium abscessus
MIKHLTRRATVALVVAAAAISPAAYPAVADTATPLNGLFALTPGSCAEGRATGSYFRMIL WP_079617725.1 hypothetical protein [Mycobacterium abscessus] Length: 330\nNumber of predicted TMHs: 1\nExp number of AAs in TMHs: 39.2571699999999\nExp number, first 60 AAs: 16.97134\nTotal prob of N-in: 0.83217\nPOSSIBLE N-term signal sequence\noutside 1 298\nTMhelix 299 321\ninside 322 330

30551 GCF_900138615.1_20034_1_15 Mycobacterium abscessus subsp. massiliense Terrabacteria group; Actinobacteria; Actinobacteria; Corynebacteriales; Mycobacteriaceae; Mycobacterium; Mycobacterium chelonae group; Mycobacterium abscessus subgroup; Mycobacterium abscessus
MIKHLTRRATVALVVAAAAISPAAYPAVADTATPLNGLFALTPGSCAEGRATGSYFRMIL WP_079617725.1 hypothetical protein [Mycobacterium abscessus] Length: 330\nNumber of predicted TMHs: 1\nExp number of AAs in TMHs: 39.2571699999999\nExp number, first 60 AAs: 16.97134\nTotal prob of N-in: 0.83217\nPOSSIBLE N-term signal sequence\noutside 1 298\nTMhelix 299 321\ninside 322 330

30552 GCF_900138795.1_10208_3_49 Mycobacterium abscessus subsp. massiliense Terrabacteria group; Actinobacteria; Actinobacteria; Corynebacteriales; Mycobacteriaceae; Mycobacterium; Mycobacterium chelonae group; Mycobacterium abscessus subgroup; Mycobacterium abscessus
MIKHLTRRATVALVVAAAAISPAAYPAVADTATPLNGLFALTPGSCAEGRATGSYFRMIL WP_005063674.1 hypothetical protein [Mycobacterium abscessus] Length: 330\nNumber of predicted TMHs: 1\nExp number of AAs in TMHs: 39.25259\nExp number, first 60 AAs: 16.98187\nTotal prob of N-in: 0.83210\nPOSSIBLE N-term signal sequence\noutside 1 298\nTMhelix 299 321\ninside 322 330

30553 GCF_900130865.1_10665_3_37 Mycobacterium abscessus subsp. massiliense Terrabacteria group; Actinobacteria; Actinobacteria; Corynebacteriales; Mycobacteriaceae; Mycobacterium; Mycobacterium chelonae group; Mycobacterium abscessus subgroup; Mycobacterium abscessus
MIKHLTRRATVALVVAAAAISPAAYPAVADTATPLNGLFALTPGSCAEGRATGSYFRMIL WP_005063674.1 hypothetical protein [Mycobacterium abscessus] Length: 330\nNumber of predicted TMHs: 1\nExp number of AAs in TMHs: 39.25259\nExp number, first 60 AAs: 16.98187\nTotal prob of N-in: 0.83210\nPOSSIBLE N-term signal sequence\noutside 1 298\nTMhelix 299 321\ninside 322 330

30554 GCF_900130635.1_10665_3_17 Mycobacterium abscessus subsp. massiliense Terrabacteria group; Actinobacteria; Actinobacteria; Corynebacteriales; Mycobacteriaceae; Mycobacterium; Mycobacterium chelonae group; Mycobacterium abscessus subgroup; Mycobacterium abscessus
MIKHLTRRATVALVVAAAAISPAAYPAVADTATPLNGLFALTPGSCAEGRATGSYFRMIL WP_005063674.1 hypothetical protein [Mycobacterium abscessus] Length: 330\nNumber of predicted TMHs: 1\nExp number of AAs in TMHs: 39.25259\nExp number, first 60 AAs: 16.98187\nTotal prob of N-in: 0.83210\nPOSSIBLE N-term signal sequence\noutside 1 298\nTMhelix 299 321\ninside 322 330

30555 GCF_900130585.1_10060_8_14 Mycobacterium abscessus subsp. massiliense Terrabacteria group; Actinobacteria; Actinobacteria; Corynebacteriales; Mycobacteriaceae; Mycobacterium; Mycobacterium chelonae group; Mycobacterium abscessus subgroup; Mycobacterium abscessus
MIKHLTRRATVALVVAAAAISPAAYPAVADTATPLNGLFALTPGSCAEGRATGSYFRMIL WP_005063674.1 hypothetical protein [Mycobacterium abscessus] Length: 330\nNumber of predicted TMHs: 1\nExp number of AAs in TMHs: 39.25259\nExp number, first 60 AAs: 16.98187\nTotal prob of N-in: 0.83210\nPOSSIBLE N-term signal sequence\noutside 1 298\nTMhelix 299 321\ninside 322 330

30556 GCF_900130665.1_10665_3_8 Mycobacterium abscessus subsp. massiliense Terrabacteria group; Actinobacteria; Actinobacteria; Corynebacteriales; Mycobacteriaceae; Mycobacterium; Mycobacterium chelonae group; Mycobacterium abscessus subgroup; Mycobacterium abscessus
MIKHLTRRATVALVVAAAAISPAAYPAVADTATPLNGLFALTPGSCAEGRATGSYFRMIL WP_005063674.1 hypothetical protein [Mycobacterium abscessus] Length: 330\nNumber of predicted TMHs: 1\nExp number of AAs in TMHs:

39.25259\nExp number, first 60 AAs: 16.98187\nTotal prob of N-in: 0.83210\nPOSSIBLE N-term signal sequence\noutside 1 298\nTMhelix 299 321\ninside 322 330

30557 GCF_900130325.1_10625_5_64 Mycobacterium abscessus subsp. massiliense Terrabacteria group; Actinobacteria; Actinobacteria; Corynebacteriales; Mycobacteriaceae; Mycobacterium; Mycobacterium chelonae group; Mycobacterium abscessus subgroup; Mycobacterium abscessus MIKHLTRRATVALVVAAAAISPAAYPAVADTATPLNGLFALTPGSCAEGRATGSYFRMIL WP_005063674.1 hypothetical protein [Mycobacterium abscessus] Length: 330\nNumber of predicted TMHs: 1\nExp number of AAs in TMHs: 39.25259\nExp number, first 60 AAs: 16.98187\nTotal prob of N-in: 0.83210\nPOSSIBLE N-term signal sequence\noutside 1 298\nTMhelix 299 321\ninside 322 330

30558 GCF_900130365.1_10625_5_69 Mycobacterium abscessus subsp. massiliense Terrabacteria group; Actinobacteria; Actinobacteria; Corynebacteriales; Mycobacteriaceae; Mycobacterium; Mycobacterium chelonae group; Mycobacterium abscessus subgroup; Mycobacterium abscessus MIKHLTRRATVALVVAAAAISPAAYPAVADTATPLNGLFALTPGSCAEGRATGSYFRMIL WP_005063674.1 hypothetical protein [Mycobacterium abscessus] Length: 330\nNumber of predicted TMHs: 1\nExp number of AAs in TMHs: 39.25259\nExp number, first 60 AAs: 16.98187\nTotal prob of N-in: 0.83210\nPOSSIBLE N-term signal sequence\noutside 1 298\nTMhelix 299 321\ninside 322 330

30559 GCF_900130345.1_10625_5_66 Mycobacterium abscessus subsp. massiliense Terrabacteria group; Actinobacteria; Actinobacteria; Corynebacteriales; Mycobacteriaceae; Mycobacterium; Mycobacterium chelonae group; Mycobacterium abscessus subgroup; Mycobacterium abscessus MIKHLTRRATVALVVAAAAISPAAYPAVADTATPLNGLFALTPGSCAEGRATGSYFRMIL WP_005063674.1 hypothetical protein [Mycobacterium abscessus] Length: 330\nNumber of predicted TMHs: 1\nExp number of AAs in TMHs: 39.25259\nExp number, first 60 AAs: 16.98187\nTotal prob of N-in: 0.83210\nPOSSIBLE N-term signal sequence\noutside 1 298\nTMhelix 299 321\ninside 322 330

30560 GCF_900130305.1_10625_5_62 Mycobacterium abscessus subsp. massiliense Terrabacteria group; Actinobacteria; Actinobacteria; Corynebacteriales; Mycobacteriaceae; Mycobacterium; Mycobacterium chelonae group; Mycobacterium abscessus subgroup; Mycobacterium abscessus MIKHLTRRATVALVVAAAAISPAAYPAVADTATPLNGLFALTPGSCAEGRATGSYFRMIL WP_005063674.1 hypothetical protein [Mycobacterium abscessus] Length: 330\nNumber of predicted TMHs: 1\nExp number of AAs in TMHs: 39.25259\nExp number, first 60 AAs: 16.98187\nTotal prob of N-in: 0.83210\nPOSSIBLE N-term signal sequence\noutside 1 298\nTMhelix 299 321\ninside 322 330

30561 GCF_900132275.1_11893_6_7 Mycobacterium abscessus subsp. massiliense Terrabacteria group; Actinobacteria; Actinobacteria; Corynebacteriales; Mycobacteriaceae; Mycobacterium; Mycobacterium chelonae group; Mycobacterium abscessus subgroup; Mycobacterium abscessus MIKHLTRRATVALVVAAAAISPAAYPAVADTATPLNGLFALTPGSCAEGRATGSYFRMIL WP_005063674.1 hypothetical protein [Mycobacterium abscessus] Length: 330\nNumber of predicted TMHs: 1\nExp number of AAs in TMHs: 39.25259\nExp number, first 60 AAs: 16.98187\nTotal prob of N-in: 0.83210\nPOSSIBLE N-term signal sequence\noutside 1 298\nTMhelix 299 321\ninside 322 330

30562 GCF_002086375.1_ASM208637v1 Mycobacterium abscessus subsp. massiliense Terrabacteria group; Actinobacteria; Actinobacteria; Corynebacteriales; Mycobacteriaceae; Mycobacterium; Mycobacterium chelonae group; Mycobacterium abscessus subgroup; Mycobacterium abscessus MIKHLTRRATVALVVAAAAISPAAYPAVADTATPLNGLFALTPGSCAEGRATGSYFRMIL WP_005063674.1 hypothetical protein [Mycobacterium abscessus] Length: 330\nNumber of predicted TMHs: 1\nExp number of AAs in TMHs: 39.25259\nExp number, first 60 AAs: 16.98187\nTotal prob of N-in: 0.83210\nPOSSIBLE N-term signal sequence\noutside 1 298\nTMhelix 299 321\ninside 322 330

30563 GCF_900134445.1_12163_1_34 Mycobacterium abscessus subsp. massiliense Terrabacteria group; Actinobacteria; Actinobacteria; Corynebacteriales; Mycobacteriaceae; Mycobacterium; Mycobacterium chelonae group; Mycobacterium abscessus subgroup; Mycobacterium abscessus MIKHLTRRATVALVVAAAAISPAAYPAVADTATPLNGLFALTPGSCAEGRATGSYFRMIL WP_005063674.1 hypothetical protein [Mycobacterium abscessus] Length: 330\nNumber of predicted TMHs: 1\nExp number of AAs in TMHs: 39.25259\nExp number, first 60 AAs: 16.98187\nTotal prob of N-in: 0.83210\nPOSSIBLE N-term signal sequence\noutside 1 298\nTMhelix 299 321\ninside 322 330

30564 GCF_900139625.1_10625_5_67 Mycobacterium abscessus subsp. massiliense Terrabacteria group; Actinobacteria; Actinobacteria; Corynebacteriales; Mycobacteriaceae; Mycobacterium; Mycobacterium chelonae group;

Mycobacterium abscessus subgroup; Mycobacterium abscessus
MIKHLTRRATVALVVAAAAISPAAYPAVADTATPLNGLFALTPGSCAEGRATGSYFRMIL WP_005063674.1 hypothetical protein [Mycobacterium abscessus] Length: 330\nNumber of predicted TMHs: 1\nExp number of AAs in TMHs: 39.25259\nExp number, first 60 AAs: 16.98187\nTotal prob of N-in: 0.83210\nPOSSIBLE N-term signal sequence\noutside 1 298\nTMhelix 299 321\ninside 322 330

30565 GCF_900139655.1_10625_5_71 Mycobacterium abscessus subsp. massiliense Terrabacteria group; Actinobacteria; Actinobacteria; Corynebacteriales; Mycobacteriaceae; Mycobacterium; Mycobacterium chelonae group; Mycobacterium abscessus subgroup; Mycobacterium abscessus
MIKHLTRRATVALVVAAAAISPAAYPAVADTATPLNGLFALTPGSCAEGRATGSYFRMIL WP_005063674.1 hypothetical protein [Mycobacterium abscessus] Length: 330\nNumber of predicted TMHs: 1\nExp number of AAs in TMHs: 39.25259\nExp number, first 60 AAs: 16.98187\nTotal prob of N-in: 0.83210\nPOSSIBLE N-term signal sequence\noutside 1 298\nTMhelix 299 321\ninside 322 330

30566 GCF_900134325.1_11893_6_35 Mycobacterium abscessus subsp. massiliense Terrabacteria group; Actinobacteria; Actinobacteria; Corynebacteriales; Mycobacteriaceae; Mycobacterium; Mycobacterium chelonae group; Mycobacterium abscessus subgroup; Mycobacterium abscessus
MIKHLTRRATVALVVAAAAISPAAYPAVADTATPLNGLFALTPGSCAEGRATGSYFRMIL WP_005063674.1 hypothetical protein [Mycobacterium abscessus] Length: 330\nNumber of predicted TMHs: 1\nExp number of AAs in TMHs: 39.25259\nExp number, first 60 AAs: 16.98187\nTotal prob of N-in: 0.83210\nPOSSIBLE N-term signal sequence\noutside 1 298\nTMhelix 299 321\ninside 322 330

30567 GCF_900131995.1_10665_4_45 Mycobacterium abscessus subsp. massiliense Terrabacteria group; Actinobacteria; Actinobacteria; Corynebacteriales; Mycobacteriaceae; Mycobacterium; Mycobacterium chelonae group; Mycobacterium abscessus subgroup; Mycobacterium abscessus
MIKHLTRRATVALVVAAAAISPAAYPAVADTATPLNGLFALTPGSCAEGRATGSYFRMIL WP_005063674.1 hypothetical protein [Mycobacterium abscessus] Length: 330\nNumber of predicted TMHs: 1\nExp number of AAs in TMHs: 39.25259\nExp number, first 60 AAs: 16.98187\nTotal prob of N-in: 0.83210\nPOSSIBLE N-term signal sequence\noutside 1 298\nTMhelix 299 321\ninside 322 330

30568 GCF_900134475.1_12163_1_36 Mycobacterium abscessus subsp. massiliense Terrabacteria group; Actinobacteria; Actinobacteria; Corynebacteriales; Mycobacteriaceae; Mycobacterium; Mycobacterium chelonae group; Mycobacterium abscessus subgroup; Mycobacterium abscessus
MIKHLTRRATVALVVAAAAISPAAYPAVADTATPLNGLFALTPGSCAEGRATGSYFRMIL WP_005063674.1 hypothetical protein [Mycobacterium abscessus] Length: 330\nNumber of predicted TMHs: 1\nExp number of AAs in TMHs: 39.25259\nExp number, first 60 AAs: 16.98187\nTotal prob of N-in: 0.83210\nPOSSIBLE N-term signal sequence\noutside 1 298\nTMhelix 299 321\ninside 322 330

30569 GCF_900134565.1_12163_1_49 Mycobacterium abscessus subsp. massiliense Terrabacteria group; Actinobacteria; Actinobacteria; Corynebacteriales; Mycobacteriaceae; Mycobacterium; Mycobacterium chelonae group; Mycobacterium abscessus subgroup; Mycobacterium abscessus
MIKHLTRRATVALVVAAAAISPAAYPAVADTATPLNGLFALTPGSCAEGRATGSYFRMIL WP_005063674.1 hypothetical protein [Mycobacterium abscessus] Length: 330\nNumber of predicted TMHs: 1\nExp number of AAs in TMHs: 39.25259\nExp number, first 60 AAs: 16.98187\nTotal prob of N-in: 0.83210\nPOSSIBLE N-term signal sequence\noutside 1 298\nTMhelix 299 321\ninside 322 330

30570 GCF_900131945.1_10665_4_35 Mycobacterium abscessus subsp. massiliense Terrabacteria group; Actinobacteria; Actinobacteria; Corynebacteriales; Mycobacteriaceae; Mycobacterium; Mycobacterium chelonae group; Mycobacterium abscessus subgroup; Mycobacterium abscessus
MIKHLTRRATVALVVAAAAISPAAYPAVADTATPLNGLFALTPGSCAEGRATGSYFRMIL WP_005063674.1 hypothetical protein [Mycobacterium abscessus] Length: 330\nNumber of predicted TMHs: 1\nExp number of AAs in TMHs: 39.25259\nExp number, first 60 AAs: 16.98187\nTotal prob of N-in: 0.83210\nPOSSIBLE N-term signal sequence\noutside 1 298\nTMhelix 299 321\ninside 322 330

30571 GCF_900134545.1_12163_1_45 Mycobacterium abscessus subsp. massiliense Terrabacteria group; Actinobacteria; Actinobacteria; Corynebacteriales; Mycobacteriaceae; Mycobacterium; Mycobacterium chelonae group; Mycobacterium abscessus subgroup; Mycobacterium abscessus
MIKHLTRRATVALVVAAAAISPAAYPAVADTATPLNGLFALTPGSCAEGRATGSYFRMIL WP_005063674.1 hypothetical protein [Mycobacterium abscessus] Length: 330\nNumber of predicted TMHs: 1\nExp number of AAs in TMHs: 39.25259\nExp number, first 60 AAs: 16.98187\nTotal prob of N-in: 0.83210\nPOSSIBLE N-term signal sequence\noutside 1 298\nTMhelix 299 321\ninside 322 330

30572 GCF_900134575.1_12163_1_47 Mycobacterium abscessus subsp. massiliense Terrabacteria group; Actinobacteria; Actinobacteria; Corynebacteriales; Mycobacteriaceae; Mycobacterium; Mycobacterium chelonae group; Mycobacterium abscessus subgroup; Mycobacterium abscessus
MIKHLTRRATVALVVAAAAISPAAYPAVADTATPLNGLFALTPGSCAEGRATGSYFRMIL WP_005063674.1 hypothetical protein [Mycobacterium abscessus] Length: 330\nNumber of predicted TMHs: 1\nExp number of AAs in TMHs: 39.25259\nExp number, first 60 AAs: 16.98187\nTotal prob of N-in: 0.83210\nPOSSIBLE N-term signal sequence\noutside 1 298\nTMhelix 299 321\ninside 322 330

30573 GCF_900134655.1_12163_2_62 Mycobacterium abscessus subsp. massiliense Terrabacteria group; Actinobacteria; Actinobacteria; Corynebacteriales; Mycobacteriaceae; Mycobacterium; Mycobacterium chelonae group; Mycobacterium abscessus subgroup; Mycobacterium abscessus
MIKHLTRRATVALVVAAAAISPAAYPAMADTATPLNGLFALTPGSCAEGRATGSYFRMIL WP_074356318.1 hypothetical protein [Mycobacterium abscessus] Length: 330\nNumber of predicted TMHs: 2\nExp number of AAs in TMHs: 39.34118\nExp number, first 60 AAs: 17.07668\nTotal prob of N-in: 0.84431\nPOSSIBLE N-term signal sequence\ninside 1 11\nTMhelix 12 29\noutside 30 298\nTMhelix 299 321\ninside 322 330

30574 GCF_900139775.1_10660_1_23 Mycobacterium abscessus subsp. massiliense Terrabacteria group; Actinobacteria; Actinobacteria; Corynebacteriales; Mycobacteriaceae; Mycobacterium; Mycobacterium chelonae group; Mycobacterium abscessus subgroup; Mycobacterium abscessus
MIKHLTRRATVALVVAAAAISPAAYPAVADTATPLNGLFALTPGSCAEGRATGSYFRMIL WP_005063674.1 hypothetical protein [Mycobacterium abscessus] Length: 330\nNumber of predicted TMHs: 1\nExp number of AAs in TMHs: 39.25259\nExp number, first 60 AAs: 16.98187\nTotal prob of N-in: 0.83210\nPOSSIBLE N-term signal sequence\noutside 1 298\nTMhelix 299 321\ninside 322 330

30575 GCF_900139795.1_10660_1_21 Mycobacterium abscessus subsp. massiliense Terrabacteria group; Actinobacteria; Actinobacteria; Corynebacteriales; Mycobacteriaceae; Mycobacterium; Mycobacterium chelonae group; Mycobacterium abscessus subgroup; Mycobacterium abscessus
MIKHLTRRATVALVVAAAAISPAAYPAVADTATPLNGLFALTPGSCAEGRATGSYFRMIL WP_005063674.1 hypothetical protein [Mycobacterium abscessus] Length: 330\nNumber of predicted TMHs: 1\nExp number of AAs in TMHs: 39.25259\nExp number, first 60 AAs: 16.98187\nTotal prob of N-in: 0.83210\nPOSSIBLE N-term signal sequence\noutside 1 298\nTMhelix 299 321\ninside 322 330

30576 GCF_900131905.1_10665_4_37 Mycobacterium abscessus subsp. massiliense Terrabacteria group; Actinobacteria; Actinobacteria; Corynebacteriales; Mycobacteriaceae; Mycobacterium; Mycobacterium chelonae group; Mycobacterium abscessus subgroup; Mycobacterium abscessus
MIKHLTRRATVALVVAAAAISPAAYPAVADTATPLNGLFALTPGSCAEGRATGSYFRMIL WP_005063674.1 hypothetical protein [Mycobacterium abscessus] Length: 330\nNumber of predicted TMHs: 1\nExp number of AAs in TMHs: 39.25259\nExp number, first 60 AAs: 16.98187\nTotal prob of N-in: 0.83210\nPOSSIBLE N-term signal sequence\noutside 1 298\nTMhelix 299 321\ninside 322 330

30577 GCF_900134835.1_10208_3_27 Mycobacterium abscessus subsp. massiliense Terrabacteria group; Actinobacteria; Actinobacteria; Corynebacteriales; Mycobacteriaceae; Mycobacterium; Mycobacterium chelonae group; Mycobacterium abscessus subgroup; Mycobacterium abscessus
MIKHLTRRATVALVVAAAAISPAAYPAVADTATPLNGLFALTPGSCAEGRATGSYFRMIL WP_005063674.1 hypothetical protein [Mycobacterium abscessus] Length: 330\nNumber of predicted TMHs: 1\nExp number of AAs in TMHs: 39.25259\nExp number, first 60 AAs: 16.98187\nTotal prob of N-in: 0.83210\nPOSSIBLE N-term signal sequence\noutside 1 298\nTMhelix 299 321\ninside 322 330

30578 GCF_900131895.1_10665_4_34 Mycobacterium abscessus subsp. massiliense Terrabacteria group; Actinobacteria; Actinobacteria; Corynebacteriales; Mycobacteriaceae; Mycobacterium; Mycobacterium chelonae group; Mycobacterium abscessus subgroup; Mycobacterium abscessus
MIKHLTRRATVALVVAAAAISPAAYPAVADTATPLNGLFALTPGSCAEGRATGSYFRMIL WP_005063674.1 hypothetical protein [Mycobacterium abscessus] Length: 330\nNumber of predicted TMHs: 1\nExp number of AAs in TMHs: 39.25259\nExp number, first 60 AAs: 16.98187\nTotal prob of N-in: 0.83210\nPOSSIBLE N-term signal sequence\noutside 1 298\nTMhelix 299 321\ninside 322 330

30579 GCF_900131815.1_11861_7_60 Mycobacterium abscessus subsp. massiliense Terrabacteria group; Actinobacteria; Actinobacteria; Corynebacteriales; Mycobacteriaceae; Mycobacterium; Mycobacterium chelonae group; Mycobacterium abscessus subgroup; Mycobacterium abscessus
MIKHLTRRATVALVVAAAAISPAAYPAVADTATPLNGLFALTPGSCAEGRATGSYFRMIL WP_005063674.1 hypothetical protein [Mycobacterium abscessus] Length: 330\nNumber of predicted TMHs: 1\nExp number of AAs in TMHs:

39.25259\nExp number, first 60 AAs:  16.98187\nTotal prob of N-in:   0.83210\nPOSSIBLE N-term signal sequence\noutside   1   298\nTMhelix   299   321\ninside   322   330

30580 GCF_900131755.1_10665_4_7 Mycobacterium abscessus subsp. massiliense Terrabacteria group; Actinobacteria; Actinobacteria; Corynebacteriales; Mycobacteriaceae; Mycobacterium; Mycobacterium chelonae group; Mycobacterium abscessus subgroup; Mycobacterium abscessus MIKHLTRRATVALVVAAAAISPAAYPAVADTATPLNGLFALTPGSCAEGRATGSYFRMIL WP_005063674.1 hypothetical protein [Mycobacterium abscessus] Length: 330\nNumber of predicted TMHs:  1\nExp number of AAs in TMHs: 39.25259\nExp number, first 60 AAs:  16.98187\nTotal prob of N-in:   0.83210\nPOSSIBLE N-term signal sequence\noutside   1   298\nTMhelix   299   321\ninside   322   330

30581 GCF_900134765.1_10208_3_18 Mycobacterium abscessus subsp. massiliense Terrabacteria group; Actinobacteria; Actinobacteria; Corynebacteriales; Mycobacteriaceae; Mycobacterium; Mycobacterium chelonae group; Mycobacterium abscessus subgroup; Mycobacterium abscessus MIKHLTRRATVALVVAAAAISPAAYPAMADTATPLNGLFALTPGSCAEGRATGSYFRMIL WP_005073213.1 hypothetical protein [Mycobacterium abscessus] Length: 330\nNumber of predicted TMHs:  2\nExp number of AAs in TMHs: 39.3416199999999\nExp number, first 60 AAs:  17.07688\nTotal prob of N-in:   0.84431\nPOSSIBLE N-term signal sequence\ninside   1   11\nTMhelix   12   29\noutside   30   298\nTMhelix   299   321\ninside   322   330

30582 GCF_900139915.1_10660_1_36 Mycobacterium abscessus subsp. massiliense Terrabacteria group; Actinobacteria; Actinobacteria; Corynebacteriales; Mycobacteriaceae; Mycobacterium; Mycobacterium chelonae group; Mycobacterium abscessus subgroup; Mycobacterium abscessus MIKHLTRRATVALVVAAAAISPAAYPAMADTATPLNGLFALTPGSCAEGRATGSYFRMIL WP_005073213.1 hypothetical protein [Mycobacterium abscessus] Length: 330\nNumber of predicted TMHs:  2\nExp number of AAs in TMHs: 39.3416199999999\nExp number, first 60 AAs:  17.07688\nTotal prob of N-in:   0.84431\nPOSSIBLE N-term signal sequence\ninside   1   11\nTMhelix   12   29\noutside   30   298\nTMhelix   299   321\ninside   322   330

30583 GCF_900131735.1_10665_4_5 Mycobacterium abscessus subsp. massiliense Terrabacteria group; Actinobacteria; Actinobacteria; Corynebacteriales; Mycobacteriaceae; Mycobacterium; Mycobacterium chelonae group; Mycobacterium abscessus subgroup; Mycobacterium abscessus MIKHLTRRATVALVVAAAAISPAAYPAVADTATPLNGLFALTPGSCAEGRATGSYFRMIL WP_005063674.1 hypothetical protein [Mycobacterium abscessus] Length: 330\nNumber of predicted TMHs:  1\nExp number of AAs in TMHs: 39.25259\nExp number, first 60 AAs:  16.98187\nTotal prob of N-in:   0.83210\nPOSSIBLE N-term signal sequence\noutside   1   298\nTMhelix   299   321\ninside   322   330

30584 GCF_900131685.1_10665_4_27 Mycobacterium abscessus subsp. massiliense Terrabacteria group; Actinobacteria; Actinobacteria; Corynebacteriales; Mycobacteriaceae; Mycobacterium; Mycobacterium chelonae group; Mycobacterium abscessus subgroup; Mycobacterium abscessus MIKHLTRRATVALVVAAAAISPAAYPAVADTATPLNGLFALTPGSCAEGRATGSYFRMIL WP_005063674.1 hypothetical protein [Mycobacterium abscessus] Length: 330\nNumber of predicted TMHs:  1\nExp number of AAs in TMHs: 39.25259\nExp number, first 60 AAs:  16.98187\nTotal prob of N-in:   0.83210\nPOSSIBLE N-term signal sequence\noutside   1   298\nTMhelix   299   321\ninside   322   330

30585 GCF_900131615.1_10665_4_21 Mycobacterium abscessus subsp. massiliense Terrabacteria group; Actinobacteria; Actinobacteria; Corynebacteriales; Mycobacteriaceae; Mycobacterium; Mycobacterium chelonae group; Mycobacterium abscessus subgroup; Mycobacterium abscessus MIKHLTRRATVALVVAAAAISPAAYPAVADTATPLNGLFALTPGSCAEGRATGSYFRMIL WP_005063674.1 hypothetical protein [Mycobacterium abscessus] Length: 330\nNumber of predicted TMHs:  1\nExp number of AAs in TMHs: 39.25259\nExp number, first 60 AAs:  16.98187\nTotal prob of N-in:   0.83210\nPOSSIBLE N-term signal sequence\noutside   1   298\nTMhelix   299   321\ninside   322   330

30586 GCF_900133695.1_12163_1_18 Mycobacterium abscessus subsp. massiliense Terrabacteria group; Actinobacteria; Actinobacteria; Corynebacteriales; Mycobacteriaceae; Mycobacterium; Mycobacterium chelonae group; Mycobacterium abscessus subgroup; Mycobacterium abscessus MIKHLTRRATVALVVAAAAISPAAYPAVADTATPLNGLFALTPGSCAEGRATGSYFRMIL WP_005063674.1 hypothetical protein [Mycobacterium abscessus] Length: 330\nNumber of predicted TMHs:  1\nExp number of AAs in TMHs: 39.25259\nExp number, first 60 AAs:  16.98187\nTotal prob of N-in:   0.83210\nPOSSIBLE N-term signal sequence\noutside   1   298\nTMhelix   299   321\ninside   322   330

30587 GCF_900135445.1_10625_4_60 Mycobacterium abscessus subsp. massiliense Terrabacteria group; Actinobacteria; Actinobacteria; Corynebacteriales; Mycobacteriaceae; Mycobacterium; Mycobacterium chelonae group;

Mycobacterium abscessus subgroup; Mycobacterium abscessus
MIKHLTRRATVALVVAAAAISPAAYPAVADTATPLNGLFALTPGSCAEGRATGSYFRMIL WP_005063674.1 hypothetical protein [Mycobacterium abscessus] Length: 330\nNumber of predicted TMHs: 1\nExp number of AAs in TMHs: 39.25259\nExp number, first 60 AAs: 16.98187\nTotal prob of N-in: 0.83210\nPOSSIBLE N-term signal sequence\noutside 1 298\nTMhelix 299 321\ninside 322 330

30588 GCF_900134025.1_10250_1_76 Mycobacterium abscessus subsp. massiliense Terrabacteria group; Actinobacteria; Actinobacteria; Corynebacteriales; Mycobacteriaceae; Mycobacterium; Mycobacterium chelonae group; Mycobacterium abscessus subgroup; Mycobacterium abscessus
MIKHLTRRATVALVVAAAAISPAAYPAVADTATPLNGLFALTPGSCAEGRATGSYFRMIL WP_005063674.1 hypothetical protein [Mycobacterium abscessus] Length: 330\nNumber of predicted TMHs: 1\nExp number of AAs in TMHs: 39.25259\nExp number, first 60 AAs: 16.98187\nTotal prob of N-in: 0.83210\nPOSSIBLE N-term signal sequence\noutside 1 298\nTMhelix 299 321\ninside 322 330

30589 GCF_900133725.1_12163_1_17 Mycobacterium abscessus subsp. massiliense Terrabacteria group; Actinobacteria; Actinobacteria; Corynebacteriales; Mycobacteriaceae; Mycobacterium; Mycobacterium chelonae group; Mycobacterium abscessus subgroup; Mycobacterium abscessus
MIKHLTRRATVALVVAAAAISPAAYPAVADTATPLNGLFALTPGSCAEGRATGSYFRMIL WP_005063674.1 hypothetical protein [Mycobacterium abscessus] Length: 330\nNumber of predicted TMHs: 1\nExp number of AAs in TMHs: 39.25259\nExp number, first 60 AAs: 16.98187\nTotal prob of N-in: 0.83210\nPOSSIBLE N-term signal sequence\noutside 1 298\nTMhelix 299 321\ninside 322 330

30590 GCF_900133195.1_12163_2_60 Mycobacterium abscessus subsp. massiliense Terrabacteria group; Actinobacteria; Actinobacteria; Corynebacteriales; Mycobacteriaceae; Mycobacterium; Mycobacterium chelonae group; Mycobacterium abscessus subgroup; Mycobacterium abscessus
MIKHLTRRATVALVVAAAAISPAAYPAMADTATPLNGLFALTPGSCAEGRATGSYFRMIL WP_074356318.1
hypothetical protein [Mycobacterium abscessus] Length: 330\nNumber of predicted TMHs: 2\nExp number of AAs in TMHs: 39.34118\nExp number, first 60 AAs: 17.07668\nTotal prob of N-in: 0.84431\nPOSSIBLE N-term signal sequence\ninside 1 11\nTMhelix 12 29\noutside 30 298\nTMhelix 299 321\ninside 322 330

30591 GCF_900140305.1_10665_2_94 Mycobacterium abscessus subsp. massiliense Terrabacteria group; Actinobacteria; Actinobacteria; Corynebacteriales; Mycobacteriaceae; Mycobacterium; Mycobacterium chelonae group; Mycobacterium abscessus subgroup; Mycobacterium abscessus
MIKHLTRRATVALVVAAAAISPAAYPAVADTATPLNGLFALTPGSCAEGRATGSYFRMIL WP_005063674.1 hypothetical protein [Mycobacterium abscessus] Length: 330\nNumber of predicted TMHs: 1\nExp number of AAs in TMHs: 39.25259\nExp number, first 60 AAs: 16.98187\nTotal prob of N-in: 0.83210\nPOSSIBLE N-term signal sequence\noutside 1 298\nTMhelix 299 321\ninside 322 330

30592 GCF_900131505.1_11893_6_24 Mycobacterium abscessus subsp. massiliense Terrabacteria group; Actinobacteria; Actinobacteria; Corynebacteriales; Mycobacteriaceae; Mycobacterium; Mycobacterium chelonae group; Mycobacterium abscessus subgroup; Mycobacterium abscessus
MIKHLTRRATVALVVAAAAISPAAYPAVADTATPLNGLFALTPGSCAEGRATGSYFRMIL WP_005063674.1 hypothetical protein [Mycobacterium abscessus] Length: 330\nNumber of predicted TMHs: 1\nExp number of AAs in TMHs: 39.25259\nExp number, first 60 AAs: 16.98187\nTotal prob of N-in: 0.83210\nPOSSIBLE N-term signal sequence\noutside 1 298\nTMhelix 299 321\ninside 322 330

30593 GCF_900133765.1_12163_1_25 Mycobacterium abscessus subsp. massiliense Terrabacteria group; Actinobacteria; Actinobacteria; Corynebacteriales; Mycobacteriaceae; Mycobacterium; Mycobacterium chelonae group; Mycobacterium abscessus subgroup; Mycobacterium abscessus
MIKHLTRRATVALVVAAAAISPAAYPAVADTATPLNGLFALTPGSCAEGRATGSYFRMIL WP_005063674.1 hypothetical protein [Mycobacterium abscessus] Length: 330\nNumber of predicted TMHs: 1\nExp number of AAs in TMHs: 39.25259\nExp number, first 60 AAs: 16.98187\nTotal prob of N-in: 0.83210\nPOSSIBLE N-term signal sequence\noutside 1 298\nTMhelix 299 321\ninside 322 330

30594 GCF_900132635.1_12163_2_77 Mycobacterium abscessus subsp. massiliense Terrabacteria group; Actinobacteria; Actinobacteria; Corynebacteriales; Mycobacteriaceae; Mycobacterium; Mycobacterium chelonae group; Mycobacterium abscessus subgroup; Mycobacterium abscessus
MIKHLTRRATVALVVAAAAISPAAYPAMADTATPLNGLFALTPGSCAEGRATGSYFRMIL WP_074356318.1
hypothetical protein [Mycobacterium abscessus] Length: 330\nNumber of predicted TMHs: 2\nExp number of AAs in TMHs: 39.34118\nExp number, first 60 AAs: 17.07668\nTotal prob of N-in: 0.84431\nPOSSIBLE N-term signal sequence\ninside 1 11\nTMhelix 12 29\noutside 30 298\nTMhelix 299 321\ninside 322 330

30595 GCF_900135965.1_12082_5_64 Mycobacterium abscessus subsp. massiliense Terrabacteria group; Actinobacteria; Actinobacteria; Corynebacteriales; Mycobacteriaceae; Mycobacterium; Mycobacterium chelonae group; Mycobacterium abscessus subgroup; Mycobacterium abscessus MIKHLTRRATVALVVAAAAISPAAYPAVADTATPLNGLFALTPGSCAEGRATGSYFRMIL WP_005063674.1 hypothetical protein [Mycobacterium abscessus] Length: 330\nNumber of predicted TMHs: 1\nExp number of AAs in TMHs: 39.25259\nExp number, first 60 AAs: 16.98187\nTotal prob of N-in: 0.83210\nPOSSIBLE N-term signal sequence\noutside 1 298\nTMhelix 299 321\ninside 322 330

30596 GCF_900135985.1_12082_5_79 Mycobacterium abscessus subsp. massiliense Terrabacteria group; Actinobacteria; Actinobacteria; Corynebacteriales; Mycobacteriaceae; Mycobacterium; Mycobacterium chelonae group; Mycobacterium abscessus subgroup; Mycobacterium abscessus MIKHLTRRATVALVVAAAAISPAAYPAVADTATPLNGLFALTPGSCAEGRATGSYFRMIL WP_005063674.1 hypothetical protein [Mycobacterium abscessus] Length: 330\nNumber of predicted TMHs: 1\nExp number of AAs in TMHs: 39.25259\nExp number, first 60 AAs: 16.98187\nTotal prob of N-in: 0.83210\nPOSSIBLE N-term signal sequence\noutside 1 298\nTMhelix 299 321\ninside 322 330

30597 GCF_900136005.1_12163_1_56 Mycobacterium abscessus subsp. massiliense Terrabacteria group; Actinobacteria; Actinobacteria; Corynebacteriales; Mycobacteriaceae; Mycobacterium; Mycobacterium chelonae group; Mycobacterium abscessus subgroup; Mycobacterium abscessus MIKHLTRRATVALVVAAAAISPAAYPAVADTATPLNGLFALTPGSCAEGRATGSYFRMIL WP_005063674.1 hypothetical protein [Mycobacterium abscessus] Length: 330\nNumber of predicted TMHs: 1\nExp number of AAs in TMHs: 39.25259\nExp number, first 60 AAs: 16.98187\nTotal prob of N-in: 0.83210\nPOSSIBLE N-term signal sequence\noutside 1 298\nTMhelix 299 321\ninside 322 330

30598 GCF_900131385.1_10665_4_1 Mycobacterium abscessus subsp. massiliense Terrabacteria group; Actinobacteria; Actinobacteria; Corynebacteriales; Mycobacteriaceae; Mycobacterium; Mycobacterium chelonae group; Mycobacterium abscessus subgroup; Mycobacterium abscessus MIKHLTRRATVALVVAAAAISPAAYPAVADTATPLNGLFALTPGSCAEGRATGSYFRMIL WP_005063674.1 hypothetical protein [Mycobacterium abscessus] Length: 330\nNumber of predicted TMHs: 1\nExp number of AAs in TMHs: 39.25259\nExp number, first 60 AAs: 16.98187\nTotal prob of N-in: 0.83210\nPOSSIBLE N-term signal sequence\noutside 1 298\nTMhelix 299 321\ninside 322 330

30599 GCF_900136195.1_12163_1_75 Mycobacterium abscessus subsp. massiliense Terrabacteria group; Actinobacteria; Actinobacteria; Corynebacteriales; Mycobacteriaceae; Mycobacterium; Mycobacterium chelonae group; Mycobacterium abscessus subgroup; Mycobacterium abscessus MIKHLTRRATVALVVAAAAISPAAYPAVADTATPLNGLFALTPGSCAEGRATGSYFRMIL WP_005063674.1 hypothetical protein [Mycobacterium abscessus] Length: 330\nNumber of predicted TMHs: 1\nExp number of AAs in TMHs: 39.25259\nExp number, first 60 AAs: 16.98187\nTotal prob of N-in: 0.83210\nPOSSIBLE N-term signal sequence\noutside 1 298\nTMhelix 299 321\ninside 322 330

30600 GCF_900136175.1_12163_1_73 Mycobacterium abscessus subsp. massiliense Terrabacteria group; Actinobacteria; Actinobacteria; Corynebacteriales; Mycobacteriaceae; Mycobacterium; Mycobacterium chelonae group; Mycobacterium abscessus subgroup; Mycobacterium abscessus MIKHLTRRATVALVVAAAAISPAAYPAVADTATPLNGLFALTPGSCAEGRATGSYFRMIL WP_005063674.1 hypothetical protein [Mycobacterium abscessus] Length: 330\nNumber of predicted TMHs: 1\nExp number of AAs in TMHs: 39.25259\nExp number, first 60 AAs: 16.98187\nTotal prob of N-in: 0.83210\nPOSSIBLE N-term signal sequence\noutside 1 298\nTMhelix 299 321\ninside 322 330

30601 GCF_900136225.1_12163_1_78 Mycobacterium abscessus subsp. massiliense Terrabacteria group; Actinobacteria; Actinobacteria; Corynebacteriales; Mycobacteriaceae; Mycobacterium; Mycobacterium chelonae group; Mycobacterium abscessus subgroup; Mycobacterium abscessus MIKHLTRRATVALVVAAAAISPAAYPAVADTATPLNGLFALTPGSCAEGRATGSYFRMIL WP_005063674.1 hypothetical protein [Mycobacterium abscessus] Length: 330\nNumber of predicted TMHs: 1\nExp number of AAs in TMHs: 39.25259\nExp number, first 60 AAs: 16.98187\nTotal prob of N-in: 0.83210\nPOSSIBLE N-term signal sequence\noutside 1 298\nTMhelix 299 321\ninside 322 330

30602 GCF_900136375.1_12163_1_89 Mycobacterium abscessus subsp. massiliense Terrabacteria group; Actinobacteria; Actinobacteria; Corynebacteriales; Mycobacteriaceae; Mycobacterium; Mycobacterium chelonae group; Mycobacterium abscessus subgroup; Mycobacterium abscessus MIKHLTRRATVALVVAAAAISPAAYPAVADTATPLNGLFALTPGSCAEGRATGSYFRMIL WP_005063674.1 hypothetical protein [Mycobacterium abscessus] Length: 330\nNumber of predicted TMHs: 1\nExp number of AAs in TMHs:

39.25259\nExp number, first 60 AAs: 16.98187\nTotal prob of N-in: 0.83210\nPOSSIBLE N-term signal sequence\noutside 1 298\nTMhelix 299 321\ninside 322 330

30603 GCF_900137295.1_12163_1_92 Mycobacterium abscessus subsp. massiliense Terrabacteria group; Actinobacteria; Actinobacteria; Corynebacteriales; Mycobacteriaceae; Mycobacterium; Mycobacterium chelonae group; Mycobacterium abscessus subgroup; Mycobacterium abscessus MIKHLTRRATVALVVAAAAISPAAYPAVADTATPLNGLFALTPGSCAEGRATGSYFRMIL WP_005063674.1 hypothetical protein [Mycobacterium abscessus] Length: 330\nNumber of predicted TMHs: 1\nExp number of AAs in TMHs: 39.25259\nExp number, first 60 AAs: 16.98187\nTotal prob of N-in: 0.83210\nPOSSIBLE N-term signal sequence\noutside 1 298\nTMhelix 299 321\ninside 322 330

30604 GCF_900131225.1_10665_3_67 Mycobacterium abscessus subsp. massiliense Terrabacteria group; Actinobacteria; Actinobacteria; Corynebacteriales; Mycobacteriaceae; Mycobacterium; Mycobacterium chelonae group; Mycobacterium abscessus subgroup; Mycobacterium abscessus MIKHLTRRATVALVVAAAAISPAAYPAVADTATPLNGLFALTPGSCAEGRATGSYFRMIL WP_005063674.1 hypothetical protein [Mycobacterium abscessus] Length: 330\nNumber of predicted TMHs: 1\nExp number of AAs in TMHs: 39.25259\nExp number, first 60 AAs: 16.98187\nTotal prob of N-in: 0.83210\nPOSSIBLE N-term signal sequence\noutside 1 298\nTMhelix 299 321\ninside 322 330

30605 GCF_900136345.1_12163_2_79 Mycobacterium abscessus subsp. massiliense Terrabacteria group; Actinobacteria; Actinobacteria; Corynebacteriales; Mycobacteriaceae; Mycobacterium; Mycobacterium chelonae group; Mycobacterium abscessus subgroup; Mycobacterium abscessus MIKHLTRRATVALVVAAAAISPAAYPAVADTATPLNGLFALTPGSCAEGRATGSYFRMIL WP_005063674.1 hypothetical protein [Mycobacterium abscessus] Length: 330\nNumber of predicted TMHs: 1\nExp number of AAs in TMHs: 39.25259\nExp number, first 60 AAs: 16.98187\nTotal prob of N-in: 0.83210\nPOSSIBLE N-term signal sequence\noutside 1 298\nTMhelix 299 321\ninside 322 330

30606 GCF_900136395.1_12163_2_85 Mycobacterium abscessus subsp. massiliense Terrabacteria group; Actinobacteria; Actinobacteria; Corynebacteriales; Mycobacteriaceae; Mycobacterium; Mycobacterium chelonae group; Mycobacterium abscessus subgroup; Mycobacterium abscessus MIKHLTRRATVALVVAAAAISPAAYPAVADTATPLNGLFALTPGSCAEGRATGSYFRMIL WP_005063674.1 hypothetical protein [Mycobacterium abscessus] Length: 330\nNumber of predicted TMHs: 1\nExp number of AAs in TMHs: 39.25259\nExp number, first 60 AAs: 16.98187\nTotal prob of N-in: 0.83210\nPOSSIBLE N-term signal sequence\noutside 1 298\nTMhelix 299 321\ninside 322 330

30607 GCF_900133025.1_11893_6_47 Mycobacterium abscessus subsp. massiliense Terrabacteria group; Actinobacteria; Actinobacteria; Corynebacteriales; Mycobacteriaceae; Mycobacterium; Mycobacterium chelonae group; Mycobacterium abscessus subgroup; Mycobacterium abscessus MIKHLTRRATVALVVAAAAISPAAYPAMADTATPLNGLFALTPGSCAEGRATGSYFRMIL WP_074356318.1 hypothetical protein [Mycobacterium abscessus] Length: 330\nNumber of predicted TMHs: 2\nExp number of AAs in TMHs: 39.34118\nExp number, first 60 AAs: 17.07668\nTotal prob of N-in: 0.84431\nPOSSIBLE N-term signal sequence\ninside 1 11\nTMhelix 12 29\noutside 30 298\nTMhelix 299 321\ninside 322 330

30608 GCF_900137335.1_12163_2_12 Mycobacterium abscessus subsp. massiliense Terrabacteria group; Actinobacteria; Actinobacteria; Corynebacteriales; Mycobacteriaceae; Mycobacterium; Mycobacterium chelonae group; Mycobacterium abscessus subgroup; Mycobacterium abscessus MIKHLTRRATVALVVAAAAISPAAYPAVADTATPLNGLFALTPGSCAEGRATGSYFRMIL WP_005063674.1 hypothetical protein [Mycobacterium abscessus] Length: 330\nNumber of predicted TMHs: 1\nExp number of AAs in TMHs: 39.25259\nExp number, first 60 AAs: 16.98187\nTotal prob of N-in: 0.83210\nPOSSIBLE N-term signal sequence\noutside 1 298\nTMhelix 299 321\ninside 322 330

30609 GCF_900141475.1_12045_8_42 Mycobacterium abscessus subsp. massiliense Terrabacteria group; Actinobacteria; Actinobacteria; Corynebacteriales; Mycobacteriaceae; Mycobacterium; Mycobacterium chelonae group; Mycobacterium abscessus subgroup; Mycobacterium abscessus MIKHLTRRATVALVVAAAAISPAAYPAVADTATPLNGLFALTPGSCAEGRATGSYFRMIL WP_005063674.1 hypothetical protein [Mycobacterium abscessus] Length: 330\nNumber of predicted TMHs: 1\nExp number of AAs in TMHs: 39.25259\nExp number, first 60 AAs: 16.98187\nTotal prob of N-in: 0.83210\nPOSSIBLE N-term signal sequence\noutside 1 298\nTMhelix 299 321\ninside 322 330

30610 GCF_900137475.1_12163_2_22 Mycobacterium abscessus subsp. massiliense Terrabacteria group; Actinobacteria; Actinobacteria; Corynebacteriales; Mycobacteriaceae; Mycobacterium; Mycobacterium chelonae group;

Mycobacterium abscessus subgroup; Mycobacterium abscessus
MIKHLTRRATVALVVAAAAISPAAYPAVADTATPLNGLFALTPGSCAEGRATGSYFRMIL WP_005063674.1 hypothetical protein [Mycobacterium abscessus] Length: 330\nNumber of predicted TMHs: 1\nExp number of AAs in TMHs: 39.25259\nExp number, first 60 AAs: 16.98187\nTotal prob of N-in: 0.83210\nPOSSIBLE N-term signal sequence\noutside 1 298\nTMhelix 299 321\ninside 322 330

30611 GCF_900133605.1_12082_5_55 Mycobacterium abscessus subsp. massiliense Terrabacteria group; Actinobacteria; Actinobacteria; Corynebacteriales; Mycobacteriaceae; Mycobacterium; Mycobacterium chelonae group; Mycobacterium abscessus subgroup; Mycobacterium abscessus
MIKHLTRRATVALVVAAAAISPAAYPAVADTATPLNGLFALTPGSCAEGRATGSYFRMIL WP_005063674.1 hypothetical protein [Mycobacterium abscessus] Length: 330\nNumber of predicted TMHs: 1\nExp number of AAs in TMHs: 39.25259\nExp number, first 60 AAs: 16.98187\nTotal prob of N-in: 0.83210\nPOSSIBLE N-term signal sequence\noutside 1 298\nTMhelix 299 321\ninside 322 330

30612 GCF_900133745.1_12163_1_16 Mycobacterium abscessus subsp. massiliense Terrabacteria group; Actinobacteria; Actinobacteria; Corynebacteriales; Mycobacteriaceae; Mycobacterium; Mycobacterium chelonae group; Mycobacterium abscessus subgroup; Mycobacterium abscessus
MIKHLTRRATVALVVAAAAISPAAYPAVADTATPLNGLFALTPGSCAEGRATGSYFRMIL WP_005063674.1 hypothetical protein [Mycobacterium abscessus] Length: 330\nNumber of predicted TMHs: 1\nExp number of AAs in TMHs: 39.25259\nExp number, first 60 AAs: 16.98187\nTotal prob of N-in: 0.83210\nPOSSIBLE N-term signal sequence\noutside 1 298\nTMhelix 299 321\ninside 322 330

30613 GCF_900137725.1_12163_2_5 Mycobacterium abscessus subsp. massiliense Terrabacteria group; Actinobacteria; Actinobacteria; Corynebacteriales; Mycobacteriaceae; Mycobacterium; Mycobacterium chelonae group; Mycobacterium abscessus subgroup; Mycobacterium abscessus
MIKHLTRRATVALVVAAAAISPAAYPAVADTATPLNGLFALTPGSCAEGRATGSYFRMIL WP_005063674.1 hypothetical protein [Mycobacterium abscessus] Length: 330\nNumber of predicted TMHs: 1\nExp number of AAs in TMHs: 39.25259\nExp number, first 60 AAs: 16.98187\nTotal prob of N-in: 0.83210\nPOSSIBLE N-term signal sequence\noutside 1 298\nTMhelix 299 321\ninside 322 330

30614 GCF_900132245.1_11893_6_30 Mycobacterium abscessus subsp. massiliense Terrabacteria group; Actinobacteria; Actinobacteria; Corynebacteriales; Mycobacteriaceae; Mycobacterium; Mycobacterium chelonae group; Mycobacterium abscessus subgroup; Mycobacterium abscessus
MIKHLTRRATVALVVAAAAISPAAYPAVADTATPLNGLFALTPGSCAEGRATGSYFRMIL WP_005063674.1 hypothetical protein [Mycobacterium abscessus] Length: 330\nNumber of predicted TMHs: 1\nExp number of AAs in TMHs: 39.25259\nExp number, first 60 AAs: 16.98187\nTotal prob of N-in: 0.83210\nPOSSIBLE N-term signal sequence\noutside 1 298\nTMhelix 299 321\ninside 322 330

30615 GCF_900130335.1_10625_5_65 Mycobacterium abscessus subsp. massiliense Terrabacteria group; Actinobacteria; Actinobacteria; Corynebacteriales; Mycobacteriaceae; Mycobacterium; Mycobacterium chelonae group; Mycobacterium abscessus subgroup; Mycobacterium abscessus
MIKHLTRRATVALVVAAAAISPAAYPAVADTATPLNGLFALTPGSCAEGRATGSYFRMIL WP_005063674.1 hypothetical protein [Mycobacterium abscessus] Length: 330\nNumber of predicted TMHs: 1\nExp number of AAs in TMHs: 39.25259\nExp number, first 60 AAs: 16.98187\nTotal prob of N-in: 0.83210\nPOSSIBLE N-term signal sequence\noutside 1 298\nTMhelix 299 321\ninside 322 330

30616 GCF_900130355.1_10625_5_68 Mycobacterium abscessus subsp. massiliense Terrabacteria group; Actinobacteria; Actinobacteria; Corynebacteriales; Mycobacteriaceae; Mycobacterium; Mycobacterium chelonae group; Mycobacterium abscessus subgroup; Mycobacterium abscessus
MIKHLTRRATVALVVAAAAISPAAYPAVADTATPLNGLFALTPGSCAEGRATGSYFRMIL WP_005063674.1 hypothetical protein [Mycobacterium abscessus] Length: 330\nNumber of predicted TMHs: 1\nExp number of AAs in TMHs: 39.25259\nExp number, first 60 AAs: 16.98187\nTotal prob of N-in: 0.83210\nPOSSIBLE N-term signal sequence\noutside 1 298\nTMhelix 299 321\ninside 322 330

30617 GCF_900130625.1_10665_3_16 Mycobacterium abscessus subsp. massiliense Terrabacteria group; Actinobacteria; Actinobacteria; Corynebacteriales; Mycobacteriaceae; Mycobacterium; Mycobacterium chelonae group; Mycobacterium abscessus subgroup; Mycobacterium abscessus
MIKHLTRRATVALVVAAAAISPAAYPAVADTATPLNGLFALTPGSCAEGRATGSYFRMIL WP_005063674.1 hypothetical protein [Mycobacterium abscessus] Length: 330\nNumber of predicted TMHs: 1\nExp number of AAs in TMHs: 39.25259\nExp number, first 60 AAs: 16.98187\nTotal prob of N-in: 0.83210\nPOSSIBLE N-term signal sequence\noutside 1 298\nTMhelix 299 321\ninside 322 330

30618 GCF_900130765.1_10665_3_27 Mycobacterium abscessus subsp. massiliense Terrabacteria group; Actinobacteria; Actinobacteria; Corynebacteriales; Mycobacteriaceae; Mycobacterium; Mycobacterium chelonae group; Mycobacterium abscessus subgroup; Mycobacterium abscessus
MIKHLTRRATVALVVAAAAISPAAYPAVADTATPLNGLFALTPGSCAEGRATGSYFRMIL WP_005063674.1 hypothetical protein [Mycobacterium abscessus] Length: 330\nNumber of predicted TMHs: 1\nExp number of AAs in TMHs: 39.25259\nExp number, first 60 AAs: 16.98187\nTotal prob of N-in: 0.83210\nPOSSIBLE N-term signal sequence\noutside 1 298\nTMhelix 299 321\ninside 322 330

30619 GCF_900130675.1_10665_3_9 Mycobacterium abscessus subsp. massiliense Terrabacteria group; Actinobacteria; Actinobacteria; Corynebacteriales; Mycobacteriaceae; Mycobacterium; Mycobacterium chelonae group; Mycobacterium abscessus subgroup; Mycobacterium abscessus
MIKHLTRRATVALVVAAAAISPAAYPAVADTATPLNGLFALTPGSCAEGRATGSYFRMIL WP_005063674.1 hypothetical protein [Mycobacterium abscessus] Length: 330\nNumber of predicted TMHs: 1\nExp number of AAs in TMHs: 39.25259\nExp number, first 60 AAs: 16.98187\nTotal prob of N-in: 0.83210\nPOSSIBLE N-term signal sequence\noutside 1 298\nTMhelix 299 321\ninside 322 330

30620 GCF_900130745.1_10665_3_22 Mycobacterium abscessus subsp. massiliense Terrabacteria group; Actinobacteria; Actinobacteria; Corynebacteriales; Mycobacteriaceae; Mycobacterium; Mycobacterium chelonae group; Mycobacterium abscessus subgroup; Mycobacterium abscessus
MIKHLTRRATVALVVAAAAISPAAYPAVADTATPLNGLFALTPGSCAEGRATGSYFRMIL WP_005063674.1 hypothetical protein [Mycobacterium abscessus] Length: 330\nNumber of predicted TMHs: 1\nExp number of AAs in TMHs: 39.25259\nExp number, first 60 AAs: 16.98187\nTotal prob of N-in: 0.83210\nPOSSIBLE N-term signal sequence\noutside 1 298\nTMhelix 299 321\ninside 322 330

30621 GCF_900130795.1_10665_3_30 Mycobacterium abscessus subsp. massiliense Terrabacteria group; Actinobacteria; Actinobacteria; Corynebacteriales; Mycobacteriaceae; Mycobacterium; Mycobacterium chelonae group; Mycobacterium abscessus subgroup; Mycobacterium abscessus
MIKHLTRRATVALVVAAAAISPAAYPAVADTATPLNGLFALTPGSCAEGRATGSYFRMIL WP_005063674.1 hypothetical protein [Mycobacterium abscessus] Length: 330\nNumber of predicted TMHs: 1\nExp number of AAs in TMHs: 39.25259\nExp number, first 60 AAs: 16.98187\nTotal prob of N-in: 0.83210\nPOSSIBLE N-term signal sequence\noutside 1 298\nTMhelix 299 321\ninside 322 330

30622 GCF_900131265.1_10665_3_71 Mycobacterium abscessus subsp. massiliense Terrabacteria group; Actinobacteria; Actinobacteria; Corynebacteriales; Mycobacteriaceae; Mycobacterium; Mycobacterium chelonae group; Mycobacterium abscessus subgroup; Mycobacterium abscessus
MIKHLTRRATVALVVAAAAISPAAYPAVADTATPLNGLFALTPGSCAEGRATGSYFRMIL WP_005063674.1 hypothetical protein [Mycobacterium abscessus] Length: 330\nNumber of predicted TMHs: 1\nExp number of AAs in TMHs: 39.25259\nExp number, first 60 AAs: 16.98187\nTotal prob of N-in: 0.83210\nPOSSIBLE N-term signal sequence\noutside 1 298\nTMhelix 299 321\ninside 322 330

30623 GCF_900131305.1_10665_3_76 Mycobacterium abscessus subsp. massiliense Terrabacteria group; Actinobacteria; Actinobacteria; Corynebacteriales; Mycobacteriaceae; Mycobacterium; Mycobacterium chelonae group; Mycobacterium abscessus subgroup; Mycobacterium abscessus
MIKHLTRRATVALVVAAAAISPAAYPAVADTATPLNGLFALTPGSCAEGRATGSYFRMIL WP_005063674.1 hypothetical protein [Mycobacterium abscessus] Length: 330\nNumber of predicted TMHs: 1\nExp number of AAs in TMHs: 39.25259\nExp number, first 60 AAs: 16.98187\nTotal prob of N-in: 0.83210\nPOSSIBLE N-term signal sequence\noutside 1 298\nTMhelix 299 321\ninside 322 330

30624 GCF_900131235.1_10665_3_68 Mycobacterium abscessus subsp. massiliense Terrabacteria group; Actinobacteria; Actinobacteria; Corynebacteriales; Mycobacteriaceae; Mycobacterium; Mycobacterium chelonae group; Mycobacterium abscessus subgroup; Mycobacterium abscessus
MIKHLTRRATVALVVAAAAISPAAYPAVADTATPLNGLFALTPGSCAEGRATGSYFRMIL WP_005063674.1 hypothetical protein [Mycobacterium abscessus] Length: 330\nNumber of predicted TMHs: 1\nExp number of AAs in TMHs: 39.25259\nExp number, first 60 AAs: 16.98187\nTotal prob of N-in: 0.83210\nPOSSIBLE N-term signal sequence\noutside 1 298\nTMhelix 299 321\ninside 322 330

30625 GCF_900131725.1_10665_4_32 Mycobacterium abscessus subsp. massiliense Terrabacteria group; Actinobacteria; Actinobacteria; Corynebacteriales; Mycobacteriaceae; Mycobacterium; Mycobacterium chelonae group; Mycobacterium abscessus subgroup; Mycobacterium abscessus
MIKHLTRRATVALVVAAAAISPAAYPAVADTATPLNGLFALTPGSCAEGRATGSYFRMIL WP_005063674.1 hypothetical protein [Mycobacterium abscessus] Length: 330\nNumber of predicted TMHs: 1\nExp number of AAs in TMHs:

39.25259\nExp number, first 60 AAs: 16.98187\nTotal prob of N-in: 0.83210\nPOSSIBLE N-term signal sequence\noutside 1 298\nTMhelix 299 321\ninside 322 330

30626 GCF_900131865.1_11893_6_6 Mycobacterium abscessus subsp. massiliense Terrabacteria group; Actinobacteria; Actinobacteria; Corynebacteriales; Mycobacteriaceae; Mycobacterium; Mycobacterium chelonae group; Mycobacterium abscessus subgroup; Mycobacterium abscessus MIKHLTRRATVALVVAAAAISPAAYPAVADTATPLNGLFALTPGSCAEGRATGSYFRMIL WP_005063674.1 hypothetical protein [Mycobacterium abscessus] Length: 330\nNumber of predicted TMHs: 1\nExp number of AAs in TMHs: 39.25259\nExp number, first 60 AAs: 16.98187\nTotal prob of N-in: 0.83210\nPOSSIBLE N-term signal sequence\noutside 1 298\nTMhelix 299 321\ninside 322 330

30627 GCF_900132225.1_11893_6_29 Mycobacterium abscessus subsp. massiliense Terrabacteria group; Actinobacteria; Actinobacteria; Corynebacteriales; Mycobacteriaceae; Mycobacterium; Mycobacterium chelonae group; Mycobacterium abscessus subgroup; Mycobacterium abscessus MIKHLTRRATVALVVAAAAISPAAYPAVADTATPLNGLFALTPGSCAEGRATGSYFRMIL WP_005063674.1 hypothetical protein [Mycobacterium abscessus] Length: 330\nNumber of predicted TMHs: 1\nExp number of AAs in TMHs: 39.25259\nExp number, first 60 AAs: 16.98187\nTotal prob of N-in: 0.83210\nPOSSIBLE N-term signal sequence\noutside 1 298\nTMhelix 299 321\ninside 322 330

30628 GCF_900132705.1_10250_1_19 Mycobacterium abscessus subsp. massiliense Terrabacteria group; Actinobacteria; Actinobacteria; Corynebacteriales; Mycobacteriaceae; Mycobacterium; Mycobacterium chelonae group; Mycobacterium abscessus subgroup; Mycobacterium abscessus MIKHLTRRATVALVVAAAAISPAAYPAVADTATPLNGLFALTPGSCAEGRATGSYFRMIL WP_005063674.1 hypothetical protein [Mycobacterium abscessus] Length: 330\nNumber of predicted TMHs: 1\nExp number of AAs in TMHs: 39.25259\nExp number, first 60 AAs: 16.98187\nTotal prob of N-in: 0.83210\nPOSSIBLE N-term signal sequence\noutside 1 298\nTMhelix 299 321\ninside 322 330

30629 GCF_900132725.1_10250_1_20 Mycobacterium abscessus subsp. massiliense Terrabacteria group; Actinobacteria; Actinobacteria; Corynebacteriales; Mycobacteriaceae; Mycobacterium; Mycobacterium chelonae group; Mycobacterium abscessus subgroup; Mycobacterium abscessus MIKHLTRRATVALVVAAAAISPAAYPAVADTATPLNGLFALTPGSCAEGRATGSYFRMIL WP_005063674.1 hypothetical protein [Mycobacterium abscessus] Length: 330\nNumber of predicted TMHs: 1\nExp number of AAs in TMHs: 39.25259\nExp number, first 60 AAs: 16.98187\nTotal prob of N-in: 0.83210\nPOSSIBLE N-term signal sequence\noutside 1 298\nTMhelix 299 321\ninside 322 330

30630 GCF_900133165.1_12163_1_4 Mycobacterium abscessus subsp. massiliense Terrabacteria group; Actinobacteria; Actinobacteria; Corynebacteriales; Mycobacteriaceae; Mycobacterium; Mycobacterium chelonae group; Mycobacterium abscessus subgroup; Mycobacterium abscessus MIKHLTRRATVALVVAAAAISPAAYPAMADTATPLNGLFALTPGSCAEGRATGSYFRMIL WP_005073213.1 hypothetical protein [Mycobacterium abscessus] Length: 330\nNumber of predicted TMHs: 2\nExp number of AAs in TMHs: 39.3416199999999\nExp number, first 60 AAs: 17.07688\nTotal prob of N-in: 0.84431\nPOSSIBLE N-term signal sequence\ninside 1 11\nTMhelix 12 29\noutside 30 298\nTMhelix 299 321\ninside 322 330

30631 GCF_900133185.1_12163_1_1 Mycobacterium abscessus subsp. massiliense Terrabacteria group; Actinobacteria; Actinobacteria; Corynebacteriales; Mycobacteriaceae; Mycobacterium; Mycobacterium chelonae group; Mycobacterium abscessus subgroup; Mycobacterium abscessus MIKHLTRRATVALVVAAAAISPAAYPAVADTATPLNGLFALTPGSCAEGRATGSYFRMIL WP_005063674.1 hypothetical protein [Mycobacterium abscessus] Length: 330\nNumber of predicted TMHs: 1\nExp number of AAs in TMHs: 39.25259\nExp number, first 60 AAs: 16.98187\nTotal prob of N-in: 0.83210\nPOSSIBLE N-term signal sequence\noutside 1 298\nTMhelix 299 321\ninside 322 330

30632 GCF_900133475.1_10625_4_76 Mycobacterium abscessus subsp. massiliense Terrabacteria group; Actinobacteria; Actinobacteria; Corynebacteriales; Mycobacteriaceae; Mycobacterium; Mycobacterium chelonae group; Mycobacterium abscessus subgroup; Mycobacterium abscessus MIKHLTRRATVALVVAAAAISPAAYPAVADTATPLNGLFALTPGSCAEGRATGSYFRMIL WP_005063674.1 hypothetical protein [Mycobacterium abscessus] Length: 330\nNumber of predicted TMHs: 1\nExp number of AAs in TMHs: 39.25259\nExp number, first 60 AAs: 16.98187\nTotal prob of N-in: 0.83210\nPOSSIBLE N-term signal sequence\noutside 1 298\nTMhelix 299 321\ninside 322 330

30633 GCF_900133865.1_10250_1_52 Mycobacterium abscessus subsp. massiliense Terrabacteria group; Actinobacteria; Actinobacteria; Corynebacteriales; Mycobacteriaceae; Mycobacterium; Mycobacterium chelonae group;

Mycobacterium abscessus subgroup; Mycobacterium abscessus
MIKHLTRRATVALVVAAAAISPAAYPAVADTATPLNGLFALTPGSCAEGRATGSYFRMIL WP_005063674.1 hypothetical protein [Mycobacterium abscessus] Length: 330\nNumber of predicted TMHs: 1\nExp number of AAs in TMHs: 39.25259\nExp number, first 60 AAs: 16.98187\nTotal prob of N-in: 0.83210\nPOSSIBLE N-term signal sequence\noutside 1 298\nTMhelix 299 321\ninside 322 330

30634 GCF_900134685.1_12163_2_78 Mycobacterium abscessus subsp. massiliense Terrabacteria group; Actinobacteria; Actinobacteria; Corynebacteriales; Mycobacteriaceae; Mycobacterium; Mycobacterium chelonae group; Mycobacterium abscessus subgroup; Mycobacterium abscessus
MIKHLTRRATVALVVAAAAISPAAYPAVADTATPLNGLFALTPGSCAEGRATGSYFRMIL WP_005063674.1 hypothetical protein [Mycobacterium abscessus] Length: 330\nNumber of predicted TMHs: 1\nExp number of AAs in TMHs: 39.25259\nExp number, first 60 AAs: 16.98187\nTotal prob of N-in: 0.83210\nPOSSIBLE N-term signal sequence\noutside 1 298\nTMhelix 299 321\ninside 322 330

30635 GCF_900134405.1_12163_1_29 Mycobacterium abscessus subsp. massiliense Terrabacteria group; Actinobacteria; Actinobacteria; Corynebacteriales; Mycobacteriaceae; Mycobacterium; Mycobacterium chelonae group; Mycobacterium abscessus subgroup; Mycobacterium abscessus
MIKHLTRRATVALVVAAAAISPAAYPAVADTATPLNGLFALTPGSCAEGRATGSYFRMIL WP_005063674.1 hypothetical protein [Mycobacterium abscessus] Length: 330\nNumber of predicted TMHs: 1\nExp number of AAs in TMHs: 39.25259\nExp number, first 60 AAs: 16.98187\nTotal prob of N-in: 0.83210\nPOSSIBLE N-term signal sequence\noutside 1 298\nTMhelix 299 321\ninside 322 330

30636 GCF_900134885.1_10208_3_3 Mycobacterium abscessus subsp. massiliense Terrabacteria group; Actinobacteria; Actinobacteria; Corynebacteriales; Mycobacteriaceae; Mycobacterium; Mycobacterium chelonae group; Mycobacterium abscessus subgroup; Mycobacterium abscessus
MIKHLTRRATVALVVAAAAISPAAYPAVADTATPLNGLFALTPGSCAEGRATGSYFRMIL WP_005063674.1 hypothetical protein [Mycobacterium abscessus] Length: 330\nNumber of predicted TMHs: 1\nExp number of AAs in TMHs: 39.25259\nExp number, first 60 AAs: 16.98187\nTotal prob of N-in: 0.83210\nPOSSIBLE N-term signal sequence\noutside 1 298\nTMhelix 299 321\ninside 322 330

30637 GCF_900135175.1_10625_4_26 Mycobacterium abscessus subsp. massiliense Terrabacteria group; Actinobacteria; Actinobacteria; Corynebacteriales; Mycobacteriaceae; Mycobacterium; Mycobacterium chelonae group; Mycobacterium abscessus subgroup; Mycobacterium abscessus
MIKHLTRRATVALVVAAAAISPAAYPAVADTATPLNGLFALTPGSCAEGRATGSYFRMIL WP_005063674.1 hypothetical protein [Mycobacterium abscessus] Length: 330\nNumber of predicted TMHs: 1\nExp number of AAs in TMHs: 39.25259\nExp number, first 60 AAs: 16.98187\nTotal prob of N-in: 0.83210\nPOSSIBLE N-term signal sequence\noutside 1 298\nTMhelix 299 321\ninside 322 330

30638 GCF_900135755.1_11861_7_77 Mycobacterium abscessus subsp. massiliense Terrabacteria group; Actinobacteria; Actinobacteria; Corynebacteriales; Mycobacteriaceae; Mycobacterium; Mycobacterium chelonae group; Mycobacterium abscessus subgroup; Mycobacterium abscessus
MIKHLTRRATVALVVAAAAISPAAYPAVADTATPLNGLFALTPGSCAEGRATGSYFRMIL WP_005063674.1 hypothetical protein [Mycobacterium abscessus] Length: 330\nNumber of predicted TMHs: 1\nExp number of AAs in TMHs: 39.25259\nExp number, first 60 AAs: 16.98187\nTotal prob of N-in: 0.83210\nPOSSIBLE N-term signal sequence\noutside 1 298\nTMhelix 299 321\ninside 322 330

30639 GCF_900136125.1_12163_1_69 Mycobacterium abscessus subsp. massiliense Terrabacteria group; Actinobacteria; Actinobacteria; Corynebacteriales; Mycobacteriaceae; Mycobacterium; Mycobacterium chelonae group; Mycobacterium abscessus subgroup; Mycobacterium abscessus
MIKHLTRRATVALVVAAAAISPAAYPAVADTATPLNGLFALTPGSCAEGRATGSYFRMIL WP_005063674.1 hypothetical protein [Mycobacterium abscessus] Length: 330\nNumber of predicted TMHs: 1\nExp number of AAs in TMHs: 39.25259\nExp number, first 60 AAs: 16.98187\nTotal prob of N-in: 0.83210\nPOSSIBLE N-term signal sequence\noutside 1 298\nTMhelix 299 321\ninside 322 330

30640 GCF_900136495.1_10208_3_74 Mycobacterium abscessus subsp. massiliense Terrabacteria group; Actinobacteria; Actinobacteria; Corynebacteriales; Mycobacteriaceae; Mycobacterium; Mycobacterium chelonae group; Mycobacterium abscessus subgroup; Mycobacterium abscessus
MIKHLTRRATVALVVAAAAISPAAYPAVADTATPLNGLFALTPGSCAEGRATGSYFRMIL WP_005063674.1 hypothetical protein [Mycobacterium abscessus] Length: 330\nNumber of predicted TMHs: 1\nExp number of AAs in TMHs: 39.25259\nExp number, first 60 AAs: 16.98187\nTotal prob of N-in: 0.83210\nPOSSIBLE N-term signal sequence\noutside 1 298\nTMhelix 299 321\ninside 322 330

30641 GCF_900136405.1_20034_1_17 Mycobacterium abscessus subsp. massiliense Terrabacteria group; Actinobacteria; Actinobacteria; Corynebacteriales; Mycobacteriaceae; Mycobacterium; Mycobacterium chelonae group; Mycobacterium abscessus subgroup; Mycobacterium abscessus MIKHLTRRATVALVVAAAAISPAAYPAVADTATPLNGLFALTPGSCAEGRATGSYFRMIL WP_079617725.1 hypothetical protein [Mycobacterium abscessus] Length: 330\nNumber of predicted TMHs: 1\nExp number of AAs in TMHs: 39.2571699999999\nExp number, first 60 AAs: 16.97134\nTotal prob of N-in: 0.83217\nPOSSIBLE N-term signal sequence\noutside 1 298\nTMhelix 299 321\ninside 322 330

30642 GCF_900136425.1_20034_1_18 Mycobacterium abscessus subsp. massiliense Terrabacteria group; Actinobacteria; Actinobacteria; Corynebacteriales; Mycobacteriaceae; Mycobacterium; Mycobacterium chelonae group; Mycobacterium abscessus subgroup; Mycobacterium abscessus MIKHLTRRATVALVVAAAAISPAAYPAVADTATPLNGLFALTPGSCAEGRATGSYFRMIL WP_079617725.1 hypothetical protein [Mycobacterium abscessus] Length: 330\nNumber of predicted TMHs: 1\nExp number of AAs in TMHs: 39.2571699999999\nExp number, first 60 AAs: 16.97134\nTotal prob of N-in: 0.83217\nPOSSIBLE N-term signal sequence\noutside 1 298\nTMhelix 299 321\ninside 322 330

30643 GCF_900138335.1_11893_7_47 Mycobacterium abscessus subsp. massiliense Terrabacteria group; Actinobacteria; Actinobacteria; Corynebacteriales; Mycobacteriaceae; Mycobacterium; Mycobacterium chelonae group; Mycobacterium abscessus subgroup; Mycobacterium abscessus MIKHLTRRATVALVVAAAAISPAAYPAVADTATPLNGLFALTPGSCAEGRATGSYFRMIL WP_005063674.1 hypothetical protein [Mycobacterium abscessus] Length: 330\nNumber of predicted TMHs: 1\nExp number of AAs in TMHs: 39.25259\nExp number, first 60 AAs: 16.98187\nTotal prob of N-in: 0.83210\nPOSSIBLE N-term signal sequence\noutside 1 298\nTMhelix 299 321\ninside 322 330

30644 GCF_900138555.1_20034_1_11 Mycobacterium abscessus subsp. massiliense Terrabacteria group; Actinobacteria; Actinobacteria; Corynebacteriales; Mycobacteriaceae; Mycobacterium; Mycobacterium chelonae group; Mycobacterium abscessus subgroup; Mycobacterium abscessus MIKHLTRRATVALVVAAAAISPAAYPAVADTATPLNGLFALTPGSCAEGRATGSYFRMIL WP_079617725.1 hypothetical protein [Mycobacterium abscessus] Length: 330\nNumber of predicted TMHs: 1\nExp number of AAs in TMHs: 39.2571699999999\nExp number, first 60 AAs: 16.97134\nTotal prob of N-in: 0.83217\nPOSSIBLE N-term signal sequence\noutside 1 298\nTMhelix 299 321\ninside 322 330

30645 GCF_900138595.1_20034_1_14 Mycobacterium abscessus subsp. massiliense Terrabacteria group; Actinobacteria; Actinobacteria; Corynebacteriales; Mycobacteriaceae; Mycobacterium; Mycobacterium chelonae group; Mycobacterium abscessus subgroup; Mycobacterium abscessus MIKHLTRRATVALVVAAAAISPAAYPAVADTATPLNGLFALTPGSCAEGRATGSYFRMIL WP_079617725.1 hypothetical protein [Mycobacterium abscessus] Length: 330\nNumber of predicted TMHs: 1\nExp number of AAs in TMHs: 39.2571699999999\nExp number, first 60 AAs: 16.97134\nTotal prob of N-in: 0.83217\nPOSSIBLE N-term signal sequence\noutside 1 298\nTMhelix 299 321\ninside 322 330

30646 GCF_900138645.1_20034_1_23 Mycobacterium abscessus subsp. massiliense Terrabacteria group; Actinobacteria; Actinobacteria; Corynebacteriales; Mycobacteriaceae; Mycobacterium; Mycobacterium chelonae group; Mycobacterium abscessus subgroup; Mycobacterium abscessus MIKHLTRRATVALVVAAAAISPAAYPAVADTATPLNGLFALTPGSCAEGRATGSYFRMIL WP_079617725.1 hypothetical protein [Mycobacterium abscessus] Length: 330\nNumber of predicted TMHs: 1\nExp number of AAs in TMHs: 39.2571699999999\nExp number, first 60 AAs: 16.97134\nTotal prob of N-in: 0.83217\nPOSSIBLE N-term signal sequence\noutside 1 298\nTMhelix 299 321\ninside 322 330

30647 GCF_900138665.1_20034_1_4 Mycobacterium abscessus subsp. massiliense Terrabacteria group; Actinobacteria; Actinobacteria; Corynebacteriales; Mycobacteriaceae; Mycobacterium; Mycobacterium chelonae group; Mycobacterium abscessus subgroup; Mycobacterium abscessus MIKHLTRRATVALVVAAAAISPAAYPAVADTATPLNGLFALTPGSCAEGRATGSYFRMIL WP_079617725.1 hypothetical protein [Mycobacterium abscessus] Length: 330\nNumber of predicted TMHs: 1\nExp number of AAs in TMHs: 39.2571699999999\nExp number, first 60 AAs: 16.97134\nTotal prob of N-in: 0.83217\nPOSSIBLE N-term signal sequence\noutside 1 298\nTMhelix 299 321\ninside 322 330

30648 GCF_900138685.1_20034_1_5 Mycobacterium abscessus subsp. massiliense Terrabacteria group; Actinobacteria; Actinobacteria; Corynebacteriales; Mycobacteriaceae; Mycobacterium; Mycobacterium chelonae group; Mycobacterium abscessus subgroup; Mycobacterium abscessus MIKHLTRRATVALVVAAAAISPAAYPAVADTATPLNGLFALTPGSCAEGRATGSYFRMIL WP_079617725.1 hypothetical protein [Mycobacterium abscessus] Length: 330\nNumber of predicted TMHs: 1\nExp number of AAs in TMHs:

39.2571699999999\nExp number, first 60 AAs: 16.97134\nTotal prob of N-in: 0.83217\nPOSSIBLE N-term signal sequence\noutside 1 298\nTMhelix 299 321\ninside 322 330

30649 GCF_900138805.1_10208_3_46 Mycobacterium abscessus subsp. massiliense Terrabacteria group; Actinobacteria; Actinobacteria; Corynebacteriales; Mycobacteriaceae; Mycobacterium; Mycobacterium chelonae group; Mycobacterium abscessus subgroup; Mycobacterium abscessus MIKHLTRRATVALVVAAAAISPAAYPAVADTATPLNGLFALTPGSCAEGRATGSYFRMIL WP_005063674.1 hypothetical protein [Mycobacterium abscessus] Length: 330\nNumber of predicted TMHs: 1\nExp number of AAs in TMHs: 39.25259\nExp number, first 60 AAs: 16.98187\nTotal prob of N-in: 0.83210\nPOSSIBLE N-term signal sequence\noutside 1 298\nTMhelix 299 321\ninside 322 330

30650 GCF_900139145.1_10465_1_60 Mycobacterium abscessus subsp. massiliense Terrabacteria group; Actinobacteria; Actinobacteria; Corynebacteriales; Mycobacteriaceae; Mycobacterium; Mycobacterium chelonae group; Mycobacterium abscessus subgroup; Mycobacterium abscessus MIKHLTRRATVALVVAAAAISPAAYPAVADTATPLNGLFALTPGSCAEGRATGSYFRMIL WP_005063674.1 hypothetical protein [Mycobacterium abscessus] Length: 330\nNumber of predicted TMHs: 1\nExp number of AAs in TMHs: 39.25259\nExp number, first 60 AAs: 16.98187\nTotal prob of N-in: 0.83210\nPOSSIBLE N-term signal sequence\noutside 1 298\nTMhelix 299 321\ninside 322 330

30651 GCF_900139445.1_10625_4_6 Mycobacterium abscessus subsp. massiliense Terrabacteria group; Actinobacteria; Actinobacteria; Corynebacteriales; Mycobacteriaceae; Mycobacterium; Mycobacterium chelonae group; Mycobacterium abscessus subgroup; Mycobacterium abscessus MIKHLTRRATVALVVAAAAISPAAYPAVADTATPLNGLFALTPGSCAEGRATGSYFRMIL WP_005063674.1 hypothetical protein [Mycobacterium abscessus] Length: 330\nNumber of predicted TMHs: 1\nExp number of AAs in TMHs: 39.25259\nExp number, first 60 AAs: 16.98187\nTotal prob of N-in: 0.83210\nPOSSIBLE N-term signal sequence\noutside 1 298\nTMhelix 299 321\ninside 322 330

30652 GCF_900140295.1_10665_2_95 Mycobacterium abscessus subsp. massiliense Terrabacteria group; Actinobacteria; Actinobacteria; Corynebacteriales; Mycobacteriaceae; Mycobacterium; Mycobacterium chelonae group; Mycobacterium abscessus subgroup; Mycobacterium abscessus MIKHLTRRATVALVVAAAAISPAAYPAVADTATPLNGLFALTPGSCAEGRATGSYFRMIL WP_005063674.1 hypothetical protein [Mycobacterium abscessus] Length: 330\nNumber of predicted TMHs: 1\nExp number of AAs in TMHs: 39.25259\nExp number, first 60 AAs: 16.98187\nTotal prob of N-in: 0.83210\nPOSSIBLE N-term signal sequence\noutside 1 298\nTMhelix 299 321\ninside 322 330

30653 GCF_900140385.1_10665_4_4 Mycobacterium abscessus subsp. massiliense Terrabacteria group; Actinobacteria; Actinobacteria; Corynebacteriales; Mycobacteriaceae; Mycobacterium; Mycobacterium chelonae group; Mycobacterium abscessus subgroup; Mycobacterium abscessus MIKHLTRRATVALVVAAAAISPAAYPAVADTATPLNGLFALTPGSCAEGRATGSYFRMIL WP_005063674.1 hypothetical protein [Mycobacterium abscessus] Length: 330\nNumber of predicted TMHs: 1\nExp number of AAs in TMHs: 39.25259\nExp number, first 60 AAs: 16.98187\nTotal prob of N-in: 0.83210\nPOSSIBLE N-term signal sequence\noutside 1 298\nTMhelix 299 321\ninside 322 330

30654 GCF_900169205.1_16933_5_1 Mycobacterium abscessus subsp. massiliense Terrabacteria group; Actinobacteria; Actinobacteria; Corynebacteriales; Mycobacteriaceae; Mycobacterium; Mycobacterium chelonae group; Mycobacterium abscessus subgroup; Mycobacterium abscessus MIKHLTRRATVALVVAAAAISPAAYPAVADTATPLNGLFALTPGSCAEGRATGSYFRMIL WP_005063674.1 hypothetical protein [Mycobacterium abscessus] Length: 330\nNumber of predicted TMHs: 1\nExp number of AAs in TMHs: 39.25259\nExp number, first 60 AAs: 16.98187\nTotal prob of N-in: 0.83210\nPOSSIBLE N-term signal sequence\noutside 1 298\nTMhelix 299 321\ninside 322 330

30655 GCF_900130945.1_10665_3_49 Mycobacterium abscessus subsp. massiliense Terrabacteria group; Actinobacteria; Actinobacteria; Corynebacteriales; Mycobacteriaceae; Mycobacterium; Mycobacterium chelonae group; Mycobacterium abscessus subgroup; Mycobacterium abscessus MIKHLTRRATVALVVAAAAISPAAYPAVADTATPLNGLFALTPGSCAEGRATGSYFRMIL WP_005063674.1 hypothetical protein [Mycobacterium abscessus] Length: 330\nNumber of predicted TMHs: 1\nExp number of AAs in TMHs: 39.25259\nExp number, first 60 AAs: 16.98187\nTotal prob of N-in: 0.83210\nPOSSIBLE N-term signal sequence\noutside 1 298\nTMhelix 299 321\ninside 322 330

30656 GCF_900136325.1_12163_1_88 Mycobacterium abscessus subsp. massiliense Terrabacteria group; Actinobacteria; Actinobacteria; Corynebacteriales; Mycobacteriaceae; Mycobacterium; Mycobacterium chelonae group;

Mycobacterium abscessus subgroup; Mycobacterium abscessus
MIKHLTRRATVALVVAAAAISPAAYPAVADTATPLNGLFALTPGSCAEGRATGSYFRMIL WP_005063674.1 hypothetical protein [Mycobacterium abscessus] Length: 330\nNumber of predicted TMHs: 1\nExp number of AAs in TMHs: 39.25259\nExp number, first 60 AAs: 16.98187\nTotal prob of N-in: 0.83210\nPOSSIBLE N-term signal sequence\noutside 1 298\nTMhelix 299 321\ninside 322 330

30657 GCF_900138915.1_10208_3_58 Mycobacterium abscessus subsp. massiliense Terrabacteria group; Actinobacteria; Actinobacteria; Corynebacteriales; Mycobacteriaceae; Mycobacterium; Mycobacterium chelonae group; Mycobacterium abscessus subgroup; Mycobacterium abscessus
MIKHLTRRATVALVVAAAAISPAAYPAVADTATPLNGLFALTPGSCAEGRATGSYFRMIL WP_005063674.1 hypothetical protein [Mycobacterium abscessus] Length: 330\nNumber of predicted TMHs: 1\nExp number of AAs in TMHs: 39.25259\nExp number, first 60 AAs: 16.98187\nTotal prob of N-in: 0.83210\nPOSSIBLE N-term signal sequence\noutside 1 298\nTMhelix 299 321\ninside 322 330

30658 GCF_900138395.1_11893_7_52 Mycobacterium abscessus subsp. massiliense Terrabacteria group; Actinobacteria; Actinobacteria; Corynebacteriales; Mycobacteriaceae; Mycobacterium; Mycobacterium chelonae group; Mycobacterium abscessus subgroup; Mycobacterium abscessus
MIKHLTRRATVALVVAAAAISPAAYPAVADTATPLNGLFALTPGSCAEGRATGSYFRMIL WP_005063674.1 hypothetical protein [Mycobacterium abscessus] Length: 330\nNumber of predicted TMHs: 1\nExp number of AAs in TMHs: 39.25259\nExp number, first 60 AAs: 16.98187\nTotal prob of N-in: 0.83210\nPOSSIBLE N-term signal sequence\noutside 1 298\nTMhelix 299 321\ninside 322 330

30659 GCF_900136635.1_10625_4_84 Mycobacterium abscessus subsp. massiliense Terrabacteria group; Actinobacteria; Actinobacteria; Corynebacteriales; Mycobacteriaceae; Mycobacterium; Mycobacterium chelonae group; Mycobacterium abscessus subgroup; Mycobacterium abscessus
MIKHLTRRATVALVVAAAAISPAAYPAVADTATPLNGLFALTPGSCAEGRATGSYFRMIL WP_005063674.1 hypothetical protein [Mycobacterium abscessus] Length: 330\nNumber of predicted TMHs: 1\nExp number of AAs in TMHs: 39.25259\nExp number, first 60 AAs: 16.98187\nTotal prob of N-in: 0.83210\nPOSSIBLE N-term signal sequence\noutside 1 298\nTMhelix 299 321\ninside 322 330

30660 GCF_900138375.1_11893_7_51 Mycobacterium abscessus subsp. massiliense Terrabacteria group; Actinobacteria; Actinobacteria; Corynebacteriales; Mycobacteriaceae; Mycobacterium; Mycobacterium chelonae group; Mycobacterium abscessus subgroup; Mycobacterium abscessus
MIKHLTRRATVALVVAAAAISPAAYPAVADTATPLNGLFALTPGSCAEGRATGSYFRMIL WP_005063674.1 hypothetical protein [Mycobacterium abscessus] Length: 330\nNumber of predicted TMHs: 1\nExp number of AAs in TMHs: 39.25259\nExp number, first 60 AAs: 16.98187\nTotal prob of N-in: 0.83210\nPOSSIBLE N-term signal sequence\noutside 1 298\nTMhelix 299 321\ninside 322 330

30661 GCF_900138355.1_11893_7_48 Mycobacterium abscessus subsp. massiliense Terrabacteria group; Actinobacteria; Actinobacteria; Corynebacteriales; Mycobacteriaceae; Mycobacterium; Mycobacterium chelonae group; Mycobacterium abscessus subgroup; Mycobacterium abscessus
MIKHLTRRATVALVVAAAAISPAAYPAVADTATPLNGLFALTPGSCAEGRATGSYFRMIL WP_005063674.1 hypothetical protein [Mycobacterium abscessus] Length: 330\nNumber of predicted TMHs: 1\nExp number of AAs in TMHs: 39.25259\nExp number, first 60 AAs: 16.98187\nTotal prob of N-in: 0.83210\nPOSSIBLE N-term signal sequence\noutside 1 298\nTMhelix 299 321\ninside 322 330

30662 GCF_900136725.1_10625_5_27 Mycobacterium abscessus subsp. massiliense Terrabacteria group; Actinobacteria; Actinobacteria; Corynebacteriales; Mycobacteriaceae; Mycobacterium; Mycobacterium chelonae group; Mycobacterium abscessus subgroup; Mycobacterium abscessus
MIKHLTRRATVALVVAAAAISPAAYPAVADTATPLNGLFALTPGSCAEGRATGSYFRMIL WP_005063674.1 hypothetical protein [Mycobacterium abscessus] Length: 330\nNumber of predicted TMHs: 1\nExp number of AAs in TMHs: 39.25259\nExp number, first 60 AAs: 16.98187\nTotal prob of N-in: 0.83210\nPOSSIBLE N-term signal sequence\noutside 1 298\nTMhelix 299 321\ninside 322 330

30663 GCF_900133735.1_12163_1_23 Mycobacterium abscessus subsp. massiliense Terrabacteria group; Actinobacteria; Actinobacteria; Corynebacteriales; Mycobacteriaceae; Mycobacterium; Mycobacterium chelonae group; Mycobacterium abscessus subgroup; Mycobacterium abscessus
MIKHLTRRATVALVVAAAAISPAAYPAVADTATPLNGLFALTPGSCAEGRATGSYFRMIL WP_005063674.1 hypothetical protein [Mycobacterium abscessus] Length: 330\nNumber of predicted TMHs: 1\nExp number of AAs in TMHs: 39.25259\nExp number, first 60 AAs: 16.98187\nTotal prob of N-in: 0.83210\nPOSSIBLE N-term signal sequence\noutside 1 298\nTMhelix 299 321\ninside 322 330

30664 GCF_900133755.1_12163_1_26 Mycobacterium abscessus subsp. massiliense Terrabacteria group; Actinobacteria; Actinobacteria; Corynebacteriales; Mycobacteriaceae; Mycobacterium; Mycobacterium chelonae group; Mycobacterium abscessus subgroup; Mycobacterium abscessus
MIKHLTRRATVALVVAAAAISPAAYPAVADTATPLNGLFALTPGSCAEGRATGSYFRMIL WP_005063674.1 hypothetical protein [Mycobacterium abscessus] Length: 330\nNumber of predicted TMHs: 1\nExp number of AAs in TMHs: 39.25259\nExp number, first 60 AAs: 16.98187\nTotal prob of N-in: 0.83210\nPOSSIBLE N-term signal sequence\noutside 1 298\nTMhelix 299 321\ninside 322 330

30665 GCF_900133825.1_12163_1_6 Mycobacterium abscessus subsp. massiliense Terrabacteria group; Actinobacteria; Actinobacteria; Corynebacteriales; Mycobacteriaceae; Mycobacterium; Mycobacterium chelonae group; Mycobacterium abscessus subgroup; Mycobacterium abscessus
MIKHLTRRATVALVVAAAAISPAAYPAVADTATPLNGLFALTPGSCAEGRATGSYFRMIL WP_005063674.1 hypothetical protein [Mycobacterium abscessus] Length: 330\nNumber of predicted TMHs: 1\nExp number of AAs in TMHs: 39.25259\nExp number, first 60 AAs: 16.98187\nTotal prob of N-in: 0.83210\nPOSSIBLE N-term signal sequence\noutside 1 298\nTMhelix 299 321\ninside 322 330

30666 GCF_900134075.1_10396_8_5 Mycobacterium abscessus subsp. massiliense Terrabacteria group; Actinobacteria; Actinobacteria; Corynebacteriales; Mycobacteriaceae; Mycobacterium; Mycobacterium chelonae group; Mycobacterium abscessus subgroup; Mycobacterium abscessus
MIKHLTRRATVALVVAAAAISPAAYPAVADTATPLNGLFALTPGSCAEGRATGSYFRMIL WP_005063674.1 hypothetical protein [Mycobacterium abscessus] Length: 330\nNumber of predicted TMHs: 1\nExp number of AAs in TMHs: 39.25259\nExp number, first 60 AAs: 16.98187\nTotal prob of N-in: 0.83210\nPOSSIBLE N-term signal sequence\noutside 1 298\nTMhelix 299 321\ninside 322 330

30667 GCF_900138315.1_11893_7_45 Mycobacterium abscessus subsp. massiliense Terrabacteria group; Actinobacteria; Actinobacteria; Corynebacteriales; Mycobacteriaceae; Mycobacterium; Mycobacterium chelonae group; Mycobacterium abscessus subgroup; Mycobacterium abscessus
MIKHLTRRATVALVVAAAAISPAAYPAVADTATPLNGLFALTPGSCAEGRATGSYFRMIL WP_005063674.1 hypothetical protein [Mycobacterium abscessus] Length: 330\nNumber of predicted TMHs: 1\nExp number of AAs in TMHs: 39.25259\nExp number, first 60 AAs: 16.98187\nTotal prob of N-in: 0.83210\nPOSSIBLE N-term signal sequence\noutside 1 298\nTMhelix 299 321\ninside 322 330

30668 GCF_900134315.1_11893_6_34 Mycobacterium abscessus subsp. massiliense Terrabacteria group; Actinobacteria; Actinobacteria; Corynebacteriales; Mycobacteriaceae; Mycobacterium; Mycobacterium chelonae group; Mycobacterium abscessus subgroup; Mycobacterium abscessus
MIKHLTRRATVALVVAAAAISPAAYPAVADTATPLNGLFALTPGSCAEGRATGSYFRMIL WP_005063674.1 hypothetical protein [Mycobacterium abscessus] Length: 330\nNumber of predicted TMHs: 1\nExp number of AAs in TMHs: 39.25259\nExp number, first 60 AAs: 16.98187\nTotal prob of N-in: 0.83210\nPOSSIBLE N-term signal sequence\noutside 1 298\nTMhelix 299 321\ninside 322 330

30669 GCF_900131585.1_10665_4_16 Mycobacterium abscessus subsp. massiliense Terrabacteria group; Actinobacteria; Actinobacteria; Corynebacteriales; Mycobacteriaceae; Mycobacterium; Mycobacterium chelonae group; Mycobacterium abscessus subgroup; Mycobacterium abscessus
MIKHLTRRATVALVVAAAAISPAAYPAVADTATPLNGLFALTPGSCAEGRATGSYFRMIL WP_005063674.1 hypothetical protein [Mycobacterium abscessus] Length: 330\nNumber of predicted TMHs: 1\nExp number of AAs in TMHs: 39.25259\nExp number, first 60 AAs: 16.98187\nTotal prob of N-in: 0.83210\nPOSSIBLE N-term signal sequence\noutside 1 298\nTMhelix 299 321\ninside 322 330

30670 GCF_900131935.1_10665_4_39 Mycobacterium abscessus subsp. massiliense Terrabacteria group; Actinobacteria; Actinobacteria; Corynebacteriales; Mycobacteriaceae; Mycobacterium; Mycobacterium chelonae group; Mycobacterium abscessus subgroup; Mycobacterium abscessus
MIKHLTRRATVALVVAAAAISPAAYPAVADTATPLNGLFALTPGSCAEGRATGSYFRMIL WP_005063674.1 hypothetical protein [Mycobacterium abscessus] Length: 330\nNumber of predicted TMHs: 1\nExp number of AAs in TMHs: 39.25259\nExp number, first 60 AAs: 16.98187\nTotal prob of N-in: 0.83210\nPOSSIBLE N-term signal sequence\noutside 1 298\nTMhelix 299 321\ninside 322 330

30671 GCF_900138445.1_11893_7_59 Mycobacterium abscessus subsp. massiliense Terrabacteria group; Actinobacteria; Actinobacteria; Corynebacteriales; Mycobacteriaceae; Mycobacterium; Mycobacterium chelonae group; Mycobacterium abscessus subgroup; Mycobacterium abscessus
MIKHLTRRATVALVVAAAAISPAAYPAVADTATPLNGLFALTPGSCAEGRATGSYFRMIL WP_005063674.1 hypothetical protein [Mycobacterium abscessus] Length: 330\nNumber of predicted TMHs: 1\nExp number of AAs in TMHs:

39.25259\nExp number, first 60 AAs: 16.98187\nTotal prob of N-in: 0.83210\nPOSSIBLE N-term signal sequence\noutside 1 298\nTMhelix 299 321\ninside 322 330

30672 GCF_900132385.1_10665_4_71 Mycobacterium abscessus subsp. massiliense Terrabacteria group; Actinobacteria; Actinobacteria; Corynebacteriales; Mycobacteriaceae; Mycobacterium; Mycobacterium chelonae group; Mycobacterium abscessus subgroup; Mycobacterium abscessus MIKHLTRRATVALVVAAAAISPAAYPAVADTATPLNGLFALTPGSCAEGRATGSYFRMIL WP_005063674.1 hypothetical protein [Mycobacterium abscessus] Length: 330\nNumber of predicted TMHs: 1\nExp number of AAs in TMHs: 39.25259\nExp number, first 60 AAs: 16.98187\nTotal prob of N-in: 0.83210\nPOSSIBLE N-term signal sequence\noutside 1 298\nTMhelix 299 321\ninside 322 330

30673 GCF_900132265.1_11893_6_46 Mycobacterium abscessus subsp. massiliense Terrabacteria group; Actinobacteria; Actinobacteria; Corynebacteriales; Mycobacteriaceae; Mycobacterium; Mycobacterium chelonae group; Mycobacterium abscessus subgroup; Mycobacterium abscessus MIKHLTRRATVALVVAAAAISPAAYPAMADTATPLNGLFALTPGSCAEGRATGSYFRMIL WP_074356318.1 hypothetical protein [Mycobacterium abscessus] Length: 330\nNumber of predicted TMHs: 2\nExp number of AAs in TMHs: 39.34118\nExp number, first 60 AAs: 17.07668\nTotal prob of N-in: 0.84431\nPOSSIBLE N-term signal sequence\ninside 1 11\nTMhelix 12 29\noutside 30 298\nTMhelix 299 321\ninside 322 330

30674 GCF_900132125.1_10665_4_57 Mycobacterium abscessus subsp. massiliense Terrabacteria group; Actinobacteria; Actinobacteria; Corynebacteriales; Mycobacteriaceae; Mycobacterium; Mycobacterium chelonae group; Mycobacterium abscessus subgroup; Mycobacterium abscessus MIKHLTRRATVALVVAAAAISPAAYPAVADTATPLNGLFALTPGSCAEGRATGSYFRMIL WP_005063674.1 hypothetical protein [Mycobacterium abscessus] Length: 330\nNumber of predicted TMHs: 1\nExp number of AAs in TMHs: 39.25259\nExp number, first 60 AAs: 16.98187\nTotal prob of N-in: 0.83210\nPOSSIBLE N-term signal sequence\noutside 1 298\nTMhelix 299 321\ninside 322 330

30675 GCF_900138465.1_11893_7_57 Mycobacterium abscessus subsp. massiliense Terrabacteria group; Actinobacteria; Actinobacteria; Corynebacteriales; Mycobacteriaceae; Mycobacterium; Mycobacterium chelonae group; Mycobacterium abscessus subgroup; Mycobacterium abscessus MIKHLTRRATVALVVAAAAISPAAYPAVADTATPLNGLFALTPGSCAEGRATGSYFRMIL WP_005063674.1 hypothetical protein [Mycobacterium abscessus] Length: 330\nNumber of predicted TMHs: 1\nExp number of AAs in TMHs: 39.25259\nExp number, first 60 AAs: 16.98187\nTotal prob of N-in: 0.83210\nPOSSIBLE N-term signal sequence\noutside 1 298\nTMhelix 299 321\ninside 322 330

30676 GCF_900132025.1_10665_4_46 Mycobacterium abscessus subsp. massiliense Terrabacteria group; Actinobacteria; Actinobacteria; Corynebacteriales; Mycobacteriaceae; Mycobacterium; Mycobacterium chelonae group; Mycobacterium abscessus subgroup; Mycobacterium abscessus MIKHLTRRATVALVVAAAAISPAAYPAVADTATPLNGLFALTPGSCAEGRATGSYFRMIL WP_005063674.1 hypothetical protein [Mycobacterium abscessus] Length: 330\nNumber of predicted TMHs: 1\nExp number of AAs in TMHs: 39.25259\nExp number, first 60 AAs: 16.98187\nTotal prob of N-in: 0.83210\nPOSSIBLE N-term signal sequence\noutside 1 298\nTMhelix 299 321\ninside 322 330

30677 GCF_900139005.1_10465_1_44 Mycobacterium abscessus subsp. massiliense Terrabacteria group; Actinobacteria; Actinobacteria; Corynebacteriales; Mycobacteriaceae; Mycobacterium; Mycobacterium chelonae group; Mycobacterium abscessus subgroup; Mycobacterium abscessus MIKHLTRRATVALVVAAAAISPAAYPAVADTATPLNGLFALTPGSCAEGRATGSYFRMIL WP_005063674.1 hypothetical protein [Mycobacterium abscessus] Length: 330\nNumber of predicted TMHs: 1\nExp number of AAs in TMHs: 39.25259\nExp number, first 60 AAs: 16.98187\nTotal prob of N-in: 0.83210\nPOSSIBLE N-term signal sequence\noutside 1 298\nTMhelix 299 321\ninside 322 330

30678 GCF_900139355.1_10625_4_12 Mycobacterium abscessus subsp. massiliense Terrabacteria group; Actinobacteria; Actinobacteria; Corynebacteriales; Mycobacteriaceae; Mycobacterium; Mycobacterium chelonae group; Mycobacterium abscessus subgroup; Mycobacterium abscessus MIKHLTRRATVALVVAAAAISPAAYPAVADTATPLNGLFALTPGSCAEGRATGSYFRMIL WP_005063674.1 hypothetical protein [Mycobacterium abscessus] Length: 330\nNumber of predicted TMHs: 1\nExp number of AAs in TMHs: 39.25259\nExp number, first 60 AAs: 16.98187\nTotal prob of N-in: 0.83210\nPOSSIBLE N-term signal sequence\noutside 1 298\nTMhelix 299 321\ninside 322 330

30679 GCF_900132895.1_11861_7_63 Mycobacterium abscessus subsp. massiliense Terrabacteria group; Actinobacteria; Actinobacteria; Corynebacteriales; Mycobacteriaceae; Mycobacterium; Mycobacterium chelonae group;

Mycobacterium abscessus subgroup; Mycobacterium abscessus
MIKHLTRRATVALVVAAAAISPAAYPAVADTATPLNGLFALTPGSCAEGRATGSYFRMIL WP_005063674.1 hypothetical protein [Mycobacterium abscessus] Length: 330\nNumber of predicted TMHs: 1\nExp number of AAs in TMHs: 39.25259\nExp number, first 60 AAs: 16.98187\nTotal prob of N-in: 0.83210\nPOSSIBLE N-term signal sequence\noutside 1 298\nTMhelix 299 321\ninside 322 330

30680 GCF_900130575.1_10060_8_12 Mycobacterium abscessus subsp. massiliense Terrabacteria group; Actinobacteria; Actinobacteria; Corynebacteriales; Mycobacteriaceae; Mycobacterium; Mycobacterium chelonae group; Mycobacterium abscessus subgroup; Mycobacterium abscessus
MIKHLTRRATVALVVAAAAISPAAYPAVADTATPLNGLFALTPGSCAEGRATGSYFRMIL WP_005063674.1 hypothetical protein [Mycobacterium abscessus] Length: 330\nNumber of predicted TMHs: 1\nExp number of AAs in TMHs: 39.25259\nExp number, first 60 AAs: 16.98187\nTotal prob of N-in: 0.83210\nPOSSIBLE N-term signal sequence\noutside 1 298\nTMhelix 299 321\ninside 322 330

30681 GCF_900137945.1_12163_2_9 Mycobacterium abscessus subsp. massiliense Terrabacteria group; Actinobacteria; Actinobacteria; Corynebacteriales; Mycobacteriaceae; Mycobacterium; Mycobacterium chelonae group; Mycobacterium abscessus subgroup; Mycobacterium abscessus
MIKHLTRRATVALVVAAAAISPAAYPAVADTATPLNGLFALTPGSCAEGRATGSYFRMIL WP_005063674.1 hypothetical protein [Mycobacterium abscessus] Length: 330\nNumber of predicted TMHs: 1\nExp number of AAs in TMHs: 39.25259\nExp number, first 60 AAs: 16.98187\nTotal prob of N-in: 0.83210\nPOSSIBLE N-term signal sequence\noutside 1 298\nTMhelix 299 321\ninside 322 330

30682 GCF_900138425.1_11893_7_53 Mycobacterium abscessus subsp. massiliense Terrabacteria group; Actinobacteria; Actinobacteria; Corynebacteriales; Mycobacteriaceae; Mycobacterium; Mycobacterium chelonae group; Mycobacterium abscessus subgroup; Mycobacterium abscessus
MIKHLTRRATVALVVAAAAISPAAYPAVADTATPLNGLFALTPGSCAEGRATGSYFRMIL WP_005063674.1 hypothetical protein [Mycobacterium abscessus] Length: 330\nNumber of predicted TMHs: 1\nExp number of AAs in TMHs: 39.25259\nExp number, first 60 AAs: 16.98187\nTotal prob of N-in: 0.83210\nPOSSIBLE N-term signal sequence\noutside 1 298\nTMhelix 299 321\ninside 322 330

30683 GCF_900130535.1_10665_3_7 Mycobacterium abscessus subsp. massiliense Terrabacteria group; Actinobacteria; Actinobacteria; Corynebacteriales; Mycobacteriaceae; Mycobacterium; Mycobacterium chelonae group; Mycobacterium abscessus subgroup; Mycobacterium abscessus
MIKHLTRRATVALVVAAAAISPAAYPAVADTATPLNGLFALTPGSCAEGRATGSYFRMIL WP_005063674.1 hypothetical protein [Mycobacterium abscessus] Length: 330\nNumber of predicted TMHs: 1\nExp number of AAs in TMHs: 39.25259\nExp number, first 60 AAs: 16.98187\nTotal prob of N-in: 0.83210\nPOSSIBLE N-term signal sequence\noutside 1 298\nTMhelix 299 321\ninside 322 330

30684 GCF_900135735.1_11861_7_85 Mycobacterium abscessus subsp. massiliense Terrabacteria group; Actinobacteria; Actinobacteria; Corynebacteriales; Mycobacteriaceae; Mycobacterium; Mycobacterium chelonae group; Mycobacterium abscessus subgroup; Mycobacterium abscessus
MIKHLTRRATVALVVAAAAISPAAYPAVADTATPLNGLFALTPGSCAEGRATGSYFRMIL WP_079617725.1 hypothetical protein [Mycobacterium abscessus] Length: 330\nNumber of predicted TMHs: 1\nExp number of AAs in TMHs: 39.2571699999999\nExp number, first 60 AAs: 16.97134\nTotal prob of N-in: 0.83217\nPOSSIBLE N-term signal sequence\noutside 1 298\nTMhelix 299 321\ninside 322 330

30685 GCF_900137555.1_12163_2_30 Mycobacterium abscessus subsp. massiliense Terrabacteria group; Actinobacteria; Actinobacteria; Corynebacteriales; Mycobacteriaceae; Mycobacterium; Mycobacterium chelonae group; Mycobacterium abscessus subgroup; Mycobacterium abscessus
MIKHLTRRATVALVVAAAAISPAAYPAVADTATPLNGLFALTPGSCAEGRATGSYFRMIL WP_005063674.1 hypothetical protein [Mycobacterium abscessus] Length: 330\nNumber of predicted TMHs: 1\nExp number of AAs in TMHs: 39.25259\nExp number, first 60 AAs: 16.98187\nTotal prob of N-in: 0.83210\nPOSSIBLE N-term signal sequence\noutside 1 298\nTMhelix 299 321\ninside 322 330

30686 GCF_900131405.1_10665_4_19 Mycobacterium abscessus subsp. massiliense Terrabacteria group; Actinobacteria; Actinobacteria; Corynebacteriales; Mycobacteriaceae; Mycobacterium; Mycobacterium chelonae group; Mycobacterium abscessus subgroup; Mycobacterium abscessus
MIKHLTRRATVALVVAAAAISPAAYPAVADTATPLNGLFALTPGSCAEGRATGSYFRMIL WP_005063674.1 hypothetical protein [Mycobacterium abscessus] Length: 330\nNumber of predicted TMHs: 1\nExp number of AAs in TMHs: 39.25259\nExp number, first 60 AAs: 16.98187\nTotal prob of N-in: 0.83210\nPOSSIBLE N-term signal sequence\noutside 1 298\nTMhelix 299 321\ninside 322 330

30687 GCF_900141425.1_11893_6_56 Mycobacterium abscessus subsp. massiliense Terrabacteria group; Actinobacteria; Actinobacteria; Corynebacteriales; Mycobacteriaceae; Mycobacterium; Mycobacterium chelonae group; Mycobacterium abscessus subgroup; Mycobacterium abscessus
MIKHLTRRATVALVVAAAAISPAAYPAVADTATPLNGLFALTPGSCAEGRATGSYFRMIL WP_005063674.1 hypothetical protein [Mycobacterium abscessus] Length: 330\nNumber of predicted TMHs: 1\nExp number of AAs in TMHs: 39.25259\nExp number, first 60 AAs: 16.98187\nTotal prob of N-in: 0.83210\nPOSSIBLE N-term signal sequence\noutside 1 298\nTMhelix 299 321\ninside 322 330

30688 GCF_900130465.1_10625_5_84 Mycobacterium abscessus subsp. massiliense Terrabacteria group; Actinobacteria; Actinobacteria; Corynebacteriales; Mycobacteriaceae; Mycobacterium; Mycobacterium chelonae group; Mycobacterium abscessus subgroup; Mycobacterium abscessus
MIKHLTRRATVALVVAAAAISPAAYPAVADTATPLNGLFALTPGSCAEGRATGSYFRMIL WP_005063674.1 hypothetical protein [Mycobacterium abscessus] Length: 330\nNumber of predicted TMHs: 1\nExp number of AAs in TMHs: 39.25259\nExp number, first 60 AAs: 16.98187\nTotal prob of N-in: 0.83210\nPOSSIBLE N-term signal sequence\noutside 1 298\nTMhelix 299 321\ninside 322 330

30689 GCF_900135425.1_10625_4_58 Mycobacterium abscessus subsp. massiliense Terrabacteria group; Actinobacteria; Actinobacteria; Corynebacteriales; Mycobacteriaceae; Mycobacterium; Mycobacterium chelonae group; Mycobacterium abscessus subgroup; Mycobacterium abscessus
MIKHLTRRATVALVVAAAAISPAAYPAVADTATPLNGLFALTPGSCAEGRATGSYFRMIL WP_005063674.1 hypothetical protein [Mycobacterium abscessus] Length: 330\nNumber of predicted TMHs: 1\nExp number of AAs in TMHs: 39.25259\nExp number, first 60 AAs: 16.98187\nTotal prob of N-in: 0.83210\nPOSSIBLE N-term signal sequence\noutside 1 298\nTMhelix 299 321\ninside 322 330

30690 GCF_900131425.1_10665_4_2 Mycobacterium abscessus subsp. massiliense Terrabacteria group; Actinobacteria; Actinobacteria; Corynebacteriales; Mycobacteriaceae; Mycobacterium; Mycobacterium chelonae group; Mycobacterium abscessus subgroup; Mycobacterium abscessus
MIKHLTRRATVALVVAAAAISPAAYPAVADTATPLNGLFALTPGSCAEGRATGSYFRMIL WP_005063674.1 hypothetical protein [Mycobacterium abscessus] Length: 330\nNumber of predicted TMHs: 1\nExp number of AAs in TMHs: 39.25259\nExp number, first 60 AAs: 16.98187\nTotal prob of N-in: 0.83210\nPOSSIBLE N-term signal sequence\noutside 1 298\nTMhelix 299 321\ninside 322 330

30691 GCF_900132815.1_10250_1_6 Mycobacterium abscessus subsp. massiliense Terrabacteria group; Actinobacteria; Actinobacteria; Corynebacteriales; Mycobacteriaceae; Mycobacterium; Mycobacterium chelonae group; Mycobacterium abscessus subgroup; Mycobacterium abscessus
MIKHLTRRATVALVVAAAAISPAAYPAVADTATPLNGLFALTPGSCAEGRATGSYFRMIL WP_005063674.1 hypothetical protein [Mycobacterium abscessus] Length: 330\nNumber of predicted TMHs: 1\nExp number of AAs in TMHs: 39.25259\nExp number, first 60 AAs: 16.98187\nTotal prob of N-in: 0.83210\nPOSSIBLE N-term signal sequence\noutside 1 298\nTMhelix 299 321\ninside 322 330

30692 GCF_900131515.1_11893_6_4 Mycobacterium abscessus subsp. massiliense Terrabacteria group; Actinobacteria; Actinobacteria; Corynebacteriales; Mycobacteriaceae; Mycobacterium; Mycobacterium chelonae group; Mycobacterium abscessus subgroup; Mycobacterium abscessus
MIKHLTRRATVALVVAAAAISPAAYPAVADTATPLNGLFALTPGSCAEGRATGSYFRMIL WP_005063674.1 hypothetical protein [Mycobacterium abscessus] Length: 330\nNumber of predicted TMHs: 1\nExp number of AAs in TMHs: 39.25259\nExp number, first 60 AAs: 16.98187\nTotal prob of N-in: 0.83210\nPOSSIBLE N-term signal sequence\noutside 1 298\nTMhelix 299 321\ninside 322 330

30693 GCF_900132795.1_10250_1_7 Mycobacterium abscessus subsp. massiliense Terrabacteria group; Actinobacteria; Actinobacteria; Corynebacteriales; Mycobacteriaceae; Mycobacterium; Mycobacterium chelonae group; Mycobacterium abscessus subgroup; Mycobacterium abscessus
MIKHLTRRATVALVVAAAAISPAAYPAMADTATPLNGLFALTPGSCAEGRATGSYFRMIL WP_005073213.1
hypothetical protein [Mycobacterium abscessus] Length: 330\nNumber of predicted TMHs: 2\nExp number of AAs in TMHs: 39.3416199999999\nExp number, first 60 AAs: 17.07688\nTotal prob of N-in: 0.84431\nPOSSIBLE N-term signal sequence\ninside 1 11\nTMhelix 12 29\noutside 30 298\nTMhelix 299 321\ninside 322 330

30694 GCF_900137485.1_12163_2_25 Mycobacterium abscessus subsp. massiliense Terrabacteria group; Actinobacteria; Actinobacteria; Corynebacteriales; Mycobacteriaceae; Mycobacterium; Mycobacterium chelonae group; Mycobacterium abscessus subgroup; Mycobacterium abscessus
MIKHLTRRATVALVVAAAAISPAAYPAVADTATPLNGLFALTPGSCAEGRATGSYFRMIL WP_005063674.1 hypothetical protein [Mycobacterium abscessus] Length: 330\nNumber of predicted TMHs: 1\nExp number of AAs in TMHs:

39.25259\nExp number, first 60 AAs: 16.98187\nTotal prob of N-in: 0.83210\nPOSSIBLE N-term signal sequence\noutside 1 298\nTMhelix 299 321\ninside 322 330

30695 GCF_900131545.1_11893_6_5 Mycobacterium abscessus subsp. massiliense Terrabacteria group; Actinobacteria; Actinobacteria; Corynebacteriales; Mycobacteriaceae; Mycobacterium; Mycobacterium chelonae group; Mycobacterium abscessus subgroup; Mycobacterium abscessus MIKHLTRRATVALVVAAAAISPAAYPAVADTATPLNGLFALTPGSCAEGRATGSYFRMIL WP_005063674.1 hypothetical protein [Mycobacterium abscessus] Length: 330\nNumber of predicted TMHs: 1\nExp number of AAs in TMHs: 39.25259\nExp number, first 60 AAs: 16.98187\nTotal prob of N-in: 0.83210\nPOSSIBLE N-term signal sequence\noutside 1 298\nTMhelix 299 321\ninside 322 330

30696 GCF_900132475.1_10665_4_80 Mycobacterium abscessus subsp. massiliense Terrabacteria group; Actinobacteria; Actinobacteria; Corynebacteriales; Mycobacteriaceae; Mycobacterium; Mycobacterium chelonae group; Mycobacterium abscessus subgroup; Mycobacterium abscessus MIKHLTRRATVALVVAAAAISPAAYPAVADTATPLNGLFALTPGSCAEGRATGSYFRMIL WP_005063674.1 hypothetical protein [Mycobacterium abscessus] Length: 330\nNumber of predicted TMHs: 1\nExp number of AAs in TMHs: 39.25259\nExp number, first 60 AAs: 16.98187\nTotal prob of N-in: 0.83210\nPOSSIBLE N-term signal sequence\noutside 1 298\nTMhelix 299 321\ninside 322 330

30697 GCF_900139645.1_10625_5_8 Mycobacterium abscessus subsp. massiliense Terrabacteria group; Actinobacteria; Actinobacteria; Corynebacteriales; Mycobacteriaceae; Mycobacterium; Mycobacterium chelonae group; Mycobacterium abscessus subgroup; Mycobacterium abscessus MIKHLTRRATVALVVAAAAISPAAYPAVADTATPLNGLFALTPGSCAEGRATGSYFRMIL WP_005063674.1 hypothetical protein [Mycobacterium abscessus] Length: 330\nNumber of predicted TMHs: 1\nExp number of AAs in TMHs: 39.25259\nExp number, first 60 AAs: 16.98187\nTotal prob of N-in: 0.83210\nPOSSIBLE N-term signal sequence\noutside 1 298\nTMhelix 299 321\ninside 322 330

30698 GCF_900130805.1_10665_3_31 Mycobacterium abscessus subsp. massiliense Terrabacteria group; Actinobacteria; Actinobacteria; Corynebacteriales; Mycobacteriaceae; Mycobacterium; Mycobacterium chelonae group; Mycobacterium abscessus subgroup; Mycobacterium abscessus MIKHLTRRATVALVVAAAAISPAAYPAVADTATPLNGLFALTPGSCAEGRATGSYFRMIL WP_005063674.1 hypothetical protein [Mycobacterium abscessus] Length: 330\nNumber of predicted TMHs: 1\nExp number of AAs in TMHs: 39.25259\nExp number, first 60 AAs: 16.98187\nTotal prob of N-in: 0.83210\nPOSSIBLE N-term signal sequence\noutside 1 298\nTMhelix 299 321\ninside 322 330

30699 GCF_900130825.1_10665_3_33 Mycobacterium abscessus subsp. massiliense Terrabacteria group; Actinobacteria; Actinobacteria; Corynebacteriales; Mycobacteriaceae; Mycobacterium; Mycobacterium chelonae group; Mycobacterium abscessus subgroup; Mycobacterium abscessus MIKHLTRRATVALVVAAAAISPAAYPAVADTATPLNGLFALTPGSCAEGRATGSYFRMIL WP_005063674.1 hypothetical protein [Mycobacterium abscessus] Length: 330\nNumber of predicted TMHs: 1\nExp number of AAs in TMHs: 39.25259\nExp number, first 60 AAs: 16.98187\nTotal prob of N-in: 0.83210\nPOSSIBLE N-term signal sequence\noutside 1 298\nTMhelix 299 321\ninside 322 330

30700 GCF_900139605.1_10625_5_54 Mycobacterium abscessus subsp. massiliense Terrabacteria group; Actinobacteria; Actinobacteria; Corynebacteriales; Mycobacteriaceae; Mycobacterium; Mycobacterium chelonae group; Mycobacterium abscessus subgroup; Mycobacterium abscessus MIKHLTRRATVALVVAAAAISPAAYPAVADTATPLNGLFALTPGSCAEGRATGSYFRMIL WP_005063674.1 hypothetical protein [Mycobacterium abscessus] Length: 330\nNumber of predicted TMHs: 1\nExp number of AAs in TMHs: 39.25259\nExp number, first 60 AAs: 16.98187\nTotal prob of N-in: 0.83210\nPOSSIBLE N-term signal sequence\noutside 1 298\nTMhelix 299 321\ninside 322 330

30701 GCF_900138085.1_11893_6_9 Mycobacterium abscessus subsp. massiliense Terrabacteria group; Actinobacteria; Actinobacteria; Corynebacteriales; Mycobacteriaceae; Mycobacterium; Mycobacterium chelonae group; Mycobacterium abscessus subgroup; Mycobacterium abscessus MIKHLTRRATVALVVAAAAISPAAYPAVADTATPLNGLFALTPGSCAEGRATGSYFRMIL WP_005063674.1 hypothetical protein [Mycobacterium abscessus] Length: 330\nNumber of predicted TMHs: 1\nExp number of AAs in TMHs: 39.25259\nExp number, first 60 AAs: 16.98187\nTotal prob of N-in: 0.83210\nPOSSIBLE N-term signal sequence\noutside 1 298\nTMhelix 299 321\ninside 322 330

30702 GCF_900136465.1_10208_3_70 Mycobacterium abscessus subsp. massiliense Terrabacteria group; Actinobacteria; Actinobacteria; Corynebacteriales; Mycobacteriaceae; Mycobacterium; Mycobacterium chelonae group;

Mycobacterium abscessus subgroup; Mycobacterium abscessus
MIKHLTRRATVALVVAAAAISPAAYPAMADTATPLNGLFALTPGSCAEGRATGSYFRMIL WP_005073213.1 hypothetical protein [Mycobacterium abscessus] Length: 330\nNumber of predicted TMHs: 2\nExp number of AAs in TMHs: 39.3416199999999\nExp number, first 60 AAs: 17.07688\nTotal prob of N-in: 0.84431\nPOSSIBLE N-term signal sequence\ninside 1 11\nTMhelix 12 29\noutside 30 298\nTMhelix 299 321\ninside 322 330

30703 GCF_900136275.1_12163_1_83 Mycobacterium abscessus subsp. massiliense Terrabacteria group; Actinobacteria; Actinobacteria; Corynebacteriales; Mycobacteriaceae; Mycobacterium; Mycobacterium chelonae group; Mycobacterium abscessus subgroup; Mycobacterium abscessus
MIKHLTRRATVALVVAAAAISPAAYPAVADTATPLNGLFALTPGSCAEGRATGSYFRMIL WP_005063674.1 hypothetical protein [Mycobacterium abscessus] Length: 330\nNumber of predicted TMHs: 1\nExp number of AAs in TMHs: 39.25259\nExp number, first 60 AAs: 16.98187\nTotal prob of N-in: 0.83210\nPOSSIBLE N-term signal sequence\noutside 1 298\nTMhelix 299 321\ninside 322 330

30704 GCF_900133595.1_11893_6_48 Mycobacterium abscessus subsp. massiliense Terrabacteria group; Actinobacteria; Actinobacteria; Corynebacteriales; Mycobacteriaceae; Mycobacterium; Mycobacterium chelonae group; Mycobacterium abscessus subgroup; Mycobacterium abscessus
MIKHLTRRATVALVVAAAAISPAAYPAMADTATPLNGLFALTPGSCAEGRATGSYFRMIL WP_074356318.1 hypothetical protein [Mycobacterium abscessus] Length: 330\nNumber of predicted TMHs: 2\nExp number of AAs in TMHs: 39.34118\nExp number, first 60 AAs: 17.07668\nTotal prob of N-in: 0.84431\nPOSSIBLE N-term signal sequence\ninside 1 11\nTMhelix 12 29\noutside 30 298\nTMhelix 299 321\ninside 322 330

30705 GCF_900139225.1_10465_1_67 Mycobacterium abscessus subsp. massiliense Terrabacteria group; Actinobacteria; Actinobacteria; Corynebacteriales; Mycobacteriaceae; Mycobacterium; Mycobacterium chelonae group; Mycobacterium abscessus subgroup; Mycobacterium abscessus
MIKHLTRRATVALVVAAAAISPAAYPAVADTATPLNGLFALTPGSCAEGRATGSYFRMIL WP_005063674.1 hypothetical protein [Mycobacterium abscessus] Length: 330\nNumber of predicted TMHs: 1\nExp number of AAs in TMHs: 39.25259\nExp number, first 60 AAs: 16.98187\nTotal prob of N-in: 0.83210\nPOSSIBLE N-term signal sequence\noutside 1 298\nTMhelix 299 321\ninside 322 330

30706 GCF_900133675.1_12163_1_21 Mycobacterium abscessus subsp. massiliense Terrabacteria group; Actinobacteria; Actinobacteria; Corynebacteriales; Mycobacteriaceae; Mycobacterium; Mycobacterium chelonae group; Mycobacterium abscessus subgroup; Mycobacterium abscessus
MIKHLTRRATVALVVAAAAISPAAYPAVADTATPLNGLFALTPGSCAEGRATGSYFRMIL WP_005063674.1 hypothetical protein [Mycobacterium abscessus] Length: 330\nNumber of predicted TMHs: 1\nExp number of AAs in TMHs: 39.25259\nExp number, first 60 AAs: 16.98187\nTotal prob of N-in: 0.83210\nPOSSIBLE N-term signal sequence\noutside 1 298\nTMhelix 299 321\ninside 322 330

30707 GCF_900136295.1_12163_1_86 Mycobacterium abscessus subsp. massiliense Terrabacteria group; Actinobacteria; Actinobacteria; Corynebacteriales; Mycobacteriaceae; Mycobacterium; Mycobacterium chelonae group; Mycobacterium abscessus subgroup; Mycobacterium abscessus
MIKHLTRRATVALVVAAAAISPAAYPAVADTATPLNGLFALTPGSCAEGRATGSYFRMIL WP_005063674.1 hypothetical protein [Mycobacterium abscessus] Length: 330\nNumber of predicted TMHs: 1\nExp number of AAs in TMHs: 39.25259\nExp number, first 60 AAs: 16.98187\nTotal prob of N-in: 0.83210\nPOSSIBLE N-term signal sequence\noutside 1 298\nTMhelix 299 321\ninside 322 330

30708 GCF_900136315.1_12163_1_85 Mycobacterium abscessus subsp. massiliense Terrabacteria group; Actinobacteria; Actinobacteria; Corynebacteriales; Mycobacteriaceae; Mycobacterium; Mycobacterium chelonae group; Mycobacterium abscessus subgroup; Mycobacterium abscessus
MIKHLTRRATVALVVAAAAISPAAYPAVADTATPLNGLFALTPGSCAEGRATGSYFRMIL WP_005063674.1 hypothetical protein [Mycobacterium abscessus] Length: 330\nNumber of predicted TMHs: 1\nExp number of AAs in TMHs: 39.25259\nExp number, first 60 AAs: 16.98187\nTotal prob of N-in: 0.83210\nPOSSIBLE N-term signal sequence\noutside 1 298\nTMhelix 299 321\ninside 322 330

30709 GCF_900137245.1_12082_5_81 Mycobacterium abscessus subsp. massiliense Terrabacteria group; Actinobacteria; Actinobacteria; Corynebacteriales; Mycobacteriaceae; Mycobacterium; Mycobacterium chelonae group; Mycobacterium abscessus subgroup; Mycobacterium abscessus
MIKHLTRRATVALVVAAAAISPAAYPAVADTATPLNGLFALTPGSCAEGRATGSYFRMIL WP_005063674.1 hypothetical protein [Mycobacterium abscessus] Length: 330\nNumber of predicted TMHs: 1\nExp number of AAs in TMHs: 39.25259\nExp number, first 60 AAs: 16.98187\nTotal prob of N-in: 0.83210\nPOSSIBLE N-term signal sequence\noutside 1 298\nTMhelix 299 321\ninside 322 330

30710 GCF_900136025.1_12163_1_58 Mycobacterium abscessus subsp. massiliense Terrabacteria group; Actinobacteria; Actinobacteria; Corynebacteriales; Mycobacteriaceae; Mycobacterium; Mycobacterium chelonae group; Mycobacterium abscessus subgroup; Mycobacterium abscessus
MIKHLTRRATVALVVAAAAISPAAYPAVADTATPLNGLFALTPGSCAEGRATGSYFRMIL WP_005063674.1 hypothetical protein [Mycobacterium abscessus] Length: 330\nNumber of predicted TMHs: 1\nExp number of AAs in TMHs: 39.25259\nExp number, first 60 AAs: 16.98187\nTotal prob of N-in: 0.83210\nPOSSIBLE N-term signal sequence\noutside 1 298\nTMhelix 299 321\ninside 322 330

30711 GCF_900135955.1_12082_5_63 Mycobacterium abscessus subsp. massiliense Terrabacteria group; Actinobacteria; Actinobacteria; Corynebacteriales; Mycobacteriaceae; Mycobacterium; Mycobacterium chelonae group; Mycobacterium abscessus subgroup; Mycobacterium abscessus
MIKHLTRRATVALVVAAAAISPAAYPAVADTATPLNGLFALTPGSCAEGRATGSYFRMIL WP_005063674.1 hypothetical protein [Mycobacterium abscessus] Length: 330\nNumber of predicted TMHs: 1\nExp number of AAs in TMHs: 39.25259\nExp number, first 60 AAs: 16.98187\nTotal prob of N-in: 0.83210\nPOSSIBLE N-term signal sequence\noutside 1 298\nTMhelix 299 321\ninside 322 330

30712 GCF_900135905.1_12078_1_69 Mycobacterium abscessus subsp. massiliense Terrabacteria group; Actinobacteria; Actinobacteria; Corynebacteriales; Mycobacteriaceae; Mycobacterium; Mycobacterium chelonae group; Mycobacterium abscessus subgroup; Mycobacterium abscessus
MIKHLTRRATVALVVAAAAISPAAYPAVADTATPLNGLFALTPGSCAEGRATGSYFRMIL WP_005063674.1 hypothetical protein [Mycobacterium abscessus] Length: 330\nNumber of predicted TMHs: 1\nExp number of AAs in TMHs: 39.25259\nExp number, first 60 AAs: 16.98187\nTotal prob of N-in: 0.83210\nPOSSIBLE N-term signal sequence\noutside 1 298\nTMhelix 299 321\ninside 322 330

30713 GCF_002140035.1_ASM214003v1 Mycobacterium abscessus subsp. massiliense Terrabacteria group; Actinobacteria; Actinobacteria; Corynebacteriales; Mycobacteriaceae; Mycobacterium; Mycobacterium chelonae group; Mycobacterium abscessus subgroup; Mycobacterium abscessus
MIKHLTRRATVALVVAAAAISPAAYPAVADTATPLNGLFALTPGSCAEGRATGSYFRMIL WP_005063674.1 hypothetical protein [Mycobacterium abscessus] Length: 330\nNumber of predicted TMHs: 1\nExp number of AAs in TMHs: 39.25259\nExp number, first 60 AAs: 16.98187\nTotal prob of N-in: 0.83210\nPOSSIBLE N-term signal sequence\noutside 1 298\nTMhelix 299 321\ninside 322 330

30714 GCF_900140365.1_10665_3_73 Mycobacterium abscessus subsp. massiliense Terrabacteria group; Actinobacteria; Actinobacteria; Corynebacteriales; Mycobacteriaceae; Mycobacterium; Mycobacterium chelonae group; Mycobacterium abscessus subgroup; Mycobacterium abscessus
MIKHLTRRATVALVVAAAAISPAAYPAVADTATPLNGLFALTPGSCAEGRATGSYFRMIL WP_005063674.1 hypothetical protein [Mycobacterium abscessus] Length: 330\nNumber of predicted TMHs: 1\nExp number of AAs in TMHs: 39.25259\nExp number, first 60 AAs: 16.98187\nTotal prob of N-in: 0.83210\nPOSSIBLE N-term signal sequence\noutside 1 298\nTMhelix 299 321\ninside 322 330

30715 GCF_900130275.1_10625_5_60 Mycobacterium abscessus subsp. massiliense Terrabacteria group; Actinobacteria; Actinobacteria; Corynebacteriales; Mycobacteriaceae; Mycobacterium; Mycobacterium chelonae group; Mycobacterium abscessus subgroup; Mycobacterium abscessus
MIKHLTRRATVALVVAAAAISPAAYPAVADTATPLNGLFALTPGSCAEGRATGSYFRMIL WP_005063674.1 hypothetical protein [Mycobacterium abscessus] Length: 330\nNumber of predicted TMHs: 1\nExp number of AAs in TMHs: 39.25259\nExp number, first 60 AAs: 16.98187\nTotal prob of N-in: 0.83210\nPOSSIBLE N-term signal sequence\noutside 1 298\nTMhelix 299 321\ninside 322 330

30716 GCF_900137695.1_12163_2_45 Mycobacterium abscessus subsp. massiliense Terrabacteria group; Actinobacteria; Actinobacteria; Corynebacteriales; Mycobacteriaceae; Mycobacterium; Mycobacterium chelonae group; Mycobacterium abscessus subgroup; Mycobacterium abscessus
MIKHLTRRATVALVVAAAAISPAAYPAVADTATPLNGLFALTPGSCAEGRATGSYFRMIL WP_005063674.1 hypothetical protein [Mycobacterium abscessus] Length: 330\nNumber of predicted TMHs: 1\nExp number of AAs in TMHs: 39.25259\nExp number, first 60 AAs: 16.98187\nTotal prob of N-in: 0.83210\nPOSSIBLE N-term signal sequence\noutside 1 298\nTMhelix 299 321\ninside 322 330

30717 GCF_900130385.1_10625_5_73 Mycobacterium abscessus subsp. massiliense Terrabacteria group; Actinobacteria; Actinobacteria; Corynebacteriales; Mycobacteriaceae; Mycobacterium; Mycobacterium chelonae group; Mycobacterium abscessus subgroup; Mycobacterium abscessus
MIKHLTRRATVALVVAAAAISPAAYPAVADTATPLNGLFALTPGSCAEGRATGSYFRMIL WP_005063674.1 hypothetical protein [Mycobacterium abscessus] Length: 330\nNumber of predicted TMHs: 1\nExp number of AAs in TMHs:

39.25259\nExp number, first 60 AAs: 16.98187\nTotal prob of N-in: 0.83210\nPOSSIBLE N-term signal sequence\noutside 1 298\nTMhelix 299 321\ninside 322 330

30718 GCF_900137835.1_12163_2_75 Mycobacterium abscessus subsp. massiliense Terrabacteria group; Actinobacteria; Actinobacteria; Corynebacteriales; Mycobacteriaceae; Mycobacterium; Mycobacterium chelonae group; Mycobacterium abscessus subgroup; Mycobacterium abscessus MIKHLTRRATVALVVAAAAISPAAYPAVADTATPLNGLFALTPGSCAEGRATGSYFRMIL WP_005063674.1 hypothetical protein [Mycobacterium abscessus] Length: 330\nNumber of predicted TMHs: 1\nExp number of AAs in TMHs: 39.25259\nExp number, first 60 AAs: 16.98187\nTotal prob of N-in: 0.83210\nPOSSIBLE N-term signal sequence\noutside 1 298\nTMhelix 299 321\ninside 322 330

30719 GCF_900139755.1_10660_1_18 Mycobacterium abscessus subsp. massiliense Terrabacteria group; Actinobacteria; Actinobacteria; Corynebacteriales; Mycobacteriaceae; Mycobacterium; Mycobacterium chelonae group; Mycobacterium abscessus subgroup; Mycobacterium abscessus MIKHLTRRATVALVVAAAAISPAAYPAVADTATPLNGLFALTPGSCAEGRATGSYFRMIL WP_005063674.1 hypothetical protein [Mycobacterium abscessus] Length: 330\nNumber of predicted TMHs: 1\nExp number of AAs in TMHs: 39.25259\nExp number, first 60 AAs: 16.98187\nTotal prob of N-in: 0.83210\nPOSSIBLE N-term signal sequence\noutside 1 298\nTMhelix 299 321\ninside 322 330

30720 GCF_900130755.1_10665_3_23 Mycobacterium abscessus subsp. massiliense Terrabacteria group; Actinobacteria; Actinobacteria; Corynebacteriales; Mycobacteriaceae; Mycobacterium; Mycobacterium chelonae group; Mycobacterium abscessus subgroup; Mycobacterium abscessus MIKHLTRRATVALVVAAAAISPAAYPAVADTATPLNGLFALTPGSCAEGRATGSYFRMIL WP_005063674.1 hypothetical protein [Mycobacterium abscessus] Length: 330\nNumber of predicted TMHs: 1\nExp number of AAs in TMHs: 39.25259\nExp number, first 60 AAs: 16.98187\nTotal prob of N-in: 0.83210\nPOSSIBLE N-term signal sequence\noutside 1 298\nTMhelix 299 321\ninside 322 330

30721 GCF_900139715.1_10660_1_16 Mycobacterium abscessus subsp. massiliense Terrabacteria group; Actinobacteria; Actinobacteria; Corynebacteriales; Mycobacteriaceae; Mycobacterium; Mycobacterium chelonae group; Mycobacterium abscessus subgroup; Mycobacterium abscessus MIKHLTRRATVALVVAAAAISPAAYPAMADTATPLNGLFALTPGSCAEGRATGSYFRMIL WP_005073213.1 hypothetical protein [Mycobacterium abscessus] Length: 330\nNumber of predicted TMHs: 2\nExp number of AAs in TMHs: 39.3416199999999\nExp number, first 60 AAs: 17.07688\nTotal prob of N-in: 0.84431\nPOSSIBLE N-term signal sequence\ninside 1 11\nTMhelix 12 29\noutside 30 298\nTMhelix 299 321\ninside 322 330

30722 GCF_900130775.1_10665_3_28 Mycobacterium abscessus subsp. massiliense Terrabacteria group; Actinobacteria; Actinobacteria; Corynebacteriales; Mycobacteriaceae; Mycobacterium; Mycobacterium chelonae group; Mycobacterium abscessus subgroup; Mycobacterium abscessus MIKHLTRRATVALVVAAAAISPAAYPAVADTATPLNGLFALTPGSCAEGRATGSYFRMIL WP_005063674.1 hypothetical protein [Mycobacterium abscessus] Length: 330\nNumber of predicted TMHs: 1\nExp number of AAs in TMHs: 39.25259\nExp number, first 60 AAs: 16.98187\nTotal prob of N-in: 0.83210\nPOSSIBLE N-term signal sequence\noutside 1 298\nTMhelix 299 321\ninside 322 330

30723 GCF_900141505.1_12078_1_64 Mycobacterium abscessus subsp. massiliense Terrabacteria group; Actinobacteria; Actinobacteria; Corynebacteriales; Mycobacteriaceae; Mycobacterium; Mycobacterium chelonae group; Mycobacterium abscessus subgroup; Mycobacterium abscessus MIKHLTRRATVALVVAAAAISPAAYPAVADTATPLNGLFALTPGSCAEGRATGSYFRMIL WP_005063674.1 hypothetical protein [Mycobacterium abscessus] Length: 330\nNumber of predicted TMHs: 1\nExp number of AAs in TMHs: 39.25259\nExp number, first 60 AAs: 16.98187\nTotal prob of N-in: 0.83210\nPOSSIBLE N-term signal sequence\noutside 1 298\nTMhelix 299 321\ninside 322 330

30724 GCF_900137875.1_12163_2_80 Mycobacterium abscessus subsp. massiliense Terrabacteria group; Actinobacteria; Actinobacteria; Corynebacteriales; Mycobacteriaceae; Mycobacterium; Mycobacterium chelonae group; Mycobacterium abscessus subgroup; Mycobacterium abscessus MIKHLTRRATVALVVAAAAISPAAYPAVADTATPLNGLFALTPGSCAEGRATGSYFRMIL WP_005063674.1 hypothetical protein [Mycobacterium abscessus] Length: 330\nNumber of predicted TMHs: 1\nExp number of AAs in TMHs: 39.25259\nExp number, first 60 AAs: 16.98187\nTotal prob of N-in: 0.83210\nPOSSIBLE N-term signal sequence\noutside 1 298\nTMhelix 299 321\ninside 322 330

30725 GCF_900134635.1_12163_2_63 Mycobacterium abscessus subsp. massiliense Terrabacteria group; Actinobacteria; Actinobacteria; Corynebacteriales; Mycobacteriaceae; Mycobacterium; Mycobacterium chelonae group;

Mycobacterium abscessus subgroup; Mycobacterium abscessus
MIKHLTRRATVALVVAAAAISPAAYPAMADTATPLNGLFALTPGSCAEGRATGSYFRMIL WP_074356318.1 hypothetical protein [Mycobacterium abscessus] Length: 330\nNumber of predicted TMHs: 2\nExp number of AAs in TMHs: 39.34118\nExp number, first 60 AAs: 17.07668\nTotal prob of N-in: 0.84431\nPOSSIBLE N-term signal sequence\ninside 1 11\nTMhelix 12 29\noutside 30 298\nTMhelix 299 321\ninside 322 330

30726 GCF_900132825.1_10625_4_75 Mycobacterium abscessus subsp. massiliense Terrabacteria group; Actinobacteria; Actinobacteria; Corynebacteriales; Mycobacteriaceae; Mycobacterium; Mycobacterium chelonae group; Mycobacterium abscessus subgroup; Mycobacterium abscessus
MIKHLTRRATVALVVAAAAISPAAYPAMADTATPLNGLFALTPGSCAEGRATGSYFRMIL WP_005073213.1 hypothetical protein [Mycobacterium abscessus] Length: 330\nNumber of predicted TMHs: 2\nExp number of AAs in TMHs: 39.3416199999999\nExp number, first 60 AAs: 17.07688\nTotal prob of N-in: 0.84431\nPOSSIBLE N-term signal sequence\ninside 1 11\nTMhelix 12 29\noutside 30 298\nTMhelix 299 321\ninside 322 330

30727 GCF_900137105.1_11893_6_36 Mycobacterium abscessus subsp. massiliense Terrabacteria group; Actinobacteria; Actinobacteria; Corynebacteriales; Mycobacteriaceae; Mycobacterium; Mycobacterium chelonae group; Mycobacterium abscessus subgroup; Mycobacterium abscessus
MIKHLTRRATVALVVAAAAISPAAYPAVADTATPLNGLFALTPGSCAEGRATGSYFRMIL WP_005063674.1 hypothetical protein [Mycobacterium abscessus] Length: 330\nNumber of predicted TMHs: 1\nExp number of AAs in TMHs: 39.25259\nExp number, first 60 AAs: 16.98187\nTotal prob of N-in: 0.83210\nPOSSIBLE N-term signal sequence\noutside 1 298\nTMhelix 299 321\ninside 322 330

30728 GCF_900138955.1_10208_3_60 Mycobacterium abscessus subsp. massiliense Terrabacteria group; Actinobacteria; Actinobacteria; Corynebacteriales; Mycobacteriaceae; Mycobacterium; Mycobacterium chelonae group; Mycobacterium abscessus subgroup; Mycobacterium abscessus
MIKHLTRRATVALVVAAAAISPAAYPAVADTATPLNGLFALTPGSCAEGRATGSYFRMIL WP_005063674.1 hypothetical protein [Mycobacterium abscessus] Length: 330\nNumber of predicted TMHs: 1\nExp number of AAs in TMHs: 39.25259\nExp number, first 60 AAs: 16.98187\nTotal prob of N-in: 0.83210\nPOSSIBLE N-term signal sequence\noutside 1 298\nTMhelix 299 321\ninside 322 330

30729 GCF_900133145.1_12082_5_54 Mycobacterium abscessus subsp. massiliense Terrabacteria group; Actinobacteria; Actinobacteria; Corynebacteriales; Mycobacteriaceae; Mycobacterium; Mycobacterium chelonae group; Mycobacterium abscessus subgroup; Mycobacterium abscessus
MIKHLTRRATVALVVAAAAISPAAYPAVADTATPLNGLFALTPGSCAEGRATGSYFRMIL WP_005063674.1 hypothetical protein [Mycobacterium abscessus] Length: 330\nNumber of predicted TMHs: 1\nExp number of AAs in TMHs: 39.25259\nExp number, first 60 AAs: 16.98187\nTotal prob of N-in: 0.83210\nPOSSIBLE N-term signal sequence\noutside 1 298\nTMhelix 299 321\ninside 322 330

30730 GCF_900133155.1_12163_1_3 Mycobacterium abscessus subsp. massiliense Terrabacteria group; Actinobacteria; Actinobacteria; Corynebacteriales; Mycobacteriaceae; Mycobacterium; Mycobacterium chelonae group; Mycobacterium abscessus subgroup; Mycobacterium abscessus
MIKHLTRRATVALVVAAAAISPAAYPAVADTATPLNGLFALTPGSCAEGRATGSYFRMIL WP_005063674.1 hypothetical protein [Mycobacterium abscessus] Length: 330\nNumber of predicted TMHs: 1\nExp number of AAs in TMHs: 39.25259\nExp number, first 60 AAs: 16.98187\nTotal prob of N-in: 0.83210\nPOSSIBLE N-term signal sequence\noutside 1 298\nTMhelix 299 321\ninside 322 330

30731 GCF_900133235.1_10250_1_28 Mycobacterium abscessus subsp. massiliense Terrabacteria group; Actinobacteria; Actinobacteria; Corynebacteriales; Mycobacteriaceae; Mycobacterium; Mycobacterium chelonae group; Mycobacterium abscessus subgroup; Mycobacterium abscessus
MIKHLTRRATVALVVAAAAISPAAYPAVADTATPLNGLFALTPGSCAEGRATGSYFRMIL WP_005063674.1 hypothetical protein [Mycobacterium abscessus] Length: 330\nNumber of predicted TMHs: 1\nExp number of AAs in TMHs: 39.25259\nExp number, first 60 AAs: 16.98187\nTotal prob of N-in: 0.83210\nPOSSIBLE N-term signal sequence\noutside 1 298\nTMhelix 299 321\ninside 322 330

30732 GCF_900138855.1_10208_3_53 Mycobacterium abscessus subsp. massiliense Terrabacteria group; Actinobacteria; Actinobacteria; Corynebacteriales; Mycobacteriaceae; Mycobacterium; Mycobacterium chelonae group; Mycobacterium abscessus subgroup; Mycobacterium abscessus
MIKHLTRRATVALVVAAAAISPAAYPAVADTATPLNGLFALTPGSCAEGRATGSYFRMIL WP_005063674.1 hypothetical protein [Mycobacterium abscessus] Length: 330\nNumber of predicted TMHs: 1\nExp number of AAs in TMHs: 39.25259\nExp number, first 60 AAs: 16.98187\nTotal prob of N-in: 0.83210\nPOSSIBLE N-term signal sequence\noutside 1 298\nTMhelix 299 321\ninside 322 330

30733 GCF_900138905.1_10208_3_57 Mycobacterium abscessus subsp. massiliense Terrabacteria group; Actinobacteria; Actinobacteria; Corynebacteriales; Mycobacteriaceae; Mycobacterium; Mycobacterium chelonae group; Mycobacterium abscessus subgroup; Mycobacterium abscessus
MIKHLTRRATVALVVAAAAISPAAYPAVADTATPLNGLFALTPGSCAEGRATGSYFRMIL WP_005063674.1 hypothetical protein [Mycobacterium abscessus] Length: 330\nNumber of predicted TMHs: 1\nExp number of AAs in TMHs: 39.25259\nExp number, first 60 AAs: 16.98187\nTotal prob of N-in: 0.83210\nPOSSIBLE N-term signal sequence\noutside 1 298\nTMhelix 299 321\ninside 322 330

30734 GCF_900133445.1_10625_4_59 Mycobacterium abscessus subsp. massiliense Terrabacteria group; Actinobacteria; Actinobacteria; Corynebacteriales; Mycobacteriaceae; Mycobacterium; Mycobacterium chelonae group; Mycobacterium abscessus subgroup; Mycobacterium abscessus
MIKHLTRRATVALVVAAAAISPAAYPAVADTATPLNGLFALTPGSCAEGRATGSYFRMIL WP_005063674.1 hypothetical protein [Mycobacterium abscessus] Length: 330\nNumber of predicted TMHs: 1\nExp number of AAs in TMHs: 39.25259\nExp number, first 60 AAs: 16.98187\nTotal prob of N-in: 0.83210\nPOSSIBLE N-term signal sequence\noutside 1 298\nTMhelix 299 321\ninside 322 330

30735 GCF_900136525.1_10208_3_71 Mycobacterium abscessus subsp. massiliense Terrabacteria group; Actinobacteria; Actinobacteria; Corynebacteriales; Mycobacteriaceae; Mycobacterium; Mycobacterium chelonae group; Mycobacterium abscessus subgroup; Mycobacterium abscessus
MIKHLTRRATVALVVAAAAISPAAYPAMADTATPLNGLFALTPGSCAEGRATGSYFRMIL WP_005073213.1
hypothetical protein [Mycobacterium abscessus] Length: 330\nNumber of predicted TMHs: 2\nExp number of AAs in TMHs: 39.3416199999999\nExp number, first 60 AAs: 17.07688\nTotal prob of N-in: 0.84431\nPOSSIBLE N-term signal sequence\ninside 1 11\nTMhelix 12 29\noutside 30 298\nTMhelix 299 321\ninside 322 330

30736 GCF_900138165.1_12082_5_85 Mycobacterium abscessus subsp. massiliense Terrabacteria group; Actinobacteria; Actinobacteria; Corynebacteriales; Mycobacteriaceae; Mycobacterium; Mycobacterium chelonae group; Mycobacterium abscessus subgroup; Mycobacterium abscessus
MIKHLTRRATVALVVAAAAISPAAYPAVADTATPLNGLFALTPGSCAEGRATGSYFRMIL WP_005063674.1 hypothetical protein [Mycobacterium abscessus] Length: 330\nNumber of predicted TMHs: 1\nExp number of AAs in TMHs: 39.25259\nExp number, first 60 AAs: 16.98187\nTotal prob of N-in: 0.83210\nPOSSIBLE N-term signal sequence\noutside 1 298\nTMhelix 299 321\ninside 322 330

30737 GCF_900131065.1_10665_3_50 Mycobacterium abscessus subsp. massiliense Terrabacteria group; Actinobacteria; Actinobacteria; Corynebacteriales; Mycobacteriaceae; Mycobacterium; Mycobacterium chelonae group; Mycobacterium abscessus subgroup; Mycobacterium abscessus
MIKHLTRRATVALVVAAAAISPAAYPAVADTATPLNGLFALTPGSCAEGRATGSYFRMIL WP_005063674.1 hypothetical protein [Mycobacterium abscessus] Length: 330\nNumber of predicted TMHs: 1\nExp number of AAs in TMHs: 39.25259\nExp number, first 60 AAs: 16.98187\nTotal prob of N-in: 0.83210\nPOSSIBLE N-term signal sequence\noutside 1 298\nTMhelix 299 321\ninside 322 330

30738 GCF_900139585.1_10625_5_47 Mycobacterium abscessus subsp. massiliense Terrabacteria group; Actinobacteria; Actinobacteria; Corynebacteriales; Mycobacteriaceae; Mycobacterium; Mycobacterium chelonae group; Mycobacterium abscessus subgroup; Mycobacterium abscessus
MIKHLTRRATVALVVAAAAISPAAYPAVADTATPLNGLFALTPGSCAEGRATGSYFRMIL WP_005063674.1 hypothetical protein [Mycobacterium abscessus] Length: 330\nNumber of predicted TMHs: 1\nExp number of AAs in TMHs: 39.25259\nExp number, first 60 AAs: 16.98187\nTotal prob of N-in: 0.83210\nPOSSIBLE N-term signal sequence\noutside 1 298\nTMhelix 299 321\ninside 322 330

30739 GCF_900139525.1_10625_5_45 Mycobacterium abscessus subsp. massiliense Terrabacteria group; Actinobacteria; Actinobacteria; Corynebacteriales; Mycobacteriaceae; Mycobacterium; Mycobacterium chelonae group; Mycobacterium abscessus subgroup; Mycobacterium abscessus
MIKHLTRRATVALVVAAAAISPAAYPAVADTATPLNGLFALTPGSCAEGRATGSYFRMIL WP_005063674.1 hypothetical protein [Mycobacterium abscessus] Length: 330\nNumber of predicted TMHs: 1\nExp number of AAs in TMHs: 39.25259\nExp number, first 60 AAs: 16.98187\nTotal prob of N-in: 0.83210\nPOSSIBLE N-term signal sequence\noutside 1 298\nTMhelix 299 321\ninside 322 330

30740 GCF_900131315.1_10665_3_77 Mycobacterium abscessus subsp. massiliense Terrabacteria group; Actinobacteria; Actinobacteria; Corynebacteriales; Mycobacteriaceae; Mycobacterium; Mycobacterium chelonae group; Mycobacterium abscessus subgroup; Mycobacterium abscessus
MIKHLTRRATVALVVAAAAISPAAYPAVADTATPLNGLFALTPGSCAEGRATGSYFRMIL WP_005063674.1 hypothetical protein [Mycobacterium abscessus] Length: 330\nNumber of predicted TMHs: 1\nExp number of AAs in TMHs:

39.25259\nExp number, first 60 AAs: 16.98187\nTotal prob of N-in: 0.83210\nPOSSIBLE N-term signal sequence\noutside 1 298\nTMhelix 299 321\ninside 322 330

30741 GCF_900131205.1_10665_3_66 Mycobacterium abscessus subsp. massiliense Terrabacteria group; Actinobacteria; Actinobacteria; Corynebacteriales; Mycobacteriaceae; Mycobacterium; Mycobacterium chelonae group; Mycobacterium abscessus subgroup; Mycobacterium abscessus MIKHLTRRATVALVVAAAAISPAAYPAVADTATPLNGLFALTPGSCAEGRATGSYFRMIL WP_005063674.1 hypothetical protein [Mycobacterium abscessus] Length: 330\nNumber of predicted TMHs: 1\nExp number of AAs in TMHs: 39.25259\nExp number, first 60 AAs: 16.98187\nTotal prob of N-in: 0.83210\nPOSSIBLE N-term signal sequence\noutside 1 298\nTMhelix 299 321\ninside 322 330

30742 GCF_900138135.1_12078_1_65 Mycobacterium abscessus subsp. massiliense Terrabacteria group; Actinobacteria; Actinobacteria; Corynebacteriales; Mycobacteriaceae; Mycobacterium; Mycobacterium chelonae group; Mycobacterium abscessus subgroup; Mycobacterium abscessus MIKHLTRRATVALVVAAAAISPAAYPAVADTATPLNGLFALTPGSCAEGRATGSYFRMIL WP_005063674.1 hypothetical protein [Mycobacterium abscessus] Length: 330\nNumber of predicted TMHs: 1\nExp number of AAs in TMHs: 39.25259\nExp number, first 60 AAs: 16.98187\nTotal prob of N-in: 0.83210\nPOSSIBLE N-term signal sequence\noutside 1 298\nTMhelix 299 321\ninside 322 330

30743 GCF_900136825.1_10625_5_39 Mycobacterium abscessus subsp. massiliense Terrabacteria group; Actinobacteria; Actinobacteria; Corynebacteriales; Mycobacteriaceae; Mycobacterium; Mycobacterium chelonae group; Mycobacterium abscessus subgroup; Mycobacterium abscessus MIKHLTRRATVALVVAAAAISPAAYPAVADTATPLNGLFALTPGSCAEGRATGSYFRMIL WP_005063674.1 hypothetical protein [Mycobacterium abscessus] Length: 330\nNumber of predicted TMHs: 1\nExp number of AAs in TMHs: 39.25259\nExp number, first 60 AAs: 16.98187\nTotal prob of N-in: 0.83210\nPOSSIBLE N-term signal sequence\noutside 1 298\nTMhelix 299 321\ninside 322 330

30744 GCF_900136755.1_10625_5_30 Mycobacterium abscessus subsp. massiliense Terrabacteria group; Actinobacteria; Actinobacteria; Corynebacteriales; Mycobacteriaceae; Mycobacterium; Mycobacterium chelonae group; Mycobacterium abscessus subgroup; Mycobacterium abscessus MIKHLTRRATVALVVAAAAISPAAYPAVADTATPLNGLFALTPGSCAEGRATGSYFRMIL WP_005063674.1 hypothetical protein [Mycobacterium abscessus] Length: 330\nNumber of predicted TMHs: 1\nExp number of AAs in TMHs: 39.25259\nExp number, first 60 AAs: 16.98187\nTotal prob of N-in: 0.83210\nPOSSIBLE N-term signal sequence\noutside 1 298\nTMhelix 299 321\ninside 322 330

30745 GCF_900131525.1_11893_6_44 Mycobacterium abscessus subsp. massiliense Terrabacteria group; Actinobacteria; Actinobacteria; Corynebacteriales; Mycobacteriaceae; Mycobacterium; Mycobacterium chelonae group; Mycobacterium abscessus subgroup; Mycobacterium abscessus MIKHLTRRATVALVVAAAAISPAAYPAMADTATPLNGLFALTPGSCAEGRATGSYFRMIL WP_074356318.1 hypothetical protein [Mycobacterium abscessus] Length: 330\nNumber of predicted TMHs: 2\nExp number of AAs in TMHs: 39.34118\nExp number, first 60 AAs: 17.07668\nTotal prob of N-in: 0.84431\nPOSSIBLE N-term signal sequence\ninside 1 11\nTMhelix 12 29\noutside 30 298\nTMhelix 299 321\ninside 322 330

30746 GCF_900131535.1_11893_6_45 Mycobacterium abscessus subsp. massiliense Terrabacteria group; Actinobacteria; Actinobacteria; Corynebacteriales; Mycobacteriaceae; Mycobacterium; Mycobacterium chelonae group; Mycobacterium abscessus subgroup; Mycobacterium abscessus MIKHLTRRATVALVVAAAAISPAAYPAMADTATPLNGLFALTPGSCAEGRATGSYFRMIL WP_074356318.1 hypothetical protein [Mycobacterium abscessus] Length: 330\nNumber of predicted TMHs: 2\nExp number of AAs in TMHs: 39.34118\nExp number, first 60 AAs: 17.07668\nTotal prob of N-in: 0.84431\nPOSSIBLE N-term signal sequence\ninside 1 11\nTMhelix 12 29\noutside 30 298\nTMhelix 299 321\ninside 322 330

30747 GCF_900136715.1_10625_5_28 Mycobacterium abscessus subsp. massiliense Terrabacteria group; Actinobacteria; Actinobacteria; Corynebacteriales; Mycobacteriaceae; Mycobacterium; Mycobacterium chelonae group; Mycobacterium abscessus subgroup; Mycobacterium abscessus MIKHLTRRATVALVVAAAAISPAAYPAVADTATPLNGLFALTPGSCAEGRATGSYFRMIL WP_005063674.1 hypothetical protein [Mycobacterium abscessus] Length: 330\nNumber of predicted TMHs: 1\nExp number of AAs in TMHs: 39.25259\nExp number, first 60 AAs: 16.98187\nTotal prob of N-in: 0.83210\nPOSSIBLE N-term signal sequence\noutside 1 298\nTMhelix 299 321\ninside 322 330

30748 GCF_900137025.1_11861_7_78 Mycobacterium abscessus subsp. massiliense Terrabacteria group; Actinobacteria; Actinobacteria; Corynebacteriales; Mycobacteriaceae; Mycobacterium; Mycobacterium chelonae group;

Mycobacterium abscessus subgroup; Mycobacterium abscessus
MIKHLTRRATVALVVAAAAISPAAYPAMADTATPLNGLFALTPGSCAEGRATGSYFRMIL WP_005073213.1 hypothetical protein [Mycobacterium abscessus] Length: 330\nNumber of predicted TMHs: 2\nExp number of AAs in TMHs: 39.3416199999999\nExp number, first 60 AAs: 17.07688\nTotal prob of N-in: 0.84431\nPOSSIBLE N-term signal sequence\ninside 1 11\nTMhelix 12 29\noutside 30 298\nTMhelix 299 321\ninside 322 330

30749 GCF_900138295.1_11893_7_41 Mycobacterium abscessus subsp. massiliense Terrabacteria group; Actinobacteria; Actinobacteria; Corynebacteriales; Mycobacteriaceae; Mycobacterium; Mycobacterium chelonae group; Mycobacterium abscessus subgroup; Mycobacterium abscessus
MIKHLTRRATVALVVAAAAISPAAYPAVADTATPLNGLFALTPGSCAEGRATGSYFRMIL WP_005063674.1 hypothetical protein [Mycobacterium abscessus] Length: 330\nNumber of predicted TMHs: 1\nExp number of AAs in TMHs: 39.25259\nExp number, first 60 AAs: 16.98187\nTotal prob of N-in: 0.83210\nPOSSIBLE N-term signal sequence\noutside 1 298\nTMhelix 299 321\ninside 322 330

30750 GCF_900138455.1_11893_7_58 Mycobacterium abscessus subsp. massiliense Terrabacteria group; Actinobacteria; Actinobacteria; Corynebacteriales; Mycobacteriaceae; Mycobacterium; Mycobacterium chelonae group; Mycobacterium abscessus subgroup; Mycobacterium abscessus
MIKHLTRRATVALVVAAAAISPAAYPAVADTATPLNGLFALTPGSCAEGRATGSYFRMIL WP_005063674.1 hypothetical protein [Mycobacterium abscessus] Length: 330\nNumber of predicted TMHs: 1\nExp number of AAs in TMHs: 39.25259\nExp number, first 60 AAs: 16.98187\nTotal prob of N-in: 0.83210\nPOSSIBLE N-term signal sequence\noutside 1 298\nTMhelix 299 321\ninside 322 330

30751 GCF_900138475.1_11893_7_60 Mycobacterium abscessus subsp. massiliense Terrabacteria group; Actinobacteria; Actinobacteria; Corynebacteriales; Mycobacteriaceae; Mycobacterium; Mycobacterium chelonae group; Mycobacterium abscessus subgroup; Mycobacterium abscessus
MIKHLTRRATVALVVAAAAISPAAYPAVADTATPLNGLFALTPGSCAEGRATGSYFRMIL WP_005063674.1 hypothetical protein [Mycobacterium abscessus] Length: 330\nNumber of predicted TMHs: 1\nExp number of AAs in TMHs: 39.25259\nExp number, first 60 AAs: 16.98187\nTotal prob of N-in: 0.83210\nPOSSIBLE N-term signal sequence\noutside 1 298\nTMhelix 299 321\ninside 322 330

30752 GCF_900131855.1_11893_6_25 Mycobacterium abscessus subsp. massiliense Terrabacteria group; Actinobacteria; Actinobacteria; Corynebacteriales; Mycobacteriaceae; Mycobacterium; Mycobacterium chelonae group; Mycobacterium abscessus subgroup; Mycobacterium abscessus
MIKHLTRRATVALVVAAAAISPAAYPAVADTATPLNGLFALTPGSCAEGRATGSYFRMIL WP_005063674.1 hypothetical protein [Mycobacterium abscessus] Length: 330\nNumber of predicted TMHs: 1\nExp number of AAs in TMHs: 39.25259\nExp number, first 60 AAs: 16.98187\nTotal prob of N-in: 0.83210\nPOSSIBLE N-term signal sequence\noutside 1 298\nTMhelix 299 321\ninside 322 330

30753 GCF_900131875.1_11893_6_26 Mycobacterium abscessus subsp. massiliense Terrabacteria group; Actinobacteria; Actinobacteria; Corynebacteriales; Mycobacteriaceae; Mycobacterium; Mycobacterium chelonae group; Mycobacterium abscessus subgroup; Mycobacterium abscessus
MIKHLTRRATVALVVAAAAISPAAYPAVADTATPLNGLFALTPGSCAEGRATGSYFRMIL WP_005063674.1 hypothetical protein [Mycobacterium abscessus] Length: 330\nNumber of predicted TMHs: 1\nExp number of AAs in TMHs: 39.25259\nExp number, first 60 AAs: 16.98187\nTotal prob of N-in: 0.83210\nPOSSIBLE N-term signal sequence\noutside 1 298\nTMhelix 299 321\ninside 322 330

30754 GCF_900138505.1_12454_1_92 Mycobacterium abscessus subsp. massiliense Terrabacteria group; Actinobacteria; Actinobacteria; Corynebacteriales; Mycobacteriaceae; Mycobacterium; Mycobacterium chelonae group; Mycobacterium abscessus subgroup; Mycobacterium abscessus
MIKHLTRRATVALVVAAAAISPAAYPAVADTATPLNGLFALTPGSCAEGRATGSYFRMIL WP_005063674.1 hypothetical protein [Mycobacterium abscessus] Length: 330\nNumber of predicted TMHs: 1\nExp number of AAs in TMHs: 39.25259\nExp number, first 60 AAs: 16.98187\nTotal prob of N-in: 0.83210\nPOSSIBLE N-term signal sequence\noutside 1 298\nTMhelix 299 321\ninside 322 330

30755 GCF_900138495.1_12454_1_93 Mycobacterium abscessus subsp. massiliense Terrabacteria group; Actinobacteria; Actinobacteria; Corynebacteriales; Mycobacteriaceae; Mycobacterium; Mycobacterium chelonae group; Mycobacterium abscessus subgroup; Mycobacterium abscessus
MIKHLTRRATVALVVAAAAISPAAYPAVADTATPLNGLFALTPGSCAEGRATGSYFRMIL WP_005063674.1 hypothetical protein [Mycobacterium abscessus] Length: 330\nNumber of predicted TMHs: 1\nExp number of AAs in TMHs: 39.25259\nExp number, first 60 AAs: 16.98187\nTotal prob of N-in: 0.83210\nPOSSIBLE N-term signal sequence\noutside 1 298\nTMhelix 299 321\ninside 322 330

30756 GCF_900132035.1_10665_4_48 Mycobacterium abscessus subsp. massiliense Terrabacteria group; Actinobacteria; Actinobacteria; Corynebacteriales; Mycobacteriaceae; Mycobacterium; Mycobacterium chelonae group; Mycobacterium abscessus subgroup; Mycobacterium abscessus
MIKHLTRRATVALVVAAAAISPAAYPAVADTATPLNGLFALTPGSCAEGRATGSYFRMIL WP_005063674.1 hypothetical protein [Mycobacterium abscessus] Length: 330\nNumber of predicted TMHs: 1\nExp number of AAs in TMHs: 39.25259\nExp number, first 60 AAs: 16.98187\nTotal prob of N-in: 0.83210\nPOSSIBLE N-term signal sequence\noutside 1 298\nTMhelix 299 321\ninside 322 330

30757 GCF_900138525.1_20034_1_19 Mycobacterium abscessus subsp. massiliense Terrabacteria group; Actinobacteria; Actinobacteria; Corynebacteriales; Mycobacteriaceae; Mycobacterium; Mycobacterium chelonae group; Mycobacterium abscessus subgroup; Mycobacterium abscessus
MIKHLTRRATVALVVAAAAISPAAYPAVADTATPLNGLFALTPGSCAEGRATGSYFRMIL WP_079617725.1 hypothetical protein [Mycobacterium abscessus] Length: 330\nNumber of predicted TMHs: 1\nExp number of AAs in TMHs: 39.2571699999999\nExp number, first 60 AAs: 16.97134\nTotal prob of N-in: 0.83217\nPOSSIBLE N-term signal sequence\noutside 1 298\nTMhelix 299 321\ninside 322 330

30758 GCF_900132075.1_10665_4_52 Mycobacterium abscessus subsp. massiliense Terrabacteria group; Actinobacteria; Actinobacteria; Corynebacteriales; Mycobacteriaceae; Mycobacterium; Mycobacterium chelonae group; Mycobacterium abscessus subgroup; Mycobacterium abscessus
MIKHLTRRATVALVVAAAAISPAAYPAVADTATPLNGLFALTPGSCAEGRATGSYFRMIL WP_005063674.1 hypothetical protein [Mycobacterium abscessus] Length: 330\nNumber of predicted TMHs: 1\nExp number of AAs in TMHs: 39.25259\nExp number, first 60 AAs: 16.98187\nTotal prob of N-in: 0.83210\nPOSSIBLE N-term signal sequence\noutside 1 298\nTMhelix 299 321\ninside 322 330

30759 GCF_900132085.1_10665_4_53 Mycobacterium abscessus subsp. massiliense Terrabacteria group; Actinobacteria; Actinobacteria; Corynebacteriales; Mycobacteriaceae; Mycobacterium; Mycobacterium chelonae group; Mycobacterium abscessus subgroup; Mycobacterium abscessus
MIKHLTRRATVALVVAAAAISPAAYPAVADTATPLNGLFALTPGSCAEGRATGSYFRMIL WP_005063674.1 hypothetical protein [Mycobacterium abscessus] Length: 330\nNumber of predicted TMHs: 1\nExp number of AAs in TMHs: 39.25259\nExp number, first 60 AAs: 16.98187\nTotal prob of N-in: 0.83210\nPOSSIBLE N-term signal sequence\noutside 1 298\nTMhelix 299 321\ninside 322 330

30760 GCF_900132215.1_11893_6_27 Mycobacterium abscessus subsp. massiliense Terrabacteria group; Actinobacteria; Actinobacteria; Corynebacteriales; Mycobacteriaceae; Mycobacterium; Mycobacterium chelonae group; Mycobacterium abscessus subgroup; Mycobacterium abscessus
MIKHLTRRATVALVVAAAAISPAAYPAVADTATPLNGLFALTPGSCAEGRATGSYFRMIL WP_005063674.1 hypothetical protein [Mycobacterium abscessus] Length: 330\nNumber of predicted TMHs: 1\nExp number of AAs in TMHs: 39.25259\nExp number, first 60 AAs: 16.98187\nTotal prob of N-in: 0.83210\nPOSSIBLE N-term signal sequence\noutside 1 298\nTMhelix 299 321\ninside 322 330

30761 GCF_900132235.1_11893_6_28 Mycobacterium abscessus subsp. massiliense Terrabacteria group; Actinobacteria; Actinobacteria; Corynebacteriales; Mycobacteriaceae; Mycobacterium; Mycobacterium chelonae group; Mycobacterium abscessus subgroup; Mycobacterium abscessus
MIKHLTRRATVALVVAAAAISPAAYPAVADTATPLNGLFALTPGSCAEGRATGSYFRMIL WP_005063674.1 hypothetical protein [Mycobacterium abscessus] Length: 330\nNumber of predicted TMHs: 1\nExp number of AAs in TMHs: 39.25259\nExp number, first 60 AAs: 16.98187\nTotal prob of N-in: 0.83210\nPOSSIBLE N-term signal sequence\noutside 1 298\nTMhelix 299 321\ninside 322 330

30762 GCF_900138575.1_20034_1_12 Mycobacterium abscessus subsp. massiliense Terrabacteria group; Actinobacteria; Actinobacteria; Corynebacteriales; Mycobacteriaceae; Mycobacterium; Mycobacterium chelonae group; Mycobacterium abscessus subgroup; Mycobacterium abscessus
MIKHLTRRATVALVVAAAAISPAAYPAVADTATPLNGLFALTPGSCAEGRATGSYFRMIL WP_079617725.1 hypothetical protein [Mycobacterium abscessus] Length: 330\nNumber of predicted TMHs: 1\nExp number of AAs in TMHs: 39.2571699999999\nExp number, first 60 AAs: 16.97134\nTotal prob of N-in: 0.83217\nPOSSIBLE N-term signal sequence\noutside 1 298\nTMhelix 299 321\ninside 322 330

30763 GCF_900138585.1_20034_1_16 Mycobacterium abscessus subsp. massiliense Terrabacteria group; Actinobacteria; Actinobacteria; Corynebacteriales; Mycobacteriaceae; Mycobacterium; Mycobacterium chelonae group; Mycobacterium abscessus subgroup; Mycobacterium abscessus
MIKHLTRRATVALVVAAAAISPAAYPAVADTATPLNGLFALTPGSCAEGRATGSYFRMIL WP_079617725.1 hypothetical protein [Mycobacterium abscessus] Length: 330\nNumber of predicted TMHs: 1\nExp number of AAs in TMHs:

39.2571699999999\nExp number, first 60 AAs: 16.97134\nTotal prob of N-in: 0.83217\nPOSSIBLE N-term signal sequence\noutside 1 298\nTMhelix 299 321\ninside 322 330

30764 GCF_900138635.1_20034_1_20 Mycobacterium abscessus subsp. massiliense Terrabacteria group; Actinobacteria; Actinobacteria; Corynebacteriales; Mycobacteriaceae; Mycobacterium; Mycobacterium chelonae group; Mycobacterium abscessus subgroup; Mycobacterium abscessus MIKHLTRRATVALVVAAAAISPAAYPAVADTATPLNGLFALTPGSCAEGRATGSYFRMIL WP_079617725.1 hypothetical protein [Mycobacterium abscessus] Length: 330\nNumber of predicted TMHs: 1\nExp number of AAs in TMHs: 39.2571699999999\nExp number, first 60 AAs: 16.97134\nTotal prob of N-in: 0.83217\nPOSSIBLE N-term signal sequence\noutside 1 298\nTMhelix 299 321\ninside 322 330

30765 GCF_900132375.1_10665_4_69 Mycobacterium abscessus subsp. massiliense Terrabacteria group; Actinobacteria; Actinobacteria; Corynebacteriales; Mycobacteriaceae; Mycobacterium; Mycobacterium chelonae group; Mycobacterium abscessus subgroup; Mycobacterium abscessus MIKHLTRRATVALVVAAAAISPAAYPAVADTATPLNGLFALTPGSCAEGRATGSYFRMIL WP_005063674.1 hypothetical protein [Mycobacterium abscessus] Length: 330\nNumber of predicted TMHs: 1\nExp number of AAs in TMHs: 39.25259\nExp number, first 60 AAs: 16.98187\nTotal prob of N-in: 0.83210\nPOSSIBLE N-term signal sequence\noutside 1 298\nTMhelix 299 321\ninside 322 330

30766 GCF_900132395.1_10665_4_70 Mycobacterium abscessus subsp. massiliense Terrabacteria group; Actinobacteria; Actinobacteria; Corynebacteriales; Mycobacteriaceae; Mycobacterium; Mycobacterium chelonae group; Mycobacterium abscessus subgroup; Mycobacterium abscessus MIKHLTRRATVALVVAAAAISPAAYPAVADTATPLNGLFALTPGSCAEGRATGSYFRMIL WP_005063674.1 hypothetical protein [Mycobacterium abscessus] Length: 330\nNumber of predicted TMHs: 1\nExp number of AAs in TMHs: 39.25259\nExp number, first 60 AAs: 16.98187\nTotal prob of N-in: 0.83210\nPOSSIBLE N-term signal sequence\noutside 1 298\nTMhelix 299 321\ninside 322 330

30767 GCF_900138655.1_20034_1_21 Mycobacterium abscessus subsp. massiliense Terrabacteria group; Actinobacteria; Actinobacteria; Corynebacteriales; Mycobacteriaceae; Mycobacterium; Mycobacterium chelonae group; Mycobacterium abscessus subgroup; Mycobacterium abscessus MIKHLTRRATVALVVAAAAISPAAYPAVADTATPLNGLFALTPGSCAEGRATGSYFRMIL WP_079617725.1 hypothetical protein [Mycobacterium abscessus] Length: 330\nNumber of predicted TMHs: 1\nExp number of AAs in TMHs: 39.2571699999999\nExp number, first 60 AAs: 16.97134\nTotal prob of N-in: 0.83217\nPOSSIBLE N-term signal sequence\noutside 1 298\nTMhelix 299 321\ninside 322 330

30768 GCF_900138695.1_20034_1_6 Mycobacterium abscessus subsp. massiliense Terrabacteria group; Actinobacteria; Actinobacteria; Corynebacteriales; Mycobacteriaceae; Mycobacterium; Mycobacterium chelonae group; Mycobacterium abscessus subgroup; Mycobacterium abscessus MIKHLTRRATVALVVAAAAISPAAYPAVADTATPLNGLFALTPGSCAEGRATGSYFRMIL WP_079617725.1 hypothetical protein [Mycobacterium abscessus] Length: 330\nNumber of predicted TMHs: 1\nExp number of AAs in TMHs: 39.2571699999999\nExp number, first 60 AAs: 16.97134\nTotal prob of N-in: 0.83217\nPOSSIBLE N-term signal sequence\noutside 1 298\nTMhelix 299 321\ninside 322 330

30769 GCF_900138675.1_20034_1_3 Mycobacterium abscessus subsp. massiliense Terrabacteria group; Actinobacteria; Actinobacteria; Corynebacteriales; Mycobacteriaceae; Mycobacterium; Mycobacterium chelonae group; Mycobacterium abscessus subgroup; Mycobacterium abscessus MIKHLTRRATVALVVAAAAISPAAYPAVADTATPLNGLFALTPGSCAEGRATGSYFRMIL WP_079617725.1 hypothetical protein [Mycobacterium abscessus] Length: 330\nNumber of predicted TMHs: 1\nExp number of AAs in TMHs: 39.2571699999999\nExp number, first 60 AAs: 16.97134\nTotal prob of N-in: 0.83217\nPOSSIBLE N-term signal sequence\noutside 1 298\nTMhelix 299 321\ninside 322 330

30770 GCF_900132695.1_10250_1_16 Mycobacterium abscessus subsp. massiliense Terrabacteria group; Actinobacteria; Actinobacteria; Corynebacteriales; Mycobacteriaceae; Mycobacterium; Mycobacterium chelonae group; Mycobacterium abscessus subgroup; Mycobacterium abscessus MIKHLTRRATVALVVAAAAISPAAYPAVADTATPLNGLFALTPGSCAEGRATGSYFRMIL WP_005063674.1 hypothetical protein [Mycobacterium abscessus] Length: 330\nNumber of predicted TMHs: 1\nExp number of AAs in TMHs: 39.25259\nExp number, first 60 AAs: 16.98187\nTotal prob of N-in: 0.83210\nPOSSIBLE N-term signal sequence\noutside 1 298\nTMhelix 299 321\ninside 322 330

30771 GCF_900138705.1_20034_1_8 Mycobacterium abscessus subsp. massiliense Terrabacteria group; Actinobacteria; Actinobacteria; Corynebacteriales; Mycobacteriaceae; Mycobacterium; Mycobacterium chelonae group;

Mycobacterium abscessus subgroup; Mycobacterium abscessus
MIKHLTRRATVALVVAAAAISPAAYPAVADTATPLNGLFALTPGSCAEGRATGSYFRMIL WP_079617725.1 hypothetical protein [Mycobacterium abscessus] Length: 330\nNumber of predicted TMHs: 1\nExp number of AAs in TMHs: 39.2571699999999\nExp number, first 60 AAs: 16.97134\nTotal prob of N-in: 0.83217\nPOSSIBLE N-term signal sequence\noutside 1 298\nTMhelix 299 321\ninside 322 330

30772 GCF_900138725.1_20034_1_7 Mycobacterium abscessus subsp. massiliense Terrabacteria group; Actinobacteria; Actinobacteria; Corynebacteriales; Mycobacteriaceae; Mycobacterium; Mycobacterium chelonae group; Mycobacterium abscessus subgroup; Mycobacterium abscessus
MIKHLTRRATVALVVAAAAISPAAYPAVADTATPLNGLFALTPGSCAEGRATGSYFRMIL WP_079617725.1 hypothetical protein [Mycobacterium abscessus] Length: 330\nNumber of predicted TMHs: 1\nExp number of AAs in TMHs: 39.2571699999999\nExp number, first 60 AAs: 16.97134\nTotal prob of N-in: 0.83217\nPOSSIBLE N-term signal sequence\noutside 1 298\nTMhelix 299 321\ninside 322 330

30773 GCF_900132925.1_11861_7_62 Mycobacterium abscessus subsp. massiliense Terrabacteria group; Actinobacteria; Actinobacteria; Corynebacteriales; Mycobacteriaceae; Mycobacterium; Mycobacterium chelonae group; Mycobacterium abscessus subgroup; Mycobacterium abscessus
MIKHLTRRATVALVVAAAAISPAAYPAVADTATPLNGLFALTPGSCAEGRATGSYFRMIL WP_005063674.1 hypothetical protein [Mycobacterium abscessus] Length: 330\nNumber of predicted TMHs: 1\nExp number of AAs in TMHs: 39.25259\nExp number, first 60 AAs: 16.98187\nTotal prob of N-in: 0.83210\nPOSSIBLE N-term signal sequence\noutside 1 298\nTMhelix 299 321\ninside 322 330

30774 GCF_900140225.1_10665_2_96 Mycobacterium abscessus subsp. massiliense Terrabacteria group; Actinobacteria; Actinobacteria; Corynebacteriales; Mycobacteriaceae; Mycobacterium; Mycobacterium chelonae group; Mycobacterium abscessus subgroup; Mycobacterium abscessus
MIKHLTRRATVALVVAAAAISPAAYPAVADTATPLNGLFALTPGSCAEGRATGSYFRMIL WP_005063674.1 hypothetical protein [Mycobacterium abscessus] Length: 330\nNumber of predicted TMHs: 1\nExp number of AAs in TMHs: 39.25259\nExp number, first 60 AAs: 16.98187\nTotal prob of N-in: 0.83210\nPOSSIBLE N-term signal sequence\noutside 1 298\nTMhelix 299 321\ninside 322 330

30775 GCF_900130375.1_10625_5_70 Mycobacterium abscessus subsp. massiliense Terrabacteria group; Actinobacteria; Actinobacteria; Corynebacteriales; Mycobacteriaceae; Mycobacterium; Mycobacterium chelonae group; Mycobacterium abscessus subgroup; Mycobacterium abscessus
MIKHLTRRATVALVVAAAAISPAAYPAVADTATPLNGLFALTPGSCAEGRATGSYFRMIL WP_005063674.1 hypothetical protein [Mycobacterium abscessus] Length: 330\nNumber of predicted TMHs: 1\nExp number of AAs in TMHs: 39.25259\nExp number, first 60 AAs: 16.98187\nTotal prob of N-in: 0.83210\nPOSSIBLE N-term signal sequence\noutside 1 298\nTMhelix 299 321\ninside 322 330

30776 GCF_900137625.1_12163_2_39 Mycobacterium abscessus subsp. massiliense Terrabacteria group; Actinobacteria; Actinobacteria; Corynebacteriales; Mycobacteriaceae; Mycobacterium; Mycobacterium chelonae group; Mycobacterium abscessus subgroup; Mycobacterium abscessus
MIKHLTRRATVALVVAAAAISPAAYPAVADTATPLNGLFALTPGSCAEGRATGSYFRMIL WP_005063674.1 hypothetical protein [Mycobacterium abscessus] Length: 330\nNumber of predicted TMHs: 1\nExp number of AAs in TMHs: 39.25259\nExp number, first 60 AAs: 16.98187\nTotal prob of N-in: 0.83210\nPOSSIBLE N-term signal sequence\noutside 1 298\nTMhelix 299 321\ninside 322 330

30777 GCF_900130285.1_10071_6_83 Mycobacterium abscessus subsp. massiliense Terrabacteria group; Actinobacteria; Actinobacteria; Corynebacteriales; Mycobacteriaceae; Mycobacterium; Mycobacterium chelonae group; Mycobacterium abscessus subgroup; Mycobacterium abscessus
MIKHLTRRATVALVVAAAAISPAAYPAVADTATPLNGLFALTPGSCAEGRATGSYFRMIL WP_005063674.1 hypothetical protein [Mycobacterium abscessus] Length: 330\nNumber of predicted TMHs: 1\nExp number of AAs in TMHs: 39.25259\nExp number, first 60 AAs: 16.98187\nTotal prob of N-in: 0.83210\nPOSSIBLE N-term signal sequence\noutside 1 298\nTMhelix 299 321\ninside 322 330

30778 GCF_900138155.1_12082_5_84 Mycobacterium abscessus subsp. massiliense Terrabacteria group; Actinobacteria; Actinobacteria; Corynebacteriales; Mycobacteriaceae; Mycobacterium; Mycobacterium chelonae group; Mycobacterium abscessus subgroup; Mycobacterium abscessus
MIKHLTRRATVALVVAAAAISPAAYPAVADTATPLNGLFALTPGSCAEGRATGSYFRMIL WP_005063674.1 hypothetical protein [Mycobacterium abscessus] Length: 330\nNumber of predicted TMHs: 1\nExp number of AAs in TMHs: 39.25259\nExp number, first 60 AAs: 16.98187\nTotal prob of N-in: 0.83210\nPOSSIBLE N-term signal sequence\noutside 1 298\nTMhelix 299 321\ninside 322 330

30779 GCF_900140755.1_10702_1_38 Mycobacterium abscessus subsp. massiliense Terrabacteria group; Actinobacteria; Actinobacteria; Corynebacteriales; Mycobacteriaceae; Mycobacterium; Mycobacterium chelonae group; Mycobacterium abscessus subgroup; Mycobacterium abscessus
MIKHLTRRATVALVVAAAAISPAAYPAMADTATPLNGLFALTPGSCAEGRATGSYFRMIL WP_005073213.1 hypothetical protein [Mycobacterium abscessus] Length: 330\nNumber of predicted TMHs: 2\nExp number of AAs in TMHs: 39.3416199999999\nExp number, first 60 AAs: 17.07688\nTotal prob of N-in: 0.84431\nPOSSIBLE N-term signal sequence\ninside 1 11\nTMhelix 12 29\noutside 30 298\nTMhelix 299 321\ninside 322 330

30780 GCF_900137985.1_11893_6_10 Mycobacterium abscessus subsp. massiliense Terrabacteria group; Actinobacteria; Actinobacteria; Corynebacteriales; Mycobacteriaceae; Mycobacterium; Mycobacterium chelonae group; Mycobacterium abscessus subgroup; Mycobacterium abscessus
MIKHLTRRATVALVVAAAAISPAAYPAVADTATPLNGLFALTPGSCAEGRATGSYFRMIL WP_005063674.1 hypothetical protein [Mycobacterium abscessus] Length: 330\nNumber of predicted TMHs: 1\nExp number of AAs in TMHs: 39.25259\nExp number, first 60 AAs: 16.98187\nTotal prob of N-in: 0.83210\nPOSSIBLE N-term signal sequence\noutside 1 298\nTMhelix 299 321\ninside 322 330

30781 GCF_900138535.1_13109_1_72 Mycobacterium abscessus subsp. massiliense Terrabacteria group; Actinobacteria; Actinobacteria; Corynebacteriales; Mycobacteriaceae; Mycobacterium; Mycobacterium chelonae group; Mycobacterium abscessus subgroup; Mycobacterium abscessus
MIKHLTRRATVALVVAAAAISPAAYPAVADTATPLNGLFALTPGSCAEGRATGSYFRMIL WP_005063674.1 hypothetical protein [Mycobacterium abscessus] Length: 330\nNumber of predicted TMHs: 1\nExp number of AAs in TMHs: 39.25259\nExp number, first 60 AAs: 16.98187\nTotal prob of N-in: 0.83210\nPOSSIBLE N-term signal sequence\noutside 1 298\nTMhelix 299 321\ninside 322 330

30782 GCF_900131635.1_10665_4_20 Mycobacterium abscessus subsp. massiliense Terrabacteria group; Actinobacteria; Actinobacteria; Corynebacteriales; Mycobacteriaceae; Mycobacterium; Mycobacterium chelonae group; Mycobacterium abscessus subgroup; Mycobacterium abscessus
MIKHLTRRATVALVVAAAAISPAAYPAVADTATPLNGLFALTPGSCAEGRATGSYFRMIL WP_005063674.1 hypothetical protein [Mycobacterium abscessus] Length: 330\nNumber of predicted TMHs: 1\nExp number of AAs in TMHs: 39.25259\nExp number, first 60 AAs: 16.98187\nTotal prob of N-in: 0.83210\nPOSSIBLE N-term signal sequence\noutside 1 298\nTMhelix 299 321\ninside 322 330

30783 GCF_900138735.1_10208_3_16 Mycobacterium abscessus subsp. massiliense Terrabacteria group; Actinobacteria; Actinobacteria; Corynebacteriales; Mycobacteriaceae; Mycobacterium; Mycobacterium chelonae group; Mycobacterium abscessus subgroup; Mycobacterium abscessus
MIKHLTRRATVALVVAAAAISPAAYPAMADTATPLNGLFALTPGSCAEGRATGSYFRMIL WP_005073213.1 hypothetical protein [Mycobacterium abscessus] Length: 330\nNumber of predicted TMHs: 2\nExp number of AAs in TMHs: 39.3416199999999\nExp number, first 60 AAs: 17.07688\nTotal prob of N-in: 0.84431\nPOSSIBLE N-term signal sequence\ninside 1 11\nTMhelix 12 29\noutside 30 298\nTMhelix 299 321\ninside 322 330

30784 GCF_900137095.1_11861_8_9 Mycobacterium abscessus subsp. massiliense Terrabacteria group; Actinobacteria; Actinobacteria; Corynebacteriales; Mycobacteriaceae; Mycobacterium; Mycobacterium chelonae group; Mycobacterium abscessus subgroup; Mycobacterium abscessus
MIKHLTRRATVALVVAAAAISPAAYPAMADTATPLNGLFALTPGSCAEGRATGSYFRMIL WP_079668796.1 hypothetical protein [Mycobacterium abscessus] Length: 320\nNumber of predicted TMHs: 2\nExp number of AAs in TMHs: 39.3533\nExp number, first 60 AAs: 17.06343\nTotal prob of N-in: 0.84669\nPOSSIBLE N-term signal sequence\ninside 1 11\nTMhelix 12 29\noutside 30 288\nTMhelix 289 311\ninside 312 320

30785 GCF_900138515.1_20034_1_1 Mycobacterium abscessus subsp. massiliense Terrabacteria group; Actinobacteria; Actinobacteria; Corynebacteriales; Mycobacteriaceae; Mycobacterium; Mycobacterium chelonae group; Mycobacterium abscessus subgroup; Mycobacterium abscessus
MIKHLTRRATVALVVAAAAISPAAYPAVADTATPLNGLFALTPGSCAEGRATGSYFRMIL WP_079617725.1 hypothetical protein [Mycobacterium abscessus] Length: 330\nNumber of predicted TMHs: 1\nExp number of AAs in TMHs: 39.2571699999999\nExp number, first 60 AAs: 16.97134\nTotal prob of N-in: 0.83217\nPOSSIBLE N-term signal sequence\noutside 1 298\nTMhelix 299 321\ninside 322 330

30786 GCF_900133015.1_11893_6_32 Mycobacterium abscessus subsp. massiliense Terrabacteria group; Actinobacteria; Actinobacteria; Corynebacteriales; Mycobacteriaceae; Mycobacterium; Mycobacterium chelonae group; Mycobacterium abscessus subgroup; Mycobacterium abscessus
MIKHLTRRATVALVVAAAAISPAAYPAVADTATPLNGLFALTPGSCAEGRATGSYFRMIL WP_005063674.1 hypothetical protein [Mycobacterium abscessus] Length: 330\nNumber of predicted TMHs: 1\nExp number of AAs in TMHs:

39.25259\nExp number, first 60 AAs:  16.98187\nTotal prob of N-in:   0.83210\nPOSSIBLE N-term signal sequence\noutside 1   298\nTMhelix   299   321\ninside   322   330

30787 GCF_900138625.1_20034_1_22 Mycobacterium abscessus subsp. massiliense Terrabacteria group; Actinobacteria; Actinobacteria; Corynebacteriales; Mycobacteriaceae; Mycobacterium; Mycobacterium chelonae group; Mycobacterium abscessus subgroup; Mycobacterium abscessus MIKHLTRRATVALVVAAAAISPAAYPAVADTATPLNGLFALTPGSCAEGRATGSYFRMIL WP_079617725.1 hypothetical protein [Mycobacterium abscessus] Length: 330\nNumber of predicted TMHs:  1\nExp number of AAs in TMHs: 39.2571699999999\nExp number, first 60 AAs:  16.97134\nTotal prob of N-in:   0.83217\nPOSSIBLE N-term signal sequence\noutside   1   298\nTMhelix   299   321\ninside   322   330

30788 GCF_900133065.1_11893_6_66 Mycobacterium abscessus subsp. massiliense Terrabacteria group; Actinobacteria; Actinobacteria; Corynebacteriales; Mycobacteriaceae; Mycobacterium; Mycobacterium chelonae group; Mycobacterium abscessus subgroup; Mycobacterium abscessus MIKHLTRRATVALVVAAAAISPAAYPAVADTATPLNGLFALTPGSCAEGRATGSYFRMIL WP_005063674.1 hypothetical protein [Mycobacterium abscessus] Length: 330\nNumber of predicted TMHs:  1\nExp number of AAs in TMHs: 39.25259\nExp number, first 60 AAs:  16.98187\nTotal prob of N-in:   0.83210\nPOSSIBLE N-term signal sequence\noutside 1   298\nTMhelix   299   321\ninside   322   330

30789 GCF_900133585.1_11893_6_33 Mycobacterium abscessus subsp. massiliense Terrabacteria group; Actinobacteria; Actinobacteria; Corynebacteriales; Mycobacteriaceae; Mycobacterium; Mycobacterium chelonae group; Mycobacterium abscessus subgroup; Mycobacterium abscessus MIKHLTRRATVALVVAAAAISPAAYPAVADTATPLNGLFALTPGSCAEGRATGSYFRMIL WP_005063674.1 hypothetical protein [Mycobacterium abscessus] Length: 330\nNumber of predicted TMHs:  1\nExp number of AAs in TMHs: 39.25259\nExp number, first 60 AAs:  16.98187\nTotal prob of N-in:   0.83210\nPOSSIBLE N-term signal sequence\noutside 1   298\nTMhelix   299   321\ninside   322   330

30790 GCF_900138715.1_20034_1_9 Mycobacterium abscessus subsp. massiliense Terrabacteria group; Actinobacteria; Actinobacteria; Corynebacteriales; Mycobacteriaceae; Mycobacterium; Mycobacterium chelonae group; Mycobacterium abscessus subgroup; Mycobacterium abscessus MIKHLTRRATVALVVAAAAISPAAYPAVADTATPLNGLFALTPGSCAEGRATGSYFRMIL WP_079617725.1 hypothetical protein [Mycobacterium abscessus] Length: 330\nNumber of predicted TMHs:  1\nExp number of AAs in TMHs: 39.2571699999999\nExp number, first 60 AAs:  16.97134\nTotal prob of N-in:   0.83217\nPOSSIBLE N-term signal sequence\noutside   1   298\nTMhelix   299   321\ninside   322   330

30791 GCF_900133615.1_12082_5_57 Mycobacterium abscessus subsp. massiliense Terrabacteria group; Actinobacteria; Actinobacteria; Corynebacteriales; Mycobacteriaceae; Mycobacterium; Mycobacterium chelonae group; Mycobacterium abscessus subgroup; Mycobacterium abscessus MIKHLTRRATVALVVAAAAISPAAYPAVADTATPLNGLFALTPGSCAEGRATGSYFRMIL WP_005063674.1 hypothetical protein [Mycobacterium abscessus] Length: 330\nNumber of predicted TMHs:  1\nExp number of AAs in TMHs: 39.25259\nExp number, first 60 AAs:  16.98187\nTotal prob of N-in:   0.83210\nPOSSIBLE N-term signal sequence\noutside 1   298\nTMhelix   299   321\ninside   322   330

30792 GCF_900138845.1_10208_3_51 Mycobacterium abscessus subsp. massiliense Terrabacteria group; Actinobacteria; Actinobacteria; Corynebacteriales; Mycobacteriaceae; Mycobacterium; Mycobacterium chelonae group; Mycobacterium abscessus subgroup; Mycobacterium abscessus MIKHLTRRATVALVVAAAAISPAAYPAVADTATPLNGLFALTPGSCAEGRATGSYFRMIL WP_005063674.1 hypothetical protein [Mycobacterium abscessus] Length: 330\nNumber of predicted TMHs:  1\nExp number of AAs in TMHs: 39.25259\nExp number, first 60 AAs:  16.98187\nTotal prob of N-in:   0.83210\nPOSSIBLE N-term signal sequence\noutside 1   298\nTMhelix   299   321\ninside   322   330

30793 GCF_900141515.1_12078_1_63 Mycobacterium abscessus subsp. massiliense Terrabacteria group; Actinobacteria; Actinobacteria; Corynebacteriales; Mycobacteriaceae; Mycobacterium; Mycobacterium chelonae group; Mycobacterium abscessus subgroup; Mycobacterium abscessus MIKHLTRRATVALVVAAAAISPAAYPAVADTATPLNGLFALTPGSCAEGRATGSYFRMIL WP_005063674.1 hypothetical protein [Mycobacterium abscessus] Length: 330\nNumber of predicted TMHs:  1\nExp number of AAs in TMHs: 39.25259\nExp number, first 60 AAs:  16.98187\nTotal prob of N-in:   0.83210\nPOSSIBLE N-term signal sequence\noutside 1   298\nTMhelix   299   321\ninside   322   330

30794 GCF_900131325.1_10665_3_79 Mycobacterium abscessus subsp. massiliense Terrabacteria group; Actinobacteria; Actinobacteria; Corynebacteriales; Mycobacteriaceae; Mycobacterium; Mycobacterium chelonae group;

Mycobacterium abscessus subgroup; Mycobacterium abscessus
MIKHLTRRATVALVVAAAAISPAAYPAVADTATPLNGLFALTPGSCAEGRATGSYFRMIL WP_005063674.1 hypothetical protein [Mycobacterium abscessus] Length: 330\nNumber of predicted TMHs: 1\nExp number of AAs in TMHs: 39.25259\nExp number, first 60 AAs: 16.98187\nTotal prob of N-in: 0.83210\nPOSSIBLE N-term signal sequence\noutside 1 298\nTMhelix 299 321\ninside 322 330

30795 GCF_900139095.1_10465_1_53 Mycobacterium abscessus subsp. massiliense Terrabacteria group; Actinobacteria; Actinobacteria; Corynebacteriales; Mycobacteriaceae; Mycobacterium; Mycobacterium chelonae group; Mycobacterium abscessus subgroup; Mycobacterium abscessus
MIKHLTRRATVALVVAAAAISPAAYPAVADTATPLNGLFALTPGSCAEGRATGSYFRMIL WP_005063674.1 hypothetical protein [Mycobacterium abscessus] Length: 330\nNumber of predicted TMHs: 1\nExp number of AAs in TMHs: 39.25259\nExp number, first 60 AAs: 16.98187\nTotal prob of N-in: 0.83210\nPOSSIBLE N-term signal sequence\noutside 1 298\nTMhelix 299 321\ninside 322 330

30796 GCF_900134495.1_12163_1_42 Mycobacterium abscessus subsp. massiliense Terrabacteria group; Actinobacteria; Actinobacteria; Corynebacteriales; Mycobacteriaceae; Mycobacterium; Mycobacterium chelonae group; Mycobacterium abscessus subgroup; Mycobacterium abscessus
MIKHLTRRATVALVVAAAAISPAAYPAVADTATPLNGLFALTPGSCAEGRATGSYFRMIL WP_005063674.1 hypothetical protein [Mycobacterium abscessus] Length: 330\nNumber of predicted TMHs: 1\nExp number of AAs in TMHs: 39.25259\nExp number, first 60 AAs: 16.98187\nTotal prob of N-in: 0.83210\nPOSSIBLE N-term signal sequence\noutside 1 298\nTMhelix 299 321\ninside 322 330

30797 GCF_900132105.1_10665_4_54 Mycobacterium abscessus subsp. massiliense Terrabacteria group; Actinobacteria; Actinobacteria; Corynebacteriales; Mycobacteriaceae; Mycobacterium; Mycobacterium chelonae group; Mycobacterium abscessus subgroup; Mycobacterium abscessus
MIKHLTRRATVALVVAAAAISPAAYPAVADTATPLNGLFALTPGSCAEGRATGSYFRMIL WP_005063674.1 hypothetical protein [Mycobacterium abscessus] Length: 330\nNumber of predicted TMHs: 1\nExp number of AAs in TMHs: 39.25259\nExp number, first 60 AAs: 16.98187\nTotal prob of N-in: 0.83210\nPOSSIBLE N-term signal sequence\noutside 1 298\nTMhelix 299 321\ninside 322 330

30798 GCF_900132065.1_10665_4_51 Mycobacterium abscessus subsp. massiliense Terrabacteria group; Actinobacteria; Actinobacteria; Corynebacteriales; Mycobacteriaceae; Mycobacterium; Mycobacterium chelonae group; Mycobacterium abscessus subgroup; Mycobacterium abscessus
MIKHLTRRATVALVVAAAAISPAAYPAVADTATPLNGLFALTPGSCAEGRATGSYFRMIL WP_005063674.1 hypothetical protein [Mycobacterium abscessus] Length: 330\nNumber of predicted TMHs: 1\nExp number of AAs in TMHs: 39.25259\nExp number, first 60 AAs: 16.98187\nTotal prob of N-in: 0.83210\nPOSSIBLE N-term signal sequence\noutside 1 298\nTMhelix 299 321\ninside 322 330

30799 GCF_900131695.1_10665_4_29 Mycobacterium abscessus subsp. massiliense Terrabacteria group; Actinobacteria; Actinobacteria; Corynebacteriales; Mycobacteriaceae; Mycobacterium; Mycobacterium chelonae group; Mycobacterium abscessus subgroup; Mycobacterium abscessus
MIKHLTRRATVALVVAAAAISPAAYPAVADTATPLNGLFALTPGSCAEGRATGSYFRMIL WP_005063674.1 hypothetical protein [Mycobacterium abscessus] Length: 330\nNumber of predicted TMHs: 1\nExp number of AAs in TMHs: 39.25259\nExp number, first 60 AAs: 16.98187\nTotal prob of N-in: 0.83210\nPOSSIBLE N-term signal sequence\noutside 1 298\nTMhelix 299 321\ninside 322 330

30800 GCF_900134865.1_10208_3_26 Mycobacterium abscessus subsp. massiliense Terrabacteria group; Actinobacteria; Actinobacteria; Corynebacteriales; Mycobacteriaceae; Mycobacterium; Mycobacterium chelonae group; Mycobacterium abscessus subgroup; Mycobacterium abscessus
MIKHLTRRATVALVVAAAAISPAAYPAVADTATPLNGLFALTPGSCAEGRATGSYFRMIL WP_005063674.1 hypothetical protein [Mycobacterium abscessus] Length: 330\nNumber of predicted TMHs: 1\nExp number of AAs in TMHs: 39.25259\nExp number, first 60 AAs: 16.98187\nTotal prob of N-in: 0.83210\nPOSSIBLE N-term signal sequence\noutside 1 298\nTMhelix 299 321\ninside 322 330

30801 GCF_900139835.1_10660_1_28 Mycobacterium abscessus subsp. massiliense Terrabacteria group; Actinobacteria; Actinobacteria; Corynebacteriales; Mycobacteriaceae; Mycobacterium; Mycobacterium chelonae group; Mycobacterium abscessus subgroup; Mycobacterium abscessus
MIKHLTRRATVALVVAAAAISPAAYPAVADTATPLNGLFALTPGSCAEGRATGSYFRMIL WP_005063674.1 hypothetical protein [Mycobacterium abscessus] Length: 330\nNumber of predicted TMHs: 1\nExp number of AAs in TMHs: 39.25259\nExp number, first 60 AAs: 16.98187\nTotal prob of N-in: 0.83210\nPOSSIBLE N-term signal sequence\noutside 1 298\nTMhelix 299 321\ninside 322 330

30802 GCF_900131655.1_10665_4_23 Mycobacterium abscessus subsp. massiliense Terrabacteria group; Actinobacteria; Actinobacteria; Corynebacteriales; Mycobacteriaceae; Mycobacterium; Mycobacterium chelonae group; Mycobacterium abscessus subgroup; Mycobacterium abscessus
MIKHLTRRATVALVVAAAAISPAAYPAVADTATPLNGLFALTPGSCAEGRATGSYFRMIL WP_005063674.1 hypothetical protein [Mycobacterium abscessus] Length: 330\nNumber of predicted TMHs: 1\nExp number of AAs in TMHs: 39.25259\nExp number, first 60 AAs: 16.98187\nTotal prob of N-in: 0.83210\nPOSSIBLE N-term signal sequence\noutside 1 298\nTMhelix 299 321\ninside 322 330

30803 GCF_900139905.1_10660_1_34 Mycobacterium abscessus subsp. massiliense Terrabacteria group; Actinobacteria; Actinobacteria; Corynebacteriales; Mycobacteriaceae; Mycobacterium; Mycobacterium chelonae group; Mycobacterium abscessus subgroup; Mycobacterium abscessus
MIKHLTRRATVALVVAAAAISPAAYPAVADTATPLNGLFALTPGSCAEGRATGSYFRMIL WP_005063674.1 hypothetical protein [Mycobacterium abscessus] Length: 330\nNumber of predicted TMHs: 1\nExp number of AAs in TMHs: 39.25259\nExp number, first 60 AAs: 16.98187\nTotal prob of N-in: 0.83210\nPOSSIBLE N-term signal sequence\noutside 1 298\nTMhelix 299 321\ninside 322 330

30804 GCF_900135835.1_11893_6_22 Mycobacterium abscessus subsp. massiliense Terrabacteria group; Actinobacteria; Actinobacteria; Corynebacteriales; Mycobacteriaceae; Mycobacterium; Mycobacterium chelonae group; Mycobacterium abscessus subgroup; Mycobacterium abscessus
MIKHLTRRATVALVVAAAAISPAAYPAMADTATPLNGLFALTPGSCAEGRATGSYFRMIL WP_074356318.1 hypothetical protein [Mycobacterium abscessus] Length: 330\nNumber of predicted TMHs: 2\nExp number of AAs in TMHs: 39.34118\nExp number, first 60 AAs: 17.07668\nTotal prob of N-in: 0.84431\nPOSSIBLE N-term signal sequence\ninside 1 11\nTMhelix 12 29\noutside 30 298\nTMhelix 299 321\ninside 322 330

30805 GCF_900140155.1_10665_2_70 Mycobacterium abscessus subsp. massiliense Terrabacteria group; Actinobacteria; Actinobacteria; Corynebacteriales; Mycobacteriaceae; Mycobacterium; Mycobacterium chelonae group; Mycobacterium abscessus subgroup; Mycobacterium abscessus
MIKHLTRRATVALVVAAAAISPAAYPAVADTATPLNGLFALTPGSCAEGRATGSYFRMIL WP_005063674.1 hypothetical protein [Mycobacterium abscessus] Length: 330\nNumber of predicted TMHs: 1\nExp number of AAs in TMHs: 39.25259\nExp number, first 60 AAs: 16.98187\nTotal prob of N-in: 0.83210\nPOSSIBLE N-term signal sequence\noutside 1 298\nTMhelix 299 321\ninside 322 330

30806 GCF_900136455.1_10208_3_67 Mycobacterium abscessus subsp. massiliense Terrabacteria group; Actinobacteria; Actinobacteria; Corynebacteriales; Mycobacteriaceae; Mycobacterium; Mycobacterium chelonae group; Mycobacterium abscessus subgroup; Mycobacterium abscessus
MIKHLTRRATVALVVAAAAISPAAYPAVADTATPLNGLFALTPGSCAEGRATGSYFRMIL WP_005063674.1 hypothetical protein [Mycobacterium abscessus] Length: 330\nNumber of predicted TMHs: 1\nExp number of AAs in TMHs: 39.25259\nExp number, first 60 AAs: 16.98187\nTotal prob of N-in: 0.83210\nPOSSIBLE N-term signal sequence\noutside 1 298\nTMhelix 299 321\ninside 322 330

30807 GCF_900130905.1_10665_3_43 Mycobacterium abscessus subsp. massiliense Terrabacteria group; Actinobacteria; Actinobacteria; Corynebacteriales; Mycobacteriaceae; Mycobacterium; Mycobacterium chelonae group; Mycobacterium abscessus subgroup; Mycobacterium abscessus
MIKHLTRRATVALVVAAAAISPAAYPAVADTATPLNGLFALTPGSCAEGRATGSYFRMIL WP_005063674.1 hypothetical protein [Mycobacterium abscessus] Length: 330\nNumber of predicted TMHs: 1\nExp number of AAs in TMHs: 39.25259\nExp number, first 60 AAs: 16.98187\nTotal prob of N-in: 0.83210\nPOSSIBLE N-term signal sequence\noutside 1 298\nTMhelix 299 321\ninside 322 330

30808 GCF_900130885.1_10665_3_41 Mycobacterium abscessus subsp. massiliense Terrabacteria group; Actinobacteria; Actinobacteria; Corynebacteriales; Mycobacteriaceae; Mycobacterium; Mycobacterium chelonae group; Mycobacterium abscessus subgroup; Mycobacterium abscessus
MIKHLTRRATVALVVAAAAISPAAYPAVADTATPLNGLFALTPGSCAEGRATGSYFRMIL WP_005063674.1 hypothetical protein [Mycobacterium abscessus] Length: 330\nNumber of predicted TMHs: 1\nExp number of AAs in TMHs: 39.25259\nExp number, first 60 AAs: 16.98187\nTotal prob of N-in: 0.83210\nPOSSIBLE N-term signal sequence\noutside 1 298\nTMhelix 299 321\ninside 322 330

30809 GCF_900130855.1_10665_3_38 Mycobacterium abscessus subsp. massiliense Terrabacteria group; Actinobacteria; Actinobacteria; Corynebacteriales; Mycobacteriaceae; Mycobacterium; Mycobacterium chelonae group; Mycobacterium abscessus subgroup; Mycobacterium abscessus
MIKHLTRRATVALVVAAAAISPAAYPAVADTATPLNGLFALTPGSCAEGRATGSYFRMIL WP_005063674.1 hypothetical protein [Mycobacterium abscessus] Length: 330\nNumber of predicted TMHs: 1\nExp number of AAs in TMHs:

39.25259\nExp number, first 60 AAs: 16.98187\nTotal prob of N-in: 0.83210\nPOSSIBLE N-term signal sequence\noutside 1 298\nTMhelix 299 321\ninside 322 330

30810 GCF_900136785.1_10625_5_38 Mycobacterium abscessus subsp. massiliense Terrabacteria group; Actinobacteria; Actinobacteria; Corynebacteriales; Mycobacteriaceae; Mycobacterium; Mycobacterium chelonae group; Mycobacterium abscessus subgroup; Mycobacterium abscessus
MIKHLTRRATVALVVAAAAISPAAYPAVADTATPLNGLFALTPGSCAEGRATGSYFRMIL WP_005063674.1 hypothetical protein [Mycobacterium abscessus] Length: 330\nNumber of predicted TMHs: 1\nExp number of AAs in TMHs: 39.25259\nExp number, first 60 AAs: 16.98187\nTotal prob of N-in: 0.83210\nPOSSIBLE N-term signal sequence\noutside 1 298\nTMhelix 299 321\ninside 322 330

30811 GCF_900130835.1_10665_3_34 Mycobacterium abscessus subsp. massiliense Terrabacteria group; Actinobacteria; Actinobacteria; Corynebacteriales; Mycobacteriaceae; Mycobacterium; Mycobacterium chelonae group; Mycobacterium abscessus subgroup; Mycobacterium abscessus
MIKHLTRRATVALVVAAAAISPAAYPAVADTATPLNGLFALTPGSCAEGRATGSYFRMIL WP_005063674.1 hypothetical protein [Mycobacterium abscessus] Length: 330\nNumber of predicted TMHs: 1\nExp number of AAs in TMHs: 39.25259\nExp number, first 60 AAs: 16.98187\nTotal prob of N-in: 0.83210\nPOSSIBLE N-term signal sequence\noutside 1 298\nTMhelix 299 321\ninside 322 330

30812 GCF_900130435.1_10625_5_79 Mycobacterium abscessus subsp. massiliense Terrabacteria group; Actinobacteria; Actinobacteria; Corynebacteriales; Mycobacteriaceae; Mycobacterium; Mycobacterium chelonae group; Mycobacterium abscessus subgroup; Mycobacterium abscessus
MIKHLTRRATVALVVAAAAISPAAYPAVADTATPLNGLFALTPGSCAEGRATGSYFRMIL WP_005063674.1 hypothetical protein [Mycobacterium abscessus] Length: 330\nNumber of predicted TMHs: 1\nExp number of AAs in TMHs: 39.25259\nExp number, first 60 AAs: 16.98187\nTotal prob of N-in: 0.83210\nPOSSIBLE N-term signal sequence\noutside 1 298\nTMhelix 299 321\ninside 322 330

30813 GCF_900137185.1_12078_1_70 Mycobacterium abscessus subsp. massiliense Terrabacteria group; Actinobacteria; Actinobacteria; Corynebacteriales; Mycobacteriaceae; Mycobacterium; Mycobacterium chelonae group; Mycobacterium abscessus subgroup; Mycobacterium abscessus
MIKHLTRRATVALVVAAAAISPAAYPAVADTATPLNGLFALTPGSCAEGRATGSYFRMIL WP_005063674.1 hypothetical protein [Mycobacterium abscessus] Length: 330\nNumber of predicted TMHs: 1\nExp number of AAs in TMHs: 39.25259\nExp number, first 60 AAs: 16.98187\nTotal prob of N-in: 0.83210\nPOSSIBLE N-term signal sequence\noutside 1 298\nTMhelix 299 321\ninside 322 330

30814 GCF_001050395.1_ASM105039v1 Mycobacterium abscessus UC22 Terrabacteria group; Actinobacteria; Actinobacteria; Corynebacteriales; Mycobacteriaceae; Mycobacterium; Mycobacterium chelonae group; Mycobacterium abscessus subgroup; Mycobacterium abscessus
MIKHLTRRATVALVVAAAAISPAAYPAMADTATPLNGLFALTPGSCAEGRATGSYFRMIL WP_049231943.1
hypothetical protein [Mycobacterium abscessus] Length: 330\nNumber of predicted TMHs: 2\nExp number of AAs in TMHs: 39.34108\nExp number, first 60 AAs: 17.07804\nTotal prob of N-in: 0.84431\nPOSSIBLE N-term signal sequence\ninside 1 11\nTMhelix 12 29\noutside 30 298\nTMhelix 299 321\ninside 322 330

30815 GCF_000443145.1_V06705 Mycobacterium abscessus V06705 Terrabacteria group; Actinobacteria; Actinobacteria; Corynebacteriales; Mycobacteriaceae; Mycobacterium; Mycobacterium chelonae group; Mycobacterium abscessus subgroup; Mycobacterium abscessus
MIKHLTRRATVALVVAAAAISPAAYPAMADTATPLNGLFALTPGSCAEGRATGSYFRMIL WP_021268995.1
hypothetical protein [Mycobacterium abscessus] Length: 330\nNumber of predicted TMHs: 2\nExp number of AAs in TMHs: 39.3406299999999\nExp number, first 60 AAs: 17.07949\nTotal prob of N-in: 0.84430\nPOSSIBLE N-term signal sequence\ninside 1 11\nTMhelix 12 29\noutside 30 298\nTMhelix 299 321\ninside 322 330

30816 GCF_000524075.1_ASM52407v1 Mycobacterium abscessus subsp. bolletii 103 Terrabacteria group; Actinobacteria; Actinobacteria; Corynebacteriales; Mycobacteriaceae; Mycobacterium; Mycobacterium chelonae group; Mycobacterium abscessus subgroup; Mycobacterium abscessus; Mycobacterium abscessus subsp. bolletii
MIKHLTRRATVALVVAAAAISPAAYPAMADTATPLNGLFALTPGSCAEGRATGSYFRMIL WP_005073213.1
hypothetical protein [Mycobacterium abscessus] Length: 330\nNumber of predicted TMHs: 2\nExp number of AAs in TMHs: 39.3416199999999\nExp number, first 60 AAs: 17.07688\nTotal prob of N-in: 0.84431\nPOSSIBLE N-term signal sequence\ninside 1 11\nTMhelix 12 29\noutside 30 298\nTMhelix 299 321\ninside 322 330

30817 GCF_000758385.1_ASM75838v1 Mycobacterium abscessus subsp. bolletii 103 Terrabacteria group; Actinobacteria; Actinobacteria; Corynebacteriales; Mycobacteriaceae; Mycobacterium; Mycobacterium chelonae group;

Mycobacterium abscessus subgroup; Mycobacterium abscessus; Mycobacterium abscessus subsp. bolletii
MIKHLTRRATVALVVAAAAISPAAYPAMADTATPLNGLFALTPGSCAEGRATGSYFRMIL WP_005073213.1 hypothetical protein [Mycobacterium abscessus] Length: 330\nNumber of predicted TMHs: 2\nExp number of AAs in TMHs: 39.3416199999999\nExp number, first 60 AAs: 17.07688\nTotal prob of N-in: 0.84431\nPOSSIBLE N-term signal sequence\ninside 1 11\nTMhelix 12 29\noutside 30 298\nTMhelix 299 321\ninside 322 330

30818 GCF_000271285.1_ASM27128v1 Mycobacterium abscessus subsp. bolletii 1S-152-0914 Terrabacteria group; Actinobacteria; Actinobacteria; Corynebacteriales; Mycobacteriaceae; Mycobacterium; Mycobacterium chelonae group; Mycobacterium abscessus subgroup; Mycobacterium abscessus; Mycobacterium abscessus subsp. bolletii
MIKHLTRRATVALVVAAAAISPAAYPAVADTATPLNGLFALTPGSCAEGRATGSYFRMIL WP_005063674.1 hypothetical protein [Mycobacterium abscessus] Length: 330\nNumber of predicted TMHs: 1\nExp number of AAs in TMHs: 39.25259\nExp number, first 60 AAs: 16.98187\nTotal prob of N-in: 0.83210\nPOSSIBLE N-term signal sequence\noutside 1 298\nTMhelix 299 321\ninside 322 330

30819 GCF_000308015.1_ASM30801v1 Mycobacterium abscessus subsp. bolletii 1S-154-0310 Terrabacteria group; Actinobacteria; Actinobacteria; Corynebacteriales; Mycobacteriaceae; Mycobacterium; Mycobacterium chelonae group; Mycobacterium abscessus subgroup; Mycobacterium abscessus; Mycobacterium abscessus subsp. bolletii
MIKHLTRRATVALVVAAAAISPAAYPAVADTATPLNGLFALTPGSCAEGRATGSYFRMIL WP_005063674.1 hypothetical protein [Mycobacterium abscessus] Length: 330\nNumber of predicted TMHs: 1\nExp number of AAs in TMHs: 39.25259\nExp number, first 60 AAs: 16.98187\nTotal prob of N-in: 0.83210\nPOSSIBLE N-term signal sequence\noutside 1 298\nTMhelix 299 321\ninside 322 330

30820 GCF_000270785.1_ASM27078v1 Mycobacterium abscessus subsp. bolletii 2B-0912-S Terrabacteria group; Actinobacteria; Actinobacteria; Corynebacteriales; Mycobacteriaceae; Mycobacterium; Mycobacterium chelonae group; Mycobacterium abscessus subgroup; Mycobacterium abscessus; Mycobacterium abscessus subsp. bolletii
MIKHLTRRATVALVVAAAAISPAAYPAVADTATPLNGLFALTPGSCAEGRATGSYFRMIL WP_005063674.1 hypothetical protein [Mycobacterium abscessus] Length: 330\nNumber of predicted TMHs: 1\nExp number of AAs in TMHs: 39.25259\nExp number, first 60 AAs: 16.98187\nTotal prob of N-in: 0.83210\nPOSSIBLE N-term signal sequence\noutside 1 298\nTMhelix 299 321\ninside 322 330

30821 GCF_000445035.1_ASM44503v1 Mycobacterium abscessus subsp. bolletii 50594 Terrabacteria group; Actinobacteria; Actinobacteria; Corynebacteriales; Mycobacteriaceae; Mycobacterium; Mycobacterium chelonae group; Mycobacterium abscessus subgroup; Mycobacterium abscessus; Mycobacterium abscessus subsp. bolletii
MIKHLTRRATVALVVAAAAISPAAYPAVADTATPLNGLFALTPGSCAEGRATGSYFRMIL WP_005063674.1 hypothetical protein [Mycobacterium abscessus] Length: 330\nNumber of predicted TMHs: 1\nExp number of AAs in TMHs: 39.25259\nExp number, first 60 AAs: 16.98187\nTotal prob of N-in: 0.83210\nPOSSIBLE N-term signal sequence\noutside 1 298\nTMhelix 299 321\ninside 322 330

30822 GCF_000239035.1_ASM23903v2 Mycobacterium abscessus subsp. bolletii BD Terrabacteria group; Actinobacteria; Actinobacteria; Corynebacteriales; Mycobacteriaceae; Mycobacterium; Mycobacterium chelonae group; Mycobacterium abscessus subgroup; Mycobacterium abscessus; Mycobacterium abscessus subsp. bolletii
MIKHLTRRATVALVVAAAAISPAAYPAMADTATPLNGLFALTPGSCAEGRATGSYFRMIL WP_005073213.1 hypothetical protein [Mycobacterium abscessus] Length: 330\nNumber of predicted TMHs: 2\nExp number of AAs in TMHs: 39.3416199999999\nExp number, first 60 AAs: 17.07688\nTotal prob of N-in: 0.84431\nPOSSIBLE N-term signal sequence\ninside 1 11\nTMhelix 12 29\noutside 30 298\nTMhelix 299 321\ninside 322 330

30823 GCF_000418575.1_CRM0020_v1.0 Mycobacterium abscessus subsp. bolletii CRM-0020 Terrabacteria group; Actinobacteria; Actinobacteria; Corynebacteriales; Mycobacteriaceae; Mycobacterium; Mycobacterium chelonae group; Mycobacterium abscessus subgroup; Mycobacterium abscessus; Mycobacterium abscessus subsp. bolletii
MIKHLTRRATVALVVAAAAISPAAYPAVADTATPLNGLFALTPGSCAEGRATGSYFRMIL WP_005063674.1 hypothetical protein [Mycobacterium abscessus] Length: 330\nNumber of predicted TMHs: 1\nExp number of AAs in TMHs: 39.25259\nExp number, first 60 AAs: 16.98187\nTotal prob of N-in: 0.83210\nPOSSIBLE N-term signal sequence\noutside 1 298\nTMhelix 299 321\ninside 322 330

30824 GCF_000363915.2_MaINCQS00594_1.0 Mycobacterium abscessus subsp. bolletii INCQS 00594 Terrabacteria group; Actinobacteria; Actinobacteria; Corynebacteriales; Mycobacteriaceae; Mycobacterium; Mycobacterium chelonae group; Mycobacterium abscessus subgroup; Mycobacterium abscessus; Mycobacterium abscessus subsp. bolletii MIKHLTRRATVALVVAAAAISPAAYPAVADTATPLNGLFALTPGSCAEGRATGSYFRMIL WP_005063674.1 hypothetical protein [Mycobacterium abscessus] Length: 330\nNumber of predicted TMHs: 1\nExp number of AAs in TMHs: 39.25259\nExp number, first 60 AAs: 16.98187\nTotal prob of N-in: 0.83210\nPOSSIBLE N-term signal sequence\noutside 1 298\nTMhelix 299 321\ninside 322 330

30825 GCF_000261085.1_ASM26108v1 Mycobacterium abscessus subsp. bolletii M18 Terrabacteria group; Actinobacteria; Actinobacteria; Corynebacteriales; Mycobacteriaceae; Mycobacterium; Mycobacterium chelonae group; Mycobacterium abscessus subgroup; Mycobacterium abscessus; Mycobacterium abscessus subsp. bolletii MIKHLTRRATVALVVAAAAISPAAYPAVADTATPLNGLFALTPGSCAEGRATGSYFRMIL WP_005063674.1 hypothetical protein [Mycobacterium abscessus] Length: 330\nNumber of predicted TMHs: 1\nExp number of AAs in TMHs: 39.25259\nExp number, first 60 AAs: 16.98187\nTotal prob of N-in: 0.83210\nPOSSIBLE N-term signal sequence\noutside 1 298\nTMhelix 299 321\ninside 322 330

30826 GCF_000239055.1_ASM23905v2 Mycobacterium abscessus subsp. massiliense CCUG 48898 = JCM 15300 Terrabacteria group; Actinobacteria; Actinobacteria; Corynebacteriales; Mycobacteriaceae; Mycobacterium; Mycobacterium chelonae group; Mycobacterium abscessus subgroup; Mycobacterium abscessus; Mycobacterium abscessus subsp. massiliense MIKHLTRRATVALVVAAAAISPAAYPAVADTATPLNGLFALTPGSCAEGRATGSYFRMIL WP_005063674.1 hypothetical protein [Mycobacterium abscessus] Length: 330\nNumber of predicted TMHs: 1\nExp number of AAs in TMHs: 39.25259\nExp number, first 60 AAs: 16.98187\nTotal prob of N-in: 0.83210\nPOSSIBLE N-term signal sequence\noutside 1 298\nTMhelix 299 321\ninside 322 330

30827 GCF_000497265.2_ASM49726v2 Mycobacterium abscessus subsp. massiliense CCUG 48898 = JCM 15300 Terrabacteria group; Actinobacteria; Actinobacteria; Corynebacteriales; Mycobacteriaceae; Mycobacterium; Mycobacterium chelonae group; Mycobacterium abscessus subgroup; Mycobacterium abscessus; Mycobacterium abscessus subsp. massiliense MIKHLTRRATVALVVAAAAISPAAYPAVADTATPLNGLFALTPGSCAEGRATGSYFRMIL WP_041043629.1 hypothetical protein [Mycobacterium abscessus] Length: 332\nNumber of predicted TMHs: 1\nExp number of AAs in TMHs: 39.25017\nExp number, first 60 AAs: 16.98962\nTotal prob of N-in: 0.83166\nPOSSIBLE N-term signal sequence\noutside 1 300\nTMhelix 301 323\ninside 324 332

30828 GCF_000277775.2_ASM27777v2 Mycobacterium abscessus subsp. massiliense str. GO 06 Terrabacteria group; Actinobacteria; Actinobacteria; Corynebacteriales; Mycobacteriaceae; Mycobacterium; Mycobacterium chelonae group; Mycobacterium abscessus subgroup; Mycobacterium abscessus; Mycobacterium abscessus subsp. massiliense MIKHLTRRATVALVVAAAAISPAAYPAVADTATPLNGLFALTPGSCAEGRATGSYFRMIL WP_005063674.1 hypothetical protein [Mycobacterium abscessus] Length: 330\nNumber of predicted TMHs: 1\nExp number of AAs in TMHs: 39.25259\nExp number, first 60 AAs: 16.98187\nTotal prob of N-in: 0.83210\nPOSSIBLE N-term signal sequence\noutside 1 298\nTMhelix 299 321\ninside 322 330

30829 GCF_001632805.1_ASM163280v1 Mycobacterium chelonae CCUG 47445 Terrabacteria group; Actinobacteria; Actinobacteria; Corynebacteriales; Mycobacteriaceae; Mycobacterium; Mycobacterium chelonae group; Mycobacterium chelonae MIKHLTRRATVALVVAAAAMSPAAYPAVADTATPLNGLFALTPGSCAEGRATGSYFRMIL WP_046252588.1 hypothetical protein [Mycobacterium chelonae] Length: 329\nNumber of predicted TMHs: 1\nExp number of AAs in TMHs: 37.06635\nExp number, first 60 AAs: 14.74582\nTotal prob of N-in: 0.73289\nPOSSIBLE N-term signal sequence\noutside 1 297\nTMhelix 298 320\ninside 321 329

30830 GCF_001570425.1_ASM157042v1 Mycobacterium brisbanense Terrabacteria group; Actinobacteria; Actinobacteria; Corynebacteriales; Mycobacteriaceae; Mycobacterium; Mycobacterium fortuitum complex MPSALPKISRRRALVSAAAVAVLGVTAAGCADPPPSPDLADLAAQLDRARSDSKLASDAA WP_062828913.1 hypothetical protein [Mycobacterium brisbanense] Length: 171\nNumber of predicted TMHs: 2\nExp number of AAs in TMHs: 33.84545\nExp number, first 60 AAs: 13.72738\nTotal prob of N-in: 0.87874\nPOSSIBLE N-term signal sequence\ninside 1 12\nTMhelix 13 31\noutside 32 146\nTMhelix 147 169\ninside 170 171

30831 GCF_001756795.1_ASM175679v1 Mycobacterium sp. (ex Dasyatis americana) Terrabacteria group; Actinobacteria; Actinobacteria; Corynebacteriales; Mycobacteriaceae; Mycobacterium; Mycobacterium fortuitum complex MSGPLGLSIGTTNLVAARVGNQPVSRRSVLTLSADRTPQVGVPVSGSGVTLSGFVERVGD WP_070189537.1 molecular chaperone, partial [Mycobacterium sp. (ex Dasyatis americana)] Length: 452\nNumber of predicted TMHs: 1\nExp number of AAs in TMHs: 24.48732\nExp number, first 60 AAs: 0.00413\nTotal prob of N-in: 0.00378\noutside 1 428\nTMhelix 429 451\ninside 452 452

30832 GCF_000016365.1_ASM1636v1 Mycobacterium gilvum PYR-GCK Terrabacteria group; Actinobacteria; Actinobacteria; Corynebacteriales; Mycobacteriaceae; Mycobacterium; Mycobacterium gilvum MPVEPERPIRTAAFFDLDKTVIAKSSTLAFSKPFFSQGLINRRAVLKSTYAQFLFSMSGA WP_041800558.1 inhibition of morphological differentiation protein [Mycobacterium gilvum] Length: 271\nNumber of predicted TMHs: 1\nExp number of AAs in TMHs: 21.74133\nExp number, first 60 AAs: 0.0141\nTotal prob of N-in: 0.03600\noutside 1 243\nTMhelix 244 266\ninside 267 271

30833 GCF_000184435.1_ASM18443v1 Mycobacterium gilvum Spyr1 Terrabacteria group; Actinobacteria; Actinobacteria; Corynebacteriales; Mycobacteriaceae; Mycobacterium; Mycobacterium gilvum MPVEPERPIRTAAFFDLDKTVIAKSSTLAFSKPFFSQGLINRRAVLKSTYAQFLFSMSGA WP_013472930.1 haloacid dehalogenase [Mycobacterium gilvum] Length: 271\nNumber of predicted TMHs: 1\nExp number of AAs in TMHs: 21.74128\nExp number, first 60 AAs: 0.0141\nTotal prob of N-in: 0.03599\noutside 1 243\nTMhelix 244 266\ninside 267 271

30834 GCF_000340435.2_ASM34043v3 Mycobacterium haemophilum DSM 44634 Terrabacteria group; Actinobacteria; Actinobacteria; Corynebacteriales; Mycobacteriaceae; Mycobacterium; Mycobacterium haemophilum MPRSGTGSKRARRGCVPRALVAVALLTSGAFAGLPPAYAISPPTIDPGAVPPDGPPGPAA WP_054880009.1 type VII secretion-associated serine protease mycosin [Mycobacterium haemophilum] Length: 591\nNumber of predicted TMHs: 2\nExp number of AAs in TMHs: 43.90598\nExp number, first 60 AAs: 20.08392\nTotal prob of N-in: 0.93656\nPOSSIBLE N-term signal sequence\ninside 1 19\nTMhelix 20 42\noutside 43 556\nTMhelix 557 579\ninside 580 591

30835 GCF_000379865.1_ASM37986v1 Mycobacterium hassiacum DSM 44199 Terrabacteria group; Actinobacteria; Actinobacteria; Corynebacteriales; Mycobacteriaceae; Mycobacterium; Mycobacterium hassiacum MTTHTTSGPARSAARSRRRRGVVRLLTVTAVLLMLAPSLIGCVRIRMSITVSPDDRVSGQ WP_081586657.1 DUF3153 domain-containing protein [Mycobacterium hassiacum] Length: 243\nNumber of predicted TMHs: 2\nExp number of AAs in TMHs: 44.45636\nExp number, first 60 AAs: 21.90125\nTotal prob of N-in: 0.99718\nPOSSIBLE N-term signal sequence\ninside 1 20\nTMhelix 21 43\noutside 44 204\nTMhelix 205 227\ninside 228 243

30836 GCF_000300375.1_ASM30037v1 Mycobacterium hassiacum DSM 44199 Terrabacteria group; Actinobacteria; Actinobacteria; Corynebacteriales; Mycobacteriaceae; Mycobacterium; Mycobacterium hassiacum MTTHTTSGPARSAARSRRRRGVVRLLTVTAVLLMLAPSLIGCVRIRMSITVSPDDRVSGQ WP_081586657.1 DUF3153 domain-containing protein [Mycobacterium hassiacum] Length: 243\nNumber of predicted TMHs: 2\nExp number of AAs in TMHs: 44.45636\nExp number, first 60 AAs: 21.90125\nTotal prob of N-in: 0.99718\nPOSSIBLE N-term signal sequence\ninside 1 20\nTMhelix 21 43\noutside 44 204\nTMhelix 205 227\ninside 228 243

30837 GCF_000612825.1_PRJEB5749_assembly_1 Mycobacterium mageritense DSM 44476 = CIP 104973 Terrabacteria group; Actinobacteria; Actinobacteria; Corynebacteriales; Mycobacteriaceae; Mycobacterium; Mycobacterium mageritense MWGTDSRRAEAVAELLHNGRDSRWRLRRCGSNSRVPRHRRSVLTHLHILHAAAEPTEQEQ WP_036439627.1 MULTISPECIES: hypothetical protein [Mycobacterium] Length: 179\nNumber of predicted TMHs: 2\nExp number of AAs in TMHs: 42.06263\nExp number, first 60 AAs: 0.00181\nTotal prob of N-in: 0.87116\ninside 1 66\nTMhelix 67 89\noutside 90 142\nTMhelix 143 162\ninside 163 179

30838 GCF_000257725.1_ASM25772v1 Mycobacterium phlei RIVM601174 Terrabacteria group; Actinobacteria; Actinobacteria; Corynebacteriales; Mycobacteriaceae; Mycobacterium; Mycobacterium phlei MEELWRARRSALAAGATCIGILLAATTFTPASAQPDEPDEAPETSVLSLPALGNDPDISL WP_040633064.1 hypothetical protein [Mycobacterium phlei] Length: 642\nNumber of predicted TMHs: 1\nExp number of AAs in TMHs: 36.34281\nExp number, first 60 AAs: 13.60653\nTotal prob of N-in: 0.65585\nPOSSIBLE N-term signal sequence\noutside 1 613\nTMhelix 614 636\ninside 637 642

30839 GCF_000257725.1_ASM25772v1 Mycobacterium phlei RIVM601174 Terrabacteria group; Actinobacteria; Actinobacteria; Corynebacteriales; Mycobacteriaceae; Mycobacterium; Mycobacterium phlei MAEGSSPAPSQPVRTAAFFDLDKTVIAKSSTLAFSKPFFDQGLINRRAVLKSSYAQFLFL WP_085980751.1 inhibition of morphological differentiation protein [Mycobacterium phlei] Length: 274\nNumber of predicted TMHs: 1\nExp number of AAs in TMHs: 21.83495\nExp number, first 60 AAs: 0.02012\nTotal prob of N-in: 0.01829\noutside 1 246\nTMhelix 247 269\ninside 270 274

30840 GCF_000230935.1_ASM23093v2 Mycobacterium rhodesiae JS60 Terrabacteria group; Actinobacteria; Actinobacteria; Corynebacteriales; Mycobacteriaceae; Mycobacterium; Mycobacterium rhodesiae MSARRRLVPYAWLGAGAITVGIGAALAHGSAVAHADDSPPGAKASSSTTSSAGKARPART WP_005140235.1 40-residue YVTN family beta-propeller repeat protein [Mycobacterium rhodesiae] Length: 536\nNumber of predicted TMHs: 2\nExp number of AAs in TMHs: 39.39001\nExp number, first 60 AAs: 21.40692\nTotal prob of N-in: 0.96699\nPOSSIBLE N-term signal sequence\ninside 1 6\nTMhelix 7 29\noutside 30 508\nTMhelix 509 531\ninside 532 536

30841 GCF_000230895.2_ASM23089v3 Mycobacterium rhodesiae NBB3 Terrabacteria group; Actinobacteria; Actinobacteria; Corynebacteriales; Mycobacteriaceae; Mycobacterium; Mycobacterium rhodesiae MVPTDNPVRTAAFFDLDKTVIAKSSTLAFSKPFFDQGLINRRTVLKSAYAQFLFLMSGAD WP_014210558.1 HAD-superfamily hydrolase [Mycobacterium rhodesiae] Length: 269\nNumber of predicted TMHs: 1\nExp number of

AAs in TMHs: 21.8978\nExp number, first 60 AAs: 0.0519\nTotal prob of N-in: 0.07159\noutside 1 241\nTMhelix 242 264\ninside 265 269

30842 GCF_002086285.1_ASM208628v1 Mycobacterium kumamotonense Terrabacteria group; Actinobacteria; Actinobacteria; Corynebacteriales; Mycobacteriaceae; Mycobacterium; Mycobacterium terrae complex MRSPKQPGSRTRTAAFLDLDHTVIAKSSALAFSKPFMDQGLINRRTVLKSSYAQFLMLLS WP_052312168.1 MULTISPECIES: inhibition of morphological differentiation protein [Mycobacterium] Length: 272\nNumber of predicted TMHs: 1\nExp number of AAs in TMHs: 21.73312\nExp number, first 60 AAs: 0.0228\nTotal prob of N-in: 0.06677\noutside 1 244\nTMhelix 245 267\ninside 268 272

30843 GCF_001679965.1_ASM167996v1 Mycobacterium kumamotonense Terrabacteria group; Actinobacteria; Actinobacteria; Corynebacteriales; Mycobacteriaceae; Mycobacterium; Mycobacterium terrae complex MRSPKQPGSRTRTAAFLDLDHTVIAKSSALAFSKPFMDQGLINRRTVLKSSYAQFLMLLS WP_052312168.1 MULTISPECIES: inhibition of morphological differentiation protein [Mycobacterium] Length: 272\nNumber of predicted TMHs: 1\nExp number of AAs in TMHs: 21.73312\nExp number, first 60 AAs: 0.0228\nTotal prob of N-in: 0.06677\noutside 1 244\nTMhelix 245 267\ninside 268 272

30844 GCF_002101775.1_ASM210177v1 Mycobacterium nonchromogenicum Terrabacteria group; Actinobacteria; Actinobacteria; Corynebacteriales; Mycobacteriaceae; Mycobacterium; Mycobacterium terrae complex MPGPLSAIDRRRVLTGAGMLALLAVAAPACGSGPTAPAVDDLEAQREAAQHDSALAAAAA WP_085137383.1 hypothetical protein [Mycobacterium nonchromogenicum]Length: 169\nNumber of predicted TMHs: 2\nExp number of AAs in TMHs: 36.65832\nExp number, first 60 AAs: 17.8686\nTotal prob of N-in: 0.83739\nPOSSIBLE N-term signal sequence\ninside 1 12\nTMhelix 13 35\noutside 36 143\nTMhelix 144 166\ninside 167 169

30845 GCF_002101885.1_ASM210188v1 Mycobacterium senuense Terrabacteria group; Actinobacteria; Actinobacteria; Corynebacteriales; Mycobacteriaceae; Mycobacterium; Mycobacterium terrae complex MSQVSGPLTAIDRRRLLSGAGVVALIALAAPACGSAPAPPAVDDLEAQRQLAQHDSELAA WP_085087113.1 hypothetical protein [Mycobacterium senuense] Length: 175\nNumber of predicted TMHs: 2\nExp number of AAs in TMHs: 34.7977\nExp number, first 60 AAs: 16.21225\nTotal prob of N-in: 0.69556\nPOSSIBLE N-term signal sequence\ninside 1 15\nTMhelix 16 38\noutside 39 150\nTMhelix 151 173\ninside 174 175

30846 GCF_000243415.2_ASM24341v3 Mycobacterium tusciae JS617 Terrabacteria group; Actinobacteria; Actinobacteria; Corynebacteriales; Mycobacteriaceae; Mycobacterium; Mycobacterium tusciae MPPANNPVRTAAFFDLDKTVIAKSSTLAFSKPFFDQGLINRRTVLKSAYAQFLFLMSGAD WP_006243858.1 haloacid dehalogenase [Mycobacterium tusciae] Length: 269\nNumber of predicted TMHs: 1\nExp number of AAs in TMHs: 22.2541\nExp number, first 60 AAs: 0.03538\nTotal prob of N-in: 0.04419\noutside 1 241\nTMhelix 242 264\ninside 265 269

30847 GCF_001655245.1_ASM165524v1 Mycobacterium vaccae 95051 Terrabacteria group; Actinobacteria; Actinobacteria; Corynebacteriales; Mycobacteriaceae; Mycobacterium; Mycobacterium vaccae MRGWQDVGVPSPPPTVSRRRVLVGAATLAALGAAVTACGTAPPPPDLDDLTTALDRARAD WP_081529171.1 hypothetical protein [Mycobacterium vaccae] Length: 177\nNumber of predicted TMHs: 2\nExp number of AAs in TMHs: 41.85195\nExp number, first 60 AAs: 20.35757\nTotal prob of N-in: 0.95179\nPOSSIBLE N-term signal sequence\ninside 1 20\nTMhelix 21 43\noutside 44 153\nTMhelix 154 176\ninside 177 177

30848 GCF_000295825.1_ASM29582v1 Mycobacterium vaccae ATCC 25954 Terrabacteria group; Actinobacteria; Actinobacteria; Corynebacteriales; Mycobacteriaceae; Mycobacterium; Mycobacterium vaccae MRGWQDVGVPSPPPTVSRRRVLVGAATLAALGAAVTACGTAPPPPDLDDLTTALDRARAD WP_081529171.1 hypothetical protein [Mycobacterium vaccae] Length: 177\nNumber of predicted TMHs: 2\nExp number of AAs in TMHs: 41.85195\nExp number, first 60 AAs: 20.35757\nTotal prob of N-in: 0.95179\nPOSSIBLE N-term signal sequence\ninside 1 20\nTMhelix 21 43\noutside 44 153\nTMhelix 154 176\ninside 177 177

30849 GCF_001552715.1_ASM155271v1 Mycobacterium vaccae NBRC 14118 = CIP 105934 Terrabacteria group; Actinobacteria; Actinobacteria; Corynebacteriales; Mycobacteriaceae; Mycobacterium; Mycobacterium vaccae MLAVAAGALIALRMVALRAAAGTGRRAVLRWSTTTGAMLLILLAAARPISGAGEETEPGR WP_060941840.1 hypothetical protein [Mycobacterium vaccae] Length: 296\nNumber of predicted TMHs: 1\nExp number of AAs in TMHs: 29.90485\nExp number, first 60 AAs: 9.19806\nTotal prob of N-in: 0.20482\noutside 1 258\nTMhelix 259 281\ninside 282 296

30850 GCF_001310275.1_ASM131027v1 Nocardia arizonensis Terrabacteria group; Actinobacteria; Actinobacteria; Corynebacteriales; Nocardiaceae; Nocardia

MIFAPMTARRAALLRLAVALSAFVAIALVHGAQCQTGMPMTMAHSAAMVGGVAGQCGGAQ WP_054811683.1 MULTISPECIES: hypothetical protein [Nocardia] Length: 156\nNumber of predicted TMHs: 2\nExp number of AAs in TMHs: 43.30523\nExp number, first 60 AAs: 21.2408\nTotal prob of N-in: 0.99795\nPOSSIBLE N-term signal sequence\ninside 1 11\nTMhelix 12 34\noutside 35 94\nTMhelix 95 117\ninside 118 156

30851 GCF_001618405.1_ASM161840v1 Nocardia arizonensis Terrabacteria group; Actinobacteria; Actinobacteria; Corynebacteriales; Nocardiaceae; Nocardia MIFAPMTARRAALLRLAVALSAFVAIALVHGAQCQTGMPMTMAHSAAMVGGVAGQCGGAQ WP_054811683.1 MULTISPECIES: hypothetical protein [Nocardia] Length: 156\nNumber of predicted TMHs: 2\nExp number of AAs in TMHs: 43.30523\nExp number, first 60 AAs: 21.2408\nTotal prob of N-in: 0.99795\nPOSSIBLE N-term signal sequence\ninside 1 11\nTMhelix 12 34\noutside 35 94\nTMhelix 95 117\ninside 118 156

30852 GCF_002209125.1_ASM220912v1 Nocardia brasiliensis Terrabacteria group; Actinobacteria; Actinobacteria; Corynebacteriales; Nocardiaceae; Nocardia MTQTFRRRAAIPLGIFVALLTILGSGTTLPLLAPPAAQAQPTGSGTVSGPKFLKLSVDSV WP_042259183.1 hypothetical protein [Nocardia brasiliensis] Length: 895\nNumber of predicted TMHs: 2\nExp number of AAs in TMHs: 43.08572\nExp number, first 60 AAs: 21.74311\nTotal prob of N-in: 0.99463\nPOSSIBLE N-term signal sequence\ninside 1 12\nTMhelix 13 35\noutside 36 826\nTMhelix 827 849\ninside 850 895

30853 GCF_002209125.1_ASM220912v1 Nocardia brasiliensis Terrabacteria group; Actinobacteria; Actinobacteria; Corynebacteriales; Nocardiaceae; Nocardia MDSPHANGRVAAFFDLDKTVIAKSSTFVFSKPFYAQGLLNRRDVLESSYAHFLFMLSGAD WP_042254975.1 inhibition of morphological differentiation protein [Nocardia brasiliensis] Length: 268\nNumber of predicted TMHs: 1\nExp number of AAs in TMHs: 22.91952\nExp number, first 60 AAs: 0.02426\nTotal prob of N-in: 0.05945\noutside 1 241\nTMhelix 242 264\ninside 265 268

30854 GCF_001689765.1_ASM168976v1 Nocardia brasiliensis Terrabacteria group; Actinobacteria; Actinobacteria; Corynebacteriales; Nocardiaceae; Nocardia MDSPHANGRVAAFFDLDKTVIAKSSTFVFSKPFYAQGLLNRRDVLESSYAHFMFMLSGAD WP_014981200.1 hydrolase [Nocardia brasiliensis] Length: 268\nNumber of predicted TMHs: 1\nExp number of AAs in TMHs: 22.91898\nExp number, first 60 AAs: 0.02348\nTotal prob of N-in: 0.05944\noutside 1 241\nTMhelix 242 264\ninside 265 268

30855 GCF_001689765.1_ASM168976v1 Nocardia brasiliensis Terrabacteria group; Actinobacteria; Actinobacteria; Corynebacteriales; Nocardiaceae; Nocardia MTQTFRRRAAIPLGIFVALLTILGSGTTLPLLAPPAAQAQPTGSGTVSGPKFLKLSVDSV WP_014989207.1 hypothetical protein [Nocardia brasiliensis] Length: 895\nNumber of predicted TMHs: 2\nExp number of AAs in TMHs: 43.08572\nExp number, first 60 AAs: 21.74309\nTotal prob of N-in: 0.99463\nPOSSIBLE N-term signal sequence\ninside 1 12\nTMhelix 13 35\noutside 36 826\nTMhelix 827 849\ninside 850 895

30856 GCF_001689765.1_ASM168976v1 Nocardia brasiliensis Terrabacteria group; Actinobacteria; Actinobacteria; Corynebacteriales; Nocardiaceae; Nocardia MKIRESRRRATLRLGSALAALAVTAGLAAVPSFAAPTSTTTTPFSTPNTDGCPQKTSPPP WP_065652221.1 penicillin-binding protein [Nocardia brasiliensis] Length: 429\nNumber of predicted TMHs: 2\nExp number of AAs in TMHs: 40.77755\nExp number, first 60 AAs: 22.37421\nTotal prob of N-in: 0.98999\nPOSSIBLE N-term signal sequence\ninside 1 11\nTMhelix 12 34\noutside 35 404\nTMhelix 405 422\ninside 423 429

30857 GCF_000717405.1_ASM71740v1 Nocardia carnea Terrabacteria group; Actinobacteria; Actinobacteria; Corynebacteriales; Nocardiaceae; Nocardia MNSRIAAFFDLDKTVIARSSTYVFSKPFFAEGLINRRAVLESSYAHFIFLLSGADHDQME WP_051157049.1 inhibition of morphological differentiation protein [Nocardia carnea] Length: 261\nNumber of predicted TMHs: 1\nExp number of AAs in TMHs: 21.14729\nExp number, first 60 AAs: 0.02759\nTotal prob of N-in: 0.02336\noutside 1 235\nTMhelix 236 258\ninside 259 261

30858 GCF_001613405.1_ASM161340v1 Nocardia crassostreae Terrabacteria group; Actinobacteria; Actinobacteria; Corynebacteriales; Nocardiaceae; Nocardia MTAESGRVAAFFDLDKTVIARSSAFVFSKPLMDQGLLSRRAVLESSYAHFLFLLSGADHD WP_067531574.1 inhibition of morphological differentiation protein [Nocardia crassostreae] Length: 266\nNumber of predicted TMHs: 1\nExp number of AAs in TMHs: 23.54072\nExp number, first 60 AAs: 0.05479\nTotal prob of N-in: 0.07338\noutside 1 239\nTMhelix 240 262\ninside 263 266

30859 GCF_000583715.2_ASM58371v2 Nocardia seriolae Terrabacteria group; Actinobacteria; Actinobacteria; Corynebacteriales; Nocardiaceae; Nocardia
MSSSPPSAPDPEGIGQATKIRKQRRTRRRLVGGLVLLMGLVGAGFAASALTPQAQVATAK WP_045436685.1
cytochrome c [Nocardia seriolae] Length: 286\nNumber of predicted TMHs: 2\nExp number of AAs in TMHs: 41.9528\nExp number, first 60 AAs: 21.62084\nTotal prob of N-in: 0.98337\nPOSSIBLE N-term signal sequence\ninside 1 29\nTMhelix 30 52\noutside 53 263\nTMhelix 264 283\ninside 284 286

30860 GCF_001885535.1_ASM188553v1 Nocardia seriolae Terrabacteria group; Actinobacteria; Actinobacteria; Corynebacteriales; Nocardiaceae; Nocardia
MSSSPPSAPDPEGIGQATKIRKQRRTRRRLVGGLVLLMGLVGAGFAASALTPQAQVATAK WP_045436685.1
cytochrome c [Nocardia seriolae] Length: 286\nNumber of predicted TMHs: 2\nExp number of AAs in TMHs: 41.9528\nExp number, first 60 AAs: 21.62084\nTotal prob of N-in: 0.98337\nPOSSIBLE N-term signal sequence\ninside 1 29\nTMhelix 30 52\noutside 53 263\nTMhelix 264 283\ninside 284 286

30861 GCF_000763165.1_ASM76316v1 Nocardia seriolae Terrabacteria group; Actinobacteria; Actinobacteria; Corynebacteriales; Nocardiaceae; Nocardia
MSSSPPSAPDPEGIGQATKIRKQRRTRRRLVGGLVLLMGLVGAGFAASALTPQAQVATAK WP_045436685.1
cytochrome c [Nocardia seriolae] Length: 286\nNumber of predicted TMHs: 2\nExp number of AAs in TMHs: 41.9528\nExp number, first 60 AAs: 21.62084\nTotal prob of N-in: 0.98337\nPOSSIBLE N-term signal sequence\ninside 1 29\nTMhelix 30 52\noutside 53 263\nTMhelix 264 283\ninside 284 286

30862 GCF_001192935.1_ASM119293v1 Nocardia seriolae Terrabacteria group; Actinobacteria; Actinobacteria; Corynebacteriales; Nocardiaceae; Nocardia
MSSSPPSAPDPEGIGQATKIRKQRRTRRRLVGGLVLLMGLVGAGFAASALTPQAQVATAK WP_045436685.1
cytochrome c [Nocardia seriolae] Length: 286\nNumber of predicted TMHs: 2\nExp number of AAs in TMHs: 41.9528\nExp number, first 60 AAs: 21.62084\nTotal prob of N-in: 0.98337\nPOSSIBLE N-term signal sequence\ninside 1 29\nTMhelix 30 52\noutside 53 263\nTMhelix 264 283\ninside 284 286

30863 GCF_002093935.1_ASM209393v1 Nocardia seriolae Terrabacteria group; Actinobacteria; Actinobacteria; Corynebacteriales; Nocardiaceae; Nocardia
MSSSPPSAPDPEGIGQATKIRKQRRTRRRLVGGLVLLMGLVGAGFAASALTPQAQVATAK WP_045436685.1
cytochrome c [Nocardia seriolae] Length: 286\nNumber of predicted TMHs: 2\nExp number of AAs in TMHs: 41.9528\nExp number, first 60 AAs: 21.62084\nTotal prob of N-in: 0.98337\nPOSSIBLE N-term signal sequence\ninside 1 29\nTMhelix 30 52\noutside 53 263\nTMhelix 264 283\ninside 284 286

30864 GCF_001865855.1_ASM186585v1 Nocardia seriolae Terrabacteria group; Actinobacteria; Actinobacteria; Corynebacteriales; Nocardiaceae; Nocardia
MSSSPPSAPDPEGIGQATKIRKQRRTRRRLVGGLVLLMGLVGAGCAASALTPQAQVATAK WP_071343934.1
cytochrome C [Nocardia seriolae] Length: 286\nNumber of predicted TMHs: 2\nExp number of AAs in TMHs: 42.46271\nExp number, first 60 AAs: 22.11907\nTotal prob of N-in: 0.97645\nPOSSIBLE N-term signal sequence\ninside 1 29\nTMhelix 30 52\noutside 53 263\nTMhelix 264 283\ninside 284 286

30865 GCF_001886715.1_ASM188671v1 Nocardia soli Terrabacteria group; Actinobacteria; Actinobacteria; Corynebacteriales; Nocardiaceae; Nocardia
MSKRRKALQGLPVAVALPAAIAFGVGPGAAQAAQVQTVTHTPDAQEPTAAVPEHAHIAPD WP_071926721.1
hypothetical protein [Nocardia soli] Length: 217\nNumber of predicted TMHs: 2\nExp number of AAs in TMHs: 58.4804\nExp number, first 60 AAs: 12.89466\nTotal prob of N-in: 0.88063\nPOSSIBLE N-term signal sequence\ninside 1 143\nTMhelix 144 166\noutside 167 175\nTMhelix 176 198\ninside 199 217

30866 GCF_001886715.1_ASM188671v1 Nocardia soli Terrabacteria group; Actinobacteria; Actinobacteria; Corynebacteriales; Nocardiaceae; Nocardia
MPGGRVAAFFDLDKTVIAKSSAFVFSKPFYAQGLINRRAVLESSYAHFLYLLSGADHDQM WP_071925899.1
inhibition of morphological differentiation protein [Nocardia soli] Length: 264\nNumber of predicted TMHs: 1\nExp number of AAs in TMHs: 22.49051\nExp number, first 60 AAs: 0.00946\nTotal prob of N-in: 0.00605\noutside 1 237\nTMhelix 238 260\ninside 261 264

30867 GCF_000482385.1_ASM48238v1 Nocardia sp. CNY236 Terrabacteria group; Actinobacteria; Actinobacteria; Corynebacteriales; Nocardiaceae; Nocardia
MPDAPHLPWRRARRGVVRAGAVVLLAALAVPMLAGCLRVQVAMGVSSNDRVSGQIVVAVV WP_051407089.1
hypothetical protein [Nocardia sp. CNY236] Length: 248\nNumber of predicted TMHs: 2\nExp number of

AAs in TMHs: 45.3810199999999\nExp number, first 60 AAs:  22.64422\nTotal prob of N-in:   0.43661\nPOSSIBLE N-term signal sequence\noutside   1   14\nTMhelix   15   37\ninside   38   197\nTMhelix   198   220\noutside   221   248

30868 GCF_000482385.1_ASM48238v1 Nocardia sp. CNY236 Terrabacteria group; Actinobacteria; Actinobacteria; Corynebacteriales; Nocardiaceae; Nocardia
MDEPSTAGRRVAAFFDLDKTVIATSSMFVFSRPFYTQGLLKRRAVLKSNYVHFLFLLSGA WP_028478144.1 inhibition of morphological differentiation protein [Nocardia sp. CNY236] Length: 271\nNumber of predicted TMHs:  1\nExp number of AAs in TMHs: 23.46341\nExp number, first 60 AAs:  0.57885\nTotal prob of N-in:   0.03511\noutside   1   242\nTMhelix   243   265\ninside   266   271

30869 GCF_001279525.1_ASM127952v1 Nocardia sp. NRRL S-836 Terrabacteria group; Actinobacteria; Actinobacteria; Corynebacteriales; Nocardiaceae; Nocardia
MMHITCTLVTGRELPLDLHVSPSKPARTRGGTGTMRHRLDTLGRRATLFTLLSAAFSWGF WP_082373817.1 hypothetical protein [Nocardia sp. NRRL S-836] Length: 441\nNumber of predicted TMHs:  2\nExp number of AAs in TMHs: 40.2827999999999\nExp number, first 60 AAs:  13.37488\nTotal prob of N-in:   0.83569\nPOSSIBLE N-term signal sequence\ninside   1   45\nTMhelix   46   68\noutside   69   409\nTMhelix   410   432\ninside   433   441

30870 GCF_001279525.1_ASM127952v1 Nocardia sp. NRRL S-836 Terrabacteria group; Actinobacteria; Actinobacteria; Corynebacteriales; Nocardiaceae; Nocardia
MSRTAAFFDLDKTVIAKSSTLAFSRPFFQEGLINRRAVLKSAYAQFVFMLAGADADQMDR WP_053737444.1 inhibition of morphological differentiation protein [Nocardia sp. NRRL S-836] Length: 265\nNumber of predicted TMHs:  1\nExp number of AAs in TMHs: 21.36907\nExp number, first 60 AAs:  0.06146\nTotal prob of N-in:   0.01498\noutside   1   235\nTMhelix   236   258\ninside   259   265

30871 GCF_001427185.1_Root136 Nocardia sp. Root136 Terrabacteria group; Actinobacteria; Actinobacteria; Corynebacteriales; Nocardiaceae; Nocardia
MSVVSDIKGSRRRIMMSRSVSYTLPAVVAVALVSAGCGTATPKPAPRAASATLTTTSKPPWP_056809982.1 hypothetical protein [Nocardia sp. Root136] Length: 172\nNumber of predicted TMHs:  2\nExp number of AAs in TMHs: 40.43904\nExp number, first 60 AAs:  22.32135\nTotal prob of N-in:   0.98353\nPOSSIBLE N-term signal sequence\ninside   1   19\nTMhelix   20   42\noutside   43   113\nTMhelix   114   131\ninside   132   172

30872 GCF_001613425.1_ASM161342v1 Nocardia vulneris Terrabacteria group; Actinobacteria; Actinobacteria; Corynebacteriales; Nocardiaceae; Nocardia
MKIRESRRRATLRLGSALAALAVTAGLAAVPSFAAPTSPTTTPFTTPNTDGCPQKTSPPP WP_052280707.1 penicillin-binding protein [Nocardia vulneris] Length: 429\nNumber of predicted TMHs:  2\nExp number of AAs in TMHs: 40.86881\nExp number, first 60 AAs:  22.38605\nTotal prob of N-in:   0.99056\nPOSSIBLE N-term signal sequence\ninside   1   11\nTMhelix   12   34\noutside   35   404\nTMhelix   405   422\ninside   423   429

30873 GCF_001613425.1_ASM161342v1 Nocardia vulneris Terrabacteria group; Actinobacteria; Actinobacteria; Corynebacteriales; Nocardiaceae; Nocardia
MDSPHANGRVAAFFDLDKTVIAKSSTFVFSKPFYAQGLLNRRDVLESSYAHFLFMLSGAD WP_043678184.1 inhibition of morphological differentiation protein [Nocardia vulneris] Length: 268\nNumber of predicted TMHs:  1\nExp number of AAs in TMHs: 22.92\nExp number, first 60 AAs:  0.02426\nTotal prob of N-in:   0.05947\noutside   1   241\nTMhelix   242   264\ninside   265   268

30874 GCF_001613425.1_ASM161342v1 Nocardia vulneris Terrabacteria group; Actinobacteria; Actinobacteria; Corynebacteriales; Nocardiaceae; Nocardia
MTQTFRRRAAIPLGIFVALLTILGSGTTLPLLAPPTAQAQPTGSGTVSGPKFLKLSVDSV WP_043676971.1 hypothetical protein [Nocardia vulneris] Length: 895\nNumber of predicted TMHs:  2\nExp number of AAs in TMHs: 43.06239\nExp number, first 60 AAs:  21.71978\nTotal prob of N-in:   0.99436\nPOSSIBLE N-term signal sequence\ninside   1   12\nTMhelix   13   35\noutside   36   826\nTMhelix   827   849\ninside   850   895

30875 GCF_000811985.1_ASM81198v1 Nocardia vulneris Terrabacteria group; Actinobacteria; Actinobacteria; Corynebacteriales; Nocardiaceae; Nocardia
MDSPHANGRVAAFFDLDKTVIAKSSTFVFSKPFYAQGLLNRRDVLESSYAHFLFMLSGAD WP_043678184.1 inhibition of morphological differentiation protein [Nocardia vulneris] Length: 268\nNumber of predicted TMHs:  1\nExp number of AAs in TMHs: 22.92\nExp number, first 60 AAs:  0.02426\nTotal prob of N-in:   0.05947\noutside   1   241\nTMhelix   242   264\ninside   265   268

30876 GCF_000811985.1_ASM81198v1 Nocardia vulneris Terrabacteria group; Actinobacteria; Actinobacteria; Corynebacteriales; Nocardiaceae; Nocardia

MKIRESRRRATLRLGSALAALAVTAGLAAVPSFAAPTSPTTTPFTTPNTDGCPQKTSPPP WP_052280707.1 penicillin-binding protein [Nocardia vulneris] Length: 429\nNumber of predicted TMHs: 2\nExp number of AAs in TMHs: 40.86881\nExp number, first 60 AAs: 22.38605\nTotal prob of N-in: 0.99056\nPOSSIBLE N-term signal sequence\ninside 1 11\nTMhelix 12 34\noutside 35 404\nTMhelix 405 422\ninside 423 429

30877 GCF_000811985.1_ASM81198v1 Nocardia vulneris Terrabacteria group; Actinobacteria; Actinobacteria; Corynebacteriales; Nocardiaceae; Nocardia MTQTFRRRAAIPLGIFVALLTILGSGTTLPLLAPPTAQAQPTGSGTVSGPKFLKLSVDSV WP_043676971.1 hypothetical protein [Nocardia vulneris] Length: 895\nNumber of predicted TMHs: 2\nExp number of AAs in TMHs: 43.06239\nExp number, first 60 AAs: 21.71978\nTotal prob of N-in: 0.99436\nPOSSIBLE N-term signal sequence\ninside 1 12\nTMhelix 13 35\noutside 36 826\nTMhelix 827 849\ninside 850 895

30878 GCF_000308455.1_ASM30845v1 Nocardia abscessus NBRC 100374 Terrabacteria group; Actinobacteria; Actinobacteria; Corynebacteriales; Nocardiaceae; Nocardia; Nocardia abscessus MPSDRHRPRLAALDRRSALRLAGGGTVGALALGALAGCTGDDVVHEPDPLAAQEVVARAD WP_043695024.1 hypothetical protein [Nocardia abscessus] Length: 182\nNumber of predicted TMHs: 1\nExp number of AAs in TMHs: 19.56143\nExp number, first 60 AAs: 4.23908\nTotal prob of N-in: 0.71894\ninside 1 157\nTMhelix 158 180\noutside 181 182

30879 GCF_000308455.1_ASM30845v1 Nocardia abscessus NBRC 100374 Terrabacteria group; Actinobacteria; Actinobacteria; Corynebacteriales; Nocardiaceae; Nocardia; Nocardia abscessus MDNSATEGGRVAAFFDLDKTVIAKSSTFVFSKPFYAQGLLNRRAVLESSYAHFLFLLSGA WP_043685756.1 inhibition of morphological differentiation protein [Nocardia abscessus] Length: 269\nNumber of predicted TMHs: 1\nExp number of AAs in TMHs: 22.51088\nExp number, first 60 AAs: 0.0738\nTotal prob of N-in: 0.06338\noutside 1 242\nTMhelix 243 265\ninside 266 269

30880 GCF_000308455.1_ASM30845v1 Nocardia abscessus NBRC 100374 Terrabacteria group; Actinobacteria; Actinobacteria; Corynebacteriales; Nocardiaceae; Nocardia; Nocardia abscessus MKITRRALLCAGAVLAVAAIGATAVPSLAEPSTTTPFTTPNTDACPQKTLPPAPIDASEV WP_043687850.1 D-alanyl-D-alanine carboxypeptidase [Nocardia abscessus] Length: 433\nNumber of predicted TMHs: 2\nExp number of AAs in TMHs: 42.54897\nExp number, first 60 AAs: 22.25475\nTotal prob of N-in: 0.99295\nPOSSIBLE N-term signal sequence\ninside 1 6\nTMhelix 7 29\noutside 30 406\nTMhelix 407 426\ninside 427 433

30881 GCF_001625085.1_ASM162508v1 Nocardia acidivorans NBRC 108247 Terrabacteria group; Actinobacteria; Actinobacteria; Corynebacteriales; Nocardiaceae; Nocardia; Nocardia acidivorans MGTVTAERGRVAAFFDLDKTVIARSSAFVFSKPFLDQGLLSRRAVLESSYAHFLFLLSGA WP_084515686.1 inhibition of morphological differentiation protein [Nocardia acidivorans] Length: 269\nNumber of predicted TMHs: 1\nExp number of AAs in TMHs: 24.04654\nExp number, first 60 AAs: 0.0463\nTotal prob of N-in: 0.13698\noutside 1 242\nTMhelix 243 265\ninside 266 269

30882 GCF_001612665.1_ASM161266v1 Nocardia alba NBRC 108234 Terrabacteria group; Actinobacteria; Actinobacteria; Corynebacteriales; Nocardiaceae; Nocardia; Nocardia alba MPGGRVAAFFDLDKTVIAKSSAFVFSKPFYAQGLINRRAVLESSYAHFLFLLSGADHDQM WP_067454360.1 inhibition of morphological differentiation protein [Nocardia alba] Length: 264\nNumber of predicted TMHs: 1\nExp number of AAs in TMHs: 22.42118\nExp number, first 60 AAs: 0.01121\nTotal prob of N-in: 0.00295\noutside 1 237\nTMhelix 238 260\ninside 261 264

30883 GCF_001612665.1_ASM161266v1 Nocardia alba NBRC 108234 Terrabacteria group; Actinobacteria; Actinobacteria; Corynebacteriales; Nocardiaceae; Nocardia; Nocardia alba MGVVSEIEGSRRRIMMSRSVSYTLPAVVAVALVSAGCGSATPKPAPRAASATLTTTSKPPWP_067458420.1 hypothetical protein [Nocardia alba] Length: 171\nNumber of predicted TMHs: 2\nExp number of AAs in TMHs: 39.9462\nExp number, first 60 AAs: 21.99767\nTotal prob of N-in: 0.96399\nPOSSIBLE N-term signal sequence\ninside 1 19\nTMhelix 20 42\noutside 43 113\nTMhelix 114 131\ninside 132 171

30884 GCF_001612665.1_ASM161266v1 Nocardia alba NBRC 108234 Terrabacteria group; Actinobacteria; Actinobacteria; Corynebacteriales; Nocardiaceae; Nocardia; Nocardia alba MTGAASRRALLRLTCAATVLGAVAGFPMPAGAIAPPRIDPGALDAALALSGRPAPLEATE WP_067448191.1 type VII secretion-associated serine protease mycosin [Nocardia alba] Length: 470\nNumber of predicted TMHs: 2\nExp number of AAs in TMHs: 42.0171\nExp number, first 60 AAs: 19.85988\nTotal prob of N-in: 0.90499\nPOSSIBLE N-term signal sequence\ninside 1 12\nTMhelix 13 35\noutside 36 434\nTMhelix 435 457\ninside 458 470

30885 GCF_001612685.1_ASM161268v1 Nocardia altamirensis NBRC 108246 Terrabacteria group; Actinobacteria; Actinobacteria; Corynebacteriales; Nocardiaceae; Nocardia; Nocardia altamirensis MDSPTADGRVAAFFDLDKTVIAKSSTFVFSRQFYAQGLLNRRDVLESSYAHFLFMLSGAD WP_069166520.1 inhibition of morphological differentiation protein [Nocardia altamirensis] Length: 268\nNumber of predicted TMHs: 1\nExp number of AAs in TMHs: 23.24781\nExp number, first 60 AAs: 0.01217\nTotal prob of N-in: 0.05189\noutside 1 241\nTMhelix 242 264\ninside 265 268

30886 GCF_001612685.1_ASM161268v1 Nocardia altamirensis NBRC 108246 Terrabacteria group; Actinobacteria; Actinobacteria; Corynebacteriales; Nocardiaceae; Nocardia; Nocardia altamirensis MKIRDSRRRATFRLGSALAVLAVSVGVAATPSFAVPPSTPPTTTPFTTPNTDSCPQKTRP WP_069163623.1 penicillin-binding protein [Nocardia altamirensis] Length: 431\nNumber of predicted TMHs: 2\nExp number of AAs in TMHs: 40.89453\nExp number, first 60 AAs: 21.94238\nTotal prob of N-in: 0.98678\nPOSSIBLE N-term signal sequence\ninside 1 11\nTMhelix 12 34\noutside 35 406\nTMhelix 407 424\ninside 425 431

30887 GCF_001612615.1_ASM161261v1 Nocardia amikacinitolerans NBRC 108937 Terrabacteria group; Actinobacteria; Actinobacteria; Corynebacteriales; Nocardiaceae; Nocardia; Nocardia amikacinitolerans MDKTVIAKSSTFVFSKPFFAQGLLNRRAVLESSYAHFLFLLSGADHDQMERMREHLTKMC WP_067790316.1 inhibition of morphological differentiation protein [Nocardia amikacinitolerans] Length: 253\nNumber of predicted TMHs: 1\nExp number of AAs in TMHs: 24.62534\nExp number, first 60 AAs: 0.13149\nTotal prob of N-in: 0.14468\noutside 1 226\nTMhelix 227 249\ninside 250 253

30888 GCF_001612725.1_ASM161272v1 Nocardia anaemiae NBRC 100462 Terrabacteria group; Actinobacteria; Actinobacteria; Corynebacteriales; Nocardiaceae; Nocardia; Nocardia anaemiae MEKAPGAVLDAPTSAAPTSFAAFVDEAGKRRDLFKMKALATGLLAFATAIYLFCRWLESR WP_062983972.1 DUF445 domain-containing protein [Nocardia anaemiae] Length: 440\nNumber of predicted TMHs: 2\nExp number of AAs in TMHs: 47.79527\nExp number, first 60 AAs: 18.29971\nTotal prob of N-in: 0.28847\nPOSSIBLE N-term signal sequence\noutside 1 37\nTMhelix 38 57\ninside 58 416\nTMhelix 417 439\noutside 440 440

30889 GCF_001612725.1_ASM161272v1 Nocardia anaemiae NBRC 100462 Terrabacteria group; Actinobacteria; Actinobacteria; Corynebacteriales; Nocardiaceae; Nocardia; Nocardia anaemiae MTATKRPGAGGERVAAFFDLDKTVIAKSSTYVFSKPFYAQGLLNRRAVLESSYAHFLFML WP_062981222.1 inhibition of morphological differentiation protein [Nocardia anaemiae] Length: 272\nNumber of predicted TMHs: 1\nExp number of AAs in TMHs: 23.22849\nExp number, first 60 AAs: 0.03248\nTotal prob of N-in: 0.06438\noutside 1 245\nTMhelix 246 268\ninside 269 272

30890 GCF_000308435.1_ASM30843v1 Nocardia araoensis NBRC 100135 Terrabacteria group; Actinobacteria; Actinobacteria; Corynebacteriales; Nocardiaceae; Nocardia; Nocardia araoensis MDHSATTGHRVAAFFDLDKTVIAKSSTYVFSKPFYAQGLLNRRAVLESSYAHFLFLLSGA WP_039794579.1 inhibition of morphological differentiation protein [Nocardia araoensis] Length: 269\nNumber of predicted TMHs: 1\nExp number of AAs in TMHs: 23.15518\nExp number, first 60 AAs: 0.08833\nTotal prob of N-in: 0.10792\noutside 1 242\nTMhelix 243 265\ninside 266 269

30891 GCF_000308435.1_ASM30843v1 Nocardia araoensis NBRC 100135 Terrabacteria group; Actinobacteria; Actinobacteria; Corynebacteriales; Nocardiaceae; Nocardia; Nocardia araoensis MKITRRALLCVGAVLAVAAFGATATPSLAAPSTTTPFTTPNTDGCPQKTSPPAPIDASEV WP_039795807.1 D-alanyl-D-alanine carboxypeptidase [Nocardia araoensis] Length: 433\nNumber of predicted TMHs: 2\nExp number of AAs in TMHs: 42.9265\nExp number, first 60 AAs: 22.09503\nTotal prob of N-in: 0.99343\nPOSSIBLE N-term signal sequence\ninside 1 6\nTMhelix 7 29\noutside 30 406\nTMhelix 407 426\ninside 427 433

30892 GCF_000308435.1_ASM30843v1 Nocardia araoensis NBRC 100135 Terrabacteria group; Actinobacteria; Actinobacteria; Corynebacteriales; Nocardiaceae; Nocardia; Nocardia araoensis MQVDAHAFKSYSAARSSRTRRRGSGSPRRRSIVSVPSFSPLSSLRGSGRTPSPSGQRFIP WP_083880810.1 magnesium transporter [Nocardia araoensis] Length: 394\nNumber of predicted TMHs: 2\nExp number of AAs in TMHs: 41.70335\nExp number, first 60 AAs: 0.00451\nTotal prob of N-in: 0.98359\ninside 1 336\nTMhelix 337 356\noutside 357 365\nTMhelix 366 388\ninside 389 394

30893 GCF_001612765.1_ASM161276v1 Nocardia arthritidis NBRC 100137 Terrabacteria group; Actinobacteria; Actinobacteria; Corynebacteriales; Nocardiaceae; Nocardia; Nocardia arthritidis MDNSAPTGGRVAAFFDLDKTVIAKSSTFVFSKPFYAQGLLNRRAVLESSYAHFLFLLSGA WP_063049436.1 inhibition of morphological differentiation protein [Nocardia arthritidis] Length: 269\nNumber of predicted TMHs: 1\nExp number of

AAs in TMHs: 22.00187\nExp number, first 60 AAs: 0.06451\nTotal prob of N-in: 0.05009\noutside 1 242\nTMhelix 243 265\ninside 266 269

30894 GCF_001612765.1_ASM161276v1 Nocardia arthritidis NBRC 100137 Terrabacteria group; Actinobacteria; Actinobacteria; Corynebacteriales; Nocardiaceae; Nocardia; Nocardia arthritidis MEVDAHAFKSYSAAASCRTRRRPCGSLRRRSIVSVPSFSPLSSLRGSGRTPAHRFIPVPT WP_084467026.1 magnesium transporter [Nocardia arthritidis] Length: 390\nNumber of predicted TMHs: 2\nExp number of AAs in TMHs: 41.72643\nExp number, first 60 AAs: 0.00825\nTotal prob of N-in: 0.97777\ninside 1 332\nTMhelix 333 352\noutside 353 361\nTMhelix 362 384\ninside 385 390

30895 GCF_001612765.1_ASM161276v1 Nocardia arthritidis NBRC 100137 Terrabacteria group; Actinobacteria; Actinobacteria; Corynebacteriales; Nocardiaceae; Nocardia; Nocardia arthritidis MKITRRALLCVGAVLAVAAIATTAAPSLAEPSTTTPFTTPNTDGCPQKTLPPAPIDASEV WP_063051558.1 penicillin-binding protein [Nocardia arthritidis] Length: 433\nNumber of predicted TMHs: 2\nExp number of AAs in TMHs: 42.50507\nExp number, first 60 AAs: 22.21406\nTotal prob of N-in: 0.99444\nPOSSIBLE N-term signal sequence\ninside 1 6\nTMhelix 7 29\noutside 30 406\nTMhelix 407 426\ninside 427 433

30896 GCF_000308415.1_ASM30841v1 Nocardia asiatica NBRC 100129 Terrabacteria group; Actinobacteria; Actinobacteria; Corynebacteriales; Nocardiaceae; Nocardia; Nocardia asiatica MDNSATAGGRVAAFFDLDKTVIAKSSTFVFSKPFYAQGLLNRRAVLESSYAHFLFLLSGA WP_043718419.1 inhibition of morphological differentiation protein [Nocardia asiatica] Length: 269\nNumber of predicted TMHs: 1\nExp number of AAs in TMHs: 22.89437\nExp number, first 60 AAs: 0.08783\nTotal prob of N-in: 0.08329\noutside 1 242\nTMhelix 243 265\ninside 266 269

30897 GCF_001612785.1_ASM161278v1 Nocardia beijingensis NBRC 16342 Terrabacteria group; Actinobacteria; Actinobacteria; Corynebacteriales; Nocardiaceae; Nocardia; Nocardia beijingensis MDKPATTGGRVAAFFDLDKTVIAKSSTFVFSKPFYAQGLLNRRAVLESSYAHFLFLLSGA WP_067811213.1 inhibition of morphological differentiation protein [Nocardia beijingensis] Length: 269\nNumber of predicted TMHs: 1\nExp number of AAs in TMHs: 22.40164\nExp number, first 60 AAs: 0.07968\nTotal prob of N-in: 0.06054\noutside 1 242\nTMhelix 243 265\ninside 266 269

30898 GCF_001612785.1_ASM161278v1 Nocardia beijingensis NBRC 16342 Terrabacteria group; Actinobacteria; Actinobacteria; Corynebacteriales; Nocardiaceae; Nocardia; Nocardia beijingensis MKITRRALLCVGAVLAVAAIGATAAPSVAAPPSTTTPFTTPNTDGCPQKTLPPTPIDTSE WP_067803507.1 penicillin-binding protein [Nocardia beijingensis] Length: 434\nNumber of predicted TMHs: 2\nExp number of AAs in TMHs: 43.1620299999999\nExp number, first 60 AAs: 22.50073\nTotal prob of N-in: 0.99625\nPOSSIBLE N-term signal sequence\ninside 1 6\nTMhelix 7 29\noutside 30 404\nTMhelix 405 427\ninside 428 434

30899 GCF_001612785.1_ASM161278v1 Nocardia beijingensis NBRC 16342 Terrabacteria group; Actinobacteria; Actinobacteria; Corynebacteriales; Nocardiaceae; Nocardia; Nocardia beijingensis MQVDAHAFKSYSAATSSRTRRRSSRRPPRRSIVSVPSFSPLSSFRGSGRTPSQPQQRFLP WP_084486325.1 magnesium transporter [Nocardia beijingensis] Length: 394\nNumber of predicted TMHs: 2\nExp number of AAs in TMHs: 41.69481\nExp number, first 60 AAs: 0.00292\nTotal prob of N-in: 0.98403\ninside 1 336\nTMhelix 337 356\noutside 357 365\nTMhelix 366 388\ninside 389 394

30900 GCF_000250675.2_ASM25067v3 Nocardia brasiliensis ATCC 700358 Terrabacteria group; Actinobacteria; Actinobacteria; Corynebacteriales; Nocardiaceae; Nocardia; Nocardia brasiliensis MKIRESRRRATLRLGSALAALAVTAGLAAVPSFAAPTSTTTTPFSTPNTDGCPQKTSPPP WP_014981882.1 D-alanyl-D-alanine carboxypeptidase [Nocardia brasiliensis] Length: 429\nNumber of predicted TMHs: 2\nExp number of AAs in TMHs: 40.77749\nExp number, first 60 AAs: 22.37433\nTotal prob of N-in: 0.98999\nPOSSIBLE N-term signal sequence\ninside 1 11\nTMhelix 12 34\noutside 35 404\nTMhelix 405 422\ninside 423 429

30901 GCF_000250675.2_ASM25067v3 Nocardia brasiliensis ATCC 700358 Terrabacteria group; Actinobacteria; Actinobacteria; Corynebacteriales; Nocardiaceae; Nocardia; Nocardia brasiliensis MTQTFRRRAAIPLGIFVALLTILGSGTTLPLLAPPAAQAQPTGSGTVSGPKFLKLSVDSV WP_014989207.1 hypothetical protein [Nocardia brasiliensis] Length: 895\nNumber of predicted TMHs: 2\nExp number of AAs in TMHs: 43.08572\nExp number, first 60 AAs: 21.74309\nTotal prob of N-in: 0.99463\nPOSSIBLE N-term signal sequence\ninside 1 12\nTMhelix 13 35\noutside 36 826\nTMhelix 827 849\ninside 850 895

30902 GCF_000250675.2_ASM25067v3 Nocardia brasiliensis ATCC 700358 Terrabacteria group; Actinobacteria; Actinobacteria; Corynebacteriales; Nocardiaceae; Nocardia; Nocardia brasiliensis

MDSPHANGRVAAFFDLDKTVIAKSSTFVFSKPFYAQGLLNRRDVLESSYAHFMFMLSGAD	WP_014981200.1	hydrolase [Nocardia brasiliensis]	Length: 268\nNumber of predicted TMHs: 1\nExp number of AAs in TMHs: 22.91898\nExp number, first 60 AAs: 0.02348\nTotal prob of N-in: 0.05944\noutside 1 241\nTMhelix 242 264\ninside 265 268

30903	GCF_000710915.1_ASM71091v1	Nocardia brasiliensis IFM 10847	Terrabacteria group; Actinobacteria; Actinobacteria; Corynebacteriales; Nocardiaceae; Nocardia; Nocardia brasiliensis	MDSPHANGRVAAFFDLDKTVIAKSSTFVFSKPFYAQGLLNRRDVLESSYAHFLFMLSGAD	WP_029893388.1	inhibition of morphological differentiation protein [Nocardia brasiliensis]	Length: 268\nNumber of predicted TMHs: 1\nExp number of AAs in TMHs: 22.69388\nExp number, first 60 AAs: 0.02318\nTotal prob of N-in: 0.04870\noutside 1 241\nTMhelix 242 264\ninside 265 268

30904	GCF_000710915.1_ASM71091v1	Nocardia brasiliensis IFM 10847	Terrabacteria group; Actinobacteria; Actinobacteria; Corynebacteriales; Nocardiaceae; Nocardia; Nocardia brasiliensis	MKIRESRRRATLRLGSALAALAVTAGLAAVPSLAAPTSTTTTPFTTPNTDGCPQKTSPPP	WP_029894928.1	penicillin-binding protein [Nocardia brasiliensis]	Length: 429\nNumber of predicted TMHs: 2\nExp number of AAs in TMHs: 40.71039\nExp number, first 60 AAs: 22.21728\nTotal prob of N-in: 0.98505\nPOSSIBLE N-term signal sequence\ninside 1 11\nTMhelix 12 34\noutside 35 404\nTMhelix 405 422\ninside 423 429

30905	GCF_000710915.1_ASM71091v1	Nocardia brasiliensis IFM 10847	Terrabacteria group; Actinobacteria; Actinobacteria; Corynebacteriales; Nocardiaceae; Nocardia; Nocardia brasiliensis	MTQTFRRRAAIPLGIFVALLTILGSGTTLPLLAPPTAQAQPTGSGTVSGPKFLKLSVDSV	WP_029897422.1	hypothetical protein [Nocardia brasiliensis]	Length: 895\nNumber of predicted TMHs: 2\nExp number of AAs in TMHs: 43.05507\nExp number, first 60 AAs: 21.71977\nTotal prob of N-in: 0.99436\nPOSSIBLE N-term signal sequence\ninside 1 12\nTMhelix 13 35\noutside 36 826\nTMhelix 827 849\ninside 850 895

30906	GCF_000308475.2_ASM30847v2	Nocardia brasiliensis NBRC 14402	Terrabacteria group; Actinobacteria; Actinobacteria; Corynebacteriales; Nocardiaceae; Nocardia; Nocardia brasiliensis	MDSPHANGRVAAFFDLDKTVIAKSSTFVFSKPFYAQGLLNRRDVLESSYAHFLFMLSGAD	WP_042254975.1	inhibition of morphological differentiation protein [Nocardia brasiliensis]	Length: 268\nNumber of predicted TMHs: 1\nExp number of AAs in TMHs: 22.91952\nExp number, first 60 AAs: 0.02426\nTotal prob of N-in: 0.05945\noutside 1 241\nTMhelix 242 264\ninside 265 268

30907	GCF_000308475.2_ASM30847v2	Nocardia brasiliensis NBRC 14402	Terrabacteria group; Actinobacteria; Actinobacteria; Corynebacteriales; Nocardiaceae; Nocardia; Nocardia brasiliensis	MTQTFRRRAAIPLGIFVALLTILGSGTTLPLLAPPAAQAQPTGSGTVSGPKFLKLSVDSV	WP_042259183.1	hypothetical protein [Nocardia brasiliensis]	Length: 895\nNumber of predicted TMHs: 2\nExp number of AAs in TMHs: 43.08572\nExp number, first 60 AAs: 21.74311\nTotal prob of N-in: 0.99463\nPOSSIBLE N-term signal sequence\ninside 1 12\nTMhelix 13 35\noutside 36 826\nTMhelix 827 849\ninside 850 895

30908	GCF_000308475.2_ASM30847v2	Nocardia brasiliensis NBRC 14402	Terrabacteria group; Actinobacteria; Actinobacteria; Corynebacteriales; Nocardiaceae; Nocardia; Nocardia brasiliensis	MKIRESRRRATLRLGSALAALAVTAGLAAVPSFAAPTSTTTTPFTTPNTDGCPQKTSPPP	WP_042263540.1	penicillin-binding protein [Nocardia brasiliensis]	Length: 429\nNumber of predicted TMHs: 2\nExp number of AAs in TMHs: 40.77805\nExp number, first 60 AAs: 22.37533\nTotal prob of N-in: 0.98999\nPOSSIBLE N-term signal sequence\ninside 1 11\nTMhelix 12 34\noutside 35 404\nTMhelix 405 422\ninside 423 429

30909	GCF_001612825.1_ASM161282v1	Nocardia caishijiensis NBRC 108228	Terrabacteria group; Actinobacteria; Actinobacteria; Corynebacteriales; Nocardiaceae; Nocardia; Nocardia caishijiensis	MAAFFDLDKTVIAKSSAFVFSKPFYAQGLLNRRAVLESSYAHFLFLLSGADHDQMERMRA	WP_067977569.1	inhibition of morphological differentiation protein [Nocardia caishijiensis]	Length: 259\nNumber of predicted TMHs: 1\nExp number of AAs in TMHs: 22.36039\nExp number, first 60 AAs: 0.02344\nTotal prob of N-in: 0.00824\noutside 1 232\nTMhelix 233 255\ninside 256 259

30910	GCF_000308515.1_ASM30851v1	Nocardia carnea NBRC 14403	Terrabacteria group; Actinobacteria; Actinobacteria; Corynebacteriales; Nocardiaceae; Nocardia; Nocardia carnea	MNSRIAAFFDLDKTVIARSSTYVFSKPFFAEGLINRRAVLESSYAHFIFLLSGADHDQME	WP_051157049.1	inhibition of morphological differentiation protein [Nocardia carnea]	Length: 261\nNumber of predicted TMHs: 1\nExp number of AAs in TMHs: 21.14729\nExp number, first 60 AAs: 0.02759\nTotal prob of N-in: 0.02336\noutside 1 235\nTMhelix 236 258\ninside 259 261

30911 GCF_000308815.1_ASM30881v1 Nocardia concava NBRC 100430 Terrabacteria group; Actinobacteria; Actinobacteria; Corynebacteriales; Nocardiaceae; Nocardia; Nocardia concava
MSSSPPSAPDPEGIGQATKIRKQRRTRRKLVGGLVLLMGLVGAGFAASALTPQAQVATAH WP_040806829.1 cytochrome c [Nocardia concava] Length: 286\nNumber of predicted TMHs: 2\nExp number of AAs in TMHs: 41.47535\nExp number, first 60 AAs: 21.27994\nTotal prob of N-in: 0.98308\nPOSSIBLE N-term signal sequence\ninside 1 29\nTMhelix 30 52\noutside 53 263\nTMhelix 264 283\ninside 284 286

30912 GCF_000308815.1_ASM30881v1 Nocardia concava NBRC 100430 Terrabacteria group; Actinobacteria; Actinobacteria; Corynebacteriales; Nocardiaceae; Nocardia; Nocardia concava
MLDHVTAERGRVAAFFDLDKTVIARSSAFVFSKPFLDQGLLSRRAVLESSYAHFLFLLQG WP_040802771.1 inhibition of morphological differentiation protein [Nocardia concava] Length: 270\nNumber of predicted TMHs: 1\nExp number of AAs in TMHs: 22.4878\nExp number, first 60 AAs: 0.02541\nTotal prob of N-in: 0.06391\noutside 1 243\nTMhelix 244 266\ninside 267 270

30913 GCF_001612865.1_ASM161286v1 Nocardia cummidelens NBRC 100378 Terrabacteria group; Actinobacteria; Actinobacteria; Corynebacteriales; Nocardiaceae; Nocardia; Nocardia cummidelens
MSVVSDIEGSRRRIMMSRSVSYTLPAVVAVALVSAGCGTATPKPAPRAASTTLTTTSKPP WP_063006426.1 hypothetical protein [Nocardia cummidelens] Length: 172\nNumber of predicted TMHs: 2\nExp number of AAs in TMHs: 40.4393200000001\nExp number, first 60 AAs: 22.30399\nTotal prob of N-in: 0.96765\nPOSSIBLE N-term signal sequence\ninside 1 19\nTMhelix 20 42\noutside 43 113\nTMhelix 114 131\ninside 132 172

30914 GCF_000284035.1_ASM28403v1 Nocardia cyriacigeorgica GUH-2 Terrabacteria group; Actinobacteria; Actinobacteria; Corynebacteriales; Nocardiaceae; Nocardia; Nocardia cyriacigeorgica
MHSLDTLKFSGRRNVLRLNEKLATIQAGTIATAAFRSELFGPHLVTGTIVRSGATGGYLL WP_048833198.1 hypothetical protein [Nocardia cyriacigeorgica] Length: 185\nNumber of predicted TMHs: 1\nExp number of AAs in TMHs: 23.64994\nExp number, first 60 AAs: 1.04401\nTotal prob of N-in: 0.18833\noutside 1 120\nTMhelix 121 143\ninside 144 185

30915 GCF_000308555.1_ASM30855v1 Nocardia cyriacigeorgica NBRC 100375 Terrabacteria group; Actinobacteria; Actinobacteria; Corynebacteriales; Nocardiaceae; Nocardia; Nocardia cyriacigeorgica
MRELRRRAAVAVAAVLTVSAVAGGPAMAEPTGTTGPTTTPFTTPNTDNCAQRTLPPPPID WP_036533687.1 D-alanyl-D-alanine carboxypeptidase [Nocardia cyriacigeorgica] Length: 427\nNumber of predicted TMHs: 1\nExp number of AAs in TMHs: 33.38256\nExp number, first 60 AAs: 13.98604\nTotal prob of N-in: 0.68477\nPOSSIBLE N-term signal sequence\noutside 1 402\nTMhelix 403 420\ninside 421 427

30916 GCF_000308575.1_ASM30857v1 Nocardia exalbida NBRC 100660 Terrabacteria group; Actinobacteria; Actinobacteria; Corynebacteriales; Nocardiaceae; Nocardia; Nocardia exalbida
MDNSAPTGGRVAAFFDLDKTVIAKSSTFVFSKPFFAQGLLNRRAVLESSYAHFLFLLSGA WP_040864499.1 MULTISPECIES: inhibition of morphological differentiation protein [Nocardia] Length: 269\nNumber of predicted TMHs: 1\nExp number of AAs in TMHs: 22.30618\nExp number, first 60 AAs: 0.08304\nTotal prob of N-in: 0.04545\noutside 1 242\nTMhelix 243 265\ninside 266 269

30917 GCF_000308575.1_ASM30857v1 Nocardia exalbida NBRC 100660 Terrabacteria group; Actinobacteria; Actinobacteria; Corynebacteriales; Nocardiaceae; Nocardia; Nocardia exalbida
MKITRRALLCVGAVLAVAAIATTAAPSLAEPSTSTPFTTPNTDGCPQKTVPPAPIDASEV WP_040865004.1 D-alanyl-D-alanine carboxypeptidase [Nocardia exalbida] Length: 433\nNumber of predicted TMHs: 2\nExp number of AAs in TMHs: 42.4731399999999\nExp number, first 60 AAs: 22.21386\nTotal prob of N-in: 0.99443\nPOSSIBLE N-term signal sequence\ninside 1 6\nTMhelix 7 29\noutside 30 406\nTMhelix 407 426\ninside 427 433

30918 GCF_001613385.1_ASM161338v1 Nocardia flavorosea NBRC 108225 Terrabacteria group; Actinobacteria; Actinobacteria; Corynebacteriales; Nocardiaceae; Nocardia; Nocardia flavorosea
MTGDRHGTNGRIAAFFDLDKTVIARSSTYVFSKPFFAEGLINRRAVLESSYAHFMFLLSG WP_062972447.1 inhibition of morphological differentiation protein [Nocardia flavorosea] Length: 268\nNumber of predicted TMHs: 1\nExp number of AAs in TMHs: 21.39264\nExp number, first 60 AAs: 0.05305\nTotal prob of N-in: 0.03922\noutside 1 242\nTMhelix 243 265\ninside 266 268

30919 GCF_001612985.1_ASM161298v1 Nocardia gamkensis NBRC 108242 Terrabacteria group; Actinobacteria; Actinobacteria; Corynebacteriales; Nocardiaceae; Nocardia; Nocardia gamkensis
MDNSAPTGGRVAAFFDLDKTVIAKSSTFVFSKPFFAQGLLNRRAVLESSYAHFLFLLSGA WP_040864499.1 MULTISPECIES: inhibition of morphological differentiation protein [Nocardia] Length: 269\nNumber of predicted TMHs: 1\nExp

number of AAs in TMHs: 22.30618\nExp number, first 60 AAs: 0.08304\nTotal prob of N-in: 0.04545\noutside 1 242\nTMhelix 243 265\ninside 266 269

30920 GCF_001613145.1_ASM161314v1 Nocardia jejuensis NBRC 103114 Terrabacteria group; Actinobacteria; Actinobacteria; Corynebacteriales; Nocardiaceae; Nocardia; Nocardia jejuensis MRQSPSRTPLGVARPDSRRRRIVIRAILAALAAATLTAAGPAPAHASPDNPAALRLDVTG WP_084479049.1 hypothetical protein [Nocardia jejuensis] Length: 214\nNumber of predicted TMHs: 2\nExp number of AAs in TMHs: 41.06655\nExp number, first 60 AAs: 21.45501\nTotal prob of N-in: 0.99133\nPOSSIBLE N-term signal sequence\ninside 1 21\nTMhelix 22 44\noutside 45 191\nTMhelix 192 209\ninside 210 214

30921 GCF_001613145.1_ASM161314v1 Nocardia jejuensis NBRC 103114 Terrabacteria group; Actinobacteria; Actinobacteria; Corynebacteriales; Nocardiaceae; Nocardia; Nocardia jejuensis MTADRGRVAAFFDLDKTVIARSSAFVFSKPFLDQGLLSRRAVLESSYAHFLFLLSGADHD WP_067703880.1 inhibition of morphological differentiation protein [Nocardia jejuensis] Length: 266\nNumber of predicted TMHs: 1\nExp number of AAs in TMHs: 23.50218\nExp number, first 60 AAs: 0.03899\nTotal prob of N-in: 0.11010\noutside 1 239\nTMhelix 240 262\ninside 263 266

30922 GCF_000308655.1_ASM30865v1 Nocardia niigatensis NBRC 100131 Terrabacteria group; Actinobacteria; Actinobacteria; Corynebacteriales; Nocardiaceae; Nocardia; Nocardia niigatensis MSIRRYPRRTARPAAAGRRRAALVIAVTLAAGLSVFAGVASGPLPGAAQAQPATTTTPFT WP_083868605.1 penicillin-binding protein [Nocardia niigatensis] Length: 447\nNumber of predicted TMHs: 2\nExp number of AAs in TMHs: 42.10192\nExp number, first 60 AAs: 21.438\nTotal prob of N-in: 0.99970\nPOSSIBLE N-term signal sequence\ninside 1 20\nTMhelix 21 43\noutside 44 417\nTMhelix 418 440\ninside 441 447

30923 GCF_000308655.1_ASM30865v1 Nocardia niigatensis NBRC 100131 Terrabacteria group; Actinobacteria; Actinobacteria; Corynebacteriales; Nocardiaceae; Nocardia; Nocardia niigatensis MLDHVTAERGRVAAFFDLDKTVIARSSAFVFSKPFLDQGLLSRRAVLESSYAHFLFLLSG WP_040854204.1 inhibition of morphological differentiation protein [Nocardia niigatensis] Length: 270\nNumber of predicted TMHs: 1\nExp number of AAs in TMHs: 23.76733\nExp number, first 60 AAs: 0.05878\nTotal prob of N-in: 0.09083\noutside 1 243\nTMhelix 244 266\ninside 267 270

30924 GCF_001613465.1_ASM161346v1 Nocardia niwae NBRC 108934 = DSM 45340 Terrabacteria group; Actinobacteria; Actinobacteria; Corynebacteriales; Nocardiaceae; Nocardia; Nocardia niwae MDNSATTGARVAAFFDLDKTVIAKSSTFVFSKPFYAQGLLNRRAVLESSYAHFLFLLSGA WP_063025198.1 inhibition of morphological differentiation protein [Nocardia niwae] Length: 269\nNumber of predicted TMHs: 1\nExp number of AAs in TMHs: 23.21611\nExp number, first 60 AAs: 0.12973\nTotal prob of N-in: 0.13078\noutside 1 242\nTMhelix 243 265\ninside 266 269

30925 GCF_001613465.1_ASM161346v1 Nocardia niwae NBRC 108934 = DSM 45340 Terrabacteria group; Actinobacteria; Actinobacteria; Corynebacteriales; Nocardiaceae; Nocardia; Nocardia niwae MKITRRALLCVGVVLAVAAIGATAAPSLAEPGTTTPFTTPNTDGCPQKTVPPAPIDASEV WP_063021406.1 penicillin-binding protein [Nocardia niwae] Length: 433\nNumber of predicted TMHs: 2\nExp number of AAs in TMHs: 42.93046\nExp number, first 60 AAs: 22.30407\nTotal prob of N-in: 0.99758\nPOSSIBLE N-term signal sequence\ninside 1 6\nTMhelix 7 29\noutside 30 406\nTMhelix 407 426\ninside 427 433

30926 GCF_001613465.1_ASM161346v1 Nocardia niwae NBRC 108934 = DSM 45340 Terrabacteria group; Actinobacteria; Actinobacteria; Corynebacteriales; Nocardiaceae; Nocardia; Nocardia niwae MQVDEHAFKSYSAAASSRTRRRSSGSLRRRSIVTVPSFSPLSSLRGSGRSPAQPEQRFIP WP_084489189.1 magnesium transporter [Nocardia niwae] Length: 394\nNumber of predicted TMHs: 2\nExp number of AAs in TMHs: 41.82922\nExp number, first 60 AAs: 0.00179\nTotal prob of N-in: 0.98019\ninside 1 336\nTMhelix 337 356\noutside 357 365\nTMhelix 366 388\ninside 389 394

30927 GCF_000308755.1_ASM30875v1 Nocardia pneumoniae NBRC 100136 Terrabacteria group; Actinobacteria; Actinobacteria; Corynebacteriales; Nocardiaceae; Nocardia; Nocardia pneumoniae MDNPPATGGRVAAFFDLDKTVIAKSSTFVFSKPFYAQGLLNRRAVLESSYAHFLFLLSGA WP_040773153.1 inhibition of morphological differentiation protein [Nocardia pneumoniae] Length: 269\nNumber of predicted TMHs: 1\nExp number of AAs in TMHs: 22.89168\nExp number, first 60 AAs: 0.05122\nTotal prob of N-in: 0.03294\noutside 1 242\nTMhelix 243 265\ninside 266 269

30928 GCF_001613105.1_ASM161310v1 Nocardia pseudobrasiliensis NBRC 108224 Terrabacteria group; Actinobacteria; Actinobacteria; Corynebacteriales; Nocardiaceae; Nocardia; Nocardia pseudobrasiliensis

MLDAVTNTRPPRTAAFFDLDKTVIAKSSAFVFSKPFFAQGLLDRRTVLESSYAHFLFLLS WP_067997993.1 inhibition of morphological differentiation protein [Nocardia pseudobrasiliensis] Length: 271\nNumber of predicted TMHs: 1\nExp number of AAs in TMHs: 22.28812\nExp number, first 60 AAs: 0.02569\nTotal prob of N-in: 0.01212\noutside 1 244\nTMhelix 245 267\ninside 268 271

30929 GCF_001613225.1_ASM161322v1 Nocardia pseudovaccinii NBRC 100343 Terrabacteria group; Actinobacteria; Actinobacteria; Corynebacteriales; Nocardiaceae; Nocardia; Nocardia pseudovaccinii
MLDASTSAPPTSFAAFVDEAGKRRDLFKMKALATGLLAFATAVYLFCRWLESRGAGDDWV WP_063046056.1 DUF445 domain-containing protein [Nocardia pseudovaccinii] Length: 433\nNumber of predicted TMHs: 2\nExp number of AAs in TMHs: 46.22302\nExp number, first 60 AAs: 18.34818\nTotal prob of N-in: 0.13710\nPOSSIBLE N-term signal sequence\noutside 1 30\nTMhelix 31 50\ninside 51 409\nTMhelix 410 432\noutside 433 433

30930 GCF_001613185.1_ASM161318v1 Nocardia puris NBRC 108233 Terrabacteria group; Actinobacteria; Actinobacteria; Corynebacteriales; Nocardiaceae; Nocardia; Nocardia puris
MIFAPMTARRAALLRLAVALSAFVAIALVHGAQCQTGMPMTMAHSAAMVGGVAGQCGGAQ WP_054811683.1 MULTISPECIES: hypothetical protein [Nocardia] Length: 156\nNumber of predicted TMHs: 2\nExp number of AAs in TMHs: 43.30523\nExp number, first 60 AAs: 21.2408\nTotal prob of N-in: 0.99795\nPOSSIBLE N-term signal sequence\ninside 1 11\nTMhelix 12 34\noutside 35 94\nTMhelix 95 117\ninside 118 156

30931 GCF_001613085.1_ASM161308v1 Nocardia salmonicida NBRC 13393 Terrabacteria group; Actinobacteria; Actinobacteria; Corynebacteriales; Nocardiaceae; Nocardia; Nocardia salmonicida
MPGGRVAAFFDLDKTVIAKSSAFVFSKPFYAQGLLNRRAVLESSYAHFLFLLSGADHDQM WP_062991432.1 inhibition of morphological differentiation protein [Nocardia salmonicida] Length: 264\nNumber of predicted TMHs: 1\nExp number of AAs in TMHs: 22.45377\nExp number, first 60 AAs: 0.01135\nTotal prob of N-in: 0.00448\noutside 1 237\nTMhelix 238 260\ninside 261 264

30932 GCF_001613285.1_ASM161328v1 Nocardia soli NBRC 100376 Terrabacteria group; Actinobacteria; Actinobacteria; Corynebacteriales; Nocardiaceae; Nocardia; Nocardia soli
MSVVSDIKGSRRRIMMSRSVSYTLPAVVAVALVSAGCGTATPKPAPRAASATLTTTSKPPWP_063055184.1 hypothetical protein [Nocardia soli] Length: 172\nNumber of predicted TMHs: 2\nExp number of AAs in TMHs: 40.4539400000001\nExp number, first 60 AAs: 22.31968\nTotal prob of N-in: 0.98309\nPOSSIBLE N-term signal sequence\ninside 1 19\nTMhelix 20 42\noutside 43 113\nTMhelix 114 131\ninside 132 172

30933 GCF_001613245.1_ASM161324v1 Nocardia speluncae NBRC 108251 Terrabacteria group; Actinobacteria; Actinobacteria; Corynebacteriales; Nocardiaceae; Nocardia; Nocardia speluncae
MTGDRHGTGSRIAAFFDLDKTVIARSSTYVFSKPFFAEGLINRRAVLESSYAHFIFLLSG WP_068046189.1 inhibition of morphological differentiation protein [Nocardia speluncae]Length: 268\nNumber of predicted TMHs: 1\nExp number of AAs in TMHs: 21.51757\nExp number, first 60 AAs: 0.05428\nTotal prob of N-in: 0.03306\noutside 1 242\nTMhelix 243 265\ninside 266 268

30934 GCF_000308715.1_ASM30871v1 Nocardia tenerifensis NBRC 101015 Terrabacteria group; Actinobacteria; Actinobacteria; Corynebacteriales; Nocardiaceae; Nocardia; Nocardia tenerifensis
MDSPTADGRVAAFFDLDKTVIAKSSTFVFSKPFYAQGLLNRRDVLESSYAHFLFMLSGAD WP_040730034.1 inhibition of morphological differentiation protein [Nocardia tenerifensis] Length: 268\nNumber of predicted TMHs: 1\nExp number of AAs in TMHs: 22.79769\nExp number, first 60 AAs: 0.02382\nTotal prob of N-in: 0.06044\noutside 1 241\nTMhelix 242 264\ninside 265 268

30935 GCF_000308795.1_ASM30879v1 Nocardia thailandica NBRC 100428 Terrabacteria group; Actinobacteria; Actinobacteria; Corynebacteriales; Nocardiaceae; Nocardia; Nocardia thailandica
MNPVWARSRRVLPASPRRGAVFLALLAAALLVLGPVLDCTVLRHESGAGHHGATGFGAPD WP_052314138.1 hypothetical protein [Nocardia thailandica] Length: 179\nNumber of predicted TMHs: 2\nExp number of AAs in TMHs: 43.83063\nExp number, first 60 AAs: 21.34293\nTotal prob of N-in: 0.99476\nPOSSIBLE N-term signal sequence\ninside 1 19\nTMhelix 20 42\noutside 43 125\nTMhelix 126 148\ninside 149 179

30936 GCF_000308795.1_ASM30879v1 Nocardia thailandica NBRC 100428 Terrabacteria group; Actinobacteria; Actinobacteria; Corynebacteriales; Nocardiaceae; Nocardia; Nocardia thailandica
MAAFFDLDKTVIAKSSAFVFSKPFYAQGLLNRRAVLESSYAHFLFLLSGADHDQMERMRA WP_052313537.1 inhibition of morphological differentiation protein [Nocardia thailandica] Length: 259\nNumber of predicted TMHs: 1\nExp number of AAs in TMHs: 22.73705\nExp number, first 60 AAs: 0.03692\nTotal prob of N-in: 0.02198\noutside 1 232\nTMhelix 233 255\ninside 256 259

30937 GCF_001613345.1_ASM161334v1 Nocardia uniformis NBRC 13702 Terrabacteria group; Actinobacteria; Actinobacteria; Corynebacteriales; Nocardiaceae; Nocardia; Nocardia uniformis
MVDGVTTEPARVAAFFDLDKTVIARSSAFAFSKPFLDQGLLSRRAVLESSYAHFLFLLSG WP_067517172.1 inhibition of morphological differentiation protein [Nocardia uniformis] Length: 270\nNumber of predicted TMHs: 1\nExp number of AAs in TMHs: 22.09622\nExp number, first 60 AAs: 0.02613\nTotal prob of N-in: 0.00922\noutside 1 243\nTMhelix 244 266\ninside 267 270

30938 GCF_000308855.1_ASM30885v1 Nocardia veterana NBRC 100344 Terrabacteria group; Actinobacteria; Actinobacteria; Corynebacteriales; Nocardiaceae; Nocardia; Nocardia veterana
MPPTRATPRIAAFFDLDKTVIAKSSAFVFSRPFFDQGLIDRRTVLESSYAHFLFLLSGAD WP_040715750.1 inhibition of morphological differentiation protein [Nocardia veterana] Length: 268\nNumber of predicted TMHs: 1\nExp number of AAs in TMHs: 23.1983\nExp number, first 60 AAs: 0.01391\nTotal prob of N-in: 0.05616\noutside 1 241\nTMhelix 242 264\ninside 265 268

30939 GCF_000308835.1_ASM30883v1 Nocardia vinacea NBRC 16497 Terrabacteria group; Actinobacteria; Actinobacteria; Corynebacteriales; Nocardiaceae; Nocardia; Nocardia vinacea
MLDAPTSAPPTSFAAFVDDAGKRRDLFKMKALATGLLAFATAVYLFCRWLESRGAGGDWV WP_051182624.1 DUF445 domain-containing protein [Nocardia vinacea] Length: 433\nNumber of predicted TMHs: 2\nExp number of AAs in TMHs: 44.2142199999999\nExp number, first 60 AAs: 18.42058\nTotal prob of N-in: 0.10627\nPOSSIBLE N-term signal sequence\noutside 1 30\nTMhelix 31 50\ninside 51 409\nTMhelix 410 432\noutside 433 433

30940 GCF_000308835.1_ASM30883v1 Nocardia vinacea NBRC 16497 Terrabacteria group; Actinobacteria; Actinobacteria; Corynebacteriales; Nocardiaceae; Nocardia; Nocardia vinacea
MTATKRPSAGGERVAAFFDLDKTVIAKSSTYVFSKPFYAQGLLNRRAVLESSYAHFLFML WP_040686119.1 inhibition of morphological differentiation protein [Nocardia vinacea] Length: 272\nNumber of predicted TMHs: 1\nExp number of AAs in TMHs: 23.47765\nExp number, first 60 AAs: 0.03971\nTotal prob of N-in: 0.07988\noutside 1 245\nTMhelix 246 268\ninside 269 272

30941 GCF_000470885.1_RhoBCP1_1.0 Rhodococcus aetherivorans Terrabacteria group; Actinobacteria; Actinobacteria; Corynebacteriales; Nocardiaceae; Rhodococcus
MRLPTSPRSRRRTLSAALLLLLVPVLAGCLRVQATMGVSADDRVSGQIVAATIPADENDK WP_006933384.1 MULTISPECIES: DUF3153 domain-containing protein [Rhodococcus] Length: 233\nNumber of predicted TMHs: 2\nExp number of AAs in TMHs: 44.90952\nExp number, first 60 AAs: 21.98985\nTotal prob of N-in: 0.95979\nPOSSIBLE N-term signal sequence\ninside 1 12\nTMhelix 13 35\noutside 36 191\nTMhelix 192 214\ninside 215 233

30942 GCF_000470885.1_RhoBCP1_1.0 Rhodococcus aetherivorans Terrabacteria group; Actinobacteria; Actinobacteria; Corynebacteriales; Nocardiaceae; Rhodococcus
MTDATVAPTQSGRIAAFFDLDKTVIAKSSALAFSKPFFAQGLLNRRAVLKSSYAQFLFML WP_006941337.1 MULTISPECIES: phosphoserine phosphatase [Rhodococcus] Length: 280\nNumber of predicted TMHs: 1\nExp number of AAs in TMHs: 22.81284\nExp number, first 60 AAs: 0.03489\nTotal prob of N-in: 0.04407\noutside 1 245\nTMhelix 246 268\ninside 269 280

30943 GCF_000982715.1_ASM98271v1 Rhodococcus aetherivorans Terrabacteria group; Actinobacteria; Actinobacteria; Corynebacteriales; Nocardiaceae; Rhodococcus
MTDATVAPTQSGRIAAFFDLDKTVIAKSSALAFSKPFFAQGLLNRRAVLKSSYAQFLFML WP_006941337.1 MULTISPECIES: phosphoserine phosphatase [Rhodococcus] Length: 280\nNumber of predicted TMHs: 1\nExp number of AAs in TMHs: 22.81284\nExp number, first 60 AAs: 0.03489\nTotal prob of N-in: 0.04407\noutside 1 245\nTMhelix 246 268\ninside 269 280

30944 GCF_000982715.1_ASM98271v1 Rhodococcus aetherivorans Terrabacteria group; Actinobacteria; Actinobacteria; Corynebacteriales; Nocardiaceae; Rhodococcus
MRLPTSPRSRRRTLSAALLLLLVPVLAGCLRVQATMGVSADDRVSGQIVAATIPADENDK WP_006933384.1 MULTISPECIES: DUF3153 domain-containing protein [Rhodococcus] Length: 233\nNumber of predicted TMHs: 2\nExp number of AAs in TMHs: 44.90952\nExp number, first 60 AAs: 21.98985\nTotal prob of N-in: 0.95979\nPOSSIBLE N-term signal sequence\ninside 1 12\nTMhelix 13 35\noutside 36 191\nTMhelix 192 214\ninside 215 233

30945 GCF_000738775.1_ASM73877v1 Rhodococcus defluvii Terrabacteria group; Actinobacteria; Actinobacteria; Corynebacteriales; Nocardiaceae; Rhodococcus
MAAHPKSTRRRILLTATASALVLGAGVAPATAVPLAGQAPLIADPGVAPTTTPPYSTPST WP_031938619.1 penicillin-binding protein [Rhodococcus defluvii] Length: 445\nNumber of predicted TMHs: 2\nExp number of AAs in TMHs:

43.3198\nExp number, first 60 AAs: 22.06467\nTotal prob of N-in: 0.98724\nPOSSIBLE N-term signal sequence\ninside 1 11\nTMhelix 12 34\noutside 35 416\nTMhelix 417 439\ninside 440 445

30946 GCF_000738775.1_ASM73877v1 Rhodococcus defluvii Terrabacteria group; Actinobacteria; Actinobacteria; Corynebacteriales; Nocardiaceae; Rhodococcus MTASPSELTERGRVAAFFDLDKTVIAKSSALAFSKPFFAQGLITRRSVLQSSYAQFLFLL WP_031940583.1 inhibition of morphological differentiation protein [Rhodococcus defluvii] Length: 272\nNumber of predicted TMHs: 1\nExp number of AAs in TMHs: 21.74784\nExp number, first 60 AAs: 0.03075\nTotal prob of N-in: 0.00696\noutside 1 244\nTMhelix 245 267\ninside 268 272

30947 GCF_900094765.1_IMG-taxon_2615840625_annotated_assembly Rhodococcus enclensis Terrabacteria group; Actinobacteria; Actinobacteria; Corynebacteriales; Nocardiaceae; Rhodococcus MSNPLGQSSNGSRRGALPTPPSGWPIGSYPTYAEAQRAVDYLSDQEFPVQDVTIVGVNLM WP_003945065.1 MULTISPECIES: hypothetical protein [Rhodococcus] Length: 164\nNumber of predicted TMHs: 2\nExp number of AAs in TMHs: 45.26265\nExp number, first 60 AAs: 0.0547\nTotal prob of N-in: 0.87268\ninside 1 73\nTMhelix 74 96\noutside 97 99\nTMhelix 100 122\ninside 123 164

30948 GCF_900094765.1_IMG-taxon_2615840625_annotated_assembly Rhodococcus enclensis Terrabacteria group; Actinobacteria; Actinobacteria; Corynebacteriales; Nocardiaceae; Rhodococcus MTDKPANGLNTGRTAAFFDLDKTIIAKSSTLAFSKPFFAQGLINRRSVLKSSYAQFLFLL WP_003946123.1 MULTISPECIES: haloacid dehalogenase [Rhodococcus] Length: 272\nNumber of predicted TMHs: 1\nExp number of AAs in TMHs: 22.87312\nExp number, first 60 AAs: 0.01889\nTotal prob of N-in: 0.03448\noutside 1 245\nTMhelix 246 268\ninside 269 272

30949 GCF_900094765.1_IMG-taxon_2615840625_annotated_assembly Rhodococcus enclensis Terrabacteria group; Actinobacteria; Actinobacteria; Corynebacteriales; Nocardiaceae; Rhodococcus MNTAKQRRALVTISRTGAMTTVAALAGSIASGRPENISWFVALRKPAFQPPPVVFPVVWT WP_058228178.1 MULTISPECIES: TspO protein [Rhodococcus] Length: 169\nNumber of predicted TMHs: 1\nExp number of AAs in TMHs: 81.87072\nExp number, first 60 AAs: 21.42694\nTotal prob of N-in: 0.96201\nPOSSIBLE N-term signal sequence\ninside 1 140\nTMhelix 141 163\noutside 164 169

30950 GCF_001456965.1_ASM145696v1 Rhodococcus enclensis Terrabacteria group; Actinobacteria; Actinobacteria; Corynebacteriales; Nocardiaceae; Rhodococcus MSNPLGQSSNGSRRGALPTPPSGWPIGSYPTYAEAQRAVDYLSDQEFPVQDVTIVGVNLM WP_003945065.1 MULTISPECIES: hypothetical protein [Rhodococcus] Length: 164\nNumber of predicted TMHs: 2\nExp number of AAs in TMHs: 45.26265\nExp number, first 60 AAs: 0.0547\nTotal prob of N-in: 0.87268\ninside 1 73\nTMhelix 74 96\noutside 97 99\nTMhelix 100 122\ninside 123 164

30951 GCF_001456965.1_ASM145696v1 Rhodococcus enclensis Terrabacteria group; Actinobacteria; Actinobacteria; Corynebacteriales; Nocardiaceae; Rhodococcus MTDKPANGLNTGRTAAFFDLDKTIIAKSSTLAFSKPFFAQGLINRRSVLKSSYAQFLFLL WP_003946123.1 MULTISPECIES: haloacid dehalogenase [Rhodococcus] Length: 272\nNumber of predicted TMHs: 1\nExp number of AAs in TMHs: 22.87312\nExp number, first 60 AAs: 0.01889\nTotal prob of N-in: 0.03448\noutside 1 245\nTMhelix 246 268\ninside 269 272

30952 GCF_001456965.1_ASM145696v1 Rhodococcus enclensis Terrabacteria group; Actinobacteria; Actinobacteria; Corynebacteriales; Nocardiaceae; Rhodococcus MNTAKQRRALVTISRTGAMTTVAALAGSIASGRPENISWFVALRKPAFQPPPVVFPVVWT WP_058228178.1 MULTISPECIES: TspO protein [Rhodococcus] Length: 169\nNumber of predicted TMHs: 1\nExp number of AAs in TMHs: 81.87072\nExp number, first 60 AAs: 21.42694\nTotal prob of N-in: 0.96201\nPOSSIBLE N-term signal sequence\ninside 1 140\nTMhelix 141 163\noutside 164 169

30953 GCF_001766885.1_ASM176688v1 Rhodococcus erythropolis Terrabacteria group; Actinobacteria; Actinobacteria; Corynebacteriales; Nocardiaceae; Rhodococcus MTDKPANGLNTGRTAAFFDLDKTIIAKSSTLAFSKPFFAQGLINRRSVLKSSYAQFLFLL WP_019746641.1 MULTISPECIES: inhibition of morphological differentiation protein [Rhodococcus] Length: 272\nNumber of predicted TMHs: 1\nExp number of AAs in TMHs: 22.87312\nExp number, first 60 AAs: 0.01889\nTotal prob of N-in: 0.03448\noutside 1 245\nTMhelix 246 268\ninside 269 272

30954 GCF_001900745.1_ASM190074v1 Rhodococcus erythropolis Terrabacteria group; Actinobacteria; Actinobacteria; Corynebacteriales; Nocardiaceae; Rhodococcus

MSNPLGQSSNGSRRGALPTPPSGWPIGSYPTYAEAQRAVDYLSDQEFPVQDVTIVGVNLM WP_003945065.1 MULTISPECIES: hypothetical protein [Rhodococcus] Length: 164\nNumber of predicted TMHs: 2\nExp number of AAs in TMHs: 45.26265\nExp number, first 60 AAs: 0.0547\nTotal prob of N-in: 0.87268\ninside 1 73\nTMhelix 74 96\noutside 97 99\nTMhelix 100 122\ninside 123 164

30955 GCF_001766885.1_ASM176688v1 Rhodococcus erythropolis Terrabacteria group; Actinobacteria; Actinobacteria; Corynebacteriales; Nocardiaceae; Rhodococcus MSNPLGQSSNGSRRGALPTPPSGWPIGSYPTYAEAQRAVDYLSDQEFPVQDVTIVGVNLM WP_003945065.1 MULTISPECIES: hypothetical protein [Rhodococcus] Length: 164\nNumber of predicted TMHs: 2\nExp number of AAs in TMHs: 45.26265\nExp number, first 60 AAs: 0.0547\nTotal prob of N-in: 0.87268\ninside 1 73\nTMhelix 74 96\noutside 97 99\nTMhelix 100 122\ninside 123 164

30956 GCF_000719985.1_ASM71998v1 Rhodococcus erythropolis Terrabacteria group; Actinobacteria; Actinobacteria; Corynebacteriales; Nocardiaceae; Rhodococcus MTDKPANGLNTGRTAAFFDLDKTIIAKSSTLAFSKPFFAQGLINRRSVLKSSYAQFLFLL WP_030535674.1 inhibition of morphological differentiation protein [Rhodococcus erythropolis] Length: 272\nNumber of predicted TMHs: 1\nExp number of AAs in TMHs: 22.86896\nExp number, first 60 AAs: 0.01898\nTotal prob of N-in: 0.03508\noutside 1 245\nTMhelix 246 268\ninside 269 272

30957 GCF_000975175.1_ASM97517v1 Rhodococcus erythropolis Terrabacteria group; Actinobacteria; Actinobacteria; Corynebacteriales; Nocardiaceae; Rhodococcus MTDKPANGLNTGRTAAFFDLDKTIIAKSSTLAFSKPFFAQGLINRRSVLKSSYAQFLFLL WP_046377845.1 inhibition of morphological differentiation protein [Rhodococcus erythropolis] Length: 272\nNumber of predicted TMHs: 1\nExp number of AAs in TMHs: 22.87381\nExp number, first 60 AAs: 0.01889\nTotal prob of N-in: 0.03451\noutside 1 245\nTMhelix 246 268\ninside 269 272

30958 GCF_001900745.1_ASM190074v1 Rhodococcus erythropolis Terrabacteria group; Actinobacteria; Actinobacteria; Corynebacteriales; Nocardiaceae; Rhodococcus MTDKPANGLKTGRTAAFFDLDKTIIAKSSTLAFSKPFFAQGLINRRSVLKSSYAQFLFLL WP_073513331.1 inhibition of morphological differentiation protein [Rhodococcus erythropolis] Length: 272\nNumber of predicted TMHs: 1\nExp number of AAs in TMHs: 22.88712\nExp number, first 60 AAs: 0.01938\nTotal prob of N-in: 0.04124\noutside 1 245\nTMhelix 246 268\ninside 269 272

30959 GCF_000747745.1_ASM74774v1 Rhodococcus erythropolis Terrabacteria group; Actinobacteria; Actinobacteria; Corynebacteriales; Nocardiaceae; Rhodococcus MTDKPANGLNTGRTAAFFDLDKTIIAKSSTLAFSKPFFAQGLINRRSVLKSSYAQFLFLL WP_003946123.1 MULTISPECIES: haloacid dehalogenase [Rhodococcus] Length: 272\nNumber of predicted TMHs: 1\nExp number of AAs in TMHs: 22.87312\nExp number, first 60 AAs: 0.01889\nTotal prob of N-in: 0.03448\noutside 1 245\nTMhelix 246 268\ninside 269 272

30960 GCF_000747745.1_ASM74774v1 Rhodococcus erythropolis Terrabacteria group; Actinobacteria; Actinobacteria; Corynebacteriales; Nocardiaceae; Rhodococcus MSNPLGQSSNGSRRGALPTPPSGWPIGSYPTYAEAQRAVDYLSDQEFPVQDVTIVGVNLM WP_003945065.1 MULTISPECIES: hypothetical protein [Rhodococcus] Length: 164\nNumber of predicted TMHs: 2\nExp number of AAs in TMHs: 45.26265\nExp number, first 60 AAs: 0.0547\nTotal prob of N-in: 0.87268\ninside 1 73\nTMhelix 74 96\noutside 97 99\nTMhelix 100 122\ninside 123 164

30961 GCF_000975175.1_ASM97517v1 Rhodococcus erythropolis Terrabacteria group; Actinobacteria; Actinobacteria; Corynebacteriales; Nocardiaceae; Rhodococcus MSNPLGQSSNGSRRGALPTPPSGWPIGSYPTYAEAQRAVDYLSDQEFPVQDVTIVGVNLM WP_003945065.1 MULTISPECIES: hypothetical protein [Rhodococcus] Length: 164\nNumber of predicted TMHs: 2\nExp number of AAs in TMHs: 45.26265\nExp number, first 60 AAs: 0.0547\nTotal prob of N-in: 0.87268\ninside 1 73\nTMhelix 74 96\noutside 97 99\nTMhelix 100 122\ninside 123 164

30962 GCF_001020225.1_ASM102022v1 Rhodococcus erythropolis Terrabacteria group; Actinobacteria; Actinobacteria; Corynebacteriales; Nocardiaceae; Rhodococcus MSNPLGQSSNGSRRGALPTPPSGWPIGSYPTYAEAQRAVDYLSDQEFPVQDVTIVGVNLM WP_003945065.1 MULTISPECIES: hypothetical protein [Rhodococcus] Length: 164\nNumber of predicted TMHs: 2\nExp number of AAs in TMHs: 45.26265\nExp number, first 60 AAs: 0.0547\nTotal prob of N-in: 0.87268\ninside 1 73\nTMhelix 74 96\noutside 97 99\nTMhelix 100 122\ninside 123 164

30963 GCF_001020225.1_ASM102022v1 Rhodococcus erythropolis Terrabacteria group; Actinobacteria; Actinobacteria; Corynebacteriales; Nocardiaceae; Rhodococcus
MTDKPANGLNTGRTAAFFDLDKTIIAKSSTLAFSKPFFAQGLINRRSVLKSSYAQFLFLL WP_003946123.1 MULTISPECIES: haloacid dehalogenase [Rhodococcus] Length: 272\nNumber of predicted TMHs: 1\nExp number of AAs in TMHs: 22.87312\nExp number, first 60 AAs: 0.01889\nTotal prob of N-in: 0.03448\noutside 1 245\nTMhelix 246 268\ninside 269 272

30964 GCF_000719985.1_ASM71998v1 Rhodococcus erythropolis Terrabacteria group; Actinobacteria; Actinobacteria; Corynebacteriales; Nocardiaceae; Rhodococcus
MSNPLGQSSNGSRRGALPTPPSGWPIGSYPTYAEAQRAVDYLSDQEFPVQDVTIVGVNLM WP_003945065.1
MULTISPECIES: hypothetical protein [Rhodococcus] Length: 164\nNumber of predicted TMHs: 2\nExp number of AAs in TMHs: 45.26265\nExp number, first 60 AAs: 0.0547\nTotal prob of N-in: 0.87268\ninside 1 73\nTMhelix 74 96\noutside 97 99\nTMhelix 100 122\ninside 123 164

30965 GCF_001715845.1_ASM171584v1 Rhodococcus erythropolis Terrabacteria group; Actinobacteria; Actinobacteria; Corynebacteriales; Nocardiaceae; Rhodococcus
MTDKPANGLNTGRTAAFFDLDKTIIAKSSTLAFSKPFFAQGLINRRSVLKSSYAQFLFLL WP_019746641.1 MULTISPECIES: inhibition of morphological differentiation protein [Rhodococcus] Length: 272\nNumber of predicted TMHs: 1\nExp number of AAs in TMHs: 22.87312\nExp number, first 60 AAs: 0.01889\nTotal prob of N-in: 0.03448\noutside 1 245\nTMhelix 246 268\ninside 269 272

30966 GCF_001831305.1_ASM183130v1 Rhodococcus erythropolis Terrabacteria group; Actinobacteria; Actinobacteria; Corynebacteriales; Nocardiaceae; Rhodococcus
MSNPLGQSSNGSRRGALPTPPSGWPIGSYPTYAEAQRAVDYLSDQEFPVQDVTIVGVNLM WP_003945065.1
MULTISPECIES: hypothetical protein [Rhodococcus] Length: 164\nNumber of predicted TMHs: 2\nExp number of AAs in TMHs: 45.26265\nExp number, first 60 AAs: 0.0547\nTotal prob of N-in: 0.87268\ninside 1 73\nTMhelix 74 96\noutside 97 99\nTMhelix 100 122\ninside 123 164

30967 GCF_001715845.1_ASM171584v1 Rhodococcus erythropolis Terrabacteria group; Actinobacteria; Actinobacteria; Corynebacteriales; Nocardiaceae; Rhodococcus
MSNPLGQSSNGSRRGALPTPPSGWPIGSYPTYAEAQRAVDYLSDQEFPVQDVTIVGVNLM WP_003945065.1
MULTISPECIES: hypothetical protein [Rhodococcus] Length: 164\nNumber of predicted TMHs: 2\nExp number of AAs in TMHs: 45.26265\nExp number, first 60 AAs: 0.0547\nTotal prob of N-in: 0.87268\ninside 1 73\nTMhelix 74 96\noutside 97 99\nTMhelix 100 122\ninside 123 164

30968 GCF_001831305.1_ASM183130v1 Rhodococcus erythropolis Terrabacteria group; Actinobacteria; Actinobacteria; Corynebacteriales; Nocardiaceae; Rhodococcus
MTDKPANGLNTGRTAAFFDLDKTIIAKSSTLAFSKPFFAQGLINRRSVLKSSYAQFLFLL WP_019746641.1 MULTISPECIES: inhibition of morphological differentiation protein [Rhodococcus] Length: 272\nNumber of predicted TMHs: 1\nExp number of AAs in TMHs: 22.87312\nExp number, first 60 AAs: 0.01889\nTotal prob of N-in: 0.03448\noutside 1 245\nTMhelix 246 268\ninside 269 272

30969 GCF_001037935.1_RhoFas1.0 Rhodococcus fascians Terrabacteria group; Actinobacteria; Actinobacteria; Corynebacteriales; Nocardiaceae; Rhodococcus
MQSTSRTSARTRRRRVLSVLALTLLVAPMLAGCLRVQVTMGVSANDRVAGQIVAATVPTS WP_045841254.1
MULTISPECIES: hypothetical protein [Rhodococcus] Length: 244\nNumber of predicted TMHs: 2\nExp number of AAs in TMHs: 45.38746\nExp number, first 60 AAs: 22.41066\nTotal prob of N-in: 0.99116\nPOSSIBLE N-term signal sequence\ninside 1 15\nTMhelix 16 38\noutside 39 197\nTMhelix 198 220\ninside 221 244

30970 GCF_001646655.1_ASM164665v1 Rhodococcus gordoniae Terrabacteria group; Actinobacteria; Actinobacteria; Corynebacteriales; Nocardiaceae; Rhodococcus
MFEFSRRALVGATTTAALMVVGAGIASAHVTVVAPGAEQGGYSVLTFRVPTESETAGTTA WP_064064187.1
nuclear export factor GLE1 [Rhodococcus gordoniae] Length: 225\nNumber of predicted TMHs: 1\nExp number of AAs in TMHs: 39.46989\nExp number, first 60 AAs: 17.37019\nTotal prob of N-in: 0.66727\nPOSSIBLE N-term signal sequence\noutside 1 197\nTMhelix 198 220\ninside 221 225

30971 GCF_002095295.1_ASM209529v1 Rhodococcus hoagii Terrabacteria group; Actinobacteria; Actinobacteria; Corynebacteriales; Nocardiaceae; Rhodococcus
MTADPSDLTRRGRTAAFFDLDKTIIAKSSALAFSKPFFAQGLITRRSVLQSSYAQFLFIL WP_005515815.1 morphological differentiation-associated protein [Rhodococcus hoagii] Length: 272\nNumber of predicted TMHs: 1\nExp number of

AAs in TMHs: 21.05758\nExp number, first 60 AAs: 0.04208\nTotal prob of N-in: 0.02567\noutside 1 244\nTMhelix 245 267\ninside 268 272

30972 GCF_002095195.1_ASM209519v1 Rhodococcus hoagii Terrabacteria group; Actinobacteria; Actinobacteria; Corynebacteriales; Nocardiaceae; Rhodococcus MAAHPKSARRRILLTATASALVLGAGGAPAAAVPLAGQSPLAADQAVAPTTTPPFSTPNT WP_022597221.1 D-alanyl-D-alanine carboxypeptidase [Rhodococcus hoagii] Length: 444\nNumber of predicted TMHs: 2\nExp number of AAs in TMHs: 42.86088\nExp number, first 60 AAs: 21.57924\nTotal prob of N-in: 0.96909\nPOSSIBLE N-term signal sequence\ninside 1 11\nTMhelix 12 34\noutside 35 412\nTMhelix 413 435\ninside 436 444

30973 GCF_002094235.1_ASM209423v1 Rhodococcus hoagii Terrabacteria group; Actinobacteria; Actinobacteria; Corynebacteriales; Nocardiaceae; Rhodococcus MTADPSDLTRRGRTAAFFDLDKTIIAKSSALAFSKPFFAQGLITRRSVLQSSYAQFLFIL WP_005515815.1 morphological differentiation-associated protein [Rhodococcus hoagii] Length: 272\nNumber of predicted TMHs: 1\nExp number of AAs in TMHs: 21.05758\nExp number, first 60 AAs: 0.04208\nTotal prob of N-in: 0.02567\noutside 1 244\nTMhelix 245 267\ninside 268 272

30974 GCF_002078625.1_ASM207862v1 Rhodococcus hoagii Terrabacteria group; Actinobacteria; Actinobacteria; Corynebacteriales; Nocardiaceae; Rhodococcus MTADPSDLTRRGRTAAFFDLDKTIIAKSSALAFSKPFFAQGLITRRSVLQSSYAQFLFIL WP_005515815.1 morphological differentiation-associated protein [Rhodococcus hoagii] Length: 272\nNumber of predicted TMHs: 1\nExp number of AAs in TMHs: 21.05758\nExp number, first 60 AAs: 0.04208\nTotal prob of N-in: 0.02567\noutside 1 244\nTMhelix 245 267\ninside 268 272

30975 GCF_001646905.1_re12 Rhodococcus hoagii Terrabacteria group; Actinobacteria; Actinobacteria; Corynebacteriales; Nocardiaceae; Rhodococcus MTADPSDLTRRGRTAAFFDLDKTIIAKSSALAFSKPFFAQGLITRRSVLQSSYAQFLFIL WP_005515815.1 morphological differentiation-associated protein [Rhodococcus hoagii] Length: 272\nNumber of predicted TMHs: 1\nExp number of AAs in TMHs: 21.05758\nExp number, first 60 AAs: 0.04208\nTotal prob of N-in: 0.02567\noutside 1 244\nTMhelix 245 267\ninside 268 272

30976 GCF_002094445.1_ASM209444v1 Rhodococcus hoagii Terrabacteria group; Actinobacteria; Actinobacteria; Corynebacteriales; Nocardiaceae; Rhodococcus MTADPSDLTRRGRTAAFFDLDKTIIAKSSALAFSKPFFAQGLITRRSVLQSSYAQFLFIL WP_005515815.1 morphological differentiation-associated protein [Rhodococcus hoagii] Length: 272\nNumber of predicted TMHs: 1\nExp number of AAs in TMHs: 21.05758\nExp number, first 60 AAs: 0.04208\nTotal prob of N-in: 0.02567\noutside 1 244\nTMhelix 245 267\ninside 268 272

30977 GCF_002095045.1_ASM209504v1 Rhodococcus hoagii Terrabacteria group; Actinobacteria; Actinobacteria; Corynebacteriales; Nocardiaceae; Rhodococcus MTADPSDLTRRGRTAAFFDLDKTIIAKSSALAFSKPFFAQGLITRRSVLQSSYAQFLFIL WP_005515815.1 morphological differentiation-associated protein [Rhodococcus hoagii] Length: 272\nNumber of predicted TMHs: 1\nExp number of AAs in TMHs: 21.05758\nExp number, first 60 AAs: 0.04208\nTotal prob of N-in: 0.02567\noutside 1 244\nTMhelix 245 267\ninside 268 272

30978 GCF_002094445.1_ASM209444v1 Rhodococcus hoagii Terrabacteria group; Actinobacteria; Actinobacteria; Corynebacteriales; Nocardiaceae; Rhodococcus MAAHPKSARRRILLTATASALVLGAGGAPAAAVPLAGQSPLAADQAVAPTTTPPFSTPNT WP_069856650.1 MULTISPECIES: D-alanyl-D-alanine carboxypeptidase [Rhodococcus] Length: 444\nNumber of predicted TMHs: 2\nExp number of AAs in TMHs: 42.7857099999999\nExp number, first 60 AAs: 21.57856\nTotal prob of N-in: 0.96909\nPOSSIBLE N-term signal sequence\ninside 1 11\nTMhelix 12 34\noutside 35 412\nTMhelix 413 435\ninside 436 444

30979 GCF_002095045.1_ASM209504v1 Rhodococcus hoagii Terrabacteria group; Actinobacteria; Actinobacteria; Corynebacteriales; Nocardiaceae; Rhodococcus MAAHPKSARRRILLTATASALVLGAGGAPAAAVPLAGQSPLAADQAVAPTTTPPFSTPNT WP_022597221.1 D-alanyl-D-alanine carboxypeptidase [Rhodococcus hoagii] Length: 444\nNumber of predicted TMHs: 2\nExp number of AAs in TMHs: 42.86088\nExp number, first 60 AAs: 21.57924\nTotal prob of N-in: 0.96909\nPOSSIBLE N-term signal sequence\ninside 1 11\nTMhelix 12 34\noutside 35 412\nTMhelix 413 435\ninside 436 444

30980	GCF_002078625.1_ASM207862v1	Rhodococcus hoagii	Terrabacteria group; Actinobacteria; Actinobacteria; Corynebacteriales; Nocardiaceae; Rhodococcus
MAAHPKSARRRILLTATASALVLGAGGAPAAAVPLAGQSPLAADQAVAPTTTPPFSTPNT	WP_005518562.1	D-alanyl-D-alanine carboxypeptidase [Rhodococcus hoagii]	Length: 444\nNumber of predicted TMHs: 2\nExp number of AAs in TMHs: 42.7856999999999\nExp number, first 60 AAs: 21.57856\nTotal prob of N-in: 0.96909\nPOSSIBLE N-term signal sequence\ninside 1 11\nTMhelix 12 34\noutside 35 412\nTMhelix 413 435\ninside 436 444

30981	GCF_002095175.1_ASM209517v1	Rhodococcus hoagii	Terrabacteria group; Actinobacteria; Actinobacteria; Corynebacteriales; Nocardiaceae; Rhodococcus
MAAHPKSARRRILLTATASALVLGAGGAPAAAVPLAGQSPLAADQAVAPTTTPPFSTPNT	WP_084866645.1	penicillin-binding protein [Rhodococcus hoagii]	Length: 444\nNumber of predicted TMHs: 2\nExp number of AAs in TMHs: 42.7847\nExp number, first 60 AAs: 21.58077\nTotal prob of N-in: 0.96910\nPOSSIBLE N-term signal sequence\ninside 1 11\nTMhelix 12 34\noutside 35 412\nTMhelix 413 435\ninside 436 444

30982	GCF_002078535.1_ASM207853v1	Rhodococcus hoagii	Terrabacteria group; Actinobacteria; Actinobacteria; Corynebacteriales; Nocardiaceae; Rhodococcus
MAAHPKSARRRILLTATASALVLGAGGAPAAAVPLAGQSPLAADQAVAPTTTPPFSTPNT	WP_081186880.1	penicillin-binding protein [Rhodococcus hoagii]	Length: 444\nNumber of predicted TMHs: 2\nExp number of AAs in TMHs: 42.77886\nExp number, first 60 AAs: 21.57747\nTotal prob of N-in: 0.96908\nPOSSIBLE N-term signal sequence\ninside 1 11\nTMhelix 12 34\noutside 35 412\nTMhelix 413 435\ninside 436 444

30983	GCF_002094235.1_ASM209423v1	Rhodococcus hoagii	Terrabacteria group; Actinobacteria; Actinobacteria; Corynebacteriales; Nocardiaceae; Rhodococcus
MAAHPKSARRRILLTATASALVLGAGGAPAAAVPLAGQSPLAADQAVAPTTTPPFSTPNT	WP_005518562.1	D-alanyl-D-alanine carboxypeptidase [Rhodococcus hoagii]	Length: 444\nNumber of predicted TMHs: 2\nExp number of AAs in TMHs: 42.7856999999999\nExp number, first 60 AAs: 21.57856\nTotal prob of N-in: 0.96909\nPOSSIBLE N-term signal sequence\ninside 1 11\nTMhelix 12 34\noutside 35 412\nTMhelix 413 435\ninside 436 444

30984	GCF_002078535.1_ASM207853v1	Rhodococcus hoagii	Terrabacteria group; Actinobacteria; Actinobacteria; Corynebacteriales; Nocardiaceae; Rhodococcus
MTADPSDLTRRGRTAAFFDLDKTIIAKSSALAFSKPFFAQGLITRRSVLQSSYAQFLFIL	WP_005515815.1	morphological differentiation-associated protein [Rhodococcus hoagii]	Length: 272\nNumber of predicted TMHs: 1\nExp number of AAs in TMHs: 21.05758\nExp number, first 60 AAs: 0.04208\nTotal prob of N-in: 0.02567\noutside 1 244\nTMhelix 245 267\ninside 268 272

30985	GCF_002094265.1_ASM209426v1	Rhodococcus hoagii	Terrabacteria group; Actinobacteria; Actinobacteria; Corynebacteriales; Nocardiaceae; Rhodococcus
MAAHPKSARRRILLTATASALVLGAGGAPAAAVPLAGQSPLAADQAVAPTTTPPFSTPNT	WP_084843816.1	penicillin-binding protein [Rhodococcus hoagii]	Length: 444\nNumber of predicted TMHs: 2\nExp number of AAs in TMHs: 42.7870899999999\nExp number, first 60 AAs: 21.57571\nTotal prob of N-in: 0.96906\nPOSSIBLE N-term signal sequence\ninside 1 11\nTMhelix 12 34\noutside 35 412\nTMhelix 413 435\ninside 436 444

30986	GCF_002095195.1_ASM209519v1	Rhodococcus hoagii	Terrabacteria group; Actinobacteria; Actinobacteria; Corynebacteriales; Nocardiaceae; Rhodococcus
MTADPSDLTRRGRTAAFFDLDKTIIAKSSALAFSKPFFAQGLITRRSVLQSSYAQFLFIL	WP_005515815.1	morphological differentiation-associated protein [Rhodococcus hoagii]	Length: 272\nNumber of predicted TMHs: 1\nExp number of AAs in TMHs: 21.05758\nExp number, first 60 AAs: 0.04208\nTotal prob of N-in: 0.02567\noutside 1 244\nTMhelix 245 267\ninside 268 272

30987	GCF_002094305.1_ASM209430v1	Rhodococcus hoagii	Terrabacteria group; Actinobacteria; Actinobacteria; Corynebacteriales; Nocardiaceae; Rhodococcus
MTADPSDLTRRGRTAAFFDLDKTIIAKSSALAFSKPFFAQGLITRRSVLQSSYAQFLFIL	WP_005515815.1	morphological differentiation-associated protein [Rhodococcus hoagii]	Length: 272\nNumber of predicted TMHs: 1\nExp number of AAs in TMHs: 21.05758\nExp number, first 60 AAs: 0.04208\nTotal prob of N-in: 0.02567\noutside 1 244\nTMhelix 245 267\ninside 268 272

30988	GCF_002094325.1_ASM209432v1	Rhodococcus hoagii	Terrabacteria group; Actinobacteria; Actinobacteria; Corynebacteriales; Nocardiaceae; Rhodococcus
MTADPSDLTRRGRTAAFFDLDKTIIAKSSALAFSKPFFAQGLITRRSVLQSSYAQFLFIL	WP_005515815.1	morphological differentiation-associated protein [Rhodococcus hoagii]	Length: 272\nNumber of predicted TMHs: 1\nExp number of

AAs in TMHs: 21.05758\nExp number, first 60 AAs: 0.04208\nTotal prob of N-in: 0.02567\noutside 1 244\nTMhelix 245 267\ninside 268 272

30989 GCF_002094325.1_ASM209432v1 Rhodococcus hoagii Terrabacteria group; Actinobacteria; Actinobacteria; Corynebacteriales; Nocardiaceae; Rhodococcus MAAHPKSARRRILLTATASALVLGAGGAPAAAVPLAGQSPLAADQAVAPTTTPPFSTPNT WP_013416807.1 D-alanyl-D-alanine carboxypeptidase [Rhodococcus hoagii] Length: 444\nNumber of predicted TMHs: 2\nExp number of AAs in TMHs: 42.7856499999999\nExp number, first 60 AAs: 21.57858\nTotal prob of N-in: 0.96909\nPOSSIBLE N-term signal sequence\ninside 1 11\nTMhelix 12 34\noutside 35 412\nTMhelix 413 435\ninside 436 444

30990 GCF_002094225.1_ASM209422v1 Rhodococcus hoagii Terrabacteria group; Actinobacteria; Actinobacteria; Corynebacteriales; Nocardiaceae; Rhodococcus MAAHPKSARRRILLTATASALVLGAGGAPAAAVPLAGQSPPAADQAVAPTTTPPFSTPNT WP_084922917.1 penicillin-binding protein [Rhodococcus hoagii] Length: 444\nNumber of predicted TMHs: 2\nExp number of AAs in TMHs: 42.41228\nExp number, first 60 AAs: 21.20376\nTotal prob of N-in: 0.95472\nPOSSIBLE N-term signal sequence\ninside 1 11\nTMhelix 12 34\noutside 35 412\nTMhelix 413 435\ninside 436 444

30991 GCF_002095035.1_ASM209503v1 Rhodococcus hoagii Terrabacteria group; Actinobacteria; Actinobacteria; Corynebacteriales; Nocardiaceae; Rhodococcus MTADPSDLTRRGRTAAFFDLDKTIIAKSSALAFSKPFFAQGLITRRSVLQSSYAQFLFIL WP_005515815.1 morphological differentiation-associated protein [Rhodococcus hoagii] Length: 272\nNumber of predicted TMHs: 1\nExp number of AAs in TMHs: 21.05758\nExp number, first 60 AAs: 0.04208\nTotal prob of N-in: 0.02567\noutside 1 244\nTMhelix 245 267\ninside 268 272

30992 GCF_002094395.1_ASM209439v1 Rhodococcus hoagii Terrabacteria group; Actinobacteria; Actinobacteria; Corynebacteriales; Nocardiaceae; Rhodococcus MTADPSDLTRRGRTAAFFDLDKTIIAKSSALAFSKPFFAQGLITRRSVLQSSYAQFLFIL WP_005515815.1 morphological differentiation-associated protein [Rhodococcus hoagii] Length: 272\nNumber of predicted TMHs: 1\nExp number of AAs in TMHs: 21.05758\nExp number, first 60 AAs: 0.04208\nTotal prob of N-in: 0.02567\noutside 1 244\nTMhelix 245 267\ninside 268 272

30993 GCF_002095235.1_ASM209523v1 Rhodococcus hoagii Terrabacteria group; Actinobacteria; Actinobacteria; Corynebacteriales; Nocardiaceae; Rhodococcus MTADPSDLTRRGRTAAFFDLDKTIIAKSSALAFSKPFFAQGLITRRSVLQSSYAQFLFIL WP_005515815.1 morphological differentiation-associated protein [Rhodococcus hoagii] Length: 272\nNumber of predicted TMHs: 1\nExp number of AAs in TMHs: 21.05758\nExp number, first 60 AAs: 0.04208\nTotal prob of N-in: 0.02567\noutside 1 244\nTMhelix 245 267\ninside 268 272

30994 GCF_002095235.1_ASM209523v1 Rhodococcus hoagii Terrabacteria group; Actinobacteria; Actinobacteria; Corynebacteriales; Nocardiaceae; Rhodococcus MAAHPKSARRRILLTATASALVLGAGGAPAAAVPLAGQSPPAADQAVAPTTTPPFSTPNT WP_084869160.1 penicillin-binding protein [Rhodococcus hoagii] Length: 445\nNumber of predicted TMHs: 2\nExp number of AAs in TMHs: 42.56153\nExp number, first 60 AAs: 21.20564\nTotal prob of N-in: 0.95473\nPOSSIBLE N-term signal sequence\ninside 1 11\nTMhelix 12 34\noutside 35 412\nTMhelix 413 435\ninside 436 445

30995 GCF_002095035.1_ASM209503v1 Rhodococcus hoagii Terrabacteria group; Actinobacteria; Actinobacteria; Corynebacteriales; Nocardiaceae; Rhodococcus MAAHPKSARRRILLTATASALVLGAGGAPAAAVPLAGQSPLAADQAVAPTTTPPFSTPNT WP_069856650.1 MULTISPECIES: D-alanyl-D-alanine carboxypeptidase [Rhodococcus] Length: 444\nNumber of predicted TMHs: 2\nExp number of AAs in TMHs: 42.7857099999999\nExp number, first 60 AAs: 21.57856\nTotal prob of N-in: 0.96909\nPOSSIBLE N-term signal sequence\ninside 1 11\nTMhelix 12 34\noutside 35 412\nTMhelix 413 435\ninside 436 444

30996 GCF_002094395.1_ASM209439v1 Rhodococcus hoagii Terrabacteria group; Actinobacteria; Actinobacteria; Corynebacteriales; Nocardiaceae; Rhodococcus MAAHPKSARRRILLTATASALVLGAGGAPAAAVPLAGQSPLAADQAVAPTTTPPFTTPNT WP_084962208.1 penicillin-binding protein [Rhodococcus hoagii] Length: 444\nNumber of predicted TMHs: 2\nExp number of AAs in TMHs: 42.7856999999999\nExp number, first 60 AAs: 21.57856\nTotal prob of N-in: 0.96909\nPOSSIBLE N-term signal sequence\ninside 1 11\nTMhelix 12 34\noutside 35 412\nTMhelix 413 435\ninside 436 444

30997 GCF_002095255.1_ASM209525v1 Rhodococcus hoagii Terrabacteria group; Actinobacteria; Actinobacteria; Corynebacteriales; Nocardiaceae; Rhodococcus
MTADPSDLTRRGRTAAFFDLDKTIIAKSSALAFSKPFFAQGLITRRSVLQSSYAQFLFIL WP_005515815.1 morphological differentiation-associated protein [Rhodococcus hoagii] Length: 272\nNumber of predicted TMHs: 1\nExp number of AAs in TMHs: 21.05758\nExp number, first 60 AAs: 0.04208\nTotal prob of N-in: 0.02567\noutside 1 244\nTMhelix 245 267\ninside 268 272

30998 GCF_001646925.1_re13 Rhodococcus hoagii Terrabacteria group; Actinobacteria; Actinobacteria; Corynebacteriales; Nocardiaceae; Rhodococcus
MARHAAKQNRWARRAVVAGALPVAIVAASAGTANAQDTGSANVPALPELPQIEVPPLYNP WP_064075989.1 hypothetical protein [Rhodococcus hoagii] Length: 174\nNumber of predicted TMHs: 2\nExp number of AAs in TMHs: 60.04882\nExp number, first 60 AAs: 16.032\nTotal prob of N-in: 0.97818\nPOSSIBLE N-term signal sequence\ninside 1 101\nTMhelix 102 124\noutside 125 133\nTMhelix 134 156\ninside 157 174

30999 GCF_001646925.1_re13 Rhodococcus hoagii Terrabacteria group; Actinobacteria; Actinobacteria; Corynebacteriales; Nocardiaceae; Rhodococcus
MAAHPKSARRRILLTATASALVLGAGGAPAAAVPLAGQSPLAADQAVAPTTTPPFSTPNT WP_064076622.1 D-alanyl-D-alanine carboxypeptidase [Rhodococcus hoagii] Length: 444\nNumber of predicted TMHs: 2\nExp number of AAs in TMHs: 42.77887\nExp number, first 60 AAs: 21.57744\nTotal prob of N-in: 0.96908\nPOSSIBLE N-term signal sequence\ninside 1 11\nTMhelix 12 34\noutside 35 412\nTMhelix 413 435\ninside 436 444

31000 GCF_002094295.1_ASM209429v1 Rhodococcus hoagii Terrabacteria group; Actinobacteria; Actinobacteria; Corynebacteriales; Nocardiaceae; Rhodococcus
MAAHPKSARRRILLTATASALVLGAGGAPAAAVPLAGQSPLAADQAVAPTTTPPFSTPNT WP_013416807.1 D-alanyl-D-alanine carboxypeptidase [Rhodococcus hoagii] Length: 444\nNumber of predicted TMHs: 2\nExp number of AAs in TMHs: 42.7856499999999\nExp number, first 60 AAs: 21.57858\nTotal prob of N-in: 0.96909\nPOSSIBLE N-term signal sequence\ninside 1 11\nTMhelix 12 34\noutside 35 412\nTMhelix 413 435\ninside 436 444

31001 GCF_001646645.1_ASM164664v1 Rhodococcus hoagii Terrabacteria group; Actinobacteria; Actinobacteria; Corynebacteriales; Nocardiaceae; Rhodococcus
MAAHPKSARRRILLTATASALVLGAGGAPAAAVPLAGQSPLAADQAVAPTTTPPFSTPNT WP_005518562.1 D-alanyl-D-alanine carboxypeptidase [Rhodococcus hoagii] Length: 444\nNumber of predicted TMHs: 2\nExp number of AAs in TMHs: 42.7856999999999\nExp number, first 60 AAs: 21.57856\nTotal prob of N-in: 0.96909\nPOSSIBLE N-term signal sequence\ninside 1 11\nTMhelix 12 34\noutside 35 412\nTMhelix 413 435\ninside 436 444

31002 GCF_001646885.1_re11 Rhodococcus hoagii Terrabacteria group; Actinobacteria; Actinobacteria; Corynebacteriales; Nocardiaceae; Rhodococcus
MAAHPKSARRRILLTATASALVLGAGGAPAAAVPLAGQSPLAADQAVAPTTTPPFSTPNT WP_005518562.1 D-alanyl-D-alanine carboxypeptidase [Rhodococcus hoagii] Length: 444\nNumber of predicted TMHs: 2\nExp number of AAs in TMHs: 42.7856999999999\nExp number, first 60 AAs: 21.57856\nTotal prob of N-in: 0.96909\nPOSSIBLE N-term signal sequence\ninside 1 11\nTMhelix 12 34\noutside 35 412\nTMhelix 413 435\ninside 436 444

31003 GCF_002095125.1_ASM209512v1 Rhodococcus hoagii Terrabacteria group; Actinobacteria; Actinobacteria; Corynebacteriales; Nocardiaceae; Rhodococcus
MTADPSDLTRRGRTAAFFDLDKTIIAKSSALAFSKPFFAQGLITRRSVLQSSYAQFLFIL WP_005515815.1 morphological differentiation-associated protein [Rhodococcus hoagii] Length: 272\nNumber of predicted TMHs: 1\nExp number of AAs in TMHs: 21.05758\nExp number, first 60 AAs: 0.04208\nTotal prob of N-in: 0.02567\noutside 1 244\nTMhelix 245 267\ninside 268 272

31004 GCF_001646925.1_re13 Rhodococcus hoagii Terrabacteria group; Actinobacteria; Actinobacteria; Corynebacteriales; Nocardiaceae; Rhodococcus
MTADPSDLTRRGRTAAFFDLDKTIIAKSSALAFSKPFFAQGLITRRSVLQSSYAQFLFIL WP_005515815.1 morphological differentiation-associated protein [Rhodococcus hoagii] Length: 272\nNumber of predicted TMHs: 1\nExp number of AAs in TMHs: 21.05758\nExp number, first 60 AAs: 0.04208\nTotal prob of N-in: 0.02567\noutside 1 244\nTMhelix 245 267\ninside 268 272

31005 GCF_002094405.1_ASM209440v1 Rhodococcus hoagii Terrabacteria group; Actinobacteria; Actinobacteria; Corynebacteriales; Nocardiaceae; Rhodococcus
MTADPSDLTRRGRTAAFFDLDKTIIAKSSALAFSKPFFAQGLITRRSVLQSSYAQFLFIL WP_005515815.1 morphological differentiation-associated protein [Rhodococcus hoagii] Length: 272\nNumber of predicted TMHs: 1\nExp number of

AAs in TMHs: 21.05758\nExp number, first 60 AAs: 0.04208\nTotal prob of N-in: 0.02567\noutside 1 244\nTMhelix 245 267\ninside 268 272

31006 GCF_001646885.1_re11 Rhodococcus hoagii Terrabacteria group; Actinobacteria; Actinobacteria; Corynebacteriales; Nocardiaceae; Rhodococcus MTADPSDLTRRGRTAAFFDLDKTIIAKSSALAFSKPFFAQGLITRRSVLQSSYAQFLFIL WP_005515815.1 morphological differentiation-associated protein [Rhodococcus hoagii] Length: 272\nNumber of predicted TMHs: 1\nExp number of AAs in TMHs: 21.05758\nExp number, first 60 AAs: 0.04208\nTotal prob of N-in: 0.02567\noutside 1 244\nTMhelix 245 267\ninside 268 272

31007 GCF_002095085.1_ASM209508v1 Rhodococcus hoagii Terrabacteria group; Actinobacteria; Actinobacteria; Corynebacteriales; Nocardiaceae; Rhodococcus MAAHPKSARRRILLTATASALVLGAGGAPAAAVPLAGQSPLAADQAVAPTTTPPFSTPNT WP_005518562.1 D-alanyl-D-alanine carboxypeptidase [Rhodococcus hoagii] Length: 444\nNumber of predicted TMHs: 2\nExp number of AAs in TMHs: 42.7856999999999\nExp number, first 60 AAs: 21.57856\nTotal prob of N-in: 0.96909\nPOSSIBLE N-term signal sequence\ninside 1 11\nTMhelix 12 34\noutside 35 412\nTMhelix 413 435\ninside 436 444

31008 GCF_002095085.1_ASM209508v1 Rhodococcus hoagii Terrabacteria group; Actinobacteria; Actinobacteria; Corynebacteriales; Nocardiaceae; Rhodococcus MTADPSDLTRRGRTAAFFDLDKTIIAKSSALAFSKPFFAQGLITRRSVLQSSYAQFLFIL WP_005515815.1 morphological differentiation-associated protein [Rhodococcus hoagii] Length: 272\nNumber of predicted TMHs: 1\nExp number of AAs in TMHs: 21.05758\nExp number, first 60 AAs: 0.04208\nTotal prob of N-in: 0.02567\noutside 1 244\nTMhelix 245 267\ninside 268 272

31009 GCF_002078545.1_ASM207854v1 Rhodococcus hoagii Terrabacteria group; Actinobacteria; Actinobacteria; Corynebacteriales; Nocardiaceae; Rhodococcus MAAHPKSARRRILLTATASALVLGAGGAPAAAVPLAGQSPPAADQAVAPTTTPPFSTPNT WP_081206222.1 penicillin-binding protein [Rhodococcus hoagii] Length: 444\nNumber of predicted TMHs: 2\nExp number of AAs in TMHs: 42.41215\nExp number, first 60 AAs: 21.20376\nTotal prob of N-in: 0.95472\nPOSSIBLE N-term signal sequence\ninside 1 11\nTMhelix 12 34\noutside 35 412\nTMhelix 413 435\ninside 436 444

31010 GCF_002095125.1_ASM209512v1 Rhodococcus hoagii Terrabacteria group; Actinobacteria; Actinobacteria; Corynebacteriales; Nocardiaceae; Rhodococcus MAAHPKSARRRILLTATASALVLGAGGAPAAAVPLAGQSPLAADQAVAPTTTPPFSTPNT WP_005518562.1 D-alanyl-D-alanine carboxypeptidase [Rhodococcus hoagii] Length: 444\nNumber of predicted TMHs: 2\nExp number of AAs in TMHs: 42.7856999999999\nExp number, first 60 AAs: 21.57856\nTotal prob of N-in: 0.96909\nPOSSIBLE N-term signal sequence\ninside 1 11\nTMhelix 12 34\noutside 35 412\nTMhelix 413 435\ninside 436 444

31011 GCF_001646645.1_ASM164664v1 Rhodococcus hoagii Terrabacteria group; Actinobacteria; Actinobacteria; Corynebacteriales; Nocardiaceae; Rhodococcus MTADPSDLTRRGRTAAFFDLDKTIIAKSSALAFSKPFFAQGLITRRSVLQSSYAQFLFIL WP_005515815.1 morphological differentiation-associated protein [Rhodococcus hoagii] Length: 272\nNumber of predicted TMHs: 1\nExp number of AAs in TMHs: 21.05758\nExp number, first 60 AAs: 0.04208\nTotal prob of N-in: 0.02567\noutside 1 244\nTMhelix 245 267\ninside 268 272

31012 GCF_002094295.1_ASM209429v1 Rhodococcus hoagii Terrabacteria group; Actinobacteria; Actinobacteria; Corynebacteriales; Nocardiaceae; Rhodococcus MTADPSDLTRRGRTAAFFDLDKTIIAKSSALAFSKPFFAQGLITRRSVLQSSYAQFLFIL WP_005515815.1 morphological differentiation-associated protein [Rhodococcus hoagii] Length: 272\nNumber of predicted TMHs: 1\nExp number of AAs in TMHs: 21.05758\nExp number, first 60 AAs: 0.04208\nTotal prob of N-in: 0.02567\noutside 1 244\nTMhelix 245 267\ninside 268 272

31013 GCF_002095295.1_ASM209529v1 Rhodococcus hoagii Terrabacteria group; Actinobacteria; Actinobacteria; Corynebacteriales; Nocardiaceae; Rhodococcus MAAHPKSARRRILLTATASALVLGAGGAPAAAVPLAGQSPLAADQAVAPTTTPPFSTPNT WP_084987279.1 penicillin-binding protein [Rhodococcus hoagii] Length: 444\nNumber of predicted TMHs: 2\nExp number of AAs in TMHs: 42.77882\nExp number, first 60 AAs: 21.57738\nTotal prob of N-in: 0.96908\nPOSSIBLE N-term signal sequence\ninside 1 11\nTMhelix 12 34\noutside 35 412\nTMhelix 413 435\ninside 436 444

31014 GCF_002094375.1_ASM209437v1 Rhodococcus hoagii Terrabacteria group; Actinobacteria; Actinobacteria; Corynebacteriales; Nocardiaceae; Rhodococcus

MTADPSDLTRRGRTAAFFDLDKTIIAKSSALAFSKPFFAQGLITRRSVLQSSYAQFLFIL WP_005515815.1 morphological differentiation-associated protein [Rhodococcus hoagii] Length: 272\nNumber of predicted TMHs: 1\nExp number of AAs in TMHs: 21.05758\nExp number, first 60 AAs: 0.04208\nTotal prob of N-in: 0.02567\noutside 1 244\nTMhelix 245 267\ninside 268 272

31015 GCF_002094265.1_ASM209426v1 Rhodococcus hoagii Terrabacteria group; Actinobacteria; Actinobacteria; Corynebacteriales; Nocardiaceae; Rhodococcus MTADPSDLTRRGRTAAFFDLDKTIIAKSSALAFSKPFFAQGLITRRSVLQSSYAQFLFIL WP_005515815.1 morphological differentiation-associated protein [Rhodococcus hoagii] Length: 272\nNumber of predicted TMHs: 1\nExp number of AAs in TMHs: 21.05758\nExp number, first 60 AAs: 0.04208\nTotal prob of N-in: 0.02567\noutside 1 244\nTMhelix 245 267\ninside 268 272

31016 GCF_002094375.1_ASM209437v1 Rhodococcus hoagii Terrabacteria group; Actinobacteria; Actinobacteria; Corynebacteriales; Nocardiaceae; Rhodococcus MAAHPKSARRRILLTATASALVLGAGGAPAAAVPLAGQSPLAADQAVAPTTTPPFSTPNT WP_005518562.1 D-alanyl-D-alanine carboxypeptidase [Rhodococcus hoagii] Length: 444\nNumber of predicted TMHs: 2\nExp number of AAs in TMHs: 42.7856999999999\nExp number, first 60 AAs: 21.57856\nTotal prob of N-in: 0.96909\nPOSSIBLE N-term signal sequence\ninside 1 11\nTMhelix 12 34\noutside 35 412\nTMhelix 413 435\ninside 436 444

31017 GCF_002094315.1_ASM209431v1 Rhodococcus hoagii Terrabacteria group; Actinobacteria; Actinobacteria; Corynebacteriales; Nocardiaceae; Rhodococcus MAAHPKSARRRILLTATASALVLGAGGAPAAAVPLAGQSPLAADQAVAPTTTPPFSTPNT WP_005518562.1 D-alanyl-D-alanine carboxypeptidase [Rhodococcus hoagii] Length: 444\nNumber of predicted TMHs: 2\nExp number of AAs in TMHs: 42.7856999999999\nExp number, first 60 AAs: 21.57856\nTotal prob of N-in: 0.96909\nPOSSIBLE N-term signal sequence\ninside 1 11\nTMhelix 12 34\noutside 35 412\nTMhelix 413 435\ninside 436 444

31018 GCF_002095115.1_ASM209511v1 Rhodococcus hoagii Terrabacteria group; Actinobacteria; Actinobacteria; Corynebacteriales; Nocardiaceae; Rhodococcus MTADPSDLTRRGRTAAFFDLDKTIIAKSSALAFSKPFFAQGLITRRSVLQSSYAQFLFIL WP_005515815.1 morphological differentiation-associated protein [Rhodococcus hoagii] Length: 272\nNumber of predicted TMHs: 1\nExp number of AAs in TMHs: 21.05758\nExp number, first 60 AAs: 0.04208\nTotal prob of N-in: 0.02567\noutside 1 244\nTMhelix 245 267\ninside 268 272

31019 GCF_002095115.1_ASM209511v1 Rhodococcus hoagii Terrabacteria group; Actinobacteria; Actinobacteria; Corynebacteriales; Nocardiaceae; Rhodococcus MAAHPKSARRRILLTATASALVLGAGGAPAAAVPLAGQSPPAADQAVAPTTTPPFSTPNT WP_084869160.1 penicillin-binding protein [Rhodococcus hoagii] Length: 445\nNumber of predicted TMHs: 2\nExp number of AAs in TMHs: 42.56153\nExp number, first 60 AAs: 21.20564\nTotal prob of N-in: 0.95473\nPOSSIBLE N-term signal sequence\ninside 1 11\nTMhelix 12 34\noutside 35 412\nTMhelix 413 435\ninside 436 445

31020 GCF_001646905.1_re12 Rhodococcus hoagii Terrabacteria group; Actinobacteria; Actinobacteria; Corynebacteriales; Nocardiaceae; Rhodococcus MAAHPKSARRRILLTATASALVLGAGGAPAAAVPLAGQSPLAADQAVAPTTTPPFSTPNT WP_022597221.1 D-alanyl-D-alanine carboxypeptidase [Rhodococcus hoagii] Length: 444\nNumber of predicted TMHs: 2\nExp number of AAs in TMHs: 42.86088\nExp number, first 60 AAs: 21.57924\nTotal prob of N-in: 0.96909\nPOSSIBLE N-term signal sequence\ninside 1 11\nTMhelix 12 34\noutside 35 412\nTMhelix 413 435\ninside 436 444

31021 GCF_002095175.1_ASM209517v1 Rhodococcus hoagii Terrabacteria group; Actinobacteria; Actinobacteria; Corynebacteriales; Nocardiaceae; Rhodococcus MTADPSDLTRRGRTAAFFDLDKTIIAKSSALAFSKPFFAQGLITRRSVLQSSYAQFLFIL WP_005515815.1 morphological differentiation-associated protein [Rhodococcus hoagii] Length: 272\nNumber of predicted TMHs: 1\nExp number of AAs in TMHs: 21.05758\nExp number, first 60 AAs: 0.04208\nTotal prob of N-in: 0.02567\noutside 1 244\nTMhelix 245 267\ninside 268 272

31022 GCF_002094315.1_ASM209431v1 Rhodococcus hoagii Terrabacteria group; Actinobacteria; Actinobacteria; Corynebacteriales; Nocardiaceae; Rhodococcus MTADPSDLTRRGRTAAFFDLDKTIIAKSSALAFSKPFFAQGLITRRSVLQSSYAQFLFIL WP_005515815.1 morphological differentiation-associated protein [Rhodococcus hoagii] Length: 272\nNumber of predicted TMHs: 1\nExp number of AAs in TMHs: 21.05758\nExp number, first 60 AAs: 0.04208\nTotal prob of N-in: 0.02567\noutside 1 244\nTMhelix 245 267\ninside 268 272

31023 GCF_002094405.1_ASM209440v1 Rhodococcus hoagii Terrabacteria group; Actinobacteria; Actinobacteria; Corynebacteriales; Nocardiaceae; Rhodococcus
MAAHPKSARRRILLTATASALVLGAGGAPAAAVPLAGQSPLAADQAVAPTTTPPFTTPNT WP_084962208.1
penicillin-binding protein [Rhodococcus hoagii] Length: 444\nNumber of predicted TMHs: 2\nExp number of AAs in TMHs: 42.7856999999999\nExp number, first 60 AAs: 21.57856\nTotal prob of N-in: 0.96909\nPOSSIBLE N-term signal sequence\ninside 1 11\nTMhelix 12 34\noutside 35 412\nTMhelix 413 435\ninside 436 444

31024 GCF_002095255.1_ASM209525v1 Rhodococcus hoagii Terrabacteria group; Actinobacteria; Actinobacteria; Corynebacteriales; Nocardiaceae; Rhodococcus
MAAHPKSARRRILLTATASALVLGAGGAPAAAVPLAGQSPLAADQAVAPTTTPPFSTPNT WP_069856650.1
MULTISPECIES: D-alanyl-D-alanine carboxypeptidase [Rhodococcus] Length: 444\nNumber of predicted TMHs: 2\nExp number of AAs in TMHs: 42.7857099999999\nExp number, first 60 AAs: 21.57856\nTotal prob of N-in: 0.96909\nPOSSIBLE N-term signal sequence\ninside 1 11\nTMhelix 12 34\noutside 35 412\nTMhelix 413 435\ninside 436 444

31025 GCF_002094305.1_ASM209430v1 Rhodococcus hoagii Terrabacteria group; Actinobacteria; Actinobacteria; Corynebacteriales; Nocardiaceae; Rhodococcus
MAAHPKSARRRILLTATASALVLGAGGAPAAAVPLAGQSPLAADQAVAPTTTPPFSTPNT WP_022597221.1 D-alanyl-D-alanine carboxypeptidase [Rhodococcus hoagii] Length: 444\nNumber of predicted TMHs: 2\nExp number of AAs in TMHs: 42.86088\nExp number, first 60 AAs: 21.57924\nTotal prob of N-in: 0.96909\nPOSSIBLE N-term signal sequence\ninside 1 11\nTMhelix 12 34\noutside 35 412\nTMhelix 413 435\ninside 436 444

31026 GCF_002095155.1_ASM209515v1 Rhodococcus hoagii Terrabacteria group; Actinobacteria; Actinobacteria; Corynebacteriales; Nocardiaceae; Rhodococcus
MAAHPKSARRRILLTATASALVLGAGGAPAAAVPLAGQSPLAADQAVAPTTTPPFSTPNT WP_013416807.1 D-alanyl-D-alanine carboxypeptidase [Rhodococcus hoagii] Length: 444\nNumber of predicted TMHs: 2\nExp number of AAs in TMHs: 42.7856499999999\nExp number, first 60 AAs: 21.57858\nTotal prob of N-in: 0.96909\nPOSSIBLE N-term signal sequence\ninside 1 11\nTMhelix 12 34\noutside 35 412\nTMhelix 413 435\ninside 436 444

31027 GCF_002095185.1_ASM209518v1 Rhodococcus hoagii Terrabacteria group; Actinobacteria; Actinobacteria; Corynebacteriales; Nocardiaceae; Rhodococcus
MTADPSDLTRRGRTAAFFDLDKTIIAKSSALAFSKPFFAQGLITRRSVLQSSYAQFLFIL WP_005515815.1 morphological differentiation-associated protein [Rhodococcus hoagii] Length: 272\nNumber of predicted TMHs: 1\nExp number of AAs in TMHs: 21.05758\nExp number, first 60 AAs: 0.04208\nTotal prob of N-in: 0.02567\noutside 1 244\nTMhelix 245 267\ninside 268 272

31028 GCF_002095155.1_ASM209515v1 Rhodococcus hoagii Terrabacteria group; Actinobacteria; Actinobacteria; Corynebacteriales; Nocardiaceae; Rhodococcus
MTADPSDLTRRGRTAAFFDLDKTIIAKSSALAFSKPFFAQGLITRRSVLQSSYAQFLFIL WP_005515815.1 morphological differentiation-associated protein [Rhodococcus hoagii] Length: 272\nNumber of predicted TMHs: 1\nExp number of AAs in TMHs: 21.05758\nExp number, first 60 AAs: 0.04208\nTotal prob of N-in: 0.02567\noutside 1 244\nTMhelix 245 267\ninside 268 272

31029 GCF_002095185.1_ASM209518v1 Rhodococcus hoagii Terrabacteria group; Actinobacteria; Actinobacteria; Corynebacteriales; Nocardiaceae; Rhodococcus
MAAHPKSARRRILLTATASALVLGAGGAPAAAVPLAGQSPLAADQAVAPTTTPPFSTPNT WP_084985101.1
penicillin-binding protein [Rhodococcus hoagii] Length: 444\nNumber of predicted TMHs: 2\nExp number of AAs in TMHs: 42.7857099999999\nExp number, first 60 AAs: 21.57856\nTotal prob of N-in: 0.96909\nPOSSIBLE N-term signal sequence\ninside 1 11\nTMhelix 12 34\noutside 35 412\nTMhelix 413 435\ninside 436 444

31030 GCF_002078515.1_ASM207851v1 Rhodococcus hoagii Terrabacteria group; Actinobacteria; Actinobacteria; Corynebacteriales; Nocardiaceae; Rhodococcus
MAAHPKSARRRILLTATASALVLGAGGAPAAAVPLAGQSPLAADQAVAPTTTPPFSTPNT WP_005518562.1 D-alanyl-D-alanine carboxypeptidase [Rhodococcus hoagii] Length: 444\nNumber of predicted TMHs: 2\nExp number of AAs in TMHs: 42.7856999999999\nExp number, first 60 AAs: 21.57856\nTotal prob of N-in: 0.96909\nPOSSIBLE N-term signal sequence\ninside 1 11\nTMhelix 12 34\noutside 35 412\nTMhelix 413 435\ninside 436 444

31031 GCF_002094225.1_ASM209422v1 Rhodococcus hoagii Terrabacteria group; Actinobacteria; Actinobacteria; Corynebacteriales; Nocardiaceae; Rhodococcus
MTADPSDLTRRGRTAAFFDLDKTIIAKSSALAFSKPFFAQGLITRRSVLQSSYAQFLFIL WP_005515815.1 morphological differentiation-associated protein [Rhodococcus hoagii] Length: 272\nNumber of predicted TMHs: 1\nExp number of

AAs in TMHs: 21.05758\nExp number, first 60 AAs: 0.04208\nTotal prob of N-in: 0.02567\noutside 1 244\nTMhelix 245 267\ninside 268 272

31032 GCF_002078545.1_ASM207854v1 Rhodococcus hoagii Terrabacteria group; Actinobacteria; Actinobacteria; Corynebacteriales; Nocardiaceae; Rhodococcus MTADPSDLTRRGRTAAFFDLDKTIIAKSSALAFSKPFFAQGLITRRSVLQSSYAQFLFIL WP_005515815.1 morphological differentiation-associated protein [Rhodococcus hoagii] Length: 272\nNumber of predicted TMHs: 1\nExp number of AAs in TMHs: 21.05758\nExp number, first 60 AAs: 0.04208\nTotal prob of N-in: 0.02567\noutside 1 244\nTMhelix 245 267\ninside 268 272

31033 GCF_002078515.1_ASM207851v1 Rhodococcus hoagii Terrabacteria group; Actinobacteria; Actinobacteria; Corynebacteriales; Nocardiaceae; Rhodococcus MTADPSDLTRRGRTAAFFDLDKTIIAKSSALAFSKPFFAQGLITRRSVLQSSYAQFLFIL WP_005515815.1 morphological differentiation-associated protein [Rhodococcus hoagii] Length: 272\nNumber of predicted TMHs: 1\nExp number of AAs in TMHs: 21.05758\nExp number, first 60 AAs: 0.04208\nTotal prob of N-in: 0.02567\noutside 1 244\nTMhelix 245 267\ninside 268 272

31034 GCF_900105375.1_IMG-taxon_2642422554_annotated_assembly Rhodococcus jostii Terrabacteria group; Actinobacteria; Actinobacteria; Corynebacteriales; Nocardiaceae; Rhodococcus MSVFASRGTHCHSRARVPPMGMSGYSGGVETAAVRRRTLVARAALLLTLAAGVLLMHSLI WP_073365742.1 hypothetical protein [Rhodococcus jostii] Length: 157\nNumber of predicted TMHs: 2\nExp number of AAs in TMHs: 43.80327\nExp number, first 60 AAs: 18.46059\nTotal prob of N-in: 0.99453\nPOSSIBLE N-term signal sequence\ninside 1 42\nTMhelix 43 65\noutside 66 94\nTMhelix 95 117\ninside 118 157

31035 GCF_001292845.1_ASM129284v1 Rhodococcus jostii Terrabacteria group; Actinobacteria; Actinobacteria; Corynebacteriales; Nocardiaceae; Rhodococcus MSVFAPRGTHCHSRVQVPPLGMSGYSGRVETATGRRRTVVARAALLIAVAAGVLLMHSLI WP_054246435.1 hypothetical protein [Rhodococcus jostii] Length: 156\nNumber of predicted TMHs: 2\nExp number of AAs in TMHs: 43.92218\nExp number, first 60 AAs: 18.72264\nTotal prob of N-in: 0.99676\nPOSSIBLE N-term signal sequence\ninside 1 44\nTMhelix 45 67\noutside 68 93\nTMhelix 94 116\ninside 117 156

31036 GCF_900105905.1_IMG-taxon_2634166362_annotated_assembly Rhodococcus koreensis Terrabacteria group; Actinobacteria; Actinobacteria; Corynebacteriales; Nocardiaceae; Rhodococcus MVVWAPAQLDVTQRMPARSRRDVFSTLRGACTNGPPVSAIALAGMQKLSSPRRRRGACVL WP_083395499.1 hypothetical protein [Rhodococcus koreensis] Length: 218\nNumber of predicted TMHs: 1\nExp number of AAs in TMHs: 19.49301\nExp number, first 60 AAs: 1.51077\nTotal prob of N-in: 0.57300\ninside 1 177\nTMhelix 178 200\noutside 201 218

31037 GCF_900111805.1_IMG-taxon_2693429870_annotated_assembly Rhodococcus kroppenstedtii Terrabacteria group; Actinobacteria; Actinobacteria; Corynebacteriales; Nocardiaceae; Rhodococcus MSSTEAPGRVAAFFDLDKTIIAKSSTLAFSKPFFDGGLINRRSVLKSSYAQFLFLVSGAD WP_068363094.1 inhibition of morphological differentiation protein [Rhodococcus kroppenstedtii] Length: 267\nNumber of predicted TMHs: 1\nExp number of AAs in TMHs: 22.38267\nExp number, first 60 AAs: 0.0098\nTotal prob of N-in: 0.00670\noutside 1 240\nTMhelix 241 263\ninside 264 267

31038 GCF_001646725.1_rkr83 Rhodococcus kroppenstedtii Terrabacteria group; Actinobacteria; Actinobacteria; Corynebacteriales; Nocardiaceae; Rhodococcus MSSTEAPGRVAAFFDLDKTIIAKSSTLAFSKPFFDGGLINRRSVLKSSYAQFLFLVSGAD WP_068363094.1 inhibition of morphological differentiation protein [Rhodococcus kroppenstedtii] Length: 267\nNumber of predicted TMHs: 1\nExp number of AAs in TMHs: 22.38267\nExp number, first 60 AAs: 0.0098\nTotal prob of N-in: 0.00670\noutside 1 240\nTMhelix 241 263\ninside 264 267

31039 GCF_001646865.1_rk21 Rhodococcus kunmingensis Terrabacteria group; Actinobacteria; Actinobacteria; Corynebacteriales; Nocardiaceae; Rhodococcus MTAQPSATTQPGRIAAFFDLDKTVIARSSTFAFSRPFFAQGLLNRRAVLKSTYAHFMFLV WP_068279521.1 inhibition of morphological differentiation protein [Rhodococcus kunmingensis] Length: 272\nNumber of predicted TMHs: 1\nExp number of AAs in TMHs: 22.77227\nExp number, first 60 AAs: 0.08432\nTotal prob of N-in: 0.02179\noutside 1 245\nTMhelix 246 268\ninside 269 272

31040 GCF_001645385.1_ASM164538v1 Rhodococcus kyotonensis Terrabacteria group; Actinobacteria; Actinobacteria; Corynebacteriales; Nocardiaceae; Rhodococcus

MQSTTREGRSRTRRRLLSALALALFVAPLLTGCLRVQVSMGISADDRVSGQIVAATVPAS WP_068420664.1 DUF3153 domain-containing protein [Rhodococcus kyotonensis] Length: 244\nNumber of predicted TMHs: 2\nExp number of AAs in TMHs: 44.7224799999999\nExp number, first 60 AAs: 22.11068\nTotal prob of N-in: 0.97753\nPOSSIBLE N-term signal sequence\ninside 1 15\nTMhelix 16 38\noutside 39 196\nTMhelix 197 219\ninside 220 244

31041 GCF_001646735.1_ASM164673v1 Rhodococcus opacus Terrabacteria group; Actinobacteria; Actinobacteria; Corynebacteriales; Nocardiaceae; Rhodococcus MSVFASRGTHCHSRVQVPPLGMSGYSGRVETATGRRRTVVARAALLIAVAAGVLLMHSLI WP_043788292.1 MULTISPECIES: hypothetical protein [Rhodococcus] Length: 156\nNumber of predicted TMHs: 2\nExp number of AAs in TMHs: 43.92061\nExp number, first 60 AAs: 18.7176\nTotal prob of N-in: 0.99776\nPOSSIBLE N-term signal sequence\ninside 1 44\nTMhelix 45 67\noutside 68 93\nTMhelix 94 116\ninside 117 156

31042 GCF_000719995.1_ASM71999v1 Rhodococcus opacus Terrabacteria group; Actinobacteria; Actinobacteria; Corynebacteriales; Nocardiaceae; Rhodococcus MSNPLGQSSNGSRRGALPTPPSGWPIGSYPTYAEAQRAVDYLSDQEFPVQDVTIVGVNLM WP_003945065.1 MULTISPECIES: hypothetical protein [Rhodococcus] Length: 164\nNumber of predicted TMHs: 2\nExp number of AAs in TMHs: 45.26265\nExp number, first 60 AAs: 0.0547\nTotal prob of N-in: 0.87268\ninside 1 73\nTMhelix 74 96\noutside 97 99\nTMhelix 100 122\ninside 123 164

31043 GCF_000719995.1_ASM71999v1 Rhodococcus opacus Terrabacteria group; Actinobacteria; Actinobacteria; Corynebacteriales; Nocardiaceae; Rhodococcus MTDKPANGLNTGRTAAFFDLDKTIIAKSSTLAFSKPFFAQGLINRRSVLKSSYAQFLFLL WP_019746641.1 MULTISPECIES: inhibition of morphological differentiation protein [Rhodococcus] Length: 272\nNumber of predicted TMHs: 1\nExp number of AAs in TMHs: 22.87312\nExp number, first 60 AAs: 0.01889\nTotal prob of N-in: 0.03448\noutside 1 245\nTMhelix 246 268\ninside 269 272

31044 GCF_001646785.1_ASM164678v1 Rhodococcus phenolicus Terrabacteria group; Actinobacteria; Actinobacteria; Corynebacteriales; Nocardiaceae; Rhodococcus MTDETQRPTEAGRIAAFFDLDKTVIAKSSILAFSRPFFDQGLLNRRTVIENTYAQFLYLM WP_068166385.1 inhibition of morphological differentiation protein [Rhodococcus phenolicus] Length: 272\nNumber of predicted TMHs: 1\nExp number of AAs in TMHs: 22.80149\nExp number, first 60 AAs: 0.02928\nTotal prob of N-in: 0.05419\noutside 1 245\nTMhelix 246 268\ninside 269 272

31045 GCF_900105195.1_IMG-taxon_2630969012_annotated_assembly Rhodococcus pyridinivorans Terrabacteria group; Actinobacteria; Actinobacteria; Corynebacteriales; Nocardiaceae; Rhodococcus MFEFSRRALVGATATAALMVAGAGIASAHVTVVAPGAEQGGYTVLTFRVPTESETAGTTA WP_006553374.1 MULTISPECIES: hypothetical protein [Rhodococcus] Length: 223\nNumber of predicted TMHs: 2\nExp number of AAs in TMHs: 40.51123\nExp number, first 60 AAs: 18.38976\nTotal prob of N-in: 0.70947\nPOSSIBLE N-term signal sequence\ninside 1 8\nTMhelix 9 31\noutside 32 195\nTMhelix 196 218\ninside 219 223

31046 GCF_001646805.1_ASM164680v1 Rhodococcus pyridinivorans Terrabacteria group; Actinobacteria; Actinobacteria; Corynebacteriales; Nocardiaceae; Rhodococcus MFEFSRRALVGATATAALMVAGAGIASAHVTVVAPGAEQGGYTVLTFRVPTESETAGTTA WP_006553374.1 MULTISPECIES: hypothetical protein [Rhodococcus] Length: 223\nNumber of predicted TMHs: 2\nExp number of AAs in TMHs: 40.51123\nExp number, first 60 AAs: 18.38976\nTotal prob of N-in: 0.70947\nPOSSIBLE N-term signal sequence\ninside 1 8\nTMhelix 9 31\noutside 32 195\nTMhelix 196 218\ninside 219 223

31047 GCF_001646745.1_ASM164674v1 Rhodococcus qingshengii Terrabacteria group; Actinobacteria; Actinobacteria; Corynebacteriales; Nocardiaceae; Rhodococcus MSNPLGQSSNGSRRGALPTPPSGWPIGSYPTYAEAQRAVDYLSDQEFPVQDVTIVGVNLM WP_003945065.1 MULTISPECIES: hypothetical protein [Rhodococcus] Length: 164\nNumber of predicted TMHs: 2\nExp number of AAs in TMHs: 45.26265\nExp number, first 60 AAs: 0.0547\nTotal prob of N-in: 0.87268\ninside 1 73\nTMhelix 74 96\noutside 97 99\nTMhelix 100 122\ninside 123 164

31048 GCF_001646745.1_ASM164674v1 Rhodococcus qingshengii Terrabacteria group; Actinobacteria; Actinobacteria; Corynebacteriales; Nocardiaceae; Rhodococcus MTDKPANGLNTGRTAAFFDLDKTIIAKSSTLAFSKPFFAQGLINRRSVLKSSYAQFLFLL WP_003946123.1 MULTISPECIES: haloacid dehalogenase [Rhodococcus] Length: 272\nNumber of predicted TMHs: 1\nExp number of AAs in TMHs: 22.87312\nExp number, first 60 AAs: 0.01889\nTotal prob of N-in: 0.03448\noutside 1 245\nTMhelix 246 268\ninside 269 272

31049 GCF_001662505.1_ASM166250v1 Rhodococcus qingshengii Terrabacteria group; Actinobacteria; Actinobacteria; Corynebacteriales; Nocardiaceae; Rhodococcus
MSNPLGQSSNGSRRGALPTPPSGWPIGSYPTYAEAQRAVDYLSDQEFPVQDVTIVGVNLM WP_003945065.1 MULTISPECIES: hypothetical protein [Rhodococcus] Length: 164\nNumber of predicted TMHs: 2\nExp number of AAs in TMHs: 45.26265\nExp number, first 60 AAs: 0.0547\nTotal prob of N-in: 0.87268\ninside 1 73\nTMhelix 74 96\noutside 97 99\nTMhelix 100 122\ninside 123 164

31050 GCF_002087025.1_ASM208702v1 Rhodococcus qingshengii Terrabacteria group; Actinobacteria; Actinobacteria; Corynebacteriales; Nocardiaceae; Rhodococcus
MTDKPANGLNTGRTAAFFDLDKTIIAKSSTLAFSKPFFAQGLINRRSVLKSSYAQFLFLL WP_003946123.1 MULTISPECIES: haloacid dehalogenase [Rhodococcus] Length: 272\nNumber of predicted TMHs: 1\nExp number of AAs in TMHs: 22.87312\nExp number, first 60 AAs: 0.01889\nTotal prob of N-in: 0.03448\noutside 1 245\nTMhelix 246 268\ninside 269 272

31051 GCF_001623435.1_ASM162343v1 Rhodococcus qingshengii Terrabacteria group; Actinobacteria; Actinobacteria; Corynebacteriales; Nocardiaceae; Rhodococcus
MSNPLGQSSNGSRRGALPTPPSGWPIGSYPTYAEAQRAVDYLSDQEFPVQDVTIVGVNLM WP_003945065.1 MULTISPECIES: hypothetical protein [Rhodococcus] Length: 164\nNumber of predicted TMHs: 2\nExp number of AAs in TMHs: 45.26265\nExp number, first 60 AAs: 0.0547\nTotal prob of N-in: 0.87268\ninside 1 73\nTMhelix 74 96\noutside 97 99\nTMhelix 100 122\ninside 123 164

31052 GCF_001623435.1_ASM162343v1 Rhodococcus qingshengii Terrabacteria group; Actinobacteria; Actinobacteria; Corynebacteriales; Nocardiaceae; Rhodococcus
MTDKPANGLNTGRTAAFFDLDKTIIAKSSTLAFSKPFFAQGLINRRSVLKSSYAQFLFLL WP_003946123.1 MULTISPECIES: haloacid dehalogenase [Rhodococcus] Length: 272\nNumber of predicted TMHs: 1\nExp number of AAs in TMHs: 22.87312\nExp number, first 60 AAs: 0.01889\nTotal prob of N-in: 0.03448\noutside 1 245\nTMhelix 246 268\ninside 269 272

31053 GCF_002087025.1_ASM208702v1 Rhodococcus qingshengii Terrabacteria group; Actinobacteria; Actinobacteria; Corynebacteriales; Nocardiaceae; Rhodococcus
MSNPLGQSSNGSRRGALPTPPSGWPIGSYPTYAEAQRAVDYLSDQEFPVQDVTIVGVNLM WP_003945065.1 MULTISPECIES: hypothetical protein [Rhodococcus] Length: 164\nNumber of predicted TMHs: 2\nExp number of AAs in TMHs: 45.26265\nExp number, first 60 AAs: 0.0547\nTotal prob of N-in: 0.87268\ninside 1 73\nTMhelix 74 96\noutside 97 99\nTMhelix 100 122\ninside 123 164

31054 GCF_001662505.1_ASM166250v1 Rhodococcus qingshengii Terrabacteria group; Actinobacteria; Actinobacteria; Corynebacteriales; Nocardiaceae; Rhodococcus
MTDKPANGLNTGRTAAFFDLDKTIIAKSSTLAFSKPFFAQGLINRRSVLKSSYAQFLFLL WP_064689112.1 inhibition of morphological differentiation protein [Rhodococcus qingshengii] Length: 272\nNumber of predicted TMHs: 1\nExp number of AAs in TMHs: 22.87302\nExp number, first 60 AAs: 0.01889\nTotal prob of N-in: 0.03449\noutside 1 245\nTMhelix 246 268\ninside 269 272

31055 GCF_001646825.1_ASM164682v1 Rhodococcus rhodochrous Terrabacteria group; Actinobacteria; Actinobacteria; Corynebacteriales; Nocardiaceae; Rhodococcus
MFEFSRRALVGATATAALMVAGAGIASAHVTVVAPGAEQGGYTVLTFRVPTESETAGTTA WP_059382055.1 nuclear export factor GLE1 [Rhodococcus rhodochrous] Length: 224\nNumber of predicted TMHs: 2\nExp number of AAs in TMHs: 40.65078\nExp number, first 60 AAs: 18.3977\nTotal prob of N-in: 0.69434\nPOSSIBLE N-term signal sequence\ninside 1 8\nTMhelix 9 31\noutside 32 196\nTMhelix 197 219\ninside 220 224

31056 GCF_000716895.1_ASM71689v1 Rhodococcus rhodochrous Terrabacteria group; Actinobacteria; Actinobacteria; Corynebacteriales; Nocardiaceae; Rhodococcus
MTDKLGNGLTTGRTAAFFDLDKTVIAKSSTLAFSRPFFAQGLINRRSVLKSSYAQFLFLL WP_033235947.1 MULTISPECIES: inhibition of morphological differentiation protein [Rhodococcus] Length: 272\nNumber of predicted TMHs: 1\nExp number of AAs in TMHs: 22.91283\nExp number, first 60 AAs: 0.00718\nTotal prob of N-in: 0.01617\noutside 1 245\nTMhelix 246 268\ninside 269 272

31057 GCF_000716895.1_ASM71689v1 Rhodococcus rhodochrous Terrabacteria group; Actinobacteria; Actinobacteria; Corynebacteriales; Nocardiaceae; Rhodococcus
MSNPLAQSPNGSRRGALPTPPSGWPIGSYPTYAEAQRAVDYLSDQEFSVQDVTIVGVNLM WP_033234426.1 MULTISPECIES: hypothetical protein [Rhodococcus] Length: 164\nNumber of predicted TMHs: 2\nExp

number of AAs in TMHs: 45.15237\nExp number, first 60 AAs: 0.04302\nTotal prob of N-in: 0.64861\ninside 1 73\nTMhelix 74 96\noutside 97 100\nTMhelix 101 123\ninside 124 164

31058 GCF_001511235.1_Rhodococcus_rhodochrous_TRN7 Rhodococcus rhodochrous Terrabacteria group; Actinobacteria; Actinobacteria; Corynebacteriales; Nocardiaceae; Rhodococcus MTDATVDPTPSGRIAAFFDLDKTVIAKSSALAFSKPFFAQGLLNRRAVLKSSYAQFLFML WP_010595022.1 MULTISPECIES: hydrolase [Rhodococcus] Length: 280\nNumber of predicted TMHs: 1\nExp number of AAs in TMHs: 23.01026\nExp number, first 60 AAs: 0.0217\nTotal prob of N-in: 0.01643\noutside 1 245\nTMhelix 246 268\ninside 269 280

31059 GCF_001051275.1_ASM105127v1 Rhodococcus ruber Terrabacteria group; Actinobacteria; Actinobacteria; Corynebacteriales; Nocardiaceae; Rhodococcus MTDATVDPTPSGRIAAFFDLDKTVIAKSSALAFSKPFFAQGLLNRRAVLKSSYAQFLFML WP_010595022.1 MULTISPECIES: hydrolase [Rhodococcus] Length: 280\nNumber of predicted TMHs: 1\nExp number of AAs in TMHs: 23.01026\nExp number, first 60 AAs: 0.0217\nTotal prob of N-in: 0.01643\noutside 1 245\nTMhelix 246 268\ninside 269 280

31060 GCF_000824945.1_BDK_PRJEB6917_v1 Rhodococcus ruber Terrabacteria group; Actinobacteria; Actinobacteria; Corynebacteriales; Nocardiaceae; Rhodococcus MTDATVDPTPSGRIAAFFDLDKTVIAKSSALAFSKPFFAQGLLNRRAVLKSSYAQFLFML WP_010595022.1 MULTISPECIES: hydrolase [Rhodococcus] Length: 280\nNumber of predicted TMHs: 1\nExp number of AAs in TMHs: 23.01026\nExp number, first 60 AAs: 0.0217\nTotal prob of N-in: 0.01643\noutside 1 245\nTMhelix 246 268\ninside 269 280

31061 GCF_001646835.1_ASM164683v1 Rhodococcus ruber Terrabacteria group; Actinobacteria; Actinobacteria; Corynebacteriales; Nocardiaceae; Rhodococcus MTDATVDPTPSGRIAAFFDLDKTVIAKSSALAFSKPFFAQGLLNRRAVLKSSYAQFLFML WP_064066535.1 inhibition of morphological differentiation protein [Rhodococcus ruber] Length: 280\nNumber of predicted TMHs: 1\nExp number of AAs in TMHs: 23.02605\nExp number, first 60 AAs: 0.02262\nTotal prob of N-in: 0.01870\noutside 1 245\nTMhelix 246 268\ninside 269 280

31062 GCF_001682295.1_ASM168229v1 Rhodococcus sp. 008 Terrabacteria group; Actinobacteria; Actinobacteria; Corynebacteriales; Nocardiaceae; Rhodococcus MNTAKQRRALVTISRTGAMTTVAALAGSIASGRPENISWFVALRKPAFQPPPVVFPVVWT WP_058228178.1 MULTISPECIES: TspO protein [Rhodococcus] Length: 169\nNumber of predicted TMHs: 1\nExp number of AAs in TMHs: 81.87072\nExp number, first 60 AAs: 21.42694\nTotal prob of N-in: 0.96201\nPOSSIBLE N-term signal sequence\ninside 1 140\nTMhelix 141 163\noutside 164 169

31063 GCF_001682295.1_ASM168229v1 Rhodococcus sp. 008 Terrabacteria group; Actinobacteria; Actinobacteria; Corynebacteriales; Nocardiaceae; Rhodococcus MSNPLGQSSNGSRRGALPTPPSGWPIGSYPTYAEAQRAVDYLSDQEFPVQDVTIVGVNLM WP_003945065.1 MULTISPECIES: hypothetical protein [Rhodococcus] Length: 164\nNumber of predicted TMHs: 2\nExp number of AAs in TMHs: 45.26265\nExp number, first 60 AAs: 0.0547\nTotal prob of N-in: 0.87268\ninside 1 73\nTMhelix 74 96\noutside 97 99\nTMhelix 100 122\ninside 123 164

31064 GCF_001682295.1_ASM168229v1 Rhodococcus sp. 008 Terrabacteria group; Actinobacteria; Actinobacteria; Corynebacteriales; Nocardiaceae; Rhodococcus MTDKPANGLNTGRTAAFFDLDKTIIAKSSTLAFSKPFFAQGLINRRSVLKSSYAQFLFLL WP_003946123.1 MULTISPECIES: haloacid dehalogenase [Rhodococcus] Length: 272\nNumber of predicted TMHs: 1\nExp number of AAs in TMHs: 22.87312\nExp number, first 60 AAs: 0.01889\nTotal prob of N-in: 0.03448\noutside 1 245\nTMhelix 246 268\ninside 269 272

31065 GCF_001757665.1_ASM175766v1 Rhodococcus sp. 1139 Terrabacteria group; Actinobacteria; Actinobacteria; Corynebacteriales; Nocardiaceae; Rhodococcus MSNPLGQSSNGSRRGALPTPPSGWPIGSYPTYAEAQRAVDYLSDQEFPVQDVTIVGVNLM WP_003945065.1 MULTISPECIES: hypothetical protein [Rhodococcus] Length: 164\nNumber of predicted TMHs: 2\nExp number of AAs in TMHs: 45.26265\nExp number, first 60 AAs: 0.0547\nTotal prob of N-in: 0.87268\ninside 1 73\nTMhelix 74 96\noutside 97 99\nTMhelix 100 122\ninside 123 164

31066 GCF_001757665.1_ASM175766v1 Rhodococcus sp. 1139 Terrabacteria group; Actinobacteria; Actinobacteria; Corynebacteriales; Nocardiaceae; Rhodococcus MTDKPANGLNTGRTAAFFDLDKTIIAKSSTLAFSKPFFAQGLINRRSVLKSSYAQFLFLL WP_003946123.1 MULTISPECIES: haloacid dehalogenase [Rhodococcus] Length: 272\nNumber of predicted TMHs: 1\nExp number of AAs in TMHs:

22.87312\nExp number, first 60 AAs: 0.01889\nTotal prob of N-in: 0.03448\noutside 1 245\nTMhelix 246 268\ninside 269 272

31067 GCF_002091935.1_ASM209193v1 Rhodococcus sp. 1159 Terrabacteria group; Actinobacteria; Actinobacteria; Corynebacteriales; Nocardiaceae; Rhodococcus MTDKPANGLNTGRTAAFFDLDKTIIAKSSTLAFSKPFFAQGLINRRSVLKSSYAQFLFLL WP_019746641.1 MULTISPECIES: inhibition of morphological differentiation protein [Rhodococcus] Length: 272\nNumber of predicted TMHs: 1\nExp number of AAs in TMHs: 22.87312\nExp number, first 60 AAs: 0.01889\nTotal prob of N-in: 0.03448\noutside 1 245\nTMhelix 246 268\ninside 269 272

31068 GCF_002091935.1_ASM209193v1 Rhodococcus sp. 1159 Terrabacteria group; Actinobacteria; Actinobacteria; Corynebacteriales; Nocardiaceae; Rhodococcus MSNPLGQSSNGSRRGALPTPPSGWPIGSYPTYAEAQRAVDYLSDQEFPVQDVTIVGVNLM WP_003945065.1 MULTISPECIES: hypothetical protein [Rhodococcus] Length: 164\nNumber of predicted TMHs: 2\nExp number of AAs in TMHs: 45.26265\nExp number, first 60 AAs: 0.0547\nTotal prob of N-in: 0.87268\ninside 1 73\nTMhelix 74 96\noutside 97 99\nTMhelix 100 122\ninside 123 164

31069 GCF_001886355.1_ASM188635v1 Rhodococcus sp. 2G Terrabacteria group; Actinobacteria; Actinobacteria; Corynebacteriales; Nocardiaceae; Rhodococcus MFEFSRRALVGATATAALMVAGAGIASAHVTVVAPGAEQGGYTVLTFRVPTESETAGTTA WP_006553374.1 MULTISPECIES: hypothetical protein [Rhodococcus] Length: 223\nNumber of predicted TMHs: 2\nExp number of AAs in TMHs: 40.51123\nExp number, first 60 AAs: 18.38976\nTotal prob of N-in: 0.70947\nPOSSIBLE N-term signal sequence\ninside 1 8\nTMhelix 9 31\noutside 32 195\nTMhelix 196 218\ninside 219 223

31070 GCF_001242945.1_Rhodococcus_sp._311R Rhodococcus sp. 311R Terrabacteria group; Actinobacteria; Actinobacteria; Corynebacteriales; Nocardiaceae; Rhodococcus MTDKPANGLNTGRTAAFFDLDKTIIAKSSTLAFSKPFFAQGLINRRSVLKSSYAQFLFLL WP_003946123.1 MULTISPECIES: haloacid dehalogenase [Rhodococcus] Length: 272\nNumber of predicted TMHs: 1\nExp number of AAs in TMHs: 22.87312\nExp number, first 60 AAs: 0.01889\nTotal prob of N-in: 0.03448\noutside 1 245\nTMhelix 246 268\ninside 269 272

31071 GCF_001242945.1_Rhodococcus_sp._311R Rhodococcus sp. 311R Terrabacteria group; Actinobacteria; Actinobacteria; Corynebacteriales; Nocardiaceae; Rhodococcus MSNPLGQSSNGSRRGALPTPPSGWPIGSYPTYAEAQRAVDYLSDQEFPVQDVTIVGVNLM WP_003945065.1 MULTISPECIES: hypothetical protein [Rhodococcus] Length: 164\nNumber of predicted TMHs: 2\nExp number of AAs in TMHs: 45.26265\nExp number, first 60 AAs: 0.0547\nTotal prob of N-in: 0.87268\ninside 1 73\nTMhelix 74 96\noutside 97 99\nTMhelix 100 122\ninside 123 164

31072 GCF_002076325.1_ASM207632v1 Rhodococcus sp. 66b Terrabacteria group; Actinobacteria; Actinobacteria; Corynebacteriales; Nocardiaceae; Rhodococcus MSNPLGQSSNGSRRGALPTPPSGWPIGSYPTYAEAQRAVDYLSDQEFPVQDVTIVGVNLM WP_003945065.1 MULTISPECIES: hypothetical protein [Rhodococcus] Length: 164\nNumber of predicted TMHs: 2\nExp number of AAs in TMHs: 45.26265\nExp number, first 60 AAs: 0.0547\nTotal prob of N-in: 0.87268\ninside 1 73\nTMhelix 74 96\noutside 97 99\nTMhelix 100 122\ninside 123 164

31073 GCF_002076325.1_ASM207632v1 Rhodococcus sp. 66b Terrabacteria group; Actinobacteria; Actinobacteria; Corynebacteriales; Nocardiaceae; Rhodococcus MTDKPANGLNTGRTAAFFDLDKTIIAKSSTLAFSKPFFAQGLINRRSVLKSSYAQFLFLL WP_080910596.1 inhibition of morphological differentiation protein [Rhodococcus sp. 66b] Length: 272\nNumber of predicted TMHs: 1\nExp number of AAs in TMHs: 22.87339\nExp number, first 60 AAs: 0.01889\nTotal prob of N-in: 0.03449\noutside 1 245\nTMhelix 246 268\ninside 269 272

31074 GCF_001665495.1_ASM166549v1 Rhodococcus sp. 852002-51564_SCH6189132-a Terrabacteria group; Actinobacteria; Actinobacteria; Corynebacteriales; Nocardiaceae; Rhodococcus MFEFSRRALVGATATAALMVAGAGIASAHVTVVAPGAEQGGYTVLTFRVPTESETAGTTA WP_024102629.1 MULTISPECIES: nuclear export factor GLE1 [Rhodococcus] Length: 223\nNumber of predicted TMHs: 2\nExp number of AAs in TMHs: 40.51123\nExp number, first 60 AAs: 18.38976\nTotal prob of N-in: 0.70947\nPOSSIBLE N-term signal sequence\ninside 1 8\nTMhelix 9 31\noutside 32 195\nTMhelix 196 218\ninside 219 223

31075 GCF_000949305.1_ASM94930v1 Rhodococcus sp. AD45 Terrabacteria group; Actinobacteria; Actinobacteria; Corynebacteriales; Nocardiaceae; Rhodococcus

MSNPLAQSPNGSRRGALPTPPSGWPIGSYPTYAEAQRAVDYLSDQEFSVQDVTIVGVNLM WP_033234426.1 MULTISPECIES: hypothetical protein [Rhodococcus] Length: 164\nNumber of predicted TMHs: 2\nExp number of AAs in TMHs: 45.15237\nExp number, first 60 AAs: 0.04302\nTotal prob of N-in: 0.64861\ninside 1 73\nTMhelix 74 96\noutside 97 100\nTMhelix 101 123\ninside 124 164

31076 GCF_000949305.1_ASM94930v1 Rhodococcus sp. AD45 Terrabacteria group; Actinobacteria; Actinobacteria; Corynebacteriales; Nocardiaceae; Rhodococcus MTDKLGNGLTTGRTAAFFDLDKTVIAKSSTLAFSRPFFAQGLINRRSVLKSSYAQFLFLL WP_033235947.1 MULTISPECIES: inhibition of morphological differentiation protein [Rhodococcus] Length: 272\nNumber of predicted TMHs: 1\nExp number of AAs in TMHs: 22.91283\nExp number, first 60 AAs: 0.00718\nTotal prob of N-in: 0.01617\noutside 1 245\nTMhelix 246 268\ninside 269 272

31077 GCF_001297885.1_ASM129788v1 Rhodococcus sp. ADH Terrabacteria group; Actinobacteria; Actinobacteria; Corynebacteriales; Nocardiaceae; Rhodococcus MSNPLGQSSNGSRRGALPTPPSGWPIGSYPTYAEAQRAVDYLSDQEFPVQDVTIVGVNLM WP_003945065.1 MULTISPECIES: hypothetical protein [Rhodococcus] Length: 164\nNumber of predicted TMHs: 2\nExp number of AAs in TMHs: 45.26265\nExp number, first 60 AAs: 0.0547\nTotal prob of N-in: 0.87268\ninside 1 73\nTMhelix 74 96\noutside 97 99\nTMhelix 100 122\ninside 123 164

31078 GCF_001297885.1_ASM129788v1 Rhodococcus sp. ADH Terrabacteria group; Actinobacteria; Actinobacteria; Corynebacteriales; Nocardiaceae; Rhodococcus MTDKPANGLNTGRTAAFFDLDKTIIAKSSTLAFSKPFFAQGLINRRSVLKSSYAQFLFLL WP_003946123.1 MULTISPECIES: haloacid dehalogenase [Rhodococcus] Length: 272\nNumber of predicted TMHs: 1\nExp number of AAs in TMHs: 22.87312\nExp number, first 60 AAs: 0.01889\nTotal prob of N-in: 0.03448\noutside 1 245\nTMhelix 246 268\ninside 269 272

31079 GCF_001652355.1_rey60 Rhodococcus sp. AJR001 Terrabacteria group; Actinobacteria; Actinobacteria; Corynebacteriales; Nocardiaceae; Rhodococcus MTDKPANGLNTGRTAAFFDLDKTIIAKSSTLAFSKPFFAQGLINRRSVLKSSYAQFLFLL WP_003946123.1 MULTISPECIES: haloacid dehalogenase [Rhodococcus] Length: 272\nNumber of predicted TMHs: 1\nExp number of AAs in TMHs: 22.87312\nExp number, first 60 AAs: 0.01889\nTotal prob of N-in: 0.03448\noutside 1 245\nTMhelix 246 268\ninside 269 272

31080 GCF_001652355.1_rey60 Rhodococcus sp. AJR001 Terrabacteria group; Actinobacteria; Actinobacteria; Corynebacteriales; Nocardiaceae; Rhodococcus MSNPLGQSSNGSRRGALPTPPSGWPIGSYPTYAEAQRAVDYLSDQEFPVQDVTIVGVNLM WP_003945065.1 MULTISPECIES: hypothetical protein [Rhodococcus] Length: 164\nNumber of predicted TMHs: 2\nExp number of AAs in TMHs: 45.26265\nExp number, first 60 AAs: 0.0547\nTotal prob of N-in: 0.87268\ninside 1 73\nTMhelix 74 96\noutside 97 99\nTMhelix 100 122\ninside 123 164

31081 GCF_001029585.1_ASM102958v1 Rhodococcus sp. ARP2 Terrabacteria group; Actinobacteria; Actinobacteria; Corynebacteriales; Nocardiaceae; Rhodococcus MTDKPANGLNTGRTAAFFDLDKTIIAKSSTLAFSKPFFAQGLINRRSVLKSSYAQFLFLL WP_019746641.1 MULTISPECIES: inhibition of morphological differentiation protein [Rhodococcus] Length: 272\nNumber of predicted TMHs: 1\nExp number of AAs in TMHs: 22.87312\nExp number, first 60 AAs: 0.01889\nTotal prob of N-in: 0.03448\noutside 1 245\nTMhelix 246 268\ninside 269 272

31082 GCF_002079265.1_ASM207926v1 Rhodococcus sp. BH4 Terrabacteria group; Actinobacteria; Actinobacteria; Corynebacteriales; Nocardiaceae; Rhodococcus MSNPLGQSSNGSRRGALPTPPSGWPIGSYPTYAEAQRAVDYLSDQEFPVQDVTIVGVNLM WP_003945065.1 MULTISPECIES: hypothetical protein [Rhodococcus] Length: 164\nNumber of predicted TMHs: 2\nExp number of AAs in TMHs: 45.26265\nExp number, first 60 AAs: 0.0547\nTotal prob of N-in: 0.87268\ninside 1 73\nTMhelix 74 96\noutside 97 99\nTMhelix 100 122\ninside 123 164

31083 GCF_002079265.1_ASM207926v1 Rhodococcus sp. BH4 Terrabacteria group; Actinobacteria; Actinobacteria; Corynebacteriales; Nocardiaceae; Rhodococcus MTDKPANGLNTGRTAAFFDLDKTIIAKSSTLAFSKPFFAQGLINRRSVLKSSYAQFLFLL WP_003946123.1 MULTISPECIES: haloacid dehalogenase [Rhodococcus] Length: 272\nNumber of predicted TMHs: 1\nExp number of AAs in TMHs: 22.87312\nExp number, first 60 AAs: 0.01889\nTotal prob of N-in: 0.03448\noutside 1 245\nTMhelix 246 268\ninside 269 272

31084 GCF_002117895.1_ASM211789v1 Rhodococcus sp. Br-6 Terrabacteria group; Actinobacteria; Actinobacteria; Corynebacteriales; Nocardiaceae; Rhodococcus
MTADPSDLTCRGRTAAFFDLDKTIIAKSSALAFSKPFFAQGLITRRSVLQSSYAQFLFIL WP_069854787.1 inhibition of morphological differentiation protein [Rhodococcus sp. Br-6] Length: 272\nNumber of predicted TMHs: 1\nExp number of AAs in TMHs: 21.09695\nExp number, first 60 AAs: 0.04002\nTotal prob of N-in: 0.01920\noutside 1 244\nTMhelix 245 267\ninside 268 272

31085 GCF_002117895.1_ASM211789v1 Rhodococcus sp. Br-6 Terrabacteria group; Actinobacteria; Actinobacteria; Corynebacteriales; Nocardiaceae; Rhodococcus
MAAHPKSARRRILLTATASALVLGAGGAPAAAVPLAGQSPLAADQAVAPTTTPPFSTPNT WP_069856650.1
MULTISPECIES: D-alanyl-D-alanine carboxypeptidase [Rhodococcus] Length: 444\nNumber of predicted TMHs: 2\nExp number of AAs in TMHs: 42.7857099999999\nExp number, first 60 AAs: 21.57856\nTotal prob of N-in: 0.96909\nPOSSIBLE N-term signal sequence\ninside 1 11\nTMhelix 12 34\noutside 35 412\nTMhelix 413 435\ninside 436 444

31086 GCF_001748445.1_ASM174844v1 Rhodococcus sp. Br-6 Terrabacteria group; Actinobacteria; Actinobacteria; Corynebacteriales; Nocardiaceae; Rhodococcus
MTADPSDLTCRGRTAAFFDLDKTIIAKSSALAFSKPFFAQGLITRRSVLQSSYAQFLFIL WP_069854787.1 inhibition of morphological differentiation protein [Rhodococcus sp. Br-6] Length: 272\nNumber of predicted TMHs: 1\nExp number of AAs in TMHs: 21.09695\nExp number, first 60 AAs: 0.04002\nTotal prob of N-in: 0.01920\noutside 1 244\nTMhelix 245 267\ninside 268 272

31087 GCF_001748445.1_ASM174844v1 Rhodococcus sp. Br-6 Terrabacteria group; Actinobacteria; Actinobacteria; Corynebacteriales; Nocardiaceae; Rhodococcus
MAAHPKSARRRILLTATASALVLGAGGAPAAAVPLAGQSPLAADQAVAPTTTPPFSTPNT WP_069856650.1
MULTISPECIES: penicillin-binding protein [Rhodococcus] Length: 444\nNumber of predicted TMHs: 2\nExp number of AAs in TMHs: 42.7857099999999\nExp number, first 60 AAs: 21.57856\nTotal prob of N-in: 0.96909\nPOSSIBLE N-term signal sequence\ninside 1 11\nTMhelix 12 34\noutside 35 412\nTMhelix 413 435\ninside 436 444

31088 GCF_000801125.1_ASM80112v1 Rhodococcus sp. Chr-9 Terrabacteria group; Actinobacteria; Actinobacteria; Corynebacteriales; Nocardiaceae; Rhodococcus
MFEFSRRALVGATATAALMVAGAGIASAHVTVVAPGAEQGGYTVLTFRVPTESETAGTTA WP_006553374.1
MULTISPECIES: hypothetical protein [Rhodococcus] Length: 223\nNumber of predicted TMHs: 2\nExp number of AAs in TMHs: 40.51123\nExp number, first 60 AAs: 18.38976\nTotal prob of N-in: 0.70947\nPOSSIBLE N-term signal sequence\ninside 1 8\nTMhelix 9 31\noutside 32 195\nTMhelix 196 218\ninside 219 223

31089 GCF_000801125.1_ASM80112v1 Rhodococcus sp. Chr-9 Terrabacteria group; Actinobacteria; Actinobacteria; Corynebacteriales; Nocardiaceae; Rhodococcus
MPRIPEPVSTPAGPIYEGRLLDRPAEDVVDQGAGFDITTLVTRRRVLGLVGAGAGALALA WP_084222307.1 3,4-dioxygenase subunit beta [Rhodococcus sp. Chr-9] Length: 435\nNumber of predicted TMHs: 1\nExp number of AAs in TMHs: 28.54246\nExp number, first 60 AAs: 4.89288\nTotal prob of N-in: 0.30466\noutside 1 362\nTMhelix 363 385\ninside 386 435

31090 GCF_001976025.1_ASM197602v1 Rhodococcus sp. D-1 Terrabacteria group; Actinobacteria; Actinobacteria; Corynebacteriales; Nocardiaceae; Rhodococcus
MSNPLGQSSNGSRRGALPTPPSGWPIGSYPTYAEAQRAVDYLSDQEFPVQDVTIVGVNLM WP_003945065.1
MULTISPECIES: hypothetical protein [Rhodococcus] Length: 164\nNumber of predicted TMHs: 2\nExp number of AAs in TMHs: 45.26265\nExp number, first 60 AAs: 0.0547\nTotal prob of N-in: 0.87268\ninside 1 73\nTMhelix 74 96\noutside 97 99\nTMhelix 100 122\ninside 123 164

31091 GCF_001976025.1_ASM197602v1 Rhodococcus sp. D-1 Terrabacteria group; Actinobacteria; Actinobacteria; Corynebacteriales; Nocardiaceae; Rhodococcus
MTDKPANGLNTGRTAAFFDLDKTIIAKSSTLAFSKPFFAQGLINRRSVLKSSYAQFLFLL WP_003946123.1 MULTISPECIES: haloacid dehalogenase [Rhodococcus] Length: 272\nNumber of predicted TMHs: 1\nExp number of AAs in TMHs: 22.87312\nExp number, first 60 AAs: 0.01889\nTotal prob of N-in: 0.03448\noutside 1 245\nTMhelix 246 268\ninside 269 272

31092 GCF_001647205.1_ASM164720v1 Rhodococcus sp. EPR-134 Terrabacteria group; Actinobacteria; Actinobacteria; Corynebacteriales; Nocardiaceae; Rhodococcus
MTDKPANGLNTGRTAAFFDLDKTIIAKSSTLAFSKPFFAQGLINRRSVLKSSYAQFLFLL WP_064112049.1 inhibition of

morphological differentiation protein [Rhodococcus sp. EPR-134]	Length: 272\nNumber of predicted TMHs: 1\nExp number of AAs in TMHs: 22.87469\nExp number, first 60 AAs: 0.0189\nTotal prob of N-in: 0.03456\noutside 1 245\nTMhelix 246 268\ninside 269 272

31093 GCF_001647205.1_ASM164720v1 Rhodococcus sp. EPR-134 Terrabacteria group; Actinobacteria; Actinobacteria; Corynebacteriales; Nocardiaceae; Rhodococcus
MSNPLGQSSNGSRRGALPTPPSGWPIGSYPTYAEAQRAVDYLSDQEFPVQDVTIVGVNLM WP_003945065.1
MULTISPECIES: hypothetical protein [Rhodococcus] Length: 164\nNumber of predicted TMHs: 2\nExp number of AAs in TMHs: 45.26265\nExp number, first 60 AAs: 0.0547\nTotal prob of N-in: 0.87268\ninside 1 73\nTMhelix 74 96\noutside 97 99\nTMhelix 100 122\ninside 123 164

31094 GCF_001647185.1_ASM164718v1 Rhodococcus sp. EPR-147 Terrabacteria group; Actinobacteria; Actinobacteria; Corynebacteriales; Nocardiaceae; Rhodococcus
MGVRSVQRRPRAVTPHRRRTVVIAAFTALSLSFDAAFGGYPARAERPVIDPFQLPAPSAP WP_082898974.1
MULTISPECIES: type VII secretion-associated serine protease mycosin [Rhodococcus] Length: 475\nNumber of predicted TMHs: 2\nExp number of AAs in TMHs: 43.84334\nExp number, first 60 AAs: 21.10889\nTotal prob of N-in: 0.98352\nPOSSIBLE N-term signal sequence\ninside 1 19\nTMhelix 20 42\noutside 43 445\nTMhelix 446 468\ninside 469 475

31095 GCF_001647175.1_ASM164717v1 Rhodococcus sp. EPR-279 Terrabacteria group; Actinobacteria; Actinobacteria; Corynebacteriales; Nocardiaceae; Rhodococcus
MQSTSRTSHRTHRRRVLSVAALALLVAPMLAGCLRVQVTMGVSANDRVAGQIVAASVPTS WP_068283778.1
DUF3153 domain-containing protein [Rhodococcus sp. EPR-279] Length: 244\nNumber of predicted TMHs: 2\nExp number of AAs in TMHs: 44.73308\nExp number, first 60 AAs: 21.88176\nTotal prob of N-in: 0.94005\nPOSSIBLE N-term signal sequence\ninside 1 15\nTMhelix 16 38\noutside 39 196\nTMhelix 197 219\ninside 220 244

31096 GCF_001647175.1_ASM164717v1 Rhodococcus sp. EPR-279 Terrabacteria group; Actinobacteria; Actinobacteria; Corynebacteriales; Nocardiaceae; Rhodococcus
MGVRSVQRRPRAVTPHRRRTVVIAAFTALSLSFDAAFGGYPARAERPVIDPFQLPAPSAP WP_082898974.1
MULTISPECIES: type VII secretion-associated serine protease mycosin [Rhodococcus] Length: 475\nNumber of predicted TMHs: 2\nExp number of AAs in TMHs: 43.84334\nExp number, first 60 AAs: 21.10889\nTotal prob of N-in: 0.98352\nPOSSIBLE N-term signal sequence\ninside 1 19\nTMhelix 20 42\noutside 43 445\nTMhelix 446 468\ninside 469 475

31097 GCF_000382865.1_ASM38286v1 Rhodococcus sp. EsD8 Terrabacteria group; Actinobacteria; Actinobacteria; Corynebacteriales; Nocardiaceae; Rhodococcus
MTDATVAPTQSGRIAAFFDLDKTVIAKSSALAFSKPFFAQGLLNRRAVLKSSYAQFLFML WP_006941337.1 MULTISPECIES: phosphoserine phosphatase [Rhodococcus] Length: 280\nNumber of predicted TMHs: 1\nExp number of AAs in TMHs: 22.81284\nExp number, first 60 AAs: 0.03489\nTotal prob of N-in: 0.04407\noutside 1 245\nTMhelix 246 268\ninside 269 280

31098 GCF_000382865.1_ASM38286v1 Rhodococcus sp. EsD8 Terrabacteria group; Actinobacteria; Actinobacteria; Corynebacteriales; Nocardiaceae; Rhodococcus
MRLPTSPRSRRRTLSAALLLLLVPVLAGCLRVQATMGVSADDRVSGQIVAATIPADENDK WP_006933384.1
MULTISPECIES: DUF3153 domain-containing protein [Rhodococcus] Length: 233\nNumber of predicted TMHs: 2\nExp number of AAs in TMHs: 44.90952\nExp number, first 60 AAs: 21.98985\nTotal prob of N-in: 0.95979\nPOSSIBLE N-term signal sequence\ninside 1 12\nTMhelix 13 35\noutside 36 191\nTMhelix 192 214\ninside 215 233

31099 GCF_001651055.1_ASM165105v1 Rhodococcus sp. HS-D2 Terrabacteria group; Actinobacteria; Actinobacteria; Corynebacteriales; Nocardiaceae; Rhodococcus
MFEFSRRALVGATATAALMVAGAGIASAHVTVVAPGAEQGGYTVLTFRVPTESETAGTTA WP_006553374.1
MULTISPECIES: hypothetical protein [Rhodococcus] Length: 223\nNumber of predicted TMHs: 2\nExp number of AAs in TMHs: 40.51123\nExp number, first 60 AAs: 18.38976\nTotal prob of N-in: 0.70947\nPOSSIBLE N-term signal sequence\ninside 1 8\nTMhelix 9 31\noutside 32 195\nTMhelix 196 218\ninside 219 223

31100 GCF_000482405.1_ASM48240v1 Rhodococcus sp. JG-3 Terrabacteria group; Actinobacteria; Actinobacteria; Corynebacteriales; Nocardiaceae; Rhodococcus
MQSTSRTSARTRRRRVLSVLALTLLVAPMLAGCLRVQVTMGVSANDRVAGQIVAATVPTS WP_045841254.1
MULTISPECIES: hypothetical protein [Rhodococcus] Length: 244\nNumber of predicted TMHs: 2\nExp number of AAs in TMHs: 45.38746\nExp number, first 60 AAs: 22.41066\nTotal prob of N-in: 0.99116\nPOSSIBLE N-term signal sequence\ninside 1 15\nTMhelix 16 38\noutside 39 197\nTMhelix 198 220\ninside 221 244

31101 GCF_001445685.1_ASM144568v1 Rhodococcus sp. KB6 Terrabacteria group; Actinobacteria; Actinobacteria; Corynebacteriales; Nocardiaceae; Rhodococcus
MTDKPANGLNTGRTAAFFDLDKTIIAKSSTLAFSKPFFAQGLINRRSVLKSSYAQFLFLL WP_003946123.1 MULTISPECIES: haloacid dehalogenase [Rhodococcus] Length: 272\nNumber of predicted TMHs: 1\nExp number of AAs in TMHs: 22.87312\nExp number, first 60 AAs: 0.01889\nTotal prob of N-in: 0.03448\noutside 1 245\nTMhelix 246 268\ninside 269 272

31102 GCF_001445685.1_ASM144568v1 Rhodococcus sp. KB6 Terrabacteria group; Actinobacteria; Actinobacteria; Corynebacteriales; Nocardiaceae; Rhodococcus
MSNPLGQSSNGSRRGALPTPPSGWPIGSYPTYAEAQRAVDYLSDQEFPVQDVTIVGVNLM WP_003945065.1 MULTISPECIES: hypothetical protein [Rhodococcus] Length: 164\nNumber of predicted TMHs: 2\nExp number of AAs in TMHs: 45.26265\nExp number, first 60 AAs: 0.0547\nTotal prob of N-in: 0.87268\ninside 1 73\nTMhelix 74 96\noutside 97 99\nTMhelix 100 122\ninside 123 164

31103 GCF_001583455.1_ASM158345v1 Rhodococcus sp. LB1 Terrabacteria group; Actinobacteria; Actinobacteria; Corynebacteriales; Nocardiaceae; Rhodococcus
MRRRTTFTSVKVIAVGLLTMLASMLGAGTALAHSVVISSTPENGAEIAAGPERVSVTFNE WP_005248910.1 MULTISPECIES: copper resistance protein C [Rhodococcus] Length: 177\nNumber of predicted TMHs: 2\nExp number of AAs in TMHs: 44.30022\nExp number, first 60 AAs: 22.13888\nTotal prob of N-in: 0.99979\nPOSSIBLE N-term signal sequence\ninside 1 12\nTMhelix 13 35\noutside 36 147\nTMhelix 148 170\ninside 171 177

31104 GCF_001426145.1_Leaf225 Rhodococcus sp. Leaf225 Terrabacteria group; Actinobacteria; Actinobacteria; Corynebacteriales; Nocardiaceae; Rhodococcus
MSTPLRPDAPGRVAAFFDLDKTIIAKSSTLAFSKPFFDGGLINRRSVLKSSYAQFLFLVS WP_056443389.1 MULTISPECIES: inhibition of morphological differentiation protein [Rhodococcus] Length: 271\nNumber of predicted TMHs: 1\nExp number of AAs in TMHs: 22.45846\nExp number, first 60 AAs: 0.00777\nTotal prob of N-in: 0.01073\noutside 1 244\nTMhelix 245 267\ninside 268 271

31105 GCF_001426065.1_Leaf258 Rhodococcus sp. Leaf258 Terrabacteria group; Actinobacteria; Actinobacteria; Corynebacteriales; Nocardiaceae; Rhodococcus
MSTPLRPDAPGRVAAFFDLDKTIIAKSSTLAFSKPFFDGGLINRRSVLKSSYAQFLFLVS WP_056443389.1 MULTISPECIES: inhibition of morphological differentiation protein [Rhodococcus] Length: 271\nNumber of predicted TMHs: 1\nExp number of AAs in TMHs: 22.45846\nExp number, first 60 AAs: 0.00777\nTotal prob of N-in: 0.01073\noutside 1 244\nTMhelix 245 267\ninside 268 271

31106 GCF_001426085.1_Leaf278 Rhodococcus sp. Leaf278 Terrabacteria group; Actinobacteria; Actinobacteria; Corynebacteriales; Nocardiaceae; Rhodococcus
MAHRRRRPHFGGQCNGHSGTKTERLSPRSHRICVREPEVSDSGTSSSDGSPGDSSGSTEP WP_082520838.1 hypothetical protein [Rhodococcus sp. Leaf278] Length: 231\nNumber of predicted TMHs: 2\nExp number of AAs in TMHs: 36.8457\nExp number, first 60 AAs: 0\nTotal prob of N-in: 0.86177\ninside 1 165\nTMhelix 166 185\noutside 186 194\nTMhelix 195 217\ninside 218 231

31107 GCF_001890475.2_ASM189047v2 Rhodococcus sp. M8 Terrabacteria group; Actinobacteria; Actinobacteria; Corynebacteriales; Nocardiaceae; Rhodococcus
MTDATVAPTQSGRIAAFFDLDKTVIAKSSALAFSKPFFAQGLLNRRAVLKSSYAQFLFML WP_072634203.1 inhibition of morphological differentiation protein [Rhodococcus sp. M8] Length: 280\nNumber of predicted TMHs: 1\nExp number of AAs in TMHs: 22.80601\nExp number, first 60 AAs: 0.03486\nTotal prob of N-in: 0.04378\noutside 1 245\nTMhelix 246 268\ninside 269 280

31108 GCF_001890475.2_ASM189047v2 Rhodococcus sp. M8 Terrabacteria group; Actinobacteria; Actinobacteria; Corynebacteriales; Nocardiaceae; Rhodococcus
MRLPTSPHSRRRTLSAALLLLLVPVLAGCLRVQATMGVSADDRVSGQIVAATIPADENDK WP_072635433.1 DUF3153 domain-containing protein [Rhodococcus sp. M8] Length: 233\nNumber of predicted TMHs: 2\nExp number of AAs in TMHs: 44.63932\nExp number, first 60 AAs: 21.71772\nTotal prob of N-in: 0.89876\nPOSSIBLE N-term signal sequence\ninside 1 12\nTMhelix 13 35\noutside 36 191\nTMhelix 192 214\ninside 215 233

31109 GCF_000256505.1_ASM25650v1 Rhodococcus sp. P14 Terrabacteria group; Actinobacteria; Actinobacteria; Corynebacteriales; Nocardiaceae; Rhodococcus
MTDATVDPTPSGRIAAFFDLDKTVIAKSSALAFSKPFFAQGLLNRRAVLKSSYAQFLFML WP_010595022.1 MULTISPECIES: hydrolase [Rhodococcus] Length: 280\nNumber of predicted TMHs: 1\nExp number of AAs in TMHs: 23.01026\nExp number, first 60 AAs: 0.0217\nTotal prob of N-in: 0.01643\noutside 1 245\nTMhelix 246 268\ninside 269 280

31110 GCF_000454285.1_RhodP27 Rhodococcus sp. P27 Terrabacteria group; Actinobacteria; Actinobacteria; Corynebacteriales; Nocardiaceae; Rhodococcus
MTDKPANGLNTGRTAAFFDLDKTIIAKSSTLAFSKPFFAQGLINRRSVLKSSYAQFLFLL WP_019746641.1 MULTISPECIES: inhibition of morphological differentiation protein [Rhodococcus] Length: 272\nNumber of predicted TMHs: 1\nExp number of AAs in TMHs: 22.87312\nExp number, first 60 AAs: 0.01889\nTotal prob of N-in: 0.03448\noutside 1 245\nTMhelix 246 268\ninside 269 272

31111 GCF_000454285.1_RhodP27 Rhodococcus sp. P27 Terrabacteria group; Actinobacteria; Actinobacteria; Corynebacteriales; Nocardiaceae; Rhodococcus
MSNPLGQSSNGSRRGALPTPPSGWPIGSYPTYAEAQRAVDYLSDQEFPVQDVTIVGVNLM WP_003945065.1 MULTISPECIES: hypothetical protein [Rhodococcus] Length: 164\nNumber of predicted TMHs: 2\nExp number of AAs in TMHs: 45.26265\nExp number, first 60 AAs: 0.0547\nTotal prob of N-in: 0.87268\ninside 1 73\nTMhelix 74 96\noutside 97 99\nTMhelix 100 122\ninside 123 164

31112 GCF_000763325.2_ASM76332v2 Rhodococcus sp. p52 Terrabacteria group; Actinobacteria; Actinobacteria; Corynebacteriales; Nocardiaceae; Rhodococcus
MFEFSRRALVGATATAALMVAGAGIASAHVTVVAPGAEQGGYTVLTFRVPTESETAGTTA WP_033097871.1 nuclear export factor GLE1 [Rhodococcus sp. p52] Length: 223\nNumber of predicted TMHs: 2\nExp number of AAs in TMHs: 40.51123\nExp number, first 60 AAs: 18.38976\nTotal prob of N-in: 0.70947\nPOSSIBLE N-term signal sequence\ninside 1 8\nTMhelix 9 31\noutside 32 195\nTMhelix 196 218\ninside 219 223

31113 GCF_001620025.1_ASM162002v1 Rhodococcus sp. PBTS 1 Terrabacteria group; Actinobacteria; Actinobacteria; Corynebacteriales; Nocardiaceae; Rhodococcus
MSSPEAPGRIAAFFDLDKTIIAKSSTLAFSKPFFDGGLINRRSVLKSSYAQFLFLVSGAD WP_068103982.1 inhibition of morphological differentiation protein [Rhodococcus sp. PBTS 1] Length: 267\nNumber of predicted TMHs: 1\nExp number of AAs in TMHs: 22.38407\nExp number, first 60 AAs: 0.00462\nTotal prob of N-in: 0.00291\noutside 1 240\nTMhelix 241 263\ninside 264 267

31114 GCF_001620005.1_ASM162000v1 Rhodococcus sp. PBTS 2 Terrabacteria group; Actinobacteria; Actinobacteria; Corynebacteriales; Nocardiaceae; Rhodococcus
MQSTSRTSARTRRRRVLSVLALTLLVAPMLAGCLRVQVTMGVSANDRVAGQIVAATVPTS WP_045841254.1 MULTISPECIES: hypothetical protein [Rhodococcus] Length: 244\nNumber of predicted TMHs: 2\nExp number of AAs in TMHs: 45.38746\nExp number, first 60 AAs: 22.41066\nTotal prob of N-in: 0.99116\nPOSSIBLE N-term signal sequence\ninside 1 15\nTMhelix 16 38\noutside 39 197\nTMhelix 198 220\ninside 221 244

31115 GCF_000963615.1_ASM96361v1 Rhodococcus sp. PML026 Terrabacteria group; Actinobacteria; Actinobacteria; Corynebacteriales; Nocardiaceae; Rhodococcus
MQSTSRTSARTRRRRVLSVLALTLLVAPMLAGCLRVQVTMGVSANDRVAGQIVAATVPTS WP_045841254.1 MULTISPECIES: hypothetical protein [Rhodococcus] Length: 244\nNumber of predicted TMHs: 2\nExp number of AAs in TMHs: 45.38746\nExp number, first 60 AAs: 22.41066\nTotal prob of N-in: 0.99116\nPOSSIBLE N-term signal sequence\ninside 1 15\nTMhelix 16 38\noutside 39 197\nTMhelix 198 220\ninside 221 244

31116 GCF_000278445.1_RhodoR1101_1.0 Rhodococcus sp. R1101 Terrabacteria group; Actinobacteria; Actinobacteria; Corynebacteriales; Nocardiaceae; Rhodococcus
MFEFSRRALVGATTTAALMVVGAGIASAHVTVVAPGAEQGGYSVLTFRVPTESETAGTTA WP_016933499.1 nuclear export factor GLE1 [Rhodococcus sp. R1101] Length: 225\nNumber of predicted TMHs: 1\nExp number of AAs in TMHs: 39.49694\nExp number, first 60 AAs: 17.38264\nTotal prob of N-in: 0.62545\nPOSSIBLE N-term signal sequence\noutside 1 197\nTMhelix 198 220\ninside 221 225

31117 GCF_001040705.1_RD6.2 Rhodococcus sp. RD6.2 Terrabacteria group; Actinobacteria; Actinobacteria; Corynebacteriales; Nocardiaceae; Rhodococcus
MTANAHDRLGEQRVAAFFDLDKTVIARSSTLAFSKPFFDQGLINRRSVLKSSYAQFLFLV WP_050063496.1 inhibition of morphological differentiation protein [Rhodococcus sp. RD6.2] Length: 272\nNumber of predicted TMHs: 1\nExp number of AAs in TMHs: 22.29849\nExp number, first 60 AAs: 0.0047\nTotal prob of N-in: 0.03428\noutside 1 245\nTMhelix 246 268\ninside 269 272

31118 GCF_001555475.1_ASM155547v1 Rhodococcus sp. SC4 Terrabacteria group; Actinobacteria; Actinobacteria; Corynebacteriales; Nocardiaceae; Rhodococcus
MSVFASRGTHCHSRAWVPPLGMSGYSGRVETATGRRRTVVARAALLIAVAAGVLLMHSLV WP_005238637.1 MULTISPECIES: hypothetical protein [Rhodococcus] Length: 157\nNumber of predicted TMHs: 2\nExp

number of AAs in TMHs: 43.88931\nExp number, first 60 AAs: 18.77006\nTotal prob of N-in: 0.99476\nPOSSIBLE N-term signal sequence\ninside 1 44\nTMhelix 45 67\noutside 68 98\nTMhelix 99 121\ninside 122 157

31119 GCF_001555475.1_ASM155547v1 Rhodococcus sp. SC4 Terrabacteria group; Actinobacteria; Actinobacteria; Corynebacteriales; Nocardiaceae; Rhodococcus MRRRTTFTSVKVIAVGLLTMLASMLGAGTALAHSVVISSTPENGAEIAAGPERVSVTFNE WP_005248910.1 MULTISPECIES: copper resistance protein C [Rhodococcus] Length: 177\nNumber of predicted TMHs: 2\nExp number of AAs in TMHs: 44.30022\nExp number, first 60 AAs: 22.13888\nTotal prob of N-in: 0.99979\nPOSSIBLE N-term signal sequence\ninside 1 12\nTMhelix 13 35\noutside 36 147\nTMhelix 148 170\ninside 171 177

31120 GCF_000686025.1_ASM68602v1 Rhodococcus sp. UNC23MFCrub1.1 Terrabacteria group; Actinobacteria; Actinobacteria; Corynebacteriales; Nocardiaceae; Rhodococcus MSTPLRPDAPGRVAAFFDLDKTIIAKSSTLAFSKPFFDGGLINRRSVLKSSYAQFLFLVS WP_027506702.1 inhibition of morphological differentiation protein [Rhodococcus sp. UNC23MFCrub1.1] Length: 271\nNumber of predicted TMHs: 1\nExp number of AAs in TMHs: 22.56564\nExp number, first 60 AAs: 0.0056\nTotal prob of N-in: 0.00523\noutside 1 244\nTMhelix 245 267\ninside 268 271

31121 GCF_000686785.1_ASM68678v1 Rhodococcus sp. UNC363MFTsu5.1 Terrabacteria group; Actinobacteria; Actinobacteria; Corynebacteriales; Nocardiaceae; Rhodococcus MTAEANRQPGNGRVAAFFDLDKTIIAKSSTLAFSKPFFDQGLINRRSVLKSSYAQFLFLL WP_027500369.1 inhibition of morphological differentiation protein [Rhodococcus sp. UNC363MFTsu5.1] Length: 272\nNumber of predicted TMHs: 1\nExp number of AAs in TMHs: 22.69072\nExp number, first 60 AAs: 0.02526\nTotal prob of N-in: 0.01820\noutside 1 245\nTMhelix 246 268\ninside 269 272

31122 GCF_001700945.1_ASM170094v1 Rhodococcus sp. WB1 Terrabacteria group; Actinobacteria; Actinobacteria; Corynebacteriales; Nocardiaceae; Rhodococcus MTDATVAPTQSGRIAAFFDLDKTVIAKSSALAFSKPFFAQGLLNRRAVLKSSYAQFLFML WP_006941337.1 MULTISPECIES: phosphoserine phosphatase [Rhodococcus] Length: 280\nNumber of predicted TMHs: 1\nExp number of AAs in TMHs: 22.81284\nExp number, first 60 AAs: 0.03489\nTotal prob of N-in: 0.04407\noutside 1 245\nTMhelix 246 268\ninside 269 280

31123 GCF_001700945.1_ASM170094v1 Rhodococcus sp. WB1 Terrabacteria group; Actinobacteria; Actinobacteria; Corynebacteriales; Nocardiaceae; Rhodococcus MRLPTSPRSRRRTLSAALLLLLVPVLAGCLRVQATMGVSADDRVSGQIVAATIPADENDK WP_006933384.1 MULTISPECIES: DUF3153 domain-containing protein [Rhodococcus] Length: 233\nNumber of predicted TMHs: 2\nExp number of AAs in TMHs: 44.90952\nExp number, first 60 AAs: 21.98985\nTotal prob of N-in: 0.95979\nPOSSIBLE N-term signal sequence\ninside 1 12\nTMhelix 13 35\noutside 36 191\nTMhelix 192 214\ninside 215 233

31124 GCF_001767395.1_ASM176739v1 Rhodococcus sp. WMMA185 Terrabacteria group; Actinobacteria; Actinobacteria; Corynebacteriales; Nocardiaceae; Rhodococcus MLKPTSTRPPTLGPIVSRRAAIRFAAASTLGVASLPLLTGCAFGEDSNAEADSLAAQSRL WP_070378241.1 hypothetical protein [Rhodococcus sp. WMMA185] Length: 177\nNumber of predicted TMHs: 1\nExp number of AAs in TMHs: 28.34379\nExp number, first 60 AAs: 8.50374999999999\nTotal prob of N-in: 0.59282\ninside 1 152\nTMhelix 153 175\noutside 176 177

31125 GCF_001653035.1_ASM165303v1 Rhodococcus sp. YH3-3 Terrabacteria group; Actinobacteria; Actinobacteria; Corynebacteriales; Nocardiaceae; Rhodococcus MSNPLGQSSNGSRRGALPTPPSGWPIGSYPTYAEAQRAVDYLSDQEFPVQDVTIVGVNLM WP_003945065.1 MULTISPECIES: hypothetical protein [Rhodococcus] Length: 164\nNumber of predicted TMHs: 2\nExp number of AAs in TMHs: 45.26265\nExp number, first 60 AAs: 0.0547\nTotal prob of N-in: 0.87268\ninside 1 73\nTMhelix 74 96\noutside 97 99\nTMhelix 100 122\ninside 123 164

31126 GCF_001653035.1_ASM165303v1 Rhodococcus sp. YH3-3 Terrabacteria group; Actinobacteria; Actinobacteria; Corynebacteriales; Nocardiaceae; Rhodococcus MTDKPANGLNTGRTAAFFDLDKTIIAKSSTLAFSKPFFAQGLINRRSVLKSSYAQFLFLL WP_003946123.1 MULTISPECIES: haloacid dehalogenase [Rhodococcus] Length: 272\nNumber of predicted TMHs: 1\nExp number of AAs in TMHs: 22.87312\nExp number, first 60 AAs: 0.01889\nTotal prob of N-in: 0.03448\noutside 1 245\nTMhelix 246 268\ninside 269 272

31127 GCF_002165495.1_ASM216549v1 Rhodococcus sp. YL-0 Terrabacteria group; Actinobacteria; Actinobacteria; Corynebacteriales; Nocardiaceae; Rhodococcus

MTDKPANGLNTGRTAAFFDLDKTIIAKSSTLAFSKPFFAQGLINRRSVLKSSYAQFLFLL WP_003946123.1 MULTISPECIES: haloacid dehalogenase [Rhodococcus] Length: 272\nNumber of predicted TMHs: 1\nExp number of AAs in TMHs: 22.87312\nExp number, first 60 AAs: 0.01889\nTotal prob of N-in: 0.03448\noutside 1 245\nTMhelix 246 268\ninside 269 272

31128 GCF_002165495.1_ASM216549v1 Rhodococcus sp. YL-0 Terrabacteria group; Actinobacteria; Actinobacteria; Corynebacteriales; Nocardiaceae; Rhodococcus MSNPLGQSSNGSRRGALPTPPSGWPIGSYPTYAEAQRAVDYLSDQEFPVQDVTIVGVNLM WP_003945065.1 MULTISPECIES: hypothetical protein [Rhodococcus] Length: 164\nNumber of predicted TMHs: 2\nExp number of AAs in TMHs: 45.26265\nExp number, first 60 AAs: 0.0547\nTotal prob of N-in: 0.87268\ninside 1 73\nTMhelix 74 96\noutside 97 99\nTMhelix 100 122\ninside 123 164

31129 GCF_001942025.1_ASM194202v1 Rhodococcus sp. YL-1 Terrabacteria group; Actinobacteria; Actinobacteria; Corynebacteriales; Nocardiaceae; Rhodococcus MSNPLGQSSNGSRRGALPTPPSGWPIGSYPTYAEAQRAVDYLSDQEFPVQDVTIVGVNLM WP_003945065.1 MULTISPECIES: hypothetical protein [Rhodococcus] Length: 164\nNumber of predicted TMHs: 2\nExp number of AAs in TMHs: 45.26265\nExp number, first 60 AAs: 0.0547\nTotal prob of N-in: 0.87268\ninside 1 73\nTMhelix 74 96\noutside 97 99\nTMhelix 100 122\ninside 123 164

31130 GCF_001942025.1_ASM194202v1 Rhodococcus sp. YL-1 Terrabacteria group; Actinobacteria; Actinobacteria; Corynebacteriales; Nocardiaceae; Rhodococcus MTDKPANGLNTGRTAAFFDLDKTIIAKSSTLAFSKPFFAQGLINRRSVLKSSYAQFLFLL WP_003946123.1 MULTISPECIES: haloacid dehalogenase [Rhodococcus] Length: 272\nNumber of predicted TMHs: 1\nExp number of AAs in TMHs: 22.87312\nExp number, first 60 AAs: 0.01889\nTotal prob of N-in: 0.03448\noutside 1 245\nTMhelix 246 268\ninside 269 272

31131 GCF_001895045.1_ASM189504v1 Rhodococcus coprophilus NBRC 100603 Terrabacteria group; Actinobacteria; Actinobacteria; Corynebacteriales; Nocardiaceae; Rhodococcus; Rhodococcus coprophilus MTGTERARRAVLTVAIALSALAVALFLGALRNDRTIEQDMGTATAEVLSAGSLRSAVAFVWP_072704387.1 DUF3592 domain-containing protein [Rhodococcus coprophilus] Length: 151\nNumber of predicted TMHs: 2\nExp number of AAs in TMHs: 43.56309\nExp number, first 60 AAs: 20.76495\nTotal prob of N-in: 0.71201\nPOSSIBLE N-term signal sequence\ninside 1 6\nTMhelix 7 29\noutside 30 106\nTMhelix 107 129\ninside 130 151

31132 GCF_000454045.1_ASM45404v1 Rhodococcus erythropolis CCM2595 Terrabacteria group; Actinobacteria; Actinobacteria; Corynebacteriales; Nocardiaceae; Rhodococcus; Rhodococcus erythropolis MSNPLGQSSNGSRRGALPTPPSGWPIGSYPTYAEAQRAVDYLSDQEFPVQDVTIVGVNLM WP_003945065.1 MULTISPECIES: hypothetical protein [Rhodococcus] Length: 164\nNumber of predicted TMHs: 2\nExp number of AAs in TMHs: 45.26265\nExp number, first 60 AAs: 0.0547\nTotal prob of N-in: 0.87268\ninside 1 73\nTMhelix 74 96\noutside 97 99\nTMhelix 100 122\ninside 123 164

31133 GCF_000454045.1_ASM45404v1 Rhodococcus erythropolis CCM2595 Terrabacteria group; Actinobacteria; Actinobacteria; Corynebacteriales; Nocardiaceae; Rhodococcus; Rhodococcus erythropolis MTDKPANGLNTGRTAAFFDLDKTIIAKSSTLAFSKPFFAQGLINRRSVLKSSYAQFLFLL WP_019746641.1 MULTISPECIES: inhibition of morphological differentiation protein [Rhodococcus] Length: 272\nNumber of predicted TMHs: 1\nExp number of AAs in TMHs: 22.87312\nExp number, first 60 AAs: 0.01889\nTotal prob of N-in: 0.03448\noutside 1 245\nTMhelix 246 268\ninside 269 272

31134 GCF_000454425.1_RodocDN1 Rhodococcus erythropolis DN1 Terrabacteria group; Actinobacteria; Actinobacteria; Corynebacteriales; Nocardiaceae; Rhodococcus; Rhodococcus erythropolis MSNPLGQSSNGSRRGALPTPPSGWPIGSYPTYAEAQRAVDYLSDQEFPVQDVTIVGVNLM WP_003945065.1 MULTISPECIES: hypothetical protein [Rhodococcus] Length: 164\nNumber of predicted TMHs: 2\nExp number of AAs in TMHs: 45.26265\nExp number, first 60 AAs: 0.0547\nTotal prob of N-in: 0.87268\ninside 1 73\nTMhelix 74 96\noutside 97 99\nTMhelix 100 122\ninside 123 164

31135 GCF_000454425.1_RodocDN1 Rhodococcus erythropolis DN1 Terrabacteria group; Actinobacteria; Actinobacteria; Corynebacteriales; Nocardiaceae; Rhodococcus; Rhodococcus erythropolis MTDKPANGLNTGRTAAFFDLDKTIIAKSSTLAFSKPFFAQGLINRRSVLKSSYAQFLFLL WP_021334243.1 inhibition of morphological differentiation protein [Rhodococcus erythropolis] Length: 272\nNumber of predicted TMHs: 1\nExp number of AAs in TMHs: 22.87334\nExp number, first 60 AAs: 0.01889\nTotal prob of N-in: 0.03449\noutside 1 245\nTMhelix 246 268\ninside 269 272

31136	GCF_001552595.1_ASM155259v1	Rhodococcus erythropolis NBRC 15567 Terrabacteria group; Actinobacteria; Actinobacteria; Corynebacteriales; Nocardiaceae; Rhodococcus; Rhodococcus erythropolis
MSNPLGQSSNGSRRGALPTPPSGWPIGSYPTYAEAQRAVDYLSDQEFPVQDVTIVGVNLM	WP_003945065.1
MULTISPECIES: hypothetical protein [Rhodococcus]	Length: 164\nNumber of predicted TMHs:  2\nExp number of AAs in TMHs: 45.26265\nExp number, first 60 AAs:  0.0547\nTotal prob of N-in:   0.87268\ninside   1   73\nTMhelix   74   96\noutside   97   99\nTMhelix   100   122\ninside   123   164

31137	GCF_001552595.1_ASM155259v1	Rhodococcus erythropolis NBRC 15567 Terrabacteria group; Actinobacteria; Actinobacteria; Corynebacteriales; Nocardiaceae; Rhodococcus; Rhodococcus erythropolis
MTDKPANGLNTGRTAAFFDLDKTIIAKSSTLAFSKPFFAQGLINRRSVLKSSYAQFLFLL	WP_060939216.1	inhibition of morphological differentiation protein [Rhodococcus erythropolis]	Length: 272\nNumber of predicted TMHs:  1\nExp number of AAs in TMHs: 23.23429\nExp number, first 60 AAs:  0.02233\nTotal prob of N-in:   0.05193\noutside   1   245\nTMhelix   246   268\ninside   269   272

31138	GCF_000010105.1_ASM1010v1	Rhodococcus erythropolis PR4	Terrabacteria group; Actinobacteria; Actinobacteria; Corynebacteriales; Nocardiaceae; Rhodococcus; Rhodococcus erythropolis
MSNPLGQSSNGSRRGALPTPPSGWPIGSYPTYAEAQRAVDYLSDQEFPVQDVTIVGVNLM	WP_003945065.1
MULTISPECIES: hypothetical protein [Rhodococcus]	Length: 164\nNumber of predicted TMHs:  2\nExp number of AAs in TMHs: 45.26265\nExp number, first 60 AAs:  0.0547\nTotal prob of N-in:   0.87268\ninside   1   73\nTMhelix   74   96\noutside   97   99\nTMhelix   100   122\ninside   123   164

31139	GCF_000010105.1_ASM1010v1	Rhodococcus erythropolis PR4	Terrabacteria group; Actinobacteria; Actinobacteria; Corynebacteriales; Nocardiaceae; Rhodococcus; Rhodococcus erythropolis
MTDKPANGLNTGRTAAFFDLDKTIIAKSSTLAFSKPFFAQGLINRRSVLKSSYAQFLFLL	WP_019746641.1	MULTISPECIES: inhibition of morphological differentiation protein [Rhodococcus]	Length: 272\nNumber of predicted TMHs:  1\nExp number of AAs in TMHs: 22.87312\nExp number, first 60 AAs:  0.01889\nTotal prob of N-in:   0.03448\noutside   1   245\nTMhelix   246   268\ninside   269   272

31140	GCF_000696675.2_ASM69667v2	Rhodococcus erythropolis R138	Terrabacteria group; Actinobacteria; Actinobacteria; Corynebacteriales; Nocardiaceae; Rhodococcus; Rhodococcus erythropolis
MTDKPANGLNTGRTAAFFDLDKTIIAKSSTLAFSKPFFAQGLINRRSVLKSSYAQFLFLL	WP_019746641.1	MULTISPECIES: inhibition of morphological differentiation protein [Rhodococcus]	Length: 272\nNumber of predicted TMHs:  1\nExp number of AAs in TMHs: 22.87312\nExp number, first 60 AAs:  0.01889\nTotal prob of N-in:   0.03448\noutside   1   245\nTMhelix   246   268\ninside   269   272

31141	GCF_000696675.2_ASM69667v2	Rhodococcus erythropolis R138	Terrabacteria group; Actinobacteria; Actinobacteria; Corynebacteriales; Nocardiaceae; Rhodococcus; Rhodococcus erythropolis
MSNPLGQSSNGSRRGALPTPPSGWPIGSYPTYAEAQRAVDYLSDQEFPVQDVTIVGVNLM	WP_003945065.1
MULTISPECIES: hypothetical protein [Rhodococcus]	Length: 164\nNumber of predicted TMHs:  2\nExp number of AAs in TMHs: 45.26265\nExp number, first 60 AAs:  0.0547\nTotal prob of N-in:   0.87268\ninside   1   73\nTMhelix   74   96\noutside   97   99\nTMhelix   100   122\ninside   123   164

31142	GCF_000174835.1_ASM17483v1	Rhodococcus erythropolis SK121	Terrabacteria group; Actinobacteria; Actinobacteria; Corynebacteriales; Nocardiaceae; Rhodococcus; Rhodococcus erythropolis
MSNPLGQSSNGSRRGALPTPPSGWPIGSYPTYAEAQRAVDYLSDQEFPVQDVTIVGVNLM	WP_003945065.1
MULTISPECIES: hypothetical protein [Rhodococcus]	Length: 164\nNumber of predicted TMHs:  2\nExp number of AAs in TMHs: 45.26265\nExp number, first 60 AAs:  0.0547\nTotal prob of N-in:   0.87268\ninside   1   73\nTMhelix   74   96\noutside   97   99\nTMhelix   100   122\ninside   123   164

31143	GCF_000174835.1_ASM17483v1	Rhodococcus erythropolis SK121	Terrabacteria group; Actinobacteria; Actinobacteria; Corynebacteriales; Nocardiaceae; Rhodococcus; Rhodococcus erythropolis
MTDKPANGLNTGRTAAFFDLDKTIIAKSSTLAFSKPFFAQGLINRRSVLKSSYAQFLFLL	WP_003946123.1	MULTISPECIES: haloacid dehalogenase [Rhodococcus]	Length: 272\nNumber of predicted TMHs:  1\nExp number of AAs in TMHs: 22.87312\nExp number, first 60 AAs:  0.01889\nTotal prob of N-in:   0.03448\noutside   1   245\nTMhelix   246   268\ninside   269   272

31144	GCF_000225665.1_ASM22566v2	Rhodococcus erythropolis XP Terrabacteria group; Actinobacteria; Actinobacteria; Corynebacteriales; Nocardiaceae; Rhodococcus; Rhodococcus erythropolis
MSNPLGQSSNGSRRGALPTPPSGWPIGSYPTYAEAQRAVDYLSDQEFPVQDVTIVGVNLM	WP_003945065.1
MULTISPECIES: hypothetical protein [Rhodococcus]	Length: 164\nNumber of predicted TMHs:  2\nExp

number of AAs in TMHs: 45.26265\nExp number, first 60 AAs: 0.0547\nTotal prob of N-in: 0.87268\ninside 1 73\nTMhelix 74 96\noutside 97 99\nTMhelix 100 122\ninside 123 164

31145 GCF_000225665.1_ASM22566v2 Rhodococcus erythropolis XP Terrabacteria group; Actinobacteria; Actinobacteria; Corynebacteriales; Nocardiaceae; Rhodococcus; Rhodococcus erythropolis MTDKPANGLNTGRTAAFFDLDKTIIAKSSTLAFSKPFFAQGLINRRSVLKSSYAQFLFLL WP_019746641.1 MULTISPECIES: inhibition of morphological differentiation protein [Rhodococcus] Length: 272\nNumber of predicted TMHs: 1\nExp number of AAs in TMHs: 22.87312\nExp number, first 60 AAs: 0.01889\nTotal prob of N-in: 0.03448\noutside 1 245\nTMhelix 246 268\ninside 269 272

31146 GCF_000760875.1_R_fas_05-561-1 Rhodococcus fascians 05-561-1 Terrabacteria group; Actinobacteria; Actinobacteria; Corynebacteriales; Nocardiaceae; Rhodococcus; Rhodococcus fascians MQSTSRTSARTRRRRVLSVLALTLLVAPMLAGCLRVQVTMGVSANDRVAGQIVAATVPTS WP_045841254.1 MULTISPECIES: hypothetical protein [Rhodococcus] Length: 244\nNumber of predicted TMHs: 2\nExp number of AAs in TMHs: 45.38746\nExp number, first 60 AAs: 22.41066\nTotal prob of N-in: 0.99116\nPOSSIBLE N-term signal sequence\ninside 1 15\nTMhelix 16 38\noutside 39 197\nTMhelix 198 220\ninside 221 244

31147 GCF_000760875.1_R_fas_05-561-1 Rhodococcus fascians 05-561-1 Terrabacteria group; Actinobacteria; Actinobacteria; Corynebacteriales; Nocardiaceae; Rhodococcus; Rhodococcus fascians MGVRSVQRRPRTVTAHRRRACVVAALTVLTLTYEALIGPSFAHAERPLIDPVQLPPPSPP WP_052057192.1 type VII secretion-associated serine protease mycosin [Rhodococcus fascians] Length: 472\nNumber of predicted TMHs: 2\nExp number of AAs in TMHs: 44.59771\nExp number, first 60 AAs: 21.4162\nTotal prob of N-in: 0.99106\nPOSSIBLE N-term signal sequence\ninside 1 19\nTMhelix 20 42\noutside 43 445\nTMhelix 446 468\ninside 469 472

31148 GCF_000760675.1_R_fas_A3b Rhodococcus fascians A3b Terrabacteria group; Actinobacteria; Actinobacteria; Corynebacteriales; Nocardiaceae; Rhodococcus; Rhodococcus fascians MQSTSRTSARTRRRRVLSVLALTLLVAPMLAGCLRVQVTMGVSANDRVAGQIVAATVPTS WP_045841254.1 MULTISPECIES: hypothetical protein [Rhodococcus] Length: 244\nNumber of predicted TMHs: 2\nExp number of AAs in TMHs: 45.38746\nExp number, first 60 AAs: 22.41066\nTotal prob of N-in: 0.99116\nPOSSIBLE N-term signal sequence\ninside 1 15\nTMhelix 16 38\noutside 39 197\nTMhelix 198 220\ninside 221 244

31149 GCF_000760775.1_R_fas_A78 Rhodococcus fascians A78 Terrabacteria group; Actinobacteria; Actinobacteria; Corynebacteriales; Nocardiaceae; Rhodococcus; Rhodococcus fascians MQSTSRTSARTRRRRVLSVLALTLLVAPMLAGCLRVQVTMGVSANDRVAGQIVAATVPTS WP_045841254.1 MULTISPECIES: hypothetical protein [Rhodococcus] Length: 244\nNumber of predicted TMHs: 2\nExp number of AAs in TMHs: 45.38746\nExp number, first 60 AAs: 22.41066\nTotal prob of N-in: 0.99116\nPOSSIBLE N-term signal sequence\ninside 1 15\nTMhelix 16 38\noutside 39 197\nTMhelix 198 220\ninside 221 244

31150 GCF_000760975.1_R_fas_D188 Rhodococcus fascians D188 Terrabacteria group; Actinobacteria; Actinobacteria; Corynebacteriales; Nocardiaceae; Rhodococcus; Rhodococcus fascians MQSTSRTSARTRRRRVLSVLALTLLVAPMLAGCLRVQVTMGVSANDRVAGQIVAATVPTS WP_045841254.1 MULTISPECIES: hypothetical protein [Rhodococcus] Length: 244\nNumber of predicted TMHs: 2\nExp number of AAs in TMHs: 45.38746\nExp number, first 60 AAs: 22.41066\nTotal prob of N-in: 0.99116\nPOSSIBLE N-term signal sequence\ninside 1 15\nTMhelix 16 38\noutside 39 197\nTMhelix 198 220\ninside 221 244

31151 GCF_001620305.1_ASM162030v1 Rhodococcus fascians D188 Terrabacteria group; Actinobacteria; Actinobacteria; Corynebacteriales; Nocardiaceae; Rhodococcus; Rhodococcus fascians MQSTSRTSARTRRRRVLSVLALTLLVAPMLAGCLRVQVTMGVSANDRVAGQIVAATVPTS WP_045841254.1 MULTISPECIES: hypothetical protein [Rhodococcus] Length: 244\nNumber of predicted TMHs: 2\nExp number of AAs in TMHs: 45.38746\nExp number, first 60 AAs: 22.41066\nTotal prob of N-in: 0.99116\nPOSSIBLE N-term signal sequence\ninside 1 15\nTMhelix 16 38\noutside 39 197\nTMhelix 198 220\ninside 221 244

31152 GCF_000760795.1_R_fas_GIC26 Rhodococcus fascians GIC26 Terrabacteria group; Actinobacteria; Actinobacteria; Corynebacteriales; Nocardiaceae; Rhodococcus; Rhodococcus fascians MQSTSRTSARTRRRRVLSVLALTLLVAPMLAGCLRVQVTMGVSANDRVAGQIVAATVPTS WP_045841254.1 MULTISPECIES: hypothetical protein [Rhodococcus] Length: 244\nNumber of predicted TMHs: 2\nExp number of AAs in TMHs: 45.38746\nExp number, first 60 AAs: 22.41066\nTotal prob of N-in: 0.99116\nPOSSIBLE N-term signal sequence\ninside 1 15\nTMhelix 16 38\noutside 39 197\nTMhelix 198 220\ninside 221 244

31153 GCF_000760815.1_R_fas_GIC36 Rhodococcus fascians GIC36 Terrabacteria group; Actinobacteria; Actinobacteria; Corynebacteriales; Nocardiaceae; Rhodococcus; Rhodococcus fascians

MQSTSRTSARTRRRRVLSVLALTLLVAPMLAGCLRVQVTMGVSANDRVAGQIVAATVPTS WP_045841254.1 MULTISPECIES: hypothetical protein [Rhodococcus] Length: 244\nNumber of predicted TMHs: 2\nExp number of AAs in TMHs: 45.38746\nExp number, first 60 AAs: 22.41066\nTotal prob of N-in: 0.99116\nPOSSIBLE N-term signal sequence\ninside 1 15\nTMhelix 16 38\noutside 39 197\nTMhelix 198 220\ninside 221 244

31154 GCF_000760995.1_R_fas_LMG3602 Rhodococcus fascians LMG 3602 Terrabacteria group; Actinobacteria; Actinobacteria; Corynebacteriales; Nocardiaceae; Rhodococcus; Rhodococcus fascians MQSTSRTSARTRRRRVLSVLALTLLVAPMLAGCLRVQVTMGVSANDRVAGQIVAATVPTS WP_045841254.1 MULTISPECIES: hypothetical protein [Rhodococcus] Length: 244\nNumber of predicted TMHs: 2\nExp number of AAs in TMHs: 45.38746\nExp number, first 60 AAs: 22.41066\nTotal prob of N-in: 0.99116\nPOSSIBLE N-term signal sequence\ninside 1 15\nTMhelix 16 38\noutside 39 197\nTMhelix 198 220\ninside 221 244

31155 GCF_000761015.1_R_fas_LMG3605 Rhodococcus fascians LMG 3605 Terrabacteria group; Actinobacteria; Actinobacteria; Corynebacteriales; Nocardiaceae; Rhodococcus; Rhodococcus fascians MQSTSRTSARTRRRRVLSVLALTLLVAPMLAGCLRVQVTMGVSANDRVAGQIVAATVPTS WP_045841254.1 MULTISPECIES: hypothetical protein [Rhodococcus] Length: 244\nNumber of predicted TMHs: 2\nExp number of AAs in TMHs: 45.38746\nExp number, first 60 AAs: 22.41066\nTotal prob of N-in: 0.99116\nPOSSIBLE N-term signal sequence\ninside 1 15\nTMhelix 16 38\noutside 39 197\nTMhelix 198 220\ninside 221 244

31156 GCF_000761035.1_R_fas_LMG3616 Rhodococcus fascians LMG 3616 Terrabacteria group; Actinobacteria; Actinobacteria; Corynebacteriales; Nocardiaceae; Rhodococcus; Rhodococcus fascians MQSTSRTSARTRRRRVLSVLALTLLVAPMLAGCLRVQVTMGVSANDRVAGQIVAATVPTS WP_045841254.1 MULTISPECIES: hypothetical protein [Rhodococcus] Length: 244\nNumber of predicted TMHs: 2\nExp number of AAs in TMHs: 45.38746\nExp number, first 60 AAs: 22.41066\nTotal prob of N-in: 0.99116\nPOSSIBLE N-term signal sequence\ninside 1 15\nTMhelix 16 38\noutside 39 197\nTMhelix 198 220\ninside 221 244

31157 GCF_000761055.1_R_fas_LMG3623 Rhodococcus fascians NBRC 12155 = LMG 3623 Terrabacteria group; Actinobacteria; Actinobacteria; Corynebacteriales; Nocardiaceae; Rhodococcus; Rhodococcus fascians MQSTSRTSARTRRRRVLSVLALTLLVAPMLAGCLRVQVTMGVSANDRVAGQIVAATVPTS WP_045841254.1 MULTISPECIES: hypothetical protein [Rhodococcus] Length: 244\nNumber of predicted TMHs: 2\nExp number of AAs in TMHs: 45.38746\nExp number, first 60 AAs: 22.41066\nTotal prob of N-in: 0.99116\nPOSSIBLE N-term signal sequence\ninside 1 15\nTMhelix 16 38\noutside 39 197\nTMhelix 198 220\ninside 221 244

31158 GCF_001894785.1_ASM189478v1 Rhodococcus fascians NBRC 12155 = LMG 3623 Terrabacteria group; Actinobacteria; Actinobacteria; Corynebacteriales; Nocardiaceae; Rhodococcus; Rhodococcus fascians MQSTSRTSARTRRRRVLSVLALTLLVAPMLAGCLRVQVTMGVSANDRVAGQIVAATVPTS WP_045841254.1 MULTISPECIES: hypothetical protein [Rhodococcus] Length: 244\nNumber of predicted TMHs: 2\nExp number of AAs in TMHs: 45.38746\nExp number, first 60 AAs: 22.41066\nTotal prob of N-in: 0.99116\nPOSSIBLE N-term signal sequence\ninside 1 15\nTMhelix 16 38\noutside 39 197\nTMhelix 198 220\ninside 221 244

31159 GCF_001894805.1_ASM189480v1 Rhodococcus globerulus NBRC 14531 Terrabacteria group; Actinobacteria; Actinobacteria; Corynebacteriales; Nocardiaceae; Rhodococcus; Rhodococcus globerulus MTDKLGNGLTTGRTAAFFDLDKTVIAKSSTLAFSRPFFAQGLINRRSVLKSSYAQFLFLL WP_033235947.1 MULTISPECIES: inhibition of morphological differentiation protein [Rhodococcus] Length: 272\nNumber of predicted TMHs: 1\nExp number of AAs in TMHs: 22.91283\nExp number, first 60 AAs: 0.00718\nTotal prob of N-in: 0.01617\noutside 1 245\nTMhelix 246 268\ninside 269 272

31160 GCF_001894805.1_ASM189480v1 Rhodococcus globerulus NBRC 14531 Terrabacteria group; Actinobacteria; Actinobacteria; Corynebacteriales; Nocardiaceae; Rhodococcus; Rhodococcus globerulus MSNPLAQSPNGSRRGALPTPPSGWPIGSYPTYAEAQRAVDYLSDQEFSVQDVTIVGVNLM WP_033234426.1 MULTISPECIES: hypothetical protein [Rhodococcus] Length: 164\nNumber of predicted TMHs: 2\nExp number of AAs in TMHs: 45.15237\nExp number, first 60 AAs: 0.04302\nTotal prob of N-in: 0.64861\ninside 1 73\nTMhelix 74 96\noutside 97 100\nTMhelix 101 123\ninside 124 164

31161 GCF_000196695.1_ASM19669v1 Rhodococcus hoagii 103S Terrabacteria group; Actinobacteria; Actinobacteria; Corynebacteriales; Nocardiaceae; Rhodococcus; Rhodococcus hoagii MAAHPKSARRRILLTATASALVLGAGGAPAAAVPLAGQSPLAADQAVAPTTTPPFSTPNT WP_013416807.1 D-alanyl-D-alanine carboxypeptidase [Rhodococcus equi] Length: 444\nNumber of predicted TMHs: 2\nExp number of AAs in TMHs: 42.7856499999999\nExp number, first 60 AAs: 21.57858\nTotal prob of N-in: 0.96909\nPOSSIBLE N-term signal sequence\ninside 1 11\nTMhelix 12 34\noutside 35 412\nTMhelix 413 435\ninside 436 444

31162 GCF_000196695.1_ASM19669v1 Rhodococcus hoagii 103S Terrabacteria group; Actinobacteria; Actinobacteria; Corynebacteriales; Nocardiaceae; Rhodococcus; Rhodococcus hoagii MTADPSDLTRRGRTAAFFDLDKTIIAKSSALAFSKPFFAQGLITRRSVLQSSYAQFLFIL WP_005515815.1 morphological differentiation-associated protein [Rhodococcus equi] Length: 272\nNumber of predicted TMHs: 1\nExp number of AAs in TMHs: 21.05758\nExp number, first 60 AAs: 0.04208\nTotal prob of N-in: 0.02567\noutside 1 244\nTMhelix 245 267\ninside 268 272

31163 GCF_000164155.2_ASM16415v2 Rhodococcus hoagii ATCC 33707 Terrabacteria group; Actinobacteria; Actinobacteria; Corynebacteriales; Nocardiaceae; Rhodococcus; Rhodococcus hoagii MAAHPKSARRRILLTATASALVLGAGGAPAAAVPLAGQSPLAADQAVAPTTTPPFSTPNT WP_005518562.1 D-alanyl-D-alanine carboxypeptidase [Rhodococcus hoagii] Length: 444\nNumber of predicted TMHs: 2\nExp number of AAs in TMHs: 42.7856999999999\nExp number, first 60 AAs: 21.57856\nTotal prob of N-in: 0.96909\nPOSSIBLE N-term signal sequence\ninside 1 11\nTMhelix 12 34\noutside 35 412\nTMhelix 413 435\ninside 436 444

31164 GCF_000164155.2_ASM16415v2 Rhodococcus hoagii ATCC 33707 Terrabacteria group; Actinobacteria; Actinobacteria; Corynebacteriales; Nocardiaceae; Rhodococcus; Rhodococcus hoagii MTADPSDLTRRGRTAAFFDLDKTIIAKSSALAFSKPFFAQGLITRRSVLQSSYAQFLFIL WP_005515815.1 morphological differentiation-associated protein [Rhodococcus hoagii] Length: 272\nNumber of predicted TMHs: 1\nExp number of AAs in TMHs: 21.05758\nExp number, first 60 AAs: 0.04208\nTotal prob of N-in: 0.02567\noutside 1 244\nTMhelix 245 267\ninside 268 272

31165 GCF_000473915.1_Prescottia_equi_Genome_sequencing Rhodococcus hoagii NBRC 101255 = C 7 Terrabacteria group; Actinobacteria; Actinobacteria; Corynebacteriales; Nocardiaceae; Rhodococcus; Rhodococcus hoagii MTADPSDLTRRGRTAAFFDLDKTIIAKSSALAFSKPFFAQGLITRRSVLQSSYAQFLFIL WP_005515815.1 morphological differentiation-associated protein [Rhodococcus hoagii] Length: 272\nNumber of predicted TMHs: 1\nExp number of AAs in TMHs: 21.05758\nExp number, first 60 AAs: 0.04208\nTotal prob of N-in: 0.02567\noutside 1 244\nTMhelix 245 267\ninside 268 272

31166 GCF_000473915.1_Prescottia_equi_Genome_sequencing Rhodococcus hoagii NBRC 101255 = C 7 Terrabacteria group; Actinobacteria; Actinobacteria; Corynebacteriales; Nocardiaceae; Rhodococcus; Rhodococcus hoagii MAAHPKSARRRILLTATASALVLGAGGAPAAAVPLAGQSPLAADQAVAPTTTPPFSTPNT WP_022597221.1 D-alanyl-D-alanine carboxypeptidase [Rhodococcus hoagii] Length: 444\nNumber of predicted TMHs: 2\nExp number of AAs in TMHs: 42.86088\nExp number, first 60 AAs: 21.57924\nTotal prob of N-in: 0.96909\nPOSSIBLE N-term signal sequence\ninside 1 11\nTMhelix 12 34\noutside 35 412\nTMhelix 413 435\ninside 436 444

31167 GCF_001552575.1_ASM155257v1 Rhodococcus hoagii NBRC 101255 = C 7 Terrabacteria group; Actinobacteria; Actinobacteria; Corynebacteriales; Nocardiaceae; Rhodococcus; Rhodococcus hoagii MTADPSDLTRRGRTAAFFDLDKTIIAKSSALAFSKPFFAQGLITRRSVLQSSYAQFLFIL WP_005515815.1 morphological differentiation-associated protein [Rhodococcus hoagii] Length: 272\nNumber of predicted TMHs: 1\nExp number of AAs in TMHs: 21.05758\nExp number, first 60 AAs: 0.04208\nTotal prob of N-in: 0.02567\noutside 1 244\nTMhelix 245 267\ninside 268 272

31168 GCF_001552575.1_ASM155257v1 Rhodococcus hoagii NBRC 101255 = C 7 Terrabacteria group; Actinobacteria; Actinobacteria; Corynebacteriales; Nocardiaceae; Rhodococcus; Rhodococcus hoagii MAAHPKSARRRILLTATASALVLGAGGAPAAAVPLAGQSPLAADQAVAPTTTPPFSTPNT WP_022597221.1 D-alanyl-D-alanine carboxypeptidase [Rhodococcus hoagii] Length: 444\nNumber of predicted TMHs: 2\nExp number of AAs in TMHs: 42.86088\nExp number, first 60 AAs: 21.57924\nTotal prob of N-in: 0.96909\nPOSSIBLE N-term signal sequence\ninside 1 11\nTMhelix 12 34\noutside 35 412\nTMhelix 413 435\ninside 436 444

31169 GCF_000260815.1_ASM26081v1 Rhodococcus imtechensis RKJ300 = JCM 13270 Terrabacteria group; Actinobacteria; Actinobacteria; Corynebacteriales; Nocardiaceae; Rhodococcus; Rhodococcus imtechensis MRRRTTFTSVKVIAVGLLTMLASMLGAGTALAHSVVISSTPENGAEIAAGPERVSVTFNE WP_039951718.1 copper resistance protein CopC [Rhodococcus imtechensis] Length: 177\nNumber of predicted TMHs: 2\nExp number of AAs in TMHs: 44.2843\nExp number, first 60 AAs: 22.13587\nTotal prob of N-in: 0.99964\nPOSSIBLE N-term signal sequence\ninside 1 12\nTMhelix 13 35\noutside 36 147\nTMhelix 148 170\ninside 171 177

31170 GCF_000260815.1_ASM26081v1 Rhodococcus imtechensis RKJ300 = JCM 13270 Terrabacteria group; Actinobacteria; Actinobacteria; Corynebacteriales; Nocardiaceae; Rhodococcus; Rhodococcus imtechensis MSVFASRGTHCHSRVQVPPLGMSGYSGRVETASGRRRTVVARAALLVAVAAGVLLMHSLI WP_039952988.1 hypothetical protein [Rhodococcus imtechensis] Length: 157\nNumber of predicted TMHs: 2\nExp number of

AAs in TMHs: 43.94831\nExp number, first 60 AAs: 18.16517\nTotal prob of N-in: 0.99763\nPOSSIBLE N-term signal sequence\ninside 1 44\nTMhelix 45 67\noutside 68 94\nTMhelix 95 117\ninside 118 157

31171 GCF_001894825.1_ASM189482v1 Rhodococcus jostii NBRC 16295 Terrabacteria group; Actinobacteria; Actinobacteria; Corynebacteriales; Nocardiaceae; Rhodococcus; Rhodococcus jostii MSVFASRGTHCHSRARVPPMGMSGYSGGVETAAVRRRTLVARAALLLTLAAGVLLMHSLI WP_073365742.1 hypothetical protein [Rhodococcus jostii] Length: 157\nNumber of predicted TMHs: 2\nExp number of AAs in TMHs: 43.80327\nExp number, first 60 AAs: 18.46059\nTotal prob of N-in: 0.99453\nPOSSIBLE N-term signal sequence\ninside 1 42\nTMhelix 43 65\noutside 66 94\nTMhelix 95 117\ninside 118 157

31172 GCF_000014565.1_ASM1456v1 Rhodococcus jostii RHA1 Terrabacteria group; Actinobacteria; Actinobacteria; Corynebacteriales; Nocardiaceae; Rhodococcus; Rhodococcus jostii MRRRTTFTSVKVIAVGLLTMLASMLGAGTAVAHSVVISSTPENGAQIAAGPERVSVTFNE WP_041811727.1 copper resistance protein CopC [Rhodococcus jostii] Length: 177\nNumber of predicted TMHs: 2\nExp number of AAs in TMHs: 44.34275\nExp number, first 60 AAs: 22.21506\nTotal prob of N-in: 0.99949\nPOSSIBLE N-term signal sequence\ninside 1 12\nTMhelix 13 35\noutside 36 147\nTMhelix 148 170\ninside 171 177

31173 GCF_001894845.1_ASM189484v1 Rhodococcus koreensis JCM 10743 = NBRC 100607 Terrabacteria group; Actinobacteria; Actinobacteria; Corynebacteriales; Nocardiaceae; Rhodococcus; Rhodococcus koreensis MVVWAPAQLDVTQRMPARSRRDVFSTLRGACTNGPPVSAIALAGMQKLSSPRRRRGACVL WP_083395499.1 hypothetical protein [Rhodococcus koreensis] Length: 218\nNumber of predicted TMHs: 1\nExp number of AAs in TMHs: 19.49301\nExp number, first 60 AAs: 1.51077\nTotal prob of N-in: 0.57300\ninside 1 177\nTMhelix 178 200\noutside 201 218

31174 GCF_001894865.1_ASM189486v1 Rhodococcus maanshanensis NBRC 100610 Terrabacteria group; Actinobacteria; Actinobacteria; Corynebacteriales; Nocardiaceae; Rhodococcus; Rhodococcus maanshanensis MTAEANRQPGNGRVAAFFDLDKTIIAKSSTLAFSKPFFDQGLINRRSVLKSSYAQFLFLL WP_072750474.1 inhibition of morphological differentiation protein [Rhodococcus maanshanensis] Length: 272\nNumber of predicted TMHs: 1\nExp number of AAs in TMHs: 22.41088\nExp number, first 60 AAs: 0.02601\nTotal prob of N-in: 0.03157\noutside 1 245\nTMhelix 246 268\ninside 269 272

31175 GCF_001894885.1_ASM189488v1 Rhodococcus marinonascens NBRC 14363 Terrabacteria group; Actinobacteria; Actinobacteria; Corynebacteriales; Nocardiaceae; Rhodococcus; Rhodococcus marinonascens MLKPPASRRLNLGPTVSRRTAIRFAAASAFGVTSIPLLTGCATADDSGAETDSLIAQSRL WP_072687607.1 hypothetical protein [Rhodococcus marinonascens] Length: 173\nNumber of predicted TMHs: 1\nExp number of AAs in TMHs: 30.25533\nExp number, first 60 AAs: 10.18519\nTotal prob of N-in: 0.67368\nPOSSIBLE N-term signal sequence\ninside 1 148\nTMhelix 149 171\noutside 172 173

31176 GCF_000010805.1_ASM1080v1 Rhodococcus opacus B4 Terrabacteria group; Actinobacteria; Actinobacteria; Corynebacteriales; Nocardiaceae; Rhodococcus; Rhodococcus opacus MRRRTTLTSVKVIAVGLLTMVASMLGAGTALAHSVVISSTPENGAEIASGPERVSVTFNE WP_012691057.1 copper resistance protein C [Rhodococcus opacus] Length: 177\nNumber of predicted TMHs: 2\nExp number of AAs in TMHs: 44.2864\nExp number, first 60 AAs: 22.13809\nTotal prob of N-in: 0.99971\nPOSSIBLE N-term signal sequence\ninside 1 12\nTMhelix 13 35\noutside 36 147\nTMhelix 148 170\ninside 171 177

31177 GCF_000264745.1_ASM26474v2 Rhodococcus opacus M213 Terrabacteria group; Actinobacteria; Actinobacteria; Corynebacteriales; Nocardiaceae; Rhodococcus; Rhodococcus opacus MRRRTTFTSVKVIAVGLLTMLASMLGAGTALAHSVVISSTPENGSEVAAGPERVSVTFNE WP_005254749.1 copper resistance protein [Rhodococcus opacus] Length: 177\nNumber of predicted TMHs: 2\nExp number of AAs in TMHs: 44.37357\nExp number, first 60 AAs: 22.2241\nTotal prob of N-in: 0.99977\nPOSSIBLE N-term signal sequence\ninside 1 12\nTMhelix 13 35\noutside 36 147\nTMhelix 148 170\ninside 171 177

31178 GCF_000599545.1_ASM59954v1 Rhodococcus opacus PD630 Terrabacteria group; Actinobacteria; Actinobacteria; Corynebacteriales; Nocardiaceae; Rhodococcus; Rhodococcus opacus MRRRTTFTSVKVIAVGLLTMLASMLGAGTALAHSVVISSTPENGAEIAAGPERVSVTFNE WP_005248910.1 MULTISPECIES: copper resistance protein C [Rhodococcus] Length: 177\nNumber of predicted TMHs: 2\nExp number of AAs in TMHs: 44.30022\nExp number, first 60 AAs: 22.13888\nTotal prob of N-in: 0.99979\nPOSSIBLE N-term signal sequence\ninside 1 12\nTMhelix 13 35\noutside 36 147\nTMhelix 148 170\ninside 171 177

31179 GCF_000234335.1_ASM23433v1 Rhodococcus opacus PD630 Terrabacteria group; Actinobacteria; Actinobacteria; Corynebacteriales; Nocardiaceae; Rhodococcus; Rhodococcus opacus

MRRRTTFTSVKVIAVGLLTMLASMLGAGTALAHSVVISSTPENGAEIAAGPERVSVTFNE WP_005248910.1 MULTISPECIES: copper resistance protein C [Rhodococcus] Length: 177\nNumber of predicted TMHs: 2\nExp number of AAs in TMHs: 44.30022\nExp number, first 60 AAs: 22.13888\nTotal prob of N-in: 0.99979\nPOSSIBLE N-term signal sequence\ninside 1 12\nTMhelix 13 35\noutside 36 147\nTMhelix 148 170\ninside 171 177

31180 GCF_000234335.1_ASM23433v1 Rhodococcus opacus PD630 Terrabacteria group; Actinobacteria; Actinobacteria; Corynebacteriales; Nocardiaceae; Rhodococcus; Rhodococcus opacus MSVFASRGTHCHSRAWVPPLGMSGYSGRVETATGRRRTVVARAALLIAVAAGVLLMHSLV WP_005238637.1 MULTISPECIES: hypothetical protein [Rhodococcus] Length: 157\nNumber of predicted TMHs: 2\nExp number of AAs in TMHs: 43.88931\nExp number, first 60 AAs: 18.77006\nTotal prob of N-in: 0.99476\nPOSSIBLE N-term signal sequence\ninside 1 44\nTMhelix 45 67\noutside 68 98\nTMhelix 99 121\ninside 122 157

31181 GCF_000599545.1_ASM59954v1 Rhodococcus opacus PD630 Terrabacteria group; Actinobacteria; Actinobacteria; Corynebacteriales; Nocardiaceae; Rhodococcus; Rhodococcus opacus MSVFASRGTHCHSRAWVPPLGMSGYSGRVETATGRRRTVVARAALLIAVAAGVLLMHSLV WP_005238637.1 MULTISPECIES: hypothetical protein [Rhodococcus] Length: 157\nNumber of predicted TMHs: 2\nExp number of AAs in TMHs: 43.88931\nExp number, first 60 AAs: 18.77006\nTotal prob of N-in: 0.99476\nPOSSIBLE N-term signal sequence\ninside 1 44\nTMhelix 45 67\noutside 68 98\nTMhelix 99 121\ninside 122 157

31182 GCF_000236965.1_ASM23696v2 Rhodococcus pyridinivorans AK37 Terrabacteria group; Actinobacteria; Actinobacteria; Corynebacteriales; Nocardiaceae; Rhodococcus; Rhodococcus pyridinivorans MFEFSRRALVGATATAALMVAGAGIASAHVTVVAPGAEQGGYTVLTFRVPTESETAGTTA WP_006553374.1 MULTISPECIES: hypothetical protein [Rhodococcus] Length: 223\nNumber of predicted TMHs: 2\nExp number of AAs in TMHs: 40.51123\nExp number, first 60 AAs: 18.38976\nTotal prob of N-in: 0.70947\nPOSSIBLE N-term signal sequence\ninside 1 8\nTMhelix 9 31\noutside 32 195\nTMhelix 196 218\ninside 219 223

31183 GCF_001894905.1_ASM189490v1 Rhodococcus pyridinivorans JCM 10940 = NBRC 100608 Terrabacteria group; Actinobacteria; Actinobacteria; Corynebacteriales; Nocardiaceae; Rhodococcus; Rhodococcus pyridinivorans MFEFSRRALVGATATAALMVAGAGIASAHVTVVAPGAEQGGYTVLTFRVPTESETAGTTA WP_006553374.1 MULTISPECIES: hypothetical protein [Rhodococcus] Length: 223\nNumber of predicted TMHs: 2\nExp number of AAs in TMHs: 40.51123\nExp number, first 60 AAs: 18.38976\nTotal prob of N-in: 0.70947\nPOSSIBLE N-term signal sequence\ninside 1 8\nTMhelix 9 31\noutside 32 195\nTMhelix 196 218\ninside 219 223

31184 GCF_001465325.1_ASM146532v1 Rhodococcus pyridinivorans KG-16 Terrabacteria group; Actinobacteria; Actinobacteria; Corynebacteriales; Nocardiaceae; Rhodococcus; Rhodococcus pyridinivorans MFEFSRRALVGATATAALMVAGAGIASAHVTVVAPGAEQGGYTVLTFRVPTESETAGTTA WP_060652333.1 nuclear export factor GLE1 [Rhodococcus pyridinivorans] Length: 223\nNumber of predicted TMHs: 1\nExp number of AAs in TMHs: 39.70445\nExp number, first 60 AAs: 17.58758\nTotal prob of N-in: 0.68701\nPOSSIBLE N-term signal sequence\noutside 1 195\nTMhelix 196 218\ninside 219 223

31185 GCF_000511305.1_ASM51130v1 Rhodococcus pyridinivorans SB3094 Terrabacteria group; Actinobacteria; Actinobacteria; Corynebacteriales; Nocardiaceae; Rhodococcus; Rhodococcus pyridinivorans MFEFSRRALVGATATAALMVAGAGIASAHVTVVAPGAEQGGYTVLTFRVPTESETAGTTA WP_024102629.1 MULTISPECIES: nuclear export factor GLE1 [Rhodococcus] Length: 223\nNumber of predicted TMHs: 2\nExp number of AAs in TMHs: 40.51123\nExp number, first 60 AAs: 18.38976\nTotal prob of N-in: 0.70947\nPOSSIBLE N-term signal sequence\ninside 1 8\nTMhelix 9 31\noutside 32 195\nTMhelix 196 218\ninside 219 223

31186 GCF_000341815.1_Whole_genome_assembly Rhodococcus qingshengii BKS 20-40 Terrabacteria group; Actinobacteria; Actinobacteria; Corynebacteriales; Nocardiaceae; Rhodococcus; Rhodococcus qingshengii MTDKPANGLNTGRTAAFFDLDKTIIAKSSTLAFSKPFFAQGLINRRSVLKSSYAQFLFLL WP_003946123.1 MULTISPECIES: haloacid dehalogenase [Rhodococcus] Length: 272\nNumber of predicted TMHs: 1\nExp number of AAs in TMHs: 22.87312\nExp number, first 60 AAs: 0.01889\nTotal prob of N-in: 0.03448\noutside 1 245\nTMhelix 246 268\ninside 269 272

31187 GCF_000341815.1_Whole_genome_assembly Rhodococcus qingshengii BKS 20-40 Terrabacteria group; Actinobacteria; Actinobacteria; Corynebacteriales; Nocardiaceae; Rhodococcus; Rhodococcus qingshengii MSNPLGQSSNGSRRGALPTPPSGWPIGSYPTYAEAQRAVDYLSDQEFPVQDVTIVGVNLM WP_003945065.1 MULTISPECIES: hypothetical protein [Rhodococcus] Length: 164\nNumber of predicted TMHs: 2\nExp number of AAs in TMHs: 45.26265\nExp number, first 60 AAs: 0.0547\nTotal prob of N-in: 0.87268\ninside 1 73\nTMhelix 74 96\noutside 97 99\nTMhelix 100 122\ninside 123 164

31188 GCF_000469645.1_ASM46964v1 Rhodococcus rhodochrous ATCC 17895 Terrabacteria group; Actinobacteria; Actinobacteria; Corynebacteriales; Nocardiaceae; Rhodococcus; Rhodococcus rhodochrous MSNPLGQSSNGSRRGALPTPPSGWPIGSYPTYAEAQRAVDYLSDQEFPVQDVTIVGVNLM WP_003945065.1 MULTISPECIES: hypothetical protein [Rhodococcus] Length: 164\nNumber of predicted TMHs: 2\nExp number of AAs in TMHs: 45.26265\nExp number, first 60 AAs: 0.0547\nTotal prob of N-in: 0.87268\ninside 1 73\nTMhelix 74 96\noutside 97 99\nTMhelix 100 122\ninside 123 164

31189 GCF_000469645.1_ASM46964v1 Rhodococcus rhodochrous ATCC 17895 Terrabacteria group; Actinobacteria; Actinobacteria; Corynebacteriales; Nocardiaceae; Rhodococcus; Rhodococcus rhodochrous MTDKPANGLNTGRTAAFFDLDKTIIAKSSTLAFSKPFFAQGLINRRSVLKSSYAQFLFLL WP_019746641.1 MULTISPECIES: inhibition of morphological differentiation protein [Rhodococcus] Length: 272\nNumber of predicted TMHs: 1\nExp number of AAs in TMHs: 22.87312\nExp number, first 60 AAs: 0.01889\nTotal prob of N-in: 0.03448\noutside 1 245\nTMhelix 246 268\ninside 269 272

31190 GCF_000517665.1_Rhodococcus_rhodochrous.strain_ATCC_21198_v1.0 Rhodococcus rhodochrous ATCC 21198 Terrabacteria group; Actinobacteria; Actinobacteria; Corynebacteriales; Nocardiaceae; Rhodococcus; Rhodococcus rhodochrous MRLPTSPRSRRRTLSAALLLLLVPVLAGCLRVQATMGVSADDRVSGQIVAATIPADENDK WP_006933384.1 MULTISPECIES: DUF3153 domain-containing protein [Rhodococcus] Length: 233\nNumber of predicted TMHs: 2\nExp number of AAs in TMHs: 44.90952\nExp number, first 60 AAs: 21.98985\nTotal prob of N-in: 0.95979\nPOSSIBLE N-term signal sequence\ninside 1 12\nTMhelix 13 35\noutside 36 191\nTMhelix 192 214\ninside 215 233

31191 GCF_000517665.1_Rhodococcus_rhodochrous.strain_ATCC_21198_v1.0 Rhodococcus rhodochrous ATCC 21198 Terrabacteria group; Actinobacteria; Actinobacteria; Corynebacteriales; Nocardiaceae; Rhodococcus; Rhodococcus rhodochrous MTDATVAPTQSGRIAAFFDLDKTVIAKSSALAFSKPFFAQGLLNRRAVLKSSYAQFLFML WP_006941337.1 MULTISPECIES: phosphoserine phosphatase [Rhodococcus] Length: 280\nNumber of predicted TMHs: 1\nExp number of AAs in TMHs: 22.81284\nExp number, first 60 AAs: 0.03489\nTotal prob of N-in: 0.04407\noutside 1 245\nTMhelix 246 268\ninside 269 280

31192 GCF_000239135.1_ASM23913v3 Rhodococcus rhodochrous BKS6-46 Terrabacteria group; Actinobacteria; Actinobacteria; Corynebacteriales; Nocardiaceae; Rhodococcus; Rhodococcus rhodochrous MFEFSRRALVGATATAALMVAGAGIASAHVTVVAPGAEQGGYTVLTFRVPTESETAGTTA WP_016695402.1 nuclear export factor GLE1 [Rhodococcus rhodochrous] Length: 224\nNumber of predicted TMHs: 2\nExp number of AAs in TMHs: 40.65107\nExp number, first 60 AAs: 18.39794\nTotal prob of N-in: 0.69434\nPOSSIBLE N-term signal sequence\ninside 1 8\nTMhelix 9 31\noutside 32 196\nTMhelix 197 219\ninside 220 224

31193 GCF_900177695.1_IMG-taxon_2527291702_annotated_assembly Rhodococcus rhodochrous J3 Terrabacteria group; Actinobacteria; Actinobacteria; Corynebacteriales; Nocardiaceae; Rhodococcus; Rhodococcus rhodochrous MFEFSRRALVGATATAALMVAGAGIASAHVTVVAPGAEQGGYTVLTFRVPTESETAGTTA WP_085468723.1 nuclear export factor GLE1 [Rhodococcus rhodochrous] Length: 224\nNumber of predicted TMHs: 1\nExp number of AAs in TMHs: 39.80087\nExp number, first 60 AAs: 17.67833\nTotal prob of N-in: 0.65314\nPOSSIBLE N-term signal sequence\noutside 1 196\nTMhelix 197 219\ninside 220 224

31194 GCF_001278665.1_ASM127866v1 Rhodococcus rhodochrous KG-21 Terrabacteria group; Actinobacteria; Actinobacteria; Corynebacteriales; Nocardiaceae; Rhodococcus; Rhodococcus rhodochrous MTDATVAPTPSGRIAAFFDLDKTVIAKSSALAFSKPFFAQGLLNRRAVLKSSYAQFLFML WP_003935262.1 MULTISPECIES: hydrolase [Rhodococcus] Length: 280\nNumber of predicted TMHs: 1\nExp number of AAs in TMHs: 22.95128\nExp number, first 60 AAs: 0.02777\nTotal prob of N-in: 0.04814\noutside 1 245\nTMhelix 246 268\ninside 269 280

31195 GCF_001047055.1_ASM104705v1 Rhodococcus rhodochrous NBRC 16069 Terrabacteria group; Actinobacteria; Actinobacteria; Corynebacteriales; Nocardiaceae; Rhodococcus; Rhodococcus rhodochrous MFEFSRRALVGATATAALMVAGAGIASAHVTVVAPGAEQGGYTVLTFRVPTESETAGTTA WP_059382055.1 nuclear export factor GLE1 [Rhodococcus rhodochrous] Length: 224\nNumber of predicted TMHs: 2\nExp number of AAs in TMHs: 40.65078\nExp number, first 60 AAs: 18.3977\nTotal prob of N-in: 0.69434\nPOSSIBLE N-term signal sequence\ninside 1 8\nTMhelix 9 31\noutside 32 196\nTMhelix 197 219\ninside 220 224

31196 GCF_000341965.1_Whole_genome_assembly Rhodococcus ruber BKS 20-38 Terrabacteria group; Actinobacteria; Actinobacteria; Corynebacteriales; Nocardiaceae; Rhodococcus; Rhodococcus ruber MTDATVAPTPSGRIAAFFDLDKTVIAKSSALAFSKPFFAQGLLNRRAVLKSSYAQFLFML WP_003935262.1 MULTISPECIES: hydrolase [Rhodococcus] Length: 280\nNumber of predicted TMHs: 1\nExp number of AAs in TMHs: 22.95128\nExp number, first 60 AAs: 0.02777\nTotal prob of N-in: 0.04814\noutside 1 245\nTMhelix 246 268\ninside 269 280

31197 GCF_000347955.2_ASM34795v2 Rhodococcus ruber Chol-4 Terrabacteria group; Actinobacteria; Actinobacteria; Corynebacteriales; Nocardiaceae; Rhodococcus; Rhodococcus ruber
MTDATVDPTPSGRIAAFFDLDKTVIAKSSALAFSKPFFAQGLLNRRAVLKSSYAQFLFML WP_010595022.1 MULTISPECIES: hydrolase [Rhodococcus] Length: 280\nNumber of predicted TMHs: 1\nExp number of AAs in TMHs: 23.01026\nExp number, first 60 AAs: 0.0217\nTotal prob of N-in: 0.01643\noutside 1 245\nTMhelix 246 268\ninside 269 280

31198 GCF_001894945.1_ASM189494v1 Rhodococcus ruber NBRC 15591 Terrabacteria group; Actinobacteria; Actinobacteria; Corynebacteriales; Nocardiaceae; Rhodococcus; Rhodococcus ruber
MTDATVDPTPSGRIAAFFDLDKTVIAKSSALAFSKPFFAQGLLNRRAVLKSSYAQFLFML WP_064066535.1 inhibition of morphological differentiation protein [Rhodococcus ruber] Length: 280\nNumber of predicted TMHs: 1\nExp number of AAs in TMHs: 23.02605\nExp number, first 60 AAs: 0.02262\nTotal prob of N-in: 0.01870\noutside 1 245\nTMhelix 246 268\ninside 269 280

31199 GCF_001894985.1_ASM189498v1 Rhodococcus tukisamuensis NBRC 100609 Terrabacteria group; Actinobacteria; Actinobacteria; Corynebacteriales; Nocardiaceae; Rhodococcus; Rhodococcus tukisamuensis
MTAAASRESEHGTVAAFFDLDKTIIAKSSTFAFSKPFFDQGLINRRSVLKSSYAHFLFML WP_072845452.1 inhibition of morphological differentiation protein [Rhodococcus tukisamuensis] Length: 272\nNumber of predicted TMHs: 1\nExp number of AAs in TMHs: 21.89604\nExp number, first 60 AAs: 0.02213\nTotal prob of N-in: 0.04408\noutside 1 245\nTMhelix 246 268\ninside 269 272

31200 GCF_000325625.1_ASM32562v1 Rhodococcus wratislaviensis IFP 2016 Terrabacteria group; Actinobacteria; Actinobacteria; Corynebacteriales; Nocardiaceae; Rhodococcus; Rhodococcus wratislaviensis
MSVFASRGTHCHSRVQVPPLGMSGYSGRVETATGRRRTVVARAALLIAVAAGVLLMHSLI WP_043788292.1
MULTISPECIES: hypothetical protein [Rhodococcus] Length: 156\nNumber of predicted TMHs: 2\nExp number of AAs in TMHs: 43.92061\nExp number, first 60 AAs: 18.7176\nTotal prob of N-in: 0.99776\nPOSSIBLE N-term signal sequence\ninside 1 44\nTMhelix 45 67\noutside 68 93\nTMhelix 94 116\ninside 117 156

31201 GCF_000583735.1_ASM58373v1 Rhodococcus wratislaviensis NBRC 100605 Terrabacteria group; Actinobacteria; Actinobacteria; Corynebacteriales; Nocardiaceae; Rhodococcus; Rhodococcus wratislaviensis
MRRRTTFTSVKVIAVGLLTMLASMLGAGTAVAHSVVISSTPENGAEIAAGPERVSVTFNE WP_037228321.1
copper resistance protein CopC [Rhodococcus wratislaviensis] Length: 177\nNumber of predicted TMHs: 2\nExp number of AAs in TMHs: 44.32039\nExp number, first 60 AAs: 22.17187\nTotal prob of N-in: 0.99970\nPOSSIBLE N-term signal sequence\ninside 1 12\nTMhelix 13 35\noutside 36 147\nTMhelix 148 170\ninside 171 177

31202 GCF_001895025.1_ASM189502v1 Rhodococcus zopfii NBRC 100606 = JCM 9919 Terrabacteria group; Actinobacteria; Actinobacteria; Corynebacteriales; Nocardiaceae; Rhodococcus; Rhodococcus zopfii
MTDETPRPTEAGRIAAFFDLDKTVIAKSSVLAFSRPFFDQGLLNRRTVIENTYAQFLYLL WP_072813620.1 inhibition of morphological differentiation protein [Rhodococcus zopfii] Length: 272\nNumber of predicted TMHs: 1\nExp number of AAs in TMHs: 22.91514\nExp number, first 60 AAs: 0.01307\nTotal prob of N-in: 0.04843\noutside 1 245\nTMhelix 246 268\ninside 269 272

31203 GCF_000185725.2_Segn_rugo_CDC_945_V2 Segniliparus rugosus ATCC BAA-974 Terrabacteria group; Actinobacteria; Actinobacteria; Corynebacteriales; Segniliparaceae; Segniliparus; Segniliparus rugosus
MSQLAAFFDLDRTLIAKSSSLAFGRPFYEQGLVGRRAVLKSSYAQFLFQLSGADHQQMDR WP_007471343.1
haloacid dehalogenase [Segniliparus rugosus] Length: 269\nNumber of predicted TMHs: 1\nExp number of AAs in TMHs: 22.34448\nExp number, first 60 AAs: 0.00297\nTotal prob of N-in: 0.03536\noutside 1 241\nTMhelix 242 264\ninside 265 269

31204 GCF_001575165.1_ASM157516v1 Tsukamurella pulmonis Terrabacteria group; Actinobacteria; Actinobacteria; Corynebacteriales; Tsukamurellaceae; Tsukamurella
MSPTAPDEAPVRRVAAFFDLDKTIIAKSSALAFSRPFFDEGLLNRRAVLKSSYAQFLMML WP_068564996.1 inhibition of morphological differentiation protein [Tsukamurella pulmonis] Length: 273\nNumber of predicted TMHs: 1\nExp number of AAs in TMHs: 22.48204\nExp number, first 60 AAs: 0.01868\nTotal prob of N-in: 0.04621\noutside 1 245\nTMhelix 246 268\ninside 269 273

31205 GCF_001575215.1_ASM157521v1 Tsukamurella pulmonis Terrabacteria group; Actinobacteria; Actinobacteria; Corynebacteriales; Tsukamurellaceae; Tsukamurella
MSPTAPDEAPVRRVAAFFDLDKTIIAKSSALAFSRPFFDEGLLNRRAVLKSSYAQFLMML WP_068532186.1 inhibition of morphological differentiation protein [Tsukamurella pulmonis] Length: 273\nNumber of predicted TMHs: 1\nExp number of AAs in TMHs: 22.48204\nExp number, first 60 AAs: 0.01868\nTotal prob of N-in: 0.04621\noutside 1 245\nTMhelix 246 268\ninside 269 273

31206	GCF_001575215.1_ASM157521v1	Tsukamurella pulmonis	Terrabacteria group; Actinobacteria; Actinobacteria; Corynebacteriales; Tsukamurellaceae; Tsukamurella
MTARTTPDDILTDDRLAAMRSAVMAEVDTDVQARRRSRRRRTMGIAAAAAIAVVAVGGGV	WP_068532605.1	hypothetical protein [Tsukamurella pulmonis]	Length: 338\nNumber of predicted TMHs: 2\nExp number of AAs in TMHs: 44.30098\nExp number, first 60 AAs: 17.75142\nTotal prob of N-in: 0.99931\nPOSSIBLE N-term signal sequence\ninside 1 42\nTMhelix 43 65\noutside 66 305\nTMhelix 306 328\ninside 329 338

31207	GCF_001575165.1_ASM157516v1	Tsukamurella pulmonis	Terrabacteria group; Actinobacteria; Actinobacteria; Corynebacteriales; Tsukamurellaceae; Tsukamurella
MTARTTPDDILTDDRLAAMRSVVMAEVDTDVQARRRSRRRRTMGIAAAAAIAVVAVGGGV	WP_068566570.1	hypothetical protein [Tsukamurella pulmonis]	Length: 338\nNumber of predicted TMHs: 2\nExp number of AAs in TMHs: 44.30098\nExp number, first 60 AAs: 17.75142\nTotal prob of N-in: 0.99931\nPOSSIBLE N-term signal sequence\ninside 1 42\nTMhelix 43 65\noutside 66 305\nTMhelix 306 328\ninside 329 338

31208	GCF_900103175.1_IMG-taxon_2634166370_annotated_assembly	Tsukamurella pulmonis	Terrabacteria group; Actinobacteria; Actinobacteria; Corynebacteriales; Tsukamurellaceae; Tsukamurella
MTARTTPDDILTDDRLAAMRSVVMAEVDTDVQARRRSRRRRTMGIAAAAAIAVVAVGGGV	WP_068566570.1	hypothetical protein [Tsukamurella pulmonis]	Length: 338\nNumber of predicted TMHs: 2\nExp number of AAs in TMHs: 44.30098\nExp number, first 60 AAs: 17.75142\nTotal prob of N-in: 0.99931\nPOSSIBLE N-term signal sequence\ninside 1 42\nTMhelix 43 65\noutside 66 305\nTMhelix 306 328\ninside 329 338

31209	GCF_900103175.1_IMG-taxon_2634166370_annotated_assembly	Tsukamurella pulmonis	Terrabacteria group; Actinobacteria; Actinobacteria; Corynebacteriales; Tsukamurellaceae; Tsukamurella
MSPTAPDEAPVRRVAAFFDLDKTIIAKSSALAFSRPFFDEGLLNRRAVLKSSYAQFLMML	WP_068564996.1	inhibition of morphological differentiation protein [Tsukamurella pulmonis]	Length: 273\nNumber of predicted TMHs: 1\nExp number of AAs in TMHs: 22.48204\nExp number, first 60 AAs: 0.01868\nTotal prob of N-in: 0.04621\noutside 1 245\nTMhelix 246 268\ninside 269 273

31210	GCF_000312385.1_ASM31238v1	Tsukamurella sp. 1534	Terrabacteria group; Actinobacteria; Actinobacteria; Corynebacteriales; Tsukamurellaceae; Tsukamurella
MSSTAPDEAPARRVAAFFDLDKTVIAKSSALAFSRPFFDEGLLNRRAVLKSSYAQFLMML	WP_019201864.1	inhibition of morphological differentiation protein [Tsukamurella sp. 1534]	Length: 273\nNumber of predicted TMHs: 1\nExp number of AAs in TMHs: 22.27175\nExp number, first 60 AAs: 0.02105\nTotal prob of N-in: 0.09422\noutside 1 245\nTMhelix 246 268\ninside 269 273

31211	GCF_000524475.1_ASM52447v1	Tomitella biformata AHU 1821	Terrabacteria group; Actinobacteria; Actinobacteria; Corynebacteriales; unclassified Corynebacteriales; Tomitella; Tomitella biformata
MVSEDNSTTTSTGRVAAFFDLDKTLIAKSSTLAFSRPFFDEGLLNRRAVLKSSYAQFLFL	WP_038170946.1	inhibition of morphological differentiation protein [Tomitella biformata]	Length: 273\nNumber of predicted TMHs: 1\nExp number of AAs in TMHs: 22.44425\nExp number, first 60 AAs: 0.06395\nTotal prob of N-in: 0.08619\noutside 1 246\nTMhelix 247 269\ninside 270 273

31212	GCF_000506245.1_Williamsia_D3	Williamsia sp. D3	Terrabacteria group; Actinobacteria; Actinobacteria; Corynebacteriales; Williamsiaceae; Williamsia
MFHADPVAHAHEHPSAPRRRSALSVVSSVIMNLLAVGGLACIVMVVLAVVFNVSLIMFKTG	WP_023955730.1	S26 family signal peptidase [Williamsia sp. D3]	Length: 208\nNumber of predicted TMHs: 2\nExp number of AAs in TMHs: 46.57551\nExp number, first 60 AAs: 23.75883\nTotal prob of N-in: 0.16230\nPOSSIBLE N-term signal sequence\noutside 1 31\nTMhelix 32 54\ninside 55 151\nTMhelix 152 174\noutside 175 208

31213	GCF_001424365.1_Leaf354	Williamsia sp. Leaf354	Terrabacteria group; Actinobacteria; Actinobacteria; Corynebacteriales; Williamsiaceae; Williamsia
MSTAQRTDPAPAPSAAPTGLSGFDAADVERRRALRKMKVVATGLLVFAAGVYLLTRWIES	WP_055787738.1	DUF445 domain-containing protein [Williamsia sp. Leaf354]	Length: 441\nNumber of predicted TMHs: 2\nExp number of AAs in TMHs: 45.03852\nExp number, first 60 AAs: 19.53741\nTotal prob of N-in: 0.97040\nPOSSIBLE N-term signal sequence\ninside 1 38\nTMhelix 39 58\noutside 59 420\nTMhelix 421 440\ninside 441 441

31214	GCF_001424365.1_Leaf354	Williamsia sp. Leaf354	Terrabacteria group; Actinobacteria; Actinobacteria; Corynebacteriales; Williamsiaceae; Williamsia
MTHSTWHDVVVPVSPVLPPHRLAPRSRTRRPAVRIAAAALLVLLGALSLTSCMERSPYVG	WP_055789914.1	hypothetical protein [Williamsia sp. Leaf354]	Length: 248\nNumber of predicted TMHs: 2\nExp number of

AAs in TMHs: 42.31824\nExp number, first 60 AAs: 20.24606\nTotal prob of N-in: 0.91228\nPOSSIBLE N-term signal sequence\ninside 1 30\nTMhelix 31 53\noutside 54 215\nTMhelix 216 238\ninside 239 248

31215 GCF_001598915.1_ASM159891v1 Williamsia muralis NBRC 105860 Terrabacteria group; Actinobacteria; Actinobacteria; Corynebacteriales; Williamsiaceae; Williamsia; Williamsia muralis MFHADPVTHAEPPSAPRRRSALSVVSSVIMNLLAVGGLACIVMVVLAVVFNVSLIMFKTG WP_084248314.1 signal peptidase I [Williamsia muralis] Length: 208\nNumber of predicted TMHs: 2\nExp number of AAs in TMHs: 46.84393\nExp number, first 60 AAs: 23.71698\nTotal prob of N-in: 0.19113\nPOSSIBLE N-term signal sequence\noutside 1 31\nTMhelix 32 54\ninside 55 151\nTMhelix 152 174\noutside 175 208

31216 GCF_900143005.1_IMG-taxon_2700988713_annotated_assembly Cryptosporangium aurantiacum Terrabacteria group; Actinobacteria; Actinobacteria; Frankiales; Cryptosporangiaceae; Cryptosporangium MPRTRRRLLAGLLATLTLFAGTVVPGAAPAAADTVRDQQWQLSFLQATSAWQYSTGRGVTV WP_084740800.1 type VII secretion-associated serine protease mycosin [Cryptosporangium aurantiacum] Length: 430\nNumber of predicted TMHs: 2\nExp number of AAs in TMHs: 42.16076\nExp number, first 60 AAs: 19.31399\nTotal prob of N-in: 0.86734\nPOSSIBLE N-term signal sequence\ninside 1 8\nTMhelix 9 31\noutside 32 380\nTMhelix 381 403\ninside 404 430

31217 GCF_900143005.1_IMG-taxon_2700988713_annotated_assembly Cryptosporangium aurantiacum Terrabacteria group; Actinobacteria; Actinobacteria; Frankiales; Cryptosporangiaceae; Cryptosporangium MLLHLPGRSASGSRARRRAVRAAAAVAAAAALTALGLPALAPGGATASSHREAPTISGFP WP_073250052.1 hypothetical protein [Cryptosporangium aurantiacum] Length: 531\nNumber of predicted TMHs: 2\nExp number of AAs in TMHs: 39.71482\nExp number, first 60 AAs: 18.91803\nTotal prob of N-in: 0.85097\nPOSSIBLE N-term signal sequence\ninside 1 19\nTMhelix 20 42\noutside 43 503\nTMhelix 504 526\ninside 527 531

31218 GCF_000013345.1_ASM1334v1 Frankia casuarinae Terrabacteria group; Actinobacteria; Actinobacteria; Frankiales; Frankiaceae; Frankia MRRSADQPGGRRGVVVVTGGSAGLGRAIVRQFARRGYDVGVLARGEDGLAGAVADVAAAG WP_011436415.1 MULTISPECIES: short-chain dehydrogenase [Frankia] Length: 346\nNumber of predicted TMHs: 1\nExp number of AAs in TMHs: 24.86933\nExp number, first 60 AAs: 4.97586\nTotal prob of N-in: 0.34375\noutside 1 316\nTMhelix 317 339\ninside 340 346

31219 GCF_000373365.1_ASM37336v1 Frankia discariae Terrabacteria group; Actinobacteria; Actinobacteria; Frankiales; Frankiaceae; Frankia MRKGMGRRAALLAMSSAATVIVTFGPGLTAAHADIFDDKGRCDDKKSEEWDIFGDKNDRD WP_026239184.1 hypothetical protein [Frankia discariae] Length: 171\nNumber of predicted TMHs: 2\nExp number of AAs in TMHs: 51.15831\nExp number, first 60 AAs: 9.70344\nTotal prob of N-in: 0.61823\ninside 1 108\nTMhelix 109 131\noutside 132 145\nTMhelix 146 163\ninside 164 171

31220 GCF_000166135.1_ASM16613v1 Frankia inefficax Terrabacteria group; Actinobacteria; Actinobacteria; Frankiales; Frankiaceae; Frankia MSRERRTARRRLLALAVGAVLAGAVAGPATAATPEARTGVPTAGARAAGRPPSFGIEPLP WP_049806950.1 hypothetical protein [Frankia inefficax] Length: 275\nNumber of predicted TMHs: 2\nExp number of AAs in TMHs: 42.9598\nExp number, first 60 AAs: 21.26499\nTotal prob of N-in: 0.99802\nPOSSIBLE N-term signal sequence\ninside 1 11\nTMhelix 12 34\noutside 35 232\nTMhelix 233 255\ninside 256 275

31221 GCF_000733325.1_ASM73332v1 Frankia sp. Allo2 Terrabacteria group; Actinobacteria; Actinobacteria; Frankiales; Frankiaceae; Frankia MRRSADQPGGRRGVVVVTGGSAGLGRAIVRQFARRGYDVGVLARGEDGLAGAVADVAAAG WP_023841236.1 MULTISPECIES: short-chain dehydrogenase of unknown substrate specificity [Frankia] Length: 346\nNumber of predicted TMHs: 1\nExp number of AAs in TMHs: 24.87487\nExp number, first 60 AAs: 4.9761\nTotal prob of N-in: 0.34378\noutside 1 316\nTMhelix 317 339\ninside 340 346

31222 GCF_000685765.2_BMG5.23v1 Frankia sp. BMG5.23 Terrabacteria group; Actinobacteria; Actinobacteria; Frankiales; Frankiaceae; Frankia MRRSADQPGGRRGVVVVTGGSAGLGRAIVRQFARRGYDVGVLARGEDGLAGAVADVAAAG WP_043591181.1 short-chain dehydrogenase [Frankia sp. BMG5.23] Length: 352\nNumber of predicted TMHs: 1\nExp number of AAs in TMHs: 28.64204\nExp number, first 60 AAs: 4.9185\nTotal prob of N-in: 0.35964\noutside 1 321\nTMhelix 322 344\ninside 345 352

31223 GCF_001636575.1_ASM163657v1 Frankia sp. BR Terrabacteria group; Actinobacteria; Actinobacteria; Frankiales; Frankiaceae; Frankia MRRSADQPGGRRGVVVVTGGSAGLGRAIVRQFARRGYDVGVLARGEDGLAGAVADVAAAG WP_063579095.1 short-chain dehydrogenase [Frankia sp. BR] Length: 346\nNumber of predicted TMHs: 1\nExp number of AAs in TMHs: 24.87201\nExp number, first 60 AAs: 4.97596\nTotal prob of N-in: 0.34376\noutside 1 316\nTMhelix 317 339\ninside 340 346

31224 GCF_001983015.1_ASM198301v1 Frankia sp. CcI156 Terrabacteria group; Actinobacteria; Actinobacteria; Frankiales; Frankiaceae; Frankia MRRSADQPGGRRGVVVVTGGSAGLGRAIVRQFARRGYDVGVLARGEDGLAGAVADVAAAG WP_076806001.1 short-chain dehydrogenase [Frankia sp. CcI156] Length: 346\nNumber of predicted TMHs: 1\nExp number of AAs in TMHs: 24.95728\nExp number, first 60 AAs: 4.97027\nTotal prob of N-in: 0.34536\noutside 1 316\nTMhelix 317 339\ninside 340 346

31225 GCF_001983215.1_ASM198321v1 Frankia sp. CcI49 Terrabacteria group; Actinobacteria; Actinobacteria; Frankiales; Frankiaceae; Frankia MSGTSTAARCVTDAGATDEHAGRRRRRLLLAGAAGCLVAGAGYLAAVDPHDPAAAMPTCP WP_083473871.1 MULTISPECIES: hypothetical protein [Frankia] Length: 153\nNumber of predicted TMHs: 2\nExp number of AAs in TMHs: 48.604\nExp number, first 60 AAs: 19.86323\nTotal prob of N-in: 0.99983\nPOSSIBLE N-term signal sequence\ninside 1 27\nTMhelix 28 47\noutside 48 86\nTMhelix 87 109\ninside 110 153

31226 GCF_000503735.2_CcI6 Frankia sp. CcI6 Terrabacteria group; Actinobacteria; Actinobacteria; Frankiales; Frankiaceae; Frankia MRRSADQPGGRRGVVVVTGGSAGLGRAIVRQFARRGYDVGVLARGEDGLAGAVADVAAAG WP_023841236.1 MULTISPECIES: short-chain dehydrogenase of unknown substrate specificity [Frankia] Length: 346\nNumber of predicted TMHs: 1\nExp number of AAs in TMHs: 24.87487\nExp number, first 60 AAs: 4.9761\nTotal prob of N-in: 0.34378\noutside 1 316\nTMhelix 317 339\ninside 340 346

31227 GCF_000732115.1_CeD Frankia sp. CeD Terrabacteria group; Actinobacteria; Actinobacteria; Frankiales; Frankiaceae; Frankia MRRSADQPGGRRGVVVVTGGSAGLGRAIVRQFARRGYDVGVLARGEDGLAGAVADVAAAG WP_035911598.1 short-chain dehydrogenase [Frankia sp. CeD] Length: 346\nNumber of predicted TMHs: 1\nExp number of AAs in TMHs: 24.85069\nExp number, first 60 AAs: 4.97505\nTotal prob of N-in: 0.34362\noutside 1 316\nTMhelix 317 339\ninside 340 346

31228 GCF_002099355.1_ASM209935v1 Frankia sp. Cg70.9 Terrabacteria group; Actinobacteria; Actinobacteria; Frankiales; Frankiaceae; Frankia MRRSADQPGGRRGVVVVTGGSAGLGRAIVRQFARRGYDVGVLARGEDGLAGAVADVAAAG WP_011436415.1 MULTISPECIES: short-chain dehydrogenase [Frankia] Length: 346\nNumber of predicted TMHs: 1\nExp number of AAs in TMHs: 24.86933\nExp number, first 60 AAs: 4.97586\nTotal prob of N-in: 0.34375\noutside 1 316\nTMhelix 317 339\ninside 340 346

31229 GCF_001756285.1_ASM175628v1 Frankia sp. CgIM4 Terrabacteria group; Actinobacteria; Actinobacteria; Frankiales; Frankiaceae; Frankia MRRSADQPGGRRGVVVVTGGSAGLGRAIVRQFARRGYDVGVLARGEDGLAGAVADVAAAG WP_070132079.1 short-chain dehydrogenase [Frankia sp. CgIM4] Length: 346\nNumber of predicted TMHs: 1\nExp number of AAs in TMHs: 24.93806\nExp number, first 60 AAs: 4.97842\nTotal prob of N-in: 0.34303\noutside 1 318\nTMhelix 319 338\ninside 339 346

31230 GCF_001854725.1_ASM185472v1 Frankia sp. CgIS1 Terrabacteria group; Actinobacteria; Actinobacteria; Frankiales; Frankiaceae; Frankia MRRSADQPGGRRGVVVVTGGSAGLGRAIVRQFARRGYDVGVLARGEDGLAGAVADVAAAG WP_011436415.1 MULTISPECIES: short-chain dehydrogenase [Frankia] Length: 346\nNumber of predicted TMHs: 1\nExp number of AAs in TMHs: 24.86933\nExp number, first 60 AAs: 4.97586\nTotal prob of N-in: 0.34375\noutside 1 316\nTMhelix 317 339\ninside 340 346

31231 GCF_000235425.2_ASM23542v3 Frankia sp. CN3 Terrabacteria group; Actinobacteria; Actinobacteria; Frankiales; Frankiaceae; Frankia MLTPFGEIPDRLARHMTMAEQHEYLRRRRPSRRAVLGSALAAVAGPTLLAGTARASVTTP WP_007511128.1 metallophosphoesterase [Frankia sp. CN3] Length: 532\nNumber of predicted TMHs: 1\nExp number of AAs in TMHs: 25.98227\nExp number, first 60 AAs: 5.34548\nTotal prob of N-in: 0.24656\noutside 1 502\nTMhelix 503 525\ninside 526 532

31232 GCF_000235425.2_ASM23542v3 Frankia sp. CN3 Terrabacteria group; Actinobacteria; Actinobacteria; Frankiales; Frankiaceae; Frankia MPPAPAHARWPRRRVLPGLLATALTGGMLLAPGLAGAALATQRGDGPWYVSALRLDEAHQ WP_007519988.1 MULTISPECIES: peptidase S8 [Frankia] Length: 449\nNumber of predicted TMHs: 1\nExp number of AAs in TMHs: 43.1342599999999\nExp number, first 60 AAs: 18.0244\nTotal prob of N-in: 0.69219\nPOSSIBLE N-term signal sequence\noutside 1 406\nTMhelix 407 429\ninside 430 449

31233 GCF_000966285.1_ASM96628v1 Frankia sp. DC12 Terrabacteria group; Actinobacteria; Actinobacteria; Frankiales; Frankiaceae; Frankia MTRRTTLYGLLGVPPSATADEIRAAYRHAARELHPDAGGSAAAFQRLTTAYHILGDPAGR WP_045877155.1 molecular chaperone DnaJ [Frankia sp. DC12] Length: 169\nNumber of predicted TMHs: 2\nExp number of AAs in TMHs: 42.99964\nExp number, first 60 AAs: 0.00179\nTotal prob of N-in: 0.99599\ninside 1 102\nTMhelix 103 122\noutside 123 126\nTMhelix 127 149\ninside 150 169

31234 GCF_000177675.1_ASM17767v1 Frankia sp. EUN1f Terrabacteria group; Actinobacteria; Actinobacteria; Frankiales; Frankiaceae; Frankia MQAVHRRSHLPLPGASLPGAALRRVAPVVALAAVLVALLGSPASAHTALTSSDPAAGATL

WP_006540442.1 copper resistance protein CopC [Frankia sp. EUN1f] Length: 224\nNumber of predicted TMHs: 2\nExp number of AAs in TMHs: 44.19722\nExp number, first 60 AAs: 20.51775\nTotal prob of N-in: 0.92132\nPOSSIBLE N-term signal sequence\ninside 1 27\nTMhelix 28 50\noutside 51 194\nTMhelix 195 217\ninside 218 224

31235 GCF_001854645.1_ASM185464v1 Frankia sp. EUN1h Terrabacteria group; Actinobacteria; Actinobacteria; Frankiales; Frankiaceae; Frankia MLTPFGEIPDRLARHMTMAEQHEYLRRRRPSRRAVLGSALAAVAGPTLLAGTXRASVTTP WP_071041290.1 hypothetical protein [Frankia sp. EUN1h] Length: 532\nNumber of predicted TMHs: 1\nExp number of AAs in TMHs: 24.88339\nExp number, first 60 AAs: 4.24714\nTotal prob of N-in: 0.19800\noutside 1 502\nTMhelix 503 525\ninside 526 532

31236 GCF_001854645.1_ASM185464v1 Frankia sp. EUN1h Terrabacteria group; Actinobacteria; Actinobacteria; Frankiales; Frankiaceae; Frankia MPPAPAHARWPRRRVLPGLLATALTGGMLLAPGLAGAALATQRGDGPWYVSALRLDEAHQ WP_007519988.1 MULTISPECIES: peptidase S8 [Frankia] Length: 449\nNumber of predicted TMHs: 1\nExp number of AAs in TMHs: 43.1342599999999\nExp number, first 60 AAs: 18.0244\nTotal prob of N-in: 0.69219\nPOSSIBLE N-term signal sequence\noutside 1 406\nTMhelix 407 429\ninside 430 449

31237 GCF_002099325.1_ASM209932v1 Frankia sp. KB5 Terrabacteria group; Actinobacteria; Actinobacteria; Frankiales; Frankiaceae; Frankia MRRSADQPGGRRGVVVVTGGSAGLGRAIVRQFARRGYDVGVLARGEDGLAGAVADVAAAG WP_085015222.1 short-chain dehydrogenase [Frankia sp. KB5] Length: 352\nNumber of predicted TMHs: 1\nExp number of AAs in TMHs: 28.63902\nExp number, first 60 AAs: 4.91833\nTotal prob of N-in: 0.35962\noutside 1 321\nTMhelix 322 344\ninside 345 352

31238 GCF_000262465.1_ASM26246v1 Frankia sp. QA3 Terrabacteria group; Actinobacteria; Actinobacteria; Frankiales; Frankiaceae; Frankia MTAATAPAPGSAGSAADAAPAGTIGRSGAYLTDGERRAVVIRGMTVPAGVTPTVGDLDTW WP_051054688.1 hypothetical protein [Frankia sp. QA3] Length: 473\nNumber of predicted TMHs: 1\nExp number of AAs in TMHs: 22.30856\nExp number, first 60 AAs: 0.05432\nTotal prob of N-in: 0.02303\noutside 1 401\nTMhelix 402 424\ninside 425 473

31239 GCF_001306465.1_ASM130646v1 Frankia sp. R43 Terrabacteria group; Actinobacteria; Actinobacteria; Frankiales; Frankiaceae; Frankia MSGTSTAARCVTDAGATDEHAGRRRRRLLLAGAAGCLVAGAGYLAAVDPHDPAAAMPTCP WP_083473871.1 MULTISPECIES: hypothetical protein [Frankia] Length: 153\nNumber of predicted TMHs: 2\nExp number of AAs in TMHs: 48.604\nExp number, first 60 AAs: 19.86323\nTotal prob of N-in: 0.99983\nPOSSIBLE N-term signal sequence\ninside 1 27\nTMhelix 28 47\noutside 48 86\nTMhelix 87 109\ninside 110 153

31240 GCF_000611815.2_ASM61181v2 Frankia sp. Thr Terrabacteria group; Actinobacteria; Actinobacteria; Frankiales; Frankiaceae; Frankia MRRSADQPGGRRGVVVVTGGSAGLGRAIVRQFARRGYDVGVLARGEDGLAGAVADVAAAG WP_011436415.1 MULTISPECIES: short-chain dehydrogenase [Frankia] Length: 346\nNumber of predicted TMHs: 1\nExp number of AAs in TMHs: 24.86933\nExp number, first 60 AAs: 4.97586\nTotal prob of N-in: 0.34375\noutside 1 316\nTMhelix 317 339\ninside 340 346

31241 GCF_900129455.1_IMG-taxon_2695420949_annotated_assembly Jatrophihabitans endophyticus Terrabacteria group; Actinobacteria; Actinobacteria; Frankiales; Frankiaceae; Jatrophihabitans MTDGSRTAAFFDLDKTIIAKSSVLAFGRPFYQGGLINRRAVLRSAYAQFMFALAGADSDQ WP_073387967.1 inhibition of morphological differentiation protein [Jatrophihabitans endophyticus] Length: 267\nNumber of predicted TMHs: 1\nExp number of AAs in TMHs: 21.19499\nExp number, first 60 AAs: 0.11085\nTotal prob of N-in: 0.02896\noutside 1 238\nTMhelix 239 261\ninside 262 267

31242 GCF_900129455.1_IMG-taxon_2695420949_annotated_assembly Jatrophihabitans endophyticus Terrabacteria group; Actinobacteria; Actinobacteria; Frankiales; Frankiaceae; Jatrophihabitans MRRAAARRRTAVVVIAVAMTTTPTPAAATGGARALLRITDPRLDEVSGLAPGHRSPGVYY WP_073385482.1 hypothetical protein [Jatrophihabitans endophyticus] Length: 375\nNumber of predicted TMHs: 2\nExp number of AAs in TMHs: 40.25121\nExp number, first 60 AAs: 19.3485\nTotal prob of N-in: 0.95622\nPOSSIBLE N-term signal sequence\ninside 1 11\nTMhelix 12 34\noutside 35 353\nTMhelix 354 371\ninside 372 375

31243 GCF_900129455.1_IMG-taxon_2695420949_annotated_assembly Jatrophihabitans endophyticus Terrabacteria group; Actinobacteria; Actinobacteria; Frankiales; Frankiaceae; Jatrophihabitans MKRRAALGAAALLVPLALAAPAAPTAAASAAAGAAAGVAAAQVFTPGGSPITGGDDAGAA WP_073390660.1 hypothetical protein [Jatrophihabitans endophyticus] Length: 438\nNumber of predicted TMHs: 2\nExp number of AAs in TMHs: 43.8383799999999\nExp number, first 60 AAs: 20.86818\nTotal prob of N-in:

0.89183\nPOSSIBLE N-term signal sequence\ninside 1 4\nTMhelix 5 27\noutside 28 403\nTMhelix 404 426\ninside 427 438

31244 GCF_000384115.1_ASM38411v1 Sporichthya polymorpha DSM 43042 Terrabacteria group; Actinobacteria; Actinobacteria; Frankiales; Sporichthyaceae; Sporichthya; Sporichthya polymorpha MSNRSVRRRRALCATGAFAILAGAPLVWSGHATAAGTFDATAAAYGFDYLMENASIAAGV WP_019873836.1 hypothetical protein [Sporichthya polymorpha] Length: 463\nNumber of predicted TMHs: 2\nExp number of AAs in TMHs: 43.23525\nExp number, first 60 AAs: 20.60924\nTotal prob of N-in: 0.95938\nPOSSIBLE N-term signal sequence\ninside 1 11\nTMhelix 12 34\noutside 35 436\nTMhelix 437 459\ninside 460 463

31245 GCF_000701365.1_ASM70136v1 Blastococcus sp. URHD0036 Terrabacteria group; Actinobacteria; Actinobacteria; Geodermatophilales; Geodermatophilaceae; Blastococcus MTRAAAFFDLDKTIIAKSSTLAFGRPFFEGGLINRRTVLKGAYAQFVFSLAGADAGQMER WP_029431055.1 inhibition of morphological differentiation protein [Blastococcus sp. URHD0036] Length: 281\nNumber of predicted TMHs: 1\nExp number of AAs in TMHs: 22.68425\nExp number, first 60 AAs: 0.0566\nTotal prob of N-in: 0.00580\noutside 1 228\nTMhelix 229 251\ninside 252 281

31246 GCF_000582785.1_B.massiliensis Candidatus Blastococcus massiliensis AP3 Terrabacteria group; Actinobacteria; Actinobacteria; Geodermatophilales; Geodermatophilaceae; Blastococcus; Candidatus Blastococcus massiliensis MTRAAAFFDLDKTIIAKSSTLAFGRPFFQGGLINRRTVLKTAYAQFVFSLAGADAQQMERWP_040337465.1 inhibition of morphological differentiation protein [Candidatus Blastococcus massiliensis] Length: 283\nNumber of predicted TMHs: 1\nExp number of AAs in TMHs: 22.334\nExp number, first 60 AAs: 0.03234\nTotal prob of N-in: 0.00669\noutside 1 233\nTMhelix 234 256\ninside 257 283

31247 GCF_900129495.1_IMG-taxon_2695420953_annotated_assembly Geodermatophilus nigrescens Terrabacteria group; Actinobacteria; Actinobacteria; Geodermatophilales; Geodermatophilaceae; Geodermatophilus MSPSSHRSPTRRPRGRRALLVTGLAAAVVAVPGVAAAAPGDAPVPLEEVGTTLEQAVTGQ WP_073418202.1 hypothetical protein [Geodermatophilus nigrescens] Length: 300\nNumber of predicted TMHs: 2\nExp number of AAs in TMHs: 38.2422\nExp number, first 60 AAs: 19.77347\nTotal prob of N-in: 0.96288\nPOSSIBLE N-term signal sequence\ninside 1 18\nTMhelix 19 38\noutside 39 266\nTMhelix 267 289\ninside 290 300

31248 GCF_900129495.1_IMG-taxon_2695420953_annotated_assembly Geodermatophilus nigrescens Terrabacteria group; Actinobacteria; Actinobacteria; Geodermatophilales; Geodermatophilaceae; Geodermatophilus MTRAAAFFDLDKTVIAKSSTLAFGRPFFAGGLINRRAVLKGAYAQFVFSLAGADAQQMER WP_073418302.1 inhibition of morphological differentiation protein [Geodermatophilus nigrescens] Length: 283\nNumber of predicted TMHs: 1\nExp number of AAs in TMHs: 22.75705\nExp number, first 60 AAs: 0.29458\nTotal prob of N-in: 0.02152\noutside 1 228\nTMhelix 229 251\ninside 252 283

31249 GCF_900115395.1_IMG-taxon_2599185183_annotated_assembly Geodermatophilus obscurus Terrabacteria group; Actinobacteria; Actinobacteria; Geodermatophilales; Geodermatophilaceae; Geodermatophilus MTRAAAFFDLDKTVIAKSSTLAFGRPFFAGGLINRRAVLKGAYAQFVFSLAGADAQQMER WP_075012953.1 inhibition of morphological differentiation protein [Geodermatophilus obscurus] Length: 289\nNumber of predicted TMHs: 1\nExp number of AAs in TMHs: 22.30849\nExp number, first 60 AAs: 0.29216\nTotal prob of N-in: 0.01990\noutside 1 232\nTMhelix 233 255\ninside 256 289

31250 GCF_900143215.1_IMG-taxon_2599185272_annotated_assembly Geodermatophilus obscurus Terrabacteria group; Actinobacteria; Actinobacteria; Geodermatophilales; Geodermatophilaceae; Geodermatophilus MRPHRNAPAPRRRPLRALAGAATALALCLTLPATAARAEEAGAPDTVVGEIVQAWPEAGP WP_072919771.1 hypothetical protein [Geodermatophilus obscurus] Length: 576\nNumber of predicted TMHs: 1\nExp number of AAs in TMHs: 35.2413\nExp number, first 60 AAs: 10.37518\nTotal prob of N-in: 0.49970\nPOSSIBLE N-term signal sequence\noutside 1 543\nTMhelix 544 566\ninside 567 576

31251 GCF_900143215.1_IMG-taxon_2599185272_annotated_assembly Geodermatophilus obscurus Terrabacteria group; Actinobacteria; Actinobacteria; Geodermatophilales; Geodermatophilaceae; Geodermatophilus MTRSAAFFDLDKTVIAKSSTLAFGRPFFQGGLINRRAVLKGAYAQFVFSLAGADAQQMER WP_072921297.1 inhibition of morphological differentiation protein [Geodermatophilus obscurus] Length: 280\nNumber of predicted TMHs: 1\nExp number of AAs in TMHs: 21.89963\nExp number, first 60 AAs: 0.05902\nTotal prob of N-in: 0.00711\noutside 1 232\nTMhelix 233 255\ninside 256 280

31252	GCF_001424455.1_Leaf369	Geodermatophilus sp. Leaf369	Terrabacteria group; Actinobacteria; Actinobacteria; Geodermatophilales; Geodermatophilaceae; Geodermatophilus
MNTNRRGLLLSGRRSCQAAALTLGALALGGVCAPAALAMQGTTDAAVPATPADASAPRIL	WP_055764009.1
hypothetical protein [Geodermatophilus sp. Leaf369]	Length: 418\nNumber of predicted TMHs: 1\nExp number of AAs in TMHs: 36.23691\nExp number, first 60 AAs: 16.75661\nTotal prob of N-in: 0.79089\nPOSSIBLE N-term signal sequence\noutside 1 389\nTMhelix 390 412\ninside 413 418

31253	GCF_001424455.1_Leaf369	Geodermatophilus sp. Leaf369	Terrabacteria group; Actinobacteria; Actinobacteria; Geodermatophilales; Geodermatophilaceae; Geodermatophilus
MTGEVGRAAAFFDLDKTVIAKSSALAFGRPFFEGGLMNRRTVLKGAYAQFVFSLAGADAA	WP_055760764.1
inhibition of morphological differentiation protein [Geodermatophilus sp. Leaf369]	Length: 281\nNumber of predicted TMHs: 1\nExp number of AAs in TMHs: 20.32803\nExp number, first 60 AAs: 0.23337\nTotal prob of N-in: 0.11257\noutside 1 239\nTMhelix 240 259\ninside 260 281

31254	GCF_000025345.1_ASM2534v1	Geodermatophilus obscurus DSM 43160	Terrabacteria group; Actinobacteria; Actinobacteria; Geodermatophilales; Geodermatophilaceae; Geodermatophilus; Geodermatophilus obscurus	MTRAAAFFDLDKTVIAKSSTLAFGRPFFQGGLINRRAVLKGAYAQFVFSLAGADAPQMER	WP_012946812.1
haloacid dehalogenase [Geodermatophilus obscurus]	Length: 280\nNumber of predicted TMHs: 1\nExp number of AAs in TMHs: 21.89401\nExp number, first 60 AAs: 0.0703\nTotal prob of N-in: 0.00681\noutside 1 232\nTMhelix 233 255\ninside 256 280

31255	GCF_000761485.1_ASM76148v1	Modestobacter caceresii	Terrabacteria group; Actinobacteria; Actinobacteria; Geodermatophilales; Geodermatophilaceae; Modestobacter
MTRAAAFFDLDKTVIAKSSTLAFGRPFFQGGLINRRAVLKGAYAQFVFSLAGADADQMER	WP_036334001.1
inhibition of morphological differentiation protein [Modestobacter caceresii]	Length: 287\nNumber of predicted TMHs: 1\nExp number of AAs in TMHs: 24.42351\nExp number, first 60 AAs: 0.13063\nTotal prob of N-in: 0.06422\noutside 1 235\nTMhelix 236 255\ninside 256 287

31256	GCF_000306785.1_ASM30678v1	Modestobacter marinus	Terrabacteria group; Actinobacteria; Actinobacteria; Geodermatophilales; Geodermatophilaceae; Modestobacter
MTCAAAFFDLDKTVIAKSSTLAFGRPFFQGGLINRRAVLKGAYAQFVFSLAGADADQMER	WP_014738740.1
inhibition of morphological differentiation protein [Modestobacter marinus]	Length: 290\nNumber of predicted TMHs: 1\nExp number of AAs in TMHs: 21.18984\nExp number, first 60 AAs: 0.06221\nTotal prob of N-in: 0.01212\noutside 1 232\nTMhelix 233 255\ninside 256 290

31257	GCF_001424485.1_Leaf380	Modestobacter sp. Leaf380	Terrabacteria group; Actinobacteria; Actinobacteria; Geodermatophilales; Geodermatophilaceae; Modestobacter
MTRSAAFFDLDKTVIAKSSTLAFGRPFFQGGLINRRAVLKGAYAQFVFSLAGADEVQMER	WP_082556774.1
inhibition of morphological differentiation protein [Modestobacter sp. Leaf380]	Length: 281\nNumber of predicted TMHs: 1\nExp number of AAs in TMHs: 20.36616\nExp number, first 60 AAs: 0.04464\nTotal prob of N-in: 0.01696\noutside 1 235\nTMhelix 236 255\ninside 256 281

31258	GCF_001424485.1_Leaf380	Modestobacter sp. Leaf380	Terrabacteria group; Actinobacteria; Actinobacteria; Geodermatophilales; Geodermatophilaceae; Modestobacter
MRTRFSENTVAAQPVTVTAARVVRPERREAFEQWATEVQALAATFPGHLGSSTLRPGPGS	WP_056291520.1
antibiotic biosynthesis monooxygenase [Modestobacter sp. Leaf380]	Length: 187\nNumber of predicted TMHs: 2\nExp number of AAs in TMHs: 42.64217\nExp number, first 60 AAs: 0.00368\nTotal prob of N-in: 0.95285\ninside 1 118\nTMhelix 119 141\noutside 142 150\nTMhelix 151 170\ninside 171 187

31259	GCF_000620205.1_ASM62020v1	Geodermatophilaceae bacterium URHA0031	Terrabacteria group; Actinobacteria; Actinobacteria; Geodermatophilales; Geodermatophilaceae; unclassified Geodermatophilaceae
MTCAAAFFDLDKTVIAKSSTLAFGRPFFQGGLINRRAVLKGAYAQFVFSLAGADADQMER	WP_034907912.1
inhibition of morphological differentiation protein [Geodermatophilaceae bacterium URHA0031]	Length: 284\nNumber of predicted TMHs: 1\nExp number of AAs in TMHs: 20.68413\nExp number, first 60 AAs: 0.04879\nTotal prob of N-in: 0.01738\noutside 1 232\nTMhelix 233 255\ninside 256 284

31260	GCF_000620185.1_ASM62018v1	Geodermatophilaceae bacterium URHB0062	Terrabacteria group; Actinobacteria; Actinobacteria; Geodermatophilales; Geodermatophilaceae; unclassified Geodermatophilaceae
MTRAAAFFDLDKTVIAKSSTLAFGRPFFHGGLINRRAVLKTAYAQLVFSFAGADAQQMER	WP_034942603.1
inhibition of morphological differentiation protein [Geodermatophilaceae bacterium URHB0062]	Length:

283\nNumber of predicted TMHs:  1\nExp number of AAs in TMHs: 21.99244\nExp number, first 60 AAs:  0.04888\nTotal prob of N-in:   0.00734\noutside   1   232\nTMhelix   233   255\ninside   256   283

31261 GCF_000620185.1_ASM62018v1 Geodermatophilaceae bacterium URHB0062 Terrabacteria group; Actinobacteria; Actinobacteria; Geodermatophilales; Geodermatophilaceae; unclassified Geodermatophilaceae MPPVRFVPAPVARRRALRSLVLAGAMAAAASLLTPPAVASAETATNPETTTTVVGQLLQT WP_026857129.1 hypothetical protein [Geodermatophilaceae bacterium URHB0062] Length: 582\nNumber of predicted TMHs: 1\nExp number of AAs in TMHs: 27.84684\nExp number, first 60 AAs:  6.43974\nTotal prob of N-in:   0.30808\noutside   1   548\nTMhelix   549   571\ninside   572   582

31262 GCF_000482705.1_ASM48270v1 Glycomyces arizonensis DSM 44726 Terrabacteria group; Actinobacteria; Actinobacteria; Glycomycetales; Glycomycetaceae; Glycomyces; Glycomyces arizonensis MTRLSRRGLIVRAGAAAVLAAAAASAQASPAVAQGGGCEGVAVVVDSGRGEPSVGCAADP WP_051393048.1 hypothetical protein [Glycomyces arizonensis] Length: 200\nNumber of predicted TMHs:  1\nExp number of AAs in TMHs: 31.19849\nExp number, first 60 AAs:  10.65187\nTotal prob of N-in:   0.52686\nPOSSIBLE N-term signal sequence\noutside   1   171\nTMhelix   172   194\ninside   195   200

31263 GCF_000427885.1_ASM42788v1 Glycomyces tenuis DSM 44171 Terrabacteria group; Actinobacteria; Actinobacteria; Glycomycetales; Glycomycetaceae; Glycomyces; Glycomyces tenuis MSKCNVSPGLPRSGNCHAYGGTVSKTNTAPRSAASARRKALAGRLGILGTAAAVTAVALA WP_081687086.1 hypothetical protein [Glycomyces tenuis] Length: 527\nNumber of predicted TMHs:  2\nExp number of AAs in TMHs: 41.19263\nExp number, first 60 AAs:  16.41886\nTotal prob of N-in:   0.83186\nPOSSIBLE N-term signal sequence\ninside   1   40\nTMhelix   41   63\noutside   64   498\nTMhelix   499   521\ninside   522   527

31264 GCF_900106035.1_IMG-taxon_2630968302_annotated_assembly Jiangella alba Terrabacteria group; Actinobacteria; Actinobacteria; Jiangellales; Jiangellaceae; Jiangella MTRRTTFGTAAALLAALALAGCGGGNDSDSGGASAADEPAPAAEAPQENAAGDGDRAAGG WP_069111596.1 hypothetical protein [Jiangella alba] Length: 312\nNumber of predicted TMHs:  1\nExp number of AAs in TMHs: 28.91282\nExp number, first 60 AAs:  4.4846\nTotal prob of N-in:   0.30520\noutside   1   268\nTMhelix   269   291\ninside   292   312

31265 GCF_900106035.1_IMG-taxon_2630968302_annotated_assembly Jiangella alba Terrabacteria group; Actinobacteria; Actinobacteria; Jiangellales; Jiangellaceae; Jiangella MSSLLVDNDDRPATAAFFDLDKTIIAKSSTLAFSKSFYQGGLINRRAVLRSAYAQFVYLV WP_069110276.1 inhibition of morphological differentiation protein [Jiangella alba] Length: 274\nNumber of predicted TMHs:  1\nExp number of AAs in TMHs: 20.5869\nExp number, first 60 AAs:  0.05075\nTotal prob of N-in:   0.01394\noutside   1   249\nTMhelix   250   269\ninside   270   274

31266 GCF_900106035.1_IMG-taxon_2630968302_annotated_assembly Jiangella alba Terrabacteria group; Actinobacteria; Actinobacteria; Jiangellales; Jiangellaceae; Jiangella MTATAKVTFSPVTSTSASSVAPRTMVSRRALVRVRSCVISLMTPQCAARLGVTMLNGRGF WP_083288753.1 hypothetical protein [Jiangella alba] Length: 920\nNumber of predicted TMHs:  2\nExp number of AAs in TMHs: 41.70473\nExp number, first 60 AAs:  0.28889\nTotal prob of N-in:   0.94129\ninside   1   110\nTMhelix   111   133\noutside   134   887\nTMhelix   888   910\ninside   911   920

31267 GCF_001708125.1_ASM170812v1 Jiangella alba Terrabacteria group; Actinobacteria; Actinobacteria; Jiangellales; Jiangellaceae; Jiangella MTATAKVTFSPVTSTSASSVAPRTMVSRRALVRVRSCVISLMTPQCAARLGVTMLNGRGF WP_083288753.1 hypothetical protein [Jiangella alba] Length: 920\nNumber of predicted TMHs:  2\nExp number of AAs in TMHs: 41.70473\nExp number, first 60 AAs:  0.28889\nTotal prob of N-in:   0.94129\ninside   1 110\nTMhelix   111   133\noutside   134   887\nTMhelix   888   910\ninside   911   920

31268 GCF_001708125.1_ASM170812v1 Jiangella alba Terrabacteria group; Actinobacteria; Actinobacteria; Jiangellales; Jiangellaceae; Jiangella MTRRTTFGTAAALLAALALAGCGGGNDSDSGGASAADEPAPAAEAPQENAAGDGDRAAGG WP_069111596.1 hypothetical protein [Jiangella alba] Length: 312\nNumber of predicted TMHs:  1\nExp number of AAs in TMHs: 28.91282\nExp number, first 60 AAs:  4.4846\nTotal prob of N-in:   0.30520\noutside   1 268\nTMhelix   269   291\ninside   292   312

31269 GCF_001708125.1_ASM170812v1 Jiangella alba Terrabacteria group; Actinobacteria; Actinobacteria; Jiangellales; Jiangellaceae; Jiangella MSSLLVDNDDRPATAAFFDLDKTIIAKSSTLAFSKSFYQGGLINRRAVLRSAYAQFVYLV WP_069110276.1 inhibition of morphological differentiation protein [Jiangella alba] Length: 274\nNumber of

predicted TMHs: 1\nExp number of AAs in TMHs: 20.5869\nExp number, first 60 AAs: 0.05075\nTotal prob of N-in: 0.01394\noutside 1 249\nTMhelix 250 269\ninside 270 274

31270 GCF_001005145.1_ASM100514v1 Jiangella alkaliphila Terrabacteria group; Actinobacteria; Actinobacteria; Jiangellales; Jiangellaceae; Jiangella MLVDNDDRPPTAAFFDLDKTIIAKSSTLAFSKSFYQGGLINRRAVLRSAYAQFVYLVGGA WP_046766506.1 inhibition of morphological differentiation protein [Jiangella alkaliphila] Length: 271\nNumber of predicted TMHs: 1\nExp number of AAs in TMHs: 20.61267\nExp number, first 60 AAs: 0.08054\nTotal prob of N-in: 0.02011\noutside 1 246\nTMhelix 247 266\ninside 267 271

31271 GCF_900105925.1_IMG-taxon_2639762615_annotated_assembly Jiangella alkaliphila Terrabacteria group; Actinobacteria; Actinobacteria; Jiangellales; Jiangellaceae; Jiangella MLVDNDDRPPTAAFFDLDKTIIAKSSTLAFSKSFYQGGLINRRAVLRSAYAQFVYLVGGA WP_046766506.1 inhibition of morphological differentiation protein [Jiangella alkaliphila] Length: 271\nNumber of predicted TMHs: 1\nExp number of AAs in TMHs: 20.61267\nExp number, first 60 AAs: 0.08054\nTotal prob of N-in: 0.02011\noutside 1 246\nTMhelix 247 266\ninside 267 271

31272 GCF_900105925.1_IMG-taxon_2639762615_annotated_assembly Jiangella alkaliphila Terrabacteria group; Actinobacteria; Actinobacteria; Jiangellales; Jiangellaceae; Jiangella MTRRTVFGTTITLLAALALAGCGGSDDSDSGSDSAGGSAADLSAPEADAPAGEGDGEAAQ WP_046770059.1 hypothetical protein [Jiangella alkaliphila] Length: 320\nNumber of predicted TMHs: 1\nExp number of AAs in TMHs: 28.00188\nExp number, first 60 AAs: 3.12434\nTotal prob of N-in: 0.18621\noutside 1 274\nTMhelix 275 297\ninside 298 320

31273 GCF_001005145.1_ASM100514v1 Jiangella alkaliphila Terrabacteria group; Actinobacteria; Actinobacteria; Jiangellales; Jiangellaceae; Jiangella MTRRTVFGTTITLLAALALAGCGGSDDSDSGSDSAGGSAADLSAPEADAPAGEGDGEAAQ WP_046770059.1 hypothetical protein [Jiangella alkaliphila] Length: 320\nNumber of predicted TMHs: 1\nExp number of AAs in TMHs: 28.00188\nExp number, first 60 AAs: 3.12434\nTotal prob of N-in: 0.18621\noutside 1 274\nTMhelix 275 297\ninside 298 320

31274 GCF_001270745.1_ASM127074v1 Jiangella muralis Terrabacteria group; Actinobacteria; Actinobacteria; Jiangellales; Jiangellaceae; Jiangella MTRRTTFGAAIALLAAVALAGCGGADDSSGGSDAAHEAPAADAPPEGGDQAPGDGDQAAG WP_053202776.1 hypothetical protein [Jiangella muralis] Length: 312\nNumber of predicted TMHs: 1\nExp number of AAs in TMHs: 36.6591\nExp number, first 60 AAs: 12.59582\nTotal prob of N-in: 0.66555\nPOSSIBLE N-term signal sequence\noutside 1 266\nTMhelix 267 289\ninside 290 312

31275 GCF_001270745.1_ASM127074v1 Jiangella muralis Terrabacteria group; Actinobacteria; Actinobacteria; Jiangellales; Jiangellaceae; Jiangella MSSLLVDNDDRPPTAAFFDLDKTIIAKSSTLAFSKSFYQGGLINRRAVLRSAYAQFVYLV WP_053206952.1 inhibition of morphological differentiation protein [Jiangella muralis] Length: 274\nNumber of predicted TMHs: 1\nExp number of AAs in TMHs: 20.58736\nExp number, first 60 AAs: 0.0512\nTotal prob of N-in: 0.01392\noutside 1 249\nTMhelix 250 269\ninside 270 274

31276 GCF_000515395.1_ASM51539v1 Jiangella gansuensis DSM 44835 Terrabacteria group; Actinobacteria; Actinobacteria; Jiangellales; Jiangellaceae; Jiangella; Jiangella gansuensis MIVDNDDRPRTAAFFDLDKTIIAKSSTLAFSRSFYQGGLINRRAVLRSAYAQFVYLVGGA WP_026874347.1 inhibition of morphological differentiation protein [Jiangella gansuensis] Length: 273\nNumber of predicted TMHs: 1\nExp number of AAs in TMHs: 20.40356\nExp number, first 60 AAs: 0.16253\nTotal prob of N-in: 0.09311\noutside 1 247\nTMhelix 248 267\ninside 268 273

31277 GCF_000515395.1_ASM51539v1 Jiangella gansuensis DSM 44835 Terrabacteria group; Actinobacteria; Actinobacteria; Jiangellales; Jiangellaceae; Jiangella; Jiangella gansuensis MRRRTAVGSIGTILAAALALGACSGGGADSATSGGADDAPAAEAGDGAAPAGGEEAADNA WP_084469651.1 hypothetical protein [Jiangella gansuensis] Length: 308\nNumber of predicted TMHs: 2\nExp number of AAs in TMHs: 42.31588\nExp number, first 60 AAs: 16.99626\nTotal prob of N-in: 0.80752\nPOSSIBLE N-term signal sequence\ninside 1 6\nTMhelix 7 29\noutside 30 269\nTMhelix 270 292\ninside 293 308

31278 GCF_000515395.1_ASM51539v1 Jiangella gansuensis DSM 44835 Terrabacteria group; Actinobacteria; Actinobacteria; Jiangellales; Jiangellaceae; Jiangella; Jiangella gansuensis MSAPLFGRRAVLALVSAAGLGIVGTLPATGLDIGSLPPQPAVRMQTFDPSAFTDAAAQLP WP_051425826.1 hypothetical protein [Jiangella gansuensis] Length: 758\nNumber of predicted TMHs: 2\nExp number of AAs in TMHs: 36.79552\nExp number, first 60 AAs: 16.67871\nTotal prob of N-in: 0.75594\nPOSSIBLE N-term signal sequence\ninside 1 11\nTMhelix 12 34\noutside 35 729\nTMhelix 730 752\ninside 753 758

31279 GCF_000017305.1_ASM1730v1 Kineococcus radiotolerans SRS30216 = ATCC BAA-149 Terrabacteria group; Actinobacteria; Actinobacteria; Kineosporiales; Kineosporiaceae; Kineococcus; Kineococcus radiotolerans MTPRSVGRRRVLAGLAAGGALLPAGAARAATPAPTATSTAGGSDLPLRVVLLDVTPSAYS WP_041292034.1 hypothetical protein [Kineococcus radiotolerans] Length: 722\nNumber of predicted TMHs: 1\nExp number of AAs in TMHs: 23.91922\nExp number, first 60 AAs: 3.11813\nTotal prob of N-in: 0.16232\noutside 1 670\nTMhelix 671 693\ninside 694 722

31280 GCF_002198675.1_ASM219867v1 Kineosporia sp. R_H_3 Terrabacteria group; Actinobacteria; Actinobacteria; Kineosporiales; Kineosporiaceae; Kineosporia MPFKEHHVGKGTIAYRGEDPEYWSRKSASWRPRRNAVTVALTVLGLLLLGFAASPLARHH WP_088318137.1 hypothetical protein [Kineosporia sp. R_H_3] Length: 168\nNumber of predicted TMHs: 2\nExp number of AAs in TMHs: 41.97158\nExp number, first 60 AAs: 21.73707\nTotal prob of N-in: 0.98211\nPOSSIBLE N-term signal sequence\ninside 1 34\nTMhelix 35 57\noutside 58 136\nTMhelix 137 156\ninside 157 168

31281 GCF_001619685.1_ASM161968v1 Brevibacterium casei Terrabacteria group; Actinobacteria; Actinobacteria; Micrococcales; Brevibacteriaceae; Brevibacterium MTTRSRSSNRRRATLGAVAAATAVGLTVLITAPGTAVPVDPDELAEAHGQLIDTDIISAD WP_063250685.1 hypothetical protein [Brevibacterium casei] Length: 1168\nNumber of predicted TMHs: 2\nExp number of AAs in TMHs: 44.26808\nExp number, first 60 AAs: 21.90715\nTotal prob of N-in: 0.99535\nPOSSIBLE N-term signal sequence\ninside 1 12\nTMhelix 13 35\noutside 36 1135\nTMhelix 1136 1158\ninside 1159 1168

31282 GCF_001584355.1_ASM158435v1 Brevibacterium ravenspurgense Terrabacteria group; Actinobacteria; Actinobacteria; Micrococcales; Brevibacteriaceae; Brevibacterium MNSSKSSTQRRRFLRAPIAALSALALTAGAFAVPSLMSGDSAAAAEFSGGIRDKSGAVEQ WP_061943284.1 cell surface protein [Brevibacterium ravenspurgense]Length: 1811\nNumber of predicted TMHs: 2\nExp number of AAs in TMHs: 44.40242\nExp number, first 60 AAs: 21.86419\nTotal prob of N-in: 0.99619\nPOSSIBLE N-term signal sequence\ninside 1 12\nTMhelix 13 35\noutside 36 1781\nTMhelix 1782 1804\ninside 1805 1811

31283 GCF_000285835.1_ASM28583v2 Brevibacterium senegalense Terrabacteria group; Actinobacteria; Actinobacteria; Micrococcales; Brevibacteriaceae; Brevibacterium MASTAATRTRRGHLGLLRGVLLGGLLVAAVITGLLAMHTLNLHGTPAAHAATAAVSVTDS WP_051013283.1 DUF2946 domain-containing protein [Brevibacterium senegalense] Length: 156\nNumber of predicted TMHs: 2\nExp number of AAs in TMHs: 43.90781\nExp number, first 60 AAs: 22.02915\nTotal prob of N-in: 0.99747\nPOSSIBLE N-term signal sequence\ninside 1 19\nTMhelix 20 42\noutside 43 98\nTMhelix 99 121\ninside 122 156

31284 GCF_900163715.1_ASM90016371v1 Brevibacterium yomogidense Terrabacteria group; Actinobacteria; Actinobacteria; Micrococcales; Brevibacteriaceae; Brevibacterium MARVNSADGFSDGHADGDAVGSSDTAGTTDPVTGIGRRTVLRTGAFGALGAAALWGLGSG WP_087002950.1 hypothetical protein [Brevibacterium yomogidense] Length: 836\nNumber of predicted TMHs: 1\nExp number of AAs in TMHs: 21.00845\nExp number, first 60 AAs: 1.21626\nTotal prob of N-in: 0.06460\noutside 1 785\nTMhelix 786 808\ninside 809 836

31285 GCF_000178455.1_ASM17845v1 Brevibacterium mcbrellneri ATCC 49030 Terrabacteria group; Actinobacteria; Actinobacteria; Micrococcales; Brevibacteriaceae; Brevibacterium; Brevibacterium mcbrellneri MHSSKTPTQRRRFLRVPIAALSALALTAGAFAVPNLMSGDSAAAATFSGGIRDKSGAVEK WP_005881676.1 cell surface protein [Brevibacterium mcbrellneri] Length: 1591\nNumber of predicted TMHs: 2\nExp number of AAs in TMHs: 43.19063\nExp number, first 60 AAs: 21.5405\nTotal prob of N-in: 0.99658\nPOSSIBLE N-term signal sequence\ninside 1 12\nTMhelix 13 35\noutside 36 1561\nTMhelix 1562 1584\ninside 1585 1591

31286 GCF_000603945.1_genome Actinotalea ferrariae CF5-4 Terrabacteria group; Actinobacteria; Actinobacteria; Micrococcales; Cellulomonadaceae; Actinotalea; Actinotalea ferrariae MVDMTSHLPRRAALALPTAALAVALLAGCVGSGAQSGDEASGGAADTSAVEESAEGGGDG WP_052023053.1 hypothetical protein [Actinotalea ferrariae] Length: 317\nNumber of predicted TMHs: 2\nExp number of AAs in TMHs: 45.22033\nExp number, first 60 AAs: 20.22254\nTotal prob of N-in: 0.88203\nPOSSIBLE N-term signal sequence\ninside 1 11\nTMhelix 12 34\noutside 35 258\nTMhelix 259 281\ninside 282 317

31287 GCF_001057055.1_ASM105705v1 Cellulomonas flavigena Terrabacteria group; Actinobacteria; Actinobacteria; Micrococcales; Cellulomonadaceae; Cellulomonas MVFRPAVNLVLLVLIVLLGAALVFLTARRATRRSFIDIFRRSMILAVVVVMGAGPSIPGE WP_048728390.1 MULTISPECIES: hypothetical protein [Micrococcales] Length: 344\nNumber of predicted TMHs: 2\nExp number of AAs in TMHs:

47.25705\nExp number, first 60 AAs: 27.33032\nTotal prob of N-in: 0.37695\nPOSSIBLE N-term signal sequence\noutside 1 4\nTMhelix 5 27\ninside 28 310\nTMhelix 311 330\noutside 331 344

31288 GCF_002165145.1_ASM216514v1 Cellulomonas iranensis Terrabacteria group; Actinobacteria; Actinobacteria; Micrococcales; Cellulomonadaceae; Cellulomonas MRDRSPGTRARRRALHATVAALGLLLVAAAPAAAAPAVDEVALTGPTAAVPAGASVDVRL WP_087510129.1 hypothetical protein [Cellulomonas iranensis] Length: 249\nNumber of predicted TMHs: 2\nExp number of AAs in TMHs: 44.50868\nExp number, first 60 AAs: 22.17952\nTotal prob of N-in: 0.99361\nPOSSIBLE N-term signal sequence\ninside 1 12\nTMhelix 13 35\noutside 36 218\nTMhelix 219 241\ninside 242 249

31289 GCF_002165145.1_ASM216514v1 Cellulomonas iranensis Terrabacteria group; Actinobacteria; Actinobacteria; Micrococcales; Cellulomonadaceae; Cellulomonas MTRPAPTARRTRAPRRALVALAAAALLGVAAGAPALAAADRTLGASGATVTVEAPAGVAA WP_087509593.1 hypothetical protein [Cellulomonas iranensis] Length: 280\nNumber of predicted TMHs: 2\nExp number of AAs in TMHs: 52.21092\nExp number, first 60 AAs: 22.48412\nTotal prob of N-in: 0.99791\nPOSSIBLE N-term signal sequence\ninside 1 16\nTMhelix 17 39\noutside 40 251\nTMhelix 252 274\ninside 275 280

31290 GCF_002165145.1_ASM216514v1 Cellulomonas iranensis Terrabacteria group; Actinobacteria; Actinobacteria; Micrococcales; Cellulomonadaceae; Cellulomonas MSRRRTALLTAPAAAAAAALTLVGAATSASAGSDAPTPYRVTPSGLELPPGDVFEVHGHV WP_087509670.1 hypothetical protein [Cellulomonas iranensis] Length: 276\nNumber of predicted TMHs: 2\nExp number of AAs in TMHs: 40.43083\nExp number, first 60 AAs: 19.82244\nTotal prob of N-in: 0.89433\nPOSSIBLE N-term signal sequence\ninside 1 6\nTMhelix 7 29\noutside 30 245\nTMhelix 246 268\ninside 269 276

31291 GCF_002165145.1_ASM216514v1 Cellulomonas iranensis Terrabacteria group; Actinobacteria; Actinobacteria; Micrococcales; Cellulomonadaceae; Cellulomonas MTTVPPTPPPGPAPLDRPAPPTRRAPRSRRRALVLLVTTALGLVGVVTLLVTWVSIGVRT WP_087508506.1 hypothetical protein [Cellulomonas iranensis] Length: 156\nNumber of predicted TMHs: 2\nExp number of AAs in TMHs: 45.81507\nExp number, first 60 AAs: 22.94356\nTotal prob of N-in: 0.87171\nPOSSIBLE N-term signal sequence\ninside 1 31\nTMhelix 32 54\noutside 55 113\nTMhelix 114 136\ninside 137 156

31292 GCF_001040865.1_ASM104086v1 Cellulomonas sp. A375-1 Terrabacteria group; Actinobacteria; Actinobacteria; Micrococcales; Cellulomonadaceae; Cellulomonas MNARRAALSVLFAVLLAAVGAAAAYAFHWYVVQPSDAELLAQARQIELPGLVADGDPAVS WP_048341338.1 hypothetical protein [Cellulomonas sp. A375-1] Length: 171\nNumber of predicted TMHs: 2\nExp number of AAs in TMHs: 44.52686\nExp number, first 60 AAs: 22.7336\nTotal prob of N-in: 0.99545\nPOSSIBLE N-term signal sequence\ninside 1 6\nTMhelix 7 29\noutside 30 142\nTMhelix 143 165\ninside 166 171

31293 GCF_001462455.1_ASM146245v1 Cellulomonas sp. B6 Terrabacteria group; Actinobacteria; Actinobacteria; Micrococcales; Cellulomonadaceae; Cellulomonas MTTVPTPPPSGPAPLGDTPPPARPAPPARPAPPARRAPRSRRRALALLVATGLGLVGVVM WP_062102724.1 hypothetical protein [Cellulomonas sp. B6] Length: 171\nNumber of predicted TMHs: 2\nExp number of AAs in TMHs: 45.9078\nExp number, first 60 AAs: 16.45666\nTotal prob of N-in: 0.95347\nPOSSIBLE N-term signal sequence\ninside 1 43\nTMhelix 44 66\noutside 67 125\nTMhelix 126 148\ninside 149 171

31294 GCF_001462455.1_ASM146245v1 Cellulomonas sp. B6 Terrabacteria group; Actinobacteria; Actinobacteria; Micrococcales; Cellulomonadaceae; Cellulomonas MVQVVRGTRRGTRADASGRGRRRAAMLAAAVVAALVAPLALVGLATPAAAAEYRPFGSVF WP_062101716.1 hypothetical protein [Cellulomonas sp. B6] Length: 1439\nNumber of predicted TMHs: 2\nExp number of AAs in TMHs: 45.4201699999999\nExp number, first 60 AAs: 22.7303\nTotal prob of N-in: 0.99972\nPOSSIBLE N-term signal sequence\ninside 1 28\nTMhelix 29 51\noutside 52 1408\nTMhelix 1409 1431\ninside 1432 1439

31295 GCF_000986825.1_ASM98682v1 Cellulomonas sp. FA1 Terrabacteria group; Actinobacteria; Actinobacteria; Micrococcales; Cellulomonadaceae; Cellulomonas MTTVPPTPPPGPAPLDCPAPPTRRAPRSRRRALALLVTTALGLVGVVTLLVTWVSIGVRT WP_046527827.1 hypothetical protein [Cellulomonas sp. FA1] Length: 156\nNumber of predicted TMHs: 2\nExp number of AAs in TMHs: 45.78671\nExp number, first 60 AAs: 22.93015\nTotal prob of N-in: 0.80655\nPOSSIBLE N-term signal sequence\ninside 1 31\nTMhelix 32 54\noutside 55 113\nTMhelix 114 136\ninside 137 156

31296 GCF_000986825.1_ASM98682v1 Cellulomonas sp. FA1 Terrabacteria group; Actinobacteria; Actinobacteria; Micrococcales; Cellulomonadaceae; Cellulomonas

MSRRRTALLTAPAAAAAAALTLVGAATSASAGSDSPTPYRVTPTGLELPAGDVFEVHGHV WP_046529179.1 hypothetical protein [Cellulomonas sp. FA1] Length: 269\nNumber of predicted TMHs: 2\nExp number of AAs in TMHs: 40.59792\nExp number, first 60 AAs: 19.93065\nTotal prob of N-in: 0.89884\nPOSSIBLE N-term signal sequence\ninside 1 6\nTMhelix 7 29\noutside 30 238\nTMhelix 239 261\ninside 262 269

31297 GCF_000708885.1_CelluHZM_1.0 Cellulomonas sp. HZM Terrabacteria group; Actinobacteria; Actinobacteria; Micrococcales; Cellulomonadaceae; Cellulomonas MDLQQAAFFDLDKTIIATSSATAFSRGFLAEGMLTRRSVLRAAYAQIAYLLGAADEDQTE WP_029289769.1 inhibition of morphological differentiation protein [Cellulomonas sp. HZM] Length: 262\nNumber of predicted TMHs: 1\nExp number of AAs in TMHs: 22.66072\nExp number, first 60 AAs: 0.05965\nTotal prob of N-in: 0.00688\noutside 1 233\nTMhelix 234 256\ninside 257 262

31298 GCF_000526515.1_ASM52651v1 Cellulomonas sp. KRMCY2 Terrabacteria group; Actinobacteria; Actinobacteria; Micrococcales; Cellulomonadaceae; Cellulomonas MSGSPSHGRASGWTCPTPAGPVEHPRPVAARHGDGHTGGPARRTVVRLSASTAPGWSGVG WP_081785062.1 hypothetical protein [Cellulomonas sp. KRMCY2] Length: 312\nNumber of predicted TMHs: 2\nExp number of AAs in TMHs: 36.38495\nExp number, first 60 AAs: 0.03038\nTotal prob of N-in: 0.70873\ninside 1 97\nTMhelix 98 120\noutside 121 281\nTMhelix 282 304\ninside 305 312

31299 GCF_001424195.1_Leaf334 Cellulomonas sp. Leaf334 Terrabacteria group; Actinobacteria; Actinobacteria; Micrococcales; Cellulomonadaceae; Cellulomonas MDEDPTQPDVLREAAFFDLDKTIIATSSATAFSRPFLAQGLLTRRSVLRTAYAQLLYLVG WP_056586227.1 inhibition of morphological differentiation protein [Cellulomonas sp. Leaf334] Length: 268\nNumber of predicted TMHs: 1\nExp number of AAs in TMHs: 21.77638\nExp number, first 60 AAs: 0.02283\nTotal prob of N-in: 0.00321\noutside 1 238\nTMhelix 239 261\ninside 262 268

31300 GCF_001424125.1_Leaf395 Cellulomonas sp. Leaf395 Terrabacteria group; Actinobacteria; Actinobacteria; Micrococcales; Cellulomonadaceae; Cellulomonas MDEDPTQPDTLREAAFFDLDKTIIATSSATAFSRPFLAQGLLTRRSVIRTAYAQLLYLVG WP_056023063.1 inhibition of morphological differentiation protein [Cellulomonas sp. Leaf395] Length: 268\nNumber of predicted TMHs: 1\nExp number of AAs in TMHs: 21.86315\nExp number, first 60 AAs: 0.0368\nTotal prob of N-in: 0.00733\noutside 1 238\nTMhelix 239 261\ninside 262 268

31301 GCF_001426725.1_Root137 Cellulomonas sp. Root137 Terrabacteria group; Actinobacteria; Actinobacteria; Micrococcales; Cellulomonadaceae; Cellulomonas MDVDPPNEDRSREAAFFDLDKTIIATSSATAFSRPFLAEGLLTRRSVVRTAYAQLLYLLG WP_056088521.1 inhibition of morphological differentiation protein [Cellulomonas sp. Root137] Length: 268\nNumber of predicted TMHs: 1\nExp number of AAs in TMHs: 21.80221\nExp number, first 60 AAs: 0.01385\nTotal prob of N-in: 0.00647\noutside 1 238\nTMhelix 239 261\ninside 262 268

31302 GCF_001426705.1_Root485 Cellulomonas sp. Root485 Terrabacteria group; Actinobacteria; Actinobacteria; Micrococcales; Cellulomonadaceae; Cellulomonas MASDRPRRRKPFDPTKLLTFCVSLAGLFAISCAFVWFGDAADHTLAVRLHTSGVTTTAHD WP_055919086.1 hypothetical protein [Cellulomonas sp. Root485] Length: 190\nNumber of predicted TMHs: 2\nExp number of AAs in TMHs: 41.47816\nExp number, first 60 AAs: 20.70247\nTotal prob of N-in: 0.99561\nPOSSIBLE N-term signal sequence\ninside 1 19\nTMhelix 20 37\noutside 38 148\nTMhelix 149 171\ninside 172 190

31303 GCF_001426705.1_Root485 Cellulomonas sp. Root485 Terrabacteria group; Actinobacteria; Actinobacteria; Micrococcales; Cellulomonadaceae; Cellulomonas MDEDLTQHDVPREAAFFDLDKTIIATSSATAFSRPFLAEGLLTRRSVLRTAYAQLLYLVG WP_055911862.1 inhibition of morphological differentiation protein [Cellulomonas sp. Root485] Length: 268\nNumber of predicted TMHs: 1\nExp number of AAs in TMHs: 21.76817\nExp number, first 60 AAs: 0.01165\nTotal prob of N-in: 0.00336\noutside 1 238\nTMhelix 239 261\ninside 262 268

31304 GCF_001429255.1_Root930 Cellulomonas sp. Root930 Terrabacteria group; Actinobacteria; Actinobacteria; Micrococcales; Cellulomonadaceae; Cellulomonas MDVDPPHEDRSREAAFFDLDKTIIATSSATAFSRPFLAEGLLTRRSVVRTAYAQLLYLLG WP_057210167.1 inhibition of morphological differentiation protein [Cellulomonas sp. Root930] Length: 268\nNumber of predicted TMHs: 1\nExp number of AAs in TMHs: 21.79446\nExp number, first 60 AAs: 0.0105\nTotal prob of N-in: 0.00376\noutside 1 238\nTMhelix 239 261\ninside 262 268

31305	GCF_000767135.1_ASM76713v1	Cellulomonas cellasea DSM 20118	Terrabacteria group; Actinobacteria; Actinobacteria; Micrococcales; Cellulomonadaceae; Cellulomonas; Cellulomonas cellasea
MLHGLRPSAHDRGAAPRRRALALTLVTGVIAASTLAAAAPASATDKPYGKPYSVAAYVYK	WP_034625822.1
hypothetical protein [Cellulomonas cellasea]	Length: 309\nNumber of predicted TMHs:  2\nExp number of AAs in TMHs: 43.36676\nExp number, first 60 AAs:  21.59788\nTotal prob of N-in:   0.97105\nPOSSIBLE N-term signal sequence\ninside   1   20\nTMhelix   21   43\noutside   44   277\nTMhelix   278   300\ninside   301   309

31306	GCF_000212695.1_ASM21269v1	Cellulomonas fimi ATCC 484	Terrabacteria group; Actinobacteria; Actinobacteria; Micrococcales; Cellulomonadaceae; Cellulomonas; Cellulomonas fimi
MVPSPATRPPRRRRWLAGTTSAALLVTAVASAATAAPLTDVHDFAGGDAQGWFSYANAGS	WP_013770863.1
glycoside hydrolase [Cellulomonas fimi]	Length: 1106\nNumber of predicted TMHs:  1\nExp number of AAs in TMHs: 32.39094\nExp number, first 60 AAs:  12.54032\nTotal prob of N-in:   0.58283\nPOSSIBLE N-term signal sequence\noutside   1  1078\nTMhelix   1079  1098\ninside   1099  1106

31307	GCF_000212695.1_ASM21269v1	Cellulomonas fimi ATCC 484	Terrabacteria group; Actinobacteria; Actinobacteria; Micrococcales; Cellulomonadaceae; Cellulomonas; Cellulomonas fimi
MKTRRAIVTAAAVAGLVLAPTAAFGYGADDYSNDGTVSDTTPAVGQAFTVTVKGPANTPV	WP_013769889.1
hypothetical protein [Cellulomonas fimi]	Length: 172\nNumber of predicted TMHs:  2\nExp number of AAs in TMHs: 46.5695\nExp number, first 60 AAs:  20.84419\nTotal prob of N-in:   0.98510\nPOSSIBLE N-term signal sequence\ninside   1   6\nTMhelix   7   26\noutside   27   140\nTMhelix   141   163\ninside   164   172

31308	GCF_001552375.1_ASM155237v1	Cellulomonas iranensis NBRC 101100 = JCM 18110	Terrabacteria group; Actinobacteria; Actinobacteria; Micrococcales; Cellulomonadaceae; Cellulomonas; Cellulomonas iranensis
MTTVPPTPPPGTPPSGPAPLDRPAPPTRRAPRSRRRALALLVTTALGLVGVVTLLVTWVSWP_070319715.1	hypothetical protein [Cellulomonas iranensis]	Length: 161\nNumber of predicted TMHs:  2\nExp number of AAs in TMHs: 45.79086\nExp number, first 60 AAs:  22.572\nTotal prob of N-in:   0.81865\nPOSSIBLE N-term signal sequence\ninside   1   36\nTMhelix   37   59\noutside   60   118\nTMhelix   119   141\ninside   142   161

31309	GCF_001552375.1_ASM155237v1	Cellulomonas iranensis NBRC 101100 = JCM 18110	Terrabacteria group; Actinobacteria; Actinobacteria; Micrococcales; Cellulomonadaceae; Cellulomonas; Cellulomonas iranensis
MSRRRTALLTAPAAAAAAALALVGAATSASAGSDSPTPYRVTPSGVELPAGDVFEVHEHV	WP_070320500.1
hypothetical protein [Cellulomonas iranensis]	Length: 276\nNumber of predicted TMHs:  2\nExp number of AAs in TMHs: 40.51817\nExp number, first 60 AAs:  20.8209\nTotal prob of N-in:   0.93304\nPOSSIBLE N-term signal sequence\ninside   1   6\nTMhelix   7   29\noutside   30   245\nTMhelix   246   268\ninside   269   276

31310	GCF_000312005.1_ASM31200v1	Cellulomonas massiliensis JC225	Terrabacteria group; Actinobacteria; Actinobacteria; Micrococcales; Cellulomonadaceae; Cellulomonas; Cellulomonas massiliensis
MPAVPAARRRRAVVAILSVLAAALLGGVVAAPAQAANIDPFVPVFSTNAHGAILMAGNTL	WP_019135425.1
hypothetical protein [Cellulomonas massiliensis]	Length: 1466\nNumber of predicted TMHs:  2\nExp number of AAs in TMHs: 48.5072299999999\nExp number, first 60 AAs:  22.63916\nTotal prob of N-in:   0.99935\nPOSSIBLE N-term signal sequence\ninside   1   12\nTMhelix   13   35\noutside   36  1440\nTMhelix   1441  1460\ninside   1461  1466

31311	GCF_001624335.1_ASM162433v1	Oerskovia enterophila	Terrabacteria group; Actinobacteria; Actinobacteria; Micrococcales; Cellulomonadaceae; Oerskovia
MLITQRGGRRRALASLLTGALVAGLAVVGVTPASAADTVMVSGGSATWNLKDSWTGYVGG	WP_068708018.1
hypothetical protein [Oerskovia enterophila]	Length: 1569\nNumber of predicted TMHs:  2\nExp number of AAs in TMHs: 44.20591\nExp number, first 60 AAs:  22.11127\nTotal prob of N-in:   0.99636\nPOSSIBLE N-term signal sequence\ninside   1   12\nTMhelix   13   35\noutside   36  1536\nTMhelix   1537  1559\ninside   1560  1569

31312	GCF_001624335.1_ASM162433v1	Oerskovia enterophila	Terrabacteria group; Actinobacteria; Actinobacteria; Micrococcales; Cellulomonadaceae; Oerskovia
MFLSTTRRTPRRTLLSALATGALATGLIGVAGMPAAVAAPLATTPLCTPSSTLPGITTGG	WP_068708877.1	hypothetical protein [Oerskovia enterophila]	Length: 467\nNumber of predicted TMHs:  2\nExp number of AAs in TMHs: 43.6508599999999\nExp number, first 60 AAs:  22.47441\nTotal prob of N-in:   0.99136\nPOSSIBLE N-term signal sequence\ninside   1   12\nTMhelix   13   35\noutside   36   436\nTMhelix   437   459\ninside   460   467

31313	GCF_001692445.1_ASM169244v1	Oerskovia enterophila	Terrabacteria group; Actinobacteria; Actinobacteria; Micrococcales; Cellulomonadaceae; Oerskovia
MLITQRGGRRRALASLLTGALVAGLAVVGVTPASAADTVMVSGGSATWNLKDSWTGYVGG	WP_068623584.1
hypothetical protein [Oerskovia enterophila]	Length: 1569\nNumber of predicted TMHs:  2\nExp number of

AAs in TMHs: 43.7230399999999\nExp number, first 60 AAs:  22.11127\nTotal prob of N-in:   0.99636\nPOSSIBLE N-term signal sequence\ninside   1   12\nTMhelix   13   35\noutside   36  1536\nTMhelix   1537  1559\ninside   1560  1569

31314 GCF_001624335.1_ASM162433v1 Oerskovia enterophila Terrabacteria group; Actinobacteria; Actinobacteria; Micrococcales; Cellulomonadaceae; Oerskovia MARRIATTDRIGRAELVAFLRANRRAVLLTRRAGGAPQMSPVAYGVDGEGRVVVSTYPDR WP_068625180.1 PPOX class F420-dependent enzyme [Oerskovia enterophila] Length: 155\nNumber of predicted TMHs: 1\nExp number of AAs in TMHs: 12.27165\nExp number, first 60 AAs:  0.00081\nTotal prob of N-in:   0.83913\ninside   1   130\nTMhelix   131   153\noutside   154   155

31315 GCF_001692445.1_ASM169244v1 Oerskovia enterophila Terrabacteria group; Actinobacteria; Actinobacteria; Micrococcales; Cellulomonadaceae; Oerskovia MFLSTTRRTPRRTLLSALATGALATGLIGVAGMPAAVAAPPATTPLCTPSSTLPGITTGG WP_068623883.1 hypothetical protein [Oerskovia enterophila] Length: 467\nNumber of predicted TMHs:  2\nExp number of AAs in TMHs: 43.37836\nExp number, first 60 AAs:  22.16176\nTotal prob of N-in:   0.98680\nPOSSIBLE N-term signal sequence\ninside   1   12\nTMhelix   13   35\noutside   36   436\nTMhelix   437   459\ninside   460   467

31316 GCF_001692445.1_ASM169244v1 Oerskovia enterophila Terrabacteria group; Actinobacteria; Actinobacteria; Micrococcales; Cellulomonadaceae; Oerskovia MARRIATTDRIGRAELVAFLRANRRAVLLTRRAGGAPQMSPVAYGVDGEGRVVVSTYPDR WP_068625180.1 PPOX class F420-dependent enzyme [Oerskovia enterophila] Length: 155\nNumber of predicted TMHs: 1\nExp number of AAs in TMHs: 12.27165\nExp number, first 60 AAs:  0.00081\nTotal prob of N-in:   0.83913\ninside   1   130\nTMhelix   131   153\noutside   154   155

31317 GCF_001429135.1_Root22 Oerskovia sp. Root22 Terrabacteria group; Actinobacteria; Actinobacteria; Micrococcales; Cellulomonadaceae; Oerskovia MTVETSRFGRAAAPTRRRLVASTSVLVLGAVMTFGAAALGPFVAPAAADQLPTCPTGDSD WP_056644573.1 MULTISPECIES: hypothetical protein [Oerskovia] Length: 781\nNumber of predicted TMHs:  2\nExp number of AAs in TMHs: 44.1633\nExp number, first 60 AAs:  22.32329\nTotal prob of N-in:   0.99620\nPOSSIBLE N-term signal sequence\ninside   1   20\nTMhelix   21   43\noutside   44   749\nTMhelix   750   772\ninside   773   781

31318 GCF_001429135.1_Root22 Oerskovia sp. Root22 Terrabacteria group; Actinobacteria; Actinobacteria; Micrococcales; Cellulomonadaceae; Oerskovia MIVAARRTWIGTAGHGHAHLSATDLAAPVLGAVLADLPASHGLGVEDVALGNCTGPGGNL WP_082618249.1 acetyl-CoA acetyltransferase [Oerskovia sp. Root22] Length: 387\nNumber of predicted TMHs:  1\nExp number of AAs in TMHs: 22.41448\nExp number, first 60 AAs:  0.11991\nTotal prob of N-in:   0.11077\noutside   1   360\nTMhelix   361   383\ninside   384   387

31319 GCF_001429135.1_Root22 Oerskovia sp. Root22 Terrabacteria group; Actinobacteria; Actinobacteria; Micrococcales; Cellulomonadaceae; Oerskovia MARRIATTDRIGRAELVAFLRANRRAVLLTRRAGGAPQMSPVAYGVDGEGRVVVSTYPDR WP_056652064.1 MULTISPECIES: pyridoxamine 5-phosphate oxidase [Oerskovia] Length: 155\nNumber of predicted TMHs: 1\nExp number of AAs in TMHs: 12.27165\nExp number, first 60 AAs:  0.00081\nTotal prob of N-in:   0.83913\ninside   1   130\nTMhelix   131   153\noutside   154   155

31320 GCF_001429135.1_Root22 Oerskovia sp. Root22 Terrabacteria group; Actinobacteria; Actinobacteria; Micrococcales; Cellulomonadaceae; Oerskovia MDSSELDSYTELTFRSSTPTPETSMFLSTTRRTTRRTLLSALATGALATGLIGVAGVPAA WP_082618215.1 hypothetical protein [Oerskovia sp. Root22] Length: 491\nNumber of predicted TMHs:  2\nExp number of AAs in TMHs: 44.1503899999999\nExp number, first 60 AAs:  21.91151\nTotal prob of N-in:   0.99031\nPOSSIBLE N-term signal sequence\ninside   1   37\nTMhelix   38   60\noutside   61   460\nTMhelix   461   483\ninside   484   491

31321 GCF_001428945.1_Root918 Oerskovia sp. Root918 Terrabacteria group; Actinobacteria; Actinobacteria; Micrococcales; Cellulomonadaceae; Oerskovia MFLSTTRRTPRRTLLSALATGALATGLIGVAGMPAAVAAPLATTPLCTPSSTLPGITTGG WP_056651555.1 hypothetical protein [Oerskovia sp. Root918] Length: 467\nNumber of predicted TMHs:  2\nExp number of AAs in TMHs: 43.79059\nExp number, first 60 AAs:  22.46878\nTotal prob of N-in:   0.99157\nPOSSIBLE N-term signal sequence\ninside   1   12\nTMhelix   13   35\noutside   36   436\nTMhelix   437   459\ninside   460   467

31322 GCF_001428945.1_Root918 Oerskovia sp. Root918 Terrabacteria group; Actinobacteria; Actinobacteria; Micrococcales; Cellulomonadaceae; Oerskovia

MVVAARRTWIGTANHGHAHLSATDLAAPVLGAVLADLPAGHGLGVEDVVLGNCTGPGGNL WP_082608743.1 acetyl-CoA acetyltransferase [Oerskovia sp. Root918] Length: 389\nNumber of predicted TMHs: 1\nExp number of AAs in TMHs: 22.42801\nExp number, first 60 AAs: 0.11572\nTotal prob of N-in: 0.11381\noutside 1 362\nTMhelix 363 385\ninside 386 389

31323 GCF_001428945.1_Root918 Oerskovia sp. Root918 Terrabacteria group; Actinobacteria; Actinobacteria; Micrococcales; Cellulomonadaceae; Oerskovia MARRIATTDRIGRAELVAFLRANRRAVLLTRRAGGAPQMSPVAYGVDGEGRVVVSTYPDR WP_056652064.1 MULTISPECIES: pyridoxamine 5-phosphate oxidase [Oerskovia] Length: 155\nNumber of predicted TMHs: 1\nExp number of AAs in TMHs: 12.27165\nExp number, first 60 AAs: 0.00081\nTotal prob of N-in: 0.83913\ninside 1 130\nTMhelix 131 153\noutside 154 155

31324 GCF_001428945.1_Root918 Oerskovia sp. Root918 Terrabacteria group; Actinobacteria; Actinobacteria; Micrococcales; Cellulomonadaceae; Oerskovia MTVETSRFGRAAAPTRRRLVASTSVLVLGAVMTFGAAALGPFVAPAAADQLPTCPTGDSD WP_056644573.1 MULTISPECIES: hypothetical protein [Oerskovia] Length: 781\nNumber of predicted TMHs: 2\nExp number of AAs in TMHs: 44.1633\nExp number, first 60 AAs: 22.32329\nTotal prob of N-in: 0.99620\nPOSSIBLE N-term signal sequence\ninside 1 20\nTMhelix 21 43\noutside 44 749\nTMhelix 750 772\ninside 773 781

31325 GCF_000718325.1_ASM71832v1 Oerskovia turbata Terrabacteria group; Actinobacteria; Actinobacteria; Micrococcales; Cellulomonadaceae; Oerskovia MTITRSPEGRTSRRRLVAATSVLVLTAALTLGAAAVGPYVAPAAAEQLPTCPDGEGDGTV WP_051702687.1 choice-of-anchor A domain-containing protein [Oerskovia turbata] Length: 777\nNumber of predicted TMHs: 2\nExp number of AAs in TMHs: 43.5078999999999\nExp number, first 60 AAs: 22.11544\nTotal prob of N-in: 0.99405\nPOSSIBLE N-term signal sequence\ninside 1 20\nTMhelix 21 43\noutside 44 745\nTMhelix 746 768\ninside 769 777

31326 GCF_000718325.1_ASM71832v1 Oerskovia turbata Terrabacteria group; Actinobacteria; Actinobacteria; Micrococcales; Cellulomonadaceae; Oerskovia MVHEAVVERPVAVRRHRGRPWTFTGDRVLITQRGGRRRALASLLTGALVAGLTVVGVAPA WP_084689956.1 hypothetical protein [Oerskovia turbata] Length: 1361\nNumber of predicted TMHs: 2\nExp number of AAs in TMHs: 42.0168599999999\nExp number, first 60 AAs: 19.85591\nTotal prob of N-in: 0.95496\nPOSSIBLE N-term signal sequence\ninside 1 39\nTMhelix 40 62\noutside 63 1331\nTMhelix 1332 1351\ninside 1352 1361

31327 GCF_000975135.1_ASM97513v1 Demequina aurantiaca Terrabacteria group; Actinobacteria; Actinobacteria; Micrococcales; Demequinaceae; Demequina MARRQTLTPPLLPGYTYVRPLGSGGFADVFLFQQNMPRRLVAVKVLLSKVQDDASARALA WP_061963199.1 serine/threonine protein kinase [Demequina aurantiaca] Length: 371\nNumber of predicted TMHs: 1\nExp number of AAs in TMHs: 24.29861\nExp number, first 60 AAs: 1.26159\nTotal prob of N-in: 0.39464\noutside 1 347\nTMhelix 348 370\ninside 371 371

31328 GCF_000975115.1_ASM97511v1 Demequina oxidasica Terrabacteria group; Actinobacteria; Actinobacteria; Micrococcales; Demequinaceae; Demequina MMALNRSVHRRAHLSPRLLAGIAVGALVALVGGSAAIATPVAVDPEPSPSASPAPEPTGT WP_062210198.1 hypothetical protein [Demequina oxidasica] Length: 680\nNumber of predicted TMHs: 2\nExp number of AAs in TMHs: 44.77132\nExp number, first 60 AAs: 22.19124\nTotal prob of N-in: 0.98176\nPOSSIBLE N-term signal sequence\ninside 1 20\nTMhelix 21 43\noutside 44 627\nTMhelix 628 650\ninside 651 680

31329 GCF_000975115.1_ASM97511v1 Demequina oxidasica Terrabacteria group; Actinobacteria; Actinobacteria; Micrococcales; Demequinaceae; Demequina MARRQTLTPPLLPGYTYVRPLGSGGFADVFLFQQNMPRRLVAVKVLLSKVQDDASARALA WP_062210432.1 serine/threonine protein kinase [Demequina oxidasica] Length: 371\nNumber of predicted TMHs: 1\nExp number of AAs in TMHs: 24.39486\nExp number, first 60 AAs: 1.42105\nTotal prob of N-in: 0.45489\noutside 1 346\nTMhelix 347 369\ninside 370 371

31330 GCF_002090095.1_ASM209009v1 Demequina sp. NBRC 110053 Terrabacteria group; Actinobacteria; Actinobacteria; Micrococcales; Demequinaceae; Demequina MRGPGHALIPGATQLFRRRVRLPTMSAMITHQPTARRSAVPAWLVAATAAIMVVLLLAAC WP_084038716.1 hypothetical protein [Demequina sp. NBRC 110053] Length: 351\nNumber of predicted TMHs: 1\nExp number of AAs in TMHs: 55.95459\nExp number, first 60 AAs: 19.93796\nTotal prob of N-in: 0.30934\nPOSSIBLE N-term signal sequence\noutside 1 297\nTMhelix 298 320\ninside 321 351

31331 GCF_002090175.1_ASM209017v1 Demequina sp. NBRC 110057 Terrabacteria group; Actinobacteria; Actinobacteria; Micrococcales; Demequinaceae; Demequina MARRQTLQPPVLPGYTYVRPLGSGGFADVFLFQQDMPRRHVAVKVLLSRIEDAQAARALA WP_084079448.1 hypothetical protein [Demequina sp. NBRC 110057] Length: 370\nNumber of predicted TMHs: 1\nExp number of AAs in TMHs: 23.85313\nExp number, first 60 AAs: 0.87613\nTotal prob of N-in: 0.47821\ninside 1 344\nTMhelix 345 367\noutside 368 370

31332 GCF_002090175.1_ASM209017v1 Demequina sp. NBRC 110057 Terrabacteria group; Actinobacteria; Actinobacteria; Micrococcales; Demequinaceae; Demequina MHRRPPIVRIALTMLIALTVSVVAIELGFWQYHRHEAKAAAIAAYDAAQVSGPAPLDDVIWP_084078503.1 hypothetical protein [Demequina sp. NBRC 110057] Length: 277\nNumber of predicted TMHs: 2\nExp number of AAs in TMHs: 41.9981399999999\nExp number, first 60 AAs: 22.17052\nTotal prob of N-in: 0.98925\nPOSSIBLE N-term signal sequence\ninside 1 6\nTMhelix 7 29\noutside 30 211\nTMhelix 212 234\ninside 235 277

31333 GCF_000971115.1_ASM97111v1 Lysinimicrobium gelatinilyticum Terrabacteria group; Actinobacteria; Actinobacteria; Micrococcales; Demequinaceae; Lysinimicrobium MSAMSTHAAARSRTTRRGAVAASILALAWALAGCTSGADESTSGGEEYGYAGTDSDAAVA WP_082096739.1 hypothetical protein [Lysinimicrobium gelatinilyticum] Length: 318\nNumber of predicted TMHs: 1\nExp number of AAs in TMHs: 33.46571\nExp number, first 60 AAs: 8.20232\nTotal prob of N-in: 0.41835\noutside 1 266\nTMhelix 267 289\ninside 290 318

31334 GCF_000971395.1_ASM97139v1 Lysinimicrobium iriomotense Terrabacteria group; Actinobacteria; Actinobacteria; Micrococcales; Demequinaceae; Lysinimicrobium MSAMTTRSAARRPLVRRGAVAAAILALAWGLAACSSGASDSAGESELGYAGGADADMEQA WP_082098982.1 hypothetical protein [Lysinimicrobium iriomotense] Length: 315\nNumber of predicted TMHs: 2\nExp number of AAs in TMHs: 44.14815\nExp number, first 60 AAs: 19.02166\nTotal prob of N-in: 0.92475\nPOSSIBLE N-term signal sequence\ninside 1 12\nTMhelix 13 32\noutside 33 264\nTMhelix 265 287\ninside 288 315

31335 GCF_000971395.1_ASM97139v1 Lysinimicrobium iriomotense Terrabacteria group; Actinobacteria; Actinobacteria; Micrococcales; Demequinaceae; Lysinimicrobium MESRVEGRGALSRRRVIAGAAWAAPVVLVAAAAPAAATSPEPTLALSPSGSSVAHKSKTF WP_062387844.1 hypothetical protein [Lysinimicrobium iriomotense] Length: 151\nNumber of predicted TMHs: 2\nExp number of AAs in TMHs: 47.06709\nExp number, first 60 AAs: 21.44236\nTotal prob of N-in: 0.98393\nPOSSIBLE N-term signal sequence\ninside 1 15\nTMhelix 16 38\noutside 39 93\nTMhelix 94 116\ninside 117 151

31336 GCF_000971235.1_ASM97123v1 Lysinimicrobium subtropicum Terrabacteria group; Actinobacteria; Actinobacteria; Micrococcales; Demequinaceae; Lysinimicrobium MSAMSIHAAARPRTTRRGAVAASILALAWALAGCTSGADESTSGGEEYGYAGADAAVAEE WP_082105535.1 hypothetical protein [Lysinimicrobium subtropicum] Length: 319\nNumber of predicted TMHs: 1\nExp number of AAs in TMHs: 29.8537\nExp number, first 60 AAs: 5.05474\nTotal prob of N-in: 0.25794\noutside 1 264\nTMhelix 265 287\ninside 288 319

31337 GCF_001807385.1_ASM180738v1 Brachybacterium sp. HMSC06H03 Terrabacteria group; Actinobacteria; Actinobacteria; Micrococcales; Dermabacteraceae; Brachybacterium MTDDRRLLDALGRGRRRLLAVALTGMLVLAQLLLGAANPAGACACGAFFDPVDQRADAGV WP_070500937.1 hypothetical protein [Brachybacterium sp. HMSC06H03] Length: 389\nNumber of predicted TMHs: 2\nExp number of AAs in TMHs: 44.18108\nExp number, first 60 AAs: 21.1817\nTotal prob of N-in: 0.96134\nPOSSIBLE N-term signal sequence\ninside 1 18\nTMhelix 19 41\noutside 42 356\nTMhelix 357 379\ninside 380 389

31338 GCF_001969445.1_ASM196944v1 Brachybacterium sp. P6-10-X1 Terrabacteria group; Actinobacteria; Actinobacteria; Micrococcales; Dermabacteraceae; Brachybacterium MKRRQLFTGVGSATLLTVLGIPAIAHADDSTDAPALDGVDRTEVPAFAGGTKHGVVETTE WP_076808965.1 hypothetical protein [Brachybacterium sp. P6-10-X1] Length: 176\nNumber of predicted TMHs: 2\nExp number of AAs in TMHs: 46.0297\nExp number, first 60 AAs: 18.00997\nTotal prob of N-in: 0.97234\nPOSSIBLE N-term signal sequence\ninside 1 6\nTMhelix 7 26\noutside 27 141\nTMhelix 142 164\ninside 165 176

31339 GCF_001807345.1_ASM180734v1 Dermabacter sp. HMSC06F07 Terrabacteria group; Actinobacteria; Actinobacteria; Micrococcales; Dermabacteraceae; Dermabacter MRLFPRRALLTGASLATASALALSAAAPALAAVNDDGSKTITVATFTDLHGHLERVPNVAWP_070435498.1 hypothetical protein [Dermabacter sp. HMSC06F07] Length: 716\nNumber of predicted TMHs: 1\nExp number of AAs in TMHs:

23.42105\nExp number, first 60 AAs: 3.11951\nTotal prob of N-in: 0.14013\noutside 1 685\nTMhelix 686 708\ninside 709 716

31340 GCF_001678905.1_ASM167890v1 Dermabacter vaginalis Terrabacteria group; Actinobacteria; Actinobacteria; Micrococcales; Dermabacteraceae; Dermabacter MRLFPRRALLTGASLATASALALSATAPALATVNDDGSKTVTVAAFTDLHGHLERVPNLA WP_065247277.1 bifunctional metallophosphatase/5-nucleotidase [Dermabacter vaginalis] Length: 704\nNumber of predicted TMHs: 1\nExp number of AAs in TMHs: 22.19933\nExp number, first 60 AAs: 1.38809\nTotal prob of N-in: 0.06273\noutside 1 673\nTMhelix 674 696\ninside 697 704

31341 GCF_900070355.1_S2_rcS3_S1_rcS3 Devriesea agamarum Terrabacteria group; Actinobacteria; Actinobacteria; Micrococcales; Dermabacteraceae; Devriesea MHRRSFFQTAGICGLALTVPSVAYAADTPVSLGFMDTPEDPTKQLETIVGRLDALPADLKWP_058235133.1 hypothetical protein [Devriesea agamarum] Length: 178\nNumber of predicted TMHs: 1\nExp number of AAs in TMHs: 28.69438\nExp number, first 60 AAs: 9.24462\nTotal prob of N-in: 0.85322\ninside 1 108\nTMhelix 109 131\noutside 132 178

31342 GCF_001403775.1_Dermabacter_indicis Helcobacillus massiliensis Terrabacteria group; Actinobacteria; Actinobacteria; Micrococcales; Dermabacteraceae; Helcobacillus MRIFSRRALLTGASLATASALAFSATAPALATVHDDGSKTVTVAAFTDLHGHLERVPNLA WP_055089149.1 bifunctional metallophosphatase/5-nucleotidase [Helcobacillus massiliensis] Length: 716\nNumber of predicted TMHs: 1\nExp number of AAs in TMHs: 24.48656\nExp number, first 60 AAs: 4.22426\nTotal prob of N-in: 0.19034\noutside 1 685\nTMhelix 686 708\ninside 709 716

31343 GCF_001556795.1_ASM155679v1 Dermacoccus sp. CCH2-D9 Terrabacteria group; Actinobacteria; Actinobacteria; Micrococcales; Dermacoccaceae; Dermacoccus MIIESMTRRAHLAAFSGLLATCLASGPPCLTNTAAASTVANMSSVAGMSAHAASASRVAT WP_062256898.1 hypothetical protein [Dermacoccus sp. CCH2-D9] Length: 809\nNumber of predicted TMHs: 1\nExp number of AAs in TMHs: 24.68595\nExp number, first 60 AAs: 3.86431\nTotal prob of N-in: 0.17052\noutside 1 768\nTMhelix 769 788\ninside 789 809

31344 GCF_001021715.1_ASM102171v1 Dermacoccus sp. PE3 Terrabacteria group; Actinobacteria; Actinobacteria; Micrococcales; Dermacoccaceae; Dermacoccus MRRRRAFFCAFPGVFVPSSLLRTRLRRTAVTSAFFPALVLGGLAISPSAAAADVTPPVVL WP_047310286.1 hypothetical protein [Dermacoccus sp. PE3] Length: 370\nNumber of predicted TMHs: 2\nExp number of AAs in TMHs: 38.00526\nExp number, first 60 AAs: 20.09698\nTotal prob of N-in: 0.91490\nPOSSIBLE N-term signal sequence\ninside 1 29\nTMhelix 30 52\noutside 53 345\nTMhelix 346 364\ninside 365 370

31345 GCF_001064185.1_ASM106418v1 Kytococcus sedentarius Terrabacteria group; Actinobacteria; Actinobacteria; Micrococcales; Dermacoccaceae; Kytococcus MVFRPAVNLVLLVLIVLLGAALVFLTARRATRRSFIDIFRRSMILAVVVVMGAGPSIPGE WP_048728390.1 MULTISPECIES: hypothetical protein [Micrococcales] Length: 344\nNumber of predicted TMHs: 2\nExp number of AAs in TMHs: 47.25705\nExp number, first 60 AAs: 27.33032\nTotal prob of N-in: 0.37695\nPOSSIBLE N-term signal sequence\noutside 1 4\nTMhelix 5 27\ninside 28 310\nTMhelix 311 330\noutside 331 344

31346 GCF_001064185.1_ASM106418v1 Kytococcus sedentarius Terrabacteria group; Actinobacteria; Actinobacteria; Micrococcales; Dermacoccaceae; Kytococcus MACAAFFDLDKTILDTSSNVALSGPFIEAGLMSRRTAVASALVHLPYLLSGADESRMQQM WP_048724847.1 hydrolase [Kytococcus sedentarius] Length: 260\nNumber of predicted TMHs: 1\nExp number of AAs in TMHs: 18.40237\nExp number, first 60 AAs: 0.03259\nTotal prob of N-in: 0.01310\noutside 1 231\nTMhelix 232 254\ninside 255 260

31347 GCF_001061885.1_ASM106188v1 Kytococcus sedentarius Terrabacteria group; Actinobacteria; Actinobacteria; Micrococcales; Dermacoccaceae; Kytococcus MARAAFFDLDKTILDTSSNVALSGPFIEAGLMSRRAAVASVLVHLPYLLSGADESRMQQM WP_048740618.1 hydrolase [Kytococcus sedentarius] Length: 260\nNumber of predicted TMHs: 1\nExp number of AAs in TMHs: 18.3244\nExp number, first 60 AAs: 0.06989\nTotal prob of N-in: 0.01564\noutside 1 231\nTMhelix 232 254\ninside 255 260

31348 GCF_001061885.1_ASM106188v1 Kytococcus sedentarius Terrabacteria group; Actinobacteria; Actinobacteria; Micrococcales; Dermacoccaceae; Kytococcus

MVFRPAVNLVLLVLIVLLGAALVFLTARRATRRSFIDILRRSMILAVVVVMGAGPSIPGE WP_009213069.1 MULTISPECIES: hypothetical protein [Actinobacteria] Length: 344\nNumber of predicted TMHs: 2\nExp number of AAs in TMHs: 46.33274\nExp number, first 60 AAs: 26.28713\nTotal prob of N-in: 0.39658\nPOSSIBLE N-term signal sequence\noutside 1 4\nTMhelix 5 27\ninside 28 310\nTMhelix 311 330\noutside 331 344

31349 GCF_001816045.1_ASM181604v1 Kytococcus sp. HMSC28H12 Terrabacteria group; Actinobacteria; Actinobacteria; Micrococcales; Dermacoccaceae; Kytococcus
MTGGQPGPYAYRPQQQPRRRGVMPLVFGGLTLLVGTPLAFLLPVAIGMGDLFGALLNPAS WP_070704842.1 hypothetical protein, partial [Kytococcus sp. HMSC28H12] Length: 219\nNumber of predicted TMHs: 2\nExp number of AAs in TMHs: 46.59389\nExp number, first 60 AAs: 23.40104\nTotal prob of N-in: 0.99731\nPOSSIBLE N-term signal sequence\ninside 1 20\nTMhelix 21 43\noutside 44 152\nTMhelix 153 175\ninside 176 219

31350 GCF_000023925.1_ASM2392v1 Kytococcus sedentarius DSM 20547 Terrabacteria group; Actinobacteria; Actinobacteria; Micrococcales; Dermacoccaceae; Kytococcus; Kytococcus sedentarius
MGSVSARRRPAPGPSGVPRASARPGRAARRAAVALSLAAVVQGLGFGGAAPAGAAPSGSV WP_081439804.1 hypothetical protein [Kytococcus sedentarius] Length: 462\nNumber of predicted TMHs: 1\nExp number of AAs in TMHs: 37.85855\nExp number, first 60 AAs: 15.25131\nTotal prob of N-in: 0.68681\nPOSSIBLE N-term signal sequence\noutside 1 436\nTMhelix 437 459\ninside 460 462

31351 GCF_001190945.1_ASM119094v1 Luteipulveratus mongoliensis Terrabacteria group; Actinobacteria; Actinobacteria; Micrococcales; Dermacoccaceae; Luteipulveratus
MSSLLIGALVLAAWCAWPRRGVLTLAPATSAVVSAASPSSSAQSSGEGPLAVAESMDLLA WP_052595558.1 hypothetical protein [Luteipulveratus mongoliensis] Length: 192\nNumber of predicted TMHs: 1\nExp number of AAs in TMHs: 30.30201\nExp number, first 60 AAs: 5.97755\nTotal prob of N-in: 0.45796\ninside 1 162\nTMhelix 163 185\noutside 186 192

31352 GCF_000298215.1_ASM29821v1 Kineosphaera limosa NBRC 100340 Terrabacteria group; Actinobacteria; Actinobacteria; Micrococcales; Dermatophilaceae; Kineosphaera; Kineosphaera limosa
MPSQTAPRHGRRSPLAFPCALVSVMLLILGLGLVGAPSAGAHAALRTSNPVDGSVVVNLP WP_006594284.1 copper resistance protein C [Kineosphaera limosa] Length: 208\nNumber of predicted TMHs: 2\nExp number of AAs in TMHs: 44.96475\nExp number, first 60 AAs: 22.76292\nTotal prob of N-in: 0.98481\nPOSSIBLE N-term signal sequence\ninside 1 12\nTMhelix 13 35\noutside 36 178\nTMhelix 179 201\ninside 202 208

31353 GCF_001637565.1_ASM163756v1 Janibacter melonis Terrabacteria group; Actinobacteria; Actinobacteria; Micrococcales; Intrasporangiaceae; Janibacter
MPGADQLATRPRETTVPLTSRRPPFATLLTGSALAAAATLALAPAAQAAEQPSATTSAAY WP_068273845.1 hypothetical protein [Janibacter melonis] Length: 456\nNumber of predicted TMHs: 1\nExp number of AAs in TMHs: 22.89185\nExp number, first 60 AAs: 2.20347\nTotal prob of N-in: 0.10173\noutside 1 425\nTMhelix 426 448\ninside 449 456

31354 GCF_000768675.1_ASM76867v1 Knoellia flava TL1 Terrabacteria group; Actinobacteria; Actinobacteria; Micrococcales; Intrasporangiaceae; Knoellia; Knoellia flava
MGRALPHRPRAERRGRPDHRGRRVVTGRRTTVVTGVLGIVAAGLVGPVATPLAAPAHAEV WP_084100611.1 hypothetical protein [Knoellia flava] Length: 430\nNumber of predicted TMHs: 2\nExp number of AAs in TMHs: 41.69295\nExp number, first 60 AAs: 20.63568\nTotal prob of N-in: 0.92946\nPOSSIBLE N-term signal sequence\ninside 1 31\nTMhelix 32 54\noutside 55 401\nTMhelix 402 424\ninside 425 430

31355 GCF_000768705.1_ASM76870v1 Knoellia sinensis KCTC 19936 Terrabacteria group; Actinobacteria; Actinobacteria; Micrococcales; Intrasporangiaceae; Knoellia; Knoellia sinensis
MVRSRDSHARTTPEWRVGRRRRRAVAGVVSILGLLAALLVGGASAQAAPAEGSAGEEKVL WP_035919084.1 ABC transporter substrate-binding protein [Knoellia sinensis] Length: 624\nNumber of predicted TMHs: 2\nExp number of AAs in TMHs: 41.68314\nExp number, first 60 AAs: 21.7536\nTotal prob of N-in: 0.99914\nPOSSIBLE N-term signal sequence\ninside 1 24\nTMhelix 25 47\noutside 48 589\nTMhelix 590 612\ninside 613 624

31356 GCF_000421185.1_ASM42118v1 Ornithinimicrobium pekingense DSM 21552 Terrabacteria group; Actinobacteria; Actinobacteria; Micrococcales; Intrasporangiaceae; Ornithinimicrobium; Ornithinimicrobium pekingense
MSTTAHRSSRPAPVPDHAPPHRRRRSAVLALVTALALLVLGACSSADLAEPGAGGSDSAG WP_022921254.1 hypothetical protein [Ornithinimicrobium pekingense] Length: 347\nNumber of predicted TMHs: 2\nExp number of AAs in TMHs: 44.6847700000001\nExp number, first 60 AAs: 20.52435\nTotal prob of N-in: 0.94411\nPOSSIBLE N-term signal sequence\ninside 1 26\nTMhelix 27 49\noutside 50 297\nTMhelix 298 320\ninside 321 347

31357 GCF_001942435.1_ASM194243v1 Serinicoccus sp. CNJ-927 Terrabacteria group; Actinobacteria; Actinobacteria; Micrococcales; Intrasporangiaceae; Serinicoccus
MTVTDPREDRALRALRRLLADLPEVPPASPAGRERRRPVGALLLAAVVAALLLVLPLLLS WP_083653165.1 hypothetical protein [Serinicoccus sp. CNJ-927] Length: 451\nNumber of predicted TMHs: 2\nExp number of AAs in TMHs: 40.72042\nExp number, first 60 AAs: 20.97923\nTotal prob of N-in: 0.99866\nPOSSIBLE N-term signal sequence\ninside 1 38\nTMhelix 39 61\noutside 62 418\nTMhelix 419 438\ninside 439 451

31358 GCF_001942225.1_ASM194222v1 Serinicoccus sp. CUA-874 Terrabacteria group; Actinobacteria; Actinobacteria; Micrococcales; Intrasporangiaceae; Serinicoccus
MAVTDPREDRAVRALRGLLADLPEVPPASPAGRERRRPVGALLVAAVVAALLLVLPLLLS WP_075814310.1 hypothetical protein [Serinicoccus sp. CUA-874] Length: 451\nNumber of predicted TMHs: 2\nExp number of AAs in TMHs: 40.70988\nExp number, first 60 AAs: 20.99628\nTotal prob of N-in: 0.99796\nPOSSIBLE N-term signal sequence\ninside 1 38\nTMhelix 39 61\noutside 62 418\nTMhelix 419 438\ninside 439 451

31359 GCF_000955875.1_ASM95587v1 Terrabacter sp. 28 Terrabacteria group; Actinobacteria; Actinobacteria; Micrococcales; Intrasporangiaceae; Terrabacter
MSRARRAPLRVVLATLVAACLAALTVAPAHAAAFRYWGFYQLTNGAWAFAQKGSDQTVPK WP_052677460.1 hypothetical protein [Terrabacter sp. 28] Length: 245\nNumber of predicted TMHs: 2\nExp number of AAs in TMHs: 43.40387\nExp number, first 60 AAs: 22.66984\nTotal prob of N-in: 0.99880\nPOSSIBLE N-term signal sequence\ninside 1 11\nTMhelix 12 34\noutside 35 215\nTMhelix 216 235\ninside 236 245

31360 GCF_001428505.1_Root181 Terrabacter sp. Root181 Terrabacteria group; Actinobacteria; Actinobacteria; Micrococcales; Intrasporangiaceae; Terrabacter
MVSDREEETVSPARRAPLRVVLAALVAACLSALTIAPAHAAAYRYWGFYQLSDGAWTFAQ WP_056758159.1 MULTISPECIES: hypothetical protein [Terrabacter] Length: 255\nNumber of predicted TMHs: 2\nExp number of AAs in TMHs: 44.09097\nExp number, first 60 AAs: 22.695\nTotal prob of N-in: 0.99107\nPOSSIBLE N-term signal sequence\ninside 1 20\nTMhelix 21 43\noutside 44 222\nTMhelix 223 245\ninside 246 255

31361 GCF_001428505.1_Root181 Terrabacter sp. Root181 Terrabacteria group; Actinobacteria; Actinobacteria; Micrococcales; Intrasporangiaceae; Terrabacter
MTATPNDTTPDTTARRALVTGASGYVGGQVVPRLLDGGWTVRVLTRRRGSMDDKPWVDRV WP_056755320.1 DNA-binding protein [Terrabacter sp. Root181] Length: 475\nNumber of predicted TMHs: 1\nExp number of AAs in TMHs: 62.08668\nExp number, first 60 AAs: 0.28089\nTotal prob of N-in: 0.57515\noutside 1 447\nTMhelix 448 470\ninside 471 475

31362 GCF_001429185.1_Root85 Terrabacter sp. Root85 Terrabacteria group; Actinobacteria; Actinobacteria; Micrococcales; Intrasporangiaceae; Terrabacter
MNPAPNPTTTDGTPRRALVTGASGYVGGQVVPRLLDEGWAVRVLTRRRASMDDKPWVDRV WP_056825897.1 DNA-binding protein [Terrabacter sp. Root85] Length: 475\nNumber of predicted TMHs: 1\nExp number of AAs in TMHs: 56.34629\nExp number, first 60 AAs: 0.05489\nTotal prob of N-in: 0.46971\noutside 1 447\nTMhelix 448 470\ninside 471 475

31363 GCF_001429645.1_Soil810 Terrabacter sp. Soil810 Terrabacteria group; Actinobacteria; Actinobacteria; Micrococcales; Intrasporangiaceae; Terrabacter
MVSDREEETVSPARRAPLRVVLAALVAACLSALTIAPAHAAAYRYWGFYQLSDGAWTFAQ WP_056758159.1 MULTISPECIES: hypothetical protein [Terrabacter] Length: 255\nNumber of predicted TMHs: 2\nExp number of AAs in TMHs: 44.09097\nExp number, first 60 AAs: 22.695\nTotal prob of N-in: 0.99107\nPOSSIBLE N-term signal sequence\ninside 1 20\nTMhelix 21 43\noutside 44 222\nTMhelix 223 245\ninside 246 255

31364 GCF_001429645.1_Soil810 Terrabacter sp. Soil810 Terrabacteria group; Actinobacteria; Actinobacteria; Micrococcales; Intrasporangiaceae; Terrabacter
MTATPNDTTPDTTARRALVTGASGYVGGQVVPRLLDGGWTVRVLTRRRGSMDDKPWVDRV WP_057587086.1 DNA-binding protein [Terrabacter sp. Soil810] Length: 475\nNumber of predicted TMHs: 1\nExp number of AAs in TMHs: 65.93886\nExp number, first 60 AAs: 0.2827\nTotal prob of N-in: 0.59287\noutside 1 447\nTMhelix 448 470\ninside 471 475

31365 GCF_001429655.1_Soil811 Terrabacter sp. Soil811 Terrabacteria group; Actinobacteria; Actinobacteria; Micrococcales; Intrasporangiaceae; Terrabacter
MNPAPNPTTTDGTPRRALVTGASGYVGGQVVPRLLDEGWAVRVLTRRRASMDDKPWVDRV WP_056774918.1 DNA-binding protein [Terrabacter sp. Soil811] Length: 475\nNumber of predicted TMHs: 1\nExp number of

AAs in TMHs: 54.62819\nExp number, first 60 AAs:  0.05238\nTotal prob of N-in:   0.44150\noutside   1   447\nTMhelix 448   470\ninside   471   475

31366 GCF_000383675.1_ASM38367v1 Terracoccus sp. 273MFTsu3.1Terrabacteria group; Actinobacteria; Actinobacteria; Micrococcales; Intrasporangiaceae; Terracoccus MNPAPNATTTDDTPRRALVTGASGYVGGQVVPRLLDEGWTVRVLTRRRASMDDKPWVDRV WP_020143569.1 hypothetical protein [Terracoccus sp. 273MFTsu3.1] Length: 475\nNumber of predicted TMHs:  1\nExp number of AAs in TMHs: 65.75881\nExp number, first 60 AAs:  0.04194\nTotal prob of N-in:   0.64818\noutside   1 447\nTMhelix   448   470\ninside   471   475

31367 GCF_000938265.1_ASM93826v1 Agreia bicolorata Terrabacteria group; Actinobacteria; Actinobacteria; Micrococcales; Microbacteriaceae; Agreia MSSPTSARRTSRSRRTPFFLRVAGAALGGLLALGGGLAFAAPAQAAAQDATFALSPVTGP WP_044442522.1 hypothetical protein [Agreia bicolorata] Length: 558\nNumber of predicted TMHs:  2\nExp number of AAs in TMHs: 68.93313\nExp number, first 60 AAs:  22.66948\nTotal prob of N-in:   0.97677\nPOSSIBLE N-term signal sequence\ninside   1   19\nTMhelix   20   42\noutside   43   525\nTMhelix   526   548\ninside   549   558

31368 GCF_000938265.1_ASM93826v1 Agreia bicolorata Terrabacteria group; Actinobacteria; Actinobacteria; Micrococcales; Microbacteriaceae; Agreia MIETVAAVDSGVRSDSPASTAHGRRRGVVREVLLNIASVLGAACIALTLAALLFNITLIM WP_044441956.1 S26 family signal peptidase [Agreia bicolorata] Length: 204\nNumber of predicted TMHs:  2\nExp number of AAs in TMHs: 54.62482\nExp number, first 60 AAs:  23.33969\nTotal prob of N-in:   0.78906\nPOSSIBLE N-term signal sequence\ninside 1   31\nTMhelix   32   54\noutside   55   159\nTMhelix   160   179\ninside   180   204

31369 GCF_001422695.1_Leaf244 Agreia sp. Leaf244 Terrabacteria group; Actinobacteria; Actinobacteria; Micrococcales; Microbacteriaceae; Agreia MQPSSPRRRPVRAAALLVSSTIALVGFAALPAAAAPGDVSIDVYTMNDFHGHLEQNLGSG WP_055945179.1 bifunctional metallophosphatase/5-nucleotidase [Agreia sp. Leaf244] Length: 739\nNumber of predicted TMHs:  2\nExp number of AAs in TMHs: 43.27834\nExp number, first 60 AAs:  22.40609\nTotal prob of N-in: 0.98818\nPOSSIBLE N-term signal sequence\ninside   1   12\nTMhelix   13   35\noutside   36   710\nTMhelix   711 730\ninside   731   739

31370 GCF_001422695.1_Leaf244 Agreia sp. Leaf244 Terrabacteria group; Actinobacteria; Actinobacteria; Micrococcales; Microbacteriaceae; Agreia MSRGPSPLARRALLAALALTLVLGGAAAPASAETGDDGTITEGTPDTSPVTVVIPKSTTP WP_055942116.1 hypothetical protein [Agreia sp. Leaf244] Length: 250\nNumber of predicted TMHs:  1\nExp number of AAs in TMHs: 45.87012\nExp number, first 60 AAs:  16.1305\nTotal prob of N-in:   0.78544\nPOSSIBLE N-term signal sequence\noutside   1 213\nTMhelix   214   233\ninside   234   250

31371 GCF_001422485.1_Leaf283 Agreia sp. Leaf283 Terrabacteria group; Actinobacteria; Actinobacteria; Micrococcales; Microbacteriaceae; Agreia MQPSSPRRRPVRAAALLVSSTIALVGFAALPASAAPGDVSIDVYTMNDFHGHLEQNLGSG WP_055926745.1 bifunctional metallophosphatase/5-nucleotidase [Agreia sp. Leaf283] Length: 740\nNumber of predicted TMHs:  2\nExp number of AAs in TMHs: 42.3759\nExp number, first 60 AAs:  21.57602\nTotal prob of N-in: 0.96913\nPOSSIBLE N-term signal sequence\ninside   1   12\nTMhelix   13   35\noutside   36   710\nTMhelix   711 730\ninside   731   740

31372 GCF_001423605.1_Leaf335 Agreia sp. Leaf335 Terrabacteria group; Actinobacteria; Actinobacteria; Micrococcales; Microbacteriaceae; Agreia MTATLTTVQDEGEADANERTASPTRRRGAVREVLLNIAAVLGAVCIALTLAALIFDITLI WP_055851701.1 signal peptidase I [Agreia sp. Leaf335] Length: 203\nNumber of predicted TMHs:  2\nExp number of AAs in TMHs: 50.16343\nExp number, first 60 AAs:  23.08773\nTotal prob of N-in:   0.97084\nPOSSIBLE N-term signal sequence\ninside 1   32\nTMhelix   33   55\noutside   56   160\nTMhelix   161   180\ninside   181   203

31373 GCF_900177765.1_IMG-taxon_2708742470_annotated_assembly Agreia sp. VKM Ac-1783 Terrabacteria group; Actinobacteria; Actinobacteria; Micrococcales; Microbacteriaceae; Agreia MHSHFFRGSVTRRRTLAGTAAIVVAASVLAPGAAFAAELPDNGGAARSEGDSTAALRASI WP_086460870.1 alkaline phosphatase [Agreia sp. VKM Ac-1783] Length: 632\nNumber of predicted TMHs:  1\nExp number of AAs in TMHs: 38.43263\nExp number, first 60 AAs:  16.534\nTotal prob of N-in:   0.79954\nPOSSIBLE N-term signal sequence\noutside   1   601\nTMhelix   602   624\ninside   625   632

31374 GCF_900177765.1_IMG-taxon_2708742470_annotated_assembly Agreia sp. VKM Ac-1783 Terrabacteria group; Actinobacteria; Actinobacteria; Micrococcales; Microbacteriaceae; Agreia
MGITPLSTPPTRRRRTLRVASVLAAATVAIAGVAVAAPAAQAAPSAANTVTVNADQAFRP WP_086459789.1 hypothetical protein [Agreia sp. VKM Ac-1783] Length: 897\nNumber of predicted TMHs: 2\nExp number of AAs in TMHs: 42.18882\nExp number, first 60 AAs: 21.55986\nTotal prob of N-in: 0.98647\nPOSSIBLE N-term signal sequence\ninside 1 19\nTMhelix 20 42\noutside 43 865\nTMhelix 866 888\ninside 889 897

31375 GCF_900167545.1_IMG-taxon_2708742401_annotated_assembly Agreia sp. VKM Ac-2052 Terrabacteria group; Actinobacteria; Actinobacteria; Micrococcales; Microbacteriaceae; Agreia
MSSPTSARRTSRSRRTPFLRVAGAALGGLLALGGGLAFAAPAQAAAQDATFALSPVTGPT WP_078713100.1 hypothetical protein [Agreia sp. VKM Ac-2052] Length: 557\nNumber of predicted TMHs: 2\nExp number of AAs in TMHs: 68.9025\nExp number, first 60 AAs: 22.27423\nTotal prob of N-in: 0.98895\nPOSSIBLE N-term signal sequence\ninside 1 20\nTMhelix 21 43\noutside 44 524\nTMhelix 525 547\ninside 548 557

31376 GCF_900167545.1_IMG-taxon_2708742401_annotated_assembly Agreia sp. VKM Ac-2052 Terrabacteria group; Actinobacteria; Actinobacteria; Micrococcales; Microbacteriaceae; Agreia
MIGTLAAADSGVRSDSPASTAYGRRRGVVREVLLNIASVLGAACIALTLAALLFNITLIM WP_078714251.1 signal peptidase I [Agreia sp. VKM Ac-2052] Length: 204\nNumber of predicted TMHs: 2\nExp number of AAs in TMHs: 55.16434\nExp number, first 60 AAs: 23.37271\nTotal prob of N-in: 0.76084\nPOSSIBLE N-term signal sequence\ninside 1 31\nTMhelix 32 54\noutside 55 159\nTMhelix 160 179\ninside 180 204

31377 GCF_001421565.1_Leaf222 Agromyces sp. Leaf222 Terrabacteria group; Actinobacteria; Actinobacteria; Micrococcales; Microbacteriaceae; Agromyces
MRASQRSNRTGVERHQHPASEVGDHRHPLARRRRRRVVAPVATAGAALLIALLAGCSAGG WP_082462074.1 hypothetical protein [Agromyces sp. Leaf222] Length: 359\nNumber of predicted TMHs: 2\nExp number of AAs in TMHs: 49.14459\nExp number, first 60 AAs: 20.95286\nTotal prob of N-in: 0.99500\nPOSSIBLE N-term signal sequence\ninside 1 36\nTMhelix 37 59\noutside 60 296\nTMhelix 297 319\ninside 320 359

31378 GCF_001620055.1_ASM162005v1 Agromyces sp. NDB4Y10 Terrabacteria group; Actinobacteria; Actinobacteria; Micrococcales; Microbacteriaceae; Agromyces
MMRRRALLAASAALTVLLLGGCGIGTSGSGSDAGDDSGGVSQPDVGVDAPAEGGEGVAPRDE WP_067945834.1 hypothetical protein [Agromyces sp. NDB4Y10] Length: 319\nNumber of predicted TMHs: 2\nExp number of AAs in TMHs: 44.09647\nExp number, first 60 AAs: 20.20496\nTotal prob of N-in: 0.90973\nPOSSIBLE N-term signal sequence\ninside 1 6\nTMhelix 7 29\noutside 30 266\nTMhelix 267 289\ninside 290 319

31379 GCF_001427265.1_Root1464 Agromyces sp. Root1464 Terrabacteria group; Actinobacteria; Actinobacteria; Micrococcales; Microbacteriaceae; Agromyces
MTTTKNATRWGRRGAVSTALLAMLAGVLAAPAPALAATEVPSDPVISTGTTTWKYLDDGS WP_056004954.1 metallophosphoesterase [Agromyces sp. Root1464] Length: 862\nNumber of predicted TMHs: 2\nExp number of AAs in TMHs: 42.61012\nExp number, first 60 AAs: 21.16485\nTotal prob of N-in: 0.97258\nPOSSIBLE N-term signal sequence\ninside 1 12\nTMhelix 13 35\noutside 36 827\nTMhelix 828 850\ninside 851 862

31380 GCF_001427265.1_Root1464 Agromyces sp. Root1464 Terrabacteria group; Actinobacteria; Actinobacteria; Micrococcales; Microbacteriaceae; Agromyces
MTDPSRLRQPQLHPQQRGSTVPDHQRPTPRRAAMLTALAIAGTVVLGGIAAAPATAAPES WP_082571447.1 hypothetical protein [Agromyces sp. Root1464] Length: 716\nNumber of predicted TMHs: 2\nExp number of AAs in TMHs: 42.2080199999999\nExp number, first 60 AAs: 20.4552\nTotal prob of N-in: 0.92042\nPOSSIBLE N-term signal sequence\ninside 1 34\nTMhelix 35 57\noutside 58 688\nTMhelix 689 708\ninside 709 716

31381 GCF_001429165.1_Root81 Agromyces sp. Root81 Terrabacteria group; Actinobacteria; Actinobacteria; Micrococcales; Microbacteriaceae; Agromyces
MTNKPYDGRAVAIERRPMLWAMFGGSILGAALLWVLWVLPIAASVANATPIPLGAAQDVQ WP_056654772.1 hypothetical protein [Agromyces sp. Root81] Length: 198\nNumber of predicted TMHs: 2\nExp number of AAs in TMHs: 45.61015\nExp number, first 60 AAs: 22.55246\nTotal prob of N-in: 0.96672\nPOSSIBLE N-term signal sequence\ninside 1 18\nTMhelix 19 41\noutside 42 169\nTMhelix 170 192\ninside 193 198

31382 GCF_001429165.1_Root81 Agromyces sp. Root81 Terrabacteria group; Actinobacteria; Actinobacteria; Micrococcales; Microbacteriaceae; Agromyces
MKNLLRALSGRPPAARHRRRQPVRAAMKAISTVAATTLILTGLTLPSFAASAEEPTGNPV WP_082600622.1 hypothetical protein [Agromyces sp. Root81] Length: 2258\nNumber of predicted TMHs: 1\nExp number of AAs in TMHs:

31.96304\nExp number, first 60 AAs: 10.54773\nTotal prob of N-in: 0.47957\nPOSSIBLE N-term signal sequence\noutside 1 2227\nTMhelix 2228 2247\ninside 2248 2258

31383 GCF_900142065.1_IMG-taxon_2698536811_annotated_assembly Agromyces cerinus subsp. cerinus Terrabacteria group; Actinobacteria; Actinobacteria; Micrococcales; Microbacteriaceae; Agromyces; Agromyces cerinus MMNVKRRLRTRSTAPATEPLGARSLRLQAGRARRRLLSFVVATALVASGGAVVMAVAPAE WP_074258532.1 hypothetical protein [Agromyces cerinus] Length: 2403\nNumber of predicted TMHs: 2\nExp number of AAs in TMHs: 45.2521\nExp number, first 60 AAs: 22.62509\nTotal prob of N-in: 0.99959\nPOSSIBLE N-term signal sequence\ninside 1 35\nTMhelix 36 58\noutside 59 2371\nTMhelix 2372 2394\ninside 2395 2403

31384 GCF_000421565.1_ASM42156v1 Agromyces subbeticus DSM 16689 Terrabacteria group; Actinobacteria; Actinobacteria; Micrococcales; Microbacteriaceae; Agromyces; Agromyces subbeticus MIDERRTVVRRIGTLGGILLVASFVVAGLATADRAAWAEETAPGENLTVDVSDGVVPPTG WP_022892658.1 hypothetical protein [Agromyces subbeticus] Length: 258\nNumber of predicted TMHs: 2\nExp number of AAs in TMHs: 44.27114\nExp number, first 60 AAs: 20.44632\nTotal prob of N-in: 0.98143\nPOSSIBLE N-term signal sequence\ninside 1 11\nTMhelix 12 31\noutside 32 222\nTMhelix 223 245\ninside 246 258

31385 GCF_000421565.1_ASM42156v1 Agromyces subbeticus DSM 16689 Terrabacteria group; Actinobacteria; Actinobacteria; Micrococcales; Microbacteriaceae; Agromyces; Agromyces subbeticus MMMMIAKRLHTRNTDPTLETRAWRLHIRRTRRKLLSTVAAVALLASGGVVAAVFAPPAPA WP_022893680.1 hypothetical protein [Agromyces subbeticus] Length: 2222\nNumber of predicted TMHs: 2\nExp number of AAs in TMHs: 44.4426399999999\nExp number, first 60 AAs: 22.5008\nTotal prob of N-in: 0.99827\nPOSSIBLE N-term signal sequence\ninside 1 33\nTMhelix 34 56\noutside 57 2189\nTMhelix 2190 2212\ninside 2213 2222

31386 GCF_000421565.1_ASM42156v1 Agromyces subbeticus DSM 16689 Terrabacteria group; Actinobacteria; Actinobacteria; Micrococcales; Microbacteriaceae; Agromyces; Agromyces subbeticus MSLPNLTGVRPRRNALDTRRVVSAALAFALVTGFVAFDGAMTPAYAAATAEVTALKTDST WP_022891046.1 rhamnosidase [Agromyces subbeticus] Length: 2036\nNumber of predicted TMHs: 2\nExp number of AAs in TMHs: 43.6075899999999\nExp number, first 60 AAs: 20.3736\nTotal prob of N-in: 0.97894\nPOSSIBLE N-term signal sequence\ninside 1 20\nTMhelix 21 40\noutside 41 1997\nTMhelix 1998 2020\ninside 2021 2036

31387 GCF_000698005.1_Cm26808.1 Clavibacter cf. michiganensis LMG 26808 Terrabacteria group; Actinobacteria; Actinobacteria; Micrococcales; Microbacteriaceae; Clavibacter MITSAPSAPRRRILRSATALVGGIALAVAVPLAASAHVRVSPDQAAAGSYSTLTFKVPTE WP_043588079.1 hypothetical protein [Clavibacter cf. michiganensis LMG 26808] Length: 254\nNumber of predicted TMHs: 2\nExp number of AAs in TMHs: 44.97655\nExp number, first 60 AAs: 22.0948\nTotal prob of N-in: 0.98294\nPOSSIBLE N-term signal sequence\ninside 1 12\nTMhelix 13 35\noutside 36 220\nTMhelix 221 243\ninside 244 254

31388 GCF_000698005.1_Cm26808.1 Clavibacter cf. michiganensis LMG 26808 Terrabacteria group; Actinobacteria; Actinobacteria; Micrococcales; Microbacteriaceae; Clavibacter MSTKTARARRTGRRSPLTTIALLAIGLLTTGGAYAMIQSGTASAEVDLKSAQTIDEGQKL WP_043586512.1 MULTISPECIES: cystathionine beta-lyase [Clavibacter] Length: 266\nNumber of predicted TMHs: 2\nExp number of AAs in TMHs: 41.67908\nExp number, first 60 AAs: 21.2051\nTotal prob of N-in: 0.99541\nPOSSIBLE N-term signal sequence\ninside 1 19\nTMhelix 20 42\noutside 43 243\nTMhelix 244 263\ninside 264 266

31389 GCF_000698005.1_Cm26808.1 Clavibacter cf. michiganensis LMG 26808 Terrabacteria group; Actinobacteria; Actinobacteria; Micrococcales; Microbacteriaceae; Clavibacter MDMSGPRRASRRRRMIGTAAGGALLAAVALGAAGVPAAVAAPEEAAQPELESAGSAPIIT WP_081840916.1 hypothetical protein [Clavibacter cf. michiganensis LMG 26808] Length: 764\nNumber of predicted TMHs: 2\nExp number of AAs in TMHs: 43.20592\nExp number, first 60 AAs: 21.71546\nTotal prob of N-in: 0.99507\nPOSSIBLE N-term signal sequence\ninside 1 19\nTMhelix 20 42\noutside 43 736\nTMhelix 737 759\ninside 760 764

31390 GCF_002151185.1_ASM215118v1 Clavibacter michiganensis Terrabacteria group; Actinobacteria; Actinobacteria; Micrococcales; Microbacteriaceae; Clavibacter MSTKTARARRTGRRSPLTTVALLAIGLLFTGGAYALIQTGTASAQVDLKSAQTIDEGQKL WP_086518190.1 cystathionine beta-lyase [Clavibacter michiganensis] Length: 266\nNumber of predicted TMHs: 2\nExp number of AAs in TMHs: 43.53364\nExp number, first 60 AAs: 21.86416\nTotal prob of N-in: 0.99450\nPOSSIBLE N-term signal sequence\ninside 1 19\nTMhelix 20 42\noutside 43 243\nTMhelix 244 263\ninside 264 266

31391 GCF_002151115.1_ASM215111v1 Clavibacter michiganensis Terrabacteria group; Actinobacteria; Actinobacteria; Micrococcales; Microbacteriaceae; Clavibacter

MSTTTARARRTGRRSPLTTVALLAIGLLFTGGAYALIQTGTASAEVDLKSAQTIDEGQKL WP_086526429.1 cystathionine beta-lyase [Clavibacter michiganensis] Length: 266\nNumber of predicted TMHs: 2\nExp number of AAs in TMHs: 42.35314\nExp number, first 60 AAs: 21.85383\nTotal prob of N-in: 0.99566\nPOSSIBLE N-term signal sequence\ninside 1 19\nTMhelix 20 42\noutside 43 243\nTMhelix 244 263\ninside 264 266

31392 GCF_002151185.1_ASM215118v1 Clavibacter michiganensis Terrabacteria group; Actinobacteria; Actinobacteria; Micrococcales; Microbacteriaceae; Clavibacter
MTTSSTPAPRRRILRSATALVGGVALAVAMPLAASAHVRVSPDQAAAGSYSTLTFKVPTE WP_086518073.1 hypothetical protein [Clavibacter michiganensis] Length: 255\nNumber of predicted TMHs: 2\nExp number of AAs in TMHs: 44.83037\nExp number, first 60 AAs: 21.90171\nTotal prob of N-in: 0.98347\nPOSSIBLE N-term signal sequence\ninside 1 12\nTMhelix 13 35\noutside 36 221\nTMhelix 222 244\ninside 245 255

31393 GCF_002151115.1_ASM215111v1 Clavibacter michiganensis Terrabacteria group; Actinobacteria; Actinobacteria; Micrococcales; Microbacteriaceae; Clavibacter
MSRRAGQPTRRGPLRRILGVLGGLATALVAVAVVAVVALSALGITRVVPVLSNSMAPGMP WP_086525499.1 signal peptidase I [Clavibacter michiganensis] Length: 200\nNumber of predicted TMHs: 2\nExp number of AAs in TMHs: 46.40822\nExp number, first 60 AAs: 24.01945\nTotal prob of N-in: 0.99546\nPOSSIBLE N-term signal sequence\ninside 1 16\nTMhelix 17 39\noutside 40 161\nTMhelix 162 184\ninside 185 200

31394 GCF_900168345.1_IMG-taxon_2708742539_annotated_assembly Clavibacter michiganensis Terrabacteria group; Actinobacteria; Actinobacteria; Micrococcales; Microbacteriaceae; Clavibacter
MSTKTARARRTGRRSPLTTIALLAIGLLTTGGAYAMIQSGTASAEVDLKSAQTIDEGQKL WP_043586512.1 MULTISPECIES: cystathionine beta-lyase [Clavibacter] Length: 266\nNumber of predicted TMHs: 2\nExp number of AAs in TMHs: 41.67908\nExp number, first 60 AAs: 21.2051\nTotal prob of N-in: 0.99541\nPOSSIBLE N-term signal sequence\ninside 1 19\nTMhelix 20 42\noutside 43 243\nTMhelix 244 263\ninside 264 266

31395 GCF_900168345.1_IMG-taxon_2708742539_annotated_assembly Clavibacter michiganensis Terrabacteria group; Actinobacteria; Actinobacteria; Micrococcales; Microbacteriaceae; Clavibacter
MTTSSPSAPRRRILRSATALVGGVALAVAVPLAASAHVRVSPDQAAAGSYSTLTFKVPTE WP_011931836.1 hypothetical protein [Clavibacter michiganensis] Length: 254\nNumber of predicted TMHs: 2\nExp number of AAs in TMHs: 44.92106\nExp number, first 60 AAs: 22.03919\nTotal prob of N-in: 0.97676\nPOSSIBLE N-term signal sequence\ninside 1 12\nTMhelix 13 35\noutside 36 220\nTMhelix 221 243\ninside 244 254

31396 GCF_002151185.1_ASM215118v1 Clavibacter michiganensis Terrabacteria group; Actinobacteria; Actinobacteria; Micrococcales; Microbacteriaceae; Clavibacter
MPRTTRLPASARPPGRRRALAATGAALGLGLALAAPLAASAHVEVDASSTAPAALSVLTFWP_086517269.1 sortase [Clavibacter michiganensis] Length: 237\nNumber of predicted TMHs: 2\nExp number of AAs in TMHs: 54.23899\nExp number, first 60 AAs: 24.44794\nTotal prob of N-in: 0.96987\nPOSSIBLE N-term signal sequence\ninside 1 19\nTMhelix 20 42\noutside 43 204\nTMhelix 205 227\ninside 228 237

31397 GCF_000785165.1_ASM78516v1 Clavibacter michiganensis Terrabacteria group; Actinobacteria; Actinobacteria; Micrococcales; Microbacteriaceae; Clavibacter
MSTKTARARRTGRRSPLTTVALLAIGLLFTGGAYALIQTGTASADVDLKSAQTIDEGQKL WP_043669791.1 cystathionine beta-lyase [Clavibacter michiganensis] Length: 266\nNumber of predicted TMHs: 2\nExp number of AAs in TMHs: 42.58177\nExp number, first 60 AAs: 21.85666\nTotal prob of N-in: 0.99680\nPOSSIBLE N-term signal sequence\ninside 1 19\nTMhelix 20 42\noutside 43 243\nTMhelix 244 263\ninside 264 266

31398 GCF_002151125.1_ASM215112v1 Clavibacter michiganensis Terrabacteria group; Actinobacteria; Actinobacteria; Micrococcales; Microbacteriaceae; Clavibacter
MTRSTLPASRRRILRSVTALVGGVALAVAVPLAASAHVRVSPDQAAAGSYSTLTFKVPTE WP_086515263.1 hypothetical protein [Clavibacter michiganensis] Length: 254\nNumber of predicted TMHs: 2\nExp number of AAs in TMHs: 45.13669\nExp number, first 60 AAs: 22.2569\nTotal prob of N-in: 0.99466\nPOSSIBLE N-term signal sequence\ninside 1 12\nTMhelix 13 35\noutside 36 220\nTMhelix 221 243\ninside 244 254

31399 GCF_002151125.1_ASM215112v1 Clavibacter michiganensis Terrabacteria group; Actinobacteria; Actinobacteria; Micrococcales; Microbacteriaceae; Clavibacter
MSTKTARARRTGRRSPLTTVALLAIGLLFTGGAYALIQSGTASAEVDLKSAQTIDEGQKL WP_086513259.1 cystathionine beta-lyase [Clavibacter michiganensis] Length: 266\nNumber of predicted TMHs: 2\nExp number of AAs in TMHs: 41.99134\nExp number, first 60 AAs: 21.49837\nTotal prob of N-in: 0.99669\nPOSSIBLE N-term signal sequence\ninside 1 19\nTMhelix 20 42\noutside 43 243\nTMhelix 244 263\ninside 264 266

31400	GCF_002151165.1_ASM215116v1	Clavibacter michiganensis	Terrabacteria group; Actinobacteria; Actinobacteria; Micrococcales; Microbacteriaceae; Clavibacter
MSTKTARARRTGRRSPLTTIALLAIGLLATGGAYSMIQSGTASAEVDLKSAQTIDEGQKL	WP_015490499.1	menaquinol-cytochrome c reductase cytochrome c subunit [Clavibacter michiganensis]	Length: 266\nNumber of predicted TMHs: 2\nExp number of AAs in TMHs: 41.68481\nExp number, first 60 AAs: 21.15138\nTotal prob of N-in: 0.99280\nPOSSIBLE N-term signal sequence\ninside 1 19\nTMhelix 20 42\noutside 43 243\nTMhelix 244 263\ninside 264 266

31401	GCF_002151165.1_ASM215116v1	Clavibacter michiganensis	Terrabacteria group; Actinobacteria; Actinobacteria; Micrococcales; Microbacteriaceae; Clavibacter
MTRSSTSAPRRRILRSATALVGGVALAVAVPLAASAHVRVSPDQAAAGSYSTLTFKVPTE WP_086521042.1	hypothetical protein [Clavibacter michiganensis]	Length: 254\nNumber of predicted TMHs: 2\nExp number of AAs in TMHs: 45.07531\nExp number, first 60 AAs: 22.14437\nTotal prob of N-in: 0.99450\nPOSSIBLE N-term signal sequence\ninside 1 12\nTMhelix 13 35\noutside 36 220\nTMhelix 221 243\ninside 244 254

31402	GCF_000785165.1_ASM78516v1	Clavibacter michiganensis	Terrabacteria group; Actinobacteria; Actinobacteria; Micrococcales; Microbacteriaceae; Clavibacter
MTPGDRIRPRDGRRPATRRGPLRRVAAALAGASLATAAVALVGLAQGSGPDGALSASAHN	WP_052129147.1
copper resistance protein CopC [Clavibacter michiganensis]	Length: 282\nNumber of predicted TMHs: 2\nExp number of AAs in TMHs: 44.6533399999999\nExp number, first 60 AAs: 21.5576\nTotal prob of N-in: 0.96824\nPOSSIBLE N-term signal sequence\ninside 1 20\nTMhelix 21 43\noutside 44 234\nTMhelix 235 257\ninside 258 282

31403	GCF_001280205.1_ASM128020v1	Clavibacter michiganensis subsp. capsici	Terrabacteria group; Actinobacteria; Actinobacteria; Micrococcales; Microbacteriaceae; Clavibacter; Clavibacter michiganensis
MTPEDRIRPRDGRRPAARRGPLRRVAAALAGASLAAAAVALVGLAQGSGPDGALSASAHN	WP_053773647.1
copper resistance protein CopC [Clavibacter michiganensis]	Length: 279\nNumber of predicted TMHs: 2\nExp number of AAs in TMHs: 45.03478\nExp number, first 60 AAs: 21.89886\nTotal prob of N-in: 0.98153\nPOSSIBLE N-term signal sequence\ninside 1 20\nTMhelix 21 43\noutside 44 231\nTMhelix 232 254\ninside 255 279

31404	GCF_001280205.1_ASM128020v1	Clavibacter michiganensis subsp. capsici	Terrabacteria group; Actinobacteria; Actinobacteria; Micrococcales; Microbacteriaceae; Clavibacter; Clavibacter michiganensis
MSRRAGRPTRRGPLRRTLGVLGGLATALVAVAVVAVVALSALGITRVVPVLSNSMAPGMP	WP_053773291.1	S26 family signal peptidase [Clavibacter michiganensis]	Length: 200\nNumber of predicted TMHs: 2\nExp number of AAs in TMHs: 46.03588\nExp number, first 60 AAs: 23.6655\nTotal prob of N-in: 0.99636\nPOSSIBLE N-term signal sequence\ninside 1 19\nTMhelix 20 42\noutside 43 161\nTMhelix 162 184\ninside 185 200

31405	GCF_001280205.1_ASM128020v1	Clavibacter michiganensis subsp. capsici	Terrabacteria group; Actinobacteria; Actinobacteria; Micrococcales; Microbacteriaceae; Clavibacter; Clavibacter michiganensis
MSTTTARARRTGRRSPLTTVALLAIGLLFTGGAYALIQTGTASAEVDLKSAQTIDEGQKL	WP_053774664.1	cystathionine beta-lyase [Clavibacter michiganensis]	Length: 266\nNumber of predicted TMHs: 2\nExp number of AAs in TMHs: 42.34952\nExp number, first 60 AAs: 21.85363\nTotal prob of N-in: 0.99581\nPOSSIBLE N-term signal sequence\ninside 1 19\nTMhelix 20 42\noutside 43 243\nTMhelix 244 263\ninside 264 266

31406	GCF_000958465.1_ASM95846v1	Clavibacter michiganensis subsp. insidiosus	Terrabacteria group; Actinobacteria; Actinobacteria; Micrococcales; Microbacteriaceae; Clavibacter; Clavibacter michiganensis
MSTKTARARRTGRRSPLTTIALLAIGLLATGGAYSMIQSGTASAEVDLKSAQTIDEGQKL	WP_015490499.1	menaquinol-cytochrome c reductase cytochrome c subunit [Clavibacter michiganensis]	Length: 266\nNumber of predicted TMHs: 2\nExp number of AAs in TMHs: 41.68481\nExp number, first 60 AAs: 21.15138\nTotal prob of N-in: 0.99280\nPOSSIBLE N-term signal sequence\ninside 1 19\nTMhelix 20 42\noutside 43 243\nTMhelix 244 263\ninside 264 266

31407	GCF_000958465.1_ASM95846v1	Clavibacter michiganensis subsp. insidiosus	Terrabacteria group; Actinobacteria; Actinobacteria; Micrococcales; Microbacteriaceae; Clavibacter; Clavibacter michiganensis
MTRSSTSAPRRRILRSATALVGGVALAVAVPLAASAHVRVSPDQAAAGSYSTLTFKVPTE WP_045526708.1	hypothetical protein [Clavibacter michiganensis]	Length: 254\nNumber of predicted TMHs: 2\nExp number of AAs in TMHs: 45.01244\nExp number, first 60 AAs: 22.14497\nTotal prob of N-in: 0.99460\nPOSSIBLE N-term signal sequence\ninside 1 12\nTMhelix 13 35\noutside 36 220\nTMhelix 221 243\ninside 244 254

31408	GCF_002159015.1_ASM215901v1	Clavibacter michiganensis subsp. michiganensis	Terrabacteria group; Actinobacteria; Actinobacteria; Micrococcales; Microbacteriaceae; Clavibacter; Clavibacter michiganensis
MSTKTARARRTGRRSPLTTIALLAIGLLTTGGAYAMIQSGTASAEVDLKSAQTIDEGQKL	WP_043586512.1	MULTISPECIES: cystathionine beta-lyase [Clavibacter]	Length: 266\nNumber of predicted TMHs: 2\nExp number of AAs in TMHs:

41.67908\nExp number, first 60 AAs: 21.2051\nTotal prob of N-in: 0.99541\nPOSSIBLE N-term signal sequence\ninside 1 19\nTMhelix 20 42\noutside 43 243\nTMhelix 244 263\ninside 264 266

31409 GCF_002151055.1_ASM215105v1 Clavibacter michiganensis subsp. michiganensis Terrabacteria group; Actinobacteria; Actinobacteria; Micrococcales; Microbacteriaceae; Clavibacter; Clavibacter michiganensis MSTKTARARRTGRRSPLTTIALLAIGLLTTGGAYAMIQSGTASAEVDLKSAQTIDEGQKL WP_043586512.1 MULTISPECIES: cystathionine beta-lyase [Clavibacter] Length: 266\nNumber of predicted TMHs: 2\nExp number of AAs in TMHs: 41.67908\nExp number, first 60 AAs: 21.2051\nTotal prob of N-in: 0.99541\nPOSSIBLE N-term signal sequence\ninside 1 19\nTMhelix 20 42\noutside 43 243\nTMhelix 244 263\ninside 264 266

31410 GCF_002151105.1_ASM215110v1 Clavibacter michiganensis subsp. michiganensis Terrabacteria group; Actinobacteria; Actinobacteria; Micrococcales; Microbacteriaceae; Clavibacter; Clavibacter michiganensis MTTSSPSAPRRRILRSATALVGGVALAVAVPLAASAHVRVSPDQAAAGSYSTLTFKVPTE WP_011931836.1 hypothetical protein [Clavibacter michiganensis] Length: 254\nNumber of predicted TMHs: 2\nExp number of AAs in TMHs: 44.92106\nExp number, first 60 AAs: 22.03919\nTotal prob of N-in: 0.97676\nPOSSIBLE N-term signal sequence\ninside 1 12\nTMhelix 13 35\noutside 36 220\nTMhelix 221 243\ninside 244 254

31411 GCF_002151005.1_ASM215100v1 Clavibacter michiganensis subsp. michiganensis Terrabacteria group; Actinobacteria; Actinobacteria; Micrococcales; Microbacteriaceae; Clavibacter; Clavibacter michiganensis MTTSSPSAPRRRILRSATALVGGVALAVAVPLAASAHVRVSPDQAAAGSYSTLTFKVPTE WP_011931836.1 hypothetical protein [Clavibacter michiganensis] Length: 254\nNumber of predicted TMHs: 2\nExp number of AAs in TMHs: 44.92106\nExp number, first 60 AAs: 22.03919\nTotal prob of N-in: 0.97676\nPOSSIBLE N-term signal sequence\ninside 1 12\nTMhelix 13 35\noutside 36 220\nTMhelix 221 243\ninside 244 254

31412 GCF_002151005.1_ASM215100v1 Clavibacter michiganensis subsp. michiganensis Terrabacteria group; Actinobacteria; Actinobacteria; Micrococcales; Microbacteriaceae; Clavibacter; Clavibacter michiganensis MSTKTARARRTGRRSPLTTIALLAIGLLTTGGAYAMIQSGTASAEVDLKSAQTIDEGQKL WP_043586512.1 MULTISPECIES: cystathionine beta-lyase [Clavibacter] Length: 266\nNumber of predicted TMHs: 2\nExp number of AAs in TMHs: 41.67908\nExp number, first 60 AAs: 21.2051\nTotal prob of N-in: 0.99541\nPOSSIBLE N-term signal sequence\ninside 1 19\nTMhelix 20 42\noutside 43 243\nTMhelix 244 263\ninside 264 266

31413 GCF_002151065.1_ASM215106v1 Clavibacter michiganensis subsp. michiganensis Terrabacteria group; Actinobacteria; Actinobacteria; Micrococcales; Microbacteriaceae; Clavibacter; Clavibacter michiganensis MSTKTARARRTGRRSPLTTIALLAIGLLTTGGAYAMIQSGTASAEVDLKSAQTIDEGQKL WP_043586512.1 MULTISPECIES: cystathionine beta-lyase [Clavibacter] Length: 266\nNumber of predicted TMHs: 2\nExp number of AAs in TMHs: 41.67908\nExp number, first 60 AAs: 21.2051\nTotal prob of N-in: 0.99541\nPOSSIBLE N-term signal sequence\ninside 1 19\nTMhelix 20 42\noutside 43 243\nTMhelix 244 263\ninside 264 266

31414 GCF_002151105.1_ASM215110v1 Clavibacter michiganensis subsp. michiganensis Terrabacteria group; Actinobacteria; Actinobacteria; Micrococcales; Microbacteriaceae; Clavibacter; Clavibacter michiganensis MSTKTARARRTGRRSPLTTIALLAIGLLTTGGAYAMIQSGTASAEVDLKSAQTIDEGQKL WP_043586512.1 MULTISPECIES: cystathionine beta-lyase [Clavibacter] Length: 266\nNumber of predicted TMHs: 2\nExp number of AAs in TMHs: 41.67908\nExp number, first 60 AAs: 21.2051\nTotal prob of N-in: 0.99541\nPOSSIBLE N-term signal sequence\ninside 1 19\nTMhelix 20 42\noutside 43 243\nTMhelix 244 263\ninside 264 266

31415 GCF_000966585.1_ASM96658v1 Clavibacter michiganensis subsp. nebraskensis Terrabacteria group; Actinobacteria; Actinobacteria; Micrococcales; Microbacteriaceae; Clavibacter; Clavibacter michiganensis MNRSSTSAPRRRILRSATALVGGVVLAVAVPLAASAHVRVSPDQAAAGSYSTLTFKVPTEWP_015489341.1 hypothetical protein [Clavibacter michiganensis] Length: 254\nNumber of predicted TMHs: 2\nExp number of AAs in TMHs: 45.06545\nExp number, first 60 AAs: 22.18461\nTotal prob of N-in: 0.99494\nPOSSIBLE N-term signal sequence\ninside 1 12\nTMhelix 13 35\noutside 36 220\nTMhelix 221 243\ninside 244 254

31416 GCF_000966585.1_ASM96658v1 Clavibacter michiganensis subsp. nebraskensis Terrabacteria group; Actinobacteria; Actinobacteria; Micrococcales; Microbacteriaceae; Clavibacter; Clavibacter michiganensis MSTKTARARRTGRRSPLTTIALLAIGLLATGGAYSMIQSGTASAEVDLKSAQTIDEGQKL WP_015490499.1 menaquinol-cytochrome c reductase cytochrome c subunit [Clavibacter michiganensis] Length: 266\nNumber of predicted TMHs: 2\nExp number of AAs in TMHs: 41.68481\nExp number, first 60 AAs: 21.15138\nTotal prob of N-in: 0.99280\nPOSSIBLE N-term signal sequence\ninside 1 19\nTMhelix 20 42\noutside 43 243\nTMhelix 244 263\ninside 264 266

31417 GCF_001643055.1_ASM164305v1 Clavibacter michiganensis subsp. nebraskensis Terrabacteria group; Actinobacteria; Actinobacteria; Micrococcales; Microbacteriaceae; Clavibacter; Clavibacter michiganensis

MSTKTARARRTGRRSPLTTIALLAIGLLATGGAYSMIQSGTASAEVDLKSAQTIDEGQKL WP_015490499.1 menaquinol-cytochrome c reductase cytochrome c subunit [Clavibacter michiganensis] Length: 266\nNumber of predicted TMHs: 2\nExp number of AAs in TMHs: 41.68481\nExp number, first 60 AAs: 21.15138\nTotal prob of N-in: 0.99280\nPOSSIBLE N-term signal sequence\ninside 1 19\nTMhelix 20 42\noutside 43 243\nTMhelix 244 263\ninside 264 266

31418 GCF_001643055.1_ASM164305v1 Clavibacter michiganensis subsp. nebraskensis Terrabacteria group; Actinobacteria; Actinobacteria; Micrococcales; Microbacteriaceae; Clavibacter; Clavibacter michiganensis MNRSSTSAPRRRILRSATALVGGVVLAVAVPLAASAHVRVSPDQAAAGSYSTLTFKVPTE WP_015489341.1 hypothetical protein [Clavibacter michiganensis] Length: 254\nNumber of predicted TMHs: 2\nExp number of AAs in TMHs: 45.06545\nExp number, first 60 AAs: 22.18461\nTotal prob of N-in: 0.99494\nPOSSIBLE N-term signal sequence\ninside 1 12\nTMhelix 13 35\noutside 36 220\nTMhelix 221 243\ninside 244 254

31419 GCF_000069225.1_ASM6922v1 Clavibacter michiganensis subsp. sepedonicus Terrabacteria group; Actinobacteria; Actinobacteria; Micrococcales; Microbacteriaceae; Clavibacter; Clavibacter michiganensis MTTSSPSRPRRRILRSATALIGGVALAVAVPLAASAHVRVSPDQAAAGSYSTLTFKVPTE WP_012300049.1 hypothetical protein [Clavibacter michiganensis] Length: 254\nNumber of predicted TMHs: 2\nExp number of AAs in TMHs: 45.0656\nExp number, first 60 AAs: 22.18545\nTotal prob of N-in: 0.99137\nPOSSIBLE N-term signal sequence\ninside 1 12\nTMhelix 13 35\noutside 36 220\nTMhelix 221 243\ninside 244 254

31420 GCF_000069225.1_ASM6922v1 Clavibacter michiganensis subsp. sepedonicus Terrabacteria group; Actinobacteria; Actinobacteria; Micrococcales; Microbacteriaceae; Clavibacter; Clavibacter michiganensis MRADAGAATSARRPPLRVLLSAAAVVALVAGLLLIGGRAAVGLMRFHDTGLPGHLTLATD WP_012300161.1 hypothetical protein [Clavibacter michiganensis] Length: 266\nNumber of predicted TMHs: 2\nExp number of AAs in TMHs: 42.54578\nExp number, first 60 AAs: 22.46043\nTotal prob of N-in: 0.94285\nPOSSIBLE N-term signal sequence\ninside 1 18\nTMhelix 19 41\noutside 42 215\nTMhelix 216 238\ninside 239 266

31421 GCF_000069225.1_ASM6922v1 Clavibacter michiganensis subsp. sepedonicus Terrabacteria group; Actinobacteria; Actinobacteria; Micrococcales; Microbacteriaceae; Clavibacter; Clavibacter michiganensis MSTKTARARRTGRRSPLTTIALLAIGLLTTGGAYAMIQSGSASAEVDLKSAQTIDEGQKL WP_012298127.1 cystathionine beta-lyase [Clavibacter michiganensis] Length: 266\nNumber of predicted TMHs: 2\nExp number of AAs in TMHs: 41.3097\nExp number, first 60 AAs: 20.78397\nTotal prob of N-in: 0.99496\nPOSSIBLE N-term signal sequence\ninside 1 19\nTMhelix 20 42\noutside 43 243\nTMhelix 244 263\ninside 264 266

31422 GCF_001618005.1_ASM161800v1 Clavibacter michiganensis subsp. tessellarius Terrabacteria group; Actinobacteria; Actinobacteria; Micrococcales; Microbacteriaceae; Clavibacter; Clavibacter michiganensis MPRTTRLPASARPPGRPRRGLVVAGAALGLGLALSAPLAASAHVELDASSTAPAALSVLT WP_063072303.1 sortase [Clavibacter michiganensis] Length: 235\nNumber of predicted TMHs: 2\nExp number of AAs in TMHs: 59.16595\nExp number, first 60 AAs: 23.87736\nTotal prob of N-in: 0.98400\nPOSSIBLE N-term signal sequence\ninside 1 19\nTMhelix 20 42\noutside 43 202\nTMhelix 203 225\ninside 226 235

31423 GCF_001618005.1_ASM161800v1 Clavibacter michiganensis subsp. tessellarius Terrabacteria group; Actinobacteria; Actinobacteria; Micrococcales; Microbacteriaceae; Clavibacter; Clavibacter michiganensis MTTSSTPAPRRRILRSATALVGGVALAVALPLAASAHVRVSPDQAAAGSYSTLTFKVPTE WP_063072393.1 hypothetical protein [Clavibacter michiganensis] Length: 254\nNumber of predicted TMHs: 2\nExp number of AAs in TMHs: 45.01187\nExp number, first 60 AAs: 22.08341\nTotal prob of N-in: 0.98921\nPOSSIBLE N-term signal sequence\ninside 1 12\nTMhelix 13 35\noutside 36 220\nTMhelix 221 243\ninside 244 254

31424 GCF_001618005.1_ASM161800v1 Clavibacter michiganensis subsp. tessellarius Terrabacteria group; Actinobacteria; Actinobacteria; Micrococcales; Microbacteriaceae; Clavibacter; Clavibacter michiganensis MSTTTARARRTGRRSPLTTVALLAIGLLFTGGAYALIQTGTASAQVDLKSAQTIDEGQKL WP_063071144.1 cystathionine beta-lyase [Clavibacter michiganensis] Length: 266\nNumber of predicted TMHs: 2\nExp number of AAs in TMHs: 42.40862\nExp number, first 60 AAs: 21.86424\nTotal prob of N-in: 0.99325\nPOSSIBLE N-term signal sequence\ninside 1 19\nTMhelix 20 42\noutside 43 243\nTMhelix 244 263\ninside 264 266

31425 GCF_001618005.1_ASM161800v1 Clavibacter michiganensis subsp. tessellarius Terrabacteria group; Actinobacteria; Actinobacteria; Micrococcales; Microbacteriaceae; Clavibacter; Clavibacter michiganensis MTPGHRIRRGPRPAGAPAAPRRRLVRVAAALSGAALATAALALVGIGQGSGPDAALPASA WP_081235111.1 copper resistance protein CopC [Clavibacter michiganensis] Length: 289\nNumber of predicted TMHs: 2\nExp number of AAs in TMHs: 44.73004\nExp number, first 60 AAs: 21.58181\nTotal prob of N-in: 0.97276\nPOSSIBLE N-term signal sequence\ninside 1 23\nTMhelix 24 46\noutside 47 233\nTMhelix 234 256\ninside 257 289

31426 GCF_000063485.1_ASM6348v1 Clavibacter michiganensis subsp. michiganensis NCPPB 382Terrabacteria group; Actinobacteria; Actinobacteria; Micrococcales; Microbacteriaceae; Clavibacter; Clavibacter michiganensis; Clavibacter michiganensis subsp. michiganensis
MSTNTARARRTGRRSPLTTIALLAIGLLTTGGAYAMIQSGTASAEVDLKSAQTIDEGQKL WP_012038522.1 cystathionine beta-lyase [Clavibacter michiganensis] Length: 266\nNumber of predicted TMHs: 2\nExp number of AAs in TMHs: 41.67438\nExp number, first 60 AAs: 21.19428\nTotal prob of N-in: 0.99382\nPOSSIBLE N-term signal sequence\ninside 1 19\nTMhelix 20 42\noutside 43 243\nTMhelix 244 263\ninside 264 266

31427 GCF_000063485.1_ASM6348v1 Clavibacter michiganensis subsp. michiganensis NCPPB 382Terrabacteria group; Actinobacteria; Actinobacteria; Micrococcales; Microbacteriaceae; Clavibacter; Clavibacter michiganensis; Clavibacter michiganensis subsp. michiganensis
MTTSSPSAPRRRILRSATALVGGVALAVAVPLAASAHVRVSPDQAAAGSYSTLTFKVPTE WP_011931836.1 hypothetical protein [Clavibacter michiganensis] Length: 254\nNumber of predicted TMHs: 2\nExp number of AAs in TMHs: 44.92106\nExp number, first 60 AAs: 22.03919\nTotal prob of N-in: 0.97676\nPOSSIBLE N-term signal sequence\ninside 1 12\nTMhelix 13 35\noutside 36 220\nTMhelix 221 243\ninside 244 254

31428 GCF_000355695.1_ASM35569v1 Clavibacter michiganensis subsp. nebraskensis NCPPB 2581
Terrabacteria group; Actinobacteria; Actinobacteria; Micrococcales; Microbacteriaceae; Clavibacter; Clavibacter michiganensis; Clavibacter michiganensis subsp. nebraskensis
MSTKTARARRTGRRSPLTTIALLAIGLLATGGAYSMIQSGTASAEVDLKSAQTIDEGQKL WP_015490499.1 menaquinol-cytochrome c reductase cytochrome c subunit [Clavibacter michiganensis] Length: 266\nNumber of predicted TMHs: 2\nExp number of AAs in TMHs: 41.68481\nExp number, first 60 AAs: 21.15138\nTotal prob of N-in: 0.99280\nPOSSIBLE N-term signal sequence\ninside 1 19\nTMhelix 20 42\noutside 43 243\nTMhelix 244 263\ninside 264 266

31429 GCF_000355695.1_ASM35569v1 Clavibacter michiganensis subsp. nebraskensis NCPPB 2581
Terrabacteria group; Actinobacteria; Actinobacteria; Micrococcales; Microbacteriaceae; Clavibacter; Clavibacter michiganensis; Clavibacter michiganensis subsp. nebraskensis
MNRSSTSAPRRRILRSATALVGGVVLAVAVPLAASAHVRVSPDQAAAGSYSTLTFKVPTEWP_015489341.1 hypothetical protein [Clavibacter michiganensis] Length: 254\nNumber of predicted TMHs: 2\nExp number of AAs in TMHs: 45.06545\nExp number, first 60 AAs: 22.18461\nTotal prob of N-in: 0.99494\nPOSSIBLE N-term signal sequence\ninside 1 12\nTMhelix 13 35\noutside 36 220\nTMhelix 221 243\ninside 244 254

31430 GCF_002096055.1_ASM209605v1 Cnuibacter physcomitrellae Terrabacteria group; Actinobacteria; Actinobacteria; Micrococcales; Microbacteriaceae; Cnuibacter
MSRRRLVIALWSAATTVLVAASPIGALPVAQADEVRSLEYWLADYGFTDAWKTTRGAGVT WP_085019923.1
peptidase S8 [Cnuibacter physcomitrellae] Length: 416\nNumber of predicted TMHs: 2\nExp number of AAs in TMHs: 44.3709899999999\nExp number, first 60 AAs: 21.49319\nTotal prob of N-in: 0.95644\nPOSSIBLE N-term signal sequence\ninside 1 6\nTMhelix 7 29\noutside 30 382\nTMhelix 383 405\ninside 406 416

31431 GCF_002096055.1_ASM209605v1 Cnuibacter physcomitrellae Terrabacteria group; Actinobacteria; Actinobacteria; Micrococcales; Microbacteriaceae; Cnuibacter
MAAHRPERARTRRRAPLRRAAALLTAAALCSPVVALTGPALAASAAPSSTAAVAGPAAYG WP_085017700.1
hypothetical protein [Cnuibacter physcomitrellae] Length: 1404\nNumber of predicted TMHs: 2\nExp number of AAs in TMHs: 45.50422\nExp number, first 60 AAs: 22.55061\nTotal prob of N-in: 0.99573\nPOSSIBLE N-term signal sequence\ninside 1 20\nTMhelix 21 43\noutside 44 1372\nTMhelix 1373 1395\ninside 1396 1404

31432 GCF_002096055.1_ASM209605v1 Cnuibacter physcomitrellae Terrabacteria group; Actinobacteria; Actinobacteria; Micrococcales; Microbacteriaceae; Cnuibacter
MTTTRDTLLGAADLERRAALRRMKALATALLVIAAVVFAVAFALEDRYAWLGYVRAAAEG WP_085018363.1
hypothetical protein [Cnuibacter physcomitrellae] Length: 426\nNumber of predicted TMHs: 2\nExp number of AAs in TMHs: 56.3035400000001\nExp number, first 60 AAs: 24.31436\nTotal prob of N-in: 0.79939\nPOSSIBLE N-term signal sequence\ninside 1 24\nTMhelix 25 44\noutside 45 402\nTMhelix 403 425\ninside 426 426

31433 GCF_002096055.1_ASM209605v1 Cnuibacter physcomitrellae Terrabacteria group; Actinobacteria; Actinobacteria; Micrococcales; Microbacteriaceae; Cnuibacter
MAFPQHPRAATRRRGLLLASAALLTAASLSIPTLASAAPGDGDLGANGGATRNDGDKSAA WP_085018623.1
alkaline phosphatase [Cnuibacter physcomitrellae] Length: 637\nNumber of predicted TMHs: 1\nExp number of AAs in TMHs: 37.72141\nExp number, first 60 AAs: 15.11706\nTotal prob of N-in: 0.67924\nPOSSIBLE N-term signal sequence\noutside 1 606\nTMhelix 607 629\ninside 630 637

31434 GCF_002096055.1_ASM209605v1 Cnuibacter physcomitrellae Terrabacteria group; Actinobacteria; Actinobacteria; Micrococcales; Microbacteriaceae; Cnuibacter
MTDPLRPRRSAVVACATHLPATLRTTAETEDRLRADNPRLTLPTGLLRRITGVESVHVAE WP_085019800.1 3-oxoacyl-ACP synthase [Cnuibacter physcomitrellae] Length: 349\nNumber of predicted TMHs: 1\nExp number of AAs in TMHs: 21.35517\nExp number, first 60 AAs: 0.00215\nTotal prob of N-in: 0.28517\noutside 1 324\nTMhelix 325 347\ninside 348 349

31435 GCF_002096055.1_ASM209605v1 Cnuibacter physcomitrellae Terrabacteria group; Actinobacteria; Actinobacteria; Micrococcales; Microbacteriaceae; Cnuibacter
MTTRRVRLRARRTPDLVKRDIVKRRTTLKKTTVLASAAAALAVAGGVLFAAPLAASAHVH WP_085021117.1 hypothetical protein [Cnuibacter physcomitrellae] Length: 275\nNumber of predicted TMHs: 2\nExp number of AAs in TMHs: 45.40013\nExp number, first 60 AAs: 22.59697\nTotal prob of N-in: 0.99968\nPOSSIBLE N-term signal sequence\ninside 1 32\nTMhelix 33 55\noutside 56 242\nTMhelix 243 265\ninside 266 275

31436 GCF_002096055.1_ASM209605v1 Cnuibacter physcomitrellae Terrabacteria group; Actinobacteria; Actinobacteria; Micrococcales; Microbacteriaceae; Cnuibacter
MRLPRRSLLALAALATIGAPVLIGATAANADPVYSEGSCVISRTVDNGPGAAQCAGADLSWP_085019858.1 hypothetical protein [Cnuibacter physcomitrellae] Length: 252\nNumber of predicted TMHs: 2\nExp number of AAs in TMHs: 54.40985\nExp number, first 60 AAs: 20.72792\nTotal prob of N-in: 0.96495\nPOSSIBLE N-term signal sequence\ninside 1 6\nTMhelix 7 29\noutside 30 221\nTMhelix 222 244\ninside 245 252

31437 GCF_001679725.1_ASM167972v1 Cryobacterium arcticum Terrabacteria group; Actinobacteria; Actinobacteria; Micrococcales; Microbacteriaceae; Cryobacterium
MNGVLTRMVNSPKRRNTVLAGGLIAALALTTLGSGIAYAAGGLAPPAVKPATIYTATAVP WP_084021289.1 hypothetical protein [Cryobacterium arcticum] Length: 755\nNumber of predicted TMHs: 2\nExp number of AAs in TMHs: 45.35746\nExp number, first 60 AAs: 22.56408\nTotal prob of N-in: 0.99679\nPOSSIBLE N-term signal sequence\ninside 1 18\nTMhelix 19 41\noutside 42 718\nTMhelix 719 741\ninside 742 755

31438 GCF_001679725.1_ASM167972v1 Cryobacterium arcticum Terrabacteria group; Actinobacteria; Actinobacteria; Micrococcales; Microbacteriaceae; Cryobacterium
MPRTQKPTAGRSRRSGRRSPLASVALIAIGLLFTGGAYAMFSTGTASAATDTASQQTVSE WP_066595335.1 cystathionine beta-lyase [Cryobacterium arcticum] Length: 270\nNumber of predicted TMHs: 2\nExp number of AAs in TMHs: 42.60328\nExp number, first 60 AAs: 21.9284\nTotal prob of N-in: 0.99703\nPOSSIBLE N-term signal sequence\ninside 1 20\nTMhelix 21 43\noutside 44 247\nTMhelix 248 267\ninside 268 270

31439 GCF_001679725.1_ASM167972v1 Cryobacterium arcticum Terrabacteria group; Actinobacteria; Actinobacteria; Micrococcales; Microbacteriaceae; Cryobacterium
MTKSSGRRRALATTVVLALGATGIGTGAVAAPAVAAPAATPLTITPNAAYQNEEFEGWGT WP_084020567.1 hypothetical protein [Cryobacterium arcticum] Length: 1431\nNumber of predicted TMHs: 2\nExp number of AAs in TMHs: 44.42273\nExp number, first 60 AAs: 22.47984\nTotal prob of N-in: 0.98389\nPOSSIBLE N-term signal sequence\ninside 1 12\nTMhelix 13 35\noutside 36 1401\nTMhelix 1402 1424\ninside 1425 1431

31440 GCF_001679725.1_ASM167972v1 Cryobacterium arcticum Terrabacteria group; Actinobacteria; Actinobacteria; Micrococcales; Microbacteriaceae; Cryobacterium
MLTSLPARRPLLAVLSALLSTVVLSTVALSGAAVGAHATEDPDGVGLTVGVVGSTPTPTP WP_066596641.1 hypothetical protein [Cryobacterium arcticum] Length: 259\nNumber of predicted TMHs: 2\nExp number of AAs in TMHs: 55.3675\nExp number, first 60 AAs: 23.48704\nTotal prob of N-in: 0.82230\nPOSSIBLE N-term signal sequence\ninside 1 11\nTMhelix 12 34\noutside 35 217\nTMhelix 218 240\ninside 241 259

31441 GCF_001679725.1_ASM167972v1 Cryobacterium arcticum Terrabacteria group; Actinobacteria; Actinobacteria; Micrococcales; Microbacteriaceae; Cryobacterium
MATTSPRRSLVRTVSGLGLTAALVFGAASSASAAVTTDGPINLGTAASYGVLAASAVTNTWP_066592494.1 hypothetical protein [Cryobacterium arcticum] Length: 442\nNumber of predicted TMHs: 2\nExp number of AAs in TMHs: 47.36896\nExp number, first 60 AAs: 20.18328\nTotal prob of N-in: 0.85273\nPOSSIBLE N-term signal sequence\ninside 1 12\nTMhelix 13 35\noutside 36 409\nTMhelix 410 432\ninside 433 442

31442 GCF_002205495.1_ASM220549v1 Cryobacterium sp. LW097 Terrabacteria group; Actinobacteria; Actinobacteria; Micrococcales; Microbacteriaceae; Cryobacterium
MPLSKTARRATLIVGATLMASCVVVAPALAATPAPTSSAFHRTATYPVYLNAPAGADAAD WP_088457652.1 alkaline phosphatase [Cryobacterium sp. LW097] Length: 943\nNumber of predicted TMHs: 2\nExp number of

AAs in TMHs: 43.79416\nExp number, first 60 AAs: 21.73046\nTotal prob of N-in: 0.98557\nPOSSIBLE N-term signal sequence\ninside 1 11\nTMhelix 12 34\noutside 35 914\nTMhelix 915 937\ninside 938 943

31443 GCF_002205495.1_ASM220549v1 Cryobacterium sp. LW097 Terrabacteria group; Actinobacteria; Actinobacteria; Micrococcales; Microbacteriaceae; Cryobacterium MPRTQKPTNGRTRRSGRRSPLASVALIAIGLLFTGGAYAMISTSTATAQTETASQQTVSE WP_088456184.1 cystathionine beta-lyase [Cryobacterium sp. LW097] Length: 270\nNumber of predicted TMHs: 2\nExp number of AAs in TMHs: 42.5859999999999\nExp number, first 60 AAs: 22.04735\nTotal prob of N-in: 0.99656\nPOSSIBLE N-term signal sequence\ninside 1 20\nTMhelix 21 43\noutside 44 247\nTMhelix 248 267\ninside 268 270

31444 GCF_000738085.1_CryRoo1.0 Cryobacterium sp. MLB-32 Terrabacteria group; Actinobacteria; Actinobacteria; Micrococcales; Microbacteriaceae; Cryobacterium MTPSIVTPRRPAIRFLLSATAGVLVALGMTVSLGVSGAWAEGSDGIAAAPSANGGVDQSR WP_051973425.1 hypothetical protein [Cryobacterium sp. MLB-32] Length: 358\nNumber of predicted TMHs: 2\nExp number of AAs in TMHs: 44.63413\nExp number, first 60 AAs: 21.82753\nTotal prob of N-in: 0.93222\nPOSSIBLE N-term signal sequence\ninside 1 12\nTMhelix 13 35\noutside 36 304\nTMhelix 305 327\ninside 328 358

31445 GCF_001475775.1_ASM147577v1 Curtobacterium citreum Terrabacteria group; Actinobacteria; Actinobacteria; Micrococcales; Microbacteriaceae; Curtobacterium MLTTTACPRRRLLLQTMATLALLLLVGFTVLMHSMAGHTAAHEHSTAAPMATAEHGHVAT WP_058741450.1 hypothetical protein [Curtobacterium citreum] Length: 151\nNumber of predicted TMHs: 2\nExp number of AAs in TMHs: 44.56636\nExp number, first 60 AAs: 21.67367\nTotal prob of N-in: 0.99368\nPOSSIBLE N-term signal sequence\ninside 1 11\nTMhelix 12 34\noutside 35 87\nTMhelix 88 110\ninside 111 151

31446 GCF_001475775.1_ASM147577v1 Curtobacterium citreum Terrabacteria group; Actinobacteria; Actinobacteria; Micrococcales; Microbacteriaceae; Curtobacterium MFNRTKSKKGRRSPMATVSLIVVGLMTTGGAYALFSSTANADDSTTTVASSQSKVNEGQK WP_058743206.1 cystathionine beta-lyase [Curtobacterium citreum] Length: 267\nNumber of predicted TMHs: 2\nExp number of AAs in TMHs: 43.51515\nExp number, first 60 AAs: 22.06861\nTotal prob of N-in: 0.91813\nPOSSIBLE N-term signal sequence\ninside 1 12\nTMhelix 13 35\noutside 36 244\nTMhelix 245 264\ninside 265 267

31447 GCF_001475775.1_ASM147577v1 Curtobacterium citreum Terrabacteria group; Actinobacteria; Actinobacteria; Micrococcales; Microbacteriaceae; Curtobacterium MIITTSPRRRLLLQVTATVALLLLVGFTVLMHTMAGHTADHEHPMTAPTAAMEHHPAAAT WP_058740988.1 hypothetical protein [Curtobacterium citreum] Length: 156\nNumber of predicted TMHs: 2\nExp number of AAs in TMHs: 50.08875\nExp number, first 60 AAs: 22.34792\nTotal prob of N-in: 0.99514\nPOSSIBLE N-term signal sequence\ninside 1 12\nTMhelix 13 35\noutside 36 92\nTMhelix 93 115\ninside 116 156

31448 GCF_000834535.1_ASM83453v1 Curtobacterium flaccumfaciens Terrabacteria group; Actinobacteria; Actinobacteria; Micrococcales; Microbacteriaceae; Curtobacterium MFNRTKSKKGRRSPMATVSLIVVGLMTTGGAYALFSSTANADDSTSTLATSSQSQVNEGE WP_042537210.1 MULTISPECIES: cystathionine beta-lyase [Curtobacterium] Length: 269\nNumber of predicted TMHs: 2\nExp number of AAs in TMHs: 43.41552\nExp number, first 60 AAs: 22.0168\nTotal prob of N-in: 0.91655\nPOSSIBLE N-term signal sequence\ninside 1 12\nTMhelix 13 35\noutside 36 246\nTMhelix 247 266\ninside 267 269

31449 GCF_001475545.1_ASM147554v1 Curtobacterium luteum Terrabacteria group; Actinobacteria; Actinobacteria; Micrococcales; Microbacteriaceae; Curtobacterium MREHVSSRRTVAAGIGERRIDGAGSRARRRAPRDRRPLLRGAVVATVTALVAATVAIQPA WP_058725996.1 hypothetical protein [Curtobacterium luteum] Length: 2532\nNumber of predicted TMHs: 2\nExp number of AAs in TMHs: 43.0098899999999\nExp number, first 60 AAs: 21.00915\nTotal prob of N-in: 0.98437\nPOSSIBLE N-term signal sequence\ninside 1 37\nTMhelix 38 60\noutside 61 2497\nTMhelix 2498 2520\ninside 2521 2532

31450 GCF_001475545.1_ASM147554v1 Curtobacterium luteum Terrabacteria group; Actinobacteria; Actinobacteria; Micrococcales; Microbacteriaceae; Curtobacterium MFNRSKSKKGRRSPLATVSLIVVGLMTTGGAYALFSSTANADDSTSTVSAASTQSKVNEG WP_058724426.1 cystathionine beta-lyase [Curtobacterium luteum] Length: 269\nNumber of predicted TMHs: 2\nExp number of AAs in TMHs: 44.03782\nExp number, first 60 AAs: 22.45253\nTotal prob of N-in: 0.88216\nPOSSIBLE N-term signal sequence\ninside 1 12\nTMhelix 13 35\noutside 36 246\nTMhelix 247 266\ninside 267 269

31451 GCF_001475745.1_ASM147574v1 Curtobacterium oceanosedimentum Terrabacteria group; Actinobacteria; Actinobacteria; Micrococcales; Microbacteriaceae; Curtobacterium

MFTRSKSKKGRRSPMATVSLIVVGLMTTGGAYALFSSTANADDSTTSAVSSQSKVNEGEK WP_058727854.1 cystathionine beta-lyase [Curtobacterium oceanosedimentum] Length: 267\nNumber of predicted TMHs: 2\nExp number of AAs in TMHs: 43.46756\nExp number, first 60 AAs: 21.99053\nTotal prob of N-in: 0.87453\nPOSSIBLE N-term signal sequence\ninside 1 12\nTMhelix 13 35\noutside 36 244\nTMhelix 245 264\ninside 265 267

31452 GCF_001476135.1_ASM147613v1 Curtobacterium oceanosedimentum Terrabacteria group; Actinobacteria; Actinobacteria; Micrococcales; Microbacteriaceae; Curtobacterium MLTTTASPRRRLLLQATATLALLLLVGFTVLMHTMAGHTDGHEHAMAAPAATAEHSHVEP WP_058728719.1 hypothetical protein [Curtobacterium oceanosedimentum] Length: 151\nNumber of predicted TMHs: 2\nExp number of AAs in TMHs: 44.97331\nExp number, first 60 AAs: 22.04768\nTotal prob of N-in: 0.99405\nPOSSIBLE N-term signal sequence\ninside 1 11\nTMhelix 12 34\noutside 35 87\nTMhelix 88 110\ninside 111 151

31453 GCF_001476135.1_ASM147613v1 Curtobacterium oceanosedimentum Terrabacteria group; Actinobacteria; Actinobacteria; Micrococcales; Microbacteriaceae; Curtobacterium MFTRSKSKKGRRSPMATVSLIVVGLMTTGGAYALFSSTANADDSTTSAVSSQSKVNEGEK WP_058727854.1 cystathionine beta-lyase [Curtobacterium oceanosedimentum] Length: 267\nNumber of predicted TMHs: 2\nExp number of AAs in TMHs: 43.46756\nExp number, first 60 AAs: 21.99053\nTotal prob of N-in: 0.87453\nPOSSIBLE N-term signal sequence\ninside 1 12\nTMhelix 13 35\noutside 36 244\nTMhelix 245 264\ninside 265 267

31454 GCF_001475745.1_ASM147574v1 Curtobacterium oceanosedimentum Terrabacteria group; Actinobacteria; Actinobacteria; Micrococcales; Microbacteriaceae; Curtobacterium MFTTTATPRRRLLLQATATLALLLLVGFTVLMHTMAGHTADHEHSMAAPAITAEHSHVAA WP_058729336.1 hypothetical protein [Curtobacterium oceanosedimentum] Length: 151\nNumber of predicted TMHs: 2\nExp number of AAs in TMHs: 44.73558\nExp number, first 60 AAs: 22.08337\nTotal prob of N-in: 0.99536\nPOSSIBLE N-term signal sequence\ninside 1 11\nTMhelix 12 34\noutside 35 87\nTMhelix 88 110\ninside 111 151

31455 GCF_001475745.1_ASM147574v1 Curtobacterium oceanosedimentum Terrabacteria group; Actinobacteria; Actinobacteria; Micrococcales; Microbacteriaceae; Curtobacterium MLTTTASPRRRLLLQATATLALLLLVGFTVLMHTMAGHTDGHEHAMAAPAATAEHSHVEP WP_058728719.1 hypothetical protein [Curtobacterium oceanosedimentum] Length: 151\nNumber of predicted TMHs: 2\nExp number of AAs in TMHs: 44.97331\nExp number, first 60 AAs: 22.04768\nTotal prob of N-in: 0.99405\nPOSSIBLE N-term signal sequence\ninside 1 11\nTMhelix 12 34\noutside 35 87\nTMhelix 88 110\ninside 111 151

31456 GCF_002025645.1_ASM202564v1 Curtobacterium pusillum Terrabacteria group; Actinobacteria; Actinobacteria; Micrococcales; Microbacteriaceae; Curtobacterium MFNRTKSKKGRRSPMATVSLIVVGLMTTGGAYALFSSTANADDSTSTVAATSQSKVNEGQ WP_079236043.1 cystathionine beta-lyase [Curtobacterium pusillum] Length: 269\nNumber of predicted TMHs: 2\nExp number of AAs in TMHs: 43.45499\nExp number, first 60 AAs: 22.03573\nTotal prob of N-in: 0.91059\nPOSSIBLE N-term signal sequence\ninside 1 12\nTMhelix 13 35\noutside 36 246\nTMhelix 247 266\ninside 267 269

31457 GCF_002025645.1_ASM202564v1 Curtobacterium pusillum Terrabacteria group; Actinobacteria; Actinobacteria; Micrococcales; Microbacteriaceae; Curtobacterium MNAHTSPRRRRRLVGSAALATAGVLVALSVPSAALAAEGDTTIDILDVNDFHGRIESEGT WP_079236843.1 hypothetical protein [Curtobacterium pusillum] Length: 780\nNumber of predicted TMHs: 2\nExp number of AAs in TMHs: 44.86784\nExp number, first 60 AAs: 22.53666\nTotal prob of N-in: 0.99575\nPOSSIBLE N-term signal sequence\ninside 1 12\nTMhelix 13 35\noutside 36 744\nTMhelix 745 767\ninside 768 780

31458 GCF_000333375.1_ASM33337v1 Curtobacterium sp. B18 Terrabacteria group; Actinobacteria; Actinobacteria; Micrococcales; Microbacteriaceae; Curtobacterium MHLTITPRRTALVAVAAAALFGSMLVAAPANAAPRSAADPLSDADTTALFAGVDTTTRTF WP_022907060.1 hypothetical protein [Curtobacterium sp. B18] Length: 203\nNumber of predicted TMHs: 2\nExp number of AAs in TMHs: 45.97309\nExp number, first 60 AAs: 21.0471\nTotal prob of N-in: 0.90398\nPOSSIBLE N-term signal sequence\ninside 1 11\nTMhelix 12 34\noutside 35 155\nTMhelix 156 178\ninside 179 203

31459 GCF_000333375.1_ASM33337v1 Curtobacterium sp. B18 Terrabacteria group; Actinobacteria; Actinobacteria; Micrococcales; Microbacteriaceae; Curtobacterium MIITTSPRRRLLLQVTATMALLLLIGFTVLMHTMAGHTADPEHPMTAPTAAMEHHPAAAT WP_022906545.1 hypothetical protein [Curtobacterium sp. B18] Length: 156\nNumber of predicted TMHs: 2\nExp number of AAs in TMHs: 51.70625\nExp number, first 60 AAs: 22.29221\nTotal prob of N-in: 0.99712\nPOSSIBLE N-term signal sequence\ninside 1 12\nTMhelix 13 35\noutside 36 92\nTMhelix 93 115\ninside 116 156

31460 GCF_000333315.1_ASM33331v1 Curtobacterium sp. B8 Terrabacteria group; Actinobacteria; Actinobacteria; Micrococcales; Microbacteriaceae; Curtobacterium
MLTTTTSPRRRLLLQATATLALLLLVGFTVLMHTMAGHTAGHEHSMATPAITAEHGHGAA WP_022903855.1
hypothetical protein [Curtobacterium sp. B8] Length: 151\nNumber of predicted TMHs: 2\nExp number of AAs in TMHs: 44.9809\nExp number, first 60 AAs: 22.09582\nTotal prob of N-in: 0.99509\nPOSSIBLE N-term signal sequence\ninside 1 11\nTMhelix 12 34\noutside 35 87\nTMhelix 88 110\ninside 111 151

31461 GCF_001806325.1_ASM180632v1 Curtobacterium sp. BH-2-1-1 Terrabacteria group; Actinobacteria; Actinobacteria; Micrococcales; Microbacteriaceae; Curtobacterium
MNAHTSPRRRRRLVGGAALATAGVLVALSVPSAAIAADGDTTIDLYDVNDFHGRIAQSIS WP_070418157.1
hypothetical protein [Curtobacterium sp. BH-2-1-1] Length: 783\nNumber of predicted TMHs: 2\nExp number of AAs in TMHs: 44.67641\nExp number, first 60 AAs: 22.79677\nTotal prob of N-in: 0.99840\nPOSSIBLE N-term signal sequence\ninside 1 12\nTMhelix 13 35\noutside 36 747\nTMhelix 748 770\ninside 771 783

31462 GCF_001806325.1_ASM180632v1 Curtobacterium sp. BH-2-1-1 Terrabacteria group; Actinobacteria; Actinobacteria; Micrococcales; Microbacteriaceae; Curtobacterium
MFNRTKSKKGRRSPMATVSLIVVGLMTTGGAYALFSSTANADDSTSTVAASSQSKVNEGQ WP_070418988.1
cystathionine beta-lyase [Curtobacterium sp. BH-2-1-1] Length: 269\nNumber of predicted TMHs: 2\nExp number of AAs in TMHs: 43.44125\nExp number, first 60 AAs: 22.03616\nTotal prob of N-in: 0.91118\nPOSSIBLE N-term signal sequence\ninside 1 12\nTMhelix 13 35\noutside 36 246\nTMhelix 247 266\ninside 267 269

31463 GCF_001742745.1_ASM174274v1 Curtobacterium sp. ER1/6 Terrabacteria group; Actinobacteria; Actinobacteria; Micrococcales; Microbacteriaceae; Curtobacterium
MFNRSKSKKGRRSPMATVSLIVVGLMTTGGAYALFSSTANADDSTTTAASSQSQVNEGEK WP_069710890.1
cystathionine beta-lyase [Curtobacterium sp. ER1/6] Length: 267\nNumber of predicted TMHs: 2\nExp number of AAs in TMHs: 43.56027\nExp number, first 60 AAs: 22.02655\nTotal prob of N-in: 0.83633\nPOSSIBLE N-term signal sequence\ninside 1 12\nTMhelix 13 35\noutside 36 244\nTMhelix 245 264\ninside 265 267

31464 GCF_001424595.1_Leaf154 Curtobacterium sp. Leaf154 Terrabacteria group; Actinobacteria; Actinobacteria; Micrococcales; Microbacteriaceae; Curtobacterium
MFNRTKSKKGRRSPMATVSLIVVGLMTTGGAYALFSSTANADDSTSTLATSSQSQVNEGE WP_042537210.1
MULTISPECIES: cystathionine beta-lyase [Curtobacterium] Length: 269\nNumber of predicted TMHs: 2\nExp number of AAs in TMHs: 43.41552\nExp number, first 60 AAs: 22.0168\nTotal prob of N-in: 0.91655\nPOSSIBLE N-term signal sequence\ninside 1 12\nTMhelix 13 35\noutside 36 246\nTMhelix 247 266\ninside 267 269

31465 GCF_001424385.1_Leaf183 Curtobacterium sp. Leaf183 Terrabacteria group; Actinobacteria; Actinobacteria; Micrococcales; Microbacteriaceae; Curtobacterium
MLTTANTTRRRLLLQATAVLALLLLVGFTVLMHSMLGHATTGDHAMTATSSMTAEHAHSE WP_056120640.1
hypothetical protein [Curtobacterium sp. Leaf183] Length: 153\nNumber of predicted TMHs: 2\nExp number of AAs in TMHs: 45.06981\nExp number, first 60 AAs: 22.07223\nTotal prob of N-in: 0.99853\nPOSSIBLE N-term signal sequence\ninside 1 11\nTMhelix 12 34\noutside 35 89\nTMhelix 90 112\ninside 113 153

31466 GCF_001424385.1_Leaf183 Curtobacterium sp. Leaf183 Terrabacteria group; Actinobacteria; Actinobacteria; Micrococcales; Microbacteriaceae; Curtobacterium
MSDDKKKGFGRRQLIGGGIAAAAVGALSTMFPELANAETRTARGVTTLNDDEWYLQEAQS WP_056124881.1
hypothetical protein [Curtobacterium sp. Leaf183] Length: 244\nNumber of predicted TMHs: 2\nExp number of AAs in TMHs: 42.4308299999999\nExp number, first 60 AAs: 18.24891\nTotal prob of N-in: 0.61614\nPOSSIBLE N-term signal sequence\ninside 1 12\nTMhelix 13 35\noutside 36 207\nTMhelix 208 230\ninside 231 244

31467 GCF_001424385.1_Leaf183 Curtobacterium sp. Leaf183 Terrabacteria group; Actinobacteria; Actinobacteria; Micrococcales; Microbacteriaceae; Curtobacterium
MFNRTKTKKGRRSPMATVSLIVVGLMTTGGAYALFSSSANADDSVSTVAASSQSQVNQGE WP_056124562.1
cystathionine beta-lyase [Curtobacterium sp. Leaf183] Length: 268\nNumber of predicted TMHs: 2\nExp number of AAs in TMHs: 43.77047\nExp number, first 60 AAs: 22.05024\nTotal prob of N-in: 0.95194\nPOSSIBLE N-term signal sequence\ninside 1 12\nTMhelix 13 35\noutside 36 245\nTMhelix 246 265\ninside 266 268

31468 GCF_001422205.1_Leaf261 Curtobacterium sp. Leaf261 Terrabacteria group; Actinobacteria; Actinobacteria; Micrococcales; Microbacteriaceae; Curtobacterium
MFTRTKTERSGAAKKGRRSPMATVSLIVVGLMTTGGAYALFSSSATADTSQSSTVAASSQ WP_055947809.1
cystathionine beta-lyase [Curtobacterium sp. Leaf261] Length: 275\nNumber of predicted TMHs: 2\nExp

number of AAs in TMHs: 41.85833\nExp number, first 60 AAs: 21.5073\nTotal prob of N-in: 0.98349\nPOSSIBLE N-term signal sequence\ninside 1 20\nTMhelix 21 43\noutside 44 252\nTMhelix 253 272\ninside 273 275

31469 GCF_001422205.1_Leaf261 Curtobacterium sp. Leaf261 Terrabacteria group; Actinobacteria; Actinobacteria; Micrococcales; Microbacteriaceae; Curtobacterium
MRHTPTRSNRRRVLAAILVAASGLGLVGVAALPAAAAAPLASQTVVSGDARFQVLSPTLIWP_082474164.1 hypothetical protein [Curtobacterium sp. Leaf261] Length: 1267\nNumber of predicted TMHs: 2\nExp number of AAs in TMHs: 44.3089699999999\nExp number, first 60 AAs: 22.61908\nTotal prob of N-in: 0.99991\nPOSSIBLE N-term signal sequence\ninside 1 12\nTMhelix 13 35\noutside 36 1238\nTMhelix 1239 1261\ninside 1262 1267

31470 GCF_001422205.1_Leaf261 Curtobacterium sp. Leaf261 Terrabacteria group; Actinobacteria; Actinobacteria; Micrococcales; Microbacteriaceae; Curtobacterium
MQQRRRLVTGVTATAGIAAAIVLGTAVAASAHISADATDTAANSYTTLTFSVPHGCDGSP WP_055952736.1 hypothetical protein [Curtobacterium sp. Leaf261] Length: 247\nNumber of predicted TMHs: 2\nExp number of AAs in TMHs: 43.82883\nExp number, first 60 AAs: 22.83295\nTotal prob of N-in: 0.99749\nPOSSIBLE N-term signal sequence\ninside 1 6\nTMhelix 7 29\noutside 30 218\nTMhelix 219 241\ninside 242 247

31471 GCF_001864835.1_ASM186483v1 Curtobacterium sp. MCBA15_001 Terrabacteria group; Actinobacteria; Actinobacteria; Micrococcales; Microbacteriaceae; Curtobacterium
MFTRTKNKKGRRSPMATVSLIVVGLMTTGGAYALFSSSANADDSVSTVAASSQSQVNEGQ WP_071300786.1 cystathionine beta-lyase [Curtobacterium sp. MCBA15_001] Length: 269\nNumber of predicted TMHs: 2\nExp number of AAs in TMHs: 43.76439\nExp number, first 60 AAs: 22.07\nTotal prob of N-in: 0.96591\nPOSSIBLE N-term signal sequence\ninside 1 12\nTMhelix 13 35\noutside 36 246\nTMhelix 247 266\ninside 267 269

31472 GCF_001864835.1_ASM186483v1 Curtobacterium sp. MCBA15_001 Terrabacteria group; Actinobacteria; Actinobacteria; Micrococcales; Microbacteriaceae; Curtobacterium
MLHSTRRTLLCATVAATLLAAPLASVTAASANTAGDGLVIQEAYLKGGSKDAFFTNKFVE WP_071299477.1 hypothetical protein [Curtobacterium sp. MCBA15_001] Length: 1023\nNumber of predicted TMHs: 1\nExp number of AAs in TMHs: 32.9593199999999\nExp number, first 60 AAs: 11.70503\nTotal prob of N-in: 0.52272\nPOSSIBLE N-term signal sequence\noutside 1 992\nTMhelix 993 1015\ninside 1016 1023

31473 GCF_001864845.1_ASM186484v1 Curtobacterium sp. MCBA15_003 Terrabacteria group; Actinobacteria; Actinobacteria; Micrococcales; Microbacteriaceae; Curtobacterium
MLPVHQGTAVPRRRPALRLVAMLGATLLAGAGLTATTAVVASEPASAVDATLDGALLCNQ WP_083404666.1 MULTISPECIES: hypothetical protein [Curtobacterium] Length: 1826\nNumber of predicted TMHs: 2\nExp number of AAs in TMHs: 43.4607999999999\nExp number, first 60 AAs: 21.23016\nTotal prob of N-in: 0.93821\nPOSSIBLE N-term signal sequence\ninside 1 18\nTMhelix 19 41\noutside 42 1784\nTMhelix 1785 1807\ninside 1808 1826

31474 GCF_001864845.1_ASM186484v1 Curtobacterium sp. MCBA15_003 Terrabacteria group; Actinobacteria; Actinobacteria; Micrococcales; Microbacteriaceae; Curtobacterium
MFNRTKSKKGRRSPMATVSLIVVGLMTTGGAYALFSSSANADDSTSTIAATSQSKVNEGQ WP_071255400.1 MULTISPECIES: cystathionine beta-lyase [Curtobacterium] Length: 269\nNumber of predicted TMHs: 2\nExp number of AAs in TMHs: 43.4648\nExp number, first 60 AAs: 22.05081\nTotal prob of N-in: 0.93134\nPOSSIBLE N-term signal sequence\ninside 1 12\nTMhelix 13 35\noutside 36 246\nTMhelix 247 266\ninside 267 269

31475 GCF_001865055.1_ASM186505v1 Curtobacterium sp. MCBA15_004 Terrabacteria group; Actinobacteria; Actinobacteria; Micrococcales; Microbacteriaceae; Curtobacterium
MGTLVPTRATERRRTTLLALTVSTLAVLLAVLVAPGPASASTTTPSDARAAVVRSIERGT WP_071246791.1 MULTISPECIES: hypothetical protein [Curtobacterium] Length: 221\nNumber of predicted TMHs: 2\nExp number of AAs in TMHs: 49.83816\nExp number, first 60 AAs: 21.52832\nTotal prob of N-in: 0.97569\nPOSSIBLE N-term signal sequence\ninside 1 15\nTMhelix 16 38\noutside 39 155\nTMhelix 156 178\ninside 179 221

31476 GCF_001865055.1_ASM186505v1 Curtobacterium sp. MCBA15_004 Terrabacteria group; Actinobacteria; Actinobacteria; Micrococcales; Microbacteriaceae; Curtobacterium
MLTTTATPRRRLLLQATATLALLLLVGFTVLMHTMAGHTADHEHPMAAPAAAAEHGHVAA WP_066651479.1 MULTISPECIES: hypothetical protein [Curtobacterium] Length: 151\nNumber of predicted TMHs: 2\nExp number of AAs in TMHs: 44.7324500000001\nExp number, first 60 AAs: 22.08752\nTotal prob of N-in: 0.99588\nPOSSIBLE N-term signal sequence\ninside 1 11\nTMhelix 12 34\noutside 35 87\nTMhelix 88 110\ninside 111 151

31477 GCF_001865055.1_ASM186505v1 Curtobacterium sp. MCBA15_004 Terrabacteria group; Actinobacteria; Actinobacteria; Micrococcales; Microbacteriaceae; Curtobacterium
MNTHTSPRRRRLVGTATLATAGVLVALSAPSAALAADGDVTIDLYNVNDFHGRIAQSISV WP_065961321.1
MULTISPECIES: hypothetical protein [Curtobacterium] Length: 775\nNumber of predicted TMHs: 2\nExp number of AAs in TMHs: 42.7160599999999\nExp number, first 60 AAs: 22.07275\nTotal prob of N-in: 0.97551\nPOSSIBLE N-term signal sequence\ninside 1 11\nTMhelix 12 34\noutside 35 738\nTMhelix 739 761\ninside 762 775

31478 GCF_001865055.1_ASM186505v1 Curtobacterium sp. MCBA15_004 Terrabacteria group; Actinobacteria; Actinobacteria; Micrococcales; Microbacteriaceae; Curtobacterium
MFTRTKTKKGRRSPMATVSLIVVGLMTTGGAYALFSSTANADDSTTATAASSQSQVNEGQ WP_065962238.1
MULTISPECIES: cystathionine beta-lyase [Curtobacterium] Length: 269\nNumber of predicted TMHs: 2\nExp number of AAs in TMHs: 43.48172\nExp number, first 60 AAs: 22.0798\nTotal prob of N-in: 0.93547\nPOSSIBLE N-term signal sequence\ninside 1 12\nTMhelix 13 35\noutside 36 246\nTMhelix 247 266\ninside 267 269

31479 GCF_001865055.1_ASM186505v1 Curtobacterium sp. MCBA15_004 Terrabacteria group; Actinobacteria; Actinobacteria; Micrococcales; Microbacteriaceae; Curtobacterium
MKPLISSPGVPSTSSPSRPGRRPPTAARLTRRRPVLSGVGLATALGLLLAPAFLASSSSA WP_071283736.1 hypothetical protein [Curtobacterium sp. MCBA15_004] Length: 666\nNumber of predicted TMHs: 2\nExp number of AAs in TMHs: 41.69364\nExp number, first 60 AAs: 19.87471\nTotal prob of N-in: 0.90577\nPOSSIBLE N-term signal sequence\ninside 1 34\nTMhelix 35 57\noutside 58 626\nTMhelix 627 649\ninside 650 666

31480 GCF_001865055.1_ASM186505v1 Curtobacterium sp. MCBA15_004 Terrabacteria group; Actinobacteria; Actinobacteria; Micrococcales; Microbacteriaceae; Curtobacterium
MNGRVPDTTMHDEGTPVPSSHRARPRRRTLRLVAGTIATTLVAGSAALLSPMTASAMANL WP_083395140.1
hypothetical protein [Curtobacterium sp. MCBA15_004] Length: 1266\nNumber of predicted TMHs: 2\nExp number of AAs in TMHs: 46.8248599999999\nExp number, first 60 AAs: 21.7179\nTotal prob of N-in: 0.99015\nPOSSIBLE N-term signal sequence\ninside 1 31\nTMhelix 32 54\noutside 55 1235\nTMhelix 1236 1258\ninside 1259 1266

31481 GCF_001864855.1_ASM186485v1 Curtobacterium sp. MCBA15_005 Terrabacteria group; Actinobacteria; Actinobacteria; Micrococcales; Microbacteriaceae; Curtobacterium
MLPLQPDDPPPLVLIEESMLTATASPRRRLLLQVTATLALLLLVGFTVLMHTMAGHTAGH WP_071244483.1
hypothetical protein [Curtobacterium sp. MCBA15_005] Length: 169\nNumber of predicted TMHs: 2\nExp number of AAs in TMHs: 44.15299\nExp number, first 60 AAs: 21.36026\nTotal prob of N-in: 0.33810\nPOSSIBLE N-term signal sequence\ninside 1 29\nTMhelix 30 52\noutside 53 105\nTMhelix 106 128\ninside 129 169

31482 GCF_001864855.1_ASM186485v1 Curtobacterium sp. MCBA15_005 Terrabacteria group; Actinobacteria; Actinobacteria; Micrococcales; Microbacteriaceae; Curtobacterium
MFNRTKSKKGRRSPMATVSLIVVGLMTTGGAYALFSSTANADDSTNTVAASSQSQVNEGE WP_071244692.1
cystathionine beta-lyase [Curtobacterium sp. MCBA15_005] Length: 269\nNumber of predicted TMHs: 2\nExp number of AAs in TMHs: 43.48948\nExp number, first 60 AAs: 22.05571\nTotal prob of N-in: 0.88824\nPOSSIBLE N-term signal sequence\ninside 1 12\nTMhelix 13 35\noutside 36 246\nTMhelix 247 266\ninside 267 269

31483 GCF_001864905.1_ASM186490v1 Curtobacterium sp. MCBA15_007 Terrabacteria group; Actinobacteria; Actinobacteria; Micrococcales; Microbacteriaceae; Curtobacterium
MFNRTKSKKGRRSPMATVSLIVVGLMTTGGAYALFSSTANADDSTSTLATSSQSQVNEGE WP_042537210.1
MULTISPECIES: cystathionine beta-lyase [Curtobacterium] Length: 269\nNumber of predicted TMHs: 2\nExp number of AAs in TMHs: 43.41552\nExp number, first 60 AAs: 22.0168\nTotal prob of N-in: 0.91655\nPOSSIBLE N-term signal sequence\ninside 1 12\nTMhelix 13 35\noutside 36 246\nTMhelix 247 266\ninside 267 269

31484 GCF_001865065.1_ASM186506v1 Curtobacterium sp. MCBA15_008 Terrabacteria group; Actinobacteria; Actinobacteria; Micrococcales; Microbacteriaceae; Curtobacterium
MFNRSKSKKGRRSPMATVSLIVVGLMTTGGAYALFSSTANADDSTSTVAASSQSKVNEGQ WP_071293026.1
cystathionine beta-lyase [Curtobacterium sp. MCBA15_008] Length: 269\nNumber of predicted TMHs: 2\nExp number of AAs in TMHs: 43.43847\nExp number, first 60 AAs: 21.98506\nTotal prob of N-in: 0.87016\nPOSSIBLE N-term signal sequence\ninside 1 12\nTMhelix 13 35\noutside 36 246\nTMhelix 247 266\ninside 267 269

31485 GCF_001864895.1_ASM186489v1 Curtobacterium sp. MCBA15_009 Terrabacteria group; Actinobacteria; Actinobacteria; Micrococcales; Microbacteriaceae; Curtobacterium
MFNRTKSKKGRRSPMATVSLIVVGLMTTGGAYALFSSSANADDSTSTIAATSQSKVNEGQ WP_071255400.1
MULTISPECIES: cystathionine beta-lyase [Curtobacterium] Length: 269\nNumber of predicted TMHs: 2\nExp

number of AAs in TMHs: 43.4648\nExp number, first 60 AAs: 22.05081\nTotal prob of N-in: 0.93134\nPOSSIBLE N-term signal sequence\ninside 1 12\nTMhelix 13 35\noutside 36 246\nTMhelix 247 266\ninside 267 269

31486 GCF_001864895.1_ASM186489v1 Curtobacterium sp. MCBA15_009 Terrabacteria group; Actinobacteria; Actinobacteria; Micrococcales; Microbacteriaceae; Curtobacterium MLPVHQGTAVPRRRPALRLVAMLGATLLAGAGLTATTAVVASEPASAVDATLDGALLCNQ WP_083420771.1 hypothetical protein [Curtobacterium sp. MCBA15_009] Length: 1826\nNumber of predicted TMHs: 2\nExp number of AAs in TMHs: 43.3669099999999\nExp number, first 60 AAs: 21.23016\nTotal prob of N-in: 0.93821\nPOSSIBLE N-term signal sequence\ninside 1 18\nTMhelix 19 41\noutside 42 1784\nTMhelix 1785 1807\ninside 1808 1826

31487 GCF_001864935.1_ASM186493v1 Curtobacterium sp. MCBA15_012 Terrabacteria group; Actinobacteria; Actinobacteria; Micrococcales; Microbacteriaceae; Curtobacterium MNTHTSPRRRRLVGTATLATAGVLVALSAPSAALAADGDVTIDLYNVNDFHGRIAQSISV WP_071247766.1 hypothetical protein [Curtobacterium sp. MCBA15_012] Length: 775\nNumber of predicted TMHs: 2\nExp number of AAs in TMHs: 43.2608599999999\nExp number, first 60 AAs: 22.07275\nTotal prob of N-in: 0.97551\nPOSSIBLE N-term signal sequence\ninside 1 11\nTMhelix 12 34\noutside 35 738\nTMhelix 739 761\ninside 762 775

31488 GCF_001864935.1_ASM186493v1 Curtobacterium sp. MCBA15_012 Terrabacteria group; Actinobacteria; Actinobacteria; Micrococcales; Microbacteriaceae; Curtobacterium MGTLVPTRATERRRTTLLALTVSTLAVLLAVLVAPGPASASTTTPSDARAAVVRSIERGT WP_071246791.1 MULTISPECIES: hypothetical protein [Curtobacterium] Length: 221\nNumber of predicted TMHs: 2\nExp number of AAs in TMHs: 49.83816\nExp number, first 60 AAs: 21.52832\nTotal prob of N-in: 0.97569\nPOSSIBLE N-term signal sequence\ninside 1 15\nTMhelix 16 38\noutside 39 155\nTMhelix 156 178\ninside 179 221

31489 GCF_001864935.1_ASM186493v1 Curtobacterium sp. MCBA15_012 Terrabacteria group; Actinobacteria; Actinobacteria; Micrococcales; Microbacteriaceae; Curtobacterium MNGRVPDTTMHDEGTPVPSSHRARPRRRTLRLVAGTIATTLVAGSAALLSPMTASAMANL WP_083394073.1 hypothetical protein [Curtobacterium sp. MCBA15_012] Length: 1263\nNumber of predicted TMHs: 2\nExp number of AAs in TMHs: 45.07833\nExp number, first 60 AAs: 21.71797\nTotal prob of N-in: 0.99016\nPOSSIBLE N-term signal sequence\ninside 1 31\nTMhelix 32 54\noutside 55 1232\nTMhelix 1233 1255\ninside 1256 1263

31490 GCF_001864935.1_ASM186493v1 Curtobacterium sp. MCBA15_012 Terrabacteria group; Actinobacteria; Actinobacteria; Micrococcales; Microbacteriaceae; Curtobacterium MKPHISSPGVPSTSSPSRPGRRPPTAARLRRRRPVLSGVGLATALGLLLAPAFLASSSSA WP_071249110.1 hypothetical protein [Curtobacterium sp. MCBA15_012] Length: 666\nNumber of predicted TMHs: 2\nExp number of AAs in TMHs: 43.1873499999999\nExp number, first 60 AAs: 21.36971\nTotal prob of N-in: 0.97401\nPOSSIBLE N-term signal sequence\ninside 1 34\nTMhelix 35 57\noutside 58 626\nTMhelix 627 649\ninside 650 666

31491 GCF_001864935.1_ASM186493v1 Curtobacterium sp. MCBA15_012 Terrabacteria group; Actinobacteria; Actinobacteria; Micrococcales; Microbacteriaceae; Curtobacterium MFTRTKTKKGRRSPMATVSLIVVGLMTTGGAYALFSSTANADDSTTATAASSQSQVNEGQ WP_065962238.1 MULTISPECIES: cystathionine beta-lyase [Curtobacterium] Length: 269\nNumber of predicted TMHs: 2\nExp number of AAs in TMHs: 43.48172\nExp number, first 60 AAs: 22.0798\nTotal prob of N-in: 0.93547\nPOSSIBLE N-term signal sequence\ninside 1 12\nTMhelix 13 35\noutside 36 246\nTMhelix 247 266\ninside 267 269

31492 GCF_001865095.1_ASM186509v1 Curtobacterium sp. MCBA15_013 Terrabacteria group; Actinobacteria; Actinobacteria; Micrococcales; Microbacteriaceae; Curtobacterium MFNRTKSKKGRRSPMATVSLIVVGLMTTGGAYALFSSTANADDSTSTVAASSQSKVNEGQ WP_071294687.1 MULTISPECIES: cystathionine beta-lyase [Curtobacterium] Length: 269\nNumber of predicted TMHs: 2\nExp number of AAs in TMHs: 43.44698\nExp number, first 60 AAs: 22.03608\nTotal prob of N-in: 0.91098\nPOSSIBLE N-term signal sequence\ninside 1 12\nTMhelix 13 35\noutside 36 246\nTMhelix 247 266\ninside 267 269

31493 GCF_001865095.1_ASM186509v1 Curtobacterium sp. MCBA15_013 Terrabacteria group; Actinobacteria; Actinobacteria; Micrococcales; Microbacteriaceae; Curtobacterium MNAHTSPRRRRRLVGGAALATAGVLVALSVPSAAIAADGDTTIDLYDVNDFHGRIAQSIS WP_071302787.1 hypothetical protein [Curtobacterium sp. MCBA15_013] Length: 781\nNumber of predicted TMHs: 2\nExp number of AAs in TMHs: 44.68179\nExp number, first 60 AAs: 22.79676\nTotal prob of N-in: 0.99840\nPOSSIBLE N-term signal sequence\ninside 1 12\nTMhelix 13 35\noutside 36 745\nTMhelix 746 768\ninside 769 781

31494 GCF_001865105.1_ASM186510v1 Curtobacterium sp. MCBA15_016 Terrabacteria group; Actinobacteria; Actinobacteria; Micrococcales; Microbacteriaceae; Curtobacterium MNAHTSPRRRRRLVGGAALATAGVLVALSVPSAAIAADGDTTIDLYDVNDFHGRIAQSIS WP_071296020.1 hypothetical protein [Curtobacterium sp. MCBA15_016] Length: 781\nNumber of predicted TMHs: 2\nExp number of AAs in TMHs: 44.68443\nExp number, first 60 AAs: 22.79676\nTotal prob of N-in: 0.99840\nPOSSIBLE N-term signal sequence\ninside 1 12\nTMhelix 13 35\noutside 36 745\nTMhelix 746 768\ninside 769 781

31495 GCF_001865105.1_ASM186510v1 Curtobacterium sp. MCBA15_016 Terrabacteria group; Actinobacteria; Actinobacteria; Micrococcales; Microbacteriaceae; Curtobacterium MLTTTASPRRRLLLQVTATLALLLLVGFTVLMHTMAGHTAGHEHPMAAPAATTEHVHAVG WP_071295101.1 hypothetical protein [Curtobacterium sp. MCBA15_016] Length: 151\nNumber of predicted TMHs: 2\nExp number of AAs in TMHs: 44.83329\nExp number, first 60 AAs: 22.15937\nTotal prob of N-in: 0.99684\nPOSSIBLE N-term signal sequence\ninside 1 11\nTMhelix 12 34\noutside 35 87\nTMhelix 88 110\ninside 111 151

31496 GCF_001865105.1_ASM186510v1 Curtobacterium sp. MCBA15_016 Terrabacteria group; Actinobacteria; Actinobacteria; Micrococcales; Microbacteriaceae; Curtobacterium MFNRTKSKKGRRSPMATVSLIVVGLMTTGGAYALFSSTANADDSTSTVAASSQSKVNEGQ WP_071294687.1 MULTISPECIES: cystathionine beta-lyase [Curtobacterium] Length: 269\nNumber of predicted TMHs: 2\nExp number of AAs in TMHs: 43.44698\nExp number, first 60 AAs: 22.03608\nTotal prob of N-in: 0.91098\nPOSSIBLE N-term signal sequence\ninside 1 12\nTMhelix 13 35\noutside 36 246\nTMhelix 247 266\ninside 267 269

31497 GCF_001864975.1_ASM186497v1 Curtobacterium sp. MMLR14_002 Terrabacteria group; Actinobacteria; Actinobacteria; Micrococcales; Microbacteriaceae; Curtobacterium MFNRTKSKKGRRSPMATVSLIVVGLMTTGGAYALFSSTANADDSTNTVAASSQSQVNEGE WP_071403834.1 MULTISPECIES: cystathionine beta-lyase [Curtobacterium] Length: 269\nNumber of predicted TMHs: 2\nExp number of AAs in TMHs: 43.48885\nExp number, first 60 AAs: 22.05574\nTotal prob of N-in: 0.88828\nPOSSIBLE N-term signal sequence\ninside 1 12\nTMhelix 13 35\noutside 36 246\nTMhelix 247 266\ninside 267 269

31498 GCF_001864985.1_ASM186498v1 Curtobacterium sp. MMLR14_006 Terrabacteria group; Actinobacteria; Actinobacteria; Micrococcales; Microbacteriaceae; Curtobacterium MFNRTKSKKGRRSPMATVSLIVVGLMTTGGAYALFSSSANADDSTSTIAATSQSKVNEGQ WP_071255400.1 MULTISPECIES: cystathionine beta-lyase [Curtobacterium] Length: 269\nNumber of predicted TMHs: 2\nExp number of AAs in TMHs: 43.4648\nExp number, first 60 AAs: 22.05081\nTotal prob of N-in: 0.93134\nPOSSIBLE N-term signal sequence\ninside 1 12\nTMhelix 13 35\noutside 36 246\nTMhelix 247 266\ninside 267 269

31499 GCF_001864985.1_ASM186498v1 Curtobacterium sp. MMLR14_006 Terrabacteria group; Actinobacteria; Actinobacteria; Micrococcales; Microbacteriaceae; Curtobacterium MLPVHQGTAVPRRRPALRLVAMLGATLLAGAGLTATTAVVASEPASAVDATLDGALLCNQ WP_083404666.1 MULTISPECIES: hypothetical protein [Curtobacterium] Length: 1826\nNumber of predicted TMHs: 2\nExp number of AAs in TMHs: 43.4607999999999\nExp number, first 60 AAs: 21.23016\nTotal prob of N-in: 0.93821\nPOSSIBLE N-term signal sequence\ninside 1 18\nTMhelix 19 41\noutside 42 1784\nTMhelix 1785 1807\ninside 1808 1826

31500 GCF_001865015.1_ASM186501v1 Curtobacterium sp. MMLR14_010 Terrabacteria group; Actinobacteria; Actinobacteria; Micrococcales; Microbacteriaceae; Curtobacterium MFNSSKSKKQGRRSPVATVSLIVVGLMTTGGAYALFSSTASADDNVSTVAASSQSQVNEG WP_071275399.1 cystathionine beta-lyase [Curtobacterium sp. MMLR14_010] Length: 270\nNumber of predicted TMHs: 2\nExp number of AAs in TMHs: 43.26206\nExp number, first 60 AAs: 21.40503\nTotal prob of N-in: 0.58219\nPOSSIBLE N-term signal sequence\noutside 1 14\nTMhelix 15 37\ninside 38 247\nTMhelix 248 267\noutside 268 270

31501 GCF_001865135.1_ASM186513v1 Curtobacterium sp. MMLR14_014 Terrabacteria group; Actinobacteria; Actinobacteria; Micrococcales; Microbacteriaceae; Curtobacterium MFNRTKSKKGRRSPMATVSLIVVGLMTTGGAYALFSSTANADDSTNTVAASSQSQVNEGE WP_071403834.1 MULTISPECIES: cystathionine beta-lyase [Curtobacterium] Length: 269\nNumber of predicted TMHs: 2\nExp number of AAs in TMHs: 43.48885\nExp number, first 60 AAs: 22.05574\nTotal prob of N-in: 0.88828\nPOSSIBLE N-term signal sequence\ninside 1 12\nTMhelix 13 35\noutside 36 246\nTMhelix 247 266\ninside 267 269

31502 GCF_000772085.1_ASM77208v1 Curtobacterium sp. MR_MD2014 Terrabacteria group; Actinobacteria; Actinobacteria; Micrococcales; Microbacteriaceae; Curtobacterium MFTRSKSKKGRRSPMATVSLIVVGLMTTGGAYALFSSTANADDSTTSAVSSQSKVNEGEK WP_066654316.1 cystathionine beta-lyase [Curtobacterium sp. MR_MD2014] Length: 267\nNumber of predicted TMHs:

2\nExp number of AAs in TMHs: 43.44778\nExp number, first 60 AAs: 21.99067\nTotal prob of N-in: 0.87547\nPOSSIBLE N-term signal sequence\ninside 1 12\nTMhelix 13 35\noutside 36 244\nTMhelix 245 264\ninside 265 267

31503 GCF_000772085.1_ASM77208v1 Curtobacterium sp. MR_MD2014 Terrabacteria group; Actinobacteria; Actinobacteria; Micrococcales; Microbacteriaceae; Curtobacterium MLTTTATPRRRLLLQATATLALLLLVGFTVLMHTMAGHTADHEHPMAAPAAAAEHGHVAA WP_066651479.1 MULTISPECIES: hypothetical protein [Curtobacterium] Length: 151\nNumber of predicted TMHs: 2\nExp number of AAs in TMHs: 44.7324500000001\nExp number, first 60 AAs: 22.08752\nTotal prob of N-in: 0.99588\nPOSSIBLE N-term signal sequence\ninside 1 11\nTMhelix 12 34\noutside 35 87\nTMhelix 88 110\ninside 111 151

31504 GCF_000772085.1_ASM77208v1 Curtobacterium sp. MR_MD2014 Terrabacteria group; Actinobacteria; Actinobacteria; Micrococcales; Microbacteriaceae; Curtobacterium MHRTNPQHVRRRRVLTTTALGLTTVVLGSTAVAAAVVPAPASASTVAEHTAKASAALRGA WP_066652216.1 hypothetical protein [Curtobacterium sp. MR_MD2014] Length: 1049\nNumber of predicted TMHs: 2\nExp number of AAs in TMHs: 44.19122\nExp number, first 60 AAs: 21.83469\nTotal prob of N-in: 0.98161\nPOSSIBLE N-term signal sequence\ninside 1 12\nTMhelix 13 35\noutside 36 1012\nTMhelix 1013 1035\ninside 1036 1049

31505 GCF_000772085.1_ASM77208v1 Curtobacterium sp. MR_MD2014 Terrabacteria group; Actinobacteria; Actinobacteria; Micrococcales; Microbacteriaceae; Curtobacterium MTGVVPVQDSAAVVSTAAARRRGVLHRLRAVVVAVAGVVGVLCVAWWVVATTTGLSLVVV WP_066656173.1 signal peptidase I [Curtobacterium sp. MR_MD2014] Length: 195\nNumber of predicted TMHs: 2\nExp number of AAs in TMHs: 49.70189\nExp number, first 60 AAs: 24.79103\nTotal prob of N-in: 0.91443\nPOSSIBLE N-term signal sequence\ninside 1 27\nTMhelix 28 50\noutside 51 159\nTMhelix 160 182\ninside 183 195

31506 GCF_001705035.1_ASM170503v1 Curtobacterium sp. UCD-KPL2560 Terrabacteria group; Actinobacteria; Actinobacteria; Micrococcales; Microbacteriaceae; Curtobacterium MLTTTASPRRRLLLQVTATLALLLLVGFTVLMHTMAGHTAGHEHPMAAPAATTEHTHAVG WP_065961555.1 hypothetical protein [Curtobacterium sp. UCD-KPL2560] Length: 151\nNumber of predicted TMHs: 2\nExp number of AAs in TMHs: 45.37742\nExp number, first 60 AAs: 22.15848\nTotal prob of N-in: 0.99656\nPOSSIBLE N-term signal sequence\ninside 1 11\nTMhelix 12 34\noutside 35 87\nTMhelix 88 110\ninside 111 151

31507 GCF_001705035.1_ASM170503v1 Curtobacterium sp. UCD-KPL2560 Terrabacteria group; Actinobacteria; Actinobacteria; Micrococcales; Microbacteriaceae; Curtobacterium MKPLISSPGVPSTSSPSRPGRRPPTAARRRRRRPVLSGVGLATALGLLLAPALLASSSSA WP_065963828.1 hypothetical protein [Curtobacterium sp. UCD-KPL2560] Length: 668\nNumber of predicted TMHs: 2\nExp number of AAs in TMHs: 43.2105899999999\nExp number, first 60 AAs: 21.41534\nTotal prob of N-in: 0.97557\nPOSSIBLE N-term signal sequence\ninside 1 34\nTMhelix 35 57\noutside 58 628\nTMhelix 629 651\ninside 652 668

31508 GCF_001705035.1_ASM170503v1 Curtobacterium sp. UCD-KPL2560 Terrabacteria group; Actinobacteria; Actinobacteria; Micrococcales; Microbacteriaceae; Curtobacterium MGTLVPTRATERRRTTLLALTVSTLAVLLAVLVAPGPASASTTTPSDARAAVVRSIERGT WP_065959450.1 hypothetical protein [Curtobacterium sp. UCD-KPL2560] Length: 221\nNumber of predicted TMHs: 2\nExp number of AAs in TMHs: 49.84262\nExp number, first 60 AAs: 21.52824\nTotal prob of N-in: 0.97562\nPOSSIBLE N-term signal sequence\ninside 1 15\nTMhelix 16 38\noutside 39 155\nTMhelix 156 178\ninside 179 221

31509 GCF_001705035.1_ASM170503v1 Curtobacterium sp. UCD-KPL2560 Terrabacteria group; Actinobacteria; Actinobacteria; Micrococcales; Microbacteriaceae; Curtobacterium MNTHTSPRRRRLVGTATLATAGVLVALSAPSAALAADGDVTIDLYNVNDFHGRIAQSISV WP_065961321.1 MULTISPECIES: hypothetical protein [Curtobacterium] Length: 775\nNumber of predicted TMHs: 2\nExp number of AAs in TMHs: 42.7160599999999\nExp number, first 60 AAs: 22.07275\nTotal prob of N-in: 0.97551\nPOSSIBLE N-term signal sequence\ninside 1 11\nTMhelix 12 34\noutside 35 738\nTMhelix 739 761\ninside 762 775

31510 GCF_001705035.1_ASM170503v1 Curtobacterium sp. UCD-KPL2560 Terrabacteria group; Actinobacteria; Actinobacteria; Micrococcales; Microbacteriaceae; Curtobacterium MSPTHRVRPRRRALRVLAATTITTIIAGGLALLSPLTASAMADLPVSGLYATVNGDSTIY WP_065961365.1 hypothetical protein [Curtobacterium sp. UCD-KPL2560] Length: 1257\nNumber of predicted TMHs: 2\nExp number of AAs in TMHs: 47.7312199999999\nExp number, first 60 AAs: 22.20368\nTotal prob of N-in: 0.99887\nPOSSIBLE N-term signal sequence\ninside 1 12\nTMhelix 13 35\noutside 36 1226\nTMhelix 1227 1249\ninside 1250 1257

31511	GCF_001705035.1_ASM170503v1	Curtobacterium sp. UCD-KPL2560	Terrabacteria group; Actinobacteria; Actinobacteria; Micrococcales; Microbacteriaceae; Curtobacterium
MFTRTKTKKGRRSPMATVSLIVVGLMTTGGAYALFSSTANADDSTTATAASSQSQVNEGQ	WP_065962238.1	MULTISPECIES: cystathionine beta-lyase [Curtobacterium]	Length: 269\nNumber of predicted TMHs: 2\nExp number of AAs in TMHs: 43.48172\nExp number, first 60 AAs: 22.0798\nTotal prob of N-in: 0.93547\nPOSSIBLE N-term signal sequence\ninside 1 12\nTMhelix 13 35\noutside 36 246\nTMhelix 247 266\ninside 267 269

31512	GCF_000686685.1_ASM68668v1	Curtobacterium sp. UNCCL17	Terrabacteria group; Actinobacteria; Actinobacteria; Micrococcales; Microbacteriaceae; Curtobacterium
MFNRTKSKKGRRSPMATVSLIVVGLMTTGGAYALFSSTANADDSTNTVAASSQSQVNEGE	WP_027465616.1	cystathionine beta-lyase [Curtobacterium sp. UNCCL17]	Length: 269\nNumber of predicted TMHs: 2\nExp number of AAs in TMHs: 43.4895\nExp number, first 60 AAs: 22.05571\nTotal prob of N-in: 0.88824\nPOSSIBLE N-term signal sequence\ninside 1 12\nTMhelix 13 35\noutside 36 246\nTMhelix 247 266\ninside 267 269

31513	GCF_000686685.1_ASM68668v1	Curtobacterium sp. UNCCL17	Terrabacteria group; Actinobacteria; Actinobacteria; Micrococcales; Microbacteriaceae; Curtobacterium
MQPDDPPPLVLIEESMLTATASPRRRLLLQVTATLALLLLVGFTVLMHTMAGHTAGHEHP	WP_027465407.1	hypothetical protein [Curtobacterium sp. UNCCL17]	Length: 166\nNumber of predicted TMHs: 2\nExp number of AAs in TMHs: 44.21329\nExp number, first 60 AAs: 21.41719\nTotal prob of N-in: 0.38614\nPOSSIBLE N-term signal sequence\ninside 1 26\nTMhelix 27 49\noutside 50 102\nTMhelix 103 125\ninside 126 166

31514	GCF_000686685.1_ASM68668v1	Curtobacterium sp. UNCCL17	Terrabacteria group; Actinobacteria; Actinobacteria; Micrococcales; Microbacteriaceae; Curtobacterium
MHHHTAAQRRRRGALTIATAMAAAAAVAGGVAAGLPASAADGYTTNASIDELHFTSETVT	WP_081827442.1	hypothetical protein [Curtobacterium sp. UNCCL17]	Length: 444\nNumber of predicted TMHs: 2\nExp number of AAs in TMHs: 44.86012\nExp number, first 60 AAs: 22.64576\nTotal prob of N-in: 0.99818\nPOSSIBLE N-term signal sequence\ninside 1 12\nTMhelix 13 35\noutside 36 414\nTMhelix 415 437\ninside 438 444

31515	GCF_001571045.1_ASM157104v1	Curtobacterium ammoniigenes NBRC 101786	Terrabacteria group; Actinobacteria; Actinobacteria; Micrococcales; Microbacteriaceae; Curtobacterium; Curtobacterium ammoniigenes
MFTRTKNDSSTGTKKGRRSPLATATLILVGLLTTGGAYALFTSSATADTAQSSAVALHSQ	WP_066518951.1	cystathionine beta-lyase [Curtobacterium ammoniigenes]	Length: 275\nNumber of predicted TMHs: 2\nExp number of AAs in TMHs: 43.39049\nExp number, first 60 AAs: 22.20931\nTotal prob of N-in: 0.92215\nPOSSIBLE N-term signal sequence\ninside 1 19\nTMhelix 20 42\noutside 43 252\nTMhelix 253 272\ninside 273 275

31516	GCF_000349565.1_Curtobacterium_flaccumfaciens_UCD-AKU	Curtobacterium flaccumfaciens UCD-AKU	Terrabacteria group; Actinobacteria; Actinobacteria; Micrococcales; Microbacteriaceae; Curtobacterium; Curtobacterium flaccumfaciens	MFNRTKSKKGRRSPMATVSLIVVGLMTTGGAYALFSSTANADDSTSTLATSSQSQVNEGE	WP_042537210.1	MULTISPECIES: cystathionine beta-lyase [Curtobacterium]	Length: 269\nNumber of predicted TMHs: 2\nExp number of AAs in TMHs: 43.41552\nExp number, first 60 AAs: 22.0168\nTotal prob of N-in: 0.91655\nPOSSIBLE N-term signal sequence\ninside 1 12\nTMhelix 13 35\noutside 36 246\nTMhelix 247 266\ninside 267 269

31517	GCF_900163665.1_ASM90016366v1	Frigoribacterium sp. JB110	Terrabacteria group; Actinobacteria; Actinobacteria; Micrococcales; Microbacteriaceae; Frigoribacterium
MHSRTTRRPLLVGGAAALGALALTFAGASAASAHVTVGSDTDEAGSYAILTFSVPHGCDG	WP_087056659.1	hypothetical protein [Frigoribacterium sp. JB110]	Length: 243\nNumber of predicted TMHs: 2\nExp number of AAs in TMHs: 43.75352\nExp number, first 60 AAs: 21.75633\nTotal prob of N-in: 0.96757\nPOSSIBLE N-term signal sequence\ninside 1 8\nTMhelix 9 31\noutside 32 214\nTMhelix 215 237\ninside 238 243

31518	GCF_900163665.1_ASM90016366v1	Frigoribacterium sp. JB110	Terrabacteria group; Actinobacteria; Actinobacteria; Micrococcales; Microbacteriaceae; Frigoribacterium
MALFKKGRRTGRRSPLATAALIGIGLILSGGIYAGASAAVAATSETAQVESALTVEDGQK	WP_087054650.1	cystathionine beta-lyase [Frigoribacterium sp. JB110]	Length: 267\nNumber of predicted TMHs: 2\nExp number of AAs in TMHs: 42.83645\nExp number, first 60 AAs: 22.50593\nTotal prob of N-in: 0.95244\nPOSSIBLE N-term signal sequence\ninside 1 20\nTMhelix 21 43\noutside 44 244\nTMhelix 245 264\ninside 265 267

31519	GCF_001423665.1_Leaf164	Frigoribacterium sp. Leaf164	Terrabacteria group; Actinobacteria; Actinobacteria; Micrococcales; Microbacteriaceae; Frigoribacterium
MRKAPVIRSSGRRRPLYPTALLVTGLVAAGAIALPATAASAADGPVVVSDSEFTDGSYVV	WP_082514234.1	hypothetical protein [Frigoribacterium sp. Leaf164]	Length: 1104\nNumber of predicted TMHs: 2\nExp number of AAs in TMHs:

42.1063\nExp number, first 60 AAs: 21.23607\nTotal prob of N-in: 0.97612\nPOSSIBLE N-term signal sequence\ninside 1 19\nTMhelix 20 42\noutside 43 1065\nTMhelix 1066 1088\ninside 1089 1104

31520 GCF_001423665.1_Leaf164 Frigoribacterium sp. Leaf164 Terrabacteria group; Actinobacteria; Actinobacteria; Micrococcales; Microbacteriaceae; Frigoribacterium
MPSSPRRRTVATTSIVTAGLLAALTGLGTVAPAAAAPTAGEVAAAQTVTEGDARFQVLSP WP_082514169.1 alpha-xylosidase [Frigoribacterium sp. Leaf164] Length: 1166\nNumber of predicted TMHs: 2\nExp number of AAs in TMHs: 39.94974\nExp number, first 60 AAs: 18.96089\nTotal prob of N-in: 0.84045\nPOSSIBLE N-term signal sequence\ninside 1 12\nTMhelix 13 35\noutside 36 1135\nTMhelix 1136 1155\ninside 1156 1166

31521 GCF_001423665.1_Leaf164 Frigoribacterium sp. Leaf164 Terrabacteria group; Actinobacteria; Actinobacteria; Micrococcales; Microbacteriaceae; Frigoribacterium
MRPRHPIAPPAETGAPRRRRRAALSLVAVATLGAGLVAAVPAQAAPTTIDILGINDFHGR WP_056051994.1 bifunctional metallophosphatase/5-nucleotidase [Frigoribacterium sp. Leaf164] Length: 757\nNumber of predicted TMHs: 2\nExp number of AAs in TMHs: 42.31268\nExp number, first 60 AAs: 21.68068\nTotal prob of N-in: 0.97543\nPOSSIBLE N-term signal sequence\ninside 1 22\nTMhelix 23 45\noutside 46 722\nTMhelix 723 745\ninside 746 757

31522 GCF_001423665.1_Leaf164 Frigoribacterium sp. Leaf164 Terrabacteria group; Actinobacteria; Actinobacteria; Micrococcales; Microbacteriaceae; Frigoribacterium
MFTRSSSSRRTAKTGRRSPVATASLLLVGLLTTGGAYALFSSTATAEESAPAAASQQSVD WP_056055014.1 cystathionine beta-lyase [Frigoribacterium sp. Leaf164] Length: 271\nNumber of predicted TMHs: 2\nExp number of AAs in TMHs: 42.02642\nExp number, first 60 AAs: 20.95733\nTotal prob of N-in: 0.97350\nPOSSIBLE N-term signal sequence\ninside 1 19\nTMhelix 20 42\noutside 43 248\nTMhelix 249 268\ninside 269 271

31523 GCF_001423665.1_Leaf164 Frigoribacterium sp. Leaf164 Terrabacteria group; Actinobacteria; Actinobacteria; Micrococcales; Microbacteriaceae; Frigoribacterium
MTALRRRTALSCASAVLVTAVVAGGIALTGAGTASAASPEPSSSSSSEPTSERLDASGVA WP_056056642.1 hypothetical protein [Frigoribacterium sp. Leaf164] Length: 648\nNumber of predicted TMHs: 2\nExp number of AAs in TMHs: 42.7046099999999\nExp number, first 60 AAs: 21.65296\nTotal prob of N-in: 0.97179\nPOSSIBLE N-term signal sequence\ninside 1 11\nTMhelix 12 34\noutside 35 587\nTMhelix 588 610\ninside 611 648

31524 GCF_001424285.1_Leaf172 Frigoribacterium sp. Leaf172 Terrabacteria group; Actinobacteria; Actinobacteria; Micrococcales; Microbacteriaceae; Frigoribacterium
MTPTTRRRRAALITGAVAISASLALAPAVASAATPSDVAENGGAARSIDDSTQVLLDSVK WP_055815126.1 alkaline phosphatase [Frigoribacterium sp. Leaf172] Length: 659\nNumber of predicted TMHs: 2\nExp number of AAs in TMHs: 40.76202\nExp number, first 60 AAs: 20.66013\nTotal prob of N-in: 0.93207\nPOSSIBLE N-term signal sequence\ninside 1 11\nTMhelix 12 34\noutside 35 620\nTMhelix 621 640\ninside 641 659

31525 GCF_001424285.1_Leaf172 Frigoribacterium sp. Leaf172 Terrabacteria group; Actinobacteria; Actinobacteria; Micrococcales; Microbacteriaceae; Frigoribacterium
MFSRTSSSRRKTGRRSPLATASLILVGLLTTGGAYALFSSTATADESAAPVASQQSIDEG WP_055813479.1 cystathionine beta-lyase [Frigoribacterium sp. Leaf172] Length: 269\nNumber of predicted TMHs: 2\nExp number of AAs in TMHs: 42.28122\nExp number, first 60 AAs: 21.12389\nTotal prob of N-in: 0.97673\nPOSSIBLE N-term signal sequence\ninside 1 20\nTMhelix 21 43\noutside 44 246\nTMhelix 247 266\ninside 267 269

31526 GCF_001423905.1_Leaf186 Frigoribacterium sp. Leaf186 Terrabacteria group; Actinobacteria; Actinobacteria; Micrococcales; Microbacteriaceae; Frigoribacterium
MSLLRASVRPRRRALSLAAFATASLLAAVAVTQAVAPAPAQAAPGLSCDQNTLYATGGNG WP_056261770.1 hypothetical protein [Frigoribacterium sp. Leaf186] Length: 1810\nNumber of predicted TMHs: 2\nExp number of AAs in TMHs: 44.2248699999999\nExp number, first 60 AAs: 20.55877\nTotal prob of N-in: 0.92131\nPOSSIBLE N-term signal sequence\ninside 1 12\nTMhelix 13 35\noutside 36 1768\nTMhelix 1769 1791\ninside 1792 1810

31527 GCF_001423905.1_Leaf186 Frigoribacterium sp. Leaf186 Terrabacteria group; Actinobacteria; Actinobacteria; Micrococcales; Microbacteriaceae; Frigoribacterium
MFSRTSSSRRTKKAGRRGPVATASLLLVGLLTTGGAYALFSSTASADESSATATSQQSID WP_056256108.1 cystathionine beta-lyase [Frigoribacterium sp. Leaf186] Length: 271\nNumber of predicted TMHs: 2\nExp number of AAs in TMHs: 42.27945\nExp number, first 60 AAs: 21.1093\nTotal prob of N-in: 0.98128\nPOSSIBLE N-term signal sequence\ninside 1 19\nTMhelix 20 42\noutside 43 248\nTMhelix 249 268\ninside 269 271

31528 GCF_001422145.1_Leaf254 Frigoribacterium sp. Leaf254 Terrabacteria group; Actinobacteria; Actinobacteria; Micrococcales; Microbacteriaceae; Frigoribacterium
MAAAAHRRDRRAGGRVRGGRTPPPGRRRDRGGAAVSGRRAAVREKGLLHALGLGLSAGLF WP_082454302.1 MULTISPECIES: signal peptidase I [Frigoribacterium] Length: 228\nNumber of predicted TMHs: 2\nExp number of AAs in TMHs: 46.26691\nExp number, first 60 AAs: 11.79273\nTotal prob of N-in: 0.99904\nPOSSIBLE N-term signal sequence\ninside 1 50\nTMhelix 51 73\noutside 74 188\nTMhelix 189 211\ninside 212 228

31529 GCF_001422145.1_Leaf254 Frigoribacterium sp. Leaf254 Terrabacteria group; Actinobacteria; Actinobacteria; Micrococcales; Microbacteriaceae; Frigoribacterium
MFTRTSSSRRTKKAGRRGPVATASLLLVGLLTTGGAYALFSSTASADESSATATSQQSID WP_054146200.1 MULTISPECIES: cystathionine beta-lyase [Frigoribacterium] Length: 271\nNumber of predicted TMHs: 2\nExp number of AAs in TMHs: 42.35339\nExp number, first 60 AAs: 21.18372\nTotal prob of N-in: 0.98724\nPOSSIBLE N-term signal sequence\ninside 1 19\nTMhelix 20 42\noutside 43 248\nTMhelix 249 268\ninside 269 271

31530 GCF_001422145.1_Leaf254 Frigoribacterium sp. Leaf254 Terrabacteria group; Actinobacteria; Actinobacteria; Micrococcales; Microbacteriaceae; Frigoribacterium
MSKNTPNRRRAALTTGVVATAAVAALALSPVAAFAATPANVAQNGGAARSIDDSTQLLLD WP_082454812.1 MULTISPECIES: alkaline phosphatase [Frigoribacterium] Length: 644\nNumber of predicted TMHs: 2\nExp number of AAs in TMHs: 41.9519399999999\nExp number, first 60 AAs: 22.56061\nTotal prob of N-in: 0.99115\nPOSSIBLE N-term signal sequence\ninside 1 12\nTMhelix 13 35\noutside 36 609\nTMhelix 610 629\ninside 630 644

31531 GCF_001422145.1_Leaf254 Frigoribacterium sp. Leaf254 Terrabacteria group; Actinobacteria; Actinobacteria; Micrococcales; Microbacteriaceae; Frigoribacterium
MSLLRASVRPRRRALSFAAFATASLLAAVAVTQAVSPAPAQAASALSCDQNTLYATGGNG WP_055974413.1 MULTISPECIES: hypothetical protein [Frigoribacterium] Length: 1812\nNumber of predicted TMHs: 2\nExp number of AAs in TMHs: 43.56745\nExp number, first 60 AAs: 20.61268\nTotal prob of N-in: 0.92525\nPOSSIBLE N-term signal sequence\ninside 1 12\nTMhelix 13 35\noutside 36 1769\nTMhelix 1770 1792\ninside 1793 1812

31532 GCF_001422285.1_Leaf263 Frigoribacterium sp. Leaf263 Terrabacteria group; Actinobacteria; Actinobacteria; Micrococcales; Microbacteriaceae; Frigoribacterium
MFSRTSSSRRKTGRRSPLATASLILVGLLTTGGAYALFSSTATADESAAPVASQQSIDEG WP_055797696.1 cystathionine beta-lyase [Frigoribacterium sp. Leaf263] Length: 269\nNumber of predicted TMHs: 2\nExp number of AAs in TMHs: 42.27812\nExp number, first 60 AAs: 21.12385\nTotal prob of N-in: 0.97673\nPOSSIBLE N-term signal sequence\ninside 1 20\nTMhelix 21 43\noutside 44 246\nTMhelix 247 266\ninside 267 269

31533 GCF_001422285.1_Leaf263 Frigoribacterium sp. Leaf263 Terrabacteria group; Actinobacteria; Actinobacteria; Micrococcales; Microbacteriaceae; Frigoribacterium
MTPTTRRRRAALITGAVAISASLALAPAVASAATPSDVAENGGAARSIDDSTQVLLDSVK WP_055795172.1 alkaline phosphatase [Frigoribacterium sp. Leaf263] Length: 659\nNumber of predicted TMHs: 2\nExp number of AAs in TMHs: 40.76202\nExp number, first 60 AAs: 20.66013\nTotal prob of N-in: 0.93207\nPOSSIBLE N-term signal sequence\ninside 1 11\nTMhelix 12 34\noutside 35 620\nTMhelix 621 640\ninside 641 659

31534 GCF_001424645.1_Leaf415 Frigoribacterium sp. Leaf415 Terrabacteria group; Actinobacteria; Actinobacteria; Micrococcales; Microbacteriaceae; Frigoribacterium
MFTRTSSSRRTKKAGRRGPVATASLLLVGLLTTGGAYALFSSTASADESSATATSQQSID WP_054146200.1 MULTISPECIES: cystathionine beta-lyase [Frigoribacterium] Length: 271\nNumber of predicted TMHs: 2\nExp number of AAs in TMHs: 42.35339\nExp number, first 60 AAs: 21.18372\nTotal prob of N-in: 0.98724\nPOSSIBLE N-term signal sequence\ninside 1 19\nTMhelix 20 42\noutside 43 248\nTMhelix 249 268\ninside 269 271

31535 GCF_001424645.1_Leaf415 Frigoribacterium sp. Leaf415 Terrabacteria group; Actinobacteria; Actinobacteria; Micrococcales; Microbacteriaceae; Frigoribacterium
MSLLRASVRPRRRALSFAAFATASLLAAVAVTQAVSPAPAQAASALSCDQNTLYATGGNG WP_055974413.1 MULTISPECIES: hypothetical protein [Frigoribacterium] Length: 1812\nNumber of predicted TMHs: 2\nExp number of AAs in TMHs: 43.56745\nExp number, first 60 AAs: 20.61268\nTotal prob of N-in: 0.92525\nPOSSIBLE N-term signal sequence\ninside 1 12\nTMhelix 13 35\noutside 36 1769\nTMhelix 1770 1792\ninside 1793 1812

31536 GCF_001424645.1_Leaf415 Frigoribacterium sp. Leaf415 Terrabacteria group; Actinobacteria; Actinobacteria; Micrococcales; Microbacteriaceae; Frigoribacterium
MAAAAHRRDRRAGGRVRGGRTPPPGRRRDRGGAAVSGRRAAVREKGLLHALGLGLSAGLF WP_082454302.1 MULTISPECIES: signal peptidase I [Frigoribacterium] Length: 228\nNumber of predicted TMHs: 2\nExp

number of AAs in TMHs: 46.26691\nExp number, first 60 AAs:  11.79273\nTotal prob of N-in:   0.99904\nPOSSIBLE N-term signal sequence\ninside   1   50\nTMhelix   51   73\noutside   74   188\nTMhelix   189   211\ninside   212   228

31537	GCF_001424645.1_Leaf415	Frigoribacterium sp. Leaf415	Terrabacteria group; Actinobacteria; Actinobacteria; Micrococcales; Microbacteriaceae; Frigoribacterium
MSKNTPNRRRAALTTGVVATAAVAALALSPVAAFAATPANVAQNGGAARSIDDSTQLLLD	WP_082454812.1
MULTISPECIES: alkaline phosphatase [Frigoribacterium]	Length: 644\nNumber of predicted TMHs:  2\nExp number of AAs in TMHs: 41.9519399999999\nExp number, first 60 AAs:  22.56061\nTotal prob of N-in:   0.99115\nPOSSIBLE N-term signal sequence\ninside   1   12\nTMhelix   13   35\noutside   36   609\nTMhelix   610   629\ninside   630   644

31538	GCF_001421865.1_Leaf44	Frigoribacterium sp. Leaf44	Terrabacteria group; Actinobacteria; Actinobacteria; Micrococcales; Microbacteriaceae; Frigoribacterium
MSKNTPNRRRAALTTGVVATAAVAALALTPVAAFAATPANVTQNGGAARSIDDSTQLLLD	WP_082452474.1
alkaline phosphatase [Frigoribacterium sp. Leaf44]	Length: 644\nNumber of predicted TMHs:  2\nExp number of AAs in TMHs: 42.44\nExp number, first 60 AAs:  22.62763\nTotal prob of N-in:   0.99445\nPOSSIBLE N-term signal sequence\ninside   1   12\nTMhelix   13   35\noutside   36   609\nTMhelix   610   629\ninside   630   644

31539	GCF_001421865.1_Leaf44	Frigoribacterium sp. Leaf44	Terrabacteria group; Actinobacteria; Actinobacteria; Micrococcales; Microbacteriaceae; Frigoribacterium
MSGRRAAVREKGLLHALGLGLSAGLFVLVLGLATVLIVVPKVAGATPLTVLTSSMEPRLP	WP_082452258.1	signal peptidase I [Frigoribacterium sp. Leaf44]	Length: 194\nNumber of predicted TMHs:  2\nExp number of AAs in TMHs: 46.66328\nExp number, first 60 AAs:  23.79138\nTotal prob of N-in:   0.99831\nPOSSIBLE N-term signal sequence\ninside   1   12\nTMhelix   13   35\noutside   36   154\nTMhelix   155   177\ninside   178   194

31540	GCF_001421865.1_Leaf44	Frigoribacterium sp. Leaf44	Terrabacteria group; Actinobacteria; Actinobacteria; Micrococcales; Microbacteriaceae; Frigoribacterium
MFTRTSSSRRNKKAGRRGPVATASLLLVGLLTTGGAYALFSSTASADESSATATSQQSID	WP_043594940.1	MULTISPECIES: cystathionine beta-lyase [Frigoribacterium]	Length: 271\nNumber of predicted TMHs:  2\nExp number of AAs in TMHs: 42.35097\nExp number, first 60 AAs:  21.18027\nTotal prob of N-in:   0.98669\nPOSSIBLE N-term signal sequence\ninside   1   19\nTMhelix   20   42\noutside   43   248\nTMhelix   249   268\ninside   269   271

31541	GCF_001421865.1_Leaf44	Frigoribacterium sp. Leaf44	Terrabacteria group; Actinobacteria; Actinobacteria; Micrococcales; Microbacteriaceae; Frigoribacterium
MSLLRASVRPRRRALSFAAFATASVLAAVAVTQAVAPAPAHAAAALSCDQNTIYATAGQG	WP_056229019.1
hypothetical protein [Frigoribacterium sp. Leaf44]	Length: 1810\nNumber of predicted TMHs:  2\nExp number of AAs in TMHs: 43.8247599999999\nExp number, first 60 AAs:  20.8631\nTotal prob of N-in:   0.93579\nPOSSIBLE N-term signal sequence\ninside   1   12\nTMhelix   13   35\noutside   36  1767\nTMhelix   1768  1790\ninside   1791  1810

31542	GCF_001421165.1_Leaf8	Frigoribacterium sp. Leaf8	Terrabacteria group; Actinobacteria; Actinobacteria; Micrococcales; Microbacteriaceae; Frigoribacterium
MSLLRASVRPRRRALSFAAFATASLLAAVAVTQAVSPAPAQAAAALSCDQNTIYATAGQG	WP_056049027.1
hypothetical protein [Frigoribacterium sp. Leaf8] Length: 1811\nNumber of predicted TMHs:  2\nExp number of AAs in TMHs: 43.61551\nExp number, first 60 AAs:  20.6871\nTotal prob of N-in:   0.92761\nPOSSIBLE N-term signal sequence\ninside   1   12\nTMhelix   13   35\noutside   36  1768\nTMhelix   1769  1791\ninside   1792  1811

31543	GCF_001421165.1_Leaf8	Frigoribacterium sp. Leaf8	Terrabacteria group; Actinobacteria; Actinobacteria; Micrococcales; Microbacteriaceae; Frigoribacterium
MFSRTSSSRRTKKAGRRGPVATASLLLVGLLTTGGAYALFSSTASADESSATATSQQSID	WP_056045057.1	cystathionine beta-lyase [Frigoribacterium sp. Leaf8]	Length: 271\nNumber of predicted TMHs:  2\nExp number of AAs in TMHs: 42.28455\nExp number, first 60 AAs:  21.10915\nTotal prob of N-in:   0.98120\nPOSSIBLE N-term signal sequence\ninside   1   19\nTMhelix   20   42\noutside   43   248\nTMhelix   249   268\ninside   269   271

31544	GCF_001421165.1_Leaf8	Frigoribacterium sp. Leaf8	Terrabacteria group; Actinobacteria; Actinobacteria; Micrococcales; Microbacteriaceae; Frigoribacterium
MLPLQPDDPPPLVLIEESMLATTASPRRRLLLQVTATLALLLLVGFTVLMHTMAGHTDGH	WP_056044687.1
hypothetical protein [Frigoribacterium sp. Leaf8] Length: 169\nNumber of predicted TMHs:  2\nExp number of AAs in TMHs: 44.55368\nExp number, first 60 AAs:  21.75008\nTotal prob of N-in:   0.70431\nPOSSIBLE N-term signal sequence\ninside   1   29\nTMhelix   30   52\noutside   53   105\nTMhelix   106   128\ninside   129   169

31545 GCF_001421165.1_Leaf8 Frigoribacterium sp. Leaf8 Terrabacteria group; Actinobacteria; Actinobacteria; Micrococcales; Microbacteriaceae; Frigoribacterium
MSGRRAAVREKGLVHALGLGLSAGLFVLVLGLATVLVVVPKVAGATPLTVLTSSMEPRLP WP_082460400.1 signal peptidase I [Frigoribacterium sp. Leaf8] Length: 194\nNumber of predicted TMHs: 2\nExp number of AAs in TMHs: 46.37938\nExp number, first 60 AAs: 23.52576\nTotal prob of N-in: 0.99838\nPOSSIBLE N-term signal sequence\ninside 1 12\nTMhelix 13 35\noutside 36 154\nTMhelix 155 177\ninside 178 194

31546 GCF_001864945.1_ASM186494v1 Frigoribacterium sp. MCBA15_019 Terrabacteria group; Actinobacteria; Actinobacteria; Micrococcales; Microbacteriaceae; Frigoribacterium
MFTRTSSSRRTKKAGRRGPVATASLLLVGLLTTGGAYALFSSTASADESSATATSQQSID WP_054146200.1 MULTISPECIES: cystathionine beta-lyase [Frigoribacterium] Length: 271\nNumber of predicted TMHs: 2\nExp number of AAs in TMHs: 42.35339\nExp number, first 60 AAs: 21.18372\nTotal prob of N-in: 0.98724\nPOSSIBLE N-term signal sequence\ninside 1 19\nTMhelix 20 42\noutside 43 248\nTMhelix 249 268\ninside 269 271

31547 GCF_001864945.1_ASM186494v1 Frigoribacterium sp. MCBA15_019 Terrabacteria group; Actinobacteria; Actinobacteria; Micrococcales; Microbacteriaceae; Frigoribacterium
MSGRRAAVREKGLLHALGLGLSAGLFVLVLGLATVLIVVPKVAGATPLTVLTSSMEPRLP WP_071254172.1 signal peptidase I [Frigoribacterium sp. MCBA15_019] Length: 202\nNumber of predicted TMHs: 2\nExp number of AAs in TMHs: 46.66883\nExp number, first 60 AAs: 23.79097\nTotal prob of N-in: 0.99828\nPOSSIBLE N-term signal sequence\ninside 1 12\nTMhelix 13 35\noutside 36 154\nTMhelix 155 177\ninside 178 202

31548 GCF_001864945.1_ASM186494v1 Frigoribacterium sp. MCBA15_019 Terrabacteria group; Actinobacteria; Actinobacteria; Micrococcales; Microbacteriaceae; Frigoribacterium
MSKNTPNRRRAALTTGVVATAAVAALALSPVAAFAATPANVAQNGGAARSIDDSTQLLLD WP_083396275.1 alkaline phosphatase [Frigoribacterium sp. MCBA15_019] Length: 644\nNumber of predicted TMHs: 2\nExp number of AAs in TMHs: 43.0790099999999\nExp number, first 60 AAs: 22.54006\nTotal prob of N-in: 0.98947\nPOSSIBLE N-term signal sequence\ninside 1 12\nTMhelix 13 35\noutside 36 609\nTMhelix 610 629\ninside 630 644

31549 GCF_000878135.1_ASM87813v1 Frigoribacterium sp. MEB024 Terrabacteria group; Actinobacteria; Actinobacteria; Micrococcales; Microbacteriaceae; Frigoribacterium
MSKNTPNRRRAALTTGVVATAAVAALALTPVAAFAATPANVAQNGGAARSIDDSTQLLLD WP_082059785.1 alkaline phosphatase [Frigoribacterium sp. MEB024] Length: 644\nNumber of predicted TMHs: 2\nExp number of AAs in TMHs: 42.22908\nExp number, first 60 AAs: 22.64461\nTotal prob of N-in: 0.99473\nPOSSIBLE N-term signal sequence\ninside 1 12\nTMhelix 13 35\noutside 36 609\nTMhelix 610 629\ninside 630 644

31550 GCF_000878135.1_ASM87813v1 Frigoribacterium sp. MEB024 Terrabacteria group; Actinobacteria; Actinobacteria; Micrococcales; Microbacteriaceae; Frigoribacterium
MSGRRAAVRDKGLLHALGLGLSAGLFVLVLGLAAVLIVVPKVAGATPLTVLTSSMEPRLP WP_082060114.1 signal peptidase I [Frigoribacterium sp. MEB024] Length: 194\nNumber of predicted TMHs: 2\nExp number of AAs in TMHs: 46.71599\nExp number, first 60 AAs: 23.83676\nTotal prob of N-in: 0.99759\nPOSSIBLE N-term signal sequence\ninside 1 12\nTMhelix 13 35\noutside 36 154\nTMhelix 155 177\ninside 178 194

31551 GCF_000878135.1_ASM87813v1 Frigoribacterium sp. MEB024 Terrabacteria group; Actinobacteria; Actinobacteria; Micrococcales; Microbacteriaceae; Frigoribacterium
MFTRTSSSRRNKKAGRRGPVATASLLLVGLLTTGGAYALFSSTASADESSATATSQQSID WP_043594940.1 MULTISPECIES: cystathionine beta-lyase [Frigoribacterium] Length: 271\nNumber of predicted TMHs: 2\nExp number of AAs in TMHs: 42.35097\nExp number, first 60 AAs: 21.18027\nTotal prob of N-in: 0.98669\nPOSSIBLE N-term signal sequence\ninside 1 19\nTMhelix 20 42\noutside 43 248\nTMhelix 249 268\ninside 269 271

31552 GCF_000878135.1_ASM87813v1 Frigoribacterium sp. MEB024 Terrabacteria group; Actinobacteria; Actinobacteria; Micrococcales; Microbacteriaceae; Frigoribacterium
MSLLRASVRPRRRALSFAAFATASVLAAVAVTQAVAPAPAHAAAALSCDQNTIYATAGQG WP_052505214.1 hypothetical protein [Frigoribacterium sp. MEB024] Length: 1811\nNumber of predicted TMHs: 2\nExp number of AAs in TMHs: 43.8271599999999\nExp number, first 60 AAs: 20.86269\nTotal prob of N-in: 0.93570\nPOSSIBLE N-term signal sequence\ninside 1 12\nTMhelix 13 35\noutside 36 1768\nTMhelix 1769 1791\ninside 1792 1811

31553 GCF_001297205.1_ASM129720v1 Frigoribacterium sp. RIT-PI-h Terrabacteria group; Actinobacteria; Actinobacteria; Micrococcales; Microbacteriaceae; Frigoribacterium
MAAAAHRGDRRAGGRVRGGRSPPPGRRRDRGGAAVSGRRAAVREKGLLHALGLGFSAGLF WP_082383249.1

signal peptidase I [Frigoribacterium sp. RIT-PI-h] Length: 228\nNumber of predicted TMHs: 2\nExp number of AAs in TMHs: 46.32862\nExp number, first 60 AAs: 11.71899\nTotal prob of N-in: 0.99706\nPOSSIBLE N-term signal sequence\ninside 1 50\nTMhelix 51 73\noutside 74 188\nTMhelix 189 211\ninside 212 228

31554 GCF_001297205.1_ASM129720v1 Frigoribacterium sp. RIT-PI-h Terrabacteria group; Actinobacteria; Actinobacteria; Micrococcales; Microbacteriaceae; Frigoribacterium MFTRTSSSRRTKKAGRRGPVATASLLLVGLLTTGGAYALFSSTASADESSATATSQQSID WP_054146200.1 MULTISPECIES: cystathionine beta-lyase [Frigoribacterium] Length: 271\nNumber of predicted TMHs: 2\nExp number of AAs in TMHs: 42.35339\nExp number, first 60 AAs: 21.18372\nTotal prob of N-in: 0.98724\nPOSSIBLE N-term signal sequence\ninside 1 19\nTMhelix 20 42\noutside 43 248\nTMhelix 249 268\ninside 269 271

31555 GCF_001423105.1_Leaf304 Frondihabitans sp. Leaf304 Terrabacteria group; Actinobacteria; Actinobacteria; Micrococcales; Microbacteriaceae; Frondihabitans MKISPSAKLASSATKGARKSVGAKSKRTGRRSPLATVSLIVVGLLTTGGAYALFTSTATA WP_084815851.1 cystathionine beta-lyase [Frondihabitans sp. Leaf304]Length: 284\nNumber of predicted TMHs: 2\nExp number of AAs in TMHs: 42.06886\nExp number, first 60 AAs: 21.51096\nTotal prob of N-in: 0.90066\nPOSSIBLE N-term signal sequence\ninside 1 32\nTMhelix 33 55\noutside 56 261\nTMhelix 262 281\ninside 282 284

31556 GCF_001423105.1_Leaf304 Frondihabitans sp. Leaf304 Terrabacteria group; Actinobacteria; Actinobacteria; Micrococcales; Microbacteriaceae; Frondihabitans MLGLSTSANNGLLWYAVAAGLALTGRTGRRAAVRGLLSLAGSSAVANLVAKPIFGGHRPV WP_084816278.1 hypothetical protein [Frondihabitans sp. Leaf304] Length: 152\nNumber of predicted TMHs: 2\nExp number of AAs in TMHs: 59.22168\nExp number, first 60 AAs: 17.24353\nTotal prob of N-in: 0.24552\nPOSSIBLE N-term signal sequence\ninside 1 90\nTMhelix 91 113\noutside 114 122\nTMhelix 123 145\ninside 146 152

31557 GCF_001577365.1_ASM157736v1 Frondihabitans sp. PAMC 28766 Terrabacteria group; Actinobacteria; Actinobacteria; Micrococcales; Microbacteriaceae; Frondihabitans MSFLSPRKRDNRRRGLLAVLAGAVVAVVVSGTALAVPAFAADQGALTVNKTVDQQNSAAL WP_066275023.1 hypothetical protein [Frondihabitans sp. PAMC 28766] Length: 2142\nNumber of predicted TMHs: 2\nExp number of AAs in TMHs: 43.7698099999999\nExp number, first 60 AAs: 22.58496\nTotal prob of N-in: 0.99749\nPOSSIBLE N-term signal sequence\ninside 1 15\nTMhelix 16 38\noutside 39 2116\nTMhelix 2117 2136\ninside 2137 2142

31558 GCF_001577365.1_ASM157736v1 Frondihabitans sp. PAMC 28766 Terrabacteria group; Actinobacteria; Actinobacteria; Micrococcales; Microbacteriaceae; Frondihabitans MFSGRRTILQLAAATVVAAVIAAMTVVPAQAATGDVSEGEGLLLSGSGLVNVDNLAELQS WP_066282195.1 hypothetical protein [Frondihabitans sp. PAMC 28766] Length: 3005\nNumber of predicted TMHs: 2\nExp number of AAs in TMHs: 46.3681399999999\nExp number, first 60 AAs: 20.65817\nTotal prob of N-in: 0.92978\nPOSSIBLE N-term signal sequence\ninside 1 6\nTMhelix 7 29\noutside 30 2971\nTMhelix 2972 2994\ninside 2995 3005

31559 GCF_001577365.1_ASM157736v1 Frondihabitans sp. PAMC 28766 Terrabacteria group; Actinobacteria; Actinobacteria; Micrococcales; Microbacteriaceae; Frondihabitans MSRRPLLRTVAAGILMSLVLTSLALVGVAAGAAAPAQAAAAPSGSAALRGQEYVALGDSY WP_066278535.1 hypothetical protein [Frondihabitans sp. PAMC 28766] Length: 396\nNumber of predicted TMHs: 2\nExp number of AAs in TMHs: 45.28481\nExp number, first 60 AAs: 22.55485\nTotal prob of N-in: 0.98496\nPOSSIBLE N-term signal sequence\ninside 1 11\nTMhelix 12 34\noutside 35 362\nTMhelix 363 385\ninside 386 396

31560 GCF_001577365.1_ASM157736v1 Frondihabitans sp. PAMC 28766 Terrabacteria group; Actinobacteria; Actinobacteria; Micrococcales; Microbacteriaceae; Frondihabitans MFSSQKSGSSAQGKKKRGRRSPLATFSLIVVGLLTTGGAYALFTSTASADSGTTATATAA WP_066279610.1 cystathionine beta-lyase [Frondihabitans sp. PAMC 28766] Length: 278\nNumber of predicted TMHs: 2\nExp number of AAs in TMHs: 43.03113\nExp number, first 60 AAs: 22.58619\nTotal prob of N-in: 0.97142\nPOSSIBLE N-term signal sequence\ninside 1 20\nTMhelix 21 43\noutside 44 255\nTMhelix 256 275\ninside 276 278

31561 GCF_000419445.1_ASM41944v1 Gryllotalpicola ginsengisoli DSM 22003 Terrabacteria group; Actinobacteria; Actinobacteria; Micrococcales; Microbacteriaceae; Gryllotalpicola; Gryllotalpicola ginsengisoli MAQRRLRVMPMPFVNRRTPVPSTAPTPVQQPARPGRRTIVDSAIYENGIRVASPGSLAET WP_027479117.1 transporter [Gryllotalpicola ginsengisoli] Length: 376\nNumber of predicted TMHs: 2\nExp number of AAs in TMHs: 42.4917\nExp number, first 60 AAs: 3e-05\nTotal prob of N-in: 0.94335\ninside 1 318\nTMhelix 319 338\noutside 339 347\nTMhelix 348 370\ninside 371 376

31562 GCF_900163695.1_ASM90016369v1 Gulosibacter sp. 10 Terrabacteria group; Actinobacteria; Actinobacteria; Micrococcales; Microbacteriaceae; Gulosibacter
MHAADPAPPPARRRALGRIALAAALAGALFAPTAAVAQPAPPADPAPAKYDIEPGSDVLI WP_087009484.1 hypothetical protein [Gulosibacter sp. 10] Length: 1324\nNumber of predicted TMHs: 1\nExp number of AAs in TMHs: 34.60242\nExp number, first 60 AAs: 13.69574\nTotal prob of N-in: 0.65652\nPOSSIBLE N-term signal sequence\noutside 1 1293\nTMhelix 1294 1316\ninside 1317 1324

31563 GCF_900163695.1_ASM90016369v1 Gulosibacter sp. 10 Terrabacteria group; Actinobacteria; Actinobacteria; Micrococcales; Microbacteriaceae; Gulosibacter
MSLATKHRPAAKKSRKRARRTPLATAALLSLGLLLTGGAYGAVDTLVANAAEDTVDVDMT WP_087007419.1 cystathionine beta-lyase [Gulosibacter sp. 10] Length: 278\nNumber of predicted TMHs: 2\nExp number of AAs in TMHs: 46.09695\nExp number, first 60 AAs: 22.49646\nTotal prob of N-in: 0.99401\nPOSSIBLE N-term signal sequence\ninside 1 20\nTMhelix 21 43\noutside 44 250\nTMhelix 251 273\ninside 274 278

31564 GCF_000425685.1_ASM42568v1 Gulosibacter molinativorax DSM 13485 Terrabacteria group; Actinobacteria; Actinobacteria; Micrococcales; Microbacteriaceae; Gulosibacter; Gulosibacter molinativorax
MSLATKSRSSAKKSRKRSRRTPLATAALLGVGLLLSGGAYGAVDTLVANAAEDAVEVDMT WP_026936019.1 cystathionine beta-lyase [Gulosibacter molinativorax] Length: 278\nNumber of predicted TMHs: 2\nExp number of AAs in TMHs: 45.25522\nExp number, first 60 AAs: 22.51137\nTotal prob of N-in: 0.99489\nPOSSIBLE N-term signal sequence\ninside 1 20\nTMhelix 21 43\noutside 44 250\nTMhelix 251 273\ninside 274 278

31565 GCF_000799285.1_ASM79928v1 Herbiconiux sp. YR403 Terrabacteria group; Actinobacteria; Actinobacteria; Micrococcales; Microbacteriaceae; Herbiconiux
MMRQSDSFASSSATKSAAARGAAGRRSAVRGVARGAAIAALALGVAVFGAAAPASAHNQV WP_047405150.1 hypothetical protein [Herbiconiux sp. YR403] Length: 230\nNumber of predicted TMHs: 2\nExp number of AAs in TMHs: 43.82641\nExp number, first 60 AAs: 21.70355\nTotal prob of N-in: 0.98335\nPOSSIBLE N-term signal sequence\ninside 1 31\nTMhelix 32 54\noutside 55 204\nTMhelix 205 227\ninside 228 230

31566 GCF_000799285.1_ASM79928v1 Herbiconiux sp. YR403 Terrabacteria group; Actinobacteria; Actinobacteria; Micrococcales; Microbacteriaceae; Herbiconiux
MHTTIRPQKRRAALMTAWAVAGATALFAGTIAAPAAAIAAPAQAEIDTVDYFADSYPGLG WP_047408652.1 hypothetical protein [Herbiconiux sp. YR403] Length: 705\nNumber of predicted TMHs: 2\nExp number of AAs in TMHs: 45.39454\nExp number, first 60 AAs: 22.52444\nTotal prob of N-in: 0.99592\nPOSSIBLE N-term signal sequence\ninside 1 12\nTMhelix 13 35\noutside 36 674\nTMhelix 675 697\ninside 698 705

31567 GCF_000799285.1_ASM79928v1 Herbiconiux sp. YR403 Terrabacteria group; Actinobacteria; Actinobacteria; Micrococcales; Microbacteriaceae; Herbiconiux
MTGASGDGRRRVVVTGMGVVSPIGVSVDSFWEALQLGRSGIATIESFDTEGLGTSFASQV WP_052175349.1 hypothetical protein [Herbiconiux sp. YR403] Length: 436\nNumber of predicted TMHs: 2\nExp number of AAs in TMHs: 42.15499\nExp number, first 60 AAs: 8.85007999999999\nTotal prob of N-in: 0.46913\noutside 1 350\nTMhelix 351 373\ninside 374 393\nTMhelix 394 416\noutside 417 436

31568 GCF_000799285.1_ASM79928v1 Herbiconiux sp. YR403 Terrabacteria group; Actinobacteria; Actinobacteria; Micrococcales; Microbacteriaceae; Herbiconiux
MMTTKNTVRWRRRAVVSTALMAVLGGVLAAPVPALAAPEVPTEPVVSTANTTWKYLDDGS WP_052176223.1 metallophosphoesterase [Herbiconiux sp. YR403]Length: 886\nNumber of predicted TMHs: 2\nExp number of AAs in TMHs: 43.83303\nExp number, first 60 AAs: 21.91886\nTotal prob of N-in: 0.99440\nPOSSIBLE N-term signal sequence\ninside 1 12\nTMhelix 13 35\noutside 36 846\nTMhelix 847 869\ninside 870 886

31569 GCF_001571005.1_ASM157100v1 Herbiconiux solani NBRC 106740 Terrabacteria group; Actinobacteria; Actinobacteria; Micrococcales; Microbacteriaceae; Herbiconiux; Herbiconiux solani
MADSRRPRRRLVAALLIPGAVLGTAGITALPAMAAPPGDLASQTVTSGDARFQVLSPTLI WP_084361599.1 alpha-xylosidase [Herbiconiux solani] Length: 1154\nNumber of predicted TMHs: 2\nExp number of AAs in TMHs: 44.53641\nExp number, first 60 AAs: 22.40454\nTotal prob of N-in: 0.99755\nPOSSIBLE N-term signal sequence\ninside 1 11\nTMhelix 12 34\noutside 35 1125\nTMhelix 1126 1148\ninside 1149 1154

31570 GCF_001571005.1_ASM157100v1 Herbiconiux solani NBRC 106740 Terrabacteria group; Actinobacteria; Actinobacteria; Micrococcales; Microbacteriaceae; Herbiconiux; Herbiconiux solani
MTVPEHARPRRRLVSTLISGAALSALLAGALAAPAAASAVGTGRDASAGTAATTSTAAAA WP_084361395.1 hypothetical protein [Herbiconiux solani] Length: 1717\nNumber of predicted TMHs: 2\nExp number of

AAs in TMHs: 45.2704499999999\nExp number, first 60 AAs: 22.32372\nTotal prob of N-in: 0.98875\nPOSSIBLE N-term signal sequence\ninside 1 12\nTMhelix 13 35\noutside 36 1688\nTMhelix 1689 1711\ninside 1712 1717

31571 GCF_001571005.1_ASM157100v1 Herbiconiux solani NBRC 106740 Terrabacteria group; Actinobacteria; Actinobacteria; Micrococcales; Microbacteriaceae; Herbiconiux; Herbiconiux solani MASHPPFRRGRRALIAAALLTAASISLPVAAFADPGDDLSVNGGATRNSGDKTAALRASI WP_066041387.1 alkaline phosphatase [Herbiconiux solani] Length: 645\nNumber of predicted TMHs: 2\nExp number of AAs in TMHs: 41.16069\nExp number, first 60 AAs: 18.85662\nTotal prob of N-in: 0.91396\nPOSSIBLE N-term signal sequence\ninside 1 12\nTMhelix 13 32\noutside 33 612\nTMhelix 613 635\ninside 636 645

31572 GCF_001571005.1_ASM157100v1 Herbiconiux solani NBRC 106740 Terrabacteria group; Actinobacteria; Actinobacteria; Micrococcales; Microbacteriaceae; Herbiconiux; Herbiconiux solani MNRRRLISAAAVLLLTAGGVAAGIAPASAAPTPGVNIHFFPEVDGGYSMGYPDGADVDLL WP_066040688.1 hypothetical protein [Herbiconiux solani] Length: 407\nNumber of predicted TMHs: 2\nExp number of AAs in TMHs: 43.5165\nExp number, first 60 AAs: 22.24262\nTotal prob of N-in: 0.99032\nPOSSIBLE N-term signal sequence\ninside 1 6\nTMhelix 7 29\noutside 30 378\nTMhelix 379 401\ninside 402 407

31573 GCF_000421825.1_ASM42182v1 Humibacter albus DSM 18994 Terrabacteria group; Actinobacteria; Actinobacteria; Micrococcales; Microbacteriaceae; Humibacter; Humibacter albus MPPRTAHAARSRRTALAGVVALAVTATVALGTAVPASADTVNTPPAGTAVDDSHYTVLDV WP_022899026.1 hypothetical protein [Humibacter albus] Length: 1022\nNumber of predicted TMHs: 2\nExp number of AAs in TMHs: 40.03831\nExp number, first 60 AAs: 20.277\nTotal prob of N-in: 0.91653\nPOSSIBLE N-term signal sequence\ninside 1 12\nTMhelix 13 35\noutside 36 994\nTMhelix 995 1014\ninside 1015 1022

31574 GCF_001984125.1_ASM198412v1 Leifsonia sp. ALI-44-B Terrabacteria group; Actinobacteria; Actinobacteria; Micrococcales; Microbacteriaceae; Leifsonia MLRKRNTSARTARKNGRRSPLATVALIAVGLLFTGGAYAAFSATTANAETPAANSQASVE WP_077053121.1 cystathionine beta-lyase [Leifsonia sp. ALI-44-B] Length: 271\nNumber of predicted TMHs: 2\nExp number of AAs in TMHs: 42.90998\nExp number, first 60 AAs: 22.26949\nTotal prob of N-in: 0.99889\nPOSSIBLE N-term signal sequence\ninside 1 20\nTMhelix 21 43\noutside 44 248\nTMhelix 249 268\ninside 269 271

31575 GCF_001422325.1_Leaf264 Leifsonia sp. Leaf264 Terrabacteria group; Actinobacteria; Actinobacteria; Micrococcales; Microbacteriaceae; Leifsonia MKLPEKLFGRAARSAAAGSHAGAVSPRRRIVRTITAIAAVVAVISGGALTAITVQAASNG WP_055907020.1 hypothetical protein [Leifsonia sp. Leaf264] Length: 1374\nNumber of predicted TMHs: 2\nExp number of AAs in TMHs: 42.9100599999999\nExp number, first 60 AAs: 20.04463\nTotal prob of N-in: 0.90734\nPOSSIBLE N-term signal sequence\ninside 1 29\nTMhelix 30 52\noutside 53 1345\nTMhelix 1346 1368\ninside 1369 1374

31576 GCF_001422325.1_Leaf264 Leifsonia sp. Leaf264 Terrabacteria group; Actinobacteria; Actinobacteria; Micrococcales; Microbacteriaceae; Leifsonia MFTRITGVPTTQNDGGQRPGSRRRSPFAVPVAMLLSAAIMLIGVPAVAIADEGVPAGETP WP_055902840.1 VWA domain-containing protein [Leifsonia sp. Leaf264] Length: 1604\nNumber of predicted TMHs: 2\nExp number of AAs in TMHs: 45.16934\nExp number, first 60 AAs: 22.39211\nTotal prob of N-in: 0.98407\nPOSSIBLE N-term signal sequence\ninside 1 26\nTMhelix 27 49\noutside 50 1573\nTMhelix 1574 1596\ninside 1597 1604

31577 GCF_002105485.1_ASM210548v1 Leifsonia sp. NCR5 Terrabacteria group; Actinobacteria; Actinobacteria; Micrococcales; Microbacteriaceae; Leifsonia MARRARRRARPRRSMLGMVGSVMLTIAAIVGVAAVLAVAASAILGIRPVVVISGSMEPVL WP_085368146.1 signal peptidase I [Leifsonia sp. NCR5] Length: 194\nNumber of predicted TMHs: 2\nExp number of AAs in TMHs: 48.89567\nExp number, first 60 AAs: 24.68755\nTotal prob of N-in: 0.99741\nPOSSIBLE N-term signal sequence\ninside 1 20\nTMhelix 21 43\noutside 44 150\nTMhelix 151 170\ninside 171 194

31578 GCF_001425085.1_Root4 Leifsonia sp. Root4 Terrabacteria group; Actinobacteria; Actinobacteria; Micrococcales; Microbacteriaceae; Leifsonia MRTHRRRPITALITCLALAVPLAGVATAATAAESSPIKINEVAQNSDVTDWVELINTGSE WP_082492136.1 hypothetical protein [Leifsonia sp. Root4] Length: 1069\nNumber of predicted TMHs: 2\nExp number of AAs in TMHs: 43.66527\nExp number, first 60 AAs: 21.43421\nTotal prob of N-in: 0.97944\nPOSSIBLE N-term signal sequence\ninside 1 8\nTMhelix 9 31\noutside 32 1040\nTMhelix 1041 1063\ninside 1064 1069

31579 GCF_001647635.1_ASM164763v1 Leifsonia xyli Terrabacteria group; Actinobacteria; Actinobacteria; Micrococcales; Microbacteriaceae; Leifsonia

MRHREDDPSRRSVVGWIAGAAAAVLLAFAIVPQSSVHAATTAFSTVTTTGSDVQNGSRAA WP_064108907.1 hypothetical protein [Leifsonia xyli] Length: 988\nNumber of predicted TMHs: 2\nExp number of AAs in TMHs: 41.7770099999999\nExp number, first 60 AAs: 20.84987\nTotal prob of N-in: 0.99929\nPOSSIBLE N-term signal sequence\ninside 1 11\nTMhelix 12 31\noutside 32 959\nTMhelix 960 982\ninside 983 988

31580 GCF_000633535.1_de_novo Leifsonia aquatica H1aii Terrabacteria group; Actinobacteria; Actinobacteria; Micrococcales; Microbacteriaceae; Leifsonia; Leifsonia aquatica MSRRAAVRALLAGAASAVALGLIASPATATTTIATTAAGASGAPLSATAPAVRLATAGGCWP_052357896.1 hypothetical protein [Leifsonia aquatica] Length: 443\nNumber of predicted TMHs: 2\nExp number of AAs in TMHs: 43.8870999999999\nExp number, first 60 AAs: 20.87734\nTotal prob of N-in: 0.91459\nPOSSIBLE N-term signal sequence\ninside 1 6\nTMhelix 7 29\noutside 30 409\nTMhelix 410 432\ninside 433 443

31581 GCF_000633535.1_de_novo Leifsonia aquatica H1aii Terrabacteria group; Actinobacteria; Actinobacteria; Micrococcales; Microbacteriaceae; Leifsonia; Leifsonia aquatica MPGALSAVAPFSLGRPFDGSRITSPDISPTRRIPVRRRRLLPATLSVLGAAATAALLTVA WP_025156969.1 hypothetical protein [Leifsonia aquatica] Length: 463\nNumber of predicted TMHs: 1\nExp number of AAs in TMHs: 86.6144599999999\nExp number, first 60 AAs: 14.9518\nTotal prob of N-in: 0.76602\nPOSSIBLE N-term signal sequence\noutside 1 431\nTMhelix 432 454\ninside 455 463

31582 GCF_000477555.1_LeRu1.0 Leifsonia rubra CMS 76R Terrabacteria group; Actinobacteria; Actinobacteria; Micrococcales; Microbacteriaceae; Leifsonia; Leifsonia rubra MSPLTAATLIGARRRDALVAQLSCTTATTATVIAPFTGQPLHELPQSSVGDVEGAAESAR WP_021809997.1 aldehyde dehydrogenase [Leifsonia rubra] Length: 520\nNumber of predicted TMHs: 1\nExp number of AAs in TMHs: 19.42317\nExp number, first 60 AAs: 0.44333\nTotal prob of N-in: 0.02457\noutside 1 499\nTMhelix 500 517\ninside 518 520

31583 GCF_900102345.1_IMG-taxon_2639762569_annotated_assembly Leucobacter chromiiresistens Terrabacteria group; Actinobacteria; Actinobacteria; Micrococcales; Microbacteriaceae; Leucobacter MTQPGSSSGLAPRRSTAPRRSAAPRRAVLVAAGALVALVGLPVAATAASAHDQLVSADPV WP_083352005.1 hypothetical protein [Leucobacter chromiiresistens] Length: 224\nNumber of predicted TMHs: 2\nExp number of AAs in TMHs: 44.78724\nExp number, first 60 AAs: 21.9749\nTotal prob of N-in: 0.96669\nPOSSIBLE N-term signal sequence\ninside 1 27\nTMhelix 28 50\noutside 51 194\nTMhelix 195 217\ninside 218 224

31584 GCF_900102345.1_IMG-taxon_2639762569_annotated_assembly Leucobacter chromiiresistens Terrabacteria group; Actinobacteria; Actinobacteria; Micrococcales; Microbacteriaceae; Leucobacter MYWRTRPRSATDPRRSALALRRCTTAVLAALLCATGAQLAPQAAIATGEAAPAESTAEAP WP_010155924.1 outer membrane protein [Leucobacter chromiiresistens] Length: 870\nNumber of predicted TMHs: 1\nExp number of AAs in TMHs: 29.64499\nExp number, first 60 AAs: 7.58866\nTotal prob of N-in: 0.35814\noutside 1 835\nTMhelix 836 858\ninside 859 870

31585 GCF_900102345.1_IMG-taxon_2639762569_annotated_assembly Leucobacter chromiiresistens Terrabacteria group; Actinobacteria; Actinobacteria; Micrococcales; Microbacteriaceae; Leucobacter MTRADTTTPARRRTSSARRRRRRTIAIAAVFVIVTGWLYLAPPLISYAGAPASCRPLAAG WP_010156067.1 hypothetical protein [Leucobacter chromiiresistens] Length: 167\nNumber of predicted TMHs: 2\nExp number of AAs in TMHs: 53.18206\nExp number, first 60 AAs: 23.21932\nTotal prob of N-in: 0.99651\nPOSSIBLE N-term signal sequence\ninside 1 24\nTMhelix 25 47\noutside 48 117\nTMhelix 118 140\ninside 141 167

31586 GCF_001477055.1_ASM147705v1 Leucobacter chromiiresistens Terrabacteria group; Actinobacteria; Actinobacteria; Micrococcales; Microbacteriaceae; Leucobacter MTLPRSSSGLAPRRGTAPRRAALAAAGALIAIAGLPAVATAASAHDQLVSADPVSGASVE WP_058594659.1 hypothetical protein [Leucobacter chromiiresistens] Length: 218\nNumber of predicted TMHs: 2\nExp number of AAs in TMHs: 45.03473\nExp number, first 60 AAs: 22.22454\nTotal prob of N-in: 0.98987\nPOSSIBLE N-term signal sequence\ninside 1 20\nTMhelix 21 43\noutside 44 188\nTMhelix 189 211\ninside 212 218

31587 GCF_000834055.1_ASM83405v1 Leucobacter komagatae Terrabacteria group; Actinobacteria; Actinobacteria; Micrococcales; Microbacteriaceae; Leucobacter MGAPMSGRHTLGARRATPRRGTVTVLVAAITVFLPVPIGPGQALIAAEWPPFSAAGSQAD WP_042545138.1 hypothetical protein [Leucobacter komagatae] Length: 514\nNumber of predicted TMHs: 2\nExp number of AAs in TMHs: 43.7433\nExp number, first 60 AAs: 21.1159\nTotal prob of N-in: 0.96604\nPOSSIBLE N-term signal sequence\ninside 1 20\nTMhelix 21 43\noutside 44 477\nTMhelix 478 500\ninside 501 514

31588 GCF_900163635.1_ASM90016363v1 Leucobacter sp. 7(1) Terrabacteria group; Actinobacteria; Actinobacteria; Micrococcales; Microbacteriaceae; Leucobacter
MRNTHTRRRTLTALVGAAALVIGSTALLAGPAFAAPGDGGTGTLTVHKLEQPEGGDFGPN WP_087014056.1 hypothetical protein [Leucobacter sp. 7(1)] Length: 492\nNumber of predicted TMHs: 2\nExp number of AAs in TMHs: 44.9324399999999\nExp number, first 60 AAs: 22.76338\nTotal prob of N-in: 0.99959\nPOSSIBLE N-term signal sequence\ninside 1 12\nTMhelix 13 35\noutside 36 460\nTMhelix 461 483\ninside 484 492

31589 GCF_000980875.1_ASM98087v1 Leucobacter sp. Ag1 Terrabacteria group; Actinobacteria; Actinobacteria; Micrococcales; Microbacteriaceae; Leucobacter
MKLTIPTRARRTALAGVAAAALAALAVSSPIGLPAAQADAAPIDGAPSIGDSLFAGIGNG WP_052961504.1 hypothetical protein [Leucobacter sp. Ag1] Length: 827\nNumber of predicted TMHs: 2\nExp number of AAs in TMHs: 44.03789\nExp number, first 60 AAs: 21.35439\nTotal prob of N-in: 0.96654\nPOSSIBLE N-term signal sequence\ninside 1 12\nTMhelix 13 35\noutside 36 776\nTMhelix 777 799\ninside 800 827

31590 GCF_000980875.1_ASM98087v1 Leucobacter sp. Ag1 Terrabacteria group; Actinobacteria; Actinobacteria; Micrococcales; Microbacteriaceae; Leucobacter
MKGPIMLGLRPPARRRRALTAGVTGTTLVAAALVGGFAAATPAAAATEVAPFAIGSLSGQ WP_052961234.1 hypothetical protein [Leucobacter sp. Ag1] Length: 685\nNumber of predicted TMHs: 2\nExp number of AAs in TMHs: 42.97104\nExp number, first 60 AAs: 20.52274\nTotal prob of N-in: 0.90440\nPOSSIBLE N-term signal sequence\ninside 1 20\nTMhelix 21 43\noutside 44 652\nTMhelix 653 675\ninside 676 685

31591 GCF_001482305.1_ASM148230v1 Leucobacter sp. G161 Terrabacteria group; Actinobacteria; Actinobacteria; Micrococcales; Microbacteriaceae; Leucobacter
MGNRMSRRGQQRKSVTPARSSRRRNLGRGFAMLVAAAVMYTGLPTAAFAAVVPPSTESSL WP_083506842.1 hypothetical protein [Leucobacter sp. G161] Length: 2984\nNumber of predicted TMHs: 2\nExp number of AAs in TMHs: 44.8867699999999\nExp number, first 60 AAs: 22.64662\nTotal prob of N-in: 0.99729\nPOSSIBLE N-term signal sequence\ninside 1 28\nTMhelix 29 51\noutside 52 2952\nTMhelix 2953 2975\ninside 2976 2984

31592 GCF_000633415.1_de_novo Leucobacter sp. PH1c Terrabacteria group; Actinobacteria; Actinobacteria; Micrococcales; Microbacteriaceae; Leucobacter
MSGCPSGHPLSSARRTHVHTPQQTSSSGSRRAIAALGAGLVGIGLLSAPAAATAAPLPATWP_081810009.1 hypothetical protein [Leucobacter sp. PH1c] Length: 312\nNumber of predicted TMHs: 1\nExp number of AAs in TMHs: 30.92665\nExp number, first 60 AAs: 8.18042\nTotal prob of N-in: 0.29591\noutside 1 275\nTMhelix 276 298\ninside 299 312

31593 GCF_000633415.1_de_novo Leucobacter sp. PH1c Terrabacteria group; Actinobacteria; Actinobacteria; Micrococcales; Microbacteriaceae; Leucobacter
MSAFPTGAPPSQQRRRALGAFGAAAIALPLVLTPGAAFAAPAPAPAVVPGGAVTAIGTGT WP_025132763.1 hypothetical protein [Leucobacter sp. PH1c] Length: 829\nNumber of predicted TMHs: 2\nExp number of AAs in TMHs: 42.5110999999999\nExp number, first 60 AAs: 20.44636\nTotal prob of N-in: 0.89507\nPOSSIBLE N-term signal sequence\ninside 1 16\nTMhelix 17 39\noutside 40 798\nTMhelix 799 821\ninside 822 829

31594 GCF_000633415.1_de_novo Leucobacter sp. PH1c Terrabacteria group; Actinobacteria; Actinobacteria; Micrococcales; Microbacteriaceae; Leucobacter
MARRRAALALGAALGIVVGGLGASAAFAAPSPEPLTVTATAPAGSTQVWIDLEGADPAEV WP_025134557.1 hypothetical protein [Leucobacter sp. PH1c] Length: 239\nNumber of predicted TMHs: 2\nExp number of AAs in TMHs: 44.7232600000001\nExp number, first 60 AAs: 22.78084\nTotal prob of N-in: 0.99939\nPOSSIBLE N-term signal sequence\ninside 1 6\nTMhelix 7 29\noutside 30 184\nTMhelix 185 207\ninside 208 239

31595 GCF_000349545.1_Leucobacter_sp._str._UCD-THU Leucobacter sp. UCD-THU Terrabacteria group; Actinobacteria; Actinobacteria; Micrococcales; Microbacteriaceae; Leucobacter
MGVGRESRRAVVRRVLSGVAAAAVVAGGLVAVSAPGATAAPGPNPYTDVRVTKKYDGTGH WP_017884253.1 hypothetical protein [Leucobacter sp. UCD-THU] Length: 2162\nNumber of predicted TMHs: 2\nExp number of AAs in TMHs: 41.3955699999999\nExp number, first 60 AAs: 20.35623\nTotal prob of N-in: 0.91174\nPOSSIBLE N-term signal sequence\ninside 1 11\nTMhelix 12 34\noutside 35 2132\nTMhelix 2133 2152\ninside 2153 2162

31596 GCF_000231305.1_ASM23130v1 Leucobacter chromiiresistens JG 31 Terrabacteria group; Actinobacteria; Actinobacteria; Micrococcales; Microbacteriaceae; Leucobacter; Leucobacter chromiiresistens
MYWRTRPRSATDPRRSALALRRCTTAVLAALLCATGAQLAPQAAIATGEAAPAESTAEAP WP_010155924.1 outer membrane protein [Leucobacter chromiiresistens] Length: 870\nNumber of predicted TMHs: 1\nExp

number of AAs in TMHs: 29.64499\nExp number, first 60 AAs: 7.58866\nTotal prob of N-in: 0.35814\noutside 1 835\nTMhelix 836 858\ninside 859 870

31597 GCF_000231305.1_ASM23130v1 Leucobacter chromiiresistens JG 31 Terrabacteria group; Actinobacteria; Actinobacteria; Micrococcales; Microbacteriaceae; Leucobacter; Leucobacter chromiiresistens MTRADTTTPARRRTSSARRRRRRTIAIAAVFVIVTGWLYLAPPLISYAGAPASCRPLAAG WP_010156067.1 hypothetical protein [Leucobacter chromiiresistens] Length: 167\nNumber of predicted TMHs: 2\nExp number of AAs in TMHs: 53.18206\nExp number, first 60 AAs: 23.21932\nTotal prob of N-in: 0.99651\nPOSSIBLE N-term signal sequence\ninside 1 24\nTMhelix 25 47\noutside 48 117\nTMhelix 118 140\ninside 141 167

31598 GCF_001273855.1_CBX130_1.1 Leucobacter musarum subsp. japonicus Terrabacteria group; Actinobacteria; Actinobacteria; Micrococcales; Microbacteriaceae; Leucobacter; Leucobacter musarum MSDRTTPSAATTPRRTRRPLLVATAVAAATLCVLPAGAAFAASAPAAPAASTSAFGAAAE WP_053387807.1 DUF4822 domain-containing protein [Leucobacter musarum] Length: 277\nNumber of predicted TMHs: 2\nExp number of AAs in TMHs: 45.1791\nExp number, first 60 AAs: 22.69626\nTotal prob of N-in: 0.99732\nPOSSIBLE N-term signal sequence\ninside 1 19\nTMhelix 20 42\noutside 43 250\nTMhelix 251 273\ninside 274 277

31599 GCF_000350525.1_Leucobacter_salsicius_M1-8T_draft_genome Leucobacter salsicius M1-8 Terrabacteria group; Actinobacteria; Actinobacteria; Micrococcales; Microbacteriaceae; Leucobacter; Leucobacter salsicius MSTTIARPSRARRTALGLATLAVAASAVVTLPLVSQPAPAQAASLPQKGCPVGSINLVIE WP_017794056.1 hypothetical protein [Leucobacter salsicius] Length: 426\nNumber of predicted TMHs: 2\nExp number of AAs in TMHs: 41.68587\nExp number, first 60 AAs: 20.2655\nTotal prob of N-in: 0.91048\nPOSSIBLE N-term signal sequence\ninside 1 12\nTMhelix 13 35\noutside 36 395\nTMhelix 396 418\ninside 419 426

31600 GCF_001854225.1_ASM185422v1 Candidatus Rhodoluna planktonica Terrabacteria group; Actinobacteria; Actinobacteria; Micrococcales; Microbacteriaceae; Luna cluster; Luna-1 subcluster; Rhodoluna MAIFKRNAKGSRRSPLAAIIVMVAGLAVTGSGYAAATAAVESVKPVSAVQSATQIEEGRK WP_070954654.1 cystathionine beta-lyase [Candidatus Rhodoluna planktonica] Length: 266\nNumber of predicted TMHs: 2\nExp number of AAs in TMHs: 43.0800799999999\nExp number, first 60 AAs: 21.78998\nTotal prob of N-in: 0.92246\nPOSSIBLE N-term signal sequence\ninside 1 12\nTMhelix 13 35\noutside 36 243\nTMhelix 244 263\ninside 264 266

31601 GCF_000699505.1_ASM69950v1 Rhodoluna lacicola Terrabacteria group; Actinobacteria; Actinobacteria; Micrococcales; Microbacteriaceae; Luna cluster; Luna-1 subcluster; Rhodoluna MSFKKNRRSPLAAIIVMAAGLALTGGGYVAATAAVEAVKPVSAVATAEQISEGRKLFLAN WP_038502532.1 cystathionine beta-lyase [Rhodoluna lacicola] Length: 262\nNumber of predicted TMHs: 2\nExp number of AAs in TMHs: 43.58605\nExp number, first 60 AAs: 22.51751\nTotal prob of N-in: 0.95258\nPOSSIBLE N-term signal sequence\ninside 1 12\nTMhelix 13 35\noutside 36 239\nTMhelix 240 259\ninside 260 262

31602 GCF_001662775.1_ASM166277v1 Microbacterium arborescens Terrabacteria group; Actinobacteria; Actinobacteria; Micrococcales; Microbacteriaceae; Microbacterium MTSPTTVTKPTKPTKPTTRAEARTSQVVTPRRRPLSRVGDAMLVLAAATGALCIALTIAA WP_064956389.1 hypothetical protein [Microbacterium arborescens] Length: 208\nNumber of predicted TMHs: 2\nExp number of AAs in TMHs: 47.87232\nExp number, first 60 AAs: 19.81629\nTotal prob of N-in: 0.95629\nPOSSIBLE N-term signal sequence\ninside 1 40\nTMhelix 41 63\noutside 64 167\nTMhelix 168 190\ninside 191 208

31603 GCF_001662775.1_ASM166277v1 Microbacterium arborescens Terrabacteria group; Actinobacteria; Actinobacteria; Micrococcales; Microbacteriaceae; Microbacterium MLRTRRAALQRPLASVLTGALVAMAAVVGVAAPATAAGPTLTTSTTVVANEKITVAITGT WP_083990418.1 hypothetical protein [Microbacterium arborescens] Length: 962\nNumber of predicted TMHs: 2\nExp number of AAs in TMHs: 41.1068399999999\nExp number, first 60 AAs: 22.33509\nTotal prob of N-in: 0.99759\nPOSSIBLE N-term signal sequence\ninside 1 12\nTMhelix 13 35\noutside 36 934\nTMhelix 935 954\ninside 955 962

31604 GCF_001662775.1_ASM166277v1 Microbacterium arborescens Terrabacteria group; Actinobacteria; Actinobacteria; Micrococcales; Microbacteriaceae; Microbacterium MPPAPSRPQSRPRRTAVPAAAVAAFTAACLTLGTAAPAGADDGELLDDGAIAPHAAPYGT WP_064955119.1 hypothetical protein [Microbacterium arborescens] Length: 702\nNumber of predicted TMHs: 1\nExp number of AAs in TMHs: 30.24407\nExp number, first 60 AAs: 8.43126000000001\nTotal prob of N-in: 0.39792\noutside 1 671\nTMhelix 672 694\ninside 695 702

31605 GCF_001974985.1_ASM197498v1 Microbacterium aurum Terrabacteria group; Actinobacteria; Actinobacteria; Micrococcales; Microbacteriaceae; Microbacterium
MTTVTDHEHPHRSGSRRRILLLSTLTTLLLAAGLLFATAAPAAAHDEIVSSSPEAGSTVS WP_076688294.1 hypothetical protein [Microbacterium aurum] Length: 238\nNumber of predicted TMHs: 2\nExp number of AAs in TMHs: 45.12157\nExp number, first 60 AAs: 22.44287\nTotal prob of N-in: 0.99979\nPOSSIBLE N-term signal sequence\ninside 1 19\nTMhelix 20 42\noutside 43 183\nTMhelix 184 206\ninside 207 238

31606 GCF_000956545.1_ASM95654v1 Microbacterium azadirachtaeTerrabacteria group; Actinobacteria; Actinobacteria; Micrococcales; Microbacteriaceae; Microbacterium
MAREKKRRTGRRSPLAAAALIGAGLMITGGVYAGASAAFAATEPASVTAAVSTEDGQKLF WP_045252334.1 cystathionine beta-lyase [Microbacterium azadirachtae] Length: 265\nNumber of predicted TMHs: 2\nExp number of AAs in TMHs: 43.14984\nExp number, first 60 AAs: 22.61095\nTotal prob of N-in: 0.99605\nPOSSIBLE N-term signal sequence\ninside 1 12\nTMhelix 13 35\noutside 36 242\nTMhelix 243 262\ninside 263 265

31607 GCF_000956505.1_ASM95650v1 Microbacterium azadirachtaeTerrabacteria group; Actinobacteria; Actinobacteria; Micrococcales; Microbacteriaceae; Microbacterium
MTNHRTAMDEEGALRSDRGWTRRRAIVAGAWSVPVIAAAVALPQASASAAITVDLLGAVD WP_045272686.1 hypothetical protein [Microbacterium azadirachtae] Length: 156\nNumber of predicted TMHs: 2\nExp number of AAs in TMHs: 44.16519\nExp number, first 60 AAs: 21.99312\nTotal prob of N-in: 0.99684\nPOSSIBLE N-term signal sequence\ninside 1 24\nTMhelix 25 47\noutside 48 122\nTMhelix 123 145\ninside 146 156

31608 GCF_000956505.1_ASM95650v1 Microbacterium azadirachtaeTerrabacteria group; Actinobacteria; Actinobacteria; Micrococcales; Microbacteriaceae; Microbacterium
MAREKKRRTGRRSPLAAAALIGAGLMITGGVYAGASAAFAATEPASVTAAVSTEDGQKLF WP_045272502.1 cystathionine beta-lyase [Microbacterium azadirachtae] Length: 265\nNumber of predicted TMHs: 2\nExp number of AAs in TMHs: 43.44151\nExp number, first 60 AAs: 22.60657\nTotal prob of N-in: 0.99528\nPOSSIBLE N-term signal sequence\ninside 1 12\nTMhelix 13 35\noutside 36 242\nTMhelix 243 262\ninside 263 265

31609 GCF_000956545.1_ASM95654v1 Microbacterium azadirachtaeTerrabacteria group; Actinobacteria; Actinobacteria; Micrococcales; Microbacteriaceae; Microbacterium
MKHIDRRAVVKGAAWSIPVVAAAAAMPLASASTPACPTCIKAGFPIIGGIISGAWTSQAV WP_045249939.1 hypothetical protein [Microbacterium azadirachtae] Length: 208\nNumber of predicted TMHs: 2\nExp number of AAs in TMHs: 83.33951\nExp number, first 60 AAs: 33.1698\nTotal prob of N-in: 0.97508\nPOSSIBLE N-term signal sequence\ninside 1 112\nTMhelix 113 135\noutside 136 181\nTMhelix 182 204\ninside 205 208

31610 GCF_000956545.1_ASM95654v1 Microbacterium azadirachtaeTerrabacteria group; Actinobacteria; Actinobacteria; Micrococcales; Microbacteriaceae; Microbacterium
MTTRNSTTDRGGAVAGDRGWTRRRALAAGAWSVPVVAAAVALPQAAASAPIVVDLVGAVD WP_045249571.1 hypothetical protein [Microbacterium azadirachtae] Length: 156\nNumber of predicted TMHs: 2\nExp number of AAs in TMHs: 43.61001\nExp number, first 60 AAs: 21.77431\nTotal prob of N-in: 0.99635\nPOSSIBLE N-term signal sequence\ninside 1 24\nTMhelix 25 47\noutside 48 122\nTMhelix 123 145\ninside 146 156

31611 GCF_001652465.1_ASM165246v1 Microbacterium chocolatum Terrabacteria group; Actinobacteria; Actinobacteria; Micrococcales; Microbacteriaceae; Microbacterium
MRNHRRSLITSLATLLLGAGLTLAVATPAQAHDELVSSSPAAESTITGSPTEITLSFSGE WP_082334822.1 hypothetical protein [Microbacterium chocolatum] Length: 223\nNumber of predicted TMHs: 2\nExp number of AAs in TMHs: 44.95384\nExp number, first 60 AAs: 22.06538\nTotal prob of N-in: 0.99691\nPOSSIBLE N-term signal sequence\ninside 1 6\nTMhelix 7 29\noutside 30 171\nTMhelix 172 194\ninside 195 223

31612 GCF_001652465.1_ASM165246v1 Microbacterium chocolatum Terrabacteria group; Actinobacteria; Actinobacteria; Micrococcales; Microbacteriaceae; Microbacterium
MSVPPRSGRRSLLATVVAAALAAAGTLAPLPAAAAEGSRVLINEVYFKGGSANAAYSSKFWP_064319272.1 multifunctional nuclease/2,3-cyclic-nucleotide 2-phosphodiesterase/5-nucleotidase/3-nucleotidase [Microbacterium chocolatum] Length: 1654\nNumber of predicted TMHs: 2\nExp number of AAs in TMHs: 42.9878299999999\nExp number, first 60 AAs: 20.99051\nTotal prob of N-in: 0.93335\nPOSSIBLE N-term signal sequence\ninside 1 11\nTMhelix 12 34\noutside 35 1626\nTMhelix 1627 1649\ninside 1650 1654

31613 GCF_001278365.1_ASM127836v1 Microbacterium chocolatum Terrabacteria group; Actinobacteria; Actinobacteria; Micrococcales; Microbacteriaceae; Microbacterium
MSVPPRSGRRSLLATVVAAALAAAGTLAPLPAAAAEGSRVLINEVYFKGGSANAAYSSKFWP_064319272.1 multifunctional nuclease/2,3-cyclic-nucleotide 2-phosphodiesterase/5-nucleotidase/3-nucleotidase

[Microbacterium chocolatum] Length: 1654\nNumber of predicted TMHs: 2\nExp number of AAs in TMHs: 42.9878299999999\nExp number, first 60 AAs: 20.99051\nTotal prob of N-in: 0.93335\nPOSSIBLE N-term signal sequence\ninside 1 11\nTMhelix 12 34\noutside 35 1626\nTMhelix 1627 1649\ninside 1650 1654

31614 GCF_001278365.1_ASM127836v1 Microbacterium chocolatum Terrabacteria group; Actinobacteria; Actinobacteria; Micrococcales; Microbacteriaceae; Microbacterium MRNHRRSLITSLATLLLGAGLTLAVATPAQAHDELVSSSPAAESTITGSPTEITLSFSGE WP_082334822.1 hypothetical protein [Microbacterium chocolatum] Length: 223\nNumber of predicted TMHs: 2\nExp number of AAs in TMHs: 44.95384\nExp number, first 60 AAs: 22.06538\nTotal prob of N-in: 0.99691\nPOSSIBLE N-term signal sequence\ninside 1 6\nTMhelix 7 29\noutside 30 171\nTMhelix 172 194\ninside 195 223

31615 GCF_001456955.1_ASM145695v1 Microbacterium enclense Terrabacteria group; Actinobacteria; Actinobacteria; Micrococcales; Microbacteriaceae; Microbacterium MCPSCAASLPRGARTTPAGAGIRVVCRAERCFVQELCGRPARRRRPLRPTAPPAGAGGGA WP_082642070.1 hypothetical protein [Microbacterium enclense] Length: 240\nNumber of predicted TMHs: 1\nExp number of AAs in TMHs: 22.79157\nExp number, first 60 AAs: 0.00099\nTotal prob of N-in: 0.25502\noutside 1 212\nTMhelix 213 235\ninside 236 240

31616 GCF_900096885.1_IMG-taxon_2616644834_annotated_assembly Microbacterium enclense Terrabacteria group; Actinobacteria; Actinobacteria; Micrococcales; Microbacteriaceae; Microbacterium MHHSPPSRRRTLAALGVIAALLGGLGIAPAAQAADRGTGFGTWAPLSRTGWHGSMRVGDV WP_058232575.1 hypothetical protein [Microbacterium enclense] Length: 501\nNumber of predicted TMHs: 2\nExp number of AAs in TMHs: 44.10168\nExp number, first 60 AAs: 21.71168\nTotal prob of N-in: 0.99421\nPOSSIBLE N-term signal sequence\ninside 1 11\nTMhelix 12 34\noutside 35 469\nTMhelix 470 492\ninside 493 501

31617 GCF_900096885.1_IMG-taxon_2616644834_annotated_assembly Microbacterium enclense Terrabacteria group; Actinobacteria; Actinobacteria; Micrococcales; Microbacteriaceae; Microbacterium MCPSCAASLPRGARTTPAGAGIRVVCRAERCFVQELCGRPARRRRPLRPTAPPAGAGGGA WP_082642070.1 hypothetical protein [Microbacterium enclense] Length: 240\nNumber of predicted TMHs: 1\nExp number of AAs in TMHs: 22.79157\nExp number, first 60 AAs: 0.00099\nTotal prob of N-in: 0.25502\noutside 1 212\nTMhelix 213 235\ninside 236 240

31618 GCF_900096885.1_IMG-taxon_2616644834_annotated_assembly Microbacterium enclense Terrabacteria group; Actinobacteria; Actinobacteria; Micrococcales; Microbacteriaceae; Microbacterium MARTPTAALSPADQQRRRALRVMKGVALGALLGMAIVFAISFSLQREVEWLQYVRAAAEG WP_058232870.1 DUF445 domain-containing protein [Microbacterium enclense] Length: 424\nNumber of predicted TMHs: 2\nExp number of AAs in TMHs: 54.6351900000001\nExp number, first 60 AAs: 22.97903\nTotal prob of N-in: 0.93544\nPOSSIBLE N-term signal sequence\ninside 1 19\nTMhelix 20 42\noutside 43 400\nTMhelix 401 423\ninside 424 424

31619 GCF_900096885.1_IMG-taxon_2616644834_annotated_assembly Microbacterium enclense Terrabacteria group; Actinobacteria; Actinobacteria; Micrococcales; Microbacteriaceae; Microbacterium MAREKKTRRASGRRSPLAAAALIGIGLLLTGGVYAGASAAMAATSSTPVSEANSATAVEE WP_058232414.1 cystathionine beta-lyase [Microbacterium enclense] Length: 270\nNumber of predicted TMHs: 2\nExp number of AAs in TMHs: 42.58573\nExp number, first 60 AAs: 22.35903\nTotal prob of N-in: 0.99105\nPOSSIBLE N-term signal sequence\ninside 1 20\nTMhelix 21 43\noutside 44 247\nTMhelix 248 267\ninside 268 270

31620 GCF_900096885.1_IMG-taxon_2616644834_annotated_assembly Microbacterium enclense Terrabacteria group; Actinobacteria; Actinobacteria; Micrococcales; Microbacteriaceae; Microbacterium MRSSQTPRRRTLPAVLTTVLLAAAALLVPALPAAAHDELIGTDPASDAVLETLPEQITLS WP_058232366.1 hypothetical protein [Microbacterium enclense] Length: 212\nNumber of predicted TMHs: 2\nExp number of AAs in TMHs: 44.63027\nExp number, first 60 AAs: 22.51034\nTotal prob of N-in: 0.99901\nPOSSIBLE N-term signal sequence\ninside 1 11\nTMhelix 12 34\noutside 35 175\nTMhelix 176 198\ninside 199 212

31621 GCF_900096885.1_IMG-taxon_2616644834_annotated_assembly Microbacterium enclense Terrabacteria group; Actinobacteria; Actinobacteria; Micrococcales; Microbacteriaceae; Microbacterium MTTPTLPPLSAERADAIESAVFGRIADERHRVRRRRRNVWTGVGAAAAVLLIAAVVGPSL WP_058231401.1 hypothetical protein [Microbacterium enclense] Length: 351\nNumber of predicted TMHs: 2\nExp number of AAs in TMHs: 44.8187\nExp number, first 60 AAs: 20.69746\nTotal prob of N-in: 0.98838\nPOSSIBLE N-term signal sequence\ninside 1 38\nTMhelix 39 61\noutside 62 313\nTMhelix 314 336\ninside 337 351

31622 GCF_001456955.1_ASM145695v1 Microbacterium enclense Terrabacteria group; Actinobacteria; Actinobacteria; Micrococcales; Microbacteriaceae; Microbacterium
MRSSQTPRRRTLPAVLTTVLLAAAALLVPALPAAAHDELIGTDPASDAVLETLPEQITLS WP_058232366.1 hypothetical protein [Microbacterium enclense] Length: 212\nNumber of predicted TMHs: 2\nExp number of AAs in TMHs: 44.63027\nExp number, first 60 AAs: 22.51034\nTotal prob of N-in: 0.99901\nPOSSIBLE N-term signal sequence\ninside 1 11\nTMhelix 12 34\noutside 35 175\nTMhelix 176 198\ninside 199 212

31623 GCF_001456955.1_ASM145695v1 Microbacterium enclense Terrabacteria group; Actinobacteria; Actinobacteria; Micrococcales; Microbacteriaceae; Microbacterium
MAREKKTRRASGRRSPLAAAALIGIGLLLTGGVYAGASAAMAATSSTPVSEANSATAVEE WP_058232414.1 cystathionine beta-lyase [Microbacterium enclense] Length: 270\nNumber of predicted TMHs: 2\nExp number of AAs in TMHs: 42.58573\nExp number, first 60 AAs: 22.35903\nTotal prob of N-in: 0.99105\nPOSSIBLE N-term signal sequence\ninside 1 20\nTMhelix 21 43\noutside 44 247\nTMhelix 248 267\ninside 268 270

31624 GCF_001456955.1_ASM145695v1 Microbacterium enclense Terrabacteria group; Actinobacteria; Actinobacteria; Micrococcales; Microbacteriaceae; Microbacterium
MHHSPPSRRRTLAALGVIAALLGGLGIAPAAQAADRGTGFGTWAPLSRTGWHGSMRVGDV WP_058232575.1 hypothetical protein [Microbacterium enclense] Length: 501\nNumber of predicted TMHs: 2\nExp number of AAs in TMHs: 44.10168\nExp number, first 60 AAs: 21.71168\nTotal prob of N-in: 0.99421\nPOSSIBLE N-term signal sequence\ninside 1 11\nTMhelix 12 34\noutside 35 469\nTMhelix 470 492\ninside 493 501

31625 GCF_001456955.1_ASM145695v1 Microbacterium enclense Terrabacteria group; Actinobacteria; Actinobacteria; Micrococcales; Microbacteriaceae; Microbacterium
MTTPTLPPLSAERADAIESAVFGRIADERHRVRRRRRNVWTGVGAAAAVLLIAAVVGPSL WP_058231401.1 hypothetical protein [Microbacterium enclense] Length: 351\nNumber of predicted TMHs: 2\nExp number of AAs in TMHs: 44.8187\nExp number, first 60 AAs: 20.69746\nTotal prob of N-in: 0.98838\nPOSSIBLE N-term signal sequence\ninside 1 38\nTMhelix 39 61\noutside 62 313\nTMhelix 314 336\ninside 337 351

31626 GCF_001456955.1_ASM145695v1 Microbacterium enclense Terrabacteria group; Actinobacteria; Actinobacteria; Micrococcales; Microbacteriaceae; Microbacterium
MARTPTAALSPADQQRRRALRVMKGVALGALLGMAIVFAISFSLQREVEWLQYVRAAAEG WP_058232870.1 DUF445 domain-containing protein [Microbacterium enclense] Length: 424\nNumber of predicted TMHs: 2\nExp number of AAs in TMHs: 54.6351900000001\nExp number, first 60 AAs: 22.97903\nTotal prob of N-in: 0.93544\nPOSSIBLE N-term signal sequence\ninside 1 19\nTMhelix 20 42\noutside 43 400\nTMhelix 401 423\ninside 424 424

31627 GCF_900163815.1_ASM90016381v1 Microbacterium esteraromaticum Terrabacteria group; Actinobacteria; Actinobacteria; Micrococcales; Microbacteriaceae; Microbacterium
MPFPPAARHRRQVVAATLAALLGGILIAPAAASADAEVEVPTAPLITTDSTWRYLDDNTD WP_087133114.1 metallophosphoesterase [Microbacterium esteraromaticum] Length: 869\nNumber of predicted TMHs: 2\nExp number of AAs in TMHs: 38.20666\nExp number, first 60 AAs: 18.7625\nTotal prob of N-in: 0.89361\nPOSSIBLE N-term signal sequence\ninside 1 12\nTMhelix 13 32\noutside 33 840\nTMhelix 841 860\ninside 861 869

31628 GCF_900163815.1_ASM90016381v1 Microbacterium esteraromaticum Terrabacteria group; Actinobacteria; Actinobacteria; Micrococcales; Microbacteriaceae; Microbacterium
MAREKKHRSRGRRSPLAAAALIGAGLLLTGGAYAGATAAIASTSDTQTSAQAELTVDDGQ WP_087129828.1 cystathionine beta-lyase [Microbacterium esteraromaticum] Length: 277\nNumber of predicted TMHs: 2\nExp number of AAs in TMHs: 42.93436\nExp number, first 60 AAs: 22.17588\nTotal prob of N-in: 0.97064\nPOSSIBLE N-term signal sequence\ninside 1 19\nTMhelix 20 42\noutside 43 254\nTMhelix 255 274\ninside 275 277

31629 GCF_002024885.1_ASM202488v1 Microbacterium foliorum Terrabacteria group; Actinobacteria; Actinobacteria; Micrococcales; Microbacteriaceae; Microbacterium
MAREKKRRSNGRRSPLAAAALIGAGLMITGAVYAGTSAAFAATDTQTTAATTLTVEDGEK WP_079113075.1 cystathionine beta-lyase [Microbacterium foliorum] Length: 267\nNumber of predicted TMHs: 2\nExp number of AAs in TMHs: 43.07809\nExp number, first 60 AAs: 22.53353\nTotal prob of N-in: 0.98246\nPOSSIBLE N-term signal sequence\ninside 1 20\nTMhelix 21 43\noutside 44 244\nTMhelix 245 264\ninside 265 267

31630 GCF_002024885.1_ASM202488v1 Microbacterium foliorum Terrabacteria group; Actinobacteria; Actinobacteria; Micrococcales; Microbacteriaceae; Microbacterium
MRRRRTALPALLVGAIALGAVLTSAPASAATLGSTVPAATAGEACTPGTRVFTPEAPPAL WP_079112183.1 hypothetical protein [Microbacterium foliorum] Length: 447\nNumber of predicted TMHs: 2\nExp number of AAs in TMHs:

43.27539\nExp number, first 60 AAs: 21.82965\nTotal prob of N-in: 0.98489\nPOSSIBLE N-term signal sequence\ninside 1 6\nTMhelix 7 29\noutside 30 389\nTMhelix 390 412\ninside 413 447

31631 GCF_000956415.1_ASM95641v1 Microbacterium foliorum Terrabacteria group; Actinobacteria; Actinobacteria; Micrococcales; Microbacteriaceae; Microbacterium MAREKKRRSNGRRSPLAAAALIGAGLMITGAVYAGTSAAFAATDTQTAASSTLTVDDGEK WP_045253754.1 MULTISPECIES: cystathionine beta-lyase [Microbacterium] Length: 267\nNumber of predicted TMHs: 2\nExp number of AAs in TMHs: 43.05922\nExp number, first 60 AAs: 22.53913\nTotal prob of N-in: 0.98687\nPOSSIBLE N-term signal sequence\ninside 1 20\nTMhelix 21 43\noutside 44 244\nTMhelix 245 264\ninside 265 267

31632 GCF_000956535.1_ASM95653v1 Microbacterium ginsengisoli Terrabacteria group; Actinobacteria; Actinobacteria; Micrococcales; Microbacteriaceae; Microbacterium MARTPKPRKAGRRSPLAAAALIGIGLLLTGGIYAGASAAVAATTDQTSTSTKASTLTVDD WP_045247257.1 cystathionine beta-lyase [Microbacterium ginsengisoli] Length: 270\nNumber of predicted TMHs: 2\nExp number of AAs in TMHs: 43.4897799999999\nExp number, first 60 AAs: 22.68752\nTotal prob of N-in: 0.90593\nPOSSIBLE N-term signal sequence\ninside 1 19\nTMhelix 20 42\noutside 43 247\nTMhelix 248 267\ninside 268 270

31633 GCF_000956535.1_ASM95653v1 Microbacterium ginsengisoli Terrabacteria group; Actinobacteria; Actinobacteria; Micrococcales; Microbacteriaceae; Microbacterium MTTVTDHEHPHRSGSRRRILLLSTLTTLLLAAGLLFATAAPAAAHDEIVSSSPEAGSTVS WP_048809232.1 MULTISPECIES: Copper resistance protein C precursor [Microbacterium] Length: 238\nNumber of predicted TMHs: 2\nExp number of AAs in TMHs: 45.12191\nExp number, first 60 AAs: 22.44287\nTotal prob of N-in: 0.99979\nPOSSIBLE N-term signal sequence\ninside 1 19\nTMhelix 20 42\noutside 43 183\nTMhelix 184 206\ninside 207 238

31634 GCF_000956535.1_ASM95653v1 Microbacterium ginsengisoli Terrabacteria group; Actinobacteria; Actinobacteria; Micrococcales; Microbacteriaceae; Microbacterium MSISQPRRRALAVIAGASLLIGIAAPGTALAASAAEPAPADVIINEAYLKGGSAGAPFTN WP_048809135.1 Endonuclease YhcR precursor [Microbacterium ginsengisoli] Length: 1638\nNumber of predicted TMHs: 2\nExp number of AAs in TMHs: 40.2516499999999\nExp number, first 60 AAs: 21.7957\nTotal prob of N-in: 0.96934\nPOSSIBLE N-term signal sequence\ninside 1 12\nTMhelix 13 35\noutside 36 1613\nTMhelix 1614 1633\ninside 1634 1638

31635 GCF_000826185.1_Microbacterium_massiliensis_G3 Microbacterium gorillae Terrabacteria group; Actinobacteria; Actinobacteria; Micrococcales; Microbacteriaceae; Microbacterium MPHTPTELLSPADQERRRALRLMKSVALGALIAMAIVFAVAFVLQREYAGWGYVRAAAEG WP_040166109.1 DUF445 domain-containing protein [Microbacterium gorillae] Length: 425\nNumber of predicted TMHs: 2\nExp number of AAs in TMHs: 51.5211\nExp number, first 60 AAs: 23.80125\nTotal prob of N-in: 0.40592\nPOSSIBLE N-term signal sequence\noutside 1 24\nTMhelix 25 44\ninside 45 400\nTMhelix 401 423\noutside 424 425

31636 GCF_000826185.1_Microbacterium_massiliensis_G3 Microbacterium gorillae Terrabacteria group; Actinobacteria; Actinobacteria; Micrococcales; Microbacteriaceae; Microbacterium MAREKKTRNNGRRSPLAAVALIGLGLLLTGGAYAGASAAVAATTDSSTSVSAAADTDTGK WP_040165408.1 cystathionine beta-lyase [Microbacterium gorillae] Length: 268\nNumber of predicted TMHs: 2\nExp number of AAs in TMHs: 43.04884\nExp number, first 60 AAs: 22.54205\nTotal prob of N-in: 0.98440\nPOSSIBLE N-term signal sequence\ninside 1 19\nTMhelix 20 42\noutside 43 245\nTMhelix 246 265\ninside 266 268

31637 GCF_000826185.1_Microbacterium_massiliensis_G3 Microbacterium gorillae Terrabacteria group; Actinobacteria; Actinobacteria; Micrococcales; Microbacteriaceae; Microbacterium MARRRLITAWAGGATVVLASVFALTPAVHAETAPSEDAPANVVVSLAWEKPESATAVPAP WP_052460806.1 hypothetical protein [Microbacterium gorillae] Length: 349\nNumber of predicted TMHs: 2\nExp number of AAs in TMHs: 41.06041\nExp number, first 60 AAs: 20.1605\nTotal prob of N-in: 0.92243\nPOSSIBLE N-term signal sequence\ninside 1 6\nTMhelix 7 29\noutside 30 315\nTMhelix 316 338\ninside 339 349

31638 GCF_000813805.1_ASM81380v1 Microbacterium hominis Terrabacteria group; Actinobacteria; Actinobacteria; Micrococcales; Microbacteriaceae; Microbacterium MSARRRRFVAVVTAVMIGLGGQAAAGAASIPAVAVAAGDSAPAALAGPSALAATAPPPGT WP_039412518.1 peptidase S8 [Microbacterium hominis] Length: 454\nNumber of predicted TMHs: 2\nExp number of AAs in TMHs: 42.9817899999999\nExp number, first 60 AAs: 20.17521\nTotal prob of N-in: 0.90527\nPOSSIBLE N-term signal sequence\ninside 1 6\nTMhelix 7 29\noutside 30 413\nTMhelix 414 436\ninside 437 454

31639 GCF_001553805.1_ASM155380v1 Microbacterium hominis Terrabacteria group; Actinobacteria; Actinobacteria; Micrococcales; Microbacteriaceae; Microbacterium

MSDQTPDAPRPTADRRKRRSVVAAITAGALAFSGLGLAALPARAAVDTGAAVLINEVYGG WP_060958951.1 cell wall protein [Microbacterium hominis] Length: 1577\nNumber of predicted TMHs: 2\nExp number of AAs in TMHs: 40.9041299999999\nExp number, first 60 AAs: 21.7961\nTotal prob of N-in: 0.99821\nPOSSIBLE N-term signal sequence\ninside 1 19\nTMhelix 20 42\noutside 43 1548\nTMhelix 1549 1571\ninside 1572 1577

31640 GCF_000813805.1_ASM81380v1 Microbacterium hominis Terrabacteria group; Actinobacteria; Actinobacteria; Micrococcales; Microbacteriaceae; Microbacterium MISHRTRRGFIDRLGASTLVAAAVVCSIGFGATSASAADAASQPPLTAKDHAITDVPGLV WP_039415654.1 cell wall protein [Microbacterium hominis] Length: 416\nNumber of predicted TMHs: 2\nExp number of AAs in TMHs: 44.46333\nExp number, first 60 AAs: 12.68979\nTotal prob of N-in: 0.73324\nPOSSIBLE N-term signal sequence\ninside 1 348\nTMhelix 349 371\noutside 372 385\nTMhelix 386 408\ninside 409 416

31641 GCF_000813805.1_ASM81380v1 Microbacterium hominis Terrabacteria group; Actinobacteria; Actinobacteria; Micrococcales; Microbacteriaceae; Microbacterium MRPTAERRNRRAIVATITAAALALSGAGFAAIPAQAAVSPNAAVLINEVYGGGGNSGATY WP_039415903.1 cell wall protein [Microbacterium hominis] Length: 1574\nNumber of predicted TMHs: 2\nExp number of AAs in TMHs: 42.44782\nExp number, first 60 AAs: 22.16313\nTotal prob of N-in: 0.99772\nPOSSIBLE N-term signal sequence\ninside 1 11\nTMhelix 12 34\noutside 35 1544\nTMhelix 1545 1564\ninside 1565 1574

31642 GCF_001553805.1_ASM155380v1 Microbacterium hominis Terrabacteria group; Actinobacteria; Actinobacteria; Micrococcales; Microbacteriaceae; Microbacterium MGTTNTAAAPARRRRALPAFVVAGLLGLIAAGSSASGAAASSPLIDATPATVTIEAPAPG WP_060958999.1 hypothetical protein [Microbacterium hominis] Length: 216\nNumber of predicted TMHs: 2\nExp number of AAs in TMHs: 44.64633\nExp number, first 60 AAs: 21.91522\nTotal prob of N-in: 0.99385\nPOSSIBLE N-term signal sequence\ninside 1 16\nTMhelix 17 39\noutside 40 177\nTMhelix 178 200\ninside 201 216

31643 GCF_001553805.1_ASM155380v1 Microbacterium hominis Terrabacteria group; Actinobacteria; Actinobacteria; Micrococcales; Microbacteriaceae; Microbacterium MAPTPTQLLSPADQERRRALRVMKGVALGALVLMAVVFVVAFVLQERYPWLEYVRAAAEG WP_060959398.1 DUF445 domain-containing protein [Microbacterium hominis] Length: 423\nNumber of predicted TMHs: 2\nExp number of AAs in TMHs: 52.34446\nExp number, first 60 AAs: 22.52241\nTotal prob of N-in: 0.78491\nPOSSIBLE N-term signal sequence\ninside 1 24\nTMhelix 25 44\noutside 45 399\nTMhelix 400 422\ninside 423 423

31644 GCF_000956475.1_ASM95647v1 Microbacterium hydrocarbonoxydans Terrabacteria group; Actinobacteria; Actinobacteria; Micrococcales; Microbacteriaceae; Microbacterium MAREKKRRSTGRRSPLAAAALIGAGLMITGAVYAGASAAFASTDTQAAAASTLSVEDGEK WP_045258269.1 cystathionine beta-lyase [Microbacterium hydrocarbonoxydans] Length: 267\nNumber of predicted TMHs: 2\nExp number of AAs in TMHs: 42.50822\nExp number, first 60 AAs: 22.33728\nTotal prob of N-in: 0.98223\nPOSSIBLE N-term signal sequence\ninside 1 20\nTMhelix 21 43\noutside 44 244\nTMhelix 245 264\ninside 265 267

31645 GCF_900105205.1_IMG-taxon_2634166363_annotated_assembly Microbacterium hydrocarbonoxydans Terrabacteria group; Actinobacteria; Actinobacteria; Micrococcales; Microbacteriaceae; Microbacterium MIGRAAEAEQHSDDVARGLPSEGGTMARRRAFIRAIVAGAAALALAIGGATTAMAADDDP WP_060928567.1 hypothetical protein [Microbacterium hydrocarbonoxydans] Length: 693\nNumber of predicted TMHs: 2\nExp number of AAs in TMHs: 43.6390999999999\nExp number, first 60 AAs: 21.83173\nTotal prob of N-in: 0.99014\nPOSSIBLE N-term signal sequence\ninside 1 30\nTMhelix 31 53\noutside 54 666\nTMhelix 667 686\ninside 687 693

31646 GCF_900105205.1_IMG-taxon_2634166363_annotated_assembly Microbacterium hydrocarbonoxydans Terrabacteria group; Actinobacteria; Actinobacteria; Micrococcales; Microbacteriaceae; Microbacterium MAREKKRRSNGRRSPLAAAALIGAGLMITGAVYAGTSAAFAASDTETTAASTLTVEDGEK WP_060926486.1 cystathionine beta-lyase [Microbacterium hydrocarbonoxydans] Length: 267\nNumber of predicted TMHs: 2\nExp number of AAs in TMHs: 42.80265\nExp number, first 60 AAs: 22.42264\nTotal prob of N-in: 0.99124\nPOSSIBLE N-term signal sequence\ninside 1 19\nTMhelix 20 42\noutside 43 244\nTMhelix 245 264\ninside 265 267

31647 GCF_000956475.1_ASM95647v1 Microbacterium hydrocarbonoxydans Terrabacteria group; Actinobacteria; Actinobacteria; Micrococcales; Microbacteriaceae; Microbacterium MTEKTTTPELPELSDDAIARIESAVFDEIASERPPTRVVERARARRRGWLTGAGIAAAFI WP_045258722.1 hypothetical protein [Microbacterium hydrocarbonoxydans] Length: 354\nNumber of predicted TMHs: 2\nExp number of AAs in TMHs: 45.85351\nExp number, first 60 AAs: 12.54678\nTotal prob of N-in: 0.99318\nPOSSIBLE N-term signal sequence\ninside 1 47\nTMhelix 48 70\noutside 71 314\nTMhelix 315 337\ninside 338 354

31648 GCF_000956575.1_ASM95657v1 Microbacterium ketosireducens Terrabacteria group; Actinobacteria; Actinobacteria; Micrococcales; Microbacteriaceae; Microbacterium
MHARRRTPMRTWLAALTAAALAVSGAVVVDVASAPPARAAYPDSFNPFSMNGGFTVYARE WP_084613500.1 hypothetical protein [Microbacterium ketosireducens] Length: 1095\nNumber of predicted TMHs: 2\nExp number of AAs in TMHs: 42.92883\nExp number, first 60 AAs: 21.80016\nTotal prob of N-in: 0.98793\nPOSSIBLE N-term signal sequence\ninside 1 11\nTMhelix 12 34\noutside 35 1066\nTMhelix 1067 1089\ninside 1090 1095

31649 GCF_001584605.1_ASM158460v1 Microbacterium laevaniformans Terrabacteria group; Actinobacteria; Actinobacteria; Micrococcales; Microbacteriaceae; Microbacterium
MITHRTRRGILGRLGATTLVAAAVVCSVGLSATSASADDSSNQPPLTAKDHAITDVPGLV WP_061683644.1 cell wall protein [Microbacterium laevaniformans] Length: 416\nNumber of predicted TMHs: 2\nExp number of AAs in TMHs: 47.61046\nExp number, first 60 AAs: 10.89251\nTotal prob of N-in: 0.70101\nPOSSIBLE N-term signal sequence\ninside 1 348\nTMhelix 349 371\noutside 372 385\nTMhelix 386 408\ninside 409 416

31650 GCF_001584605.1_ASM158460v1 Microbacterium laevaniformans Terrabacteria group; Actinobacteria; Actinobacteria; Micrococcales; Microbacteriaceae; Microbacterium
MTCIPVADRASRRRSPTAGIHRRSFLLPGLAALLLTAGLLFAVAAPAAAHDEIVSSSPEA WP_082784215.1 hypothetical protein [Microbacterium laevaniformans] Length: 245\nNumber of predicted TMHs: 2\nExp number of AAs in TMHs: 45.28546\nExp number, first 60 AAs: 22.61402\nTotal prob of N-in: 0.99745\nPOSSIBLE N-term signal sequence\ninside 1 23\nTMhelix 24 46\noutside 47 188\nTMhelix 189 211\ninside 212 245

31651 GCF_001584605.1_ASM158460v1 Microbacterium laevaniformans Terrabacteria group; Actinobacteria; Actinobacteria; Micrococcales; Microbacteriaceae; Microbacterium
MSDPTRDATRPTAERRNRRAIVATITAGALALSGAGFATIPAQAAVSPNAAVLINEVYGG WP_061683200.1 cell wall protein [Microbacterium laevaniformans] Length: 1582\nNumber of predicted TMHs: 2\nExp number of AAs in TMHs: 42.4232499999999\nExp number, first 60 AAs: 22.06233\nTotal prob of N-in: 0.99607\nPOSSIBLE N-term signal sequence\ninside 1 19\nTMhelix 20 42\noutside 43 1552\nTMhelix 1553 1572\ninside 1573 1582

31652 GCF_000802305.1_ASM80230v1 Microbacterium mangrovi Terrabacteria group; Actinobacteria; Actinobacteria; Micrococcales; Microbacteriaceae; Microbacterium
MAREKKRRNSGRRSPLAAAVLIGVGLLLTGGVYAGASAAMAATNDTTQQSSNLTAEDGKK WP_039400505.1 cystathionine beta-lyase [Microbacterium mangrovi] Length: 267\nNumber of predicted TMHs: 2\nExp number of AAs in TMHs: 43.01344\nExp number, first 60 AAs: 22.52295\nTotal prob of N-in: 0.97888\nPOSSIBLE N-term signal sequence\ninside 1 19\nTMhelix 20 42\noutside 43 244\nTMhelix 245 264\ninside 265 267

31653 GCF_000802305.1_ASM80230v1 Microbacterium mangrovi Terrabacteria group; Actinobacteria; Actinobacteria; Micrococcales; Microbacteriaceae; Microbacterium
MNGNTLKTSPSRARRTLLAAAGVTAGCALALAVPLAASAHVEVSPATAPAGGTTRLTFQF WP_052226469.1 hypothetical protein [Microbacterium mangrovi] Length: 238\nNumber of predicted TMHs: 2\nExp number of AAs in TMHs: 43.74791\nExp number, first 60 AAs: 21.92788\nTotal prob of N-in: 0.98185\nPOSSIBLE N-term signal sequence\ninside 1 16\nTMhelix 17 39\noutside 40 211\nTMhelix 212 234\ninside 235 238

31654 GCF_000632065.1_Willow_Endophyte_RIT293 Microbacterium oleivorans Terrabacteria group; Actinobacteria; Actinobacteria; Micrococcales; Microbacteriaceae; Microbacterium
MTTRSARRSALTLSGLALGAALVLAAPLAASAHVHVHSDDAAAGATSRLEFSFSHGCDGA WP_036311780.1 nuclear export factor GLE1 [Microbacterium oleivorans] Length: 232\nNumber of predicted TMHs: 2\nExp number of AAs in TMHs: 44.16283\nExp number, first 60 AAs: 21.69563\nTotal prob of N-in: 0.99301\nPOSSIBLE N-term signal sequence\ninside 1 12\nTMhelix 13 35\noutside 36 204\nTMhelix 205 227\ninside 228 232

31655 GCF_001975955.1_ASM197595v1 Microbacterium oleivorans Terrabacteria group; Actinobacteria; Actinobacteria; Micrococcales; Microbacteriaceae; Microbacterium
MTTRSARRSALTLSGLALGAALVLAAPLAASAHVHVHSDDAAAGATSRLEFSFSHGCDGA WP_036311780.1 nuclear export factor GLE1 [Microbacterium oleivorans] Length: 232\nNumber of predicted TMHs: 2\nExp number of AAs in TMHs: 44.16283\nExp number, first 60 AAs: 21.69563\nTotal prob of N-in: 0.99301\nPOSSIBLE N-term signal sequence\ninside 1 12\nTMhelix 13 35\noutside 36 204\nTMhelix 205 227\ninside 228 232

31656 GCF_001644225.1_ASM164422v1 Microbacterium oleivorans Terrabacteria group; Actinobacteria; Actinobacteria; Micrococcales; Microbacteriaceae; Microbacterium
MTTRSPRRSALTLTGIALGGALVLAAPLAASAHVHVHAEGAAAGATSRLDFSFSHGCDGS WP_064001498.1 nuclear export factor GLE1 [Microbacterium oleivorans] Length: 225\nNumber of predicted TMHs: 2\nExp

number of AAs in TMHs: 44.07702\nExp number, first 60 AAs:  21.55133\nTotal prob of N-in:   0.98012\nPOSSIBLE N-term signal sequence\ninside   1   8\nTMhelix   9   31\noutside   32   198\nTMhelix   199   221\ninside   222   225

31657 GCF_001691565.1_ASM169156v1 Microbacterium oleivorans Terrabacteria group; Actinobacteria; Actinobacteria; Micrococcales; Microbacteriaceae; Microbacterium
MTTRSARRSALTLTGLALGGALVLAAPLAASAHVHVHADGAAAGATSRLEFSFSHGCDGA WP_065569994.1 nuclear export factor GLE1 [Microbacterium oleivorans] Length: 228\nNumber of predicted TMHs:  2\nExp number of AAs in TMHs: 44.4092199999999\nExp number, first 60 AAs:  21.77766\nTotal prob of N-in:   0.99150\nPOSSIBLE N-term signal sequence\ninside   1   12\nTMhelix   13   35\noutside   36   200\nTMhelix   201   223\ninside   224   228

31658 GCF_000956405.1_ASM95640v1 Microbacterium oxydans Terrabacteria group; Actinobacteria; Actinobacteria; Micrococcales; Microbacteriaceae; Microbacterium
MPRLTSASRRRLIATLPITALLAGSLLATSLPAAAAADPIDGAHTSGDPMFPNIGNGGY WP_045262964.1 hypothetical protein [Microbacterium oxydans] Length: 786\nNumber of predicted TMHs:  2\nExp number of AAs in TMHs: 41.98049\nExp number, first 60 AAs:  21.09708\nTotal prob of N-in:   0.94119\nPOSSIBLE N-term signal sequence\ninside 1   11\nTMhelix   12   34\noutside   35   753\nTMhelix   754   776\ninside   777   786

31659 GCF_000956525.1_ASM95652v1 Microbacterium oxydans Terrabacteria group; Actinobacteria; Actinobacteria; Micrococcales; Microbacteriaceae; Microbacterium
MAREKKRRSNGRRSPLAAVALIGAGLMITGAVYAGASAAFAATDTQSAATTQLTVEDGEK WP_045279983.1 cystathionine beta-lyase [Microbacterium oxydans] Length: 267\nNumber of predicted TMHs:  2\nExp number of AAs in TMHs: 43.12583\nExp number, first 60 AAs:  22.56227\nTotal prob of N-in:   0.98350\nPOSSIBLE N-term signal sequence\ninside   1   20\nTMhelix   21   43\noutside   44   244\nTMhelix   245   264\ninside   265   267

31660 GCF_000956525.1_ASM95652v1 Microbacterium oxydans Terrabacteria group; Actinobacteria; Actinobacteria; Micrococcales; Microbacteriaceae; Microbacterium
MLRLTSPARRRLLATLPVTTLLAGSLLVTSIPASAADGPIDGPRTAGDAMFPNVGNGGYD WP_045277607.1 hypothetical protein [Microbacterium oxydans] Length: 786\nNumber of predicted TMHs:  2\nExp number of AAs in TMHs: 43.1874399999999\nExp number, first 60 AAs:  18.68148\nTotal prob of N-in:   0.83911\nPOSSIBLE N-term signal sequence\ninside   1   11\nTMhelix   12   34\noutside   35   757\nTMhelix   758   777\ninside   778   786

31661 GCF_000956405.1_ASM95640v1 Microbacterium oxydans Terrabacteria group; Actinobacteria; Actinobacteria; Micrococcales; Microbacteriaceae; Microbacterium
MRDAEEGRRPAVVRGVIVAAVVSTALILIGALPAAAIYEPDPVETPRDGFITEVTITGSQY WP_045264519.1 hypothetical protein [Microbacterium oxydans] Length: 490\nNumber of predicted TMHs:  2\nExp number of AAs in TMHs: 40.1086299999999\nExp number, first 60 AAs:  22.48553\nTotal prob of N-in:   0.99426\nPOSSIBLE N-term signal sequence\ninside   1   11\nTMhelix   12   34\noutside   35   460\nTMhelix   461   478\ninside   479   490

31662 GCF_000956405.1_ASM95640v1 Microbacterium oxydans Terrabacteria group; Actinobacteria; Actinobacteria; Micrococcales; Microbacteriaceae; Microbacterium
MKMTRRRGMLRAVLGGMAAFALALGGATAGAAAEDPVVTGSISGTVTADAGGAALEGIQV WP_082066551.1 hypothetical protein [Microbacterium oxydans] Length: 676\nNumber of predicted TMHs:  2\nExp number of AAs in TMHs: 42.61554\nExp number, first 60 AAs:  20.90439\nTotal prob of N-in:   0.98281\nPOSSIBLE N-term signal sequence\ninside   1   11\nTMhelix   12   34\noutside   35   647\nTMhelix   648   670\ninside   671   676

31663 GCF_001476635.1_ASM147663v1 Microbacterium oxydans Terrabacteria group; Actinobacteria; Actinobacteria; Micrococcales; Microbacteriaceae; Microbacterium
MAREKKRRSSGRRSPLAAAALIGAGLMITGAVYAGATAAFAATDTQSAATSQLTVEDGKK WP_058632814.1 cystathionine beta-lyase [Microbacterium oxydans] Length: 267\nNumber of predicted TMHs:  2\nExp number of AAs in TMHs: 43.1932\nExp number, first 60 AAs:  22.66658\nTotal prob of N-in:   0.98686\nPOSSIBLE N-term signal sequence\ninside   1   20\nTMhelix   21   43\noutside   44   244\nTMhelix   245   264\ninside   265   267

31664 GCF_000956405.1_ASM95640v1 Microbacterium oxydans Terrabacteria group; Actinobacteria; Actinobacteria; Micrococcales; Microbacteriaceae; Microbacterium
MRRRAALATLVLGAAVFGAPTSALASTSAPASIVSAATGACEPGNIVFTPQAPAALAALG WP_052674706.1 hypothetical protein [Microbacterium oxydans] Length: 442\nNumber of predicted TMHs:  2\nExp number of AAs in TMHs: 39.3946699999999\nExp number, first 60 AAs:  17.96849\nTotal prob of N-in:   0.82510\nPOSSIBLE N-term signal sequence\ninside   1   6\nTMhelix   7   29\noutside   30   385\nTMhelix   386   408\ninside   409   442

31665 GCF_001887285.1_ASM188728v1 Microbacterium paludicola Terrabacteria group; Actinobacteria; Actinobacteria; Micrococcales; Microbacteriaceae; Microbacterium
MTSPTTVTKPTKPATRAEARTARVVTPRRRPLSRVGDAMLVLAAATGALCIALTIAAAVF WP_071918655.1 hypothetical protein [Microbacterium paludicola] Length: 205\nNumber of predicted TMHs: 2\nExp number of AAs in TMHs: 47.81219\nExp number, first 60 AAs: 22.60801\nTotal prob of N-in: 0.93513\nPOSSIBLE N-term signal sequence\ninside 1 37\nTMhelix 38 60\noutside 61 164\nTMhelix 165 187\ninside 188 205

31666 GCF_001887285.1_ASM188728v1 Microbacterium paludicola Terrabacteria group; Actinobacteria; Actinobacteria; Micrococcales; Microbacteriaceae; Microbacterium
MLRTRRAALQRPLASVLTGALVAMAAVVGVAAPATAAGPTLTTSTTVVANEKITVAITGT WP_084670347.1 hypothetical protein [Microbacterium paludicola] Length: 962\nNumber of predicted TMHs: 2\nExp number of AAs in TMHs: 41.5393199999999\nExp number, first 60 AAs: 22.33509\nTotal prob of N-in: 0.99758\nPOSSIBLE N-term signal sequence\ninside 1 12\nTMhelix 13 35\noutside 36 934\nTMhelix 935 954\ninside 955 962

31667 GCF_001887285.1_ASM188728v1 Microbacterium paludicola Terrabacteria group; Actinobacteria; Actinobacteria; Micrococcales; Microbacteriaceae; Microbacterium
MPPAPSRPQSRPRRTAVPAAAVAAFTAACLTLGTAAPAGAADGELLDDGAIAPHAAPYGT WP_071919170.1 hypothetical protein [Microbacterium paludicola] Length: 702\nNumber of predicted TMHs: 1\nExp number of AAs in TMHs: 34.41766\nExp number, first 60 AAs: 12.4098\nTotal prob of N-in: 0.56985\nPOSSIBLE N-term signal sequence\noutside 1 671\nTMhelix 672 694\ninside 695 702

31668 GCF_900105335.1_IMG-taxon_2634166897_annotated_assembly Microbacterium paraoxydansTerrabacteria group; Actinobacteria; Actinobacteria; Micrococcales; Microbacteriaceae; Microbacterium
MTRRRMLRGAVALVAVAASVLLLGSTATPTPTPPPVADDPADPVRAAEYWLDGAGIRDAW WP_060921984.1 peptidase S8 [Microbacterium paraoxydans] Length: 439\nNumber of predicted TMHs: 2\nExp number of AAs in TMHs: 41.47033\nExp number, first 60 AAs: 19.80703\nTotal prob of N-in: 0.93228\nPOSSIBLE N-term signal sequence\ninside 1 6\nTMhelix 7 29\noutside 30 397\nTMhelix 398 420\ninside 421 439

31669 GCF_900105335.1_IMG-taxon_2634166897_annotated_assembly Microbacterium paraoxydansTerrabacteria group; Actinobacteria; Actinobacteria; Micrococcales; Microbacteriaceae; Microbacterium
MAREKKRRSSGRRSPLAAAALIGAGLMITGAVYAGATAAFAANDAPTASTQLTVEDGEKL WP_025105058.1 MULTISPECIES: cystathionine beta-lyase [Microbacterium] Length: 266\nNumber of predicted TMHs: 2\nExp number of AAs in TMHs: 43.19361\nExp number, first 60 AAs: 22.60029\nTotal prob of N-in: 0.99024\nPOSSIBLE N-term signal sequence\ninside 1 19\nTMhelix 20 42\noutside 43 243\nTMhelix 244 263\ninside 264 266

31670 GCF_900105335.1_IMG-taxon_2634166897_annotated_assembly Microbacterium paraoxydansTerrabacteria group; Actinobacteria; Actinobacteria; Micrococcales; Microbacteriaceae; Microbacterium
MSNTTIRPRRPRRTLLGATGIVGGLALALAVPAMAGAHVSVSPDELAAGDHGVLTFSFSH WP_060922111.1 hypothetical protein [Microbacterium paraoxydans] Length: 224\nNumber of predicted TMHs: 2\nExp number of AAs in TMHs: 45.2104\nExp number, first 60 AAs: 22.3675\nTotal prob of N-in: 0.99874\nPOSSIBLE N-term signal sequence\ninside 1 12\nTMhelix 13 35\noutside 36 197\nTMhelix 198 220\ninside 221 224

31671 GCF_900105335.1_IMG-taxon_2634166897_annotated_assembly Microbacterium paraoxydansTerrabacteria group; Actinobacteria; Actinobacteria; Micrococcales; Microbacteriaceae; Microbacterium
MRRSSTFGEGIAAGSAALEVDRDSIGLRRPPPEGGDMARRRAFFRSIMGGAAALALALGG WP_082750119.1 hypothetical protein [Microbacterium paraoxydans] Length: 706\nNumber of predicted TMHs: 2\nExp number of AAs in TMHs: 39.42056\nExp number, first 60 AAs: 15.85922\nTotal prob of N-in: 0.96446\nPOSSIBLE N-term signal sequence\ninside 1 42\nTMhelix 43 65\noutside 66 679\nTMhelix 680 699\ninside 700 706

31672 GCF_900105335.1_IMG-taxon_2634166897_annotated_assembly Microbacterium paraoxydansTerrabacteria group; Actinobacteria; Actinobacteria; Micrococcales; Microbacteriaceae; Microbacterium
MRRRAALATLVVGAAVWGTPAAAVAAPTAPTTIVSMATGACEPGTVVFAPEAPSALAALG WP_060923224.1 hypothetical protein [Microbacterium paraoxydans] Length: 436\nNumber of predicted TMHs: 2\nExp number of AAs in TMHs: 43.05017\nExp number, first 60 AAs: 21.16204\nTotal prob of N-in: 0.94802\nPOSSIBLE N-term signal sequence\ninside 1 6\nTMhelix 7 29\noutside 30 386\nTMhelix 387 409\ninside 410 436

31673 GCF_000763375.1_ASM76337v1 Microbacterium profundi Terrabacteria group; Actinobacteria; Actinobacteria; Micrococcales; Microbacteriaceae; Microbacterium
MVREKKRRSNGRRSPLAAAALIGAGLLITGGIYAGASAAIAATDTRTSASTMSVEDGQKL WP_033107511.1 cystathionine beta-lyase [Microbacterium profundi] Length: 266\nNumber of predicted TMHs: 2\nExp number of AAs in TMHs:

43.77471\nExp number, first 60 AAs:  22.82329\nTotal prob of N-in:   0.95662\nPOSSIBLE N-term signal sequence\ninside   1   19\nTMhelix   20   42\noutside   43   243\nTMhelix   244   263\ninside   264   266

31674 GCF_000763375.1_ASM76337v1 Microbacterium profundi Terrabacteria group; Actinobacteria; Actinobacteria; Micrococcales; Microbacteriaceae; Microbacterium
MLVSRERRRGTLLAVGITALMAATLMTPTIAMAADDGATDITENGGATRNTEDRTAAVRE WP_084595551.1
alkaline phosphatase [Microbacterium profundi] Length: 631\nNumber of predicted TMHs:  2\nExp number of AAs in TMHs: 44.99075\nExp number, first 60 AAs:  22.00918\nTotal prob of N-in:   0.99224\nPOSSIBLE N-term signal sequence\ninside   1   11\nTMhelix   12   34\noutside   35   600\nTMhelix   601   623\ninside   624   631

31675 GCF_001689915.1_ASM168991v1 Microbacterium sediminis Terrabacteria group; Actinobacteria; Actinobacteria; Micrococcales; Microbacteriaceae; Microbacterium
MARTPTELLSPADQDRRRALAGMKAVAIGALLATAALFVIGFVGQERIPALAYVRAAGEG WP_067025400.1
DUF445 domain-containing protein [Microbacterium sediminis] Length: 423\nNumber of predicted TMHs: 2\nExp number of AAs in TMHs: 50.64566\nExp number, first 60 AAs:  25.01051\nTotal prob of N-in:   0.72683\nPOSSIBLE N-term signal sequence\ninside   1   19\nTMhelix   20   42\noutside   43   399\nTMhelix   400   422\ninside   423   423

31676 GCF_001689915.1_ASM168991v1 Microbacterium sediminis Terrabacteria group; Actinobacteria; Actinobacteria; Micrococcales; Microbacteriaceae; Microbacterium
MAREKKMRRSGGRRSPLATAALIGIGLLITGGGYAAASAAVASTAETSQVQTALTAEDGE WP_067023990.1
cystathionine beta-lyase [Microbacterium sediminis] Length: 268\nNumber of predicted TMHs:  2\nExp number of AAs in TMHs: 42.89546\nExp number, first 60 AAs:  22.36437\nTotal prob of N-in:   0.97996\nPOSSIBLE N-term signal sequence\ninside   1   19\nTMhelix   20   42\noutside   43   245\nTMhelix   246   265\ninside   266   268

31677 GCF_000383475.1_ASM38347v1 Microbacterium sp. 11MF Terrabacteria group; Actinobacteria; Actinobacteria; Micrococcales; Microbacteriaceae; Microbacterium
MTLRPARRSALTLTGLALGGALVLAAPLAASAHVHVHSDDAAAGATSRLEFSFSHGCDGA WP_020096836.1
hypothetical protein [Microbacterium sp. 11MF] Length: 228\nNumber of predicted TMHs:  2\nExp number of AAs in TMHs: 43.8300999999999\nExp number, first 60 AAs:  21.21718\nTotal prob of N-in:   0.96486\nPOSSIBLE N-term signal sequence\ninside   1   12\nTMhelix   13   35\noutside   36   201\nTMhelix   202   224\ninside   225   228

31678 GCF_001889265.1_ASM188926v1 Microbacterium sp. 1.5R Terrabacteria group; Actinobacteria; Actinobacteria; Micrococcales; Microbacteriaceae; Microbacterium
MAQRRALIHAVLGGVAAIALALGGATAAGAVEGDPTIDPGATTGEVIATTDDIAADPLEG WP_083569401.1
hypothetical protein [Microbacterium sp. 1.5R] Length: 669\nNumber of predicted TMHs:  2\nExp number of AAs in TMHs: 43.3673\nExp number, first 60 AAs:  22.46287\nTotal prob of N-in:   0.99339\nPOSSIBLE N-term signal sequence\ninside   1   6\nTMhelix   7   29\noutside   30   639\nTMhelix   640   662\ninside   663   669

31679 GCF_001889265.1_ASM188926v1 Microbacterium sp. 1.5R Terrabacteria group; Actinobacteria; Actinobacteria; Micrococcales; Microbacteriaceae; Microbacterium
MERRRRFLAGLTVLALAAASLAFGAAPAFAVAAFSIQKSAVEAGPFGPGDTVTYQITVNC WP_072591806.1
hypothetical protein [Microbacterium sp. 1.5R] Length: 2989\nNumber of predicted TMHs:  2\nExp number of AAs in TMHs: 43.32463\nExp number, first 60 AAs:  22.92081\nTotal prob of N-in:   0.99960\nPOSSIBLE N-term signal sequence\ninside   1   6\nTMhelix   7   29\noutside   30  2958\nTMhelix   2959  2978\ninside   2979  2989

31680 GCF_001889265.1_ASM188926v1 Microbacterium sp. 1.5R Terrabacteria group; Actinobacteria; Actinobacteria; Micrococcales; Microbacteriaceae; Microbacterium
MAREKKRRSNGRRSPLAAAALIGAGLMITGAVYAGTSAAFAATDTQTTAASTLTVEDGEK WP_053096702.1
MULTISPECIES: cystathionine beta-lyase [Microbacterium] Length: 267\nNumber of predicted TMHs:  2\nExp number of AAs in TMHs: 43.06753\nExp number, first 60 AAs:  22.53635\nTotal prob of N-in:   0.98491\nPOSSIBLE N-term signal sequence\ninside   1   20\nTMhelix   21   43\noutside   44   244\nTMhelix   245   264\ninside   265   267

31681 GCF_001889265.1_ASM188926v1 Microbacterium sp. 1.5R Terrabacteria group; Actinobacteria; Actinobacteria; Micrococcales; Microbacteriaceae; Microbacterium
MRRTRRGVIAAVGAMALAAAIAVPSSATAAAASGLWWFDRGKVQEAHDAGFDGAGIKIAV WP_072590666.1
hypothetical protein [Microbacterium sp. 1.5R] Length: 445\nNumber of predicted TMHs:  2\nExp number of AAs in TMHs: 46.17889\nExp number, first 60 AAs:  23.29353\nTotal prob of N-in:   0.99582\nPOSSIBLE N-term signal sequence\ninside   1   6\nTMhelix   7   29\noutside   30   405\nTMhelix   406   428\ninside   429   445

31682 GCF_001889265.1_ASM188926v1 Microbacterium sp. 1.5R Terrabacteria group; Actinobacteria; Actinobacteria; Micrococcales; Microbacteriaceae; Microbacterium

MNDTSTEPLPELTDAAVERIERAVFEDIAAELPRQSPAEKRGRVRRRRWLTGTGIAAAFV WP_072591188.1 hypothetical protein [Microbacterium sp. 1.5R] Length: 359\nNumber of predicted TMHs: 2\nExp number of AAs in TMHs: 45.75742\nExp number, first 60 AAs: 11.19729\nTotal prob of N-in: 0.99168\nPOSSIBLE N-term signal sequence\ninside 1 48\nTMhelix 49 71\noutside 72 317\nTMhelix 318 340\ninside 341 359

31683 GCF_000380605.1_ASM38060v1 Microbacterium sp. 292MF Terrabacteria group; Actinobacteria; Actinobacteria; Micrococcales; Microbacteriaceae; Microbacterium MTVTSPVGLSPADQERRRALRRMKGVALGALLFMAALFAVSFAFQQEVPALAYLRAAGEG WP_026305687.1 DUF445 domain-containing protein [Microbacterium sp. 292MF] Length: 423\nNumber of predicted TMHs: 2\nExp number of AAs in TMHs: 48.13551\nExp number, first 60 AAs: 22.0785\nTotal prob of N-in: 0.91130\nPOSSIBLE N-term signal sequence\ninside 1 24\nTMhelix 25 44\noutside 45 399\nTMhelix 400 422\ninside 423 423

31684 GCF_000380605.1_ASM38060v1 Microbacterium sp. 292MF Terrabacteria group; Actinobacteria; Actinobacteria; Micrococcales; Microbacteriaceae; Microbacterium MQRRRGLVAGLTAMVLAAASVFVGVAPAALAEDTAAFSIQKTTVEEGPFSPGDTVTYQII WP_081634708.1 hypothetical protein [Microbacterium sp. 292MF] Length: 2990\nNumber of predicted TMHs: 2\nExp number of AAs in TMHs: 46.1530099999999\nExp number, first 60 AAs: 22.78922\nTotal prob of N-in: 0.99912\nPOSSIBLE N-term signal sequence\ninside 1 6\nTMhelix 7 29\noutside 30 2959\nTMhelix 2960 2982\ninside 2983 2990

31685 GCF_001005635.1_ASM100563v1 Microbacterium sp. Ag1 Terrabacteria group; Actinobacteria; Actinobacteria; Micrococcales; Microbacteriaceae; Microbacterium MAREKKRRSSGRRSPLAAAALIGAGLMITGAVYAGATAAFAATDTQSAATSQLTVEDGKK WP_046749294.1 cystathionine beta-lyase [Microbacterium sp. Ag1] Length: 267\nNumber of predicted TMHs: 2\nExp number of AAs in TMHs: 43.19301\nExp number, first 60 AAs: 22.66657\nTotal prob of N-in: 0.98686\nPOSSIBLE N-term signal sequence\ninside 1 20\nTMhelix 21 43\noutside 44 244\nTMhelix 245 264\ninside 265 267

31686 GCF_002204305.1_ASM220430v1 Microbacterium sp. AISO3 Terrabacteria group; Actinobacteria; Actinobacteria; Micrococcales; Microbacteriaceae; Microbacterium MPPAPSRPQSRPRRTAVPAAAVAAFTAACLTLGTAAPAGAADGELLDDGAIAPHAAPYGT WP_023955709.1 putative phospholipid phosphatase [Microbacterium sp. TS-1] Length: 698\nNumber of predicted TMHs: 1\nExp number of AAs in TMHs: 34.2210699999999\nExp number, first 60 AAs: 12.40938\nTotal prob of N-in: 0.56987\nPOSSIBLE N-term signal sequence\noutside 1 667\nTMhelix 668 690\ninside 691 698

31687 GCF_002204305.1_ASM220430v1 Microbacterium sp. AISO3 Terrabacteria group; Actinobacteria; Actinobacteria; Micrococcales; Microbacteriaceae; Microbacterium MLRTRRAALQRPLASVLTGALVAMAAVVGVAAPATAAGPTLTTSTTVVANEKITVAITGT WP_088379655.1 hypothetical protein [Microbacterium sp. AISO3] Length: 962\nNumber of predicted TMHs: 2\nExp number of AAs in TMHs: 41.3927899999999\nExp number, first 60 AAs: 22.33509\nTotal prob of N-in: 0.99759\nPOSSIBLE N-term signal sequence\ninside 1 12\nTMhelix 13 35\noutside 36 934\nTMhelix 935 954\ninside 955 962

31688 GCF_001878835.1_ASM187883v1 Microbacterium sp. AR7-10 Terrabacteria group; Actinobacteria; Actinobacteria; Micrococcales; Microbacteriaceae; Microbacterium MAREKKHRSRGRRSPLAAAALIGAGLLLTGGVYAGASAAMASTTGTQSSATAELTVEDGQ WP_071643062.1 cystathionine beta-lyase [Microbacterium sp. AR7-10] Length: 277\nNumber of predicted TMHs: 2\nExp number of AAs in TMHs: 42.96614\nExp number, first 60 AAs: 22.16823\nTotal prob of N-in: 0.95757\nPOSSIBLE N-term signal sequence\ninside 1 12\nTMhelix 13 35\noutside 36 254\nTMhelix 255 274\ninside 275 277

31689 GCF_000333395.1_ASM33339v1 Microbacterium sp. B19 Terrabacteria group; Actinobacteria; Actinobacteria; Micrococcales; Microbacteriaceae; Microbacterium MSRRRTIALRVVVGVFLVVAAAGATAPAVASDSPEDGTPLSVTITDGSTPTPSASPSRST WP_022877795.1 hypothetical protein [Microbacterium sp. B19] Length: 261\nNumber of predicted TMHs: 2\nExp number of AAs in TMHs: 46.46331\nExp number, first 60 AAs: 21.84175\nTotal prob of N-in: 0.99751\nPOSSIBLE N-term signal sequence\ninside 1 6\nTMhelix 7 29\noutside 30 222\nTMhelix 223 245\ninside 246 261

31690 GCF_000333395.1_ASM33339v1 Microbacterium sp. B19 Terrabacteria group; Actinobacteria; Actinobacteria; Micrococcales; Microbacteriaceae; Microbacterium MAREKKPRRSNGRRSPLAAAALIGIGLLLTGGIYAGASAAMAATDTPTSASNSASTVDEG WP_022878542.1 MULTISPECIES: cystathionine beta-lyase [Microbacterium] Length: 269\nNumber of predicted TMHs: 2\nExp number of AAs in TMHs: 42.73548\nExp number, first 60 AAs: 22.46923\nTotal prob of N-in: 0.98833\nPOSSIBLE N-term signal sequence\ninside 1 20\nTMhelix 21 43\noutside 44 246\nTMhelix 247 266\ninside 267 269

31691 GCF_000333415.1_ASM33341v1 Microbacterium sp. B24 Terrabacteria group; Actinobacteria; Actinobacteria; Micrococcales; Microbacteriaceae; Microbacterium
MLPRTRRRTWFASGLTGLLTAGAVFALVAGLGVTIAPTGARAADSSQVTVTAKAQDRDLA WP_036316991.1
hypothetical protein [Microbacterium sp. B24] Length: 843\nNumber of predicted TMHs: 2\nExp number of AAs in TMHs: 45.3895\nExp number, first 60 AAs: 22.80755\nTotal prob of N-in: 0.99593\nPOSSIBLE N-term signal sequence\ninside 1 12\nTMhelix 13 35\noutside 36 809\nTMhelix 810 832\ninside 833 843

31692 GCF_001792815.1_ASM179281v1 Microbacterium sp. BH-3-3-3 Terrabacteria group; Actinobacteria; Actinobacteria; Micrococcales; Microbacteriaceae; Microbacterium
MARTPSTLLSPADQERRRALRVMKGVALGALLGMAVIFAISFSLQREIEWMQYVRAAAEG WP_070409598.1
DUF445 domain-containing protein [Microbacterium sp. BH-3-3-3] Length: 424\nNumber of predicted TMHs: 2\nExp number of AAs in TMHs: 49.5427699999999\nExp number, first 60 AAs: 22.08609\nTotal prob of N-in: 0.70994\nPOSSIBLE N-term signal sequence\ninside 1 19\nTMhelix 20 42\noutside 43 400\nTMhelix 401 423\ninside 424 424

31693 GCF_001792815.1_ASM179281v1 Microbacterium sp. BH-3-3-3 Terrabacteria group; Actinobacteria; Actinobacteria; Micrococcales; Microbacteriaceae; Microbacterium
MAREKKTRRASGRRSPLAAAALIGIGLLLTGGVYAGASAAMAATNATPTSEANSATAVEE WP_055835609.1
MULTISPECIES: cystathionine beta-lyase [Microbacterium] Length: 270\nNumber of predicted TMHs: 2\nExp number of AAs in TMHs: 42.56802\nExp number, first 60 AAs: 22.36159\nTotal prob of N-in: 0.98585\nPOSSIBLE N-term signal sequence\ninside 1 20\nTMhelix 21 43\noutside 44 247\nTMhelix 248 267\ninside 268 270

31694 GCF_001792815.1_ASM179281v1 Microbacterium sp. BH-3-3-3 Terrabacteria group; Actinobacteria; Actinobacteria; Micrococcales; Microbacteriaceae; Microbacterium
MARRQDAVPTGVPVPSPTSSRRAALIALIALVGLSVPTTAQAAARTDCRPVPTPAADILR WP_083285711.1 hypothetical protein [Microbacterium sp. BH-3-3-3] Length: 613\nNumber of predicted TMHs: 1\nExp number of AAs in TMHs: 29.39309\nExp number, first 60 AAs: 7.46704\nTotal prob of N-in: 0.37223\noutside 1 572\nTMhelix 573 595\ninside 596 613

31695 GCF_000582705.1_MIC448 Microbacterium sp. C448 Terrabacteria group; Actinobacteria; Actinobacteria; Micrococcales; Microbacteriaceae; Microbacterium
MPREKKRRSGRRSPLAAAGLIGLGLVLTGGLYAGASAAMASTTDTAATQTLTVEDGKKLF WP_036297118.1
cystathionine beta-lyase [Microbacterium sp. C448] Length: 265\nNumber of predicted TMHs: 2\nExp number of AAs in TMHs: 43.08387\nExp number, first 60 AAs: 22.53811\nTotal prob of N-in: 0.96806\nPOSSIBLE N-term signal sequence\ninside 1 12\nTMhelix 13 35\noutside 36 242\nTMhelix 243 262\ninside 263 265

31696 GCF_000799385.1_ASM79938v1 Microbacterium sp. CF335 Terrabacteria group; Actinobacteria; Actinobacteria; Micrococcales; Microbacteriaceae; Microbacterium
MVSGGRRARRRRLIPLAAMITLGAVVGFGSPAAATTSEPDGQWWYGAYGVEQVHAEGWTG WP_081993675.1
hypothetical protein [Microbacterium sp. CF335] Length: 429\nNumber of predicted TMHs: 2\nExp number of AAs in TMHs: 45.43474\nExp number, first 60 AAs: 22.38606\nTotal prob of N-in: 0.99488\nPOSSIBLE N-term signal sequence\ninside 1 12\nTMhelix 13 35\noutside 36 394\nTMhelix 395 417\ninside 418 429

31697 GCF_000799385.1_ASM79938v1 Microbacterium sp. CF335 Terrabacteria group; Actinobacteria; Actinobacteria; Micrococcales; Microbacteriaceae; Microbacterium
MPRHPHRARRPGIAHAPRRRRALIATAIAALVITPLSVTAVATAVSLNADTTPTATLSAT WP_047543190.1 hypothetical protein [Microbacterium sp. CF335] Length: 411\nNumber of predicted TMHs: 2\nExp number of AAs in TMHs: 44.48992\nExp number, first 60 AAs: 22.55518\nTotal prob of N-in: 0.99370\nPOSSIBLE N-term signal sequence\ninside 1 22\nTMhelix 23 45\noutside 46 379\nTMhelix 380 402\ninside 403 411

31698 GCF_000799385.1_ASM79938v1 Microbacterium sp. CF335 Terrabacteria group; Actinobacteria; Actinobacteria; Micrococcales; Microbacteriaceae; Microbacterium
MTREKKRRPSGRRSPLAAAALIGIGLLLTGGIYAGASAAMAATTDTTPAASSLTAEDGQK WP_047545539.1 cystathionine beta-lyase [Microbacterium sp. CF335] Length: 267\nNumber of predicted TMHs: 2\nExp number of AAs in TMHs: 43.06272\nExp number, first 60 AAs: 22.5685\nTotal prob of N-in: 0.96454\nPOSSIBLE N-term signal sequence\ninside 1 20\nTMhelix 21 43\noutside 44 244\nTMhelix 245 264\ninside 265 267

31699 GCF_000799385.1_ASM79938v1 Microbacterium sp. CF335 Terrabacteria group; Actinobacteria; Actinobacteria; Micrococcales; Microbacteriaceae; Microbacterium
MRAARHRRRTTVGFGVGVGVALAITVGIAPAVAVDPAELEDRVTVAAVRGHLQALQDIAT WP_081993913.1
hypothetical protein [Microbacterium sp. CF335] Length: 551\nNumber of predicted TMHs: 2\nExp number of

AAs in TMHs: 44.23741\nExp number, first 60 AAs: 22.74723\nTotal prob of N-in: 0.99985\nPOSSIBLE N-term signal sequence\ninside 1 11\nTMhelix 12 34\noutside 35 522\nTMhelix 523 545\ninside 546 551

31700 GCF_000799385.1_ASM79938v1 Microbacterium sp. CF335 Terrabacteria group; Actinobacteria; Actinobacteria; Micrococcales; Microbacteriaceae; Microbacterium
MPRPRRRALRVALVALVAGALAAGAAVPAQAATYVAIAGSGSTWSQNALDQWRRNVASNY WP_052174493.1
phosphate ABC transporter substrate-binding protein PstS [Microbacterium sp. CF335] Length: 486\nNumber of predicted TMHs: 2\nExp number of AAs in TMHs: 44.48261\nExp number, first 60 AAs: 22.04107\nTotal prob of N-in: 0.98322\nPOSSIBLE N-term signal sequence\ninside 1 11\nTMhelix 12 34\noutside 35 456\nTMhelix 457 479\ninside 480 486

31701 GCF_000799385.1_ASM79938v1 Microbacterium sp. CF335 Terrabacteria group; Actinobacteria; Actinobacteria; Micrococcales; Microbacteriaceae; Microbacterium
MNITASARRRLLNGAAGATLLTAGLVLAAPLAASAHVTVTPDQSASAGGHGVLTFAFSHG WP_047542555.1
hypothetical protein [Microbacterium sp. CF335] Length: 236\nNumber of predicted TMHs: 2\nExp number of AAs in TMHs: 44.88236\nExp number, first 60 AAs: 22.09932\nTotal prob of N-in: 0.98634\nPOSSIBLE N-term signal sequence\ninside 1 11\nTMhelix 12 34\noutside 35 208\nTMhelix 209 231\ninside 232 236

31702 GCF_001266755.1_ASM126675v1 Microbacterium sp. CGR1 Terrabacteria group; Actinobacteria; Actinobacteria; Micrococcales; Microbacteriaceae; Microbacterium
MNDTSTEPLPELTDAAVERIERAVFEDIAAELPRQSPAEKRGRVRRRRWLTGTGIAAAFV WP_053095880.1 hypothetical protein [Microbacterium sp. CGR1] Length: 359\nNumber of predicted TMHs: 2\nExp number of AAs in TMHs: 45.7576\nExp number, first 60 AAs: 11.19729\nTotal prob of N-in: 0.99168\nPOSSIBLE N-term signal sequence\ninside 1 48\nTMhelix 49 71\noutside 72 317\nTMhelix 318 340\ninside 341 359

31703 GCF_001266755.1_ASM126675v1 Microbacterium sp. CGR1 Terrabacteria group; Actinobacteria; Actinobacteria; Micrococcales; Microbacteriaceae; Microbacterium
MRRTRRGVIAAVGAMALAAAIAVPSSATAAAASGLWWFDRGKVQEAHDAGFDGAGIKIAV WP_082297971.1
hypothetical protein [Microbacterium sp. CGR1] Length: 445\nNumber of predicted TMHs: 2\nExp number of AAs in TMHs: 46.21086\nExp number, first 60 AAs: 23.29353\nTotal prob of N-in: 0.99582\nPOSSIBLE N-term signal sequence\ninside 1 6\nTMhelix 7 29\noutside 30 405\nTMhelix 406 428\ninside 429 445

31704 GCF_001266755.1_ASM126675v1 Microbacterium sp. CGR1 Terrabacteria group; Actinobacteria; Actinobacteria; Micrococcales; Microbacteriaceae; Microbacterium
MAREKKRRSNGRRSPLAAAALIGAGLMITGAVYAGTSAAFAATDTQTTAASTLTVEDGEK WP_053096702.1
MULTISPECIES: cystathionine beta-lyase [Microbacterium] Length: 267\nNumber of predicted TMHs: 2\nExp number of AAs in TMHs: 43.06753\nExp number, first 60 AAs: 22.53635\nTotal prob of N-in: 0.98491\nPOSSIBLE N-term signal sequence\ninside 1 20\nTMhelix 21 43\noutside 44 244\nTMhelix 245 264\ninside 265 267

31705 GCF_001266755.1_ASM126675v1 Microbacterium sp. CGR1 Terrabacteria group; Actinobacteria; Actinobacteria; Micrococcales; Microbacteriaceae; Microbacterium
MEFLEGKTLRRRRAIGGGIAGVTVGAMILAGALVPTAANALDDANPNDPSVAQGQIIQLP WP_053098651.1
hypothetical protein [Microbacterium sp. CGR1] Length: 1899\nNumber of predicted TMHs: 2\nExp number of AAs in TMHs: 41.3709599999999\nExp number, first 60 AAs: 21.67795\nTotal prob of N-in: 0.96849\nPOSSIBLE N-term signal sequence\ninside 1 12\nTMhelix 13 35\noutside 36 1866\nTMhelix 1867 1889\ninside 1890 1899

31706 GCF_001595495.1_ASM159549v1 Microbacterium sp. CH1 Terrabacteria group; Actinobacteria; Actinobacteria; Micrococcales; Microbacteriaceae; Microbacterium
MSNTTIRPRRPRRTLLGATGVVGGLALALAVPAMAGAHVSVSPDELAAGDHGVLTFSFSH WP_025103163.1
MULTISPECIES: hypothetical protein [Microbacterium] Length: 224\nNumber of predicted TMHs: 2\nExp number of AAs in TMHs: 45.16071\nExp number, first 60 AAs: 22.31743\nTotal prob of N-in: 0.99849\nPOSSIBLE N-term signal sequence\ninside 1 12\nTMhelix 13 35\noutside 36 197\nTMhelix 198 220\ninside 221 224

31707 GCF_001595495.1_ASM159549v1 Microbacterium sp. CH1 Terrabacteria group; Actinobacteria; Actinobacteria; Micrococcales; Microbacteriaceae; Microbacterium
MRRSSTFGEGVAAGSAELEVDRDSTGLRRPPPEGGDMARRRAFFRSIMGGAAALALALGG WP_082802980.1
hypothetical protein [Microbacterium sp. CH1] Length: 710\nNumber of predicted TMHs: 2\nExp number of AAs in TMHs: 41.50673\nExp number, first 60 AAs: 15.28926\nTotal prob of N-in: 0.95913\nPOSSIBLE N-term signal sequence\ninside 1 42\nTMhelix 43 65\noutside 66 683\nTMhelix 684 703\ninside 704 710

31708 GCF_001595495.1_ASM159549v1 Microbacterium sp. CH1 Terrabacteria group; Actinobacteria; Actinobacteria; Micrococcales; Microbacteriaceae; Microbacterium
MHKNLPRRTAIVLSAVLLSGAVVVATPTVAVADDNFDSSYPVADSTISGSPDEISLMFTG WP_029266231.1 MULTISPECIES: hypothetical protein [Microbacterium] Length: 230\nNumber of predicted TMHs: 2\nExp number of AAs in TMHs: 44.08665\nExp number, first 60 AAs: 21.4021\nTotal prob of N-in: 0.99476\nPOSSIBLE N-term signal sequence\ninside 1 8\nTMhelix 9 31\noutside 32 181\nTMhelix 182 204\ninside 205 230

31709 GCF_001595495.1_ASM159549v1 Microbacterium sp. CH1 Terrabacteria group; Actinobacteria; Actinobacteria; Micrococcales; Microbacteriaceae; Microbacterium
MTRRRMLRGAVALVAVAASVLLLGSTATPTPTPPPVADDPADPVRAAEYWLDGARIRDAW WP_062633540.1 peptidase S8 [Microbacterium sp. CH1] Length: 439\nNumber of predicted TMHs: 2\nExp number of AAs in TMHs: 41.4776299999999\nExp number, first 60 AAs: 19.80315\nTotal prob of N-in: 0.93229\nPOSSIBLE N-term signal sequence\ninside 1 6\nTMhelix 7 29\noutside 30 397\nTMhelix 398 420\ninside 421 439

31710 GCF_001595495.1_ASM159549v1 Microbacterium sp. CH1 Terrabacteria group; Actinobacteria; Actinobacteria; Micrococcales; Microbacteriaceae; Microbacterium
MAREKKRRSSGRRSPLAAAALIGAGLMITGAVYAGATAAFAANDAPTASTQLTVEDGEKL WP_062634020.1 cystathionine beta-lyase [Microbacterium sp. CH1] Length: 266\nNumber of predicted TMHs: 2\nExp number of AAs in TMHs: 43.19357\nExp number, first 60 AAs: 22.60028\nTotal prob of N-in: 0.99024\nPOSSIBLE N-term signal sequence\ninside 1 19\nTMhelix 20 42\noutside 43 243\nTMhelix 244 263\ninside 264 266

31711 GCF_001595495.1_ASM159549v1 Microbacterium sp. CH1 Terrabacteria group; Actinobacteria; Actinobacteria; Micrococcales; Microbacteriaceae; Microbacterium
MRRRAALATLVVGAAVWGTPAAAVAAPTAPTTIVSMATGACEPGTVVLAPEAPSALAALG WP_062635393.1 hypothetical protein [Microbacterium sp. CH1] Length: 436\nNumber of predicted TMHs: 2\nExp number of AAs in TMHs: 43.0398599999999\nExp number, first 60 AAs: 21.15577\nTotal prob of N-in: 0.94801\nPOSSIBLE N-term signal sequence\ninside 1 6\nTMhelix 7 29\noutside 30 386\nTMhelix 387 409\ninside 410 436

31712 GCF_000685355.1_Microbacteriumx4 Microbacterium sp. CH12i Terrabacteria group; Actinobacteria; Actinobacteria; Micrococcales; Microbacteriaceae; Microbacterium
MAREKKRRSNGRRSPLAAAALIGAGLLITGGIYAGASAAIAATDTQTTASTQLTVDDGEK WP_036278033.1 cystathionine beta-lyase [Microbacterium sp. CH12i] Length: 267\nNumber of predicted TMHs: 2\nExp number of AAs in TMHs: 43.39724\nExp number, first 60 AAs: 22.67147\nTotal prob of N-in: 0.97172\nPOSSIBLE N-term signal sequence\ninside 1 20\nTMhelix 21 43\noutside 44 244\nTMhelix 245 264\ninside 265 267

31713 GCF_000708305.1_Cr-K1W Microbacterium sp. Cr-K1W Terrabacteria group; Actinobacteria; Actinobacteria; Micrococcales; Microbacteriaceae; Microbacterium
MRASRRATIAGTIVLIAVLVGVLPASAATVASWATWQPLVGVGGAFTTSVEVAAQPAISA WP_029264221.1 MULTISPECIES: hypothetical protein [Microbacterium] Length: 652\nNumber of predicted TMHs: 2\nExp number of AAs in TMHs: 44.4223399999999\nExp number, first 60 AAs: 23.03257\nTotal prob of N-in: 0.99909\nPOSSIBLE N-term signal sequence\ninside 1 12\nTMhelix 13 35\noutside 36 623\nTMhelix 624 646\ninside 647 652

31714 GCF_000708305.1_Cr-K1W Microbacterium sp. Cr-K1W Terrabacteria group; Actinobacteria; Actinobacteria; Micrococcales; Microbacteriaceae; Microbacterium
MAREKKRRSSGRRSPLAAAALIGAGLMITGAVYAGATAAFAASDTPTASTQLTVEDGEKL WP_029261592.1 MULTISPECIES: cystathionine beta-lyase [Microbacterium] Length: 266\nNumber of predicted TMHs: 2\nExp number of AAs in TMHs: 43.19358\nExp number, first 60 AAs: 22.65324\nTotal prob of N-in: 0.99358\nPOSSIBLE N-term signal sequence\ninside 1 19\nTMhelix 20 42\noutside 43 243\nTMhelix 244 263\ninside 264 266

31715 GCF_000708305.1_Cr-K1W Microbacterium sp. Cr-K1W Terrabacteria group; Actinobacteria; Actinobacteria; Micrococcales; Microbacteriaceae; Microbacterium
MTEEIQKNKGISRRTIVKGAAWSVPVIAAAVATPLAAASGDVEVGAFSVDGDCGTLGLLF WP_029261182.1 MULTISPECIES: hypothetical protein [Microbacterium] Length: 168\nNumber of predicted TMHs: 2\nExp number of AAs in TMHs: 44.6481\nExp number, first 60 AAs: 19.70819\nTotal prob of N-in: 0.97105\nPOSSIBLE N-term signal sequence\ninside 1 19\nTMhelix 20 42\noutside 43 126\nTMhelix 127 149\ninside 150 168

31716 GCF_000708305.1_Cr-K1W Microbacterium sp. Cr-K1W Terrabacteria group; Actinobacteria; Actinobacteria; Micrococcales; Microbacteriaceae; Microbacterium
MLPSDRRLFEVERDGTVPRATSLRRVGNMARRRAFIQAVIGGVAALALAFGGATSAMAAD WP_029262819.1 MULTISPECIES: hypothetical protein [Microbacterium] Length: 667\nNumber of predicted TMHs: 2\nExp

number of AAs in TMHs: 50.6035399999999\nExp number, first 60 AAs:  21.40123\nTotal prob of N-in: 0.94756\nPOSSIBLE N-term signal sequence\ninside   1   33\nTMhelix   34   56\noutside   57   640\nTMhelix   641 660\ninside   661   667

31717 GCF_000708325.1_Cr-K20 Microbacterium sp. Cr-K20 Terrabacteria group; Actinobacteria; Actinobacteria; Micrococcales; Microbacteriaceae; Microbacterium
MTEEIQKNKGISRRTIVKGAAWSVPVIAAAVATPLAAASGDVEVGAFSVDGDCGTLGLLF WP_029261182.1
MULTISPECIES: hypothetical protein [Microbacterium] Length: 168\nNumber of predicted TMHs:  2\nExp number of AAs in TMHs: 44.6481\nExp number, first 60 AAs:  19.70819\nTotal prob of N-in:   0.97105\nPOSSIBLE N-term signal sequence\ninside   1   19\nTMhelix   20   42\noutside   43   126\nTMhelix   127   149\ninside   150   168

31718 GCF_000708325.1_Cr-K20 Microbacterium sp. Cr-K20 Terrabacteria group; Actinobacteria; Actinobacteria; Micrococcales; Microbacteriaceae; Microbacterium
MAREKKRRSSGRRSPLAAAALIGAGLMITGAVYAGATAAFAASDTPTASTQLTVEDGEKL WP_029261592.1
MULTISPECIES: cystathionine beta-lyase [Microbacterium] Length: 266\nNumber of predicted TMHs:  2\nExp number of AAs in TMHs: 43.19358\nExp number, first 60 AAs:  22.65324\nTotal prob of N-in:   0.99358\nPOSSIBLE N-term signal sequence\ninside   1   19\nTMhelix   20   42\noutside   43   243\nTMhelix   244   263\ninside   264   266

31719 GCF_000708325.1_Cr-K20 Microbacterium sp. Cr-K20 Terrabacteria group; Actinobacteria; Actinobacteria; Micrococcales; Microbacteriaceae; Microbacterium
MRASRRATIAGTIVLIAVLVGVLPASAATVASWATWQPLVGVGGAFTTSVEVAAQPAISA WP_029264221.1
MULTISPECIES: hypothetical protein [Microbacterium] Length: 652\nNumber of predicted TMHs:  2\nExp number of AAs in TMHs: 44.4223399999999\nExp number, first 60 AAs:  23.03257\nTotal prob of N-in: 0.99909\nPOSSIBLE N-term signal sequence\ninside   1   12\nTMhelix   13   35\noutside   36   623\nTMhelix   624 646\ninside   647   652

31720 GCF_000708325.1_Cr-K20 Microbacterium sp. Cr-K20 Terrabacteria group; Actinobacteria; Actinobacteria; Micrococcales; Microbacteriaceae; Microbacterium
MLPSDRRLFEVERDGTVPRATSLRRVGNMARRRAFIQAVIGGVAALALAFGGATSAMAAD WP_029262819.1
MULTISPECIES: hypothetical protein [Microbacterium] Length: 667\nNumber of predicted TMHs:  2\nExp number of AAs in TMHs: 50.6035399999999\nExp number, first 60 AAs:  21.40123\nTotal prob of N-in: 0.94756\nPOSSIBLE N-term signal sequence\ninside   1   33\nTMhelix   34   56\noutside   57   640\nTMhelix   641 660\ninside   661   667

31721 GCF_000708365.1_Cr-K29 Microbacterium sp. Cr-K29 Terrabacteria group; Actinobacteria; Actinobacteria; Micrococcales; Microbacteriaceae; Microbacterium
MNRDCTRRWDHRANGLPRKVRRMTRRRGMLRAVLGGMAAIAIALGGATAVVAAEDPAPTG WP_081860119.1
hypothetical protein [Microbacterium sp. Cr-K29] Length: 697\nNumber of predicted TMHs:  2\nExp number of AAs in TMHs: 44.46897\nExp number, first 60 AAs:  22.81605\nTotal prob of N-in:   0.99997\nPOSSIBLE N-term signal sequence\ninside   1   29\nTMhelix   30   52\noutside   53   667\nTMhelix   668   690\ninside   691   697

31722 GCF_000708365.1_Cr-K29 Microbacterium sp. Cr-K29 Terrabacteria group; Actinobacteria; Actinobacteria; Micrococcales; Microbacteriaceae; Microbacterium
MAREKKRRSSGRRSPLAAAALIGAGLMITGAVYAGATAAFAATDTQSAATSQLTVEDGKK WP_029258331.1
MULTISPECIES: cystathionine beta-lyase [Microbacterium] Length: 267\nNumber of predicted TMHs:  2\nExp number of AAs in TMHs: 43.18585\nExp number, first 60 AAs:  22.66682\nTotal prob of N-in:   0.98705\nPOSSIBLE N-term signal sequence\ninside   1   20\nTMhelix   21   43\noutside   44   244\nTMhelix   245   264\ninside   265   267

31723 GCF_000708345.1_Cr-K32 Microbacterium sp. Cr-K32 Terrabacteria group; Actinobacteria; Actinobacteria; Micrococcales; Microbacteriaceae; Microbacterium
MELQERTLRRRRAVGGGIAGVTVGAVILASALVPTAANALDAANPNDPSVAQGQIIQLPA WP_051667978.1
hypothetical protein [Microbacterium sp. Cr-K32] Length: 1733\nNumber of predicted TMHs:  2\nExp number of AAs in TMHs: 43.70005\nExp number, first 60 AAs:  22.32675\nTotal prob of N-in:   0.99668\nPOSSIBLE N-term signal sequence\ninside   1   12\nTMhelix   13   35\noutside   36  1700\nTMhelix   1701  1723\ninside   1724  1733

31724 GCF_000708345.1_Cr-K32 Microbacterium sp. Cr-K32 Terrabacteria group; Actinobacteria; Actinobacteria; Micrococcales; Microbacteriaceae; Microbacterium
MAREKKRRSSGRRSPLAAAALIGAGLMITGAVYAGATAAFAATDTQSAATSQLTVEDGKK WP_029258331.1
MULTISPECIES: cystathionine beta-lyase [Microbacterium] Length: 267\nNumber of predicted TMHs:  2\nExp number of AAs in TMHs: 43.18585\nExp number, first 60 AAs:  22.66682\nTotal prob of N-in:   0.98705\nPOSSIBLE N-term signal sequence\ninside   1   20\nTMhelix   21   43\noutside   44   244\nTMhelix   245   264\ninside   265   267

31725 GCF_000708345.1_Cr-K32 Microbacterium sp. Cr-K32 Terrabacteria group; Actinobacteria; Actinobacteria; Micrococcales; Microbacteriaceae; Microbacterium MHKNLPRRTAIVLSAVLLSGAVVVATPTVAVADDNFDSSYPVADSTISGSPDEISLMFTG WP_029266231.1 MULTISPECIES: hypothetical protein [Microbacterium] Length: 230\nNumber of predicted TMHs: 2\nExp number of AAs in TMHs: 44.08665\nExp number, first 60 AAs: 21.4021\nTotal prob of N-in: 0.99476\nPOSSIBLE N-term signal sequence\ninside 1 8\nTMhelix 9 31\noutside 32 181\nTMhelix 182 204\ninside 205 230

31726 GCF_000708345.1_Cr-K32 Microbacterium sp. Cr-K32 Terrabacteria group; Actinobacteria; Actinobacteria; Micrococcales; Microbacteriaceae; Microbacterium MNRDCTRRWDHRANGLPRKVRRMTRRRGMLRAVLGGMAAIAIALGGATAVVAAEDPAPTG WP_081859623.1 hypothetical protein [Microbacterium sp. Cr-K32] Length: 697\nNumber of predicted TMHs: 2\nExp number of AAs in TMHs: 45.28162\nExp number, first 60 AAs: 22.81605\nTotal prob of N-in: 0.99997\nPOSSIBLE N-term signal sequence\ninside 1 29\nTMhelix 30 52\noutside 53 667\nTMhelix 668 690\ninside 691 697

31727 GCF_001984105.1_ASM198410v1 Microbacterium sp. CSI-V Terrabacteria group; Actinobacteria; Actinobacteria; Micrococcales; Microbacteriaceae; Microbacterium MSDFRERSDAEAPATPSRRRLRIRRAHADAANRGARRTVLPLAAGLVAGVTVFALVSGGV WP_077052164.1 hypothetical protein [Microbacterium sp. CSI-V] Length: 877\nNumber of predicted TMHs: 2\nExp number of AAs in TMHs: 44.0403099999999\nExp number, first 60 AAs: 21.03564\nTotal prob of N-in: 0.97485\nPOSSIBLE N-term signal sequence\ninside 1 38\nTMhelix 39 61\noutside 62 846\nTMhelix 847 869\ninside 870 877

31728 GCF_001984105.1_ASM198410v1 Microbacterium sp. CSI-V Terrabacteria group; Actinobacteria; Actinobacteria; Micrococcales; Microbacteriaceae; Microbacterium MIESPRPKGHSMHHPPPPRRRLLRTALTLALIAVGVATVPVLATAPAHAAEQGTGFGTWA WP_077052147.1 hypothetical protein [Microbacterium sp. CSI-V] Length: 514\nNumber of predicted TMHs: 2\nExp number of AAs in TMHs: 42.95366\nExp number, first 60 AAs: 20.42082\nTotal prob of N-in: 0.91679\nPOSSIBLE N-term signal sequence\ninside 1 20\nTMhelix 21 43\noutside 44 483\nTMhelix 484 506\ninside 507 514

31729 GCF_001984105.1_ASM198410v1 Microbacterium sp. CSI-V Terrabacteria group; Actinobacteria; Actinobacteria; Micrococcales; Microbacteriaceae; Microbacterium MTTPSLPPLRAERVDAIESAVFARIADERAARRRRRRNIWTGVGAAAAVVVVAAVVGPAL WP_077051665.1 hypothetical protein [Microbacterium sp. CSI-V] Length: 361\nNumber of predicted TMHs: 2\nExp number of AAs in TMHs: 45.9534599999999\nExp number, first 60 AAs: 21.43306\nTotal prob of N-in: 0.99602\nPOSSIBLE N-term signal sequence\ninside 1 38\nTMhelix 39 61\noutside 62 323\nTMhelix 324 346\ninside 347 361

31730 GCF_001984105.1_ASM198410v1 Microbacterium sp. CSI-V Terrabacteria group; Actinobacteria; Actinobacteria; Micrococcales; Microbacteriaceae; Microbacterium MAREKKPRRSNGRRSPLAAAALIGIGLLLTGGIYAGASAAMAATDTPTSASNSASTVEEG WP_077049912.1 cystathionine beta-lyase [Microbacterium sp. CSI-V] Length: 269\nNumber of predicted TMHs: 2\nExp number of AAs in TMHs: 42.94385\nExp number, first 60 AAs: 22.46061\nTotal prob of N-in: 0.98943\nPOSSIBLE N-term signal sequence\ninside 1 20\nTMhelix 21 43\noutside 44 246\nTMhelix 247 266\ninside 267 269

31731 GCF_001262495.1_GCS4v1 Microbacterium sp. GCS4 Terrabacteria group; Actinobacteria; Actinobacteria; Micrococcales; Microbacteriaceae; Microbacterium MRTFSTRPSTARRTVVAGLVTGAVLALAVPAMASAHVTVSPDELAAGDHGTLTFAFSHGC WP_050721264.1 hypothetical protein [Microbacterium sp. GCS4] Length: 234\nNumber of predicted TMHs: 2\nExp number of AAs in TMHs: 45.15322\nExp number, first 60 AAs: 21.95005\nTotal prob of N-in: 0.97825\nPOSSIBLE N-term signal sequence\ninside 1 12\nTMhelix 13 35\noutside 36 207\nTMhelix 208 230\ninside 231 234

31732 GCF_001262495.1_GCS4v1 Microbacterium sp. GCS4 Terrabacteria group; Actinobacteria; Actinobacteria; Micrococcales; Microbacteriaceae; Microbacterium MTASIPENGLRARLLRLARRSVVVATVLGLCAAGAAGATTATAATDADDPSVELTVSAGT WP_050723562.1 hypothetical protein [Microbacterium sp. GCS4] Length: 711\nNumber of predicted TMHs: 2\nExp number of AAs in TMHs: 42.8751999999999\nExp number, first 60 AAs: 21.2912\nTotal prob of N-in: 0.95406\nPOSSIBLE N-term signal sequence\ninside 1 20\nTMhelix 21 43\noutside 44 661\nTMhelix 662 684\ninside 685 711

31733 GCF_001262495.1_GCS4v1 Microbacterium sp. GCS4 Terrabacteria group; Actinobacteria; Actinobacteria; Micrococcales; Microbacteriaceae; Microbacterium MAREKKRRSNGRRSPLAAAALIGAGLMITGAVYAGASAAFAATDTQTAASSTLTVEDGEK WP_050723763.1 cystathionine beta-lyase [Microbacterium sp. GCS4] Length: 267\nNumber of predicted TMHs: 2\nExp

number of AAs in TMHs: 43.26712\nExp number, first 60 AAs: 22.61188\nTotal prob of N-in: 0.98268\nPOSSIBLE N-term signal sequence\ninside 1 20\nTMhelix 21 43\noutside 44 244\nTMhelix 245 264\ninside 265 267

31734 GCF_001650405.1_ASM165040v1 Microbacterium sp. H83 Terrabacteria group; Actinobacteria; Actinobacteria; Micrococcales; Microbacteriaceae; Microbacterium
MNHQTTDELPDLPDTSITRIEKAVFDEIADERPRTPASGSRARSRRRGWLTGVGVAAAFA WP_067114872.1
hypothetical protein [Microbacterium sp. H83] Length: 367\nNumber of predicted TMHs: 2\nExp number of AAs in TMHs: 45.62708\nExp number, first 60 AAs: 12.08431\nTotal prob of N-in: 0.98053\nPOSSIBLE N-term signal sequence\ninside 1 47\nTMhelix 48 70\noutside 71 323\nTMhelix 324 346\ninside 347 367

31735 GCF_001650405.1_ASM165040v1 Microbacterium sp. H83 Terrabacteria group; Actinobacteria; Actinobacteria; Micrococcales; Microbacteriaceae; Microbacterium
MMRRRLRGRRAVVRSTTALLAACLLAAATTAPASAAGTTPSAIAPATSVSMATQGQACEP WP_082914201.1
hypothetical protein [Microbacterium sp. H83] Length: 470\nNumber of predicted TMHs: 2\nExp number of AAs in TMHs: 40.75132\nExp number, first 60 AAs: 19.1788\nTotal prob of N-in: 0.87523\nPOSSIBLE N-term signal sequence\ninside 1 11\nTMhelix 12 34\noutside 35 403\nTMhelix 404 426\ninside 427 470

31736 GCF_001650405.1_ASM165040v1 Microbacterium sp. H83 Terrabacteria group; Actinobacteria; Actinobacteria; Micrococcales; Microbacteriaceae; Microbacterium
MHRDETTHRSGVEGATMRVHRPSTVRSARRSILAAGVAASVGVVMLTPVAATAAAAETTF WP_082914497.1
hypothetical protein [Microbacterium sp. H83] Length: 2023\nNumber of predicted TMHs: 2\nExp number of AAs in TMHs: 43.99839\nExp number, first 60 AAs: 22.48521\nTotal prob of N-in: 0.99010\nPOSSIBLE N-term signal sequence\ninside 1 31\nTMhelix 32 54\noutside 55 1993\nTMhelix 1994 2016\ninside 2017 2023

31737 GCF_001650405.1_ASM165040v1 Microbacterium sp. H83 Terrabacteria group; Actinobacteria; Actinobacteria; Micrococcales; Microbacteriaceae; Microbacterium
MTATTLENGPRARLHRLARRSVVFAIVLGTCGSGAGSAFAADGAVADTTPVELHVTAGVR WP_067122045.1
hypothetical protein [Microbacterium sp. H83] Length: 718\nNumber of predicted TMHs: 2\nExp number of AAs in TMHs: 40.97314\nExp number, first 60 AAs: 20.23419\nTotal prob of N-in: 0.95402\nPOSSIBLE N-term signal sequence\ninside 1 20\nTMhelix 21 40\noutside 41 653\nTMhelix 654 673\ninside 674 718

31738 GCF_001650405.1_ASM165040v1 Microbacterium sp. H83 Terrabacteria group; Actinobacteria; Actinobacteria; Micrococcales; Microbacteriaceae; Microbacterium
MAREKKRRSNGRRSPLAAAALIGAGLLITGAVYAGATAAIASTDTRSSAASTLTAEDGAK WP_067120488.1 cystathionine beta-lyase [Microbacterium sp. H83] Length: 267\nNumber of predicted TMHs: 2\nExp number of AAs in TMHs: 43.11477\nExp number, first 60 AAs: 22.51533\nTotal prob of N-in: 0.98281\nPOSSIBLE N-term signal sequence\ninside 1 20\nTMhelix 21 43\noutside 44 244\nTMhelix 245 264\ninside 265 267

31739 GCF_001639925.1_Microbacterium_sp._strain_HM58-_2_assembly01 Microbacterium sp. HM58-2 Terrabacteria group; Actinobacteria; Actinobacteria; Micrococcales; Microbacteriaceae; Microbacterium
MAREKKRRSNGRRSPLAAAALIGAGLMITGAVYAGASAAFAATDTQTAASTTLTVEDGEK WP_067352906.1
cystathionine beta-lyase [Microbacterium sp. HM58-2] Length: 267\nNumber of predicted TMHs: 2\nExp number of AAs in TMHs: 43.03307\nExp number, first 60 AAs: 22.61035\nTotal prob of N-in: 0.98110\nPOSSIBLE N-term signal sequence\ninside 1 20\nTMhelix 21 43\noutside 44 244\nTMhelix 245 264\ninside 265 267

31740 GCF_001866135.1_ASM186613v1 Microbacterium sp. LCT-H2 Terrabacteria group; Actinobacteria; Actinobacteria; Micrococcales; Microbacteriaceae; Microbacterium
MAREKKRRSSGRRSPLAAAALIGAGLMITGAVYAGATAAFAANDAPTASTQLTVEDGEKL WP_071329602.1
cystathionine beta-lyase [Microbacterium sp. LCT-H2] Length: 266\nNumber of predicted TMHs: 2\nExp number of AAs in TMHs: 43.1952\nExp number, first 60 AAs: 22.60029\nTotal prob of N-in: 0.99025\nPOSSIBLE N-term signal sequence\ninside 1 19\nTMhelix 20 42\noutside 43 243\nTMhelix 244 263\ninside 264 266

31741 GCF_001866135.1_ASM186613v1 Microbacterium sp. LCT-H2 Terrabacteria group; Actinobacteria; Actinobacteria; Micrococcales; Microbacteriaceae; Microbacterium
MRRRAALATLVVGAAVCGTPAAAVAAPTAPTTIVSMATGACEPGTVVFAPEAPSALAALG WP_071327739.1
hypothetical protein [Microbacterium sp. LCT-H2] Length: 436\nNumber of predicted TMHs: 2\nExp number of AAs in TMHs: 43.27242\nExp number, first 60 AAs: 21.39091\nTotal prob of N-in: 0.95808\nPOSSIBLE N-term signal sequence\ninside 1 6\nTMhelix 7 29\noutside 30 386\nTMhelix 387 409\ninside 410 436

31742 GCF_001866135.1_ASM186613v1 Microbacterium sp. LCT-H2 Terrabacteria group; Actinobacteria; Actinobacteria; Micrococcales; Microbacteriaceae; Microbacterium

MTRRRMLRSAVALVAVAASVLLLGSTPSPTPTPPVADDPADPVRAAEYWLDGAGIRDAWQ WP_071328144.1 peptidase S8 [Microbacterium sp. LCT-H2] Length: 438\nNumber of predicted TMHs: 2\nExp number of AAs in TMHs: 37.53461\nExp number, first 60 AAs: 15.82295\nTotal prob of N-in: 0.79826\nPOSSIBLE N-term signal sequence\ninside 1 6\nTMhelix 7 26\noutside 27 396\nTMhelix 397 419\ninside 420 438

31743 GCF_001866135.1_ASM186613v1 Microbacterium sp. LCT-H2 Terrabacteria group; Actinobacteria; Actinobacteria; Micrococcales; Microbacteriaceae; Microbacterium MSNTTIRPRRPRRAFLGATGIVGGLALALAVPAMAGAHVSVSPDELAAGDHGVLTFSFSH WP_071328382.1 hypothetical protein [Microbacterium sp. LCT-H2] Length: 224\nNumber of predicted TMHs: 2\nExp number of AAs in TMHs: 45.4029\nExp number, first 60 AAs: 22.56265\nTotal prob of N-in: 0.99942\nPOSSIBLE N-term signal sequence\ninside 1 12\nTMhelix 13 35\noutside 36 197\nTMhelix 198 220\ninside 221 224

31744 GCF_001866135.1_ASM186613v1 Microbacterium sp. LCT-H2 Terrabacteria group; Actinobacteria; Actinobacteria; Micrococcales; Microbacteriaceae; Microbacterium MLFSQQSSPQRDEPQQDRRRRRRLLSPRRPWAVATMLALVLGSLVAPVLPATAAPGDPAQ WP_083391816.1 hypothetical protein [Microbacterium sp. LCT-H2] Length: 2129\nNumber of predicted TMHs: 2\nExp number of AAs in TMHs: 43.85489\nExp number, first 60 AAs: 20.73873\nTotal prob of N-in: 0.91695\nPOSSIBLE N-term signal sequence\ninside 1 30\nTMhelix 31 53\noutside 54 2092\nTMhelix 2093 2115\ninside 2116 2129

31745 GCF_001866135.1_ASM186613v1 Microbacterium sp. LCT-H2 Terrabacteria group; Actinobacteria; Actinobacteria; Micrococcales; Microbacteriaceae; Microbacterium MHKNLPRRTAIVLSAVLLSGAVVVATPTVAVADDNFDSSYPVADSTISGSPDEISLMFTG WP_029266231.1 MULTISPECIES: hypothetical protein [Microbacterium] Length: 230\nNumber of predicted TMHs: 2\nExp number of AAs in TMHs: 44.08665\nExp number, first 60 AAs: 21.4021\nTotal prob of N-in: 0.99476\nPOSSIBLE N-term signal sequence\ninside 1 8\nTMhelix 9 31\noutside 32 181\nTMhelix 182 204\ninside 205 230

31746 GCF_001424225.1_Leaf151 Microbacterium sp. Leaf151 Terrabacteria group; Actinobacteria; Actinobacteria; Micrococcales; Microbacteriaceae; Microbacterium MTTPALPSLSAERVDAMEFAVLGRIADERHRVRRRRRNVWTGVGAAAAVVVVAAVVGPAL WP_055954023.1 hypothetical protein [Microbacterium sp. Leaf151] Length: 352\nNumber of predicted TMHs: 2\nExp number of AAs in TMHs: 45.85276\nExp number, first 60 AAs: 21.54922\nTotal prob of N-in: 0.99541\nPOSSIBLE N-term signal sequence\ninside 1 38\nTMhelix 39 61\noutside 62 314\nTMhelix 315 337\ninside 338 352

31747 GCF_001424225.1_Leaf151 Microbacterium sp. Leaf151 Terrabacteria group; Actinobacteria; Actinobacteria; Micrococcales; Microbacteriaceae; Microbacterium MAREKKTRRSSGRRSPLAAAALIGIGLLLTGGAYAGASAAMAATTTTPVSEANSATAVEEWP_055957397.1 cystathionine beta-lyase [Microbacterium sp. Leaf151] Length: 270\nNumber of predicted TMHs: 2\nExp number of AAs in TMHs: 42.51091\nExp number, first 60 AAs: 22.28922\nTotal prob of N-in: 0.97978\nPOSSIBLE N-term signal sequence\ninside 1 20\nTMhelix 21 43\noutside 44 247\nTMhelix 248 267\ninside 268 270

31748 GCF_001424225.1_Leaf151 Microbacterium sp. Leaf151 Terrabacteria group; Actinobacteria; Actinobacteria; Micrococcales; Microbacteriaceae; Microbacterium MRSSQAPRRRTLPAVLAAAFLAAAALLAPALPAAAHDELIATDPTADAVLESPPAQITLS WP_055957548.1 hypothetical protein [Microbacterium sp. Leaf151] Length: 213\nNumber of predicted TMHs: 2\nExp number of AAs in TMHs: 44.49909\nExp number, first 60 AAs: 22.42343\nTotal prob of N-in: 0.99885\nPOSSIBLE N-term signal sequence\ninside 1 11\nTMhelix 12 34\noutside 35 176\nTMhelix 177 199\ninside 200 213

31749 GCF_001424225.1_Leaf151 Microbacterium sp. Leaf151 Terrabacteria group; Actinobacteria; Actinobacteria; Micrococcales; Microbacteriaceae; Microbacterium MARTSTAALSPADQERRRALRVMKGVALGALLGMAIVFAISFSLQREVEWLQYVRAAAEG WP_055957235.1 DUF445 domain-containing protein [Microbacterium sp. Leaf151] Length: 424\nNumber of predicted TMHs: 2\nExp number of AAs in TMHs: 51.31844\nExp number, first 60 AAs: 22.33409\nTotal prob of N-in: 0.94723\nPOSSIBLE N-term signal sequence\ninside 1 19\nTMhelix 20 42\noutside 43 400\nTMhelix 401 423\ninside 424 424

31750 GCF_001423615.1_Leaf159 Microbacterium sp. Leaf159 Terrabacteria group; Actinobacteria; Actinobacteria; Micrococcales; Microbacteriaceae; Microbacterium MAREKKRRSNGRRSPLAAVALIGAGLMITGAVYAGTSAAFAATDTQTTAASTLTVEDGEK WP_056307037.1 cystathionine beta-lyase [Microbacterium sp. Leaf159] Length: 267\nNumber of predicted TMHs: 2\nExp number of AAs in TMHs: 42.99658\nExp number, first 60 AAs: 22.47532\nTotal prob of N-in: 0.98505\nPOSSIBLE N-term signal sequence\ninside 1 12\nTMhelix 13 35\noutside 36 244\nTMhelix 245 264\ninside 265 267

31751 GCF_001423615.1_Leaf159 Microbacterium sp. Leaf159 Terrabacteria group; Actinobacteria; Actinobacteria; Micrococcales; Microbacteriaceae; Microbacterium
MARRRAFIRAAVSGAVAFALALGGATAAVAEDADPPAAAGTISGTVTRGDDGSAVAGVGV WP_082488321.1 hypothetical protein [Microbacterium sp. Leaf159] Length: 577\nNumber of predicted TMHs: 2\nExp number of AAs in TMHs: 40.97074\nExp number, first 60 AAs: 21.86404\nTotal prob of N-in: 0.96841\nPOSSIBLE N-term signal sequence\ninside 1 6\nTMhelix 7 29\noutside 30 550\nTMhelix 551 570\ninside 571 577

31752 GCF_001423645.1_Leaf161 Microbacterium sp. Leaf161 Terrabacteria group; Actinobacteria; Actinobacteria; Micrococcales; Microbacteriaceae; Microbacterium
MRIIEGKTLRRRRAIGGGIAGVTVGAVVLAGALVPTAANALDDANPNDPSVAQGQIVQLP WP_056375501.1 hypothetical protein [Microbacterium sp. Leaf161] Length: 1955\nNumber of predicted TMHs: 2\nExp number of AAs in TMHs: 41.5828\nExp number, first 60 AAs: 22.24161\nTotal prob of N-in: 0.99211\nPOSSIBLE N-term signal sequence\ninside 1 12\nTMhelix 13 35\noutside 36 1922\nTMhelix 1923 1945\ninside 1946 1955

31753 GCF_001423645.1_Leaf161 Microbacterium sp. Leaf161 Terrabacteria group; Actinobacteria; Actinobacteria; Micrococcales; Microbacteriaceae; Microbacterium
MRRRRTALPALLVGAIALGAVLTSAPASAATLGSTVAAATAGEACTPGTRVFTPEAPPAL WP_056375517.1 hypothetical protein [Microbacterium sp. Leaf161] Length: 446\nNumber of predicted TMHs: 2\nExp number of AAs in TMHs: 43.35153\nExp number, first 60 AAs: 21.90715\nTotal prob of N-in: 0.98474\nPOSSIBLE N-term signal sequence\ninside 1 6\nTMhelix 7 29\noutside 30 389\nTMhelix 390 412\ninside 413 446

31754 GCF_001423645.1_Leaf161 Microbacterium sp. Leaf161 Terrabacteria group; Actinobacteria; Actinobacteria; Micrococcales; Microbacteriaceae; Microbacterium
MAREKKRRSNGRRSPLAAAALIGAGLMITGAVYAGTSAAFAATDTQTTAASTLTVEDGEK WP_056377205.1 cystathionine beta-lyase [Microbacterium sp. Leaf161] Length: 267\nNumber of predicted TMHs: 2\nExp number of AAs in TMHs: 43.07155\nExp number, first 60 AAs: 22.53642\nTotal prob of N-in: 0.98490\nPOSSIBLE N-term signal sequence\ninside 1 20\nTMhelix 21 43\noutside 44 244\nTMhelix 245 264\ninside 265 267

31755 GCF_001423785.1_Leaf179 Microbacterium sp. Leaf179 Terrabacteria group; Actinobacteria; Actinobacteria; Micrococcales; Microbacteriaceae; Microbacterium
MARRQDAVPTGVPVPSPTSSRRAALIALIALVGLSVPTTAQAAARTDCRPVPAPAADILR WP_082509289.1 hypothetical protein [Microbacterium sp. Leaf179] Length: 607\nNumber of predicted TMHs: 1\nExp number of AAs in TMHs: 29.06673\nExp number, first 60 AAs: 7.6261\nTotal prob of N-in: 0.37682\noutside 1 566\nTMhelix 567 589\ninside 590 607

31756 GCF_001423785.1_Leaf179 Microbacterium sp. Leaf179 Terrabacteria group; Actinobacteria; Actinobacteria; Micrococcales; Microbacteriaceae; Microbacterium
MARTPSTLLSPADQERRRALRVMKGVALGALLGMAVIFAISFSLQREIEWMQYVRAAAEG WP_055937988.1 DUF445 domain-containing protein [Microbacterium sp. Leaf179] Length: 424\nNumber of predicted TMHs: 2\nExp number of AAs in TMHs: 49.54129\nExp number, first 60 AAs: 22.08605\nTotal prob of N-in: 0.70994\nPOSSIBLE N-term signal sequence\ninside 1 19\nTMhelix 20 42\noutside 43 400\nTMhelix 401 423\ninside 424 424

31757 GCF_001423785.1_Leaf179 Microbacterium sp. Leaf179 Terrabacteria group; Actinobacteria; Actinobacteria; Micrococcales; Microbacteriaceae; Microbacterium
MAREKKTRRASGRRSPLAAAALIGIGLLLTGGVYAGASAAMAATNATPTSEANSATAVEE WP_055835609.1 MULTISPECIES: cystathionine beta-lyase [Microbacterium] Length: 270\nNumber of predicted TMHs: 2\nExp number of AAs in TMHs: 42.56802\nExp number, first 60 AAs: 22.36159\nTotal prob of N-in: 0.98585\nPOSSIBLE N-term signal sequence\ninside 1 20\nTMhelix 21 43\noutside 44 247\nTMhelix 248 267\ninside 268 270

31758 GCF_001421275.1_Leaf203 Microbacterium sp. Leaf203 Terrabacteria group; Actinobacteria; Actinobacteria; Micrococcales; Microbacteriaceae; Microbacterium
MARTPTRALSPADQERRRALRVMKGVALGALIAMALVFAVSFSLQREVEWLQYVRAAAEG WP_056229005.1 MULTISPECIES: DUF445 domain-containing protein [Microbacterium] Length: 424\nNumber of predicted TMHs: 2\nExp number of AAs in TMHs: 51.3979999999999\nExp number, first 60 AAs: 22.34727\nTotal prob of N-in: 0.95486\nPOSSIBLE N-term signal sequence\ninside 1 19\nTMhelix 20 42\noutside 43 400\nTMhelix 401 423\ninside 424 424

31759 GCF_001421275.1_Leaf203 Microbacterium sp. Leaf203 Terrabacteria group; Actinobacteria; Actinobacteria; Micrococcales; Microbacteriaceae; Microbacterium
MTPRRRLLPAVLAAGLLAAAALFVPASPAAAHDELVSTDPAAGATLEALPAQITFSYSAD WP_056227666.1 hypothetical protein [Microbacterium sp. Leaf203] Length: 209\nNumber of predicted TMHs: 2\nExp number of AAs in TMHs:

43.74896\nExp number, first 60 AAs:  21.65191\nTotal prob of N-in:   0.99736\nPOSSIBLE N-term signal sequence\ninside 1 6\nTMhelix 7 29\noutside 30 171\nTMhelix 172 194\ninside 195 209

31760 GCF_001421275.1_Leaf203 Microbacterium sp. Leaf203 Terrabacteria group; Actinobacteria; Actinobacteria; Micrococcales; Microbacteriaceae; Microbacterium
MVLSRPVSEDRSRRGIVRRAGSLVAAAAVVLSGLVPAVAASAAPLPTPLASYDFAATTGT WP_082450279.1 hypothetical protein [Microbacterium sp. Leaf203] Length: 1835\nNumber of predicted TMHs:  2\nExp number of AAs in TMHs: 45.02031\nExp number, first 60 AAs:  22.36042\nTotal prob of N-in:   0.99619\nPOSSIBLE N-term signal sequence\ninside 1 20\nTMhelix 21 43\noutside 44 1801\nTMhelix 1802 1824\ninside 1825 1835

31761 GCF_001421275.1_Leaf203 Microbacterium sp. Leaf203 Terrabacteria group; Actinobacteria; Actinobacteria; Micrococcales; Microbacteriaceae; Microbacterium
MAREKKTRRASGRRSPLAAAALIGIGLLLTGGVYAGASAAMAATSGTPTSEANSATAVEE WP_056227819.1 MULTISPECIES: cystathionine beta-lyase [Microbacterium] Length: 270\nNumber of predicted TMHs:  2\nExp number of AAs in TMHs: 42.60447\nExp number, first 60 AAs:  22.3786\nTotal prob of N-in:   0.99243\nPOSSIBLE N-term signal sequence\ninside 1 20\nTMhelix 21 43\noutside 44 247\nTMhelix 248 267\ninside 268 270

31762 GCF_001423485.1_Leaf320 Microbacterium sp. Leaf320 Terrabacteria group; Actinobacteria; Actinobacteria; Micrococcales; Microbacteriaceae; Microbacterium
MAREKKRRSNGRRSPLAAVALIGAGLMITGAVYAGTSAAFAATDTQTTAESTLTVEDGEK WP_056516548.1 cystathionine beta-lyase [Microbacterium sp. Leaf320] Length: 267\nNumber of predicted TMHs:  2\nExp number of AAs in TMHs: 42.92656\nExp number, first 60 AAs:  22.40765\nTotal prob of N-in:   0.98499\nPOSSIBLE N-term signal sequence\ninside 1 12\nTMhelix 13 35\noutside 36 244\nTMhelix 245 264\ninside 265 267

31763 GCF_001423805.1_Leaf347 Microbacterium sp. Leaf347 Terrabacteria group; Actinobacteria; Actinobacteria; Micrococcales; Microbacteriaceae; Microbacterium
MARTPKPRKAGRRSPLAAAALIGIGLLLTGGIYAGASAAVAATTDQTSTSAKASTLTVDD WP_056209929.1 MULTISPECIES: cystathionine beta-lyase [Microbacterium] Length: 270\nNumber of predicted TMHs:  2\nExp number of AAs in TMHs: 43.4441299999999\nExp number, first 60 AAs:  22.68386\nTotal prob of N-in:   0.91165\nPOSSIBLE N-term signal sequence\ninside 1 19\nTMhelix 20 42\noutside 43 247\nTMhelix 248 267\ninside 268 270

31764 GCF_001423805.1_Leaf347 Microbacterium sp. Leaf347 Terrabacteria group; Actinobacteria; Actinobacteria; Micrococcales; Microbacteriaceae; Microbacterium
MPRTPLAALSPADQARRRALRRMKAVALAALLAMAVLFAISFPLQATVPALAYVRAAAEG WP_056210180.1 MULTISPECIES: DUF445 domain-containing protein [Microbacterium] Length: 424\nNumber of predicted TMHs:  2\nExp number of AAs in TMHs: 56.0630900000002\nExp number, first 60 AAs:  26.19731\nTotal prob of N-in:   0.96574\nPOSSIBLE N-term signal sequence\ninside 1 25\nTMhelix 26 48\noutside 49 400\nTMhelix 401 423\ninside 424 424

31765 GCF_001423805.1_Leaf347 Microbacterium sp. Leaf347 Terrabacteria group; Actinobacteria; Actinobacteria; Micrococcales; Microbacteriaceae; Microbacterium
MSISQPRRRALAIIAGASLLIGIAAPGTALAASAAEPAPADVIINEAYLKGGSAGAPFTN WP_082483085.1 MULTISPECIES: multifunctional nuclease/2,3-cyclic-nucleotide 2-phosphodiesterase/5-nucleotidase/3-nucleotidase [Microbacterium] Length: 1638\nNumber of predicted TMHs:  2\nExp number of AAs in TMHs: 40.3707599999999\nExp number, first 60 AAs:  21.9023\nTotal prob of N-in:   0.97390\nPOSSIBLE N-term signal sequence\ninside 1 12\nTMhelix 13 35\noutside 36 1613\nTMhelix 1614 1633\ninside 1634 1638

31766 GCF_001423825.1_Leaf351 Microbacterium sp. Leaf351 Terrabacteria group; Actinobacteria; Actinobacteria; Micrococcales; Microbacteriaceae; Microbacterium
MPRTPLAALSPADQARRRALRRMKAVALAALLAMAVLFAISFPLQATVPALAYVRAAAEG WP_056210180.1 MULTISPECIES: DUF445 domain-containing protein [Microbacterium] Length: 424\nNumber of predicted TMHs:  2\nExp number of AAs in TMHs: 56.0630900000002\nExp number, first 60 AAs:  26.19731\nTotal prob of N-in:   0.96574\nPOSSIBLE N-term signal sequence\ninside 1 25\nTMhelix 26 48\noutside 49 400\nTMhelix 401 423\ninside 424 424

31767 GCF_001423825.1_Leaf351 Microbacterium sp. Leaf351 Terrabacteria group; Actinobacteria; Actinobacteria; Micrococcales; Microbacteriaceae; Microbacterium
MARTPKPRKAGRRSPLAAAALIGIGLLLTGGIYAGASAAVAATTDQTSTSAKASTLTVDD WP_056209929.1 MULTISPECIES: cystathionine beta-lyase [Microbacterium] Length: 270\nNumber of predicted TMHs:  2\nExp number of AAs in TMHs: 43.4441299999999\nExp number, first 60 AAs:  22.68386\nTotal prob of N-in:   0.91165\nPOSSIBLE N-term signal sequence\ninside 1 19\nTMhelix 20 42\noutside 43 247\nTMhelix 248 267\ninside 268 270

31768 GCF_001423825.1_Leaf351 Microbacterium sp. Leaf351 Terrabacteria group; Actinobacteria; Actinobacteria; Micrococcales; Microbacteriaceae; Microbacterium
MSISQPRRRALAIIAGASLLIGIAAPGTALAASAAEPAPADVIINEAYLKGGSAGAPFTN WP_082483085.1 MULTISPECIES: multifunctional nuclease/2,3-cyclic-nucleotide 2-phosphodiesterase/5-nucleotidase/3-nucleotidase [Microbacterium]
Length: 1638\nNumber of predicted TMHs: 2\nExp number of AAs in TMHs: 40.3707599999999\nExp number, first 60 AAs: 21.9023\nTotal prob of N-in: 0.97390\nPOSSIBLE N-term signal sequence\ninside 1 12\nTMhelix 13 35\noutside 36 1613\nTMhelix 1614 1633\ninside 1634 1638

31769 GCF_001424715.1_Leaf436 Microbacterium sp. Leaf436 Terrabacteria group; Actinobacteria; Actinobacteria; Micrococcales; Microbacteriaceae; Microbacterium
MTSPVAPRRRTLPALLATALLAAAALLVPALPAAAHDELVSTDPSADAVLETLPAEITFT WP_082516518.1 hypothetical protein [Microbacterium sp. Leaf436] Length: 220\nNumber of predicted TMHs: 2\nExp number of AAs in TMHs: 44.99039\nExp number, first 60 AAs: 22.54572\nTotal prob of N-in: 0.99787\nPOSSIBLE N-term signal sequence\ninside 1 11\nTMhelix 12 34\noutside 35 180\nTMhelix 181 203\ninside 204 220

31770 GCF_001424715.1_Leaf436 Microbacterium sp. Leaf436 Terrabacteria group; Actinobacteria; Actinobacteria; Micrococcales; Microbacteriaceae; Microbacterium
MAREKKTRRASGRRSPLAAAALIGIGLLLTGGVYAGASAAMAATNATPTSEANSATAVEE WP_055835609.1
MULTISPECIES: cystathionine beta-lyase [Microbacterium] Length: 270\nNumber of predicted TMHs: 2\nExp number of AAs in TMHs: 42.56802\nExp number, first 60 AAs: 22.36159\nTotal prob of N-in: 0.98585\nPOSSIBLE N-term signal sequence\ninside 1 20\nTMhelix 21 43\noutside 44 247\nTMhelix 248 267\ninside 268 270

31771 GCF_001424715.1_Leaf436 Microbacterium sp. Leaf436 Terrabacteria group; Actinobacteria; Actinobacteria; Micrococcales; Microbacteriaceae; Microbacterium
MARTPSTLLSPADQERRRALRVMKGVALGALLGMAVIFAISFSLQREIEWMQYVRAAAEG WP_055838053.1
DUF445 domain-containing protein [Microbacterium sp. Leaf436] Length: 424\nNumber of predicted TMHs: 2\nExp number of AAs in TMHs: 49.13359\nExp number, first 60 AAs: 22.06226\nTotal prob of N-in: 0.72081\nPOSSIBLE N-term signal sequence\ninside 1 19\nTMhelix 20 42\noutside 43 400\nTMhelix 401 423\ninside 424 424

31772 GCF_002114135.1_ASM211413v1 Microbacterium sp. LEMMJ01 Terrabacteria group; Actinobacteria; Actinobacteria; Micrococcales; Microbacteriaceae; Microbacterium
MAREKKRRSSGRRSPLAAAALIGAGLMITGAVYAGATAAFAANDAPTASTQLTVEDGEKL WP_025105058.1
MULTISPECIES: cystathionine beta-lyase [Microbacterium] Length: 266\nNumber of predicted TMHs: 2\nExp number of AAs in TMHs: 43.19361\nExp number, first 60 AAs: 22.60029\nTotal prob of N-in: 0.99024\nPOSSIBLE N-term signal sequence\ninside 1 19\nTMhelix 20 42\noutside 43 243\nTMhelix 244 263\ninside 264 266

31773 GCF_002114135.1_ASM211413v1 Microbacterium sp. LEMMJ01 Terrabacteria group; Actinobacteria; Actinobacteria; Micrococcales; Microbacteriaceae; Microbacterium
MRRRAALATLVVGAAVWGAPAAAVAAPTAPTTIVSMATGACEPGTVVFAPEAPSPLAALG WP_085604923.1
hypothetical protein [Microbacterium sp. LEMMJ01] Length: 436\nNumber of predicted TMHs: 2\nExp number of AAs in TMHs: 43.5319099999999\nExp number, first 60 AAs: 21.62528\nTotal prob of N-in: 0.96876\nPOSSIBLE N-term signal sequence\ninside 1 6\nTMhelix 7 29\noutside 30 386\nTMhelix 387 409\ninside 410 436

31774 GCF_002114135.1_ASM211413v1 Microbacterium sp. LEMMJ01 Terrabacteria group; Actinobacteria; Actinobacteria; Micrococcales; Microbacteriaceae; Microbacterium
MFMQFSLHTNSEREEKKDRRTRRRLLSPRRPWAVATMLALVLGSLVAPVLPATAAPGDPA WP_085604174.1
hypothetical protein [Microbacterium sp. LEMMJ01] Length: 2245\nNumber of predicted TMHs: 2\nExp number of AAs in TMHs: 43.3115499999999\nExp number, first 60 AAs: 20.6902\nTotal prob of N-in: 0.91391\nPOSSIBLE N-term signal sequence\ninside 1 31\nTMhelix 32 54\noutside 55 2209\nTMhelix 2210 2232\ninside 2233 2245

31775 GCF_002114135.1_ASM211413v1 Microbacterium sp. LEMMJ01 Terrabacteria group; Actinobacteria; Actinobacteria; Micrococcales; Microbacteriaceae; Microbacterium
MTRRRMLRGAVALVAVAASVLLLGSTATPTPTPPPVADDPADPVRAAEYWLDGARIRDAW WP_036292654.1
MULTISPECIES: peptidase S8 [Microbacterium] Length: 439\nNumber of predicted TMHs: 2\nExp number of AAs in TMHs: 41.4776299999999\nExp number, first 60 AAs: 19.80316\nTotal prob of N-in: 0.93229\nPOSSIBLE N-term signal sequence\ninside 1 6\nTMhelix 7 29\noutside 30 397\nTMhelix 398 420\ninside 421 439

31776 GCF_002114135.1_ASM211413v1 Microbacterium sp. LEMMJ01 Terrabacteria group; Actinobacteria; Actinobacteria; Micrococcales; Microbacteriaceae; Microbacterium
MSNTTIRPRRPRRTLLGATGVVGGLALALAVPAMAGAHVSVSPDELAAGDHGVLTFSFSH WP_085607271.1

hypothetical protein [Microbacterium sp. LEMMJ01] Length: 224\nNumber of predicted TMHs: 2\nExp number of AAs in TMHs: 45.15794\nExp number, first 60 AAs: 22.31633\nTotal prob of N-in: 0.99851\nPOSSIBLE N-term signal sequence\ninside 1 12\nTMhelix 13 35\noutside 36 197\nTMhelix 198 220\ninside 221 224

31777 GCF_002114135.1_ASM211413v1 Microbacterium sp. LEMMJ01 Terrabacteria group; Actinobacteria; Actinobacteria; Micrococcales; Microbacteriaceae; Microbacterium MRRSSTFGEGVAAGSAELEVDRDSTGLRRPPPEGGDMARRRAFFRSIMGGAAALALALGG WP_085605654.1 hypothetical protein [Microbacterium sp. LEMMJ01] Length: 710\nNumber of predicted TMHs: 2\nExp number of AAs in TMHs: 38.7195899999999\nExp number, first 60 AAs: 14.85922\nTotal prob of N-in: 0.85732\nPOSSIBLE N-term signal sequence\ninside 1 41\nTMhelix 42 64\noutside 65 685\nTMhelix 686 703\ninside 704 710

31778 GCF_000834175.1_ASM83417v1 Microbacterium sp. MEJ108Y Terrabacteria group; Actinobacteria; Actinobacteria; Micrococcales; Microbacteriaceae; Microbacterium MAREKKRRSNGRRSPLAAAALIGAGLMITGAVYAGTSAAFAATDTQTTAANTLTVEDGEK WP_042538393.1 cystathionine beta-lyase [Microbacterium sp. MEJ108Y] Length: 267\nNumber of predicted TMHs: 2\nExp number of AAs in TMHs: 43.08096\nExp number, first 60 AAs: 22.53588\nTotal prob of N-in: 0.98271\nPOSSIBLE N-term signal sequence\ninside 1 20\nTMhelix 21 43\noutside 44 244\nTMhelix 245 264\ninside 265 267

31779 GCF_000834175.1_ASM83417v1 Microbacterium sp. MEJ108Y Terrabacteria group; Actinobacteria; Actinobacteria; Micrococcales; Microbacteriaceae; Microbacterium MRRRRTALPALLVGAIALGAVLTSAPASAATLGSTVPAATAGEACTPGTVVFTPEAPPAL WP_052493031.1 hypothetical protein [Microbacterium sp. MEJ108Y] Length: 447\nNumber of predicted TMHs: 2\nExp number of AAs in TMHs: 43.27129\nExp number, first 60 AAs: 21.82331\nTotal prob of N-in: 0.98487\nPOSSIBLE N-term signal sequence\ninside 1 6\nTMhelix 7 29\noutside 30 389\nTMhelix 390 412\ninside 413 447

31780 GCF_000834175.1_ASM83417v1 Microbacterium sp. MEJ108Y Terrabacteria group; Actinobacteria; Actinobacteria; Micrococcales; Microbacteriaceae; Microbacterium MAQRRALIHAVLGGVAAIALALGGATAAGAVEDDPPTGAVADVVAAAAGETADPLEGVAD WP_082041131.1 hypothetical protein [Microbacterium sp. MEJ108Y] Length: 645\nNumber of predicted TMHs: 2\nExp number of AAs in TMHs: 42.7227199999999\nExp number, first 60 AAs: 22.41885\nTotal prob of N-in: 0.99140\nPOSSIBLE N-term signal sequence\ninside 1 6\nTMhelix 7 29\noutside 30 618\nTMhelix 619 638\ninside 639 645

31781 GCF_000585455.1_Denovo_assembly Microbacterium sp. MRS-1 Terrabacteria group; Actinobacteria; Actinobacteria; Micrococcales; Microbacteriaceae; Microbacterium MRPTAERRKRRSFVAAITAGALALTGAGLAALPARAAVSADAAVLINEVYGGGGNSKATY WP_036284994.1 cell wall protein [Microbacterium sp. MRS-1] Length: 1578\nNumber of predicted TMHs: 2\nExp number of AAs in TMHs: 42.09439\nExp number, first 60 AAs: 21.69899\nTotal prob of N-in: 0.99859\nPOSSIBLE N-term signal sequence\ninside 1 11\nTMhelix 12 34\noutside 35 1548\nTMhelix 1549 1568\ninside 1569 1578

31782 GCF_001314225.1_ASM131422v1 Microbacterium sp. No. 7 Terrabacteria group; Actinobacteria; Actinobacteria; Micrococcales; Microbacteriaceae; Microbacterium MTTVTDHEHPHRSGSRRRILLLSTLTTLLLAAGLLFATAAPAAAHDEIVSSSPEAGSTVS WP_054678997.1 hypothetical protein [Microbacterium sp. No. 7] Length: 238\nNumber of predicted TMHs: 2\nExp number of AAs in TMHs: 45.15401\nExp number, first 60 AAs: 22.45081\nTotal prob of N-in: 0.99983\nPOSSIBLE N-term signal sequence\ninside 1 19\nTMhelix 20 42\noutside 43 183\nTMhelix 184 206\ninside 207 238

31783 GCF_001314225.1_ASM131422v1 Microbacterium sp. No. 7 Terrabacteria group; Actinobacteria; Actinobacteria; Micrococcales; Microbacteriaceae; Microbacterium MPTTTRRPALRSGRALLASGLALGVCALGVGVVPAYADAAHPDHVALFGEFGTYWQAEAF WP_082405844.1 hypothetical protein [Microbacterium sp. No. 7] Length: 1004\nNumber of predicted TMHs: 2\nExp number of AAs in TMHs: 44.1389099999999\nExp number, first 60 AAs: 21.45908\nTotal prob of N-in: 0.95846\nPOSSIBLE N-term signal sequence\ninside 1 12\nTMhelix 13 35\noutside 36 974\nTMhelix 975 997\ninside 998 1004

31784 GCF_001558975.1_ASM155897v1 Microbacterium sp. PAMC 28756 Terrabacteria group; Actinobacteria; Actinobacteria; Micrococcales; Microbacteriaceae; Microbacterium MAREKKRRSSGRRSPLAAAALIGAGLMITGAVYAGATAAFAANDAPTASTQLTVEDGEKL WP_025105058.1 MULTISPECIES: cystathionine beta-lyase [Microbacterium] Length: 266\nNumber of predicted TMHs: 2\nExp number of AAs in TMHs: 43.19361\nExp number, first 60 AAs: 22.60029\nTotal prob of N-in: 0.99024\nPOSSIBLE N-term signal sequence\ninside 1 19\nTMhelix 20 42\noutside 43 243\nTMhelix 244 263\ninside 264 266

31785 GCF_001558975.1_ASM155897v1 Microbacterium sp. PAMC 28756 Terrabacteria group; Actinobacteria; Actinobacteria; Micrococcales; Microbacteriaceae; Microbacterium
MRRRAALATLVVGAAVWGTPAAVVAAPTAPTTIVSMATGACEPGTVVFAPEAPSALAALG WP_025104647.1
MULTISPECIES: hypothetical protein [Microbacterium] Length: 436\nNumber of predicted TMHs: 2\nExp number of AAs in TMHs: 43.5666099999999\nExp number, first 60 AAs: 21.6829\nTotal prob of N-in: 0.96731\nPOSSIBLE N-term signal sequence\ninside 1 6\nTMhelix 7 29\noutside 30 386\nTMhelix 387 409\ninside 410 436

31786 GCF_001558975.1_ASM155897v1 Microbacterium sp. PAMC 28756 Terrabacteria group; Actinobacteria; Actinobacteria; Micrococcales; Microbacteriaceae; Microbacterium
MSNTTIRPRRPRRTLLGATGVVGGLALALAVPAMAGAHVSVSPDELAAGDHGVLTFSFSH WP_025103163.1
MULTISPECIES: hypothetical protein [Microbacterium] Length: 224\nNumber of predicted TMHs: 2\nExp number of AAs in TMHs: 45.16071\nExp number, first 60 AAs: 22.31743\nTotal prob of N-in: 0.99849\nPOSSIBLE N-term signal sequence\ninside 1 12\nTMhelix 13 35\noutside 36 197\nTMhelix 198 220\ninside 221 224

31787 GCF_001558975.1_ASM155897v1 Microbacterium sp. PAMC 28756 Terrabacteria group; Actinobacteria; Actinobacteria; Micrococcales; Microbacteriaceae; Microbacterium
MTRRRMLRGAVALVAVAASVLLLGSTATPTPTPPPVADDPADPVRAAEYWLDGARIRDAW WP_036292654.1
MULTISPECIES: peptidase S8 [Microbacterium] Length: 439\nNumber of predicted TMHs: 2\nExp number of AAs in TMHs: 41.4776299999999\nExp number, first 60 AAs: 19.80316\nTotal prob of N-in: 0.93229\nPOSSIBLE N-term signal sequence\ninside 1 6\nTMhelix 7 29\noutside 30 397\nTMhelix 398 420\ninside 421 439

31788 GCF_001426835.1_Root1433D1 Microbacterium sp. Root1433D1 Terrabacteria group; Actinobacteria; Actinobacteria; Micrococcales; Microbacteriaceae; Microbacterium
MAREKKRRSGGRRSPLAAAALIGAGLMITGAVYAGASAAFAASDTQSAATTQLTVEDGEK WP_055867422.1
cystathionine beta-lyase [Microbacterium sp. Root1433D1]Length: 267\nNumber of predicted TMHs: 2\nExp number of AAs in TMHs: 42.96638\nExp number, first 60 AAs: 22.51273\nTotal prob of N-in: 0.98149\nPOSSIBLE N-term signal sequence\ninside 1 19\nTMhelix 20 42\noutside 43 244\nTMhelix 245 264\ninside 265 267

31789 GCF_001426835.1_Root1433D1 Microbacterium sp. Root1433D1 Terrabacteria group; Actinobacteria; Actinobacteria; Micrococcales; Microbacteriaceae; Microbacterium
MARRRAFIRAVLGAAAALALALGGVTAAAAEDDPVVTGSISGTVTSTVGGTALAGIEVRA WP_055866250.1
hypothetical protein [Microbacterium sp. Root1433D1] Length: 682\nNumber of predicted TMHs: 2\nExp number of AAs in TMHs: 66.9881300000003\nExp number, first 60 AAs: 32.48513\nTotal prob of N-in: 0.99227\nPOSSIBLE N-term signal sequence\ninside 1 6\nTMhelix 7 29\noutside 30 652\nTMhelix 653 675\ninside 676 682

31790 GCF_001426835.1_Root1433D1 Microbacterium sp. Root1433D1 Terrabacteria group; Actinobacteria; Actinobacteria; Micrococcales; Microbacteriaceae; Microbacterium
MPRLTSPARRRLIATLPITALLAGSLLATSLPAAAADDSIAGPRTSGDPMFPNIGNGGYD WP_082531423.1 hypothetical protein [Microbacterium sp. Root1433D1] Length: 792\nNumber of predicted TMHs: 2\nExp number of AAs in TMHs: 42.0447899999999\nExp number, first 60 AAs: 21.07478\nTotal prob of N-in: 0.93172\nPOSSIBLE N-term signal sequence\ninside 1 11\nTMhelix 12 34\noutside 35 759\nTMhelix 760 782\ninside 783 792

31791 GCF_001426835.1_Root1433D1 Microbacterium sp. Root1433D1 Terrabacteria group; Actinobacteria; Actinobacteria; Micrococcales; Microbacteriaceae; Microbacterium
MKTHPRPEPARRRALGGGVIAAAAALTLLLPALPAAAIAPMADPASLPATEPMTPQRIEA WP_055869169.1
hypothetical protein [Microbacterium sp. Root1433D1] Length: 1004\nNumber of predicted TMHs: 2\nExp number of AAs in TMHs: 44.18681\nExp number, first 60 AAs: 22.42461\nTotal prob of N-in: 0.99416\nPOSSIBLE N-term signal sequence\ninside 1 12\nTMhelix 13 35\noutside 36 976\nTMhelix 977 999\ninside 1000 1004

31792 GCF_001427145.1_Root166 Microbacterium sp. Root166 Terrabacteria group; Actinobacteria; Actinobacteria; Micrococcales; Microbacteriaceae; Microbacterium
MTSSPDRRRSAVAALAIAAGLVLAPVAASPAHAGAAASAVLHSTDAAAQPALHVGVRQAA WP_055960901.1
hypothetical protein [Microbacterium sp. Root166] Length: 189\nNumber of predicted TMHs: 2\nExp number of AAs in TMHs: 45.01594\nExp number, first 60 AAs: 21.95234\nTotal prob of N-in: 0.99068\nPOSSIBLE N-term signal sequence\ninside 1 11\nTMhelix 12 34\noutside 35 158\nTMhelix 159 181\ninside 182 189

31793 GCF_001427145.1_Root166 Microbacterium sp. Root166 Terrabacteria group; Actinobacteria; Actinobacteria; Micrococcales; Microbacteriaceae; Microbacterium
MTAQLDSRPRRRRTLAAVAAAVVVAALSPLGGVLATPAYAAGDVCDICSLNFDVTACEKI WP_055962946.1
hypothetical protein [Microbacterium sp. Root166] Length: 288\nNumber of predicted TMHs: 2\nExp

number of AAs in TMHs: 45.0209099999999\nExp number, first 60 AAs: 22.33645\nTotal prob of N-in: 0.97980\nPOSSIBLE N-term signal sequence\ninside 1 12\nTMhelix 13 35\noutside 36 253\nTMhelix 254 276\ninside 277 288

31794 GCF_001428975.1_Root280D1 Microbacterium sp. Root280D1 Terrabacteria group; Actinobacteria; Actinobacteria; Micrococcales; Microbacteriaceae; Microbacterium MAREKKRRSNGRRSPLAAAALIGAGLMITGAVYAGTSAAFAATDTQTTAASTLTVEDGEK WP_053096702.1 MULTISPECIES: cystathionine beta-lyase [Microbacterium] Length: 267\nNumber of predicted TMHs: 2\nExp number of AAs in TMHs: 43.06753\nExp number, first 60 AAs: 22.53635\nTotal prob of N-in: 0.98491\nPOSSIBLE N-term signal sequence\ninside 1 20\nTMhelix 21 43\noutside 44 244\nTMhelix 245 264\ninside 265 267

31795 GCF_001428975.1_Root280D1 Microbacterium sp. Root280D1 Terrabacteria group; Actinobacteria; Actinobacteria; Micrococcales; Microbacteriaceae; Microbacterium MRRTRRGVIAAVGAMALAAAIAVPSSATAAAASGLWWFDRGKVQEAHDAGFDGAGIKIAV WP_082586491.1 hypothetical protein [Microbacterium sp. Root280D1] Length: 445\nNumber of predicted TMHs: 2\nExp number of AAs in TMHs: 46.179\nExp number, first 60 AAs: 23.29353\nTotal prob of N-in: 0.99582\nPOSSIBLE N-term signal sequence\ninside 1 6\nTMhelix 7 29\noutside 30 405\nTMhelix 406 428\ninside 429 445

31796 GCF_001425645.1_Root322 Microbacterium sp. Root322 Terrabacteria group; Actinobacteria; Actinobacteria; Micrococcales; Microbacteriaceae; Microbacterium MPRLTSPARRRLIATLPITALLAGSLLATSLPAAAADDSVAGPRTSGDPMFPNIGNGGYD WP_082524328.1 hypothetical protein [Microbacterium sp. Root322] Length: 792\nNumber of predicted TMHs: 2\nExp number of AAs in TMHs: 41.5665899999999\nExp number, first 60 AAs: 20.59012\nTotal prob of N-in: 0.91171\nPOSSIBLE N-term signal sequence\ninside 1 11\nTMhelix 12 34\noutside 35 759\nTMhelix 760 782\ninside 783 792

31797 GCF_001425645.1_Root322 Microbacterium sp. Root322 Terrabacteria group; Actinobacteria; Actinobacteria; Micrococcales; Microbacteriaceae; Microbacterium MARRRALIRAVLGAAAALALALGGVTPAAAEDDPVVTGSISGTVTSTVGGTTLAGIEVRAWP_055874771.1 hypothetical protein [Microbacterium sp. Root322] Length: 679\nNumber of predicted TMHs: 2\nExp number of AAs in TMHs: 52.3772799999999\nExp number, first 60 AAs: 25.52116\nTotal prob of N-in: 0.96715\nPOSSIBLE N-term signal sequence\ninside 1 6\nTMhelix 7 29\noutside 30 649\nTMhelix 650 672\ninside 673 679

31798 GCF_001425645.1_Root322 Microbacterium sp. Root322 Terrabacteria group; Actinobacteria; Actinobacteria; Micrococcales; Microbacteriaceae; Microbacterium MKTRPRPGPARRRALGGGVIAAAAALTLLLPALPAAAIAPMADPASLPATEPMTPQRIEA WP_055873157.1 hypothetical protein [Microbacterium sp. Root322] Length: 1004\nNumber of predicted TMHs: 2\nExp number of AAs in TMHs: 44.54294\nExp number, first 60 AAs: 22.45496\nTotal prob of N-in: 0.99551\nPOSSIBLE N-term signal sequence\ninside 1 12\nTMhelix 13 35\noutside 36 976\nTMhelix 977 999\ninside 1000 1004

31799 GCF_001425645.1_Root322 Microbacterium sp. Root322 Terrabacteria group; Actinobacteria; Actinobacteria; Micrococcales; Microbacteriaceae; Microbacterium MAREKKRRSGGRRSPLAAAALIGAGLMITGAVYAGATAAFAASDTQSAATSQLTVEDGEK WP_055872347.1 cystathionine beta-lyase [Microbacterium sp. Root322] Length: 267\nNumber of predicted TMHs: 2\nExp number of AAs in TMHs: 43.03254\nExp number, first 60 AAs: 22.59443\nTotal prob of N-in: 0.98774\nPOSSIBLE N-term signal sequence\ninside 1 19\nTMhelix 20 42\noutside 43 244\nTMhelix 245 264\ninside 265 267

31800 GCF_001426925.1_Root53 Microbacterium sp. Root53 Terrabacteria group; Actinobacteria; Actinobacteria; Micrococcales; Microbacteriaceae; Microbacterium MAREKKRRSNGRRSPLAAAALIGIGLLITGGAYAGASAAVAASTETTAASTTLTVEDGEK WP_055991805.1 cystathionine beta-lyase [Microbacterium sp. Root53] Length: 267\nNumber of predicted TMHs: 2\nExp number of AAs in TMHs: 43.46469\nExp number, first 60 AAs: 22.50179\nTotal prob of N-in: 0.97459\nPOSSIBLE N-term signal sequence\ninside 1 19\nTMhelix 20 42\noutside 43 244\nTMhelix 245 264\ninside 265 267

31801 GCF_001426995.1_Root553 Microbacterium sp. Root553 Terrabacteria group; Actinobacteria; Actinobacteria; Micrococcales; Microbacteriaceae; Microbacterium MAREKKRRSNGRRSPLAAAALIGAGLMITGAVYAGTSAAFAATDTQTAASSTLTVDDGEK WP_045253754.1 MULTISPECIES: cystathionine beta-lyase [Microbacterium] Length: 267\nNumber of predicted TMHs: 2\nExp number of AAs in TMHs: 43.05922\nExp number, first 60 AAs: 22.53913\nTotal prob of N-in: 0.98687\nPOSSIBLE N-term signal sequence\ninside 1 20\nTMhelix 21 43\noutside 44 244\nTMhelix 245 264\ninside 265 267

31802 GCF_001427525.1_Root61 Microbacterium sp. Root61 Terrabacteria group; Actinobacteria; Actinobacteria; Micrococcales; Microbacteriaceae; Microbacterium
MAQEKKRRSTGRRSPLAAAALIGIGLLLTGGIYAGASAAMAATADTTATTALTVDDGKKL WP_056217059.1 cystathionine beta-lyase [Microbacterium sp. Root61] Length: 266\nNumber of predicted TMHs: 2\nExp number of AAs in TMHs: 43.04937\nExp number, first 60 AAs: 22.61911\nTotal prob of N-in: 0.97058\nPOSSIBLE N-term signal sequence\ninside 1 19\nTMhelix 20 42\noutside 43 243\nTMhelix 244 263\ninside 264 266

31803 GCF_001427525.1_Root61 Microbacterium sp. Root61 Terrabacteria group; Actinobacteria; Actinobacteria; Micrococcales; Microbacteriaceae; Microbacterium
MARTPTALLSPADQERRRALQLMKGVALGALLFMAVVFVVAFAWQESVPALAYVRAAAEG WP_056219696.1 DUF445 domain-containing protein [Microbacterium sp. Root61] Length: 423\nNumber of predicted TMHs: 2\nExp number of AAs in TMHs: 53.3644600000003\nExp number, first 60 AAs: 24.04806\nTotal prob of N-in: 0.90893\nPOSSIBLE N-term signal sequence\ninside 1 24\nTMhelix 25 44\noutside 45 399\nTMhelix 400 422\ninside 423 423

31804 GCF_900156435.1_IMG-taxon_2710264749_annotated_assembly Microbacterium sp. RU1D Terrabacteria group; Actinobacteria; Actinobacteria; Micrococcales; Microbacteriaceae; Microbacterium
MARTPTAALSPADQERRRALHVMKGVALGALLAMALVFAVSFSLQREVEWLQYVRAAAEG WP_076676667.1 DUF445 domain-containing protein [Microbacterium sp. RU1D] Length: 424\nNumber of predicted TMHs: 2\nExp number of AAs in TMHs: 53.0642799999999\nExp number, first 60 AAs: 23.12903\nTotal prob of N-in: 0.85509\nPOSSIBLE N-term signal sequence\ninside 1 19\nTMhelix 20 42\noutside 43 400\nTMhelix 401 423\ninside 424 424

31805 GCF_900156435.1_IMG-taxon_2710264749_annotated_assembly Microbacterium sp. RU1D Terrabacteria group; Actinobacteria; Actinobacteria; Micrococcales; Microbacteriaceae; Microbacterium
MTSSLAPRRRTLPAVLASALLAAAALLAPASPAAAHDELVSTDPSADAVLDALPPQITLT WP_076674855.1 hypothetical protein [Microbacterium sp. RU1D] Length: 213\nNumber of predicted TMHs: 2\nExp number of AAs in TMHs: 43.44635\nExp number, first 60 AAs: 21.42596\nTotal prob of N-in: 0.96678\nPOSSIBLE N-term signal sequence\ninside 1 11\nTMhelix 12 34\noutside 35 176\nTMhelix 177 199\ninside 200 213

31806 GCF_900156435.1_IMG-taxon_2710264749_annotated_assembly Microbacterium sp. RU1D Terrabacteria group; Actinobacteria; Actinobacteria; Micrococcales; Microbacteriaceae; Microbacterium
MAREKKTRRASGRRSPLAAAALIGIGLLLTGGVYAGASAAMAATTTVPESEVNSATAVEE WP_076674935.1 cystathionine beta-lyase [Microbacterium sp. RU1D] Length: 270\nNumber of predicted TMHs: 2\nExp number of AAs in TMHs: 42.49875\nExp number, first 60 AAs: 22.27213\nTotal prob of N-in: 0.98422\nPOSSIBLE N-term signal sequence\ninside 1 20\nTMhelix 21 43\noutside 44 247\nTMhelix 248 267\ninside 268 270

31807 GCF_900155915.1_IMG-taxon_2708742473_annotated_assembly Microbacterium sp. RU33B Terrabacteria group; Actinobacteria; Actinobacteria; Micrococcales; Microbacteriaceae; Microbacterium
MARTPTSQLSPADQERRRALRVMKGVALGALIGMAVVFVFAFAFQQQVPWLAYVRAAAEG WP_076680123.1 DUF445 domain-containing protein [Microbacterium sp. RU33B] Length: 423\nNumber of predicted TMHs: 2\nExp number of AAs in TMHs: 55.9879900000001\nExp number, first 60 AAs: 24.55661\nTotal prob of N-in: 0.95585\nPOSSIBLE N-term signal sequence\ninside 1 24\nTMhelix 25 44\noutside 45 399\nTMhelix 400 422\ninside 423 423

31808 GCF_900156455.1_IMG-taxon_2710264756_annotated_assembly Microbacterium sp. RURRCA19A Terrabacteria group; Actinobacteria; Actinobacteria; Micrococcales; Microbacteriaceae; Microbacterium
MAREKKPRRSNGRRSPLAAAALIGIGLLLTGGIYAGASAAMAATDTPTSASNSASTVEEG WP_076490678.1 cystathionine beta-lyase [Microbacterium sp. RURRCA19A] Length: 269\nNumber of predicted TMHs: 2\nExp number of AAs in TMHs: 42.94594\nExp number, first 60 AAs: 22.46062\nTotal prob of N-in: 0.98945\nPOSSIBLE N-term signal sequence\ninside 1 20\nTMhelix 21 43\noutside 44 246\nTMhelix 247 266\ninside 267 269

31809 GCF_900156455.1_IMG-taxon_2710264756_annotated_assembly Microbacterium sp. RURRCA19A Terrabacteria group; Actinobacteria; Actinobacteria; Micrococcales; Microbacteriaceae; Microbacterium
MSDFRERSDAEGSATPSRRRRRVGRAQSDAATRSARRAVLPLAAGLVAGVTVFALVSGGI WP_076494745.1 hypothetical protein [Microbacterium sp. RURRCA19A] Length: 876\nNumber of predicted TMHs: 2\nExp number of AAs in TMHs: 43.99054\nExp number, first 60 AAs: 21.30587\nTotal prob of N-in: 0.98961\nPOSSIBLE N-term signal sequence\ninside 1 38\nTMhelix 39 61\noutside 62 845\nTMhelix 846 868\ninside 869 876

31810 GCF_900156455.1_IMG-taxon_2710264756_annotated_assembly Microbacterium sp. RURRCA19A Terrabacteria group; Actinobacteria; Actinobacteria; Micrococcales; Microbacteriaceae; Microbacterium

MIESPRPKGHSMHHPPPRRRRPLLTALTLALVAVVGVPALAAAPAHAAERGTGFGTWAPL WP_076494321.1 hypothetical protein [Microbacterium sp. RURRCA19A] Length: 516\nNumber of predicted TMHs: 2\nExp number of AAs in TMHs: 44.63364\nExp number, first 60 AAs: 21.82122\nTotal prob of N-in: 0.96720\nPOSSIBLE N-term signal sequence\ninside 1 22\nTMhelix 23 45\noutside 46 485\nTMhelix 486 508\ninside 509 516

31811 GCF_000967865.1_ASM96786v1 Microbacterium sp. SA39 Terrabacteria group; Actinobacteria; Actinobacteria; Micrococcales; Microbacteriaceae; Microbacterium MPVVPLDSIRCRIIGRSSRPSGTTRSLRARPPSRGGTVTRRRGMMRAVLGGAVAFALALG WP_082087440.1 hypothetical protein [Microbacterium sp. SA39] Length: 612\nNumber of predicted TMHs: 1\nExp number of AAs in TMHs: 86.9416200000002\nExp number, first 60 AAs: 14.31186\nTotal prob of N-in: 0.96547\nPOSSIBLE N-term signal sequence\noutside 1 585\nTMhelix 586 605\ninside 606 612

31812 GCF_000967865.1_ASM96786v1 Microbacterium sp. SA39 Terrabacteria group; Actinobacteria; Actinobacteria; Micrococcales; Microbacteriaceae; Microbacterium MARDKKRRSGGRRSPLAAAALIGAGLMITGAVYAGASAAFAATDTQTVATELTVEDGEKL WP_046012606.1 cystathionine beta-lyase [Microbacterium sp. SA39] Length: 266\nNumber of predicted TMHs: 2\nExp number of AAs in TMHs: 43.97477\nExp number, first 60 AAs: 22.54577\nTotal prob of N-in: 0.93410\nPOSSIBLE N-term signal sequence\ninside 1 19\nTMhelix 20 42\noutside 43 243\nTMhelix 244 263\ninside 264 266

31813 GCF_000967865.1_ASM96786v1 Microbacterium sp. SA39 Terrabacteria group; Actinobacteria; Actinobacteria; Micrococcales; Microbacteriaceae; Microbacterium MSVKTPETGFRARLQQLARRTAVFAIALTLACVASPGAVSAAETPPADDEQSVELYVSAG WP_046014689.1 hypothetical protein [Microbacterium sp. SA39] Length: 724\nNumber of predicted TMHs: 1\nExp number of AAs in TMHs: 50.41966\nExp number, first 60 AAs: 18.79271\nTotal prob of N-in: 0.83868\nPOSSIBLE N-term signal sequence\noutside 1 666\nTMhelix 667 686\ninside 687 724

31814 GCF_001619615.1_ASM161961v1 Microbacterium sp. T32 Terrabacteria group; Actinobacteria; Actinobacteria; Micrococcales; Microbacteriaceae; Microbacterium MAREKKPRRSNGRRSPLAAAALIGIGLLLTGGIYAGASAAMAATDTPTSASNSASTVDEG WP_022878542.1 MULTISPECIES: cystathionine beta-lyase [Microbacterium] Length: 269\nNumber of predicted TMHs: 2\nExp number of AAs in TMHs: 42.73548\nExp number, first 60 AAs: 22.46923\nTotal prob of N-in: 0.98833\nPOSSIBLE N-term signal sequence\ninside 1 20\nTMhelix 21 43\noutside 44 246\nTMhelix 247 266\ninside 267 269

31815 GCF_001620065.1_ASM162006v1 Microbacterium sp. TNHR37B Terrabacteria group; Actinobacteria; Actinobacteria; Micrococcales; Microbacteriaceae; Microbacterium MTTVTDHEHPHRSGSRRRILLLSTLTTLLLAAGLLFATAAPAAAHDEIVSSSPEAGSTVS WP_048809232.1 MULTISPECIES: Copper resistance protein C precursor [Microbacterium] Length: 238\nNumber of predicted TMHs: 2\nExp number of AAs in TMHs: 45.12191\nExp number, first 60 AAs: 22.44287\nTotal prob of N-in: 0.99979\nPOSSIBLE N-term signal sequence\ninside 1 19\nTMhelix 20 42\noutside 43 183\nTMhelix 184 206\ninside 207 238

31816 GCF_001620065.1_ASM162006v1 Microbacterium sp. TNHR37B Terrabacteria group; Actinobacteria; Actinobacteria; Micrococcales; Microbacteriaceae; Microbacterium MTALARRREEVDVKGPPVARSMMRVPARHLRDDGRRRGGRRDGGRRGAVLVTVGALVASL WP_082829233.1 hypothetical protein [Microbacterium sp. TNHR37B] Length: 2127\nNumber of predicted TMHs: 2\nExp number of AAs in TMHs: 44.51793\nExp number, first 60 AAs: 13.68431\nTotal prob of N-in: 0.97850\nPOSSIBLE N-term signal sequence\ninside 1 46\nTMhelix 47 69\noutside 70 2096\nTMhelix 2097 2119\ninside 2120 2127

31817 GCF_001620065.1_ASM162006v1 Microbacterium sp. TNHR37B Terrabacteria group; Actinobacteria; Actinobacteria; Micrococcales; Microbacteriaceae; Microbacterium MNADDTELPPLSDARVREIEESLLATIHEEGRRDSRPKAHPVRRRRRWVAGATAAAAVVV WP_067162731.1 hypothetical protein [Microbacterium sp. TNHR37B] Length: 366\nNumber of predicted TMHs: 2\nExp number of AAs in TMHs: 45.12764\nExp number, first 60 AAs: 12.77597\nTotal prob of N-in: 0.98889\nPOSSIBLE N-term signal sequence\ninside 1 47\nTMhelix 48 70\noutside 71 324\nTMhelix 325 347\ninside 348 366

31818 GCF_000509385.1_ASM50938v1 Microbacterium sp. TS-1 Terrabacteria group; Actinobacteria; Actinobacteria; Micrococcales; Microbacteriaceae; Microbacterium MLRTRRAALQRPLASVLTGALVAMAAVVGVAAPATAAGPTLTTSTTVVANEKITVAITGT WP_023953107.1 hypothetical protein [Microbacterium sp. TS-1] Length: 962\nNumber of predicted TMHs: 2\nExp number of AAs in TMHs: 41.0879599999999\nExp number, first 60 AAs: 22.33509\nTotal prob of N-in: 0.99759\nPOSSIBLE N-term signal sequence\ninside 1 12\nTMhelix 13 35\noutside 36 934\nTMhelix 935 954\ninside 955 962

31819 GCF_000340625.1_Microbacterium_sp._UCD-TDU Microbacterium sp. UCD-TDU Terrabacteria group; Actinobacteria; Actinobacteria; Micrococcales; Microbacteriaceae; Microbacterium MAREKKRRSGGRRSPLAAAALIGAGLMITGAVYAGASAAFAASDTQSAATSQLTVEDGKK WP_017830256.1 cystathionine beta-lyase [Microbacterium sp. UCD-TDU] Length: 267\nNumber of predicted TMHs: 2\nExp number of AAs in TMHs: 43.04812\nExp number, first 60 AAs: 22.514\nTotal prob of N-in: 0.98345\nPOSSIBLE N-term signal sequence\ninside 1 19\nTMhelix 20 42\noutside 43 244\nTMhelix 245 264\ninside 265 267

31820 GCF_000340625.1_Microbacterium_sp._UCD-TDU Microbacterium sp. UCD-TDU Terrabacteria group; Actinobacteria; Actinobacteria; Micrococcales; Microbacteriaceae; Microbacterium MTTRPRPEPARRRALGGGVIAAAAALTLLLPALPAAAIAPLADPASLPASEPMTPQRIEA WP_017830792.1 hypothetical protein [Microbacterium sp. UCD-TDU] Length: 1004\nNumber of predicted TMHs: 2\nExp number of AAs in TMHs: 44.194\nExp number, first 60 AAs: 22.39902\nTotal prob of N-in: 0.99318\nPOSSIBLE N-term signal sequence\ninside 1 12\nTMhelix 13 35\noutside 36 976\nTMhelix 977 999\ninside 1000 1004

31821 GCF_000340625.1_Microbacterium_sp._UCD-TDU Microbacterium sp. UCD-TDU Terrabacteria group; Actinobacteria; Actinobacteria; Micrococcales; Microbacteriaceae; Microbacterium MARRRAFIRVVLGAAAALALALGGVTAAAAEDDPVVTGSISGTVTSTVGGTALAGIEVRA WP_020380574.1 hypothetical protein [Microbacterium sp. UCD-TDU] Length: 680\nNumber of predicted TMHs: 2\nExp number of AAs in TMHs: 61.6964300000003\nExp number, first 60 AAs: 31.71806\nTotal prob of N-in: 0.99602\nPOSSIBLE N-term signal sequence\ninside 1 6\nTMhelix 7 29\noutside 30 650\nTMhelix 651 673\ninside 674 680

31822 GCF_000340625.1_Microbacterium_sp._UCD-TDU Microbacterium sp. UCD-TDU Terrabacteria group; Actinobacteria; Actinobacteria; Micrococcales; Microbacteriaceae; Microbacterium MRRRAALATLVLGAAVFGAPASALASTSAPASIVSAATGACEPGNVVFTPEAPAALAALG WP_051056635.1 hypothetical protein [Microbacterium sp. UCD-TDU] Length: 436\nNumber of predicted TMHs: 2\nExp number of AAs in TMHs: 40.44281\nExp number, first 60 AAs: 19.34712\nTotal prob of N-in: 0.88927\nPOSSIBLE N-term signal sequence\ninside 1 6\nTMhelix 7 29\noutside 30 385\nTMhelix 386 408\ninside 409 436

31823 GCF_000620445.1_ASM62044v1 Microbacterium sp. UNC423CL45Tsu Terrabacteria group; Actinobacteria; Actinobacteria; Micrococcales; Microbacteriaceae; Microbacterium MPSPRRSLITTLATLLLGAGLTLAVATPAQAHDELVSSYPAADSTVSDSPAEITLSFSGE WP_081782667.1 MULTISPECIES: hypothetical protein [Microbacterium] Length: 225\nNumber of predicted TMHs: 2\nExp number of AAs in TMHs: 44.44474\nExp number, first 60 AAs: 21.72587\nTotal prob of N-in: 0.97817\nPOSSIBLE N-term signal sequence\ninside 1 6\nTMhelix 7 29\noutside 30 173\nTMhelix 174 196\ninside 197 225

31824 GCF_000746195.1_ASM74619v1 Microbacterium sp. UNCCL10Terrabacteria group; Actinobacteria; Actinobacteria; Micrococcales; Microbacteriaceae; Microbacterium MPSPRRSLITTLATLLLGAGLTLAVATPAQAHDELVSSYPAADSTVSDSPAEITLSFSGE WP_081782667.1 MULTISPECIES: hypothetical protein [Microbacterium] Length: 225\nNumber of predicted TMHs: 2\nExp number of AAs in TMHs: 44.44474\nExp number, first 60 AAs: 21.72587\nTotal prob of N-in: 0.97817\nPOSSIBLE N-term signal sequence\ninside 1 6\nTMhelix 7 29\noutside 30 173\nTMhelix 174 196\ninside 197 225

31825 GCF_000425985.1_ASM42598v1 Microbacterium sp. URHA0036 Terrabacteria group; Actinobacteria; Actinobacteria; Micrococcales; Microbacteriaceae; Microbacterium MAREKKRRSSGRRSPLAAAALIGAGLMITGAVYAGASAAFAASDTQSATTQLTVEDGEKL WP_028501544.1 cystathionine beta-lyase [Microbacterium sp. URHA0036] Length: 266\nNumber of predicted TMHs: 2\nExp number of AAs in TMHs: 43.0957\nExp number, first 60 AAs: 22.57321\nTotal prob of N-in: 0.98600\nPOSSIBLE N-term signal sequence\ninside 1 19\nTMhelix 20 42\noutside 43 243\nTMhelix 244 263\ninside 264 266

31826 GCF_001513675.1_ASM151367v1 Microbacterium sp. XT11 Terrabacteria group; Actinobacteria; Actinobacteria; Micrococcales; Microbacteriaceae; Microbacterium MAREKKRRSHGRRSPLAAAALIGAGLMITGAVYAGASAAFAATDTQASASSTLTVEDGEK WP_067200502.1 cystathionine beta-lyase [Microbacterium sp. XT11] Length: 267\nNumber of predicted TMHs: 2\nExp number of AAs in TMHs: 43.08953\nExp number, first 60 AAs: 22.61395\nTotal prob of N-in: 0.98501\nPOSSIBLE N-term signal sequence\ninside 1 20\nTMhelix 21 43\noutside 44 244\nTMhelix 245 264\ninside 265 267

31827 GCF_001513675.1_ASM151367v1 Microbacterium sp. XT11 Terrabacteria group; Actinobacteria; Actinobacteria; Micrococcales; Microbacteriaceae; Microbacterium MTKLEGQTLRRRRAIGGGIATATAGAVILASALVPTAASALEGADPSDPSAASGRILNIG WP_067196734.1 hypothetical protein [Microbacterium sp. XT11] Length: 1242\nNumber of predicted TMHs: 2\nExp number of AAs in TMHs:

43.6094999999999\nExp number, first 60 AAs: 21.45777\nTotal prob of N-in: 0.95554\nPOSSIBLE N-term signal sequence\ninside 1 12\nTMhelix 13 35\noutside 36 1209\nTMhelix 1210 1232\ninside 1233 1242

31828 GCF_000800925.1_ZOR0019.1 Microbacterium sp. ZOR0019 Terrabacteria group; Actinobacteria; Actinobacteria; Micrococcales; Microbacteriaceae; Microbacterium MAREKKRRSSGRRSPLAAAALIGAGLMITGAVYAGATAAFAASDTQAATTQLTVEDGEKL WP_047524047.1 cystathionine beta-lyase [Microbacterium sp. ZOR0019] Length: 266\nNumber of predicted TMHs: 2\nExp number of AAs in TMHs: 43.08837\nExp number, first 60 AAs: 22.64303\nTotal prob of N-in: 0.98567\nPOSSIBLE N-term signal sequence\ninside 1 19\nTMhelix 20 42\noutside 43 243\nTMhelix 244 263\ninside 264 266

31829 GCF_000800925.1_ZOR0019.1 Microbacterium sp. ZOR0019 Terrabacteria group; Actinobacteria; Actinobacteria; Micrococcales; Microbacteriaceae; Microbacterium MNTHPGRTRARRTTITAIAATALTAVMTFVGAGAATAAEYDYLAAIDAASITVTTTNGGS WP_047519257.1 hypothetical protein [Microbacterium sp. ZOR0019] Length: 523\nNumber of predicted TMHs: 2\nExp number of AAs in TMHs: 47.80756\nExp number, first 60 AAs: 23.14771\nTotal prob of N-in: 0.94933\nPOSSIBLE N-term signal sequence\ninside 1 12\nTMhelix 13 35\noutside 36 480\nTMhelix 481 503\ninside 504 523

31830 GCF_001476285.1_ASM147628v1 Microbacterium testaceum Terrabacteria group; Actinobacteria; Actinobacteria; Micrococcales; Microbacteriaceae; Microbacterium MARTPTALLSPADQERRRALRRMKGVALGALIGMAVIFAVAFAFQRDVTWLQYVRAAAEG WP_058595107.1 DUF445 domain-containing protein [Microbacterium testaceum] Length: 423\nNumber of predicted TMHs: 2\nExp number of AAs in TMHs: 52.81835\nExp number, first 60 AAs: 21.83984\nTotal prob of N-in: 0.92687\nPOSSIBLE N-term signal sequence\ninside 1 24\nTMhelix 25 44\noutside 45 399\nTMhelix 400 422\ninside 423 423

31831 GCF_001476655.1_ASM147665v1 Microbacterium testaceum Terrabacteria group; Actinobacteria; Actinobacteria; Micrococcales; Microbacteriaceae; Microbacterium MSRIGRASRSRARRGPVGLAAPRVAATGVALALLVLAGSTGASASTPSPEDSGTLSVTIT WP_058622210.1 hypothetical protein [Microbacterium testaceum] Length: 275\nNumber of predicted TMHs: 1\nExp number of AAs in TMHs: 40.86125\nExp number, first 60 AAs: 18.27352\nTotal prob of N-in: 0.84916\nPOSSIBLE N-term signal sequence\noutside 1 234\nTMhelix 235 257\ninside 258 275

31832 GCF_001476655.1_ASM147665v1 Microbacterium testaceum Terrabacteria group; Actinobacteria; Actinobacteria; Micrococcales; Microbacteriaceae; Microbacterium MAREKKPRRASGRRSPLAAAALIGIGLLLTGGVYAGASAAMAATDTPTSNASSATAVEEG WP_058622312.1 cystathionine beta-lyase [Microbacterium testaceum] Length: 269\nNumber of predicted TMHs: 2\nExp number of AAs in TMHs: 42.86904\nExp number, first 60 AAs: 22.45657\nTotal prob of N-in: 0.98783\nPOSSIBLE N-term signal sequence\ninside 1 20\nTMhelix 21 43\noutside 44 246\nTMhelix 247 266\ninside 267 269

31833 GCF_001476285.1_ASM147628v1 Microbacterium testaceum Terrabacteria group; Actinobacteria; Actinobacteria; Micrococcales; Microbacteriaceae; Microbacterium MAREKKPRRASGRRSPLAAAALIGIGLLLTGGVYAGASAAMAATDTPTSSASSATAVEEG WP_058595114.1 cystathionine beta-lyase [Microbacterium testaceum] Length: 269\nNumber of predicted TMHs: 2\nExp number of AAs in TMHs: 42.85479\nExp number, first 60 AAs: 22.45656\nTotal prob of N-in: 0.98860\nPOSSIBLE N-term signal sequence\ninside 1 20\nTMhelix 21 43\noutside 44 246\nTMhelix 247 266\ninside 267 269

31834 GCF_001476275.1_ASM147627v1 Microbacterium testaceum Terrabacteria group; Actinobacteria; Actinobacteria; Micrococcales; Microbacteriaceae; Microbacterium MAREKKPRRASGRRSPLAAAALIGIGLLLTGGVYAGASAAMAATDTPTSSASSATAVEEG WP_058595114.1 cystathionine beta-lyase [Microbacterium testaceum] Length: 269\nNumber of predicted TMHs: 2\nExp number of AAs in TMHs: 42.85479\nExp number, first 60 AAs: 22.45656\nTotal prob of N-in: 0.98860\nPOSSIBLE N-term signal sequence\ninside 1 20\nTMhelix 21 43\noutside 44 246\nTMhelix 247 266\ninside 267 269

31835 GCF_001476665.1_ASM147666v1 Microbacterium testaceum Terrabacteria group; Actinobacteria; Actinobacteria; Micrococcales; Microbacteriaceae; Microbacterium MARTPTALLSPADQERRRALRRMKGVALGALIGMAVIFAVAFAFQRDVTWLQYVRAAAEG WP_058613477.1 DUF445 domain-containing protein [Microbacterium testaceum] Length: 423\nNumber of predicted TMHs: 2\nExp number of AAs in TMHs: 52.82263\nExp number, first 60 AAs: 21.83919\nTotal prob of N-in: 0.92688\nPOSSIBLE N-term signal sequence\ninside 1 24\nTMhelix 25 44\noutside 45 399\nTMhelix 400 422\ninside 423 423

31836 GCF_001477115.1_ASM147711v1 Microbacterium testaceum Terrabacteria group; Actinobacteria; Actinobacteria; Micrococcales; Microbacteriaceae; Microbacterium

MARTPTALLSPADQERRRALRRMKGVALGALIGMAVIFAVAFAFQRDVTWLQYVRAAAEG WP_058595107.1 DUF445 domain-containing protein [Microbacterium testaceum] Length: 423\nNumber of predicted TMHs: 2\nExp number of AAs in TMHs: 52.81835\nExp number, first 60 AAs: 21.83984\nTotal prob of N-in: 0.92687\nPOSSIBLE N-term signal sequence\ninside 1 24\nTMhelix 25 44\noutside 45 399\nTMhelix 400 422\ninside 423 423

31837 GCF_001477115.1_ASM147711v1 Microbacterium testaceum Terrabacteria group; Actinobacteria; Actinobacteria; Micrococcales; Microbacteriaceae; Microbacterium MAREKKPRRASGRRSPLAAAALIGIGLLLTGGVYAGASAAMAATDTPTSSASSATAVEEG WP_058595114.1 cystathionine beta-lyase [Microbacterium testaceum] Length: 269\nNumber of predicted TMHs: 2\nExp number of AAs in TMHs: 42.85479\nExp number, first 60 AAs: 22.45656\nTotal prob of N-in: 0.98860\nPOSSIBLE N-term signal sequence\ninside 1 20\nTMhelix 21 43\noutside 44 246\nTMhelix 247 266\ninside 267 269

31838 GCF_001476655.1_ASM147665v1 Microbacterium testaceum Terrabacteria group; Actinobacteria; Actinobacteria; Micrococcales; Microbacteriaceae; Microbacterium MARTPTAQLSPADQERRRALARMKGVALGALIGMAVIFAVAFAFQRDVTWLQYVRAAAEG WP_058623401.1 DUF445 domain-containing protein [Microbacterium testaceum] Length: 423\nNumber of predicted TMHs: 2\nExp number of AAs in TMHs: 53.07996\nExp number, first 60 AAs: 22.13391\nTotal prob of N-in: 0.93413\nPOSSIBLE N-term signal sequence\ninside 1 24\nTMhelix 25 44\noutside 45 399\nTMhelix 400 422\ninside 423 423

31839 GCF_001476665.1_ASM147666v1 Microbacterium testaceum Terrabacteria group; Actinobacteria; Actinobacteria; Micrococcales; Microbacteriaceae; Microbacterium MAREKKPRRASGRRSPLAAAALIGIGLLLTGGVYAGASAAMAATDTPTSSASSATAVEEG WP_058595114.1 cystathionine beta-lyase [Microbacterium testaceum] Length: 269\nNumber of predicted TMHs: 2\nExp number of AAs in TMHs: 42.85479\nExp number, first 60 AAs: 22.45656\nTotal prob of N-in: 0.98860\nPOSSIBLE N-term signal sequence\ninside 1 20\nTMhelix 21 43\noutside 44 246\nTMhelix 247 266\ninside 267 269

31840 GCF_001476275.1_ASM147627v1 Microbacterium testaceum Terrabacteria group; Actinobacteria; Actinobacteria; Micrococcales; Microbacteriaceae; Microbacterium MARTPTALLSPADQERRRALRRMKGVALGSLIGMAVVFAVAFAFQRDVTWLQYVRAAAEG WP_058629783.1 DUF445 domain-containing protein [Microbacterium testaceum] Length: 423\nNumber of predicted TMHs: 2\nExp number of AAs in TMHs: 52.79642\nExp number, first 60 AAs: 21.75557\nTotal prob of N-in: 0.93661\nPOSSIBLE N-term signal sequence\ninside 1 24\nTMhelix 25 44\noutside 45 399\nTMhelix 400 422\ninside 423 423

31841 GCF_000299315.2_ASM29931v2 Microbacterium barkeri 2011-R4 Terrabacteria group; Actinobacteria; Actinobacteria; Micrococcales; Microbacteriaceae; Microbacterium; Microbacterium barkeri MNSTPSPRDVRRTHRVRSSSRQGRRRRTTVTQVVLLTSALLFTVTTATMLPPQAQAEPLP WP_017204160.1 hypothetical protein [Microbacterium barkeri] Length: 215\nNumber of predicted TMHs: 2\nExp number of AAs in TMHs: 41.26402\nExp number, first 60 AAs: 20.41219\nTotal prob of N-in: 0.97530\nPOSSIBLE N-term signal sequence\ninside 1 29\nTMhelix 30 52\noutside 53 187\nTMhelix 188 210\ninside 211 215

31842 GCF_000299315.2_ASM29931v2 Microbacterium barkeri 2011-R4 Terrabacteria group; Actinobacteria; Actinobacteria; Micrococcales; Microbacteriaceae; Microbacterium; Microbacterium barkeri MAREKKHRSRGRRSPLAAAALIGAGLLITGAVYAGASAAFAASDTQSASTQLTVEDGEKL WP_026049807.1 cystathionine beta-lyase [Microbacterium barkeri] Length: 266\nNumber of predicted TMHs: 2\nExp number of AAs in TMHs: 43.13969\nExp number, first 60 AAs: 22.58772\nTotal prob of N-in: 0.98833\nPOSSIBLE N-term signal sequence\ninside 1 19\nTMhelix 20 42\noutside 43 243\nTMhelix 244 263\ninside 264 266

31843 GCF_000299315.2_ASM29931v2 Microbacterium barkeri 2011-R4 Terrabacteria group; Actinobacteria; Actinobacteria; Micrococcales; Microbacteriaceae; Microbacterium; Microbacterium barkeri MSRNTSTRTRRTAIASAGILGGAALVLAVPAMAGAHVGVSPDELVAGDHGVLTFSFSHGC WP_017203113.1 hypothetical protein [Microbacterium barkeri] Length: 224\nNumber of predicted TMHs: 2\nExp number of AAs in TMHs: 45.43315\nExp number, first 60 AAs: 22.59807\nTotal prob of N-in: 0.99990\nPOSSIBLE N-term signal sequence\ninside 1 12\nTMhelix 13 35\noutside 36 195\nTMhelix 196 218\ninside 219 224

31844 GCF_000422745.1_ASM42274v1 Microbacterium gubbeenense DSM 15944 Terrabacteria group; Actinobacteria; Actinobacteria; Micrococcales; Microbacteriaceae; Microbacterium; Microbacterium gubbeenense MARRQWIQKAAAAVAAAALAVMAAPAAAAAEPYSFTTDVTEVADQRIDIAIDGTGFGDVR WP_029153468.1 hypothetical protein [Microbacterium gubbeenense] Length: 1051\nNumber of predicted TMHs: 2\nExp number of AAs in TMHs: 39.12765\nExp number, first 60 AAs: 18.04687\nTotal prob of N-in: 0.89175\nPOSSIBLE N-term signal sequence\ninside 1 6\nTMhelix 7 29\noutside 30 1018\nTMhelix 1019 1041\ninside 1042 1051

31845 GCF_000422745.1_ASM42274v1 Microbacterium gubbeenense DSM 15944 Terrabacteria group; Actinobacteria; Actinobacteria; Micrococcales; Microbacteriaceae; Microbacterium; Microbacterium gubbeenense MAREKLKRRARGRRSPLAAGILIGVGLLMTGGIYAGASAAVAATDEATPITASVDEGEKL WP_029152180.1 cystathionine beta-lyase [Microbacterium gubbeenense] Length: 266\nNumber of predicted TMHs: 2\nExp number of AAs in TMHs: 43.76456\nExp number, first 60 AAs: 22.69181\nTotal prob of N-in: 0.99743\nPOSSIBLE N-term signal sequence\ninside 1 20\nTMhelix 21 43\noutside 44 243\nTMhelix 244 263\ninside 264 266

31846 GCF_000422745.1_ASM42274v1 Microbacterium gubbeenense DSM 15944 Terrabacteria group; Actinobacteria; Actinobacteria; Micrococcales; Microbacteriaceae; Microbacterium; Microbacterium gubbeenense MMSEIERDLPELSDESVLRIERAVMDDIGRERQTARTRAKKRRGWVGAGLAAAAVVAGAI WP_029154188.1 hypothetical protein [Microbacterium gubbeenense] Length: 327\nNumber of predicted TMHs: 2\nExp number of AAs in TMHs: 50.56976\nExp number, first 60 AAs: 16.73724\nTotal prob of N-in: 0.98451\nPOSSIBLE N-term signal sequence\ninside 1 43\nTMhelix 44 66\noutside 67 289\nTMhelix 290 312\ninside 313 327

31847 GCF_001592125.1_ASM159212v1 Microbacterium hominis NBRC 15708 Terrabacteria group; Actinobacteria; Actinobacteria; Micrococcales; Microbacteriaceae; Microbacterium; Microbacterium hominis MLRPRRRTLLASTASGFVTAALVFTLVAGANVALGSPAQAATPASSAVTVTAAEQDRDLT WP_061782355.1 hypothetical protein [Microbacterium hominis] Length: 840\nNumber of predicted TMHs: 2\nExp number of AAs in TMHs: 43.6755899999999\nExp number, first 60 AAs: 21.13056\nTotal prob of N-in: 0.94339\nPOSSIBLE N-term signal sequence\ninside 1 12\nTMhelix 13 35\noutside 36 809\nTMhelix 810 832\ninside 833 840

31848 GCF_001592125.1_ASM159212v1 Microbacterium hominis NBRC 15708 Terrabacteria group; Actinobacteria; Actinobacteria; Micrococcales; Microbacteriaceae; Microbacterium; Microbacterium hominis MAPTPTQLLSPADQERRRALRVMKGVALGALVLMAVVFVVAFVLQERYPWLEYVRAAAEG WP_060959398.1 DUF445 domain-containing protein [Microbacterium hominis] Length: 423\nNumber of predicted TMHs: 2\nExp number of AAs in TMHs: 52.34446\nExp number, first 60 AAs: 22.52241\nTotal prob of N-in: 0.78491\nPOSSIBLE N-term signal sequence\ninside 1 24\nTMhelix 25 44\noutside 45 399\nTMhelix 400 422\ninside 423 423

31849 GCF_001592125.1_ASM159212v1 Microbacterium hominis NBRC 15708 Terrabacteria group; Actinobacteria; Actinobacteria; Micrococcales; Microbacteriaceae; Microbacterium; Microbacterium hominis MGTTNTAAAPARRRRALPAFVVAGLLGLIAAGSSASGAAASSPLIDATPATVTIEAPAPG WP_061782010.1 hypothetical protein [Microbacterium hominis] Length: 216\nNumber of predicted TMHs: 2\nExp number of AAs in TMHs: 44.64644\nExp number, first 60 AAs: 21.9152\nTotal prob of N-in: 0.99385\nPOSSIBLE N-term signal sequence\ninside 1 16\nTMhelix 17 39\noutside 40 177\nTMhelix 178 200\ninside 201 216

31850 GCF_001592125.1_ASM159212v1 Microbacterium hominis NBRC 15708 Terrabacteria group; Actinobacteria; Actinobacteria; Micrococcales; Microbacteriaceae; Microbacterium; Microbacterium hominis MSDQTPDAPRPTADRRKRRSVVAAITAGALAFSGLGLAALPARAAVDTGAAVLINEVYGG WP_061781777.1 cell wall protein [Microbacterium hominis] Length: 1578\nNumber of predicted TMHs: 2\nExp number of AAs in TMHs: 42.4667499999999\nExp number, first 60 AAs: 21.7961\nTotal prob of N-in: 0.99821\nPOSSIBLE N-term signal sequence\ninside 1 19\nTMhelix 20 42\noutside 43 1548\nTMhelix 1549 1571\ninside 1572 1578

31851 GCF_001552455.1_ASM155245v1 Microbacterium hydrocarbonoxydans NBRC 103074 Terrabacteria group; Actinobacteria; Actinobacteria; Micrococcales; Microbacteriaceae; Microbacterium; Microbacterium hydrocarbonoxydans MIGRAAEAEQHSDDVARGLPSEGGTMARRRAFIRAIVAGAAALALAIGGATTAMAADDDP WP_060928567.1 hypothetical protein [Microbacterium hydrocarbonoxydans] Length: 693\nNumber of predicted TMHs: 2\nExp number of AAs in TMHs: 43.6390999999999\nExp number, first 60 AAs: 21.83173\nTotal prob of N-in: 0.99014\nPOSSIBLE N-term signal sequence\ninside 1 30\nTMhelix 31 53\noutside 54 666\nTMhelix 667 686\ninside 687 693

31852 GCF_001552455.1_ASM155245v1 Microbacterium hydrocarbonoxydans NBRC 103074 Terrabacteria group; Actinobacteria; Actinobacteria; Micrococcales; Microbacteriaceae; Microbacterium; Microbacterium hydrocarbonoxydans MAREKKRRSNGRRSPLAAAALIGAGLMITGAVYAGTSAAFAASDTETTAASTLTVEDGEK WP_060926486.1 cystathionine beta-lyase [Microbacterium hydrocarbonoxydans] Length: 267\nNumber of predicted TMHs: 2\nExp number of AAs in TMHs: 42.80265\nExp number, first 60 AAs: 22.42264\nTotal prob of N-in: 0.99124\nPOSSIBLE N-term signal sequence\ninside 1 19\nTMhelix 20 42\noutside 43 244\nTMhelix 245 264\ninside 265 267

31853 GCF_000422385.1_ASM42238v1 Microbacterium indicum DSM 19969 Terrabacteria group; Actinobacteria; Actinobacteria; Micrococcales; Microbacteriaceae; Microbacterium; Microbacterium indicum MTKKRKAGGRRSPLAAAALILIGLVATGGVYAGATAAVAATNDDSAVSTSHASAEDGEKL WP_084502677.1

cystathionine beta-lyase [Microbacterium indicum] Length: 266\nNumber of predicted TMHs: 2\nExp number of AAs in TMHs: 42.9940099999999\nExp number, first 60 AAs: 22.62661\nTotal prob of N-in: 0.98724\nPOSSIBLE N-term signal sequence\ninside 1 12\nTMhelix 13 35\noutside 36 243\nTMhelix 244 263\ninside 264 266

31854 GCF_000255595.1_OR221_CLC Microbacterium laevaniformans OR221Terrabacteria group; Actinobacteria; Actinobacteria; Micrococcales; Microbacteriaceae; Microbacterium; Microbacterium laevaniformans MNRRRTLASLLLVALLSVGSALAAPASAEEGAMSWSVQPSTPSGPDGRSDLSYQVGPGAV WP_005054386.1 hypothetical protein [Microbacterium laevaniformans] Length: 370\nNumber of predicted TMHs: 2\nExp number of AAs in TMHs: 50.19953\nExp number, first 60 AAs: 19.48038\nTotal prob of N-in: 0.97033\nPOSSIBLE N-term signal sequence\ninside 1 6\nTMhelix 7 26\noutside 27 312\nTMhelix 313 335\ninside 336 370

31855 GCF_000255595.1_OR221_CLC Microbacterium laevaniformans OR221Terrabacteria group; Actinobacteria; Actinobacteria; Micrococcales; Microbacteriaceae; Microbacterium; Microbacterium laevaniformans MRPTAERRKRRSFVAAITAGALALTGAGLAALPARAAVSADAAVLINEVYGGGGNSKATY WP_043340214.1 cell wall protein [Microbacterium laevaniformans] Length: 1578\nNumber of predicted TMHs: 2\nExp number of AAs in TMHs: 42.09309\nExp number, first 60 AAs: 21.69911\nTotal prob of N-in: 0.99858\nPOSSIBLE N-term signal sequence\ninside 1 11\nTMhelix 12 34\noutside 35 1548\nTMhelix 1549 1568\ninside 1569 1578

31856 GCF_000422405.1_ASM42240v1 Microbacterium luticocti DSM 19459 Terrabacteria group; Actinobacteria; Actinobacteria; Micrococcales; Microbacteriaceae; Microbacterium; Microbacterium luticocti MAREKKRRASGRRSPLAAVVLIGVGLLMTGGIYAGASAAMAATGDTSTSTALTVEDGKKL WP_029145108.1 cystathionine beta-lyase [Microbacterium luticocti] Length: 266\nNumber of predicted TMHs: 2\nExp number of AAs in TMHs: 43.48118\nExp number, first 60 AAs: 22.52528\nTotal prob of N-in: 0.99051\nPOSSIBLE N-term signal sequence\ninside 1 19\nTMhelix 20 42\noutside 43 243\nTMhelix 244 263\ninside 264 266

31857 GCF_000455825.1_ASM45582v1 Microbacterium maritypicum MF109 Terrabacteria group; Actinobacteria; Actinobacteria; Micrococcales; Microbacteriaceae; Microbacterium; Microbacterium maritypicum MAREKKRRSSGRRSPLAAAALIGAGLMITGAVYAGATAAFAATDTQSAATSQLTVEDGKK WP_029258331.1 MULTISPECIES: cystathionine beta-lyase [Microbacterium] Length: 267\nNumber of predicted TMHs: 2\nExp number of AAs in TMHs: 43.18585\nExp number, first 60 AAs: 22.66682\nTotal prob of N-in: 0.98705\nPOSSIBLE N-term signal sequence\ninside 1 20\nTMhelix 21 43\noutside 44 244\nTMhelix 245 264\ninside 265 267

31858 GCF_000455825.1_ASM45582v1 Microbacterium maritypicum MF109 Terrabacteria group; Actinobacteria; Actinobacteria; Micrococcales; Microbacteriaceae; Microbacterium; Microbacterium maritypicum MNRDCTRRWDHRANGLPRKVRRMTRRRGMLRAVLGGMAAIAIALGGATAVVAAEDPAPTG WP_084487221.1 hypothetical protein [Microbacterium maritypicum] Length: 697\nNumber of predicted TMHs: 2\nExp number of AAs in TMHs: 44.46894\nExp number, first 60 AAs: 22.81605\nTotal prob of N-in: 0.99997\nPOSSIBLE N-term signal sequence\ninside 1 29\nTMhelix 30 52\noutside 53 667\nTMhelix 668 690\ninside 691 697

31859 GCF_000455825.1_ASM45582v1 Microbacterium maritypicum MF109 Terrabacteria group; Actinobacteria; Actinobacteria; Micrococcales; Microbacteriaceae; Microbacterium; Microbacterium maritypicum MELQERTLRRRRAVGGGIAGVTVGAVILASALVPTAANALDAANPNDPSVAQGQIIQLPA WP_021198471.1 hypothetical protein [Microbacterium maritypicum] Length: 1733\nNumber of predicted TMHs: 2\nExp number of AAs in TMHs: 43.71797\nExp number, first 60 AAs: 22.32964\nTotal prob of N-in: 0.99668\nPOSSIBLE N-term signal sequence\ninside 1 12\nTMhelix 13 35\noutside 36 1700\nTMhelix 1701 1723\ninside 1724 1733

31860 GCF_001552475.1_ASM155247v1 Microbacterium oleivorans NBRC 103075 Terrabacteria group; Actinobacteria; Actinobacteria; Micrococcales; Microbacteriaceae; Microbacterium; Microbacterium oleivorans MTTRSARRSALTLSGLALGGALVLAAPLAASAHVHVHSDDAAAGATSRLEFSFSHGCDGA WP_060915309.1 nuclear export factor GLE1 [Microbacterium oleivorans] Length: 232\nNumber of predicted TMHs: 2\nExp number of AAs in TMHs: 44.1002899999999\nExp number, first 60 AAs: 21.6335\nTotal prob of N-in: 0.99061\nPOSSIBLE N-term signal sequence\ninside 1 12\nTMhelix 13 35\noutside 36 204\nTMhelix 205 227\ninside 228 232

31861 GCF_000380465.1_ASM38046v1 Microbacterium paraoxydans 77MFTsu3.2 Terrabacteria group; Actinobacteria; Actinobacteria; Micrococcales; Microbacteriaceae; Microbacterium; Microbacterium paraoxydans MTLRPARRSALTLTGLALGGALVLAAPLAASAHVHVHSDDASAGATSRLEFSFSHGCDGA WP_018188612.1 hypothetical protein [Microbacterium paraoxydans] Length: 228\nNumber of predicted TMHs: 2\nExp number of AAs in TMHs: 43.89919\nExp number, first 60 AAs: 21.28469\nTotal prob of N-in: 0.96719\nPOSSIBLE N-term signal sequence\ninside 1 12\nTMhelix 13 35\noutside 36 201\nTMhelix 202 224\ninside 225 228

31862 GCF_000633215.1_de_novo Microbacterium paraoxydans DH1b Terrabacteria group; Actinobacteria; Actinobacteria; Micrococcales; Microbacteriaceae; Microbacterium; Microbacterium paraoxydans
MRRRAALATLVVGAAVWGTPAAVVAAPTAPTTIVSMATGACEPGTVVFAPEAPSALAALG WP_025104647.1
MULTISPECIES: hypothetical protein [Microbacterium] Length: 436\nNumber of predicted TMHs: 2\nExp number of AAs in TMHs: 43.5666099999999\nExp number, first 60 AAs: 21.6829\nTotal prob of N-in: 0.96731\nPOSSIBLE N-term signal sequence\ninside 1 6\nTMhelix 7 29\noutside 30 386\nTMhelix 387 409\ninside 410 436

31863 GCF_000633215.1_de_novo Microbacterium paraoxydans DH1b Terrabacteria group; Actinobacteria; Actinobacteria; Micrococcales; Microbacteriaceae; Microbacterium; Microbacterium paraoxydans
MSNTTIRPRRPRRTLLGATGVVGGLALALAVPAMAGAHVSVSPDELAAGDHGVLTFSFSH WP_025103163.1
MULTISPECIES: hypothetical protein [Microbacterium] Length: 224\nNumber of predicted TMHs: 2\nExp number of AAs in TMHs: 45.16071\nExp number, first 60 AAs: 22.31743\nTotal prob of N-in: 0.99849\nPOSSIBLE N-term signal sequence\ninside 1 12\nTMhelix 13 35\noutside 36 197\nTMhelix 198 220\ninside 221 224

31864 GCF_000633215.1_de_novo Microbacterium paraoxydans DH1b Terrabacteria group; Actinobacteria; Actinobacteria; Micrococcales; Microbacteriaceae; Microbacterium; Microbacterium paraoxydans
MTRRRMLRGAVALVAVAASVLLLGSTATPTPTPPPVADDPADPVRAAEYWLDGARIRDAW WP_036292654.1
MULTISPECIES: peptidase S8 [Microbacterium] Length: 439\nNumber of predicted TMHs: 2\nExp number of AAs in TMHs: 41.4776299999999\nExp number, first 60 AAs: 19.80316\nTotal prob of N-in: 0.93229\nPOSSIBLE N-term signal sequence\ninside 1 6\nTMhelix 7 29\noutside 30 397\nTMhelix 398 420\ninside 421 439

31865 GCF_000633215.1_de_novo Microbacterium paraoxydans DH1b Terrabacteria group; Actinobacteria; Actinobacteria; Micrococcales; Microbacteriaceae; Microbacterium; Microbacterium paraoxydans
MAREKKRRSSGRRSPLAAAALIGAGLMITGAVYAGATAAFAANDAPTASTQLTVEDGEKL WP_025105058.1
MULTISPECIES: cystathionine beta-lyase [Microbacterium] Length: 266\nNumber of predicted TMHs: 2\nExp number of AAs in TMHs: 43.19361\nExp number, first 60 AAs: 22.60029\nTotal prob of N-in: 0.99024\nPOSSIBLE N-term signal sequence\ninside 1 19\nTMhelix 20 42\noutside 43 243\nTMhelix 244 263\ninside 264 266

31866 GCF_001552495.1_ASM155249v1 Microbacterium paraoxydans NBRC 103076 Terrabacteria group; Actinobacteria; Actinobacteria; Micrococcales; Microbacteriaceae; Microbacterium; Microbacterium paraoxydans
MTRRRMLRGAVALVAVAASVLLLGSTATPTPTPPPVADDPADPVRAAEYWLDGAGIRDAW WP_060921984.1
peptidase S8 [Microbacterium paraoxydans] Length: 439\nNumber of predicted TMHs: 2\nExp number of AAs in TMHs: 41.47033\nExp number, first 60 AAs: 19.80703\nTotal prob of N-in: 0.93228\nPOSSIBLE N-term signal sequence\ninside 1 6\nTMhelix 7 29\noutside 30 397\nTMhelix 398 420\ninside 421 439

31867 GCF_001552495.1_ASM155249v1 Microbacterium paraoxydans NBRC 103076 Terrabacteria group; Actinobacteria; Actinobacteria; Micrococcales; Microbacteriaceae; Microbacterium; Microbacterium paraoxydans
MSNTTIRPRRPRRTLLGATGIVGGLALALAVPAMAGAHVSVSPDELAAGDHGVLTFSFSH WP_060922111.1
hypothetical protein [Microbacterium paraoxydans] Length: 224\nNumber of predicted TMHs: 2\nExp number of AAs in TMHs: 45.2104\nExp number, first 60 AAs: 22.3675\nTotal prob of N-in: 0.99874\nPOSSIBLE N-term signal sequence\ninside 1 12\nTMhelix 13 35\noutside 36 197\nTMhelix 198 220\ninside 221 224

31868 GCF_001552495.1_ASM155249v1 Microbacterium paraoxydans NBRC 103076 Terrabacteria group; Actinobacteria; Actinobacteria; Micrococcales; Microbacteriaceae; Microbacterium; Microbacterium paraoxydans
MRRRAALATLVVGAAVWGTPAAAVAAPTAPTTIVSMATGACEPGTVVFAPEAPSALAALG WP_060923224.1
hypothetical protein [Microbacterium paraoxydans] Length: 436\nNumber of predicted TMHs: 2\nExp number of AAs in TMHs: 43.05017\nExp number, first 60 AAs: 21.16204\nTotal prob of N-in: 0.94802\nPOSSIBLE N-term signal sequence\ninside 1 6\nTMhelix 7 29\noutside 30 386\nTMhelix 387 409\ninside 410 436

31869 GCF_001552495.1_ASM155249v1 Microbacterium paraoxydans NBRC 103076 Terrabacteria group; Actinobacteria; Actinobacteria; Micrococcales; Microbacteriaceae; Microbacterium; Microbacterium paraoxydans
MRRSSTFGEGIAAGSAALEVDRDSIGLRRPPPEGGDMARRRAFFRSIMGGAAALALALGG WP_082750119.1
hypothetical protein [Microbacterium paraoxydans] Length: 706\nNumber of predicted TMHs: 2\nExp number of AAs in TMHs: 39.42056\nExp number, first 60 AAs: 15.85922\nTotal prob of N-in: 0.96446\nPOSSIBLE N-term signal sequence\ninside 1 42\nTMhelix 43 65\noutside 66 679\nTMhelix 680 699\ninside 700 706

31870 GCF_001552495.1_ASM155249v1 Microbacterium paraoxydans NBRC 103076 Terrabacteria group; Actinobacteria; Actinobacteria; Micrococcales; Microbacteriaceae; Microbacterium; Microbacterium paraoxydans
MAREKKRRSSGRRSPLAAAALIGAGLMITGAVYAGATAAFAANDAPTASTQLTVEDGEKL WP_025105058.1
MULTISPECIES: cystathionine beta-lyase [Microbacterium] Length: 266\nNumber of predicted TMHs: 2\nExp

number of AAs in TMHs: 43.19361\nExp number, first 60 AAs: 22.60029\nTotal prob of N-in: 0.99024\nPOSSIBLE N-term signal sequence\ninside 1 19\nTMhelix 20 42\noutside 43 243\nTMhelix 244 263\ninside 264 266

31871 GCF_001552355.1_ASM155235v1 Microbacterium resistens NBRC 103078 Terrabacteria group; Actinobacteria; Actinobacteria; Micrococcales; Microbacteriaceae; Microbacterium; Microbacterium resistens MAREKKRRSGGRRSPLAAAALIGAGLMITGGIYAGASAAFAASEPTAATAQVSVEEGEKL WP_067244188.1 cystathionine beta-lyase [Microbacterium resistens] Length: 266\nNumber of predicted TMHs: 2\nExp number of AAs in TMHs: 43.06691\nExp number, first 60 AAs: 22.43844\nTotal prob of N-in: 0.98597\nPOSSIBLE N-term signal sequence\ninside 1 19\nTMhelix 20 42\noutside 43 243\nTMhelix 244 263\ninside 264 266

31872 GCF_001552355.1_ASM155235v1 Microbacterium resistens NBRC 103078 Terrabacteria group; Actinobacteria; Actinobacteria; Micrococcales; Microbacteriaceae; Microbacterium; Microbacterium resistens MNVIHPSPSARAARPARRRSVVRTALVGVVVALAAVLGTALPAAAHDELVDSTPALTDDG WP_084344888.1 hypothetical protein [Microbacterium resistens] Length: 234\nNumber of predicted TMHs: 2\nExp number of AAs in TMHs: 45.34935\nExp number, first 60 AAs: 22.58333\nTotal prob of N-in: 0.99933\nPOSSIBLE N-term signal sequence\ninside 1 20\nTMhelix 21 43\noutside 44 194\nTMhelix 195 217\ninside 218 234

31873 GCF_000411455.1_Micr_sp_oral_taxon_186_F0373_V1 Microbacterium sp. oral taxon 186 str. F0373 Terrabacteria group; Actinobacteria; Actinobacteria; Micrococcales; Microbacteriaceae; Microbacterium; Microbacterium sp. oral taxon 186 MITHRTRRGILGRLGATTLVAAAVVCSVGLSATSASADDSSNQPPLTAKDHAITDVPGLV WP_016464758.1 LPXTG-domain-containing protein cell wall anchor domain [Microbacterium sp. oral taxon 186] Length: 416\nNumber of predicted TMHs: 2\nExp number of AAs in TMHs: 48.72466\nExp number, first 60 AAs: 10.14989\nTotal prob of N-in: 0.69404\nPOSSIBLE N-term signal sequence\ninside 1 348\nTMhelix 349 371\noutside 372 385\nTMhelix 386 408\ninside 409 416

31874 GCF_000411455.1_Micr_sp_oral_taxon_186_F0373_V1 Microbacterium sp. oral taxon 186 str. F0373 Terrabacteria group; Actinobacteria; Actinobacteria; Micrococcales; Microbacteriaceae; Microbacterium; Microbacterium sp. oral taxon 186 MRPTAERRKRRSFVAAITAGALALTGAGLAALPARAAVSADAAVLINEVYGGGGNSKATY WP_016465141.1 hypothetical protein [Microbacterium sp. oral taxon 186] Length: 1578\nNumber of predicted TMHs: 2\nExp number of AAs in TMHs: 42.1566799999999\nExp number, first 60 AAs: 21.69899\nTotal prob of N-in: 0.99859\nPOSSIBLE N-term signal sequence\ninside 1 11\nTMhelix 12 34\noutside 35 1548\nTMhelix 1549 1568\ninside 1569 1578

31875 GCF_000411455.1_Micr_sp_oral_taxon_186_F0373_V1 Microbacterium sp. oral taxon 186 str. F0373 Terrabacteria group; Actinobacteria; Actinobacteria; Micrococcales; Microbacteriaceae; Microbacterium; Microbacterium sp. oral taxon 186 MRNPRRTLIATLAALVLGTGLTLAVASPAQAHDELVSSYPEADTTVPESPAEITLSFSGE WP_016463500.1 hypothetical protein [Microbacterium sp. oral taxon 186] Length: 225\nNumber of predicted TMHs: 2\nExp number of AAs in TMHs: 44.61069\nExp number, first 60 AAs: 21.90325\nTotal prob of N-in: 0.99568\nPOSSIBLE N-term signal sequence\ninside 1 6\nTMhelix 7 29\noutside 30 173\nTMhelix 174 196\ninside 197 225

31876 GCF_900104345.1_IMG-taxon_2623620614_annotated_assembly Microbacterium testaceum StLB037 Terrabacteria group; Actinobacteria; Actinobacteria; Micrococcales; Microbacteriaceae; Microbacterium; Microbacterium testaceum MTPRRRLLPAVLAAGLLAAAALFVPASPAAAHDELVSTDPAAGATLEALPAQITFSYSAD WP_081349655.1 hypothetical protein [Microbacterium testaceum] Length: 209\nNumber of predicted TMHs: 2\nExp number of AAs in TMHs: 43.74902\nExp number, first 60 AAs: 21.65195\nTotal prob of N-in: 0.99736\nPOSSIBLE N-term signal sequence\ninside 1 6\nTMhelix 7 29\noutside 30 171\nTMhelix 172 194\ninside 195 209

31877 GCF_900104345.1_IMG-taxon_2623620614_annotated_assembly Microbacterium testaceum StLB037 Terrabacteria group; Actinobacteria; Actinobacteria; Micrococcales; Microbacteriaceae; Microbacterium; Microbacterium testaceum MVLSRPVSEDRSRRGIVRRAGSLVAAAAVVLSGLVPAVAASAAPLPTPLASYDFAATTGT WP_081349694.1 hypothetical protein [Microbacterium testaceum] Length: 1835\nNumber of predicted TMHs: 2\nExp number of AAs in TMHs: 45.08561\nExp number, first 60 AAs: 22.36042\nTotal prob of N-in: 0.99619\nPOSSIBLE N-term signal sequence\ninside 1 20\nTMhelix 21 43\noutside 44 1801\nTMhelix 1802 1824\ninside 1825 1835

31878 GCF_000202635.1_ASM20263v1 Microbacterium testaceum StLB037 Terrabacteria group; Actinobacteria; Actinobacteria; Micrococcales; Microbacteriaceae; Microbacterium; Microbacterium testaceum MAREKKPRRASGRRSPLAAAALIGIGLLLTGGVYAGASAAMAATDTPTSSASSATAVEEG WP_013583167.1 cystathionine beta-lyase [Microbacterium testaceum] Length: 269\nNumber of predicted TMHs: 2\nExp

number of AAs in TMHs: 42.85483\nExp number, first 60 AAs: 22.45656\nTotal prob of N-in: 0.98860\nPOSSIBLE N-term signal sequence\ninside 1 20\nTMhelix 21 43\noutside 44 246\nTMhelix 247 266\ninside 267 269

31879 GCF_900104345.1_IMG-taxon_2623620614_annotated_assembly Microbacterium testaceum StLB037 Terrabacteria group; Actinobacteria; Actinobacteria; Micrococcales; Microbacteriaceae; Microbacterium; Microbacterium testaceum MARTPTRALSPADQERRRALRVMKGVALGALIAMALVFAVSFSLQREVEWLQYVRAAAEG WP_056229005.1 MULTISPECIES: DUF445 domain-containing protein [Microbacterium] Length: 424\nNumber of predicted TMHs: 2\nExp number of AAs in TMHs: 51.3979999999999\nExp number, first 60 AAs: 22.34727\nTotal prob of N-in: 0.95486\nPOSSIBLE N-term signal sequence\ninside 1 19\nTMhelix 20 42\noutside 43 400\nTMhelix 401 423\ninside 424 424

31880 GCF_900104345.1_IMG-taxon_2623620614_annotated_assembly Microbacterium testaceum StLB037 Terrabacteria group; Actinobacteria; Actinobacteria; Micrococcales; Microbacteriaceae; Microbacterium; Microbacterium testaceum MAREKKTRRASGRRSPLAAAALIGIGLLLTGGVYAGASAAMAATSGTPTSEANSATAVEE WP_056227819.1 MULTISPECIES: cystathionine beta-lyase [Microbacterium] Length: 270\nNumber of predicted TMHs: 2\nExp number of AAs in TMHs: 42.60447\nExp number, first 60 AAs: 22.3786\nTotal prob of N-in: 0.99243\nPOSSIBLE N-term signal sequence\ninside 1 20\nTMhelix 21 43\noutside 44 247\nTMhelix 248 267\ninside 268 270

31881 GCF_000202635.1_ASM20263v1 Microbacterium testaceum StLB037 Terrabacteria group; Actinobacteria; Actinobacteria; Micrococcales; Microbacteriaceae; Microbacterium; Microbacterium testaceum MHHSRSSRPRRRRRRALVAVLTLVAAAIALFAPPATAADRGTGFGTWAPLSSTGWHGSMRV WP_013585746.1 hypothetical protein [Microbacterium testaceum] Length: 505\nNumber of predicted TMHs: 2\nExp number of AAs in TMHs: 43.26249\nExp number, first 60 AAs: 21.27046\nTotal prob of N-in: 0.99865\nPOSSIBLE N-term signal sequence\ninside 1 12\nTMhelix 13 35\noutside 36 474\nTMhelix 475 497\ninside 498 505

31882 GCF_000202635.1_ASM20263v1 Microbacterium testaceum StLB037 Terrabacteria group; Actinobacteria; Actinobacteria; Micrococcales; Microbacteriaceae; Microbacterium; Microbacterium testaceum MARTPTAQLSPADQERRRALARMKGVALGALIGMAVIFAVAFAFQRDVTWLQYVRAAAEG WP_013585924.1 DUF445 domain-containing protein [Microbacterium testaceum] Length: 423\nNumber of predicted TMHs: 2\nExp number of AAs in TMHs: 53.07995\nExp number, first 60 AAs: 22.13391\nTotal prob of N-in: 0.93413\nPOSSIBLE N-term signal sequence\ninside 1 24\nTMhelix 25 44\noutside 45 399\nTMhelix 400 422\ninside 423 423

31883 GCF_900104895.1_IMG-taxon_2634166341_annotated_assembly Microterricola viridarii Terrabacteria group; Actinobacteria; Actinobacteria; Micrococcales; Microbacteriaceae; Microterricola MKLHPRRTAVRGAASALAILGIATGLAVAGASAGHAAEIPIPVALDFTELQPNTPVSKTI WP_083364995.1 hypothetical protein [Microterricola viridarii] Length: 201\nNumber of predicted TMHs: 2\nExp number of AAs in TMHs: 43.76457\nExp number, first 60 AAs: 22.13874\nTotal prob of N-in: 0.99513\nPOSSIBLE N-term signal sequence\ninside 1 11\nTMhelix 12 34\noutside 35 157\nTMhelix 158 180\ninside 181 201

31884 GCF_900163835.1_ASM90016383v1 Mycetocola reblochoni REB411 Terrabacteria group; Actinobacteria; Actinobacteria; Micrococcales; Microbacteriaceae; Mycetocola; Mycetocola reblochoni MPSPTPRTRRHRRRRSLLIAATALTATALAPATAASAAPAVLGTPVLTAPATAEVGDVIT WP_087136771.1 hypothetical protein [Mycetocola reblochoni] Length: 269\nNumber of predicted TMHs: 2\nExp number of AAs in TMHs: 44.16315\nExp number, first 60 AAs: 21.77628\nTotal prob of N-in: 0.96999\nPOSSIBLE N-term signal sequence\ninside 1 15\nTMhelix 16 38\noutside 39 240\nTMhelix 241 263\ninside 264 269

31885 GCF_900163835.1_ASM90016383v1 Mycetocola reblochoni REB411 Terrabacteria group; Actinobacteria; Actinobacteria; Micrococcales; Microbacteriaceae; Mycetocola; Mycetocola reblochoni MPAGSASRRPRARARRAAVVGIVCAALTAGALVAAPASAAPAVAAAEQQHPDAVVTEPGT WP_087136226.1 hypothetical protein [Mycetocola reblochoni] Length: 1103\nNumber of predicted TMHs: 2\nExp number of AAs in TMHs: 56.2800499999998\nExp number, first 60 AAs: 21.67412\nTotal prob of N-in: 0.98459\nPOSSIBLE N-term signal sequence\ninside 1 16\nTMhelix 17 39\noutside 40 1047\nTMhelix 1048 1070\ninside 1071 1103

31886 GCF_900163835.1_ASM90016383v1 Mycetocola reblochoni REB411 Terrabacteria group; Actinobacteria; Actinobacteria; Micrococcales; Microbacteriaceae; Mycetocola; Mycetocola reblochoni MRGRNRPHTRGRRALLAVGLVTAVTATAVVVPSAPPAEAASTLSAFDAVNQFIGTEMNDG WP_087136031.1 hypothetical protein [Mycetocola reblochoni] Length: 1883\nNumber of predicted TMHs: 2\nExp number of AAs in TMHs: 43.54419\nExp number, first 60 AAs: 21.18792\nTotal prob of N-in: 0.97522\nPOSSIBLE N-term signal sequence\ninside 1 12\nTMhelix 13 35\noutside 36 1853\nTMhelix 1854 1876\ninside 1877 1883

31887 GCF_900163835.1_ASM90016383v1 Mycetocola reblochoni REB411 Terrabacteria group; Actinobacteria; Actinobacteria; Micrococcales; Microbacteriaceae; Mycetocola; Mycetocola reblochoni
MPNPHTPRRRSRRPALVSALLTVGLAAGTLVAVAPAAHALTPSAGWAELEGRIDAIVDEA WP_087137159.1 hypothetical protein [Mycetocola reblochoni] Length: 703\nNumber of predicted TMHs: 2\nExp number of AAs in TMHs: 43.50312\nExp number, first 60 AAs: 21.53939\nTotal prob of N-in: 0.96718\nPOSSIBLE N-term signal sequence\ninside 1 12\nTMhelix 13 35\noutside 36 673\nTMhelix 674 696\ninside 697 703

31888 GCF_900167575.1_IMG-taxon_2708742390_annotated_assembly Okibacterium fritillariae Terrabacteria group; Actinobacteria; Actinobacteria; Micrococcales; Microbacteriaceae; Okibacterium
MLRKRNTSARTARKNGRRSPLATVALIAVGLLFTGGAYAAFSATTANAETPTANTQASVE WP_079727669.1 cystathionine beta-lyase [Okibacterium fritillariae] Length: 271\nNumber of predicted TMHs: 2\nExp number of AAs in TMHs: 42.9109499999999\nExp number, first 60 AAs: 22.26181\nTotal prob of N-in: 0.99868\nPOSSIBLE N-term signal sequence\ninside 1 20\nTMhelix 21 43\noutside 44 248\nTMhelix 249 268\ninside 269 271

31889 GCF_900167175.1_IMG-taxon_2708742383_annotated_assembly Plantibacter cousiniae Terrabacteria group; Actinobacteria; Actinobacteria; Micrococcales; Microbacteriaceae; Plantibacter
MTVSRRRRLVGTAATAFLAVAAVLAVAQPASAHNSIISTTPAEGSTITEQPEQLVITTND WP_079705109.1 hypothetical protein [Plantibacter cousiniae] Length: 218\nNumber of predicted TMHs: 2\nExp number of AAs in TMHs: 42.26288\nExp number, first 60 AAs: 20.71984\nTotal prob of N-in: 0.98447\nPOSSIBLE N-term signal sequence\ninside 1 8\nTMhelix 9 31\noutside 32 179\nTMhelix 180 199\ninside 200 218

31890 GCF_900167175.1_IMG-taxon_2708742383_annotated_assembly Plantibacter cousiniae Terrabacteria group; Actinobacteria; Actinobacteria; Micrococcales; Microbacteriaceae; Plantibacter
MKTDPTQTTTTKASKKAARATRKSGRRSPFASAALLAIGLLVTGGAYAAVSVSTASAEPV WP_064296976.1 MULTISPECIES: cystathionine beta-lyase [Plantibacter] Length: 281\nNumber of predicted TMHs: 2\nExp number of AAs in TMHs: 43.19064\nExp number, first 60 AAs: 22.44082\nTotal prob of N-in: 0.97415\nPOSSIBLE N-term signal sequence\ninside 1 27\nTMhelix 28 50\noutside 51 258\nTMhelix 259 278\ninside 279 281

31891 GCF_900167175.1_IMG-taxon_2708742383_annotated_assembly Plantibacter cousiniae Terrabacteria group; Actinobacteria; Actinobacteria; Micrococcales; Microbacteriaceae; Plantibacter
MRGRPGGAGIGPARDGGRLWGSDGPRRGDRRRPRGRRGALSALSAVLLAFLLVGTQALPA WP_079705475.1 peptidase S8 [Plantibacter cousiniae] Length: 457\nNumber of predicted TMHs: 2\nExp number of AAs in TMHs: 44.01078\nExp number, first 60 AAs: 21.11893\nTotal prob of N-in: 0.97494\nPOSSIBLE N-term signal sequence\ninside 1 37\nTMhelix 38 60\noutside 61 424\nTMhelix 425 447\ninside 448 457

31892 GCF_900177795.1_IMG-taxon_2708742472_annotated_assembly Plantibacter elymi Terrabacteria group; Actinobacteria; Actinobacteria; Micrococcales; Microbacteriaceae; Plantibacter
MSMGNSRRSLLHGKHRYGVIVLVLAGASVLAPAGLSPVSATARSVVDVPPVTPEEVAPAA WP_086474051.1 hypothetical protein [Plantibacter elymi] Length: 298\nNumber of predicted TMHs: 2\nExp number of AAs in TMHs: 41.01093\nExp number, first 60 AAs: 19.27561\nTotal prob of N-in: 0.88061\nPOSSIBLE N-term signal sequence\ninside 1 19\nTMhelix 20 42\noutside 43 262\nTMhelix 263 285\ninside 286 298

31893 GCF_900177795.1_IMG-taxon_2708742472_annotated_assembly Plantibacter elymi Terrabacteria group; Actinobacteria; Actinobacteria; Micrococcales; Microbacteriaceae; Plantibacter
MTVSRRRRLVGTAATAFLAVAAVLAVAQPASAHNSIISTTPAEGSTITEQPEQLVITTND WP_086474591.1 hypothetical protein [Plantibacter elymi] Length: 218\nNumber of predicted TMHs: 2\nExp number of AAs in TMHs: 42.24965\nExp number, first 60 AAs: 20.70954\nTotal prob of N-in: 0.98424\nPOSSIBLE N-term signal sequence\ninside 1 8\nTMhelix 9 31\noutside 32 179\nTMhelix 180 199\ninside 200 218

31894 GCF_900177795.1_IMG-taxon_2708742472_annotated_assembly Plantibacter elymi Terrabacteria group; Actinobacteria; Actinobacteria; Micrococcales; Microbacteriaceae; Plantibacter
MKTDPTQTTTTKASKKAARATRKSGRRSPFASAALLAIGLLVTGGAYAAVSVSTASAEPV WP_064296976.1 MULTISPECIES: cystathionine beta-lyase [Plantibacter] Length: 281\nNumber of predicted TMHs: 2\nExp number of AAs in TMHs: 43.19064\nExp number, first 60 AAs: 22.44082\nTotal prob of N-in: 0.97415\nPOSSIBLE N-term signal sequence\ninside 1 27\nTMhelix 28 50\noutside 51 258\nTMhelix 259 278\ninside 279 281

31895 GCF_900177615.1_IMG-taxon_2708742471_annotated_assembly Plantibacter flavus Terrabacteria group; Actinobacteria; Actinobacteria; Micrococcales; Microbacteriaceae; Plantibacter
MKTDPTQTTTTKASKKAARATRKSGRRSPFASAALLAIGLLVTGGAYAAVSVSTASAEPV WP_064296976.1 MULTISPECIES: cystathionine beta-lyase [Plantibacter] Length: 281\nNumber of predicted TMHs: 2\nExp number of AAs in TMHs:

43.19064\nExp number, first 60 AAs:  22.44082\nTotal prob of N-in:   0.97415\nPOSSIBLE N-term signal sequence\ninside 1   27\nTMhelix   28   50\noutside   51   258\nTMhelix   259   278\ninside   279   281

31896 GCF_900177615.1_IMG-taxon_2708742471_annotated_assembly Plantibacter flavus Terrabacteria group; Actinobacteria; Actinobacteria; Micrococcales; Microbacteriaceae; Plantibacter
MTVSRRRRLVGTAATAFLAVAAVLAVAQPASAHNSIISTTPAEGSTISEQPEQLVITTND WP_085511799.1 hypothetical protein [Plantibacter flavus] Length: 218\nNumber of predicted TMHs:  2\nExp number of AAs in TMHs: 42.39928\nExp number, first 60 AAs:  20.5969\nTotal prob of N-in:   0.98181\nPOSSIBLE N-term signal sequence\ninside   1   8\nTMhelix 9   31\noutside   32   176\nTMhelix   177   199\ninside   200   218

31897 GCF_002024505.1_ASM202450v1 Plantibacter flavus Terrabacteria group; Actinobacteria; Actinobacteria; Micrococcales; Microbacteriaceae; Plantibacter
MKTDPTQTTTTKASKKAARATRKSGRRSPFASAALLAIGLLVTGGAYAAVSVSTASAEPV WP_064296976.1 MULTISPECIES: cystathionine beta-lyase [Plantibacter] Length: 281\nNumber of predicted TMHs:  2\nExp number of AAs in TMHs: 43.19064\nExp number, first 60 AAs:  22.44082\nTotal prob of N-in:   0.97415\nPOSSIBLE N-term signal sequence\ninside 1   27\nTMhelix   28   50\noutside   51   258\nTMhelix   259   278\ninside   279   281

31898 GCF_002024505.1_ASM202450v1 Plantibacter flavus Terrabacteria group; Actinobacteria; Actinobacteria; Micrococcales; Microbacteriaceae; Plantibacter
MTVSRRRRLVGTAATAFLAVAAVLAVAQPASAHNSIISTTPAEGSTITEQPEQLVITTND WP_079001510.1 hypothetical protein [Plantibacter flavus] Length: 219\nNumber of predicted TMHs:  2\nExp number of AAs in TMHs: 42.59737\nExp number, first 60 AAs:  20.72258\nTotal prob of N-in:   0.98601\nPOSSIBLE N-term signal sequence\ninside   1   8\nTMhelix 9   31\noutside   32   177\nTMhelix   178   200\ninside   201   219

31899 GCF_001650455.1_ASM165045v1 Plantibacter sp. H53 Terrabacteria group; Actinobacteria; Actinobacteria; Micrococcales; Microbacteriaceae; Plantibacter
MKTDPTQTTTTKASKKAARATRKSGRRSPFASAALLAIGLLVTGGAYAAVSVSTASAEPV WP_064296976.1 MULTISPECIES: cystathionine beta-lyase [Plantibacter] Length: 281\nNumber of predicted TMHs:  2\nExp number of AAs in TMHs: 43.19064\nExp number, first 60 AAs:  22.44082\nTotal prob of N-in:   0.97415\nPOSSIBLE N-term signal sequence\ninside 1   27\nTMhelix   28   50\noutside   51   258\nTMhelix   259   278\ninside   279   281

31900 GCF_001650455.1_ASM165045v1 Plantibacter sp. H53 Terrabacteria group; Actinobacteria; Actinobacteria; Micrococcales; Microbacteriaceae; Plantibacter
MTVSRRRRLVGTAATAFLAVAAVLAVAQPASAHNSIISTTPAEGSTITEQPEQLVITTND WP_064294246.1 hypothetical protein [Plantibacter sp. H53] Length: 219\nNumber of predicted TMHs:  2\nExp number of AAs in TMHs: 42.57156\nExp number, first 60 AAs:  20.68928\nTotal prob of N-in:   0.98524\nPOSSIBLE N-term signal sequence\ninside 1   8\nTMhelix   9   31\noutside   32   177\nTMhelix   178   200\ninside   201   219

31901 GCF_001421315.1_ASM142131v1 Plantibacter sp. Leaf1 Terrabacteria group; Actinobacteria; Actinobacteria; Micrococcales; Microbacteriaceae; Plantibacter
MKTDPTQPTATKSSKKTARANRKSGRRSPFASAALLAIGLLVTGGAYAAVSVSTASAEPV WP_056005492.1 MULTISPECIES: cystathionine beta-lyase [Plantibacter] Length: 281\nNumber of predicted TMHs:  2\nExp number of AAs in TMHs: 42.69846\nExp number, first 60 AAs:  22.33873\nTotal prob of N-in:   0.94518\nPOSSIBLE N-term signal sequence\ninside 1   27\nTMhelix   28   50\noutside   51   258\nTMhelix   259   278\ninside   279   281

31902 GCF_001423725.1_Leaf171 Plantibacter sp. Leaf171 Terrabacteria group; Actinobacteria; Actinobacteria; Micrococcales; Microbacteriaceae; Plantibacter
MKTDPTQPTATKSSKKTARANRKSGRRSPFASAALLAIGLLVTGGAYAAVSVSTASAEPV WP_056005492.1 MULTISPECIES: cystathionine beta-lyase [Plantibacter] Length: 281\nNumber of predicted TMHs:  2\nExp number of AAs in TMHs: 42.69846\nExp number, first 60 AAs:  22.33873\nTotal prob of N-in:   0.94518\nPOSSIBLE N-term signal sequence\ninside 1   27\nTMhelix   28   50\noutside   51   258\nTMhelix   259   278\ninside   279   281

31903 GCF_001423725.1_Leaf171 Plantibacter sp. Leaf171 Terrabacteria group; Actinobacteria; Actinobacteria; Micrococcales; Microbacteriaceae; Plantibacter
MTTARRRRLVGSVATAFLAAAAVFAVAGPAAAHNTIVSTSPAEGSTVTEQPEQVVITTND WP_056779050.1
hypothetical protein [Plantibacter sp. Leaf171] Length: 228\nNumber of predicted TMHs:  2\nExp number of AAs in TMHs: 43.8018\nExp number, first 60 AAs:  22.16838\nTotal prob of N-in:   0.99827\nPOSSIBLE N-term signal sequence\ninside   1   8\nTMhelix   9   31\noutside   32   180\nTMhelix   181   203\ninside   204   228

31904 GCF_001423185.1_Leaf314 Plantibacter sp. Leaf314 Terrabacteria group; Actinobacteria; Actinobacteria; Micrococcales; Microbacteriaceae; Plantibacter

MKTDPTQPTATKSSKKTARANRKSGRRSPFASAALLAIGLLVTGGAYAAVSVSTASAEPV WP_056005492.1 MULTISPECIES: cystathionine beta-lyase [Plantibacter] Length: 281\nNumber of predicted TMHs: 2\nExp number of AAs in TMHs: 42.69846\nExp number, first 60 AAs: 22.33873\nTotal prob of N-in: 0.94518\nPOSSIBLE N-term signal sequence\ninside 1 27\nTMhelix 28 50\noutside 51 258\nTMhelix 259 278\ninside 279 281

31905 GCF_001423185.1_Leaf314 Plantibacter sp. Leaf314 Terrabacteria group; Actinobacteria; Actinobacteria; Micrococcales; Microbacteriaceae; Plantibacter
MTTARRRRLVGSVATAFLAAAAVFAMAGPAAAHNSIVSTSPAEGSTVTEQPEQVVITTND WP_056008599.1 hypothetical protein [Plantibacter sp. Leaf314] Length: 228\nNumber of predicted TMHs: 2\nExp number of AAs in TMHs: 43.43222\nExp number, first 60 AAs: 21.97531\nTotal prob of N-in: 0.99825\nPOSSIBLE N-term signal sequence\ninside 1 8\nTMhelix 9 31\noutside 32 180\nTMhelix 181 203\ninside 204 228

31906 GCF_001865025.1_ASM186502v1 Plantibacter sp. MMLR14_011 Terrabacteria group; Actinobacteria; Actinobacteria; Micrococcales; Microbacteriaceae; Plantibacter
MTVSRRRRLVGTAATAFLAVAAVLAVAQPASAHNSIISTTPAEGSTITEQPEQLVITTND WP_071259775.1 hypothetical protein [Plantibacter sp. MMLR14_011] Length: 219\nNumber of predicted TMHs: 2\nExp number of AAs in TMHs: 42.5715\nExp number, first 60 AAs: 20.68929\nTotal prob of N-in: 0.98524\nPOSSIBLE N-term signal sequence\ninside 1 8\nTMhelix 9 31\noutside 32 177\nTMhelix 178 200\ninside 201 219

31907 GCF_001865025.1_ASM186502v1 Plantibacter sp. MMLR14_011 Terrabacteria group; Actinobacteria; Actinobacteria; Micrococcales; Microbacteriaceae; Plantibacter
MKTDPTQTTTTKASKKAARATRKSGRRSPFASAALLAIGLLVTGGAYAAVSVSTASAEPV WP_064296976.1 MULTISPECIES: cystathionine beta-lyase [Plantibacter] Length: 281\nNumber of predicted TMHs: 2\nExp number of AAs in TMHs: 43.19064\nExp number, first 60 AAs: 22.44082\nTotal prob of N-in: 0.97415\nPOSSIBLE N-term signal sequence\ninside 1 27\nTMhelix 28 50\noutside 51 258\nTMhelix 259 278\ninside 279 281

31908 GCF_900014995.1_Draft_genome_of_Pseudoclavibacter_helvolus_G8 Pseudoclavibacter helvolus Terrabacteria group; Actinobacteria; Actinobacteria; Micrococcales; Microbacteriaceae; Pseudoclavibacter
MFRTTKPRTAKSATRKTARRRTPLATAALLGAGLLLTGGAYTAIDSATTAAAAEEAAPAS WP_068476337.1 cystathionine beta-lyase [Pseudoclavibacter helvolus] Length: 277\nNumber of predicted TMHs: 2\nExp number of AAs in TMHs: 42.03393\nExp number, first 60 AAs: 21.8436\nTotal prob of N-in: 0.98952\nPOSSIBLE N-term signal sequence\ninside 1 20\nTMhelix 21 43\noutside 44 253\nTMhelix 254 273\ninside 274 277

31909 GCF_900016305.1_Draft_genome_of_Kocuria_palustris_strain_W4 Pseudoclavibacter helvolus Terrabacteria group; Actinobacteria; Actinobacteria; Micrococcales; Microbacteriaceae; Pseudoclavibacter
MSRTPARTSREGRRRAALPLATALAATALTVGGGIATAGAAEADFTYPAAIDPATITIKT WP_068478771.1 hypothetical protein [Pseudoclavibacter helvolus] Length: 400\nNumber of predicted TMHs: 2\nExp number of AAs in TMHs: 41.26959\nExp number, first 60 AAs: 21.48122\nTotal prob of N-in: 0.96808\nPOSSIBLE N-term signal sequence\ninside 1 16\nTMhelix 17 39\noutside 40 372\nTMhelix 373 392\ninside 393 400

31910 GCF_900016305.1_Draft_genome_of_Kocuria_palustris_strain_W4 Pseudoclavibacter helvolus Terrabacteria group; Actinobacteria; Actinobacteria; Micrococcales; Microbacteriaceae; Pseudoclavibacter
MFRTTKPRTAKSATRKTARRRTPLATAALLGAGLLLTGGAYTAIDSATTAAAAEEAAPAS WP_068476337.1 cystathionine beta-lyase [Pseudoclavibacter helvolus] Length: 277\nNumber of predicted TMHs: 2\nExp number of AAs in TMHs: 42.03393\nExp number, first 60 AAs: 21.8436\nTotal prob of N-in: 0.98952\nPOSSIBLE N-term signal sequence\ninside 1 20\nTMhelix 21 43\noutside 44 253\nTMhelix 254 273\ninside 274 277

31911 GCF_001570905.1_ASM157090v1 Zimmermannella alba NBRC 15616 Terrabacteria group; Actinobacteria; Actinobacteria; Micrococcales; Microbacteriaceae; Pseudoclavibacter; Pseudoclavibacter alba
MFKKPRQTRKSAAARRRSPLATTVLLGLALLVTGGSYAALDSATSAQAETDLSSPQMISE WP_066080782.1 cystathionine beta-lyase [Pseudoclavibacter alba] Length: 271\nNumber of predicted TMHs: 2\nExp number of AAs in TMHs: 39.32759\nExp number, first 60 AAs: 19.97975\nTotal prob of N-in: 0.97797\nPOSSIBLE N-term signal sequence\ninside 1 19\nTMhelix 20 39\noutside 40 247\nTMhelix 248 267\ninside 268 271

31912 GCF_001570925.1_ASM157092v1 Zimmermannella bifida NBRC 103089 Terrabacteria group; Actinobacteria; Actinobacteria; Micrococcales; Microbacteriaceae; Pseudoclavibacter; Pseudoclavibacter bifida
MSLTTTNRSAAKKSRKRGRRSPLATAGLLGLGLMLTGGAYGAIDSFVENAAANESQVDMN WP_066055322.1 cystathionine beta-lyase [Pseudoclavibacter bifida] Length: 278\nNumber of predicted TMHs: 2\nExp number of AAs in TMHs: 43.22055\nExp number, first 60 AAs: 22.70406\nTotal prob of N-in: 0.99784\nPOSSIBLE N-term signal sequence\ninside 1 20\nTMhelix 21 43\noutside 44 250\nTMhelix 251 273\ninside 274 278

31913 GCF_000381765.1_ASM38176v1 Zimmermannella faecalis ATCC 13722 Terrabacteria group; Actinobacteria; Actinobacteria; Micrococcales; Microbacteriaceae; Pseudoclavibacter; Pseudoclavibacter faecalis MSLVTQHRSTAKKSRKRGRRSPLATAALLGVGLMLTGGAYGAVDTLVANAADGTTEVDMN WP_019617431.1 hypothetical protein [Pseudoclavibacter faecalis] Length: 278\nNumber of predicted TMHs: 2\nExp number of AAs in TMHs: 44.36339\nExp number, first 60 AAs: 22.65176\nTotal prob of N-in: 0.99664\nPOSSIBLE N-term signal sequence\ninside 1 20\nTMhelix 21 43\noutside 44 250\nTMhelix 251 273\ninside 274 278

31914 GCF_000423505.1_ASM42350v1 Pseudoclavibacter soli DSM 23366 Terrabacteria group; Actinobacteria; Actinobacteria; Micrococcales; Microbacteriaceae; Pseudoclavibacter; Pseudoclavibacter soli MEMNTHTKRRGIFASVGALLTAAALALTGAGLATAAPTIPDSGATANLHITKLTTPRGGA WP_084626871.1 hypothetical protein [Pseudoclavibacter soli] Length: 544\nNumber of predicted TMHs: 2\nExp number of AAs in TMHs: 44.9396199999999\nExp number, first 60 AAs: 22.60229\nTotal prob of N-in: 0.99826\nPOSSIBLE N-term signal sequence\ninside 1 11\nTMhelix 12 34\noutside 35 510\nTMhelix 511 533\ninside 534 544

31915 GCF_000423505.1_ASM42350v1 Pseudoclavibacter soli DSM 23366 Terrabacteria group; Actinobacteria; Actinobacteria; Micrococcales; Microbacteriaceae; Pseudoclavibacter; Pseudoclavibacter soli MKIRTTKKTSTTKRRARRRGPLATVGLLAVGLAVTGGGYAAVAQIGANDTEATATVTQST WP_051223931.1 cystathionine beta-lyase [Pseudoclavibacter soli] Length: 276\nNumber of predicted TMHs: 2\nExp number of AAs in TMHs: 44.07323\nExp number, first 60 AAs: 22.78652\nTotal prob of N-in: 0.99988\nPOSSIBLE N-term signal sequence\ninside 1 20\nTMhelix 21 43\noutside 44 249\nTMhelix 250 272\ninside 273 276

31916 GCF_001423885.1_Leaf185 Rathayibacter sp. Leaf185 Terrabacteria group; Actinobacteria; Actinobacteria; Micrococcales; Microbacteriaceae; Rathayibacter MPHMSERHPSDRASERRRLVHRARGAVAATAGIAVLAVGFIVGSPAQAATKPDLGTAGAF WP_082498971.1 MULTISPECIES: hypothetical protein [Rathayibacter] Length: 590\nNumber of predicted TMHs: 2\nExp number of AAs in TMHs: 46.98245\nExp number, first 60 AAs: 21.8541\nTotal prob of N-in: 0.97872\nPOSSIBLE N-term signal sequence\ninside 1 19\nTMhelix 20 42\noutside 43 556\nTMhelix 557 579\ninside 580 590

31917 GCF_001423885.1_Leaf185 Rathayibacter sp. Leaf185 Terrabacteria group; Actinobacteria; Actinobacteria; Micrococcales; Microbacteriaceae; Rathayibacter MHRRVLTPPERPRRRRAVPQLLRSTGAAALSIALIALATPAHAETPEATPQASSSSVTAT WP_056039821.1 MULTISPECIES: hypothetical protein [Rathayibacter] Length: 737\nNumber of predicted TMHs: 2\nExp number of AAs in TMHs: 39.78478\nExp number, first 60 AAs: 19.06627\nTotal prob of N-in: 0.87139\nPOSSIBLE N-term signal sequence\ninside 1 20\nTMhelix 21 43\noutside 44 680\nTMhelix 681 700\ninside 701 737

31918 GCF_001423005.1_Leaf294 Rathayibacter sp. Leaf294 Terrabacteria group; Actinobacteria; Actinobacteria; Micrococcales; Microbacteriaceae; Rathayibacter MHRRVLTPPERPRRRRAVPQLLRSTGAAALSIALIALATPAHAETPEATPQASSSSVTAT WP_056039821.1 MULTISPECIES: hypothetical protein [Rathayibacter] Length: 737\nNumber of predicted TMHs: 2\nExp number of AAs in TMHs: 39.78478\nExp number, first 60 AAs: 19.06627\nTotal prob of N-in: 0.87139\nPOSSIBLE N-term signal sequence\ninside 1 20\nTMhelix 21 43\noutside 44 680\nTMhelix 681 700\ninside 701 737

31919 GCF_001423005.1_Leaf294 Rathayibacter sp. Leaf294 Terrabacteria group; Actinobacteria; Actinobacteria; Micrococcales; Microbacteriaceae; Rathayibacter MPHMSERHPSDRASERRRLVHRARGAVAATAGIAVLAVGFIVGSPAQAATKPDLGTAGAF WP_082498971.1 MULTISPECIES: hypothetical protein [Rathayibacter] Length: 590\nNumber of predicted TMHs: 2\nExp number of AAs in TMHs: 46.98245\nExp number, first 60 AAs: 21.8541\nTotal prob of N-in: 0.97872\nPOSSIBLE N-term signal sequence\ninside 1 19\nTMhelix 20 42\noutside 43 556\nTMhelix 557 579\ninside 580 590

31920 GCF_001423045.1_Leaf296 Rathayibacter sp. Leaf296 Terrabacteria group; Actinobacteria; Actinobacteria; Micrococcales; Microbacteriaceae; Rathayibacter MTETPDLLPPLDEHRVSAIEARVFRSLTAARAARAARRRRALSIGGAAAGLLVLAVVVGP WP_056867345.1 hypothetical protein [Rathayibacter sp. Leaf296] Length: 367\nNumber of predicted TMHs: 2\nExp number of AAs in TMHs: 46.26065\nExp number, first 60 AAs: 19.2423\nTotal prob of N-in: 0.97563\nPOSSIBLE N-term signal sequence\ninside 1 40\nTMhelix 41 63\noutside 64 332\nTMhelix 333 355\ninside 356 367

31921 GCF_001423045.1_Leaf296 Rathayibacter sp. Leaf296 Terrabacteria group; Actinobacteria; Actinobacteria; Micrococcales; Microbacteriaceae; Rathayibacter MRSTTPGRRRGLIRTLSVTAVGAVAGAAMLGAAPAMAAEGDIAIDVLSINDFHGRIEANG WP_082485567.1 hypothetical protein [Rathayibacter sp. Leaf296] Length: 746\nNumber of predicted TMHs: 2\nExp number of

AAs in TMHs: 42.7094\nExp number, first 60 AAs:  21.74271\nTotal prob of N-in:   0.97872\nPOSSIBLE N-term signal sequence\ninside   1   11\nTMhelix   12   34\noutside   35   716\nTMhelix   717   739\ninside   740   746

31922 GCF_001423055.1_Leaf299 Rathayibacter sp. Leaf299 Terrabacteria group; Actinobacteria; Actinobacteria; Micrococcales; Microbacteriaceae; Rathayibacter
MLSSSLRRRALVSTVAPALAVALVFGSASTAAAADFVDPVTGAVVQAAATDAALTVTAGA WP_055794040.1 hypothetical protein [Rathayibacter sp. Leaf299] Length: 680\nNumber of predicted TMHs:  2\nExp number of AAs in TMHs: 36.6034399999999\nExp number, first 60 AAs:  14.42743\nTotal prob of N-in:   0.62173\nPOSSIBLE N-term signal sequence\ninside   1   11\nTMhelix   12   34\noutside   35   619\nTMhelix   620   642\ninside   643   680

31923 GCF_001423055.1_Leaf299 Rathayibacter sp. Leaf299 Terrabacteria group; Actinobacteria; Actinobacteria; Micrococcales; Microbacteriaceae; Rathayibacter
MTETPDLLPPLDEHRVSAIEARVFRSLAAARAARAAHRRRALSIGGAAAGLLVLAVVVGP WP_055794768.1 hypothetical protein [Rathayibacter sp. Leaf299] Length: 360\nNumber of predicted TMHs:  2\nExp number of AAs in TMHs: 45.82839\nExp number, first 60 AAs:  18.83219\nTotal prob of N-in:   0.94378\nPOSSIBLE N-term signal sequence\ninside 1   40\nTMhelix   41   63\noutside   64   325\nTMhelix   326   348\ninside   349   360

31924 GCF_001423055.1_Leaf299 Rathayibacter sp. Leaf299 Terrabacteria group; Actinobacteria; Actinobacteria; Micrococcales; Microbacteriaceae; Rathayibacter
MRSTTPGRRRGLIRTLSVTAVGAVAGAAMLGAAPAMAAEGDIAIDVLSINDFHGRIEANG WP_082481119.1 hypothetical protein [Rathayibacter sp. Leaf299] Length: 746\nNumber of predicted TMHs:  2\nExp number of AAs in TMHs: 42.71421\nExp number, first 60 AAs:  21.74273\nTotal prob of N-in:   0.97872\nPOSSIBLE N-term signal sequence\ninside   1   11\nTMhelix   12   34\noutside   35   716\nTMhelix   717   739\ninside   740   746

31925 GCF_900177245.1_IMG-taxon_2708742461_annotated_assembly Rathayibacter sp. VKM Ac-2121 Terrabacteria group; Actinobacteria; Actinobacteria; Micrococcales; Microbacteriaceae; Rathayibacter
MHRRVLTLPRRRRRRAVPQLLRLVGAAVLSFALVGVASPATASTGTLEPSDPASGTVTATWP_085477620.1 hypothetical protein [Rathayibacter sp. VKM Ac-2121] Length: 740\nNumber of predicted TMHs:  2\nExp number of AAs in TMHs: 42.6194799999999\nExp number, first 60 AAs:  21.96524\nTotal prob of N-in:   0.99784\nPOSSIBLE N-term signal sequence\ninside   1   19\nTMhelix   20   42\noutside   43   683\nTMhelix   684   703\ninside   704   740

31926 GCF_001995175.1_ASM199517v1 Rathayibacter sp. VKM Ac-2630 Terrabacteria group; Actinobacteria; Actinobacteria; Micrococcales; Microbacteriaceae; Rathayibacter
MLPTTRSSRRRGVLRAAVVLSAASALAALGAAPASAAEQETGWVRVAHLSPDTKAVDVTL WP_077223961.1 hypothetical protein [Rathayibacter sp. VKM Ac-2630] Length: 300\nNumber of predicted TMHs:  2\nExp number of AAs in TMHs: 42.96486\nExp number, first 60 AAs:  20.1673\nTotal prob of N-in:   0.93256\nPOSSIBLE N-term signal sequence\ninside   1   11\nTMhelix   12   34\noutside   35   267\nTMhelix   268   290\ninside   291   300

31927 GCF_001641005.1_ASM164100v1 Rathayibacter tritici Terrabacteria group; Actinobacteria; Actinobacteria; Micrococcales; Microbacteriaceae; Rathayibacter
MRSTSPGRRGLFRLLSATAVGAVAGAALLGATPALAAEGDVAIDILSINDFHGRLAADRS WP_068255954.1 hypothetical protein [Rathayibacter tritici] Length: 742\nNumber of predicted TMHs:  2\nExp number of AAs in TMHs: 41.4490299999999\nExp number, first 60 AAs:  20.34798\nTotal prob of N-in:   0.91308\nPOSSIBLE N-term signal sequence\ninside   1   11\nTMhelix   12   34\noutside   35   712\nTMhelix   713   735\ninside   736   742

31928 GCF_900098805.1_Microbacterium_sp._strain_Alg239_V18 Microbacteriaceae bacteriumTerrabacteria group; Actinobacteria; Actinobacteria; Micrococcales; Microbacteriaceae; unclassified Microbacteriaceae
MPNTSSTGPERTTDLTTVTDHEHPHRSGSRRRILILSSLTSLLLTAGLLFATAAPAAAHD WP_083336454.1 hypothetical protein [Microbacteriaceae bacterium] Length: 240\nNumber of predicted TMHs:  2\nExp number of AAs in TMHs: 44.6558099999999\nExp number, first 60 AAs:  21.96101\nTotal prob of N-in:   0.97904\nPOSSIBLE N-term signal sequence\ninside   1   32\nTMhelix   33   55\noutside   56   197\nTMhelix   198   220\ninside   221   240

31929 GCF_002157505.1_ASM215750v1 Arthrobacter agilis Terrabacteria group; Actinobacteria; Actinobacteria; Micrococcales; Micrococcaceae; Arthrobacter
MTMELSLPQRVAHSVGEDPLSERVAVTQEVIYGPLLGWARRSPFHTGVLGHSVHPLLTDL WP_087075781.1 hypothetical protein [Arthrobacter agilis] Length: 192\nNumber of predicted TMHs:  2\nExp number of AAs in TMHs: 55.4806\nExp number, first 60 AAs:  0.73489\nTotal prob of N-in:   0.61273\ninside   1   114\nTMhelix   115 136\noutside   137   145\nTMhelix   146   168\ninside   169   192

31930 GCF_001445575.1_ASM144557v1 Arthrobacter alpinus Terrabacteria group; Actinobacteria; Actinobacteria; Micrococcales; Micrococcaceae; Arthrobacter

MAWGSAVNGATAAEAPTQGRRRTFKNGEKTGVLWWAGQIFSWLLLFTVLAVAAVMIVIPR	WP_082634382.1	signal peptidase I [Arthrobacter alpinus]	Length: 204\nNumber of predicted TMHs: 2\nExp number of AAs in TMHs: 51.98143\nExp number, first 60 AAs: 23.14846\nTotal prob of N-in: 0.59708\nPOSSIBLE N-term signal sequence\ninside 1 38\nTMhelix 39 61\noutside 62 171\nTMhelix 172 194\ninside 195 204

31931	GCF_001445575.1_ASM144557v1	Arthrobacter alpinus	Terrabacteria group; Actinobacteria; Actinobacteria; Micrococcales; Micrococcaceae; Arthrobacter	MKIRHRTGRRAVVAALTAAALTLASAAIALPGANADIKPALAPAVLAAAAPSGAVPLTPNWP_062285750.1	hypothetical protein [Arthrobacter alpinus]	Length: 1187\nNumber of predicted TMHs: 2\nExp number of AAs in TMHs: 42.0274699999999\nExp number, first 60 AAs: 21.46542\nTotal prob of N-in: 0.99474\nPOSSIBLE N-term signal sequence\ninside 1 11\nTMhelix 12 34\noutside 35 1157\nTMhelix 1158 1177\ninside 1178 1187

31932	GCF_001457025.1_ASM145702v1	Arthrobacter enclensis	Terrabacteria group; Actinobacteria; Actinobacteria; Micrococcales; Micrococcaceae; Arthrobacter	MTAARRAHLVLGAATGILLSATLASPATADDGSSTAPAPIQSAPAQPAPVQSAPVQSTSG	WP_058266926.1	hypothetical protein [Arthrobacter enclensis]	Length: 673\nNumber of predicted TMHs: 2\nExp number of AAs in TMHs: 40.55989\nExp number, first 60 AAs: 17.88816\nTotal prob of N-in: 0.81555\nPOSSIBLE N-term signal sequence\ninside 1 6\nTMhelix 7 29\noutside 30 642\nTMhelix 643 665\ninside 666 673

31933	GCF_900094675.1_IMG-taxon_2615840696_annotated_assembly	Arthrobacter enclensis	Terrabacteria group; Actinobacteria; Actinobacteria; Micrococcales; Micrococcaceae; Arthrobacter	MTAARRAHLVLGAATGILLSATLASPATADDGSSTAPAPIQSAPAQPAPVQSAPVQSTSG	WP_058266926.1	hypothetical protein [Arthrobacter enclensis]	Length: 673\nNumber of predicted TMHs: 2\nExp number of AAs in TMHs: 40.55989\nExp number, first 60 AAs: 17.88816\nTotal prob of N-in: 0.81555\nPOSSIBLE N-term signal sequence\ninside 1 6\nTMhelix 7 29\noutside 30 642\nTMhelix 643 665\ninside 666 673

31934	GCF_002189495.1_ASM218949v1	Arthrobacter globiformis	Terrabacteria group; Actinobacteria; Actinobacteria; Micrococcales; Micrococcaceae; Arthrobacter	MPQRVARAVGSSPVSEKLATAQELLYGPIPAWARRSPFHTGVLGHSAHPPLTDVTIGCWT	WP_087873679.1	hypothetical protein [Arthrobacter globiformis]	Length: 181\nNumber of predicted TMHs: 2\nExp number of AAs in TMHs: 50.95853\nExp number, first 60 AAs: 0.20239\nTotal prob of N-in: 0.81578\ninside 1 108\nTMhelix 109 131\noutside 132 140\nTMhelix 141 163\ninside 164 181

31935	GCF_000482545.1_ASM48254v1	Arthrobacter sp. 35W	Terrabacteria group; Actinobacteria; Actinobacteria; Micrococcales; Micrococcaceae; Arthrobacter	MSPTRPRRRLMGMGVAGALLAGTVIPLAMAAPALAVPGIATITSANLSVAVSTAFPQATS	WP_081709824.1	hypothetical protein [Arthrobacter sp. 35W]	Length: 1943\nNumber of predicted TMHs: 2\nExp number of AAs in TMHs: 45.0607099999999\nExp number, first 60 AAs: 22.72862\nTotal prob of N-in: 0.98964\nPOSSIBLE N-term signal sequence\ninside 1 12\nTMhelix 13 35\noutside 36 1912\nTMhelix 1913 1935\ninside 1936 1943

31936	GCF_000755585.2_ASM75558v2	Arthrobacter sp. A3Terrabacteria group; Actinobacteria; Actinobacteria; Micrococcales; Micrococcaceae; Arthrobacter	MQVSRTPLGVAGVHPRRRGLVSLFTLFLCVAFVTLGLSAVRILGSLTPASANPGTPGVTQ WP_083508733.1	hypothetical protein [Arthrobacter sp. A3] Length: 917\nNumber of predicted TMHs: 2\nExp number of AAs in TMHs: 46.68582\nExp number, first 60 AAs: 22.03027\nTotal prob of N-in: 0.99894\nPOSSIBLE N-term signal sequence\ninside 1 19\nTMhelix 20 42\noutside 43 883\nTMhelix 884 906\ninside 907 917

31937	GCF_001512285.1_ASM151228v1	Arthrobacter sp. EpRS71	Terrabacteria group; Actinobacteria; Actinobacteria; Micrococcales; Micrococcaceae; Arthrobacter	MSVLPSWGRFGRSEKMPGVRRDSSARPRIPPSRRRSILQIPAAALAVAVSAGSLMLGGTP	WP_062072668.1	hypothetical protein [Arthrobacter sp. EpRS71]	Length: 1337\nNumber of predicted TMHs: 1\nExp number of AAs in TMHs: 33.78389\nExp number, first 60 AAs: 10.09686\nTotal prob of N-in: 0.50185\nPOSSIBLE N-term signal sequence\noutside 1 1308\nTMhelix 1309 1331\ninside 1332 1337

31938	GCF_001263615.1_ASM126361v1	Arthrobacter sp. Hiyo1	Terrabacteria group; Actinobacteria; Actinobacteria; Micrococcales; Micrococcaceae; Arthrobacter	MGSAFAAIAVSLVLVFAAILGLAYQGRTRRRSILRTLRGPLGSNGPMNNTSRGEAAGTLR WP_059389499.1	secretion system protein [Arthrobacter sp. Hiyo1]	Length: 210\nNumber of predicted TMHs: 2\nExp number of AAs in TMHs: 53.62104\nExp number, first 60 AAs: 21.5588\nTotal prob of N-in: 0.02546\nPOSSIBLE N-term signal sequence\noutside 1 3\nTMhelix 4 26\ninside 27 180\nTMhelix 181 203\noutside 204 210

31939 GCF_001807395.1_ASM180739v1 Arthrobacter sp. HMSC06H05 Terrabacteria group; Actinobacteria; Actinobacteria; Micrococcales; Micrococcaceae; Arthrobacter
MKALSQRRRSPLALLALLVLGLVVTGGVYTVAGSVTSANAEAQATAEGDVENGSKLFSAN WP_035755245.1
MULTISPECIES: cystathionine beta-lyase [Micrococcaceae] Length: 266\nNumber of predicted TMHs: 2\nExp number of AAs in TMHs: 43.13616\nExp number, first 60 AAs: 21.97996\nTotal prob of N-in: 0.99736\nPOSSIBLE N-term signal sequence\ninside 1 12\nTMhelix 13 35\noutside 36 238\nTMhelix 239 261\ninside 262 266

31940 GCF_001807575.1_ASM180757v1 Arthrobacter sp. HMSC08H08 Terrabacteria group; Actinobacteria; Actinobacteria; Micrococcales; Micrococcaceae; Arthrobacter
MKALSQRRRSPLALLALLVLGLVVTGGVYTVAGSVTSANAEAQATAEGDVENGSKLFSAN WP_035755245.1
MULTISPECIES: cystathionine beta-lyase [Micrococcaceae] Length: 266\nNumber of predicted TMHs: 2\nExp number of AAs in TMHs: 43.13616\nExp number, first 60 AAs: 21.97996\nTotal prob of N-in: 0.99736\nPOSSIBLE N-term signal sequence\ninside 1 12\nTMhelix 13 35\noutside 36 238\nTMhelix 239 261\ninside 262 266

31941 GCF_000950575.1_ASM95057v1 Arthrobacter sp. IHBB 11108 Terrabacteria group; Actinobacteria; Actinobacteria; Micrococcales; Micrococcaceae; Arthrobacter
MATTTRRSTLRVTTAMALSVLSLLGVLSVASVTPASADEMRDNQYWLDEYGITKAWESTK WP_052663444.1
peptidase S8 [Arthrobacter sp. IHBB 11108] Length: 439\nNumber of predicted TMHs: 2\nExp number of AAs in TMHs: 44.42909\nExp number, first 60 AAs: 21.11609\nTotal prob of N-in: 0.94984\nPOSSIBLE N-term signal sequence\ninside 1 11\nTMhelix 12 34\noutside 35 402\nTMhelix 403 425\ninside 426 439

31942 GCF_002008315.1_ASM200831v1 Arthrobacter sp. KBS0703 Terrabacteria group; Actinobacteria; Actinobacteria; Micrococcales; Micrococcaceae; Arthrobacter
MTGAILSAAITAMVLAAAAALAFTGHSRTRRRLLSGNGLILNTGTHGGASGRPAPTGQGT WP_078026535.1
secretion system protein [Arthrobacter sp. KBS0703] Length: 204\nNumber of predicted TMHs: 2\nExp number of AAs in TMHs: 59.79683\nExp number, first 60 AAs: 22.52593\nTotal prob of N-in: 0.40974\nPOSSIBLE N-term signal sequence\noutside 1 3\nTMhelix 4 23\ninside 24 174\nTMhelix 175 197\noutside 198 204

31943 GCF_001422665.1_Leaf234 Arthrobacter sp. Leaf234 Terrabacteria group; Actinobacteria; Actinobacteria; Micrococcales; Micrococcaceae; Arthrobacter
MPSPTSTSRITARRAAMHRTAARAAAVLAAVIVGGAGLLALPASGAVVPAVSSSAQAAVL WP_055769734.1
hypothetical protein [Arthrobacter sp. Leaf234] Length: 163\nNumber of predicted TMHs: 2\nExp number of AAs in TMHs: 45.19227\nExp number, first 60 AAs: 22.71823\nTotal prob of N-in: 0.99974\nPOSSIBLE N-term signal sequence\ninside 1 20\nTMhelix 21 43\noutside 44 129\nTMhelix 130 152\ninside 153 163

31944 GCF_000281065.1_ASM28106v1 Arthrobacter sp. M2012083 Terrabacteria group; Actinobacteria; Actinobacteria; Micrococcales; Micrococcaceae; Arthrobacter
MHHSSVRRNPGTPSLGSARRRTTVAVGAASGVLLLAGAIMVQQSPAFAATPVNLGTATSY WP_026005431.1
sortase [Arthrobacter sp. M2012083] Length: 434\nNumber of predicted TMHs: 2\nExp number of AAs in TMHs: 55.5954900000001\nExp number, first 60 AAs: 21.14515\nTotal prob of N-in: 0.92508\nPOSSIBLE N-term signal sequence\ninside 1 23\nTMhelix 24 41\noutside 42 401\nTMhelix 402 424\ninside 425 434

31945 GCF_001969265.1_ASM196926v1 Arthrobacter sp. QXT-31 Terrabacteria group; Actinobacteria; Actinobacteria; Micrococcales; Micrococcaceae; Arthrobacter
MPQRVARAVGSSPVSEKLAAAQELLYGPIPAWARRSLFHTGVLGHSAHPPLTDVTIGCWT WP_076799932.1
hypothetical protein [Arthrobacter sp. QXT-31] Length: 181\nNumber of predicted TMHs: 2\nExp number of AAs in TMHs: 51.09886\nExp number, first 60 AAs: 0.27352\nTotal prob of N-in: 0.81629\ninside 1 108\nTMhelix 109 131\noutside 132 140\nTMhelix 141 163\ninside 164 181

31946 GCF_001969265.1_ASM196926v1 Arthrobacter sp. QXT-31 Terrabacteria group; Actinobacteria; Actinobacteria; Micrococcales; Micrococcaceae; Arthrobacter
MPYDAGRMRARRGTVLLALLLGLAATVIPTAPASAHTYLESSDPAPDAQLSAAPDRVSLI WP_083706190.1 hypothetical protein [Arthrobacter sp. QXT-31] Length: 208\nNumber of predicted TMHs: 2\nExp number of AAs in TMHs: 44.40544\nExp number, first 60 AAs: 22.02418\nTotal prob of N-in: 0.99884\nPOSSIBLE N-term signal sequence\ninside 1 12\nTMhelix 13 35\noutside 36 175\nTMhelix 176 198\ninside 199 208

31947 GCF_000294695.2_ASM29469v2 Arthrobacter sp. Rue61a Terrabacteria group; Actinobacteria; Actinobacteria; Micrococcales; Micrococcaceae; Arthrobacter
MHHSSARKDSRTPWMGSRRRRRTVSVGAASGILLLAGAIILQQSPAFAATPVNLGTAASY WP_014920658.1
hypothetical protein [Arthrobacter sp. Rue61a] Length: 390\nNumber of predicted TMHs: 2\nExp number of

AAs in TMHs: 64.01804\nExp number, first 60 AAs: 24.51191\nTotal prob of N-in: 0.99583\nPOSSIBLE N-term signal sequence\ninside 1 23\nTMhelix 24 46\noutside 47 357\nTMhelix 358 380\ninside 381 390

31948 GCF_001428005.1_Soil736 Arthrobacter sp. Soil736 Terrabacteria group; Actinobacteria; Actinobacteria; Micrococcales; Micrococcaceae; Arthrobacter
MTGPILSAALVAMVLAAAAALAFTGHTRTRRRLLGRSGLIPNAATHGGASGSAPAGHGTG WP_056634648.1
secretion system protein [Arthrobacter sp. Soil736] Length: 203\nNumber of predicted TMHs: 2\nExp number of AAs in TMHs: 58.47437\nExp number, first 60 AAs: 23.12341\nTotal prob of N-in: 0.28475\nPOSSIBLE N-term signal sequence\noutside 1 4\nTMhelix 5 27\ninside 28 173\nTMhelix 174 196\noutside 197 203

31949 GCF_002009585.1_ASM200958v1 Arthrobacter sp. SRS-W-1-2016 Terrabacteria group; Actinobacteria; Actinobacteria; Micrococcales; Micrococcaceae; Arthrobacter
MLQTRSSLRWVGVERRRGFLGEKVKIWSLAVVAATAFAYLGAVPAMAAPSDVLDVSGVPV WP_078106369.1
hypothetical protein [Arthrobacter sp. SRS-W-1-2016] Length: 416\nNumber of predicted TMHs: 2\nExp number of AAs in TMHs: 45.78508\nExp number, first 60 AAs: 21.8762\nTotal prob of N-in: 0.97862\nPOSSIBLE N-term signal sequence\ninside 1 25\nTMhelix 26 48\noutside 49 373\nTMhelix 374 396\ninside 397 416

31950 GCF_002009585.1_ASM200958v1 Arthrobacter sp. SRS-W-1-2016 Terrabacteria group; Actinobacteria; Actinobacteria; Micrococcales; Micrococcaceae; Arthrobacter
MKSSRRRTLKTASAPMMAAGLLAFGVSAASAHVNVNPDDPAAGGYTHLTFNVPNESATAK WP_078105861.1
nuclear export factor GLE1 [Arthrobacter sp. SRS-W-1-2016] Length: 240\nNumber of predicted TMHs: 2\nExp number of AAs in TMHs: 40.09707\nExp number, first 60 AAs: 17.10686\nTotal prob of N-in: 0.90299\nPOSSIBLE N-term signal sequence\ninside 1 12\nTMhelix 13 35\noutside 36 209\nTMhelix 210 232\ninside 233 240

31951 GCF_001750145.1_ASM175014v1 Arthrobacter sp. U41 Terrabacteria group; Actinobacteria; Actinobacteria; Micrococcales; Micrococcaceae; Arthrobacter
MVSSAPSHAQSRVESGDGSGTRRPRVRRRTAVGTAMGASALLAWALVGQSPAYAADAPVG WP_083266569.1
hypothetical protein [Arthrobacter sp. U41] Length: 341\nNumber of predicted TMHs: 1\nExp number of AAs in TMHs: 36.70894\nExp number, first 60 AAs: 14.55491\nTotal prob of N-in: 0.69458\nPOSSIBLE N-term signal sequence\noutside 1 309\nTMhelix 310 332\ninside 333 341

31952 GCF_900105345.1_IMG-taxon_2639762519_annotated_assembly Arthrobacter woluwensis Terrabacteria group; Actinobacteria; Actinobacteria; Micrococcales; Micrococcaceae; Arthrobacter
MLLRTLPSVLRRRAFIAAVTMASLAAVSLPLTIPADAQAAPPPSNGNAWGPDGTFDSDSV WP_066212419.1
hypothetical protein [Arthrobacter woluwensis] Length: 927\nNumber of predicted TMHs: 1\nExp number of AAs in TMHs: 34.71802\nExp number, first 60 AAs: 12.2401\nTotal prob of N-in: 0.55122\nPOSSIBLE N-term signal sequence\noutside 1 895\nTMhelix 896 918\ninside 919 927

31953 GCF_001552095.1_ASM155209v1 Arthrobacter woluwensis NBRC 107840 Terrabacteria group; Actinobacteria; Actinobacteria; Micrococcales; Micrococcaceae; Arthrobacter; Arthrobacter woluwensis
MLLRTLPSVLRRRAFIAAVTMASLAAVSLPLTIPADAQAAPPPSNGNAWGPDGTFDSDSV WP_066212419.1
hypothetical protein [Arthrobacter woluwensis] Length: 927\nNumber of predicted TMHs: 1\nExp number of AAs in TMHs: 34.71802\nExp number, first 60 AAs: 12.2401\nTotal prob of N-in: 0.55122\nPOSSIBLE N-term signal sequence\noutside 1 895\nTMhelix 896 918\ninside 919 927

31954 GCF_000688395.1_ASM68839v1 Haematomicrobium sanguinis DSM 21259 Terrabacteria group; Actinobacteria; Actinobacteria; Micrococcales; Micrococcaceae; Haematomicrobium; Haematomicrobium sanguinis
MKYESVPSSRASAKRRLGRRGIVAAATSILTTLALLPGIAIAAPSDLGPDPAQDTASTSS WP_026529730.1 hypothetical protein [Haematomicrobium sanguinis]Length: 881\nNumber of predicted TMHs: 2\nExp number of AAs in TMHs: 44.64672\nExp number, first 60 AAs: 22.73491\nTotal prob of N-in: 0.99889\nPOSSIBLE N-term signal sequence\ninside 1 20\nTMhelix 21 43\noutside 44 851\nTMhelix 852 874\ninside 875 881

31955 GCF_001887245.1_ASM188724v1 Neomicrococcus aestuarii Terrabacteria group; Actinobacteria; Actinobacteria; Micrococcales; Micrococcaceae; Neomicrococcus
MSHHDSAEISRNSKPSRRRVVSAAAWSAPVIAAAGVAPFSAASTPAGLTVGWTSQYSENL WP_071894446.1
hypothetical protein [Neomicrococcus aestuarii] Length: 232\nNumber of predicted TMHs: 2\nExp number of AAs in TMHs: 45.05142\nExp number, first 60 AAs: 22.19879\nTotal prob of N-in: 0.97894\nPOSSIBLE N-term signal sequence\ninside 1 19\nTMhelix 20 42\noutside 43 172\nTMhelix 173 195\ninside 196 232

31956 GCF_001483765.1_ASM148376v1 Nesterenkonia jeotgali Terrabacteria group; Actinobacteria; Actinobacteria; Micrococcales; Micrococcaceae; Nesterenkonia

MTTSRRRTIGAGATLGAAVLLLSPTMAHASTEAGSEVEWPEEFTSAYTVMATPDEVINSD WP_058888770.1 hypothetical protein [Nesterenkonia jeotgali] Length: 238\nNumber of predicted TMHs: 1\nExp number of AAs in TMHs: 31.18607\nExp number, first 60 AAs: 9.17231\nTotal prob of N-in: 0.63113\noutside 1 209\nTMhelix 210 232\ninside 233 238

31957 GCF_000582475.1_NestAN1.1 Nesterenkonia sp. AN1 Terrabacteria group; Actinobacteria; Actinobacteria; Micrococcales; Micrococcaceae; Nesterenkonia MMKQRRRTIGAGATLGAAVLLLSPTMAQASTEAGTEVEWPEEFTSAYTVMATPDEVINSD WP_036474851.1 hypothetical protein [Nesterenkonia sp. AN1] Length: 238\nNumber of predicted TMHs: 1\nExp number of AAs in TMHs: 29.44918\nExp number, first 60 AAs: 8.23727\nTotal prob of N-in: 0.61699\noutside 1 209\nTMhelix 210 232\ninside 233 238

31958 GCF_000220985.1_ASM22098v2 Nesterenkonia sp. F Terrabacteria group; Actinobacteria; Actinobacteria; Micrococcales; Micrococcaceae; Nesterenkonia MTSWTASPSRWPPVASPRSPVDRRTSQTMSPMPPQSPEPPRRRRPLRLLRSTAVVGALVP WP_083830309.1 hypothetical protein [Nesterenkonia sp. F] Length: 537\nNumber of predicted TMHs: 1\nExp number of AAs in TMHs: 25.86672\nExp number, first 60 AAs: 2.02461\nTotal prob of N-in: 0.17255\noutside 1 506\nTMhelix 507 529\ninside 530 537

31959 GCF_000220985.1_ASM22098v2 Nesterenkonia sp. F Terrabacteria group; Actinobacteria; Actinobacteria; Micrococcales; Micrococcaceae; Nesterenkonia MDHHTPDTTPRRRGPRRAALAGLAGASALALGTVTAAPAWAHDELISATPESGQILTEIP WP_010525312.1 hypothetical protein [Nesterenkonia sp. F] Length: 267\nNumber of predicted TMHs: 2\nExp number of AAs in TMHs: 44.34944\nExp number, first 60 AAs: 22.05365\nTotal prob of N-in: 0.98899\nPOSSIBLE N-term signal sequence\ninside 1 19\nTMhelix 20 42\noutside 43 218\nTMhelix 219 241\ninside 242 267

31960 GCF_000220985.1_ASM22098v2 Nesterenkonia sp. F Terrabacteria group; Actinobacteria; Actinobacteria; Micrococcales; Micrococcaceae; Nesterenkonia MSTFDTQPRRTALAVTGGLSLVVGLVLLSPAAAVATTTATPTATASPSSTPTADGSESPS WP_010524920.1 hypothetical protein [Nesterenkonia sp. F] Length: 438\nNumber of predicted TMHs: 2\nExp number of AAs in TMHs: 44.01765\nExp number, first 60 AAs: 22.52268\nTotal prob of N-in: 0.98559\nPOSSIBLE N-term signal sequence\ninside 1 12\nTMhelix 13 35\noutside 36 410\nTMhelix 411 433\ninside 434 438

31961 GCF_000514015.1_ASM51401v1 Paenarthrobacter nicotinovorans 231Sha2.1M6 Terrabacteria group; Actinobacteria; Actinobacteria; Micrococcales; Micrococcaceae; Paenarthrobacter; Paenarthrobacter nicotinovorans MSIPDGGTRHVLDSNRWFSFRVERRAMSDEVTVSKPHSRRAVVKGAAWSVPVIAAAVAAP WP_051421565.1 hypothetical protein [Paenarthrobacter nicotinovorans] Length: 239\nNumber of predicted TMHs: 1\nExp number of AAs in TMHs: 52.79679\nExp number, first 60 AAs: 12.04975\nTotal prob of N-in: 0.58689\nPOSSIBLE N-term signal sequence\noutside 1 196\nTMhelix 197 219\ninside 220 239

31962 GCF_000758985.1_04_NF40_HMP2128v01 Pseudoglutamicibacter albus DNF00011 Terrabacteria group; Actinobacteria; Actinobacteria; Micrococcales; Micrococcaceae; Pseudoglutamicibacter; Pseudoglutamicibacter albus MKALSQRRRSPLALLALLVLGLVVTGGVYTVAGSVTSANAEAQATAEGDVENGSKLFSAN WP_035755245.1 MULTISPECIES: cystathionine beta-lyase [Micrococcaceae] Length: 266\nNumber of predicted TMHs: 2\nExp number of AAs in TMHs: 43.13616\nExp number, first 60 AAs: 21.97996\nTotal prob of N-in: 0.99736\nPOSSIBLE N-term signal sequence\ninside 1 12\nTMhelix 13 35\noutside 36 238\nTMhelix 239 261\ninside 262 266

31963 GCF_001065485.1_ASM106548v1 Rothia mucilaginosa Terrabacteria group; Actinobacteria; Actinobacteria; Micrococcales; Micrococcaceae; Rothia MQRRKVLITSLTLAAAAGAISCTAATAAETPTQSGSITIRDNGQPNRDQTRDLTVALIIK WP_049347585.1 hypothetical protein [Rothia mucilaginosa] Length: 386\nNumber of predicted TMHs: 2\nExp number of AAs in TMHs: 36.37783\nExp number, first 60 AAs: 15.53159\nTotal prob of N-in: 0.70919\nPOSSIBLE N-term signal sequence\ninside 1 6\nTMhelix 7 29\noutside 30 357\nTMhelix 358 380\ninside 381 386

31964 GCF_001061655.1_ASM106165v1 Rothia mucilaginosa Terrabacteria group; Actinobacteria; Actinobacteria; Micrococcales; Micrococcaceae; Rothia MQRRKVLITSLTLAAAAGAISCTAATAAETPTQSGSITIRDNGQPNRDQTRSLTVALITK WP_049350406.1 hypothetical protein [Rothia mucilaginosa] Length: 385\nNumber of predicted TMHs: 2\nExp number of AAs in TMHs: 36.09891\nExp number, first 60 AAs: 15.55229\nTotal prob of N-in: 0.70904\nPOSSIBLE N-term signal sequence\ninside 1 6\nTMhelix 7 29\noutside 30 358\nTMhelix 359 378\ninside 379 385

31965 GCF_002087015.1_ASM208701v1 Rothia nasimurium Terrabacteria group; Actinobacteria; Actinobacteria; Micrococcales; Micrococcaceae; Rothia MERRKFLVGATVATAIAVTSSKANSAEISVIDPDIKLFIRENPELENSVTELLRKGEISQ WP_083090608.1 hypothetical protein [Rothia nasimurium] Length: 169\nNumber of predicted TMHs: 2\nExp number of AAs in TMHs: 37.77653\nExp number, first 60 AAs: 0.09024\nTotal prob of N-in: 0.90379\ninside 1 100\nTMhelix 101 123\noutside 124 132\nTMhelix 133 155\ninside 156 169

31966 GCF_001814485.1_ASM181448v1 Rothia sp. HMSC066G07 Terrabacteria group; Actinobacteria; Actinobacteria; Micrococcales; Micrococcaceae; Rothia MQRRKVLITSLTLAAAAGAISCTAATAAETPTQSGSITIRDNGQPNRDQTRYLTVALITK WP_070690408.1 hypothetical protein [Rothia sp. HMSC066G07] Length: 368\nNumber of predicted TMHs: 2\nExp number of AAs in TMHs: 35.6763\nExp number, first 60 AAs: 15.47275\nTotal prob of N-in: 0.71087\nPOSSIBLE N-term signal sequence\ninside 1 6\nTMhelix 7 29\noutside 30 339\nTMhelix 340 362\ninside 363 368

31967 GCF_001813725.1_ASM181372v1 Rothia sp. HMSC072B04 Terrabacteria group; Actinobacteria; Actinobacteria; Micrococcales; Micrococcaceae; Rothia MSATMNRRTAFRALGVGTATIALGCAAAPAAQAQQMAHPQKAMTADQQLKLILRRLEQVP WP_070688290.1 hypothetical protein [Rothia sp. HMSC072B04] Length: 173\nNumber of predicted TMHs: 2\nExp number of AAs in TMHs: 41.5834\nExp number, first 60 AAs: 19.54519\nTotal prob of N-in: 0.96972\nPOSSIBLE N-term signal sequence\ninside 1 12\nTMhelix 13 35\noutside 36 116\nTMhelix 117 139\ninside 140 173

31968 GCF_001722485.1_ASM172248v1 Cellulosimicrobium cellulans Terrabacteria group; Actinobacteria; Actinobacteria; Micrococcales; Promicromonosporaceae; Cellulosimicrobium MTHATTRRRARRRAALAGGLALAASLLAVPAAHAAPTVAQIDVIAPASVDAGAPIEVTVA WP_069385866.1 hypothetical protein [Cellulosimicrobium cellulans] Length: 252\nNumber of predicted TMHs: 2\nExp number of AAs in TMHs: 45.48942\nExp number, first 60 AAs: 21.98818\nTotal prob of N-in: 0.99329\nPOSSIBLE N-term signal sequence\ninside 1 12\nTMhelix 13 35\noutside 36 221\nTMhelix 222 244\ninside 245 252

31969 GCF_002162335.1_ASM216233v1 Cellulosimicrobium cellulans Terrabacteria group; Actinobacteria; Actinobacteria; Micrococcales; Promicromonosporaceae; Cellulosimicrobium MSTTTGRRTLVRATTAAALAASAALLAGPAAHAAEDDTYVPSQPSMRLTTLVSACEQDAA WP_087470543.1 hypothetical protein [Cellulosimicrobium cellulans] Length: 224\nNumber of predicted TMHs: 1\nExp number of AAs in TMHs: 36.74159\nExp number, first 60 AAs: 14.2507\nTotal prob of N-in: 0.67474\nPOSSIBLE N-term signal sequence\noutside 1 195\nTMhelix 196 218\ninside 219 224

31970 GCF_002162335.1_ASM216233v1 Cellulosimicrobium cellulans Terrabacteria group; Actinobacteria; Actinobacteria; Micrococcales; Promicromonosporaceae; Cellulosimicrobium MTRSPSRRRLGRRAALVGALAVAATALTVPAHAAPSASDATLDAQSTAEVGTSLTVTVAT WP_087472132.1 hypothetical protein [Cellulosimicrobium cellulans] Length: 247\nNumber of predicted TMHs: 2\nExp number of AAs in TMHs: 42.64075\nExp number, first 60 AAs: 20.10343\nTotal prob of N-in: 0.91703\nPOSSIBLE N-term signal sequence\ninside 1 12\nTMhelix 13 35\noutside 36 216\nTMhelix 217 239\ninside 240 247

31971 GCF_001722485.1_ASM172248v1 Cellulosimicrobium cellulans Terrabacteria group; Actinobacteria; Actinobacteria; Micrococcales; Promicromonosporaceae; Cellulosimicrobium MPAQMRARASLLTMRRSRLGKTGPLSHRRPEENTMNIRRRAVVALAAAAGALALAAGPAA WP_083259649.1 hypothetical protein [Cellulosimicrobium cellulans] Length: 252\nNumber of predicted TMHs: 2\nExp number of AAs in TMHs: 43.27904\nExp number, first 60 AAs: 19.34825\nTotal prob of N-in: 0.99564\nPOSSIBLE N-term signal sequence\ninside 1 40\nTMhelix 41 63\noutside 64 223\nTMhelix 224 246\ninside 247 252

31972 GCF_001019615.1_ASM101961v1 Cellulosimicrobium funkei Terrabacteria group; Actinobacteria; Actinobacteria; Micrococcales; Promicromonosporaceae; Cellulosimicrobium MSTTTGRRTLVRATTAAALAASAALLAGPAAHAAEDDTYVPSQPSMRLTTLVSACEQDAA WP_047233459.1 hypothetical protein [Cellulosimicrobium funkei] Length: 224\nNumber of predicted TMHs: 1\nExp number of AAs in TMHs: 36.66786\nExp number, first 60 AAs: 14.17403\nTotal prob of N-in: 0.67653\nPOSSIBLE N-term signal sequence\noutside 1 195\nTMhelix 196 218\ninside 219 224

31973 GCF_001019615.1_ASM101961v1 Cellulosimicrobium funkei Terrabacteria group; Actinobacteria; Actinobacteria; Micrococcales; Promicromonosporaceae; Cellulosimicrobium MTPSPSRRRLGRRAALVGGLAVAATALTAPAHAAPSVSAVALDAPSSVEVGAPITVTVAA WP_047234617.1 hypothetical protein [Cellulosimicrobium funkei] Length: 248\nNumber of predicted TMHs: 1\nExp number of AAs in TMHs: 46.85658\nExp number, first 60 AAs: 17.47946\nTotal prob of N-in: 0.70962\nPOSSIBLE N-term signal sequence\noutside 1 217\nTMhelix 218 240\ninside 241 248

31974 GCF_001647255.1_ASM164725v1 Cellulosimicrobium sp. I38E Terrabacteria group; Actinobacteria; Actinobacteria; Micrococcales; Promicromonosporaceae; Cellulosimicrobium
MTSTTAVAHGRRRPSTRPDRRRLLAGLVAALLLGVVLVPVAPPPTASAATLEPSGLQTGM WP_064316121.1 hypothetical protein [Cellulosimicrobium sp. I38E] Length: 1489\nNumber of predicted TMHs: 2\nExp number of AAs in TMHs: 42.82173\nExp number, first 60 AAs: 21.35008\nTotal prob of N-in: 0.99743\nPOSSIBLE N-term signal sequence\ninside 1 20\nTMhelix 21 43\noutside 44 1460\nTMhelix 1461 1483\ninside 1484 1489

31975 GCF_001647255.1_ASM164725v1 Cellulosimicrobium sp. I38E Terrabacteria group; Actinobacteria; Actinobacteria; Micrococcales; Promicromonosporaceae; Cellulosimicrobium
MTPSPSRRRLGRRAALVGGLAVAATALTAPAHAAPSVSAVALDAPSSVEVGAPITVTVAA WP_064315323.1 hypothetical protein [Cellulosimicrobium sp. I38E] Length: 248\nNumber of predicted TMHs: 1\nExp number of AAs in TMHs: 42.42507\nExp number, first 60 AAs: 15.5747\nTotal prob of N-in: 0.67461\nPOSSIBLE N-term signal sequence\noutside 1 217\nTMhelix 218 240\ninside 241 248

31976 GCF_001647255.1_ASM164725v1 Cellulosimicrobium sp. I38E Terrabacteria group; Actinobacteria; Actinobacteria; Micrococcales; Promicromonosporaceae; Cellulosimicrobium
MSTTTGRRTLVRATTAAALAASAALLAGPAAHAAEDDTYVPSQPSMRLTTLVSACEQDAA WP_064317010.1 hypothetical protein [Cellulosimicrobium sp. I38E] Length: 224\nNumber of predicted TMHs: 1\nExp number of AAs in TMHs: 36.76089\nExp number, first 60 AAs: 14.26765\nTotal prob of N-in: 0.67330\nPOSSIBLE N-term signal sequence\noutside 1 195\nTMhelix 196 218\ninside 219 224

31977 GCF_001274015.1_ASM127401v1 Cellulosimicrobium cellulans F16 Terrabacteria group; Actinobacteria; Actinobacteria; Micrococcales; Promicromonosporaceae; Cellulosimicrobium; Cellulosimicrobium cellulans
MTPSPSRRRLGRRAALVGALAVAATTLTAPAHAAPSVSAVALDAPSSVEVGAPITVTVAA WP_053370785.1 hypothetical protein [Cellulosimicrobium cellulans] Length: 248\nNumber of predicted TMHs: 1\nExp number of AAs in TMHs: 37.4679\nExp number, first 60 AAs: 13.54758\nTotal prob of N-in: 0.59979\nPOSSIBLE N-term signal sequence\noutside 1 217\nTMhelix 218 240\ninside 241 248

31978 GCF_001274015.1_ASM127401v1 Cellulosimicrobium cellulans F16 Terrabacteria group; Actinobacteria; Actinobacteria; Micrococcales; Promicromonosporaceae; Cellulosimicrobium; Cellulosimicrobium cellulans
MSTTTGRRTLVRATPAAALAASAALLAGPAAHAAEDDTYVPSQPSMRLTTLVSACEQDAA WP_053369103.1 hypothetical protein [Cellulosimicrobium cellulans] Length: 224\nNumber of predicted TMHs: 1\nExp number of AAs in TMHs: 31.86243\nExp number, first 60 AAs: 9.38835\nTotal prob of N-in: 0.47629\noutside 1 195\nTMhelix 196 218\ninside 219 224

31979 GCF_001750215.1_ASM175021v1 Isoptericola variabilis Terrabacteria group; Actinobacteria; Actinobacteria; Micrococcales; Promicromonosporaceae; Isoptericola
MNSTARRSAVIVGAAGLVALGALAAPASADTAENASLSVLHGVPDTPVDVWVNDERTLDD WP_069984382.1 hypothetical protein [Isoptericola variabilis] Length: 298\nNumber of predicted TMHs: 2\nExp number of AAs in TMHs: 42.69521\nExp number, first 60 AAs: 19.90674\nTotal prob of N-in: 0.92479\nPOSSIBLE N-term signal sequence\ninside 1 6\nTMhelix 7 29\noutside 30 250\nTMhelix 251 273\ninside 274 298

31980 GCF_001636295.1_ASM163629v1 Isoptericola dokdonensis DS-3 Terrabacteria group; Actinobacteria; Actinobacteria; Micrococcales; Promicromonosporaceae; Isoptericola; Isoptericola dokdonensis
MEADRRRRRAGARVTRTRRQWVTLGLGAVVLAALCVTAAFWQWHRYTDREAQIALVEANY WP_083973524.1 hypothetical protein [Isoptericola dokdonensis] Length: 303\nNumber of predicted TMHs: 2\nExp number of AAs in TMHs: 42.74295\nExp number, first 60 AAs: 22.71384\nTotal prob of N-in: 1.00000\nPOSSIBLE N-term signal sequence\ninside 1 20\nTMhelix 21 43\noutside 44 234\nTMhelix 235 254\ninside 255 303

31981 GCF_000215105.1_ASM21510v1 Isoptericola variabilis 225 Terrabacteria group; Actinobacteria; Actinobacteria; Micrococcales; Promicromonosporaceae; Isoptericola; Isoptericola variabilis
MNHARRRRTLASTLVGILVAAFAAVGLSTAPVSAQPAERGPVERITYCHATGSETNPYVL WP_013840046.1 hypothetical protein [Isoptericola variabilis] Length: 299\nNumber of predicted TMHs: 2\nExp number of AAs in TMHs: 44.17863\nExp number, first 60 AAs: 21.80007\nTotal prob of N-in: 0.99261\nPOSSIBLE N-term signal sequence\ninside 1 11\nTMhelix 12 34\noutside 35 268\nTMhelix 269 291\ninside 292 299

31982 GCF_000215105.1_ASM21510v1 Isoptericola variabilis 225 Terrabacteria group; Actinobacteria; Actinobacteria; Micrococcales; Promicromonosporaceae; Isoptericola; Isoptericola variabilis
MTSSSGVRRRPWLAAPLAVAAALALSGCMKMDLALTLTEGDTASGTFVVAFSDELAEAMG WP_013837494.1 hypothetical protein [Isoptericola variabilis] Length: 257\nNumber of predicted TMHs: 1\nExp number of

AAs in TMHs: 36.25591\nExp number, first 60 AAs: 13.53188\nTotal prob of N-in: 0.65373\nPOSSIBLE N-term signal sequence\noutside 1 218\nTMhelix 219 241\ninside 242 257

31983 GCF_900167525.1_IMG-taxon_2596583702_annotated_assembly Krasilnikoviella flava Terrabacteria group; Actinobacteria; Actinobacteria; Micrococcales; Promicromonosporaceae; Krasilnikoviella MTENSASTRASSVPPVREAEIDEATHRRNPVPTQAGTGAVHLGQVPAPPAPPTVAPASSS WP_079574122.1 hypothetical protein [Krasilnikoviella flava] Length: 285\nNumber of predicted TMHs: 2\nExp number of AAs in TMHs: 51.92438\nExp number, first 60 AAs: 0.00165\nTotal prob of N-in: 0.67711\ninside 1 185\nTMhelix 186 208\noutside 209 245\nTMhelix 246 268\ninside 269 285

31984 GCF_900167525.1_IMG-taxon_2596583702_annotated_assembly Krasilnikoviella flava Terrabacteria group; Actinobacteria; Actinobacteria; Micrococcales; Promicromonosporaceae; Krasilnikoviella MHPQTARRRRALVALLAVALAPVLGVAVAQPASAHDRLLSSDPADGASVDGAPTEIALVF WP_079573477.1 hypothetical protein [Krasilnikoviella flava] Length: 256\nNumber of predicted TMHs: 2\nExp number of AAs in TMHs: 44.3943\nExp number, first 60 AAs: 21.723\nTotal prob of N-in: 0.99965\nPOSSIBLE N-term signal sequence\ninside 1 11\nTMhelix 12 34\noutside 35 185\nTMhelix 186 208\ninside 209 256

31985 GCF_000385135.1_ASM38513v1 Promicromonospora sukumoe 327MFSha3.1 Terrabacteria group; Actinobacteria; Actinobacteria; Micrococcales; Promicromonosporaceae; Promicromonospora; Promicromonospora sukumoe MRVISLLRSAFAGPRRGPVGALAILTVIVGVVTMHSMSGSPTSHGHAVPHETSVAAVEQA WP_020014906.1 hypothetical protein [Promicromonospora sukumoe] Length: 164\nNumber of predicted TMHs: 2\nExp number of AAs in TMHs: 41.9375\nExp number, first 60 AAs: 20.57821\nTotal prob of N-in: 0.99268\nPOSSIBLE N-term signal sequence\ninside 1 16\nTMhelix 17 39\noutside 40 106\nTMhelix 107 126\ninside 127 164

31986 GCF_000307245.1_ASM30724v1 Promicromonosporaceae bacterium W15 Terrabacteria group; Actinobacteria; Actinobacteria; Micrococcales; Promicromonosporaceae; unclassified Promicromonosporaceae MTGRRRLVRTWATGAVVALVMSSAVAVSGAGPASADPPTGGSTIVDETFTNPTVPDDAWQ WP_036947581.1 hypothetical protein [Promicromonosporaceae bacterium W15] Length: 1535\nNumber of predicted TMHs: 2\nExp number of AAs in TMHs: 42.19932\nExp number, first 60 AAs: 20.21362\nTotal prob of N-in: 0.91077\nPOSSIBLE N-term signal sequence\ninside 1 6\nTMhelix 7 29\noutside 30 1503\nTMhelix 1504 1526\ninside 1527 1535

31987 GCF_000421225.1_ASM42122v1 Ruania albidiflava DSM 18029 Terrabacteria group; Actinobacteria; Actinobacteria; Micrococcales; Ruaniaceae; Ruania; Ruania albidiflava MRTQRTRRSALVAGGTLAVAAGLVLGGAVAANAHVSIAEDAVEAGAYEILTVSVPHGCEG WP_022919134.1 nuclear export factor GLE1 [Ruania albidiflava] Length: 244\nNumber of predicted TMHs: 2\nExp number of AAs in TMHs: 44.36057\nExp number, first 60 AAs: 22.24853\nTotal prob of N-in: 0.99250\nPOSSIBLE N-term signal sequence\ninside 1 8\nTMhelix 9 31\noutside 32 215\nTMhelix 216 238\ninside 239 244

31988 GCF_001425965.1_Leaf3 Sanguibacter sp. Leaf3 Terrabacteria group; Actinobacteria; Actinobacteria; Micrococcales; Sanguibacteraceae; Sanguibacter MLEHPLKARSRRRTPILVACGLSLGLTVLAASPASAEPTVTSTTLTATPSVTVGDAVTVD WP_056132372.1 hypothetical protein [Sanguibacter sp. Leaf3] Length: 261\nNumber of predicted TMHs: 1\nExp number of AAs in TMHs: 31.60388\nExp number, first 60 AAs: 9.71767\nTotal prob of N-in: 0.44816\noutside 1 230\nTMhelix 231 253\ninside 254 261

31989 GCF_001552735.1_ASM155273v1 Sanguibacter suarezii NBRC 16159 Terrabacteria group; Actinobacteria; Actinobacteria; Micrococcales; Sanguibacteraceae; Sanguibacter; Sanguibacter suarezii MNARPQARRTMFGALALTAATTLAGLTMIPAAQAAEGDAQLSVLHAVPGLTVDVWVNGER WP_066464177.1 hypothetical protein [Sanguibacter suarezii] Length: 284\nNumber of predicted TMHs: 2\nExp number of AAs in TMHs: 41.86019\nExp number, first 60 AAs: 22.07655\nTotal prob of N-in: 0.99248\nPOSSIBLE N-term signal sequence\ninside 1 11\nTMhelix 12 34\noutside 35 254\nTMhelix 255 274\ninside 275 284

31990 GCF_001552735.1_ASM155273v1 Sanguibacter suarezii NBRC 16159 Terrabacteria group; Actinobacteria; Actinobacteria; Micrococcales; Sanguibacteraceae; Sanguibacter; Sanguibacter suarezii MSQLTPRRARRGVVSLALFALLGAVIAAPTSAVAAPAAPPAAPPAVPTEPVVSTATTTWR WP_066462875.1 metallophosphoesterase [Sanguibacter suarezii] Length: 928\nNumber of predicted TMHs: 2\nExp number of AAs in TMHs: 42.80955\nExp number, first 60 AAs: 22.63209\nTotal prob of N-in: 0.99881\nPOSSIBLE N-term signal sequence\ninside 1 12\nTMhelix 13 35\noutside 36 888\nTMhelix 889 911\ninside 912 928

31991 GCF_000962825.1_ASM96282v1 Actinoplanes rectilineatus Terrabacteria group; Actinobacteria; Actinobacteria; Micromonosporales; Micromonosporaceae; Actinoplanes

MRIGRRATVVALALLVTVMTGTPASAAKRLSVSKTENLSRSGETVTVTGSGYNVEKGIYV WP_052720950.1 hypothetical protein [Actinoplanes rectilineatus] Length: 261\nNumber of predicted TMHs: 2\nExp number of AAs in TMHs: 38.93242\nExp number, first 60 AAs: 17.56845\nTotal prob of N-in: 0.90546\nPOSSIBLE N-term signal sequence\ninside 1 6\nTMhelix 7 26\noutside 27 229\nTMhelix 230 252\ninside 253 261

31992 GCF_001553785.1_ASM155378v1 Actinoplanes sp. TFC3 Terrabacteria group; Actinobacteria; Actinobacteria; Micromonosporales; Micromonosporaceae; Actinoplanes MTRLEKPRRAAILAATVAAGVLGLAAPAVAGVTVNPPSAPQGSGQDLAFRVTNTASTPIT WP_067500999.1 hypothetical protein [Actinoplanes sp. TFC3] Length: 281\nNumber of predicted TMHs: 2\nExp number of AAs in TMHs: 44.17818\nExp number, first 60 AAs: 22.40249\nTotal prob of N-in: 0.97658\nPOSSIBLE N-term signal sequence\ninside 1 11\nTMhelix 12 34\noutside 35 210\nTMhelix 211 233\ninside 234 281

31993 GCF_000785985.1_ASM78598v1 Actinoplanes utahensis Terrabacteria group; Actinobacteria; Actinobacteria; Micromonosporales; Micromonosporaceae; Actinoplanes MAVPLDSRTMSPAAVNRRPAISEWSRKGFAIIISVVTAGVLAAAPATAAPGQNCAGPGEA WP_083999025.1 hypothetical protein [Actinoplanes utahensis] Length: 422\nNumber of predicted TMHs: 1\nExp number of AAs in TMHs: 51.34336\nExp number, first 60 AAs: 14.14252\nTotal prob of N-in: 0.64462\nPOSSIBLE N-term signal sequence\noutside 1 366\nTMhelix 367 389\ninside 390 422

31994 GCF_000785985.1_ASM78598v1 Actinoplanes utahensis Terrabacteria group; Actinobacteria; Actinobacteria; Micromonosporales; Micromonosporaceae; Actinoplanes MRLVPLLRRRRTAFAAGAVALVAAVTAALFLNTGPAAATLTSLNGTPTCDCVVKDSSTGK WP_043530441.1 hypothetical protein [Actinoplanes utahensis] Length: 181\nNumber of predicted TMHs: 2\nExp number of AAs in TMHs: 42.48647\nExp number, first 60 AAs: 21.39897\nTotal prob of N-in: 0.99943\nPOSSIBLE N-term signal sequence\ninside 1 12\nTMhelix 13 35\noutside 36 152\nTMhelix 153 172\ninside 173 181

31995 GCF_000785985.1_ASM78598v1 Actinoplanes utahensis Terrabacteria group; Actinobacteria; Actinobacteria; Micromonosporales; Micromonosporaceae; Actinoplanes MPYTTARRARRRVIVASGLLAGALVITPASPAAAYDPAAGREVLGAGRHVDAIYAEVTGG WP_043523714.1 anch_rpt_wall multi-domain protein [Actinoplanes utahensis] Length: 576\nNumber of predicted TMHs: 2\nExp number of AAs in TMHs: 44.08588\nExp number, first 60 AAs: 22.67969\nTotal prob of N-in: 0.99853\nPOSSIBLE N-term signal sequence\ninside 1 12\nTMhelix 13 35\noutside 36 544\nTMhelix 545 567\ninside 568 576

31996 GCF_000494755.1_ASM49475v1 Actinoplanes friuliensis DSM 7358 Terrabacteria group; Actinobacteria; Actinobacteria; Micromonosporales; Micromonosporaceae; Actinoplanes; Actinoplanes friuliensis MNDCGAVFPRRKAVRDVKRLTTIGVLAGLGVLVLAGPAAAHVEVSADKTRAGATDVTLTF WP_052359452.1 hypothetical protein [Actinoplanes friuliensis] Length: 242\nNumber of predicted TMHs: 2\nExp number of AAs in TMHs: 43.03242\nExp number, first 60 AAs: 22.16854\nTotal prob of N-in: 0.99102\nPOSSIBLE N-term signal sequence\ninside 1 19\nTMhelix 20 42\noutside 43 215\nTMhelix 216 235\ninside 236 242

31997 GCF_000284295.1_ASM28429v1 Actinoplanes missouriensis 431 Terrabacteria group; Actinobacteria; Actinobacteria; Micromonosporales; Micromonosporaceae; Actinoplanes; Actinoplanes missouriensis MRLVPQSLRRRRALLAASGAALVAAIATVLVLNSTPAAATWQTFSGTPTCDCVVQDSKTN WP_014445984.1 hypothetical protein [Actinoplanes missouriensis] Length: 182\nNumber of predicted TMHs: 2\nExp number of AAs in TMHs: 42.04348\nExp number, first 60 AAs: 21.51419\nTotal prob of N-in: 0.99953\nPOSSIBLE N-term signal sequence\ninside 1 12\nTMhelix 13 35\noutside 36 153\nTMhelix 154 173\ninside 174 182

31998 GCF_000379685.1_ASM37968v1 Catelliglobosispora koreensis DSM 44566 Terrabacteria group; Actinobacteria; Actinobacteria; Micromonosporales; Micromonosporaceae; Catelliglobosispora; Catelliglobosispora koreensis MVSGSRTRRKVVSALRMLAALMLAGGIYTLISPGAAANNAPLSAAAQEGKELYASCISCH WP_020521338.1 hypothetical protein [Catelliglobosispora koreensis] Length: 259\nNumber of predicted TMHs: 2\nExp number of AAs in TMHs: 41.87999\nExp number, first 60 AAs: 21.04823\nTotal prob of N-in: 0.97917\nPOSSIBLE N-term signal sequence\ninside 1 11\nTMhelix 12 34\noutside 35 237\nTMhelix 238 257\ninside 258 259

31999 GCF_000716715.1_ASM71671v1 Dactylosporangium aurantiacum Terrabacteria group; Actinobacteria; Actinobacteria; Micromonosporales; Micromonosporaceae; Dactylosporangium MRPTTERHVQRHRIGRGTIVQTPATAGRRTAVVAVLALVLSLVAPAGPAAAAGHARITGT WP_081970237.1 hypothetical protein [Dactylosporangium aurantiacum] Length: 571\nNumber of predicted TMHs: 2\nExp number of AAs in TMHs: 44.40076\nExp number, first 60 AAs: 21.20055\nTotal prob of N-in: 0.95368\nPOSSIBLE N-term signal sequence\ninside 1 30\nTMhelix 31 53\noutside 54 541\nTMhelix 542 564\ninside 565 571

32000 GCF_000428945.1_ASM42894v1 Hamadaea tsunoensis DSM 44101 Terrabacteria group; Actinobacteria; Actinobacteria; Micromonosporales; Micromonosporaceae; Hamadaea; Hamadaea tsunoensis
MGPGQPVGGRRGRRPGGRGAGGVRPHRSGPGDRRTLVMLRRTLVAAAVAVVAVFAATPAY WP_084557071.1 type VII secretion-associated serine protease mycosin [Hamadaea tsunoensis] Length: 425\nNumber of predicted TMHs: 2\nExp number of AAs in TMHs: 39.88088\nExp number, first 60 AAs: 15.99667\nTotal prob of N-in: 0.77020\nPOSSIBLE N-term signal sequence\ninside 1 41\nTMhelix 42 61\noutside 62 380\nTMhelix 381 403\ninside 404 425

32001 GCF_000379825.1_ASM37982v1 Longispora albida DSM 44784 Terrabacteria group; Actinobacteria; Actinobacteria; Micromonosporales; Micromonosporaceae; Longispora; Longispora albida
MTHHRTGRRRTALAGSLTGLLIAAVFLGGPASPVYAGPAEDRQRVQILSETGTPSIRAAA WP_018353464.1 hypothetical protein [Longispora albida] Length: 389\nNumber of predicted TMHs: 2\nExp number of AAs in TMHs: 43.41256\nExp number, first 60 AAs: 22.33153\nTotal prob of N-in: 0.99646\nPOSSIBLE N-term signal sequence\ninside 1 12\nTMhelix 13 35\noutside 36 359\nTMhelix 360 382\ninside 383 389

32002 GCF_900156065.1_IMG-taxon_2681813563_annotated_assembly Micromonospora avicenniae Terrabacteria group; Actinobacteria; Actinobacteria; Micromonosporales; Micromonosporaceae; Micromonospora
MASVDVPAGRARRRRAVPALGVTVTLCVLTALAVMPGPAAPGDPTASPVAVLTDPDEQGG WP_076470653.1
hypothetical protein [Micromonospora avicenniae] Length: 465\nNumber of predicted TMHs: 2\nExp number of AAs in TMHs: 44.6044699999999\nExp number, first 60 AAs: 20.35565\nTotal prob of N-in: 0.93675\nPOSSIBLE N-term signal sequence\ninside 1 16\nTMhelix 17 39\noutside 40 424\nTMhelix 425 447\ninside 448 465

32003 GCF_900091535.1_IMG-taxon_2622736592_annotated_assembly Micromonospora carbonaceaTerrabacteria group; Actinobacteria; Actinobacteria; Micromonosporales; Micromonosporaceae; Micromonospora
MARVDVRPVRRAADRRRAALLAASVAVGLLVSAAGPPGWATAAPTPARAAAAPADPPAGP WP_074477036.1
hypothetical protein [Micromonospora carbonacea] Length: 471\nNumber of predicted TMHs: 2\nExp number of AAs in TMHs: 45.22939\nExp number, first 60 AAs: 22.24869\nTotal prob of N-in: 0.99353\nPOSSIBLE N-term signal sequence\ninside 1 19\nTMhelix 20 42\noutside 43 437\nTMhelix 438 460\ninside 461 471

32004 GCF_900091535.1_IMG-taxon_2622736592_annotated_assembly Micromonospora carbonaceaTerrabacteria group; Actinobacteria; Actinobacteria; Micromonosporales; Micromonosporaceae; Micromonospora
MIRPTPVHPSAAVPRRPRRTPFVAVLAGLALAAWPSAALARPAPAPSAPAPSAPTPSAPT WP_074479339.1
hypothetical protein [Micromonospora carbonacea] Length: 483\nNumber of predicted TMHs: 1\nExp number of AAs in TMHs: 37.17079\nExp number, first 60 AAs: 15.57873\nTotal prob of N-in: 0.79026\nPOSSIBLE N-term signal sequence\noutside 1 455\nTMhelix 456 478\ninside 479 483

32005 GCF_900091535.1_IMG-taxon_2622736592_annotated_assembly Micromonospora carbonaceaTerrabacteria group; Actinobacteria; Actinobacteria; Micromonosporales; Micromonosporaceae; Micromonospora
MIRPKLSPRRPLVVAAAALAGLATAFAVSSPALATSPSPTATASPTPTAAATPPANEGPA WP_074473800.1 hypothetical protein [Micromonospora carbonacea] Length: 423\nNumber of predicted TMHs: 2\nExp number of AAs in TMHs: 43.44825\nExp number, first 60 AAs: 20.81217\nTotal prob of N-in: 0.93277\nPOSSIBLE N-term signal sequence\ninside 1 11\nTMhelix 12 34\noutside 35 390\nTMhelix 391 413\ninside 414 423

32006 GCF_000876225.1_ASM87622v1 Micromonospora carbonaceaTerrabacteria group; Actinobacteria; Actinobacteria; Micromonosporales; Micromonosporaceae; Micromonospora
MIRPKLSPRRPLVVAAAALAGLATAFAASSPALATSPSPSASATATASPSPTVTPTSPAP WP_052503667.1 hypothetical protein [Micromonospora carbonacea] Length: 426\nNumber of predicted TMHs: 2\nExp number of AAs in TMHs: 42.38952\nExp number, first 60 AAs: 19.77049\nTotal prob of N-in: 0.89142\nPOSSIBLE N-term signal sequence\ninside 1 11\nTMhelix 12 34\noutside 35 392\nTMhelix 393 415\ninside 416 426

32007 GCF_000718555.1_ASM71855v1 Micromonospora chokoriensis Terrabacteria group; Actinobacteria; Actinobacteria; Micromonosporales; Micromonosporaceae; Micromonospora
MTTMGAGMDGQVGRRRSALLAATALAATLVLTGCGAGDSGSKDAAGSAAEAPANGADREG WP_084597727.1
hypothetical protein [Micromonospora chokoriensis] Length: 334\nNumber of predicted TMHs: 1\nExp number of AAs in TMHs: 34.09278\nExp number, first 60 AAs: 5.71551\nTotal prob of N-in: 0.28121\noutside 1 285\nTMhelix 286 308\ninside 309 334

32008 GCF_900143515.1_IMG-taxon_2630968262_annotated_assembly Micromonospora cremea Terrabacteria group; Actinobacteria; Actinobacteria; Micromonosporales; Micromonosporaceae; Micromonospora
MDGRTGRRRSALLAATALAAALALTACGAGDSGARDAADTAAEVPGANNGGADKDSAAGA WP_074316730.1
hypothetical protein [Micromonospora cremea] Length: 313\nNumber of predicted TMHs: 1\nExp number of

AAs in TMHs: 41.21812\nExp number, first 60 AAs: 11.60182\nTotal prob of N-in: 0.65638\nPOSSIBLE N-term signal sequence\noutside 1 264\nTMhelix 265 287\ninside 288 313

32009 GCF_000717585.1_ASM71758v1 Micromonospora globosa Terrabacteria group; Actinobacteria; Actinobacteria; Micromonosporales; Micromonosporaceae; Micromonospora
MRARRRTILTTVLLTAAVVATPAPAHAGTATGSGGQKLTVTPSTGLSRSGATVSVSGRGY WP_013476452.1
MULTISPECIES: hypothetical protein [Micromonospora] Length: 275\nNumber of predicted TMHs: 2\nExp number of AAs in TMHs: 39.10581\nExp number, first 60 AAs: 12.54633\nTotal prob of N-in: 0.82084\nPOSSIBLE N-term signal sequence\ninside 1 6\nTMhelix 7 29\noutside 30 245\nTMhelix 246 265\ninside 266 275

32010 GCF_000718515.1_ASM71851v1 Micromonospora parva Terrabacteria group; Actinobacteria; Actinobacteria; Micromonosporales; Micromonosporaceae; Micromonospora
MTTMGAGMDGRVGRRRGALLAATALAAALALTGCGAGDSGAQDTAGSAVEAPPAKGGEVP WP_081883711.1
hypothetical protein [Micromonospora parva] Length: 320\nNumber of predicted TMHs: 1\nExp number of AAs in TMHs: 37.30537\nExp number, first 60 AAs: 10.09046\nTotal prob of N-in: 0.48179\nPOSSIBLE N-term signal sequence\noutside 1 271\nTMhelix 272 294\ninside 295 320

32011 GCF_000718515.1_ASM71851v1 Micromonospora parva Terrabacteria group; Actinobacteria; Actinobacteria; Micromonosporales; Micromonosporaceae; Micromonospora
MARVKAPPGNRAEGRRRTPVLALALGLFAALALGPAGASAATPDTEPVLSSAGPVNEPGD WP_030329257.1
hypothetical protein [Micromonospora parva] Length: 494\nNumber of predicted TMHs: 2\nExp number of AAs in TMHs: 43.0195\nExp number, first 60 AAs: 21.98391\nTotal prob of N-in: 0.98991\nPOSSIBLE N-term signal sequence\ninside 1 19\nTMhelix 20 42\noutside 43 453\nTMhelix 454 473\ninside 474 494

32012 GCF_000721115.1_ASM72111v1 Micromonospora purpureochromogenes Terrabacteria group; Actinobacteria; Actinobacteria; Micromonosporales; Micromonosporaceae; Micromonospora
MRARRRTIWTTVLLTAAVVATPAPAHAGTATGSGGQKLTVTPSTGLSRSGATVSVSGRGY WP_030499203.1
hypothetical protein [Micromonospora purpureochromogenes] Length: 275\nNumber of predicted TMHs: 2\nExp number of AAs in TMHs: 39.5272\nExp number, first 60 AAs: 15.33016\nTotal prob of N-in: 0.85520\nPOSSIBLE N-term signal sequence\ninside 1 6\nTMhelix 7 29\noutside 30 242\nTMhelix 243 265\ninside 266 275

32013 GCF_000721115.1_ASM72111v1 Micromonospora purpureochromogenes Terrabacteria group; Actinobacteria; Actinobacteria; Micromonosporales; Micromonosporaceae; Micromonospora
MPGGPPRRLARRALLAVASALVVVAGPLTPPAQAAPSVPVAAPQRQWAAPDRVLRGDHVR WP_052165692.1 type VII secretion-associated serine protease mycosin [Micromonospora purpureochromogenes] Length: 432\nNumber of predicted TMHs: 1\nExp number of AAs in TMHs: 31.64164\nExp number, first 60 AAs: 12.60653\nTotal prob of N-in: 0.58336\nPOSSIBLE N-term signal sequence\noutside 1 405\nTMhelix 406 424\ninside 425 432

32014 GCF_000721115.1_ASM72111v1 Micromonospora purpureochromogenes Terrabacteria group; Actinobacteria; Actinobacteria; Micromonosporales; Micromonosporaceae; Micromonospora
MINKRRSTIVLFGLLAAAALIGPSPAGADDETTEPVAEPPKVELVLDVSGSMRARDIDGR WP_030501147.1 inter-alpha-trypsin inhibitor heavy chain H5 [Micromonospora purpureochromogenes] Length: 430\nNumber of predicted TMHs: 1\nExp number of AAs in TMHs: 37.80531\nExp number, first 60 AAs: 15.13224\nTotal prob of N-in: 0.78730\nPOSSIBLE N-term signal sequence\noutside 1 397\nTMhelix 398 420\ninside 421 430

32015 GCF_001542325.1_ASM154232v1 Micromonospora rifamycinica Terrabacteria group; Actinobacteria; Actinobacteria; Micromonosporales; Micromonosporaceae; Micromonospora
MITPSRRSAVVRLAAALLAAGLALLVAATPARAHNSLTSASPAREAMLPSAPTEITLEFL WP_067311181.1 copper resistance protein CopC [Micromonospora rifamycinica] Length: 190\nNumber of predicted TMHs: 2\nExp number of AAs in TMHs: 43.2758\nExp number, first 60 AAs: 21.50342\nTotal prob of N-in: 0.99270\nPOSSIBLE N-term signal sequence\ninside 1 8\nTMhelix 9 31\noutside 32 160\nTMhelix 161 183\ninside 184 190

32016 GCF_001567585.1_ASM156758v1 Micromonospora rosaria Terrabacteria group; Actinobacteria; Actinobacteria; Micromonosporales; Micromonosporaceae; Micromonospora
MVRELSPRPGGHRRTALAAAVAAGLVALVLTPGPARAQGDLVAADPPAGATLTAAPAEVA WP_067359874.1
hypothetical protein [Micromonospora rosaria] Length: 201\nNumber of predicted TMHs: 1\nExp number of AAs in TMHs: 39.57859\nExp number, first 60 AAs: 16.81517\nTotal prob of N-in: 0.59519\nPOSSIBLE N-term signal sequence\noutside 1 160\nTMhelix 161 183\ninside 184 201

32017 GCF_001567585.1_ASM156758v1 Micromonospora rosaria Terrabacteria group; Actinobacteria; Actinobacteria; Micromonosporales; Micromonosporaceae; Micromonospora

MIIKRRSAVVLVGLLAATVTTGPAPALAAAEPVAEPPKVELVLDVSGSMAARDIDGRSRI WP_067365922.1 alpha-1-antitrypsin [Micromonospora rosaria] Length: 427\nNumber of predicted TMHs: 2\nExp number of AAs in TMHs: 43.70339\nExp number, first 60 AAs: 21.11102\nTotal prob of N-in: 0.94314\nPOSSIBLE N-term signal sequence\ninside 1 6\nTMhelix 7 29\noutside 30 394\nTMhelix 395 417\ninside 418 427

32018 GCF_000158815.1_ASM15881v1 Micromonospora sp. ATCC 39149 Terrabacteria group; Actinobacteria; Actinobacteria; Micromonosporales; Micromonosporaceae; Micromonospora MDGHARRRLLTAVGLVALLAAGGCSADSDGAGEPSAAVAPEQAAPADGGKAEQGADAGGT WP_007074548.1 DUF4349 domain-containing protein [Micromonospora sp. ATCC 39149] Length: 313\nNumber of predicted TMHs: 1\nExp number of AAs in TMHs: 29.21801\nExp number, first 60 AAs: 0.52334\nTotal prob of N-in: 0.08631\noutside 1 265\nTMhelix 266 288\ninside 289 313

32019 GCF_001905095.1_ASM190509v1 Micromonospora sp. CB01531 Terrabacteria group; Actinobacteria; Actinobacteria; Micromonosporales; Micromonosporaceae; Micromonospora MSERIISTATVGASWRRRAQRGDGVNAERGRRRAVSLAVVGLAVVLAAGACGADSGQNDQ WP_083673252.1 hypothetical protein [Micromonospora sp. CB01531] Length: 357\nNumber of predicted TMHs: 1\nExp number of AAs in TMHs: 40.92784\nExp number, first 60 AAs: 12.80116\nTotal prob of N-in: 0.66890\nPOSSIBLE N-term signal sequence\noutside 1 289\nTMhelix 290 312\ninside 313 357

32020 GCF_001905095.1_ASM190509v1 Micromonospora sp. CB01531 Terrabacteria group; Actinobacteria; Actinobacteria; Micromonosporales; Micromonosporaceae; Micromonospora MVSVTRRGAMLIVAAAACAAPLLGAVPALAAPPPGALVAGPTDPPDQGQQQPEPVDLVFV WP_073839607.1 hypothetical protein [Micromonospora sp. CB01531] Length: 197\nNumber of predicted TMHs: 2\nExp number of AAs in TMHs: 45.67794\nExp number, first 60 AAs: 22.00485\nTotal prob of N-in: 0.98601\nPOSSIBLE N-term signal sequence\ninside 1 8\nTMhelix 9 31\noutside 32 163\nTMhelix 164 186\ninside 187 197

32021 GCF_001905095.1_ASM190509v1 Micromonospora sp. CB01531 Terrabacteria group; Actinobacteria; Actinobacteria; Micromonosporales; Micromonosporaceae; Micromonospora MGVDSGRRAILRAAVAGGVGTGLAATGLAAPAGAAPGEATPSGRGPHIAILLYDGLTALD WP_073838665.1 hypothetical protein [Micromonospora sp. CB01531] Length: 242\nNumber of predicted TMHs: 2\nExp number of AAs in TMHs: 65.65506\nExp number, first 60 AAs: 18.90379\nTotal prob of N-in: 0.86565\nPOSSIBLE N-term signal sequence\ninside 1 141\nTMhelix 142 164\noutside 165 195\nTMhelix 196 218\ninside 219 242

32022 GCF_001709195.1_ASM170919v1 Micromonospora sp. II Terrabacteria group; Actinobacteria; Actinobacteria; Micromonosporales; Micromonosporaceae; Micromonospora MRARRRTIWTTVLLTAAVVATPAPAHAGTATGSGGQKLTVTPSTGLSRSGATVSVSGRGY WP_069088473.1 hypothetical protein [Micromonospora sp. II] Length: 275\nNumber of predicted TMHs: 2\nExp number of AAs in TMHs: 38.98833\nExp number, first 60 AAs: 16.92498\nTotal prob of N-in: 0.84015\nPOSSIBLE N-term signal sequence\ninside 1 6\nTMhelix 7 29\noutside 30 242\nTMhelix 243 265\ninside 266 275

32023 GCF_001709195.1_ASM170919v1 Micromonospora sp. II Terrabacteria group; Actinobacteria; Actinobacteria; Micromonosporales; Micromonosporaceae; Micromonospora MTRDRQGLPGGPPRRVARRALLAVASALVVVAGPLTPPAQAAPSVPVAASQRQLAAPDRV WP_082930992.1 MULTISPECIES: type VII secretion-associated serine protease mycosin [Micromonospora] Length: 439\nNumber of predicted TMHs: 2\nExp number of AAs in TMHs: 35.35577\nExp number, first 60 AAs: 16.29188\nTotal prob of N-in: 0.76062\nPOSSIBLE N-term signal sequence\ninside 1 19\nTMhelix 20 42\noutside 43 412\nTMhelix 413 431\ninside 432 439

32024 GCF_001709195.1_ASM170919v1 Micromonospora sp. II Terrabacteria group; Actinobacteria; Actinobacteria; Micromonosporales; Micromonosporaceae; Micromonospora MINKRRSTIVLFGLLTAAALIGPSPAGADDETSEPVAEPPKVELVLDVSGSMRARDIDGR WP_043326826.1 MULTISPECIES: inter-alpha-trypsin inhibitor heavy chain H5 [Micromonospora] Length: 431\nNumber of predicted TMHs: 1\nExp number of AAs in TMHs: 35.67832\nExp number, first 60 AAs: 12.95828\nTotal prob of N-in: 0.68956\nPOSSIBLE N-term signal sequence\noutside 1 398\nTMhelix 399 421\ninside 422 431

32025 GCF_000177655.2_ASM17765v2 Micromonospora sp. L5 Terrabacteria group; Actinobacteria; Actinobacteria; Micromonosporales; Micromonosporaceae; Micromonospora MRARRRTILTTVLLTAAVVATPAPAHAGTATGSGGQKLTVTPSTGLSRSGATVSVSGRGY WP_013476452.1 MULTISPECIES: hypothetical protein [Micromonospora] Length: 275\nNumber of predicted TMHs: 2\nExp number of AAs in TMHs: 39.10581\nExp number, first 60 AAs: 12.54633\nTotal prob of N-in: 0.82084\nPOSSIBLE N-term signal sequence\ninside 1 6\nTMhelix 7 29\noutside 30 245\nTMhelix 246 265\ninside 266 275

32026 GCF_000568915.1_ASM56891v1 Micromonospora sp. M42 Terrabacteria group; Actinobacteria; Actinobacteria; Micromonosporales; Micromonosporaceae; Micromonospora
MRARRRTIWTTVLLTAAVVATPAPAHAGTATGSGGQKLTVTPSTGLSRSGATVSVSGRGY WP_043323399.1
hypothetical protein [Micromonospora sp. M42] Length: 275\nNumber of predicted TMHs: 2\nExp number of AAs in TMHs: 39.55871\nExp number, first 60 AAs: 15.22973\nTotal prob of N-in: 0.85614\nPOSSIBLE N-term signal sequence\ninside 1 6\nTMhelix 7 29\noutside 30 242\nTMhelix 243 265\ninside 266 275

32027 GCF_000568915.1_ASM56891v1 Micromonospora sp. M42 Terrabacteria group; Actinobacteria; Actinobacteria; Micromonosporales; Micromonosporaceae; Micromonospora
MINKRRSTIVLFGLLTAAALIGPSPAGADDETSEPVAEPPKVELVLDVSGSMRARDIDGR WP_043326826.1 MULTISPECIES: inter-alpha-trypsin inhibitor heavy chain H5 [Micromonospora] Length: 431\nNumber of predicted TMHs: 1\nExp number of AAs in TMHs: 35.67832\nExp number, first 60 AAs: 12.95828\nTotal prob of N-in: 0.68956\nPOSSIBLE N-term signal sequence\noutside 1 398\nTMhelix 399 421\ninside 422 431

32028 GCF_000568915.1_ASM56891v1 Micromonospora sp. M42 Terrabacteria group; Actinobacteria; Actinobacteria; Micromonosporales; Micromonosporaceae; Micromonospora
MTRDRQGLPGGPPRRVARRALLAVASQLVVVAGPLTPPAQAGGVSGPVAASQRQLAAPDR WP_043327879.1 type VII secretion-associated serine protease [Micromonospora sp. M42] Length: 440\nNumber of predicted TMHs: 1\nExp number of AAs in TMHs: 22.51666\nExp number, first 60 AAs: 3.35902\nTotal prob of N-in: 0.16870\noutside 1 413\nTMhelix 414 432\ninside 433 440

32029 GCF_001652995.1_ASM165299v1 Micromonospora sp. NBRC 110037 Terrabacteria group; Actinobacteria; Actinobacteria; Micromonosporales; Micromonosporaceae; Micromonospora
MINKRRSTIVLFGLLTAAALIGPSPAGADDETSEPVAEPPKVELVLDVSGSMRARDIDGR WP_043326826.1 MULTISPECIES: inter-alpha-trypsin inhibitor heavy chain H5 [Micromonospora] Length: 431\nNumber of predicted TMHs: 1\nExp number of AAs in TMHs: 35.67832\nExp number, first 60 AAs: 12.95828\nTotal prob of N-in: 0.68956\nPOSSIBLE N-term signal sequence\noutside 1 398\nTMhelix 399 421\ninside 422 431

32030 GCF_001652995.1_ASM165299v1 Micromonospora sp. NBRC 110037 Terrabacteria group; Actinobacteria; Actinobacteria; Micromonosporales; Micromonosporaceae; Micromonospora
MRARRRTIWTTVLLTAAVVATPAPAHAGTATGSGGQKLTVTPSTGLSRSGATVSVSGRGY WP_064445663.1
hypothetical protein [Micromonospora sp. NBRC 110037] Length: 275\nNumber of predicted TMHs: 2\nExp number of AAs in TMHs: 39.04872\nExp number, first 60 AAs: 16.74466\nTotal prob of N-in: 0.84186\nPOSSIBLE N-term signal sequence\ninside 1 6\nTMhelix 7 29\noutside 30 242\nTMhelix 243 265\ninside 266 275

32031 GCF_001652995.1_ASM165299v1 Micromonospora sp. NBRC 110037 Terrabacteria group; Actinobacteria; Actinobacteria; Micromonosporales; Micromonosporaceae; Micromonospora
MNRRSFLIRLACVLLVAGGGALLWLPFHTQMELDDWAVTLRERGTPARAVVVDQVTESGG WP_064448876.1
hypothetical protein [Micromonospora sp. NBRC 110037] Length: 183\nNumber of predicted TMHs: 2\nExp number of AAs in TMHs: 44.64915\nExp number, first 60 AAs: 22.25385\nTotal prob of N-in: 0.99905\nPOSSIBLE N-term signal sequence\ninside 1 4\nTMhelix 5 27\noutside 28 123\nTMhelix 124 146\ninside 147 183

32032 GCF_001652995.1_ASM165299v1 Micromonospora sp. NBRC 110037 Terrabacteria group; Actinobacteria; Actinobacteria; Micromonosporales; Micromonosporaceae; Micromonospora
MTRDRQGLPGGPPRRVARRALLAVASALVVVAGPLTPPAQAAPSVPVAASQRQLAAPDRV WP_082930992.1
MULTISPECIES: type VII secretion-associated serine protease mycosin [Micromonospora] Length: 439\nNumber of predicted TMHs: 2\nExp number of AAs in TMHs: 35.35577\nExp number, first 60 AAs: 16.29188\nTotal prob of N-in: 0.76062\nPOSSIBLE N-term signal sequence\ninside 1 19\nTMhelix 20 42\noutside 43 412\nTMhelix 413 431\ninside 432 439

32033 GCF_001279555.1_ASM127955v1 Micromonospora sp. NRRL B-16802 Terrabacteria group; Actinobacteria; Actinobacteria; Micromonosporales; Micromonosporaceae; Micromonospora
MDGQAGRRRGALLTAMALTATLAMAGCGADDSGARDSAASAPERAVAEGGAAEKNSGTDA WP_053659339.1
hypothetical protein [Micromonospora sp. NRRL B-16802] Length: 313\nNumber of predicted TMHs: 1\nExp number of AAs in TMHs: 28.98588\nExp number, first 60 AAs: 2.02856\nTotal prob of N-in: 0.16443\noutside 1 264\nTMhelix 265 287\ninside 288 313

32034 GCF_002006875.1_ASM200687v1 Micromonospora sp. Rc5 Terrabacteria group; Actinobacteria; Actinobacteria; Micromonosporales; Micromonosporaceae; Micromonospora
MIRPKLSPRRPLVVAAAALAGLATAFAASSPALATSPSPSASATATASPSPTVTPTSPAP WP_077936608.1 hypothetical protein [Micromonospora sp. Rc5] Length: 426\nNumber of predicted TMHs: 2\nExp number of AAs in TMHs:

42.38952\nExp number, first 60 AAs: 19.77049\nTotal prob of N-in: 0.89142\nPOSSIBLE N-term signal sequence\ninside 1 11\nTMhelix 12 34\noutside 35 392\nTMhelix 393 415\ninside 416 426

32035 GCF_001029515.1_ASM102951v1 Micromonospora sp. RV43 Terrabacteria group; Actinobacteria; Actinobacteria; Micromonosporales; Micromonosporaceae; Micromonospora MRARRRTILTTVLLTAAVVATPAPAHAGTATGSGGQKLTVTPSTGLSRSGATVSVSGRGY WP_047890386.1 hypothetical protein [Micromonospora sp. RV43] Length: 275\nNumber of predicted TMHs: 2\nExp number of AAs in TMHs: 38.97815\nExp number, first 60 AAs: 12.81101\nTotal prob of N-in: 0.81704\nPOSSIBLE N-term signal sequence\ninside 1 6\nTMhelix 7 29\noutside 30 245\nTMhelix 246 265\ninside 266 275

32036 GCF_001905625.1_ASM190562v1 Micromonospora sp. TSRI0369 Terrabacteria group; Actinobacteria; Actinobacteria; Micromonosporales; Micromonosporaceae; Micromonospora MRARRRTIPTTVLLTVAVVAATPAPAHAGTATGSGGQKLTVTPSTGLSRSGATVSVSGRG WP_073828243.1 hypothetical protein [Micromonospora sp. TSRI0369] Length: 274\nNumber of predicted TMHs: 1\nExp number of AAs in TMHs: 37.48081\nExp number, first 60 AAs: 15.53907\nTotal prob of N-in: 0.79053\nPOSSIBLE N-term signal sequence\noutside 1 241\nTMhelix 242 264\ninside 265 274

32037 GCF_001905625.1_ASM190562v1 Micromonospora sp. TSRI0369 Terrabacteria group; Actinobacteria; Actinobacteria; Micromonosporales; Micromonosporaceae; Micromonospora MPGGPPRLARRALLAVASALVVVAGPLTLPAQAAPSVPVAAQQQRAAPDRALRGDHVRDE WP_083664749.1 type VII secretion-associated serine protease mycosin [Micromonospora sp. TSRI0369] Length: 430\nNumber of predicted TMHs: 2\nExp number of AAs in TMHs: 34.8715\nExp number, first 60 AAs: 15.94115\nTotal prob of N-in: 0.73138\nPOSSIBLE N-term signal sequence\ninside 1 11\nTMhelix 12 34\noutside 35 403\nTMhelix 404 422\ninside 423 430

32038 GCF_001905625.1_ASM190562v1 Micromonospora sp. TSRI0369 Terrabacteria group; Actinobacteria; Actinobacteria; Micromonosporales; Micromonosporaceae; Micromonospora MINKRRSTIVLFGLLAAAALIGPSPAGADDETTEPVAEPPKVELVLDVSGSMRARDIDGR WP_073825409.1 alpha-1-antitrypsin [Micromonospora sp. TSRI0369] Length: 430\nNumber of predicted TMHs: 1\nExp number of AAs in TMHs: 37.80533\nExp number, first 60 AAs: 15.13209\nTotal prob of N-in: 0.78730\nPOSSIBLE N-term signal sequence\noutside 1 397\nTMhelix 398 420\ninside 421 430

32039 GCF_001855515.1_ASM185551v1 Micromonospora sp. WMMB235 Terrabacteria group; Actinobacteria; Actinobacteria; Micromonosporales; Micromonosporaceae; Micromonospora MRARRRTILTTVLLTAAVVATPAPAHAGTATGSGGQKLTVTPSTGLSRSGATVSVSGRGY WP_071094599.1 hypothetical protein [Micromonospora sp. WMMB235] Length: 275\nNumber of predicted TMHs: 2\nExp number of AAs in TMHs: 38.87866\nExp number, first 60 AAs: 12.96444\nTotal prob of N-in: 0.81250\nPOSSIBLE N-term signal sequence\ninside 1 6\nTMhelix 7 29\noutside 30 245\nTMhelix 246 268\ninside 269 275

32040 GCF_000145235.1_ASM14523v1 Micromonospora aurantiaca ATCC 27029 Terrabacteria group; Actinobacteria; Actinobacteria; Micromonosporales; Micromonosporaceae; Micromonospora; Micromonospora aurantiaca MRARRRTILTTVLLTAAVVATPAPAHAGTATGSGGQKLTVTPSTGLSRSGATVSVSGRGY WP_013286804.1 hypothetical protein [Micromonospora aurantiaca] Length: 275\nNumber of predicted TMHs: 2\nExp number of AAs in TMHs: 39.15805\nExp number, first 60 AAs: 12.43647\nTotal prob of N-in: 0.82242\nPOSSIBLE N-term signal sequence\ninside 1 6\nTMhelix 7 29\noutside 30 245\nTMhelix 246 265\ninside 266 275

32041 GCF_000297395.2_ASM29739v2 Micromonospora lupini str. Lupac 08 Terrabacteria group; Actinobacteria; Actinobacteria; Micromonosporales; Micromonosporaceae; Micromonospora; Micromonospora lupini MDGRTGRRRSALLAAGTLSVALALAGCSAGDSGSADSAGPAAKAPLAEAGAADKGSGADA WP_007455506.1 hypothetical protein [Micromonospora lupini] Length: 313\nNumber of predicted TMHs: 1\nExp number of AAs in TMHs: 35.99004\nExp number, first 60 AAs: 3.93929\nTotal prob of N-in: 0.60355\noutside 1 263\nTMhelix 264 286\ninside 287 313

32042 GCF_000297395.2_ASM29739v2 Micromonospora lupini str. Lupac 08 Terrabacteria group; Actinobacteria; Actinobacteria; Micromonosporales; Micromonosporaceae; Micromonospora; Micromonospora lupini MDADRTARLQGNQIAAYERRGPFWGRLTQLATAVALAAALVGSSLVLDTVASFYPGTGVV WP_007457857.1 hypothetical protein [Micromonospora lupini] Length: 198\nNumber of predicted TMHs: 2\nExp number of AAs in TMHs: 44.20402\nExp number, first 60 AAs: 19.88605\nTotal prob of N-in: 0.89301\nPOSSIBLE N-term signal sequence\ninside 1 22\nTMhelix 23 45\noutside 46 158\nTMhelix 159 181\ninside 182 198

32043 GCF_000514555.1_ASM51455v1 Salinispora arenicola CNX482 Terrabacteria group; Actinobacteria; Actinobacteria; Micromonosporales; Micromonosporaceae; Salinispora; Salinispora arenicola
MNRRRRRVVVVGAGLAGSLAALYLARQGHEVDIFERRPDPRSALAGPEGRSINLGLSARG WP_028190038.1
kynurenine 3-monooxygenase [Salinispora arenicola] Length: 454\nNumber of predicted TMHs: 2\nExp number of AAs in TMHs: 38.62593\nExp number, first 60 AAs: 18.79111\nTotal prob of N-in: 0.97305\nPOSSIBLE N-term signal sequence\ninside 1 6\nTMhelix 7 24\noutside 25 426\nTMhelix 427 446\ninside 447 454

32044 GCF_000514935.1_ASM51493v1 Salinispora arenicola CNX891 Terrabacteria group; Actinobacteria; Actinobacteria; Micromonosporales; Micromonosporaceae; Salinispora; Salinispora arenicola
MNRRRRRVVVVGAGLAGSLAALYLARQGHEVDIFERRPDPRSALAGPEGRSINLGLSARG WP_028191314.1
kynurenine 3-monooxygenase [Salinispora arenicola] Length: 454\nNumber of predicted TMHs: 2\nExp number of AAs in TMHs: 39.02761\nExp number, first 60 AAs: 18.81275\nTotal prob of N-in: 0.97302\nPOSSIBLE N-term signal sequence\ninside 1 6\nTMhelix 7 24\noutside 25 426\nTMhelix 427 446\ninside 447 454

32045 GCF_000375225.1_ASM37522v1 Salinispora pacifica CNQ768 Terrabacteria group; Actinobacteria; Actinobacteria; Micromonosporales; Micromonosporaceae; Salinispora; Salinispora pacifica
MSSGWARRRTHILGLLPGWALVVLIALTTVPTAAAAEEPSPGLGQSIAADQPITTGRAEL WP_026185219.1 hypothetical protein [Salinispora pacifica] Length: 326\nNumber of predicted TMHs: 2\nExp number of AAs in TMHs: 45.48688\nExp number, first 60 AAs: 22.57131\nTotal prob of N-in: 0.99344\nPOSSIBLE N-term signal sequence\ninside 1 11\nTMhelix 12 34\noutside 35 281\nTMhelix 282 304\ninside 305 326

32046 GCF_000514775.1_ASM51477v1 Salinispora pacifica CNR114 Terrabacteria group; Actinobacteria; Actinobacteria; Micromonosporales; Micromonosporaceae; Salinispora; Salinispora pacifica
MSSGWARRRTHILGLLSGWALVVLIALTTVPTAAAAEEPSPGLGQSIAADQPITTGRAEL WP_028193122.1 hypothetical protein [Salinispora pacifica] Length: 326\nNumber of predicted TMHs: 2\nExp number of AAs in TMHs: 45.50858\nExp number, first 60 AAs: 22.64756\nTotal prob of N-in: 0.99571\nPOSSIBLE N-term signal sequence\ninside 1 11\nTMhelix 12 34\noutside 35 281\nTMhelix 282 304\ninside 305 326

32047 GCF_000514755.1_ASM51475v1 Salinispora pacifica CNR894 Terrabacteria group; Actinobacteria; Actinobacteria; Micromonosporales; Micromonosporaceae; Salinispora; Salinispora pacifica
MSSGWARRRTHILGLLPGWALVVLIALTTVPTAAAAEEPSPGLGQSIAADQPITTGRAEL WP_026185219.1 hypothetical protein [Salinispora pacifica] Length: 326\nNumber of predicted TMHs: 2\nExp number of AAs in TMHs: 45.48688\nExp number, first 60 AAs: 22.57131\nTotal prob of N-in: 0.99344\nPOSSIBLE N-term signal sequence\ninside 1 11\nTMhelix 12 34\noutside 35 281\nTMhelix 282 304\ninside 305 326

32048 GCF_000514755.1_ASM51475v1 Salinispora pacifica CNR894 Terrabacteria group; Actinobacteria; Actinobacteria; Micromonosporales; Micromonosporaceae; Salinispora; Salinispora pacifica
MAAQPGREGAVVKGTRRGLVATIAVAVVGPLLAVVVGLLLAVGAANATAGPASAQVVSAG WP_080678665.1
hypothetical protein [Salinispora pacifica] Length: 234\nNumber of predicted TMHs: 2\nExp number of AAs in TMHs: 45.06626\nExp number, first 60 AAs: 23.07635\nTotal prob of N-in: 0.99097\nPOSSIBLE N-term signal sequence\ninside 1 19\nTMhelix 20 42\noutside 43 202\nTMhelix 203 225\ninside 226 234

32049 GCF_000514795.1_ASM51479v1 Salinispora pacifica CNS103 Terrabacteria group; Actinobacteria; Actinobacteria; Micromonosporales; Micromonosporaceae; Salinispora; Salinispora pacifica
MSSGWARRRTHILGLLPGWALVVLIALTTVPTAAAAEEPSPGLGQSIAADQPITTGRAEL WP_026185219.1 hypothetical protein [Salinispora pacifica] Length: 326\nNumber of predicted TMHs: 2\nExp number of AAs in TMHs: 45.48688\nExp number, first 60 AAs: 22.57131\nTotal prob of N-in: 0.99344\nPOSSIBLE N-term signal sequence\ninside 1 11\nTMhelix 12 34\noutside 35 281\nTMhelix 282 304\ninside 305 326

32050 GCF_000424905.1_ASM42490v1 Salinispora pacifica CNS237 Terrabacteria group; Actinobacteria; Actinobacteria; Micromonosporales; Micromonosporaceae; Salinispora; Salinispora pacifica
MSSGRARRRTHVLGLLPGWALVVLIALTAVPTAAAAEEPTPGLSQSIAADQPIATGRAEVWP_027653432.1 hypothetical protein [Salinispora pacifica] Length: 326\nNumber of predicted TMHs: 2\nExp number of AAs in TMHs: 45.33208\nExp number, first 60 AAs: 22.6361\nTotal prob of N-in: 0.99806\nPOSSIBLE N-term signal sequence\ninside 1 11\nTMhelix 12 34\noutside 35 281\nTMhelix 282 304\ninside 305 326

32051 GCF_000514535.1_ASM51453v1 Salinispora pacifica CNT001 Terrabacteria group; Actinobacteria; Actinobacteria; Micromonosporales; Micromonosporaceae; Salinispora; Salinispora pacifica
MSSGWARRRTHILGLLPGWALVVLIALTTVPTAAAAEEPSPGLGQSIAADQPITTGRAEL WP_026185219.1 hypothetical protein [Salinispora pacifica] Length: 326\nNumber of predicted TMHs: 2\nExp number of AAs in TMHs: 45.48688\nExp

number, first 60 AAs: 22.57131\nTotal prob of N-in: 0.99344\nPOSSIBLE N-term signal sequence\ninside 1 11\nTMhelix 12 34\noutside 35 281\nTMhelix 282 304\ninside 305 326

32052 GCF_000514575.1_ASM51457v1 Salinispora pacifica CNT003 Terrabacteria group; Actinobacteria; Actinobacteria; Micromonosporales; Micromonosporaceae; Salinispora; Salinispora pacifica
MSSGWARRRTHILGLLPGWALVVLIALTTVPTAAAAEEPSPGLGQSIAADQPITTGRAEL WP_026185219.1 hypothetical protein [Salinispora pacifica] Length: 326\nNumber of predicted TMHs: 2\nExp number of AAs in TMHs: 45.48688\nExp number, first 60 AAs: 22.57131\nTotal prob of N-in: 0.99344\nPOSSIBLE N-term signal sequence\ninside 1 11\nTMhelix 12 34\noutside 35 281\nTMhelix 282 304\ninside 305 326

32053 GCF_000514475.1_ASM51447v1 Salinispora pacifica CNT131 Terrabacteria group; Actinobacteria; Actinobacteria; Micromonosporales; Micromonosporaceae; Salinispora; Salinispora pacifica
MSSGWARRRTHILGLLPGWALVVLIALTTVPTAAAAEEPSPGLGQSIAADQPITTGRAEL WP_027658311.1 hypothetical protein [Salinispora pacifica] Length: 326\nNumber of predicted TMHs: 2\nExp number of AAs in TMHs: 45.48608\nExp number, first 60 AAs: 22.57149\nTotal prob of N-in: 0.99345\nPOSSIBLE N-term signal sequence\ninside 1 11\nTMhelix 12 34\noutside 35 281\nTMhelix 282 304\ninside 305 326

32054 GCF_000514415.1_ASM51441v1 Salinispora pacifica CNT603 Terrabacteria group; Actinobacteria; Actinobacteria; Micromonosporales; Micromonosporaceae; Salinispora; Salinispora pacifica
MSSGWARRRTHILGLLPGWALVVLIALTTVPTAAAAEEPSPGLGQSIAADQPITTGRAEL WP_027658311.1 hypothetical protein [Salinispora pacifica] Length: 326\nNumber of predicted TMHs: 2\nExp number of AAs in TMHs: 45.48608\nExp number, first 60 AAs: 22.57149\nTotal prob of N-in: 0.99345\nPOSSIBLE N-term signal sequence\ninside 1 11\nTMhelix 12 34\noutside 35 281\nTMhelix 282 304\ninside 305 326

32055 GCF_000514735.1_ASM51473v1 Salinispora pacifica CNT796 Terrabacteria group; Actinobacteria; Actinobacteria; Micromonosporales; Micromonosporaceae; Salinispora; Salinispora pacifica
MSSGWARRRTHILGLLPGWALVVLIALTTVPTAAAAEEPSPGLGQSIAADQPITTGRAEL WP_026273477.1 hypothetical protein [Salinispora pacifica] Length: 326\nNumber of predicted TMHs: 2\nExp number of AAs in TMHs: 45.4815299999999\nExp number, first 60 AAs: 22.57031\nTotal prob of N-in: 0.99339\nPOSSIBLE N-term signal sequence\ninside 1 11\nTMhelix 12 34\noutside 35 281\nTMhelix 282 304\ninside 305 326

32056 GCF_000375285.1_ASM37528v1 Salinispora pacifica CNT851 Terrabacteria group; Actinobacteria; Actinobacteria; Micromonosporales; Micromonosporaceae; Salinispora; Salinispora pacifica
MSSGWARRRTHILGLLPGWALVVLIALTTVPTAAAAEEPSPGLGQSIAADQPITTGRAEL WP_026273477.1 hypothetical protein [Salinispora pacifica] Length: 326\nNumber of predicted TMHs: 2\nExp number of AAs in TMHs: 45.4815299999999\nExp number, first 60 AAs: 22.57031\nTotal prob of N-in: 0.99339\nPOSSIBLE N-term signal sequence\ninside 1 11\nTMhelix 12 34\noutside 35 281\nTMhelix 282 304\ninside 305 326

32057 GCF_000424925.1_ASM42492v1 Salinispora pacifica CNY239 Terrabacteria group; Actinobacteria; Actinobacteria; Micromonosporales; Micromonosporaceae; Salinispora; Salinispora pacifica
MSSGWARRRTHILGLLPGWALVVLIALTTVPTAAAAEEPSPGLGQSIAADQPITTGRAEL WP_026185219.1 hypothetical protein [Salinispora pacifica] Length: 326\nNumber of predicted TMHs: 2\nExp number of AAs in TMHs: 45.48688\nExp number, first 60 AAs: 22.57131\nTotal prob of N-in: 0.99344\nPOSSIBLE N-term signal sequence\ninside 1 11\nTMhelix 12 34\noutside 35 281\nTMhelix 282 304\ninside 305 326

32058 GCF_000378845.1_ASM37884v1 Salinispora pacifica DSM 45544 Terrabacteria group; Actinobacteria; Actinobacteria; Micromonosporales; Micromonosporaceae; Salinispora; Salinispora pacifica
MSSGWARRRTHILGLLPGWALVVLIALTTVPTAAAAEEPSPGLGQSIAADQPITTGRAEL WP_026185219.1 hypothetical protein [Salinispora pacifica] Length: 326\nNumber of predicted TMHs: 2\nExp number of AAs in TMHs: 45.48688\nExp number, first 60 AAs: 22.57131\nTotal prob of N-in: 0.99344\nPOSSIBLE N-term signal sequence\ninside 1 11\nTMhelix 12 34\noutside 35 281\nTMhelix 282 304\ninside 305 326

32059 GCF_000426645.1_ASM42664v1 Nakamurella lactea DSM 19367 Terrabacteria group; Actinobacteria; Actinobacteria; Nakamurellales; Nakamurellaceae; Nakamurella; Nakamurella lactea
MRAGRLAAVIALVLALAAPLLFAGLPAAAAGTTVGAVDRRSAVVPPAAAGTPGTTADRGS WP_029136656.1 hypothetical protein [Nakamurella lactea] Length: 870\nNumber of predicted TMHs: 2\nExp number of AAs in TMHs: 45.96285\nExp number, first 60 AAs: 22.742\nTotal prob of N-in: 0.99641\nPOSSIBLE N-term signal sequence\ninside 1 6\nTMhelix 7 29\noutside 30 805\nTMhelix 806 828\ninside 829 870

32060 GCF_000024365.1_ASM2436v1 Nakamurella multipartita DSM 44233 Terrabacteria group; Actinobacteria; Actinobacteria; Nakamurellales; Nakamurellaceae; Nakamurella; Nakamurella multipartita

MSERRVNHQVDPSAPGGPAAVSGAPRRPHVVVLGGGFGGLTAVRALGKADVDITLVDRHT WP_015747587.1 NADH dehydrogenase family protein [Nakamurella multipartita] Length: 445\nNumber of predicted TMHs: 1\nExp number of AAs in TMHs: 22.49257\nExp number, first 60 AAs: 1.39595\nTotal prob of N-in: 0.18378\noutside 1 384\nTMhelix 385 407\ninside 408 445

32061 GCF_000373925.1_ASM37392v1 Actinopolymorpha alba DSM 45243 Terrabacteria group; Actinobacteria; Actinobacteria; Propionibacteriales; Nocardioidaceae; Actinopolymorpha; Actinopolymorpha alba MSATDGSRPNAAPTAVFFDLDKTILARSSTLAFSRHFYQGGLINRRTVIRSAYAQFVYLL WP_020577100.1 hypothetical protein [Actinopolymorpha alba] Length: 298\nNumber of predicted TMHs: 1\nExp number of AAs in TMHs: 18.35794\nExp number, first 60 AAs: 0.07397\nTotal prob of N-in: 0.09392\noutside 1 246\nTMhelix 247 264\ninside 265 298

32062 GCF_900167475.1_IMG-taxon_2708742384_annotated_assembly Aeromicrobium choanae Terrabacteria group; Actinobacteria; Actinobacteria; Propionibacteriales; Nocardioidaceae; Aeromicrobium MATPAGRPAAFFDLDKTIIARSSTLAFSKSLFAEGLLSRRAVLRSAYAQLVFSTGGADHD WP_078699469.1 inhibition of morphological differentiation protein [Aeromicrobium choanae] Length: 271\nNumber of predicted TMHs: 1\nExp number of AAs in TMHs: 17.96964\nExp number, first 60 AAs: 0.02577\nTotal prob of N-in: 0.08850\noutside 1 241\nTMhelix 242 264\ninside 265 271

32063 GCF_001509405.1_ASM150940v1 Aeromicrobium erythreum Terrabacteria group; Actinobacteria; Actinobacteria; Propionibacteriales; Nocardioidaceae; Aeromicrobium MRERAYPGRVSPRSAAFFDLDKTIIARSSALAFSKPFFEGGLLTRRAVLRSAYANFMFGL WP_083516460.1 inhibition of morphological differentiation protein [Aeromicrobium erythreum] Length: 283\nNumber of predicted TMHs: 1\nExp number of AAs in TMHs: 19.80946\nExp number, first 60 AAs: 0.03481\nTotal prob of N-in: 0.04686\noutside 1 247\nTMhelix 248 265\ninside 266 283

32064 GCF_001422765.1_Leaf245 Aeromicrobium sp. Leaf245 Terrabacteria group; Actinobacteria; Actinobacteria; Propionibacteriales; Nocardioidaceae; Aeromicrobium MSPRSAAFFDLDKTIIARSSTLAFSKPFFEGGLLTRRAVLRSAYANFLFGLNGADHDQLE WP_056552788.1 inhibition of morphological differentiation protein [Aeromicrobium sp. Leaf245] Length: 275\nNumber of predicted TMHs: 1\nExp number of AAs in TMHs: 16.60561\nExp number, first 60 AAs: 0.02376\nTotal prob of N-in: 0.00666\noutside 1 238\nTMhelix 239 256\ninside 257 275

32065 GCF_001422865.1_Leaf272 Aeromicrobium sp. Leaf272 Terrabacteria group; Actinobacteria; Actinobacteria; Propionibacteriales; Nocardioidaceae; Aeromicrobium MSPRSAAFFDLDKTIIARSSTLAFSKPFFEGGLLTRRAVLRSAYANFLFGLNGADHDQLE WP_055969644.1 MULTISPECIES: inhibition of morphological differentiation protein [Aeromicrobium] Length: 275\nNumber of predicted TMHs: 1\nExp number of AAs in TMHs: 16.02606\nExp number, first 60 AAs: 0.01897\nTotal prob of N-in: 0.00480\noutside 1 238\nTMhelix 239 256\ninside 257 275

32066 GCF_001423335.1_Leaf289 Aeromicrobium sp. Leaf289 Terrabacteria group; Actinobacteria; Actinobacteria; Propionibacteriales; Nocardioidaceae; Aeromicrobium MSPRTAAFFDLDKTIIARSSTLAFSKPFFEGGLLTRRAVLRSAYANFLFGLNGADHDQLE WP_082506551.1 inhibition of morphological differentiation protein [Aeromicrobium sp. Leaf289] Length: 275\nNumber of predicted TMHs: 1\nExp number of AAs in TMHs: 16.016\nExp number, first 60 AAs: 0.02153\nTotal prob of N-in: 0.00697\noutside 1 238\nTMhelix 239 256\ninside 257 275

32067 GCF_001423365.1_Leaf291 Aeromicrobium sp. Leaf291 Terrabacteria group; Actinobacteria; Actinobacteria; Propionibacteriales; Nocardioidaceae; Aeromicrobium MSPRSAAFFDLDKTIIARSSTLAFSKPFFEGGLLTRRAVLRSAYANFLFGLNGADHDQLE WP_055969644.1 MULTISPECIES: inhibition of morphological differentiation protein [Aeromicrobium] Length: 275\nNumber of predicted TMHs: 1\nExp number of AAs in TMHs: 16.02606\nExp number, first 60 AAs: 0.01897\nTotal prob of N-in: 0.00480\noutside 1 238\nTMhelix 239 256\ninside 257 275

32068 GCF_002174305.1_ASM217430v1 Aeromicrobium sp. PE09-221 Terrabacteria group; Actinobacteria; Actinobacteria; Propionibacteriales; Nocardioidaceae; Aeromicrobium MGRTAAFFDLDKTIIARSSTLAFSRPLFAEGLLSRRAVLRSAYAQFVFAANGADHEQMEA WP_087624051.1 inhibition of morphological differentiation protein [Aeromicrobium sp. PE09-221] Length: 263\nNumber of predicted TMHs: 1\nExp number of AAs in TMHs: 21.205\nExp number, first 60 AAs: 0.01831\nTotal prob of N-in: 0.02058\noutside 1 235\nTMhelix 236 258\ninside 259 263

32069 GCF_002174305.1_ASM217430v1 Aeromicrobium sp. PE09-221 Terrabacteria group; Actinobacteria; Actinobacteria; Propionibacteriales; Nocardioidaceae; Aeromicrobium
MLSQRRAAVLASAAVLMALVPSVATAAPAEPPPPPSIELPPPPPAAPLSGVLVSEVASGG WP_087622105.1 hypothetical protein [Aeromicrobium sp. PE09-221] Length: 1389\nNumber of predicted TMHs: 2\nExp number of AAs in TMHs: 41.06627\nExp number, first 60 AAs: 20.17522\nTotal prob of N-in: 0.90312\nPOSSIBLE N-term signal sequence\ninside 1 6\nTMhelix 7 29\noutside 30 1361\nTMhelix 1362 1384\ninside 1385 1389

32070 GCF_002174305.1_ASM217430v1 Aeromicrobium sp. PE09-221 Terrabacteria group; Actinobacteria; Actinobacteria; Propionibacteriales; Nocardioidaceae; Aeromicrobium
MKRSPRSRDVLRRSRRAAVVATAVSALGVATLAPAAAEQTQAVPLSNGAERTLTVTPDAG WP_087625019.1 hypothetical protein [Aeromicrobium sp. PE09-221] Length: 907\nNumber of predicted TMHs: 2\nExp number of AAs in TMHs: 40.1580499999999\nExp number, first 60 AAs: 17.75215\nTotal prob of N-in: 0.87941\nPOSSIBLE N-term signal sequence\ninside 1 16\nTMhelix 17 36\noutside 37 878\nTMhelix 879 901\ninside 902 907

32071 GCF_002174305.1_ASM217430v1 Aeromicrobium sp. PE09-221 Terrabacteria group; Actinobacteria; Actinobacteria; Propionibacteriales; Nocardioidaceae; Aeromicrobium
MRRQRRAAVATLAAMSGCLVLATQAPMVASAVATGVKGDVSITNTETVDAKLSATGRVSS WP_087624756.1 hypothetical protein [Aeromicrobium sp. PE09-221] Length: 788\nNumber of predicted TMHs: 2\nExp number of AAs in TMHs: 37.1836799999999\nExp number, first 60 AAs: 20.72207\nTotal prob of N-in: 0.93703\nPOSSIBLE N-term signal sequence\ninside 1 6\nTMhelix 7 29\noutside 30 763\nTMhelix 764 783\ninside 784 788

32072 GCF_001428805.1_Root236 Aeromicrobium sp. Root236 Terrabacteria group; Actinobacteria; Actinobacteria; Propionibacteriales; Nocardioidaceae; Aeromicrobium
MSPTRSAAFFDLDKTIIAKSSTLAFSKPFFQGGLLSRRAVLRSAYANFMFALNGADHDKL WP_056401745.1 inhibition of morphological differentiation protein [Aeromicrobium sp. Root236] Length: 269\nNumber of predicted TMHs: 1\nExp number of AAs in TMHs: 20.69758\nExp number, first 60 AAs: 0.03514\nTotal prob of N-in: 0.02611\noutside 1 236\nTMhelix 237 259\ninside 260 269

32073 GCF_001425755.1_Root344 Aeromicrobium sp. Root344 Terrabacteria group; Actinobacteria; Actinobacteria; Propionibacteriales; Nocardioidaceae; Aeromicrobium
MLDSRRRSLEGEPVSSLLSAPRRRRRAALAALLSIVAVPLAAASTADAADACDPAQACLI WP_082530627.1 hypothetical protein [Aeromicrobium sp. Root344] Length: 442\nNumber of predicted TMHs: 1\nExp number of AAs in TMHs: 29.01192\nExp number, first 60 AAs: 11.56069\nTotal prob of N-in: 0.56881\nPOSSIBLE N-term signal sequence\noutside 1 405\nTMhelix 406 424\ninside 425 442

32074 GCF_001426755.1_Root495 Aeromicrobium sp. Root495 Terrabacteria group; Actinobacteria; Actinobacteria; Propionibacteriales; Nocardioidaceae; Aeromicrobium
MTTAGTRSAAFFDLDKTIIAKSSTLAFSKPFFQGGLLTRRAVLRSAYANFMFSLQGADHD WP_056288184.1 inhibition of morphological differentiation protein [Aeromicrobium sp. Root495] Length: 268\nNumber of predicted TMHs: 1\nExp number of AAs in TMHs: 17.15077\nExp number, first 60 AAs: 0.05563\nTotal prob of N-in: 0.11879\noutside 1 241\nTMhelix 242 260\ninside 261 268

32075 GCF_000160775.2_ASM16077v2 Aeromicrobium marinum DSM 15272 Terrabacteria group; Actinobacteria; Actinobacteria; Propionibacteriales; Nocardioidaceae; Aeromicrobium; Aeromicrobium marinum
MSRAAAFFDLDKTIIAKSSTLAFSRPFYDGGLLSRRAVLRSAYAQFVFAISGADHDQLEK WP_040320320.1 inhibition of morphological differentiation protein [Aeromicrobium marinum] Length: 266\nNumber of predicted TMHs: 1\nExp number of AAs in TMHs: 22.13249\nExp number, first 60 AAs: 0.00927\nTotal prob of N-in: 0.00213\noutside 1 234\nTMhelix 235 257\ninside 258 266

32076 GCF_000312105.1_ASM31210v1 Aeromicrobium massiliense JC14 Terrabacteria group; Actinobacteria; Actinobacteria; Propionibacteriales; Nocardioidaceae; Aeromicrobium; Aeromicrobium massiliense
MSIPPFQPVPPPWPHRRRDPLGSWGWLVLAVTALGLSVVSLLPPDDADVDAWRSHGTATI WP_019146500.1 hypothetical protein [Aeromicrobium massiliense] Length: 161\nNumber of predicted TMHs: 2\nExp number of AAs in TMHs: 44.90146\nExp number, first 60 AAs: 22.1202\nTotal prob of N-in: 0.95469\nPOSSIBLE N-term signal sequence\ninside 1 20\nTMhelix 21 43\noutside 44 123\nTMhelix 124 146\ninside 147 161

32077 GCF_000312105.1_ASM31210v1 Aeromicrobium massiliense JC14 Terrabacteria group; Actinobacteria; Actinobacteria; Propionibacteriales; Nocardioidaceae; Aeromicrobium; Aeromicrobium massiliense
MTRSAAFFDLDKTIIAKSSTLAFSRPFFDGGLLNRRAVLRSAYAQFVFAVSGADHDQLEK WP_019144452.1 hypothetical

protein [Aeromicrobium massiliense]	Length: 261\nNumber of predicted TMHs:  1\nExp number of AAs in TMHs: 21.36485\nExp number, first 60 AAs:  0.01492\nTotal prob of N-in:   0.03437\noutside   1   236\nTMhelix   237   256\ninside   257   261

32078	GCF_000312105.1_ASM31210v1	Aeromicrobium massiliense JC14	Terrabacteria group; Actinobacteria; Actinobacteria; Propionibacteriales; Nocardioidaceae; Aeromicrobium; Aeromicrobium massiliense	MNPHETDRADLPELGDDRVDAMLSAITTRIDAEDRERSRRRRTTLLAAAAAVVVGVGAGT	WP_019144836.1	hypothetical protein [Aeromicrobium massiliense]	Length: 283\nNumber of predicted TMHs:  2\nExp number of AAs in TMHs: 42.88053\nExp number, first 60 AAs:  16.40129\nTotal prob of N-in:   0.98606\nPOSSIBLE N-term signal sequence\ninside   1   43\nTMhelix   44   66\noutside   67   253\nTMhelix   254   276\ninside   277   283

32079	GCF_001984195.1_ASM198419v1	Kribbella sp. ALI-6-A	Terrabacteria group; Actinobacteria; Actinobacteria; Propionibacteriales; Nocardioidaceae; Kribbella	MEHLRTPRPAAFFDLDKTIIARSSTLAFSRPFYAGGLINRRTVLRSAYAQFVYLLGGADH	WP_077017852.1	inhibition of morphological differentiation protein [Kribbella sp. ALI-6-A]	Length: 272\nNumber of predicted TMHs:  1\nExp number of AAs in TMHs: 20.82366\nExp number, first 60 AAs:  0.05961\nTotal prob of N-in:   0.02121\noutside   1   243\nTMhelix   244   263\ninside   264   272

32080	GCF_000372465.1_ASM37246v1	Kribbella catacumbae DSM 19601	Terrabacteria group; Actinobacteria; Actinobacteria; Propionibacteriales; Nocardioidaceae; Kribbella; Kribbella catacumbae	MSAPQKRRKLLRAVIVVGLLGGIGLAWMQSASGSPDASPTPTPKTTPSAVKPGSKPTPQP	WP_020392913.1	hypothetical protein [Kribbella catacumbae]	Length: 676\nNumber of predicted TMHs:  2\nExp number of AAs in TMHs: 38.06768\nExp number, first 60 AAs:  18.7707\nTotal prob of N-in:   0.96829\nPOSSIBLE N-term signal sequence\ninside   1   12\nTMhelix   13   31\noutside   32   615\nTMhelix   616   635\ninside   636   676

32081	GCF_000372465.1_ASM37246v1	Kribbella catacumbae DSM 19601	Terrabacteria group; Actinobacteria; Actinobacteria; Propionibacteriales; Nocardioidaceae; Kribbella; Kribbella catacumbae	MLVPMEPPETTRSAAFFDLDKTIIARSSTLAFSRPFYAGGLINRRTVLRSAYAQFMYLLG	WP_020392791.1	inhibition of morphological differentiation protein [Kribbella catacumbae]	Length: 276\nNumber of predicted TMHs:  1\nExp number of AAs in TMHs: 20.47998\nExp number, first 60 AAs:  0.13115\nTotal prob of N-in:   0.01398\noutside   1   246\nTMhelix   247   266\ninside   267   276

32082	GCF_000372465.1_ASM37246v1	Kribbella catacumbae DSM 19601	Terrabacteria group; Actinobacteria; Actinobacteria; Propionibacteriales; Nocardioidaceae; Kribbella; Kribbella catacumbae	MAVHGARVRRRAVVAVRIMVASVLLSGTTLMAFPQQAQAATVRDFEPLFKAQVNGANLIT	WP_020386861.1	hypothetical protein [Kribbella catacumbae]	Length: 1857\nNumber of predicted TMHs:  2\nExp number of AAs in TMHs: 44.4150399999999\nExp number, first 60 AAs:  21.87008\nTotal prob of N-in:   0.99523\nPOSSIBLE N-term signal sequence\ninside   1   12\nTMhelix   13   35\noutside   36  1828\nTMhelix   1829  1851\ninside   1852  1857

32083	GCF_000024345.1_ASM2434v1	Kribbella flavida DSM 17836	Terrabacteria group; Actinobacteria; Actinobacteria; Propionibacteriales; Nocardioidaceae; Kribbella; Kribbella flavida	MLDLMEHPRTPRPAAFFDLDKTIIARSSTLAFSRPFYAGGLINRRTVLRSAYAQFVYLLG	WP_012918225.1	haloacid dehalogenase [Kribbella flavida]	Length: 276\nNumber of predicted TMHs:  1\nExp number of AAs in TMHs: 20.00954\nExp number, first 60 AAs:  0.3088\nTotal prob of N-in:   0.15228\noutside   1   247\nTMhelix   248   267\ninside   268   276

32084	GCF_000720335.1_ASM72033v1	Marmoricola aequoreus	Terrabacteria group; Actinobacteria; Actinobacteria; Propionibacteriales; Nocardioidaceae; Marmoricola	MTRRRPLLLRLAAPTLLLTLALPLAACAGSTSTSDSGASGSSASDSGGADSGSEPAPAPD	WP_030484094.1	hypothetical protein, partial [Marmoricola aequoreus]	Length: 317\nNumber of predicted TMHs:  2\nExp number of AAs in TMHs: 41.80833\nExp number, first 60 AAs:  16.24769\nTotal prob of N-in:   0.75174\nPOSSIBLE N-term signal sequence\ninside   1   6\nTMhelix   7   29\noutside   30   279\nTMhelix   280   302\ninside   303   317

32085	GCF_000389985.1_1.0	Nocardioides sp. CF8	Terrabacteria group; Actinobacteria; Actinobacteria; Propionibacteriales; Nocardioidaceae; Nocardioides	MPRPTAAFFDLDKTIIAKSSTLAFSKPFQAGGLISRRAMLRSTYAQFVYLVGGADHDQME	WP_010831317.1	haloacid dehalogenase [Nocardioides sp. CF8]	Length: 265\nNumber of predicted TMHs:  1\nExp number of AAs in TMHs: 18.81233\nExp number, first 60 AAs:  0.02625\nTotal prob of N-in:   0.00963\noutside   1   238\nTMhelix   239   258\ninside   259   265

32086 GCF_000364605.1_ASM36460v1 Nocardioides sp. Iso805N Terrabacteria group; Actinobacteria; Actinobacteria; Propionibacteriales; Nocardioidaceae; Nocardioides
MRLLNRSAGRLSRHRRGPIAGLVLLLLGLALSGGLYAAFSPSSSADTAEDQAAMAAKGRQ WP_017933931.1
hypothetical protein [Nocardioides sp. Iso805N] Length: 291\nNumber of predicted TMHs: 2\nExp number of AAs in TMHs: 42.43096\nExp number, first 60 AAs: 20.91183\nTotal prob of N-in: 0.99625\nPOSSIBLE N-term signal sequence\ninside 1 16\nTMhelix 17 39\noutside 40 258\nTMhelix 259 281\ninside 282 291

32087 GCF_000364605.1_ASM36460v1 Nocardioides sp. Iso805N Terrabacteria group; Actinobacteria; Actinobacteria; Propionibacteriales; Nocardioidaceae; Nocardioides
MIDRHRDDAAAPSRRTVVAAVLLRALGFALIAFVLGAGVLGRDWYATRHRPVEHAVVVAV WP_017935992.1
hypothetical protein [Nocardioides sp. Iso805N] Length: 179\nNumber of predicted TMHs: 2\nExp number of AAs in TMHs: 44.53623\nExp number, first 60 AAs: 21.46345\nTotal prob of N-in: 0.94700\nPOSSIBLE N-term signal sequence\ninside 1 19\nTMhelix 20 42\noutside 43 139\nTMhelix 140 162\ninside 163 179

32088 GCF_000519005.1_ASM51900v1 Nocardioides sp. J54 Terrabacteria group; Actinobacteria; Actinobacteria; Propionibacteriales; Nocardioidaceae; Nocardioides
MRLLNRSAGRLSRHRRGPLAGLVVLVLGLLLTGGLYAALAPAQADTDADQEALVQEGREL WP_028655215.1
cytochrome c [Nocardioides sp. J54] Length: 300\nNumber of predicted TMHs: 2\nExp number of AAs in TMHs: 42.59238\nExp number, first 60 AAs: 21.62165\nTotal prob of N-in: 0.99539\nPOSSIBLE N-term signal sequence\ninside 1 19\nTMhelix 20 42\noutside 43 266\nTMhelix 267 286\ninside 287 300

32089 GCF_000519005.1_ASM51900v1 Nocardioides sp. J54 Terrabacteria group; Actinobacteria; Actinobacteria; Propionibacteriales; Nocardioidaceae; Nocardioides
MARPGAAPTAAFFDLDKTIIAKSSVLAFSKPFQAGGLISRRAVLRSAYAQFVFMVGGADH WP_028655975.1
inhibition of morphological differentiation protein [Nocardioides sp. J54] Length: 269\nNumber of predicted TMHs: 1\nExp number of AAs in TMHs: 20.12636\nExp number, first 60 AAs: 0.04489\nTotal prob of N-in: 0.01621\noutside 1 242\nTMhelix 243 262\ninside 263 269

32090 GCF_000015265.1_ASM1526v1 Nocardioides sp. JS614 Terrabacteria group; Actinobacteria; Actinobacteria; Propionibacteriales; Nocardioidaceae; Nocardioides
MATRSTAAFFDLDKTIIAKSSTLAFSKPFQAGGLISRRAVLRSAYAQFVYLVGGADHDQM WP_041546100.1 inhibition of morphological differentiation protein [Nocardioides sp. JS614] Length: 266\nNumber of predicted TMHs: 1\nExp number of AAs in TMHs: 19.43393\nExp number, first 60 AAs: 0.05928\nTotal prob of N-in: 0.02059\noutside 1 237\nTMhelix 238 257\ninside 258 266

32091 GCF_000015265.1_ASM1526v1 Nocardioides sp. JS614 Terrabacteria group; Actinobacteria; Actinobacteria; Propionibacteriales; Nocardioidaceae; Nocardioides
MRLLNRSAGRLSRHRRGPLAGLAVVLLGLLMTGGLYTVFSPAQAGSAAASEEQIAKGREL WP_011756571.1
cystathionine beta-lyase [Nocardioides sp. JS614] Length: 289\nNumber of predicted TMHs: 2\nExp number of AAs in TMHs: 42.1921699999999\nExp number, first 60 AAs: 21.37698\nTotal prob of N-in: 0.99636\nPOSSIBLE N-term signal sequence\ninside 1 16\nTMhelix 17 39\noutside 40 257\nTMhelix 258 277\ninside 278 289

32092 GCF_001422955.1_Leaf285 Nocardioides sp. Leaf285 Terrabacteria group; Actinobacteria; Actinobacteria; Propionibacteriales; Nocardioidaceae; Nocardioides
MTPWPPGDPGAPVAAGAARCGTARRAVVAAVPVAAVHLAAVLVAAVLTAAVLTAVVPPAA WP_056865087.1
hypothetical protein [Nocardioides sp. Leaf285] Length: 494\nNumber of predicted TMHs: 2\nExp number of AAs in TMHs: 44.6266499999999\nExp number, first 60 AAs: 22.36414\nTotal prob of N-in: 0.87683\nPOSSIBLE N-term signal sequence\ninside 1 26\nTMhelix 27 49\noutside 50 465\nTMhelix 466 488\ninside 489 494

32093 GCF_001422955.1_Leaf285 Nocardioides sp. Leaf285 Terrabacteria group; Actinobacteria; Actinobacteria; Propionibacteriales; Nocardioidaceae; Nocardioides
MPGAAFFDLDKTIIAKSSAMAFTKPFQAGGLISRRAVVRSAYGQFVYLTGGADHDQMEKM WP_056863190.1
MULTISPECIES: inhibition of morphological differentiation protein [Nocardioides] Length: 262\nNumber of predicted TMHs: 1\nExp number of AAs in TMHs: 20.29868\nExp number, first 60 AAs: 0.01087\nTotal prob of N-in: 0.00484\noutside 1 233\nTMhelix 234 253\ninside 254 262

32094 GCF_001422955.1_Leaf285 Nocardioides sp. Leaf285 Terrabacteria group; Actinobacteria; Actinobacteria; Propionibacteriales; Nocardioidaceae; Nocardioides
MSDDLRETMSLLADRADPSGAPPAPDELWARGAAYRRRRRAVGAAGVAAVVVLLAALGGL WP_056862610.1
hypothetical protein [Nocardioides sp. Leaf285] Length: 455\nNumber of predicted TMHs: 2\nExp number of

AAs in TMHs: 41.8414\nExp number, first 60 AAs: 19.92345\nTotal prob of N-in: 0.99904\nPOSSIBLE N-term signal sequence\ninside 1 39\nTMhelix 40 62\noutside 63 429\nTMhelix 430 449\ninside 450 455

32095 GCF_001423135.1_Leaf307 Nocardioides sp. Leaf307 Terrabacteria group; Actinobacteria; Actinobacteria; Propionibacteriales; Nocardioidaceae; Nocardioides MPGAAFFDLDKTIIAKSSAMAFTKPFQAGGLISRRAVVRSAYGQFVYLTGGADHDQMEKM WP_056863190.1 MULTISPECIES: inhibition of morphological differentiation protein [Nocardioides] Length: 262\nNumber of predicted TMHs: 1\nExp number of AAs in TMHs: 20.29868\nExp number, first 60 AAs: 0.01087\nTotal prob of N-in: 0.00484\noutside 1 233\nTMhelix 234 253\ninside 254 262

32096 GCF_001423135.1_Leaf307 Nocardioides sp. Leaf307 Terrabacteria group; Actinobacteria; Actinobacteria; Propionibacteriales; Nocardioidaceae; Nocardioides MSDDLRETMSLLADRADPSGAPPAPDELWARGAAYRRRRRAVGAAGVAAVLVLLAALGGL WP_056904455.1 hypothetical protein [Nocardioides sp. Leaf307] Length: 455\nNumber of predicted TMHs: 2\nExp number of AAs in TMHs: 41.84581\nExp number, first 60 AAs: 19.92363\nTotal prob of N-in: 0.99901\nPOSSIBLE N-term signal sequence\ninside 1 39\nTMhelix 40 62\noutside 63 429\nTMhelix 430 449\ninside 450 455

32097 GCF_002117955.1_ASM211795v1 Nocardioides sp. PD653 Terrabacteria group; Actinobacteria; Actinobacteria; Propionibacteriales; Nocardioidaceae; Nocardioides MAPRPTAAFFDLDKTIIAKSSTLAFSKPFQAGGLISRRAVLRSAYAQFVYLVGGADHDQMWP_085872748.1 MULTISPECIES: inhibition of morphological differentiation protein [Nocardioides] Length: 266\nNumber of predicted TMHs: 1\nExp number of AAs in TMHs: 20.29345\nExp number, first 60 AAs: 0.02143\nTotal prob of N-in: 0.00640\noutside 1 239\nTMhelix 240 259\ninside 260 266

32098 GCF_002117935.1_ASM211793v1 Nocardioides sp. PD653-B2 Terrabacteria group; Actinobacteria; Actinobacteria; Propionibacteriales; Nocardioidaceae; Nocardioides MAPRPTAAFFDLDKTIIAKSSTLAFSKPFQAGGLISRRAVLRSAYAQFVYLVGGADHDQMWP_085872748.1 MULTISPECIES: inhibition of morphological differentiation protein [Nocardioides] Length: 266\nNumber of predicted TMHs: 1\nExp number of AAs in TMHs: 20.29345\nExp number, first 60 AAs: 0.02143\nTotal prob of N-in: 0.00640\noutside 1 239\nTMhelix 240 259\ninside 260 266

32099 GCF_001425025.1_Root122 Nocardioides sp. Root122 Terrabacteria group; Actinobacteria; Actinobacteria; Propionibacteriales; Nocardioidaceae; Nocardioides MRVLNRTAGRLSRHRRGPLAGLVVLLLGLLISGSLYTALAPAQADSQASETEQVAQGKELWP_056906450.1 cytochrome c [Nocardioides sp. Root122] Length: 290\nNumber of predicted TMHs: 2\nExp number of AAs in TMHs: 42.25996\nExp number, first 60 AAs: 21.48839\nTotal prob of N-in: 0.99779\nPOSSIBLE N-term signal sequence\ninside 1 19\nTMhelix 20 42\noutside 43 257\nTMhelix 258 277\ninside 278 290

32100 GCF_001425025.1_Root122 Nocardioides sp. Root122 Terrabacteria group; Actinobacteria; Actinobacteria; Propionibacteriales; Nocardioidaceae; Nocardioides MPRPTAAFFDLDKTIIAKSSTLAFSREFQAGGLISRRAMLRSAYAQFLFFTGGADHDQMD WP_056909410.1 inhibition of morphological differentiation protein [Nocardioides sp. Root122] Length: 265\nNumber of predicted TMHs: 1\nExp number of AAs in TMHs: 19.58224\nExp number, first 60 AAs: 0.01888\nTotal prob of N-in: 0.00539\noutside 1 236\nTMhelix 237 256\ninside 257 265

32101 GCF_001425175.1_Root1257 Nocardioides sp. Root1257 Terrabacteria group; Actinobacteria; Actinobacteria; Propionibacteriales; Nocardioidaceae; Nocardioides MAHTAAFFDLDKTIIAKSSTLAFSKPFQAGGLITRRAVLRSAYAQFVYLVGGADHDQMEK WP_056157122.1 MULTISPECIES: inhibition of morphological differentiation protein [Nocardioides] Length: 264\nNumber of predicted TMHs: 1\nExp number of AAs in TMHs: 22.07499\nExp number, first 60 AAs: 0.03256\nTotal prob of N-in: 0.00278\noutside 1 233\nTMhelix 234 256\ninside 257 264

32102 GCF_001425175.1_Root1257 Nocardioides sp. Root1257 Terrabacteria group; Actinobacteria; Actinobacteria; Propionibacteriales; Nocardioidaceae; Nocardioides MRLLNRSAGRLSRHRRGPLASVLVIIAGLLLTGGIFTAVVNTTAQADSTSSNEELVAQGR WP_056148929.1 MULTISPECIES: cytochrome c [Nocardioides] Length: 291\nNumber of predicted TMHs: 2\nExp number of AAs in TMHs: 43.76053\nExp number, first 60 AAs: 21.84605\nTotal prob of N-in: 0.99652\nPOSSIBLE N-term signal sequence\ninside 1 19\nTMhelix 20 42\noutside 43 255\nTMhelix 256 278\ninside 279 291

32103 GCF_001428565.1_Root190 Nocardioides sp. Root190 Terrabacteria group; Actinobacteria; Actinobacteria; Propionibacteriales; Nocardioidaceae; Nocardioides

MRLLNRSAGRLSRHRRGPIAGLVVLLLGLLLTGGLYAAFAPAQADTDQNNEELVKEGREL WP_082606064.1 cytochrome C [Nocardioides sp. Root190] Length: 300\nNumber of predicted TMHs: 2\nExp number of AAs in TMHs: 42.46426\nExp number, first 60 AAs: 21.53049\nTotal prob of N-in: 0.99599\nPOSSIBLE N-term signal sequence\ninside 1 20\nTMhelix 21 40\noutside 41 266\nTMhelix 267 286\ninside 287 300

32104 GCF_001428565.1_Root190 Nocardioides sp. Root190 Terrabacteria group; Actinobacteria; Actinobacteria; Propionibacteriales; Nocardioidaceae; Nocardioides
MRPAPRRRPVALAALVAAAAMLSLPVAAATTTAASAATTGPGGTPVRTSSIPAACGMLDL WP_056749018.1 hypothetical protein [Nocardioides sp. Root190] Length: 224\nNumber of predicted TMHs: 2\nExp number of AAs in TMHs: 43.85165\nExp number, first 60 AAs: 22.31303\nTotal prob of N-in: 0.98741\nPOSSIBLE N-term signal sequence\ninside 1 11\nTMhelix 12 34\noutside 35 197\nTMhelix 198 220\ninside 221 224

32105 GCF_001428565.1_Root190 Nocardioides sp. Root190 Terrabacteria group; Actinobacteria; Actinobacteria; Propionibacteriales; Nocardioidaceae; Nocardioides
MAQLARPRTAPTAAFFDLDKTIIAKSSVLAFSKPFQAGGLISRRAVLRSAYAQFVFMVGG WP_056754805.1 inhibition of morphological differentiation protein [Nocardioides sp. Root190] Length: 272\nNumber of predicted TMHs: 1\nExp number of AAs in TMHs: 19.69792\nExp number, first 60 AAs: 0.0934\nTotal prob of N-in: 0.05934\noutside 1 245\nTMhelix 246 265\ninside 266 272

32106 GCF_001428725.1_Root224 Nocardioides sp. Root224 Terrabacteria group; Actinobacteria; Actinobacteria; Propionibacteriales; Nocardioidaceae; Nocardioides
MRLLNRSAGRLSRHRRGPLASVLVIIAGLLLTGGIFTAVVNTTAQADSTSSNEELVAQGR WP_056148929.1 MULTISPECIES: cytochrome c [Nocardioides] Length: 291\nNumber of predicted TMHs: 2\nExp number of AAs in TMHs: 43.76053\nExp number, first 60 AAs: 21.84605\nTotal prob of N-in: 0.99652\nPOSSIBLE N-term signal sequence\ninside 1 19\nTMhelix 20 42\noutside 43 255\nTMhelix 256 278\ninside 279 291

32107 GCF_001428725.1_Root224 Nocardioides sp. Root224 Terrabacteria group; Actinobacteria; Actinobacteria; Propionibacteriales; Nocardioidaceae; Nocardioides
MAHTAAFFDLDKTIIAKSSTLAFSKPFQAGGLITRRAVLRSAYAQFVYLVGGADHDQMEK WP_056157122.1 MULTISPECIES: inhibition of morphological differentiation protein [Nocardioides] Length: 264\nNumber of predicted TMHs: 1\nExp number of AAs in TMHs: 22.07499\nExp number, first 60 AAs: 0.03256\nTotal prob of N-in: 0.00278\noutside 1 233\nTMhelix 234 256\ninside 257 264

32108 GCF_001428825.1_Root240 Nocardioides sp. Root240 Terrabacteria group; Actinobacteria; Actinobacteria; Propionibacteriales; Nocardioidaceae; Nocardioides
MARPGTSLTAAFFDLDKTIIAKSSVLAFSKPFQAGGLISRRAVLRSAYAQFVFMIGGADH WP_056894020.1 MULTISPECIES: inhibition of morphological differentiation protein [Nocardioides] Length: 269\nNumber of predicted TMHs: 1\nExp number of AAs in TMHs: 19.67064\nExp number, first 60 AAs: 0.1355\nTotal prob of N-in: 0.02101\noutside 1 242\nTMhelix 243 262\ninside 263 269

32109 GCF_001428825.1_Root240 Nocardioides sp. Root240 Terrabacteria group; Actinobacteria; Actinobacteria; Propionibacteriales; Nocardioidaceae; Nocardioides
MRLLNRSAGRLSRHRRGPIAGLLVLALGLLLTGGLYAAFAPAQADSSQSQDELVKEGREL WP_082612055.1 MULTISPECIES: cytochrome C [Nocardioides] Length: 300\nNumber of predicted TMHs: 2\nExp number of AAs in TMHs: 42.38029\nExp number, first 60 AAs: 21.5931\nTotal prob of N-in: 0.99844\nPOSSIBLE N-term signal sequence\ninside 1 20\nTMhelix 21 40\noutside 41 266\nTMhelix 267 286\ninside 287 300

32110 GCF_001428765.1_Root79 Nocardioides sp. Root79 Terrabacteria group; Actinobacteria; Actinobacteria; Propionibacteriales; Nocardioidaceae; Nocardioides
MARPGTSLTAAFFDLDKTIIAKSSVLAFSKPFQAGGLISRRAVLRSAYAQFVFMIGGADH WP_056894020.1 MULTISPECIES: inhibition of morphological differentiation protein [Nocardioides] Length: 269\nNumber of predicted TMHs: 1\nExp number of AAs in TMHs: 19.67064\nExp number, first 60 AAs: 0.1355\nTotal prob of N-in: 0.02101\noutside 1 242\nTMhelix 243 262\ninside 263 269

32111 GCF_001428765.1_Root79 Nocardioides sp. Root79 Terrabacteria group; Actinobacteria; Actinobacteria; Propionibacteriales; Nocardioidaceae; Nocardioides
MRLLNRSAGRLSRHRRGPIAGLLVLALGLLLTGGLYAAFAPAQADSSQSQDELVKEGREL WP_082612055.1 MULTISPECIES: cytochrome C [Nocardioides] Length: 300\nNumber of predicted TMHs: 2\nExp number of AAs in TMHs: 42.38029\nExp number, first 60 AAs: 21.5931\nTotal prob of N-in: 0.99844\nPOSSIBLE N-term signal sequence\ninside 1 20\nTMhelix 21 40\noutside 41 266\nTMhelix 267 286\ninside 287 300

32112 GCF_001428125.1_Soil774 Nocardioides sp. Soil774 Terrabacteria group; Actinobacteria; Actinobacteria; Propionibacteriales; Nocardioidaceae; Nocardioides
MRLLNRTAGRLSRHRRGPLAGLVVLLLGLLISGSLYTALAPAQAQNQASETDQVAQGKEL WP_056602383.1 cytochrome c [Nocardioides sp. Soil774] Length: 290\nNumber of predicted TMHs: 2\nExp number of AAs in TMHs: 42.25789\nExp number, first 60 AAs: 21.51654\nTotal prob of N-in: 0.99503\nPOSSIBLE N-term signal sequence\ninside 1 19\nTMhelix 20 42\noutside 43 257\nTMhelix 258 277\ninside 278 290

32113 GCF_001428125.1_Soil774 Nocardioides sp. Soil774 Terrabacteria group; Actinobacteria; Actinobacteria; Propionibacteriales; Nocardioidaceae; Nocardioides
MPRPTAAFFDLDKTIIAKSSTLAFSREFQAGGLISRRAMLRSAYAQFVFFTGGADHDQMD WP_056601961.1 inhibition of morphological differentiation protein [Nocardioides sp. Soil774] Length: 263\nNumber of predicted TMHs: 1\nExp number of AAs in TMHs: 19.27285\nExp number, first 60 AAs: 0.01828\nTotal prob of N-in: 0.00612\noutside 1 236\nTMhelix 237 256\ninside 257 263

32114 GCF_001428425.1_Soil777 Nocardioides sp. Soil777 Terrabacteria group; Actinobacteria; Actinobacteria; Propionibacteriales; Nocardioidaceae; Nocardioides
MADMSRPTAAFFDLDKTIIAKSSTLAFSKPFQAGGLISRRAMLRSTYAQFVYLVGGADHD WP_056888206.1 inhibition of morphological differentiation protein [Nocardioides sp. Soil777] Length: 268\nNumber of predicted TMHs: 1\nExp number of AAs in TMHs: 19.86779\nExp number, first 60 AAs: 0.03455\nTotal prob of N-in: 0.01321\noutside 1 239\nTMhelix 240 259\ninside 260 268

32115 GCF_001428425.1_Soil777 Nocardioides sp. Soil777 Terrabacteria group; Actinobacteria; Actinobacteria; Propionibacteriales; Nocardioidaceae; Nocardioides
MRLLNRSAGRISRHRRGPLAGLVVLLLGLLMTGGLYTAFSPAQAESTSDEELIAEGRELF WP_082599763.1 cytochrome C [Nocardioides sp. Soil777] Length: 291\nNumber of predicted TMHs: 2\nExp number of AAs in TMHs: 42.28088\nExp number, first 60 AAs: 21.26528\nTotal prob of N-in: 0.99711\nPOSSIBLE N-term signal sequence\ninside 1 16\nTMhelix 17 39\noutside 40 256\nTMhelix 257 276\ninside 277 291

32116 GCF_001429555.1_Soil797 Nocardioides sp. Soil797 Terrabacteria group; Actinobacteria; Actinobacteria; Propionibacteriales; Nocardioidaceae; Nocardioides
MSPAHRQAAFFDLDKTIIAKSSTLAFSKPFQAGGLISRRAVLRSAYAQFVYLVGGADHDQ WP_057324237.1 inhibition of morphological differentiation protein [Nocardioides sp. Soil797] Length: 266\nNumber of predicted TMHs: 1\nExp number of AAs in TMHs: 20.71939\nExp number, first 60 AAs: 0.07694\nTotal prob of N-in: 0.12153\noutside 1 237\nTMhelix 238 260\ninside 261 266

32117 GCF_001429625.1_Soil805 Nocardioides sp. Soil805 Terrabacteria group; Actinobacteria; Actinobacteria; Propionibacteriales; Nocardioidaceae; Nocardioides
MPRPTAAFFDLDKTIIAKSSTLAFSKPFQAGGLISRRAMLRSTYAQFVYLVGGADHDQME WP_082624094.1 inhibition of morphological differentiation protein [Nocardioides sp. Soil805] Length: 265\nNumber of predicted TMHs: 1\nExp number of AAs in TMHs: 20.95992\nExp number, first 60 AAs: 0.01164\nTotal prob of N-in: 0.00346\noutside 1 236\nTMhelix 237 256\ninside 257 265

32118 GCF_001429625.1_Soil805 Nocardioides sp. Soil805 Terrabacteria group; Actinobacteria; Actinobacteria; Propionibacteriales; Nocardioidaceae; Nocardioides
MADRRRRLLTVLAGAAVLAPFAVGWAMGAEPDDGRVVFSFADPEIAESSGLVAAGDRVWT WP_056925535.1 hypothetical protein [Nocardioides sp. Soil805] Length: 331\nNumber of predicted TMHs: 2\nExp number of AAs in TMHs: 42.82907\nExp number, first 60 AAs: 22.29667\nTotal prob of N-in: 0.99305\nPOSSIBLE N-term signal sequence\ninside 1 6\nTMhelix 7 29\noutside 30 305\nTMhelix 306 328\ninside 329 331

32119 GCF_000620705.1_ASM62070v1 Nocardioides sp. URHA0020 Terrabacteria group; Actinobacteria; Actinobacteria; Propionibacteriales; Nocardioidaceae; Nocardioides
MARTAAFFDLDKTIIAKSSTLAFSKPFQAGGLISRRAVLRSAYAQFVYLVGGADHDQMEKWP_036507088.1 inhibition of morphological differentiation protein [Nocardioides sp. URHA0020] Length: 264\nNumber of predicted TMHs: 1\nExp number of AAs in TMHs: 22.00715\nExp number, first 60 AAs: 0.03637\nTotal prob of N-in: 0.00393\noutside 1 233\nTMhelix 234 256\ninside 257 264

32120 GCF_000620645.1_ASM62064v1 Nocardioides sp. URHA0032 Terrabacteria group; Actinobacteria; Actinobacteria; Propionibacteriales; Nocardioidaceae; Nocardioides
MAFRPTAAFFDLDKTVIAKSSTLAFSKPFQAGGLISRRAVLRSAYAQFVYLVGGADHDQM WP_028638696.1 inhibition of morphological differentiation protein [Nocardioides sp. URHA0032] Length: 266\nNumber of

predicted TMHs: 1\nExp number of AAs in TMHs: 22.02439\nExp number, first 60 AAs: 0.01651\nTotal prob of N-in: 0.00246\noutside 1 235\nTMhelix 236 258\ninside 259 266

32121 GCF_002165235.1_ASM216523v1 Nocardioides sp. XL1 Terrabacteria group; Actinobacteria; Actinobacteria; Propionibacteriales; Nocardioidaceae; Nocardioides MATRSTAAFFDLDKTIIAKSSTLAFSKPFQAGGLISRRAVLRSAYAQFVYLVGGADHDQM WP_041546100.1 inhibition of morphological differentiation protein [Nocardioides sp. JS614] Length: 266\nNumber of predicted TMHs: 1\nExp number of AAs in TMHs: 19.43393\nExp number, first 60 AAs: 0.05928\nTotal prob of N-in: 0.02059\noutside 1 237\nTMhelix 238 257\ninside 258 266

32122 GCF_002165235.1_ASM216523v1 Nocardioides sp. XL1 Terrabacteria group; Actinobacteria; Actinobacteria; Propionibacteriales; Nocardioidaceae; Nocardioides MRLLNRSAGRLSRHRRGPLAGLAVVLLGLLMTGGLYTVFSPAQAGSAAASEEQIAKGREL WP_011756571.1 cystathionine beta-lyase [Nocardioides sp. JS614] Length: 289\nNumber of predicted TMHs: 2\nExp number of AAs in TMHs: 42.1921699999999\nExp number, first 60 AAs: 21.37698\nTotal prob of N-in: 0.99636\nPOSSIBLE N-term signal sequence\ninside 1 16\nTMhelix 17 39\noutside 40 257\nTMhelix 258 277\ninside 278 289

32123 GCF_000426525.1_ASM42652v1 Nocardioides alkalitolerans DSM 16699 Terrabacteria group; Actinobacteria; Actinobacteria; Propionibacteriales; Nocardioidaceae; Nocardioides; Nocardioides alkalitolerans MRFLNRSAGRISRHRRGPVAGLLVLLLGLALTGTAYAAFAPAGQAEQAVADQELIDEGRE WP_084540955.1 cytochrome C [Nocardioides alkalitolerans] Length: 293\nNumber of predicted TMHs: 2\nExp number of AAs in TMHs: 42.6639899999999\nExp number, first 60 AAs: 21.67154\nTotal prob of N-in: 0.99841\nPOSSIBLE N-term signal sequence\ninside 1 19\nTMhelix 20 42\noutside 43 261\nTMhelix 262 281\ninside 282 293

32124 GCF_000426525.1_ASM42652v1 Nocardioides alkalitolerans DSM 16699 Terrabacteria group; Actinobacteria; Actinobacteria; Propionibacteriales; Nocardioidaceae; Nocardioides; Nocardioides alkalitolerans MLEPDASPPRAARRPSRRSAVLRSVLAITVVVVGLALMLGSPPAPSGSVAGTATAPALDP WP_028474384.1 hypothetical protein [Nocardioides alkalitolerans] Length: 797\nNumber of predicted TMHs: 2\nExp number of AAs in TMHs: 42.76385\nExp number, first 60 AAs: 20.94882\nTotal prob of N-in: 0.99501\nPOSSIBLE N-term signal sequence\ninside 1 19\nTMhelix 20 42\noutside 43 760\nTMhelix 761 783\ninside 784 797

32125 GCF_000426525.1_ASM42652v1 Nocardioides alkalitolerans DSM 16699 Terrabacteria group; Actinobacteria; Actinobacteria; Propionibacteriales; Nocardioidaceae; Nocardioides; Nocardioides alkalitolerans MPGSAPAGRSAAFFDLDKTIIAKSSTLAFSKPFQAGGLISRRAVLRSAYAQFVYLVGGAD WP_028474727.1 inhibition of morphological differentiation protein [Nocardioides alkalitolerans] Length: 268\nNumber of predicted TMHs: 1\nExp number of AAs in TMHs: 22.06708\nExp number, first 60 AAs: 0.01362\nTotal prob of N-in: 0.00142\noutside 1 236\nTMhelix 237 259\ninside 260 268

32126 GCF_000426525.1_ASM42652v1 Nocardioides alkalitolerans DSM 16699 Terrabacteria group; Actinobacteria; Actinobacteria; Propionibacteriales; Nocardioidaceae; Nocardioides; Nocardioides alkalitolerans MPRLSALSPSTDHSRRAPVLGLLVALVAATLVPLLAAAPAQAASRVSVANPDGDAVVDAT WP_052336245.1 hypothetical protein [Nocardioides alkalitolerans] Length: 382\nNumber of predicted TMHs: 2\nExp number of AAs in TMHs: 43.4149399999999\nExp number, first 60 AAs: 20.41703\nTotal prob of N-in: 0.93549\nPOSSIBLE N-term signal sequence\ninside 1 19\nTMhelix 20 42\noutside 43 353\nTMhelix 354 376\ninside 377 382

32127 GCF_001653335.1_ASM165333v1 Nocardioides dokdonensis FR1436 Terrabacteria group; Actinobacteria; Actinobacteria; Propionibacteriales; Nocardioidaceae; Nocardioides; Nocardioides dokdonensis MRFVNRSAGRLSRHRRGPLAGLVLLLVGLVTAGGLYTALAPAQAQDSDTLAIEVEKGREL WP_068110452.1 cytochrome C [Nocardioides dokdonensis] Length: 289\nNumber of predicted TMHs: 2\nExp number of AAs in TMHs: 42.99969\nExp number, first 60 AAs: 21.84417\nTotal prob of N-in: 0.99872\nPOSSIBLE N-term signal sequence\ninside 1 19\nTMhelix 20 42\noutside 43 257\nTMhelix 258 277\ninside 278 289

32128 GCF_001653335.1_ASM165333v1 Nocardioides dokdonensis FR1436 Terrabacteria group; Actinobacteria; Actinobacteria; Propionibacteriales; Nocardioidaceae; Nocardioides; Nocardioides dokdonensis MSESTDLQRRLDALAERAVREAGPARPDDAWSRGRAWQRRRRHVTSAVVGVAVLVVLVVG WP_068112638.1 hypothetical protein [Nocardioides dokdonensis] Length: 452\nNumber of predicted TMHs: 2\nExp number of AAs in TMHs: 41.36426\nExp number, first 60 AAs: 16.62165\nTotal prob of N-in: 0.99930\nPOSSIBLE N-term signal sequence\ninside 1 43\nTMhelix 44 66\noutside 67 426\nTMhelix 427 446\ninside 447 452

32129 GCF_001653335.1_ASM165333v1 Nocardioides dokdonensis FR1436 Terrabacteria group; Actinobacteria; Actinobacteria; Propionibacteriales; Nocardioidaceae; Nocardioides; Nocardioides dokdonensis MTASAAFFDLDKTIIAKSSTLAFSKPFQAGGLISRRAVLRSAYAQFVYLVGGADHDQMEK WP_068107955.1 inhibition of morphological differentiation protein [Nocardioides dokdonensis] Length: 261\nNumber of predicted TMHs: 1\nExp number of AAs in TMHs: 20.65475\nExp number, first 60 AAs: 0.04711\nTotal prob of N-in: 0.05759\noutside 1 234\nTMhelix 235 254\ninside 255 261

32130 GCF_000422825.1_ASM42282v1 Nocardioides insulae DSM 17944 Terrabacteria group; Actinobacteria; Actinobacteria; Propionibacteriales; Nocardioidaceae; Nocardioides; Nocardioides insulae MNARNSPSRPRRGREERRGLLRPLPALLVALLLGGALGTASAVVPGHHPATTASDYPSDQ WP_028659604.1 hypothetical protein [Nocardioides insulae] Length: 252\nNumber of predicted TMHs: 2\nExp number of AAs in TMHs: 45.52471\nExp number, first 60 AAs: 22.49084\nTotal prob of N-in: 0.99566\nPOSSIBLE N-term signal sequence\ninside 1 19\nTMhelix 20 42\noutside 43 220\nTMhelix 221 243\ninside 244 252

32131 GCF_001552535.1_ASM155253v1 Nocardioides jensenii JCM 1364 = NBRC 14755 Terrabacteria group; Actinobacteria; Actinobacteria; Propionibacteriales; Nocardioidaceae; Nocardioides; Nocardioides jensenii MRISTLVARRSRRATIGMLFAALGSFALLAPAPATPALPSSAADVRSEQWWLDVLGVEKIWP_082748224.1 hypothetical protein [Nocardioides jensenii] Length: 430\nNumber of predicted TMHs: 2\nExp number of AAs in TMHs: 42.49509\nExp number, first 60 AAs: 20.37693\nTotal prob of N-in: 0.94965\nPOSSIBLE N-term signal sequence\ninside 1 12\nTMhelix 13 35\noutside 36 396\nTMhelix 397 419\ninside 420 430

32132 GCF_000785495.1_ASM78549v1 Pimelobacter simplex Terrabacteria group; Actinobacteria; Actinobacteria; Propionibacteriales; Nocardioidaceae; Pimelobacter MSRRRSALVGAAVAGVLSFALLTPGSGTAVAQTADSAPVRASAPAKDGFRTTKSIDRTFV WP_038677511.1 hypothetical protein [Pimelobacter simplex] Length: 864\nNumber of predicted TMHs: 2\nExp number of AAs in TMHs: 41.86391\nExp number, first 60 AAs: 19.98799\nTotal prob of N-in: 0.91705\nPOSSIBLE N-term signal sequence\ninside 1 6\nTMhelix 7 29\noutside 30 830\nTMhelix 831 853\ninside 854 864

32133 GCF_900114845.1_IMG-taxon_2687453782_annotated_assembly Pimelobacter simplex Terrabacteria group; Actinobacteria; Actinobacteria; Propionibacteriales; Nocardioidaceae; Pimelobacter MSRRRSALVGAAVAGVLSFALLTPGSGTAVAQTADSAPVRASAPAKDGFRTTKSIDRTFV WP_038677511.1 hypothetical protein [Pimelobacter simplex] Length: 864\nNumber of predicted TMHs: 2\nExp number of AAs in TMHs: 41.86391\nExp number, first 60 AAs: 19.98799\nTotal prob of N-in: 0.91705\nPOSSIBLE N-term signal sequence\ninside 1 6\nTMhelix 7 29\noutside 30 830\nTMhelix 831 853\ninside 854 864

32134 GCF_900114845.1_IMG-taxon_2687453782_annotated_assembly Pimelobacter simplex Terrabacteria group; Actinobacteria; Actinobacteria; Propionibacteriales; Nocardioidaceae; Pimelobacter MGPMAPAGTPPTAAFFDLDKTIIAKSSVLAFSKPFQAGGLISRRAVLRSAYAQFVFMVGG WP_038676148.1 inhibition of morphological differentiation protein [Pimelobacter simplex] Length: 272\nNumber of predicted TMHs: 1\nExp number of AAs in TMHs: 20.18839\nExp number, first 60 AAs: 0.01403\nTotal prob of N-in: 0.00386\noutside 1 245\nTMhelix 246 265\ninside 266 272

32135 GCF_000785495.1_ASM78549v1 Pimelobacter simplex Terrabacteria group; Actinobacteria; Actinobacteria; Propionibacteriales; Nocardioidaceae; Pimelobacter MGPMAPAGTPPTAAFFDLDKTIIAKSSVLAFSKPFQAGGLISRRAVLRSAYAQFVFMVGG WP_038676148.1 inhibition of morphological differentiation protein [Pimelobacter simplex] Length: 272\nNumber of predicted TMHs: 1\nExp number of AAs in TMHs: 20.18839\nExp number, first 60 AAs: 0.01403\nTotal prob of N-in: 0.00386\noutside 1 245\nTMhelix 246 265\ninside 266 272

32136 GCF_000423465.1_ASM42346v1 Propionicicella superfundia DSM 22317Terrabacteria group; Actinobacteria; Actinobacteria; Propionibacteriales; Nocardioidaceae; Propionicicella; Propionicicella superfundia MTHADRPARTRALRRRRPVLGVLGIVLAGAALLAPSAAWAETAPPDDPNLKQTLSPDLAI WP_084634577.1 hypothetical protein [Propionicicella superfundia] Length: 377\nNumber of predicted TMHs: 2\nExp number of AAs in TMHs: 44.40327\nExp number, first 60 AAs: 21.60842\nTotal prob of N-in: 0.99919\nPOSSIBLE N-term signal sequence\ninside 1 19\nTMhelix 20 42\noutside 43 306\nTMhelix 307 329\ninside 330 377

32137 GCF_000145455.1_ASM14545v1 Propionibacterium acnes HL027PA1 Terrabacteria group; Actinobacteria; Actinobacteria; Propionibacteriales; Propionibacteriaceae; Cutibacterium; Cutibacterium acnes MMANNYGVSRRTLLGVGGLMLGVGLVGCSDSEEGWESVTVDRITVSHPTEWVEKPPENEN WP_002529663.1 hypothetical protein [Cutibacterium acnes] Length: 174\nNumber of predicted TMHs: 1\nExp number of

AAs in TMHs: 25.993\nExp number, first 60 AAs: 6.91708\nTotal prob of N-in: 0.85026\ninside 1 133\nTMhelix 134 156\noutside 157 174

32138 GCF_000270245.1_ASM27024v1 Microlunatus phosphovorus NM-1 Terrabacteria group; Actinobacteria; Actinobacteria; Propionibacteriales; Propionibacteriaceae; Microlunatus; Microlunatus phosphovorus MLLANQLPAGRTVRRRPTWRRRALLTAAGLTATSLTLLPITAEAHVRVISDGANSGGYSAWP_049804801.1 hypothetical protein [Microlunatus phosphovorus] Length: 263\nNumber of predicted TMHs: 1\nExp number of AAs in TMHs: 29.0719\nExp number, first 60 AAs: 8.81589\nTotal prob of N-in: 0.43893\noutside 1 234\nTMhelix 235 254\ninside 255 263

32139 GCF_000413335.1_Prop_sp_HGH0353_V1 Propionibacterium sp. HGH0353 Terrabacteria group; Actinobacteria; Actinobacteria; Propionibacteriales; Propionibacteriaceae; Propionibacterium MTAGDVMSRRSVLAGGALGGLTIGAGLVGCSSSQNADAQGDSSWVKAEVGRVSVRHPAEW WP_016666890.1 hypothetical protein [Propionibacterium sp. HGH0353] Length: 184\nNumber of predicted TMHs: 1\nExp number of AAs in TMHs: 29.00499\nExp number, first 60 AAs: 9.15704\nTotal prob of N-in: 0.67196\ninside 1 142\nTMhelix 143 165\noutside 166 184

32140 GCF_001997345.1_ASM199734v1 Tessaracoccus sp. NSG39 Terrabacteria group; Actinobacteria; Actinobacteria; Propionibacteriales; Propionibacteriaceae; Tessaracoccus MSTNDSLGSAPRRRRSRLAAREGRRRALAIGIVSALSASLVFASGLAYANDPTSPPEPVE WP_077686904.1 hypothetical protein [Tessaracoccus sp. NSG39] Length: 1433\nNumber of predicted TMHs: 2\nExp number of AAs in TMHs: 39.90628\nExp number, first 60 AAs: 20.99707\nTotal prob of N-in: 0.93262\nPOSSIBLE N-term signal sequence\ninside 1 26\nTMhelix 27 49\noutside 50 1407\nTMhelix 1408 1425\ninside 1426 1433

32141 GCF_001997345.1_ASM199734v1 Tessaracoccus sp. NSG39 Terrabacteria group; Actinobacteria; Actinobacteria; Propionibacteriales; Propionibacteriaceae; Tessaracoccus MSLTESTRTTARRGVLARLTALLALTFMALMAALPAPAHADGADNEVGACLSADKVWLLI WP_077686118.1 hypothetical protein [Tessaracoccus sp. NSG39] Length: 249\nNumber of predicted TMHs: 2\nExp number of AAs in TMHs: 43.21645\nExp number, first 60 AAs: 20.69734\nTotal prob of N-in: 0.98251\nPOSSIBLE N-term signal sequence\ninside 1 12\nTMhelix 13 35\noutside 36 213\nTMhelix 214 236\ninside 237 249

32142 GCF_001956895.1_ASM195689v1 Tessaracoccus sp. ZS01 Terrabacteria group; Actinobacteria; Actinobacteria; Propionibacteriales; Propionibacteriaceae; Tessaracoccus MRRKKRLKIGWSPLVALIGALVPVAALAPRRPLVSGPHTVAALAQQCRETGATGRALADE WP_083665280.1 hypothetical protein [Tessaracoccus sp. ZS01] Length: 214\nNumber of predicted TMHs: 2\nExp number of AAs in TMHs: 38.50518\nExp number, first 60 AAs: 18.80592\nTotal prob of N-in: 0.98413\nPOSSIBLE N-term signal sequence\ninside 1 6\nTMhelix 7 29\noutside 30 169\nTMhelix 170 192\ninside 193 214

32143 GCF_000619965.1_ASM61996v1 Propionibacteriaceae bacterium P6A17 Terrabacteria group; Actinobacteria; Actinobacteria; Propionibacteriales; Propionibacteriaceae; unclassified Propionibacteriaceae MAPTTAAQIRRGRRPLLRSALLGGLLVAAVIAGLLAMHTFNLHGALPTREAVAAPLGHSG WP_028705928.1 hypothetical protein [Propionibacteriaceae bacterium P6A17] Length: 156\nNumber of predicted TMHs: 2\nExp number of AAs in TMHs: 41.65076\nExp number, first 60 AAs: 21.14987\nTotal prob of N-in: 0.99803\nPOSSIBLE N-term signal sequence\ninside 1 19\nTMhelix 20 42\noutside 43 93\nTMhelix 94 116\ninside 117 156

32144 GCF_000716095.1_ASM71609v1 Actinoalloteichus cyanogriseus Terrabacteria group; Actinobacteria; Actinobacteria; Pseudonocardiales; Pseudonocardiaceae; Actinoalloteichus MTKASPRRGLLTAVLAAGLVAAGSALGATTAAAQVEEEPAGETLLDVAAEALPTEMRIAV WP_026419747.1 hypothetical protein [Actinoalloteichus cyanogriseus] Length: 695\nNumber of predicted TMHs: 2\nExp number of AAs in TMHs: 40.91056\nExp number, first 60 AAs: 19.00368\nTotal prob of N-in: 0.84396\nPOSSIBLE N-term signal sequence\ninside 1 12\nTMhelix 13 35\noutside 36 659\nTMhelix 660 682\ninside 683 695

32145 GCF_000718005.1_ASM71800v1 Actinoalloteichus cyanogriseus Terrabacteria group; Actinobacteria; Actinobacteria; Pseudonocardiales; Pseudonocardiaceae; Actinoalloteichus MTKASPRRGLLTAVLAAGLVAAGSALGATTAAAQVEEEPAGETLLDVAAEALPTEMRIAV WP_026419747.1 hypothetical protein [Actinoalloteichus cyanogriseus] Length: 695\nNumber of predicted TMHs: 2\nExp number of AAs in TMHs: 40.91056\nExp number, first 60 AAs: 19.00368\nTotal prob of N-in: 0.84396\nPOSSIBLE N-term signal sequence\ninside 1 12\nTMhelix 13 35\noutside 36 659\nTMhelix 660 682\ninside 683 695

32146 GCF_001747425.1_ASM174742v1 Actinoalloteichus hymeniacidonis Terrabacteria group; Actinobacteria; Actinobacteria; Pseudonocardiales; Pseudonocardiaceae; Actinoalloteichus

MIFSMGRTRRAVVAVATGLLAVSIPLTATPALAQSEETEEAITDIARYAPADPANLPQNV WP_084643428.1 type VII secretion-associated serine protease mycosin [Actinoalloteichus hymeniacidonis] Length: 476\nNumber of predicted TMHs: 1\nExp number of AAs in TMHs: 36.93763\nExp number, first 60 AAs: 14.16061\nTotal prob of N-in: 0.66206\nPOSSIBLE N-term signal sequence\noutside 1 440\nTMhelix 441 463\ninside 464 476

32147 GCF_000429185.1_ASM42918v1 Actinoalloteichus cyanogriseus DSM 43889 Terrabacteria group; Actinobacteria; Actinobacteria; Pseudonocardiales; Pseudonocardiaceae; Actinoalloteichus; Actinoalloteichus cyanogriseus MTKASPRRGLLTAVLAAGLVAAGSALGATTAAAQVEEEPAGETLLDVAAEALPTEMRIAV WP_026419747.1 hypothetical protein [Actinoalloteichus cyanogriseus] Length: 695\nNumber of predicted TMHs: 2\nExp number of AAs in TMHs: 40.91056\nExp number, first 60 AAs: 19.00368\nTotal prob of N-in: 0.84396\nPOSSIBLE N-term signal sequence\ninside 1 12\nTMhelix 13 35\noutside 36 659\nTMhelix 660 682\ninside 683 695

32148 GCF_000239155.1_ASM23915v2 Actinoalloteichus spitiensis RMV-1378 Terrabacteria group; Actinobacteria; Actinobacteria; Pseudonocardiales; Pseudonocardiaceae; Actinoalloteichus; Actinoalloteichus spitiensis MTKASPRRGLLTAVLAAGLVAAGSALGATTAAAQAEEEPTGETLLEVAAEALPTEMRIAV WP_016699903.1 hypothetical protein [Actinoalloteichus spitiensis] Length: 695\nNumber of predicted TMHs: 2\nExp number of AAs in TMHs: 40.59508\nExp number, first 60 AAs: 18.41822\nTotal prob of N-in: 0.82190\nPOSSIBLE N-term signal sequence\ninside 1 12\nTMhelix 13 35\noutside 36 661\nTMhelix 662 684\ninside 685 695

32149 GCF_001940455.1_ASM194045v1 Actinokineospora bangkokensis Terrabacteria group; Actinobacteria; Actinobacteria; Pseudonocardiales; Pseudonocardiaceae; Actinokineospora MISQKTRRGAVIGALTAAVLMAGAAPASAAPGDGSAYVAAGTVTLLGVPAVTIPPQAATS WP_075978649.1 hypothetical protein [Actinokineospora bangkokensis] Length: 246\nNumber of predicted TMHs: 2\nExp number of AAs in TMHs: 81.49001\nExp number, first 60 AAs: 31.12588\nTotal prob of N-in: 0.90852\nPOSSIBLE N-term signal sequence\ninside 1 8\nTMhelix 9 31\noutside 32 216\nTMhelix 217 239\ninside 240 246

32150 GCF_001940455.1_ASM194045v1 Actinokineospora bangkokensis Terrabacteria group; Actinobacteria; Actinobacteria; Pseudonocardiales; Pseudonocardiaceae; Actinokineospora MRARRSALAAVAVLTATVVVPGLGSPATAQQGNTLRIAVTQEVDSLNPFLSSTRTGTDILWP_075974846.1 peptide ABC transporter substrate-binding protein [Actinokineospora bangkokensis] Length: 595\nNumber of predicted TMHs: 2\nExp number of AAs in TMHs: 43.7075\nExp number, first 60 AAs: 21.58158\nTotal prob of N-in: 0.96291\nPOSSIBLE N-term signal sequence\ninside 1 6\nTMhelix 7 29\noutside 30 561\nTMhelix 562 584\ninside 585 595

32151 GCF_000482865.1_ASM48286v1 Actinokineospora inagensis DSM 44258 Terrabacteria group; Actinobacteria; Actinobacteria; Pseudonocardiales; Pseudonocardiaceae; Actinokineospora; Actinokineospora inagensis MLKRWPRRACLVGAVVTTAVLAGALPASAAPGDGSAVVVTADIALLGRSTANVGPLAPSS WP_026421352.1 hypothetical protein [Actinokineospora inagensis] Length: 243\nNumber of predicted TMHs: 1\nExp number of AAs in TMHs: 39.59695\nExp number, first 60 AAs: 17.18125\nTotal prob of N-in: 0.82066\nPOSSIBLE N-term signal sequence\noutside 1 212\nTMhelix 213 235\ninside 236 243

32152 GCF_000379625.1_ASM37962v1 Actinomycetospora chiangmaiensis DSM 45062 Terrabacteria group; Actinobacteria; Actinobacteria; Pseudonocardiales; Pseudonocardiaceae; Actinomycetospora; Actinomycetospora chiangmaiensis MTVLQPKRRAAVERSASAVSAASAVSAAATVAAPAARRSWRVSSAPQPAVGHLVVDASHS WP_018334839.1 hypothetical protein [Actinomycetospora chiangmaiensis] Length: 174\nNumber of predicted TMHs: 1\nExp number of AAs in TMHs: 24.6358\nExp number, first 60 AAs: 1.21094\nTotal prob of N-in: 0.89513\ninside 1 147\nTMhelix 148 170\noutside 171 174

32153 GCF_001921215.1_ASM192121v1 Actinophytocola xinjiangensis Terrabacteria group; Actinobacteria; Actinobacteria; Pseudonocardiales; Pseudonocardiaceae; Actinophytocola MARRTVMALFARACAVALGLSLGAVAGVAAADTTTGTVDPAGDRPGFTVDVGDGVFAALD WP_075135817.1 hypothetical protein [Actinophytocola xinjiangensis] Length: 364\nNumber of predicted TMHs: 2\nExp number of AAs in TMHs: 42.20061\nExp number, first 60 AAs: 19.387\nTotal prob of N-in: 0.86937\nPOSSIBLE N-term signal sequence\ninside 1 8\nTMhelix 9 31\noutside 32 334\nTMhelix 335 357\ninside 358 364

32154 GCF_001984155.1_ASM198415v1 Actinosynnema sp. ALI-1.44 Terrabacteria group; Actinobacteria; Actinobacteria; Pseudonocardiales; Pseudonocardiaceae; Actinosynnema MTDAPKRVAAFFDLDKTVIAKSSTLAFSRPFFHGGLINRRAVLKSAYAQFVFMQSGADAD WP_076990428.1 inhibition of morphological differentiation protein [Actinosynnema sp. ALI-1.44] Length: 267\nNumber of predicted TMHs: 1\nExp number of AAs in TMHs: 22.63859\nExp number, first 60 AAs: 0.01898\nTotal prob of N-in: 0.01217\noutside 1 238\nTMhelix 239 261\ninside 262 267

32155 GCF_000023245.1_ASM2324v1 Actinosynnema mirum DSM 43827 Terrabacteria group; Actinobacteria; Actinobacteria; Pseudonocardiales; Pseudonocardiaceae; Actinosynnema; Actinosynnema mirum MTDPAAEGTRVAAFFDLDKTVIAKSSTLAFSRPFFQEGLINRRAVLKSAYAQFVFMLAGA WP_012782906.1 haloacid dehalogenase [Actinosynnema mirum] Length: 270\nNumber of predicted TMHs: 1\nExp number of AAs in TMHs: 22.90977\nExp number, first 60 AAs: 0.06825\nTotal prob of N-in: 0.02678\noutside 1 241\nTMhelix 242 264\ninside 265 270

32156 GCF_900119165.1_IMG-taxon_2642422517_annotated_assembly Amycolatopsis australiensis Terrabacteria group; Actinobacteria; Actinobacteria; Pseudonocardiales; Pseudonocardiaceae; Amycolatopsis MSRRALVLAVLTGAALLMTAPVSGAAQAAGISSADAAAAHAAAVRLQDTLVGFFAQAPDA WP_072476476.1 hypothetical protein [Amycolatopsis australiensis] Length: 252\nNumber of predicted TMHs: 2\nExp number of AAs in TMHs: 43.29093\nExp number, first 60 AAs: 21.07751\nTotal prob of N-in: 0.95492\nPOSSIBLE N-term signal sequence\ninside 1 6\nTMhelix 7 29\noutside 30 224\nTMhelix 225 247\ninside 248 252

32157 GCF_001953865.1_ASM195386v1 Amycolatopsis coloradensis Terrabacteria group; Actinobacteria; Actinobacteria; Pseudonocardiales; Pseudonocardiaceae; Amycolatopsis MAQPSTTPSPVAAFFDLDKTIIASSSALAFSKPLLREGLINRRAALRSAYAQLVFSLAGA WP_076155971.1 inhibition of morphological differentiation protein [Amycolatopsis coloradensis] Length: 265\nNumber of predicted TMHs: 1\nExp number of AAs in TMHs: 20.34481\nExp number, first 60 AAs: 0.05492\nTotal prob of N-in: 0.01546\noutside 1 235\nTMhelix 236 258\ninside 259 265

32158 GCF_001953865.1_ASM195386v1 Amycolatopsis coloradensis Terrabacteria group; Actinobacteria; Actinobacteria; Pseudonocardiales; Pseudonocardiaceae; Amycolatopsis MKSAKTTRRTVLTGVAAAAAFGLATGPAQAETKGLRIGILLYDGFSLLDPTGPAEVLSRL WP_076168824.1 thiamine biosynthesis protein ThiJ [Amycolatopsis coloradensis] Length: 247\nNumber of predicted TMHs: 2\nExp number of AAs in TMHs: 33.4458\nExp number, first 60 AAs: 4.99306\nTotal prob of N-in: 0.91037\ninside 1 130\nTMhelix 131 148\noutside 149 180\nTMhelix 181 203\ninside 204 247

32159 GCF_000732925.1_ASM73292v1 Amycolatopsis japonica Terrabacteria group; Actinobacteria; Actinobacteria; Pseudonocardiales; Pseudonocardiaceae; Amycolatopsis MDRPSTTPSQVAAFFDLDKTIIASSSALAFSKPLLREGLINRRAALRSAYAQLVFSLAGA WP_016337846.1 MULTISPECIES: haloacid dehalogenase [Amycolatopsis] Length: 265\nNumber of predicted TMHs: 1\nExp number of AAs in TMHs: 21.25757\nExp number, first 60 AAs: 0.07089\nTotal prob of N-in: 0.02314\noutside 1 235\nTMhelix 236 258\ninside 259 265

32160 GCF_002155975.1_ASM215597v1 Amycolatopsis kentuckyensis Terrabacteria group; Actinobacteria; Actinobacteria; Pseudonocardiales; Pseudonocardiaceae; Amycolatopsis MSRRAIVLAVLTGAALLMTAPVSGAAQAAGISSADASAAHGAAAGFQDVLVRFFAQAPGA WP_086837935.1 hypothetical protein [Amycolatopsis kentuckyensis] Length: 249\nNumber of predicted TMHs: 2\nExp number of AAs in TMHs: 43.97093\nExp number, first 60 AAs: 21.5551\nTotal prob of N-in: 0.97252\nPOSSIBLE N-term signal sequence\ninside 1 6\nTMhelix 7 29\noutside 30 221\nTMhelix 222 244\ninside 245 249

32161 GCF_002155975.1_ASM215597v1 Amycolatopsis kentuckyensis Terrabacteria group; Actinobacteria; Actinobacteria; Pseudonocardiales; Pseudonocardiaceae; Amycolatopsis MAHRRKPLARSLFALLTAVLGVLALAPPAAAQVSIVPDRVDGGTHTFAFRLANGRADTKS WP_086840305.1 hypothetical protein [Amycolatopsis kentuckyensis] Length: 251\nNumber of predicted TMHs: 2\nExp number of AAs in TMHs: 42.8345899999999\nExp number, first 60 AAs: 21.74991\nTotal prob of N-in: 0.99474\nPOSSIBLE N-term signal sequence\ninside 1 12\nTMhelix 13 35\noutside 36 207\nTMhelix 208 227\ninside 228 251

32162 GCF_900105855.1_IMG-taxon_2634166275_annotated_assembly Amycolatopsis keratiniphila Terrabacteria group; Actinobacteria; Actinobacteria; Pseudonocardiales; Pseudonocardiaceae; Amycolatopsis MDRPSTTPSQVAAFFDLDKTIIASSSALAFSKPLLREGLINRRAALRSAYAQLVFSLAGA WP_072029633.1 inhibition of morphological differentiation protein [Amycolatopsis keratiniphila] Length: 265\nNumber of predicted TMHs: 1\nExp number of AAs in TMHs: 21.15989\nExp number, first 60 AAs: 0.06604\nTotal prob of N-in: 0.01470\noutside 1 235\nTMhelix 236 258\ninside 259 265

32163 GCF_000700945.1_ASM70094v1 Amycolatopsis mediterranei Terrabacteria group; Actinobacteria; Actinobacteria; Pseudonocardiales; Pseudonocardiaceae; Amycolatopsis MSRRALVLAVLTGAALLMTAPVSGAAQAAGISSADASAAHDAAAGFQDVLVRFFAQAPGA WP_013229737.1 hypothetical protein [Amycolatopsis mediterranei] Length: 251\nNumber of predicted TMHs: 2\nExp

number of AAs in TMHs: 43.69728\nExp number, first 60 AAs: 21.22417\nTotal prob of N-in: 0.96054\nPOSSIBLE N-term signal sequence\ninside 1 6\nTMhelix 7 29\noutside 30 223\nTMhelix 224 246\ninside 247 251

32164 GCF_000696405.1_ASM69640v1 Amycolatopsis mediterranei Terrabacteria group; Actinobacteria; Actinobacteria; Pseudonocardiales; Pseudonocardiaceae; Amycolatopsis MSRRALVLAVLTGAALLMTAPVSGAAQAAGISSADASAAHDAAAGFQDVLVRFFAQAPGA WP_013229737.1 hypothetical protein [Amycolatopsis mediterranei] Length: 251\nNumber of predicted TMHs: 2\nExp number of AAs in TMHs: 43.69728\nExp number, first 60 AAs: 21.22417\nTotal prob of N-in: 0.96054\nPOSSIBLE N-term signal sequence\ninside 1 6\nTMhelix 7 29\noutside 30 223\nTMhelix 224 246\ninside 247 251

32165 GCF_001742805.1_ASM174280v1 Amycolatopsis mediterranei Terrabacteria group; Actinobacteria; Actinobacteria; Pseudonocardiales; Pseudonocardiaceae; Amycolatopsis MSRRALVLAVLTGAALLMTAPVSGAAQAAGISSADASAAHDAAAGFQDVLVRFFAQAPGA WP_013229737.1 hypothetical protein [Amycolatopsis mediterranei] Length: 251\nNumber of predicted TMHs: 2\nExp number of AAs in TMHs: 43.69728\nExp number, first 60 AAs: 21.22417\nTotal prob of N-in: 0.96054\nPOSSIBLE N-term signal sequence\ninside 1 6\nTMhelix 7 29\noutside 30 223\nTMhelix 224 246\ninside 247 251

32166 GCF_000943515.2_ASM94351v2 Amycolatopsis orientalis Terrabacteria group; Actinobacteria; Actinobacteria; Pseudonocardiales; Pseudonocardiaceae; Amycolatopsis MTPSPDRQDPPPAASPTTITQYQILATRRQAFDTLMWQVPALSLTAQSFLLSLAYGSQST WP_065912755.1 hypothetical protein [Amycolatopsis orientalis] Length: 155\nNumber of predicted TMHs: 2\nExp number of AAs in TMHs: 53.26599\nExp number, first 60 AAs: 9.62812\nTotal prob of N-in: 0.40727\noutside 1 60\nTMhelix 61 83\ninside 84 130\nTMhelix 131 153\noutside 154 155

32167 GCF_000943515.2_ASM94351v2 Amycolatopsis orientalis Terrabacteria group; Actinobacteria; Actinobacteria; Pseudonocardiales; Pseudonocardiaceae; Amycolatopsis MAQPSTTPSPVAAFFDLDKTIIASSSALAFSKPLLREGLINRRAALRSAYAQLVFSLAGA WP_044853684.1 inhibition of morphological differentiation protein [Amycolatopsis orientalis] Length: 265\nNumber of predicted TMHs: 1\nExp number of AAs in TMHs: 21.22898\nExp number, first 60 AAs: 0.0406\nTotal prob of N-in: 0.00848\noutside 1 235\nTMhelix 236 258\ninside 259 265

32168 GCF_001613935.1_ASM161393v1 Amycolatopsis regifaucium Terrabacteria group; Actinobacteria; Actinobacteria; Pseudonocardiales; Pseudonocardiaceae; Amycolatopsis MAQPSTTPSPVAAFFDLDKTIIASSSALAFSKPLLREGLINRRAALRSAYAQLVFSLAGA WP_061985788.1 inhibition of morphological differentiation protein [Amycolatopsis regifaucium] Length: 265\nNumber of predicted TMHs: 1\nExp number of AAs in TMHs: 20.37475\nExp number, first 60 AAs: 0.04932\nTotal prob of N-in: 0.01281\noutside 1 235\nTMhelix 236 258\ninside 259 265

32169 GCF_001558125.2_ASM155812v2 Amycolatopsis regifaucium Terrabacteria group; Actinobacteria; Actinobacteria; Pseudonocardiales; Pseudonocardiaceae; Amycolatopsis MAQPSTTPSPVAAFFDLDKTIIASSSALAFSKPLLREGLINRRAALRSAYAQLVFSLAGA WP_061985788.1 inhibition of morphological differentiation protein [Amycolatopsis regifaucium] Length: 265\nNumber of predicted TMHs: 1\nExp number of AAs in TMHs: 20.37475\nExp number, first 60 AAs: 0.04932\nTotal prob of N-in: 0.01281\noutside 1 235\nTMhelix 236 258\ninside 259 265

32170 GCF_900113515.1_IMG-taxon_2634166327_annotated_assembly Amycolatopsis regifaucium Terrabacteria group; Actinobacteria; Actinobacteria; Pseudonocardiales; Pseudonocardiaceae; Amycolatopsis MAQPSTTPSPVAAFFDLDKTIIASSSALAFSKPLLREGLINRRAALRSAYAQLVFSLAGA WP_061985788.1 inhibition of morphological differentiation protein [Amycolatopsis regifaucium] Length: 265\nNumber of predicted TMHs: 1\nExp number of AAs in TMHs: 20.37475\nExp number, first 60 AAs: 0.04932\nTotal prob of N-in: 0.01281\noutside 1 235\nTMhelix 236 258\ninside 259 265

32171 GCF_000695625.1_ASM69562v1 Amycolatopsis rifamycinica Terrabacteria group; Actinobacteria; Actinobacteria; Pseudonocardiales; Pseudonocardiaceae; Amycolatopsis MSRRALVLAVLTGAALLMTAPVSGAAQAAGISSADAFAAHSVAAGYQDVLVRFFAQAPGA WP_043786267.1 hypothetical protein [Amycolatopsis rifamycinica] Length: 249\nNumber of predicted TMHs: 2\nExp number of AAs in TMHs: 44.28505\nExp number, first 60 AAs: 21.88341\nTotal prob of N-in: 0.97372\nPOSSIBLE N-term signal sequence\ninside 1 6\nTMhelix 7 29\noutside 30 221\nTMhelix 222 244\ninside 245 249

32172 GCF_001905755.1_ASM190575v1 Amycolatopsis sp. CB00013 Terrabacteria group; Actinobacteria; Actinobacteria; Pseudonocardiales; Pseudonocardiaceae; Amycolatopsis

MDRPSTTPSQVAAFFDLDKTIIASSSALAFSKPLLREGLINRRAALRSAYAQLVFSLAGA WP_016337846.1 MULTISPECIES: haloacid dehalogenase [Amycolatopsis] Length: 265\nNumber of predicted TMHs: 1\nExp number of AAs in TMHs: 21.25757\nExp number, first 60 AAs: 0.07089\nTotal prob of N-in: 0.02314\noutside 1 235\nTMhelix 236 258\ninside 259 265

32173 GCF_001651785.1_ASM165178v1 Amycolatopsis sp. M39 Terrabacteria group; Actinobacteria; Actinobacteria; Pseudonocardiales; Pseudonocardiaceae; Amycolatopsis MAEPRSEEAPRRVAAFFDLDKTIIASSSALAFSKPLLREGLINRRAALRSAYAQLVFSLA WP_067576983.1 inhibition of morphological differentiation protein [Amycolatopsis sp. M39] Length: 273\nNumber of predicted TMHs: 1\nExp number of AAs in TMHs: 20.58369\nExp number, first 60 AAs: 0.16447\nTotal prob of N-in: 0.02965\noutside 1 242\nTMhelix 243 262\ninside 263 273

32174 GCF_000754115.1_ASM75411v1 Amycolatopsis sp. MJM2582 Terrabacteria group; Actinobacteria; Actinobacteria; Pseudonocardiales; Pseudonocardiaceae; Amycolatopsis MDRPSTTPSQVAAFFDLDKTIIASSSALAFSKPLLREGLINRRAALRSAYAQLVFSLAGA WP_016337846.1 MULTISPECIES: haloacid dehalogenase [Amycolatopsis] Length: 265\nNumber of predicted TMHs: 1\nExp number of AAs in TMHs: 21.25757\nExp number, first 60 AAs: 0.07089\nTotal prob of N-in: 0.02314\noutside 1 235\nTMhelix 236 258\ninside 259 265

32175 GCF_000716785.1_ASM71678v1 Amycolatopsis vancoresmycina Terrabacteria group; Actinobacteria; Actinobacteria; Pseudonocardiales; Pseudonocardiaceae; Amycolatopsis MSRRALVLAVLTGAALLMTAPVSGAAQAAGISSADAVAAHTAAAGLQDTLVRFFAAAPDA WP_033261304.1 hypothetical protein [Amycolatopsis vancoresmycina] Length: 249\nNumber of predicted TMHs: 2\nExp number of AAs in TMHs: 43.5249799999999\nExp number, first 60 AAs: 21.10003\nTotal prob of N-in: 0.95576\nPOSSIBLE N-term signal sequence\ninside 1 6\nTMhelix 7 29\noutside 30 221\nTMhelix 222 244\ninside 245 249

32176 GCF_000716785.1_ASM71678v1 Amycolatopsis vancoresmycina Terrabacteria group; Actinobacteria; Actinobacteria; Pseudonocardiales; Pseudonocardiaceae; Amycolatopsis MLSRTTPDTPRRTVLVRKLLTVAALAAFVTGGAFLSTGTAAATTTLVSNSCTGSVTGQMG WP_033261107.1 hypothetical protein [Amycolatopsis vancoresmycina] Length: 304\nNumber of predicted TMHs: 2\nExp number of AAs in TMHs: 44.4226\nExp number, first 60 AAs: 22.43038\nTotal prob of N-in: 0.96923\nPOSSIBLE N-term signal sequence\ninside 1 19\nTMhelix 20 42\noutside 43 277\nTMhelix 278 300\ninside 301 304

32177 GCF_000384215.1_ASM38421v1 Amycolatopsis alba DSM 44262 Terrabacteria group; Actinobacteria; Actinobacteria; Pseudonocardiales; Pseudonocardiaceae; Amycolatopsis; Amycolatopsis alba MDRPSTTPSPVAAFFDLDKTIIASSSALAFSKPLLREGLINRRAALRSAYAQLVFSLAGA WP_020632106.1 inhibition of morphological differentiation protein [Amycolatopsis alba] Length: 265\nNumber of predicted TMHs: 1\nExp number of AAs in TMHs: 21.22987\nExp number, first 60 AAs: 0.05\nTotal prob of N-in: 0.01159\noutside 1 235\nTMhelix 236 258\ninside 259 265

32178 GCF_000340415.1_Amycolatopsis_azurea Amycolatopsis azurea DSM 43854 Terrabacteria group; Actinobacteria; Actinobacteria; Pseudonocardiales; Pseudonocardiaceae; Amycolatopsis; Amycolatopsis azurea MAQPSTTPSPVAAFFDLDKTIIASSSALAFSKPLLREGLINRRAALRSAYAQLVFSLAGA WP_005168538.1 phosphoserine phosphatase [Amycolatopsis azurea] Length: 265\nNumber of predicted TMHs: 1\nExp number of AAs in TMHs: 21.3345\nExp number, first 60 AAs: 0.04125\nTotal prob of N-in: 0.00874\noutside 1 235\nTMhelix 236 258\ninside 259 265

32179 GCF_001995215.1_ASM199521v1 Amycolatopsis azurea DSM 43854 Terrabacteria group; Actinobacteria; Actinobacteria; Pseudonocardiales; Pseudonocardiaceae; Amycolatopsis; Amycolatopsis azurea MAQPSTTPSPVAAFFDLDKTIIASSSALAFSKPLLREGLINRRAALRSAYAQLVFSLAGA WP_005168538.1 phosphoserine phosphatase [Amycolatopsis azurea] Length: 265\nNumber of predicted TMHs: 1\nExp number of AAs in TMHs: 21.3345\nExp number, first 60 AAs: 0.04125\nTotal prob of N-in: 0.00874\noutside 1 235\nTMhelix 236 258\ninside 259 265

32180 GCF_000383915.1_ASM38391v1 Amycolatopsis benzoatilytica AK 16/65 Terrabacteria group; Actinobacteria; Actinobacteria; Pseudonocardiales; Pseudonocardiaceae; Amycolatopsis; Amycolatopsis benzoatilytica MAELRSEEAPRRVAAFFDLDKTIIASSSALAFSKPLLREGLINRRAALRSAYAQLVFSLA WP_020664196.1 inhibition of morphological differentiation protein [Amycolatopsis benzoatilytica] Length: 273\nNumber of predicted TMHs: 1\nExp number of AAs in TMHs: 20.73735\nExp number, first 60 AAs: 0.14615\nTotal prob of N-in: 0.01653\noutside 1 242\nTMhelix 243 262\ninside 263 273

32181 GCF_000342005.1_Whole_genome_assembly Amycolatopsis decaplanina DSM 44594Terrabacteria group; Actinobacteria; Actinobacteria; Pseudonocardiales; Pseudonocardiaceae; Amycolatopsis; Amycolatopsis decaplanina MDRPSTTPTPVAAFFDLDKTIIASSSALAFSKPLLREGLINRRAALRSAYAQLVFSLAGA WP_007028480.1 haloacid dehalogenase [Amycolatopsis decaplanina] Length: 265\nNumber of predicted TMHs: 1\nExp number of AAs in TMHs: 21.22489\nExp number, first 60 AAs: 0.05973\nTotal prob of N-in: 0.01661\noutside 1 235\nTMhelix 236 258\ninside 259 265

32182 GCF_001620365.2_ASM162036v2 Amycolatopsis keratiniphila subsp. keratiniphila Terrabacteria group; Actinobacteria; Actinobacteria; Pseudonocardiales; Pseudonocardiaceae; Amycolatopsis; Amycolatopsis keratiniphila MDRPSTTPSQVAAFFDLDKTIIASSSALAFSKPLLREGLINRRAALRSAYAQLVFSLAGA WP_016337846.1 MULTISPECIES: haloacid dehalogenase [Amycolatopsis] Length: 265\nNumber of predicted TMHs: 1\nExp number of AAs in TMHs: 21.25757\nExp number, first 60 AAs: 0.07089\nTotal prob of N-in: 0.02314\noutside 1 235\nTMhelix 236 258\ninside 259 265

32183 GCF_001953855.1_ASM195385v1 Amycolatopsis keratiniphila subsp. nogabecina Terrabacteria group; Actinobacteria; Actinobacteria; Pseudonocardiales; Pseudonocardiaceae; Amycolatopsis; Amycolatopsis keratiniphila MDRPSTTPSQVAAFFDLDKTIIASSSALAFSKPLLREGLINRRAALRSAYAQLVFSLAGA WP_072029633.1 inhibition of morphological differentiation protein [Amycolatopsis keratiniphila] Length: 265\nNumber of predicted TMHs: 1\nExp number of AAs in TMHs: 21.15989\nExp number, first 60 AAs: 0.06604\nTotal prob of N-in: 0.01470\noutside 1 235\nTMhelix 236 258\ninside 259 265

32184 GCF_000454025.1_ASM45402v1 Amycolatopsis mediterranei RB Terrabacteria group; Actinobacteria; Actinobacteria; Pseudonocardiales; Pseudonocardiaceae; Amycolatopsis; Amycolatopsis mediterranei MSRRALVLAVLTGAALLMTAPVSGAAQAAGISSADASAAHDAAAGFQDVLVRFFAQAPGA WP_013229737.1 hypothetical protein [Amycolatopsis mediterranei] Length: 251\nNumber of predicted TMHs: 2\nExp number of AAs in TMHs: 43.69728\nExp number, first 60 AAs: 21.22417\nTotal prob of N-in: 0.96054\nPOSSIBLE N-term signal sequence\ninside 1 6\nTMhelix 7 29\noutside 30 223\nTMhelix 224 246\ninside 247 251

32185 GCF_000220945.1_ASM22094v1 Amycolatopsis mediterranei S699 Terrabacteria group; Actinobacteria; Actinobacteria; Pseudonocardiales; Pseudonocardiaceae; Amycolatopsis; Amycolatopsis mediterranei MSRRALVLAVLTGAALLMTAPVSGAAQAAGISSADASAAHDAAAGFQDVLVRFFAQAPGA WP_013229737.1 hypothetical protein [Amycolatopsis mediterranei] Length: 251\nNumber of predicted TMHs: 2\nExp number of AAs in TMHs: 43.69728\nExp number, first 60 AAs: 21.22417\nTotal prob of N-in: 0.96054\nPOSSIBLE N-term signal sequence\ninside 1 6\nTMhelix 7 29\noutside 30 223\nTMhelix 224 246\ninside 247 251

32186 GCF_000282715.1_ASM28271v1 Amycolatopsis mediterranei S699 Terrabacteria group; Actinobacteria; Actinobacteria; Pseudonocardiales; Pseudonocardiaceae; Amycolatopsis; Amycolatopsis mediterranei MSRRALVLAVLTGAALLMTAPVSGAAQAAGISSADASAAHDAAAGFQDVLVRFFAQAPGA WP_013229737.1 hypothetical protein [Amycolatopsis mediterranei] Length: 251\nNumber of predicted TMHs: 2\nExp number of AAs in TMHs: 43.69728\nExp number, first 60 AAs: 21.22417\nTotal prob of N-in: 0.96054\nPOSSIBLE N-term signal sequence\ninside 1 6\nTMhelix 7 29\noutside 30 223\nTMhelix 224 246\ninside 247 251

32187 GCF_000196835.1_ASM19683v1 Amycolatopsis mediterranei U32 Terrabacteria group; Actinobacteria; Actinobacteria; Pseudonocardiales; Pseudonocardiaceae; Amycolatopsis; Amycolatopsis mediterranei MSRRALVLAVLTGAALLMTAPVSGAAQAAGISSADASAAHDAAAGFQDVLVRFFAQAPGA YP_003770106.1 hypothetical protein AMED_8000 [Amycolatopsis mediterranei U32] Length: 251\nNumber of predicted TMHs: 2\nExp number of AAs in TMHs: 43.69728\nExp number, first 60 AAs: 21.22417\nTotal prob of N-in: 0.96054\nPOSSIBLE N-term signal sequence\ninside 1 6\nTMhelix 7 29\noutside 30 223\nTMhelix 224 246\ninside 247 251

32188 GCF_000478275.1_AO9412_01 Amycolatopsis orientalis DSM 40040 = KCTC 9412 Terrabacteria group; Actinobacteria; Actinobacteria; Pseudonocardiales; Pseudonocardiaceae; Amycolatopsis; Amycolatopsis orientalis MAQPSTTPSPVAAFFDLDKTIIASSSALAFSKPLLREGLINRRAALRSAYAQLVFSLAGA WP_037318469.1 inhibition of morphological differentiation protein [Amycolatopsis orientalis] Length: 265\nNumber of predicted TMHs: 1\nExp number of AAs in TMHs: 21.2288\nExp number, first 60 AAs: 0.0406\nTotal prob of N-in: 0.00848\noutside 1 235\nTMhelix 236 258\ninside 259 265

32189 GCF_000411975.1_AO43388_01 Amycolatopsis orientalis DSM 43388 Terrabacteria group; Actinobacteria; Actinobacteria; Pseudonocardiales; Pseudonocardiaceae; Amycolatopsis; Amycolatopsis orientalis MAEPRSEEAPRRVAAFFDLDKTIIASSSALAFSKPLLREGLINRRAALRSAYAQLVFSLA WP_037369238.1 inhibition of morphological differentiation protein [Amycolatopsis orientalis] Length: 273\nNumber of predicted TMHs: 1\nExp

number of AAs in TMHs: 20.79733\nExp number, first 60 AAs: 0.15987\nTotal prob of N-in: 0.02100\noutside 1 242\nTMhelix 243 262\ninside 263 273

32190 GCF_000411995.1_AO46075_01 Amycolatopsis orientalis DSM 46075 Terrabacteria group; Actinobacteria; Actinobacteria; Pseudonocardiales; Pseudonocardiaceae; Amycolatopsis; Amycolatopsis orientalis MTDDAHPATSPDPEDQAFPGRSRKTRARRTALWIGLGSAAVFVAGLVFLLLPVTALYYLD WP_043837707.1 hypothetical protein [Amycolatopsis orientalis] Length: 178\nNumber of predicted TMHs: 2\nExp number of AAs in TMHs: 43.86203\nExp number, first 60 AAs: 23.11593\nTotal prob of N-in: 0.99804\nPOSSIBLE N-term signal sequence\ninside 1 30\nTMhelix 31 53\noutside 54 138\nTMhelix 139 158\ninside 159 178

32191 GCF_000411995.1_AO46075_01 Amycolatopsis orientalis DSM 46075 Terrabacteria group; Actinobacteria; Actinobacteria; Pseudonocardiales; Pseudonocardiaceae; Amycolatopsis; Amycolatopsis orientalis MAEPRSEEAPRRVAAFFDLDKTIIASSSALAFSKPLLREGLINRRAALRSAYAQLVFSLA WP_043827107.1 inhibition of morphological differentiation protein [Amycolatopsis orientalis] Length: 273\nNumber of predicted TMHs: 1\nExp number of AAs in TMHs: 20.68178\nExp number, first 60 AAs: 0.16236\nTotal prob of N-in: 0.02572\noutside 1 242\nTMhelix 243 262\ninside 263 273

32192 GCF_000400635.2_ASM40063v2 Amycolatopsis orientalis HCCB10007 Terrabacteria group; Actinobacteria; Actinobacteria; Pseudonocardiales; Pseudonocardiaceae; Amycolatopsis; Amycolatopsis orientalis MDRPSTTPSQVAAFFDLDKTIIASSSALAFSKPLLREGLINRRAALRSAYAQLVFSLAGA WP_016337846.1 MULTISPECIES: haloacid dehalogenase [Amycolatopsis] Length: 265\nNumber of predicted TMHs: 1\nExp number of AAs in TMHs: 21.25757\nExp number, first 60 AAs: 0.07089\nTotal prob of N-in: 0.02314\noutside 1 235\nTMhelix 236 258\ninside 259 265

32193 GCF_000519205.1_ASM51920v1 Amycolatopsis taiwanensis DSM 45107 Terrabacteria group; Actinobacteria; Actinobacteria; Pseudonocardiales; Pseudonocardiaceae; Amycolatopsis; Amycolatopsis taiwanensis MAEPSGSPATLPNRVAAFFDLDKTIIASSSALAFSKPMLRHGLISRRAALKSAYAQLVFS WP_027942832.1 inhibition of morphological differentiation protein [Amycolatopsis taiwanensis] Length: 273\nNumber of predicted TMHs: 1\nExp number of AAs in TMHs: 22.83208\nExp number, first 60 AAs: 0.04134\nTotal prob of N-in: 0.00725\noutside 1 244\nTMhelix 245 267\ninside 268 273

32194 GCF_000388135.1_Whole_genome_assembly Amycolatopsis vancoresmycina DSM 44592 Terrabacteria group; Actinobacteria; Actinobacteria; Pseudonocardiales; Pseudonocardiaceae; Amycolatopsis; Amycolatopsis vancoresmycina MLSRTTPDTPRRTVLVRKLLTVAALAAFVTGGAFLSTGTAAATTTLVSNSCTGSVTGQMG WP_033261107.1 hypothetical protein [Amycolatopsis vancoresmycina] Length: 304\nNumber of predicted TMHs: 2\nExp number of AAs in TMHs: 44.4226\nExp number, first 60 AAs: 22.43038\nTotal prob of N-in: 0.96923\nPOSSIBLE N-term signal sequence\ninside 1 19\nTMhelix 20 42\noutside 43 277\nTMhelix 278 300\ninside 301 304

32195 GCF_000388135.1_Whole_genome_assembly Amycolatopsis vancoresmycina DSM 44592 Terrabacteria group; Actinobacteria; Actinobacteria; Pseudonocardiales; Pseudonocardiaceae; Amycolatopsis; Amycolatopsis vancoresmycina MSRRALVLAVLTGAALLMTAPVSGAAQAAGISSADAVAAHTAAAGLQDTLVRFFAAAPDA WP_033261304.1 hypothetical protein [Amycolatopsis vancoresmycina] Length: 249\nNumber of predicted TMHs: 2\nExp number of AAs in TMHs: 43.5249799999999\nExp number, first 60 AAs: 21.10003\nTotal prob of N-in: 0.95576\nPOSSIBLE N-term signal sequence\ninside 1 6\nTMhelix 7 29\noutside 30 221\nTMhelix 222 244\ninside 245 249

32196 GCF_002155995.1_ASM215599v1 Crossiella equi Terrabacteria group; Actinobacteria; Actinobacteria; Pseudonocardiales; Pseudonocardiaceae; Crossiella MAGKRRALVTVCALCALLAAPVAQAQQCASPQGTYGGPVSWAQRLVGAERVWPLTDGAGQ WP_086782709.1 hypothetical protein [Crossiella equi] Length: 368\nNumber of predicted TMHs: 1\nExp number of AAs in TMHs: 70.29935\nExp number, first 60 AAs: 9.13696\nTotal prob of N-in: 0.75600\noutside 1 334\nTMhelix 335 357\ninside 358 368

32197 GCF_002155995.1_ASM215599v1 Crossiella equi Terrabacteria group; Actinobacteria; Actinobacteria; Pseudonocardiales; Pseudonocardiaceae; Crossiella MLTMEGVAAQGKQGRTMGIRRRRALAAVPGAVLLAQLFGGGVAAAHVTAHVHGEQPERGG WP_086788397.1 hypothetical protein [Crossiella equi] Length: 236\nNumber of predicted TMHs: 2\nExp number of AAs in TMHs: 44.12745\nExp number, first 60 AAs: 21.32943\nTotal prob of N-in: 0.94654\nPOSSIBLE N-term signal sequence\ninside 1 24\nTMhelix 25 47\noutside 48 206\nTMhelix 207 229\ninside 230 236

32198 GCF_900176515.1_IMG-taxon_2595698251_annotated_assembly Kibdelosporangium aridum Terrabacteria group; Actinobacteria; Actinobacteria; Pseudonocardiales; Pseudonocardiaceae; Kibdelosporangium MNALRGRRKPLAMAGALLIGSALAGFIPGSASASSHREAPLIAADPTVDNTDVYAFVSPD WP_033387677.1 hypothetical protein [Kibdelosporangium aridum] Length: 520\nNumber of predicted TMHs: 2\nExp number of AAs in TMHs: 39.2692899999999\nExp number, first 60 AAs: 18.01693\nTotal prob of N-in: 0.83690\nPOSSIBLE N-term signal sequence\ninside 1 11\nTMhelix 12 34\noutside 35 489\nTMhelix 490 512\ninside 513 520

32199 GCF_900176515.1_IMG-taxon_2595698251_annotated_assembly Kibdelosporangium aridum Terrabacteria group; Actinobacteria; Actinobacteria; Pseudonocardiales; Pseudonocardiaceae; Kibdelosporangium MTEATKRVAAFFDLDKTVIAKSSTLAFSRPFFQGGLINRRAVLKSAYAQFVFMQSGADAD WP_033389965.1 inhibition of morphological differentiation protein [Kibdelosporangium aridum] Length: 267\nNumber of predicted TMHs: 1\nExp number of AAs in TMHs: 22.49583\nExp number, first 60 AAs: 0.02533\nTotal prob of N-in: 0.03976\noutside 1 237\nTMhelix 238 260\ninside 261 267

32200 GCF_001302585.1_ASM130258v1 Kibdelosporangium phytohabitans Terrabacteria group; Actinobacteria; Actinobacteria; Pseudonocardiales; Pseudonocardiaceae; Kibdelosporangium MNSLRGRRKPLAMAGALLIGTAFAGFVPGSATASSHREAPLIAADPTVDNTDVYAFVSPD WP_054293102.1 hypothetical protein [Kibdelosporangium phytohabitans] Length: 519\nNumber of predicted TMHs: 2\nExp number of AAs in TMHs: 41.67126\nExp number, first 60 AAs: 19.85895\nTotal prob of N-in: 0.90829\nPOSSIBLE N-term signal sequence\ninside 1 11\nTMhelix 12 34\noutside 35 489\nTMhelix 490 512\ninside 513 519

32201 GCF_001302585.1_ASM130258v1 Kibdelosporangium phytohabitans Terrabacteria group; Actinobacteria; Actinobacteria; Pseudonocardiales; Pseudonocardiaceae; Kibdelosporangium MSPPPAMRRRRMVVALVATGSTVLSAVAVGLVPGSAVASSHREAPLIAADPPVDNTDVYA WP_054296410.1 hypothetical protein [Kibdelosporangium phytohabitans] Length: 516\nNumber of predicted TMHs: 2\nExp number of AAs in TMHs: 38.54706\nExp number, first 60 AAs: 21.66288\nTotal prob of N-in: 0.98695\nPOSSIBLE N-term signal sequence\ninside 1 12\nTMhelix 13 35\noutside 36 485\nTMhelix 486 505\ninside 506 516

32202 GCF_001302585.1_ASM130258v1 Kibdelosporangium phytohabitans Terrabacteria group; Actinobacteria; Actinobacteria; Pseudonocardiales; Pseudonocardiaceae; Kibdelosporangium MTDATKRVAAFFDLDKTVIAKSSTLAFSRPFFHGGLINRRAVLKSAYAQFVFMQSGADAD WP_054287803.1 inhibition of morphological differentiation protein [Kibdelosporangium phytohabitans] Length: 267\nNumber of predicted TMHs: 1\nExp number of AAs in TMHs: 22.61759\nExp number, first 60 AAs: 0.04039\nTotal prob of N-in: 0.02841\noutside 1 238\nTMhelix 239 261\ninside 262 267

32203 GCF_000826545.1_High_quaKibdelosporangium_sp._MJ126-NF4 Kibdelosporangium sp. MJ126-NF4 Terrabacteria group; Actinobacteria; Actinobacteria; Pseudonocardiales; Pseudonocardiaceae; Kibdelosporangium MIVLRGRRKPLTMVVALLAGIALAGIVPGSATASSHREAPLIAADPTVDNTDVYAFVSPD WP_042196000.1 hypothetical protein [Kibdelosporangium sp. MJ126-NF4] Length: 519\nNumber of predicted TMHs: 2\nExp number of AAs in TMHs: 43.07474\nExp number, first 60 AAs: 21.20824\nTotal prob of N-in: 0.97823\nPOSSIBLE N-term signal sequence\ninside 1 12\nTMhelix 13 35\noutside 36 489\nTMhelix 490 512\ninside 513 519

32204 GCF_000826545.1_High_quaKibdelosporangium_sp._MJ126-NF4 Kibdelosporangium sp. MJ126-NF4 Terrabacteria group; Actinobacteria; Actinobacteria; Pseudonocardiales; Pseudonocardiaceae; Kibdelosporangium MTDATKRVAAFFDLDKTVIAKSSTLAFSRPFFHGGLINRRAVLKSAYAQFVFMQSGADAD WP_042198105.1 inhibition of morphological differentiation protein [Kibdelosporangium sp. MJ126-NF4] Length: 267\nNumber of predicted TMHs: 1\nExp number of AAs in TMHs: 22.73851\nExp number, first 60 AAs: 0.04089\nTotal prob of N-in: 0.01855\noutside 1 238\nTMhelix 239 261\ninside 262 267

32205 GCF_000568255.1_ASM56825v1 Kutzneria sp. 744 Terrabacteria group; Actinobacteria; Actinobacteria; Pseudonocardiales; Pseudonocardiaceae; Kutzneria MDPSTGARVAAFFDLDKTVIAKSSLLAFSRPFFQGGLINRRAVLKSAYAQFVFMQSGADA WP_043720026.1 inhibition of morphological differentiation protein [Kutzneria sp. 744] Length: 268\nNumber of predicted TMHs: 1\nExp number of AAs in TMHs: 22.76032\nExp number, first 60 AAs: 0.02307\nTotal prob of N-in: 0.00672\noutside 1 239\nTMhelix 240 262\ninside 263 268

32206 GCF_000525635.1_ASM52563v1 Kutzneria albida DSM 43870 Terrabacteria group; Actinobacteria; Actinobacteria; Pseudonocardiales; Pseudonocardiaceae; Kutzneria; Kutzneria albida MDPSTGARTAAFFDLDKTVIAKSSLLAFSRPFFQEGLINRRAVLKSAYAQFVFMQAGADA WP_025353981.1 inhibition of morphological differentiation protein [Kutzneria albida] Length: 268\nNumber of predicted TMHs:

1\nExp number of AAs in TMHs: 22.75379\nExp number, first 60 AAs: 0.01047\nTotal prob of N-in: 0.00845\noutside 1 239\nTMhelix 240 262\ninside 263 268

32207 GCF_000525635.1_ASM52563v1 Kutzneria albida DSM 43870 Terrabacteria group; Actinobacteria; Actinobacteria; Pseudonocardiales; Pseudonocardiaceae; Kutzneria; Kutzneria albida
MITAAMRRRAAMLVVGLGVLSAPLAGCASGTTQVSNTANTGNPLDVGTEVTQTQWGPLGP WP_081789339.1
hypothetical protein [Kutzneria albida] Length: 262\nNumber of predicted TMHs: 2\nExp number of AAs in TMHs: 37.80024\nExp number, first 60 AAs: 16.67183\nTotal prob of N-in: 0.90978\nPOSSIBLE N-term signal sequence\ninside 1 12\nTMhelix 13 35\noutside 36 228\nTMhelix 229 248\ninside 249 262

32208 GCF_000955955.1_ASM95595v1 Lechevalieria aerocolonigenes Terrabacteria group; Actinobacteria; Actinobacteria; Pseudonocardiales; Pseudonocardiaceae; Lechevalieria
MSERPARRGLLRCTALVTAAVLAFGGTSTALAQEDPTSTSPTPTPTETVTPPPTETVTPP WP_045310765.1 hypothetical protein [Lechevalieria aerocolonigenes] Length: 1606\nNumber of predicted TMHs: 1\nExp number of AAs in TMHs: 32.30244\nExp number, first 60 AAs: 9.86259\nTotal prob of N-in: 0.46179\noutside 1 1577\nTMhelix 1578 1600\ninside 1601 1606

32209 GCF_000719205.1_ASM71920v1 Lechevalieria aerocolonigenes Terrabacteria group; Actinobacteria; Actinobacteria; Pseudonocardiales; Pseudonocardiaceae; Lechevalieria
MTRSQRRRRRHFSLAALVWAVGATALLGLGAGIATAPSSWLVEDIPQSVVTEQRTRTALA WP_030471366.1
HAMP domain-containing protein [Lechevalieria aerocolonigenes] Length: 347\nNumber of predicted TMHs: 2\nExp number of AAs in TMHs: 45.73118\nExp number, first 60 AAs: 22.73572\nTotal prob of N-in: 0.99960\nPOSSIBLE N-term signal sequence\ninside 1 11\nTMhelix 12 34\noutside 35 276\nTMhelix 277 299\ninside 300 347

32210 GCF_000719205.1_ASM71920v1 Lechevalieria aerocolonigenes Terrabacteria group; Actinobacteria; Actinobacteria; Pseudonocardiales; Pseudonocardiaceae; Lechevalieria
MSDRPARRGLLRFTAIATAALLAFGGATTGALAQEPETPTSTVEPTPTSEPSAPPSTTDP WP_030465437.1 hypothetical protein [Lechevalieria aerocolonigenes] Length: 1055\nNumber of predicted TMHs: 2\nExp number of AAs in TMHs: 41.71412\nExp number, first 60 AAs: 19.2151\nTotal prob of N-in: 0.88494\nPOSSIBLE N-term signal sequence\ninside 1 8\nTMhelix 9 31\noutside 32 1025\nTMhelix 1026 1048\ninside 1049 1055

32211 GCF_000719205.1_ASM71920v1 Lechevalieria aerocolonigenes Terrabacteria group; Actinobacteria; Actinobacteria; Pseudonocardiales; Pseudonocardiaceae; Lechevalieria
MSRIAAFFDLDKTVIAKSSTLAFSRPFFQEGLINRRAVLKSAYAQFVFMLAGADADQMDR WP_051783973.1
inhibition of morphological differentiation protein [Lechevalieria aerocolonigenes] Length: 264\nNumber of predicted TMHs: 1\nExp number of AAs in TMHs: 21.87786\nExp number, first 60 AAs: 0.04144\nTotal prob of N-in: 0.01017\noutside 1 235\nTMhelix 236 258\ninside 259 264

32212 GCF_000974445.1_ASM97444v1 Lechevalieria aerocolonigenes Terrabacteria group; Actinobacteria; Actinobacteria; Pseudonocardiales; Pseudonocardiaceae; Lechevalieria
MTRSQRRRRRHFSLAALVWAVGATALLGLGAGIATAPSSWLVEDIPQSVVTEQRTRTALA WP_030471366.1
HAMP domain-containing protein [Lechevalieria aerocolonigenes] Length: 347\nNumber of predicted TMHs: 2\nExp number of AAs in TMHs: 45.73118\nExp number, first 60 AAs: 22.73572\nTotal prob of N-in: 0.99960\nPOSSIBLE N-term signal sequence\ninside 1 11\nTMhelix 12 34\noutside 35 276\nTMhelix 277 299\ninside 300 347

32213 GCF_000974445.1_ASM97444v1 Lechevalieria aerocolonigenes Terrabacteria group; Actinobacteria; Actinobacteria; Pseudonocardiales; Pseudonocardiaceae; Lechevalieria
MSRIAAFFDLDKTVIAKSSTLAFSRPFFQEGLINRRAVLKSAYAQFVFMLAGADADQMDR WP_051783973.1
inhibition of morphological differentiation protein [Lechevalieria aerocolonigenes] Length: 264\nNumber of predicted TMHs: 1\nExp number of AAs in TMHs: 21.87786\nExp number, first 60 AAs: 0.04144\nTotal prob of N-in: 0.01017\noutside 1 235\nTMhelix 236 258\ninside 259 264

32214 GCF_000974445.1_ASM97444v1 Lechevalieria aerocolonigenes Terrabacteria group; Actinobacteria; Actinobacteria; Pseudonocardiales; Pseudonocardiaceae; Lechevalieria
MSDRPARRGLLRFTAIATAALLAFGGATTGALAQEPETPTSTVEPTPTSEPSAPPSTTDP WP_030465437.1 hypothetical protein [Lechevalieria aerocolonigenes] Length: 1055\nNumber of predicted TMHs: 2\nExp number of AAs in TMHs: 41.71412\nExp number, first 60 AAs: 19.2151\nTotal prob of N-in: 0.88494\nPOSSIBLE N-term signal sequence\ninside 1 8\nTMhelix 9 31\noutside 32 1025\nTMhelix 1026 1048\ninside 1049 1055

32215 GCF_000955955.1_ASM95595v1 Lechevalieria aerocolonigenes Terrabacteria group; Actinobacteria; Actinobacteria; Pseudonocardiales; Pseudonocardiaceae; Lechevalieria

MSIAESFERVQRAYADGERRPLLGYLASLCSYGALVGVATAVGRQRGTRLPERFSAGDTV WP_045313515.1 hypothetical protein [Lechevalieria aerocolonigenes] Length: 165\nNumber of predicted TMHs: 2\nExp number of AAs in TMHs: 40.15631\nExp number, first 60 AAs: 22.53531\nTotal prob of N-in: 0.97257\nPOSSIBLE N-term signal sequence\ninside 1 20\nTMhelix 21 43\noutside 44 113\nTMhelix 114 136\ninside 137 165

32216 GCF_000955955.1_ASM95595v1 Lechevalieria aerocolonigenes Terrabacteria group; Actinobacteria; Actinobacteria; Pseudonocardiales; Pseudonocardiaceae; Lechevalieria MSRIAAFFDLDKTVIAKSSTLAFSRPFFQEGLINRRAVLKSAYAQLVFMLAGADADQMDR WP_082114596.1 inhibition of morphological differentiation protein [Lechevalieria aerocolonigenes] Length: 264\nNumber of predicted TMHs: 1\nExp number of AAs in TMHs: 21.5353\nExp number, first 60 AAs: 0.03134\nTotal prob of N-in: 0.00683\noutside 1 235\nTMhelix 236 258\ninside 259 264

32217 GCF_900176525.1_IMG-taxon_2595698253_annotated_assembly Lentzea albidocapillata Terrabacteria group; Actinobacteria; Actinobacteria; Pseudonocardiales; Pseudonocardiaceae; Lentzea MSAPPPARRRRRAAVSLIAVGSMVLAAGLIGGSGPAGASSHREAPLIAGDPAVDNTDVYA WP_030476464.1 hypothetical protein [Lentzea albidocapillata] Length: 529\nNumber of predicted TMHs: 2\nExp number of AAs in TMHs: 41.30154\nExp number, first 60 AAs: 21.09152\nTotal prob of N-in: 0.99141\nPOSSIBLE N-term signal sequence\ninside 1 12\nTMhelix 13 35\noutside 36 496\nTMhelix 497 516\ninside 517 529

32218 GCF_000719115.1_ASM71911v1 Lentzea albidocapillata Terrabacteria group; Actinobacteria; Actinobacteria; Pseudonocardiales; Pseudonocardiaceae; Lentzea MSIAESFERAQLAYADGERRPLFGYLASLCSYGALVGVATAVGRHRGTRLPQRFTPDDII WP_030481134.1 hypothetical protein [Lentzea albidocapillata] Length: 165\nNumber of predicted TMHs: 2\nExp number of AAs in TMHs: 41.96153\nExp number, first 60 AAs: 22.47874\nTotal prob of N-in: 0.92341\nPOSSIBLE N-term signal sequence\ninside 1 20\nTMhelix 21 43\noutside 44 113\nTMhelix 114 136\ninside 137 165

32219 GCF_000719115.1_ASM71911v1 Lentzea albidocapillata Terrabacteria group; Actinobacteria; Actinobacteria; Pseudonocardiales; Pseudonocardiaceae; Lentzea MSAPPPARRRRRAAVSLIAVGSMVLAAGLIGGSGPAGASSHREAPLIAGDPAVDNTDVYA WP_030476464.1 hypothetical protein [Lentzea albidocapillata] Length: 529\nNumber of predicted TMHs: 2\nExp number of AAs in TMHs: 41.30154\nExp number, first 60 AAs: 21.09152\nTotal prob of N-in: 0.99141\nPOSSIBLE N-term signal sequence\ninside 1 12\nTMhelix 13 35\noutside 36 496\nTMhelix 497 516\ninside 517 529

32220 GCF_000719115.1_ASM71911v1 Lentzea albidocapillata Terrabacteria group; Actinobacteria; Actinobacteria; Pseudonocardiales; Pseudonocardiaceae; Lentzea MSRIAAFFDLDKTVIAKSSTLAFSRPFFQEGLINRRAVLKSAYAQFVFMLAGADADQMDR WP_051770057.1 inhibition of morphological differentiation protein [Lentzea albidocapillata] Length: 264\nNumber of predicted TMHs: 1\nExp number of AAs in TMHs: 21.93782\nExp number, first 60 AAs: 0.04436\nTotal prob of N-in: 0.01309\noutside 1 235\nTMhelix 236 258\ninside 259 264

32221 GCF_900176525.1_IMG-taxon_2595698253_annotated_assembly Lentzea albidocapillata Terrabacteria group; Actinobacteria; Actinobacteria; Pseudonocardiales; Pseudonocardiaceae; Lentzea MSRIAAFFDLDKTVIAKSSTLAFSRPFFQEGLINRRAVLKSAYAQFVFMLAGADADQMDR WP_051770057.1 inhibition of morphological differentiation protein [Lentzea albidocapillata] Length: 264\nNumber of predicted TMHs: 1\nExp number of AAs in TMHs: 21.93782\nExp number, first 60 AAs: 0.04436\nTotal prob of N-in: 0.01309\noutside 1 235\nTMhelix 236 258\ninside 259 264

32222 GCF_900176525.1_IMG-taxon_2595698253_annotated_assembly Lentzea albidocapillata Terrabacteria group; Actinobacteria; Actinobacteria; Pseudonocardiales; Pseudonocardiaceae; Lentzea MSIAESFERAQLAYADGERRPLFGYLASLCSYGALVGVATAVGRHRGTRLPQRFTPDDII WP_030481134.1 hypothetical protein [Lentzea albidocapillata] Length: 165\nNumber of predicted TMHs: 2\nExp number of AAs in TMHs: 41.96153\nExp number, first 60 AAs: 22.47874\nTotal prob of N-in: 0.92341\nPOSSIBLE N-term signal sequence\ninside 1 20\nTMhelix 21 43\noutside 44 113\nTMhelix 114 136\ninside 137 165

32223 GCF_001701025.1_ASM170102v1 Lentzea guizhouensis Terrabacteria group; Actinobacteria; Actinobacteria; Pseudonocardiales; Pseudonocardiaceae; Lentzea MSRTAAFFDLDKTVIAKSSTLAFSRPFFQEGLINRRAVLKSAYAQFVFMLAGADADQMDR WP_065915214.1 inhibition of morphological differentiation protein [Lentzea guizhouensis] Length: 264\nNumber of predicted TMHs: 1\nExp number of AAs in TMHs: 21.44533\nExp number, first 60 AAs: 0.06163\nTotal prob of N-in: 0.01520\noutside 1 235\nTMhelix 236 258\ninside 259 264

32224 GCF_001701025.1_ASM170102v1 Lentzea guizhouensis Terrabacteria group; Actinobacteria; Actinobacteria; Pseudonocardiales; Pseudonocardiaceae; Lentzea
MSAPPPARRRRRAAVSLIAVGSTVLAAGLIGGSGPAGASSHREAPLIAGDPAVDNTDVYA WP_065919765.1
hypothetical protein [Lentzea guizhouensis] Length: 528\nNumber of predicted TMHs: 2\nExp number of AAs in TMHs: 38.99584\nExp number, first 60 AAs: 20.86561\nTotal prob of N-in: 0.98119\nPOSSIBLE N-term signal sequence\ninside 1 12\nTMhelix 13 35\noutside 36 496\nTMhelix 497 516\ninside 517 528

32225 GCF_002150765.1_ASM215076v1 Lentzea kentuckyensis Terrabacteria group; Actinobacteria; Actinobacteria; Pseudonocardiales; Pseudonocardiaceae; Lentzea
MSRIAAFFDLDKTVIAKSSTLAFSRPFFQEGLINRRAVLKSAYAQFVFMLAGADADQMDR WP_086663159.1
inhibition of morphological differentiation protein [Lentzea kentuckyensis] Length: 264\nNumber of predicted TMHs: 1\nExp number of AAs in TMHs: 21.84425\nExp number, first 60 AAs: 0.04423\nTotal prob of N-in: 0.01350\noutside 1 235\nTMhelix 236 258\ninside 259 264

32226 GCF_000717175.1_ASM71717v1 Pseudonocardia autotrophica Terrabacteria group; Actinobacteria; Actinobacteria; Pseudonocardiales; Pseudonocardiaceae; Pseudonocardia
MFTRTRRTALRATAVTAAAGGLLLAGTVPAFAHVTAQPSTAEQGGYAVISMRVPNESDTA WP_081923875.1
nuclear export factor GLE1 [Pseudonocardia autotrophica] Length: 256\nNumber of predicted TMHs: 1\nExp number of AAs in TMHs: 36.46455\nExp number, first 60 AAs: 16.90456\nTotal prob of N-in: 0.90503\nPOSSIBLE N-term signal sequence\ninside 1 213\nTMhelix 214 236\noutside 237 256

32227 GCF_001294605.1_ASM129460v1 Pseudonocardia sp. AL041005-10 Terrabacteria group; Actinobacteria; Actinobacteria; Pseudonocardiales; Pseudonocardiaceae; Pseudonocardia
MTAGLPAGAAAGRRGPVTGRPARGSGSNGYATCSASVSPTRTSASPCISRVAACCSTADR WP_083474024.1
hypothetical protein [Pseudonocardia sp. AL041005-10] Length: 404\nNumber of predicted TMHs: 1\nExp number of AAs in TMHs: 31.83868\nExp number, first 60 AAs: 1e-05\nTotal prob of N-in: 0.05806\noutside 1 361\nTMhelix 362 384\ninside 385 404

32228 GCF_001942185.1_ASM194218v1 Pseudonocardia sp. CNS-004 Terrabacteria group; Actinobacteria; Actinobacteria; Pseudonocardiales; Pseudonocardiaceae; Pseudonocardia
MSGALPPQLPARSGAAAFFDLDKTIIAGSSALAFSRPFRRQGLISRRAALRSGYAQLLLV WP_075952708.1 inhibition of morphological differentiation protein [Pseudonocardia sp. CNS-004] Length: 272\nNumber of predicted TMHs: 1\nExp number of AAs in TMHs: 21.37559\nExp number, first 60 AAs: 0.18455\nTotal prob of N-in: 0.00671\noutside 1 244\nTMhelix 245 267\ninside 268 272

32229 GCF_000620785.1_ASM62078v1 Pseudonocardia acaciae DSM 45401 Terrabacteria group; Actinobacteria; Actinobacteria; Pseudonocardiales; Pseudonocardiaceae; Pseudonocardia; Pseudonocardia acaciae
MLAGVPGPAPIAAFFDLDKTIIAGSSTLAFSGPLRDRGLIGRRALLRSGYAQLLLMVAGA WP_037069421.1 inhibition of morphological differentiation protein [Pseudonocardia acaciae] Length: 282\nNumber of predicted TMHs: 1\nExp number of AAs in TMHs: 22.29246\nExp number, first 60 AAs: 0.2437\nTotal prob of N-in: 0.00742\noutside 1 250\nTMhelix 251 273\ninside 274 282

32230 GCF_000620785.1_ASM62078v1 Pseudonocardia acaciae DSM 45401 Terrabacteria group; Actinobacteria; Actinobacteria; Pseudonocardiales; Pseudonocardiaceae; Pseudonocardia; Pseudonocardia acaciae
MLGAVQRNLRQPTVVKAARGLSHFGEHAAGWLAVGVLGAALNRSRRREWLYAAGGVAVAH WP_051580762.1
phosphatase PAP2 family protein [Pseudonocardia acaciae] Length: 168\nNumber of predicted TMHs: 1\nExp number of AAs in TMHs: 42.7501\nExp number, first 60 AAs: 14.01996\nTotal prob of N-in: 0.54342\nPOSSIBLE N-term signal sequence\noutside 1 133\nTMhelix 134 156\ninside 157 168

32231 GCF_000620785.1_ASM62078v1 Pseudonocardia acaciae DSM 45401 Terrabacteria group; Actinobacteria; Actinobacteria; Pseudonocardiales; Pseudonocardiaceae; Pseudonocardia; Pseudonocardia acaciae
MTSLLLLAAALVIAPSSSPGRSRLLLLYPPDTPPGPTPQTRRPLLAAGCGLAIALLTAST WP_037069415.1 hypothetical protein [Pseudonocardia acaciae] Length: 238\nNumber of predicted TMHs: 2\nExp number of AAs in TMHs: 52.27851\nExp number, first 60 AAs: 10.56311\nTotal prob of N-in: 0.16146\nPOSSIBLE N-term signal sequence\noutside 1 57\nTMhelix 58 80\ninside 81 209\nTMhelix 210 232\noutside 233 238

32232 GCF_000196675.1_ASM19667v1 Pseudonocardia dioxanivorans CB1190 Terrabacteria group; Actinobacteria; Actinobacteria; Pseudonocardiales; Pseudonocardiaceae; Pseudonocardia; Pseudonocardia dioxanivorans
MQSPPVTGVRSPRRRALSAVLMAVLVLGALSGCARVRTALALQPDDTVTGEVVIATPQKT WP_013674910.1
hypothetical protein [Pseudonocardia dioxanivorans] Length: 230\nNumber of predicted TMHs: 2\nExp

number of AAs in TMHs: 42.73709\nExp number, first 60 AAs: 20.28515\nTotal prob of N-in: 0.86357\nPOSSIBLE N-term signal sequence\ninside 1 16\nTMhelix 17 39\noutside 40 194\nTMhelix 195 217\ninside 218 230

32233 GCF_000196675.1_ASM19667v1 Pseudonocardia dioxanivorans CB1190 Terrabacteria group; Actinobacteria; Actinobacteria; Pseudonocardiales; Pseudonocardiaceae; Pseudonocardia; Pseudonocardia dioxanivorans MTTTETPVPQAPPVRTPPPPPTAPPPPAAPSSRGRARRRTAFRVAGVVLAALVLGVLLAA WP_013677670.1 DUF4349 domain-containing protein [Pseudonocardia dioxanivorans] Length: 362\nNumber of predicted TMHs: 2\nExp number of AAs in TMHs: 45.56278\nExp number, first 60 AAs: 18.04262\nTotal prob of N-in: 0.99979\nPOSSIBLE N-term signal sequence\ninside 1 41\nTMhelix 42 64\noutside 65 306\nTMhelix 307 329\ninside 330 362

32234 GCF_000196675.1_ASM19667v1 Pseudonocardia dioxanivorans CB1190 Terrabacteria group; Actinobacteria; Actinobacteria; Pseudonocardiales; Pseudonocardiaceae; Pseudonocardia; Pseudonocardia dioxanivorans MPGPPPHVPTSAAFFDLDKTIIAGSSALAFSRPFRSHGLIGRRAVLRSVYAQMMVLASGA WP_013672618.1 haloacid dehalogenase [Pseudonocardia dioxanivorans] Length: 270\nNumber of predicted TMHs: 1\nExp number of AAs in TMHs: 21.02065\nExp number, first 60 AAs: 0.05078\nTotal prob of N-in: 0.00189\noutside 1 241\nTMhelix 242 261\ninside 262 270

32235 GCF_000429025.1_ASM42902v1 Pseudonocardia spinosispora DSM 44797 Terrabacteria group; Actinobacteria; Actinobacteria; Pseudonocardiales; Pseudonocardiaceae; Pseudonocardia; Pseudonocardia spinosispora MSTSRSTRRALVRAAATVAAAAALTLAGTGIASAHVTAQPGTAEQGSYAKVAFRVPSELP WP_028938488.1 hypothetical protein [Pseudonocardia spinosispora] Length: 259\nNumber of predicted TMHs: 2\nExp number of AAs in TMHs: 41.58702\nExp number, first 60 AAs: 19.9374\nTotal prob of N-in: 0.89743\nPOSSIBLE N-term signal sequence\ninside 1 11\nTMhelix 12 34\noutside 35 226\nTMhelix 227 249\ninside 250 259

32236 GCF_000527075.1_ASM52707v1 Saccharomonospora sp. CNQ490 Terrabacteria group; Actinobacteria; Actinobacteria; Pseudonocardiales; Pseudonocardiaceae; Saccharomonospora MPARDDAPTAAFFDLDKTIIASSSALAFSRPFLQQGLINRRAALKSAYAQLLFALSGADA WP_024875433.1 MULTISPECIES: inhibition of morphological differentiation protein [Saccharomonospora] Length: 271\nNumber of predicted TMHs: 1\nExp number of AAs in TMHs: 21.63397\nExp number, first 60 AAs: 0.07887\nTotal prob of N-in: 0.05335\noutside 1 236\nTMhelix 237 259\ninside 260 271

32237 GCF_001942305.1_ASM194230v1 Saccharomonospora sp. CUA-673 Terrabacteria group; Actinobacteria; Actinobacteria; Pseudonocardiales; Pseudonocardiaceae; Saccharomonospora MSTYPSRRRAYLVRGAAVTAALGLTLSAGAGLASAHVTANVYGDQPEQGGFGAVSLRVPN WP_075849736.1 nuclear export factor GLE1 [Saccharomonospora sp. CUA-673] Length: 243\nNumber of predicted TMHs: 2\nExp number of AAs in TMHs: 41.93023\nExp number, first 60 AAs: 21.06631\nTotal prob of N-in: 0.80446\nPOSSIBLE N-term signal sequence\ninside 1 11\nTMhelix 12 34\noutside 35 210\nTMhelix 211 233\ninside 234 243

32238 GCF_002077655.1_ASM207765v1 Saccharomonospora sp. LRS4.154 Terrabacteria group; Actinobacteria; Actinobacteria; Pseudonocardiales; Pseudonocardiaceae; Saccharomonospora MPARDDAPTAAFFDLDKTIIASSSALAFSRPFLQQGLINRRAALKSAYAQLLFALSGADA WP_024875433.1 MULTISPECIES: inhibition of morphological differentiation protein [Saccharomonospora] Length: 271\nNumber of predicted TMHs: 1\nExp number of AAs in TMHs: 21.63397\nExp number, first 60 AAs: 0.07887\nTotal prob of N-in: 0.05335\noutside 1 236\nTMhelix 237 259\ninside 260 271

32239 GCF_000430445.1_ASM43044v1 [Actinopolyspora] iraqiensis IQ-H1 Terrabacteria group; Actinobacteria; Actinobacteria; Pseudonocardiales; Pseudonocardiaceae; Saccharomonospora; [Actinopolyspora] iraqiensis MTAADTASPHRTAAFFDLDKTIIASSSALAFSRTFLRQGLINRRAALRSAYAQLVFSLSG WP_026453218.1 inhibition of morphological differentiation protein [[Actinopolyspora] iraqiensis] Length: 272\nNumber of predicted TMHs: 1\nExp number of AAs in TMHs: 23.0432\nExp number, first 60 AAs: 0.33537\nTotal prob of N-in: 0.02336\noutside 1 240\nTMhelix 241 263\ninside 264 272

32240 GCF_000231055.2_ASM23105v3 Saccharomonospora azurea NA-128 Terrabacteria group; Actinobacteria; Actinobacteria; Pseudonocardiales; Pseudonocardiaceae; Saccharomonospora; Saccharomonospora azurea MPAPDKTTTATSAAFFDLDKTIIASSSALAFSKPFLRQGLINRRAALKSAYAQLMFALSG WP_005441733.1 haloacid dehalogenase [Saccharomonospora azurea] Length: 274\nNumber of predicted TMHs: 1\nExp number of AAs in TMHs: 22.1932\nExp number, first 60 AAs: 0.14492\nTotal prob of N-in: 0.09047\noutside 1 240\nTMhelix 241 263\ninside 264 274

32241 GCF_000231055.2_ASM23105v3 Saccharomonospora azurea NA-128 Terrabacteria group; Actinobacteria; Actinobacteria; Pseudonocardiales; Pseudonocardiaceae; Saccharomonospora; Saccharomonospora azurea MRRKRHRRAHVFALPTAASLLVLLAAPAAAQTVPTTPPPVPAEEGQPPISSETKPPGRIM WP_005441710.1 hypothetical protein [Saccharomonospora azurea] Length: 561\nNumber of predicted TMHs: 2\nExp number of AAs in TMHs: 40.67922\nExp number, first 60 AAs: 20.01727\nTotal prob of N-in: 0.95768\nPOSSIBLE N-term signal sequence\ninside 1 8\nTMhelix 9 31\noutside 32 520\nTMhelix 521 543\ninside 544 561

32242 GCF_000236985.1_ASM23698v2 Saccharomonospora azurea SZMC 14600 Terrabacteria group; Actinobacteria; Actinobacteria; Pseudonocardiales; Pseudonocardiaceae; Saccharomonospora; Saccharomonospora azurea MPAPDKTTTATSAAFFDLDKTIIASSSALAFSKPFLRQGLINRRAALKSAYAQLMFALSG WP_005441733.1 haloacid dehalogenase [Saccharomonospora azurea] Length: 274\nNumber of predicted TMHs: 1\nExp number of AAs in TMHs: 22.1932\nExp number, first 60 AAs: 0.14492\nTotal prob of N-in: 0.09047\noutside 1 240\nTMhelix 241 263\ninside 264 274

32243 GCF_000236985.1_ASM23698v2 Saccharomonospora azurea SZMC 14600 Terrabacteria group; Actinobacteria; Actinobacteria; Pseudonocardiales; Pseudonocardiaceae; Saccharomonospora; Saccharomonospora azurea MRRKRHRRAHVFALPTAASLLVLLAAPAAAQTVPTTPPPVPAEEGQPPISSETKPPGRIM WP_005449049.1 hypothetical protein [Saccharomonospora azurea] Length: 561\nNumber of predicted TMHs: 2\nExp number of AAs in TMHs: 40.67922\nExp number, first 60 AAs: 20.01727\nTotal prob of N-in: 0.95768\nPOSSIBLE N-term signal sequence\ninside 1 8\nTMhelix 9 31\noutside 32 520\nTMhelix 521 543\ninside 544 561

32244 GCF_000243395.2_ASM24339v3 Saccharomonospora glauca K62 Terrabacteria group; Actinobacteria; Actinobacteria; Pseudonocardiales; Pseudonocardiaceae; Saccharomonospora; Saccharomonospora glauca MRTVGPERAGEAALQARREHVLRLGSWGVLGAASVITLLGVVLVMGAFRNDQLIAENHGT WP_085978096.1 hypothetical protein [Saccharomonospora glauca] Length: 166\nNumber of predicted TMHs: 2\nExp number of AAs in TMHs: 45.54583\nExp number, first 60 AAs: 22.81884\nTotal prob of N-in: 0.92945\nPOSSIBLE N-term signal sequence\ninside 1 20\nTMhelix 21 43\noutside 44 124\nTMhelix 125 147\ninside 148 166

32245 GCF_000243395.2_ASM24339v3 Saccharomonospora glauca K62 Terrabacteria group; Actinobacteria; Actinobacteria; Pseudonocardiales; Pseudonocardiaceae; Saccharomonospora; Saccharomonospora glauca MRSARTMKRRRALALAATIGIGGAVLAPTPAWAQDDAASQETVGYATPPPVDPGQRPNDD WP_005461404.1 type VII secretion-associated serine protease [Saccharomonospora glauca] Length: 505\nNumber of predicted TMHs: 2\nExp number of AAs in TMHs: 43.96032\nExp number, first 60 AAs: 21.13641\nTotal prob of N-in: 0.98805\nPOSSIBLE N-term signal sequence\ninside 1 11\nTMhelix 12 34\noutside 35 466\nTMhelix 467 489\ninside 490 505

32246 GCF_000231035.2_ASM23103v3 Saccharomonospora paurometabolica YIM 90007 Terrabacteria group; Actinobacteria; Actinobacteria; Pseudonocardiales; Pseudonocardiaceae; Saccharomonospora; Saccharomonospora paurometabolica MAAADTAAPHRTAAFFDLDKTIIASSSALAFSRTFLRQGLINRRAALRSAYAQLVFSLSG WP_028662618.1 inhibition of morphological differentiation protein [Saccharomonospora paurometabolica] Length: 272\nNumber of predicted TMHs: 1\nExp number of AAs in TMHs: 23.01682\nExp number, first 60 AAs: 0.29506\nTotal prob of N-in: 0.01910\noutside 1 240\nTMhelix 241 263\ninside 264 272

32247 GCF_000383795.1_ASM38379v1 Saccharomonospora saliphila YIM 90502 Terrabacteria group; Actinobacteria; Actinobacteria; Pseudonocardiales; Pseudonocardiaceae; Saccharomonospora; Saccharomonospora saliphila MLTPRRSTVARAVSLATVPLVTASLSAVPAADAVSPADASLKGAAPVGAPRQPPAVGAEP WP_019816430.1 hypothetical protein [Saccharomonospora saliphila] Length: 378\nNumber of predicted TMHs: 1\nExp number of AAs in TMHs: 27.6647\nExp number, first 60 AAs: 6.98592\nTotal prob of N-in: 0.33515\noutside 1 353\nTMhelix 354 373\ninside 374 378

32248 GCF_000194155.1_ASM19415v1 Saccharopolyspora spinosa NRRL 18395 Terrabacteria group; Actinobacteria; Actinobacteria; Pseudonocardiales; Pseudonocardiaceae; Saccharopolyspora; Saccharopolyspora spinosa MTEPANPSAPTGRRVAAFFDLDKTVIAKSSTLAFSRPFFQEGLINRRAVLKSAYAQFVFM WP_029535547.1 inhibition of morphological differentiation protein [Saccharopolyspora spinosa] Length: 275\nNumber of predicted TMHs: 1\nExp number of AAs in TMHs: 22.81093\nExp number, first 60 AAs: 0.03198\nTotal prob of N-in: 0.00601\noutside 1 245\nTMhelix 246 268\ninside 269 275

32249 GCF_001984175.1_ASM198417v1 Saccharothrix sp. ALI-22-I Terrabacteria group; Actinobacteria; Actinobacteria; Pseudonocardiales; Pseudonocardiaceae; Saccharothrix MRVRMRRRAAVAGLVAVAAVVVGAAPASAALGDASAYGAKLNLSLLGGHAVSADPFAADP WP_077008428.1 hypothetical protein [Saccharothrix sp. ALI-22-I] Length: 267\nNumber of predicted TMHs: 2\nExp number of

AAs in TMHs: 50.81643\nExp number, first 60 AAs:  20.09516\nTotal prob of N-in:   0.88264\nPOSSIBLE N-term signal sequence\ninside   1   8\nTMhelix   9   31\noutside   32   237\nTMhelix   238   260\ninside   261   267

32250 GCF_001984175.1_ASM198417v1 Saccharothrix sp. ALI-22-I Terrabacteria group; Actinobacteria; Actinobacteria; Pseudonocardiales; Pseudonocardiaceae; Saccharothrix MTEHANEGSRVAAFFDLDKTVIAKSSTLAFSRPFFQEGLINRRAVLKSAYAQFVFMLAGA WP_077010340.1 inhibition of morphological differentiation protein [Saccharothrix sp. ALI-22-I] Length: 270\nNumber of predicted TMHs:  1\nExp number of AAs in TMHs: 22.93301\nExp number, first 60 AAs:  0.15877\nTotal prob of N-in:   0.02369\noutside   1   241\nTMhelix   242   264\ninside   265   270

32251 GCF_001905065.1_ASM190506v1 Saccharothrix sp. CB00851 Terrabacteria group; Actinobacteria; Actinobacteria; Pseudonocardiales; Pseudonocardiaceae; Saccharothrix MRLRMTRRAAVAGLVAVAAVVVGAPPASAAPGDASAHGARLDLSLLGRDAVSAGPFAAST WP_073889111.1 hypothetical protein [Saccharothrix sp. CB00851] Length: 257\nNumber of predicted TMHs:  2\nExp number of AAs in TMHs: 45.27483\nExp number, first 60 AAs:  6.39191\nTotal prob of N-in:   0.94283\ninside   1   188\nTMhelix   189   211\noutside   212   225\nTMhelix   226   248\ninside   249   257

32252 GCF_001905065.1_ASM190506v1 Saccharothrix sp. CB00851 Terrabacteria group; Actinobacteria; Actinobacteria; Pseudonocardiales; Pseudonocardiaceae; Saccharothrix MTEHATDGSRVAAFFDLDKTVIAKSSTLAFSRPFFQEGLINRRAVLKSAYAQFVFMLAGA WP_073886865.1 inhibition of morphological differentiation protein [Saccharothrix sp. CB00851] Length: 270\nNumber of predicted TMHs:  1\nExp number of AAs in TMHs: 22.97153\nExp number, first 60 AAs:  0.19109\nTotal prob of N-in:   0.02849\noutside   1   241\nTMhelix   242   264\ninside   265   270

32253 GCF_000716595.1_ASM71659v1 Saccharothrix sp. NRRL B-16314 Terrabacteria group; Actinobacteria; Actinobacteria; Pseudonocardiales; Pseudonocardiaceae; Saccharothrix MTEHATEGSRVAAFFDLDKTVIAKSSTLAFSRPFFQEGLINRRAVLKSAYAQFVFMLAGA WP_033441918.1 inhibition of morphological differentiation protein [Saccharothrix sp. NRRL B-16314] Length: 270\nNumber of predicted TMHs:  1\nExp number of AAs in TMHs: 22.92102\nExp number, first 60 AAs:  0.15866\nTotal prob of N-in:   0.02316\noutside   1   241\nTMhelix   242   264\ninside   265   270

32254 GCF_000716595.1_ASM71659v1 Saccharothrix sp. NRRL B-16314 Terrabacteria group; Actinobacteria; Actinobacteria; Pseudonocardiales; Pseudonocardiaceae; Saccharothrix MRVRMTRRAAVAGLVAVAALVVGAAPASAAPGDASAYGVKLDLSLLGSDAVSAGPFAAAD WP_033438454.1 hypothetical protein [Saccharothrix sp. NRRL B-16314] Length: 257\nNumber of predicted TMHs:  2\nExp number of AAs in TMHs: 45.45304\nExp number, first 60 AAs:  16.83429\nTotal prob of N-in:   0.95461\nPOSSIBLE N-term signal sequence\ninside   1   8\nTMhelix   9   31\noutside   32   225\nTMhelix   226   248\ninside   249   257

32255 GCF_001280085.1_ASM128008v1 Saccharothrix sp. NRRL B-16348 Terrabacteria group; Actinobacteria; Actinobacteria; Pseudonocardiales; Pseudonocardiaceae; Saccharothrix MTEHATEGSRVAAFFDLDKTVIAKSSTLAFSRPFFQEGLINRRAVLKSAYAQFVFMLAGA WP_053715797.1 inhibition of morphological differentiation protein [Saccharothrix sp. NRRL B-16348] Length: 270\nNumber of predicted TMHs:  1\nExp number of AAs in TMHs: 22.92484\nExp number, first 60 AAs:  0.15865\nTotal prob of N-in:   0.02339\noutside   1   241\nTMhelix   242   264\ninside   265   270

32256 GCF_000955975.1_ASM95597v1 Saccharothrix sp. ST-888 Terrabacteria group; Actinobacteria; Actinobacteria; Pseudonocardiales; Pseudonocardiaceae; Saccharothrix MSAESRTHRPSRRTRRAAVAATATALVGALAIAPAAARPAAPTLRYTCVFPTIGGQAITA WP_052682257.1 hypothetical protein [Saccharothrix sp. ST-888] Length: 317\nNumber of predicted TMHs:  2\nExp number of AAs in TMHs: 44.2412499999999\nExp number, first 60 AAs:  20.28416\nTotal prob of N-in:   0.97917\nPOSSIBLE N-term signal sequence\ninside   1   16\nTMhelix   17   36\noutside   37   280\nTMhelix   281   303\ninside   304   317

32257 GCF_000716755.1_ASM71675v1 Saccharothrix syringae Terrabacteria group; Actinobacteria; Actinobacteria; Pseudonocardiales; Pseudonocardiaceae; Saccharothrix MTDRATEGSRVAAFFDLDKTVIAKSSTLAFSRPFFQEGLINRRAVLKSAYAQFVFMLAGA WP_033433487.1 inhibition of morphological differentiation protein [Saccharothrix syringae] Length: 270\nNumber of predicted TMHs:  1\nExp number of AAs in TMHs: 23.07606\nExp number, first 60 AAs:  0.29584\nTotal prob of N-in:   0.04408\noutside   1   241\nTMhelix   242   264\ninside   265   270

32258 GCF_000328705.1_ASM32870v1 Saccharothrix espanaensis DSM 44229 Terrabacteria group; Actinobacteria; Actinobacteria; Pseudonocardiales; Pseudonocardiaceae; Saccharothrix; Saccharothrix espanaensis

MTEHATEGSRVAAFFDLDKTVIAKSSTLAFSRPFFQEGLINRRAVLKSAYAQFVFMLAGA WP_015097763.1 hypothetical protein [Saccharothrix espanaensis] Length: 270\nNumber of predicted TMHs: 1\nExp number of AAs in TMHs: 22.84433\nExp number, first 60 AAs: 0.15873\nTotal prob of N-in: 0.02044\noutside 1 241\nTMhelix 242 264\ninside 265 270

32259 GCF_000427905.1_ASM42790v1 Thermocrispum agreste DSM 44070 Terrabacteria group; Actinobacteria; Actinobacteria; Pseudonocardiales; Pseudonocardiaceae; Thermocrispum; Thermocrispum agreste MGRRVATRRTAMAGAVALTFLFGTIGSGLAPVPAAAQQEEKPPWAPQGPISSSVPKDTGR WP_084609345.1 hypothetical protein [Thermocrispum agreste] Length: 580\nNumber of predicted TMHs: 2\nExp number of AAs in TMHs: 44.9239299999999\nExp number, first 60 AAs: 22.16668\nTotal prob of N-in: 0.99176\nPOSSIBLE N-term signal sequence\ninside 1 12\nTMhelix 13 35\noutside 36 544\nTMhelix 545 567\ninside 568 580

32260 GCF_000427905.1_ASM42790v1 Thermocrispum agreste DSM 44070 Terrabacteria group; Actinobacteria; Actinobacteria; Pseudonocardiales; Pseudonocardiaceae; Thermocrispum; Thermocrispum agreste MESAGPVAAFFDLDKTIIASSSALAFSKPLLRQGLINRRAALRSAYAQLMFSLAGADEQT WP_028848535.1 inhibition of morphological differentiation protein [Thermocrispum agreste] Length: 276\nNumber of predicted TMHs: 1\nExp number of AAs in TMHs: 22.78352\nExp number, first 60 AAs: 0.02832\nTotal prob of N-in: 0.00372\noutside 1 236\nTMhelix 237 259\ninside 260 276

32261 GCF_000427825.1_ASM42782v1 Thermocrispum municipale DSM 44069 Terrabacteria group; Actinobacteria; Actinobacteria; Pseudonocardiales; Pseudonocardiaceae; Thermocrispum; Thermocrispum municipale MESAGAVAAFFDLDKTIIASSSALAFSKPLLRQGLINRRAALRSAYAQLMFSLAGADEQT WP_028849764.1 inhibition of morphological differentiation protein [Thermocrispum municipale] Length: 269\nNumber of predicted TMHs: 1\nExp number of AAs in TMHs: 22.8648\nExp number, first 60 AAs: 0.06839\nTotal prob of N-in: 0.00914\noutside 1 236\nTMhelix 237 259\ninside 260 269

32262 GCF_000717275.1_ASM71727v1 Kitasatospora arboriphila Terrabacteria group; Actinobacteria; Actinobacteria; Streptomycetales; Streptomycetaceae; Kitasatospora MFHADPVAHAEPPSAPRRRSALSVVSSVIMNLLAVGGLACIVMVVLAVVFNVSLIMFKTG WP_081906626.1 MULTISPECIES: signal peptidase I [Actinobacteria] Length: 208\nNumber of predicted TMHs: 2\nExp number of AAs in TMHs: 46.44187\nExp number, first 60 AAs: 23.63275\nTotal prob of N-in: 0.15209\nPOSSIBLE N-term signal sequence\noutside 1 31\nTMhelix 32 54\ninside 55 151\nTMhelix 152 174\noutside 175 208

32263 GCF_000717185.1_ASM71718v1 Kitasatospora phosalacinea Terrabacteria group; Actinobacteria; Actinobacteria; Streptomycetales; Streptomycetaceae; Kitasatospora MPRTAAFFDLDKTIIAKSSALAFSRPFYQGGLINRRSVVKSAYAQFVFLVGGADHDQMEK WP_033251405.1 inhibition of morphological differentiation protein [Kitasatospora phosalacinea] Length: 267\nNumber of predicted TMHs: 1\nExp number of AAs in TMHs: 21.17715\nExp number, first 60 AAs: 0.01564\nTotal prob of N-in: 0.00561\noutside 1 236\nTMhelix 237 259\ninside 260 267

32264 GCF_000717185.1_ASM71718v1 Kitasatospora phosalacinea Terrabacteria group; Actinobacteria; Actinobacteria; Streptomycetales; Streptomycetaceae; Kitasatospora MQRSRAPRRSLVPAALALGLVLGAAGVAAAHVEVDSPTAQALAVDARVAFSAEGESSTAG WP_033251316.1 hypothetical protein [Kitasatospora phosalacinea] Length: 247\nNumber of predicted TMHs: 2\nExp number of AAs in TMHs: 43.42674\nExp number, first 60 AAs: 20.80472\nTotal prob of N-in: 0.97082\nPOSSIBLE N-term signal sequence\ninside 1 11\nTMhelix 12 34\noutside 35 216\nTMhelix 217 239\ninside 240 247

32265 GCF_000716545.1_ASM71654v1 Kitasatospora phosalacinea Terrabacteria group; Actinobacteria; Actinobacteria; Streptomycetales; Streptomycetaceae; Kitasatospora MPRTAAFFDLDKTIIAKSSALAFSRPFYQGGLINRRSVVKSAYAQFVFLVGGADHDQMEK WP_033218537.1 inhibition of morphological differentiation protein [Kitasatospora phosalacinea] Length: 267\nNumber of predicted TMHs: 1\nExp number of AAs in TMHs: 21.34419\nExp number, first 60 AAs: 0.01642\nTotal prob of N-in: 0.00759\noutside 1 236\nTMhelix 237 259\ninside 260 267

32266 GCF_000716545.1_ASM71654v1 Kitasatospora phosalacinea Terrabacteria group; Actinobacteria; Actinobacteria; Streptomycetales; Streptomycetaceae; Kitasatospora MLARRRTALPAQAVPAVPVRVTAARPPALRRWAVLGGAALPLLLAAPALAADPAAGPAAG WP_033214217.1 hypothetical protein [Kitasatospora phosalacinea] Length: 392\nNumber of predicted TMHs: 1\nExp number of AAs in TMHs: 29.41171\nExp number, first 60 AAs: 10.77125\nTotal prob of N-in: 0.51669\nPOSSIBLE N-term signal sequence\noutside 1 369\nTMhelix 370 389\ninside 390 392

32267 GCF_000716545.1_ASM71654v1 Kitasatospora phosalacinea Terrabacteria group; Actinobacteria; Actinobacteria; Streptomycetales; Streptomycetaceae; Kitasatospora
MPHPRRPSVRLLTRRRLDRRQDAHHRRGRGARCNPLRRGVDRARRRAVALACCALGAAVA WP_033218839.1
hypothetical protein [Kitasatospora phosalacinea] Length: 154\nNumber of predicted TMHs: 2\nExp number of AAs in TMHs: 45.73448\nExp number, first 60 AAs: 13.96155\nTotal prob of N-in: 0.99999\nPOSSIBLE N-term signal sequence\ninside 1 46\nTMhelix 47 69\noutside 70 83\nTMhelix 84 106\ninside 107 154

32268 GCF_000716545.1_ASM71654v1 Kitasatospora phosalacinea Terrabacteria group; Actinobacteria; Actinobacteria; Streptomycetales; Streptomycetaceae; Kitasatospora
MQRHPRVTRRSAVPAALALGLLLGTAGTAAAHVEVESSTAQALAVDAQVAFSAEGESSTA WP_033220623.1
hypothetical protein [Kitasatospora phosalacinea] Length: 252\nNumber of predicted TMHs: 2\nExp number of AAs in TMHs: 42.45338\nExp number, first 60 AAs: 19.66423\nTotal prob of N-in: 0.92613\nPOSSIBLE N-term signal sequence\ninside 1 12\nTMhelix 13 35\noutside 36 223\nTMhelix 224 246\ninside 247 252

32269 GCF_000717185.1_ASM71718v1 Kitasatospora phosalacinea Terrabacteria group; Actinobacteria; Actinobacteria; Streptomycetales; Streptomycetaceae; Kitasatospora
MAGGRGDRRLWREARPCRPRRARTGTGGPAPHVHPPVRRSGRRGLMDWANTAAGTAACCA WP_081973614.1
hypothetical protein [Kitasatospora phosalacinea] Length: 203\nNumber of predicted TMHs: 2\nExp number of AAs in TMHs: 45.72429\nExp number, first 60 AAs: 9.80847\nTotal prob of N-in: 0.01172\noutside 1 50\nTMhelix 51 73\ninside 74 172\nTMhelix 173 195\noutside 196 203

32270 GCF_000716965.1_ASM71696v1 Kitasatospora setaeTerrabacteria group; Actinobacteria; Actinobacteria; Streptomycetales; Streptomycetaceae; Kitasatospora
MTPRHRPAGTGTPLPSTRPAALPLSLLAARTTLLGPTRAPRRRPAVAGTLLLSLLAACTT WP_014135556.1 hypothetical protein [Kitasatospora setae] Length: 262\nNumber of predicted TMHs: 2\nExp number of AAs in TMHs: 42.60646\nExp number, first 60 AAs: 14.17945\nTotal prob of N-in: 0.95716\nPOSSIBLE N-term signal sequence\ninside 1 45\nTMhelix 46 68\noutside 69 235\nTMhelix 236 254\ninside 255 262

32271 GCF_001905465.1_ASM190546v1 Kitasatospora sp. CB01950 Terrabacteria group; Actinobacteria; Actinobacteria; Streptomycetales; Streptomycetaceae; Kitasatospora
MPRTAAFFDLDKTIIAKSSALAFSRPFYQGGLINRRSVVKSAYAQFVFLVGGADHDQMEK WP_073808310.1
inhibition of morphological differentiation protein [Kitasatospora sp. CB01950]Length: 266\nNumber of predicted TMHs: 1\nExp number of AAs in TMHs: 21.1912\nExp number, first 60 AAs: 0.01533\nTotal prob of N-in: 0.00668\noutside 1 236\nTMhelix 237 259\ninside 260 266

32272 GCF_000710465.2_KspMBT63_1.0 Kitasatospora sp. MBT63 Terrabacteria group; Actinobacteria; Actinobacteria; Streptomycetales; Streptomycetaceae; Kitasatospora
MTRRGLILALAIPLSTVTGAGVAHADVCTTMTSGTYRIDGVYGAVPKVRTGGSRQTDNTL WP_033826143.1
hypothetical protein [Kitasatospora sp. MBT63] Length: 325\nNumber of predicted TMHs: 1\nExp number of AAs in TMHs: 31.1159\nExp number, first 60 AAs: 11.5177\nTotal prob of N-in: 0.53193\nPOSSIBLE N-term signal sequence\noutside 1 283\nTMhelix 284 306\ninside 307 325

32273 GCF_001182915.1_ASM118291v1 Kitasatospora sp. MY 5-36 Terrabacteria group; Actinobacteria; Actinobacteria; Streptomycetales; Streptomycetaceae; Kitasatospora
MRTAAFFDLDKTIIAKSSALAFSRPFYQGGLINRRAVLRSAYAQFVFLVGGADHDQMEKM WP_049660070.1
inhibition of morphological differentiation protein [Kitasatospora sp. MY 5-36] Length: 265\nNumber of predicted TMHs: 1\nExp number of AAs in TMHs: 22.66962\nExp number, first 60 AAs: 0.11451\nTotal prob of N-in: 0.01940\noutside 1 235\nTMhelix 236 258\ninside 259 265

32274 GCF_001182915.1_ASM118291v1 Kitasatospora sp. MY 5-36 Terrabacteria group; Actinobacteria; Actinobacteria; Streptomycetales; Streptomycetaceae; Kitasatospora
MGQATGSPDHRTTAVRTPATRRRRAVRVAVATALTALPLAAVLTVGSAAPLAPPLGACSG WP_082345927.1
hypothetical protein [Kitasatospora sp. MY 5-36] Length: 501\nNumber of predicted TMHs: 2\nExp number of AAs in TMHs: 43.72521\nExp number, first 60 AAs: 21.66527\nTotal prob of N-in: 0.96948\nPOSSIBLE N-term signal sequence\ninside 1 25\nTMhelix 26 48\noutside 49 471\nTMhelix 472 494\ninside 495 501

32275 GCF_001182915.1_ASM118291v1 Kitasatospora sp. MY 5-36 Terrabacteria group; Actinobacteria; Actinobacteria; Streptomycetales; Streptomycetaceae; Kitasatospora
MPGGGARARRRTALAALLLPLLGAALLLASAAPAQAAGYRYWSFWRGAEGGGWTYQQQGP WP_063780413.1
hypothetical protein [Kitasatospora sp. MY 5-36] Length: 252\nNumber of predicted TMHs: 2\nExp number of

AAs in TMHs: 40.6461\nExp number, first 60 AAs: 22.27916\nTotal prob of N-in: 0.99516\nPOSSIBLE N-term signal sequence\ninside 1 12\nTMhelix 13 35\noutside 36 225\nTMhelix 226 244\ninside 245 252

32276 GCF_000721345.1_ASM72134v1 Kitasatospora sp. NRRL B-11411 Terrabacteria group; Actinobacteria; Actinobacteria; Streptomycetales; Streptomycetaceae; Kitasatospora MFISDKVCLLPPDPTPSARRHVERRGHVKGPNSLATQLASATALAIAVLLAGAPAAPAAD WP_081967024.1 hypothetical protein [Kitasatospora sp. NRRL B-11411] Length: 419\nNumber of predicted TMHs: 1\nExp number of AAs in TMHs: 23.13073\nExp number, first 60 AAs: 1.21759\nTotal prob of N-in: 0.05516\noutside 1 388\nTMhelix 389 411\ninside 412 419

32277 GCF_000721345.1_ASM72134v1 Kitasatospora sp. NRRL B-11411 Terrabacteria group; Actinobacteria; Actinobacteria; Streptomycetales; Streptomycetaceae; Kitasatospora MIAKSSALAFSRPFYQGGLINRRSVVKSAYTQFIFLVGGADHDQMEKMRAYLSALTRGWN WP_051817180.1 inhibition of morphological differentiation protein [Kitasatospora sp. NRRL B-11411] Length: 254\nNumber of predicted TMHs: 1\nExp number of AAs in TMHs: 21.22202\nExp number, first 60 AAs: 0.03004\nTotal prob of N-in: 0.00980\noutside 1 223\nTMhelix 224 246\ninside 247 254

32278 GCF_001424875.1_Root107 Kitasatospora sp. Root107 Terrabacteria group; Actinobacteria; Actinobacteria; Streptomycetales; Streptomycetaceae; Kitasatospora MTGTGYGRRRAALAVAGAATAAAVLLSGCGAGRDSGGRATAADAVAPQAQTQKMAEGAAA WP_057229896.1 hypothetical protein [Kitasatospora sp. Root107] Length: 326\nNumber of predicted TMHs: 1\nExp number of AAs in TMHs: 37.59259\nExp number, first 60 AAs: 13.84188\nTotal prob of N-in: 0.66844\nPOSSIBLE N-term signal sequence\noutside 1 284\nTMhelix 285 307\ninside 308 326

32279 GCF_001424875.1_Root107 Kitasatospora sp. Root107 Terrabacteria group; Actinobacteria; Actinobacteria; Streptomycetales; Streptomycetaceae; Kitasatospora MQTMRRTRRTAVRAALLATTVLAAALCGTASAQAAAGDGIAGYQSAQQILRSDQLRDTVS WP_057231034.1 MULTISPECIES: hypothetical protein [Kitasatospora] Length: 313\nNumber of predicted TMHs: 2\nExp number of AAs in TMHs: 38.37448\nExp number, first 60 AAs: 21.80767\nTotal prob of N-in: 0.97566\nPOSSIBLE N-term signal sequence\ninside 1 11\nTMhelix 12 34\noutside 35 285\nTMhelix 286 303\ninside 304 313

32280 GCF_001424875.1_Root107 Kitasatospora sp. Root107 Terrabacteria group; Actinobacteria; Actinobacteria; Streptomycetales; Streptomycetaceae; Kitasatospora MSAPTLDRPVGTTEPAPPPRATRPPVSAARRRVVQAGWATSSLAGLVLGFAVYLPALSGL WP_063857240.1 MULTISPECIES: sortase [Kitasatospora] Length: 297\nNumber of predicted TMHs: 2\nExp number of AAs in TMHs: 56.64354\nExp number, first 60 AAs: 16.43276\nTotal prob of N-in: 0.57515\nPOSSIBLE N-term signal sequence\noutside 1 242\nTMhelix 243 265\ninside 266 269\nTMhelix 270 292\noutside 293 297

32281 GCF_001429805.1_Root187 Kitasatospora sp. Root187 Terrabacteria group; Actinobacteria; Actinobacteria; Streptomycetales; Streptomycetaceae; Kitasatospora MQTMRRTRRTAVRAALLATTVLAAALCGTASAQAAAGDGIAGYQSAQQILRSDQLRDTVS WP_057231034.1 MULTISPECIES: hypothetical protein [Kitasatospora] Length: 313\nNumber of predicted TMHs: 2\nExp number of AAs in TMHs: 38.37448\nExp number, first 60 AAs: 21.80767\nTotal prob of N-in: 0.97566\nPOSSIBLE N-term signal sequence\ninside 1 11\nTMhelix 12 34\noutside 35 285\nTMhelix 286 303\ninside 304 313

32282 GCF_001429805.1_Root187 Kitasatospora sp. Root187 Terrabacteria group; Actinobacteria; Actinobacteria; Streptomycetales; Streptomycetaceae; Kitasatospora MSAPTLDRPVGTTEPAPPPRATRPPVSAARRRVVQAGWATSSLAGLVLGFAVYLPALSGL WP_063857240.1 MULTISPECIES: sortase [Kitasatospora] Length: 297\nNumber of predicted TMHs: 2\nExp number of AAs in TMHs: 56.64354\nExp number, first 60 AAs: 16.43276\nTotal prob of N-in: 0.57515\nPOSSIBLE N-term signal sequence\noutside 1 242\nTMhelix 243 265\ninside 266 269\nTMhelix 270 292\noutside 293 297

32283 GCF_000744785.1_ASM74478v1 Kitasatospora azatica KCTC 9699 Terrabacteria group; Actinobacteria; Actinobacteria; Streptomycetales; Streptomycetaceae; Kitasatospora; Kitasatospora azatica MFSTRTRRAAIGTAAAGLLAAAATAVPAHAAGISLQLRLPNSLQIPLTPPADPASDSNSL WP_035847823.1 hypothetical protein [Kitasatospora azatica] Length: 565\nNumber of predicted TMHs: 2\nExp number of AAs in TMHs: 43.25635\nExp number, first 60 AAs: 21.15119\nTotal prob of N-in: 0.94543\nPOSSIBLE N-term signal sequence\ninside 1 8\nTMhelix 9 31\noutside 32 533\nTMhelix 534 556\ninside 557 565

32284 GCF_000696185.1_ASM69618v1 Kitasatospora cheerisanensis KCTC 2395 Terrabacteria group; Actinobacteria; Actinobacteria; Streptomycetales; Streptomycetaceae; Kitasatospora; Kitasatospora cheerisanensis

MTTVSTRRRVLAGAALAAGLFCTTAPAFAALTSPEATAYVVSADALNGAAAVPPQPLSTY WP_035863006.1 hypothetical protein [Kitasatospora cheerisanensis] Length: 245\nNumber of predicted TMHs: 2\nExp number of AAs in TMHs: 59.43337\nExp number, first 60 AAs: 16.57021\nTotal prob of N-in: 0.67205\nPOSSIBLE N-term signal sequence\ninside 1 179\nTMhelix 180 202\noutside 203 216\nTMhelix 217 239\ninside 240 245

32285 GCF_000744225.1_ASM74422v1 Kitasatospora mediocidica KCTC 9733 Terrabacteria group; Actinobacteria; Actinobacteria; Streptomycetales; Streptomycetaceae; Kitasatospora; Kitasatospora mediocidica MRPFAAPARTAAFFDLDKTIIAKSSALAFSRPFYQGGLINRRAVLKSAYAQFVFLVGGAD WP_035802465.1 inhibition of morphological differentiation protein [Kitasatospora mediocidica] Length: 272\nNumber of predicted TMHs: 1\nExp number of AAs in TMHs: 20.66367\nExp number, first 60 AAs: 0.09388\nTotal prob of N-in: 0.09658\noutside 1 242\nTMhelix 243 265\ninside 266 272

32286 GCF_000269985.1_ASM26998v1 Kitasatospora setae KM-6054 Terrabacteria group; Actinobacteria; Actinobacteria; Streptomycetales; Streptomycetaceae; Kitasatospora; Kitasatospora setae MTPRHRPAGTGTPLPSTRPAALPLSLLAARTTLLGPTRAPRRRPAVAGTLLLSLLAACTT WP_014135556.1 hypothetical protein [Kitasatospora setae] Length: 262\nNumber of predicted TMHs: 2\nExp number of AAs in TMHs: 42.60646\nExp number, first 60 AAs: 14.17945\nTotal prob of N-in: 0.95716\nPOSSIBLE N-term signal sequence\ninside 1 45\nTMhelix 46 68\noutside 69 235\nTMhelix 236 254\ninside 255 262

32287 GCF_000787755.1_ASM78775v1 Streptacidiphilus albus Terrabacteria group; Actinobacteria; Actinobacteria; Streptomycetales; Streptomycetaceae; Streptacidiphilus MSGRRRTAVLGLLSGALLLPAVPAAAAPASGSGSLPGIAQTVSDSQQGCVKASSAQVGRT WP_034087229.1 hypothetical protein [Streptacidiphilus albus] Length: 409\nNumber of predicted TMHs: 2\nExp number of AAs in TMHs: 43.94332\nExp number, first 60 AAs: 20.12928\nTotal prob of N-in: 0.92482\nPOSSIBLE N-term signal sequence\ninside 1 6\nTMhelix 7 29\noutside 30 370\nTMhelix 371 393\ninside 394 409

32288 GCF_000787755.1_ASM78775v1 Streptacidiphilus albus Terrabacteria group; Actinobacteria; Actinobacteria; Streptomycetales; Streptomycetaceae; Streptacidiphilus MRTAAFFDLDKTVIAKSSALAFSRPLYRGGLINRRAVLKSAYAQFVFLAGGADHDQMEKM WP_034093582.1 inhibition of morphological differentiation protein [Streptacidiphilus albus] Length: 266\nNumber of predicted TMHs: 1\nExp number of AAs in TMHs: 20.19992\nExp number, first 60 AAs: 0.06516\nTotal prob of N-in: 0.03615\noutside 1 238\nTMhelix 239 258\ninside 259 266

32289 GCF_000787755.1_ASM78775v1 Streptacidiphilus albus Terrabacteria group; Actinobacteria; Actinobacteria; Streptomycetales; Streptomycetaceae; Streptacidiphilus MPARRRALTATATAVVLAATTAVLAAGAANAAPVPATSNVTVTLGTPAPAGALTPGGGTE WP_052069070.1 hypothetical protein [Streptacidiphilus albus] Length: 393\nNumber of predicted TMHs: 2\nExp number of AAs in TMHs: 42.61644\nExp number, first 60 AAs: 21.65551\nTotal prob of N-in: 0.95551\nPOSSIBLE N-term signal sequence\ninside 1 6\nTMhelix 7 29\noutside 30 366\nTMhelix 367 386\ninside 387 393

32290 GCF_000787775.1_ASM78777v1 Streptacidiphilus carbonis Terrabacteria group; Actinobacteria; Actinobacteria; Streptomycetales; Streptomycetaceae; Streptacidiphilus MRQAARRGLLTVMATGSVLASTAGYAYADAGAVGSASDSPGVGSGNAVQVPVDVPVNVCG WP_042405542.1 hypothetical protein [Streptacidiphilus carbonis] Length: 283\nNumber of predicted TMHs: 1\nExp number of AAs in TMHs: 23.59323\nExp number, first 60 AAs: 6.70588\nTotal prob of N-in: 0.33346\noutside 1 257\nTMhelix 258 275\ninside 276 283

32291 GCF_000787775.1_ASM78777v1 Streptacidiphilus carbonis Terrabacteria group; Actinobacteria; Actinobacteria; Streptomycetales; Streptomycetaceae; Streptacidiphilus MRTAAFFDLDKTVIAKSSALAFSRPLYRGGLINRRAVLKSAYAQFVFLAGGADHDQMEKM WP_042407587.1 inhibition of morphological differentiation protein [Streptacidiphilus carbonis] Length: 265\nNumber of predicted TMHs: 1\nExp number of AAs in TMHs: 20.81682\nExp number, first 60 AAs: 0.0509\nTotal prob of N-in: 0.02879\noutside 1 237\nTMhelix 238 257\ninside 258 265

32292 GCF_000787775.1_ASM78777v1 Streptacidiphilus carbonis Terrabacteria group; Actinobacteria; Actinobacteria; Streptomycetales; Streptomycetaceae; Streptacidiphilus MDTAGDRTAVSRHHNRRRALPALLVAAALAATAPGPASALTHPPRTALASDGQCVFPAAS WP_052433307.1 type VII secretion-associated serine protease mycosin [Streptacidiphilus carbonis] Length: 418\nNumber of predicted TMHs: 2\nExp number of AAs in TMHs: 43.36405\nExp number, first 60 AAs: 20.01915\nTotal prob of N-in: 0.94301\nPOSSIBLE N-term signal sequence\ninside 1 19\nTMhelix 20 42\noutside 43 386\nTMhelix 387 409\ninside 410 418

32293 GCF_000719095.1_ASM71909v1 Streptacidiphilus jeojiense Terrabacteria group; Actinobacteria; Actinobacteria; Streptomycetales; Streptomycetaceae; Streptacidiphilus
MRQAARRGLLTVMATGSVLASTAGYAYADAGAAGSASDSPGVGSGNAVQVPVDVPVNVCG WP_030256123.1
hypothetical protein [Streptacidiphilus jeojiense] Length: 267\nNumber of predicted TMHs: 1\nExp number of AAs in TMHs: 25.81152\nExp number, first 60 AAs: 7.70261\nTotal prob of N-in: 0.37552\noutside 1 241\nTMhelix 242 259\ninside 260 267

32294 GCF_000719095.1_ASM71909v1 Streptacidiphilus jeojiense Terrabacteria group; Actinobacteria; Actinobacteria; Streptomycetales; Streptomycetaceae; Streptacidiphilus
MRTAAFFDLDKTVIAKSSALAFSRPLYRGGLINRRAVLKSAYAQFVFLAGGADHDQMEKM WP_030253917.1
inhibition of morphological differentiation protein [Streptacidiphilus jeojiense] Length: 265\nNumber of predicted TMHs: 1\nExp number of AAs in TMHs: 20.82989\nExp number, first 60 AAs: 0.04899\nTotal prob of N-in: 0.02103\noutside 1 237\nTMhelix 238 257\ninside 258 265

32295 GCF_900109465.1_IMG-taxon_2675903135_annotated_assembly Streptacidiphilus jiangxiensis Terrabacteria group; Actinobacteria; Actinobacteria; Streptomycetales; Streptomycetaceae; Streptacidiphilus
MTAVQPITPHEGEVAVAPKRRRWVFRLALVLLGLGTVLTVTGGARTLGRVHDYAVPTTSM WP_082015435.1
signal peptidase I [Streptacidiphilus jiangxiensis] Length: 248\nNumber of predicted TMHs: 2\nExp number of AAs in TMHs: 47.86134\nExp number, first 60 AAs: 21.22851\nTotal prob of N-in: 0.97854\nPOSSIBLE N-term signal sequence\ninside 1 22\nTMhelix 23 45\noutside 46 198\nTMhelix 199 230\ninside 231 248

32296 GCF_000787795.1_ASM78779v1 Streptacidiphilus jiangxiensis Terrabacteria group; Actinobacteria; Actinobacteria; Streptomycetales; Streptomycetaceae; Streptacidiphilus
MTAVQPITPHEGEVAVAPKRRRWVFRLALVLLGLGTVLTVTGGARTLGRVHDYAVPTTSM WP_082015435.1
signal peptidase I [Streptacidiphilus jiangxiensis] Length: 248\nNumber of predicted TMHs: 2\nExp number of AAs in TMHs: 47.86134\nExp number, first 60 AAs: 21.22851\nTotal prob of N-in: 0.97854\nPOSSIBLE N-term signal sequence\ninside 1 22\nTMhelix 23 45\noutside 46 198\nTMhelix 199 230\ninside 231 248

32297 GCF_000787815.1_ASM78781v1 Streptacidiphilus neutrinimicus Terrabacteria group; Actinobacteria; Actinobacteria; Streptomycetales; Streptomycetaceae; Streptacidiphilus
MLQRRGAVVFGAVGAFGVVFGAIGLLFAAIGTAVGVVIVRRWATARQTVGTGLVAEGVVL WP_084729695.1
hypothetical protein [Streptacidiphilus neutrinimicus] Length: 167\nNumber of predicted TMHs: 2\nExp number of AAs in TMHs: 51.65032\nExp number, first 60 AAs: 25.71152\nTotal prob of N-in: 0.07634\nPOSSIBLE N-term signal sequence\noutside 1 14\nTMhelix 15 37\ninside 38 131\nTMhelix 132 154\noutside 155 167

32298 GCF_000744705.1_ASM74470v1 Streptacidiphilus albus JL83 Terrabacteria group; Actinobacteria; Actinobacteria; Streptomycetales; Streptomycetaceae; Streptacidiphilus; Streptacidiphilus albus
MSGRRRTAVLGLLSGALLLPAVPAAAAPASGSGSLPGIAQTVSDSQQGCVKASSAQVGRT WP_034087229.1
hypothetical protein [Streptacidiphilus albus] Length: 409\nNumber of predicted TMHs: 2\nExp number of AAs in TMHs: 43.94332\nExp number, first 60 AAs: 20.12928\nTotal prob of N-in: 0.92482\nPOSSIBLE N-term signal sequence\ninside 1 6\nTMhelix 7 29\noutside 30 370\nTMhelix 371 393\ninside 394 409

32299 GCF_000744705.1_ASM74470v1 Streptacidiphilus albus JL83 Terrabacteria group; Actinobacteria; Actinobacteria; Streptomycetales; Streptomycetaceae; Streptacidiphilus; Streptacidiphilus albus
MRTAAFFDLDKTVIAKSSALAFSRPLYRGGLINRRAVLKSAYAQFVFLAGGADHDQMEKM WP_034093582.1
inhibition of morphological differentiation protein [Streptacidiphilus albus] Length: 266\nNumber of predicted TMHs: 1\nExp number of AAs in TMHs: 20.19992\nExp number, first 60 AAs: 0.06516\nTotal prob of N-in: 0.03615\noutside 1 238\nTMhelix 239 258\ninside 259 266

32300 GCF_000744705.1_ASM74470v1 Streptacidiphilus albus JL83 Terrabacteria group; Actinobacteria; Actinobacteria; Streptomycetales; Streptomycetaceae; Streptacidiphilus; Streptacidiphilus albus
MPARRRALTATATAVVLAATTAVLAAGAANAAPVPATSNVTVTLGTPAPAGALTPGGGTE WP_052069070.1
hypothetical protein [Streptacidiphilus albus] Length: 393\nNumber of predicted TMHs: 2\nExp number of AAs in TMHs: 42.61644\nExp number, first 60 AAs: 21.65551\nTotal prob of N-in: 0.95551\nPOSSIBLE N-term signal sequence\ninside 1 6\nTMhelix 7 29\noutside 30 366\nTMhelix 367 386\ninside 387 393

32301 GCF_000744815.1_ASM74481v1 Streptacidiphilus oryzae TH49 Terrabacteria group; Actinobacteria; Actinobacteria; Streptomycetales; Streptomycetaceae; Streptacidiphilus; Streptacidiphilus oryzae
MSTVIARRAAVVGALAGAAVLAAAGTASAHVTVNPQSAAKGSYTEVSFRAPNEEDAADTT WP_051940138.1
hypothetical protein [Streptacidiphilus oryzae] Length: 250\nNumber of predicted TMHs: 2\nExp number of

AAs in TMHs: 40.9261\nExp number, first 60 AAs: 19.37032\nTotal prob of N-in: 0.73558\nPOSSIBLE N-term signal sequence\ninside 1 8\nTMhelix 9 31\noutside 32 214\nTMhelix 215 237\ninside 238 250

32302 GCF_000744815.1_ASM74481v1 Streptacidiphilus oryzae TH49 Terrabacteria group; Actinobacteria; Actinobacteria; Streptomycetales; Streptomycetaceae; Streptacidiphilus; Streptacidiphilus oryzae MRTLRRFAGAEHNPLLRTFDRRRSALALLTAVLLCGAVAAGVLAGLIGSRLEQGLADRQV WP_037568650.1 hypothetical protein [Streptacidiphilus oryzae] Length: 199\nNumber of predicted TMHs: 2\nExp number of AAs in TMHs: 45.32153\nExp number, first 60 AAs: 22.86974\nTotal prob of N-in: 0.99871\nPOSSIBLE N-term signal sequence\ninside 1 24\nTMhelix 25 47\noutside 48 151\nTMhelix 152 174\ninside 175 199

32303 GCF_000744815.1_ASM74481v1 Streptacidiphilus oryzae TH49 Terrabacteria group; Actinobacteria; Actinobacteria; Streptomycetales; Streptomycetaceae; Streptacidiphilus; Streptacidiphilus oryzae MIGRRALVTAAIGGAGTVLAAAGGAFAAEPSAAAAGQPADFQQATVACAAVAHGGQAAIT WP_037573464.1 hypothetical protein [Streptacidiphilus oryzae] Length: 197\nNumber of predicted TMHs: 2\nExp number of AAs in TMHs: 46.13492\nExp number, first 60 AAs: 21.00848\nTotal prob of N-in: 0.93893\nPOSSIBLE N-term signal sequence\ninside 1 6\nTMhelix 7 29\noutside 30 167\nTMhelix 168 190\ninside 191 197

32304 GCF_000744655.1_ASM74465v1 Streptacidiphilus rugosus AM-16 Terrabacteria group; Actinobacteria; Actinobacteria; Streptomycetales; Streptomycetaceae; Streptacidiphilus; Streptacidiphilus rugosus MVFARARRTALLGAVAAAAVISGGPTALPAFAAGAAPAGGGRQFTTVVTSPEVTLLPQSG WP_037602190.1 hypothetical protein [Streptacidiphilus rugosus] Length: 564\nNumber of predicted TMHs: 2\nExp number of AAs in TMHs: 41.4276499999999\nExp number, first 60 AAs: 21.88828\nTotal prob of N-in: 0.95949\nPOSSIBLE N-term signal sequence\ninside 1 11\nTMhelix 12 34\noutside 35 531\nTMhelix 532 554\ninside 555 564

32305 GCF_000721375.1_ASM72137v1 Actinosporangium sp. NRRL B-3428 Terrabacteria group; Actinobacteria; Actinobacteria; Streptomycetales; Streptomycetaceae; Streptomyces MLDHVETHSLPRTAAFFDLDKTVIAKSSTLTFSKSFYQGGLINRRAVLRTAYTQFIFLAG WP_052581388.1 inhibition of morphological differentiation protein [Actinosporangium sp. NRRL B-3428] Length: 277\nNumber of predicted TMHs: 1\nExp number of AAs in TMHs: 21.48722\nExp number, first 60 AAs: 0.04977\nTotal prob of N-in: 0.01602\noutside 1 247\nTMhelix 248 270\ninside 271 277

32306 GCF_000721375.1_ASM72137v1 Actinosporangium sp. NRRL B-3428 Terrabacteria group; Actinobacteria; Actinobacteria; Streptomycetales; Streptomycetaceae; Streptomyces MTHGLRLGFGARKRLPQRRSLVGALASTAAVTLAIGGAPAPAMAADTKSQQWYLDAMHAD WP_052582755.1 serine protease [Actinosporangium sp. NRRL B-3428] Length: 425\nNumber of predicted TMHs: 2\nExp number of AAs in TMHs: 43.74782\nExp number, first 60 AAs: 21.10372\nTotal prob of N-in: 0.95547\nPOSSIBLE N-term signal sequence\ninside 1 20\nTMhelix 21 43\noutside 44 396\nTMhelix 397 419\ninside 420 425

32307 GCF_000721375.1_ASM72137v1 Actinosporangium sp. NRRL B-3428 Terrabacteria group; Actinobacteria; Actinobacteria; Streptomycetales; Streptomycetaceae; Streptomyces MSDPPPPARRRRRYLVASGVAAFALTAGLAAAPAAPAAPAPADSAAGQTNAQAPQAKAKGKT WP_052582802.1 ABC transporter substrate-binding protein [Actinosporangium sp. NRRL B-3428] Length: 633\nNumber of predicted TMHs: 2\nExp number of AAs in TMHs: 43.37375\nExp number, first 60 AAs: 21.26464\nTotal prob of N-in: 0.95081\nPOSSIBLE N-term signal sequence\ninside 1 12\nTMhelix 13 35\noutside 36 599\nTMhelix 600 622\ninside 623 633

32308 GCF_000717245.1_ASM71724v1 [Kitasatospora] papulosa Terrabacteria group; Actinobacteria; Actinobacteria; Streptomycetales; Streptomycetaceae; Streptomyces MRTTRRNTLRRGGVGALLAAVTLLTSAGAAVADDPSPSASADGGTAGPTEAGTTFRTATA WP_030124501.1 hypothetical protein [[Kitasatospora] papulosa] Length: 274\nNumber of predicted TMHs: 2\nExp number of AAs in TMHs: 40.1724199999999\nExp number, first 60 AAs: 19.30552\nTotal prob of N-in: 0.99359\nPOSSIBLE N-term signal sequence\ninside 1 12\nTMhelix 13 32\noutside 33 237\nTMhelix 238 260\ninside 261 274

32309 GCF_000717245.1_ASM71724v1 [Kitasatospora] papulosa Terrabacteria group; Actinobacteria; Actinobacteria; Streptomycetales; Streptomycetaceae; Streptomyces MTTHPSPRRRLLTAAAATAVLLVTLPVLPAAAEDSTQCTFPSKKYAGRPWSLQRVLMDEL WP_030125141.1 type VII secretion-associated serine protease [[Kitasatospora] papulosa] Length: 414\nNumber of predicted TMHs: 1\nExp number of AAs in TMHs: 36.07069\nExp number, first 60 AAs: 13.50637\nTotal prob of N-in: 0.65550\nPOSSIBLE N-term signal sequence\noutside 1 378\nTMhelix 379 401\ninside 402 414

32310 GCF_000717245.1_ASM71724v1 [Kitasatospora] papulosa Terrabacteria group; Actinobacteria; Actinobacteria; Streptomycetales; Streptomycetaceae; Streptomyces

MLTFVENCFSPRTAAFFDLDKTVIAKSSTLTFSKSFYQGGLINRRAVLRTAYTQFVFLAG WP_014155461.1 MULTISPECIES: haloacid dehalogenase [Streptomyces] Length: 279\nNumber of predicted TMHs: 1\nExp number of AAs in TMHs: 21.10966\nExp number, first 60 AAs: 0.01497\nTotal prob of N-in: 0.00234\noutside 1 248\nTMhelix 249 268\ninside 269 279

32311 GCF_001751365.1_ASM175136v1 Streptomyces abyssalis Terrabacteria group; Actinobacteria; Actinobacteria; Streptomycetales; Streptomycetaceae; Streptomyces MITHVLSSRARRSVLLSAATVMVGGAMVSVPAHAASADGHSDSTGNATAVAARADLQVSV WP_070013327.1 MULTISPECIES: hypothetical protein [Streptomyces] Length: 306\nNumber of predicted TMHs: 2\nExp number of AAs in TMHs: 42.74517\nExp number, first 60 AAs: 21.24237\nTotal prob of N-in: 0.95367\nPOSSIBLE N-term signal sequence\ninside 1 12\nTMhelix 13 35\noutside 36 274\nTMhelix 275 297\ninside 298 306

32312 GCF_001751365.1_ASM175136v1 Streptomyces abyssalis Terrabacteria group; Actinobacteria; Actinobacteria; Streptomycetales; Streptomycetaceae; Streptomyces MTQQRRATMTKWALIVGNVTFGLFLAGYALFGLDGVGRTYLTGEKTTAKVQNCIPPSSSK WP_070011590.1 MULTISPECIES: hypothetical protein [Streptomyces] Length: 156\nNumber of predicted TMHs: 2\nExp number of AAs in TMHs: 43.6246\nExp number, first 60 AAs: 20.89944\nTotal prob of N-in: 0.99825\nPOSSIBLE N-term signal sequence\ninside 1 11\nTMhelix 12 31\noutside 32 108\nTMhelix 109 131\ninside 132 156

32313 GCF_001751365.1_ASM175136v1 Streptomyces abyssalis Terrabacteria group; Actinobacteria; Actinobacteria; Streptomycetales; Streptomycetaceae; Streptomyces MLGFVENLAMRRTAAFFDLDKTVIAKSSTLAFGRSFYQGGLINRRAVLRTAYAQFVYLLG WP_070009382.1 MULTISPECIES: inhibition of morphological differentiation protein [Streptomyces] Length: 280\nNumber of predicted TMHs: 1\nExp number of AAs in TMHs: 22.1319\nExp number, first 60 AAs: 0.01219\nTotal prob of N-in: 0.00185\noutside 1 246\nTMhelix 247 269\ninside 270 280

32314 GCF_001751345.1_ASM175134v1 Streptomyces abyssalis Terrabacteria group; Actinobacteria; Actinobacteria; Streptomycetales; Streptomycetaceae; Streptomyces MTQQRRATMTKWALIVGNVTFGLFLAGYALFGLDGVGRTYLTGEKTTAKVQNCIPPSSSK WP_070011590.1 MULTISPECIES: hypothetical protein [Streptomyces] Length: 156\nNumber of predicted TMHs: 2\nExp number of AAs in TMHs: 43.6246\nExp number, first 60 AAs: 20.89944\nTotal prob of N-in: 0.99825\nPOSSIBLE N-term signal sequence\ninside 1 11\nTMhelix 12 31\noutside 32 108\nTMhelix 109 131\ninside 132 156

32315 GCF_001751345.1_ASM175134v1 Streptomyces abyssalis Terrabacteria group; Actinobacteria; Actinobacteria; Streptomycetales; Streptomycetaceae; Streptomyces MLGFVENLAMRRTAAFFDLDKTVIAKSSTLAFGRSFYQGGLINRRAVLRTAYAQFVYLLG WP_070009382.1 MULTISPECIES: inhibition of morphological differentiation protein [Streptomyces] Length: 280\nNumber of predicted TMHs: 1\nExp number of AAs in TMHs: 22.1319\nExp number, first 60 AAs: 0.01219\nTotal prob of N-in: 0.00185\noutside 1 246\nTMhelix 247 269\ninside 270 280

32316 GCF_001751345.1_ASM175134v1 Streptomyces abyssalis Terrabacteria group; Actinobacteria; Actinobacteria; Streptomycetales; Streptomycetaceae; Streptomyces MITHVLSSRARRSVLLSAATVMVGGAMVSVPAHAASADGHSDSTGNATAVAARADLQVSV WP_070013327.1 MULTISPECIES: hypothetical protein [Streptomyces] Length: 306\nNumber of predicted TMHs: 2\nExp number of AAs in TMHs: 42.74517\nExp number, first 60 AAs: 21.24237\nTotal prob of N-in: 0.95367\nPOSSIBLE N-term signal sequence\ninside 1 12\nTMhelix 13 35\noutside 36 274\nTMhelix 275 297\ninside 298 306

32317 GCF_001941265.1_ASM194126v1 Streptomyces acidiscabies Terrabacteria group; Actinobacteria; Actinobacteria; Streptomycetales; Streptomycetaceae; Streptomyces MLRDVENHPSPRTAAFFDLDKTVIAKSSTLTFGKSFYQGGLINRRAALRTAYAQFVFLVG WP_010359176.1 morphological differentiation-associated protein [Streptomyces acidiscabies] Length: 275\nNumber of predicted TMHs: 1\nExp number of AAs in TMHs: 21.76485\nExp number, first 60 AAs: 0.08203\nTotal prob of N-in: 0.06750\noutside 1 245\nTMhelix 246 268\ninside 269 275

32318 GCF_001941265.1_ASM194126v1 Streptomyces acidiscabies Terrabacteria group; Actinobacteria; Actinobacteria; Streptomycetales; Streptomycetaceae; Streptomyces MTENTSRRPALLTRAKALPPWTLLAAGALAGGLAGGAYGVLKPPVYTATAYVVAVPAEKS WP_010355677.1 hypothetical protein [Streptomyces acidiscabies] Length: 232\nNumber of predicted TMHs: 2\nExp number of AAs in TMHs: 43.18943\nExp number, first 60 AAs: 21.07852\nTotal prob of N-in: 0.99398\nPOSSIBLE N-term signal sequence\ninside 1 20\nTMhelix 21 40\noutside 41 164\nTMhelix 165 187\ninside 188 232

32319 GCF_001941325.1_ASM194132v1 Streptomyces acidiscabies Terrabacteria group; Actinobacteria; Actinobacteria; Streptomycetales; Streptomycetaceae; Streptomyces
MTENTSRRPALLTRAKALPPWTLLAAGALAGGLAGGAYGVLKPPVYTATAYVVAVPAEKS WP_010355677.1
hypothetical protein [Streptomyces acidiscabies] Length: 232\nNumber of predicted TMHs: 2\nExp number of AAs in TMHs: 43.18943\nExp number, first 60 AAs: 21.07852\nTotal prob of N-in: 0.99398\nPOSSIBLE N-term signal sequence\ninside 1 20\nTMhelix 21 40\noutside 41 164\nTMhelix 165 187\ninside 188 232

32320 GCF_001941285.1_ASM194128v1 Streptomyces acidiscabies Terrabacteria group; Actinobacteria; Actinobacteria; Streptomycetales; Streptomycetaceae; Streptomyces
MTENTSRRPALLTRAKALPPWTLLAAGALAGGLAGGAYGVLKPPVYTATAYVVAVPAEKS WP_010355677.1
hypothetical protein [Streptomyces acidiscabies] Length: 232\nNumber of predicted TMHs: 2\nExp number of AAs in TMHs: 43.18943\nExp number, first 60 AAs: 21.07852\nTotal prob of N-in: 0.99398\nPOSSIBLE N-term signal sequence\ninside 1 20\nTMhelix 21 40\noutside 41 164\nTMhelix 165 187\ninside 188 232

32321 GCF_001949645.1_ASM194964v1 Streptomyces acidiscabies Terrabacteria group; Actinobacteria; Actinobacteria; Streptomycetales; Streptomycetaceae; Streptomyces
MTENTSRRPALLTRAKALPPWTLLAAGALAGGLAGGAYGVLKPPVYTATAYVVAVPAEKS WP_010355677.1
hypothetical protein [Streptomyces acidiscabies] Length: 232\nNumber of predicted TMHs: 2\nExp number of AAs in TMHs: 43.18943\nExp number, first 60 AAs: 21.07852\nTotal prob of N-in: 0.99398\nPOSSIBLE N-term signal sequence\ninside 1 20\nTMhelix 21 40\noutside 41 164\nTMhelix 165 187\ninside 188 232

32322 GCF_001485105.1_ASM148510v1 Streptomyces acidiscabies Terrabacteria group; Actinobacteria; Actinobacteria; Streptomycetales; Streptomycetaceae; Streptomyces
MLRDVENHPSPRTAAFFDLDKTVIAKSSTLTFGKSFYQGGLINRRAALRTAYAQFVFLVG WP_050371489.1 inhibition of morphological differentiation protein [Streptomyces acidiscabies] Length: 275\nNumber of predicted TMHs: 1\nExp number of AAs in TMHs: 21.76486\nExp number, first 60 AAs: 0.08203\nTotal prob of N-in: 0.06750\noutside 1 245\nTMhelix 246 268\ninside 269 275

32323 GCF_001949645.1_ASM194964v1 Streptomyces acidiscabies Terrabacteria group; Actinobacteria; Actinobacteria; Streptomycetales; Streptomycetaceae; Streptomyces
MLRDVENHPSPRTAAFFDLDKTVIAKSSTLTFGKSFYQGGLINRRAALRTAYAQFVFLVG WP_050371489.1 inhibition of morphological differentiation protein [Streptomyces acidiscabies] Length: 275\nNumber of predicted TMHs: 1\nExp number of AAs in TMHs: 21.76486\nExp number, first 60 AAs: 0.08203\nTotal prob of N-in: 0.06750\noutside 1 245\nTMhelix 246 268\ninside 269 275

32324 GCF_001485105.1_ASM148510v1 Streptomyces acidiscabies Terrabacteria group; Actinobacteria; Actinobacteria; Streptomycetales; Streptomycetaceae; Streptomyces
MTENTSRRPALLTRAKALPPWTLLAAGALAGGLAGGAYGVLKPPVYTATAYVVAVPAEKS WP_010355677.1
hypothetical protein [Streptomyces acidiscabies] Length: 232\nNumber of predicted TMHs: 2\nExp number of AAs in TMHs: 43.18943\nExp number, first 60 AAs: 21.07852\nTotal prob of N-in: 0.99398\nPOSSIBLE N-term signal sequence\ninside 1 20\nTMhelix 21 40\noutside 41 164\nTMhelix 165 187\ninside 188 232

32325 GCF_001941275.1_ASM194127v1 Streptomyces acidiscabies Terrabacteria group; Actinobacteria; Actinobacteria; Streptomycetales; Streptomycetaceae; Streptomyces
MPARRSALALAAFLAAVALTLAGCGANDDTGSAAQDKSAAKEGAVKGDADTAPGASGSGP WP_075732192.1
hypothetical protein [Streptomyces acidiscabies] Length: 312\nNumber of predicted TMHs: 1\nExp number of AAs in TMHs: 36.52187\nExp number, first 60 AAs: 13.37346\nTotal prob of N-in: 0.62648\nPOSSIBLE N-term signal sequence\noutside 1 261\nTMhelix 262 284\ninside 285 312

32326 GCF_001941275.1_ASM194127v1 Streptomyces acidiscabies Terrabacteria group; Actinobacteria; Actinobacteria; Streptomycetales; Streptomycetaceae; Streptomyces
MTENTSRRPALLTRAKALPPWTLLAAGALAGGLAGGAYGVLKPPVYTATAYVVAVPAEKS WP_010355677.1
hypothetical protein [Streptomyces acidiscabies] Length: 232\nNumber of predicted TMHs: 2\nExp number of AAs in TMHs: 43.18943\nExp number, first 60 AAs: 21.07852\nTotal prob of N-in: 0.99398\nPOSSIBLE N-term signal sequence\ninside 1 20\nTMhelix 21 40\noutside 41 164\nTMhelix 165 187\ninside 188 232

32327 GCF_001941275.1_ASM194127v1 Streptomyces acidiscabies Terrabacteria group; Actinobacteria; Actinobacteria; Streptomycetales; Streptomycetaceae; Streptomyces
MLRDVENHPPPRTAAFFDLDKTVIAKSSTLTFGKSFYQGGLINRRAALRTAYAQFVFLVG WP_075737737.1 inhibition of morphological differentiation protein [Streptomyces acidiscabies] Length: 275\nNumber of predicted TMHs: 1\nExp

number of AAs in TMHs: 21.74548\nExp number, first 60 AAs: 0.07328\nTotal prob of N-in: 0.04595\noutside 1 245\nTMhelix 246 268\ninside 269 275

32328 GCF_001189015.1_ASM118901v1 Streptomyces acidiscabies Terrabacteria group; Actinobacteria; Actinobacteria; Streptomycetales; Streptomycetaceae; Streptomyces MTENTSRRPALLTRAKALPPWTLLAAGALAGGLAGGAYGALKPPVYTATAYVVAVPAEKS WP_050371013.1 hypothetical protein [Streptomyces acidiscabies] Length: 232\nNumber of predicted TMHs: 2\nExp number of AAs in TMHs: 43.09223\nExp number, first 60 AAs: 20.98625\nTotal prob of N-in: 0.99336\nPOSSIBLE N-term signal sequence\ninside 1 20\nTMhelix 21 40\noutside 41 164\nTMhelix 165 187\ninside 188 232

32329 GCF_001189015.1_ASM118901v1 Streptomyces acidiscabies Terrabacteria group; Actinobacteria; Actinobacteria; Streptomycetales; Streptomycetaceae; Streptomyces MHLTRRTLLKTTALTGAASALALPTTTAAAAPTAYTHPGLLHTAADLTRMATKVKANSAP WP_050369614.1 Tat pathway signal sequence domain protein [Streptomyces acidiscabies] Length: 512\nNumber of predicted TMHs: 1\nExp number of AAs in TMHs: 26.16091\nExp number, first 60 AAs: 7.54418\nTotal prob of N-in: 0.34221\noutside 1 463\nTMhelix 464 483\ninside 484 512

32330 GCF_001189015.1_ASM118901v1 Streptomyces acidiscabies Terrabacteria group; Actinobacteria; Actinobacteria; Streptomycetales; Streptomycetaceae; Streptomyces MLRDVENHPSPRTAAFFDLDKTVIAKSSTLTFGKSFYQGGLINRRAALRTAYAQFVFLVG WP_050371489.1 inhibition of morphological differentiation protein [Streptomyces acidiscabies] Length: 275\nNumber of predicted TMHs: 1\nExp number of AAs in TMHs: 21.76486\nExp number, first 60 AAs: 0.08203\nTotal prob of N-in: 0.06750\noutside 1 245\nTMhelix 246 268\ninside 269 275

32331 GCF_001941325.1_ASM194132v1 Streptomyces acidiscabies Terrabacteria group; Actinobacteria; Actinobacteria; Streptomycetales; Streptomycetaceae; Streptomyces MLRDVENHPSPRTAAFFDLDKTVIAKSSTLTFGKSFYQGGLINRRAALRTAYAQFVFLVG WP_075779868.1 inhibition of morphological differentiation protein [Streptomyces acidiscabies] Length: 275\nNumber of predicted TMHs: 1\nExp number of AAs in TMHs: 21.76503\nExp number, first 60 AAs: 0.08203\nTotal prob of N-in: 0.06750\noutside 1 245\nTMhelix 246 268\ninside 269 275

32332 GCF_001941285.1_ASM194128v1 Streptomyces acidiscabies Terrabacteria group; Actinobacteria; Actinobacteria; Streptomycetales; Streptomycetaceae; Streptomyces MLRDVENHPSPRTAAFFDLDKTVIAKSSTLTFGKSFYQGGLINRRAALRTAYAQFVFLVG WP_010359176.1 morphological differentiation-associated protein [Streptomyces acidiscabies] Length: 275\nNumber of predicted TMHs: 1\nExp number of AAs in TMHs: 21.76485\nExp number, first 60 AAs: 0.08203\nTotal prob of N-in: 0.06750\noutside 1 245\nTMhelix 246 268\ninside 269 275

32333 GCF_002150735.1_ASM215073v1 Streptomyces africanus Terrabacteria group; Actinobacteria; Actinobacteria; Streptomycetales; Streptomycetaceae; Streptomyces MLKGVENHLSPRTAAFFDLDKTVIAKSSTLTFSKSFYQGGLINRRAVLRTAYAQFVFLAG WP_086565263.1 inhibition of morphological differentiation protein [Streptomyces africanus] Length: 277\nNumber of predicted TMHs: 1\nExp number of AAs in TMHs: 21.48796\nExp number, first 60 AAs: 0.03993\nTotal prob of N-in: 0.00320\noutside 1 247\nTMhelix 248 270\ninside 271 277

32334 GCF_002150735.1_ASM215073v1 Streptomyces africanus Terrabacteria group; Actinobacteria; Actinobacteria; Streptomycetales; Streptomycetaceae; Streptomyces MRHARRRVVRRVTRLAAVGGLLLGGAMVTNAVASESPDPSIGVPYSARPATSMGTDLVSR WP_086565138.1 serine protease [Streptomyces africanus] Length: 448\nNumber of predicted TMHs: 2\nExp number of AAs in TMHs: 40.85027\nExp number, first 60 AAs: 21.03078\nTotal prob of N-in: 0.95813\nPOSSIBLE N-term signal sequence\ninside 1 11\nTMhelix 12 34\noutside 35 409\nTMhelix 410 432\ninside 433 448

32335 GCF_002150735.1_ASM215073v1 Streptomyces africanus Terrabacteria group; Actinobacteria; Actinobacteria; Streptomycetales; Streptomycetaceae; Streptomyces MHTRRPALHPVPALTALLLAAALALTGCSGAGDDSAGSSAADRGAAAPQSDAKAGANEAA WP_086565734.1 hypothetical protein, partial [Streptomyces africanus] Length: 291\nNumber of predicted TMHs: 1\nExp number of AAs in TMHs: 33.65901\nExp number, first 60 AAs: 10.76661\nTotal prob of N-in: 0.45880\nPOSSIBLE N-term signal sequence\noutside 1 265\nTMhelix 266 288\ninside 289 291

32336 GCF_002192455.1_ASM219245v1 Streptomyces albireticuli Terrabacteria group; Actinobacteria; Actinobacteria; Streptomycetales; Streptomycetaceae; Streptomyces

MTVTSRTSPAARRARTTRRGALAVLATATALGATAFAVPAHAADAGASSAPAPKAAPKFD WP_087928343.1 hypothetical protein [Streptomyces albireticuli] Length: 546\nNumber of predicted TMHs: 2\nExp number of AAs in TMHs: 42.42347\nExp number, first 60 AAs: 22.11952\nTotal prob of N-in: 0.99903\nPOSSIBLE N-term signal sequence\ninside 1 20\nTMhelix 21 43\noutside 44 514\nTMhelix 515 537\ninside 538 546

32337 GCF_002192455.1_ASM219245v1 Streptomyces albireticuli Terrabacteria group; Actinobacteria; Actinobacteria; Streptomycetales; Streptomycetaceae; Streptomyces MIVYMMGVGFMGRRRTVRRAAVSIAVPGAVGLALAAAPAWAGAPPHADYQAVAQPVVGRV WP_087926919.1 hypothetical protein [Streptomyces albireticuli] Length: 226\nNumber of predicted TMHs: 2\nExp number of AAs in TMHs: 40.67562\nExp number, first 60 AAs: 22.53173\nTotal prob of N-in: 0.99853\nPOSSIBLE N-term signal sequence\ninside 1 20\nTMhelix 21 43\noutside 44 201\nTMhelix 202 221\ninside 222 226

32338 GCF_002192455.1_ASM219245v1 Streptomyces albireticuli Terrabacteria group; Actinobacteria; Actinobacteria; Streptomycetales; Streptomycetaceae; Streptomyces MLGRVENHSLSRTAAFFDLDKTVIAKSSTLTFSKSFYQGGLINRRAVLRTAYAQFVFLAG WP_087927003.1 inhibition of morphological differentiation protein [Streptomyces albireticuli] Length: 281\nNumber of predicted TMHs: 1\nExp number of AAs in TMHs: 21.72502\nExp number, first 60 AAs: 0.03502\nTotal prob of N-in: 0.00485\noutside 1 246\nTMhelix 247 269\ninside 270 281

32339 GCF_000716675.1_ASM71667v1 Streptomyces alboflavus Terrabacteria group; Actinobacteria; Actinobacteria; Streptomycetales; Streptomycetaceae; Streptomyces MLNLVENHSSPRTAAFFDLDKTVIAKSSTLTFSKSFYQGGLINRRAVLRTAYAQFVFLAG WP_030359659.1 MULTISPECIES: inhibition of morphological differentiation protein [Streptomyces] Length: 274\nNumber of predicted TMHs: 1\nExp number of AAs in TMHs: 21.81923\nExp number, first 60 AAs: 0.095\nTotal prob of N-in: 0.03042\noutside 1 244\nTMhelix 245 267\ninside 268 274

32340 GCF_002189675.1_ASM218967v1 Streptomyces alboflavus Terrabacteria group; Actinobacteria; Actinobacteria; Streptomycetales; Streptomycetaceae; Streptomyces MLNLVENHSSPRTAAFFDLDKTVIAKSSTLTFSKSFYQGGLINRRAVLRTAYAQFVFLAG WP_030359659.1 MULTISPECIES: inhibition of morphological differentiation protein [Streptomyces] Length: 274\nNumber of predicted TMHs: 1\nExp number of AAs in TMHs: 21.81923\nExp number, first 60 AAs: 0.095\nTotal prob of N-in: 0.03042\noutside 1 244\nTMhelix 245 267\ninside 268 274

32341 GCF_002082585.1_ASM208258v1 Streptomyces albolongus Terrabacteria group; Actinobacteria; Actinobacteria; Streptomycetales; Streptomycetaceae; Streptomyces MLSVVENCFSPRTAAFFDLDKTVIAKSSTLTFSKSFYQGGLINRRAVLRTAYTQFVFLAG WP_084747213.1 inhibition of morphological differentiation protein [Streptomyces albolongus] Length: 281\nNumber of predicted TMHs: 1\nExp number of AAs in TMHs: 21.7246\nExp number, first 60 AAs: 0.01851\nTotal prob of N-in: 0.00390\noutside 1 247\nTMhelix 248 270\ninside 271 281

32342 GCF_002082585.1_ASM208258v1 Streptomyces albolongus Terrabacteria group; Actinobacteria; Actinobacteria; Streptomycetales; Streptomycetaceae; Streptomyces MGIAVSGRRTPLSGGAVLASAALIALTATAPAQAAGTGPDLGVGALTPVTGIAPGGTFGL WP_084749454.1 peptidase [Streptomyces albolongus] Length: 443\nNumber of predicted TMHs: 1\nExp number of AAs in TMHs: 36.2296899999999\nExp number, first 60 AAs: 15.22843\nTotal prob of N-in: 0.65905\nPOSSIBLE N-term signal sequence\noutside 1 411\nTMhelix 412 434\ninside 435 443

32343 GCF_002150845.1_ASM215084v1 Streptomyces alboverticillatus Terrabacteria group; Actinobacteria; Actinobacteria; Streptomycetales; Streptomycetaceae; Streptomyces MRLTRRAALAATTALTAALLAPAAAATPASAPGGSTATVLNGSGECDYPAKPIKGTPWAL WP_086573719.1 type VII secretion-associated serine protease mycosin [Streptomyces alboverticillatus] Length: 415\nNumber of predicted TMHs: 2\nExp number of AAs in TMHs: 39.57527\nExp number, first 60 AAs: 17.18095\nTotal prob of N-in: 0.79776\nPOSSIBLE N-term signal sequence\ninside 1 6\nTMhelix 7 29\noutside 30 383\nTMhelix 384 406\ninside 407 415

32344 GCF_001646665.1_ASM164666v1 Streptomyces albulus Terrabacteria group; Actinobacteria; Actinobacteria; Streptomycetales; Streptomycetaceae; Streptomyces MNSTTSRPSARRRTIAAAVAAFAAGPALLTAPAAHATGGEGRASAVVLRTDLDVALLNKA WP_064073155.1 hypothetical protein [Streptomyces albulus] Length: 314\nNumber of predicted TMHs: 2\nExp number of AAs in TMHs: 39.82809\nExp number, first 60 AAs: 19.23574\nTotal prob of N-in: 0.87259\nPOSSIBLE N-term signal sequence\ninside 1 12\nTMhelix 13 35\noutside 36 280\nTMhelix 281 303\ninside 304 314

32345 GCF_001646665.1_ASM164666v1 Streptomyces albulus Terrabacteria group; Actinobacteria; Actinobacteria; Streptomycetales; Streptomycetaceae; Streptomyces
MTNRRRRTAVTAALAATAFAVLPATPARADGIRAQQWGLEALHTETAWRTTKGAGVTVAV WP_064068125.1 type VII secretion-associated serine protease mycosin [Streptomyces albulus] Length: 392\nNumber of predicted TMHs: 1\nExp number of AAs in TMHs: 39.1337\nExp number, first 60 AAs: 6.68851999999999\nTotal prob of N-in: 0.82296\noutside 1 361\nTMhelix 362 384\ninside 385 392

32346 GCF_001646665.1_ASM164666v1 Streptomyces albulus Terrabacteria group; Actinobacteria; Actinobacteria; Streptomycetales; Streptomycetaceae; Streptomyces
MLRLVENHSLPRTAAFFDLDKTVIAKSSTLTFSKSFYQGGLINRRAVLRTAYIQFVFLAG WP_064067283.1 inhibition of morphological differentiation protein [Streptomyces albulus] Length: 276\nNumber of predicted TMHs: 1\nExp number of AAs in TMHs: 21.88051\nExp number, first 60 AAs: 0.05896\nTotal prob of N-in: 0.00539\noutside 1 247\nTMhelix 248 270\ninside 271 276

32347 GCF_000695235.1_ASM69523v1 Streptomyces albulus Terrabacteria group; Actinobacteria; Actinobacteria; Streptomycetales; Streptomycetaceae; Streptomyces
MTLMRPTGPPRPTGRRRRATLAATAALVLAGAGVVTAAGTAQAADVAYKTECQPPPISGL WP_038519829.1 MULTISPECIES: hypothetical protein [Streptomyces] Length: 479\nNumber of predicted TMHs: 2\nExp number of AAs in TMHs: 43.80161\nExp number, first 60 AAs: 21.18149\nTotal prob of N-in: 0.97217\nPOSSIBLE N-term signal sequence\ninside 1 20\nTMhelix 21 43\noutside 44 450\nTMhelix 451 473\ninside 474 479

32348 GCF_001632865.1_ASM163286v1 Streptomyces ambofaciens Terrabacteria group; Actinobacteria; Actinobacteria; Streptomycetales; Streptomycetaceae; Streptomyces
MPRTAAFFDLDKTVIAKSSTLTFSKSFYQGGLINRRAVLRTAYAQFVFLAGGADHDQMER WP_063482640.1 inhibition of morphological differentiation protein [Streptomyces ambofaciens] Length: 268\nNumber of predicted TMHs: 1\nExp number of AAs in TMHs: 21.53677\nExp number, first 60 AAs: 0.04893\nTotal prob of N-in: 0.00773\noutside 1 238\nTMhelix 239 261\ninside 262 268

32349 GCF_001632865.1_ASM163286v1 Streptomyces ambofaciens Terrabacteria group; Actinobacteria; Actinobacteria; Streptomycetales; Streptomycetaceae; Streptomyces
MRHARRRIVRRVARLAAVGGLLLGGTMVTRAVASEPPGPPTGPHATVRVASGTGAGLVSR WP_053140927.1 protease [Streptomyces ambofaciens] Length: 455\nNumber of predicted TMHs: 2\nExp number of AAs in TMHs: 41.9456\nExp number, first 60 AAs: 21.00104\nTotal prob of N-in: 0.96796\nPOSSIBLE N-term signal sequence\ninside 1 11\nTMhelix 12 34\noutside 35 416\nTMhelix 417 439\ninside 440 455

32350 GCF_001632865.1_ASM163286v1 Streptomyces ambofaciens Terrabacteria group; Actinobacteria; Actinobacteria; Streptomycetales; Streptomycetaceae; Streptomyces
MGHNRRRRPTGARRATFGACALILGGGGLVAVNVFASATEAGDPAVPLGSSGVAATVDCP WP_079155936.1 hypothetical protein [Streptomyces ambofaciens] Length: 732\nNumber of predicted TMHs: 2\nExp number of AAs in TMHs: 42.81686\nExp number, first 60 AAs: 21.63504\nTotal prob of N-in: 0.98648\nPOSSIBLE N-term signal sequence\ninside 1 16\nTMhelix 17 39\noutside 40 704\nTMhelix 705 727\ninside 728 732

32351 GCF_001953875.1_ASM195387v1 Streptomyces amritsarensis Terrabacteria group; Actinobacteria; Actinobacteria; Streptomycetales; Streptomycetaceae; Streptomyces
MSPVRRRTLVRAAAVTACAGSLLALPTAAALAEGVPAVSSAHSSAPQRTLVKSLSLADGV WP_060176777.1 MULTISPECIES: hypothetical protein [Streptomyces] Length: 210\nNumber of predicted TMHs: 2\nExp number of AAs in TMHs: 42.68199\nExp number, first 60 AAs: 20.014\nTotal prob of N-in: 0.93067\nPOSSIBLE N-term signal sequence\ninside 1 8\nTMhelix 9 31\noutside 32 180\nTMhelix 181 203\ninside 204 210

32352 GCF_001953875.1_ASM195387v1 Streptomyces amritsarensis Terrabacteria group; Actinobacteria; Actinobacteria; Streptomycetales; Streptomycetaceae; Streptomyces
MPARRRTVTPIRIAAAGLAPLALAAYAAGPAAAHGSMTDPVSRVAACYAEGPESPKSAAC WP_060179978.1 MULTISPECIES: chitin-binding protein [Streptomyces] Length: 355\nNumber of predicted TMHs: 2\nExp number of AAs in TMHs: 41.76409\nExp number, first 60 AAs: 20.71548\nTotal prob of N-in: 0.94609\nPOSSIBLE N-term signal sequence\ninside 1 12\nTMhelix 13 35\noutside 36 320\nTMhelix 321 343\ninside 344 355

32353 GCF_001953875.1_ASM195387v1 Streptomyces amritsarensis Terrabacteria group; Actinobacteria; Actinobacteria; Streptomycetales; Streptomycetaceae; Streptomyces
MPQNRRPRRPLLAGAFALTLAAGAAATAGAAADAATTAAAAPHPPYALRLDGAGECTFPM WP_079250908.1 type VII secretion-associated serine protease mycosin [Streptomyces amritsarensis] Length: 424\nNumber of predicted TMHs:

2\nExp number of AAs in TMHs: 41.33732\nExp number, first 60 AAs: 19.12426\nTotal prob of N-in: 0.87967\nPOSSIBLE N-term signal sequence\ninside 1 11\nTMhelix 12 34\noutside 35 390\nTMhelix 391 413\ninside 414 424

32354 GCF_002154415.1_ASM215441v1 Streptomyces angustmyceticus Terrabacteria group; Actinobacteria; Actinobacteria; Streptomycetales; Streptomycetaceae; Streptomyces MRQRRRAAVAPVAALVLAGAGIVATAGTAQAAEVSYKTECLPPPISGLPPVEGTTKVAVT WP_086719786.1 hypothetical protein [Streptomyces angustmyceticus] Length: 485\nNumber of predicted TMHs: 2\nExp number of AAs in TMHs: 44.92096\nExp number, first 60 AAs: 22.23913\nTotal prob of N-in: 0.99553\nPOSSIBLE N-term signal sequence\ninside 1 6\nTMhelix 7 29\noutside 30 454\nTMhelix 455 477\ninside 478 485

32355 GCF_001270635.1_ASM127063v1 Streptomyces antibioticus Terrabacteria group; Actinobacteria; Actinobacteria; Streptomycetales; Streptomycetaceae; Streptomyces MLRGVENHSLPRAAAFFDLDKTVIAKSSTLTFSKSFYQGGLINRRAALRTAYAQFVFLVG WP_030648527.1 MULTISPECIES: inhibition of morphological differentiation protein [Streptomyces] Length: 276\nNumber of predicted TMHs: 1\nExp number of AAs in TMHs: 21.19627\nExp number, first 60 AAs: 0.04374\nTotal prob of N-in: 0.00486\noutside 1 248\nTMhelix 249 268\ninside 269 276

32356 GCF_001514065.1_ASM151406v1 Streptomyces antibioticus Terrabacteria group; Actinobacteria; Actinobacteria; Streptomycetales; Streptomycetaceae; Streptomyces MRGRARRRLLAVVLALLATQLGSLAAPAYACGCGAMVPDTARQVAVAREVSVVRWDGRRE WP_079083022.1 hypothetical protein [Streptomyces antibioticus] Length: 377\nNumber of predicted TMHs: 2\nExp number of AAs in TMHs: 41.67793\nExp number, first 60 AAs: 19.09345\nTotal prob of N-in: 0.87800\nPOSSIBLE N-term signal sequence\ninside 1 8\nTMhelix 9 31\noutside 32 330\nTMhelix 331 353\ninside 354 377

32357 GCF_001514065.1_ASM151406v1 Streptomyces antibioticus Terrabacteria group; Actinobacteria; Actinobacteria; Streptomycetales; Streptomycetaceae; Streptomyces MLGVVENHSLPRTAAFFDLDKTVIAKSSTLTFSKSFYQGGLINRRAAVRTAYAQFVFLVG WP_059194055.1 inhibition of morphological differentiation protein [Streptomyces antibioticus] Length: 277\nNumber of predicted TMHs: 1\nExp number of AAs in TMHs: 21.81716\nExp number, first 60 AAs: 0.01292\nTotal prob of N-in: 0.00208\noutside 1 247\nTMhelix 248 270\ninside 271 277

32358 GCF_002019855.1_ASM201985v1 Streptomyces antibioticus Terrabacteria group; Actinobacteria; Actinobacteria; Streptomycetales; Streptomycetaceae; Streptomyces MLMLVENHSLPRTAAFFDLDKTVIAKSSTLTFSKSFYQGGLINRRAALRTAYAQFVFLVG WP_030795901.1 MULTISPECIES: inhibition of morphological differentiation protein [Streptomyces] Length: 277\nNumber of predicted TMHs: 1\nExp number of AAs in TMHs: 21.68626\nExp number, first 60 AAs: 0.01173\nTotal prob of N-in: 0.00195\noutside 1 247\nTMhelix 248 270\ninside 271 277

32359 GCF_001514065.1_ASM151406v1 Streptomyces antibioticus Terrabacteria group; Actinobacteria; Actinobacteria; Streptomycetales; Streptomycetaceae; Streptomyces MSKRRRAVPDRARVVLALCGAGVLIGVAVAPAQAAEVSYATHCVPPAGVGLDPVDGTTKV WP_059196190.1 hypothetical protein [Streptomyces antibioticus] Length: 422\nNumber of predicted TMHs: 2\nExp number of AAs in TMHs: 42.0056399999999\nExp number, first 60 AAs: 21.65802\nTotal prob of N-in: 0.99530\nPOSSIBLE N-term signal sequence\ninside 1 12\nTMhelix 13 35\noutside 36 396\nTMhelix 397 416\ninside 417 422

32360 GCF_001270635.1_ASM127063v1 Streptomyces antibioticus Terrabacteria group; Actinobacteria; Actinobacteria; Streptomycetales; Streptomycetaceae; Streptomyces MRHARRRVVRRVTRLAAVGGLLLGGAMVTQAAMASETPPASARPLAATGDTGAALVARLG WP_030640851.1 MULTISPECIES: protease [Streptomyces] Length: 458\nNumber of predicted TMHs: 2\nExp number of AAs in TMHs: 43.16453\nExp number, first 60 AAs: 22.25986\nTotal prob of N-in: 0.99239\nPOSSIBLE N-term signal sequence\ninside 1 11\nTMhelix 12 34\noutside 35 419\nTMhelix 420 442\ninside 443 458

32361 GCF_001270635.1_ASM127063v1 Streptomyces antibioticus Terrabacteria group; Actinobacteria; Actinobacteria; Streptomycetales; Streptomycetaceae; Streptomyces MRTPHTARRRRALAGVLLAACLALAGCGGADDMGGSSKAASDSKAEAAGAPGGSARNGRA WP_030645059.1 MULTISPECIES: hypothetical protein [Streptomyces] Length: 324\nNumber of predicted TMHs: 2\nExp number of AAs in TMHs: 42.21572\nExp number, first 60 AAs: 19.28007\nTotal prob of N-in: 0.97332\nPOSSIBLE N-term signal sequence\ninside 1 12\nTMhelix 13 30\noutside 31 259\nTMhelix 260 282\ninside 283 324

32362 GCF_001514065.1_ASM151406v1 Streptomyces antibioticus Terrabacteria group; Actinobacteria; Actinobacteria; Streptomycetales; Streptomycetaceae; Streptomyces

MRHARRRMVRRVTRLAAVGGLLLGGTMVTRAVASEPTGPSALPRTTAAAAAQTGTELVAR WP_059191146.1 protease [Streptomyces antibioticus] Length: 459\nNumber of predicted TMHs: 2\nExp number of AAs in TMHs: 40.982\nExp number, first 60 AAs: 21.18109\nTotal prob of N-in: 0.95964\nPOSSIBLE N-term signal sequence\ninside 1 11\nTMhelix 12 34\noutside 35 420\nTMhelix 421 443\ninside 444 459

32363 GCF_001514065.1_ASM151406v1 Streptomyces antibioticus Terrabacteria group; Actinobacteria; Actinobacteria; Streptomycetales; Streptomycetaceae; Streptomyces MTENPTGPRRPAVLTRAKTLPAWSLLAAGAVAGGLLGGGYGLAKTPAYTATSYVIAVPNE WP_059191885.1 hypothetical protein [Streptomyces antibioticus] Length: 217\nNumber of predicted TMHs: 2\nExp number of AAs in TMHs: 42.66442\nExp number, first 60 AAs: 22.59281\nTotal prob of N-in: 0.93940\nPOSSIBLE N-term signal sequence\ninside 1 20\nTMhelix 21 43\noutside 44 166\nTMhelix 167 189\ninside 190 217

32364 GCF_000968685.2_ASM96868v2 Streptomyces antioxidans Terrabacteria group; Actinobacteria; Actinobacteria; Streptomycetales; Streptomycetaceae; Streptomyces MSSPDREVRAARNDTARTARRPVVAVTGAASGPGALLAQRLAESEEVKQVLAIDERRGEV WP_046090413.1 NAD-dependent dehydratase [Streptomyces antioxidans] Length: 369\nNumber of predicted TMHs: 1\nExp number of AAs in TMHs: 20.17761\nExp number, first 60 AAs: 0.63413\nTotal prob of N-in: 0.04297\noutside 1 339\nTMhelix 340 359\ninside 360 369

32365 GCF_900119365.1_IMG-taxon_2616644941_annotated_assembly Streptomyces atratus Terrabacteria group; Actinobacteria; Actinobacteria; Streptomycetales; Streptomycetaceae; Streptomyces MLCLVENRFLPRTAAFFDLDKTVIAKSSTLTFSKSFYQGGLINRRAVLRTAYAQFVFLAG WP_072487140.1 inhibition of morphological differentiation protein [Streptomyces atratus] Length: 279\nNumber of predicted TMHs: 1\nExp number of AAs in TMHs: 21.15546\nExp number, first 60 AAs: 0.01931\nTotal prob of N-in: 0.00209\noutside 1 247\nTMhelix 248 270\ninside 271 279

32366 GCF_000745675.1_ASM74567v1 Streptomyces atratus Terrabacteria group; Actinobacteria; Actinobacteria; Streptomycetales; Streptomycetaceae; Streptomyces MLCLVENHSLPRTAAFFDLDKTVIAKSSTLTFSKSFYQGGLINRRAVLRTAYAQFVFLAG WP_037690843.1 inhibition of morphological differentiation protein [Streptomyces atratus] Length: 279\nNumber of predicted TMHs: 1\nExp number of AAs in TMHs: 22.28174\nExp number, first 60 AAs: 0.02138\nTotal prob of N-in: 0.00289\noutside 1 245\nTMhelix 246 268\ninside 269 279

32367 GCF_000745675.1_ASM74567v1 Streptomyces atratus Terrabacteria group; Actinobacteria; Actinobacteria; Streptomycetales; Streptomycetaceae; Streptomyces MSPIGRRRWLHVSCGGSTILNSNTFRLAALAVAAAPVALLVAVPAHATGSTTTTGGGKAS WP_078839971.1 hypothetical protein [Streptomyces atratus] Length: 336\nNumber of predicted TMHs: 1\nExp number of AAs in TMHs: 35.22148\nExp number, first 60 AAs: 14.59425\nTotal prob of N-in: 0.65371\nPOSSIBLE N-term signal sequence\noutside 1 307\nTMhelix 308 330\ninside 331 336

32368 GCF_001418325.1_ASM141832v1 Streptomyces atriruber Terrabacteria group; Actinobacteria; Actinobacteria; Streptomycetales; Streptomycetaceae; Streptomyces MLGLVENHSLPRTAAFFDLDKTVIAKSSTLTFSKSFYQGGLINRRAVLRTAYAQFVFLAG WP_055569268.1 inhibition of morphological differentiation protein [Streptomyces atriruber] Length: 274\nNumber of predicted TMHs: 1\nExp number of AAs in TMHs: 21.74977\nExp number, first 60 AAs: 0.02371\nTotal prob of N-in: 0.00378\noutside 1 244\nTMhelix 245 267\ninside 268 274

32369 GCF_001418335.1_ASM141833v1 Streptomyces aurantiacus Terrabacteria group; Actinobacteria; Actinobacteria; Streptomycetales; Streptomycetaceae; Streptomyces MAANRRRPIALAAACATAVTLGATALAATSASAAEVPLTGYELTWGIKESYRNYVTGMAA WP_055512896.1 hypothetical protein [Streptomyces aurantiacus] Length: 501\nNumber of predicted TMHs: 2\nExp number of AAs in TMHs: 42.21002\nExp number, first 60 AAs: 20.24774\nTotal prob of N-in: 0.90253\nPOSSIBLE N-term signal sequence\ninside 1 8\nTMhelix 9 31\noutside 32 463\nTMhelix 464 486\ninside 487 501

32370 GCF_001418335.1_ASM141833v1 Streptomyces aurantiacus Terrabacteria group; Actinobacteria; Actinobacteria; Streptomycetales; Streptomycetaceae; Streptomyces MPAPAPPHRRTYVTLLCAAVLVALLPAGQANAASRTVQGGRLDWGIKSSFQSYVTGPIAN WP_055512898.1 Htaa domain protein [Streptomyces aurantiacus] Length: 501\nNumber of predicted TMHs: 1\nExp number of AAs in TMHs: 40.17084\nExp number, first 60 AAs: 16.34543\nTotal prob of N-in: 0.63817\nPOSSIBLE N-term signal sequence\noutside 1 450\nTMhelix 451 473\ninside 474 501

32371 GCF_001418335.1_ASM141833v1 Streptomyces aurantiacus Terrabacteria group; Actinobacteria; Actinobacteria; Streptomycetales; Streptomycetaceae; Streptomyces
MFVRRRAAVLAATAVIGTAFAPAALADESPSASPSPSTSPAAPATPAGLYGSTDPTYDGVWP_055508106.1 hypothetical protein [Streptomyces aurantiacus] Length: 436\nNumber of predicted TMHs: 2\nExp number of AAs in TMHs: 38.08454\nExp number, first 60 AAs: 15.83519\nTotal prob of N-in: 0.80928\nPOSSIBLE N-term signal sequence\ninside 1 6\nTMhelix 7 26\noutside 27 405\nTMhelix 406 428\ninside 429 436

32372 GCF_000725745.1_Doro.v1.0Streptomyces aureocirculatus Terrabacteria group; Actinobacteria; Actinobacteria; Streptomycetales; Streptomycetaceae; Streptomyces
MSRHARRPQRRTHLRAVAVATAVAGGVLAAPTASAFASDATPKPKPTPPTSAPAPEEVAE WP_030565856.1 hypothetical protein [Streptomyces aureocirculatus] Length: 166\nNumber of predicted TMHs: 2\nExp number of AAs in TMHs: 42.45076\nExp number, first 60 AAs: 22.28303\nTotal prob of N-in: 0.99896\nPOSSIBLE N-term signal sequence\ninside 1 12\nTMhelix 13 35\noutside 36 136\nTMhelix 137 156\ninside 157 166

32373 GCF_000725745.1_Doro.v1.0Streptomyces aureocirculatus Terrabacteria group; Actinobacteria; Actinobacteria; Streptomycetales; Streptomycetaceae; Streptomyces
MLDLVENHPLPRTAAFFDLDKTVIAKSSTLTFSKSFYQGGLINRRAVLRTAYAQFVFLAG WP_030561212.1 inhibition of morphological differentiation protein [Streptomyces aureocirculatus] Length: 274\nNumber of predicted TMHs: 1\nExp number of AAs in TMHs: 21.90269\nExp number, first 60 AAs: 0.02723\nTotal prob of N-in: 0.00377\noutside 1 244\nTMhelix 245 267\ninside 268 274

32374 GCF_000720475.1_ASM72047v1 Streptomyces aureocirculatus Terrabacteria group; Actinobacteria; Actinobacteria; Streptomycetales; Streptomycetaceae; Streptomyces
MSRHARRPQRRTHLRAVAVATAVAGGVLAAPTASAFASDATPKPKPTPPTSAPAPEEVAE WP_030565856.1 hypothetical protein [Streptomyces aureocirculatus] Length: 166\nNumber of predicted TMHs: 2\nExp number of AAs in TMHs: 42.45076\nExp number, first 60 AAs: 22.28303\nTotal prob of N-in: 0.99896\nPOSSIBLE N-term signal sequence\ninside 1 12\nTMhelix 13 35\noutside 36 136\nTMhelix 137 156\ninside 157 166

32375 GCF_000720475.1_ASM72047v1 Streptomyces aureocirculatus Terrabacteria group; Actinobacteria; Actinobacteria; Streptomycetales; Streptomycetaceae; Streptomyces
MLDLVENHPLPRTAAFFDLDKTVIAKSSTLTFSKSFYQGGLINRRAVLRTAYAQFVFLAG WP_030561212.1 inhibition of morphological differentiation protein [Streptomyces aureocirculatus] Length: 274\nNumber of predicted TMHs: 1\nExp number of AAs in TMHs: 21.90269\nExp number, first 60 AAs: 0.02723\nTotal prob of N-in: 0.00377\noutside 1 244\nTMhelix 245 267\ninside 268 274

32376 GCF_002082605.1_ASM208260v1 Streptomyces aureofaciens Terrabacteria group; Actinobacteria; Actinobacteria; Streptomycetales; Streptomycetaceae; Streptomyces
MRTAAFFDLDKTIIAKSSALAFSRSFYQGGLINRRAVLRSAYAQFVFLVGGADHDQMEKM WP_030288083.1 MULTISPECIES: inhibition of morphological differentiation protein [Streptomyces] Length: 272\nNumber of predicted TMHs: 1\nExp number of AAs in TMHs: 22.68364\nExp number, first 60 AAs: 0.10845\nTotal prob of N-in: 0.00817\noutside 1 235\nTMhelix 236 258\ninside 259 272

32377 GCF_001188955.3_ASM118895v3 Streptomyces aureofaciens Terrabacteria group; Actinobacteria; Actinobacteria; Streptomycetales; Streptomycetaceae; Streptomyces
MRTAAFFDLDKTIIAKSSALAFSRSFYQGGLINRRAVLRSAYAQFVFLVGGADHDQMEKM WP_030288083.1 MULTISPECIES: inhibition of morphological differentiation protein [Streptomyces] Length: 272\nNumber of predicted TMHs: 1\nExp number of AAs in TMHs: 22.68364\nExp number, first 60 AAs: 0.10845\nTotal prob of N-in: 0.00817\noutside 1 235\nTMhelix 236 258\ninside 259 272

32378 GCF_000719175.1_ASM71917v1 Streptomyces aureofaciens Terrabacteria group; Actinobacteria; Actinobacteria; Streptomycetales; Streptomycetaceae; Streptomyces
MRTAAFFDLDKTIIAKSSALAFSRSFYQGGLINRRAVLRSAYAQFVFLVGGADHDQMEKM WP_030288083.1 MULTISPECIES: inhibition of morphological differentiation protein [Streptomyces] Length: 272\nNumber of predicted TMHs: 1\nExp number of AAs in TMHs: 22.68364\nExp number, first 60 AAs: 0.10845\nTotal prob of N-in: 0.00817\noutside 1 235\nTMhelix 236 258\ninside 259 272

32379 GCF_001270665.1_ASM127066v1 Streptomyces aureofaciens Terrabacteria group; Actinobacteria; Actinobacteria; Streptomycetales; Streptomycetaceae; Streptomyces
MESRQPVGPRANGHRAAHRRRAAVALAGAVCAVGLTATTAAPALADTNVPLNNGQCNFGA WP_078586913.1 MULTISPECIES: type VII secretion-associated serine protease mycosin [Streptomyces] Length: 427\nNumber of predicted TMHs: 2\nExp number of AAs in TMHs: 43.8948\nExp number, first 60 AAs: 21.08952\nTotal prob of N-in:

0.94565\nPOSSIBLE N-term signal sequence\ninside 1 20\nTMhelix 21 43\noutside 44 389\nTMhelix 390 412\ninside 413 427

32380 GCF_000978515.1_ASM97851v1 Streptomyces aureofaciens Terrabacteria group; Actinobacteria; Actinobacteria; Streptomycetales; Streptomycetaceae; Streptomyces MRTAAFFDLDKTIIAKSSALAFSRSFYQGGLINRRAVLRSAYAQFVFLVGGADHDQMEKM WP_030288083.1 MULTISPECIES: inhibition of morphological differentiation protein [Streptomyces] Length: 272\nNumber of predicted TMHs: 1\nExp number of AAs in TMHs: 22.68364\nExp number, first 60 AAs: 0.10845\nTotal prob of N-in: 0.00817\noutside 1 235\nTMhelix 236 258\ninside 259 272

32381 GCF_001270665.1_ASM127066v1 Streptomyces aureofaciens Terrabacteria group; Actinobacteria; Actinobacteria; Streptomycetales; Streptomycetaceae; Streptomyces MTERSAPAASATGPGTPARRTAVRAAATGTLAAALLGLAPQSALAATPAPPPAPAISAVT WP_030180617.1 MULTISPECIES: hypothetical protein [Streptomyces] Length: 293\nNumber of predicted TMHs: 1\nExp number of AAs in TMHs: 24.94793\nExp number, first 60 AAs: 4.55383\nTotal prob of N-in: 0.22746\noutside 1 264\nTMhelix 265 287\ninside 288 293

32382 GCF_000720845.1_ASM72084v1 Streptomyces aureofaciens Terrabacteria group; Actinobacteria; Actinobacteria; Streptomycetales; Streptomycetaceae; Streptomyces MRHARRRMVRRVARLAAVGGLLLGGTMVTRAVASEPPDASAAPPTLARTASGDGAGLVSR WP_052841124.1 protease [Streptomyces aureofaciens] Length: 447\nNumber of predicted TMHs: 2\nExp number of AAs in TMHs: 42.10606\nExp number, first 60 AAs: 20.72333\nTotal prob of N-in: 0.95955\nPOSSIBLE N-term signal sequence\ninside 1 11\nTMhelix 12 34\noutside 35 408\nTMhelix 409 431\ninside 432 447

32383 GCF_000716885.1_ASM71688v1 Streptomyces aureofaciens Terrabacteria group; Actinobacteria; Actinobacteria; Streptomycetales; Streptomycetaceae; Streptomyces MTEFSKARRTAVTALLAAGLALVTAQAAHADPAAHADTSAQGAAPAQGAPEPVPAADLAR WP_033347631.1 hypothetical protein [Streptomyces aureofaciens] Length: 308\nNumber of predicted TMHs: 1\nExp number of AAs in TMHs: 28.54559\nExp number, first 60 AAs: 6.15238\nTotal prob of N-in: 0.33326\noutside 1 278\nTMhelix 279 301\ninside 302 308

32384 GCF_000716885.1_ASM71688v1 Streptomyces aureofaciens Terrabacteria group; Actinobacteria; Actinobacteria; Streptomycetales; Streptomycetaceae; Streptomyces MRTAAFFDLDKTIIAKSSALAFSRSFYQGGLINRRAVLRSAYAQFVFLVGGADHDQMEKM WP_033347125.1 inhibition of morphological differentiation protein [Streptomyces aureofaciens] Length: 272\nNumber of predicted TMHs: 1\nExp number of AAs in TMHs: 22.68211\nExp number, first 60 AAs: 0.1088\nTotal prob of N-in: 0.00869\noutside 1 235\nTMhelix 236 258\ninside 259 272

32385 GCF_000725495.1_Doro.v1.0 Streptomyces aureus Terrabacteria group; Actinobacteria; Actinobacteria; Streptomycetales; Streptomycetaceae; Streptomyces MAAHRRRPLALAAAVAVALGAGALAVTSASAAEAPLKDYELTWGIKQSYRSYVTGMAAGS WP_037623895.1 hypothetical protein [Streptomyces aureus] Length: 514\nNumber of predicted TMHs: 2\nExp number of AAs in TMHs: 43.61986\nExp number, first 60 AAs: 21.54559\nTotal prob of N-in: 0.96800\nPOSSIBLE N-term signal sequence\ninside 1 8\nTMhelix 9 31\noutside 32 484\nTMhelix 485 507\ninside 508 514

32386 GCF_000725495.1_Doro.v1.0 Streptomyces aureus Terrabacteria group; Actinobacteria; Actinobacteria; Streptomycetales; Streptomycetaceae; Streptomyces MRHARRRIVRRGARLAAVGGLLCGGFMVTRAMAAEPSDASRTTASSAQAAADTGAGLVSR WP_037617620.1 protease [Streptomyces aureus] Length: 461\nNumber of predicted TMHs: 2\nExp number of AAs in TMHs: 39.60243\nExp number, first 60 AAs: 21.46355\nTotal prob of N-in: 0.98155\nPOSSIBLE N-term signal sequence\ninside 1 11\nTMhelix 12 34\noutside 35 422\nTMhelix 423 445\ninside 446 461

32387 GCF_000725495.1_Doro.v1.0 Streptomyces aureus Terrabacteria group; Actinobacteria; Actinobacteria; Streptomycetales; Streptomycetaceae; Streptomyces MLDLVENHSLPRTAAFFDLDKTVIAKSSTLTFSKSFYQGGLISRRAALRTAYIQFVFLAG WP_037625437.1 inhibition of morphological differentiation protein [Streptomyces aureus] Length: 277\nNumber of predicted TMHs: 1\nExp number of AAs in TMHs: 21.73262\nExp number, first 60 AAs: 0.0479\nTotal prob of N-in: 0.00310\noutside 1 247\nTMhelix 248 270\ninside 271 277

32388 GCF_001418135.1_ASM141813v1 Streptomyces aureus Terrabacteria group; Actinobacteria; Actinobacteria; Streptomycetales; Streptomycetaceae; Streptomyces

MLRLVENHSLPRTAAFFDLDKTVIAKSSTLTFSKSFYRGGLISRRAALRTAYIQFVFLAG WP_055602495.1 inhibition of morphological differentiation protein [Streptomyces aureus] Length: 288\nNumber of predicted TMHs: 1\nExp number of AAs in TMHs: 22.10255\nExp number, first 60 AAs: 0.0542\nTotal prob of N-in: 0.00438\noutside 1 247\nTMhelix 248 270\ninside 271 288

32389 GCF_001418135.1_ASM141813v1 Streptomyces aureus Terrabacteria group; Actinobacteria; Actinobacteria; Streptomycetales; Streptomycetaceae; Streptomyces
MTGRRRLVGRVLGAAGLTAAVGAIVLLGTGYTTATVAGDSMDPTYPRGARVFFERIDGGE WP_055602302.1 signal peptidase I [Streptomyces aureus] Length: 217\nNumber of predicted TMHs: 2\nExp number of AAs in TMHs: 47.30141\nExp number, first 60 AAs: 23.05586\nTotal prob of N-in: 0.99139\nPOSSIBLE N-term signal sequence\ninside 1 6\nTMhelix 7 29\noutside 30 170\nTMhelix 171 193\ninside 194 217

32390 GCF_000725495.1_Doro.v1.0 Streptomyces aureus Terrabacteria group; Actinobacteria; Actinobacteria; Streptomycetales; Streptomycetaceae; Streptomyces
MRDSSGLSRRAALGLAAALPLAAATATTADATSTIGGERLARAGIQVSGASGLPKHLTAR WP_051901016.1 D-alanyl-D-alanine carboxypeptidase [Streptomyces aureus] Length: 406\nNumber of predicted TMHs: 1\nExp number of AAs in TMHs: 30.0237\nExp number, first 60 AAs: 7.83751\nTotal prob of N-in: 0.36098\noutside 1 349\nTMhelix 350 372\ninside 373 406

32391 GCF_001983975.1_ASM198397v1 Streptomyces autolyticus Terrabacteria group; Actinobacteria; Actinobacteria; Streptomycetales; Streptomycetaceae; Streptomyces
MLGVVENHSLPRTAAFFDLDKTVIAKSSALTFSKSFYQGGLINRRAVLRTAYAQFVFLAG WP_079258151.1 inhibition of morphological differentiation protein [Streptomyces autolyticus] Length: 278\nNumber of predicted TMHs: 1\nExp number of AAs in TMHs: 21.25435\nExp number, first 60 AAs: 0.02058\nTotal prob of N-in: 0.00296\noutside 1 247\nTMhelix 248 267\ninside 268 278

32392 GCF_000721255.1_ASM72125v1 Streptomyces avellaneus Terrabacteria group; Actinobacteria; Actinobacteria; Streptomycetales; Streptomycetaceae; Streptomyces
MRTAAFFDLDKTIIAKSSALAFSRSFYQGGLINRRAVLRSAYAQFVFLVGGADHDQMEKM WP_030288083.1 MULTISPECIES: inhibition of morphological differentiation protein [Streptomyces] Length: 272\nNumber of predicted TMHs: 1\nExp number of AAs in TMHs: 22.68364\nExp number, first 60 AAs: 0.10845\nTotal prob of N-in: 0.00817\noutside 1 235\nTMhelix 236 258\ninside 259 272

32393 GCF_002017835.1_ASM201783v1 Streptomyces avermitilis Terrabacteria group; Actinobacteria; Actinobacteria; Streptomycetales; Streptomycetaceae; Streptomyces
MLGPVENHSLPRTAAFFDLDKTVIAKSSTLTFSKSFYQGGLINRRAVLRTAYAQFVFLAG WP_010986028.1 morphological differentiation-associated protein [Streptomyces avermitilis] Length: 277\nNumber of predicted TMHs: 1\nExp number of AAs in TMHs: 21.0967\nExp number, first 60 AAs: 0.01343\nTotal prob of N-in: 0.00115\noutside 1 248\nTMhelix 249 268\ninside 269 277

32394 GCF_002017835.1_ASM201783v1 Streptomyces avermitilis Terrabacteria group; Actinobacteria; Actinobacteria; Streptomycetales; Streptomycetaceae; Streptomyces
MMSTPMPYVTEATLRGTFARRAALAATTTAVLAATTVLLPAAAPAAVADSTTGQCTFPSK WP_010983976.1 type VII secretion-associated serine protease mycosin [Streptomyces avermitilis] Length: 432\nNumber of predicted TMHs: 1\nExp number of AAs in TMHs: 37.55284\nExp number, first 60 AAs: 15.0823\nTotal prob of N-in: 0.66729\nPOSSIBLE N-term signal sequence\noutside 1 395\nTMhelix 396 418\ninside 419 432

32395 GCF_002017835.1_ASM201783v1 Streptomyces avermitilis Terrabacteria group; Actinobacteria; Actinobacteria; Streptomycetales; Streptomycetaceae; Streptomyces
MGRNTRRRPTGPRRATFAAVALILGGGGLLAANVYASATEGGLGSEPVRQTDSDGNAWGR WP_078234756.1 hypothetical protein [Streptomyces avermitilis] Length: 675\nNumber of predicted TMHs: 2\nExp number of AAs in TMHs: 39.46572\nExp number, first 60 AAs: 21.74729\nTotal prob of N-in: 0.99097\nPOSSIBLE N-term signal sequence\ninside 1 16\nTMhelix 17 39\noutside 40 649\nTMhelix 650 667\ninside 668 675

32396 GCF_002017835.1_ASM201783v1 Streptomyces avermitilis Terrabacteria group; Actinobacteria; Actinobacteria; Streptomycetales; Streptomycetaceae; Streptomyces
MTPPLPCLCHPAVGRHGLTLYGPRMRDSSRLSRRAVLGLAAAAPLAAAAPASASAATVIG WP_010987315.1 D-alanyl-D-alanine carboxypeptidase [Streptomyces avermitilis] Length: 446\nNumber of predicted TMHs: 1\nExp number of AAs in TMHs: 28.11838\nExp number, first 60 AAs: 6.58154\nTotal prob of N-in: 0.28816\noutside 1 373\nTMhelix 374 396\ninside 397 446

32397	GCF_002017835.1_ASM201783v1	Streptomyces avermitilis	Terrabacteria group; Actinobacteria; Actinobacteria; Streptomycetales; Streptomycetaceae; Streptomyces
MRTDRTLRSQPVNQDSSGRSISRREALRGAGAVAVAGAGAAVLGGASTAVAADAFAHPGL	WP_010988599.1	hypothetical protein [Streptomyces avermitilis]	Length: 506\nNumber of predicted TMHs: 2\nExp number of AAs in TMHs: 45.1170199999999\nExp number, first 60 AAs: 22.25086\nTotal prob of N-in: 0.97947\nPOSSIBLE N-term signal sequence\ninside 1 28\nTMhelix 29 51\noutside 52 474\nTMhelix 475 497\ninside 498 506

32398	GCF_001514135.1_ASM151413v1	Streptomyces avermitilis	Terrabacteria group; Actinobacteria; Actinobacteria; Streptomycetales; Streptomycetaceae; Streptomyces
MLGPVENHSLPRTAAFFDLDKTVIAKSSTLTFSKSFYQGGLINRRAVLRTAYAQFVFLAG	WP_010986028.1	morphological differentiation-associated protein [Streptomyces avermitilis]	Length: 277\nNumber of predicted TMHs: 1\nExp number of AAs in TMHs: 21.0967\nExp number, first 60 AAs: 0.01343\nTotal prob of N-in: 0.00115\noutside 1 248\nTMhelix 249 268\ninside 269 277

32399	GCF_001514135.1_ASM151413v1	Streptomyces avermitilis	Terrabacteria group; Actinobacteria; Actinobacteria; Streptomycetales; Streptomycetaceae; Streptomyces
MTPPLPCLCHPAVGRHGLTLYGPRMRDSSRLSRRAVLGLAAAAPLAAAAPASASAATVIG	WP_010987315.1	D-alanyl-D-alanine carboxypeptidase [Streptomyces avermitilis]	Length: 446\nNumber of predicted TMHs: 1\nExp number of AAs in TMHs: 28.11838\nExp number, first 60 AAs: 6.58154\nTotal prob of N-in: 0.28816\noutside 1 373\nTMhelix 374 396\ninside 397 446

32400	GCF_001514135.1_ASM151413v1	Streptomyces avermitilis	Terrabacteria group; Actinobacteria; Actinobacteria; Streptomycetales; Streptomycetaceae; Streptomyces
MMSTPMPYVTEATLRGTFARRAALAATTTAVLAATTVLLPAAAPAAVADSTTGQCTFPSK	WP_010983976.1	type VII secretion-associated serine protease mycosin [Streptomyces avermitilis]	Length: 432\nNumber of predicted TMHs: 1\nExp number of AAs in TMHs: 37.55284\nExp number, first 60 AAs: 15.0823\nTotal prob of N-in: 0.66729\nPOSSIBLE N-term signal sequence\noutside 1 395\nTMhelix 396 418\ninside 419 432

32401	GCF_001514135.1_ASM151413v1	Streptomyces avermitilis	Terrabacteria group; Actinobacteria; Actinobacteria; Streptomycetales; Streptomycetaceae; Streptomyces
MRTDRTLRSQPVNQDSSGRSISRREALRGAGAVAVAGAGAAVLGGASTAVAADAFAHPGL	WP_010988599.1	hypothetical protein [Streptomyces avermitilis]	Length: 506\nNumber of predicted TMHs: 2\nExp number of AAs in TMHs: 45.1170199999999\nExp number, first 60 AAs: 22.25086\nTotal prob of N-in: 0.97947\nPOSSIBLE N-term signal sequence\ninside 1 28\nTMhelix 29 51\noutside 52 474\nTMhelix 475 497\ninside 498 506

32402	GCF_000719135.1_ASM71913v1	Streptomyces avicenniae	Terrabacteria group; Actinobacteria; Actinobacteria; Streptomycetales; Streptomycetaceae; Streptomyces
MTRTTGTARRAALAAVPLLAALLLAGCTGSDGGGGGSNADSDMAVPEASADERGASEAED	WP_052851228.1	hypothetical protein [Streptomyces avicenniae]	Length: 315\nNumber of predicted TMHs: 2\nExp number of AAs in TMHs: 42.86087\nExp number, first 60 AAs: 18.913\nTotal prob of N-in: 0.88460\nPOSSIBLE N-term signal sequence\ninside 1 11\nTMhelix 12 34\noutside 35 263\nTMhelix 264 286\ninside 287 315

32403	GCF_001270025.1_Sazu_1.0	Streptomyces azureus	Terrabacteria group; Actinobacteria; Actinobacteria; Streptomycetales; Streptomycetaceae; Streptomyces
MPAPKKTARRSLLVTSAVLSSLALAAPAAVAAPSPSGSPSVSPSGSPSATPPASMSSVGG	WP_059415102.1	D-alanyl-D-alanine carboxypeptidase [Streptomyces azureus]	Length: 439\nNumber of predicted TMHs: 1\nExp number of AAs in TMHs: 38.8486199999999\nExp number, first 60 AAs: 16.15992\nTotal prob of N-in: 0.77257\nPOSSIBLE N-term signal sequence\noutside 1 400\nTMhelix 401 423\ninside 424 439

32404	GCF_001270025.1_Sazu_1.0	Streptomyces azureus	Terrabacteria group; Actinobacteria; Actinobacteria; Streptomycetales; Streptomycetaceae; Streptomyces
MLKGVENHSLPRTAAFFDLDKTVIAKSSTLTFSKSFYQGGLINRRAVLRTAYAQFVFLAG	WP_059419055.1	inhibition of morphological differentiation protein [Streptomyces azureus]	Length: 277\nNumber of predicted TMHs: 1\nExp number of AAs in TMHs: 21.60114\nExp number, first 60 AAs: 0.04042\nTotal prob of N-in: 0.00395\noutside 1 247\nTMhelix 248 270\ninside 271 277

32405	GCF_001270025.1_Sazu_1.0	Streptomyces azureus	Terrabacteria group; Actinobacteria; Actinobacteria; Streptomycetales; Streptomycetaceae; Streptomyces
MRHARRRVVRRVTRLAAVGGLLLGGAMVTRAVASEPPDPTIGVPYSARPATSMGTDLVSR	WP_059423739.1	protease [Streptomyces azureus]	Length: 452\nNumber of predicted TMHs: 2\nExp number of AAs in

TMHs: 41.02725\nExp number, first 60 AAs: 20.65724\nTotal prob of N-in: 0.95252\nPOSSIBLE N-term signal sequence\ninside 1 11\nTMhelix 12 34\noutside 35 413\nTMhelix 414 436\ninside 437 452

32406 GCF_001418165.1_ASM141816v1 Streptomyces bambergiensis Terrabacteria group; Actinobacteria; Actinobacteria; Streptomycetales; Streptomycetaceae; Streptomyces MNSTDSRASARRRAVVATATVLAAGPVALAGAGAAQATGEHGSASAAVLRTGLDVALLDR WP_055606274.1 hypothetical protein [Streptomyces bambergiensis] Length: 339\nNumber of predicted TMHs: 2\nExp number of AAs in TMHs: 42.3989099999999\nExp number, first 60 AAs: 21.44381\nTotal prob of N-in: 0.95532\nPOSSIBLE N-term signal sequence\ninside 1 12\nTMhelix 13 35\noutside 36 310\nTMhelix 311 333\ninside 334 339

32407 GCF_001418165.1_ASM141816v1 Streptomyces bambergiensis Terrabacteria group; Actinobacteria; Actinobacteria; Streptomycetales; Streptomycetaceae; Streptomyces MPARRKAVVAAVGIAPLALTGLASSPAAAHGSMGDPVSRVAQCFAEGPESPKSAACRAAV WP_055609296.1 chitin-binding protein [Streptomyces bambergiensis] Length: 365\nNumber of predicted TMHs: 2\nExp number of AAs in TMHs: 40.61378\nExp number, first 60 AAs: 21.64187\nTotal prob of N-in: 0.96324\nPOSSIBLE N-term signal sequence\ninside 1 6\nTMhelix 7 29\noutside 30 331\nTMhelix 332 354\ninside 355 365

32408 GCF_000719285.1_ASM71928v1 Streptomyces bicolor Terrabacteria group; Actinobacteria; Actinobacteria; Streptomycetales; Streptomycetaceae; Streptomyces MRHARRRLVRRVTRLAAVGGLLLGGTMVTRAVASEPPDASAVPRTYAASAADTGSDLVSS WP_031487939.1 protease [Streptomyces bicolor] Length: 454\nNumber of predicted TMHs: 2\nExp number of AAs in TMHs: 39.8857\nExp number, first 60 AAs: 19.14966\nTotal prob of N-in: 0.87506\nPOSSIBLE N-term signal sequence\ninside 1 11\nTMhelix 12 34\noutside 35 415\nTMhelix 416 438\ninside 439 454

32409 GCF_000719285.1_ASM71928v1 Streptomyces bicolor Terrabacteria group; Actinobacteria; Actinobacteria; Streptomycetales; Streptomycetaceae; Streptomyces MNPGTTNVVTTPRRTFMSRRTAPLAVLLAAGLALLPPTAAHADSIRARQWALDAMHTQEA WP_051828326.1 type VII secretion-associated serine protease mycosin [Streptomyces bicolor] Length: 403\nNumber of predicted TMHs: 2\nExp number of AAs in TMHs: 43.21414\nExp number, first 60 AAs: 19.62061\nTotal prob of N-in: 0.92026\nPOSSIBLE N-term signal sequence\ninside 1 20\nTMhelix 21 40\noutside 41 371\nTMhelix 372 394\ninside 395 403

32410 GCF_000725465.1_Doro.v1.0Streptomyces bicolor Terrabacteria group; Actinobacteria; Actinobacteria; Streptomycetales; Streptomycetaceae; Streptomyces MRHARRRLVRRVTRLAAVGGLLLGGTMVTRAVASEPPDASAVPRTYAASAADTGSDLVSS WP_031487939.1 protease [Streptomyces bicolor] Length: 454\nNumber of predicted TMHs: 2\nExp number of AAs in TMHs: 39.8857\nExp number, first 60 AAs: 19.14966\nTotal prob of N-in: 0.87506\nPOSSIBLE N-term signal sequence\ninside 1 11\nTMhelix 12 34\noutside 35 415\nTMhelix 416 438\ninside 439 454

32411 GCF_000725465.1_Doro.v1.0Streptomyces bicolor Terrabacteria group; Actinobacteria; Actinobacteria; Streptomycetales; Streptomycetaceae; Streptomyces MNPGTTNVVTTPRRTFMSRRTAPLAVLLAAGLALLPPTAAHADSIRARQWALDAMHTQEA WP_051828326.1 type VII secretion-associated serine protease mycosin [Streptomyces bicolor] Length: 403\nNumber of predicted TMHs: 2\nExp number of AAs in TMHs: 43.21414\nExp number, first 60 AAs: 19.62061\nTotal prob of N-in: 0.92026\nPOSSIBLE N-term signal sequence\ninside 1 20\nTMhelix 21 40\noutside 41 371\nTMhelix 372 394\ninside 395 403

32412 GCF_000716465.1_ASM71646v1 Streptomyces bikiniensis Terrabacteria group; Actinobacteria; Actinobacteria; Streptomycetales; Streptomycetaceae; Streptomyces MRARTPAHRLSSPSRRALVVAATTALCALGLPFSTPAVADDGSQCTYPGKTYEGRPWALQ WP_030217787.1 type VII secretion-associated serine protease [Streptomyces bikiniensis] Length: 421\nNumber of predicted TMHs: 1\nExp number of AAs in TMHs: 33.34379\nExp number, first 60 AAs: 10.74408\nTotal prob of N-in: 0.49773\nPOSSIBLE N-term signal sequence\noutside 1 385\nTMhelix 386 408\ninside 409 421

32413 GCF_000718595.1_ASM71859v1 Streptomyces bikiniensis Terrabacteria group; Actinobacteria; Actinobacteria; Streptomycetales; Streptomycetaceae; Streptomyces MRARTPAHRLSSPSRRALVVAATTALCALGLPFSTPAVADDGSQCTYPGKTYEGRPWALQ WP_030217787.1 type VII secretion-associated serine protease [Streptomyces bikiniensis] Length: 421\nNumber of predicted TMHs: 1\nExp number of AAs in TMHs: 33.34379\nExp number, first 60 AAs: 10.74408\nTotal prob of N-in: 0.49773\nPOSSIBLE N-term signal sequence\noutside 1 385\nTMhelix 386 408\ninside 409 421

32414 GCF_000716625.1_ASM71662v1 Streptomyces bikiniensis Terrabacteria group; Actinobacteria; Actinobacteria; Streptomycetales; Streptomycetaceae; Streptomyces
MRARTPAHRLSSPSRRALVVAATTALCALGLPFSTPAVADDGSQCTYPGKTYEGRPWALQ WP_030217787.1 type VII secretion-associated serine protease [Streptomyces bikiniensis] Length: 421\nNumber of predicted TMHs: 1\nExp number of AAs in TMHs: 33.34379\nExp number, first 60 AAs: 10.74408\nTotal prob of N-in: 0.49773\nPOSSIBLE N-term signal sequence\noutside 1 385\nTMhelix 386 408\ninside 409 421

32415 GCF_000716625.1_ASM71662v1 Streptomyces bikiniensis Terrabacteria group; Actinobacteria; Actinobacteria; Streptomycetales; Streptomycetaceae; Streptomyces
MGTEQQARRWSRARRRLVSVLTVSFLALGAGLTHLTAASPASADPIAKCTRTKGAIVAVD WP_030211287.1 peptidase [Streptomyces bikiniensis] Length: 939\nNumber of predicted TMHs: 2\nExp number of AAs in TMHs: 42.43235\nExp number, first 60 AAs: 21.42616\nTotal prob of N-in: 0.99165\nPOSSIBLE N-term signal sequence\ninside 1 16\nTMhelix 17 39\noutside 40 899\nTMhelix 900 922\ninside 923 939

32416 GCF_000716465.1_ASM71646v1 Streptomyces bikiniensis Terrabacteria group; Actinobacteria; Actinobacteria; Streptomycetales; Streptomycetaceae; Streptomyces
MGTEQQARRWSRARRRLVSVLTVSFLALGAGLTHLTAASPASADPIAKCTRTKGAIVAVD WP_030211287.1 peptidase [Streptomyces bikiniensis] Length: 939\nNumber of predicted TMHs: 2\nExp number of AAs in TMHs: 42.43235\nExp number, first 60 AAs: 21.42616\nTotal prob of N-in: 0.99165\nPOSSIBLE N-term signal sequence\ninside 1 16\nTMhelix 17 39\noutside 40 899\nTMhelix 900 922\ninside 923 939

32417 GCF_000716625.1_ASM71662v1 Streptomyces bikiniensis Terrabacteria group; Actinobacteria; Actinobacteria; Streptomycetales; Streptomycetaceae; Streptomyces
MLGLVENHSLPRTAAFFDLDKTVIAKSSTLTFSKSFYRGGLISRRAALRTAYIQFVFLAG WP_030205982.1 inhibition of morphological differentiation protein [Streptomyces bikiniensis] Length: 289\nNumber of predicted TMHs: 1\nExp number of AAs in TMHs: 22.11305\nExp number, first 60 AAs: 0.03412\nTotal prob of N-in: 0.00287\noutside 1 247\nTMhelix 248 270\ninside 271 289

32418 GCF_000718595.1_ASM71859v1 Streptomyces bikiniensis Terrabacteria group; Actinobacteria; Actinobacteria; Streptomycetales; Streptomycetaceae; Streptomyces
MGTEQQARRWSRARRRLVSVLTVSFLALGAGLTHLTAASPASADPIAKCTRTKGAIVAVD WP_030211287.1 peptidase [Streptomyces bikiniensis] Length: 939\nNumber of predicted TMHs: 2\nExp number of AAs in TMHs: 42.43235\nExp number, first 60 AAs: 21.42616\nTotal prob of N-in: 0.99165\nPOSSIBLE N-term signal sequence\ninside 1 16\nTMhelix 17 39\noutside 40 899\nTMhelix 900 922\ninside 923 939

32419 GCF_000718595.1_ASM71859v1 Streptomyces bikiniensis Terrabacteria group; Actinobacteria; Actinobacteria; Streptomycetales; Streptomycetaceae; Streptomyces
MLGLVENHSLPRTAAFFDLDKTVIAKSSTLTFSKSFYRGGLISRRAALRTAYIQFVFLAG WP_030205982.1 inhibition of morphological differentiation protein [Streptomyces bikiniensis] Length: 289\nNumber of predicted TMHs: 1\nExp number of AAs in TMHs: 22.11305\nExp number, first 60 AAs: 0.03412\nTotal prob of N-in: 0.00287\noutside 1 247\nTMhelix 248 270\ninside 271 289

32420 GCF_000716465.1_ASM71646v1 Streptomyces bikiniensis Terrabacteria group; Actinobacteria; Actinobacteria; Streptomycetales; Streptomycetaceae; Streptomyces
MLGLVENHSLPRTAAFFDLDKTVIAKSSTLTFSKSFYRGGLISRRAALRTAYIQFVFLAG WP_030205982.1 inhibition of morphological differentiation protein [Streptomyces bikiniensis] Length: 289\nNumber of predicted TMHs: 1\nExp number of AAs in TMHs: 22.11305\nExp number, first 60 AAs: 0.03412\nTotal prob of N-in: 0.00287\noutside 1 247\nTMhelix 248 270\ninside 271 289

32421 GCF_002154575.1_ASM215457v1 Streptomyces bobili Terrabacteria group; Actinobacteria; Actinobacteria; Streptomycetales; Streptomycetaceae; Streptomyces
MLKGVENHSLPRTAAFFDLDKTVIAKSSTLTFSKSFYQGGLINRRAALRTAYAQFVFLVG WP_086765648.1 inhibition of morphological differentiation protein [Streptomyces bobili] Length: 278\nNumber of predicted TMHs: 1\nExp number of AAs in TMHs: 21.53853\nExp number, first 60 AAs: 0.01954\nTotal prob of N-in: 0.00254\noutside 1 247\nTMhelix 248 270\ninside 271 278

32422 GCF_002154575.1_ASM215457v1 Streptomyces bobili Terrabacteria group; Actinobacteria; Actinobacteria; Streptomycetales; Streptomycetaceae; Streptomyces
MRPVSPPGVRFHDGGGVGDGTSRRLPGTFSVVPAPLKQTARRTLLVTSATLASLALTAPV WP_086765287.1 D-alanyl-D-alanine carboxypeptidase [Streptomyces bobili] Length: 484\nNumber of predicted TMHs: 1\nExp number of

AAs in TMHs: 24.93345\nExp number, first 60 AAs: 1.76955\nTotal prob of N-in: 0.10409\noutside 1 447\nTMhelix 448 470\ninside 471 484

32423 GCF_002154575.1_ASM215457v1 Streptomyces bobili Terrabacteria group; Actinobacteria; Actinobacteria; Streptomycetales; Streptomycetaceae; Streptomyces MSERERAVPRGRRRAAVAAVAVLAGGSAVLLGPAALQAQAAEVSYATRCIPPAGIGLPDV WP_086771392.1 hypothetical protein [Streptomyces bobili] Length: 443\nNumber of predicted TMHs: 2\nExp number of AAs in TMHs: 41.17668\nExp number, first 60 AAs: 20.56493\nTotal prob of N-in: 0.94233\nPOSSIBLE N-term signal sequence\ninside 1 15\nTMhelix 16 38\noutside 39 416\nTMhelix 417 436\ninside 437 443

32424 GCF_000717015.1_ASM71701v1 Streptomyces brasiliensis Terrabacteria group; Actinobacteria; Actinobacteria; Streptomycetales; Streptomycetaceae; Streptomyces MLSVVENCFSPRTAAFFDLDKTVIAKSSTLTFSKSFYQGGLINRRAVLRTAYTQFVFMAG WP_030113359.1 MULTISPECIES: inhibition of morphological differentiation protein [Streptomyces] Length: 280\nNumber of predicted TMHs: 1\nExp number of AAs in TMHs: 21.68695\nExp number, first 60 AAs: 0.01801\nTotal prob of N-in: 0.00793\noutside 1 247\nTMhelix 248 270\ninside 271 280

32425 GCF_000717015.1_ASM71701v1 Streptomyces brasiliensis Terrabacteria group; Actinobacteria; Actinobacteria; Streptomycetales; Streptomycetaceae; Streptomyces MGSSWNERRLERRVRPGDGKPLKPFRWWQLTRRSLLSLALPVGGGLSATYTVEVKHGGDA WP_030113438.1 MULTISPECIES: hypothetical protein [Streptomyces] Length: 218\nNumber of predicted TMHs: 2\nExp number of AAs in TMHs: 41.40199\nExp number, first 60 AAs: 0.36812\nTotal prob of N-in: 0.97576\ninside 1 135\nTMhelix 136 158\noutside 159 182\nTMhelix 183 200\ninside 201 218

32426 GCF_001514215.1_ASM151421v1 Streptomyces bungoensis Terrabacteria group; Actinobacteria; Actinobacteria; Streptomycetales; Streptomycetaceae; Streptomyces MLRDVENHSFPRAAAFFDLDKTVIAKSSTLTFGKSFYQGGLISRRAALRTAYAQFVFLAG WP_061927642.1 inhibition of morphological differentiation protein [Streptomyces bungoensis] Length: 277\nNumber of predicted TMHs: 1\nExp number of AAs in TMHs: 21.18261\nExp number, first 60 AAs: 0.03636\nTotal prob of N-in: 0.00551\noutside 1 247\nTMhelix 248 270\ninside 271 277

32427 GCF_001514215.1_ASM151421v1 Streptomyces bungoensis Terrabacteria group; Actinobacteria; Actinobacteria; Streptomycetales; Streptomycetaceae; Streptomyces MRHARRRVVRRVTRLAAVGGLLLGGAMVTQAAMASETPPASARALSSPGTATGTGAALVA WP_061927802.1 protease [Streptomyces bungoensis] Length: 459\nNumber of predicted TMHs: 2\nExp number of AAs in TMHs: 43.1389\nExp number, first 60 AAs: 22.2645\nTotal prob of N-in: 0.99225\nPOSSIBLE N-term signal sequence\ninside 1 11\nTMhelix 12 34\noutside 35 420\nTMhelix 421 443\ninside 444 459

32428 GCF_001187435.1_ASM118743v1 Streptomyces caatingaensis Terrabacteria group; Actinobacteria; Actinobacteria; Streptomycetales; Streptomycetaceae; Streptomyces MLGGVENHSLRRTAAFFDLDKTVIAKSSTLTFGKSFYQGGLINRRAALRTAYTQFVFLLG WP_049718970.1 inhibition of morphological differentiation protein [Streptomyces caatingaensis] Length: 285\nNumber of predicted TMHs: 1\nExp number of AAs in TMHs: 21.95901\nExp number, first 60 AAs: 0.0119\nTotal prob of N-in: 0.00089\noutside 1 246\nTMhelix 247 269\ninside 270 285

32429 GCF_001279005.1_ASM127900v1 Streptomyces caelestis Terrabacteria group; Actinobacteria; Actinobacteria; Streptomycetales; Streptomycetaceae; Streptomyces MLRDVENHSMPRTAAFFDLDKTVIAKSSALTFSKSFYQGGLINRRAVLRTAYSQFVFLAG WP_030827973.1 MULTISPECIES: inhibition of morphological differentiation protein [Streptomyces] Length: 277\nNumber of predicted TMHs: 1\nExp number of AAs in TMHs: 21.88014\nExp number, first 60 AAs: 0.04028\nTotal prob of N-in: 0.00638\noutside 1 247\nTMhelix 248 270\ninside 271 277

32430 GCF_001279005.1_ASM127900v1 Streptomyces caelestis Terrabacteria group; Actinobacteria; Actinobacteria; Streptomycetales; Streptomycetaceae; Streptomyces MRHARRRVVRRVTRLAAVGGLLLGATMVTHAVASEPRAPDAVPFATAEGTVSTGPGADLV WP_030821297.1 MULTISPECIES: protease [Streptomyces] Length: 459\nNumber of predicted TMHs: 2\nExp number of AAs in TMHs: 42.46692\nExp number, first 60 AAs: 22.12504\nTotal prob of N-in: 0.98832\nPOSSIBLE N-term signal sequence\ninside 1 11\nTMhelix 12 34\noutside 35 420\nTMhelix 421 443\ninside 444 459

32431 GCF_001514235.1_ASM151423v1 Streptomyces caeruleatus Terrabacteria group; Actinobacteria; Actinobacteria; Streptomycetales; Streptomycetaceae; Streptomyces

MRHARRRLVRRVTRLAAVGGLLLGGTMVTRAVASEPPDASSVPRTYAASAASTGGDLVAR WP_062720468.1 protease [Streptomyces caeruleatus] Length: 456\nNumber of predicted TMHs: 2\nExp number of AAs in TMHs: 39.61108\nExp number, first 60 AAs: 19.03424\nTotal prob of N-in: 0.87092\nPOSSIBLE N-term signal sequence\ninside 1 11\nTMhelix 12 34\noutside 35 417\nTMhelix 418 440\ninside 441 456

32432 GCF_001517125.1_ASM151712v1 Streptomyces canus Terrabacteria group; Actinobacteria; Actinobacteria; Streptomycetales; Streptomycetaceae; Streptomyces MRHARRRIVRRVARLAAVGGLLIGGTMVTRAVASEPPDDSAAPRTLAQHASGVGADLVSR WP_059296777.1 protease [Streptomyces canus] Length: 447\nNumber of predicted TMHs: 2\nExp number of AAs in TMHs: 41.76525\nExp number, first 60 AAs: 21.26659\nTotal prob of N-in: 0.97026\nPOSSIBLE N-term signal sequence\ninside 1 11\nTMhelix 12 34\noutside 35 408\nTMhelix 409 431\ninside 432 447

32433 GCF_001517125.1_ASM151712v1 Streptomyces canus Terrabacteria group; Actinobacteria; Actinobacteria; Streptomycetales; Streptomycetaceae; Streptomyces MGHNRRRRPTGARRATFGAVALILGGGGLVAVNVFASATESGNAAVPLGSSGVAATVDCP WP_079032704.1 hypothetical protein [Streptomyces canus] Length: 686\nNumber of predicted TMHs: 2\nExp number of AAs in TMHs: 43.24499\nExp number, first 60 AAs: 21.94699\nTotal prob of N-in: 0.99510\nPOSSIBLE N-term signal sequence\ninside 1 16\nTMhelix 17 39\noutside 40 658\nTMhelix 659 681\ninside 682 686

32434 GCF_001514145.1_ASM151414v1 Streptomyces canus Terrabacteria group; Actinobacteria; Actinobacteria; Streptomycetales; Streptomycetaceae; Streptomyces MLGVVENHSLPRTAAFFDLDKTVIAKSSTLTFSKSFYQGGLINRRAALRTAYAQFVFLVG WP_059209118.1 MULTISPECIES: inhibition of morphological differentiation protein [Streptomyces] Length: 277\nNumber of predicted TMHs: 1\nExp number of AAs in TMHs: 21.66888\nExp number, first 60 AAs: 0.01246\nTotal prob of N-in: 0.00150\noutside 1 247\nTMhelix 248 270\ninside 271 277

32435 GCF_001514145.1_ASM151414v1 Streptomyces canus Terrabacteria group; Actinobacteria; Actinobacteria; Streptomycetales; Streptomycetaceae; Streptomyces MTRRALLLSLLAGLLLATASPAHAAGYRYWSFWDRTGSTWTYATEGPSTAVPSDGDVQGF WP_059205742.1 hypothetical protein [Streptomyces canus] Length: 209\nNumber of predicted TMHs: 2\nExp number of AAs in TMHs: 41.63641\nExp number, first 60 AAs: 21.76576\nTotal prob of N-in: 0.96796\nPOSSIBLE N-term signal sequence\ninside 1 4\nTMhelix 5 27\noutside 28 181\nTMhelix 182 201\ninside 202 209

32436 GCF_001514145.1_ASM151414v1 Streptomyces canus Terrabacteria group; Actinobacteria; Actinobacteria; Streptomycetales; Streptomycetaceae; Streptomyces MRHARRRVVRRVTRLAAVGGLLLGGAMVTRAVASETPLITAVPHTYAMEAGQTGADLVSR WP_059204190.1 protease [Streptomyces canus] Length: 457\nNumber of predicted TMHs: 2\nExp number of AAs in TMHs: 39.14878\nExp number, first 60 AAs: 20.93747\nTotal prob of N-in: 0.97106\nPOSSIBLE N-term signal sequence\ninside 1 11\nTMhelix 12 34\noutside 35 418\nTMhelix 419 441\ninside 442 457

32437 GCF_000716845.1_ASM71684v1 Streptomyces capuensis Terrabacteria group; Actinobacteria; Actinobacteria; Streptomycetales; Streptomycetaceae; Streptomyces MTERSAPAASATGPGTPARRTAVRAAATGTLAAALLGLAPQSALAATPAPPPAPAISAVT WP_030180617.1 MULTISPECIES: hypothetical protein [Streptomyces] Length: 293\nNumber of predicted TMHs: 1\nExp number of AAs in TMHs: 24.94793\nExp number, first 60 AAs: 4.55383\nTotal prob of N-in: 0.22746\noutside 1 264\nTMhelix 265 287\ninside 288 293

32438 GCF_000716845.1_ASM71684v1 Streptomyces capuensis Terrabacteria group; Actinobacteria; Actinobacteria; Streptomycetales; Streptomycetaceae; Streptomyces MTRRPLLRAAGVALSAAALTVLPALPAHADGIRAQEWALKAMNTQDAWETTKGEGITVAV WP_003987438.1 MULTISPECIES: type VII secretion-associated serine protease [Streptomyces] Length: 386\nNumber of predicted TMHs: 1\nExp number of AAs in TMHs: 34.80558\nExp number, first 60 AAs: 10.27095\nTotal prob of N-in: 0.55118\nPOSSIBLE N-term signal sequence\noutside 1 359\nTMhelix 360 382\ninside 383 386

32439 GCF_000716845.1_ASM71684v1 Streptomyces capuensis Terrabacteria group; Actinobacteria; Actinobacteria; Streptomycetales; Streptomycetaceae; Streptomyces MESRQPVGPRANGHRAAHRRRAAVALAGAVCAVGLTATTAAPALADTNVPLNNGQCNFGA WP_078586913.1 MULTISPECIES: type VII secretion-associated serine protease mycosin [Streptomyces] Length: 427\nNumber of predicted TMHs: 2\nExp number of AAs in TMHs: 43.8948\nExp number, first 60 AAs: 21.08952\nTotal prob of N-in: 0.94565\nPOSSIBLE N-term signal sequence\ninside 1 20\nTMhelix 21 43\noutside 44 389\nTMhelix 390 412\ninside 413 427

32440 GCF_000718295.1_ASM71829v1 Streptomyces capuensis Terrabacteria group; Actinobacteria; Actinobacteria; Streptomycetales; Streptomycetaceae; Streptomyces
MESRQPVGPRANGHRAAHRRRAAVALAGAVCAVGLTATTAAPALADTNVPLNNGQCNFGA WP_078586913.1
MULTISPECIES: type VII secretion-associated serine protease mycosin [Streptomyces] Length: 427\nNumber of predicted TMHs: 2\nExp number of AAs in TMHs: 43.8948\nExp number, first 60 AAs: 21.08952\nTotal prob of N-in: 0.94565\nPOSSIBLE N-term signal sequence\ninside 1 20\nTMhelix 21 43\noutside 44 389\nTMhelix 390 412\ninside 413 427

32441 GCF_000718295.1_ASM71829v1 Streptomyces capuensis Terrabacteria group; Actinobacteria; Actinobacteria; Streptomycetales; Streptomycetaceae; Streptomyces
MTRRPLLRAAGVALSAAALTVLPALPAHADGIRAQEWALKAMNTQDAWETTKGEGITVAV WP_003987438.1
MULTISPECIES: type VII secretion-associated serine protease [Streptomyces] Length: 386\nNumber of predicted TMHs: 1\nExp number of AAs in TMHs: 34.80558\nExp number, first 60 AAs: 10.27095\nTotal prob of N-in: 0.55118\nPOSSIBLE N-term signal sequence\noutside 1 359\nTMhelix 360 382\ninside 383 386

32442 GCF_000718295.1_ASM71829v1 Streptomyces capuensis Terrabacteria group; Actinobacteria; Actinobacteria; Streptomycetales; Streptomycetaceae; Streptomyces
MTERSAPAASATGPGTPARRTAVRAAATGTLAAALLGLAPQSALAATPAPPPAPAISAVT WP_030180617.1
MULTISPECIES: hypothetical protein [Streptomyces] Length: 293\nNumber of predicted TMHs: 1\nExp number of AAs in TMHs: 24.94793\nExp number, first 60 AAs: 4.55383\nTotal prob of N-in: 0.22746\noutside 1 264\nTMhelix 265 287\ninside 288 293

32443 GCF_002154505.1_ASM215450v1 Streptomyces carpinensis Terrabacteria group; Actinobacteria; Actinobacteria; Streptomycetales; Streptomycetaceae; Streptomyces
MASRGRRATIRSLGVLAGAAALTVLGGNAPGWAAAPTGQHNHGSTATPIEHVVVLFDENI WP_086728543.1
phospholipase [Streptomyces carpinensis] Length: 614\nNumber of predicted TMHs: 2\nExp number of AAs in TMHs: 43.20534\nExp number, first 60 AAs: 22.01548\nTotal prob of N-in: 0.97258\nPOSSIBLE N-term signal sequence\ninside 1 12\nTMhelix 13 35\noutside 36 581\nTMhelix 582 604\ninside 605 614

32444 GCF_002154505.1_ASM215450v1 Streptomyces carpinensis Terrabacteria group; Actinobacteria; Actinobacteria; Streptomycetales; Streptomycetaceae; Streptomyces
MLKDVENHSLPRTAAFFDLDKTVIAKSSTLTFSKSFYQGGLINRRAVLRTAYAQFVFLAG WP_086727571.1 inhibition of morphological differentiation protein [Streptomyces carpinensis] Length: 277\nNumber of predicted TMHs: 1\nExp number of AAs in TMHs: 21.69758\nExp number, first 60 AAs: 0.05984\nTotal prob of N-in: 0.00614\noutside 1 247\nTMhelix 248 270\ninside 271 277

32445 GCF_002154275.1_ASM215427v1 Streptomyces castelarensis Terrabacteria group; Actinobacteria; Actinobacteria; Streptomycetales; Streptomycetaceae; Streptomyces
MLGVVENHSLPRTAAFFDLDKTVIAKSSAFTFSKSFYQGGLINRRAVLRTAYAQFVFLAG WP_086706467.1 inhibition of morphological differentiation protein [Streptomyces castelarensis] Length: 278\nNumber of predicted TMHs: 1\nExp number of AAs in TMHs: 21.26173\nExp number, first 60 AAs: 0.02696\nTotal prob of N-in: 0.00325\noutside 1 247\nTMhelix 248 267\ninside 268 278

32446 GCF_000718015.1_ASM71801v1 Streptomyces catenulae Terrabacteria group; Actinobacteria; Actinobacteria; Streptomycetales; Streptomycetaceae; Streptomyces
MSSPDPTVRAARNADAPTPQTGRGTEEPAPRPGRRPVIAVTGAATGVGALLTRALAASDE WP_030280942.1
NAD-dependent dehydratase [Streptomyces catenulae] Length: 383\nNumber of predicted TMHs: 1\nExp number of AAs in TMHs: 21.01945\nExp number, first 60 AAs: 0.78385\nTotal prob of N-in: 0.03213\noutside 1 353\nTMhelix 354 373\ninside 374 383

32447 GCF_000718015.1_ASM71801v1 Streptomyces catenulae Terrabacteria group; Actinobacteria; Actinobacteria; Streptomycetales; Streptomycetaceae; Streptomyces
MRKRRSVLRRRGGAGVAAGAALLLAGAGLVATAPTAQAADTAYRTECTPPPISGLPPVEG WP_078653899.1
hypothetical protein [Streptomyces catenulae] Length: 475\nNumber of predicted TMHs: 2\nExp number of AAs in TMHs: 44.73993\nExp number, first 60 AAs: 22.35721\nTotal prob of N-in: 0.99693\nPOSSIBLE N-term signal sequence\ninside 1 12\nTMhelix 13 35\noutside 36 444\nTMhelix 445 467\ninside 468 475

32448 GCF_001513965.1_ASM151396v1 Streptomyces cellostaticus Terrabacteria group; Actinobacteria; Actinobacteria; Streptomycetales; Streptomycetaceae; Streptomyces
MRHARRRVVRQVTRLAAVGGLLLGGAMVAQAAMASEPPAASAKPLSTAGTTGDPGAALVA WP_066994743.1
protease [Streptomyces cellostaticus] Length: 460\nNumber of predicted TMHs: 2\nExp number of AAs in

TMHs: 43.64958\nExp number, first 60 AAs: 22.14405\nTotal prob of N-in: 0.98997\nPOSSIBLE N-term signal sequence\ninside 1 11\nTMhelix 12 34\noutside 35 421\nTMhelix 422 444\ninside 445 460

32449 GCF_001513965.1_ASM151396v1 Streptomyces cellostaticus Terrabacteria group; Actinobacteria; Actinobacteria; Streptomycetales; Streptomycetaceae; Streptomyces MLRGVENHSLPRAAAFFDLDKTVIAKSSTLTFSKSFYQGGLINRRAALRTAYAQFVFLVG WP_067006145.1 inhibition of morphological differentiation protein [Streptomyces cellostaticus] Length: 277\nNumber of predicted TMHs: 1\nExp number of AAs in TMHs: 21.63795\nExp number, first 60 AAs: 0.04799\nTotal prob of N-in: 0.00701\noutside 1 247\nTMhelix 248 270\ninside 271 277

32450 GCF_001513965.1_ASM151396v1 Streptomyces cellostaticus Terrabacteria group; Actinobacteria; Actinobacteria; Streptomycetales; Streptomycetaceae; Streptomyces MASRGRRAAVRSLGALAGAAALTVLGSNAPGWAAVTSGHASTATPVKHVVVLFDENISFD WP_066994399.1 phospholipase [Streptomyces cellostaticus] Length: 605\nNumber of predicted TMHs: 2\nExp number of AAs in TMHs: 40.3053\nExp number, first 60 AAs: 20.18901\nTotal prob of N-in: 0.89012\nPOSSIBLE N-term signal sequence\ninside 1 12\nTMhelix 13 35\noutside 36 576\nTMhelix 577 599\ninside 600 605

32451 GCF_001513965.1_ASM151396v1 Streptomyces cellostaticus Terrabacteria group; Actinobacteria; Actinobacteria; Streptomycetales; Streptomycetaceae; Streptomyces MHVTSQTLSRRSLIKAAGIATTATALGTSTTPAAADTAFAHPGLLHTRTDLARMAAKVKA WP_067001745.1 Tat pathway signal sequence domain protein [Streptomyces cellostaticus] Length: 483\nNumber of predicted TMHs: 1\nExp number of AAs in TMHs: 28.79489\nExp number, first 60 AAs: 9.14217\nTotal prob of N-in: 0.40684\noutside 1 452\nTMhelix 453 472\ninside 473 483

32452 GCF_001513965.1_ASM151396v1 Streptomyces cellostaticus Terrabacteria group; Actinobacteria; Actinobacteria; Streptomycetales; Streptomycetaceae; Streptomyces MPAPKKTGRRSLLVTSATLLTLSVTAPAALAAPTPQADPTATPPAHMSQVGGVRLGQSGT WP_066996028.1 D-alanyl-D-alanine carboxypeptidase [Streptomyces cellostaticus] Length: 423\nNumber of predicted TMHs: 1\nExp number of AAs in TMHs: 36.09038\nExp number, first 60 AAs: 13.38395\nTotal prob of N-in: 0.62579\nPOSSIBLE N-term signal sequence\noutside 1 384\nTMhelix 385 407\ninside 408 423

32453 GCF_000720995.1_ASM72099v1 Streptomyces celluloflavus Terrabacteria group; Actinobacteria; Actinobacteria; Streptomycetales; Streptomycetaceae; Streptomyces MSSPDPTVRAARNTAAKTEQTDGTTDRSRRRPVVAVTGAASGVGALLTRALVASDEVKRV WP_052860708.1 NAD-dependent dehydratase [Streptomyces celluloflavus] Length: 379\nNumber of predicted TMHs: 1\nExp number of AAs in TMHs: 23.70715\nExp number, first 60 AAs: 3.64488\nTotal prob of N-in: 0.17065\noutside 1 349\nTMhelix 350 369\ninside 370 379

32454 GCF_000721105.1_ASM72110v1 Streptomyces cellulosae Terrabacteria group; Actinobacteria; Actinobacteria; Streptomycetales; Streptomycetaceae; Streptomyces MRHARRRVVRRVTRLAAVGGLLLGGTMVTRAMASETPAVPAPRTFAQQAQQAGTMGEGLV WP_030675198.1 protease [Streptomyces cellulosae] Length: 461\nNumber of predicted TMHs: 2\nExp number of AAs in TMHs: 40.79684\nExp number, first 60 AAs: 20.32731\nTotal prob of N-in: 0.95276\nPOSSIBLE N-term signal sequence\ninside 1 11\nTMhelix 12 34\noutside 35 422\nTMhelix 423 445\ninside 446 461

32455 GCF_000721105.1_ASM72110v1 Streptomyces cellulosae Terrabacteria group; Actinobacteria; Actinobacteria; Streptomycetales; Streptomycetaceae; Streptomyces MLELVENHSLPRTAAFFDLDKTVIAKSSTLTFGKSFYQGGLINRRAALRTAYAQFVFLVG WP_030668062.1 inhibition of morphological differentiation protein [Streptomyces cellulosae] Length: 277\nNumber of predicted TMHs: 1\nExp number of AAs in TMHs: 21.70055\nExp number, first 60 AAs: 0.01184\nTotal prob of N-in: 0.00190\noutside 1 247\nTMhelix 248 270\ninside 271 277

32456 GCF_000721105.1_ASM72110v1 Streptomyces cellulosae Terrabacteria group; Actinobacteria; Actinobacteria; Streptomycetales; Streptomycetaceae; Streptomyces MSISRRTAVRTARVLGVASASAALALSAAGSALACNISEFSAEAKCDGTKGVIIVTDKDA WP_030667206.1 membrane protein [Streptomyces cellulosae] Length: 238\nNumber of predicted TMHs: 2\nExp number of AAs in TMHs: 40.21389\nExp number, first 60 AAs: 18.17553\nTotal prob of N-in: 0.85697\nPOSSIBLE N-term signal sequence\ninside 1 11\nTMhelix 12 34\noutside 35 206\nTMhelix 207 229\ninside 230 238

32457 GCF_001294335.1_ASM129433v1 Streptomyces chattanoogensis Terrabacteria group; Actinobacteria; Actinobacteria; Streptomycetales; Streptomycetaceae; Streptomyces

MTARISHTRRGLVTVGSALGALTLTLSGTASAAAAEDVQSKQWYLDALYAEKIWKKTTGK WP_053925821.1 hypothetical protein [Streptomyces chattanoogensis] Length: 412\nNumber of predicted TMHs: 2\nExp number of AAs in TMHs: 43.9498\nExp number, first 60 AAs: 8.07250999999999\nTotal prob of N-in: 0.91814\ninside 1 262\nTMhelix 263 285\noutside 286 383\nTMhelix 384 406\ninside 407 412

32458 GCF_001294335.1_ASM129433v1 Streptomyces chattanoogensis Terrabacteria group; Actinobacteria; Actinobacteria; Streptomycetales; Streptomycetaceae; Streptomyces MARRHRRGVVALLGGLLLTLAAAPLQTTPAAANAPAAPTTEGTGMVMVLDSSGSMAEKDG WP_053922847.1 hypothetical protein [Streptomyces chattanoogensis] Length: 644\nNumber of predicted TMHs: 2\nExp number of AAs in TMHs: 40.76939\nExp number, first 60 AAs: 18.58987\nTotal prob of N-in: 0.90690\nPOSSIBLE N-term signal sequence\ninside 1 6\nTMhelix 7 29\noutside 30 600\nTMhelix 601 623\ninside 624 644

32459 GCF_001514045.1_ASM151404v1 Streptomyces ciscaucasicus Terrabacteria group; Actinobacteria; Actinobacteria; Streptomycetales; Streptomycetaceae; Streptomyces MLGVVENHSLPRTAAFFDLDKTVIAKSSTLTFSKSFYQGGLINRRAALRTAYAQFVFLVG WP_059209118.1 MULTISPECIES: inhibition of morphological differentiation protein [Streptomyces] Length: 277\nNumber of predicted TMHs: 1\nExp number of AAs in TMHs: 21.66888\nExp number, first 60 AAs: 0.01246\nTotal prob of N-in: 0.00150\noutside 1 247\nTMhelix 248 270\ninside 271 277

32460 GCF_001514045.1_ASM151404v1 Streptomyces ciscaucasicus Terrabacteria group; Actinobacteria; Actinobacteria; Streptomycetales; Streptomycetaceae; Streptomyces MSSPDPQVRAARNQSTNPTARRPVVAVTGAASGIGALLTERLAASEEIKQVIALDERRGE WP_062046107.1 NAD-dependent dehydratase [Streptomyces ciscaucasicus] Length: 371\nNumber of predicted TMHs: 1\nExp number of AAs in TMHs: 20.46582\nExp number, first 60 AAs: 0.45237\nTotal prob of N-in: 0.03872\noutside 1 341\nTMhelix 342 361\ninside 362 371

32461 GCF_001514045.1_ASM151404v1 Streptomyces ciscaucasicus Terrabacteria group; Actinobacteria; Actinobacteria; Streptomycetales; Streptomycetaceae; Streptomyces MTRRALLLSLLAGLLLATASPAHAAGYRYWSFWDRTGSTWTYATQGPSTAVPSDGDVQGF WP_062042357.1 hypothetical protein [Streptomyces ciscaucasicus] Length: 210\nNumber of predicted TMHs: 2\nExp number of AAs in TMHs: 42.05573\nExp number, first 60 AAs: 21.72101\nTotal prob of N-in: 0.97201\nPOSSIBLE N-term signal sequence\ninside 1 4\nTMhelix 5 27\noutside 28 182\nTMhelix 183 202\ninside 203 210

32462 GCF_001514045.1_ASM151404v1 Streptomyces ciscaucasicus Terrabacteria group; Actinobacteria; Actinobacteria; Streptomycetales; Streptomycetaceae; Streptomyces MRHARRRVVRRVTRLAAVGGLLLGGAMVTRAVASETPPITAVPHTYAMESGQTGADLVSR WP_062039979.1 protease [Streptomyces ciscaucasicus] Length: 457\nNumber of predicted TMHs: 2\nExp number of AAs in TMHs: 38.7961099999999\nExp number, first 60 AAs: 20.58524\nTotal prob of N-in: 0.95681\nPOSSIBLE N-term signal sequence\ninside 1 11\nTMhelix 12 34\noutside 35 418\nTMhelix 419 441\ninside 442 457

32463 GCF_001693675.1_ASM169367v1 Streptomyces clavuligerus Terrabacteria group; Actinobacteria; Actinobacteria; Streptomycetales; Streptomycetaceae; Streptomyces MRIRTPPTPPGEGRARRPGPRHSGPPTRRGLMRIRRTLVIAAATAVISPAALWVAPAASA WP_003961538.1 hypothetical protein [Streptomyces clavuligerus] Length: 440\nNumber of predicted TMHs: 2\nExp number of AAs in TMHs: 42.46109\nExp number, first 60 AAs: 20.42926\nTotal prob of N-in: 0.95678\nPOSSIBLE N-term signal sequence\ninside 1 37\nTMhelix 38 60\noutside 61 408\nTMhelix 409 431\ninside 432 440

32464 GCF_001514055.1_ASM151405v1 Streptomyces corchorusii Terrabacteria group; Actinobacteria; Actinobacteria; Streptomycetales; Streptomycetaceae; Streptomyces MRHARRRVVRRVTRLAAVGGLLLGGAMVAQAAMASETPSAPARPLAATGDTGAALVARLG WP_059262504.1 protease [Streptomyces corchorusii] Length: 458\nNumber of predicted TMHs: 2\nExp number of AAs in TMHs: 43.91143\nExp number, first 60 AAs: 22.34125\nTotal prob of N-in: 0.99551\nPOSSIBLE N-term signal sequence\ninside 1 11\nTMhelix 12 34\noutside 35 419\nTMhelix 420 442\ninside 443 458

32465 GCF_001514055.1_ASM151405v1 Streptomyces corchorusii Terrabacteria group; Actinobacteria; Actinobacteria; Streptomycetales; Streptomycetaceae; Streptomyces MLRDVENHSLPRAAAFFDLDKTVIAKSSTLTFSKSFYQGGLINRRAALRTAYAQFVFLAG WP_014674034.1 MULTISPECIES: morphological differentiation-associated protein [Streptomyces] Length: 276\nNumber of predicted TMHs: 1\nExp number of AAs in TMHs: 21.37944\nExp number, first 60 AAs: 0.04579\nTotal prob of N-in: 0.01060\noutside 1 247\nTMhelix 248 270\ninside 271 276

32466 GCF_001514055.1_ASM151405v1 Streptomyces corchorusii Terrabacteria group; Actinobacteria; Actinobacteria; Streptomycetales; Streptomycetaceae; Streptomyces
MRRRAVLARGAVIGLLLGLSAALATPTPAPAVPAGTAADARPGSYAFTPGARTVAGTKGT WP_059263647.1 hypothetical protein [Streptomyces corchorusii] Length: 444\nNumber of predicted TMHs: 2\nExp number of AAs in TMHs: 42.90228\nExp number, first 60 AAs: 20.35668\nTotal prob of N-in: 0.94457\nPOSSIBLE N-term signal sequence\ninside 1 4\nTMhelix 5 27\noutside 28 406\nTMhelix 407 429\ninside 430 444

32467 GCF_001513975.1_ASM151397v1 Streptomyces curacoi Terrabacteria group; Actinobacteria; Actinobacteria; Streptomycetales; Streptomycetaceae; Streptomyces
MLRVVENHSLPRTAAFFDLDKTVIAKSSTLTFSKSFYQGGLINRRAVLRTAYAQFVFLAG WP_062146253.1 inhibition of morphological differentiation protein [Streptomyces curacoi] Length: 277\nNumber of predicted TMHs: 1\nExp number of AAs in TMHs: 21.73787\nExp number, first 60 AAs: 0.05602\nTotal prob of N-in: 0.00650\noutside 1 247\nTMhelix 248 270\ninside 271 277

32468 GCF_001513975.1_ASM151397v1 Streptomyces curacoi Terrabacteria group; Actinobacteria; Actinobacteria; Streptomycetales; Streptomycetaceae; Streptomyces
MLGFLRSGSPHLTPAPARRRVLAVLLTLLALQAGSLVAPAWACGCGAMVAGGAGRVSVGR WP_062143793.1 hypothetical protein [Streptomyces curacoi] Length: 392\nNumber of predicted TMHs: 2\nExp number of AAs in TMHs: 36.88197\nExp number, first 60 AAs: 18.0377\nTotal prob of N-in: 0.77625\nPOSSIBLE N-term signal sequence\ninside 1 20\nTMhelix 21 43\noutside 44 340\nTMhelix 341 363\ninside 364 392

32469 GCF_001513975.1_ASM151397v1 Streptomyces curacoi Terrabacteria group; Actinobacteria; Actinobacteria; Streptomycetales; Streptomycetaceae; Streptomyces
MSTVGNRDENRRRPLVTRRTGLLGGLLAASIALVPPTTAHADAIRARQWALDAMRTQEAW WP_062154544.1 type VII secretion-associated serine protease mycosin [Streptomyces curacoi] Length: 402\nNumber of predicted TMHs: 2\nExp number of AAs in TMHs: 43.05532\nExp number, first 60 AAs: 19.80685\nTotal prob of N-in: 0.95778\nPOSSIBLE N-term signal sequence\ninside 1 20\nTMhelix 21 43\noutside 44 370\nTMhelix 371 393\ninside 394 402

32470 GCF_001418175.1_ASM141817v1 Streptomyces cyanoalbus Terrabacteria group; Actinobacteria; Actinobacteria; Streptomycetales; Streptomycetaceae; Streptomyces
MPARRRRSTVIAAAVATAAALGATALTGATSAAAAEVPLTGYELTWGIKESYRTYVATYA WP_055631020.1 hypothetical protein [Streptomyces cyanoalbus] Length: 529\nNumber of predicted TMHs: 2\nExp number of AAs in TMHs: 43.95837\nExp number, first 60 AAs: 21.56657\nTotal prob of N-in: 0.95274\nPOSSIBLE N-term signal sequence\ninside 1 12\nTMhelix 13 35\noutside 36 498\nTMhelix 499 521\ninside 522 529

32471 GCF_001418175.1_ASM141817v1 Streptomyces cyanoalbus Terrabacteria group; Actinobacteria; Actinobacteria; Streptomycetales; Streptomycetaceae; Streptomyces
MRHARRRVVRRVTRLAAVGGLLLGATMVTRAVASEPPAPDPVPYAAAEEDTGPGSALVAR WP_079047754.1 serine protease [Streptomyces cyanoalbus] Length: 454\nNumber of predicted TMHs: 2\nExp number of AAs in TMHs: 40.86658\nExp number, first 60 AAs: 20.38102\nTotal prob of N-in: 0.94285\nPOSSIBLE N-term signal sequence\ninside 1 11\nTMhelix 12 34\noutside 35 415\nTMhelix 416 438\ninside 439 454

32472 GCF_002154435.1_ASM215443v1 Streptomyces daghestanicus Terrabacteria group; Actinobacteria; Actinobacteria; Streptomycetales; Streptomycetaceae; Streptomyces
MPAPKKTARRSLLVTSALLSSLALAAPAAVAAPSPSGSPSASPSGSPSATPPASMSTVGG WP_086838583.1 D-alanyl-D-alanine carboxypeptidase [Streptomyces daghestanicus] Length: 439\nNumber of predicted TMHs: 2\nExp number of AAs in TMHs: 39.1078599999999\nExp number, first 60 AAs: 16.47026\nTotal prob of N-in: 0.78478\nPOSSIBLE N-term signal sequence\ninside 1 11\nTMhelix 12 34\noutside 35 400\nTMhelix 401 423\ninside 424 439

32473 GCF_002154435.1_ASM215443v1 Streptomyces daghestanicus Terrabacteria group; Actinobacteria; Actinobacteria; Streptomycetales; Streptomycetaceae; Streptomyces
MPARRTAVTVAALGALPLALTGLTATPAAAHGSMGDPVSRVSQCYAEGPESPTSAACKAA WP_086855645.1 chitin-binding protein [Streptomyces daghestanicus] Length: 343\nNumber of predicted TMHs: 2\nExp number of AAs in TMHs: 41.08624\nExp number, first 60 AAs: 20.95487\nTotal prob of N-in: 0.94439\nPOSSIBLE N-term signal sequence\ninside 1 6\nTMhelix 7 29\noutside 30 308\nTMhelix 309 331\ninside 332 343

32474 GCF_002154435.1_ASM215443v1 Streptomyces daghestanicus Terrabacteria group; Actinobacteria; Actinobacteria; Streptomycetales; Streptomycetaceae; Streptomyces
MLKGVENHSLPRTAAFFDLDKTVIAKSSTLTFSKSFYQGGLINRRAVLRTAYAQFVFLAG WP_086848378.1 inhibition of morphological differentiation protein [Streptomyces daghestanicus] Length: 277\nNumber of predicted TMHs: 1\nExp

number of AAs in TMHs: 21.59438\nExp number, first 60 AAs: 0.04109\nTotal prob of N-in: 0.00443\noutside 1 247\nTMhelix 248 270\ninside 271 277

32475 GCF_002154435.1_ASM215443v1 Streptomyces daghestanicus Terrabacteria group; Actinobacteria; Actinobacteria; Streptomycetales; Streptomycetaceae; Streptomyces MRHARRRVVRRVTRLAAVGGLLLGGAMVTRAVASEPPDPTIGVPYSAQPATSMGTDLVSR WP_086855524.1 serine protease [Streptomyces daghestanicus] Length: 449\nNumber of predicted TMHs: 2\nExp number of AAs in TMHs: 41.02523\nExp number, first 60 AAs: 20.65468\nTotal prob of N-in: 0.95252\nPOSSIBLE N-term signal sequence\ninside 1 11\nTMhelix 12 34\noutside 35 410\nTMhelix 411 433\ninside 434 449

32476 GCF_001270575.1_ASM127057v1 Streptomyces decoyicus Terrabacteria group; Actinobacteria; Actinobacteria; Streptomycetales; Streptomycetaceae; Streptomyces MRQRRRAAVASAAALVLAGAGIVAAAGTAQAAEVSYKTECLPPPISGLPPIQGTTKVAVS WP_053209708.1 hypothetical protein [Streptomyces decoyicus] Length: 476\nNumber of predicted TMHs: 2\nExp number of AAs in TMHs: 45.04223\nExp number, first 60 AAs: 22.36683\nTotal prob of N-in: 0.99805\nPOSSIBLE N-term signal sequence\ninside 1 6\nTMhelix 7 29\noutside 30 445\nTMhelix 446 468\ninside 469 476

32477 GCF_001418405.1_ASM141840v1 Streptomyces diastatochromogenes Terrabacteria group; Actinobacteria; Actinobacteria; Streptomycetales; Streptomycetaceae; Streptomyces MVTVRPAHPPAERPRQGEDALQMPDRRRRAALLRPIPATTGWGLAPVTSPGAVLLPRERT WP_079073274.1 hypothetical protein [Streptomyces diastatochromogenes] Length: 332\nNumber of predicted TMHs: 2\nExp number of AAs in TMHs: 48.47014\nExp number, first 60 AAs: 0.91583\nTotal prob of N-in: 0.58613\ninside 1 188\nTMhelix 189 211\noutside 212 266\nTMhelix 267 289\ninside 290 332

32478 GCF_001418405.1_ASM141840v1 Streptomyces diastatochromogenes Terrabacteria group; Actinobacteria; Actinobacteria; Streptomycetales; Streptomycetaceae; Streptomyces MTLMRPTGPPRPTGRRRRATLAATAALVLAGAGVVTAAGTAQAADVAYKTECQPPPISGL WP_038519829.1 MULTISPECIES: hypothetical protein [Streptomyces] Length: 479\nNumber of predicted TMHs: 2\nExp number of AAs in TMHs: 43.80161\nExp number, first 60 AAs: 21.18149\nTotal prob of N-in: 0.97217\nPOSSIBLE N-term signal sequence\ninside 1 20\nTMhelix 21 43\noutside 44 450\nTMhelix 451 473\ninside 474 479

32479 GCF_001550305.1_ASM155030v1 Streptomyces diastatochromogenes Terrabacteria group; Actinobacteria; Actinobacteria; Streptomycetales; Streptomycetaceae; Streptomyces MLGLVENHSLPRTAAFFDLDKTVIAKSSTLTFSKSFYQGGLINRRAVLRTAYAQFVFMAG WP_054237412.1 inhibition of morphological differentiation protein [Streptomyces diastatochromogenes] Length: 277\nNumber of predicted TMHs: 1\nExp number of AAs in TMHs: 21.57763\nExp number, first 60 AAs: 0.0148\nTotal prob of N-in: 0.00176\noutside 1 247\nTMhelix 248 270\ninside 271 277

32480 GCF_001550305.1_ASM155030v1 Streptomyces diastatochromogenes Terrabacteria group; Actinobacteria; Actinobacteria; Streptomycetales; Streptomycetaceae; Streptomyces MTAMSRSRRRVVAAVSALLAALVLALLPLPSAATSAAAADGAGEDSAVTVTGKQGRYDDF WP_060897625.1 hypothetical protein [Streptomyces diastatochromogenes] Length: 811\nNumber of predicted TMHs: 2\nExp number of AAs in TMHs: 44.40947\nExp number, first 60 AAs: 21.56022\nTotal prob of N-in: 0.99281\nPOSSIBLE N-term signal sequence\ninside 1 11\nTMhelix 12 34\noutside 35 782\nTMhelix 783 805\ninside 806 811

32481 GCF_001550305.1_ASM155030v1 Streptomyces diastatochromogenes Terrabacteria group; Actinobacteria; Actinobacteria; Streptomycetales; Streptomycetaceae; Streptomyces MRHARRRIVRRGTRLAAVGGVLCGGLMVTHAMASEPSAASRAPQSATLAAAGKGAGLVST WP_054236526.1 protease [Streptomyces diastatochromogenes] Length: 460\nNumber of predicted TMHs: 2\nExp number of AAs in TMHs: 42.1892999999999\nExp number, first 60 AAs: 20.80533\nTotal prob of N-in: 0.96509\nPOSSIBLE N-term signal sequence\ninside 1 11\nTMhelix 12 34\noutside 35 421\nTMhelix 422 444\ninside 445 460

32482 GCF_000725475.1_Doro.v1.0Streptomyces durhamensis Terrabacteria group; Actinobacteria; Actinobacteria; Streptomycetales; Streptomycetaceae; Streptomyces MRHARRRIVRRVTRLAAVGGLLLGGAMVAQAAMASETPPAARPFASATGGADTGAALVAR WP_031170051.1 protease [Streptomyces durhamensis] Length: 460\nNumber of predicted TMHs: 2\nExp number of AAs in TMHs: 43.52477\nExp number, first 60 AAs: 22.60958\nTotal prob of N-in: 0.99750\nPOSSIBLE N-term signal sequence\ninside 1 11\nTMhelix 12 34\noutside 35 421\nTMhelix 422 444\ninside 445 460

32483 GCF_000725475.1_Doro.v1.0Streptomyces durhamensis Terrabacteria group; Actinobacteria; Actinobacteria; Streptomycetales; Streptomycetaceae; Streptomyces

MASRGRRTAMRSIGALAGAAALTMLGGNTPGWAAAPSGAQSGHPSTATPIKHVVVLFDEN WP_031168597.1 phospholipase C [Streptomyces durhamensis] Length: 611\nNumber of predicted TMHs: 2\nExp number of AAs in TMHs: 41.23741\nExp number, first 60 AAs: 20.65073\nTotal prob of N-in: 0.91300\nPOSSIBLE N-term signal sequence\ninside 1 12\nTMhelix 13 35\noutside 36 580\nTMhelix 581 603\ninside 604 611

32484 GCF_000725475.1_Doro.v1.0Streptomyces durhamensis Terrabacteria group; Actinobacteria; Actinobacteria; Streptomycetales; Streptomycetaceae; Streptomyces MLRGVENHSLPRAAAFFDLDKTVIAKSSTLTFSKSFYQGGLINRRAALRTAYAQFVFLAG WP_031164946.1 inhibition of morphological differentiation protein [Streptomyces durhamensis] Length: 276\nNumber of predicted TMHs: 1\nExp number of AAs in TMHs: 21.75907\nExp number, first 60 AAs: 0.02927\nTotal prob of N-in: 0.00464\noutside 1 247\nTMhelix 248 270\ninside 271 276

32485 GCF_001418395.1_ASM141839v1 Streptomyces emeiensis Terrabacteria group; Actinobacteria; Actinobacteria; Streptomycetales; Streptomycetaceae; Streptomyces MPRTAAFFDLDKTVIAKSSTLTFSKSFYQGGLINRRAVLRTAYAQFVFLAGGADHDQMER WP_074995091.1 inhibition of morphological differentiation protein [Streptomyces emeiensis] Length: 268\nNumber of predicted TMHs: 1\nExp number of AAs in TMHs: 21.15656\nExp number, first 60 AAs: 0.05988\nTotal prob of N-in: 0.00562\noutside 1 238\nTMhelix 239 261\ninside 262 268

32486 GCF_001418395.1_ASM141839v1 Streptomyces emeiensis Terrabacteria group; Actinobacteria; Actinobacteria; Streptomycetales; Streptomycetaceae; Streptomyces MNSTNSRVLARRRTVVAAAAVLAAGSAALAGAGTAQATGEPGSASAAVLRTGLDVALLDE WP_055573436.1 hypothetical protein [Streptomyces emeiensis] Length: 331\nNumber of predicted TMHs: 2\nExp number of AAs in TMHs: 42.56422\nExp number, first 60 AAs: 20.58738\nTotal prob of N-in: 0.92706\nPOSSIBLE N-term signal sequence\ninside 1 12\nTMhelix 13 35\noutside 36 302\nTMhelix 303 325\ninside 326 331

32487 GCF_900101585.1_IMG-taxon_2663762722_annotated_assembly Streptomyces emeiensis Terrabacteria group; Actinobacteria; Actinobacteria; Streptomycetales; Streptomycetaceae; Streptomyces MPRTAAFFDLDKTVIAKSSTLTFSKSFYQGGLINRRAVLRTAYAQFVFLAGGADHDQMER WP_074995091.1 inhibition of morphological differentiation protein [Streptomyces emeiensis] Length: 268\nNumber of predicted TMHs: 1\nExp number of AAs in TMHs: 21.15656\nExp number, first 60 AAs: 0.05988\nTotal prob of N-in: 0.00562\noutside 1 238\nTMhelix 239 261\ninside 262 268

32488 GCF_900101585.1_IMG-taxon_2663762722_annotated_assembly Streptomyces emeiensis Terrabacteria group; Actinobacteria; Actinobacteria; Streptomycetales; Streptomycetaceae; Streptomyces MRHARRRVVRRVTRLAAVGGLLLGATMVTRAVAGEPPAPDPVPYAGAQEDTGPGSALVAR WP_055573085.1 MULTISPECIES: protease [Streptomyces] Length: 453\nNumber of predicted TMHs: 2\nExp number of AAs in TMHs: 41.2757\nExp number, first 60 AAs: 21.21396\nTotal prob of N-in: 0.96361\nPOSSIBLE N-term signal sequence\ninside 1 11\nTMhelix 12 34\noutside 35 414\nTMhelix 415 437\ninside 438 453

32489 GCF_900101585.1_IMG-taxon_2663762722_annotated_assembly Streptomyces emeiensis Terrabacteria group; Actinobacteria; Actinobacteria; Streptomycetales; Streptomycetaceae; Streptomyces MNSTNSRVLARRRTVVAAAAVLAAGSAALAGAGTAQATGEPGSASAAVLRTGLDVALLDE WP_055573436.1 hypothetical protein [Streptomyces emeiensis] Length: 331\nNumber of predicted TMHs: 2\nExp number of AAs in TMHs: 42.56422\nExp number, first 60 AAs: 20.58738\nTotal prob of N-in: 0.92706\nPOSSIBLE N-term signal sequence\ninside 1 12\nTMhelix 13 35\noutside 36 302\nTMhelix 303 325\ninside 326 331

32490 GCF_001418395.1_ASM141839v1 Streptomyces emeiensis Terrabacteria group; Actinobacteria; Actinobacteria; Streptomycetales; Streptomycetaceae; Streptomyces MRHARRRVVRRVTRLAAVGGLLLGATMVTRAVAGEPPAPDPVPYAGAQEDTGPGSALVAR WP_055573085.1 MULTISPECIES: protease [Streptomyces] Length: 453\nNumber of predicted TMHs: 2\nExp number of AAs in TMHs: 41.2757\nExp number, first 60 AAs: 21.21396\nTotal prob of N-in: 0.96361\nPOSSIBLE N-term signal sequence\ninside 1 11\nTMhelix 12 34\noutside 35 414\nTMhelix 415 437\ninside 438 453

32491 GCF_001553475.1_ASM155347v1 Streptomyces endus Terrabacteria group; Actinobacteria; Actinobacteria; Streptomycetales; Streptomycetaceae; Streptomyces MLSVVENHSLPRTAAFFDLDKTVIAKSSALTFSKSFYQGGLINRRAVLRTAYAQFVFLAG WP_067076293.1 inhibition of morphological differentiation protein [Streptomyces endus] Length: 281\nNumber of predicted TMHs: 1\nExp number of AAs in TMHs: 21.26898\nExp number, first 60 AAs: 0.02452\nTotal prob of N-in: 0.00328\noutside 1 246\nTMhelix 247 269\ninside 270 281

32492 GCF_000725555.1_Doro.v1.0 Streptomyces erythrochromogenes Terrabacteria group; Actinobacteria; Actinobacteria; Streptomycetales; Streptomycetaceae; Streptomyces MPARRRTVTPIRIAAAGLAPLALAAYAAVPAAAHGSMTDPVSRVAACYAEGPESPKSAAC WP_031154014.1 chitin-binding protein [Streptomyces erythrochromogenes] Length: 360\nNumber of predicted TMHs: 2\nExp number of AAs in TMHs: 43.21648\nExp number, first 60 AAs: 21.4915\nTotal prob of N-in: 0.97649\nPOSSIBLE N-term signal sequence\ninside 1 12\nTMhelix 13 35\noutside 36 325\nTMhelix 326 348\ninside 349 360

32493 GCF_000725555.1_Doro.v1.0 Streptomyces erythrochromogenes Terrabacteria group; Actinobacteria; Actinobacteria; Streptomycetales; Streptomycetaceae; Streptomyces MSTVRRRTLVRAAAVTACAGSLLALPAAAALAEGVPAASSAHASAPQRTLVKSLSLADGV WP_031148216.1 hypothetical protein [Streptomyces erythrochromogenes] Length: 210\nNumber of predicted TMHs: 2\nExp number of AAs in TMHs: 43.71999\nExp number, first 60 AAs: 21.22463\nTotal prob of N-in: 0.97855\nPOSSIBLE N-term signal sequence\ninside 1 8\nTMhelix 9 31\noutside 32 180\nTMhelix 181 203\ninside 204 210

32494 GCF_000725555.1_Doro.v1.0 Streptomyces erythrochromogenes Terrabacteria group; Actinobacteria; Actinobacteria; Streptomycetales; Streptomycetaceae; Streptomyces MALDVIRSGPPDKRPRRNLFKRGVLVLLPAALLAAATLVPNLAHALEAPVNLGTATNYAV WP_031153501.1 hypothetical protein [Streptomyces erythrochromogenes] Length: 530\nNumber of predicted TMHs: 2\nExp number of AAs in TMHs: 43.1695999999999\nExp number, first 60 AAs: 20.83129\nTotal prob of N-in: 0.94544\nPOSSIBLE N-term signal sequence\ninside 1 22\nTMhelix 23 45\noutside 46 495\nTMhelix 496 518\ninside 519 530

32495 GCF_001550325.1_ASM155032v1 Streptomyces europaeiscabiei Terrabacteria group; Actinobacteria; Actinobacteria; Streptomycetales; Streptomycetaceae; Streptomyces MLGPVENHSLPRTAAFFDLDKTVIAKSSTLTFSKSFYQGGLINRRAALRTAYTQFVFLAG WP_045563048.1 MULTISPECIES: inhibition of morphological differentiation protein [Streptomyces] Length: 274\nNumber of predicted TMHs: 1\nExp number of AAs in TMHs: 21.74861\nExp number, first 60 AAs: 0.04228\nTotal prob of N-in: 0.01050\noutside 1 244\nTMhelix 245 267\ninside 268 274

32496 GCF_001550325.1_ASM155032v1 Streptomyces europaeiscabiei Terrabacteria group; Actinobacteria; Actinobacteria; Streptomycetales; Streptomycetaceae; Streptomyces MTAEISRRSTLKFAGIAAGTAGPGIGTGLAAAPAFAAGGGLAHPGLLHTGADLARMAAKV WP_046705711.1 Tat pathway signal sequence domain protein [Streptomyces europaeiscabiei] Length: 495\nNumber of predicted TMHs: 2\nExp number of AAs in TMHs: 40.19305\nExp number, first 60 AAs: 22.08887\nTotal prob of N-in: 0.96313\nPOSSIBLE N-term signal sequence\ninside 1 12\nTMhelix 13 35\noutside 36 463\nTMhelix 464 483\ninside 484 495

32497 GCF_001189025.1_ASM118902v1 Streptomyces europaeiscabiei Terrabacteria group; Actinobacteria; Actinobacteria; Streptomycetales; Streptomycetaceae; Streptomyces MLSVVENCFSPRTAAFFDLDKTVIAKSSTLTFSKSFYQGGLINRRAVLRTAYAQFVFLAG WP_050362505.1 inhibition of morphological differentiation protein [Streptomyces europaeiscabiei] Length: 281\nNumber of predicted TMHs: 1\nExp number of AAs in TMHs: 21.45203\nExp number, first 60 AAs: 0.03283\nTotal prob of N-in: 0.00694\noutside 1 247\nTMhelix 248 270\ninside 271 281

32498 GCF_000988945.1_ASM98894v1 Streptomyces europaeiscabiei Terrabacteria group; Actinobacteria; Actinobacteria; Streptomycetales; Streptomycetaceae; Streptomyces MLGPVENHSLPRTAAFFDLDKTVIAKSSTLTFSKSFYQGGLINRRAALRTAYTQFVFLAG WP_045563048.1 MULTISPECIES: inhibition of morphological differentiation protein [Streptomyces] Length: 274\nNumber of predicted TMHs: 1\nExp number of AAs in TMHs: 21.74861\nExp number, first 60 AAs: 0.04228\nTotal prob of N-in: 0.01050\noutside 1 244\nTMhelix 245 267\ninside 268 274

32499 GCF_001550375.1_ASM155037v1 Streptomyces europaeiscabiei Terrabacteria group; Actinobacteria; Actinobacteria; Streptomycetales; Streptomycetaceae; Streptomyces MLGPVENHSLPRTAAFFDLDKTVIAKSSTLTFSKSFYQGGLINRRAALRTAYTQFVFLAG WP_045563048.1 MULTISPECIES: inhibition of morphological differentiation protein [Streptomyces] Length: 274\nNumber of predicted TMHs: 1\nExp number of AAs in TMHs: 21.74861\nExp number, first 60 AAs: 0.04228\nTotal prob of N-in: 0.01050\noutside 1 244\nTMhelix 245 267\ninside 268 274

32500 GCF_000988945.1_ASM98894v1 Streptomyces europaeiscabiei Terrabacteria group; Actinobacteria; Actinobacteria; Streptomycetales; Streptomycetaceae; Streptomyces MTAEISRRSTLKFAGIAAGTAGPGIGTGLAAAPAFAAGGGLAHPGLLHTGADLARMAAKV WP_046705711.1 Tat pathway signal sequence domain protein [Streptomyces europaeiscabiei] Length: 495\nNumber of predicted TMHs:

2\nExp number of AAs in TMHs: 40.19305\nExp number, first 60 AAs: 22.08887\nTotal prob of N-in: 0.96313\nPOSSIBLE N-term signal sequence\ninside 1 12\nTMhelix 13 35\noutside 36 463\nTMhelix 464 483\ninside 484 495

32501 GCF_000718175.1_ASM71817v1 Streptomyces exfoliatus Terrabacteria group; Actinobacteria; Actinobacteria; Streptomycetales; Streptomycetaceae; Streptomyces MLSLVENHSLPRTAAFFDLDKTVIAKSSTLTFSKSFYRGGLISRRAALRTAYIQFVFLAG WP_030554423.1 inhibition of morphological differentiation protein [Streptomyces exfoliatus] Length: 288\nNumber of predicted TMHs: 1\nExp number of AAs in TMHs: 21.7579\nExp number, first 60 AAs: 0.0305\nTotal prob of N-in: 0.00210\noutside 1 247\nTMhelix 248 270\ninside 271 288

32502 GCF_000725545.1_Doro.v1.0Streptomyces exfoliatus Terrabacteria group; Actinobacteria; Actinobacteria; Streptomycetales; Streptomycetaceae; Streptomyces MTSRPTSRRTAVAVTAVLAFSGLASVPASAHGSMTDPVSRVSACYAEGPESPRSAACKAA WP_030551490.1 chitin-binding protein [Streptomyces exfoliatus] Length: 338\nNumber of predicted TMHs: 1\nExp number of AAs in TMHs: 37.40431\nExp number, first 60 AAs: 15.07628\nTotal prob of N-in: 0.72301\nPOSSIBLE N-term signal sequence\noutside 1 308\nTMhelix 309 331\ninside 332 338

32503 GCF_000718175.1_ASM71817v1 Streptomyces exfoliatus Terrabacteria group; Actinobacteria; Actinobacteria; Streptomycetales; Streptomycetaceae; Streptomyces MTSRPTSRRTAVAVTAVLAFSGLASVPASAHGSMTDPVSRVSACYAEGPESPRSAACKAA WP_030551490.1 chitin-binding protein [Streptomyces exfoliatus] Length: 338\nNumber of predicted TMHs: 1\nExp number of AAs in TMHs: 37.40431\nExp number, first 60 AAs: 15.07628\nTotal prob of N-in: 0.72301\nPOSSIBLE N-term signal sequence\noutside 1 308\nTMhelix 309 331\ninside 332 338

32504 GCF_000725545.1_Doro.v1.0Streptomyces exfoliatus Terrabacteria group; Actinobacteria; Actinobacteria; Streptomycetales; Streptomycetaceae; Streptomyces MLSLVENHSLPRTAAFFDLDKTVIAKSSTLTFSKSFYRGGLISRRAALRTAYIQFVFLAG WP_030554423.1 inhibition of morphological differentiation protein [Streptomyces exfoliatus] Length: 288\nNumber of predicted TMHs: 1\nExp number of AAs in TMHs: 21.7579\nExp number, first 60 AAs: 0.0305\nTotal prob of N-in: 0.00210\noutside 1 247\nTMhelix 248 270\ninside 271 288

32505 GCF_000716535.1_ASM71653v1 Streptomyces flaveolus Terrabacteria group; Actinobacteria; Actinobacteria; Streptomycetales; Streptomycetaceae; Streptomyces MRTPHTARRRRALAGVLLAACLALAGCGGADDMGGSSKAASDSKAEAAGAPGGSARNGRA WP_030645059.1 MULTISPECIES: hypothetical protein [Streptomyces] Length: 324\nNumber of predicted TMHs: 2\nExp number of AAs in TMHs: 42.21572\nExp number, first 60 AAs: 19.28007\nTotal prob of N-in: 0.97332\nPOSSIBLE N-term signal sequence\ninside 1 12\nTMhelix 13 30\noutside 31 259\nTMhelix 260 282\ninside 283 324

32506 GCF_000716535.1_ASM71653v1 Streptomyces flaveolus Terrabacteria group; Actinobacteria; Actinobacteria; Streptomycetales; Streptomycetaceae; Streptomyces MLRGVENHSLPRAAAFFDLDKTVIAKSSTLTFSKSFYQGGLINRRAALRTAYAQFVFLVG WP_030648527.1 MULTISPECIES: inhibition of morphological differentiation protein [Streptomyces] Length: 276\nNumber of predicted TMHs: 1\nExp number of AAs in TMHs: 21.19627\nExp number, first 60 AAs: 0.04374\nTotal prob of N-in: 0.00486\noutside 1 248\nTMhelix 249 268\ninside 269 276

32507 GCF_000716535.1_ASM71653v1 Streptomyces flaveolus Terrabacteria group; Actinobacteria; Actinobacteria; Streptomycetales; Streptomycetaceae; Streptomyces MRHARRRVVRRVTRLAAVGGLLLGGAMVTQAAMASETPPASARPLAATGDTGAALVARLG WP_030640851.1 MULTISPECIES: protease [Streptomyces] Length: 458\nNumber of predicted TMHs: 2\nExp number of AAs in TMHs: 43.16453\nExp number, first 60 AAs: 22.25986\nTotal prob of N-in: 0.99239\nPOSSIBLE N-term signal sequence\ninside 1 11\nTMhelix 12 34\noutside 35 419\nTMhelix 420 442\ninside 443 458

32508 GCF_000720755.1_ASM72075v1 Streptomyces flaveus Terrabacteria group; Actinobacteria; Actinobacteria; Streptomycetales; Streptomycetaceae; Streptomyces MRHARRRVVRRVTRLAAVGGLLLGGAMVTQAAMASETPPASARPLAATGDTGAALVARLG WP_030640851.1 MULTISPECIES: protease [Streptomyces] Length: 458\nNumber of predicted TMHs: 2\nExp number of AAs in TMHs: 43.16453\nExp number, first 60 AAs: 22.25986\nTotal prob of N-in: 0.99239\nPOSSIBLE N-term signal sequence\ninside 1 11\nTMhelix 12 34\noutside 35 419\nTMhelix 420 442\ninside 443 458

32509 GCF_000720755.1_ASM72075v1 Streptomyces flaveus Terrabacteria group; Actinobacteria; Actinobacteria; Streptomycetales; Streptomycetaceae; Streptomyces

MRTPHTARRRRALAGVLLAACLALAGCGGADDMGGSSKAASDSKAEAAGAPGGSARNGRA WP_030645059.1 MULTISPECIES: hypothetical protein [Streptomyces] Length: 324\nNumber of predicted TMHs: 2\nExp number of AAs in TMHs: 42.21572\nExp number, first 60 AAs: 19.28007\nTotal prob of N-in: 0.97332\nPOSSIBLE N-term signal sequence\ninside 1 12\nTMhelix 13 30\noutside 31 259\nTMhelix 260 282\ninside 283 324

32510 GCF_000720755.1_ASM72075v1 Streptomyces flaveus Terrabacteria group; Actinobacteria; Actinobacteria; Streptomycetales; Streptomycetaceae; Streptomyces MLRGVENHSLPRAAAFFDLDKTVIAKSSTLTFSKSFYQGGLINRRAALRTAYAQFVFLVG WP_030648527.1 MULTISPECIES: inhibition of morphological differentiation protein [Streptomyces] Length: 276\nNumber of predicted TMHs: 1\nExp number of AAs in TMHs: 21.19627\nExp number, first 60 AAs: 0.04374\nTotal prob of N-in: 0.00486\noutside 1 248\nTMhelix 249 268\ninside 269 276

32511 GCF_000717595.1_ASM71759v1 Streptomyces flavochromogenes Terrabacteria group; Actinobacteria; Actinobacteria; Streptomycetales; Streptomycetaceae; Streptomyces MLSLVENHSLPRTAAFFDLDKTVIAKSSTLTFSKSFYRGGLISRRAALRTAYIQFVFLAG WP_030323443.1 inhibition of morphological differentiation protein [Streptomyces flavochromogenes] Length: 288\nNumber of predicted TMHs: 1\nExp number of AAs in TMHs: 22.10428\nExp number, first 60 AAs: 0.03256\nTotal prob of N-in: 0.00239\noutside 1 247\nTMhelix 248 270\ninside 271 288

32512 GCF_000725785.1_Doro.v1.0Streptomyces flavovariabilis Terrabacteria group; Actinobacteria; Actinobacteria; Streptomycetales; Streptomycetaceae; Streptomyces MRHARRRVVRRVTRLAAVGGLLLGGAMVTNAVASEPPAPSIGVPFTAPPATGEGTDLVSR WP_031133743.1 MULTISPECIES: protease [Streptomyces] Length: 448\nNumber of predicted TMHs: 2\nExp number of AAs in TMHs: 41.67951\nExp number, first 60 AAs: 21.32501\nTotal prob of N-in: 0.96855\nPOSSIBLE N-term signal sequence\ninside 1 11\nTMhelix 12 34\noutside 35 409\nTMhelix 410 432\ninside 433 448

32513 GCF_000725785.1_Doro.v1.0Streptomyces flavovariabilis Terrabacteria group; Actinobacteria; Actinobacteria; Streptomycetales; Streptomycetaceae; Streptomyces MLWGVENHSLPRTAAFFDLDKTVIAKSSTLTFSKSFYQGGLINRRAVLRTAYAQFVFLAG WP_031140406.1 MULTISPECIES: inhibition of morphological differentiation protein [Streptomyces] Length: 277\nNumber of predicted TMHs: 1\nExp number of AAs in TMHs: 21.58162\nExp number, first 60 AAs: 0.0297\nTotal prob of N-in: 0.00292\noutside 1 247\nTMhelix 248 270\ninside 271 277

32514 GCF_001190165.1_ASM119016v1 Streptomyces fradiae Terrabacteria group; Actinobacteria; Actinobacteria; Streptomycetales; Streptomycetaceae; Streptomyces MENPSLPRTAAFFDLDKTVIAKSSALAFSKSFYQGGLINRRAVLRTAYAQFVYLLGGADH WP_078649556.1 inhibition of morphological differentiation protein [Streptomyces fradiae] Length: 284\nNumber of predicted TMHs: 1\nExp number of AAs in TMHs: 20.93248\nExp number, first 60 AAs: 0.03949\nTotal prob of N-in: 0.02447\noutside 1 251\nTMhelix 252 274\ninside 275 284

32515 GCF_002119225.1_ASM211922v1 Streptomyces fradiae Terrabacteria group; Actinobacteria; Actinobacteria; Streptomycetales; Streptomycetaceae; Streptomyces MRTVRSRRTAVTAALLTGAALLGGLAAPGAALAASPTPSPAATDGEPPAPTEAGTSFRSA WP_031129975.1 hypothetical protein [Streptomyces sp. NRRL WC-3719] Length: 266\nNumber of predicted TMHs: 2\nExp number of AAs in TMHs: 41.1869\nExp number, first 60 AAs: 21.9617\nTotal prob of N-in: 0.99139\nPOSSIBLE N-term signal sequence\ninside 1 12\nTMhelix 13 35\noutside 36 229\nTMhelix 230 252\ninside 253 266

32516 GCF_001190165.1_ASM119016v1 Streptomyces fradiae Terrabacteria group; Actinobacteria; Actinobacteria; Streptomycetales; Streptomycetaceae; Streptomyces MRRNTRRRPTGARRRTFAAVALILGGGGLVAANTYASAGEGDSGGRVQDQSGGDQAATIS WP_043464200.1 hypothetical protein [Streptomyces fradiae] Length: 702\nNumber of predicted TMHs: 2\nExp number of AAs in TMHs: 41.46852\nExp number, first 60 AAs: 21.48105\nTotal prob of N-in: 0.99637\nPOSSIBLE N-term signal sequence\ninside 1 16\nTMhelix 17 39\noutside 40 673\nTMhelix 674 693\ninside 694 702

32517 GCF_002119225.1_ASM211922v1 Streptomyces fradiae Terrabacteria group; Actinobacteria; Actinobacteria; Streptomycetales; Streptomycetaceae; Streptomyces MLWLVENHSSPRTAAFFDLDKTVIAKSSTLTFGKSFYQGGLINRRDALRTAYTQFVFLVG WP_051838844.1 inhibition of morphological differentiation protein [Streptomyces sp. NRRL WC-3719] Length: 299\nNumber of predicted TMHs: 1\nExp number of AAs in TMHs: 21.32356\nExp number, first 60 AAs: 0.00575\nTotal prob of N-in: 0.00344\noutside 1 247\nTMhelix 248 270\ninside 271 299

32518 GCF_001756925.1_ASM175692v1 Streptomyces fradiae Terrabacteria group; Actinobacteria; Actinobacteria; Streptomycetales; Streptomycetaceae; Streptomyces
MRRNTRRRPTGARRRTFAAVALILGGGGLVAANTYASAGEGDSGGRVQDQSGGDQAATIS WP_043464200.1 hypothetical protein [Streptomyces fradiae] Length: 702\nNumber of predicted TMHs: 2\nExp number of AAs in TMHs: 41.46852\nExp number, first 60 AAs: 21.48105\nTotal prob of N-in: 0.99637\nPOSSIBLE N-term signal sequence\ninside 1 16\nTMhelix 17 39\noutside 40 673\nTMhelix 674 693\ninside 694 702

32519 GCF_002154445.1_ASM215444v1 Streptomyces fradiae Terrabacteria group; Actinobacteria; Actinobacteria; Streptomycetales; Streptomycetaceae; Streptomyces
MLWLVENHSSPRTAAFFDLDKTVIAKSSTLTFGKSFYQGGLINRRDALRTAYTQFVFLVG WP_051838844.1 MULTISPECIES: inhibition of morphological differentiation protein [Streptomyces] Length: 299\nNumber of predicted TMHs: 1\nExp number of AAs in TMHs: 21.32356\nExp number, first 60 AAs: 0.00575\nTotal prob of N-in: 0.00344\noutside 1 247\nTMhelix 248 270\ninside 271 299

32520 GCF_001756925.1_ASM175692v1 Streptomyces fradiae Terrabacteria group; Actinobacteria; Actinobacteria; Streptomycetales; Streptomycetaceae; Streptomyces
MENPSLPRTAAFFDLDKTVIAKSSALAFSKSFYQGGLINRRAVLRTAYAQFVYLLGGADH WP_078649556.1 inhibition of morphological differentiation protein [Streptomyces fradiae] Length: 284\nNumber of predicted TMHs: 1\nExp number of AAs in TMHs: 20.93248\nExp number, first 60 AAs: 0.03949\nTotal prob of N-in: 0.02447\noutside 1 251\nTMhelix 252 274\ninside 275 284

32521 GCF_000700005.1_ASM70000v1 Streptomyces fradiae Terrabacteria group; Actinobacteria; Actinobacteria; Streptomycetales; Streptomycetaceae; Streptomyces
MENPSLPRTAAFFDLDKTVIAKSSALAFSKSFYQGGLINRRAVLRTAYAQFVYLLGGADH WP_078649556.1 inhibition of morphological differentiation protein [Streptomyces fradiae] Length: 284\nNumber of predicted TMHs: 1\nExp number of AAs in TMHs: 20.93248\nExp number, first 60 AAs: 0.03949\nTotal prob of N-in: 0.02447\noutside 1 251\nTMhelix 252 274\ninside 275 284

32522 GCF_000700005.1_ASM70000v1 Streptomyces fradiae Terrabacteria group; Actinobacteria; Actinobacteria; Streptomycetales; Streptomycetaceae; Streptomyces
MRRNTRRRPTGARRRTFAAVALILGGGGLVAANTYASAGEGDSGGRVQDQSGGDQAATIS WP_043464200.1 hypothetical protein [Streptomyces fradiae] Length: 702\nNumber of predicted TMHs: 2\nExp number of AAs in TMHs: 41.46852\nExp number, first 60 AAs: 21.48105\nTotal prob of N-in: 0.99637\nPOSSIBLE N-term signal sequence\ninside 1 16\nTMhelix 17 39\noutside 40 673\nTMhelix 674 693\ninside 694 702

32523 GCF_002154445.1_ASM215444v1 Streptomyces fradiae Terrabacteria group; Actinobacteria; Actinobacteria; Streptomycetales; Streptomycetaceae; Streptomyces
MRTVRSRRTAVTAALLTGAALLGGLAAPGAALAASPTPSPAATDGEPPAPTEAGTSFRSA WP_031129975.1 MULTISPECIES: hypothetical protein [Streptomyces] Length: 266\nNumber of predicted TMHs: 2\nExp number of AAs in TMHs: 41.1869\nExp number, first 60 AAs: 21.9617\nTotal prob of N-in: 0.99139\nPOSSIBLE N-term signal sequence\ninside 1 12\nTMhelix 13 35\noutside 36 229\nTMhelix 230 252\ninside 253 266

32524 GCF_000718165.1_ASM71816v1 Streptomyces fulvoviolaceus Terrabacteria group; Actinobacteria; Actinobacteria; Streptomycetales; Streptomycetaceae; Streptomyces
MSRRTAVAAALVAPFLLTVFAAAPAQAHGAPTDPVSRVYACSPDGGSPSGSAACRAAVAA WP_030602996.1 chitin-binding protein [Streptomyces fulvoviolaceus] Length: 306\nNumber of predicted TMHs: 2\nExp number of AAs in TMHs: 41.90152\nExp number, first 60 AAs: 20.92942\nTotal prob of N-in: 0.97440\nPOSSIBLE N-term signal sequence\ninside 1 6\nTMhelix 7 29\noutside 30 278\nTMhelix 279 301\ninside 302 306

32525 GCF_000718165.1_ASM71816v1 Streptomyces fulvoviolaceus Terrabacteria group; Actinobacteria; Actinobacteria; Streptomycetales; Streptomycetaceae; Streptomyces
MPAPKKTARRSLLVTSASVLSSLALTAAPAFAAPSPSPSSSSSASSSSASPSTSPSATP WP_030597466.1 D-alanyl-D-alanine carboxypeptidase [Streptomyces fulvoviolaceus] Length: 441\nNumber of predicted TMHs: 1\nExp number of AAs in TMHs: 37.22161\nExp number, first 60 AAs: 14.52911\nTotal prob of N-in: 0.67659\nPOSSIBLE N-term signal sequence\noutside 1 402\nTMhelix 403 425\ninside 426 441

32526 GCF_000718165.1_ASM71816v1 Streptomyces fulvoviolaceus Terrabacteria group; Actinobacteria; Actinobacteria; Streptomycetales; Streptomycetaceae; Streptomyces
MLWVVENHSLPRTAAFFDLDKTVIAKSSTLTFSKSFYQGGLINRRAALRTAYAQFVFLVG WP_030599799.1 inhibition of morphological differentiation protein [Streptomyces fulvoviolaceus] Length: 277\nNumber of predicted TMHs: 1\nExp

number of AAs in TMHs: 21.8633\nExp number, first 60 AAs: 0.01796\nTotal prob of N-in: 0.00225\noutside 1 247\nTMhelix 248 270\ninside 271 277

32527 GCF_000718165.1_ASM71816v1 Streptomyces fulvoviolaceus Terrabacteria group; Actinobacteria; Actinobacteria; Streptomycetales; Streptomycetaceae; Streptomyces MPATPRRRRPVALAAAVATAAALGATALATVGASTASAAEVPLTGYELTWGIKQSYRTYV WP_030601925.1 hypothetical protein [Streptomyces fulvoviolaceus] Length: 497\nNumber of predicted TMHs: 2\nExp number of AAs in TMHs: 43.44062\nExp number, first 60 AAs: 21.40581\nTotal prob of N-in: 0.95286\nPOSSIBLE N-term signal sequence\ninside 1 12\nTMhelix 13 35\noutside 36 462\nTMhelix 463 485\ninside 486 497

32528 GCF_000718165.1_ASM71816v1 Streptomyces fulvoviolaceus Terrabacteria group; Actinobacteria; Actinobacteria; Streptomycetales; Streptomycetaceae; Streptomyces MRHARRRVVRRVTRLAAVGGLLLGGTMVTRAVASETPGDSALTRTYAQEAGDTGAGLVSR WP_030605201.1 protease [Streptomyces fulvoviolaceus] Length: 458\nNumber of predicted TMHs: 2\nExp number of AAs in TMHs: 41.96268\nExp number, first 60 AAs: 20.78979\nTotal prob of N-in: 0.94489\nPOSSIBLE N-term signal sequence\ninside 1 11\nTMhelix 12 34\noutside 35 419\nTMhelix 420 442\ninside 443 458

32529 GCF_000717065.1_ASM71706v1 Streptomyces fulvoviridis Terrabacteria group; Actinobacteria; Actinobacteria; Streptomycetales; Streptomycetaceae; Streptomyces MTTHPSPRRRLLTAAAATAVLLVTLPVLPAAAEDSTQCTFPSKKYAGRPWSLQRVLMDEL WP_033249645.1 MULTISPECIES: type VII secretion-associated serine protease [Streptomyces] Length: 414\nNumber of predicted TMHs: 1\nExp number of AAs in TMHs: 36.07361\nExp number, first 60 AAs: 13.5063\nTotal prob of N-in: 0.65550\nPOSSIBLE N-term signal sequence\noutside 1 378\nTMhelix 379 401\ninside 402 414

32530 GCF_000717065.1_ASM71706v1 Streptomyces fulvoviridis Terrabacteria group; Actinobacteria; Actinobacteria; Streptomycetales; Streptomycetaceae; Streptomyces MRTTRRNTLRRGGVGALLAAVTLLTSAGAAVADDPSPSASAAGGTAGPTEAGTTFRTATA WP_033249102.1 MULTISPECIES: hypothetical protein [Streptomyces] Length: 274\nNumber of predicted TMHs: 2\nExp number of AAs in TMHs: 39.9840299999999\nExp number, first 60 AAs: 19.24596\nTotal prob of N-in: 0.99309\nPOSSIBLE N-term signal sequence\ninside 1 12\nTMhelix 13 32\noutside 33 237\nTMhelix 238 260\ninside 261 274

32531 GCF_000717065.1_ASM71706v1 Streptomyces fulvoviridis Terrabacteria group; Actinobacteria; Actinobacteria; Streptomycetales; Streptomycetaceae; Streptomyces MLTFVENCFSPRTAAFFDLDKTVIAKSSTLTFSKSFYQGGLINRRAVLRTAYTQFVFLAG WP_014155461.1 MULTISPECIES: haloacid dehalogenase [Streptomyces] Length: 279\nNumber of predicted TMHs: 1\nExp number of AAs in TMHs: 21.10966\nExp number, first 60 AAs: 0.01497\nTotal prob of N-in: 0.00234\noutside 1 248\nTMhelix 249 268\ninside 269 279

32532 GCF_000772895.1_ASM77289v1 Streptomyces galbus Terrabacteria group; Actinobacteria; Actinobacteria; Streptomycetales; Streptomycetaceae; Streptomyces MYRLSARRATARRTTGCRADGRRGTGRRAVVTTAAATVVAPLLLTVGAGGPAWAHGAPTD WP_033529488.1 chitin-binding protein [Streptomyces galbus] Length: 354\nNumber of predicted TMHs: 2\nExp number of AAs in TMHs: 38.1615199999999\nExp number, first 60 AAs: 20.31808\nTotal prob of N-in: 0.91591\nPOSSIBLE N-term signal sequence\ninside 1 30\nTMhelix 31 53\noutside 54 320\nTMhelix 321 343\ninside 344 354

32533 GCF_001879105.1_ASM187910v1 Streptomyces gilvigriseus Terrabacteria group; Actinobacteria; Actinobacteria; Streptomycetales; Streptomycetaceae; Streptomyces MSTRRRALRGAALAAAGAVALSAAALPAHAVGSASPAVPKPPAKMSAVGGARLAKPGVQV WP_071657364.1 D-alanyl-D-alanine carboxypeptidase [Streptomyces gilvigriseus] Length: 409\nNumber of predicted TMHs: 2\nExp number of AAs in TMHs: 41.1843599999999\nExp number, first 60 AAs: 19.0888\nTotal prob of N-in: 0.88023\nPOSSIBLE N-term signal sequence\ninside 1 6\nTMhelix 7 29\noutside 30 374\nTMhelix 375 397\ninside 398 409

32534 GCF_001879105.1_ASM187910v1 Streptomyces gilvigriseus Terrabacteria group; Actinobacteria; Actinobacteria; Streptomycetales; Streptomycetaceae; Streptomyces MSMMRDLRSAVRPARRRAALPASSPSAALPVAVRPEPPSAIVDCAVYHEGRRTTGLLAPE WP_071657115.1 magnesium transporter CorA [Streptomyces gilvigriseus] Length: 375\nNumber of predicted TMHs: 2\nExp number of AAs in TMHs: 39.31863\nExp number, first 60 AAs: 9e-05\nTotal prob of N-in: 0.92186\ninside 1 317\nTMhelix 318 337\noutside 338 346\nTMhelix 347 364\ninside 365 375

32535 GCF_002082195.1_ASM208219v1 Streptomyces gilvosporeus Terrabacteria group; Actinobacteria; Actinobacteria; Streptomycetales; Streptomycetaceae; Streptomyces
MTDRRRRAALPPLPATTPGAFLLREHKPSPSGAPQASGGPDRSGTDKDSRESSSGSPQAP WP_083107039.1 hypothetical protein [Streptomyces gilvosporeus] Length: 318\nNumber of predicted TMHs: 2\nExp number of AAs in TMHs: 46.4947\nExp number, first 60 AAs: 0.00035\nTotal prob of N-in: 0.92953\ninside 1 185\nTMhelix 186 208\noutside 209 256\nTMhelix 257 279\ninside 280 318

32536 GCF_002082195.1_ASM208219v1 Streptomyces gilvosporeus Terrabacteria group; Actinobacteria; Actinobacteria; Streptomycetales; Streptomycetaceae; Streptomyces
MARRRRRGAVALLGGLLLALVATPLHTTPAVAAPAAPSAEGSGMVMVLDSSGSMAEKDGS WP_083106339.1 hypothetical protein [Streptomyces gilvosporeus] Length: 641\nNumber of predicted TMHs: 2\nExp number of AAs in TMHs: 43.74221\nExp number, first 60 AAs: 20.99402\nTotal prob of N-in: 0.96790\nPOSSIBLE N-term signal sequence\ninside 1 11\nTMhelix 12 34\noutside 35 597\nTMhelix 598 620\ninside 621 641

32537 GCF_002154375.1_ASM215437v1 Streptomyces glaucescens Terrabacteria group; Actinobacteria; Actinobacteria; Streptomycetales; Streptomycetaceae; Streptomyces
MLRDVENHSLPRTAAFFDLDKTVIAKSSTLTFSKSFYQGGLINRRAALRTAYAQFVFLVG WP_086736002.1 inhibition of morphological differentiation protein [Streptomyces glaucescens] Length: 277\nNumber of predicted TMHs: 1\nExp number of AAs in TMHs: 21.69454\nExp number, first 60 AAs: 0.03199\nTotal prob of N-in: 0.00646\noutside 1 247\nTMhelix 248 270\ninside 271 277

32538 GCF_000761215.1_ASM76121v1 Streptomyces glaucescens Terrabacteria group; Actinobacteria; Actinobacteria; Streptomycetales; Streptomycetaceae; Streptomyces
MTRRRTTRRATLLGVLLAASLTLVPPTVAHADGIRAKQWGLDALHTQEAWRTTKGAGITV WP_043499168.1 type VII secretion-associated serine protease [Streptomyces glaucescens] Length: 390\nNumber of predicted TMHs: 2\nExp number of AAs in TMHs: 42.0802199999999\nExp number, first 60 AAs: 19.1798\nTotal prob of N-in: 0.98813\nPOSSIBLE N-term signal sequence\ninside 1 11\nTMhelix 12 34\noutside 35 360\nTMhelix 361 383\ninside 384 390

32539 GCF_000761215.1_ASM76121v1 Streptomyces glaucescens Terrabacteria group; Actinobacteria; Actinobacteria; Streptomycetales; Streptomycetaceae; Streptomyces
MLWVVENHSLPRTAAFFDLDKTVIAKSSTLTFSKSFYQGGLINRRAALRTAYAQFVFLVG WP_043502838.1 inhibition of morphological differentiation protein [Streptomyces glaucescens] Length: 277\nNumber of predicted TMHs: 1\nExp number of AAs in TMHs: 21.65085\nExp number, first 60 AAs: 0.02185\nTotal prob of N-in: 0.00394\noutside 1 247\nTMhelix 248 270\ninside 271 277

32540 GCF_001418565.1_ASM141856v1 Streptomyces graminilatus Terrabacteria group; Actinobacteria; Actinobacteria; Streptomycetales; Streptomycetaceae; Streptomyces
MRHARRRIVRRATRLAAVGGVILGGVMVTQAVASEPPPANLPFSSAQAAGTTGNELVTEL WP_055528030.1 protease [Streptomyces graminilatus] Length: 452\nNumber of predicted TMHs: 2\nExp number of AAs in TMHs: 43.72565\nExp number, first 60 AAs: 21.92901\nTotal prob of N-in: 0.99058\nPOSSIBLE N-term signal sequence\ninside 1 11\nTMhelix 12 34\noutside 35 413\nTMhelix 414 436\ninside 437 452

32541 GCF_001418565.1_ASM141856v1 Streptomyces graminilatus Terrabacteria group; Actinobacteria; Actinobacteria; Streptomycetales; Streptomycetaceae; Streptomyces
MLTLVENHSLPRTAAFFDLDKTVIAKSSTLTFSKSFYQGGLINRRAVLRTAYAQFVFLAG WP_055531585.1 inhibition of morphological differentiation protein [Streptomyces graminilatus] Length: 277\nNumber of predicted TMHs: 1\nExp number of AAs in TMHs: 21.5636\nExp number, first 60 AAs: 0.02401\nTotal prob of N-in: 0.00358\noutside 1 247\nTMhelix 248 270\ninside 271 277

32542 GCF_000716525.1_ASM71652v1 Streptomyces graminis Terrabacteria group; Actinobacteria; Actinobacteria; Streptomycetales; Streptomycetaceae; Streptomyces
MSPRRRRATVLAAATVLCAAAAPVAAAAPSGTAAAGPVPAVKPAKLPAGLYGDADPKYDG WP_016468754.1 MULTISPECIES: hypothetical protein [Streptomyces] Length: 457\nNumber of predicted TMHs: 2\nExp number of AAs in TMHs: 40.9820199999999\nExp number, first 60 AAs: 18.73808\nTotal prob of N-in: 0.84735\nPOSSIBLE N-term signal sequence\ninside 1 6\nTMhelix 7 29\noutside 30 427\nTMhelix 428 450\ninside 451 457

32543 GCF_000716525.1_ASM71652v1 Streptomyces graminis Terrabacteria group; Actinobacteria; Actinobacteria; Streptomycetales; Streptomycetaceae; Streptomyces
MLSGVENLAMPRTAAFFDLDKTVIAKSSTLTFGKSFYQGGLINRRAVLRTAYAQFVYLLG WP_016473027.1 MULTISPECIES: haloacid dehalogenase [Streptomyces] Length: 284\nNumber of predicted TMHs: 1\nExp number of AAs in TMHs:

21.94508\nExp number, first 60 AAs: 0.01548\nTotal prob of N-in: 0.00204\noutside 1 246\nTMhelix 247 269\ninside 270 284

32544 GCF_001542625.1_ASM154262v2 Streptomyces griseochromogenes Terrabacteria group; Actinobacteria; Actinobacteria; Streptomycetales; Streptomycetaceae; Streptomyces MASRGRRAAMRSLGALAGAAALTVLGSNAPGWAAAPAGAPGGHSSTTTPIKHVVVLFDEN WP_067305609.1 phospholipase [Streptomyces griseochromogenes] Length: 609\nNumber of predicted TMHs: 2\nExp number of AAs in TMHs: 41.72401\nExp number, first 60 AAs: 20.45814\nTotal prob of N-in: 0.90429\nPOSSIBLE N-term signal sequence\ninside 1 12\nTMhelix 13 35\noutside 36 578\nTMhelix 579 601\ninside 602 609

32545 GCF_001542625.1_ASM154262v2 Streptomyces griseochromogenes Terrabacteria group; Actinobacteria; Actinobacteria; Streptomycetales; Streptomycetaceae; Streptomyces MRHARRRVVRRVTRLAAVGGLLLGGAMVTQAAMASEPPAASAKPLSTAEAAGDSGAALVA WP_067305341.1 protease [Streptomyces griseochromogenes] Length: 458\nNumber of predicted TMHs: 2\nExp number of AAs in TMHs: 43.28472\nExp number, first 60 AAs: 22.19106\nTotal prob of N-in: 0.99185\nPOSSIBLE N-term signal sequence\ninside 1 11\nTMhelix 12 34\noutside 35 419\nTMhelix 420 442\ninside 443 458

32546 GCF_001542625.1_ASM154262v2 Streptomyces griseochromogenes Terrabacteria group; Actinobacteria; Actinobacteria; Streptomycetales; Streptomycetaceae; Streptomyces MPAPMKTGRRSVLVTSATLLSLSVTAPAALAAPTPSTSPTATPPAHMSQVGGARLGQPGT WP_067299969.1 D-alanyl-D-alanine carboxypeptidase [Streptomyces griseochromogenes] Length: 422\nNumber of predicted TMHs: 1\nExp number of AAs in TMHs: 30.4099199999999\nExp number, first 60 AAs: 8.21715999999999\nTotal prob of N-in: 0.38774\noutside 1 383\nTMhelix 384 406\ninside 407 422

32547 GCF_001542625.1_ASM154262v2 Streptomyces griseochromogenes Terrabacteria group; Actinobacteria; Actinobacteria; Streptomycetales; Streptomycetaceae; Streptomyces MLRGVENHSLPRAAAFFDLDKTVIAKSSTLTFSKSFYQGGLINRRAALRTAYAQFVFLVG WP_067299551.1 inhibition of morphological differentiation protein [Streptomyces griseochromogenes] Length: 277\nNumber of predicted TMHs: 1\nExp number of AAs in TMHs: 21.57481\nExp number, first 60 AAs: 0.04272\nTotal prob of N-in: 0.00434\noutside 1 247\nTMhelix 248 270\ninside 271 277

32548 GCF_001270585.1_ASM127058v1 Streptomyces griseoflavus Terrabacteria group; Actinobacteria; Actinobacteria; Streptomycetales; Streptomycetaceae; Streptomyces MTLTRRRHLVRSTAVAAVAGSALLLPAAAAFADSPQPTTAGSPGPAGDQQQKDDRHKDDQ WP_030603358.1 MULTISPECIES: hypothetical protein [Streptomyces] Length: 239\nNumber of predicted TMHs: 2\nExp number of AAs in TMHs: 41.4211\nExp number, first 60 AAs: 20.09485\nTotal prob of N-in: 0.95720\nPOSSIBLE N-term signal sequence\ninside 1 12\nTMhelix 13 32\noutside 33 207\nTMhelix 208 230\ninside 231 239

32549 GCF_001270585.1_ASM127058v1 Streptomyces griseoflavus Terrabacteria group; Actinobacteria; Actinobacteria; Streptomycetales; Streptomycetaceae; Streptomyces MTERSAPAASATAPGTPARRTAVRAAATGTLAAALLGLAPQSALAATPAPPPAPAISAVT WP_030603830.1 MULTISPECIES: hypothetical protein [Streptomyces] Length: 293\nNumber of predicted TMHs: 1\nExp number of AAs in TMHs: 24.61251\nExp number, first 60 AAs: 5.37395\nTotal prob of N-in: 0.26790\noutside 1 264\nTMhelix 265 287\ninside 288 293

32550 GCF_001270585.1_ASM127058v1 Streptomyces griseoflavus Terrabacteria group; Actinobacteria; Actinobacteria; Streptomycetales; Streptomycetaceae; Streptomyces MTRRPLLRAAGVALSAAALTVLPALPAHADGIRAQEWALKAMNTQDAWETTKGEGITVAV WP_030595365.1 MULTISPECIES: type VII secretion-associated serine protease [Streptomyces] Length: 386\nNumber of predicted TMHs: 1\nExp number of AAs in TMHs: 37.57611\nExp number, first 60 AAs: 9.6597\nTotal prob of N-in: 0.57980\noutside 1 359\nTMhelix 360 382\ninside 383 386

32551 GCF_000718315.1_ASM71831v1 Streptomyces griseofuscus Terrabacteria group; Actinobacteria; Actinobacteria; Streptomycetales; Streptomycetaceae; Streptomyces MRESTRLSRRAVLGLTAAALPTSAVLPLTAAAPARAASPATAIGGERLARGGLQVDGAAA WP_078967395.1 D-alanyl-D-alanine carboxypeptidase [Streptomyces griseofuscus] Length: 422\nNumber of predicted TMHs: 2\nExp number of AAs in TMHs: 33.32176\nExp number, first 60 AAs: 17.42176\nTotal prob of N-in: 0.79075\nPOSSIBLE N-term signal sequence\ninside 1 11\nTMhelix 12 34\noutside 35 354\nTMhelix 355 377\ninside 378 422

32552 GCF_000718315.1_ASM71831v1 Streptomyces griseofuscus Terrabacteria group; Actinobacteria; Actinobacteria; Streptomycetales; Streptomycetaceae; Streptomyces

MLGDVENHSLPRAAAFFDLDKTVIAKSSTLTFSKSFYQGGLINRRAALRTAYAQFVFLAG WP_037663065.1 inhibition of morphological differentiation protein [Streptomyces griseofuscus] Length: 276\nNumber of predicted TMHs: 1\nExp number of AAs in TMHs: 21.51155\nExp number, first 60 AAs: 0.02847\nTotal prob of N-in: 0.00286\noutside 1 247\nTMhelix 248 270\ninside 271 276

32553 GCF_000718315.1_ASM71831v1 Streptomyces griseofuscus Terrabacteria group; Actinobacteria; Actinobacteria; Streptomycetales; Streptomycetaceae; Streptomyces MPAPKNTARRSLLISAATLLSLAVTAPAALAAPTADKAPAATPPARMSTLGGARLGLPGTWP_037656023.1 D-alanyl-D-alanine carboxypeptidase [Streptomyces griseofuscus] Length: 407\nNumber of predicted TMHs: 2\nExp number of AAs in TMHs: 43.0967799999999\nExp number, first 60 AAs: 20.50033\nTotal prob of N-in: 0.94110\nPOSSIBLE N-term signal sequence\ninside 1 11\nTMhelix 12 34\noutside 35 368\nTMhelix 369 391\ninside 392 407

32554 GCF_000718315.1_ASM71831v1 Streptomyces griseofuscus Terrabacteria group; Actinobacteria; Actinobacteria; Streptomycetales; Streptomycetaceae; Streptomyces MASRGRRAAMRSLGALVGVAALTALGSNAPGWAATSAHGHGASSTATPIKHVVVLFDENI WP_037653933.1 phospholipase C [Streptomyces griseofuscus] Length: 606\nNumber of predicted TMHs: 2\nExp number of AAs in TMHs: 42.02161\nExp number, first 60 AAs: 20.61423\nTotal prob of N-in: 0.91707\nPOSSIBLE N-term signal sequence\ninside 1 12\nTMhelix 13 35\noutside 36 575\nTMhelix 576 598\ninside 599 606

32555 GCF_002114065.1_ASM211406v1 Streptomyces griseofuscus Terrabacteria group; Actinobacteria; Actinobacteria; Streptomycetales; Streptomycetaceae; Streptomyces MLGDVENHSLPRAAAFFDLDKTVIAKSSTLTFSKSFYQGGLINRRAALRTAYAQFVFLAG WP_085565851.1 inhibition of morphological differentiation protein [Streptomyces griseofuscus] Length: 276\nNumber of predicted TMHs: 1\nExp number of AAs in TMHs: 21.51155\nExp number, first 60 AAs: 0.02847\nTotal prob of N-in: 0.00286\noutside 1 247\nTMhelix 248 270\ninside 271 276

32556 GCF_002114065.1_ASM211406v1 Streptomyces griseofuscus Terrabacteria group; Actinobacteria; Actinobacteria; Streptomycetales; Streptomycetaceae; Streptomyces MPAPMNTARRSLLISAATLLSLAVTAPAALAAPATGKAPAATPPAHMSTLGGARLGLPGT WP_085566213.1 D-alanyl-D-alanine carboxypeptidase [Streptomyces griseofuscus] Length: 407\nNumber of predicted TMHs: 2\nExp number of AAs in TMHs: 43.3236\nExp number, first 60 AAs: 20.78286\nTotal prob of N-in: 0.94284\nPOSSIBLE N-term signal sequence\ninside 1 11\nTMhelix 12 34\noutside 35 368\nTMhelix 369 391\ninside 392 407

32557 GCF_002114065.1_ASM211406v1 Streptomyces griseofuscus Terrabacteria group; Actinobacteria; Actinobacteria; Streptomycetales; Streptomycetaceae; Streptomyces MASRGRRAAMRSLGALVGVAALTALGSNAPGWAATSAHGHGASSTATPIKHVVVLFDENI WP_085563424.1 phospholipase [Streptomyces griseofuscus] Length: 607\nNumber of predicted TMHs: 2\nExp number of AAs in TMHs: 41.65184\nExp number, first 60 AAs: 20.61429\nTotal prob of N-in: 0.91707\nPOSSIBLE N-term signal sequence\ninside 1 12\nTMhelix 13 35\noutside 36 576\nTMhelix 577 599\ninside 600 607

32558 GCF_002114065.1_ASM211406v1 Streptomyces griseofuscus Terrabacteria group; Actinobacteria; Actinobacteria; Streptomycetales; Streptomycetaceae; Streptomyces MRESTRLSRRAVLGLTAAALPTSAVLPLTAAAPAHAASPATAIGGERLARGGLQVDGAAA WP_085568484.1 D-alanyl-D-alanine carboxypeptidase [Streptomyces griseofuscus] Length: 427\nNumber of predicted TMHs: 2\nExp number of AAs in TMHs: 32.04232\nExp number, first 60 AAs: 16.15686\nTotal prob of N-in: 0.73007\nPOSSIBLE N-term signal sequence\ninside 1 11\nTMhelix 12 34\noutside 35 359\nTMhelix 360 382\ninside 383 427

32559 GCF_000721185.1_ASM72118v1 Streptomyces griseolus Terrabacteria group; Actinobacteria; Actinobacteria; Streptomycetales; Streptomycetaceae; Streptomyces MTAGMSQTWKRRAALGSRGRLLGTVAVAAAWTVGMAGAAPSAVAMDVQSKQWYLSAMHAE WP_030630441.1 MULTISPECIES: serine protease [Streptomyces] Length: 420\nNumber of predicted TMHs: 2\nExp number of AAs in TMHs: 44.93486\nExp number, first 60 AAs: 22.25757\nTotal prob of N-in: 0.98803\nPOSSIBLE N-term signal sequence\ninside 1 20\nTMhelix 21 43\noutside 44 391\nTMhelix 392 414\ninside 415 420

32560 GCF_000721185.1_ASM72118v1 Streptomyces griseolus Terrabacteria group; Actinobacteria; Actinobacteria; Streptomycetales; Streptomycetaceae; Streptomyces MLSFVENCFSPRTAAFFDLDKTVIAKSSTLAFSKSFYQGGLINRRAVLRTAYTQFVFLAG WP_030629408.1 MULTISPECIES: inhibition of morphological differentiation protein [Streptomyces] Length: 279\nNumber of predicted TMHs: 1\nExp number of AAs in TMHs: 21.81622\nExp number, first 60 AAs: 0.01771\nTotal prob of N-in: 0.00802\noutside 1 247\nTMhelix 248 270\ninside 271 279

32561 GCF_000717735.1_ASM71773v1 Streptomyces griseoluteus Terrabacteria group; Actinobacteria; Actinobacteria; Streptomycetales; Streptomycetaceae; Streptomyces
MTSHPTSRPVSRRTAVAVTAVLAFSGLASVPASAHGSMTDPVSRVSACYAEGPESPRSAA WP_030221273.1 chitin-binding protein [Streptomyces griseoluteus] Length: 342\nNumber of predicted TMHs: 1\nExp number of AAs in TMHs: 32.96856\nExp number, first 60 AAs: 10.69885\nTotal prob of N-in: 0.48496\nPOSSIBLE N-term signal sequence\noutside 1 312\nTMhelix 313 335\ninside 336 342

32562 GCF_000717735.1_ASM71773v1 Streptomyces griseoluteus Terrabacteria group; Actinobacteria; Actinobacteria; Streptomycetales; Streptomycetaceae; Streptomyces
MLRLVENHSLPRTAAFFDLDKTVIAKSSTLTFSKSFYRGGLISRRAALRTAYIQFVFLAG WP_030209056.1 inhibition of morphological differentiation protein [Streptomyces griseoluteus] Length: 288\nNumber of predicted TMHs: 1\nExp number of AAs in TMHs: 21.78436\nExp number, first 60 AAs: 0.05317\nTotal prob of N-in: 0.00392\noutside 1 247\nTMhelix 248 270\ninside 271 288

32563 GCF_000717735.1_ASM71773v1 Streptomyces griseoluteus Terrabacteria group; Actinobacteria; Actinobacteria; Streptomycetales; Streptomycetaceae; Streptomyces
MKAAAGRRRAALAAGAAAGALLTLTSVAAAPAQAAGYRYWSFWESSGGKPWAYATQGPAT WP_030207910.1 hypothetical protein [Streptomyces griseoluteus] Length: 232\nNumber of predicted TMHs: 2\nExp number of AAs in TMHs: 41.8024\nExp number, first 60 AAs: 21.39395\nTotal prob of N-in: 0.95836\nPOSSIBLE N-term signal sequence\ninside 1 11\nTMhelix 12 34\noutside 35 203\nTMhelix 204 226\ninside 227 232

32564 GCF_000709915.1_JSD Streptomyces griseorubens Terrabacteria group; Actinobacteria; Actinobacteria; Streptomycetales; Streptomycetaceae; Streptomyces
MRHARRRVVRRVTRLAAVGGLLLGGTMVTHAVASEPGTPDAVPFSTASPADASGPGAGLV WP_037643279.1 MULTISPECIES: protease [Streptomyces] Length: 457\nNumber of predicted TMHs: 2\nExp number of AAs in TMHs: 38.82844\nExp number, first 60 AAs: 21.8309\nTotal prob of N-in: 0.97535\nPOSSIBLE N-term signal sequence\ninside 1 11\nTMhelix 12 34\noutside 35 418\nTMhelix 419 441\ninside 442 457

32565 GCF_000709915.1_JSD Streptomyces griseorubens Terrabacteria group; Actinobacteria; Actinobacteria; Streptomycetales; Streptomycetaceae; Streptomyces
MLRGVENHLSPRTAAFFDLDKTVIAKSSTLTFSKSFYQGGLINRRAVLRTAYSQFVFLAG WP_033274879.1 MULTISPECIES: inhibition of morphological differentiation protein [Actinobacteria] Length: 277\nNumber of predicted TMHs: 1\nExp number of AAs in TMHs: 21.81221\nExp number, first 60 AAs: 0.02078\nTotal prob of N-in: 0.00383\noutside 1 247\nTMhelix 248 270\ninside 271 277

32566 GCF_001514205.1_ASM151420v1 Streptomyces griseoruber Terrabacteria group; Actinobacteria; Actinobacteria; Streptomycetales; Streptomycetaceae; Streptomyces
MRHARRRIVRRVTRLAAVGGLLLGGTMVTQAAMATETPATSATPRTTPESVVTNPGTALV WP_055632075.1 protease [Streptomyces griseoruber] Length: 465\nNumber of predicted TMHs: 2\nExp number of AAs in TMHs: 42.36765\nExp number, first 60 AAs: 21.77359\nTotal prob of N-in: 0.97964\nPOSSIBLE N-term signal sequence\ninside 1 11\nTMhelix 12 34\noutside 35 424\nTMhelix 425 447\ninside 448 465

32567 GCF_001514205.1_ASM151420v1 Streptomyces griseoruber Terrabacteria group; Actinobacteria; Actinobacteria; Streptomycetales; Streptomycetaceae; Streptomyces
MLLVVENHSLPRTAAFFDLDKTVIAKSSTLTFSKSFYQGGLINRRAALRTAYAQFVFLVG WP_055638307.1 inhibition of morphological differentiation protein [Streptomyces griseoruber] Length: 276\nNumber of predicted TMHs: 1\nExp number of AAs in TMHs: 22.07321\nExp number, first 60 AAs: 0.01445\nTotal prob of N-in: 0.00182\noutside 1 246\nTMhelix 247 269\ninside 270 276

32568 GCF_001418585.1_ASM141858v1 Streptomyces griseoruber Terrabacteria group; Actinobacteria; Actinobacteria; Streptomycetales; Streptomycetaceae; Streptomyces
MLLVVENHSLPRTAAFFDLDKTVIAKSSTLTFSKSFYQGGLINRRAALRTAYAQFVFLVG WP_055638307.1 inhibition of morphological differentiation protein [Streptomyces griseoruber] Length: 276\nNumber of predicted TMHs: 1\nExp number of AAs in TMHs: 22.07321\nExp number, first 60 AAs: 0.01445\nTotal prob of N-in: 0.00182\noutside 1 246\nTMhelix 247 269\ninside 270 276

32569 GCF_001418585.1_ASM141858v1 Streptomyces griseoruber Terrabacteria group; Actinobacteria; Actinobacteria; Streptomycetales; Streptomycetaceae; Streptomyces
MRHARRRIVRRVTRLAAVGGLLLGGTMVTQAAMATETPATSATPRTTPESVVTNPGTALV WP_055632075.1 protease [Streptomyces griseoruber] Length: 465\nNumber of predicted TMHs: 2\nExp number of AAs in

TMHs: 42.36765\nExp number, first 60 AAs: 21.77359\nTotal prob of N-in: 0.97964\nPOSSIBLE N-term signal sequence\ninside 1 11\nTMhelix 12 34\noutside 35 424\nTMhelix 425 447\ninside 448 465

32570 GCF_001514195.1_ASM151419v1 Streptomyces griseorubiginosus Terrabacteria group; Actinobacteria; Actinobacteria; Streptomycetales; Streptomycetaceae; Streptomyces MLGVVDNHSLPRTAAFFDLDKTVIAKSSTLTFSKSFYQGGLINRRAALRTAYAQFVFLVG WP_062026023.1 MULTISPECIES: inhibition of morphological differentiation protein [Streptomyces] Length: 277\nNumber of predicted TMHs: 1\nExp number of AAs in TMHs: 21.69291\nExp number, first 60 AAs: 0.0377\nTotal prob of N-in: 0.00335\noutside 1 247\nTMhelix 248 270\ninside 271 277

32571 GCF_001514195.1_ASM151419v1 Streptomyces griseorubiginosus Terrabacteria group; Actinobacteria; Actinobacteria; Streptomycetales; Streptomycetaceae; Streptomyces MRHARRRVVRRVTQLAAVGGLLLGGTMVTRAVASEPPATAPPRTFAQQAGGTGADIVARL WP_062239856.1 protease [Streptomyces griseorubiginosus] Length: 454\nNumber of predicted TMHs: 2\nExp number of AAs in TMHs: 39.03657\nExp number, first 60 AAs: 18.87788\nTotal prob of N-in: 0.86917\nPOSSIBLE N-term signal sequence\ninside 1 11\nTMhelix 12 34\noutside 35 415\nTMhelix 416 438\ninside 439 454

32572 GCF_000376565.1_ASM37656v1 Streptomyces hokutonensis Terrabacteria group; Actinobacteria; Actinobacteria; Streptomycetales; Streptomycetaceae; Streptomyces MRHARRRVVRRVTRLTAVGGLLLGGAMVTQAAMASDTPARDAATLSAPQTPADKGSALVA WP_043259400.1 S1 family peptidase [Streptomyces hokutonensis] Length: 456\nNumber of predicted TMHs: 2\nExp number of AAs in TMHs: 40.76231\nExp number, first 60 AAs: 21.69878\nTotal prob of N-in: 0.98031\nPOSSIBLE N-term signal sequence\ninside 1 11\nTMhelix 12 34\noutside 35 417\nTMhelix 418 440\ninside 441 456

32573 GCF_000376565.1_ASM37656v1 Streptomyces hokutonensis Terrabacteria group; Actinobacteria; Actinobacteria; Streptomycetales; Streptomycetaceae; Streptomyces MLWVVENHSLPRTAAFFDLDKTVIAKSSTLTFGKSFYQGGLINRRAALRTAYAQFVFLAG WP_019070283.1 inhibition of morphological differentiation protein [Streptomyces hokutonensis] Length: 280\nNumber of predicted TMHs: 1\nExp number of AAs in TMHs: 21.59912\nExp number, first 60 AAs: 0.03892\nTotal prob of N-in: 0.00614\noutside 1 247\nTMhelix 248 270\ninside 271 280

32574 GCF_001005085.2_ASM100508v2 Streptomyces humi Terrabacteria group; Actinobacteria; Actinobacteria; Streptomycetales; Streptomycetaceae; Streptomyces MLRGVENHSLPRTAAFFDLDKTVIAKSSTLTFSKSFYQGGLINRRAALRNAYSQFVFLVG WP_071659674.1 inhibition of morphological differentiation protein [Streptomyces humi] Length: 274\nNumber of predicted TMHs: 1\nExp number of AAs in TMHs: 21.62121\nExp number, first 60 AAs: 0.00913\nTotal prob of N-in: 0.02319\noutside 1 244\nTMhelix 245 267\ninside 268 274

32575 GCF_001005085.2_ASM100508v2 Streptomyces humi Terrabacteria group; Actinobacteria; Actinobacteria; Streptomycetales; Streptomycetaceae; Streptomyces MRHARRRVVRRGTQLAAVGGLLLGGAMVTTAMASETPGTSAVPFSSASAAGDTGAALVAK WP_046733394.1 protease [Streptomyces humi] Length: 457\nNumber of predicted TMHs: 2\nExp number of AAs in TMHs: 40.7149299999999\nExp number, first 60 AAs: 21.1553\nTotal prob of N-in: 0.96518\nPOSSIBLE N-term signal sequence\ninside 1 11\nTMhelix 12 34\noutside 35 418\nTMhelix 419 441\ninside 442 457

32576 GCF_001005085.2_ASM100508v2 Streptomyces humi Terrabacteria group; Actinobacteria; Actinobacteria; Streptomycetales; Streptomycetaceae; Streptomyces MASTGRRTAMRSLGALAGAAALTVLGGTAPSWAAVPAGHTGGSSSTATPIKHVVVLFDEN WP_046734512.1 phospholipase C [Streptomyces humi] Length: 611\nNumber of predicted TMHs: 2\nExp number of AAs in TMHs: 42.94398\nExp number, first 60 AAs: 22.22243\nTotal prob of N-in: 0.97827\nPOSSIBLE N-term signal sequence\ninside 1 12\nTMhelix 13 35\noutside 36 580\nTMhelix 581 603\ninside 604 611

32577 GCF_002021875.1_ASM202187v1 Streptomyces hygroscopicus Terrabacteria group; Actinobacteria; Actinobacteria; Streptomycetales; Streptomycetaceae; Streptomyces MLGVVENHSLPRTAAFFDLDKTVIAKSSAFTFSKSFYQGGLINRRAVLRTAYAQFVFLAG WP_078642258.1 inhibition of morphological differentiation protein [Streptomyces hygroscopicus] Length: 278\nNumber of predicted TMHs: 1\nExp number of AAs in TMHs: 21.25954\nExp number, first 60 AAs: 0.03117\nTotal prob of N-in: 0.00369\noutside 1 247\nTMhelix 248 267\ninside 268 278

32578 GCF_002021875.1_ASM202187v1 Streptomyces hygroscopicus Terrabacteria group; Actinobacteria; Actinobacteria; Streptomycetales; Streptomycetaceae; Streptomyces

MSSPDREVRAARNDTARTARRPVVAVTGAASGPGALLTQRLAESEEVKQVLAIDERRGEV WP_078643469.1 NAD-dependent dehydratase [Streptomyces hygroscopicus] Length: 369\nNumber of predicted TMHs: 1\nExp number of AAs in TMHs: 19.99839\nExp number, first 60 AAs: 0.38218\nTotal prob of N-in: 0.03041\noutside 1 339\nTMhelix 340 359\ninside 360 369

32579 GCF_000717055.1_ASM71705v1 Streptomyces iakyrus Terrabacteria group; Actinobacteria; Actinobacteria; Streptomycetales; Streptomycetaceae; Streptomyces MLWGVENHSLPRTAAFFDLDKTVIAKSSTLTFSKSFYQGGLINRRAVLRTAYAQFVFLAG WP_033310194.1 inhibition of morphological differentiation protein [Streptomyces iakyrus] Length: 277\nNumber of predicted TMHs: 1\nExp number of AAs in TMHs: 21.58162\nExp number, first 60 AAs: 0.0297\nTotal prob of N-in: 0.00292\noutside 1 247\nTMhelix 248 270\ninside 271 277

32580 GCF_000717055.1_ASM71705v1 Streptomyces iakyrus Terrabacteria group; Actinobacteria; Actinobacteria; Streptomycetales; Streptomycetaceae; Streptomyces MTGSRRSGRSPPVSTRRTTVHLRPPAAVLAAAVIAVAGPAVVAPLAHAEEGAPELVVSAL WP_051814589.1 LPXTG cell wall anchor domain-containing protein [Streptomyces iakyrus] Length: 425\nNumber of predicted TMHs: 2\nExp number of AAs in TMHs: 41.66329\nExp number, first 60 AAs: 20.51333\nTotal prob of N-in: 0.92235\nPOSSIBLE N-term signal sequence\ninside 1 25\nTMhelix 26 48\noutside 49 393\nTMhelix 394 416\ninside 417 425

32581 GCF_000717055.1_ASM71705v1 Streptomyces iakyrus Terrabacteria group; Actinobacteria; Actinobacteria; Streptomycetales; Streptomycetaceae; Streptomyces MRHARRRVVRRVTRLAAVGGLLLGGAMVTTAVASEPSAPVIGVPFTAPPATGEGTDLVTQ WP_033312445.1 protease [Streptomyces iakyrus] Length: 448\nNumber of predicted TMHs: 2\nExp number of AAs in TMHs: 42.38013\nExp number, first 60 AAs: 22.02692\nTotal prob of N-in: 0.99052\nPOSSIBLE N-term signal sequence\ninside 1 11\nTMhelix 12 34\noutside 35 409\nTMhelix 410 432\ninside 433 448

32582 GCF_000717055.1_ASM71705v1 Streptomyces iakyrus Terrabacteria group; Actinobacteria; Actinobacteria; Streptomycetales; Streptomycetaceae; Streptomyces MNVIHRRAAVRTARSLVVPTAVLAVLAVSAPRAAAHTELDTGSPGANTTLAGLPPQATLT WP_051814695.1 hypothetical protein [Streptomyces iakyrus] Length: 199\nNumber of predicted TMHs: 2\nExp number of AAs in TMHs: 42.6511\nExp number, first 60 AAs: 21.26567\nTotal prob of N-in: 0.96546\nPOSSIBLE N-term signal sequence\ninside 1 12\nTMhelix 13 35\noutside 36 167\nTMhelix 168 190\ninside 191 199

32583 GCF_000938975.1_Siranensis Streptomyces iranensis Terrabacteria group; Actinobacteria; Actinobacteria; Streptomycetales; Streptomycetaceae; Streptomyces MSSPDREVRAARNDTARTTRRPVVAVTGAASGPGALLTQRLAESEEVKQVLAIDERRGEV WP_044570187.1 NAD-dependent dehydratase [Streptomyces iranensis] Length: 369\nNumber of predicted TMHs: 1\nExp number of AAs in TMHs: 20.05676\nExp number, first 60 AAs: 0.45079\nTotal prob of N-in: 0.03345\noutside 1 339\nTMhelix 340 359\ninside 360 369

32584 GCF_001642995.1_ASM164299v1 Streptomyces jeddahensis Terrabacteria group; Actinobacteria; Actinobacteria; Streptomycetales; Streptomycetaceae; Streptomyces MRRRRLLTGTVLAAAAAGLSVAPAHASDANALDLYPSTTAPGATVSANTKACGAGETATG WP_067285055.1 hypothetical protein [Streptomyces jeddahensis] Length: 172\nNumber of predicted TMHs: 2\nExp number of AAs in TMHs: 39.30265\nExp number, first 60 AAs: 18.99445\nTotal prob of N-in: 0.93512\nPOSSIBLE N-term signal sequence\ninside 1 6\nTMhelix 7 26\noutside 27 141\nTMhelix 142 164\ninside 165 172

32585 GCF_001642995.1_ASM164299v1 Streptomyces jeddahensis Terrabacteria group; Actinobacteria; Actinobacteria; Streptomycetales; Streptomycetaceae; Streptomyces MFSAFSVHRRGLVRLATAALAPGLVAAGAIVAAAPAVADDVPQHQGGATATLTGLKIHGE WP_078066793.1 peptidase [Streptomyces jeddahensis] Length: 468\nNumber of predicted TMHs: 2\nExp number of AAs in TMHs: 41.71253\nExp number, first 60 AAs: 19.62129\nTotal prob of N-in: 0.86202\nPOSSIBLE N-term signal sequence\ninside 1 11\nTMhelix 12 34\noutside 35 436\nTMhelix 437 459\ninside 460 468

32586 GCF_001642995.1_ASM164299v1 Streptomyces jeddahensis Terrabacteria group; Actinobacteria; Actinobacteria; Streptomycetales; Streptomycetaceae; Streptomyces MLWLVENHSSPRTAAFFDLDKTVIAKSSTLTFSKSFYQGGLINRRAVLRTAYAQFVFLAG WP_067282842.1 inhibition of morphological differentiation protein [Streptomyces jeddahensis] Length: 277\nNumber of predicted TMHs: 1\nExp number of AAs in TMHs: 21.57952\nExp number, first 60 AAs: 0.03476\nTotal prob of N-in: 0.00602\noutside 1 247\nTMhelix 248 270\ninside 271 277

32587 GCF_001642995.1_ASM164299v1 Streptomyces jeddahensis Terrabacteria group; Actinobacteria; Actinobacteria; Streptomycetales; Streptomycetaceae; Streptomyces
MDTPAQHDRHGRGRTSRRRPLRLLLAAAAAAFTLTAGLATPLNPAAEQAKAADDGKKVLT WP_067284647.1 ABC transporter substrate-binding protein [Streptomyces jeddahensis] Length: 629\nNumber of predicted TMHs: 2\nExp number of AAs in TMHs: 43.55049\nExp number, first 60 AAs: 21.28953\nTotal prob of N-in: 0.99735\nPOSSIBLE N-term signal sequence\ninside 1 20\nTMhelix 21 40\noutside 41 595\nTMhelix 596 618\ninside 619 629

32588 GCF_001418465.1_ASM141846v1 Streptomyces kanamyceticus Terrabacteria group; Actinobacteria; Actinobacteria; Streptomycetales; Streptomycetaceae; Streptomyces
MLGLVENHSLPRTAAFFDLDKTVIAKSSTLTFSKSFYQGGLINRRAVLRTAYAQFVFLAG WP_055556308.1 inhibition of morphological differentiation protein [Streptomyces kanamyceticus] Length: 274\nNumber of predicted TMHs: 1\nExp number of AAs in TMHs: 21.49094\nExp number, first 60 AAs: 0.02161\nTotal prob of N-in: 0.00426\noutside 1 244\nTMhelix 245 267\ninside 268 274

32589 GCF_001418465.1_ASM141846v1 Streptomyces kanamyceticus Terrabacteria group; Actinobacteria; Actinobacteria; Streptomycetales; Streptomycetaceae; Streptomyces
MNLLLLGRRTAMGSLTALLLATGVWASWDTARLLAGGPERGVLTVTGCGSDECAGWYVSA WP_055548827.1 hypothetical protein [Streptomyces kanamyceticus] Length: 155\nNumber of predicted TMHs: 2\nExp number of AAs in TMHs: 55.12288\nExp number, first 60 AAs: 13.06454\nTotal prob of N-in: 0.60784\nPOSSIBLE N-term signal sequence\noutside 1 106\nTMhelix 107 129\ninside 130 133\nTMhelix 134 153\noutside 154 155

32590 GCF_001484625.1_ASM148462v1 Streptomyces kanasensis Terrabacteria group; Actinobacteria; Actinobacteria; Streptomycetales; Streptomycetaceae; Streptomyces
MLGLVENHSLPRTAAFFDLDKTVIAKSSTLTFSRSFYQGGLINRRAALRTAYTQFVFLAG WP_058940847.1 inhibition of morphological differentiation protein [Streptomyces kanasensis] Length: 301\nNumber of predicted TMHs: 1\nExp number of AAs in TMHs: 22.51971\nExp number, first 60 AAs: 0.00595\nTotal prob of N-in: 0.00126\noutside 1 247\nTMhelix 248 270\ninside 271 301

32591 GCF_000717655.1_ASM71765v1 Streptomyces katrae Terrabacteria group; Actinobacteria; Actinobacteria; Streptomycetales; Streptomycetaceae; Streptomyces
MSTVNRRTLVRAAAVTVCAGTLLGLPAAAALAEGVPAASAAHSAGAPRTLVKSLALADGV WP_030295329.1 hypothetical protein [Streptomyces katrae] Length: 200\nNumber of predicted TMHs: 2\nExp number of AAs in TMHs: 44.29223\nExp number, first 60 AAs: 21.67235\nTotal prob of N-in: 0.99186\nPOSSIBLE N-term signal sequence\ninside 1 8\nTMhelix 9 31\noutside 32 170\nTMhelix 171 193\ninside 194 200

32592 GCF_000717655.1_ASM71765v1 Streptomyces katrae Terrabacteria group; Actinobacteria; Actinobacteria; Streptomycetales; Streptomycetaceae; Streptomyces
MPARRRTVTRIAAAGLAPLAVAAYAAGPAAAHGSMTDPVSRVAACYAEGPESPKSAACKA WP_030302478.1 chitin-binding protein [Streptomyces katrae] Length: 347\nNumber of predicted TMHs: 2\nExp number of AAs in TMHs: 41.43052\nExp number, first 60 AAs: 19.16065\nTotal prob of N-in: 0.90165\nPOSSIBLE N-term signal sequence\ninside 1 11\nTMhelix 12 31\noutside 32 312\nTMhelix 313 335\ninside 336 347

32593 GCF_002028425.1_ASM202842v1 Streptomyces katrae Terrabacteria group; Actinobacteria; Actinobacteria; Streptomycetales; Streptomycetaceae; Streptomyces
MSSPDPQARAPHDAENRPEHGDSAPGVRQPRNEAAPRRRSPVIAVTGAASGVGAALVGRL WP_079429679.1 NAD-dependent dehydratase [Streptomyces katrae] Length: 388\nNumber of predicted TMHs: 1\nExp number of AAs in TMHs: 20.67061\nExp number, first 60 AAs: 0.72514\nTotal prob of N-in: 0.05093\noutside 1 359\nTMhelix 360 379\ninside 380 388

32594 GCF_002028425.1_ASM202842v1 Streptomyces katrae Terrabacteria group; Actinobacteria; Actinobacteria; Streptomycetales; Streptomycetaceae; Streptomyces
MNTVSRRTIVRATAVTACAAALLALPTAAALADGVPAASAAGAQRTLVSSLSLADGQSTA WP_079429764.1 hypothetical protein [Streptomyces katrae] Length: 200\nNumber of predicted TMHs: 2\nExp number of AAs in TMHs: 43.8425899999999\nExp number, first 60 AAs: 21.19636\nTotal prob of N-in: 0.98514\nPOSSIBLE N-term signal sequence\ninside 1 8\nTMhelix 9 31\noutside 32 170\nTMhelix 171 193\ninside 194 200

32595 GCF_001906585.1_ASM190658v1 Streptomyces kebangsaanensis Terrabacteria group; Actinobacteria; Actinobacteria; Streptomycetales; Streptomycetaceae; Streptomyces
MPATATATTTRRRPLVFAAAVATATALGATVLATAPSASAASVPLTGYELSWGIKESYRT WP_073951852.1 hypothetical protein [Streptomyces kebangsaanensis] Length: 525\nNumber of predicted TMHs: 2\nExp number of AAs in

TMHs: 44.76629\nExp number, first 60 AAs: 22.37971\nTotal prob of N-in: 0.98711\nPOSSIBLE N-term signal sequence\ninside 1 12\nTMhelix 13 35\noutside 36 493\nTMhelix 494 516\ninside 517 525

32596 GCF_000715625.1_ASM71562v1 Streptomyces lavendulae Terrabacteria group; Actinobacteria; Actinobacteria; Streptomycetales; Streptomycetaceae; Streptomyces MTQSHAPRRALLATAFVLTLASGGTGTAAAAGAAAPQPPYALLLDGAGECTFPMKKQIED WP_030009522.1 type VII secretion-associated serine protease [Streptomyces lavendulae] Length: 418\nNumber of predicted TMHs: 2\nExp number of AAs in TMHs: 44.27734\nExp number, first 60 AAs: 21.97445\nTotal prob of N-in: 0.96130\nPOSSIBLE N-term signal sequence\ninside 1 11\nTMhelix 12 34\noutside 35 384\nTMhelix 385 407\ninside 408 418

32597 GCF_000715625.1_ASM71562v1 Streptomyces lavendulae Terrabacteria group; Actinobacteria; Actinobacteria; Streptomycetales; Streptomycetaceae; Streptomyces MAEAADNQGASPAPARRRWLRRAIVLLAGTPVLAALIYSPAPRAEAAQASAVDVQLTSMA WP_030008397.1 hypothetical protein [Streptomyces lavendulae] Length: 754\nNumber of predicted TMHs: 2\nExp number of AAs in TMHs: 39.91177\nExp number, first 60 AAs: 18.26857\nTotal prob of N-in: 0.89446\nPOSSIBLE N-term signal sequence\ninside 1 18\nTMhelix 19 41\noutside 42 687\nTMhelix 688 710\ninside 711 754

32598 GCF_000718625.1_ASM71862v1 Streptomyces lavenduligriseus Terrabacteria group; Actinobacteria; Actinobacteria; Streptomycetales; Streptomycetaceae; Streptomyces MLVAVYGPRVRECSRLSRRAVLGLTAAALPLSTATDATAATAATTVVGGERLARTGVQVH WP_037702946.1 D-alanyl-D-alanine carboxypeptidase [Streptomyces lavenduligriseus] Length: 409\nNumber of predicted TMHs: 1\nExp number of AAs in TMHs: 24.67588\nExp number, first 60 AAs: 3.7446\nTotal prob of N-in: 0.16601\noutside 1 363\nTMhelix 364 386\ninside 387 409

32599 GCF_000718625.1_ASM71862v1 Streptomyces lavenduligriseus Terrabacteria group; Actinobacteria; Actinobacteria; Streptomycetales; Streptomycetaceae; Streptomyces MRHARRRVVRRVTRLAAACGLLLGGTMVARAAGADQPPAAGPGRPGDTGAALVARLGTAR WP_030779875.1 protease [Streptomyces lavenduligriseus] Length: 438\nNumber of predicted TMHs: 2\nExp number of AAs in TMHs: 43.00694\nExp number, first 60 AAs: 21.36633\nTotal prob of N-in: 0.96357\nPOSSIBLE N-term signal sequence\ninside 1 11\nTMhelix 12 34\noutside 35 399\nTMhelix 400 422\ninside 423 438

32600 GCF_000718625.1_ASM71862v1 Streptomyces lavenduligriseus Terrabacteria group; Actinobacteria; Actinobacteria; Streptomycetales; Streptomycetaceae; Streptomyces MLKGVENHSLPRAAAFFDLDKTVIAKSSTLTFSKSFYQGGLINRRAALRTAYAQFVFLAG WP_030781867.1 inhibition of morphological differentiation protein [Streptomyces lavenduligriseus] Length: 276\nNumber of predicted TMHs: 1\nExp number of AAs in TMHs: 21.83016\nExp number, first 60 AAs: 0.02674\nTotal prob of N-in: 0.00322\noutside 1 247\nTMhelix 248 270\ninside 271 276

32601 GCF_000718625.1_ASM71862v1 Streptomyces lavenduligriseus Terrabacteria group; Actinobacteria; Actinobacteria; Streptomycetales; Streptomycetaceae; Streptomyces MTRPTGRRRALAAVASLLLTGSLALLPAAPAHADGLRAQQWALTALHLEEAWRTTEGRGV WP_030781235.1 type VII secretion-associated serine protease [Streptomyces lavenduligriseus] Length: 392\nNumber of predicted TMHs: 1\nExp number of AAs in TMHs: 40.99946\nExp number, first 60 AAs: 14.66354\nTotal prob of N-in: 0.85201\nPOSSIBLE N-term signal sequence\noutside 1 364\nTMhelix 365 387\ninside 388 392

32602 GCF_000696595.1_StrepLeeu1.0 Streptomyces leeuwenhoekii Terrabacteria group; Actinobacteria; Actinobacteria; Streptomycetales; Streptomycetaceae; Streptomyces MGRRGASGRVPARRRAVTGRRAAPVLMATALAVLCTTAAAPARAVPSAGPPGGYAFAEDA WP_029383025.1 hypothetical protein [Streptomyces leeuwenhoekii] Length: 442\nNumber of predicted TMHs: 2\nExp number of AAs in TMHs: 43.87924\nExp number, first 60 AAs: 19.04459\nTotal prob of N-in: 0.90873\nPOSSIBLE N-term signal sequence\ninside 1 22\nTMhelix 23 45\noutside 46 408\nTMhelix 409 431\ninside 432 442

32603 GCF_001047315.1_ASM104731v1 Streptomyces leeuwenhoekii Terrabacteria group; Actinobacteria; Actinobacteria; Streptomycetales; Streptomycetaceae; Streptomyces MLGVVENHSLPRTAAFFDLDKTVIAKSSTLTFSKSFYQGGLINRRAVLRTAYAQFVFLAG WP_029382559.1 inhibition of morphological differentiation protein [Streptomyces leeuwenhoekii] Length: 277\nNumber of predicted TMHs: 1\nExp number of AAs in TMHs: 21.5732\nExp number, first 60 AAs: 0.02604\nTotal prob of N-in: 0.00255\noutside 1 247\nTMhelix 248 270\ninside 271 277

32604 GCF_000696595.1_StrepLeeu1.0 Streptomyces leeuwenhoekii Terrabacteria group; Actinobacteria; Actinobacteria; Streptomycetales; Streptomycetaceae; Streptomyces

MLGVVENHSLPRTAAFFDLDKTVIAKSSTLTFSKSFYQGGLINRRAVLRTAYAQFVFLAG WP_029382559.1 inhibition of morphological differentiation protein [Streptomyces leeuwenhoekii] Length: 277\nNumber of predicted TMHs: 1\nExp number of AAs in TMHs: 21.5732\nExp number, first 60 AAs: 0.02604\nTotal prob of N-in: 0.00255\noutside 1 247\nTMhelix 248 270\ninside 271 277

32605 GCF_000696595.1_StrepLeeu1.0 Streptomyces leeuwenhoekii Terrabacteria group; Actinobacteria; Actinobacteria; Streptomycetales; Streptomycetaceae; Streptomyces MAVLSFLSRPASRRGLIRPARVLGVAAASAALAAGAAGTALACDIGEFTAAAKCDGGKGV WP_029385521.1 hypothetical protein [Streptomyces leeuwenhoekii] Length: 244\nNumber of predicted TMHs: 1\nExp number of AAs in TMHs: 37.65256\nExp number, first 60 AAs: 15.54812\nTotal prob of N-in: 0.69301\nPOSSIBLE N-term signal sequence\noutside 1 212\nTMhelix 213 235\ninside 236 244

32606 GCF_001047315.1_ASM104731v1 Streptomyces leeuwenhoekii Terrabacteria group; Actinobacteria; Actinobacteria; Streptomycetales; Streptomycetaceae; Streptomyces MAVLSFLSRPASRRGLIRPARVLGVAAASAALAAGAAGTALACNIGEFTAAAKCDGGKGV WP_048572334.1 hypothetical protein [Streptomyces leeuwenhoekii] Length: 244\nNumber of predicted TMHs: 1\nExp number of AAs in TMHs: 37.76783\nExp number, first 60 AAs: 15.54081\nTotal prob of N-in: 0.68422\nPOSSIBLE N-term signal sequence\noutside 1 212\nTMhelix 213 235\ninside 236 244

32607 GCF_001047315.1_ASM104731v1 Streptomyces leeuwenhoekii Terrabacteria group; Actinobacteria; Actinobacteria; Streptomycetales; Streptomycetaceae; Streptomyces MGRRGASGRVPARRRAVTGRRAAPVLMATALAVLCTTAAAPARAVPSAGPPGGYAFAEDA WP_048571945.1 hypothetical protein [Streptomyces leeuwenhoekii] Length: 442\nNumber of predicted TMHs: 2\nExp number of AAs in TMHs: 43.88002\nExp number, first 60 AAs: 19.04463\nTotal prob of N-in: 0.90873\nPOSSIBLE N-term signal sequence\ninside 1 22\nTMhelix 23 45\noutside 46 408\nTMhelix 409 431\ninside 432 442

32608 GCF_000696595.1_StrepLeeu1.0 Streptomyces leeuwenhoekii Terrabacteria group; Actinobacteria; Actinobacteria; Streptomycetales; Streptomycetaceae; Streptomyces MFGIRRRPTMRLSAALLVAGLTAGAAVATAGTAAAEDGHNPAGGVTATLDGLSVYGDAVL WP_029382662.1 TQXA domain-containing protein [Streptomyces leeuwenhoekii] Length: 463\nNumber of predicted TMHs: 2\nExp number of AAs in TMHs: 44.27486\nExp number, first 60 AAs: 21.99306\nTotal prob of N-in: 0.97202\nPOSSIBLE N-term signal sequence\ninside 1 11\nTMhelix 12 34\noutside 35 434\nTMhelix 435 457\ninside 458 463

32609 GCF_001013905.1_sleC34 Streptomyces leeuwenhoekii Terrabacteria group; Actinobacteria; Actinobacteria; Streptomycetales; Streptomycetaceae; Streptomyces MAVLSFLSRPASRRGLIRPARVLGVAAASAALAAGAAGTALACDIGEFTAAAKCDGGKGV WP_029385521.1 hypothetical protein [Streptomyces leeuwenhoekii] Length: 244\nNumber of predicted TMHs: 1\nExp number of AAs in TMHs: 37.65256\nExp number, first 60 AAs: 15.54812\nTotal prob of N-in: 0.69301\nPOSSIBLE N-term signal sequence\noutside 1 212\nTMhelix 213 235\ninside 236 244

32610 GCF_001013905.1_sleC34 Streptomyces leeuwenhoekii Terrabacteria group; Actinobacteria; Actinobacteria; Streptomycetales; Streptomycetaceae; Streptomyces MFGIRRRPTMRLSAALLVAGLTAGAAVATAGTAAAEDGHNPAGGVTATLDGLSVYGDAVL WP_029382662.1 TQXA domain-containing protein [Streptomyces leeuwenhoekii] Length: 463\nNumber of predicted TMHs: 2\nExp number of AAs in TMHs: 44.27486\nExp number, first 60 AAs: 21.99306\nTotal prob of N-in: 0.97202\nPOSSIBLE N-term signal sequence\ninside 1 11\nTMhelix 12 34\noutside 35 434\nTMhelix 435 457\ninside 458 463

32611 GCF_001013905.1_sleC34 Streptomyces leeuwenhoekii Terrabacteria group; Actinobacteria; Actinobacteria; Streptomycetales; Streptomycetaceae; Streptomyces MLGVVENHSLPRTAAFFDLDKTVIAKSSTLTFSKSFYQGGLINRRAVLRTAYAQFVFLAG WP_029382559.1 inhibition of morphological differentiation protein [Streptomyces leeuwenhoekii] Length: 277\nNumber of predicted TMHs: 1\nExp number of AAs in TMHs: 21.5732\nExp number, first 60 AAs: 0.02604\nTotal prob of N-in: 0.00255\noutside 1 247\nTMhelix 248 270\ninside 271 277

32612 GCF_001013905.1_sleC34 Streptomyces leeuwenhoekii Terrabacteria group; Actinobacteria; Actinobacteria; Streptomycetales; Streptomycetaceae; Streptomyces MGRRGASGRVPARRRAVTGRRAAPVLMATALAVLCTTAAAPARAVPSAGPPGGYAFAEDA WP_029383025.1 hypothetical protein [Streptomyces leeuwenhoekii] Length: 442\nNumber of predicted TMHs: 2\nExp number of AAs in TMHs: 43.87924\nExp number, first 60 AAs: 19.04459\nTotal prob of N-in: 0.90873\nPOSSIBLE N-term signal sequence\ninside 1 22\nTMhelix 23 45\noutside 46 408\nTMhelix 409 431\ninside 432 442

32613 GCF_001685355.1_ASM168535v1 Streptomyces lincolnensis Terrabacteria group; Actinobacteria; Actinobacteria; Streptomycetales; Streptomycetaceae; Streptomyces
MRHARRRVVRRVTRLAAVGGLLLGGTMVTRAMASETPAPIPRTYAQTAGDMGADLVTRLG WP_067443535.1 protease [Streptomyces lincolnensis] Length: 457\nNumber of predicted TMHs: 2\nExp number of AAs in TMHs: 42.3592499999999\nExp number, first 60 AAs: 20.89921\nTotal prob of N-in: 0.96739\nPOSSIBLE N-term signal sequence\ninside 1 11\nTMhelix 12 34\noutside 35 418\nTMhelix 419 441\ninside 442 457

32614 GCF_001685355.1_ASM168535v1 Streptomyces lincolnensis Terrabacteria group; Actinobacteria; Actinobacteria; Streptomycetales; Streptomycetaceae; Streptomyces
MGRRDFHARTSETRRSTSVGRNKRRRPTGARRATFAAVALMLGGSGLVAANVYASATEDG WP_079164990.1 hypothetical protein [Streptomyces lincolnensis] Length: 716\nNumber of predicted TMHs: 2\nExp number of AAs in TMHs: 40.32839\nExp number, first 60 AAs: 21.71651\nTotal prob of N-in: 0.97454\nPOSSIBLE N-term signal sequence\ninside 1 32\nTMhelix 33 55\noutside 56 689\nTMhelix 690 707\ninside 708 716

32615 GCF_001685355.1_ASM168535v1 Streptomyces lincolnensis Terrabacteria group; Actinobacteria; Actinobacteria; Streptomycetales; Streptomycetaceae; Streptomyces
MLGVVENHSLPRTAAFFDLDKTVIAKSSTLTFGKSFYQGGLINRRAALRTAYAQFVFLVG WP_067437309.1 inhibition of morphological differentiation protein [Streptomyces lincolnensis] Length: 277\nNumber of predicted TMHs: 1\nExp number of AAs in TMHs: 21.91031\nExp number, first 60 AAs: 0.01971\nTotal prob of N-in: 0.00412\noutside 1 247\nTMhelix 248 270\ninside 271 277

32616 GCF_001514125.1_ASM151412v1 Streptomyces longwoodensis Terrabacteria group; Actinobacteria; Actinobacteria; Streptomycetales; Streptomycetaceae; Streptomyces
MSSPDPQVRAARNHTTPAGGRRPVVAVTGAATGVGALLTERLAASEEIKQVVAVDERRGE WP_067236815.1 NAD-dependent dehydratase [Streptomyces longwoodensis] Length: 369\nNumber of predicted TMHs: 1\nExp number of AAs in TMHs: 20.29062\nExp number, first 60 AAs: 0.23799\nTotal prob of N-in: 0.02875\noutside 1 340\nTMhelix 341 360\ninside 361 369

32617 GCF_001514125.1_ASM151412v1 Streptomyces longwoodensis Terrabacteria group; Actinobacteria; Actinobacteria; Streptomycetales; Streptomycetaceae; Streptomyces
MLRGVENHSLPRTAAFFDLDKTVIAKSSTLTFSKSFYQGGLINRRAALRTAYAQFVFLVG WP_067240848.1 inhibition of morphological differentiation protein [Streptomyces longwoodensis] Length: 278\nNumber of predicted TMHs: 1\nExp number of AAs in TMHs: 22.01271\nExp number, first 60 AAs: 0.02652\nTotal prob of N-in: 0.01528\noutside 1 247\nTMhelix 248 270\ninside 271 278

32618 GCF_001700515.1_ASM170051v1 Streptomyces lushanensis Terrabacteria group; Actinobacteria; Actinobacteria; Streptomycetales; Streptomycetaceae; Streptomyces
MLGLVENHSLPRTAAFFDLDKTVIAKSSTLTFSKSFYQGGLINRRAVLRTAYAQFVFLAG WP_066950817.1 inhibition of morphological differentiation protein [Streptomyces lushanensis] Length: 278\nNumber of predicted TMHs: 1\nExp number of AAs in TMHs: 21.75453\nExp number, first 60 AAs: 0.01299\nTotal prob of N-in: 0.00142\noutside 1 247\nTMhelix 248 270\ninside 271 278

32619 GCF_001748305.1_ASM174830v1 Streptomyces luteocolor Terrabacteria group; Actinobacteria; Actinobacteria; Streptomycetales; Streptomycetaceae; Streptomyces
MSTHRRTALLRTTAVSALVGGALLIPAAGAFADSPAPEPTASGTQAPDTTGTPSQKPTQK WP_069885441.1 hypothetical protein [Streptomyces luteocolor] Length: 206\nNumber of predicted TMHs: 2\nExp number of AAs in TMHs: 42.91622\nExp number, first 60 AAs: 21.21089\nTotal prob of N-in: 0.98746\nPOSSIBLE N-term signal sequence\ninside 1 6\nTMhelix 7 29\noutside 30 176\nTMhelix 177 199\ninside 200 206

32620 GCF_001748305.1_ASM174830v1 Streptomyces luteocolor Terrabacteria group; Actinobacteria; Actinobacteria; Streptomycetales; Streptomycetaceae; Streptomyces
MLGLVENHSLPRTAAFFDLDKTVIAKSSTLTFSKSFYQGGLINRRAVLRTAYAQFVFLAG WP_055699300.1 MULTISPECIES: inhibition of morphological differentiation protein [Streptomyces] Length: 288\nNumber of predicted TMHs: 1\nExp number of AAs in TMHs: 21.95172\nExp number, first 60 AAs: 0.01273\nTotal prob of N-in: 0.00126\noutside 1 247\nTMhelix 248 270\ninside 271 288

32621 GCF_000717345.1_ASM71734v1 Streptomyces lydicus Terrabacteria group; Actinobacteria; Actinobacteria; Streptomycetales; Streptomycetaceae; Streptomyces
MAHRPRRGAVALLGGLLLALTAAPFPAIAAAPAAAPPADGSGMVMVLDSSGSMAGSDGSG WP_051732961.1 hypothetical protein [Streptomyces lydicus] Length: 629\nNumber of predicted TMHs: 2\nExp number of

AAs in TMHs: 44.84918\nExp number, first 60 AAs: 22.66877\nTotal prob of N-in: 0.99410\nPOSSIBLE N-term signal sequence\ninside 1 12\nTMhelix 13 35\noutside 36 599\nTMhelix 600 622\ninside 623 629

32622 GCF_000717345.1_ASM71734v1 Streptomyces lydicus Terrabacteria group; Actinobacteria; Actinobacteria; Streptomycetales; Streptomycetaceae; Streptomyces MRQRRRAAVASAAALVLAGAGVLTAAGTARAAEVSYKTECLPPPISGLPPIEGTTKVDVS WP_033266588.1 hypothetical protein [Streptomyces lydicus] Length: 473\nNumber of predicted TMHs: 2\nExp number of AAs in TMHs: 45.01544\nExp number, first 60 AAs: 22.34257\nTotal prob of N-in: 0.99681\nPOSSIBLE N-term signal sequence\ninside 1 6\nTMhelix 7 29\noutside 30 442\nTMhelix 443 465\ninside 466 473

32623 GCF_001984445.1_ASM198444v1 Streptomyces lydicus Terrabacteria group; Actinobacteria; Actinobacteria; Streptomycetales; Streptomycetaceae; Streptomyces MIPEHTSRRAALAALPAALALTVALASPAAADGNGAYQINLAPLNNSGSKGVAMLSLKGN WP_033270775.1 hypothetical protein [Streptomyces lydicus] Length: 277\nNumber of predicted TMHs: 1\nExp number of AAs in TMHs: 33.97362\nExp number, first 60 AAs: 13.34373\nTotal prob of N-in: 0.64960\nPOSSIBLE N-term signal sequence\noutside 1 235\nTMhelix 236 258\ninside 259 277

32624 GCF_000717345.1_ASM71734v1 Streptomyces lydicus Terrabacteria group; Actinobacteria; Actinobacteria; Streptomycetales; Streptomycetaceae; Streptomyces MIPEHTSRRAALAALPAALALTVALASPAAADGNGAYQINLAPLNNSGSKGVAMLSLKGN WP_033270775.1 hypothetical protein [Streptomyces lydicus] Length: 277\nNumber of predicted TMHs: 1\nExp number of AAs in TMHs: 33.97362\nExp number, first 60 AAs: 13.34373\nTotal prob of N-in: 0.64960\nPOSSIBLE N-term signal sequence\noutside 1 235\nTMhelix 236 258\ninside 259 277

32625 GCF_000717345.1_ASM71734v1 Streptomyces lydicus Terrabacteria group; Actinobacteria; Actinobacteria; Streptomycetales; Streptomycetaceae; Streptomyces MAPLAPMAQQAPSTHPVAFLARRAAMALAAAAVLGGAAAPAAYAVDGADGAPAAGSPKKL WP_033271713.1 hypothetical protein [Streptomyces lydicus] Length: 444\nNumber of predicted TMHs: 1\nExp number of AAs in TMHs: 27.03128\nExp number, first 60 AAs: 5.01738\nTotal prob of N-in: 0.21802\noutside 1 414\nTMhelix 415 437\ninside 438 444

32626 GCF_001984445.1_ASM198444v1 Streptomyces lydicus Terrabacteria group; Actinobacteria; Actinobacteria; Streptomycetales; Streptomycetaceae; Streptomyces MRQRRRAAVASAAALVLAGAGVLTAAGTARAAEVSYKTECLPPPISGLPPIEGTTKVDVS WP_077192611.1 hypothetical protein [Streptomyces lydicus] Length: 477\nNumber of predicted TMHs: 2\nExp number of AAs in TMHs: 45.01601\nExp number, first 60 AAs: 22.34219\nTotal prob of N-in: 0.99680\nPOSSIBLE N-term signal sequence\ninside 1 6\nTMhelix 7 29\noutside 30 446\nTMhelix 447 469\ninside 470 477

32627 GCF_000952035.2_ASM95203v2 Streptomyces lydicus Terrabacteria group; Actinobacteria; Actinobacteria; Streptomycetales; Streptomycetaceae; Streptomyces MARRHRRGVVALLGGLLLTLAAAPLQTTPAAANAPAAPTTEGTGMVMVLDSSGSMAEKDS WP_046927080.1 VWA domain-containing protein [Streptomyces lydicus] Length: 644\nNumber of predicted TMHs: 2\nExp number of AAs in TMHs: 41.22218\nExp number, first 60 AAs: 18.59059\nTotal prob of N-in: 0.90690\nPOSSIBLE N-term signal sequence\ninside 1 6\nTMhelix 7 29\noutside 30 600\nTMhelix 601 623\ninside 624 644

32628 GCF_001729485.1_ASM172948v1 Streptomyces lydicus Terrabacteria group; Actinobacteria; Actinobacteria; Streptomycetales; Streptomycetaceae; Streptomyces MIPEHTSRRAALAALPAALALTVALASPATADGNGAYQINLAQLNNSGSKGVAMLSLKGN WP_069568061.1 hypothetical protein [Streptomyces lydicus] Length: 278\nNumber of predicted TMHs: 1\nExp number of AAs in TMHs: 31.812\nExp number, first 60 AAs: 10.84054\nTotal prob of N-in: 0.53293\nPOSSIBLE N-term signal sequence\noutside 1 236\nTMhelix 237 259\ninside 260 278

32629 GCF_001729485.1_ASM172948v1 Streptomyces lydicus Terrabacteria group; Actinobacteria; Actinobacteria; Streptomycetales; Streptomycetaceae; Streptomyces MRQRRRAAVASAAALVLAGAGVLTAAGTARAAEVSYKTECLPPPISGLPPIEGTTKVDVS WP_069567401.1 hypothetical protein [Streptomyces lydicus] Length: 476\nNumber of predicted TMHs: 2\nExp number of AAs in TMHs: 45.01597\nExp number, first 60 AAs: 22.34219\nTotal prob of N-in: 0.99680\nPOSSIBLE N-term signal sequence\ninside 1 6\nTMhelix 7 29\noutside 30 445\nTMhelix 446 468\ninside 469 476

32630 GCF_001984445.1_ASM198444v1 Streptomyces lydicus Terrabacteria group; Actinobacteria; Actinobacteria; Streptomycetales; Streptomycetaceae; Streptomyces

MAHRPRRGAVALLGGLLLALTAAPFPAIAAAPAAAPPAAGSGMVMVLDSSGSMAGSDGSG WP_077192932.1 hypothetical protein [Streptomyces lydicus] Length: 629\nNumber of predicted TMHs: 2\nExp number of AAs in TMHs: 45.0108\nExp number, first 60 AAs: 22.75228\nTotal prob of N-in: 0.99516\nPOSSIBLE N-term signal sequence\ninside 1 12\nTMhelix 13 35\noutside 36 599\nTMhelix 600 622\ninside 623 629

32631 GCF_001984445.1_ASM198444v1 Streptomyces lydicus Terrabacteria group; Actinobacteria; Actinobacteria; Streptomycetales; Streptomycetaceae; Streptomyces MAPLAPMAQQAPSTHPVAFLARRAAMALAAAAVLGGAAAPAAYAVDGADGAPAAGSPKKL WP_077194704.1 hypothetical protein [Streptomyces lydicus] Length: 443\nNumber of predicted TMHs: 1\nExp number of AAs in TMHs: 27.11779\nExp number, first 60 AAs: 4.86684\nTotal prob of N-in: 0.21746\noutside 1 413\nTMhelix 414 436\ninside 437 443

32632 GCF_001729485.1_ASM172948v1 Streptomyces lydicus Terrabacteria group; Actinobacteria; Actinobacteria; Streptomycetales; Streptomycetaceae; Streptomyces MAPMAPMAQQAPSTHPVAFLARRAAMALAAAAVLGGAAAPAAYAADGADSAPAGSAKKLP WP_079145617.1 hypothetical protein [Streptomyces lydicus] Length: 447\nNumber of predicted TMHs: 1\nExp number of AAs in TMHs: 28.18162\nExp number, first 60 AAs: 6.19741\nTotal prob of N-in: 0.28683\noutside 1 417\nTMhelix 418 440\ninside 441 447

32633 GCF_000980885.2_ASM98088v2 Streptomyces malaysiense Terrabacteria group; Actinobacteria; Actinobacteria; Streptomycetales; Streptomycetaceae; Streptomyces MLGDVENHSLPRAAAFFDLDKTVIAKSSTLTFSKSFYQGGLINRRAALRTAYAQFVFLVG WP_046419038.1 inhibition of morphological differentiation protein [Streptomyces malaysiense] Length: 276\nNumber of predicted TMHs: 1\nExp number of AAs in TMHs: 21.5294\nExp number, first 60 AAs: 0.04752\nTotal prob of N-in: 0.00365\noutside 1 247\nTMhelix 248 270\ninside 271 276

32634 GCF_000980885.2_ASM98088v2 Streptomyces malaysiense Terrabacteria group; Actinobacteria; Actinobacteria; Streptomycetales; Streptomycetaceae; Streptomyces MPAPMNTARRSLLAGAATLLSLAVTAPAALAAPAAGRTPAATPPAHMSTLGGARLGRPGN WP_046417906.1 D-alanyl-D-alanine carboxypeptidase [Streptomyces malaysiense] Length: 407\nNumber of predicted TMHs: 2\nExp number of AAs in TMHs: 43.10384\nExp number, first 60 AAs: 20.55731\nTotal prob of N-in: 0.92768\nPOSSIBLE N-term signal sequence\ninside 1 11\nTMhelix 12 34\noutside 35 368\nTMhelix 369 391\ninside 392 407

32635 GCF_000980885.2_ASM98088v2 Streptomyces malaysiense Terrabacteria group; Actinobacteria; Actinobacteria; Streptomycetales; Streptomycetaceae; Streptomyces MASRGRRAAMRSLGALAGVAALTALGSNAPGWAATPAHVHGASSTATPIKHVVVLFDENI WP_046423759.1 phospholipase C [Streptomyces malaysiense] Length: 607\nNumber of predicted TMHs: 2\nExp number of AAs in TMHs: 40.3177\nExp number, first 60 AAs: 18.91286\nTotal prob of N-in: 0.84132\nPOSSIBLE N-term signal sequence\ninside 1 12\nTMhelix 13 35\noutside 36 576\nTMhelix 577 599\ninside 600 607

32636 GCF_000974985.2_ASM97498v2 Streptomyces mangrovisoli Terrabacteria group; Actinobacteria; Actinobacteria; Streptomycetales; Streptomycetaceae; Streptomyces MRAGARRAAVLAAATALTATTSVLALPAAVAHADDGGECTFPSKNYSGRPWALQRVNLDE WP_046584733.1 type VII secretion-associated serine protease mycosin [Streptomyces mangrovisoli] Length: 416\nNumber of predicted TMHs: 2\nExp number of AAs in TMHs: 41.69002\nExp number, first 60 AAs: 19.20462\nTotal prob of N-in: 0.85232\nPOSSIBLE N-term signal sequence\ninside 1 8\nTMhelix 9 31\noutside 32 379\nTMhelix 380 402\ninside 403 416

32637 GCF_000974985.2_ASM97498v2 Streptomyces mangrovisoli Terrabacteria group; Actinobacteria; Actinobacteria; Streptomycetales; Streptomycetaceae; Streptomyces MTPLFSKGGAGRRSLVTLVCGALAAGGLTAAGVAALQPEAASASSHREAPLISGTPQYDN WP_046584143.1 hypothetical protein [Streptomyces mangrovisoli] Length: 513\nNumber of predicted TMHs: 1\nExp number of AAs in TMHs: 34.8929\nExp number, first 60 AAs: 12.17704\nTotal prob of N-in: 0.51354\nPOSSIBLE N-term signal sequence\noutside 1 482\nTMhelix 483 505\ninside 506 513

32638 GCF_000974985.2_ASM97498v2 Streptomyces mangrovisoli Terrabacteria group; Actinobacteria; Actinobacteria; Streptomycetales; Streptomycetaceae; Streptomyces MRHARRRVVRRVTRLAAVGGILLGGAMAAQAAMADEAPRTGATVLRGSTGSAEDPGASLV WP_046587340.1 protease [Streptomyces mangrovisoli] Length: 461\nNumber of predicted TMHs: 2\nExp number of AAs in TMHs: 43.03206\nExp number, first 60 AAs: 21.91509\nTotal prob of N-in: 0.99388\nPOSSIBLE N-term signal sequence\ninside 1 11\nTMhelix 12 34\noutside 35 422\nTMhelix 423 445\ninside 446 461

32639 GCF_000974985.2_ASM97498v2 Streptomyces mangrovisoli Terrabacteria group; Actinobacteria; Actinobacteria; Streptomycetales; Streptomycetaceae; Streptomyces
MLGGVENHSLPRTAAFFDLDKTVIAKSSTLTFSKSFYQGGLINRRAVLRTAYTQFVFLVG WP_046581547.1 inhibition of morphological differentiation protein [Streptomyces mangrovisoli] Length: 277\nNumber of predicted TMHs: 1\nExp number of AAs in TMHs: 21.96343\nExp number, first 60 AAs: 0.01239\nTotal prob of N-in: 0.00106\noutside 1 247\nTMhelix 248 270\ninside 271 277

32640 GCF_000718985.1_ASM71898v1 Streptomyces megasporus Terrabacteria group; Actinobacteria; Actinobacteria; Streptomycetales; Streptomycetaceae; Streptomyces
MLTLVENHSLPRTAAFFDLDKTVIAKSSTLTFGKSFYQGGLINRRDALRTAYTQFVYLVG WP_031509229.1 inhibition of morphological differentiation protein [Streptomyces megasporus] Length: 278\nNumber of predicted TMHs: 1\nExp number of AAs in TMHs: 21.73447\nExp number, first 60 AAs: 0.00183\nTotal prob of N-in: 0.00182\noutside 1 246\nTMhelix 247 269\ninside 270 278

32641 GCF_000718985.1_ASM71898v1 Streptomyces megasporus Terrabacteria group; Actinobacteria; Actinobacteria; Streptomycetales; Streptomycetaceae; Streptomyces
MTAGHNHGPSGMPPRRTTRRRLLAVLAAAGAWTTCLVGPAPMAHADDVRSQQWYLDAMQA WP_051717410.1 serine protease [Streptomyces megasporus] Length: 415\nNumber of predicted TMHs: 2\nExp number of AAs in TMHs: 43.26764\nExp number, first 60 AAs: 21.10168\nTotal prob of N-in: 0.94825\nPOSSIBLE N-term signal sequence\ninside 1 20\nTMhelix 21 43\noutside 44 386\nTMhelix 387 409\ninside 410 415

32642 GCF_002154605.1_ASM215460v1 Streptomyces milbemycinicus Terrabacteria group; Actinobacteria; Actinobacteria; Streptomycetales; Streptomycetaceae; Streptomyces
MGRRAALLCAVLLSTLVSVVALGASPAQAAGYRYWSFWQRERGAGGWTYATQGPATARPG WP_014180138.1 hypothetical protein [Streptomyces bingchenggensis] Length: 224\nNumber of predicted TMHs: 2\nExp number of AAs in TMHs: 43.52954\nExp number, first 60 AAs: 22.06599\nTotal prob of N-in: 0.99215\nPOSSIBLE N-term signal sequence\ninside 1 6\nTMhelix 7 29\noutside 30 193\nTMhelix 194 216\ninside 217 224

32643 GCF_002154605.1_ASM215460v1 Streptomyces milbemycinicus Terrabacteria group; Actinobacteria; Actinobacteria; Streptomycetales; Streptomycetaceae; Streptomyces
MLGVVENHSLPRTAAFFDLDKTVIAKSSTLTFSKSFYQGGLINRRAVLRTAYAQFVFLAG WP_014178415.1 morphological differentiation-associated protein [Streptomyces bingchenggensis] Length: 278\nNumber of predicted TMHs: 1\nExp number of AAs in TMHs: 21.89798\nExp number, first 60 AAs: 0.02259\nTotal prob of N-in: 0.00436\noutside 1 246\nTMhelix 247 269\ninside 270 278

32644 GCF_002154605.1_ASM215460v1 Streptomyces milbemycinicus Terrabacteria group; Actinobacteria; Actinobacteria; Streptomycetales; Streptomycetaceae; Streptomyces
MSAADDGGNRSLGQLVATATTELSALVHDEIALAKAEFRDSARRTLFGSGAAIVAGVLLL WP_014178408.1 phage holin family protein [Streptomyces bingchenggensis] Length: 153\nNumber of predicted TMHs: 2\nExp number of AAs in TMHs: 46.5707000000001\nExp number, first 60 AAs: 13.62843\nTotal prob of N-in: 0.95447\nPOSSIBLE N-term signal sequence\ninside 1 45\nTMhelix 46 68\noutside 69 82\nTMhelix 83 105\ninside 106 153

32645 GCF_000746395.1_ASM74639v1 Streptomyces mirabilis Terrabacteria group; Actinobacteria; Actinobacteria; Streptomycetales; Streptomycetaceae; Streptomyces
MRHARRRVVRRVTRLAAVGGLLLGGAMVTQAAMASDSSGTSAATLKSSQTPADEGAALVS WP_037734823.1 protease [Streptomyces mirabilis] Length: 460\nNumber of predicted TMHs: 2\nExp number of AAs in TMHs: 43.20789\nExp number, first 60 AAs: 21.93902\nTotal prob of N-in: 0.98870\nPOSSIBLE N-term signal sequence\ninside 1 11\nTMhelix 12 34\noutside 35 421\nTMhelix 422 444\ninside 445 460

32646 GCF_000746395.1_ASM74639v1 Streptomyces mirabilis Terrabacteria group; Actinobacteria; Actinobacteria; Streptomycetales; Streptomycetaceae; Streptomyces
MPTSTHAPARAGRRAALAGAAALTTTLVLAAPAAAHAEVEADRPQALAQNVTLSFVSEAE WP_037734597.1 hypothetical protein [Streptomyces mirabilis] Length: 247\nNumber of predicted TMHs: 1\nExp number of AAs in TMHs: 37.19047\nExp number, first 60 AAs: 15.10737\nTotal prob of N-in: 0.47070\nPOSSIBLE N-term signal sequence\noutside 1 215\nTMhelix 216 238\ninside 239 247

32647 GCF_900112965.1_IMG-taxon_2616644814_annotated_assembly Streptomyces mirabilis Terrabacteria group; Actinobacteria; Actinobacteria; Streptomycetales; Streptomycetaceae; Streptomyces
MLGLVENHSLPRTAAFFDLDKTVIAKSSTLTFSKSFYQGGLINRRAVLRTAYAQFVFMAG WP_075033560.1 inhibition of morphological differentiation protein [Streptomyces mirabilis] Length: 277\nNumber of predicted TMHs: 1\nExp

number of AAs in TMHs: 21.53459\nExp number, first 60 AAs: 0.01511\nTotal prob of N-in: 0.00191\noutside 1 247\nTMhelix 248 270\ninside 271 277

32648 GCF_000745715.1_ASM74571v1 Streptomyces mirabilis Terrabacteria group; Actinobacteria; Actinobacteria; Streptomycetales; Streptomycetaceae; Streptomyces MRHARRRVVRRVTRLAAVGGLLLGGVMVTRAVASEPPAATTVPHTYAMKAGQTGAELVSR WP_037714072.1 protease [Streptomyces mirabilis] Length: 455\nNumber of predicted TMHs: 2\nExp number of AAs in TMHs: 39.1832299999999\nExp number, first 60 AAs: 20.91059\nTotal prob of N-in: 0.96789\nPOSSIBLE N-term signal sequence\ninside 1 11\nTMhelix 12 34\noutside 35 416\nTMhelix 417 439\ninside 440 455

32649 GCF_000745715.1_ASM74571v1 Streptomyces mirabilis Terrabacteria group; Actinobacteria; Actinobacteria; Streptomycetales; Streptomycetaceae; Streptomyces MLGVVENHSLPRTAAFFDLDKTVIAKSSTLTFSKSFYQGGLINRRAALRTAYAQFVFLVG WP_037717716.1 inhibition of morphological differentiation protein [Streptomyces mirabilis] Length: 277\nNumber of predicted TMHs: 1\nExp number of AAs in TMHs: 21.65845\nExp number, first 60 AAs: 0.01246\nTotal prob of N-in: 0.00153\noutside 1 247\nTMhelix 248 270\ninside 271 277

32650 GCF_000745715.1_ASM74571v1 Streptomyces mirabilis Terrabacteria group; Actinobacteria; Actinobacteria; Streptomycetales; Streptomycetaceae; Streptomyces MSARRKAVVAAVGLTPLALTALAAAPASAHGSMGDPVSRVAQCYAEGPESPKSNACRAAV WP_037715556.1 chitin-binding protein [Streptomyces mirabilis] Length: 325\nNumber of predicted TMHs: 2\nExp number of AAs in TMHs: 42.7439499999999\nExp number, first 60 AAs: 21.88936\nTotal prob of N-in: 0.98743\nPOSSIBLE N-term signal sequence\ninside 1 6\nTMhelix 7 29\noutside 30 289\nTMhelix 290 312\ninside 313 325

32651 GCF_900112965.1_IMG-taxon_2616644814_annotated_assembly Streptomyces mirabilis Terrabacteria group; Actinobacteria; Actinobacteria; Streptomycetales; Streptomycetaceae; Streptomyces MRDSSPVPPRAARLSRRAALGLAAAVPLAAASPASAATPVIGGERLARAGVQVRGAAGLP WP_075026534.1 D-alanyl-D-alanine carboxypeptidase [Streptomyces mirabilis] Length: 405\nNumber of predicted TMHs: 1\nExp number of AAs in TMHs: 32.83518\nExp number, first 60 AAs: 13.08694\nTotal prob of N-in: 0.58076\nPOSSIBLE N-term signal sequence\noutside 1 355\nTMhelix 356 378\ninside 379 405

32652 GCF_900112965.1_IMG-taxon_2616644814_annotated_assembly Streptomyces mirabilis Terrabacteria group; Actinobacteria; Actinobacteria; Streptomycetales; Streptomycetaceae; Streptomyces MRHARRRIVRRGTRLAAVGGVLCGGLMVTHAMASEPSAASRAPQSATLAAAGKGAGLVST WP_075028213.1 serine protease [Streptomyces mirabilis] Length: 460\nNumber of predicted TMHs: 2\nExp number of AAs in TMHs: 42.1919299999999\nExp number, first 60 AAs: 20.81044\nTotal prob of N-in: 0.96509\nPOSSIBLE N-term signal sequence\ninside 1 11\nTMhelix 12 34\noutside 35 421\nTMhelix 422 444\ninside 445 460

32653 GCF_900104815.1_IMG-taxon_2634166373_annotated_assembly Streptomyces misionensis Terrabacteria group; Actinobacteria; Actinobacteria; Streptomycetales; Streptomycetaceae; Streptomyces MASRGRRAAMRSLGALAGVAALTALGSTAPGWAATPVHAHGKPSSTATPIQHVVVLFDEN WP_074990644.1 phospholipase [Streptomyces misionensis] Length: 608\nNumber of predicted TMHs: 2\nExp number of AAs in TMHs: 43.20074\nExp number, first 60 AAs: 21.90049\nTotal prob of N-in: 0.96888\nPOSSIBLE N-term signal sequence\ninside 1 12\nTMhelix 13 35\noutside 36 579\nTMhelix 580 602\ninside 603 608

32654 GCF_900104815.1_IMG-taxon_2634166373_annotated_assembly Streptomyces misionensis Terrabacteria group; Actinobacteria; Actinobacteria; Streptomycetales; Streptomycetaceae; Streptomyces MVRNPLLGAVYRPGVRESSRLSRRAVLGLTAAALPLSAAAPASAEPTAVGGARLARRGVQ WP_079172327.1 MULTISPECIES: D-alanyl-D-alanine carboxypeptidase [Streptomyces] Length: 432\nNumber of predicted TMHs: 1\nExp number of AAs in TMHs: 17.37505\nExp number, first 60 AAs: 0.69826\nTotal prob of N-in: 0.03361\noutside 1 367\nTMhelix 368 387\ninside 388 432

32655 GCF_000715845.1_ASM71584v1 Streptomyces monomycini Terrabacteria group; Actinobacteria; Actinobacteria; Streptomycetales; Streptomycetaceae; Streptomyces MFSSKPSRRRTVRTAAVGVLTAVACATVMAGSAGAIVNGANSTESYPFMATIPESAPKDG WP_030019685.1 trypsin [Streptomyces monomycini] Length: 321\nNumber of predicted TMHs: 2\nExp number of AAs in TMHs: 41.77674\nExp number, first 60 AAs: 20.83157\nTotal prob of N-in: 0.91811\nPOSSIBLE N-term signal sequence\ninside 1 16\nTMhelix 17 39\noutside 40 289\nTMhelix 290 312\ninside 313 321

32656 GCF_000715845.1_ASM71584v1 Streptomyces monomycini Terrabacteria group; Actinobacteria; Actinobacteria; Streptomycetales; Streptomycetaceae; Streptomyces

MTPSTSRRNMLRGSVATAALAVGPALAGTAHAAPSPGTPHVGILLYDGYSLLDPTGPAEI WP_030019312.1 thiamine biosynthesis protein ThiJ [Streptomyces monomycini] Length: 252\nNumber of predicted TMHs: 2\nExp number of AAs in TMHs: 37.27536\nExp number, first 60 AAs: 5.06298\nTotal prob of N-in: 0.86430\ninside 1 132\nTMhelix 133 152\noutside 153 185\nTMhelix 186 208\ninside 209 252

32657 GCF_000715845.1_ASM71584v1 Streptomyces monomycini Terrabacteria group; Actinobacteria; Actinobacteria; Streptomycetales; Streptomycetaceae; Streptomyces MTRRPLLRAAGVALSAAALTVLPVLPAHADGIRTQEWALNALHAEKAWETTKGDGITVAV WP_030018074.1 type VII secretion-associated serine protease [Streptomyces monomycini] Length: 386\nNumber of predicted TMHs: 1\nExp number of AAs in TMHs: 34.08011\nExp number, first 60 AAs: 12.74997\nTotal prob of N-in: 0.61092\nPOSSIBLE N-term signal sequence\noutside 1 359\nTMhelix 360 382\ninside 383 386

32658 GCF_002154555.1_ASM215455v1 Streptomyces murinus Terrabacteria group; Actinobacteria; Actinobacteria; Streptomycetales; Streptomycetaceae; Streptomyces MASRGRRAAMRSLGALVGVAALTALGSNAPGWAATSAHGHGASSTATPIKHVVVLFDENI WP_086811000.1 phospholipase [Streptomyces murinus] Length: 607\nNumber of predicted TMHs: 2\nExp number of AAs in TMHs: 41.50147\nExp number, first 60 AAs: 20.61396\nTotal prob of N-in: 0.91708\nPOSSIBLE N-term signal sequence\ninside 1 12\nTMhelix 13 35\noutside 36 576\nTMhelix 577 599\ninside 600 607

32659 GCF_002154555.1_ASM215455v1 Streptomyces murinus Terrabacteria group; Actinobacteria; Actinobacteria; Streptomycetales; Streptomycetaceae; Streptomyces MLGDVENHSLPRAAAFFDLDKTVIAKSSTLTFSKSFYQGGLINRRAALRTAYAQFVFLAG WP_086812318.1 inhibition of morphological differentiation protein [Streptomyces murinus] Length: 276\nNumber of predicted TMHs: 1\nExp number of AAs in TMHs: 21.51155\nExp number, first 60 AAs: 0.02847\nTotal prob of N-in: 0.00286\noutside 1 247\nTMhelix 248 270\ninside 271 276

32660 GCF_002154555.1_ASM215455v1 Streptomyces murinus Terrabacteria group; Actinobacteria; Actinobacteria; Streptomycetales; Streptomycetaceae; Streptomyces MPAPKNTARRSLLISAATLLSLAVTAPAALAAPAAGKAPAATPPAHMSTLGGARLGLPGT WP_086806635.1 D-alanyl-D-alanine carboxypeptidase [Streptomyces murinus] Length: 407\nNumber of predicted TMHs: 2\nExp number of AAs in TMHs: 44.0074\nExp number, first 60 AAs: 21.46482\nTotal prob of N-in: 0.96880\nPOSSIBLE N-term signal sequence\ninside 1 11\nTMhelix 12 34\noutside 35 368\nTMhelix 369 391\ninside 392 407

32661 GCF_001700505.1_ASM170050v1 Streptomyces mutomycini Terrabacteria group; Actinobacteria; Actinobacteria; Streptomycetales; Streptomycetaceae; Streptomyces MLTFVENCFSPRTAAFFDLDKTVIAKSSTLTFSKSFYQGGLINRRAVLRTAYTQFVFLAG WP_031090886.1 MULTISPECIES: inhibition of morphological differentiation protein [Streptomyces] Length: 279\nNumber of predicted TMHs: 1\nExp number of AAs in TMHs: 21.53635\nExp number, first 60 AAs: 0.01474\nTotal prob of N-in: 0.00168\noutside 1 247\nTMhelix 248 270\ninside 271 279

32662 GCF_001700505.1_ASM170050v1 Streptomyces mutomycini Terrabacteria group; Actinobacteria; Actinobacteria; Streptomycetales; Streptomycetaceae; Streptomyces MRSSRRNTVRRGCVGALLAAVTLLTSAGGAVADDPSASASPADGESPGPTEAGTTFRTAT WP_031100002.1 MULTISPECIES: hypothetical protein [Streptomyces] Length: 275\nNumber of predicted TMHs: 2\nExp number of AAs in TMHs: 37.98301\nExp number, first 60 AAs: 18.54979\nTotal prob of N-in: 0.98601\nPOSSIBLE N-term signal sequence\ninside 1 12\nTMhelix 13 32\noutside 33 239\nTMhelix 240 262\ninside 263 275

32663 GCF_001753705.1_ASM175370v1 Streptomyces nanshensis Terrabacteria group; Actinobacteria; Actinobacteria; Streptomycetales; Streptomycetaceae; Streptomyces MSPRRRRATVLTAATVLSAAAAPVASAAAPAAPVTSVAASPARTTASGLPEGLYGTADPK WP_069991284.1 MULTISPECIES: hypothetical protein [Streptomyces] Length: 443\nNumber of predicted TMHs: 1\nExp number of AAs in TMHs: 34.01561\nExp number, first 60 AAs: 11.7064\nTotal prob of N-in: 0.52682\nPOSSIBLE N-term signal sequence\noutside 1 413\nTMhelix 414 436\ninside 437 443

32664 GCF_001751255.1_ASM175125v1 Streptomyces nanshensis Terrabacteria group; Actinobacteria; Actinobacteria; Streptomycetales; Streptomycetaceae; Streptomyces MLAFVENLAMRRTAAFFDLDKTVIAKSSTLAFGRSFYQGGLINRRAVLRTAYAQFVYLLG WP_070017769.1 inhibition of morphological differentiation protein [Streptomyces nanshensis] Length: 280\nNumber of predicted TMHs: 1\nExp number of AAs in TMHs: 22.09504\nExp number, first 60 AAs: 0.01684\nTotal prob of N-in: 0.00311\noutside 1 246\nTMhelix 247 269\ninside 270 280

32665	GCF_001753705.1_ASM175370v1	Streptomyces nanshensis	Terrabacteria group; Actinobacteria; Actinobacteria; Streptomycetales; Streptomycetaceae; Streptomyces
MLGAVKNLAMPRTAAFFDLDKTVIAKSSTLTFGKSFYQGGLINRRAVLRTAYAQFVYLLG WP_027760567.1 MULTISPECIES: inhibition of morphological differentiation protein [Streptomyces] Length: 281\nNumber of predicted TMHs: 1\nExp number of AAs in TMHs: 22.28973\nExp number, first 60 AAs: 0.03324\nTotal prob of N-in: 0.01080\noutside 1 246\nTMhelix 247 269\ninside 270 281

32666	GCF_001753705.1_ASM175370v1	Streptomyces nanshensis	Terrabacteria group; Actinobacteria; Actinobacteria; Streptomycetales; Streptomycetaceae; Streptomyces
MITRRRLVTGACALLMALPLGLLPTGTASAEPASDEDAPKVELTLDVSGSMRKRDIDGRS WP_026005050.1 MULTISPECIES: inter-alpha-trypsin inhibitor heavy chain H5 [Streptomyces] Length: 421\nNumber of predicted TMHs: 2\nExp number of AAs in TMHs: 38.90213\nExp number, first 60 AAs: 16.54504\nTotal prob of N-in: 0.77694\nPOSSIBLE N-term signal sequence\ninside 1 6\nTMhelix 7 29\noutside 30 388\nTMhelix 389 411\ninside 412 421

32667	GCF_001753705.1_ASM175370v1	Streptomyces nanshensis	Terrabacteria group; Actinobacteria; Actinobacteria; Streptomycetales; Streptomycetaceae; Streptomyces
MIFRMRAFWGMRIDDPWHDALATGGAEVHGTPVDTPPRRAAPPPRREPVPSAVHGVYPTV WP_079163992.1 hypothetical protein [Streptomyces nanshensis] Length: 177\nNumber of predicted TMHs: 2\nExp number of AAs in TMHs: 45.16849\nExp number, first 60 AAs: 0.00025\nTotal prob of N-in: 0.99463\ninside 1 75\nTMhelix 76 98\noutside 99 107\nTMhelix 108 130\ninside 131 177

32668	GCF_001751255.1_ASM175125v1	Streptomyces nanshensis	Terrabacteria group; Actinobacteria; Actinobacteria; Streptomycetales; Streptomycetaceae; Streptomyces
MITHVSGSRARRSALLGAAAVLAGGTFLATPAHAAPAHGHSGSKGKAGATAARADLQVSV WP_070014799.1 hypothetical protein [Streptomyces nanshensis] Length: 308\nNumber of predicted TMHs: 2\nExp number of AAs in TMHs: 43.21283\nExp number, first 60 AAs: 21.89459\nTotal prob of N-in: 0.97828\nPOSSIBLE N-term signal sequence\ninside 1 12\nTMhelix 13 35\noutside 36 277\nTMhelix 278 300\ninside 301 308

32669	GCF_001751305.1_ASM175130v1	Streptomyces nanshensis	Terrabacteria group; Actinobacteria; Actinobacteria; Streptomycetales; Streptomycetaceae; Streptomyces
MLSVVENCFSPRTAAFFDLDKTVIAKSSTLTFSKSFYQGGLINRRAVLRTAYTQFVFLAG WP_070204432.1 inhibition of morphological differentiation protein [Streptomyces nanshensis] Length: 282\nNumber of predicted TMHs: 1\nExp number of AAs in TMHs: 21.55264\nExp number, first 60 AAs: 0.02024\nTotal prob of N-in: 0.00656\noutside 1 247\nTMhelix 248 270\ninside 271 282

32670	GCF_001418645.1_ASM141864v1	Streptomyces neyagawaensis	Terrabacteria group; Actinobacteria; Actinobacteria; Streptomycetales; Streptomycetaceae; Streptomyces
MRRPSRRPFLAGLAVLAGALLAAVPAVPAAADDGDFTVKDPRITESSGLAASRQHPGVYW WP_055541462.1 hypothetical protein [Streptomyces neyagawaensis] Length: 338\nNumber of predicted TMHs: 2\nExp number of AAs in TMHs: 40.41527\nExp number, first 60 AAs: 21.69848\nTotal prob of N-in: 0.97051\nPOSSIBLE N-term signal sequence\ninside 1 8\nTMhelix 9 31\noutside 32 307\nTMhelix 308 330\ninside 331 338

32671	GCF_000718305.1_ASM71830v1	Streptomyces niger	Terrabacteria group; Actinobacteria; Actinobacteria; Streptomycetales; Streptomycetaceae; Streptomyces
MSDTTLSSTAPVATPATTPSRRRKALRRAPLVGGAAAGAVLLVAGPAFAHVTVQPGQAAK WP_052866162.1 hypothetical protein [Streptomyces niger] Length: 264\nNumber of predicted TMHs: 2\nExp number of AAs in TMHs: 44.36695\nExp number, first 60 AAs: 22.51893\nTotal prob of N-in: 0.97697\nPOSSIBLE N-term signal sequence\ninside 1 30\nTMhelix 31 53\noutside 54 236\nTMhelix 237 259\ninside 260 264

32672	GCF_000718305.1_ASM71830v1	Streptomyces niger	Terrabacteria group; Actinobacteria; Actinobacteria; Streptomycetales; Streptomycetaceae; Streptomyces
MTDRRRRAALLLLPMPATAPGVALLPREHKPSSSSEGEDRGREDRGREGRDGDQDNPFAA WP_052863743.1 hypothetical protein [Streptomyces niger] Length: 316\nNumber of predicted TMHs: 2\nExp number of AAs in TMHs: 47.49879\nExp number, first 60 AAs: 0.73676\nTotal prob of N-in: 0.88056\ninside 1 160\nTMhelix 161 183\noutside 184 254\nTMhelix 255 277\ninside 278 316

32673	GCF_001419795.1_ASM141979v1	Streptomyces niveiscabiei	Terrabacteria group; Actinobacteria; Actinobacteria; Streptomycetales; Streptomycetaceae; Streptomyces
MLRDVENHSSPRTAAFFDLDKTVIAKSSTLTFGKSFYQGGLINRRAALRTAYAQFVFLVG WP_055724198.1 inhibition of morphological differentiation protein [Streptomyces niveiscabiei] Length: 275\nNumber of predicted TMHs: 1\nExp

number of AAs in TMHs: 21.82896\nExp number, first 60 AAs: 0.09472\nTotal prob of N-in: 0.10028\noutside 1 245\nTMhelix 246 268\ninside 269 275

32674 GCF_001723055.1_ASM172305v1 Streptomyces niveus Terrabacteria group; Actinobacteria; Actinobacteria; Streptomycetales; Streptomycetaceae; Streptomyces MNASRTNTSRRGVSRPSTVGRTRTARRAAVVGGVAASAVVLMAGPALAHVGVQPVGEAAK WP_079127887.1 hypothetical protein [Streptomyces niveus] Length: 281\nNumber of predicted TMHs: 2\nExp number of AAs in TMHs: 43.33022\nExp number, first 60 AAs: 22.41672\nTotal prob of N-in: 0.99726\nPOSSIBLE N-term signal sequence\ninside 1 29\nTMhelix 30 52\noutside 53 244\nTMhelix 245 267\ninside 268 281

32675 GCF_001723055.1_ASM172305v1 Streptomyces niveus Terrabacteria group; Actinobacteria; Actinobacteria; Streptomycetales; Streptomycetaceae; Streptomyces MLGIVENHSLPRTAAFFDLDKTVIAKSSTLTFSKSFYQGGLINRRAVLRTAYTQFVFLAG WP_023539673.1 inhibition of morphological differentiation protein [Streptomyces niveus] Length: 278\nNumber of predicted TMHs: 1\nExp number of AAs in TMHs: 21.70268\nExp number, first 60 AAs: 0.01114\nTotal prob of N-in: 0.00185\noutside 1 247\nTMhelix 248 270\ninside 271 278

32676 GCF_002009175.1_ASM200917v1 Streptomyces niveus Terrabacteria group; Actinobacteria; Actinobacteria; Streptomycetales; Streptomycetaceae; Streptomyces MLDFVENHSLPRTAAFFDLDKTVIAKSSTLTFSKSFYQGGLINRRAVLRTAYTQFVFLAG WP_078075838.1 inhibition of morphological differentiation protein [Streptomyces niveus] Length: 278\nNumber of predicted TMHs: 1\nExp number of AAs in TMHs: 21.73904\nExp number, first 60 AAs: 0.01087\nTotal prob of N-in: 0.00142\noutside 1 247\nTMhelix 248 270\ninside 271 278

32677 GCF_002009175.1_ASM200917v1 Streptomyces niveus Terrabacteria group; Actinobacteria; Actinobacteria; Streptomycetales; Streptomycetaceae; Streptomyces MNASRTNTSRRGVPQPSTAGRTRTARRAAVVGGVAASAVVLLAGPALAHVGVQPVGEAAK WP_078076280.1 hypothetical protein [Streptomyces niveus] Length: 284\nNumber of predicted TMHs: 2\nExp number of AAs in TMHs: 42.74269\nExp number, first 60 AAs: 22.46781\nTotal prob of N-in: 0.99780\nPOSSIBLE N-term signal sequence\ninside 1 29\nTMhelix 30 52\noutside 53 245\nTMhelix 246 268\ninside 269 284

32678 GCF_001723075.1_ASM172307v1 Streptomyces niveus Terrabacteria group; Actinobacteria; Actinobacteria; Streptomycetales; Streptomycetaceae; Streptomyces MLGIVENHSLPRTAAFFDLDKTVIAKSSTLTFSKSFYQGGLINRRAVLRTAYTQFVFLAG WP_023539673.1 inhibition of morphological differentiation protein [Streptomyces niveus] Length: 278\nNumber of predicted TMHs: 1\nExp number of AAs in TMHs: 21.70268\nExp number, first 60 AAs: 0.01114\nTotal prob of N-in: 0.00185\noutside 1 247\nTMhelix 248 270\ninside 271 278

32679 GCF_001723075.1_ASM172307v1 Streptomyces niveus Terrabacteria group; Actinobacteria; Actinobacteria; Streptomycetales; Streptomycetaceae; Streptomyces MNASRTNTSRRGVSRPSTVGRTRTARRAAVVGGVAASAVVLMAGPALAHVGVQPVGEAAK WP_023540332.1 hypothetical protein [Streptomyces niveus] Length: 281\nNumber of predicted TMHs: 2\nExp number of AAs in TMHs: 43.33007\nExp number, first 60 AAs: 22.41673\nTotal prob of N-in: 0.99726\nPOSSIBLE N-term signal sequence\ninside 1 29\nTMhelix 30 52\noutside 53 244\nTMhelix 245 267\ninside 268 281

32680 GCF_000819545.1_ASM81954v1 Streptomyces nodosus Terrabacteria group; Actinobacteria; Actinobacteria; Streptomycetales; Streptomycetaceae; Streptomyces MDKTVIAKSSTLTFSKSFYQGGLINRRAALRTAYTQFVFLAGGADHDQMERMREYLSAMC WP_043448926.1 inhibition of morphological differentiation protein [Streptomyces nodosus] Length: 256\nNumber of predicted TMHs: 1\nExp number of AAs in TMHs: 21.54796\nExp number, first 60 AAs: 0.10931\nTotal prob of N-in: 0.04477\noutside 1 226\nTMhelix 227 249\ninside 250 256

32681 GCF_000819545.1_ASM81954v1 Streptomyces nodosus Terrabacteria group; Actinobacteria; Actinobacteria; Streptomycetales; Streptomycetaceae; Streptomyces MGHARRRIVRRGTRIAAIGGVLLGGLMVTQALASEPSGGRGSGDSSARAASRTAADLVSK WP_043446940.1 protease [Streptomyces nodosus] Length: 459\nNumber of predicted TMHs: 2\nExp number of AAs in TMHs: 39.26784\nExp number, first 60 AAs: 20.76196\nTotal prob of N-in: 0.96272\nPOSSIBLE N-term signal sequence\ninside 1 11\nTMhelix 12 34\noutside 35 420\nTMhelix 421 443\ninside 444 459

32682 GCF_000819545.1_ASM81954v1 Streptomyces nodosus Terrabacteria group; Actinobacteria; Actinobacteria; Streptomycetales; Streptomycetaceae; Streptomyces

MTRALPCLCDPAVCRRQLPLYGPRVRESSPLPPAARMSRRAVLGLGAAVPLTLAAPAAAS WP_052454075.1 D-alanyl-D-alanine carboxypeptidase [Streptomyces nodosus] Length: 429\nNumber of predicted TMHs: 1\nExp number of AAs in TMHs: 23.23766\nExp number, first 60 AAs: 2.18008\nTotal prob of N-in: 0.11792\noutside 1 390\nTMhelix 391 413\ninside 414 429

32683 GCF_000819545.1_ASM81954v1 Streptomyces nodosus Terrabacteria group; Actinobacteria; Actinobacteria; Streptomycetales; Streptomycetaceae; Streptomyces MKRNSRRRPTGARRATFAAVALIVGGGGLIAANVYASASEVGWGGANDGEQTVSSETVTI WP_043437661.1 hypothetical protein [Streptomyces nodosus] Length: 676\nNumber of predicted TMHs: 2\nExp number of AAs in TMHs: 39.54792\nExp number, first 60 AAs: 21.53649\nTotal prob of N-in: 0.99443\nPOSSIBLE N-term signal sequence\ninside 1 16\nTMhelix 17 39\noutside 40 653\nTMhelix 654 671\ninside 672 676

32684 GCF_000716875.1_ASM71687v1 Streptomyces novaecaesareae Terrabacteria group; Actinobacteria; Actinobacteria; Streptomycetales; Streptomycetaceae; Streptomyces MRTAAFFDLDKTIIAKSSALAFSRPFYQGGLINRRAVLRSAYAQFVFLVGGADHDQMEKM WP_030259263.1 MULTISPECIES: inhibition of morphological differentiation protein [Streptomyces] Length: 265\nNumber of predicted TMHs: 1\nExp number of AAs in TMHs: 22.6499\nExp number, first 60 AAs: 0.08965\nTotal prob of N-in: 0.01242\noutside 1 235\nTMhelix 236 258\ninside 259 265

32685 GCF_001751245.1_ASM175124v1 Streptomyces oceani Terrabacteria group; Actinobacteria; Actinobacteria; Streptomycetales; Streptomycetaceae; Streptomyces MLALVEIPSLPRTAAFFDLDKTVIAKSSTLTFGKSFYRGGLINRRAVLRTAYAQFVYLAG WP_070198729.1 inhibition of morphological differentiation protein [Streptomyces oceani] Length: 274\nNumber of predicted TMHs: 1\nExp number of AAs in TMHs: 22.51975\nExp number, first 60 AAs: 0.01\nTotal prob of N-in: 0.00192\noutside 1 247\nTMhelix 248 270\ninside 271 274

32686 GCF_000720485.1_ASM72048v1 Streptomyces ochraceiscleroticus Terrabacteria group; Actinobacteria; Actinobacteria; Streptomycetales; Streptomycetaceae; Streptomyces MRAFSHSPCPSCLRSVLTMSHNTAVSGPSRRRTVLRRLPVVGGAAAGTVLLLAGPAFAHV WP_078648520.1 hypothetical protein [Streptomyces ochraceiscleroticus] Length: 273\nNumber of predicted TMHs: 2\nExp number of AAs in TMHs: 43.78067\nExp number, first 60 AAs: 20.61193\nTotal prob of N-in: 0.92273\nPOSSIBLE N-term signal sequence\ninside 1 39\nTMhelix 40 62\noutside 63 245\nTMhelix 246 268\ninside 269 273

32687 GCF_000696115.1_SoWt_assembly Streptomyces olindensis Terrabacteria group; Actinobacteria; Actinobacteria; Streptomycetales; Streptomycetaceae; Streptomyces MPAPKKTARRSLLVTSAALSSLALAAPAAVAAPSPSPSPSGSPSASPSATPPASMSTVGG WP_031119869.1 MULTISPECIES: D-alanyl-D-alanine carboxypeptidase [Streptomyces] Length: 439\nNumber of predicted TMHs: 1\nExp number of AAs in TMHs: 36.6566799999999\nExp number, first 60 AAs: 14.09768\nTotal prob of N-in: 0.67577\nPOSSIBLE N-term signal sequence\noutside 1 400\nTMhelix 401 423\ninside 424 439

32688 GCF_000696115.1_SoWt_assembly Streptomyces olindensis Terrabacteria group; Actinobacteria; Actinobacteria; Streptomycetales; Streptomycetaceae; Streptomyces MAVLSVSRRTTRRRTVARSVRILGVASASAALALGAAGNALACNINEFSAEAKCDGDKGV WP_051648926.1 hypothetical protein [Streptomyces olindensis] Length: 237\nNumber of predicted TMHs: 2\nExp number of AAs in TMHs: 43.85284\nExp number, first 60 AAs: 21.83045\nTotal prob of N-in: 0.98574\nPOSSIBLE N-term signal sequence\ninside 1 19\nTMhelix 20 42\noutside 43 205\nTMhelix 206 228\ninside 229 237

32689 GCF_000696115.1_SoWt_assembly Streptomyces olindensis Terrabacteria group; Actinobacteria; Actinobacteria; Streptomycetales; Streptomycetaceae; Streptomyces MLKGVENHSLPRTAAFFDLDKTVIAKSSTLTFSKSFYQGGLINRRAVLRTAYAQFVFLAG WP_031115976.1 MULTISPECIES: inhibition of morphological differentiation protein [Streptomyces] Length: 277\nNumber of predicted TMHs: 1\nExp number of AAs in TMHs: 21.58823\nExp number, first 60 AAs: 0.04125\nTotal prob of N-in: 0.00423\noutside 1 247\nTMhelix 248 270\ninside 271 277

32690 GCF_000696115.1_SoWt_assembly Streptomyces olindensis Terrabacteria group; Actinobacteria; Actinobacteria; Streptomycetales; Streptomycetaceae; Streptomyces MRHARRRVVRRVTRLAAVGGLLLGGAMVTRAVASEPPDPSIGVPYSARPATGMGTDLVSR WP_031120420.1 MULTISPECIES: protease [Streptomyces] Length: 450\nNumber of predicted TMHs: 2\nExp number of AAs in TMHs: 41.20976\nExp number, first 60 AAs: 20.68383\nTotal prob of N-in: 0.95239\nPOSSIBLE N-term signal sequence\ninside 1 11\nTMhelix 12 34\noutside 35 411\nTMhelix 412 434\ninside 435 450

32691 GCF_000716815.1_ASM71681v1 Streptomyces olivaceus Terrabacteria group; Actinobacteria; Actinobacteria; Streptomycetales; Streptomycetaceae; Streptomyces
MLTFVENCFSPRTAAFFDLDKTVIAKSSTLTFSKSFYQGGLINRRAVLRTAYTQFVFLAG WP_014155461.1 MULTISPECIES: haloacid dehalogenase [Streptomyces] Length: 279\nNumber of predicted TMHs: 1\nExp number of AAs in TMHs: 21.10966\nExp number, first 60 AAs: 0.01497\nTotal prob of N-in: 0.00234\noutside 1 248\nTMhelix 249 268\ninside 269 279

32692 GCF_000721235.1_ASM72123v1 Streptomyces olivaceus Terrabacteria group; Actinobacteria; Actinobacteria; Streptomycetales; Streptomycetaceae; Streptomyces
MRHARRRIVRRAARLAAVGGLLIGGTMVTRAVASEPADAPAAPHTLARSASGAGAGLVER WP_031031837.1 protease [Streptomyces olivaceus] Length: 453\nNumber of predicted TMHs: 2\nExp number of AAs in TMHs: 39.99145\nExp number, first 60 AAs: 20.05262\nTotal prob of N-in: 0.94476\nPOSSIBLE N-term signal sequence\ninside 1 11\nTMhelix 12 34\noutside 35 414\nTMhelix 415 437\ninside 438 453

32693 GCF_000721235.1_ASM72123v1 Streptomyces olivaceus Terrabacteria group; Actinobacteria; Actinobacteria; Streptomycetales; Streptomycetaceae; Streptomyces
MSESVTTQEHVLGHNRRRRPTGARRATFGAVALILGGGGLVAVNVFASATESGNTAVPLG WP_078892164.1 hypothetical protein [Streptomyces olivaceus] Length: 700\nNumber of predicted TMHs: 2\nExp number of AAs in TMHs: 43.48821\nExp number, first 60 AAs: 22.40841\nTotal prob of N-in: 0.99501\nPOSSIBLE N-term signal sequence\ninside 1 25\nTMhelix 26 48\noutside 49 672\nTMhelix 673 695\ninside 696 700

32694 GCF_000716815.1_ASM71681v1 Streptomyces olivaceus Terrabacteria group; Actinobacteria; Actinobacteria; Streptomycetales; Streptomycetaceae; Streptomyces
MRTTRRNTLRRGGVGALLAAVTLLTSAGAAVADDPSPSASAAGGTAGPTEAGTTFRTATA WP_033249102.1 MULTISPECIES: hypothetical protein [Streptomyces] Length: 274\nNumber of predicted TMHs: 2\nExp number of AAs in TMHs: 39.9840299999999\nExp number, first 60 AAs: 19.24596\nTotal prob of N-in: 0.99309\nPOSSIBLE N-term signal sequence\ninside 1 12\nTMhelix 13 32\noutside 33 237\nTMhelix 238 260\ninside 261 274

32695 GCF_000716815.1_ASM71681v1 Streptomyces olivaceus Terrabacteria group; Actinobacteria; Actinobacteria; Streptomycetales; Streptomycetaceae; Streptomyces
MTTHPSPRRRLLTAAAATAVLLVTLPVLPAAAEDSTQCTFPSKKYAGRPWSLQRVLMDEL WP_033249645.1 MULTISPECIES: type VII secretion-associated serine protease [Streptomyces] Length: 414\nNumber of predicted TMHs: 1\nExp number of AAs in TMHs: 36.07361\nExp number, first 60 AAs: 13.5063\nTotal prob of N-in: 0.65550\nPOSSIBLE N-term signal sequence\noutside 1 378\nTMhelix 379 401\ninside 402 414

32696 GCF_001514115.1_ASM151411v1 Streptomyces olivochromogenes Terrabacteria group; Actinobacteria; Actinobacteria; Streptomycetales; Streptomycetaceae; Streptomyces
MRDSSPVPPRAAQLSRRAALGLAAAVPLAAASPASAATPVIGGERLARAGVQVGAAAGLP WP_079064839.1 D-alanyl-D-alanine carboxypeptidase [Streptomyces olivochromogenes] Length: 426\nNumber of predicted TMHs: 1\nExp number of AAs in TMHs: 27.56408\nExp number, first 60 AAs: 8.95229999999999\nTotal prob of N-in: 0.39774\noutside 1 355\nTMhelix 356 378\ninside 379 426

32697 GCF_001514115.1_ASM151411v1 Streptomyces olivochromogenes Terrabacteria group; Actinobacteria; Actinobacteria; Streptomycetales; Streptomycetaceae; Streptomyces
MLGLVENHSLPRTAAFFDLDKTVIAKSSTLTFSKSFYQGGLINRRAVLRTAYAQFVFMAG WP_067373858.1 inhibition of morphological differentiation protein [Streptomyces olivochromogenes] Length: 277\nNumber of predicted TMHs: 1\nExp number of AAs in TMHs: 21.53459\nExp number, first 60 AAs: 0.01511\nTotal prob of N-in: 0.00191\noutside 1 247\nTMhelix 248 270\ninside 271 277

32698 GCF_001514115.1_ASM151411v1 Streptomyces olivochromogenes Terrabacteria group; Actinobacteria; Actinobacteria; Streptomycetales; Streptomycetaceae; Streptomyces
MAANRRRPLALAAAVAVALGAAALAVPHASAAEAPLKDYELTWGIKQSYRAYVTGMAAGT WP_067363291.1 hypothetical protein [Streptomyces olivochromogenes] Length: 495\nNumber of predicted TMHs: 2\nExp number of AAs in TMHs: 43.31557\nExp number, first 60 AAs: 21.23139\nTotal prob of N-in: 0.96724\nPOSSIBLE N-term signal sequence\ninside 1 8\nTMhelix 9 31\noutside 32 466\nTMhelix 467 489\ninside 490 495

32699 GCF_001514115.1_ASM151411v1 Streptomyces olivochromogenes Terrabacteria group; Actinobacteria; Actinobacteria; Streptomycetales; Streptomycetaceae; Streptomyces
MRHARRRIVRRGARLAAVGGVLCGGLMVTHAMASEPSAASRAPQSATLAAAGKGAGLVST WP_067360011.1 protease [Streptomyces olivochromogenes] Length: 460\nNumber of predicted TMHs: 2\nExp number of

AAs in TMHs: 42.6832299999999\nExp number, first 60 AAs: 21.30253\nTotal prob of N-in: 0.98480\nPOSSIBLE N-term signal sequence\ninside 1 11\nTMhelix 12 34\noutside 35 421\nTMhelix 422 444\ninside 445 460

32700 GCF_001418475.1_ASM141847v1 Streptomyces ossamyceticus Terrabacteria group; Actinobacteria; Actinobacteria; Streptomycetales; Streptomycetaceae; Streptomyces MRRPSRRPLLAGLAVLAGASLAAVPAVPAAADDGNFTVKDPRITESSGLAASRQHPGVYW WP_055522229.1 hypothetical protein [Streptomyces ossamyceticus] Length: 339\nNumber of predicted TMHs: 2\nExp number of AAs in TMHs: 37.08029\nExp number, first 60 AAs: 19.53523\nTotal prob of N-in: 0.87747\nPOSSIBLE N-term signal sequence\ninside 1 8\nTMhelix 9 31\noutside 32 308\nTMhelix 309 331\ninside 332 339

32701 GCF_001418475.1_ASM141847v1 Streptomyces ossamyceticus Terrabacteria group; Actinobacteria; Actinobacteria; Streptomycetales; Streptomycetaceae; Streptomyces MRHARRRVVKRVARLAAVGGLLCGGLMVTQAAMATESTTSPPATRSSVLAASGTGTGLVS WP_055520900.1 protease [Streptomyces ossamyceticus] Length: 457\nNumber of predicted TMHs: 2\nExp number of AAs in TMHs: 38.19913\nExp number, first 60 AAs: 22.21382\nTotal prob of N-in: 0.99347\nPOSSIBLE N-term signal sequence\ninside 1 11\nTMhelix 12 34\noutside 35 418\nTMhelix 419 441\ninside 442 457

32702 GCF_001418475.1_ASM141847v1 Streptomyces ossamyceticus Terrabacteria group; Actinobacteria; Actinobacteria; Streptomycetales; Streptomycetaceae; Streptomyces MTATVVTAGDPAPASSAPARTRRARRHRRTALLALPLAFSLALLPATAAHADGIRDQQWA WP_055521889.1 type VII secretion-associated serine protease mycosin [Streptomyces ossamyceticus] Length: 415\nNumber of predicted TMHs: 2\nExp number of AAs in TMHs: 42.37886\nExp number, first 60 AAs: 20.80408\nTotal prob of N-in: 0.98754\nPOSSIBLE N-term signal sequence\ninside 1 31\nTMhelix 32 54\noutside 55 380\nTMhelix 381 403\ninside 404 415

32703 GCF_001414545.1_ASM141454v1 Streptomyces pactum Terrabacteria group; Actinobacteria; Actinobacteria; Streptomycetales; Streptomycetaceae; Streptomyces MGHNRRRRPTGARRATFGAFALILGGGGLVAVNVFASATEAGDTAVPLGSSGVAATVDCP WP_079160679.1 hypothetical protein [Streptomyces pactum] Length: 716\nNumber of predicted TMHs: 2\nExp number of AAs in TMHs: 43.25514\nExp number, first 60 AAs: 22.02209\nTotal prob of N-in: 0.99646\nPOSSIBLE N-term signal sequence\ninside 1 16\nTMhelix 17 39\noutside 40 688\nTMhelix 689 711\ninside 712 716

32704 GCF_001414545.1_ASM141454v1 Streptomyces pactum Terrabacteria group; Actinobacteria; Actinobacteria; Streptomycetales; Streptomycetaceae; Streptomyces MRHARRRRIVRRVARLAAVGGLLVGGTMVTRAVASEPPDTSAAPHTSARPASGTGTGAGL WP_055421240.1 protease [Streptomyces pactum] Length: 457\nNumber of predicted TMHs: 2\nExp number of AAs in TMHs: 42.76973\nExp number, first 60 AAs: 21.51108\nTotal prob of N-in: 0.98654\nPOSSIBLE N-term signal sequence\ninside 1 12\nTMhelix 13 35\noutside 36 418\nTMhelix 419 441\ninside 442 457

32705 GCF_001767375.1_ASM176737v1 Streptomyces pactum Terrabacteria group; Actinobacteria; Actinobacteria; Streptomycetales; Streptomycetaceae; Streptomyces MSESVTTQEHVLGHNRRRRPTGARRATFGAVALILGGGGLVAVNVFASATESGNTAVPLG WP_079155135.1 hypothetical protein [Streptomyces pactum] Length: 700\nNumber of predicted TMHs: 2\nExp number of AAs in TMHs: 43.48821\nExp number, first 60 AAs: 22.40841\nTotal prob of N-in: 0.99501\nPOSSIBLE N-term signal sequence\ninside 1 25\nTMhelix 26 48\noutside 49 672\nTMhelix 673 695\ninside 696 700

32706 GCF_001767375.1_ASM176737v1 Streptomyces pactum Terrabacteria group; Actinobacteria; Actinobacteria; Streptomycetales; Streptomycetaceae; Streptomyces MRHARRRIVRRAARLAAVGGLLIGGTMVTRAVASEPADAPAAPHTLARSASGAGAGLVER WP_070389912.1 serine protease [Streptomyces pactum] Length: 453\nNumber of predicted TMHs: 2\nExp number of AAs in TMHs: 39.99153\nExp number, first 60 AAs: 20.05268\nTotal prob of N-in: 0.94476\nPOSSIBLE N-term signal sequence\ninside 1 11\nTMhelix 12 34\noutside 35 414\nTMhelix 415 437\ninside 438 453

32707 GCF_001414545.1_ASM141454v1 Streptomyces pactum Terrabacteria group; Actinobacteria; Actinobacteria; Streptomycetales; Streptomycetaceae; Streptomyces MPHSKPRTAAFFDLDKTVIAKSSTLTFSKSFYQGGLINRRAVLRTAYAQFVFLAGGADHD WP_055418670.1 inhibition of morphological differentiation protein [Streptomyces pactum] Length: 272\nNumber of predicted TMHs: 1\nExp number of AAs in TMHs: 21.31692\nExp number, first 60 AAs: 0.1342\nTotal prob of N-in: 0.03488\noutside 1 242\nTMhelix 243 265\ninside 266 272

32708 GCF_002005225.1_ASM200522v1 Streptomyces pactum Terrabacteria group; Actinobacteria; Actinobacteria; Streptomycetales; Streptomycetaceae; Streptomyces
MGHNRRRRPTGARRATFGAFALILGGGGLVAVNVFASATEAGDTAVPLGSSGVAATVDCP WP_079160679.1 hypothetical protein [Streptomyces pactum] Length: 716\nNumber of predicted TMHs: 2\nExp number of AAs in TMHs: 43.25514\nExp number, first 60 AAs: 22.02209\nTotal prob of N-in: 0.99646\nPOSSIBLE N-term signal sequence\ninside 1 16\nTMhelix 17 39\noutside 40 688\nTMhelix 689 711\ninside 712 716

32709 GCF_002005225.1_ASM200522v1 Streptomyces pactum Terrabacteria group; Actinobacteria; Actinobacteria; Streptomycetales; Streptomycetaceae; Streptomyces
MRHARRRRIVRRVARLAAVGGLLVGGTMVTRAVASEPPDTSAAPHTSARPASGTGTGAGL WP_055421240.1 protease [Streptomyces pactum] Length: 457\nNumber of predicted TMHs: 2\nExp number of AAs in TMHs: 42.76973\nExp number, first 60 AAs: 21.51108\nTotal prob of N-in: 0.98654\nPOSSIBLE N-term signal sequence\ninside 1 12\nTMhelix 13 35\noutside 36 418\nTMhelix 419 441\ninside 442 457

32710 GCF_002005225.1_ASM200522v1 Streptomyces pactum Terrabacteria group; Actinobacteria; Actinobacteria; Streptomycetales; Streptomycetaceae; Streptomyces
MPHSKPRTAAFFDLDKTVIAKSSTLTFSKSFYQGGLINRRAVLRTAYAQFVFLAGGADHD WP_055418670.1 inhibition of morphological differentiation protein [Streptomyces pactum] Length: 272\nNumber of predicted TMHs: 1\nExp number of AAs in TMHs: 21.31692\nExp number, first 60 AAs: 0.1342\nTotal prob of N-in: 0.03488\noutside 1 242\nTMhelix 243 265\ninside 266 272

32711 GCF_001660045.1_ASM166004v1 Streptomyces parvulus Terrabacteria group; Actinobacteria; Actinobacteria; Streptomycetales; Streptomycetaceae; Streptomyces
MRHARRRIVRRVARLAAVGGLLVGGTMVTRAVASEPPDASAAPHTLSRTASGAGLVAELG WP_064731184.1 protease [Streptomyces parvulus] Length: 446\nNumber of predicted TMHs: 2\nExp number of AAs in TMHs: 40.95714\nExp number, first 60 AAs: 20.49447\nTotal prob of N-in: 0.95321\nPOSSIBLE N-term signal sequence\ninside 1 11\nTMhelix 12 34\noutside 35 407\nTMhelix 408 430\ninside 431 446

32712 GCF_001660045.1_ASM166004v1 Streptomyces parvulus Terrabacteria group; Actinobacteria; Actinobacteria; Streptomycetales; Streptomycetaceae; Streptomyces
MPRTAAFFDLDKTVIAKSSTLTFSKSFYQGGLINRRAVLRTAYAQFVFLAGGADHDQMER WP_064729057.1 inhibition of morphological differentiation protein [Streptomyces parvulus] Length: 268\nNumber of predicted TMHs: 1\nExp number of AAs in TMHs: 21.5878\nExp number, first 60 AAs: 0.03456\nTotal prob of N-in: 0.00383\noutside 1 238\nTMhelix 239 261\ninside 262 268

32713 GCF_001418495.1_ASM141849v1 Streptomyces pathocidini Terrabacteria group; Actinobacteria; Actinobacteria; Streptomycetales; Streptomycetaceae; Streptomyces
MSSPDREVRAARNGAAAGGRRPIVAVTGAATGIGDLLTRRLAESEEIKQVVAIDERRGEV WP_055470700.1 NAD-dependent dehydratase [Streptomyces pathocidini] Length: 369\nNumber of predicted TMHs: 1\nExp number of AAs in TMHs: 19.6922\nExp number, first 60 AAs: 0.11172\nTotal prob of N-in: 0.01626\noutside 1 339\nTMhelix 340 359\ninside 360 369

32714 GCF_001418495.1_ASM141849v1 Streptomyces pathocidini Terrabacteria group; Actinobacteria; Actinobacteria; Streptomycetales; Streptomycetaceae; Streptomyces
MLASVEKHSLPRTAAFFDLDKTVIAKSSTLTFSKSFYQGGLINRRGVLRSAYAQFVYLVG WP_055474294.1 inhibition of morphological differentiation protein [Streptomyces pathocidini] Length: 278\nNumber of predicted TMHs: 1\nExp number of AAs in TMHs: 21.36097\nExp number, first 60 AAs: 0.01012\nTotal prob of N-in: 0.00173\noutside 1 246\nTMhelix 247 269\ninside 270 278

32715 GCF_001418495.1_ASM141849v1 Streptomyces pathocidini Terrabacteria group; Actinobacteria; Actinobacteria; Streptomycetales; Streptomycetaceae; Streptomyces
MSHCNFSDVSAFLAASPDASPSAGFGRSASGSGRSASVPARRRPVPRYHGAVLTIAAAAL WP_079101010.1 D-alanyl-D-alanine carboxypeptidase [Streptomyces pathocidini] Length: 469\nNumber of predicted TMHs: 2\nExp number of AAs in TMHs: 40.4948999999999\nExp number, first 60 AAs: 9.34577\nTotal prob of N-in: 0.75874\ninside 1 49\nTMhelix 50 72\noutside 73 425\nTMhelix 426 448\ninside 449 469

32716 GCF_900142575.1_IMG-taxon_2667527448_annotated_assembly Streptomyces paucisporeus Terrabacteria group; Actinobacteria; Actinobacteria; Streptomycetales; Streptomycetaceae; Streptomyces
MPLRRRAALVAAAATTLTALTAPIALADTTAPAATTAAAPSRGDLYGTADPTYDGVWRQS WP_073494280.1 hypothetical protein [Streptomyces paucisporeus] Length: 419\nNumber of predicted TMHs: 1\nExp

number of AAs in TMHs: 36.38326\nExp number, first 60 AAs: 14.19238\nTotal prob of N-in: 0.68180\nPOSSIBLE N-term signal sequence\noutside 1 390\nTMhelix 391 413\ninside 414 419

32717 GCF_900142575.1_IMG-taxon_2667527448_annotated_assembly Streptomyces paucisporeus Terrabacteria group; Actinobacteria; Actinobacteria; Streptomycetales; Streptomycetaceae; Streptomyces MGRRRPPPRRRPLPRPAAGLLIAALTGLAAAALLPATAAWAAPPVAGSQDGSLIMVLDSS WP_079189651.1 hypothetical protein [Streptomyces paucisporeus] Length: 662\nNumber of predicted TMHs: 2\nExp number of AAs in TMHs: 42.97249\nExp number, first 60 AAs: 22.50413\nTotal prob of N-in: 0.98881\nPOSSIBLE N-term signal sequence\ninside 1 19\nTMhelix 20 42\noutside 43 629\nTMhelix 630 649\ninside 650 662

32718 GCF_900142575.1_IMG-taxon_2667527448_annotated_assembly Streptomyces paucisporeus Terrabacteria group; Actinobacteria; Actinobacteria; Streptomycetales; Streptomycetaceae; Streptomyces MSINRRTAIAVRSIGVASAAAALALGVAGSAFACSITEFTPSASCDDSHHGVIQVLDQDG WP_073497979.1 hypothetical protein [Streptomyces paucisporeus] Length: 217\nNumber of predicted TMHs: 2\nExp number of AAs in TMHs: 42.95629\nExp number, first 60 AAs: 21.54418\nTotal prob of N-in: 0.98236\nPOSSIBLE N-term signal sequence\ninside 1 12\nTMhelix 13 35\noutside 36 185\nTMhelix 186 208\ninside 209 217

32719 GCF_000725565.1_Doro.v1.0Streptomyces peucetius Terrabacteria group; Actinobacteria; Actinobacteria; Streptomycetales; Streptomycetaceae; Streptomyces MTERSAPAASATGPGTPARRTAVRAAATGTLAAALLGLAPQSALAATPAPPPAPAISAVA WP_032915072.1 MULTISPECIES: hypothetical protein [Streptomyces] Length: 293\nNumber of predicted TMHs: 1\nExp number of AAs in TMHs: 23.67473\nExp number, first 60 AAs: 4.47659\nTotal prob of N-in: 0.22667\noutside 1 264\nTMhelix 265 287\ninside 288 293

32720 GCF_000725565.1_Doro.v1.0Streptomyces peucetius Terrabacteria group; Actinobacteria; Actinobacteria; Streptomycetales; Streptomycetaceae; Streptomyces MTLTRRRHLVRSTAVAAVAGSALLLPAAAAFADSPQPTTAGSPGAAGDQQQKDDRQKDDQ WP_031191693.1 MULTISPECIES: hypothetical protein [Streptomyces] Length: 250\nNumber of predicted TMHs: 2\nExp number of AAs in TMHs: 40.95506\nExp number, first 60 AAs: 19.80107\nTotal prob of N-in: 0.94361\nPOSSIBLE N-term signal sequence\ninside 1 12\nTMhelix 13 32\noutside 33 218\nTMhelix 219 241\ninside 242 250

32721 GCF_000725565.1_Doro.v1.0Streptomyces peucetius Terrabacteria group; Actinobacteria; Actinobacteria; Streptomycetales; Streptomycetaceae; Streptomyces MESRQPVGPRANGHRAAHRRRAAVALAGAVCAVGLTATTAAPALADTNVPLNNGQCNFGA WP_078586913.1 MULTISPECIES: type VII secretion-associated serine protease mycosin [Streptomyces] Length: 427\nNumber of predicted TMHs: 2\nExp number of AAs in TMHs: 43.8948\nExp number, first 60 AAs: 21.08952\nTotal prob of N-in: 0.94565\nPOSSIBLE N-term signal sequence\ninside 1 20\nTMhelix 21 43\noutside 44 389\nTMhelix 390 412\ninside 413 427

32722 GCF_000725565.1_Doro.v1.0Streptomyces peucetius Terrabacteria group; Actinobacteria; Actinobacteria; Streptomycetales; Streptomycetaceae; Streptomyces MTRRPLLRAAGVALSAAALTVLPALPAHADGIRAQEWALKAMNTQDAWETTKGEGITVAV WP_031187823.1 MULTISPECIES: type VII secretion-associated serine protease [Streptomyces] Length: 386\nNumber of predicted TMHs: 1\nExp number of AAs in TMHs: 36.26455\nExp number, first 60 AAs: 9.94874\nTotal prob of N-in: 0.56610\noutside 1 359\nTMhelix 360 382\ninside 383 386

32723 GCF_001418655.1_ASM141865v1 Streptomyces phaeochromogenes Terrabacteria group; Actinobacteria; Actinobacteria; Streptomycetales; Streptomycetaceae; Streptomyces MLDPVENHSLPRTAAFFDLDKTVIAKSSTLTFSKSFYQGGLINRRAVLRTAYAQFVFLAG WP_055613406.1 inhibition of morphological differentiation protein [Streptomyces phaeochromogenes] Length: 274\nNumber of predicted TMHs: 1\nExp number of AAs in TMHs: 21.51672\nExp number, first 60 AAs: 0.04416\nTotal prob of N-in: 0.00900\noutside 1 244\nTMhelix 245 267\ninside 268 274

32724 GCF_001418655.1_ASM141865v1 Streptomyces phaeochromogenes Terrabacteria group; Actinobacteria; Actinobacteria; Streptomycetales; Streptomycetaceae; Streptomyces MAANRRRPIALAAACATAVTLGATALAATSASAAEVPLKGYEFTWGIKQSYRTYVTGMAA WP_055612260.1 hypothetical protein [Streptomyces phaeochromogenes] Length: 491\nNumber of predicted TMHs: 2\nExp number of AAs in TMHs: 43.1126299999999\nExp number, first 60 AAs: 20.95836\nTotal prob of N-in: 0.92905\nPOSSIBLE N-term signal sequence\ninside 1 8\nTMhelix 9 31\noutside 32 458\nTMhelix 459 481\ninside 482 491

32725 GCF_001896135.1_ASM189613v1 Streptomyces phaeoluteigriseus Terrabacteria group; Actinobacteria; Actinobacteria; Streptomycetales; Streptomycetaceae; Streptomyces
MSRITLPRGTRRTPLAVAAATTAVLLAAVPAAAHVEVESPGAQALAENVRLTFAAESESA WP_073499707.1 hypothetical protein [Streptomyces phaeoluteigriseus] Length: 247\nNumber of predicted TMHs: 2\nExp number of AAs in TMHs: 42.36018\nExp number, first 60 AAs: 20.79144\nTotal prob of N-in: 0.92652\nPOSSIBLE N-term signal sequence\ninside 1 12\nTMhelix 13 35\noutside 36 216\nTMhelix 217 239\ninside 240 247

32726 GCF_001896135.1_ASM189613v1 Streptomyces phaeoluteigriseus Terrabacteria group; Actinobacteria; Actinobacteria; Streptomycetales; Streptomycetaceae; Streptomyces
MLKGVENHSLPRTAAFFDLDKTVIAKSSTLTFSKSFYQGGLINRRAALRTAYAQFVFLVG WP_073492691.1 inhibition of morphological differentiation protein [Streptomyces phaeoluteigriseus] Length: 278\nNumber of predicted TMHs: 1\nExp number of AAs in TMHs: 21.54977\nExp number, first 60 AAs: 0.01951\nTotal prob of N-in: 0.00272\noutside 1 247\nTMhelix 248 270\ninside 271 278

32727 GCF_001513985.1_ASM151398v1 Streptomyces phaeopurpureus Terrabacteria group; Actinobacteria; Actinobacteria; Streptomycetales; Streptomycetaceae; Streptomyces
MLGVVDNHSLPRTAAFFDLDKTVIAKSSTLTFSKSFYQGGLINRRAALRTAYAQFVFLVG WP_062026023.1 MULTISPECIES: inhibition of morphological differentiation protein [Streptomyces] Length: 277\nNumber of predicted TMHs: 1\nExp number of AAs in TMHs: 21.69291\nExp number, first 60 AAs: 0.0377\nTotal prob of N-in: 0.00335\noutside 1 247\nTMhelix 248 270\ninside 271 277

32728 GCF_001513985.1_ASM151398v1 Streptomyces phaeopurpureus Terrabacteria group; Actinobacteria; Actinobacteria; Streptomycetales; Streptomycetaceae; Streptomyces
MRHARRRVVRRVTQLAAVGGLLLGGTMVTRAVASEPPATAPPRTFAQQAGETGADIVARL WP_062020264.1 protease [Streptomyces phaeopurpureus] Length: 454\nNumber of predicted TMHs: 2\nExp number of AAs in TMHs: 39.0222299999999\nExp number, first 60 AAs: 18.86541\nTotal prob of N-in: 0.86908\nPOSSIBLE N-term signal sequence\ninside 1 11\nTMhelix 12 34\noutside 35 415\nTMhelix 416 438\ninside 439 454

32729 GCF_002119195.1_ASM211919v1 Streptomyces platensis Terrabacteria group; Actinobacteria; Actinobacteria; Streptomycetales; Streptomycetaceae; Streptomyces
MRQRRRAAVAATAALVLAGAGIMTAAGTAQAAEVSYKTECLPPPISGLPPIQGTTKVAVS WP_085926357.1 hypothetical protein [Streptomyces platensis] Length: 472\nNumber of predicted TMHs: 2\nExp number of AAs in TMHs: 44.90961\nExp number, first 60 AAs: 22.22636\nTotal prob of N-in: 0.99685\nPOSSIBLE N-term signal sequence\ninside 1 6\nTMhelix 7 29\noutside 30 441\nTMhelix 442 464\ninside 465 472

32730 GCF_002119195.1_ASM211919v1 Streptomyces platensis Terrabacteria group; Actinobacteria; Actinobacteria; Streptomycetales; Streptomycetaceae; Streptomyces
MTDRRRRAALLLLLPATAPGAVLLPRERKSSSSSEPTTPDSPDTERDNPFAPPPAGTPD WP_085927641.1 hypothetical protein [Streptomyces platensis] Length: 293\nNumber of predicted TMHs: 2\nExp number of AAs in TMHs: 47.79611\nExp number, first 60 AAs: 0.40235\nTotal prob of N-in: 0.97240\ninside 1 154\nTMhelix 155 177\noutside 178 233\nTMhelix 234 256\ninside 257 293

32731 GCF_002119195.1_ASM211919v1 Streptomyces platensis Terrabacteria group; Actinobacteria; Actinobacteria; Streptomycetales; Streptomycetaceae; Streptomyces
MNSASPAPDSSCRAARPARSGARPPLTRRRPAVVLALLAGLLAAAVAPAAAAGRSTGDTT WP_085928192.1 hypothetical protein [Streptomyces platensis] Length: 160\nNumber of predicted TMHs: 2\nExp number of AAs in TMHs: 44.26559\nExp number, first 60 AAs: 21.30785\nTotal prob of N-in: 0.88625\nPOSSIBLE N-term signal sequence\ninside 1 30\nTMhelix 31 53\noutside 54 125\nTMhelix 126 148\ninside 149 160

32732 GCF_000816465.2_ASM81646v3 Streptomyces pluripotens Terrabacteria group; Actinobacteria; Actinobacteria; Streptomycetales; Streptomycetaceae; Streptomyces
MLKDVENHSLPRAAAFFDLDKTVIAKSSTLTFSKSFYQGGLINRRAALRTAYAQFVFLAG WP_039651231.1 MULTISPECIES: inhibition of morphological differentiation protein [Streptomyces] Length: 275\nNumber of predicted TMHs: 1\nExp number of AAs in TMHs: 21.73004\nExp number, first 60 AAs: 0.0183\nTotal prob of N-in: 0.00576\noutside 1 247\nTMhelix 248 270\ninside 271 275

32733 GCF_000816465.2_ASM81646v3 Streptomyces pluripotens Terrabacteria group; Actinobacteria; Actinobacteria; Streptomycetales; Streptomycetaceae; Streptomyces
MASRGRRTAMRSLGALAGAAALTALGSSAPGWAAAPSGAHHGHSSTRTPIKHVVVLFDEN WP_039652300.1 MULTISPECIES: phospholipase C [Streptomyces] Length: 609\nNumber of predicted TMHs: 2\nExp number of

AAs in TMHs: 42.3825\nExp number, first 60 AAs: 21.14153\nTotal prob of N-in: 0.93320\nPOSSIBLE N-term signal sequence\ninside 1 12\nTMhelix 13 35\noutside 36 578\nTMhelix 579 601\ninside 602 609

32734 GCF_000802245.2_ASM80224v2 Streptomyces pluripotens Terrabacteria group; Actinobacteria; Actinobacteria; Streptomycetales; Streptomycetaceae; Streptomyces MLKDVENHSLPRAAAFFDLDKTVIAKSSTLTFSKSFYQGGLINRRAALRTAYAQFVFLAG WP_039651231.1 MULTISPECIES: inhibition of morphological differentiation protein [Streptomyces] Length: 275\nNumber of predicted TMHs: 1\nExp number of AAs in TMHs: 21.73004\nExp number, first 60 AAs: 0.0183\nTotal prob of N-in: 0.00576\noutside 1 247\nTMhelix 248 270\ninside 271 275

32735 GCF_000802245.2_ASM80224v2 Streptomyces pluripotens Terrabacteria group; Actinobacteria; Actinobacteria; Streptomycetales; Streptomycetaceae; Streptomyces MRHARRRVVRRLTRLAAVGGLLLGCTMVARAAVASEPPPASARSLSSAGNTGAAWVARLG WP_043435093.1 S1 family peptidase [Streptomyces pluripotens] Length: 457\nNumber of predicted TMHs: 2\nExp number of AAs in TMHs: 42.37495\nExp number, first 60 AAs: 22.37635\nTotal prob of N-in: 0.98715\nPOSSIBLE N-term signal sequence\ninside 1 11\nTMhelix 12 34\noutside 35 418\nTMhelix 419 441\ninside 442 457

32736 GCF_000802245.2_ASM80224v2 Streptomyces pluripotens Terrabacteria group; Actinobacteria; Actinobacteria; Streptomycetales; Streptomycetaceae; Streptomyces MLDAVYGPRVRECSRLSRRAALGLTAAALPLSVATPAAAATVIGGERLARSGVQVRGASG WP_039657545.1 MULTISPECIES: D-alanyl-D-alanine carboxypeptidase [Streptomyces] Length: 408\nNumber of predicted TMHs: 2\nExp number of AAs in TMHs: 39.3505\nExp number, first 60 AAs: 21.43868\nTotal prob of N-in: 0.93948\nPOSSIBLE N-term signal sequence\ninside 1 20\nTMhelix 21 43\noutside 44 357\nTMhelix 358 380\ninside 381 408

32737 GCF_000802245.2_ASM80224v2 Streptomyces pluripotens Terrabacteria group; Actinobacteria; Actinobacteria; Streptomycetales; Streptomycetaceae; Streptomyces MASRGRRTAMRSLGALAGAAALTALGSSAPGWAAAPSGAHHGHSSTRTPIKHVVVLFDEN WP_039652300.1 MULTISPECIES: phospholipase C [Streptomyces] Length: 609\nNumber of predicted TMHs: 2\nExp number of AAs in TMHs: 42.3825\nExp number, first 60 AAs: 21.14153\nTotal prob of N-in: 0.93320\nPOSSIBLE N-term signal sequence\ninside 1 12\nTMhelix 13 35\noutside 36 578\nTMhelix 579 601\ninside 602 609

32738 GCF_000816465.2_ASM81646v3 Streptomyces pluripotens Terrabacteria group; Actinobacteria; Actinobacteria; Streptomycetales; Streptomycetaceae; Streptomyces MLDAVYGPRVRECSRLSRRAALGLTAAALPLSVATPAAAATVIGGERLARSGVQVRGASG WP_039657545.1 MULTISPECIES: D-alanyl-D-alanine carboxypeptidase [Streptomyces] Length: 408\nNumber of predicted TMHs: 2\nExp number of AAs in TMHs: 39.3505\nExp number, first 60 AAs: 21.43868\nTotal prob of N-in: 0.93948\nPOSSIBLE N-term signal sequence\ninside 1 20\nTMhelix 21 43\noutside 44 357\nTMhelix 358 380\ninside 381 408

32739 GCF_000816465.2_ASM81646v3 Streptomyces pluripotens Terrabacteria group; Actinobacteria; Actinobacteria; Streptomycetales; Streptomycetaceae; Streptomyces MRHARRRVVRRLTRLAAVGGLLLGCTMVARAAVASEPPPASARSLSSAGNTGAAWVARLG WP_043405418.1 protease [Streptomyces pluripotens] Length: 457\nNumber of predicted TMHs: 2\nExp number of AAs in TMHs: 42.37515\nExp number, first 60 AAs: 22.37634\nTotal prob of N-in: 0.98715\nPOSSIBLE N-term signal sequence\ninside 1 11\nTMhelix 12 34\noutside 35 418\nTMhelix 419 441\ninside 442 457

32740 GCF_001419695.1_ASM141969v1 Streptomyces prasinopilosus Terrabacteria group; Actinobacteria; Actinobacteria; Streptomycetales; Streptomycetaceae; Streptomyces MNSTNSRVPARRRTVVAAAAVLAAGSAALAGAGTAQATGEPGSASAAVLRTGLDVALLDG WP_055694013.1 hypothetical protein [Streptomyces prasinopilosus] Length: 331\nNumber of predicted TMHs: 2\nExp number of AAs in TMHs: 42.6664\nExp number, first 60 AAs: 20.68706\nTotal prob of N-in: 0.93125\nPOSSIBLE N-term signal sequence\ninside 1 12\nTMhelix 13 35\noutside 36 302\nTMhelix 303 325\ninside 326 331

32741 GCF_001419695.1_ASM141969v1 Streptomyces prasinopilosus Terrabacteria group; Actinobacteria; Actinobacteria; Streptomycetales; Streptomycetaceae; Streptomyces MRHARRRVVRRVTRLAAVGGLLLGATMVTRAVAGEPPAPDPVPYAGAQEDTGPGSALVAR WP_055573085.1 MULTISPECIES: protease [Streptomyces] Length: 453\nNumber of predicted TMHs: 2\nExp number of AAs in TMHs: 41.2757\nExp number, first 60 AAs: 21.21396\nTotal prob of N-in: 0.96361\nPOSSIBLE N-term signal sequence\ninside 1 11\nTMhelix 12 34\noutside 35 414\nTMhelix 415 437\ninside 438 453

32742 GCF_001419705.1_ASM141970v1 Streptomyces prasinus Terrabacteria group; Actinobacteria; Actinobacteria; Streptomycetales; Streptomycetaceae; Streptomyces

MPARRKAVVAAVGIVPLALTGLASSPAAAHGSMGDPVSRVAQCFAEGPESPKSAACRAAV WP_055688801.1 chitin-binding protein [Streptomyces prasinus] Length: 362\nNumber of predicted TMHs: 2\nExp number of AAs in TMHs: 40.96719\nExp number, first 60 AAs: 21.99065\nTotal prob of N-in: 0.97860\nPOSSIBLE N-term signal sequence\ninside 1 6\nTMhelix 7 29\noutside 30 328\nTMhelix 329 351\ninside 352 362

32743 GCF_001419705.1_ASM141970v1 Streptomyces prasinus Terrabacteria group; Actinobacteria; Actinobacteria; Streptomycetales; Streptomycetaceae; Streptomyces MRHARRRVVRRVTRLAAVGGVLLGATMVTRAVASEPPAPAPVPYAATEEDAGPGSALVAR WP_079040672.1 serine protease [Streptomyces prasinus] Length: 454\nNumber of predicted TMHs: 2\nExp number of AAs in TMHs: 41.17037\nExp number, first 60 AAs: 20.65859\nTotal prob of N-in: 0.95061\nPOSSIBLE N-term signal sequence\ninside 1 11\nTMhelix 12 34\noutside 35 415\nTMhelix 416 438\ninside 439 454

32744 GCF_001419705.1_ASM141970v1 Streptomyces prasinus Terrabacteria group; Actinobacteria; Actinobacteria; Streptomycetales; Streptomycetaceae; Streptomyces MNSTDSRASARRRAVVATVTVLAAGPVALAGAGAAQATGEHGSASAAVLRTGLDVALLDR WP_055688174.1 hypothetical protein [Streptomyces prasinus] Length: 338\nNumber of predicted TMHs: 2\nExp number of AAs in TMHs: 43.2681599999999\nExp number, first 60 AAs: 21.89484\nTotal prob of N-in: 0.97359\nPOSSIBLE N-term signal sequence\ninside 1 12\nTMhelix 13 35\noutside 36 309\nTMhelix 310 332\ninside 333 338

32745 GCF_001278075.1_ASM127807v1 Streptomyces pristinaespiralis Terrabacteria group; Actinobacteria; Actinobacteria; Streptomycetales; Streptomycetaceae; Streptomyces MTGRRRTPVAAAVAALTGAVPLLLAAGAPALAHGTPTDPVSRAAVCGLDGAQRASGACRA WP_005307481.1 secreted cellulose-binding protein [Streptomyces pristinaespiralis] Length: 325\nNumber of predicted TMHs: 2\nExp number of AAs in TMHs: 42.9823\nExp number, first 60 AAs: 21.39181\nTotal prob of N-in: 0.96758\nPOSSIBLE N-term signal sequence\ninside 1 8\nTMhelix 9 31\noutside 32 296\nTMhelix 297 319\ninside 320 325

32746 GCF_001278075.1_ASM127807v1 Streptomyces pristinaespiralis Terrabacteria group; Actinobacteria; Actinobacteria; Streptomycetales; Streptomycetaceae; Streptomyces MLCLVENHSLPRTAAFFDLDKTVIAKSSTLTFSKSFYQGGLINRRAVLRTAYAQFVFLAG WP_005314378.1 morphological differentiation-associated protein [Streptomyces pristinaespiralis] Length: 282\nNumber of predicted TMHs: 1\nExp number of AAs in TMHs: 21.64546\nExp number, first 60 AAs: 0.02262\nTotal prob of N-in: 0.00298\noutside 1 247\nTMhelix 248 270\ninside 271 282

32747 GCF_002154585.1_ASM215458v1 Streptomyces pseudogriseolus Terrabacteria group; Actinobacteria; Actinobacteria; Streptomycetales; Streptomycetaceae; Streptomyces MLRNVENHLLPRTAAFFDLDKTVIAKSSTLTFSKSFYQGGLINRRAVLRTAYSQFVFLAG WP_031020047.1 MULTISPECIES: inhibition of morphological differentiation protein [Streptomyces] Length: 277\nNumber of predicted TMHs: 1\nExp number of AAs in TMHs: 21.81477\nExp number, first 60 AAs: 0.02315\nTotal prob of N-in: 0.00436\noutside 1 247\nTMhelix 248 270\ninside 271 277

32748 GCF_001513955.1_ASM151395v1 Streptomyces pseudovenezuelae Terrabacteria group; Actinobacteria; Actinobacteria; Streptomycetales; Streptomycetaceae; Streptomyces MRHARRRVVRRVTRLAAVGGLLLGGAMVTRAVASEPPATTAVPRTYAMKAGQTGTDLVSQ WP_031038994.1 MULTISPECIES: protease [Streptomyces] Length: 455\nNumber of predicted TMHs: 2\nExp number of AAs in TMHs: 39.2163499999999\nExp number, first 60 AAs: 20.94931\nTotal prob of N-in: 0.95945\nPOSSIBLE N-term signal sequence\ninside 1 11\nTMhelix 12 34\noutside 35 416\nTMhelix 417 439\ninside 440 455

32749 GCF_001513955.1_ASM151395v1 Streptomyces pseudovenezuelae Terrabacteria group; Actinobacteria; Actinobacteria; Streptomycetales; Streptomycetaceae; Streptomyces MVTMHLTMSRRQGRTCGWRLRRNPLRRRSYFVEAWLRLATGILAMTVAIVTGVLTARAVE WP_062230060.1 hypothetical protein [Streptomyces pseudovenezuelae] Length: 199\nNumber of predicted TMHs: 2\nExp number of AAs in TMHs: 44.98971\nExp number, first 60 AAs: 22.5146\nTotal prob of N-in: 0.99810\nPOSSIBLE N-term signal sequence\ninside 1 33\nTMhelix 34 56\noutside 57 148\nTMhelix 149 171\ninside 172 199

32750 GCF_001513955.1_ASM151395v1 Streptomyces pseudovenezuelae Terrabacteria group; Actinobacteria; Actinobacteria; Streptomycetales; Streptomycetaceae; Streptomyces MLGVVENHSLPRTAAFFDLDKTVIAKSSTLTFSKSFYQGGLINRRAALRTAYAQFVFLVG WP_031044737.1 MULTISPECIES: inhibition of morphological differentiation protein [Streptomyces] Length: 277\nNumber of predicted TMHs: 1\nExp number of AAs in TMHs: 21.65845\nExp number, first 60 AAs: 0.01246\nTotal prob of N-in: 0.00153\noutside 1 247\nTMhelix 248 270\ninside 271 277

32751 GCF_001513955.1_ASM151395v1 Streptomyces pseudovenezuelae Terrabacteria group; Actinobacteria; Actinobacteria; Streptomycetales; Streptomycetaceae; Streptomyces
MTRRAVVLSLLTGLFLTAMSPAHATGYRYWSFWDRTAGTWTYATQGPSTTVPSDGDVQGF WP_031060443.1 MULTISPECIES: hypothetical protein [Streptomyces] Length: 213\nNumber of predicted TMHs: 2\nExp number of AAs in TMHs: 41.14384\nExp number, first 60 AAs: 21.27398\nTotal prob of N-in: 0.95944\nPOSSIBLE N-term signal sequence\ninside 1 4\nTMhelix 5 27\noutside 28 184\nTMhelix 185 204\ninside 205 213

32752 GCF_001735805.1_ASM173580v1 Streptomyces puniciscabiei Terrabacteria group; Actinobacteria; Actinobacteria; Streptomycetales; Streptomycetaceae; Streptomyces
MPARKNTGRRSLLVTSAALLSLSATAPAALAAPAPSTGPTATPPARMSTVGGARLGQPGT WP_069779106.1 D-alanyl-D-alanine carboxypeptidase [Streptomyces puniciscabiei] Length: 403\nNumber of predicted TMHs: 2\nExp number of AAs in TMHs: 39.48881\nExp number, first 60 AAs: 17.03209\nTotal prob of N-in: 0.80110\nPOSSIBLE N-term signal sequence\ninside 1 11\nTMhelix 12 34\noutside 35 364\nTMhelix 365 387\ninside 388 403

32753 GCF_001735805.1_ASM173580v1 Streptomyces puniciscabiei Terrabacteria group; Actinobacteria; Actinobacteria; Streptomycetales; Streptomycetaceae; Streptomyces
MLGAVYGPGVRECSRPSRRALLGLTAAALPLSTATEAAATTVIGGERLARNGVQVRGASG WP_079161534.1 D-alanyl-D-alanine carboxypeptidase [Streptomyces puniciscabiei] Length: 415\nNumber of predicted TMHs: 1\nExp number of AAs in TMHs: 20.28335\nExp number, first 60 AAs: 4.4026\nTotal prob of N-in: 0.19431\noutside 1 361\nTMhelix 362 381\ninside 382 415

32754 GCF_001735805.1_ASM173580v1 Streptomyces puniciscabiei Terrabacteria group; Actinobacteria; Actinobacteria; Streptomycetales; Streptomycetaceae; Streptomyces
MITGFSRRSVVSACSLCVAGVLALAPAAASAAPRAGEPAPPGPRATMPGPSLLYRSGTQV WP_079161659.1 D-alanyl-D-alanine carboxypeptidase [Streptomyces puniciscabiei] Length: 395\nNumber of predicted TMHs: 1\nExp number of AAs in TMHs: 30.57726\nExp number, first 60 AAs: 12.06682\nTotal prob of N-in: 0.55983\nPOSSIBLE N-term signal sequence\noutside 1 362\nTMhelix 363 385\ninside 386 395

32755 GCF_001735805.1_ASM173580v1 Streptomyces puniciscabiei Terrabacteria group; Actinobacteria; Actinobacteria; Streptomycetales; Streptomycetaceae; Streptomyces
MSARARRAAVVVAVLGLPPVALTVLAAAPASAHGSMGDPVSRVSQCYAEGPESPKSGACK WP_069781738.1 chitin-binding protein [Streptomyces puniciscabiei] Length: 320\nNumber of predicted TMHs: 2\nExp number of AAs in TMHs: 44.24057\nExp number, first 60 AAs: 21.67843\nTotal prob of N-in: 0.98334\nPOSSIBLE N-term signal sequence\ninside 1 6\nTMhelix 7 29\noutside 30 286\nTMhelix 287 309\ninside 310 320

32756 GCF_001419685.1_ASM141968v1 Streptomyces puniciscabiei Terrabacteria group; Actinobacteria; Actinobacteria; Streptomycetales; Streptomycetaceae; Streptomyces
MLWGVENHSLPRAAAFFDLDKTVIAKSSTLTFGKSFYQGGLINRRAALRTAYAQFVFLAGWP_055704759.1 inhibition of morphological differentiation protein [Streptomyces puniciscabiei] Length: 276\nNumber of predicted TMHs: 1\nExp number of AAs in TMHs: 21.81896\nExp number, first 60 AAs: 0.02104\nTotal prob of N-in: 0.00237\noutside 1 247\nTMhelix 248 270\ninside 271 276

32757 GCF_001419685.1_ASM141968v1 Streptomyces puniciscabiei Terrabacteria group; Actinobacteria; Actinobacteria; Streptomycetales; Streptomycetaceae; Streptomyces
MASRGRRAAMRSLGALAGAAALTVLGSNAPGWAAAPAAAQGGNSSTVTPIKHVVVLFDEN WP_055709373.1 phospholipase [Streptomyces puniciscabiei] Length: 613\nNumber of predicted TMHs: 2\nExp number of AAs in TMHs: 41.76488\nExp number, first 60 AAs: 20.12741\nTotal prob of N-in: 0.89027\nPOSSIBLE N-term signal sequence\ninside 1 12\nTMhelix 13 35\noutside 36 584\nTMhelix 585 607\ninside 608 613

32758 GCF_001419685.1_ASM141968v1 Streptomyces puniciscabiei Terrabacteria group; Actinobacteria; Actinobacteria; Streptomycetales; Streptomycetaceae; Streptomyces
MRHARRRIVRRVTRLAAVGGLLLGGAMVAQAAMASETPPAAARPLAASGASGAGDTGAAL WP_055709441.1 protease [Streptomyces puniciscabiei] Length: 463\nNumber of predicted TMHs: 2\nExp number of AAs in TMHs: 43.7115699999999\nExp number, first 60 AAs: 22.37425\nTotal prob of N-in: 0.99546\nPOSSIBLE N-term signal sequence\ninside 1 11\nTMhelix 12 34\noutside 35 424\nTMhelix 425 447\ninside 448 463

32759 GCF_001735805.1_ASM173580v1 Streptomyces puniciscabiei Terrabacteria group; Actinobacteria; Actinobacteria; Streptomycetales; Streptomycetaceae; Streptomyces
MRRCGRRAVFGVGAVRTALLPGLCALLVLVTGAAPASATPAGDAQAAPYAFAPGARTVAG WP_069779816.1 hypothetical protein [Streptomyces puniciscabiei] Length: 449\nNumber of predicted TMHs: 2\nExp number of AAs in TMHs: 44.2264699999999\nExp number, first 60 AAs: 21.98102\nTotal prob of N-in:

0.96913\nPOSSIBLE N-term signal sequence\ninside 1 11\nTMhelix 12 34\noutside 35 412\nTMhelix 413 435\ninside 436 449

32760 GCF_001735805.1_ASM173580v1 Streptomyces puniciscabiei Terrabacteria group; Actinobacteria; Actinobacteria; Streptomycetales; Streptomycetaceae; Streptomyces MASRGRRAAIRSLGALTGAAALTVLGSNAPGWAAVSAGQSSTATPIKHVVVLFDENISFD WP_069776808.1 phospholipase [Streptomyces puniciscabiei] Length: 609\nNumber of predicted TMHs: 2\nExp number of AAs in TMHs: 38.75078\nExp number, first 60 AAs: 17.68294\nTotal prob of N-in: 0.78031\nPOSSIBLE N-term signal sequence\ninside 1 12\nTMhelix 13 35\noutside 36 580\nTMhelix 581 603\ninside 604 609

32761 GCF_001735805.1_ASM173580v1 Streptomyces puniciscabiei Terrabacteria group; Actinobacteria; Actinobacteria; Streptomycetales; Streptomycetaceae; Streptomyces MRHARRRIVRRVTRLAAVGGLLLGGAMVTQAAMASETPPAAARTLSSAAGTPDTGAALVS WP_069776897.1 protease [Streptomyces puniciscabiei] Length: 461\nNumber of predicted TMHs: 2\nExp number of AAs in TMHs: 43.1269299999999\nExp number, first 60 AAs: 22.22211\nTotal prob of N-in: 0.99185\nPOSSIBLE N-term signal sequence\ninside 1 11\nTMhelix 12 34\noutside 35 422\nTMhelix 423 445\ninside 446 461

32762 GCF_001735805.1_ASM173580v1 Streptomyces puniciscabiei Terrabacteria group; Actinobacteria; Actinobacteria; Streptomycetales; Streptomycetaceae; Streptomyces MLMDVENHSLPRAAAFFDLDKTVIAKSSTLTFSKSFYQGGLINRRAALRTAYAQFVFLVG WP_069779543.1 inhibition of morphological differentiation protein [Streptomyces puniciscabiei] Length: 276\nNumber of predicted TMHs: 1\nExp number of AAs in TMHs: 21.85301\nExp number, first 60 AAs: 0.03885\nTotal prob of N-in: 0.00344\noutside 1 247\nTMhelix 248 270\ninside 271 276

32763 GCF_000717965.1_ASM71796v1 Streptomyces purpeochromogenes Terrabacteria group; Actinobacteria; Actinobacteria; Streptomycetales; Streptomycetaceae; Streptomyces MGSSWNERRLERRVRPGDGKPLKPFRWWQLTRRSLLSLALPVGGGLSATYTVEVKHGGDA WP_030113438.1 MULTISPECIES: hypothetical protein [Streptomyces] Length: 218\nNumber of predicted TMHs: 2\nExp number of AAs in TMHs: 41.40199\nExp number, first 60 AAs: 0.36812\nTotal prob of N-in: 0.97576\ninside 1 135\nTMhelix 136 158\noutside 159 182\nTMhelix 183 200\ninside 201 218

32764 GCF_000717965.1_ASM71796v1 Streptomyces purpeochromogenes Terrabacteria group; Actinobacteria; Actinobacteria; Streptomycetales; Streptomycetaceae; Streptomyces MLSVVENCFSPRTAAFFDLDKTVIAKSSTLTFSKSFYQGGLINRRAVLRTAYTQFVFMAG WP_030113359.1 MULTISPECIES: inhibition of morphological differentiation protein [Streptomyces] Length: 280\nNumber of predicted TMHs: 1\nExp number of AAs in TMHs: 21.68695\nExp number, first 60 AAs: 0.01801\nTotal prob of N-in: 0.00793\noutside 1 247\nTMhelix 248 270\ninside 271 280

32765 GCF_000718615.1_ASM71861v1 Streptomyces purpeochromogenes Terrabacteria group; Actinobacteria; Actinobacteria; Streptomycetales; Streptomycetaceae; Streptomyces MGSSWNERRLERRVRPGDGKPLKPFRWWQLTRRSLLSLALPVGGGLSATYTVEVKHGGDA WP_030113438.1 MULTISPECIES: hypothetical protein [Streptomyces] Length: 218\nNumber of predicted TMHs: 2\nExp number of AAs in TMHs: 41.40199\nExp number, first 60 AAs: 0.36812\nTotal prob of N-in: 0.97576\ninside 1 135\nTMhelix 136 158\noutside 159 182\nTMhelix 183 200\ninside 201 218

32766 GCF_000718615.1_ASM71861v1 Streptomyces purpeochromogenes Terrabacteria group; Actinobacteria; Actinobacteria; Streptomycetales; Streptomycetaceae; Streptomyces MLSVVENCFSPRTAAFFDLDKTVIAKSSTLTFSKSFYQGGLINRRAVLRTAYTQFVFMAG WP_030113359.1 MULTISPECIES: inhibition of morphological differentiation protein [Streptomyces] Length: 280\nNumber of predicted TMHs: 1\nExp number of AAs in TMHs: 21.68695\nExp number, first 60 AAs: 0.01801\nTotal prob of N-in: 0.00793\noutside 1 247\nTMhelix 248 270\ninside 271 280

32767 GCF_000717725.1_ASM71772v1 Streptomyces pyridomyceticus Terrabacteria group; Actinobacteria; Actinobacteria; Streptomycetales; Streptomycetaceae; Streptomyces MRTAAFFDLDKTIIAKSSALAFSRPFYQGGLINRRAVLRSAYAQFVFLVGGADHDQMEKM WP_030259263.1 MULTISPECIES: inhibition of morphological differentiation protein [Streptomyces] Length: 265\nNumber of predicted TMHs: 1\nExp number of AAs in TMHs: 22.6499\nExp number, first 60 AAs: 0.08965\nTotal prob of N-in: 0.01242\noutside 1 235\nTMhelix 236 258\ninside 259 265

32768 GCF_001751375.1_ASM175137v1 Streptomyces qinglanensis Terrabacteria group; Actinobacteria; Actinobacteria; Streptomycetales; Streptomycetaceae; Streptomyces

MLGAVKNLAMPRTAAFFDLDKTVIAKSSTLTFGKSFYQGGLINRRAVLRTAYAQFVYLLG WP_027760567.1 MULTISPECIES: inhibition of morphological differentiation protein [Streptomyces] Length: 281\nNumber of predicted TMHs: 1\nExp number of AAs in TMHs: 22.28973\nExp number, first 60 AAs: 0.03324\nTotal prob of N-in: 0.01080\noutside 1 246\nTMhelix 247 269\ninside 270 281

32769 GCF_001751375.1_ASM175137v1 Streptomyces qinglanensis Terrabacteria group; Actinobacteria; Actinobacteria; Streptomycetales; Streptomycetaceae; Streptomyces MIFRMRAFWGMRIDDPWHDALATGGAEVHGTPVDTPPRRTAPPPRREPVPSAVHGVYPTV WP_079135177.1 hypothetical protein [Streptomyces qinglanensis]Length: 177\nNumber of predicted TMHs: 2\nExp number of AAs in TMHs: 45.1685\nExp number, first 60 AAs: 0.00025\nTotal prob of N-in: 0.99463\ninside 1 75\nTMhelix 76 98\noutside 99 107\nTMhelix 108 130\ninside 131 177

32770 GCF_001751375.1_ASM175137v1 Streptomyces qinglanensis Terrabacteria group; Actinobacteria; Actinobacteria; Streptomycetales; Streptomycetaceae; Streptomyces MITRRRLVTGACALLMALPLGLLPTGTASAEPASDEDAPKVELTLDVSGSMRKRDIDGRS WP_026005050.1 MULTISPECIES: inter-alpha-trypsin inhibitor heavy chain H5 [Streptomyces] Length: 421\nNumber of predicted TMHs: 2\nExp number of AAs in TMHs: 38.90213\nExp number, first 60 AAs: 16.54504\nTotal prob of N-in: 0.77694\nPOSSIBLE N-term signal sequence\ninside 1 6\nTMhelix 7 29\noutside 30 388\nTMhelix 389 411\ninside 412 421

32771 GCF_900111245.1_IMG-taxon_2675903055_annotated_assembly Streptomyces qinglanensis Terrabacteria group; Actinobacteria; Actinobacteria; Streptomycetales; Streptomycetaceae; Streptomyces MSPRRRRATVLAAATVLSAAAAPVASAAAPAATATSVAASPARTTASELPDGLYGTADPK WP_074998544.1 hypothetical protein [Streptomyces qinglanensis]Length: 443\nNumber of predicted TMHs: 1\nExp number of AAs in TMHs: 37.1860499999999\nExp number, first 60 AAs: 14.86289\nTotal prob of N-in: 0.66546\nPOSSIBLE N-term signal sequence\noutside 1 413\nTMhelix 414 436\ninside 437 443

32772 GCF_900111245.1_IMG-taxon_2675903055_annotated_assembly Streptomyces qinglanensis Terrabacteria group; Actinobacteria; Actinobacteria; Streptomycetales; Streptomycetaceae; Streptomyces MITRRRLVTGACALLMALPLGLLPTGTASAEPASDENAPKVELTLDVSGSMRKRDIDGRS WP_075001037.1 alpha-1-antitrypsin [Streptomyces qinglanensis] Length: 421\nNumber of predicted TMHs: 2\nExp number of AAs in TMHs: 39.33688\nExp number, first 60 AAs: 16.97811\nTotal prob of N-in: 0.79271\nPOSSIBLE N-term signal sequence\ninside 1 6\nTMhelix 7 29\noutside 30 388\nTMhelix 389 411\ninside 412 421

32773 GCF_900111245.1_IMG-taxon_2675903055_annotated_assembly Streptomyces qinglanensis Terrabacteria group; Actinobacteria; Actinobacteria; Streptomycetales; Streptomycetaceae; Streptomyces MLGAVKNLAMPRTAAFFDLDKTVIAKSSTLTFGKSFYQGGLINRRAVLRTAYAQFVYLLG WP_075002780.1 inhibition of morphological differentiation protein [Streptomyces qinglanensis] Length: 281\nNumber of predicted TMHs: 1\nExp number of AAs in TMHs: 22.33775\nExp number, first 60 AAs: 0.03383\nTotal prob of N-in: 0.01169\noutside 1 247\nTMhelix 248 270\ninside 271 281

32774 GCF_001751375.1_ASM175137v1 Streptomyces qinglanensis Terrabacteria group; Actinobacteria; Actinobacteria; Streptomycetales; Streptomycetaceae; Streptomyces MSPRRRRATVLTAATVLSAAAAPVASAAAPAAPVTSVAASPARTTASGLPEGLYGTADPK WP_069991284.1 MULTISPECIES: hypothetical protein [Streptomyces] Length: 443\nNumber of predicted TMHs: 1\nExp number of AAs in TMHs: 34.01561\nExp number, first 60 AAs: 11.7064\nTotal prob of N-in: 0.52682\nPOSSIBLE N-term signal sequence\noutside 1 413\nTMhelix 414 436\ninside 437 443

32775 GCF_001509475.1_ASM150947v1 Streptomyces regalis Terrabacteria group; Actinobacteria; Actinobacteria; Streptomycetales; Streptomycetaceae; Streptomyces MRHARRRLVRRVTRLAAVGGLLLGGTMVTRAVASEPPDGSTVPRTYAASAADTGSDLVAR WP_062714790.1 protease [Streptomyces regalis] Length: 459\nNumber of predicted TMHs: 2\nExp number of AAs in TMHs: 40.55491\nExp number, first 60 AAs: 19.21127\nTotal prob of N-in: 0.87902\nPOSSIBLE N-term signal sequence\ninside 1 11\nTMhelix 12 34\noutside 35 420\nTMhelix 421 443\ninside 444 459

32776 GCF_001509475.1_ASM150947v1 Streptomyces regalis Terrabacteria group; Actinobacteria; Actinobacteria; Streptomycetales; Streptomycetaceae; Streptomyces MPAPSKKTARRALLVTSAVVSSLALTAPVSQAAPKPSPSTSPSATPPPNMSTVGGERLGQ WP_062702374.1 D-alanyl-D-alanine carboxypeptidase [Streptomyces regalis] Length: 429\nNumber of predicted TMHs: 1\nExp number of AAs in TMHs: 31.80528\nExp number, first 60 AAs: 9.06174\nTotal prob of N-in: 0.46468\noutside 1 390\nTMhelix 391 413\ninside 414 429

32777 GCF_001270505.1_ASM127050v1 Streptomyces resistomycificus Terrabacteria group; Actinobacteria; Actinobacteria; Streptomycetales; Streptomycetaceae; Streptomyces MRHARRRVVRRVTRLAAVGGLLLGGTMVTRAVASEPSEASAVPRTYAEQSADTGTGLVSR WP_053190400.1 protease [Streptomyces resistomycificus] Length: 456\nNumber of predicted TMHs: 2\nExp number of AAs in TMHs: 41.0919099999999\nExp number, first 60 AAs: 19.88525\nTotal prob of N-in: 0.91112\nPOSSIBLE N-term signal sequence\ninside 1 11\nTMhelix 12 34\noutside 35 417\nTMhelix 418 440\ninside 441 456

32778 GCF_001270505.1_ASM127050v1 Streptomyces resistomycificus Terrabacteria group; Actinobacteria; Actinobacteria; Streptomycetales; Streptomycetaceae; Streptomyces MLMVVENHSLPRTAAFFDLDKTVIAKSSTLTFSKSFYQGGLINRRAALRTAYAQFVFLVG WP_030043278.1 inhibition of morphological differentiation protein [Streptomyces resistomycificus] Length: 277\nNumber of predicted TMHs: 1\nExp number of AAs in TMHs: 21.85426\nExp number, first 60 AAs: 0.01664\nTotal prob of N-in: 0.00264\noutside 1 247\nTMhelix 248 270\ninside 271 277

32779 GCF_001514265.1_ASM151426v1 Streptomyces resistomycificus Terrabacteria group; Actinobacteria; Actinobacteria; Streptomycetales; Streptomycetaceae; Streptomyces MRHARRRVVRRVTRLAAVGGLLLGGTMVTRAVASEPSEASAVPRTYAEQSADTGTGLVSR WP_053190400.1 protease [Streptomyces resistomycificus] Length: 456\nNumber of predicted TMHs: 2\nExp number of AAs in TMHs: 41.0919099999999\nExp number, first 60 AAs: 19.88525\nTotal prob of N-in: 0.91112\nPOSSIBLE N-term signal sequence\ninside 1 11\nTMhelix 12 34\noutside 35 417\nTMhelix 418 440\ninside 441 456

32780 GCF_001514265.1_ASM151426v1 Streptomyces resistomycificus Terrabacteria group; Actinobacteria; Actinobacteria; Streptomycetales; Streptomycetaceae; Streptomyces MLMVVENHSLPRTAAFFDLDKTVIAKSSTLTFSKSFYQGGLINRRAALRTAYAQFVFLVG WP_030043278.1 inhibition of morphological differentiation protein [Streptomyces resistomycificus] Length: 277\nNumber of predicted TMHs: 1\nExp number of AAs in TMHs: 21.85426\nExp number, first 60 AAs: 0.01664\nTotal prob of N-in: 0.00264\noutside 1 247\nTMhelix 248 270\ninside 271 277

32781 GCF_000716215.1_ASM71621v1 Streptomyces resistomycificus Terrabacteria group; Actinobacteria; Actinobacteria; Streptomycetales; Streptomycetaceae; Streptomyces MLMVVENHSLPRTAAFFDLDKTVIAKSSTLTFSKSFYQGGLINRRAALRTAYAQFVFLVG WP_030043278.1 inhibition of morphological differentiation protein [Streptomyces resistomycificus] Length: 277\nNumber of predicted TMHs: 1\nExp number of AAs in TMHs: 21.85426\nExp number, first 60 AAs: 0.01664\nTotal prob of N-in: 0.00264\noutside 1 247\nTMhelix 248 270\ninside 271 277

32782 GCF_001511815.1_TUE45 Streptomyces reticuli Terrabacteria group; Actinobacteria; Actinobacteria; Streptomycetales; Streptomycetaceae; Streptomyces MLRDVENHSLPRAAAFFDLDKTVIAKSSTLTFSKSFYQGGLINRRAALRTAYAQFVFLAG WP_059251463.1 inhibition of morphological differentiation protein [Streptomyces reticuli] Length: 276\nNumber of predicted TMHs: 1\nExp number of AAs in TMHs: 21.75336\nExp number, first 60 AAs: 0.04333\nTotal prob of N-in: 0.00752\noutside 1 247\nTMhelix 248 270\ninside 271 276

32783 GCF_001511815.1_TUE45 Streptomyces reticuli Terrabacteria group; Actinobacteria; Actinobacteria; Streptomycetales; Streptomycetaceae; Streptomyces MAWQRPVRFGRPLANPLFDAVYRPGVRESSRLSRRAVLGLTAAALPLSTATEAAAATAVV WP_059249036.1 D-alanyl-D-alanine carboxypeptidase [Streptomyces reticuli] Length: 429\nNumber of predicted TMHs: 1\nExp number of AAs in TMHs: 22.00794\nExp number, first 60 AAs: 2.04588\nTotal prob of N-in: 0.09347\noutside 1 374\nTMhelix 375 397\ninside 398 429

32784 GCF_001511815.1_TUE45 Streptomyces reticuli Terrabacteria group; Actinobacteria; Actinobacteria; Streptomycetales; Streptomycetaceae; Streptomyces MPAPKPAGRRSLLVTSAALLSLSVTAPTALAAPTPSTSPSATPPAQMSTLGGARLGMPGT WP_059252215.1 D-alanyl-D-alanine carboxypeptidase [Streptomyces reticuli] Length: 413\nNumber of predicted TMHs: 1\nExp number of AAs in TMHs: 27.80819\nExp number, first 60 AAs: 5.26239\nTotal prob of N-in: 0.24702\noutside 1 374\nTMhelix 375 397\ninside 398 413

32785 GCF_001511815.1_TUE45 Streptomyces reticuli Terrabacteria group; Actinobacteria; Actinobacteria; Streptomycetales; Streptomycetaceae; Streptomyces MRHARRRVVRRVTRLAAVGGLLLGGAMVAQAAVAGETPPASARALTAADDPGATLVARLG WP_059246863.1 protease [Streptomyces reticuli] Length: 457\nNumber of predicted TMHs: 2\nExp number of AAs in

TMHs: 43.8404599999999\nExp number, first 60 AAs: 22.54822\nTotal prob of N-in: 0.99526\nPOSSIBLE N-term signal sequence\ninside 1 11\nTMhelix 12 34\noutside 35 418\nTMhelix 419 441\ninside 442 457

32786 GCF_002154675.1_ASM215467v1 Streptomyces reticuliscabiei Terrabacteria group; Actinobacteria; Actinobacteria; Streptomycetales; Streptomycetaceae; Streptomyces MRHARRRIVRRATRLAAVGGVILGGLMVTQAVASEPPEVNLPFSSAQAAGTTGSELVTEL WP_006380203.1 S1 family peptidase [Streptomyces turgidiscabies] Length: 452\nNumber of predicted TMHs: 2\nExp number of AAs in TMHs: 42.58327\nExp number, first 60 AAs: 21.83569\nTotal prob of N-in: 0.98683\nPOSSIBLE N-term signal sequence\ninside 1 11\nTMhelix 12 34\noutside 35 413\nTMhelix 414 436\ninside 437 452

32787 GCF_002154675.1_ASM215467v1 Streptomyces reticuliscabiei Terrabacteria group; Actinobacteria; Actinobacteria; Streptomycetales; Streptomycetaceae; Streptomyces MPARRRPAVVTTVTALGIAALALGATPASAHGSMGDPVSRVAQCYAEGPESPTSAACRAA WP_006376680.1 chitin-binding protein [Streptomyces turgidiscabies] Length: 353\nNumber of predicted TMHs: 2\nExp number of AAs in TMHs: 41.16593\nExp number, first 60 AAs: 20.07244\nTotal prob of N-in: 0.93537\nPOSSIBLE N-term signal sequence\ninside 1 6\nTMhelix 7 29\noutside 30 318\nTMhelix 319 341\ninside 342 353

32788 GCF_002154675.1_ASM215467v1 Streptomyces reticuliscabiei Terrabacteria group; Actinobacteria; Actinobacteria; Streptomycetales; Streptomycetaceae; Streptomyces MLSLVENHSLPRAAAFFDLDKTVIAKSSTLTFSKSFYQGGLINRRAVLRTAYAQFVFLAG WP_044473232.1 inhibition of morphological differentiation protein [Streptomyces turgidiscabies] Length: 277\nNumber of predicted TMHs: 1\nExp number of AAs in TMHs: 21.55643\nExp number, first 60 AAs: 0.01804\nTotal prob of N-in: 0.00246\noutside 1 247\nTMhelix 248 270\ninside 271 277

32789 GCF_002155885.1_ASM215588v1 Streptomyces rhizosphaericus Terrabacteria group; Actinobacteria; Actinobacteria; Streptomycetales; Streptomycetaceae; Streptomyces MSSPDREVRAARNDTARTARRPVVAVTGAASGPGALLTQRLAESEEVKQVLAIDERRGEV WP_086886330.1 NAD-dependent dehydratase [Streptomyces rhizosphaericus] Length: 369\nNumber of predicted TMHs: 1\nExp number of AAs in TMHs: 19.99751\nExp number, first 60 AAs: 0.38218\nTotal prob of N-in: 0.03037\noutside 1 339\nTMhelix 340 359\ninside 360 369

32790 GCF_000707965.1_rev-51 Streptomyces rimosus Terrabacteria group; Actinobacteria; Actinobacteria; Streptomycetales; Streptomycetaceae; Streptomyces MESRQPVGPRANGHRAAHRRRAAVALAGAVCAVGLTATTAAPALADTNVPLNNGQCNFGA WP_078586913.1 MULTISPECIES: type VII secretion-associated serine protease mycosin [Streptomyces] Length: 427\nNumber of predicted TMHs: 2\nExp number of AAs in TMHs: 43.8948\nExp number, first 60 AAs: 21.08952\nTotal prob of N-in: 0.94565\nPOSSIBLE N-term signal sequence\ninside 1 20\nTMhelix 21 43\noutside 44 389\nTMhelix 390 412\ninside 413 427

32791 GCF_000707965.1_rev-51 Streptomyces rimosus Terrabacteria group; Actinobacteria; Actinobacteria; Streptomycetales; Streptomycetaceae; Streptomyces MTERSAPAASATGPGTPARRTAVRAAATGTLAAALLGLAPQSALAATPAPPPAPAISAVT WP_030180617.1 MULTISPECIES: hypothetical protein [Streptomyces] Length: 293\nNumber of predicted TMHs: 1\nExp number of AAs in TMHs: 24.94793\nExp number, first 60 AAs: 4.55383\nTotal prob of N-in: 0.22746\noutside 1 264\nTMhelix 265 287\ninside 288 293

32792 GCF_000707945.1_mv9-51 Streptomyces rimosus Terrabacteria group; Actinobacteria; Actinobacteria; Streptomycetales; Streptomycetaceae; Streptomyces MTRRPLLRAAGVALSAAALTVLPALPAHADGIRAQEWALKAMNTQDAWETTKGEGITVAV WP_003987438.1 MULTISPECIES: type VII secretion-associated serine protease [Streptomyces] Length: 386\nNumber of predicted TMHs: 1\nExp number of AAs in TMHs: 34.80558\nExp number, first 60 AAs: 10.27095\nTotal prob of N-in: 0.55118\nPOSSIBLE N-term signal sequence\noutside 1 359\nTMhelix 360 382\ninside 383 386

32793 GCF_000707945.1_mv9-51 Streptomyces rimosus Terrabacteria group; Actinobacteria; Actinobacteria; Streptomycetales; Streptomycetaceae; Streptomyces MESRQPVGPRANGHRAAHRRRAAVALAGAVCAVGLTATTAAPALADTNVPLNNGQCNFGA WP_078586913.1 MULTISPECIES: type VII secretion-associated serine protease mycosin [Streptomyces] Length: 427\nNumber of predicted TMHs: 2\nExp number of AAs in TMHs: 43.8948\nExp number, first 60 AAs: 21.08952\nTotal prob of N-in: 0.94565\nPOSSIBLE N-term signal sequence\ninside 1 20\nTMhelix 21 43\noutside 44 389\nTMhelix 390 412\ninside 413 427

32794 GCF_000707965.1_rev-51 Streptomyces rimosus Terrabacteria group; Actinobacteria; Actinobacteria; Streptomycetales; Streptomycetaceae; Streptomyces
MTRRPLLRAAGVALSAAALTVLPALPAHADGIRAQEWALKAMNTQDAWETTKGEGITVAV WP_003987438.1
MULTISPECIES: type VII secretion-associated serine protease [Streptomyces] Length: 386\nNumber of predicted TMHs: 1\nExp number of AAs in TMHs: 34.80558\nExp number, first 60 AAs: 10.27095\nTotal prob of N-in: 0.55118\nPOSSIBLE N-term signal sequence\noutside 1 359\nTMhelix 360 382\ninside 383 386

32795 GCF_000719265.1_ASM71926v1 Streptomyces roseoverticillatus Terrabacteria group; Actinobacteria; Actinobacteria; Streptomycetales; Streptomycetaceae; Streptomyces
MDDERQAGAAAPVVRRRWQRARRAAVVAALALGASLLVSPASAQGTEQARPAVVTAVAER WP_052392303.1
hypothetical protein [Streptomyces roseoverticillatus] Length: 151\nNumber of predicted TMHs: 2\nExp number of AAs in TMHs: 43.64169\nExp number, first 60 AAs: 20.43026\nTotal prob of N-in: 0.98742\nPOSSIBLE N-term signal sequence\ninside 1 23\nTMhelix 24 46\noutside 47 116\nTMhelix 117 139\ninside 140 151

32796 GCF_000719265.1_ASM71926v1 Streptomyces roseoverticillatus Terrabacteria group; Actinobacteria; Actinobacteria; Streptomycetales; Streptomycetaceae; Streptomyces
MLGIVENHSLPRTAAFFDLDKTVIAKSSTLTFSKSFYQGGLINRRAVLRTAYAQFVFLAG WP_030367291.1 inhibition of morphological differentiation protein [Streptomyces roseoverticillatus] Length: 281\nNumber of predicted TMHs: 1\nExp number of AAs in TMHs: 21.75531\nExp number, first 60 AAs: 0.02274\nTotal prob of N-in: 0.00463\noutside 1 246\nTMhelix 247 269\ninside 270 281

32797 GCF_000719015.1_ASM71901v1 Streptomyces ruber Terrabacteria group; Actinobacteria; Actinobacteria; Streptomycetales; Streptomycetaceae; Streptomyces
MLNLVENHSSPRTAAFFDLDKTVIAKSSTLTFSKSFYQGGLINRRAVLRTAYAQFVFLAG WP_030359659.1 MULTISPECIES: inhibition of morphological differentiation protein [Streptomyces] Length: 274\nNumber of predicted TMHs: 1\nExp number of AAs in TMHs: 21.81923\nExp number, first 60 AAs: 0.095\nTotal prob of N-in: 0.03042\noutside 1 244\nTMhelix 245 267\ninside 268 274

32798 GCF_000725575.1_Doro.v1.0Streptomyces ruber Terrabacteria group; Actinobacteria; Actinobacteria; Streptomycetales; Streptomycetaceae; Streptomyces
MLNLVENHSSPRTAAFFDLDKTVIAKSSTLTFSKSFYQGGLINRRAVLRTAYAQFVFLAG WP_030359659.1 MULTISPECIES: inhibition of morphological differentiation protein [Streptomyces] Length: 274\nNumber of predicted TMHs: 1\nExp number of AAs in TMHs: 21.81923\nExp number, first 60 AAs: 0.095\nTotal prob of N-in: 0.03042\noutside 1 244\nTMhelix 245 267\ninside 268 274

32799 GCF_001723065.1_ASM172306v1 Streptomyces rubidus Terrabacteria group; Actinobacteria; Actinobacteria; Streptomycetales; Streptomycetaceae; Streptomyces
MSTNRRTAIAVRTIGVASAAAALALGVAGTALACSITEFTPSAGCDDSGHGYIQVVDKDG WP_069464636.1
hypothetical protein [Streptomyces rubidus] Length: 224\nNumber of predicted TMHs: 2\nExp number of AAs in TMHs: 43.13601\nExp number, first 60 AAs: 21.73054\nTotal prob of N-in: 0.98724\nPOSSIBLE N-term signal sequence\ninside 1 12\nTMhelix 13 35\noutside 36 193\nTMhelix 194 216\ninside 217 224

32800 GCF_900110255.1_IMG-taxon_2671180039_annotated_assembly Streptomyces rubidus Terrabacteria group; Actinobacteria; Actinobacteria; Streptomycetales; Streptomycetaceae; Streptomyces
MSTNRRTAIAVRTIGVASAAAALALGVAGTALACSITEFTPSAGCDDSGHGYIQVVDKDG WP_069464636.1
hypothetical protein [Streptomyces rubidus] Length: 224\nNumber of predicted TMHs: 2\nExp number of AAs in TMHs: 43.13601\nExp number, first 60 AAs: 21.73054\nTotal prob of N-in: 0.98724\nPOSSIBLE N-term signal sequence\ninside 1 12\nTMhelix 13 35\noutside 36 193\nTMhelix 194 216\ninside 217 224

32801 GCF_001750785.1_ASM175078v1 Streptomyces rubrolavendulae Terrabacteria group; Actinobacteria; Actinobacteria; Streptomycetales; Streptomycetaceae; Streptomyces
MLWLVENHSSPRTAAFFDLDKTVIAKSSTLTFGKSFYQGGLINRRDALRTAYTQFVFLVG WP_069977397.1 inhibition of morphological differentiation protein [Streptomyces rubrolavendulae] Length: 299\nNumber of predicted TMHs: 1\nExp number of AAs in TMHs: 21.33343\nExp number, first 60 AAs: 0.00541\nTotal prob of N-in: 0.00286\noutside 1 247\nTMhelix 248 270\ninside 271 299

32802 GCF_000738715.1_ASM73871v1 Streptomyces scabiei Terrabacteria group; Actinobacteria; Actinobacteria; Streptomycetales; Streptomycetaceae; Streptomyces
MYRLSARRTGAHPAGAHRPGAHHDSAHRDSGRGDGGRRAAVATAAAVVVAPLLLTAGTAT WP_037724552.1
chitin-binding protein, partial [Streptomyces scabiei] Length: 349\nNumber of predicted TMHs: 1\nExp

number of AAs in TMHs: 37.01665\nExp number, first 60 AAs: 14.51724\nTotal prob of N-in: 0.69707\nPOSSIBLE N-term signal sequence\noutside 1 318\nTMhelix 319 341\ninside 342 349

32803 GCF_002155725.1_ASM215572v1 Streptomyces scabiei Terrabacteria group; Actinobacteria; Actinobacteria; Streptomycetales; Streptomycetaceae; Streptomyces MTAQISRRSTLKIAGIAAGAAGLGIGTGLAAAPAAAAGAAFAHPGLLHTTADLARMAAKV WP_086800221.1 Tat pathway signal sequence domain protein [Streptomyces scabiei] Length: 505\nNumber of predicted TMHs: 2\nExp number of AAs in TMHs: 37.55328\nExp number, first 60 AAs: 23.08815\nTotal prob of N-in: 0.99584\nPOSSIBLE N-term signal sequence\ninside 1 12\nTMhelix 13 35\noutside 36 474\nTMhelix 475 492\ninside 493 505

32804 GCF_002155765.1_ASM215576v1 Streptomyces scabiei Terrabacteria group; Actinobacteria; Actinobacteria; Streptomycetales; Streptomycetaceae; Streptomyces MHRLSARRATARRAVVTTAAATVXLLTVGAAGPASAHGAPTDPVSRVSACSPEGGSQDSA WP_086786502.1 chitin-binding protein [Streptomyces scabiei] Length: 342\nNumber of predicted TMHs: 2\nExp number of AAs in TMHs: 37.69395\nExp number, first 60 AAs: 16.59818\nTotal prob of N-in: 0.79066\nPOSSIBLE N-term signal sequence\ninside 1 12\nTMhelix 13 35\noutside 36 306\nTMhelix 307 329\ninside 330 342

32805 GCF_002155725.1_ASM215572v1 Streptomyces scabiei Terrabacteria group; Actinobacteria; Actinobacteria; Streptomycetales; Streptomycetaceae; Streptomyces MSSHRRAAIAGSLLTASFTSVLILGFTASADDEGGLSNHGGKAVGAAPSDVKLSTLLPKE WP_086804664.1 cell wall protein [Streptomyces scabiei] Length: 260\nNumber of predicted TMHs: 2\nExp number of AAs in TMHs: 40.35984\nExp number, first 60 AAs: 21.42036\nTotal prob of N-in: 0.96588\nPOSSIBLE N-term signal sequence\ninside 1 6\nTMhelix 7 29\noutside 30 229\nTMhelix 230 252\ninside 253 260

32806 GCF_002155725.1_ASM215572v1 Streptomyces scabiei Terrabacteria group; Actinobacteria; Actinobacteria; Streptomycetales; Streptomycetaceae; Streptomyces MLGPVENHSLPRTAAFFDLDKTVIAKSSTLTFSKSFYQGGLINRRAALRTAYTQFVFLAG WP_086802367.1 inhibition of morphological differentiation protein [Streptomyces scabiei] Length: 274\nNumber of predicted TMHs: 1\nExp number of AAs in TMHs: 21.74271\nExp number, first 60 AAs: 0.04858\nTotal prob of N-in: 0.01223\noutside 1 244\nTMhelix 245 267\ninside 268 274

32807 GCF_000738695.1_ASM73869v1 Streptomyces scabiei Terrabacteria group; Actinobacteria; Actinobacteria; Streptomycetales; Streptomycetaceae; Streptomyces MLGTSHDTYPRPEDPHVTAEISRRSTLKIAGIAAGTAGLGIGTGLAATPASAAGGALAHP WP_079024514.1 Tat pathway signal sequence domain protein [Streptomyces scabiei] Length: 513\nNumber of predicted TMHs: 1\nExp number of AAs in TMHs: 27.33045\nExp number, first 60 AAs: 10.00799\nTotal prob of N-in: 0.44423\nPOSSIBLE N-term signal sequence\noutside 1 481\nTMhelix 482 501\ninside 502 513

32808 GCF_000738695.1_ASM73869v1 Streptomyces scabiei Terrabacteria group; Actinobacteria; Actinobacteria; Streptomycetales; Streptomycetaceae; Streptomyces MNYSSLSRRGALAAVAGSTVALAAPGAAGAATRASAPSAGSTVRAASVNQLLSELSEGQK WP_037703295.1 hypothetical protein [Streptomyces scabiei] Length: 548\nNumber of predicted TMHs: 2\nExp number of AAs in TMHs: 48.2866999999999\nExp number, first 60 AAs: 8.37457\nTotal prob of N-in: 0.39524\ninside 1 446\nTMhelix 447 469\noutside 470 478\nTMhelix 479 501\ninside 502 548

32809 GCF_001579685.1_ASM157968v1 Streptomyces scabiei Terrabacteria group; Actinobacteria; Actinobacteria; Streptomycetales; Streptomycetaceae; Streptomyces MLGVVENHSLPRTAAFFDLDKTVIAKSSTLTFSKSFYQGGLINRRAALRTAYIQFVFLAG WP_013001810.1 morphological differentiation protein [Streptomyces scabiei] Length: 274\nNumber of predicted TMHs: 1\nExp number of AAs in TMHs: 21.78293\nExp number, first 60 AAs: 0.08751\nTotal prob of N-in: 0.00802\noutside 1 244\nTMhelix 245 267\ninside 268 274

32810 GCF_002155725.1_ASM215572v1 Streptomyces scabiei Terrabacteria group; Actinobacteria; Actinobacteria; Streptomycetales; Streptomycetaceae; Streptomyces MSKAHARALSARIRRSTETWGLRSGWDRRPTVRAAATALIATTSVLALPATTAWADAGQC WP_086798046.1 type VII secretion-associated serine protease mycosin [Streptomyces scabiei] Length: 442\nNumber of predicted TMHs: 1\nExp number of AAs in TMHs: 30.01981\nExp number, first 60 AAs: 7.69319\nTotal prob of N-in: 0.35249\noutside 1 400\nTMhelix 401 423\ninside 424 442

32811 GCF_002155765.1_ASM215576v1 Streptomyces scabiei Terrabacteria group; Actinobacteria; Actinobacteria; Streptomycetales; Streptomycetaceae; Streptomyces

MLGPVENHSLPRTAAFFDLDKTVIAKSSTLTFSKSFYQGGLINRRAALRTAYTQFVFLAG WP_045563048.1 MULTISPECIES: inhibition of morphological differentiation protein [Streptomyces] Length: 274\nNumber of predicted TMHs: 1\nExp number of AAs in TMHs: 21.74861\nExp number, first 60 AAs: 0.04228\nTotal prob of N-in: 0.01050\noutside 1 244\nTMhelix 245 267\ninside 268 274

32812 GCF_001550215.1_ASM155021v1 Streptomyces scabiei Terrabacteria group; Actinobacteria; Actinobacteria; Streptomycetales; Streptomycetaceae; Streptomyces MLGVVENHSLPRTAAFFDLDKTVIAKSSTLTFSKSFYQGGLINRRAALRTAYIQFVFLAG WP_013001810.1 morphological differentiation protein [Streptomyces scabiei] Length: 274\nNumber of predicted TMHs: 1\nExp number of AAs in TMHs: 21.78293\nExp number, first 60 AAs: 0.08751\nTotal prob of N-in: 0.00802\noutside 1 244\nTMhelix 245 267\ninside 268 274

32813 GCF_001550295.1_ASM155029v1 Streptomyces scabiei Terrabacteria group; Actinobacteria; Actinobacteria; Streptomycetales; Streptomycetaceae; Streptomyces MLGVVENHSLPRTAAFFDLDKTVIAKSSTLTFSKSFYQGGLINRRAALRTAYIQFVFLAG WP_013001810.1 morphological differentiation protein [Streptomyces scabiei] Length: 274\nNumber of predicted TMHs: 1\nExp number of AAs in TMHs: 21.78293\nExp number, first 60 AAs: 0.08751\nTotal prob of N-in: 0.00802\noutside 1 244\nTMhelix 245 267\ninside 268 274

32814 GCF_001550235.1_ASM155023v1 Streptomyces scabiei Terrabacteria group; Actinobacteria; Actinobacteria; Streptomycetales; Streptomycetaceae; Streptomyces MLGVVENHSLPRTAAFFDLDKTVIAKSSTLTFSKSFYQGGLINRRAALRTAYIQFVFLAG WP_013001810.1 morphological differentiation protein [Streptomyces scabiei] Length: 274\nNumber of predicted TMHs: 1\nExp number of AAs in TMHs: 21.78293\nExp number, first 60 AAs: 0.08751\nTotal prob of N-in: 0.00802\noutside 1 244\nTMhelix 245 267\ninside 268 274

32815 GCF_002155735.1_ASM215573v1 Streptomyces scabiei Terrabacteria group; Actinobacteria; Actinobacteria; Streptomycetales; Streptomycetaceae; Streptomyces MTAEISRRSTLKIAGIAAGAAGLGVGTGVAATPASAAGGALAHPGLLHTGADLARMAAKV WP_086747795.1 Tat pathway signal sequence domain protein [Streptomyces scabiei] Length: 496\nNumber of predicted TMHs: 2\nExp number of AAs in TMHs: 38.92429\nExp number, first 60 AAs: 21.36239\nTotal prob of N-in: 0.95780\nPOSSIBLE N-term signal sequence\ninside 1 12\nTMhelix 13 35\noutside 36 464\nTMhelix 465 484\ninside 485 496

32816 GCF_002156075.1_ASM215607v1 Streptomyces scabiei Terrabacteria group; Actinobacteria; Actinobacteria; Streptomycetales; Streptomycetaceae; Streptomyces MCSPPGACPPQAAVPRSEDPPVTAEISRRSTLKFAGVAGAAASAAGLGIGTGLTATPASA WP_086754741.1 Tat pathway signal sequence domain protein [Streptomyces scabiei] Length: 533\nNumber of predicted TMHs: 1\nExp number of AAs in TMHs: 20.37128\nExp number, first 60 AAs: 4.13294\nTotal prob of N-in: 0.18329\noutside 1 494\nTMhelix 495 514\ninside 515 533

32817 GCF_001485125.1_ASM148512v1 Streptomyces scabiei Terrabacteria group; Actinobacteria; Actinobacteria; Streptomycetales; Streptomycetaceae; Streptomyces MLGVVENHSLPRTAAFFDLDKTVIAKSSTLTFSKSFYQGGLINRRAALRTAYIQFVFLAG WP_013001810.1 morphological differentiation protein [Streptomyces scabiei] Length: 274\nNumber of predicted TMHs: 1\nExp number of AAs in TMHs: 21.78293\nExp number, first 60 AAs: 0.08751\nTotal prob of N-in: 0.00802\noutside 1 244\nTMhelix 245 267\ninside 268 274

32818 GCF_001005405.1_ASM100540v1 Streptomyces scabiei Terrabacteria group; Actinobacteria; Actinobacteria; Streptomycetales; Streptomycetaceae; Streptomyces MYRLSARRTGAHPAGAHRPGAHHDSAHRDSGRGDGGRRAAVATAAAVVVAPLLLTAGTAT WP_046773940.1 chitin-binding protein [Streptomyces scabiei] Length: 352\nNumber of predicted TMHs: 1\nExp number of AAs in TMHs: 36.84987\nExp number, first 60 AAs: 14.48401\nTotal prob of N-in: 0.69739\nPOSSIBLE N-term signal sequence\noutside 1 318\nTMhelix 319 341\ninside 342 352

32819 GCF_001550225.1_ASM155022v1 Streptomyces scabiei Terrabacteria group; Actinobacteria; Actinobacteria; Streptomycetales; Streptomycetaceae; Streptomyces MLGVVENHSLPRTAAFFDLDKTVIAKSSTLTFSKSFYQGGLINRRAALRTAYIQFVFLAG WP_013001810.1 morphological differentiation protein [Streptomyces scabiei] Length: 274\nNumber of predicted TMHs: 1\nExp number of AAs in TMHs: 21.78293\nExp number, first 60 AAs: 0.08751\nTotal prob of N-in: 0.00802\noutside 1 244\nTMhelix 245 267\ninside 268 274

32820 GCF_001550245.1_ASM155024v1 Streptomyces scabiei Terrabacteria group; Actinobacteria; Actinobacteria; Streptomycetales; Streptomycetaceae; Streptomyces
MLGVVENHSLPRTAAFFDLDKTVIAKSSTLTFSKSFYQGGLINRRAALRTAYIQFVFLAG WP_013001810.1 morphological differentiation protein [Streptomyces scabiei] Length: 274\nNumber of predicted TMHs: 1\nExp number of AAs in TMHs: 21.78293\nExp number, first 60 AAs: 0.08751\nTotal prob of N-in: 0.00802\noutside 1 244\nTMhelix 245 267\ninside 268 274

32821 GCF_001005405.1_ASM100540v1 Streptomyces scabiei Terrabacteria group; Actinobacteria; Actinobacteria; Streptomycetales; Streptomycetaceae; Streptomyces
MLGVVENHSLPRTAAFFDLDKTVIAKSSTLTFSKSFYQGGLINRRAALRTAYIQFVFLAG WP_013001810.1 morphological differentiation protein [Streptomyces scabiei] Length: 274\nNumber of predicted TMHs: 1\nExp number of AAs in TMHs: 21.78293\nExp number, first 60 AAs: 0.08751\nTotal prob of N-in: 0.00802\noutside 1 244\nTMhelix 245 267\ninside 268 274

32822 GCF_001572115.1_ASM157211v1 Streptomyces scabiei Terrabacteria group; Actinobacteria; Actinobacteria; Streptomycetales; Streptomycetaceae; Streptomyces
MLGVVENHSLPRTAAFFDLDKTVIAKSSTLTFSKSFYQGGLINRRAALRTAYIQFVFLAG WP_013001810.1 morphological differentiation protein [Streptomyces scabiei] Length: 274\nNumber of predicted TMHs: 1\nExp number of AAs in TMHs: 21.78293\nExp number, first 60 AAs: 0.08751\nTotal prob of N-in: 0.00802\noutside 1 244\nTMhelix 245 267\ninside 268 274

32823 GCF_000738715.1_ASM73871v1 Streptomyces scabiei Terrabacteria group; Actinobacteria; Actinobacteria; Streptomycetales; Streptomycetaceae; Streptomyces
MLGVVENHSLPRTAAFFDLDKTVIAKSSTLTFSKSFYQGGLINRRAALRTAYIQFVFLAG WP_013001810.1 morphological differentiation protein [Streptomyces scabiei] Length: 274\nNumber of predicted TMHs: 1\nExp number of AAs in TMHs: 21.78293\nExp number, first 60 AAs: 0.08751\nTotal prob of N-in: 0.00802\noutside 1 244\nTMhelix 245 267\ninside 268 274

32824 GCF_001485125.1_ASM148512v1 Streptomyces scabiei Terrabacteria group; Actinobacteria; Actinobacteria; Streptomycetales; Streptomycetaceae; Streptomyces
MYRLSARRTGAHPAGAHRPGAHRDAALRDTGRGDGGRRAAVATAAAVVVAPLLLTAGTAT WP_059083597.1 chitin-binding protein [Streptomyces scabiei] Length: 351\nNumber of predicted TMHs: 1\nExp number of AAs in TMHs: 38.3310799999999\nExp number, first 60 AAs: 15.89555\nTotal prob of N-in: 0.75912\nPOSSIBLE N-term signal sequence\noutside 1 317\nTMhelix 318 340\ninside 341 351

32825 GCF_002155735.1_ASM215573v1 Streptomyces scabiei Terrabacteria group; Actinobacteria; Actinobacteria; Streptomycetales; Streptomycetaceae; Streptomyces
MLGLVENHSLPRTAAFFDLDKTVIAKSSTLTFSKSFYQGGLINRRAALRTAYTQFVFLAG WP_086753873.1 inhibition of morphological differentiation protein [Streptomyces scabiei] Length: 274\nNumber of predicted TMHs: 1\nExp number of AAs in TMHs: 21.73907\nExp number, first 60 AAs: 0.03386\nTotal prob of N-in: 0.00976\noutside 1 244\nTMhelix 245 267\ninside 268 274

32826 GCF_002156075.1_ASM215607v1 Streptomyces scabiei Terrabacteria group; Actinobacteria; Actinobacteria; Streptomycetales; Streptomycetaceae; Streptomyces
MPPPPRPSVRRRAPVLPAALSAAALIGTVLGTAAPAYADPAAEVTPYSVAPGGTLTFSVS WP_086763368.1 hypothetical protein [Streptomyces scabiei] Length: 235\nNumber of predicted TMHs: 2\nExp number of AAs in TMHs: 36.65116\nExp number, first 60 AAs: 16.5969\nTotal prob of N-in: 0.71909\nPOSSIBLE N-term signal sequence\ninside 1 12\nTMhelix 13 35\noutside 36 204\nTMhelix 205 227\ninside 228 235

32827 GCF_002156075.1_ASM215607v1 Streptomyces scabiei Terrabacteria group; Actinobacteria; Actinobacteria; Streptomycetales; Streptomycetaceae; Streptomyces
MLGLVENHSLPRTAAFFDLDKTVIAKSSTLTFSKSFYQGGLINRRDALRTAYTQFVFLAG WP_086764495.1 inhibition of morphological differentiation protein [Streptomyces scabiei] Length: 274\nNumber of predicted TMHs: 1\nExp number of AAs in TMHs: 21.70759\nExp number, first 60 AAs: 0.0064\nTotal prob of N-in: 0.00952\noutside 1 244\nTMhelix 245 267\ninside 268 274

32828 GCF_002156075.1_ASM215607v1 Streptomyces scabiei Terrabacteria group; Actinobacteria; Actinobacteria; Streptomycetales; Streptomycetaceae; Streptomyces
MSVAGPARLPPARNSPVSTAFRTPRVSTYVRPRAGRRAALVGAAALTAVLALAGPAAAHA WP_086765003.1 hypothetical protein [Streptomyces scabiei] Length: 276\nNumber of predicted TMHs: 2\nExp number of

AAs in TMHs: 39.77576\nExp number, first 60 AAs:  17.61956\nTotal prob of N-in:   0.80151\nPOSSIBLE N-term signal sequence\ninside   1   37\nTMhelix   38   60\noutside   61   244\nTMhelix   245   267\ninside   268   276

32829 GCF_002024165.1_SSNF3.0 Streptomyces scabrisporus Terrabacteria group; Actinobacteria; Actinobacteria; Streptomycetales; Streptomycetaceae; Streptomyces
METQAPKRTAAFFDLDKTIIARSSALAFSRSFYHGGLINRRSVLKSAYAQFVFLVGGADH WP_078976509.1 inhibition of morphological differentiation protein [Streptomyces scabrisporus] Length: 268\nNumber of predicted TMHs:  1\nExp number of AAs in TMHs: 21.11556\nExp number, first 60 AAs:  0.18413\nTotal prob of N-in:   0.03661\noutside   1   243\nTMhelix   244   263\ninside   264   268

32830 GCF_000719105.1_ASM71910v1 Streptomyces sclerotialus Terrabacteria group; Actinobacteria; Actinobacteria; Streptomycetales; Streptomycetaceae; Streptomyces
MRRRRPFPLRQLLVGLAGFLLTAAVAAPAVAADAAEFTIEDPRITESSGLAASRLHPGVY WP_030576332.1 hypothetical protein [Streptomyces sclerotialus] Length: 334\nNumber of predicted TMHs:  2\nExp number of AAs in TMHs: 37.20215\nExp number, first 60 AAs:  21.38525\nTotal prob of N-in:   0.99205\nPOSSIBLE N-term signal sequence\ninside   1   11\nTMhelix   12   34\noutside   35   308\nTMhelix   309   326\ninside   327   334

32831 GCF_000720555.1_ASM72055v1 Streptomyces sclerotialus Terrabacteria group; Actinobacteria; Actinobacteria; Streptomycetales; Streptomycetaceae; Streptomyces
MSDNTLSRNTRSHSTLTKAPATAPATALSAGPAERGPSAAPSRRRTALRRLPLVGGAAAG WP_078888588.1 hypothetical protein [Streptomyces sclerotialus] Length: 290\nNumber of predicted TMHs:  2\nExp number of AAs in TMHs: 44.45611\nExp number, first 60 AAs:  8.62243\nTotal prob of N-in:   0.99238\ninside   1   52\nTMhelix   53   75\noutside   76   262\nTMhelix   263   285\ninside   286   290

32832 GCF_000720555.1_ASM72055v1 Streptomyces sclerotialus Terrabacteria group; Actinobacteria; Actinobacteria; Streptomycetales; Streptomycetaceae; Streptomyces
MTDRRRRAALLLLPPPATAGWYPQPATAPGTVLLSREHKPSSPSEGQQGAGTGGTGDPDG WP_030609692.1 hypothetical protein [Streptomyces sclerotialus] Length: 332\nNumber of predicted TMHs:  2\nExp number of AAs in TMHs: 46.7054\nExp number, first 60 AAs:  0.03164\nTotal prob of N-in:   0.86932\ninside   1   179\nTMhelix   180   202\noutside   203   270\nTMhelix   271   293\ninside   294   332

32833 GCF_000719105.1_ASM71910v1 Streptomyces sclerotialus Terrabacteria group; Actinobacteria; Actinobacteria; Streptomycetales; Streptomycetaceae; Streptomyces
MLGLVENHSLPRTAAFFDLDKTVIAKSSTLTFSKSFYQGGLINRRAVLRTAYAQFVFLAG WP_030573624.1 inhibition of morphological differentiation protein [Streptomyces sclerotialus] Length: 274\nNumber of predicted TMHs:  1\nExp number of AAs in TMHs: 21.6491\nExp number, first 60 AAs:  0.01489\nTotal prob of N-in:   0.00202\noutside   1   244\nTMhelix   245   267\ninside   268   274

32834 GCF_000725795.1_Doro.v1.0Streptomyces seoulensis Terrabacteria group; Actinobacteria; Actinobacteria; Streptomycetales; Streptomycetaceae; Streptomyces
MRHARRRVVRRVTRLAAVGGLLLGGGMVAQAAMASETPAAPATTFSAAGTAGTRGAALAE WP_031179362.1 protease [Streptomyces seoulensis] Length: 457\nNumber of predicted TMHs:  2\nExp number of AAs in TMHs: 42.9601599999999\nExp number, first 60 AAs:  22.0795\nTotal prob of N-in:   0.99055\nPOSSIBLE N-term signal sequence\ninside   1   11\nTMhelix   12   34\noutside   35   418\nTMhelix   419   441\ninside   442   457

32835 GCF_000725795.1_Doro.v1.0Streptomyces seoulensis Terrabacteria group; Actinobacteria; Actinobacteria; Streptomycetales; Streptomycetaceae; Streptomyces
MLSDVENHSLPRAAAFFDLDKTVIAKSSTLTFSKSFYQGGLINRRAALRTAYAQFVFLAG WP_031182723.1 inhibition of morphological differentiation protein [Streptomyces seoulensis] Length: 276\nNumber of predicted TMHs:  1\nExp number of AAs in TMHs: 21.55439\nExp number, first 60 AAs:  0.0355\nTotal prob of N-in:   0.00654\noutside   1   246\nTMhelix   247   269\ninside   270   276

32836 GCF_001419745.1_ASM141974v1 Streptomyces silaceus Terrabacteria group; Actinobacteria; Actinobacteria; Streptomycetales; Streptomycetaceae; Streptomyces
MLGLVENHSLPRTAAFFDLDKTVIAKSSTLTFSKSFYQGGLINRRAVLRTAYAQFVFLAG WP_055699300.1 MULTISPECIES: inhibition of morphological differentiation protein [Streptomyces] Length: 288\nNumber of predicted TMHs:  1\nExp number of AAs in TMHs: 21.95172\nExp number, first 60 AAs:  0.01273\nTotal prob of N-in:   0.00126\noutside   1   247\nTMhelix   248   270\ninside   271   288

32837 GCF_001975025.1_ASM197502v1 Streptomyces silaceus Terrabacteria group; Actinobacteria; Actinobacteria; Streptomycetales; Streptomycetaceae; Streptomyces

MRITGPRRGRRRARRALVTALAATLVLAPAATARADDGIRVQQWGLGAMRTDQVWQTTKG WP_076687056.1 type VII secretion-associated serine protease mycosin [Streptomyces silaceus] Length: 404\nNumber of predicted TMHs: 2\nExp number of AAs in TMHs: 39.2287199999999\nExp number, first 60 AAs: 16.36883\nTotal prob of N-in: 0.89406\nPOSSIBLE N-term signal sequence\ninside 1 12\nTMhelix 13 35\noutside 36 365\nTMhelix 366 388\ninside 389 404

32838 GCF_001975025.1_ASM197502v1 Streptomyces silaceus Terrabacteria group; Actinobacteria; Actinobacteria; Streptomycetales; Streptomycetaceae; Streptomyces
MLSIVENHSLPRTAAFFDLDKTVIAKSSTLTFSKSFYQGGLINRRAVLRTAYTQFVFLAG WP_062774786.1 MULTISPECIES: inhibition of morphological differentiation protein [Streptomyces] Length: 274\nNumber of predicted TMHs: 1\nExp number of AAs in TMHs: 21.79397\nExp number, first 60 AAs: 0.03036\nTotal prob of N-in: 0.02122\noutside 1 244\nTMhelix 245 267\ninside 268 274

32839 GCF_001482415.1_ASM148241v1 Streptomyces silvensis Terrabacteria group; Actinobacteria; Actinobacteria; Streptomycetales; Streptomycetaceae; Streptomyces
MSRHARRPQRRSPLRAVAVATAVAGGVLAAPAASAFADDASPKPSKAPTSAPSPEDVAKK WP_058848772.1 hypothetical protein [Streptomyces silvensis] Length: 195\nNumber of predicted TMHs: 2\nExp number of AAs in TMHs: 42.62847\nExp number, first 60 AAs: 22.213\nTotal prob of N-in: 0.99607\nPOSSIBLE N-term signal sequence\ninside 1 12\nTMhelix 13 35\noutside 36 165\nTMhelix 166 185\ninside 186 195

32840 GCF_001482415.1_ASM148241v1 Streptomyces silvensis Terrabacteria group; Actinobacteria; Actinobacteria; Streptomycetales; Streptomycetaceae; Streptomyces
MLDLVENHSLPRTAAFFDLDKTVIAKSSTLTFSKSFYQGGLINRRAVLRTAYAQFVFLAG WP_058850224.1 inhibition of morphological differentiation protein [Streptomyces silvensis] Length: 274\nNumber of predicted TMHs: 1\nExp number of AAs in TMHs: 21.95083\nExp number, first 60 AAs: 0.07871\nTotal prob of N-in: 0.01708\noutside 1 244\nTMhelix 245 267\ninside 268 274

32841 GCF_000424945.1_ASM42494v1 Streptomyces sp. 142MFCol3.1 Terrabacteria group; Actinobacteria; Actinobacteria; Streptomycetales; Streptomycetaceae; Streptomyces
MRHARRRIVRRGARLAAVGGLLCGGLMVTQAMASEPSDTSRAAGSRAQAAVDTGTALVAR WP_028804190.1 protease [Streptomyces sp. 142MFCol3.1] Length: 464\nNumber of predicted TMHs: 2\nExp number of AAs in TMHs: 40.67548\nExp number, first 60 AAs: 20.98584\nTotal prob of N-in: 0.97228\nPOSSIBLE N-term signal sequence\ninside 1 11\nTMhelix 12 34\noutside 35 425\nTMhelix 426 448\ninside 449 464

32842 GCF_000424945.1_ASM42494v1 Streptomyces sp. 142MFCol3.1 Terrabacteria group; Actinobacteria; Actinobacteria; Streptomycetales; Streptomycetaceae; Streptomyces
MLRLVENHSLPRTAAFFDLDKTVIAKSSTLTFSKSFYQGGLINRRAALRTAYAQFVFLAG WP_028799143.1 inhibition of morphological differentiation protein [Streptomyces sp. 142MFCol3.1] Length: 277\nNumber of predicted TMHs: 1\nExp number of AAs in TMHs: 21.68878\nExp number, first 60 AAs: 0.02514\nTotal prob of N-in: 0.00307\noutside 1 247\nTMhelix 248 270\ninside 271 277

32843 GCF_000424945.1_ASM42494v1 Streptomyces sp. 142MFCol3.1 Terrabacteria group; Actinobacteria; Actinobacteria; Streptomycetales; Streptomycetaceae; Streptomyces
MRDSSPRTPRSAPPSPRAAAVPAARSAVLSRRAALGLAAAVPLAASGVPAASAATTIGGE WP_078629026.1 D-alanyl-D-alanine carboxypeptidase [Streptomyces sp. 142MFCol3.1] Length: 415\nNumber of predicted TMHs: 1\nExp number of AAs in TMHs: 35.6859499999999\nExp number, first 60 AAs: 13.98427\nTotal prob of N-in: 0.62036\nPOSSIBLE N-term signal sequence\noutside 1 371\nTMhelix 372 394\ninside 395 415

32844 GCF_000818195.1_Streptomyces_species_150FB_genome_v1 Streptomyces sp. 150FB Terrabacteria group; Actinobacteria; Actinobacteria; Streptomycetales; Streptomycetaceae; Streptomyces
MPDRRRRTALHLPQPAGLVLLPRERQSSSTGSSGSSGSSGSSGSPSGSEGSSGSSGSDGA WP_052489045.1 hypothetical protein [Streptomyces sp. 150FB] Length: 362\nNumber of predicted TMHs: 2\nExp number of AAs in TMHs: 46.05649\nExp number, first 60 AAs: 0\nTotal prob of N-in: 0.91266\ninside 1 198\nTMhelix 199 221\noutside 222 300\nTMhelix 301 323\ninside 324 362

32845 GCF_000818195.1_Streptomyces_species_150FB_genome_v1 Streptomyces sp. 150FB Terrabacteria group; Actinobacteria; Actinobacteria; Streptomycetales; Streptomycetaceae; Streptomyces
MHAHTTRRRTAVTALVRTAVLALLVVMATAGLGAGPALADDGDVTWTVRTAANGYGADRS WP_040026619.1 dihydroorotate oxidase [Streptomyces sp. 150FB] Length: 359\nNumber of predicted TMHs: 2\nExp number of AAs in TMHs: 45.76482\nExp number, first 60 AAs: 21.44055\nTotal prob of N-in: 0.98507\nPOSSIBLE N-term signal sequence\ninside 1 12\nTMhelix 13 35\noutside 36 303\nTMhelix 304 326\ninside 327 359

32846 GCF_000818195.1_Streptomyces_species_150FB_genome_v1 Streptomyces sp. 150FB Terrabacteria group; Actinobacteria; Actinobacteria; Streptomycetales; Streptomycetaceae; Streptomyces MLDVVENHYLPRTAAFFDLDKTVIAKSSTLTFSRSFYQGGLINRRAVLRTAYAQFVFLAG WP_040022593.1 inhibition of morphological differentiation protein [Streptomyces sp. 150FB] Length: 278\nNumber of predicted TMHs: 1\nExp number of AAs in TMHs: 21.75156\nExp number, first 60 AAs: 0.04103\nTotal prob of N-in: 0.00491\noutside 1 247\nTMhelix 248 270\ninside 271 278

32847 GCF_000383635.1_ASM38363v1 Streptomyces sp. 303MFCol5.2 Terrabacteria group; Actinobacteria; Actinobacteria; Streptomycetales; Streptomycetaceae; Streptomyces MKKLSAARRPTLATAVAATAVLLTAGPASAHVEVESESAQALARNVEIAFDAESESATAGWP_020129099.1 hypothetical protein [Streptomyces sp. 303MFCol5.2] Length: 237\nNumber of predicted TMHs: 1\nExp number of AAs in TMHs: 29.62578\nExp number, first 60 AAs: 7.79251\nTotal prob of N-in: 0.43716\noutside 1 205\nTMhelix 206 228\ninside 229 237

32848 GCF_000383635.1_ASM38363v1 Streptomyces sp. 303MFCol5.2 Terrabacteria group; Actinobacteria; Actinobacteria; Streptomycetales; Streptomycetaceae; Streptomyces MHAFSRPSPRLSRRSLLALAATAATAPLAAAPLAATPAAAASDVVVGGARLVADGVQVNG WP_020126801.1 hypothetical protein [Streptomyces sp. 303MFCol5.2] Length: 397\nNumber of predicted TMHs: 1\nExp number of AAs in TMHs: 22.15408\nExp number, first 60 AAs: 3.31406\nTotal prob of N-in: 0.15043\noutside 1 360\nTMhelix 361 383\ninside 384 397

32849 GCF_000383635.1_ASM38363v1 Streptomyces sp. 303MFCol5.2 Terrabacteria group; Actinobacteria; Actinobacteria; Streptomycetales; Streptomycetaceae; Streptomyces MLMVVENHSLPRTAAFFDLDKTVIAKSSTLTFSKSFYQGGLINRRAALRTAYAQFVFLVG WP_028806757.1 inhibition of morphological differentiation protein [Streptomyces sp. 303MFCol5.2] Length: 277\nNumber of predicted TMHs: 1\nExp number of AAs in TMHs: 21.39246\nExp number, first 60 AAs: 0.01616\nTotal prob of N-in: 0.00222\noutside 1 247\nTMhelix 248 270\ninside 271 277

32850 GCF_900129855.1_IMG-taxon_2695421014_annotated_assembly Streptomyces sp. 3124.6 Terrabacteria group; Actinobacteria; Actinobacteria; Streptomycetales; Streptomycetaceae; Streptomyces MLLLVENHSLPRTAAFFDLDKTVIAKSSTLTFSKSFYQGGLINRRAALRTAYAQFVFLVG WP_079660780.1 inhibition of morphological differentiation protein [Streptomyces sp. 3124.6] Length: 279\nNumber of predicted TMHs: 1\nExp number of AAs in TMHs: 21.45206\nExp number, first 60 AAs: 0.01527\nTotal prob of N-in: 0.00337\noutside 1 247\nTMhelix 248 270\ninside 271 279

32851 GCF_900129855.1_IMG-taxon_2695421014_annotated_assembly Streptomyces sp. 3124.6 Terrabacteria group; Actinobacteria; Actinobacteria; Streptomycetales; Streptomycetaceae; Streptomyces MHAFSRSSPRLSRRSLLAFAAAAPLGAPSVSPTSRPPLADGVQVDGGSPLPEKLTARAWL WP_079662723.1 D-alanyl-D-alanine carboxypeptidase [Streptomyces sp. 3124.6] Length: 393\nNumber of predicted TMHs: 1\nExp number of AAs in TMHs: 22.44026\nExp number, first 60 AAs: 0.08359\nTotal prob of N-in: 0.00542\noutside 1 347\nTMhelix 348 370\ninside 371 393

32852 GCF_002028385.1_ASM202838v1 Streptomyces sp. 3211 Terrabacteria group; Actinobacteria; Actinobacteria; Streptomycetales; Streptomycetaceae; Streptomyces MSTVSRRTLVRAAAVTACAGSLLALPTAAALAEGVPAASSARSAAPRLSLVKSLSLADGT WP_079406108.1 hypothetical protein [Streptomyces sp. 3211] Length: 217\nNumber of predicted TMHs: 2\nExp number of AAs in TMHs: 41.61766\nExp number, first 60 AAs: 19.00963\nTotal prob of N-in: 0.88905\nPOSSIBLE N-term signal sequence\ninside 1 8\nTMhelix 9 31\noutside 32 187\nTMhelix 188 210\ninside 211 217

32853 GCF_002028385.1_ASM202838v1 Streptomyces sp. 3211 Terrabacteria group; Actinobacteria; Actinobacteria; Streptomycetales; Streptomycetaceae; Streptomyces MPARRRTAVALTRIASAGLAPLALAAYAAAPAVAHGSMTDPVSRVAACYAEGPESPKTAA WP_079404501.1 chitin-binding protein [Streptomyces sp. 3211] Length: 362\nNumber of predicted TMHs: 2\nExp number of AAs in TMHs: 42.23849\nExp number, first 60 AAs: 20.35407\nTotal prob of N-in: 0.90789\nPOSSIBLE N-term signal sequence\ninside 1 12\nTMhelix 13 35\noutside 36 325\nTMhelix 326 348\ninside 349 362

32854 GCF_002028385.1_ASM202838v1 Streptomyces sp. 3211 Terrabacteria group; Actinobacteria; Actinobacteria; Streptomycetales; Streptomycetaceae; Streptomyces MNRTHAPRRALLAAVFALTLASAGAGAGAASAAEAAPAPQAPYAQDPYGLRLDGAGECTF WP_079406304.1 type VII secretion-associated serine protease mycosin [Streptomyces sp. 3211] Length: 426\nNumber of predicted TMHs:

2\nExp number of AAs in TMHs: 42.51495\nExp number, first 60 AAs: 20.21837\nTotal prob of N-in: 0.92838\nPOSSIBLE N-term signal sequence\ninside 1 10\nTMhelix 11 33\noutside 34 392\nTMhelix 393 415\ninside 416 426

32855 GCF_002028385.1_ASM202838v1 Streptomyces sp. 3211 Terrabacteria group; Actinobacteria; Actinobacteria; Streptomycetales; Streptomycetaceae; Streptomyces
MRRRSPVIAVTGAASGVGAALVSRLAASDEVKQVVAIDERRGDCAAAQWHILDVRDPAIA WP_051779201.1
MULTISPECIES: NAD-dependent dehydratase [Streptomyces] Length: 353\nNumber of predicted TMHs: 1\nExp number of AAs in TMHs: 30.20524\nExp number, first 60 AAs: 7.40836\nTotal prob of N-in: 0.39674\noutside 1 324\nTMhelix 325 347\ninside 348 353

32856 GCF_000383655.1_ASM38365v1 Streptomyces sp. 351MFTsu5.1 Terrabacteria group; Actinobacteria; Actinobacteria; Streptomycetales; Streptomycetaceae; Streptomyces
MLGVVDNHSLPRTAAFFDLDKTVIAKSSTLTFSKSFYQGGLINRRAALRTAYAQFVFLVG WP_028809122.1 inhibition of morphological differentiation protein [Streptomyces sp. 351MFTsu5.1] Length: 277\nNumber of predicted TMHs: 1\nExp number of AAs in TMHs: 21.69284\nExp number, first 60 AAs: 0.0377\nTotal prob of N-in: 0.00335\noutside 1 247\nTMhelix 248 270\ninside 271 277

32857 GCF_000383655.1_ASM38365v1 Streptomyces sp. 351MFTsu5.1 Terrabacteria group; Actinobacteria; Actinobacteria; Streptomycetales; Streptomycetaceae; Streptomyces
MRHARRRVVRRVTRLAAVGGLLLGGTMVTRAVASEPPATAPPRTFAQQAGETGADIVARL WP_028809158.1
protease [Streptomyces sp. 351MFTsu5.1] Length: 454\nNumber of predicted TMHs: 2\nExp number of AAs in TMHs: 40.2269799999999\nExp number, first 60 AAs: 20.5339\nTotal prob of N-in: 0.94038\nPOSSIBLE N-term signal sequence\ninside 1 11\nTMhelix 12 34\noutside 35 415\nTMhelix 416 438\ninside 439 454

32858 GCF_001484705.1_ASM148470v1 Streptomyces sp. 4F Terrabacteria group; Actinobacteria; Actinobacteria; Streptomycetales; Streptomycetaceae; Streptomyces
MLRGVENHLSPRTAAFFDLDKTVIAKSSTLTFSKSFYQGGLINRRAVLRTAYSQFVFLAG WP_033274879.1 MULTISPECIES: inhibition of morphological differentiation protein [Actinobacteria] Length: 277\nNumber of predicted TMHs: 1\nExp number of AAs in TMHs: 21.81221\nExp number, first 60 AAs: 0.02078\nTotal prob of N-in: 0.00383\noutside 1 247\nTMhelix 248 270\ninside 271 277

32859 GCF_001484705.1_ASM148470v1 Streptomyces sp. 4F Terrabacteria group; Actinobacteria; Actinobacteria; Streptomycetales; Streptomycetaceae; Streptomyces
MRHARRRVVRRVTRLAAVGGLLLGGTMVTHAVASEPGTPDAVPFSTASPADASGPGAGLV WP_037643279.1
MULTISPECIES: protease [Streptomyces] Length: 457\nNumber of predicted TMHs: 2\nExp number of AAs in TMHs: 38.82844\nExp number, first 60 AAs: 21.8309\nTotal prob of N-in: 0.97535\nPOSSIBLE N-term signal sequence\ninside 1 11\nTMhelix 12 34\noutside 35 418\nTMhelix 419 441\ninside 442 457

32860 GCF_000816025.1_ASM81602v1 Streptomyces sp. 769 Terrabacteria group; Actinobacteria; Actinobacteria; Streptomycetales; Streptomycetaceae; Streptomyces
MTLMRLTRPTRPTGQRRRAVTAAAAALMLAGAGVVATAGTAQAAEVAYLTECQPPPISGL WP_078875879.1
hypothetical protein [Streptomyces sp. 769] Length: 482\nNumber of predicted TMHs: 2\nExp number of AAs in TMHs: 43.73957\nExp number, first 60 AAs: 21.57355\nTotal prob of N-in: 0.98375\nPOSSIBLE N-term signal sequence\ninside 1 18\nTMhelix 19 41\noutside 42 452\nTMhelix 453 475\ninside 476 482

32861 GCF_000297635.1_ASM29763v1 Streptomyces sp. AA0539 Terrabacteria group; Actinobacteria; Actinobacteria; Streptomycetales; Streptomycetaceae; Streptomyces
MLGQVENLSQPRTAAFFDLDKTVIAKSSTVTFSRSFYQGGLINRRAVLRTAYAQFLYLLG WP_019433990.1 inhibition of morphological differentiation protein [Streptomyces sp. AA0539] Length: 280\nNumber of predicted TMHs: 1\nExp number of AAs in TMHs: 22.36259\nExp number, first 60 AAs: 0.00852\nTotal prob of N-in: 0.00102\noutside 1 246\nTMhelix 247 269\ninside 270 280

32862 GCF_000297635.1_ASM29763v1 Streptomyces sp. AA0539 Terrabacteria group; Actinobacteria; Actinobacteria; Streptomycetales; Streptomycetaceae; Streptomyces
MTPTRRALVTTLALTAALTTTPTAQAADTPQHPWYFDTYRIEDVWEHTTGEGITVAVIDS WP_078575958.1
hypothetical protein [Streptomyces sp. AA0539] Length: 395\nNumber of predicted TMHs: 1\nExp number of AAs in TMHs: 23.25517\nExp number, first 60 AAs: 1.93018\nTotal prob of N-in: 0.10747\noutside 1 368\nTMhelix 369 391\ninside 392 395

32863 GCF_000297635.1_ASM29763v1 Streptomyces sp. AA0539 Terrabacteria group; Actinobacteria; Actinobacteria; Streptomycetales; Streptomycetaceae; Streptomyces

MHLTRPRRGALALAGAAAIVFATPVGAVAAPGSPDDALPEALFGAADPSYDGVWRQSIAL WP_019433066.1 hypothetical protein [Streptomyces sp. AA0539] Length: 468\nNumber of predicted TMHs: 2\nExp number of AAs in TMHs: 42.42524\nExp number, first 60 AAs: 20.20913\nTotal prob of N-in: 0.92223\nPOSSIBLE N-term signal sequence\ninside 1 8\nTMhelix 9 31\noutside 32 415\nTMhelix 416 438\ninside 439 468

32864 GCF_000297635.1_ASM29763v1 Streptomyces sp. AA0539 Terrabacteria group; Actinobacteria; Actinobacteria; Streptomycetales; Streptomycetaceae; Streptomyces MTPTRRALVTTLALTATLTTAPITTAQAADTPQHPWYFDTYRIDEVWEHTTGEGITVAVI WP_019433298.1 hypothetical protein [Streptomyces sp. AA0539] Length: 407\nNumber of predicted TMHs: 1\nExp number of AAs in TMHs: 32.05254\nExp number, first 60 AAs: 10.6765\nTotal prob of N-in: 0.48444\nPOSSIBLE N-term signal sequence\noutside 1 375\nTMhelix 376 398\ninside 399 407

32865 GCF_000280905.1_ASM28090v1 Streptomyces sp. AA1529 Terrabacteria group; Actinobacteria; Actinobacteria; Streptomycetales; Streptomycetaceae; Streptomyces MRIDDPWHDALATGGAEVHGTPVDTPPRRTAPPPRREPVPSAVHGVYPTVQRSAAFRQVR WP_037708340.1 DUF485 domain-containing protein [Streptomyces sp. AA1529] Length: 167\nNumber of predicted TMHs: 2\nExp number of AAs in TMHs: 45.2832\nExp number, first 60 AAs: 0.00094\nTotal prob of N-in: 0.99724\ninside 1 65\nTMhelix 66 88\noutside 89 97\nTMhelix 98 120\ninside 121 167

32866 GCF_000280905.1_ASM28090v1 Streptomyces sp. AA1529 Terrabacteria group; Actinobacteria; Actinobacteria; Streptomycetales; Streptomycetaceae; Streptomyces MITRRRLVTGACALLMALPLGLLPTGTASAEPASDEDAPKVELTLDVSGSMRKRDIDGRS WP_026005050.1 MULTISPECIES: inter-alpha-trypsin inhibitor heavy chain H5 [Streptomyces] Length: 421\nNumber of predicted TMHs: 2\nExp number of AAs in TMHs: 38.90213\nExp number, first 60 AAs: 16.54504\nTotal prob of N-in: 0.77694\nPOSSIBLE N-term signal sequence\ninside 1 6\nTMhelix 7 29\noutside 30 388\nTMhelix 389 411\ninside 412 421

32867 GCF_000280905.1_ASM28090v1 Streptomyces sp. AA1529 Terrabacteria group; Actinobacteria; Actinobacteria; Streptomycetales; Streptomycetaceae; Streptomyces MSPRRRRATVLTAATVLSAAAAPVASAAAPAAPVTSVAASPARTTTSGLPEGLYGTADPK WP_019357822.1 hypothetical protein [Streptomyces sp. AA1529] Length: 443\nNumber of predicted TMHs: 1\nExp number of AAs in TMHs: 34.00137\nExp number, first 60 AAs: 11.69215\nTotal prob of N-in: 0.52626\nPOSSIBLE N-term signal sequence\noutside 1 413\nTMhelix 414 436\ninside 437 443

32868 GCF_000280905.1_ASM28090v1 Streptomyces sp. AA1529 Terrabacteria group; Actinobacteria; Actinobacteria; Streptomycetales; Streptomycetaceae; Streptomyces MLGAVKNLAMPRTAAFFDLDKTVIAKSSTLTFGKSFYQGGLINRRAVLRTAYAQFVYLLG WP_019355563.1 inhibition of morphological differentiation protein [Streptomyces sp. AA1529] Length: 281\nNumber of predicted TMHs: 1\nExp number of AAs in TMHs: 22.26414\nExp number, first 60 AAs: 0.03425\nTotal prob of N-in: 0.01227\noutside 1 246\nTMhelix 247 269\ninside 270 281

32869 GCF_000158875.1_ASM15887v1 Streptomyces sp. AA4 Terrabacteria group; Actinobacteria; Actinobacteria; Streptomycetales; Streptomycetaceae; Streptomyces MAEPRSEEAPRRVAAFFDLDKTIIASSSALAFSKPLLREGLINRRAALRSAYAQLVFSLA WP_009084385.1 haloacid dehalogenase [Streptomyces sp. AA4] Length: 273\nNumber of predicted TMHs: 1\nExp number of AAs in TMHs: 20.42196\nExp number, first 60 AAs: 0.16961\nTotal prob of N-in: 0.03962\noutside 1 242\nTMhelix 243 262\ninside 263 273

32870 GCF_000818175.1_ASM81817v1 Streptomyces sp. AcH 505 Terrabacteria group; Actinobacteria; Actinobacteria; Streptomycetales; Streptomycetaceae; Streptomyces MPDRRRRTALHLPPPSGPVLLQRERQSSSTGPTGSSGSSGPAEPGDPASGSGSGSGSGSA WP_041985421.1 hypothetical protein [Streptomyces sp. AcH 505] Length: 345\nNumber of predicted TMHs: 2\nExp number of AAs in TMHs: 46.9565600000001\nExp number, first 60 AAs: 0\nTotal prob of N-in: 0.82331\ninside 1 191\nTMhelix 192 214\noutside 215 283\nTMhelix 284 306\ninside 307 345

32871 GCF_000818175.1_ASM81817v1 Streptomyces sp. AcH 505 Terrabacteria group; Actinobacteria; Actinobacteria; Streptomycetales; Streptomycetaceae; Streptomyces MHSADAPTRQDRRRAVRAAALTAAVALVAFGTGAAPAFADAGQHKAGQHKPKDNSAKTAT WP_041984158.1 phospholipase C [Streptomyces sp. AcH 505] Length: 628\nNumber of predicted TMHs: 2\nExp number of AAs in TMHs: 42.58161\nExp number, first 60 AAs: 21.0542\nTotal prob of N-in: 0.98913\nPOSSIBLE N-term signal sequence\ninside 1 19\nTMhelix 20 42\noutside 43 597\nTMhelix 598 620\ninside 621 628

32872 GCF_000177175.2_ASM17717v2 Streptomyces sp. ACT-1 Terrabacteria group; Actinobacteria; Actinobacteria; Streptomycetales; Streptomycetaceae; Streptomyces
MGTEQQARRGPRARRGLLSALTASLLTLGAALTHLTTASPASADPIAKCTRTKGAIVAVD WP_003965687.1 peptidase [Streptomyces sp. ACT-1] Length: 931\nNumber of predicted TMHs: 2\nExp number of AAs in TMHs: 40.26699\nExp number, first 60 AAs: 18.69314\nTotal prob of N-in: 0.84908\nPOSSIBLE N-term signal sequence\ninside 1 15\nTMhelix 16 38\noutside 39 893\nTMhelix 894 916\ninside 917 931

32873 GCF_000177175.2_ASM17717v2 Streptomyces sp. ACT-1 Terrabacteria group; Actinobacteria; Actinobacteria; Streptomycetales; Streptomycetaceae; Streptomyces
MLSVVENCFSPRTAAFFDLDKTVIAKSSTLTFSKSFYQGGLINRRAVLRTAYAQFVFLAG WP_003967438.1 MULTISPECIES: haloacid dehalogenase [Streptomyces] Length: 281\nNumber of predicted TMHs: 1\nExp number of AAs in TMHs: 21.47765\nExp number, first 60 AAs: 0.03233\nTotal prob of N-in: 0.00650\noutside 1 247\nTMhelix 248 270\ninside 271 281

32874 GCF_900177255.1_IMG-taxon_2524614730_annotated_assembly Streptomyces sp. Amel2xC10 Terrabacteria group; Actinobacteria; Actinobacteria; Streptomycetales; Streptomycetaceae; Streptomyces
MLMLVENHSLPRTAAFFDLDKTVIAKSSTLTFSKSFYQGGLINRRAALRTAYAQFVFLVG WP_030795901.1 inhibition of morphological differentiation protein [Streptomyces antibioticus] Length: 277\nNumber of predicted TMHs: 1\nExp number of AAs in TMHs: 21.68626\nExp number, first 60 AAs: 0.01173\nTotal prob of N-in: 0.00195\noutside 1 247\nTMhelix 248 270\ninside 271 277

32875 GCF_000383935.1_ASM38393v1 Streptomyces sp. Amel2xE9 Terrabacteria group; Actinobacteria; Actinobacteria; Streptomycetales; Streptomycetaceae; Streptomyces
MLRVVENHSLPRTAAFFDLDKTVIAKSSTLTFSKSFYQGGLINRRAVLRTAYAQFVFLAG WP_009189832.1 MULTISPECIES: haloacid dehalogenase [Streptomyces] Length: 277\nNumber of predicted TMHs: 1\nExp number of AAs in TMHs: 21.60611\nExp number, first 60 AAs: 0.04466\nTotal prob of N-in: 0.00565\noutside 1 247\nTMhelix 248 270\ninside 271 277

32876 GCF_000383935.1_ASM38393v1 Streptomyces sp. Amel2xE9 Terrabacteria group; Actinobacteria; Actinobacteria; Streptomycetales; Streptomycetaceae; Streptomyces
MAGRGRRTTLRSLGALTGAVALTVLGPAAPGRAAGRPAPPATPIEHVVVLFDENISFDHY WP_019983617.1 hypothetical protein [Streptomyces sp. Amel2xE9] Length: 608\nNumber of predicted TMHs: 1\nExp number of AAs in TMHs: 38.82952\nExp number, first 60 AAs: 16.93737\nTotal prob of N-in: 0.82198\nPOSSIBLE N-term signal sequence\noutside 1 576\nTMhelix 577 599\ninside 600 608

32877 GCF_000383935.1_ASM38393v1 Streptomyces sp. Amel2xE9 Terrabacteria group; Actinobacteria; Actinobacteria; Streptomycetales; Streptomycetaceae; Streptomyces
MPATTGDRSRRRPLAFAAAVATAAAIGAASLAAAPTTAAAGGIPLKGYELTWGIKETYRS WP_019982084.1 hypothetical protein [Streptomyces sp. Amel2xE9] Length: 538\nNumber of predicted TMHs: 2\nExp number of AAs in TMHs: 45.1150299999999\nExp number, first 60 AAs: 22.65991\nTotal prob of N-in: 0.99239\nPOSSIBLE N-term signal sequence\ninside 1 12\nTMhelix 13 35\noutside 36 504\nTMhelix 505 527\ninside 528 538

32878 GCF_000383935.1_ASM38393v1 Streptomyces sp. Amel2xE9 Terrabacteria group; Actinobacteria; Actinobacteria; Streptomycetales; Streptomycetaceae; Streptomyces
MRHARRRVVRRAARLTAVAGLLLGTAMVAQSALAGEPSGTRASPRSSAGSPAAPGSGLVA WP_037725443.1 protease [Streptomyces sp. Amel2xE9] Length: 469\nNumber of predicted TMHs: 2\nExp number of AAs in TMHs: 43.0612199999999\nExp number, first 60 AAs: 21.83139\nTotal prob of N-in: 0.98424\nPOSSIBLE N-term signal sequence\ninside 1 11\nTMhelix 12 34\noutside 35 430\nTMhelix 431 453\ninside 454 469

32879 GCF_000383935.1_ASM38393v1 Streptomyces sp. Amel2xE9 Terrabacteria group; Actinobacteria; Actinobacteria; Streptomycetales; Streptomycetaceae; Streptomyces
MPAPHKTARRTLLVASAALLSVSLTAPAALAAPSPKTSRPAAPPARMSTVGGARLGQAGT WP_019981628.1 D-alanyl-D-alanine carboxypeptidase [Streptomyces sp. Amel2xE9] Length: 427\nNumber of predicted TMHs: 2\nExp number of AAs in TMHs: 41.66102\nExp number, first 60 AAs: 19.31271\nTotal prob of N-in: 0.91456\nPOSSIBLE N-term signal sequence\ninside 1 11\nTMhelix 12 34\noutside 35 388\nTMhelix 389 411\ninside 412 427

32880 GCF_001704635.1_ASM170463v1 Streptomyces sparsogenes Terrabacteria group; Actinobacteria; Actinobacteria; Streptomycetales; Streptomycetaceae; Streptomyces
MVTGEVSEARRKAVGLGSGACHALGLMVLAITEWVRADLKDATSLASHSYLKDMLRLAAD WP_065963662.1 hypothetical protein [Streptomyces sparsogenes] Length: 152\nNumber of predicted TMHs: 2\nExp

number of AAs in TMHs: 71.82401\nExp number, first 60 AAs: 22.34642\nTotal prob of N-in: 0.99607\nPOSSIBLE N-term signal sequence\ninside 1 12\nTMhelix 13 35\noutside 36 127\nTMhelix 128 150\ninside 151 152

32881 GCF_001704635.1_ASM170463v1 Streptomyces sparsogenes Terrabacteria group; Actinobacteria; Actinobacteria; Streptomycetales; Streptomycetaceae; Streptomyces MLGVVENHSLPRTAAFFDLDKTVIAKSSTLTFSKSFYQGGLINRRAVLRTAYAQFVFLAG WP_065961517.1 inhibition of morphological differentiation protein [Streptomyces sparsogenes] Length: 278\nNumber of predicted TMHs: 1\nExp number of AAs in TMHs: 20.71292\nExp number, first 60 AAs: 0.02768\nTotal prob of N-in: 0.00716\noutside 1 246\nTMhelix 247 269\ninside 270 278

32882 GCF_001704635.1_ASM170463v1 Streptomyces sparsogenes Terrabacteria group; Actinobacteria; Actinobacteria; Streptomycetales; Streptomycetaceae; Streptomyces MSAADDDGNRSLGQLVATATTELSALVHDEIALAKAEFRDSARRTLLGSGAAVSAGVLVL WP_065961531.1 transporter [Streptomyces sparsogenes] Length: 153\nNumber of predicted TMHs: 2\nExp number of AAs in TMHs: 46.4376800000001\nExp number, first 60 AAs: 13.14988\nTotal prob of N-in: 0.92902\nPOSSIBLE N-term signal sequence\ninside 1 45\nTMhelix 46 68\noutside 69 82\nTMhelix 83 105\ninside 106 153

32883 GCF_001704635.1_ASM170463v1 Streptomyces sparsogenes Terrabacteria group; Actinobacteria; Actinobacteria; Streptomycetales; Streptomycetaceae; Streptomyces MSSPDREVRAARNDTPRTARRPVVAVTGAASGAGALLTQRLAESDEIKHVLAIDERRGDV WP_065964212.1 NAD-dependent dehydratase [Streptomyces sparsogenes] Length: 369\nNumber of predicted TMHs: 1\nExp number of AAs in TMHs: 21.31707\nExp number, first 60 AAs: 1.44367\nTotal prob of N-in: 0.08113\noutside 1 339\nTMhelix 340 359\ninside 360 369

32884 GCF_001704635.1_ASM170463v1 Streptomyces sparsogenes Terrabacteria group; Actinobacteria; Actinobacteria; Streptomycetales; Streptomycetaceae; Streptomyces MQLPSLSVRRRALLALAGAVLVSAAGFAPSAEAQDIRSQQWYLDAMKADAIWKVSRGQGI WP_076970707.1 serine protease [Streptomyces sparsogenes] Length: 413\nNumber of predicted TMHs: 1\nExp number of AAs in TMHs: 36.9750299999999\nExp number, first 60 AAs: 14.81189\nTotal prob of N-in: 0.75311\nPOSSIBLE N-term signal sequence\noutside 1 384\nTMhelix 385 407\ninside 408 413

32885 GCF_001279985.1_ASM127998v1 Streptomyces sp. AS58 Terrabacteria group; Actinobacteria; Actinobacteria; Streptomycetales; Streptomycetaceae; Streptomyces MPMSRSIEPRFRPRRAYVMAAAVTGALLTAAPAAADGTPTAPDGGGAANGKPTVVLVHGA WP_053756300.1 alpha/beta hydrolase [Streptomyces sp. AS58] Length: 327\nNumber of predicted TMHs: 1\nExp number of AAs in TMHs: 31.62069\nExp number, first 60 AAs: 9.41823\nTotal prob of N-in: 0.45769\noutside 1 295\nTMhelix 296 318\ninside 319 327

32886 GCF_001279985.1_ASM127998v1 Streptomyces sp. AS58 Terrabacteria group; Actinobacteria; Actinobacteria; Streptomycetales; Streptomycetaceae; Streptomyces MSSPDPQVRAARNHSTPSSRRAPVVAVTGAATGVGALLTARLAASEEIKQVIAIDERRGE WP_053757466.1 NAD-dependent dehydratase [Streptomyces sp. AS58] Length: 370\nNumber of predicted TMHs: 1\nExp number of AAs in TMHs: 27.96977\nExp number, first 60 AAs: 7.23733\nTotal prob of N-in: 0.30067\noutside 1 340\nTMhelix 341 360\ninside 361 370

32887 GCF_001279985.1_ASM127998v1 Streptomyces sp. AS58 Terrabacteria group; Actinobacteria; Actinobacteria; Streptomycetales; Streptomycetaceae; Streptomyces MLRGVENHSLPRTAAFFDLDKTVIAKSSTLTFSKSFYQGGLINRRAVLRTAYTQFVFLAG WP_053756520.1 inhibition of morphological differentiation protein [Streptomyces sp. AS58] Length: 277\nNumber of predicted TMHs: 1\nExp number of AAs in TMHs: 21.63851\nExp number, first 60 AAs: 0.0252\nTotal prob of N-in: 0.00356\noutside 1 247\nTMhelix 248 270\ninside 271 277

32888 GCF_001279985.1_ASM127998v1 Streptomyces sp. AS58 Terrabacteria group; Actinobacteria; Actinobacteria; Streptomycetales; Streptomycetaceae; Streptomyces MRHARRRIVRRATRLAAVGGLLLGAGMVTQAVASEPSGSAGVPNASVRSAAGTGEDLVER WP_053757042.1 protease [Streptomyces sp. AS58] Length: 457\nNumber of predicted TMHs: 2\nExp number of AAs in TMHs: 40.1675\nExp number, first 60 AAs: 20.42459\nTotal prob of N-in: 0.94222\nPOSSIBLE N-term signal sequence\ninside 1 11\nTMhelix 12 34\noutside 35 416\nTMhelix 417 439\ninside 440 457

32889 GCF_000373645.1_ASM37364v1 Streptomyces sp. ATexAB-D23 Terrabacteria group; Actinobacteria; Actinobacteria; Streptomycetales; Streptomycetaceae; Streptomyces

MTRTLARRATFSLAALALTLGPAAPAHTSETLPPGAGSDAAPRTGTGRKVIGDGHVDLGP WP_018552017.1 hypothetical protein [Streptomyces sp. ATexAB-D23] Length: 308\nNumber of predicted TMHs: 1\nExp number of AAs in TMHs: 22.08837\nExp number, first 60 AAs: 1.95287\nTotal prob of N-in: 0.18249\noutside 1 265\nTMhelix 266 285\ninside 286 308

32890 GCF_000373645.1_ASM37364v1 Streptomyces sp. ATexAB-D23 Terrabacteria group; Actinobacteria; Actinobacteria; Streptomycetales; Streptomycetaceae; Streptomyces MLCLVENRFLPRTAAFFDLDKTVIAKSSTLTFSKSFYHGGLINRRAVLRTAYVQFVFLAG WP_026249521.1 inhibition of morphological differentiation protein [Streptomyces sp. ATexAB-D23] Length: 279\nNumber of predicted TMHs: 1\nExp number of AAs in TMHs: 21.74932\nExp number, first 60 AAs: 0.03032\nTotal prob of N-in: 0.00322\noutside 1 247\nTMhelix 248 270\ninside 271 279

32891 GCF_000373645.1_ASM37364v1 Streptomyces sp. ATexAB-D23 Terrabacteria group; Actinobacteria; Actinobacteria; Streptomycetales; Streptomycetaceae; Streptomyces MTRRARRRFLGALCAATAFALLPAAPAAADTIRAQQWGLDALHTDRAWQTTRGKGVTVAV WP_018556187.1 type VII secretion-associated serine protease [Streptomyces sp. ATexAB-D23] Length: 396\nNumber of predicted TMHs: 2\nExp number of AAs in TMHs: 41.73659\nExp number, first 60 AAs: 16.3824\nTotal prob of N-in: 0.90889\nPOSSIBLE N-term signal sequence\ninside 1 6\nTMhelix 7 29\noutside 30 360\nTMhelix 361 383\ninside 384 396

32892 GCF_001469455.2_ASM146945v2 Streptomyces sp. AVP053U2 Terrabacteria group; Actinobacteria; Actinobacteria; Streptomycetales; Streptomycetaceae; Streptomyces MLRDVENHSLPRTAAFFDLDKTVIAKSSTLTFSKSFYQGGLINRRAVLRTAYAQFVFLAG WP_042163730.1 MULTISPECIES: inhibition of morphological differentiation protein [Streptomyces] Length: 277\nNumber of predicted TMHs: 1\nExp number of AAs in TMHs: 20.8169\nExp number, first 60 AAs: 0.10818\nTotal prob of N-in: 0.02276\noutside 1 247\nTMhelix 248 270\ninside 271 277

32893 GCF_001469455.2_ASM146945v2 Streptomyces sp. AVP053U2 Terrabacteria group; Actinobacteria; Actinobacteria; Streptomycetales; Streptomycetaceae; Streptomyces MRHARRRVVRRVTRLAAVGGLLLGATMVTRAVASEPPAPAPVPYAASEEDTGPGSALVSR WP_078878315.1 MULTISPECIES: serine protease [Streptomyces] Length: 454\nNumber of predicted TMHs: 2\nExp number of AAs in TMHs: 37.19435\nExp number, first 60 AAs: 20.65846\nTotal prob of N-in: 0.95154\nPOSSIBLE N-term signal sequence\ninside 1 11\nTMhelix 12 34\noutside 35 415\nTMhelix 416 438\ninside 439 454

32894 GCF_000470535.1_STREPTOMYCES1 Streptomyces sp. AW19M42 Terrabacteria group; Actinobacteria; Actinobacteria; Streptomycetales; Streptomycetaceae; Streptomyces MLGLVENCFLPRTAAFFDLDKTVIAKSSTLTFSKSFYQGGLINRRAVLRTAYAQFVFLAG WP_024493800.1 inhibition of morphological differentiation protein [Streptomyces sp. AW19M42] Length: 279\nNumber of predicted TMHs: 1\nExp number of AAs in TMHs: 21.57305\nExp number, first 60 AAs: 0.00893\nTotal prob of N-in: 0.00091\noutside 1 247\nTMhelix 248 270\ninside 271 279

32895 GCF_002081495.1_ASM208149v1 Streptomyces sp. B9173 Terrabacteria group; Actinobacteria; Actinobacteria; Streptomycetales; Streptomycetaceae; Streptomyces MLMAVENHSLPRTAAFFDLDKTVIAKSSTLTFSKSFYQGGLINRRAALRTAYAQFVFLVG WP_081547920.1 inhibition of morphological differentiation protein [Streptomyces sp. B9173] Length: 277\nNumber of predicted TMHs: 1\nExp number of AAs in TMHs: 21.51658\nExp number, first 60 AAs: 0.01718\nTotal prob of N-in: 0.00189\noutside 1 247\nTMhelix 248 270\ninside 271 277

32896 GCF_002081495.1_ASM208149v1 Streptomyces sp. B9173 Terrabacteria group; Actinobacteria; Actinobacteria; Streptomycetales; Streptomycetaceae; Streptomyces MHAFSRPFPGSSPRPSRRSLLTLAATAPLTVAPLAATPAAAASDVVVGGARLVADGVQVN WP_081546686.1 D-alanyl-D-alanine carboxypeptidase [Streptomyces sp. B9173] Length: 398\nNumber of predicted TMHs: 1\nExp number of AAs in TMHs: 21.77012\nExp number, first 60 AAs: 2.71293\nTotal prob of N-in: 0.11889\noutside 1 361\nTMhelix 362 384\ninside 385 398

32897 GCF_000373665.1_ASM37366v1 Streptomyces sp. BoleA5 Terrabacteria group; Actinobacteria; Actinobacteria; Streptomycetales; Streptomycetaceae; Streptomyces MLVLVENHLPPRTAAFFDLDKTVIAKSSTLAFSRPFYQGGLINRRAVLRTAYAQFVYLLG WP_018562178.1 inhibition of morphological differentiation protein [Streptomyces sp. BoleA5] Length: 276\nNumber of predicted TMHs: 1\nExp number of AAs in TMHs: 21.99676\nExp number, first 60 AAs: 0.01532\nTotal prob of N-in: 0.00363\noutside 1 245\nTMhelix 246 268\ninside 269 276

32898 GCF_001905575.1_ASM190557v1 Streptomyces sp. CB00072 Terrabacteria group; Actinobacteria; Actinobacteria; Streptomycetales; Streptomycetaceae; Streptomyces
MLSVVENCFSPRTAAFFDLDKTVIAKSSTLTFSKSFYQGGLINRRAVLRTAYAQFVFLAG WP_003967438.1 MULTISPECIES: haloacid dehalogenase [Streptomyces] Length: 281\nNumber of predicted TMHs: 1\nExp number of AAs in TMHs: 21.47765\nExp number, first 60 AAs: 0.03233\nTotal prob of N-in: 0.00650\noutside 1 247\nTMhelix 248 270\ninside 271 281

32899 GCF_001905595.1_ASM190559v1 Streptomyces sp. CB00316 Terrabacteria group; Actinobacteria; Actinobacteria; Streptomycetales; Streptomycetaceae; Streptomyces
MLSVVENCFSPRTAAFFDLDKTVIAKSSTLTFSKSFYQGGLINRRAVLRTAYTQFVFLAG WP_073874550.1 inhibition of morphological differentiation protein [Streptomyces sp. CB00316] Length: 281\nNumber of predicted TMHs: 1\nExp number of AAs in TMHs: 21.75071\nExp number, first 60 AAs: 0.02133\nTotal prob of N-in: 0.00734\noutside 1 247\nTMhelix 248 270\ninside 271 281

32900 GCF_001905445.1_ASM190544v1 Streptomyces sp. CB00455 Terrabacteria group; Actinobacteria; Actinobacteria; Streptomycetales; Streptomycetaceae; Streptomyces
MRRRTAFPALLFPALMAALGLTLLAPATAHAHGDTVKVAVTGQRDGHVTADVTWQNDGDA WP_073918172.1 hypothetical protein [Streptomyces sp. CB00455]Length: 227\nNumber of predicted TMHs: 2\nExp number of AAs in TMHs: 45.25023\nExp number, first 60 AAs: 22.48068\nTotal prob of N-in: 0.99688\nPOSSIBLE N-term signal sequence\ninside 1 6\nTMhelix 7 29\noutside 30 186\nTMhelix 187 209\ninside 210 227

32901 GCF_001905445.1_ASM190544v1 Streptomyces sp. CB00455 Terrabacteria group; Actinobacteria; Actinobacteria; Streptomycetales; Streptomycetaceae; Streptomyces
MTQSPGRRRALPAALALVALCATPAAASAAVPDTDTAPTASGHATPPQRPYALRPDGAGE WP_073910812.1 type VII secretion-associated serine protease mycosin [Streptomyces sp. CB00455] Length: 429\nNumber of predicted TMHs: 1\nExp number of AAs in TMHs: 27.93184\nExp number, first 60 AAs: 5.46891\nTotal prob of N-in: 0.28498\noutside 1 395\nTMhelix 396 418\ninside 419 429

32902 GCF_001905445.1_ASM190544v1 Streptomyces sp. CB00455 Terrabacteria group; Actinobacteria; Actinobacteria; Streptomycetales; Streptomycetaceae; Streptomyces
MPARRRTVTLTRIASAGLAPVAVAAYAAAPAVAHGSMTDPVSRVAACYAEGPESPKSAAC WP_073920678.1 chitin-binding protein [Streptomyces sp. CB00455] Length: 367\nNumber of predicted TMHs: 2\nExp number of AAs in TMHs: 43.24238\nExp number, first 60 AAs: 20.68387\nTotal prob of N-in: 0.94137\nPOSSIBLE N-term signal sequence\ninside 1 12\nTMhelix 13 35\noutside 36 332\nTMhelix 333 355\ninside 356 367

32903 GCF_001905735.1_ASM190573v1 Streptomyces sp. CB01249 Terrabacteria group; Actinobacteria; Actinobacteria; Streptomycetales; Streptomycetaceae; Streptomyces
MLCRVENRFLPRTAAFFDLDKTVIAKSSTLTFSKSFYHGGLINRRAVLRTAYTQFVFLAG WP_073864053.1 inhibition of morphological differentiation protein [Streptomyces sp. CB01249] Length: 279\nNumber of predicted TMHs: 1\nExp number of AAs in TMHs: 21.97754\nExp number, first 60 AAs: 0.03012\nTotal prob of N-in: 0.00683\noutside 1 247\nTMhelix 248 270\ninside 271 279

32904 GCF_001905795.1_ASM190579v1 Streptomyces sp. CB01580 Terrabacteria group; Actinobacteria; Actinobacteria; Streptomycetales; Streptomycetaceae; Streptomyces
MSRRARRRVLGAVCAATAFALLPAVPARADAIRDQQWGLEALHADQAWRTTKGRGITVAV WP_073787791.1 type VII secretion-associated serine protease mycosin [Streptomyces sp. CB01580] Length: 399\nNumber of predicted TMHs: 1\nExp number of AAs in TMHs: 37.14287\nExp number, first 60 AAs: 12.34034\nTotal prob of N-in: 0.80528\nPOSSIBLE N-term signal sequence\noutside 1 365\nTMhelix 366 388\ninside 389 399

32905 GCF_001905795.1_ASM190579v1 Streptomyces sp. CB01580 Terrabacteria group; Actinobacteria; Actinobacteria; Streptomycetales; Streptomycetaceae; Streptomyces
MLCVVENRFLPRTAAFFDLDKTVIAKSSTLTFSKSFYHGGLINRRAVLRTAYAQFVFLAG WP_073788300.1 inhibition of morphological differentiation protein [Streptomyces sp. CB01580] Length: 279\nNumber of predicted TMHs: 1\nExp number of AAs in TMHs: 21.2254\nExp number, first 60 AAs: 0.02935\nTotal prob of N-in: 0.00590\noutside 1 247\nTMhelix 248 270\ninside 271 279

32906 GCF_001905795.1_ASM190579v1 Streptomyces sp. CB01580 Terrabacteria group; Actinobacteria; Actinobacteria; Streptomycetales; Streptomycetaceae; Streptomyces
MAVARRPLLTATAAGTLLAALWFVPSANATDERTDGTGGTTTSVTGPTGRSTGSADRGTD WP_073781669.1 hypothetical protein [Streptomyces sp. CB01580]Length: 153\nNumber of predicted TMHs: 2\nExp number of

AAs in TMHs: 40.79269\nExp number, first 60 AAs: 19.68684\nTotal prob of N-in: 0.97398\nPOSSIBLE N-term signal sequence\ninside 1 6\nTMhelix 7 24\noutside 25 122\nTMhelix 123 145\ninside 146 153

32907 GCF_001905855.1_ASM190585v1 Streptomyces sp. CB01883 Terrabacteria group; Actinobacteria; Actinobacteria; Streptomycetales; Streptomycetaceae; Streptomyces MISVVHGPEVTRRAPVARALALLTLTWTLVGCPVGSASADACAYASTGPEGTTAVAYAGG WP_073891173.1 hypothetical protein [Streptomyces sp. CB01883]Length: 205\nNumber of predicted TMHs: 1\nExp number of AAs in TMHs: 27.40561\nExp number, first 60 AAs: 8.64203\nTotal prob of N-in: 0.34089\noutside 1 179\nTMhelix 180 202\ninside 203 205

32908 GCF_001905855.1_ASM190585v1 Streptomyces sp. CB01883 Terrabacteria group; Actinobacteria; Actinobacteria; Streptomycetales; Streptomycetaceae; Streptomyces MLRGVENHSLPRAAAFFDLDKTVIAKSSTLTFSKSFYQGGLINRRAALRTAYAQFVFLAG WP_073901248.1 inhibition of morphological differentiation protein [Streptomyces sp. CB01883] Length: 276\nNumber of predicted TMHs: 1\nExp number of AAs in TMHs: 21.7541\nExp number, first 60 AAs: 0.03332\nTotal prob of N-in: 0.00354\noutside 1 247\nTMhelix 248 270\ninside 271 276

32909 GCF_001905855.1_ASM190585v1 Streptomyces sp. CB01883 Terrabacteria group; Actinobacteria; Actinobacteria; Streptomycetales; Streptomycetaceae; Streptomyces MFAPQKTGRRSLLVTSATVLTLSVTAPAALAAPTPPASPSATPPAGMSTLGGARLGMAGT WP_079192310.1 D-alanyl-D-alanine carboxypeptidase [Streptomyces sp. CB01883] Length: 411\nNumber of predicted TMHs: 1\nExp number of AAs in TMHs: 35.47565\nExp number, first 60 AAs: 12.92169\nTotal prob of N-in: 0.60506\nPOSSIBLE N-term signal sequence\noutside 1 372\nTMhelix 373 395\ninside 396 411

32910 GCF_001905855.1_ASM190585v1 Streptomyces sp. CB01883 Terrabacteria group; Actinobacteria; Actinobacteria; Streptomycetales; Streptomycetaceae; Streptomyces MRHARRRVVRRVTRLAAVGGLLLGGAMVTQAAMASETPPATTGALRSAAGAGDPGDALVA WP_073890878.1 serine protease [Streptomyces sp. CB01883] Length: 461\nNumber of predicted TMHs: 2\nExp number of AAs in TMHs: 41.2035499999999\nExp number, first 60 AAs: 21.99116\nTotal prob of N-in: 0.98918\nPOSSIBLE N-term signal sequence\ninside 1 11\nTMhelix 12 34\noutside 35 422\nTMhelix 423 445\ninside 446 461

32911 GCF_001905825.1_ASM190582v1 Streptomyces sp. CB02009 Terrabacteria group; Actinobacteria; Actinobacteria; Streptomycetales; Streptomycetaceae; Streptomyces MLRLVENHSLPRTAAFFDLDKTVIAKSSTLTFSKSFYRGGLISRRAALRTAYIQFVFLAG WP_073908070.1 inhibition of morphological differentiation protein [Streptomyces sp. CB02009] Length: 288\nNumber of predicted TMHs: 1\nExp number of AAs in TMHs: 22.11365\nExp number, first 60 AAs: 0.05482\nTotal prob of N-in: 0.00466\noutside 1 247\nTMhelix 248 270\ninside 271 288

32912 GCF_001905545.1_ASM190554v1 Streptomyces sp. CB02056 Terrabacteria group; Actinobacteria; Actinobacteria; Streptomycetales; Streptomycetaceae; Streptomyces MRTAAFFDLDKTIIAKSSALAFSRPFYQGGLINRRAVLRSAYAQFVFLVGGADHDQMEKM WP_074002134.1 inhibition of morphological differentiation protein [Streptomyces sp. CB02056] Length: 265\nNumber of predicted TMHs: 1\nExp number of AAs in TMHs: 22.66915\nExp number, first 60 AAs: 0.1145\nTotal prob of N-in: 0.01938\noutside 1 235\nTMhelix 236 258\ninside 259 265

32913 GCF_001905545.1_ASM190554v1 Streptomyces sp. CB02056 Terrabacteria group; Actinobacteria; Actinobacteria; Streptomycetales; Streptomycetaceae; Streptomyces MGQATGSPDHRTTAVRTPATRRRRAVRVAVATALTALPLAAVLTVGSAAPLAPPLGACSG WP_079271739.1 hypothetical protein [Streptomyces sp. CB02056]Length: 496\nNumber of predicted TMHs: 2\nExp number of AAs in TMHs: 43.72653\nExp number, first 60 AAs: 21.66519\nTotal prob of N-in: 0.96948\nPOSSIBLE N-term signal sequence\ninside 1 25\nTMhelix 26 48\noutside 49 466\nTMhelix 467 489\ninside 490 496

32914 GCF_001905505.1_ASM190550v1 Streptomyces sp. CB02058 Terrabacteria group; Actinobacteria; Actinobacteria; Streptomycetales; Streptomycetaceae; Streptomyces MLTFVENCFSPRTAAFFDLDKTVIAKSSTLTFSKSFYQGGLINRRAVLRTAYAQFVFLAG WP_073750303.1 inhibition of morphological differentiation protein [Streptomyces sp. CB02058] Length: 279\nNumber of predicted TMHs: 1\nExp number of AAs in TMHs: 21.58956\nExp number, first 60 AAs: 0.02381\nTotal prob of N-in: 0.00185\noutside 1 247\nTMhelix 248 270\ninside 271 279

32915 GCF_001905645.1_ASM190564v1 Streptomyces sp. CB02115 Terrabacteria group; Actinobacteria; Actinobacteria; Streptomycetales; Streptomycetaceae; Streptomyces

MLSVVENCFSPRTAAFFDLDKTVIAKSSTLTFSKSFYQGGLINRRAVLRTAYAQFVFLAG WP_032791512.1 MULTISPECIES: inhibition of morphological differentiation protein [Streptomyces] Length: 281\nNumber of predicted TMHs: 1\nExp number of AAs in TMHs: 21.51851\nExp number, first 60 AAs: 0.0316\nTotal prob of N-in: 0.00586\noutside 1 247\nTMhelix 248 270\ninside 271 281

32916 GCF_001905785.1_ASM190578v1 Streptomyces sp. CB02130 Terrabacteria group; Actinobacteria; Actinobacteria; Streptomycetales; Streptomycetaceae; Streptomyces MLSVVENCFSPRTAAFFDLDKTVIAKSSTLTFSKSFYQGGLINRRAVLRTAYTQFVFLAG WP_073826592.1 inhibition of morphological differentiation protein [Streptomyces sp. CB02130] Length: 281\nNumber of predicted TMHs: 1\nExp number of AAs in TMHs: 21.75194\nExp number, first 60 AAs: 0.0182\nTotal prob of N-in: 0.00408\noutside 1 247\nTMhelix 248 270\ninside 271 281

32917 GCF_001905785.1_ASM190578v1 Streptomyces sp. CB02130 Terrabacteria group; Actinobacteria; Actinobacteria; Streptomycetales; Streptomycetaceae; Streptomyces MGLSWNDRRIERRVKPGDGKPLKPFRWWQMTRRSLLSIALPVGGGLSATYTVEVKHGGDA WP_073822692.1 hypothetical protein [Streptomyces sp. CB02130]Length: 218\nNumber of predicted TMHs: 2\nExp number of AAs in TMHs: 40.89884\nExp number, first 60 AAs: 0.90638\nTotal prob of N-in: 0.96439\ninside 1 135\nTMhelix 136 158\noutside 159 182\nTMhelix 183 200\ninside 201 218

32918 GCF_001905785.1_ASM190578v1 Streptomyces sp. CB02130 Terrabacteria group; Actinobacteria; Actinobacteria; Streptomycetales; Streptomycetaceae; Streptomyces MSPPTPRRRALPVLTAALAALLCGLVQPAAADDPPPDWTARPAGGTDRDARPYVYLEGTP WP_010059884.1 MULTISPECIES: hypothetical protein [Streptomyces] Length: 322\nNumber of predicted TMHs: 1\nExp number of AAs in TMHs: 28.26039\nExp number, first 60 AAs: 6.03507\nTotal prob of N-in: 0.31029\noutside 1 249\nTMhelix 250 272\ninside 273 322

32919 GCF_001905665.1_ASM190566v1 Streptomyces sp. CB02261 Terrabacteria group; Actinobacteria; Actinobacteria; Streptomycetales; Streptomycetaceae; Streptomyces MLRLVENHSLPRTAAFFDLDKTVIAKSSTLTFSKSFYRGGLISRRAALRTAYIQFVFLAG WP_073811089.1 inhibition of morphological differentiation protein [Streptomyces sp. CB02261] Length: 288\nNumber of predicted TMHs: 1\nExp number of AAs in TMHs: 21.86782\nExp number, first 60 AAs: 0.05475\nTotal prob of N-in: 0.00459\noutside 1 247\nTMhelix 248 270\ninside 271 288

32920 GCF_001905905.1_ASM190590v1 Streptomyces sp. CB02366 Terrabacteria group; Actinobacteria; Actinobacteria; Streptomycetales; Streptomycetaceae; Streptomyces MLSVVENCFSPRTAAFFDLDKTVIAKSSTLTFSKSFYQGGLINRRAVLRTAYTQFVFMAG WP_073765520.1 inhibition of morphological differentiation protein [Streptomyces sp. CB02366] Length: 281\nNumber of predicted TMHs: 1\nExp number of AAs in TMHs: 21.48553\nExp number, first 60 AAs: 0.01642\nTotal prob of N-in: 0.00370\noutside 1 247\nTMhelix 248 270\ninside 271 281

32921 GCF_001905725.1_ASM190572v1 Streptomyces sp. CB02400 Terrabacteria group; Actinobacteria; Actinobacteria; Streptomycetales; Streptomycetaceae; Streptomyces MRHARRRVVRRVTRLAAVGGLLLGATMVTHAVASEPRAPDAVPFATAEETAAGPGAGLVS WP_073934166.1 serine protease [Streptomyces sp. CB02400] Length: 458\nNumber of predicted TMHs: 2\nExp number of AAs in TMHs: 41.57111\nExp number, first 60 AAs: 22.12305\nTotal prob of N-in: 0.98832\nPOSSIBLE N-term signal sequence\ninside 1 11\nTMhelix 12 34\noutside 35 419\nTMhelix 420 442\ninside 443 458

32922 GCF_001905725.1_ASM190572v1 Streptomyces sp. CB02400 Terrabacteria group; Actinobacteria; Actinobacteria; Streptomycetales; Streptomycetaceae; Streptomyces MLKDVENHSMPRTAAFFDLDKTVIAKSSTLTFSKSFYQGGLINRRAVLRTAYSQFVFLAG WP_073936685.1 inhibition of morphological differentiation protein [Streptomyces sp. CB02400] Length: 277\nNumber of predicted TMHs: 1\nExp number of AAs in TMHs: 21.77688\nExp number, first 60 AAs: 0.04819\nTotal prob of N-in: 0.00665\noutside 1 247\nTMhelix 248 270\ninside 271 277

32923 GCF_001905385.1_ASM190538v1 Streptomyces sp. CB02414 Terrabacteria group; Actinobacteria; Actinobacteria; Streptomycetales; Streptomycetaceae; Streptomyces MPRTAAFFDLDKTVIAKSSTLTFSKSFYQGGLINRRAVLRTAYAQFVFLAGGADHDQMER WP_073726846.1 inhibition of morphological differentiation protein [Streptomyces sp. CB02414] Length: 269\nNumber of predicted TMHs: 1\nExp number of AAs in TMHs: 21.719\nExp number, first 60 AAs: 0.06255\nTotal prob of N-in: 0.00712\noutside 1 238\nTMhelix 239 261\ninside 262 269

32924 GCF_001905385.1_ASM190538v1 Streptomyces sp. CB02414 Terrabacteria group; Actinobacteria; Actinobacteria; Streptomycetales; Streptomycetaceae; Streptomyces
MRHARRRIVRRVARLAAVGGLLLGGTMVTRAVATEPPDAAVPHTLARSASGTGAGLVTRL WP_073728708.1 serine protease [Streptomyces sp. CB02414] Length: 438\nNumber of predicted TMHs: 2\nExp number of AAs in TMHs: 41.8399\nExp number, first 60 AAs: 21.23316\nTotal prob of N-in: 0.96605\nPOSSIBLE N-term signal sequence\ninside 1 11\nTMhelix 12 34\noutside 35 399\nTMhelix 400 422\ninside 423 438

32925 GCF_001905385.1_ASM190538v1 Streptomyces sp. CB02414 Terrabacteria group; Actinobacteria; Actinobacteria; Streptomycetales; Streptomycetaceae; Streptomyces
MGHNRRRRPTGARRATFGAFALILGGGGLVAVNVFASATEAGDTAVPLGSSGVAATIDCP WP_079187565.1 hypothetical protein [Streptomyces sp. CB02414]Length: 693\nNumber of predicted TMHs: 2\nExp number of AAs in TMHs: 43.08151\nExp number, first 60 AAs: 22.02227\nTotal prob of N-in: 0.99647\nPOSSIBLE N-term signal sequence\ninside 1 16\nTMhelix 17 39\noutside 40 666\nTMhelix 667 686\ninside 687 693

32926 GCF_001905705.1_ASM190570v1 Streptomyces sp. CB02460 Terrabacteria group; Actinobacteria; Actinobacteria; Streptomycetales; Streptomycetaceae; Streptomyces
MLCRVENRFLPRTAAFFDLDKTVIAKSSTLTFSKSFYHGGLINRRAVLRTAYTQFVFLAG WP_073966429.1 inhibition of morphological differentiation protein [Streptomyces sp. CB02460] Length: 279\nNumber of predicted TMHs: 1\nExp number of AAs in TMHs: 21.7716\nExp number, first 60 AAs: 0.03\nTotal prob of N-in: 0.00612\noutside 1 247\nTMhelix 248 270\ninside 271 279

32927 GCF_001905365.1_ASM190536v1 Streptomyces sp. CB02488 Terrabacteria group; Actinobacteria; Actinobacteria; Streptomycetales; Streptomycetaceae; Streptomyces
MLCLVENCFLPRTAAFFDLDKTVIAKSSTLTFSKSFYQGGLINRRAVLRTAYAQFVFLAG WP_073733604.1 inhibition of morphological differentiation protein [Streptomyces sp. CB02488] Length: 279\nNumber of predicted TMHs: 1\nExp number of AAs in TMHs: 21.22998\nExp number, first 60 AAs: 0.01456\nTotal prob of N-in: 0.00151\noutside 1 247\nTMhelix 248 270\ninside 271 279

32928 GCF_001905365.1_ASM190536v1 Streptomyces sp. CB02488 Terrabacteria group; Actinobacteria; Actinobacteria; Streptomycetales; Streptomycetaceae; Streptomyces
MFNPSPTPLARTGARRATLTALVVASSAGLAISPLGASGAVAAEKKHPITIAVTTDPARP WP_073738813.1 hypothetical protein [Streptomyces sp. CB02488] Length: 479\nNumber of predicted TMHs: 2\nExp number of AAs in TMHs: 35.78177\nExp number, first 60 AAs: 14.70374\nTotal prob of N-in: 0.64135\nPOSSIBLE N-term signal sequence\ninside 1 20\nTMhelix 21 43\noutside 44 447\nTMhelix 448 470\ninside 471 479

32929 GCF_001905005.1_ASM190500v1 Streptomyces sp. CB02923 Terrabacteria group; Actinobacteria; Actinobacteria; Streptomycetales; Streptomycetaceae; Streptomyces
MGSRANGHRAGHRRRATVALAGAASAVALSCATAAPASADPNVPLNSGQCTFGTDDIKET WP_073760718.1 type VII secretion-associated serine protease mycosin [Streptomyces sp. CB02923] Length: 424\nNumber of predicted TMHs: 1\nExp number of AAs in TMHs: 31.75765\nExp number, first 60 AAs: 9.17505\nTotal prob of N-in: 0.49082\noutside 1 383\nTMhelix 384 406\ninside 407 424

32930 GCF_001905005.1_ASM190500v1 Streptomyces sp. CB02923 Terrabacteria group; Actinobacteria; Actinobacteria; Streptomycetales; Streptomycetaceae; Streptomyces
MALTRRRTLVRSAAVAAVTGSALLLPAAAAFADSPQPTGRSSENSAEKSTEKSTEKPSTE WP_073758166.1 hypothetical protein [Streptomyces sp. CB02923] Length: 231\nNumber of predicted TMHs: 2\nExp number of AAs in TMHs: 38.59901\nExp number, first 60 AAs: 20.17497\nTotal prob of N-in: 0.96895\nPOSSIBLE N-term signal sequence\ninside 1 12\nTMhelix 13 32\noutside 33 199\nTMhelix 200 222\ninside 223 231

32931 GCF_001905005.1_ASM190500v1 Streptomyces sp. CB02923 Terrabacteria group; Actinobacteria; Actinobacteria; Streptomycetales; Streptomycetaceae; Streptomyces
MTEHRAPAVSATAAGRGTPARRAAIRAAATGALTAALLGLAPQAALAATPAPPAAPAASA WP_073760183.1 hypothetical protein [Streptomyces sp. CB02923]Length: 302\nNumber of predicted TMHs: 1\nExp number of AAs in TMHs: 27.48486\nExp number, first 60 AAs: 8.84691\nTotal prob of N-in: 0.41327\noutside 1 273\nTMhelix 274 296\ninside 297 302

32932 GCF_001905885.1_ASM190588v1 Streptomyces sp. CB03234 Terrabacteria group; Actinobacteria; Actinobacteria; Streptomycetales; Streptomycetaceae; Streptomyces
MTRNRRTALAAALAATAFAVLPATPAHADAIRAQQWGLEAMRTADAWRTTKGEGITVAVL WP_073758261.1 type VII secretion-associated serine protease mycosin [Streptomyces sp. CB03234] Length: 391\nNumber of predicted TMHs: 2\nExp number of AAs in TMHs: 39.5888199999999\nExp number, first 60 AAs: 16.5639\nTotal prob of N-in:

0.85887\nPOSSIBLE N-term signal sequence\ninside 1 6\nTMhelix 7 26\noutside 27 360\nTMhelix 361 383\ninside 384 391

32933 GCF_001905885.1_ASM190588v1 Streptomyces sp. CB03234 Terrabacteria group; Actinobacteria; Actinobacteria; Streptomycetales; Streptomycetaceae; Streptomyces MLGLVENHSLPRTAAFFDLDKTVIAKSSTLTFSKSFYQGGLINRRAVLRTAYIQFVFLAG WP_073756918.1 inhibition of morphological differentiation protein [Streptomyces sp. CB03234] Length: 276\nNumber of predicted TMHs: 1\nExp number of AAs in TMHs: 21.83497\nExp number, first 60 AAs: 0.03078\nTotal prob of N-in: 0.00268\noutside 1 247\nTMhelix 248 270\ninside 271 276

32934 GCF_001905885.1_ASM190588v1 Streptomyces sp. CB03234 Terrabacteria group; Actinobacteria; Actinobacteria; Streptomycetales; Streptomycetaceae; Streptomyces MNSNTVRLPARLPAARRRALSTAAAAVALAAGPALLAAPMAHATGGGEGRASAVVLRTDL WP_073752756.1 hypothetical protein [Streptomyces sp. CB03234]Length: 313\nNumber of predicted TMHs: 2\nExp number of AAs in TMHs: 41.38363\nExp number, first 60 AAs: 21.12161\nTotal prob of N-in: 0.95598\nPOSSIBLE N-term signal sequence\ninside 1 19\nTMhelix 20 42\noutside 43 279\nTMhelix 280 302\ninside 303 313

32935 GCF_002099185.1_ASM209918v1 Streptomyces sp. CB03238 Terrabacteria group; Actinobacteria; Actinobacteria; Streptomycetales; Streptomycetaceae; Streptomyces MTHRRRTAFAAALAATALAVLPATPAHADAIRAQQWGLEAMHTTEAWRTTQGEGITVAVL WP_084897512.1 type VII secretion-associated serine protease mycosin [Streptomyces sp. CB03238] Length: 391\nNumber of predicted TMHs: 1\nExp number of AAs in TMHs: 38.72386\nExp number, first 60 AAs: 15.13582\nTotal prob of N-in: 0.80329\nPOSSIBLE N-term signal sequence\noutside 1 360\nTMhelix 361 383\ninside 384 391

32936 GCF_002099185.1_ASM209918v1 Streptomyces sp. CB03238 Terrabacteria group; Actinobacteria; Actinobacteria; Streptomycetales; Streptomycetaceae; Streptomyces MLGLVENHSLPRTAAFFDLDKTVIAKSSTLTFSKSFYQGGLINRRAVLRTAYIQFVFLAG WP_084898257.1 inhibition of morphological differentiation protein [Streptomyces sp. CB03238] Length: 276\nNumber of predicted TMHs: 1\nExp number of AAs in TMHs: 21.83497\nExp number, first 60 AAs: 0.03078\nTotal prob of N-in: 0.00268\noutside 1 247\nTMhelix 248 270\ninside 271 276

32937 GCF_001905025.1_ASM190502v1 Streptomyces sp. CB03578 Terrabacteria group; Actinobacteria; Actinobacteria; Streptomycetales; Streptomycetaceae; Streptomyces MRPEHGDGAQSVRQPRNPAGPARRGPVIAVTGAAGGVGAALVSRLVASEEVKAVVAIDER WP_037799844.1 MULTISPECIES: NAD-dependent dehydratase [Streptomyces] Length: 373\nNumber of predicted TMHs: 1\nExp number of AAs in TMHs: 23.21663\nExp number, first 60 AAs: 2.48036\nTotal prob of N-in: 0.08776\noutside 1 344\nTMhelix 345 364\ninside 365 373

32938 GCF_001905025.1_ASM190502v1 Streptomyces sp. CB03578 Terrabacteria group; Actinobacteria; Actinobacteria; Streptomycetales; Streptomycetaceae; Streptomyces MSTVSRRTLVRAAAVTACAGSLLALPAAAALAEGLPASAASGAPAVTASQRVLLKSLSLA WP_073798262.1 hypothetical protein [Streptomyces sp. CB03578] Length: 230\nNumber of predicted TMHs: 2\nExp number of AAs in TMHs: 43.05456\nExp number, first 60 AAs: 20.57051\nTotal prob of N-in: 0.95017\nPOSSIBLE N-term signal sequence\ninside 1 8\nTMhelix 9 31\noutside 32 200\nTMhelix 201 223\ninside 224 230

32939 GCF_001905025.1_ASM190502v1 Streptomyces sp. CB03578 Terrabacteria group; Actinobacteria; Actinobacteria; Streptomycetales; Streptomycetaceae; Streptomyces MAEAADNQGAVPAPARRRWLRRAVVLLAGTPVLAALVYAPAPPAEAADAPVDVQLDTLAP WP_079196203.1 hypothetical protein [Streptomyces sp. CB03578]Length: 753\nNumber of predicted TMHs: 2\nExp number of AAs in TMHs: 39.77324\nExp number, first 60 AAs: 18.12819\nTotal prob of N-in: 0.87708\nPOSSIBLE N-term signal sequence\ninside 1 18\nTMhelix 19 41\noutside 42 686\nTMhelix 687 709\ninside 710 753

32940 GCF_001905045.1_ASM190504v1 Streptomyces sp. CB03911 Terrabacteria group; Actinobacteria; Actinobacteria; Streptomycetales; Streptomycetaceae; Streptomyces MRSSAISRFTKNVPSLSRRSALTVGATALVLGAGLPLLPGAGAAWACGAPDDAPVTAPAA WP_073925819.1 hypothetical protein [Streptomyces sp. CB03911]Length: 343\nNumber of predicted TMHs: 2\nExp number of AAs in TMHs: 39.8085\nExp number, first 60 AAs: 19.13928\nTotal prob of N-in: 0.86246\nPOSSIBLE N-term signal sequence\ninside 1 20\nTMhelix 21 43\noutside 44 314\nTMhelix 315 337\ninside 338 343

32941 GCF_001905045.1_ASM190504v1 Streptomyces sp. CB03911 Terrabacteria group; Actinobacteria; Actinobacteria; Streptomycetales; Streptomycetaceae; Streptomyces

MLRPHDAGPRTAAFFDLDKTIIAKSSALAFSRPFYQGGLINRRAVLKSAYAQFVFLVGGA WP_073924785.1 inhibition of morphological differentiation protein [Streptomyces sp. CB03911] Length: 273\nNumber of predicted TMHs: 1\nExp number of AAs in TMHs: 21.19493\nExp number, first 60 AAs: 0.04855\nTotal prob of N-in: 0.01928\noutside 1 245\nTMhelix 246 265\ninside 266 273

32942 GCF_001865245.1_ASM186524v1 Streptomyces sp. CC53 Terrabacteria group; Actinobacteria; Actinobacteria; Streptomycetales; Streptomycetaceae; Streptomyces MLGFVENHSSPRTAAFFDLDKTVIAKSSALTFSKSFYQGGLISRRAALRTAYTQFVFLAG WP_071281018.1 inhibition of morphological differentiation protein [Streptomyces sp. CC53] Length: 292\nNumber of predicted TMHs: 1\nExp number of AAs in TMHs: 20.18777\nExp number, first 60 AAs: 0.01132\nTotal prob of N-in: 0.00139\noutside 1 248\nTMhelix 249 268\ninside 269 292

32943 GCF_001595515.1_ASM159551v1 Streptomyces sp. CC71 Terrabacteria group; Actinobacteria; Actinobacteria; Streptomycetales; Streptomycetaceae; Streptomyces MRHARRRMVRRVARLAAVGGLLLGGTMVTRAVASEPPDAPAVPYTSARAASGTGAGLVSR WP_062132366.1 protease [Streptomyces sp. CC71] Length: 458\nNumber of predicted TMHs: 2\nExp number of AAs in TMHs: 39.93875\nExp number, first 60 AAs: 21.01585\nTotal prob of N-in: 0.96415\nPOSSIBLE N-term signal sequence\ninside 1 11\nTMhelix 12 34\noutside 35 419\nTMhelix 420 442\ninside 443 458

32944 GCF_001865255.1_ASM186525v1 Streptomyces sp. CC77 Terrabacteria group; Actinobacteria; Actinobacteria; Streptomycetales; Streptomycetaceae; Streptomyces MLGFVENHSSPRTAAFFDLDKTVIAKSSALTFSKSFYQGGLISRRAALRTAYTQFVFLAG WP_071274060.1 inhibition of morphological differentiation protein [Streptomyces sp. CC77] Length: 292\nNumber of predicted TMHs: 1\nExp number of AAs in TMHs: 20.33548\nExp number, first 60 AAs: 0.00997\nTotal prob of N-in: 0.00095\noutside 1 248\nTMhelix 249 268\ninside 269 292

32945 GCF_000373305.1_ASM37330v1 Streptomyces sp. CcalMP-8WTerrabacteria group; Actinobacteria; Actinobacteria; Streptomycetales; Streptomycetaceae; Streptomyces MLSVVENCFSPRTAAFFDLDKTVIAKSSTLTFSKSFYQGGLINRRAVLRTAYTQFVFLAG WP_018489460.1 inhibition of morphological differentiation protein [Streptomyces sp. CcalMP-8W]Length: 280\nNumber of predicted TMHs: 1\nExp number of AAs in TMHs: 21.37533\nExp number, first 60 AAs: 0.02224\nTotal prob of N-in: 0.00962\noutside 1 245\nTMhelix 246 268\ninside 269 280

32946 GCF_000772045.1_ASM77204v1 Streptomyces sp. CCM_MD2014 Terrabacteria group; Actinobacteria; Actinobacteria; Streptomycetales; Streptomycetaceae; Streptomyces MRHARRRIVRRVARLAAVGGLLLGGTMVTRAVASEPPGPSAAPRALAQTASGAGAALVAR WP_061440805.1 protease [Streptomyces sp. CCM_MD2014] Length: 447\nNumber of predicted TMHs: 2\nExp number of AAs in TMHs: 41.75804\nExp number, first 60 AAs: 21.4444\nTotal prob of N-in: 0.97402\nPOSSIBLE N-term signal sequence\ninside 1 11\nTMhelix 12 34\noutside 35 408\nTMhelix 409 431\ninside 432 447

32947 GCF_000772045.1_ASM77204v1 Streptomyces sp. CCM_MD2014 Terrabacteria group; Actinobacteria; Actinobacteria; Streptomycetales; Streptomycetaceae; Streptomyces MTQEHALGHNRRRRPTGARRATFAAVALILGGGGLVAANVFASATESGNAAVPLGSSGVA WP_078884480.1 hypothetical protein [Streptomyces sp. CCM_MD2014] Length: 693\nNumber of predicted TMHs: 2\nExp number of AAs in TMHs: 43.59072\nExp number, first 60 AAs: 22.35935\nTotal prob of N-in: 0.99893\nPOSSIBLE N-term signal sequence\ninside 1 20\nTMhelix 21 43\noutside 44 665\nTMhelix 666 688\ninside 689 693

32948 GCF_001484565.1_ASM148456v1 Streptomyces sp. CdTB01 Terrabacteria group; Actinobacteria; Actinobacteria; Streptomycetales; Streptomycetaceae; Streptomyces MLGVVENQSLPRTAAFFDLDKTVIAKSSTLTFGKSFYQGGLINRRAALRTAYAQFVFLLG WP_058924086.1 inhibition of morphological differentiation protein [Streptomyces sp. CdTB01] Length: 277\nNumber of predicted TMHs: 1\nExp number of AAs in TMHs: 21.69975\nExp number, first 60 AAs: 0.02187\nTotal prob of N-in: 0.00243\noutside 1 247\nTMhelix 248 270\ninside 271 277

32949 GCF_001484565.1_ASM148456v1 Streptomyces sp. CdTB01 Terrabacteria group; Actinobacteria; Actinobacteria; Streptomycetales; Streptomycetaceae; Streptomyces MTRRAVLVLTTLFLTLGLAAPAGATGYRYWSFWTYDSGRWTYATQGPATARPSDGDVQGF WP_058925582.1 hypothetical protein [Streptomyces sp. CdTB01] Length: 207\nNumber of predicted TMHs: 2\nExp number of AAs in TMHs: 43.39459\nExp number, first 60 AAs: 22.38515\nTotal prob of N-in: 0.99467\nPOSSIBLE N-term signal sequence\ninside 1 4\nTMhelix 5 27\noutside 28 179\nTMhelix 180 199\ninside 200 207

32950 GCF_001278095.1_ASM127809v1 Streptomyces sp. CFMR 7 Terrabacteria group; Actinobacteria; Actinobacteria; Streptomycetales; Streptomycetaceae; Streptomyces
MLSVVENCFSPRTAAFFDLDKTVIAKSSTLTFSKSFYQGGLINRRAVLRTAYTQFVFMAG WP_053560604.1 inhibition of morphological differentiation protein [Streptomyces sp. CFMR 7] Length: 282\nNumber of predicted TMHs: 1\nExp number of AAs in TMHs: 21.47617\nExp number, first 60 AAs: 0.01835\nTotal prob of N-in: 0.00838\noutside 1 247\nTMhelix 248 270\ninside 271 282

32951 GCF_000377965.1_ASM37796v1 Streptomyces sp. CNB091 Terrabacteria group; Actinobacteria; Actinobacteria; Streptomycetales; Streptomycetaceae; Streptomyces
MLSVVENCFSPRTAAFFDLDKTVIAKSSTLTFSKSFYQGGLINRRAVLRTAYAQFVFLAG WP_018958840.1 inhibition of morphological differentiation protein [Streptomyces sp. CNB091] Length: 281\nNumber of predicted TMHs: 1\nExp number of AAs in TMHs: 21.43674\nExp number, first 60 AAs: 0.03371\nTotal prob of N-in: 0.00768\noutside 1 247\nTMhelix 248 270\ninside 271 281

32952 GCF_000514715.1_ASM51471v1 Streptomyces sp. CNH099 Terrabacteria group; Actinobacteria; Actinobacteria; Streptomycetales; Streptomycetaceae; Streptomyces
MLALVENHSLPRTAAFFDLDKTVIAKSSTLTFSRSFYHGGLINRRAVLRSAYSQFIYLLG WP_027753026.1 inhibition of morphological differentiation protein [Streptomyces sp. CNH099] Length: 276\nNumber of predicted TMHs: 1\nExp number of AAs in TMHs: 21.845\nExp number, first 60 AAs: 0.00301\nTotal prob of N-in: 0.00136\noutside 1 246\nTMhelix 247 269\ninside 270 276

32953 GCF_000527195.1_ASM52719v1 Streptomyces sp. CNH189 Terrabacteria group; Actinobacteria; Actinobacteria; Streptomycetales; Streptomycetaceae; Streptomyces
MLRDVENHLLPRTAAFFDLDKTVIAKSSTLTFSKSFYQGGLINRRAVLRTAYAQFVFLAG WP_078540984.1 inhibition of morphological differentiation protein [Streptomyces sp. CNH189] Length: 276\nNumber of predicted TMHs: 1\nExp number of AAs in TMHs: 21.9161\nExp number, first 60 AAs: 0.05243\nTotal prob of N-in: 0.00490\noutside 1 247\nTMhelix 248 270\ninside 271 276

32954 GCF_000527195.1_ASM52719v1 Streptomyces sp. CNH189 Terrabacteria group; Actinobacteria; Actinobacteria; Streptomycetales; Streptomycetaceae; Streptomyces
MPGRHRRRSVLAAMAAAPLAGALGACSGGADDAPSGSSTDSKRAVRVTFWSALRGSQEVV WP_078541208.1 hypothetical protein [Streptomyces sp. CNH189] Length: 220\nNumber of predicted TMHs: 1\nExp number of AAs in TMHs: 30.05139\nExp number, first 60 AAs: 5.81512999999999\nTotal prob of N-in: 0.76659\ninside 1 183\nTMhelix 184 206\noutside 207 220

32955 GCF_000527195.1_ASM52719v1 Streptomyces sp. CNH189 Terrabacteria group; Actinobacteria; Actinobacteria; Streptomycetales; Streptomycetaceae; Streptomyces
MRHARRRVVRRVTRLAAVGGLLLGATMVTQAVAGEPGTPDAVPFSTAAPTDKAAGPGAAL WP_037722459.1 protease [Streptomyces sp. CNH189] Length: 460\nNumber of predicted TMHs: 2\nExp number of AAs in TMHs: 41.46399\nExp number, first 60 AAs: 22.18037\nTotal prob of N-in: 0.98470\nPOSSIBLE N-term signal sequence\ninside 1 11\nTMhelix 12 34\noutside 35 421\nTMhelix 422 444\ninside 445 460

32956 GCF_000424825.1_ASM42482v1 Streptomyces sp. CNH287 Terrabacteria group; Actinobacteria; Actinobacteria; Streptomycetales; Streptomycetaceae; Streptomyces
MARTPLHHARRRPLTAGGLTAAVLAVCTAAFGALPAAAAPGPGSVPAAPVPAAVAGPEGD WP_051261987.1 hypothetical protein [Streptomyces sp. CNH287] Length: 236\nNumber of predicted TMHs: 2\nExp number of AAs in TMHs: 42.0568\nExp number, first 60 AAs: 22.21009\nTotal prob of N-in: 0.98908\nPOSSIBLE N-term signal sequence\ninside 1 12\nTMhelix 13 35\noutside 36 202\nTMhelix 203 225\ninside 226 236

32957 GCF_000424825.1_ASM42482v1 Streptomyces sp. CNH287 Terrabacteria group; Actinobacteria; Actinobacteria; Streptomycetales; Streptomycetaceae; Streptomyces
MLAVVEIQSLPRTAAFFDLDKTVIAKSSTLTFSKSFYHGGLINRRAVLRTAYAQFVFLLG WP_027748344.1 inhibition of morphological differentiation protein [Streptomyces sp. CNH287] Length: 279\nNumber of predicted TMHs: 1\nExp number of AAs in TMHs: 22.36057\nExp number, first 60 AAs: 0.03175\nTotal prob of N-in: 0.00298\noutside 1 247\nTMhelix 248 270\ninside 271 279

32958 GCF_000482585.1_ASM48258v1 Streptomyces sp. CNQ329 Terrabacteria group; Actinobacteria; Actinobacteria; Streptomycetales; Streptomycetaceae; Streptomyces
MLALVENHSLPRTAAFFDLDKTVIAKSSTLTFSRSFYAGGLINRRAVLRSAYSQFLYLLG WP_027773270.1 inhibition of morphological differentiation protein [Streptomyces sp. CNQ329] Length: 275\nNumber of predicted TMHs: 1\nExp

number of AAs in TMHs: 21.16452\nExp number, first 60 AAs: 0.0049\nTotal prob of N-in: 0.00132\noutside 1 246\nTMhelix 247 269\ninside 270 275

32959 GCF_000797385.1_ASM79738v1 Streptomyces sp. CNQ431 Terrabacteria group; Actinobacteria; Actinobacteria; Streptomycetales; Streptomycetaceae; Streptomyces MNPNRRPVVLAAAVATALGLGATALTLPAVAAEAPRAGAPVMDLTDGTLNWGVKESFRRY WP_033948155.1 membrane protein [Streptomyces sp. CNQ431] Length: 521\nNumber of predicted TMHs: 2\nExp number of AAs in TMHs: 42.7912\nExp number, first 60 AAs: 22.03081\nTotal prob of N-in: 0.97262\nPOSSIBLE N-term signal sequence\ninside 1 6\nTMhelix 7 29\noutside 30 486\nTMhelix 487 509\ninside 510 521

32960 GCF_000797385.1_ASM79738v1 Streptomyces sp. CNQ431 Terrabacteria group; Actinobacteria; Actinobacteria; Streptomycetales; Streptomycetaceae; Streptomyces MASPHRRRAAVAVRALVALTALLCATGLAPAATSGPAARAAVAEVPPPTATTSVVTVRTG WP_033949764.1 VWA domain-containing protein [Streptomyces sp. CNQ431] Length: 835\nNumber of predicted TMHs: 2\nExp number of AAs in TMHs: 40.56285\nExp number, first 60 AAs: 20.3579\nTotal prob of N-in: 0.93764\nPOSSIBLE N-term signal sequence\ninside 1 11\nTMhelix 12 34\noutside 35 796\nTMhelix 797 819\ninside 820 835

32961 GCF_000797385.1_ASM79738v1 Streptomyces sp. CNQ431 Terrabacteria group; Actinobacteria; Actinobacteria; Streptomycetales; Streptomycetaceae; Streptomyces MGGRGTTAQARGRGRLPGRPSDPAPTGGNDVITGVNLRMPARRTALPALLLTVTALLAVP WP_033950195.1 MULTISPECIES: hypothetical protein [Streptomyces] Length: 301\nNumber of predicted TMHs: 2\nExp number of AAs in TMHs: 41.7448\nExp number, first 60 AAs: 13.52141\nTotal prob of N-in: 0.87370\nPOSSIBLE N-term signal sequence\ninside 1 44\nTMhelix 45 67\noutside 68 273\nTMhelix 274 296\ninside 297 301

32962 GCF_000797385.1_ASM79738v1 Streptomyces sp. CNQ431 Terrabacteria group; Actinobacteria; Actinobacteria; Streptomycetales; Streptomycetaceae; Streptomyces MPDRRSAVSLRLPLPTAAPGFGGLALPGAAATAPDAGDVVAVRESSEKERKSSSDPEGRS WP_033947955.1 hypothetical protein [Streptomyces sp. CNQ431] Length: 292\nNumber of predicted TMHs: 2\nExp number of AAs in TMHs: 49.52916\nExp number, first 60 AAs: 3.9366\nTotal prob of N-in: 0.40255\noutside 1 164\nTMhelix 165 187\ninside 188 229\nTMhelix 230 252\noutside 253 292

32963 GCF_000797385.1_ASM79738v1 Streptomyces sp. CNQ431 Terrabacteria group; Actinobacteria; Actinobacteria; Streptomycetales; Streptomycetaceae; Streptomyces MLDPVENRSLPRTAAFFDLDKTVIAKSSTLTFSKSFYQGGLINRRAVLRTAYTQFVFLAG WP_023422160.1 MULTISPECIES: SsgB protein [Streptomyces] Length: 279\nNumber of predicted TMHs: 1\nExp number of AAs in TMHs: 21.73892\nExp number, first 60 AAs: 0.02846\nTotal prob of N-in: 0.01117\noutside 1 247\nTMhelix 248 270\ninside 271 279

32964 GCF_001011035.1_ASM101103v1 Streptomyces sp. CNQ-509 Terrabacteria group; Actinobacteria; Actinobacteria; Streptomycetales; Streptomycetaceae; Streptomyces MLALVENHSLPRTAAFFDLDKTVIAKSSTLTFTRSFYHGGLMNRRAVLRSAYSQFIYLLG WP_047016536.1 inhibition of morphological differentiation protein [Streptomyces sp. CNQ-509] Length: 276\nNumber of predicted TMHs: 1\nExp number of AAs in TMHs: 22.31919\nExp number, first 60 AAs: 0.00562\nTotal prob of N-in: 0.00145\noutside 1 246\nTMhelix 247 269\ninside 270 276

32965 GCF_000701285.1_ASM70128v1 Streptomyces sp. CNQ-525 Terrabacteria group; Actinobacteria; Actinobacteria; Streptomycetales; Streptomycetaceae; Streptomyces MLALVENHSLPRTAAFFDLDKTVIAKSSTLTFSRSFYHGGLISRRAVLRSAYSQFIYLLG WP_037753578.1 inhibition of morphological differentiation protein [Streptomyces sp. CNQ-525] Length: 276\nNumber of predicted TMHs: 1\nExp number of AAs in TMHs: 22.14218\nExp number, first 60 AAs: 0.00323\nTotal prob of N-in: 0.00173\noutside 1 246\nTMhelix 247 269\ninside 270 276

32966 GCF_000377105.1_ASM37710v1 Streptomyces sp. CNQ766 Terrabacteria group; Actinobacteria; Actinobacteria; Streptomycetales; Streptomycetaceae; Streptomyces MLALVENHSLPRTAAFFDLDKTVIAKSSTLTFSRSFYHGGLINRRAVLRSAYSQFIYLLG WP_018835865.1 MULTISPECIES: inhibition of morphological differentiation protein [Streptomyces] Length: 276\nNumber of predicted TMHs: 1\nExp number of AAs in TMHs: 22.3164\nExp number, first 60 AAs: 0.00298\nTotal prob of N-in: 0.00131\noutside 1 246\nTMhelix 247 269\ninside 270 276

32967 GCF_000377105.1_ASM37710v1 Streptomyces sp. CNQ766 Terrabacteria group; Actinobacteria; Actinobacteria; Streptomycetales; Streptomycetaceae; Streptomyces MLRRLRALAATLLPRRRVVAVAAATLLTSTLAVGTDLAGAPPAHAAARTVDGGRLDWGVK WP_018838279.1

hypothetical protein [Streptomyces sp. CNQ766] Length: 519\nNumber of predicted TMHs: 1\nExp number of AAs in TMHs: 26.88059\nExp number, first 60 AAs: 6.44244\nTotal prob of N-in: 0.29990\noutside 1 459\nTMhelix 460 482\ninside 483 519

32968 GCF_000424805.1_ASM42480v1 Streptomyces sp. CNQ865 Terrabacteria group; Actinobacteria; Actinobacteria; Streptomycetales; Streptomycetaceae; Streptomyces MLALVENHSLPRTAAFFDLDKTVIAKSSTLTFSRSFYHGGLINRRAVLRSAYSQFIYLLG WP_018835865.1 MULTISPECIES: inhibition of morphological differentiation protein [Streptomyces] Length: 276\nNumber of predicted TMHs: 1\nExp number of AAs in TMHs: 22.3164\nExp number, first 60 AAs: 0.00298\nTotal prob of N-in: 0.00131\noutside 1 246\nTMhelix 247 269\ninside 270 276

32969 GCF_000424805.1_ASM42480v1 Streptomyces sp. CNQ865 Terrabacteria group; Actinobacteria; Actinobacteria; Streptomycetales; Streptomycetaceae; Streptomyces MLRRLRALAATLLPRRRVVAVAAATLLTSTLAVGADLAGAPPAHAAARTVDGGRLDWGIK WP_027767851.1 hypothetical protein [Streptomyces sp. CNQ865] Length: 516\nNumber of predicted TMHs: 1\nExp number of AAs in TMHs: 29.39726\nExp number, first 60 AAs: 8.96069\nTotal prob of N-in: 0.41577\noutside 1 456\nTMhelix 457 479\ninside 480 516

32970 GCF_000515055.1_ASM51505v1 Streptomyces sp. CNR698 Terrabacteria group; Actinobacteria; Actinobacteria; Streptomycetales; Streptomycetaceae; Streptomyces MLCVVENHSLPRTAAFFDLDKTVIAKSSALTFSKSFYHGGLINRRAVLRTAYTQFVFLAG WP_027733739.1 inhibition of morphological differentiation protein [Streptomyces sp. CNR698] Length: 281\nNumber of predicted TMHs: 1\nExp number of AAs in TMHs: 21.90816\nExp number, first 60 AAs: 0.03275\nTotal prob of N-in: 0.00432\noutside 1 247\nTMhelix 248 270\ninside 271 281

32971 GCF_000515055.1_ASM51505v1 Streptomyces sp. CNR698 Terrabacteria group; Actinobacteria; Actinobacteria; Streptomycetales; Streptomycetaceae; Streptomyces MPVARRPLLTATAAGSLLCALWFVPSANATADGESGHPDRARSGTSASGVQDVEGAERVE WP_027731726.1 hypothetical protein [Streptomyces sp. CNR698] Length: 151\nNumber of predicted TMHs: 2\nExp number of AAs in TMHs: 38.12757\nExp number, first 60 AAs: 17.80846\nTotal prob of N-in: 0.89036\nPOSSIBLE N-term signal sequence\ninside 1 6\nTMhelix 7 29\noutside 30 120\nTMhelix 121 143\ninside 144 151

32972 GCF_000377125.1_ASM37712v1 Streptomyces sp. CNS335 Terrabacteria group; Actinobacteria; Actinobacteria; Streptomycetales; Streptomycetaceae; Streptomyces MLALVENHSLPRTAAFFDLDKTVIAKSSTLTFSRSFYHGGLINRRAVLRSAYSQFIYLLG WP_018841337.1 inhibition of morphological differentiation protein [Streptomyces sp. CNS335] Length: 276\nNumber of predicted TMHs: 1\nExp number of AAs in TMHs: 22.33174\nExp number, first 60 AAs: 0.01514\nTotal prob of N-in: 0.00199\noutside 1 246\nTMhelix 247 269\ninside 270 276

32973 GCF_000377125.1_ASM37712v1 Streptomyces sp. CNS335 Terrabacteria group; Actinobacteria; Actinobacteria; Streptomycetales; Streptomycetaceae; Streptomyces MLRRLRALAATLLPRRRVVAVAAATLLTSTLAVGTDLAGAPPAHAAARTVDGGRLDWGIK WP_018844154.1 hypothetical protein [Streptomyces sp. CNS335] Length: 516\nNumber of predicted TMHs: 1\nExp number of AAs in TMHs: 26.57291\nExp number, first 60 AAs: 6.44234\nTotal prob of N-in: 0.29995\noutside 1 455\nTMhelix 456 478\ninside 479 516

32974 GCF_000426325.1_ASM42632v1 Streptomyces sp. CNS606 Terrabacteria group; Actinobacteria; Actinobacteria; Streptomycetales; Streptomycetaceae; Streptomyces MPRRAGGGSGPRRRAPLRALVAAVLGAGMLVSAAVPAQAAGYRYWSFWERTDSAWTYARQ WP_051307977.1 hypothetical protein [Streptomyces sp. CNS606] Length: 235\nNumber of predicted TMHs: 2\nExp number of AAs in TMHs: 44.33942\nExp number, first 60 AAs: 22.14611\nTotal prob of N-in: 0.98655\nPOSSIBLE N-term signal sequence\ninside 1 19\nTMhelix 20 42\noutside 43 205\nTMhelix 206 228\ninside 229 235

32975 GCF_000426325.1_ASM42632v1 Streptomyces sp. CNS606 Terrabacteria group; Actinobacteria; Actinobacteria; Streptomycetales; Streptomycetaceae; Streptomyces MRRDARPGRRRSALVGLTVAGLLATTTAGVAQAEDTDEPGATVPVSFTGPESYALSLQAQ WP_078529637.1 hypothetical protein [Streptomyces sp. CNS606] Length: 570\nNumber of predicted TMHs: 2\nExp number of AAs in TMHs: 40.15291\nExp number, first 60 AAs: 17.95323\nTotal prob of N-in: 0.90287\nPOSSIBLE N-term signal sequence\ninside 1 11\nTMhelix 12 31\noutside 32 530\nTMhelix 531 553\ninside 554 570

32976 GCF_000365385.1_ASM36538v1 Streptomyces sp. CNS615 Terrabacteria group; Actinobacteria; Actinobacteria; Streptomycetales; Streptomycetaceae; Streptomyces
MPVARRPLLTATAAGSLLCALWFVPSANATADGESGHPDRARSGASASGVQDADGAEGVQ WP_017946382.1 hypothetical protein [Streptomyces sp. CNS615] Length: 160\nNumber of predicted TMHs: 2\nExp number of AAs in TMHs: 37.10394\nExp number, first 60 AAs: 17.06003\nTotal prob of N-in: 0.85767\nPOSSIBLE N-term signal sequence\ninside 1 6\nTMhelix 7 29\noutside 30 129\nTMhelix 130 152\ninside 153 160

32977 GCF_000365385.1_ASM36538v1 Streptomyces sp. CNS615 Terrabacteria group; Actinobacteria; Actinobacteria; Streptomycetales; Streptomycetaceae; Streptomyces
MLCVVENHSLPRTAAFFDLDKTVIAKSSALTFSKSFYHGGLINRRAVLRTAYTQFVFLAG WP_017947803.1 MULTISPECIES: inhibition of morphological differentiation protein [Streptomyces] Length: 281\nNumber of predicted TMHs: 1\nExp number of AAs in TMHs: 21.92185\nExp number, first 60 AAs: 0.03178\nTotal prob of N-in: 0.00389\noutside 1 247\nTMhelix 248 270\ninside 271 281

32978 GCF_000702365.1_ASM70236v1 Streptomyces sp. CNS654 Terrabacteria group; Actinobacteria; Actinobacteria; Streptomycetales; Streptomycetaceae; Streptomyces
MGLSWNDRRLERRVKPGDGKPLKPFRWWQMTRRSLLSIALPLAGGGGLSATYTVEVKHGG WP_032767146.1 hypothetical protein [Streptomyces sp. CNS654] Length: 220\nNumber of predicted TMHs: 2\nExp number of AAs in TMHs: 41.55178\nExp number, first 60 AAs: 6.12906\nTotal prob of N-in: 0.89949\ninside 1 137\nTMhelix 138 160\noutside 161 184\nTMhelix 185 202\ninside 203 220

32979 GCF_000702365.1_ASM70236v1 Streptomyces sp. CNS654 Terrabacteria group; Actinobacteria; Actinobacteria; Streptomycetales; Streptomycetaceae; Streptomyces
MLSVVENCFSPRTAAFFDLDKTVIAKSSTLTFSKSFYQGGLINRRAVLRTAYTQFVFLAG WP_032769244.1 MULTISPECIES: inhibition of morphological differentiation protein [Streptomyces] Length: 281\nNumber of predicted TMHs: 1\nExp number of AAs in TMHs: 21.46267\nExp number, first 60 AAs: 0.01959\nTotal prob of N-in: 0.00583\noutside 1 247\nTMhelix 248 270\ninside 271 281

32980 GCF_000377525.1_ASM37752v1 Streptomyces sp. CNT302 Terrabacteria group; Actinobacteria; Actinobacteria; Streptomycetales; Streptomycetaceae; Streptomyces
MLCVVENHSLPRTAAFFDLDKTVIAKSSALTFSKSFYHGGLINRRAVLRTAYTQFVFLAG WP_017947803.1 MULTISPECIES: inhibition of morphological differentiation protein [Streptomyces] Length: 281\nNumber of predicted TMHs: 1\nExp number of AAs in TMHs: 21.92185\nExp number, first 60 AAs: 0.03178\nTotal prob of N-in: 0.00389\noutside 1 247\nTMhelix 248 270\ninside 271 281

32981 GCF_000424765.1_ASM42476v1 Streptomyces sp. CNT318 Terrabacteria group; Actinobacteria; Actinobacteria; Streptomycetales; Streptomycetaceae; Streptomyces
MITRRRLVTGACALLMALPLGLLPTGTASAEPASDEDAPKVELTLDVSGSMRKRDIDGRS WP_026005050.1 MULTISPECIES: inter-alpha-trypsin inhibitor heavy chain H5 [Streptomyces] Length: 421\nNumber of predicted TMHs: 2\nExp number of AAs in TMHs: 38.90213\nExp number, first 60 AAs: 16.54504\nTotal prob of N-in: 0.77694\nPOSSIBLE N-term signal sequence\ninside 1 6\nTMhelix 7 29\noutside 30 388\nTMhelix 389 411\ninside 412 421

32982 GCF_000424765.1_ASM42476v1 Streptomyces sp. CNT318 Terrabacteria group; Actinobacteria; Actinobacteria; Streptomycetales; Streptomycetaceae; Streptomyces
MSPRRRRATVLTAATVLSAAAAPVASAAAPAAPVTSVAASPARTTASGLPEGLYGTADPK WP_027759771.1 hypothetical protein [Streptomyces sp. CNT318] Length: 443\nNumber of predicted TMHs: 1\nExp number of AAs in TMHs: 34.01542\nExp number, first 60 AAs: 11.70667\nTotal prob of N-in: 0.52681\nPOSSIBLE N-term signal sequence\noutside 1 413\nTMhelix 414 436\ninside 437 443

32983 GCF_000424765.1_ASM42476v1 Streptomyces sp. CNT318 Terrabacteria group; Actinobacteria; Actinobacteria; Streptomycetales; Streptomycetaceae; Streptomyces
MLGAVKNLAMPRTAAFFDLDKTVIAKSSTLTFGKSFYQGGLINRRAVLRTAYAQFVYLLG WP_027760567.1 MULTISPECIES: inhibition of morphological differentiation protein [Streptomyces] Length: 281\nNumber of predicted TMHs: 1\nExp number of AAs in TMHs: 22.28973\nExp number, first 60 AAs: 0.03324\nTotal prob of N-in: 0.01080\noutside 1 246\nTMhelix 247 269\ninside 270 281

32984 GCF_000424765.1_ASM42476v1 Streptomyces sp. CNT318 Terrabacteria group; Actinobacteria; Actinobacteria; Streptomycetales; Streptomycetaceae; Streptomyces
MIFRMRAFWGMRIDDPWHDALATGGAEVHGTPVDTPPRRTAPPPRREPVPSAVHGVYPTV WP_027760894.1 hypothetical protein [Streptomyces sp. CNT318] Length: 177\nNumber of predicted TMHs: 2\nExp number of

AAs in TMHs: 44.64462\nExp number, first 60 AAs:  0.00024\nTotal prob of N-in:   0.99631\ninside   1   75\nTMhelix   76   98\noutside   99   107\nTMhelix   108   130\ninside   131   177

32985 GCF_000424785.1_ASM42478v1 Streptomyces sp. CNT360 Terrabacteria group; Actinobacteria; Actinobacteria; Streptomycetales; Streptomycetaceae; Streptomyces MLSGVENLALPRTAAFFDLDKTVIAKSSTLTFGKSFYQGGLINRRAVLRTAYAQFVYLLG WP_027739687.1 inhibition of morphological differentiation protein [Streptomyces sp. CNT360] Length: 283\nNumber of predicted TMHs:  1\nExp number of AAs in TMHs: 21.96145\nExp number, first 60 AAs:  0.01027\nTotal prob of N-in:   0.00078\noutside   1   246\nTMhelix   247   269\ninside   270   283

32986 GCF_000424785.1_ASM42478v1 Streptomyces sp. CNT360 Terrabacteria group; Actinobacteria; Actinobacteria; Streptomycetales; Streptomycetaceae; Streptomyces MSPRRRRATVLTAATMLCAAAAPMASAAAPQAAPRTVPHAASQPASAVAAGAAKPKLPAG WP_027740156.1 hypothetical protein [Streptomyces sp. CNT360] Length: 473\nNumber of predicted TMHs:  1\nExp number of AAs in TMHs: 35.14682\nExp number, first 60 AAs:  12.81027\nTotal prob of N-in:   0.58893\nPOSSIBLE N-term signal sequence\noutside   1   443\nTMhelix   444   466\ninside   467   473

32987 GCF_000514995.1_ASM51499v1 Streptomyces sp. CNT371 Terrabacteria group; Actinobacteria; Actinobacteria; Streptomycetales; Streptomycetaceae; Streptomyces MLRRLRALAATLLPRRRVVAVAAATLLTSTLAVGTDLAGAPPAHAAAGTVDGGRLDWGIK WP_027743790.1 hypothetical protein [Streptomyces sp. CNT371] Length: 513\nNumber of predicted TMHs:  1\nExp number of AAs in TMHs: 25.62012\nExp number, first 60 AAs:  5.36475\nTotal prob of N-in:   0.25046\noutside   1   453\nTMhelix   454   476\ninside   477   513

32988 GCF_000514995.1_ASM51499v1 Streptomyces sp. CNT371 Terrabacteria group; Actinobacteria; Actinobacteria; Streptomycetales; Streptomycetaceae; Streptomyces MLALVENHSLPRTAAFFDLDKTVIAKSSTLTFSRSFYHGGLINRRAVLRSAYSQFIYLLG WP_018835865.1 MULTISPECIES: inhibition of morphological differentiation protein [Streptomyces] Length: 276\nNumber of predicted TMHs:  1\nExp number of AAs in TMHs: 22.3164\nExp number, first 60 AAs:  0.00298\nTotal prob of N-in:   0.00131\noutside   1   246\nTMhelix   247   269\ninside   270   276

32989 GCF_000377545.1_ASM37754v1 Streptomyces sp. CNY228 Terrabacteria group; Actinobacteria; Actinobacteria; Streptomycetales; Streptomycetaceae; Streptomyces MLDPVENRSLPRTAAFFDLDKTVIAKSSTLTFSKSFYQGGLINRRAVLRTAYTQFVFLAG WP_010643895.1 MULTISPECIES: morphological differentiation-associated protein [Streptomyces] Length: 290\nNumber of predicted TMHs:  1\nExp number of AAs in TMHs: 21.93803\nExp number, first 60 AAs:  0.02636\nTotal prob of N-in:   0.00751\noutside   1   247\nTMhelix   248   270\ninside   271   290

32990 GCF_000377545.1_ASM37754v1 Streptomyces sp. CNY228 Terrabacteria group; Actinobacteria; Actinobacteria; Streptomycetales; Streptomycetaceae; Streptomyces MGGRGTTAQARERGRLPGRPSDPAPTGGNDVITGVNLRMPARRTALPALLLTATALLAVP WP_026282568.1 hypothetical protein [Streptomyces sp. CNY228] Length: 304\nNumber of predicted TMHs:  2\nExp number of AAs in TMHs: 39.02666\nExp number, first 60 AAs:  11.63983\nTotal prob of N-in:   0.75391\nPOSSIBLE N-term signal sequence\ninside   1   44\nTMhelix   45   67\noutside   68   276\nTMhelix   277   299\ninside   300   304

32991 GCF_000377545.1_ASM37754v1 Streptomyces sp. CNY228 Terrabacteria group; Actinobacteria; Actinobacteria; Streptomycetales; Streptomycetaceae; Streptomyces MSRRTRTARRRPVAAAVLAAGAVLLPAVPASAASGADQDFVIEDPRITESSGLAASKRHP WP_026282589.1 hypothetical protein [Streptomyces sp. CNY228] Length: 352\nNumber of predicted TMHs:  2\nExp number of AAs in TMHs: 39.5043\nExp number, first 60 AAs:  21.11165\nTotal prob of N-in:   0.99083\nPOSSIBLE N-term signal sequence\ninside   1   12\nTMhelix   13   35\noutside   36   324\nTMhelix   325   344\ninside   345   352

32992 GCF_000377545.1_ASM37754v1 Streptomyces sp. CNY228 Terrabacteria group; Actinobacteria; Actinobacteria; Streptomycetales; Streptomycetaceae; Streptomyces MPDRRSAVSLRLPLPTAAPGFGGLALPGAAATAPDAWGVVAVRESSEKERKSSSEPEGRS WP_018894266.1 hypothetical protein [Streptomyces sp. CNY228] Length: 292\nNumber of predicted TMHs:  2\nExp number of AAs in TMHs: 51.76527\nExp number, first 60 AAs:  6.06558\nTotal prob of N-in:   0.33682\noutside   1   164\nTMhelix   165   187\ninside   188   229\nTMhelix   230   252\noutside   253   292

32993 GCF_000377545.1_ASM37754v1 Streptomyces sp. CNY228 Terrabacteria group; Actinobacteria; Actinobacteria; Streptomycetales; Streptomycetaceae; Streptomyces

MSRMRGSSGGRIAGRRAAVVLAVAVLGLAGCSAGPDSGASSAGDRAVAPSAADGAGGGQA WP_018894652.1 hypothetical protein [Streptomyces sp. CNY228] Length: 334\nNumber of predicted TMHs: 1\nExp number of AAs in TMHs: 36.39009\nExp number, first 60 AAs: 13.60232\nTotal prob of N-in: 0.64387\nPOSSIBLE N-term signal sequence\noutside 1 269\nTMhelix 270 292\ninside 293 334

32994 GCF_000377545.1_ASM37754v1 Streptomyces sp. CNY228 Terrabacteria group; Actinobacteria; Actinobacteria; Streptomycetales; Streptomycetaceae; Streptomyces MTPNRRPVVLAAAVATALGLGATALTLPAVAAEAPRAGAPVMDLTDGTLDWGVKESFRRY WP_018896317.1 membrane protein [Streptomyces sp. CNY228] Length: 511\nNumber of predicted TMHs: 2\nExp number of AAs in TMHs: 42.8566599999999\nExp number, first 60 AAs: 22.03084\nTotal prob of N-in: 0.97272\nPOSSIBLE N-term signal sequence\ninside 1 6\nTMhelix 7 29\noutside 30 476\nTMhelix 477 499\ninside 500 511

32995 GCF_000377165.1_ASM37716v1 Streptomyces sp. CNY243 Terrabacteria group; Actinobacteria; Actinobacteria; Streptomycetales; Streptomycetaceae; Streptomyces MLRRLRALAATLLPRRRVVAVAAATLLTSTLAVGTDLAGAPPAHAAARTVDGGRLDWGIK WP_018853632.1 hypothetical protein [Streptomyces sp. CNY243] Length: 515\nNumber of predicted TMHs: 1\nExp number of AAs in TMHs: 26.63686\nExp number, first 60 AAs: 6.44153\nTotal prob of N-in: 0.30013\noutside 1 458\nTMhelix 459 481\ninside 482 515

32996 GCF_000377165.1_ASM37716v1 Streptomyces sp. CNY243 Terrabacteria group; Actinobacteria; Actinobacteria; Streptomycetales; Streptomycetaceae; Streptomyces MLALVENHSLPRTAAFFDLDKTVIAKSSTLTFSRSFYHGGLINRRAVLRSAYSQFIYLLG WP_018835865.1 MULTISPECIES: inhibition of morphological differentiation protein [Streptomyces] Length: 276\nNumber of predicted TMHs: 1\nExp number of AAs in TMHs: 22.3164\nExp number, first 60 AAs: 0.00298\nTotal prob of N-in: 0.00131\noutside 1 246\nTMhelix 247 269\ninside 270 276

32997 GCF_002188345.1_ASM218834v1 Streptomyces sp. CS057 Terrabacteria group; Actinobacteria; Actinobacteria; Streptomycetales; Streptomycetaceae; Streptomyces MLSVVENRFSPRTAAFFDLDKTVIAKSSTLTFSKSFYQGGLINRRAVLRTAYAQFVFLAG WP_087765561.1 inhibition of morphological differentiation protein [Streptomyces sp. CS057] Length: 281\nNumber of predicted TMHs: 1\nExp number of AAs in TMHs: 21.46406\nExp number, first 60 AAs: 0.04078\nTotal prob of N-in: 0.00982\noutside 1 247\nTMhelix 248 270\ninside 271 281

32998 GCF_002188345.1_ASM218834v1 Streptomyces sp. CS057 Terrabacteria group; Actinobacteria; Actinobacteria; Streptomycetales; Streptomycetaceae; Streptomyces MGTEQQARRWPRARRGLLSALTASLLTLGASLTHLTAASPASADPIAKCTRTKGAIVAVDWP_087764645.1 peptidase [Streptomyces sp. CS057] Length: 927\nNumber of predicted TMHs: 2\nExp number of AAs in TMHs: 38.42564\nExp number, first 60 AAs: 18.04221\nTotal prob of N-in: 0.81398\nPOSSIBLE N-term signal sequence\ninside 1 15\nTMhelix 16 38\noutside 39 889\nTMhelix 890 912\ninside 913 927

32999 GCF_002188365.1_ASM218836v1 Streptomyces sp. CS113 Terrabacteria group; Actinobacteria; Actinobacteria; Streptomycetales; Streptomycetaceae; Streptomyces MGHNRRRRPTGARRATFGAVALILGGSGLVAVNVFASATESGNAAVPLGSSGVAATVDCP WP_087811701.1 hypothetical protein [Streptomyces sp. CS113] Length: 707\nNumber of predicted TMHs: 2\nExp number of AAs in TMHs: 43.11227\nExp number, first 60 AAs: 21.83889\nTotal prob of N-in: 0.99023\nPOSSIBLE N-term signal sequence\ninside 1 16\nTMhelix 17 39\noutside 40 679\nTMhelix 680 702\ninside 703 707

33000 GCF_002188365.1_ASM218836v1 Streptomyces sp. CS113 Terrabacteria group; Actinobacteria; Actinobacteria; Streptomycetales; Streptomycetaceae; Streptomyces MPRTAAFFDLDKTVIAKSSTLTFSKSFYQGGLINRRAVLRTAYAQFVFLAGGADHDQMER WP_087808038.1 inhibition of morphological differentiation protein [Streptomyces sp. CS113] Length: 272\nNumber of predicted TMHs: 1\nExp number of AAs in TMHs: 21.78654\nExp number, first 60 AAs: 0.03303\nTotal prob of N-in: 0.00253\noutside 1 238\nTMhelix 239 261\ninside 262 272

33001 GCF_002188365.1_ASM218836v1 Streptomyces sp. CS113 Terrabacteria group; Actinobacteria; Actinobacteria; Streptomycetales; Streptomycetaceae; Streptomyces MRHARRRIVRRVARLAAVGGLLAGSAMVAHAGASEPPGASAAPRTLARTASGAGAGLVSR WP_087804880.1 serine protease [Streptomyces sp. CS113] Length: 447\nNumber of predicted TMHs: 2\nExp number of AAs in TMHs: 41.70534\nExp number, first 60 AAs: 21.2728\nTotal prob of N-in: 0.97452\nPOSSIBLE N-term signal sequence\ninside 1 11\nTMhelix 12 34\noutside 35 408\nTMhelix 409 431\ninside 432 447

33002 GCF_002188335.1_ASM218833v1 Streptomyces sp. CS159 Terrabacteria group; Actinobacteria; Actinobacteria; Streptomycetales; Streptomycetaceae; Streptomyces
MGHNRRRRPTGARRATFGAVALILGGSGLVAVNVFASATESGNAAVPLGSSGVAATVDCP WP_087792726.1
hypothetical protein [Streptomyces sp. CS159] Length: 702\nNumber of predicted TMHs: 2\nExp number of AAs in TMHs: 43.21345\nExp number, first 60 AAs: 21.83862\nTotal prob of N-in: 0.99025\nPOSSIBLE N-term signal sequence\ninside 1 16\nTMhelix 17 39\noutside 40 674\nTMhelix 675 697\ninside 698 702

33003 GCF_002188335.1_ASM218833v1 Streptomyces sp. CS159 Terrabacteria group; Actinobacteria; Actinobacteria; Streptomycetales; Streptomycetaceae; Streptomyces
MRHARRRIVRRAARLAAVGGLLLGGTMVTRAVAGEPPGPSAAPRTLAQTASGAGAGLVAR WP_087788440.1
serine protease [Streptomyces sp. CS159] Length: 452\nNumber of predicted TMHs: 2\nExp number of AAs in TMHs: 42.02165\nExp number, first 60 AAs: 21.46147\nTotal prob of N-in: 0.97024\nPOSSIBLE N-term signal sequence\ninside 1 11\nTMhelix 12 34\noutside 35 413\nTMhelix 414 436\ninside 437 452

33004 GCF_002188335.1_ASM218833v1 Streptomyces sp. CS159 Terrabacteria group; Actinobacteria; Actinobacteria; Streptomycetales; Streptomycetaceae; Streptomyces
MPRTAAFFDLDKTVIAKSSTLTFSKSFYQGGLINRRAVLRTAYAQFVFLAGGADHDQMER WP_042826664.1
inhibition of morphological differentiation protein [Streptomyces coelicoflavus] Length: 268\nNumber of predicted TMHs: 1\nExp number of AAs in TMHs: 21.70608\nExp number, first 60 AAs: 0.03423\nTotal prob of N-in: 0.00318\noutside 1 238\nTMhelix 239 261\ninside 262 268

33005 GCF_002188355.1_ASM218835v1 Streptomyces sp. CS227 Terrabacteria group; Actinobacteria; Actinobacteria; Streptomycetales; Streptomycetaceae; Streptomyces
MSRMRGTSGGRIVGRRAAVVLAVAVLGLAGCSAGSDSGASSAGDRAAAPSAADGAGGGQA WP_087772526.1
hypothetical protein [Streptomyces sp. CS227] Length: 337\nNumber of predicted TMHs: 2\nExp number of AAs in TMHs: 40.26754\nExp number, first 60 AAs: 17.48296\nTotal prob of N-in: 0.82273\nPOSSIBLE N-term signal sequence\ninside 1 11\nTMhelix 12 34\noutside 35 269\nTMhelix 270 292\ninside 293 337

33006 GCF_002188355.1_ASM218835v1 Streptomyces sp. CS227 Terrabacteria group; Actinobacteria; Actinobacteria; Streptomycetales; Streptomycetaceae; Streptomyces
MSRRTRTARRRPVAAAVLAAGAVLLPVVPASAASGADQDFVIEDPRITESSGLAASKRHP WP_087773605.1
hypothetical protein [Streptomyces sp. CS227] Length: 348\nNumber of predicted TMHs: 2\nExp number of AAs in TMHs: 36.16596\nExp number, first 60 AAs: 21.28342\nTotal prob of N-in: 0.99454\nPOSSIBLE N-term signal sequence\ninside 1 12\nTMhelix 13 35\noutside 36 322\nTMhelix 323 340\ninside 341 348

33007 GCF_002188355.1_ASM218835v1 Streptomyces sp. CS227 Terrabacteria group; Actinobacteria; Actinobacteria; Streptomycetales; Streptomycetaceae; Streptomyces
MTPNRRPVVLAAAVATALGLGATALTLPAVAAEAPRAGAPVMDLTDGTLDWGVKESFRRY WP_087776164.1
hypothetical protein [Streptomyces sp. CS227] Length: 518\nNumber of predicted TMHs: 2\nExp number of AAs in TMHs: 43.22224\nExp number, first 60 AAs: 22.03093\nTotal prob of N-in: 0.97254\nPOSSIBLE N-term signal sequence\ninside 1 6\nTMhelix 7 29\noutside 30 483\nTMhelix 484 506\ninside 507 518

33008 GCF_002188355.1_ASM218835v1 Streptomyces sp. CS227 Terrabacteria group; Actinobacteria; Actinobacteria; Streptomycetales; Streptomycetaceae; Streptomyces
MAPPHRRRAAVVVRALAALTALLCATGLAPAAPSGLAARAVVAEVPPPTATTSVVTVRTG WP_087773882.1
hypothetical protein [Streptomyces sp. CS227] Length: 829\nNumber of predicted TMHs: 2\nExp number of AAs in TMHs: 39.58431\nExp number, first 60 AAs: 19.37531\nTotal prob of N-in: 0.90443\nPOSSIBLE N-term signal sequence\ninside 1 11\nTMhelix 12 34\noutside 35 790\nTMhelix 791 813\ninside 814 829

33009 GCF_002188355.1_ASM218835v1 Streptomyces sp. CS227 Terrabacteria group; Actinobacteria; Actinobacteria; Streptomycetales; Streptomycetaceae; Streptomyces
MGGRGTTARARERGRLPGRPCDPAPTGGNDVITGVNLRMPARRTALPALLLTATALLAVP WP_087774025.1
hypothetical protein [Streptomyces sp. CS227] Length: 305\nNumber of predicted TMHs: 2\nExp number of AAs in TMHs: 40.09898\nExp number, first 60 AAs: 12.40746\nTotal prob of N-in: 0.80261\nPOSSIBLE N-term signal sequence\ninside 1 44\nTMhelix 45 67\noutside 68 277\nTMhelix 278 300\ninside 301 305

33010 GCF_002188355.1_ASM218835v1 Streptomyces sp. CS227 Terrabacteria group; Actinobacteria; Actinobacteria; Streptomycetales; Streptomycetaceae; Streptomyces
MLDPVENRSLPRTAAFFDLDKTVIAKSSTLTFSKSFYQGGLINRRAVLRTAYTQFVFLAG WP_087774259.1 inhibition of morphological differentiation protein [Streptomyces sp. CS227] Length: 279\nNumber of predicted TMHs: 1\nExp

number of AAs in TMHs: 21.72379\nExp number, first 60 AAs: 0.0316\nTotal prob of N-in: 0.01274\noutside 1 247\nTMhelix 248 270\ninside 271 279

33011 GCF_000805335.1_ASM80533v1 Streptomyces sp. CT34 Terrabacteria group; Actinobacteria; Actinobacteria; Streptomycetales; Streptomycetaceae; Streptomyces
MPDRRRRAALLRPTPATAGWGFQPATAPGAVLLPRERKSSSSPDSPTPDSPSSGRPDPAG WP_043265717.1 hypothetical protein [Streptomyces sp. CT34] Length: 312\nNumber of predicted TMHs: 2\nExp number of AAs in TMHs: 46.16194\nExp number, first 60 AAs: 0.01197\nTotal prob of N-in: 0.92218\ninside 1 175\nTMhelix 176 198\noutside 199 249\nTMhelix 250 272\ninside 273 312

33012 GCF_000805335.1_ASM80533v1 Streptomyces sp. CT34 Terrabacteria group; Actinobacteria; Actinobacteria; Streptomycetales; Streptomycetaceae; Streptomyces
MRQRRRAAVASMAALVLAGAGTVAVAGTAQAAEVSYKTECLPPPISGLPPIEGTTKVDIG WP_043264623.1 hypothetical protein [Streptomyces sp. CT34] Length: 489\nNumber of predicted TMHs: 2\nExp number of AAs in TMHs: 45.06744\nExp number, first 60 AAs: 22.26146\nTotal prob of N-in: 0.99593\nPOSSIBLE N-term signal sequence\ninside 1 6\nTMhelix 7 29\noutside 30 458\nTMhelix 459 481\ninside 482 489

33013 GCF_000514055.1_ASM51405v1 Streptomyces sp. DpondAA-B6 Terrabacteria group; Actinobacteria; Actinobacteria; Streptomycetales; Streptomycetaceae; Streptomyces
MLSFVENCFSPRTAAFFDLDKTVIAKSSTLAFSKSFYQGGLINRRAVLRTAYTQFVFLAG WP_028442100.1 inhibition of morphological differentiation protein [Streptomyces sp. DpondAA-B6] Length: 279\nNumber of predicted TMHs: 1\nExp number of AAs in TMHs: 21.81622\nExp number, first 60 AAs: 0.01771\nTotal prob of N-in: 0.00802\noutside 1 247\nTMhelix 248 270\ninside 271 279

33014 GCF_001514285.1_ASM151428v1 Streptomyces sp. DSM 15324 Terrabacteria group; Actinobacteria; Actinobacteria; Streptomycetales; Streptomycetaceae; Streptomyces
MRHARRRIVRRVTRLAAVGGLLLGGTMVTQAAMATETPATSATPRTAPESVVTNPGAALV WP_067254820.1 protease [Streptomyces sp. DSM 15324] Length: 464\nNumber of predicted TMHs: 2\nExp number of AAs in TMHs: 42.67542\nExp number, first 60 AAs: 21.78052\nTotal prob of N-in: 0.97963\nPOSSIBLE N-term signal sequence\ninside 1 11\nTMhelix 12 34\noutside 35 425\nTMhelix 426 448\ninside 449 464

33015 GCF_000382745.1_ASM38274v1 Streptomyces sp. DvalAA-83 Terrabacteria group; Actinobacteria; Actinobacteria; Streptomycetales; Streptomycetaceae; Streptomyces
MLTFVENCFSPRTAAFFDLDKTVIAKSSTLTFSKSFYQGGLINRRAVLRTAYTQFVFLAG WP_014046857.1 MULTISPECIES: haloacid dehalogenase [Streptomyces] Length: 279\nNumber of predicted TMHs: 1\nExp number of AAs in TMHs: 21.69088\nExp number, first 60 AAs: 0.01488\nTotal prob of N-in: 0.00190\noutside 1 247\nTMhelix 248 270\ninside 271 279

33016 GCF_000162775.1_ASM16277v1 Streptomyces sp. e14 Terrabacteria group; Actinobacteria; Actinobacteria; Streptomycetales; Streptomycetaceae; Streptomyces
MLRVVENHSLPRTAAFFDLDKTVIAKSSTLTFSKSFYQGGLINRRAVLRTAYAQFVFLAG WP_009189832.1 MULTISPECIES: haloacid dehalogenase [Streptomyces] Length: 277\nNumber of predicted TMHs: 1\nExp number of AAs in TMHs: 21.60611\nExp number, first 60 AAs: 0.04466\nTotal prob of N-in: 0.00565\noutside 1 247\nTMhelix 248 270\ninside 271 277

33017 GCF_000162775.1_ASM16277v1 Streptomyces sp. e14 Terrabacteria group; Actinobacteria; Actinobacteria; Streptomycetales; Streptomycetaceae; Streptomyces
MPAPHKTARRTLLVASAALLSVSLTAPAALAAPSPKTSKPATPPARMSTVGGARLGQAGT WP_043261201.1 D-alanyl-D-alanine carboxypeptidase [Streptomyces sp. e14] Length: 427\nNumber of predicted TMHs: 2\nExp number of AAs in TMHs: 41.37581\nExp number, first 60 AAs: 19.02121\nTotal prob of N-in: 0.90216\nPOSSIBLE N-term signal sequence\ninside 1 11\nTMhelix 12 34\noutside 35 388\nTMhelix 389 411\ninside 412 427

33018 GCF_001493375.1_Streptomyces_specialis Streptomyces specialis Terrabacteria group; Actinobacteria; Actinobacteria; Streptomycetales; Streptomycetaceae; Streptomyces
MPPRRRALPAAVARGALLAPGPPLGAAPAHADSIRDQQQWALDAINAPEAWETTRGAGVT WP_059008513.1 type VII secretion-associated serine protease mycosin [Streptomyces specialis] Length: 408\nNumber of predicted TMHs: 1\nExp number of AAs in TMHs: 24.03844\nExp number, first 60 AAs: 0.04025\nTotal prob of N-in: 0.09609\noutside 1 365\nTMhelix 366 388\ninside 389 408

33019 GCF_001493375.1_Streptomyces_specialis Streptomyces specialis Terrabacteria group; Actinobacteria; Actinobacteria; Streptomycetales; Streptomycetaceae; Streptomyces

MVKILAQPRTAAFFDLDKTVIAKSSTLTFSRSFYQGGLINRRGVLRSAYAQFVYLLGGAD WP_059010571.1 inhibition of morphological differentiation protein [Streptomyces specialis] Length: 284\nNumber of predicted TMHs: 1\nExp number of AAs in TMHs: 20.33255\nExp number, first 60 AAs: 0.05465\nTotal prob of N-in: 0.04360\noutside 1 243\nTMhelix 244 266\ninside 267 284

33020 GCF_001493375.1_Streptomyces_specialis Streptomyces specialis Terrabacteria group; Actinobacteria; Actinobacteria; Streptomycetales; Streptomycetaceae; Streptomyces MPTWTPTWIRPRRAAVAAALLLPLAAGSPAAAADEPAVFTLDDPRIVESSGLQASDRHPG WP_059006136.1 hypothetical protein [Streptomyces specialis] Length: 362\nNumber of predicted TMHs: 1\nExp number of AAs in TMHs: 22.54572\nExp number, first 60 AAs: 3.0151\nTotal prob of N-in: 0.16749\noutside 1 333\nTMhelix 334 356\ninside 357 362

33021 GCF_001493375.1_Streptomyces_specialis Streptomyces specialis Terrabacteria group; Actinobacteria; Actinobacteria; Streptomycetales; Streptomycetaceae; Streptomyces MPLSGRSGDVKPYQGVPHVYNSAYRRLGSRRSAVLVATTAALLGSVSPALAAEGDQDERG WP_079032150.1 hypothetical protein [Streptomyces specialis] Length: 366\nNumber of predicted TMHs: 1\nExp number of AAs in TMHs: 22.04808\nExp number, first 60 AAs: 1.40544\nTotal prob of N-in: 0.06972\noutside 1 335\nTMhelix 336 355\ninside 356 366

33022 GCF_001746285.1_ASM174628v1 Streptomyces sp. EN16 Terrabacteria group; Actinobacteria; Actinobacteria; Streptomycetales; Streptomycetaceae; Streptomyces MLSVVENCFSPRTAAFFDLDKTVIAKSSTLTFSKSFYQGGLINRRAVLRTAYAQFVFLAG WP_069752100.1 inhibition of morphological differentiation protein [Streptomyces sp. EN16] Length: 281\nNumber of predicted TMHs: 1\nExp number of AAs in TMHs: 21.45971\nExp number, first 60 AAs: 0.03431\nTotal prob of N-in: 0.00824\noutside 1 247\nTMhelix 248 270\ninside 271 281

33023 GCF_001746305.1_ASM174630v1 Streptomyces sp. EN23 Terrabacteria group; Actinobacteria; Actinobacteria; Streptomycetales; Streptomycetaceae; Streptomyces MLSVVENCFSPRTAAFFDLDKTVIAKSSTLTFSKSFYQGGLINRRAVLRTAYTQFVFLAG WP_069740854.1 inhibition of morphological differentiation protein [Streptomyces sp. EN23] Length: 281\nNumber of predicted TMHs: 1\nExp number of AAs in TMHs: 21.35068\nExp number, first 60 AAs: 0.0254\nTotal prob of N-in: 0.01352\noutside 1 247\nTMhelix 248 270\ninside 271 281

33024 GCF_001746315.1_ASM174631v1 Streptomyces sp. EN27 Terrabacteria group; Actinobacteria; Actinobacteria; Streptomycetales; Streptomycetaceae; Streptomyces MLSVVENCFSPRTAAFFDLDKTVIAKSSTLTFSKSFYQGGLINRRAVLRTAYTQFVFLAG WP_069735505.1 inhibition of morphological differentiation protein [Streptomyces sp. EN27] Length: 281\nNumber of predicted TMHs: 1\nExp number of AAs in TMHs: 21.41509\nExp number, first 60 AAs: 0.02225\nTotal prob of N-in: 0.00934\noutside 1 247\nTMhelix 248 270\ninside 271 281

33025 GCF_001746315.1_ASM174631v1 Streptomyces sp. EN27 Terrabacteria group; Actinobacteria; Actinobacteria; Streptomycetales; Streptomycetaceae; Streptomyces MGLSWNDRRLERRVKPGDGKPLKPFRWWQMTRRSLLSLPLPVGGGISATYTVEVKHGGDP WP_069735562.1 hypothetical protein [Streptomyces sp. EN27] Length: 219\nNumber of predicted TMHs: 2\nExp number of AAs in TMHs: 40.11862\nExp number, first 60 AAs: 0.2816\nTotal prob of N-in: 0.96088\ninside 1 136\nTMhelix 137 159\noutside 160 183\nTMhelix 184 201\ninside 202 219

33026 GCF_001653515.1_ASM165351v1 Streptomyces sp. ERV7 Terrabacteria group; Actinobacteria; Actinobacteria; Streptomycetales; Streptomycetaceae; Streptomyces MLGGLFAAARSGGRRGVVLLAVALLAGAQTAQAAVTVAYGDRERLHRLDSYPAWGGAHDA WP_067159911.1 hypothetical protein [Streptomyces sp. ERV7] Length: 390\nNumber of predicted TMHs: 1\nExp number of AAs in TMHs: 28.95502\nExp number, first 60 AAs: 8.85697\nTotal prob of N-in: 0.36665\noutside 1 359\nTMhelix 360 379\ninside 380 390

33027 GCF_001653515.1_ASM165351v1 Streptomyces sp. ERV7 Terrabacteria group; Actinobacteria; Actinobacteria; Streptomycetales; Streptomycetaceae; Streptomyces MLAFVENHSLPRTAAFFDLDKTVIAKSSTLTFSKSFYQGGLINRRAVLRTAYAQFVFLAG WP_067161478.1 inhibition of morphological differentiation protein [Streptomyces sp. ERV7] Length: 278\nNumber of predicted TMHs: 1\nExp number of AAs in TMHs: 21.6764\nExp number, first 60 AAs: 0.0177\nTotal prob of N-in: 0.00216\noutside 1 247\nTMhelix 248 270\ninside 271 278

33028 GCF_900079405.2_Streptomyces_sp_F1_v3 Streptomyces sp. F-1 Terrabacteria group; Actinobacteria; Actinobacteria; Streptomycetales; Streptomycetaceae; Streptomyces
MVRNPLLGAVYRPGVRESSRLSRRAVLGLTAAALPLSAAAPASAEPTAVGGARLARRGVQ WP_079172327.1
MULTISPECIES: D-alanyl-D-alanine carboxypeptidase [Streptomyces] Length: 432\nNumber of predicted TMHs: 1\nExp number of AAs in TMHs: 17.37505\nExp number, first 60 AAs: 0.69826\nTotal prob of N-in: 0.03361\noutside 1 367\nTMhelix 368 387\ninside 388 432

33029 GCF_900079405.2_Streptomyces_sp_F1_v3 Streptomyces sp. F-1 Terrabacteria group; Actinobacteria; Actinobacteria; Streptomycetales; Streptomycetaceae; Streptomyces
MASRGRRAAMRSLGALAGVAALTALGSTAPGWAATPVHAHGKPSSTATPIQHVVVLFDEN WP_070024923.1
phospholipase [Streptomyces sp. F-1] Length: 608\nNumber of predicted TMHs: 2\nExp number of AAs in TMHs: 43.20869\nExp number, first 60 AAs: 21.90049\nTotal prob of N-in: 0.96888\nPOSSIBLE N-term signal sequence\ninside 1 12\nTMhelix 13 35\noutside 36 579\nTMhelix 580 602\ninside 603 608

33030 GCF_001653115.1_ASM165311v1 Streptomyces sp. F-3 Terrabacteria group; Actinobacteria; Actinobacteria; Streptomycetales; Streptomycetaceae; Streptomyces
MLATPRRRPLALVAAMATATALGATVLVAAPTAAAAETPLSGYELTWGIKESYRTYVVRY WP_067392033.1
hypothetical protein [Streptomyces sp. F-3] Length: 507\nNumber of predicted TMHs: 2\nExp number of AAs in TMHs: 43.58403\nExp number, first 60 AAs: 21.17274\nTotal prob of N-in: 0.93360\nPOSSIBLE N-term signal sequence\ninside 1 11\nTMhelix 12 34\noutside 35 476\nTMhelix 477 499\ninside 500 507

33031 GCF_001653115.1_ASM165311v1 Streptomyces sp. F-3 Terrabacteria group; Actinobacteria; Actinobacteria; Streptomycetales; Streptomycetaceae; Streptomyces
MLGGVENHSLPRTAAFFDLDKTVIAKSSTLTFSKSFYQGGLINRRAVLRTAYAQFVFLAG WP_067393794.1 inhibition of morphological differentiation protein [Streptomyces sp. F-3] Length: 276\nNumber of predicted TMHs: 1\nExp number of AAs in TMHs: 20.62704\nExp number, first 60 AAs: 0.01656\nTotal prob of N-in: 0.00242\noutside 1 248\nTMhelix 249 268\ninside 269 276

33032 GCF_002007685.1_ASM200768v1 Streptomyces sp. fd1-xmd Terrabacteria group; Actinobacteria; Actinobacteria; Streptomycetales; Streptomycetaceae; Streptomyces
MPARRRTVTPIRIAAAGLAPLALAAYAAGPAAAHGSMTDPVSRVAACYAEGPESPKSAAC WP_078093932.1
chitin-binding protein [Streptomyces sp. fd1-xmd] Length: 355\nNumber of predicted TMHs: 2\nExp number of AAs in TMHs: 41.76409\nExp number, first 60 AAs: 20.71548\nTotal prob of N-in: 0.94609\nPOSSIBLE N-term signal sequence\ninside 1 12\nTMhelix 13 35\noutside 36 320\nTMhelix 321 343\ninside 344 355

33033 GCF_002007685.1_ASM200768v1 Streptomyces sp. fd1-xmd Terrabacteria group; Actinobacteria; Actinobacteria; Streptomycetales; Streptomycetaceae; Streptomyces
MSPVRRRTLVRAAAVTACAGSLLALPTAAALAEGVPAVSSAHSSAPQRTLVKSLSLADGV WP_078095079.1
hypothetical protein [Streptomyces sp. fd1-xmd] Length: 210\nNumber of predicted TMHs: 2\nExp number of AAs in TMHs: 42.682\nExp number, first 60 AAs: 20.01401\nTotal prob of N-in: 0.93067\nPOSSIBLE N-term signal sequence\ninside 1 8\nTMhelix 9 31\noutside 32 180\nTMhelix 181 203\ninside 204 210

33034 GCF_002007685.1_ASM200768v1 Streptomyces sp. fd1-xmd Terrabacteria group; Actinobacteria; Actinobacteria; Streptomycetales; Streptomycetaceae; Streptomyces
MKNLKRVPLAGRRTAVAAAAAATAVALAGPAYAVSGGPATPSPSASVPASASPSADGPRP WP_078095829.1
hypothetical protein [Streptomyces sp. fd1-xmd] Length: 155\nNumber of predicted TMHs: 2\nExp number of AAs in TMHs: 44.05631\nExp number, first 60 AAs: 21.72286\nTotal prob of N-in: 0.98134\nPOSSIBLE N-term signal sequence\ninside 1 12\nTMhelix 13 35\noutside 36 126\nTMhelix 127 149\ninside 150 155

33035 GCF_002007685.1_ASM200768v1 Streptomyces sp. fd1-xmd Terrabacteria group; Actinobacteria; Actinobacteria; Streptomycetales; Streptomycetaceae; Streptomyces
MPQNRRPRRPLLAGAFALTLAAGAAATAGAAADAATTAAAAPHPPYALRLDGAGECTFPM WP_078095181.1
MULTISPECIES: type VII secretion-associated serine protease mycosin [Streptomyces] Length: 424\nNumber of predicted TMHs: 2\nExp number of AAs in TMHs: 41.33936\nExp number, first 60 AAs: 19.1446\nTotal prob of N-in: 0.88160\nPOSSIBLE N-term signal sequence\ninside 1 11\nTMhelix 12 34\noutside 35 390\nTMhelix 391 413\ninside 414 424

33036 GCF_001431765.1_ASM143176v1 Streptomyces sp. FR-008 Terrabacteria group; Actinobacteria; Actinobacteria; Streptomycetales; Streptomycetaceae; Streptomyces
MSAAAPEAAPRTARPAPRAGALLRRLLWWCVLTLTGGVERRGALPPGGCVVVANHTSHAD WP_015507057.1
MULTISPECIES: 1-acyl-sn-glycerol-3-phosphate acyltransferase [Streptomyces] Length: 401\nNumber of

predicted TMHs:  2\nExp number of AAs in TMHs: 83.99314\nExp number, first 60 AAs:  1.08255\nTotal prob of N-in:   0.25169\noutside   1   353\nTMhelix   354   371\ninside   372   377\nTMhelix   378   400\noutside   401   401

33037 GCF_001431765.1_ASM143176v1 Streptomyces sp. FR-008 Terrabacteria group; Actinobacteria; Actinobacteria; Streptomycetales; Streptomycetaceae; Streptomyces
MTPNRRPVVLAAAVATALGLGATALTLPAVAAEAPRAGAPVMDLTDGTLDWGVKESFRRY WP_075987369.1
hypothetical protein [Streptomyces sp. FR-008] Length: 514\nNumber of predicted TMHs:  2\nExp number of AAs in TMHs: 42.8564199999999\nExp number, first 60 AAs:  22.03086\nTotal prob of N-in:   0.97276\nPOSSIBLE N-term signal sequence\ninside   1   6\nTMhelix   7   29\noutside   30   479\nTMhelix   480   502\ninside   503   514

33038 GCF_001431765.1_ASM143176v1 Streptomyces sp. FR-008 Terrabacteria group; Actinobacteria; Actinobacteria; Streptomycetales; Streptomycetaceae; Streptomyces
MGGRGTTAQARERGRLPGRPSDPAPTGGNDVITGVNLRMPARRTALPALLLTATALLAVP WP_075986836.1
hypothetical protein [Streptomyces sp. FR-008] Length: 303\nNumber of predicted TMHs:  2\nExp number of AAs in TMHs: 38.94004\nExp number, first 60 AAs:  11.595\nTotal prob of N-in:   0.75054\nPOSSIBLE N-term signal sequence\ninside   1   44\nTMhelix   45   67\noutside   68   275\nTMhelix   276   298\ninside   299   303

33039 GCF_001431765.1_ASM143176v1 Streptomyces sp. FR-008 Terrabacteria group; Actinobacteria; Actinobacteria; Streptomycetales; Streptomycetaceae; Streptomyces
MSRRTRTARRRPVAAAVLAAGAVLLPAVPASAASGADQDFVIEDPRITESSGLAASKRHP WP_075986595.1
hypothetical protein [Streptomyces sp. FR-008] Length: 350\nNumber of predicted TMHs:  2\nExp number of AAs in TMHs: 39.44236\nExp number, first 60 AAs:  21.11619\nTotal prob of N-in:   0.99084\nPOSSIBLE N-term signal sequence\ninside   1   12\nTMhelix   13   35\noutside   36   322\nTMhelix   323   342\ninside   343   350

33040 GCF_001431765.1_ASM143176v1 Streptomyces sp. FR-008 Terrabacteria group; Actinobacteria; Actinobacteria; Streptomycetales; Streptomycetaceae; Streptomyces
MSRMRGSSGGRIAGRRAAVVLAVAVLGLAGCSAGPDSGASSAGDRAVAPSAADGAGGGQA WP_075986046.1
hypothetical protein [Streptomyces sp. FR-008] Length: 334\nNumber of predicted TMHs:  1\nExp number of AAs in TMHs: 36.36626\nExp number, first 60 AAs:  13.57659\nTotal prob of N-in:   0.64436\nPOSSIBLE N-term signal sequence\noutside   1   269\nTMhelix   270   292\ninside   293   334

33041 GCF_001431765.1_ASM143176v1 Streptomyces sp. FR-008 Terrabacteria group; Actinobacteria; Actinobacteria; Streptomycetales; Streptomycetaceae; Streptomyces
MLDPVENRSLPRTAAFFDLDKTVIAKSSTLTFSKSFYQGGLINRRAVLRTAYTQFVFLAG WP_010643895.1 MULTISPECIES: morphological differentiation-associated protein [Streptomyces] Length: 290\nNumber of predicted TMHs:  1\nExp number of AAs in TMHs: 21.93803\nExp number, first 60 AAs:  0.02636\nTotal prob of N-in:   0.00751\noutside   1   247\nTMhelix   248   270\ninside   271   290

33042 GCF_000958545.1_ASM95854v1 Streptomyces sp. FxanaA7 Terrabacteria group; Actinobacteria; Actinobacteria; Streptomycetales; Streptomycetaceae; Streptomyces
MLGPVENHSLPRTAAFFDLDKTVIAKSSTLTFSKSFYQGGLINRRAALRTAYTQFVFLAG WP_045563048.1 MULTISPECIES: inhibition of morphological differentiation protein [Streptomyces] Length: 274\nNumber of predicted TMHs:  1\nExp number of AAs in TMHs: 21.74861\nExp number, first 60 AAs:  0.04228\nTotal prob of N-in:   0.01050\noutside   1   244\nTMhelix   245   267\ninside   268   274

33043 GCF_000958545.1_ASM95854v1 Streptomyces sp. FxanaA7 Terrabacteria group; Actinobacteria; Actinobacteria; Streptomycetales; Streptomycetaceae; Streptomyces
MHRLSARRATARRAVVTTAAATVAAPLLLTVGAAGPASAHGAPTDPVSRVSACSPEGGSQ WP_045559812.1
chitin-binding protein [Streptomyces sp. FxanaA7] Length: 345\nNumber of predicted TMHs:  2\nExp number of AAs in TMHs: 41.27369\nExp number, first 60 AAs:  20.17854\nTotal prob of N-in:   0.90795\nPOSSIBLE N-term signal sequence\ninside   1   12\nTMhelix   13   35\noutside   36   309\nTMhelix   310   332\ninside   333   345

33044 GCF_000375625.1_ASM37562v1 Streptomyces sp. FxanaC1 Terrabacteria group; Actinobacteria; Actinobacteria; Streptomycetales; Streptomycetaceae; Streptomyces
MRQRRRAAVASMAALMLAGAGIMTAAGTAQAAEVSYKTECLPPPISGLPPIQGTTKVAVS WP_026170271.1
hypothetical protein [Streptomyces sp. FxanaC1] Length: 468\nNumber of predicted TMHs:  2\nExp number of AAs in TMHs: 44.90721\nExp number, first 60 AAs:  22.21535\nTotal prob of N-in:   0.99649\nPOSSIBLE N-term signal sequence\ninside   1   6\nTMhelix   7   29\noutside   30   437\nTMhelix   438   460\ninside   461   468

33045 GCF_000375625.1_ASM37562v1 Streptomyces sp. FxanaC1 Terrabacteria group; Actinobacteria; Actinobacteria; Streptomycetales; Streptomycetaceae; Streptomyces

MTDRRRRAALLLPLPATSPGVVLLPRERKSSSSSPEPTAPDSPGTEQDNPFAPPPAGAPD WP_018089553.1 hypothetical protein [Streptomyces sp. FxanaC1] Length: 293\nNumber of predicted TMHs: 2\nExp number of AAs in TMHs: 47.07369\nExp number, first 60 AAs: 0.0236\nTotal prob of N-in: 0.97534\ninside 1 154\nTMhelix 155 177\noutside 178 231\nTMhelix 232 254\ninside 255 293

33046 GCF_000375625.1_ASM37562v1 Streptomyces sp. FxanaC1 Terrabacteria group; Actinobacteria; Actinobacteria; Streptomycetales; Streptomycetaceae; Streptomyces MNSASPAPDSSHRAARPARSGPGSPLTRRRPAVVLALLAGLLAATVAPAAAAGRSTGDTA WP_018089851.1 hypothetical protein [Streptomyces sp. FxanaC1] Length: 161\nNumber of predicted TMHs: 2\nExp number of AAs in TMHs: 43.6816\nExp number, first 60 AAs: 20.71649\nTotal prob of N-in: 0.78139\nPOSSIBLE N-term signal sequence\ninside 1 32\nTMhelix 33 52\noutside 53 125\nTMhelix 126 148\ninside 149 161

33047 GCF_000373465.1_ASM37346v1 Streptomyces sp. FxanaD5 Terrabacteria group; Actinobacteria; Actinobacteria; Streptomycetales; Streptomycetaceae; Streptomyces MLKDVENHLSPRTAAFFDLDKTVIAKSSTLTFSKSFYQGGLINRRAVLRTAYSQFVFLAG WP_019527085.1 inhibition of morphological differentiation protein [Streptomyces sp. FxanaD5] Length: 277\nNumber of predicted TMHs: 1\nExp number of AAs in TMHs: 21.90135\nExp number, first 60 AAs: 0.03207\nTotal prob of N-in: 0.00818\noutside 1 247\nTMhelix 248 270\ninside 271 277

33048 GCF_001636945.1_ASM163694v1 Streptomyces sp. FXJ1.172 Terrabacteria group; Actinobacteria; Actinobacteria; Streptomycetales; Streptomycetaceae; Streptomyces MASRGRRAAMRSLGALAGAAALTVLGSNAPGWAAAPEAHGGHSSTATPIKHVVVLFDENI WP_067044735.1 phospholipase [Streptomyces sp. FXJ1.172] Length: 608\nNumber of predicted TMHs: 2\nExp number of AAs in TMHs: 40.91259\nExp number, first 60 AAs: 19.41356\nTotal prob of N-in: 0.85933\nPOSSIBLE N-term signal sequence\ninside 1 12\nTMhelix 13 35\noutside 36 577\nTMhelix 578 600\ninside 601 608

33049 GCF_001636945.1_ASM163694v1 Streptomyces sp. FXJ1.172 Terrabacteria group; Actinobacteria; Actinobacteria; Streptomycetales; Streptomycetaceae; Streptomyces MLGDVENHSLPRAAAFFDLDKTVIAKSSTLTFSKSFYQGGLINRRAALRTAYAQFVFLAG WP_067046700.1 inhibition of morphological differentiation protein [Streptomyces sp. FXJ1.172] Length: 276\nNumber of predicted TMHs: 1\nExp number of AAs in TMHs: 21.70658\nExp number, first 60 AAs: 0.02142\nTotal prob of N-in: 0.00255\noutside 1 247\nTMhelix 248 270\ninside 271 276

33050 GCF_001636945.1_ASM163694v1 Streptomyces sp. FXJ1.172 Terrabacteria group; Actinobacteria; Actinobacteria; Streptomycetales; Streptomycetaceae; Streptomyces MSTAKKSARRSLLVTSVVLPFLAATAPPVLAVFWPSASPTATPPARMSTVGGERLGRPGT WP_067039987.1 D-alanyl-D-alanine carboxypeptidase [Streptomyces sp. FXJ1.172] Length: 380\nNumber of predicted TMHs: 2\nExp number of AAs in TMHs: 44.68628\nExp number, first 60 AAs: 22.7497\nTotal prob of N-in: 0.99507\nPOSSIBLE N-term signal sequence\ninside 1 11\nTMhelix 12 34\noutside 35 353\nTMhelix 354 376\ninside 377 380

33051 GCF_001636945.1_ASM163694v1 Streptomyces sp. FXJ1.172 Terrabacteria group; Actinobacteria; Actinobacteria; Streptomycetales; Streptomycetaceae; Streptomyces MTPLVSRSGGPGRGTGRRGLVALLCGALAAGGLAAAGVATLAPGAAFASSHREAPLISGT WP_067052849.1 hypothetical protein [Streptomyces sp. FXJ1.172] Length: 517\nNumber of predicted TMHs: 2\nExp number of AAs in TMHs: 45.27927\nExp number, first 60 AAs: 22.64952\nTotal prob of N-in: 0.99382\nPOSSIBLE N-term signal sequence\ninside 1 19\nTMhelix 20 42\noutside 43 486\nTMhelix 487 509\ninside 510 517

33052 GCF_001636945.1_ASM163694v1 Streptomyces sp. FXJ1.172 Terrabacteria group; Actinobacteria; Actinobacteria; Streptomycetales; Streptomycetaceae; Streptomyces MRHARRRIVRRVTRLAAVGGLLLGGAMVTQAAMASQTPPAATRTLSAASGAQDTGAALVA WP_067044048.1 protease [Streptomyces sp. FXJ1.172] Length: 461\nNumber of predicted TMHs: 2\nExp number of AAs in TMHs: 43.54154\nExp number, first 60 AAs: 22.24375\nTotal prob of N-in: 0.99339\nPOSSIBLE N-term signal sequence\ninside 1 11\nTMhelix 12 34\noutside 35 422\nTMhelix 423 445\ninside 446 461

33053 GCF_001636945.1_ASM163694v1 Streptomyces sp. FXJ1.172 Terrabacteria group; Actinobacteria; Actinobacteria; Streptomycetales; Streptomycetaceae; Streptomyces MSTRRTAVVAALGGAVLALCAAPASAHGSMGDPVSRVSQCYAEGPESPKSAACKAAVAAG WP_067052820.1 chitin-binding protein [Streptomyces sp. FXJ1.172] Length: 329\nNumber of predicted TMHs: 2\nExp number of AAs in TMHs: 42.77393\nExp number, first 60 AAs: 20.51311\nTotal prob of N-in: 0.97221\nPOSSIBLE N-term signal sequence\ninside 1 6\nTMhelix 7 26\noutside 27 298\nTMhelix 299 321\ninside 322 329

33054 GCF_000404005.1_Streptomyces_sp._FXJ7.023 Streptomyces sp. FXJ7.023 Terrabacteria group; Actinobacteria; Actinobacteria; Streptomycetales; Streptomycetaceae; Streptomyces
MRHARRRIVRRAARLAAVGGLLIGGTMVTRAVASEPADAPAAPHTLARSASGAGAGLVER WP_037766766.1 protease [Streptomyces sp. FXJ7.023] Length: 443\nNumber of predicted TMHs: 2\nExp number of AAs in TMHs: 39.96207\nExp number, first 60 AAs: 20.02097\nTotal prob of N-in: 0.94485\nPOSSIBLE N-term signal sequence\ninside 1 11\nTMhelix 12 34\noutside 35 404\nTMhelix 405 427\ninside 428 443

33055 GCF_000404005.1_Streptomyces_sp._FXJ7.023 Streptomyces sp. FXJ7.023 Terrabacteria group; Actinobacteria; Actinobacteria; Streptomycetales; Streptomycetaceae; Streptomyces
MSESVTTQEHVLGHNRRRRPTGARRATFGAVALILGGGGLVAVNVFASATESGNTAVPLG WP_078535903.1 hypothetical protein [Streptomyces sp. FXJ7.023] Length: 700\nNumber of predicted TMHs: 2\nExp number of AAs in TMHs: 43.48821\nExp number, first 60 AAs: 22.40841\nTotal prob of N-in: 0.99501\nPOSSIBLE N-term signal sequence\ninside 1 25\nTMhelix 26 48\noutside 49 672\nTMhelix 673 695\ninside 696 700

33056 GCF_000495635.1_GBA94-10.20 Streptomyces sp. GBA 94-10 Terrabacteria group; Actinobacteria; Actinobacteria; Streptomycetales; Streptomycetaceae; Streptomyces
MTPNRRPVVLAAAVATALGLGATALTLPAVAAEAPRAGAPVMDLTDGTLNWGVKESFRRY WP_023419704.1 MULTISPECIES: membrane protein [Streptomyces] Length: 518\nNumber of predicted TMHs: 2\nExp number of AAs in TMHs: 42.79336\nExp number, first 60 AAs: 22.03121\nTotal prob of N-in: 0.97269\nPOSSIBLE N-term signal sequence\ninside 1 6\nTMhelix 7 29\noutside 30 483\nTMhelix 484 506\ninside 507 518

33057 GCF_000495635.1_GBA94-10.20 Streptomyces sp. GBA 94-10 Terrabacteria group; Actinobacteria; Actinobacteria; Streptomycetales; Streptomycetaceae; Streptomyces
MSRRTRTARRRPVAAAVLAAGAVLLPVVPASAASGADQDFVIEDPRITESSGLAASRKHP WP_023421988.1 hypothetical protein [Streptomyces sp. GBA 94-10] Length: 348\nNumber of predicted TMHs: 2\nExp number of AAs in TMHs: 37.75573\nExp number, first 60 AAs: 21.29767\nTotal prob of N-in: 0.99446\nPOSSIBLE N-term signal sequence\ninside 1 12\nTMhelix 13 35\noutside 36 320\nTMhelix 321 340\ninside 341 348

33058 GCF_000495635.1_GBA94-10.20 Streptomyces sp. GBA 94-10 Terrabacteria group; Actinobacteria; Actinobacteria; Streptomycetales; Streptomycetaceae; Streptomyces
MGGRGTTAQARGRGRLPGRPSDPAPTGGNDVITGVNLRMPARRTALPALLLTVTALLAVP WP_037787275.1 hypothetical protein [Streptomyces sp. GBA 94-10] Length: 301\nNumber of predicted TMHs: 2\nExp number of AAs in TMHs: 41.72499\nExp number, first 60 AAs: 13.51624\nTotal prob of N-in: 0.87299\nPOSSIBLE N-term signal sequence\ninside 1 44\nTMhelix 45 67\noutside 68 273\nTMhelix 274 296\ninside 297 301

33059 GCF_000495635.1_GBA94-10.20 Streptomyces sp. GBA 94-10 Terrabacteria group; Actinobacteria; Actinobacteria; Streptomycetales; Streptomycetaceae; Streptomyces
MLDPVENRSLPRTAAFFDLDKTVIAKSSTLTFSKSFYQGGLINRRAVLRTAYTQFVFLAG WP_023422160.1 MULTISPECIES: SsgB protein [Streptomyces] Length: 279\nNumber of predicted TMHs: 1\nExp number of AAs in TMHs: 21.73892\nExp number, first 60 AAs: 0.02846\nTotal prob of N-in: 0.01117\noutside 1 247\nTMhelix 248 270\ninside 271 279

33060 GCF_000495635.1_GBA94-10.20 Streptomyces sp. GBA 94-10 Terrabacteria group; Actinobacteria; Actinobacteria; Streptomycetales; Streptomycetaceae; Streptomyces
MASPHRRRAAVAVRALVALTALLCATGLAPAATSGPAARAAVAEVPPPTATTSVVTVRTG WP_037787307.1 VWA domain-containing protein [Streptomyces sp. GBA 94-10] Length: 833\nNumber of predicted TMHs: 2\nExp number of AAs in TMHs: 40.56284\nExp number, first 60 AAs: 20.3579\nTotal prob of N-in: 0.93764\nPOSSIBLE N-term signal sequence\ninside 1 11\nTMhelix 12 34\noutside 35 794\nTMhelix 795 817\ninside 818 833

33061 GCF_000495635.1_GBA94-10.20 Streptomyces sp. GBA 94-10 Terrabacteria group; Actinobacteria; Actinobacteria; Streptomycetales; Streptomycetaceae; Streptomyces
MPDRRSAVSLRLPLPTAAPGFGGLALPGAAATAPDAGDVVAVRESSEKERKSSSDPEGRS WP_023419047.1 MULTISPECIES: hypothetical protein [Streptomyces] Length: 292\nNumber of predicted TMHs: 2\nExp number of AAs in TMHs: 49.52916\nExp number, first 60 AAs: 3.9366\nTotal prob of N-in: 0.40255\noutside 1 164\nTMhelix 165 187\ninside 188 229\nTMhelix 230 252\noutside 253 292

33062 GCF_002027195.1_ASM202719v1 Streptomyces sp. GKU 895 Terrabacteria group; Actinobacteria; Actinobacteria; Streptomycetales; Streptomycetaceae; Streptomyces
MRHARRRMVRRVTRLAAVGGLLLGGTMVTRAVASEPEGPSAFPRTTAASAAQTGTELVSR WP_079308928.1 serine protease [Streptomyces sp. GKU 895] Length: 459\nNumber of predicted TMHs: 2\nExp number of AAs in TMHs: 41.09761\nExp number, first 60 AAs: 21.29899\nTotal prob of N-in: 0.96043\nPOSSIBLE N-term signal sequence\ninside 1 11\nTMhelix 12 34\noutside 35 420\nTMhelix 421 443\ninside 444 459

33063 GCF_002027195.1_ASM202719v1 Streptomyces sp. GKU 895 Terrabacteria group; Actinobacteria; Actinobacteria; Streptomycetales; Streptomycetaceae; Streptomyces
MSKRRRAVPDRARVALALCGAGVLIGVAVAPAQAAEVSNATHCVPPAGVGLDPVDGTTKV WP_079311137.1
hypothetical protein [Streptomyces sp. GKU 895] Length: 432\nNumber of predicted TMHs: 2\nExp number of AAs in TMHs: 41.80138\nExp number, first 60 AAs: 21.45749\nTotal prob of N-in: 0.98738\nPOSSIBLE N-term signal sequence\ninside 1 12\nTMhelix 13 35\noutside 36 406\nTMhelix 407 426\ninside 427 432

33064 GCF_002027195.1_ASM202719v1 Streptomyces sp. GKU 895 Terrabacteria group; Actinobacteria; Actinobacteria; Streptomycetales; Streptomycetaceae; Streptomyces
MLGVVENHSLPRTAAFFDLDKTVIAKSSTLTFSKSFYQGGLINRRAAVRTAYAQFVFLVG WP_079309325.1 inhibition of morphological differentiation protein [Streptomyces sp. GKU 895] Length: 277\nNumber of predicted TMHs: 1\nExp number of AAs in TMHs: 21.774\nExp number, first 60 AAs: 0.01464\nTotal prob of N-in: 0.00283\noutside 1 247\nTMhelix 248 270\ninside 271 277

33065 GCF_001279495.1_ASM127949v1 Streptomyces sp. H021 Terrabacteria group; Actinobacteria; Actinobacteria; Streptomycetales; Streptomycetaceae; Streptomyces
MALDVIRSGPPDKRPRRNLFKRGVLVLLPAALLAAATLVPNLAHALEAPVNLGTATNYAV WP_053631548.1
MULTISPECIES: hypothetical protein [Streptomyces] Length: 546\nNumber of predicted TMHs: 2\nExp number of AAs in TMHs: 43.12563\nExp number, first 60 AAs: 20.82835\nTotal prob of N-in: 0.94536\nPOSSIBLE N-term signal sequence\ninside 1 22\nTMhelix 23 45\noutside 46 511\nTMhelix 512 534\ninside 535 546

33066 GCF_001279495.1_ASM127949v1 Streptomyces sp. H021 Terrabacteria group; Actinobacteria; Actinobacteria; Streptomycetales; Streptomycetaceae; Streptomyces
MSTVRRRTLVRAAAVTACAGSLLALPTAAALAEGVPAASSAHSAAPQRTLVKSLSLADGV WP_053629362.1
hypothetical protein [Streptomyces sp. H021] Length: 218\nNumber of predicted TMHs: 2\nExp number of AAs in TMHs: 42.9309\nExp number, first 60 AAs: 20.3393\nTotal prob of N-in: 0.95701\nPOSSIBLE N-term signal sequence\ninside 1 8\nTMhelix 9 31\noutside 32 188\nTMhelix 189 211\ninside 212 218

33067 GCF_001279495.1_ASM127949v1 Streptomyces sp. H021 Terrabacteria group; Actinobacteria; Actinobacteria; Streptomycetales; Streptomycetaceae; Streptomyces
MSARRRTVMPIRIAAAGAAPLALAVYAAVPAVAHGSMTDPVSRVAACYAEGPESPKSAAC WP_053632541.1
MULTISPECIES: chitin-binding protein [Streptomyces] Length: 345\nNumber of predicted TMHs: 2\nExp number of AAs in TMHs: 43.49631\nExp number, first 60 AAs: 21.79151\nTotal prob of N-in: 0.98454\nPOSSIBLE N-term signal sequence\ninside 1 12\nTMhelix 13 35\noutside 36 310\nTMhelix 311 333\ninside 334 345

33068 GCF_001279495.1_ASM127949v1 Streptomyces sp. H021 Terrabacteria group; Actinobacteria; Actinobacteria; Streptomycetales; Streptomycetaceae; Streptomyces
MPPYRDARRALLAAAFALTLTAAGATAATVAQHPPYALRLDGAGECTFPMKKQIADRPWA WP_053630392.1 type VII secretion-associated serine protease mycosin [Streptomyces sp. H021] Length: 414\nNumber of predicted TMHs: 2\nExp number of AAs in TMHs: 42.6394099999999\nExp number, first 60 AAs: 20.30353\nTotal prob of N-in: 0.94987\nPOSSIBLE N-term signal sequence\ninside 1 11\nTMhelix 12 31\noutside 32 380\nTMhelix 381 403\ninside 404 414

33069 GCF_001279925.1_ASM127992v1 Streptomyces sp. H036 Terrabacteria group; Actinobacteria; Actinobacteria; Streptomycetales; Streptomycetaceae; Streptomyces
MSTVRRRTLVRAAAVTACAGSLLALPTAAALAEGVPAASSAHSAAPQRTLVKSLSLADGV WP_053611851.1
MULTISPECIES: hypothetical protein [Streptomyces] Length: 218\nNumber of predicted TMHs: 2\nExp number of AAs in TMHs: 42.93081\nExp number, first 60 AAs: 20.33935\nTotal prob of N-in: 0.95701\nPOSSIBLE N-term signal sequence\ninside 1 8\nTMhelix 9 31\noutside 32 188\nTMhelix 189 211\ninside 212 218

33070 GCF_001279925.1_ASM127992v1 Streptomyces sp. H036 Terrabacteria group; Actinobacteria; Actinobacteria; Streptomycetales; Streptomycetaceae; Streptomyces
MPQYRDARRALLATAFALTLTAAGATVATAAPHPPYALRLDGAGECTFPMKKQIADRPWA WP_030651182.1
MULTISPECIES: type VII secretion-associated serine protease [Streptomyces] Length: 414\nNumber of predicted TMHs: 2\nExp number of AAs in TMHs: 43.048\nExp number, first 60 AAs: 20.71014\nTotal prob of N-in: 0.96189\nPOSSIBLE N-term signal sequence\ninside 1 8\nTMhelix 9 31\noutside 32 380\nTMhelix 381 403\ninside 404 414

33071 GCF_000498935.1_HCCB10043 Streptomyces sp. HCCB10043Terrabacteria group; Actinobacteria; Actinobacteria; Streptomycetales; Streptomycetaceae; Streptomyces
MGLSWNDRRLERRVKPGDGKPLKPFRWWQMTRRSLLSIALPVGGGGGLSATYTVEVKHGG WP_023608351.1

MULTISPECIES: hypothetical protein [Streptomyces] Length: 220\nNumber of predicted TMHs: 2\nExp number of AAs in TMHs: 41.83877\nExp number, first 60 AAs: 2.68896\nTotal prob of N-in: 0.92352\ninside 1 137\nTMhelix 138 160\noutside 161 184\nTMhelix 185 202\ninside 203 220

33072 GCF_000498935.1_HCCB10043 Streptomyces sp. HCCB10043Terrabacteria group; Actinobacteria; Actinobacteria; Streptomycetales; Streptomycetaceae; Streptomyces MTRQPGPEAPRTLLRPTRPARRPHLTALLALLALLPALLVLTAPGARAADRDPAVSLSKK WP_032797602.1 hypothetical protein, partial [Streptomyces sp. HCCB10043] Length: 316\nNumber of predicted TMHs: 2\nExp number of AAs in TMHs: 42.1432\nExp number, first 60 AAs: 20.29596\nTotal prob of N-in: 0.79693\nPOSSIBLE N-term signal sequence\ninside 1 23\nTMhelix 24 46\noutside 47 277\nTMhelix 278 300\ninside 301 316

33073 GCF_000498935.1_HCCB10043 Streptomyces sp. HCCB10043Terrabacteria group; Actinobacteria; Actinobacteria; Streptomycetales; Streptomycetaceae; Streptomyces MLSVVENCFSPRTAAFFDLDKTVIAKSSTLTFSKSFYQGGLINRRAVLRTAYTQFVFLAG WP_006126413.1 MULTISPECIES: morphological differentiation protein inhibitor [Streptomyces] Length: 281\nNumber of predicted TMHs: 1\nExp number of AAs in TMHs: 21.46267\nExp number, first 60 AAs: 0.01959\nTotal prob of N-in: 0.00583\noutside 1 247\nTMhelix 248 270\ninside 271 281

33074 GCF_000411315.1_Stre_sp_HGB0020_V1 Streptomyces sp. HGB0020 Terrabacteria group; Actinobacteria; Actinobacteria; Streptomycetales; Streptomycetaceae; Streptomyces MGRNTRRRPTGPRRATFAAFALMLGGGGLVAANVYASATEDGSGTDAQTRSAAAATIDCP WP_016430591.1 hypothetical protein [Streptomyces sp. HGB0020] Length: 703\nNumber of predicted TMHs: 2\nExp number of AAs in TMHs: 41.21164\nExp number, first 60 AAs: 21.17959\nTotal prob of N-in: 0.97075\nPOSSIBLE N-term signal sequence\ninside 1 16\nTMhelix 17 39\noutside 40 676\nTMhelix 677 696\ninside 697 703

33075 GCF_000411315.1_Stre_sp_HGB0020_V1 Streptomyces sp. HGB0020 Terrabacteria group; Actinobacteria; Actinobacteria; Streptomycetales; Streptomycetaceae; Streptomyces MLGVVENHSLHRTAAFFDLDKTVIAKSSTLTFSKSFYQGGLINRRAALRTAYAQFVFLVG WP_016437318.1 haloacid dehalogenase [Streptomyces sp. HGB0020] Length: 277\nNumber of predicted TMHs: 1\nExp number of AAs in TMHs: 21.41547\nExp number, first 60 AAs: 0.01468\nTotal prob of N-in: 0.00145\noutside 1 247\nTMhelix 248 270\ninside 271 277

33076 GCF_000411315.1_Stre_sp_HGB0020_V1 Streptomyces sp. HGB0020 Terrabacteria group; Actinobacteria; Actinobacteria; Streptomycetales; Streptomycetaceae; Streptomyces MRHARRRMVRRVTRLAAVGGLLLGGAMVTQAAMASETPGTSALPFSASGTSDSPGAALVS WP_037779956.1 protease [Streptomyces sp. HGB0020] Length: 459\nNumber of predicted TMHs: 2\nExp number of AAs in TMHs: 41.67733\nExp number, first 60 AAs: 22.27934\nTotal prob of N-in: 0.99303\nPOSSIBLE N-term signal sequence\ninside 1 11\nTMhelix 12 34\noutside 35 420\nTMhelix 421 443\ninside 444 459

33077 GCF_001672315.1_ASM167231v1 Streptomyces sp. H-KF8 Terrabacteria group; Actinobacteria; Actinobacteria; Streptomycetales; Streptomycetaceae; Streptomyces MNSTDSRASARRRAVVATVTVLAAGPAALAGAGAAQATGEHGSASAAVLRTGLDVALLDR WP_065002969.1 hypothetical protein [Streptomyces sp. H-KF8] Length: 339\nNumber of predicted TMHs: 2\nExp number of AAs in TMHs: 42.8206799999999\nExp number, first 60 AAs: 21.44943\nTotal prob of N-in: 0.95557\nPOSSIBLE N-term signal sequence\ninside 1 12\nTMhelix 13 35\noutside 36 310\nTMhelix 311 333\ninside 334 339

33078 GCF_001672315.1_ASM167231v1 Streptomyces sp. H-KF8 Terrabacteria group; Actinobacteria; Actinobacteria; Streptomycetales; Streptomycetaceae; Streptomyces MPARRKAVVAAVGIVPLALTGLASSPAAAHGSMGDPVSRVAQCFAEGPESPKSAACRAAV WP_065002899.1 chitin-binding protein [Streptomyces sp. H-KF8] Length: 365\nNumber of predicted TMHs: 2\nExp number of AAs in TMHs: 40.96373\nExp number, first 60 AAs: 21.99051\nTotal prob of N-in: 0.97857\nPOSSIBLE N-term signal sequence\ninside 1 6\nTMhelix 7 29\noutside 30 331\nTMhelix 332 354\ninside 355 365

33079 GCF_001672315.1_ASM167231v1 Streptomyces sp. H-KF8 Terrabacteria group; Actinobacteria; Actinobacteria; Streptomycetales; Streptomycetaceae; Streptomyces MRHARRRVVRRVTRLAAVGGVLLGATMVTRAVASEPPAPAPVPYAAAEDDAGPGSALVAR WP_079130289.1 serine protease [Streptomyces sp. H-KF8] Length: 454\nNumber of predicted TMHs: 2\nExp number of AAs in TMHs: 41.17048\nExp number, first 60 AAs: 20.65848\nTotal prob of N-in: 0.95062\nPOSSIBLE N-term signal sequence\ninside 1 11\nTMhelix 12 34\noutside 35 415\nTMhelix 416 438\ninside 439 454

33080 GCF_001044185.1_ASM104418v1 Streptomyces sp. HNS054 Terrabacteria group; Actinobacteria; Actinobacteria; Streptomycetales; Streptomycetaceae; Streptomyces
MLKGVENHLSPRTAAFFDLDKTVIAKSSTLTFSKSFYQGGLINRRAVLRTAYSQFVFLAG WP_048456974.1 inhibition of morphological differentiation protein [Streptomyces sp. HNS054] Length: 277\nNumber of predicted TMHs: 1\nExp number of AAs in TMHs: 21.81151\nExp number, first 60 AAs: 0.02023\nTotal prob of N-in: 0.00357\noutside 1 247\nTMhelix 248 270\ninside 271 277

33081 GCF_001044185.1_ASM104418v1 Streptomyces sp. HNS054 Terrabacteria group; Actinobacteria; Actinobacteria; Streptomycetales; Streptomycetaceae; Streptomyces
MRHARRRVVRRVTRLAAVGGLLLGGTMVTHAVASEPGTPDAVPFSTASPADAHASGPGAD WP_048454886.1 protease [Streptomyces sp. HNS054] Length: 459\nNumber of predicted TMHs: 2\nExp number of AAs in TMHs: 39.4274\nExp number, first 60 AAs: 21.83298\nTotal prob of N-in: 0.97534\nPOSSIBLE N-term signal sequence\ninside 1 11\nTMhelix 12 34\noutside 35 420\nTMhelix 421 443\ninside 444 459

33082 GCF_000411495.1_Stre_sp_HPH0547_V1 Streptomyces sp. HPH0547 Terrabacteria group; Actinobacteria; Actinobacteria; Streptomycetales; Streptomycetaceae; Streptomyces
MSPRRRRATVLAAATVLCAAAAPVAAAAPSGTAAAGPVPAVKPAKLPAGLYGDADPKYDG WP_016468754.1 MULTISPECIES: hypothetical protein [Streptomyces] Length: 457\nNumber of predicted TMHs: 2\nExp number of AAs in TMHs: 40.9820199999999\nExp number, first 60 AAs: 18.73808\nTotal prob of N-in: 0.84735\nPOSSIBLE N-term signal sequence\ninside 1 6\nTMhelix 7 29\noutside 30 427\nTMhelix 428 450\ninside 451 457

33083 GCF_000411495.1_Stre_sp_HPH0547_V1 Streptomyces sp. HPH0547 Terrabacteria group; Actinobacteria; Actinobacteria; Streptomycetales; Streptomycetaceae; Streptomyces
MLSGVENLAMPRTAAFFDLDKTVIAKSSTLTFGKSFYQGGLINRRAVLRTAYAQFVYLLG WP_016473027.1 MULTISPECIES: haloacid dehalogenase [Streptomyces] Length: 284\nNumber of predicted TMHs: 1\nExp number of AAs in TMHs: 21.94508\nExp number, first 60 AAs: 0.01548\nTotal prob of N-in: 0.00204\noutside 1 246\nTMhelix 247 269\ninside 270 284

33084 GCF_001983595.1_ASM198359v1 Streptomyces sp. IB2014 011-1 Terrabacteria group; Actinobacteria; Actinobacteria; Streptomycetales; Streptomycetaceae; Streptomyces
MGIAASGRRTLLSATAVSATAALIALGAAPAQADAIKPDLGVRALAPVTGIAAGSGFGLP WP_076967992.1 peptidase [Streptomyces sp. IB2014 011-1] Length: 446\nNumber of predicted TMHs: 1\nExp number of AAs in TMHs: 36.96806\nExp number, first 60 AAs: 16.71434\nTotal prob of N-in: 0.43098\nPOSSIBLE N-term signal sequence\noutside 1 414\nTMhelix 415 437\ninside 438 446

33085 GCF_001983595.1_ASM198359v1 Streptomyces sp. IB2014 011-1 Terrabacteria group; Actinobacteria; Actinobacteria; Streptomycetales; Streptomycetaceae; Streptomyces
MLSVVENCFSPRTAAFFDLDKTVIAKSSTLTFSKSFYQGGLINRRAVLRTAYTQFVFLAG WP_076967101.1 inhibition of morphological differentiation protein [Streptomyces sp. IB2014 011-1] Length: 281\nNumber of predicted TMHs: 1\nExp number of AAs in TMHs: 21.4543\nExp number, first 60 AAs: 0.02143\nTotal prob of N-in: 0.00744\noutside 1 247\nTMhelix 248 270\ninside 271 281

33086 GCF_001279935.1_ASM127993v1 Streptomyces sp. IGB124 Terrabacteria group; Actinobacteria; Actinobacteria; Streptomycetales; Streptomycetaceae; Streptomyces
MPPYRDARRALLAAAFALTLTAAGATAATAAQHPPYALRLDGAGECTFPMKKQIADRPWA WP_053634588.1 MULTISPECIES: type VII secretion-associated serine protease mycosin [Streptomyces] Length: 414\nNumber of predicted TMHs: 2\nExp number of AAs in TMHs: 42.2832099999999\nExp number, first 60 AAs: 19.94733\nTotal prob of N-in: 0.93627\nPOSSIBLE N-term signal sequence\ninside 1 11\nTMhelix 12 31\noutside 32 380\nTMhelix 381 403\ninside 404 414

33087 GCF_001279935.1_ASM127993v1 Streptomyces sp. IGB124 Terrabacteria group; Actinobacteria; Actinobacteria; Streptomycetales; Streptomycetaceae; Streptomyces
MSTVRRRTLVRAAAVTACAGSLLALPTAAALAEGVPAASSAHSAAPQRTLVKSLSLADGV WP_053677522.1 MULTISPECIES: hypothetical protein [Streptomyces] Length: 218\nNumber of predicted TMHs: 2\nExp number of AAs in TMHs: 42.93092\nExp number, first 60 AAs: 20.3393\nTotal prob of N-in: 0.95701\nPOSSIBLE N-term signal sequence\ninside 1 8\nTMhelix 9 31\noutside 32 188\nTMhelix 189 211\ninside 212 218

33088 GCF_001279935.1_ASM127993v1 Streptomyces sp. IGB124 Terrabacteria group; Actinobacteria; Actinobacteria; Streptomycetales; Streptomycetaceae; Streptomyces
MALDVIRSGPPDKRPRRNLFKRGVLVLLPAALLAAATLVPNLAHALEAPVNLGTATNYAV WP_053686608.1

hypothetical protein [Streptomyces sp. IGB124] Length: 543\nNumber of predicted TMHs: 2\nExp number of AAs in TMHs: 43.1289199999999\nExp number, first 60 AAs: 20.82859\nTotal prob of N-in: 0.94536\nPOSSIBLE N-term signal sequence\ninside 1 22\nTMhelix 23 45\noutside 46 508\nTMhelix 509 531\ninside 532 543

33089 GCF_001279935.1_ASM127993v1 Streptomyces sp. IGB124 Terrabacteria group; Actinobacteria; Actinobacteria; Streptomycetales; Streptomycetaceae; Streptomyces MSARRRTVMPIRIAAAGAAPLALAVYAAVPAVAHGSMTDPVSRVAACYAEGPESPKSAAC WP_053687452.1 chitin-binding protein [Streptomyces sp. IGB124] Length: 345\nNumber of predicted TMHs: 2\nExp number of AAs in TMHs: 43.49631\nExp number, first 60 AAs: 21.79151\nTotal prob of N-in: 0.98454\nPOSSIBLE N-term signal sequence\ninside 1 12\nTMhelix 13 35\noutside 36 310\nTMhelix 311 333\ninside 334 345

33090 GCF_001531145.1_ASM153114v1 Streptomyces sp. IMTB 1903 Terrabacteria group; Actinobacteria; Actinobacteria; Streptomycetales; Streptomycetaceae; Streptomyces MSPVRRRTLVRAAAVTACAGSLLALPTAAALAEGVPAVSSAHSSAPQRTLVKSLSLADGV WP_060176777.1 MULTISPECIES: hypothetical protein [Streptomyces] Length: 210\nNumber of predicted TMHs: 2\nExp number of AAs in TMHs: 42.68199\nExp number, first 60 AAs: 20.014\nTotal prob of N-in: 0.93067\nPOSSIBLE N-term signal sequence\ninside 1 8\nTMhelix 9 31\noutside 32 180\nTMhelix 181 203\ninside 204 210

33091 GCF_001531145.1_ASM153114v1 Streptomyces sp. IMTB 1903 Terrabacteria group; Actinobacteria; Actinobacteria; Streptomycetales; Streptomycetaceae; Streptomyces MKNLKRVPLAGRRTAVAAAAAATAVALAGPAYAVSGGPAAPSPSASVPASASPSADGPRP WP_060178439.1 hypothetical protein [Streptomyces sp. IMTB 1903] Length: 155\nNumber of predicted TMHs: 2\nExp number of AAs in TMHs: 44.03601\nExp number, first 60 AAs: 21.70359\nTotal prob of N-in: 0.98055\nPOSSIBLE N-term signal sequence\ninside 1 12\nTMhelix 13 35\noutside 36 126\nTMhelix 127 149\ninside 150 155

33092 GCF_001531145.1_ASM153114v1 Streptomyces sp. IMTB 1903 Terrabacteria group; Actinobacteria; Actinobacteria; Streptomycetales; Streptomycetaceae; Streptomyces MPARRRTVTPIRIAAAGLAPLALAAYAAGPAAAHGSMTDPVSRVAACYAEGPESPKSAAC WP_060179978.1 MULTISPECIES: chitin-binding protein [Streptomyces] Length: 355\nNumber of predicted TMHs: 2\nExp number of AAs in TMHs: 41.76409\nExp number, first 60 AAs: 20.71548\nTotal prob of N-in: 0.94609\nPOSSIBLE N-term signal sequence\ninside 1 12\nTMhelix 13 35\noutside 36 320\nTMhelix 321 343\ninside 344 355

33093 GCF_001953885.1_ASM195388v1 Streptomyces sp. IMTB 2501 Terrabacteria group; Actinobacteria; Actinobacteria; Streptomycetales; Streptomycetaceae; Streptomyces MRGRGRRTAWRWREREGRFGRRAVFGAGALRTVLLSGLCALLVLPAAGTATAPAAAAEDG WP_076089841.1 hypothetical protein [Streptomyces sp. IMTB 2501] Length: 467\nNumber of predicted TMHs: 2\nExp number of AAs in TMHs: 44.33798\nExp number, first 60 AAs: 22.05899\nTotal prob of N-in: 0.96379\nPOSSIBLE N-term signal sequence\ninside 1 22\nTMhelix 23 45\noutside 46 429\nTMhelix 430 452\ninside 453 467

33094 GCF_001953885.1_ASM195388v1 Streptomyces sp. IMTB 2501 Terrabacteria group; Actinobacteria; Actinobacteria; Streptomycetales; Streptomycetaceae; Streptomyces MRHARRRIVRRVTRLAAVGGLLLGGAMVTQAAMASETPPAARTLAASDGTGGTGAALVAR WP_076082650.1 serine protease [Streptomyces sp. IMTB 2501] Length: 458\nNumber of predicted TMHs: 2\nExp number of AAs in TMHs: 43.51889\nExp number, first 60 AAs: 22.42186\nTotal prob of N-in: 0.99423\nPOSSIBLE N-term signal sequence\ninside 1 11\nTMhelix 12 34\noutside 35 419\nTMhelix 420 442\ninside 443 458

33095 GCF_001953885.1_ASM195388v1 Streptomyces sp. IMTB 2501 Terrabacteria group; Actinobacteria; Actinobacteria; Streptomycetales; Streptomycetaceae; Streptomyces MLGGVENHSLPRAAAFFDLDKTVIAKSSTLTFSKSFYQGGLINRRAALRTAYAQFVFLVG WP_076094070.1 inhibition of morphological differentiation protein [Streptomyces sp. IMTB 2501] Length: 276\nNumber of predicted TMHs: 1\nExp number of AAs in TMHs: 21.52324\nExp number, first 60 AAs: 0.02215\nTotal prob of N-in: 0.00182\noutside 1 247\nTMhelix 248 270\ninside 271 276

33096 GCF_001953885.1_ASM195388v1 Streptomyces sp. IMTB 2501 Terrabacteria group; Actinobacteria; Actinobacteria; Streptomycetales; Streptomycetaceae; Streptomyces MASRGRRAAMRSLGALAGAAALTVLGSNAPGWAAAPAAAHGGHSSTATPIKHVVVLFDEN WP_076082929.1 phospholipase [Streptomyces sp. IMTB 2501] Length: 609\nNumber of predicted TMHs: 2\nExp number of AAs in TMHs: 41.9199\nExp number, first 60 AAs: 20.22142\nTotal prob of N-in: 0.89354\nPOSSIBLE N-term signal sequence\ninside 1 12\nTMhelix 13 35\noutside 36 578\nTMhelix 579 601\ninside 602 609

33097 GCF_001417695.1_ASM141769v1 Streptomyces sp. JHA19 Terrabacteria group; Actinobacteria; Actinobacteria; Streptomycetales; Streptomycetaceae; Streptomyces
MTRRRPLATITPVSVLPLVFTSGWASGVNAYAVVLLLGIFGATGLSDDVPEALQRPDVLI WP_079033850.1 hypothetical protein [Streptomyces sp. JHA19] Length: 217\nNumber of predicted TMHs: 2\nExp number of AAs in TMHs: 63.47\nExp number, first 60 AAs: 24.31847\nTotal prob of N-in: 0.90067\nPOSSIBLE N-term signal sequence\ninside 1 12\nTMhelix 13 35\noutside 36 166\nTMhelix 167 189\ninside 190 217

33098 GCF_001417695.1_ASM141769v1 Streptomyces sp. JHA19 Terrabacteria group; Actinobacteria; Actinobacteria; Streptomycetales; Streptomycetaceae; Streptomyces
MSTSEQHGRARRRLVAATGAAALTLAAGLVPQQAAAADGDKVLTVAVAQSVDSLSPFLAQ WP_055621846.1 ABC transporter substrate-binding protein [Streptomyces sp. JHA19] Length: 607\nNumber of predicted TMHs: 2\nExp number of AAs in TMHs: 39.49026\nExp number, first 60 AAs: 17.02282\nTotal prob of N-in: 0.79169\nPOSSIBLE N-term signal sequence\ninside 1 12\nTMhelix 13 35\noutside 36 574\nTMhelix 575 597\ninside 598 607

33099 GCF_001417695.1_ASM141769v1 Streptomyces sp. JHA19 Terrabacteria group; Actinobacteria; Actinobacteria; Streptomycetales; Streptomycetaceae; Streptomyces
MRHARRRLVRRVTRLAAVGGLLLGGTMVTRAVASESAGPSAVPRTFAESATPGPGGELVA WP_055624432.1 protease [Streptomyces sp. JHA19] Length: 454\nNumber of predicted TMHs: 2\nExp number of AAs in TMHs: 40.33519\nExp number, first 60 AAs: 19.63278\nTotal prob of N-in: 0.89666\nPOSSIBLE N-term signal sequence\ninside 1 11\nTMhelix 12 34\noutside 35 415\nTMhelix 416 438\ninside 439 454

33100 GCF_002005565.1_ASM200556v1 Streptomyces sp. JHA26 Terrabacteria group; Actinobacteria; Actinobacteria; Streptomycetales; Streptomycetaceae; Streptomyces
MLGVVENHSSPRTAAFFDLDKTVIAKSSTLTFSKSFYQGGLINRRAALRTAYTQFVFLAG WP_077796161.1 inhibition of morphological differentiation protein [Streptomyces sp. JHA26] Length: 277\nNumber of predicted TMHs: 1\nExp number of AAs in TMHs: 21.59354\nExp number, first 60 AAs: 0.01136\nTotal prob of N-in: 0.00306\noutside 1 247\nTMhelix 248 270\ninside 271 277

33101 GCF_002005565.1_ASM200556v1 Streptomyces sp. JHA26 Terrabacteria group; Actinobacteria; Actinobacteria; Streptomycetales; Streptomycetaceae; Streptomyces
MRHARRRIVRRVARLAAVGGLLLGTAMVTRAVAGEPPGAPAVPHTKALPVSDVGSALVSR WP_077797205.1 serine protease [Streptomyces sp. JHA26] Length: 453\nNumber of predicted TMHs: 2\nExp number of AAs in TMHs: 41.64787\nExp number, first 60 AAs: 21.61669\nTotal prob of N-in: 0.98443\nPOSSIBLE N-term signal sequence\ninside 1 11\nTMhelix 12 34\noutside 35 414\nTMhelix 415 437\ninside 438 453

33102 GCF_000743295.1_ASM74329v1 Streptomyces sp. JS01 Terrabacteria group; Actinobacteria; Actinobacteria; Streptomycetales; Streptomycetaceae; Streptomyces
MLSVVENCFSPRTAAFFDLDKTVIAKSSTLTFSKSFYQGGLINRRAVLRTAYTQFVFLAG WP_032782321.1 inhibition of morphological differentiation protein [Streptomyces sp. JS01] Length: 281\nNumber of predicted TMHs: 1\nExp number of AAs in TMHs: 21.42564\nExp number, first 60 AAs: 0.02005\nTotal prob of N-in: 0.00656\noutside 1 247\nTMhelix 248 270\ninside 271 281

33103 GCF_000743295.1_ASM74329v1 Streptomyces sp. JS01 Terrabacteria group; Actinobacteria; Actinobacteria; Streptomycetales; Streptomycetaceae; Streptomyces
MGLGWNDRRLERRVKPGDGKPLKPFRWWQMTRRSLLSIALPVGGGLSATYTVEVKHGGDA WP_032784043.1 hypothetical protein [Streptomyces sp. JS01] Length: 218\nNumber of predicted TMHs: 2\nExp number of AAs in TMHs: 40.51382\nExp number, first 60 AAs: 1.33243\nTotal prob of N-in: 0.92476\ninside 1 135\nTMhelix 136 158\noutside 159 182\nTMhelix 183 200\ninside 201 218

33104 GCF_001014595.1_ASM101459v1 Streptomyces sp. KE1 Terrabacteria group; Actinobacteria; Actinobacteria; Streptomycetales; Streptomycetaceae; Streptomyces
MSRRTRTARRRPVAAAVLAAGVVLLPTVPAAAAPGADQDFVIEDPRITESSGLAASKRHP WP_047142306.1 hypothetical protein [Streptomyces sp. KE1] Length: 349\nNumber of predicted TMHs: 2\nExp number of AAs in TMHs: 37.65137\nExp number, first 60 AAs: 21.48597\nTotal prob of N-in: 0.99654\nPOSSIBLE N-term signal sequence\ninside 1 12\nTMhelix 13 35\noutside 36 320\nTMhelix 321 340\ninside 341 349

33105 GCF_001014595.1_ASM101459v1 Streptomyces sp. KE1 Terrabacteria group; Actinobacteria; Actinobacteria; Streptomycetales; Streptomycetaceae; Streptomyces
MPDRRSAVSLRLPLPTAAPGFGGLALPGAAATAPEAWGVVAVRESSEKERKSSSEPEGRS WP_047138782.1 hypothetical protein [Streptomyces sp. KE1] Length: 292\nNumber of predicted TMHs: 2\nExp number of

AAs in TMHs: 51.41526\nExp number, first 60 AAs: 5.73689\nTotal prob of N-in: 0.32906\noutside 1 164\nTMhelix 165 187\ninside 188 229\nTMhelix 230 252\noutside 253 292

33106 GCF_001014595.1_ASM101459v1 Streptomyces sp. KE1 Terrabacteria group; Actinobacteria; Actinobacteria; Streptomycetales; Streptomycetaceae; Streptomyces MTPNRRPVVLAAAVATALGLGATALTLPAVAAEAPRAGAPVMDLTDGTLNWGVKESFRRY WP_047138664.1 membrane protein [Streptomyces sp. KE1] Length: 518\nNumber of predicted TMHs: 2\nExp number of AAs in TMHs: 42.9854299999999\nExp number, first 60 AAs: 22.03093\nTotal prob of N-in: 0.97245\nPOSSIBLE N-term signal sequence\ninside 1 6\nTMhelix 7 29\noutside 30 483\nTMhelix 484 506\ninside 507 518

33107 GCF_001014595.1_ASM101459v1 Streptomyces sp. KE1 Terrabacteria group; Actinobacteria; Actinobacteria; Streptomycetales; Streptomycetaceae; Streptomyces MLDPVENRSLPRTAAFFDLDKTVIAKSSTLTFSKSFYQGGLINRRAVLRTAYTQFVFLAG WP_030698235.1 MULTISPECIES: inhibition of morphological differentiation protein [Streptomyces] Length: 279\nNumber of predicted TMHs: 1\nExp number of AAs in TMHs: 21.72379\nExp number, first 60 AAs: 0.0316\nTotal prob of N-in: 0.01274\noutside 1 247\nTMhelix 248 270\ninside 271 279

33108 GCF_001014595.1_ASM101459v1 Streptomyces sp. KE1 Terrabacteria group; Actinobacteria; Actinobacteria; Streptomycetales; Streptomycetaceae; Streptomyces MGGRGTTARARERGRLPGRPSDPAPTGGNDVITGVNLRMPARRTALPALLLTATALLAVP WP_047139237.1 hypothetical protein [Streptomyces sp. KE1] Length: 305\nNumber of predicted TMHs: 2\nExp number of AAs in TMHs: 40.14857\nExp number, first 60 AAs: 12.43541\nTotal prob of N-in: 0.80614\nPOSSIBLE N-term signal sequence\ninside 1 44\nTMhelix 45 67\noutside 68 277\nTMhelix 278 300\ninside 301 305

33109 GCF_001014595.1_ASM101459v1 Streptomyces sp. KE1 Terrabacteria group; Actinobacteria; Actinobacteria; Streptomycetales; Streptomycetaceae; Streptomyces MSAAAPEAAPRTKRPVPRAGALLRRLLWWCVLTLTGGVERRGALPPGGCVVVANHTSHAD WP_047143257.1 1-acyl-sn-glycerol-3-phosphate acyltransferase [Streptomyces sp. KE1] Length: 401\nNumber of predicted TMHs: 1\nExp number of AAs in TMHs: 79.71362\nExp number, first 60 AAs: 1.92175\nTotal prob of N-in: 0.18877\noutside 1 377\nTMhelix 378 400\ninside 401 401

33110 GCF_000373625.1_ASM37362v1 Streptomyces sp. LaPpAH-108 Terrabacteria group; Actinobacteria; Actinobacteria; Streptomycetales; Streptomycetaceae; Streptomyces MRHARRRVVRRVTRLAAVGGLLLGGAMVTQAAMASEPPQAPATPLSTAGTAGETGSALTA WP_037801963.1 S1 family peptidase [Streptomyces sp. LaPpAH-108] Length: 460\nNumber of predicted TMHs: 2\nExp number of AAs in TMHs: 42.6041399999999\nExp number, first 60 AAs: 21.99418\nTotal prob of N-in: 0.98948\nPOSSIBLE N-term signal sequence\ninside 1 11\nTMhelix 12 34\noutside 35 421\nTMhelix 422 444\ninside 445 460

33111 GCF_000373625.1_ASM37362v1 Streptomyces sp. LaPpAH-108 Terrabacteria group; Actinobacteria; Actinobacteria; Streptomycetales; Streptomycetaceae; Streptomyces MSTQSIRRRGPLVRCALVGAVLASGVVVGAGPASAAPVQVSDASFAWGLSGEQGGGAFFG WP_018546292.1 hypothetical protein [Streptomyces sp. LaPpAH-108] Length: 614\nNumber of predicted TMHs: 2\nExp number of AAs in TMHs: 42.2604\nExp number, first 60 AAs: 20.05305\nTotal prob of N-in: 0.90578\nPOSSIBLE N-term signal sequence\ninside 1 12\nTMhelix 13 35\noutside 36 580\nTMhelix 581 603\ninside 604 614

33112 GCF_000373625.1_ASM37362v1 Streptomyces sp. LaPpAH-108 Terrabacteria group; Actinobacteria; Actinobacteria; Streptomycetales; Streptomycetaceae; Streptomyces MITGTVGRVHHRGMRRRATVTLAAVAASLTLVAPASPAPQDDGGLRIEVPEVYYTQTTDP WP_018545442.1 hypothetical protein [Streptomyces sp. LaPpAH-108] Length: 508\nNumber of predicted TMHs: 1\nExp number of AAs in TMHs: 36.6571\nExp number, first 60 AAs: 13.85657\nTotal prob of N-in: 0.72490\nPOSSIBLE N-term signal sequence\noutside 1 460\nTMhelix 461 483\ninside 484 508

33113 GCF_000373625.1_ASM37362v1 Streptomyces sp. LaPpAH-108 Terrabacteria group; Actinobacteria; Actinobacteria; Streptomycetales; Streptomycetaceae; Streptomyces MLRDVENHSLPRAAAFFDLDKTVIAKSSTLTFSKSFYQGGLINRRAALRTAYAQFVFLAG WP_018547951.1 inhibition of morphological differentiation protein [Streptomyces sp. LaPpAH-108] Length: 277\nNumber of predicted TMHs: 1\nExp number of AAs in TMHs: 21.4517\nExp number, first 60 AAs: 0.0456\nTotal prob of N-in: 0.00444\noutside 1 247\nTMhelix 248 270\ninside 271 277

33114 GCF_000373625.1_ASM37362v1 Streptomyces sp. LaPpAH-108 Terrabacteria group; Actinobacteria; Actinobacteria; Streptomycetales; Streptomycetaceae; Streptomyces

MSTGFSRRATVSICSLCAAGVLALAPAASALGHDAGEPAPPAPRAAVPRPSLLYGSGPQV WP_037801489.1 D-alanyl-D-alanine carboxypeptidase [Streptomyces sp. LaPpAH-108] Length: 394\nNumber of predicted TMHs: 1\nExp number of AAs in TMHs: 31.29536\nExp number, first 60 AAs: 10.40065\nTotal prob of N-in: 0.49744\nPOSSIBLE N-term signal sequence\noutside 1 360\nTMhelix 361 383\ninside 384 394

33115 GCF_000373525.1_ASM37352v1 Streptomyces sp. LaPpAH-165 Terrabacteria group; Actinobacteria; Actinobacteria; Streptomycetales; Streptomycetaceae; Streptomyces MLCRVENRFLPRTAAFFDLDKTVIAKSSTLTFSKSFYHGGLINRRAVLRTAYTQFVFLAG WP_020207266.1 MULTISPECIES: inhibition of morphological differentiation protein [Streptomyces] Length: 279\nNumber of predicted TMHs: 1\nExp number of AAs in TMHs: 21.97754\nExp number, first 60 AAs: 0.03012\nTotal prob of N-in: 0.00683\noutside 1 247\nTMhelix 248 270\ninside 271 279

33116 GCF_000373225.1_ASM37322v1 Streptomyces sp. LaPpAH-202 Terrabacteria group; Actinobacteria; Actinobacteria; Streptomycetales; Streptomycetaceae; Streptomyces MLDPVENRSLPRTAAFFDLDKTVIAKSSTLTFSKSFYQGGLINRRAVLRTAYTQFVFLAG WP_018470348.1 MULTISPECIES: inhibition of morphological differentiation protein [Streptomyces] Length: 290\nNumber of predicted TMHs: 1\nExp number of AAs in TMHs: 21.82254\nExp number, first 60 AAs: 0.02718\nTotal prob of N-in: 0.00834\noutside 1 247\nTMhelix 248 270\ninside 271 290

33117 GCF_000373225.1_ASM37322v1 Streptomyces sp. LaPpAH-202 Terrabacteria group; Actinobacteria; Actinobacteria; Streptomycetales; Streptomycetaceae; Streptomyces MPDRRSAVSLRLPLPTAAPGFGGLALPGAAATAPDAWGVVAVRESSEKERKSSSEPEGRS WP_018468438.1 hypothetical protein [Streptomyces sp. LaPpAH-202] Length: 292\nNumber of predicted TMHs: 2\nExp number of AAs in TMHs: 51.77674\nExp number, first 60 AAs: 6.07612\nTotal prob of N-in: 0.33780\noutside 1 164\nTMhelix 165 187\ninside 188 229\nTMhelix 230 252\noutside 253 292

33118 GCF_000373225.1_ASM37322v1 Streptomyces sp. LaPpAH-202 Terrabacteria group; Actinobacteria; Actinobacteria; Streptomycetales; Streptomycetaceae; Streptomyces MSRMRGSSGGRIAGRRAAVVLAVAVLGLAGCSAGPDSGASSAGDRAVAPSAADGAGGGQA WP_018469475.1 DUF4349 domain-containing protein [Streptomyces sp. LaPpAH-202] Length: 334\nNumber of predicted TMHs: 1\nExp number of AAs in TMHs: 36.36626\nExp number, first 60 AAs: 13.57659\nTotal prob of N-in: 0.64436\nPOSSIBLE N-term signal sequence\noutside 1 269\nTMhelix 270 292\ninside 293 334

33119 GCF_000373225.1_ASM37322v1 Streptomyces sp. LaPpAH-202 Terrabacteria group; Actinobacteria; Actinobacteria; Streptomycetales; Streptomycetaceae; Streptomyces MGGRGTTAQARERGRLPGRPSDPAPTGGNDVITGVNLRMPARRTALPALLLTATALLAVP WP_026235269.1 hypothetical protein [Streptomyces sp. LaPpAH-202] Length: 304\nNumber of predicted TMHs: 2\nExp number of AAs in TMHs: 39.18452\nExp number, first 60 AAs: 11.73851\nTotal prob of N-in: 0.76139\nPOSSIBLE N-term signal sequence\ninside 1 44\nTMhelix 45 67\noutside 68 276\nTMhelix 277 299\ninside 300 304

33120 GCF_000373225.1_ASM37322v1 Streptomyces sp. LaPpAH-202 Terrabacteria group; Actinobacteria; Actinobacteria; Streptomycetales; Streptomycetaceae; Streptomyces MTPNRRPVVLAAAVATALGLGATALTLPAVAAEAPRAGAPVMDLTDGTLDWGVKESFRRY WP_018470706.1 membrane protein [Streptomyces sp. LaPpAH-202] Length: 514\nNumber of predicted TMHs: 2\nExp number of AAs in TMHs: 42.8538999999999\nExp number, first 60 AAs: 22.03085\nTotal prob of N-in: 0.97272\nPOSSIBLE N-term signal sequence\ninside 1 6\nTMhelix 7 29\noutside 30 479\nTMhelix 480 502\ninside 503 514

33121 GCF_000375725.1_ASM37572v1 Streptomyces sp. LaPpAH-95 Terrabacteria group; Actinobacteria; Actinobacteria; Streptomycetales; Streptomycetaceae; Streptomyces MLCRVENRFLPRTAAFFDLDKTVIAKSSTLTFSKSFYHGGLINRRAVLRTAYTQFVFLAG WP_020207266.1 MULTISPECIES: inhibition of morphological differentiation protein [Streptomyces] Length: 279\nNumber of predicted TMHs: 1\nExp number of AAs in TMHs: 21.97754\nExp number, first 60 AAs: 0.03012\nTotal prob of N-in: 0.00683\noutside 1 247\nTMhelix 248 270\ninside 271 279

33122 GCF_001746365.1_ASM174636v1 Streptomyces sp. LUP30 Terrabacteria group; Actinobacteria; Actinobacteria; Streptomycetales; Streptomycetaceae; Streptomyces MLLVVENHSLPRTAAFFDLDKTVIAKSSTLTFGKSFYQGGLINRRAALRTAYAQFVFLVG WP_069768418.1 inhibition of morphological differentiation protein [Streptomyces sp. LUP30] Length: 277\nNumber of predicted TMHs: 1\nExp number of AAs in TMHs: 21.35815\nExp number, first 60 AAs: 0.01081\nTotal prob of N-in: 0.00139\noutside 1 247\nTMhelix 248 270\ninside 271 277

33123 GCF_001746295.1_ASM174629v1 Streptomyces sp. LUP47B Terrabacteria group; Actinobacteria; Actinobacteria; Streptomycetales; Streptomycetaceae; Streptomyces
MLGVVENHSLPRTAAFFDLDKTVIAKSSTLTFSKSFYQGGLINRRAALRTAYAQFVFLVG WP_069764576.1 inhibition of morphological differentiation protein [Streptomyces sp. LUP47B] Length: 277\nNumber of predicted TMHs: 1\nExp number of AAs in TMHs: 21.6355\nExp number, first 60 AAs: 0.01559\nTotal prob of N-in: 0.00302\noutside 1 247\nTMhelix 248 270\ninside 271 277

33124 GCF_001746295.1_ASM174629v1 Streptomyces sp. LUP47B Terrabacteria group; Actinobacteria; Actinobacteria; Streptomycetales; Streptomycetaceae; Streptomyces
MTRRALLLSLLAGLLLASASPAHAAGYRYWSFWDRTGSTWTYATEGPSTAVPSDGDVQGF WP_069759320.1 hypothetical protein [Streptomyces sp. LUP47B] Length: 209\nNumber of predicted TMHs: 2\nExp number of AAs in TMHs: 41.71883\nExp number, first 60 AAs: 21.44531\nTotal prob of N-in: 0.96360\nPOSSIBLE N-term signal sequence\ninside 1 4\nTMhelix 5 27\noutside 28 181\nTMhelix 182 201\ninside 202 209

33125 GCF_001746295.1_ASM174629v1 Streptomyces sp. LUP47B Terrabacteria group; Actinobacteria; Actinobacteria; Streptomycetales; Streptomycetaceae; Streptomyces
MRHARRRVVRRVTRLAAVGGLLLGGAMVTRAVASEIPAVPAVPHTYAMKAGQTGADLVSR WP_069759747.1 protease [Streptomyces sp. LUP47B] Length: 456\nNumber of predicted TMHs: 2\nExp number of AAs in TMHs: 38.9149699999999\nExp number, first 60 AAs: 20.70057\nTotal prob of N-in: 0.95923\nPOSSIBLE N-term signal sequence\ninside 1 11\nTMhelix 12 34\noutside 35 417\nTMhelix 418 440\ninside 441 456

33126 GCF_001746295.1_ASM174629v1 Streptomyces sp. LUP47B Terrabacteria group; Actinobacteria; Actinobacteria; Streptomycetales; Streptomycetaceae; Streptomyces
MSSPDPQVRAARNQSTNPAARRPVVAVTGAASGIGALLTERLAASEEIKQVIALDERRGE WP_069765881.1 NAD-dependent dehydratase [Streptomyces sp. LUP47B] Length: 371\nNumber of predicted TMHs: 1\nExp number of AAs in TMHs: 20.4092\nExp number, first 60 AAs: 0.39598\nTotal prob of N-in: 0.03604\noutside 1 341\nTMhelix 342 361\ninside 362 371

33127 GCF_000800535.1_ASM80053v1 Streptomyces sp. M10 Terrabacteria group; Actinobacteria; Actinobacteria; Streptomycetales; Streptomycetaceae; Streptomyces
MTPNRRPVVLAAAVATALGLGATALTLPAVAAEAPRAGAPVMDLTDGTLNWGVKESFRRY WP_047471042.1 membrane protein [Streptomyces sp. M10] Length: 518\nNumber of predicted TMHs: 2\nExp number of AAs in TMHs: 43.04364\nExp number, first 60 AAs: 22.03099\nTotal prob of N-in: 0.97252\nPOSSIBLE N-term signal sequence\ninside 1 6\nTMhelix 7 29\noutside 30 483\nTMhelix 484 506\ninside 507 518

33128 GCF_000800535.1_ASM80053v1 Streptomyces sp. M10 Terrabacteria group; Actinobacteria; Actinobacteria; Streptomycetales; Streptomycetaceae; Streptomyces
MLDPVENRSLPRTAAFFDLDKTVIAKSSTLTFSKSFYQGGLINRRAVLRTAYTQFVFLAG WP_030698235.1 MULTISPECIES: inhibition of morphological differentiation protein [Streptomyces] Length: 279\nNumber of predicted TMHs: 1\nExp number of AAs in TMHs: 21.72379\nExp number, first 60 AAs: 0.0316\nTotal prob of N-in: 0.01274\noutside 1 247\nTMhelix 248 270\ninside 271 279

33129 GCF_000800535.1_ASM80053v1 Streptomyces sp. M10 Terrabacteria group; Actinobacteria; Actinobacteria; Streptomycetales; Streptomycetaceae; Streptomyces
MGGRGTTARARERGRLPGRPSDPAPTGGNDVITGVNLRMPARRTALPALLLTATALLAVP WP_047466284.1 hypothetical protein [Streptomyces sp. M10] Length: 305\nNumber of predicted TMHs: 2\nExp number of AAs in TMHs: 39.93611\nExp number, first 60 AAs: 12.29671\nTotal prob of N-in: 0.79704\nPOSSIBLE N-term signal sequence\ninside 1 44\nTMhelix 45 67\noutside 68 277\nTMhelix 278 300\ninside 301 305

33130 GCF_000800535.1_ASM80053v1 Streptomyces sp. M10 Terrabacteria group; Actinobacteria; Actinobacteria; Streptomycetales; Streptomycetaceae; Streptomyces
MAPPHRRRAAVAVRAFVALTALLCATGLAPAPATAPVAVAEVPAPTPTTSVVTVRTGGDR WP_030696185.1 MULTISPECIES: VWA domain-containing protein [Streptomyces] Length: 823\nNumber of predicted TMHs: 2\nExp number of AAs in TMHs: 39.18881\nExp number, first 60 AAs: 18.98212\nTotal prob of N-in: 0.86306\nPOSSIBLE N-term signal sequence\ninside 1 12\nTMhelix 13 35\noutside 36 784\nTMhelix 785 807\ninside 808 823

33131 GCF_000800535.1_ASM80053v1 Streptomyces sp. M10 Terrabacteria group; Actinobacteria; Actinobacteria; Streptomycetales; Streptomycetaceae; Streptomyces
MSRRTRTARRRPVAAAVLAAGAVLLPVVPASAASGADQDFVIEDPRITESSGLAASKRHP WP_047471185.1 hypothetical protein [Streptomyces sp. M10] Length: 348\nNumber of predicted TMHs: 2\nExp number of

AAs in TMHs: 38.4095\nExp number, first 60 AAs: 21.30138\nTotal prob of N-in: 0.99458\nPOSSIBLE N-term signal sequence\ninside 1 12\nTMhelix 13 35\noutside 36 322\nTMhelix 323 340\ninside 341 348

33132 GCF_001970135.1_ASM197013v1 Streptomyces sp. M1013 Terrabacteria group; Actinobacteria; Actinobacteria; Streptomycetales; Streptomycetaceae; Streptomyces MRHARRRIVRRVARLAAVGGLLIGGTMVTRAVASEPPDDSAAPRTLAQRASGVGADLVSR WP_076972742.1 serine protease [Streptomyces sp. M1013] Length: 447\nNumber of predicted TMHs: 2\nExp number of AAs in TMHs: 41.76533\nExp number, first 60 AAs: 21.2658\nTotal prob of N-in: 0.97026\nPOSSIBLE N-term signal sequence\ninside 1 11\nTMhelix 12 34\noutside 35 408\nTMhelix 409 431\ninside 432 447

33133 GCF_001970135.1_ASM197013v1 Streptomyces sp. M1013 Terrabacteria group; Actinobacteria; Actinobacteria; Streptomycetales; Streptomycetaceae; Streptomyces MGHNRRRRPTGARRATFGAVALILGGGGLVAVNVFASATESGNAAVPLGSSGVAATVDCP WP_076972671.1 hypothetical protein [Streptomyces sp. M1013] Length: 681\nNumber of predicted TMHs: 2\nExp number of AAs in TMHs: 43.24502\nExp number, first 60 AAs: 21.947\nTotal prob of N-in: 0.99510\nPOSSIBLE N-term signal sequence\ninside 1 16\nTMhelix 17 39\noutside 40 653\nTMhelix 654 676\ninside 677 681

33134 GCF_002078175.1_ASM207817v1 Streptomyces sp. M41(2017) Terrabacteria group; Actinobacteria; Actinobacteria; Streptomycetales; Streptomycetaceae; Streptomyces MRHARRRIVRRGARFAAVGGLLCGGLMVTHAMASEPSDSSRATGSSVRSAVTTGAGLVSR WP_081220707.1 serine protease [Streptomyces sp. M41(2017)] Length: 464\nNumber of predicted TMHs: 2\nExp number of AAs in TMHs: 42.19651\nExp number, first 60 AAs: 21.34978\nTotal prob of N-in: 0.98621\nPOSSIBLE N-term signal sequence\ninside 1 11\nTMhelix 12 34\noutside 35 425\nTMhelix 426 448\ninside 449 464

33135 GCF_002078175.1_ASM207817v1 Streptomyces sp. M41(2017) Terrabacteria group; Actinobacteria; Actinobacteria; Streptomycetales; Streptomycetaceae; Streptomyces MTRGHGPMKTPEQPRRRLVRPRLVAAAGVAALTLSAGLATPLNPAPQQAQAKEGKKVLTV WP_081219734.1 ABC transporter substrate-binding protein [Streptomyces sp. M41(2017)] Length: 621\nNumber of predicted TMHs: 1\nExp number of AAs in TMHs: 37.67234\nExp number, first 60 AAs: 15.36834\nTotal prob of N-in: 0.73340\nPOSSIBLE N-term signal sequence\noutside 1 587\nTMhelix 588 610\ninside 611 621

33136 GCF_002078175.1_ASM207817v1 Streptomyces sp. M41(2017) Terrabacteria group; Actinobacteria; Actinobacteria; Streptomycetales; Streptomycetaceae; Streptomyces MLSLVENHSLPRTAAFFDLDKTVIAKSSTLTFSKSFYQGGLINRRAALRTAYIQFVFLAG WP_081218531.1 inhibition of morphological differentiation protein [Streptomyces sp. M41(2017)] Length: 277\nNumber of predicted TMHs: 1\nExp number of AAs in TMHs: 21.60746\nExp number, first 60 AAs: 0.04754\nTotal prob of N-in: 0.00355\noutside 1 247\nTMhelix 248 270\ninside 271 277

33137 GCF_002078175.1_ASM207817v1 Streptomyces sp. M41(2017) Terrabacteria group; Actinobacteria; Actinobacteria; Streptomycetales; Streptomycetaceae; Streptomyces MRDSSGLSRRAALGLAAAAPLAAATAATASAAGTIGGARLGRTGVQASGGTALPKHLTAR WP_081216934.1 D-alanyl-D-alanine carboxypeptidase [Streptomyces sp. M41(2017)] Length: 396\nNumber of predicted TMHs: 2\nExp number of AAs in TMHs: 42.1599799999999\nExp number, first 60 AAs: 20.90609\nTotal prob of N-in: 0.92223\nPOSSIBLE N-term signal sequence\ninside 1 12\nTMhelix 13 35\noutside 36 348\nTMhelix 349 371\ninside 372 396

33138 GCF_000972435.1_StrepMBT28_1.0 Streptomyces sp. MBT28 Terrabacteria group; Actinobacteria; Actinobacteria; Streptomycetales; Streptomycetaceae; Streptomyces MRHARRRMVRRVARLAAVGGLLLGGTMVTRAVASEPPDAPAVPYTSARAASGTGAGLVSR WP_046248529.1 protease [Streptomyces sp. MBT28] Length: 456\nNumber of predicted TMHs: 2\nExp number of AAs in TMHs: 39.86986\nExp number, first 60 AAs: 20.99795\nTotal prob of N-in: 0.96422\nPOSSIBLE N-term signal sequence\ninside 1 11\nTMhelix 12 34\noutside 35 417\nTMhelix 418 440\ninside 441 456

33139 GCF_001445655.1_ASM144565v1 Streptomyces sp. MBT76 Terrabacteria group; Actinobacteria; Actinobacteria; Streptomycetales; Streptomycetaceae; Streptomyces MLGTVENHSLPRTAAFFDLDKTVIAKSSTLTFSKSFYQGGLINRRAVLRTAYAQFVFLAG WP_058045554.1 inhibition of morphological differentiation protein [Streptomyces sp. MBT76] Length: 283\nNumber of predicted TMHs: 1\nExp number of AAs in TMHs: 21.63718\nExp number, first 60 AAs: 0.03478\nTotal prob of N-in: 0.00731\noutside 1 246\nTMhelix 247 269\ninside 270 283

33140 GCF_001445655.1_ASM144565v1 Streptomyces sp. MBT76 Terrabacteria group; Actinobacteria; Actinobacteria; Streptomycetales; Streptomycetaceae; Streptomyces
MMILPGIAALGAVVAAVRVRRRRTVRRRIAALFGGASATAAKGQRLRALTGRLPGIRDGL WP_058047046.1 type II secretion protein F [Streptomyces sp. MBT76] Length: 255\nNumber of predicted TMHs: 2\nExp number of AAs in TMHs: 54.18488\nExp number, first 60 AAs: 6.35407\nTotal prob of N-in: 0.27470\noutside 1 65\nTMhelix 66 88\ninside 89 224\nTMhelix 225 247\noutside 248 255

33141 GCF_000412265.2_ASM41226v2 Streptomyces sp. Mg1 Terrabacteria group; Actinobacteria; Actinobacteria; Streptomycetales; Streptomycetaceae; Streptomyces
MRPEHGDGAQSVRQPRNPAGPARRGPVIAVTGAAGGVGAALVSRLVASEEVKAVVAIDER WP_037799844.1 MULTISPECIES: NAD-dependent dehydratase [Streptomyces] Length: 373\nNumber of predicted TMHs: 1\nExp number of AAs in TMHs: 23.21663\nExp number, first 60 AAs: 2.48036\nTotal prob of N-in: 0.08776\noutside 1 344\nTMhelix 345 364\ninside 365 373

33142 GCF_000412265.2_ASM41226v2 Streptomyces sp. Mg1 Terrabacteria group; Actinobacteria; Actinobacteria; Streptomycetales; Streptomycetaceae; Streptomyces
MAEAADNQGAVPAPARRRWLRRAVVLLAGTPVLAALVYAPAPPAEAADAPVDVQLDTLAP WP_078489131.1 hypothetical protein [Streptomyces sp. Mg1] Length: 753\nNumber of predicted TMHs: 2\nExp number of AAs in TMHs: 39.77324\nExp number, first 60 AAs: 18.12819\nTotal prob of N-in: 0.87708\nPOSSIBLE N-term signal sequence\ninside 1 18\nTMhelix 19 41\noutside 42 686\nTMhelix 687 709\ninside 710 753

33143 GCF_000412265.2_ASM41226v2 Streptomyces sp. Mg1 Terrabacteria group; Actinobacteria; Actinobacteria; Streptomycetales; Streptomycetaceae; Streptomyces
MSTVSRRPLVRAAAVSACAGSLLALPAAAALAEGLPAAAASGAPAVTASQRVLLKSLSLA WP_047960688.1 hypothetical protein [Streptomyces sp. Mg1] Length: 227\nNumber of predicted TMHs: 2\nExp number of AAs in TMHs: 41.75249\nExp number, first 60 AAs: 19.15078\nTotal prob of N-in: 0.89121\nPOSSIBLE N-term signal sequence\ninside 1 12\nTMhelix 13 35\noutside 36 197\nTMhelix 198 220\ninside 221 227

33144 GCF_000154885.1_ASM15488v1 Streptomyces sp. Mg1 Terrabacteria group; Actinobacteria; Actinobacteria; Streptomycetales; Streptomycetaceae; Streptomyces
MAEAADNQGAVPAPARRRWLRRAVVLLAGTPVLAALVYAPAPPAEAADAPVDVQLDTLAP WP_078489131.1 hypothetical protein [Streptomyces sp. Mg1] Length: 753\nNumber of predicted TMHs: 2\nExp number of AAs in TMHs: 39.77324\nExp number, first 60 AAs: 18.12819\nTotal prob of N-in: 0.87708\nPOSSIBLE N-term signal sequence\ninside 1 18\nTMhelix 19 41\noutside 42 686\nTMhelix 687 709\ninside 710 753

33145 GCF_001905425.1_ASM190542v1 Streptomyces sp. MJM1172 Terrabacteria group; Actinobacteria; Actinobacteria; Streptomycetales; Streptomycetaceae; Streptomyces
MSTVSRRTLVRAAAVTACAGSLLALAAAAALAEGLPAAAASGAPAAAASQRVLLKSLSLA WP_073773078.1 hypothetical protein [Streptomyces sp. MJM1172] Length: 238\nNumber of predicted TMHs: 2\nExp number of AAs in TMHs: 44.17143\nExp number, first 60 AAs: 21.56477\nTotal prob of N-in: 0.98409\nPOSSIBLE N-term signal sequence\ninside 1 8\nTMhelix 9 31\noutside 32 208\nTMhelix 209 231\ninside 232 238

33146 GCF_001905425.1_ASM190542v1 Streptomyces sp. MJM1172 Terrabacteria group; Actinobacteria; Actinobacteria; Streptomycetales; Streptomycetaceae; Streptomyces
MAEAADNQGAVPAPARRRWLRRAVVLLAGTPVLAALVYAPAPPAEAADAPVDVQLDTLAP WP_079248511.1 hypothetical protein [Streptomyces sp. MJM1172] Length: 753\nNumber of predicted TMHs: 2\nExp number of AAs in TMHs: 39.77324\nExp number, first 60 AAs: 18.12819\nTotal prob of N-in: 0.87708\nPOSSIBLE N-term signal sequence\ninside 1 18\nTMhelix 19 41\noutside 42 686\nTMhelix 687 709\ninside 710 753

33147 GCF_001905425.1_ASM190542v1 Streptomyces sp. MJM1172 Terrabacteria group; Actinobacteria; Actinobacteria; Streptomycetales; Streptomycetaceae; Streptomyces
MRPEHGDGAQSVRQPRNPAGPARRGPVIAVTGAAGGVGAALVSRLVASEEVKAVVAIDER WP_073773307.1 NAD-dependent dehydratase [Streptomyces sp. MJM1172]Length: 373\nNumber of predicted TMHs: 1\nExp number of AAs in TMHs: 22.55459\nExp number, first 60 AAs: 2.2306\nTotal prob of N-in: 0.08403\noutside 1 344\nTMhelix 345 364\ninside 365 373

33148 GCF_001625365.1_ASM162536v1 Streptomyces sp. MJM8645 Terrabacteria group; Actinobacteria; Actinobacteria; Streptomycetales; Streptomycetaceae; Streptomyces
MTDLGKARRAAVTALLAAGLGLALAPAAHADGPAGAPVPVPAADLANARGVVQDQAVLDK WP_063348001.1 hypothetical protein [Streptomyces sp. MJM8645] Length: 282\nNumber of predicted TMHs: 1\nExp

number of AAs in TMHs: 31.63308\nExp number, first 60 AAs: 10.38181\nTotal prob of N-in: 0.54563\nPOSSIBLE N-term signal sequence\noutside 1 250\nTMhelix 251 273\ninside 274 282

33149 GCF_001625365.1_ASM162536v1 Streptomyces sp. MJM8645 Terrabacteria group; Actinobacteria; Actinobacteria; Streptomycetales; Streptomycetaceae; Streptomyces MRACSFSYARRNVSMGQGTGTPDHRTTAVRREPATRRRRTVRVAVATVVAALPLAAVLTV WP_079135562.1 hypothetical protein [Streptomyces sp. MJM8645] Length: 505\nNumber of predicted TMHs: 2\nExp number of AAs in TMHs: 44.2843899999999\nExp number, first 60 AAs: 18.56552\nTotal prob of N-in: 0.99633\nPOSSIBLE N-term signal sequence\ninside 1 40\nTMhelix 41 63\noutside 64 475\nTMhelix 476 498\ninside 499 505

33150 GCF_001279965.1_ASM127996v1 Streptomyces sp. MMG1064 Terrabacteria group; Actinobacteria; Actinobacteria; Streptomycetales; Streptomycetaceae; Streptomyces MSARRRTVMPIRIAAAGAAPLALAVYAAVPAVAHGSMTDPVSRVAACYAEGPESPKSAAC WP_053633676.1 MULTISPECIES: chitin-binding protein [Streptomyces] Length: 347\nNumber of predicted TMHs: 2\nExp number of AAs in TMHs: 43.49607\nExp number, first 60 AAs: 21.79131\nTotal prob of N-in: 0.98451\nPOSSIBLE N-term signal sequence\ninside 1 12\nTMhelix 13 35\noutside 36 312\nTMhelix 313 335\ninside 336 347

33151 GCF_001279965.1_ASM127996v1 Streptomyces sp. MMG1064 Terrabacteria group; Actinobacteria; Actinobacteria; Streptomycetales; Streptomycetaceae; Streptomyces MTSIEGLPVSTVRRRTLVRAAAVTACAGSLLALPTAAALAEGVPAASSAHSAAPQRTLVK WP_053687614.1 hypothetical protein [Streptomyces sp. MMG1064] Length: 226\nNumber of predicted TMHs: 2\nExp number of AAs in TMHs: 41.09152\nExp number, first 60 AAs: 18.51524\nTotal prob of N-in: 0.85066\nPOSSIBLE N-term signal sequence\ninside 1 16\nTMhelix 17 39\noutside 40 196\nTMhelix 197 219\ninside 220 226

33152 GCF_001279965.1_ASM127996v1 Streptomyces sp. MMG1064 Terrabacteria group; Actinobacteria; Actinobacteria; Streptomycetales; Streptomycetaceae; Streptomyces MPPYRDARRALLAAAFALTLTAAGATAATAAQHPPYALRLDGAGECTFPMKKQIADRPWA WP_053634588.1 MULTISPECIES: type VII secretion-associated serine protease mycosin [Streptomyces] Length: 414\nNumber of predicted TMHs: 2\nExp number of AAs in TMHs: 42.2832099999999\nExp number, first 60 AAs: 19.94733\nTotal prob of N-in: 0.93627\nPOSSIBLE N-term signal sequence\ninside 1 11\nTMhelix 12 31\noutside 32 380\nTMhelix 381 403\ninside 404 414

33153 GCF_001279965.1_ASM127996v1 Streptomyces sp. MMG1064 Terrabacteria group; Actinobacteria; Actinobacteria; Streptomycetales; Streptomycetaceae; Streptomyces MALDVIRSGPPDKRPRRNLFKRGVLVLLPAALLAAATLVPNLAHALEAPVNLGTATNYAV WP_053631548.1 MULTISPECIES: hypothetical protein [Streptomyces] Length: 546\nNumber of predicted TMHs: 2\nExp number of AAs in TMHs: 43.12563\nExp number, first 60 AAs: 20.82835\nTotal prob of N-in: 0.94536\nPOSSIBLE N-term signal sequence\ninside 1 22\nTMhelix 23 45\noutside 46 511\nTMhelix 512 534\ninside 535 546

33154 GCF_001280015.1_ASM128001v1 Streptomyces sp. MMG1121 Terrabacteria group; Actinobacteria; Actinobacteria; Streptomycetales; Streptomycetaceae; Streptomyces MRHARRRIVRRVTRLAAVGGLLLGGAMVAQAAMASETPPAAGAGDTGAALVARLGTGHTA WP_053660334.1 protease [Streptomyces sp. MMG1121] Length: 449\nNumber of predicted TMHs: 2\nExp number of AAs in TMHs: 43.5307\nExp number, first 60 AAs: 22.15096\nTotal prob of N-in: 0.99390\nPOSSIBLE N-term signal sequence\ninside 1 11\nTMhelix 12 34\noutside 35 410\nTMhelix 411 433\ninside 434 449

33155 GCF_001280015.1_ASM128001v1 Streptomyces sp. MMG1121 Terrabacteria group; Actinobacteria; Actinobacteria; Streptomycetales; Streptomycetaceae; Streptomyces MPAPKKTGRRSLLVTSATLLSLSVTAPAALAAPTPSTSPTATPPAGMSTVGGARLGQPGT WP_053657310.1 D-alanyl-D-alanine carboxypeptidase [Streptomyces sp. MMG1121] Length: 406\nNumber of predicted TMHs: 1\nExp number of AAs in TMHs: 33.48115\nExp number, first 60 AAs: 11.09308\nTotal prob of N-in: 0.52676\nPOSSIBLE N-term signal sequence\noutside 1 368\nTMhelix 369 391\ninside 392 406

33156 GCF_001280015.1_ASM128001v1 Streptomyces sp. MMG1121 Terrabacteria group; Actinobacteria; Actinobacteria; Streptomycetales; Streptomycetaceae; Streptomyces MASRGRRAAMRSLGALAGAAALTVLGSNAPGWAAVPAHGHGGRSSTATPIQHVVVLFDEN WP_053667439.1 phospholipase [Streptomyces sp. MMG1121] Length: 609\nNumber of predicted TMHs: 2\nExp number of AAs in TMHs: 42.98489\nExp number, first 60 AAs: 21.46149\nTotal prob of N-in: 0.94171\nPOSSIBLE N-term signal sequence\ninside 1 12\nTMhelix 13 35\noutside 36 578\nTMhelix 579 601\ninside 602 609

33157 GCF_001280015.1_ASM128001v1 Streptomyces sp. MMG1121 Terrabacteria group; Actinobacteria; Actinobacteria; Streptomycetales; Streptomycetaceae; Streptomyces
MLRGVENHSLPRAAAFFDLDKTVIAKSSTLTFSKSFYQGGLINRRAALRTAYAQFVFLAG WP_053656312.1 inhibition of morphological differentiation protein [Streptomyces sp. MMG1121] Length: 276\nNumber of predicted TMHs: 1\nExp number of AAs in TMHs: 21.38531\nExp number, first 60 AAs: 0.03078\nTotal prob of N-in: 0.00641\noutside 1 247\nTMhelix 248 270\ninside 271 276

33158 GCF_001280015.1_ASM128001v1 Streptomyces sp. MMG1121 Terrabacteria group; Actinobacteria; Actinobacteria; Streptomycetales; Streptomycetaceae; Streptomyces
MSTRRKVVVAALAGTALALCAAPASAHGSMGDPVSRVSQCYAEGPESPRSAACEAAVAAG WP_053654655.1 chitin-binding protein [Streptomyces sp. MMG1121] Length: 323\nNumber of predicted TMHs: 2\nExp number of AAs in TMHs: 42.4140599999999\nExp number, first 60 AAs: 20.60892\nTotal prob of N-in: 0.97805\nPOSSIBLE N-term signal sequence\ninside 1 6\nTMhelix 7 26\noutside 27 289\nTMhelix 290 312\ninside 313 323

33159 GCF_001279735.1_ASM127973v1 Streptomyces sp. MMG1522 Terrabacteria group; Actinobacteria; Actinobacteria; Streptomycetales; Streptomycetaceae; Streptomyces
MGSSWNERRLERRVRPGDGKPLKPFRWWQLTRRSLLSLALPVGGGLSATYTVEVKHGGDA WP_030113438.1 MULTISPECIES: hypothetical protein [Streptomyces] Length: 218\nNumber of predicted TMHs: 2\nExp number of AAs in TMHs: 41.40199\nExp number, first 60 AAs: 0.36812\nTotal prob of N-in: 0.97576\ninside 1 135\nTMhelix 136 158\noutside 159 182\nTMhelix 183 200\ninside 201 218

33160 GCF_001279735.1_ASM127973v1 Streptomyces sp. MMG1522 Terrabacteria group; Actinobacteria; Actinobacteria; Streptomycetales; Streptomycetaceae; Streptomyces
MLSVVENCFSPRTAAFFDLDKTVIAKSSTLTFSKSFYQGGLINRRAVLRTAYTQFVFMAG WP_030189579.1 MULTISPECIES: inhibition of morphological differentiation protein [Streptomyces] Length: 280\nNumber of predicted TMHs: 1\nExp number of AAs in TMHs: 21.68695\nExp number, first 60 AAs: 0.01801\nTotal prob of N-in: 0.00793\noutside 1 247\nTMhelix 248 270\ninside 271 280

33161 GCF_001279775.1_ASM127977v1 Streptomyces sp. MMG1533 Terrabacteria group; Actinobacteria; Actinobacteria; Streptomycetales; Streptomycetaceae; Streptomyces
MRHARRRVLRRVTRLAAVGGLLLGGTMVTRAMASEPPDGSAVPRTYASQPADTGTALVSR WP_053748266.1 protease [Streptomyces sp. MMG1533] Length: 456\nNumber of predicted TMHs: 2\nExp number of AAs in TMHs: 40.97544\nExp number, first 60 AAs: 19.44062\nTotal prob of N-in: 0.91448\nPOSSIBLE N-term signal sequence\ninside 1 11\nTMhelix 12 34\noutside 35 417\nTMhelix 418 440\ninside 441 456

33162 GCF_001279775.1_ASM127977v1 Streptomyces sp. MMG1533 Terrabacteria group; Actinobacteria; Actinobacteria; Streptomycetales; Streptomycetaceae; Streptomyces
MLGAVENHSLPRTAAFFDLDKTVIAKSSTLTFSKSFYQGGLINRRAALRTAYAQFVFLVG WP_053750078.1 inhibition of morphological differentiation protein [Streptomyces sp. MMG1533] Length: 277\nNumber of predicted TMHs: 1\nExp number of AAs in TMHs: 21.77724\nExp number, first 60 AAs: 0.02207\nTotal prob of N-in: 0.00595\noutside 1 247\nTMhelix 248 270\ninside 271 277

33163 GCF_000935135.2_ASM93513v2 Streptomyces sp. MNU77 Terrabacteria group; Actinobacteria; Actinobacteria; Streptomycetales; Streptomycetaceae; Streptomyces
MLSVVENCFSPRTAAFFDLDKTVIAKSSTLTFSKSFYQGGLINRRAVLRTAYAQFVFLAG WP_047177942.1 inhibition of morphological differentiation protein [Streptomyces sp. MNU77] Length: 281\nNumber of predicted TMHs: 1\nExp number of AAs in TMHs: 21.45456\nExp number, first 60 AAs: 0.03333\nTotal prob of N-in: 0.00732\noutside 1 247\nTMhelix 248 270\ninside 271 281

33164 GCF_002090335.1_ASM209033v1 Streptomyces sp. MOE7 Terrabacteria group; Actinobacteria; Actinobacteria; Streptomycetales; Streptomycetaceae; Streptomyces
MRQRRRAAVASAAALVLAGAGVLTAAGTARAAEVSYKTECLPPPISGLPPIEGTTKVDVS WP_084771524.1 hypothetical protein [Streptomyces sp. MOE7] Length: 477\nNumber of predicted TMHs: 2\nExp number of AAs in TMHs: 45.01602\nExp number, first 60 AAs: 22.34219\nTotal prob of N-in: 0.99680\nPOSSIBLE N-term signal sequence\ninside 1 6\nTMhelix 7 29\noutside 30 446\nTMhelix 447 469\ninside 470 477

33165 GCF_002090335.1_ASM209033v1 Streptomyces sp. MOE7 Terrabacteria group; Actinobacteria; Actinobacteria; Streptomycetales; Streptomycetaceae; Streptomyces
MAPLAPMAQQAPSTHPVAFLARRAAMALAAAAVLGGAAAPAAYAVDGADAAPAAGSTEKL WP_084776136.1 hypothetical protein [Streptomyces sp. MOE7] Length: 444\nNumber of predicted TMHs: 1\nExp number of

AAs in TMHs: 26.66057\nExp number, first 60 AAs: 4.59087\nTotal prob of N-in: 0.20017\noutside 1 414\nTMhelix 415 437\ninside 438 444

33166 GCF_001984575.1_ASM198457v1 Streptomyces sp. MP131-18 Terrabacteria group; Actinobacteria; Actinobacteria; Streptomycetales; Streptomycetaceae; Streptomyces MKILAQPRTAAFFDLDKTVIAKSSTLTFSRSFYQGGLINRRAVLRSAYAQFVYLLGGADH WP_077060362.1 inhibition of morphological differentiation protein [Streptomyces sp. MP131-18] Length: 280\nNumber of predicted TMHs: 1\nExp number of AAs in TMHs: 21.81284\nExp number, first 60 AAs: 0.03189\nTotal prob of N-in: 0.00429\noutside 1 242\nTMhelix 243 265\ninside 266 280

33167 GCF_001866665.1_ASM186666v1 Streptomyces sp. MUSC 1 Terrabacteria group; Actinobacteria; Actinobacteria; Streptomycetales; Streptomycetaceae; Streptomyces MASRGRRAAMRSLGALAGAAALTVLGSNAPGWAAAPETHGGHSSTATPIKHVVVLFDENI WP_071381436.1 phospholipase [Streptomyces sp. MUSC 1] Length: 608\nNumber of predicted TMHs: 2\nExp number of AAs in TMHs: 40.30999\nExp number, first 60 AAs: 19.29947\nTotal prob of N-in: 0.85415\nPOSSIBLE N-term signal sequence\ninside 1 12\nTMhelix 13 35\noutside 36 579\nTMhelix 580 602\ninside 603 608

33168 GCF_001866665.1_ASM186666v1 Streptomyces sp. MUSC 1 Terrabacteria group; Actinobacteria; Actinobacteria; Streptomycetales; Streptomycetaceae; Streptomyces MRHARRRIVRRVTRLAAVGGLLLGGVMVTQAAMASETPRAATRTLSSAGGTGDPGAALVA WP_071381920.1 serine protease [Streptomyces sp. MUSC 1] Length: 466\nNumber of predicted TMHs: 2\nExp number of AAs in TMHs: 43.2614\nExp number, first 60 AAs: 22.36782\nTotal prob of N-in: 0.99669\nPOSSIBLE N-term signal sequence\ninside 1 11\nTMhelix 12 34\noutside 35 427\nTMhelix 428 450\ninside 451 466

33169 GCF_001866665.1_ASM186666v1 Streptomyces sp. MUSC 1 Terrabacteria group; Actinobacteria; Actinobacteria; Streptomycetales; Streptomycetaceae; Streptomyces MLRGVENHSSPRAAAFFDLDKTVIAKSSTLTFSKSFYQGGLINRRAALRTAYAQFVFLAG WP_071383617.1 inhibition of morphological differentiation protein [Streptomyces sp. MUSC 1] Length: 276\nNumber of predicted TMHs: 1\nExp number of AAs in TMHs: 21.72108\nExp number, first 60 AAs: 0.0471\nTotal prob of N-in: 0.00633\noutside 1 247\nTMhelix 248 270\ninside 271 276

33170 GCF_000816485.1_ASM81648v1 Streptomyces sp. MUSC 125 Terrabacteria group; Actinobacteria; Actinobacteria; Streptomycetales; Streptomycetaceae; Streptomyces MLDAVYGPRVRECSRLSRRAALGLTAAALPLSVATPAAAATVIGGERLARSGVQVRGASG WP_039657545.1 MULTISPECIES: D-alanyl-D-alanine carboxypeptidase [Streptomyces] Length: 408\nNumber of predicted TMHs: 2\nExp number of AAs in TMHs: 39.3505\nExp number, first 60 AAs: 21.43868\nTotal prob of N-in: 0.93948\nPOSSIBLE N-term signal sequence\ninside 1 20\nTMhelix 21 43\noutside 44 357\nTMhelix 358 380\ninside 381 408

33171 GCF_000816485.1_ASM81648v1 Streptomyces sp. MUSC 125 Terrabacteria group; Actinobacteria; Actinobacteria; Streptomycetales; Streptomycetaceae; Streptomyces MLKDVENHSLPRAAAFFDLDKTVIAKSSTLTFSKSFYQGGLINRRAALRTAYAQFVFLAG WP_039651231.1 MULTISPECIES: inhibition of morphological differentiation protein [Streptomyces] Length: 275\nNumber of predicted TMHs: 1\nExp number of AAs in TMHs: 21.73004\nExp number, first 60 AAs: 0.0183\nTotal prob of N-in: 0.00576\noutside 1 247\nTMhelix 248 270\ninside 271 275

33172 GCF_000816485.1_ASM81648v1 Streptomyces sp. MUSC 125 Terrabacteria group; Actinobacteria; Actinobacteria; Streptomycetales; Streptomycetaceae; Streptomyces MASRGRRTAMRSLGALAGAAALTALGSSAPGWAAAPSGAHHGHSSTRTPIKHVVVLFDEN WP_039652300.1 MULTISPECIES: phospholipase C [Streptomyces] Length: 609\nNumber of predicted TMHs: 2\nExp number of AAs in TMHs: 42.3825\nExp number, first 60 AAs: 21.14153\nTotal prob of N-in: 0.93320\nPOSSIBLE N-term signal sequence\ninside 1 12\nTMhelix 13 35\noutside 36 578\nTMhelix 579 601\ninside 602 609

33173 GCF_000816485.1_ASM81648v1 Streptomyces sp. MUSC 125 Terrabacteria group; Actinobacteria; Actinobacteria; Streptomycetales; Streptomycetaceae; Streptomyces MRHARRRVVRRLTRLAAVGGLLLGCTMVARAAVASEPPPASARSLSSAGNTGAAWVARLG WP_039650162.1 protease [Streptomyces sp. MUSC 125] Length: 457\nNumber of predicted TMHs: 2\nExp number of AAs in TMHs: 42.37515\nExp number, first 60 AAs: 22.37634\nTotal prob of N-in: 0.98715\nPOSSIBLE N-term signal sequence\ninside 1 11\nTMhelix 12 34\noutside 35 418\nTMhelix 419 441\ninside 442 457

33174 GCF_001866675.1_ASM186667v1 Streptomyces sp. MUSC 14 Terrabacteria group; Actinobacteria; Actinobacteria; Streptomycetales; Streptomycetaceae; Streptomyces

MLRGVENHSSPRAAAFFDLDKTVIAKSSTLTFSKSFYQGGLINRRAALRTAYAQFVFLAG WP_071373222.1 inhibition of morphological differentiation protein [Streptomyces sp. MUSC 14] Length: 276\nNumber of predicted TMHs: 1\nExp number of AAs in TMHs: 21.72126\nExp number, first 60 AAs: 0.0471\nTotal prob of N-in: 0.00633\noutside 1 247\nTMhelix 248 270\ninside 271 276

33175 GCF_001866675.1_ASM186667v1 Streptomyces sp. MUSC 14 Terrabacteria group; Actinobacteria; Actinobacteria; Streptomycetales; Streptomycetaceae; Streptomyces MARTSRSRSRRGNRAHPSGARRTTLAAAVLLLGGAALVVAGVYASAAEGRGAGQATAGQG WP_079176493.1 hypothetical protein [Streptomyces sp. MUSC 14] Length: 476\nNumber of predicted TMHs: 2\nExp number of AAs in TMHs: 43.99795\nExp number, first 60 AAs: 22.27229\nTotal prob of N-in: 0.99840\nPOSSIBLE N-term signal sequence\ninside 1 24\nTMhelix 25 47\noutside 48 446\nTMhelix 447 469\ninside 470 476

33176 GCF_001866675.1_ASM186667v1 Streptomyces sp. MUSC 14 Terrabacteria group; Actinobacteria; Actinobacteria; Streptomycetales; Streptomycetaceae; Streptomyces MRHARRRIVRRVTRLAAVGGLLLGGVMVTQAAMASETPRAATRTLSSAGGTADTGAALVA WP_071371441.1 serine protease [Streptomyces sp. MUSC 14] Length: 460\nNumber of predicted TMHs: 2\nExp number of AAs in TMHs: 43.37445\nExp number, first 60 AAs: 22.3736\nTotal prob of N-in: 0.99669\nPOSSIBLE N-term signal sequence\ninside 1 11\nTMhelix 12 34\noutside 35 421\nTMhelix 422 444\ninside 445 460

33177 GCF_001866675.1_ASM186667v1 Streptomyces sp. MUSC 14 Terrabacteria group; Actinobacteria; Actinobacteria; Streptomycetales; Streptomycetaceae; Streptomyces MASRGRRAAMRSLGALAGAAALTVLGSNAPGWAAAPETHGGHSSTATPIKHVVVLFDENI WP_071371529.1 phospholipase [Streptomyces sp. MUSC 14] Length: 608\nNumber of predicted TMHs: 2\nExp number of AAs in TMHs: 40.3054\nExp number, first 60 AAs: 19.30094\nTotal prob of N-in: 0.85414\nPOSSIBLE N-term signal sequence\ninside 1 12\nTMhelix 13 35\noutside 36 579\nTMhelix 580 602\ninside 603 608

33178 GCF_001866645.1_ASM186664v1 Streptomyces sp. MUSC 93 Terrabacteria group; Actinobacteria; Actinobacteria; Streptomycetales; Streptomycetaceae; Streptomyces MLRPVENHSLPRTAAFFDLDKTVIAKSSTLTFSKSFYQGGLINRRAVLRTAYAQFVFLTG WP_071367571.1 inhibition of morphological differentiation protein [Streptomyces sp. MUSC 93] Length: 274\nNumber of predicted TMHs: 1\nExp number of AAs in TMHs: 21.64139\nExp number, first 60 AAs: 0.02336\nTotal prob of N-in: 0.01613\noutside 1 244\nTMhelix 245 267\ninside 268 274

33179 GCF_001866645.1_ASM186664v1 Streptomyces sp. MUSC 93 Terrabacteria group; Actinobacteria; Actinobacteria; Streptomycetales; Streptomycetaceae; Streptomyces MRDSFLPSPCAVQMSRRAVLGLAAAAPLTLAAPASASPELSAAAPLSTSLGREGAPMAVG WP_079168661.1 D-alanyl-D-alanine carboxypeptidase [Streptomyces sp. MUSC 93] Length: 412\nNumber of predicted TMHs: 1\nExp number of AAs in TMHs: 22.3454\nExp number, first 60 AAs: 0.67304\nTotal prob of N-in: 0.03413\noutside 1 372\nTMhelix 373 395\ninside 396 412

33180 GCF_001652895.1_ASM165289v1 Streptomyces sp. NBRC 109436 Terrabacteria group; Actinobacteria; Actinobacteria; Streptomycetales; Streptomycetaceae; Streptomyces MLSVVENHSLPRTAAFFDLDKTVIAKSSALTFSKSFYQGGLINRRAVLRTAYAQFVFLAG WP_060951846.1 MULTISPECIES: inhibition of morphological differentiation protein [Streptomyces] Length: 281\nNumber of predicted TMHs: 1\nExp number of AAs in TMHs: 21.2919\nExp number, first 60 AAs: 0.02346\nTotal prob of N-in: 0.00265\noutside 1 246\nTMhelix 247 269\ninside 270 281

33181 GCF_000974485.1_ASM97448v1 Streptomyces sp. NBRC 109706 Terrabacteria group; Actinobacteria; Actinobacteria; Streptomycetales; Streptomycetaceae; Streptomyces MPQLRKPPGPSRRRAVPVTALLAVGLLALAAPAVHADENADEGDDLEQTVAPEEENATGR WP_062204714.1 hypothetical protein [Streptomyces sp. NBRC 109706] Length: 335\nNumber of predicted TMHs: 1\nExp number of AAs in TMHs: 25.65752\nExp number, first 60 AAs: 3.62585\nTotal prob of N-in: 0.15397\noutside 1 287\nTMhelix 288 310\ninside 311 335

33182 GCF_000829715.2_ASM82971v2 Streptomyces sp. NBRC 110027 Terrabacteria group; Actinobacteria; Actinobacteria; Streptomycetales; Streptomycetaceae; Streptomyces MRQRRRAAVAASAALVLAGAGIMTAAGTAQAAEVSYKTECLPPPISGLPPIQGTTKVAVS WP_042156490.1 hypothetical protein [Streptomyces sp. NBRC 110027] Length: 473\nNumber of predicted TMHs: 2\nExp number of AAs in TMHs: 44.82676\nExp number, first 60 AAs: 22.14478\nTotal prob of N-in: 0.99489\nPOSSIBLE N-term signal sequence\ninside 1 6\nTMhelix 7 29\noutside 30 442\nTMhelix 443 465\ninside 466 473

33183	GCF_000829715.2_ASM82971v2	Streptomyces sp. NBRC 110027	Terrabacteria group; Actinobacteria; Actinobacteria; Streptomycetales; Streptomycetaceae; Streptomyces
MNSASPAPDSSHRAARPARSGARSPLTRRRPAVVLALLAGLLAAAVAPAAAAGRGAGDTA	WP_042151785.1	hypothetical protein [Streptomyces sp. NBRC 110027]	Length: 160\nNumber of predicted TMHs: 2\nExp number of AAs in TMHs: 44.36876\nExp number, first 60 AAs: 21.40893\nTotal prob of N-in: 0.93763\nPOSSIBLE N-term signal sequence\ninside 1 30\nTMhelix 31 53\noutside 54 125\nTMhelix 126 148\ninside 149 160

33184	GCF_000829715.2_ASM82971v2	Streptomyces sp. NBRC 110027	Terrabacteria group; Actinobacteria; Actinobacteria; Streptomycetales; Streptomycetaceae; Streptomyces
MTDRRRRAALLLPLPATAPGVVLLPRERKSSSSSSSSSEPTAPDSPDTERDNPFAPPPAG	WP_042147472.1	hypothetical protein [Streptomyces sp. NBRC 110027]	Length: 293\nNumber of predicted TMHs: 2\nExp number of AAs in TMHs: 47.1297\nExp number, first 60 AAs: 0.06538\nTotal prob of N-in: 0.97696\ninside 1 154\nTMhelix 155 177\noutside 178 233\nTMhelix 234 256\ninside 257 293

33185	GCF_001417735.1_ASM141773v1	Streptomyces sp. NBRC 110028	Terrabacteria group; Actinobacteria; Actinobacteria; Streptomycetales; Streptomycetaceae; Streptomyces
MSSPDREVRAARNDTARTARRPVVAVTGAASGPGALLTQRLAESEEIKQVLALDERRGEV	WP_055547987.1	NAD-dependent dehydratase [Streptomyces sp. NBRC 110028]	Length: 369\nNumber of predicted TMHs: 1\nExp number of AAs in TMHs: 20.11208\nExp number, first 60 AAs: 0.51851\nTotal prob of N-in: 0.03619\noutside 1 339\nTMhelix 340 359\ninside 360 369

33186	GCF_001417675.1_ASM141767v1	Streptomyces sp. NBRC 110030	Terrabacteria group; Actinobacteria; Actinobacteria; Streptomycetales; Streptomycetaceae; Streptomyces
MPRTAAFFDLDKTVIAKSSTLTFSKSFYQGGLINRRAVLRTAYAQFVFLAGGADHDQMER	WP_037798454.1	MULTISPECIES: inhibition of morphological differentiation protein [Streptomyces]	Length: 268\nNumber of predicted TMHs: 1\nExp number of AAs in TMHs: 21.72001\nExp number, first 60 AAs: 0.03441\nTotal prob of N-in: 0.00339\noutside 1 238\nTMhelix 239 261\ninside 262 268

33187	GCF_001417675.1_ASM141767v1	Streptomyces sp. NBRC 110030	Terrabacteria group; Actinobacteria; Actinobacteria; Streptomycetales; Streptomycetaceae; Streptomyces
MRHARRRIVRRVARLAAVGGLLLGATMVTRAVAGEPPGPSAAPRTLAQSASGTGAALVAR	WP_031084860.1	MULTISPECIES: protease [Streptomyces]	Length: 447\nNumber of predicted TMHs: 2\nExp number of AAs in TMHs: 40.53891\nExp number, first 60 AAs: 22.02945\nTotal prob of N-in: 0.98901\nPOSSIBLE N-term signal sequence\ninside 1 11\nTMhelix 12 34\noutside 35 408\nTMhelix 409 431\ninside 432 447

33188	GCF_000829695.1_ASM82969v1	Streptomyces sp. NBRC 110035	Terrabacteria group; Actinobacteria; Actinobacteria; Streptomycetales; Streptomycetaceae; Streptomyces
MRHARRRVVRRVTRLAAVGGLLLGATMVTRAVASEPPAPAPVPYAASEEDTGPGSALVSR	WP_078878315.1	MULTISPECIES: serine protease [Streptomyces]	Length: 454\nNumber of predicted TMHs: 2\nExp number of AAs in TMHs: 37.19435\nExp number, first 60 AAs: 20.65846\nTotal prob of N-in: 0.95154\nPOSSIBLE N-term signal sequence\ninside 1 11\nTMhelix 12 34\noutside 35 415\nTMhelix 416 438\ninside 439 454

33189	GCF_000829695.1_ASM82969v1	Streptomyces sp. NBRC 110035	Terrabacteria group; Actinobacteria; Actinobacteria; Streptomycetales; Streptomycetaceae; Streptomyces
MLRDVENHSLPRTAAFFDLDKTVIAKSSTLTFSKSFYQGGLINRRAVLRTAYAQFVFLAG	WP_042163730.1	MULTISPECIES: inhibition of morphological differentiation protein [Streptomyces]	Length: 277\nNumber of predicted TMHs: 1\nExp number of AAs in TMHs: 20.8169\nExp number, first 60 AAs: 0.10818\nTotal prob of N-in: 0.02276\noutside 1 247\nTMhelix 248 270\ninside 271 277

33190	GCF_001895105.1_ASM189510v1	Streptomyces sp. NBRC 110465	Terrabacteria group; Actinobacteria; Actinobacteria; Streptomycetales; Streptomycetaceae; Streptomyces
MLSVVENCFSPRTAAFFDLDKTVIAKSSTLTFSKSFYQGGLINRRAVLRTAYTQFVFLAG	WP_073219259.1	inhibition of morphological differentiation protein [Streptomyces sp. NBRC 110465]	Length: 281\nNumber of predicted TMHs: 1\nExp number of AAs in TMHs: 21.4716\nExp number, first 60 AAs: 0.01878\nTotal prob of N-in: 0.00472\noutside 1 247\nTMhelix 248 270\ninside 271 281

33191	GCF_001184025.1_ASM118402v1	Streptomyces sp. NBRC 110468	Terrabacteria group; Actinobacteria; Actinobacteria; Streptomycetales; Streptomycetaceae; Streptomyces
MHAFSRPFPGSSPRPFPGSSPRPSRRSLLALAATAPLTVAPLAATPAAAASDVVVGGARL	WP_078890368.1	D-alanyl-D-alanine carboxypeptidase [Streptomyces sp. NBRC 110468]Length: 406\nNumber of predicted TMHs: 1\nExp number of

AAs in TMHs: 19.01573\nExp number, first 60 AAs: 0.51468\nTotal prob of N-in: 0.02294\noutside 1 369\nTMhelix 370 392\ninside 393 406

33192 GCF_001184025.1_ASM118402v1 Streptomyces sp. NBRC 110468 Terrabacteria group; Actinobacteria; Actinobacteria; Streptomycetales; Streptomycetaceae; Streptomyces MRRRGWIPGATAVALVAASAVTRFTVYPALHQVPAGSETTFRLQGTATLLNSPALSGGSG WP_078890523.1 hypothetical protein [Streptomyces sp. NBRC 110468] Length: 344\nNumber of predicted TMHs: 2\nExp number of AAs in TMHs: 37.75726\nExp number, first 60 AAs: 18.1194\nTotal prob of N-in: 0.81194\nPOSSIBLE N-term signal sequence\ninside 1 4\nTMhelix 5 27\noutside 28 302\nTMhelix 303 320\ninside 321 344

33193 GCF_001184025.1_ASM118402v1 Streptomyces sp. NBRC 110468 Terrabacteria group; Actinobacteria; Actinobacteria; Streptomycetales; Streptomycetaceae; Streptomyces MLMAVENHSLPRTAAFFDLDKTVIAKSSTLTFSKSFYQGGLINRRAALRTAYAQFVFLVG WP_062651410.1 inhibition of morphological differentiation protein [Streptomyces sp. NBRC 110468] Length: 277\nNumber of predicted TMHs: 1\nExp number of AAs in TMHs: 21.51658\nExp number, first 60 AAs: 0.01718\nTotal prob of N-in: 0.00189\noutside 1 247\nTMhelix 248 270\ninside 271 277

33194 GCF_001590865.1_ASM159086v1 Streptomyces sp. NBRC 110611 Terrabacteria group; Actinobacteria; Actinobacteria; Streptomycetales; Streptomycetaceae; Streptomyces MTDRRRRAALLLPLPATSPGAVLLRDRKSPSSSPEPSSPPSGSPSGPSGSSPSPAPGSGQ WP_066925780.1 hypothetical protein [Streptomyces sp. NBRC 110611] Length: 336\nNumber of predicted TMHs: 2\nExp number of AAs in TMHs: 48.53292\nExp number, first 60 AAs: 0.00755\nTotal prob of N-in: 0.97671\ninside 1 196\nTMhelix 197 219\noutside 220 272\nTMhelix 273 295\ninside 296 336

33195 GCF_001590865.1_ASM159086v1 Streptomyces sp. NBRC 110611 Terrabacteria group; Actinobacteria; Actinobacteria; Streptomycetales; Streptomycetaceae; Streptomyces MRQRRRAAVASAAALVLAGAGVMTMAGPARAAEVSYRTECLPPPISGLPPIAGTTKVAVD WP_066932926.1 hypothetical protein [Streptomyces sp. NBRC 110611] Length: 440\nNumber of predicted TMHs: 2\nExp number of AAs in TMHs: 44.26048\nExp number, first 60 AAs: 21.81646\nTotal prob of N-in: 0.99657\nPOSSIBLE N-term signal sequence\ninside 1 6\nTMhelix 7 29\noutside 30 409\nTMhelix 410 432\ninside 433 440

33196 GCF_001590865.1_ASM159086v1 Streptomyces sp. NBRC 110611 Terrabacteria group; Actinobacteria; Actinobacteria; Streptomycetales; Streptomycetaceae; Streptomyces MTARRPLLTAATAGSVLFALWFVPSANAIVGIDGAKDSSDSSLSGGSGLSSSGLSDGSG WP_066926162.1 hypothetical protein [Streptomyces sp. NBRC 110611] Length: 158\nNumber of predicted TMHs: 2\nExp number of AAs in TMHs: 43.52933\nExp number, first 60 AAs: 21.91233\nTotal prob of N-in: 0.99501\nPOSSIBLE N-term signal sequence\ninside 1 6\nTMhelix 7 29\noutside 30 127\nTMhelix 128 150\ninside 151 158

33197 GCF_000716455.1_ASM71645v1 Streptomyces sp. NRRL B-11253 Terrabacteria group; Actinobacteria; Actinobacteria; Streptomycetales; Streptomycetaceae; Streptomyces MTLTRRRHLVRSTAVAAVAGSALLLPAAAAFADSPQPTTAGSPGAAGDQQQKDDRQKDDQ WP_031191693.1 MULTISPECIES: hypothetical protein [Streptomyces] Length: 250\nNumber of predicted TMHs: 2\nExp number of AAs in TMHs: 40.95506\nExp number, first 60 AAs: 19.80107\nTotal prob of N-in: 0.94361\nPOSSIBLE N-term signal sequence\ninside 1 12\nTMhelix 13 32\noutside 33 218\nTMhelix 219 241\ninside 242 250

33198 GCF_000716455.1_ASM71645v1 Streptomyces sp. NRRL B-11253 Terrabacteria group; Actinobacteria; Actinobacteria; Streptomycetales; Streptomycetaceae; Streptomyces MESRQPVGPRANGHRAAHRRRAAVALAGAVCAVGLTATTAAPALADTNVPLNNGQCNFGA WP_078586913.1 MULTISPECIES: type VII secretion-associated serine protease mycosin [Streptomyces] Length: 427\nNumber of predicted TMHs: 2\nExp number of AAs in TMHs: 43.8948\nExp number, first 60 AAs: 21.08952\nTotal prob of N-in: 0.94565\nPOSSIBLE N-term signal sequence\ninside 1 20\nTMhelix 21 43\noutside 44 389\nTMhelix 390 412\ninside 413 427

33199 GCF_000716455.1_ASM71645v1 Streptomyces sp. NRRL B-11253 Terrabacteria group; Actinobacteria; Actinobacteria; Streptomycetales; Streptomycetaceae; Streptomyces MTERSAPAASATGPGTPARRTAVRAAATGTLAAALLGLAPQSALAATPAPPPAPAISAVA WP_032915072.1 MULTISPECIES: hypothetical protein [Streptomyces] Length: 293\nNumber of predicted TMHs: 1\nExp number of AAs in TMHs: 23.67473\nExp number, first 60 AAs: 4.47659\nTotal prob of N-in: 0.22667\noutside 1 264\nTMhelix 265 287\ninside 288 293

33200 GCF_000716455.1_ASM71645v1 Streptomyces sp. NRRL B-11253 Terrabacteria group; Actinobacteria; Actinobacteria; Streptomycetales; Streptomycetaceae; Streptomyces
MTRRPLLRAAGVALSAAALTVLPALPAHADGIRAQEWALKAMNTQDAWETTKGEGITVAV WP_031187823.1
MULTISPECIES: type VII secretion-associated serine protease [Streptomyces] Length: 386\nNumber of predicted TMHs: 1\nExp number of AAs in TMHs: 36.26455\nExp number, first 60 AAs: 9.94874\nTotal prob of N-in: 0.56610\noutside 1 359\nTMhelix 360 382\ninside 383 386

33201 GCF_001280065.1_ASM128006v1 Streptomyces sp. NRRL B-1140 Terrabacteria group; Actinobacteria; Actinobacteria; Streptomycetales; Streptomycetaceae; Streptomyces
MLWGVENHSLPRTAAFFDLDKTVIAKSSTLTFSKSFYQGGLINRRAVLRTAYAQFVFLAG WP_053669683.1 inhibition of morphological differentiation protein [Streptomyces sp. NRRL B-1140] Length: 277\nNumber of predicted TMHs: 1\nExp number of AAs in TMHs: 21.58162\nExp number, first 60 AAs: 0.0297\nTotal prob of N-in: 0.00292\noutside 1 247\nTMhelix 248 270\ninside 271 277

33202 GCF_001280065.1_ASM128006v1 Streptomyces sp. NRRL B-1140 Terrabacteria group; Actinobacteria; Actinobacteria; Streptomycetales; Streptomycetaceae; Streptomyces
MRHARRRVVRRVTRLAAVGGLLLGGAMVTNAVASEPSAPVIGVPFTAPPATGEGADLVSR WP_053673401.1
protease [Streptomyces sp. NRRL B-1140] Length: 448\nNumber of predicted TMHs: 2\nExp number of AAs in TMHs: 41.65134\nExp number, first 60 AAs: 21.29759\nTotal prob of N-in: 0.96649\nPOSSIBLE N-term signal sequence\ninside 1 11\nTMhelix 12 34\noutside 35 409\nTMhelix 410 432\ninside 433 448

33203 GCF_000717635.1_ASM71763v1 Streptomyces sp. NRRL B-12105 Terrabacteria group; Actinobacteria; Actinobacteria; Streptomycetales; Streptomycetaceae; Streptomyces
MLGPVEKHSSPRTAAFFDLDKTVIAKSSTLTFSKSFYQGGLINRRAVLRTAYAQFVFLAG WP_030378643.1 MULTISPECIES: inhibition of morphological differentiation protein [Streptomyces] Length: 277\nNumber of predicted TMHs: 1\nExp number of AAs in TMHs: 21.67091\nExp number, first 60 AAs: 0.03021\nTotal prob of N-in: 0.00251\noutside 1 247\nTMhelix 248 270\ninside 271 277

33204 GCF_000717635.1_ASM71763v1 Streptomyces sp. NRRL B-12105 Terrabacteria group; Actinobacteria; Actinobacteria; Streptomycetales; Streptomycetaceae; Streptomyces
MRDSSPLSRRALLGLTAAAPLAAAVPAATATSAYAATAIGGELLGRSGLQVRGGVDVPAK WP_051707667.1
MULTISPECIES: D-alanyl-D-alanine carboxypeptidase [Streptomyces] Length: 403\nNumber of predicted TMHs: 2\nExp number of AAs in TMHs: 37.68999\nExp number, first 60 AAs: 17.47041\nTotal prob of N-in: 0.76165\nPOSSIBLE N-term signal sequence\ninside 1 11\nTMhelix 12 34\noutside 35 355\nTMhelix 356 378\ninside 379 403

33205 GCF_000717635.1_ASM71763v1 Streptomyces sp. NRRL B-12105 Terrabacteria group; Actinobacteria; Actinobacteria; Streptomycetales; Streptomycetaceae; Streptomyces
MPASHAKNGRRRAPVAAVALLSLPALPFAVPGTAFAAPMPPSPGASASTDAAHTPGAGAS WP_030994093.1
MULTISPECIES: D-alanyl-D-alanine carboxypeptidase [Streptomyces] Length: 465\nNumber of predicted TMHs: 2\nExp number of AAs in TMHs: 43.7500699999999\nExp number, first 60 AAs: 21.56521\nTotal prob of N-in: 0.96490\nPOSSIBLE N-term signal sequence\ninside 1 12\nTMhelix 13 35\noutside 36 426\nTMhelix 427 449\ninside 450 465

33206 GCF_000719625.1_ASM71962v1 Streptomyces sp. NRRL B-1322 Terrabacteria group; Actinobacteria; Actinobacteria; Streptomycetales; Streptomycetaceae; Streptomyces
MSTVRRRTLVRAAAVTACAGSLLALPTAAALAEGVPAASSAHSAAPQRTLVKSLSLADGV WP_030660217.1
MULTISPECIES: hypothetical protein [Streptomyces] Length: 218\nNumber of predicted TMHs: 2\nExp number of AAs in TMHs: 42.00955\nExp number, first 60 AAs: 19.35386\nTotal prob of N-in: 0.95630\nPOSSIBLE N-term signal sequence\ninside 1 8\nTMhelix 9 31\noutside 32 188\nTMhelix 189 211\ninside 212 218

33207 GCF_000719625.1_ASM71962v1 Streptomyces sp. NRRL B-1322 Terrabacteria group; Actinobacteria; Actinobacteria; Streptomycetales; Streptomycetaceae; Streptomyces
MPQYRDARRALLATAFALTLTAAGATVATAAPHPPYALRLDGAGECTFPMKKQIADRPWA WP_030651182.1
MULTISPECIES: type VII secretion-associated serine protease [Streptomyces] Length: 414\nNumber of predicted TMHs: 2\nExp number of AAs in TMHs: 43.048\nExp number, first 60 AAs: 20.71014\nTotal prob of N-in: 0.96189\nPOSSIBLE N-term signal sequence\ninside 1 8\nTMhelix 9 31\noutside 32 380\nTMhelix 381 403\ninside 404 414

33208 GCF_000720325.1_ASM72032v1 Streptomyces sp. NRRL B-1347 Terrabacteria group; Actinobacteria; Actinobacteria; Streptomycetales; Streptomycetaceae; Streptomyces
MAAPRRPIVRAAAVATAVALGASALTLPAFAADEPAAEPAAGSPKAPPVYSVVDGTLNWG WP_030679056.1

hypothetical protein [Streptomyces sp. NRRL B-1347] Length: 510\nNumber of predicted TMHs: 1\nExp number of AAs in TMHs: 33.75407\nExp number, first 60 AAs: 14.25199\nTotal prob of N-in: 0.66566\nPOSSIBLE N-term signal sequence\noutside 1 475\nTMhelix 476 498\ninside 499 510

33209 GCF_000720325.1_ASM72032v1 Streptomyces sp. NRRL B-1347 Terrabacteria group; Actinobacteria; Actinobacteria; Streptomycetales; Streptomycetaceae; Streptomyces MTTTAPAPRRTAPRRRRRLLGAALLFLTGVFVLTFFAYPGHTLVTRMYKQGVAVDATTVA WP_030675263.1 hypothetical protein [Streptomyces sp. NRRL B-1347] Length: 158\nNumber of predicted TMHs: 2\nExp number of AAs in TMHs: 42.44253\nExp number, first 60 AAs: 21.30732\nTotal prob of N-in: 0.99904\nPOSSIBLE N-term signal sequence\ninside 1 19\nTMhelix 20 42\noutside 43 128\nTMhelix 129 151\ninside 152 158

33210 GCF_000720325.1_ASM72032v1 Streptomyces sp. NRRL B-1347 Terrabacteria group; Actinobacteria; Actinobacteria; Streptomycetales; Streptomycetaceae; Streptomyces MLGLVENHSLPRTAAFFDLDKTVIAKSSTLTFSKSFYQGGLINRRAVLRTAYAQFVFLAG WP_030677863.1 inhibition of morphological differentiation protein [Streptomyces sp. NRRL B-1347] Length: 275\nNumber of predicted TMHs: 1\nExp number of AAs in TMHs: 21.53775\nExp number, first 60 AAs: 0.02051\nTotal prob of N-in: 0.00391\noutside 1 244\nTMhelix 245 267\ninside 268 275

33211 GCF_000719655.1_ASM71965v1 Streptomyces sp. NRRL B-1381 Terrabacteria group; Actinobacteria; Actinobacteria; Streptomycetales; Streptomycetaceae; Streptomyces MLSVVENCFSPRTAAFFDLDKTVIAKSSTLTFSKSFYQGGLINRRAVLRTAYTQFVFMAG WP_030334371.1 inhibition of morphological differentiation protein [Streptomyces sp. NRRL B-1381] Length: 280\nNumber of predicted TMHs: 1\nExp number of AAs in TMHs: 21.68695\nExp number, first 60 AAs: 0.01801\nTotal prob of N-in: 0.00793\noutside 1 247\nTMhelix 248 270\ninside 271 280

33212 GCF_000719655.1_ASM71965v1 Streptomyces sp. NRRL B-1381 Terrabacteria group; Actinobacteria; Actinobacteria; Streptomycetales; Streptomycetaceae; Streptomyces MGSSWNERRLERRVRPGDGKPLKPFRWWQLTRRSLLSLALPVGGGLSATYTVEVKHGGDA WP_030189408.1 MULTISPECIES: hypothetical protein [Streptomyces] Length: 218\nNumber of predicted TMHs: 2\nExp number of AAs in TMHs: 41.32127\nExp number, first 60 AAs: 0.37701\nTotal prob of N-in: 0.97413\ninside 1 135\nTMhelix 136 158\noutside 159 182\nTMhelix 183 200\ninside 201 218

33213 GCF_000720405.1_ASM72040v1 Streptomyces sp. NRRL B-24051 Terrabacteria group; Actinobacteria; Actinobacteria; Streptomycetales; Streptomycetaceae; Streptomyces MLTFVENCFSPRTAAFFDLDKTVIAKSSTLTFSKSFYQGGLINRRAVLRTAYTQFVFLAG WP_014155461.1 MULTISPECIES: haloacid dehalogenase [Streptomyces] Length: 279\nNumber of predicted TMHs: 1\nExp number of AAs in TMHs: 21.10966\nExp number, first 60 AAs: 0.01497\nTotal prob of N-in: 0.00234\noutside 1 248\nTMhelix 249 268\ninside 269 279

33214 GCF_000720405.1_ASM72040v1 Streptomyces sp. NRRL B-24051 Terrabacteria group; Actinobacteria; Actinobacteria; Streptomycetales; Streptomycetaceae; Streptomyces MTTHPSPRRRLLTAAAATAVLLVTLPVLPAAAEDSTQCTFPSKKYAGRPWSLQRVLMDEL WP_014153729.1 MULTISPECIES: type VII secretion-associated serine protease [Streptomyces] Length: 414\nNumber of predicted TMHs: 1\nExp number of AAs in TMHs: 36.07361\nExp number, first 60 AAs: 13.5063\nTotal prob of N-in: 0.65550\nPOSSIBLE N-term signal sequence\noutside 1 378\nTMhelix 379 401\ninside 402 414

33215 GCF_000720405.1_ASM72040v1 Streptomyces sp. NRRL B-24051 Terrabacteria group; Actinobacteria; Actinobacteria; Streptomycetales; Streptomycetaceae; Streptomyces MRTTRRNTLRRGGVGALLAAVTLLTSAGAAVADDPSPSASADGGTAGPTEAGTTFRTATA WP_030879423.1 hypothetical protein [Streptomyces sp. NRRL B-24051] Length: 274\nNumber of predicted TMHs: 2\nExp number of AAs in TMHs: 40.1726899999999\nExp number, first 60 AAs: 19.30585\nTotal prob of N-in: 0.99359\nPOSSIBLE N-term signal sequence\ninside 1 12\nTMhelix 13 32\noutside 33 237\nTMhelix 238 260\ninside 261 274

33216 GCF_001282115.1_ASM128211v1 Streptomyces sp. NRRL B-24085 Terrabacteria group; Actinobacteria; Actinobacteria; Streptomycetales; Streptomycetaceae; Streptomyces MRHARRRVVRRVTRLAAVGGLLLGGVMVTRAVASEPPATTAVPHTYAVKAGRTGAELVSR WP_053849945.1 protease [Streptomyces sp. NRRL B-24085] Length: 455\nNumber of predicted TMHs: 2\nExp number of AAs in TMHs: 39.1350199999999\nExp number, first 60 AAs: 20.92997\nTotal prob of N-in: 0.96760\nPOSSIBLE N-term signal sequence\ninside 1 11\nTMhelix 12 34\noutside 35 416\nTMhelix 417 439\ninside 440 455

33217	GCF_001282115.1_ASM128211v1	Streptomyces sp. NRRL B-24085	Terrabacteria group; Actinobacteria; Actinobacteria; Streptomycetales; Streptomycetaceae; Streptomyces
MLGVVENHSLPRTAAFFDLDKTVIAKSSTLTFSKSFYQGGLINRRAALRTAYAQFVFLVG	WP_053848156.1	inhibition of morphological differentiation protein [Streptomyces sp. NRRL B-24085]	Length: 277\nNumber of predicted TMHs: 1\nExp number of AAs in TMHs: 21.65845\nExp number, first 60 AAs: 0.01246\nTotal prob of N-in: 0.00153\noutside 1 247\nTMhelix 248 270\ninside 271 277

33218	GCF_001282115.1_ASM128211v1	Streptomyces sp. NRRL B-24085	Terrabacteria group; Actinobacteria; Actinobacteria; Streptomycetales; Streptomycetaceae; Streptomyces
MTRRTVLLSLLTGLLLAGVSPAHATGYRYWSFWDRTGTAWAYATQGPSTAVPSDGDVQGF	WP_053849849.1	hypothetical protein [Streptomyces sp. NRRL B-24085]	Length: 210\nNumber of predicted TMHs: 2\nExp number of AAs in TMHs: 41.42069\nExp number, first 60 AAs: 20.82539\nTotal prob of N-in: 0.96457\nPOSSIBLE N-term signal sequence\ninside 1 4\nTMhelix 5 27\noutside 28 178\nTMhelix 179 201\ninside 202 210

33219	GCF_002156095.1_ASM215609v1	Streptomyces sp. NRRL B-24085	Terrabacteria group; Actinobacteria; Actinobacteria; Streptomycetales; Streptomycetaceae; Streptomyces
MLGVVENHSLPRTAAFFDLDKTVIAKSSTLTFSKSFYQGGLINRRAALRTAYAQFVFLVG	WP_053848156.1	inhibition of morphological differentiation protein [Streptomyces sp. NRRL B-24085]	Length: 277\nNumber of predicted TMHs: 1\nExp number of AAs in TMHs: 21.65845\nExp number, first 60 AAs: 0.01246\nTotal prob of N-in: 0.00153\noutside 1 247\nTMhelix 248 270\ninside 271 277

33220	GCF_002156095.1_ASM215609v1	Streptomyces sp. NRRL B-24085	Terrabacteria group; Actinobacteria; Actinobacteria; Streptomycetales; Streptomycetaceae; Streptomyces
MHARTADTRRSSSVGRSNRRRPTGARRATFAAVALMLGGGGLVAANVYASATESGTPQQT	WP_086724782.1	hypothetical protein [Streptomyces sp. NRRL B-24085]	Length: 714\nNumber of predicted TMHs: 2\nExp number of AAs in TMHs: 41.32272\nExp number, first 60 AAs: 22.15581\nTotal prob of N-in: 0.99572\nPOSSIBLE N-term signal sequence\ninside 1 27\nTMhelix 28 50\noutside 51 688\nTMhelix 689 706\ninside 707 714

33221	GCF_002156095.1_ASM215609v1	Streptomyces sp. NRRL B-24085	Terrabacteria group; Actinobacteria; Actinobacteria; Streptomycetales; Streptomycetaceae; Streptomyces
MRHARRRVVRRVTRLAAVGGLLLGGVMVTRAVASEPPATTAVPHTYAVKAGRTGAELVSR	WP_053849945.1	S1 family peptidase [Streptomyces sp. NRRL B-24085]	Length: 455\nNumber of predicted TMHs: 2\nExp number of AAs in TMHs: 39.1350199999999\nExp number, first 60 AAs: 20.92997\nTotal prob of N-in: 0.96760\nPOSSIBLE N-term signal sequence\ninside 1 11\nTMhelix 12 34\noutside 35 416\nTMhelix 417 439\ninside 440 455

33222	GCF_002156095.1_ASM215609v1	Streptomyces sp. NRRL B-24085	Terrabacteria group; Actinobacteria; Actinobacteria; Streptomycetales; Streptomycetaceae; Streptomyces
MTRRTVLLSLLTGLLLAGVSPAHATGYRYWSFWDRTGTAWAYATQGPSTAVPSDGDVQGF	WP_053849849.1	hypothetical protein [Streptomyces sp. NRRL B-24085]	Length: 210\nNumber of predicted TMHs: 2\nExp number of AAs in TMHs: 41.42069\nExp number, first 60 AAs: 20.82539\nTotal prob of N-in: 0.96457\nPOSSIBLE N-term signal sequence\ninside 1 4\nTMhelix 5 27\noutside 28 178\nTMhelix 179 201\ninside 202 210

33223	GCF_000717715.1_ASM71771v1	Streptomyces sp. NRRL B-24484	Terrabacteria group; Actinobacteria; Actinobacteria; Streptomycetales; Streptomycetaceae; Streptomyces
MPGRPAARRAAVPAQAGPSAPAAGAREASAPAVPAQGTADAAGVPGAADAAEVYRAVQSS	WP_078661650.1	hypothetical protein [Streptomyces sp. NRRL B-24484]	Length: 152\nNumber of predicted TMHs: 2\nExp number of AAs in TMHs: 43.70467\nExp number, first 60 AAs: 0.00016\nTotal prob of N-in: 0.99889\ninside 1 71\nTMhelix 72 94\noutside 95 103\nTMhelix 104 126\ninside 127 152

33224	GCF_000717715.1_ASM71771v1	Streptomyces sp. NRRL B-24484	Terrabacteria group; Actinobacteria; Actinobacteria; Streptomycetales; Streptomycetaceae; Streptomyces
MRTTPARRTALAAAVAGAAVVVLAGPAAAHVTVQPGSVPQGGYTAVAFRVPDESDTASTV	WP_030271099.1	membrane protein [Streptomyces sp. NRRL B-24484]	Length: 242\nNumber of predicted TMHs: 2\nExp number of AAs in TMHs: 40.98593\nExp number, first 60 AAs: 19.9375\nTotal prob of N-in: 0.94789\nPOSSIBLE N-term signal sequence\ninside 1 11\nTMhelix 12 34\noutside 35 211\nTMhelix 212 234\ninside 235 242

33225	GCF_002156145.1_ASM215614v1	Streptomyces sp. NRRL B-24572	Terrabacteria group; Actinobacteria; Actinobacteria; Streptomycetales; Streptomycetaceae; Streptomyces
MLGLVENHSLPRTAAFFDLDKTVIAKSSTLTFSKSFYRGGLISRRAALRTAYIQFVFLAG	WP_086832055.1	inhibition of morphological differentiation protein [Streptomyces sp. NRRL B-24572]	Length: 288\nNumber of predicted TMHs:

1\nExp number of AAs in TMHs: 22.01996\nExp number, first 60 AAs: 0.02671\nTotal prob of N-in: 0.00194\noutside 1 247\nTMhelix 248 270\ninside 271 288

33226 GCF_000721715.1_ASM72171v1 Streptomyces sp. NRRL B-24720 Terrabacteria group; Actinobacteria; Actinobacteria; Streptomycetales; Streptomycetaceae; Streptomyces MLCLVENHSLPRTAAFFDLDKTVIAKSSTLTFSKSFYQGGLINRRAVLRTAYAQFVFLAG WP_030925136.1 inhibition of morphological differentiation protein [Streptomyces sp. NRRL B-24720] Length: 279\nNumber of predicted TMHs: 1\nExp number of AAs in TMHs: 21.31113\nExp number, first 60 AAs: 0.02048\nTotal prob of N-in: 0.00231\noutside 1 247\nTMhelix 248 270\ninside 271 279

33227 GCF_000981895.1_ASM98189v1 Streptomyces sp. NRRL B-24891 Terrabacteria group; Actinobacteria; Actinobacteria; Streptomycetales; Streptomycetaceae; Streptomyces MTLFSGSWRRRGALITVATAGAIVGLSAVPAVAHTPNWSVTCSEVSVDLTAYGNSDQNTV WP_046502582.1 LPXTG-motif cell wall anchor domain protein [Streptomyces sp. NRRL B-24891]Length: 216\nNumber of predicted TMHs: 2\nExp number of AAs in TMHs: 43.74146\nExp number, first 60 AAs: 21.59256\nTotal prob of N-in: 0.98317\nPOSSIBLE N-term signal sequence\ninside 1 11\nTMhelix 12 34\noutside 35 185\nTMhelix 186 208\ninside 209 216

33228 GCF_000981895.1_ASM98189v1 Streptomyces sp. NRRL B-24891 Terrabacteria group; Actinobacteria; Actinobacteria; Streptomycetales; Streptomycetaceae; Streptomyces MLMPVENHSLPRTAAFFDLDKTVIAKSSTLTFSKSFYQGGLINRRAVLRTAYAQFVFLAG WP_046496584.1 inhibition of morphological differentiation protein [Streptomyces sp. NRRL B-24891] Length: 278\nNumber of predicted TMHs: 1\nExp number of AAs in TMHs: 21.62982\nExp number, first 60 AAs: 0.01894\nTotal prob of N-in: 0.00256\noutside 1 247\nTMhelix 248 270\ninside 271 278

33229 GCF_000981895.1_ASM98189v1 Streptomyces sp. NRRL B-24891 Terrabacteria group; Actinobacteria; Actinobacteria; Streptomycetales; Streptomycetaceae; Streptomyces MNQSHQSSPPPRRAARRGPVAARLRALPVWGIVPAAALAGALAGGAYGLLRTPEYSATSY WP_052744599.1 hypothetical protein, partial [Streptomyces sp. NRRL B-24891] Length: 195\nNumber of predicted TMHs: 2\nExp number of AAs in TMHs: 43.83057\nExp number, first 60 AAs: 22.46533\nTotal prob of N-in: 0.24093\nPOSSIBLE N-term signal sequence\noutside 1 27\nTMhelix 28 50\ninside 51 170\nTMhelix 171 193\noutside 194 195

33230 GCF_000981895.1_ASM98189v1 Streptomyces sp. NRRL B-24891 Terrabacteria group; Actinobacteria; Actinobacteria; Streptomycetales; Streptomycetaceae; Streptomyces MQDRRRRTALHLPPPAGPVLLPRERQSSSTGPSGTPGPSGSPGASGSADSEQDRAPAGGP WP_079023587.1 hypothetical protein [Streptomyces sp. NRRL B-24891] Length: 364\nNumber of predicted TMHs: 2\nExp number of AAs in TMHs: 45.9187\nExp number, first 60 AAs: 0.0002\nTotal prob of N-in: 0.73554\ninside 1 205\nTMhelix 206 228\noutside 229 302\nTMhelix 303 325\ninside 326 364

33231 GCF_000981895.1_ASM98189v1 Streptomyces sp. NRRL B-24891 Terrabacteria group; Actinobacteria; Actinobacteria; Streptomycetales; Streptomycetaceae; Streptomyces MHASVPRRKSDPRRQPAPDRQPAPLRKTAPRRRTAVAGLVRTAALALLMVVAVAGLSAGP WP_079023627.1 DUF916 domain-containing protein [Streptomyces sp. NRRL B-24891] Length: 383\nNumber of predicted TMHs: 2\nExp number of AAs in TMHs: 45.27448\nExp number, first 60 AAs: 21.53128\nTotal prob of N-in: 0.97452\nPOSSIBLE N-term signal sequence\ninside 1 36\nTMhelix 37 59\noutside 60 327\nTMhelix 328 350\ninside 351 383

33232 GCF_000719505.1_ASM71950v1 Streptomyces sp. NRRL B-3229 Terrabacteria group; Actinobacteria; Actinobacteria; Streptomycetales; Streptomycetaceae; Streptomyces MRHARRRVVRRVTRLAAVGGLLLGGTMVTRAVASEPPATAPPRTFAQQAGETGTDIVARL WP_030323245.1 protease [Streptomyces sp. NRRL B-3229] Length: 454\nNumber of predicted TMHs: 2\nExp number of AAs in TMHs: 40.6905799999999\nExp number, first 60 AAs: 20.53903\nTotal prob of N-in: 0.94036\nPOSSIBLE N-term signal sequence\ninside 1 11\nTMhelix 12 34\noutside 35 415\nTMhelix 416 438\ninside 439 454

33233 GCF_000719505.1_ASM71950v1 Streptomyces sp. NRRL B-3229 Terrabacteria group; Actinobacteria; Actinobacteria; Streptomycetales; Streptomycetaceae; Streptomyces MLGVVDNHSLPRTAAFFDLDKTVIAKSSTLTFSKSFYQGGLINRRAALRTAYAQFVFLVG WP_030324035.1 inhibition of morphological differentiation protein [Streptomyces sp. NRRL B-3229] Length: 277\nNumber of predicted TMHs: 1\nExp number of AAs in TMHs: 21.69258\nExp number, first 60 AAs: 0.03764\nTotal prob of N-in: 0.00334\noutside 1 247\nTMhelix 248 270\ninside 271 277

33234 GCF_000725635.1_Doro.v1.0 Streptomyces sp. NRRL B-3253 Terrabacteria group; Actinobacteria; Actinobacteria; Streptomycetales; Streptomycetaceae; Streptomyces
MAPPHRRRAAVAVRALVALTALLCTTGLAPAATSGPAARAAVAEVPPPTATTSVVTVRTG WP_008409019.1 MULTISPECIES: VWA domain-containing protein [Streptomyces] Length: 835\nNumber of predicted TMHs: 2\nExp number of AAs in TMHs: 37.77539\nExp number, first 60 AAs: 18.55298\nTotal prob of N-in: 0.85432\nPOSSIBLE N-term signal sequence\ninside 1 11\nTMhelix 12 34\noutside 35 796\nTMhelix 797 819\ninside 820 835

33235 GCF_000725635.1_Doro.v1.0 Streptomyces sp. NRRL B-3253 Terrabacteria group; Actinobacteria; Actinobacteria; Streptomycetales; Streptomycetaceae; Streptomyces
MPDRRSAVSLRLPLPTAAPGFGGLALPGAAATAPDAWGVVAVRESSEKERKSSSEPEGRS WP_015507753.1 MULTISPECIES: hypothetical protein [Streptomyces] Length: 292\nNumber of predicted TMHs: 2\nExp number of AAs in TMHs: 51.77675\nExp number, first 60 AAs: 6.07612\nTotal prob of N-in: 0.33780\noutside 1 164\nTMhelix 165 187\ninside 188 229\nTMhelix 230 252\noutside 253 292

33236 GCF_000725635.1_Doro.v1.0 Streptomyces sp. NRRL B-3253 Terrabacteria group; Actinobacteria; Actinobacteria; Streptomycetales; Streptomycetaceae; Streptomyces
MLDPVENRSLPRTAAFFDLDKTVIAKSSTLTFSKSFYQGGLINRRAVLRTAYTQFVFLAG WP_018470348.1 MULTISPECIES: inhibition of morphological differentiation protein [Streptomyces] Length: 290\nNumber of predicted TMHs: 1\nExp number of AAs in TMHs: 21.82254\nExp number, first 60 AAs: 0.02718\nTotal prob of N-in: 0.00834\noutside 1 247\nTMhelix 248 270\ninside 271 290

33237 GCF_000725635.1_Doro.v1.0 Streptomyces sp. NRRL B-3253 Terrabacteria group; Actinobacteria; Actinobacteria; Streptomycetales; Streptomycetaceae; Streptomyces
MSRRTRTARRRPVAAAVLAAGAVLLPAVPASAASGADQDFVIEDPRITESSGLAASKRHP WP_030309758.1 MULTISPECIES: hypothetical protein [Streptomyces] Length: 348\nNumber of predicted TMHs: 2\nExp number of AAs in TMHs: 39.44414\nExp number, first 60 AAs: 21.11647\nTotal prob of N-in: 0.99086\nPOSSIBLE N-term signal sequence\ninside 1 12\nTMhelix 13 35\noutside 36 320\nTMhelix 321 340\ninside 341 348

33238 GCF_000725635.1_Doro.v1.0 Streptomyces sp. NRRL B-3253 Terrabacteria group; Actinobacteria; Actinobacteria; Streptomycetales; Streptomycetaceae; Streptomyces
MSAAAPEAAPRTARPLPRAGALLRRLLWWCVLTLTGGVERRGALPPGGCVVVANHTSHAD WP_031177737.1 1-acyl-sn-glycerol-3-phosphate acyltransferase [Streptomyces sp. NRRL B-3253] Length: 401\nNumber of predicted TMHs: 2\nExp number of AAs in TMHs: 82.13307\nExp number, first 60 AAs: 0.94183\nTotal prob of N-in: 0.15922\noutside 1 353\nTMhelix 354 371\ninside 372 377\nTMhelix 378 400\noutside 401 401

33239 GCF_000725635.1_Doro.v1.0 Streptomyces sp. NRRL B-3253 Terrabacteria group; Actinobacteria; Actinobacteria; Streptomycetales; Streptomycetaceae; Streptomyces
MTPNRRPVVLAAAVATALGLGATALTLPAVAAEAPRAGAPVMDLTDGTLDWGVKESFRRY WP_031178127.1 membrane protein [Streptomyces sp. NRRL B-3253] Length: 514\nNumber of predicted TMHs: 2\nExp number of AAs in TMHs: 42.8567099999999\nExp number, first 60 AAs: 22.03087\nTotal prob of N-in: 0.97276\nPOSSIBLE N-term signal sequence\ninside 1 6\nTMhelix 7 29\noutside 30 479\nTMhelix 480 502\ninside 503 514

33240 GCF_000725635.1_Doro.v1.0 Streptomyces sp. NRRL B-3253 Terrabacteria group; Actinobacteria; Actinobacteria; Streptomycetales; Streptomycetaceae; Streptomyces
MGGRGTTAQARERGRLPGRPSDPAPTGGNDVITGVNLRMPARRTALPALLLTATALLAVP WP_031178864.1 hypothetical protein [Streptomyces sp. NRRL B-3253] Length: 304\nNumber of predicted TMHs: 2\nExp number of AAs in TMHs: 39.19634\nExp number, first 60 AAs: 11.7572\nTotal prob of N-in: 0.76169\nPOSSIBLE N-term signal sequence\ninside 1 44\nTMhelix 45 67\noutside 68 276\nTMhelix 277 299\ninside 300 304

33241 GCF_001279545.1_ASM127954v1 Streptomyces sp. NRRL B-3648 Terrabacteria group; Actinobacteria; Actinobacteria; Streptomycetales; Streptomycetaceae; Streptomyces
MPAPKTTGRRSLLVTSATLLSLTVTAPAALAAQAPSPSPTATPPAGMSTLGGARLGQPGT WP_053707807.1 D-alanyl-D-alanine carboxypeptidase [Streptomyces sp. NRRL B-3648] Length: 421\nNumber of predicted TMHs: 1\nExp number of AAs in TMHs: 36.32953\nExp number, first 60 AAs: 13.85229\nTotal prob of N-in: 0.64632\nPOSSIBLE N-term signal sequence\noutside 1 382\nTMhelix 383 405\ninside 406 421

33242 GCF_001279545.1_ASM127954v1 Streptomyces sp. NRRL B-3648 Terrabacteria group; Actinobacteria; Actinobacteria; Streptomycetales; Streptomycetaceae; Streptomyces
MLRGVENHSLPRAAAFFDLDKTVIAKSSTLTFSKSFYQGGLINRRAALRTAYAQFVFLAG WP_053708104.1 inhibition of morphological differentiation protein [Streptomyces sp. NRRL B-3648] Length: 276\nNumber of predicted TMHs:

1\nExp number of AAs in TMHs: 21.32665\nExp number, first 60 AAs: 0.02846\nTotal prob of N-in: 0.00390\noutside 1 247\nTMhelix 248 270\ninside 271 276

33243 GCF_001279545.1_ASM127954v1 Streptomyces sp. NRRL B-3648 Terrabacteria group; Actinobacteria; Actinobacteria; Streptomycetales; Streptomycetaceae; Streptomyces MRHARRRVVRRVTRLAAVGGLLLGGAMVAQAAMASEPPPASARPFAATDDPGAALVARLG WP_053711347.1 protease [Streptomyces sp. NRRL B-3648] Length: 458\nNumber of predicted TMHs: 2\nExp number of AAs in TMHs: 43.9295\nExp number, first 60 AAs: 22.34314\nTotal prob of N-in: 0.99589\nPOSSIBLE N-term signal sequence\ninside 1 11\nTMhelix 12 34\noutside 35 419\nTMhelix 420 442\ninside 443 458

33244 GCF_000719415.1_ASM71941v1 Streptomyces sp. NRRL B-5680 Terrabacteria group; Actinobacteria; Actinobacteria; Streptomycetales; Streptomycetaceae; Streptomyces MRTAAFFDLDKTIIAKSSALAFSRSFYQGGLINRRAVLRSAYAQFVFLVGGADHDQMEKM WP_030288083.1 MULTISPECIES: inhibition of morphological differentiation protein [Streptomyces] Length: 272\nNumber of predicted TMHs: 1\nExp number of AAs in TMHs: 22.68364\nExp number, first 60 AAs: 0.10845\nTotal prob of N-in: 0.00817\noutside 1 235\nTMhelix 236 258\ninside 259 272

33245 GCF_000719925.1_ASM71992v1 Streptomyces sp. NRRL F-2202 Terrabacteria group; Actinobacteria; Actinobacteria; Streptomycetales; Streptomycetaceae; Streptomyces MLSVVENCFSPRTAAFFDLDKTVIAKSSTLTFSKSFYQGGLINRRAVLRTAYTQFVFMAG WP_030189579.1 MULTISPECIES: inhibition of morphological differentiation protein [Streptomyces] Length: 280\nNumber of predicted TMHs: 1\nExp number of AAs in TMHs: 21.68695\nExp number, first 60 AAs: 0.01801\nTotal prob of N-in: 0.00793\noutside 1 247\nTMhelix 248 270\ninside 271 280

33246 GCF_000719925.1_ASM71992v1 Streptomyces sp. NRRL F-2202 Terrabacteria group; Actinobacteria; Actinobacteria; Streptomycetales; Streptomycetaceae; Streptomyces MGSSWNERWLERCVRPGDGKPLKPFRWWQLTRRSLLSLALPVGGGLSATYTVEVKHGGDA WP_032773289.1 hypothetical protein [Streptomyces sp. NRRL F-2202] Length: 218\nNumber of predicted TMHs: 2\nExp number of AAs in TMHs: 41.82036\nExp number, first 60 AAs: 0.95179\nTotal prob of N-in: 0.93795\ninside 1 135\nTMhelix 136 158\noutside 159 182\nTMhelix 183 200\ninside 201 218

33247 GCF_001279425.1_ASM127942v1 Streptomyces sp. NRRL F-2295 Terrabacteria group; Actinobacteria; Actinobacteria; Streptomycetales; Streptomycetaceae; Streptomyces MLSVVENCFSPRTAAFFDLDKTVIAKSSTLTFSKSFYQGGLINRRAVLRTAYTQFVFMAG WP_030189579.1 MULTISPECIES: inhibition of morphological differentiation protein [Streptomyces] Length: 280\nNumber of predicted TMHs: 1\nExp number of AAs in TMHs: 21.68695\nExp number, first 60 AAs: 0.01801\nTotal prob of N-in: 0.00793\noutside 1 247\nTMhelix 248 270\ninside 271 280

33248 GCF_001279425.1_ASM127942v1 Streptomyces sp. NRRL F-2295 Terrabacteria group; Actinobacteria; Actinobacteria; Streptomycetales; Streptomycetaceae; Streptomyces MGSSWNERWLERRVRPGDGKPLKPFRWWQLTRRSLLSLALPVGGGLSATYTVEVKHGGDA WP_030733765.1 MULTISPECIES: hypothetical protein [Streptomyces] Length: 218\nNumber of predicted TMHs: 2\nExp number of AAs in TMHs: 41.82036\nExp number, first 60 AAs: 0.95179\nTotal prob of N-in: 0.93795\ninside 1 135\nTMhelix 136 158\noutside 159 182\nTMhelix 183 200\ninside 201 218

33249 GCF_000718375.1_ASM71837v1 Streptomyces sp. NRRL F-2305 Terrabacteria group; Actinobacteria; Actinobacteria; Streptomycetales; Streptomycetaceae; Streptomyces MLRDVENHSMPRTAAFFDLDKTVIAKSSALTFSKSFYQGGLINRRAVLRTAYSQFVFLAG WP_030827973.1 MULTISPECIES: inhibition of morphological differentiation protein [Streptomyces] Length: 277\nNumber of predicted TMHs: 1\nExp number of AAs in TMHs: 21.88014\nExp number, first 60 AAs: 0.04028\nTotal prob of N-in: 0.00638\noutside 1 247\nTMhelix 248 270\ninside 271 277

33250 GCF_000718375.1_ASM71837v1 Streptomyces sp. NRRL F-2305 Terrabacteria group; Actinobacteria; Actinobacteria; Streptomycetales; Streptomycetaceae; Streptomyces MRHARRRVVRRVTRLAAVGGLLLGATMVTHAVASEPRAPDAVPFATAEGTVSTGPGADLV WP_030821297.1 MULTISPECIES: protease [Streptomyces] Length: 459\nNumber of predicted TMHs: 2\nExp number of AAs in TMHs: 42.46692\nExp number, first 60 AAs: 22.12504\nTotal prob of N-in: 0.98832\nPOSSIBLE N-term signal sequence\ninside 1 11\nTMhelix 12 34\noutside 35 420\nTMhelix 421 443\ninside 444 459

33251 GCF_000720115.1_ASM72011v1 Streptomyces sp. NRRL F-2580 Terrabacteria group; Actinobacteria; Actinobacteria; Streptomycetales; Streptomycetaceae; Streptomyces

MSTVSRRTLVRAAAVTACAASLLALPTAAALAEGVPAASSAHSAAPQRTLVKSLSLADGA WP_030708058.1 hypothetical protein [Streptomyces sp. NRRL F-2580] Length: 210\nNumber of predicted TMHs: 2\nExp number of AAs in TMHs: 42.8229899999999\nExp number, first 60 AAs: 20.21275\nTotal prob of N-in: 0.93851\nPOSSIBLE N-term signal sequence\ninside 1 8\nTMhelix 9 31\noutside 32 180\nTMhelix 181 203\ninside 204 210

33252 GCF_000720115.1_ASM72011v1 Streptomyces sp. NRRL F-2580 Terrabacteria group; Actinobacteria; Actinobacteria; Streptomycetales; Streptomycetaceae; Streptomyces MPARRRTVTLARIASASFAPLALAVYAAAPAVAHGSMTDPVSRVAACYAEGPEAPKSAAC WP_030720207.1 chitin-binding protein [Streptomyces sp. NRRL F-2580] Length: 355\nNumber of predicted TMHs: 2\nExp number of AAs in TMHs: 43.3557\nExp number, first 60 AAs: 20.91603\nTotal prob of N-in: 0.95106\nPOSSIBLE N-term signal sequence\ninside 1 12\nTMhelix 13 35\noutside 36 319\nTMhelix 320 342\ninside 343 355

33253 GCF_000720115.1_ASM72011v1 Streptomyces sp. NRRL F-2580 Terrabacteria group; Actinobacteria; Actinobacteria; Streptomycetales; Streptomycetaceae; Streptomyces MRRRRPALALALALALGSVLAVLAAAPALAAGYRYWSFWDGSGGGQWSYATQGPSMARPK WP_030722950.1 hypothetical protein [Streptomyces sp. NRRL F-2580] Length: 219\nNumber of predicted TMHs: 2\nExp number of AAs in TMHs: 43.66634\nExp number, first 60 AAs: 22.9582\nTotal prob of N-in: 0.99979\nPOSSIBLE N-term signal sequence\ninside 1 6\nTMhelix 7 29\noutside 30 189\nTMhelix 190 212\ninside 213 219

33254 GCF_000719345.1_ASM71934v1 Streptomyces sp. NRRL F-2664 Terrabacteria group; Actinobacteria; Actinobacteria; Streptomycetales; Streptomycetaceae; Streptomyces MSTVRRRTLVRAAAVTACAGSLLALPTAAALAEGVPAASSARSAAPQRTLVKTLSLADGV WP_030768766.1 hypothetical protein [Streptomyces sp. NRRL F-2664] Length: 210\nNumber of predicted TMHs: 2\nExp number of AAs in TMHs: 43.1572\nExp number, first 60 AAs: 20.54025\nTotal prob of N-in: 0.96296\nPOSSIBLE N-term signal sequence\ninside 1 8\nTMhelix 9 31\noutside 32 180\nTMhelix 181 203\ninside 204 210

33255 GCF_000719345.1_ASM71934v1 Streptomyces sp. NRRL F-2664 Terrabacteria group; Actinobacteria; Actinobacteria; Streptomycetales; Streptomycetaceae; Streptomyces MSARRRTVMPVRIAAAGLAPLALAAYAAVPAAAHGSMTDPVSRVAACYAEGPESPESAAC WP_030770144.1 chitin-binding protein [Streptomyces sp. NRRL F-2664] Length: 360\nNumber of predicted TMHs: 2\nExp number of AAs in TMHs: 42.76177\nExp number, first 60 AAs: 21.45112\nTotal prob of N-in: 0.97434\nPOSSIBLE N-term signal sequence\ninside 1 12\nTMhelix 13 35\noutside 36 325\nTMhelix 326 348\ninside 349 360

33256 GCF_000719345.1_ASM71934v1 Streptomyces sp. NRRL F-2664 Terrabacteria group; Actinobacteria; Actinobacteria; Streptomycetales; Streptomycetaceae; Streptomyces MPTSPYRAPRRALLVAASALVLTAGGGAAAAGPAGAPHPPYTVHLDGAGECTFPMKKQIE WP_030767314.1 type VII secretion-associated serine protease [Streptomyces sp. NRRL F-2664] Length: 419\nNumber of predicted TMHs: 2\nExp number of AAs in TMHs: 43.80117\nExp number, first 60 AAs: 21.50089\nTotal prob of N-in: 0.97473\nPOSSIBLE N-term signal sequence\ninside 1 12\nTMhelix 13 35\noutside 36 385\nTMhelix 386 408\ninside 409 419

33257 GCF_000720145.1_ASM72014v1 Streptomyces sp. NRRL F-2747 Terrabacteria group; Actinobacteria; Actinobacteria; Streptomycetales; Streptomycetaceae; Streptomyces MAEAADIQGGPLAPARRRWLRRAVVLLAGTPVLAALVYSPAPQAAQAAEAGAVDVQLDGM WP_030857734.1 hypothetical protein [Streptomyces sp. NRRL F-2747] Length: 755\nNumber of predicted TMHs: 1\nExp number of AAs in TMHs: 35.29064\nExp number, first 60 AAs: 13.58287\nTotal prob of N-in: 0.65390\nPOSSIBLE N-term signal sequence\noutside 1 688\nTMhelix 689 711\ninside 712 755

33258 GCF_000720145.1_ASM72014v1 Streptomyces sp. NRRL F-2747 Terrabacteria group; Actinobacteria; Actinobacteria; Streptomycetales; Streptomycetaceae; Streptomyces MSTVDRRTLVRAAAVTVCAGTLLALPAAAALAEGVPAASAAHSAGAPRTLLKSLALADGV WP_030859193.1 hypothetical protein [Streptomyces sp. NRRL F-2747] Length: 200\nNumber of predicted TMHs: 2\nExp number of AAs in TMHs: 44.2935899999999\nExp number, first 60 AAs: 21.67581\nTotal prob of N-in: 0.99046\nPOSSIBLE N-term signal sequence\ninside 1 8\nTMhelix 9 31\noutside 32 170\nTMhelix 171 193\ninside 194 200

33259 GCF_000720145.1_ASM72014v1 Streptomyces sp. NRRL F-2747 Terrabacteria group; Actinobacteria; Actinobacteria; Streptomycetales; Streptomycetaceae; Streptomyces MRSTTRRTTLRTAAVFAGAAAVLALPVGSAFADSPAGPEQEVLPGVEQPGGEKQAEQKPG WP_030870946.1 hypothetical protein [Streptomyces sp. NRRL F-2747] Length: 356\nNumber of predicted TMHs: 2\nExp

number of AAs in TMHs: 40.82592\nExp number, first 60 AAs:  19.87782\nTotal prob of N-in:   0.98004\nPOSSIBLE N-term signal sequence\ninside   1   12\nTMhelix   13   32\noutside   33   325\nTMhelix   326   348\ninside   349   356

33260 GCF_000720145.1_ASM72014v1 Streptomyces sp. NRRL F-2747 Terrabacteria group; Actinobacteria; Actinobacteria; Streptomycetales; Streptomycetaceae; Streptomyces MPARRRRTVTRIAAVGLAPLAVAAYAAGPAAAHGSMTDPVSRVAACYAEGPESPKSAACK WP_030870554.1 chitin-binding protein [Streptomyces sp. NRRL F-2747] Length: 349\nNumber of predicted TMHs:  2\nExp number of AAs in TMHs: 43.67489\nExp number, first 60 AAs:  21.3688\nTotal prob of N-in:   0.98815\nPOSSIBLE N-term signal sequence\ninside   1   11\nTMhelix   12   34\noutside   35   313\nTMhelix   314   336\ninside   337   349

33261 GCF_000719895.1_ASM71989v1 Streptomyces sp. NRRL F-2799 Terrabacteria group; Actinobacteria; Actinobacteria; Streptomycetales; Streptomycetaceae; Streptomyces MRHARRRVVRRVTRLAAVGGLLLGGGMVTQAAMASETPAAPATTFSAAGTAGTKGAALAE WP_030805059.1 protease [Streptomyces sp. NRRL F-2799] Length: 455\nNumber of predicted TMHs:  2\nExp number of AAs in TMHs: 42.77147\nExp number, first 60 AAs:  21.9037\nTotal prob of N-in:   0.98398\nPOSSIBLE N-term signal sequence\ninside   1   11\nTMhelix   12   34\noutside   35   416\nTMhelix   417   439\ninside   440   455

33262 GCF_000719895.1_ASM71989v1 Streptomyces sp. NRRL F-2799 Terrabacteria group; Actinobacteria; Actinobacteria; Streptomycetales; Streptomycetaceae; Streptomyces MLMDVENHSLPRAAAFFDLDKTVIAKSSTLTFSKSFYQGGLINRRAALRTAYAQFVFLAG WP_030811411.1 inhibition of morphological differentiation protein [Streptomyces sp. NRRL F-2799] Length: 276\nNumber of predicted TMHs: 1\nExp number of AAs in TMHs: 21.42076\nExp number, first 60 AAs:  0.03674\nTotal prob of N-in:   0.00932\noutside   1   246\nTMhelix   247   269\ninside   270   276

33263 GCF_000719915.1_ASM71991v1 Streptomyces sp. NRRL F-2890 Terrabacteria group; Actinobacteria; Actinobacteria; Streptomycetales; Streptomycetaceae; Streptomyces MLGQVENLSQPRTAAFFDLDKTVIAKSSTVTFSRSFYQGGLINRRAVLRTAYAQFLYLVG WP_046723933.1 MULTISPECIES: inhibition of morphological differentiation protein [Streptomyces] Length: 290\nNumber of predicted TMHs:  1\nExp number of AAs in TMHs: 22.31044\nExp number, first 60 AAs:  0.0082\nTotal prob of N-in:   0.00135\noutside   1   246\nTMhelix   247   269\ninside   270   290

33264 GCF_000719915.1_ASM71991v1 Streptomyces sp. NRRL F-2890 Terrabacteria group; Actinobacteria; Actinobacteria; Streptomycetales; Streptomycetaceae; Streptomyces MRTPARAPVLSTARRSVLLVLLLLATQLVAVAWPAYACGCGGMVTGPGQRLTVEEETSAV WP_046722871.1 MULTISPECIES: hypothetical protein [Streptomyces] Length: 373\nNumber of predicted TMHs:  2\nExp number of AAs in TMHs: 43.02627\nExp number, first 60 AAs:  20.69791\nTotal prob of N-in:   0.97846\nPOSSIBLE N-term signal sequence\ninside   1   16\nTMhelix   17   36\noutside   37   338\nTMhelix   339   361\ninside   362   373

33265 GCF_000720075.1_ASM72007v1 Streptomyces sp. NRRL F-3213 Terrabacteria group; Actinobacteria; Actinobacteria; Streptomycetales; Streptomycetaceae; Streptomyces MAEPRSEEAPWRVAAFFDLDKTIIASSSALAFSKPLLREGLINRRAALRSAYAQLVFSLA WP_037808354.1 inhibition of morphological differentiation protein [Streptomyces sp. NRRL F-3213] Length: 273\nNumber of predicted TMHs: 1\nExp number of AAs in TMHs: 20.5984\nExp number, first 60 AAs:  0.07863\nTotal prob of N-in:   0.02144\noutside   1   242\nTMhelix   243   262\ninside   263   273

33266 GCF_000720075.1_ASM72007v1 Streptomyces sp. NRRL F-3213 Terrabacteria group; Actinobacteria; Actinobacteria; Streptomycetales; Streptomycetaceae; Streptomyces MTDDPRPATSPGPDDQAFPGRPRKPLARRPALWIGLASAACFITGIVFFLLPVTALYYLD WP_037807269.1 hypothetical protein [Streptomyces sp. NRRL F-3213] Length: 172\nNumber of predicted TMHs:  2\nExp number of AAs in TMHs: 44.96873\nExp number, first 60 AAs:  22.96129\nTotal prob of N-in:   0.91147\nPOSSIBLE N-term signal sequence\ninside   1   30\nTMhelix   31   53\noutside   54   131\nTMhelix   132   154\ninside   155   172

33267 GCF_000720055.1_ASM72005v1 Streptomyces sp. NRRL F-3218 Terrabacteria group; Actinobacteria; Actinobacteria; Streptomycetales; Streptomycetaceae; Streptomyces MLSVVENCFSPRTAAFFDLDKTVIAKSSTLTFSKSFYQGGLINRRAVLRTAYTQFVFMAG WP_030113359.1 MULTISPECIES: inhibition of morphological differentiation protein [Streptomyces] Length: 280\nNumber of predicted TMHs:  1\nExp number of AAs in TMHs: 21.68695\nExp number, first 60 AAs:  0.01801\nTotal prob of N-in:   0.00793\noutside   1   247\nTMhelix   248   270\ninside   271   280

33268 GCF_000720055.1_ASM72005v1 Streptomyces sp. NRRL F-3218 Terrabacteria group; Actinobacteria; Actinobacteria; Streptomycetales; Streptomycetaceae; Streptomyces

MGSSWNERRLERRVRPGDGKPLKPFRWWQLTRRSLLSLALPVGGGLSATYTVEVKHGGDA WP_030113438.1 MULTISPECIES: hypothetical protein [Streptomyces] Length: 218\nNumber of predicted TMHs: 2\nExp number of AAs in TMHs: 41.40199\nExp number, first 60 AAs: 0.36812\nTotal prob of N-in: 0.97576\ninside 1 135\nTMhelix 136 158\noutside 159 182\nTMhelix 183 200\ninside 201 218

33269 GCF_000720065.1_ASM72006v1 Streptomyces sp. NRRL F-3273 Terrabacteria group; Actinobacteria; Actinobacteria; Streptomycetales; Streptomycetaceae; Streptomyces MGSSWNERRLERRVRPGDGKPLKPFRWWQLTRRSLLSLALPVGGGLSATYTVEVKHGGDA WP_030113438.1 MULTISPECIES: hypothetical protein [Streptomyces] Length: 218\nNumber of predicted TMHs: 2\nExp number of AAs in TMHs: 41.40199\nExp number, first 60 AAs: 0.36812\nTotal prob of N-in: 0.97576\ninside 1 135\nTMhelix 136 158\noutside 159 182\nTMhelix 183 200\ninside 201 218

33270 GCF_000720065.1_ASM72006v1 Streptomyces sp. NRRL F-3273 Terrabacteria group; Actinobacteria; Actinobacteria; Streptomycetales; Streptomycetaceae; Streptomyces MLSVVENCFSPRTAAFFDLDKTVIAKSSTLTFSKSFYQGGLINRRAVLRTAYTQFVFMAG WP_030113359.1 MULTISPECIES: inhibition of morphological differentiation protein [Streptomyces] Length: 280\nNumber of predicted TMHs: 1\nExp number of AAs in TMHs: 21.68695\nExp number, first 60 AAs: 0.01801\nTotal prob of N-in: 0.00793\noutside 1 247\nTMhelix 248 270\ninside 271 280

33271 GCF_000721525.1_ASM72152v1 Streptomyces sp. NRRL F-3307 Terrabacteria group; Actinobacteria; Actinobacteria; Streptomycetales; Streptomycetaceae; Streptomyces MLRGVENHSLPRAAAFFDLDKTVIAKSSTLTFSKSFYQGGLINRRAALRTAYAQFVFLVG WP_030648527.1 MULTISPECIES: inhibition of morphological differentiation protein [Streptomyces] Length: 276\nNumber of predicted TMHs: 1\nExp number of AAs in TMHs: 21.19627\nExp number, first 60 AAs: 0.04374\nTotal prob of N-in: 0.00486\noutside 1 248\nTMhelix 249 268\ninside 269 276

33272 GCF_000721525.1_ASM72152v1 Streptomyces sp. NRRL F-3307 Terrabacteria group; Actinobacteria; Actinobacteria; Streptomycetales; Streptomycetaceae; Streptomyces MRHARRRVVRRVTRLAAVGGLLLGGAMVTQAAMASETPPASARPLAATGDTGAALVARLG WP_030640851.1 MULTISPECIES: protease [Streptomyces] Length: 458\nNumber of predicted TMHs: 2\nExp number of AAs in TMHs: 43.16453\nExp number, first 60 AAs: 22.25986\nTotal prob of N-in: 0.99239\nPOSSIBLE N-term signal sequence\ninside 1 11\nTMhelix 12 34\noutside 35 419\nTMhelix 420 442\ninside 443 458

33273 GCF_000721525.1_ASM72152v1 Streptomyces sp. NRRL F-3307 Terrabacteria group; Actinobacteria; Actinobacteria; Streptomycetales; Streptomycetaceae; Streptomyces MRTPHTARRRRALAGVLLAACLALAGCGGADDMGGSSKAASDSKAEAAGAPGGSARNGRA WP_030645059.1 MULTISPECIES: hypothetical protein [Streptomyces] Length: 324\nNumber of predicted TMHs: 2\nExp number of AAs in TMHs: 42.21572\nExp number, first 60 AAs: 19.28007\nTotal prob of N-in: 0.97332\nPOSSIBLE N-term signal sequence\ninside 1 12\nTMhelix 13 30\noutside 31 259\nTMhelix 260 282\ninside 283 324

33274 GCF_000718395.1_ASM71839v1 Streptomyces sp. NRRL F-4335 Terrabacteria group; Actinobacteria; Actinobacteria; Streptomycetales; Streptomycetaceae; Streptomyces MSTVSRRTLVRAAAVTACAGSLLALPTAAALAEGVPAASSARSAAPQPILVKSLSLADGA WP_052873759.1 hypothetical protein [Streptomyces sp. NRRL F-4335] Length: 217\nNumber of predicted TMHs: 2\nExp number of AAs in TMHs: 41.60117\nExp number, first 60 AAs: 18.98833\nTotal prob of N-in: 0.88870\nPOSSIBLE N-term signal sequence\ninside 1 8\nTMhelix 9 31\noutside 32 187\nTMhelix 188 210\ninside 211 217

33275 GCF_000718395.1_ASM71839v1 Streptomyces sp. NRRL F-4335 Terrabacteria group; Actinobacteria; Actinobacteria; Streptomycetales; Streptomycetaceae; Streptomyces MRRRSPVIAVTGAASGVGAALVSRLAASDEVKQVVAIDERRGDCAAAQWHILDVRDPAIA WP_051779201.1 MULTISPECIES: NAD-dependent dehydratase [Streptomyces] Length: 353\nNumber of predicted TMHs: 1\nExp number of AAs in TMHs: 30.20524\nExp number, first 60 AAs: 7.40836\nTotal prob of N-in: 0.39674\noutside 1 324\nTMhelix 325 347\ninside 348 353

33276 GCF_000718395.1_ASM71839v1 Streptomyces sp. NRRL F-4335 Terrabacteria group; Actinobacteria; Actinobacteria; Streptomycetales; Streptomycetaceae; Streptomyces MNRTHAPRRALLAAAFALTLASAGTGAGAGTASAAEAAPAPQPPYALRLDGAGECTFPMK WP_052872212.1 type VII secretion-associated serine protease mycosin [Streptomyces sp. NRRL F-4335] Length: 423\nNumber of predicted TMHs: 2\nExp number of AAs in TMHs: 41.2103499999999\nExp number, first 60 AAs: 18.90391\nTotal prob of N-in: 0.85156\nPOSSIBLE N-term signal sequence\ninside 1 11\nTMhelix 12 34\noutside 35 389\nTMhelix 390 412\ninside 413 423

33277	GCF_000718395.1_ASM71839v1	Streptomyces sp. NRRL F-4335	Terrabacteria group; Actinobacteria; Actinobacteria; Streptomycetales; Streptomycetaceae; Streptomyces
	MPARRRTAVTLARIASAGLAPLALAAYAAVPAVAHGSMTDPVSRVAACYAEGPESPKSAA	WP_052875733.1
	chitin-binding protein [Streptomyces sp. NRRL F-4335]	Length: 361\nNumber of predicted TMHs:  2\nExp number of AAs in TMHs: 43.29585\nExp number, first 60 AAs:  21.37525\nTotal prob of N-in:   0.95060\nPOSSIBLE N-term signal sequence\ninside   1   11\nTMhelix   12   34\noutside   35   324\nTMhelix   325   347\ninside   348   361

33278	GCF_000956015.1_ASM95601v1	Streptomyces sp. NRRL F-4428	Terrabacteria group; Actinobacteria; Actinobacteria; Streptomycetales; Streptomycetaceae; Streptomyces
	MSARRRTVTPIRIAAAGFAPLALAVYAAVPAAAHGSMTDPVSRVAACYAEGPESPKSAAC	WP_045324290.1
	chitin-binding protein [Streptomyces sp. NRRL F-4428]	Length: 366\nNumber of predicted TMHs:  2\nExp number of AAs in TMHs: 44.00081\nExp number, first 60 AAs:  21.86126\nTotal prob of N-in:   0.99272\nPOSSIBLE N-term signal sequence\ninside   1   12\nTMhelix   13   35\noutside   36   331\nTMhelix   332   354\ninside   355   366

33279	GCF_000719855.1_ASM71985v1	Streptomyces sp. NRRL F-4474	Terrabacteria group; Actinobacteria; Actinobacteria; Streptomycetales; Streptomycetaceae; Streptomyces
	MPARRRTVTPIRIAAAGLAPLALAAYAAGPAAAHGSMTDPVSRVAACYAEGPESPKSAAC	WP_030855280.1
	chitin-binding protein [Streptomyces sp. NRRL F-4474]	Length: 353\nNumber of predicted TMHs:  2\nExp number of AAs in TMHs: 41.7659\nExp number, first 60 AAs:  20.71651\nTotal prob of N-in:   0.94616\nPOSSIBLE N-term signal sequence\ninside   1   12\nTMhelix   13   35\noutside   36   318\nTMhelix   319   341\ninside   342   353

33280	GCF_000719855.1_ASM71985v1	Streptomyces sp. NRRL F-4474	Terrabacteria group; Actinobacteria; Actinobacteria; Streptomycetales; Streptomycetaceae; Streptomyces
	MSPVRRRTLVRAAAVTACAGSLLALPTAAALAEGVPAVSSAHSSAPQRTLVKSLSLADGV	WP_030853527.1
	hypothetical protein [Streptomyces sp. NRRL F-4474]	Length: 210\nNumber of predicted TMHs:  2\nExp number of AAs in TMHs: 42.68223\nExp number, first 60 AAs:  20.01388\nTotal prob of N-in:   0.93067\nPOSSIBLE N-term signal sequence\ninside   1   8\nTMhelix   9   31\noutside   32   180\nTMhelix   181   203\ninside   204   210

33281	GCF_000719855.1_ASM71985v1	Streptomyces sp. NRRL F-4474	Terrabacteria group; Actinobacteria; Actinobacteria; Streptomycetales; Streptomycetaceae; Streptomyces
	MPQNRRPRRPLLAGAFALTLAAGAAATAGAAADAATTAAAAPHPPYALRLDGAGECTFPM	WP_078095181.1
	MULTISPECIES: type VII secretion-associated serine protease mycosin [Streptomyces]	Length: 424\nNumber of predicted TMHs:  2\nExp number of AAs in TMHs: 41.33936\nExp number, first 60 AAs:  19.1446\nTotal prob of N-in:   0.88160\nPOSSIBLE N-term signal sequence\ninside   1   11\nTMhelix   12   34\noutside   35   390\nTMhelix   391   413\ninside   414   424

33282	GCF_001509485.1_ASM150948v1	Streptomyces sp. NRRL F-4489	Terrabacteria group; Actinobacteria; Actinobacteria; Streptomycetales; Streptomycetaceae; Streptomyces
	MPDRRRRAALLTSPAATAGWYPQPVTAPGAVLLPRERSSSSAPDSAPAGSGRGGRDDRDD	WP_066983488.1
	hypothetical protein [Streptomyces sp. NRRL F-4489]	Length: 309\nNumber of predicted TMHs:  2\nExp number of AAs in TMHs: 46.6180000000001\nExp number, first 60 AAs:  0.01304\nTotal prob of N-in:   0.92193\ninside   1   169\nTMhelix   170   192\noutside   193   246\nTMhelix   247   269\ninside   270   309

33283	GCF_001509485.1_ASM150948v1	Streptomyces sp. NRRL F-4489	Terrabacteria group; Actinobacteria; Actinobacteria; Streptomycetales; Streptomycetaceae; Streptomyces
	MSSFLSRRRRALRAAATSAAVAAAVLGPAAGAFAAPAAGPGGGAASVAGPTGGASSVAGH	WP_066980116.1
	hypothetical protein [Streptomyces sp. NRRL F-4489]	Length: 391\nNumber of predicted TMHs:  2\nExp number of AAs in TMHs: 39.04605\nExp number, first 60 AAs:  18.00095\nTotal prob of N-in:   0.80827\nPOSSIBLE N-term signal sequence\ninside   1   11\nTMhelix   12   34\noutside   35   358\nTMhelix   359   381\ninside   382   391

33284	GCF_001279605.1_ASM127960v1	Streptomyces sp. NRRL F-4707	Terrabacteria group; Actinobacteria; Actinobacteria; Streptomycetales; Streptomycetaceae; Streptomyces
	MPRTAAFFDLDKTVIAKSSTLTFSKSFYQGGLINRRAVLRTAYAQFVFLAGGADHDQMER	WP_037825525.1
	MULTISPECIES: inhibition of morphological differentiation protein [Streptomyces]	Length: 268\nNumber of predicted TMHs:  1\nExp number of AAs in TMHs: 21.72001\nExp number, first 60 AAs:  0.03441\nTotal prob of N-in:   0.00339\noutside   1   238\nTMhelix   239   261\ninside   262   268

33285	GCF_001279605.1_ASM127960v1	Streptomyces sp. NRRL F-4707	Terrabacteria group; Actinobacteria; Actinobacteria; Streptomycetales; Streptomycetaceae; Streptomyces
	MRHARRRIVRRVARLAAVGGLLLGATMVTRAVAGEPPGPSAAPRTLAQSASGTGAALVAR	WP_031084860.1
	MULTISPECIES: protease [Streptomyces]	Length: 447\nNumber of predicted TMHs:  2\nExp number of

AAs in TMHs: 40.53891\nExp number, first 60 AAs: 22.02945\nTotal prob of N-in: 0.98901\nPOSSIBLE N-term signal sequence\ninside 1 11\nTMhelix 12 34\noutside 35 408\nTMhelix 409 431\ninside 432 447

33286 GCF_001279355.1_ASM127935v1 Streptomyces sp. NRRL F-4711 Terrabacteria group; Actinobacteria; Actinobacteria; Streptomycetales; Streptomycetaceae; Streptomyces MPRTAAFFDLDKTVIAKSSTLTFSKSFYQGGLINRRAVLRTAYAQFVFLAGGADHDQMER WP_037825525.1 MULTISPECIES: inhibition of morphological differentiation protein [Streptomyces] Length: 268\nNumber of predicted TMHs: 1\nExp number of AAs in TMHs: 21.72001\nExp number, first 60 AAs: 0.03441\nTotal prob of N-in: 0.00339\noutside 1 238\nTMhelix 239 261\ninside 262 268

33287 GCF_001279355.1_ASM127935v1 Streptomyces sp. NRRL F-4711 Terrabacteria group; Actinobacteria; Actinobacteria; Streptomycetales; Streptomycetaceae; Streptomyces MRHARRRIVRRVARLAAVGGLLLGATMVTRAVAGEPPGPSAAPRTLAQSASGTGAALVAR WP_030402928.1 MULTISPECIES: protease [Streptomyces] Length: 447\nNumber of predicted TMHs: 2\nExp number of AAs in TMHs: 40.5389\nExp number, first 60 AAs: 22.02944\nTotal prob of N-in: 0.98901\nPOSSIBLE N-term signal sequence\ninside 1 11\nTMhelix 12 34\noutside 35 408\nTMhelix 409 431\ninside 432 447

33288 GCF_000721445.1_ASM72144v1 Streptomyces sp. NRRL F-5053 Terrabacteria group; Actinobacteria; Actinobacteria; Streptomycetales; Streptomycetaceae; Streptomyces MLGPVENVPMPRTAAFFDLDKTVIAKSSTLTFGKPFYQGGLINRRAVLRTAYAQFVYLLG WP_030886561.1 inhibition of morphological differentiation protein [Streptomyces sp. NRRL F-5053] Length: 285\nNumber of predicted TMHs: 1\nExp number of AAs in TMHs: 21.98752\nExp number, first 60 AAs: 0.01823\nTotal prob of N-in: 0.00265\noutside 1 246\nTMhelix 247 269\ninside 270 285

33289 GCF_000719815.1_ASM71981v1 Streptomyces sp. NRRL F-5065 Terrabacteria group; Actinobacteria; Actinobacteria; Streptomycetales; Streptomycetaceae; Streptomyces MNTTKARRTAVPSLFLAAALALVPAPAAHADGIRAQQWGLDALRTQEVWRTTKGAGVTVA WP_037857213.1 type VII secretion-associated serine protease [Streptomyces sp. NRRL F-5065] Length: 392\nNumber of predicted TMHs: 2\nExp number of AAs in TMHs: 38.21496\nExp number, first 60 AAs: 13.22144\nTotal prob of N-in: 0.83256\nPOSSIBLE N-term signal sequence\ninside 1 12\nTMhelix 13 30\noutside 31 359\nTMhelix 360 382\ninside 383 392

33290 GCF_000719815.1_ASM71981v1 Streptomyces sp. NRRL F-5065 Terrabacteria group; Actinobacteria; Actinobacteria; Streptomycetales; Streptomycetaceae; Streptomyces MPRTAAFFDLDKTVIAKSSTLTFSKSFYQGGLINRRAVLRTAYAQFIFLAGGADHDQMER WP_078856197.1 MULTISPECIES: inhibition of morphological differentiation protein [Streptomyces] Length: 268\nNumber of predicted TMHs: 1\nExp number of AAs in TMHs: 21.65373\nExp number, first 60 AAs: 0.08568\nTotal prob of N-in: 0.00693\noutside 1 238\nTMhelix 239 261\ninside 262 268

33291 GCF_000719815.1_ASM71981v1 Streptomyces sp. NRRL F-5065 Terrabacteria group; Actinobacteria; Actinobacteria; Streptomycetales; Streptomycetaceae; Streptomyces MGHNRRRRPTGARRATFGAFALILGGGGLVAANVIASATETSNAAVPLGGSGIAATVDCP WP_078855939.1 hypothetical protein [Streptomyces sp. NRRL F-5065] Length: 671\nNumber of predicted TMHs: 2\nExp number of AAs in TMHs: 43.52923\nExp number, first 60 AAs: 21.90243\nTotal prob of N-in: 0.99338\nPOSSIBLE N-term signal sequence\ninside 1 16\nTMhelix 17 39\noutside 40 641\nTMhelix 642 664\ninside 665 671

33292 GCF_000719815.1_ASM71981v1 Streptomyces sp. NRRL F-5065 Terrabacteria group; Actinobacteria; Actinobacteria; Streptomycetales; Streptomycetaceae; Streptomyces MGHARRRIVRRVARLAAVGGLLLGGTMVTRAVASEPSGARAPAAASASAAGAALVERLGT WP_030420451.1 MULTISPECIES: protease [Streptomyces] Length: 451\nNumber of predicted TMHs: 2\nExp number of AAs in TMHs: 40.90346\nExp number, first 60 AAs: 20.62799\nTotal prob of N-in: 0.94294\nPOSSIBLE N-term signal sequence\ninside 1 11\nTMhelix 12 34\noutside 35 412\nTMhelix 413 435\ninside 436 451

33293 GCF_001507435.1_ASM150743v1 Streptomyces sp. NRRL F-5122 Terrabacteria group; Actinobacteria; Actinobacteria; Streptomycetales; Streptomycetaceae; Streptomyces MRHARRRIVRRGARIAAVGGVLLGGLMVTQALAAEPSGAPSPGGASAQGTSRTVAGLMSQ WP_059128045.1 protease [Streptomyces sp. NRRL F-5122] Length: 465\nNumber of predicted TMHs: 2\nExp number of AAs in TMHs: 38.85787\nExp number, first 60 AAs: 22.19197\nTotal prob of N-in: 0.99349\nPOSSIBLE N-term signal sequence\ninside 1 11\nTMhelix 12 34\noutside 35 426\nTMhelix 427 449\ninside 450 465

33294 GCF_001507435.1_ASM150743v1 Streptomyces sp. NRRL F-5122 Terrabacteria group; Actinobacteria; Actinobacteria; Streptomycetales; Streptomycetaceae; Streptomyces

MSIARRATVRRLLMTGAAALTLTAVTGGAAWAGGTPGGDGWNKGGRGYASGKGPGTVTQA WP_059130410.1 hypothetical protein [Streptomyces sp. NRRL F-5122] Length: 308\nNumber of predicted TMHs: 2\nExp number of AAs in TMHs: 43.80996\nExp number, first 60 AAs: 22.38991\nTotal prob of N-in: 0.99286\nPOSSIBLE N-term signal sequence\ninside 1 11\nTMhelix 12 34\noutside 35 275\nTMhelix 276 298\ninside 299 308

33295 GCF_001507435.1_ASM150743v1 Streptomyces sp. NRRL F-5122 Terrabacteria group; Actinobacteria; Actinobacteria; Streptomycetales; Streptomycetaceae; Streptomyces MGRNSRRRPTGARRATFAAVALILGGGGLVAANVYASASESWGGGANQNGTDQVKSGTVT WP_059129719.1 hypothetical protein [Streptomyces sp. NRRL F-5122] Length: 692\nNumber of predicted TMHs: 2\nExp number of AAs in TMHs: 41.163\nExp number, first 60 AAs: 21.27247\nTotal prob of N-in: 0.98327\nPOSSIBLE N-term signal sequence\ninside 1 16\nTMhelix 17 39\noutside 40 664\nTMhelix 665 684\ninside 685 692

33296 GCF_001507435.1_ASM150743v1 Streptomyces sp. NRRL F-5122 Terrabacteria group; Actinobacteria; Actinobacteria; Streptomycetales; Streptomycetaceae; Streptomyces MASRGRRATMRGLGALLGAAALTALGANVPAWAAPPAGHGHGQSQSSTATPIQHVVVLFD WP_059127900.1 phospholipase [Streptomyces sp. NRRL F-5122] Length: 617\nNumber of predicted TMHs: 2\nExp number of AAs in TMHs: 44.11818\nExp number, first 60 AAs: 22.26799\nTotal prob of N-in: 0.98714\nPOSSIBLE N-term signal sequence\ninside 1 11\nTMhelix 12 34\noutside 35 584\nTMhelix 585 607\ninside 608 617

33297 GCF_001507435.1_ASM150743v1 Streptomyces sp. NRRL F-5122 Terrabacteria group; Actinobacteria; Actinobacteria; Streptomycetales; Streptomycetaceae; Streptomyces MRDFSPPSPHAARMSRRAVLGLTAAAPLALAAPASALTTVGGERLGRSGVQVRQPAGLPK WP_079050130.1 D-alanyl-D-alanine carboxypeptidase [Streptomyces sp. NRRL F-5122] Length: 389\nNumber of predicted TMHs: 1\nExp number of AAs in TMHs: 36.59458\nExp number, first 60 AAs: 15.07249\nTotal prob of N-in: 0.64894\nPOSSIBLE N-term signal sequence\noutside 1 352\nTMhelix 353 375\ninside 376 389

33298 GCF_001507435.1_ASM150743v1 Streptomyces sp. NRRL F-5122 Terrabacteria group; Actinobacteria; Actinobacteria; Streptomycetales; Streptomycetaceae; Streptomyces MLGPVENHSLPRTAAFFDLDKTVIAKSSTLTFSKSFYQGGLINRRAALRTAYAQFVFLAG WP_059133608.1 inhibition of morphological differentiation protein [Streptomyces sp. NRRL F-5122] Length: 274\nNumber of predicted TMHs: 1\nExp number of AAs in TMHs: 21.09341\nExp number, first 60 AAs: 0.01808\nTotal prob of N-in: 0.00500\noutside 1 244\nTMhelix 245 267\ninside 268 274

33299 GCF_000721495.1_ASM72149v1 Streptomyces sp. NRRL F-5123 Terrabacteria group; Actinobacteria; Actinobacteria; Streptomycetales; Streptomycetaceae; Streptomyces MSLRRRAALLVTAASTLTALAAPAALADTTPPAAPSTPPVTRADLYGAGDPTYDGVWRQG WP_031517753.1 hypothetical protein [Streptomyces sp. NRRL F-5123] Length: 420\nNumber of predicted TMHs: 1\nExp number of AAs in TMHs: 34.06812\nExp number, first 60 AAs: 11.50031\nTotal prob of N-in: 0.55852\nPOSSIBLE N-term signal sequence\noutside 1 391\nTMhelix 392 414\ninside 415 420

33300 GCF_000721495.1_ASM72149v1 Streptomyces sp. NRRL F-5123 Terrabacteria group; Actinobacteria; Actinobacteria; Streptomycetales; Streptomycetaceae; Streptomyces MTARTRTRRTALATAVVLAAGAAALGMPAAASAAGSKTLPLSSGTFDWSVKRSLWSYVKD WP_031524926.1 hypothetical protein [Streptomyces sp. NRRL F-5123] Length: 460\nNumber of predicted TMHs: 2\nExp number of AAs in TMHs: 43.5218099999999\nExp number, first 60 AAs: 22.11719\nTotal prob of N-in: 0.99749\nPOSSIBLE N-term signal sequence\ninside 1 11\nTMhelix 12 34\noutside 35 433\nTMhelix 434 456\ninside 457 460

33301 GCF_000721495.1_ASM72149v1 Streptomyces sp. NRRL F-5123 Terrabacteria group; Actinobacteria; Actinobacteria; Streptomycetales; Streptomycetaceae; Streptomyces MSAHDTTGRRRTLLVRSGGLLTALLAAIGLVLLPAPAGGQARAAAGTGDSSVTKSGTKGP WP_031516122.1 hypothetical protein [Streptomyces sp. NRRL F-5123] Length: 841\nNumber of predicted TMHs: 2\nExp number of AAs in TMHs: 42.41046\nExp number, first 60 AAs: 22.49828\nTotal prob of N-in: 0.99956\nPOSSIBLE N-term signal sequence\ninside 1 12\nTMhelix 13 35\noutside 36 786\nTMhelix 787 809\ninside 810 841

33302 GCF_000719775.1_ASM71977v1 Streptomyces sp. NRRL F-5135 Terrabacteria group; Actinobacteria; Actinobacteria; Streptomycetales; Streptomycetaceae; Streptomyces MIIRRRSALALCGVLTTLAVGLLPPAASAAPTADEPAGAAAPKVDLVLDVSGSMRTRDID WP_030752169.1 inter-alpha-trypsin inhibitor heavy chain H5 [Streptomyces sp. NRRL F-5135] Length: 431\nNumber of predicted TMHs: 2\nExp number of AAs in TMHs: 40.94925\nExp number, first 60 AAs: 18.70743\nTotal prob of N-in: 0.85617\nPOSSIBLE N-term signal sequence\ninside 1 6\nTMhelix 7 29\noutside 30 399\nTMhelix 400 422\ninside 423 431

33303 GCF_000719775.1_ASM71977v1 Streptomyces sp. NRRL F-5135 Terrabacteria group; Actinobacteria; Actinobacteria; Streptomycetales; Streptomycetaceae; Streptomyces
MLGPVENHSLPRTAAFFDLDKTVIAKSSTLTFSKSFYQGGLINRRAVLRTAYAQFVFLAG WP_030751128.1 inhibition of morphological differentiation protein [Streptomyces sp. NRRL F-5135] Length: 281\nNumber of predicted TMHs: 1\nExp number of AAs in TMHs: 21.86781\nExp number, first 60 AAs: 0.01989\nTotal prob of N-in: 0.00302\noutside 1 247\nTMhelix 248 270\ninside 271 281

33304 GCF_000721515.1_ASM72151v1 Streptomyces sp. NRRL F-5193 Terrabacteria group; Actinobacteria; Actinobacteria; Streptomycetales; Streptomycetaceae; Streptomyces
MSKKPNRRRTAVVLTAAVAVWAPAGLAAAPAAHAEVVDVTYECATKIGDRGAVSPVDITA WP_043232110.1 hypothetical protein [Streptomyces sp. NRRL F-5193] Length: 238\nNumber of predicted TMHs: 2\nExp number of AAs in TMHs: 42.55779\nExp number, first 60 AAs: 20.04651\nTotal prob of N-in: 0.94607\nPOSSIBLE N-term signal sequence\ninside 1 11\nTMhelix 12 34\noutside 35 206\nTMhelix 207 229\ninside 230 238

33305 GCF_000721515.1_ASM72151v1 Streptomyces sp. NRRL F-5193 Terrabacteria group; Actinobacteria; Actinobacteria; Streptomycetales; Streptomycetaceae; Streptomyces
MLSVVENHSLPRTAAFFDLDKTVIAKSSTLTFSKSFYRGGLISRRAALRTAYIQFVFLVG WP_043221120.1 inhibition of morphological differentiation protein [Streptomyces sp. NRRL F-5193] Length: 284\nNumber of predicted TMHs: 1\nExp number of AAs in TMHs: 20.94703\nExp number, first 60 AAs: 0.07763\nTotal prob of N-in: 0.00508\noutside 1 248\nTMhelix 249 268\ninside 269 284

33306 GCF_000716605.1_ASM71660v1 Streptomyces sp. NRRL F-525 Terrabacteria group; Actinobacteria; Actinobacteria; Streptomycetales; Streptomycetaceae; Streptomyces
MRHARRRVIRRVTRLAAVGGLLLGGAMVTQAAMASDSSGTSAATLSAPQTPADKGSALVA WP_033280326.1 protease [Streptomyces sp. NRRL F-525] Length: 455\nNumber of predicted TMHs: 2\nExp number of AAs in TMHs: 40.9905299999999\nExp number, first 60 AAs: 21.90209\nTotal prob of N-in: 0.98806\nPOSSIBLE N-term signal sequence\ninside 1 11\nTMhelix 12 34\noutside 35 416\nTMhelix 417 439\ninside 440 455

33307 GCF_000716605.1_ASM71660v1 Streptomyces sp. NRRL F-525 Terrabacteria group; Actinobacteria; Actinobacteria; Streptomycetales; Streptomycetaceae; Streptomyces
MRDSSRLTRRAALGLTAAALPLAAATPASAATAIGGERLARDGIQAGDASTLPKKLTARA WP_033281943.1 D-alanyl-D-alanine carboxypeptidase [Streptomyces sp. NRRL F-525] Length: 391\nNumber of predicted TMHs: 2\nExp number of AAs in TMHs: 37.9809699999999\nExp number, first 60 AAs: 20.85497\nTotal prob of N-in: 0.92021\nPOSSIBLE N-term signal sequence\ninside 1 11\nTMhelix 12 34\noutside 35 350\nTMhelix 351 373\ninside 374 391

33308 GCF_000716605.1_ASM71660v1 Streptomyces sp. NRRL F-525 Terrabacteria group; Actinobacteria; Actinobacteria; Streptomycetales; Streptomycetaceae; Streptomyces
MFISSTRRGRRELVASAVLGLLLVGATDVPAHADSTRRQQWFLDAMKAEQMWRTSTGEGV WP_033280011.1 type VII secretion-associated serine protease [Streptomyces sp. NRRL F-525] Length: 412\nNumber of predicted TMHs: 1\nExp number of AAs in TMHs: 35.0579899999999\nExp number, first 60 AAs: 12.17362\nTotal prob of N-in: 0.60981\nPOSSIBLE N-term signal sequence\noutside 1 383\nTMhelix 384 406\ninside 407 412

33309 GCF_000716605.1_ASM71660v1 Streptomyces sp. NRRL F-525 Terrabacteria group; Actinobacteria; Actinobacteria; Streptomycetales; Streptomycetaceae; Streptomyces
MLWVVENHSLPRTAAFFDLDKTVIAKSSTLTFGKSFYQGGLINRRAALRTAYAQFVFLTG WP_033286987.1 inhibition of morphological differentiation protein [Streptomyces sp. NRRL F-525] Length: 280\nNumber of predicted TMHs: 1\nExp number of AAs in TMHs: 21.57218\nExp number, first 60 AAs: 0.01999\nTotal prob of N-in: 0.00434\noutside 1 247\nTMhelix 248 270\ninside 271 280

33310 GCF_000721435.1_ASM72143v1 Streptomyces sp. NRRL F-5527 Terrabacteria group; Actinobacteria; Actinobacteria; Streptomycetales; Streptomycetaceae; Streptomyces
MRHARRRVVRRVTRLAAVGGLLLGGTMVTHAVASEPGTPDAVPFSTASPGDGAAAGPGAA WP_031063202.1 protease [Streptomyces sp. NRRL F-5527] Length: 461\nNumber of predicted TMHs: 2\nExp number of AAs in TMHs: 39.18369\nExp number, first 60 AAs: 21.83654\nTotal prob of N-in: 0.97534\nPOSSIBLE N-term signal sequence\ninside 1 11\nTMhelix 12 34\noutside 35 422\nTMhelix 423 445\ninside 446 461

33311 GCF_000721435.1_ASM72143v1 Streptomyces sp. NRRL F-5527 Terrabacteria group; Actinobacteria; Actinobacteria; Streptomycetales; Streptomycetaceae; Streptomyces
MLRNVENHLLPRTAAFFDLDKTVIAKSSTLTFSKSFYQGGLINRRAVLRTAYSQFVFLAG WP_031020047.1 MULTISPECIES: inhibition of morphological differentiation protein [Streptomyces] Length: 277\nNumber of predicted TMHs: 1\nExp

number of AAs in TMHs: 21.81477\nExp number, first 60 AAs: 0.02315\nTotal prob of N-in: 0.00436\noutside 1 247\nTMhelix 248 270\ninside 271 277

33312 GCF_000719665.1_ASM71966v1 Streptomyces sp. NRRL F-5555 Terrabacteria group; Actinobacteria; Actinobacteria; Streptomycetales; Streptomycetaceae; Streptomyces MRHARRRIVRRVARLAAVGGLLLGATMVTRAVAGEPPGPSAAPRTLAQSASGTGAALVAR WP_030402928.1 MULTISPECIES: protease [Streptomyces] Length: 447\nNumber of predicted TMHs: 2\nExp number of AAs in TMHs: 40.5389\nExp number, first 60 AAs: 22.02944\nTotal prob of N-in: 0.98901\nPOSSIBLE N-term signal sequence\ninside 1 11\nTMhelix 12 34\noutside 35 408\nTMhelix 409 431\ninside 432 447

33313 GCF_000719665.1_ASM71966v1 Streptomyces sp. NRRL F-5555 Terrabacteria group; Actinobacteria; Actinobacteria; Streptomycetales; Streptomycetaceae; Streptomyces MPRTAAFFDLDKTVIAKSSTLTFSKSFYQGGLINRRAVLRTAYAQFVFLAGGADHDQMER WP_037825525.1 MULTISPECIES: inhibition of morphological differentiation protein [Streptomyces] Length: 268\nNumber of predicted TMHs: 1\nExp number of AAs in TMHs: 21.72001\nExp number, first 60 AAs: 0.03441\nTotal prob of N-in: 0.00339\noutside 1 238\nTMhelix 239 261\ninside 262 268

33314 GCF_000719475.1_ASM71947v1 Streptomyces sp. NRRL F-5630 Terrabacteria group; Actinobacteria; Actinobacteria; Streptomycetales; Streptomycetaceae; Streptomyces MLGLVENHSLPRTAAFFDLDKTVIAKSSTLTFSKSFYQGGLINRRAALRTAYTQFVFLVG WP_030999187.1 inhibition of morphological differentiation protein [Streptomyces sp. NRRL F-5630] Length: 275\nNumber of predicted TMHs: 1\nExp number of AAs in TMHs: 21.765\nExp number, first 60 AAs: 0.01534\nTotal prob of N-in: 0.00351\noutside 1 245\nTMhelix 246 268\ninside 269 275

33315 GCF_000725645.1_Doro.v1.0Streptomyces sp. NRRL F-5635 Terrabacteria group; Actinobacteria; Actinobacteria; Streptomycetales; Streptomycetaceae; Streptomyces MRHARRRIVRRVARLAAVGGLLLGATMVTRAVAGEPPGPSAAPRTLAQSAPGTGAALVAR WP_031185450.1 MULTISPECIES: protease [Streptomyces] Length: 447\nNumber of predicted TMHs: 2\nExp number of AAs in TMHs: 40.6709\nExp number, first 60 AAs: 22.00678\nTotal prob of N-in: 0.98904\nPOSSIBLE N-term signal sequence\ninside 1 11\nTMhelix 12 34\noutside 35 408\nTMhelix 409 431\ninside 432 447

33316 GCF_000725645.1_Doro.v1.0Streptomyces sp. NRRL F-5635 Terrabacteria group; Actinobacteria; Actinobacteria; Streptomycetales; Streptomycetaceae; Streptomyces MPRTAAFFDLDKTVIAKSSTLTFSKSFYQGGLINRRAVLRTAYAQFVFLAGGADHDQMER WP_037798454.1 MULTISPECIES: inhibition of morphological differentiation protein [Streptomyces] Length: 268\nNumber of predicted TMHs: 1\nExp number of AAs in TMHs: 21.72001\nExp number, first 60 AAs: 0.03441\nTotal prob of N-in: 0.00339\noutside 1 238\nTMhelix 239 261\ninside 262 268

33317 GCF_000719425.1_ASM71942v1 Streptomyces sp. NRRL F-5639 Terrabacteria group; Actinobacteria; Actinobacteria; Streptomycetales; Streptomycetaceae; Streptomyces MSPRRRRATVLAAATVLCAAAAPVAAAAPSGTAAAGPVPAVKPAKLPAGLYGDADPKYDG WP_031028222.1 hypothetical protein [Streptomyces sp. NRRL F-5639] Length: 457\nNumber of predicted TMHs: 2\nExp number of AAs in TMHs: 40.9820499999999\nExp number, first 60 AAs: 18.73809\nTotal prob of N-in: 0.84735\nPOSSIBLE N-term signal sequence\ninside 1 6\nTMhelix 7 29\noutside 30 427\nTMhelix 428 450\ninside 451 457

33318 GCF_000719425.1_ASM71942v1 Streptomyces sp. NRRL F-5639 Terrabacteria group; Actinobacteria; Actinobacteria; Streptomycetales; Streptomycetaceae; Streptomyces MLSGVENLAMPRTAAFFDLDKTVIAKSSTLTFGKSFYQGGLINRRAVLRTAYAQFVYLLG WP_016473027.1 MULTISPECIES: haloacid dehalogenase [Streptomyces] Length: 284\nNumber of predicted TMHs: 1\nExp number of AAs in TMHs: 21.94508\nExp number, first 60 AAs: 0.01548\nTotal prob of N-in: 0.00204\noutside 1 246\nTMhelix 247 269\ninside 270 284

33319 GCF_000721365.1_ASM72136v1 Streptomyces sp. NRRL F-5650 Terrabacteria group; Actinobacteria; Actinobacteria; Streptomycetales; Streptomycetaceae; Streptomyces MPRTAAFFDLDKTVIAKSSTLTFSKSFYQGGLINRRAVLRTAYAQFVFLAGGADHDQMER WP_037836119.1 inhibition of morphological differentiation protein [Streptomyces sp. NRRL F-5650] Length: 268\nNumber of predicted TMHs: 1\nExp number of AAs in TMHs: 21.72001\nExp number, first 60 AAs: 0.03441\nTotal prob of N-in: 0.00339\noutside 1 238\nTMhelix 239 261\ninside 262 268

33320 GCF_000719585.1_ASM71958v1 Streptomyces sp. NRRL F-5681 Terrabacteria group; Actinobacteria; Actinobacteria; Streptomycetales; Streptomycetaceae; Streptomyces
MLSVVENCFSPRTAAFFDLDKTVIAKSSTLTFSKSFYQGGLINRRAVLRTAYTQFVFMAG WP_030189579.1 MULTISPECIES: inhibition of morphological differentiation protein [Streptomyces] Length: 280\nNumber of predicted TMHs: 1\nExp number of AAs in TMHs: 21.68695\nExp number, first 60 AAs: 0.01801\nTotal prob of N-in: 0.00793\noutside 1 247\nTMhelix 248 270\ninside 271 280

33321 GCF_000719585.1_ASM71958v1 Streptomyces sp. NRRL F-5681 Terrabacteria group; Actinobacteria; Actinobacteria; Streptomycetales; Streptomycetaceae; Streptomyces
MGSSWNERRLERRVRPGDGKPLKPFRWWQLTRRSLLSLALPVGGGLSATYTVEVKHGGDA WP_030189408.1 MULTISPECIES: hypothetical protein [Streptomyces] Length: 218\nNumber of predicted TMHs: 2\nExp number of AAs in TMHs: 41.32127\nExp number, first 60 AAs: 0.37701\nTotal prob of N-in: 0.97413\ninside 1 135\nTMhelix 136 158\noutside 159 182\nTMhelix 183 200\ninside 201 218

33322 GCF_000721415.1_ASM72141v1 Streptomyces sp. NRRL F-5702 Terrabacteria group; Actinobacteria; Actinobacteria; Streptomycetales; Streptomycetaceae; Streptomyces
MLSVVENCFSPRTAAFFDLDKTVIAKSSTLTFSKSFYQGGLINRRAVLRTAYTQFVFMAG WP_030113359.1 MULTISPECIES: inhibition of morphological differentiation protein [Streptomyces] Length: 280\nNumber of predicted TMHs: 1\nExp number of AAs in TMHs: 21.68695\nExp number, first 60 AAs: 0.01801\nTotal prob of N-in: 0.00793\noutside 1 247\nTMhelix 248 270\ninside 271 280

33323 GCF_000721415.1_ASM72141v1 Streptomyces sp. NRRL F-5702 Terrabacteria group; Actinobacteria; Actinobacteria; Streptomycetales; Streptomycetaceae; Streptomyces
MGPSWNERRLERRVRPGDGKPLKPFRWWQLTRRSLLSLALPVGGGLSATYTVEVKHGGDA WP_031052890.1 hypothetical protein [Streptomyces sp. NRRL F-5702] Length: 218\nNumber of predicted TMHs: 2\nExp number of AAs in TMHs: 41.37841\nExp number, first 60 AAs: 0.49329\nTotal prob of N-in: 0.96232\ninside 1 135\nTMhelix 136 158\noutside 159 182\nTMhelix 183 200\ninside 201 218

33324 GCF_000719555.1_ASM71955v1 Streptomyces sp. NRRL F-5727 Terrabacteria group; Actinobacteria; Actinobacteria; Streptomycetales; Streptomycetaceae; Streptomyces
MLSVVENHSLPRTAAFFDLDKTVIAKSSTLTFSKSFYRGGLISRRAALRTAYIQFVFLVG WP_031006574.1 inhibition of morphological differentiation protein [Streptomyces sp. NRRL F-5727] Length: 285\nNumber of predicted TMHs: 1\nExp number of AAs in TMHs: 21.571\nExp number, first 60 AAs: 0.07589\nTotal prob of N-in: 0.00491\noutside 1 247\nTMhelix 248 270\ninside 271 285

33325 GCF_001279365.1_ASM127936v1 Streptomyces sp. NRRL F-5755 Terrabacteria group; Actinobacteria; Actinobacteria; Streptomycetales; Streptomycetaceae; Streptomyces
MTERSAPAASATAPGTPARRTAVRAAATGTLAAALLGLAPQSALAATPAPPPAPAISAVT WP_053696604.1 hypothetical protein [Streptomyces sp. NRRL F-5755] Length: 293\nNumber of predicted TMHs: 1\nExp number of AAs in TMHs: 24.62822\nExp number, first 60 AAs: 5.3697\nTotal prob of N-in: 0.26855\noutside 1 264\nTMhelix 265 287\ninside 288 293

33326 GCF_001279365.1_ASM127936v1 Streptomyces sp. NRRL F-5755 Terrabacteria group; Actinobacteria; Actinobacteria; Streptomycetales; Streptomycetaceae; Streptomyces
MTLTRRRHLVRSTAVAAVAGSALLLPAAAAFADTPQPTTAVSPGADGGASRTPTEKPSQN WP_053695080.1 hypothetical protein [Streptomyces sp. NRRL F-5755] Length: 243\nNumber of predicted TMHs: 2\nExp number of AAs in TMHs: 41.24606\nExp number, first 60 AAs: 20.00589\nTotal prob of N-in: 0.94877\nPOSSIBLE N-term signal sequence\ninside 1 12\nTMhelix 13 32\noutside 33 211\nTMhelix 212 234\ninside 235 243

33327 GCF_000719735.1_ASM71973v1 Streptomyces sp. NRRL F-5917 Terrabacteria group; Actinobacteria; Actinobacteria; Streptomycetales; Streptomycetaceae; Streptomyces
MSPRRRRATVLAAATVLCAAAAPVAAAAPSGTAAAGPVPAVKPAKLPAGLYGDADPKYDG WP_030406608.1 hypothetical protein [Streptomyces sp. NRRL F-5917] Length: 444\nNumber of predicted TMHs: 2\nExp number of AAs in TMHs: 40.97887\nExp number, first 60 AAs: 18.73362\nTotal prob of N-in: 0.84743\nPOSSIBLE N-term signal sequence\ninside 1 6\nTMhelix 7 29\noutside 30 414\nTMhelix 415 437\ninside 438 444

33328 GCF_000719735.1_ASM71973v1 Streptomyces sp. NRRL F-5917 Terrabacteria group; Actinobacteria; Actinobacteria; Streptomycetales; Streptomycetaceae; Streptomyces
MLSGVENLAMPRTAAFFDLDKTVIAKSSTLTFGKSFYQGGLINRRAVLRTAYAQFVYLLG WP_016473027.1 MULTISPECIES: haloacid dehalogenase [Streptomyces] Length: 284\nNumber of predicted TMHs: 1\nExp number of AAs in TMHs:

21.94508\nExp number, first 60 AAs: 0.01548\nTotal prob of N-in: 0.00204\noutside 1 246\nTMhelix 247 269\ninside 270 284

33329 GCF_001280095.1_ASM128009v1 Streptomyces sp. NRRL F-6491 Terrabacteria group; Actinobacteria; Actinobacteria; Streptomycetales; Streptomycetaceae; Streptomyces MSAAVAARRAVVGGSAAVCLVLAGGSGTWAHGAPGGDGWEKRNAGSYKAADSVPSQDTSG WP_053642605.1 MULTISPECIES: hypothetical protein [Streptomyces] Length: 328\nNumber of predicted TMHs: 1\nExp number of AAs in TMHs: 35.08404\nExp number, first 60 AAs: 13.59263\nTotal prob of N-in: 0.62726\nPOSSIBLE N-term signal sequence\noutside 1 295\nTMhelix 296 318\ninside 319 328

33330 GCF_001280095.1_ASM128009v1 Streptomyces sp. NRRL F-6491 Terrabacteria group; Actinobacteria; Actinobacteria; Streptomycetales; Streptomycetaceae; Streptomyces MLGLVENHSLPRTAAFFDLDKTVIAKSSTLTFSKSFYRGGLISRRAALRTAYIQFVFLAG WP_053641382.1 MULTISPECIES: inhibition of morphological differentiation protein [Streptomyces] Length: 289\nNumber of predicted TMHs: 1\nExp number of AAs in TMHs: 22.10367\nExp number, first 60 AAs: 0.03062\nTotal prob of N-in: 0.00243\noutside 1 247\nTMhelix 248 270\ninside 271 289

33331 GCF_001280125.1_ASM128012v1 Streptomyces sp. NRRL F-6492 Terrabacteria group; Actinobacteria; Actinobacteria; Streptomycetales; Streptomycetaceae; Streptomyces MSAAVAARRAVVGGSAAVCLVLAGGSGTWAHGAPGGDGWEKRNAGSYKAADSVPSQDTSG WP_053642605.1 MULTISPECIES: hypothetical protein [Streptomyces] Length: 328\nNumber of predicted TMHs: 1\nExp number of AAs in TMHs: 35.08404\nExp number, first 60 AAs: 13.59263\nTotal prob of N-in: 0.62726\nPOSSIBLE N-term signal sequence\noutside 1 295\nTMhelix 296 318\ninside 319 328

33332 GCF_001280125.1_ASM128012v1 Streptomyces sp. NRRL F-6492 Terrabacteria group; Actinobacteria; Actinobacteria; Streptomycetales; Streptomycetaceae; Streptomyces MLGLVENHSLPRTAAFFDLDKTVIAKSSTLTFSKSFYRGGLISRRAALRTAYIQFVFLAG WP_053641382.1 MULTISPECIES: inhibition of morphological differentiation protein [Streptomyces] Length: 289\nNumber of predicted TMHs: 1\nExp number of AAs in TMHs: 22.10367\nExp number, first 60 AAs: 0.03062\nTotal prob of N-in: 0.00243\noutside 1 247\nTMhelix 248 270\ninside 271 289

33333 GCF_000721455.1_ASM72145v1 Streptomyces sp. NRRL F-6628 Terrabacteria group; Actinobacteria; Actinobacteria; Streptomycetales; Streptomycetaceae; Streptomyces MTPNRRPVVLAAAVATALGLGATALTLPAVAAEAPRAGAPVMDLTDGTLDWGVKESFRRY WP_037844263.1 membrane protein [Streptomyces sp. NRRL F-6628] Length: 514\nNumber of predicted TMHs: 2\nExp number of AAs in TMHs: 42.8565999999999\nExp number, first 60 AAs: 22.03087\nTotal prob of N-in: 0.97276\nPOSSIBLE N-term signal sequence\ninside 1 6\nTMhelix 7 29\noutside 30 479\nTMhelix 480 502\ninside 503 514

33334 GCF_000721455.1_ASM72145v1 Streptomyces sp. NRRL F-6628 Terrabacteria group; Actinobacteria; Actinobacteria; Streptomycetales; Streptomycetaceae; Streptomyces MPDRRSAVSLRLPLPTAAPGFGGLALPGAAATAPDAWGVVAVRESSEKERKSSSEPEGRS WP_037843057.1 MULTISPECIES: hypothetical protein [Streptomyces] Length: 292\nNumber of predicted TMHs: 2\nExp number of AAs in TMHs: 51.76527\nExp number, first 60 AAs: 6.06558\nTotal prob of N-in: 0.33682\noutside 1 164\nTMhelix 165 187\ninside 188 229\nTMhelix 230 252\noutside 253 292

33335 GCF_000721455.1_ASM72145v1 Streptomyces sp. NRRL F-6628 Terrabacteria group; Actinobacteria; Actinobacteria; Streptomycetales; Streptomycetaceae; Streptomyces MSRRTRTARRRPVAAAVLAAGAVLLPAVPASAASGADQDFVIEDPRITESSGLAASKRHP WP_030771374.1 MULTISPECIES: hypothetical protein [Streptomyces] Length: 350\nNumber of predicted TMHs: 2\nExp number of AAs in TMHs: 39.44236\nExp number, first 60 AAs: 21.1162\nTotal prob of N-in: 0.99084\nPOSSIBLE N-term signal sequence\ninside 1 12\nTMhelix 13 35\noutside 36 322\nTMhelix 323 342\ninside 343 350

33336 GCF_000721455.1_ASM72145v1 Streptomyces sp. NRRL F-6628 Terrabacteria group; Actinobacteria; Actinobacteria; Streptomycetales; Streptomycetaceae; Streptomyces MLDPVENRSLPRTAAFFDLDKTVIAKSSTLTFSKSFYQGGLINRRAVLRTAYTQFVFLAG WP_018470348.1 MULTISPECIES: inhibition of morphological differentiation protein [Streptomyces] Length: 290\nNumber of predicted TMHs: 1\nExp number of AAs in TMHs: 21.82254\nExp number, first 60 AAs: 0.02718\nTotal prob of N-in: 0.00834\noutside 1 247\nTMhelix 248 270\ninside 271 290

33337 GCF_000721455.1_ASM72145v1 Streptomyces sp. NRRL F-6628 Terrabacteria group; Actinobacteria; Actinobacteria; Streptomycetales; Streptomycetaceae; Streptomyces
MGGRGTTAQARERGRLPGRPSDPAPTGGNDVITGVNLRMPARRTALPALLLTATALLAVP WP_037845628.1
hypothetical protein [Streptomyces sp. NRRL F-6628] Length: 304\nNumber of predicted TMHs: 2\nExp number of AAs in TMHs: 39.19493\nExp number, first 60 AAs: 11.75707\nTotal prob of N-in: 0.76164\nPOSSIBLE N-term signal sequence\ninside 1 44\nTMhelix 45 67\noutside 68 276\nTMhelix 277 299\ninside 300 304

33338 GCF_000721455.1_ASM72145v1 Streptomyces sp. NRRL F-6628 Terrabacteria group; Actinobacteria; Actinobacteria; Streptomycetales; Streptomycetaceae; Streptomyces
MSRMRGSSGGRIAGRRAAVVLAVAVLGLAGCSAGPDSGASSAGDRAVAPSAADGAGGGQA WP_037844815.1
lipoprotein [Streptomyces sp. NRRL F-6628] Length: 334\nNumber of predicted TMHs: 1\nExp number of AAs in TMHs: 36.36626\nExp number, first 60 AAs: 13.57659\nTotal prob of N-in: 0.64436\nPOSSIBLE N-term signal sequence\noutside 1 269\nTMhelix 270 292\ninside 293 334

33339 GCF_000716615.1_ASM71661v1 Streptomyces sp. NRRL F-6674 Terrabacteria group; Actinobacteria; Actinobacteria; Streptomycetales; Streptomycetaceae; Streptomyces
MRDSSPLSRRALLGLTAAAPLAAAVPAATATSAYAATAIGGELLGRSGLQVRGGVDVPAK WP_051707667.1
MULTISPECIES: D-alanyl-D-alanine carboxypeptidase [Streptomyces] Length: 403\nNumber of predicted TMHs: 2\nExp number of AAs in TMHs: 37.68999\nExp number, first 60 AAs: 17.47041\nTotal prob of N-in: 0.76165\nPOSSIBLE N-term signal sequence\ninside 1 11\nTMhelix 12 34\noutside 35 355\nTMhelix 356 378\ninside 379 403

33340 GCF_000716615.1_ASM71661v1 Streptomyces sp. NRRL F-6674 Terrabacteria group; Actinobacteria; Actinobacteria; Streptomycetales; Streptomycetaceae; Streptomyces
MPASHAKNGRRRAPVAAVALLSLPALPFAVPGTAFAAPMPPSPGASASTDAAHTPGAGAS WP_030994093.1
MULTISPECIES: D-alanyl-D-alanine carboxypeptidase [Streptomyces] Length: 465\nNumber of predicted TMHs: 2\nExp number of AAs in TMHs: 43.7500699999999\nExp number, first 60 AAs: 21.56521\nTotal prob of N-in: 0.96490\nPOSSIBLE N-term signal sequence\ninside 1 12\nTMhelix 13 35\noutside 36 426\nTMhelix 427 449\ninside 450 465

33341 GCF_000716615.1_ASM71661v1 Streptomyces sp. NRRL F-6674 Terrabacteria group; Actinobacteria; Actinobacteria; Streptomycetales; Streptomycetaceae; Streptomyces
MLGPVEKHSSPRTAAFFDLDKTVIAKSSTLTFSKSFYQGGLINRRAVLRTAYAQFVFLAG WP_030378643.1 MULTISPECIES: inhibition of morphological differentiation protein [Streptomyces] Length: 277\nNumber of predicted TMHs: 1\nExp number of AAs in TMHs: 21.67091\nExp number, first 60 AAs: 0.03021\nTotal prob of N-in: 0.00251\noutside 1 247\nTMhelix 248 270\ninside 271 277

33342 GCF_000725625.1_Doro.v1.0Streptomyces sp. NRRL F-6676 Terrabacteria group; Actinobacteria; Actinobacteria; Streptomycetales; Streptomycetaceae; Streptomyces
MPASHAKNGRRRAPVAAVALLSLPALPFAVPGTAFAAPMPPSPGASASTDAAHTPGAGAS WP_030994093.1
MULTISPECIES: D-alanyl-D-alanine carboxypeptidase [Streptomyces] Length: 465\nNumber of predicted TMHs: 2\nExp number of AAs in TMHs: 43.7500699999999\nExp number, first 60 AAs: 21.56521\nTotal prob of N-in: 0.96490\nPOSSIBLE N-term signal sequence\ninside 1 12\nTMhelix 13 35\noutside 36 426\nTMhelix 427 449\ninside 450 465

33343 GCF_000725625.1_Doro.v1.0Streptomyces sp. NRRL F-6676 Terrabacteria group; Actinobacteria; Actinobacteria; Streptomycetales; Streptomycetaceae; Streptomyces
MRDSSPLSRRALLGLTAAAPLAAAVPAATATSAYAATAIGGELLGRSGLQVRGGVDVPAK WP_051707667.1
MULTISPECIES: D-alanyl-D-alanine carboxypeptidase [Streptomyces] Length: 403\nNumber of predicted TMHs: 2\nExp number of AAs in TMHs: 37.68999\nExp number, first 60 AAs: 17.47041\nTotal prob of N-in: 0.76165\nPOSSIBLE N-term signal sequence\ninside 1 11\nTMhelix 12 34\noutside 35 355\nTMhelix 356 378\ninside 379 403

33344 GCF_000725625.1_Doro.v1.0Streptomyces sp. NRRL F-6676 Terrabacteria group; Actinobacteria; Actinobacteria; Streptomycetales; Streptomycetaceae; Streptomyces
MLGPVEKHSSPRTAAFFDLDKTVIAKSSTLTFSKSFYQGGLINRRAVLRTAYAQFVFLAG WP_030378643.1 MULTISPECIES: inhibition of morphological differentiation protein [Streptomyces] Length: 277\nNumber of predicted TMHs: 1\nExp number of AAs in TMHs: 21.67091\nExp number, first 60 AAs: 0.03021\nTotal prob of N-in: 0.00251\noutside 1 247\nTMhelix 248 270\ninside 271 277

33345 GCF_000719745.1_ASM71974v1 Streptomyces sp. NRRL F-6677 Terrabacteria group; Actinobacteria; Actinobacteria; Streptomycetales; Streptomycetaceae; Streptomyces
MRDSSPLSRRALLGLTAAAPLAAAVPAATATSAYAATAIGGELLGRSGLQVRGGVDVPAK WP_051707667.1

MULTISPECIES: D-alanyl-D-alanine carboxypeptidase [Streptomyces] Length: 403\nNumber of predicted TMHs: 2\nExp number of AAs in TMHs: 37.68999\nExp number, first 60 AAs: 17.47041\nTotal prob of N-in: 0.76165\nPOSSIBLE N-term signal sequence\ninside 1 11\nTMhelix 12 34\noutside 35 355\nTMhelix 356 378\ninside 379 403

33346 GCF_000719745.1_ASM71974v1 Streptomyces sp. NRRL F-6677 Terrabacteria group; Actinobacteria; Actinobacteria; Streptomycetales; Streptomycetaceae; Streptomyces
MPASHAKNGRRRAPVAAVALLSLPALPFAVPGTAFAAPMPPSPGASASTDAAHTPGAGAS WP_030994093.1
MULTISPECIES: D-alanyl-D-alanine carboxypeptidase [Streptomyces] Length: 465\nNumber of predicted TMHs: 2\nExp number of AAs in TMHs: 43.7500699999999\nExp number, first 60 AAs: 21.56521\nTotal prob of N-in: 0.96490\nPOSSIBLE N-term signal sequence\ninside 1 12\nTMhelix 13 35\noutside 36 426\nTMhelix 427 449\ninside 450 465

33347 GCF_000719745.1_ASM71974v1 Streptomyces sp. NRRL F-6677 Terrabacteria group; Actinobacteria; Actinobacteria; Streptomycetales; Streptomycetaceae; Streptomyces
MLGPVEKHSSPRTAAFFDLDKTVIAKSSTLTFSKSFYQGGLINRRAVLRTAYAQFVFLAG WP_030378643.1 MULTISPECIES: inhibition of morphological differentiation protein [Streptomyces] Length: 277\nNumber of predicted TMHs: 1\nExp number of AAs in TMHs: 21.67091\nExp number, first 60 AAs: 0.03021\nTotal prob of N-in: 0.00251\noutside 1 247\nTMhelix 248 270\ninside 271 277

33348 GCF_001279615.1_ASM127961v1 Streptomyces sp. NRRL F-7442 Terrabacteria group; Actinobacteria; Actinobacteria; Streptomycetales; Streptomycetaceae; Streptomyces
MPRTAAFFDLDKTVIAKSSTLTFSKSFYQGGLINRRAVLRTAYAQFVFLAGGADHDQMER WP_037798454.1
MULTISPECIES: inhibition of morphological differentiation protein [Streptomyces] Length: 268\nNumber of predicted TMHs: 1\nExp number of AAs in TMHs: 21.72001\nExp number, first 60 AAs: 0.03441\nTotal prob of N-in: 0.00339\noutside 1 238\nTMhelix 239 261\ninside 262 268

33349 GCF_001279615.1_ASM127961v1 Streptomyces sp. NRRL F-7442 Terrabacteria group; Actinobacteria; Actinobacteria; Streptomycetales; Streptomycetaceae; Streptomyces
MRHARRRIVRRVARLAAVGGLLLGATMVTRAVAGEPPGPSAAPRTLAQSAPGTGAALVAR WP_031185450.1
MULTISPECIES: protease [Streptomyces] Length: 447\nNumber of predicted TMHs: 2\nExp number of AAs in TMHs: 40.6709\nExp number, first 60 AAs: 22.00678\nTotal prob of N-in: 0.98904\nPOSSIBLE N-term signal sequence\ninside 1 11\nTMhelix 12 34\noutside 35 408\nTMhelix 409 431\ninside 432 447

33350 GCF_000720135.1_ASM72013v1 Streptomyces sp. NRRL S-1022 Terrabacteria group; Actinobacteria; Actinobacteria; Streptomycetales; Streptomycetaceae; Streptomyces
MLRGVENHSLPRAAAFFDLDKTVIAKSSTLTFSKSFYQGGLINRRAALRTAYAQFVFLAG WP_030342607.1 inhibition of morphological differentiation protein [Streptomyces sp. NRRL S-1022] Length: 276\nNumber of predicted TMHs: 1\nExp number of AAs in TMHs: 21.50956\nExp number, first 60 AAs: 0.02808\nTotal prob of N-in: 0.00355\noutside 1 247\nTMhelix 248 270\ninside 271 276

33351 GCF_000720135.1_ASM72013v1 Streptomyces sp. NRRL S-1022 Terrabacteria group; Actinobacteria; Actinobacteria; Streptomycetales; Streptomycetaceae; Streptomyces
MPAPKTTGRRSLLVTSATLLSLTVTAPAALAAQAPSPSPTATPPAGMSTLGGARLGQPGT WP_030345652.1 D-alanyl-D-alanine carboxypeptidase [Streptomyces sp. NRRL S-1022] Length: 421\nNumber of predicted TMHs: 1\nExp number of AAs in TMHs: 36.40568\nExp number, first 60 AAs: 13.8689\nTotal prob of N-in: 0.64582\nPOSSIBLE N-term signal sequence\noutside 1 382\nTMhelix 383 405\ninside 406 421

33352 GCF_000720135.1_ASM72013v1 Streptomyces sp. NRRL S-1022 Terrabacteria group; Actinobacteria; Actinobacteria; Streptomycetales; Streptomycetaceae; Streptomyces
MRHARRRVVRRVTRLAAVGGLLLGGAMVAQAAMASEPPPASARPFAATDDPGAALVARLG WP_030348401.1
protease [Streptomyces sp. NRRL S-1022] Length: 458\nNumber of predicted TMHs: 2\nExp number of AAs in TMHs: 43.92907\nExp number, first 60 AAs: 22.34314\nTotal prob of N-in: 0.99589\nPOSSIBLE N-term signal sequence\ninside 1 11\nTMhelix 12 34\noutside 35 419\nTMhelix 420 442\ninside 443 458

33353 GCF_000966965.1_ASM96696v1 Streptomyces sp. NRRL S-104 Terrabacteria group; Actinobacteria; Actinobacteria; Streptomycetales; Streptomycetaceae; Streptomyces
MSTVRRRTLVRAAAVTACAGSLLALPTAAALAEGVPAASSAHSAAPQRTLVKSLSLADGV WP_030834094.1
MULTISPECIES: hypothetical protein [Streptomyces] Length: 218\nNumber of predicted TMHs: 2\nExp number of AAs in TMHs: 42.93094\nExp number, first 60 AAs: 20.33933\nTotal prob of N-in: 0.95701\nPOSSIBLE N-term signal sequence\ninside 1 8\nTMhelix 9 31\noutside 32 188\nTMhelix 189 211\ninside 212 218

33354 GCF_000966965.1_ASM96696v1 Streptomyces sp. NRRL S-104 Terrabacteria group; Actinobacteria; Actinobacteria; Streptomycetales; Streptomycetaceae; Streptomyces
MPPYRDARRALLAAAFALTLTAAGATAATAAQHPPYALRLDGAGECTFPMKKQIADRPWA WP_030823957.1
MULTISPECIES: type VII secretion-associated serine protease [Streptomyces] Length: 414\nNumber of predicted TMHs: 2\nExp number of AAs in TMHs: 42.2832099999999\nExp number, first 60 AAs: 19.94733\nTotal prob of N-in: 0.93627\nPOSSIBLE N-term signal sequence\ninside 1 11\nTMhelix 12 31\noutside 32 380\nTMhelix 381 403\ninside 404 414

33355 GCF_000716335.1_ASM71633v1 Streptomyces sp. NRRL S-118 Terrabacteria group; Actinobacteria; Actinobacteria; Streptomycetales; Streptomycetaceae; Streptomyces
MTPVMSQVPKRSTSTARRGLLCAVAAAGAWAVGISGSATAATITDPQSKQWYLNAMRADE WP_031074839.1
serine protease [Streptomyces sp. NRRL S-118] Length: 431\nNumber of predicted TMHs: 2\nExp number of AAs in TMHs: 43.10018\nExp number, first 60 AAs: 21.90514\nTotal prob of N-in: 0.96063\nPOSSIBLE N-term signal sequence\ninside 1 20\nTMhelix 21 43\noutside 44 405\nTMhelix 406 425\ninside 426 431

33356 GCF_000716335.1_ASM71633v1 Streptomyces sp. NRRL S-118 Terrabacteria group; Actinobacteria; Actinobacteria; Streptomycetales; Streptomycetaceae; Streptomyces
MSSPDPQVRAARNHAAPSAGRRPVVAVTGAASGVGDLLIRRLVASEEIKQVIAIDERRGD WP_031076788.1
NAD-dependent dehydratase [Streptomyces sp. NRRL S-118] Length: 369\nNumber of predicted TMHs: 1\nExp number of AAs in TMHs: 19.35758\nExp number, first 60 AAs: 0.04815\nTotal prob of N-in: 0.02239\noutside 1 340\nTMhelix 341 360\ninside 361 369

33357 GCF_000716335.1_ASM71633v1 Streptomyces sp. NRRL S-118 Terrabacteria group; Actinobacteria; Actinobacteria; Streptomycetales; Streptomycetaceae; Streptomyces
MLRLVENHSLPRTAAFFDLDKTVIAKSSTLTFSKSFYQGGLINRRAVLRTAYIQFVFLAG WP_031077574.1 inhibition of morphological differentiation protein [Streptomyces sp. NRRL S-118] Length: 276\nNumber of predicted TMHs: 1\nExp number of AAs in TMHs: 21.68141\nExp number, first 60 AAs: 0.08198\nTotal prob of N-in: 0.00677\noutside 1 247\nTMhelix 248 270\ninside 271 276

33358 GCF_000719785.1_ASM71978v1 Streptomyces sp. NRRL S-1314 Terrabacteria group; Actinobacteria; Actinobacteria; Streptomycetales; Streptomycetaceae; Streptomyces
MRHARRRVVRRVTRLAAVGGLLLGGTMVTHAVASEPGTPDAVPFSTASPGDGAAAGPGAA WP_031020792.1
protease [Streptomyces sp. NRRL S-1314] Length: 461\nNumber of predicted TMHs: 2\nExp number of AAs in TMHs: 39.18364\nExp number, first 60 AAs: 21.83653\nTotal prob of N-in: 0.97534\nPOSSIBLE N-term signal sequence\ninside 1 11\nTMhelix 12 34\noutside 35 422\nTMhelix 423 445\ninside 446 461

33359 GCF_000719785.1_ASM71978v1 Streptomyces sp. NRRL S-1314 Terrabacteria group; Actinobacteria; Actinobacteria; Streptomycetales; Streptomycetaceae; Streptomyces
MLRNVENHLLPRTAAFFDLDKTVIAKSSTLTFSKSFYQGGLINRRAVLRTAYSQFVFLAG WP_031020047.1 MULTISPECIES: inhibition of morphological differentiation protein [Streptomyces] Length: 277\nNumber of predicted TMHs: 1\nExp number of AAs in TMHs: 21.81477\nExp number, first 60 AAs: 0.02315\nTotal prob of N-in: 0.00436\noutside 1 247\nTMhelix 248 270\ninside 271 277

33360 GCF_000719365.1_ASM71936v1 Streptomyces sp. NRRL S-1448 Terrabacteria group; Actinobacteria; Actinobacteria; Streptomycetales; Streptomycetaceae; Streptomyces
MAHRPRRGAVALLGGLMLAVAAVAAPFQATAAGMPKAAPAAAGSGLVMVLDSSGSMSGSD WP_078865984.1
hypothetical protein [Streptomyces sp. NRRL S-1448] Length: 645\nNumber of predicted TMHs: 2\nExp number of AAs in TMHs: 44.02953\nExp number, first 60 AAs: 21.78753\nTotal prob of N-in: 0.99472\nPOSSIBLE N-term signal sequence\ninside 1 8\nTMhelix 9 31\noutside 32 601\nTMhelix 602 624\ninside 625 645

33361 GCF_000719365.1_ASM71936v1 Streptomyces sp. NRRL S-1448 Terrabacteria group; Actinobacteria; Actinobacteria; Streptomycetales; Streptomycetaceae; Streptomyces
MENHSVPHSTPRTAAFFDLDKTVIAKSSTLTFSKSFYQGGLINRRAVLRTAYAQFVFLAG WP_030412601.1 inhibition of morphological differentiation protein [Streptomyces sp. NRRL S-1448] Length: 294\nNumber of predicted TMHs: 1\nExp number of AAs in TMHs: 21.76226\nExp number, first 60 AAs: 0.06756\nTotal prob of N-in: 0.00677\noutside 1 247\nTMhelix 248 270\ninside 271 294

33362 GCF_000719365.1_ASM71936v1 Streptomyces sp. NRRL S-1448 Terrabacteria group; Actinobacteria; Actinobacteria; Streptomycetales; Streptomycetaceae; Streptomyces
MASRGRRPAIRTLGALVGAAALTVLGTTAPSWAAGHQPTPSRTATPIKHVVVLFDENISF WP_030412678.1
phospholipase C [Streptomyces sp. NRRL S-1448] Length: 612\nNumber of predicted TMHs: 2\nExp number of

AAs in TMHs: 43.17234\nExp number, first 60 AAs: 21.95766\nTotal prob of N-in: 0.97589\nPOSSIBLE N-term signal sequence\ninside 1 12\nTMhelix 13 35\noutside 36 580\nTMhelix 581 603\ninside 604 612

33363 GCF_000719365.1_ASM71936v1 Streptomyces sp. NRRL S-1448 Terrabacteria group; Actinobacteria; Actinobacteria; Streptomycetales; Streptomycetaceae; Streptomyces MPTSTSTSTPVNSARSGSGSPRPSAARRRTAVVLALLAGLLAAVVAPATAAVRTAPETARWP_030417060.1 hypothetical protein [Streptomyces sp. NRRL S-1448] Length: 174\nNumber of predicted TMHs: 2\nExp number of AAs in TMHs: 45.25934\nExp number, first 60 AAs: 22.10905\nTotal prob of N-in: 0.58843\nPOSSIBLE N-term signal sequence\noutside 1 29\nTMhelix 30 52\ninside 53 137\nTMhelix 138 160\noutside 161 174

33364 GCF_000720505.1_ASM72050v1 Streptomyces sp. NRRL S-146 Terrabacteria group; Actinobacteria; Actinobacteria; Streptomycetales; Streptomycetaceae; Streptomyces MLWGVENHSMPRTAAFFDLDKTVIAKSSTLTFSKSFYQGGLINRRAVLRTAYAQFVFLAG WP_031111211.1 inhibition of morphological differentiation protein [Streptomyces sp. NRRL S-146] Length: 277\nNumber of predicted TMHs: 1\nExp number of AAs in TMHs: 21.61697\nExp number, first 60 AAs: 0.04777\nTotal prob of N-in: 0.00483\noutside 1 247\nTMhelix 248 270\ninside 271 277

33365 GCF_000720505.1_ASM72050v1 Streptomyces sp. NRRL S-146 Terrabacteria group; Actinobacteria; Actinobacteria; Streptomycetales; Streptomycetaceae; Streptomyces MLRSSCLSGPTKNPTRRTTVRLRPSAAVLAAAVIAVAGPAVAAPVAQAEEGGPELVVSAL WP_051853789.1 hypothetical protein [Streptomyces sp. NRRL S-146] Length: 423\nNumber of predicted TMHs: 1\nExp number of AAs in TMHs: 37.5811\nExp number, first 60 AAs: 16.49931\nTotal prob of N-in: 0.74084\nPOSSIBLE N-term signal sequence\noutside 1 391\nTMhelix 392 414\ninside 415 423

33366 GCF_000722185.1_ASM72218v1 Streptomyces sp. NRRL S-15 Terrabacteria group; Actinobacteria; Actinobacteria; Streptomycetales; Streptomycetaceae; Streptomyces MLTFVENCFSPRTAAFFDLDKTVIAKSSTLTFSKSFYQGGLINRRAVLRTAYTQFVFLAG WP_031090886.1 MULTISPECIES: inhibition of morphological differentiation protein [Streptomyces] Length: 279\nNumber of predicted TMHs: 1\nExp number of AAs in TMHs: 21.53635\nExp number, first 60 AAs: 0.01474\nTotal prob of N-in: 0.00168\noutside 1 247\nTMhelix 248 270\ninside 271 279

33367 GCF_000722185.1_ASM72218v1 Streptomyces sp. NRRL S-15 Terrabacteria group; Actinobacteria; Actinobacteria; Streptomycetales; Streptomycetaceae; Streptomyces MRSSRRNTVRRGCVGALLAAVTLLTSAGGAVADDPSASASPADGESPGPTEAGTTFRTAT WP_031100002.1 MULTISPECIES: hypothetical protein [Streptomyces] Length: 275\nNumber of predicted TMHs: 2\nExp number of AAs in TMHs: 37.98301\nExp number, first 60 AAs: 18.54979\nTotal prob of N-in: 0.98601\nPOSSIBLE N-term signal sequence\ninside 1 12\nTMhelix 13 32\noutside 33 239\nTMhelix 240 262\ninside 263 275

33368 GCF_001509505.1_ASM150950v1 Streptomyces sp. NRRL S-1521 Terrabacteria group; Actinobacteria; Actinobacteria; Streptomycetales; Streptomycetaceae; Streptomyces MRITGPRRGRRRARRALVTALAATLVLALAATARADDGIRVQQWGLSAMRTGQVWQTTKG WP_062779360.1 type VII secretion-associated serine protease mycosin [Streptomyces sp. NRRL S-1521] Length: 404\nNumber of predicted TMHs: 2\nExp number of AAs in TMHs: 40.4223\nExp number, first 60 AAs: 19.87331\nTotal prob of N-in: 0.97818\nPOSSIBLE N-term signal sequence\ninside 1 12\nTMhelix 13 35\noutside 36 365\nTMhelix 366 388\ninside 389 404

33369 GCF_001509505.1_ASM150950v1 Streptomyces sp. NRRL S-1521 Terrabacteria group; Actinobacteria; Actinobacteria; Streptomycetales; Streptomycetaceae; Streptomyces MLSIVENHSLPRTAAFFDLDKTVIAKSSTLTFSKSFYQGGLINRRAVLRTAYTQFVFLAG WP_062774786.1 MULTISPECIES: inhibition of morphological differentiation protein [Streptomyces] Length: 274\nNumber of predicted TMHs: 1\nExp number of AAs in TMHs: 21.79397\nExp number, first 60 AAs: 0.03036\nTotal prob of N-in: 0.02122\noutside 1 244\nTMhelix 245 267\ninside 268 274

33370 GCF_000719605.1_ASM71960v1 Streptomyces sp. NRRL S-1813 Terrabacteria group; Actinobacteria; Actinobacteria; Streptomycetales; Streptomycetaceae; Streptomyces MRQRRRAAVASAAALVLAGAGIVAAAGTARAAEVAYKTECLPPPISGLPPIQGTTKVAVS WP_037839904.1 hypothetical protein [Streptomyces sp. NRRL S-1813] Length: 472\nNumber of predicted TMHs: 2\nExp number of AAs in TMHs: 45.0969499999999\nExp number, first 60 AAs: 22.39331\nTotal prob of N-in: 0.99870\nPOSSIBLE N-term signal sequence\ninside 1 6\nTMhelix 7 29\noutside 30 441\nTMhelix 442 464\ninside 465 472

33371 GCF_000719575.1_ASM71957v1 Streptomyces sp. NRRL S-1824 Terrabacteria group; Actinobacteria; Actinobacteria; Streptomycetales; Streptomycetaceae; Streptomyces MSPIGRRRWLHVSCGGSTILNSNTFRLAALAVAAAPVALLVAAPAHATGATTTGGGKAS WP_078852820.1 hypothetical protein [Streptomyces sp. NRRL S-1824] Length: 336\nNumber of predicted TMHs: 1\nExp number of AAs in TMHs: 33.92281\nExp number, first 60 AAs: 13.21814\nTotal prob of N-in: 0.59095\nPOSSIBLE N-term signal sequence\noutside 1 306\nTMhelix 307 329\ninside 330 336

33372 GCF_000719575.1_ASM71957v1 Streptomyces sp. NRRL S-1824 Terrabacteria group; Actinobacteria; Actinobacteria; Streptomycetales; Streptomycetaceae; Streptomyces MLCLVENHSLPRTAAFFDLDKTVIAKSSTLTFSKSFYQGGLINRRAVLRTAYAQFVFLAG WP_030972213.1 inhibition of morphological differentiation protein [Streptomyces sp. NRRL S-1824] Length: 279\nNumber of predicted TMHs: 1\nExp number of AAs in TMHs: 21.3107\nExp number, first 60 AAs: 0.02047\nTotal prob of N-in: 0.00230\noutside 1 247\nTMhelix 248 270\ninside 271 279

33373 GCF_000721335.1_ASM72133v1 Streptomyces sp. NRRL S-1831 Terrabacteria group; Actinobacteria; Actinobacteria; Streptomycetales; Streptomycetaceae; Streptomyces MPRTAAFFDLDKTVIAKSSTLTFSKSFYQGGLINRRAVLRTAYAQFVFLAGGADHDQMER WP_037832651.1 inhibition of morphological differentiation protein [Streptomyces sp. NRRL S-1831] Length: 267\nNumber of predicted TMHs: 1\nExp number of AAs in TMHs: 20.58486\nExp number, first 60 AAs: 0.03487\nTotal prob of N-in: 0.00369\noutside 1 239\nTMhelix 240 259\ninside 260 267

33374 GCF_000721335.1_ASM72133v1 Streptomyces sp. NRRL S-1831 Terrabacteria group; Actinobacteria; Actinobacteria; Streptomycetales; Streptomycetaceae; Streptomyces MRHARRRIVRRVARLAAVGGLLLGATMVTRAVAGEPPGPSAAPRTLAQSASGTGAALVAR WP_031084860.1 MULTISPECIES: protease [Streptomyces] Length: 447\nNumber of predicted TMHs: 2\nExp number of AAs in TMHs: 40.53891\nExp number, first 60 AAs: 22.02945\nTotal prob of N-in: 0.98901\nPOSSIBLE N-term signal sequence\ninside 1 11\nTMhelix 12 34\noutside 35 408\nTMhelix 409 431\ninside 432 447

33375 GCF_000721305.1_ASM72130v1 Streptomyces sp. NRRL S-1868 Terrabacteria group; Actinobacteria; Actinobacteria; Streptomycetales; Streptomycetaceae; Streptomyces MLGPVENVPMPRTAAFFDLDKTVIAKSSTLTFGKPFYQGGLINRRAVLRTAYAQFVYLLG WP_030883110.1 inhibition of morphological differentiation protein [Streptomyces sp. NRRL S-1868] Length: 285\nNumber of predicted TMHs: 1\nExp number of AAs in TMHs: 21.98851\nExp number, first 60 AAs: 0.01836\nTotal prob of N-in: 0.00267\noutside 1 246\nTMhelix 247 269\ninside 270 285

33376 GCF_000725655.1_Doro.v1.0 Streptomyces sp. NRRL S-1896 Terrabacteria group; Actinobacteria; Actinobacteria; Streptomycetales; Streptomycetaceae; Streptomyces MRTPHTARRRRALAGVLLAACLALAGCGGADDMGGSSKAASDSKAEAAGAPGGSARNGRA WP_031095397.1 MULTISPECIES: hypothetical protein [Streptomyces] Length: 324\nNumber of predicted TMHs: 2\nExp number of AAs in TMHs: 42.45736\nExp number, first 60 AAs: 19.4804\nTotal prob of N-in: 0.97333\nPOSSIBLE N-term signal sequence\ninside 1 12\nTMhelix 13 30\noutside 31 259\nTMhelix 260 282\ninside 283 324

33377 GCF_000725655.1_Doro.v1.0 Streptomyces sp. NRRL S-1896 Terrabacteria group; Actinobacteria; Actinobacteria; Streptomycetales; Streptomycetaceae; Streptomyces MRHARRRVVRRVTRLAAVGGLLLGGAMVTQAAMASETPPASARPLAATGDTGAALVARLG WP_031092578.1 MULTISPECIES: protease [Streptomyces] Length: 457\nNumber of predicted TMHs: 2\nExp number of AAs in TMHs: 43.16462\nExp number, first 60 AAs: 22.25978\nTotal prob of N-in: 0.99239\nPOSSIBLE N-term signal sequence\ninside 1 11\nTMhelix 12 34\noutside 35 418\nTMhelix 419 441\ninside 442 457

33378 GCF_000725655.1_Doro.v1.0 Streptomyces sp. NRRL S-1896 Terrabacteria group; Actinobacteria; Actinobacteria; Streptomycetales; Streptomycetaceae; Streptomyces MLRGVENHSLPRAAAFFDLDKTVIAKSSTLTFSKSFYQGGLINRRAALRTAYAQFVFLVG WP_031092052.1 MULTISPECIES: inhibition of morphological differentiation protein [Streptomyces] Length: 276\nNumber of predicted TMHs: 1\nExp number of AAs in TMHs: 21.19627\nExp number, first 60 AAs: 0.04374\nTotal prob of N-in: 0.00486\noutside 1 248\nTMhelix 249 268\ninside 269 276

33379 GCF_000718935.1_ASM71893v1 Streptomyces sp. NRRL S-237 Terrabacteria group; Actinobacteria; Actinobacteria; Streptomycetales; Streptomycetaceae; Streptomyces MPARRRRTAVTLTRIAAAGLAPLAVAAYAAAPAVAHGSMTDPVSRVAACYAEGPESPKSA WP_030723165.1 chitin-binding protein [Streptomyces sp. NRRL S-237] Length: 359\nNumber of predicted TMHs: 2\nExp

number of AAs in TMHs: 43.41228\nExp number, first 60 AAs:  21.5805\nTotal prob of N-in:   0.96980\nPOSSIBLE N-term signal sequence\ninside   1   11\nTMhelix   12   34\noutside   35   322\nTMhelix   323   345\ninside   346   359

33380 GCF_000718935.1_ASM71893v1 Streptomyces sp. NRRL S-237 Terrabacteria group; Actinobacteria; Actinobacteria; Streptomycetales; Streptomycetaceae; Streptomyces MSSPDPQFRAMPDDENSSEHGESAPDVRRPRNPEGMRRRSPVIAVTGAASGVGAALVSRL WP_030711692.1 NAD-dependent dehydratase [Streptomyces sp. NRRL S-237] Length: 388\nNumber of predicted TMHs: 1\nExp number of AAs in TMHs: 23.52632\nExp number, first 60 AAs:  1.26426\nTotal prob of N-in:   0.06307\noutside   1   359\nTMhelix   360   382\ninside   383   388

33381 GCF_000718935.1_ASM71893v1 Streptomyces sp. NRRL S-237 Terrabacteria group; Actinobacteria; Actinobacteria; Streptomycetales; Streptomycetaceae; Streptomyces MTQTQASRRALLAAAFALTLASGGTGTAAAATGAAPQPPYALRLDGAGECTFPMKKQIED WP_030718710.1 type VII secretion-associated serine protease mycosin [Streptomyces sp. NRRL S-237] Length: 418\nNumber of predicted TMHs:  2\nExp number of AAs in TMHs: 44.65104\nExp number, first 60 AAs:  22.33517\nTotal prob of N-in: 0.98107\nPOSSIBLE N-term signal sequence\ninside   1   11\nTMhelix   12   34\noutside   35   384\nTMhelix   385   407\ninside   408   418

33382 GCF_000718935.1_ASM71893v1 Streptomyces sp. NRRL S-237 Terrabacteria group; Actinobacteria; Actinobacteria; Streptomycetales; Streptomycetaceae; Streptomyces MSTVSRRTLVRAAAVTACAGSLLALPTAAALADGVPAASSAHSAAPQSILVKSLALADGVWP_030721008.1 hypothetical protein [Streptomyces sp. NRRL S-237] Length: 218\nNumber of predicted TMHs:  2\nExp number of AAs in TMHs: 42.759\nExp number, first 60 AAs:  20.14498\nTotal prob of N-in:   0.93966\nPOSSIBLE N-term signal sequence\ninside   1   8\nTMhelix   9   31\noutside   32   188\nTMhelix   189   211\ninside   212   218

33383 GCF_000717925.1_ASM71792v1 Streptomyces sp. NRRL S-241 Terrabacteria group; Actinobacteria; Actinobacteria; Streptomycetales; Streptomycetaceae; Streptomyces MSTVSRRTLVRAAAVTACAGSLLALPTAAALAEGVPAASSARSAAPQLSLVKSLSLADGT WP_030386983.1 hypothetical protein [Streptomyces sp. NRRL S-241] Length: 217\nNumber of predicted TMHs:  2\nExp number of AAs in TMHs: 41.60696\nExp number, first 60 AAs:  18.99598\nTotal prob of N-in:   0.88897\nPOSSIBLE N-term signal sequence\ninside   1   8\nTMhelix   9   31\noutside   32   187\nTMhelix   188   210\ninside   211   217

33384 GCF_000717925.1_ASM71792v1 Streptomyces sp. NRRL S-241 Terrabacteria group; Actinobacteria; Actinobacteria; Streptomycetales; Streptomycetaceae; Streptomyces MRRRSPVIAVTGAASGVGAALVSRLAASDEVKQVVAIDERRGDCAAAQWHILDVRDPAIA WP_051779201.1 MULTISPECIES: NAD-dependent dehydratase [Streptomyces] Length: 353\nNumber of predicted TMHs: 1\nExp number of AAs in TMHs: 30.20524\nExp number, first 60 AAs:  7.40836\nTotal prob of N-in:   0.39674\noutside   1   324\nTMhelix   325   347\ninside   348   353

33385 GCF_000717925.1_ASM71792v1 Streptomyces sp. NRRL S-241 Terrabacteria group; Actinobacteria; Actinobacteria; Streptomycetales; Streptomycetaceae; Streptomyces MNRTHAPRRPLLAAAFALTFTFTLASAGAGAGAASAAEAAPAPQPPYGQAPYGQAPYGLR WP_078626310.1 type VII secretion-associated serine protease mycosin [Streptomyces sp. NRRL S-241] Length: 435\nNumber of predicted TMHs:  2\nExp number of AAs in TMHs: 44.30371\nExp number, first 60 AAs:  22.00943\nTotal prob of N-in: 0.97370\nPOSSIBLE N-term signal sequence\ninside   1   11\nTMhelix   12   34\noutside   35   401\nTMhelix   402   424\ninside   425   435

33386 GCF_000717925.1_ASM71792v1 Streptomyces sp. NRRL S-241 Terrabacteria group; Actinobacteria; Actinobacteria; Streptomycetales; Streptomycetaceae; Streptomyces MPARRRTAVVLTRIASAGLAPLALAAYAAAPAVAHGSMTDPVSRVAACYAEGPESPKTAA WP_030388762.1 chitin-binding protein [Streptomyces sp. NRRL S-241] Length: 359\nNumber of predicted TMHs:  2\nExp number of AAs in TMHs: 42.99055\nExp number, first 60 AAs:  21.0956\nTotal prob of N-in:   0.93766\nPOSSIBLE N-term signal sequence\ninside   1   8\nTMhelix   9   31\noutside   32   322\nTMhelix   323   345\ninside   346   359

33387 GCF_000717915.1_ASM71791v1 Streptomyces sp. NRRL S-244 Terrabacteria group; Actinobacteria; Actinobacteria; Streptomycetales; Streptomycetaceae; Streptomyces MTHDPRPTTYEYEKSPHMRSTTRRTTLRTAAVSAGAAAVLALPVGSAFADSSAGPEQEVL WP_030152910.1 hypothetical protein [Streptomyces sp. NRRL S-244] Length: 375\nNumber of predicted TMHs:  1\nExp number of AAs in TMHs: 35.30013\nExp number, first 60 AAs:  15.08312\nTotal prob of N-in:   0.70707\nPOSSIBLE N-term signal sequence\noutside   1   344\nTMhelix   345   367\ninside   368   375

33388 GCF_000717915.1_ASM71791v1 Streptomyces sp. NRRL S-244 Terrabacteria group; Actinobacteria; Actinobacteria; Streptomycetales; Streptomycetaceae; Streptomyces
MAEAADIQGAPPAPARRRWLRRAVVLLAGTPVLAALVYSPAPQAAQAADAGTVDVQLDGM WP_030153186.1 hypothetical protein [Streptomyces sp. NRRL S-244] Length: 755\nNumber of predicted TMHs: 1\nExp number of AAs in TMHs: 36.36037\nExp number, first 60 AAs: 14.65303\nTotal prob of N-in: 0.70793\nPOSSIBLE N-term signal sequence\noutside 1 688\nTMhelix 689 711\ninside 712 755

33389 GCF_000717915.1_ASM71791v1 Streptomyces sp. NRRL S-244 Terrabacteria group; Actinobacteria; Actinobacteria; Streptomycetales; Streptomycetaceae; Streptomyces
MSTVSRRTLVRAAAVTVCAGTLLALPAAAALAEGLPAASAARSAGAPRTLVKSLALADGV WP_030153679.1 hypothetical protein [Streptomyces sp. NRRL S-244] Length: 200\nNumber of predicted TMHs: 2\nExp number of AAs in TMHs: 44.19665\nExp number, first 60 AAs: 21.60089\nTotal prob of N-in: 0.98812\nPOSSIBLE N-term signal sequence\ninside 1 8\nTMhelix 9 31\noutside 32 170\nTMhelix 171 193\ninside 194 200

33390 GCF_000718775.1_ASM71877v1 Streptomyces sp. NRRL S-31 Terrabacteria group; Actinobacteria; Actinobacteria; Streptomycetales; Streptomycetaceae; Streptomyces
MLRGVENHSLPRAAAFFDLDKTVIAKSSTLTFSKSFYQGGLINRRAALRTAYAQFVFLAG WP_030734150.1 inhibition of morphological differentiation protein [Streptomyces sp. NRRL S-31] Length: 276\nNumber of predicted TMHs: 1\nExp number of AAs in TMHs: 21.78494\nExp number, first 60 AAs: 0.02923\nTotal prob of N-in: 0.00487\noutside 1 247\nTMhelix 248 270\ninside 271 276

33391 GCF_000718775.1_ASM71877v1 Streptomyces sp. NRRL S-31 Terrabacteria group; Actinobacteria; Actinobacteria; Streptomycetales; Streptomycetaceae; Streptomyces
MTRSTGRRRRAVASGAAVSLLLAGSLALLPSTEARADGLRGQQWGLSALHLDEVWQTTKG WP_030744629.1 type VII secretion-associated serine protease [Streptomyces sp. NRRL S-31] Length: 395\nNumber of predicted TMHs: 2\nExp number of AAs in TMHs: 39.9869699999999\nExp number, first 60 AAs: 16.34886\nTotal prob of N-in: 0.93100\nPOSSIBLE N-term signal sequence\ninside 1 11\nTMhelix 12 31\noutside 32 365\nTMhelix 366 388\ninside 389 395

33392 GCF_000718775.1_ASM71877v1 Streptomyces sp. NRRL S-31 Terrabacteria group; Actinobacteria; Actinobacteria; Streptomycetales; Streptomycetaceae; Streptomyces
MRHARRRVVRRVTRLAAIGGLLLGGAMVTQAAMASGTPPVSAAAPGSAGGTGSTGAALVA WP_030749214.1 protease [Streptomyces sp. NRRL S-31] Length: 459\nNumber of predicted TMHs: 2\nExp number of AAs in TMHs: 43.5458599999999\nExp number, first 60 AAs: 22.2712\nTotal prob of N-in: 0.99429\nPOSSIBLE N-term signal sequence\ninside 1 11\nTMhelix 12 34\noutside 35 420\nTMhelix 421 443\ninside 444 459

33393 GCF_000721595.1_ASM72159v1 Streptomyces sp. NRRL S-325 Terrabacteria group; Actinobacteria; Actinobacteria; Streptomycetales; Streptomycetaceae; Streptomyces
MTTHPSPRRRLLTAAAATAVLLVTLPVLPAAAEDSTQCTFPSKKYAGRPWSLQRVLMDEL WP_014153729.1 MULTISPECIES: type VII secretion-associated serine protease [Streptomyces] Length: 414\nNumber of predicted TMHs: 1\nExp number of AAs in TMHs: 36.07361\nExp number, first 60 AAs: 13.5063\nTotal prob of N-in: 0.65550\nPOSSIBLE N-term signal sequence\noutside 1 378\nTMhelix 379 401\ninside 402 414

33394 GCF_000721595.1_ASM72159v1 Streptomyces sp. NRRL S-325 Terrabacteria group; Actinobacteria; Actinobacteria; Streptomycetales; Streptomycetaceae; Streptomyces
MRTTRRNTLRRGGVGALLAAVTLLTSAGAAVADDPSPSASADGGAAGPTEAGTTFRTATA WP_014152884.1 MULTISPECIES: hypothetical protein [Streptomyces] Length: 274\nNumber of predicted TMHs: 2\nExp number of AAs in TMHs: 40.1724799999999\nExp number, first 60 AAs: 19.30637\nTotal prob of N-in: 0.99359\nPOSSIBLE N-term signal sequence\ninside 1 12\nTMhelix 13 32\noutside 33 237\nTMhelix 238 260\ninside 261 274

33395 GCF_000721595.1_ASM72159v1 Streptomyces sp. NRRL S-325 Terrabacteria group; Actinobacteria; Actinobacteria; Streptomycetales; Streptomycetaceae; Streptomyces
MLTFVENCFSPRTAAFFDLDKTVIAKSSTLTFSKSFYQGGLINRRAVLRTAYTQFVFLAG WP_037829548.1 inhibition of morphological differentiation protein [Streptomyces sp. NRRL S-325] Length: 279\nNumber of predicted TMHs: 1\nExp number of AAs in TMHs: 21.11388\nExp number, first 60 AAs: 0.0147\nTotal prob of N-in: 0.00170\noutside 1 248\nTMhelix 249 268\ninside 269 279

33396 GCF_000720175.1_ASM72017v1 Streptomyces sp. NRRL S-337 Terrabacteria group; Actinobacteria; Actinobacteria; Streptomycetales; Streptomycetaceae; Streptomyces
MENHSVPHSTPRTAAFFDLDKTVIAKSSTLTFSKSFYQGGLINRRAVLRTAYAQFVFLAG WP_030801458.1 inhibition of

morphological differentiation protein [Streptomyces sp. NRRL S-337] Length: 294\nNumber of predicted TMHs: 1\nExp number of AAs in TMHs: 21.76682\nExp number, first 60 AAs: 0.06748\nTotal prob of N-in: 0.00668\noutside 1 247\nTMhelix 248 270\ninside 271 294

33397 GCF_000720175.1_ASM72017v1 Streptomyces sp. NRRL S-337 Terrabacteria group; Actinobacteria; Actinobacteria; Streptomycetales; Streptomycetaceae; Streptomyces MASRGRRPAIRTLGALAGAAALTVLGTTAPSWAAGHQPTPSRTATPIKHVVVLFDENISF WP_030804226.1 phospholipase C [Streptomyces sp. NRRL S-337] Length: 612\nNumber of predicted TMHs: 2\nExp number of AAs in TMHs: 41.38115\nExp number, first 60 AAs: 21.4618\nTotal prob of N-in: 0.95333\nPOSSIBLE N-term signal sequence\ninside 1 12\nTMhelix 13 35\noutside 36 580\nTMhelix 581 603\ninside 604 612

33398 GCF_000720175.1_ASM72017v1 Streptomyces sp. NRRL S-337 Terrabacteria group; Actinobacteria; Actinobacteria; Streptomycetales; Streptomycetaceae; Streptomyces MPVNSVRSGSGSPRPSAARRRTAVVLALLAGLLAAAVAPATAAVRTAPETVRTATVRTES WP_030814550.1 hypothetical protein [Streptomyces sp. NRRL S-337] Length: 177\nNumber of predicted TMHs: 2\nExp number of AAs in TMHs: 45.14851\nExp number, first 60 AAs: 22.0508\nTotal prob of N-in: 0.66642\nPOSSIBLE N-term signal sequence\noutside 1 21\nTMhelix 22 44\ninside 45 140\nTMhelix 141 163\noutside 164 177

33399 GCF_000720185.1_ASM72018v1 Streptomyces sp. NRRL S-340 Terrabacteria group; Actinobacteria; Actinobacteria; Streptomycetales; Streptomycetaceae; Streptomyces MLDVVENHSLPRTAAFFDLDKTVIAKSSTLTFSKSFYQGGLINRRAVLRTAYAQFVFLAG WP_037850574.1 inhibition of morphological differentiation protein [Streptomyces sp. NRRL S-340] Length: 277\nNumber of predicted TMHs: 1\nExp number of AAs in TMHs: 21.55629\nExp number, first 60 AAs: 0.03599\nTotal prob of N-in: 0.00601\noutside 1 247\nTMhelix 248 270\ninside 271 277

33400 GCF_000720185.1_ASM72018v1 Streptomyces sp. NRRL S-340 Terrabacteria group; Actinobacteria; Actinobacteria; Streptomycetales; Streptomycetaceae; Streptomyces MPAAFRTTRRSLLVTSAALLSVSLTAPAALAAPTAPKTPSATPPAHMSGVGGARLGLPGT WP_037861009.1 D-alanyl-D-alanine carboxypeptidase [Streptomyces sp. NRRL S-340] Length: 422\nNumber of predicted TMHs: 2\nExp number of AAs in TMHs: 39.5465499999999\nExp number, first 60 AAs: 17.21371\nTotal prob of N-in: 0.79451\nPOSSIBLE N-term signal sequence\ninside 1 11\nTMhelix 12 34\noutside 35 383\nTMhelix 384 406\ninside 407 422

33401 GCF_000719705.1_ASM71970v1 Streptomyces sp. NRRL S-350 Terrabacteria group; Actinobacteria; Actinobacteria; Streptomycetales; Streptomycetaceae; Streptomyces MHGKSTSGARADLRRSRRATVLVLGAAALLSLAGPVHAAPAKDPLPADIPDYQVALDLVK WP_030231344.1 hypothetical protein [Streptomyces sp. NRRL S-350] Length: 289\nNumber of predicted TMHs: 2\nExp number of AAs in TMHs: 40.88755\nExp number, first 60 AAs: 19.77948\nTotal prob of N-in: 0.98649\nPOSSIBLE N-term signal sequence\ninside 1 19\nTMhelix 20 39\noutside 40 258\nTMhelix 259 281\ninside 282 289

33402 GCF_000719705.1_ASM71970v1 Streptomyces sp. NRRL S-350 Terrabacteria group; Actinobacteria; Actinobacteria; Streptomycetales; Streptomycetaceae; Streptomyces MRTAAFFDLDKTIIAKSSALAFSRPFYQGGLINRRAVLRSAYAQFVFLVGGADHDQMEKM WP_030240935.1 inhibition of morphological differentiation protein [Streptomyces sp. NRRL S-350] Length: 265\nNumber of predicted TMHs: 1\nExp number of AAs in TMHs: 22.65233\nExp number, first 60 AAs: 0.09228\nTotal prob of N-in: 0.01601\noutside 1 235\nTMhelix 236 258\ninside 259 265

33403 GCF_000721605.1_ASM72160v1 Streptomyces sp. NRRL S-37 Terrabacteria group; Actinobacteria; Actinobacteria; Streptomycetales; Streptomycetaceae; Streptomyces MLKDVENHSMPRTAAFFDLDKTVIAKSSTLTFSKSFYQGGLINRRAVLRTAYSQFVFLAG WP_030863277.1 inhibition of morphological differentiation protein [Streptomyces sp. NRRL S-37] Length: 277\nNumber of predicted TMHs: 1\nExp number of AAs in TMHs: 21.92108\nExp number, first 60 AAs: 0.04798\nTotal prob of N-in: 0.00652\noutside 1 247\nTMhelix 248 270\ninside 271 277

33404 GCF_000721605.1_ASM72160v1 Streptomyces sp. NRRL S-37 Terrabacteria group; Actinobacteria; Actinobacteria; Streptomycetales; Streptomycetaceae; Streptomyces MRHARRRVVRRVTRLAAVGGLLLGATMVTHAVAGEPRAPGAVPFATAGEAAAGPGAALVA WP_030863718.1 protease [Streptomyces sp. NRRL S-37] Length: 458\nNumber of predicted TMHs: 2\nExp number of AAs in TMHs: 41.52598\nExp number, first 60 AAs: 22.51222\nTotal prob of N-in: 0.99487\nPOSSIBLE N-term signal sequence\ninside 1 11\nTMhelix 12 34\noutside 35 419\nTMhelix 420 442\ninside 443 458

33405 GCF_000720645.1_ASM72064v1 Streptomyces sp. NRRL S-378 Terrabacteria group; Actinobacteria; Actinobacteria; Streptomycetales; Streptomycetaceae; Streptomyces
MSARRRTVTPTRIAAAGVAPLALVAYAAVPAAAHGSMTDPVSRVAACYAEGPESPKSAAC WP_030963406.1 chitin-binding protein [Streptomyces sp. NRRL S-378] Length: 357\nNumber of predicted TMHs: 2\nExp number of AAs in TMHs: 43.46732\nExp number, first 60 AAs: 21.78466\nTotal prob of N-in: 0.99137\nPOSSIBLE N-term signal sequence\ninside 1 12\nTMhelix 13 35\noutside 36 322\nTMhelix 323 345\ninside 346 357

33406 GCF_000720645.1_ASM72064v1 Streptomyces sp. NRRL S-378 Terrabacteria group; Actinobacteria; Actinobacteria; Streptomycetales; Streptomycetaceae; Streptomyces
MSTVRRRTLVRAAAVTACAGSLLALPTAAALAEGVPAASSAHSTASQRTLVKSLALADGV WP_030964283.1 hypothetical protein [Streptomyces sp. NRRL S-378] Length: 210\nNumber of predicted TMHs: 2\nExp number of AAs in TMHs: 43.31967\nExp number, first 60 AAs: 20.82626\nTotal prob of N-in: 0.96498\nPOSSIBLE N-term signal sequence\ninside 1 8\nTMhelix 9 31\noutside 32 180\nTMhelix 181 203\ninside 204 210

33407 GCF_000720645.1_ASM72064v1 Streptomyces sp. NRRL S-378 Terrabacteria group; Actinobacteria; Actinobacteria; Streptomycetales; Streptomycetaceae; Streptomyces
MPRRAVLAGAFVLALTAGGAAGTAGAATGAAPAAATAPHPPYALRLDGAGECTFPMKKQI WP_030958662.1 type VII secretion-associated serine protease mycosin [Streptomyces sp. NRRL S-378] Length: 420\nNumber of predicted TMHs: 2\nExp number of AAs in TMHs: 44.35622\nExp number, first 60 AAs: 22.02457\nTotal prob of N-in: 0.96003\nPOSSIBLE N-term signal sequence\ninside 1 4\nTMhelix 5 27\noutside 28 386\nTMhelix 387 409\ninside 410 420

33408 GCF_000721625.1_ASM72162v1 Streptomyces sp. NRRL S-384 Terrabacteria group; Actinobacteria; Actinobacteria; Streptomycetales; Streptomycetaceae; Streptomyces
MIAKSSALAFSRPFYQGGLINRRSVVKSAYTQFIFLVGGADHDQMEKMRAYLSALTRGWN WP_051827913.1 inhibition of morphological differentiation protein [Streptomyces sp. NRRL S-384] Length: 254\nNumber of predicted TMHs: 1\nExp number of AAs in TMHs: 21.22202\nExp number, first 60 AAs: 0.03004\nTotal prob of N-in: 0.00980\noutside 1 223\nTMhelix 224 246\ninside 247 254

33409 GCF_001294345.1_ASM129434v1 Streptomyces sp. NRRL S-4 Terrabacteria group; Actinobacteria; Actinobacteria; Streptomycetales; Streptomycetaceae; Streptomyces
MLTFVENCFSPRTAAFFDLDKTVIAKSSTLTFSKSFYQGGLINRRAVLRTAYTQFVFLAG WP_031090886.1 MULTISPECIES: inhibition of morphological differentiation protein [Streptomyces] Length: 279\nNumber of predicted TMHs: 1\nExp number of AAs in TMHs: 21.53635\nExp number, first 60 AAs: 0.01474\nTotal prob of N-in: 0.00168\noutside 1 247\nTMhelix 248 270\ninside 271 279

33410 GCF_001294345.1_ASM129434v1 Streptomyces sp. NRRL S-4 Terrabacteria group; Actinobacteria; Actinobacteria; Streptomycetales; Streptomycetaceae; Streptomyces
MRSSRRNTVRRGCVGALLAAVTLLTSAGGAVADDPSASASPADGESPGPTEAGTTFRTAT WP_031100002.1 MULTISPECIES: hypothetical protein [Streptomyces] Length: 275\nNumber of predicted TMHs: 2\nExp number of AAs in TMHs: 37.98301\nExp number, first 60 AAs: 18.54979\nTotal prob of N-in: 0.98601\nPOSSIBLE N-term signal sequence\ninside 1 12\nTMhelix 13 32\noutside 33 239\nTMhelix 240 262\ninside 263 275

33411 GCF_000717875.1_ASM71787v1 Streptomyces sp. NRRL S-455 Terrabacteria group; Actinobacteria; Actinobacteria; Streptomycetales; Streptomycetaceae; Streptomyces
MLRGVENHSLPRTAAFFDLDKTVIAKSSTLTFSKSFYQGGLINRRAVLRTAYAQFVFLAG WP_030235813.1 MULTISPECIES: inhibition of morphological differentiation protein [Streptomyces] Length: 277\nNumber of predicted TMHs: 1\nExp number of AAs in TMHs: 21.66866\nExp number, first 60 AAs: 0.03762\nTotal prob of N-in: 0.00406\noutside 1 247\nTMhelix 248 270\ninside 271 277

33412 GCF_000717875.1_ASM71787v1 Streptomyces sp. NRRL S-455 Terrabacteria group; Actinobacteria; Actinobacteria; Streptomycetales; Streptomycetaceae; Streptomyces
MRHARRRVVRRVTRLAAVGGLLLGGAMVTTAVASEPPDPSIGVPYSARPATSMGADLVSR WP_030244608.1 MULTISPECIES: protease [Streptomyces] Length: 445\nNumber of predicted TMHs: 2\nExp number of AAs in TMHs: 41.14419\nExp number, first 60 AAs: 21.93617\nTotal prob of N-in: 0.98872\nPOSSIBLE N-term signal sequence\ninside 1 11\nTMhelix 12 34\noutside 35 406\nTMhelix 407 429\ninside 430 445

33413 GCF_000717875.1_ASM71787v1 Streptomyces sp. NRRL S-455 Terrabacteria group; Actinobacteria; Actinobacteria; Streptomycetales; Streptomycetaceae; Streptomyces
MPARRKAVTVAALGILPLALTAAPAAAHGSMGDPVSRVSQCYAEGPESPKSAACGAAVAA WP_030247876.1 MULTISPECIES: chitin-binding protein [Streptomyces] Length: 335\nNumber of predicted TMHs: 2\nExp

number of AAs in TMHs: 41.5621\nExp number, first 60 AAs: 21.08464\nTotal prob of N-in: 0.96521\nPOSSIBLE N-term signal sequence\ninside 1 6\nTMhelix 7 29\noutside 30 300\nTMhelix 301 323\ninside 324 335

33414 GCF_000722015.1_ASM72201v1 Streptomyces sp. NRRL S-474 Terrabacteria group; Actinobacteria; Actinobacteria; Streptomycetales; Streptomycetaceae; Streptomyces MRHARRRVVRRVTRLAAVGGLLLGGAMVTTAVASEPPDPSIGVPYSARPATSMGADLVSR WP_030244608.1 MULTISPECIES: protease [Streptomyces] Length: 445\nNumber of predicted TMHs: 2\nExp number of AAs in TMHs: 41.14419\nExp number, first 60 AAs: 21.93617\nTotal prob of N-in: 0.98872\nPOSSIBLE N-term signal sequence\ninside 1 11\nTMhelix 12 34\noutside 35 406\nTMhelix 407 429\ninside 430 445

33415 GCF_000722015.1_ASM72201v1 Streptomyces sp. NRRL S-474 Terrabacteria group; Actinobacteria; Actinobacteria; Streptomycetales; Streptomycetaceae; Streptomyces MPARRKAVTVAALGILPLALTAAPAAAHGSMGDPVSRVSQCYAEGPESPKSAACGAAVAA WP_030247876.1 MULTISPECIES: chitin-binding protein [Streptomyces] Length: 335\nNumber of predicted TMHs: 2\nExp number of AAs in TMHs: 41.5621\nExp number, first 60 AAs: 21.08464\nTotal prob of N-in: 0.96521\nPOSSIBLE N-term signal sequence\ninside 1 6\nTMhelix 7 29\noutside 30 300\nTMhelix 301 323\ninside 324 335

33416 GCF_000722015.1_ASM72201v1 Streptomyces sp. NRRL S-474 Terrabacteria group; Actinobacteria; Actinobacteria; Streptomycetales; Streptomycetaceae; Streptomyces MLRGVENHSLPRTAAFFDLDKTVIAKSSTLTFSKSFYQGGLINRRAVLRTAYAQFVFLAG WP_030235813.1 MULTISPECIES: inhibition of morphological differentiation protein [Streptomyces] Length: 277\nNumber of predicted TMHs: 1\nExp number of AAs in TMHs: 21.66866\nExp number, first 60 AAs: 0.03762\nTotal prob of N-in: 0.00406\noutside 1 247\nTMhelix 248 270\ninside 271 277

33417 GCF_000720215.1_ASM72021v1 Streptomyces sp. NRRL S-475 Terrabacteria group; Actinobacteria; Actinobacteria; Streptomycetales; Streptomycetaceae; Streptomyces MRHARRRVVRRVTRLAAVGGLLLGGAMVTNAVASEPAAPSIGVPFTAPPATGEGTDLVSR WP_030847554.1 protease [Streptomyces sp. NRRL S-475] Length: 448\nNumber of predicted TMHs: 2\nExp number of AAs in TMHs: 41.0893\nExp number, first 60 AAs: 21.34656\nTotal prob of N-in: 0.96868\nPOSSIBLE N-term signal sequence\ninside 1 11\nTMhelix 12 34\noutside 35 409\nTMhelix 410 432\ninside 433 448

33418 GCF_000720215.1_ASM72021v1 Streptomyces sp. NRRL S-475 Terrabacteria group; Actinobacteria; Actinobacteria; Streptomycetales; Streptomycetaceae; Streptomyces MLWDVENHSMPRTAAFFDLDKTVIAKSSTLTFSKSFYQGGLINRRAVLRTAYAQFVFLAG WP_030846111.1 inhibition of morphological differentiation protein [Streptomyces sp. NRRL S-475] Length: 277\nNumber of predicted TMHs: 1\nExp number of AAs in TMHs: 21.58227\nExp number, first 60 AAs: 0.06522\nTotal prob of N-in: 0.00812\noutside 1 247\nTMhelix 248 270\ninside 271 277

33419 GCF_000720215.1_ASM72021v1 Streptomyces sp. NRRL S-475 Terrabacteria group; Actinobacteria; Actinobacteria; Streptomycetales; Streptomycetaceae; Streptomyces MPAPKQTTRRSLLVTSALLSFLALAAPAAVAAPKPSDSPSASASPSATPPASMSSVGGAR WP_030842128.1 D-alanyl-D-alanine carboxypeptidase [Streptomyces sp. NRRL S-475] Length: 435\nNumber of predicted TMHs: 2\nExp number of AAs in TMHs: 41.64801\nExp number, first 60 AAs: 19.03343\nTotal prob of N-in: 0.94385\nPOSSIBLE N-term signal sequence\ninside 1 11\nTMhelix 12 31\noutside 32 396\nTMhelix 397 419\ninside 420 435

33420 GCF_000721955.1_ASM72195v1 Streptomyces sp. NRRL S-481 Terrabacteria group; Actinobacteria; Actinobacteria; Streptomycetales; Streptomycetaceae; Streptomyces MRHARRRVVRRVTRLAAVGGLLLGGAMVTTAVASEPPDPSIGVPYSARPATSMGADLVSR WP_030244608.1 MULTISPECIES: protease [Streptomyces] Length: 445\nNumber of predicted TMHs: 2\nExp number of AAs in TMHs: 41.14419\nExp number, first 60 AAs: 21.93617\nTotal prob of N-in: 0.98872\nPOSSIBLE N-term signal sequence\ninside 1 11\nTMhelix 12 34\noutside 35 406\nTMhelix 407 429\ninside 430 445

33421 GCF_000721955.1_ASM72195v1 Streptomyces sp. NRRL S-481 Terrabacteria group; Actinobacteria; Actinobacteria; Streptomycetales; Streptomycetaceae; Streptomyces MLRGVENHSLPRTAAFFDLDKTVIAKSSTLTFSKSFYQGGLINRRAVLRTAYAQFVFLAG WP_030235813.1 MULTISPECIES: inhibition of morphological differentiation protein [Streptomyces] Length: 277\nNumber of predicted TMHs: 1\nExp number of AAs in TMHs: 21.66866\nExp number, first 60 AAs: 0.03762\nTotal prob of N-in: 0.00406\noutside 1 247\nTMhelix 248 270\ninside 271 277

33422 GCF_000721955.1_ASM72195v1 Streptomyces sp. NRRL S-481 Terrabacteria group; Actinobacteria; Actinobacteria; Streptomycetales; Streptomycetaceae; Streptomyces

MPARRKAVTVVALGILPLALTAAPAAAHGSMGDPVSRVSQCYAEGPESPKSAACGAAVAV WP_030953727.1 chitin-binding protein [Streptomyces sp. NRRL S-481] Length: 335\nNumber of predicted TMHs: 2\nExp number of AAs in TMHs: 41.98733\nExp number, first 60 AAs: 21.50433\nTotal prob of N-in: 0.98290\nPOSSIBLE N-term signal sequence\ninside 1 6\nTMhelix 7 29\noutside 30 300\nTMhelix 301 323\ninside 324 335

33423 GCF_000720795.1_ASM72079v1 Streptomyces sp. NRRL S-515 Terrabacteria group; Actinobacteria; Actinobacteria; Streptomycetales; Streptomycetaceae; Streptomyces MSSPDPQFRAMPDDENSSEHGESAPGVRRPRNPDGIRRRSPVIAVTGAASGVGAALVSRL WP_030764580.1 MULTISPECIES: NAD-dependent dehydratase [Streptomyces] Length: 388\nNumber of predicted TMHs: 1\nExp number of AAs in TMHs: 23.07357\nExp number, first 60 AAs: 0.86549\nTotal prob of N-in: 0.04440\noutside 1 359\nTMhelix 360 382\ninside 383 388

33424 GCF_000720795.1_ASM72079v1 Streptomyces sp. NRRL S-515 Terrabacteria group; Actinobacteria; Actinobacteria; Streptomycetales; Streptomycetaceae; Streptomyces MSTVSRRTLVRAAAVTACAGSLLALPTAAALADGVPAASSAHSAAPRAILVKSLALADGVWP_030771538.1 MULTISPECIES: hypothetical protein [Streptomyces] Length: 222\nNumber of predicted TMHs: 2\nExp number of AAs in TMHs: 42.55344\nExp number, first 60 AAs: 19.94438\nTotal prob of N-in: 0.93139\nPOSSIBLE N-term signal sequence\ninside 1 8\nTMhelix 9 31\noutside 32 192\nTMhelix 193 215\ninside 216 222

33425 GCF_000720795.1_ASM72079v1 Streptomyces sp. NRRL S-515 Terrabacteria group; Actinobacteria; Actinobacteria; Streptomycetales; Streptomycetaceae; Streptomyces MNPTHAPRRTLLVAAFALTLASGGTGTAAAVTTAAPQPPYALRLDGAGECTFPMKKQIED WP_051782042.1 MULTISPECIES: type VII secretion-associated serine protease mycosin [Streptomyces] Length: 418\nNumber of predicted TMHs: 2\nExp number of AAs in TMHs: 44.64396\nExp number, first 60 AAs: 22.32784\nTotal prob of N-in: 0.97809\nPOSSIBLE N-term signal sequence\ninside 1 11\nTMhelix 12 34\noutside 35 384\nTMhelix 385 407\ninside 408 418

33426 GCF_000720795.1_ASM72079v1 Streptomyces sp. NRRL S-515 Terrabacteria group; Actinobacteria; Actinobacteria; Streptomycetales; Streptomycetaceae; Streptomyces MPARRRTAVTLTRIAAAGLAPLALAAYAAAPAVAHGSMTDPVSRVAACYAEGPESPKSAA WP_030773238.1 MULTISPECIES: chitin-binding protein [Streptomyces] Length: 362\nNumber of predicted TMHs: 2\nExp number of AAs in TMHs: 43.34514\nExp number, first 60 AAs: 21.32926\nTotal prob of N-in: 0.95286\nPOSSIBLE N-term signal sequence\ninside 1 12\nTMhelix 13 35\noutside 36 324\nTMhelix 325 347\ninside 348 362

33427 GCF_000718955.1_ASM71895v1 Streptomyces sp. NRRL S-575 Terrabacteria group; Actinobacteria; Actinobacteria; Streptomycetales; Streptomycetaceae; Streptomyces MPARRRTAVTLTRIAAAGLAPLALAAYAAAPAVAHGSMTDPVSRVAACYAEGPESPKSAA WP_030773238.1 MULTISPECIES: chitin-binding protein [Streptomyces] Length: 362\nNumber of predicted TMHs: 2\nExp number of AAs in TMHs: 43.34514\nExp number, first 60 AAs: 21.32926\nTotal prob of N-in: 0.95286\nPOSSIBLE N-term signal sequence\ninside 1 12\nTMhelix 13 35\noutside 36 324\nTMhelix 325 347\ninside 348 362

33428 GCF_000718955.1_ASM71895v1 Streptomyces sp. NRRL S-575 Terrabacteria group; Actinobacteria; Actinobacteria; Streptomycetales; Streptomycetaceae; Streptomyces MNPTHAPRRTLLVAAFALTLASGGTGTAAAVTTAAPQPPYALRLDGAGECTFPMKKQIED WP_051782042.1 MULTISPECIES: type VII secretion-associated serine protease mycosin [Streptomyces] Length: 418\nNumber of predicted TMHs: 2\nExp number of AAs in TMHs: 44.64396\nExp number, first 60 AAs: 22.32784\nTotal prob of N-in: 0.97809\nPOSSIBLE N-term signal sequence\ninside 1 11\nTMhelix 12 34\noutside 35 384\nTMhelix 385 407\ninside 408 418

33429 GCF_000718955.1_ASM71895v1 Streptomyces sp. NRRL S-575 Terrabacteria group; Actinobacteria; Actinobacteria; Streptomycetales; Streptomycetaceae; Streptomyces MSSPDPQFRAMPDDENSSEHGESAPGVRRPRNPDGIRRRSPVIAVTGAASGVGAALVSRL WP_030764580.1 MULTISPECIES: NAD-dependent dehydratase [Streptomyces] Length: 388\nNumber of predicted TMHs: 1\nExp number of AAs in TMHs: 23.07357\nExp number, first 60 AAs: 0.86549\nTotal prob of N-in: 0.04440\noutside 1 359\nTMhelix 360 382\ninside 383 388

33430 GCF_000718955.1_ASM71895v1 Streptomyces sp. NRRL S-575 Terrabacteria group; Actinobacteria; Actinobacteria; Streptomycetales; Streptomycetaceae; Streptomyces MSTVSRRTLVRAAAVTACAGSLLALPTAAALADGVPAASSAHSAAPRAILVKSLALADGVWP_030771538.1 MULTISPECIES: hypothetical protein [Streptomyces] Length: 222\nNumber of predicted TMHs: 2\nExp number of AAs in TMHs:

42.55344\nExp number, first 60 AAs: 19.94438\nTotal prob of N-in: 0.93139\nPOSSIBLE N-term signal sequence\ninside 1 8\nTMhelix 9 31\noutside 32 192\nTMhelix 193 215\ninside 216 222

33431 GCF_000725705.1_Doro.v1.0 Streptomyces sp. NRRL S-623 Terrabacteria group; Actinobacteria; Actinobacteria; Streptomycetales; Streptomycetaceae; Streptomyces MLSVVENCFSPRTAAFFDLDKTVIAKSSTLTFSKSFYQGGLINRRAVLRTAYTQFVFMAG WP_031123945.1 MULTISPECIES: inhibition of morphological differentiation protein [Streptomyces] Length: 281\nNumber of predicted TMHs: 1\nExp number of AAs in TMHs: 21.79856\nExp number, first 60 AAs: 0.01652\nTotal prob of N-in: 0.00400\noutside 1 247\nTMhelix 248 270\ninside 271 281

33432 GCF_000725705.1_Doro.v1.0 Streptomyces sp. NRRL S-623 Terrabacteria group; Actinobacteria; Actinobacteria; Streptomycetales; Streptomycetaceae; Streptomyces MGNAVSGRRTLLSGTAVLAAVALIALGSAPAQADETEPDLGVRGLTPVTGITPGSGFGLP WP_031122887.1 peptidase [Streptomyces sp. NRRL S-623] Length: 434\nNumber of predicted TMHs: 1\nExp number of AAs in TMHs: 34.96425\nExp number, first 60 AAs: 14.18722\nTotal prob of N-in: 0.62194\nPOSSIBLE N-term signal sequence\noutside 1 402\nTMhelix 403 425\ninside 426 434

33433 GCF_000725705.1_Doro.v1.0 Streptomyces sp. NRRL S-623 Terrabacteria group; Actinobacteria; Actinobacteria; Streptomycetales; Streptomycetaceae; Streptomyces MFGRTGRRSAVTVLAAVVLWGPTGQDTAVADSRHPVIETLPEQEVRAAVAAARTPVAVNL WP_031124694.1 hypothetical protein [Streptomyces sp. NRRL S-623] Length: 259\nNumber of predicted TMHs: 1\nExp number of AAs in TMHs: 18.48871\nExp number, first 60 AAs: 0.18238\nTotal prob of N-in: 0.12868\noutside 1 229\nTMhelix 230 247\ninside 248 259

33434 GCF_000720765.1_ASM72076v1 Streptomyces sp. NRRL S-646 Terrabacteria group; Actinobacteria; Actinobacteria; Streptomycetales; Streptomycetaceae; Streptomyces MTVIAVWRRLEAVEESRKREESDPMSMLIVGQEQVVAARPVVPRGRRRYLRYRPEHVLVG WP_030939315.1 hypothetical protein [Streptomyces sp. NRRL S-646] Length: 173\nNumber of predicted TMHs: 2\nExp number of AAs in TMHs: 66.17058\nExp number, first 60 AAs: 5.09928\nTotal prob of N-in: 0.98882\ninside 1 55\nTMhelix 56 78\noutside 79 142\nTMhelix 143 165\ninside 166 173

33435 GCF_000720765.1_ASM72076v1 Streptomyces sp. NRRL S-646 Terrabacteria group; Actinobacteria; Actinobacteria; Streptomycetales; Streptomycetaceae; Streptomyces MRHARRRIVRRVTRLAAVGGLLLGGAMVTQAAMASETPGTSALPLKASGTSDNPGAALVS WP_030937896.1 protease [Streptomyces sp. NRRL S-646] Length: 460\nNumber of predicted TMHs: 2\nExp number of AAs in TMHs: 43.74712\nExp number, first 60 AAs: 22.24674\nTotal prob of N-in: 0.99204\nPOSSIBLE N-term signal sequence\ninside 1 11\nTMhelix 12 34\noutside 35 421\nTMhelix 422 444\ninside 445 460

33436 GCF_000720765.1_ASM72076v1 Streptomyces sp. NRRL S-646 Terrabacteria group; Actinobacteria; Actinobacteria; Streptomycetales; Streptomycetaceae; Streptomyces MFSSVSAPSVRRRGAVRLAAATLVSGLVAVGVLTGAGTAAAEETTQSQGGATATMGGLKT WP_078912459.1 peptidase [Streptomyces sp. NRRL S-646] Length: 469\nNumber of predicted TMHs: 1\nExp number of AAs in TMHs: 40.1295899999999\nExp number, first 60 AAs: 17.94219\nTotal prob of N-in: 0.76745\nPOSSIBLE N-term signal sequence\noutside 1 438\nTMhelix 439 461\ninside 462 469

33437 GCF_000718945.1_ASM71894v1 Streptomyces sp. NRRL S-813 Terrabacteria group; Actinobacteria; Actinobacteria; Streptomycetales; Streptomycetaceae; Streptomyces MRHARRRVVRRVTRLTAVAGLLLGGAMVAQAAMADEAPATAGAAPGSVRSAAATGAATAA WP_030172626.1 protease [Streptomyces sp. NRRL S-813] Length: 468\nNumber of predicted TMHs: 2\nExp number of AAs in TMHs: 41.9386399999999\nExp number, first 60 AAs: 21.93043\nTotal prob of N-in: 0.98777\nPOSSIBLE N-term signal sequence\ninside 1 11\nTMhelix 12 34\noutside 35 429\nTMhelix 430 452\ninside 453 468

33438 GCF_000718945.1_ASM71894v1 Streptomyces sp. NRRL S-813 Terrabacteria group; Actinobacteria; Actinobacteria; Streptomycetales; Streptomycetaceae; Streptomyces MLRDVENHSLPRTAAFFDLDKTVIAKSSTLTFSKSFYQGGLINRRAVLRTAYAQFVFLAG WP_030177246.1 inhibition of morphological differentiation protein [Streptomyces sp. NRRL S-813] Length: 277\nNumber of predicted TMHs: 1\nExp number of AAs in TMHs: 21.70113\nExp number, first 60 AAs: 0.06486\nTotal prob of N-in: 0.00788\noutside 1 247\nTMhelix 248 270\ninside 271 277

33439 GCF_000718945.1_ASM71894v1 Streptomyces sp. NRRL S-813 Terrabacteria group; Actinobacteria; Actinobacteria; Streptomycetales; Streptomycetaceae; Streptomyces

MPGPVTTTSRSTFVRRTSRRRPTAPRRATFAAVALILGGGGLMVGNVYASATEPDRNQDV WP_078917162.1 hypothetical protein [Streptomyces sp. NRRL S-813] Length: 571\nNumber of predicted TMHs: 2\nExp number of AAs in TMHs: 40.71623\nExp number, first 60 AAs: 21.61366\nTotal prob of N-in: 0.96642\nPOSSIBLE N-term signal sequence\ninside 1 29\nTMhelix 30 52\noutside 53 545\nTMhelix 546 563\ninside 564 571

33440 GCF_000718945.1_ASM71894v1 Streptomyces sp. NRRL S-813 Terrabacteria group; Actinobacteria; Actinobacteria; Streptomycetales; Streptomycetaceae; Streptomyces MADSARRATMRTLGALAGAAALTVLGGTAPTWAAIPAGHHGHPATATPIEHVVVLFDENV WP_037870844.1 phospholipase C [Streptomyces sp. NRRL S-813] Length: 614\nNumber of predicted TMHs: 2\nExp number of AAs in TMHs: 44.04396\nExp number, first 60 AAs: 22.65784\nTotal prob of N-in: 0.99437\nPOSSIBLE N-term signal sequence\ninside 1 12\nTMhelix 13 35\noutside 36 583\nTMhelix 584 606\ninside 607 614

33441 GCF_000718945.1_ASM71894v1 Streptomyces sp. NRRL S-813 Terrabacteria group; Actinobacteria; Actinobacteria; Streptomycetales; Streptomycetaceae; Streptomyces MASRGRRATIRSLGALAGAAALTVLGAAVPGWAAAPTGHHGHGSTATPIKHVVVLFDENI WP_030179510.1 phospholipase C [Streptomyces sp. NRRL S-813] Length: 614\nNumber of predicted TMHs: 2\nExp number of AAs in TMHs: 43.69554\nExp number, first 60 AAs: 22.65866\nTotal prob of N-in: 0.99704\nPOSSIBLE N-term signal sequence\ninside 1 12\nTMhelix 13 35\noutside 36 581\nTMhelix 582 604\ninside 605 614

33442 GCF_000718945.1_ASM71894v1 Streptomyces sp. NRRL S-813 Terrabacteria group; Actinobacteria; Actinobacteria; Streptomycetales; Streptomycetaceae; Streptomyces MLNVRPFRKAQEARTAGTAGTAGTAGTAGRRAALITAATALTAATSAITLPAPAAHADDGQCT WP_078917309.1 type VII secretion-associated serine protease mycosin [Streptomyces sp. NRRL S-813] Length: 443\nNumber of predicted TMHs: 1\nExp number of AAs in TMHs: 25.31296\nExp number, first 60 AAs: 3.0718\nTotal prob of N-in: 0.14969\noutside 1 399\nTMhelix 400 422\ninside 423 443

33443 GCF_000719335.1_ASM71933v1 Streptomyces sp. NRRL S-87 Terrabacteria group; Actinobacteria; Actinobacteria; Streptomycetales; Streptomycetaceae; Streptomyces MNALRWGRRGLVVAVAGAVLAAGGGGAAVADGIRDRQWGLAALHARQAWRTTQGAGVTVA WP_037887152.1 type VII secretion-associated serine protease [Streptomyces sp. NRRL S-87] Length: 397\nNumber of predicted TMHs: 2\nExp number of AAs in TMHs: 39.7278\nExp number, first 60 AAs: 20.13167\nTotal prob of N-in: 0.93584\nPOSSIBLE N-term signal sequence\ninside 1 10\nTMhelix 11 33\noutside 34 363\nTMhelix 364 383\ninside 384 397

33444 GCF_000719335.1_ASM71933v1 Streptomyces sp. NRRL S-87 Terrabacteria group; Actinobacteria; Actinobacteria; Streptomycetales; Streptomycetaceae; Streptomyces MQTTQTLRRARRAAVRAAVLTTATLAAALCGSVSAHGAEGDGIAGYPSAQQTLRDDQLRD WP_037887801.1 hypothetical protein [Streptomyces sp. NRRL S-87] Length: 314\nNumber of predicted TMHs: 2\nExp number of AAs in TMHs: 40.85955\nExp number, first 60 AAs: 21.88326\nTotal prob of N-in: 0.96272\nPOSSIBLE N-term signal sequence\ninside 1 12\nTMhelix 13 35\noutside 36 283\nTMhelix 284 306\ninside 307 314

33445 GCF_000719335.1_ASM71933v1 Streptomyces sp. NRRL S-87 Terrabacteria group; Actinobacteria; Actinobacteria; Streptomycetales; Streptomycetaceae; Streptomyces MNRRTTVRAAAALALAGLLAGPAALAEAAPTTARAATAAVAPGSAAAAGAISVAGTAAAT WP_051795396.1 hypothetical protein [Streptomyces sp. NRRL S-87] Length: 155\nNumber of predicted TMHs: 2\nExp number of AAs in TMHs: 48.39985\nExp number, first 60 AAs: 25.02824\nTotal prob of N-in: 0.91257\nPOSSIBLE N-term signal sequence\ninside 1 6\nTMhelix 7 29\noutside 30 112\nTMhelix 113 135\ninside 136 155

33446 GCF_000717945.1_ASM71794v1 Streptomyces sp. NRRL S-920 Terrabacteria group; Actinobacteria; Actinobacteria; Streptomycetales; Streptomycetaceae; Streptomyces MLKPVENHSLPRAAAFFDLDKTVIAKSSTLTFSKSFYQGGLINRRAVLRTAYAQFVFLAG WP_030782118.1 inhibition of morphological differentiation protein [Streptomyces sp. NRRL S-920] Length: 274\nNumber of predicted TMHs: 1\nExp number of AAs in TMHs: 21.75286\nExp number, first 60 AAs: 0.03562\nTotal prob of N-in: 0.00868\noutside 1 244\nTMhelix 245 267\ninside 268 274

33447 GCF_000717945.1_ASM71794v1 Streptomyces sp. NRRL S-920 Terrabacteria group; Actinobacteria; Actinobacteria; Streptomycetales; Streptomycetaceae; Streptomyces MPARRKAVAIAAAGVAPLALTALSVTPAAAHGSMTDPVSRVSACFQEGPEAPKSAACKAA WP_030778982.1 chitin-binding protein [Streptomyces sp. NRRL S-920] Length: 342\nNumber of predicted TMHs: 2\nExp number of AAs in TMHs: 39.9235\nExp number, first 60 AAs: 21.37424\nTotal prob of N-in: 0.95986\nPOSSIBLE N-term signal sequence\ninside 1 6\nTMhelix 7 29\noutside 30 309\nTMhelix 310 332\ninside 333 342

33448 GCF_000717945.1_ASM71794v1 Streptomyces sp. NRRL S-920 Terrabacteria group; Actinobacteria; Actinobacteria; Streptomycetales; Streptomycetaceae; Streptomyces
MNSFTFRMPACRRHRRRLVVVATATALAAGPAALAGAGSARATGGDHDRGGRADAVVLRT WP_030791129.1
hypothetical protein [Streptomyces sp. NRRL S-920] Length: 315\nNumber of predicted TMHs: 2\nExp number of AAs in TMHs: 39.47007\nExp number, first 60 AAs: 19.2274\nTotal prob of N-in: 0.89121\nPOSSIBLE N-term signal sequence\ninside 1 16\nTMhelix 17 39\noutside 40 285\nTMhelix 286 308\ninside 309 315

33449 GCF_000720235.1_ASM72023v1 Streptomyces sp. NRRL S-98 Terrabacteria group; Actinobacteria; Actinobacteria; Streptomycetales; Streptomycetaceae; Streptomyces
MSTVRRRTLVRAAAVTACAGSLLALPTAAALAEGVPAASSAHSAAPQRTLVKSLSLADGV WP_030834094.1
MULTISPECIES: hypothetical protein [Streptomyces] Length: 218\nNumber of predicted TMHs: 2\nExp number of AAs in TMHs: 42.93094\nExp number, first 60 AAs: 20.33933\nTotal prob of N-in: 0.95701\nPOSSIBLE N-term signal sequence\ninside 1 8\nTMhelix 9 31\noutside 32 188\nTMhelix 189 211\ninside 212 218

33450 GCF_000720235.1_ASM72023v1 Streptomyces sp. NRRL S-98 Terrabacteria group; Actinobacteria; Actinobacteria; Streptomycetales; Streptomycetaceae; Streptomyces
MPPYRDARRALLAAAFALTLTAAGATAATAAQHPPYALRLDGAGECTFPMKKQIADRPWA WP_030823957.1
MULTISPECIES: type VII secretion-associated serine protease [Streptomyces] Length: 414\nNumber of predicted TMHs: 2\nExp number of AAs in TMHs: 42.2832099999999\nExp number, first 60 AAs: 19.94733\nTotal prob of N-in: 0.93627\nPOSSIBLE N-term signal sequence\ninside 1 11\nTMhelix 12 31\noutside 32 380\nTMhelix 381 403\ninside 404 414

33451 GCF_000720235.1_ASM72023v1 Streptomyces sp. NRRL S-98 Terrabacteria group; Actinobacteria; Actinobacteria; Streptomycetales; Streptomycetaceae; Streptomyces
MALDVIRSGPPDKRPRRNLFKRGVLVLLPAALLAAATLVPNLAHALEAPVNLGTATNYAV WP_030831911.1
MULTISPECIES: hypothetical protein [Streptomyces] Length: 518\nNumber of predicted TMHs: 2\nExp number of AAs in TMHs: 43.15964\nExp number, first 60 AAs: 20.83032\nTotal prob of N-in: 0.94539\nPOSSIBLE N-term signal sequence\ninside 1 22\nTMhelix 23 45\noutside 46 483\nTMhelix 484 506\ninside 507 518

33452 GCF_000720235.1_ASM72023v1 Streptomyces sp. NRRL S-98 Terrabacteria group; Actinobacteria; Actinobacteria; Streptomycetales; Streptomycetaceae; Streptomyces
MSARRRTVMPIRIAAAGAAPLALAVYAAVPAVAHGSMTDPVSRVAACYAEGPESPKSAAC WP_030831682.1
chitin-binding protein [Streptomyces sp. NRRL S-98] Length: 373\nNumber of predicted TMHs: 2\nExp number of AAs in TMHs: 43.4943\nExp number, first 60 AAs: 21.78981\nTotal prob of N-in: 0.98431\nPOSSIBLE N-term signal sequence\ninside 1 12\nTMhelix 13 35\noutside 36 338\nTMhelix 339 361\ninside 362 373

33453 GCF_000718915.1_ASM71891v1 Streptomyces sp. NRRL WC-3540 Terrabacteria group; Actinobacteria; Actinobacteria; Streptomycetales; Streptomycetaceae; Streptomyces
MGSSWNERRLERRVRPGDGKPLKPFRWWQLTRRSLLSLALPVGGGLSATYTVEVKHGGDA WP_030189408.1
MULTISPECIES: hypothetical protein [Streptomyces] Length: 218\nNumber of predicted TMHs: 2\nExp number of AAs in TMHs: 41.32127\nExp number, first 60 AAs: 0.37701\nTotal prob of N-in: 0.97413\ninside 1 135\nTMhelix 136 158\noutside 159 182\nTMhelix 183 200\ninside 201 218

33454 GCF_000718915.1_ASM71891v1 Streptomyces sp. NRRL WC-3540 Terrabacteria group; Actinobacteria; Actinobacteria; Streptomycetales; Streptomycetaceae; Streptomyces
MLSVVENCFSPRTAAFFDLDKTVIAKSSTLTFSKSFYQGGLINRRAVLRTAYTQFVFMAG WP_030113359.1 MULTISPECIES: inhibition of morphological differentiation protein [Streptomyces] Length: 280\nNumber of predicted TMHs: 1\nExp number of AAs in TMHs: 21.68695\nExp number, first 60 AAs: 0.01801\nTotal prob of N-in: 0.00793\noutside 1 247\nTMhelix 248 270\ninside 271 280

33455 GCF_000721935.1_ASM72193v1 Streptomyces sp. NRRL WC-3549 Terrabacteria group; Actinobacteria; Actinobacteria; Streptomycetales; Streptomycetaceae; Streptomyces
MLTFVENCFSPRTAAFFDLDKTVIAKSSTLAFSKSFYQGGLINRRAVLRTAYTQFVFLAG WP_031080970.1 inhibition of morphological differentiation protein [Streptomyces sp. NRRL WC-3549] Length: 279\nNumber of predicted TMHs: 1\nExp number of AAs in TMHs: 22.11621\nExp number, first 60 AAs: 0.01488\nTotal prob of N-in: 0.00196\noutside 1 247\nTMhelix 248 270\ninside 271 279

33456 GCF_001509785.1_ASM150978v1 Streptomyces sp. NRRL WC-3604 Terrabacteria group; Actinobacteria; Actinobacteria; Streptomycetales; Streptomycetaceae; Streptomyces
MRHARRRLVRRVTRLAAVGGLLLGGTMVTRAVASEPPGPSAVPRTFAESATPGPGGELVA WP_062670020.1
MULTISPECIES: protease [Streptomyces] Length: 454\nNumber of predicted TMHs: 2\nExp number of

AAs in TMHs: 40.90396\nExp number, first 60 AAs: 19.80239\nTotal prob of N-in: 0.90419\nPOSSIBLE N-term signal sequence\ninside 1 11\nTMhelix 12 34\noutside 35 415\nTMhelix 416 438\ninside 439 454

33457 GCF_001509795.1_ASM150979v1 Streptomyces sp. NRRL WC-3605 Terrabacteria group; Actinobacteria; Actinobacteria; Streptomycetales; Streptomycetaceae; Streptomyces MRHARRRLVRRVTRLAAVGGLLLGGTMVTRAVASEPPGPSAVPRTFAESATPGPGGELVA WP_062670020.1 MULTISPECIES: protease [Streptomyces] Length: 454\nNumber of predicted TMHs: 2\nExp number of AAs in TMHs: 40.90396\nExp number, first 60 AAs: 19.80239\nTotal prob of N-in: 0.90419\nPOSSIBLE N-term signal sequence\ninside 1 11\nTMhelix 12 34\noutside 35 415\nTMhelix 416 438\ninside 439 454

33458 GCF_001280005.1_ASM128000v1 Streptomyces sp. NRRL WC-3618 Terrabacteria group; Actinobacteria; Actinobacteria; Streptomycetales; Streptomycetaceae; Streptomyces MLSLVESHSLPRTAAFFDLDKTVIAKSSTLTFSKSFYQGGLINRRAVLRTAYAQFVFLAG WP_053745792.1 inhibition of morphological differentiation protein [Streptomyces sp. NRRL WC-3618] Length: 277\nNumber of predicted TMHs: 1\nExp number of AAs in TMHs: 21.51056\nExp number, first 60 AAs: 0.01412\nTotal prob of N-in: 0.00181\noutside 1 247\nTMhelix 248 270\ninside 271 277

33459 GCF_001280005.1_ASM128000v1 Streptomyces sp. NRRL WC-3618 Terrabacteria group; Actinobacteria; Actinobacteria; Streptomycetales; Streptomycetaceae; Streptomyces MRHARRRIVRRATRLAAVGGVILGGVMVTQAVASEPPPTNLTFSSARAAGTTGSDLVTEL WP_053742850.1 protease [Streptomyces sp. NRRL WC-3618] Length: 452\nNumber of predicted TMHs: 2\nExp number of AAs in TMHs: 43.5821\nExp number, first 60 AAs: 21.89352\nTotal prob of N-in: 0.98959\nPOSSIBLE N-term signal sequence\ninside 1 11\nTMhelix 12 34\noutside 35 413\nTMhelix 414 436\ninside 437 452

33460 GCF_000718785.1_ASM71878v1 Streptomyces sp. NRRL WC-3626 Terrabacteria group; Actinobacteria; Actinobacteria; Streptomycetales; Streptomycetaceae; Streptomyces MPAPKNTARRPLLIASAVLLSTALTSPVALAAPKPSTSPTATPPAKMSVVGGARLAQPGT WP_030228728.1 D-alanyl-D-alanine carboxypeptidase [Streptomyces sp. NRRL WC-3626] Length: 425\nNumber of predicted TMHs: 1\nExp number of AAs in TMHs: 36.69477\nExp number, first 60 AAs: 14.45435\nTotal prob of N-in: 0.72822\nPOSSIBLE N-term signal sequence\noutside 1 386\nTMhelix 387 409\ninside 410 425

33461 GCF_000718785.1_ASM71878v1 Streptomyces sp. NRRL WC-3626 Terrabacteria group; Actinobacteria; Actinobacteria; Streptomycetales; Streptomycetaceae; Streptomyces MLRDVENHSMPRTAAFFDLDKTVIAKSSTLTFSKSFYQGGLINRRAVLRTAYSQFVFLAG WP_030225070.1 inhibition of morphological differentiation protein [Streptomyces sp. NRRL WC-3626] Length: 277\nNumber of predicted TMHs: 1\nExp number of AAs in TMHs: 21.83865\nExp number, first 60 AAs: 0.05095\nTotal prob of N-in: 0.00781\noutside 1 247\nTMhelix 248 270\ninside 271 277

33462 GCF_000718785.1_ASM71878v1 Streptomyces sp. NRRL WC-3626 Terrabacteria group; Actinobacteria; Actinobacteria; Streptomycetales; Streptomycetaceae; Streptomyces MRHARRRVVRRVTRLAAVGGLLLGATMVTHAVASEPRAPGAVPLSTAGEKASAGPGDELV WP_030218994.1 protease [Streptomyces sp. NRRL WC-3626] Length: 459\nNumber of predicted TMHs: 2\nExp number of AAs in TMHs: 40.21826\nExp number, first 60 AAs: 22.04274\nTotal prob of N-in: 0.98888\nPOSSIBLE N-term signal sequence\ninside 1 11\nTMhelix 12 34\noutside 35 420\nTMhelix 421 443\ninside 444 459

33463 GCF_001279025.1_ASM127902v1 Streptomyces sp. NRRL WC-3701 Terrabacteria group; Actinobacteria; Actinobacteria; Streptomycetales; Streptomycetaceae; Streptomyces MTRRPLLRAAGVALSAAALTVLPALPAHADGIRAQEWALKAMNTQDAWETTKGEGITVAV WP_003987438.1 MULTISPECIES: type VII secretion-associated serine protease [Streptomyces] Length: 386\nNumber of predicted TMHs: 1\nExp number of AAs in TMHs: 34.80558\nExp number, first 60 AAs: 10.27095\nTotal prob of N-in: 0.55118\nPOSSIBLE N-term signal sequence\noutside 1 359\nTMhelix 360 382\ninside 383 386

33464 GCF_001279025.1_ASM127902v1 Streptomyces sp. NRRL WC-3701 Terrabacteria group; Actinobacteria; Actinobacteria; Streptomycetales; Streptomycetaceae; Streptomyces MTERSAPAASATGPGTPARRTAVRAAATGTLAAALLGLAPQSALAATPAPPPAPAISAVT WP_030180617.1 MULTISPECIES: hypothetical protein [Streptomyces] Length: 293\nNumber of predicted TMHs: 1\nExp number of AAs in TMHs: 24.94793\nExp number, first 60 AAs: 4.55383\nTotal prob of N-in: 0.22746\noutside 1 264\nTMhelix 265 287\ninside 288 293

33465 GCF_001279025.1_ASM127902v1 Streptomyces sp. NRRL WC-3701 Terrabacteria group; Actinobacteria; Actinobacteria; Streptomycetales; Streptomycetaceae; Streptomyces

MESRQPVGPRANGHRAAHRRRAAVALAGAVCAVGLTATTAAPALADTNVPLNNGQCNFGA WP_078586913.1 MULTISPECIES: type VII secretion-associated serine protease mycosin [Streptomyces] Length: 427\nNumber of predicted TMHs: 2\nExp number of AAs in TMHs: 43.8948\nExp number, first 60 AAs: 21.08952\nTotal prob of N-in: 0.94565\nPOSSIBLE N-term signal sequence\ninside 1 20\nTMhelix 21 43\noutside 44 389\nTMhelix 390 412\ninside 413 427

33466 GCF_000717805.1_ASM71780v1 Streptomyces sp. NRRL WC-3702 Terrabacteria group; Actinobacteria; Actinobacteria; Streptomycetales; Streptomycetaceae; Streptomyces MTRRPLLRAAGVALSAAALTVLPALPAHADGIRAQEWALKAMNTQDAWETTKGEGITVAV WP_003987438.1 MULTISPECIES: type VII secretion-associated serine protease [Streptomyces] Length: 386\nNumber of predicted TMHs: 1\nExp number of AAs in TMHs: 34.80558\nExp number, first 60 AAs: 10.27095\nTotal prob of N-in: 0.55118\nPOSSIBLE N-term signal sequence\noutside 1 359\nTMhelix 360 382\ninside 383 386

33467 GCF_000717805.1_ASM71780v1 Streptomyces sp. NRRL WC-3702 Terrabacteria group; Actinobacteria; Actinobacteria; Streptomycetales; Streptomycetaceae; Streptomyces MESRQPVGPRANGHRAAHRRRAAVALAGAVCAVGLTATTAAPALADTNVPLNNGQCNFGA WP_078586913.1 MULTISPECIES: type VII secretion-associated serine protease mycosin [Streptomyces] Length: 427\nNumber of predicted TMHs: 2\nExp number of AAs in TMHs: 43.8948\nExp number, first 60 AAs: 21.08952\nTotal prob of N-in: 0.94565\nPOSSIBLE N-term signal sequence\ninside 1 20\nTMhelix 21 43\noutside 44 389\nTMhelix 390 412\ninside 413 427

33468 GCF_000718815.1_ASM71881v1 Streptomyces sp. NRRL WC-3703 Terrabacteria group; Actinobacteria; Actinobacteria; Streptomycetales; Streptomycetaceae; Streptomyces MESRQPVGPRANGHRAAHRRRAAVALAGAVCAVGLTATTAAPALADTNVPLNNGQCNFGA WP_078586913.1 MULTISPECIES: type VII secretion-associated serine protease mycosin [Streptomyces] Length: 427\nNumber of predicted TMHs: 2\nExp number of AAs in TMHs: 43.8948\nExp number, first 60 AAs: 21.08952\nTotal prob of N-in: 0.94565\nPOSSIBLE N-term signal sequence\ninside 1 20\nTMhelix 21 43\noutside 44 389\nTMhelix 390 412\ninside 413 427

33469 GCF_000718815.1_ASM71881v1 Streptomyces sp. NRRL WC-3703 Terrabacteria group; Actinobacteria; Actinobacteria; Streptomycetales; Streptomycetaceae; Streptomyces MTRRPLLRAAGVALSAAALTVLPALPAHADGIRAQEWALKAMNTQDAWETTKGEGITVAV WP_003987438.1 MULTISPECIES: type VII secretion-associated serine protease [Streptomyces] Length: 386\nNumber of predicted TMHs: 1\nExp number of AAs in TMHs: 34.80558\nExp number, first 60 AAs: 10.27095\nTotal prob of N-in: 0.55118\nPOSSIBLE N-term signal sequence\noutside 1 359\nTMhelix 360 382\ninside 383 386

33470 GCF_000718815.1_ASM71881v1 Streptomyces sp. NRRL WC-3703 Terrabacteria group; Actinobacteria; Actinobacteria; Streptomycetales; Streptomycetaceae; Streptomyces MTERSAPAASATGPGTPARRTAVRAAATGTLAAALLGLAPQSALAATPAPPPAPAISAVT WP_030180617.1 MULTISPECIES: hypothetical protein [Streptomyces] Length: 293\nNumber of predicted TMHs: 1\nExp number of AAs in TMHs: 24.94793\nExp number, first 60 AAs: 4.55383\nTotal prob of N-in: 0.22746\noutside 1 264\nTMhelix 265 287\ninside 288 293

33471 GCF_000725715.1_Doro.v1.0Streptomyces sp. NRRL WC-3719 Terrabacteria group; Actinobacteria; Actinobacteria; Streptomycetales; Streptomycetaceae; Streptomyces MRTVRSRRTAVTAALLTGAALLGGLAAPGAALAASPTPSPAATDGEPPAPTEAGTSFRSA WP_031129975.1 hypothetical protein [Streptomyces sp. NRRL WC-3719] Length: 266\nNumber of predicted TMHs: 2\nExp number of AAs in TMHs: 41.1869\nExp number, first 60 AAs: 21.9617\nTotal prob of N-in: 0.99139\nPOSSIBLE N-term signal sequence\ninside 1 12\nTMhelix 13 35\noutside 36 229\nTMhelix 230 252\ninside 253 266

33472 GCF_000725715.1_Doro.v1.0Streptomyces sp. NRRL WC-3719 Terrabacteria group; Actinobacteria; Actinobacteria; Streptomycetales; Streptomycetaceae; Streptomyces MLWLVENHSSPRTAAFFDLDKTVIAKSSTLTFGKSFYQGGLINRRDALRTAYTQFVFLVG WP_051838844.1 inhibition of morphological differentiation protein [Streptomyces sp. NRRL WC-3719] Length: 299\nNumber of predicted TMHs: 1\nExp number of AAs in TMHs: 21.32356\nExp number, first 60 AAs: 0.00575\nTotal prob of N-in: 0.00344\noutside 1 247\nTMhelix 248 270\ninside 271 299

33473 GCF_001280045.1_ASM128004v1 Streptomyces sp. NRRL WC-3723 Terrabacteria group; Actinobacteria; Actinobacteria; Streptomycetales; Streptomycetaceae; Streptomyces MRHARRRVVRRVTRLAAVGGLLLGGAMVTQAAMASETPPASARPLAATGDTGAALVARLG WP_031092578.1 MULTISPECIES: protease [Streptomyces] Length: 457\nNumber of predicted TMHs: 2\nExp number of

AAs in TMHs: 43.16462\nExp number, first 60 AAs: 22.25978\nTotal prob of N-in: 0.99239\nPOSSIBLE N-term signal sequence\ninside 1 11\nTMhelix 12 34\noutside 35 418\nTMhelix 419 441\ninside 442 457

33474 GCF_001280045.1_ASM128004v1 Streptomyces sp. NRRL WC-3723 Terrabacteria group; Actinobacteria; Actinobacteria; Streptomycetales; Streptomycetaceae; Streptomyces MRTPHTARRRRALAGVLLAACLALAGCGGADDMGGSSKAASDSKAEAAGAPGGSARNGRA WP_031095397.1 MULTISPECIES: hypothetical protein [Streptomyces] Length: 324\nNumber of predicted TMHs: 2\nExp number of AAs in TMHs: 42.45736\nExp number, first 60 AAs: 19.4804\nTotal prob of N-in: 0.97333\nPOSSIBLE N-term signal sequence\ninside 1 12\nTMhelix 13 30\noutside 31 259\nTMhelix 260 282\ninside 283 324

33475 GCF_001280045.1_ASM128004v1 Streptomyces sp. NRRL WC-3723 Terrabacteria group; Actinobacteria; Actinobacteria; Streptomycetales; Streptomycetaceae; Streptomyces MLRGVENHSLPRAAAFFDLDKTVIAKSSTLTFSKSFYQGGLINRRAALRTAYAQFVFLVG WP_031092052.1 MULTISPECIES: inhibition of morphological differentiation protein [Streptomyces] Length: 276\nNumber of predicted TMHs: 1\nExp number of AAs in TMHs: 21.19627\nExp number, first 60 AAs: 0.04374\nTotal prob of N-in: 0.00486\noutside 1 248\nTMhelix 249 268\ninside 269 276

33476 GCF_000720295.1_ASM72029v1 Streptomyces sp. NRRL WC-3725 Terrabacteria group; Actinobacteria; Actinobacteria; Streptomycetales; Streptomycetaceae; Streptomyces MLRGVENHSLPRAAAFFDLDKTVIAKSSTLTFSKSFYQGGLINRRAALRTAYAQFVFLVG WP_030648527.1 MULTISPECIES: inhibition of morphological differentiation protein [Streptomyces] Length: 276\nNumber of predicted TMHs: 1\nExp number of AAs in TMHs: 21.19627\nExp number, first 60 AAs: 0.04374\nTotal prob of N-in: 0.00486\noutside 1 248\nTMhelix 249 268\ninside 269 276

33477 GCF_000720295.1_ASM72029v1 Streptomyces sp. NRRL WC-3725 Terrabacteria group; Actinobacteria; Actinobacteria; Streptomycetales; Streptomycetaceae; Streptomyces MRHARRRVVRRVTRLAAVGGLLLGGAMVTQAAMASETPPASARPLAATGDTGAALVARLG WP_030984557.1 MULTISPECIES: protease [Streptomyces] Length: 458\nNumber of predicted TMHs: 2\nExp number of AAs in TMHs: 43.16635\nExp number, first 60 AAs: 22.26001\nTotal prob of N-in: 0.99239\nPOSSIBLE N-term signal sequence\ninside 1 11\nTMhelix 12 34\noutside 35 419\nTMhelix 420 442\ninside 443 458

33478 GCF_000720295.1_ASM72029v1 Streptomyces sp. NRRL WC-3725 Terrabacteria group; Actinobacteria; Actinobacteria; Streptomycetales; Streptomycetaceae; Streptomyces MRTPHTARRRRALAGVLLAACLALAGCGGADDMGGSSKAASDSKAEAAGAPGGSARNGRA WP_031023561.1 hypothetical protein [Streptomyces sp. NRRL WC-3725] Length: 324\nNumber of predicted TMHs: 2\nExp number of AAs in TMHs: 42.45736\nExp number, first 60 AAs: 19.48055\nTotal prob of N-in: 0.97333\nPOSSIBLE N-term signal sequence\ninside 1 12\nTMhelix 13 30\noutside 31 259\nTMhelix 260 282\ninside 283 324

33479 GCF_000717095.1_ASM71709v1 Streptomyces sp. NRRL WC-3744 Terrabacteria group; Actinobacteria; Actinobacteria; Streptomycetales; Streptomycetaceae; Streptomyces MRTPHTARRRRALAGVLLAACLALAGCGGADDMGGSSKAASDSKAEAAGAPGGSARNGRA WP_030991215.1 hypothetical protein [Streptomyces sp. NRRL WC-3744] Length: 324\nNumber of predicted TMHs: 2\nExp number of AAs in TMHs: 42.47004\nExp number, first 60 AAs: 19.49302\nTotal prob of N-in: 0.97332\nPOSSIBLE N-term signal sequence\ninside 1 12\nTMhelix 13 30\noutside 31 259\nTMhelix 260 282\ninside 283 324

33480 GCF_000717895.1_ASM71789v1 Streptomyces sp. NRRL WC-3744 Terrabacteria group; Actinobacteria; Actinobacteria; Streptomycetales; Streptomycetaceae; Streptomyces MRHARRRVVRRVTRLAAVGGLLLGGAMVTQAAMASETPPASARPLAATGDTGAALVARLG WP_030984557.1 MULTISPECIES: protease [Streptomyces] Length: 458\nNumber of predicted TMHs: 2\nExp number of AAs in TMHs: 43.16635\nExp number, first 60 AAs: 22.26001\nTotal prob of N-in: 0.99239\nPOSSIBLE N-term signal sequence\ninside 1 11\nTMhelix 12 34\noutside 35 419\nTMhelix 420 442\ninside 443 458

33481 GCF_000717895.1_ASM71789v1 Streptomyces sp. NRRL WC-3744 Terrabacteria group; Actinobacteria; Actinobacteria; Streptomycetales; Streptomycetaceae; Streptomyces MRTPHTARRRRALAGVLLAACLALAGCGGADDMGGSSKAASDSKAEAAGAPGGSARNGRA WP_030991215.1 hypothetical protein [Streptomyces sp. NRRL WC-3744] Length: 324\nNumber of predicted TMHs: 2\nExp number of AAs in TMHs: 42.47004\nExp number, first 60 AAs: 19.49302\nTotal prob of N-in: 0.97332\nPOSSIBLE N-term signal sequence\ninside 1 12\nTMhelix 13 30\noutside 31 259\nTMhelix 260 282\ninside 283 324

33482 GCF_000717895.1_ASM71789v1 Streptomyces sp. NRRL WC-3744 Terrabacteria group; Actinobacteria; Actinobacteria; Streptomycetales; Streptomycetaceae; Streptomyces

MLRGVENHSLPRAAAFFDLDKTVIAKSSTLTFSKSFYQGGLINRRAALRTAYAQFVFLVG WP_030981928.1 inhibition of morphological differentiation protein [Streptomyces sp. NRRL WC-3744] Length: 276\nNumber of predicted TMHs: 1\nExp number of AAs in TMHs: 21.26121\nExp number, first 60 AAs: 0.04391\nTotal prob of N-in: 0.00495\noutside 1 248\nTMhelix 249 268\ninside 269 276

33483 GCF_000717095.1_ASM71709v1 Streptomyces sp. NRRL WC-3744 Terrabacteria group; Actinobacteria; Actinobacteria; Streptomycetales; Streptomycetaceae; Streptomyces MRHARRRVVRRVTRLAAVGGLLLGGAMVTQAAMASETPPASARPLAATGDTGAALVARLG WP_030984557.1 MULTISPECIES: protease [Streptomyces] Length: 458\nNumber of predicted TMHs: 2\nExp number of AAs in TMHs: 43.16635\nExp number, first 60 AAs: 22.26001\nTotal prob of N-in: 0.99239\nPOSSIBLE N-term signal sequence\ninside 1 11\nTMhelix 12 34\noutside 35 419\nTMhelix 420 442\ninside 443 458

33484 GCF_000717095.1_ASM71709v1 Streptomyces sp. NRRL WC-3744 Terrabacteria group; Actinobacteria; Actinobacteria; Streptomycetales; Streptomycetaceae; Streptomyces MLRGVENHSLPRAAAFFDLDKTVIAKSSTLTFSKSFYQGGLINRRAALRTAYAQFVFLVG WP_030981928.1 inhibition of morphological differentiation protein [Streptomyces sp. NRRL WC-3744] Length: 276\nNumber of predicted TMHs: 1\nExp number of AAs in TMHs: 21.26121\nExp number, first 60 AAs: 0.04391\nTotal prob of N-in: 0.00495\noutside 1 248\nTMhelix 249 268\ninside 269 276

33485 GCF_000721945.1_ASM72194v1 Streptomyces sp. NRRL WC-3774 Terrabacteria group; Actinobacteria; Actinobacteria; Streptomycetales; Streptomycetaceae; Streptomyces MTRRAVVLSLLTGLFLTAMSPAHATGYRYWSFWDRTAGTWTYATQGPSTTVPSDGDVQGF WP_031060443.1 MULTISPECIES: hypothetical protein [Streptomyces] Length: 213\nNumber of predicted TMHs: 2\nExp number of AAs in TMHs: 41.14384\nExp number, first 60 AAs: 21.27398\nTotal prob of N-in: 0.95944\nPOSSIBLE N-term signal sequence\ninside 1 4\nTMhelix 5 27\noutside 28 184\nTMhelix 185 204\ninside 205 213

33486 GCF_000721945.1_ASM72194v1 Streptomyces sp. NRRL WC-3774 Terrabacteria group; Actinobacteria; Actinobacteria; Streptomycetales; Streptomycetaceae; Streptomyces MRHARRRVVRRVTRLAAVGGLLLGGAMVTRAVASEPPATTAVPRTYAMKAGQTGTDLVSQ WP_031038994.1 MULTISPECIES: protease [Streptomyces] Length: 455\nNumber of predicted TMHs: 2\nExp number of AAs in TMHs: 39.2163499999999\nExp number, first 60 AAs: 20.94931\nTotal prob of N-in: 0.95945\nPOSSIBLE N-term signal sequence\ninside 1 11\nTMhelix 12 34\noutside 35 416\nTMhelix 417 439\ninside 440 455

33487 GCF_000721945.1_ASM72194v1 Streptomyces sp. NRRL WC-3774 Terrabacteria group; Actinobacteria; Actinobacteria; Streptomycetales; Streptomycetaceae; Streptomyces MHLTMSRRQGRTCGWRLRRNPLRRRSYFVEAWLRLATGILAMTVAIVTGVLTARAVEGTL WP_051833035.1 hypothetical protein [Streptomyces sp. NRRL WC-3774] Length: 196\nNumber of predicted TMHs: 2\nExp number of AAs in TMHs: 44.99961\nExp number, first 60 AAs: 22.5245\nTotal prob of N-in: 0.99861\nPOSSIBLE N-term signal sequence\ninside 1 30\nTMhelix 31 53\noutside 54 145\nTMhelix 146 168\ninside 169 196

33488 GCF_000721945.1_ASM72194v1 Streptomyces sp. NRRL WC-3774 Terrabacteria group; Actinobacteria; Actinobacteria; Streptomycetales; Streptomycetaceae; Streptomyces MLGVVENHSLPRTAAFFDLDKTVIAKSSTLTFSKSFYQGGLINRRAALRTAYAQFVFLVG WP_031044737.1 MULTISPECIES: inhibition of morphological differentiation protein [Streptomyces] Length: 277\nNumber of predicted TMHs: 1\nExp number of AAs in TMHs: 21.65845\nExp number, first 60 AAs: 0.01246\nTotal prob of N-in: 0.00153\noutside 1 247\nTMhelix 248 270\ninside 271 277

33489 GCF_000720675.1_ASM72067v1 Streptomyces sp. NRRL WC-3795 Terrabacteria group; Actinobacteria; Actinobacteria; Streptomycetales; Streptomycetaceae; Streptomyces MRHARRRMVRRVARLAAVGGLLLGGTMVTRAVASEPPDAPAVPYTSARAASGTGAGLVSR WP_031015861.1 protease [Streptomyces sp. NRRL WC-3795] Length: 459\nNumber of predicted TMHs: 2\nExp number of AAs in TMHs: 39.87129\nExp number, first 60 AAs: 20.99973\nTotal prob of N-in: 0.96422\nPOSSIBLE N-term signal sequence\ninside 1 11\nTMhelix 12 34\noutside 35 420\nTMhelix 421 443\ninside 444 459

33490 GCF_000698495.1_NTK937 Streptomyces sp. NTK 937 Terrabacteria group; Actinobacteria; Actinobacteria; Streptomycetales; Streptomycetaceae; Streptomyces MLSFVENCFSPRTAAFFDLDKTVIAKSSTLAFSKSFYQGGLINRRAVLRTAYTQFVFLAG WP_037877868.1 inhibition of morphological differentiation protein [Streptomyces sp. NTK 937] Length: 279\nNumber of predicted TMHs: 1\nExp number of AAs in TMHs: 21.58503\nExp number, first 60 AAs: 0.0171\nTotal prob of N-in: 0.00744\noutside 1 247\nTMhelix 248 270\ninside 271 279

33491	GCF_001705785.1_ASM170578v1	Streptomyces sporocinereus	Terrabacteria group; Actinobacteria; Actinobacteria; Streptomycetales; Streptomycetaceae; Streptomyces
	MLSVVENHSLPRTAAFFDLDKTVIAKSSALTFSKSFYQGGLINRRAVLRTAYAQFVFLAG	WP_066027256.1	inhibition of morphological differentiation protein [Streptomyces sporocinereus]	Length: 281\nNumber of predicted TMHs:  1\nExp number of AAs in TMHs: 21.29191\nExp number, first 60 AAs:  0.02347\nTotal prob of N-in:   0.00265\noutside   1   246\nTMhelix   247   269\ninside   270   281

33492	GCF_001570645.1_ASM157064v1	Streptomyces sporocinereus	Terrabacteria group; Actinobacteria; Actinobacteria; Streptomycetales; Streptomycetaceae; Streptomyces
	MLSVVENHSLPRTAAFFDLDKTVIAKSSALTFSKSFYQGGLINRRAVLRTAYAQFVFLAG	WP_060951846.1	MULTISPECIES: inhibition of morphological differentiation protein [Streptomyces]	Length: 281\nNumber of predicted TMHs:  1\nExp number of AAs in TMHs: 21.2919\nExp number, first 60 AAs:  0.02346\nTotal prob of N-in:   0.00265\noutside   1   246\nTMhelix   247   269\ninside   270   281

33493	GCF_000364805.1_ASM36480v1	Streptomyces sp. PAMC 26508	Terrabacteria group; Actinobacteria; Actinobacteria; Streptomycetales; Streptomycetaceae; Streptomyces
	MLTFVENCFSPRTAAFFDLDKTVIAKSSTLTFSKSFYQGGLINRRAVLRTAYTQFVFLAG	WP_014155461.1	MULTISPECIES: haloacid dehalogenase [Streptomyces]	Length: 279\nNumber of predicted TMHs:  1\nExp number of AAs in TMHs: 21.10966\nExp number, first 60 AAs:  0.01497\nTotal prob of N-in:   0.00234\noutside   1   248\nTMhelix   249   268\ninside   269   279

33494	GCF_000364805.1_ASM36480v1	Streptomyces sp. PAMC 26508	Terrabacteria group; Actinobacteria; Actinobacteria; Streptomycetales; Streptomycetaceae; Streptomyces
	MTTHPSPRRRLLTAAAATAVLLVTLPVLPAAAEDSTQCTFPSKKYAGRPWSLQRVLMDEL	WP_014153729.1
	MULTISPECIES: type VII secretion-associated serine protease [Streptomyces]	Length: 414\nNumber of predicted TMHs:  1\nExp number of AAs in TMHs: 36.07361\nExp number, first 60 AAs:  13.5063\nTotal prob of N-in:   0.65550\nPOSSIBLE N-term signal sequence\noutside   1   378\nTMhelix   379   401\ninside   402   414

33495	GCF_000364805.1_ASM36480v1	Streptomyces sp. PAMC 26508	Terrabacteria group; Actinobacteria; Actinobacteria; Streptomycetales; Streptomycetaceae; Streptomyces
	MRTTRRNTLRRGGVGALLAAVTLLTSAGAAVADDPSPSASADGGAAGPTEAGTTFRTATA	WP_014152884.1
	MULTISPECIES: hypothetical protein [Streptomyces]	Length: 274\nNumber of predicted TMHs:  2\nExp number of AAs in TMHs: 40.1724799999999\nExp number, first 60 AAs:  19.30637\nTotal prob of N-in:   0.99359\nPOSSIBLE N-term signal sequence\ninside   1   12\nTMhelix   13   32\noutside   33   237\nTMhelix   238   260\ninside   261   274

33496	GCF_000612545.1_P1-1	Streptomyces sp. PCS3-D2	Terrabacteria group; Actinobacteria; Actinobacteria; Streptomycetales; Streptomycetaceae; Streptomyces
	MPARRRTVTPIRIAAAGLAPLALAAYAAVPAAAHGSMTDPVSRVAACYAEGPESPKSAAC	WP_037925675.1
	chitin-binding protein [Streptomyces sp. PCS3-D2]	Length: 356\nNumber of predicted TMHs:  2\nExp number of AAs in TMHs: 43.27536\nExp number, first 60 AAs:  21.49429\nTotal prob of N-in:   0.97664\nPOSSIBLE N-term signal sequence\ninside   1   12\nTMhelix   13   35\noutside   36   321\nTMhelix   322   344\ninside   345   356

33497	GCF_000612545.1_P1-1	Streptomyces sp. PCS3-D2	Terrabacteria group; Actinobacteria; Actinobacteria; Streptomycetales; Streptomycetaceae; Streptomyces
	MSTARRRTLVRATAVTACAGSLLALPTAAALAEGVPAAASAHSSAAQRTLVKSLELADGV	WP_037927858.1
	hypothetical protein [Streptomyces sp. PCS3-D2]	Length: 210\nNumber of predicted TMHs:  2\nExp number of AAs in TMHs: 41.43362\nExp number, first 60 AAs:  18.49721\nTotal prob of N-in:   0.91294\nPOSSIBLE N-term signal sequence\ninside   1   8\nTMhelix   9   31\noutside   32   180\nTMhelix   181   203\ninside   204   210

33498	GCF_000612545.1_P1-1	Streptomyces sp. PCS3-D2	Terrabacteria group; Actinobacteria; Actinobacteria; Streptomycetales; Streptomycetaceae; Streptomyces
	MPQNRKPRRALLIAACALTLTSASASASASASASAAGAAPTRTAPPHAPYGLRLDGAGEC	WP_078557024.1	type VII secretion-associated serine protease mycosin [Streptomyces sp. PCS3-D2]	Length: 428\nNumber of predicted TMHs:  1\nExp number of AAs in TMHs: 27.32096\nExp number, first 60 AAs:  4.97605\nTotal prob of N-in:   0.28887\noutside   1   394\nTMhelix   395   417\ninside   418   428

33499	GCF_900187925.1_IMG-taxon_2524614673_annotated_assembly	Streptomyces sp. PgraA7	Terrabacteria group; Actinobacteria; Actinobacteria; Streptomycetales; Streptomycetaceae; Streptomyces
	MGLSWNDRRLERRVKPGDGKPLKPFRWWQMTRRSLLSIALPVGGGGGLSATYTVEVKHGG	WP_088577171.1
	hypothetical protein [Streptomyces sp. PgraA7]	Length: 220\nNumber of predicted TMHs:  2\nExp number of

AAs in TMHs: 42.07118\nExp number, first 60 AAs: 2.58869\nTotal prob of N-in: 0.93135\ninside 1 137\nTMhelix 138 160\noutside 161 184\nTMhelix 185 202\ninside 203 220

33500 GCF_900187925.1_IMG-taxon_2524614673_annotated_assembly Streptomyces sp. PgraA7 Terrabacteria group; Actinobacteria; Actinobacteria; Streptomycetales; Streptomycetaceae; Streptomyces MLSVVENCFSPRTAAFFDLDKTVIAKSSTLTFSKSFYQGGLINRRAVLRTAYTQFVFLAG WP_088576745.1 inhibition of morphological differentiation protein [Streptomyces sp. PgraA7] Length: 281\nNumber of predicted TMHs: 1\nExp number of AAs in TMHs: 21.47468\nExp number, first 60 AAs: 0.01919\nTotal prob of N-in: 0.00538\noutside 1 247\nTMhelix 248 270\ninside 271 281

33501 GCF_000590515.1_Streptomyces_sp._PRh5_version1.0 Streptomyces sp. PRh5 Terrabacteria group; Actinobacteria; Actinobacteria; Streptomycetales; Streptomycetaceae; Streptomyces MSSPDREVRAARNDTARTARRPVVAVTGAASGPGALLTQRLAESEEVKQVLAIDERRGEV WP_037952333.1 NAD-dependent dehydratase [Streptomyces sp. PRh5] Length: 369\nNumber of predicted TMHs: 1\nExp number of AAs in TMHs: 19.99765\nExp number, first 60 AAs: 0.38218\nTotal prob of N-in: 0.03037\noutside 1 339\nTMhelix 340 359\ninside 360 369

33502 GCF_000373685.1_ASM37368v1 Streptomyces sp. PsTaAH-124 Terrabacteria group; Actinobacteria; Actinobacteria; Streptomycetales; Streptomycetaceae; Streptomyces MSRTTGGRSRRRPLALAAAVATVTATVASVGAAALAGAPGAAAADRPLSGYELTWGIKDS WP_018570515.1 hypothetical protein [Streptomyces sp. PsTaAH-124] Length: 538\nNumber of predicted TMHs: 2\nExp number of AAs in TMHs: 45.55135\nExp number, first 60 AAs: 22.95714\nTotal prob of N-in: 0.99303\nPOSSIBLE N-term signal sequence\ninside 1 12\nTMhelix 13 35\noutside 36 506\nTMhelix 507 529\ninside 530 538

33503 GCF_000373685.1_ASM37368v1 Streptomyces sp. PsTaAH-124 Terrabacteria group; Actinobacteria; Actinobacteria; Streptomycetales; Streptomycetaceae; Streptomyces MASRGRRATMRNLGALAGAAALTVLGGAAPGWAASPTPDGHRPPATPIEHVVVLFDENIS WP_026252868.1 MULTISPECIES: phospholipase C [Streptomyces] Length: 613\nNumber of predicted TMHs: 2\nExp number of AAs in TMHs: 44.60387\nExp number, first 60 AAs: 22.3165\nTotal prob of N-in: 0.99261\nPOSSIBLE N-term signal sequence\ninside 1 12\nTMhelix 13 35\noutside 36 582\nTMhelix 583 605\ninside 606 613

33504 GCF_000373685.1_ASM37368v1 Streptomyces sp. PsTaAH-124 Terrabacteria group; Actinobacteria; Actinobacteria; Streptomycetales; Streptomycetaceae; Streptomyces MLDLVENHSLPRTAAFFDLDKTVIAKSSTLTFSKSFYQGGLINRRAVLRTAYAQFVFLAG WP_078531172.1 inhibition of morphological differentiation protein [Streptomyces sp. PsTaAH-124] Length: 276\nNumber of predicted TMHs: 1\nExp number of AAs in TMHs: 21.81496\nExp number, first 60 AAs: 0.03069\nTotal prob of N-in: 0.00631\noutside 1 247\nTMhelix 248 270\ninside 271 276

33505 GCF_001687325.1_ASM168732v1 Streptomyces sp. PTY087I2 Terrabacteria group; Actinobacteria; Actinobacteria; Streptomycetales; Streptomycetaceae; Streptomyces MLSVVENCFSPRTAAFFDLDKTVIAKSSTLTFSKSFYQGGLINRRAVLRTAYTQFVFLAG WP_065487922.1 inhibition of morphological differentiation protein [Streptomyces sp. PTY087I2] Length: 281\nNumber of predicted TMHs: 1\nExp number of AAs in TMHs: 21.50792\nExp number, first 60 AAs: 0.01798\nTotal prob of N-in: 0.00376\noutside 1 247\nTMhelix 248 270\ninside 271 281

33506 GCF_000495755.1_PVA94-07.23 Streptomyces sp. PVA 94-07 Terrabacteria group; Actinobacteria; Actinobacteria; Streptomycetales; Streptomycetaceae; Streptomyces MGGRGTTAQARGRGRLPGRPSDPAPTGGNDVITGVNLRMPARRTALPALLLTVTALLAVP WP_033950195.1 MULTISPECIES: hypothetical protein [Streptomyces] Length: 301\nNumber of predicted TMHs: 2\nExp number of AAs in TMHs: 41.7448\nExp number, first 60 AAs: 13.52141\nTotal prob of N-in: 0.87370\nPOSSIBLE N-term signal sequence\ninside 1 44\nTMhelix 45 67\noutside 68 273\nTMhelix 274 296\ninside 297 301

33507 GCF_000495755.1_PVA94-07.23 Streptomyces sp. PVA 94-07 Terrabacteria group; Actinobacteria; Actinobacteria; Streptomycetales; Streptomycetaceae; Streptomyces MLDPVENRSLPRTAAFFDLDKTVIAKSSTLTFSKSFYQGGLINRRAVLRTAYTQFVFLAG WP_023418091.1 SsgB protein [Streptomyces sp. PVA 94-07] Length: 279\nNumber of predicted TMHs: 1\nExp number of AAs in TMHs: 21.68246\nExp number, first 60 AAs: 0.03234\nTotal prob of N-in: 0.01551\noutside 1 247\nTMhelix 248 270\ninside 271 279

33508 GCF_000495755.1_PVA94-07.23 Streptomyces sp. PVA 94-07 Terrabacteria group; Actinobacteria; Actinobacteria; Streptomycetales; Streptomycetaceae; Streptomyces

MSRRTRTARRRPVAAAVLAAGAVLLPVVPASAASGADQDFVIEDPRITESSGLAASRKHP WP_023417760.1 hypothetical protein [Streptomyces sp. PVA 94-07] Length: 344\nNumber of predicted TMHs: 2\nExp number of AAs in TMHs: 38.78147\nExp number, first 60 AAs: 21.30202\nTotal prob of N-in: 0.99448\nPOSSIBLE N-term signal sequence\ninside 1 12\nTMhelix 13 35\noutside 36 316\nTMhelix 317 336\ninside 337 344

33509 GCF_000495755.1_PVA94-07.23 Streptomyces sp. PVA 94-07 Terrabacteria group; Actinobacteria; Actinobacteria; Streptomycetales; Streptomycetaceae; Streptomyces MTPNRRPVVLAAAVATALGLGATALTLPAVAAEAPRAGAPVMDLTDGTLNWGVKESFRRY WP_023419704.1 MULTISPECIES: membrane protein [Streptomyces] Length: 518\nNumber of predicted TMHs: 2\nExp number of AAs in TMHs: 42.79336\nExp number, first 60 AAs: 22.03121\nTotal prob of N-in: 0.97269\nPOSSIBLE N-term signal sequence\ninside 1 6\nTMhelix 7 29\noutside 30 483\nTMhelix 484 506\ninside 507 518

33510 GCF_000495755.1_PVA94-07.23 Streptomyces sp. PVA 94-07 Terrabacteria group; Actinobacteria; Actinobacteria; Streptomycetales; Streptomycetaceae; Streptomyces MPDRRSAVSLRLPLPTAAPGFGGLALPGAAATAPDAGDVVAVRESSEKERKSSSDPEGRS WP_023419047.1 MULTISPECIES: hypothetical protein [Streptomyces] Length: 292\nNumber of predicted TMHs: 2\nExp number of AAs in TMHs: 49.52916\nExp number, first 60 AAs: 3.9366\nTotal prob of N-in: 0.40255\noutside 1 164\nTMhelix 165 187\ninside 188 229\nTMhelix 230 252\noutside 253 292

33511 GCF_001426325.1_Root1295 Streptomyces sp. Root1295 Terrabacteria group; Actinobacteria; Actinobacteria; Streptomycetales; Streptomycetaceae; Streptomyces MLSVVENCFSPRTAAFFDLDKTVIAKSSTLTFSKSFYQGGLINRRAVLRTAYAQFVFLAG WP_032791512.1 MULTISPECIES: inhibition of morphological differentiation protein [Streptomyces] Length: 281\nNumber of predicted TMHs: 1\nExp number of AAs in TMHs: 21.51851\nExp number, first 60 AAs: 0.0316\nTotal prob of N-in: 0.00586\noutside 1 247\nTMhelix 248 270\ninside 271 281

33512 GCF_001426405.1_Root1304 Streptomyces sp. Root1304 Terrabacteria group; Actinobacteria; Actinobacteria; Streptomycetales; Streptomycetaceae; Streptomyces MLRLVENHSLPRTAAFFDLDKTVIAKSSTLTFSKSFYRGGLISRRAALRTAYIQFVFLAG WP_056569321.1 MULTISPECIES: inhibition of morphological differentiation protein [Streptomyces] Length: 288\nNumber of predicted TMHs: 1\nExp number of AAs in TMHs: 21.90302\nExp number, first 60 AAs: 0.0542\nTotal prob of N-in: 0.00429\noutside 1 247\nTMhelix 248 270\ninside 271 288

33513 GCF_001426405.1_Root1304 Streptomyces sp. Root1304 Terrabacteria group; Actinobacteria; Actinobacteria; Streptomycetales; Streptomycetaceae; Streptomyces MKPARIARRRRALRNSLMALACAVVVTVTGQPVTGVAEAAPAPKLKPGKRCDQMYGVKNV WP_056564030.1 MULTISPECIES: hypothetical protein [Streptomyces] Length: 774\nNumber of predicted TMHs: 2\nExp number of AAs in TMHs: 42.9008\nExp number, first 60 AAs: 20.98442\nTotal prob of N-in: 0.94780\nPOSSIBLE N-term signal sequence\ninside 1 12\nTMhelix 13 35\noutside 36 742\nTMhelix 743 765\ninside 766 774

33514 GCF_001426405.1_Root1304 Streptomyces sp. Root1304 Terrabacteria group; Actinobacteria; Actinobacteria; Streptomycetales; Streptomycetaceae; Streptomyces MRTDTALRPEPAPAATAPDETVAAPASRRSPLARIAGRAPASPAPAAAAHNIGAVTTALL WP_056564148.1 MULTISPECIES: HPP family protein+B94 [Streptomyces] Length: 195\nNumber of predicted TMHs: 2\nExp number of AAs in TMHs: 79.81699\nExp number, first 60 AAs: 10.80285\nTotal prob of N-in: 0.92446\nPOSSIBLE N-term signal sequence\ninside 1 120\nTMhelix 121 143\noutside 144 157\nTMhelix 158 180\ninside 181 195

33515 GCF_001426425.1_Root1310 Streptomyces sp. Root1310 Terrabacteria group; Actinobacteria; Actinobacteria; Streptomycetales; Streptomycetaceae; Streptomyces MLMVVENHSLPRTAAFFDLDKTVIAKSSTLTFSKSFYQGGLINRRAALRTAYAQFVFLVG WP_054241799.1 inhibition of morphological differentiation protein [Streptomyces sp. Root1310] Length: 277\nNumber of predicted TMHs: 1\nExp number of AAs in TMHs: 21.36377\nExp number, first 60 AAs: 0.01645\nTotal prob of N-in: 0.00243\noutside 1 247\nTMhelix 248 270\ninside 271 277

33516 GCF_001426425.1_Root1310 Streptomyces sp. Root1310 Terrabacteria group; Actinobacteria; Actinobacteria; Streptomycetales; Streptomycetaceae; Streptomyces MPLYGPAVHAFSRSSPRLSRRSLLALAATAATAPLTAVPLAATPAAAASDVVVGGPRLVAWP_057600964.1 D-alanyl-D-alanine carboxypeptidase [Streptomyces sp. Root1310] Length: 405\nNumber of predicted TMHs: 1\nExp number of AAs in TMHs: 23.76841\nExp number, first 60 AAs: 2.86039\nTotal prob of N-in: 0.12811\noutside 1 368\nTMhelix 369 391\ninside 392 405

33517 GCF_001426425.1_Root1310 Streptomyces sp. Root1310 Terrabacteria group; Actinobacteria; Actinobacteria; Streptomycetales; Streptomycetaceae; Streptomyces
MRRRAFFATLPGWAALLFPPGLGAAPAQAADPVAPYVELTPNTGQLASGGAPLTVDGLFH WP_057601763.1 hypothetical protein [Streptomyces sp. Root1310] Length: 304\nNumber of predicted TMHs: 1\nExp number of AAs in TMHs: 25.24305\nExp number, first 60 AAs: 4.52925\nTotal prob of N-in: 0.21891\noutside 1 275\nTMhelix 276 295\ninside 296 304

33518 GCF_001426615.1_Root1319 Streptomyces sp. Root1319 Terrabacteria group; Actinobacteria; Actinobacteria; Streptomycetales; Streptomycetaceae; Streptomyces
MLTIVENCFSPRTAAFFDLDKTVIAKSSTLTFSKSFYQGGLINRRAVLRTAYTQFVFLAG WP_056793391.1 MULTISPECIES: inhibition of morphological differentiation protein [Streptomyces] Length: 279\nNumber of predicted TMHs: 1\nExp number of AAs in TMHs: 21.77146\nExp number, first 60 AAs: 0.02229\nTotal prob of N-in: 0.00403\noutside 1 247\nTMhelix 248 270\ninside 271 279

33519 GCF_001428885.1_Root264 Streptomyces sp. Root264 Terrabacteria group; Actinobacteria; Actinobacteria; Streptomycetales; Streptomycetaceae; Streptomyces
MLLVVENHSLPRTAAFFDLDKTVIAKSSTLTFSKSFYQGGLINRRAALRTAYAQFVFLVG WP_057577822.1 inhibition of morphological differentiation protein [Streptomyces sp. Root264] Length: 277\nNumber of predicted TMHs: 1\nExp number of AAs in TMHs: 21.51622\nExp number, first 60 AAs: 0.01088\nTotal prob of N-in: 0.00161\noutside 1 247\nTMhelix 248 270\ninside 271 277

33520 GCF_001428885.1_Root264 Streptomyces sp. Root264 Terrabacteria group; Actinobacteria; Actinobacteria; Streptomycetales; Streptomycetaceae; Streptomyces
MPRRATVAGIAAAVLTVALLSGVAHAQDLNCSDFTYQEDAQAVFDQDPSDPNRLDEDRGP WP_057581109.1 excalibur calcium-binding protein [Streptomyces sp. Root264] Length: 155\nNumber of predicted TMHs: 2\nExp number of AAs in TMHs: 41.55882\nExp number, first 60 AAs: 20.60869\nTotal prob of N-in: 0.99301\nPOSSIBLE N-term signal sequence\ninside 1 6\nTMhelix 7 26\noutside 27 125\nTMhelix 126 148\ninside 149 155

33521 GCF_001428885.1_Root264 Streptomyces sp. Root264 Terrabacteria group; Actinobacteria; Actinobacteria; Streptomycetales; Streptomycetaceae; Streptomyces
MHAFSGPSPHQPPQRSPHKPAHQPSAPPSAAPFSRRSLLAAAAAAPLAAAPILAAPAAAA WP_079041571.1 D-alanyl-D-alanine carboxypeptidase [Streptomyces sp. Root264] Length: 413\nNumber of predicted TMHs: 1\nExp number of AAs in TMHs: 23.74998\nExp number, first 60 AAs: 1.68044\nTotal prob of N-in: 0.07248\noutside 1 378\nTMhelix 379 401\ninside 402 413

33522 GCF_001425805.1_Root369 Streptomyces sp. Root369 Terrabacteria group; Actinobacteria; Actinobacteria; Streptomycetales; Streptomycetaceae; Streptomyces
MTRRALLLSLLAGLLLATASPAHAAGYRYWSFWDRTGSTWTYATEGPSTAVPSDGDVQGF WP_057615152.1 hypothetical protein [Streptomyces sp. Root369] Length: 209\nNumber of predicted TMHs: 2\nExp number of AAs in TMHs: 41.9244\nExp number, first 60 AAs: 21.52429\nTotal prob of N-in: 0.94703\nPOSSIBLE N-term signal sequence\ninside 1 4\nTMhelix 5 27\noutside 28 181\nTMhelix 182 201\ninside 202 209

33523 GCF_001425805.1_Root369 Streptomyces sp. Root369 Terrabacteria group; Actinobacteria; Actinobacteria; Streptomycetales; Streptomycetaceae; Streptomyces
MSSPDPQVRAARNQSTNPAARRPVVAVTGAASGIGALLTERLAASEEIKQVIALDERRGE WP_057612235.1 NAD-dependent dehydratase [Streptomyces sp. Root369] Length: 371\nNumber of predicted TMHs: 1\nExp number of AAs in TMHs: 20.4092\nExp number, first 60 AAs: 0.39598\nTotal prob of N-in: 0.03604\noutside 1 341\nTMhelix 342 361\ninside 362 371

33524 GCF_001425805.1_Root369 Streptomyces sp. Root369 Terrabacteria group; Actinobacteria; Actinobacteria; Streptomycetales; Streptomycetaceae; Streptomyces
MRTSRSRRSVLPARALAGLLLAAGLALTGCSGGADSSSDGGSALSDDKAAQSQAEGAAGA WP_057611173.1 hypothetical protein [Streptomyces sp. Root369] Length: 329\nNumber of predicted TMHs: 1\nExp number of AAs in TMHs: 32.1141\nExp number, first 60 AAs: 9.03512\nTotal prob of N-in: 0.66853\noutside 1 264\nTMhelix 265 287\ninside 288 329

33525 GCF_001425805.1_Root369 Streptomyces sp. Root369 Terrabacteria group; Actinobacteria; Actinobacteria; Streptomycetales; Streptomycetaceae; Streptomyces
MRHARRRVVRRVTRLAAVGGLLLGGAMVTRAVASETPPITAVPHTYAMEAGQTGADLVSR WP_057613027.1 protease [Streptomyces sp. Root369] Length: 457\nNumber of predicted TMHs: 2\nExp number of AAs in

TMHs: 38.7970699999999\nExp number, first 60 AAs:  20.58462\nTotal prob of N-in:   0.95681\nPOSSIBLE N-term signal sequence\ninside   1   11\nTMhelix   12   34\noutside   35   418\nTMhelix   419   441\ninside   442   457

33526 GCF_001425805.1_Root369 Streptomyces sp. Root369 Terrabacteria group; Actinobacteria; Actinobacteria; Streptomycetales; Streptomycetaceae; Streptomyces
MLGVVENHSLPRTAAFFDLDKTVIAKSSTLTFSKSFYQGGLINRRAALRTAYAQFVFLVG WP_057612289.1 inhibition of morphological differentiation protein [Streptomyces sp. Root369] Length: 277\nNumber of predicted TMHs:  1\nExp number of AAs in TMHs: 21.66888\nExp number, first 60 AAs:  0.01246\nTotal prob of N-in:   0.00150\noutside   1   247\nTMhelix   248   270\ninside   271   277

33527 GCF_001426585.1_Root431 Streptomyces sp. Root431 Terrabacteria group; Actinobacteria; Actinobacteria; Streptomycetales; Streptomycetaceae; Streptomyces
MLRLVENHSLPRTAAFFDLDKTVIAKSSTLTFSKSFYRGGLISRRAALRTAYIQFVFLVG WP_056652902.1 inhibition of morphological differentiation protein [Streptomyces sp. Root431] Length: 287\nNumber of predicted TMHs:  1\nExp number of AAs in TMHs: 20.91039\nExp number, first 60 AAs:  0.08487\nTotal prob of N-in:   0.00641\noutside   1   248\nTMhelix   249   268\ninside   269   287

33528 GCF_001427245.1_Root55 Streptomyces sp. Root55 Terrabacteria group; Actinobacteria; Actinobacteria; Streptomycetales; Streptomycetaceae; Streptomyces
MLTIVENCFSPRTAAFFDLDKTVIAKSSTLTFSKSFYQGGLINRRAVLRTAYTQFVFLAG WP_056793391.1 MULTISPECIES: inhibition of morphological differentiation protein [Streptomyces] Length: 279\nNumber of predicted TMHs:  1\nExp number of AAs in TMHs: 21.77146\nExp number, first 60 AAs:  0.02229\nTotal prob of N-in:   0.00403\noutside   1   247\nTMhelix   248   270\ninside   271   279

33529 GCF_001427565.1_Root63 Streptomyces sp. Root63 Terrabacteria group; Actinobacteria; Actinobacteria; Streptomycetales; Streptomycetaceae; Streptomyces
MLSVVENCFSPRTAAFFDLDKTVIAKSSTLTFSKSFYQGGLINRRAVLRTAYAQFVFLAG WP_032791512.1 MULTISPECIES: inhibition of morphological differentiation protein [Streptomyces] Length: 281\nNumber of predicted TMHs:  1\nExp number of AAs in TMHs: 21.51851\nExp number, first 60 AAs:  0.0316\nTotal prob of N-in:   0.00586\noutside   1   247\nTMhelix   248   270\ninside   271   281

33530 GCF_001427745.1_Root66D1 Streptomyces sp. Root66D1 Terrabacteria group; Actinobacteria; Actinobacteria; Streptomycetales; Streptomycetaceae; Streptomyces
MLRLVENHSLPRTAAFFDLDKTVIAKSSTLTFSKSFYRGGLISRRAALRTAYIQFVFLAG WP_056569321.1 MULTISPECIES: inhibition of morphological differentiation protein [Streptomyces] Length: 288\nNumber of predicted TMHs:  1\nExp number of AAs in TMHs: 21.90302\nExp number, first 60 AAs:  0.0542\nTotal prob of N-in:   0.00429\noutside   1   247\nTMhelix   248   270\ninside   271   288

33531 GCF_001427745.1_Root66D1 Streptomyces sp. Root66D1 Terrabacteria group; Actinobacteria; Actinobacteria; Streptomycetales; Streptomycetaceae; Streptomyces
MKPARIARRRRALRNSLMALACAVVVTVTGQPVTGVAEAAPAPKLKPGKRCDQMYGVKNV WP_056564030.1
MULTISPECIES: hypothetical protein [Streptomyces] Length: 774\nNumber of predicted TMHs:  2\nExp number of AAs in TMHs: 42.9008\nExp number, first 60 AAs:  20.98442\nTotal prob of N-in:   0.94780\nPOSSIBLE N-term signal sequence\ninside   1   12\nTMhelix   13   35\noutside   36   742\nTMhelix   743   765\ninside   766   774

33532 GCF_001427745.1_Root66D1 Streptomyces sp. Root66D1 Terrabacteria group; Actinobacteria; Actinobacteria; Streptomycetales; Streptomycetaceae; Streptomyces
MRTDTALRPEPAPAATAPDETVAAPASRRSPLARIAGRAPASPAPAAAAHNIGAVTTALL WP_056564148.1
MULTISPECIES: HPP family protein+B94 [Streptomyces] Length: 195\nNumber of predicted TMHs:  2\nExp number of AAs in TMHs: 79.81699\nExp number, first 60 AAs:  10.80285\nTotal prob of N-in:   0.92446\nPOSSIBLE N-term signal sequence\ninside   1   120\nTMhelix   121   143\noutside   144   157\nTMhelix   158   180\ninside   181   195

33533 GCF_001640565.1_ASM164056v1 Streptomyces sp. RTd22 Terrabacteria group; Actinobacteria; Actinobacteria; Streptomycetales; Streptomycetaceae; Streptomyces
MSSPDREVRAARNDTARTARRPVVAVTGAASGPGALLTQRLAESEEIKQVLALDERRGEV WP_009715199.1
MULTISPECIES: NAD-dependent dehydratase [Streptomyces] Length: 369\nNumber of predicted TMHs:  1\nExp number of AAs in TMHs: 20.51064\nExp number, first 60 AAs:  0.50973\nTotal prob of N-in:   0.05336\noutside   1   339\nTMhelix   340   359\ninside   360   369

33534 GCF_001650215.1_ASM165021v1 Streptomyces sp. RTd22 Terrabacteria group; Actinobacteria; Actinobacteria; Streptomycetales; Streptomycetaceae; Streptomyces

MSSPDREVRAARNDTARTARRPVVAVTGAASGPGALLTQRLAESEEIKQVLALDERRGEV WP_009715199.1 MULTISPECIES: NAD-dependent dehydratase [Streptomyces] Length: 369\nNumber of predicted TMHs: 1\nExp number of AAs in TMHs: 20.51064\nExp number, first 60 AAs: 0.50973\nTotal prob of N-in: 0.05336\noutside 1 339\nTMhelix 340 359\ninside 360 369

33535 GCF_001514305.1_ASM151430v1 Streptomyces sp. RV15 Terrabacteria group; Actinobacteria; Actinobacteria; Streptomycetales; Streptomycetaceae; Streptomyces MRHARRQVVRRVTRLAAVGGLLLGGTMVTRAVASEPPDASAVPRTTAASAADMGSGLVTR WP_067026487.1 protease [Streptomyces sp. RV15] Length: 459\nNumber of predicted TMHs: 2\nExp number of AAs in TMHs: 40.66974\nExp number, first 60 AAs: 19.65471\nTotal prob of N-in: 0.89774\nPOSSIBLE N-term signal sequence\ninside 1 11\nTMhelix 12 34\noutside 35 420\nTMhelix 421 443\ninside 444 459

33536 GCF_001514305.1_ASM151430v1 Streptomyces sp. RV15 Terrabacteria group; Actinobacteria; Actinobacteria; Streptomycetales; Streptomycetaceae; Streptomyces MLKPVENHSLPRTAAFFDLDKTVIAKSSTLTFSKSFYQGGLINRRAVLRTAYAQFVFLAG WP_067033297.1 inhibition of morphological differentiation protein [Streptomyces sp. RV15] Length: 277\nNumber of predicted TMHs: 1\nExp number of AAs in TMHs: 21.61285\nExp number, first 60 AAs: 0.02941\nTotal prob of N-in: 0.00415\noutside 1 247\nTMhelix 248 270\ninside 271 277

33537 GCF_001611795.1_ASM161179v1 Streptomyces sp. S10(2016) Terrabacteria group; Actinobacteria; Actinobacteria; Streptomycetales; Streptomycetaceae; Streptomyces MRHARRRVVRRVTRLAAVGGLLLGGAMVTNAVASEPSAPVIGVPFTAPPATGEGTDLVSR WP_062930905.1 protease [Streptomyces sp. S10(2016)] Length: 448\nNumber of predicted TMHs: 2\nExp number of AAs in TMHs: 40.77223\nExp number, first 60 AAs: 21.28837\nTotal prob of N-in: 0.96650\nPOSSIBLE N-term signal sequence\ninside 1 11\nTMhelix 12 34\noutside 35 409\nTMhelix 410 432\ninside 433 448

33538 GCF_001611795.1_ASM161179v1 Streptomyces sp. S10(2016) Terrabacteria group; Actinobacteria; Actinobacteria; Streptomycetales; Streptomycetaceae; Streptomyces MLWGVENHSLPRTAAFFDLDKTVIAKSSTLTFSKSFYQGGLINRRAVLRTAYAQFVFLAG WP_062928299.1 inhibition of morphological differentiation protein [Streptomyces sp. S10(2016)] Length: 277\nNumber of predicted TMHs: 1\nExp number of AAs in TMHs: 21.5954\nExp number, first 60 AAs: 0.02912\nTotal prob of N-in: 0.00274\noutside 1 247\nTMhelix 248 270\ninside 271 277

33539 GCF_001611795.1_ASM161179v1 Streptomyces sp. S10(2016) Terrabacteria group; Actinobacteria; Actinobacteria; Streptomycetales; Streptomycetaceae; Streptomyces MTGGPRSGRSPPVSTRRTTVHLRPPAAVLAAAVIAVAGPAVAAPVAHAEEGAPELVVSAL WP_062927141.1 hypothetical protein [Streptomyces sp. S10(2016)] Length: 427\nNumber of predicted TMHs: 1\nExp number of AAs in TMHs: 35.43163\nExp number, first 60 AAs: 14.13528\nTotal prob of N-in: 0.63588\nPOSSIBLE N-term signal sequence\noutside 1 395\nTMhelix 396 418\ninside 419 427

33540 GCF_000297715.1_ASM29771v1 Streptomyces sp. S4 Terrabacteria group; Actinobacteria; Actinobacteria; Streptomycetales; Streptomycetaceae; Streptomyces MTPNRRPVVLAAAVATALGLGATALTLPAVAAEAPRAGAPVMDLTDGTLDWGVKESFRRY WP_010637519.1 LPXTG-motif cell wall anchor domain-containing protein, partial [Streptomyces sp. S4] Length: 503\nNumber of predicted TMHs: 2\nExp number of AAs in TMHs: 37.73818\nExp number, first 60 AAs: 22.02851\nTotal prob of N-in: 0.96635\nPOSSIBLE N-term signal sequence\ninside 1 6\nTMhelix 7 29\noutside 30 479\nTMhelix 480 502\ninside 503 503

33541 GCF_000297715.1_ASM29771v1 Streptomyces sp. S4 Terrabacteria group; Actinobacteria; Actinobacteria; Streptomycetales; Streptomycetaceae; Streptomyces MLDPVENRSLPRTAAFFDLDKTVIAKSSTLTFSKSFYQGGLINRRAVLRTAYTQFVFLAG WP_010643895.1 MULTISPECIES: morphological differentiation-associated protein [Streptomyces] Length: 290\nNumber of predicted TMHs: 1\nExp number of AAs in TMHs: 21.93803\nExp number, first 60 AAs: 0.02636\nTotal prob of N-in: 0.00751\noutside 1 247\nTMhelix 248 270\ninside 271 290

33542 GCF_000297715.1_ASM29771v1 Streptomyces sp. S4 Terrabacteria group; Actinobacteria; Actinobacteria; Streptomycetales; Streptomycetaceae; Streptomyces MSRRTRTARRRPVAAAVLAAGAVLLPAVPASAASGADQDFVIEDPRITESSGLAASKRHP WP_010643297.1 hypothetical protein [Streptomyces sp. S4] Length: 348\nNumber of predicted TMHs: 2\nExp number of AAs in TMHs: 39.45061\nExp number, first 60 AAs: 21.10527\nTotal prob of N-in: 0.99085\nPOSSIBLE N-term signal sequence\ninside 1 12\nTMhelix 13 35\noutside 36 320\nTMhelix 321 340\ninside 341 348

33543 GCF_000297715.1_ASM29771v1 Streptomyces sp. S4 Terrabacteria group; Actinobacteria; Actinobacteria; Streptomycetales; Streptomycetaceae; Streptomyces
MAPPHRRRAAVAVRALVALTALLCTTGLAPAATSGPAARAAVAEVPPPTATSSVVTVRTG WP_026048504.1 VWA domain-containing protein [Streptomyces sp. S4] Length: 946\nNumber of predicted TMHs: 2\nExp number of AAs in TMHs: 38.76119\nExp number, first 60 AAs: 18.55273\nTotal prob of N-in: 0.85430\nPOSSIBLE N-term signal sequence\ninside 1 11\nTMhelix 12 34\noutside 35 907\nTMhelix 908 930\ninside 931 946

33544 GCF_000297715.1_ASM29771v1 Streptomyces sp. S4 Terrabacteria group; Actinobacteria; Actinobacteria; Streptomycetales; Streptomycetaceae; Streptomyces
MSRMRGSSGDRIAGRRAAVVLAVAVLGLAGCSAGPDSGASSAGDRAVAPSAADGAGGGQA WP_010641263.1 hypothetical protein [Streptomyces sp. S4] Length: 334\nNumber of predicted TMHs: 1\nExp number of AAs in TMHs: 38.13059\nExp number, first 60 AAs: 15.33908\nTotal prob of N-in: 0.72816\nPOSSIBLE N-term signal sequence\noutside 1 269\nTMhelix 270 292\ninside 293 334

33545 GCF_000297715.1_ASM29771v1 Streptomyces sp. S4 Terrabacteria group; Actinobacteria; Actinobacteria; Streptomycetales; Streptomycetaceae; Streptomyces
MPDRRSAVSLRLPLPTAAPGFGGLALPGAAATAPDAWGVVAVRESSEKERKSSSEPEGRS WP_010638641.1 hypothetical protein [Streptomyces sp. S4] Length: 292\nNumber of predicted TMHs: 2\nExp number of AAs in TMHs: 51.76528\nExp number, first 60 AAs: 6.06558\nTotal prob of N-in: 0.33682\noutside 1 164\nTMhelix 165 187\ninside 188 229\nTMhelix 230 252\noutside 253 292

33546 GCF_002094995.1_ASM209499v1 Streptomyces sp. S8 Terrabacteria group; Actinobacteria; Actinobacteria; Streptomycetales; Streptomycetaceae; Streptomyces
MSGHTARRAAVTVLSAALWSPAAQAAAVADPPRPVVERLPEQDVRVAVAAARTPAAVNLA WP_084993274.1 hypothetical protein [Streptomyces sp. S8] Length: 257\nNumber of predicted TMHs: 1\nExp number of AAs in TMHs: 18.92101\nExp number, first 60 AAs: 0.44389\nTotal prob of N-in: 0.23765\noutside 1 228\nTMhelix 229 246\ninside 247 257

33547 GCF_002094995.1_ASM209499v1 Streptomyces sp. S8 Terrabacteria group; Actinobacteria; Actinobacteria; Streptomycetales; Streptomycetaceae; Streptomyces
MLSVVENCFSPRTAAFFDLDKTVIAKSSTLTFSKSFYQGGLINRRAVLRTAYTQFVFLAG WP_084994192.1 inhibition of morphological differentiation protein [Streptomyces sp. S8] Length: 282\nNumber of predicted TMHs: 1\nExp number of AAs in TMHs: 21.56954\nExp number, first 60 AAs: 0.01975\nTotal prob of N-in: 0.00580\noutside 1 247\nTMhelix 248 270\ninside 271 282

33548 GCF_000179215.1_ASM17921v1 Streptomyces sp. SA3_actF Terrabacteria group; Actinobacteria; Actinobacteria; Streptomycetales; Streptomycetaceae; Streptomyces
MLGPVENHSLPRTAAFFDLDKTVIAKSSTLTFSKSFYQGGLINRRAALRTAYTQFVFLVG WP_007823897.1 MULTISPECIES: morphological differentiation-associated protein [Streptomyces] Length: 276\nNumber of predicted TMHs: 1\nExp number of AAs in TMHs: 21.70944\nExp number, first 60 AAs: 0.02049\nTotal prob of N-in: 0.00432\noutside 1 244\nTMhelix 245 267\ninside 268 276

33549 GCF_000179195.1_ASM17919v1 Streptomyces sp. SA3_actG Terrabacteria group; Actinobacteria; Actinobacteria; Streptomycetales; Streptomycetaceae; Streptomyces
MLGPVENHSLPRTAAFFDLDKTVIAKSSTLTFSKSFYQGGLINRRAALRTAYTQFVFLVG WP_007823897.1 MULTISPECIES: morphological differentiation-associated protein [Streptomyces] Length: 276\nNumber of predicted TMHs: 1\nExp number of AAs in TMHs: 21.70944\nExp number, first 60 AAs: 0.02049\nTotal prob of N-in: 0.00432\noutside 1 244\nTMhelix 245 267\ninside 268 276

33550 GCF_001654495.1_ASM165449v1 Streptomyces sp. SAT1 Terrabacteria group; Actinobacteria; Actinobacteria; Streptomycetales; Streptomycetaceae; Streptomyces
MSCSVLRKGRRSTVRAVLVGAAVAAAVLGPAAATFAATTVPGAAAPSATAPAKTARDAVP WP_064534583.1 hypothetical protein [Streptomyces sp. SAT1] Length: 328\nNumber of predicted TMHs: 2\nExp number of AAs in TMHs: 42.51994\nExp number, first 60 AAs: 22.39194\nTotal prob of N-in: 0.98704\nPOSSIBLE N-term signal sequence\ninside 1 12\nTMhelix 13 35\noutside 36 295\nTMhelix 296 318\ninside 319 328

33551 GCF_001654495.1_ASM165449v1 Streptomyces sp. SAT1 Terrabacteria group; Actinobacteria; Actinobacteria; Streptomycetales; Streptomycetaceae; Streptomyces
MLDLVENHSLPRTAAFFDLDKTVIAKSSTLTFSKSFYQGGLINRRAVLRTAYAQFVFLAG WP_079159013.1 inhibition of morphological differentiation protein [Streptomyces sp. SAT1] Length: 276\nNumber of predicted TMHs: 1\nExp

number of AAs in TMHs: 21.79835\nExp number, first 60 AAs: 0.035\nTotal prob of N-in: 0.00783\noutside 1 247\nTMhelix 248 270\ninside 271 276

33552 GCF_001654495.1_ASM165449v1 Streptomyces sp. SAT1 Terrabacteria group; Actinobacteria; Actinobacteria; Streptomycetales; Streptomycetaceae; Streptomyces MSRTTGGRSRRRPLALAAAVATVTATVASVGAAALAGAPGAAAADRPLSGYELTWGIKDS WP_064532817.1 hypothetical protein [Streptomyces sp. SAT1] Length: 538\nNumber of predicted TMHs: 2\nExp number of AAs in TMHs: 45.48526\nExp number, first 60 AAs: 22.95721\nTotal prob of N-in: 0.99305\nPOSSIBLE N-term signal sequence\ninside 1 12\nTMhelix 13 35\noutside 36 506\nTMhelix 507 529\ninside 530 538

33553 GCF_001654495.1_ASM165449v1 Streptomyces sp. SAT1 Terrabacteria group; Actinobacteria; Actinobacteria; Streptomycetales; Streptomycetaceae; Streptomyces MASRGRRATMRNLGALAGAAALTVLGGAAPGLAAPAAPDGHRLTATPIEHVVVLFDENIS WP_064536857.1 phospholipase [Streptomyces sp. SAT1]Length: 613\nNumber of predicted TMHs: 2\nExp number of AAs in TMHs: 44.58795\nExp number, first 60 AAs: 22.34964\nTotal prob of N-in: 0.99302\nPOSSIBLE N-term signal sequence\ninside 1 11\nTMhelix 12 34\noutside 35 582\nTMhelix 583 605\ninside 606 613

33554 GCF_000373405.1_ASM37340v1 Streptomyces sp. ScaeMP-e10 Terrabacteria group; Actinobacteria; Actinobacteria; Streptomycetales; Streptomycetaceae; Streptomyces MGIAASGRRTLLSATAVSATAALIALGAAPAQADATEPDLGVRALAPVAGIAAGSGFGLP WP_018509850.1 hypothetical protein [Streptomyces sp. ScaeMP-e10] Length: 444\nNumber of predicted TMHs: 1\nExp number of AAs in TMHs: 35.0191\nExp number, first 60 AAs: 13.00487\nTotal prob of N-in: 0.29251\nPOSSIBLE N-term signal sequence\noutside 1 412\nTMhelix 413 435\ninside 436 444

33555 GCF_000373405.1_ASM37340v1 Streptomyces sp. ScaeMP-e10 Terrabacteria group; Actinobacteria; Actinobacteria; Streptomycetales; Streptomycetaceae; Streptomyces MLSVVENCFSPRTAAFFDLDKTVIAKSSTLTFSKSFYQGGLINRRAVLRTAYTQFVFLAG WP_018514217.1 inhibition of morphological differentiation protein [Streptomyces sp. ScaeMP-e10] Length: 281\nNumber of predicted TMHs: 1\nExp number of AAs in TMHs: 21.4543\nExp number, first 60 AAs: 0.02143\nTotal prob of N-in: 0.00744\noutside 1 247\nTMhelix 248 270\ninside 271 281

33556 GCF_002128305.1_ASM212830v1 Streptomyces sp. SCSIO 03032 Terrabacteria group; Actinobacteria; Actinobacteria; Streptomycetales; Streptomycetaceae; Streptomyces MEILAQPRTAAFFDLDKTVIAKSSALTFSRSFYQGGLINRRAVLRSAYAQFVYLLGAADH WP_086159819.1 inhibition of morphological differentiation protein [Streptomyces sp. SCSIO 03032] Length: 280\nNumber of predicted TMHs: 1\nExp number of AAs in TMHs: 20.94557\nExp number, first 60 AAs: 0.02721\nTotal prob of N-in: 0.00848\noutside 1 242\nTMhelix 243 265\ninside 266 280

33557 GCF_002128305.1_ASM212830v1 Streptomyces sp. SCSIO 03032 Terrabacteria group; Actinobacteria; Actinobacteria; Streptomycetales; Streptomycetaceae; Streptomyces MSVIRGFRRLPGRARESFRTRPAPGPGTAAERVDLAHPRRSPLGTAVVNCAVYRDGVRRE WP_086158898.1 magnesium transporter CorA [Streptomyces sp. SCSIO 03032] Length: 379\nNumber of predicted TMHs: 2\nExp number of AAs in TMHs: 41.19053\nExp number, first 60 AAs: 0.00018\nTotal prob of N-in: 0.96035\ninside 1 321\nTMhelix 322 341\noutside 342 350\nTMhelix 351 373\ninside 374 379

33558 GCF_002080455.1_ASM208045v1 Streptomyces sp. Sge12 Terrabacteria group; Actinobacteria; Actinobacteria; Streptomycetales; Streptomycetaceae; Streptomyces MNRIHAPRRPLLAAAFALTFTLASAGAGAGAASAAEAAPAPQAPYAQAPYGLRLDGAGEC WP_081522290.1 type VII secretion-associated serine protease mycosin [Streptomyces sp. Sge12] Length: 428\nNumber of predicted TMHs: 2\nExp number of AAs in TMHs: 42.86554\nExp number, first 60 AAs: 20.57064\nTotal prob of N-in: 0.91715\nPOSSIBLE N-term signal sequence\ninside 1 11\nTMhelix 12 34\noutside 35 394\nTMhelix 395 417\ninside 418 428

33559 GCF_002080455.1_ASM208045v1 Streptomyces sp. Sge12 Terrabacteria group; Actinobacteria; Actinobacteria; Streptomycetales; Streptomycetaceae; Streptomyces MSTVSRRTLVRAAAVTACAGSLLALPTAAALAEGVPAASSARSAAPQLSLVKSLSLADGT WP_081522449.1 hypothetical protein [Streptomyces sp. Sge12] Length: 217\nNumber of predicted TMHs: 2\nExp number of AAs in TMHs: 41.6065599999999\nExp number, first 60 AAs: 18.99766\nTotal prob of N-in: 0.88899\nPOSSIBLE N-term signal sequence\ninside 1 8\nTMhelix 9 31\noutside 32 187\nTMhelix 188 210\ninside 211 217

33560 GCF_002080455.1_ASM208045v1 Streptomyces sp. Sge12 Terrabacteria group; Actinobacteria; Actinobacteria; Streptomycetales; Streptomycetaceae; Streptomyces

MPARRRTAVALTRIASAGLAPLALAAYAAAPAVAHGSMTDPVSRVAACYAEGPESPKTAA WP_081523924.1 chitin-binding protein [Streptomyces sp. Sge12] Length: 359\nNumber of predicted TMHs: 2\nExp number of AAs in TMHs: 42.24546\nExp number, first 60 AAs: 20.34508\nTotal prob of N-in: 0.90797\nPOSSIBLE N-term signal sequence\ninside 1 12\nTMhelix 13 35\noutside 36 322\nTMhelix 323 345\ninside 346 359

33561 GCF_002080455.1_ASM208045v1 Streptomyces sp. Sge12 Terrabacteria group; Actinobacteria; Actinobacteria; Streptomycetales; Streptomycetaceae; Streptomyces MRRRSPVIAVTGAASGVGAALVSRLAASDEVKQVVAIDERRGDCAAAQWHILDVRDPAIA WP_051779201.1 MULTISPECIES: NAD-dependent dehydratase [Streptomyces] Length: 353\nNumber of predicted TMHs: 1\nExp number of AAs in TMHs: 30.20524\nExp number, first 60 AAs: 7.40836\nTotal prob of N-in: 0.39674\noutside 1 324\nTMhelix 325 347\ninside 348 353

33562 GCF_000177195.2_ASM17719v2 Streptomyces sp. SirexAA-E Terrabacteria group; Actinobacteria; Actinobacteria; Streptomycetales; Streptomycetaceae; Streptomyces MLTFVENCFSPRTAAFFDLDKTVIAKSSTLTFSKSFYQGGLINRRAVLRTAYTQFVFLAG WP_014046857.1 MULTISPECIES: haloacid dehalogenase [Streptomyces] Length: 279\nNumber of predicted TMHs: 1\nExp number of AAs in TMHs: 21.69088\nExp number, first 60 AAs: 0.01488\nTotal prob of N-in: 0.00190\noutside 1 247\nTMhelix 248 270\ninside 271 279

33563 GCF_000647875.1_ASM64787v1 Streptomyces sp. SolWspMP-sol2th Terrabacteria group; Actinobacteria; Actinobacteria; Streptomycetales; Streptomycetaceae; Streptomyces MSGHTARRAAVTVLSAAALWSPAAQAAAVADPPRPVVERLPEQDVRVAVAAARTPAAVNL WP_028416940.1 hypothetical protein [Streptomyces sp. SolWspMP-sol2th] Length: 258\nNumber of predicted TMHs: 1\nExp number of AAs in TMHs: 19.40294\nExp number, first 60 AAs: 0.92809\nTotal prob of N-in: 0.25360\noutside 1 229\nTMhelix 230 247\ninside 248 258

33564 GCF_000647875.1_ASM64787v1 Streptomyces sp. SolWspMP-sol2th Terrabacteria group; Actinobacteria; Actinobacteria; Streptomycetales; Streptomycetaceae; Streptomyces MLSVVENCFSPRTAAFFDLDKTVIAKSSTLTFSKSFYQGGLINRRAVLRTAYTQFVFLAG WP_028416068.1 inhibition of morphological differentiation protein [Streptomyces sp. SolWspMP-sol2th] Length: 279\nNumber of predicted TMHs: 1\nExp number of AAs in TMHs: 21.58339\nExp number, first 60 AAs: 0.02018\nTotal prob of N-in: 0.00632\noutside 1 247\nTMhelix 248 270\ninside 271 279

33565 GCF_000154905.1_ASM15490v1 Streptomyces sp. SPB074 Terrabacteria group; Actinobacteria; Actinobacteria; Streptomycetales; Streptomycetaceae; Streptomyces MLRPVENHSLPRTAAFFDLDKTVIAKSSTLTFSKSFYQGGLINRRAALRTAYTQFVFLVG WP_008746524.1 haloacid dehalogenase [Streptomyces sp. SPB074] Length: 275\nNumber of predicted TMHs: 1\nExp number of AAs in TMHs: 21.76861\nExp number, first 60 AAs: 0.02556\nTotal prob of N-in: 0.01167\noutside 1 245\nTMhelix 246 268\ninside 269 275

33566 GCF_000158855.1_ASM15885v1 Streptomyces sp. SPB78 Terrabacteria group; Actinobacteria; Actinobacteria; Streptomycetales; Streptomycetaceae; Streptomyces MLGPVENHSLPRTAAFFDLDKTVIAKSSTLTFSKSFYQGGLINRRAALRTAYTQFVFLVG WP_007823897.1 MULTISPECIES: morphological differentiation-associated protein [Streptomyces] Length: 276\nNumber of predicted TMHs: 1\nExp number of AAs in TMHs: 21.70944\nExp number, first 60 AAs: 0.02049\nTotal prob of N-in: 0.00432\noutside 1 244\nTMhelix 245 267\ninside 268 276

33567 GCF_001748165.1_ASM174816v1 Streptomyces sp. SPMA113 Terrabacteria group; Actinobacteria; Actinobacteria; Streptomycetales; Streptomycetaceae; Streptomyces MLGVVENHSLPRTAAFFDLDKTVIAKSSALTFSKSFYQGGLINRRAVLRTAYAQFVFLAG WP_069860967.1 inhibition of morphological differentiation protein [Streptomyces sp. SPMA113] Length: 278\nNumber of predicted TMHs: 1\nExp number of AAs in TMHs: 21.25438\nExp number, first 60 AAs: 0.02058\nTotal prob of N-in: 0.00296\noutside 1 247\nTMhelix 248 267\ninside 268 278

33568 GCF_000424845.1_ASM42484v1 Streptomyces sp. TAA040 Terrabacteria group; Actinobacteria; Actinobacteria; Streptomycetales; Streptomycetaceae; Streptomyces MFPRRSTVALAAAAVLCTTWAPAAHADESPSAAPLPGGLHGTSDPQFDGVFRQSLALLAQ WP_028425756.1 hypothetical protein [Streptomyces sp. TAA040] Length: 441\nNumber of predicted TMHs: 1\nExp number of AAs in TMHs: 23.75768\nExp number, first 60 AAs: 1.7266\nTotal prob of N-in: 0.09208\noutside 1 410\nTMhelix 411 433\ninside 434 441

33569 GCF_000424845.1_ASM42484v1 Streptomyces sp. TAA040 Terrabacteria group; Actinobacteria; Actinobacteria; Streptomycetales; Streptomycetaceae; Streptomyces
MTSFKTSSFARRAAVATTLTAAAVLVGGGPALAHVTVDPEEVPAGGYATINFKVPNERDN WP_028427408.1
membrane protein [Streptomyces sp. TAA040] Length: 247\nNumber of predicted TMHs: 2\nExp number of AAs in TMHs: 40.02701\nExp number, first 60 AAs: 19.28689\nTotal prob of N-in: 0.85125\nPOSSIBLE N-term signal sequence\ninside 1 12\nTMhelix 13 35\noutside 36 219\nTMhelix 220 242\ninside 243 247

33570 GCF_000424965.1_ASM42496v1 Streptomyces sp. TAA204 Terrabacteria group; Actinobacteria; Actinobacteria; Streptomycetales; Streptomycetaceae; Streptomyces
MLALVEKLAMRRTAAFFDLDKTVIAKSSTLTFGRSFYQGGLINRRAVLRTAYAQFVYLLG WP_028430715.1 inhibition of morphological differentiation protein [Streptomyces sp. TAA204] Length: 280\nNumber of predicted TMHs: 1\nExp number of AAs in TMHs: 22.09776\nExp number, first 60 AAs: 0.01841\nTotal prob of N-in: 0.00375\noutside 1 246\nTMhelix 247 269\ninside 270 280

33571 GCF_000426165.1_ASM42616v1 Streptomyces sp. TAA486 Terrabacteria group; Actinobacteria; Actinobacteria; Streptomycetales; Streptomycetaceae; Streptomyces
MLGFVENLAMRRTAAFFDLDKTVIAKSSTLAFGRSFYQGGLINRRAVLRTAYAQFVYLLG WP_028438492.1 inhibition of morphological differentiation protein [Streptomyces sp. TAA486] Length: 280\nNumber of predicted TMHs: 1\nExp number of AAs in TMHs: 21.83069\nExp number, first 60 AAs: 0.01347\nTotal prob of N-in: 0.00232\noutside 1 247\nTMhelix 248 270\ninside 271 280

33572 GCF_000426165.1_ASM42616v1 Streptomyces sp. TAA486 Terrabacteria group; Actinobacteria; Actinobacteria; Streptomycetales; Streptomycetaceae; Streptomyces
MIMHVSSRRARRAALLGTATIVVGGAMISVPAHAASTDGHSGSTGKATAVAARADLNVSV WP_051345639.1
hypothetical protein [Streptomyces sp. TAA486] Length: 297\nNumber of predicted TMHs: 2\nExp number of AAs in TMHs: 43.92749\nExp number, first 60 AAs: 22.17403\nTotal prob of N-in: 0.99314\nPOSSIBLE N-term signal sequence\ninside 1 12\nTMhelix 13 35\noutside 36 266\nTMhelix 267 289\ninside 290 297

33573 GCF_001941845.1_ASM194184v1 Streptomyces sp. TN58 Terrabacteria group; Actinobacteria; Actinobacteria; Streptomycetales; Streptomycetaceae; Streptomyces
MSARRRTVTPIRIAAAGFAPLALAVYAAVPAAAHGSMTDPVSRVAACYAEGPESPKSAAC WP_075970044.1
chitin-binding protein [Streptomyces sp. TN58] Length: 366\nNumber of predicted TMHs: 2\nExp number of AAs in TMHs: 44.00081\nExp number, first 60 AAs: 21.86126\nTotal prob of N-in: 0.99272\nPOSSIBLE N-term signal sequence\ninside 1 12\nTMhelix 13 35\noutside 36 331\nTMhelix 332 354\ninside 355 366

33574 GCF_000259895.1_ASM25989v1 Streptomyces sp. TOR3209 Terrabacteria group; Actinobacteria; Actinobacteria; Streptomycetales; Streptomycetaceae; Streptomyces
MRHARRRMVRRVARLAAVGGLLLGGTMVTRAVASEPPDAPAVPYTSARAASGTGAGLVSR WP_029393839.1 S1 family peptidase [Streptomyces sp. TOR3209] Length: 456\nNumber of predicted TMHs: 2\nExp number of AAs in TMHs: 37.7479\nExp number, first 60 AAs: 20.97699\nTotal prob of N-in: 0.96434\nPOSSIBLE N-term signal sequence\ninside 1 11\nTMhelix 12 34\noutside 35 417\nTMhelix 418 440\ninside 441 456

33575 GCF_000259895.1_ASM25989v1 Streptomyces sp. TOR3209 Terrabacteria group; Actinobacteria; Actinobacteria; Streptomycetales; Streptomycetaceae; Streptomyces
MGHNRRRRPTGARRATFGACALILGGGGLVAVNVFASATEGGNTAVPLGSSGVAATIDCP WP_086024792.1
hypothetical protein [Streptomyces sp. TOR3209] Length: 672\nNumber of predicted TMHs: 2\nExp number of AAs in TMHs: 42.96643\nExp number, first 60 AAs: 21.64952\nTotal prob of N-in: 0.98800\nPOSSIBLE N-term signal sequence\ninside 1 16\nTMhelix 17 39\noutside 40 644\nTMhelix 645 667\ninside 668 672

33576 GCF_001417775.1_ASM141777v1 Streptomyces sp. TP-A0356 Terrabacteria group; Actinobacteria; Actinobacteria; Streptomycetales; Streptomycetaceae; Streptomyces
MRRNTRRRPTGARRATFGAVALILGGGGLVAANVYASASEGWGGGPQSSQSGDQVLTAGT WP_055490993.1
hypothetical protein [Streptomyces sp. TP-A0356] Length: 715\nNumber of predicted TMHs: 2\nExp number of AAs in TMHs: 41.62086\nExp number, first 60 AAs: 21.55327\nTotal prob of N-in: 0.99442\nPOSSIBLE N-term signal sequence\ninside 1 16\nTMhelix 17 39\noutside 40 687\nTMhelix 688 707\ninside 708 715

33577 GCF_001417775.1_ASM141777v1 Streptomyces sp. TP-A0356 Terrabacteria group; Actinobacteria; Actinobacteria; Streptomycetales; Streptomycetaceae; Streptomyces
MPVRRRAVVAAVGAAPLLMAGPAAVPAVAHGSMGGPVSRVSQCSAEGPENPRSNACRAAV WP_055492640.1
chitin-binding protein [Streptomyces sp. TP-A0356] Length: 325\nNumber of predicted TMHs: 2\nExp number of AAs in TMHs: 43.9448999999999\nExp number, first 60 AAs: 22.34253\nTotal prob of N-in:

0.98443\nPOSSIBLE N-term signal sequence\ninside 1 6\nTMhelix 7 29\noutside 30 290\nTMhelix 291 313\ninside 314 325

33578 GCF_001417775.1_ASM141777v1 Streptomyces sp. TP-A0356 Terrabacteria group; Actinobacteria; Actinobacteria; Streptomycetales; Streptomycetaceae; Streptomyces MLRPVENHSLLRTAAFFDLDKTVIAKSSTLTFGKSFYQGGLINRRAALRTAYAQFVFLAG WP_055492069.1 inhibition of morphological differentiation protein [Streptomyces sp. TP-A0356] Length: 274\nNumber of predicted TMHs: 1\nExp number of AAs in TMHs: 21.58081\nExp number, first 60 AAs: 0.02616\nTotal prob of N-in: 0.01471\noutside 1 244\nTMhelix 245 267\ninside 268 274

33579 GCF_001417775.1_ASM141777v1 Streptomyces sp. TP-A0356 Terrabacteria group; Actinobacteria; Actinobacteria; Streptomycetales; Streptomycetaceae; Streptomyces MRHARRRIVRRGARFAAVGGLLLGGLMVTQALASEPSGAQTPDGTSAQSAAGTGSDLVSR WP_055489236.1 protease [Streptomyces sp. TP-A0356] Length: 466\nNumber of predicted TMHs: 2\nExp number of AAs in TMHs: 40.3840199999999\nExp number, first 60 AAs: 21.23621\nTotal prob of N-in: 0.98607\nPOSSIBLE N-term signal sequence\ninside 1 11\nTMhelix 12 34\noutside 35 427\nTMhelix 428 450\ninside 451 466

33580 GCF_001748085.1_ASM174808v1 Streptomyces sp. TP-A0874 Terrabacteria group; Actinobacteria; Actinobacteria; Streptomycetales; Streptomycetaceae; Streptomyces MSSSDPNVRAARNATSTAGRRPVVAVTGAASGVGALLTERLAASEEAERVVAIDDRPGAE WP_069815624.1 NAD-dependent dehydratase [Streptomyces sp. TP-A0874] Length: 369\nNumber of predicted TMHs: 1\nExp number of AAs in TMHs: 20.35623\nExp number, first 60 AAs: 0.58777\nTotal prob of N-in: 0.03516\noutside 1 339\nTMhelix 340 359\ninside 360 369

33581 GCF_001748085.1_ASM174808v1 Streptomyces sp. TP-A0874 Terrabacteria group; Actinobacteria; Actinobacteria; Streptomycetales; Streptomycetaceae; Streptomyces MLSDVENHSLPRTAAFFDLDKTVIAKSSTLAFSKSFYQGGLINRRAVLRTAYAQFVYLLG WP_069813307.1 inhibition of morphological differentiation protein [Streptomyces sp. TP-A0874] Length: 277\nNumber of predicted TMHs: 1\nExp number of AAs in TMHs: 20.52786\nExp number, first 60 AAs: 0.04515\nTotal prob of N-in: 0.02746\noutside 1 247\nTMhelix 248 267\ninside 268 277

33582 GCF_001293595.1_ASM129359v1 Streptomyces sp. TP-A0875 Terrabacteria group; Actinobacteria; Actinobacteria; Streptomycetales; Streptomycetaceae; Streptomyces MGHARRRIVRRVARLAAVGGLLLGGTMVTRAVASEPSGARAPAAASASAAGAALVERLGT WP_030420451.1 MULTISPECIES: protease [Streptomyces] Length: 451\nNumber of predicted TMHs: 2\nExp number of AAs in TMHs: 40.90346\nExp number, first 60 AAs: 20.62799\nTotal prob of N-in: 0.94294\nPOSSIBLE N-term signal sequence\ninside 1 11\nTMhelix 12 34\noutside 35 412\nTMhelix 413 435\ninside 436 451

33583 GCF_001293595.1_ASM129359v1 Streptomyces sp. TP-A0875 Terrabacteria group; Actinobacteria; Actinobacteria; Streptomycetales; Streptomycetaceae; Streptomyces MNTTKARRTAVPSLFLAAALALVPAPAAHADGIRAQQWGLDALRTQEVWRTTKGADVTVA WP_053913500.1 type VII secretion-associated serine protease mycosin [Streptomyces sp. TP-A0875] Length: 392\nNumber of predicted TMHs: 2\nExp number of AAs in TMHs: 38.21452\nExp number, first 60 AAs: 13.22131\nTotal prob of N-in: 0.83256\nPOSSIBLE N-term signal sequence\ninside 1 12\nTMhelix 13 30\noutside 31 359\nTMhelix 360 382\ninside 383 392

33584 GCF_001293595.1_ASM129359v1 Streptomyces sp. TP-A0875 Terrabacteria group; Actinobacteria; Actinobacteria; Streptomycetales; Streptomycetaceae; Streptomyces MPRTAAFFDLDKTVIAKSSTLTFSKSFYQGGLINRRAVLRTAYAQFIFLAGGADHDQMER WP_078856197.1 MULTISPECIES: inhibition of morphological differentiation protein [Streptomyces] Length: 268\nNumber of predicted TMHs: 1\nExp number of AAs in TMHs: 21.65373\nExp number, first 60 AAs: 0.08568\nTotal prob of N-in: 0.00693\noutside 1 238\nTMhelix 239 261\ninside 262 268

33585 GCF_001293595.1_ASM129359v1 Streptomyces sp. TP-A0875 Terrabacteria group; Actinobacteria; Actinobacteria; Streptomycetales; Streptomycetaceae; Streptomyces MGHNRRKRPTGARRATFGAFALILGGGGLVAANVIASATETSNAAVPLGGSGIAATVDCP WP_078962511.1 hypothetical protein [Streptomyces sp. TP-A0875] Length: 671\nNumber of predicted TMHs: 2\nExp number of AAs in TMHs: 43.52668\nExp number, first 60 AAs: 21.89339\nTotal prob of N-in: 0.99283\nPOSSIBLE N-term signal sequence\ninside 1 16\nTMhelix 17 39\noutside 40 641\nTMhelix 642 664\ninside 665 671

33586 GCF_001905845.1_ASM190584v1 Streptomyces sp. TSRI0107 Terrabacteria group; Actinobacteria; Actinobacteria; Streptomycetales; Streptomycetaceae; Streptomyces

MLCLVENHSLPRTAAFFDLDKTVIAKSSTLTFSKSFYQGGLINRRAALRTAYAQFVFLVG WP_073943235.1 inhibition of morphological differentiation protein [Streptomyces sp. TSRI0107] Length: 277\nNumber of predicted TMHs: 1\nExp number of AAs in TMHs: 21.54348\nExp number, first 60 AAs: 0.01371\nTotal prob of N-in: 0.00226\noutside 1 247\nTMhelix 248 270\ninside 271 277

33587 GCF_001905845.1_ASM190584v1 Streptomyces sp. TSRI0107 Terrabacteria group; Actinobacteria; Actinobacteria; Streptomycetales; Streptomycetaceae; Streptomyces
MSSRRRAAIAGSLVAASFSAVVILSTAAGAKDEGPGSSEGGKAVEQAPAGVELTTLLPER WP_073938500.1 hypothetical protein [Streptomyces sp. TSRI0107] Length: 239\nNumber of predicted TMHs: 2\nExp number of AAs in TMHs: 42.00761\nExp number, first 60 AAs: 20.99545\nTotal prob of N-in: 0.97184\nPOSSIBLE N-term signal sequence\ninside 1 6\nTMhelix 7 29\noutside 30 209\nTMhelix 210 232\ninside 233 239

33588 GCF_001905485.1_ASM190548v1 Streptomyces sp. TSRI0261 Terrabacteria group; Actinobacteria; Actinobacteria; Streptomycetales; Streptomycetaceae; Streptomyces
MLSVVENRFSPRTAAFFDLDKTVIAKSSTLTFSKSFYQGGLINRRAVLRTAYAQFVFLAG WP_073800253.1 inhibition of morphological differentiation protein [Streptomyces sp. TSRI0261] Length: 281\nNumber of predicted TMHs: 1\nExp number of AAs in TMHs: 21.48735\nExp number, first 60 AAs: 0.04008\nTotal prob of N-in: 0.00871\noutside 1 247\nTMhelix 248 270\ninside 271 281

33589 GCF_001905125.1_ASM190512v1 Streptomyces sp. TSRI0281 Terrabacteria group; Actinobacteria; Actinobacteria; Streptomycetales; Streptomycetaceae; Streptomyces
MLTFVENCFSPRTAAFFDLDKTVIAKSSTLTFSKSFYQGGLINRRAVLRTAYAQFVFLAG WP_073719540.1 inhibition of morphological differentiation protein [Streptomyces sp. TSRI0281] Length: 280\nNumber of predicted TMHs: 1\nExp number of AAs in TMHs: 21.47833\nExp number, first 60 AAs: 0.02539\nTotal prob of N-in: 0.00347\noutside 1 247\nTMhelix 248 270\ninside 271 280

33590 GCF_001905125.1_ASM190512v1 Streptomyces sp. TSRI0281 Terrabacteria group; Actinobacteria; Actinobacteria; Streptomycetales; Streptomycetaceae; Streptomyces
MRTSSSRKVVARRGTVASTALVLLLVGAAPASADSVRGDQWHLDAMKAEEMWAVSTGADV WP_073723783.1 type VII secretion-associated serine protease mycosin [Streptomyces sp. TSRI0281] Length: 452\nNumber of predicted TMHs: 2\nExp number of AAs in TMHs: 37.64808\nExp number, first 60 AAs: 15.09368\nTotal prob of N-in: 0.71157\nPOSSIBLE N-term signal sequence\ninside 1 8\nTMhelix 9 31\noutside 32 381\nTMhelix 382 404\ninside 405 452

33591 GCF_001905125.1_ASM190512v1 Streptomyces sp. TSRI0281 Terrabacteria group; Actinobacteria; Actinobacteria; Streptomycetales; Streptomycetaceae; Streptomyces
MPDRRRRTAFRLARSSGAPGSSGAPGSSGPSDSAGPSGPSDPSGSSGPSDPSGSSGSSDD WP_073718266.1
hypothetical protein [Streptomyces sp. TSRI0281] Length: 318\nNumber of predicted TMHs: 2\nExp number of AAs in TMHs: 46.96774\nExp number, first 60 AAs: 0.00015\nTotal prob of N-in: 0.91207\ninside 1 168\nTMhelix 169 191\noutside 192 256\nTMhelix 257 279\ninside 280 318

33592 GCF_001905525.1_ASM190552v1 Streptomyces sp. TSRI0395 Terrabacteria group; Actinobacteria; Actinobacteria; Streptomycetales; Streptomycetaceae; Streptomyces
MLSVVENCFSPRTAAFFDLDKTVIAKSSTLTFSKSFYQGGLINRRAVLRTAYAQFVFLAG WP_073958104.1 inhibition of morphological differentiation protein [Streptomyces sp. TSRI0395] Length: 281\nNumber of predicted TMHs: 1\nExp number of AAs in TMHs: 21.43834\nExp number, first 60 AAs: 0.03377\nTotal prob of N-in: 0.00778\noutside 1 247\nTMhelix 248 270\ninside 271 281

33593 GCF_001905405.1_ASM190540v1 Streptomyces sp. TSRI0445 Terrabacteria group; Actinobacteria; Actinobacteria; Streptomycetales; Streptomycetaceae; Streptomyces
MLSVVENCFSPRTAAFFDLDKTVIAKSSTLTFSKSFYQGGLINRRAVLRTAYTQFVFLAG WP_030813057.1 MULTISPECIES: inhibition of morphological differentiation protein [Streptomyces] Length: 281\nNumber of predicted TMHs: 1\nExp number of AAs in TMHs: 21.46267\nExp number, first 60 AAs: 0.01959\nTotal prob of N-in: 0.00583\noutside 1 247\nTMhelix 248 270\ninside 271 281

33594 GCF_001905405.1_ASM190540v1 Streptomyces sp. TSRI0445 Terrabacteria group; Actinobacteria; Actinobacteria; Streptomycetales; Streptomycetaceae; Streptomyces
MGLSWNDRRLERRVKPGDGKPLKPFRWWQMTRRSLLSIALPVGGGLSATYTVEVKHGGDP WP_073775700.1
hypothetical protein [Streptomyces sp. TSRI0445] Length: 218\nNumber of predicted TMHs: 2\nExp number of AAs in TMHs: 39.57631\nExp number, first 60 AAs: 1.48435\nTotal prob of N-in: 0.93375\ninside 1 135\nTMhelix 136 158\noutside 159 182\nTMhelix 183 200\ninside 201 218

33595 GCF_001905405.1_ASM190540v1 Streptomyces sp. TSRI0445 Terrabacteria group; Actinobacteria; Actinobacteria; Streptomycetales; Streptomycetaceae; Streptomyces
MSPPTPRRRRALPVLTAALAALLCGLAPPAAADGPPDWAARPTGGTDRDARPYVYLEGTP WP_073779743.1
hypothetical protein [Streptomyces sp. TSRI0445] Length: 322\nNumber of predicted TMHs: 1\nExp number of AAs in TMHs: 32.99085\nExp number, first 60 AAs: 11.21825\nTotal prob of N-in: 0.56076\nPOSSIBLE N-term signal sequence\noutside 1 249\nTMhelix 250 272\ninside 273 322

33596 GCF_000213055.1_ASM21305v1 Streptomyces sp. Tu6071 Terrabacteria group; Actinobacteria; Actinobacteria; Streptomycetales; Streptomycetaceae; Streptomyces
MLGPVENHSLPRTAAFFDLDKTVIAKSSTLTFSKSFYQGGLINRRAALRTAYTQFVFLVG WP_007823897.1 MULTISPECIES: morphological differentiation-associated protein [Streptomyces] Length: 276\nNumber of predicted TMHs: 1\nExp number of AAs in TMHs: 21.70944\nExp number, first 60 AAs: 0.02049\nTotal prob of N-in: 0.00432\noutside 1 244\nTMhelix 245 267\ninside 268 276

33597 GCF_000611795.1_Tu61761.0 Streptomyces sp. Tu 6176 Terrabacteria group; Actinobacteria; Actinobacteria; Streptomycetales; Streptomycetaceae; Streptomyces
MASRGRRATMRNLGALAGAAALTVLGGAAPGWAASPTPDGHRPPATPIEHVVVLFDENIS WP_026252868.1
MULTISPECIES: phospholipase C [Streptomyces] Length: 613\nNumber of predicted TMHs: 2\nExp number of AAs in TMHs: 44.60387\nExp number, first 60 AAs: 22.3165\nTotal prob of N-in: 0.99261\nPOSSIBLE N-term signal sequence\ninside 1 12\nTMhelix 13 35\noutside 36 582\nTMhelix 583 605\ninside 606 613

33598 GCF_000611795.1_Tu61761.0 Streptomyces sp. Tu 6176 Terrabacteria group; Actinobacteria; Actinobacteria; Streptomycetales; Streptomycetaceae; Streptomyces
MLDLVENHSLPRTAAFFDLDKTVIAKSSTLTFSKSFYQGGLINRRAVLRTAYAQFVFLAG WP_078510011.1 inhibition of morphological differentiation protein [Streptomyces sp. Tu 6176] Length: 276\nNumber of predicted TMHs: 1\nExp number of AAs in TMHs: 21.81496\nExp number, first 60 AAs: 0.03069\nTotal prob of N-in: 0.00630\noutside 1 247\nTMhelix 248 270\ninside 271 276

33599 GCF_000611795.1_Tu61761.0 Streptomyces sp. Tu 6176 Terrabacteria group; Actinobacteria; Actinobacteria; Streptomycetales; Streptomycetaceae; Streptomyces
MSRTTGGRSRRRPLALAAAVATVTATVASVGAAALAGAPGAAAADRPLSGYELTWGIKDS WP_037893396.1
hypothetical protein [Streptomyces sp. Tu 6176] Length: 538\nNumber of predicted TMHs: 2\nExp number of AAs in TMHs: 45.50227\nExp number, first 60 AAs: 22.95749\nTotal prob of N-in: 0.99312\nPOSSIBLE N-term signal sequence\ninside 1 12\nTMhelix 13 35\noutside 36 506\nTMhelix 507 529\ninside 530 538

33600 GCF_001931635.1_ASM193163v1 Streptomyces sp. Tue 6075 Terrabacteria group; Actinobacteria; Actinobacteria; Streptomycetales; Streptomycetaceae; Streptomyces
MLSVVENCFSPRTAAFFDLDKTVIAKSSTLTFSKSFYQGGLINRRAVLRTAYTQFVFLAG WP_032769244.1 MULTISPECIES: inhibition of morphological differentiation protein [Streptomyces] Length: 281\nNumber of predicted TMHs: 1\nExp number of AAs in TMHs: 21.46267\nExp number, first 60 AAs: 0.01959\nTotal prob of N-in: 0.00583\noutside 1 247\nTMhelix 248 270\ninside 271 281

33601 GCF_001931635.1_ASM193163v1 Streptomyces sp. Tue 6075 Terrabacteria group; Actinobacteria; Actinobacteria; Streptomycetales; Streptomycetaceae; Streptomyces
MGLSWNDRRLERRVKPGDGKPLKPFRWWQMTRRSLLSIALPVGGGLSATYTVEVKHGGDP WP_075265106.1
hypothetical protein [Streptomyces sp. Tue 6075] Length: 219\nNumber of predicted TMHs: 2\nExp number of AAs in TMHs: 40.67373\nExp number, first 60 AAs: 1.11343\nTotal prob of N-in: 0.94509\ninside 1 136\nTMhelix 137 159\noutside 160 183\nTMhelix 184 201\ninside 202 219

33602 GCF_000686765.1_ASM68676v1 Streptomyces sp. UNC401CLCol Terrabacteria group; Actinobacteria; Actinobacteria; Streptomycetales; Streptomycetaceae; Streptomyces
MRHARRRVVRRVTRLAAVGGLLLGGTMVTHAVASEPGTPDAVPFSTASPADASGPGAGLV WP_037953023.1
protease [Streptomyces sp. UNC401CLCol] Length: 457\nNumber of predicted TMHs: 2\nExp number of AAs in TMHs: 41.25029\nExp number, first 60 AAs: 21.83601\nTotal prob of N-in: 0.97534\nPOSSIBLE N-term signal sequence\ninside 1 11\nTMhelix 12 34\noutside 35 418\nTMhelix 419 441\ninside 442 457

33603 GCF_000686765.1_ASM68676v1 Streptomyces sp. UNC401CLCol Terrabacteria group; Actinobacteria; Actinobacteria; Streptomycetales; Streptomycetaceae; Streptomyces
MLKDVENHLSPRTAAFFDLDKTVIAKSSTLTFSKSFYQGGLINRRAVLRTAYSQFVFLAG WP_028959472.1 inhibition of morphological differentiation protein [Streptomyces sp. UNC401CLCol] Length: 277\nNumber of predicted TMHs:

1\nExp number of AAs in TMHs: 21.90146\nExp number, first 60 AAs: 0.03208\nTotal prob of N-in: 0.00818\noutside 1 247\nTMhelix 248 270\ninside 271 277

33604 GCF_000701325.1_ASM70132v1 Streptomyces sp. URHA0041 Terrabacteria group; Actinobacteria; Actinobacteria; Streptomycetales; Streptomycetaceae; Streptomyces MPLRRRAALVVAAATTLTALTAPVALADTTAPAAATTAAPSRGDLYGAADPTYDGVWRQS WP_033174746.1 hypothetical protein [Streptomyces sp. URHA0041] Length: 418\nNumber of predicted TMHs: 2\nExp number of AAs in TMHs: 40.81218\nExp number, first 60 AAs: 15.78383\nTotal prob of N-in: 0.76681\nPOSSIBLE N-term signal sequence\ninside 1 6\nTMhelix 7 26\noutside 27 389\nTMhelix 390 412\ninside 413 418

33605 GCF_000701325.1_ASM70132v1 Streptomyces sp. URHA0041 Terrabacteria group; Actinobacteria; Actinobacteria; Streptomycetales; Streptomycetaceae; Streptomyces MSINRRTAIAVRSIGVASAAAALALGVAGSAFACSITEFTPSASCDDSNHGVIKVLDQDG WP_051674631.1 hypothetical protein [Streptomyces sp. URHA0041] Length: 225\nNumber of predicted TMHs: 2\nExp number of AAs in TMHs: 42.82562\nExp number, first 60 AAs: 21.45845\nTotal prob of N-in: 0.97925\nPOSSIBLE N-term signal sequence\ninside 1 12\nTMhelix 13 35\noutside 36 193\nTMhelix 194 216\ninside 217 225

33606 GCF_000239075.1_ASM23907v2 Streptomyces sp. W007 Terrabacteria group; Actinobacteria; Actinobacteria; Streptomycetales; Streptomycetaceae; Streptomyces MLSVVENCFSPRTAAFFDLDKTVIAKSSTLTFSKSFYQGGLINRRAVLRTAYAQFVFLAG WP_032791512.1 MULTISPECIES: inhibition of morphological differentiation protein [Streptomyces] Length: 281\nNumber of predicted TMHs: 1\nExp number of AAs in TMHs: 21.51851\nExp number, first 60 AAs: 0.0316\nTotal prob of N-in: 0.00586\noutside 1 247\nTMhelix 248 270\ninside 271 281

33607 GCF_001592685.1_ASM159268v1 Streptomyces sp. WAC04657 Terrabacteria group; Actinobacteria; Actinobacteria; Streptomycetales; Streptomycetaceae; Streptomyces MLGLVENHSLPRAAAFFDLDKTVIAKSSTLTFSKSFYRGGLISRRAALRTAYIQFVFLAG WP_062754201.1 inhibition of morphological differentiation protein [Streptomyces sp. WAC04657] Length: 289\nNumber of predicted TMHs: 1\nExp number of AAs in TMHs: 22.01861\nExp number, first 60 AAs: 0.03663\nTotal prob of N-in: 0.00318\noutside 1 247\nTMhelix 248 270\ninside 271 289

33608 GCF_001279725.1_ASM127972v1 Streptomyces sp. WM4235 Terrabacteria group; Actinobacteria; Actinobacteria; Streptomycetales; Streptomycetaceae; Streptomyces MSTVSRRSLVRAAAVTACAGSLLALPTAAALADGVPAASTGQRTLVKSLSLADGESLARI WP_053676489.1 hypothetical protein [Streptomyces sp. WM4235] Length: 201\nNumber of predicted TMHs: 2\nExp number of AAs in TMHs: 42.82591\nExp number, first 60 AAs: 20.26699\nTotal prob of N-in: 0.94037\nPOSSIBLE N-term signal sequence\ninside 1 8\nTMhelix 9 31\noutside 32 171\nTMhelix 172 194\ninside 195 201

33609 GCF_001279725.1_ASM127972v1 Streptomyces sp. WM4235 Terrabacteria group; Actinobacteria; Actinobacteria; Streptomycetales; Streptomycetaceae; Streptomyces MAEAAEIQGAVPAPARRRWLRRAVVMLAGTPVLAALVYAPAPPAQAAESPVDVQLSEVAP WP_053683954.1 hypothetical protein [Streptomyces sp. WM4235] Length: 755\nNumber of predicted TMHs: 1\nExp number of AAs in TMHs: 37.60415\nExp number, first 60 AAs: 15.93969\nTotal prob of N-in: 0.75553\nPOSSIBLE N-term signal sequence\noutside 1 688\nTMhelix 689 711\ninside 712 755

33610 GCF_001279725.1_ASM127972v1 Streptomyces sp. WM4235 Terrabacteria group; Actinobacteria; Actinobacteria; Streptomycetales; Streptomycetaceae; Streptomyces MSTYTRSRRRTVLRAAVATAALAGALLAPSTAAFAADSPKAADSPKATDASKTAGAPKAA WP_053682322.1 hypothetical protein [Streptomyces sp. WM4235] Length: 363\nNumber of predicted TMHs: 2\nExp number of AAs in TMHs: 43.94944\nExp number, first 60 AAs: 22.18969\nTotal prob of N-in: 0.98579\nPOSSIBLE N-term signal sequence\ninside 1 11\nTMhelix 12 34\noutside 35 332\nTMhelix 333 355\ninside 356 363

33611 GCF_001279445.1_ASM127944v1 Streptomyces sp. WM6349 Terrabacteria group; Actinobacteria; Actinobacteria; Streptomycetales; Streptomycetaceae; Streptomyces MPQYRDARRALLATAFALTLTAAGATVATAAPHPPYALRLDGAGECTFPMKKQIADRPWA WP_030651182.1 MULTISPECIES: type VII secretion-associated serine protease [Streptomyces] Length: 414\nNumber of predicted TMHs: 2\nExp number of AAs in TMHs: 43.048\nExp number, first 60 AAs: 20.71014\nTotal prob of N-in: 0.96189\nPOSSIBLE N-term signal sequence\ninside 1 8\nTMhelix 9 31\noutside 32 380\nTMhelix 381 403\ninside 404 414

33612 GCF_001279445.1_ASM127944v1 Streptomyces sp. WM6349 Terrabacteria group; Actinobacteria; Actinobacteria; Streptomycetales; Streptomycetaceae; Streptomyces
MSTVRRRTLVRAAAVTACAGSLLALPTAAALAEGVPAASSAHSAAPQRTLVKSLSLADGV WP_053611851.1
MULTISPECIES: hypothetical protein [Streptomyces] Length: 218\nNumber of predicted TMHs: 2\nExp number of AAs in TMHs: 42.93081\nExp number, first 60 AAs: 20.33935\nTotal prob of N-in: 0.95701\nPOSSIBLE N-term signal sequence\ninside 1 8\nTMhelix 9 31\noutside 32 188\nTMhelix 189 211\ninside 212 218

33613 GCF_001279625.1_ASM127962v1 Streptomyces sp. WM6368 Terrabacteria group; Actinobacteria; Actinobacteria; Streptomycetales; Streptomycetaceae; Streptomyces
MRRRSPVIAVTGAASGVGAALVSRLAASDEVKQVVAIDERRGDCAAAQWHILDVRDPAIA WP_051779201.1
MULTISPECIES: NAD-dependent dehydratase [Streptomyces] Length: 353\nNumber of predicted TMHs: 1\nExp number of AAs in TMHs: 30.20524\nExp number, first 60 AAs: 7.40836\nTotal prob of N-in: 0.39674\noutside 1 324\nTMhelix 325 347\ninside 348 353

33614 GCF_001279625.1_ASM127962v1 Streptomyces sp. WM6368 Terrabacteria group; Actinobacteria; Actinobacteria; Streptomycetales; Streptomycetaceae; Streptomyces
MPARRRTAVALTRIASAGLAPLALAAYAAAPAVAHGSMTDPVSRVAACYAEGPESPKTAA WP_053704554.1
chitin-binding protein [Streptomyces sp. WM6368] Length: 355\nNumber of predicted TMHs: 2\nExp number of AAs in TMHs: 42.25019\nExp number, first 60 AAs: 20.34906\nTotal prob of N-in: 0.90818\nPOSSIBLE N-term signal sequence\ninside 1 12\nTMhelix 13 35\noutside 36 318\nTMhelix 319 341\ninside 342 355

33615 GCF_001279625.1_ASM127962v1 Streptomyces sp. WM6368 Terrabacteria group; Actinobacteria; Actinobacteria; Streptomycetales; Streptomycetaceae; Streptomyces
MNRTHAPRRPLLAAAFALTLTFTLASAGVGAGAASAAESAPAPQPPYGQAPSAQAPYGLR WP_078972394.1 type VII secretion-associated serine protease mycosin [Streptomyces sp. WM6368] Length: 435\nNumber of predicted TMHs: 2\nExp number of AAs in TMHs: 44.48013\nExp number, first 60 AAs: 22.18583\nTotal prob of N-in: 0.97823\nPOSSIBLE N-term signal sequence\ninside 1 11\nTMhelix 12 34\noutside 35 401\nTMhelix 402 424\ninside 425 435

33616 GCF_001279625.1_ASM127962v1 Streptomyces sp. WM6368 Terrabacteria group; Actinobacteria; Actinobacteria; Streptomycetales; Streptomycetaceae; Streptomyces
MSTVSRRTLVRAAAVTACAGSLLALPTAAALAEGVPAASSARSAAPQLSLVKSLSLADGT WP_053169188.1 MULTISPECIES: hypothetical protein [Streptomyces] Length: 217\nNumber of predicted TMHs: 2\nExp number of AAs in TMHs: 41.6059599999999\nExp number, first 60 AAs: 18.99757\nTotal prob of N-in: 0.88899\nPOSSIBLE N-term signal sequence\ninside 1 8\nTMhelix 9 31\noutside 32 187\nTMhelix 188 210\ninside 211 217

33617 GCF_001279655.1_ASM127965v1 Streptomyces sp. WM6372 Terrabacteria group; Actinobacteria; Actinobacteria; Streptomycetales; Streptomycetaceae; Streptomyces
MSTVDRRTLVRAAAVTACAGTLLALPAAAALAEGVPAASAARSAGAPRTLVKSLALADGV WP_053689381.1
hypothetical protein [Streptomyces sp. WM6372] Length: 200\nNumber of predicted TMHs: 2\nExp number of AAs in TMHs: 44.08263\nExp number, first 60 AAs: 21.47969\nTotal prob of N-in: 0.97870\nPOSSIBLE N-term signal sequence\ninside 1 8\nTMhelix 9 31\noutside 32 170\nTMhelix 171 193\ninside 194 200

33618 GCF_001279655.1_ASM127965v1 Streptomyces sp. WM6372 Terrabacteria group; Actinobacteria; Actinobacteria; Streptomycetales; Streptomycetaceae; Streptomyces
MPARRRRTVTRIAAAGLAPLAVAAYAAGPAAAHGSMTDPVSRVAACYAEGPESPKSAACK WP_053691541.1
chitin-binding protein [Streptomyces sp. WM6372] Length: 345\nNumber of predicted TMHs: 2\nExp number of AAs in TMHs: 43.47605\nExp number, first 60 AAs: 21.10113\nTotal prob of N-in: 0.97582\nPOSSIBLE N-term signal sequence\ninside 1 11\nTMhelix 12 34\noutside 35 310\nTMhelix 311 333\ninside 334 345

33619 GCF_001279655.1_ASM127965v1 Streptomyces sp. WM6372 Terrabacteria group; Actinobacteria; Actinobacteria; Streptomycetales; Streptomycetaceae; Streptomyces
MGTEQQAKRQSGTRRRFLSLVTASFLTLGAGAAFLAAPGSASADPIEQCTPTTGALVAVD WP_053690890.1
peptidase [Streptomyces sp. WM6372] Length: 931\nNumber of predicted TMHs: 2\nExp number of AAs in TMHs: 42.56464\nExp number, first 60 AAs: 21.37149\nTotal prob of N-in: 0.99387\nPOSSIBLE N-term signal sequence\ninside 1 16\nTMhelix 17 39\noutside 40 893\nTMhelix 894 916\ninside 917 931

33620 GCF_001279685.1_ASM127968v1 Streptomyces sp. WM6373 Terrabacteria group; Actinobacteria; Actinobacteria; Streptomycetales; Streptomycetaceae; Streptomyces
MSARRRTVMPIRIAAAGAAPLALAVYAAVPAVAHGSMTDPVSRVAACYAEGPESPKSAAC WP_053633676.1
MULTISPECIES: chitin-binding protein [Streptomyces] Length: 347\nNumber of predicted TMHs: 2\nExp

number of AAs in TMHs: 43.49607\nExp number, first 60 AAs: 21.79131\nTotal prob of N-in: 0.98451\nPOSSIBLE N-term signal sequence\ninside 1 12\nTMhelix 13 35\noutside 36 312\nTMhelix 313 335\ninside 336 347

33621 GCF_001279685.1_ASM127968v1 Streptomyces sp. WM6373 Terrabacteria group; Actinobacteria; Actinobacteria; Streptomycetales; Streptomycetaceae; Streptomyces MALDVIRSGPPDKRPRRNLFKRGVLVLLPAALLAAATLVPNLAHALEAPVNLGTATNYAV WP_053706276.1 hypothetical protein [Streptomyces sp. WM6373] Length: 540\nNumber of predicted TMHs: 2\nExp number of AAs in TMHs: 43.13228\nExp number, first 60 AAs: 20.82873\nTotal prob of N-in: 0.94536\nPOSSIBLE N-term signal sequence\ninside 1 22\nTMhelix 23 45\noutside 46 505\nTMhelix 506 528\ninside 529 540

33622 GCF_001279685.1_ASM127968v1 Streptomyces sp. WM6373 Terrabacteria group; Actinobacteria; Actinobacteria; Streptomycetales; Streptomycetaceae; Streptomyces MPPYRDARRALLAAAFALTLTAAGATAATVAQHPPYALRLDGAGECTFPMKKQIADRPWA WP_053706793.1 type VII secretion-associated serine protease mycosin [Streptomyces sp. WM6373] Length: 414\nNumber of predicted TMHs: 2\nExp number of AAs in TMHs: 42.6403399999999\nExp number, first 60 AAs: 20.30353\nTotal prob of N-in: 0.94987\nPOSSIBLE N-term signal sequence\ninside 1 11\nTMhelix 12 31\noutside 32 380\nTMhelix 381 403\ninside 404 414

33623 GCF_001279685.1_ASM127968v1 Streptomyces sp. WM6373 Terrabacteria group; Actinobacteria; Actinobacteria; Streptomycetales; Streptomycetaceae; Streptomyces MSTVRRRTLVRAAAVTACAGSLLALPTAAALAEGVPAASSAHSAAPQRTLVKSLSLADGV WP_053677522.1 MULTISPECIES: hypothetical protein [Streptomyces] Length: 218\nNumber of predicted TMHs: 2\nExp number of AAs in TMHs: 42.93092\nExp number, first 60 AAs: 20.3393\nTotal prob of N-in: 0.95701\nPOSSIBLE N-term signal sequence\ninside 1 8\nTMhelix 9 31\noutside 32 188\nTMhelix 189 211\ninside 212 218

33624 GCF_001279695.1_ASM127969v1 Streptomyces sp. WM6378 Terrabacteria group; Actinobacteria; Actinobacteria; Streptomycetales; Streptomycetaceae; Streptomyces MTFPKLSTAFEQRGPRRRRAAVVAALGFTPLALTGLAAGPAAAHGSMTDPVSRVSACYAE WP_053723408.1 chitin-binding protein [Streptomyces sp. WM6378] Length: 342\nNumber of predicted TMHs: 2\nExp number of AAs in TMHs: 44.0137\nExp number, first 60 AAs: 22.24178\nTotal prob of N-in: 0.98909\nPOSSIBLE N-term signal sequence\ninside 1 20\nTMhelix 21 43\noutside 44 307\nTMhelix 308 330\ninside 331 342

33625 GCF_001279695.1_ASM127969v1 Streptomyces sp. WM6378 Terrabacteria group; Actinobacteria; Actinobacteria; Streptomycetales; Streptomycetaceae; Streptomyces MPANGPATRRPAVRRSAAVAACVALAAGPLLLAAPARATGVEGDGGGRASAVVLRTGLDV WP_053729412.1 hypothetical protein [Streptomyces sp. WM6378] Length: 316\nNumber of predicted TMHs: 1\nExp number of AAs in TMHs: 35.80277\nExp number, first 60 AAs: 14.34937\nTotal prob of N-in: 0.66227\nPOSSIBLE N-term signal sequence\noutside 1 284\nTMhelix 285 307\ninside 308 316

33626 GCF_001279695.1_ASM127969v1 Streptomyces sp. WM6378 Terrabacteria group; Actinobacteria; Actinobacteria; Streptomycetales; Streptomycetaceae; Streptomyces MLGLVENHSLPRTAAFFDLDKTVIAKSSTLTFSKSFYQGGLINRRAVLRTAYAQFVFLAG WP_053728105.1 inhibition of morphological differentiation protein [Streptomyces sp. WM6378] Length: 278\nNumber of predicted TMHs: 1\nExp number of AAs in TMHs: 21.82136\nExp number, first 60 AAs: 0.03955\nTotal prob of N-in: 0.01019\noutside 1 247\nTMhelix 248 270\ninside 271 278

33627 GCF_001279695.1_ASM127969v1 Streptomyces sp. WM6378 Terrabacteria group; Actinobacteria; Actinobacteria; Streptomycetales; Streptomycetaceae; Streptomyces MARTVPSRKTLRPWRPGRRPAIGAAALALTVSAVLPAQGAAADDAKGGEKVLTVAVSQSV WP_053728959.1 ABC transporter substrate-binding protein [Streptomyces sp. WM6378] Length: 615\nNumber of predicted TMHs: 2\nExp number of AAs in TMHs: 41.73499\nExp number, first 60 AAs: 18.87831\nTotal prob of N-in: 0.86472\nPOSSIBLE N-term signal sequence\ninside 1 19\nTMhelix 20 42\noutside 43 582\nTMhelix 583 605\ninside 606 615

33628 GCF_000972385.1_ASM97238v1 Streptomyces sp. WM6386 Terrabacteria group; Actinobacteria; Actinobacteria; Streptomycetales; Streptomycetaceae; Streptomyces MRQRRRVLPGARVALALGCAGALIGVAAGPAQAAEVSFATHCIPPVGDPVDGTTKVEITA WP_046257517.1 hypothetical protein [Streptomyces sp. WM6386] Length: 422\nNumber of predicted TMHs: 2\nExp number of AAs in TMHs: 40.3796699999999\nExp number, first 60 AAs: 20.56619\nTotal prob of N-in: 0.96484\nPOSSIBLE N-term signal sequence\ninside 1 6\nTMhelix 7 29\noutside 30 395\nTMhelix 396 415\ninside 416 422

33629 GCF_000972385.1_ASM97238v1 Streptomyces sp. WM6386 Terrabacteria group; Actinobacteria; Actinobacteria; Streptomycetales; Streptomycetaceae; Streptomyces MSSRRRTTIAGSLVTASFSAVLILSTAAGADEGPGSSKGGKAVDEAPAGVKLTTLLPEQI WP_046259605.1 hypothetical protein [Streptomyces sp. WM6386] Length: 237\nNumber of predicted TMHs: 2\nExp number of AAs in TMHs: 36.51247\nExp number, first 60 AAs: 18.70908\nTotal prob of N-in: 0.92819\nPOSSIBLE N-term signal sequence\ninside 1 6\nTMhelix 7 29\noutside 30 209\nTMhelix 210 232\ninside 233 237

33630 GCF_000972385.1_ASM97238v1 Streptomyces sp. WM6386 Terrabacteria group; Actinobacteria; Actinobacteria; Streptomycetales; Streptomycetaceae; Streptomyces MSAPQKTARRSLLVTSAALTSFALAAAPAFAAPGPTPSSSASSSASASTSPSATPPANMS WP_046261706.1 D-alanyl-D-alanine carboxypeptidase [Streptomyces sp. WM6386] Length: 435\nNumber of predicted TMHs: 2\nExp number of AAs in TMHs: 42.1077699999999\nExp number, first 60 AAs: 19.43792\nTotal prob of N-in: 0.91768\nPOSSIBLE N-term signal sequence\ninside 1 11\nTMhelix 12 34\noutside 35 398\nTMhelix 399 421\ninside 422 435

33631 GCF_000972385.1_ASM97238v1 Streptomyces sp. WM6386 Terrabacteria group; Actinobacteria; Actinobacteria; Streptomycetales; Streptomycetaceae; Streptomyces MLRVVENHSLPRTAAFFDLDKTVIAKSSTLTFSKSFYQGGLINRRAALRTAYAQFVFLVG WP_046260303.1 inhibition of morphological differentiation protein [Streptomyces sp. WM6386] Length: 277\nNumber of predicted TMHs: 1\nExp number of AAs in TMHs: 21.6983\nExp number, first 60 AAs: 0.0288\nTotal prob of N-in: 0.00837\noutside 1 247\nTMhelix 248 270\ninside 271 277

33632 GCF_000972385.1_ASM97238v1 Streptomyces sp. WM6386 Terrabacteria group; Actinobacteria; Actinobacteria; Streptomycetales; Streptomycetaceae; Streptomyces MRHARRRMVRRVTRLAAVGGLLLGSVMVTRAMASESSDASALPRTTAQSASAMGSDLVSR WP_046258199.1 protease [Streptomyces sp. WM6386] Length: 460\nNumber of predicted TMHs: 2\nExp number of AAs in TMHs: 40.83897\nExp number, first 60 AAs: 21.0256\nTotal prob of N-in: 0.96912\nPOSSIBLE N-term signal sequence\ninside 1 11\nTMhelix 12 34\noutside 35 421\nTMhelix 422 444\ninside 445 460

33633 GCF_000972395.1_ASM97239v1 Streptomyces sp. WM6391 Terrabacteria group; Actinobacteria; Actinobacteria; Streptomycetales; Streptomycetaceae; Streptomyces MPRTAAFFDLDKTVIAKSSTLTFSKSFYQGGLINRRAVLRTAYAQFVFLAGGADHDQMER WP_003975378.1 MULTISPECIES: morphological differentiation-associated protein [Streptomyces] Length: 268\nNumber of predicted TMHs: 1\nExp number of AAs in TMHs: 21.51645\nExp number, first 60 AAs: 0.03737\nTotal prob of N-in: 0.00617\noutside 1 238\nTMhelix 239 261\ninside 262 268

33634 GCF_000972395.1_ASM97239v1 Streptomyces sp. WM6391 Terrabacteria group; Actinobacteria; Actinobacteria; Streptomycetales; Streptomycetaceae; Streptomyces MRHARRRIVRRVARLAAVGGLLLGGTMVTRAVASEPPDASAAPRTFAQTPSGAGGDLVSR WP_037666917.1 MULTISPECIES: protease [Streptomyces] Length: 454\nNumber of predicted TMHs: 2\nExp number of AAs in TMHs: 40.7848\nExp number, first 60 AAs: 21.15262\nTotal prob of N-in: 0.96513\nPOSSIBLE N-term signal sequence\ninside 1 11\nTMhelix 12 34\noutside 35 415\nTMhelix 416 438\ninside 439 454

33635 GCF_900090145.1_IMG-taxon_2522125135_annotated_assembly Streptomyces sp. WMMB 322 Terrabacteria group; Actinobacteria; Actinobacteria; Streptomycetales; Streptomycetaceae; Streptomyces MLGFVENLAMRRTAAFFDLDKTVIAKSSTLAFGRSFYQGGLINRRAVLRTAYAQFVYLLG WP_055487677.1 inhibition of morphological differentiation protein [Streptomyces sp. WMMB 322] Length: 280\nNumber of predicted TMHs: 1\nExp number of AAs in TMHs: 21.76891\nExp number, first 60 AAs: 0.01082\nTotal prob of N-in: 0.00113\noutside 1 246\nTMhelix 247 269\ninside 270 280

33636 GCF_001418285.1_ASM141828v1 Streptomyces sp. WMMB 322 Terrabacteria group; Actinobacteria; Actinobacteria; Streptomycetales; Streptomycetaceae; Streptomyces MLGFVENLAMRRTAAFFDLDKTVIAKSSTLAFGRSFYQGGLINRRAVLRTAYAQFVYLLG WP_055487677.1 inhibition of morphological differentiation protein [Streptomyces sp. WMMB 322] Length: 280\nNumber of predicted TMHs: 1\nExp number of AAs in TMHs: 21.76891\nExp number, first 60 AAs: 0.01082\nTotal prob of N-in: 0.00113\noutside 1 246\nTMhelix 247 269\ninside 270 280

33637 GCF_000964305.1_ASM96430v1 Streptomyces sp. WMMB 714 Terrabacteria group; Actinobacteria; Actinobacteria; Streptomycetales; Streptomycetaceae; Streptomyces MLAFVENLAMRRTAAFFDLDKTVIAKSSTLTFGRSFYQGGLINRRAVLRTAYAQFVYLLG WP_045865951.1 inhibition of morphological differentiation protein [Streptomyces sp. WMMB 714] Length: 279\nNumber of predicted TMHs:

1\nExp number of AAs in TMHs: 21.88895\nExp number, first 60 AAs: 0.01585\nTotal prob of N-in: 0.00266\noutside 1 247\nTMhelix 248 270\ninside 271 279

33638 GCF_900090125.1_IMG-taxon_2522125136_annotated_assembly Streptomyces sp. WMMB 714 Terrabacteria group; Actinobacteria; Actinobacteria; Streptomycetales; Streptomycetaceae; Streptomyces MLAFVENLAMRRTAAFFDLDKTVIAKSSTLTFGRSFYQGGLINRRAVLRTAYAQFVYLLG WP_045865951.1 inhibition of morphological differentiation protein [Streptomyces sp. WMMB 714] Length: 279\nNumber of predicted TMHs: 1\nExp number of AAs in TMHs: 21.88895\nExp number, first 60 AAs: 0.01585\nTotal prob of N-in: 0.00266\noutside 1 247\nTMhelix 248 270\ninside 271 279

33639 GCF_001279455.1_ASM127945v1 Streptomyces sp. XY152 Terrabacteria group; Actinobacteria; Actinobacteria; Streptomycetales; Streptomycetaceae; Streptomyces MLRDVENHSMPRTAAFFDLDKTVIAKSSALTFSKSFYQGGLINRRAVLRTAYSQFVFLAG WP_053637387.1 inhibition of morphological differentiation protein [Streptomyces sp. XY152] Length: 277\nNumber of predicted TMHs: 1\nExp number of AAs in TMHs: 21.88019\nExp number, first 60 AAs: 0.04028\nTotal prob of N-in: 0.00638\noutside 1 247\nTMhelix 248 270\ninside 271 277

33640 GCF_001279455.1_ASM127945v1 Streptomyces sp. XY152 Terrabacteria group; Actinobacteria; Actinobacteria; Streptomycetales; Streptomycetaceae; Streptomyces MRHARRRVVRRVTRLAAVGGLLLGATMVTHAVASEPRAPDAVPFATAEGTVSTGPGADLV WP_030821297.1 MULTISPECIES: protease [Streptomyces] Length: 459\nNumber of predicted TMHs: 2\nExp number of AAs in TMHs: 42.46692\nExp number, first 60 AAs: 22.12504\nTotal prob of N-in: 0.98832\nPOSSIBLE N-term signal sequence\ninside 1 11\nTMhelix 12 34\noutside 35 420\nTMhelix 421 443\ninside 444 459

33641 GCF_001280885.1_ASM128088v1 Streptomyces sp. XY332 Terrabacteria group; Actinobacteria; Actinobacteria; Streptomycetales; Streptomycetaceae; Streptomyces MSTVNRRTLVRAAAVTVCAGTLLGLPAAAALAEGVPAASAAHSAGAPRTLVKSLALADGM WP_053784674.1 hypothetical protein [Streptomyces sp. XY332] Length: 200\nNumber of predicted TMHs: 2\nExp number of AAs in TMHs: 44.25577\nExp number, first 60 AAs: 21.62733\nTotal prob of N-in: 0.99073\nPOSSIBLE N-term signal sequence\ninside 1 8\nTMhelix 9 31\noutside 32 170\nTMhelix 171 193\ninside 194 200

33642 GCF_001280885.1_ASM128088v1 Streptomyces sp. XY332 Terrabacteria group; Actinobacteria; Actinobacteria; Streptomycetales; Streptomycetaceae; Streptomyces MPARRRTVTRIAAVGLAPLAMAAYAAAPAAAHGSMTDPVSRVAACYAEGPESPKSAACKA WP_053785886.1 chitin-binding protein [Streptomyces sp. XY332] Length: 347\nNumber of predicted TMHs: 2\nExp number of AAs in TMHs: 42.5924\nExp number, first 60 AAs: 20.31231\nTotal prob of N-in: 0.95534\nPOSSIBLE N-term signal sequence\ninside 1 11\nTMhelix 12 31\noutside 32 312\nTMhelix 313 335\ninside 336 347

33643 GCF_001279815.1_ASM127981v1 Streptomyces sp. XY37 Terrabacteria group; Actinobacteria; Actinobacteria; Streptomycetales; Streptomycetaceae; Streptomyces MSARRRTVMPIRIAAAGAAPLALAVYAAVPAVAHGSMTDPVSRVAACYAEGPESPKSAAC WP_053633676.1 MULTISPECIES: chitin-binding protein [Streptomyces] Length: 347\nNumber of predicted TMHs: 2\nExp number of AAs in TMHs: 43.49607\nExp number, first 60 AAs: 21.79131\nTotal prob of N-in: 0.98451\nPOSSIBLE N-term signal sequence\ninside 1 12\nTMhelix 13 35\noutside 36 312\nTMhelix 313 335\ninside 336 347

33644 GCF_001279815.1_ASM127981v1 Streptomyces sp. XY37 Terrabacteria group; Actinobacteria; Actinobacteria; Streptomycetales; Streptomycetaceae; Streptomyces MPPYRDARRALLAAAFALTLTAAGATAATAAQHPPYALRLDGAGECTFPMKKQIADRPWA WP_053634588.1 MULTISPECIES: type VII secretion-associated serine protease mycosin [Streptomyces] Length: 414\nNumber of predicted TMHs: 2\nExp number of AAs in TMHs: 42.2832099999999\nExp number, first 60 AAs: 19.94733\nTotal prob of N-in: 0.93627\nPOSSIBLE N-term signal sequence\ninside 1 11\nTMhelix 12 31\noutside 32 380\nTMhelix 381 403\ninside 404 414

33645 GCF_001279815.1_ASM127981v1 Streptomyces sp. XY37 Terrabacteria group; Actinobacteria; Actinobacteria; Streptomycetales; Streptomycetaceae; Streptomyces MSTVRRRTLVRAAAVTACAGSLLALPTAAALAEGVPAASSAHSAAPQRTLVKSLSLADGV WP_030834094.1 MULTISPECIES: hypothetical protein [Streptomyces] Length: 218\nNumber of predicted TMHs: 2\nExp number of AAs in TMHs: 42.93094\nExp number, first 60 AAs: 20.33933\nTotal prob of N-in: 0.95701\nPOSSIBLE N-term signal sequence\ninside 1 8\nTMhelix 9 31\noutside 32 188\nTMhelix 189 211\ninside 212 218

33646 GCF_001279765.1_ASM127976v1 Streptomyces sp. XY413 Terrabacteria group; Actinobacteria; Actinobacteria; Streptomycetales; Streptomycetaceae; Streptomyces
MPPYRDARRALLAAAFALTLTAAGATAATAAQHPPYALRLDGAGECTFPMKKQIADRPWA WP_053634588.1
MULTISPECIES: type VII secretion-associated serine protease mycosin [Streptomyces] Length: 414\nNumber of predicted TMHs: 2\nExp number of AAs in TMHs: 42.2832099999999\nExp number, first 60 AAs: 19.94733\nTotal prob of N-in: 0.93627\nPOSSIBLE N-term signal sequence\ninside 1 11\nTMhelix 12 31\noutside 32 380\nTMhelix 381 403\ninside 404 414

33647 GCF_001279765.1_ASM127976v1 Streptomyces sp. XY413 Terrabacteria group; Actinobacteria; Actinobacteria; Streptomycetales; Streptomycetaceae; Streptomyces
MALDVIRSGPPDKRPRRNLFKRGVLVLLPAALLAAATLVPNLAHALEAPVNLGTATNYAV WP_053713884.1
hypothetical protein [Streptomyces sp. XY413] Length: 533\nNumber of predicted TMHs: 2\nExp number of AAs in TMHs: 43.1408699999999\nExp number, first 60 AAs: 20.8292\nTotal prob of N-in: 0.94537\nPOSSIBLE N-term signal sequence\ninside 1 22\nTMhelix 23 45\noutside 46 498\nTMhelix 499 521\ninside 522 533

33648 GCF_001279765.1_ASM127976v1 Streptomyces sp. XY413 Terrabacteria group; Actinobacteria; Actinobacteria; Streptomycetales; Streptomycetaceae; Streptomyces
MSARRRTVMPIRIAAAGAAPLALAVYAAVPAVAHGSMTDPVSRVAACYAEGPESPKSAAC WP_053713597.1
chitin-binding protein [Streptomyces sp. XY413] Length: 345\nNumber of predicted TMHs: 2\nExp number of AAs in TMHs: 43.49631\nExp number, first 60 AAs: 21.79151\nTotal prob of N-in: 0.98454\nPOSSIBLE N-term signal sequence\ninside 1 12\nTMhelix 13 35\noutside 36 310\nTMhelix 311 333\ninside 334 345

33649 GCF_001279765.1_ASM127976v1 Streptomyces sp. XY413 Terrabacteria group; Actinobacteria; Actinobacteria; Streptomycetales; Streptomycetaceae; Streptomyces
MSTVRRRTLVRAAAVTACAGSLLALPTAAALAEGVPAASSAHSAAPQRTLVKSLSLADGV WP_030834094.1
MULTISPECIES: hypothetical protein [Streptomyces] Length: 218\nNumber of predicted TMHs: 2\nExp number of AAs in TMHs: 42.93094\nExp number, first 60 AAs: 20.33933\nTotal prob of N-in: 0.95701\nPOSSIBLE N-term signal sequence\ninside 1 8\nTMhelix 9 31\noutside 32 188\nTMhelix 189 211\ninside 212 218

33650 GCF_001279485.1_ASM127948v1 Streptomyces sp. XY511 Terrabacteria group; Actinobacteria; Actinobacteria; Streptomycetales; Streptomycetaceae; Streptomyces
MPQYRDARRALLATAFALTLTAAGATAATAAPHPPYALRLDGAGECTFPMKKQIADRPWA WP_053611783.1
MULTISPECIES: type VII secretion-associated serine protease mycosin [Streptomyces] Length: 414\nNumber of predicted TMHs: 2\nExp number of AAs in TMHs: 42.58933\nExp number, first 60 AAs: 20.25125\nTotal prob of N-in: 0.94555\nPOSSIBLE N-term signal sequence\ninside 1 8\nTMhelix 9 31\noutside 32 380\nTMhelix 381 403\ninside 404 414

33651 GCF_001279485.1_ASM127948v1 Streptomyces sp. XY511 Terrabacteria group; Actinobacteria; Actinobacteria; Streptomycetales; Streptomycetaceae; Streptomyces
MSTVRRRTLVRAAAVTACAGSLLALPTAAALAEGVPAASSAHSAAPQRTLVKSLSLADGV WP_053611851.1
MULTISPECIES: hypothetical protein [Streptomyces] Length: 218\nNumber of predicted TMHs: 2\nExp number of AAs in TMHs: 42.93081\nExp number, first 60 AAs: 20.33935\nTotal prob of N-in: 0.95701\nPOSSIBLE N-term signal sequence\ninside 1 8\nTMhelix 9 31\noutside 32 188\nTMhelix 189 211\ninside 212 218

33652 GCF_001279845.1_ASM127984v1 Streptomyces sp. XY533 Terrabacteria group; Actinobacteria; Actinobacteria; Streptomycetales; Streptomycetaceae; Streptomyces
MSTVRRRTLVRAAAVTACAGSLLALPTAAALAEGVPAASSAHSAAPQRTLVKSLSLADGV WP_053611851.1
MULTISPECIES: hypothetical protein [Streptomyces] Length: 218\nNumber of predicted TMHs: 2\nExp number of AAs in TMHs: 42.93081\nExp number, first 60 AAs: 20.33935\nTotal prob of N-in: 0.95701\nPOSSIBLE N-term signal sequence\ninside 1 8\nTMhelix 9 31\noutside 32 188\nTMhelix 189 211\ninside 212 218

33653 GCF_001279845.1_ASM127984v1 Streptomyces sp. XY533 Terrabacteria group; Actinobacteria; Actinobacteria; Streptomycetales; Streptomycetaceae; Streptomyces
MPQYRDARRALLATAFALTLTAAGATAATAAPHPPYALRLDGAGECTFPMKKQIADRPWA WP_053611783.1
MULTISPECIES: type VII secretion-associated serine protease mycosin [Streptomyces] Length: 414\nNumber of predicted TMHs: 2\nExp number of AAs in TMHs: 42.58933\nExp number, first 60 AAs: 20.25125\nTotal prob of N-in: 0.94555\nPOSSIBLE N-term signal sequence\ninside 1 8\nTMhelix 9 31\noutside 32 380\nTMhelix 381 403\ninside 404 414

33654 GCF_001279855.1_ASM127985v1 Streptomyces sp. XY58 Terrabacteria group; Actinobacteria; Actinobacteria; Streptomycetales; Streptomycetaceae; Streptomyces

MSARRRTVMPIRIAAAGAAPLALAVYAAVPAVAHGSMTDPVSRVAACYAEGPESPKSAAC WP_053633676.1 MULTISPECIES: chitin-binding protein [Streptomyces] Length: 347\nNumber of predicted TMHs: 2\nExp number of AAs in TMHs: 43.49607\nExp number, first 60 AAs: 21.79131\nTotal prob of N-in: 0.98451\nPOSSIBLE N-term signal sequence\ninside 1 12\nTMhelix 13 35\noutside 36 312\nTMhelix 313 335\ninside 336 347

33655 GCF_001279855.1_ASM127985v1 Streptomyces sp. XY58 Terrabacteria group; Actinobacteria; Actinobacteria; Streptomycetales; Streptomycetaceae; Streptomyces MSTVRRRTLVRAAAVTACAGSLLALPTAAALAEGVPAASSAHSAAPQRTLVKSLSLADGV WP_030834094.1 MULTISPECIES: hypothetical protein [Streptomyces] Length: 218\nNumber of predicted TMHs: 2\nExp number of AAs in TMHs: 42.93094\nExp number, first 60 AAs: 20.33933\nTotal prob of N-in: 0.95701\nPOSSIBLE N-term signal sequence\ninside 1 8\nTMhelix 9 31\noutside 32 188\nTMhelix 189 211\ninside 212 218

33656 GCF_001279855.1_ASM127985v1 Streptomyces sp. XY58 Terrabacteria group; Actinobacteria; Actinobacteria; Streptomycetales; Streptomycetaceae; Streptomyces MALDVIRSGPPDKRPRRNLFKRGVLVLLPAALLAAATLVPNLAHALEAPVNLGTATNYAV WP_053667212.1 hypothetical protein [Streptomyces sp. XY58] Length: 525\nNumber of predicted TMHs: 2\nExp number of AAs in TMHs: 43.1518499999999\nExp number, first 60 AAs: 20.82977\nTotal prob of N-in: 0.94538\nPOSSIBLE N-term signal sequence\ninside 1 22\nTMhelix 23 45\noutside 46 490\nTMhelix 491 513\ninside 514 525

33657 GCF_001279855.1_ASM127985v1 Streptomyces sp. XY58 Terrabacteria group; Actinobacteria; Actinobacteria; Streptomycetales; Streptomycetaceae; Streptomyces MPPYRDARRALLAAAFALTLTAAGATAATAAQHPPYALRLDGAGECTFPMKKQIADRPWA WP_053634588.1 MULTISPECIES: type VII secretion-associated serine protease mycosin [Streptomyces] Length: 414\nNumber of predicted TMHs: 2\nExp number of AAs in TMHs: 42.2832099999999\nExp number, first 60 AAs: 19.94733\nTotal prob of N-in: 0.93627\nPOSSIBLE N-term signal sequence\ninside 1 11\nTMhelix 12 31\noutside 32 380\nTMhelix 381 403\ninside 404 414

33658 GCF_001279885.1_ASM127988v1 Streptomyces sp. XY593 Terrabacteria group; Actinobacteria; Actinobacteria; Streptomycetales; Streptomycetaceae; Streptomyces MPQYRDARRALLATAFALTLTAAGATVATAAPHPPYALRLDGAGECTFPMKKQIADRPWA WP_053685516.1 type VII secretion-associated serine protease mycosin [Streptomyces sp. XY593] Length: 414\nNumber of predicted TMHs: 2\nExp number of AAs in TMHs: 43.0473099999999\nExp number, first 60 AAs: 20.71013\nTotal prob of N-in: 0.96189\nPOSSIBLE N-term signal sequence\ninside 1 8\nTMhelix 9 31\noutside 32 380\nTMhelix 381 403\ninside 404 414

33659 GCF_001279885.1_ASM127988v1 Streptomyces sp. XY593 Terrabacteria group; Actinobacteria; Actinobacteria; Streptomycetales; Streptomycetaceae; Streptomyces MSTVRRRTLVRAAAVTACAGSLLALPTAAALAEGVPAASSAHSAAPQRTLVKSLSLADGV WP_053611851.1 MULTISPECIES: hypothetical protein [Streptomyces] Length: 218\nNumber of predicted TMHs: 2\nExp number of AAs in TMHs: 42.93081\nExp number, first 60 AAs: 20.33935\nTotal prob of N-in: 0.95701\nPOSSIBLE N-term signal sequence\ninside 1 8\nTMhelix 9 31\noutside 32 188\nTMhelix 189 211\ninside 212 218

33660 GCF_001279905.1_ASM127990v1 Streptomyces sp. XY66 Terrabacteria group; Actinobacteria; Actinobacteria; Streptomycetales; Streptomycetaceae; Streptomyces MSARRRTVMPIRIAAAGAAPLALAVYAAVPAVAHGSMTDPVSRVAACYAEGPESPKSAAC WP_053632541.1 MULTISPECIES: chitin-binding protein [Streptomyces] Length: 345\nNumber of predicted TMHs: 2\nExp number of AAs in TMHs: 43.49631\nExp number, first 60 AAs: 21.79151\nTotal prob of N-in: 0.98454\nPOSSIBLE N-term signal sequence\ninside 1 12\nTMhelix 13 35\noutside 36 310\nTMhelix 311 333\ninside 334 345

33661 GCF_001279905.1_ASM127990v1 Streptomyces sp. XY66 Terrabacteria group; Actinobacteria; Actinobacteria; Streptomycetales; Streptomycetaceae; Streptomyces MALDVIRSGPPDKRPRRNLFKRGVLVLLPAALLAAATLVPNLAHALEAPVNLGTATNYAV WP_053679816.1 hypothetical protein [Streptomyces sp. XY66] Length: 546\nNumber of predicted TMHs: 2\nExp number of AAs in TMHs: 43.12563\nExp number, first 60 AAs: 20.82835\nTotal prob of N-in: 0.94536\nPOSSIBLE N-term signal sequence\ninside 1 22\nTMhelix 23 45\noutside 46 511\nTMhelix 512 534\ninside 535 546

33662 GCF_001279905.1_ASM127990v1 Streptomyces sp. XY66 Terrabacteria group; Actinobacteria; Actinobacteria; Streptomycetales; Streptomycetaceae; Streptomyces MPPYRDARRALLAAAFALTLTAAGATAATAAQHPPYALRLDGAGECTFPMKKQIADRPWA WP_053678064.1 type VII secretion-associated serine protease mycosin [Streptomyces sp. XY66] Length: 414\nNumber of predicted TMHs: 2\nExp number of AAs in TMHs: 42.2832099999999\nExp number, first 60 AAs: 19.94733\nTotal prob of N-in:

0.93627\nPOSSIBLE N-term signal sequence\ninside 1 11\nTMhelix 12 31\noutside 32 380\nTMhelix 381 403\ninside 404 414

33663 GCF_001279905.1_ASM127990v1 Streptomyces sp. XY66 Terrabacteria group; Actinobacteria; Actinobacteria; Streptomycetales; Streptomycetaceae; Streptomyces MSTVRRRTLVRAAAVTACAGSLLALPTAAALAEGVPAASSAHSAAPQRTLVKSLSLADGV WP_053677522.1 MULTISPECIES: hypothetical protein [Streptomyces] Length: 218\nNumber of predicted TMHs: 2\nExp number of AAs in TMHs: 42.93092\nExp number, first 60 AAs: 20.3393\nTotal prob of N-in: 0.95701\nPOSSIBLE N-term signal sequence\ninside 1 8\nTMhelix 9 31\noutside 32 188\nTMhelix 189 211\ninside 212 218

33664 GCF_001746425.1_ASM174642v1 Streptomyces subrutilus Terrabacteria group; Actinobacteria; Actinobacteria; Streptomycetales; Streptomycetaceae; Streptomyces MSARRRTVTLLRIASAGLAPLAVAAYAAAPAVAHGSMTDPVSRVAACYAEGPESPRSGAC WP_069919632.1 chitin-binding protein [Streptomyces subrutilus] Length: 346\nNumber of predicted TMHs: 2\nExp number of AAs in TMHs: 43.41134\nExp number, first 60 AAs: 21.10185\nTotal prob of N-in: 0.95546\nPOSSIBLE N-term signal sequence\ninside 1 12\nTMhelix 13 35\noutside 36 311\nTMhelix 312 334\ninside 335 346

33665 GCF_001746425.1_ASM174642v1 Streptomyces subrutilus Terrabacteria group; Actinobacteria; Actinobacteria; Streptomycetales; Streptomycetaceae; Streptomyces MTQLPPARRAPLAAALALLGVCAVAPAAVPPPAAASARPVAPEAPYALRLDGAGECTFPM WP_069922204.1 type VII secretion-associated serine protease mycosin [Streptomyces subrutilus] Length: 424\nNumber of predicted TMHs: 1\nExp number of AAs in TMHs: 37.93343\nExp number, first 60 AAs: 15.62416\nTotal prob of N-in: 0.74279\nPOSSIBLE N-term signal sequence\noutside 1 390\nTMhelix 391 413\ninside 414 424

33666 GCF_001746425.1_ASM174642v1 Streptomyces subrutilus Terrabacteria group; Actinobacteria; Actinobacteria; Streptomycetales; Streptomycetaceae; Streptomyces MSTVSRRTLVRAAAVTACAGSLLALPAAAALAEGVPAASSAHSAAPQAVLVKSLTLADGT WP_069921941.1 hypothetical protein [Streptomyces subrutilus] Length: 210\nNumber of predicted TMHs: 2\nExp number of AAs in TMHs: 43.42767\nExp number, first 60 AAs: 20.77799\nTotal prob of N-in: 0.95628\nPOSSIBLE N-term signal sequence\ninside 1 8\nTMhelix 9 31\noutside 32 180\nTMhelix 181 203\ninside 204 210

33667 GCF_001746425.1_ASM174642v1 Streptomyces subrutilus Terrabacteria group; Actinobacteria; Actinobacteria; Streptomycetales; Streptomycetaceae; Streptomyces MRAEHGEDVPGVRQPRNEDGLPRRSPVIAVTGAASGVGAALVSRLAASDEVKQVVAIDER WP_079154649.1 NAD-dependent dehydratase [Streptomyces subrutilus] Length: 373\nNumber of predicted TMHs: 1\nExp number of AAs in TMHs: 21.14969\nExp number, first 60 AAs: 0.94191\nTotal prob of N-in: 0.04218\noutside 1 344\nTMhelix 345 364\ninside 365 373

33668 GCF_002148965.1_ASM214896v1 Streptomyces swartbergensisTerrabacteria group; Actinobacteria; Actinobacteria; Streptomycetales; Streptomycetaceae; Streptomyces MLKGVENHSLPRTAAFFDLDKTVIAKSSTLTFSKSFYQGGLINRRAVLRTAYAQFVFLAG WP_086605494.1 inhibition of morphological differentiation protein [Streptomyces swartbergensis]Length: 277\nNumber of predicted TMHs: 1\nExp number of AAs in TMHs: 21.5893\nExp number, first 60 AAs: 0.0407\nTotal prob of N-in: 0.00409\noutside 1 247\nTMhelix 248 270\ninside 271 277

33669 GCF_002148965.1_ASM214896v1 Streptomyces swartbergensisTerrabacteria group; Actinobacteria; Actinobacteria; Streptomycetales; Streptomycetaceae; Streptomyces MRHARRRVVRRVTRLAAVGGLLLGGAMVTRAVASEPPDPTIGVPYSAQPATSMGTALVSR WP_086602166.1 serine protease [Streptomyces swartbergensis] Length: 449\nNumber of predicted TMHs: 2\nExp number of AAs in TMHs: 41.04413\nExp number, first 60 AAs: 20.67218\nTotal prob of N-in: 0.95256\nPOSSIBLE N-term signal sequence\ninside 1 11\nTMhelix 12 34\noutside 35 410\nTMhelix 411 433\ninside 434 449

33670 GCF_001543925.1_ASM154392v1 Streptomyces thermoautotrophicus Terrabacteria group; Actinobacteria; Actinobacteria; Streptomycetales; Streptomycetaceae; Streptomyces MLISVTHQVPPRTAAFFDLDKTIIAKSSALAFGKPLYQGGLINRRAVLKIAYAQFVYLIG WP_066883711.1 inhibition of morphological differentiation protein [Streptomyces thermoautotrophicus] Length: 277\nNumber of predicted TMHs: 1\nExp number of AAs in TMHs: 18.96456\nExp number, first 60 AAs: 0.14481\nTotal prob of N-in: 0.01611\noutside 1 247\nTMhelix 248 267\ninside 268 277

33671 GCF_001543895.1_ASM154389v1 Streptomyces thermoautotrophicus Terrabacteria group; Actinobacteria; Actinobacteria; Streptomycetales; Streptomycetaceae; Streptomyces

MLISVTHQVPPRTAAFFDLDKTIIAKSSALAFGKPLYQGGLINRRAVLKIAYAQFVYLIG WP_066883711.1 inhibition of morphological differentiation protein [Streptomyces thermoautotrophicus] Length: 277\nNumber of predicted TMHs: 1\nExp number of AAs in TMHs: 18.96456\nExp number, first 60 AAs: 0.14481\nTotal prob of N-in: 0.01611\noutside 1 247\nTMhelix 248 267\ninside 268 277

33672 GCF_002155915.1_ASM215591v1 Streptomyces thermovulgarisTerrabacteria group; Actinobacteria; Actinobacteria; Streptomycetales; Streptomycetaceae; Streptomyces
MLATPRRRPLALVAAMATATALGATVLVAAPTAAAAETPLSGYELTWGIKESYRTYVVRY WP_067392033.1 hypothetical protein [Streptomyces sp. F-3] Length: 507\nNumber of predicted TMHs: 2\nExp number of AAs in TMHs: 43.58403\nExp number, first 60 AAs: 21.17274\nTotal prob of N-in: 0.93360\nPOSSIBLE N-term signal sequence\ninside 1 11\nTMhelix 12 34\noutside 35 476\nTMhelix 477 499\ninside 500 507

33673 GCF_001419765.1_ASM141976v1 Streptomyces torulosus Terrabacteria group; Actinobacteria; Actinobacteria; Streptomycetales; Streptomycetaceae; Streptomyces
MLSPVENHSLPRTAAFFDLDKTVIAKSSTLTFSKSFYHGGLINRRAALRTAYTQFVFLAG WP_055712420.1 inhibition of morphological differentiation protein [Streptomyces torulosus] Length: 274\nNumber of predicted TMHs: 1\nExp number of AAs in TMHs: 21.79227\nExp number, first 60 AAs: 0.06364\nTotal prob of N-in: 0.01330\noutside 1 244\nTMhelix 245 267\ninside 268 274

33674 GCF_001419765.1_ASM141976v1 Streptomyces torulosus Terrabacteria group; Actinobacteria; Actinobacteria; Streptomycetales; Streptomycetaceae; Streptomyces
MRRPSRRPLLAGLAVLAGALLAAVPAVPAAADDGNFTVKDPRITESSGLAASRRHPGVYW WP_055717924.1 hypothetical protein [Streptomyces torulosus] Length: 335\nNumber of predicted TMHs: 2\nExp number of AAs in TMHs: 39.1840199999999\nExp number, first 60 AAs: 21.73077\nTotal prob of N-in: 0.97179\nPOSSIBLE N-term signal sequence\ninside 1 8\nTMhelix 9 31\noutside 32 304\nTMhelix 305 327\ninside 328 335

33675 GCF_000725125.1_ASM72512v1 Streptomyces toyocaensis Terrabacteria group; Actinobacteria; Actinobacteria; Streptomycetales; Streptomycetaceae; Streptomyces
MRRPGSDEGWRVRGWACTARGRAGSRARRRPVAGCVGVAVVLGLGGVLPGQTAAAAAGPA WP_078903354.1 hypothetical protein [Streptomyces toyocaensis] Length: 457\nNumber of predicted TMHs: 2\nExp number of AAs in TMHs: 41.1417799999999\nExp number, first 60 AAs: 19.13161\nTotal prob of N-in: 0.86600\nPOSSIBLE N-term signal sequence\ninside 1 31\nTMhelix 32 54\noutside 55 427\nTMhelix 428 450\ninside 451 457

33676 GCF_000725125.1_ASM72512v1 Streptomyces toyocaensis Terrabacteria group; Actinobacteria; Actinobacteria; Streptomycetales; Streptomycetaceae; Streptomyces
MPRTAAFFDLDKTVIAKSSTLTFSKSFYQGGLINRRAVLRTAYAQFVFLAGGADHDQMER WP_037933720.1 inhibition of morphological differentiation protein [Streptomyces toyocaensis] Length: 268\nNumber of predicted TMHs: 1\nExp number of AAs in TMHs: 21.80961\nExp number, first 60 AAs: 0.06012\nTotal prob of N-in: 0.00598\noutside 1 238\nTMhelix 239 261\ninside 262 268

33677 GCF_000725125.1_ASM72512v1 Streptomyces toyocaensis Terrabacteria group; Actinobacteria; Actinobacteria; Streptomycetales; Streptomycetaceae; Streptomyces
MRHARRRVVRRVTRLAAVGGLLLGATMVTHAVAGEPPRVPEAVPFSTAEEPAVGPGAGLV WP_037928336.1 protease [Streptomyces toyocaensis] Length: 459\nNumber of predicted TMHs: 2\nExp number of AAs in TMHs: 41.76359\nExp number, first 60 AAs: 21.99118\nTotal prob of N-in: 0.98691\nPOSSIBLE N-term signal sequence\ninside 1 11\nTMhelix 12 34\noutside 35 420\nTMhelix 421 443\ninside 444 459

33678 GCF_002155905.1_ASM215590v1 Streptomyces tricolor Terrabacteria group; Actinobacteria; Actinobacteria; Streptomycetales; Streptomycetaceae; Streptomyces
MLRDVENHSLPRAAAFFDLDKTVIAKSSTLTFSKSFYQGGLINRRAALRTAYAQFVFLAG WP_059251463.1 inhibition of morphological differentiation protein [Streptomyces reticuli] Length: 276\nNumber of predicted TMHs: 1\nExp number of AAs in TMHs: 21.75336\nExp number, first 60 AAs: 0.04333\nTotal prob of N-in: 0.00752\noutside 1 247\nTMhelix 248 270\ninside 271 276

33679 GCF_002155905.1_ASM215590v1 Streptomyces tricolor Terrabacteria group; Actinobacteria; Actinobacteria; Streptomycetales; Streptomycetaceae; Streptomyces
MRHARRRVVRRVTRLAAVGGLVLGGAMVAQAAVAGETPPASARALTAADDPGATLVARLG WP_086698134.1 serine protease [Streptomyces tricolor]Length: 457\nNumber of predicted TMHs: 2\nExp number of AAs in TMHs: 43.8388699999999\nExp number, first 60 AAs: 22.54664\nTotal prob of N-in: 0.99517\nPOSSIBLE N-term signal sequence\ninside 1 11\nTMhelix 12 34\noutside 35 418\nTMhelix 419 441\ninside 442 457

33680 GCF_002155905.1_ASM215590v1 Streptomyces tricolor Terrabacteria group; Actinobacteria; Actinobacteria; Streptomycetales; Streptomycetaceae; Streptomyces
MAWQRPVRFGRPLANPLFDAVYRPGVRESSRLSRRAVLGLTAAALPLSTATEAAAATAVV WP_086697472.1 D-alanyl-D-alanine carboxypeptidase [Streptomyces tricolor] Length: 429\nNumber of predicted TMHs: 1\nExp number of AAs in TMHs: 20.98436\nExp number, first 60 AAs: 2.04585\nTotal prob of N-in: 0.09333\noutside 1 374\nTMhelix 375 397\ninside 398 429

33681 GCF_002007125.1_ASM200712v1 Streptomyces tsukubensis Terrabacteria group; Actinobacteria; Actinobacteria; Streptomycetales; Streptomycetaceae; Streptomyces
MLGHVENHSSPRTAAFFDLDKTVIAKSSTLTFSKSFYQGGLINRRAALRTAYTQFVFLVG WP_077966662.1 inhibition of morphological differentiation protein [Streptomyces tsukubensis] Length: 278\nNumber of predicted TMHs: 1\nExp number of AAs in TMHs: 20.00748\nExp number, first 60 AAs: 0.02589\nTotal prob of N-in: 0.00485\noutside 1 247\nTMhelix 248 267\ninside 268 278

33682 GCF_002007125.1_ASM200712v1 Streptomyces tsukubensis Terrabacteria group; Actinobacteria; Actinobacteria; Streptomycetales; Streptomycetaceae; Streptomyces
MARDERSRKRERRRAVRLGALAAAAGLAVLGGAWPSSAVAPTGGKAPAHSRTATPVKHLV WP_077973964.1 phospholipase [Streptomyces tsukubensis] Length: 625\nNumber of predicted TMHs: 2\nExp number of AAs in TMHs: 43.80785\nExp number, first 60 AAs: 21.44345\nTotal prob of N-in: 0.98712\nPOSSIBLE N-term signal sequence\ninside 1 19\nTMhelix 20 42\noutside 43 595\nTMhelix 596 618\ninside 619 625

33683 GCF_001485145.1_ASM148514v1 Streptomyces turgidiscabies Terrabacteria group; Actinobacteria; Actinobacteria; Streptomycetales; Streptomycetaceae; Streptomyces
MLSLVENHSLPRAAAFFDLDKTVIAKSSTLTFSKSFYQGGLINRRAVLRTAYAQFVFLAG WP_044473232.1 inhibition of morphological differentiation protein [Streptomyces turgidiscabies] Length: 277\nNumber of predicted TMHs: 1\nExp number of AAs in TMHs: 21.55643\nExp number, first 60 AAs: 0.01804\nTotal prob of N-in: 0.00246\noutside 1 247\nTMhelix 248 270\ninside 271 277

33684 GCF_001485145.1_ASM148514v1 Streptomyces turgidiscabies Terrabacteria group; Actinobacteria; Actinobacteria; Streptomycetales; Streptomycetaceae; Streptomyces
MRHARRRIVRRATRLAAVGGVILGGLMVTQAVASEPPEVNLPFSSAQAAGTTGSELVTEL WP_006380203.1 alpha-lytic protease PRO domain protein [Streptomyces turgidiscabies] Length: 452\nNumber of predicted TMHs: 2\nExp number of AAs in TMHs: 42.58327\nExp number, first 60 AAs: 21.83569\nTotal prob of N-in: 0.98683\nPOSSIBLE N-term signal sequence\ninside 1 11\nTMhelix 12 34\noutside 35 413\nTMhelix 414 436\ninside 437 452

33685 GCF_001485145.1_ASM148514v1 Streptomyces turgidiscabies Terrabacteria group; Actinobacteria; Actinobacteria; Streptomycetales; Streptomycetaceae; Streptomyces
MPARRRPAVVTTVTALGIAALALGATPASAHGSMGDPVSRVAQCYAEGPESPTSAACRAA WP_006376680.1 chitin-binding protein [Streptomyces turgidiscabies] Length: 353\nNumber of predicted TMHs: 2\nExp number of AAs in TMHs: 41.16593\nExp number, first 60 AAs: 20.07244\nTotal prob of N-in: 0.93537\nPOSSIBLE N-term signal sequence\ninside 1 6\nTMhelix 7 29\noutside 30 318\nTMhelix 319 341\ninside 342 353

33686 GCF_001905345.1_ASM190534v1 Streptomyces uncialis Terrabacteria group; Actinobacteria; Actinobacteria; Streptomycetales; Streptomycetaceae; Streptomyces
MTRSPAPAARRRTLRRWAALTAAAATTATVTVLPAASPALADTGQCTFPSKAYPGRPWAL WP_073793385.1 type VII secretion-associated serine protease mycosin [Streptomyces uncialis] Length: 425\nNumber of predicted TMHs: 2\nExp number of AAs in TMHs: 40.8704\nExp number, first 60 AAs: 18.56764\nTotal prob of N-in: 0.84256\nPOSSIBLE N-term signal sequence\ninside 1 16\nTMhelix 17 39\noutside 40 386\nTMhelix 387 409\ninside 410 425

33687 GCF_001905345.1_ASM190534v1 Streptomyces uncialis Terrabacteria group; Actinobacteria; Actinobacteria; Streptomycetales; Streptomycetaceae; Streptomyces
MLGLVENHSLPRTAAFFDLDKTVIAKSSALTFSKSFYHGGLINRRAVLRTAYAQFVFLAG WP_073795406.1 inhibition of morphological differentiation protein [Streptomyces uncialis] Length: 277\nNumber of predicted TMHs: 1\nExp number of AAs in TMHs: 21.17723\nExp number, first 60 AAs: 0.03744\nTotal prob of N-in: 0.00640\noutside 1 247\nTMhelix 248 270\ninside 271 277

33688 GCF_001905345.1_ASM190534v1 Streptomyces uncialis Terrabacteria group; Actinobacteria; Actinobacteria; Streptomycetales; Streptomycetaceae; Streptomyces
MSATDGTGGSDAGPRTASGPDVGPPPPGAPRARRGAMLAGAVVALGCVLFLGGFVWGAVL WP_073793324.1 signal peptidase I [Streptomyces uncialis] Length: 267\nNumber of predicted TMHs: 2\nExp number of

AAs in TMHs: 45.33235\nExp number, first 60 AAs: 22.21897\nTotal prob of N-in: 0.93057\nPOSSIBLE N-term signal sequence\ninside 1 37\nTMhelix 38 60\noutside 61 224\nTMhelix 225 247\ninside 248 267

33689 GCF_000955965.1_ASM95596v1 Streptomyces variegatus Terrabacteria group; Actinobacteria; Actinobacteria; Streptomycetales; Streptomycetaceae; Streptomyces MLWGVENHSLPRTAAFFDLDKTVIAKSSTLTFSKSFYQGGLINRRAVLRTAYAQFVFLAG WP_031140406.1 MULTISPECIES: inhibition of morphological differentiation protein [Streptomyces] Length: 277\nNumber of predicted TMHs: 1\nExp number of AAs in TMHs: 21.58162\nExp number, first 60 AAs: 0.0297\nTotal prob of N-in: 0.00292\noutside 1 247\nTMhelix 248 270\ninside 271 277

33690 GCF_000955965.1_ASM95596v1 Streptomyces variegatus Terrabacteria group; Actinobacteria; Actinobacteria; Streptomycetales; Streptomycetaceae; Streptomyces MRHARRRVVRRVTRLAAVGGLLLGGAMVTNAVASEPPAPSIGVPFTAPPATGEGTDLVSR WP_031133743.1 MULTISPECIES: protease [Streptomyces] Length: 448\nNumber of predicted TMHs: 2\nExp number of AAs in TMHs: 41.67951\nExp number, first 60 AAs: 21.32501\nTotal prob of N-in: 0.96855\nPOSSIBLE N-term signal sequence\ninside 1 11\nTMhelix 12 34\noutside 35 409\nTMhelix 410 432\ninside 433 448

33691 GCF_000718635.1_ASM71863v1 Streptomyces varsoviensis Terrabacteria group; Actinobacteria; Actinobacteria; Streptomycetales; Streptomycetaceae; Streptomyces MSSPDPHVRAARNGTAHGGAKSGRRPVVAVTGAAAGIGALLAQRLAESEEIKQVIAIDER WP_030878525.1 NAD-dependent dehydratase [Streptomyces varsoviensis] Length: 374\nNumber of predicted TMHs: 1\nExp number of AAs in TMHs: 20.91377\nExp number, first 60 AAs: 0.48118\nTotal prob of N-in: 0.03030\noutside 1 344\nTMhelix 345 364\ninside 365 374

33692 GCF_000718635.1_ASM71863v1 Streptomyces varsoviensis Terrabacteria group; Actinobacteria; Actinobacteria; Streptomycetales; Streptomycetaceae; Streptomyces MLSLVENHSLPRTAAFFDLDKTVIAKSSTLTFSKSFYQGGLINRRAVLRTAYAQFVFLAG WP_030880801.1 inhibition of morphological differentiation protein [Streptomyces varsoviensis] Length: 281\nNumber of predicted TMHs: 1\nExp number of AAs in TMHs: 21.23966\nExp number, first 60 AAs: 0.02346\nTotal prob of N-in: 0.00446\noutside 1 246\nTMhelix 247 269\ninside 270 281

33693 GCF_000719255.1_ASM71925v1 Streptomyces varsoviensis Terrabacteria group; Actinobacteria; Actinobacteria; Streptomycetales; Streptomycetaceae; Streptomyces MLSLVENHSLPRTAAFFDLDKTVIAKSSTLTFSKSFYQGGLINRRAVLRTAYAQFVFLAG WP_030880801.1 inhibition of morphological differentiation protein [Streptomyces varsoviensis] Length: 281\nNumber of predicted TMHs: 1\nExp number of AAs in TMHs: 21.23966\nExp number, first 60 AAs: 0.02346\nTotal prob of N-in: 0.00446\noutside 1 246\nTMhelix 247 269\ninside 270 281

33694 GCF_000719255.1_ASM71925v1 Streptomyces varsoviensis Terrabacteria group; Actinobacteria; Actinobacteria; Streptomycetales; Streptomycetaceae; Streptomyces MSSPDPHVRAARNGTAHGGAKSGRRPVVAVTGAAAGIGALLAQRLAESEEIKQVIAIDER WP_030878525.1 NAD-dependent dehydratase [Streptomyces varsoviensis] Length: 374\nNumber of predicted TMHs: 1\nExp number of AAs in TMHs: 20.91377\nExp number, first 60 AAs: 0.48118\nTotal prob of N-in: 0.03030\noutside 1 344\nTMhelix 345 364\ninside 365 374

33695 GCF_001406115.1_Streptomyces_venezuelae_ATCC_15439 Streptomyces venezuelae Terrabacteria group; Actinobacteria; Actinobacteria; Streptomycetales; Streptomycetaceae; Streptomyces MLWLVENHSLPRTAAFFDLDKTVIAKSSTLTFSKSFYRGGLISRRAALRTAYIQFVFLVG WP_055642706.1 inhibition of morphological differentiation protein [Streptomyces venezuelae] Length: 288\nNumber of predicted TMHs: 1\nExp number of AAs in TMHs: 22.08564\nExp number, first 60 AAs: 0.06209\nTotal prob of N-in: 0.00395\noutside 1 247\nTMhelix 248 270\ninside 271 288

33696 GCF_001886595.1_ASM188659v1 Streptomyces venezuelae Terrabacteria group; Actinobacteria; Actinobacteria; Streptomycetales; Streptomycetaceae; Streptomyces MLRLVENHSLPRTAAFFDLDKTVIAKSSTLTFSKSFYRGGLISRRAALRTAYIQFVFLAG WP_015034529.1 phosphoserine phosphatase [Streptomyces venezuelae] Length: 288\nNumber of predicted TMHs: 1\nExp number of AAs in TMHs: 22.12812\nExp number, first 60 AAs: 0.0544\nTotal prob of N-in: 0.00444\noutside 1 247\nTMhelix 248 270\ninside 271 288

33697 GCF_001443625.1_ASM144362v1 Streptomyces venezuelae Terrabacteria group; Actinobacteria; Actinobacteria; Streptomycetales; Streptomycetaceae; Streptomyces

MYPRPRRRRVLAAVSLLLLGALLSPAATASASPAAPGTPEPARMDLVLDLSGSMNKSDAG WP_079040852.1 hypothetical protein [Streptomyces venezuelae] Length: 627\nNumber of predicted TMHs: 2\nExp number of AAs in TMHs: 41.77417\nExp number, first 60 AAs: 20.47276\nTotal prob of N-in: 0.92014\nPOSSIBLE N-term signal sequence\ninside 1 12\nTMhelix 13 35\noutside 36 600\nTMhelix 601 623\ninside 624 627

33698 GCF_001443625.1_ASM144362v1 Streptomyces venezuelae Terrabacteria group; Actinobacteria; Actinobacteria; Streptomycetales; Streptomycetaceae; Streptomyces MLWLVENHSLPRTAAFFDLDKTVIAKSSTLTFSKSFYRGGLISRRAALRTAYIQFVFLVG WP_055642706.1 inhibition of morphological differentiation protein [Streptomyces venezuelae] Length: 288\nNumber of predicted TMHs: 1\nExp number of AAs in TMHs: 22.08564\nExp number, first 60 AAs: 0.06209\nTotal prob of N-in: 0.00395\noutside 1 247\nTMhelix 248 270\ninside 271 288

33699 GCF_000830005.1_ASM83000v1 Streptomyces vietnamensis Terrabacteria group; Actinobacteria; Actinobacteria; Streptomycetales; Streptomycetaceae; Streptomyces MLGLVENHSLPRTAAFFDLDKTVIAKSSTLTFSKSFYRGGLISRRAALRTAYIQFVFLAG WP_041129907.1 inhibition of morphological differentiation protein [Streptomyces vietnamensis] Length: 288\nNumber of predicted TMHs: 1\nExp number of AAs in TMHs: 22.01399\nExp number, first 60 AAs: 0.02671\nTotal prob of N-in: 0.00194\noutside 1 247\nTMhelix 248 270\ninside 271 288

33700 GCF_002154465.1_ASM215446v1 Streptomyces vinaceus Terrabacteria group; Actinobacteria; Actinobacteria; Streptomycetales; Streptomycetaceae; Streptomyces MLKGVENHLSPRTAAFFDLDKTVIAKSSTLTFSKSFYQGGLINRRAVLRTAYSQFVFLAG WP_086704966.1 inhibition of morphological differentiation protein [Streptomyces vinaceus] Length: 277\nNumber of predicted TMHs: 1\nExp number of AAs in TMHs: 21.81151\nExp number, first 60 AAs: 0.02023\nTotal prob of N-in: 0.00358\noutside 1 247\nTMhelix 248 270\ninside 271 277

33701 GCF_000717215.1_ASM71721v1 Streptomyces vinaceus Terrabacteria group; Actinobacteria; Actinobacteria; Streptomycetales; Streptomycetaceae; Streptomyces MLSVVENCFSPRTAAFFDLDKTVIAKSSTLTFSKSFYQGGLINRRAVLRTAYTQFVFMAG WP_043247809.1 inhibition of morphological differentiation protein [Streptomyces vinaceus] Length: 280\nNumber of predicted TMHs: 1\nExp number of AAs in TMHs: 21.68705\nExp number, first 60 AAs: 0.01801\nTotal prob of N-in: 0.00793\noutside 1 247\nTMhelix 248 270\ninside 271 280

33702 GCF_000717215.1_ASM71721v1 Streptomyces vinaceus Terrabacteria group; Actinobacteria; Actinobacteria; Streptomycetales; Streptomycetaceae; Streptomyces MGSSWNERRLERRVRPGGGKPLKPFRWWQLTRRSLLSLALPVGGGLSATYTVEVKHGGDA WP_043247767.1 hypothetical protein [Streptomyces vinaceus] Length: 218\nNumber of predicted TMHs: 2\nExp number of AAs in TMHs: 41.32\nExp number, first 60 AAs: 0.37575\nTotal prob of N-in: 0.97418\ninside 1 135\nTMhelix 136 158\noutside 159 182\nTMhelix 183 200\ninside 201 218

33703 GCF_000717845.1_ASM71784v1 Streptomyces violaceoruber Terrabacteria group; Actinobacteria; Actinobacteria; Streptomycetales; Streptomycetaceae; Streptomyces MPRTAAFFDLDKTVIAKSSTLTFSKSFYQGGLINRRAVLRTAYAQFVFLAGGADHDQMER WP_003975378.1 MULTISPECIES: morphological differentiation-associated protein [Streptomyces] Length: 268\nNumber of predicted TMHs: 1\nExp number of AAs in TMHs: 21.51645\nExp number, first 60 AAs: 0.03737\nTotal prob of N-in: 0.00617\noutside 1 238\nTMhelix 239 261\ninside 262 268

33704 GCF_000717845.1_ASM71784v1 Streptomyces violaceoruber Terrabacteria group; Actinobacteria; Actinobacteria; Streptomycetales; Streptomycetaceae; Streptomyces MRHARRRIVRRVARLAAVGGLLLGGTMVTRAVASEPPDASAAPRTFAQTPSGAGGDLVSR WP_030872231.1 protease [Streptomyces violaceoruber] Length: 454\nNumber of predicted TMHs: 2\nExp number of AAs in TMHs: 40.78158\nExp number, first 60 AAs: 21.15281\nTotal prob of N-in: 0.96513\nPOSSIBLE N-term signal sequence\ninside 1 11\nTMhelix 12 34\noutside 35 415\nTMhelix 416 438\ninside 439 454

33705 GCF_000717845.1_ASM71784v1 Streptomyces violaceoruber Terrabacteria group; Actinobacteria; Actinobacteria; Streptomycetales; Streptomycetaceae; Streptomyces MGHNRRRRPTGARRATFGAVALILGGSGLVAVNVFASATESGNPAVPLGSSGIAATVDCP WP_078652983.1 hypothetical protein [Streptomyces violaceoruber] Length: 691\nNumber of predicted TMHs: 2\nExp number of AAs in TMHs: 43.09161\nExp number, first 60 AAs: 21.84277\nTotal prob of N-in: 0.99047\nPOSSIBLE N-term signal sequence\ninside 1 16\nTMhelix 17 39\noutside 40 663\nTMhelix 664 686\ninside 687 691

33706 GCF_002082175.1_ASM208217v1 Streptomyces violaceoruber Terrabacteria group; Actinobacteria; Actinobacteria; Streptomycetales; Streptomycetaceae; Streptomyces
MLSVVENCFSPRTAAFFDLDKTVIAKSSTLTFSKSFYQGGLINRRAVLRTAYTQFVFMAG WP_030113359.1 MULTISPECIES: inhibition of morphological differentiation protein [Streptomyces] Length: 280\nNumber of predicted TMHs: 1\nExp number of AAs in TMHs: 21.68695\nExp number, first 60 AAs: 0.01801\nTotal prob of N-in: 0.00793\noutside 1 247\nTMhelix 248 270\ninside 271 280

33707 GCF_000720805.1_ASM72080v1 Streptomyces violaceoruber Terrabacteria group; Actinobacteria; Actinobacteria; Streptomycetales; Streptomycetaceae; Streptomyces
MSISRRTAVRTARILGVASASAALALSAAGSALACNISEFSAEAKCDGTKGVIIVTDKDA WP_030942286.1 membrane protein [Streptomyces violaceoruber] Length: 239\nNumber of predicted TMHs: 2\nExp number of AAs in TMHs: 40.30742\nExp number, first 60 AAs: 18.27777\nTotal prob of N-in: 0.87593\nPOSSIBLE N-term signal sequence\ninside 1 11\nTMhelix 12 34\noutside 35 207\nTMhelix 208 230\ninside 231 239

33708 GCF_000720805.1_ASM72080v1 Streptomyces violaceoruber Terrabacteria group; Actinobacteria; Actinobacteria; Streptomycetales; Streptomycetaceae; Streptomyces
MLGLVENHSLPRTAAFFDLDKTVIAKSSTLTFGKSFYQGGLINRRAALRTAYAQFVFLVG WP_030939271.1 inhibition of morphological differentiation protein [Streptomyces violaceoruber] Length: 277\nNumber of predicted TMHs: 1\nExp number of AAs in TMHs: 21.69946\nExp number, first 60 AAs: 0.01081\nTotal prob of N-in: 0.00155\noutside 1 247\nTMhelix 248 270\ninside 271 277

33709 GCF_000720805.1_ASM72080v1 Streptomyces violaceoruber Terrabacteria group; Actinobacteria; Actinobacteria; Streptomycetales; Streptomycetaceae; Streptomyces
MRHARRRVVRRVTRLAAVGGLLLGGTMVTRAMASESPAVPAPRTFAQQAQQAGTTGEGLV WP_030931144.1
protease [Streptomyces violaceoruber] Length: 461\nNumber of predicted TMHs: 2\nExp number of AAs in TMHs: 40.79187\nExp number, first 60 AAs: 20.32233\nTotal prob of N-in: 0.95261\nPOSSIBLE N-term signal sequence\ninside 1 11\nTMhelix 12 34\noutside 35 422\nTMhelix 423 445\ninside 446 461

33710 GCF_000717995.1_ASM71799v1 Streptomyces violaceorubidus Terrabacteria group; Actinobacteria; Actinobacteria; Streptomycetales; Streptomycetaceae; Streptomyces
MGHNRRRRPTGARRATFGAVALILGGSGLVAVNVFASATESGDAAVPLGSSGVAATVDCP WP_078606519.1
hypothetical protein [Streptomyces violaceorubidus] Length: 694\nNumber of predicted TMHs: 2\nExp number of AAs in TMHs: 43.11842\nExp number, first 60 AAs: 21.79934\nTotal prob of N-in: 0.98877\nPOSSIBLE N-term signal sequence\ninside 1 16\nTMhelix 17 39\noutside 40 666\nTMhelix 667 689\ninside 690 694

33711 GCF_000717995.1_ASM71799v1 Streptomyces violaceorubidus Terrabacteria group; Actinobacteria; Actinobacteria; Streptomycetales; Streptomycetaceae; Streptomyces
MPTWTASATSAGAAPIPASARRSGPAPRPCVRTTEREGPMAARRSTVLLVTLAAVLVAAA WP_078606556.1
hypothetical protein [Streptomyces violaceorubidus] Length: 389\nNumber of predicted TMHs: 2\nExp number of AAs in TMHs: 49.13536\nExp number, first 60 AAs: 15.01435\nTotal prob of N-in: 0.75322\nPOSSIBLE N-term signal sequence\ninside 1 45\nTMhelix 46 68\noutside 69 339\nTMhelix 340 362\ninside 363 389

33712 GCF_001509775.1_ASM150977v1 Streptomyces violaceusniger Terrabacteria group; Actinobacteria; Actinobacteria; Streptomycetales; Streptomycetaceae; Streptomyces
MLGVVENHSLPRTAAFFDLDKTVIAKSSAFTFSKSFYQGGLINRRAVLRTAYAQFVFLAG WP_059142394.1 inhibition of morphological differentiation protein [Streptomyces violaceusniger] Length: 278\nNumber of predicted TMHs: 1\nExp number of AAs in TMHs: 21.26535\nExp number, first 60 AAs: 0.03012\nTotal prob of N-in: 0.00337\noutside 1 247\nTMhelix 248 267\ninside 268 278

33713 GCF_001509775.1_ASM150977v1 Streptomyces violaceusniger Terrabacteria group; Actinobacteria; Actinobacteria; Streptomycetales; Streptomycetaceae; Streptomyces
MSSPDREVRAARNDTARTARRPVVAVTGAASGPGALLTQRLAESEEVKQVLAIDERRGEV WP_059145201.1
NAD-dependent dehydratase [Streptomyces violaceusniger] Length: 369\nNumber of predicted TMHs: 1\nExp number of AAs in TMHs: 19.99644\nExp number, first 60 AAs: 0.3822\nTotal prob of N-in: 0.03032\noutside 1 339\nTMhelix 340 359\ninside 360 369

33714 GCF_000717745.1_ASM71774v1 Streptomyces violens Terrabacteria group; Actinobacteria; Actinobacteria; Streptomycetales; Streptomycetaceae; Streptomyces
MSNNMSDHTLPSDSPAAAPSRRRTVLRRLPVVGGAAAGAVLLLAGPAFAHVTIQPGQAAK WP_078600540.1
hypothetical protein [Streptomyces violens] Length: 267\nNumber of predicted TMHs: 2\nExp number of

AAs in TMHs: 44.288\nExp number, first 60 AAs:  22.53514\nTotal prob of N-in:   0.95194\nPOSSIBLE N-term signal sequence\ninside   1   30\nTMhelix   31   53\noutside   54   239\nTMhelix   240   262\ninside   263   267

33715 GCF_000717745.1_ASM71774v1 Streptomyces violens Terrabacteria group; Actinobacteria; Actinobacteria; Streptomycetales; Streptomycetaceae; Streptomyces MTDRRRRAALLLALPATAPGSAVLLPRERKPSPSSPEGASGDGRGAGAGGQDGRDRDRDN WP_030254192.1 hypothetical protein [Streptomyces violens] Length: 327\nNumber of predicted TMHs:  2\nExp number of AAs in TMHs: 48.15729\nExp number, first 60 AAs:  1.50789\nTotal prob of N-in:   0.77755\ninside   1   175\nTMhelix   176   198\noutside   199   265\nTMhelix   266   288\ninside   289   327

33716 GCF_000720455.1_ASM72045v1 Streptomyces virginiae Terrabacteria group; Actinobacteria; Actinobacteria; Streptomycetales; Streptomycetaceae; Streptomyces MSARRRTVMPIRIAAAGAAPLALAVYAAVPAVAHGSMTDPVSRVAACYAEGPESPKSAAC WP_030901979.1 chitin-binding protein [Streptomyces virginiae] Length: 359\nNumber of predicted TMHs:  2\nExp number of AAs in TMHs: 43.495\nExp number, first 60 AAs:  21.79041\nTotal prob of N-in:   0.98439\nPOSSIBLE N-term signal sequence\ninside   1   12\nTMhelix   13   35\noutside   36   324\nTMhelix   325   347\ninside   348   359

33717 GCF_000716685.1_ASM71668v1 Streptomyces virginiae Terrabacteria group; Actinobacteria; Actinobacteria; Streptomycetales; Streptomycetaceae; Streptomyces MPARRRTAVTLTRIAAAGLAPLAVAAYAAAPAVAHGSMTDPVSRVAACYAEGPESPKSAA WP_033223868.1 chitin-binding protein [Streptomyces virginiae] Length: 354\nNumber of predicted TMHs:  2\nExp number of AAs in TMHs: 43.28025\nExp number, first 60 AAs:  21.31605\nTotal prob of N-in:   0.95206\nPOSSIBLE N-term signal sequence\ninside   1   12\nTMhelix   13   35\noutside   36   317\nTMhelix   318   340\ninside   341   354

33718 GCF_000716685.1_ASM71668v1 Streptomyces virginiae Terrabacteria group; Actinobacteria; Actinobacteria; Streptomycetales; Streptomycetaceae; Streptomyces MRRRNPVIAVTGAASGVGAALVSRLAASEEVKQVVAIDERRGDCAAAQWHILDVRDPAIA WP_051763101.1 NAD-dependent dehydratase [Streptomyces virginiae] Length: 353\nNumber of predicted TMHs:  1\nExp number of AAs in TMHs: 30.65483\nExp number, first 60 AAs:  7.65343\nTotal prob of N-in:   0.43524\noutside   1   324\nTMhelix   325   347\ninside   348   353

33719 GCF_000716685.1_ASM71668v1 Streptomyces virginiae Terrabacteria group; Actinobacteria; Actinobacteria; Streptomycetales; Streptomycetaceae; Streptomyces MTTASRRTLVRAAAVTACAGSLLALPTAVALADGVPAASSAHSAAPRSILVKSLALADGV WP_033218789.1 hypothetical protein [Streptomyces virginiae] Length: 218\nNumber of predicted TMHs:  2\nExp number of AAs in TMHs: 43.61745\nExp number, first 60 AAs:  21.00356\nTotal prob of N-in:   0.97242\nPOSSIBLE N-term signal sequence\ninside   1   8\nTMhelix   9   31\noutside   32   188\nTMhelix   189   211\ninside   212   218

33720 GCF_000716685.1_ASM71668v1 Streptomyces virginiae Terrabacteria group; Actinobacteria; Actinobacteria; Streptomycetales; Streptomycetaceae; Streptomyces MPAKPPAAPPTAARRRLVTVAAVTLIAAAAAPATVAYAIPSATVSPTAVAPGGRVALNVA WP_033226880.1 hypothetical protein [Streptomyces virginiae] Length: 170\nNumber of predicted TMHs:  2\nExp number of AAs in TMHs: 43.22489\nExp number, first 60 AAs:  22.49358\nTotal prob of N-in:   0.99114\nPOSSIBLE N-term signal sequence\ninside   1   16\nTMhelix   17   39\noutside   40   139\nTMhelix   140   162\ninside   163   170

33721 GCF_001270565.1_ASM127056v1 Streptomyces virginiae Terrabacteria group; Actinobacteria; Actinobacteria; Streptomycetales; Streptomycetaceae; Streptomyces MPARRRTAVALTRIASAGLAPLALAAYAAAPAVAHGSMTDPVSRVAACYAEGPESPKTAA WP_053168276.1 chitin-binding protein [Streptomyces virginiae] Length: 357\nNumber of predicted TMHs:  2\nExp number of AAs in TMHs: 42.24451\nExp number, first 60 AAs:  20.35603\nTotal prob of N-in:   0.90812\nPOSSIBLE N-term signal sequence\ninside   1   12\nTMhelix   13   35\noutside   36   320\nTMhelix   321   343\ninside   344   357

33722 GCF_001270565.1_ASM127056v1 Streptomyces virginiae Terrabacteria group; Actinobacteria; Actinobacteria; Streptomycetales; Streptomycetaceae; Streptomyces MSTVSRRTLVRAAAVTACAGSLLALPTAAALAEGVPAASSARSAAPQLSLVKSLSLADGT WP_053169188.1 MULTISPECIES: hypothetical protein [Streptomyces] Length: 217\nNumber of predicted TMHs:  2\nExp number of AAs in TMHs: 41.6059599999999\nExp number, first 60 AAs:  18.99757\nTotal prob of N-in:   0.88899\nPOSSIBLE N-term signal sequence\ninside   1   8\nTMhelix   9   31\noutside   32   187\nTMhelix   188   210\ninside   211   217

33723 GCF_001270565.1_ASM127056v1 Streptomyces virginiae Terrabacteria group; Actinobacteria; Actinobacteria; Streptomycetales; Streptomycetaceae; Streptomyces

MNRTHAPRRALLAAAFALTLASAGAGAGAASAAEAAPAPQPPYGQAPYAQAPYGLRLNGA WP_053177305.1 type VII secretion-associated serine protease mycosin [Streptomyces virginiae] Length: 431\nNumber of predicted TMHs: 2\nExp number of AAs in TMHs: 41.2144\nExp number, first 60 AAs: 18.91654\nTotal prob of N-in: 0.87140\nPOSSIBLE N-term signal sequence\ninside 1 10\nTMhelix 11 33\noutside 34 397\nTMhelix 398 420\ninside 421 431

33724 GCF_000716685.1_ASM71668v1 Streptomyces virginiae Terrabacteria group; Actinobacteria; Actinobacteria; Streptomycetales; Streptomycetaceae; Streptomyces MTQTHASRRALLVAAFALTLASGGTGTAAAATGAAPQPPYALRLDGAGECTFPMKKQIED WP_051763475.1 type VII secretion-associated serine protease mycosin [Streptomyces virginiae] Length: 418\nNumber of predicted TMHs: 2\nExp number of AAs in TMHs: 44.79427\nExp number, first 60 AAs: 22.50766\nTotal prob of N-in: 0.98852\nPOSSIBLE N-term signal sequence\ninside 1 11\nTMhelix 12 34\noutside 35 384\nTMhelix 385 407\ninside 408 418

33725 GCF_001270565.1_ASM127056v1 Streptomyces virginiae Terrabacteria group; Actinobacteria; Actinobacteria; Streptomycetales; Streptomycetaceae; Streptomyces MRRRSPVIAVTGAASGVGAALVSRLAASDEVKQVVAIDERRGDCAAAQWHILDVRDPAIA WP_053167991.1 NAD-dependent dehydratase [Streptomyces virginiae] Length: 353\nNumber of predicted TMHs: 1\nExp number of AAs in TMHs: 30.20328\nExp number, first 60 AAs: 7.40817\nTotal prob of N-in: 0.39678\noutside 1 324\nTMhelix 325 347\ninside 348 353

33726 GCF_000720455.1_ASM72045v1 Streptomyces virginiae Terrabacteria group; Actinobacteria; Actinobacteria; Streptomycetales; Streptomycetaceae; Streptomyces MSTVRRRTLVRAAAVTACAGSLLALPTAAALAEGVPAASSAHSAAPQRTLVKSLSLADGV WP_030834094.1 MULTISPECIES: hypothetical protein [Streptomyces] Length: 218\nNumber of predicted TMHs: 2\nExp number of AAs in TMHs: 42.93094\nExp number, first 60 AAs: 20.33933\nTotal prob of N-in: 0.95701\nPOSSIBLE N-term signal sequence\ninside 1 8\nTMhelix 9 31\noutside 32 188\nTMhelix 189 211\ninside 212 218

33727 GCF_000720455.1_ASM72045v1 Streptomyces virginiae Terrabacteria group; Actinobacteria; Actinobacteria; Streptomycetales; Streptomycetaceae; Streptomyces MALDVIRSGPPDKRPRRNLFKRGVLVLLPAALLAAATLVPNLAHALEAPVNLGTATNYAV WP_030831911.1 MULTISPECIES: hypothetical protein [Streptomyces] Length: 518\nNumber of predicted TMHs: 2\nExp number of AAs in TMHs: 43.15964\nExp number, first 60 AAs: 20.83032\nTotal prob of N-in: 0.94539\nPOSSIBLE N-term signal sequence\ninside 1 22\nTMhelix 23 45\noutside 46 483\nTMhelix 484 506\ninside 507 518

33728 GCF_000720455.1_ASM72045v1 Streptomyces virginiae Terrabacteria group; Actinobacteria; Actinobacteria; Streptomycetales; Streptomycetaceae; Streptomyces MPPYRDARRALLAAAFALTLTAAGATAATAAQHPPYALRLDGAGECTFPMKKQIADRPWA WP_030823957.1 MULTISPECIES: type VII secretion-associated serine protease [Streptomyces] Length: 414\nNumber of predicted TMHs: 2\nExp number of AAs in TMHs: 42.2832099999999\nExp number, first 60 AAs: 19.94733\nTotal prob of N-in: 0.93627\nPOSSIBLE N-term signal sequence\ninside 1 11\nTMhelix 12 31\noutside 32 380\nTMhelix 381 403\ninside 404 414

33729 GCF_001888885.1_ASM188888v1 Streptomyces viridifaciens Terrabacteria group; Actinobacteria; Actinobacteria; Streptomycetales; Streptomycetaceae; Streptomyces MSSTTRIRRSRRAVVAAATLGVLALGGSLALPAQTAFAGALHDAYHGALSVQAPAAVGFA WP_072653681.1 hypothetical protein [Streptomyces viridifaciens] Length: 438\nNumber of predicted TMHs: 2\nExp number of AAs in TMHs: 38.5871\nExp number, first 60 AAs: 21.63021\nTotal prob of N-in: 0.99806\nPOSSIBLE N-term signal sequence\ninside 1 12\nTMhelix 13 35\noutside 36 412\nTMhelix 413 431\ninside 432 438

33730 GCF_001888885.1_ASM188888v1 Streptomyces viridifaciens Terrabacteria group; Actinobacteria; Actinobacteria; Streptomycetales; Streptomycetaceae; Streptomyces MRTAAFFDLDKTIIAKSSALAFSRSFYQGGLINRRAVLRSAYAQFVFLVGGADHDQMEKM WP_030288083.1 MULTISPECIES: inhibition of morphological differentiation protein [Streptomyces] Length: 272\nNumber of predicted TMHs: 1\nExp number of AAs in TMHs: 22.68364\nExp number, first 60 AAs: 0.10845\nTotal prob of N-in: 0.00817\noutside 1 235\nTMhelix 236 258\ninside 259 272

33731 GCF_002156055.1_ASM215605v1 Streptomyces viridochromogenes Terrabacteria group; Actinobacteria; Actinobacteria; Streptomycetales; Streptomycetaceae; Streptomyces MRHARRRLVRRVTRLAAVGGLLLGGTMVTRAVASEPPGPSAVPRTFAESATPGPGGELVA WP_086867759.1 serine protease [Streptomyces viridochromogenes] Length: 454\nNumber of predicted TMHs: 2\nExp number of AAs in TMHs: 40.46713\nExp number, first 60 AAs: 19.79816\nTotal prob of N-in: 0.90421\nPOSSIBLE N-term signal sequence\ninside 1 11\nTMhelix 12 34\noutside 35 415\nTMhelix 416 438\ninside 439 454

33732	GCF_001047325.1_ASM104732v1	Streptomyces viridochromogenes	Terrabacteria group; Actinobacteria; Actinobacteria; Streptomycetales; Streptomycetaceae; Streptomyces
	MRHARRRLVRRVTRLAAVGGLLLGGTMVTRAVASEPPDASAVPRTYAASAAGTGSDLVAR	WP_048578925.1
	protease [Streptomyces viridochromogenes]	Length: 457\nNumber of predicted TMHs:  2\nExp number of AAs in TMHs: 40.19694\nExp number, first 60 AAs:  19.15121\nTotal prob of N-in:   0.87507\nPOSSIBLE N-term signal sequence\ninside   1   11\nTMhelix   12   34\noutside   35   418\nTMhelix   419   441\ninside   442   457

33733	GCF_001270495.1_P009IDBAStreptomyces viridochromogenes	Terrabacteria group; Actinobacteria; Actinobacteria; Streptomycetales; Streptomycetaceae; Streptomyces
	MRHARRRLVRRVTRLAAVGGLLLGGTMVTRAVASEPPDASAVPRTYAASVAGTGSDLVAR	WP_053195972.1
	protease [Streptomyces viridochromogenes]	Length: 457\nNumber of predicted TMHs:  2\nExp number of AAs in TMHs: 40.19809\nExp number, first 60 AAs:  19.1523\nTotal prob of N-in:   0.87507\nPOSSIBLE N-term signal sequence\ninside   1   11\nTMhelix   12   34\noutside   35   418\nTMhelix   419   441\ninside   442   457

33734	GCF_001270495.1_P009IDBAStreptomyces viridochromogenes	Terrabacteria group; Actinobacteria; Actinobacteria; Streptomycetales; Streptomycetaceae; Streptomyces
	MPAPKKTARRALLVTSAVVSSLALAAPVSYAAPKPSPSTTPSATPPANMSTVGGERLGQP	WP_053196823.1	D-alanyl-D-alanine carboxypeptidase [Streptomyces viridochromogenes]	Length: 428\nNumber of predicted TMHs: 1\nExp number of AAs in TMHs: 35.69963\nExp number, first 60 AAs:  13.54643\nTotal prob of N-in:   0.67039\nPOSSIBLE N-term signal sequence\noutside   1   389\nTMhelix   390   412\ninside   413   428

33735	GCF_001047325.1_ASM104732v1	Streptomyces viridochromogenes	Terrabacteria group; Actinobacteria; Actinobacteria; Streptomycetales; Streptomycetaceae; Streptomyces
	MPAPKKTARRALLVTSAVVSSLALAAPVSYAAPKPSPSTTPSATPPANMSTVGGERLGQP	WP_048587024.1	D-alanyl-D-alanine carboxypeptidase [Streptomyces viridochromogenes]	Length: 428\nNumber of predicted TMHs: 1\nExp number of AAs in TMHs: 35.69944\nExp number, first 60 AAs:  13.54643\nTotal prob of N-in:   0.67039\nPOSSIBLE N-term signal sequence\noutside   1   389\nTMhelix   390   412\ninside   413   428

33736	GCF_001270515.1_P002IDBA1	Streptomyces viridochromogenes	Terrabacteria group; Actinobacteria; Actinobacteria; Streptomycetales; Streptomycetaceae; Streptomyces
	MLGLVENHSLPRTAAFFDLDKTVIAKSSTLTFSKSFYRGGLISRRAALRTAYIQFVFLAG	WP_033202868.1	MULTISPECIES: inhibition of morphological differentiation protein [Streptomyces]	Length: 288\nNumber of predicted TMHs:  1\nExp number of AAs in TMHs: 21.84873\nExp number, first 60 AAs:  0.0299\nTotal prob of N-in:   0.00233\noutside   1   247\nTMhelix   248   270\ninside   271   288

33737	GCF_001270485.1_P006IDBAStreptomyces viridochromogenes	Terrabacteria group; Actinobacteria; Actinobacteria; Streptomycetales; Streptomycetaceae; Streptomyces
	MRHARRRLVRRVTRLAAVGGLLLGGTMVTRAVASEPPDASAVPRTYAASVAGTGSDLVAR	WP_053195972.1
	protease [Streptomyces viridochromogenes]	Length: 457\nNumber of predicted TMHs:  2\nExp number of AAs in TMHs: 40.19809\nExp number, first 60 AAs:  19.1523\nTotal prob of N-in:   0.87507\nPOSSIBLE N-term signal sequence\ninside   1   11\nTMhelix   12   34\noutside   35   418\nTMhelix   419   441\ninside   442   457

33738	GCF_001270485.1_P006IDBAStreptomyces viridochromogenes	Terrabacteria group; Actinobacteria; Actinobacteria; Streptomycetales; Streptomycetaceae; Streptomyces
	MPAPKKTARRALLVTSAVVSSLALAAPVSYAAPKPSPSTTPSATPPANMSTVGGERLGQP	WP_053196823.1	D-alanyl-D-alanine carboxypeptidase [Streptomyces viridochromogenes]	Length: 428\nNumber of predicted TMHs: 1\nExp number of AAs in TMHs: 35.69963\nExp number, first 60 AAs:  13.54643\nTotal prob of N-in:   0.67039\nPOSSIBLE N-term signal sequence\noutside   1   389\nTMhelix   390   412\ninside   413   428

33739	GCF_002078235.1_ASM207823v1	Streptomyces viridosporus	Terrabacteria group; Actinobacteria; Actinobacteria; Streptomycetales; Streptomycetaceae; Streptomyces
	MRHARRRVVRRVTRLAAVGGLLLGATMVTRAVASEPAPPNAVPHTAAEQSADPGTDLASR	WP_081238835.1
	serine protease [Streptomyces viridosporus]	Length: 457\nNumber of predicted TMHs:  2\nExp number of AAs in TMHs: 41.29878\nExp number, first 60 AAs:  20.23638\nTotal prob of N-in:   0.94065\nPOSSIBLE N-term signal sequence\ninside   1   11\nTMhelix   12   34\noutside   35   418\nTMhelix   419   441\ninside   442   457

33740	GCF_002078235.1_ASM207823v1	Streptomyces viridosporus	Terrabacteria group; Actinobacteria; Actinobacteria; Streptomycetales; Streptomycetaceae; Streptomyces
	MLKGVENHSSPRTAAFFDLDKTVIAKSSTLTFSKSFYQGGLINRRAVLRTAYAQFVFLAG	WP_004985180.1	MULTISPECIES: morphological differentiation-associated protein [Streptomyces]	Length: 277\nNumber of predicted TMHs:  1\nExp

number of AAs in TMHs: 21.98535\nExp number, first 60 AAs: 0.08708\nTotal prob of N-in: 0.00829\noutside 1 247\nTMhelix 248 270\ninside 271 277

33741 GCF_001445835.1_ASM144583v1 Streptomyces vitaminophilus Terrabacteria group; Actinobacteria; Actinobacteria; Streptomycetales; Streptomycetaceae; Streptomyces MLIPVENHPQPRAAAFFDLDKTVIAKSSALAFGRTFYRGGLINRRAVLRTAYAQFVFLAG WP_040912223.1 inhibition of morphological differentiation protein [Streptomyces vitaminophilus] Length: 275\nNumber of predicted TMHs: 1\nExp number of AAs in TMHs: 22.67849\nExp number, first 60 AAs: 0.03519\nTotal prob of N-in: 0.00631\noutside 1 243\nTMhelix 244 266\ninside 267 275

33742 GCF_001445835.1_ASM144583v1 Streptomyces vitaminophilus Terrabacteria group; Actinobacteria; Actinobacteria; Streptomycetales; Streptomycetaceae; Streptomyces MRAGRRALLGALVGFLLGPVLVAGATTPASAETIEEGQWGLDAIHARDVWPTTRGEGTVV WP_018382267.1 type VII secretion-associated serine protease [Streptomyces vitaminophilus] Length: 398\nNumber of predicted TMHs: 2\nExp number of AAs in TMHs: 43.13129\nExp number, first 60 AAs: 21.19527\nTotal prob of N-in: 0.97230\nPOSSIBLE N-term signal sequence\ninside 1 6\nTMhelix 7 29\noutside 30 367\nTMhelix 368 390\ninside 391 398

33743 GCF_000698945.1_LGO_A23_AS7 Streptomyces wadayamensis Terrabacteria group; Actinobacteria; Actinobacteria; Streptomycetales; Streptomycetaceae; Streptomyces MPDRRSAVSLRLPLPTAAPGFGGLALPGAAATAPDAWGVVAVRESSETERKSSSEPEGRS WP_049978295.1 hypothetical protein [Streptomyces wadayamensis] Length: 292\nNumber of predicted TMHs: 2\nExp number of AAs in TMHs: 51.14145\nExp number, first 60 AAs: 5.46126\nTotal prob of N-in: 0.34800\noutside 1 164\nTMhelix 165 187\ninside 188 229\nTMhelix 230 252\noutside 253 292

33744 GCF_000698945.1_LGO_A23_AS7 Streptomyces wadayamensis Terrabacteria group; Actinobacteria; Actinobacteria; Streptomycetales; Streptomycetaceae; Streptomyces MSRMRGSSGGRIAGRRAAVVLAVAVLGLAGCSAGPDSGASSAGDRAVAPSAADGAGGGQA WP_049978979.1 MULTISPECIES: hypothetical protein [Streptomyces] Length: 334\nNumber of predicted TMHs: 1\nExp number of AAs in TMHs: 36.36536\nExp number, first 60 AAs: 13.57637\nTotal prob of N-in: 0.64435\nPOSSIBLE N-term signal sequence\noutside 1 269\nTMhelix 270 292\ninside 293 334

33745 GCF_000698945.1_LGO_A23_AS7 Streptomyces wadayamensis Terrabacteria group; Actinobacteria; Actinobacteria; Streptomycetales; Streptomycetaceae; Streptomyces MGGRGTTAQARERGRLPGRPSDPAPTGGNDVITGVNLRMPARRTALPALLLTATALLAVP WP_049978783.1 hypothetical protein [Streptomyces wadayamensis] Length: 304\nNumber of predicted TMHs: 2\nExp number of AAs in TMHs: 40.0518\nExp number, first 60 AAs: 12.23688\nTotal prob of N-in: 0.79325\nPOSSIBLE N-term signal sequence\ninside 1 44\nTMhelix 45 67\noutside 68 276\nTMhelix 277 299\ninside 300 304

33746 GCF_000698945.1_LGO_A23_AS7 Streptomyces wadayamensis Terrabacteria group; Actinobacteria; Actinobacteria; Streptomycetales; Streptomycetaceae; Streptomyces MTPNRRPVVLAAAVATALGLGATALTLPAVAAEAPRAGAPVMDLTDGTLDWGVKESFRRY WP_049977564.1 hypothetical protein [Streptomyces wadayamensis] Length: 514\nNumber of predicted TMHs: 2\nExp number of AAs in TMHs: 42.8566099999999\nExp number, first 60 AAs: 22.03087\nTotal prob of N-in: 0.97276\nPOSSIBLE N-term signal sequence\ninside 1 6\nTMhelix 7 29\noutside 30 479\nTMhelix 480 502\ninside 503 514

33747 GCF_000698945.1_LGO_A23_AS7 Streptomyces wadayamensis Terrabacteria group; Actinobacteria; Actinobacteria; Streptomycetales; Streptomycetaceae; Streptomyces MLDPVENRSLPRTAAFFDLDKTVIAKSSTLTFSKSFYQGGLINRRAVLRTAYTQFVFLAG WP_010643895.1 MULTISPECIES: morphological differentiation-associated protein [Streptomyces] Length: 290\nNumber of predicted TMHs: 1\nExp number of AAs in TMHs: 21.93803\nExp number, first 60 AAs: 0.02636\nTotal prob of N-in: 0.00751\noutside 1 247\nTMhelix 248 270\ninside 271 290

33748 GCF_000698945.1_LGO_A23_AS7 Streptomyces wadayamensis Terrabacteria group; Actinobacteria; Actinobacteria; Streptomycetales; Streptomycetaceae; Streptomyces MSRRTRTARRRPVAAAVLAAGAVLLPAVPASAASGADQDFVIEDPRITESSGLAASKRHP WP_049978038.1 hypothetical protein [Streptomyces wadayamensis] Length: 352\nNumber of predicted TMHs: 2\nExp number of AAs in TMHs: 39.5043\nExp number, first 60 AAs: 21.11165\nTotal prob of N-in: 0.99083\nPOSSIBLE N-term signal sequence\ninside 1 12\nTMhelix 13 35\noutside 36 324\nTMhelix 325 344\ninside 345 352

33749 GCF_000698945.1_LGO_A23_AS7 Streptomyces wadayamensis Terrabacteria group; Actinobacteria; Actinobacteria; Streptomycetales; Streptomycetaceae; Streptomyces
MSAAAPEAAPRTARPAPRAGALLRRLLWWCVLTLTGGVERRGALPPGGCVVVANHTSHAD WP_049978612.1 1-acyl-sn-glycerol-3-phosphate acyltransferase [Streptomyces wadayamensis] Length: 401\nNumber of predicted TMHs: 2\nExp number of AAs in TMHs: 82.23516\nExp number, first 60 AAs: 1.119\nTotal prob of N-in: 0.15176\noutside 1 353\nTMhelix 354 371\ninside 372 377\nTMhelix 378 400\noutside 401 401

33750 GCF_000716445.1_ASM71644v1 Streptomyces wedmorensis Terrabacteria group; Actinobacteria; Actinobacteria; Streptomycetales; Streptomycetaceae; Streptomyces
MLGLVENHSLPRTAAFFDLDKTVIAKSSTLTFSKSFYRGGLISRRAALRTAYIQFVFLAG WP_033202868.1 MULTISPECIES: inhibition of morphological differentiation protein [Streptomyces] Length: 288\nNumber of predicted TMHs: 1\nExp number of AAs in TMHs: 21.84873\nExp number, first 60 AAs: 0.0299\nTotal prob of N-in: 0.00233\noutside 1 247\nTMhelix 248 270\ninside 271 288

33751 GCF_000725725.1_Doro.v1.0Streptomyces xanthophaeus Terrabacteria group; Actinobacteria; Actinobacteria; Streptomycetales; Streptomycetaceae; Streptomyces
MRSPRRPRRTTLRTVVVVAGASAVLALPVGSAFADSPAVPDPQVLPGIDQPQVDQPQVDR WP_031154496.1
hypothetical protein [Streptomyces xanthophaeus] Length: 359\nNumber of predicted TMHs: 2\nExp number of AAs in TMHs: 38.8735899999999\nExp number, first 60 AAs: 21.05183\nTotal prob of N-in: 0.95583\nPOSSIBLE N-term signal sequence\ninside 1 11\nTMhelix 12 34\noutside 35 327\nTMhelix 328 350\ninside 351 359

33752 GCF_000725725.1_Doro.v1.0Streptomyces xanthophaeus Terrabacteria group; Actinobacteria; Actinobacteria; Streptomycetales; Streptomycetaceae; Streptomyces
MSTVSRRTLVRAAAVTACAGSLLALPAAAALAEGVPAASSAHSAAPQRTLVKSLSLADGV WP_031150581.1
hypothetical protein [Streptomyces xanthophaeus] Length: 224\nNumber of predicted TMHs: 2\nExp number of AAs in TMHs: 42.34141\nExp number, first 60 AAs: 19.79868\nTotal prob of N-in: 0.91699\nPOSSIBLE N-term signal sequence\ninside 1 8\nTMhelix 9 31\noutside 32 194\nTMhelix 195 217\ninside 218 224

33753 GCF_000725805.1_Doro.v1.0Streptomyces xanthophaeus Terrabacteria group; Actinobacteria; Actinobacteria; Streptomycetales; Streptomycetaceae; Streptomyces
MSTVSRRTLVRAAAVTACAGSLLALPAAAALAEGVPAASSAHSAAPQRTLVKSLSLADGV WP_031150581.1
hypothetical protein [Streptomyces xanthophaeus] Length: 224\nNumber of predicted TMHs: 2\nExp number of AAs in TMHs: 42.34141\nExp number, first 60 AAs: 19.79868\nTotal prob of N-in: 0.91699\nPOSSIBLE N-term signal sequence\ninside 1 8\nTMhelix 9 31\noutside 32 194\nTMhelix 195 217\ninside 218 224

33754 GCF_000725805.1_Doro.v1.0Streptomyces xanthophaeus Terrabacteria group; Actinobacteria; Actinobacteria; Streptomycetales; Streptomycetaceae; Streptomyces
MRSPRRPRRTTLRTAVVVAGASAVLALPVGSAFADSPAVPDPQVLPGIDQPQVDQPQVDQ WP_031140980.1
hypothetical protein [Streptomyces xanthophaeus] Length: 364\nNumber of predicted TMHs: 2\nExp number of AAs in TMHs: 37.71054\nExp number, first 60 AAs: 19.87823\nTotal prob of N-in: 0.91453\nPOSSIBLE N-term signal sequence\ninside 1 11\nTMhelix 12 34\noutside 35 332\nTMhelix 333 355\ninside 356 364

33755 GCF_000993785.2_ASM99378v2 Streptomyces xiamenensis Terrabacteria group; Actinobacteria; Actinobacteria; Streptomycetales; Streptomycetaceae; Streptomyces
MRTPARAPVLSTARRSVLLVLLLLATQLVAVAWPAYACGCGGMVTGPGQRLTVEEETSAV WP_046722871.1
MULTISPECIES: hypothetical protein [Streptomyces] Length: 373\nNumber of predicted TMHs: 2\nExp number of AAs in TMHs: 43.02627\nExp number, first 60 AAs: 20.69791\nTotal prob of N-in: 0.97846\nPOSSIBLE N-term signal sequence\ninside 1 16\nTMhelix 17 36\noutside 37 338\nTMhelix 339 361\ninside 362 373

33756 GCF_000993785.2_ASM99378v2 Streptomyces xiamenensis Terrabacteria group; Actinobacteria; Actinobacteria; Streptomycetales; Streptomycetaceae; Streptomyces
MLGQVENLSQPRTAAFFDLDKTVIAKSSTVTFSRSFYQGGLINRRAVLRTAYAQFLYLVG WP_046723933.1 MULTISPECIES: inhibition of morphological differentiation protein [Streptomyces] Length: 290\nNumber of predicted TMHs: 1\nExp number of AAs in TMHs: 22.31044\nExp number, first 60 AAs: 0.0082\nTotal prob of N-in: 0.00135\noutside 1 246\nTMhelix 247 269\ninside 270 290

33757 GCF_000262345.1_Streptomyces_sulphureus_L180_version_1.0 Streptomyces xiaopingdaonensis
Terrabacteria group; Actinobacteria; Actinobacteria; Streptomycetales; Streptomycetaceae; Streptomyces
MLGSVENVSLPRTAAFFDLDKTVIAKSSTLTFGRSFYQGGLINRRAVLRTAYAQFVYLLG WP_016906398.1 inhibition of morphological differentiation protein [Streptomyces xiaopingdaonensis] Length: 285\nNumber of predicted TMHs:

1\nExp number of AAs in TMHs: 21.86514\nExp number, first 60 AAs: 0.0427\nTotal prob of N-in: 0.01166\noutside 1 246\nTMhelix 247 269\ninside 270 285

33758 GCF_000262345.1_Streptomyces_sulphureus_L180_version_1.0 Streptomyces xiaopingdaonensis Terrabacteria group; Actinobacteria; Actinobacteria; Streptomycetales; Streptomycetaceae; Streptomyces MASAHAAVPRPRSSASGRRRSPLVRVLLVGVLALLGTGLVPPSNAVGDEGSDAPGLEAPD WP_020372371.1 hypothetical protein [Streptomyces xiaopingdaonensis] Length: 519\nNumber of predicted TMHs: 2\nExp number of AAs in TMHs: 40.4465\nExp number, first 60 AAs: 18.3316\nTotal prob of N-in: 0.88773\nPOSSIBLE N-term signal sequence\ninside 1 20\nTMhelix 21 40\noutside 41 489\nTMhelix 490 512\ninside 513 519

33759 GCF_000716435.1_ASM71643v1 Streptomyces xylophagus Terrabacteria group; Actinobacteria; Actinobacteria; Streptomycetales; Streptomycetaceae; Streptomyces MRHARRRVIRRVTRLAAVGGLLLGGAMVTQAAMASDSSGTSAATLSAPQTPADKGSALVA WP_043662306.1 protease [Streptomyces xylophagus] Length: 455\nNumber of predicted TMHs: 2\nExp number of AAs in TMHs: 41.0119299999999\nExp number, first 60 AAs: 21.9039\nTotal prob of N-in: 0.98807\nPOSSIBLE N-term signal sequence\ninside 1 11\nTMhelix 12 34\noutside 35 416\nTMhelix 417 439\ninside 440 455

33760 GCF_000716435.1_ASM71643v1 Streptomyces xylophagus Terrabacteria group; Actinobacteria; Actinobacteria; Streptomycetales; Streptomycetaceae; Streptomyces MLWVVESHSLPRTAAFFDLDKTVIAKSSTLTFGKSFYQGGLINRRAALRTAYAQFVFLTG WP_043667555.1 inhibition of morphological differentiation protein [Streptomyces xylophagus] Length: 277\nNumber of predicted TMHs: 1\nExp number of AAs in TMHs: 21.73689\nExp number, first 60 AAs: 0.01233\nTotal prob of N-in: 0.00169\noutside 1 247\nTMhelix 248 270\ninside 271 277

33761 GCF_001005295.1_ASM100529v1 Streptomyces yangpuensis Terrabacteria group; Actinobacteria; Actinobacteria; Streptomycetales; Streptomycetaceae; Streptomyces MSTVRRRTLVRAAAVTACAGSLLALPTAAALAEGVPAASSAHSAATQRTLVKSLALADGV WP_046776149.1 hypothetical protein [Streptomyces yangpuensis]Length: 210\nNumber of predicted TMHs: 2\nExp number of AAs in TMHs: 43.2821399999999\nExp number, first 60 AAs: 20.78902\nTotal prob of N-in: 0.96435\nPOSSIBLE N-term signal sequence\ninside 1 8\nTMhelix 9 31\noutside 32 180\nTMhelix 181 203\ninside 204 210

33762 GCF_001005295.1_ASM100529v1 Streptomyces yangpuensis Terrabacteria group; Actinobacteria; Actinobacteria; Streptomycetales; Streptomycetaceae; Streptomyces MPRRAILAGAFALTLTAGGGTGTAAAATAAGAAPAAAVPHPPYALRLDGAGECTFPMKKQ WP_079025685.1 type VII secretion-associated serine protease mycosin [Streptomyces yangpuensis] Length: 421\nNumber of predicted TMHs: 2\nExp number of AAs in TMHs: 42.68606\nExp number, first 60 AAs: 20.35078\nTotal prob of N-in: 0.86814\nPOSSIBLE N-term signal sequence\ninside 1 6\nTMhelix 7 29\noutside 30 387\nTMhelix 388 410\ninside 411 421

33763 GCF_001005295.1_ASM100529v1 Streptomyces yangpuensis Terrabacteria group; Actinobacteria; Actinobacteria; Streptomycetales; Streptomycetaceae; Streptomyces MSARRRTVTPTRIAAAGLAPLALVAYAAVPAAAHGSMTDPVSRVAACYAEGPESPKSAAC WP_046780570.1 chitin-binding protein [Streptomyces yangpuensis] Length: 358\nNumber of predicted TMHs: 2\nExp number of AAs in TMHs: 43.51279\nExp number, first 60 AAs: 21.78799\nTotal prob of N-in: 0.99153\nPOSSIBLE N-term signal sequence\ninside 1 12\nTMhelix 13 35\noutside 36 323\nTMhelix 324 346\ninside 347 358

33764 GCF_000716805.1_ASM71680v1 Streptomyces yerevanensis Terrabacteria group; Actinobacteria; Actinobacteria; Streptomycetales; Streptomycetaceae; Streptomyces MLGLVENHSLPRTAAFFDLDKTVIAKSSTLTFSKSFYQGGLINRRAVLRTAYAQFVFLAG WP_033321745.1 inhibition of morphological differentiation protein [Streptomyces yerevanensis] Length: 274\nNumber of predicted TMHs: 1\nExp number of AAs in TMHs: 21.39259\nExp number, first 60 AAs: 0.02617\nTotal prob of N-in: 0.00572\noutside 1 244\nTMhelix 245 267\ninside 268 274

33765 GCF_000716805.1_ASM71680v1 Streptomyces yerevanensis Terrabacteria group; Actinobacteria; Actinobacteria; Streptomycetales; Streptomycetaceae; Streptomyces MAADRRRPIALAGAIATALAFGATALVATTASAAEVPLKDYELTWGIKESYRTYVTGMAA WP_033325580.1 hypothetical protein [Streptomyces yerevanensis] Length: 497\nNumber of predicted TMHs: 2\nExp number of AAs in TMHs: 44.63803\nExp number, first 60 AAs: 22.38021\nTotal prob of N-in: 0.98907\nPOSSIBLE N-term signal sequence\ninside 1 8\nTMhelix 9 31\noutside 32 465\nTMhelix 466 488\ninside 489 497

33766 GCF_000716805.1_ASM71680v1 Streptomyces yerevanensis Terrabacteria group; Actinobacteria; Actinobacteria; Streptomycetales; Streptomycetaceae; Streptomyces

MGAGDCHARPPPNRRSTPVRRNTRRRPTGARRTTFAAVALMLGGGALVAANVYATATESG WP_078491660.1 hypothetical protein [Streptomyces yerevanensis] Length: 711\nNumber of predicted TMHs: 2\nExp number of AAs in TMHs: 40.95203\nExp number, first 60 AAs: 21.96458\nTotal prob of N-in: 0.97581\nPOSSIBLE N-term signal sequence\ninside 1 33\nTMhelix 34 56\noutside 57 685\nTMhelix 686 703\ninside 704 711

33767 GCF_001514035.1_ASM151403v1 Streptomyces yokosukanensis Terrabacteria group; Actinobacteria; Actinobacteria; Streptomycetales; Streptomycetaceae; Streptomyces MASRGRRAAMRSLGALAGAAALTVLGSNAPGWAATPSGHGGHSSTVTPIKHVVVLFDENI WP_067117413.1 phospholipase [Streptomyces yokosukanensis] Length: 607\nNumber of predicted TMHs: 2\nExp number of AAs in TMHs: 40.42894\nExp number, first 60 AAs: 19.37379\nTotal prob of N-in: 0.86159\nPOSSIBLE N-term signal sequence\ninside 1 12\nTMhelix 13 35\noutside 36 576\nTMhelix 577 599\ninside 600 607

33768 GCF_001514035.1_ASM151403v1 Streptomyces yokosukanensis Terrabacteria group; Actinobacteria; Actinobacteria; Streptomycetales; Streptomycetaceae; Streptomyces MLRGVENHSLPRAAAFFDLDKTVIAKSSTLTFSKSFYQGGLINRRAALRTAYAQFVFLAG WP_067116570.1 inhibition of morphological differentiation protein [Streptomyces yokosukanensis] Length: 276\nNumber of predicted TMHs: 1\nExp number of AAs in TMHs: 21.62516\nExp number, first 60 AAs: 0.02774\nTotal prob of N-in: 0.00306\noutside 1 247\nTMhelix 248 270\ninside 271 276

33769 GCF_001514035.1_ASM151403v1 Streptomyces yokosukanensis Terrabacteria group; Actinobacteria; Actinobacteria; Streptomycetales; Streptomycetaceae; Streptomyces MRHARRRIVRRVTRLAAVGGLLLGGAMVTQAAMASQTPPAAGRTLSSAAGAGDTGGALVA WP_067117143.1 protease [Streptomyces yokosukanensis] Length: 459\nNumber of predicted TMHs: 2\nExp number of AAs in TMHs: 43.76628\nExp number, first 60 AAs: 22.30576\nTotal prob of N-in: 0.99405\nPOSSIBLE N-term signal sequence\ninside 1 11\nTMhelix 12 34\noutside 35 420\nTMhelix 421 443\ninside 444 459

33770 GCF_001514035.1_ASM151403v1 Streptomyces yokosukanensis Terrabacteria group; Actinobacteria; Actinobacteria; Streptomycetales; Streptomycetaceae; Streptomyces MPAPKPTGRRSLLVTSATLLSLSATAPAALAAPPPGASPTATPPARMSTVGGERLGQPGT WP_067119775.1 D-alanyl-D-alanine carboxypeptidase [Streptomyces yokosukanensis] Length: 408\nNumber of predicted TMHs: 1\nExp number of AAs in TMHs: 26.38776\nExp number, first 60 AAs: 3.77921\nTotal prob of N-in: 0.19103\noutside 1 369\nTMhelix 370 392\ninside 393 408

33771 GCF_900142595.1_IMG-taxon_2667527450_annotated_assembly Streptomyces yunnanensis Terrabacteria group; Actinobacteria; Actinobacteria; Streptomycetales; Streptomycetaceae; Streptomyces MTLMRLTRPTRPTGQRRRAVTAAAAALLLAGAGVVATAGTAQAAEVAYLTECQPPPISGL WP_079181755.1 hypothetical protein [Streptomyces yunnanensis] Length: 483\nNumber of predicted TMHs: 2\nExp number of AAs in TMHs: 43.81733\nExp number, first 60 AAs: 21.64556\nTotal prob of N-in: 0.98720\nPOSSIBLE N-term signal sequence\ninside 1 18\nTMhelix 19 41\noutside 42 453\nTMhelix 454 476\ninside 477 483

33772 GCF_000720835.1_ASM72083v1 Streptomyces achromogenes subsp. achromogenes Terrabacteria group; Actinobacteria; Actinobacteria; Streptomycetales; Streptomycetaceae; Streptomyces; Streptomyces achromogenes MYGPRVRECSPLSRRAVLGLTAAALPLSTATDATAATAATTVVGGGLLSRTGVQVRGASG WP_051754093.1 D-alanyl-D-alanine carboxypeptidase [Streptomyces achromogenes] Length: 414\nNumber of predicted TMHs: 1\nExp number of AAs in TMHs: 22.42124\nExp number, first 60 AAs: 1.07314\nTotal prob of N-in: 0.04929\noutside 1 358\nTMhelix 359 381\ninside 382 414

33773 GCF_000720835.1_ASM72083v1 Streptomyces achromogenes subsp. achromogenes Terrabacteria group; Actinobacteria; Actinobacteria; Streptomycetales; Streptomycetaceae; Streptomyces; Streptomyces achromogenes MRHARRRVVRRVTRLAAAVGLLLGGAMVARAAGADQPPAAPPGRPGDTGAALVARLGTAR WP_030607634.1 protease [Streptomyces achromogenes] Length: 438\nNumber of predicted TMHs: 2\nExp number of AAs in TMHs: 43.6146899999999\nExp number, first 60 AAs: 21.90131\nTotal prob of N-in: 0.98898\nPOSSIBLE N-term signal sequence\ninside 1 11\nTMhelix 12 34\noutside 35 399\nTMhelix 400 422\ninside 423 438

33774 GCF_000720835.1_ASM72083v1 Streptomyces achromogenes subsp. achromogenes Terrabacteria group; Actinobacteria; Actinobacteria; Streptomycetales; Streptomycetaceae; Streptomyces; Streptomyces achromogenes MLRGVENHSLPRAAAFFDLDKTVIAKSSTLTFSKSFYQGGLINRRAALRTAYAQFVFLAG WP_030606360.1 inhibition of morphological differentiation protein [Streptomyces achromogenes] Length: 276\nNumber of predicted TMHs: 1\nExp number of AAs in TMHs: 21.83157\nExp number, first 60 AAs: 0.02782\nTotal prob of N-in: 0.00345\noutside 1 247\nTMhelix 248 270\ninside 271 276

33775 GCF_000242715.1_ASM24271v2 Streptomyces acidiscabies 84-104 Terrabacteria group; Actinobacteria; Actinobacteria; Streptomycetales; Streptomycetaceae; Streptomyces; Streptomyces acidiscabies
MTENTSRRPALLTRAKALPPWTLLAAGALAGGLAGGAYGVLKPPVYTATAYVVAVPAEKS WP_010355677.1
hypothetical protein [Streptomyces acidiscabies] Length: 232\nNumber of predicted TMHs: 2\nExp number of AAs in TMHs: 43.18943\nExp number, first 60 AAs: 21.07852\nTotal prob of N-in: 0.99398\nPOSSIBLE N-term signal sequence\ninside 1 20\nTMhelix 21 40\noutside 41 164\nTMhelix 165 187\ninside 188 232

33776 GCF_000242715.1_ASM24271v2 Streptomyces acidiscabies 84-104 Terrabacteria group; Actinobacteria; Actinobacteria; Streptomycetales; Streptomycetaceae; Streptomyces; Streptomyces acidiscabies
MLRDVENHPSPRTAAFFDLDKTVIAKSSTLTFGKSFYQGGLINRRAALRTAYAQFVFLVG WP_010359176.1 morphological differentiation-associated protein [Streptomyces acidiscabies] Length: 275\nNumber of predicted TMHs: 1\nExp number of AAs in TMHs: 21.76485\nExp number, first 60 AAs: 0.08203\nTotal prob of N-in: 0.06750\noutside 1 245\nTMhelix 246 268\ninside 269 275

33777 GCF_000242715.1_ASM24271v2 Streptomyces acidiscabies 84-104 Terrabacteria group; Actinobacteria; Actinobacteria; Streptomycetales; Streptomycetaceae; Streptomyces; Streptomyces acidiscabies
MHLTRRTLLKTTALTGAASALALPTTTAAAAPTAYAHPGLLHTAADLTRMATKVKANSAP WP_010356313.1
hypothetical protein [Streptomyces acidiscabies] Length: 523\nNumber of predicted TMHs: 1\nExp number of AAs in TMHs: 26.06153\nExp number, first 60 AAs: 7.92269\nTotal prob of N-in: 0.35771\noutside 1 471\nTMhelix 472 491\ninside 492 523

33778 GCF_000415505.1_STAFG Streptomyces afghaniensis 772 Terrabacteria group; Actinobacteria; Actinobacteria; Streptomycetales; Streptomycetaceae; Streptomyces; Streptomyces afghaniensis
MPAPKKTARRSLLVTSAVLSSLALAAPAAVAAPSPSGSPSASPSGSPSATPPASMSTVGG WP_020273774.1 D-alanyl-D-alanine carboxypeptidase [Streptomyces afghaniensis] Length: 439\nNumber of predicted TMHs: 2\nExp number of AAs in TMHs: 39.0843799999999\nExp number, first 60 AAs: 16.45469\nTotal prob of N-in: 0.78103\nPOSSIBLE N-term signal sequence\ninside 1 11\nTMhelix 12 34\noutside 35 400\nTMhelix 401 423\ninside 424 439

33779 GCF_000415505.1_STAFG Streptomyces afghaniensis 772 Terrabacteria group; Actinobacteria; Actinobacteria; Streptomycetales; Streptomycetaceae; Streptomyces; Streptomyces afghaniensis
MPARRTAVTVAALGALPLALTATPAAAHGSMGDPVSRVSQCYAEGPESPKSAACKAAVGA WP_020274847.1
chitin-binding protein [Streptomyces afghaniensis] Length: 331\nNumber of predicted TMHs: 2\nExp number of AAs in TMHs: 38.9520299999999\nExp number, first 60 AAs: 18.48862\nTotal prob of N-in: 0.84671\nPOSSIBLE N-term signal sequence\ninside 1 6\nTMhelix 7 29\noutside 30 299\nTMhelix 300 322\ninside 323 331

33780 GCF_000415505.1_STAFG Streptomyces afghaniensis 772 Terrabacteria group; Actinobacteria; Actinobacteria; Streptomycetales; Streptomycetaceae; Streptomyces; Streptomyces afghaniensis
MLKGVENHSSPRTAAFFDLDKTVIAKSSTLTFSKSFYQGGLINRRAVLRTAYAQFVFLAG WP_020271824.1 inhibition of morphological differentiation protein [Streptomyces afghaniensis] Length: 277\nNumber of predicted TMHs: 1\nExp number of AAs in TMHs: 21.5561\nExp number, first 60 AAs: 0.06768\nTotal prob of N-in: 0.00889\noutside 1 247\nTMhelix 248 270\ninside 271 277

33781 GCF_000415505.1_STAFG Streptomyces afghaniensis 772 Terrabacteria group; Actinobacteria; Actinobacteria; Streptomycetales; Streptomycetaceae; Streptomyces; Streptomyces afghaniensis
MRHARRRVVRRVTRLAAVGGLLLGGAMVTNAVASEPPDPSIGVPYSARPATSMGTDLVSR WP_037667955.1
protease [Streptomyces afghaniensis] Length: 448\nNumber of predicted TMHs: 2\nExp number of AAs in TMHs: 40.90959\nExp number, first 60 AAs: 21.09066\nTotal prob of N-in: 0.96065\nPOSSIBLE N-term signal sequence\ninside 1 11\nTMhelix 12 34\noutside 35 409\nTMhelix 410 432\ninside 433 448

33782 GCF_000935185.2_ASM93518v2 Streptomyces ahygroscopicus subsp. wuyiensis Terrabacteria group; Actinobacteria; Actinobacteria; Streptomycetales; Streptomycetaceae; Streptomyces; Streptomyces ahygroscopicus
MVTVRPAHPPAERPRQGEDALQMPDRRRRAALLRPIQATTGWGLAPVTSPGAVLLPRERT WP_078948375.1
hypothetical protein [Streptomyces ahygroscopicus] Length: 331\nNumber of predicted TMHs: 2\nExp number of AAs in TMHs: 48.0519499999999\nExp number, first 60 AAs: 0.49785\nTotal prob of N-in: 0.59124\ninside 1 187\nTMhelix 188 210\noutside 211 265\nTMhelix 266 288\ninside 289 331

33783 GCF_000935185.2_ASM93518v2 Streptomyces ahygroscopicus subsp. wuyiensis Terrabacteria group; Actinobacteria; Actinobacteria; Streptomycetales; Streptomycetaceae; Streptomyces; Streptomyces ahygroscopicus
MTLMRPTGPPRPTGRRRRATLAATAALVLAGAGVVTAAGTAQAADVAYKTECQPPPISGL WP_037635128.1
MULTISPECIES: hypothetical protein [Streptomyces] Length: 479\nNumber of predicted TMHs: 2\nExp

number of AAs in TMHs: 43.80161\nExp number, first 60 AAs:  21.18149\nTotal prob of N-in:   0.97217\nPOSSIBLE N-term signal sequence\ninside   1   20\nTMhelix   21   43\noutside   44   450\nTMhelix   451   473\ninside   474   479

33784	GCF_900171555.1_Salb	Streptomyces albidoflavus	Terrabacteria group; Actinobacteria; Actinobacteria; Streptomycetales; Streptomycetaceae; Streptomyces; Streptomyces albidoflavus group
	MLDPVENRSLPRTAAFFDLDKTVIAKSSTLTFSKSFYQGGLINRRAVLRTAYTQFVFLAG	WP_003949650.1	MULTISPECIES: morphological differentiation-associated protein [Streptomyces]	Length: 290\nNumber of predicted TMHs:  1\nExp number of AAs in TMHs: 21.6224\nExp number, first 60 AAs:  0.02852\nTotal prob of N-in:   0.00967\noutside   1   247\nTMhelix   248   270\ninside   271   290

33785	GCF_000719955.1_ASM71995v1	Streptomyces albidoflavus	Terrabacteria group; Actinobacteria; Actinobacteria; Streptomycetales; Streptomycetaceae; Streptomyces; Streptomyces albidoflavus group
	MLDPVENRSLPRTAAFFDLDKTVIAKSSTLTFSKSFYQGGLINRRAVLRTAYTQFVFLAG	WP_018470348.1	MULTISPECIES: inhibition of morphological differentiation protein [Streptomyces]	Length: 290\nNumber of predicted TMHs:  1\nExp number of AAs in TMHs: 21.82254\nExp number, first 60 AAs:  0.02718\nTotal prob of N-in:   0.00834\noutside   1   247\nTMhelix   248   270\ninside   271   290

33786	GCF_000719955.1_ASM71995v1	Streptomyces albidoflavus	Terrabacteria group; Actinobacteria; Actinobacteria; Streptomycetales; Streptomycetaceae; Streptomyces; Streptomyces albidoflavus group
	MPDRRSAVSLRLPLPTAAPGFGGLALPGAAATAPDAWGVVAVRESSETERKSSSEPEGRS	WP_030304860.1
	hypothetical protein [Streptomyces albidoflavus]	Length: 292\nNumber of predicted TMHs:  2\nExp number of AAs in TMHs: 51.2115\nExp number, first 60 AAs:  5.54647\nTotal prob of N-in:   0.33599\noutside   1   164\nTMhelix   165   187\ninside   188   229\nTMhelix   230   252\noutside   253   292

33787	GCF_900171555.1_Salb	Streptomyces albidoflavus	Terrabacteria group; Actinobacteria; Actinobacteria; Streptomycetales; Streptomycetaceae; Streptomyces; Streptomyces albidoflavus group
	MPDRRSAVSLRLPLPTAAPGFGGLALPGAAATAPDAWGVVAVRESSEKERKSSSEPEGRS	WP_085478814.1
	hypothetical protein [Streptomyces albidoflavus]	Length: 292\nNumber of predicted TMHs:  2\nExp number of AAs in TMHs: 51.79394\nExp number, first 60 AAs:  6.09103\nTotal prob of N-in:   0.33998\noutside   1   164\nTMhelix   165   187\ninside   188   229\nTMhelix   230   252\noutside   253   292

33788	GCF_900171555.1_Salb	Streptomyces albidoflavus	Terrabacteria group; Actinobacteria; Actinobacteria; Streptomycetales; Streptomycetaceae; Streptomyces; Streptomyces albidoflavus group
	MGGRGTTAQARERGRLPGRPSDPAPTGGNDVITGVNLRMPARRTALPALLLTATALLAVP	WP_003949316.1
	MULTISPECIES: hypothetical protein [Streptomyces]	Length: 304\nNumber of predicted TMHs:  2\nExp number of AAs in TMHs: 39.19493\nExp number, first 60 AAs:  11.75707\nTotal prob of N-in:   0.76164\nPOSSIBLE N-term signal sequence\ninside   1   44\nTMhelix   45   67\noutside   68   276\nTMhelix   277   299\ninside   300   304

33789	GCF_001865795.1_ASM186579v1	Streptomyces albidoflavus	Terrabacteria group; Actinobacteria; Actinobacteria; Streptomycetales; Streptomycetaceae; Streptomyces; Streptomyces albidoflavus group
	MSRRTRTARRRPVAAAVLAAGAVLLPAVPASAASGADQDFVIEDPRITESSGLAASKRHP	WP_033240369.1
	MULTISPECIES: hypothetical protein [Streptomyces]	Length: 350\nNumber of predicted TMHs:  2\nExp number of AAs in TMHs: 39.44481\nExp number, first 60 AAs:  21.11089\nTotal prob of N-in:   0.99076\nPOSSIBLE N-term signal sequence\ninside   1   12\nTMhelix   13   35\noutside   36   322\nTMhelix   323   342\ninside   343   350

33790	GCF_900171555.1_Salb	Streptomyces albidoflavus	Terrabacteria group; Actinobacteria; Actinobacteria; Streptomycetales; Streptomycetaceae; Streptomyces; Streptomyces albidoflavus group
	MSRRTRTARRRPVAAAVLAAGAVLLPAVPASAASGADQDFVIEDPRITESSGLAASKRHP	WP_030309758.1
	MULTISPECIES: hypothetical protein [Streptomyces]	Length: 348\nNumber of predicted TMHs:  2\nExp number of AAs in TMHs: 39.44414\nExp number, first 60 AAs:  21.11647\nTotal prob of N-in:   0.99086\nPOSSIBLE N-term signal sequence\ninside   1   12\nTMhelix   13   35\noutside   36   320\nTMhelix   321   340\ninside   341   348

33791	GCF_900171555.1_Salb	Streptomyces albidoflavus	Terrabacteria group; Actinobacteria; Actinobacteria; Streptomycetales; Streptomycetaceae; Streptomyces; Streptomyces albidoflavus group
	MSAAEPEAAPRTARPAPRAGALLRRLLWWCVLTLTGGVERRGALPPGGCVVVANHTSHAD	WP_085478689.1	1-acyl-sn-glycerol-3-phosphate acyltransferase [Streptomyces albidoflavus]	Length: 401\nNumber of predicted TMHs:  2\nExp number of AAs in TMHs: 84.11332\nExp number, first 60 AAs:  0.96241\nTotal prob of N-in:   0.22265\noutside   1   353\nTMhelix   354   371\ninside   372   377\nTMhelix   378   400\noutside   401   401

33792	GCF_900171555.1_Salb	Streptomyces albidoflavus	Terrabacteria group; Actinobacteria; Actinobacteria; Streptomycetales; Streptomycetaceae; Streptomyces; Streptomyces albidoflavus group

MSRMRGSSGGRIAGRRAAVVLAVAVLGLAGCSAGPDSGASSAGDRAVAPSAADGAGGGQA WP_049978979.1 MULTISPECIES: hypothetical protein [Streptomyces] Length: 334\nNumber of predicted TMHs: 1\nExp number of AAs in TMHs: 36.36536\nExp number, first 60 AAs: 13.57637\nTotal prob of N-in: 0.64435\nPOSSIBLE N-term signal sequence\noutside 1 269\nTMhelix 270 292\ninside 293 334

33793 GCF_000719955.1_ASM71995v1 Streptomyces albidoflavus Terrabacteria group; Actinobacteria; Actinobacteria; Streptomycetales; Streptomycetaceae; Streptomyces; Streptomyces albidoflavus group MGGRGTTAQARERGRLPGRPSDPAPTGGNDVITGVNLRMPARRTALPALLLTATALLAVP WP_030310179.1 hypothetical protein [Streptomyces albidoflavus] Length: 304\nNumber of predicted TMHs: 2\nExp number of AAs in TMHs: 40.02442\nExp number, first 60 AAs: 12.22539\nTotal prob of N-in: 0.79211\nPOSSIBLE N-term signal sequence\ninside 1 44\nTMhelix 45 67\noutside 68 276\nTMhelix 277 299\ninside 300 304

33794 GCF_000719955.1_ASM71995v1 Streptomyces albidoflavus Terrabacteria group; Actinobacteria; Actinobacteria; Streptomycetales; Streptomycetaceae; Streptomyces; Streptomyces albidoflavus group MSRRTRTARRRPVAAAVLAAGAVLLPAVPASAASGADQDFVIEDPRITESSGLAASKRHP WP_030309758.1 MULTISPECIES: hypothetical protein [Streptomyces] Length: 348\nNumber of predicted TMHs: 2\nExp number of AAs in TMHs: 39.44414\nExp number, first 60 AAs: 21.11647\nTotal prob of N-in: 0.99086\nPOSSIBLE N-term signal sequence\ninside 1 12\nTMhelix 13 35\noutside 36 320\nTMhelix 321 340\ninside 341 348

33795 GCF_001865795.1_ASM186579v1 Streptomyces albidoflavus Terrabacteria group; Actinobacteria; Actinobacteria; Streptomycetales; Streptomycetaceae; Streptomyces; Streptomyces albidoflavus group MLDPVENRSLPRTAAFFDLDKTVIAKSSTLTFSKSFYQGGLINRRAVLRTAYTQFVFLAG WP_018470348.1 MULTISPECIES: inhibition of morphological differentiation protein [Streptomyces] Length: 290\nNumber of predicted TMHs: 1\nExp number of AAs in TMHs: 21.82254\nExp number, first 60 AAs: 0.02718\nTotal prob of N-in: 0.00834\noutside 1 247\nTMhelix 248 270\ninside 271 290

33796 GCF_001865795.1_ASM186579v1 Streptomyces albidoflavus Terrabacteria group; Actinobacteria; Actinobacteria; Streptomycetales; Streptomycetaceae; Streptomyces; Streptomyces albidoflavus group MPDRRSAVSLRLPLPTAAPGFGGLALPGAAATAPDAWGVVAVRESSEKERKSSSEPEGRS WP_033237593.1 MULTISPECIES: hypothetical protein [Streptomyces] Length: 292\nNumber of predicted TMHs: 2\nExp number of AAs in TMHs: 51.77677\nExp number, first 60 AAs: 6.07612\nTotal prob of N-in: 0.33780\noutside 1 164\nTMhelix 165 187\ninside 188 229\nTMhelix 230 252\noutside 253 292

33797 GCF_001865795.1_ASM186579v1 Streptomyces albidoflavus Terrabacteria group; Actinobacteria; Actinobacteria; Streptomycetales; Streptomycetaceae; Streptomyces; Streptomyces albidoflavus group MTPNRRPVVLAAAVATALGLGATALTLPAVAAEAPRAGAPVMDLTDGTLDWGVKESFRRY WP_071338956.1 hypothetical protein [Streptomyces albidoflavus] Length: 514\nNumber of predicted TMHs: 2\nExp number of AAs in TMHs: 42.8607399999999\nExp number, first 60 AAs: 22.03087\nTotal prob of N-in: 0.97279\nPOSSIBLE N-term signal sequence\ninside 1 6\nTMhelix 7 29\noutside 30 479\nTMhelix 480 502\ninside 503 514

33798 GCF_001865795.1_ASM186579v1 Streptomyces albidoflavus Terrabacteria group; Actinobacteria; Actinobacteria; Streptomycetales; Streptomycetaceae; Streptomyces; Streptomyces albidoflavus group MSRMRGSSGGRIAGRRAAVVLAVAVLGLAGCSAGPDSGASSAGDRAVAPSAADGAGGGQA WP_071338394.1 hypothetical protein [Streptomyces albidoflavus] Length: 334\nNumber of predicted TMHs: 1\nExp number of AAs in TMHs: 36.36695\nExp number, first 60 AAs: 13.57728\nTotal prob of N-in: 0.64434\nPOSSIBLE N-term signal sequence\noutside 1 269\nTMhelix 270 292\ninside 293 334

33799 GCF_000719955.1_ASM71995v1 Streptomyces albidoflavus Terrabacteria group; Actinobacteria; Actinobacteria; Streptomycetales; Streptomycetaceae; Streptomyces; Streptomyces albidoflavus group MSRMRGSSGGRIAGRRAAVVLAVAVLGLAGCSAGPDSGASSAADRAVAPSAADGAGGGQA WP_030307403.1 MULTISPECIES: hypothetical protein [Streptomyces] Length: 334\nNumber of predicted TMHs: 1\nExp number of AAs in TMHs: 36.07114\nExp number, first 60 AAs: 13.28349\nTotal prob of N-in: 0.63206\nPOSSIBLE N-term signal sequence\noutside 1 269\nTMhelix 270 292\ninside 293 334

33800 GCF_000719955.1_ASM71995v1 Streptomyces albidoflavus Terrabacteria group; Actinobacteria; Actinobacteria; Streptomycetales; Streptomycetaceae; Streptomyces; Streptomyces albidoflavus group MTPNRRPVVLAAAVATALGLGATALTLPAVAAEAPRAGAPVMDLTDGTLDWGVKESFRRY WP_030306677.1 membrane protein [Streptomyces albidoflavus] Length: 514\nNumber of predicted TMHs: 2\nExp number of AAs in TMHs: 42.8566699999999\nExp number, first 60 AAs: 22.03087\nTotal prob of N-in: 0.97276\nPOSSIBLE N-term signal sequence\ninside 1 6\nTMhelix 7 29\noutside 30 479\nTMhelix 480 502\ninside 503 514

33801	GCF_001865795.1_ASM186579v1	Streptomyces albidoflavus	Terrabacteria group; Actinobacteria; Actinobacteria; Streptomycetales; Streptomycetaceae; Streptomyces; Streptomyces albidoflavus group
MGGRGTTAQARERGRLPGRPSDPAPTGGNDVITGVNLRMPARRTALPALLLTATALLAVP	WP_071337319.1
hypothetical protein [Streptomyces albidoflavus] Length: 304\nNumber of predicted TMHs: 2\nExp number of AAs in TMHs: 38.94121\nExp number, first 60 AAs: 11.57756\nTotal prob of N-in: 0.75058\nPOSSIBLE N-term signal sequence\ninside 1 44\nTMhelix 45 67\noutside 68 276\nTMhelix 277 299\ninside 300 304

33802	GCF_001865795.1_ASM186579v1	Streptomyces albidoflavus	Terrabacteria group; Actinobacteria; Actinobacteria; Streptomycetales; Streptomycetaceae; Streptomyces; Streptomyces albidoflavus group
MAPPHRRRAAVAVRALIALTALLCTTGLAPAATSGPAARAAVAEVPPPTATTSVVTVRTG	WP_071339460.1
hypothetical protein [Streptomyces albidoflavus] Length: 835\nNumber of predicted TMHs: 2\nExp number of AAs in TMHs: 39.53793\nExp number, first 60 AAs: 19.33093\nTotal prob of N-in: 0.88869\nPOSSIBLE N-term signal sequence\ninside 1 11\nTMhelix 12 34\noutside 35 796\nTMhelix 797 819\ninside 820 835

33803	GCF_001865795.1_ASM186579v1	Streptomyces albidoflavus	Terrabacteria group; Actinobacteria; Actinobacteria; Streptomycetales; Streptomycetaceae; Streptomyces; Streptomyces albidoflavus group
MSAAAPEAAPRTARPAPRAGALLRRLLWWCVLTLTGGVERRGALPPGGCVVVANHTSHAD	WP_071336819.1	1-acyl-sn-glycerol-3-phosphate acyltransferase [Streptomyces albidoflavus]	Length: 401\nNumber of predicted TMHs: 2\nExp number of AAs in TMHs: 84.15533\nExp number, first 60 AAs: 1.12621\nTotal prob of N-in: 0.13765\noutside 1 353\nTMhelix 354 371\ninside 372 377\nTMhelix 378 400\noutside 401 401

33804	GCF_001704195.1_ASM170419v1	Streptomyces sampsonii	Terrabacteria group; Actinobacteria; Actinobacteria; Streptomycetales; Streptomycetaceae; Streptomyces; Streptomyces albidoflavus group
MGGRGTTAQARERGRLPGRPSDPAPTGGNDVITGVNLRVPARRTALPALLLTATALLAVP	WP_067413686.1
hypothetical protein [Streptomyces sampsonii]	Length: 304\nNumber of predicted TMHs: 1\nExp number of AAs in TMHs: 33.72431\nExp number, first 60 AAs: 7.93262\nTotal prob of N-in: 0.52185\noutside 1 276\nTMhelix 277 299\ninside 300 304

33805	GCF_001704195.1_ASM170419v1	Streptomyces sampsonii	Terrabacteria group; Actinobacteria; Actinobacteria; Streptomycetales; Streptomycetaceae; Streptomyces; Streptomyces albidoflavus group
MSRRTRTARRRPVAAAVLAAGAVLLPAVPASAASGADQDFVIEDPRITESSGLAASKRHP	WP_067414640.1
hypothetical protein [Streptomyces sampsonii]	Length: 350\nNumber of predicted TMHs: 2\nExp number of AAs in TMHs: 39.44236\nExp number, first 60 AAs: 21.1162\nTotal prob of N-in: 0.99084\nPOSSIBLE N-term signal sequence\ninside 1 12\nTMhelix 13 35\noutside 36 322\nTMhelix 323 342\ninside 343 350

33806	GCF_001704195.1_ASM170419v1	Streptomyces sampsonii	Terrabacteria group; Actinobacteria; Actinobacteria; Streptomycetales; Streptomycetaceae; Streptomyces; Streptomyces albidoflavus group
MLDPVENRSLPRTAAFFDLDKTVIAKSSTLTFSKSFYQGGLINRRAVLRTAYTQFVFLAG	WP_067414143.1	inhibition of morphological differentiation protein [Streptomyces sampsonii]	Length: 290\nNumber of predicted TMHs: 1\nExp number of AAs in TMHs: 21.62258\nExp number, first 60 AAs: 0.02853\nTotal prob of N-in: 0.00967\noutside 1 247\nTMhelix 248 270\ninside 271 290

33807	GCF_001865315.1_ASM186531v1	Streptomyces sampsonii	Terrabacteria group; Actinobacteria; Actinobacteria; Streptomycetales; Streptomycetaceae; Streptomyces; Streptomyces albidoflavus group
MLDPVENRSLPRTAAFFDLDKTVIAKSSTLTFSKSFYQGGLINRRAVLRTAYTQFVFLAG	WP_003949650.1	MULTISPECIES: morphological differentiation-associated protein [Streptomyces]	Length: 290\nNumber of predicted TMHs: 1\nExp number of AAs in TMHs: 21.6224\nExp number, first 60 AAs: 0.02852\nTotal prob of N-in: 0.00967\noutside 1 247\nTMhelix 248 270\ninside 271 290

33808	GCF_001865315.1_ASM186531v1	Streptomyces sampsonii	Terrabacteria group; Actinobacteria; Actinobacteria; Streptomycetales; Streptomycetaceae; Streptomyces; Streptomyces albidoflavus group
MPDRRSAVSLRLPLPTAAPGFGGLALPGAAATAPDAWGVVAVRESSEKERKSSSEPEGRS	WP_015507753.1
MULTISPECIES: hypothetical protein [Streptomyces]	Length: 292\nNumber of predicted TMHs: 2\nExp number of AAs in TMHs: 51.77675\nExp number, first 60 AAs: 6.07612\nTotal prob of N-in: 0.33780\noutside 1 164\nTMhelix 165 187\ninside 188 229\nTMhelix 230 252\noutside 253 292

33809	GCF_001704195.1_ASM170419v1	Streptomyces sampsonii	Terrabacteria group; Actinobacteria; Actinobacteria; Streptomycetales; Streptomycetaceae; Streptomyces; Streptomyces albidoflavus group
MTPNRRPVVLAAAVATALGLGATALTLPAVAAEAPRAGAPVMDLTDGTLDWGVKESFRRY	WP_067411621.1
hypothetical protein [Streptomyces sampsonii]	Length: 514\nNumber of predicted TMHs: 2\nExp number of

AAs in TMHs: 42.8565699999999\nExp number, first 60 AAs: 22.03087\nTotal prob of N-in: 0.97276\nPOSSIBLE N-term signal sequence\ninside 1 6\nTMhelix 7 29\noutside 30 479\nTMhelix 480 502\ninside 503 514

33810 GCF_001704195.1_ASM170419v1 Streptomyces sampsonii Terrabacteria group; Actinobacteria; Actinobacteria; Streptomycetales; Streptomycetaceae; Streptomyces; Streptomyces albidoflavus group MSRMRGSSGGRIAGRRTAVVLAVAVLGLAGCSAGPDSGASSAGDRAVAPSAADGAGGGQA WP_067416561.1 hypothetical protein [Streptomyces sampsonii] Length: 334\nNumber of predicted TMHs: 1\nExp number of AAs in TMHs: 33.41361\nExp number, first 60 AAs: 10.62695\nTotal prob of N-in: 0.50969\nPOSSIBLE N-term signal sequence\noutside 1 269\nTMhelix 270 292\ninside 293 334

33811 GCF_001704195.1_ASM170419v1 Streptomyces sampsonii Terrabacteria group; Actinobacteria; Actinobacteria; Streptomycetales; Streptomycetaceae; Streptomyces; Streptomyces albidoflavus group MPDRRSAVSLRLPLPTAAPGFGGLALPGAAATAPDAWGVVAVRESSEKERKSSSEPEGRS WP_015507753.1 MULTISPECIES: hypothetical protein [Streptomyces] Length: 292\nNumber of predicted TMHs: 2\nExp number of AAs in TMHs: 51.77675\nExp number, first 60 AAs: 6.07612\nTotal prob of N-in: 0.33780\noutside 1 164\nTMhelix 165 187\ninside 188 229\nTMhelix 230 252\noutside 253 292

33812 GCF_001865315.1_ASM186531v1 Streptomyces sampsonii Terrabacteria group; Actinobacteria; Actinobacteria; Streptomycetales; Streptomycetaceae; Streptomyces; Streptomyces albidoflavus group MSRMRGSSGGRIAGRRTAVVLAVAVLGLAGCSAGPDSGASSAGDRAVAPSAADGAGGGQA WP_067416561.1 hypothetical protein [Streptomyces sampsonii] Length: 334\nNumber of predicted TMHs: 1\nExp number of AAs in TMHs: 33.41361\nExp number, first 60 AAs: 10.62695\nTotal prob of N-in: 0.50969\nPOSSIBLE N-term signal sequence\noutside 1 269\nTMhelix 270 292\ninside 293 334

33813 GCF_001865315.1_ASM186531v1 Streptomyces sampsonii Terrabacteria group; Actinobacteria; Actinobacteria; Streptomycetales; Streptomycetaceae; Streptomyces; Streptomyces albidoflavus group MGGRGTTAQARERGRLPGRPSDPAPTGGNDVITGVNLRVPARRTALPALLLTATALLAVP WP_067413686.1 hypothetical protein [Streptomyces sampsonii] Length: 304\nNumber of predicted TMHs: 1\nExp number of AAs in TMHs: 33.72431\nExp number, first 60 AAs: 7.93262\nTotal prob of N-in: 0.52185\noutside 1 276\nTMhelix 277 299\ninside 300 304

33814 GCF_001865315.1_ASM186531v1 Streptomyces sampsonii Terrabacteria group; Actinobacteria; Actinobacteria; Streptomycetales; Streptomycetaceae; Streptomyces; Streptomyces albidoflavus group MTPNRRPVVLAAAVATALGLGATALTLPAVAAEAPRAGAPVMDLTDGTLDWGVKESFRRY WP_067411621.1 hypothetical protein [Streptomyces sampsonii] Length: 514\nNumber of predicted TMHs: 2\nExp number of AAs in TMHs: 42.8565699999999\nExp number, first 60 AAs: 22.03087\nTotal prob of N-in: 0.97276\nPOSSIBLE N-term signal sequence\ninside 1 6\nTMhelix 7 29\noutside 30 479\nTMhelix 480 502\ninside 503 514

33815 GCF_001865315.1_ASM186531v1 Streptomyces sampsonii Terrabacteria group; Actinobacteria; Actinobacteria; Streptomycetales; Streptomycetaceae; Streptomyces; Streptomyces albidoflavus group MSRRTRTARRRPVAAAVLAAGAVLLPAVPASAASGADQDFVIEDPRITESSGLAASKRHP WP_067414640.1 hypothetical protein [Streptomyces sampsonii] Length: 350\nNumber of predicted TMHs: 2\nExp number of AAs in TMHs: 39.44236\nExp number, first 60 AAs: 21.1162\nTotal prob of N-in: 0.99084\nPOSSIBLE N-term signal sequence\ninside 1 12\nTMhelix 13 35\noutside 36 322\nTMhelix 323 342\ninside 343 350

33816 GCF_000203835.1_ASM20383v1 Streptomyces coelicolor A3(2) Terrabacteria group; Actinobacteria; Actinobacteria; Streptomycetales; Streptomycetaceae; Streptomyces; Streptomyces albidoflavus group; Streptomyces coelicolor MPRTAAFFDLDKTVIAKSSTLTFSKSFYQGGLINRRAVLRTAYAQFVFLAGGADHDQMER NP_627756.1 morphological differentiation-associated protein [Streptomyces coelicolor A3(2)] Length: 268\nNumber of predicted TMHs: 1\nExp number of AAs in TMHs: 21.51645\nExp number, first 60 AAs: 0.03737\nTotal prob of N-in: 0.00617\noutside 1 238\nTMhelix 239 261\ninside 262 268

33817 GCF_000203835.1_ASM20383v1 Streptomyces coelicolor A3(2) Terrabacteria group; Actinobacteria; Actinobacteria; Streptomycetales; Streptomycetaceae; Streptomyces; Streptomyces albidoflavus group; Streptomyces coelicolor MGHNRRRRPTGARRATFGAVALILGGSGLVAVNVFASATESGNPAVPLGSSGVAATVDCP NP_624953.1 hypothetical protein SCO0644 [Streptomyces coelicolor A3(2)] Length: 718\nNumber of predicted TMHs: 2\nExp number of AAs in TMHs: 43.09057\nExp number, first 60 AAs: 21.84171\nTotal prob of N-in: 0.99047\nPOSSIBLE N-term signal sequence\ninside 1 16\nTMhelix 17 39\noutside 40 690\nTMhelix 691 713\ninside 714 718

33818 GCF_000203835.1_ASM20383v1 Streptomyces coelicolor A3(2) Terrabacteria group; Actinobacteria; Actinobacteria; Streptomycetales; Streptomycetaceae; Streptomyces; Streptomyces albidoflavus group; Streptomyces coelicolor MRHARRRIVRRVARLAAVGGLLLGGTMVTRAVASEPPDASTAPRTFAQTPSGAGGDLVSR NP_625036.1 protease [Streptomyces coelicolor A3(2)] Length: 454\nNumber of predicted TMHs: 2\nExp number of AAs in TMHs: 40.7318399999999\nExp number, first 60 AAs: 21.1031\nTotal prob of N-in: 0.96387\nPOSSIBLE N-term signal sequence\ninside 1 11\nTMhelix 12 34\noutside 35 415\nTMhelix 416 438\ninside 439 454

33819 GCF_000403765.2_SAlb2 Streptomyces albulus CCRC 11814 Terrabacteria group; Actinobacteria; Actinobacteria; Streptomycetales; Streptomycetaceae; Streptomyces; Streptomyces albulus MVTVRPAHPPAERPRQGEDALQMPDRRRRAALLRPIPATTGWGLAPVTSPGAVLLPRERT WP_016570515.1 hypothetical protein [Streptomyces albulus] Length: 331\nNumber of predicted TMHs: 2\nExp number of AAs in TMHs: 48.4885399999999\nExp number, first 60 AAs: 0.90344\nTotal prob of N-in: 0.52918\ninside 1 187\nTMhelix 188 210\noutside 211 265\nTMhelix 266 288\ninside 289 331

33820 GCF_000403765.2_SAlb2 Streptomyces albulus CCRC 11814 Terrabacteria group; Actinobacteria; Actinobacteria; Streptomycetales; Streptomycetaceae; Streptomyces; Streptomyces albulus MTLMRPTGPPRPTGRRRRATLAATAALVLAGAGVVTAAGTAQAADVAYKTECQPPPISGL WP_037635128.1 MULTISPECIES: hypothetical protein [Streptomyces] Length: 479\nNumber of predicted TMHs: 2\nExp number of AAs in TMHs: 43.80161\nExp number, first 60 AAs: 21.18149\nTotal prob of N-in: 0.97217\nPOSSIBLE N-term signal sequence\ninside 1 20\nTMhelix 21 43\noutside 44 450\nTMhelix 451 473\ninside 474 479

33821 GCF_000504065.1_Velvet_for_version_02_of_Streptomyces_albulus_PD-1 Streptomyces albulus PD-1 Terrabacteria group; Actinobacteria; Actinobacteria; Streptomycetales; Streptomycetaceae; Streptomyces; Streptomyces albulus MVTVRPAHPPAERPRQGEDALQMPDRRRRAALLRPIPATTGWGLAPVTSPGAVLLPRERT WP_016570515.1 hypothetical protein [Streptomyces albulus] Length: 331\nNumber of predicted TMHs: 2\nExp number of AAs in TMHs: 48.4885399999999\nExp number, first 60 AAs: 0.90344\nTotal prob of N-in: 0.52918\ninside 1 187\nTMhelix 188 210\noutside 211 265\nTMhelix 266 288\ninside 289 331

33822 GCF_000504065.1_Velvet_for_version_02_of_Streptomyces_albulus_PD-1 Streptomyces albulus PD-1 Terrabacteria group; Actinobacteria; Actinobacteria; Streptomycetales; Streptomycetaceae; Streptomyces; Streptomyces albulus MTLMRPTGPPRPTGRRRRATLAATAALVLAGAGVVTAAGTAQAADVAYKTECQPPPISGL WP_037635128.1 MULTISPECIES: hypothetical protein [Streptomyces] Length: 479\nNumber of predicted TMHs: 2\nExp number of AAs in TMHs: 43.80161\nExp number, first 60 AAs: 21.18149\nTotal prob of N-in: 0.97217\nPOSSIBLE N-term signal sequence\ninside 1 20\nTMhelix 21 43\noutside 44 450\nTMhelix 451 473\ninside 474 479

33823 GCF_000963515.1_ASM96351v1 Streptomyces albulus ZPM Terrabacteria group; Actinobacteria; Actinobacteria; Streptomycetales; Streptomycetaceae; Streptomyces; Streptomyces albulus MTLMRPTGPPRPTGRRRRATLAATAALVLAGAGVVTAAGTAQAADVAYKTECQPPPISGL WP_037635128.1 MULTISPECIES: hypothetical protein [Streptomyces] Length: 479\nNumber of predicted TMHs: 2\nExp number of AAs in TMHs: 43.80161\nExp number, first 60 AAs: 21.18149\nTotal prob of N-in: 0.97217\nPOSSIBLE N-term signal sequence\ninside 1 20\nTMhelix 21 43\noutside 44 450\nTMhelix 451 473\ninside 474 479

33824 GCF_000963515.1_ASM96351v1 Streptomyces albulus ZPM Terrabacteria group; Actinobacteria; Actinobacteria; Streptomycetales; Streptomycetaceae; Streptomyces; Streptomyces albulus MVTVRPAHPPAERPRQGEDALQMPDRRRRAALLRPIPATTGWGLAPVTSPGAVLLPRERT WP_016570515.1 hypothetical protein [Streptomyces albulus] Length: 331\nNumber of predicted TMHs: 2\nExp number of AAs in TMHs: 48.4885399999999\nExp number, first 60 AAs: 0.90344\nTotal prob of N-in: 0.52918\ninside 1 187\nTMhelix 188 210\noutside 211 265\nTMhelix 266 288\ninside 289 331

33825 GCF_000813365.1_ASM81336v1 Streptomyces albusTerrabacteria group; Actinobacteria; Actinobacteria; Streptomycetales; Streptomycetaceae; Streptomyces; Streptomyces albus group MLSGVENLAMPRTAAFFDLDKTVIAKSSTLTFGKSFYQGGLINRRAVLRTAYAQFVYLLG WP_016473027.1 MULTISPECIES: haloacid dehalogenase [Streptomyces] Length: 284\nNumber of predicted TMHs: 1\nExp number of AAs in TMHs: 21.94508\nExp number, first 60 AAs: 0.01548\nTotal prob of N-in: 0.00204\noutside 1 246\nTMhelix 247 269\ninside 270 284

33826 GCF_001753425.1_ASM175342v1 Streptomyces albusTerrabacteria group; Actinobacteria; Actinobacteria; Streptomycetales; Streptomycetaceae; Streptomyces; Streptomyces albus group MRRRPARRSPLSLLLLGTGTFLLVLAPLLAWYVEPQARRTPLDVDVTTVFEGSGSYFDTA WP_040251858.1 hypothetical protein [Streptomyces albus] Length: 336\nNumber of predicted TMHs:  2\nExp number of AAs in TMHs: 43.0981\nExp number, first 60 AAs:  21.80498\nTotal prob of N-in:   0.99690\nPOSSIBLE N-term signal sequence\ninside   1   10\nTMhelix 11   33\noutside   34   289\nTMhelix   290   312\ninside   313   336

33827 GCF_001577385.1_ASM157738v1 Streptomyces albusTerrabacteria group; Actinobacteria; Actinobacteria; Streptomycetales; Streptomycetaceae; Streptomyces; Streptomyces albus group MSRMRGSSGGRIAGRRAAVVLAVAVLGLAGCSAGPDSGASSAGDRAVAPSAADGAGGGQA WP_061403993.1 hypothetical protein [Streptomyces albus] Length: 334\nNumber of predicted TMHs:  1\nExp number of AAs in TMHs: 36.36643\nExp number, first 60 AAs:  13.57798\nTotal prob of N-in:   0.64438\nPOSSIBLE N-term signal sequence\noutside   1   269\nTMhelix   270   292\ninside   293   334

33828 GCF_001577385.1_ASM157738v1 Streptomyces albusTerrabacteria group; Actinobacteria; Actinobacteria; Streptomycetales; Streptomycetaceae; Streptomyces; Streptomyces albus group MPDRRSAVSLRLPLPTAAPGFGGLALPGAAATAPDAWGVVAVRESSEKERKSSSEPEGRS WP_015507753.1 MULTISPECIES: hypothetical protein [Streptomyces] Length: 292\nNumber of predicted TMHs:  2\nExp number of AAs in TMHs: 51.77675\nExp number, first 60 AAs:  6.07612\nTotal prob of N-in:   0.33780\noutside   1 164\nTMhelix   165   187\ninside   188   229\nTMhelix   230   252\noutside   253   292

33829 GCF_001577385.1_ASM157738v1 Streptomyces albusTerrabacteria group; Actinobacteria; Actinobacteria; Streptomycetales; Streptomycetaceae; Streptomyces; Streptomyces albus group MSRRTRTARRRPVAAAVLAAGAVLLPAVPASAASGADQDFVIEDPRITESSGLAASKRHP WP_061404434.1 hypothetical protein [Streptomyces albus] Length: 345\nNumber of predicted TMHs:  2\nExp number of AAs in TMHs: 39.70691\nExp number, first 60 AAs:  21.11813\nTotal prob of N-in:   0.99086\nPOSSIBLE N-term signal sequence\ninside   1   12\nTMhelix   13   35\noutside   36   317\nTMhelix   318   337\ninside   338   345

33830 GCF_000827005.1_ASM82700v1 Streptomyces albusTerrabacteria group; Actinobacteria; Actinobacteria; Streptomycetales; Streptomycetaceae; Streptomyces; Streptomyces albus group MRRRPARRSPLSLLLLGTGTFLLVLAPLLAWYVEPQARRTPLDVDVTTVFEGSGSYFDTA WP_040251858.1 hypothetical protein [Streptomyces albus] Length: 336\nNumber of predicted TMHs:  2\nExp number of AAs in TMHs: 43.0981\nExp number, first 60 AAs:  21.80498\nTotal prob of N-in:   0.99690\nPOSSIBLE N-term signal sequence\ninside   1   10\nTMhelix 11   33\noutside   34   289\nTMhelix   290   312\ninside   313   336

33831 GCF_001577385.1_ASM157738v1 Streptomyces albusTerrabacteria group; Actinobacteria; Actinobacteria; Streptomycetales; Streptomycetaceae; Streptomyces; Streptomyces albus group MSAAAPEAAPRTARPAPRAGALLRRLLWWCVLTLTGGVERRGALPPGGCVVVANHTSHAD WP_015507057.1 MULTISPECIES: 1-acyl-sn-glycerol-3-phosphate acyltransferase [Streptomyces] Length: 401\nNumber of predicted TMHs:  2\nExp number of AAs in TMHs: 83.99314\nExp number, first 60 AAs:  1.08255\nTotal prob of N-in: 0.25169\noutside   1   353\nTMhelix   354   371\ninside   372   377\nTMhelix   378   400\noutside   401   401

33832 GCF_001577385.1_ASM157738v1 Streptomyces albusTerrabacteria group; Actinobacteria; Actinobacteria; Streptomycetales; Streptomycetaceae; Streptomyces; Streptomyces albus group MAPPHRRRAAVAVRALVALTALLCTTGLAPAATSGPAARAAVAEVPPPTATTSVVTVRTG WP_008409019.1 MULTISPECIES: VWA domain-containing protein [Streptomyces] Length: 835\nNumber of predicted TMHs: 2\nExp number of AAs in TMHs: 37.77539\nExp number, first 60 AAs:  18.55298\nTotal prob of N-in:   0.85432\nPOSSIBLE N-term signal sequence\ninside   1   11\nTMhelix   12   34\noutside   35   796\nTMhelix   797   819\ninside   820   835

33833 GCF_001577385.1_ASM157738v1 Streptomyces albusTerrabacteria group; Actinobacteria; Actinobacteria; Streptomycetales; Streptomycetaceae; Streptomyces; Streptomyces albus group MLDPVENRSLPRTAAFFDLDKTVIAKSSTLTFSKSFYQGGLINRRAVLRTAYTQFVFLAG WP_003949650.1 MULTISPECIES: morphological differentiation-associated protein [Streptomyces] Length: 290\nNumber of predicted TMHs:  1\nExp number of AAs in TMHs: 21.6224\nExp number, first 60 AAs:  0.02852\nTotal prob of N-in:   0.00967\noutside   1 247\nTMhelix   248   270\ninside   271   290

33834 GCF_000813365.1_ASM81336v1 Streptomyces albusTerrabacteria group; Actinobacteria; Actinobacteria; Streptomycetales; Streptomycetaceae; Streptomyces; Streptomyces albus group MSPRRRRATVLAAATVLCAAAAPVAAAAPSGTAAAGPVPAVKPAKLPAGLYGDADPKYDG WP_016468754.1 MULTISPECIES: hypothetical protein [Streptomyces] Length: 457\nNumber of predicted TMHs:  2\nExp number of AAs in TMHs: 40.9820199999999\nExp number, first 60 AAs:  18.73808\nTotal prob of N-in: 0.84735\nPOSSIBLE N-term signal sequence\ninside   1   6\nTMhelix   7   29\noutside   30   427\nTMhelix   428 450\ninside   451   457

33835 GCF_001753425.1_ASM175342v1 Streptomyces albusTerrabacteria group; Actinobacteria; Actinobacteria; Streptomycetales; Streptomycetaceae; Streptomyces; Streptomyces albus group
MAVIRRRTAFAASVAAALTLGAAGSVALPAVAAPGEGAPAAKAIELKDGTLDWGFKESFR WP_040254138.1 membrane protein [Streptomyces albus] Length: 500\nNumber of predicted TMHs: 2\nExp number of AAs in TMHs: 41.82416\nExp number, first 60 AAs: 20.29992\nTotal prob of N-in: 0.90699\nPOSSIBLE N-term signal sequence\ninside 1 8\nTMhelix 9 31\noutside 32 467\nTMhelix 468 490\ninside 491 500

33836 GCF_000827005.1_ASM82700v1 Streptomyces albusTerrabacteria group; Actinobacteria; Actinobacteria; Streptomycetales; Streptomycetaceae; Streptomyces; Streptomyces albus group
MAVIRRRTAFAASVAAALTLGAAGSVALPAVAAPGEGAPAAKAIELKDGTLDWGFKESFR WP_040254138.1 membrane protein [Streptomyces albus] Length: 500\nNumber of predicted TMHs: 2\nExp number of AAs in TMHs: 41.82416\nExp number, first 60 AAs: 20.29992\nTotal prob of N-in: 0.90699\nPOSSIBLE N-term signal sequence\ninside 1 8\nTMhelix 9 31\noutside 32 467\nTMhelix 468 490\ninside 491 500

33837 GCF_001577385.1_ASM157738v1 Streptomyces albusTerrabacteria group; Actinobacteria; Actinobacteria; Streptomycetales; Streptomycetaceae; Streptomyces; Streptomyces albus group
MTPNRRPVVLAAAVATALGLGATALTLPAVAAEAPRAGAPVMDLTDGTLDWGVKESFRRY WP_061405035.1 hypothetical protein [Streptomyces albus] Length: 518\nNumber of predicted TMHs: 2\nExp number of AAs in TMHs: 42.8565299999999\nExp number, first 60 AAs: 22.03089\nTotal prob of N-in: 0.97282\nPOSSIBLE N-term signal sequence\ninside 1 6\nTMhelix 7 29\noutside 30 483\nTMhelix 484 506\ninside 507 518

33838 GCF_001418125.1_ASM141812v1 Streptomyces almquistii Terrabacteria group; Actinobacteria; Actinobacteria; Streptomycetales; Streptomycetaceae; Streptomyces; Streptomyces albus group
MLSGVENLAMPRTAAFFDLDKTVIAKSSTLTFGKSFYQGGLINRRAVLRTAYAQFVYLLG WP_016473027.1 MULTISPECIES: haloacid dehalogenase [Streptomyces] Length: 284\nNumber of predicted TMHs: 1\nExp number of AAs in TMHs: 21.94508\nExp number, first 60 AAs: 0.01548\nTotal prob of N-in: 0.00204\noutside 1 246\nTMhelix 247 269\ninside 270 284

33839 GCF_001418125.1_ASM141812v1 Streptomyces almquistii Terrabacteria group; Actinobacteria; Actinobacteria; Streptomycetales; Streptomycetaceae; Streptomyces; Streptomyces albus group
MSPRRRRATVLAAATVLCAAAAPVAAAAPSGTAAAGPVPAVKPAKLPAGLYGDADPKYDG WP_016468754.1 MULTISPECIES: hypothetical protein [Streptomyces] Length: 457\nNumber of predicted TMHs: 2\nExp number of AAs in TMHs: 40.9820199999999\nExp number, first 60 AAs: 18.73808\nTotal prob of N-in: 0.84735\nPOSSIBLE N-term signal sequence\ninside 1 6\nTMhelix 7 29\noutside 30 427\nTMhelix 428 450\ninside 451 457

33840 GCF_001418505.1_ASM141850v1 Streptomyces flocculus Terrabacteria group; Actinobacteria; Actinobacteria; Streptomycetales; Streptomycetaceae; Streptomyces; Streptomyces albus group
MLSGVENLAMPRTAAFFDLDKTVIAKSSTLTFGKSFYQGGLINRRAVLRTAYAQFVYLLG WP_055499288.1 inhibition of morphological differentiation protein [Streptomyces flocculus] Length: 284\nNumber of predicted TMHs: 1\nExp number of AAs in TMHs: 21.94508\nExp number, first 60 AAs: 0.01548\nTotal prob of N-in: 0.00204\noutside 1 246\nTMhelix 247 269\ninside 270 284

33841 GCF_001418505.1_ASM141850v1 Streptomyces flocculus Terrabacteria group; Actinobacteria; Actinobacteria; Streptomycetales; Streptomycetaceae; Streptomyces; Streptomyces albus group
MSPRRRRATVLAAATVLCAAAAPVAAAAPSGTAAAGPVPAVKPAKLPAGLYGDADPKYDG WP_055498263.1 hypothetical protein [Streptomyces flocculus] Length: 459\nNumber of predicted TMHs: 2\nExp number of AAs in TMHs: 41.01463\nExp number, first 60 AAs: 18.72899\nTotal prob of N-in: 0.84741\nPOSSIBLE N-term signal sequence\ninside 1 6\nTMhelix 7 29\noutside 30 429\nTMhelix 430 452\ninside 453 459

33842 GCF_001418545.1_ASM141854v1 Streptomyces gibsonii Terrabacteria group; Actinobacteria; Actinobacteria; Streptomycetales; Streptomycetaceae; Streptomyces; Streptomyces albus group
MSPRRRRATVLAAATVLCAAAAPVAAAAPSGTAAAGPVPAVKPAKLPAGLYGDADPKYDG WP_016468754.1 MULTISPECIES: hypothetical protein [Streptomyces] Length: 457\nNumber of predicted TMHs: 2\nExp number of AAs in TMHs: 40.9820199999999\nExp number, first 60 AAs: 18.73808\nTotal prob of N-in: 0.84735\nPOSSIBLE N-term signal sequence\ninside 1 6\nTMhelix 7 29\noutside 30 427\nTMhelix 428 450\ninside 451 457

33843 GCF_001418545.1_ASM141854v1 Streptomyces gibsonii Terrabacteria group; Actinobacteria; Actinobacteria; Streptomycetales; Streptomycetaceae; Streptomyces; Streptomyces albus group
MLSGVENLAMPRTAAFFDLDKTVIAKSSTLTFGKSFYQGGLINRRAVLRTAYAQFVYLLG WP_016473027.1 MULTISPECIES:

haloacid dehalogenase [Streptomyces] Length: 284\nNumber of predicted TMHs: 1\nExp number of AAs in TMHs: 21.94508\nExp number, first 60 AAs: 0.01548\nTotal prob of N-in: 0.00204\noutside 1 246\nTMhelix 247 269\ninside 270 284

33844 GCF_001661615.1_ASM166161v1 Streptomyces rangoonensis Terrabacteria group; Actinobacteria; Actinobacteria; Streptomycetales; Streptomycetaceae; Streptomyces; Streptomyces albus group MLSGVENLAMPRTAAFFDLDKTVIAKSSTLTFGKSFYQGGLINRRAVLRTAYAQFVYLLG WP_016473027.1 MULTISPECIES: haloacid dehalogenase [Streptomyces] Length: 284\nNumber of predicted TMHs: 1\nExp number of AAs in TMHs: 21.94508\nExp number, first 60 AAs: 0.01548\nTotal prob of N-in: 0.00204\noutside 1 246\nTMhelix 247 269\ninside 270 284

33845 GCF_001661615.1_ASM166161v1 Streptomyces rangoonensis Terrabacteria group; Actinobacteria; Actinobacteria; Streptomycetales; Streptomycetaceae; Streptomyces; Streptomyces albus group MSPRRRRATVLAAATVLCAAAAPVAAAAPSGTAAAGPVPAVKPAKLPAGLYGDADPKYDG WP_016468754.1 MULTISPECIES: hypothetical protein [Streptomyces] Length: 457\nNumber of predicted TMHs: 2\nExp number of AAs in TMHs: 40.9820199999999\nExp number, first 60 AAs: 18.73808\nTotal prob of N-in: 0.84735\nPOSSIBLE N-term signal sequence\ninside 1 6\nTMhelix 7 29\noutside 30 427\nTMhelix 428 450\ninside 451 457

33846 GCF_000156475.1_ASM15647v1 Streptomyces albus J1074 Terrabacteria group; Actinobacteria; Actinobacteria; Streptomycetales; Streptomycetaceae; Streptomyces; Streptomyces albus group; Streptomyces albus MSAAAPEAAPRTARPAPRAGALLRRLLWWCVLTLTGGVERRGALPPGGCVVVANHTSHAD WP_015507057.1 MULTISPECIES: 1-acyl-sn-glycerol-3-phosphate acyltransferase [Streptomyces] Length: 401\nNumber of predicted TMHs: 2\nExp number of AAs in TMHs: 83.99314\nExp number, first 60 AAs: 1.08255\nTotal prob of N-in: 0.25169\noutside 1 353\nTMhelix 354 371\ninside 372 377\nTMhelix 378 400\noutside 401 401

33847 GCF_000359525.1_ASM35952v1 Streptomyces albus J1074 Terrabacteria group; Actinobacteria; Actinobacteria; Streptomycetales; Streptomycetaceae; Streptomyces; Streptomyces albus group; Streptomyces albus MSAAAPEAAPRTARPAPRAGALLRRLLWWCVLTLTGGVERRGALPPGGCVVVANHTSHAD WP_015507057.1 MULTISPECIES: 1-acyl-sn-glycerol-3-phosphate acyltransferase [Streptomyces] Length: 401\nNumber of predicted TMHs: 2\nExp number of AAs in TMHs: 83.99314\nExp number, first 60 AAs: 1.08255\nTotal prob of N-in: 0.25169\noutside 1 353\nTMhelix 354 371\ninside 372 377\nTMhelix 378 400\noutside 401 401

33848 GCF_000359525.1_ASM35952v1 Streptomyces albus J1074 Terrabacteria group; Actinobacteria; Actinobacteria; Streptomycetales; Streptomycetaceae; Streptomyces; Streptomyces albus group; Streptomyces albus MPDRRSAVSLRLPLPTAAPGFGGLALPGAAATAPDAWGVVAVRESSEKERKSSSEPEGRS WP_015507753.1 MULTISPECIES: hypothetical protein [Streptomyces] Length: 292\nNumber of predicted TMHs: 2\nExp number of AAs in TMHs: 51.77675\nExp number, first 60 AAs: 6.07612\nTotal prob of N-in: 0.33780\noutside 1 164\nTMhelix 165 187\ninside 188 229\nTMhelix 230 252\noutside 253 292

33849 GCF_000359525.1_ASM35952v1 Streptomyces albus J1074 Terrabacteria group; Actinobacteria; Actinobacteria; Streptomycetales; Streptomycetaceae; Streptomyces; Streptomyces albus group; Streptomyces albus MAPPHRRRAAVAVRALVALTALLCTTGLAPAATSGPAARAAVAEVPPPTATTSVVTVRTG WP_008409019.1 MULTISPECIES: VWA domain-containing protein [Streptomyces] Length: 835\nNumber of predicted TMHs: 2\nExp number of AAs in TMHs: 37.77539\nExp number, first 60 AAs: 18.55298\nTotal prob of N-in: 0.85432\nPOSSIBLE N-term signal sequence\ninside 1 11\nTMhelix 12 34\noutside 35 796\nTMhelix 797 819\ninside 820 835

33850 GCF_000359525.1_ASM35952v1 Streptomyces albus J1074 Terrabacteria group; Actinobacteria; Actinobacteria; Streptomycetales; Streptomycetaceae; Streptomyces; Streptomyces albus group; Streptomyces albus MSRMRGSSGGRIAGRRAAVVLAVAVLGLAGCSAGPDSGASSAGDRAVAPSAADGAGGGQA WP_015506895.1 MULTISPECIES: DUF4349 domain-containing protein [Streptomyces] Length: 334\nNumber of predicted TMHs: 1\nExp number of AAs in TMHs: 36.36573\nExp number, first 60 AAs: 13.5773\nTotal prob of N-in: 0.64439\nPOSSIBLE N-term signal sequence\noutside 1 269\nTMhelix 270 292\ninside 293 334

33851 GCF_000156475.1_ASM15647v1 Streptomyces albus J1074 Terrabacteria group; Actinobacteria; Actinobacteria; Streptomycetales; Streptomycetaceae; Streptomyces; Streptomyces albus group; Streptomyces albus MPDRRSAVSLRLPLPTAAPGFGGLALPGAAATAPDAWGVVAVRESSEKERKSSSEPEGRS WP_015507753.1 MULTISPECIES: hypothetical protein [Streptomyces] Length: 292\nNumber of predicted TMHs: 2\nExp number of AAs in TMHs: 51.77675\nExp number, first 60 AAs: 6.07612\nTotal prob of N-in: 0.33780\noutside 1 164\nTMhelix 165 187\ninside 188 229\nTMhelix 230 252\noutside 253 292

33852 GCF_000359525.1_ASM35952v1 Streptomyces albus J1074 Terrabacteria group; Actinobacteria; Actinobacteria; Streptomycetales; Streptomycetaceae; Streptomyces; Streptomyces albus group; Streptomyces albus MTPNRRPVVLAAAVATALGLGATALTLPAVAAEAPRAGAPVMDLTDGTLDWGVKESFRRY WP_015507978.1 MULTISPECIES: membrane protein [Streptomyces] Length: 514\nNumber of predicted TMHs: 2\nExp number of AAs in TMHs: 42.8566699999999\nExp number, first 60 AAs: 22.03087\nTotal prob of N-in: 0.97276\nPOSSIBLE N-term signal sequence\ninside 1 6\nTMhelix 7 29\noutside 30 479\nTMhelix 480 502\ninside 503 514

33853 GCF_000359525.1_ASM35952v1 Streptomyces albus J1074 Terrabacteria group; Actinobacteria; Actinobacteria; Streptomycetales; Streptomycetaceae; Streptomyces; Streptomyces albus group; Streptomyces albus MLDPVENRSLPRTAAFFDLDKTVIAKSSTLTFSKSFYQGGLINRRAVLRTAYTQFVFLAG WP_003949650.1 MULTISPECIES: morphological differentiation-associated protein [Streptomyces] Length: 290\nNumber of predicted TMHs: 1\nExp number of AAs in TMHs: 21.6224\nExp number, first 60 AAs: 0.02852\nTotal prob of N-in: 0.00967\noutside 1 247\nTMhelix 248 270\ninside 271 290

33854 GCF_000359525.1_ASM35952v1 Streptomyces albus J1074 Terrabacteria group; Actinobacteria; Actinobacteria; Streptomycetales; Streptomycetaceae; Streptomyces; Streptomyces albus group; Streptomyces albus MGGRGTTAQARERGRLPGRPSDPAPTGGNDVITGVNLRMPARRTALPALLLTATALLAVP WP_003949316.1 MULTISPECIES: hypothetical protein [Streptomyces] Length: 304\nNumber of predicted TMHs: 2\nExp number of AAs in TMHs: 39.19493\nExp number, first 60 AAs: 11.75707\nTotal prob of N-in: 0.76164\nPOSSIBLE N-term signal sequence\ninside 1 44\nTMhelix 45 67\noutside 68 276\nTMhelix 277 299\ninside 300 304

33855 GCF_000359525.1_ASM35952v1 Streptomyces albus J1074 Terrabacteria group; Actinobacteria; Actinobacteria; Streptomycetales; Streptomycetaceae; Streptomyces; Streptomyces albus group; Streptomyces albus MSRRTRTARRRPVAAAVLAAGAVLLPAVPASAASGADQDFVIEDPRITESSGLAASKRHP WP_003950051.1 hypothetical protein [Streptomyces albus] Length: 348\nNumber of predicted TMHs: 2\nExp number of AAs in TMHs: 39.50741\nExp number, first 60 AAs: 21.11214\nTotal prob of N-in: 0.99087\nPOSSIBLE N-term signal sequence\ninside 1 12\nTMhelix 13 35\noutside 36 320\nTMhelix 321 340\ninside 341 348

33856 GCF_000156475.1_ASM15647v1 Streptomyces albus J1074 Terrabacteria group; Actinobacteria; Actinobacteria; Streptomycetales; Streptomycetaceae; Streptomyces; Streptomyces albus group; Streptomyces albus MLDPVENRSLPRTAAFFDLDKTVIAKSSTLTFSKSFYQGGLINRRAVLRTAYTQFVFLAG WP_003949650.1 MULTISPECIES: morphological differentiation-associated protein [Streptomyces] Length: 290\nNumber of predicted TMHs: 1\nExp number of AAs in TMHs: 21.6224\nExp number, first 60 AAs: 0.02852\nTotal prob of N-in: 0.00967\noutside 1 247\nTMhelix 248 270\ninside 271 290

33857 GCF_000156475.1_ASM15647v1 Streptomyces albus J1074 Terrabacteria group; Actinobacteria; Actinobacteria; Streptomycetales; Streptomycetaceae; Streptomyces; Streptomyces albus group; Streptomyces albus MGGRGTTAQARERGRLPGRPSDPAPTGGNDVITGVNLRMPARRTALPALLLTATALLAVP WP_003949316.1 MULTISPECIES: hypothetical protein [Streptomyces] Length: 304\nNumber of predicted TMHs: 2\nExp number of AAs in TMHs: 39.19493\nExp number, first 60 AAs: 11.75707\nTotal prob of N-in: 0.76164\nPOSSIBLE N-term signal sequence\ninside 1 44\nTMhelix 45 67\noutside 68 276\nTMhelix 277 299\ninside 300 304

33858 GCF_000156475.1_ASM15647v1 Streptomyces albus J1074 Terrabacteria group; Actinobacteria; Actinobacteria; Streptomycetales; Streptomycetaceae; Streptomyces; Streptomyces albus group; Streptomyces albus MSRRTRTARRRPVAAAVLAAGAVLLPAVPASAASGADQDFVIEDPRITESSGLAASKRHP WP_003950051.1 hypothetical protein [Streptomyces albus] Length: 348\nNumber of predicted TMHs: 2\nExp number of AAs in TMHs: 39.50741\nExp number, first 60 AAs: 21.11214\nTotal prob of N-in: 0.99087\nPOSSIBLE N-term signal sequence\ninside 1 12\nTMhelix 13 35\noutside 36 320\nTMhelix 321 340\ninside 341 348

33859 GCF_000716945.1_ASM71694v1 Streptomyces albus subsp. albus Terrabacteria group; Actinobacteria; Actinobacteria; Streptomycetales; Streptomycetaceae; Streptomyces; Streptomyces albus group; Streptomyces albusMLSVVENCFSPRTAAFFDLDKTVIAKSSTLTFSKSFYQGGLINRRAVLRTAYTQFVFLAG WP_032776898.1 inhibition of morphological differentiation protein [Streptomyces albus] Length: 281\nNumber of predicted TMHs: 1\nExp number of AAs in TMHs: 21.46265\nExp number, first 60 AAs: 0.01958\nTotal prob of N-in: 0.00583\noutside 1 247\nTMhelix 248 270\ninside 271 281

33860 GCF_000716945.1_ASM71694v1 Streptomyces albus subsp. albus Terrabacteria group; Actinobacteria; Actinobacteria; Streptomycetales; Streptomycetaceae; Streptomyces; Streptomyces albus group; Streptomyces albusMGLSWNDRRLERRVKPGDGKPLKPFRWWQMTRRSLLSIALPVGGGGGLSATYTVEVKHGG WP_032776788.1 hypothetical protein [Streptomyces albus] Length: 220\nNumber of predicted TMHs:

2\nExp number of AAs in TMHs: 42.07089\nExp number, first 60 AAs: 2.58855\nTotal prob of N-in: 0.93135\ninside 1 137\nTMhelix 138 160\noutside 161 184\nTMhelix 185 202\ninside 203 220

33861 GCF_001541145.1_ASM154114v1 Streptomyces albus subsp. albus Terrabacteria group; Actinobacteria; Actinobacteria; Streptomycetales; Streptomycetaceae; Streptomyces; Streptomyces albus group; Streptomyces albusMTERSAPAASATAPGTPARRTAVRAAATGTLAAALLGLAPQSALAATPAPPPAPAISAVT WP_030603830.1 MULTISPECIES: hypothetical protein [Streptomyces] Length: 293\nNumber of predicted TMHs: 1\nExp number of AAs in TMHs: 24.61251\nExp number, first 60 AAs: 5.37395\nTotal prob of N-in: 0.26790\noutside 1 264\nTMhelix 265 287\ninside 288 293

33862 GCF_000720915.1_ASM72091v1 Streptomyces albus subsp. albus Terrabacteria group; Actinobacteria; Actinobacteria; Streptomycetales; Streptomycetaceae; Streptomyces; Streptomyces albus group; Streptomyces albusMRITPSRRRTLRGSTAAVAATALVALGAAPALADEPEPDLGVGTIAPIKGVQAGAAFSTP WP_030626375.1 hypothetical protein [Streptomyces albus] Length: 431\nNumber of predicted TMHs: 1\nExp number of AAs in TMHs: 34.30376\nExp number, first 60 AAs: 14.19651\nTotal prob of N-in: 0.69603\nPOSSIBLE N-term signal sequence\noutside 1 399\nTMhelix 400 422\ninside 423 431

33863 GCF_000720915.1_ASM72091v1 Streptomyces albus subsp. albus Terrabacteria group; Actinobacteria; Actinobacteria; Streptomycetales; Streptomycetaceae; Streptomyces; Streptomyces albus group; Streptomyces albusMLSVVENCFSPRTAAFFDLDKTVIAKSSTLTFSKSFYQGGLINRRAVLRTAYAQFVFLAG WP_030634155.1 inhibition of morphological differentiation protein [Streptomyces albus] Length: 282\nNumber of predicted TMHs: 1\nExp number of AAs in TMHs: 21.69171\nExp number, first 60 AAs: 0.02994\nTotal prob of N-in: 0.00448\noutside 1 247\nTMhelix 248 270\ninside 271 282

33864 GCF_000725885.1_ASM72588v1 Streptomyces albus subsp. albus Terrabacteria group; Actinobacteria; Actinobacteria; Streptomycetales; Streptomycetaceae; Streptomyces; Streptomyces albus group; Streptomyces albusMLSGVENLAMPRTAAFFDLDKTVIAKSSTLTFGKSFYQGGLINRRAVLRTAYAQFVYLLG WP_016473027.1 MULTISPECIES: haloacid dehalogenase [Streptomyces] Length: 284\nNumber of predicted TMHs: 1\nExp number of AAs in TMHs: 21.94508\nExp number, first 60 AAs: 0.01548\nTotal prob of N-in: 0.00204\noutside 1 246\nTMhelix 247 269\ninside 270 284

33865 GCF_000720885.1_ASM72088v1 Streptomyces albus subsp. albus Terrabacteria group; Actinobacteria; Actinobacteria; Streptomycetales; Streptomycetaceae; Streptomyces; Streptomyces albus group; Streptomyces albusMSPRRRRATVLAAATVLCAAAAPVAAAAPSGTAAAGPVPAVKPAQLPAGLYGDADPKYDG WP_037611286.1 hypothetical protein [Streptomyces albus] Length: 457\nNumber of predicted TMHs: 2\nExp number of AAs in TMHs: 40.71176\nExp number, first 60 AAs: 18.46969\nTotal prob of N-in: 0.83862\nPOSSIBLE N-term signal sequence\ninside 1 6\nTMhelix 7 29\noutside 30 427\nTMhelix 428 450\ninside 451 457

33866 GCF_000720885.1_ASM72088v1 Streptomyces albus subsp. albus Terrabacteria group; Actinobacteria; Actinobacteria; Streptomycetales; Streptomycetaceae; Streptomyces; Streptomyces albus group; Streptomyces albusMLSGVENLAMPRTAAFFDLDKTVIAKSSTLTFGKSFYQGGLINRRAVLRTAYAQFVYLLG WP_016473027.1 MULTISPECIES: haloacid dehalogenase [Streptomyces] Length: 284\nNumber of predicted TMHs: 1\nExp number of AAs in TMHs: 21.94508\nExp number, first 60 AAs: 0.01548\nTotal prob of N-in: 0.00204\noutside 1 246\nTMhelix 247 269\ninside 270 284

33867 GCF_000725885.1_ASM72588v1 Streptomyces albus subsp. albus Terrabacteria group; Actinobacteria; Actinobacteria; Streptomycetales; Streptomycetaceae; Streptomyces; Streptomyces albus group; Streptomyces albusMSPRRRRATVLAAATVLCAAAAPVAAAAPSGTAAAGPVPAVKPAKLPAGLYGDADPKYDG WP_016468754.1 MULTISPECIES: hypothetical protein [Streptomyces] Length: 457\nNumber of predicted TMHs: 2\nExp number of AAs in TMHs: 40.9820199999999\nExp number, first 60 AAs: 18.73808\nTotal prob of N-in: 0.84735\nPOSSIBLE N-term signal sequence\ninside 1 6\nTMhelix 7 29\noutside 30 427\nTMhelix 428 450\ninside 451 457

33868 GCF_000719865.1_ASM71986v1 Streptomyces albus subsp. albus Terrabacteria group; Actinobacteria; Actinobacteria; Streptomycetales; Streptomycetaceae; Streptomyces; Streptomyces albus group; Streptomyces albusMRRNKRRRPTGARRSTFAAFALILGGGGLVAVNAVASATGGDSGGSRVQGQSGGDPGAAT WP_037680772.1 hypothetical protein [Streptomyces albus] Length: 704\nNumber of predicted TMHs: 2\nExp number of AAs in TMHs: 43.74659\nExp number, first 60 AAs: 21.94326\nTotal prob of N-in: 0.99862\nPOSSIBLE N-term signal sequence\ninside 1 16\nTMhelix 17 39\noutside 40 672\nTMhelix 673 695\ninside 696 704

33869 GCF_000720915.1_ASM72091v1 Streptomyces albus subsp. albus Terrabacteria group; Actinobacteria; Actinobacteria; Streptomycetales; Streptomycetaceae; Streptomyces; Streptomyces albus group; Streptomyces albusMDPVTDGESRSRRSFVRARAALVAGALGLLLAVGGCGASGDASDSGAKAADGKAAPREGF WP_030631798.1 hypothetical protein [Streptomyces albus] Length: 332\nNumber of predicted TMHs: 2\nExp number of AAs in TMHs: 43.52123\nExp number, first 60 AAs: 20.52541\nTotal prob of N-in: 0.93684\nPOSSIBLE N-term signal sequence\ninside 1 19\nTMhelix 20 42\noutside 43 279\nTMhelix 280 302\ninside 303 332

33870 GCF_001267885.1_ASM126788v1 Streptomyces ambofaciens ATCC 23877 Terrabacteria group; Actinobacteria; Actinobacteria; Streptomycetales; Streptomycetaceae; Streptomyces; Streptomyces ambofaciens MGHNRRRRPTGARRATFGACALILGGGGLVAVNVFASATEAGDPAVPLGSSGVAATVDCP WP_079030850.1 hypothetical protein [Streptomyces ambofaciens] Length: 740\nNumber of predicted TMHs: 2\nExp number of AAs in TMHs: 42.92424\nExp number, first 60 AAs: 21.63506\nTotal prob of N-in: 0.98648\nPOSSIBLE N-term signal sequence\ninside 1 16\nTMhelix 17 39\noutside 40 712\nTMhelix 713 735\ninside 736 740

33871 GCF_001267885.1_ASM126788v1 Streptomyces ambofaciens ATCC 23877 Terrabacteria group; Actinobacteria; Actinobacteria; Streptomycetales; Streptomycetaceae; Streptomyces; Streptomyces ambofaciens MPRTAAFFDLDKTVIAKSSTLTFSKSFYQGGLINRRAVLRTAYAQFVFLAGGADHDQMER WP_063482640.1 inhibition of morphological differentiation protein [Streptomyces ambofaciens] Length: 268\nNumber of predicted TMHs: 1\nExp number of AAs in TMHs: 21.53677\nExp number, first 60 AAs: 0.04893\nTotal prob of N-in: 0.00773\noutside 1 238\nTMhelix 239 261\ninside 262 268

33872 GCF_001267885.1_ASM126788v1 Streptomyces ambofaciens ATCC 23877 Terrabacteria group; Actinobacteria; Actinobacteria; Streptomycetales; Streptomycetaceae; Streptomyces; Streptomyces ambofaciens MNADEQQCEEDTRVDRRRGLIQRVAVVCGLVGALMAVGPLTARADTGPVDAAVALELDRP WP_053141924.1 hypothetical protein [Streptomyces ambofaciens] Length: 367\nNumber of predicted TMHs: 2\nExp number of AAs in TMHs: 42.35848\nExp number, first 60 AAs: 21.59679\nTotal prob of N-in: 0.98989\nPOSSIBLE N-term signal sequence\ninside 1 19\nTMhelix 20 42\noutside 43 337\nTMhelix 338 360\ninside 361 367

33873 GCF_001267885.1_ASM126788v1 Streptomyces ambofaciens ATCC 23877 Terrabacteria group; Actinobacteria; Actinobacteria; Streptomycetales; Streptomycetaceae; Streptomyces; Streptomyces ambofaciens MRHARRRIVRRVARLAAVGGLLLGGTMVTRAVASEPPGPPTGPHATVRVASGTGAGLVSR WP_053140927.1 protease [Streptomyces ambofaciens] Length: 455\nNumber of predicted TMHs: 2\nExp number of AAs in TMHs: 41.9456\nExp number, first 60 AAs: 21.00104\nTotal prob of N-in: 0.96796\nPOSSIBLE N-term signal sequence\ninside 1 11\nTMhelix 12 34\noutside 35 416\nTMhelix 417 439\ninside 440 455

33874 GCF_000414115.1_STRAU Streptomyces aurantiacus JA 4570 Terrabacteria group; Actinobacteria; Actinobacteria; Streptomycetales; Streptomycetaceae; Streptomyces; Streptomyces aurantiacus MLKLVENHSLPRTAAFFDLDKTVIAKSSTLTFSKSFYQGGLINRRAVLRTAYSQFVFLAG WP_016642743.1 inhibition of morphological differentiation protein [Streptomyces aurantiacus] Length: 274\nNumber of predicted TMHs: 1\nExp number of AAs in TMHs: 21.87751\nExp number, first 60 AAs: 0.01641\nTotal prob of N-in: 0.00977\noutside 1 244\nTMhelix 245 267\ninside 268 274

33875 GCF_000280865.1_ASM28086v1 Streptomyces auratus AGR0001 Terrabacteria group; Actinobacteria; Actinobacteria; Streptomycetales; Streptomycetaceae; Streptomyces; Streptomyces auratus MSMNSAQSRPDSSRANARPALSRTLRRRPAVVLALLAGLLAAAIAPAAAAGRAVPDTARV WP_006604260.1 hypothetical protein [Streptomyces auratus] Length: 153\nNumber of predicted TMHs: 2\nExp number of AAs in TMHs: 44.59393\nExp number, first 60 AAs: 21.62249\nTotal prob of N-in: 0.98957\nPOSSIBLE N-term signal sequence\ninside 1 30\nTMhelix 31 53\noutside 54 117\nTMhelix 118 140\ninside 141 153

33876 GCF_000280865.1_ASM28086v1 Streptomyces auratus AGR0001 Terrabacteria group; Actinobacteria; Actinobacteria; Streptomycetales; Streptomycetaceae; Streptomyces; Streptomyces auratus MASRGRRATIRSLGALTGAAALAALGGGAPAWAAAGAPDSSHTATPIKHVVVLFDENISF WP_040901191.1 phospholipase C [Streptomyces auratus] Length: 614\nNumber of predicted TMHs: 2\nExp number of AAs in TMHs: 44.85567\nExp number, first 60 AAs: 22.28923\nTotal prob of N-in: 0.98038\nPOSSIBLE N-term signal sequence\ninside 1 12\nTMhelix 13 35\noutside 36 581\nTMhelix 582 604\ninside 605 614

33877 GCF_000280865.1_ASM28086v1 Streptomyces auratus AGR0001 Terrabacteria group; Actinobacteria; Actinobacteria; Streptomycetales; Streptomycetaceae; Streptomyces; Streptomyces auratus MRQRRRAAVASAAALMLAGAGIMTAAGTAQAAEVSYKTECLPPPISGLPPIEGTTKVAVS WP_040899100.1 hypothetical protein [Streptomyces auratus] Length: 483\nNumber of predicted TMHs: 2\nExp number of

AAs in TMHs: 44.7211\nExp number, first 60 AAs: 22.19544\nTotal prob of N-in: 0.99510\nPOSSIBLE N-term signal sequence\ninside 1 6\nTMhelix 7 29\noutside 30 452\nTMhelix 453 475\ninside 476 483

33878 GCF_000280865.1_ASM28086v1 Streptomyces auratus AGR0001 Terrabacteria group; Actinobacteria; Actinobacteria; Streptomycetales; Streptomycetaceae; Streptomyces; Streptomyces auratus MPPHNRHVSDPVAHLAPPAPRRRALPRRLAAPLGALTAALAAFAYVGAVDPHQPGHYPVC WP_006606205.1 membrane protein [Streptomyces auratus] Length: 155\nNumber of predicted TMHs: 2\nExp number of AAs in TMHs: 62.88645\nExp number, first 60 AAs: 20.27938\nTotal prob of N-in: 0.98670\nPOSSIBLE N-term signal sequence\ninside 1 28\nTMhelix 29 48\noutside 49 89\nTMhelix 90 112\ninside 113 155

33879 GCF_000968255.1_ASM96825v1 Streptomyces avermitilis MA-4680 = NBRC 14893 Terrabacteria group; Actinobacteria; Actinobacteria; Streptomycetales; Streptomycetaceae; Streptomyces; Streptomyces avermitilis MLGPVENHSLPRTAAFFDLDKTVIAKSSTLTFSKSFYQGGLINRRAVLRTAYAQFVFLAG WP_010986028.1 morphological differentiation-associated protein [Streptomyces avermitilis] Length: 277\nNumber of predicted TMHs: 1\nExp number of AAs in TMHs: 21.0967\nExp number, first 60 AAs: 0.01343\nTotal prob of N-in: 0.00115\noutside 1 248\nTMhelix 249 268\ninside 269 277

33880 GCF_000968255.1_ASM96825v1 Streptomyces avermitilis MA-4680 = NBRC 14893 Terrabacteria group; Actinobacteria; Actinobacteria; Streptomycetales; Streptomycetaceae; Streptomyces; Streptomyces avermitilis MGRNTRRRPTGPRRATFAAVALILGGGGLLAANVYASATEGGLGSEPVRQTDSDGNAWGR WP_078234756.1 hypothetical protein [Streptomyces avermitilis] Length: 675\nNumber of predicted TMHs: 2\nExp number of AAs in TMHs: 39.46572\nExp number, first 60 AAs: 21.74729\nTotal prob of N-in: 0.99097\nPOSSIBLE N-term signal sequence\ninside 1 16\nTMhelix 17 39\noutside 40 649\nTMhelix 650 667\ninside 668 675

33881 GCF_000968255.1_ASM96825v1 Streptomyces avermitilis MA-4680 = NBRC 14893 Terrabacteria group; Actinobacteria; Actinobacteria; Streptomycetales; Streptomycetaceae; Streptomyces; Streptomyces avermitilis MRHARRRIVRRGARFAAVGGVLLGGLMVTQALASEPSSASPATTSSTETAAGTGSGLVSR WP_037649294.1 protease [Streptomyces avermitilis] Length: 459\nNumber of predicted TMHs: 2\nExp number of AAs in TMHs: 40.08158\nExp number, first 60 AAs: 21.26163\nTotal prob of N-in: 0.98513\nPOSSIBLE N-term signal sequence\ninside 1 11\nTMhelix 12 34\noutside 35 420\nTMhelix 421 443\ninside 444 459

33882 GCF_000009765.2_ASM976v2 Streptomyces avermitilis MA-4680 = NBRC 14893 Terrabacteria group; Actinobacteria; Actinobacteria; Streptomycetales; Streptomycetaceae; Streptomyces; Streptomyces avermitilis MMSTPMPYVTEATLRGTFARRAALAATTTAVLAATTVLLPAAAPAAVADSTTGQCTFPSK WP_010983976.1 type VII secretion-associated serine protease mycosin [Streptomyces avermitilis] Length: 432\nNumber of predicted TMHs: 1\nExp number of AAs in TMHs: 37.55284\nExp number, first 60 AAs: 15.0823\nTotal prob of N-in: 0.66729\nPOSSIBLE N-term signal sequence\noutside 1 395\nTMhelix 396 418\ninside 419 432

33883 GCF_000009765.2_ASM976v2 Streptomyces avermitilis MA-4680 = NBRC 14893 Terrabacteria group; Actinobacteria; Actinobacteria; Streptomycetales; Streptomycetaceae; Streptomyces; Streptomyces avermitilis MLGPVENHSLPRTAAFFDLDKTVIAKSSTLTFSKSFYQGGLINRRAVLRTAYAQFVFLAG WP_010986028.1 morphological differentiation-associated protein [Streptomyces avermitilis] Length: 277\nNumber of predicted TMHs: 1\nExp number of AAs in TMHs: 21.0967\nExp number, first 60 AAs: 0.01343\nTotal prob of N-in: 0.00115\noutside 1 248\nTMhelix 249 268\ninside 269 277

33884 GCF_000968255.1_ASM96825v1 Streptomyces avermitilis MA-4680 = NBRC 14893 Terrabacteria group; Actinobacteria; Actinobacteria; Streptomycetales; Streptomycetaceae; Streptomyces; Streptomyces avermitilis MTPPLPCLCHPAVGRHGLTLYGPRMRDSSRLSRRAVLGLAAAAPLAAAAPASASAATVIG WP_010987315.1 D-alanyl-D-alanine carboxypeptidase [Streptomyces avermitilis] Length: 446\nNumber of predicted TMHs: 1\nExp number of AAs in TMHs: 28.11838\nExp number, first 60 AAs: 6.58154\nTotal prob of N-in: 0.28816\noutside 1 373\nTMhelix 374 396\ninside 397 446

33885 GCF_000009765.2_ASM976v2 Streptomyces avermitilis MA-4680 = NBRC 14893 Terrabacteria group; Actinobacteria; Actinobacteria; Streptomycetales; Streptomycetaceae; Streptomyces; Streptomyces avermitilis MTPPLPCLCHPAVGRHGLTLYGPRMRDSSRLSRRAVLGLAAAAPLAAAAPASASAATVIG WP_010987315.1 D-alanyl-D-alanine carboxypeptidase [Streptomyces avermitilis] Length: 446\nNumber of predicted TMHs: 1\nExp number of AAs in TMHs: 28.11838\nExp number, first 60 AAs: 6.58154\nTotal prob of N-in: 0.28816\noutside 1 373\nTMhelix 374 396\ninside 397 446

33886 GCF_000968255.1_ASM96825v1 Streptomyces avermitilis MA-4680 = NBRC 14893 Terrabacteria group; Actinobacteria; Actinobacteria; Streptomycetales; Streptomycetaceae; Streptomyces; Streptomyces avermitilis

MRTDRTLRSQPVNQDSSGRSISRREALRGAGAVAVAGAGAAVLGGASTAVAADAFAHPGL WP_010988599.1 hypothetical protein [Streptomyces avermitilis] Length: 506\nNumber of predicted TMHs: 2\nExp number of AAs in TMHs: 45.1170199999999\nExp number, first 60 AAs: 22.25086\nTotal prob of N-in: 0.97947\nPOSSIBLE N-term signal sequence\ninside 1 28\nTMhelix 29 51\noutside 52 474\nTMhelix 475 497\ninside 498 506

33887 GCF_000009765.2_ASM976v2 Streptomyces avermitilis MA-4680 = NBRC 14893 Terrabacteria group; Actinobacteria; Actinobacteria; Streptomycetales; Streptomycetaceae; Streptomyces; Streptomyces avermitilis MRTDRTLRSQPVNQDSSGRSISRREALRGAGAVAVAGAGAAVLGGASTAVAADAFAHPGL WP_010988599.1 hypothetical protein [Streptomyces avermitilis] Length: 506\nNumber of predicted TMHs: 2\nExp number of AAs in TMHs: 45.1170199999999\nExp number, first 60 AAs: 22.25086\nTotal prob of N-in: 0.97947\nPOSSIBLE N-term signal sequence\ninside 1 28\nTMhelix 29 51\noutside 52 474\nTMhelix 475 497\ninside 498 506

33888 GCF_000968255.1_ASM96825v1 Streptomyces avermitilis MA-4680 = NBRC 14893 Terrabacteria group; Actinobacteria; Actinobacteria; Streptomycetales; Streptomycetaceae; Streptomyces; Streptomyces avermitilis MMSTPMPYVTEATLRGTFARRAALAATTTAVLAATTVLLPAAAPAAVADSTTGQCTFPSK WP_010983976.1 type VII secretion-associated serine protease mycosin [Streptomyces avermitilis] Length: 432\nNumber of predicted TMHs: 1\nExp number of AAs in TMHs: 37.55284\nExp number, first 60 AAs: 15.0823\nTotal prob of N-in: 0.66729\nPOSSIBLE N-term signal sequence\noutside 1 395\nTMhelix 396 418\ninside 419 432

33889 GCF_000764715.1_ASM76471v1 Streptomyces avermitilis MA-4680 = NBRC 14893 Terrabacteria group; Actinobacteria; Actinobacteria; Streptomycetales; Streptomycetaceae; Streptomyces; Streptomyces avermitilis MLGPVENHSLPRTAAFFDLDKTVIAKSSTLTFSKSFYQGGLINRRAVLRTAYAQFVFLAG WP_010986028.1 morphological differentiation-associated protein [Streptomyces avermitilis] Length: 277\nNumber of predicted TMHs: 1\nExp number of AAs in TMHs: 21.0967\nExp number, first 60 AAs: 0.01343\nTotal prob of N-in: 0.00115\noutside 1 248\nTMhelix 249 268\ninside 269 277

33890 GCF_000764715.1_ASM76471v1 Streptomyces avermitilis MA-4680 = NBRC 14893 Terrabacteria group; Actinobacteria; Actinobacteria; Streptomycetales; Streptomycetaceae; Streptomyces; Streptomyces avermitilis MGRNTRRRPTGPRRATFAAVALILGGGGLLAANVYASATEGGLGSEPVRQTDSDGNAWGR WP_078234756.1 hypothetical protein [Streptomyces avermitilis] Length: 675\nNumber of predicted TMHs: 2\nExp number of AAs in TMHs: 39.46572\nExp number, first 60 AAs: 21.74729\nTotal prob of N-in: 0.99097\nPOSSIBLE N-term signal sequence\ninside 1 16\nTMhelix 17 39\noutside 40 649\nTMhelix 650 667\ninside 668 675

33891 GCF_000764715.1_ASM76471v1 Streptomyces avermitilis MA-4680 = NBRC 14893 Terrabacteria group; Actinobacteria; Actinobacteria; Streptomycetales; Streptomycetaceae; Streptomyces; Streptomyces avermitilis MTPPLPCLCHPAVGRHGLTLYGPRMRDSSRLSRRAVLGLAAAAPLAAAAPASASAATVIG WP_010987315.1 D-alanyl-D-alanine carboxypeptidase [Streptomyces avermitilis] Length: 446\nNumber of predicted TMHs: 1\nExp number of AAs in TMHs: 28.11838\nExp number, first 60 AAs: 6.58154\nTotal prob of N-in: 0.28816\noutside 1 373\nTMhelix 374 396\ninside 397 446

33892 GCF_000764715.1_ASM76471v1 Streptomyces avermitilis MA-4680 = NBRC 14893 Terrabacteria group; Actinobacteria; Actinobacteria; Streptomycetales; Streptomycetaceae; Streptomyces; Streptomyces avermitilis MRTDRTLRSQPVNQDSSGRSISRREALRGAGAVAVAGAGAAVLGGASTAVAADAFAHPGL WP_010988599.1 hypothetical protein [Streptomyces avermitilis] Length: 506\nNumber of predicted TMHs: 2\nExp number of AAs in TMHs: 45.1170199999999\nExp number, first 60 AAs: 22.25086\nTotal prob of N-in: 0.97947\nPOSSIBLE N-term signal sequence\ninside 1 28\nTMhelix 29 51\noutside 52 474\nTMhelix 475 497\ninside 498 506

33893 GCF_000764715.1_ASM76471v1 Streptomyces avermitilis MA-4680 = NBRC 14893 Terrabacteria group; Actinobacteria; Actinobacteria; Streptomycetales; Streptomycetaceae; Streptomyces; Streptomyces avermitilis MRHARRRIVRRGARFAAVGGVLLGGLMVTQALASEPSSASPATTSSTETAAGTGSGLVSR WP_037649294.1 protease [Streptomyces avermitilis] Length: 459\nNumber of predicted TMHs: 2\nExp number of AAs in TMHs: 40.08158\nExp number, first 60 AAs: 21.26163\nTotal prob of N-in: 0.98513\nPOSSIBLE N-term signal sequence\ninside 1 11\nTMhelix 12 34\noutside 35 420\nTMhelix 421 443\ninside 444 459

33894 GCF_000764715.1_ASM76471v1 Streptomyces avermitilis MA-4680 = NBRC 14893 Terrabacteria group; Actinobacteria; Actinobacteria; Streptomycetales; Streptomycetaceae; Streptomyces; Streptomyces avermitilis MMSTPMPYVTEATLRGTFARRAALAATTTAVLAATTVLLPAAAPAAVADSTTGQCTFPSK WP_010983976.1 type VII secretion-associated serine protease mycosin [Streptomyces avermitilis] Length: 432\nNumber of predicted TMHs: 1\nExp number of AAs in TMHs: 37.55284\nExp number, first 60 AAs: 15.0823\nTotal prob of N-in: 0.66729\nPOSSIBLE N-term signal sequence\noutside 1 395\nTMhelix 396 418\ninside 419 432

33895 GCF_000092385.1_ASM9238v1 Streptomyces bingchenggensis BCW-1 Terrabacteria group; Actinobacteria; Actinobacteria; Streptomycetales; Streptomycetaceae; Streptomyces; Streptomyces bingchenggensis MGRRAALLCAVLLSTLVSVVALGASPAQAAGYRYWSFWQRERGAGGWTYATQGPATARPG WP_014180138.1 hypothetical protein [Streptomyces bingchenggensis] Length: 224\nNumber of predicted TMHs: 2\nExp number of AAs in TMHs: 43.52954\nExp number, first 60 AAs: 22.06599\nTotal prob of N-in: 0.99215\nPOSSIBLE N-term signal sequence\ninside 1 6\nTMhelix 7 29\noutside 30 193\nTMhelix 194 216\ninside 217 224

33896 GCF_000092385.1_ASM9238v1 Streptomyces bingchenggensis BCW-1 Terrabacteria group; Actinobacteria; Actinobacteria; Streptomycetales; Streptomycetaceae; Streptomyces; Streptomyces bingchenggensis MLGVVENHSLPRTAAFFDLDKTVIAKSSTLTFSKSFYQGGLINRRAVLRTAYAQFVFLAG WP_014178415.1 morphological differentiation-associated protein [Streptomyces bingchenggensis] Length: 278\nNumber of predicted TMHs: 1\nExp number of AAs in TMHs: 21.89798\nExp number, first 60 AAs: 0.02259\nTotal prob of N-in: 0.00436\noutside 1 246\nTMhelix 247 269\ninside 270 278

33897 GCF_000092385.1_ASM9238v1 Streptomyces bingchenggensis BCW-1 Terrabacteria group; Actinobacteria; Actinobacteria; Streptomycetales; Streptomycetaceae; Streptomyces; Streptomyces bingchenggensis MSAADDGGNRSLGQLVATATTELSALVHDEIALAKAEFRDSARRTLFGSGAAIVAGVLLL WP_014178408.1 phage holin family protein [Streptomyces bingchenggensis] Length: 153\nNumber of predicted TMHs: 2\nExp number of AAs in TMHs: 46.5707000000001\nExp number, first 60 AAs: 13.62843\nTotal prob of N-in: 0.95447\nPOSSIBLE N-term signal sequence\ninside 1 45\nTMhelix 46 68\noutside 69 82\nTMhelix 83 105\ninside 106 153

33898 GCF_000092385.1_ASM9238v1 Streptomyces bingchenggensis BCW-1 Terrabacteria group; Actinobacteria; Actinobacteria; Streptomycetales; Streptomycetaceae; Streptomyces; Streptomyces bingchenggensis MSSPDREVRAARNDPPRTARRPVVAVTGAASGAGALLTQRLAESDETKHVLAIDERRGDV WP_014176621.1 NAD-dependent dehydratase [Streptomyces bingchenggensis] Length: 369\nNumber of predicted TMHs: 1\nExp number of AAs in TMHs: 20.16076\nExp number, first 60 AAs: 0.56792\nTotal prob of N-in: 0.03858\noutside 1 339\nTMhelix 340 359\ninside 360 369

33899 GCF_000383595.1_ASM38359v1 Streptomyces bottropensis ATCC 25435 Terrabacteria group; Actinobacteria; Actinobacteria; Streptomycetales; Streptomycetaceae; Streptomyces; Streptomyces bottropensis MLGPVENHSLPRTAAFFDLDKTVIAKSSTLTFSKSFYQGGLINRRAALRTAYTQFVFLAG WP_005475982.1 morphological differentiation protein [Streptomyces bottropensis] Length: 274\nNumber of predicted TMHs: 1\nExp number of AAs in TMHs: 21.70339\nExp number, first 60 AAs: 0.01912\nTotal prob of N-in: 0.00405\noutside 1 244\nTMhelix 245 267\ninside 268 274

33900 GCF_000340335.1_ASM34033v2 Streptomyces bottropensis ATCC 25435 Terrabacteria group; Actinobacteria; Actinobacteria; Streptomycetales; Streptomycetaceae; Streptomyces; Streptomyces bottropensis MLGPVENHSLPRTAAFFDLDKTVIAKSSTLTFSKSFYQGGLINRRAALRTAYTQFVFLAG WP_005475982.1 morphological differentiation protein [Streptomyces bottropensis] Length: 274\nNumber of predicted TMHs: 1\nExp number of AAs in TMHs: 21.70339\nExp number, first 60 AAs: 0.01912\nTotal prob of N-in: 0.00405\noutside 1 244\nTMhelix 245 267\ninside 268 274

33901 GCF_002154495.1_ASM215449v1 Streptomyces cacaoi subsp. cacaoi Terrabacteria group; Actinobacteria; Actinobacteria; Streptomycetales; Streptomycetaceae; Streptomyces; Streptomyces cacaoi MLGPVENVPMPRTAAFFDLDKTVIAKSSTLTFGKPFYQGGLINRRAVLRTAYAQFVYLLG WP_030883110.1 inhibition of morphological differentiation protein [Streptomyces sp. NRRL S-1868] Length: 285\nNumber of predicted TMHs: 1\nExp number of AAs in TMHs: 21.98851\nExp number, first 60 AAs: 0.01836\nTotal prob of N-in: 0.00267\noutside 1 246\nTMhelix 247 269\ninside 270 285

33902 GCF_000383615.1_ASM38361v1 Streptomyces canus 299MFChir4.1 Terrabacteria group; Actinobacteria; Actinobacteria; Streptomycetales; Streptomycetaceae; Streptomyces; Streptomyces canus MLGVVENHSLPRTAAFFDLDKTVIAKSSTLTFSKSFYQGGLINRRAALRTAYAQFVFLVG WP_020118792.1 inhibition of morphological differentiation protein [Streptomyces canus] Length: 277\nNumber of predicted TMHs: 1\nExp number of AAs in TMHs: 21.68342\nExp number, first 60 AAs: 0.03177\nTotal prob of N-in: 0.00309\noutside 1 247\nTMhelix 248 270\ninside 271 277

33903 GCF_000383615.1_ASM38361v1 Streptomyces canus 299MFChir4.1 Terrabacteria group; Actinobacteria; Actinobacteria; Streptomycetales; Streptomycetaceae; Streptomyces; Streptomyces canus MRHARRRVVRRVTRLAAVGGLLLGGAMVTRAVASEPPATTTAVPHTYSTKAGQTGTDLVA WP_033415245.1 protease [Streptomyces canus] Length: 456\nNumber of predicted TMHs: 2\nExp number of AAs in

TMHs: 39.0290599999999\nExp number, first 60 AAs: 20.3092\nTotal prob of N-in: 0.94569\nPOSSIBLE N-term signal sequence\ninside 1 11\nTMhelix 12 34\noutside 35 417\nTMhelix 418 440\ninside 441 456

33904 GCF_000240165.1_ASM24016v1 Streptomyces cattleya NRRL 8057 = DSM 46488 Terrabacteria group; Actinobacteria; Actinobacteria; Streptomycetales; Streptomycetaceae; Streptomyces; Streptomyces cattleya MLVTVENHSLPRTAAFFDLDKTVIAKSSTLAFSKSFYHGGLINRRAVLRTAYAQFVYLLG WP_014143607.1 inhibition of morphological differentiation protein [Streptomyces cattleya] Length: 289\nNumber of predicted TMHs: 1\nExp number of AAs in TMHs: 22.05528\nExp number, first 60 AAs: 0.08979\nTotal prob of N-in: 0.03204\noutside 1 244\nTMhelix 245 267\ninside 268 289

33905 GCF_000240165.1_ASM24016v1 Streptomyces cattleya NRRL 8057 = DSM 46488 Terrabacteria group; Actinobacteria; Actinobacteria; Streptomycetales; Streptomycetaceae; Streptomyces; Streptomyces cattleya MRRRSALVVLALAVLCATLSPLLRWYAFPRLARVPAGQYQVVVLEARNATLLDYTRMKAV WP_014141891.1 DUF3068 domain-containing protein [Streptomyces cattleya] Length: 338\nNumber of predicted TMHs: 2\nExp number of AAs in TMHs: 43.8990299999999\nExp number, first 60 AAs: 21.88274\nTotal prob of N-in: 0.99194\nPOSSIBLE N-term signal sequence\ninside 1 4\nTMhelix 5 27\noutside 28 282\nTMhelix 283 305\ninside 306 338

33906 GCF_000237305.1_ASM23730v1 Streptomyces cattleya NRRL 8057 = DSM 46488 Terrabacteria group; Actinobacteria; Actinobacteria; Streptomycetales; Streptomycetaceae; Streptomyces; Streptomyces cattleya MRRRSALVVLALAVLCATLSPLLRWYAFPRLARVPAGQYQVVVLEARNATLLDYTRMKAV WP_014141891.1 DUF3068 domain-containing protein [Streptomyces cattleya] Length: 338\nNumber of predicted TMHs: 2\nExp number of AAs in TMHs: 43.8990299999999\nExp number, first 60 AAs: 21.88274\nTotal prob of N-in: 0.99194\nPOSSIBLE N-term signal sequence\ninside 1 4\nTMhelix 5 27\noutside 28 282\nTMhelix 283 305\ninside 306 338

33907 GCF_000237305.1_ASM23730v1 Streptomyces cattleya NRRL 8057 = DSM 46488 Terrabacteria group; Actinobacteria; Actinobacteria; Streptomycetales; Streptomycetaceae; Streptomyces; Streptomyces cattleya MLVTVENHSLPRTAAFFDLDKTVIAKSSTLAFSKSFYHGGLINRRAVLRTAYAQFVYLLG WP_014143607.1 inhibition of morphological differentiation protein [Streptomyces cattleya] Length: 289\nNumber of predicted TMHs: 1\nExp number of AAs in TMHs: 22.05528\nExp number, first 60 AAs: 0.08979\nTotal prob of N-in: 0.03204\noutside 1 244\nTMhelix 245 267\ninside 268 289

33908 GCF_000226455.1_ASM22645v1 Streptomyces chartreusis NRRL 12338 Terrabacteria group; Actinobacteria; Actinobacteria; Streptomycetales; Streptomycetaceae; Streptomyces; Streptomyces chartreusis MLKGVENHSLPRTAAFFDLDKTVIAKSSTLTFSKSFYQGGLINRRAVLRTAYAQFVFLAG WP_010031922.1 morphological differentiation-associated protein [Streptomyces chartreusis] Length: 277\nNumber of predicted TMHs: 1\nExp number of AAs in TMHs: 21.58881\nExp number, first 60 AAs: 0.03583\nTotal prob of N-in: 0.00417\noutside 1 247\nTMhelix 248 270\ninside 271 277

33909 GCF_000226455.1_ASM22645v1 Streptomyces chartreusis NRRL 12338 Terrabacteria group; Actinobacteria; Actinobacteria; Streptomycetales; Streptomycetaceae; Streptomyces; Streptomyces chartreusis MPARRTAVTVTALGALPLALTGPAVTPAAAHGSMGDPVSRVSQCYAEGPESPKSAACEAA WP_010046142.1 chitin-binding protein [Streptomyces chartreusis]Length: 333\nNumber of predicted TMHs: 2\nExp number of AAs in TMHs: 34.5491\nExp number, first 60 AAs: 15.76315\nTotal prob of N-in: 0.72655\nPOSSIBLE N-term signal sequence\ninside 1 6\nTMhelix 7 29\noutside 30 298\nTMhelix 299 321\ninside 322 333

33910 GCF_000226455.1_ASM22645v1 Streptomyces chartreusis NRRL 12338 Terrabacteria group; Actinobacteria; Actinobacteria; Streptomycetales; Streptomycetaceae; Streptomyces; Streptomyces chartreusis MTDTPPRQRRSTLARLRGLPLWSVLAAGTLVGGLLGGAYGLLTPSTYTATSYVIAVPTEK WP_029181655.1 hypothetical protein [Streptomyces chartreusis] Length: 218\nNumber of predicted TMHs: 2\nExp number of AAs in TMHs: 42.82398\nExp number, first 60 AAs: 22.79704\nTotal prob of N-in: 0.99407\nPOSSIBLE N-term signal sequence\ninside 1 20\nTMhelix 21 43\noutside 44 164\nTMhelix 165 187\ninside 188 218

33911 GCF_000226455.1_ASM22645v1 Streptomyces chartreusis NRRL 12338 Terrabacteria group; Actinobacteria; Actinobacteria; Streptomycetales; Streptomycetaceae; Streptomyces; Streptomyces chartreusis MRHARRRVVRRVTRLAAVGGLLLGGAMVTRAVASEPPDPSIGVPYAARPATSMGTDLVSR WP_040902940.1 S1 family peptidase [Streptomyces chartreusis] Length: 449\nNumber of predicted TMHs: 2\nExp number of AAs in TMHs: 41.03581\nExp number, first 60 AAs: 20.67043\nTotal prob of N-in: 0.95243\nPOSSIBLE N-term signal sequence\ninside 1 11\nTMhelix 12 34\noutside 35 410\nTMhelix 411 433\ninside 434 449

33912 GCF_001885705.1_ASM188570v1 Streptomyces cinnamoneus Terrabacteria group; Actinobacteria; Actinobacteria; Streptomycetales; Streptomycetaceae; Streptomyces; Streptomyces cinnamoneus group
MLGGVENHSLPRTAAFFDLDKTVIAKSSTLTFSKSFYQGGLINRRAALRTAYTQFVFLAG WP_079273997.1 inhibition of morphological differentiation protein [Streptomyces cinnamoneus] Length: 299\nNumber of predicted TMHs: 1\nExp number of AAs in TMHs: 20.67927\nExp number, first 60 AAs: 0.01369\nTotal prob of N-in: 0.00468\noutside 1 246\nTMhelix 247 269\ninside 270 299

33913 GCF_000154925.1_ASM15492v1 Streptomyces clavuligerus ATCC 27064 Terrabacteria group; Actinobacteria; Actinobacteria; Streptomycetales; Streptomycetaceae; Streptomyces; Streptomyces clavuligerus
MRIRTPPTPPGEGRARRPGPRHSGPPTRRGLMRIRRTLVIAAATAVISPAALWVAPAASA WP_003961538.1
hypothetical protein [Streptomyces clavuligerus] Length: 440\nNumber of predicted TMHs: 2\nExp number of AAs in TMHs: 42.46109\nExp number, first 60 AAs: 20.42926\nTotal prob of N-in: 0.95678\nPOSSIBLE N-term signal sequence\ninside 1 37\nTMhelix 38 60\noutside 61 408\nTMhelix 409 431\ninside 432 440

33914 GCF_000148465.1_Scla_1.0 Streptomyces clavuligerus ATCC 27064 Terrabacteria group; Actinobacteria; Actinobacteria; Streptomycetales; Streptomycetaceae; Streptomyces; Streptomyces clavuligerus
MRIRTPPTPPGEGRARRPGPRHSGPPTRRGLMRIRRTLVIAAATAVISPAALWVAPAASA WP_003961538.1
hypothetical protein [Streptomyces clavuligerus] Length: 440\nNumber of predicted TMHs: 2\nExp number of AAs in TMHs: 42.46109\nExp number, first 60 AAs: 20.42926\nTotal prob of N-in: 0.95678\nPOSSIBLE N-term signal sequence\ninside 1 37\nTMhelix 38 60\noutside 61 408\nTMhelix 409 431\ninside 432 440

33915 GCF_000163875.1_Strep_clav_ATCC27064 Streptomyces clavuligerus ATCC 27064 Terrabacteria group; Actinobacteria; Actinobacteria; Streptomycetales; Streptomycetaceae; Streptomyces; Streptomyces clavuligerus
MRIRTPPTPPGEGRARRPGPRHSGPPTRRGLMRIRRTLVIAAATAVISPAALWVAPAASA WP_003961538.1
hypothetical protein [Streptomyces clavuligerus] Length: 440\nNumber of predicted TMHs: 2\nExp number of AAs in TMHs: 42.46109\nExp number, first 60 AAs: 20.42926\nTotal prob of N-in: 0.95678\nPOSSIBLE N-term signal sequence\ninside 1 37\nTMhelix 38 60\noutside 61 408\nTMhelix 409 431\ninside 432 440

33916 GCF_000241835.1_ASM24183v2 Streptomyces coelicoflavus ZG0656 Terrabacteria group; Actinobacteria; Actinobacteria; Streptomycetales; Streptomycetaceae; Streptomyces; Streptomyces coelicoflavus
MPRTAAFFDLDKTVIAKSSTLTFSKSFYQGGLINRRAVLRTAYAQFVFLAGGADHDQMER WP_042826664.1
inhibition of morphological differentiation protein [Streptomyces coelicoflavus] Length: 268\nNumber of predicted TMHs: 1\nExp number of AAs in TMHs: 21.70608\nExp number, first 60 AAs: 0.03423\nTotal prob of N-in: 0.00318\noutside 1 238\nTMhelix 239 261\ninside 262 268

33917 GCF_000931445.1_ASM93144v1 Streptomyces cyaneogriseus subsp. noncyanogenus Terrabacteria group; Actinobacteria; Actinobacteria; Streptomycetales; Streptomycetaceae; Streptomyces; Streptomyces cyaneogriseus
MAVLSFLSRPASRRGPIRPARVLAVAAASAALAAGAAGTALACDISEFTAAAKCDGGKGV WP_044384405.1
hypothetical protein [Streptomyces cyaneogriseus] Length: 242\nNumber of predicted TMHs: 1\nExp number of AAs in TMHs: 37.47191\nExp number, first 60 AAs: 15.22936\nTotal prob of N-in: 0.67851\nPOSSIBLE N-term signal sequence\noutside 1 210\nTMhelix 211 233\ninside 234 242

33918 GCF_000931445.1_ASM93144v1 Streptomyces cyaneogriseus subsp. noncyanogenus Terrabacteria group; Actinobacteria; Actinobacteria; Streptomycetales; Streptomycetaceae; Streptomyces; Streptomyces cyaneogriseus
MLGVVENHSLPRTAAFFDLDKTVIAKSSTLTFSKSFYQGGLINRRAVLRTAYAQFVFLAG WP_044382412.1 inhibition of morphological differentiation protein [Streptomyces cyaneogriseus] Length: 277\nNumber of predicted TMHs: 1\nExp number of AAs in TMHs: 21.68981\nExp number, first 60 AAs: 0.03348\nTotal prob of N-in: 0.00488\noutside 1 247\nTMhelix 248 270\ninside 271 277

33919 GCF_000349325.1_ASM34932v1 Streptomyces davawensis JCM 4913 Terrabacteria group; Actinobacteria; Actinobacteria; Streptomycetales; Streptomycetaceae; Streptomyces; Streptomyces davawensis
MSKRRRAIPVVAAALLLGVAASPARAAEVSYATECVPPSISGLPPVHGTTKVEITAPETA WP_015659944.1 hypothetical protein [Streptomyces davawensis] Length: 414\nNumber of predicted TMHs: 1\nExp number of AAs in TMHs: 32.21683\nExp number, first 60 AAs: 11.97423\nTotal prob of N-in: 0.63649\nPOSSIBLE N-term signal sequence\noutside 1 388\nTMhelix 389 408\ninside 409 414

33920 GCF_000349325.1_ASM34932v1 Streptomyces davawensis JCM 4913 Terrabacteria group; Actinobacteria; Actinobacteria; Streptomycetales; Streptomycetaceae; Streptomyces; Streptomyces davawensis
MRRNTRRRPTGPRRATFAAIALILGGSGLVATNIYASATEGDGTSAQDGGATIDCPDVGA WP_015655538.1
secreted protein [Streptomyces davawensis] Length: 626\nNumber of predicted TMHs: 2\nExp number of

AAs in TMHs: 39.75959\nExp number, first 60 AAs: 21.36683\nTotal prob of N-in: 0.98308\nPOSSIBLE N-term signal sequence\ninside 1 16\nTMhelix 17 39\noutside 40 601\nTMhelix 602 619\ninside 620 626

33921 GCF_000349325.1_ASM34932v1 Streptomyces davawensis JCM 4913 Terrabacteria group; Actinobacteria; Actinobacteria; Streptomycetales; Streptomycetaceae; Streptomyces; Streptomyces davawensis MRHARRRIVRRVTRLAAVGGLLLGSAMVTRAVASEPSDTAPGASVRSAVDTGAGLVERLG WP_015655752.1 S1 family peptidase [Streptomyces davawensis] Length: 460\nNumber of predicted TMHs: 2\nExp number of AAs in TMHs: 40.25728\nExp number, first 60 AAs: 18.95942\nTotal prob of N-in: 0.87961\nPOSSIBLE N-term signal sequence\ninside 1 11\nTMhelix 12 34\noutside 35 421\nTMhelix 422 444\ninside 445 460

33922 GCF_000518045.1_StrExf1.0 Streptomyces exfoliatus DSM 41693 Terrabacteria group; Actinobacteria; Actinobacteria; Streptomycetales; Streptomycetaceae; Streptomyces; Streptomyces exfoliatus MYPRPRRRRVLAAVSALMLGALLSPAATASASPAAPGAPEPARIDLVLDLSGSMNKNDAG WP_078626837.1 hypothetical protein [Streptomyces exfoliatus] Length: 634\nNumber of predicted TMHs: 1\nExp number of AAs in TMHs: 37.43801\nExp number, first 60 AAs: 15.28626\nTotal prob of N-in: 0.74523\nPOSSIBLE N-term signal sequence\noutside 1 606\nTMhelix 607 629\ninside 630 634

33923 GCF_000518045.1_StrExf1.0 Streptomyces exfoliatus DSM 41693 Terrabacteria group; Actinobacteria; Actinobacteria; Streptomycetales; Streptomycetaceae; Streptomyces; Streptomyces exfoliatus MLRLVENHSLPRTAAFFDLDKTVIAKSSTLTFSKSFYRGGLISRRAALRTAYIQFVFLVG WP_024760195.1 inhibition of morphological differentiation protein [Streptomyces exfoliatus] Length: 288\nNumber of predicted TMHs: 1\nExp number of AAs in TMHs: 22.1127\nExp number, first 60 AAs: 0.08459\nTotal prob of N-in: 0.00642\noutside 1 247\nTMhelix 248 270\ninside 271 288

33924 GCF_000156695.2_ASM15669v2 Streptomyces roseosporus NRRL 11379Terrabacteria group; Actinobacteria; Actinobacteria; Streptomycetales; Streptomycetaceae; Streptomyces; Streptomyces filamentosus MGLSWNDRRLERRVKPGDGKPLKPFRWWQMTRRSLLSIALPVGGGGGLSATYTVEVKHGG WP_023608351.1 MULTISPECIES: hypothetical protein [Streptomyces] Length: 220\nNumber of predicted TMHs: 2\nExp number of AAs in TMHs: 41.83877\nExp number, first 60 AAs: 2.68896\nTotal prob of N-in: 0.92352\ninside 1 137\nTMhelix 138 160\noutside 161 184\nTMhelix 185 202\ninside 203 220

33925 GCF_000156695.2_ASM15669v2 Streptomyces roseosporus NRRL 11379Terrabacteria group; Actinobacteria; Actinobacteria; Streptomycetales; Streptomycetaceae; Streptomyces; Streptomyces filamentosus MSPPTSRRRRALPVLTAALAALLYGLAPPAAADDGPADWTARPAGTGTGRDARPYVYLEG WP_010070991.1 hypothetical protein [Streptomyces filamentosus] Length: 308\nNumber of predicted TMHs: 1\nExp number of AAs in TMHs: 33.67054\nExp number, first 60 AAs: 11.54547\nTotal prob of N-in: 0.58310\nPOSSIBLE N-term signal sequence\noutside 1 251\nTMhelix 252 274\ninside 275 308

33926 GCF_000156695.2_ASM15669v2 Streptomyces roseosporus NRRL 11379Terrabacteria group; Actinobacteria; Actinobacteria; Streptomycetales; Streptomycetaceae; Streptomyces; Streptomyces filamentosus MSGTVSSPRRRAVASGAPAAGAPVPSAPRIPRPIRLAAHAAALTPVPSGLWRIAIALGWD WP_006124616.1 hypothetical protein [Streptomyces filamentosus] Length: 194\nNumber of predicted TMHs: 2\nExp number of AAs in TMHs: 76.26624\nExp number, first 60 AAs: 12.45245\nTotal prob of N-in: 0.74830\nPOSSIBLE N-term signal sequence\ninside 1 122\nTMhelix 123 145\noutside 146 159\nTMhelix 160 182\ninside 183 194

33927 GCF_000156695.2_ASM15669v2 Streptomyces roseosporus NRRL 11379Terrabacteria group; Actinobacteria; Actinobacteria; Streptomycetales; Streptomycetaceae; Streptomyces; Streptomyces filamentosus MLSVVENCFSPRTAAFFDLDKTVIAKSSTLTFSKSFYQGGLINRRAVLRTAYTQFVFLAG WP_006126413.1 MULTISPECIES: morphological differentiation protein inhibitor [Streptomyces] Length: 281\nNumber of predicted TMHs: 1\nExp number of AAs in TMHs: 21.46267\nExp number, first 60 AAs: 0.01959\nTotal prob of N-in: 0.00583\noutside 1 247\nTMhelix 248 270\ninside 271 281

33928 GCF_000156455.1_ASM15645v1 Streptomyces roseosporus NRRL 15998Terrabacteria group; Actinobacteria; Actinobacteria; Streptomycetales; Streptomycetaceae; Streptomyces; Streptomyces filamentosus MGLSWNDRRLERRVKPGDGKPLKPFRWWQMTRRSLLSIALPVGGGGGLSATYTVEVKHGG WP_023608351.1 MULTISPECIES: hypothetical protein [Streptomyces] Length: 220\nNumber of predicted TMHs: 2\nExp number of AAs in TMHs: 41.83877\nExp number, first 60 AAs: 2.68896\nTotal prob of N-in: 0.92352\ninside 1 137\nTMhelix 138 160\noutside 161 184\nTMhelix 185 202\ninside 203 220

33929 GCF_000156455.1_ASM15645v1 Streptomyces roseosporus NRRL 15998Terrabacteria group; Actinobacteria; Actinobacteria; Streptomycetales; Streptomycetaceae; Streptomyces; Streptomyces filamentosus

MSPPTSRRRRALPVLTAALAALLYGLAPPAAADDGPADWTARPAGTGTGRDARPYVYLEG WP_010070991.1 hypothetical protein [Streptomyces filamentosus] Length: 308\nNumber of predicted TMHs: 1\nExp number of AAs in TMHs: 33.67054\nExp number, first 60 AAs: 11.54547\nTotal prob of N-in: 0.58310\nPOSSIBLE N-term signal sequence\noutside 1 251\nTMhelix 252 274\ninside 275 308

33930 GCF_000156455.1_ASM15645v1 Streptomyces roseosporus NRRL 15998Terrabacteria group; Actinobacteria; Actinobacteria; Streptomycetales; Streptomycetaceae; Streptomyces; Streptomyces filamentosus MLSVVENCFSPRTAAFFDLDKTVIAKSSTLTFSKSFYQGGLINRRAVLRTAYTQFVFLAG WP_006126413.1 MULTISPECIES: morphological differentiation protein inhibitor [Streptomyces] Length: 281\nNumber of predicted TMHs: 1\nExp number of AAs in TMHs: 21.46267\nExp number, first 60 AAs: 0.01959\nTotal prob of N-in: 0.00583\noutside 1 247\nTMhelix 248 270\ninside 271 281

33931 GCF_000156455.1_ASM15645v1 Streptomyces roseosporus NRRL 15998Terrabacteria group; Actinobacteria; Actinobacteria; Streptomycetales; Streptomycetaceae; Streptomyces; Streptomyces filamentosus MSGTVSSPRRRAVASGAPAAGAPVPSAPRIPRPIRLAAHAAALTPVPSGLWRIAIALGWD WP_006124616.1 hypothetical protein [Streptomyces filamentosus] Length: 194\nNumber of predicted TMHs: 2\nExp number of AAs in TMHs: 76.26624\nExp number, first 60 AAs: 12.45245\nTotal prob of N-in: 0.74830\nPOSSIBLE N-term signal sequence\ninside 1 122\nTMhelix 123 145\noutside 146 159\nTMhelix 160 182\ninside 183 194

33932 GCF_000385945.1_ASM38594v1 Streptomyces fulvissimus DSM 40593 Terrabacteria group; Actinobacteria; Actinobacteria; Streptomycetales; Streptomycetaceae; Streptomyces; Streptomyces fulvissimus MLSVVENCFSPRTAAFFDLDKTVIAKSSTLTFSKSFYQGGLINRRAVLRTAYTQFVFMAG WP_015609440.1 morphological differentiation protein inhibitor [Streptomyces fulvissimus]Length: 281\nNumber of predicted TMHs: 1\nExp number of AAs in TMHs: 21.79852\nExp number, first 60 AAs: 0.01652\nTotal prob of N-in: 0.00400\noutside 1 247\nTMhelix 248 270\ninside 271 281

33933 GCF_000385945.1_ASM38594v1 Streptomyces fulvissimus DSM 40593 Terrabacteria group; Actinobacteria; Actinobacteria; Streptomycetales; Streptomycetaceae; Streptomyces; Streptomyces fulvissimus MGNAVSGRRTLLSGTAVLAAVALIALGSAPAQADETEPDLGVRGLTPVTGIAPGSGFGLP WP_015611441.1 LPXTG-motif cell wall anchor domain protein [Streptomyces fulvissimus] Length: 434\nNumber of predicted TMHs: 1\nExp number of AAs in TMHs: 35.23171\nExp number, first 60 AAs: 14.6668\nTotal prob of N-in: 0.61603\nPOSSIBLE N-term signal sequence\noutside 1 402\nTMhelix 403 425\ninside 426 434

33934 GCF_000342345.1_Whole_genome_assembly Streptomyces gancidicus BKS 13-15 Terrabacteria group; Actinobacteria; Actinobacteria; Streptomycetales; Streptomycetaceae; Streptomyces; Streptomyces gancidicus MLRNVENHLLPRTAAFFDLDKTVIAKSSTLTFSKSFYQGGLINRRAVLRTAYSQFVFLAG WP_031020047.1 MULTISPECIES: inhibition of morphological differentiation protein [Streptomyces] Length: 277\nNumber of predicted TMHs: 1\nExp number of AAs in TMHs: 21.81477\nExp number, first 60 AAs: 0.02315\nTotal prob of N-in: 0.00436\noutside 1 247\nTMhelix 248 270\ninside 271 277

33935 GCF_000156435.1_ASM15643v1 Streptomyces ghanaensis ATCC 14672 Terrabacteria group; Actinobacteria; Actinobacteria; Streptomycetales; Streptomycetaceae; Streptomyces; Streptomyces ghanaensis MMVVCCFGLVGYGFGSVADGLAGRKERRPAVLRGTAALAGACALAVYAWGLLGVAGAWME WP_086015626.1 hypothetical protein [Streptomyces ghanaensis] Length: 151\nNumber of predicted TMHs: 2\nExp number of AAs in TMHs: 49.28399\nExp number, first 60 AAs: 27.19793\nTotal prob of N-in: 0.77805\nPOSSIBLE N-term signal sequence\ninside 1 36\nTMhelix 37 59\noutside 60 118\nTMhelix 119 141\ninside 142 151

33936 GCF_000156435.1_ASM15643v1 Streptomyces ghanaensis ATCC 14672 Terrabacteria group; Actinobacteria; Actinobacteria; Streptomycetales; Streptomycetaceae; Streptomyces; Streptomyces ghanaensis MRHARRRVVRRVTRLAAVGGLLLGATMVTRAVASEPAPPNAVPHTAAGQSAGPGTDLASR WP_039830567.1 MULTISPECIES: S1 family peptidase [Streptomyces] Length: 457\nNumber of predicted TMHs: 2\nExp number of AAs in TMHs: 41.29905\nExp number, first 60 AAs: 20.23659\nTotal prob of N-in: 0.94066\nPOSSIBLE N-term signal sequence\ninside 1 11\nTMhelix 12 34\noutside 35 418\nTMhelix 419 441\ninside 442 457

33937 GCF_000156435.1_ASM15643v1 Streptomyces ghanaensis ATCC 14672 Terrabacteria group; Actinobacteria; Actinobacteria; Streptomycetales; Streptomycetaceae; Streptomyces; Streptomyces ghanaensis MLKGVENHSSPRTAAFFDLDKTVIAKSSTLTFSKSFYQGGLINRRAVLRTAYAQFVFLAG WP_004985180.1 MULTISPECIES: morphological differentiation-associated protein [Streptomyces] Length: 277\nNumber of predicted TMHs: 1\nExp number of AAs in TMHs: 21.98535\nExp number, first 60 AAs: 0.08708\nTotal prob of N-in: 0.00829\noutside 1 247\nTMhelix 248 270\ninside 271 277

33938 GCF_000204605.1_ASM20460v1 Streptomyces griseoaurantiacus M045 Terrabacteria group; Actinobacteria; Actinobacteria; Streptomycetales; Streptomycetaceae; Streptomyces; Streptomyces griseoaurantiacus MLGFVEKHPSPRTAAFFDLDKTVIAKSSTLTFSKSFYQGGLINRRAVLRTAYAQFVFLAG WP_006142037.1 morphological differentiation-associated protein [Streptomyces griseoaurantiacus] Length: 277\nNumber of predicted TMHs: 1\nExp number of AAs in TMHs: 21.65846\nExp number, first 60 AAs: 0.01915\nTotal prob of N-in: 0.00156\noutside 1 247\nTMhelix 248 270\ninside 271 277

33939 GCF_000158975.1_ASM15897v1 Streptomyces griseoflavus Tu4000 Terrabacteria group; Actinobacteria; Actinobacteria; Streptomycetales; Streptomycetaceae; Streptomyces; Streptomyces griseoflavus MLSTVENHSMPRTAAFFDLDKTVIAKSSTLTFSKSFYQGGLINRRAVLRTAYSQFVFLAG WP_004928262.1 haloacid dehalogenase [Streptomyces griseoflavus] Length: 277\nNumber of predicted TMHs: 1\nExp number of AAs in TMHs: 21.80519\nExp number, first 60 AAs: 0.03597\nTotal prob of N-in: 0.00690\noutside 1 247\nTMhelix 248 270\ninside 271 277

33940 GCF_000718135.1_ASM71813v1 Streptomyces cyaneofuscatus Terrabacteria group; Actinobacteria; Actinobacteria; Streptomycetales; Streptomycetaceae; Streptomyces; Streptomyces griseus group MGIAASGRRTLLSATAVSATAALIALGAAPAQADAIKPDLGVRALAPVTGIAAGSGFGLP WP_030568954.1 peptidase [Streptomyces cyaneofuscatus] Length: 446\nNumber of predicted TMHs: 1\nExp number of AAs in TMHs: 37.10562\nExp number, first 60 AAs: 16.77727\nTotal prob of N-in: 0.43256\nPOSSIBLE N-term signal sequence\noutside 1 414\nTMhelix 415 437\ninside 438 446

33941 GCF_000718135.1_ASM71813v1 Streptomyces cyaneofuscatus Terrabacteria group; Actinobacteria; Actinobacteria; Streptomycetales; Streptomycetaceae; Streptomyces; Streptomyces griseus group MLSLVENCFSPRTAAFFDLDKTVIAKSSTLTFSKSFYQGGLINRRAVLRTAYTQFVFLAG WP_030565022.1 inhibition of morphological differentiation protein [Streptomyces cyaneofuscatus] Length: 281\nNumber of predicted TMHs: 1\nExp number of AAs in TMHs: 21.44434\nExp number, first 60 AAs: 0.01381\nTotal prob of N-in: 0.00446\noutside 1 247\nTMhelix 248 270\ninside 271 281

33942 GCF_002154385.1_ASM215438v1 Streptomyces albovinaceus Terrabacteria group; Actinobacteria; Actinobacteria; Streptomycetales; Streptomycetaceae; Streptomyces; Streptomyces griseus group; Streptomyces albovinaceus subgroup MGLSWNDRRIERRVKPGDGKPLKPFRWWQMTRRSLLSIALPVGGGPSATYTVEVKHGGDP WP_086673651.1 hypothetical protein [Streptomyces albovinaceus] Length: 218\nNumber of predicted TMHs: 2\nExp number of AAs in TMHs: 39.13586\nExp number, first 60 AAs: 0.17243\nTotal prob of N-in: 0.96472\ninside 1 135\nTMhelix 136 158\noutside 159 182\nTMhelix 183 200\ninside 201 218

33943 GCF_002154385.1_ASM215438v1 Streptomyces albovinaceus Terrabacteria group; Actinobacteria; Actinobacteria; Streptomycetales; Streptomycetaceae; Streptomyces; Streptomyces griseus group; Streptomyces albovinaceus subgroup MSPPTPRRRRALPVLTAALAALLCGLAPPAAADGPPDWAARPTGGTDRDARPYVYLEGTP WP_030803089.1 hypothetical protein [Streptomyces mediolani] Length: 322\nNumber of predicted TMHs: 1\nExp number of AAs in TMHs: 32.99085\nExp number, first 60 AAs: 11.21825\nTotal prob of N-in: 0.56076\nPOSSIBLE N-term signal sequence\noutside 1 249\nTMhelix 250 272\ninside 273 322

33944 GCF_000721685.1_ASM72168v1 Streptomyces mediolani Terrabacteria group; Actinobacteria; Actinobacteria; Streptomycetales; Streptomycetaceae; Streptomyces; Streptomyces griseus group; Streptomyces albovinaceus subgroup MGLSWNDRRLERRVKPGDGKPLKPFRWWQMTRRSLLSIALPVGGGPSATYTVEVKHGGDP WP_030812892.1 hypothetical protein [Streptomyces mediolani] Length: 218\nNumber of predicted TMHs: 2\nExp number of AAs in TMHs: 39.98254\nExp number, first 60 AAs: 0.09302\nTotal prob of N-in: 0.97329\ninside 1 135\nTMhelix 136 158\noutside 159 182\nTMhelix 183 200\ninside 201 218

33945 GCF_000721685.1_ASM72168v1 Streptomyces mediolani Terrabacteria group; Actinobacteria; Actinobacteria; Streptomycetales; Streptomycetaceae; Streptomyces; Streptomyces griseus group; Streptomyces albovinaceus subgroup MSPPTPRRRRALPVLTAALAALLCGLAPPAAADGPPDWAARPTGGTDRDARPYVYLEGTP WP_030803089.1 hypothetical protein [Streptomyces mediolani] Length: 322\nNumber of predicted TMHs: 1\nExp number of AAs in TMHs: 32.99085\nExp number, first 60 AAs: 11.21825\nTotal prob of N-in: 0.56076\nPOSSIBLE N-term signal sequence\noutside 1 249\nTMhelix 250 272\ninside 273 322

33946 GCF_000721685.1_ASM72168v1 Streptomyces mediolani Terrabacteria group; Actinobacteria; Actinobacteria; Streptomycetales; Streptomycetaceae; Streptomyces; Streptomyces griseus group; Streptomyces albovinaceus subgroup MLSVVENCFSPRTAAFFDLDKTVIAKSSTLTFSKSFYQGGLINRRAVLRTAYTQFVFLAG WP_030813057.1 MULTISPECIES: inhibition of morphological differentiation protein [Streptomyces]

Length: 281\nNumber of predicted TMHs: 1\nExp number of AAs in TMHs: 21.46267\nExp number, first 60 AAs: 0.01959\nTotal prob of N-in: 0.00583\noutside 1 247\nTMhelix 248 270\ninside 271 281

33947 GCF_000261345.2_ASM26134v2 Streptomyces globisporus C-1027 Terrabacteria group; Actinobacteria; Actinobacteria; Streptomycetales; Streptomycetaceae; Streptomyces; Streptomyces griseus group; Streptomyces albovinaceus subgroup; Streptomyces globisporus MSPPTPRRRALPVLTAALAALLCGLVQPAAADDPPPDWTARPAGGTDRDARPYVYLEGTP WP_010059884.1 MULTISPECIES: hypothetical protein [Streptomyces] Length: 322\nNumber of predicted TMHs: 1\nExp number of AAs in TMHs: 28.26039\nExp number, first 60 AAs: 6.03507\nTotal prob of N-in: 0.31029\noutside 1 249\nTMhelix 250 272\ninside 273 322

33948 GCF_000261345.2_ASM26134v2 Streptomyces globisporus C-1027 Terrabacteria group; Actinobacteria; Actinobacteria; Streptomycetales; Streptomycetaceae; Streptomyces; Streptomyces griseus group; Streptomyces albovinaceus subgroup; Streptomyces globisporus MLSVVENCFSPRTAAFFDLDKTVIAKSSTLTFSKSFYQGGLINRRAVLRTAYTQFVFLAG WP_010056081.1 morphological differentiation-associated protein [Streptomyces globisporus] Length: 281\nNumber of predicted TMHs: 1\nExp number of AAs in TMHs: 21.80054\nExp number, first 60 AAs: 0.01819\nTotal prob of N-in: 0.00405\noutside 1 247\nTMhelix 248 270\ninside 271 281

33949 GCF_000261345.2_ASM26134v2 Streptomyces globisporus C-1027 Terrabacteria group; Actinobacteria; Actinobacteria; Streptomycetales; Streptomycetaceae; Streptomyces; Streptomyces griseus group; Streptomyces albovinaceus subgroup; Streptomyces globisporus MGLSWNDRRLERRVKPGDGKPLKPFRWWQMTRRSLLSIALPVGGGLSATYTVEVKHGGDA WP_029182569.1 hypothetical protein [Streptomyces globisporus] Length: 218\nNumber of predicted TMHs: 2\nExp number of AAs in TMHs: 41.12263\nExp number, first 60 AAs: 1.21859\nTotal prob of N-in: 0.93922\ninside 1 135\nTMhelix 136 158\noutside 159 182\nTMhelix 183 200\ninside 201 218

33950 GCF_000718455.1_ASM71845v1 Streptomyces globisporus subsp. globisporus Terrabacteria group; Actinobacteria; Actinobacteria; Streptomycetales; Streptomycetaceae; Streptomyces; Streptomyces griseus group; Streptomyces albovinaceus subgroup; Streptomyces globisporus MLSVVENCFSPRTAAFFDLDKTVIAKSSTLTFSKSFYQGGLINRRAVLRTAYTQFVFLAG WP_030590952.1 MULTISPECIES: inhibition of morphological differentiation protein [Streptomyces griseus group] Length: 281\nNumber of predicted TMHs: 1\nExp number of AAs in TMHs: 21.42564\nExp number, first 60 AAs: 0.02005\nTotal prob of N-in: 0.00656\noutside 1 247\nTMhelix 248 270\ninside 271 281

33951 GCF_000718455.1_ASM71845v1 Streptomyces globisporus subsp. globisporus Terrabacteria group; Actinobacteria; Actinobacteria; Streptomycetales; Streptomycetaceae; Streptomyces; Streptomyces griseus group; Streptomyces albovinaceus subgroup; Streptomyces globisporus MRSENRSRRSFVRARAALVAGTLGVLLAVTGCGASGDSSDSGAKADVKSAPREGFADGEA WP_030592787.1 hypothetical protein [Streptomyces globisporus] Length: 326\nNumber of predicted TMHs: 2\nExp number of AAs in TMHs: 41.41561\nExp number, first 60 AAs: 18.44381\nTotal prob of N-in: 0.85973\nPOSSIBLE N-term signal sequence\ninside 1 11\nTMhelix 12 34\noutside 35 274\nTMhelix 275 297\ninside 298 326

33952 GCF_000720875.1_ASM72087v1 Streptomyces globisporus subsp. globisporus Terrabacteria group; Actinobacteria; Actinobacteria; Streptomycetales; Streptomycetaceae; Streptomyces; Streptomyces griseus group; Streptomyces albovinaceus subgroup; Streptomyces globisporus MLGLVENHSLPRTAAFFDLDKTVIAKSSTLTFSKSFYRGGLISRRAALRTAYIQFVFLAG WP_030692543.1 inhibition of morphological differentiation protein [Streptomyces globisporus] Length: 288\nNumber of predicted TMHs: 1\nExp number of AAs in TMHs: 22.13469\nExp number, first 60 AAs: 0.02672\nTotal prob of N-in: 0.00192\noutside 1 247\nTMhelix 248 270\ninside 271 288

33953 GCF_000720875.1_ASM72087v1 Streptomyces globisporus subsp. globisporus Terrabacteria group; Actinobacteria; Actinobacteria; Streptomycetales; Streptomycetaceae; Streptomyces; Streptomyces griseus group; Streptomyces albovinaceus subgroup; Streptomyces globisporus MKPARIARRRRAFRNGLTALVCATVVTLSGQTGAGVAEAAPPKLKPGKRCDELYGVKNVP WP_030689701.1 hypothetical protein [Streptomyces globisporus] Length: 774\nNumber of predicted TMHs: 2\nExp number of AAs in TMHs: 42.83008\nExp number, first 60 AAs: 20.42349\nTotal prob of N-in: 0.91237\nPOSSIBLE N-term signal sequence\ninside 1 12\nTMhelix 13 35\noutside 36 743\nTMhelix 744 766\ninside 767 774

33954 GCF_001434355.1_ASM143435v1 Streptomyces anulatus Terrabacteria group; Actinobacteria; Actinobacteria; Streptomycetales; Streptomycetaceae; Streptomyces; Streptomyces griseus group; Streptomyces anulatus subgroup MLSVVENCFSPRTAAFFDLDKTVIAKSSTLTFSKSFYQGGLINRRAVLRTAYAQFVFLAG WP_032791512.1 MULTISPECIES: inhibition of morphological differentiation protein [Streptomyces] Length: 281\nNumber of predicted TMHs: 1\nExp number of AAs in TMHs: 21.51851\nExp number, first 60 AAs: 0.0316\nTotal prob of N-in: 0.00586\noutside 1 247\nTMhelix 248 270\ninside 271 281

33955 GCF_000717025.1_ASM71702v1 Streptomyces atroolivaceus Terrabacteria group; Actinobacteria; Actinobacteria; Streptomycetales; Streptomycetaceae; Streptomyces; Streptomyces griseus group; Streptomyces atroolivaceus subgroup MTAGMSQTRKRRTPLELRGRLLGTVAMAAAWTVGLTAAAPSAFSADVQSKQWYLEAMQAE WP_033304178.1 serine protease [Streptomyces atroolivaceus] Length: 417\nNumber of predicted TMHs: 2\nExp number of AAs in TMHs: 44.0090299999999\nExp number, first 60 AAs: 22.04121\nTotal prob of N-in: 0.98829\nPOSSIBLE N-term signal sequence\ninside 1 20\nTMhelix 21 43\noutside 44 388\nTMhelix 389 411\ninside 412 417

33956 GCF_000717025.1_ASM71702v1 Streptomyces atroolivaceus Terrabacteria group; Actinobacteria; Actinobacteria; Streptomycetales; Streptomycetaceae; Streptomyces; Streptomyces griseus group; Streptomyces atroolivaceus subgroup MLTFVENCFSPRTAAFFDLDKTVIAKSSTLTFSKSFYQGGLINRRAVLRTAYTQFVFLAG WP_033296553.1 inhibition of morphological differentiation protein [Streptomyces atroolivaceus] Length: 279\nNumber of predicted TMHs: 1\nExp number of AAs in TMHs: 21.59235\nExp number, first 60 AAs: 0.01473\nTotal prob of N-in: 0.00189\noutside 1 247\nTMhelix 248 270\ninside 271 279

33957 GCF_000717025.1_ASM71702v1 Streptomyces atroolivaceus Terrabacteria group; Actinobacteria; Actinobacteria; Streptomycetales; Streptomycetaceae; Streptomyces; Streptomyces griseus group; Streptomyces atroolivaceus subgroup MSPIGRRRWLHVSCGGSTTVNSNTFRAAALAAAAVPVALLVAVPAQAATATTPATPAGGH WP_051710137.1 hypothetical protein [Streptomyces atroolivaceus] Length: 329\nNumber of predicted TMHs: 1\nExp number of AAs in TMHs: 31.73212\nExp number, first 60 AAs: 12.30362\nTotal prob of N-in: 0.55528\nPOSSIBLE N-term signal sequence\noutside 1 300\nTMhelix 301 323\ninside 324 329

33958 GCF_000716025.1_ASM71602v1 Streptomyces baarnensis Terrabacteria group; Actinobacteria; Actinobacteria; Streptomycetales; Streptomycetaceae; Streptomyces; Streptomyces griseus group; Streptomyces fimicarius subgroup MLSVVENCFSPRTAAFFDLDKTVIAKSSTLTFSKSFYQGGLINRRAVLRTAYAQFVFLAG WP_030087051.1 inhibition of morphological differentiation protein [Streptomyces baarnensis] Length: 282\nNumber of predicted TMHs: 1\nExp number of AAs in TMHs: 21.69171\nExp number, first 60 AAs: 0.02994\nTotal prob of N-in: 0.00448\noutside 1 247\nTMhelix 248 270\ninside 271 282

33959 GCF_000716025.1_ASM71602v1 Streptomyces baarnensis Terrabacteria group; Actinobacteria; Actinobacteria; Streptomycetales; Streptomycetaceae; Streptomyces; Streptomyces griseus group; Streptomyces fimicarius subgroup MDPVTDGESGSRRSFVRARAALVAGALGLLLAVGGCGASGDASDDGAKAADGKAAPREGF WP_030079718.1 hypothetical protein [Streptomyces baarnensis] Length: 332\nNumber of predicted TMHs: 2\nExp number of AAs in TMHs: 42.13704\nExp number, first 60 AAs: 19.16768\nTotal prob of N-in: 0.77796\nPOSSIBLE N-term signal sequence\ninside 1 19\nTMhelix 20 42\noutside 43 279\nTMhelix 280 302\ninside 303 332

33960 GCF_000716025.1_ASM71602v1 Streptomyces baarnensis Terrabacteria group; Actinobacteria; Actinobacteria; Streptomycetales; Streptomycetaceae; Streptomyces; Streptomyces griseus group; Streptomyces fimicarius subgroup MRITPSRRRTLRGSTAAVAATALVALGAAPALADEPEPDLGVGTIAPIKGVQAGAAFSTP WP_030079189.1 hypothetical protein [Streptomyces baarnensis] Length: 431\nNumber of predicted TMHs: 1\nExp number of AAs in TMHs: 34.30376\nExp number, first 60 AAs: 14.19651\nTotal prob of N-in: 0.69603\nPOSSIBLE N-term signal sequence\noutside 1 399\nTMhelix 400 422\ninside 423 431

33961 GCF_002154345.1_ASM215434v1 Streptomyces fimicarius Terrabacteria group; Actinobacteria; Actinobacteria; Streptomycetales; Streptomycetaceae; Streptomyces; Streptomyces griseus group; Streptomyces fimicarius subgroup MDPVTDGESRSRRSFVRARAALVAGALGLLLAVGGCGASGDASDSGAKAADGKAAPREGF WP_086778779.1 hypothetical protein [Streptomyces fimicarius] Length: 332\nNumber of predicted TMHs: 2\nExp number of AAs in TMHs: 43.52131\nExp number, first 60 AAs: 20.52533\nTotal prob of N-in: 0.93696\nPOSSIBLE N-term signal sequence\ninside 1 19\nTMhelix 20 42\noutside 43 279\nTMhelix 280 302\ninside 303 332

33962 GCF_002154345.1_ASM215434v1 Streptomyces fimicarius Terrabacteria group; Actinobacteria; Actinobacteria; Streptomycetales; Streptomycetaceae; Streptomyces; Streptomyces griseus group; Streptomyces fimicarius subgroup MLSVVENCFSPRTAAFFDLDKTVIAKSSTLTFSKSFYQGGLINRRAVLRTAYAQFVFLAG WP_030087051.1 inhibition of morphological differentiation protein [Streptomyces baarnensis] Length: 282\nNumber of predicted TMHs: 1\nExp number of AAs in TMHs: 21.69171\nExp number, first 60 AAs: 0.02994\nTotal prob of N-in: 0.00448\noutside 1 247\nTMhelix 248 270\ninside 271 282

33963 GCF_002154345.1_ASM215434v1 Streptomyces fimicarius Terrabacteria group; Actinobacteria; Actinobacteria; Streptomycetales; Streptomycetaceae; Streptomyces; Streptomyces griseus group; Streptomyces fimicarius subgroup MRITPSRRRTLRGSTAAVAATALVALGAAPALADEPEPDLGVGTIAPIKGVQAGSAFSTP WP_086778847.1 peptidase [Streptomyces fimicarius] Length: 431\nNumber of predicted TMHs: 1\nExp number of AAs in TMHs: 34.90588\nExp number, first 60 AAs: 14.18986\nTotal prob of N-in: 0.69576\nPOSSIBLE N-term signal sequence\noutside 1 399\nTMhelix 400 422\ninside 423 431

33964 GCF_000720395.1_ASM72039v1 Streptomyces flavovirens Terrabacteria group; Actinobacteria; Actinobacteria; Streptomycetales; Streptomycetaceae; Streptomyces; Streptomyces griseus group; Streptomyces flavovirens subgroup MRTTRRNTLRRGGVGALLAAVTLLTSAGAAVADDPSPSASADGGTAGPTEAGTTFRTATA WP_030637428.1 hypothetical protein [Streptomyces flavovirens] Length: 274\nNumber of predicted TMHs: 2\nExp number of AAs in TMHs: 40.1726399999999\nExp number, first 60 AAs: 19.30599\nTotal prob of N-in: 0.99359\nPOSSIBLE N-term signal sequence\ninside 1 12\nTMhelix 13 32\noutside 33 237\nTMhelix 238 260\ninside 261 274

33965 GCF_000720395.1_ASM72039v1 Streptomyces flavovirens Terrabacteria group; Actinobacteria; Actinobacteria; Streptomycetales; Streptomycetaceae; Streptomyces; Streptomyces griseus group; Streptomyces flavovirens subgroup MTTHPSPRRRLLTAAAATAVLLVTLPVLPAAAEDSTQCTFPSKKYAGRPWSLQRVLMDEL WP_014153729.1 MULTISPECIES: type VII secretion-associated serine protease [Streptomyces] Length: 414\nNumber of predicted TMHs: 1\nExp number of AAs in TMHs: 36.07361\nExp number, first 60 AAs: 13.5063\nTotal prob of N-in: 0.65550\nPOSSIBLE N-term signal sequence\noutside 1 378\nTMhelix 379 401\ninside 402 414

33966 GCF_000720395.1_ASM72039v1 Streptomyces flavovirens Terrabacteria group; Actinobacteria; Actinobacteria; Streptomycetales; Streptomycetaceae; Streptomyces; Streptomyces griseus group; Streptomyces flavovirens subgroup MLTFVENCFSPRTAAFFDLDKTVIAKSSTLTFSKSFYQGGLINRRAVLRTAYTQFVFLAG WP_014155461.1 MULTISPECIES: haloacid dehalogenase [Streptomyces] Length: 279\nNumber of predicted TMHs: 1\nExp number of AAs in TMHs: 21.10966\nExp number, first 60 AAs: 0.01497\nTotal prob of N-in: 0.00234\noutside 1 248\nTMhelix 249 268\ninside 269 279

33967 GCF_001715295.1_ASM171529v1 Streptomyces griseus Terrabacteria group; Actinobacteria; Actinobacteria; Streptomycetales; Streptomycetaceae; Streptomyces; Streptomyces griseus group; Streptomyces griseus subgroup MLTFVENCFSPRTAAFFDLDKTVIAKSSTLTFSKSFYQGGLINRRAVLRTAYVQFVFLAG WP_069171894.1 inhibition of morphological differentiation protein [Streptomyces griseus] Length: 279\nNumber of predicted TMHs: 1\nExp number of AAs in TMHs: 21.66536\nExp number, first 60 AAs: 0.05636\nTotal prob of N-in: 0.00555\noutside 1 247\nTMhelix 248 270\ninside 271 279

33968 GCF_000932225.1_ASM93222v1 Streptomyces griseus Terrabacteria group; Actinobacteria; Actinobacteria; Streptomycetales; Streptomycetaceae; Streptomyces; Streptomyces griseus group; Streptomyces griseus subgroup MLSVVENCFSPRTAAFFDLDKTVIAKSSTLTFSKSFYQGGLINRRAVLRTAYTQFVFLAG WP_030813057.1 MULTISPECIES: inhibition of morphological differentiation protein [Streptomyces] Length: 281\nNumber of predicted TMHs: 1\nExp number of AAs in TMHs: 21.46267\nExp number, first 60 AAs: 0.01959\nTotal prob of N-in: 0.00583\noutside 1 247\nTMhelix 248 270\ninside 271 281

33969 GCF_000932225.1_ASM93222v1 Streptomyces griseus Terrabacteria group; Actinobacteria; Actinobacteria; Streptomycetales; Streptomycetaceae; Streptomyces; Streptomyces griseus group; Streptomyces griseus subgroup MGLSWNDRRIERRVKPGDGKPLKPFRWWQMTRRSLLSIALPVGGGLSATYTVEVKHGGDP WP_044374596.1 hypothetical protein [Streptomyces griseus] Length: 218\nNumber of predicted TMHs: 2\nExp number of AAs in TMHs: 39.66996\nExp number, first 60 AAs: 1.35906\nTotal prob of N-in: 0.95905\ninside 1 135\nTMhelix 136 158\noutside 159 182\nTMhelix 183 200\ninside 201 218

33970 GCF_001723115.1_ASM172311v1 Streptomyces griseus Terrabacteria group; Actinobacteria; Actinobacteria; Streptomycetales; Streptomycetaceae; Streptomyces; Streptomyces griseus group; Streptomyces griseus subgroup MGTEQQARRGPRARRGLLSALTASLLTLGAALTHLTTASPASADPIAKCTRTKGAIVAVD WP_012378693.1 peptidase [Streptomyces griseus] Length: 931\nNumber of predicted TMHs: 2\nExp number of AAs in TMHs: 39.57519\nExp number, first 60 AAs: 18.69314\nTotal prob of N-in: 0.84908\nPOSSIBLE N-term signal sequence\ninside 1 15\nTMhelix 16 38\noutside 39 893\nTMhelix 894 916\ninside 917 931

33971 GCF_001723115.1_ASM172311v1 Streptomyces griseus Terrabacteria group; Actinobacteria; Actinobacteria; Streptomycetales; Streptomycetaceae; Streptomyces; Streptomyces griseus group; Streptomyces griseus subgroup MLSVVENCFSPRTAAFFDLDKTVIAKSSTLTFSKSFYQGGLINRRAVLRTAYAQFVFLAG WP_012379811.1 morphological differentiation-associated protein [Streptomyces griseus] Length: 281\nNumber of predicted TMHs: 1\nExp number of

AAs in TMHs: 21.467\nExp number, first 60 AAs: 0.03606\nTotal prob of N-in: 0.00978\noutside 1 247\nTMhelix 248 270\ninside 271 281

33972 GCF_900105705.1_IMG-taxon_2634166340_annotated_assembly Streptomyces griseus Terrabacteria group; Actinobacteria; Actinobacteria; Streptomycetales; Streptomycetaceae; Streptomyces; Streptomyces griseus group; Streptomyces griseus subgroup MLSVVENCFSPRTAAFFDLDKTVIAKSSTLTFSKSFYQGGLINRRAVLRTAYAQFVFLAG WP_012379811.1 morphological differentiation-associated protein [Streptomyces griseus] Length: 281\nNumber of predicted TMHs: 1\nExp number of AAs in TMHs: 21.467\nExp number, first 60 AAs: 0.03606\nTotal prob of N-in: 0.00978\noutside 1 247\nTMhelix 248 270\ninside 271 281

33973 GCF_900105705.1_IMG-taxon_2634166340_annotated_assembly Streptomyces griseus Terrabacteria group; Actinobacteria; Actinobacteria; Streptomycetales; Streptomycetaceae; Streptomyces; Streptomyces griseus group; Streptomyces griseus subgroup MGTEQQARRGPRARRGLLSALTASLLTLGAALTHLTTASPASADPIAKCTRTKGAIVAVD WP_012378693.1 peptidase [Streptomyces griseus] Length: 931\nNumber of predicted TMHs: 2\nExp number of AAs in TMHs: 39.57519\nExp number, first 60 AAs: 18.69314\nTotal prob of N-in: 0.84908\nPOSSIBLE N-term signal sequence\ninside 1 15\nTMhelix 16 38\noutside 39 893\nTMhelix 894 916\ninside 917 931

33974 GCF_001715295.1_ASM171529v1 Streptomyces griseus Terrabacteria group; Actinobacteria; Actinobacteria; Streptomycetales; Streptomycetaceae; Streptomyces; Streptomyces griseus group; Streptomyces griseus subgroup MHNDRHTDSRPTDRPGRRRRRAVLTAGLLALLLSTAACGAGDGGGSTDRSVAEDRKAAAP WP_069169392.1 hypothetical protein [Streptomyces griseus] Length: 347\nNumber of predicted TMHs: 2\nExp number of AAs in TMHs: 43.51586\nExp number, first 60 AAs: 20.50112\nTotal prob of N-in: 0.96490\nPOSSIBLE N-term signal sequence\ninside 1 20\nTMhelix 21 43\noutside 44 279\nTMhelix 280 302\ninside 303 347

33975 GCF_000719495.1_ASM71949v1 Streptomyces griseus subsp. griseus Terrabacteria group; Actinobacteria; Actinobacteria; Streptomycetales; Streptomycetaceae; Streptomyces; Streptomyces griseus group; Streptomyces griseus subgroup; Streptomyces griseus MTPNRRPVVLAAAVATALGLGATALTLPAVAAEAPRAGAPVMDLTDGTLNWGVKESFRRY WP_030699100.1 membrane protein [Streptomyces griseus] Length: 519\nNumber of predicted TMHs: 2\nExp number of AAs in TMHs: 42.17467\nExp number, first 60 AAs: 22.03072\nTotal prob of N-in: 0.97139\nPOSSIBLE N-term signal sequence\ninside 1 6\nTMhelix 7 29\noutside 30 484\nTMhelix 485 507\ninside 508 519

33976 GCF_000721035.1_ASM72103v1 Streptomyces griseus subsp. griseus Terrabacteria group; Actinobacteria; Actinobacteria; Streptomycetales; Streptomycetaceae; Streptomyces; Streptomyces griseus group; Streptomyces griseus subgroup; Streptomyces griseus MLSVVENHSLPRAAAFFDLDKTVIAKSSTLTFSKSFYRGGLISRRAALRTAYIQFVFLVG WP_030749261.1 inhibition of morphological differentiation protein [Streptomyces griseus] Length: 284\nNumber of predicted TMHs: 1\nExp number of AAs in TMHs: 21.36258\nExp number, first 60 AAs: 0.07194\nTotal prob of N-in: 0.00449\noutside 1 247\nTMhelix 248 270\ninside 271 284

33977 GCF_000721035.1_ASM72103v1 Streptomyces griseus subsp. griseus Terrabacteria group; Actinobacteria; Actinobacteria; Streptomycetales; Streptomycetaceae; Streptomyces; Streptomyces griseus group; Streptomyces griseus subgroup; Streptomyces griseus MSKKPNRRRTAVVLTAAVAAWGPAGLAAAPAARADVVDVSYACRTKIGDRSAVSPVDVTA WP_030754832.1 hypothetical protein [Streptomyces griseus] Length: 239\nNumber of predicted TMHs: 2\nExp number of AAs in TMHs: 42.77698\nExp number, first 60 AAs: 15.49111\nTotal prob of N-in: 0.90685\nPOSSIBLE N-term signal sequence\ninside 1 11\nTMhelix 12 34\noutside 35 202\nTMhelix 203 225\ninside 226 239

33978 GCF_000719495.1_ASM71949v1 Streptomyces griseus subsp. griseus Terrabacteria group; Actinobacteria; Actinobacteria; Streptomycetales; Streptomycetaceae; Streptomyces; Streptomyces griseus group; Streptomyces griseus subgroup; Streptomyces griseus MAPPHRRRAAVAVRAFVALTALLCATGLAPAPATAPVAVAEVPAPTPTTSVVTVRTGGDR WP_030696185.1 MULTISPECIES: VWA domain-containing protein [Streptomyces] Length: 823\nNumber of predicted TMHs: 2\nExp number of AAs in TMHs: 39.18881\nExp number, first 60 AAs: 18.98212\nTotal prob of N-in: 0.86306\nPOSSIBLE N-term signal sequence\ninside 1 12\nTMhelix 13 35\noutside 36 784\nTMhelix 785 807\ninside 808 823

33979 GCF_000719355.1_ASM71935v1 Streptomyces griseus subsp. griseus Terrabacteria group; Actinobacteria; Actinobacteria; Streptomycetales; Streptomycetaceae; Streptomyces; Streptomyces griseus group; Streptomyces griseus subgroup; Streptomyces griseus MSPPTPRRRRALPVLTAALAALLCGLAPPAAADDGPPDWTARPAGGTDRDARPYVYLEGT WP_030710305.1 hypothetical protein [Streptomyces griseus] Length: 323\nNumber of predicted TMHs: 1\nExp number of

AAs in TMHs: 30.27003\nExp number, first 60 AAs:  9.06873\nTotal prob of N-in:   0.48624\noutside   1   250\nTMhelix   251   273\ninside   274   323

33980 GCF_000719355.1_ASM71935v1 Streptomyces griseus subsp. griseus Terrabacteria group; Actinobacteria; Actinobacteria; Streptomycetales; Streptomycetaceae; Streptomyces; Streptomyces griseus group; Streptomyces griseus subgroup; Streptomyces griseus
MGLSWNDRRLERRVKPGDGKPLKPFRWWQMTRRSLLSIALPVGGGLSATYTVEVKHGGDP WP_030722038.1 hypothetical protein [Streptomyces griseus] Length: 218\nNumber of predicted TMHs:  2\nExp number of AAs in TMHs: 41.12382\nExp number, first 60 AAs:  0.82594\nTotal prob of N-in:   0.96139\ninside   1   135\nTMhelix   136   158\noutside   159   182\nTMhelix   183   200\ninside   201   218

33981 GCF_000719435.1_ASM71943v1 Streptomyces griseus subsp. griseus Terrabacteria group; Actinobacteria; Actinobacteria; Streptomycetales; Streptomycetaceae; Streptomyces; Streptomyces griseus group; Streptomyces griseus subgroup; Streptomyces griseus
MPDRRSAVSLRLPLPTAAPGFGGLALPGAAATAPDAWGVVAVRESSEKERKSSSEPEGRS WP_015507753.1 MULTISPECIES: hypothetical protein [Streptomyces] Length: 292\nNumber of predicted TMHs:  2\nExp number of AAs in TMHs: 51.77675\nExp number, first 60 AAs:  6.07612\nTotal prob of N-in:   0.33780\noutside   1   164\nTMhelix   165   187\ninside   188   229\nTMhelix   230   252\noutside   253   292

33982 GCF_000719435.1_ASM71943v1 Streptomyces griseus subsp. griseus Terrabacteria group; Actinobacteria; Actinobacteria; Streptomycetales; Streptomycetaceae; Streptomyces; Streptomyces griseus group; Streptomyces griseus subgroup; Streptomyces griseus
MSRRTRTARRRPVAAAVLAAGAVLLPAVPASAASGADQDFVIEDPRITESSGLAASKRHP WP_030771374.1 MULTISPECIES: hypothetical protein [Streptomyces] Length: 350\nNumber of predicted TMHs:  2\nExp number of AAs in TMHs: 39.44236\nExp number, first 60 AAs:  21.1162\nTotal prob of N-in:   0.99084\nPOSSIBLE N-term signal sequence\ninside   1   12\nTMhelix   13   35\noutside   36   322\nTMhelix   323   342\ninside   343   350

33983 GCF_000719435.1_ASM71943v1 Streptomyces griseus subsp. griseus Terrabacteria group; Actinobacteria; Actinobacteria; Streptomycetales; Streptomycetaceae; Streptomyces; Streptomyces griseus group; Streptomyces griseus subgroup; Streptomyces griseus
MGGRGTTAQARERGRLPGRPSDPAPTGGNDVITGVNLRVPARRTALPALLLTATALLAVP WP_030772467.1 hypothetical protein [Streptomyces griseus] Length: 304\nNumber of predicted TMHs:  1\nExp number of AAs in TMHs: 33.74215\nExp number, first 60 AAs:  7.95715\nTotal prob of N-in:   0.52144\noutside   1   276\nTMhelix   277   299\ninside   300   304

33984 GCF_000720255.1_ASM72025v1 Streptomyces griseus subsp. griseus Terrabacteria group; Actinobacteria; Actinobacteria; Streptomycetales; Streptomycetaceae; Streptomyces; Streptomyces griseus group; Streptomyces griseus subgroup; Streptomyces griseus
MSPTTTRRNVLRGTLATAALATTAATTTDPARAATPTRPDPAGSGPHIGILLYDGYSLLD WP_037673990.1 thiamine biosynthesis protein ThiJ [Streptomyces griseus] Length: 260\nNumber of predicted TMHs:  2\nExp number of AAs in TMHs: 31.69359\nExp number, first 60 AAs:  0.1859\nTotal prob of N-in:   0.82385\ninside   1   141\nTMhelix   142   159\noutside   160   192\nTMhelix   193   215\ninside   216   260

33985 GCF_000719435.1_ASM71943v1 Streptomyces griseus subsp. griseus Terrabacteria group; Actinobacteria; Actinobacteria; Streptomycetales; Streptomycetaceae; Streptomyces; Streptomyces griseus group; Streptomyces griseus subgroup; Streptomyces griseus
MSAAAPEAAPRTARPAPRAGALLRRLLWWCVLTLTGGVERRGALPPGGCVVVANHTSHAD WP_015507057.1 MULTISPECIES: 1-acyl-sn-glycerol-3-phosphate acyltransferase [Streptomyces] Length: 401\nNumber of predicted TMHs:  2\nExp number of AAs in TMHs: 83.99314\nExp number, first 60 AAs:  1.08255\nTotal prob of N-in:   0.25169\noutside   1   353\nTMhelix   354   371\ninside   372   377\nTMhelix   378   400\noutside   401   401

33986 GCF_000719435.1_ASM71943v1 Streptomyces griseus subsp. griseus Terrabacteria group; Actinobacteria; Actinobacteria; Streptomycetales; Streptomycetaceae; Streptomyces; Streptomyces griseus group; Streptomyces griseus subgroup; Streptomyces griseus
MTPNRRPVVLAAAVATALGLGATALTLPAVAAEAPRAGAPVMDLTDGTLDWGVKESFRRY WP_030769190.1 membrane protein [Streptomyces griseus] Length: 514\nNumber of predicted TMHs:  2\nExp number of AAs in TMHs: 42.8566699999999\nExp number, first 60 AAs:  22.03087\nTotal prob of N-in:   0.97276\nPOSSIBLE N-term signal sequence\ninside   1   6\nTMhelix   7   29\noutside   30   479\nTMhelix   480   502\ninside   503   514

33987 GCF_000717375.1_ASM71737v1 Streptomyces griseus subsp. griseus Terrabacteria group; Actinobacteria; Actinobacteria; Streptomycetales; Streptomycetaceae; Streptomyces; Streptomyces griseus group;

Streptomyces griseus subgroup; Streptomyces griseus
MGGRGTTAQARERGRLPGRPSDPAPTGGNDVITGVNLRMPARRTALPALLLTATALLVVP WP_033240522.1 hypothetical protein [Streptomyces griseus] Length: 304\nNumber of predicted TMHs: 2\nExp number of AAs in TMHs: 41.13193\nExp number, first 60 AAs: 13.08129\nTotal prob of N-in: 0.84550\nPOSSIBLE N-term signal sequence\ninside 1 44\nTMhelix 45 67\noutside 68 276\nTMhelix 277 299\ninside 300 304

33988 GCF_000717795.1_ASM71779v1 Streptomyces griseus subsp. griseus Terrabacteria group; Actinobacteria; Actinobacteria; Streptomycetales; Streptomycetaceae; Streptomyces; Streptomyces griseus group; Streptomyces griseus subgroup; Streptomyces griseus
MLSVVENCFSPRTAAFFDLDKTVIAKSSTLTFSKPFYQGGLINRRAVLRTAYAQFVFLAG WP_030701076.1 inhibition of morphological differentiation protein [Streptomyces griseus] Length: 281\nNumber of predicted TMHs: 1\nExp number of AAs in TMHs: 21.47312\nExp number, first 60 AAs: 0.02825\nTotal prob of N-in: 0.00628\noutside 1 247\nTMhelix 248 270\ninside 271 281

33989 GCF_000719355.1_ASM71935v1 Streptomyces griseus subsp. griseus Terrabacteria group; Actinobacteria; Actinobacteria; Streptomycetales; Streptomycetaceae; Streptomyces; Streptomyces griseus group; Streptomyces griseus subgroup; Streptomyces griseus
MLSVVENCFSPRTAAFFDLDKTVIAKSSTLTFSKSFYQGGLINRRAVLRTAYTQFVFLAG WP_030590952.1 MULTISPECIES: inhibition of morphological differentiation protein [Streptomyces griseus group] Length: 281\nNumber of predicted TMHs: 1\nExp number of AAs in TMHs: 21.42564\nExp number, first 60 AAs: 0.02005\nTotal prob of N-in: 0.00656\noutside 1 247\nTMhelix 248 270\ninside 271 281

33990 GCF_000717375.1_ASM71737v1 Streptomyces griseus subsp. griseus Terrabacteria group; Actinobacteria; Actinobacteria; Streptomycetales; Streptomycetaceae; Streptomyces; Streptomyces griseus group; Streptomyces griseus subgroup; Streptomyces griseus
MSAAAPEAAPRTARPLPRAGALLRRLLWWCVLTLTGGVERRGALPPGGCVVVANHTSHAD WP_033239148.1 1-acyl-sn-glycerol-3-phosphate acyltransferase [Streptomyces griseus] Length: 401\nNumber of predicted TMHs: 2\nExp number of AAs in TMHs: 81.71127\nExp number, first 60 AAs: 0.949709999999999\nTotal prob of N-in: 0.13673\noutside 1 353\nTMhelix 354 371\ninside 372 377\nTMhelix 378 400\noutside 401 401

33991 GCF_000717375.1_ASM71737v1 Streptomyces griseus subsp. griseus Terrabacteria group; Actinobacteria; Actinobacteria; Streptomycetales; Streptomycetaceae; Streptomyces; Streptomyces griseus group; Streptomyces griseus subgroup; Streptomyces griseus
MAPPHRRRAAVAVRALIALTALLCTTGLAPAATSGPAARAAVAEVPPPTATTSVVTVRTG WP_033239281.1 VWA domain-containing protein [Streptomyces griseus] Length: 832\nNumber of predicted TMHs: 2\nExp number of AAs in TMHs: 39.56407\nExp number, first 60 AAs: 19.33092\nTotal prob of N-in: 0.88869\nPOSSIBLE N-term signal sequence\ninside 1 11\nTMhelix 12 34\noutside 35 793\nTMhelix 794 816\ninside 817 832

33992 GCF_000717375.1_ASM71737v1 Streptomyces griseus subsp. griseus Terrabacteria group; Actinobacteria; Actinobacteria; Streptomycetales; Streptomycetaceae; Streptomyces; Streptomyces griseus group; Streptomyces griseus subgroup; Streptomyces griseus
MTPNRRPVVLAAAVATALGLGATALTLPAVAAEAPRAGAPVMDLTDGTLDWGVKESFRRY WP_033239537.1 membrane protein [Streptomyces griseus] Length: 511\nNumber of predicted TMHs: 2\nExp number of AAs in TMHs: 42.8611299999999\nExp number, first 60 AAs: 22.03087\nTotal prob of N-in: 0.97274\nPOSSIBLE N-term signal sequence\ninside 1 6\nTMhelix 7 29\noutside 30 476\nTMhelix 477 499\ninside 500 511

33993 GCF_000717375.1_ASM71737v1 Streptomyces griseus subsp. griseus Terrabacteria group; Actinobacteria; Actinobacteria; Streptomycetales; Streptomycetaceae; Streptomyces; Streptomyces griseus group; Streptomyces griseus subgroup; Streptomyces griseus
MSRRTRTARRRPVAAAVLAAGAVLLPAVPASAASGADQDFVIEDPRITESSGLAASKRHP WP_033240369.1 MULTISPECIES: hypothetical protein [Streptomyces] Length: 350\nNumber of predicted TMHs: 2\nExp number of AAs in TMHs: 39.44481\nExp number, first 60 AAs: 21.11089\nTotal prob of N-in: 0.99076\nPOSSIBLE N-term signal sequence\ninside 1 12\nTMhelix 13 35\noutside 36 322\nTMhelix 323 342\ninside 343 350

33994 GCF_000717795.1_ASM71779v1 Streptomyces griseus subsp. griseus Terrabacteria group; Actinobacteria; Actinobacteria; Streptomycetales; Streptomycetaceae; Streptomyces; Streptomyces griseus group; Streptomyces griseus subgroup; Streptomyces griseus
MGTEQQARRGPRARRGLLSALTASLLTLGAALTHLTTASPASADPIAKCTRTKGAIVAVD WP_030703601.1 peptidase [Streptomyces griseus] Length: 927\nNumber of predicted TMHs: 2\nExp number of AAs in TMHs: 40.34487\nExp number, first 60 AAs: 18.69268\nTotal prob of N-in: 0.84908\nPOSSIBLE N-term signal sequence\ninside 1 15\nTMhelix 16 38\noutside 39 889\nTMhelix 890 912\ninside 913 927

33995 GCF_000721575.1_ASM72157v1 Streptomyces griseus subsp. griseus Terrabacteria group; Actinobacteria; Actinobacteria; Streptomycetales; Streptomycetaceae; Streptomyces; Streptomyces griseus group; Streptomyces griseus subgroup; Streptomyces griseus
MGSSWNERRLERRVRPGDGKPLKPFRWWQLTRRSLLSLSLPVGGGLSATYTVEVKHGGDA WP_030847770.1 hypothetical protein [Streptomyces griseus] Length: 218\nNumber of predicted TMHs: 2\nExp number of AAs in TMHs: 41.10642\nExp number, first 60 AAs: 0.13743\nTotal prob of N-in: 0.98269\ninside 1 135\nTMhelix 136 158\noutside 159 182\nTMhelix 183 200\ninside 201 218

33996 GCF_000721575.1_ASM72157v1 Streptomyces griseus subsp. griseus Terrabacteria group; Actinobacteria; Actinobacteria; Streptomycetales; Streptomycetaceae; Streptomyces; Streptomyces griseus group; Streptomyces griseus subgroup; Streptomyces griseus
MLSVVENCFSPRTAAFFDLDKTVIAKSSTLTFSKSFYQGGLINRRAVLRTAYTQFVFMAG WP_030851795.1 inhibition of morphological differentiation protein [Streptomyces griseus] Length: 280\nNumber of predicted TMHs: 1\nExp number of AAs in TMHs: 21.70106\nExp number, first 60 AAs: 0.01764\nTotal prob of N-in: 0.00692\noutside 1 247\nTMhelix 248 270\ninside 271 280

33997 GCF_000717375.1_ASM71737v1 Streptomyces griseus subsp. griseus Terrabacteria group; Actinobacteria; Actinobacteria; Streptomycetales; Streptomycetaceae; Streptomyces; Streptomyces griseus group; Streptomyces griseus subgroup; Streptomyces griseus
MSRMRGSSGGRIAGRRAAVVLAVAVLGLAGCSAGPDSGASWAGDRAVAPSAADGAGGGQA WP_033240774.1 lipoprotein [Streptomyces griseus] Length: 334\nNumber of predicted TMHs: 1\nExp number of AAs in TMHs: 36.98141\nExp number, first 60 AAs: 14.18969\nTotal prob of N-in: 0.66733\nPOSSIBLE N-term signal sequence\noutside 1 269\nTMhelix 270 292\ninside 293 334

33998 GCF_001509565.1_ASM150956v1 Streptomyces griseus subsp. griseus Terrabacteria group; Actinobacteria; Actinobacteria; Streptomycetales; Streptomycetaceae; Streptomyces; Streptomyces griseus group; Streptomyces griseus subgroup; Streptomyces griseus
MSRMRGSSGGRIAGRRAAVVLAVAVLGLAGCSAGPDSGASSAADRAVAPSAADGAGGGQA WP_030307403.1 MULTISPECIES: hypothetical protein [Streptomyces] Length: 334\nNumber of predicted TMHs: 1\nExp number of AAs in TMHs: 36.07114\nExp number, first 60 AAs: 13.28349\nTotal prob of N-in: 0.63206\nPOSSIBLE N-term signal sequence\noutside 1 269\nTMhelix 270 292\ninside 293 334

33999 GCF_001509565.1_ASM150956v1 Streptomyces griseus subsp. griseus Terrabacteria group; Actinobacteria; Actinobacteria; Streptomycetales; Streptomycetaceae; Streptomyces; Streptomyces griseus group; Streptomyces griseus subgroup; Streptomyces griseus
MLDPVENRSLPRTAAFFDLDKTVIAKSSTLTFSKSFYQGGLINRRAVLRTAYTQFVFLAG WP_018470348.1 MULTISPECIES: inhibition of morphological differentiation protein [Streptomyces] Length: 290\nNumber of predicted TMHs: 1\nExp number of AAs in TMHs: 21.82254\nExp number, first 60 AAs: 0.02718\nTotal prob of N-in: 0.00834\noutside 1 247\nTMhelix 248 270\ninside 271 290

34000 GCF_001509565.1_ASM150956v1 Streptomyces griseus subsp. griseus Terrabacteria group; Actinobacteria; Actinobacteria; Streptomycetales; Streptomycetaceae; Streptomyces; Streptomyces griseus group; Streptomyces griseus subgroup; Streptomyces griseus
MTPNRRPVVLAAAVATALGLGATALTLPAVAAEAPRAGAPVMDLTDGTLDWGVKESFRRY WP_015507978.1 MULTISPECIES: membrane protein [Streptomyces] Length: 514\nNumber of predicted TMHs: 2\nExp number of AAs in TMHs: 42.8566699999999\nExp number, first 60 AAs: 22.03087\nTotal prob of N-in: 0.97276\nPOSSIBLE N-term signal sequence\ninside 1 6\nTMhelix 7 29\noutside 30 479\nTMhelix 480 502\ninside 503 514

34001 GCF_000718525.1_ASM71852v1 Streptomyces griseus subsp. griseus Terrabacteria group; Actinobacteria; Actinobacteria; Streptomycetales; Streptomycetaceae; Streptomyces; Streptomyces griseus group; Streptomyces griseus subgroup; Streptomyces griseus
MSKKPNRRRTAVVLTAAVAAWGPAGLAAAPAARADVVDVSYACRTKIGDRSAVSPVDVTA WP_030754832.1 hypothetical protein [Streptomyces griseus] Length: 239\nNumber of predicted TMHs: 2\nExp number of AAs in TMHs: 42.77698\nExp number, first 60 AAs: 15.49111\nTotal prob of N-in: 0.90685\nPOSSIBLE N-term signal sequence\ninside 1 11\nTMhelix 12 34\noutside 35 202\nTMhelix 203 225\ninside 226 239

34002 GCF_000719435.1_ASM71943v1 Streptomyces griseus subsp. griseus Terrabacteria group; Actinobacteria; Actinobacteria; Streptomycetales; Streptomycetaceae; Streptomyces; Streptomyces griseus group; Streptomyces griseus subgroup; Streptomyces griseus
MSRMRGSSGGRIAGRRAAVVLAVAVLGLAGCSAGPDSGASSAGDRAVAPSAADGAGGGQA WP_015506895.1

MULTISPECIES: lipoprotein [Streptomyces] Length: 334\nNumber of predicted TMHs: 1\nExp number of AAs in TMHs: 36.36573\nExp number, first 60 AAs: 13.5773\nTotal prob of N-in: 0.64439\nPOSSIBLE N-term signal sequence\noutside 1 269\nTMhelix 270 292\ninside 293 334

34003 GCF_000719435.1_ASM71943v1 Streptomyces griseus subsp. griseus Terrabacteria group; Actinobacteria; Actinobacteria; Streptomycetales; Streptomycetaceae; Streptomyces; Streptomyces griseus group; Streptomyces griseus subgroup; Streptomyces griseus
MLDPVENRSLPRTAAFFDLDKTVIAKSSTLTFSKSFYQGGLINRRAVLRTAYTQFVFLAG WP_010643895.1 MULTISPECIES: morphological differentiation-associated protein [Streptomyces] Length: 290\nNumber of predicted TMHs: 1\nExp number of AAs in TMHs: 21.93803\nExp number, first 60 AAs: 0.02636\nTotal prob of N-in: 0.00751\noutside 1 247\nTMhelix 248 270\ninside 271 290

34004 GCF_001509565.1_ASM150956v1 Streptomyces griseus subsp. griseus Terrabacteria group; Actinobacteria; Actinobacteria; Streptomycetales; Streptomycetaceae; Streptomyces; Streptomyces griseus group; Streptomyces griseus subgroup; Streptomyces griseus
MSRRTRTARRRPVAAAVLAAGAVLLPAVPASAASGADQDFVIEDPRITESSGLAASKRHP WP_030309758.1
MULTISPECIES: hypothetical protein [Streptomyces] Length: 348\nNumber of predicted TMHs: 2\nExp number of AAs in TMHs: 39.44414\nExp number, first 60 AAs: 21.11647\nTotal prob of N-in: 0.99086\nPOSSIBLE N-term signal sequence\ninside 1 12\nTMhelix 13 35\noutside 36 320\nTMhelix 321 340\ninside 341 348

34005 GCF_001509565.1_ASM150956v1 Streptomyces griseus subsp. griseus Terrabacteria group; Actinobacteria; Actinobacteria; Streptomycetales; Streptomycetaceae; Streptomyces; Streptomyces griseus group; Streptomyces griseus subgroup; Streptomyces griseus
MPDRRSAVSLRLPLPTAAPGFGGLALPGAAATAPDAWGVVAVRESSEKERKSSSEPEGRS WP_037843057.1
MULTISPECIES: hypothetical protein [Streptomyces] Length: 292\nNumber of predicted TMHs: 2\nExp number of AAs in TMHs: 51.76527\nExp number, first 60 AAs: 6.06558\nTotal prob of N-in: 0.33682\noutside 1 164\nTMhelix 165 187\ninside 188 229\nTMhelix 230 252\noutside 253 292

34006 GCF_000718525.1_ASM71852v1 Streptomyces griseus subsp. griseus Terrabacteria group; Actinobacteria; Actinobacteria; Streptomycetales; Streptomycetaceae; Streptomyces; Streptomyces griseus group; Streptomyces griseus subgroup; Streptomyces griseus
MLSVVENHSLPRAAAFFDLDKTVIAKSSTLTFSKSFYRGGLISRRAALRTAYIQFVFLVG WP_030749261.1 inhibition of morphological differentiation protein [Streptomyces griseus] Length: 284\nNumber of predicted TMHs: 1\nExp number of AAs in TMHs: 21.36258\nExp number, first 60 AAs: 0.07194\nTotal prob of N-in: 0.00449\noutside 1 247\nTMhelix 248 270\ninside 271 284

34007 GCF_001509565.1_ASM150956v1 Streptomyces griseus subsp. griseus Terrabacteria group; Actinobacteria; Actinobacteria; Streptomycetales; Streptomycetaceae; Streptomyces; Streptomyces griseus group; Streptomyces griseus subgroup; Streptomyces griseus
MGGRGTTAQARERGRLPGRPSDPAPTGGNDVITGVNLRMPARRTALPALLLTATALLAVP WP_059214191.1
hypothetical protein [Streptomyces griseus] Length: 304\nNumber of predicted TMHs: 2\nExp number of AAs in TMHs: 38.95199\nExp number, first 60 AAs: 11.59641\nTotal prob of N-in: 0.75090\nPOSSIBLE N-term signal sequence\ninside 1 44\nTMhelix 45 67\noutside 68 276\nTMhelix 277 299\ninside 300 304

34008 GCF_000720255.1_ASM72025v1 Streptomyces griseus subsp. griseus Terrabacteria group; Actinobacteria; Actinobacteria; Streptomycetales; Streptomycetaceae; Streptomyces; Streptomyces griseus group; Streptomyces griseus subgroup; Streptomyces griseus
MLKGVENHSLPRTAAFFDLDKTVIAKSSTLTFSKSFYQGGLINRRAALRTAYAQFVFLVG WP_037676937.1 inhibition of morphological differentiation protein [Streptomyces griseus] Length: 277\nNumber of predicted TMHs: 1\nExp number of AAs in TMHs: 21.40362\nExp number, first 60 AAs: 0.04209\nTotal prob of N-in: 0.00426\noutside 1 247\nTMhelix 248 270\ninside 271 277

34009 GCF_001509565.1_ASM150956v1 Streptomyces griseus subsp. griseus Terrabacteria group; Actinobacteria; Actinobacteria; Streptomycetales; Streptomycetaceae; Streptomyces; Streptomyces griseus group; Streptomyces griseus subgroup; Streptomyces griseus
MSAAAPEAAPRTARPAPRAGALLRRLLWWCVLTLTGGVERRGALPPGGCVVVANHTSHAD WP_015507057.1
MULTISPECIES: 1-acyl-sn-glycerol-3-phosphate acyltransferase [Streptomyces] Length: 401\nNumber of predicted TMHs: 2\nExp number of AAs in TMHs: 83.99314\nExp number, first 60 AAs: 1.08255\nTotal prob of N-in: 0.25169\noutside 1 353\nTMhelix 354 371\ninside 372 377\nTMhelix 378 400\noutside 401 401

34010 GCF_000719495.1_ASM71949v1 Streptomyces griseus subsp. griseus Terrabacteria group; Actinobacteria; Actinobacteria; Streptomycetales; Streptomycetaceae; Streptomyces; Streptomyces griseus group; Streptomyces griseus subgroup; Streptomyces griseus
MGGRGTTARARERGRLPGRPSDPAPTGGNDVITGVNLRMPARRTALPALLLTATALLAVP WP_030696962.1 hypothetical protein [Streptomyces griseus] Length: 305\nNumber of predicted TMHs: 2\nExp number of AAs in TMHs: 40.14857\nExp number, first 60 AAs: 12.43541\nTotal prob of N-in: 0.80614\nPOSSIBLE N-term signal sequence\ninside 1 44\nTMhelix 45 67\noutside 68 277\nTMhelix 278 300\ninside 301 305

34011 GCF_000719495.1_ASM71949v1 Streptomyces griseus subsp. griseus Terrabacteria group; Actinobacteria; Actinobacteria; Streptomycetales; Streptomycetaceae; Streptomyces; Streptomyces griseus group; Streptomyces griseus subgroup; Streptomyces griseus
MSRRTRTARRRPVAAAVLAAGAVLLPVVPASAASGADQDFVIEDPRITESSGLAASKRHP WP_030698069.1 hypothetical protein [Streptomyces griseus] Length: 348\nNumber of predicted TMHs: 2\nExp number of AAs in TMHs: 38.26588\nExp number, first 60 AAs: 21.30028\nTotal prob of N-in: 0.99451\nPOSSIBLE N-term signal sequence\ninside 1 12\nTMhelix 13 35\noutside 36 322\nTMhelix 323 340\ninside 341 348

34012 GCF_000719495.1_ASM71949v1 Streptomyces griseus subsp. griseus Terrabacteria group; Actinobacteria; Actinobacteria; Streptomycetales; Streptomycetaceae; Streptomyces; Streptomyces griseus group; Streptomyces griseus subgroup; Streptomyces griseus
MLDPVENRSLPRTAAFFDLDKTVIAKSSTLTFSKSFYQGGLINRRAVLRTAYTQFVFLAG WP_030698235.1 MULTISPECIES: inhibition of morphological differentiation protein [Streptomyces] Length: 279\nNumber of predicted TMHs: 1\nExp number of AAs in TMHs: 21.72379\nExp number, first 60 AAs: 0.0316\nTotal prob of N-in: 0.01274\noutside 1 247\nTMhelix 248 270\ninside 271 279

34013 GCF_001723125.1_ASM172312v1 Streptomyces griseus subsp. griseus Terrabacteria group; Actinobacteria; Actinobacteria; Streptomycetales; Streptomycetaceae; Streptomyces; Streptomyces griseus group; Streptomyces griseus subgroup; Streptomyces griseus
MLSVVENCFSPRTAAFFDLDKTVIAKSSTLTFSKSFYQGGLINRRAVLRTAYAQFVFLAG WP_012379811.1 morphological differentiation-associated protein [Streptomyces griseus] Length: 281\nNumber of predicted TMHs: 1\nExp number of AAs in TMHs: 21.467\nExp number, first 60 AAs: 0.03606\nTotal prob of N-in: 0.00978\noutside 1 247\nTMhelix 248 270\ninside 271 281

34014 GCF_000717375.1_ASM71737v1 Streptomyces griseus subsp. griseus Terrabacteria group; Actinobacteria; Actinobacteria; Streptomycetales; Streptomycetaceae; Streptomyces; Streptomyces griseus group; Streptomyces griseus subgroup; Streptomyces griseus
MLDPVENRSLPRTAAFFDLDKTVIAKSSTLTFSKSFYQGGLINRRAVLRTAYTQFVFLAG WP_018470348.1 MULTISPECIES: inhibition of morphological differentiation protein [Streptomyces] Length: 290\nNumber of predicted TMHs: 1\nExp number of AAs in TMHs: 21.82254\nExp number, first 60 AAs: 0.02718\nTotal prob of N-in: 0.00834\noutside 1 247\nTMhelix 248 270\ninside 271 290

34015 GCF_000717375.1_ASM71737v1 Streptomyces griseus subsp. griseus Terrabacteria group; Actinobacteria; Actinobacteria; Streptomycetales; Streptomycetaceae; Streptomyces; Streptomyces griseus group; Streptomyces griseus subgroup; Streptomyces griseus
MPDRRSAVSLRLPLPTAAPGFGGLALPGAAATAPDAWGVVAVRESSEKERKSSSEPEGRS WP_033237593.1 MULTISPECIES: hypothetical protein [Streptomyces] Length: 292\nNumber of predicted TMHs: 2\nExp number of AAs in TMHs: 51.77677\nExp number, first 60 AAs: 6.07612\nTotal prob of N-in: 0.33780\noutside 1 164\nTMhelix 165 187\ninside 188 229\nTMhelix 230 252\noutside 253 292

34016 GCF_001270675.1_ASM127067v1 Streptomyces griseus subsp. rhodochrous Terrabacteria group; Actinobacteria; Actinobacteria; Streptomycetales; Streptomycetaceae; Streptomyces; Streptomyces griseus group; Streptomyces griseus subgroup; Streptomyces griseus
MGSSWNERRLERRVRPGDGKPLKPFRWWQLTRRSLLSLALPVGGGLSATYTVEVKHGGDA WP_030189408.1 MULTISPECIES: hypothetical protein [Streptomyces] Length: 218\nNumber of predicted TMHs: 2\nExp number of AAs in TMHs: 41.32127\nExp number, first 60 AAs: 0.37701\nTotal prob of N-in: 0.97413\ninside 1 135\nTMhelix 136 158\noutside 159 182\nTMhelix 183 200\ninside 201 218

34017 GCF_001270675.1_ASM127067v1 Streptomyces griseus subsp. rhodochrous Terrabacteria group; Actinobacteria; Actinobacteria; Streptomycetales; Streptomycetaceae; Streptomyces; Streptomyces griseus group; Streptomyces griseus subgroup; Streptomyces griseus
MLSVVENCFSPRTAAFFDLDKTVIAKSSTLTFSKSFYQGGLINRRAVLRTAYTQFVFMAG WP_030113359.1 MULTISPECIES: inhibition of morphological differentiation protein [Streptomyces] Length: 280\nNumber of predicted TMHs: 1\nExp

number of AAs in TMHs: 21.68695\nExp number, first 60 AAs: 0.01801\nTotal prob of N-in: 0.00793\noutside 1 247\nTMhelix 248 270\ninside 271 280

34018 GCF_000718205.1_ASM71820v1 Streptomyces griseus subsp. rhodochrous Terrabacteria group; Actinobacteria; Actinobacteria; Streptomycetales; Streptomycetaceae; Streptomyces; Streptomyces griseus group; Streptomyces griseus subgroup; Streptomyces griseus MGSSWNERRLERRVRPGDGKPLKPFRWWQLTRRSLLSLALPVGGGLSATYTVEVKHGGDA WP_030189408.1 MULTISPECIES: hypothetical protein [Streptomyces] Length: 218\nNumber of predicted TMHs: 2\nExp number of AAs in TMHs: 41.32127\nExp number, first 60 AAs: 0.37701\nTotal prob of N-in: 0.97413\ninside 1 135\nTMhelix 136 158\noutside 159 182\nTMhelix 183 200\ninside 201 218

34019 GCF_000719035.1_ASM71903v1 Streptomyces griseus subsp. rhodochrous Terrabacteria group; Actinobacteria; Actinobacteria; Streptomycetales; Streptomycetaceae; Streptomyces; Streptomyces griseus group; Streptomyces griseus subgroup; Streptomyces griseus MGSSWNERRLERRVRPGDGKPLKPFRWWQLTRRSLLSLALPVGGGLSATYTVEVKHGGDA WP_030189408.1 MULTISPECIES: hypothetical protein [Streptomyces] Length: 218\nNumber of predicted TMHs: 2\nExp number of AAs in TMHs: 41.32127\nExp number, first 60 AAs: 0.37701\nTotal prob of N-in: 0.97413\ninside 1 135\nTMhelix 136 158\noutside 159 182\nTMhelix 183 200\ninside 201 218

34020 GCF_000718205.1_ASM71820v1 Streptomyces griseus subsp. rhodochrous Terrabacteria group; Actinobacteria; Actinobacteria; Streptomycetales; Streptomycetaceae; Streptomyces; Streptomyces griseus group; Streptomyces griseus subgroup; Streptomyces griseus MLSVVENCFSPRTAAFFDLDKTVIAKSSTLTFSKSFYQGGLINRRAVLRTAYTQFVFMAG WP_030113359.1 MULTISPECIES: inhibition of morphological differentiation protein [Streptomyces] Length: 280\nNumber of predicted TMHs: 1\nExp number of AAs in TMHs: 21.68695\nExp number, first 60 AAs: 0.01801\nTotal prob of N-in: 0.00793\noutside 1 247\nTMhelix 248 270\ninside 271 280

34021 GCF_000719035.1_ASM71903v1 Streptomyces griseus subsp. rhodochrous Terrabacteria group; Actinobacteria; Actinobacteria; Streptomycetales; Streptomycetaceae; Streptomyces; Streptomyces griseus group; Streptomyces griseus subgroup; Streptomyces griseus MLSVVENCFSPRTAAFFDLDKTVIAKSSTLTFSKSFYQGGLINRRAVLRTAYTQFVFMAG WP_030113359.1 MULTISPECIES: inhibition of morphological differentiation protein [Streptomyces] Length: 280\nNumber of predicted TMHs: 1\nExp number of AAs in TMHs: 21.68695\nExp number, first 60 AAs: 0.01801\nTotal prob of N-in: 0.00793\noutside 1 247\nTMhelix 248 270\ninside 271 280

34022 GCF_000721205.1_ASM72120v1 Streptomyces griseus subsp. rhodochrous Terrabacteria group; Actinobacteria; Actinobacteria; Streptomycetales; Streptomycetaceae; Streptomyces; Streptomyces griseus group; Streptomyces griseus subgroup; Streptomyces griseus MGSSWNERRLERRVRPGDGKPLKPFRWWQLTRRSLLSLALPVGGGLSATYTVEVKHGGDA WP_030189408.1 MULTISPECIES: hypothetical protein [Streptomyces] Length: 218\nNumber of predicted TMHs: 2\nExp number of AAs in TMHs: 41.32127\nExp number, first 60 AAs: 0.37701\nTotal prob of N-in: 0.97413\ninside 1 135\nTMhelix 136 158\noutside 159 182\nTMhelix 183 200\ninside 201 218

34023 GCF_000721205.1_ASM72120v1 Streptomyces griseus subsp. rhodochrous Terrabacteria group; Actinobacteria; Actinobacteria; Streptomycetales; Streptomycetaceae; Streptomyces; Streptomyces griseus group; Streptomyces griseus subgroup; Streptomyces griseus MLSVVENCFSPRTAAFFDLDKTVIAKSSTLTFSKSFYQGGLINRRAVLRTAYTQFVFMAG WP_030113359.1 MULTISPECIES: inhibition of morphological differentiation protein [Streptomyces] Length: 280\nNumber of predicted TMHs: 1\nExp number of AAs in TMHs: 21.68695\nExp number, first 60 AAs: 0.01801\nTotal prob of N-in: 0.00793\noutside 1 247\nTMhelix 248 270\ninside 271 280

34024 GCF_000718235.1_ASM71823v1 Streptomyces griseus subsp. rhodochrous Terrabacteria group; Actinobacteria; Actinobacteria; Streptomycetales; Streptomycetaceae; Streptomyces; Streptomyces griseus group; Streptomyces griseus subgroup; Streptomyces griseus MGSSWNERWLERRVRPGDGKPLKPFRWWQLTRRSLLSLALPVGGGLSATYTVEVKHGGDA WP_030733765.1 MULTISPECIES: hypothetical protein [Streptomyces] Length: 218\nNumber of predicted TMHs: 2\nExp number of AAs in TMHs: 41.82036\nExp number, first 60 AAs: 0.95179\nTotal prob of N-in: 0.93795\ninside 1 135\nTMhelix 136 158\noutside 159 182\nTMhelix 183 200\ninside 201 218

34025 GCF_000718235.1_ASM71823v1 Streptomyces griseus subsp. rhodochrous Terrabacteria group; Actinobacteria; Actinobacteria; Streptomycetales; Streptomycetaceae; Streptomyces; Streptomyces griseus group;

Streptomyces griseus subgroup; Streptomyces griseus
MLSVVENCFSPRTAAFFDLDKTVIAKSSTLTFSKSFYQGGLINRRAVLRTAYTQFVFMAG WP_030113359.1 MULTISPECIES: inhibition of morphological differentiation protein [Streptomyces] Length: 280\nNumber of predicted TMHs: 1\nExp number of AAs in TMHs: 21.68695\nExp number, first 60 AAs: 0.01801\nTotal prob of N-in: 0.00793\noutside 1 247\nTMhelix 248 270\ninside 271 280

34026 GCF_000010605.1_ASM1060v1 Streptomyces griseus subsp. griseus NBRC 13350 Terrabacteria group; Actinobacteria; Actinobacteria; Streptomycetales; Streptomycetaceae; Streptomyces; Streptomyces griseus group; Streptomyces griseus subgroup; Streptomyces griseus; Streptomyces griseus subsp. griseus
MLSVVENCFSPRTAAFFDLDKTVIAKSSTLTFSKSFYQGGLINRRAVLRTAYAQFVFLAG WP_012379811.1 morphological differentiation-associated protein [Streptomyces griseus] Length: 281\nNumber of predicted TMHs: 1\nExp number of AAs in TMHs: 21.467\nExp number, first 60 AAs: 0.03606\nTotal prob of N-in: 0.00978\noutside 1 247\nTMhelix 248 270\ninside 271 281

34027 GCF_000010605.1_ASM1060v1 Streptomyces griseus subsp. griseus NBRC 13350 Terrabacteria group; Actinobacteria; Actinobacteria; Streptomycetales; Streptomycetaceae; Streptomyces; Streptomyces griseus group; Streptomyces griseus subgroup; Streptomyces griseus; Streptomyces griseus subsp. griseus
MGTEQQARRGPRARRGLLSALTASLLTLGAALTHLTTASPASADPIAKCTRTKGAIVAVD WP_012378693.1 peptidase [Streptomyces griseus] Length: 931\nNumber of predicted TMHs: 2\nExp number of AAs in TMHs: 39.57519\nExp number, first 60 AAs: 18.69314\nTotal prob of N-in: 0.84908\nPOSSIBLE N-term signal sequence\ninside 1 15\nTMhelix 16 38\noutside 39 893\nTMhelix 894 916\ninside 917 931

34028 GCF_000720535.1_ASM72053v1 Streptomyces halstedii Terrabacteria group; Actinobacteria; Actinobacteria; Streptomycetales; Streptomycetaceae; Streptomyces; Streptomyces griseus group; Streptomyces halstedii subgroup MLSFVENCFSPRTAAFFDLDKTVIAKSSTLAFSKSFYQGGLINRRAVLRTAYTQFVFLAG WP_030629408.1 MULTISPECIES: inhibition of morphological differentiation protein [Streptomyces] Length: 279\nNumber of predicted TMHs: 1\nExp number of AAs in TMHs: 21.81622\nExp number, first 60 AAs: 0.01771\nTotal prob of N-in: 0.00802\noutside 1 247\nTMhelix 248 270\ninside 271 279

34029 GCF_000720535.1_ASM72053v1 Streptomyces halstedii Terrabacteria group; Actinobacteria; Actinobacteria; Streptomycetales; Streptomycetaceae; Streptomyces; Streptomyces griseus group; Streptomyces halstedii subgroup MTAGMSQTWKRRAALGSRGRLLGTVAVAAAWTVGMAGAAPSAVAMDVQSKQWYLSAMHAE WP_030630441.1
MULTISPECIES: serine protease [Streptomyces] Length: 420\nNumber of predicted TMHs: 2\nExp number of AAs in TMHs: 44.93486\nExp number, first 60 AAs: 22.25757\nTotal prob of N-in: 0.98803\nPOSSIBLE N-term signal sequence\ninside 1 20\nTMhelix 21 43\noutside 44 391\nTMhelix 392 414\ninside 415 420

34030 GCF_000716935.1_ASM71693v1 Streptomyces alboviridis Terrabacteria group; Actinobacteria; Actinobacteria; Streptomycetales; Streptomycetaceae; Streptomyces; Streptomyces griseus group; Streptomyces microflavus subgroup MLSVVENCFSPRTAAFFDLDKTVIAKSSTLTFSKSFYQGGLINRRAVLRTAYTQFVFMAG
WP_032758724.1 inhibition of morphological differentiation protein [Streptomyces alboviridis] Length: 281\nNumber of predicted TMHs: 1\nExp number of AAs in TMHs: 21.79849\nExp number, first 60 AAs: 0.01651\nTotal prob of N-in: 0.00400\noutside 1 247\nTMhelix 248 270\ninside 271 281

34031 GCF_000716935.1_ASM71693v1 Streptomyces alboviridis Terrabacteria group; Actinobacteria; Actinobacteria; Streptomycetales; Streptomycetaceae; Streptomyces; Streptomyces griseus group; Streptomyces microflavus subgroup MGNAVSGRRTLLSGTAVLAAVALIALGSAPAQADETEPDLGVRGLTPVTGIAPGSGFGLP
WP_032758621.1 peptidase [Streptomyces alboviridis] Length: 434\nNumber of predicted TMHs: 1\nExp number of AAs in TMHs: 35.2262\nExp number, first 60 AAs: 14.69427\nTotal prob of N-in: 0.61645\nPOSSIBLE N-term signal sequence\noutside 1 402\nTMhelix 403 425\ninside 426 434

34032 GCF_001418625.1_ASM141862v1 Streptomyces luridiscabiei Terrabacteria group; Actinobacteria; Actinobacteria; Streptomycetales; Streptomycetaceae; Streptomyces; Streptomyces griseus group; Streptomyces microflavus subgroup MGNAVSGRRTLLSGTAVLAAVALIALGSAPAQADETEPDLGVRGLTPVTGITPGSGFGLP
WP_055558699.1 peptidase [Streptomyces luridiscabiei] Length: 434\nNumber of predicted TMHs: 1\nExp number of AAs in TMHs: 34.64005\nExp number, first 60 AAs: 14.18972\nTotal prob of N-in: 0.62197\nPOSSIBLE N-term signal sequence\noutside 1 402\nTMhelix 403 425\ninside 426 434

34033 GCF_001418625.1_ASM141862v1 Streptomyces luridiscabiei Terrabacteria group; Actinobacteria; Actinobacteria; Streptomycetales; Streptomycetaceae; Streptomyces; Streptomyces griseus group; Streptomyces microflavus subgroup MLSVVENCFSPRTAAFFDLDKTVIAKSSTLTFSKSFYQGGLINRRAVLRTAYTQFVFMAG
WP_031123945.1 MULTISPECIES: inhibition of morphological differentiation protein [Streptomyces]

Length: 281\nNumber of predicted TMHs: 1\nExp number of AAs in TMHs: 21.79856\nExp number, first 60 AAs: 0.01652\nTotal prob of N-in: 0.00400\noutside 1 247\nTMhelix 248 270\ninside 271 281

34034 GCF_001418625.1_ASM141862v1 Streptomyces luridiscabiei Terrabacteria group; Actinobacteria; Actinobacteria; Streptomycetales; Streptomycetaceae; Streptomyces; Streptomyces griseus group; Streptomyces microflavus subgroup MFGRTGRRSAVTVLAAVVLWGPTGQDTAVADSRHPVIETLPEQEVRAAVAAARTPVAVNL WP_055561060.1 hypothetical protein [Streptomyces luridiscabiei] Length: 259\nNumber of predicted TMHs: 1\nExp number of AAs in TMHs: 16.50156\nExp number, first 60 AAs: 0.17582\nTotal prob of N-in: 0.11936\noutside 1 228\nTMhelix 229 247\ninside 248 259

34035 GCF_000717645.1_ASM71764v1 Streptomyces californicus Terrabacteria group; Actinobacteria; Actinobacteria; Streptomycetales; Streptomycetaceae; Streptomyces; Streptomyces griseus group; Streptomyces puniceus subgroup MGSSWNERRLERRVRPGDGKPLKPFRWWQLTRRSLLSLALPVGGGLSATYTVEVKHGGDA WP_030113438.1 MULTISPECIES: hypothetical protein [Streptomyces] Length: 218\nNumber of predicted TMHs: 2\nExp number of AAs in TMHs: 41.40199\nExp number, first 60 AAs: 0.36812\nTotal prob of N-in: 0.97576\ninside 1 135\nTMhelix 136 158\noutside 159 182\nTMhelix 183 200\ninside 201 218

34036 GCF_000717645.1_ASM71764v1 Streptomyces californicus Terrabacteria group; Actinobacteria; Actinobacteria; Streptomycetales; Streptomycetaceae; Streptomyces; Streptomyces griseus group; Streptomyces puniceus subgroup MLSVVENCFSPRTAAFFDLDKTVIAKSSTLTFSKSFYQGGLINRRAVLRTAYTQFVFMAG WP_030113359.1 MULTISPECIES: inhibition of morphological differentiation protein [Streptomyces] Length: 280\nNumber of predicted TMHs: 1\nExp number of AAs in TMHs: 21.68695\nExp number, first 60 AAs: 0.01801\nTotal prob of N-in: 0.00793\noutside 1 247\nTMhelix 248 270\ninside 271 280

34037 GCF_000718245.1_ASM71824v1 Streptomyces californicus Terrabacteria group; Actinobacteria; Actinobacteria; Streptomycetales; Streptomycetaceae; Streptomyces; Streptomyces griseus group; Streptomyces puniceus subgroup MLSVVENCFSPRTAAFFDLDKTVIAKSSTLTFSKSFYQGGLINRRAVLRTAYTQFVFMAG WP_030113359.1 MULTISPECIES: inhibition of morphological differentiation protein [Streptomyces] Length: 280\nNumber of predicted TMHs: 1\nExp number of AAs in TMHs: 21.68695\nExp number, first 60 AAs: 0.01801\nTotal prob of N-in: 0.00793\noutside 1 247\nTMhelix 248 270\ninside 271 280

34038 GCF_000718245.1_ASM71824v1 Streptomyces californicus Terrabacteria group; Actinobacteria; Actinobacteria; Streptomycetales; Streptomycetaceae; Streptomyces; Streptomyces griseus group; Streptomyces puniceus subgroup MGSSWNERRLERRVRPGDGKPLKPFRWWQLTRRSLLSLALPVGGGLSATYTVEVKHGGDA WP_030113438.1 MULTISPECIES: hypothetical protein [Streptomyces] Length: 218\nNumber of predicted TMHs: 2\nExp number of AAs in TMHs: 41.40199\nExp number, first 60 AAs: 0.36812\nTotal prob of N-in: 0.97576\ninside 1 135\nTMhelix 136 158\noutside 159 182\nTMhelix 183 200\ninside 201 218

34039 GCF_000717665.1_ASM71766v1 Streptomyces floridae Terrabacteria group; Actinobacteria; Actinobacteria; Streptomycetales; Streptomycetaceae; Streptomyces; Streptomyces griseus group; Streptomyces puniceus subgroup MLSVVENCFSPRTAAFFDLDKTVIAKSSTLTFSKSFYQGGLINRRAVLRTAYTQFVFMAG WP_030189579.1 MULTISPECIES: inhibition of morphological differentiation protein [Streptomyces] Length: 280\nNumber of predicted TMHs: 1\nExp number of AAs in TMHs: 21.68695\nExp number, first 60 AAs: 0.01801\nTotal prob of N-in: 0.00793\noutside 1 247\nTMhelix 248 270\ninside 271 280

34040 GCF_000717665.1_ASM71766v1 Streptomyces floridae Terrabacteria group; Actinobacteria; Actinobacteria; Streptomycetales; Streptomycetaceae; Streptomyces; Streptomyces griseus group; Streptomyces puniceus subgroup MGSSWNERRLERRVRPGDGKPLKPFRWWQLTRRSLLSLALPVGGGLSATYTVEVKHGGDA WP_030189408.1 MULTISPECIES: hypothetical protein [Streptomyces] Length: 218\nNumber of predicted TMHs: 2\nExp number of AAs in TMHs: 41.32127\nExp number, first 60 AAs: 0.37701\nTotal prob of N-in: 0.97413\ninside 1 135\nTMhelix 136 158\noutside 159 182\nTMhelix 183 200\ninside 201 218

34041 GCF_000718695.1_ASM71869v1 Streptomyces puniceus Terrabacteria group; Actinobacteria; Actinobacteria; Streptomycetales; Streptomycetaceae; Streptomyces; Streptomyces griseus group; Streptomyces puniceus subgroup MLSVVENCFSPRTAAFFDLDKTVIAKSSTLTFSKSFYQGGLINRRAVLRTAYTQFVFMAG WP_030189579.1 MULTISPECIES: inhibition of morphological differentiation protein [Streptomyces] Length: 280\nNumber of predicted TMHs: 1\nExp number of AAs in TMHs: 21.68695\nExp number, first 60 AAs: 0.01801\nTotal prob of N-in: 0.00793\noutside 1 247\nTMhelix 248 270\ninside 271 280

34042 GCF_000718695.1_ASM71869v1 Streptomyces puniceus Terrabacteria group; Actinobacteria; Actinobacteria; Streptomycetales; Streptomycetaceae; Streptomyces; Streptomyces griseus group; Streptomyces puniceus subgroup MGSSWNERRLERRVRPGDGKPLKPFRWWQLTRRSLLSLALPVGGGLSATYTVEVKHGGDA WP_030189408.1 MULTISPECIES: hypothetical protein [Streptomyces] Length: 218\nNumber of predicted TMHs: 2\nExp number of AAs in TMHs: 41.32127\nExp number, first 60 AAs: 0.37701\nTotal prob of N-in: 0.97413\ninside 1 135\nTMhelix 136 158\noutside 159 182\nTMhelix 183 200\ninside 201 218

34043 GCF_000719195.1_ASM71919v1 Streptomyces puniceus Terrabacteria group; Actinobacteria; Actinobacteria; Streptomycetales; Streptomycetaceae; Streptomyces; Streptomyces griseus group; Streptomyces puniceus subgroup MLSVVENCFSPRTAAFFDLDKTVIAKSSTLTFSKSFYQGGLINRRAVLRTAYTQFVFMAG WP_030189579.1 MULTISPECIES: inhibition of morphological differentiation protein [Streptomyces] Length: 280\nNumber of predicted TMHs: 1\nExp number of AAs in TMHs: 21.68695\nExp number, first 60 AAs: 0.01801\nTotal prob of N-in: 0.00793\noutside 1 247\nTMhelix 248 270\ninside 271 280

34044 GCF_000719195.1_ASM71919v1 Streptomyces puniceus Terrabacteria group; Actinobacteria; Actinobacteria; Streptomycetales; Streptomycetaceae; Streptomyces; Streptomyces griseus group; Streptomyces puniceus subgroup MGSSWNERRLERRVRPGDGKPLKPFRWWQLTRRSLLSLALPVGGGLSATYTVEVKHGGDA WP_030189408.1 MULTISPECIES: hypothetical protein [Streptomyces] Length: 218\nNumber of predicted TMHs: 2\nExp number of AAs in TMHs: 41.32127\nExp number, first 60 AAs: 0.37701\nTotal prob of N-in: 0.97413\ninside 1 135\nTMhelix 136 158\noutside 159 182\nTMhelix 183 200\ninside 201 218

34045 GCF_000158915.1_ASM15891v1 Streptomyces himastatinicus ATCC 53653 Terrabacteria group; Actinobacteria; Actinobacteria; Streptomycetales; Streptomycetaceae; Streptomyces; Streptomyces himastatinicus MSSPDREVRAARNDTARTARRPVVAVTGAASGPGALLTQRLAESEEIKQVLALDERRGEV WP_009715199.1 MULTISPECIES: NAD-dependent dehydratase [Streptomyces] Length: 369\nNumber of predicted TMHs: 1\nExp number of AAs in TMHs: 20.51064\nExp number, first 60 AAs: 0.50973\nTotal prob of N-in: 0.05336\noutside 1 339\nTMhelix 340 359\ninside 360 369

34046 GCF_000721535.1_ASM72153v1 Streptomyces hygroscopicus subsp. hygroscopicus Terrabacteria group; Actinobacteria; Actinobacteria; Streptomycetales; Streptomycetaceae; Streptomyces; Streptomyces hygroscopicus MLSVVENHSLPRTAAFFDLDKTVIAKSSALTFSKSFYQGGLINRRAVLRTAYAQFVFLAG WP_030838032.1 inhibition of morphological differentiation protein [Streptomyces hygroscopicus] Length: 281\nNumber of predicted TMHs: 1\nExp number of AAs in TMHs: 21.24913\nExp number, first 60 AAs: 0.0235\nTotal prob of N-in: 0.00267\noutside 1 246\nTMhelix 247 269\ninside 270 281

34047 GCF_001553435.1_ASM155343v1 Streptomyces hygroscopicus subsp. hygroscopicus Terrabacteria group; Actinobacteria; Actinobacteria; Streptomycetales; Streptomycetaceae; Streptomyces; Streptomyces hygroscopicus MLSVVENHSLPRTAAFFDLDKTVIAKSSALTFSKSFYQGGLINRRAVLRTAYAQFVFLAG WP_060951846.1 MULTISPECIES: inhibition of morphological differentiation protein [Streptomyces] Length: 281\nNumber of predicted TMHs: 1\nExp number of AAs in TMHs: 21.2919\nExp number, first 60 AAs: 0.02346\nTotal prob of N-in: 0.00265\noutside 1 246\nTMhelix 247 269\ninside 270 281

34048 GCF_001553455.1_ASM155345v1 Streptomyces hygroscopicus subsp. hygroscopicus Terrabacteria group; Actinobacteria; Actinobacteria; Streptomycetales; Streptomycetaceae; Streptomyces; Streptomyces hygroscopicus MLSVVENHSLPRTAAFFDLDKTVIAKSSALTFSKSFYQGGLINRRAVLRTAYAQFVFLAG WP_060942826.1 inhibition of morphological differentiation protein [Streptomyces hygroscopicus] Length: 281\nNumber of predicted TMHs: 1\nExp number of AAs in TMHs: 21.25421\nExp number, first 60 AAs: 0.02347\nTotal prob of N-in: 0.00267\noutside 1 246\nTMhelix 247 269\ninside 270 281

34049 GCF_001447075.1_ASM144707v1 Streptomyces hygroscopicus subsp. limoneus Terrabacteria group; Actinobacteria; Actinobacteria; Streptomycetales; Streptomycetaceae; Streptomyces; Streptomyces hygroscopicus MLRDVENHSLPRAAAFFDLDKTVIAKSSTLTFGKSFYQGGLINRRAALRTAYAQFVFLAG WP_058081378.1 inhibition of morphological differentiation protein [Streptomyces hygroscopicus] Length: 276\nNumber of predicted TMHs: 1\nExp number of AAs in TMHs: 21.38024\nExp number, first 60 AAs: 0.04668\nTotal prob of N-in: 0.01061\noutside 1 247\nTMhelix 248 270\ninside 271 276

34050 GCF_001447075.1_ASM144707v1 Streptomyces hygroscopicus subsp. limoneus Terrabacteria group; Actinobacteria; Actinobacteria; Streptomycetales; Streptomycetaceae; Streptomyces; Streptomyces hygroscopicus MRHARRRVVRRVTRLAAVGGLLLGGAMVTQAAMASETPSAPARPLAATGDTGAALVARLG WP_058083229.1 protease [Streptomyces hygroscopicus] Length: 458\nNumber of predicted TMHs: 2\nExp number of AAs in TMHs: 43.8397399999999\nExp number, first 60 AAs: 22.26878\nTotal prob of N-in: 0.99236\nPOSSIBLE N-term signal sequence\ninside 1 11\nTMhelix 12 34\noutside 35 419\nTMhelix 420 442\ninside 443 458

34051 GCF_000245355.1_ASM24535v1 Streptomyces hygroscopicus subsp. jinggangensis 5008 Terrabacteria group; Actinobacteria; Actinobacteria; Streptomycetales; Streptomycetaceae; Streptomyces; Streptomyces hygroscopicus; Streptomyces hygroscopicus subsp. jinggangensis
MRHARRRVVRRVTRLAAVGGLLLGGAMVTQAAMASETPSAPARPLAATGDTGAALVARLG WP_014676716.1 S1 family peptidase [Streptomyces hygroscopicus] Length: 458\nNumber of predicted TMHs: 2\nExp number of AAs in TMHs: 43.5516799999999\nExp number, first 60 AAs: 22.26814\nTotal prob of N-in: 0.99236\nPOSSIBLE N-term signal sequence\ninside 1 11\nTMhelix 12 34\noutside 35 419\nTMhelix 420 442\ninside 443 458

34052 GCF_000245355.1_ASM24535v1 Streptomyces hygroscopicus subsp. jinggangensis 5008 Terrabacteria group; Actinobacteria; Actinobacteria; Streptomycetales; Streptomycetaceae; Streptomyces; Streptomyces hygroscopicus; Streptomyces hygroscopicus subsp. jinggangensis
MLRDVENHSLPRAAAFFDLDKTVIAKSSTLTFSKSFYQGGLINRRAALRTAYAQFVFLAG WP_014674034.1 MULTISPECIES: morphological differentiation-associated protein [Streptomyces] Length: 276\nNumber of predicted TMHs: 1\nExp number of AAs in TMHs: 21.37944\nExp number, first 60 AAs: 0.04579\nTotal prob of N-in: 0.01060\noutside 1 247\nTMhelix 248 270\ninside 271 276

34053 GCF_000245355.1_ASM24535v1 Streptomyces hygroscopicus subsp. jinggangensis 5008 Terrabacteria group; Actinobacteria; Actinobacteria; Streptomycetales; Streptomycetaceae; Streptomyces; Streptomyces hygroscopicus; Streptomyces hygroscopicus subsp. jinggangensis
MRRRAVLARGAVIGLLLGLSAALATPTPAPAVPAGAGTAADARPGSYAFTPGARTVAGTK WP_014673788.1 hypothetical protein [Streptomyces hygroscopicus] Length: 446\nNumber of predicted TMHs: 2\nExp number of AAs in TMHs: 43.09951\nExp number, first 60 AAs: 20.54675\nTotal prob of N-in: 0.94830\nPOSSIBLE N-term signal sequence\ninside 1 4\nTMhelix 5 27\noutside 28 408\nTMhelix 409 431\ninside 432 446

34054 GCF_000340845.1_ASM34084v1 Streptomyces hygroscopicus subsp. jinggangensis TL01 Terrabacteria group; Actinobacteria; Actinobacteria; Streptomycetales; Streptomycetaceae; Streptomyces; Streptomyces hygroscopicus; Streptomyces hygroscopicus subsp. jinggangensis
MRHARRRVVRRVTRLAAVGGLLLGGAMVTQAAMASETPSAPARPLAATGDTGAALVARLG WP_014676716.1 S1 family peptidase [Streptomyces hygroscopicus] Length: 458\nNumber of predicted TMHs: 2\nExp number of AAs in TMHs: 43.5516799999999\nExp number, first 60 AAs: 22.26814\nTotal prob of N-in: 0.99236\nPOSSIBLE N-term signal sequence\ninside 1 11\nTMhelix 12 34\noutside 35 419\nTMhelix 420 442\ninside 443 458

34055 GCF_000340845.1_ASM34084v1 Streptomyces hygroscopicus subsp. jinggangensis TL01 Terrabacteria group; Actinobacteria; Actinobacteria; Streptomycetales; Streptomycetaceae; Streptomyces; Streptomyces hygroscopicus; Streptomyces hygroscopicus subsp. jinggangensis
MRRRAVLARGAVIGLLLGLSAALATPTPAPAVPAGAGTAADARPGSYAFTPGARTVAGTK WP_014673788.1 hypothetical protein [Streptomyces hygroscopicus] Length: 446\nNumber of predicted TMHs: 2\nExp number of AAs in TMHs: 43.09951\nExp number, first 60 AAs: 20.54675\nTotal prob of N-in: 0.94830\nPOSSIBLE N-term signal sequence\ninside 1 4\nTMhelix 5 27\noutside 28 408\nTMhelix 409 431\ninside 432 446

34056 GCF_000340845.1_ASM34084v1 Streptomyces hygroscopicus subsp. jinggangensis TL01 Terrabacteria group; Actinobacteria; Actinobacteria; Streptomycetales; Streptomycetaceae; Streptomyces; Streptomyces hygroscopicus; Streptomyces hygroscopicus subsp. jinggangensis
MLRDVENHSLPRAAAFFDLDKTVIAKSSTLTFSKSFYQGGLINRRAALRTAYAQFVFLAG WP_014674034.1 MULTISPECIES: morphological differentiation-associated protein [Streptomyces] Length: 276\nNumber of predicted TMHs: 1\nExp number of AAs in TMHs: 21.37944\nExp number, first 60 AAs: 0.04579\nTotal prob of N-in: 0.01060\noutside 1 247\nTMhelix 248 270\ninside 271 276

34057 GCF_000317595.1_Sipom9103v1.0 Streptomyces ipomoeae 91-03 Terrabacteria group; Actinobacteria; Actinobacteria; Streptomycetales; Streptomycetaceae; Streptomyces; Streptomyces ipomoeae
MLGPVENHSLPRTAAFFDLDKTVIAKSSTLTFSKSFYQGGLINRRAALRTAYTQFLFLAG WP_048821724.1 inhibition of morphological differentiation protein [Streptomyces ipomoeae] Length: 274\nNumber of predicted TMHs: 1\nExp number of AAs in TMHs: 21.72054\nExp number, first 60 AAs: 0.01569\nTotal prob of N-in: 0.00306\noutside 1 244\nTMhelix 245 267\ninside 268 274

34058 GCF_000718705.1_ASM71870v1 Streptomyces lavendulae subsp. lavendulae Terrabacteria group; Actinobacteria; Actinobacteria; Streptomycetales; Streptomycetaceae; Streptomyces; Streptomyces lavendulae
MPQYRDARRALLATAFALTLTAAGATVATAAPHPPYALRLDGAGECTFPMKKQIADRPWA WP_030651182.1 MULTISPECIES: type VII secretion-associated serine protease [Streptomyces] Length: 414\nNumber of predicted TMHs: 2\nExp number of AAs in TMHs: 43.048\nExp number, first 60 AAs: 20.71014\nTotal prob of N-in:

0.96189\nPOSSIBLE N-term signal sequence\ninside 1 8\nTMhelix 9 31\noutside 32 380\nTMhelix 381 403\ninside 404 414

34059 GCF_000718705.1_ASM71870v1 Streptomyces lavendulae subsp. lavendulae Terrabacteria group; Actinobacteria; Actinobacteria; Streptomycetales; Streptomycetaceae; Streptomyces; Streptomyces lavendulae MSTVRRRTLVRAAAVTACAGSLLALPTAAALAEGVPAASSAHSAAPQRTLVKSLSLADGV WP_030660217.1 MULTISPECIES: hypothetical protein [Streptomyces] Length: 218\nNumber of predicted TMHs: 2\nExp number of AAs in TMHs: 42.00955\nExp number, first 60 AAs: 19.35386\nTotal prob of N-in: 0.95630\nPOSSIBLE N-term signal sequence\ninside 1 8\nTMhelix 9 31\noutside 32 188\nTMhelix 189 211\ninside 212 218

34060 GCF_000718155.1_ASM71815v1 Streptomyces lavendulae subsp. lavendulae Terrabacteria group; Actinobacteria; Actinobacteria; Streptomycetales; Streptomycetaceae; Streptomyces; Streptomyces lavendulae MSTVSRRSTVRAAAVTACVAGLLALPAATAFAEGRPASGAQRTLVKSIFLADGRSTAKVYWP_030235708.1 hypothetical protein [Streptomyces lavendulae] Length: 212\nNumber of predicted TMHs: 2\nExp number of AAs in TMHs: 43.22327\nExp number, first 60 AAs: 20.6263\nTotal prob of N-in: 0.96743\nPOSSIBLE N-term signal sequence\ninside 1 12\nTMhelix 13 32\noutside 33 182\nTMhelix 183 205\ninside 206 212

34061 GCF_000718155.1_ASM71815v1 Streptomyces lavendulae subsp. lavendulae Terrabacteria group; Actinobacteria; Actinobacteria; Streptomycetales; Streptomycetaceae; Streptomyces; Streptomyces lavendulae MSSHRRPAFLAAAVLTAATLTAAAFVIPATAADAVPAAGAPAAAAPVKVIGGTLDWGLLA WP_030238604.1 hypothetical protein [Streptomyces lavendulae] Length: 495\nNumber of predicted TMHs: 2\nExp number of AAs in TMHs: 44.67959\nExp number, first 60 AAs: 22.31854\nTotal prob of N-in: 0.97027\nPOSSIBLE N-term signal sequence\ninside 1 6\nTMhelix 7 29\noutside 30 463\nTMhelix 464 486\ninside 487 495

34062 GCF_000721155.1_ASM72115v1 Streptomyces lavendulae subsp. lavendulae Terrabacteria group; Actinobacteria; Actinobacteria; Streptomycetales; Streptomycetaceae; Streptomyces; Streptomyces lavendulae MTRRPLLRAAGVALSAAALTVLPALPAHADGIRAQEWALKAMNTQDAWETTKGEGITVAV WP_003987438.1 MULTISPECIES: type VII secretion-associated serine protease [Streptomyces] Length: 386\nNumber of predicted TMHs: 1\nExp number of AAs in TMHs: 34.80558\nExp number, first 60 AAs: 10.27095\nTotal prob of N-in: 0.55118\nPOSSIBLE N-term signal sequence\noutside 1 359\nTMhelix 360 382\ninside 383 386

34063 GCF_000721155.1_ASM72115v1 Streptomyces lavendulae subsp. lavendulae Terrabacteria group; Actinobacteria; Actinobacteria; Streptomycetales; Streptomycetaceae; Streptomyces; Streptomyces lavendulae MTERSAPAASATGPGTPARRTAVRAAATGTLAAALLGLAPQSALAATPAPPPAPAISAVT WP_030180617.1 MULTISPECIES: hypothetical protein [Streptomyces] Length: 293\nNumber of predicted TMHs: 1\nExp number of AAs in TMHs: 24.94793\nExp number, first 60 AAs: 4.55383\nTotal prob of N-in: 0.22746\noutside 1 264\nTMhelix 265 287\ninside 288 293

34064 GCF_000718855.1_ASM71885v1 Streptomyces lavendulae subsp. lavendulae Terrabacteria group; Actinobacteria; Actinobacteria; Streptomycetales; Streptomycetaceae; Streptomyces; Streptomyces lavendulae MESRQPVGPRANGHRAAHRRRAAVALAGAVCAVGLTATTAAPALADTNVPLNNGQCNFGA WP_078586913.1 MULTISPECIES: type VII secretion-associated serine protease mycosin [Streptomyces] Length: 427\nNumber of predicted TMHs: 2\nExp number of AAs in TMHs: 43.8948\nExp number, first 60 AAs: 21.08952\nTotal prob of N-in: 0.94565\nPOSSIBLE N-term signal sequence\ninside 1 20\nTMhelix 21 43\noutside 44 389\nTMhelix 390 412\ninside 413 427

34065 GCF_000718855.1_ASM71885v1 Streptomyces lavendulae subsp. lavendulae Terrabacteria group; Actinobacteria; Actinobacteria; Streptomycetales; Streptomycetaceae; Streptomyces; Streptomyces lavendulae MTERSAPAASATGPGTPARRTAVRAAATGTLAAALLGLAPQSALAATPAPPPAPAISAVT WP_030180617.1 MULTISPECIES: hypothetical protein [Streptomyces] Length: 293\nNumber of predicted TMHs: 1\nExp number of AAs in TMHs: 24.94793\nExp number, first 60 AAs: 4.55383\nTotal prob of N-in: 0.22746\noutside 1 264\nTMhelix 265 287\ninside 288 293

34066 GCF_000718855.1_ASM71885v1 Streptomyces lavendulae subsp. lavendulae Terrabacteria group; Actinobacteria; Actinobacteria; Streptomycetales; Streptomycetaceae; Streptomyces; Streptomyces lavendulae MTRRPLLRAAGVALSAAALTVLPALPAHADGIRAQEWALKAMNTQDAWETTKGEGITVAV WP_003987438.1 MULTISPECIES: type VII secretion-associated serine protease [Streptomyces] Length: 386\nNumber of predicted TMHs: 1\nExp number of AAs in TMHs: 34.80558\nExp number, first 60 AAs: 10.27095\nTotal prob of N-in: 0.55118\nPOSSIBLE N-term signal sequence\noutside 1 359\nTMhelix 360 382\ninside 383 386

34067 GCF_000718155.1_ASM71815v1 Streptomyces lavendulae subsp. lavendulae Terrabacteria group; Actinobacteria; Actinobacteria; Streptomycetales; Streptomycetaceae; Streptomyces; Streptomyces lavendulae
MAEAADIQGAAPAPARRRWLRRAVVLLTGTPVLAALIYAPAPPAQAADGSVDVQLTEVSP WP_030225214.1
hypothetical protein [Streptomyces lavendulae] Length: 755\nNumber of predicted TMHs: 2\nExp number of AAs in TMHs: 39.64079\nExp number, first 60 AAs: 18.06334\nTotal prob of N-in: 0.86085\nPOSSIBLE N-term signal sequence\ninside 1 18\nTMhelix 19 41\noutside 42 688\nTMhelix 689 711\ninside 712 755

34068 GCF_000718155.1_ASM71815v1 Streptomyces lavendulae subsp. lavendulae Terrabacteria group; Actinobacteria; Actinobacteria; Streptomycetales; Streptomycetaceae; Streptomyces; Streptomyces lavendulae
MSSPDPQARAPHDAENRPEHGDSAPGVRQSRNQAEPRRRSPVIAVTGAASGVGAALVARL WP_030225883.1
NAD-dependent dehydratase [Streptomyces lavendulae] Length: 388\nNumber of predicted TMHs: 1\nExp number of AAs in TMHs: 21.30752\nExp number, first 60 AAs: 1.20635\nTotal prob of N-in: 0.07392\noutside 1 359\nTMhelix 360 379\ninside 380 388

34069 GCF_000718155.1_ASM71815v1 Streptomyces lavendulae subsp. lavendulae Terrabacteria group; Actinobacteria; Actinobacteria; Streptomycetales; Streptomycetaceae; Streptomyces; Streptomyces lavendulae
MKNLKRVPLSARRAAVVAAAGAVVALGGPAFASEGASSGAAAPSVTTSAAPSPVPSTPGG WP_051840821.1
LPXTG cell wall anchor domain-containing protein [Streptomyces lavendulae] Length: 160\nNumber of predicted TMHs: 2\nExp number of AAs in TMHs: 42.45248\nExp number, first 60 AAs: 20.2554\nTotal prob of N-in: 0.97700\nPOSSIBLE N-term signal sequence\ninside 1 12\nTMhelix 13 32\noutside 33 131\nTMhelix 132 154\ninside 155 160

34070 GCF_000721155.1_ASM72115v1 Streptomyces lavendulae subsp. lavendulae Terrabacteria group; Actinobacteria; Actinobacteria; Streptomycetales; Streptomycetaceae; Streptomyces; Streptomyces lavendulae
MESRQPVGPRANGHRAAHRRRAAVALAGAVCAVGLTATTAAPALADTNVPLNNGQCNFGA WP_078586913.1
MULTISPECIES: type VII secretion-associated serine protease mycosin [Streptomyces] Length: 427\nNumber of predicted TMHs: 2\nExp number of AAs in TMHs: 43.8948\nExp number, first 60 AAs: 21.08952\nTotal prob of N-in: 0.94565\nPOSSIBLE N-term signal sequence\ninside 1 20\nTMhelix 21 43\noutside 44 389\nTMhelix 390 412\ninside 413 427

34071 GCF_000403665.1_Streptomyces_lividans_1326 Streptomyces lividans 1326 Terrabacteria group; Actinobacteria; Actinobacteria; Streptomycetales; Streptomycetaceae; Streptomyces; Streptomyces lividans
MRHARRRIVRRVARLAAVGGLLLGGTMVTRAVASEPPDASAAPRTFAQTPSGAGGDLVSR WP_016325237.1
hypothetical protein [Streptomyces lividans] Length: 454\nNumber of predicted TMHs: 2\nExp number of AAs in TMHs: 40.7848\nExp number, first 60 AAs: 21.15262\nTotal prob of N-in: 0.96513\nPOSSIBLE N-term signal sequence\ninside 1 11\nTMhelix 12 34\noutside 35 415\nTMhelix 416 438\ninside 439 454

34072 GCF_000403665.1_Streptomyces_lividans_1326 Streptomyces lividans 1326 Terrabacteria group; Actinobacteria; Actinobacteria; Streptomycetales; Streptomycetaceae; Streptomyces; Streptomyces lividans
MPRTAAFFDLDKTVIAKSSTLTFSKSFYQGGLINRRAVLRTAYAQFVFLAGGADHDQMER WP_003975378.1
MULTISPECIES: morphological differentiation-associated protein [Streptomyces] Length: 268\nNumber of predicted TMHs: 1\nExp number of AAs in TMHs: 21.51645\nExp number, first 60 AAs: 0.03737\nTotal prob of N-in: 0.00617\noutside 1 238\nTMhelix 239 261\ninside 262 268

34073 GCF_000403665.1_Streptomyces_lividans_1326 Streptomyces lividans 1326 Terrabacteria group; Actinobacteria; Actinobacteria; Streptomycetales; Streptomycetaceae; Streptomyces; Streptomyces lividans
MGHNRRRRPTGARRATFGAVALILGGSGLVAVNVFASATESGNPAVPLGSSGIAATVDCP WP_079021358.1
hypothetical protein [Streptomyces lividans] Length: 700\nNumber of predicted TMHs: 2\nExp number of AAs in TMHs: 43.09162\nExp number, first 60 AAs: 21.84277\nTotal prob of N-in: 0.99047\nPOSSIBLE N-term signal sequence\ninside 1 16\nTMhelix 17 39\noutside 40 672\nTMhelix 673 695\ninside 696 700

34074 GCF_000739105.1_ASM73910v1 Streptomyces lividans TK24 Terrabacteria group; Actinobacteria; Actinobacteria; Streptomycetales; Streptomycetaceae; Streptomyces; Streptomyces lividans
MRHARRRIVRRVARLAAVGGLLLGGTMVTRAVASEPPDASAAPRTFAQTPSGAGGDLVSR WP_037666917.1
MULTISPECIES: protease [Streptomyces] Length: 454\nNumber of predicted TMHs: 2\nExp number of AAs in TMHs: 40.7848\nExp number, first 60 AAs: 21.15262\nTotal prob of N-in: 0.96513\nPOSSIBLE N-term signal sequence\ninside 1 11\nTMhelix 12 34\noutside 35 415\nTMhelix 416 438\ninside 439 454

34075 GCF_000739105.1_ASM73910v1 Streptomyces lividans TK24 Terrabacteria group; Actinobacteria; Actinobacteria; Streptomycetales; Streptomycetaceae; Streptomyces; Streptomyces lividans
MPRTAAFFDLDKTVIAKSSTLTFSKSFYQGGLINRRAVLRTAYAQFVFLAGGADHDQMER WP_003975378.1

MULTISPECIES: morphological differentiation-associated protein [Streptomyces] Length: 268\nNumber of predicted TMHs: 1\nExp number of AAs in TMHs: 21.51645\nExp number, first 60 AAs: 0.03737\nTotal prob of N-in: 0.00617\noutside 1 238\nTMhelix 239 261\ninside 262 268

34076 GCF_000158935.1_ASM15893v1 Streptomyces lividans TK24 Terrabacteria group; Actinobacteria; Actinobacteria; Streptomycetales; Streptomycetaceae; Streptomyces; Streptomyces lividans MPRTAAFFDLDKTVIAKSSTLTFSKSFYQGGLINRRAVLRTAYAQFVFLAGGADHDQMER WP_003975378.1 MULTISPECIES: morphological differentiation-associated protein [Streptomyces] Length: 268\nNumber of predicted TMHs: 1\nExp number of AAs in TMHs: 21.51645\nExp number, first 60 AAs: 0.03737\nTotal prob of N-in: 0.00617\noutside 1 238\nTMhelix 239 261\ninside 262 268

34077 GCF_000158935.1_ASM15893v1 Streptomyces lividans TK24 Terrabacteria group; Actinobacteria; Actinobacteria; Streptomycetales; Streptomycetaceae; Streptomyces; Streptomyces lividans MRHARRRIVRRVARLAAVGGLLLGGTMVTRAVASEPPDASAAPRTFAQTPSGAGGDLVSR WP_037666917.1 MULTISPECIES: S1 family peptidase [Streptomyces] Length: 454\nNumber of predicted TMHs: 2\nExp number of AAs in TMHs: 40.7848\nExp number, first 60 AAs: 21.15262\nTotal prob of N-in: 0.96513\nPOSSIBLE N-term signal sequence\ninside 1 11\nTMhelix 12 34\noutside 35 415\nTMhelix 416 438\ninside 439 454

34078 GCF_000739105.1_ASM73910v1 Streptomyces lividans TK24 Terrabacteria group; Actinobacteria; Actinobacteria; Streptomycetales; Streptomycetaceae; Streptomyces; Streptomyces lividans MGHNRRRRPTGARRATFGAVALILGGSGLVAVNVFASATESGNPAVPLGSSGIAATVDCP WP_079021358.1 hypothetical protein [Streptomyces lividans] Length: 700\nNumber of predicted TMHs: 2\nExp number of AAs in TMHs: 43.09162\nExp number, first 60 AAs: 21.84277\nTotal prob of N-in: 0.99047\nPOSSIBLE N-term signal sequence\ninside 1 16\nTMhelix 17 39\noutside 40 672\nTMhelix 673 695\ninside 696 700

34079 GCF_000158935.1_ASM15893v1 Streptomyces lividans TK24 Terrabacteria group; Actinobacteria; Actinobacteria; Streptomycetales; Streptomycetaceae; Streptomyces; Streptomyces lividans MGHNRRRRPTGARRATFGAVALILGGSGLVAVNVFASATESGNPAVPLGSSGIAATVDCP WP_079021358.1 hypothetical protein [Streptomyces lividans] Length: 700\nNumber of predicted TMHs: 2\nExp number of AAs in TMHs: 43.09162\nExp number, first 60 AAs: 21.84277\nTotal prob of N-in: 0.99047\nPOSSIBLE N-term signal sequence\ninside 1 16\nTMhelix 17 39\noutside 40 672\nTMhelix 673 695\ninside 696 700

34080 GCF_000342125.1_S.mobaraensisn_1.0 Streptomyces mobaraensis NBRC 13819 = DSM 40847 Terrabacteria group; Actinobacteria; Actinobacteria; Streptomycetales; Streptomycetaceae; Streptomyces; Streptomyces mobaraensis MLGDVENHSSRRTAAFFDLDKTVIAKSSTLTFGKSFYQGGLINRRAALRTAYIQFVFLLG WP_004939755.1 phosphoserine phosphatase [Streptomyces mobaraensis] Length: 282\nNumber of predicted TMHs: 1\nExp number of AAs in TMHs: 22.41627\nExp number, first 60 AAs: 0.70135\nTotal prob of N-in: 0.04383\noutside 1 246\nTMhelix 247 269\ninside 270 282

34081 GCF_000342125.1_S.mobaraensisn_1.0 Streptomyces mobaraensis NBRC 13819 = DSM 40847 Terrabacteria group; Actinobacteria; Actinobacteria; Streptomycetales; Streptomycetaceae; Streptomyces; Streptomyces mobaraensis MSSTTRSSFGMPVSRRAVVPSAAALAAALTALSASPASAAPATGDHRTGAADAVVLRAGL WP_004943253.1 hypothetical protein [Streptomyces mobaraensis] Length: 327\nNumber of predicted TMHs: 1\nExp number of AAs in TMHs: 25.45905\nExp number, first 60 AAs: 3.51694\nTotal prob of N-in: 0.16298\noutside 1 294\nTMhelix 295 317\ninside 318 327

34082 GCF_000935125.1_SNA_1.0 Streptomyces natalensis ATCC 27448 Terrabacteria group; Actinobacteria; Actinobacteria; Streptomycetales; Streptomycetaceae; Streptomyces; Streptomyces natalensis MENHSVPHSTPRTAAFFDLDKTVIAKSSTLTFSKSFYQGGLINRRAVLRTAYAQFVFLAG WP_044362437.1 inhibition of morphological differentiation protein [Streptomyces natalensis] Length: 285\nNumber of predicted TMHs: 1\nExp number of AAs in TMHs: 21.90974\nExp number, first 60 AAs: 0.06659\nTotal prob of N-in: 0.00615\noutside 1 247\nTMhelix 248 270\ninside 271 285

34083 GCF_000935125.1_SNA_1.0 Streptomyces natalensis ATCC 27448 Terrabacteria group; Actinobacteria; Actinobacteria; Streptomycetales; Streptomycetaceae; Streptomyces; Streptomyces natalensis MRQGEDTLQMTDRRRRAALLPLPATTPGAFLLREHKPSPSGASQASGGPDRSGTDKDSPD WP_078970402.1 hypothetical protein [Streptomyces natalensis] Length: 330\nNumber of predicted TMHs: 2\nExp number of AAs in TMHs: 46.75151\nExp number, first 60 AAs: 0.01844\nTotal prob of N-in: 0.75813\ninside 1 196\nTMhelix 197 219\noutside 220 268\nTMhelix 269 291\ninside 292 330

34084 GCF_000497425.1_S.niveus_v1 Streptomyces niveus NCIMB 11891 Terrabacteria group; Actinobacteria; Actinobacteria; Streptomycetales; Streptomycetaceae; Streptomyces; Streptomyces niveus MNASRTNTSRRGVSRPSTVGRTRTARRAAVVGGVAASAVVLMAGPALAHVGVQPVGEAAK WP_023540332.1 hypothetical protein [Streptomyces niveus] Length: 281\nNumber of predicted TMHs: 2\nExp number of AAs in TMHs: 43.33007\nExp number, first 60 AAs: 22.41673\nTotal prob of N-in: 0.99726\nPOSSIBLE N-term signal sequence\ninside 1 29\nTMhelix 30 52\noutside 53 244\nTMhelix 245 267\ninside 268 281

34085 GCF_000497425.1_S.niveus_v1 Streptomyces niveus NCIMB 11891 Terrabacteria group; Actinobacteria; Actinobacteria; Streptomycetales; Streptomycetaceae; Streptomyces; Streptomyces niveus MLGIVENHSLPRTAAFFDLDKTVIAKSSTLTFSKSFYQGGLINRRAVLRTAYTQFVFLAG WP_023539673.1 inhibition of morphological differentiation protein [Streptomyces niveus] Length: 278\nNumber of predicted TMHs: 1\nExp number of AAs in TMHs: 21.70268\nExp number, first 60 AAs: 0.01114\nTotal prob of N-in: 0.00185\noutside 1 247\nTMhelix 248 270\ninside 271 278

34086 GCF_001704275.1_ASM170427v1 Streptomyces noursei ATCC 11455 Terrabacteria group; Actinobacteria; Actinobacteria; Streptomycetales; Streptomycetaceae; Streptomyces; Streptomyces noursei MTLMRLTRPTRPTGQRRRAVTAAAAALMLAGAGVVTTAGTAQAAEVAYLTECQPPPISGL WP_079142857.1 hypothetical protein [Streptomyces noursei] Length: 480\nNumber of predicted TMHs: 2\nExp number of AAs in TMHs: 43.51111\nExp number, first 60 AAs: 21.33744\nTotal prob of N-in: 0.97258\nPOSSIBLE N-term signal sequence\ninside 1 18\nTMhelix 19 41\noutside 42 450\nTMhelix 451 473\ninside 474 480

34087 GCF_002128465.1_ASM212846v1 Streptomyces pharetrae CZA14 Terrabacteria group; Actinobacteria; Actinobacteria; Streptomycetales; Streptomycetaceae; Streptomyces; Streptomyces pharetrae MLRDVENHSLPRTAAFFDLDKTVIAKSSTLTFSKSFYQGGLINRRAALRTAYAQFVFLVG WP_086168507.1 inhibition of morphological differentiation protein [Streptomyces pharetrae] Length: 277\nNumber of predicted TMHs: 1\nExp number of AAs in TMHs: 21.69454\nExp number, first 60 AAs: 0.03199\nTotal prob of N-in: 0.00646\noutside 1 247\nTMhelix 248 270\ninside 271 277

34088 GCF_000176115.2_ASM17611v2 Streptomyces pratensis ATCC 33331 Terrabacteria group; Actinobacteria; Actinobacteria; Streptomycetales; Streptomycetaceae; Streptomyces; Streptomyces pratensis MLTFVENCFSPRTAAFFDLDKTVIAKSSTLTFSKSFYQGGLINRRAVLRTAYTQFVFLAG WP_014155461.1 MULTISPECIES: haloacid dehalogenase [Streptomyces] Length: 279\nNumber of predicted TMHs: 1\nExp number of AAs in TMHs: 21.10966\nExp number, first 60 AAs: 0.01497\nTotal prob of N-in: 0.00234\noutside 1 248\nTMhelix 249 268\ninside 269 279

34089 GCF_000176115.2_ASM17611v2 Streptomyces pratensis ATCC 33331 Terrabacteria group; Actinobacteria; Actinobacteria; Streptomycetales; Streptomycetaceae; Streptomyces; Streptomyces pratensis MTTHPSPRRRLLTAAAATAVLLVTLPVLPAAAEDSTQCTFPSKKYAGRPWSLQRVLMDEL WP_014153729.1 MULTISPECIES: type VII secretion-associated serine protease [Streptomyces] Length: 414\nNumber of predicted TMHs: 1\nExp number of AAs in TMHs: 36.07361\nExp number, first 60 AAs: 13.5063\nTotal prob of N-in: 0.65550\nPOSSIBLE N-term signal sequence\noutside 1 378\nTMhelix 379 401\ninside 402 414

34090 GCF_000176115.2_ASM17611v2 Streptomyces pratensis ATCC 33331 Terrabacteria group; Actinobacteria; Actinobacteria; Streptomycetales; Streptomycetaceae; Streptomyces; Streptomyces pratensis MRTTRRNTLRRGGVGALLAAVTLLTSAGAAVADDPSPSASADGGAAGPTEAGTTFRTATA WP_014152884.1 MULTISPECIES: hypothetical protein [Streptomyces] Length: 274\nNumber of predicted TMHs: 2\nExp number of AAs in TMHs: 40.1724799999999\nExp number, first 60 AAs: 19.30637\nTotal prob of N-in: 0.99359\nPOSSIBLE N-term signal sequence\ninside 1 12\nTMhelix 13 32\noutside 33 237\nTMhelix 238 260\ninside 261 274

34091 GCF_000154945.1_ASM15494v1 Streptomyces pristinaespiralis ATCC 25486 Terrabacteria group; Actinobacteria; Actinobacteria; Streptomycetales; Streptomycetaceae; Streptomyces; Streptomyces pristinaespiralis MLCLVENHSLPRTAAFFDLDKTVIAKSSTLTFSKSFYQGGLINRRAVLRTAYAQFVFLAG WP_005314378.1 morphological differentiation-associated protein [Streptomyces pristinaespiralis] Length: 282\nNumber of predicted TMHs: 1\nExp number of AAs in TMHs: 21.64546\nExp number, first 60 AAs: 0.02262\nTotal prob of N-in: 0.00298\noutside 1 247\nTMhelix 248 270\ninside 271 282

34092 GCF_000154945.1_ASM15494v1 Streptomyces pristinaespiralis ATCC 25486 Terrabacteria group; Actinobacteria; Actinobacteria; Streptomycetales; Streptomycetaceae; Streptomyces; Streptomyces pristinaespiralis MTGRRRTPVAAAVAALTGAVPLLLAAGAPALAHGTPTDPVSRAAVCGLDGAQRASGACRA WP_005307481.1 secreted cellulose-binding protein [Streptomyces pristinaespiralis] Length: 325\nNumber of predicted TMHs:

2\nExp number of AAs in TMHs: 42.9823\nExp number, first 60 AAs: 21.39181\nTotal prob of N-in: 0.96758\nPOSSIBLE N-term signal sequence\ninside 1 8\nTMhelix 9 31\noutside 32 296\nTMhelix 297 319\ninside 320 325

34093 GCF_000367365.1_ASM36736v1 Streptomyces prunicolor NBRC 13075 Terrabacteria group; Actinobacteria; Actinobacteria; Streptomycetales; Streptomycetaceae; Streptomyces; Streptomyces prunicolor MLSAFPALSLRGRRGAVRLVAATWTCGLVAAGVLAGAGPAVADETPQTQGGATATIGGLK WP_019056443.1 TQXA domain-containing protein [Streptomyces prunicolor] Length: 468\nNumber of predicted TMHs: 1\nExp number of AAs in TMHs: 34.54359\nExp number, first 60 AAs: 12.31738\nTotal prob of N-in: 0.53676\nPOSSIBLE N-term signal sequence\noutside 1 437\nTMhelix 438 460\ninside 461 468

34094 GCF_000367365.1_ASM36736v1 Streptomyces prunicolor NBRC 13075 Terrabacteria group; Actinobacteria; Actinobacteria; Streptomycetales; Streptomycetaceae; Streptomyces; Streptomyces prunicolor MRHARRRVIRRVTRLAAVGGLLLGGAMVTQAAMASDSSGTSAATLSAPQTPADKGSALAA WP_026151595.1 S1 family peptidase [Streptomyces prunicolor] Length: 455\nNumber of predicted TMHs: 2\nExp number of AAs in TMHs: 40.97196\nExp number, first 60 AAs: 21.90254\nTotal prob of N-in: 0.98805\nPOSSIBLE N-term signal sequence\ninside 1 11\nTMhelix 12 34\noutside 35 416\nTMhelix 417 439\ninside 440 455

34095 GCF_000367365.1_ASM36736v1 Streptomyces prunicolor NBRC 13075 Terrabacteria group; Actinobacteria; Actinobacteria; Streptomycetales; Streptomycetaceae; Streptomyces; Streptomyces prunicolor MLWVVENHSLPRTAAFFDLDKTVIAKSSTLTFGKSFYQGGLINRRAALRTAYAQFVFLTG WP_019063766.1 inhibition of morphological differentiation protein [Streptomyces prunicolor] Length: 280\nNumber of predicted TMHs: 1\nExp number of AAs in TMHs: 21.61017\nExp number, first 60 AAs: 0.01811\nTotal prob of N-in: 0.00326\noutside 1 247\nTMhelix 248 270\ninside 271 280

34096 GCF_000384175.1_ASM38417v1 Streptomyces purpureus KA281 Terrabacteria group; Actinobacteria; Actinobacteria; Streptomycetales; Streptomycetaceae; Streptomyces; Streptomyces purpureus MLGLVENHSLPRTAAFFDLDKTVIAKSSTLTFSKSFYQGGLINRRAVLRTAYIQFVFLVG WP_019888290.1 inhibition of morphological differentiation protein [Streptomyces purpureus] Length: 278\nNumber of predicted TMHs: 1\nExp number of AAs in TMHs: 21.53038\nExp number, first 60 AAs: 0.0529\nTotal prob of N-in: 0.00335\noutside 1 247\nTMhelix 248 270\ninside 271 278

34097 GCF_000707925.1_r6_51 Streptomyces rimosus R6-500 Terrabacteria group; Actinobacteria; Actinobacteria; Streptomycetales; Streptomycetaceae; Streptomyces; Streptomyces rimosus MTRRPLLRAAGVALSAAALTVLPALPAHADGIRAQEWALKAMNTQDAWETTKGEGITVAV WP_003987438.1 MULTISPECIES: type VII secretion-associated serine protease [Streptomyces] Length: 386\nNumber of predicted TMHs: 1\nExp number of AAs in TMHs: 34.80558\nExp number, first 60 AAs: 10.27095\nTotal prob of N-in: 0.55118\nPOSSIBLE N-term signal sequence\noutside 1 359\nTMhelix 360 382\ninside 383 386

34098 GCF_001279345.1_ASM127934v1 Streptomyces rimosus subsp. pseudoverticillatus Terrabacteria group; Actinobacteria; Actinobacteria; Streptomycetales; Streptomycetaceae; Streptomyces; Streptomyces rimosus MTERSAPAASATGPGTPARRTAVRAAATGTLAAALLGLAPQSALAATPAPPPAPAISAVT WP_053800621.1 hypothetical protein [Streptomyces rimosus] Length: 293\nNumber of predicted TMHs: 1\nExp number of AAs in TMHs: 24.94731\nExp number, first 60 AAs: 4.55388\nTotal prob of N-in: 0.22744\noutside 1 264\nTMhelix 265 287\ninside 288 293

34099 GCF_000721045.1_ASM72104v1 Streptomyces rimosus subsp. rimosus Terrabacteria group; Actinobacteria; Actinobacteria; Streptomycetales; Streptomycetaceae; Streptomyces; Streptomyces rimosus MTRRPLLRAAGVALSAAALTVLPALPAHADGIRAQEWALKAMNTQDAWETTKGEGITVAV WP_003987438.1 MULTISPECIES: type VII secretion-associated serine protease [Streptomyces] Length: 386\nNumber of predicted TMHs: 1\nExp number of AAs in TMHs: 34.80558\nExp number, first 60 AAs: 10.27095\nTotal prob of N-in: 0.55118\nPOSSIBLE N-term signal sequence\noutside 1 359\nTMhelix 360 382\ninside 383 386

34100 GCF_001279095.1_ASM127909v1 Streptomyces rimosus subsp. rimosus Terrabacteria group; Actinobacteria; Actinobacteria; Streptomycetales; Streptomycetaceae; Streptomyces; Streptomyces rimosus MTERSAPAASATGPGTPARRTAVRAAATGTLAAALLGLAPQSALAATPAPPPAPAISAVT WP_030180617.1 MULTISPECIES: hypothetical protein [Streptomyces] Length: 293\nNumber of predicted TMHs: 1\nExp number of AAs in TMHs: 24.94793\nExp number, first 60 AAs: 4.55383\nTotal prob of N-in: 0.22746\noutside 1 264\nTMhelix 265 287\ninside 288 293

34101 GCF_000720595.1_ASM72059v1 Streptomyces rimosus subsp. rimosus Terrabacteria group; Actinobacteria; Actinobacteria; Streptomycetales; Streptomycetaceae; Streptomyces; Streptomyces rimosus

MESRQPVGPRANGHRAAHRRRAAVALAGAVCAVGLTATTAAPALADTNVPLNNGQCNFGA WP_078586913.1 MULTISPECIES: type VII secretion-associated serine protease mycosin [Streptomyces] Length: 427\nNumber of predicted TMHs: 2\nExp number of AAs in TMHs: 43.8948\nExp number, first 60 AAs: 21.08952\nTotal prob of N-in: 0.94565\nPOSSIBLE N-term signal sequence\ninside 1 20\nTMhelix 21 43\noutside 44 389\nTMhelix 390 412\ninside 413 427

34102 GCF_000718865.1_ASM71886v1 Streptomyces rimosus subsp. rimosus Terrabacteria group; Actinobacteria; Actinobacteria; Streptomycetales; Streptomycetaceae; Streptomyces; Streptomyces rimosus MESRQPVGPRANGHRAAHRRRAAVALAGAVCAVGLTATTAAPALADTNVPLNNGQCNFGA WP_078586913.1 MULTISPECIES: type VII secretion-associated serine protease mycosin [Streptomyces] Length: 427\nNumber of predicted TMHs: 2\nExp number of AAs in TMHs: 43.8948\nExp number, first 60 AAs: 21.08952\nTotal prob of N-in: 0.94565\nPOSSIBLE N-term signal sequence\ninside 1 20\nTMhelix 21 43\noutside 44 389\nTMhelix 390 412\ninside 413 427

34103 GCF_000718755.1_ASM71875v1 Streptomyces rimosus subsp. rimosus Terrabacteria group; Actinobacteria; Actinobacteria; Streptomycetales; Streptomycetaceae; Streptomyces; Streptomyces rimosus MTLTRRRHLVRSTAVAAVAGSALLLPAAAAFADSPQPTTAGSPGPAGDQQQKDDRHKDDQ WP_030603358.1 MULTISPECIES: hypothetical protein [Streptomyces] Length: 239\nNumber of predicted TMHs: 2\nExp number of AAs in TMHs: 41.4211\nExp number, first 60 AAs: 20.09485\nTotal prob of N-in: 0.95720\nPOSSIBLE N-term signal sequence\ninside 1 12\nTMhelix 13 32\noutside 33 207\nTMhelix 208 230\ninside 231 239

34104 GCF_000716515.1_ASM71651v1 Streptomyces rimosus subsp. rimosus Terrabacteria group; Actinobacteria; Actinobacteria; Streptomycetales; Streptomycetaceae; Streptomyces; Streptomyces rimosus MESRQPVGPRANGHRAAHRRRAAVALAGAVCAVGLTATTAAPALADTNVPLNNGQCNFGA WP_078586913.1 MULTISPECIES: type VII secretion-associated serine protease mycosin [Streptomyces] Length: 427\nNumber of predicted TMHs: 2\nExp number of AAs in TMHs: 43.8948\nExp number, first 60 AAs: 21.08952\nTotal prob of N-in: 0.94565\nPOSSIBLE N-term signal sequence\ninside 1 20\nTMhelix 21 43\noutside 44 389\nTMhelix 390 412\ninside 413 427

34105 GCF_000720715.1_ASM72071v1 Streptomyces rimosus subsp. rimosus Terrabacteria group; Actinobacteria; Actinobacteria; Streptomycetales; Streptomycetaceae; Streptomyces; Streptomyces rimosus MTERSAPAASATGPGTPARRTAVRAAATGTLAAALLGLAPQSALAATPAPPPAPAISAVT WP_030180617.1 MULTISPECIES: hypothetical protein [Streptomyces] Length: 293\nNumber of predicted TMHs: 1\nExp number of AAs in TMHs: 24.94793\nExp number, first 60 AAs: 4.55383\nTotal prob of N-in: 0.22746\noutside 1 264\nTMhelix 265 287\ninside 288 293

34106 GCF_000720715.1_ASM72071v1 Streptomyces rimosus subsp. rimosus Terrabacteria group; Actinobacteria; Actinobacteria; Streptomycetales; Streptomycetaceae; Streptomyces; Streptomyces rimosus MTRRPLLRAAGVALSAAALTVLPALPAHADGIRAQEWALKAMNTQDAWETTKGEGITVAV WP_003987438.1 MULTISPECIES: type VII secretion-associated serine protease [Streptomyces] Length: 386\nNumber of predicted TMHs: 1\nExp number of AAs in TMHs: 34.80558\nExp number, first 60 AAs: 10.27095\nTotal prob of N-in: 0.55118\nPOSSIBLE N-term signal sequence\noutside 1 359\nTMhelix 360 382\ninside 383 386

34107 GCF_000718675.1_ASM71867v1 Streptomyces rimosus subsp. rimosus Terrabacteria group; Actinobacteria; Actinobacteria; Streptomycetales; Streptomycetaceae; Streptomyces; Streptomyces rimosus MTRRPLLRAAGVALSAAALTVLPALPAHADGIRAQEWALKAMNTQDAWETTKGEGITVAV WP_030373662.1 type VII secretion-associated serine protease [Streptomyces rimosus] Length: 386\nNumber of predicted TMHs: 1\nExp number of AAs in TMHs: 36.27224\nExp number, first 60 AAs: 9.95056\nTotal prob of N-in: 0.56617\noutside 1 359\nTMhelix 360 382\ninside 383 386

34108 GCF_000721665.1_ASM72166v1 Streptomyces rimosus subsp. rimosus Terrabacteria group; Actinobacteria; Actinobacteria; Streptomycetales; Streptomycetaceae; Streptomyces; Streptomyces rimosus MTLTRRRHLVRSTAVAAVAGSALLLPAAAAFADSPQPTTAGSPGAAGDQQQKDDRQKDDQ WP_030654335.1 hypothetical protein [Streptomyces rimosus] Length: 245\nNumber of predicted TMHs: 2\nExp number of AAs in TMHs: 41.16745\nExp number, first 60 AAs: 19.93579\nTotal prob of N-in: 0.94992\nPOSSIBLE N-term signal sequence\ninside 1 12\nTMhelix 13 32\noutside 33 213\nTMhelix 214 236\ninside 237 245

34109 GCF_000717285.1_ASM71728v1 Streptomyces rimosus subsp. rimosus Terrabacteria group; Actinobacteria; Actinobacteria; Streptomycetales; Streptomycetaceae; Streptomyces; Streptomyces rimosus MTRRPLLRAAGVALSAAALTVLPALPAHADGIRAQEWALKAMNTQDAWETTKGEGITVAV WP_003987438.1 MULTISPECIES: type VII secretion-associated serine protease [Streptomyces] Length: 386\nNumber of

predicted TMHs: 1\nExp number of AAs in TMHs: 34.80558\nExp number, first 60 AAs: 10.27095\nTotal prob of N-in: 0.55118\nPOSSIBLE N-term signal sequence\noutside 1 359\nTMhelix 360 382\ninside 383 386

34110 GCF_000717285.1_ASM71728v1 Streptomyces rimosus subsp. rimosus Terrabacteria group; Actinobacteria; Actinobacteria; Streptomycetales; Streptomycetaceae; Streptomyces; Streptomyces rimosus MTERSAPAASATGPGTPARRTAVRAAATGTLAAALLGLAPQSALAATPAPPPAPAISAVT WP_030180617.1 MULTISPECIES: hypothetical protein [Streptomyces] Length: 293\nNumber of predicted TMHs: 1\nExp number of AAs in TMHs: 24.94793\nExp number, first 60 AAs: 4.55383\nTotal prob of N-in: 0.22746\noutside 1 264\nTMhelix 265 287\ninside 288 293

34111 GCF_000721665.1_ASM72166v1 Streptomyces rimosus subsp. rimosus Terrabacteria group; Actinobacteria; Actinobacteria; Streptomycetales; Streptomycetaceae; Streptomyces; Streptomyces rimosus MTERSAPAASATGPGTPARRTAVRAAATGTLAAALLGLAPQSALAATPAPPPAPAISAVT WP_030651718.1 hypothetical protein [Streptomyces rimosus] Length: 293\nNumber of predicted TMHs: 1\nExp number of AAs in TMHs: 23.70867\nExp number, first 60 AAs: 4.49823\nTotal prob of N-in: 0.22752\noutside 1 264\nTMhelix 265 287\ninside 288 293

34112 GCF_000720595.1_ASM72059v1 Streptomyces rimosus subsp. rimosus Terrabacteria group; Actinobacteria; Actinobacteria; Streptomycetales; Streptomycetaceae; Streptomyces; Streptomyces rimosus MTERSAPAASATGPGTPARRTAVRAAATGTLAAALLGLAPQSALAATPAPPPAPAISAVT WP_030634038.1 hypothetical protein [Streptomyces rimosus] Length: 293\nNumber of predicted TMHs: 1\nExp number of AAs in TMHs: 23.67352\nExp number, first 60 AAs: 4.47531\nTotal prob of N-in: 0.22663\noutside 1 264\nTMhelix 265 287\ninside 288 293

34113 GCF_000720605.1_ASM72060v1 Streptomyces rimosus subsp. rimosus Terrabacteria group; Actinobacteria; Actinobacteria; Streptomycetales; Streptomycetaceae; Streptomyces; Streptomyces rimosus MESRQPVGPRANGHRAAHRRRAAVALAGAVCAVGLTATTAAPALADTNVPLNNGQCNFGA WP_078586913.1 MULTISPECIES: type VII secretion-associated serine protease mycosin [Streptomyces] Length: 427\nNumber of predicted TMHs: 2\nExp number of AAs in TMHs: 43.8948\nExp number, first 60 AAs: 21.08952\nTotal prob of N-in: 0.94565\nPOSSIBLE N-term signal sequence\ninside 1 20\nTMhelix 21 43\noutside 44 389\nTMhelix 390 412\ninside 413 427

34114 GCF_000720725.1_ASM72072v1 Streptomyces rimosus subsp. rimosus Terrabacteria group; Actinobacteria; Actinobacteria; Streptomycetales; Streptomycetaceae; Streptomyces; Streptomyces rimosus MTRRPLLRAAGVALSAAALTVLPALPAHADGIRAQEWALKAMNAQDAWETTKGDGITVAV WP_030669572.1 type VII secretion-associated serine protease [Streptomyces rimosus] Length: 386\nNumber of predicted TMHs: 1\nExp number of AAs in TMHs: 37.36955\nExp number, first 60 AAs: 9.67184\nTotal prob of N-in: 0.57940\noutside 1 359\nTMhelix 360 382\ninside 383 386

34115 GCF_000720725.1_ASM72072v1 Streptomyces rimosus subsp. rimosus Terrabacteria group; Actinobacteria; Actinobacteria; Streptomycetales; Streptomycetaceae; Streptomyces; Streptomyces rimosus MTITRRRKLVRTTAVAAVAGSAMLLPAAAAFADSPQPTTAGSPGAEGRQQKDDRQKDDRQ WP_030672730.1 hypothetical protein [Streptomyces rimosus] Length: 242\nNumber of predicted TMHs: 2\nExp number of AAs in TMHs: 41.51356\nExp number, first 60 AAs: 20.19999\nTotal prob of N-in: 0.97716\nPOSSIBLE N-term signal sequence\ninside 1 12\nTMhelix 13 32\noutside 33 210\nTMhelix 211 233\ninside 234 242

34116 GCF_000717815.1_ASM71781v1 Streptomyces rimosus subsp. rimosus Terrabacteria group; Actinobacteria; Actinobacteria; Streptomycetales; Streptomycetaceae; Streptomyces; Streptomyces rimosus MTLTRRRHLVRSTAVAAVAGSALLLPAAAAFADSPQPTTAGSPGAAGDQQQKDDRQKDDQ WP_030376124.1 hypothetical protein [Streptomyces rimosus] Length: 250\nNumber of predicted TMHs: 2\nExp number of AAs in TMHs: 40.95491\nExp number, first 60 AAs: 19.80137\nTotal prob of N-in: 0.94361\nPOSSIBLE N-term signal sequence\ninside 1 12\nTMhelix 13 32\noutside 33 218\nTMhelix 219 241\ninside 242 250

34117 GCF_000717815.1_ASM71781v1 Streptomyces rimosus subsp. rimosus Terrabacteria group; Actinobacteria; Actinobacteria; Streptomycetales; Streptomycetaceae; Streptomyces; Streptomyces rimosus MTERSAPAASATGPGTPARRTAVRAAATGTLAAALLGLAPQSALAATPAPPPAPAISAVT WP_033026720.1 hypothetical protein [Streptomyces rimosus] Length: 293\nNumber of predicted TMHs: 1\nExp number of AAs in TMHs: 23.67277\nExp number, first 60 AAs: 4.47541\nTotal prob of N-in: 0.22660\noutside 1 264\nTMhelix 265 287\ninside 288 293

34118 GCF_000717815.1_ASM71781v1 Streptomyces rimosus subsp. rimosus Terrabacteria group; Actinobacteria; Actinobacteria; Streptomycetales; Streptomycetaceae; Streptomyces; Streptomyces rimosus MESRQPVGPRANGHRAAHRRRAAVALAGAVCAVGLTATTAAPALADTNVPLNNGQCNFGA WP_078586913.1 MULTISPECIES: type VII secretion-associated serine protease mycosin [Streptomyces] Length: 427\nNumber of predicted TMHs: 2\nExp number of AAs in TMHs: 43.8948\nExp number, first 60 AAs: 21.08952\nTotal prob of N-in: 0.94565\nPOSSIBLE N-term signal sequence\ninside 1 20\nTMhelix 21 43\noutside 44 389\nTMhelix 390 412\ninside 413 427

34119 GCF_000718895.1_ASM71889v1 Streptomyces rimosus subsp. rimosus Terrabacteria group; Actinobacteria; Actinobacteria; Streptomycetales; Streptomycetaceae; Streptomyces; Streptomyces rimosus MTERSAPAASATGPGTPARRTAVRAAATGTLAAALLGLAPQSALAATPAPPPAPAISAVT WP_030180617.1 MULTISPECIES: hypothetical protein [Streptomyces] Length: 293\nNumber of predicted TMHs: 1\nExp number of AAs in TMHs: 24.94793\nExp number, first 60 AAs: 4.55383\nTotal prob of N-in: 0.22746\noutside 1 264\nTMhelix 265 287\ninside 288 293

34120 GCF_000718895.1_ASM71889v1 Streptomyces rimosus subsp. rimosus Terrabacteria group; Actinobacteria; Actinobacteria; Streptomycetales; Streptomycetaceae; Streptomyces; Streptomyces rimosus MESRQPVGPRANGHRAAHRRRAAVALAGAVCAVGLTATTAAPALADTNVPLNNGQCNFGA WP_078586913.1 MULTISPECIES: type VII secretion-associated serine protease mycosin [Streptomyces] Length: 427\nNumber of predicted TMHs: 2\nExp number of AAs in TMHs: 43.8948\nExp number, first 60 AAs: 21.08952\nTotal prob of N-in: 0.94565\nPOSSIBLE N-term signal sequence\ninside 1 20\nTMhelix 21 43\noutside 44 389\nTMhelix 390 412\ninside 413 427

34121 GCF_000716745.1_ASM71674v1 Streptomyces rimosus subsp. rimosus Terrabacteria group; Actinobacteria; Actinobacteria; Streptomycetales; Streptomycetaceae; Streptomyces; Streptomyces rimosus MESRQPVGPRANGHRAAHRRRAAVALAGAVCAVGLTATTAAPALADTNVPLNNGQCNFGA WP_078586913.1 MULTISPECIES: type VII secretion-associated serine protease mycosin [Streptomyces] Length: 427\nNumber of predicted TMHs: 2\nExp number of AAs in TMHs: 43.8948\nExp number, first 60 AAs: 21.08952\nTotal prob of N-in: 0.94565\nPOSSIBLE N-term signal sequence\ninside 1 20\nTMhelix 21 43\noutside 44 389\nTMhelix 390 412\ninside 413 427

34122 GCF_000721085.1_ASM72108v1 Streptomyces rimosus subsp. rimosus Terrabacteria group; Actinobacteria; Actinobacteria; Streptomycetales; Streptomycetaceae; Streptomyces; Streptomyces rimosus MTRRPLLRAAGVALSAAALTVLPALPAHADGIRAQEWALKAMNTQDAWETTKGEGITVAV WP_003987438.1 MULTISPECIES: type VII secretion-associated serine protease [Streptomyces] Length: 386\nNumber of predicted TMHs: 1\nExp number of AAs in TMHs: 34.80558\nExp number, first 60 AAs: 10.27095\nTotal prob of N-in: 0.55118\nPOSSIBLE N-term signal sequence\noutside 1 359\nTMhelix 360 382\ninside 383 386

34123 GCF_000721085.1_ASM72108v1 Streptomyces rimosus subsp. rimosus Terrabacteria group; Actinobacteria; Actinobacteria; Streptomycetales; Streptomycetaceae; Streptomyces; Streptomyces rimosus MTERSAPAASATGPGTPARRTAVRAAATGTLAAALLGLAPQSALAATPAPPPAPAISAVT WP_030180617.1 MULTISPECIES: hypothetical protein [Streptomyces] Length: 293\nNumber of predicted TMHs: 1\nExp number of AAs in TMHs: 24.94793\nExp number, first 60 AAs: 4.55383\nTotal prob of N-in: 0.22746\noutside 1 264\nTMhelix 265 287\ninside 288 293

34124 GCF_000721085.1_ASM72108v1 Streptomyces rimosus subsp. rimosus Terrabacteria group; Actinobacteria; Actinobacteria; Streptomycetales; Streptomycetaceae; Streptomyces; Streptomyces rimosus MESRQPVGPRANGHRAAHRRRAAVALAGAVCAVGLTATTAAPALADTNVPLNNGQCNFGA WP_078586913.1 MULTISPECIES: type VII secretion-associated serine protease mycosin [Streptomyces] Length: 427\nNumber of predicted TMHs: 2\nExp number of AAs in TMHs: 43.8948\nExp number, first 60 AAs: 21.08952\nTotal prob of N-in: 0.94565\nPOSSIBLE N-term signal sequence\ninside 1 20\nTMhelix 21 43\noutside 44 389\nTMhelix 390 412\ninside 413 427

34125 GCF_000720605.1_ASM72060v1 Streptomyces rimosus subsp. rimosus Terrabacteria group; Actinobacteria; Actinobacteria; Streptomycetales; Streptomycetaceae; Streptomyces; Streptomyces rimosus MTERSAPAASATGPGTPARRTAVRAAATGTLAAALLGLAPQSALAATPAPPPAPAISAVT WP_030180617.1 MULTISPECIES: hypothetical protein [Streptomyces] Length: 293\nNumber of predicted TMHs: 1\nExp number of AAs in TMHs: 24.94793\nExp number, first 60 AAs: 4.55383\nTotal prob of N-in: 0.22746\noutside 1 264\nTMhelix 265 287\ninside 288 293

34126 GCF_000720725.1_ASM72072v1 Streptomyces rimosus subsp. rimosus Terrabacteria group; Actinobacteria; Actinobacteria; Streptomycetales; Streptomycetaceae; Streptomyces; Streptomyces rimosus MTERSAPAASATGRGTPARRTAVRAAATGTLAAALLGLAPQSALAATPAPPPAPAAPAVT WP_030680149.1 hypothetical protein [Streptomyces rimosus] Length: 293\nNumber of predicted TMHs: 1\nExp number of AAs in TMHs: 29.887\nExp number, first 60 AAs: 10.47538\nTotal prob of N-in: 0.50317\nPOSSIBLE N-term signal sequence\noutside 1 264\nTMhelix 265 287\ninside 288 293

34127 GCF_001279105.1_ASM127910v1 Streptomyces rimosus subsp. rimosus Terrabacteria group; Actinobacteria; Actinobacteria; Streptomycetales; Streptomycetaceae; Streptomyces; Streptomyces rimosus MTERSAPAASATGPGTPARRTAVRAAATGTLAAALLGLAPQSALAATPAPPPAPAISAVT WP_030180617.1 MULTISPECIES: hypothetical protein [Streptomyces] Length: 293\nNumber of predicted TMHs: 1\nExp number of AAs in TMHs: 24.94793\nExp number, first 60 AAs: 4.55383\nTotal prob of N-in: 0.22746\noutside 1 264\nTMhelix 265 287\ninside 288 293

34128 GCF_001279105.1_ASM127910v1 Streptomyces rimosus subsp. rimosus Terrabacteria group; Actinobacteria; Actinobacteria; Streptomycetales; Streptomycetaceae; Streptomyces; Streptomyces rimosus MESRQPVGPRANGHRAAHRRRAAVALAGAVCAVGLTATTAAPALADTNVPLNNGQCNFGA WP_078586913.1 MULTISPECIES: type VII secretion-associated serine protease mycosin [Streptomyces] Length: 427\nNumber of predicted TMHs: 2\nExp number of AAs in TMHs: 43.8948\nExp number, first 60 AAs: 21.08952\nTotal prob of N-in: 0.94565\nPOSSIBLE N-term signal sequence\ninside 1 20\nTMhelix 21 43\noutside 44 389\nTMhelix 390 412\ninside 413 427

34129 GCF_001507395.1_ASM150739v1 Streptomyces rimosus subsp. rimosus Terrabacteria group; Actinobacteria; Actinobacteria; Streptomycetales; Streptomycetaceae; Streptomyces; Streptomyces rimosus MTERSAPAASATGPGTPARRTAVRAAATGTLAAALLGLAPQSALAATPAPPPAPAISAVT WP_030180617.1 MULTISPECIES: hypothetical protein [Streptomyces] Length: 293\nNumber of predicted TMHs: 1\nExp number of AAs in TMHs: 24.94793\nExp number, first 60 AAs: 4.55383\nTotal prob of N-in: 0.22746\noutside 1 264\nTMhelix 265 287\ninside 288 293

34130 GCF_001507395.1_ASM150739v1 Streptomyces rimosus subsp. rimosus Terrabacteria group; Actinobacteria; Actinobacteria; Streptomycetales; Streptomycetaceae; Streptomyces; Streptomyces rimosus MESRQPVGPRANGHRAAHRRRAAVALAGAVCAVGLTATTAAPALADTNVPLNNGQCNFGA WP_078586913.1 MULTISPECIES: type VII secretion-associated serine protease mycosin [Streptomyces] Length: 427\nNumber of predicted TMHs: 2\nExp number of AAs in TMHs: 43.8948\nExp number, first 60 AAs: 21.08952\nTotal prob of N-in: 0.94565\nPOSSIBLE N-term signal sequence\ninside 1 20\nTMhelix 21 43\noutside 44 389\nTMhelix 390 412\ninside 413 427

34131 GCF_000716515.1_ASM71651v1 Streptomyces rimosus subsp. rimosus Terrabacteria group; Actinobacteria; Actinobacteria; Streptomycetales; Streptomycetaceae; Streptomyces; Streptomyces rimosus MTRRPLLRAAGVALSAAALTVLPALPAHADGIRAQEWALKAMNTQDAWETTKGEGITVAV WP_033033885.1 type VII secretion-associated serine protease [Streptomyces rimosus] Length: 386\nNumber of predicted TMHs: 1\nExp number of AAs in TMHs: 34.66295\nExp number, first 60 AAs: 10.30287\nTotal prob of N-in: 0.54968\nPOSSIBLE N-term signal sequence\noutside 1 359\nTMhelix 360 382\ninside 383 386

34132 GCF_000718755.1_ASM71875v1 Streptomyces rimosus subsp. rimosus Terrabacteria group; Actinobacteria; Actinobacteria; Streptomycetales; Streptomycetaceae; Streptomyces; Streptomyces rimosus MTRRPLLRAAGVALSAAALTVLPALPAHADGIRAQEWALKAMNTQDAWETTKGEGITVAV WP_030595365.1 MULTISPECIES: type VII secretion-associated serine protease [Streptomyces] Length: 386\nNumber of predicted TMHs: 1\nExp number of AAs in TMHs: 37.57611\nExp number, first 60 AAs: 9.6597\nTotal prob of N-in: 0.57980\noutside 1 359\nTMhelix 360 382\ninside 383 386

34133 GCF_000720595.1_ASM72059v1 Streptomyces rimosus subsp. rimosus Terrabacteria group; Actinobacteria; Actinobacteria; Streptomycetales; Streptomycetaceae; Streptomyces; Streptomyces rimosus MTRRPLLRAAGVALSAAALTVLPALPAHADGIRAQEWALKAMNTQDAWETTKGEGITVAV WP_030626968.1 type VII secretion-associated serine protease [Streptomyces rimosus] Length: 386\nNumber of predicted TMHs: 1\nExp number of AAs in TMHs: 36.54747\nExp number, first 60 AAs: 9.94287\nTotal prob of N-in: 0.56654\noutside 1 359\nTMhelix 360 382\ninside 383 386

34134 GCF_000718675.1_ASM71867v1 Streptomyces rimosus subsp. rimosus Terrabacteria group; Actinobacteria; Actinobacteria; Streptomycetales; Streptomycetaceae; Streptomyces; Streptomyces rimosus MTERSAPAASATGPGTPARRTAVRAAATGTLAAALLGLAPQSALAATPAPPPAPAISAVT WP_030651718.1

hypothetical protein [Streptomyces rimosus] Length: 293\nNumber of predicted TMHs: 1\nExp number of AAs in TMHs: 23.70867\nExp number, first 60 AAs: 4.49823\nTotal prob of N-in: 0.22752\noutside 1 264\nTMhelix 265 287\ninside 288 293

34135 GCF_000720595.1_ASM72059v1 Streptomyces rimosus subsp. rimosus Terrabacteria group; Actinobacteria; Actinobacteria; Streptomycetales; Streptomycetaceae; Streptomyces; Streptomyces rimosus MTLTRRRHLVRSTAVAAVAGSALLLPAAAAFADSPQPTTAGSPGAAGDQQQKDDRQKDDQ WP_030628046.1 hypothetical protein [Streptomyces rimosus] Length: 245\nNumber of predicted TMHs: 2\nExp number of AAs in TMHs: 41.16899\nExp number, first 60 AAs: 19.93162\nTotal prob of N-in: 0.94990\nPOSSIBLE N-term signal sequence\ninside 1 12\nTMhelix 13 32\noutside 33 213\nTMhelix 214 236\ninside 237 245

34136 GCF_000721665.1_ASM72166v1 Streptomyces rimosus subsp. rimosus Terrabacteria group; Actinobacteria; Actinobacteria; Streptomycetales; Streptomycetaceae; Streptomyces; Streptomyces rimosus MTRRPLLRAAGVALSAAALTVLPALPAHADGIRAQEWALKAMNTQDAWETTKGEGITVAV WP_030373662.1 type VII secretion-associated serine protease [Streptomyces rimosus] Length: 386\nNumber of predicted TMHs: 1\nExp number of AAs in TMHs: 36.27224\nExp number, first 60 AAs: 9.95056\nTotal prob of N-in: 0.56617\noutside 1 359\nTMhelix 360 382\ninside 383 386

34137 GCF_000717285.1_ASM71728v1 Streptomyces rimosus subsp. rimosus Terrabacteria group; Actinobacteria; Actinobacteria; Streptomycetales; Streptomycetaceae; Streptomyces; Streptomyces rimosus MESRQPVGPRANGHRAAHRRRAAVALAGAVCAVGLTATTAAPALADTNVPLNNGQCNFGA WP_078586913.1 MULTISPECIES: type VII secretion-associated serine protease mycosin [Streptomyces] Length: 427\nNumber of predicted TMHs: 2\nExp number of AAs in TMHs: 43.8948\nExp number, first 60 AAs: 21.08952\nTotal prob of N-in: 0.94565\nPOSSIBLE N-term signal sequence\ninside 1 20\nTMhelix 21 43\noutside 44 389\nTMhelix 390 412\ninside 413 427

34138 GCF_001279065.1_ASM127906v1 Streptomyces rimosus subsp. rimosus Terrabacteria group; Actinobacteria; Actinobacteria; Streptomycetales; Streptomycetaceae; Streptomyces; Streptomyces rimosus MTRRPLLRAAGVALSAAALTVLPALPAHADGIRAQEWALKAMNTQDAWETTKGEGITVAV WP_031187823.1 MULTISPECIES: type VII secretion-associated serine protease [Streptomyces] Length: 386\nNumber of predicted TMHs: 1\nExp number of AAs in TMHs: 36.26455\nExp number, first 60 AAs: 9.94874\nTotal prob of N-in: 0.56610\noutside 1 359\nTMhelix 360 382\ninside 383 386

34139 GCF_001279065.1_ASM127906v1 Streptomyces rimosus subsp. rimosus Terrabacteria group; Actinobacteria; Actinobacteria; Streptomycetales; Streptomycetaceae; Streptomyces; Streptomyces rimosus MTLTRRRHLVRSTAVAAVAGSALLLPAAAAFADSPQPTTAGSPGAAGDQQQKDDRQKDDQ WP_031191693.1 MULTISPECIES: hypothetical protein [Streptomyces] Length: 250\nNumber of predicted TMHs: 2\nExp number of AAs in TMHs: 40.95506\nExp number, first 60 AAs: 19.80107\nTotal prob of N-in: 0.94361\nPOSSIBLE N-term signal sequence\ninside 1 12\nTMhelix 13 32\noutside 33 218\nTMhelix 219 241\ninside 242 250

34140 GCF_001279065.1_ASM127906v1 Streptomyces rimosus subsp. rimosus Terrabacteria group; Actinobacteria; Actinobacteria; Streptomycetales; Streptomycetaceae; Streptomyces; Streptomyces rimosus MTERSAPAASATGPGTPARRTAVRAAATGTLAAALLGLAPQSALAATPAPPPAPAISAVA WP_032915072.1 MULTISPECIES: hypothetical protein [Streptomyces] Length: 293\nNumber of predicted TMHs: 1\nExp number of AAs in TMHs: 23.67473\nExp number, first 60 AAs: 4.47659\nTotal prob of N-in: 0.22667\noutside 1 264\nTMhelix 265 287\ninside 288 293

34141 GCF_001279065.1_ASM127906v1 Streptomyces rimosus subsp. rimosus Terrabacteria group; Actinobacteria; Actinobacteria; Streptomycetales; Streptomycetaceae; Streptomyces; Streptomyces rimosus MESRQPVGPRANGHRAAHRRRAAVALAGAVCAVGLTATTAAPALADTNVPLNNGQCNFGA WP_078586913.1 MULTISPECIES: type VII secretion-associated serine protease mycosin [Streptomyces] Length: 427\nNumber of predicted TMHs: 2\nExp number of AAs in TMHs: 43.8948\nExp number, first 60 AAs: 21.08952\nTotal prob of N-in: 0.94565\nPOSSIBLE N-term signal sequence\ninside 1 20\nTMhelix 21 43\noutside 44 389\nTMhelix 390 412\ninside 413 427

34142 GCF_000718865.1_ASM71886v1 Streptomyces rimosus subsp. rimosus Terrabacteria group; Actinobacteria; Actinobacteria; Streptomycetales; Streptomycetaceae; Streptomyces; Streptomyces rimosus MTERSAPAASATGPGTPARRTAVRAAATGTLAAALLGLAPQSALAATPAPPPAPAISAVT WP_030180617.1 MULTISPECIES: hypothetical protein [Streptomyces] Length: 293\nNumber of predicted TMHs: 1\nExp number of AAs in TMHs: 24.94793\nExp number, first 60 AAs: 4.55383\nTotal prob of N-in: 0.22746\noutside 1 264\nTMhelix 265 287\ninside 288 293

34143 GCF_000720715.1_ASM72071v1 Streptomyces rimosus subsp. rimosus Terrabacteria group; Actinobacteria; Actinobacteria; Streptomycetales; Streptomycetaceae; Streptomyces; Streptomyces rimosus MESRQPVGPRANGHRAAHRRRAAVALAGAVCAVGLTATTAAPALADTNVPLNNGQCNFGA WP_078586913.1 MULTISPECIES: type VII secretion-associated serine protease mycosin [Streptomyces] Length: 427\nNumber of predicted TMHs: 2\nExp number of AAs in TMHs: 43.8948\nExp number, first 60 AAs: 21.08952\nTotal prob of N-in: 0.94565\nPOSSIBLE N-term signal sequence\ninside 1 20\nTMhelix 21 43\noutside 44 389\nTMhelix 390 412\ninside 413 427

34144 GCF_000720605.1_ASM72060v1 Streptomyces rimosus subsp. rimosus Terrabacteria group; Actinobacteria; Actinobacteria; Streptomycetales; Streptomycetaceae; Streptomyces; Streptomyces rimosus MTRRPLLRAAGVALSAAALTVLPALPAHADGIRAQEWALKAMNTQDAWETTKGEGITVAV WP_003987438.1 MULTISPECIES: type VII secretion-associated serine protease [Streptomyces] Length: 386\nNumber of predicted TMHs: 1\nExp number of AAs in TMHs: 34.80558\nExp number, first 60 AAs: 10.27095\nTotal prob of N-in: 0.55118\nPOSSIBLE N-term signal sequence\noutside 1 359\nTMhelix 360 382\ninside 383 386

34145 GCF_000720685.1_ASM72068v1 Streptomyces rimosus subsp. rimosus Terrabacteria group; Actinobacteria; Actinobacteria; Streptomycetales; Streptomycetaceae; Streptomyces; Streptomyces rimosus MTERSAPAASATGPGTPARRTAVRAAATGTLAAALLGLAPQSALAATPAPPPAPAISAVT WP_030180617.1 MULTISPECIES: hypothetical protein [Streptomyces] Length: 293\nNumber of predicted TMHs: 1\nExp number of AAs in TMHs: 24.94793\nExp number, first 60 AAs: 4.55383\nTotal prob of N-in: 0.22746\noutside 1 264\nTMhelix 265 287\ninside 288 293

34146 GCF_000720565.1_ASM72056v1 Streptomyces rimosus subsp. rimosus Terrabacteria group; Actinobacteria; Actinobacteria; Streptomycetales; Streptomycetaceae; Streptomyces; Streptomyces rimosus MTERSAPAASATGPGTPARRTAVRAAATGTLAAALLGLAPQSALAATPAPPPAPAISAVT WP_030180617.1 MULTISPECIES: hypothetical protein [Streptomyces] Length: 293\nNumber of predicted TMHs: 1\nExp number of AAs in TMHs: 24.94793\nExp number, first 60 AAs: 4.55383\nTotal prob of N-in: 0.22746\noutside 1 264\nTMhelix 265 287\ninside 288 293

34147 GCF_000720685.1_ASM72068v1 Streptomyces rimosus subsp. rimosus Terrabacteria group; Actinobacteria; Actinobacteria; Streptomycetales; Streptomycetaceae; Streptomyces; Streptomyces rimosus MESRQPVGPRANGHRAAHRRRAAVALAGAVCAVGLTATTAAPALADTNVPLNNGQCNFGA WP_078586913.1 MULTISPECIES: type VII secretion-associated serine protease mycosin [Streptomyces] Length: 427\nNumber of predicted TMHs: 2\nExp number of AAs in TMHs: 43.8948\nExp number, first 60 AAs: 21.08952\nTotal prob of N-in: 0.94565\nPOSSIBLE N-term signal sequence\ninside 1 20\nTMhelix 21 43\noutside 44 389\nTMhelix 390 412\ninside 413 427

34148 GCF_000720565.1_ASM72056v1 Streptomyces rimosus subsp. rimosus Terrabacteria group; Actinobacteria; Actinobacteria; Streptomycetales; Streptomycetaceae; Streptomyces; Streptomyces rimosus MESRQPVGPRANGHRAAHRRRAAVALAGAVCAVGLTATTAAPALADTNVPLNNGQCNFGA WP_078586913.1 MULTISPECIES: type VII secretion-associated serine protease mycosin [Streptomyces] Length: 427\nNumber of predicted TMHs: 2\nExp number of AAs in TMHs: 43.8948\nExp number, first 60 AAs: 21.08952\nTotal prob of N-in: 0.94565\nPOSSIBLE N-term signal sequence\ninside 1 20\nTMhelix 21 43\noutside 44 389\nTMhelix 390 412\ninside 413 427

34149 GCF_000720685.1_ASM72068v1 Streptomyces rimosus subsp. rimosus Terrabacteria group; Actinobacteria; Actinobacteria; Streptomycetales; Streptomycetaceae; Streptomyces; Streptomyces rimosus MTRRPLLRAAGVALSAAALTVLPALPAHADGIRAQEWALKAMNTQDAWETTKGEGITVAV WP_003987438.1 MULTISPECIES: type VII secretion-associated serine protease [Streptomyces] Length: 386\nNumber of predicted TMHs: 1\nExp number of AAs in TMHs: 34.80558\nExp number, first 60 AAs: 10.27095\nTotal prob of N-in: 0.55118\nPOSSIBLE N-term signal sequence\noutside 1 359\nTMhelix 360 382\ninside 383 386

34150 GCF_000720565.1_ASM72056v1 Streptomyces rimosus subsp. rimosus Terrabacteria group; Actinobacteria; Actinobacteria; Streptomycetales; Streptomycetaceae; Streptomyces; Streptomyces rimosus MTRRPLLRAAGVALSAAALTVLPALPAHADGIRAQEWALKAMNTQDAWETTKGEGITVAV WP_003987438.1 MULTISPECIES: type VII secretion-associated serine protease [Streptomyces] Length: 386\nNumber of predicted TMHs: 1\nExp number of AAs in TMHs: 34.80558\nExp number, first 60 AAs: 10.27095\nTotal prob of N-in: 0.55118\nPOSSIBLE N-term signal sequence\noutside 1 359\nTMhelix 360 382\ninside 383 386

34151 GCF_000719185.1_ASM71918v1 Streptomyces rimosus subsp. rimosus Terrabacteria group; Actinobacteria; Actinobacteria; Streptomycetales; Streptomycetaceae; Streptomyces; Streptomyces rimosus

MTERSAPAASATAPGTPARRTAVRAAATGTLAAALLGLAPQSALAATPAPPPAPAISAVT WP_030603830.1 MULTISPECIES: hypothetical protein [Streptomyces] Length: 293\nNumber of predicted TMHs: 1\nExp number of AAs in TMHs: 24.61251\nExp number, first 60 AAs: 5.37395\nTotal prob of N-in: 0.26790\noutside 1 264\nTMhelix 265 287\ninside 288 293

34152 GCF_000721045.1_ASM72104v1 Streptomyces rimosus subsp. rimosus Terrabacteria group; Actinobacteria; Actinobacteria; Streptomycetales; Streptomycetaceae; Streptomyces; Streptomyces rimosus MESRQPVGPRANGHRAAHRRRAAVALAGAVCAVGLTATTAAPALADTNVPLNNGQCNFGA WP_078586913.1 MULTISPECIES: type VII secretion-associated serine protease mycosin [Streptomyces] Length: 427\nNumber of predicted TMHs: 2\nExp number of AAs in TMHs: 43.8948\nExp number, first 60 AAs: 21.08952\nTotal prob of N-in: 0.94565\nPOSSIBLE N-term signal sequence\ninside 1 20\nTMhelix 21 43\noutside 44 389\nTMhelix 390 412\ninside 413 427

34153 GCF_000721045.1_ASM72104v1 Streptomyces rimosus subsp. rimosus Terrabacteria group; Actinobacteria; Actinobacteria; Streptomycetales; Streptomycetaceae; Streptomyces; Streptomyces rimosus MTERSAPAASATGPGTPARRTAVRAAATGTLAAALLGLAPQSALAATPAPPPAPAISAVT WP_030180617.1 MULTISPECIES: hypothetical protein [Streptomyces] Length: 293\nNumber of predicted TMHs: 1\nExp number of AAs in TMHs: 24.94793\nExp number, first 60 AAs: 4.55383\nTotal prob of N-in: 0.22746\noutside 1 264\nTMhelix 265 287\ninside 288 293

34154 GCF_001279375.1_ASM127937v1 Streptomyces rimosus subsp. rimosus Terrabacteria group; Actinobacteria; Actinobacteria; Streptomycetales; Streptomycetaceae; Streptomyces; Streptomyces rimosus MESRQPVGPRANGHRAAHRRRAAVALAGAVCAVGLTATTAAPALADTNVPLNNGQCNFGA WP_078586913.1 MULTISPECIES: type VII secretion-associated serine protease mycosin [Streptomyces] Length: 427\nNumber of predicted TMHs: 2\nExp number of AAs in TMHs: 43.8948\nExp number, first 60 AAs: 21.08952\nTotal prob of N-in: 0.94565\nPOSSIBLE N-term signal sequence\ninside 1 20\nTMhelix 21 43\noutside 44 389\nTMhelix 390 412\ninside 413 427

34155 GCF_001279095.1_ASM127909v1 Streptomyces rimosus subsp. rimosus Terrabacteria group; Actinobacteria; Actinobacteria; Streptomycetales; Streptomycetaceae; Streptomyces; Streptomyces rimosus MESRQPVGPRANGHRAAHRRRAAVALAGAVCAVGLTATTAAPALADTNVPLNNGQCNFGA WP_078586913.1 MULTISPECIES: type VII secretion-associated serine protease mycosin [Streptomyces] Length: 427\nNumber of predicted TMHs: 2\nExp number of AAs in TMHs: 43.8948\nExp number, first 60 AAs: 21.08952\nTotal prob of N-in: 0.94565\nPOSSIBLE N-term signal sequence\ninside 1 20\nTMhelix 21 43\noutside 44 389\nTMhelix 390 412\ninside 413 427

34156 GCF_001279375.1_ASM127937v1 Streptomyces rimosus subsp. rimosus Terrabacteria group; Actinobacteria; Actinobacteria; Streptomycetales; Streptomycetaceae; Streptomyces; Streptomyces rimosus MTERSAPAASATGPGTPARRTAVRAAATGTLAAALLGLAPQSALAATPAPPPAPAISAVT WP_030180617.1 MULTISPECIES: hypothetical protein [Streptomyces] Length: 293\nNumber of predicted TMHs: 1\nExp number of AAs in TMHs: 24.94793\nExp number, first 60 AAs: 4.55383\nTotal prob of N-in: 0.22746\noutside 1 264\nTMhelix 265 287\ninside 288 293

34157 GCF_000719185.1_ASM71918v1 Streptomyces rimosus subsp. rimosus Terrabacteria group; Actinobacteria; Actinobacteria; Streptomycetales; Streptomycetaceae; Streptomyces; Streptomyces rimosus MTLTRRRHLVRSTAVAAVAGSALLLPAAAAFADSPQPTTAGSPGPAGDQQQKDDRHKDDQ WP_030603358.1 MULTISPECIES: hypothetical protein [Streptomyces] Length: 239\nNumber of predicted TMHs: 2\nExp number of AAs in TMHs: 41.4211\nExp number, first 60 AAs: 20.09485\nTotal prob of N-in: 0.95720\nPOSSIBLE N-term signal sequence\ninside 1 12\nTMhelix 13 32\noutside 33 207\nTMhelix 208 230\ninside 231 239

34158 GCF_000719185.1_ASM71918v1 Streptomyces rimosus subsp. rimosus Terrabacteria group; Actinobacteria; Actinobacteria; Streptomycetales; Streptomycetaceae; Streptomyces; Streptomyces rimosus MTRRPLLRAAGVALSAAALTVLPALPAHADGIRAQEWALKAMNTQDAWETTKGEGITVAV WP_030595365.1 MULTISPECIES: type VII secretion-associated serine protease [Streptomyces] Length: 386\nNumber of predicted TMHs: 1\nExp number of AAs in TMHs: 37.57611\nExp number, first 60 AAs: 9.6597\nTotal prob of N-in: 0.57980\noutside 1 359\nTMhelix 360 382\ninside 383 386

34159 GCF_000718865.1_ASM71886v1 Streptomyces rimosus subsp. rimosus Terrabacteria group; Actinobacteria; Actinobacteria; Streptomycetales; Streptomycetaceae; Streptomyces; Streptomyces rimosus MTRRPLLRAAGVALSAAALTVLPALPAHADGIRAQEWALKAMNTQDAWETTKGEGITVAV WP_003987438.1 MULTISPECIES: type VII secretion-associated serine protease [Streptomyces] Length: 386\nNumber of

predicted TMHs: 1\nExp number of AAs in TMHs: 34.80558\nExp number, first 60 AAs: 10.27095\nTotal prob of N-in: 0.55118\nPOSSIBLE N-term signal sequence\noutside 1 359\nTMhelix 360 382\ninside 383 386

34160 GCF_000716515.1_ASM71651v1 Streptomyces rimosus subsp. rimosus Terrabacteria group; Actinobacteria; Actinobacteria; Streptomycetales; Streptomycetaceae; Streptomyces; Streptomyces rimosus MTERSAPAASATGPGTPARRTAVRAAATGTLAAALLGLAPQSALAATPAPPPAPAISAVT WP_030180617.1 MULTISPECIES: hypothetical protein [Streptomyces] Length: 293\nNumber of predicted TMHs: 1\nExp number of AAs in TMHs: 24.94793\nExp number, first 60 AAs: 4.55383\nTotal prob of N-in: 0.22746\noutside 1 264\nTMhelix 265 287\ninside 288 293

34161 GCF_000718675.1_ASM71867v1 Streptomyces rimosus subsp. rimosus Terrabacteria group; Actinobacteria; Actinobacteria; Streptomycetales; Streptomycetaceae; Streptomyces; Streptomyces rimosus MTLTRRRHLVRSTAVAAVAGSALLLPAAAAFADSPQPTTAGSPGAAGDQQQKDDRQKDDQ WP_030654335.1 hypothetical protein [Streptomyces rimosus] Length: 245\nNumber of predicted TMHs: 2\nExp number of AAs in TMHs: 41.16745\nExp number, first 60 AAs: 19.93579\nTotal prob of N-in: 0.94992\nPOSSIBLE N-term signal sequence\ninside 1 12\nTMhelix 13 32\noutside 33 213\nTMhelix 214 236\ninside 237 245

34162 GCF_001279015.1_ASM127901v1 Streptomyces rimosus subsp. rimosus Terrabacteria group; Actinobacteria; Actinobacteria; Streptomycetales; Streptomycetaceae; Streptomyces; Streptomyces rimosus MESRQPVGPRANGHRAAHRRRAAVALAGAVCAVGLTATTAAPALADTNVPLNNGQCNFGA WP_078586913.1 MULTISPECIES: type VII secretion-associated serine protease mycosin [Streptomyces] Length: 427\nNumber of predicted TMHs: 2\nExp number of AAs in TMHs: 43.8948\nExp number, first 60 AAs: 21.08952\nTotal prob of N-in: 0.94565\nPOSSIBLE N-term signal sequence\ninside 1 20\nTMhelix 21 43\noutside 44 389\nTMhelix 390 412\ninside 413 427

34163 GCF_000717815.1_ASM71781v1 Streptomyces rimosus subsp. rimosus Terrabacteria group; Actinobacteria; Actinobacteria; Streptomycetales; Streptomycetaceae; Streptomyces; Streptomyces rimosus MTRRPLLRAAGVALSAAALTVLPALPAHADGIRAQEWALKAMNTQDAWETTKGEGITVAV WP_030373662.1 type VII secretion-associated serine protease [Streptomyces rimosus] Length: 386\nNumber of predicted TMHs: 1\nExp number of AAs in TMHs: 36.27224\nExp number, first 60 AAs: 9.95056\nTotal prob of N-in: 0.56617\noutside 1 359\nTMhelix 360 382\ninside 383 386

34164 GCF_000716745.1_ASM71674v1 Streptomyces rimosus subsp. rimosus Terrabacteria group; Actinobacteria; Actinobacteria; Streptomycetales; Streptomycetaceae; Streptomyces; Streptomyces rimosus MTRRPLLRAAGVALSAAALTVLPALPAHADGIRAQEWALKAMNTQDAWETTKGEGITVAV WP_003987438.1 MULTISPECIES: type VII secretion-associated serine protease [Streptomyces] Length: 386\nNumber of predicted TMHs: 1\nExp number of AAs in TMHs: 34.80558\nExp number, first 60 AAs: 10.27095\nTotal prob of N-in: 0.55118\nPOSSIBLE N-term signal sequence\noutside 1 359\nTMhelix 360 382\ninside 383 386

34165 GCF_000716745.1_ASM71674v1 Streptomyces rimosus subsp. rimosus Terrabacteria group; Actinobacteria; Actinobacteria; Streptomycetales; Streptomycetaceae; Streptomyces; Streptomyces rimosus MTERSAPAASATGPGTPARRTAVRAAATGTLAAALLGLAPQSALAATPAPPPAPAISAVT WP_030180617.1 MULTISPECIES: hypothetical protein [Streptomyces] Length: 293\nNumber of predicted TMHs: 1\nExp number of AAs in TMHs: 24.94793\nExp number, first 60 AAs: 4.55383\nTotal prob of N-in: 0.22746\noutside 1 264\nTMhelix 265 287\ninside 288 293

34166 GCF_000718835.1_ASM71883v1 Streptomyces rimosus subsp. rimosus Terrabacteria group; Actinobacteria; Actinobacteria; Streptomycetales; Streptomycetaceae; Streptomyces; Streptomyces rimosus MTERSAPAASATGPGTPARRTAVRAAATGTLAAALLGLAPQSALAATPAPPPAPAISAVT WP_030180617.1 MULTISPECIES: hypothetical protein [Streptomyces] Length: 293\nNumber of predicted TMHs: 1\nExp number of AAs in TMHs: 24.94793\nExp number, first 60 AAs: 4.55383\nTotal prob of N-in: 0.22746\noutside 1 264\nTMhelix 265 287\ninside 288 293

34167 GCF_000718835.1_ASM71883v1 Streptomyces rimosus subsp. rimosus Terrabacteria group; Actinobacteria; Actinobacteria; Streptomycetales; Streptomycetaceae; Streptomyces; Streptomyces rimosus MTRRPLLRAAGVALSAAALTVLPALPAHADGIRAQEWALKAMNTQDAWETTKGEGITVAV WP_031024350.1 type VII secretion-associated serine protease [Streptomyces rimosus] Length: 386\nNumber of predicted TMHs: 1\nExp number of AAs in TMHs: 34.58477\nExp number, first 60 AAs: 10.28663\nTotal prob of N-in: 0.54981\nPOSSIBLE N-term signal sequence\noutside 1 359\nTMhelix 360 382\ninside 383 386

34168	GCF_000718835.1_ASM71883v1	Streptomyces rimosus subsp. rimosus	Terrabacteria group; Actinobacteria; Actinobacteria; Streptomycetales; Streptomycetaceae; Streptomyces; Streptomyces rimosus
MESRQPVGPRANGHRAAHRRRAAVALAGAVCAVGLTATTAAPALADTNVPLNNGQCNFGA	WP_078586913.1
MULTISPECIES: type VII secretion-associated serine protease mycosin [Streptomyces]	Length: 427\nNumber of predicted TMHs:  2\nExp number of AAs in TMHs: 43.8948\nExp number, first 60 AAs:  21.08952\nTotal prob of N-in:  0.94565\nPOSSIBLE N-term signal sequence\ninside   1   20\nTMhelix   21   43\noutside   44   389\nTMhelix   390   412\ninside   413   427

34169	GCF_000718715.1_ASM71871v1	Streptomyces rimosus subsp. rimosus	Terrabacteria group; Actinobacteria; Actinobacteria; Streptomycetales; Streptomycetaceae; Streptomyces; Streptomyces rimosus
MTRRPLLRAAGVALSAAALTVLPALPAHADGIRAQEWALKAMNTQDAWETTKGEGITVAV	WP_003987438.1
MULTISPECIES: type VII secretion-associated serine protease [Streptomyces]	Length: 386\nNumber of predicted TMHs:  1\nExp number of AAs in TMHs: 34.80558\nExp number, first 60 AAs:  10.27095\nTotal prob of N-in:  0.55118\nPOSSIBLE N-term signal sequence\noutside   1   359\nTMhelix   360   382\ninside   383   386

34170	GCF_000718715.1_ASM71871v1	Streptomyces rimosus subsp. rimosus	Terrabacteria group; Actinobacteria; Actinobacteria; Streptomycetales; Streptomycetaceae; Streptomyces; Streptomyces rimosus
MTERSAPAASATGPGTPARRTAVRAAATGTLAAALLGLAPQSALAATPAPPPAPAISAVT	WP_030180617.1
MULTISPECIES: hypothetical protein [Streptomyces]	Length: 293\nNumber of predicted TMHs:  1\nExp number of AAs in TMHs: 24.94793\nExp number, first 60 AAs:  4.55383\nTotal prob of N-in:   0.22746\noutside   1   264\nTMhelix   265   287\ninside   288   293

34171	GCF_000718715.1_ASM71871v1	Streptomyces rimosus subsp. rimosus	Terrabacteria group; Actinobacteria; Actinobacteria; Streptomycetales; Streptomycetaceae; Streptomyces; Streptomyces rimosus
MESRQPVGPRANGHRAAHRRRAAVALAGAVCAVGLTATTAAPALADTNVPLNNGQCNFGA	WP_078586913.1
MULTISPECIES: type VII secretion-associated serine protease mycosin [Streptomyces]	Length: 427\nNumber of predicted TMHs:  2\nExp number of AAs in TMHs: 43.8948\nExp number, first 60 AAs:  21.08952\nTotal prob of N-in:  0.94565\nPOSSIBLE N-term signal sequence\ninside   1   20\nTMhelix   21   43\noutside   44   389\nTMhelix   390   412\ninside   413   427

34172	GCF_000718895.1_ASM71889v1	Streptomyces rimosus subsp. rimosus	Terrabacteria group; Actinobacteria; Actinobacteria; Streptomycetales; Streptomycetaceae; Streptomyces; Streptomyces rimosus
MTRRPLLRAAGVALSAAALTVLPALPAHADGIRAQEWALKAMNTQDAWETTKGEGITVAV	WP_003987438.1
MULTISPECIES: type VII secretion-associated serine protease [Streptomyces]	Length: 386\nNumber of predicted TMHs:  1\nExp number of AAs in TMHs: 34.80558\nExp number, first 60 AAs:  10.27095\nTotal prob of N-in:  0.55118\nPOSSIBLE N-term signal sequence\noutside   1   359\nTMhelix   360   382\ninside   383   386

34173	GCF_001279015.1_ASM127901v1	Streptomyces rimosus subsp. rimosus	Terrabacteria group; Actinobacteria; Actinobacteria; Streptomycetales; Streptomycetaceae; Streptomyces; Streptomyces rimosus
MTERSAPAASATGPGTPARRTAVRAAATGTLAAALLGLAPQSALAATPAPPPAPAISAVT	WP_030180617.1
MULTISPECIES: hypothetical protein [Streptomyces]	Length: 293\nNumber of predicted TMHs:  1\nExp number of AAs in TMHs: 24.94793\nExp number, first 60 AAs:  4.55383\nTotal prob of N-in:   0.22746\noutside   1   264\nTMhelix   265   287\ninside   288   293

34174	GCF_001279075.1_ASM127907v1	Streptomyces rimosus subsp. rimosus	Terrabacteria group; Actinobacteria; Actinobacteria; Streptomycetales; Streptomycetaceae; Streptomyces; Streptomyces rimosus
MTRRPLLRAAGVALSAAALTVLPALPAHADGIRAQEWALKAMNTQDAWETTKGEGITVAV	WP_003987438.1
MULTISPECIES: type VII secretion-associated serine protease [Streptomyces]	Length: 386\nNumber of predicted TMHs:  1\nExp number of AAs in TMHs: 34.80558\nExp number, first 60 AAs:  10.27095\nTotal prob of N-in:  0.55118\nPOSSIBLE N-term signal sequence\noutside   1   359\nTMhelix   360   382\ninside   383   386

34175	GCF_001279075.1_ASM127907v1	Streptomyces rimosus subsp. rimosus	Terrabacteria group; Actinobacteria; Actinobacteria; Streptomycetales; Streptomycetaceae; Streptomyces; Streptomyces rimosus
MTERSAPAASATGPGTPARRTAVRAAATGTLAAALLGLAPQSALAATPAPPPAPAISAVT	WP_030180617.1
MULTISPECIES: hypothetical protein [Streptomyces]	Length: 293\nNumber of predicted TMHs:  1\nExp number of AAs in TMHs: 24.94793\nExp number, first 60 AAs:  4.55383\nTotal prob of N-in:   0.22746\noutside   1   264\nTMhelix   265   287\ninside   288   293

34176	GCF_001279075.1_ASM127907v1	Streptomyces rimosus subsp. rimosus	Terrabacteria group; Actinobacteria; Actinobacteria; Streptomycetales; Streptomycetaceae; Streptomyces; Streptomyces rimosus
MESRQPVGPRANGHRAAHRRRAAVALAGAVCAVGLTATTAAPALADTNVPLNNGQCNFGA	WP_078586913.1

MULTISPECIES: type VII secretion-associated serine protease mycosin [Streptomyces]	Length: 427\nNumber of predicted TMHs:  2\nExp number of AAs in TMHs: 43.8948\nExp number, first 60 AAs:  21.08952\nTotal prob of N-in:   0.94565\nPOSSIBLE N-term signal sequence\ninside   1   20\nTMhelix   21   43\noutside   44   389\nTMhelix   390   412\ninside   413   427

34177	GCF_000331185.1_ASM33118v1	Streptomyces rimosus subsp. rimosus ATCC 10970	Terrabacteria group; Actinobacteria; Actinobacteria; Streptomycetales; Streptomycetaceae; Streptomyces; Streptomyces rimosus; Streptomyces rimosus subsp. rimosus	MESRQPVGPRANGHRAAHRRRAAVALAGAVCAVGLTATTAAPALADTNVPLNNGQCNFGA	WP_078586913.1	MULTISPECIES: type VII secretion-associated serine protease mycosin [Streptomyces]	Length: 427\nNumber of predicted TMHs:  2\nExp number of AAs in TMHs: 43.8948\nExp number, first 60 AAs:  21.08952\nTotal prob of N-in:   0.94565\nPOSSIBLE N-term signal sequence\ninside   1   20\nTMhelix   21   43\noutside   44   389\nTMhelix   390   412\ninside   413   427

34178	GCF_000331185.1_ASM33118v1	Streptomyces rimosus subsp. rimosus ATCC 10970	Terrabacteria group; Actinobacteria; Actinobacteria; Streptomycetales; Streptomycetaceae; Streptomyces; Streptomyces rimosus; Streptomyces rimosus subsp. rimosus	MTRRPLLRAAGVALSAAALTVLPALPAHADGIRAQEWALKAMNTQDAWETTKGEGITVAV	WP_003987438.1	MULTISPECIES: type VII secretion-associated serine protease [Streptomyces]	Length: 386\nNumber of predicted TMHs:  1\nExp number of AAs in TMHs: 34.80558\nExp number, first 60 AAs:  10.27095\nTotal prob of N-in:   0.55118\nPOSSIBLE N-term signal sequence\noutside   1   359\nTMhelix   360   382\ninside   383   386

34179	GCF_000497445.1_S.roseochromogenes_v1	Streptomyces roseochromogenus subsp. oscitans DS 12.976	Terrabacteria group; Actinobacteria; Actinobacteria; Streptomycetales; Streptomycetaceae; Streptomyces; Streptomyces roseochromogenus; Streptomyces roseochromogenus subsp. oscitans	MPQESVAGRAGRRTPVLAILVGLTLMLVAHVVSCALHHLDGAPHIAARTVSMTSEAQGAG	WP_023544957.1	hypothetical protein [Streptomyces roseochromogenus]	Length: 152\nNumber of predicted TMHs:  2\nExp number of AAs in TMHs: 37.04417\nExp number, first 60 AAs:  21.57796\nTotal prob of N-in:   0.69171\nPOSSIBLE N-term signal sequence\ninside   1   16\nTMhelix   17   39\noutside   40   96\nTMhelix   97   119\ninside   120   152

34180	GCF_000497445.1_S.roseochromogenes_v1	Streptomyces roseochromogenus subsp. oscitans DS 12.976	Terrabacteria group; Actinobacteria; Actinobacteria; Streptomycetales; Streptomycetaceae; Streptomyces; Streptomyces roseochromogenus; Streptomyces roseochromogenus subsp. oscitans	MLGGVENHSLPRAAAFFDLDKTVIAKSSTLTFSKSFYQGGLINRRAALRTAYAQFVFLAG	WP_023547759.1	inhibition of morphological differentiation protein [Streptomyces roseochromogenus]	Length: 276\nNumber of predicted TMHs:  1\nExp number of AAs in TMHs: 21.79414\nExp number, first 60 AAs:  0.01539\nTotal prob of N-in:   0.00192\noutside   1   247\nTMhelix   248   270\ninside   271   276

34181	GCF_000497445.1_S.roseochromogenes_v1	Streptomyces roseochromogenus subsp. oscitans DS 12.976	Terrabacteria group; Actinobacteria; Actinobacteria; Streptomycetales; Streptomycetaceae; Streptomyces; Streptomyces roseochromogenus; Streptomyces roseochromogenus subsp. oscitans	MRGRGRRTAWRWREGRLGRRAVFGAGAVRTVVLSGLCALLVLPTVGTATAPAAAAEDGRS	WP_051430100.1	hypothetical protein [Streptomyces roseochromogenus]	Length: 465\nNumber of predicted TMHs:  2\nExp number of AAs in TMHs: 45.2062\nExp number, first 60 AAs:  23.04524\nTotal prob of N-in:   0.99192\nPOSSIBLE N-term signal sequence\ninside   1   20\nTMhelix   21   43\noutside   44   427\nTMhelix   428   450\ninside   451   465

34182	GCF_000497445.1_S.roseochromogenes_v1	Streptomyces roseochromogenus subsp. oscitans DS 12.976	Terrabacteria group; Actinobacteria; Actinobacteria; Streptomycetales; Streptomycetaceae; Streptomyces; Streptomyces roseochromogenus; Streptomyces roseochromogenus subsp. oscitans	MRHARRRIVRRVTRLAAVGGLLLGGAMVTQAAMASETPPAARTLAASEGTGGTGAALVAR	WP_031224006.1	protease [Streptomyces roseochromogenus]	Length: 458\nNumber of predicted TMHs:  2\nExp number of AAs in TMHs: 43.3893299999999\nExp number, first 60 AAs:  22.42564\nTotal prob of N-in:   0.99423\nPOSSIBLE N-term signal sequence\ninside   1   11\nTMhelix   12   34\noutside   35   419\nTMhelix   420   442\ninside   443   458

34183	GCF_000091305.1_ASM9130v1	Streptomyces scabiei 87.22	Terrabacteria group; Actinobacteria; Actinobacteria; Streptomycetales; Streptomycetaceae; Streptomyces; Streptomyces scabiei	MYRLSARRTGAHPAGAHRPGAHHDSAHRDSGRGDGGRRAAVATAAAVVVAPLLLTAGTAT	WP_012999856.1	chitin-binding protein [Streptomyces scabiei]	Length: 352\nNumber of predicted TMHs:  1\nExp number of AAs in TMHs: 36.84986\nExp number, first 60 AAs:  14.48401\nTotal prob of N-in:   0.69739\nPOSSIBLE N-term signal sequence\noutside   1   318\nTMhelix   319   341\ninside   342   352

34184	GCF_000091305.1_ASM9130v1	Streptomyces scabiei 87.22	Terrabacteria group; Actinobacteria; Actinobacteria; Streptomycetales; Streptomycetaceae; Streptomyces; Streptomyces scabiei

MLGVVENHSLPRTAAFFDLDKTVIAKSSTLTFSKSFYQGGLINRRAALRTAYIQFVFLAG WP_013001810.1 morphological differentiation protein [Streptomyces scabiei] Length: 274\nNumber of predicted TMHs: 1\nExp number of AAs in TMHs: 21.78293\nExp number, first 60 AAs: 0.08751\nTotal prob of N-in: 0.00802\noutside 1 244\nTMhelix 245 267\ninside 268 274

34185 GCF_000372745.1_ASM37274v1 Streptomyces scabrisporus DSM 41855 Terrabacteria group; Actinobacteria; Actinobacteria; Streptomycetales; Streptomycetaceae; Streptomyces; Streptomyces scabrisporus METQAPKRTAAFFDLDKTIIARSSALAFSRSFYHGGLINRRSVLKSAYAQFVFLVGGADH WP_020554935.1 inhibition of morphological differentiation protein [Streptomyces scabrisporus] Length: 268\nNumber of predicted TMHs: 1\nExp number of AAs in TMHs: 20.94084\nExp number, first 60 AAs: 0.18415\nTotal prob of N-in: 0.04150\noutside 1 243\nTMhelix 244 263\ninside 264 268

34186 GCF_000718095.1_ASM71809v1 Streptomyces scopuliridis RB72 Terrabacteria group; Actinobacteria; Actinobacteria; Streptomycetales; Streptomycetaceae; Streptomyces; Streptomyces scopuliridis MLGLVENHSLPRTAAFFDLDKTVIAKSSTLTFSKSFYQGGLINRRAVLRTAYAQFVFLAG WP_030353265.1 inhibition of morphological differentiation protein [Streptomyces scopuliridis] Length: 278\nNumber of predicted TMHs: 1\nExp number of AAs in TMHs: 21.73001\nExp number, first 60 AAs: 0.01336\nTotal prob of N-in: 0.00155\noutside 1 247\nTMhelix 248 270\ninside 271 278

34187 GCF_000718095.1_ASM71809v1 Streptomyces scopuliridis RB72 Terrabacteria group; Actinobacteria; Actinobacteria; Streptomycetales; Streptomycetaceae; Streptomyces; Streptomyces scopuliridis MPDRRRRTALHLPPPSGPVLLPRERQSSSTGPSETSEHSGTGAPGTSGTPGTAGTPGAES WP_030351526.1 hypothetical protein [Streptomyces scopuliridis] Length: 322\nNumber of predicted TMHs: 2\nExp number of AAs in TMHs: 46.22906\nExp number, first 60 AAs: 0\nTotal prob of N-in: 0.48471\noutside 1 174\nTMhelix 175 197\ninside 198 263\nTMhelix 264 286\noutside 287 322

34188 GCF_000258595.1_Ssomaliensis1 Streptomyces somaliensis DSM 40738 Terrabacteria group; Actinobacteria; Actinobacteria; Streptomycetales; Streptomycetaceae; Streptomyces; Streptomyces somaliensis MRSTNRRRANRRRTAVLAAAAFLGITLPAPPAAADAADTPGQCTYPAKPYEGRPWALQRV WP_010471532.1 type VII secretion-associated serine protease [Streptomyces somaliensis] Length: 422\nNumber of predicted TMHs: 2\nExp number of AAs in TMHs: 41.11469\nExp number, first 60 AAs: 18.54862\nTotal prob of N-in: 0.92149\nPOSSIBLE N-term signal sequence\ninside 1 12\nTMhelix 13 32\noutside 33 383\nTMhelix 384 406\ninside 407 422

34189 GCF_000258595.1_Ssomaliensis1 Streptomyces somaliensis DSM 40738 Terrabacteria group; Actinobacteria; Actinobacteria; Streptomycetales; Streptomycetaceae; Streptomyces; Streptomyces somaliensis MLGLVENHSSPRTAAFFDLDKTVIAKSSTLTFGKSFYQGGLINRRDALRTAYTQFVFLAG WP_010473267.1 haloacid dehalogenase [Streptomyces somaliensis] Length: 287\nNumber of predicted TMHs: 1\nExp number of AAs in TMHs: 20.79632\nExp number, first 60 AAs: 0.00263\nTotal prob of N-in: 0.00140\noutside 1 248\nTMhelix 249 268\ninside 269 287

34190 GCF_001969965.1_Sparso1.0Streptomyces sparsogenes DSM 40356 Terrabacteria group; Actinobacteria; Actinobacteria; Streptomycetales; Streptomycetaceae; Streptomyces; Streptomyces sparsogenes MSSPDREVRAARNDTPRTARRPVVAVTGAASGAGALLTQRLAESDEIKHVLAIDERRGDV WP_065964212.1 NAD-dependent dehydratase [Streptomyces sparsogenes] Length: 369\nNumber of predicted TMHs: 1\nExp number of AAs in TMHs: 21.31707\nExp number, first 60 AAs: 1.44367\nTotal prob of N-in: 0.08113\noutside 1 339\nTMhelix 340 359\ninside 360 369

34191 GCF_001969965.1_Sparso1.0Streptomyces sparsogenes DSM 40356 Terrabacteria group; Actinobacteria; Actinobacteria; Streptomycetales; Streptomycetaceae; Streptomyces; Streptomyces sparsogenes MLGVVENHSLPRTAAFFDLDKTVIAKSSTLTFSKSFYQGGLINRRAVLRTAYAQFVFLAG WP_065961517.1 inhibition of morphological differentiation protein [Streptomyces sparsogenes] Length: 278\nNumber of predicted TMHs: 1\nExp number of AAs in TMHs: 20.71292\nExp number, first 60 AAs: 0.02768\nTotal prob of N-in: 0.00716\noutside 1 246\nTMhelix 247 269\ninside 270 278

34192 GCF_001969965.1_Sparso1.0Streptomyces sparsogenes DSM 40356 Terrabacteria group; Actinobacteria; Actinobacteria; Streptomycetales; Streptomycetaceae; Streptomyces; Streptomyces sparsogenes MSAADDDGNRSLGQLVATATTELSALVHDEIALAKAEFRDSARRTLLGSGAAVSAGVLVL WP_065961531.1 transporter [Streptomyces sparsogenes] Length: 153\nNumber of predicted TMHs: 2\nExp number of AAs in TMHs: 46.4376800000001\nExp number, first 60 AAs: 13.14988\nTotal prob of N-in: 0.92902\nPOSSIBLE N-term signal sequence\ninside 1 45\nTMhelix 46 68\noutside 69 82\nTMhelix 83 105\ninside 106 153

34193 GCF_001969965.1_Sparso1.0Streptomyces sparsogenes DSM 40356 Terrabacteria group; Actinobacteria; Actinobacteria; Streptomycetales; Streptomycetaceae; Streptomyces; Streptomyces sparsogenes MVTGEVSEARRKAVGLGSGACHALGLMVLAITEWVRADLKDATSLASHSYLKDMLRLAAD WP_065963662.1 hypothetical protein [Streptomyces sparsogenes] Length: 152\nNumber of predicted TMHs: 2\nExp number of AAs in TMHs: 71.82401\nExp number, first 60 AAs: 22.34642\nTotal prob of N-in: 0.99607\nPOSSIBLE N-term signal sequence\ninside 1 12\nTMhelix 13 35\noutside 36 127\nTMhelix 128 150\ninside 151 152

34194 GCF_001969965.1_Sparso1.0Streptomyces sparsogenes DSM 40356 Terrabacteria group; Actinobacteria; Actinobacteria; Streptomycetales; Streptomycetaceae; Streptomyces; Streptomyces sparsogenes MQLPSLSVRRRALLALAGAVLVSAAGFAPSAEAQDIRSQQWYLDAMKADAIWKVSRGQGI WP_076970707.1 serine protease [Streptomyces sparsogenes] Length: 413\nNumber of predicted TMHs: 1\nExp number of AAs in TMHs: 36.9750299999999\nExp number, first 60 AAs: 14.81189\nTotal prob of N-in: 0.75311\nPOSSIBLE N-term signal sequence\noutside 1 384\nTMhelix 385 407\ninside 408 413

34195 GCF_000381025.1_ASM38102v1 Streptomyces sulphureus DSM 40104 Terrabacteria group; Actinobacteria; Actinobacteria; Streptomycetales; Streptomycetaceae; Streptomyces; Streptomyces sulphureus MLGSVENVSLPRTAAFFDLDKTVIAKSSTLTFGRSFYQGGLINRRAVLRTAYAQFVYLLG WP_019547849.1 inhibition of morphological differentiation protein [Streptomyces sulphureus] Length: 285\nNumber of predicted TMHs: 1\nExp number of AAs in TMHs: 21.86514\nExp number, first 60 AAs: 0.0427\nTotal prob of N-in: 0.01166\noutside 1 246\nTMhelix 247 269\ninside 270 285

34196 GCF_000154965.1_ASM15496v1 Streptomyces sviceus ATCC 29083 Terrabacteria group; Actinobacteria; Actinobacteria; Streptomycetales; Streptomycetaceae; Streptomyces; Streptomyces sviceus MLGVVENHSLPRTAAFFDLDKTVIAKSSTLTFSKSFYQGGLINRRAALRTAYAQFVFLVG WP_007383682.1 haloacid dehalogenase [Streptomyces sviceus] Length: 277\nNumber of predicted TMHs: 1\nExp number of AAs in TMHs: 21.65845\nExp number, first 60 AAs: 0.01246\nTotal prob of N-in: 0.00153\noutside 1 247\nTMhelix 248 270\ninside 271 277

34197 GCF_000154965.1_ASM15496v1 Streptomyces sviceus ATCC 29083 Terrabacteria group; Actinobacteria; Actinobacteria; Streptomycetales; Streptomycetaceae; Streptomyces; Streptomyces sviceus MRHARRRVVRRVTRLAAVGGLLLGGAMVTRAVASEPPATTAVPRTYAMEAGRTGEDLVSR WP_007386666.1 S1 family peptidase [Streptomyces sviceus] Length: 456\nNumber of predicted TMHs: 2\nExp number of AAs in TMHs: 39.6745299999999\nExp number, first 60 AAs: 20.95884\nTotal prob of N-in: 0.95939\nPOSSIBLE N-term signal sequence\ninside 1 11\nTMhelix 12 34\noutside 35 417\nTMhelix 418 440\ninside 441 456

34198 GCF_000478605.2_ASM47860v2 Streptomyces thermolilacinus SPC6 Terrabacteria group; Actinobacteria; Actinobacteria; Streptomycetales; Streptomycetaceae; Streptomyces; Streptomyces thermolilacinus MLWLVENHSSPRTAAFFDLDKTVIAKSSTLTFSKSFYQGGLINRRDALRTAYTQFVFLVG WP_023587342.1 inhibition of morphological differentiation protein [Streptomyces thermolilacinus] Length: 284\nNumber of predicted TMHs: 1\nExp number of AAs in TMHs: 21.42315\nExp number, first 60 AAs: 0.00544\nTotal prob of N-in: 0.00549\noutside 1 247\nTMhelix 248 270\ninside 271 284

34199 GCF_000297155.2_ASM29715v2 Streptomyces tsukubensis NRRL18488 Terrabacteria group; Actinobacteria; Actinobacteria; Streptomycetales; Streptomycetaceae; Streptomyces; Streptomyces tsukubensis MTNATTARRAAVALAVSAVLGTALAPAATAATTPPPAAPLPSGLYGTTDPAYDGVWRQSL WP_006349525.1 hypothetical protein [Streptomyces tsukubensis] Length: 420\nNumber of predicted TMHs: 2\nExp number of AAs in TMHs: 41.76333\nExp number, first 60 AAs: 20.89463\nTotal prob of N-in: 0.96876\nPOSSIBLE N-term signal sequence\ninside 1 11\nTMhelix 12 34\noutside 35 391\nTMhelix 392 414\ninside 415 420

34200 GCF_000297155.2_ASM29715v2 Streptomyces tsukubensis NRRL18488 Terrabacteria group; Actinobacteria; Actinobacteria; Streptomycetales; Streptomycetaceae; Streptomyces; Streptomyces tsukubensis MITNPRQPRRTPLPLRAAAPAALACGLALVFGTAQPAAAHAGVTASDPRALAKNVTVTLT WP_006345762.1 hypothetical protein [Streptomyces tsukubensis] Length: 272\nNumber of predicted TMHs: 2\nExp number of AAs in TMHs: 38.93806\nExp number, first 60 AAs: 17.84283\nTotal prob of N-in: 0.86155\nPOSSIBLE N-term signal sequence\ninside 1 12\nTMhelix 13 35\noutside 36 245\nTMhelix 246 265\ninside 266 272

34201 GCF_000297155.2_ASM29715v2 Streptomyces tsukubensis NRRL18488 Terrabacteria group; Actinobacteria; Actinobacteria; Streptomycetales; Streptomycetaceae; Streptomyces; Streptomyces tsukubensis MPVIRRIRHAVRRAYRPKVDLSHPRRSPLGSSVINAVVYEDGRRRPCDGTAADALRQVRK WP_006347905.1 magnesium transporter CorA [Streptomyces tsukubensis] Length: 362\nNumber of predicted TMHs: 2\nExp number of AAs in TMHs: 40.76808\nExp number, first 60 AAs: 0\nTotal prob of N-in: 0.94191\ninside 1 304\nTMhelix 305 324\noutside 325 333\nTMhelix 334 356\ninside 357 362

34202 GCF_000331005.1_Sturgcar8v1.0 Streptomyces turgidiscabies Car8 Terrabacteria group; Actinobacteria; Actinobacteria; Streptomycetales; Streptomycetaceae; Streptomyces; Streptomyces turgidiscabies MPARRRPAVVTTVTALGIAALALGATPASAHGSMGDPVSRVAQCYAEGPESPTSAACRAA WP_006376680.1 chitin-binding protein [Streptomyces turgidiscabies] Length: 353\nNumber of predicted TMHs: 2\nExp number of AAs in TMHs: 41.16593\nExp number, first 60 AAs: 20.07244\nTotal prob of N-in: 0.93537\nPOSSIBLE N-term signal sequence\ninside 1 6\nTMhelix 7 29\noutside 30 318\nTMhelix 319 341\ninside 342 353

34203 GCF_000331005.1_Sturgcar8v1.0 Streptomyces turgidiscabies Car8 Terrabacteria group; Actinobacteria; Actinobacteria; Streptomycetales; Streptomycetaceae; Streptomyces; Streptomyces turgidiscabies MLSLVENHSLPRAAAFFDLDKTVIAKSSTLTFSKSFYQGGLINRRAVLRTAYAQFVFLAG WP_044473232.1 inhibition of morphological differentiation protein [Streptomyces turgidiscabies] Length: 277\nNumber of predicted TMHs: 1\nExp number of AAs in TMHs: 21.55643\nExp number, first 60 AAs: 0.01804\nTotal prob of N-in: 0.00246\noutside 1 247\nTMhelix 248 270\ninside 271 277

34204 GCF_000331005.1_Sturgcar8v1.0 Streptomyces turgidiscabies Car8 Terrabacteria group; Actinobacteria; Actinobacteria; Streptomycetales; Streptomycetaceae; Streptomyces; Streptomyces turgidiscabies MRHARRRIVRRATRLAAVGGVILGGLMVTQAVASEPPEVNLPFSSAQAAGTTGSELVTEL WP_006380203.1 S1 family peptidase [Streptomyces turgidiscabies] Length: 452\nNumber of predicted TMHs: 2\nExp number of AAs in TMHs: 42.58327\nExp number, first 60 AAs: 21.83569\nTotal prob of N-in: 0.98683\nPOSSIBLE N-term signal sequence\ninside 1 11\nTMhelix 12 34\noutside 35 413\nTMhelix 414 436\ninside 437 452

34205 GCF_000253235.1_ASM25323v1 Streptomyces venezuelae ATCC 10712 Terrabacteria group; Actinobacteria; Actinobacteria; Streptomycetales; Streptomycetaceae; Streptomyces; Streptomyces venezuelae MLRLVENHSLPRTAAFFDLDKTVIAKSSTLTFSKSFYRGGLISRRAALRTAYIQFVFLAG WP_015034529.1 phosphoserine phosphatase [Streptomyces venezuelae] Length: 288\nNumber of predicted TMHs: 1\nExp number of AAs in TMHs: 22.12812\nExp number, first 60 AAs: 0.0544\nTotal prob of N-in: 0.00444\noutside 1 247\nTMhelix 248 270\ninside 271 288

34206 GCF_000147815.2_ASM14781v3 Streptomyces violaceusniger Tu 4113 Terrabacteria group; Actinobacteria; Actinobacteria; Streptomycetales; Streptomycetaceae; Streptomyces; Streptomyces violaceusniger MSSPDREVRAARNDTARAARRPVVAVTGAASGPGALLTQRLAESEEVKQVLAIDERRGEV WP_014054769.1 NAD-dependent dehydratase [Streptomyces violaceusniger] Length: 371\nNumber of predicted TMHs: 1\nExp number of AAs in TMHs: 21.65842\nExp number, first 60 AAs: 0.31702\nTotal prob of N-in: 0.02575\noutside 1 339\nTMhelix 340 362\ninside 363 371

34207 GCF_000147815.2_ASM14781v3 Streptomyces violaceusniger Tu 4113 Terrabacteria group; Actinobacteria; Actinobacteria; Streptomycetales; Streptomycetaceae; Streptomyces; Streptomyces violaceusniger MPFKKYGARRRALSALAMLGTALLAVGGTAPAAAAQDISSQQWYLKDMQADDMWAVSKGE WP_014055290.1 serine protease [Streptomyces violaceusniger] Length: 418\nNumber of predicted TMHs: 2\nExp number of AAs in TMHs: 45.48173\nExp number, first 60 AAs: 22.56387\nTotal prob of N-in: 0.99121\nPOSSIBLE N-term signal sequence\ninside 1 12\nTMhelix 13 35\noutside 36 385\nTMhelix 386 408\ninside 409 418

34208 GCF_000147815.2_ASM14781v3 Streptomyces violaceusniger Tu 4113 Terrabacteria group; Actinobacteria; Actinobacteria; Streptomycetales; Streptomycetaceae; Streptomyces; Streptomyces violaceusniger MLGVVENHSLPRTAAFFDLDKTVIAKSSAFTFSKSFYQGGLINRRAVLRTAYAQFVFLAG WP_043237453.1 inhibition of morphological differentiation protein [Streptomyces violaceusniger] Length: 278\nNumber of predicted TMHs: 1\nExp number of AAs in TMHs: 21.24863\nExp number, first 60 AAs: 0.01593\nTotal prob of N-in: 0.00269\noutside 1 247\nTMhelix 248 267\ninside 268 278

34209 GCF_000158955.1_ASM15895v1 Streptomyces viridochromogenes DSM 40736 Terrabacteria group; Actinobacteria; Actinobacteria; Streptomycetales; Streptomycetaceae; Streptomyces; Streptomyces viridochromogenes MRHARRRVVRRVTRLAAVGGLLLGGAMVTQAVASEPPDPTIGLPYSAGAATSKGTDLVSS WP_003994962.1 S1 family peptidase [Streptomyces viridochromogenes] Length: 446\nNumber of predicted TMHs: 2\nExp number of AAs in TMHs: 42.17679\nExp number, first 60 AAs: 21.82103\nTotal prob of N-in: 0.97984\nPOSSIBLE N-term signal sequence\ninside 1 11\nTMhelix 12 34\noutside 35 407\nTMhelix 408 430\ninside 431 446

34210 GCF_000158955.1_ASM15895v1 Streptomyces viridochromogenes DSM 40736 Terrabacteria group; Actinobacteria; Actinobacteria; Streptomycetales; Streptomycetaceae; Streptomyces; Streptomyces viridochromogenes

MLRGVENHSLPRTAAFFDLDKTVIAKSSTLTFSKSFYQGGLINRRAVLRTAYAQFVFLAG WP_003991891.1 morphological differentiation-associated protein [Streptomyces viridochromogenes] Length: 277\nNumber of predicted TMHs: 1\nExp number of AAs in TMHs: 21.87124\nExp number, first 60 AAs: 0.04195\nTotal prob of N-in: 0.00411\noutside 1 247\nTMhelix 248 270\ninside 271 277

34211 GCF_000158955.1_ASM15895v1 Streptomyces viridochromogenes DSM 40736 Terrabacteria group; Actinobacteria; Actinobacteria; Streptomycetales; Streptomycetaceae; Streptomyces; Streptomyces viridochromogenes MTGSRRSGRRAFGPTRIPTRRNPVHLRPSAAVLAAAVLAVAGPAVAAPMAQAEEGEPALV WP_003990586.1 hypothetical protein [Streptomyces viridochromogenes] Length: 429\nNumber of predicted TMHs: 2\nExp number of AAs in TMHs: 40.57541\nExp number, first 60 AAs: 19.12042\nTotal prob of N-in: 0.87646\nPOSSIBLE N-term signal sequence\ninside 1 29\nTMhelix 30 52\noutside 53 397\nTMhelix 398 420\ninside 421 429

34212 GCF_000332625.1_STVIR_1.0 Streptomyces viridochromogenes Tue57 Terrabacteria group; Actinobacteria; Actinobacteria; Streptomycetales; Streptomycetaceae; Streptomyces; Streptomyces viridochromogenes MRHARRRLVRRVTRLAAVGGLLLGATMVTRAVASEPPDASAAPRTYAASAAGTGGELVSR WP_037885301.1 S1 family peptidase [Streptomyces viridochromogenes] Length: 457\nNumber of predicted TMHs: 2\nExp number of AAs in TMHs: 41.4791\nExp number, first 60 AAs: 20.02476\nTotal prob of N-in: 0.91471\nPOSSIBLE N-term signal sequence\ninside 1 11\nTMhelix 12 34\noutside 35 418\nTMhelix 419 441\ninside 442 457

34213 GCF_000332625.1_STVIR_1.0 Streptomyces viridochromogenes Tue57 Terrabacteria group; Actinobacteria; Actinobacteria; Streptomycetales; Streptomycetaceae; Streptomyces; Streptomyces viridochromogenes MLRGVENHSLPRTAAFFDLDKTVIAKSSTLTFSKSFYQGGLINRRAVLRTAYAQFVFLAG WP_037887166.1 inhibition of morphological differentiation protein [Streptomyces viridochromogenes] Length: 277\nNumber of predicted TMHs: 1\nExp number of AAs in TMHs: 21.71387\nExp number, first 60 AAs: 0.03511\nTotal prob of N-in: 0.00450\noutside 1 247\nTMhelix 248 270\ninside 271 277

34214 GCF_000332625.1_STVIR_1.0 Streptomyces viridochromogenes Tue57 Terrabacteria group; Actinobacteria; Actinobacteria; Streptomycetales; Streptomycetaceae; Streptomyces; Streptomyces viridochromogenes MSTVGNRDENRRRSLVTRRTGLLGGLLAASLALVPPTTAHADAIRARQWALDAMRTQEAW WP_004002572.1 type VII secretion-associated serine protease [Streptomyces viridochromogenes] Length: 402\nNumber of predicted TMHs: 2\nExp number of AAs in TMHs: 42.63925\nExp number, first 60 AAs: 19.86206\nTotal prob of N-in: 0.96585\nPOSSIBLE N-term signal sequence\ninside 1 20\nTMhelix 21 43\noutside 44 370\nTMhelix 371 393\ninside 394 402

34215 GCF_000316095.1_ASM31609v1 Streptomyces viridosporus T7A Terrabacteria group; Actinobacteria; Actinobacteria; Streptomycetales; Streptomycetaceae; Streptomyces; Streptomyces viridosporus MLKGVENHSSPRTAAFFDLDKTVIAKSSTLTFSKSFYQGGLINRRAVLRTAYAQFVFLAG WP_004985180.1 MULTISPECIES: morphological differentiation-associated protein [Streptomyces] Length: 277\nNumber of predicted TMHs: 1\nExp number of AAs in TMHs: 21.98535\nExp number, first 60 AAs: 0.08708\nTotal prob of N-in: 0.00829\noutside 1 247\nTMhelix 248 270\ninside 271 277

34216 GCF_000316095.1_ASM31609v1 Streptomyces viridosporus T7A Terrabacteria group; Actinobacteria; Actinobacteria; Streptomycetales; Streptomycetaceae; Streptomyces; Streptomyces viridosporus MRHARRRVVRRVTRLAAVGGLLLGATMVTRAVASEPAPPNAVPHTAAGQSAGPGTDLASR WP_039830567.1 MULTISPECIES: S1 family peptidase [Streptomyces] Length: 457\nNumber of predicted TMHs: 2\nExp number of AAs in TMHs: 41.29905\nExp number, first 60 AAs: 20.23659\nTotal prob of N-in: 0.94066\nPOSSIBLE N-term signal sequence\ninside 1 11\nTMhelix 12 34\noutside 35 418\nTMhelix 419 441\ninside 442 457

34217 GCF_000380165.1_ASM38016v1 Streptomyces vitaminophilus DSM 41686 Terrabacteria group; Actinobacteria; Actinobacteria; Streptomycetales; Streptomycetaceae; Streptomyces; Streptomyces vitaminophilus MLIPVENHPQPRAAAFFDLDKTVIAKSSALAFGRTFYRGGLINRRAVLRTAYAQFVFLAG WP_040912223.1 inhibition of morphological differentiation protein [Streptomyces vitaminophilus] Length: 275\nNumber of predicted TMHs: 1\nExp number of AAs in TMHs: 22.67849\nExp number, first 60 AAs: 0.03519\nTotal prob of N-in: 0.00631\noutside 1 243\nTMhelix 244 266\ninside 267 275

34218 GCF_000380165.1_ASM38016v1 Streptomyces vitaminophilus DSM 41686 Terrabacteria group; Actinobacteria; Actinobacteria; Streptomycetales; Streptomycetaceae; Streptomyces; Streptomyces vitaminophilus MRAGRRALLGALVGFLLGPVLVAGATTPASAETIEEGQWGLDAIHARDVWPTTRGEGTVV WP_018382267.1 type VII secretion-associated serine protease [Streptomyces vitaminophilus] Length: 398\nNumber of predicted TMHs: 2\nExp number of AAs in TMHs: 43.13129\nExp number, first 60 AAs: 21.19527\nTotal prob of N-in: 0.97230\nPOSSIBLE N-term signal sequence\ninside 1 6\nTMhelix 7 29\noutside 30 367\nTMhelix 368 390\ninside 391 398

34219 GCF_000745345.1_ASM74534v1 Streptomyces yeochonensis CN732 Terrabacteria group; Actinobacteria; Actinobacteria; Streptomycetales; Streptomycetaceae; Streptomyces; Streptomyces yeochonensis MPLRRRAALLLAALAAPLLGAAPLLGAAPAGAATPPAGLYGTADPTYDGVWRQSYAFLAL WP_037906505.1 hypothetical protein [Streptomyces yeochonensis] Length: 423\nNumber of predicted TMHs: 2\nExp number of AAs in TMHs: 49.66418\nExp number, first 60 AAs: 23.04819\nTotal prob of N-in: 0.96316\nPOSSIBLE N-term signal sequence\ninside 1 6\nTMhelix 7 29\noutside 30 394\nTMhelix 395 417\ninside 418 423

34220 GCF_000225525.1_ASM22552v1 Streptomyces zinciresistens K42 Terrabacteria group; Actinobacteria; Actinobacteria; Streptomycetales; Streptomycetaceae; Streptomyces; Streptomyces zinciresistens MRHARRRLVRRVTRLAAVGGLLLGGTMVTRAVASEPPAATGVPRTFAASAARTGGELVAR WP_043733548.1 S1 family peptidase [Streptomyces zinciresistens] Length: 456\nNumber of predicted TMHs: 2\nExp number of AAs in TMHs: 36.6973\nExp number, first 60 AAs: 19.74247\nTotal prob of N-in: 0.90206\nPOSSIBLE N-term signal sequence\ninside 1 11\nTMhelix 12 34\noutside 35 417\nTMhelix 418 440\ninside 441 456

34221 GCF_000497405.1_v1 Streptomycetaceae bacterium MP113-05 Terrabacteria group; Actinobacteria; Actinobacteria; Streptomycetales; Streptomycetaceae; unclassified Streptomycetaceae GRRRGAVRASVAVALAAVPCAAGYAVPVPTGAADTVDIAVVQGNVQQPGMDFLGRPMLIL WP_023531630.1 apolipoprotein N-acyltransferase, partial [Streptomycetaceae bacterium MP113-05] Length: 339\nNumber of predicted TMHs: 1\nExp number of AAs in TMHs: 32.69704\nExp number, first 60 AAs: 15.21755\nTotal prob of N-in: 0.75892\nPOSSIBLE N-term signal sequence\noutside 1 300\nTMhelix 301 323\ninside 324 339

34222 GCF_900167435.1_IMG-taxon_2524614583_annotated_assembly Marinactinospora thermotolerans DSM 45154 Terrabacteria group; Actinobacteria; Actinobacteria; Streptosporangiales; Nocardiopsaceae; Marinactinospora; Marinactinospora thermotolerans MNLCVGALHGRRRAVALACLTLATALLAGCSESRSNLGTASVPDAAMEAAPPEGGADGAA WP_078762545.1 hypothetical protein [Marinactinospora thermotolerans] Length: 317\nNumber of predicted TMHs: 1\nExp number of AAs in TMHs: 25.02454\nExp number, first 60 AAs: 2.06727\nTotal prob of N-in: 0.11464\noutside 1 261\nTMhelix 262 284\ninside 285 317

34223 GCF_001877055.1_ASM187705v1 Nocardiopsis dassonvillei Terrabacteria group; Actinobacteria; Actinobacteria; Streptosporangiales; Nocardiopsaceae; Nocardiopsis MPTNAARPPLTPGRRRTAIALGAVFVGSSLVAAMSPAHAEPSPSASASPAPSTPPSSPEP WP_071622919.1 hypothetical protein [Nocardiopsis dassonvillei] Length: 354\nNumber of predicted TMHs: 1\nExp number of AAs in TMHs: 28.85488\nExp number, first 60 AAs: 7.1312\nTotal prob of N-in: 0.34980\noutside 1 315\nTMhelix 316 338\ninside 339 354

34224 GCF_001877055.1_ASM187705v1 Nocardiopsis dassonvillei Terrabacteria group; Actinobacteria; Actinobacteria; Streptosporangiales; Nocardiopsaceae; Nocardiopsis MQAALSLRPRNGRRRGLVPPLVLTAALVLSAPAAALAQSVPDPSPDRSPGQDPARPRVTT WP_071621346.1 cell wall protein [Nocardiopsis dassonvillei] Length: 507\nNumber of predicted TMHs: 1\nExp number of AAs in TMHs: 30.7224\nExp number, first 60 AAs: 8.54114\nTotal prob of N-in: 0.40719\noutside 1 471\nTMhelix 472 494\ninside 495 507

34225 GCF_900141985.1_IMG-taxon_2675903133_annotated_assembly Nocardiopsis flavescens Terrabacteria group; Actinobacteria; Actinobacteria; Streptosporangiales; Nocardiopsaceae; Nocardiopsis MDHPRAPHGGGRRRVLACGATALALVLAPVAAAAQPADPSPAPPAADDTAVPQVLDVSDA WP_073383984.1 hypothetical protein [Nocardiopsis flavescens] Length: 420\nNumber of predicted TMHs: 1\nExp number of AAs in TMHs: 30.32888\nExp number, first 60 AAs: 8.74748\nTotal prob of N-in: 0.40370\noutside 1 377\nTMhelix 378 400\ninside 401 420

34226 GCF_900141985.1_IMG-taxon_2675903133_annotated_assembly Nocardiopsis flavescens Terrabacteria group; Actinobacteria; Actinobacteria; Streptosporangiales; Nocardiopsaceae; Nocardiopsis MRSTPAPGAPRRRAVGRAAAALLLALGPVAGPAAAAADTAADEPPPPLPQVFEFKGERQA WP_073384391.1 peptidase S8 and S53 subtilisin kexin sedolisin [Nocardiopsis flavescens] Length: 427\nNumber of predicted TMHs: 1\nExp number of AAs in TMHs: 33.55704\nExp number, first 60 AAs: 9.70598\nTotal prob of N-in: 0.46390\noutside 1 372\nTMhelix 373 395\ninside 396 427

34227 GCF_900141985.1_IMG-taxon_2675903133_annotated_assembly Nocardiopsis flavescens Terrabacteria group; Actinobacteria; Actinobacteria; Streptosporangiales; Nocardiopsaceae; Nocardiopsis MPPRRTPLLALLGTAALGAALLSPASAGAQDAAHADAAVSSARWLAGELQEDGSLPGPFG WP_073378631.1 hypothetical protein [Nocardiopsis flavescens] Length: 493\nNumber of predicted TMHs: 1\nExp number of

AAs in TMHs: 30.03589\nExp number, first 60 AAs: 7.31642\nTotal prob of N-in: 0.33238\noutside 1 460\nTMhelix 461 483\ninside 484 493

34228 GCF_001942255.1_ASM194225v1 Nocardiopsis sp. CNR-923 Terrabacteria group; Actinobacteria; Actinobacteria; Streptosporangiales; Nocardiopsaceae; Nocardiopsis
MPGPTLGAGADGTSGASGVPDPEGRGAPEDLPPRRGAFLVALGAFLLTVAALLPLYVYDR WP_083678488.1
hypothetical protein [Nocardiopsis sp. CNR-923] Length: 358\nNumber of predicted TMHs: 2\nExp number of AAs in TMHs: 43.49522\nExp number, first 60 AAs: 21.10819\nTotal prob of N-in: 0.80769\nPOSSIBLE N-term signal sequence\ninside 1 35\nTMhelix 36 58\noutside 59 305\nTMhelix 306 328\ninside 329 358

34229 GCF_000381685.1_ASM38168v1 Nocardiopsis sp. CNS-639 Terrabacteria group; Actinobacteria; Actinobacteria; Streptosporangiales; Nocardiopsaceae; Nocardiopsis
MQAALSLRPRNGRRRGLVPPLVLTAALVLSAPAAALAQPVPAPSPGQSPGQDPVRPRVTT WP_020722865.1
hypothetical protein [Nocardiopsis sp. CNS-639] Length: 506\nNumber of predicted TMHs: 1\nExp number of AAs in TMHs: 32.43198\nExp number, first 60 AAs: 10.01934\nTotal prob of N-in: 0.47458\nPOSSIBLE N-term signal sequence\noutside 1 470\nTMhelix 471 493\ninside 494 506

34230 GCF_001083815.1_ASM108381v1 Nocardiopsis sp. SBT366 Terrabacteria group; Actinobacteria; Actinobacteria; Streptosporangiales; Nocardiopsaceae; Nocardiopsis
MTRVACGTDRRGAFGRRAGGIGAGLLAAVLCSAPAVADSGGFGDADAVRLSLSGAAESPL WP_049577979.1
hypothetical protein [Nocardiopsis sp. SBT366] Length: 206\nNumber of predicted TMHs: 2\nExp number of AAs in TMHs: 39.20395\nExp number, first 60 AAs: 20.1823\nTotal prob of N-in: 0.98092\nPOSSIBLE N-term signal sequence\ninside 1 19\nTMhelix 20 42\noutside 43 179\nTMhelix 180 197\ninside 198 206

34231 GCF_000341205.1_v1.06 Nocardiopsis baichengensis YIM 90130 Terrabacteria group; Actinobacteria; Actinobacteria; Streptosporangiales; Nocardiopsaceae; Nocardiopsis; Nocardiopsis baichengensis
MARRRSASRRSLLPKGVCAFGTAFFALLTLACAALTFDVSSTAVEHTAAAQGRAGEVGTV WP_017557735.1
hypothetical protein [Nocardiopsis baichengensis] Length: 186\nNumber of predicted TMHs: 2\nExp number of AAs in TMHs: 45.2014\nExp number, first 60 AAs: 22.1432\nTotal prob of N-in: 0.99726\nPOSSIBLE N-term signal sequence\ninside 1 11\nTMhelix 12 34\noutside 35 145\nTMhelix 146 168\ninside 169 186

34232 GCF_000341205.1_v1.06 Nocardiopsis baichengensis YIM 90130 Terrabacteria group; Actinobacteria; Actinobacteria; Streptosporangiales; Nocardiopsaceae; Nocardiopsis; Nocardiopsis baichengensis
MGTNLDVASDRKRRSTGRRAVVLLGTGAAGAAVLAGALALSSVAATLPGGLGPCVPGDCP WP_017559785.1
choice-of-anchor A domain-containing protein [Nocardiopsis baichengensis] Length: 536\nNumber of predicted TMHs: 2\nExp number of AAs in TMHs: 44.77822\nExp number, first 60 AAs: 22.52366\nTotal prob of N-in: 0.99986\nPOSSIBLE N-term signal sequence\ninside 1 20\nTMhelix 21 43\noutside 44 503\nTMhelix 504 526\ninside 527 536

34233 GCF_000341205.1_v1.06 Nocardiopsis baichengensis YIM 90130 Terrabacteria group; Actinobacteria; Actinobacteria; Streptosporangiales; Nocardiopsaceae; Nocardiopsis; Nocardiopsis baichengensis
MPIAFPPSPTRSLRPARRPGRPVGPRRTALRGTAAAAALALAAGAGAAPAAADGGADERT WP_017559909.1
hypothetical protein [Nocardiopsis baichengensis] Length: 625\nNumber of predicted TMHs: 1\nExp number of AAs in TMHs: 23.15967\nExp number, first 60 AAs: 0.34004\nTotal prob of N-in: 0.01565\noutside 1 591\nTMhelix 592 614\ninside 615 625

34234 GCF_000341185.1_SOAPdenovo_v1.06Nocardiopsis chromatogenes YIM 90109 Terrabacteria group; Actinobacteria; Actinobacteria; Streptosporangiales; Nocardiopsaceae; Nocardiopsis; Nocardiopsis chromatogenes
MPTAFPPIPTRSPRPSRRPGGPGRPRRGALRAAAAAAALALAAGTAAPAAADSAAPEGDT WP_017622416.1
hypothetical protein [Nocardiopsis chromatogenes] Length: 624\nNumber of predicted TMHs: 1\nExp number of AAs in TMHs: 22.89151\nExp number, first 60 AAs: 0.41007\nTotal prob of N-in: 0.01904\noutside 1 590\nTMhelix 591 613\ninside 614 624

34235 GCF_001552695.1_ASM155269v1 Nocardiopsis dassonvillei subsp. albirubida NBRC 13392 Terrabacteria group; Actinobacteria; Actinobacteria; Streptosporangiales; Nocardiopsaceae; Nocardiopsis; Nocardiopsis dassonvillei; Nocardiopsis dassonvillei subsp. albirubida
MPTNAARPPLTPGRRRTAVALGAVFVGSSLVAVMSPAHAEPSPSASASPSVPPSPPPSAS WP_061081165.1
hypothetical protein [Nocardiopsis dassonvillei] Length: 354\nNumber of predicted TMHs: 1\nExp number of AAs in TMHs: 29.52771\nExp number, first 60 AAs: 8.48553\nTotal prob of N-in: 0.41384\noutside 1 315\nTMhelix 316 338\ninside 339 354

34236 GCF_000092985.1_ASM9298v1 Nocardiopsis dassonvillei subsp. dassonvillei DSM 43111 Terrabacteria group; Actinobacteria; Actinobacteria; Streptosporangiales; Nocardiopsaceae; Nocardiopsis; Nocardiopsis dassonvillei; Nocardiopsis dassonvillei subsp. dassonvillei
MPTNAARPPLTPGRRRTAIALGAVFVGSSLVAAMSPAHAEPSPSASASPAPSTPPSSPEP WP_013155373.1 hypothetical protein [Nocardiopsis dassonvillei] Length: 354\nNumber of predicted TMHs: 1\nExp number of AAs in TMHs: 28.85412\nExp number, first 60 AAs: 7.13127\nTotal prob of N-in: 0.34979\noutside 1 315\nTMhelix 316 338\ninside 339 354

34237 GCF_000092985.1_ASM9298v1 Nocardiopsis dassonvillei subsp. dassonvillei DSM 43111 Terrabacteria group; Actinobacteria; Actinobacteria; Streptosporangiales; Nocardiopsaceae; Nocardiopsis; Nocardiopsis dassonvillei; Nocardiopsis dassonvillei subsp. dassonvillei
MQAALSLRPRNGRRRGLVPPLVLTAALVLSAPAAALAQSVPDPSPDRSPGQDPARPRVTT WP_013151425.1 LPXTG-motif cell wall anchor domain-containing protein [Nocardiopsis dassonvillei] Length: 515\nNumber of predicted TMHs: 1\nExp number of AAs in TMHs: 30.72264\nExp number, first 60 AAs: 8.54136\nTotal prob of N-in: 0.40715\noutside 1 479\nTMhelix 480 502\ninside 503 515

34238 GCF_000341245.1_SOAPdenovo_v1.06 Nocardiopsis halophila DSM 44494 Terrabacteria group; Actinobacteria; Actinobacteria; Streptosporangiales; Nocardiopsaceae; Nocardiopsis; Nocardiopsis halophila
MPIAFPPSPTRSLRPARRPGRPAGPRRTALRGAAAAAALALAAGAGAAPAAADGGADERT WP_017541123.1 hypothetical protein [Nocardiopsis halophila] Length: 625\nNumber of predicted TMHs: 1\nExp number of AAs in TMHs: 23.52021\nExp number, first 60 AAs: 0.74144\nTotal prob of N-in: 0.03361\noutside 1 591\nTMhelix 592 614\ninside 615 625

34239 GCF_000341245.1_SOAPdenovo_v1.06 Nocardiopsis halophila DSM 44494 Terrabacteria group; Actinobacteria; Actinobacteria; Streptosporangiales; Nocardiopsaceae; Nocardiopsis; Nocardiopsis halophila
MGTNLDVASDRKRRSTGRRAVVLLGTGAAGAAVLAGALALSSVAATLPGGLGPCVPGDCP WP_017539431.1 choice-of-anchor A domain-containing protein [Nocardiopsis halophila] Length: 517\nNumber of predicted TMHs: 2\nExp number of AAs in TMHs: 44.72432\nExp number, first 60 AAs: 22.52366\nTotal prob of N-in: 0.99986\nPOSSIBLE N-term signal sequence\ninside 1 20\nTMhelix 21 43\noutside 44 484\nTMhelix 485 507\ninside 508 517

34240 GCF_000341065.1_SOAPdenovo_v1.06 Nocardiopsis halotolerans DSM 44410 Terrabacteria group; Actinobacteria; Actinobacteria; Streptosporangiales; Nocardiopsaceae; Nocardiopsis; Nocardiopsis halotolerans
MPTDAARSPLTPGHRRRAIALGAVFVGSSLIAVMSPAYAEPSPSGTPSGSAPPPTSSASP WP_026122751.1 hypothetical protein [Nocardiopsis halotolerans] Length: 396\nNumber of predicted TMHs: 1\nExp number of AAs in TMHs: 36.80587\nExp number, first 60 AAs: 15.36291\nTotal prob of N-in: 0.75784\nPOSSIBLE N-term signal sequence\noutside 1 357\nTMhelix 358 380\ninside 381 396

34241 GCF_000341265.1_SOAPdenovo_1.06 Nocardiopsis prasina DSM 43845 Terrabacteria group; Actinobacteria; Actinobacteria; Streptosporangiales; Nocardiopsaceae; Nocardiopsis; Nocardiopsis prasina
MTRVACGTDRRGTFGRRAGGVVAGLVAAVLCAAPAVADSGGFGDADAVRLSLSGVDEPPL WP_017546715.1 hypothetical protein [Nocardiopsis prasina] Length: 206\nNumber of predicted TMHs: 2\nExp number of AAs in TMHs: 39.92781\nExp number, first 60 AAs: 20.77635\nTotal prob of N-in: 0.99503\nPOSSIBLE N-term signal sequence\ninside 1 19\nTMhelix 20 42\noutside 43 179\nTMhelix 180 197\ninside 198 206

34242 GCF_000340945.1_SOAPdenovo1.06 Nocardiopsis synnemataformans DSM 44143 Terrabacteria group; Actinobacteria; Actinobacteria; Streptosporangiales; Nocardiopsaceae; Nocardiopsis; Nocardiopsis synnemataformans
MPTNAARPPLAPGRRRTAVALGAVFVGSSLVAVMPPAYAEPSPSASASPSPSTPPSSSPE WP_026114600.1 hypothetical protein [Nocardiopsis synnemataformans] Length: 361\nNumber of predicted TMHs: 1\nExp number of AAs in TMHs: 35.00192\nExp number, first 60 AAs: 13.3071\nTotal prob of N-in: 0.63411\nPOSSIBLE N-term signal sequence\noutside 1 322\nTMhelix 323 345\ninside 346 361

34243 GCF_000340945.1_SOAPdenovo1.06 Nocardiopsis synnemataformans DSM 44143 Terrabacteria group; Actinobacteria; Actinobacteria; Streptosporangiales; Nocardiopsaceae; Nocardiopsis; Nocardiopsis synnemataformans
MQAALSPRPRNGRRRGLVPPLVLTAAVVLSAPAALAQPVPDPPPGRDPAHPRVTTVDRTR WP_020480233.1 hypothetical protein [Nocardiopsis synnemataformans] Length: 509\nNumber of predicted TMHs: 1\nExp number of AAs in TMHs: 30.34609\nExp number, first 60 AAs: 8.11285\nTotal prob of N-in: 0.38902\noutside 1 473\nTMhelix 474 496\ninside 497 509

34244 GCF_000826685.1_ASM82668v1 Streptomonospora alba Terrabacteria group; Actinobacteria; Actinobacteria; Streptosporangiales; Nocardiopsaceae; Streptomonospora

MTLATPALPDPALQRRRPSRARRPAVAAAALLGSLALATATAPAALAHDTLIGSSPEDGD WP_040269875.1 hypothetical protein [Streptomonospora alba] Length: 222\nNumber of predicted TMHs: 2\nExp number of AAs in TMHs: 43.59587\nExp number, first 60 AAs: 21.88108\nTotal prob of N-in: 0.96455\nPOSSIBLE N-term signal sequence\ninside 1 24\nTMhelix 25 47\noutside 48 185\nTMhelix 186 208\ninside 209 222

34245 GCF_001592045.1_ASM159204v1 Thermobifida fusca NBRC 14071 = JCM 3263 Terrabacteria group; Actinobacteria; Actinobacteria; Streptosporangiales; Nocardiopsaceae; Thermobifida; Thermobifida fusca MPPGMDKARVSRPRRRAWLGVPVGAAFIGCLLVPTLPASADATLDLDLSSRSALPGTAFS WP_011293161.1 hypothetical protein [Thermobifida fusca] Length: 347\nNumber of predicted TMHs: 2\nExp number of AAs in TMHs: 42.74049\nExp number, first 60 AAs: 20.59314\nTotal prob of N-in: 0.95099\nPOSSIBLE N-term signal sequence\ninside 1 16\nTMhelix 17 39\noutside 40 308\nTMhelix 309 331\ninside 332 347

34246 GCF_000401915.1_ASM40191v1 Thermobifida fusca TM51 Terrabacteria group; Actinobacteria; Actinobacteria; Streptosporangiales; Nocardiopsaceae; Thermobifida; Thermobifida fusca MPPGMDKARVSRPRRRAWLGVPVGAAFIGCLLVPTLPASADATLDLDLSSRSALPGTAFS WP_011293161.1 hypothetical protein [Thermobifida fusca] Length: 347\nNumber of predicted TMHs: 2\nExp number of AAs in TMHs: 42.74049\nExp number, first 60 AAs: 20.59314\nTotal prob of N-in: 0.95099\nPOSSIBLE N-term signal sequence\ninside 1 16\nTMhelix 17 39\noutside 40 308\nTMhelix 309 331\ninside 332 347

34247 GCF_000012405.1_ASM1240v1 Thermobifida fusca YX Terrabacteria group; Actinobacteria; Actinobacteria; Streptosporangiales; Nocardiopsaceae; Thermobifida; Thermobifida fusca MPPGMDKARVSRPRRRAWLGVPVGAAFIGCLLVPTLPASADATLDLDLSSRSALPGTAFS WP_011293161.1 hypothetical protein [Thermobifida fusca] Length: 347\nNumber of predicted TMHs: 2\nExp number of AAs in TMHs: 42.74049\nExp number, first 60 AAs: 20.59314\nTotal prob of N-in: 0.95099\nPOSSIBLE N-term signal sequence\ninside 1 16\nTMhelix 17 39\noutside 40 308\nTMhelix 309 331\ninside 332 347

34248 GCF_000719065.1_ASM71906v1 Herbidospora cretacea Terrabacteria group; Actinobacteria; Actinobacteria; Streptosporangiales; Streptosporangiaceae; Herbidospora MNLEARRALLALAAFATVGGATIALPSAAHAAPTWTATTTADDDDRGNDDDDRRGGDDDD WP_030450198.1 hypothetical protein [Herbidospora cretacea] Length: 159\nNumber of predicted TMHs: 2\nExp number of AAs in TMHs: 44.35671\nExp number, first 60 AAs: 21.93388\nTotal prob of N-in: 0.98615\nPOSSIBLE N-term signal sequence\ninside 1 6\nTMhelix 7 29\noutside 30 128\nTMhelix 129 151\ninside 152 159

34249 GCF_001570585.1_ASM157058v1 Herbidospora daliensis Terrabacteria group; Actinobacteria; Actinobacteria; Streptosporangiales; Streptosporangiaceae; Herbidospora MNLEARRALLALAAFTTVGGAATVALPAAAQAAPTWTATATDDDDDDRGDDDDDRPRGGVE WP_062432348.1 hypothetical protein [Herbidospora daliensis] Length: 248\nNumber of predicted TMHs: 2\nExp number of AAs in TMHs: 41.6921\nExp number, first 60 AAs: 19.52846\nTotal prob of N-in: 0.86989\nPOSSIBLE N-term signal sequence\ninside 1 6\nTMhelix 7 29\noutside 30 217\nTMhelix 218 240\ninside 241 248

34250 GCF_001570545.1_ASM157054v1 Herbidospora mongoliensis Terrabacteria group; Actinobacteria; Actinobacteria; Streptosporangiales; Streptosporangiaceae; Herbidospora MRRRTLLSSAALAFALTGAVPAHAEPAPLTQVKPAWRQDRQIADLKIKESSGLYASPLHP WP_066361022.1 hypothetical protein [Herbidospora mongoliensis] Length: 338\nNumber of predicted TMHs: 1\nExp number of AAs in TMHs: 25.23812\nExp number, first 60 AAs: 5.1397\nTotal prob of N-in: 0.27149\noutside 1 313\nTMhelix 314 333\ninside 334 338

34251 GCF_001570545.1_ASM157054v1 Herbidospora mongoliensis Terrabacteria group; Actinobacteria; Actinobacteria; Streptosporangiales; Streptosporangiaceae; Herbidospora MNLETRRALLAIAAFATVGGASVALPAAAHTAPMWSATVADDDDDRGGDDDDDRRGGDDN WP_066361959.1 hypothetical protein [Herbidospora mongoliensis] Length: 221\nNumber of predicted TMHs: 2\nExp number of AAs in TMHs: 43.65931\nExp number, first 60 AAs: 21.82106\nTotal prob of N-in: 0.97029\nPOSSIBLE N-term signal sequence\ninside 1 6\nTMhelix 7 29\noutside 30 190\nTMhelix 191 213\ninside 214 221

34252 GCF_001570525.1_ASM157052v1 Herbidospora sakaeratensis Terrabacteria group; Actinobacteria; Actinobacteria; Streptosporangiales; Streptosporangiaceae; Herbidospora MNLEARRALLALAAFATVGGATAALPAAAHASPTWTATVADDDSRDDDDDDRQGGDDDDD WP_062330275.1 hypothetical protein [Herbidospora sakaeratensis] Length: 201\nNumber of predicted TMHs: 2\nExp number of AAs in TMHs: 43.69996\nExp number, first 60 AAs: 21.39805\nTotal prob of N-in: 0.95389\nPOSSIBLE N-term signal sequence\ninside 1 6\nTMhelix 7 29\noutside 30 170\nTMhelix 171 193\ninside 194 201

34253 GCF_001570565.1_ASM157056v1 Herbidospora yilanensis Terrabacteria group; Actinobacteria; Actinobacteria; Streptosporangiales; Streptosporangiaceae; Herbidospora
MNLEARRALLALAAFATVGCTTAALPSTAHAAPVWTTTATDDDDDRGGADDDERPSGGVE WP_062355275.1
hypothetical protein [Herbidospora yilanensis] Length: 157\nNumber of predicted TMHs: 2\nExp number of AAs in TMHs: 43.33212\nExp number, first 60 AAs: 20.90349\nTotal prob of N-in: 0.94636\nPOSSIBLE N-term signal sequence\ninside 1 6\nTMhelix 7 29\noutside 30 126\nTMhelix 127 149\ninside 150 157

34254 GCF_000718545.1_ASM71854v1 Microbispora sp. NRRL B-24597 Terrabacteria group; Actinobacteria; Actinobacteria; Streptosporangiales; Streptosporangiaceae; Microbispora
MFHADPVAHAEPPSAPRRRSALSVVSSVIMNLLAVGGLACIVMVVLAVVFNVSLIMFKTG WP_081906626.1
MULTISPECIES: signal peptidase I [Actinobacteria] Length: 208\nNumber of predicted TMHs: 2\nExp number of AAs in TMHs: 46.44187\nExp number, first 60 AAs: 23.63275\nTotal prob of N-in: 0.15209\nPOSSIBLE N-term signal sequence\noutside 1 31\nTMhelix 32 54\ninside 55 151\nTMhelix 152 174\noutside 175 208

34255 GCF_000721275.1_ASM72127v1 Microtetraspora glauca Terrabacteria group; Actinobacteria; Actinobacteria; Streptosporangiales; Streptosporangiaceae; Microtetraspora
MKATSTASGRRRMLDHSLLVLGSGALATAMAVLALGPGVSSASSHREAPLIAGDPRADNT WP_030497999.1
hypothetical protein [Microtetraspora glauca] Length: 516\nNumber of predicted TMHs: 2\nExp number of AAs in TMHs: 36.4547199999999\nExp number, first 60 AAs: 21.9514\nTotal prob of N-in: 0.98836\nPOSSIBLE N-term signal sequence\ninside 1 16\nTMhelix 17 39\noutside 40 490\nTMhelix 491 508\ninside 509 516

34256 GCF_000721275.1_ASM72127v1 Microtetraspora glauca Terrabacteria group; Actinobacteria; Actinobacteria; Streptosporangiales; Streptosporangiaceae; Microtetraspora
MSAAVAARRAVVGGMAAVCLTLGGGTAAWAQGTPGGEGWEGRGARAYTAEDAVPAQDLSG WP_030493088.1
hypothetical protein [Microtetraspora glauca] Length: 341\nNumber of predicted TMHs: 1\nExp number of AAs in TMHs: 36.5315\nExp number, first 60 AAs: 15.04339\nTotal prob of N-in: 0.69897\nPOSSIBLE N-term signal sequence\noutside 1 309\nTMhelix 310 332\ninside 333 341

34257 GCF_000721275.1_ASM72127v1 Microtetraspora glauca Terrabacteria group; Actinobacteria; Actinobacteria; Streptosporangiales; Streptosporangiaceae; Microtetraspora
MLSVVENHSLPRAAAFFDLDKTVIAKSSTLTFSKSFYRGGLISRRAALRTAYIQFVFLVG WP_030495816.1 inhibition of morphological differentiation protein [Microtetraspora glauca] Length: 285\nNumber of predicted TMHs: 1\nExp number of AAs in TMHs: 21.46708\nExp number, first 60 AAs: 0.07804\nTotal prob of N-in: 0.00510\noutside 1 247\nTMhelix 248 270\ninside 271 285

34258 GCF_001570405.1_ASM157040v1 Microtetraspora niveoalba Terrabacteria group; Actinobacteria; Actinobacteria; Streptosporangiales; Streptosporangiaceae; Microtetraspora
MTGIRSGRHSTDGELFEPPARRDHVWLANIYATPWHLPEGVPSEPPGTYLRLGRLERHLA WP_067184399.1
hypothetical protein [Microtetraspora niveoalba]Length: 438\nNumber of predicted TMHs: 2\nExp number of AAs in TMHs: 47.86408\nExp number, first 60 AAs: 0.00101\nTotal prob of N-in: 0.85759\ninside 1 356\nTMhelix 357 379\noutside 380 398\nTMhelix 399 421\ninside 422 438

34259 GCF_001570405.1_ASM157040v1 Microtetraspora niveoalba Terrabacteria group; Actinobacteria; Actinobacteria; Streptosporangiales; Streptosporangiaceae; Microtetraspora
MDDPRAHVFIVDHVNPGTTFTRRLRVNSSSPRRQHVELYAAAAGIAHGTFTFAPGRTAND WP_084516786.1
hypothetical protein [Microtetraspora niveoalba]Length: 289\nNumber of predicted TMHs: 1\nExp number of AAs in TMHs: 22.98081\nExp number, first 60 AAs: 0.23566\nTotal prob of N-in: 0.01440\noutside 1 256\nTMhelix 257 279\ninside 280 289

34260 GCF_001696485.1_ASM169648v1 Planobispora rosea Terrabacteria group; Actinobacteria; Actinobacteria; Streptosporangiales; Streptosporangiaceae; Planobispora
MARRTALIPLLALACAGLLSPAAHADTHSVLPQPPEPLFHVLGNTINEASGLAVSADGKR WP_068920822.1 hypothetical protein [Planobispora rosea] Length: 376\nNumber of predicted TMHs: 1\nExp number of AAs in TMHs: 27.30617\nExp number, first 60 AAs: 5.05648\nTotal prob of N-in: 0.26283\noutside 1 333\nTMhelix 334 356\ninside 357 376

34261 GCF_900115095.1_IMG-taxon_2634166331_annotated_assembly Actinomadura madurae Terrabacteria group; Actinobacteria; Actinobacteria; Streptosporangiales; Thermomonosporaceae; Actinomadura
MERAGRRLFLGSAGLGALPVWLEALPAMPRRAVLIPTAANPLPSAPFVDALADLLRAEGL WP_075022990.1
hypothetical protein [Actinomadura madurae] Length: 203\nNumber of predicted TMHs: 2\nExp number of AAs in TMHs: 43.81103\nExp number, first 60 AAs: 3.15956\nTotal prob of N-in: 0.85445\ninside 1 106\nTMhelix 107 129\noutside 130 143\nTMhelix 144 161\ninside 162 203

34262 GCF_001552155.1_ASM155215v1 Actinomadura formosensis NBRC 14204 Terrabacteria group; Actinobacteria; Actinobacteria; Streptosporangiales; Thermomonosporaceae; Actinomadura; Actinomadura formosensis MMPQGDANVRRPRRSLITFLWRFGLVVVLVLTGSLGGLGYTLLAPATYTATAYVLVVNGG WP_067793415.1 hypothetical protein [Actinomadura formosensis] Length: 233\nNumber of predicted TMHs: 2\nExp number of AAs in TMHs: 43.09319\nExp number, first 60 AAs: 23.28707\nTotal prob of N-in: 0.99340\nPOSSIBLE N-term signal sequence\ninside 1 20\nTMhelix 21 43\noutside 44 169\nTMhelix 170 192\ninside 193 233

34263 GCF_001552175.1_ASM155217v1 Actinomadura kijaniata NBRC 14229 Terrabacteria group; Actinobacteria; Actinobacteria; Streptosporangiales; Thermomonosporaceae; Actinomadura; Actinomadura kijaniata MMTKRLPSRRPALLAGLLAVFLAALGGLFAAPASAHTRLISSTPGKDATAANVTEVAPVF WP_067825074.1 hypothetical protein [Actinomadura kijaniata] Length: 200\nNumber of predicted TMHs: 2\nExp number of AAs in TMHs: 44.87965\nExp number, first 60 AAs: 22.1267\nTotal prob of N-in: 0.99867\nPOSSIBLE N-term signal sequence\ninside 1 12\nTMhelix 13 35\noutside 36 164\nTMhelix 165 187\ninside 188 200

34264 GCF_001552175.1_ASM155217v1 Actinomadura kijaniata NBRC 14229 Terrabacteria group; Actinobacteria; Actinobacteria; Streptosporangiales; Thermomonosporaceae; Actinomadura; Actinomadura kijaniata MTSAHRHRRRFLLGLPPLAGVLLILLGFVLGPPAGQWARERAAGIGTPVGPGEPITLVRD WP_067818420.1 hypothetical protein [Actinomadura kijaniata] Length: 176\nNumber of predicted TMHs: 2\nExp number of AAs in TMHs: 44.2229\nExp number, first 60 AAs: 21.48054\nTotal prob of N-in: 0.98212\nPOSSIBLE N-term signal sequence\ninside 1 11\nTMhelix 12 34\noutside 35 141\nTMhelix 142 164\ninside 165 176

34265 GCF_001552215.1_ASM155221v1 Actinomadura macra NBRC 14102 Terrabacteria group; Actinobacteria; Actinobacteria; Streptosporangiales; Thermomonosporaceae; Actinomadura; Actinomadura macra MSLRPSRRRRAVRAAAALVLLGTASAFPAPASASPAGSAAGSVPGGLAADARAPKVAFTI WP_084265062.1 hypothetical protein [Actinomadura macra] Length: 360\nNumber of predicted TMHs: 1\nExp number of AAs in TMHs: 25.96788\nExp number, first 60 AAs: 8.61988\nTotal prob of N-in: 0.44905\noutside 1 334\nTMhelix 335 352\ninside 353 360

34266 GCF_000468475.2_Amad2 Actinomadura madurae LIID-AJ290 Terrabacteria group; Actinobacteria; Actinobacteria; Streptosporangiales; Thermomonosporaceae; Actinomadura; Actinomadura madurae MERAGRRLFLGSAGLGALPAWLEALPAMPRRAVLIPTAANPLPSAPFVDALADLLRAEGL WP_021595392.1 hypothetical protein [Actinomadura madurae] Length: 203\nNumber of predicted TMHs: 2\nExp number of AAs in TMHs: 41.3253600000001\nExp number, first 60 AAs: 1.96999\nTotal prob of N-in: 0.84336\ninside 1 106\nTMhelix 107 129\noutside 130 143\nTMhelix 144 161\ninside 162 203

34267 GCF_000425065.1_ASM42506v1 Actinomadura rifamycini DSM 43936 Terrabacteria group; Actinobacteria; Actinobacteria; Streptosporangiales; Thermomonosporaceae; Actinomadura; Actinomadura rifamycini MSLIPHARRTAVVAVLGAAGLVGLATAASAHVTVNPDAAEQGSWAKVSFRVPNERDDAAT WP_026403212.1 membrane protein [Actinomadura rifamycini] Length: 237\nNumber of predicted TMHs: 2\nExp number of AAs in TMHs: 44.28757\nExp number, first 60 AAs: 22.14357\nTotal prob of N-in: 0.96225\nPOSSIBLE N-term signal sequence\ninside 1 11\nTMhelix 12 34\noutside 35 207\nTMhelix 208 230\ninside 231 237

34268 GCF_000718255.1_ASM71825v1 Spirillospora albida Terrabacteria group; Actinobacteria; Actinobacteria; Streptosporangiales; Thermomonosporaceae; Spirillospora MHSLPRGCAPAHSLLPRYGIEGSPPHRRQALRKFIGHALTGAGAFLLVGGALVRFQVAPA WP_084219429.1 hypothetical protein [Spirillospora albida] Length: 348\nNumber of predicted TMHs: 2\nExp number of AAs in TMHs: 34.84906\nExp number, first 60 AAs: 17.29037\nTotal prob of N-in: 0.75368\nPOSSIBLE N-term signal sequence\ninside 1 33\nTMhelix 34 53\noutside 54 312\nTMhelix 313 332\ninside 333 348

34269 GCF_001298565.1_Actinobacteria.strain_OK006_v1.0 Actinobacteria bacterium OK006 Terrabacteria group; Actinobacteria; Actinobacteria; unclassified Actinobacteria; unclassified Actinobacteria (class) (miscellaneous) MRHARRRIVRRGTRLAAVGGVLCGGLMVTHAMASEPSAASRAPQSATLAAAGKGAGLVST WP_054236526.1 protease [Streptomyces diastatochromogenes] Length: 460\nNumber of predicted TMHs: 2\nExp number of AAs in TMHs: 42.1892999999999\nExp number, first 60 AAs: 20.80533\nTotal prob of N-in: 0.96509\nPOSSIBLE N-term signal sequence\ninside 1 11\nTMhelix 12 34\noutside 35 421\nTMhelix 422 444\ninside 445 460

34270 GCF_001298565.1_Actinobacteria.strain_OK006_v1.0 Actinobacteria bacterium OK006 Terrabacteria group; Actinobacteria; Actinobacteria; unclassified Actinobacteria; unclassified Actinobacteria (class) (miscellaneous) MCETPSTRKSAPKTPDPAAGRRRSALAGNPSGHQHQGRPRPPGRSGPAPQARTNEALTVE WP_082417500.1 hypothetical protein [Actinobacteria bacterium OK006] Length: 163\nNumber of predicted TMHs: 2\nExp number of AAs in TMHs: 44.21018\nExp number, first 60 AAs: 0\nTotal prob of N-in: 0.99247\ninside 1 101\nTMhelix 102 124\noutside 125 127\nTMhelix 128 150\ninside 151 163

34271 GCF_001298565.1_Actinobacteria.strain_OK006_v1.0 Actinobacteria bacterium OK006 Terrabacteria group; Actinobacteria; Actinobacteria; unclassified Actinobacteria; unclassified Actinobacteria (class) (miscellaneous) MLGLVENHSLPRTAAFFDLDKTVIAKSSTLTFSKSFYQGGLINRRAVLRTAYAQFVFMAG WP_054237412.1 inhibition of morphological differentiation protein [Streptomyces diastatochromogenes] Length: 277\nNumber of predicted TMHs: 1\nExp number of AAs in TMHs: 21.57763\nExp number, first 60 AAs: 0.0148\nTotal prob of N-in: 0.00176\noutside 1 247\nTMhelix 248 270\ninside 271 277

34272 GCF_001298565.1_Actinobacteria.strain_OK006_v1.0 Actinobacteria bacterium OK006 Terrabacteria group; Actinobacteria; Actinobacteria; unclassified Actinobacteria; unclassified Actinobacteria (class) (miscellaneous) MRDSSPVPPRAARLSRRAALGLAGGLPLAAASPASAATPVIGGERLARAGVQVRGAAGLP WP_054230871.1 D-alanyl-D-alanine carboxypeptidase [Actinobacteria bacterium OK006] Length: 408\nNumber of predicted TMHs: 1\nExp number of AAs in TMHs: 27.92832\nExp number, first 60 AAs: 9.40055\nTotal prob of N-in: 0.41854\noutside 1 355\nTMhelix 356 378\ninside 379 408

34273 GCF_001298575.1_Actinobacteria.strain_OK074_v1.0 Actinobacteria bacterium OK074 Terrabacteria group; Actinobacteria; Actinobacteria; unclassified Actinobacteria; unclassified Actinobacteria (class) (miscellaneous) MLKLVENHSLPRTAAFFDLDKTVIAKSSTLTFGKSFYQGGLINRRAVLRTAYAQFVFLAG WP_054215940.1 inhibition of morphological differentiation protein [Actinobacteria bacterium OK074] Length: 277\nNumber of predicted TMHs: 1\nExp number of AAs in TMHs: 21.44628\nExp number, first 60 AAs: 0.02615\nTotal prob of N-in: 0.00273\noutside 1 247\nTMhelix 248 270\ninside 271 277

34274 GCF_001298575.1_Actinobacteria.strain_OK074_v1.0 Actinobacteria bacterium OK074 Terrabacteria group; Actinobacteria; Actinobacteria; unclassified Actinobacteria; unclassified Actinobacteria (class) (miscellaneous) MALVPLASRVPSPPFRTRRRPVRPVQALAATLLVAALALTGCSGADDSGSSGSAADKAAP WP_054216528.1 hypothetical protein [Actinobacteria bacterium OK074] Length: 329\nNumber of predicted TMHs: 1\nExp number of AAs in TMHs: 24.11859\nExp number, first 60 AAs: 0.99861\nTotal prob of N-in: 0.05873\noutside 1 284\nTMhelix 285 307\ninside 308 329

34275 GCF_001298575.1_Actinobacteria.strain_OK074_v1.0 Actinobacteria bacterium OK074 Terrabacteria group; Actinobacteria; Actinobacteria; unclassified Actinobacteria; unclassified Actinobacteria (class) (miscellaneous) MRDSSRLSRRAVLGFAAAAPLAAAVPTPAAATSAAVVGGSRLGRSGAQVRGGSALPKGLS WP_054220105.1 D-alanyl-D-alanine carboxypeptidase [Actinobacteria bacterium OK074] Length: 402\nNumber of predicted TMHs: 2\nExp number of AAs in TMHs: 41.00387\nExp number, first 60 AAs: 19.31233\nTotal prob of N-in: 0.85284\nPOSSIBLE N-term signal sequence\ninside 1 11\nTMhelix 12 34\noutside 35 362\nTMhelix 363 385\ninside 386 402

34276 GCF_001298575.1_Actinobacteria.strain_OK074_v1.0 Actinobacteria bacterium OK074 Terrabacteria group; Actinobacteria; Actinobacteria; unclassified Actinobacteria; unclassified Actinobacteria (class) (miscellaneous) MTPISRSGTGRRSLVTLICGALAAGGLTAAGVSVLEPGAASASSHREAPLISGTPQYDNT WP_054214097.1 hypothetical protein [Actinobacteria bacterium OK074] Length: 509\nNumber of predicted TMHs: 2\nExp number of AAs in TMHs: 44.81981\nExp number, first 60 AAs: 22.10773\nTotal prob of N-in: 0.98112\nPOSSIBLE N-term signal sequence\ninside 1 12\nTMhelix 13 35\noutside 36 477\nTMhelix 478 500\ninside 501 509

34277 GCF_001298545.1_Actinobacteria.strain_OV320_v1.0 Actinobacteria bacterium OV320 Terrabacteria group; Actinobacteria; Actinobacteria; unclassified Actinobacteria; unclassified Actinobacteria (class) (miscellaneous) MKKLSAARRPTLATAVAATAVLLTAGPASAHVEVESESAQALAQNVEIAFDAESESATAG WP_054238826.1 hypothetical protein [Actinobacteria bacterium OV320] Length: 238\nNumber of predicted TMHs: 1\nExp number of AAs in TMHs: 27.98071\nExp number, first 60 AAs: 6.34107\nTotal prob of N-in: 0.36008\noutside 1 206\nTMhelix 207 229\ninside 230 238

34278 GCF_001298545.1_Actinobacteria.strain_OV320_v1.0 Actinobacteria bacterium OV320 Terrabacteria group; Actinobacteria; Actinobacteria; unclassified Actinobacteria; unclassified Actinobacteria (class) (miscellaneous) MLMVVENHSLPRTAAFFDLDKTVIAKSSTLTFSKSFYQGGLINRRAALRTAYAQFVFLVG WP_054241799.1 inhibition of morphological differentiation protein [Streptomyces sp. Root1310] Length: 277\nNumber of predicted TMHs: 1\nExp number of AAs in TMHs: 21.36377\nExp number, first 60 AAs: 0.01645\nTotal prob of N-in: 0.00243\noutside 1 247\nTMhelix 248 270\ninside 271 277

34279 GCF_001298545.1_Actinobacteria.strain_OV320_v1.0 Actinobacteria bacterium OV320 Terrabacteria group; Actinobacteria; Actinobacteria; unclassified Actinobacteria; unclassified Actinobacteria (class) (miscellaneous)

MYGPAVHAFSRPSPRLSRRSLLALVATAVTAPLTAAPLAATPAAAASDVVVGGARLVADG WP_054238031.1 D-alanyl-D-alanine carboxypeptidase [Actinobacteria bacterium OV320] Length: 402\nNumber of predicted TMHs: 1\nExp number of AAs in TMHs: 34.6896\nExp number, first 60 AAs: 15.10847\nTotal prob of N-in: 0.68558\nPOSSIBLE N-term signal sequence\noutside 1 365\nTMhelix 366 388\ninside 389 402

34280 GCF_001298555.1_Actinobacteria.strain_OV450_v1.0 Actinobacteria bacterium OV450 Terrabacteria group; Actinobacteria; Actinobacteria; unclassified Actinobacteria; unclassified Actinobacteria (class) (miscellaneous)
MPARRRTVTRIATAALAPLAVAAYAAAPAAAHGSMTDPVSRVAACYAEGPESPKSAACKA WP_054225684.1 chitin-binding protein [Actinobacteria bacterium OV450] Length: 341\nNumber of predicted TMHs: 2\nExp number of AAs in TMHs: 41.64412\nExp number, first 60 AAs: 19.09323\nTotal prob of N-in: 0.89549\nPOSSIBLE N-term signal sequence\ninside 1 6\nTMhelix 7 29\noutside 30 305\nTMhelix 306 328\ninside 329 341

34281 GCF_000471045.1_LLX17V1 actinobacterium LLX17 Terrabacteria group; Actinobacteria; Actinobacteria; unclassified Actinobacteria; unclassified Actinobacteria (class) (miscellaneous)
MLGTMDDRPPARAAAFFDLDKTIIAKSSTLAFSRPFFAEGLLTRRAVLRGAYAQFVFGVS WP_023647230.1 inhibition of morphological differentiation protein [actinobacterium LLX17] Length: 274\nNumber of predicted TMHs: 1\nExp number of AAs in TMHs: 21.74625\nExp number, first 60 AAs: 0.02409\nTotal prob of N-in: 0.02539\noutside 1 246\nTMhelix 247 268\ninside 269 274

34282 GCF_001552785.1_ASM155278v1 Atopobium deltae Terrabacteria group; Actinobacteria; Coriobacteriia; Coriobacteriales; Atopobiaceae; Atopobium
MNERFNLRVLNAKAHIRAWSRRRTLIALASAALAMLLASPFTAQALQTATKTVSALPAAT WP_066304274.1 hypothetical protein [Atopobium deltae] Length: 863\nNumber of predicted TMHs: 2\nExp number of AAs in TMHs: 40.8792\nExp number, first 60 AAs: 19.87057\nTotal prob of N-in: 0.91545\nPOSSIBLE N-term signal sequence\ninside 1 23\nTMhelix 24 46\noutside 47 835\nTMhelix 836 858\ninside 859 863

34283 GCF_001438885.1_ASM143888v1 Atopobium rimae Terrabacteria group; Actinobacteria; Coriobacteriia; Coriobacteriales; Atopobiaceae; Atopobium
MTFDQDARTSQDGIQNKAISRREVLKLFVGAGAAAAVTASLSLRPFTALATTQSEIDAAQ WP_003150348.1 hypothetical protein [Atopobium rimae] Length: 431\nNumber of predicted TMHs: 1\nExp number of AAs in TMHs: 50.58803\nExp number, first 60 AAs: 13.09674\nTotal prob of N-in: 0.71580\nPOSSIBLE N-term signal sequence\noutside 1 366\nTMhelix 367 389\ninside 390 431

34284 GCF_001813205.1_ASM181320v1 Atopobium sp. HMSC064B08 Terrabacteria group; Actinobacteria; Coriobacteriia; Coriobacteriales; Atopobiaceae; Atopobium
MTFDQDARTSQDGIQNKAISRREVLKLFVGAGAAAAVTASLSLRPFTALATTQSEIDAAQ WP_070699546.1 Tat pathway signal protein [Atopobium sp. HMSC064B08] Length: 431\nNumber of predicted TMHs: 1\nExp number of AAs in TMHs: 48.59403\nExp number, first 60 AAs: 12.742\nTotal prob of N-in: 0.72134\nPOSSIBLE N-term signal sequence\noutside 1 366\nTMhelix 367 389\ninside 390 431

34285 GCF_000174015.1_ASM17401v1 Atopobium rimae ATCC 49626 Terrabacteria group; Actinobacteria; Coriobacteriia; Coriobacteriales; Atopobiaceae; Atopobium; Atopobium rimae
MTFDQDARTSQDGIQNKAISRREVLKLFVGAGAAAAVTASLSLRPFTALATTQSEIDAAQ WP_003150348.1 hypothetical protein [Atopobium rimae] Length: 431\nNumber of predicted TMHs: 1\nExp number of AAs in TMHs: 50.58803\nExp number, first 60 AAs: 13.09674\nTotal prob of N-in: 0.71580\nPOSSIBLE N-term signal sequence\noutside 1 366\nTMhelix 367 389\ninside 390 431

34286 GCF_001494635.1_ASM149463v1 Olsenella scatoligenes Terrabacteria group; Actinobacteria; Coriobacteriia; Coriobacteriales; Atopobiaceae; Olsenella
MAKNTTRRKFIGSTIASVGLLGVTAAANVAANKFSSLLDHYVGGRPATVEKVEGSEGWDA WP_059052747.1 hypothetical protein [Olsenella scatoligenes] Length: 983\nNumber of predicted TMHs: 2\nExp number of AAs in TMHs: 39.3749399999999\nExp number, first 60 AAs: 17.87244\nTotal prob of N-in: 0.85023\nPOSSIBLE N-term signal sequence\ninside 1 8\nTMhelix 9 31\noutside 32 924\nTMhelix 925 947\ninside 948 983

34287 GCF_002160255.1_ASM216025v1 Olsenella sp. An188Terrabacteria group; Actinobacteria; Coriobacteriia; Coriobacteriales; Atopobiaceae; Olsenella
MADKPKKQPKKMTRRGFVASGVGAGALVGVTVAANVGTNMFKSALDHYVGGGETTITNAP WP_087351888.1 hypothetical protein [Olsenella sp. An188] Length: 982\nNumber of predicted TMHs: 2\nExp number of AAs in TMHs: 41.7690799999999\nExp number, first 60 AAs: 21.73505\nTotal prob of N-in: 0.99645\nPOSSIBLE N-term signal sequence\ninside 1 15\nTMhelix 16 38\noutside 39 939\nTMhelix 940 962\ninside 963 982

34288 GCF_002160255.1_ASM216025v1 Olsenella sp. An188Terrabacteria group; Actinobacteria; Coriobacteriia; Coriobacteriales; Atopobiaceae; Olsenella
MSRTVTDAAQAASELSERFSRARRAVFFGGAGVSTASGIPDFRSADGLYHQHFRYPPETM WP_087353490.1
NAD-dependent protein deacylase [Olsenella sp. An188] Length: 245\nNumber of predicted TMHs: 1\nExp number of AAs in TMHs: 18.16924\nExp number, first 60 AAs: 0.30951\nTotal prob of N-in: 0.80026\ninside 1 191\nTMhelix 192 214\noutside 215 245

34289 GCF_002159735.1_ASM215973v1 Olsenella sp. An270Terrabacteria group; Actinobacteria; Coriobacteriia; Coriobacteriales; Atopobiaceae; Olsenella
MSRRRTFPPALLALLVTLAVALVAAPAARAGTEGVTKLADPDTSASWQEVFRSDAGALST WP_087227073.1
hypothetical protein [Olsenella sp. An270] Length: 1019\nNumber of predicted TMHs: 2\nExp number of AAs in TMHs: 42.65049\nExp number, first 60 AAs: 21.13004\nTotal prob of N-in: 0.98406\nPOSSIBLE N-term signal sequence\ninside 1 6\nTMhelix 7 29\noutside 30 992\nTMhelix 993 1015\ninside 1016 1019

34290 GCF_900119625.1_IMG-taxon_2606217754_annotated_assembly Olsenella sp. kh2p3 Terrabacteria group; Actinobacteria; Coriobacteriia; Coriobacteriales; Atopobiaceae; Olsenella
MAKSTTRRKFIGSTIASVGLLGVTAAANVAANKFSSLLDHYVGGRPATVEKVQGSEDWDA WP_072354479.1
hypothetical protein [Olsenella sp. kh2p3] Length: 987\nNumber of predicted TMHs: 2\nExp number of AAs in TMHs: 38.2375599999999\nExp number, first 60 AAs: 16.73644\nTotal prob of N-in: 0.79599\nPOSSIBLE N-term signal sequence\ninside 1 8\nTMhelix 9 31\noutside 32 929\nTMhelix 930 952\ninside 953 987

34291 GCF_900128445.1_PRJEB18031 Olsenella sp. Marseille-P2912Terrabacteria group; Actinobacteria; Coriobacteriia; Coriobacteriales; Atopobiaceae; Olsenella
MSNEARPKTLVTRRGLFGLGALGASVAVGAGGVMEQFASTLDTNLGTQSEKLTSESTENE WP_083662161.1 beta-glucosidase [Olsenella sp. Marseille-P2912] Length: 1081\nNumber of predicted TMHs: 1\nExp number of AAs in TMHs: 39.10533\nExp number, first 60 AAs: 16.75668\nTotal prob of N-in: 0.77868\nPOSSIBLE N-term signal sequence\noutside 1 1051\nTMhelix 1052 1074\ninside 1075 1081

34292 GCF_900128445.1_PRJEB18031 Olsenella sp. Marseille-P2912Terrabacteria group; Actinobacteria; Coriobacteriia; Coriobacteriales; Atopobiaceae; Olsenella
MHRTNRRAHVISALVAICATLAFSLAPLNAQALSTDSFYTGTKSGGGSRVNAEPVTAETGWP_075279198.1 hypothetical protein [Olsenella sp. Marseille-P2912] Length: 670\nNumber of predicted TMHs: 2\nExp number of AAs in TMHs: 42.2108199999999\nExp number, first 60 AAs: 21.10187\nTotal prob of N-in: 0.98056\nPOSSIBLE N-term signal sequence\ninside 1 12\nTMhelix 13 35\noutside 36 642\nTMhelix 643 665\ninside 666 670

34293 GCF_900128445.1_PRJEB18031 Olsenella sp. Marseille-P2912Terrabacteria group; Actinobacteria; Coriobacteriia; Coriobacteriales; Atopobiaceae; Olsenella
MADAKKPRKKMTRRGFIASGATAAGLVGVTVAANVGTNMFKSAIDHYIGGGETTITNAPG WP_075279362.1
hypothetical protein [Olsenella sp. Marseille-P2912] Length: 979\nNumber of predicted TMHs: 2\nExp number of AAs in TMHs: 38.48892\nExp number, first 60 AAs: 21.38514\nTotal prob of N-in: 0.98663\nPOSSIBLE N-term signal sequence\ninside 1 12\nTMhelix 13 35\noutside 36 939\nTMhelix 940 962\ninside 963 979

34294 GCF_900155635.1_Olsenella_urinainfantis Olsenella sp. Marseille-P3197Terrabacteria group; Actinobacteria; Coriobacteriia; Coriobacteriales; Atopobiaceae; Olsenella
MKDITRRRFLVGGTAVGGAAIVGANVAANKFSSLLDHYLGGRPYTVETVAGSEDWDSEYY WP_077598534.1
hypothetical protein [Olsenella sp. Marseille-P3197] Length: 986\nNumber of predicted TMHs: 2\nExp number of AAs in TMHs: 36.18712\nExp number, first 60 AAs: 17.79808\nTotal prob of N-in: 0.87002\nPOSSIBLE N-term signal sequence\ninside 1 8\nTMhelix 9 28\noutside 29 932\nTMhelix 933 955\ninside 956 986

34295 GCF_000333815.1_ASM33381v1 Collinsella massiliensis Terrabacteria group; Actinobacteria; Coriobacteriia; Coriobacteriales; Coriobacteriaceae; Collinsella
MSEVEPTPHAPLALNRRSFLKLVIAASGTAAAASALPATASAADAPTSYPQKFTFAIMSD WP_019239921.1 MULTISPECIES: hypothetical protein [Coriobacteriaceae] Length: 906\nNumber of predicted TMHs: 1\nExp number of AAs in TMHs: 22.17985\nExp number, first 60 AAs: 1.99651\nTotal prob of N-in: 0.08978\noutside 1 872\nTMhelix 873 895\ninside 896 906

34296 GCF_000333815.1_ASM33381v1 Collinsella massiliensis Terrabacteria group; Actinobacteria; Coriobacteriia; Coriobacteriales; Coriobacteriaceae; Collinsella
MDLFIETPRQVSERRERELRAQGDRRDALQGVPDVASASSGCGAASISAGGRVGGEACAC WP_019239402.1
MULTISPECIES: hypothetical protein [Coriobacteriaceae] Length: 170\nNumber of predicted TMHs: 1\nExp

number of AAs in TMHs: 22.82928\nExp number, first 60 AAs: 0.0111\nTotal prob of N-in: 0.97202\ninside 1 112\nTMhelix 113 135\noutside 136 170

34297 GCF_000333815.1_ASM33381v1 Collinsella massiliensis Terrabacteria group; Actinobacteria; Coriobacteriia; Coriobacteriales; Coriobacteriaceae; Collinsella MRPRRAIITAGLAFALACAPVVASTAFADSDDPAVAYDKADFYEESAASAATRSASSARS WP_019238747.1 MULTISPECIES: hypothetical protein [Coriobacteriaceae] Length: 538\nNumber of predicted TMHs: 2\nExp number of AAs in TMHs: 40.93725\nExp number, first 60 AAs: 19.46599\nTotal prob of N-in: 0.89735\nPOSSIBLE N-term signal sequence\ninside 1 6\nTMhelix 7 29\noutside 30 499\nTMhelix 500 522\ninside 523 538

34298 GCF_900119895.1_PRJEB18019 Collinsella phocaeensis Terrabacteria group; Actinobacteria; Coriobacteriia; Coriobacteriales; Coriobacteriaceae; Collinsella MGRAVAVEQRCISRRGALAGTALLAGAAIAGFPATAAEGLPDEPYTAWPGYASTAPVSVV WP_072414687.1 hypothetical protein [Collinsella phocaeensis] Length: 529\nNumber of predicted TMHs: 1\nExp number of AAs in TMHs: 38.7520000000001\nExp number, first 60 AAs: 16.31125\nTotal prob of N-in: 0.74959\nPOSSIBLE N-term signal sequence\noutside 1 470\nTMhelix 471 493\ninside 494 529

34299 GCF_000321165.1_ASM32116v1 Enorma timonensis Terrabacteria group; Actinobacteria; Coriobacteriia; Coriobacteriales; Coriobacteriaceae; Enorma MDLFIETPRQVSERRERELRAQGDRRDALQGVPDVASASSGCGAASISAGGRVGGEACAC WP_019239402.1 MULTISPECIES: hypothetical protein [Coriobacteriaceae] Length: 170\nNumber of predicted TMHs: 1\nExp number of AAs in TMHs: 22.82928\nExp number, first 60 AAs: 0.0111\nTotal prob of N-in: 0.97202\ninside 1 112\nTMhelix 113 135\noutside 136 170

34300 GCF_000321165.1_ASM32116v1 Enorma timonensis Terrabacteria group; Actinobacteria; Coriobacteriia; Coriobacteriales; Coriobacteriaceae; Enorma MRPRRAIITAGLAFALACAPVVASTAFADSDDPAVAYDKADFYEESAASAATRSASSARS WP_019238747.1 MULTISPECIES: hypothetical protein [Coriobacteriaceae] Length: 538\nNumber of predicted TMHs: 2\nExp number of AAs in TMHs: 40.93725\nExp number, first 60 AAs: 19.46599\nTotal prob of N-in: 0.89735\nPOSSIBLE N-term signal sequence\ninside 1 6\nTMhelix 7 29\noutside 30 499\nTMhelix 500 522\ninside 523 538

34301 GCF_000321165.1_ASM32116v1 Enorma timonensis Terrabacteria group; Actinobacteria; Coriobacteriia; Coriobacteriales; Coriobacteriaceae; Enorma MSEVEPTPHAPLALNRRSFLKLVIAASGTAAAASALPATASAADAPTSYPQKFTFAIMSD WP_019239921.1 MULTISPECIES: hypothetical protein [Coriobacteriaceae] Length: 906\nNumber of predicted TMHs: 1\nExp number of AAs in TMHs: 22.17985\nExp number, first 60 AAs: 1.99651\nTotal prob of N-in: 0.08978\noutside 1 872\nTMhelix 873 895\ninside 896 906

34302 GCF_000191845.1_ASM19184v1 Eggerthella sp. HGA1 Terrabacteria group; Actinobacteria; Coriobacteriia; Eggerthellales; Eggerthellaceae; Eggerthella MGQVLKREHGGRRAAFAALLTTGMLAALLALTGIAAPAPAYAADQLKGTGTADDPVLIYT WP_009608731.1 fibronectin type III domain protein [Eggerthella sp. HGA1] Length: 3833\nNumber of predicted TMHs: 2\nExp number of AAs in TMHs: 45.2765399999999\nExp number, first 60 AAs: 22.60569\nTotal prob of N-in: 0.99010\nPOSSIBLE N-term signal sequence\ninside 1 12\nTMhelix 13 35\noutside 36 3803\nTMhelix 3804 3826\ninside 3827 3833

34303 GCF_000270285.1_ASM27028v1 Eggerthella sp. YY7918 Terrabacteria group; Actinobacteria; Coriobacteriia; Eggerthellales; Eggerthellaceae; Eggerthella MFEREKSMQRENINTQNNPAPRGGIDRRSFLKGTAALGAVAALGTLAGCASKSADETAPT WP_083833034.1 hypothetical protein [Eggerthella sp. YY7918] Length: 592\nNumber of predicted TMHs: 1\nExp number of AAs in TMHs: 19.44139\nExp number, first 60 AAs: 0.91333\nTotal prob of N-in: 0.08330\noutside 1 563\nTMhelix 564 586\ninside 587 592

34304 GCF_000270285.1_ASM27028v1 Eggerthella sp. YY7918 Terrabacteria group; Actinobacteria; Coriobacteriia; Eggerthellales; Eggerthellaceae; Eggerthella MESISRRSFVSGAAATGILATFGGLAGCAPQAVDSKANSNSEGILDANSLEQKWSFEVPP WP_013980000.1 3-oxosteroid 1-dehydrogenase [Eggerthella sp. YY7918] Length: 582\nNumber of predicted TMHs: 1\nExp number of AAs in TMHs: 26.30906\nExp number, first 60 AAs: 6.63779\nTotal prob of N-in: 0.31312\noutside 1 553\nTMhelix 554 576\ninside 577 582

34305 GCF_000763035.1_Egge_lent_1_1_60AFAA_V1 Eggerthella lenta 1_1_60AFAA Terrabacteria group; Actinobacteria; Coriobacteriia; Eggerthellales; Eggerthellaceae; Eggerthella; Eggerthella lenta MGQVLKREHGGRRAAFAALLTTGMLAALLALTGIAAPAPAYAADQLKGTGTADDPVLIYT WP_035586485.1 hypothetical protein [Eggerthella lenta] Length: 3834\nNumber of predicted TMHs: 2\nExp number of AAs in TMHs: 45.20712\nExp number, first 60 AAs: 22.60558\nTotal prob of N-in: 0.99081\nPOSSIBLE N-term signal sequence\ninside 1 12\nTMhelix 13 35\noutside 36 3804\nTMhelix 3805 3827\ninside 3828 3834

34306 GCF_000763035.1_Egge_lent_1_1_60AFAA_V1 Eggerthella lenta 1_1_60AFAA Terrabacteria group; Actinobacteria; Coriobacteriia; Eggerthellales; Eggerthellaceae; Eggerthella; Eggerthella lenta MMTRRRSRRGALALAAAALAVAAALAMPMFVRPAAAYAAPAEPPTVVYDSAAREFSVRNA WP_052106175.1 hypothetical protein [Eggerthella lenta] Length: 254\nNumber of predicted TMHs: 2\nExp number of AAs in TMHs: 42.98868\nExp number, first 60 AAs: 20.72679\nTotal prob of N-in: 0.99910\nPOSSIBLE N-term signal sequence\ninside 1 11\nTMhelix 12 31\noutside 32 224\nTMhelix 225 247\ninside 248 254

34307 GCF_002159895.1_ASM215989v1 Gordonibacter urolithinfaciens Terrabacteria group; Actinobacteria; Coriobacteriia; Eggerthellales; Eggerthellaceae; Gordonibacter MPAETLVPPRFRAPEPQPRRRALTLAIVVCALATVLFALAPGQADAAPAGPFAVEGPADA WP_087192245.1 hypothetical protein [Gordonibacter urolithinfaciens] Length: 410\nNumber of predicted TMHs: 2\nExp number of AAs in TMHs: 54.1608000000001\nExp number, first 60 AAs: 20.83232\nTotal prob of N-in: 0.93540\nPOSSIBLE N-term signal sequence\ninside 1 20\nTMhelix 21 43\noutside 44 382\nTMhelix 383 405\ninside 406 410

34308 GCF_001029505.1_ASM102950v1 Rubrobacter aplysinae Terrabacteria group; Actinobacteria; Rubrobacteria; Rubrobacterales; Rubrobacteraceae; Rubrobacter MFGMPDTTTRTPRFARRSNLLLVLAAISVTFMLFYPHIVSAQEEADELDDSDANAQSCPG WP_047866510.1 hypothetical protein [Rubrobacter aplysinae] Length: 213\nNumber of predicted TMHs: 2\nExp number of AAs in TMHs: 41.99417\nExp number, first 60 AAs: 20.71388\nTotal prob of N-in: 0.99929\nPOSSIBLE N-term signal sequence\ninside 1 19\nTMhelix 20 39\noutside 40 186\nTMhelix 187 209\ninside 210 213

34309 GCF_000420025.1_ASM42002v1 Patulibacter americanus DSM 16676 Terrabacteria group; Actinobacteria; Thermoleophilia; Solirubrobacterales; Patulibacteraceae; Patulibacter; Patulibacter americanus MTRRALIAVALTAVAGGATFALAQDPAPEAPQTWRTRPEVPGVATTLVPFGVEATADGAR WP_022929851.1 hypothetical protein [Patulibacter americanus] Length: 1668\nNumber of predicted TMHs: 1\nExp number of AAs in TMHs: 36.45228\nExp number, first 60 AAs: 15.99847\nTotal prob of N-in: 0.80724\nPOSSIBLE N-term signal sequence\noutside 1 1630\nTMhelix 1631 1653\ninside 1654 1668

34310 GCF_000519325.1_ASM51932v1 Patulibacter minatonensis DSM 18081 Terrabacteria group; Actinobacteria; Thermoleophilia; Solirubrobacterales; Patulibacteraceae; Patulibacter; Patulibacter minatonensis MSRRAALAAAIAVVVPLAGAGTAGAHVQLQPSTVAPDDPVEFTVLIPSEREAHTTKVDLK WP_026912667.1 hypothetical protein [Patulibacter minatonensis] Length: 223\nNumber of predicted TMHs: 2\nExp number of AAs in TMHs: 42.03005\nExp number, first 60 AAs: 22.14916\nTotal prob of N-in: 0.98324\nPOSSIBLE N-term signal sequence\ninside 1 6\nTMhelix 7 29\noutside 30 197\nTMhelix 198 217\ninside 218 223

34311 GCF_000519325.1_ASM51932v1 Patulibacter minatonensis DSM 18081 Terrabacteria group; Actinobacteria; Thermoleophilia; Solirubrobacterales; Patulibacteraceae; Patulibacter; Patulibacter minatonensis MTSRRSTTGTGPDRTGRPAGPRRRALATVLTAALFAATAGPAAAQSTTAPTTQLPGSGAT WP_026910841.1 hypothetical protein [Patulibacter minatonensis] Length: 303\nNumber of predicted TMHs: 1\nExp number of AAs in TMHs: 35.70474\nExp number, first 60 AAs: 13.22617\nTotal prob of N-in: 0.77779\nPOSSIBLE N-term signal sequence\noutside 1 254\nTMhelix 255 277\ninside 278 303

34312 GCF_000425945.1_ASM42594v1 Solirubrobacter sp. URHD0082 Terrabacteria group; Actinobacteria; Thermoleophilia; Solirubrobacterales; Solirubrobacteraceae; Solirubrobacter MTRRTLMRPRTLAAALATAALALPATAQAHVTLQPSSVPAGGDDTLLDVRVPNERSDAST WP_081685795.1 hypothetical protein [Solirubrobacter sp. URHD0082] Length: 232\nNumber of predicted TMHs: 2\nExp number of AAs in TMHs: 37.06208\nExp number, first 60 AAs: 14.7597\nTotal prob of N-in: 0.84482\nPOSSIBLE N-term signal sequence\ninside 1 11\nTMhelix 12 34\noutside 35 201\nTMhelix 202 224\ninside 225 232

34313 GCF_000423665.1_ASM42366v1 Solirubrobacter soli DSM 22325 Terrabacteria group; Actinobacteria; Thermoleophilia; Solirubrobacterales; Solirubrobacteraceae; Solirubrobacter; Solirubrobacter soli MLAADPMTRRRRGWVRVLIGLAMVGFGALGVVVGIAGAVNARSDIEDDAVARAPLGQPTS WP_028065148.1 hypothetical protein [Solirubrobacter soli] Length: 175\nNumber of predicted TMHs: 2\nExp number of

AAs in TMHs: 45.93275\nExp number, first 60 AAs: 22.99823\nTotal prob of N-in: 0.98811\nPOSSIBLE N-term signal sequence\ninside 1 12\nTMhelix 13 35\noutside 36 146\nTMhelix 147 169\ninside 170 175

34314 GCF_001458035.1_TKA4.10 Chthonomonas calidirosea Terrabacteria group; Armatimonadetes; Chthonomonadetes; Chthonomonadales; Chthonomonadaceae; Chthonomonas MSPPVGGNPPQPSARSRGIARRQFLTSALASLGAAVFAGASLESAQSAQALSTGSPPLPP WP_016484140.1 Rhs family protein [Chthonomonas calidirosea] Length: 1076\nNumber of predicted TMHs: 2\nExp number of AAs in TMHs: 45.49587\nExp number, first 60 AAs: 0.61125\nTotal prob of N-in: 0.03049\noutside 1 1006\nTMhelix 1007 1029\ninside 1030 1035\nTMhelix 1036 1058\noutside 1059 1076

34315 GCF_000427095.1_T49 Chthonomonas calidirosea T49 Terrabacteria group; Armatimonadetes; Chthonomonadetes; Chthonomonadales; Chthonomonadaceae; Chthonomonas; Chthonomonas calidirosea MSPPVGGNPPQPSARSRGIARRQFLTSALASLGAAVFAGASLESAQSAQALSTGSPPLPP WP_016484140.1 Rhs family protein [Chthonomonas calidirosea] Length: 1076\nNumber of predicted TMHs: 2\nExp number of AAs in TMHs: 45.49587\nExp number, first 60 AAs: 0.61125\nTotal prob of N-in: 0.03049\noutside 1 1006\nTMhelix 1007 1029\ninside 1030 1035\nTMhelix 1036 1058\noutside 1059 1076

34316 GCF_001536245.1_IMG-taxon_2616645014_annotated_assembly Armatimonadetes bacterium DC Terrabacteria group; Armatimonadetes; unclassified Armatimonadetes MELSGLVKPLFISCALEELHTLIERRNHIMKNRWMTWLTGCVCALSLTSVARVEAQLTYGWP_082704683.1 hypothetical protein [Armatimonadetes bacterium DC] Length: 327\nNumber of predicted TMHs: 1\nExp number of AAs in TMHs: 22.27468\nExp number, first 60 AAs: 1.3326\nTotal prob of N-in: 0.04371\noutside 1 299\nTMhelix 300 322\ninside 323 327

34317 GCF_001306095.1_ASM130609v1 Leptolinea tardivitalis Terrabacteria group; Chloroflexi; Anaerolineae; Anaerolineales; Anaerolineaceae; Leptolinea MNKFVRILLVILAALIVAAGGFWSGMQFSRRQAVSLPGPAAITTDTANGITPGDRSSRNR WP_062423052.1 zinc ribbon domain-containing protein [Leptolinea tardivitalis] Length: 206\nNumber of predicted TMHs: 2\nExp number of AAs in TMHs: 44.38525\nExp number, first 60 AAs: 20.93556\nTotal prob of N-in: 0.94679\nPOSSIBLE N-term signal sequence\ninside 1 6\nTMhelix 7 26\noutside 27 144\nTMhelix 145 167\ninside 168 206

34318 GCF_001050275.1_ASM105027v1 Leptolinea tardivitalis Terrabacteria group; Chloroflexi; Anaerolineae; Anaerolineales; Anaerolineaceae; Leptolinea MNKFVRILLVILAALIVAAGGFWSGMQFSRRQAVSLPGPAAITTDTANGITPGDRSSRNR WP_062423052.1 zinc ribbon domain-containing protein [Leptolinea tardivitalis] Length: 206\nNumber of predicted TMHs: 2\nExp number of AAs in TMHs: 44.38525\nExp number, first 60 AAs: 20.93556\nTotal prob of N-in: 0.94679\nPOSSIBLE N-term signal sequence\ninside 1 6\nTMhelix 7 26\noutside 27 144\nTMhelix 145 167\ninside 168 206

34319 GCF_000281175.1_ASM28117v1 Caldilinea aerophila DSM 14535 = NBRC 104270 Terrabacteria group; Chloroflexi; Caldilineae; Caldilineales; Caldilineaceae; Caldilinea; Caldilinea aerophila MTGRRNFFLRIVNGIVAIALTFSAVPAAAAPTLQRAEILRDCSVVSEESLQEELNAVVQG WP_014433859.1 hypothetical protein [Caldilinea aerophila] Length: 623\nNumber of predicted TMHs: 2\nExp number of AAs in TMHs: 53.98834\nExp number, first 60 AAs: 22.53257\nTotal prob of N-in: 0.98840\nPOSSIBLE N-term signal sequence\ninside 1 6\nTMhelix 7 29\noutside 30 589\nTMhelix 590 612\ninside 613 623

34320 GCF_000022185.1_ASM2218v1 Chloroflexus sp. Y-400-fl Terrabacteria group; Chloroflexi; Chloroflexia; Chloroflexales; Chloroflexineae; Chloroflexaceae; Chloroflexus MISLERRQIVSSPHIISNQPAQSRRPRVVIVGAGFGGLAAVRTLAQAPVDVLLINRTNYH WP_012259196.1 MULTISPECIES: NADH dehydrogenase family protein [Chloroflexus] Length: 455\nNumber of predicted TMHs: 1\nExp number of AAs in TMHs: 31.19511\nExp number, first 60 AAs: 6.45419\nTotal prob of N-in: 0.37882\noutside 1 391\nTMhelix 392 409\ninside 410 455

34321 GCF_000018865.1_ASM1886v1 Chloroflexus aurantiacus J-10-fl Terrabacteria group; Chloroflexi; Chloroflexia; Chloroflexales; Chloroflexineae; Chloroflexaceae; Chloroflexus; Chloroflexus aurantiacus MISLERRQIVSSPHIISNQPAQSRRPRVVIVGAGFGGLAAVRTLAQAPVDVLLINRTNYH YP_001636932.1 FAD-dependent pyridine nucleotide-disulfide oxidoreductase [Chloroflexus aurantiacus J-10-fl] Length: 455\nNumber of predicted TMHs: 1\nExp number of AAs in TMHs: 31.19511\nExp number, first 60 AAs: 6.45419\nTotal prob of N-in: 0.37882\noutside 1 391\nTMhelix 392 409\ninside 410 455

34322 GCF_002007905.1_ASM200790v1 Dehalococcoides mccartyi Terrabacteria group; Chloroflexi; Dehalococcoidia; Dehalococcoidales; Dehalococcoidaceae; Dehalococcoides

MFNTKLTRRDFVQLAAGSTAALSLGALKLPEFEKMFAEALKEIPVIWLQGAGCNGCTIST WP_058292395.1 oxidoreductase [Dehalococcoides mccartyi] Length: 354\nNumber of predicted TMHs: 1\nExp number of AAs in TMHs: 24.35363\nExp number, first 60 AAs: 0.46096\nTotal prob of N-in: 0.29153\noutside 1 323\nTMhelix 324 343\ninside 344 354

34323 GCF_002024205.1_ASM202420v1 Dehalococcoides mccartyi Terrabacteria group; Chloroflexi; Dehalococcoidia; Dehalococcoidales; Dehalococcoidaceae; Dehalococcoides MFNTKLTRRDFVQLAAGSTAALSLGALKLPEFEKMFAEALKEIPVIWLQGAGCNGCTIST WP_011308746.1 oxidoreductase [Dehalococcoides mccartyi] Length: 354\nNumber of predicted TMHs: 1\nExp number of AAs in TMHs: 20.9917\nExp number, first 60 AAs: 0.48583\nTotal prob of N-in: 0.16076\noutside 1 323\nTMhelix 324 343\ninside 344 354

34324 GCF_002028285.1_ASM202828v1 Dehalococcoides mccartyi Terrabacteria group; Chloroflexi; Dehalococcoidia; Dehalococcoidales; Dehalococcoidaceae; Dehalococcoides MFNTKLTRRDFVQLAAGSTAALSLGALKLPEFEKMFAEALKEIPVIWLQGAGCNGCTIST WP_011308746.1 oxidoreductase [Dehalococcoides mccartyi] Length: 354\nNumber of predicted TMHs: 1\nExp number of AAs in TMHs: 20.9917\nExp number, first 60 AAs: 0.48583\nTotal prob of N-in: 0.16076\noutside 1 323\nTMhelix 324 343\ninside 344 354

34325 GCF_002007845.1_ASM200784v1 Dehalococcoides mccartyi Terrabacteria group; Chloroflexi; Dehalococcoidia; Dehalococcoidales; Dehalococcoidaceae; Dehalococcoides MFNTKLTRRDFVQLAAGSTAALSLGALKLPEFEKMFAEALKEIPVIWLQGAGCNGCTIST WP_011308746.1 oxidoreductase [Dehalococcoides mccartyi] Length: 354\nNumber of predicted TMHs: 1\nExp number of AAs in TMHs: 20.9917\nExp number, first 60 AAs: 0.48583\nTotal prob of N-in: 0.16076\noutside 1 323\nTMhelix 324 343\ninside 344 354

34326 GCF_000741845.1_ASM74184v1 Dehalococcoides mccartyi Terrabacteria group; Chloroflexi; Dehalococcoidia; Dehalococcoidales; Dehalococcoidaceae; Dehalococcoides MFNTKLTRRDFVQLAAGSTAALSLGALKLPEFEKMFAEALKEIPVIWLQGAGCNGCTIST WP_012881470.1 MULTISPECIES: oxidoreductase [Dehalococcoides] Length: 354\nNumber of predicted TMHs: 1\nExp number of AAs in TMHs: 20.49086\nExp number, first 60 AAs: 0.55141\nTotal prob of N-in: 0.13926\noutside 1 323\nTMhelix 324 343\ninside 344 354

34327 GCF_001889305.1_ASM188930v1 Dehalococcoides mccartyi Terrabacteria group; Chloroflexi; Dehalococcoidia; Dehalococcoidales; Dehalococcoidaceae; Dehalococcoides MFNTKLTRRDFVQLAAGSTAALSLGALKLPEFEKMFAEALKEIPVIWLQGAGCNGCTIST WP_012881470.1 MULTISPECIES: oxidoreductase [Dehalococcoides] Length: 354\nNumber of predicted TMHs: 1\nExp number of AAs in TMHs: 20.49086\nExp number, first 60 AAs: 0.55141\nTotal prob of N-in: 0.13926\noutside 1 323\nTMhelix 324 343\ninside 344 354

34328 GCF_002021775.1_ASM202177v1 Dehalococcoides mccartyi Terrabacteria group; Chloroflexi; Dehalococcoidia; Dehalococcoidales; Dehalococcoidaceae; Dehalococcoides MFNTKLTRRDFVQLAAGSTAALSLGALKLPEFEKMFAEALKEIPVIWLQGAGCNGCTIST WP_011308746.1 oxidoreductase [Dehalococcoides mccartyi] Length: 354\nNumber of predicted TMHs: 1\nExp number of AAs in TMHs: 20.9917\nExp number, first 60 AAs: 0.48583\nTotal prob of N-in: 0.16076\noutside 1 323\nTMhelix 324 343\ninside 344 354

34329 GCF_002024225.1_ASM202422v1 Dehalococcoides mccartyi Terrabacteria group; Chloroflexi; Dehalococcoidia; Dehalococcoidales; Dehalococcoidaceae; Dehalococcoides MFNTKLTRRDFVQLAAGSTAALSLGALKLPEFEKMFAEALKEIPVIWLQGAGCNGCTIST WP_011308746.1 oxidoreductase [Dehalococcoides mccartyi] Length: 354\nNumber of predicted TMHs: 1\nExp number of AAs in TMHs: 20.9917\nExp number, first 60 AAs: 0.48583\nTotal prob of N-in: 0.16076\noutside 1 323\nTMhelix 324 343\ninside 344 354

34330 GCF_001459095.1_ASM145909v1 Dehalococcoides mccartyi Terrabacteria group; Chloroflexi; Dehalococcoidia; Dehalococcoidales; Dehalococcoidaceae; Dehalococcoides MFNTKLTRRDFVQLAAGSTAALSLGALKLPEFEKMFAEALKEIPVIWLQGAGCNGCTIST WP_058292395.1 oxidoreductase [Dehalococcoides mccartyi] Length: 354\nNumber of predicted TMHs: 1\nExp number of AAs in TMHs: 24.35363\nExp number, first 60 AAs: 0.46096\nTotal prob of N-in: 0.29153\noutside 1 323\nTMhelix 324 343\ninside 344 354

34331 GCF_001460955.1_ASM146095v1 Dehalococcoides mccartyi Terrabacteria group; Chloroflexi; Dehalococcoidia; Dehalococcoidales; Dehalococcoidaceae; Dehalococcoides
MFNTKLTRRDFVQLAAGSTAALSLGALKLPEFEKMFAEALKEIPVIWLQGAGCNGCTIST WP_011308746.1
oxidoreductase [Dehalococcoides mccartyi] Length: 354\nNumber of predicted TMHs: 1\nExp number of AAs in TMHs: 20.9917\nExp number, first 60 AAs: 0.48583\nTotal prob of N-in: 0.16076\noutside 1 323\nTMhelix 324 343\ninside 344 354

34332 GCF_002007825.1_ASM200782v1 Dehalococcoides mccartyi Terrabacteria group; Chloroflexi; Dehalococcoidia; Dehalococcoidales; Dehalococcoidaceae; Dehalococcoides
MFNTKLTRRDFVQLAAGSTAALSLGALKLPEFEKMFAEALKEIPVIWLQGAGCNGCTIST WP_058292395.1
oxidoreductase [Dehalococcoides mccartyi] Length: 354\nNumber of predicted TMHs: 1\nExp number of AAs in TMHs: 24.35363\nExp number, first 60 AAs: 0.46096\nTotal prob of N-in: 0.29153\noutside 1 323\nTMhelix 324 343\ninside 344 354

34333 GCF_002007865.1_ASM200786v1 Dehalococcoides mccartyi Terrabacteria group; Chloroflexi; Dehalococcoidia; Dehalococcoidales; Dehalococcoidaceae; Dehalococcoides
MFNTKLTRRDFVQLAAGSTAALSLGALKLPEFEKMFAEALKEIPVIWLQGAGCNGCTIST WP_011308746.1
oxidoreductase [Dehalococcoides mccartyi] Length: 354\nNumber of predicted TMHs: 1\nExp number of AAs in TMHs: 20.9917\nExp number, first 60 AAs: 0.48583\nTotal prob of N-in: 0.16076\noutside 1 323\nTMhelix 324 343\ninside 344 354

34334 GCF_001515295.1_ASM151529v1 Dehalococcoides mccartyi Terrabacteria group; Chloroflexi; Dehalococcoidia; Dehalococcoidales; Dehalococcoidaceae; Dehalococcoides
MFNTKLTRRDFVQLAAGSTAALSLGALKLPEFEKMFAEALKEIPVIWLQGAGCNGCTIST WP_011308746.1
oxidoreductase [Dehalococcoides mccartyi] Length: 354\nNumber of predicted TMHs: 1\nExp number of AAs in TMHs: 20.9917\nExp number, first 60 AAs: 0.48583\nTotal prob of N-in: 0.16076\noutside 1 323\nTMhelix 324 343\ninside 344 354

34335 GCF_001610775.1_ASM161077v1 Dehalococcoides mccartyi Terrabacteria group; Chloroflexi; Dehalococcoidia; Dehalococcoidales; Dehalococcoidaceae; Dehalococcoides
MFNTKLTRRDFVQLAAGSTAALSLGALKLPEFEKMFAEALKEIPVIWLQGAGCNGCTIST WP_011308746.1
oxidoreductase [Dehalococcoides mccartyi] Length: 354\nNumber of predicted TMHs: 1\nExp number of AAs in TMHs: 20.9917\nExp number, first 60 AAs: 0.48583\nTotal prob of N-in: 0.16076\noutside 1 323\nTMhelix 324 343\ninside 344 354

34336 GCF_001010485.1_ASM101048v1 Dehalococcoides sp. UCH007 Terrabacteria group; Chloroflexi; Dehalococcoidia; Dehalococcoidales; Dehalococcoidaceae; Dehalococcoides
MFNTKLTRRDFVQLAAGSTAALSLGALKLPEFEKMFAEALKEIPVIWLQGAGCNGCTIST WP_012881470.1 MULTISPECIES: oxidoreductase [Dehalococcoides] Length: 354\nNumber of predicted TMHs: 1\nExp number of AAs in TMHs: 20.49086\nExp number, first 60 AAs: 0.55141\nTotal prob of N-in: 0.13926\noutside 1 323\nTMhelix 324 343\ninside 344 354

34337 GCF_000011905.1_ASM1190v1 Dehalococcoides mccartyi 195 Terrabacteria group; Chloroflexi; Dehalococcoidia; Dehalococcoidales; Dehalococcoidaceae; Dehalococcoides; Dehalococcoides mccartyi
MFNTKLTRRDFVQLAAGSTAALSLGALKLPEFEKMFAEALKEIPVIWLQGAGCNGCTIST WP_010935917.1
oxidoreductase [Dehalococcoides mccartyi] Length: 354\nNumber of predicted TMHs: 1\nExp number of AAs in TMHs: 25.92917\nExp number, first 60 AAs: 0.51518\nTotal prob of N-in: 0.35146\noutside 1 323\nTMhelix 324 343\ninside 344 354

34338 GCF_000016705.1_ASM1670v1 Dehalococcoides mccartyi BAV1 Terrabacteria group; Chloroflexi; Dehalococcoidia; Dehalococcoidales; Dehalococcoidaceae; Dehalococcoides; Dehalococcoides mccartyi
MFNTKLTRRDFVQLAAGSTAALSLGALKLPEFEKMFAEALKEIPVIWLQGAGCNGCTIST WP_011308746.1
oxidoreductase [Dehalococcoides mccartyi] Length: 354\nNumber of predicted TMHs: 1\nExp number of AAs in TMHs: 20.9917\nExp number, first 60 AAs: 0.48583\nTotal prob of N-in: 0.16076\noutside 1 323\nTMhelix 324 343\ninside 344 354

34339 GCF_000341695.1_ASM34169v1 Dehalococcoides mccartyi BTF08 Terrabacteria group; Chloroflexi; Dehalococcoidia; Dehalococcoidales; Dehalococcoidaceae; Dehalococcoides; Dehalococcoides mccartyi
MFNTKLTRRDFVQLAAGSTAALSLGALKLPEFEKMFAEALKEIPVIWLQGAGCNGCTIST WP_011308746.1
oxidoreductase [Dehalococcoides mccartyi] Length: 354\nNumber of predicted TMHs: 1\nExp number of

AAs in TMHs: 20.9917\nExp number, first 60 AAs:  0.48583\nTotal prob of N-in:   0.16076\noutside   1   323\nTMhelix   324   343\ninside   344   354

34340 GCF_000009025.1_ASM902v1 Dehalococcoides mccartyi CBDB1 Terrabacteria group; Chloroflexi; Dehalococcoidia; Dehalococcoidales; Dehalococcoidaceae; Dehalococcoides; Dehalococcoides mccartyi MFNTKLTRRDFVQLAAGSTAALSLGALKLPEFEKMFAEALKEIPVIWLQGAGCNGCTIST WP_011308746.1 oxidoreductase [Dehalococcoides mccartyi] Length: 354\nNumber of predicted TMHs:  1\nExp number of AAs in TMHs: 20.9917\nExp number, first 60 AAs:  0.48583\nTotal prob of N-in:   0.16076\noutside   1   323\nTMhelix   324   343\ninside   344   354

34341 GCF_000830925.1_ASM83092v1 Dehalococcoides mccartyi CG1 Terrabacteria group; Chloroflexi; Dehalococcoidia; Dehalococcoidales; Dehalococcoidaceae; Dehalococcoides; Dehalococcoides mccartyi MFNTKLTRRDFVQLAAGSTAALSLGALKLPEFEKMFAEALKEIPVIWLQGAGCNGCTIST WP_012881470.1 MULTISPECIES: oxidoreductase [Dehalococcoides] Length: 354\nNumber of predicted TMHs:  1\nExp number of AAs in TMHs: 20.49086\nExp number, first 60 AAs:  0.55141\nTotal prob of N-in:   0.13926\noutside   1   323\nTMhelix   324   343\ninside   344   354

34342 GCF_000830905.1_ASM83090v1 Dehalococcoides mccartyi CG4 Terrabacteria group; Chloroflexi; Dehalococcoidia; Dehalococcoidales; Dehalococcoidaceae; Dehalococcoides; Dehalococcoides mccartyi MFNTKLTRRDFVQLAAGSTAALSLGALKLPEFEKMFAEALKEIPVIWLQGAGCNGCTIST WP_041340943.1 oxidoreductase [Dehalococcoides mccartyi] Length: 354\nNumber of predicted TMHs:  1\nExp number of AAs in TMHs: 25.50402\nExp number, first 60 AAs:  0.51172\nTotal prob of N-in:   0.33453\noutside   1   323\nTMhelix   324   343\ninside   344   354

34343 GCF_000830885.1_ASM83088v1 Dehalococcoides mccartyi CG5 Terrabacteria group; Chloroflexi; Dehalococcoidia; Dehalococcoidales; Dehalococcoidaceae; Dehalococcoides; Dehalococcoides mccartyi MFNTKLTRRDFVQLAAGSTAALSLGALKLPEFEKMFAEALKEIPVIWLQGAGCNGCTIST WP_011308746.1 oxidoreductase [Dehalococcoides mccartyi] Length: 354\nNumber of predicted TMHs:  1\nExp number of AAs in TMHs: 20.9917\nExp number, first 60 AAs:  0.48583\nTotal prob of N-in:   0.16076\noutside   1   323\nTMhelix   324   343\ninside   344   354

34344 GCF_000341655.1_ASM34165v1 Dehalococcoides mccartyi DCMB5 Terrabacteria group; Chloroflexi; Dehalococcoidia; Dehalococcoidales; Dehalococcoidaceae; Dehalococcoides; Dehalococcoides mccartyi MFNTKLTRRDFVQLAAGSTAALSLGALKLPEFEKMFAEALKEIPVIWLQGAGCNGCTIST WP_011308746.1 oxidoreductase [Dehalococcoides mccartyi] Length: 354\nNumber of predicted TMHs:  1\nExp number of AAs in TMHs: 20.9917\nExp number, first 60 AAs:  0.48583\nTotal prob of N-in:   0.16076\noutside   1   323\nTMhelix   324   343\ninside   344   354

34345 GCF_000025585.1_ASM2558v1 Dehalococcoides mccartyi GT Terrabacteria group; Chloroflexi; Dehalococcoidia; Dehalococcoidales; Dehalococcoidaceae; Dehalococcoides; Dehalococcoides mccartyi MFNTKLTRRDFVQLAAGSTAALSLGALKLPEFEKMFAEALKEIPVIWLQGAGCNGCTIST WP_011308746.1 oxidoreductase [Dehalococcoides mccartyi] Length: 354\nNumber of predicted TMHs:  1\nExp number of AAs in TMHs: 20.9917\nExp number, first 60 AAs:  0.48583\nTotal prob of N-in:   0.16076\noutside   1   323\nTMhelix   324   343\ninside   344   354

34346 GCF_000499365.1_ASM49936v1 Dehalococcoides mccartyi GY50 Terrabacteria group; Chloroflexi; Dehalococcoidia; Dehalococcoidales; Dehalococcoidaceae; Dehalococcoides; Dehalococcoides mccartyi MFNTKLTRRDFVQLAAGSTAALSLGALKLPEFEKMFAEALKEIPVIWLQGAGCNGCTIST WP_012881470.1 MULTISPECIES: oxidoreductase [Dehalococcoides] Length: 354\nNumber of predicted TMHs:  1\nExp number of AAs in TMHs: 20.49086\nExp number, first 60 AAs:  0.55141\nTotal prob of N-in:   0.13926\noutside   1   323\nTMhelix   324   343\ninside   344   354

34347 GCF_001547795.1_ASM154779v1 Dehalococcoides mccartyi IBARAKI Terrabacteria group; Chloroflexi; Dehalococcoidia; Dehalococcoidales; Dehalococcoidaceae; Dehalococcoides; Dehalococcoides mccartyi MFNTKLTRRDFVQLAAGSTAALSLGALKLPEFEKMFAEALKEIPVIWLQGAGCNGCTIST WP_011308746.1 oxidoreductase [Dehalococcoides mccartyi] Length: 354\nNumber of predicted TMHs:  1\nExp number of AAs in TMHs: 20.9917\nExp number, first 60 AAs:  0.48583\nTotal prob of N-in:   0.16076\noutside   1   323\nTMhelix   324   343\ninside   344   354

34348 GCF_000025025.1_ASM2502v1 Dehalococcoides mccartyi VS Terrabacteria group; Chloroflexi; Dehalococcoidia; Dehalococcoidales; Dehalococcoidaceae; Dehalococcoides; Dehalococcoides mccartyi

MFNTKLTRRDFVQLAAGSTAALSLGALKLPEFEKMFAEALKEIPVIWLQGAGCNGCTIST WP_012881470.1 MULTISPECIES: oxidoreductase [Dehalococcoides] Length: 354\nNumber of predicted TMHs: 1\nExp number of AAs in TMHs: 20.49086\nExp number, first 60 AAs: 0.55141\nTotal prob of N-in: 0.13926\noutside 1 323\nTMhelix 324 343\ninside 344 354

34349 GCF_000178855.1_ASM17885v1 Ktedonobacter racemifer DSM 44963 Terrabacteria group; Chloroflexi; Ktedonobacteria; Ktedonobacterales; Ktedonobacteraceae; Ktedonobacter; Ktedonobacter racemifer MVHSRRRLLYPTLLLTLTLMGLLLMGSSPRLAYADGGAPNLAYVAGASQGLSIIDIATQK WP_007911811.1 hypothetical protein [Ktedonobacter racemifer] Length: 390\nNumber of predicted TMHs: 2\nExp number of AAs in TMHs: 47.00837\nExp number, first 60 AAs: 21.79767\nTotal prob of N-in: 0.95351\nPOSSIBLE N-term signal sequence\ninside 1 6\nTMhelix 7 29\noutside 30 355\nTMhelix 356 378\ninside 379 390

34350 GCF_900187885.1_IMG-taxon_2140918011_annotated_assembly Thermoflexus hugenholtzii JAD2 Terrabacteria group; Chloroflexi; Thermoflexia; Thermoflexales; Thermoflexaceae; Thermoflexus; Thermoflexus hugenholtzii MGRRGLLWAVAILTFAADQGSKAWVRENLALYESWAPIPALARYFTFTHVTNTGIAFGLF WP_088571893.1 signal peptidase II [Thermoflexus hugenholtzii] Length: 167\nNumber of predicted TMHs: 2\nExp number of AAs in TMHs: 60.17038\nExp number, first 60 AAs: 5.99476\nTotal prob of N-in: 0.43388\ninside 1 58\nTMhelix 59 81\noutside 82 123\nTMhelix 124 146\ninside 147 167

34351 GCF_001870225.1_ASM187022v1 Gloeomargarita lithophora Alchichica-D10 Terrabacteria group; Cyanobacteria/Melainabacteria group; Cyanobacteria; Gloeoemargaritales; Gloeomargaritaceae; Gloeomargarita; Gloeomargarita lithophora MAAEQVWEQPEFYDPTPQDFNAYGGGRRRKAVLVLSLLWGGTLALHWVSWGTWVVWALTT WP_084111478.1 hypothetical protein [Gloeomargarita lithophora]Length: 443\nNumber of predicted TMHs: 2\nExp number of AAs in TMHs: 45.8449\nExp number, first 60 AAs: 22.51878\nTotal prob of N-in: 0.87567\nPOSSIBLE N-term signal sequence\ninside 1 31\nTMhelix 32 54\noutside 55 408\nTMhelix 409 426\ninside 427 443

34352 GCF_000346485.2_ASM34648v2 Scytonema hofmannii PCC 7110 Terrabacteria group; Cyanobacteria/Melainabacteria group; Cyanobacteria; Nostocales; Scytonemataceae; Scytonema; Scytonema hofmannii MSRYALEPNTINELVQLLQPFLEDERDRRPFLVLALGHSLPVLERLIWSGSVATFIPDMI WP_017741353.1 hypothetical protein [Scytonema hofmannii] Length: 468\nNumber of predicted TMHs: 1\nExp number of AAs in TMHs: 19.96811\nExp number, first 60 AAs: 0.13249\nTotal prob of N-in: 0.04627\noutside 1 430\nTMhelix 431 450\ninside 451 468

34353 GCF_000817735.2_ASM81773v2 Scytonema millei VB511283 Terrabacteria group; Cyanobacteria/Melainabacteria group; Cyanobacteria; Nostocales; Scytonemataceae; Scytonema; Scytonema millei MSPTQTNTTVKLLSPTLHLYHYVLRNSINERPESLNQRRAFFSKNLHQLTSHLTSSSGKN WP_069349982.1 hypothetical protein [Scytonema millei] Length: 481\nNumber of predicted TMHs: 2\nExp number of AAs in TMHs: 45.18616\nExp number, first 60 AAs: 0.01433\nTotal prob of N-in: 0.82753\ninside 1 404\nTMhelix 405 427\noutside 428 446\nTMhelix 447 469\ninside 470 481

34354 GCF_000828075.2_ASM82807v2 Tolypothrix campylonemoides VB511288 Terrabacteria group; Cyanobacteria/Melainabacteria group; Cyanobacteria; Nostocales; Tolypothrichaceae; Tolypothrix; Tolypothrix campylonemoides MSRTAPLPGRARRGAFVALAIALAVAFAGFVALGAWQLQRMQWKHDLIARVEARVGAPPV WP_071838795.1 hypothetical protein [Tolypothrix campylonemoides] Length: 255\nNumber of predicted TMHs: 2\nExp number of AAs in TMHs: 42.83017\nExp number, first 60 AAs: 22.46216\nTotal prob of N-in: 0.79093\nPOSSIBLE N-term signal sequence\ninside 1 12\nTMhelix 13 35\noutside 36 216\nTMhelix 217 236\ninside 237 255

34355 GCF_000312205.1_ASM31220v1 Microcystis aeruginosa PCC 7941 Terrabacteria group; Cyanobacteria/Melainabacteria group; Cyanobacteria; Oscillatoriophycideae; Chroococcales; Microcystaceae; Microcystis; Microcystis aeruginosa MSDELIQKTFDEVSSITRSFDAARRKLIDYQMQVMAGMNNISLMPDVITLAIDAVDSSPD WP_002778173.1 hypothetical protein [Microcystis aeruginosa] Length: 294\nNumber of predicted TMHs: 2\nExp number of AAs in TMHs: 47.91357\nExp number, first 60 AAs: 1.34309\nTotal prob of N-in: 0.90188\ninside 1 210\nTMhelix 211 230\noutside 231 239\nTMhelix 240 262\ninside 263 294

34356 GCF_000317045.1_ASM31704v1 Geitlerinema sp. PCC 7407 Terrabacteria group; Cyanobacteria/Melainabacteria group; Cyanobacteria; Oscillatoriophycideae; Oscillatoriales; Coleofasciculaceae; Geitlerinema MSVHRRPLIVSSWVLASLLAIGAPQRAIAHAGHGNEFQSAPAASSSAIAVDETTAERLGI WP_015172123.1 RND family efflux transporter MFP subunit [Geitlerinema sp. PCC 7407] Length: 523\nNumber of predicted TMHs: 1\nExp number of AAs in TMHs: 25.55953\nExp number, first 60 AAs: 7.84302\nTotal prob of N-in: 0.36697\noutside 1 475\nTMhelix 476 493\ninside 494 523

34357 GCF_000317495.1_ASM31749v1 Crinalium epipsammum PCC 9333 Terrabacteria group; Cyanobacteria/Melainabacteria group; Cyanobacteria; Oscillatoriophycideae; Oscillatoriales; Gomontiellaceae; Crinalium; Crinalium epipsammum MKYSRFNSRRNFIWMVGISSLGAALGGLSRKSLAQRPPIVNPISPNPRSFRLAPNTQVLL WP_015202080.1 hypothetical protein [Crinalium epipsammum] Length: 310\nNumber of predicted TMHs: 2\nExp number of AAs in TMHs: 39.87992\nExp number, first 60 AAs: 12.53794\nTotal prob of N-in: 0.86668\nPOSSIBLE N-term signal sequence\ninside 1 11\nTMhelix 12 29\noutside 30 256\nTMhelix 257 279\ninside 280 310

34358 GCF_000210375.1_ASM21037v1 Arthrospira platensis NIES-39 Terrabacteria group; Cyanobacteria/Melainabacteria group; Cyanobacteria; Oscillatoriophycideae; Oscillatoriales; Microcoleaceae; Arthrospira; Arthrospira platensis MTIDEEERRELMNGYQSNNPINQATAKDLFNTFGDLKTLGSNQNNELRLNENLYFVKVRP WP_006619571.1 HAMP domain-containing protein [Arthrospira platensis] Length: 153\nNumber of predicted TMHs: 1\nExp number of AAs in TMHs: 20.54881\nExp number, first 60 AAs: 1e-05\nTotal prob of N-in: 0.95634\ninside 1 94\nTMhelix 95 114\noutside 115 153

34359 GCF_000175415.3_ASM17541v3 Arthrospira platensis str. Paraca Terrabacteria group; Cyanobacteria/Melainabacteria group; Cyanobacteria; Oscillatoriophycideae; Oscillatoriales; Microcoleaceae; Arthrospira; Arthrospira platensis MTIDEEERRELMNGYQSNNPINQATAKDLFNTFGDLKTLGSNQNNELRLNENLYFVKVRP WP_006619571.1 HAMP domain-containing protein [Arthrospira platensis] Length: 153\nNumber of predicted TMHs: 1\nExp number of AAs in TMHs: 20.54881\nExp number, first 60 AAs: 1e-05\nTotal prob of N-in: 0.95634\ninside 1 94\nTMhelix 95 114\noutside 115 153

34360 GCF_001611905.1_ASM161190v1 Arthrospira platensis YZ Terrabacteria group; Cyanobacteria/Melainabacteria group; Cyanobacteria; Oscillatoriophycideae; Oscillatoriales; Microcoleaceae; Arthrospira; Arthrospira platensis MTIDEEERRELMNGYQSNNPINQATAKDLFNTFGDLKTLGSNQNNELRLNENLYFVKVRP WP_006619571.1 HAMP domain-containing protein [Arthrospira platensis] Length: 153\nNumber of predicted TMHs: 1\nExp number of AAs in TMHs: 20.54881\nExp number, first 60 AAs: 1e-05\nTotal prob of N-in: 0.95634\ninside 1 94\nTMhelix 95 114\noutside 115 153

34361 GCF_000180455.1_ASM18045v1 [Oscillatoria] sp. PCC 6506 Terrabacteria group; Cyanobacteria/Melainabacteria group; Cyanobacteria; Oscillatoriophycideae; Oscillatoriales; Microcoleaceae; Kamptonema MQQTQISTTVQVLSPTLHLYHYVLRNGINESFETLEKRRETFSQQLQQFTSQLTVTSGKK WP_007352828.1 MULTISPECIES: hypothetical protein [Kamptonema] Length: 476\nNumber of predicted TMHs: 1\nExp number of AAs in TMHs: 28.47874\nExp number, first 60 AAs: 0.03459\nTotal prob of N-in: 0.39413\noutside 1 441\nTMhelix 442 464\ninside 465 476

34362 GCF_000332155.1_ASM33215v1 Kamptonema formosum PCC 6407 Terrabacteria group; Cyanobacteria/Melainabacteria group; Cyanobacteria; Oscillatoriophycideae; Oscillatoriales; Microcoleaceae; Kamptonema; Kamptonema formosum MQQTQISTTVQVLSPTLHLYHYVLRNGINESFETLEKRRETFSQQLQQFTSQLTVTSGKK WP_007352828.1 MULTISPECIES: hypothetical protein [Kamptonema] Length: 476\nNumber of predicted TMHs: 1\nExp number of AAs in TMHs: 28.47874\nExp number, first 60 AAs: 0.03459\nTotal prob of N-in: 0.39413\noutside 1 441\nTMhelix 442 464\ninside 465 476

34363 GCF_000332335.1_ASM33233v1 Oscillatoria sp. PCC 10802 Terrabacteria group; Cyanobacteria/Melainabacteria group; Cyanobacteria; Oscillatoriophycideae; Oscillatoriales; Oscillatoriaceae; Oscillatoria MSRVLEIPQTQTSTAIEARRSWLTSGWLYLILGILLNGIIWGGAILAMKKQAPNYTAQWSLH WP_017719097.1 hypothetical protein [Oscillatoria sp. PCC 10802] Length: 471\nNumber of predicted TMHs: 2\nExp number of AAs in TMHs: 44.13668\nExp number, first 60 AAs: 22.65796\nTotal prob of N-in: 0.93554\nPOSSIBLE N-term signal sequence\ninside 1 20\nTMhelix 21 43\noutside 44 436\nTMhelix 437 459\ninside 460 471

34364 GCF_001939115.1_ASM193911v1 Oscillatoriales cyanobacterium hensonii Terrabacteria group; Cyanobacteria/Melainabacteria group; Cyanobacteria; Oscillatoriophycideae; Oscillatoriales; unclassified Oscillatoriales; unclassified Oscillatoriales (miscellaneous) MSVLPNDSLKLQRPAESGPYRNDRSVEKTYIDKTYRWNRERYSRRRRTFDIWSFVLQLLA WP_075596774.1 hypothetical protein [Oscillatoriales cyanobacterium hensonii] Length: 587\nNumber of predicted TMHs: 1\nExp number of AAs in TMHs: 43.99264\nExp number, first 60 AAs: 1.62205\nTotal prob of N-in: 0.30408\noutside 1 554\nTMhelix 555 577\ninside 578 587

34365 GCF_001482745.1_ASM148274v1 Oscillatoriales cyanobacterium MTP1 Terrabacteria group; Cyanobacteria/Melainabacteria group; Cyanobacteria; Oscillatoriophycideae; Oscillatoriales; unclassified Oscillatoriales; unclassified Oscillatoriales (miscellaneous) MTQSASRRKHIVVVGAGFGGVAFCQSFPEGLADITLVDRNNYHLFQPLLYQVATADLSPA WP_058867355.1

pyridine nucleotide-disulfide oxidoreductase [Chloracidobacterium thermophilum] Length: 429\nNumber of predicted TMHs: 1\nExp number of AAs in TMHs: 27.77766\nExp number, first 60 AAs: 5.33717\nTotal prob of N-in: 0.41544\noutside 1 376\nTMhelix 377 399\ninside 400 429

34366 GCF_000332055.1_ASM33205v1 Xenococcus sp. PCC 7305 Terrabacteria group; Cyanobacteria/Melainabacteria group; Cyanobacteria; Pleurocapsales; Xenococcaceae; Xenococcus MKQKNKFYSSKILKKIKRRRLFDYSYETPGSLPGTLQISQDAIFPPIDLIDYNSEQVRLH WP_006508745.1 magnesium and cobalt transport protein CorA [Xenococcus sp. PCC 7305] Length: 386\nNumber of predicted TMHs: 2\nExp number of AAs in TMHs: 45.50272\nExp number, first 60 AAs: 0.00164\nTotal prob of N-in: 0.99570\ninside 1 311\nTMhelix 312 334\noutside 335 348\nTMhelix 349 371\ninside 372 386

34367 GCF_000482245.1_LepHIscaffolds Leptolyngbya sp. Heron Island J Terrabacteria group; Cyanobacteria/Melainabacteria group; Cyanobacteria; Synechococcales; Leptolyngbyaceae; Leptolyngbya MNFIRNVIDLNIQYLWAATRRGIVPATSAFVLTVLGAGAVASMTVPTYLAEGKLLFKRDR WP_023070472.1 capsular exopolysaccharide biosynthesis protein [Leptolyngbya sp. Heron Island J] Length: 496\nNumber of predicted TMHs: 2\nExp number of AAs in TMHs: 44.25028\nExp number, first 60 AAs: 22.07535\nTotal prob of N-in: 0.93478\nPOSSIBLE N-term signal sequence\ninside 1 27\nTMhelix 28 50\noutside 51 453\nTMhelix 454 476\ninside 477 496

34368 GCF_001485215.1_ASM148521v1 Leptolyngbya sp. NIES-2104 Terrabacteria group; Cyanobacteria/Melainabacteria group; Cyanobacteria; Synechococcales; Leptolyngbyaceae; Leptolyngbya MLAVKLQCDSSFGSRHSKSIVIDGDRMSRFTTTNIHQQPPQCPASRRNVLIVGGGPAGLA WP_082690100.1 FAD-binding monooxygenase [Leptolyngbya sp. NIES-2104] Length: 528\nNumber of predicted TMHs: 1\nExp number of AAs in TMHs: 27.76976\nExp number, first 60 AAs: 0.81366\nTotal prob of N-in: 0.07951\noutside 1 503\nTMhelix 504 526\ninside 527 528

34369 GCF_000316115.1_ASM31611v1 Leptolyngbya sp. PCC 7375 Terrabacteria group; Cyanobacteria/Melainabacteria group; Cyanobacteria; Synechococcales; Leptolyngbyaceae; Leptolyngbya MNFIRNVIDLNLQYLFAATRRGIVPATTAFVLTVLGAAAVAKLQKPTFLSEGKLLFKPDR WP_006519775.1 exopolysaccharide biosynthesis protein [Leptolyngbya sp. PCC 7375] Length: 472\nNumber of predicted TMHs: 2\nExp number of AAs in TMHs: 41.1002199999999\nExp number, first 60 AAs: 18.39113\nTotal prob of N-in: 0.86565\nPOSSIBLE N-term signal sequence\ninside 1 20\nTMhelix 21 40\noutside 41 428\nTMhelix 429 451\ninside 452 472

34370 GCF_000309385.1_ASM30938v1 Nodosilinea nodulosa PCC 7104 Terrabacteria group; Cyanobacteria/Melainabacteria group; Cyanobacteria; Synechococcales; Leptolyngbyaceae; Nodosilinea; Nodosilinea nodulosa MTRSAFPLLRNRRAALLAFALGATLVGGCAEAPNLTLNQDAMTTDAEVAMGGAAAGAENQ WP_017297372.1 hypothetical protein [Nodosilinea nodulosa] Length: 321\nNumber of predicted TMHs: 1\nExp number of AAs in TMHs: 27.60554\nExp number, first 60 AAs: 5.94837\nTotal prob of N-in: 0.24561\noutside 1 280\nTMhelix 281 303\ninside 304 321

34371 GCF_000478825.2_ASM47882v2 Synechocystis sp. PCC 6714 Terrabacteria group; Cyanobacteria/Melainabacteria group; Cyanobacteria; Synechococcales; Merismopediaceae; Synechocystis MNAPSSPRRPHVVIVGGGFAGLYTAKHLRRSPVEVTLIDKRNFHLFQPLLYQVATGSLSP WP_028947702.1 pyridine nucleotide-disulfide oxidoreductase [Synechocystis sp. PCC 6714] Length: 445\nNumber of predicted TMHs: 1\nExp number of AAs in TMHs: 20.5934\nExp number, first 60 AAs: 0.18576\nTotal prob of N-in: 0.05214\noutside 1 374\nTMhelix 375 397\ninside 398 445

34372 GCF_001318385.1_ASM131838v1 Synechocystis sp. PCC 6803 Terrabacteria group; Cyanobacteria/Melainabacteria group; Cyanobacteria; Synechococcales; Merismopediaceae; Synechocystis MNSPTSPRRPHVVIVGGGFAGLYTAKNLRRSPVDITLIDKRNFHLFQPLLYQVATGSLSP WP_010872417.1 pyridine nucleotide-disulfide oxidoreductase [Synechocystis sp. PCC 6803] Length: 445\nNumber of predicted TMHs: 1\nExp number of AAs in TMHs: 21.054\nExp number, first 60 AAs: 0.32765\nTotal prob of N-in: 0.07004\noutside 1 376\nTMhelix 377 399\ninside 400 445

34373 GCF_000270265.1_ASM27026v1 Synechocystis sp. PCC 6803 Terrabacteria group; Cyanobacteria/Melainabacteria group; Cyanobacteria; Synechococcales; Merismopediaceae; Synechocystis MNSPTSPRRPHVVIVGGGFAGLYTAKNLRRSPVDITLIDKRNFHLFQPLLYQVATGSLSP WP_010872417.1 pyridine nucleotide-disulfide oxidoreductase [Synechocystis sp. PCC 6803] Length: 445\nNumber of predicted TMHs: 1\nExp number of AAs in TMHs: 21.054\nExp number, first 60 AAs: 0.32765\nTotal prob of N-in: 0.07004\noutside 1 376\nTMhelix 377 399\ninside 400 445

34374	GCF_000340785.1_ASM34078v1	Synechocystis sp. PCC 6803	Terrabacteria group; Cyanobacteria/Melainabacteria group; Cyanobacteria; Synechococcales; Merismopediaceae; Synechocystis
	MNSPTSPRRPHVVIVGGGFAGLYTAKNLRRSPVDITLIDKRNFHLFQPLLYQVATGSLSP	WP_010872417.1	pyridine nucleotide-disulfide oxidoreductase [Synechocystis sp. PCC 6803]	Length: 445\nNumber of predicted TMHs: 1\nExp number of AAs in TMHs: 21.054\nExp number, first 60 AAs: 0.32765\nTotal prob of N-in: 0.07004\noutside 1 376\nTMhelix 377 399\ninside 400 445

34375	GCF_000009725.1_ASM972v1	Synechocystis sp. PCC 6803	Terrabacteria group; Cyanobacteria/Melainabacteria group; Cyanobacteria; Synechococcales; Merismopediaceae; Synechocystis
	MNSPTSPRRPHVVIVGGGFAGLYTAKNLRRSPVDITLIDKRNFHLFQPLLYQVATGSLSP	WP_010872417.1	pyridine nucleotide-disulfide oxidoreductase [Synechocystis sp. PCC 6803]	Length: 445\nNumber of predicted TMHs: 1\nExp number of AAs in TMHs: 21.054\nExp number, first 60 AAs: 0.32765\nTotal prob of N-in: 0.07004\noutside 1 376\nTMhelix 377 399\ninside 400 445

34376	GCF_000284135.1_ASM28413v1	Synechocystis sp. PCC 6803 substr. GT-ITerrabacteria group; Cyanobacteria/Melainabacteria group; Cyanobacteria; Synechococcales; Merismopediaceae; Synechocystis; Synechocystis sp. PCC 6803	MNSPTSPRRPHVVIVGGGFAGLYTAKNLRRSPVDITLIDKRNFHLFQPLLYQVATGSLSP	WP_010872417.1
	pyridine nucleotide-disulfide oxidoreductase [Synechocystis sp. PCC 6803]	Length: 445\nNumber of predicted TMHs: 1\nExp number of AAs in TMHs: 21.054\nExp number, first 60 AAs: 0.32765\nTotal prob of N-in: 0.07004\noutside 1 376\nTMhelix 377 399\ninside 400 445

34377	GCF_000284215.1_ASM28421v1	Synechocystis sp. PCC 6803 substr. PCC-N	Terrabacteria group; Cyanobacteria/Melainabacteria group; Cyanobacteria; Synechococcales; Merismopediaceae; Synechocystis; Synechocystis sp. PCC 6803	MNSPTSPRRPHVVIVGGGFAGLYTAKNLRRSPVDITLIDKRNFHLFQPLLYQVATGSLSP	WP_010872417.1
	pyridine nucleotide-disulfide oxidoreductase [Synechocystis sp. PCC 6803]	Length: 445\nNumber of predicted TMHs: 1\nExp number of AAs in TMHs: 21.054\nExp number, first 60 AAs: 0.32765\nTotal prob of N-in: 0.07004\noutside 1 376\nTMhelix 377 399\ninside 400 445

34378	GCF_000284455.1_ASM28445v1	Synechocystis sp. PCC 6803 substr. PCC-P	Terrabacteria group; Cyanobacteria/Melainabacteria group; Cyanobacteria; Synechococcales; Merismopediaceae; Synechocystis; Synechocystis sp. PCC 6803	MNSPTSPRRPHVVIVGGGFAGLYTAKNLRRSPVDITLIDKRNFHLFQPLLYQVATGSLSP	WP_010872417.1
	pyridine nucleotide-disulfide oxidoreductase [Synechocystis sp. PCC 6803]	Length: 445\nNumber of predicted TMHs: 1\nExp number of AAs in TMHs: 21.054\nExp number, first 60 AAs: 0.32765\nTotal prob of N-in: 0.07004\noutside 1 376\nTMhelix 377 399\ninside 400 445

34379	GCF_000332175.1_ASM33217v1	Pseudanabaena sp. PCC 6802Terrabacteria group; Cyanobacteria/Melainabacteria group; Cyanobacteria; Synechococcales; Pseudanabaenaceae; Pseudanabaena
	MGNNSRRRIIKSIFWSVCGACLAIALLGIFSGFPESALALKRNPAIDPTSSVHLTMGNPS	WP_019502533.1	hypothetical protein [Pseudanabaena sp. PCC 6802] Length: 324\nNumber of predicted TMHs: 2\nExp number of AAs in TMHs: 41.28395\nExp number, first 60 AAs: 21.57464\nTotal prob of N-in: 0.99884\nPOSSIBLE N-term signal sequence\ninside 1 11\nTMhelix 12 34\noutside 35 297\nTMhelix 298 316\ninside 317 324

34380	GCF_002198095.1_ASM219809v1	Deinococcus indicus	Terrabacteria group; Deinococcus-Thermus; Deinococci; Deinococcales; Deinococcaceae; Deinococcus
	MTAPGSGVRRRRALRRLGVTGLWLAALLTVGAWLGVWLTLAPARAALTGAGQSLATLDRQ	WP_088246978.1
	hypothetical protein [Deinococcus indicus]	Length: 190\nNumber of predicted TMHs: 2\nExp number of AAs in TMHs: 44.90066\nExp number, first 60 AAs: 22.1987\nTotal prob of N-in: 0.69128\nPOSSIBLE N-term signal sequence\ninside 1 18\nTMhelix 19 41\noutside 42 155\nTMhelix 156 178\ninside 179 190

34381	GCF_002017875.1_ASM201787v1	Deinococcus sp. LM3	Terrabacteria group; Deinococcus-Thermus; Deinococci; Deinococcales; Deinococcaceae; Deinococcus
	MTAPGSRVRRRRALRRLGVTGLWLAALLTVGAWLGVFLALAPARAALNGAAQSLGTLDRQ	WP_078302060.1
	hypothetical protein [Deinococcus sp. LM3]	Length: 190\nNumber of predicted TMHs: 2\nExp number of AAs in TMHs: 44.40162\nExp number, first 60 AAs: 22.50818\nTotal prob of N-in: 0.90888\nPOSSIBLE N-term signal sequence\ninside 1 20\nTMhelix 21 43\noutside 44 150\nTMhelix 151 173\ninside 174 190

34382	GCF_000701425.1_ASM70142v1	Deinococcus frigens DSM 12807	Terrabacteria group; Deinococcus-Thermus; Deinococci; Deinococcales; Deinococcaceae; Deinococcus; Deinococcus frigens
	MRGGVGCRLSAVEANSNRLTVKPSKRRKHLASCLLTSVFCLSLAHATAPATLPLTLPPAQ	WP_084147912.1	hypothetical protein [Deinococcus frigens]Length: 188\nNumber of predicted TMHs: 2\nExp number of AAs in TMHs: 40.06831\nExp

number, first 60 AAs: 18.31091\nTotal prob of N-in: 0.92647\nPOSSIBLE N-term signal sequence\ninside 1 29\nTMhelix 30 52\noutside 53 131\nTMhelix 132 154\ninside 155 188

34383 GCF_000373205.1_ASM37320v1 Meiothermus timidus DSM 17022 Terrabacteria group; Deinococcus-Thermus; Deinococci; Thermales; Thermaceae; Meiothermus; Meiothermus timidus MSMEPHTQSDELSLRDLYLVLKRRQNLILGLTLGAAILVFAVSQVWPKTYSSKVVLSLSF WP_018466007.1 hypothetical protein [Meiothermus timidus] Length: 319\nNumber of predicted TMHs: 2\nExp number of AAs in TMHs: 42.15524\nExp number, first 60 AAs: 20.02973\nTotal prob of N-in: 0.46664\nPOSSIBLE N-term signal sequence\noutside 1 26\nTMhelix 27 46\ninside 47 273\nTMhelix 274 296\noutside 297 319

34384 GCF_001399675.1_ASM139967v1 Alicyclobacillus ferrooxydans Terrabacteria group; Firmicutes; Bacilli; Bacillales; Alicyclobacillaceae; Alicyclobacillus MLWERRRCMISVILVTYNSMSVLPDCLASLEINPIASELEVVVADNASSDGTLRWLMEYQ WP_083486416.1 hypothetical protein [Alicyclobacillus ferrooxydans] Length: 306\nNumber of predicted TMHs: 1\nExp number of AAs in TMHs: 30.87849\nExp number, first 60 AAs: 0.0351\nTotal prob of N-in: 0.73747\ninside 1 272\nTMhelix 273 295\noutside 296 306

34385 GCF_000714935.1_GST4V1 Tumebacillus flagellatus Terrabacteria group; Firmicutes; Bacilli; Bacillales; Alicyclobacillaceae; Tumebacillus MTERRKRRRRMIRFFTSVALCISLCLNAGEAFAVAQPLPASESLQNTFLTSTNQTLTPVT WP_081857365.1 hypothetical protein [Tumebacillus flagellatus] Length: 738\nNumber of predicted TMHs: 2\nExp number of AAs in TMHs: 45.14873\nExp number, first 60 AAs: 22.59571\nTotal prob of N-in: 0.99278\nPOSSIBLE N-term signal sequence\ninside 1 11\nTMhelix 12 34\noutside 35 699\nTMhelix 700 722\ninside 723 738

34386 GCF_001629795.1_ASM162979v1 Aeribacillus pallidus Terrabacteria group; Firmicutes; Bacilli; Bacillales; Bacillaceae; Aeribacillus MKKRKRRSMLAGASAVLLVLVSVLTPIHTYATATVAGDAEQQMEEQTKSTDKPNQEQPLH WP_063387033.1 hypothetical protein [Aeribacillus pallidus] Length: 1811\nNumber of predicted TMHs: 2\nExp number of AAs in TMHs: 41.3549\nExp number, first 60 AAs: 21.58793\nTotal prob of N-in: 0.97527\nPOSSIBLE N-term signal sequence\ninside 1 12\nTMhelix 13 35\noutside 36 1782\nTMhelix 1783 1801\ninside 1802 1811

34387 GCF_001865995.1_ASM186599v1 Anaerobacillus arseniciselenatis Terrabacteria group; Firmicutes; Bacilli; Bacillales; Bacillaceae; Anaerobacillus MTEKNKLDRRTFLKRSAQATVTAGVVAATSNSVFAERDEDKVNGYVASIIDLTKCDGCAN WP_071313372.1 oxidoreductase [Anaerobacillus arseniciselenatis] Length: 356\nNumber of predicted TMHs: 1\nExp number of AAs in TMHs: 20.05239\nExp number, first 60 AAs: 0.22205\nTotal prob of N-in: 0.45976\noutside 1 311\nTMhelix 312 334\ninside 335 356

34388 GCF_001418025.1_ASM141802v1 Anoxybacillus suryakundensis Terrabacteria group; Firmicutes; Bacilli; Bacillales; Bacillaceae; Anoxybacillus MSEKISRRTFLKRSTVAGATAGMLLATKTPGYAQTDKSDENTLIGSLIDLTKCDGCAAYD WP_032100996.1 MULTISPECIES: oxidoreductase [Anoxybacillus] Length: 350\nNumber of predicted TMHs: 1\nExp number of AAs in TMHs: 20.31145\nExp number, first 60 AAs: 0.0764\nTotal prob of N-in: 0.07656\noutside 1 310\nTMhelix 311 333\ninside 334 350

34389 GCF_001517225.1_JGI_assembly Anoxybacillus suryakundensis Terrabacteria group; Firmicutes; Bacilli; Bacillales; Bacillaceae; Anoxybacillus MSEKISRRTFLKRSTVAGATAGMLLATKTPGYAQTDKSDENTLIGSLIDLTKCDGCAAYD WP_032100996.1 MULTISPECIES: oxidoreductase [Anoxybacillus] Length: 350\nNumber of predicted TMHs: 1\nExp number of AAs in TMHs: 20.31145\nExp number, first 60 AAs: 0.0764\nTotal prob of N-in: 0.07656\noutside 1 310\nTMhelix 311 333\ninside 334 350

34390 GCF_000353425.1_Anoxybacillus_flavithermus_AK1 Anoxybacillus flavithermus AK1 Terrabacteria group; Firmicutes; Bacilli; Bacillales; Bacillaceae; Anoxybacillus; Anoxybacillus flavithermus MSEKISRRTFLKRSTVAGATAGILLATKTPGYAQTDKSDENTLIGSLIDLTKCDGCAAYD WP_003398576.1 4Fe-4S ferredoxin iron-sulfur-binding domain-containing protein [Anoxybacillus flavithermus] Length: 350\nNumber of predicted TMHs: 1\nExp number of AAs in TMHs: 19.94586\nExp number, first 60 AAs: 0.12703\nTotal prob of N-in: 0.10184\noutside 1 310\nTMhelix 311 333\ninside 334 350

34391 GCF_000753835.1_ASM75383v1 Anoxybacillus flavithermus subsp. yunnanensis str. E13 Terrabacteria group; Firmicutes; Bacilli; Bacillales; Bacillaceae; Anoxybacillus; Anoxybacillus flavithermus; Anoxybacillus flavithermus subsp. yunnanensis MSEKISRRTFLKRSTVAGATAGMLLATKTPGYAQTDKSDENTLIGSLIDLTKCDGCAAYD WP_032100996.1 MULTISPECIES: oxidoreductase [Anoxybacillus] Length: 350\nNumber of predicted TMHs: 1\nExp number of

AAs in TMHs: 20.31145\nExp number, first 60 AAs:  0.0764\nTotal prob of N-in:   0.07656\noutside   1   310\nTMhelix   311   333\ninside   334   350

34392 GCF_001307105.1_ASM130710v1 Bacillus australimaris Terrabacteria group; Firmicutes; Bacilli; Bacillales; Bacillaceae; Bacillus MKKQTHRSKRRKNLSILLTSIMMISLILPAVPTHAAIENGNAVKVSILATTDVHANMMNY WP_060699953.1 2,3-cyclic-nucleotide 2-phosphodiesterase [Bacillus australimaris] Length: 1429\nNumber of predicted TMHs:  2\nExp number of AAs in TMHs: 44.32614\nExp number, first 60 AAs:  21.76433\nTotal prob of N-in:   0.99871\nPOSSIBLE N-term signal sequence\ninside   1   12\nTMhelix   13   35\noutside   36  1395\nTMhelix   1396   1418\ninside   1419  1429

34393 GCF_001721685.1_ASM172168v1 Bacillus beveridgei Terrabacteria group; Firmicutes; Bacilli; Bacillales; Bacillaceae; Bacillus MDRRNFLKRASAATAAAAVMTTSNKWVSASAIDEDKQIGSIIDLSKCDGCEYQDTPLCVA WP_069364808.1 oxidoreductase [Bacillus beveridgei] Length: 338\nNumber of predicted TMHs:  1\nExp number of AAs in TMHs: 19.95188\nExp number, first 60 AAs:  0.0244\nTotal prob of N-in:   0.23680\noutside   1   297\nTMhelix   298   320\ninside   321   338

34394 GCF_001038565.1_ASM103856v1 Bacillus pseudalcaliphilus Terrabacteria group; Firmicutes; Bacilli; Bacillales; Bacillaceae; Bacillus MNVFFNTSRAFHICLLYYSLYLKRRRSNMLKRITQSSIIVLVVMGLMVTQGFAHTHLESS WP_083991168.1 hypothetical protein [Bacillus pseudalcaliphilus] Length: 225\nNumber of predicted TMHs:  2\nExp number of AAs in TMHs: 47.68901\nExp number, first 60 AAs:  25.52687\nTotal prob of N-in:   0.73273\nPOSSIBLE N-term signal sequence\ninside   1   32\nTMhelix   33   55\noutside   56   198\nTMhelix   199   221\ninside   222   225

34395 GCF_002174275.1_ASM217427v1 Bacillus pumilus Terrabacteria group; Firmicutes; Bacilli; Bacillales; Bacillaceae; Bacillus MKKQAHRSKRRKNLSILLTSIMMISLILPAVPIHAETESGKAVKVSILATTDVHANMMNY WP_088000724.1 2,3-cyclic-nucleotide 2-phosphodiesterase [Bacillus pumilus] Length: 1429\nNumber of predicted TMHs:  2\nExp number of AAs in TMHs: 44.9065399999999\nExp number, first 60 AAs:  22.32205\nTotal prob of N-in:   0.99930\nPOSSIBLE N-term signal sequence\ninside   1   12\nTMhelix   13   35\noutside   36  1395\nTMhelix   1396  1418\ninside   1419  1429

34396 GCF_000714495.2_ASM71449v2 Bacillus pumilus Terrabacteria group; Firmicutes; Bacilli; Bacillales; Bacillaceae; Bacillus MKKQRRKNLSILLTSIMMISLILPAVPTYAETENGNAVKVSILATTDVHANMMNYDYYSD WP_034665156.1 2,3-cyclic-nucleotide 2-phosphodiesterase [Bacillus pumilus] Length: 1424\nNumber of predicted TMHs:  2\nExp number of AAs in TMHs: 43.0923\nExp number, first 60 AAs:  20.52829\nTotal prob of N-in:   0.97431\nPOSSIBLE N-term signal sequence\ninside   1   8\nTMhelix   9   31\noutside   32  1390\nTMhelix   1391   1413\ninside   1414  1424

34397 GCF_001431785.1_ASM143178v1 Bacillus pumilus Terrabacteria group; Firmicutes; Bacilli; Bacillales; Bacillaceae; Bacillus MKKQAHRSKRRKNLSILLTSMMLISLILPAVPTHAETESGSAVKVSILATTDVHANMMNY WP_057078327.1 2,3-cyclic-nucleotide 2-phosphodiesterase [Bacillus pumilus] Length: 1426\nNumber of predicted TMHs:  2\nExp number of AAs in TMHs: 44.27896\nExp number, first 60 AAs:  21.58314\nTotal prob of N-in:   0.99584\nPOSSIBLE N-term signal sequence\ninside   1   12\nTMhelix   13   35\noutside   36  1392\nTMhelix   1393   1415\ninside   1416  1426

34398 GCF_001038765.1_ASM103876v1 Bacillus pumilus Terrabacteria group; Firmicutes; Bacilli; Bacillales; Bacillaceae; Bacillus MKKQAHRSKRRKNLSILLTSIMMISLILPAVPTHAAIENGDAVKVSILATTDVHANMMNY WP_048240120.1 2,3-cyclic-nucleotide 2-phosphodiesterase [Bacillus pumilus] Length: 1430\nNumber of predicted TMHs:  2\nExp number of AAs in TMHs: 44.27925\nExp number, first 60 AAs:  21.71641\nTotal prob of N-in:   0.99812\nPOSSIBLE N-term signal sequence\ninside   1   12\nTMhelix   13   35\noutside   36  1396\nTMhelix   1397   1419\ninside   1420  1430

34399 GCF_001675655.1_ASM167565v1 Bacillus pumilus Terrabacteria group; Firmicutes; Bacilli; Bacillales; Bacillaceae; Bacillus MKKQAHRSKRRKNLSILLTSIMMISLILPAVPTHAETENGKAVKVSILATTDVHANMMNY WP_065097475.1 2,3-cyclic-nucleotide 2-phosphodiesterase [Bacillus pumilus] Length: 1429\nNumber of predicted TMHs:  2\nExp number of AAs in TMHs: 44.26709\nExp number, first 60 AAs:  21.69265\nTotal prob of N-in:   0.99788\nPOSSIBLE N-term signal sequence\ninside   1   12\nTMhelix   13   35\noutside   36  1395\nTMhelix   1396   1418\ninside   1419  1429

34400 GCF_001704975.1_ASM170497v1 Bacillus pumilus Terrabacteria group; Firmicutes; Bacilli; Bacillales; Bacillaceae; Bacillus MKKQTHRSKRRKNLSILLTSIMMISLILPAVPTHAETENGKAVKVSILATTDVHANMMNY WP_066031168.1 2,3-cyclic-nucleotide 2-phosphodiesterase [Bacillus pumilus] Length: 1429\nNumber of predicted TMHs:  2\nExp number of AAs in TMHs: 44.27703\nExp number, first 60 AAs:  21.70262\nTotal prob of N-in:

0.99834\nPOSSIBLE N-term signal sequence\ninside 1 12\nTMhelix 13 35\noutside 36 1395\nTMhelix 1396 1418\ninside 1419 1429

34401 GCF_001687085.1_ASM168708v1 Bacillus pumilus Terrabacteria group; Firmicutes; Bacilli; Bacillales; Bacillaceae; Bacillus MKKQAHRSKRRKNLSILLTSMMLISLILPAVPTHAETESGSAVKVSILATTDVHANMMNY WP_065460006.1 2,3-cyclic-nucleotide 2-phosphodiesterase [Bacillus pumilus] Length: 1426\nNumber of predicted TMHs: 2\nExp number of AAs in TMHs: 44.13563\nExp number, first 60 AAs: 21.58314\nTotal prob of N-in: 0.99584\nPOSSIBLE N-term signal sequence\ninside 1 12\nTMhelix 13 35\noutside 36 1392\nTMhelix 1393 1415\ninside 1416 1426

34402 GCF_000828375.1_ASM82837v1 Bacillus pumilus Terrabacteria group; Firmicutes; Bacilli; Bacillales; Bacillaceae; Bacillus MKKQTNRSKRRKNLSILLTSIMMISLILPAVPTHAAIENGNAVKVSILATTDVHANMMNY WP_041087244.1 2,3-cyclic-nucleotide 2-phosphodiesterase [Bacillus pumilus] Length: 1430\nNumber of predicted TMHs: 2\nExp number of AAs in TMHs: 44.32819\nExp number, first 60 AAs: 21.76977\nTotal prob of N-in: 0.99896\nPOSSIBLE N-term signal sequence\ninside 1 12\nTMhelix 13 35\noutside 36 1396\nTMhelix 1397 1419\ninside 1420 1430

34403 GCF_000828455.1_ASM82845v1 Bacillus pumilus Terrabacteria group; Firmicutes; Bacilli; Bacillales; Bacillaceae; Bacillus MKKQAHRSKRRKNLSILLTSIMLISLILPAVPTHAETESGSAVKVSILATTDVHANMMNY WP_041090692.1 2,3-cyclic-nucleotide 2-phosphodiesterase [Bacillus pumilus] Length: 1426\nNumber of predicted TMHs: 2\nExp number of AAs in TMHs: 44.19855\nExp number, first 60 AAs: 21.62366\nTotal prob of N-in: 0.99772\nPOSSIBLE N-term signal sequence\ninside 1 12\nTMhelix 13 35\noutside 36 1392\nTMhelix 1393 1415\ninside 1416 1426

34404 GCF_000691485.2_ASM69148v2 Bacillus pumilus Terrabacteria group; Firmicutes; Bacilli; Bacillales; Bacillaceae; Bacillus MKKQRRKNLSILLTSIMMISLILPAVPTYAETENGNAVKVSILATTDVHANMMNYDYYSD WP_034665156.1 2,3-cyclic-nucleotide 2-phosphodiesterase [Bacillus pumilus] Length: 1424\nNumber of predicted TMHs: 2\nExp number of AAs in TMHs: 43.0923\nExp number, first 60 AAs: 20.52829\nTotal prob of N-in: 0.97431\nPOSSIBLE N-term signal sequence\ninside 1 8\nTMhelix 9 31\noutside 32 1390\nTMhelix 1391 1413\ninside 1414 1424

34405 GCF_001017485.1_BpV2.0 Bacillus pumilus Terrabacteria group; Firmicutes; Bacilli; Bacillales; Bacillaceae; Bacillus MKKQTHRSKRRKNLSILLTSIMMISLILPAVPTHAAIENGDAVKVSILATTDVHANMMNY WP_047203589.1 2,3-cyclic-nucleotide 2-phosphodiesterase [Bacillus pumilus] Length: 1430\nNumber of predicted TMHs: 2\nExp number of AAs in TMHs: 44.2879\nExp number, first 60 AAs: 21.72523\nTotal prob of N-in: 0.99853\nPOSSIBLE N-term signal sequence\ninside 1 12\nTMhelix 13 35\noutside 36 1396\nTMhelix 1397 1419\ninside 1420 1430

34406 GCF_001908475.1_ASM190847v1 Bacillus pumilus Terrabacteria group; Firmicutes; Bacilli; Bacillales; Bacillaceae; Bacillus MKKQAHRSKRRKNLSILLTSMMLISLILPAVPTHAETESGSAVKVSILATTDVHANMMNY WP_060832163.1 2,3-cyclic-nucleotide 2-phosphodiesterase [Bacillus pumilus] Length: 1426\nNumber of predicted TMHs: 2\nExp number of AAs in TMHs: 44.29415\nExp number, first 60 AAs: 21.58314\nTotal prob of N-in: 0.99584\nPOSSIBLE N-term signal sequence\ninside 1 12\nTMhelix 13 35\noutside 36 1392\nTMhelix 1393 1415\ninside 1416 1426

34407 GCF_001700735.1_ASM170073v1 Bacillus pumilus Terrabacteria group; Firmicutes; Bacilli; Bacillales; Bacillaceae; Bacillus MKKQAHRSKRRKNLSILLTSMMLISLILPAVPTHAETESGSAVKVSILATTDVHANMMNY WP_057078327.1 2,3-cyclic-nucleotide 2-phosphodiesterase [Bacillus pumilus] Length: 1426\nNumber of predicted TMHs: 2\nExp number of AAs in TMHs: 44.27896\nExp number, first 60 AAs: 21.58314\nTotal prob of N-in: 0.99584\nPOSSIBLE N-term signal sequence\ninside 1 12\nTMhelix 13 35\noutside 36 1392\nTMhelix 1393 1415\ninside 1416 1426

34408 GCF_001191605.1_ASM119160v1 Bacillus pumilus Terrabacteria group; Firmicutes; Bacilli; Bacillales; Bacillaceae; Bacillus MKKQAHRSKRRKNLSILLTSMMLISLILPAVPTHAETESGSAVKVSILATTDVHANMMNY WP_050827057.1 2,3-cyclic-nucleotide 2-phosphodiesterase [Bacillus pumilus] Length: 1426\nNumber of predicted TMHs: 2\nExp number of AAs in TMHs: 44.2942\nExp number, first 60 AAs: 21.58314\nTotal prob of N-in: 0.99584\nPOSSIBLE N-term signal sequence\ninside 1 12\nTMhelix 13 35\noutside 36 1392\nTMhelix 1393 1415\ninside 1416 1426

34409 GCF_001543165.1_ASM154316v1 Bacillus pumilus Terrabacteria group; Firmicutes; Bacilli; Bacillales; Bacillaceae; Bacillus MKKQTHRSKRRKNLSILLTSIMLISLILPAVPTHAETESGSAVKVSILATTDVHANMMNY WP_060780877.1 2,3-cyclic-nucleotide 2-phosphodiesterase [Bacillus pumilus] Length: 1426\nNumber of predicted TMHs:  2\nExp number of AAs in TMHs: 44.32978\nExp number, first 60 AAs:  21.63439\nTotal prob of N-in: 0.99822\nPOSSIBLE N-term signal sequence\ninside   1   12\nTMhelix   13   35\noutside   36  1392\nTMhelix   1393 1415\ninside   1416  1426

34410 GCF_001431145.1_ASM143114v1 Bacillus pumilus Terrabacteria group; Firmicutes; Bacilli; Bacillales; Bacillaceae; Bacillus MKKQAHRSKRRKNLSILLTSMMLISLILPAVPTHAETESGSAVKVSILATTDVHANMMNY WP_057078327.1 2,3-cyclic-nucleotide 2-phosphodiesterase [Bacillus pumilus] Length: 1426\nNumber of predicted TMHs:  2\nExp number of AAs in TMHs: 44.27896\nExp number, first 60 AAs:  21.58314\nTotal prob of N-in: 0.99584\nPOSSIBLE N-term signal sequence\ninside   1   12\nTMhelix   13   35\noutside   36  1392\nTMhelix   1393 1415\ninside   1416  1426

34411 GCF_001578205.1_ASM157820v1 Bacillus pumilus Terrabacteria group; Firmicutes; Bacilli; Bacillales; Bacillaceae; Bacillus MKKQAHRSKRRKNLSILLTSIMMISLILPAVPTHAETENGKAVKVSILATTDVHANMMNY WP_058015724.1 2,3-cyclic-nucleotide 2-phosphodiesterase [Bacillus pumilus] Length: 1427\nNumber of predicted TMHs:  2\nExp number of AAs in TMHs: 44.26714\nExp number, first 60 AAs:  21.69265\nTotal prob of N-in: 0.99788\nPOSSIBLE N-term signal sequence\ninside   1   12\nTMhelix   13   35\noutside   36  1393\nTMhelix   1394 1416\ninside   1417  1427

34412 GCF_000604385.1_B.pum_Fairview_1.0 Bacillus pumilus Terrabacteria group; Firmicutes; Bacilli; Bacillales; Bacillaceae; Bacillus MKKQAHRSKRRKNLSILLTSIMMISLILPAVPTHAAIENGDAVKVSILATTDVHANMMNY WP_025092693.1 2,3-cyclic-nucleotide 2-phosphodiesterase [Bacillus pumilus] Length: 1430\nNumber of predicted TMHs:  2\nExp number of AAs in TMHs: 44.28052\nExp number, first 60 AAs:  21.7164\nTotal prob of N-in: 0.99812\nPOSSIBLE N-term signal sequence\ninside   1   12\nTMhelix   13   35\noutside   36  1396\nTMhelix   1397 1419\ninside   1420  1430

34413 GCF_000828345.1_ASM82834v1 Bacillus pumilus Terrabacteria group; Firmicutes; Bacilli; Bacillales; Bacillaceae; Bacillus MKKQAHRSKRRKNLSILLTSIMMISLILPAVPTHAETENGKAVKVSILATTDVHANMMNY WP_044139739.1 2,3-cyclic-nucleotide 2-phosphodiesterase [Bacillus pumilus] Length: 1429\nNumber of predicted TMHs:  2\nExp number of AAs in TMHs: 44.26709\nExp number, first 60 AAs:  21.69265\nTotal prob of N-in: 0.99788\nPOSSIBLE N-term signal sequence\ninside   1   12\nTMhelix   13   35\noutside   36  1395\nTMhelix   1396 1418\ninside   1419  1429

34414 GCF_000828425.1_ASM82842v1 Bacillus pumilus Terrabacteria group; Firmicutes; Bacilli; Bacillales; Bacillaceae; Bacillus MKKQAHRSKRRKNLSILLTSIMMISLILPAVPTHAAIENGDAVKVSILATTDVHANMMNY WP_041116297.1 2,3-cyclic-nucleotide 2-phosphodiesterase [Bacillus pumilus] Length: 1430\nNumber of predicted TMHs:  2\nExp number of AAs in TMHs: 44.27946\nExp number, first 60 AAs:  21.7164\nTotal prob of N-in: 0.99812\nPOSSIBLE N-term signal sequence\ninside   1   12\nTMhelix   13   35\noutside   36  1396\nTMhelix   1397 1419\ninside   1420  1430

34415 GCF_001038905.1_ASM103890v1 Bacillus pumilus Terrabacteria group; Firmicutes; Bacilli; Bacillales; Bacillaceae; Bacillus MKKQAHRSKRRKNLSILLTSIMMISLILPAVPTHAAIENGDAVKVSILATTDVHANMMNY WP_048240120.1 2,3-cyclic-nucleotide 2-phosphodiesterase [Bacillus pumilus] Length: 1430\nNumber of predicted TMHs:  2\nExp number of AAs in TMHs: 44.27925\nExp number, first 60 AAs:  21.71641\nTotal prob of N-in: 0.99812\nPOSSIBLE N-term signal sequence\ninside   1   12\nTMhelix   13   35\noutside   36  1396\nTMhelix   1397 1419\ninside   1420  1430

34416 GCF_000972685.1_ASM97268v1 Bacillus pumilus Terrabacteria group; Firmicutes; Bacilli; Bacillales; Bacillaceae; Bacillus MKKQAHRSKRRKNLSILLTSIMLISLILPAVPTHAETESGSAVKVSILATTDVHANMMNY WP_046342947.1 2,3-cyclic-nucleotide 2-phosphodiesterase [Bacillus pumilus] Length: 1426\nNumber of predicted TMHs:  2\nExp number of AAs in TMHs: 44.33421\nExp number, first 60 AAs:  21.62366\nTotal prob of N-in: 0.99772\nPOSSIBLE N-term signal sequence\ninside   1   12\nTMhelix   13   35\noutside   36  1392\nTMhelix   1393 1415\ninside   1416  1426

34417 GCF_000828395.1_ASM82839v1 Bacillus pumilus Terrabacteria group; Firmicutes; Bacilli; Bacillales; Bacillaceae; Bacillus MKKQAHRSKRRKNLSILLTSIMMISLILPAVPTHAAIENGDAVKVSILATTDVHANMMNY WP_041106302.1 2,3-cyclic-nucleotide 2-phosphodiesterase [Bacillus pumilus] Length: 1430\nNumber of predicted TMHs:  2\nExp number of AAs in TMHs: 44.279\nExp number, first 60 AAs:  21.71641\nTotal prob of N-in: 0.99812\nPOSSIBLE N-term signal sequence\ninside   1   12\nTMhelix   13   35\noutside   36  1396\nTMhelix   1397 1419\ninside   1420  1430

34418 GCF_001578165.1_ASM157816v1 Bacillus pumilus Terrabacteria group; Firmicutes; Bacilli; Bacillales; Bacillaceae; Bacillus MKKQAHRSKRRKNLSILLTSIMLISLILPAVPTHAETESGSAVKVSILATTDVHANMMNY WP_061418357.1 2,3-cyclic-nucleotide 2-phosphodiesterase [Bacillus pumilus] Length: 1426\nNumber of predicted TMHs: 2\nExp number of AAs in TMHs: 44.33421\nExp number, first 60 AAs: 21.62366\nTotal prob of N-in: 0.99772\nPOSSIBLE N-term signal sequence\ninside 1 12\nTMhelix 13 35\noutside 36 1392\nTMhelix 1393 1415\ninside 1416 1426

34419 GCF_001548215.1_ASM154821v1 Bacillus pumilus Terrabacteria group; Firmicutes; Bacilli; Bacillales; Bacillaceae; Bacillus MKKQAHRSKRRKNLSILLTSMMLISLILPAVPTHAETESGSAVKVSILATTDVHANMMNY WP_060832163.1 2,3-cyclic-nucleotide 2-phosphodiesterase [Bacillus pumilus] Length: 1426\nNumber of predicted TMHs: 2\nExp number of AAs in TMHs: 44.29415\nExp number, first 60 AAs: 21.58314\nTotal prob of N-in: 0.99584\nPOSSIBLE N-term signal sequence\ninside 1 12\nTMhelix 13 35\noutside 36 1392\nTMhelix 1393 1415\ninside 1416 1426

34420 GCF_001444515.1_ASM144451v1 Bacillus pumilus Terrabacteria group; Firmicutes; Bacilli; Bacillales; Bacillaceae; Bacillus MKKQAHRSKRRKNLSILLTSIMMISLILPAVPTHAETENGKAVKVSILATTDVHANMMNY WP_058015724.1 2,3-cyclic-nucleotide 2-phosphodiesterase [Bacillus pumilus] Length: 1427\nNumber of predicted TMHs: 2\nExp number of AAs in TMHs: 44.26714\nExp number, first 60 AAs: 21.69265\nTotal prob of N-in: 0.99788\nPOSSIBLE N-term signal sequence\ninside 1 12\nTMhelix 13 35\noutside 36 1393\nTMhelix 1394 1416\ninside 1417 1427

34421 GCF_001286985.1_Bacillus_JRS3 Bacillus pumilus Terrabacteria group; Firmicutes; Bacilli; Bacillales; Bacillaceae; Bacillus MKKQTNRSKRRKNLSILLTSIMMISLILPAVPTHAAIENGNAVKVSILATTDVHANMMNY WP_073978587.1 2,3-cyclic-nucleotide 2-phosphodiesterase [Bacillus pumilus] Length: 1428\nNumber of predicted TMHs: 2\nExp number of AAs in TMHs: 44.3329\nExp number, first 60 AAs: 21.76945\nTotal prob of N-in: 0.99895\nPOSSIBLE N-term signal sequence\ninside 1 12\nTMhelix 13 35\noutside 36 1394\nTMhelix 1395 1417\ninside 1418 1428

34422 GCF_001043695.1_ASM104369v1 Bacillus pumilus Terrabacteria group; Firmicutes; Bacilli; Bacillales; Bacillaceae; Bacillus MKKQAHRSKRRKNLSILLTSIMMISLILPAVPTHAAIENGDAVKVSILATTDVHANMMNY WP_048240120.1 2,3-cyclic-nucleotide 2-phosphodiesterase [Bacillus pumilus] Length: 1430\nNumber of predicted TMHs: 2\nExp number of AAs in TMHs: 44.27925\nExp number, first 60 AAs: 21.71641\nTotal prob of N-in: 0.99812\nPOSSIBLE N-term signal sequence\ninside 1 12\nTMhelix 13 35\noutside 36 1396\nTMhelix 1397 1419\ninside 1420 1430

34423 GCF_000590455.1_ASM59045v1 Bacillus pumilus Terrabacteria group; Firmicutes; Bacilli; Bacillales; Bacillaceae; Bacillus MKKQAHRSKRRKNLSILLTSIMLISLILPAVPTHAETESGSAVKVSILATTDVHANMMNY WP_025206927.1 2,3-cyclic-nucleotide 2-phosphodiesterase [Bacillus pumilus] Length: 1426\nNumber of predicted TMHs: 2\nExp number of AAs in TMHs: 44.31905\nExp number, first 60 AAs: 21.62366\nTotal prob of N-in: 0.99772\nPOSSIBLE N-term signal sequence\ninside 1 12\nTMhelix 13 35\noutside 36 1392\nTMhelix 1393 1415\ninside 1416 1426

34424 GCF_001183525.1_ASM118352v1 Bacillus pumilus Terrabacteria group; Firmicutes; Bacilli; Bacillales; Bacillaceae; Bacillus MKKQAHRSKRRKNLSILLTSIMMISLILPAVPTHAETENGKAVKVSILATTDVHANMMNY WP_050944794.1 2,3-cyclic-nucleotide 2-phosphodiesterase [Bacillus pumilus] Length: 1429\nNumber of predicted TMHs: 2\nExp number of AAs in TMHs: 44.26709\nExp number, first 60 AAs: 21.69265\nTotal prob of N-in: 0.99788\nPOSSIBLE N-term signal sequence\ninside 1 12\nTMhelix 13 35\noutside 36 1395\nTMhelix 1396 1418\ninside 1419 1429

34425 GCF_000715185.1_ASM71518v1 Bacillus safensis Terrabacteria group; Firmicutes; Bacilli; Bacillales; Bacillaceae; Bacillus MMKQTHRSKRRKNLSILLTSIMMISLILPAVPTHAAIENGDAVKVSILATTDVHANMMNY WP_034623151.1 2,3-cyclic-nucleotide 2-phosphodiesterase [Bacillus safensis] Length: 1430\nNumber of predicted TMHs: 2\nExp number of AAs in TMHs: 44.27021\nExp number, first 60 AAs: 21.70607\nTotal prob of N-in: 0.99764\nPOSSIBLE N-term signal sequence\ninside 1 12\nTMhelix 13 35\noutside 36 1396\nTMhelix 1397 1419\ninside 1420 1430

34426 GCF_002077215.1_ASM207721v1 Bacillus safensis Terrabacteria group; Firmicutes; Bacilli; Bacillales; Bacillaceae; Bacillus MKKQAHRSKRRKNLSILLTSIMMISLILPAVPTHAAIENGDAVKVSILATTDVHANMMNY WP_081125245.1 2,3-cyclic-nucleotide 2-phosphodiesterase [Bacillus safensis] Length: 1430\nNumber of predicted TMHs: 2\nExp number of AAs in TMHs: 44.27946\nExp number, first 60 AAs: 21.7164\nTotal prob of N-in:

0.99812\nPOSSIBLE N-term signal sequence\ninside 1 12\nTMhelix 13 35\noutside 36 1396\nTMhelix 1397 1419\ninside 1420 1430

34427 GCF_002151195.1_ASM215119v1 Bacillus safensis Terrabacteria group; Firmicutes; Bacilli; Bacillales; Bacillaceae; Bacillus MKKQAHRSKRRKNLSILLTSIMMISLILPAVPTHAAIENGDAVKVSILATTDVHANMMNY WP_087976431.1 2,3-cyclic-nucleotide 2-phosphodiesterase [Bacillus safensis] Length: 1430\nNumber of predicted TMHs: 2\nExp number of AAs in TMHs: 44.22071\nExp number, first 60 AAs: 21.71641\nTotal prob of N-in: 0.99812\nPOSSIBLE N-term signal sequence\ninside 1 12\nTMhelix 13 35\noutside 36 1396\nTMhelix 1397 1419\ninside 1420 1430

34428 GCF_000935315.1_Bac_saf_RIT372_v1 Bacillus safensis Terrabacteria group; Firmicutes; Bacilli; Bacillales; Bacillaceae; Bacillus MKKQAHRSKRRKNLSILLTSIMMISLILPAVPTHAAIENGDAVKVSILATTDVHANMMNY WP_044334648.1 2,3-cyclic-nucleotide 2-phosphodiesterase [Bacillus safensis] Length: 1430\nNumber of predicted TMHs: 2\nExp number of AAs in TMHs: 44.28052\nExp number, first 60 AAs: 21.7164\nTotal prob of N-in: 0.99812\nPOSSIBLE N-term signal sequence\ninside 1 12\nTMhelix 13 35\noutside 36 1396\nTMhelix 1397 1419\ninside 1420 1430

34429 GCF_001895885.1_ASM189588v1 Bacillus safensis Terrabacteria group; Firmicutes; Bacilli; Bacillales; Bacillaceae; Bacillus MKKQAHRSKRRKNLSILLTSIMMISLILPAVPTHAAIENGDAVKVSILATTDVHANMMNY WP_073204289.1 2,3-cyclic-nucleotide 2-phosphodiesterase [Bacillus safensis] Length: 1430\nNumber of predicted TMHs: 2\nExp number of AAs in TMHs: 44.2803\nExp number, first 60 AAs: 21.71641\nTotal prob of N-in: 0.99812\nPOSSIBLE N-term signal sequence\ninside 1 12\nTMhelix 13 35\noutside 36 1396\nTMhelix 1397 1419\ninside 1420 1430

34430 GCF_001653905.1_ASM165390v1 Bacillus safensis Terrabacteria group; Firmicutes; Bacilli; Bacillales; Bacillaceae; Bacillus MKKQTHRSKRRKNLSILLTSIMMISLILPAVPTHAAIENGDAVKVSILATTDVHANMMNY WP_064497724.1 2,3-cyclic-nucleotide 2-phosphodiesterase [Bacillus safensis] Length: 1430\nNumber of predicted TMHs: 2\nExp number of AAs in TMHs: 44.2879\nExp number, first 60 AAs: 21.72523\nTotal prob of N-in: 0.99853\nPOSSIBLE N-term signal sequence\ninside 1 12\nTMhelix 13 35\noutside 36 1396\nTMhelix 1397 1419\ninside 1420 1430

34431 GCF_001938705.1_ASM193870v1 Bacillus safensis Terrabacteria group; Firmicutes; Bacilli; Bacillales; Bacillaceae; Bacillus MKKQTNRSKRRKNLSILLTSIMMISLILPAVPTHATIENGNAVKVSILATTDVHANMMNY WP_075611027.1 2,3-cyclic-nucleotide 2-phosphodiesterase [Bacillus safensis] Length: 1430\nNumber of predicted TMHs: 2\nExp number of AAs in TMHs: 44.28146\nExp number, first 60 AAs: 21.71761\nTotal prob of N-in: 0.99889\nPOSSIBLE N-term signal sequence\ninside 1 12\nTMhelix 13 35\noutside 36 1396\nTMhelix 1397 1419\ninside 1420 1430

34432 GCF_002155005.1_ASM215500v1 Bacillus safensis Terrabacteria group; Firmicutes; Bacilli; Bacillales; Bacillaceae; Bacillus MKKQAHRSKRRKNLSILLTSIMMISLILPAVPTHAAIENGDAVKVSILATTDVHANMMNY WP_087977862.1 2,3-cyclic-nucleotide 2-phosphodiesterase [Bacillus safensis] Length: 1430\nNumber of predicted TMHs: 2\nExp number of AAs in TMHs: 44.28052\nExp number, first 60 AAs: 21.7164\nTotal prob of N-in: 0.99812\nPOSSIBLE N-term signal sequence\ninside 1 12\nTMhelix 13 35\noutside 36 1396\nTMhelix 1397 1419\ninside 1420 1430

34433 GCF_001677975.1_ASM167797v1 Bacillus safensis Terrabacteria group; Firmicutes; Bacilli; Bacillales; Bacillaceae; Bacillus MKKQAHRSKRRKNLSILLTSIMMISLILPAVPTHAAIENGDAVKVSILATTDVHANMMNY WP_065215161.1 2,3-cyclic-nucleotide 2-phosphodiesterase [Bacillus safensis] Length: 1430\nNumber of predicted TMHs: 2\nExp number of AAs in TMHs: 44.27946\nExp number, first 60 AAs: 21.7164\nTotal prob of N-in: 0.99812\nPOSSIBLE N-term signal sequence\ninside 1 12\nTMhelix 13 35\noutside 36 1396\nTMhelix 1397 1419\ninside 1420 1430

34434 GCF_000972825.1_ASM97282v1 Bacillus safensis Terrabacteria group; Firmicutes; Bacilli; Bacillales; Bacillaceae; Bacillus MKKQAHRSKRRKNLSILLTSIMMISLILPAVPTHAAIENGDAVKVSILATTDVHANMMNY WP_046314257.1 2,3-cyclic-nucleotide 2-phosphodiesterase [Bacillus safensis] Length: 1430\nNumber of predicted TMHs: 2\nExp number of AAs in TMHs: 44.27946\nExp number, first 60 AAs: 21.7164\nTotal prob of N-in: 0.99812\nPOSSIBLE N-term signal sequence\ninside 1 12\nTMhelix 13 35\noutside 36 1396\nTMhelix 1397 1419\ninside 1420 1430

34435 GCF_001938685.1_ASM193868v1 Bacillus safensis Terrabacteria group; Firmicutes; Bacilli; Bacillales; Bacillaceae; Bacillus MKKQTNRSKRRKNLSILLTSIMMISLILPAVPTHATIENGNAVKVSILATTDVHANMMNY

WP_075611027.1 2,3-cyclic-nucleotide 2-phosphodiesterase [Bacillus safensis] Length: 1430\nNumber of predicted TMHs: 2\nExp number of AAs in TMHs: 44.28146\nExp number, first 60 AAs: 21.71761\nTotal prob of N-in: 0.99889\nPOSSIBLE N-term signal sequence\ninside 1 12\nTMhelix 13 35\noutside 36 1396\nTMhelix 1397 1419\ninside 1420 1430

34436 GCF_001938665.1_ASM193866v1 Bacillus safensis Terrabacteria group; Firmicutes; Bacilli; Bacillales; Bacillaceae; Bacillus MKKQAHRSKRRKNLSILLTSIMMISLILPAVPTHAAIENGDAVKVSILATTDIHANMMNY WP_075622551.1 2,3-cyclic-nucleotide 2-phosphodiesterase [Bacillus safensis] Length: 1430\nNumber of predicted TMHs: 2\nExp number of AAs in TMHs: 44.27533\nExp number, first 60 AAs: 21.71648\nTotal prob of N-in: 0.99812\nPOSSIBLE N-term signal sequence\ninside 1 12\nTMhelix 13 35\noutside 36 1396\nTMhelix 1397 1419\ninside 1420 1430

34437 GCF_001766395.1_ASM176639v1 Bacillus safensis Terrabacteria group; Firmicutes; Bacilli; Bacillales; Bacillaceae; Bacillus MKKQAHRSKRRKNLSILLTSIMMISLILPAVPTHAAIENGDAVKVSILATTDVHANMMNY WP_070326779.1 2,3-cyclic-nucleotide 2-phosphodiesterase [Bacillus safensis] Length: 1430\nNumber of predicted TMHs: 2\nExp number of AAs in TMHs: 44.27946\nExp number, first 60 AAs: 21.7164\nTotal prob of N-in: 0.99812\nPOSSIBLE N-term signal sequence\ninside 1 12\nTMhelix 13 35\noutside 36 1396\nTMhelix 1397 1419\ninside 1420 1430

34438 GCF_000242895.2_ASM24289v3 Bacillus sp. 1NLA3E Terrabacteria group; Firmicutes; Bacilli; Bacillales; Bacillaceae; Bacillus MNKQFFKKKTAKRRSFMAMKVIALLATTTLALAPINTSASLLGGGLTSELTGTVSQVTGA WP_015592261.1 hypothetical protein [Bacillus sp. 1NLA3E] Length: 445\nNumber of predicted TMHs: 2\nExp number of AAs in TMHs: 45.8628599999999\nExp number, first 60 AAs: 22.56259\nTotal prob of N-in: 0.99851\nPOSSIBLE N-term signal sequence\ninside 1 20\nTMhelix 21 43\noutside 44 417\nTMhelix 418 440\ninside 441 445

34439 GCF_001518755.1_ASM151875v1 Bacillus sp. AM 13(2015) Terrabacteria group; Firmicutes; Bacilli; Bacillales; Bacillaceae; Bacillus MKKQTHRSKRRKNLSILLTSIMMISLILPAVPTHAAIENGDAVKVSILATTDVHANMMNY WP_059377708.1 2,3-cyclic-nucleotide 2-phosphodiesterase [Bacillus sp. AM 13(2015)]Length: 1430\nNumber of predicted TMHs: 2\nExp number of AAs in TMHs: 44.28365\nExp number, first 60 AAs: 21.72523\nTotal prob of N-in: 0.99853\nPOSSIBLE N-term signal sequence\ninside 1 12\nTMhelix 13 35\noutside 36 1396\nTMhelix 1397 1419\ninside 1420 1430

34440 GCF_001420655.1_ASM142065v1 Bacillus sp. FJAT-21955 Terrabacteria group; Firmicutes; Bacilli; Bacillales; Bacillaceae; Bacillus MKKQAHRSKRRKNLSILLTSMMLISLILPAVPTHAETESGSAVKVSILATTDVHANMMNY WP_045034474.1 MULTISPECIES: 2,3-cyclic-nucleotide 2-phosphodiesterase [Bacillus] Length: 1426\nNumber of predicted TMHs: 2\nExp number of AAs in TMHs: 44.15856\nExp number, first 60 AAs: 21.58314\nTotal prob of N-in: 0.99584\nPOSSIBLE N-term signal sequence\ninside 1 12\nTMhelix 13 35\noutside 36 1392\nTMhelix 1393 1415\ninside 1416 1426

34441 GCF_001439625.1_ASM143962v1 Bacillus sp. FJAT-25496 Terrabacteria group; Firmicutes; Bacilli; Bacillales; Bacillaceae; Bacillus MRTISRRKLIKYSSVLLVSFVFLQLGINVNTISAKSTYTYNSKEFHSVINKLDMQGHADH WP_057772245.1 hypothetical protein [Bacillus sp. FJAT-25496] Length: 170\nNumber of predicted TMHs: 2\nExp number of AAs in TMHs: 44.41077\nExp number, first 60 AAs: 21.95605\nTotal prob of N-in: 0.99849\nPOSSIBLE N-term signal sequence\ninside 1 11\nTMhelix 12 34\noutside 35 109\nTMhelix 110 132\ninside 133 170

34442 GCF_001482475.1_ASM148247v1 Bacillus sp. G1(2015b) Terrabacteria group; Firmicutes; Bacilli; Bacillales; Bacillaceae; Bacillus MKKQTHRSKRRKNLSILLTSIMMISLILPAVPTHAAIENGDAVKVSILATTDVHANMMNY WP_058837479.1 2,3-cyclic-nucleotide 2-phosphodiesterase [Bacillus sp. G1(2015b)] Length: 1430\nNumber of predicted TMHs: 2\nExp number of AAs in TMHs: 44.35236\nExp number, first 60 AAs: 21.72523\nTotal prob of N-in: 0.99853\nPOSSIBLE N-term signal sequence\ninside 1 12\nTMhelix 13 35\noutside 36 1395\nTMhelix 1396 1418\ninside 1419 1430

34443 GCF_001974945.1_ASM197494v1 Bacillus sp. I-2 Terrabacteria group; Firmicutes; Bacilli; Bacillales; Bacillaceae; Bacillus MKKQAHRSKRRKNLSILLTSIMMISLILPAVPTHAAIENGDAVKVSILATTDVHANMMNY WP_076839111.1 2,3-cyclic-nucleotide 2-phosphodiesterase [Bacillus sp. I-2] Length: 1428\nNumber of predicted TMHs: 2\nExp number of AAs in TMHs: 44.33928\nExp number, first 60 AAs: 21.7164\nTotal prob of N-in: 0.99812\nPOSSIBLE N-term signal sequence\ninside 1 12\nTMhelix 13 35\noutside 36 1393\nTMhelix 1394 1416\ninside 1417 1428

34444 GCF_000986655.1_ASM98665v1 Bacillus sp. L_1B0_12 Terrabacteria group; Firmicutes; Bacilli; Bacillales; Bacillaceae; Bacillus MKKQAHRSKRRKNLSILLTSIMLISLILPAVPTHAETESGSAVKVSILATTDVHANMMNY WP_046527355.1 2,3-cyclic-nucleotide 2-phosphodiesterase [Bacillus sp. L_1B0_12] Length: 1426\nNumber of predicted TMHs: 2\nExp number of AAs in TMHs: 44.31901\nExp number, first 60 AAs: 21.62366\nTotal prob of N-in: 0.99772\nPOSSIBLE N-term signal sequence\ninside 1 12\nTMhelix 13 35\noutside 36 1392\nTMhelix 1393 1415\ninside 1416 1426

34445 GCF_001426125.1_Leaf49 Bacillus sp. Leaf49 Terrabacteria group; Firmicutes; Bacilli; Bacillales; Bacillaceae; Bacillus MKKQAHRSKRRKNLSILLTSIMLISLILPAVPTHAETESGSAVKVSILATTDVHANMMNY WP_056703320.1 2,3-cyclic-nucleotide 2-phosphodiesterase [Bacillus sp. Leaf49] Length: 1426\nNumber of predicted TMHs: 2\nExp number of AAs in TMHs: 44.33503\nExp number, first 60 AAs: 21.62365\nTotal prob of N-in: 0.99772\nPOSSIBLE N-term signal sequence\ninside 1 12\nTMhelix 13 35\noutside 36 1392\nTMhelix 1393 1415\ninside 1416 1426

34446 GCF_001043795.1_ASM104379v1 Bacillus sp. LK10 Terrabacteria group; Firmicutes; Bacilli; Bacillales; Bacillaceae; Bacillus MKKQAHRSKRRKNLSILLTSIMLISLILPAVPTHAETESSSAVKVSILATTDVHANMMNY WP_048002088.1 MULTISPECIES: 2,3-cyclic-nucleotide 2-phosphodiesterase [Bacillus] Length: 1426\nNumber of predicted TMHs: 2\nExp number of AAs in TMHs: 44.32494\nExp number, first 60 AAs: 21.61406\nTotal prob of N-in: 0.99744\nPOSSIBLE N-term signal sequence\ninside 1 12\nTMhelix 13 35\noutside 36 1392\nTMhelix 1393 1415\ninside 1416 1426

34447 GCF_000264255.1_ASM26425v1 Bacillus sp. M 2-6 Terrabacteria group; Firmicutes; Bacilli; Bacillales; Bacillaceae; Bacillus MKKQAHRSKRRKNLSILLTSMMLISLILPAVPTHAETESGSAVKVSILATTDVHANMMNY WP_008341758.1 2,3-cyclic-nucleotide 2-phosphodiesterase [Bacillus sp. M 2-6] Length: 1426\nNumber of predicted TMHs: 2\nExp number of AAs in TMHs: 44.13563\nExp number, first 60 AAs: 21.58314\nTotal prob of N-in: 0.99584\nPOSSIBLE N-term signal sequence\ninside 1 12\nTMhelix 13 35\noutside 36 1392\nTMhelix 1393 1415\ninside 1416 1426

34448 GCF_001429445.1_Root920 Bacillus sp. Root920 Terrabacteria group; Firmicutes; Bacilli; Bacillales; Bacillaceae; Bacillus MKKQAHRSKRRKNLSILLTSIMMISLILPAVPTHAAIENGDAVKVSILATTDVHANMMNY WP_056767698.1 2,3-cyclic-nucleotide 2-phosphodiesterase [Bacillus sp. Root920] Length: 1430\nNumber of predicted TMHs: 2\nExp number of AAs in TMHs: 44.27688\nExp number, first 60 AAs: 21.7164\nTotal prob of N-in: 0.99812\nPOSSIBLE N-term signal sequence\ninside 1 12\nTMhelix 13 35\noutside 36 1396\nTMhelix 1397 1419\ninside 1420 1430

34449 GCF_900156275.1_IMG-taxon_2681812863_annotated_assembly Bacillus sp. RRD69 Terrabacteria group; Firmicutes; Bacilli; Bacillales; Bacillaceae; Bacillus MKKQAHRSKRRKNLSILLTSIMLISLILPAVPTHAETESGSAVKVSILATTDVHANMMNY WP_076822846.1 2,3-cyclic-nucleotide 2-phosphodiesterase [Bacillus sp. RRD69] Length: 1426\nNumber of predicted TMHs: 2\nExp number of AAs in TMHs: 44.3342\nExp number, first 60 AAs: 21.62366\nTotal prob of N-in: 0.99772\nPOSSIBLE N-term signal sequence\ninside 1 12\nTMhelix 13 35\noutside 36 1392\nTMhelix 1393 1415\ninside 1416 1426

34450 GCF_900119345.1_IMG-taxon_2606217744_annotated_assembly Bacillus sp. ru9509.4 Terrabacteria group; Firmicutes; Bacilli; Bacillales; Bacillaceae; Bacillus MKKQAHRSKRRKNLSILLTSIMLISLILPAVPTHAETESGSAVKVSILATTDVHANMMNY WP_072367755.1 2,3-cyclic-nucleotide 2-phosphodiesterase [Bacillus sp. ru9509.4] Length: 1426\nNumber of predicted TMHs: 2\nExp number of AAs in TMHs: 44.19855\nExp number, first 60 AAs: 21.62366\nTotal prob of N-in: 0.99772\nPOSSIBLE N-term signal sequence\ninside 1 12\nTMhelix 13 35\noutside 36 1392\nTMhelix 1393 1415\ninside 1416 1426

34451 GCF_001444735.1_ASM144473v1 Bacillus sp. TH007 Terrabacteria group; Firmicutes; Bacilli; Bacillales; Bacillaceae; Bacillus MKKQAHRSKRRKNLSILLTSMMLISLILPAVPTHAETESGSAVKVSILATTDVHANMMNY WP_058336529.1 2,3-cyclic-nucleotide 2-phosphodiesterase [Bacillus sp. TH007] Length: 1426\nNumber of predicted TMHs: 2\nExp number of AAs in TMHs: 44.279\nExp number, first 60 AAs: 21.58314\nTotal prob of N-in: 0.99584\nPOSSIBLE N-term signal sequence\ninside 1 12\nTMhelix 13 35\noutside 36 1392\nTMhelix 1393 1415\ninside 1416 1426

34452 GCF_000701305.1_ASM70130v1 Bacillus sp. UNC125MFCrub1.1 Terrabacteria group; Firmicutes; Bacilli; Bacillales; Bacillaceae; Bacillus MKKQAHRSKRRKNLSILLTSMMLISLILPAVPTYAETESGSAVKVSILATTDVHANMMNY WP_035391349.1 2,3-cyclic-nucleotide 2-phosphodiesterase [Bacillus sp. UNC125MFCrub1.1] Length: 1426\nNumber of predicted TMHs: 2\nExp number of AAs in TMHs: 45.15201\nExp number, first 60 AAs: 22.44158\nTotal prob of N-in: 0.99878\nPOSSIBLE N-term signal sequence\ninside 1 12\nTMhelix 13 35\noutside 36 1392\nTMhelix 1393 1415\ninside 1416 1426

34453 GCF_000800825.1_ASM80082v1 Bacillus sp. WP8 Terrabacteria group; Firmicutes; Bacilli; Bacillales; Bacillaceae; Bacillus MKKQTHRSKRRKNLSILLTSIMMISLILPAVPTHAAIENGDAVKVSILATTDVHANMMNY WP_039177137.1 2,3-cyclic-nucleotide 2-phosphodiesterase [Bacillus sp. WP8] Length: 1430\nNumber of predicted TMHs: 2\nExp number of AAs in TMHs: 44.28789\nExp number, first 60 AAs: 21.72523\nTotal prob of N-in: 0.99853\nPOSSIBLE N-term signal sequence\ninside 1 12\nTMhelix 13 35\noutside 36 1396\nTMhelix 1397 1419\ninside 1420 1430

34454 GCF_001857925.1_ASM185792v1 Bacillus xiamenensis Terrabacteria group; Firmicutes; Bacilli; Bacillales; Bacillaceae; Bacillus MKKQAHRSKRRKNLSILLTSMMMISLILPAVPTHAETESGSAVKVSILATTDVHANMMNY WP_071167907.1 2,3-cyclic-nucleotide 2-phosphodiesterase [Bacillus xiamenensis] Length: 1422\nNumber of predicted TMHs: 2\nExp number of AAs in TMHs: 44.17744\nExp number, first 60 AAs: 21.55676\nTotal prob of N-in: 0.99473\nPOSSIBLE N-term signal sequence\ninside 1 12\nTMhelix 13 35\noutside 36 1388\nTMhelix 1389 1411\ninside 1412 1422

34455 GCF_000300535.1_ASM30053v1 Bacillus xiamenensis Terrabacteria group; Firmicutes; Bacilli; Bacillales; Bacillaceae; Bacillus MKKQAHRSKRRKNLSILLTSMMMISLILPAVPTHAETESGSAVKVSILATTDVHANMMNY WP_008357702.1 2,3-cyclic-nucleotide 2-phosphodiesterase [Bacillus xiamenensis] Length: 1422\nNumber of predicted TMHs: 2\nExp number of AAs in TMHs: 44.17744\nExp number, first 60 AAs: 21.55676\nTotal prob of N-in: 0.99473\nPOSSIBLE N-term signal sequence\ninside 1 12\nTMhelix 13 35\noutside 36 1388\nTMhelix 1389 1411\ninside 1412 1422

34456 GCF_000715205.1_SOAPdenovo_v1.05 Bacillus zhangzhouensis Terrabacteria group; Firmicutes; Bacilli; Bacillales; Bacillaceae; Bacillus MKKQAHRSKRRKNLSILLTSIMMISLILPAVPTHAETENGKAVKVSILATTDVHANMMNY WP_034323498.1 2,3-cyclic-nucleotide 2-phosphodiesterase [Bacillus zhangzhouensis] Length: 1426\nNumber of predicted TMHs: 2\nExp number of AAs in TMHs: 44.26725\nExp number, first 60 AAs: 21.69267\nTotal prob of N-in: 0.99788\nPOSSIBLE N-term signal sequence\ninside 1 12\nTMhelix 13 35\noutside 36 1392\nTMhelix 1393 1415\ninside 1416 1426

34457 GCF_000831655.1_ASM83165v1 Bacillus aerophilus Terrabacteria group; Firmicutes; Bacilli; Bacillales; Bacillaceae; Bacillus; Bacillus altitudinis complex MKKQAHRSKRRKNLSILLTSIMLISLILPAVPTHAETESGSAVKVSILATTDVHANMRNY WP_041506603.1 2,3-cyclic-nucleotide 2-phosphodiesterase [Bacillus aerophilus] Length: 1426\nNumber of predicted TMHs: 2\nExp number of AAs in TMHs: 44.17567\nExp number, first 60 AAs: 21.6235\nTotal prob of N-in: 0.99772\nPOSSIBLE N-term signal sequence\ninside 1 12\nTMhelix 13 35\noutside 36 1392\nTMhelix 1393 1415\ninside 1416 1426

34458 GCF_000789425.2_ASM78942v2 Bacillus altitudinis Terrabacteria group; Firmicutes; Bacilli; Bacillales; Bacillaceae; Bacillus; Bacillus altitudinis complex MKKQAHRSKRRKNLSILLTSIMLISLILPAVPTHAETESGSAVKVSILATTDVHANMMNY WP_039168035.1 2,3-cyclic-nucleotide 2-phosphodiesterase [Bacillus altitudinis] Length: 1426\nNumber of predicted TMHs: 2\nExp number of AAs in TMHs: 44.33504\nExp number, first 60 AAs: 21.62365\nTotal prob of N-in: 0.99772\nPOSSIBLE N-term signal sequence\ninside 1 12\nTMhelix 13 35\noutside 36 1392\nTMhelix 1393 1415\ninside 1416 1426

34459 GCF_001029865.1_RIT380_V1 Bacillus altitudinis Terrabacteria group; Firmicutes; Bacilli; Bacillales; Bacillaceae; Bacillus; Bacillus altitudinis complex MKKQAHRSKRRKNLSILLTSIMLISLILPAVPTHAETESGSAVKVSILATTDVHANMMNY WP_047946510.1 2,3-cyclic-nucleotide 2-phosphodiesterase [Bacillus altitudinis] Length: 1426\nNumber of predicted TMHs: 2\nExp number of AAs in TMHs: 44.33421\nExp number, first 60 AAs: 21.62366\nTotal prob of N-in: 0.99772\nPOSSIBLE N-term signal sequence\ninside 1 12\nTMhelix 13 35\noutside 36 1392\nTMhelix 1393 1415\ninside 1416 1426

34460 GCF_002042895.1_ASM204289v1 Bacillus altitudinis Terrabacteria group; Firmicutes; Bacilli; Bacillales; Bacillaceae; Bacillus; Bacillus altitudinis complex MKKQAHRSKRRKNLSILLTSIMLISLILPAVPTHAETESGSAVKVSILATTDVHANMMNY WP_079919181.1 2,3-cyclic-nucleotide 2-phosphodiesterase [Bacillus altitudinis] Length: 1426\nNumber of predicted TMHs: 2\nExp number of AAs in TMHs: 44.31905\nExp number, first 60 AAs: 21.62366\nTotal prob of N-in: 0.99772\nPOSSIBLE N-term signal sequence\ninside 1 12\nTMhelix 13 35\noutside 36 1392\nTMhelix 1393 1415\ninside 1416 1426

34461 GCF_000949525.1_ASM94952v1 Bacillus altitudinis Terrabacteria group; Firmicutes; Bacilli; Bacillales; Bacillaceae; Bacillus; Bacillus altitudinis complex MKKQAHRSKRRKNLSILLTSMMLISLILPAVPTHAETESGSAVKVSILATTDVHANMMNY WP_045034474.1 MULTISPECIES: 2,3-cyclic-nucleotide 2-phosphodiesterase [Bacillus] Length: 1426\nNumber of predicted TMHs:

2\nExp number of AAs in TMHs: 44.15856\nExp number, first 60 AAs:  21.58314\nTotal prob of N-in:   0.99584\nPOSSIBLE N-term signal sequence\ninside   1   12\nTMhelix   13   35\noutside   36  1392\nTMhelix   1393  1415\ninside   1416  1426

34462 GCF_001896335.1_ASM189633v1 Bacillus altitudinis Terrabacteria group; Firmicutes; Bacilli; Bacillales; Bacillaceae; Bacillus; Bacillus altitudinis complex
MKKQAHRSKRRKNLSILLTSMMLISLILPAVPTHAETESGSAVKVSILATTDVHANMMNY WP_073415363.1 2,3-cyclic-nucleotide 2-phosphodiesterase [Bacillus altitudinis] Length: 1426\nNumber of predicted TMHs:  2\nExp number of AAs in TMHs: 44.13812\nExp number, first 60 AAs:  21.58314\nTotal prob of N-in:   0.99584\nPOSSIBLE N-term signal sequence\ninside   1   12\nTMhelix   13   35\noutside   36  1392\nTMhelix   1393  1415\ninside   1416  1426

34463 GCF_001457015.1_ASM145701v1 Bacillus cellulasensis Terrabacteria group; Firmicutes; Bacilli; Bacillales; Bacillaceae; Bacillus; Bacillus altitudinis complex
MKKQAHRSKRRKNLSILLTSMMLISLILPAVPTHAETESGSAVKVSILATTDVHANMMNY WP_058214343.1 2,3-cyclic-nucleotide 2-phosphodiesterase [Bacillus cellulasensis] Length: 1426\nNumber of predicted TMHs:  2\nExp number of AAs in TMHs: 44.15851\nExp number, first 60 AAs:  21.58314\nTotal prob of N-in:   0.99584\nPOSSIBLE N-term signal sequence\ninside   1   12\nTMhelix   13   35\noutside   36  1392\nTMhelix   1393  1415\ninside   1416  1426

34464 GCF_001038775.1_ASM103877v1 Bacillus stratosphericus Terrabacteria group; Firmicutes; Bacilli; Bacillales; Bacillaceae; Bacillus; Bacillus altitudinis complex
MKKQAHRSKRRKNLSILLTSIMLISLILPAVPTHAETESSSAVKVSILATTDVHANMMNY WP_048002088.1 MULTISPECIES: 2,3-cyclic-nucleotide 2-phosphodiesterase [Bacillus] Length: 1426\nNumber of predicted TMHs:  2\nExp number of AAs in TMHs: 44.32494\nExp number, first 60 AAs:  21.61406\nTotal prob of N-in:   0.99744\nPOSSIBLE N-term signal sequence\ninside   1   12\nTMhelix   13   35\noutside   36  1392\nTMhelix   1393  1415\ninside   1416  1426

34465 GCF_001038915.1_ASM103891v1 Bacillus stratosphericus Terrabacteria group; Firmicutes; Bacilli; Bacillales; Bacillaceae; Bacillus; Bacillus altitudinis complex
MKKQAHRSKRRKNLSILLTSIMLISLILPAVPTHAETESSSAVKVSILATTDVHANMMNY WP_048002088.1 MULTISPECIES: 2,3-cyclic-nucleotide 2-phosphodiesterase [Bacillus] Length: 1426\nNumber of predicted TMHs:  2\nExp number of AAs in TMHs: 44.32494\nExp number, first 60 AAs:  21.61406\nTotal prob of N-in:   0.99744\nPOSSIBLE N-term signal sequence\ninside   1   12\nTMhelix   13   35\noutside   36  1392\nTMhelix   1393  1415\ninside   1416  1426

34466 GCF_001043535.1_ASM104353v1 Bacillus stratosphericus Terrabacteria group; Firmicutes; Bacilli; Bacillales; Bacillaceae; Bacillus; Bacillus altitudinis complex
MKKQAHRSKRRKNLSILLTSIMLISLILPAVPTHAETESSSAVKVSILATTDVHANMMNY WP_048002088.1 MULTISPECIES: 2,3-cyclic-nucleotide 2-phosphodiesterase [Bacillus] Length: 1426\nNumber of predicted TMHs:  2\nExp number of AAs in TMHs: 44.32494\nExp number, first 60 AAs:  21.61406\nTotal prob of N-in:   0.99744\nPOSSIBLE N-term signal sequence\ninside   1   12\nTMhelix   13   35\noutside   36  1392\nTMhelix   1393  1415\ninside   1416  1426

34467 GCF_002077245.1_ASM207724v1 Bacillus stratosphericus Terrabacteria group; Firmicutes; Bacilli; Bacillales; Bacillaceae; Bacillus; Bacillus altitudinis complex
MKKQAHRSKRRKNLSILLTSIMLISLILPAVPTHAETESGSAVKVSILATTDVHANMMNY WP_081114445.1 2,3-cyclic-nucleotide 2-phosphodiesterase [Bacillus stratosphericus] Length: 1426\nNumber of predicted TMHs:  2\nExp number of AAs in TMHs: 44.3342\nExp number, first 60 AAs:  21.62366\nTotal prob of N-in:   0.99772\nPOSSIBLE N-term signal sequence\ninside   1   12\nTMhelix   13   35\noutside   36  1392\nTMhelix   1393  1415\ninside   1416  1426

34468 GCF_001038845.1_ASM103884v1 Bacillus stratosphericus Terrabacteria group; Firmicutes; Bacilli; Bacillales; Bacillaceae; Bacillus; Bacillus altitudinis complex
MKKQAHRSKRRKNLSILLTSMMLISLILPAVPTHAETESGSAVKVSILATTDVHANMMNY WP_048002721.1 2,3-cyclic-nucleotide 2-phosphodiesterase [Bacillus stratosphericus] Length: 1426\nNumber of predicted TMHs:  2\nExp number of AAs in TMHs: 44.27902\nExp number, first 60 AAs:  21.58314\nTotal prob of N-in:   0.99584\nPOSSIBLE N-term signal sequence\ninside   1   12\nTMhelix   13   35\noutside   36  1392\nTMhelix   1393  1415\ninside   1416  1426

34469 GCF_001039015.1_ASM103901v1 Bacillus stratosphericus Terrabacteria group; Firmicutes; Bacilli; Bacillales; Bacillaceae; Bacillus; Bacillus altitudinis complex
MKKQAHRSKRRKNLSILLTSMMLISLILPAVPTHAETESGSAVKVSILATTDVHANMMNY WP_048002721.1 2,3-cyclic-nucleotide 2-phosphodiesterase [Bacillus stratosphericus] Length: 1426\nNumber of predicted TMHs:  2\nExp number of AAs in TMHs: 44.27902\nExp number, first 60 AAs:  21.58314\nTotal prob of N-in:   0.99584\nPOSSIBLE N-term signal sequence\ninside   1   12\nTMhelix   13   35\noutside   36  1392\nTMhelix   1393  1415\ninside   1416  1426

34470 GCF_001265125.1_ASM126512v1 Bacillus stratosphericus Terrabacteria group; Firmicutes; Bacilli; Bacillales; Bacillaceae; Bacillus; Bacillus altitudinis complex

MKKQAHRSKRRKNLSILLTSIMLISLILPAVPTHAETESGSAVKVSILATTDVHANMMNY WP_052321015.1 2,3-cyclic-nucleotide 2-phosphodiesterase [Bacillus stratosphericus] Length: 1426\nNumber of predicted TMHs: 2\nExp number of AAs in TMHs: 44.33433\nExp number, first 60 AAs: 21.62366\nTotal prob of N-in: 0.99772\nPOSSIBLE N-term signal sequence\ninside 1 12\nTMhelix 13 35\noutside 36 1392\nTMhelix 1393 1415\ninside 1416 1426

34471 GCF_000691145.1_ASM69114v1 Bacillus altitudinis 41KF2b Terrabacteria group; Firmicutes; Bacilli; Bacillales; Bacillaceae; Bacillus; Bacillus altitudinis complex; Bacillus altitudinis MKKQAHRSKRRKNLSILLTSMMLISLILPAVPTHAETESGSAVKVSILATTDVHANMMNY WP_035703674.1 2,3-cyclic-nucleotide 2-phosphodiesterase [Bacillus altitudinis] Length: 1426\nNumber of predicted TMHs: 2\nExp number of AAs in TMHs: 44.1588\nExp number, first 60 AAs: 21.58314\nTotal prob of N-in: 0.99584\nPOSSIBLE N-term signal sequence\ninside 1 12\nTMhelix 13 35\noutside 36 1392\nTMhelix 1393 1415\ninside 1416 1426

34472 GCF_000353245.1_BStr_LAMA585_V1 Bacillus stratosphericus LAMA 585 Terrabacteria group; Firmicutes; Bacilli; Bacillales; Bacillaceae; Bacillus; Bacillus altitudinis complex; Bacillus stratosphericus MKKQAHRSKRRKNLSILLTSMMLISLILPAVPTYAETESGSTVKVSILATTDVHANMMNY WP_039962570.1 2,3-cyclic-nucleotide 2-phosphodiesterase [Bacillus stratosphericus] Length: 1426\nNumber of predicted TMHs: 2\nExp number of AAs in TMHs: 45.13845\nExp number, first 60 AAs: 22.43047\nTotal prob of N-in: 0.99873\nPOSSIBLE N-term signal sequence\ninside 1 12\nTMhelix 13 35\noutside 36 1392\nTMhelix 1393 1415\ninside 1416 1426

34473 GCF_002158195.1_ASM215819v1 Bacillus pseudomycoides Terrabacteria group; Firmicutes; Bacilli; Bacillales; Bacillaceae; Bacillus; Bacillus cereus group MKNKKRRATVALTTGLALTAVTPYGVGHAEETDQLQVQIQEDSFRTGELTQPSQKAPENV WP_088094647.1 peptidase M4 [Bacillus pseudomycoides] Length: 818\nNumber of predicted TMHs: 1\nExp number of AAs in TMHs: 27.75035\nExp number, first 60 AAs: 8.24655\nTotal prob of N-in: 0.39630\noutside 1 787\nTMhelix 788 810\ninside 811 818

34474 GCF_000293685.1_Baci_cere_HD73_V1 Bacillus cereus HD73 Terrabacteria group; Firmicutes; Bacilli; Bacillales; Bacillaceae; Bacillus; Bacillus cereus group; Bacillus cereus MVNTKQRVAKRRKALNEIQNLVGNASNLLIIHYSCESFYDIKDGRTPRITSIAVRFFNTG WP_000248334.1 MULTISPECIES: hypothetical protein [Bacillus cereus group] Length: 262\nNumber of predicted TMHs: 1\nExp number of AAs in TMHs: 18.81871\nExp number, first 60 AAs: 0.0768900000000001\nTotal prob of N-in: 0.93351\ninside 1 240\nTMhelix 241 258\noutside 259 262

34475 GCF_000399485.1_Baci_cere_VD136_V1 Bacillus cereus VD136 Terrabacteria group; Firmicutes; Bacilli; Bacillales; Bacillaceae; Bacillus; Bacillus cereus group; Bacillus cereus MKNKKRRATVALTTGLALTAVTPYGVGHAEETDQLQVQIQEDSFRTGELTQPSQKAPENV WP_016115629.1 LPXTG-domain-containing protein cell wall anchor domain [Bacillus cereus] Length: 818\nNumber of predicted TMHs: 1\nExp number of AAs in TMHs: 27.17531\nExp number, first 60 AAs: 8.25431\nTotal prob of N-in: 0.39573\noutside 1 792\nTMhelix 793 810\ninside 811 818

34476 GCF_000290975.2_Baci_cere_VD142_V2 Bacillus cereus VD142 Terrabacteria group; Firmicutes; Bacilli; Bacillales; Bacillaceae; Bacillus; Bacillus cereus group; Bacillus cereus MVNTKQRVAKRRKALNDIQNLVGNASNLLIIHYSCESFYDIKDGRTPRITSIAVRFFNTG WP_002198638.1 hypothetical protein [Bacillus cereus] Length: 262\nNumber of predicted TMHs: 1\nExp number of AAs in TMHs: 18.96033\nExp number, first 60 AAs: 0.18244\nTotal prob of N-in: 0.86909\ninside 1 240\nTMhelix 241 258\noutside 259 262

34477 GCF_000399585.1_Baci_cere_VDM006_V1 Bacillus cereus VDM006 Terrabacteria group; Firmicutes; Bacilli; Bacillales; Bacillaceae; Bacillus; Bacillus cereus group; Bacillus cereus MKNKKRRATVALTTGLALTAVTPYGVGHAEETDQLQVQIQEDSFRTGELTQPSQKAPENV WP_016115629.1 LPXTG-domain-containing protein cell wall anchor domain [Bacillus cereus] Length: 818\nNumber of predicted TMHs: 1\nExp number of AAs in TMHs: 27.17531\nExp number, first 60 AAs: 8.25431\nTotal prob of N-in: 0.39573\noutside 1 792\nTMhelix 793 810\ninside 811 818

34478 GCF_000399605.1_Baci_cere_VDM021_V1 Bacillus cereus VDM021 Terrabacteria group; Firmicutes; Bacilli; Bacillales; Bacillaceae; Bacillus; Bacillus cereus group; Bacillus cereus MKNKKRRATVALTTGLALTAVTPYGVGHAEETDQLQVQIQEDSFRTGELTQPSQKAPENV WP_016133109.1 LPXTG-domain-containing protein cell wall anchor domain [Bacillus cereus] Length: 818\nNumber of predicted TMHs: 1\nExp number of AAs in TMHs: 27.75035\nExp number, first 60 AAs: 8.24655\nTotal prob of N-in: 0.39630\noutside 1 787\nTMhelix 788 810\ninside 811 818

34479 GCF_002146845.1_ASM214684v1 Bacillus thuringiensis serovar alesti Terrabacteria group; Firmicutes; Bacilli; Bacillales; Bacillaceae; Bacillus; Bacillus cereus group; Bacillus thuringiensis MVNTKQRVAKRRKALNEIQNLVGNASNLLIIHYSCESFYDIKDGRTPRITSIAVRFFNTG WP_000248334.1 MULTISPECIES: hypothetical protein [Bacillus cereus group] Length: 262\nNumber of predicted TMHs: 1\nExp number of AAs in TMHs: 18.81871\nExp number, first 60 AAs: 0.0768900000000001\nTotal prob of N-in: 0.93351\ninside 1 240\nTMhelix 241 258\noutside 259 262

34480 GCF_001640965.1_ASM164096v1 Bacillus thuringiensis serovar alesti Terrabacteria group; Firmicutes; Bacilli; Bacillales; Bacillaceae; Bacillus; Bacillus cereus group; Bacillus thuringiensis MVNTKQRVAKRRKALNEIQNLVGNASNLLIIHYSCESFYDIKDGRTPRITSIAVRFFNTG WP_000248334.1 MULTISPECIES: hypothetical protein [Bacillus cereus group] Length: 262\nNumber of predicted TMHs: 1\nExp number of AAs in TMHs: 18.81871\nExp number, first 60 AAs: 0.0768900000000001\nTotal prob of N-in: 0.93351\ninside 1 240\nTMhelix 241 258\noutside 259 262

34481 GCF_000338755.1_ASM33875v1 Bacillus thuringiensis serovar kurstaki str. HD73 Terrabacteria group; Firmicutes; Bacilli; Bacillales; Bacillaceae; Bacillus; Bacillus cereus group; Bacillus thuringiensis; Bacillus thuringiensis serovar kurstaki MVNTKQRVAKRRKALNEIQNLVGNASNLLIIHYSCESFYDIKDGRTPRITSIAVRFFNTG WP_000248334.1 MULTISPECIES: hypothetical protein [Bacillus cereus group] Length: 262\nNumber of predicted TMHs: 1\nExp number of AAs in TMHs: 18.81871\nExp number, first 60 AAs: 0.0768900000000001\nTotal prob of N-in: 0.93351\ninside 1 240\nTMhelix 241 258\noutside 259 262

34482 GCF_000172815.1_ASM17281v1 Bacillus pumilus ATCC 7061 Terrabacteria group; Firmicutes; Bacilli; Bacillales; Bacillaceae; Bacillus; Bacillus pumilus MKKQAHRSKRRKNLSILLTSIMMISLILPAVPTHAETENGKAVKVSILATTDVHANMMNY WP_003214062.1 2,3-cyclic-nucleotide 2-phosphodiesterase [Bacillus pumilus] Length: 1429\nNumber of predicted TMHs: 2\nExp number of AAs in TMHs: 44.26687\nExp number, first 60 AAs: 21.69267\nTotal prob of N-in: 0.99788\nPOSSIBLE N-term signal sequence\ninside 1 12\nTMhelix 13 35\noutside 36 1395\nTMhelix 1396 1418\ninside 1419 1429

34483 GCF_000299555.1_BA06_V1.0 Bacillus pumilus BA06 Terrabacteria group; Firmicutes; Bacilli; Bacillales; Bacillaceae; Bacillus; Bacillus pumilus MKKQAHRSKRRKNLSILLTSMMLISLILPAVPTYAETESGSAVKVSILATTDVHANMMNY WP_017366822.1 2,3-cyclic-nucleotide 2-phosphodiesterase [Bacillus pumilus] Length: 1426\nNumber of predicted TMHs: 2\nExp number of AAs in TMHs: 45.13672\nExp number, first 60 AAs: 22.44158\nTotal prob of N-in: 0.99878\nPOSSIBLE N-term signal sequence\ninside 1 12\nTMhelix 13 35\noutside 36 1392\nTMhelix 1393 1415\ninside 1416 1426

34484 GCF_000299555.1_BA06_V1.0 Bacillus pumilus BA06 Terrabacteria group; Firmicutes; Bacilli; Bacillales; Bacillaceae; Bacillus; Bacillus pumilus MVNTRQRLNKRRKTLNEIQNLVNNAANLLIIHYSCESFYHIKDGRTPRITSIAVRYFHTG WP_017366477.1 hypothetical protein [Bacillus pumilus] Length: 266\nNumber of predicted TMHs: 1\nExp number of AAs in TMHs: 23.64345\nExp number, first 60 AAs: 0.80473\nTotal prob of N-in: 0.85783\ninside 1 240\nTMhelix 241 263\noutside 264 266

34485 GCF_000444805.1_B._pumilus Bacillus pumilus CCMA-560 Terrabacteria group; Firmicutes; Bacilli; Bacillales; Bacillaceae; Bacillus; Bacillus pumilus MKKQAHRSKRRKNLSILLTSIMMISLILPAVPTHAAIENGDAVKVSILATTDVHANMMNY WP_024425852.1 2,3-cyclic-nucleotide 2-phosphodiesterase [Bacillus pumilus] Length: 1430\nNumber of predicted TMHs: 2\nExp number of AAs in TMHs: 44.27506\nExp number, first 60 AAs: 21.7164\nTotal prob of N-in: 0.99812\nPOSSIBLE N-term signal sequence\ninside 1 12\nTMhelix 13 35\noutside 36 1396\nTMhelix 1397 1419\ninside 1420 1430

34486 GCF_000508145.1_INR7_01 Bacillus pumilus INR7 Terrabacteria group; Firmicutes; Bacilli; Bacillales; Bacillaceae; Bacillus; Bacillus pumilus MKKQAHRSKRRKNLSILLTSIMLISLILPAVPTHAETESGSAVKVSILATTDVHANMMNY WP_024720241.1 2,3-cyclic-nucleotide 2-phosphodiesterase [Bacillus pumilus] Length: 1426\nNumber of predicted TMHs: 2\nExp number of AAs in TMHs: 44.20037\nExp number, first 60 AAs: 21.62364\nTotal prob of N-in: 0.99772\nPOSSIBLE N-term signal sequence\ninside 1 12\nTMhelix 13 35\noutside 36 1392\nTMhelix 1393 1415\ninside 1416 1426

34487 GCF_000225935.1_ASM22593v1 Bacillus pumilus S-1 Terrabacteria group; Firmicutes; Bacilli; Bacillales; Bacillaceae; Bacillus; Bacillus pumilus MKKQAHRSKRRKNLSILLTSIMLISLILPAVPTHAETESGSAVKVSILATTDVHANMMNY WP_029575436.1 2,3-cyclic-nucleotide 2-phosphodiesterase [Bacillus pumilus] Length: 1426\nNumber of predicted TMHs: 2\nExp number of AAs in TMHs: 44.33433\nExp number, first 60 AAs: 21.62366\nTotal prob of N-in: 0.99772\nPOSSIBLE N-term signal sequence\ninside 1 12\nTMhelix 13 35\noutside 36 1392\nTMhelix 1393 1415\ninside 1416 1426

34488 GCF_000017885.4_ASM1788v4 Bacillus pumilus SAFR-032 Terrabacteria group; Firmicutes; Bacilli; Bacillales; Bacillaceae; Bacillus; Bacillus pumilus
MKKQAHRSKRRKNLSILLTSIMMISLILPAVPIHAETESGKAVKVSILATTDVHANMMNY WP_041815277.1 2,3-cyclic-nucleotide 2-phosphodiesterase [Bacillus pumilus] Length: 1429\nNumber of predicted TMHs: 2\nExp number of AAs in TMHs: 44.89176\nExp number, first 60 AAs: 22.32206\nTotal prob of N-in: 0.99930\nPOSSIBLE N-term signal sequence\ninside 1 12\nTMhelix 13 35\noutside 36 1395\nTMhelix 1396 1418\ninside 1419 1429

34489 GCF_000691165.1_ASM69116v1 Bacillus safensis FO-36b Terrabacteria group; Firmicutes; Bacilli; Bacillales; Bacillaceae; Bacillus; Bacillus safensis
MKKQAHRSKRRKNLSILLTSIMMISLILPAVPTHAAIENGDAVKVSILATTDVHANMMNY WP_034283690.1 2,3-cyclic-nucleotide 2-phosphodiesterase [Bacillus safensis] Length: 1430\nNumber of predicted TMHs: 2\nExp number of AAs in TMHs: 44.27946\nExp number, first 60 AAs: 21.7164\nTotal prob of N-in: 0.99812\nPOSSIBLE N-term signal sequence\ninside 1 12\nTMhelix 13 35\noutside 36 1396\nTMhelix 1397 1419\ninside 1420 1430

34490 GCF_001624605.1_ASM162460v1 Geobacillus sp. 8 Terrabacteria group; Firmicutes; Bacilli; Bacillales; Bacillaceae; Geobacillus MKKRKRRSMLAGASAVLLVLVSVLTPIHTYATATVAGNAEQQMEKQTKSTDKPNQEQPLH WP_066251562.1 hypothetical protein [Geobacillus sp. 8]Length: 1822\nNumber of predicted TMHs: 2\nExp number of AAs in TMHs: 41.99153\nExp number, first 60 AAs: 21.92775\nTotal prob of N-in: 0.98227\nPOSSIBLE N-term signal sequence\ninside 1 12\nTMhelix 13 35\noutside 36 1793\nTMhelix 1794 1812\ninside 1813 1822

34491 GCF_001049895.1_Rubeoparvulum_massiliensis Rubeoparvulum massiliense Terrabacteria group; Firmicutes; Bacilli; Bacillales; Bacillaceae; Rubeoparvulum
MNVSLYLRGYIFVKDLSRRAFLKRTVAAGAAGAVTLATSKTKVFATSEKEESMGTLIDLT WP_048602062.1 oxidoreductase [Rubeoparvulum massiliense] Length: 348\nNumber of predicted TMHs: 1\nExp number of AAs in TMHs: 22.31818\nExp number, first 60 AAs: 1.65087\nTotal prob of N-in: 0.09153\noutside 1 315\nTMhelix 316 338\ninside 339 348

34492 GCF_000411915.1_ExiS17_1.0 Exiguobacterium sp. S17 Terrabacteria group; Firmicutes; Bacilli; Bacillales; Bacillales incertae sedis; Bacillales Family XII. Incertae Sedis; Exiguobacterium
MALDRRQMLQLGTLIAGGWAVSSVLKRTAPIGRDVANPEALAAIFDDRRSPASGPPTASL WP_035386738.1 MULTISPECIES: DSBA oxidoreductase [Bacteria] Length: 214\nNumber of predicted TMHs: 1\nExp number of AAs in TMHs: 21.55582\nExp number, first 60 AAs: 0.17312\nTotal prob of N-in: 0.48486\ninside 1 174\nTMhelix 175 197\noutside 198 214

34493 GCF_001187725.1_ASM118772v1 Brevibacillus reuszeri Terrabacteria group; Firmicutes; Bacilli; Bacillales; Paenibacillaceae; Brevibacillus
MLQRRNVFAVLFAFFVSSSFFFQGIPAHACSCSTPSAQEAIQQSPAIFSGKVTQVKETND WP_084766093.1 hypothetical protein [Brevibacillus reuszeri] Length: 194\nNumber of predicted TMHs: 2\nExp number of AAs in TMHs: 42.73371\nExp number, first 60 AAs: 20.76477\nTotal prob of N-in: 0.99468\nPOSSIBLE N-term signal sequence\ninside 1 6\nTMhelix 7 29\noutside 30 164\nTMhelix 165 187\ninside 188 194

34494 GCF_000829465.1_ASM82946v1 Cohnella kolymensis Terrabacteria group; Firmicutes; Bacilli; Bacillales; Paenibacillaceae; Cohnella MEQSDSKIVYVRMRRRAVVTPGSLITLGVVARLVADPDTEAKLKQLPLHLVSEKDGNLLL WP_041066153.1 stage V sporulation protein AA [Cohnella kolymensis] Length: 214\nNumber of predicted TMHs: 2\nExp number of AAs in TMHs: 50.67284\nExp number, first 60 AAs: 7.07635\nTotal prob of N-in: 0.88301\ninside 1 101\nTMhelix 102 124\noutside 125 143\nTMhelix 144 166\ninside 167 214

34495 GCF_000214295.1_ASM21429v1 Paenibacillus sp. HGF7 Terrabacteria group; Firmicutes; Bacilli; Bacillales; Paenibacillaceae; Paenibacillus
MKHNHPVRGRKQERDGDGKRDSFRIARRGLIVALLIGASLTGCSAADKGASTADQKSGVA WP_009671820.1 MULTISPECIES: DUF4349 domain-containing protein [Paenibacillus] Length: 364\nNumber of predicted TMHs: 1\nExp number of AAs in TMHs: 29.68081\nExp number, first 60 AAs: 6.12147\nTotal prob of N-in: 0.32286\noutside 1 296\nTMhelix 297 319\ninside 320 364

34496 GCF_000411255.1_Paen_sp_HGH0039_V1 Paenibacillus sp. HGH0039 Terrabacteria group; Firmicutes; Bacilli; Bacillales; Paenibacillaceae; Paenibacillus
MKHNHPVRGRKQERDGDGKRDSFRIARRGLIVALLIGASLTGCSAADKGASTADQKSGVA WP_009671820.1 MULTISPECIES: hypothetical protein [Paenibacillus] Length: 364\nNumber of predicted TMHs: 1\nExp number of AAs in TMHs: 29.68081\nExp number, first 60 AAs: 6.12147\nTotal prob of N-in: 0.32286\noutside 1 296\nTMhelix 297 319\ninside 320 364

34497 GCF_000093085.1_ASM9308v1 [Bacillus] selenitireducens MLS10 Terrabacteria group; Firmicutes; Bacilli; Bacillales; Sporolactobacillaceae; unclassified Sporolactobacillaceae; [Bacillus] selenitireducens MMNRRNFLKRASAATATAAVMTTSQKWVSASGIDDDKQIGSIIDLSKCDGCEYENTPLCV WP_013173107.1 oxidoreductase [[Bacillus] selenitireducens] Length: 337\nNumber of predicted TMHs: 1\nExp number of AAs in TMHs: 21.65097\nExp number, first 60 AAs: 0.01788\nTotal prob of N-in: 0.23533\noutside 1 298\nTMhelix 299 321\ninside 322 337

34498 GCF_000330705.1_Crm Salinicoccus carnicancri Crm Terrabacteria group; Firmicutes; Bacilli; Bacillales; Staphylococcaceae; Salinicoccus; Salinicoccus carnicancri MDKVNFTMKRRNLLTAALALALLMPQHDALATTEVPSEENASEEFTSEDNGSSQNPSTEE WP_017548268.1 hypothetical protein [Salinicoccus carnicancri] Length: 549\nNumber of predicted TMHs: 1\nExp number of AAs in TMHs: 22.91357\nExp number, first 60 AAs: 0.86101\nTotal prob of N-in: 0.04679\noutside 1 522\nTMhelix 523 545\ninside 546 549

34499 GCF_002005165.1_ASM200516v1 Novibacillus thermophilus Terrabacteria group; Firmicutes; Bacilli; Bacillales; Thermoactinomycetaceae; Novibacillus MWKRRAVITSLCMTLVIGLVVAAVPTAHAAVGFSLYTPFSGISATPGETIDYAVEVLNDT WP_077721366.1 hypothetical protein [Novibacillus thermophilus] Length: 387\nNumber of predicted TMHs: 2\nExp number of AAs in TMHs: 45.3509\nExp number, first 60 AAs: 22.72953\nTotal prob of N-in: 0.99920\nPOSSIBLE N-term signal sequence\ninside 1 6\nTMhelix 7 29\noutside 30 358\nTMhelix 359 381\ninside 382 387

34500 GCF_001742305.1_ASM174230v1 Desulfuribacillus stibiiarsenatis Terrabacteria group; Firmicutes; Bacilli; Bacillales; unclassified Bacillales; Desulfuribacillus MNAKVNRRTFLDRSIKAVGITSIAIAGLPKNIEASTTHQGKQYGTMIDLTLCDGCPHLDT WP_069701376.1 oxidoreductase [Desulfuribacillus stibiiarsenatis] Length: 337\nNumber of predicted TMHs: 1\nExp number of AAs in TMHs: 19.4641\nExp number, first 60 AAs: 0.04791\nTotal prob of N-in: 0.74671\ninside 1 305\nTMhelix 306 325\noutside 326 337

34501 GCF_000698125.1_ASM69812v1 Geomicrobium sp. JCM 19038 Terrabacteria group; Firmicutes; Bacilli; Bacillales; unclassified Bacillales; Geomicrobium MMTMILKEMKDTYRDRRTVFFTTAIPMISVFALVFFFSFFNTSDVHIGIDENIDAGLIES WP_042416226.1 hypothetical protein [Geomicrobium sp. JCM 19038]Length: 217\nNumber of predicted TMHs: 2\nExp number of AAs in TMHs: 43.94395\nExp number, first 60 AAs: 21.63641\nTotal prob of N-in: 0.99512\nPOSSIBLE N-term signal sequence\ninside 1 19\nTMhelix 20 42\noutside 43 156\nTMhelix 157 179\ninside 180 217

34502 GCF_001885915.1_ASM188591v1 Enterococcus gilvusTerrabacteria group; Firmicutes; Bacilli; Lactobacillales; Enterococcaceae; Enterococcus MKRRALFLSVLLLSLFSPLAVVFADETKKEQPLPFYVQPVLPENQLDSSVGYFYLPSTPN WP_010782219.1 hypothetical protein [Enterococcus gilvus] Length: 343\nNumber of predicted TMHs: 2\nExp number of AAs in TMHs: 41.39353\nExp number, first 60 AAs: 19.01837\nTotal prob of N-in: 0.93656\nPOSSIBLE N-term signal sequence\ninside 1 4\nTMhelix 5 24\noutside 25 311\nTMhelix 312 334\ninside 335 343

34503 GCF_001990645.1_ASM199064v1 Enterococcus mundtii Terrabacteria group; Firmicutes; Bacilli; Lactobacillales; Enterococcaceae; Enterococcus MRTRRKLLNSALILGLTLGNLNGTVVQAVTVLNNLGMTQNRPVSNKQKKDEAKSVIENTK WP_077151302.1 hypothetical protein [Enterococcus mundtii] Length: 1727\nNumber of predicted TMHs: 2\nExp number of AAs in TMHs: 40.73523\nExp number, first 60 AAs: 21.52341\nTotal prob of N-in: 0.95423\nPOSSIBLE N-term signal sequence\ninside 1 6\nTMhelix 7 29\noutside 30 1693\nTMhelix 1694 1713\ninside 1714 1727

34504 GCF_001705215.1_ASM170521v1 Enterococcus mundtii Terrabacteria group; Firmicutes; Bacilli; Lactobacillales; Enterococcaceae; Enterococcus MNQTKKFFSDCLIKGGNNLRTRRKLLNSALILGLTLGNLNGTVVQAVTVLNNLGMTQNRP WP_081308284.1 hypothetical protein [Enterococcus mundtii] Length: 1745\nNumber of predicted TMHs: 2\nExp number of AAs in TMHs: 38.99239\nExp number, first 60 AAs: 20.09499\nTotal prob of N-in: 0.89525\nPOSSIBLE N-term signal sequence\ninside 1 24\nTMhelix 25 47\noutside 48 1711\nTMhelix 1712 1731\ninside 1732 1745

34505 GCF_001601555.1_ASM160155v1 Enterococcus pernyi Terrabacteria group; Firmicutes; Bacilli; Lactobacillales; Enterococcaceae; Enterococcus MNQTKKFFSDCLIKGGNNLRTRRKLLNSALILGLTLGNLNGTVVQAVTVLNNLGMTQNRP WP_084812831.1 hypothetical protein [Enterococcus pernyi] Length: 1745\nNumber of predicted TMHs: 2\nExp number of

AAs in TMHs: 39.30765\nExp number, first 60 AAs: 20.09552\nTotal prob of N-in: 0.89527\nPOSSIBLE N-term signal sequence\ninside 1 24\nTMhelix 25 47\noutside 48 1711\nTMhelix 1712 1731\ninside 1732 1745

34506 GCF_000394615.1_Ente_gilf_BAA-350_PQ1_CCUG_45553_V1 Enterococcus gilvus ATCC BAA-350 Terrabacteria group; Firmicutes; Bacilli; Lactobacillales; Enterococcaceae; Enterococcus; Enterococcus gilvus MKRRALFLSVLLLSLFSPLAVVFADETKKEQPLPFYVQPVLPENQLDSSVGYFYLPSTPN WP_010782219.1 hypothetical protein [Enterococcus gilvus] Length: 343\nNumber of predicted TMHs: 2\nExp number of AAs in TMHs: 41.39353\nExp number, first 60 AAs: 19.01837\nTotal prob of N-in: 0.93656\nPOSSIBLE N-term signal sequence\ninside 1 4\nTMhelix 5 24\noutside 25 311\nTMhelix 312 334\ninside 335 343

34507 GCF_000407545.1_Ente_gilf_BAA-350_PQ1_CCUG_45553_V2 Enterococcus gilvus ATCC BAA-350 Terrabacteria group; Firmicutes; Bacilli; Lactobacillales; Enterococcaceae; Enterococcus; Enterococcus gilvus MKRRALFLSVLLLSLFSPLAVVFADETKKEQPLPFYVQPVLPENQLDSSVGYFYLPSTPN WP_010782219.1 hypothetical protein [Enterococcus gilvus] Length: 343\nNumber of predicted TMHs: 2\nExp number of AAs in TMHs: 41.39353\nExp number, first 60 AAs: 19.01837\nTotal prob of N-in: 0.93656\nPOSSIBLE N-term signal sequence\ninside 1 4\nTMhelix 5 24\noutside 25 311\nTMhelix 312 334\ninside 335 343

34508 GCF_001064985.1_ASM106498v1 Lactobacillus gasseri Terrabacteria group; Firmicutes; Bacilli; Lactobacillales; Lactobacillaceae; Lactobacillus MKRRTTMLMSALFVAGAFAVSGNVAKADTVQANNSNNVNSTEVKQNNTNASANQVATKTP WP_049151215.1 hypothetical protein [Lactobacillus gasseri] Length: 503\nNumber of predicted TMHs: 1\nExp number of AAs in TMHs: 21.04352\nExp number, first 60 AAs: 3.29631\nTotal prob of N-in: 0.18457\noutside 1 474\nTMhelix 475 494\ninside 495 503

34509 GCF_002007185.1_ASM200718v1 Lactobacillus gasseri Terrabacteria group; Firmicutes; Bacilli; Lactobacillales; Lactobacillaceae; Lactobacillus MKRRTTMLMSALFVAGAFAVSGNVVKADTVQANNSNNVNSTEVKQNNTNANANQVATKTP WP_077959486.1 hypothetical protein [Lactobacillus gasseri] Length: 513\nNumber of predicted TMHs: 1\nExp number of AAs in TMHs: 21.65843\nExp number, first 60 AAs: 3.89648\nTotal prob of N-in: 0.21086\noutside 1 484\nTMhelix 485 504\ninside 505 513

34510 GCF_001437125.1_ASM143712v1 Lactobacillus paucivorans Terrabacteria group; Firmicutes; Bacilli; Lactobacillales; Lactobacillaceae; Lactobacillus MSVDQQQAKNVQVSHRRQWLFVGVTAAVLSTELLVVGVTSAQAATGDQTPTAVSGEKPAE WP_057878938.1 hypothetical protein [Lactobacillus paucivorans] Length: 771\nNumber of predicted TMHs: 2\nExp number of AAs in TMHs: 39.06259\nExp number, first 60 AAs: 20.85698\nTotal prob of N-in: 0.94689\nPOSSIBLE N-term signal sequence\ninside 1 18\nTMhelix 19 41\noutside 42 746\nTMhelix 747 764\ninside 765 771

34511 GCF_000227195.1_Lactobacillus_sp_7_1_47FAA_V1 Lactobacillus sp. 7_1_47FAA Terrabacteria group; Firmicutes; Bacilli; Lactobacillales; Lactobacillaceae; Lactobacillus MVFNRRRSNLCYGIRKLAIGVVSMVISVAFLTEISVAQAKENTSNYQEHLVDKKASVADL WP_009310390.1 hypothetical protein [Lactobacillus sp. 7_1_47FAA] Length: 902\nNumber of predicted TMHs: 2\nExp number of AAs in TMHs: 40.02553\nExp number, first 60 AAs: 19.49231\nTotal prob of N-in: 0.91958\nPOSSIBLE N-term signal sequence\ninside 1 11\nTMhelix 12 34\noutside 35 875\nTMhelix 876 898\ninside 899 902

34512 GCF_000814185.1_ASM81418v1 Lactobacillus sp. FMNP02 Terrabacteria group; Firmicutes; Bacilli; Lactobacillales; Lactobacillaceae; Lactobacillus MMKAARRTPIFAALIMLFAQLFVPASIAFAASMTNIKVADWHNTWHLYLFNGLHWTDNGM WP_039639977.1 MULTISPECIES: adhesin [Lactobacillus] Length: 1270\nNumber of predicted TMHs: 2\nExp number of AAs in TMHs: 45.1134199999999\nExp number, first 60 AAs: 22.15672\nTotal prob of N-in: 0.99247\nPOSSIBLE N-term signal sequence\ninside 1 8\nTMhelix 9 31\noutside 32 1241\nTMhelix 1242 1264\ninside 1265 1270

34513 GCF_002027415.1_ASM202741v1 Lactobacillus casei Terrabacteria group; Firmicutes; Bacilli; Lactobacillales; Lactobacillaceae; Lactobacillus; Lactobacillus casei group MMKTARRSAVIVALVSLFVQLFIPASMAIAASMTNTKVADWRNTWHLHLFNGLHWTDTGM WP_079322621.1 adhesin [Lactobacillus casei] Length: 1270\nNumber of predicted TMHs: 2\nExp number of AAs in TMHs: 45.2706099999999\nExp number, first 60 AAs: 22.36632\nTotal prob of N-in: 0.99670\nPOSSIBLE N-term signal sequence\ninside 1 8\nTMhelix 9 31\noutside 32 1241\nTMhelix 1242 1264\ninside 1265 1270

34514 GCF_000309565.2_ASM30956v2 Lactobacillus casei 12A Terrabacteria group; Firmicutes; Bacilli; Lactobacillales; Lactobacillaceae; Lactobacillus; Lactobacillus casei group; Lactobacillus casei

MRQRKHYEMYKSRRPLMTCIIAISSFAIMGPLTNKTTVRADAITVKNTILPSSNESSENN WP_003563282.1 mucin-binding protein [Lactobacillus casei] Length: 761\nNumber of predicted TMHs: 1\nExp number of AAs in TMHs: 33.56858\nExp number, first 60 AAs: 15.25573\nTotal prob of N-in: 0.77310\nPOSSIBLE N-term signal sequence\noutside 1 736\nTMhelix 737 754\ninside 755 761

34515 GCF_000472345.1_ASM47234v1 Lactobacillus casei 12A Terrabacteria group; Firmicutes; Bacilli; Lactobacillales; Lactobacillaceae; Lactobacillus; Lactobacillus casei group; Lactobacillus casei MRQRKHYEMYKSRRPLMTCIIAISSFAIMGPLTNKTTVRADAITVKNTILPSSNESSENN WP_003563282.1 mucin-binding protein [Lactobacillus casei] Length: 761\nNumber of predicted TMHs: 1\nExp number of AAs in TMHs: 33.56858\nExp number, first 60 AAs: 15.25573\nTotal prob of N-in: 0.77310\nPOSSIBLE N-term signal sequence\noutside 1 736\nTMhelix 737 754\ninside 755 761

34516 GCF_000309585.1_ASM30958v1 Lactobacillus casei 21/1 Terrabacteria group; Firmicutes; Bacilli; Lactobacillales; Lactobacillaceae; Lactobacillus; Lactobacillus casei group; Lactobacillus casei MRQRKHYEMYKSRRPLMTCIIAISSFAIMGPLTNKTTVRADAITVKNTILPSSNESSENN WP_003563282.1 mucin-binding protein [Lactobacillus casei] Length: 761\nNumber of predicted TMHs: 1\nExp number of AAs in TMHs: 33.56858\nExp number, first 60 AAs: 15.25573\nTotal prob of N-in: 0.77310\nPOSSIBLE N-term signal sequence\noutside 1 736\nTMhelix 737 754\ninside 755 761

34517 GCF_000829055.1_ASM82905v1 Lactobacillus casei subsp. casei ATCC 393 Terrabacteria group; Firmicutes; Bacilli; Lactobacillales; Lactobacillaceae; Lactobacillus; Lactobacillus casei group; Lactobacillus casei MMKAARRTPIFAALIMLFAQLFVPASIAFAASMTNIKVADWHNTWHLYLFNGLHWTDNGM WP_039639977.1 MULTISPECIES: adhesin [Lactobacillus] Length: 1270\nNumber of predicted TMHs: 2\nExp number of AAs in TMHs: 45.1134199999999\nExp number, first 60 AAs: 22.15672\nTotal prob of N-in: 0.99247\nPOSSIBLE N-term signal sequence\ninside 1 8\nTMhelix 9 31\noutside 32 1241\nTMhelix 1242 1264\ninside 1265 1270

34518 GCF_000409875.1_Lpp46.v.1 Lactobacillus paracasei subsp. paracasei Lpp46 Terrabacteria group; Firmicutes; Bacilli; Lactobacillales; Lactobacillaceae; Lactobacillus; Lactobacillus casei group; Lactobacillus paracasei; Lactobacillus paracasei subsp. paracasei MRQRKHYEMYKSRRPLMTCIIAISSFAIMGPLTNKTTVRADAITVKNTILPSSNESSENN WP_003573464.1 mucin-binding protein [Lactobacillus paracasei] Length: 761\nNumber of predicted TMHs: 1\nExp number of AAs in TMHs: 33.56858\nExp number, first 60 AAs: 15.25573\nTotal prob of N-in: 0.77310\nPOSSIBLE N-term signal sequence\noutside 1 736\nTMhelix 737 754\ninside 755 761

34519 GCF_000185405.1_ASM18540v1 Lactobacillus iners ATCC 55195 Terrabacteria group; Firmicutes; Bacilli; Lactobacillales; Lactobacillaceae; Lactobacillus; Lactobacillus iners MVFNRRRSNLCYGIRKLAIGVVSMVISVAFLTEISVAQAKENTSNYQEHLVDKKASVADL WP_006736993.1 hypothetical protein [Lactobacillus iners] Length: 1021\nNumber of predicted TMHs: 2\nExp number of AAs in TMHs: 38.46488\nExp number, first 60 AAs: 19.49238\nTotal prob of N-in: 0.91958\nPOSSIBLE N-term signal sequence\ninside 1 11\nTMhelix 12 34\noutside 35 993\nTMhelix 994 1016\ninside 1017 1021

34520 GCF_001436505.1_ASM143650v1 Lactobacillus rennini DSM 20253 Terrabacteria group; Firmicutes; Bacilli; Lactobacillales; Lactobacillaceae; Lactobacillus; Lactobacillus rennini MNYSRRPVLKNTLYASGALLLLVGSSVPASAAVVTGAVQTSSATARTGLADMSILENTSL WP_057873001.1 hypothetical protein [Lactobacillus rennini] Length: 787\nNumber of predicted TMHs: 2\nExp number of AAs in TMHs: 41.22311\nExp number, first 60 AAs: 21.23285\nTotal prob of N-in: 0.92641\nPOSSIBLE N-term signal sequence\ninside 1 12\nTMhelix 13 35\noutside 36 761\nTMhelix 762 781\ninside 782 787

34521 GCF_000233755.1_ASM23375v1 Lactobacillus rhamnosus ATCC 8530 Terrabacteria group; Firmicutes; Bacilli; Lactobacillales; Lactobacillaceae; Lactobacillus; Lactobacillus rhamnosus MMKTARRSAVIVALVSLFVQLFIPASMAIAASMTNTKVADWRNTWHLHLFNGMNWTENNI WP_005691205.1 adhesin [Lactobacillus rhamnosus] Length: 1270\nNumber of predicted TMHs: 2\nExp number of AAs in TMHs: 45.05551\nExp number, first 60 AAs: 22.35575\nTotal prob of N-in: 0.99750\nPOSSIBLE N-term signal sequence\ninside 1 8\nTMhelix 9 31\noutside 32 1241\nTMhelix 1242 1264\ninside 1265 1270

34522 GCF_000160175.1_ASM16017v1 Lactobacillus rhamnosus LMS2-1 Terrabacteria group; Firmicutes; Bacilli; Lactobacillales; Lactobacillaceae; Lactobacillus; Lactobacillus rhamnosus MMKTARRSAVIVALVSLFVQLFIPASMAIAASMTNTKVADWRNTWHLHLFNGMNWTENNI WP_005691205.1 adhesin [Lactobacillus rhamnosus] Length: 1270\nNumber of predicted TMHs: 2\nExp number of AAs in TMHs: 45.05551\nExp number, first 60 AAs: 22.35575\nTotal prob of N-in: 0.99750\nPOSSIBLE N-term signal sequence\ninside 1 8\nTMhelix 9 31\noutside 32 1241\nTMhelix 1242 1264\ninside 1265 1270

34523 GCF_001436695.1_ASM143669v1 Lactobacillus taiwanensis DSM 21401 Terrabacteria group; Firmicutes; Bacilli; Lactobacillales; Lactobacillaceae; Lactobacillus; Lactobacillus taiwanensis
MKRRTTMLMSALFVAGAFAVSGNVAKADAVQANNSNNVNSTEVKQNNTNASANQVATKTP WP_057718161.1 hypothetical protein [Lactobacillus taiwanensis] Length: 499\nNumber of predicted TMHs: 1\nExp number of AAs in TMHs: 22.14244\nExp number, first 60 AAs: 3.50611\nTotal prob of N-in: 0.19958\noutside 1 470\nTMhelix 471 492\ninside 493 499

34524 GCF_001884305.1_ASM188430v1 Weissella confusa Terrabacteria group; Firmicutes; Bacilli; Lactobacillales; Leuconostocaceae; Weissella
MYNRRRIMTKLKSLLVMALTVIGVGLFMPNVHAQSSMDFVVSPKLPDTQVDKKAGFFNFQ WP_071707692.1 cell surface protein [Weissella confusa] Length: 375\nNumber of predicted TMHs: 2\nExp number of AAs in TMHs: 44.1175999999999\nExp number, first 60 AAs: 20.84065\nTotal prob of N-in: 0.99775\nPOSSIBLE N-term signal sequence\ninside 1 12\nTMhelix 13 35\noutside 36 330\nTMhelix 331 353\ninside 354 375

34525 GCF_000239955.1_ASM23995v2 Weissella confusa LBAE C39-2 Terrabacteria group; Firmicutes; Bacilli; Lactobacillales; Leuconostocaceae; Weissella; Weissella confusa
MYNRRRIMTKLKSLLVMALTVIGVGLFMPNVHAQSSMDFVVSPKLPDTQVDKKAGFFNFQ WP_003609186.1 cell surface protein [Weissella confusa] Length: 375\nNumber of predicted TMHs: 2\nExp number of AAs in TMHs: 44.0810699999999\nExp number, first 60 AAs: 20.83119\nTotal prob of N-in: 0.99767\nPOSSIBLE N-term signal sequence\ninside 1 12\nTMhelix 13 35\noutside 36 330\nTMhelix 331 353\ninside 354 375

34526 GCF_000959895.1_ASM95989v1 Streptococcus cristatus Terrabacteria group; Firmicutes; Bacilli; Lactobacillales; Streptococcaceae; Streptococcus
MEAIFLTHRSRRKLFSLGICSFLLGLSVSVTPMAVVGADEVQQAESQLEQIADSSMEEDD WP_005592214.1 hypothetical protein [Streptococcus cristatus] Length: 161\nNumber of predicted TMHs: 2\nExp number of AAs in TMHs: 42.26278\nExp number, first 60 AAs: 22.26992\nTotal prob of N-in: 0.93359\nPOSSIBLE N-term signal sequence\ninside 1 12\nTMhelix 13 35\noutside 36 135\nTMhelix 136 155\ninside 156 161

34527 GCF_001069775.1_ASM106977v1 Streptococcus cristatus Terrabacteria group; Firmicutes; Bacilli; Lactobacillales; Streptococcaceae; Streptococcus
MEANFLTHRSRRKLFSLSICSFLLGLSISVTPMAVVGADEVQQAESQPEQIADSSVEEDD WP_048791748.1 hypothetical protein [Streptococcus cristatus] Length: 161\nNumber of predicted TMHs: 2\nExp number of AAs in TMHs: 41.8938\nExp number, first 60 AAs: 22.13637\nTotal prob of N-in: 0.92154\nPOSSIBLE N-term signal sequence\ninside 1 12\nTMhelix 13 35\noutside 36 135\nTMhelix 136 155\ninside 156 161

34528 GCF_001578775.1_ASM157877v1 Streptococcus cristatus Terrabacteria group; Firmicutes; Bacilli; Lactobacillales; Streptococcaceae; Streptococcus
MEANFLTHRSRRKLFSLGICSFLLGLSISVTPMAVVGADEVQQAESQLEQIADSSVEEDD WP_061423038.1 hypothetical protein [Streptococcus cristatus] Length: 161\nNumber of predicted TMHs: 2\nExp number of AAs in TMHs: 42.12914\nExp number, first 60 AAs: 22.36688\nTotal prob of N-in: 0.92557\nPOSSIBLE N-term signal sequence\ninside 1 12\nTMhelix 13 35\noutside 36 135\nTMhelix 136 155\ninside 156 161

34529 GCF_001588855.1_ASM158885v1 Streptococcus cristatus Terrabacteria group; Firmicutes; Bacilli; Lactobacillales; Streptococcaceae; Streptococcus
MEANFLTHRSRRKLFSLGICSFLLGLSVSVTPMAVVGADEVQQAESQLEQIADSSMEEDD WP_045496817.1 hypothetical protein [Streptococcus cristatus] Length: 161\nNumber of predicted TMHs: 2\nExp number of AAs in TMHs: 42.03119\nExp number, first 60 AAs: 22.2855\nTotal prob of N-in: 0.88185\nPOSSIBLE N-term signal sequence\ninside 1 12\nTMhelix 13 35\noutside 36 135\nTMhelix 136 155\ninside 156 161

34530 GCF_001589115.1_ASM158911v1 Streptococcus cristatus Terrabacteria group; Firmicutes; Bacilli; Lactobacillales; Streptococcaceae; Streptococcus
MEANFLTHRSRRKLFSLGICSFLLGLSVSVAPMAVVGADEVQQAEPQLEQIADSSVEEDD WP_061910244.1 hypothetical protein [Streptococcus cristatus] Length: 161\nNumber of predicted TMHs: 2\nExp number of AAs in TMHs: 42.16932\nExp number, first 60 AAs: 22.43962\nTotal prob of N-in: 0.93881\nPOSSIBLE N-term signal sequence\ninside 1 12\nTMhelix 13 35\noutside 36 135\nTMhelix 136 155\ninside 156 161

34531 GCF_000959885.1_ASM95988v1 Streptococcus cristatus Terrabacteria group; Firmicutes; Bacilli; Lactobacillales; Streptococcaceae; Streptococcus
MEANFLTHRSRRKLFSLGICSFLLGLSVSVTPMAVVGADEVQQAESQLEQIADSSMEEDD WP_045496817.1 hypothetical protein [Streptococcus cristatus] Length: 161\nNumber of predicted TMHs: 2\nExp number of

AAs in TMHs: 42.03119\nExp number, first 60 AAs: 22.2855\nTotal prob of N-in: 0.88185\nPOSSIBLE N-term signal sequence\ninside 1 12\nTMhelix 13 35\noutside 36 135\nTMhelix 136 155\ninside 156 161

34532 GCF_000724645.1_ASM72464v1 Streptococcus parasanguinis Terrabacteria group; Firmicutes; Bacilli; Lactobacillales; Streptococcaceae; Streptococcus
MQIRWTLKTVICHSKNKRRKTMNKKVILTTVATIAAIATAGGVKADEFNGDLAKDSIGLT WP_080713446.1 hypothetical protein [Streptococcus parasanguinis] Length: 571\nNumber of predicted TMHs: 2\nExp number of AAs in TMHs: 39.45348\nExp number, first 60 AAs: 18.89972\nTotal prob of N-in: 0.97933\nPOSSIBLE N-term signal sequence\ninside 1 25\nTMhelix 26 43\noutside 44 543\nTMhelix 544 566\ninside 567 571

34533 GCF_000385925.1_ASM38592v1 Streptococcus cristatus AS 1.3089 Terrabacteria group; Firmicutes; Bacilli; Lactobacillales; Streptococcaceae; Streptococcus; Streptococcus cristatus
MEANFLTHRSRRKLFSLGICSCLLGLSVSVTPMAVVGADEVQQAESQLEQIADSSVEEDDWP_015604698.1 hypothetical protein [Streptococcus cristatus] Length: 161\nNumber of predicted TMHs: 2\nExp number of AAs in TMHs: 42.20583\nExp number, first 60 AAs: 21.9882\nTotal prob of N-in: 0.89896\nPOSSIBLE N-term signal sequence\ninside 1 12\nTMhelix 13 35\noutside 36 135\nTMhelix 136 155\ninside 156 161

34534 GCF_000187855.1_ASM18785v1 Streptococcus cristatus ATCC 51100 Terrabacteria group; Firmicutes; Bacilli; Lactobacillales; Streptococcaceae; Streptococcus; Streptococcus cristatus
MEAIFLTHRSRRKLFSLGICSFLLGLSVSVTPMAVVGADEVQQAESQLEQIADSSMEEDD WP_005592214.1 hypothetical protein [Streptococcus cristatus] Length: 161\nNumber of predicted TMHs: 2\nExp number of AAs in TMHs: 42.26278\nExp number, first 60 AAs: 22.26992\nTotal prob of N-in: 0.93359\nPOSSIBLE N-term signal sequence\ninside 1 12\nTMhelix 13 35\noutside 36 135\nTMhelix 136 155\ninside 156 161

34535 GCF_000222765.1_ASM22276v2 Streptococcus cristatus ATCC 51100 Terrabacteria group; Firmicutes; Bacilli; Lactobacillales; Streptococcaceae; Streptococcus; Streptococcus cristatus
MEAIFLTHRSRRKLFSLGICSFLLGLSVSVTPMAVVGADEVQQAESQLEQIADSSMEEDD WP_005592214.1 hypothetical protein [Streptococcus cristatus] Length: 161\nNumber of predicted TMHs: 2\nExp number of AAs in TMHs: 42.26278\nExp number, first 60 AAs: 22.26992\nTotal prob of N-in: 0.93359\nPOSSIBLE N-term signal sequence\ninside 1 12\nTMhelix 13 35\noutside 36 135\nTMhelix 136 155\ninside 156 161

34536 GCF_000706805.1_ASM70680v1 Streptococcus equi subsp. ruminatorum CECT 5772 Terrabacteria group; Firmicutes; Bacilli; Lactobacillales; Streptococcaceae; Streptococcus; Streptococcus dysgalactiae group; Streptococcus equi; Streptococcus equi subsp. ruminatorum
MSIIKGERNEQKKSAKRRRKNLITKLAMTSALTLGVGAAATLAGQTEVRAEVLTLNMKDK WP_037580257.1
hypothetical protein [Streptococcus equi] Length: 299\nNumber of predicted TMHs: 2\nExp number of AAs in TMHs: 37.31466\nExp number, first 60 AAs: 22.47846\nTotal prob of N-in: 0.99106\nPOSSIBLE N-term signal sequence\ninside 1 20\nTMhelix 21 43\noutside 44 274\nTMhelix 275 292\ninside 293 299

34537 GCF_000146585.1_ASM14658v1 Streptococcus mitis ATCC 6249 Terrabacteria group; Firmicutes; Bacilli; Lactobacillales; Streptococcaceae; Streptococcus; Streptococcus mitis
MGYIFFIHSINFVHKSLHYWTRDFFKNNRRSLLMKTKSAINLFATAILAIATIGSNTIYA WP_004240092.1 cell wall surface anchor signal protein [Streptococcus mitis] Length: 527\nNumber of predicted TMHs: 2\nExp number of AAs in TMHs: 36.96706\nExp number, first 60 AAs: 19.10654\nTotal prob of N-in: 0.78464\nPOSSIBLE N-term signal sequence\ninside 1 37\nTMhelix 38 60\noutside 61 504\nTMhelix 505 522\ninside 523 527

34538 GCF_000441095.1_version1 Streptococcus suis YS21 Terrabacteria group; Firmicutes; Bacilli; Lactobacillales; Streptococcaceae; Streptococcus; Streptococcus suis
MKNRRTFMKKLTKLFSVFATLLTVLGTLVSLAPVAHAATKETKVIVHKIVMDEKDFNKFT WP_079270589.1 hypothetical protein [Streptococcus suis] Length: 656\nNumber of predicted TMHs: 2\nExp number of AAs in TMHs: 44.92098\nExp number, first 60 AAs: 22.56802\nTotal prob of N-in: 0.99400\nPOSSIBLE N-term signal sequence\ninside 1 12\nTMhelix 13 35\noutside 36 624\nTMhelix 625 647\ninside 648 656

34539 GCF_001579785.1_ASM157978v1 Clostridium perfringens Terrabacteria group; Firmicutes; Clostridia; Clostridiales; Clostridiaceae; Clostridium
MNFKKRRGVLILLLSFLLTSMLTSNVFAKPSSGNFSKNPSESKKSSSFSSSRKSFNSSKS WP_061429810.1 hypothetical protein [Clostridium perfringens] Length: 172\nNumber of predicted TMHs: 2\nExp number of AAs in TMHs: 42.36995\nExp number, first 60 AAs: 20.07702\nTotal prob of N-in: 0.97486\nPOSSIBLE N-term signal sequence\ninside 1 8\nTMhelix 9 28\noutside 29 147\nTMhelix 148 170\ninside 171 172

34540 GCF_002018235.1_ASM201823v1 Clostridium perfringens Terrabacteria group; Firmicutes; Clostridia; Clostridiales; Clostridiaceae; Clostridium
MTNKQKMRKRRSALLCVFAAFAFSFNPLGKIAYATPKQDTSVVNQVQTQEETSNSIQNQP WP_078233233.1 endo-beta-N-acetylglucosaminidase [Clostridium perfringens] Length: 1135\nNumber of predicted TMHs: 2\nExp number of AAs in TMHs: 40.08135\nExp number, first 60 AAs: 21.87491\nTotal prob of N-in: 0.99025\nPOSSIBLE N-term signal sequence\ninside 1 12\nTMhelix 13 35\noutside 36 1107\nTMhelix 1108 1130\ninside 1131 1135

34541 GCF_001517625.1_Clostridium_bouchedurhonense Clostridium sp. AT4 Terrabacteria group; Firmicutes; Clostridia; Clostridiales; Clostridiaceae; Clostridium
MNEYKNRRQLLCLAVTVGMMFTAVNVSSKTVWAAAEQVIEYDPNQEEKTHPGSIILKQNL WP_066550529.1 hypothetical protein [Clostridium sp. AT4] Length: 332\nNumber of predicted TMHs: 1\nExp number of AAs in TMHs: 33.05552\nExp number, first 60 AAs: 11.31482\nTotal prob of N-in: 0.52662\nPOSSIBLE N-term signal sequence\noutside 1 295\nTMhelix 296 318\ninside 319 332

34542 GCF_000270305.1_ASM27030v1 Clostridium sp. SY8519 Terrabacteria group; Firmicutes; Clostridia; Clostridiales; Clostridiaceae; Clostridium
MLRQNFLSRHHQRKAQRQNRRPIMRKRNLLAALLAAMLCCGLCYTTAFAQTNEPEAAPAE WP_083834998.1 hypothetical protein [Clostridium sp. SY8519] Length: 202\nNumber of predicted TMHs: 2\nExp number of AAs in TMHs: 43.66027\nExp number, first 60 AAs: 21.77469\nTotal prob of N-in: 0.99909\nPOSSIBLE N-term signal sequence\ninside 1 28\nTMhelix 29 48\noutside 49 135\nTMhelix 136 158\ninside 159 202

34543 GCF_000320405.1_ASM32040v1 Clostridium celatum DSM 1785 Terrabacteria group; Firmicutes; Clostridia; Clostridiales; Clostridiaceae; Clostridium; Clostridium celatum
MRRRNVMNKNISLVIAIAVTLGQIPVSALANTGNGAEAIIREAKAEKAEGVFNLHLSIAI WP_005209628.1 hypothetical protein [Clostridium celatum] Length: 408\nNumber of predicted TMHs: 2\nExp number of AAs in TMHs: 39.85173\nExp number, first 60 AAs: 18.57792\nTotal prob of N-in: 0.90834\nPOSSIBLE N-term signal sequence\ninside 1 12\nTMhelix 13 35\noutside 36 380\nTMhelix 381 403\ninside 404 408

34544 GCF_000768585.1_4570_1.0 Clostridium novyi A str. 4570 Terrabacteria group; Firmicutes; Clostridia; Clostridiales; Clostridiaceae; Clostridium; Clostridium novyi; Clostridium novyi A
MQRRKIFKKFLRAFVVGLCMVQVLQINTAKGLDRSSKLDTSITVLTGTKPFDKKRDYLYP WP_039249807.1 hypothetical protein [Clostridium novyi] Length: 806\nNumber of predicted TMHs: 1\nExp number of AAs in TMHs: 21.71743\nExp number, first 60 AAs: 3.46746\nTotal prob of N-in: 0.19039\noutside 1 779\nTMhelix 780 799\ninside 800 806

34545 GCF_001405675.1_13414_6_34 Hungatella hathewayi Terrabacteria group; Firmicutes; Clostridia; Clostridiales; Clostridiaceae; Hungatella
MTAKRRQYLWSAAAVLYVLFIFSNSMKTADLSSADSGAVLKLVQQVLTAGGVDSTIITEH WP_002602281.1 VanZ family protein [Hungatella hathewayi] Length: 157\nNumber of predicted TMHs: 2\nExp number of AAs in TMHs: 70.28314\nExp number, first 60 AAs: 17.9558\nTotal prob of N-in: 0.97824\nPOSSIBLE N-term signal sequence\ninside 1 6\nTMhelix 7 26\noutside 27 123\nTMhelix 124 146\ninside 147 157

34546 GCF_000371445.1_Clos_hath_12489931_V1 Hungatella hathewayi 12489931 Terrabacteria group; Firmicutes; Clostridia; Clostridiales; Clostridiaceae; Hungatella; Hungatella hathewayi
MTAKRRQYLWSAAAVLYVLFIFSNSMKTADLSSADSGAVLKLVQQVLTAGGVDSTIITEH WP_002602281.1 VanZ family protein [Hungatella hathewayi] Length: 157\nNumber of predicted TMHs: 2\nExp number of AAs in TMHs: 70.28314\nExp number, first 60 AAs: 17.9558\nTotal prob of N-in: 0.97824\nPOSSIBLE N-term signal sequence\ninside 1 6\nTMhelix 7 26\noutside 27 123\nTMhelix 124 146\ninside 147 157

34547 GCF_000508905.1_ASM50890v1 Hungatella hathewayi VE202-04 Terrabacteria group; Firmicutes; Clostridia; Clostridiales; Clostridiaceae; Hungatella; Hungatella hathewayi
MTAKRRQYLWSAAAVLYVLFIFSNSMKTADLSSADSGAVLKLVQQVLTAGGVDSTIITEH WP_002602281.1 VanZ family protein [Hungatella hathewayi] Length: 157\nNumber of predicted TMHs: 2\nExp number of AAs in TMHs: 70.28314\nExp number, first 60 AAs: 17.9558\nTotal prob of N-in: 0.97824\nPOSSIBLE N-term signal sequence\ninside 1 6\nTMhelix 7 26\noutside 27 123\nTMhelix 124 146\ninside 147 157

34548 GCF_000471805.1_ASM47180v1 Hungatella hathewayi VE202-11 Terrabacteria group; Firmicutes; Clostridia; Clostridiales; Clostridiaceae; Hungatella; Hungatella hathewayi
MTAKRRQYLWSAAAVLYVLFIFSNSMKTADLSSADSGAVLKLVQQVLTAGGVDSTIITEH WP_002602281.1 VanZ family protein [Hungatella hathewayi] Length: 157\nNumber of predicted TMHs: 2\nExp number of AAs in TMHs: 70.28314\nExp number, first 60 AAs: 17.9558\nTotal prob of N-in: 0.97824\nPOSSIBLE N-term signal sequence\ninside 1 6\nTMhelix 7 26\noutside 27 123\nTMhelix 124 146\ninside 147 157

34549 GCF_900095865.1_PRJEB15310 Massilioclostridium coli Terrabacteria group; Firmicutes; Clostridia; Clostridiales; Clostridiaceae; Massilioclostridium
MKRRRLLSLGLVLAMSTSILIGNFSTATAEGASSDFYNQLVNHYTTPDIENRTEVRWWMA WP_069988814.1 hypothetical protein [Massilioclostridium coli] Length: 1501\nNumber of predicted TMHs: 2\nExp number of AAs in TMHs: 40.54755\nExp number, first 60 AAs: 20.78959\nTotal prob of N-in: 0.92625\nPOSSIBLE N-term signal sequence\ninside 1 6\nTMhelix 7 29\noutside 30 1475\nTMhelix 1476 1495\ninside 1496 1501

34550 GCF_000184705.1_ASM18470v1 Thermaerobacter marianensis DSM 12885 Terrabacteria group; Firmicutes; Clostridia; Clostridiales; Clostridiales incertae sedis; Clostridiales Family XVII. Incertae Sedis; Thermaerobacter; Thermaerobacter marianensis MADRADAHRRRVLWRGTNTAVLTVAVLALLVLANVFAGRYSWRYDATQQKIYSLSPSTYE WP_013494899.1 ABC transporter [Thermaerobacter marianensis] Length: 503\nNumber of predicted TMHs: 2\nExp number of AAs in TMHs: 45.09534\nExp number, first 60 AAs: 22.11191\nTotal prob of N-in: 0.99265\nPOSSIBLE N-term signal sequence\ninside 1 19\nTMhelix 20 42\noutside 43 475\nTMhelix 476 498\ninside 499 503

34551 GCF_000183545.2_ASM18354v3 Thermaerobacter subterraneus DSM 13965 Terrabacteria group; Firmicutes; Clostridia; Clostridiales; Clostridiales incertae sedis; Clostridiales Family XVII. Incertae Sedis; Thermaerobacter; Thermaerobacter subterraneus MAERADAHRRRVLWRGTNTAVLTVAVLALLVLANVFAARYSWRYDATAQKIYSLSPSTHQ WP_006904371.1 ABC transporter [Thermaerobacter subterraneus] Length: 506\nNumber of predicted TMHs: 2\nExp number of AAs in TMHs: 46.07585\nExp number, first 60 AAs: 22.14956\nTotal prob of N-in: 0.99809\nPOSSIBLE N-term signal sequence\ninside 1 19\nTMhelix 20 42\noutside 43 478\nTMhelix 479 501\ninside 502 506

34552 GCF_001405175.1_13470_2_54 [Eubacterium] halliiTerrabacteria group; Firmicutes; Clostridia; Clostridiales; Eubacteriaceae; Eubacterium
MVPYLENFIKHNRRSFMFRSLKKIASVVMATAMAVTTAVPGTAIVANAETTTIKTRVKNS WP_082425579.1 hypothetical protein [[Eubacterium] hallii] Length: 2577\nNumber of predicted TMHs: 2\nExp number of AAs in TMHs: 42.50799\nExp number, first 60 AAs: 20.45845\nTotal prob of N-in: 0.89275\nPOSSIBLE N-term signal sequence\ninside 1 23\nTMhelix 24 46\noutside 47 2548\nTMhelix 2549 2571\ninside 2572 2577

34553 GCF_000019165.1_ASM1916v1 Heliobacterium modesticaldum Ice1 Terrabacteria group; Firmicutes; Clostridia; Clostridiales; Heliobacteriaceae; Heliobacterium; Heliobacterium modesticaldum
MTDVSRRTFFRRAIASGVAGAGVLTLGPKGPAFADADAVGTMIDLTRCDGCAGEQTPRCV WP_012282812.1 oxidoreductase [Heliobacterium modesticaldum] Length: 328\nNumber of predicted TMHs: 1\nExp number of AAs in TMHs: 22.41058\nExp number, first 60 AAs: 3.60189\nTotal prob of N-in: 0.32794\noutside 1 301\nTMhelix 302 321\ninside 322 328

34554 GCF_000019165.1_ASM1916v1 Heliobacterium modesticaldum Ice1 Terrabacteria group; Firmicutes; Clostridia; Clostridiales; Heliobacteriaceae; Heliobacterium; Heliobacterium modesticaldum
MGKLVGAALIVGAFSAGGFMTARDLRRRRQILASLQSALAMLATEVAFRATHLPEALVQV WP_012281658.1 stage iii sporulation protein spoab [Heliobacterium modesticaldum] Length: 172\nNumber of predicted TMHs: 2\nExp number of AAs in TMHs: 34.69903\nExp number, first 60 AAs: 18.39161\nTotal prob of N-in: 0.01531\nPOSSIBLE N-term signal sequence\noutside 1 3\nTMhelix 4 21\ninside 22 154\nTMhelix 155 171\noutside 172 172

34555 GCF_000019165.1_ASM1916v1 Heliobacterium modesticaldum Ice1 Terrabacteria group; Firmicutes; Clostridia; Clostridiales; Heliobacteriaceae; Heliobacterium; Heliobacterium modesticaldum
MAKEETYYQYLKRRGVSRRDFLKFCTVMAASLGLSAGSVPKIAEALETKKRIPVIWRHLV WP_012282566.1 uptake [ni/fe] hydrogenase small subunit [Heliobacterium modesticaldum] Length: 372\nNumber of predicted TMHs: 2\nExp number of AAs in TMHs: 37.2974399999999\nExp number, first 60 AAs: 18.50811\nTotal prob of N-in: 0.88581\nPOSSIBLE N-term signal sequence\ninside 1 20\nTMhelix 21 43\noutside 44 329\nTMhelix 330 352\ninside 353 372

34556 GCF_001404655.1_13414_6_14 Anaerostipes hadrus Terrabacteria group; Firmicutes; Clostridia; Clostridiales; Lachnospiraceae; Anaerostipes
MKKARRTLMALGVVVFSLSAALTAKIKEKSVLAQLNCSKAKPITSSWGDDKHIFSAFDDQWP_055198347.1 peptidase [Anaerostipes hadrus] Length: 1228\nNumber of predicted TMHs: 2\nExp number of AAs in TMHs: 31.83459\nExp number, first 60 AAs: 14.19329\nTotal prob of N-in: 0.77497\nPOSSIBLE N-term signal sequence\ninside 1 6\nTMhelix 7 24\noutside 25 1206\nTMhelix 1207 1224\ninside 1225 1228

34557 GCF_001405315.1_13470_2_60 [Ruminococcus] torques Terrabacteria group; Firmicutes; Clostridia; Clostridiales; Lachnospiraceae; Blautia MRVKRRKTMALLAAVSIVCAQSTVSDVFAAGAEEVAAQENNAVSIEYLTLQPGERDDSVN WP_009319933.1 hypothetical protein [[Ruminococcus] torques] Length: 1161\nNumber of predicted TMHs:

1\nExp number of AAs in TMHs: 23.64632\nExp number, first 60 AAs: 1.33478\nTotal prob of N-in: 0.10565\noutside 1 1127\nTMhelix 1128 1150\ninside 1151 1161

34558 GCF_000424505.1_ASM42450v1 Butyrivibrio sp. XBB1001 Terrabacteria group; Firmicutes; Clostridia; Clostridiales; Lachnospiraceae; Butyrivibrio MSRRKIIALVRVFLSLLVIAAMGSSRVIAIASDYDQTLISVDESEQAVYEKGDFEALGDE WP_026488219.1 hypothetical protein [Butyrivibrio sp. XBB1001] Length: 603\nNumber of predicted TMHs: 2\nExp number of AAs in TMHs: 40.64136\nExp number, first 60 AAs: 22.02793\nTotal prob of N-in: 0.99476\nPOSSIBLE N-term signal sequence\ninside 1 6\nTMhelix 7 29\noutside 30 574\nTMhelix 575 597\ninside 598 603

34559 GCF_900101295.1_IMG-taxon_2606217764_annotated_assembly [Clostridium] aminophilum Terrabacteria group; Firmicutes; Clostridia; Clostridiales; Lachnospiraceae; Lachnoclostridium MKIPKAKVGIDRRQLIAAGAGILLGAMMCVVSSAEDHSGIRSFHRNLYGGSGRDEEVIVS WP_051684734.1 hypothetical protein [[Clostridium] aminophilum] Length: 407\nNumber of predicted TMHs: 2\nExp number of AAs in TMHs: 54.5417\nExp number, first 60 AAs: 15.55639\nTotal prob of N-in: 0.53835\nPOSSIBLE N-term signal sequence\noutside 1 225\nTMhelix 226 243\ninside 244 380\nTMhelix 381 403\noutside 404 407

34560 GCF_900112885.1_IMG-taxon_2593339256_annotated_assembly [Clostridium] aminophilum Terrabacteria group; Firmicutes; Clostridia; Clostridiales; Lachnospiraceae; Lachnoclostridium MKIPKAKVGIDRRQLIAAGAGILLGAMMCVVSSAEDHSGIRSFHRNLYGGSGRDEEVIVS WP_051684734.1 hypothetical protein [[Clostridium] aminophilum] Length: 407\nNumber of predicted TMHs: 2\nExp number of AAs in TMHs: 54.5417\nExp number, first 60 AAs: 15.55639\nTotal prob of N-in: 0.53835\nPOSSIBLE N-term signal sequence\noutside 1 225\nTMhelix 226 243\ninside 244 380\nTMhelix 381 403\noutside 404 407

34561 GCF_002160535.1_ASM216053v1 [Clostridium] saccharolyticum Terrabacteria group; Firmicutes; Clostridia; Clostridiales; Lachnospiraceae; Lachnoclostridium MFIYKRRAAVTAAAAMLTGFLALYPGGNMEVYADLEPFYNLSVSVSSELDRPLNEKIEVY WP_087383135.1 hypothetical protein [[Clostridium] saccharolyticum] Length: 286\nNumber of predicted TMHs: 1\nExp number of AAs in TMHs: 34.68186\nExp number, first 60 AAs: 15.59199\nTotal prob of N-in: 0.78703\nPOSSIBLE N-term signal sequence\noutside 1 263\nTMhelix 264 281\ninside 282 286

34562 GCF_900120345.1_PRJEB18024 Lachnoclostridium phocaeense Terrabacteria group; Firmicutes; Clostridia; Clostridiales; Lachnospiraceae; Lachnoclostridium MRGRRRRILAFVLSAALACSMMPFSASTVSAEEPAQEQDAEILTETGVIVPEDPGLPDNEWP_076779723.1 hypothetical protein [Lachnoclostridium phocaeense] Length: 1732\nNumber of predicted TMHs: 2\nExp number of AAs in TMHs: 38.1541699999999\nExp number, first 60 AAs: 16.22973\nTotal prob of N-in: 0.76635\nPOSSIBLE N-term signal sequence\ninside 1 6\nTMhelix 7 29\noutside 30 1703\nTMhelix 1704 1726\ninside 1727 1732

34563 GCF_002160765.1_ASM216076v1 Lachnoclostridium sp. An138 Terrabacteria group; Firmicutes; Clostridia; Clostridiales; Lachnospiraceae; Lachnoclostridium MKKRKRRPAVLLCNAAGTAVLVLLILACLPLTLPRLAGYRLYTVVSGSMEPEIPVGSLVM WP_087308291.1 signal peptidase I [Lachnoclostridium sp. An138] Length: 184\nNumber of predicted TMHs: 2\nExp number of AAs in TMHs: 44.17425\nExp number, first 60 AAs: 22.73543\nTotal prob of N-in: 0.99823\nPOSSIBLE N-term signal sequence\ninside 1 11\nTMhelix 12 34\noutside 35 142\nTMhelix 143 165\ninside 166 184

34564 GCF_000711825.1_ASM71182v1 [Clostridium] aminophilum DSM 10710 Terrabacteria group; Firmicutes; Clostridia; Clostridiales; Lachnospiraceae; Lachnoclostridium; [Clostridium] aminophilum MKIPKAKVGIDRRQLIAAGAGILLGAMMCVVSSAEDHSGIRSFHRNLYGGSGRDEEVIVS WP_051684734.1 hypothetical protein [[Clostridium] aminophilum] Length: 407\nNumber of predicted TMHs: 2\nExp number of AAs in TMHs: 54.5417\nExp number, first 60 AAs: 15.55639\nTotal prob of N-in: 0.53835\nPOSSIBLE N-term signal sequence\noutside 1 225\nTMhelix 226 243\ninside 244 380\nTMhelix 381 403\noutside 404 407

34565 GCF_000471845.1_ASM47184v1 [Clostridium] scindens VE202-05 Terrabacteria group; Firmicutes; Clostridia; Clostridiales; Lachnospiraceae; Lachnoclostridium; [Clostridium] scindens MGKRRRALLFAGAVTVLAFVFILFFVLSGQISENLEESSYDYLYGSTEIIRRQIDNGVRE WP_029467300.1 hypothetical protein [[Clostridium] scindens] Length: 337\nNumber of predicted TMHs: 2\nExp number of AAs in TMHs: 44.3669\nExp number, first 60 AAs: 22.06633\nTotal prob of N-in: 0.99955\nPOSSIBLE N-term signal sequence\ninside 1 6\nTMhelix 7 29\noutside 30 284\nTMhelix 285 307\ninside 308 337

34566 GCF_001406555.1_14207_7_9 Lachnospira pectinoschiza Terrabacteria group; Firmicutes; Clostridia; Clostridiales; Lachnospiraceae; Lachnospira

MKSRRRREWFRRSSAMFMALALIISGLSLPEGLFSITAKAAEKASAEWTVSSTMYADGDN WP_055306235.1 hypothetical protein [Lachnospira pectinoschiza] Length: 2411\nNumber of predicted TMHs: 2\nExp number of AAs in TMHs: 41.8215\nExp number, first 60 AAs: 20.67592\nTotal prob of N-in: 0.98428\nPOSSIBLE N-term signal sequence\ninside 1 12\nTMhelix 13 35\noutside 36 2382\nTMhelix 2383 2405\ninside 2406 2411

34567 GCF_001405835.1_13470_2_55 Lachnospira pectinoschiza Terrabacteria group; Firmicutes; Clostridia; Clostridiales; Lachnospiraceae; Lachnospira MKSRRRREWFRRSSAMFMALALIISGLSLPEGLFSITAKAAEKASAEWTVSSTMYADGDN WP_055173989.1 hypothetical protein [Lachnospira pectinoschiza] Length: 2403\nNumber of predicted TMHs: 2\nExp number of AAs in TMHs: 41.81899\nExp number, first 60 AAs: 20.67683\nTotal prob of N-in: 0.98440\nPOSSIBLE N-term signal sequence\ninside 1 12\nTMhelix 13 35\noutside 36 2374\nTMhelix 2375 2397\ninside 2398 2403

34568 GCF_001940225.1_ASM194022v1 Roseburia sp. 499 Terrabacteria group; Firmicutes; Clostridia; Clostridiales; Lachnospiraceae; Roseburia MALKGSRRQAVIVAVSGCLLAGLLALTDGKNKMDDSVERQEPGKGSQEQKYILNAEEVLE WP_075721826.1 hypothetical protein [Roseburia sp. 499] Length: 404\nNumber of predicted TMHs: 2\nExp number of AAs in TMHs: 63.2481\nExp number, first 60 AAs: 13.75664\nTotal prob of N-in: 0.32018\nPOSSIBLE N-term signal sequence\noutside 1 218\nTMhelix 219 236\ninside 237 378\nTMhelix 379 401\noutside 402 404

34569 GCF_900104635.1_PRJEB16356 Tyzzerella sp. Marseille-P3062 Terrabacteria group; Firmicutes; Clostridia; Clostridiales; Lachnospiraceae; Tyzzerella MQKRRRIIAIVLTAALSGTMLPANSLTMYAADKLSPAKSIRADVDKTKFTHKEWTGTDYTWP_071144471.1 glycoside hydrolase family 2 [Tyzzerella sp. Marseille-P3062] Length: 2606\nNumber of predicted TMHs: 2\nExp number of AAs in TMHs: 41.19167\nExp number, first 60 AAs: 22.03444\nTotal prob of N-in: 0.98620\nPOSSIBLE N-term signal sequence\ninside 1 6\nTMhelix 7 29\noutside 30 2581\nTMhelix 2582 2601\ninside 2602 2606

34570 GCF_000156035.2_ASM15603v2 Tyzzerella nexilis DSM 1787 Terrabacteria group; Firmicutes; Clostridia; Clostridiales; Lachnospiraceae; Tyzzerella; Tyzzerella nexilis MQKRRRIIAIVLTAALSGTMLPANSLTMYAADKLSPAKSIRADVDKTKFTHKEWTGTDYTWP_004614900.1 glycoside hydrolase family 2 [Tyzzerella nexilis] Length: 2606\nNumber of predicted TMHs: 2\nExp number of AAs in TMHs: 41.19167\nExp number, first 60 AAs: 22.03444\nTotal prob of N-in: 0.98620\nPOSSIBLE N-term signal sequence\ninside 1 6\nTMhelix 7 29\noutside 30 2581\nTMhelix 2582 2601\ninside 2602 2606

34571 GCF_000218445.1_Lach_bact_1_1_57FAA_V1 Lachnospiraceae bacterium 1_1_57FAA Terrabacteria group; Firmicutes; Clostridia; Clostridiales; Lachnospiraceae; unclassified Lachnospiraceae MRVKRRKTMALLAAVSIVCAQSTVSDVFAAGAEEVAAQENNAVSIEYLTLQPGERDDSVN WP_009243371.1 MULTISPECIES: hypothetical protein [unclassified Lachnospiraceae] Length: 1161\nNumber of predicted TMHs: 1\nExp number of AAs in TMHs: 23.64508\nExp number, first 60 AAs: 1.33479\nTotal prob of N-in: 0.10563\noutside 1 1127\nTMhelix 1128 1150\ninside 1151 1161

34572 GCF_000209405.1_Lach_bact_3_1_46FAA_V1 Lachnospiraceae bacterium 3_1_46FAA Terrabacteria group; Firmicutes; Clostridia; Clostridiales; Lachnospiraceae; unclassified Lachnospiraceae MRVKRRKTMALLAAVSIVCAQSTVSDVFAAGAEEVAAQENNAVSIEYLTLQPGERDDSVN WP_009319933.1 hypothetical protein [[Ruminococcus] torques] Length: 1161\nNumber of predicted TMHs: 1\nExp number of AAs in TMHs: 23.64632\nExp number, first 60 AAs: 1.33478\nTotal prob of N-in: 0.10565\noutside 1 1127\nTMhelix 1128 1150\ninside 1151 1161

34573 GCF_000403475.2_Lach_bact_3-2_V1 Lachnospiraceae bacterium 3-2 Terrabacteria group; Firmicutes; Clostridia; Clostridiales; Lachnospiraceae; unclassified Lachnospiraceae MNRRNLIRKVTAVVVLFAMAFNFVPFFVMYETKAASHNITESYVGHYDQYHNGVHQMKYQ WP_016223196.1 hypothetical protein [Lachnospiraceae bacterium 3-2] Length: 795\nNumber of predicted TMHs: 2\nExp number of AAs in TMHs: 40.3876\nExp number, first 60 AAs: 19.19422\nTotal prob of N-in: 0.99886\nPOSSIBLE N-term signal sequence\ninside 1 12\nTMhelix 13 30\noutside 31 769\nTMhelix 770 792\ninside 793 795

34574 GCF_000185545.1_ASM18554v1 Lachnospiraceae bacterium 8_1_57FAA Terrabacteria group; Firmicutes; Clostridia; Clostridiales; Lachnospiraceae; unclassified Lachnospiraceae MRVKRRKTMALLAAVSIVCAQSTVSDVFAAGAEEVAAQENNAVSIEYLTLQPGERDDSVN WP_009243371.1 MULTISPECIES: hypothetical protein [unclassified Lachnospiraceae] Length: 1161\nNumber of predicted TMHs: 1\nExp number of AAs in TMHs: 23.64508\nExp number, first 60 AAs: 1.33479\nTotal prob of N-in: 0.10563\noutside 1 1127\nTMhelix 1128 1150\ninside 1151 1161

34575 GCF_000702145.1_ASM70214v1 Lachnospiraceae bacterium FD2005 Terrabacteria group; Firmicutes; Clostridia; Clostridiales; Lachnospiraceae; unclassified Lachnospiraceae
MRRTICRATHQSLQQHMKQRRKLMKNKIRLITVAAALMIGSMPLTAFAQVPDNVDTSVID WP_081826815.1
hypothetical protein [Lachnospiraceae bacterium FD2005] Length: 239\nNumber of predicted TMHs: 2\nExp number of AAs in TMHs: 43.69764\nExp number, first 60 AAs: 21.87742\nTotal prob of N-in: 0.99982\nPOSSIBLE N-term signal sequence\ninside 1 27\nTMhelix 28 50\noutside 51 183\nTMhelix 184 203\ninside 204 239

34576 GCF_000469425.1_ASM46942v1 Oscillibacter sp. KLE 1728 Terrabacteria group; Firmicutes; Clostridia; Clostridiales; Oscillospiraceae; Oscillibacter
MEHRDWKNSLQETAGRHASRRGAFSAGLTALAVAAVLVFNLLAAQLPEQWAQIDLTGSGI WP_021751121.1
MULTISPECIES: hypothetical protein [Oscillibacter] Length: 472\nNumber of predicted TMHs: 2\nExp number of AAs in TMHs: 44.9401199999999\nExp number, first 60 AAs: 22.34504\nTotal prob of N-in: 0.99114\nPOSSIBLE N-term signal sequence\ninside 1 20\nTMhelix 21 43\noutside 44 441\nTMhelix 442 464\ninside 465 472

34577 GCF_000469445.2_ASM46944v2 Oscillibacter sp. KLE 1745 Terrabacteria group; Firmicutes; Clostridia; Clostridiales; Oscillospiraceae; Oscillibacter
MEHRDWKNSLQETAGRHASRRGAFSAGLTALAVAAVLVFNLLAAQLPEQWAQIDLTGSGI WP_021751121.1
MULTISPECIES: hypothetical protein [Oscillibacter] Length: 472\nNumber of predicted TMHs: 2\nExp number of AAs in TMHs: 44.9401199999999\nExp number, first 60 AAs: 22.34504\nTotal prob of N-in: 0.99114\nPOSSIBLE N-term signal sequence\ninside 1 20\nTMhelix 21 43\noutside 44 441\nTMhelix 442 464\ninside 465 472

34578 GCF_000471765.1_ASM47176v1 Oscillospiraceae bacterium VE202-24 Terrabacteria group; Firmicutes; Clostridia; Clostridiales; Oscillospiraceae; unclassified Oscillospiraceae
MEHRDWKNSLQETAGRHASRRGAFSAGLTALAVAAVLVFNLLAAQLPEQWAQIDLTGSGI WP_025544707.1
hypothetical protein [Oscillospiraceae bacterium VE202-24] Length: 472\nNumber of predicted TMHs: 2\nExp number of AAs in TMHs: 44.9401299999999\nExp number, first 60 AAs: 22.34504\nTotal prob of N-in: 0.99114\nPOSSIBLE N-term signal sequence\ninside 1 20\nTMhelix 21 43\noutside 44 441\nTMhelix 442 464\ninside 465 472

34579 GCF_000620305.1_ASM62030v1 Desulfitibacter alkalitolerans DSM 16504 Terrabacteria group; Firmicutes; Clostridia; Clostridiales; Peptococcaceae; Desulfitibacter; Desulfitibacter alkalitolerans
MVINMERRRFIKTSLTAGFAVAINPLSAASVLADNNKEAILMDNVKCVNCQRCVKACSDL WP_051534117.1
hypothetical protein [Desulfitibacter alkalitolerans] Length: 278\nNumber of predicted TMHs: 2\nExp number of AAs in TMHs: 42.46421\nExp number, first 60 AAs: 2.21901\nTotal prob of N-in: 0.87286\ninside 1 194\nTMhelix 195 212\noutside 213 236\nTMhelix 237 259\ninside 260 278

34580 GCF_001468785.1_ASM146878v1 Desulfitobacterium hafniense Terrabacteria group; Firmicutes; Clostridia; Clostridiales; Peptococcaceae; Desulfitobacterium
MLNRREFLKLVVKGAILGNFISLVTPELEKALAQGEINKLPIIMVETGTCTGDSISLDNI WP_005808601.1
oxidoreductase [Desulfitobacterium hafniense] Length: 375\nNumber of predicted TMHs: 1\nExp number of AAs in TMHs: 18.35233\nExp number, first 60 AAs: 0.599149999999999\nTotal prob of N-in: 0.06532\noutside 1 314\nTMhelix 315 337\ninside 338 375

34581 GCF_000384015.1_ASM38401v1 Desulfitobacterium sp. PCE1 Terrabacteria group; Firmicutes; Clostridia; Clostridiales; Peptococcaceae; Desulfitobacterium
MLNRREFLKLVVKGAILGNFISLVTPELEKALAQGEIKKLPIVMVETGTCTGDSISLDNI WP_019849464.1
oxidoreductase [Desulfitobacterium sp. PCE1] Length: 374\nNumber of predicted TMHs: 1\nExp number of AAs in TMHs: 18.94958\nExp number, first 60 AAs: 0.617569999999999\nTotal prob of N-in: 0.08603\noutside 1 314\nTMhelix 315 337\ninside 338 374

34582 GCF_000243155.2_ASM24315v3 Desulfitobacterium dehalogenans ATCC 51507 Terrabacteria group; Firmicutes; Clostridia; Clostridiales; Peptococcaceae; Desulfitobacterium; Desulfitobacterium dehalogenans
MLNRRKFLKLVVKGAILGNFISLVTPELEKALAQGEIKKLPIVMVETGTCTGDSISLDNI WP_014792615.1
oxidoreductase [Desulfitobacterium dehalogenans] Length: 374\nNumber of predicted TMHs: 1\nExp number of AAs in TMHs: 19.42614\nExp number, first 60 AAs: 1.03837\nTotal prob of N-in: 0.14511\noutside 1 314\nTMhelix 315 337\ninside 338 374

34583 GCF_000021925.1_ASM2192v1 Desulfitobacterium hafniense DCB-2 Terrabacteria group; Firmicutes; Clostridia; Clostridiales; Peptococcaceae; Desulfitobacterium; Desulfitobacterium hafniense

MLNRREFLKLVVKGAILGNFISLVTPELEKALAQGEINKLPIIMVETGTCTGDSISLDNI WP_005808601.1 oxidoreductase [Desulfitobacterium hafniense] Length: 375\nNumber of predicted TMHs: 1\nExp number of AAs in TMHs: 18.35233\nExp number, first 60 AAs: 0.599149999999999\nTotal prob of N-in: 0.06532\noutside 1 314\nTMhelix 315 337\ninside 338 375

34584 GCF_000238035.1_ASM23803v1 Desulfitobacterium hafniense DP7 Terrabacteria group; Firmicutes; Clostridia; Clostridiales; Peptococcaceae; Desulfitobacterium; Desulfitobacterium hafniense MLNRREFLKLVVKGAILGNFISLVTPELEKALAQGEINKLPIIMVETGTCTGDSISLDNI WP_005808601.1 oxidoreductase [Desulfitobacterium hafniense] Length: 375\nNumber of predicted TMHs: 1\nExp number of AAs in TMHs: 18.35233\nExp number, first 60 AAs: 0.599149999999999\nTotal prob of N-in: 0.06532\noutside 1 314\nTMhelix 315 337\ninside 338 375

34585 GCF_000379505.1_ASM37950v1 Desulfitobacterium hafniense PCP-1 Terrabacteria group; Firmicutes; Clostridia; Clostridiales; Peptococcaceae; Desulfitobacterium; Desulfitobacterium hafniense MLNRREFLKLVVKGAILGNFISLVTPELEKALAQGEINKLPIIMVETGTCTGDSISLDNI WP_018214190.1 oxidoreductase [Desulfitobacterium hafniense] Length: 375\nNumber of predicted TMHs: 1\nExp number of AAs in TMHs: 17.99108\nExp number, first 60 AAs: 0.597589999999999\nTotal prob of N-in: 0.06788\noutside 1 314\nTMhelix 315 337\ninside 338 375

34586 GCF_000378805.1_ASM37880v1 Desulfitobacterium hafniense TCP-A Terrabacteria group; Firmicutes; Clostridia; Clostridiales; Peptococcaceae; Desulfitobacterium; Desulfitobacterium hafniense MLNRREFLKLVVKGAILGNFISLVTPELEKALAQGEINKLPIIMVETGTCTGDSISLDNI WP_018214190.1 oxidoreductase [Desulfitobacterium hafniense] Length: 375\nNumber of predicted TMHs: 1\nExp number of AAs in TMHs: 17.99108\nExp number, first 60 AAs: 0.597589999999999\nTotal prob of N-in: 0.06788\noutside 1 314\nTMhelix 315 337\ninside 338 375

34587 GCF_000010045.1_ASM1004v1 Desulfitobacterium hafniense Y51 Terrabacteria group; Firmicutes; Clostridia; Clostridiales; Peptococcaceae; Desulfitobacterium; Desulfitobacterium hafniense MLNRREFLKLVVKGAILGNFISLVTPELEKALAQGEINKLPIIMVETGTCTGDSISLDNI WP_011459323.1 oxidoreductase [Desulfitobacterium hafniense] Length: 374\nNumber of predicted TMHs: 1\nExp number of AAs in TMHs: 17.3859\nExp number, first 60 AAs: 0.603089999999999\nTotal prob of N-in: 0.05869\noutside 1 314\nTMhelix 315 337\ninside 338 374

34588 GCF_001029285.1_ASM102928v1 Desulfosporosinus acididurans Terrabacteria group; Firmicutes; Clostridia; Clostridiales; Peptococcaceae; Desulfosporosinus MLDRREFLKLIVKGAILGNFYQLITPHLSRAIAAGDIKKKLPVIMIETGTCTGDSISFDN WP_047808671.1 oxidoreductase [Desulfosporosinus acididurans] Length: 377\nNumber of predicted TMHs: 1\nExp number of AAs in TMHs: 19.46481\nExp number, first 60 AAs: 0.75492\nTotal prob of N-in: 0.08730\noutside 1 315\nTMhelix 316 338\ninside 339 377

34589 GCF_002196785.1_ASM219678v1 Desulfosporosinus sp. FKA Terrabacteria group; Firmicutes; Clostridia; Clostridiales; Peptococcaceae; Desulfosporosinus MLDRREFLKLIVKGAILGNFYQLITPHLSRAIAAGDIKKKLPVIMIETGTCTGDSISFDN WP_088187428.1 oxidoreductase [Desulfosporosinus sp. FKA] Length: 377\nNumber of predicted TMHs: 1\nExp number of AAs in TMHs: 19.4649\nExp number, first 60 AAs: 0.755\nTotal prob of N-in: 0.08730\noutside 1 315\nTMhelix 316 338\ninside 339 377

34590 GCF_002196705.1_ASM219670v1 Desulfosporosinus sp. FKB Terrabacteria group; Firmicutes; Clostridia; Clostridiales; Peptococcaceae; Desulfosporosinus MLDRREFLKLIVKGAILGNFYQLITPHLSQAIAAGDIKKKLPVIMIETGTCTGDSISFDN WP_088228118.1 oxidoreductase [Desulfosporosinus sp. FKB] Length: 377\nNumber of predicted TMHs: 1\nExp number of AAs in TMHs: 20.27115\nExp number, first 60 AAs: 0.53681\nTotal prob of N-in: 0.04459\noutside 1 315\nTMhelix 316 338\ninside 339 377

34591 GCF_001936615.1_ASM193661v1 Desulfosporosinus sp. OL Terrabacteria group; Firmicutes; Clostridia; Clostridiales; Peptococcaceae; Desulfosporosinus MLSRRDFLKLVVKGAVLGNFYQLITPPLAQAVAAGEVKKLPVVMIETGTCTGDSISLDNI WP_075364354.1 oxidoreductase [Desulfosporosinus sp. OL] Length: 372\nNumber of predicted TMHs: 1\nExp number of AAs in TMHs: 16.71122\nExp number, first 60 AAs: 0.40232\nTotal prob of N-in: 0.03906\noutside 1 314\nTMhelix 315 337\ninside 338 372

34592 GCF_900129935.1_IMG-taxon_2585428067_annotated_assembly Desulfosporosinus lacus DSM 15449 Terrabacteria group; Firmicutes; Clostridia; Clostridiales; Peptococcaceae; Desulfosporosinus; Desulfosporosinus lacus MKKDDFISRRTFFKRTTAAGLLGALAATGLNKPVQAAGLSNGKLGTLIDLSKCDGCKSYG WP_073030090.1 oxidoreductase [Desulfosporosinus lacus] Length: 358\nNumber of predicted TMHs: 1\nExp number of AAs in TMHs: 23.53875\nExp number, first 60 AAs: 3.4523\nTotal prob of N-in: 0.24910\noutside 1 309\nTMhelix 310 332\ninside 333 358

34593 GCF_000235605.1_ASM23560v1 Desulfosporosinus orientis DSM 765 Terrabacteria group; Firmicutes; Clostridia; Clostridiales; Peptococcaceae; Desulfosporosinus; Desulfosporosinus orientis MLSRRDFFKLVIKGAVLGNFCQLISPPLAEAVTAGEVNKLPVVMIETGSCTGDSISLDNI WP_014182846.1 oxidoreductase [Desulfosporosinus orientis] Length: 372\nNumber of predicted TMHs: 1\nExp number of AAs in TMHs: 14.35115\nExp number, first 60 AAs: 0.09804\nTotal prob of N-in: 0.02966\noutside 1 314\nTMhelix 315 337\ninside 338 372

34594 GCF_000244895.1_ASM24489v1 Desulfosporosinus youngiae DSM 17734 Terrabacteria group; Firmicutes; Clostridia; Clostridiales; Peptococcaceae; Desulfosporosinus; Desulfosporosinus youngiae MNDNISRRTFFKRTTVAGLLGALAATGLDKPVQAAGLSSGKLGTLIDLTKCDGCKDFETP WP_007782756.1 oxidoreductase [Desulfosporosinus youngiae] Length: 353\nNumber of predicted TMHs: 1\nExp number of AAs in TMHs: 20.99045\nExp number, first 60 AAs: 0.79929\nTotal prob of N-in: 0.12586\noutside 1 307\nTMhelix 308 327\ninside 328 353

34595 GCF_900129895.1_IMG-taxon_2585428063_annotated_assembly Desulfotomaculum hydrothermale Lam5 = DSM 18033 Terrabacteria group; Firmicutes; Clostridia; Clostridiales; Peptococcaceae; Desulfotomaculum; Desulfotomaculum hydrothermale MAGLSRREFIKRSVAGGVAGSAVLYGFHKPAAAAAKGVEGSVGTFIDLSQCNGCAGQSIP WP_008409599.1 4Fe-4S ferredoxin iron-sulfur-binding domain-containing protein [Desulfotomaculum hydrothermale] Length: 338\nNumber of predicted TMHs: 1\nExp number of AAs in TMHs: 21.13587\nExp number, first 60 AAs: 0.88594\nTotal prob of N-in: 0.14567\noutside 1 306\nTMhelix 307 329\ninside 330 338

34596 GCF_000315365.1_ASM31536v1 Desulfotomaculum hydrothermale Lam5 = DSM 18033 Terrabacteria group; Firmicutes; Clostridia; Clostridiales; Peptococcaceae; Desulfotomaculum; Desulfotomaculum hydrothermale MAGLSRREFIKRSVAGGVAGSAVLYGFHKPAAAAAKGVEGSVGTFIDLSQCNGCAGQSIP WP_008409599.1 4Fe-4S ferredoxin iron-sulfur-binding domain-containing protein [Desulfotomaculum hydrothermale] Length: 338\nNumber of predicted TMHs: 1\nExp number of AAs in TMHs: 21.13587\nExp number, first 60 AAs: 0.88594\nTotal prob of N-in: 0.14567\noutside 1 306\nTMhelix 307 329\ninside 330 338

34597 GCF_900129195.1_IMG-taxon_2582580748_annotated_assembly Desulfotomaculum putei DSM 12395 Terrabacteria group; Firmicutes; Clostridia; Clostridiales; Peptococcaceae; Desulfotomaculum; Desulfotomaculum putei MSGLSRREFLKRSLATGIAGGAVLYGGNKTAGASTKSLTGPVGTLIDLTKCNGCAGEKVA WP_073234829.1 oxidoreductase [Desulfotomaculum putei] Length: 337\nNumber of predicted TMHs: 1\nExp number of AAs in TMHs: 20.76383\nExp number, first 60 AAs: 0.25623\nTotal prob of N-in: 0.04094\noutside 1 305\nTMhelix 306 328\ninside 329 337

34598 GCF_000451365.2_ASM45136v2 Clostridioides difficile Y165 Terrabacteria group; Firmicutes; Clostridia; Clostridiales; Peptostreptococcaceae; Clostridioides; Clostridioides difficile TRRGYTGEQTARRVLDSNGLYDVKIEMVRGHLSDHYDPRRKAVRLSEDVYYGTSITSVAV WP_021412057.1 neutral zinc metallopeptidase, partial [Clostridioides difficile] Length: 190\nNumber of predicted TMHs: 2\nExp number of AAs in TMHs: 66.8346\nExp number, first 60 AAs: 2.71054\nTotal prob of N-in: 0.81079\ninside 1 84\nTMhelix 85 107\noutside 108 161\nTMhelix 162 184\ninside 185 190

34599 GCF_000147675.1_ASM14767v1 Peptostreptococcus stomatis DSM 17678 Terrabacteria group; Firmicutes; Clostridia; Clostridiales; Peptostreptococcaceae; Peptostreptococcus; Peptostreptococcus stomatis MRGKKDERRPPLNAVNTVITHFSITDQDGNPVTELAKYKKFKLKISWDASSYGAGNLKAG WP_007790986.1 LPXTG-motif cell wall anchor domain protein [Peptostreptococcus stomatis] Length: 705\nNumber of predicted TMHs: 1\nExp number of AAs in TMHs: 22.37313\nExp number, first 60 AAs: 0.00324\nTotal prob of N-in: 0.00346\noutside 1 675\nTMhelix 676 698\ninside 699 705

34600 GCF_002160175.1_ASM216017v1 Faecalibacterium sp. An192 Terrabacteria group; Firmicutes; Clostridia; Clostridiales; Ruminococcaceae; Faecalibacterium MKKISRRSFLKVAGAAAATMGLAACGGSQSTTAASGGAAAGSELSGELVFTIWDNNLMEY WP_087364128.1 sugar ABC transporter substrate-binding protein [Faecalibacterium sp. An192] Length: 447\nNumber of

predicted TMHs: 1\nExp number of AAs in TMHs: 23.74691\nExp number, first 60 AAs: 1.91765\nTotal prob of N-in: 0.15000\noutside 1 410\nTMhelix 411 433\ninside 434 447

34601 GCF_900167555.1_IMG-taxon_2590828806_annotated_assembly Gemmiger formicilis Terrabacteria group; Firmicutes; Clostridia; Clostridiales; Ruminococcaceae; Gemmiger
MKALKNWFSRRGALPLTAYLLALAVWLVLGAVHFGSDAAARAQGRLAEETMAVTDWQLVG WP_078784464.1
hypothetical protein [Gemmiger formicilis] Length: 213\nNumber of predicted TMHs: 2\nExp number of AAs in TMHs: 45.33855\nExp number, first 60 AAs: 22.47451\nTotal prob of N-in: 0.97164\nPOSSIBLE N-term signal sequence\ninside 1 11\nTMhelix 12 34\noutside 35 182\nTMhelix 183 205\ninside 206 213

34602 GCF_000154345.1_ASM15434v1 [Clostridium] leptum DSM 753 Terrabacteria group; Firmicutes; Clostridia; Clostridiales; Ruminococcaceae; Ruminiclostridium; [Clostridium] leptum
MKRRKLLSLFSVLAFVLSVQFSVLGVSGPGAEDAPADMSGTALNAQEREAFHRNVADDEA WP_003528239.1
hypothetical protein [[Clostridium] leptum] Length: 372\nNumber of predicted TMHs: 2\nExp number of AAs in TMHs: 42.21352\nExp number, first 60 AAs: 19.68002\nTotal prob of N-in: 0.91055\nPOSSIBLE N-term signal sequence\ninside 1 6\nTMhelix 7 29\noutside 30 342\nTMhelix 343 365\ninside 366 372

34603 GCF_000157955.1_ASM15795v1 Subdoligranulum variabile DSM 15176 Terrabacteria group; Firmicutes; Clostridia; Clostridiales; Ruminococcaceae; Subdoligranulum; Subdoligranulum variabile
MKTLQRWFSRRGALPLTCYLAAAALWLVLGAVHWGSDALARSAGTMYAAEIPVTDWQLAG WP_007047833.1
hypothetical protein [Subdoligranulum variabile] Length: 221\nNumber of predicted TMHs: 2\nExp number of AAs in TMHs: 45.06312\nExp number, first 60 AAs: 22.59627\nTotal prob of N-in: 0.99809\nPOSSIBLE N-term signal sequence\ninside 1 11\nTMhelix 12 34\noutside 35 186\nTMhelix 187 209\ninside 210 221

34604 GCF_002159455.1_ASM215945v1 Flavonifractor sp. An306 Terrabacteria group; Firmicutes; Clostridia; Clostridiales; unclassified Clostridiales; Flavonifractor
MKRRRLFSLLTALALLAALLPARAAEEVFSIATPQDLARFAQLCTKDTWSQGVTAELTAD WP_087347104.1 hypothetical protein [Flavonifractor sp. An306] Length: 1053\nNumber of predicted TMHs: 1\nExp number of AAs in TMHs: 32.0665799999999\nExp number, first 60 AAs: 9.69669\nTotal prob of N-in: 0.51333\noutside 1 1014\nTMhelix 1015 1037\ninside 1038 1053

34605 GCF_002160435.1_ASM216043v1 Pseudoflavonifractor sp. An176 Terrabacteria group; Firmicutes; Clostridia; Clostridiales; unclassified Clostridiales; Pseudoflavonifractor
MLHLSCVFVQRGVLRPSARRTGFLPGMKRRRNIMFQKKRLLAGALALVMGANLVTVGAFA WP_087232861.1
hypothetical protein [Pseudoflavonifractor sp. An176] Length: 1895\nNumber of predicted TMHs: 2\nExp number of AAs in TMHs: 42.84195\nExp number, first 60 AAs: 20.18837\nTotal prob of N-in: 0.96887\nPOSSIBLE N-term signal sequence\ninside 1 39\nTMhelix 40 62\noutside 63 1867\nTMhelix 1868 1890\ninside 1891 1895

34606 GCF_002160275.1_ASM216027v1 Pseudoflavonifractor sp. An187 Terrabacteria group; Firmicutes; Clostridia; Clostridiales; unclassified Clostridiales; Pseudoflavonifractor
MLHLSCVLVQRGVLRPSARRTGFLPGMKRRRNIMFQKKRLLAGALALVMGANLVTVGAFA WP_087392496.1
hypothetical protein [Pseudoflavonifractor sp. An187] Length: 1896\nNumber of predicted TMHs: 2\nExp number of AAs in TMHs: 42.84116\nExp number, first 60 AAs: 20.18494\nTotal prob of N-in: 0.96860\nPOSSIBLE N-term signal sequence\ninside 1 39\nTMhelix 40 62\noutside 63 1868\nTMhelix 1869 1891\ninside 1892 1896

34607 GCF_002161675.1_ASM216167v1 Pseudoflavonifractor sp. An44 Terrabacteria group; Firmicutes; Clostridia; Clostridiales; unclassified Clostridiales; Pseudoflavonifractor
MLHLSCVFVQRGVLRPSARRTGFLPGMKRRRNIMFQKKRLLAGALALVMGANLVTVGAFA WP_087370191.1
hypothetical protein [Pseudoflavonifractor sp. An44] Length: 1906\nNumber of predicted TMHs: 2\nExp number of AAs in TMHs: 42.84195\nExp number, first 60 AAs: 20.18837\nTotal prob of N-in: 0.96887\nPOSSIBLE N-term signal sequence\ninside 1 39\nTMhelix 40 62\noutside 63 1878\nTMhelix 1879 1901\ninside 1902 1906

34608 GCF_000020005.1_ASM2000v1 Natranaerobius thermophilus JW/NM-WN-LF Terrabacteria group; Firmicutes; Clostridia; Natranaerobiales; Natranaerobiaceae; Natranaerobius; Natranaerobius thermophilus
MKENKGFKLKRRQFIQSMAGTGLLAGLSLSAPRIAEAGENNGSEYAMMVDKTECVGCNAC WP_012447020.1 4Fe-4S ferredoxin [Natranaerobius thermophilus] Length: 284\nNumber of predicted TMHs: 1\nExp number of AAs in TMHs: 21.56893\nExp number, first 60 AAs: 1.28\nTotal prob of N-in: 0.12581\noutside 1 247\nTMhelix 248 267\ninside 268 284

34609 GCF_000189775.2_ASM18977v3 Thermoanaerobacterium xylanolyticum LX-11 Terrabacteria group; Firmicutes; Clostridia; Thermoanaerobacterales; Thermoanaerobacterales Family III. Incertae Sedis;

Thermoanaerobacterium; Thermoanaerobacterium xylanolyticum
MLENGYVLTGIHNRRKYVVEKKIGHGGVANVYLVHDEENLKYALKISEDLISITREHKVL WP_013787072.1 serine/threonine protein kinase [Thermoanaerobacterium xylanolyticum] Length: 279\nNumber of predicted TMHs: 1\nExp number of AAs in TMHs: 28.25075\nExp number, first 60 AAs: 0.00048\nTotal prob of N-in: 0.28266\noutside 1 255\nTMhelix 256 278\ninside 279 279

34610 GCF_000157995.1_ASM15799v1 Holdemania filiformis DSM 12042 Terrabacteria group; Firmicutes; Erysipelotrichia; Erysipelotrichales; Erysipelotrichaceae; Holdemania; Holdemania filiformis
MEKNKTKGITRRDFIKGAAAGAAGLVGANLLSGCQNTASVPASAKIFNEPEKWDYECDVV WP_006057305.1 FAD binding domain-containing protein [Holdemania filiformis] Length: 559\nNumber of predicted TMHs: 1\nExp number of AAs in TMHs: 21.50808\nExp number, first 60 AAs: 2.52214\nTotal prob of N-in: 0.14289\noutside 1 529\nTMhelix 530 552\ninside 553 559

34611 GCF_000327285.1_ASM32728v1 Holdemania massiliensis AP2 Terrabacteria group; Firmicutes; Erysipelotrichia; Erysipelotrichales; Erysipelotrichaceae; Holdemania; Holdemania massiliensis
MQRRDIMKKKRGHSILIAGLSLGMFFALQPVAAQTPSTEQKDQASIHFADPEPLNWNTNL WP_081587929.1 hypothetical protein [Holdemania massiliensis] Length: 963\nNumber of predicted TMHs: 2\nExp number of AAs in TMHs: 43.15417\nExp number, first 60 AAs: 20.42133\nTotal prob of N-in: 0.97859\nPOSSIBLE N-term signal sequence\ninside 1 12\nTMhelix 13 32\noutside 33 931\nTMhelix 932 954\ninside 955 963

34612 GCF_001544015.1_ASM154401v1 Limnochorda pilosaTerrabacteria group; Firmicutes; Limnochordia; Limnochordales; Limnochordaceae; Limnochorda
MGTRFGVEGEPGMAQEAPLPTSYDDEIDLRALVEVLLRRRSLILGLVAVAVATAAALSYF WP_082726110.1 hypothetical protein [Limnochorda pilosa] Length: 288\nNumber of predicted TMHs: 2\nExp number of AAs in TMHs: 43.63042\nExp number, first 60 AAs: 19.07369\nTotal prob of N-in: 0.14542\nPOSSIBLE N-term signal sequence\noutside 1 40\nTMhelix 41 63\ninside 64 252\nTMhelix 253 275\noutside 276 288

34613 GCF_002159575.1_ASM215957v1 Megamonas hypermegale Terrabacteria group; Firmicutes; Negativicutes; Selenomonadales; Selenomonadaceae; Megamonas
MEKYSSIWDMYQKKGMSRRSFIKACTAMAAMLGIAPTMLSNVVEAAEKRLPVVVWLHGHE WP_027889017.1 uptake hydrogenase small subunit [Megamonas hypermegale] Length: 366\nNumber of predicted TMHs: 2\nExp number of AAs in TMHs: 39.99358\nExp number, first 60 AAs: 21.28602\nTotal prob of N-in: 0.97452\nPOSSIBLE N-term signal sequence\ninside 1 20\nTMhelix 21 43\noutside 44 326\nTMhelix 327 349\ninside 350 366

34614 GCF_000423385.1_ASM42338v1 Megamonas hypermegale DSM 1672 Terrabacteria group; Firmicutes; Negativicutes; Selenomonadales; Selenomonadaceae; Megamonas; Megamonas hypermegale
MEKYSSIWDMYQKKGMSRRSFIKACTAMAAMLGIAPTMLSNVVEAAEKRLPVVVWLHGHE WP_027889017.1 uptake hydrogenase small subunit [Megamonas hypermegale] Length: 366\nNumber of predicted TMHs: 2\nExp number of AAs in TMHs: 39.99358\nExp number, first 60 AAs: 21.28602\nTotal prob of N-in: 0.97452\nPOSSIBLE N-term signal sequence\ninside 1 20\nTMhelix 21 43\noutside 44 326\nTMhelix 327 349\ninside 350 366

34615 GCF_900116485.1_IMG-taxon_2595698242_annotated_assembly Selenomonas ruminantium Terrabacteria group; Firmicutes; Negativicutes; Selenomonadales; Selenomonadaceae; Selenomonas
MENEESIDLGRLMQIMLERRKVVGGIIAGCTIVAMGAALMMPKQYESTTLVQTRSAGKDI WP_075429010.1 chain-length determining protein [Selenomonas ruminantium] Length: 451\nNumber of predicted TMHs: 2\nExp number of AAs in TMHs: 54.56223\nExp number, first 60 AAs: 20.32235\nTotal prob of N-in: 0.89546\nPOSSIBLE N-term signal sequence\ninside 1 20\nTMhelix 21 40\noutside 41 424\nTMhelix 425 444\ninside 445 451

34616 GCF_900113715.1_IMG-taxon_2623620514_annotated_assembly Selenomonas ruminantium Terrabacteria group; Firmicutes; Negativicutes; Selenomonadales; Selenomonadaceae; Selenomonas
MEHEESIDLERLMQIMLERRKVVGGIIAGCTALAIGTAFILPKQYESMTMVQTRTAGNDL WP_075444307.1 chain-length determining protein [Selenomonas ruminantium] Length: 449\nNumber of predicted TMHs: 2\nExp number of AAs in TMHs: 48.00994\nExp number, first 60 AAs: 20.79607\nTotal prob of N-in: 0.94732\nPOSSIBLE N-term signal sequence\ninside 1 20\nTMhelix 21 40\noutside 41 422\nTMhelix 423 445\ninside 446 449

34617 GCF_000686945.1_ASM68694v1 Selenomonas sp. AE3005 Terrabacteria group; Firmicutes; Negativicutes; Selenomonadales; Selenomonadaceae; Selenomonas
MENSQEEAIDLGRLMQIIYDRRKIVGGIVGGCTALALIVSLVLPKTYESTTLVQTRSAGK WP_028128640.1 chain-length determining protein [Selenomonas sp. AE3005] Length: 453\nNumber of predicted TMHs: 2\nExp number of AAs in TMHs: 52.4383700000001\nExp number, first 60 AAs: 20.0795\nTotal prob of N-in: 0.62989\nPOSSIBLE N-term signal sequence\ninside 1 23\nTMhelix 24 43\noutside 44 426\nTMhelix 427 446\ninside 447 453

34618	GCF_000424065.1_ASM42406v1	Selenomonas ruminantium subsp. ruminantium ATCC 12561	Terrabacteria group; Firmicutes; Negativicutes; Selenomonadales; Selenomonadaceae; Selenomonas; Selenomonas ruminantium; Selenomonas ruminantium subsp. ruminantium	MENNVQKEESIDLGKLMQIMYNRRKTVGTIVGGCTALALIVSFALPKTYESTTLVQTRSN WP_026765944.1	chain-length determining protein [Selenomonas ruminantium]	Length: 448\nNumber of predicted TMHs: 2\nExp number of AAs in TMHs: 51.78854\nExp number, first 60 AAs: 20.21123\nTotal prob of N-in: 0.90705\nPOSSIBLE N-term signal sequence\ninside 1 26\nTMhelix 27 46\noutside 47 421\nTMhelix 422 441\ninside 442 448

34619	GCF_000219125.1_ASM21912v1	Acetonema longum DSM 6540	Terrabacteria group; Firmicutes; Negativicutes; Selenomonadales; Sporomusaceae; Acetonema; Acetonema longum	MTTGTSLWDLFEQKSMSRRSFLKTCTALTALMGLPAAMVRQVVEAAEKKALPVVIWLHGH	WP_004573251.1	hydrogenase (NiFe) small subunit HydA [Acetonema longum]	Length: 367\nNumber of predicted TMHs: 1\nExp number of AAs in TMHs: 24.23215\nExp number, first 60 AAs: 9.35975\nTotal prob of N-in: 0.44268\noutside 1 325\nTMhelix 326 348\ninside 349 367

34620	GCF_001546845.1_ASM154684v1	Veillonella atypica	Terrabacteria group; Firmicutes; Negativicutes; Veillonellales; Veillonellaceae; Veillonella	MEQIIDIKDIAKILVRKRRTIINVTGICILLGGAYLIVTPSTYQSTAMLRIKQTQGLGNS	WP_060807115.1	chain-length determining protein [Veillonella atypica]	Length: 479\nNumber of predicted TMHs: 2\nExp number of AAs in TMHs: 44.52906\nExp number, first 60 AAs: 22.22581\nTotal prob of N-in: 0.99992\nPOSSIBLE N-term signal sequence\ninside 1 20\nTMhelix 21 43\noutside 44 409\nTMhelix 410 432\ninside 433 479

34621	GCF_000163715.1_ASM16371v1	Veillonella sp. 3_1_44	Terrabacteria group; Firmicutes; Negativicutes; Veillonellales; Veillonellaceae; Veillonella	MEQIIDLKEVGRVLIKKRRKIINITVACMVLGGAYAFLAPSTYQSTSMLRIKQAQGLSNS	WP_008713663.1	chain-length determining protein [Veillonella sp. 3_1_44]	Length: 481\nNumber of predicted TMHs: 2\nExp number of AAs in TMHs: 44.24382\nExp number, first 60 AAs: 22.16752\nTotal prob of N-in: 0.99962\nPOSSIBLE N-term signal sequence\ninside 1 20\nTMhelix 21 43\noutside 44 411\nTMhelix 412 434\ninside 435 481

34622	GCF_000286635.1_VspACP1v1.0	Veillonella sp. ACP1Terrabacteria group; Firmicutes; Negativicutes; Veillonellales; Veillonellaceae; Veillonella	MEQIIDIKDIAKILVRKRRTIINVTGICILLGGAYLIVTPSTYQSTAMLRIKQTQGLGNS	WP_009660956.1	chain-length determining protein [Veillonella sp. ACP1]	Length: 479\nNumber of predicted TMHs: 2\nExp number of AAs in TMHs: 44.52906\nExp number, first 60 AAs: 22.22581\nTotal prob of N-in: 0.99992\nPOSSIBLE N-term signal sequence\ninside 1 20\nTMhelix 21 43\noutside 44 409\nTMhelix 410 432\ninside 433 479

34623	GCF_000411535.1_Veil_sp_HPA0037_V1	Veillonella sp. HPA0037	Terrabacteria group; Firmicutes; Negativicutes; Veillonellales; Veillonellaceae; Veillonella	MEQIIDIKDIAKILVRKRRTIINVTGICILLGGAYLIVTPSTYQSTAMLRIKQTQGLGNS	WP_005380966.1	MULTISPECIES: chain-length determining protein [Veillonella]	Length: 479\nNumber of predicted TMHs: 2\nExp number of AAs in TMHs: 44.52515\nExp number, first 60 AAs: 22.22581\nTotal prob of N-in: 0.99992\nPOSSIBLE N-term signal sequence\ninside 1 20\nTMhelix 21 43\noutside 44 409\nTMhelix 410 432\ninside 433 479

34624	GCF_000524355.1_VspICM51av1.0	Veillonella sp. ICM51a	Terrabacteria group; Firmicutes; Negativicutes; Veillonellales; Veillonellaceae; Veillonella	MEQIIDIKDIAKILVRKRRTIINVTGICILLGGAYLIVTPSTYQSTAMLRIKQTQGLGNS	WP_038125040.1	chain-length determining protein [Veillonella sp. ICM51a]	Length: 479\nNumber of predicted TMHs: 2\nExp number of AAs in TMHs: 44.52906\nExp number, first 60 AAs: 22.22581\nTotal prob of N-in: 0.99992\nPOSSIBLE N-term signal sequence\ninside 1 20\nTMhelix 21 43\noutside 44 409\nTMhelix 410 432\ninside 433 479

34625	GCF_000179735.1_ASM17973v1	Veillonella atypica ACS-049-V-Sch6	Terrabacteria group; Firmicutes; Negativicutes; Veillonellales; Veillonellaceae; Veillonella; Veillonella atypica	MEQIIDIKDIAKILVRKRRTIINVTGICILLGGAYLIVTPSTYQSTAMLRIKQTQGLGNS	WP_005376845.1	chain-length determining protein [Veillonella atypica]	Length: 479\nNumber of predicted TMHs: 2\nExp number of AAs in TMHs: 43.66089\nExp number, first 60 AAs: 22.22578\nTotal prob of N-in: 0.99990\nPOSSIBLE N-term signal sequence\ninside 1 20\nTMhelix 21 43\noutside 44 409\nTMhelix 410 432\ninside 433 479

34626	GCF_000179755.1_ASM17975v1	Veillonella atypica ACS-134-V-Col7a	Terrabacteria group; Firmicutes; Negativicutes; Veillonellales; Veillonellaceae; Veillonella; Veillonella atypica	MEQIIDIKDIAKILVRKRRTIINVTGICILLGGAYLIVTPSTYQSTAMLRIKQTQGLGNS	WP_005380966.1	MULTISPECIES: chain-length determining protein [Veillonella]	Length: 479\nNumber of predicted TMHs: 2\nExp number of AAs in

TMHs: 44.52515\nExp number, first 60 AAs: 22.22581\nTotal prob of N-in: 0.99992\nPOSSIBLE N-term signal sequence\ninside 1 20\nTMhelix 21 43\noutside 44 409\nTMhelix 410 432\ninside 433 479

34627 GCF_000318355.2_ASM31835v2 Veillonella atypica KON Terrabacteria group; Firmicutes; Negativicutes; Veillonellales; Veillonellaceae; Veillonella; Veillonella atypica MEQIIDIKDIAKILVRKRRTIINVTGICILLGGAYLIVTPSTYQSTAMLRIKQTQGLGNS WP_005384244.1 chain length determinant protein [Veillonella atypica] Length: 479\nNumber of predicted TMHs: 2\nExp number of AAs in TMHs: 44.46596\nExp number, first 60 AAs: 22.2258\nTotal prob of N-in: 0.99992\nPOSSIBLE N-term signal sequence\ninside 1 20\nTMhelix 21 43\noutside 44 409\nTMhelix 410 432\ninside 433 479

34628 GCF_000024945.1_ASM2494v1 Veillonella parvula DSM 2008 Terrabacteria group; Firmicutes; Negativicutes; Veillonellales; Veillonellaceae; Veillonella; Veillonella parvula MEQIIDLKEVGRVLIKKRRKIINITVACMVLGGAYAFLAPSTYQSTSMLRIKQAQSLSNS WP_012864172.1 chain-length determining protein [Veillonella parvula] Length: 481\nNumber of predicted TMHs: 2\nExp number of AAs in TMHs: 43.80378\nExp number, first 60 AAs: 22.16678\nTotal prob of N-in: 0.99966\nPOSSIBLE N-term signal sequence\ninside 1 20\nTMhelix 21 43\noutside 44 410\nTMhelix 411 433\ninside 434 481

34629 GCF_000163295.1_ASM16329v1 Anaerococcus vaginalis ATCC 51170 Terrabacteria group; Firmicutes; Tissierellia; Tissierellales; Peptoniphilaceae; Anaerococcus; Anaerococcus vaginalis MFSERIFKKINKRRKFMNKNKKIISASLALALGVSLLIPNAKANEDKTVEKALKEPVQTS WP_004838290.1 hypothetical protein [Anaerococcus vaginalis] Length: 342\nNumber of predicted TMHs: 1\nExp number of AAs in TMHs: 29.26514\nExp number, first 60 AAs: 14.29408\nTotal prob of N-in: 0.75631\nPOSSIBLE N-term signal sequence\noutside 1 314\nTMhelix 315 337\ninside 338 342

34630 GCF_900088125.1_PRJEB14116 Peptoniphilus sp. KHD5 Terrabacteria group; Firmicutes; Tissierellia; Tissierellales; Peptoniphilaceae; Peptoniphilus MTSNKEIRDVENFSAIPLSNESIKKRRNNLLKSVIGASVIATGLTMMPQASFAQVAEQPP WP_068455654.1 hypothetical protein [Peptoniphilus sp. KHD5] Length: 1907\nNumber of predicted TMHs: 1\nExp number of AAs in TMHs: 31.88943\nExp number, first 60 AAs: 16.22517\nTotal prob of N-in: 0.76828\nPOSSIBLE N-term signal sequence\noutside 1 1880\nTMhelix 1881 1898\ninside 1899 1907

34631 GCF_900120375.1_PRJEB18029 Ndongobacter massiliensis Terrabacteria group; Firmicutes; unclassified Firmicutes sensu stricto; Ndongobacter MRDGRQFRRRRREILFSLLQSGILPLLRTPRWNEELPETQERLLHERRAELGRRLKKWLE WP_072515150.1 hypothetical protein [Ndongobacter massiliensis] Length: 552\nNumber of predicted TMHs: 2\nExp number of AAs in TMHs: 46.27031\nExp number, first 60 AAs: 0.01037\nTotal prob of N-in: 0.95615\ninside 1 491\nTMhelix 492 514\noutside 515 523\nTMhelix 524 546\ninside 547 552

34632 GCF_001652585.1_ASM165258v1 Thermosulfurimonas dismutans Thermodesulfobacteria; Thermodesulfobacteria; Thermodesulfobacteriales; Thermodesulfobacteriaceae; Thermosulfurimonas MWGYFHILERDTHTKARLGRLVTRRGIIETPVFMPVATAGSIKAVPPEVVTGLGTRVILA WP_068669105.1 tRNA guanosine(34) transglycosylase Tgt [Thermosulfurimonas dismutans] Length: 376\nNumber of predicted TMHs: 1\nExp number of AAs in TMHs: 20.74017\nExp number, first 60 AAs: 0.09835\nTotal prob of N-in: 0.07762\noutside 1 321\nTMhelix 322 344\ninside 345 376

34633 GCF_000753055.1_Cuture_of_bacteria_from_human_gut bacterium OL-1 unclassified Bacteria; unclassified Bacteria (miscellaneous) MAKNTTRRKFIGSTIASVGLLGVTAAANVAANKFSSLLDHYVGGRPATVEKVEGSEDWDA WP_052009812.1 hypothetical protein [bacterium OL-1] Length: 983\nNumber of predicted TMHs: 2\nExp number of AAs in TMHs: 38.8245999999999\nExp number, first 60 AAs: 17.87023\nTotal prob of N-in: 0.85013\nPOSSIBLE N-term signal sequence\ninside 1 8\nTMhelix 9 31\noutside 32 924\nTMhelix 925 947\ninside 948 983

34634 GCF_000752675.1_Jeddahella_massiliensis bacterium OL-1 unclassified Bacteria; unclassified Bacteria (miscellaneous) MAKNTTRRKFIGSTIASVGLLGVTAAANVAANKFSSLLDHYVGGRPATVEKVEGSEDWDA WP_052009812.1 hypothetical protein [bacterium OL-1] Length: 983\nNumber of predicted TMHs: 2\nExp number of AAs in TMHs: 38.8245999999999\nExp number, first 60 AAs: 17.87023\nTotal prob of N-in: 0.85013\nPOSSIBLE N-term signal sequence\ninside 1 8\nTMhelix 9 31\noutside 32 924\nTMhelix 925 947\ninside 948 983

**Dataset S2**

| | |
|---|---|
| WP_048847868.1 | hypothetical protein |
| WP_086613831.1 | hypothetical protein |
| WP_006226706.1 | hypothetical protein |
| WP_088590309.1 | hypothetical protein |
| WP_081322336.1 | cytochrome c-type biogenesis protein CcmH |
| WP_080969392.1 | cytochrome c-type biogenesis protein CcmH |
| WP_080982266.1 | cytochrome c-type biogenesis protein CcmH |
| WP_081247826.1 | cytochrome c-type biogenesis protein CcmH |
| WP_029313089.1 | DUF302 domain-containing protein |
| WP_073494280.1 | terpene cyclase/mutase family protein |
| WP_069464636.1 | LAETG motif-containing sortase-dependent surface protein |
| WP_037906505.1 | terpene cyclase/mutase family protein |
| WP_026419747.1 | Ig-like domain-containing protein |
| WP_016699903.1 | S1 family peptidase |
| WP_075978649.1 | choice-of-anchor P family protein |
| WP_026421352.1 | choice-of-anchor P family protein |
| WP_067818420.1 | hypothetical protein |
| WP_026410447.1 | hypothetical protein |
| WP_073453776.1 | GDSL-type esterase/lipase family protein |
| WP_058237475.1 | HAD family hydrolase |
| WP_021607699.1 | CshA/CshB family fibrillar adhesin-related protein |
| WP_021608244.1 | CshA/CshB family fibrillar adhesin-related protein |
| WP_008733546.1 | hypothetical protein |
| WP_043534360.1 | hypothetical protein |
| WP_067940768.1 | hypothetical protein |
| WP_070725291.1 | HAD family hydrolase |
| WP_050793000.1 | hypothetical protein |
| WP_009213069.1 | VWA domain-containing protein |
| WP_009409131.1 | hypothetical protein |
| WP_009233255.1 | CshA/CshB family fibrillar adhesin-related protein |
| WP_017177339.1 | hypothetical protein |
| WP_067783620.1 | LPXTG cell wall anchor domain-containing protein |
| WP_018334839.1 | hypothetical protein |
| WP_014445984.1 | hypothetical protein |
| WP_067500999.1 | DUF1775 domain-containing protein |
| WP_043530441.1 | hypothetical protein |
| WP_034263843.1 | hypothetical protein |
| WP_016442126.1 | DUF11 domain-containing protein |
| WP_063387033.1 | lamin tail domain-containing protein |
| WP_019144452.1 | HAD family hydrolase |
| WP_087622105.1 | lamin tail domain-containing protein |
| WP_019196192.1 | GumC family protein |

| | |
|---|---|
| WP_044442522.1 | Ig-like domain-containing protein |
| WP_078713100.1 | Ig-like domain-containing protein |
| WP_055942116.1 | hypothetical protein |
| WP_086459789.1 | RICIN domain-containing protein |
| WP_082462074.1 | DUF4349 domain-containing protein |
| WP_067945834.1 | DUF4349 domain-containing protein |
| WP_082600622.1 | DUF11 domain-containing protein |
| WP_022892658.1 | hypothetical protein |
| WP_063508475.1 | SURF1 family protein |
| WP_062817430.1 | SURF1 family protein |
| WP_018698701.1 | TIGR02186 family protein |
| WP_072476476.1 | hypothetical protein |
| WP_086837935.1 | hypothetical protein |
| WP_013229737.1 | hypothetical protein |
| WP_065912755.1 | hypothetical protein |
| WP_043786267.1 | hypothetical protein |
| WP_033261304.1 | hypothetical protein |
| WP_012632768.1 | DUF4091 domain-containing protein |
| WP_011985708.1 | DUF4091 domain-containing protein |
| WP_012525440.1 | DUF4091 domain-containing protein |
| WP_059439524.1 | DUF4091 domain-containing protein |
| WP_035022923.1 | hypothetical protein |
| WP_087075781.1 | hypothetical protein |
| WP_062285750.1 | family 20 glycosylhydrolase |
| WP_081709824.1 | endo-alpha-N-acetylgalactosaminidase family protein |
| WP_055769734.1 | hypothetical protein |
| WP_083706190.1 | copper resistance protein CopC |
| WP_083266569.1 | ice-binding family protein |
| WP_066212419.1 | hypothetical protein |
| WP_066304274.1 | C1 family peptidase |
| WP_048884997.1 | MotA/TolQ/ExbB proton channel family protein |
| WP_047093296.1 | serine hydrolase domain-containing protein |
| WP_055888441.1 | TIGR02186 family protein |
| WP_017366477.1 | hypothetical protein |
| WP_002198638.1 | hypothetical protein |
| WP_057772245.1 | cytochrome c-type biogenesis protein CcmH |
| WP_015592261.1 | hypothetical protein |
| WP_000248334.1 | hypothetical protein |
| WP_085346658.1 | endo-1,4-beta-xylanase |
| WP_055427006.1 | transporter substrate-binding domain-containing protein |
| WP_052826618.1 | hypothetical protein |
| WP_080788544.1 | endo-1,4-beta-xylanase |
| WP_033501893.1 | endo-1,4-beta-xylanase |
| WP_006294973.1 | Cna B-type domain-containing protein |

| | |
|---|---|
| WP_081277489.1 | type II secretion system F family protein |
| WP_033496003.1 | metallophosphoesterase |
| WP_082617556.1 | DUF1134 domain-containing protein |
| WP_054144656.1 | DUF1134 domain-containing protein |
| WP_069879344.1 | DsrE family protein |
| WP_082540354.1 | DsbA family protein |
| WP_038359302.1 | DUF1134 domain-containing protein |
| WP_055729429.1 | DUF1134 domain-containing protein |
| WP_071164834.1 | SpaA isopeptide-forming pilin-related protein |
| WP_070500937.1 | DUF2330 domain-containing protein |
| WP_076808965.1 | hypothetical protein |
| WP_027521434.1 | SURF1 family cytochrome oxidase biogenesis protein |
| WP_061024826.1 | hypothetical protein |
| WP_063250685.1 | choice-of-anchor G family protein |
| WP_084542572.1 | VWA domain-containing protein |
| WP_043182909.1 | hypothetical protein |
| WP_011659356.1 | hypothetical protein |
| WP_012366622.1 | hypothetical protein |
| WP_006752991.1 | hypothetical protein |
| WP_006761406.1 | hypothetical protein |
| WP_059585348.1 | hypothetical protein |
| WP_059236981.1 | hypothetical protein |
| WP_059239508.1 | hypothetical protein |
| WP_077021267.1 | hypothetical protein |
| WP_034201760.1 | hypothetical protein |
| WP_006483583.1 | hypothetical protein |
| WP_048986526.1 | hypothetical protein |
| WP_062884514.1 | hypothetical protein |
| WP_050012404.1 | hypothetical protein |
| WP_040131189.1 | hypothetical protein |
| WP_060268096.1 | hypothetical protein |
| WP_027787499.1 | hypothetical protein |
| WP_027787377.1 | hypothetical protein |
| WP_060086226.1 | hypothetical protein |
| WP_059488505.1 | hypothetical protein |
| WP_059485162.1 | hypothetical protein |
| WP_059700133.1 | hypothetical protein |
| WP_060050342.1 | hypothetical protein |
| WP_059677781.1 | hypothetical protein |
| WP_060069881.1 | hypothetical protein |
| WP_060280409.1 | hypothetical protein |
| WP_059700302.1 | hypothetical protein |
| WP_059621060.1 | hypothetical protein |
| WP_059496097.1 | hypothetical protein |

| | |
|---|---|
| WP_048243592.1 | hypothetical protein |
| WP_059525490.1 | hypothetical protein |
| WP_060221786.1 | hypothetical protein |
| WP_060084896.1 | hypothetical protein |
| WP_059682727.1 | hypothetical protein |
| WP_059526018.1 | hypothetical protein |
| WP_060172271.1 | hypothetical protein |
| WP_059861788.1 | hypothetical protein |
| WP_059904810.1 | hypothetical protein |
| WP_060343570.1 | hypothetical protein |
| WP_060308855.1 | hypothetical protein |
| WP_059588290.1 | hypothetical protein |
| WP_027781201.1 | hypothetical protein |
| WP_041490491.1 | hypothetical protein |
| WP_039366460.1 | hypothetical protein |
| WP_046548430.1 | hypothetical protein |
| WP_046545623.1 | hypothetical protein |
| WP_071332150.1 | hypothetical protein |
| WP_047850963.1 | hypothetical protein |
| WP_046545747.1 | hypothetical protein |
| WP_011352018.1 | hypothetical protein |
| WP_069247875.1 | hypothetical protein |
| WP_069260495.1 | hypothetical protein |
| WP_006410297.1 | PepSY-associated TM helix domain-containing protein |
| WP_059479327.1 | hypothetical protein |
| WP_060254451.1 | hypothetical protein |
| WP_060299571.1 | hypothetical protein |
| WP_040127356.1 | hypothetical protein |
| WP_059514641.1 | hypothetical protein |
| WP_047900707.1 | hypothetical protein |
| WP_034180619.1 | hypothetical protein |
| WP_072441791.1 | hypothetical protein |
| WP_034189316.1 | hypothetical protein |
| WP_059557670.1 | hypothetical protein |
| WP_069617352.1 | hypothetical protein |
| WP_021159606.1 | hypothetical protein |
| WP_059588168.1 | hypothetical protein |
| WP_059789976.1 | DUF3999 domain-containing protein |
| WP_060030825.1 | DUF3999 domain-containing protein |
| WP_060353807.1 | DUF3999 domain-containing protein |
| WP_059823442.1 | DUF3999 domain-containing protein |
| WP_059835798.1 | DUF3999 domain-containing protein |
| WP_059781113.1 | DUF3999 domain-containing protein |
| WP_069748857.1 | hypothetical protein |

| | |
|---|---|
| WP_069745444.1 | hypothetical protein |
| WP_047902623.1 | hypothetical protein |
| WP_059884723.1 | hypothetical protein |
| WP_059801629.1 | hypothetical protein |
| WP_060157732.1 | hypothetical protein |
| WP_059921586.1 | hypothetical protein |
| WP_060291916.1 | hypothetical protein |
| WP_060363596.1 | hypothetical protein |
| WP_059542378.1 | hypothetical protein |
| WP_064558476.1 | anti-virulence regulator CigR family protein |
| WP_026488219.1 | hypothetical protein |
| WP_021688355.1 | SURF1 family protein |
| WP_014433859.1 | hypothetical protein |
| WP_021091342.1 | hydrogenase small subunit |
| WP_048669916.1 | DUF2062 domain-containing protein |
| WP_043684137.1 | GumC family protein |
| WP_020521338.1 | c-type cytochrome |
| WP_034625822.1 | LPXTG cell wall anchor domain-containing protein |
| WP_013769889.1 | hypothetical protein |
| WP_048728390.1 | VWA domain-containing protein |
| WP_087509670.1 | LPXTG cell wall anchor domain-containing protein |
| WP_070320500.1 | LPXTG cell wall anchor domain-containing protein |
| WP_019135425.1 | Ig-like domain-containing protein |
| WP_048341338.1 | hypothetical protein |
| WP_062101716.1 | Ig-like domain-containing protein |
| WP_046529179.1 | LPXTG cell wall anchor domain-containing protein |
| WP_055919086.1 | hypothetical protein |
| WP_069385866.1 | cohesin domain-containing protein |
| WP_087472132.1 | cohesin domain-containing protein |
| WP_053370785.1 | cohesin domain-containing protein |
| WP_047233459.1 | hypothetical protein |
| WP_064315323.1 | cohesin domain-containing protein |
| WP_011910314.1 | hydrogenase small subunit |
| WP_014099504.1 | LEA type 2 family protein |
| WP_052374220.1 | hypothetical protein |
| WP_086521042.1 | YcnI family protein |
| WP_086518073.1 | YcnI family protein |
| WP_086515263.1 | YcnI family protein |
| WP_011931836.1 | YcnI family protein |
| WP_045526708.1 | YcnI family protein |
| WP_015489341.1 | YcnI family protein |
| WP_012300161.1 | hypothetical protein |
| WP_012300049.1 | YcnI family protein |
| WP_063072393.1 | YcnI family protein |

| | |
|---|---|
| WP_043588079.1 | YcnI family protein |
| WP_087364264.1 | Wzz/FepE/Etk N-terminal domain-containing protein |
| WP_005209628.1 | LPXTG cell wall anchor domain-containing protein |
| WP_039249807.1 | hypothetical protein |
| WP_061429810.1 | hypothetical protein |
| WP_066550529.1 | hypothetical protein |
| WP_085018363.1 | DUF445 domain-containing protein |
| WP_085019858.1 | pentapeptide repeat-containing protein |
| WP_072414687.1 | hypothetical protein |
| WP_082855643.1 | hypothetical protein |
| WP_075727961.1 | hypothetical protein |
| WP_051866792.1 | copper resistance protein CopC |
| WP_066523952.1 | hypothetical protein |
| WP_083292036.1 | DUF4185 domain-containing protein |
| WP_070568623.1 | DUF4185 domain-containing protein |
| WP_003856086.1 | exo-alpha-sialidase |
| WP_018023019.1 | copper resistance protein CopC |
| WP_006062599.1 | hypothetical protein |
| WP_025403387.1 | hypothetical protein |
| WP_075664640.1 | porin PorA family protein |
| WP_074508171.1 | exo-alpha-sialidase |
| WP_077312225.1 | exo-alpha-sialidase |
| WP_044027174.1 | exo-alpha-sialidase |
| WP_081803761.1 | S8 family serine peptidase |
| WP_051106395.1 | hypothetical protein |
| WP_051904793.1 | DUF4185 domain-containing protein |
| WP_080971943.1 | Rib/alpha-like domain-containing protein |
| WP_035002726.1 | Rib/alpha-like domain-containing protein |
| WP_051879522.1 | Rib/alpha-like domain-containing protein |
| WP_043014821.1 | Rib/alpha-like domain-containing protein |
| WP_035003110.1 | hypothetical protein |
| WP_051894060.1 | Rib/alpha-like domain-containing protein |
| WP_051894215.1 | Rib/alpha-like domain-containing protein |
| WP_055174723.1 | hypothetical protein |
| WP_047261182.1 | HtaA domain-containing protein |
| WP_055122566.1 | hypothetical protein |
| WP_066490205.1 | hypothetical protein |
| WP_018121428.1 | D-alanyl-D-alanine carboxypeptidase family protein |
| WP_023019056.1 | D-alanyl-D-alanine carboxypeptidase family protein |
| WP_070844356.1 | hypothetical protein |
| WP_070816867.1 | DUF4185 domain-containing protein |
| WP_070819865.1 | DUF4185 domain-containing protein |
| WP_070769551.1 | lamin tail domain-containing protein |
| WP_070614106.1 | DUF4185 domain-containing protein |

| | |
|---|---|
| WP_070508929.1 | HtaA domain-containing protein |
| WP_070537220.1 | hypothetical protein |
| WP_070420471.1 | HtaA domain-containing protein |
| WP_070763091.1 | DUF4185 domain-containing protein |
| WP_070791085.1 | DUF4185 domain-containing protein |
| WP_083316703.1 | DUF4185 domain-containing protein |
| WP_070452682.1 | DUF4185 domain-containing protein |
| WP_070433763.1 | TIGR03773 family transporter-associated surface protein |
| WP_070975208.1 | Rib/alpha-like domain-containing protein |
| WP_071573543.1 | Rib/alpha-like domain-containing protein |
| WP_083584111.1 | Rib/alpha-like domain-containing protein |
| WP_071567683.1 | Rib/alpha-like domain-containing protein |
| WP_070362059.1 | Rib/alpha-like domain-containing protein |
| WP_049151632.1 | HtaA domain-containing protein |
| WP_046646201.1 | HtaA domain-containing protein |
| WP_049146658.1 | HtaA domain-containing protein |
| WP_049064233.1 | HtaA domain-containing protein |
| WP_049160753.1 | HtaA domain-containing protein |
| WP_005531760.1 | HtaA domain-containing protein |
| WP_019193688.1 | HtaA domain-containing protein |
| WP_054314932.1 | DUF3068 domain-containing protein |
| WP_052844124.1 | choice-of-anchor M domain-containing protein |
| WP_015202080.1 | hypothetical protein |
| WP_086782709.1 | S8 family serine peptidase |
| WP_084021289.1 | fibronectin type III domain-containing protein |
| WP_051973425.1 | DUF916 domain-containing protein |
| WP_073250052.1 | DUF4331 domain-containing protein |
| WP_059410969.1 | hypothetical protein |
| WP_006159236.1 | hypothetical protein |
| WP_011517311.1 | hypothetical protein |
| WP_062796007.1 | hypothetical protein |
| WP_011616288.1 | hypothetical protein |
| WP_078198379.1 | hypothetical protein |
| WP_042883642.1 | hypothetical protein |
| WP_018004969.1 | hypothetical protein |
| WP_063239409.1 | tripartite tricarboxylate transporter substrate binding protein |
| WP_010811364.1 | hypothetical protein |
| WP_020206616.1 | hypothetical protein |
| WP_029046437.1 | hypothetical protein |
| WP_012354378.1 | hypothetical protein |
| WP_025585582.1 | hypothetical protein |
| WP_058740988.1 | hypothetical protein |
| WP_058725996.1 | choice-of-anchor G family protein |
| WP_058729336.1 | hypothetical protein |

| | |
|---|---|
| WP_058728719.1 | hypothetical protein |
| WP_066651479.1 | hypothetical protein |
| WP_070418157.1 | bifunctional UDP-sugar hydrolase/5'-nucleotidase |
| WP_056120640.1 | hypothetical protein |
| WP_055952736.1 | YcnI family protein |
| WP_071299477.1 | ExeM/NucH family extracellular endonuclease |
| WP_065961321.1 | bifunctional UDP-sugar hydrolase/5'-nucleotidase |
| WP_071244483.1 | hypothetical protein |
| WP_071249110.1 | DUF11 domain-containing protein |
| WP_071302787.1 | bifunctional UDP-sugar hydrolase/5'-nucleotidase |
| WP_071296020.1 | bifunctional UDP-sugar hydrolase/5'-nucleotidase |
| WP_066652216.1 | hypothetical protein |
| WP_065959450.1 | hypothetical protein |
| WP_027465407.1 | hypothetical protein |
| WP_002529663.1 | hypothetical protein |
| WP_088246978.1 | hypothetical protein |
| WP_078302060.1 | hypothetical protein |
| WP_062387844.1 | hypothetical protein |
| WP_084078503.1 | SURF1 family protein |
| WP_070435498.1 | 5'-nucleotidase C-terminal domain-containing protein |
| WP_028311354.1 | hypothetical protein |
| WP_085318242.1 | hypothetical protein |
| WP_051534117.1 | 4Fe-4S dicluster domain-containing protein |
| WP_028318203.1 | 4Fe-4S dicluster domain-containing protein |
| WP_005988823.1 | hypothetical protein |
| WP_028580723.1 | 4Fe-4S dicluster domain-containing protein |
| WP_035247752.1 | 4Fe-4S dicluster domain-containing protein |
| WP_084604463.1 | cellulose biosynthesis cyclic di-GMP-binding regulatory protein BcsB |
| WP_066728980.1 | hydrogenase 2 operon protein HybA |
| WP_058235133.1 | hypothetical protein |
| WP_020438466.1 | YaaW family protein |
| WP_061228443.1 | hypothetical protein |
| WP_007630985.1 | hypothetical protein |
| WP_084765547.1 | hypothetical protein |
| WP_025544707.1 | Gldg family protein |
| WP_035353858.1 | twin-arginine translocation signal domain-containing protein |
| WP_052106175.1 | hypothetical protein |
| WP_019238747.1 | hypothetical protein |
| WP_010782219.1 | DUF916 domain-containing protein |
| WP_077151302.1 | putative mucin/carbohydrate-binding domain-containing protein |
| WP_001515206.1 | YaaW family protein |
| WP_028116374.1 | hypothetical protein |
| WP_087347104.1 | hypothetical protein |

| | |
|---|---|
| WP_056044687.1 | hypothetical protein |
| WP_056056642.1 | right-handed parallel beta-helix repeat-containing protein |
| WP_056261770.1 | Ig-like domain-containing protein |
| WP_055974413.1 | Ig-like domain-containing protein |
| WP_056049027.1 | Ig-like domain-containing protein |
| WP_052505214.1 | Ig-like domain-containing protein |
| WP_066278535.1 | SGNH/GDSL hydrolase family protein |
| WP_083445970.1 | AhpA/YtjB family protein |
| WP_004128748.1 | hypothetical protein |
| WP_004136222.1 | hypothetical protein |
| WP_026850174.1 | hypothetical protein |
| WP_078784464.1 | hypothetical protein |
| WP_040101249.1 | hydrogenase 2 operon protein HybA |
| WP_040201369.1 | hydrogenase 2 operon protein HybA |
| WP_072919771.1 | hypothetical protein |
| WP_055764009.1 | SpaA isopeptide-forming pilin-related protein |
| WP_042416226.1 | ABC transporter permease subunit |
| WP_020676593.1 | 4Fe-4S dicluster domain-containing protein |
| WP_062015704.1 | sugar dehydrogenase complex small subunit |
| WP_082785218.1 | sugar dehydrogenase complex small subunit |
| WP_052327572.1 | superinfection immunity protein |
| WP_062446384.1 | superinfection immunity protein |
| WP_062455509.1 | superinfection immunity protein |
| WP_062495313.1 | superinfection immunity protein |
| WP_051393048.1 | hypothetical protein |
| WP_007318184.1 | HAD-IB family hydrolase |
| WP_035726910.1 | YcnI family protein |
| WP_026919933.1 | YcnI family protein |
| WP_014361013.1 | YcnI family protein |
| WP_006339621.1 | hypothetical protein |
| WP_007618975.1 | HAD family hydrolase |
| WP_079930690.1 | YcnI family protein |
| WP_087192245.1 | hypothetical protein |
| WP_026529730.1 | SpaA isopeptide-forming pilin-related protein |
| WP_011396790.1 | hypothetical protein |
| WP_004088007.1 | 4Fe-4S dicluster domain-containing protein |
| WP_050465065.1 | Wzz/FepE/Etk N-terminal domain-containing protein |
| WP_053198831.1 | Wzz/FepE/Etk N-terminal domain-containing protein |
| WP_016834044.1 | Wzz/FepE/Etk N-terminal domain-containing protein |
| WP_050476765.1 | Wzz/FepE/Etk N-terminal domain-containing protein |
| WP_034291961.1 | Wzz/FepE/Etk N-terminal domain-containing protein |
| WP_066040688.1 | hypothetical protein |
| WP_030450198.1 | hypothetical protein |
| WP_062432348.1 | hypothetical protein |

| | |
|---|---|
| WP_066361022.1 | hypothetical protein |
| WP_062330275.1 | hypothetical protein |
| WP_062355275.1 | hypothetical protein |
| WP_022899026.1 | discoidin domain-containing protein |
| WP_035564895.1 | hypothetical protein |
| WP_069984382.1 | DUF4397 domain-containing protein |
| WP_013837494.1 | hypothetical protein |
| WP_038497525.1 | hypothetical protein |
| WP_035821865.1 | hypothetical protein |
| WP_073390660.1 | hypothetical protein |
| WP_069111596.1 | DUF4349 domain-containing protein |
| WP_046770059.1 | DUF4349 domain-containing protein |
| WP_084469651.1 | DUF4349 domain-containing protein |
| WP_053202776.1 | DUF4349 domain-containing protein |
| WP_007352828.1 | hypothetical protein |
| WP_033387677.1 | DUF4331 domain-containing protein |
| WP_054296410.1 | DUF4331 domain-containing protein |
| WP_054293102.1 | DUF4331 domain-containing protein |
| WP_042196000.1 | DUF4331 domain-containing protein |
| WP_041292034.1 | DUF6049 family protein |
| WP_088318137.1 | hypothetical protein |
| WP_033347631.1 | hypothetical protein |
| WP_035847823.1 | hypothetical protein |
| WP_035863006.1 | choice-of-anchor P family protein |
| WP_033251316.1 | DUF1775 domain-containing protein |
| WP_033214217.1 | hypothetical protein |
| WP_014135556.1 | SCO2322 family protein |
| WP_033826143.1 | hypothetical protein |
| WP_057229896.1 | DUF4349 domain-containing protein |
| WP_063444545.1 | YaaW family protein |
| WP_079574122.1 | DUF3566 domain-containing protein |
| WP_020392913.1 | right-handed parallel beta-helix repeat-containing protein |
| WP_007911811.1 | YncE family protein |
| WP_081789339.1 | DUF4142 domain-containing protein |
| WP_076779723.1 | Ig-like domain-containing protein |
| WP_055173989.1 | pectinesterase family protein |
| WP_055306235.1 | pectinesterase family protein |
| WP_077959486.1 | hypothetical protein |
| WP_049151215.1 | hypothetical protein |
| WP_009310390.1 | alpha/beta hydrolase |
| WP_006736993.1 | DUF2974 domain-containing protein |
| WP_057718161.1 | hypothetical protein |
| WP_026199800.1 | hypothetical protein |
| WP_003150348.1 | CHAP domain-containing protein |

| | |
|---|---|
| WP_058527013.1 | hypothetical protein |
| WP_052357896.1 | S8 family serine peptidase |
| WP_055907020.1 | hypothetical protein |
| WP_030465437.1 | SdrD B-like domain-containing protein |
| WP_045310765.1 | SdrD B-like domain-containing protein |
| WP_030481134.1 | DUF1360 domain-containing protein |
| WP_004786587.1 | hypothetical protein |
| WP_035983057.1 | hypothetical protein |
| WP_002985403.1 | hypothetical protein |
| WP_015681490.1 | hypothetical protein |
| WP_083352005.1 | copper resistance protein CopC |
| WP_058594659.1 | copper resistance protein CopC |
| WP_010156067.1 | hypothetical protein |
| WP_017794056.1 | hypothetical protein |
| WP_087014056.1 | SpaH/EbpB family LPXTG-anchored major pilin |
| WP_052961504.1 | M1 family aminopeptidase |
| WP_025134557.1 | hypothetical protein |
| WP_057878938.1 | leucine-rich repeat protein |
| WP_057873001.1 | LPXTG cell wall anchor domain-containing protein |
| WP_018353464.1 | ALF repeat-containing protein |
| WP_058835255.1 | SURF1 family protein |
| WP_052595558.1 | type II secretion system F family protein |
| WP_031371299.1 | SURF1 family protein |
| WP_057920519.1 | SURF1 family protein |
| WP_046659779.1 | SURF1 family protein |
| WP_036112499.1 | SURF1 family protein |
| WP_036194832.1 | SURF1 family protein |
| WP_074872556.1 | SURF1 family protein |
| WP_057945932.1 | SURF1 family protein |
| WP_055909325.1 | SURF1 family protein |
| WP_056178414.1 | SURF1 family protein |
| WP_056100861.1 | SURF1 family protein |
| WP_056306103.1 | SURF1 family protein |
| WP_057164304.1 | SURF1 family protein |
| WP_057161099.1 | SURF1 family protein |
| WP_078758809.1 | SURF1 family protein |
| WP_072908093.1 | 4Fe-4S dicluster domain-containing protein |
| WP_014871804.1 | Wzz/FepE/Etk N-terminal domain-containing protein |
| WP_067213518.1 | IS4 family transposase |
| WP_030484094.1 | DUF4349 domain-containing protein |
| WP_036240134.1 | hypothetical protein |
| WP_069988814.1 | glycosyl hydrolase |
| WP_020157341.1 | hypothetical protein |
| WP_060849731.1 | Rieske 2Fe-2S domain-containing protein |

| | |
|---|---|
| WP_066924169.1 | DUF4349 domain-containing protein |
| WP_048435058.1 | hypothetical protein |
| WP_026610324.1 | hypothetical protein |
| WP_052700213.1 | FAD/NAD(P)-binding oxidoreductase |
| WP_026598801.1 | hypothetical protein |
| WP_022948069.1 | SURF1 family protein |
| WP_083960596.1 | TIGR03758 family integrating conjugative element protein |
| WP_020186848.1 | NAD(P)/FAD-dependent oxidoreductase |
| WP_003597644.1 | DUF1134 domain-containing protein |
| WP_015822348.1 | DUF1134 domain-containing protein |
| WP_012453627.1 | DUF1134 domain-containing protein |
| WP_017485181.1 | DUF1134 domain-containing protein |
| WP_060769419.1 | DUF1134 domain-containing protein |
| WP_064031969.1 | hypothetical protein |
| WP_017203113.1 | YcnI family protein |
| WP_064956389.1 | hypothetical protein |
| WP_082334822.1 | copper resistance protein CopC |
| WP_076688294.1 | copper resistance protein CopC |
| WP_045272686.1 | hypothetical protein |
| WP_045249571.1 | hypothetical protein |
| WP_058232366.1 | copper resistance protein CopC |
| WP_079112183.1 | S8 family serine peptidase |
| WP_052460806.1 | hypothetical protein |
| WP_029153468.1 | HtaA domain-containing protein |
| WP_061782355.1 | hypothetical protein |
| WP_045258722.1 | DUF4349 domain-containing protein |
| WP_084613500.1 | DUF5979 domain-containing protein |
| WP_082784215.1 | copper resistance protein CopC |
| WP_052226469.1 | DUF1775 domain-containing protein |
| WP_021198471.1 | choice-of-anchor G family protein |
| WP_052674706.1 | S8 family serine peptidase |
| WP_045277607.1 | M1 family metallopeptidase |
| WP_071919170.1 | phosphatase PAP2 family protein |
| WP_025103163.1 | YcnI family protein |
| WP_060922111.1 | YcnI family protein |
| WP_025104647.1 | S8 family serine peptidase |
| WP_084344888.1 | copper resistance protein CopC |
| WP_083569401.1 | carboxypeptidase-like regulatory domain-containing protein |
| WP_020096836.1 | DUF1775 domain-containing protein |
| WP_082586491.1 | S8 family serine peptidase |
| WP_018188612.1 | DUF1775 domain-containing protein |
| WP_088379655.1 | HtaA domain-containing protein |
| WP_029261182.1 | hypothetical protein |
| WP_036316991.1 | hypothetical protein |

| | |
|---|---|
| WP_082297971.1 | S8 family serine peptidase |
| WP_077052164.1 | hypothetical protein |
| WP_029264221.1 | carboxypeptidase regulatory-like domain-containing protein |
| WP_029266231.1 | copper resistance protein CopC |
| WP_050721264.1 | YcnI family protein |
| WP_067114872.1 | DUF4349 domain-containing protein |
| WP_087056659.1 | YcnI family protein |
| WP_071327739.1 | S8 family serine peptidase |
| WP_085604923.1 | S8 family serine peptidase |
| WP_055954023.1 | DUF4349 domain-containing protein |
| WP_082488321.1 | carboxypeptidase-like regulatory domain-containing protein |
| WP_056375517.1 | S8 family serine peptidase |
| WP_082450279.1 | alpha-L-arabinofuranosidase C-terminal domain-containing protein |
| WP_082516518.1 | copper resistance protein CopC |
| WP_052493031.1 | S8 family serine peptidase |
| WP_054678997.1 | copper resistance protein CopC |
| WP_076674855.1 | copper resistance protein CopC |
| WP_076494745.1 | hypothetical protein |
| WP_055866250.1 | carboxypeptidase-like regulatory domain-containing protein |
| WP_055962946.1 | fibronectin type III domain-containing protein |
| WP_082524328.1 | M1 family aminopeptidase |
| WP_046014689.1 | DUF6049 family protein |
| WP_067162731.1 | DUF4349 domain-containing protein |
| WP_023953107.1 | HtaA domain-containing protein |
| WP_020380574.1 | carboxypeptidase-like regulatory domain-containing protein |
| WP_067196734.1 | choice-of-anchor G family protein |
| WP_047519257.1 | DUF5979 domain-containing protein |
| WP_016463500.1 | copper resistance protein CopC |
| WP_058622210.1 | hypothetical protein |
| WP_013585746.1 | hypothetical protein |
| WP_002778173.1 | hypothetical protein |
| WP_013286804.1 | neocarzinostatin apoprotein domain-containing protein |
| WP_076470653.1 | hypothetical protein |
| WP_074473800.1 | hypothetical protein |
| WP_064445663.1 | neocarzinostatin apoprotein domain-containing protein |
| WP_069088473.1 | neocarzinostatin apoprotein domain-containing protein |
| WP_030499203.1 | neocarzinostatin apoprotein domain-containing protein |
| WP_084597727.1 | DUF4349 domain-containing protein |
| WP_074316730.1 | DUF4349 domain-containing protein |
| WP_013476452.1 | neocarzinostatin apoprotein domain-containing protein |
| WP_052503667.1 | hypothetical protein |
| WP_007457857.1 | DUF6346 domain-containing protein |
| WP_030329257.1 | hypothetical protein |

| | |
|---|---|
| WP_067359874.1 | copper resistance protein CopC |
| WP_073839607.1 | hypothetical protein |
| WP_077936608.1 | hypothetical protein |
| WP_043323399.1 | neocarzinostatin apoprotein domain-containing protein |
| WP_047890386.1 | neocarzinostatin apoprotein domain-containing protein |
| WP_073828243.1 | hypothetical protein |
| WP_071094599.1 | neocarzinostatin apoprotein domain-containing protein |
| WP_083364995.1 | hypothetical protein |
| WP_067184399.1 | hypothetical protein |
| WP_084452563.1 | SURF1 family protein |
| WP_075217005.1 | SURF1 family protein |
| WP_087136771.1 | cohesin domain-containing protein |
| WP_087136226.1 | hypothetical protein |
| WP_065015780.1 | hypothetical protein |
| WP_083087896.1 | hypothetical protein |
| WP_066975116.1 | hypothetical protein |
| WP_067958698.1 | hypothetical protein |
| WP_005063674.1 | hypothetical protein |
| WP_005073213.1 | hypothetical protein |
| WP_074319467.1 | hypothetical protein |
| WP_052618647.1 | hypothetical protein |
| WP_049231943.1 | hypothetical protein |
| WP_021268995.1 | hypothetical protein |
| WP_074356318.1 | hypothetical protein |
| WP_074250269.1 | hypothetical protein |
| WP_079642101.1 | hypothetical protein |
| WP_074258304.1 | hypothetical protein |
| WP_074340136.1 | hypothetical protein |
| WP_074343781.1 | hypothetical protein |
| WP_079583063.1 | hypothetical protein |
| WP_079655361.1 | hypothetical protein |
| WP_074254038.1 | hypothetical protein |
| WP_074330522.1 | hypothetical protein |
| WP_079668796.1 | hypothetical protein |
| WP_079617725.1 | hypothetical protein |
| WP_041043629.1 | hypothetical protein |
| WP_030094292.1 | hypothetical protein |
| WP_070949002.1 | hypothetical protein |
| WP_070941957.1 | hypothetical protein |
| WP_078321647.1 | hypothetical protein |
| WP_046252588.1 | hypothetical protein |
| WP_070936631.1 | hypothetical protein |
| WP_078334177.1 | hypothetical protein |
| WP_078282577.1 | hypothetical protein |

| | |
|---|---|
| WP_043079207.1 | hypothetical protein |
| WP_064632136.1 | hypothetical protein |
| WP_078359203.1 | DUF4349 domain-containing protein |
| WP_078324820.1 | hypothetical protein |
| WP_070913086.1 | hypothetical protein |
| WP_085130694.1 | hypothetical protein |
| WP_047318566.1 | hypothetical protein |
| WP_085087113.1 | hypothetical protein |
| WP_065064343.1 | DUF4349 domain-containing protein |
| WP_064961509.1 | DUF4349 domain-containing protein |
| WP_064899274.1 | DUF4349 domain-containing protein |
| WP_085140301.1 | DUF4349 domain-containing protein |
| WP_076216568.1 | DUF4349 domain-containing protein |
| WP_047037357.1 | DUF4349 domain-containing protein |
| WP_036393700.1 | DUF4349 domain-containing protein |
| WP_083145723.1 | twin-arginine translocation signal domain-containing protein |
| WP_029136656.1 | DUF6049 family protein |
| WP_072515150.1 | AarF/UbiB family protein |
| WP_071894446.1 | hypothetical protein |
| WP_058888770.1 | CHRD domain-containing protein |
| WP_036474851.1 | CHRD domain-containing protein |
| WP_084183403.1 | protease pro-enzyme activation domain-containing protein |
| WP_071675328.1 | hypothetical protein |
| WP_036481955.1 | SURF1 family protein |
| WP_053378551.1 | carboxypeptidase-like regulatory domain-containing protein |
| WP_043695024.1 | hypothetical protein |
| WP_054811683.1 | hypothetical protein |
| WP_042259183.1 | hypothetical protein |
| WP_014989207.1 | hypothetical protein |
| WP_029897422.1 | hypothetical protein |
| WP_063055184.1 | hypothetical protein |
| WP_063006426.1 | hypothetical protein |
| WP_051407089.1 | hypothetical protein |
| WP_056809982.1 | hypothetical protein |
| WP_052314138.1 | hypothetical protein |
| WP_043676971.1 | hypothetical protein |
| WP_028474384.1 | DUF6049 family protein |
| WP_068112638.1 | hypothetical protein |
| WP_028659604.1 | hypothetical protein |
| WP_082748224.1 | S8 family serine peptidase |
| WP_017935992.1 | hypothetical protein |
| WP_056862610.1 | hypothetical protein |
| WP_056904455.1 | hypothetical protein |
| WP_056749018.1 | hypothetical protein |

| | |
|---|---|
| WP_056925535.1 | hypothetical protein |
| WP_061081165.1 | hypothetical protein |
| WP_017557735.1 | hypothetical protein |
| WP_017622416.1 | ABC transporter substrate-binding protein |
| WP_013155373.1 | hypothetical protein |
| WP_071622919.1 | hypothetical protein |
| WP_073378631.1 | prenyltransferase/squalene oxidase repeat-containing protein |
| WP_017541123.1 | ABC transporter substrate-binding protein |
| WP_026122751.1 | hypothetical protein |
| WP_017546715.1 | hypothetical protein |
| WP_020722865.1 | hypothetical protein |
| WP_049577979.1 | hypothetical protein |
| WP_017297372.1 | DUF4349 domain-containing protein |
| WP_068086363.1 | nuclear transport factor 2 family protein |
| WP_061552847.1 | sugar dehydrogenase complex small subunit |
| WP_068623584.1 | HtaA domain-containing protein |
| WP_068708877.1 | choice-of-anchor A family protein |
| WP_056644573.1 | choice-of-anchor A family protein |
| WP_087351888.1 | glycoside hydrolase family 3 protein |
| WP_087227073.1 | hypothetical protein |
| WP_077598534.1 | glycoside hydrolase family 3 C-terminal domain-containing protein |
| WP_022921254.1 | DUF4349 domain-containing protein |
| WP_017719097.1 | hypothetical protein |
| WP_028604672.1 | glycosyltransferase |
| WP_009671820.1 | DUF4349 domain-containing protein |
| WP_050472495.1 | DUF1134 domain-containing protein |
| WP_026239184.1 | hypothetical protein |
| WP_022929851.1 | hypothetical protein |
| WP_026912667.1 | DUF1775 domain-containing protein |
| WP_058720172.1 | hypothetical protein |
| WP_029915213.1 | hydrogenase 2 operon protein HybA |
| WP_057298407.1 | hypothetical protein |
| WP_068455654.1 | G5 domain-containing protein |
| WP_051940138.1 | YcnI family protein |
| WP_019208887.1 | Wzz/FepE/Etk N-terminal domain-containing protein |
| WP_038677511.1 | hypothetical protein |
| WP_079705109.1 | copper resistance protein CopC |
| WP_079001510.1 | copper resistance protein CopC |
| WP_085511799.1 | copper resistance protein CopC |
| WP_064294246.1 | copper resistance protein CopC |
| WP_056779050.1 | copper resistance protein CopC |
| WP_056008599.1 | copper resistance protein CopC |
| WP_071259775.1 | copper resistance protein CopC |

| | |
|---|---|
| WP_086474591.1 | copper resistance protein CopC |
| WP_064075989.1 | hypothetical protein |
| WP_018362232.1 | hypothetical protein |
| WP_084634577.1 | choice-of-anchor M domain-containing protein |
| WP_019502533.1 | DNA/RNA non-specific endonuclease |
| WP_058266926.1 | S1 family peptidase |
| WP_068478771.1 | Ig-like domain-containing protein |
| WP_071041290.1 | metallophosphoesterase family protein |
| WP_077934522.1 | hypothetical protein |
| WP_041119189.1 | hypothetical protein |
| WP_053152798.1 | hypothetical protein |
| WP_060840229.1 | hypothetical protein |
| WP_029978019.1 | hypothetical protein |
| WP_060844157.1 | hypothetical protein |
| WP_037069415.1 | type II secretion system F family protein |
| WP_013674910.1 | DUF3153 domain-containing protein |
| WP_028938488.1 | YcnI family protein |
| WP_057657592.1 | SURF1 family protein |
| WP_019717437.1 | hypothetical protein |
| WP_023470008.1 | hypothetical protein |
| WP_064051878.1 | ABC transporter permease |
| WP_087451541.1 | hypothetical protein |
| WP_085477620.1 | DUF6049 family protein |
| WP_056039821.1 | DUF6049 family protein |
| WP_056867345.1 | DUF4349 domain-containing protein |
| WP_055794040.1 | hypothetical protein |
| WP_077223961.1 | DUF4397 domain-containing protein |
| WP_068255954.1 | bifunctional UDP-sugar hydrolase/5'-nucleotidase |
| WP_007608427.1 | hypothetical protein |
| WP_043292041.1 | hypothetical protein |
| WP_015448274.1 | hypothetical protein |
| WP_027490105.1 | hypothetical protein |
| WP_077554286.1 | DUF2167 domain-containing protein |
| WP_027484742.1 | hypothetical protein |
| WP_056388158.1 | DUF3999 family protein |
| WP_003945065.1 | hypothetical protein |
| WP_045841254.1 | DUF3153 domain-containing protein |
| WP_033234426.1 | hypothetical protein |
| WP_072687607.1 | hypothetical protein |
| WP_006553374.1 | YcnI family protein |
| WP_070378241.1 | hypothetical protein |
| WP_054163953.1 | DUF1134 domain-containing protein |
| WP_011504069.1 | DUF1134 domain-containing protein |
| WP_009814865.1 | hypothetical protein |

| | |
|---|---|
| WP_049350406.1 | LPXTG cell wall anchor domain-containing protein |
| WP_049347585.1 | hypothetical protein |
| WP_083090608.1 | twin-arginine translocation signal domain-containing protein |
| WP_070690408.1 | hypothetical protein |
| WP_070688290.1 | hypothetical protein |
| WP_013628474.1 | hypothetical protein |
| WP_047866510.1 | hypothetical protein |
| WP_005441710.1 | hypothetical protein |
| WP_005449049.1 | hypothetical protein |
| WP_085978096.1 | hypothetical protein |
| WP_015097763.1 | HAD-IB family hydrolase |
| WP_077008428.1 | choice-of-anchor P family protein |
| WP_073889111.1 | choice-of-anchor P family protein |
| WP_033438454.1 | choice-of-anchor P family protein |
| WP_052682257.1 | hypothetical protein |
| WP_013061775.1 | Gldg family protein |
| WP_017548268.1 | hypothetical protein |
| WP_024269372.1 | hypothetical protein |
| WP_023226644.1 | hypothetical protein |
| WP_023206755.1 | anti-virulence regulatory inner membrane protein CigR |
| WP_053234517.1 | hypothetical protein |
| WP_056132372.1 | cohesin domain-containing protein |
| WP_051259476.1 | TIGR03773 family transporter-associated surface protein |
| WP_010585528.1 | hypothetical protein |
| WP_017741353.1 | tetratricopeptide repeat protein |
| WP_083653165.1 | hypothetical protein |
| WP_075814310.1 | hypothetical protein |
| WP_061795424.1 | anti-virulence regulator CigR family protein |
| WP_061807603.1 | anti-virulence regulator CigR family protein |
| WP_063989634.1 | anti-virulence regulator CigR family protein |
| WP_087763533.1 | anti-virulence regulator CigR family protein |
| WP_004941297.1 | anti-virulence regulator CigR family protein |
| WP_060427691.1 | anti-virulence regulator CigR family protein |
| WP_060419303.1 | anti-virulence regulator CigR family protein |
| WP_060432922.1 | anti-virulence regulator CigR family protein |
| WP_060424250.1 | anti-virulence regulator CigR family protein |
| WP_055316240.1 | anti-virulence regulator CigR family protein |
| WP_086579703.1 | anti-virulence regulator CigR family protein |
| WP_079451327.1 | anti-virulence regulator CigR family protein |
| WP_055312290.1 | anti-virulence regulator CigR family protein |
| WP_061872057.1 | anti-virulence regulator CigR family protein |
| WP_082997693.1 | anti-virulence regulator CigR family protein |
| WP_077039006.1 | anti-virulence regulator CigR family protein |
| WP_074307161.1 | hypothetical protein |

| | |
|---|---|
| WP_012237712.1 | hypothetical protein |
| WP_049949682.1 | hypothetical protein |
| WP_037451034.1 | TIGR02186 family protein |
| WP_013846095.1 | TIGR02186 family protein |
| WP_046763191.1 | TIGR02186 family protein |
| WP_066606327.1 | DUF2585 domain-containing protein |
| WP_025550228.1 | TIGR02186 family protein |
| WP_007684696.1 | TIGR02186 family protein |
| WP_013039593.1 | TIGR02186 family protein |
| WP_061939000.1 | TIGR02186 family protein |
| WP_030541714.1 | DUF2585 domain-containing protein |
| WP_056427251.1 | flagellar basal-body MS-ring/collar protein FliF |
| WP_037541996.1 | DUF2585 domain-containing protein |
| WP_077132096.1 | hypothetical protein |
| WP_046574408.1 | hypothetical protein |
| WP_019873836.1 | hypothetical protein |
| WP_005412559.1 | gluconate 2-dehydrogenase subunit 3 family protein |
| WP_005415638.1 | gluconate 2-dehydrogenase subunit 3 family protein |
| WP_061480680.1 | DUF4349 domain-containing protein |
| WP_065186376.1 | DUF4349 domain-containing protein |
| WP_005415830.1 | DUF4349 domain-containing protein |
| WP_088496929.1 | DUF4349 domain-containing protein |
| WP_065177445.1 | DUF4349 domain-containing protein |
| WP_053497069.1 | DUF4349 domain-containing protein |
| WP_034087229.1 | S8 family serine peptidase |
| WP_042405542.1 | chaplin |
| WP_030256123.1 | chaplin |
| WP_037602190.1 | LPXTG cell wall anchor domain-containing protein |
| WP_048791748.1 | hypothetical protein |
| WP_045496817.1 | hypothetical protein |
| WP_005592214.1 | hypothetical protein |
| WP_061423038.1 | hypothetical protein |
| WP_061910244.1 | hypothetical protein |
| WP_015604698.1 | hypothetical protein |
| WP_037580257.1 | LPXTG cell wall anchor domain-containing protein |
| WP_040269875.1 | copper resistance protein CopC |
| WP_070013327.1 | SCO1860 family LAETG-anchored protein |
| WP_070011590.1 | hypothetical protein |
| WP_010355677.1 | hypothetical protein |
| WP_050371013.1 | hypothetical protein |
| WP_010356313.1 | alginate lyase family protein |
| WP_086565734.1 | DUF4349 domain-containing protein |
| WP_015507753.1 | hypothetical protein |
| WP_030307403.1 | DUF4349 domain-containing protein |

| | |
|---|---|
| WP_061404434.1 | hypothetical protein |
| WP_010638641.1 | hypothetical protein |
| WP_085478814.1 | hypothetical protein |
| WP_030309758.1 | hypothetical protein |
| WP_049978979.1 | DUF4349 domain-containing protein |
| WP_049977564.1 | HtaA domain-containing protein |
| WP_030304860.1 | hypothetical protein |
| WP_033237593.1 | hypothetical protein |
| WP_030698069.1 | hypothetical protein |
| WP_037843057.1 | hypothetical protein |
| WP_071338394.1 | DUF4349 domain-containing protein |
| WP_061405035.1 | HtaA domain-containing protein |
| WP_047471185.1 | hypothetical protein |
| WP_087928343.1 | HtaA domain-containing protein |
| WP_030803089.1 | hypothetical protein |
| WP_040251858.1 | DUF3068 domain-containing protein |
| WP_016468754.1 | prenyltransferase/squalene oxidase repeat-containing protein |
| WP_055498263.1 | prenyltransferase/squalene oxidase repeat-containing protein |
| WP_032776788.1 | hypothetical protein |
| WP_037611286.1 | prenyltransferase/squalene oxidase repeat-containing protein |
| WP_030631798.1 | DUF4349 domain-containing protein |
| WP_030603830.1 | hypothetical protein |
| WP_037680772.1 | DUF1996 domain-containing protein |
| WP_053141924.1 | hypothetical protein |
| WP_060176777.1 | hypothetical protein |
| WP_086719786.1 | hypothetical protein |
| WP_059191885.1 | hypothetical protein |
| WP_030645059.1 | DUF4349 domain-containing protein |
| WP_055508106.1 | terpene cyclase/mutase family protein |
| WP_040899100.1 | hypothetical protein |
| WP_030565856.1 | hypothetical protein |
| WP_037623895.1 | HtaA domain-containing protein |
| WP_078234756.1 | DUF1996 domain-containing protein |
| WP_052851228.1 | DUF4349 domain-containing protein |
| WP_014152884.1 | hypothetical protein |
| WP_030079189.1 | LAETG motif-containing sortase-dependent surface protein |
| WP_014180138.1 | SCO2322 family protein |
| WP_030113438.1 | hypothetical protein |
| WP_086763368.1 | hypothetical protein |
| WP_030189408.1 | hypothetical protein |
| WP_059205742.1 | SCO2322 family protein |
| WP_062042357.1 | SCO2322 family protein |
| WP_030180617.1 | hypothetical protein |
| WP_029181655.1 | hypothetical protein |

| | |
|---|---|
| WP_053925821.1 | S8 family serine peptidase |
| WP_059263647.1 | hypothetical protein |
| WP_062143793.1 | DUF2330 domain-containing protein |
| WP_044384405.1 | LAETG motif-containing sortase-dependent surface protein |
| WP_015659944.1 | hypothetical protein |
| WP_053209708.1 | hypothetical protein |
| WP_031148216.1 | hypothetical protein |
| WP_006124616.1 | hypothetical protein |
| WP_010070991.1 | hypothetical protein |
| WP_030637428.1 | hypothetical protein |
| WP_031129975.1 | hypothetical protein |
| WP_043464200.1 | DUF1996 domain-containing protein |
| WP_030601925.1 | HtaA domain-containing protein |
| WP_079040852.1 | VWA domain-containing protein |
| WP_083107039.1 | hypothetical protein |
| WP_010059884.1 | hypothetical protein |
| WP_030689701.1 | hypothetical protein |
| WP_030592787.1 | DUF4349 domain-containing protein |
| WP_078489131.1 | DUF6049 family protein |
| WP_079196203.1 | DUF6049 family protein |
| WP_069169392.1 | DUF4349 domain-containing protein |
| WP_030754832.1 | hypothetical protein |
| WP_030847770.1 | hypothetical protein |
| WP_030710305.1 | hypothetical protein |
| WP_030733765.1 | hypothetical protein |
| WP_055631020.1 | HtaA domain-containing protein |
| WP_051814695.1 | copper resistance protein CopC |
| WP_067285055.1 | hypothetical protein |
| WP_055548827.1 | hypothetical protein |
| WP_030295329.1 | hypothetical protein |
| WP_079429764.1 | hypothetical protein |
| WP_073951852.1 | HtaA domain-containing protein |
| WP_030238604.1 | HtaA domain-containing protein |
| WP_030225214.1 | DUF6049 family protein |
| WP_030660217.1 | hypothetical protein |
| WP_029385521.1 | LAETG motif-containing sortase-dependent surface protein |
| WP_048572334.1 | LAETG motif-containing sortase-dependent surface protein |
| WP_016325237.1 | S1 family peptidase |
| WP_079021358.1 | DUF1996 domain-containing protein |
| WP_069885441.1 | hypothetical protein |
| WP_042156490.1 | hypothetical protein |
| WP_051732961.1 | VWA domain-containing protein |
| WP_077192932.1 | VWA domain-containing protein |
| WP_033266588.1 | hypothetical protein |

| | |
|---|---|
| WP_046584143.1 | DUF4331 domain-containing protein |
| WP_030812892.1 | hypothetical protein |
| WP_004943253.1 | SCO1860 family LAETG-anchored protein |
| WP_031100002.1 | hypothetical protein |
| WP_070014799.1 | SCO1860 family LAETG-anchored protein |
| WP_055541462.1 | hypothetical protein |
| WP_052863743.1 | hypothetical protein |
| WP_078076280.1 | YcnI family protein |
| WP_064073155.1 | SCO1860 family LAETG-anchored protein |
| WP_037635128.1 | hypothetical protein |
| WP_038519829.1 | hypothetical protein |
| WP_079023587.1 | hypothetical protein |
| WP_078892164.1 | DUF1996 domain-containing protein |
| WP_078535903.1 | DUF1996 domain-containing protein |
| WP_079155135.1 | DUF1996 domain-containing protein |
| WP_067363291.1 | HtaA domain-containing protein |
| WP_055522229.1 | hypothetical protein |
| WP_032915072.1 | hypothetical protein |
| WP_055612260.1 | HtaA domain-containing protein |
| WP_073499707.1 | DUF1775 domain-containing protein |
| WP_085927641.1 | hypothetical protein |
| WP_085926357.1 | hypothetical protein |
| WP_055573436.1 | SCO1860 family LAETG-anchored protein |
| WP_055694013.1 | SCO1860 family LAETG-anchored protein |
| WP_055606274.1 | SCO1860 family LAETG-anchored protein |
| WP_055688174.1 | SCO1860 family LAETG-anchored protein |
| WP_031060443.1 | SCO2322 family protein |
| WP_074998544.1 | prenyltransferase/squalene oxidase repeat-containing protein |
| WP_069991284.1 | prenyltransferase/squalene oxidase repeat-containing protein |
| WP_053800621.1 | hypothetical protein |
| WP_030603358.1 | hypothetical protein |
| WP_030672730.1 | hypothetical protein |
| WP_033026720.1 | hypothetical protein |
| WP_030634038.1 | hypothetical protein |
| WP_030651718.1 | hypothetical protein |
| WP_051430100.1 | hypothetical protein |
| WP_067416561.1 | DUF4349 domain-containing protein |
| WP_037703295.1 | TULIP family P47-like protein |
| WP_030351526.1 | hypothetical protein |
| WP_058848772.1 | hypothetical protein |
| WP_052489045.1 | hypothetical protein |
| WP_020129099.1 | DUF1775 domain-containing protein |
| WP_079406108.1 | hypothetical protein |
| WP_033947955.1 | hypothetical protein |

| | |
|---|---|
| WP_078875879.1 | hypothetical protein |
| WP_019433298.1 | S8 family serine peptidase |
| WP_019357822.1 | prenyltransferase/squalene oxidase repeat-containing protein |
| WP_023419047.1 | hypothetical protein |
| WP_018552017.1 | TIGR03773 family transporter-associated surface protein |
| WP_019983617.1 | alkaline phosphatase family protein |
| WP_033240369.1 | hypothetical protein |
| WP_073918172.1 | hypothetical protein |
| WP_073781669.1 | hypothetical protein |
| WP_073738813.1 | hypothetical protein |
| WP_073758166.1 | hypothetical protein |
| WP_073752756.1 | SCO1860 family LAETG-anchored protein |
| WP_087776164.1 | HtaA domain-containing protein |
| WP_043265717.1 | hypothetical protein |
| WP_058925582.1 | SCO2322 family protein |
| WP_061403993.1 | DUF4349 domain-containing protein |
| WP_067159911.1 | hypothetical protein |
| WP_067392033.1 | HtaA domain-containing protein |
| WP_075987369.1 | HtaA domain-containing protein |
| WP_067052849.1 | DUF4331 domain-containing protein |
| WP_018089851.1 | hypothetical protein |
| WP_075986595.1 | hypothetical protein |
| WP_065002969.1 | SCO1860 family LAETG-anchored protein |
| WP_053629362.1 | hypothetical protein |
| WP_053677522.1 | hypothetical protein |
| WP_060178439.1 | LPXTG cell wall anchor domain-containing protein |
| WP_076089841.1 | hypothetical protein |
| WP_053611851.1 | hypothetical protein |
| WP_032773289.1 | hypothetical protein |
| WP_047138782.1 | hypothetical protein |
| WP_069759320.1 | SCO2322 family protein |
| WP_018546292.1 | HtaA domain-containing protein |
| WP_018468438.1 | hypothetical protein |
| WP_076972671.1 | DUF1996 domain-containing protein |
| WP_030991215.1 | DUF4349 domain-containing protein |
| WP_073773078.1 | hypothetical protein |
| WP_063348001.1 | hypothetical protein |
| WP_030834094.1 | hypothetical protein |
| WP_084771524.1 | hypothetical protein |
| WP_079176493.1 | DUF1996 domain-containing protein |
| WP_047960688.1 | hypothetical protein |
| WP_062204714.1 | choice-of-anchor M domain-containing protein |
| WP_078890523.1 | porin PorA family protein |
| WP_066926162.1 | hypothetical protein |

| | |
|---|---|
| WP_071338956.1 | HtaA domain-containing protein |
| WP_030679056.1 | HtaA domain-containing protein |
| WP_030879423.1 | hypothetical protein |
| WP_053849849.1 | SCO2322 family protein |
| WP_030722950.1 | SCO2322 family protein |
| WP_030768766.1 | hypothetical protein |
| WP_030857734.1 | DUF6049 family protein |
| WP_046722871.1 | DUF2330 domain-containing protein |
| WP_030853527.1 | hypothetical protein |
| WP_066980116.1 | hypothetical protein |
| WP_078855939.1 | DUF1996 domain-containing protein |
| WP_059130410.1 | LAETG motif-containing sortase-dependent surface protein |
| WP_031517753.1 | terpene cyclase/mutase family protein |
| WP_043232110.1 | hypothetical protein |
| WP_031028222.1 | prenyltransferase/squalene oxidase repeat-containing protein |
| WP_031052890.1 | hypothetical protein |
| WP_053696604.1 | hypothetical protein |
| WP_030406608.1 | prenyltransferase/squalene oxidase repeat-containing protein |
| WP_053642605.1 | LAETG motif-containing sortase-dependent surface protein |
| WP_030771374.1 | hypothetical protein |
| WP_078865984.1 | VWA domain-containing protein |
| WP_037839904.1 | hypothetical protein |
| WP_078852820.1 | SCO1860 family LAETG-anchored protein |
| WP_031095397.1 | DUF4349 domain-containing protein |
| WP_030721008.1 | hypothetical protein |
| WP_030386983.1 | hypothetical protein |
| WP_030153186.1 | DUF6049 family protein |
| WP_030231344.1 | LPXTG cell wall anchor domain-containing protein |
| WP_030964283.1 | hypothetical protein |
| WP_030771538.1 | hypothetical protein |
| WP_030939315.1 | hypothetical protein |
| WP_051795396.1 | hypothetical protein |
| WP_030791129.1 | SCO1860 family LAETG-anchored protein |
| WP_031023561.1 | DUF4349 domain-containing protein |
| WP_037927858.1 | hypothetical protein |
| WP_088577171.1 | hypothetical protein |
| WP_018570515.1 | HtaA domain-containing protein |
| WP_073775700.1 | hypothetical protein |
| WP_057601763.1 | hypothetical protein |
| WP_057611173.1 | DUF4349 domain-containing protein |
| WP_084993274.1 | hypothetical protein |
| WP_064534583.1 | hypothetical protein |
| WP_023608351.1 | hypothetical protein |
| WP_006345762.1 | DUF1775 domain-containing protein |

| | |
|---|---|
| WP_018509850.1 | LPXTG cell wall anchor domain-containing protein |
| WP_081522449.1 | hypothetical protein |
| WP_028416940.1 | hypothetical protein |
| WP_055490993.1 | DUF1996 domain-containing protein |
| WP_078962511.1 | DUF1996 domain-containing protein |
| WP_073938500.1 | hypothetical protein |
| WP_073718266.1 | hypothetical protein |
| WP_037893396.1 | HtaA domain-containing protein |
| WP_075265106.1 | hypothetical protein |
| WP_053676489.1 | hypothetical protein |
| WP_053169188.1 | hypothetical protein |
| WP_053689381.1 | hypothetical protein |
| WP_046259605.1 | LPXTG cell wall anchor domain-containing protein |
| WP_053784674.1 | hypothetical protein |
| WP_078095079.1 | hypothetical protein |
| WP_065963662.1 | hypothetical protein |
| WP_059006136.1 | hypothetical protein |
| WP_069921941.1 | hypothetical protein |
| WP_055717924.1 | hypothetical protein |
| WP_006349525.1 | terpene cyclase/mutase family protein |
| WP_043247767.1 | hypothetical protein |
| WP_078606519.1 | DUF1996 domain-containing protein |
| WP_078600540.1 | YcnI family protein |
| WP_033218789.1 | hypothetical protein |
| WP_049978038.1 | hypothetical protein |
| WP_031150581.1 | hypothetical protein |
| WP_020372371.1 | hypothetical protein |
| WP_046776149.1 | hypothetical protein |
| WP_033325580.1 | HtaA domain-containing protein |
| WP_003298078.1 | hypothetical protein |
| WP_007047833.1 | hypothetical protein |
| WP_087488285.1 | sugar dehydrogenase complex small subunit |
| WP_038020607.1 | sugar dehydrogenase complex small subunit |
| WP_020143569.1 | tryptophan-rich sensory protein |
| WP_077686904.1 | SpaA isopeptide-forming pilin-related protein |
| WP_083665280.1 | hypothetical protein |
| WP_053334878.1 | 4Fe-4S dicluster domain-containing protein |
| WP_011293161.1 | hypothetical protein |
| WP_075279362.1 | glycoside hydrolase family 3 protein |
| WP_015258557.1 | ethylbenzene dehydrogenase-related protein |
| WP_007194980.1 | hypothetical protein |
| WP_041066896.1 | hypothetical protein |
| WP_059052747.1 | glycoside hydrolase family 3 C-terminal domain-containing protein |

| | |
|---|---|
| WP_080999149.1 | leucine-rich repeat domain-containing protein |
| WP_080712419.1 | leucine-rich repeat domain-containing protein |
| WP_080689132.1 | leucine-rich repeat domain-containing protein |
| WP_080754043.1 | leucine-rich repeat domain-containing protein |
| WP_068566570.1 | DUF4349 domain-containing protein |
| WP_068532605.1 | DUF4349 domain-containing protein |
| WP_081857365.1 | cellulose biosynthesis cyclic di-GMP-binding regulatory protein BcsB |
| WP_052706166.1 | hypothetical protein |
| WP_055789914.1 | DUF3153 domain-containing protein |
| WP_038217501.1 | SURF1 family protein |
| WP_051684734.1 | type II secretion system F family protein |
| WP_030124501.1 | hypothetical protein |
| WP_009319933.1 | choice-of-anchor I family protein |
| WP_009243371.1 | choice-of-anchor I family protein |
| WP_019617431.1 | cytochrome c |
